# Symmetry as a Fundamental Principle in Defining Gene Expression and Phenotypic Traits

**DOI:** 10.1101/2025.01.27.634930

**Authors:** Cheng Zhang, Cristina Correia, Taylor Weiskittel, Shyang-Hong Tan, Zhuofei Zhang, Kok-Siong Yeo, Shizhen Zhu, Choong-Yong Ung, Hu Li

## Abstract

Symmetry refers to properties that remain invariant upon mathematical transformations. The principles of symmetry have guided numerous important discoveries in physics and chemistry but not in biology and medicine. Here, we aim to explore the presence of symmetry relationships at the gene expression level as a mean to distinguish between healthy and disease states. We deployed Learning-Based Invariant Feature Engineering - LIFE, a hybrid machine learning approach implemented with two symmetric invariant feature functions (IFFs) to identify Invariant Feature Genes (IFGs), which are gene pairs whose IFF single-value outputs remain invariant across individual samples in a given biological phenotype. Our multiclass classification results across the transcriptomes of 25 normal organs, 25 cancer types, and blood samples obtained from 4 different types of neurodegenerative diseases revealed the presence of unique phenotype-specific IFGs. We constructed networks using these IFGs (IF-Nets) and intriguingly, we demonstrated that the hubs could serve as information encoders, capable of reconstructing sample-wise expression values in relation to their counterpart genes. More importantly, we found that hubs of cancer IF-Nets were enriched with both approved and clinical trial drugs, highlighting “symmetry breaking” as a novel approach for treating diseases.

## INTRODUCTION

Symmetry is the main fingerprint of natural laws. Our daily sense of aesthetics is shaped by the symmetric appearance of objects, such as the bilateral symmetry of a human face or the radial symmetry of a sunflower. In mathematics, symmetry is a kind of invariance where the property of a mathematical object or a mathematical function remains unchanged when subjected to a given transformation (reflection, rotation, and translation). The transformation that preserves an invariance property of an object or a system is called symmetry transformation. For decades, physicists used symmetry principles as roadmaps to make novel discoveries^1,2^. For instance, physicists use the principle of symmetry to build Standard Model to explain what made our universe^3,4^. Chemists similarly use the principle of symmetry to design reaction paths to synthesize novel molecules^5,6^. While mathematicians admire the beauty of bilateral and radial symmetries in various biological constructs including icosahedral makeups of viral particles, the symmetry nature of gene activities that underpin the phenotypic property of a biological state has not been demonstrated. The concept of symmetry is rarely invoked as a guiding principle in biology and medicine and is often underutilized as a tool for discovery. Often the concept of symmetry in biology is confined at the esthetic aspects of symmetrical organization of biological structures, such as the icosahedral arrangement of proteins in viral particles. Beyond the structural considerations, the presence of symmetry in biological processes, such as gene expression remains largely enigmatic.

Compared to the activity of individual genes, biological phenotypes - such as a tissue type and or disease states, are relatively invariant (i.e., stable) despite the underlying genetic heterogeneity. Thus, we set out to explore whether biological phenotypes can be considered symmetric with respect to certain mathematical transformations, with a focus on gene expression patterns. We first hypothesize that there are relationships between gene expressions that remain invariant across individuals displaying the same biological phenotype. Our Gene Expression Symmetry Hypothesis (GESH) posits that the invariant nature of phenotypic traits in cells is defined by a set of genes exhibiting specific symmetric expression relationships. In this study, we sought to explore the validity of GESH and investigate the symmetry nature of gene activities in living systems.

We first devised a novel machine learning approach, combined with mathematical functions that describe different modes of symmetric relationships between expressed genes, and tested GESH on transcriptomes derived from various normal tissues, disease types and blood. We then employed systems-based analyses to characterize the biological properties of these symmetrically expressed genes pairs, exploring their roles in shaping phenotypic traits and investigating the pharmacologic potential of symmetrically expressed genes in diseases.

## RESULTS

### The Design of Invariant Feature Functions and Learning-Based Invariant Feature Engineering

Our first step was to devise mathematic functions that describe specific modes of symmetric relationships among expressed genes. Since there is no single gene whose expression level is invariant under any circumstance, the simplest scenario is to look for gene pairs whose expression levels after subjected to a mathematical function display symmetric expression relations across individuals.

Gene pairs that satisfied this criterion are defined as Invariant Feature Genes (IFGs). The corresponding symmetry transformations are called Invariant Feature Functions (IFFs). Hence, IFGs are gene pairs whose expression levels, after subjected to a given IFF, yield single-value outputs that are invariant (i.e. unchanged) across individuals assigned to the same biological phenotype, regardless of the heterogeneity of single-gene expression activities (**Figure 1A**).

**Figure 1.**
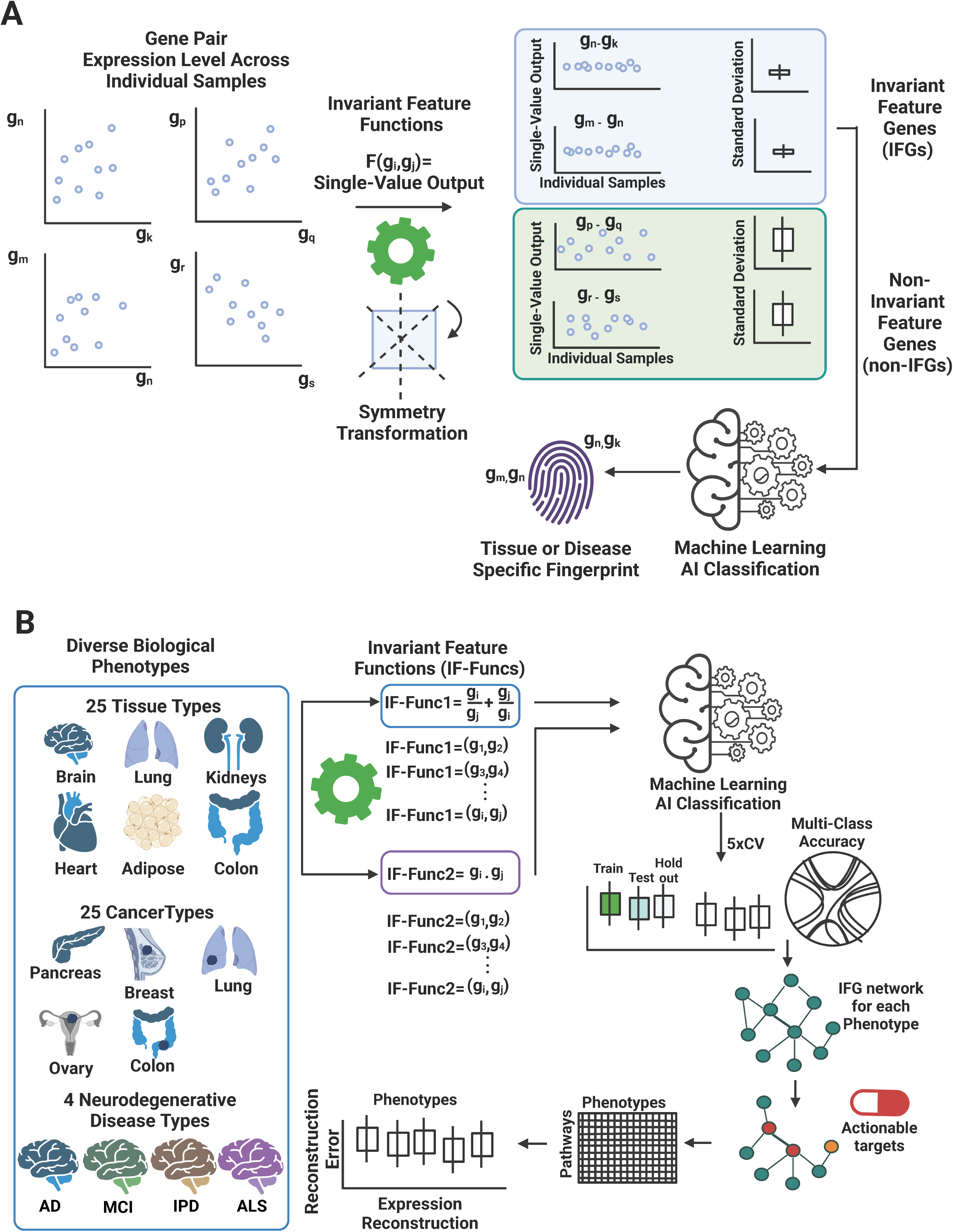
Symmetric gene expression relationships define a phenotypic trait. (**A**) Key concepts for detecting symmetric gene expression relationships across individual samples under given a biological phenotype. Here, the expression values of gene pair serve as input for an invariant feature function (IFF1 and IFF2), a mathematical function that describes a symmetrical relationship between a gene pair. If a pair of genes exhibit symmetric gene expression relationship, its single-value outputs across samples obtained from different individuals belonging to the same biological phenotype shows small variability (i.e., a small standard deviation), in contrast to a gene pair that do not display symmetric expression relations. Gene pairs that exhibit symmetric expression relationships in a biological phenotype are called Invariant Feature Genes (IFGs) and they can serve as fingerprints to define a specific biological phenotype. (**B**) Overall design of Machine Learning-Based Invariant Feature Engineering (LIFE). Two different invariant feature functions, IFF1 and IFF2, are incorporated into LIFE. Transcriptomics data from 25 normal organs and 25 cancer types, and blood transcriptomics across four neurological disorders were used in this proof-of-principle studies to explore IFGs, and their power to classify phenotypes. Multi-class classification with cross-validation were used to assess IFGs as phenotype-specific fingerprints that can robustly assign individual samples to their respective biological phenotypes. In addition, disease-specific networks constructed from IFGs (IF-Nets) are a valuable platform for drug target discovery. AD: Alzheimer’s disease; MCI: mild cognitive impair; IPD: idiopathic Parkinson’s disease; and ALS: amyotrophic lateral sclerosis.

Building upon this idea, we devised two IFFs that describe two different modes of symmetric expression relationships among gene pairs as followed: 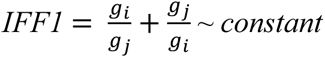 and *IFF2* = 𝑔*_i_* · 𝑔*_j_* ∼ *constant* where 𝑔*_i_* and 𝑔*_j_* are expression levels of gene-*i* and gene-*j*, respectively in a sample for a given biological phenotype. Both *IFF1* and *IFF2* describe symmetric expression relations between genes 𝑔*_i_* and 𝑔*_j_* since the subscripts *i* and *j* can be interchanged without altering the computation results. Pairs of gene-*i* and gene-*j* that exhibit invariant IFF single-value outputs across samples with respect to a biological phenotype are considered IFG candidates. Ideally, IFGs should produce invariant (i.e., constant) IFF single-value outputs across individual samples within the same biological phenotypes. However, in practice, the observed gene expression levels are subjected to both biological and experimental noise, meaning that the computed IFF single-value outputs are not perfectly invariant but instead fluctuate within a small spread of values. As such we defined IFGs as exhibiting low variability, if their standard deviations (σ) for computed IFF single-value outputs were small, compared to gene pairs that did not show such stability in their expression values (**Figure 1A**).

Next, we implemented *IFF1* and *IFF2* into a machine learning-based strategy called Learning-Based Invariant Feature Engineering (LIFE) to identify IFGs with respect to each biological phenotype (**Figure 1B**). Because IFGs are specific to a phenotype and their IFF single-value outputs are invariant across individuals for a given phenotype, they can serve as “fingerprints” to characterize molecular properties underpinning a phenotypic state. Next, we employed multiclass classification learning strategies to assess the performance of IFGs in classifying samples into their respective phenotypes. We used transcriptomics data derived from different biological conditions as proof-of-concept studies: (i) 25 types of normal organs extracted from the Genotype-Tissue Expression (GTEx) Portal, (ii) 25 cancer types from The Cancer Genome Atlas Program (TCGA), (iii) blood-based transcriptomics in neurological disorders (**Supplementary Data 1)**.

### IFGs are Phenotype-Specific Fingerprints

We computed single-value outputs with respect to *IFF1* and *IFF2* for all possible gene pair combinations for genes whose expression levels exhibit normal distribution in each phenotype (0.48% to 32% of genes after filtering, out of 20,272 genes), as determined by the Shapiro-Wilk test with an FDR cutoff of 0.05. For each biological phenotype, we selected top 1000 gene pairs with smallest standard deviations (SDs) with respect to *IFF1* (IF1_1000_) (**Supplementary Data 2**) and *IFF2* (IF2_1000_) (**Supplementary Data 3**) as IFG candidates.

We first inspected the gene expression patterns for the top 20 IFG candidates across normal organs and cancers. We found IFG candidates derived from *IFF1* (**Supplementary Data 4** for organs and **Supplementary Data 6** for cancers) exhibited higher linearity than IFG candidates derived from *IFF2* (**Supplementary Data 5** for organs and **Supplementary Data 7** for cancers). Yet, many IFG candidates derived from *IFF1* also displayed poor linearity in their expression patterns, such as JUNB-UBA52 (**Supplementary Data 4**) in blood and MYL12B-ARF1 from breast cancer (**Supplementary Data 6**). This suggests that linearity of gene pair expression is not the determining factor gene expression symmetry. Instead, it is a function to either *IFF1* or *IFF2*, that produces invariant single-value outputs, regardless of the expression heterogeneity among individuals.

We next tested IF1_1000_ and IF2_1000_ for their capabilities in acting as phenotype-specific fingerprints through multi-class classification, employing a 5-fold cross-validation and hold-out strategies on 25 types of normal organs and 25 types of cancers, respectively. Averaged results of 5-fold cross-validations presented in the chord diagrams show good multi-class classification performance (**Figure 2**) with IF1_1000_ (**Figure 2A** and **Figure 2C**) showing a better classification performance than IF2_1000_ (**Figure 2B** and **Figure 2D**) for both normal organ and cancer types. Plots for prediction accuracies are given in **Supplementary Figures 1-4** and confusion matrices are provided in **Supplementary Figures 5-8**. General good multiclass classification performance (>70% overall classification accuracies) for both IF1_1000_ and IF2_1000_ pairs indicates that IFG candidates are phenotype-specific fingerprints. This strongly suggests that IFGs can be used to define phenotypic states across a variety of tissues and cancer types.

**Figure 2.**
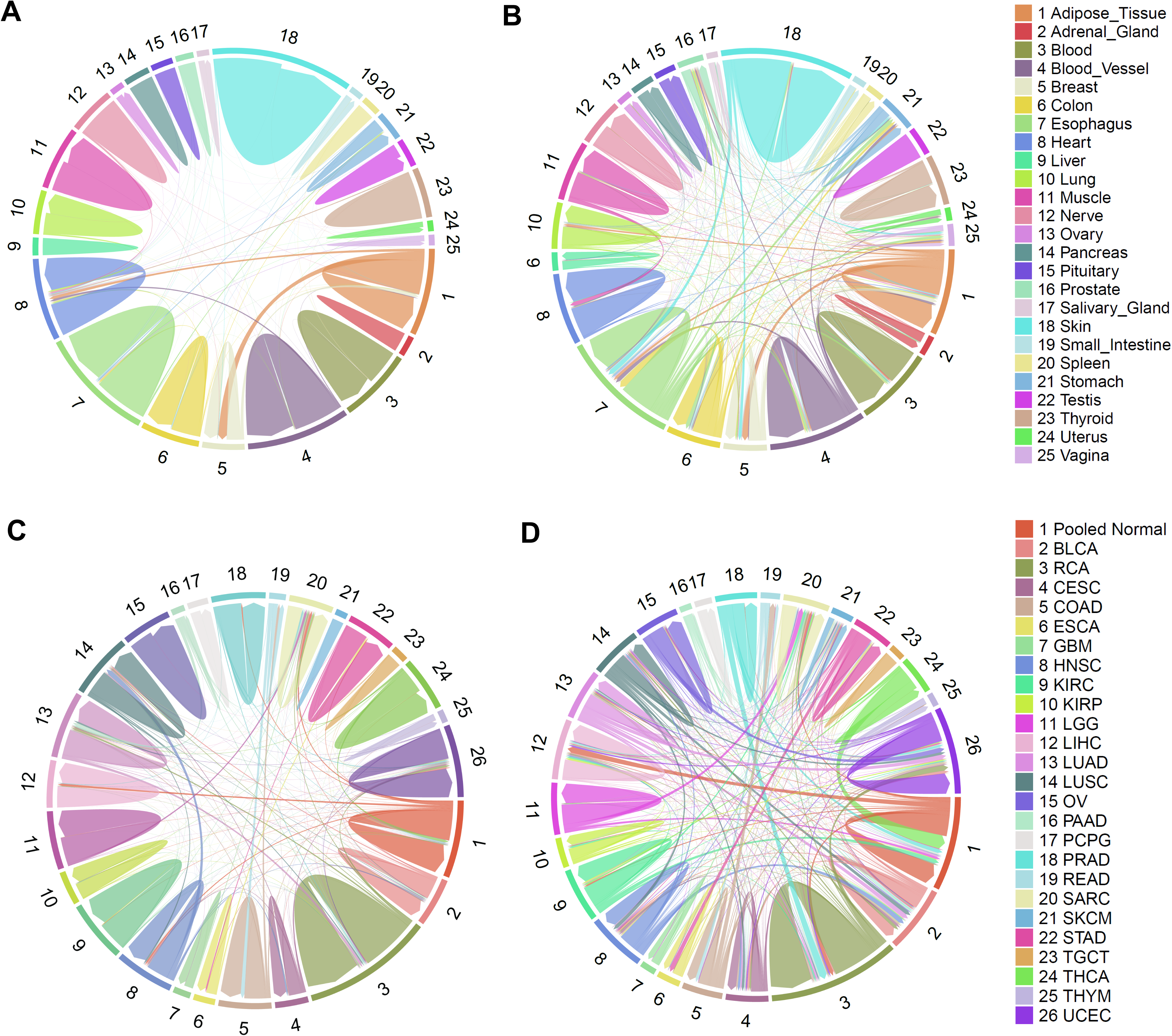
Multi-class classification performance for top 1000 Invariant Feature Genes (IFGs) derived with IFF1 (IF1_1000_) and IFF2 (IF2_1000_) across diverse normal organs and cancer types. Chord diagrams for the averaged results of multiclass classification performance derived from the 5-fold cross-validation with top 1000 most stable genes pairs (with smallest standard deviation) computed on IFF single-value outputs across samples for a given phenotype. These chord diagrams show the number of samples that are correctly classified (self-looping connections) and misclassified (cross-state connections) in **A**. 25 normal organs on IF1_1000_; **B**. 25 normal organs on IF2_1000_; **C**. 25 cancer types on IF1_1000_; and **D**. 25 cancer types on IF2_1000_.

### Blood-Borne IFGs Inform on Disease Types

Next, we explored whether patient-derived blood transcriptomes could be used to identify blood-borne IFGs that define disease types. This would be particularly valuable for longitudinal monitoring of patients using a minimally invasive approach. We are especially interested in identifying blood-borne IFGs that can reliably inform pathological incidents at distant sites such as the brain. The blood can capture a broad range of molecular information and pathological activity at distinct sites, to a certain extent, that can be extracted from blood transcriptomics to provide a representative view of the disease^7^.

To test the validity of this approach we used blood transcriptomics data corresponding to 4 different neurodegenerative diseases: Alzheimer’s disease (AD), mild cognitive impair (MCI), idiopathic Parkinson’s disease (IPD), and amyotrophic lateral sclerosis (ALS). All possible gene pairs combinations of individual genes exhibiting expression patterns that follow a normal distribution were used to compute single-value outputs for *IFF1* and *IFF2*. IF1_1000_ (**Supplementary Data 8**) and IF2_1000_ (**Supplementary Data 9**) corresponding to each neurodegenerative disease were identified and subjected to multi-class classification using a 5-fold cross-validation and hold-out strategies, like previously used, to evaluate the classification performance of IF1_1000_ and IF2_1000_ against normal blood samples.

Both blood-borne IF1_1000_ and IF2_1000_ exhibit > 75% classification accuracies for either 5-fold cross validations and/or hold-out tests across multiple cases for AD and MCI (**Supplementary Figure 9** for IF1_1000_ and **Supplementary Figure 10** for IF2_1000_), IPD (**Supplementary Figure 11** for IF1_1000_ and **Supplementary Figure 12** for IF2_1000_) and ALS (**Supplementary Figure 13** for IF1_1000_ and **Supplementary Figure 14** for IF2_1000_). However, we found that blood-borne symmetric gene expression relationships showed a lower classification performance compared to IF1_1000_ and IF2_1000_ identified from normal organs and cancers. This is not surprising as blood contains signaling molecules released from multiple organs throughout the body. Nonetheless, our results demonstrate that symmetric gene expression relationships can be extracted from blood transcriptomes and used inform on the presence of pathology at distinct body sites, in this case in the brain. Blood-borne IFGs offer unique advantages over conventional biomarkers, such as genetic alterations or over-expressed genes, because their single-value outputs are more stable (i.e., quasi-invariant) across individuals, making them more reliable for disease diagnosis and monitoring. In contrast traditional biomarkers often fluctuate among individuals, reducing their consistency and reliability.

### IFG-Enriched Pathways Underpin the Characteristics of Phenotypic Traits

We next asked which biological pathways are enriched with IFGs using as input IF1_1000_ and IF2_1000_ from normal organs and cancer types. These IFG-enriched pathways can provide insights into which biological processes are governed by symmetric gene expression relationships and help elucidate their roles in defining a biological state. Results of the over-representation analyses for enriched pathways in normal organs (**Supplementary Data 10** for IF1_1000_ and **Supplementary Data 11** for IF2_1000_) and cancers (**Supplementary Data 12** for IF1_1000_ and **Supplementary Data 13** for IF2_1000_) are provided.

Of particular interest, are the biological pathways constrained by symmetric gene expression relationships across multiple organs and cancer types. This generic IFG-constrained pathways, can elucidate which disease-driven pathways are constrained in pathological contexts but not in most normal conditions. Heatmaps for pathways enriched in at least four normal organs (**Supplementary Figure 15** for IF1_1000_ and **Supplementary Figure 16** for IF2_1000_) or cancers (**Supplementary Figure 17** for IF1_1000_ and **Supplementary Figure 18** for IF2_1000_) are shown. Intriguingly, only a few IFG-enriched pathways were shared between normal organs and cancers (**Supplementary Figure 19** and **Supplementary Data 14**), indicating that distinct pathways are confined by symmetric gene expression relationships between in heathy and cancer types. Interestingly, IF2_1000_ captures tRNA related pathways and in cancers tRNA modification appears to be altered. Additional work on these pathways is needed to fully realize their clinical importance in the near future.

### Hubs of Networks Constructed from IFGs are Information Encoders

Based on symmetric expression relationships of IFGs as depicted by *IFF1* and *IFF2* and their quasi-invariant single-value outputs for a given biological phenotype, we reasoned that knowing the expression value of a gene in an IFG pair will allow us to compute the symmetric expression value of its counterpart. This so called “information encoding” property of IFGs holds both biological and clinical importance, in particular for genes acting as hubs within an Invariant Feature Network (IF-Net), i.e., a network constructed from IFGs for a given biological phenotype. This encoding property is a consequence of the expression of IFGs being “locked” by a set of hub genes in IF-Net according to a specific mode of gene expression symmetry relationship (e.g. symmetry mode depicted in *IFF1*). Hence, we envisaged hubs of IF-Nets are information encoders that their expression values can reconstruct expression values of most counterpart IFGs corresponding to each sample under the same biological phenotype.

We tested this idea by focusing on IF-Nets derived from both IF1_1000_ and IF2_1000_ from 25 cancer types. We defined hubs as genes in the top 10% of the most connected genes. We employed mean relative error (MRE) and correlation coefficient (R) as metrics to measure errors in reconstructed gene expression values from gene hubs. **Figure 3** shows small reconstruction errors and high correlations to true expression values, indicating that IF-Net hubs are indeed information encoders for gene expression. We found that hubs of IF-Nets from *IFF1* (**Figure 3A**) provide a better reconstruction performance than IF-Net hubs from *IFF2* (**Figure 3B**) across all individual samples for a given cancer type. This suggests that IFGs that exhibit *IFF1* symmetric expression relations are subjected to tighter biological constraints. However, the nature of these biological constraints and the underlying mechanisms remained to be elucidated.

**Figure 3.**
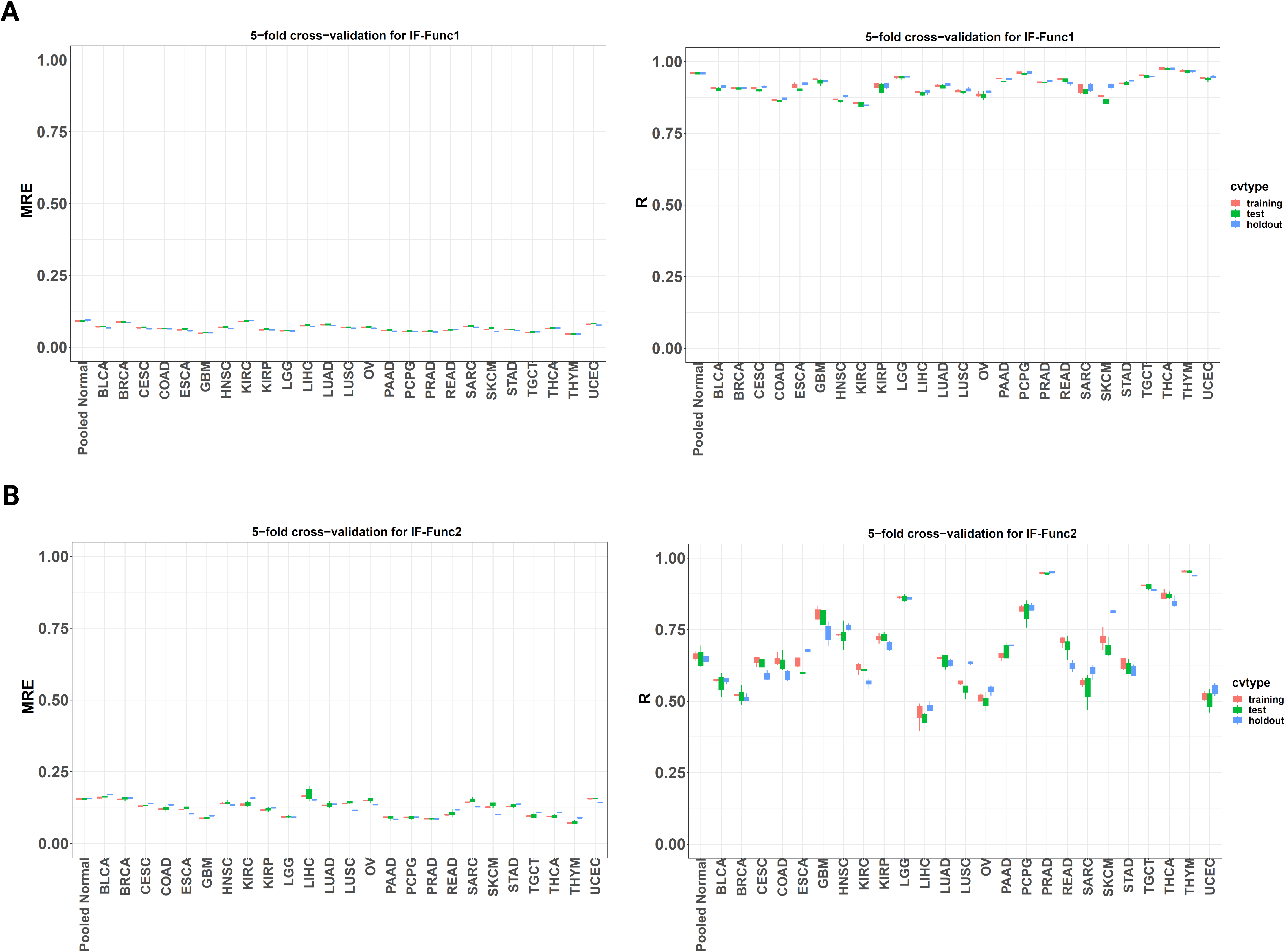
Hubs of IF-Nets are information encoders that can be used to reconstruct expression values of their counterpart IFGs. The reconstruction of gene expression performance was calculated based on reconstruction errors of gene expression and the robustness of the reconstruction results were assessed using a 5-fold cross-validation, using the mean relative error (MRE) and correlation coefficient (R) metrics. (**A**) Reconstruction expression errors for hubs in *IFF1-*derived IF-Nets across 25 cancer types. (**B**) Reconstruction expression errors for hubs in *IFF2*-derived IF-Nets across 25 cancer types.

The fact that IF-Net hubs are information encoders of gene expression also indicate that the expression of IFGs in an individual are modulated as a whole, with each IFG directly or indirectly imposes its symmetric constraint to other IFGs, forming a coordinated gene expression web whose collective activities defined the property of a biological phenotype. This finding is important in two major aspects. First, it is possible just to measure the expression values of hub genes, especially for hubs that also present in multiple cancer types where expression values of their corresponding associated invariant feature gene counterparts can be reliably reconstructed. Second, expression values of IF-Net hubs and selected counterpart IFGs can serve as a “reduced” set of genes that encode most information of gene expression that can be used as molecular fingerprints of a given biological phenotype. Hence, information encoding properties of IF-Net hubs can be used as a new type of diagnosis, where a sample can be assigned to a given disease type by evaluating the reconstruction error of gene expression values correspond to a disease type.

### IF-Nets are Pharmacological Discovery Platforms

Since the symmetric expression relationships of IFGs defined the identity and activities of a given biological phenotype and IFG connected in IF-Net acted as a whole, we next questioned whether IFGs, especially IF-Net hubs are promising intervention targets for disease treatment. We tested this idea using drugs with known targets to explore how many of these targets constitute IF-Nets with respect to IF1_1000_ and IF2_1000_ across 25 cancer types. We extracted information from Drug Repurposing Hub^8^ to map drugs with known targets and clinical phases onto IF-Nets across 25 cancer types. Additionally, we also mapped drug targets onto IF-Nets across 25 normal organs, with the purpose to contrast pharmacological interventions in cancer tissues against normal organs as a potential approach to prevent adverse drug responses. The interactive drug-IF-Net results can be explored at life.hulilab.org.

Intriguingly, IF-Net hubs, especially *IFF1-*derived IF-Nets corresponding to cancers and normal organs are enriched with drug targets. **Figure 4** provides drug target-enriched results for breast and colon cancers as illustrative examples, showing that especially *IFF1-*derived IF-Nets are valuable pharmacological discovery platforms. For instance, our results suggests that COX7C, which involves in mitochondrial electron transport chain is a target of launched clinical drugs enriched in breast cancer IF-Net (**Figure 4A**) and colon cancer IF-Net (**Figure 4B**). The role of COX7C in breast cancer etiology is unknown but is associated with venous thromboembolism in colon cancer patients^9^. Another mitochondrial gene IARS2, is known to involve in the development of some cancer types^10,11^, is a breast cancer IF-Net hub that can be targeted by approved clinical drugs (**Figure 4A**). Interestingly, PARP1 which is essential for DNA repair and the progression of several types of cancers, including breast cancer, is the drug target-enriched hub in breast cancer IF-Net. On contrary, a number of ribosomal genes (e.g., RPL19, RPL37, RPL23A) are enriched as drug target in breast IF-Net (**Figure 4A**), suggesting that drugs targeting ribosomal genes should be used with caution to avoid potential unwanted side effects in breast cancer patients^12^.

**Figure 4.**
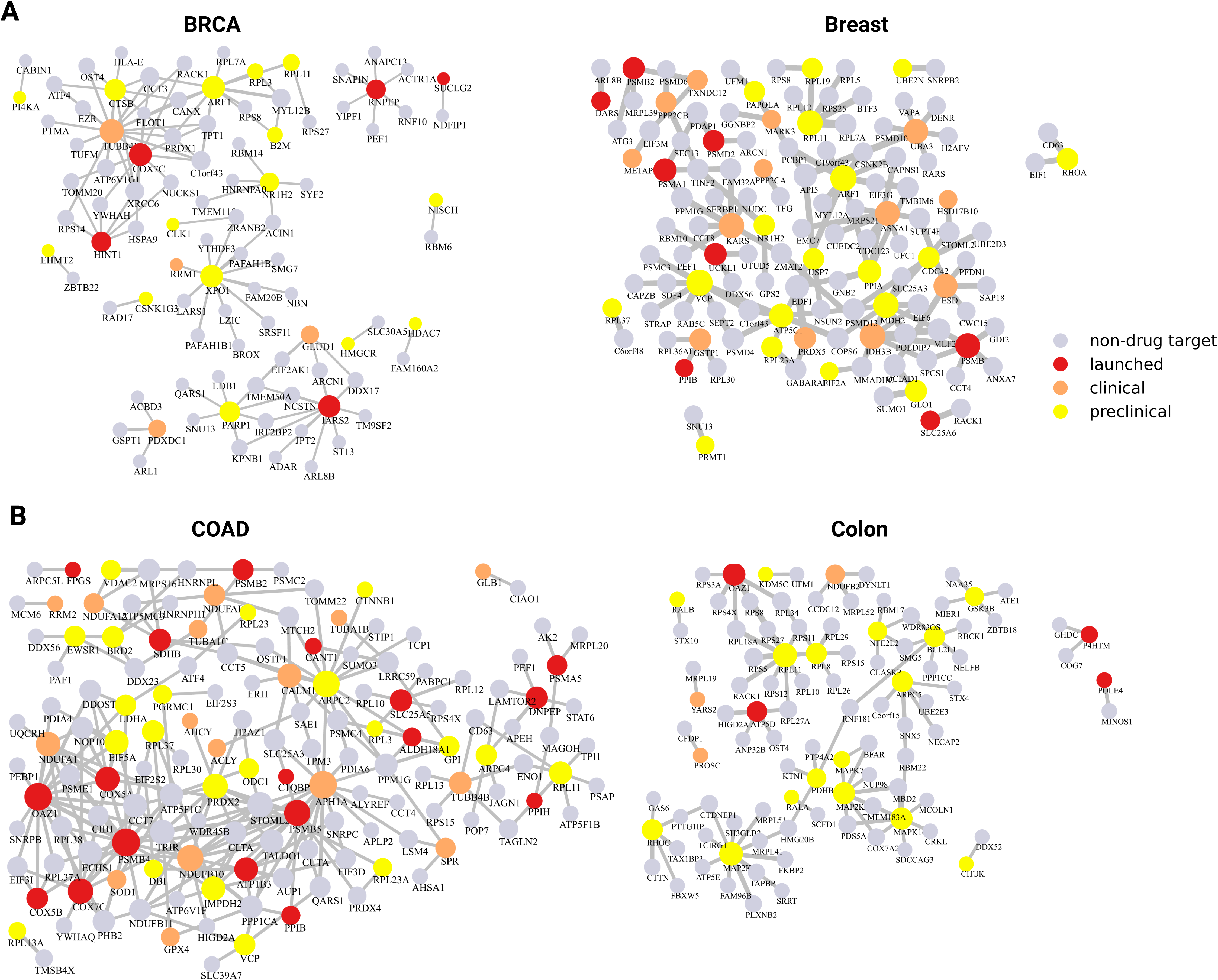
IFGs Known drug targets within IF-Nets for two selected cancer types with respect to organs of cancer origin constructed from IF1_1000_: (**A**) BRCA and breast, (**B**) COAD and colon. Nodes indicate IFGs present in IF-Nets and edges represent symmetric expressed gene pairs as described by *IFF1*. Red nodes denote targets of FDA approved drugs, orange nodes denote targets whose drugs are in Phase 3 trials, yellow nodes denote targets under preclinical trials, and grey nodes denote genes without available drug information.

In summary, IF-Nets are discovery platforms not only to uncover symmetric gene expression relationships that underpin disease traits but more importantly they provide novel clues to identify actionable targets and enable the repurposing of clinically approved drugs for the treatment of new diseases.

## DISCUSSION

The concept of symmetry has been widely used as a guiding principle in physics and chemistry to drive new discoveries. However, the concept of symmetry is rarely invoked in biological sciences to understand how a cell behaves and how a disease develops. In mathematical terms, symmetry refers to a property (structural or behavioral) that remains invariant (i.e., unchanged) upon a transformation (or mapping) by a given mathematical operation (e.g., rotation). In biology, we can perceive genetic variability as a form of transformation imposed on a living organism. A biological phenomenon that satisfies this criterion of symmetry is for example the constrained by the total daily energy expenditure of an individual^13,14^ or the energetic constraints at the cellular level^15,16^ that restrict the expression of certain genes whose collective activities are needed to maintain the stability of a phenotypic property. All these phenomena inspired us to formulate and investigate symmetric relationships among expressed genes from transcriptomics data of known phenotypic traits. In this context, we can reason that the stability of biological phenotypes, such as cell types and disease states in the face of genetic variations aligns the mathematical definition of symmetry

We proposed our Gene Expression Symmetry Hypothesis (GESH) which predicts that a set of genes pairs whose expression are confined by certain modes of symmetric relationships in defining a biological phenotype. We devised two Invariant Feature Functions, *IFF1* and *IFF2*, that describe two different modes of symmetric relationship of gene expression. We implemented these functions into the design of Machine Learning-Based Invariant Feature Engineering (LIFE) platform to detect Invariant Feature Genes (IFGs), gene pairs whose expressions are confined by symmetric modes as described by *IFF1* or *IFF2*. We tested our idea using transcriptomics data derived from 25 normal organs and 25 cancer types and blood across 4 neurologic diseases.

Robust multi-class classification results identified symmetric gene expression relationship across diverse biological phenotypes, at least in 25 normal organs and 25 cancer types investigated in this work. The fact that these symmetric gene expression relationships define the identity of biological phenotypes, regardless of their variable genetic backgrounds and heterogeneity of gene expression among individuals opens a new window for understanding underlying pathways or processes that help sustain their identity in health and disease.

We next investigated whether IFGs also can be detected in patients’ blood transcriptomes. This is clinically important because blood-based diagnosis is minimally invasive, rapid, and has a low associated cost. Biomarkers derived from blood transcriptomes have been routinely used in diagnosis^17,18^. However, many of these blood-borne biomarkers are based on expression profiles of single genes which are sensitive to genetic variations and gene expression heterogeneity among individuals. Invariant disease-specific blood-borne gene pairs, can serve as robust genetic fingerprints to indicate the presence of diseases because these are not dependent on genetic variations and gene expression heterogeneity in patients.

In this study, we used blood transcriptomics data derived from four types of neurodegenerative disorders as case studies. Our choice of the brain is based on the fact that it is a remote organ, and generating of omics data for diagnosing and monitoring progression is prohibited in living individuals. We identified blood-borne disease-specific IFGs specific to distinct neurogenerative disorders when compared with matched controls. Our work therefore holds great promise as a new kind of non-invasive fingerprints for disease detection.

Our study also suggests that IFGs, directly or indirectly, function as a coordinated whole within a network that we named Invariant Feature Network (IF-Net). The coordinated nature of IFGs within an IF-Net implies that the expression activities of IFGs can be encoded by hub genes in these networks. In other words, given expression values of IF-Net hub genes, we can reconstruct the expression values of the corresponding IFGs in a network. Our study demonstrates that the expression of IF-Net hub genes enables the reconstruction expression of their counterpart IFGs with small reconstruction errors, indicating that IF-Net hub genes act as information encoders. Such highly coordinated networking behavior is not found in other experimental or data-driven biological networks such as protein-protein interaction (PPI) network and correlative-based reverse engineered gene-gene association networks, like those inferred from Pearson correlation and mutual information. This is because PPI networks or a correlative-based reverse engineered networks cannot be used to reconstruct gene expression values with high precision since they do tend to not behave as a coordinated whole. For instance, PPI network is constructed from various experimental results derived from different biological contexts^19,20^.

We tested the pharmacological value of IF-Nets by mapping the targets of known FDA approved clinical drug onto IF-Nets for each normal organs and cancer types. Importantly, the coordinated whole properties of IF-Nets immediately suggest their pharmacological values in target and drug discovery. Selected drug targets were enriched in hubs within cancer IF-Nets derived with *IFF1*, indicating that disease IF-Nets offer a novel pharmacological discovery platform for drug repurposing, as well as a valuable resource to identify new actionable targets and potential drugs.

Equally important, we found that hubs of *IFF1-*derived IF-Nets with respect to normal organs are also enriched with clinical drugs. We reason that these are targets that should be avoided to reduce unwanted adverse reactions in a targeted organ (e.g., targeted therapeutics for breast cancer IFGs should not intervene IFGs in normal breast tissues). In contrast, *IFF2-* derived cancer IF-Nets are poorly enriched in known clinical drugs, suggesting that hubs in these networks are either under-explored targets or undruggable. The pharmacological values of *IFF2-* derived IF-Nets remains to be further investigated.

In sum, we demonstrated that gene expression symmetric relationships hold a unique value in identifying biological phenotypes. Additionally, we highlighted the pharmacological and clinical potential for disease-specific IFGs and IF-Nets. However, the discovery of the symmetry of gene expression relationships also adds up new complexities in our understanding of how biological systems behave. New questions remain to be answered. For instance, beside the symmetric relations described in *IFF1* and *IFF2* in this work, are there other modes of gene expression symmetry? What kind of regulatory mechanisms are involved to keep symmetric gene expression relationships in check? Are there inter-organ symmetric relationships between expressed genes that may confer memory-like “locked” states in disease states^21^? Unlocking the answers for these questions will undoubtedly provide a new paradigm shift in medicine.

## MATERIALS AND METHODS

### Dataset and Data Processing

Transcriptomics data for a total of 25 organ types from The Genotype-Tissue Expression (GTEx) project (https://gtexportal.org/) and 25 cancer types from The Cancer Genome Atlas (TCGA) program (https://www.cancer.gov/ccg/research/genome-sequencing/tcga) were used in this work (**Supplementary Data 1**). Downloaded gene-level raw counts were converted to rpkm for all protein-code genes using R package edgeR^22^. The rpkm values were log-transformed using formula log2(v+2) where v is rpkm value for each gene in each sample to make the transformed value no less than 1. Samples were filtered to exclude FFPE (formalin-fixed paraffin-embedded) samples as the RNA molecules in these samples can be highly degraded. The TCGA 25 pooled normal samples correspond to the pooled non-cancer samples identified from each of the 25 cancer types and were used as control (i.e., normal) for comparison with the cancer samples. Blood microarray data for case/control studies with at least 100 samples per group corresponding from four neurodegenerative disorders, Alzheimer’s disease (AD) (GEO Accession: GSE63060), mild cognitive impairment (MCI) (GEO Accession: GSE63060), idiopathic Parkinson’s disease (IPD) (GEO Accession: GSE99039), and amyotrophic lateral sclerosis (ALS) (GEO Accession: GSE112676) were downloaded to identity blood-based invariant feature genes (IFGs) that can reliably inform on pathology at distinct sites. Processed gene intensity data from GEO were used for the analysis after log2 transformation and shifting to ensure all intensity values were not smaller than 1. Additional description on these datasets is provided in Supplementary Data 1.

### Learning-Based Invariant Feature Engineering (LIFE) and the Identification of Invariant Feature Gene Pair Candidates

The Learning-Based Invariant Feature Engineering (LIFE) approach is designed based on two hypothetical scenarios indicated in **Figure 1**. We designed two Invariant Feature Functions, IFF1 and IFF2, to describe different symmetric gene expression relationships. Here,

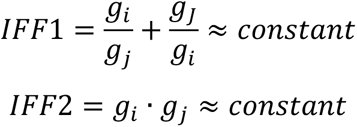

where g_i_ and g_j_ are expression levels of gene-i and gene-j, respectively. Note that interchange of subscript *i* and *j* will not change the computed results, hence both IFF1 and IFF2 capture different symmetric expression relationships between genes pairs. Single-output values for both IFF1 and IFF2 were computed for all possible combinations of gene pairs across 25 normal organs and 25 cancer types, respectively. Mean and standard deviation (σ) for each gene pair were computed. Top 1000 gene pairs with smallest SDs identified from IFF1 (IF1_1000_) and IFF2 (IF2_1000_) were deemed candidates of invariant feature genes (**Figure 1**).

### Accessing the Performance of Invariant Feature Candidates in Assigning Samples to Correct Phenotypic Class

To test the capability of IF1_1000_ and IF2_1000_ as phenotype-specific fingerprints, we employed a 5-fold cross-validation strategy to assess the robustness of IF1_1000_ and IF2_1000_ in assigning individual samples to their respective biological phenotypes. Here, samples from each phenotype were randomly split into 5 portions with approximate equal sample size, with four portions used for training and one portion for testing. Taking IFF1 as an example, top 1000 gene pairs with the smallest standard deviation (σ) computed for IFF1 in a given phenotype were identified and used to testing their ability to classify samples in test sets (the remaining one portion was not used for training) to their respective biological state according to the phenotype score:

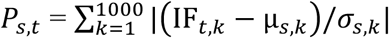

where 𝑃*_s,t_*_,’_ is the phenotype score of state *s* for test sample *t* from the test set; (𝐼𝐹*_t_*, 𝑝𝑎𝑖𝑟*_k_*) is the IFF value of pair *k* of sample *t*, µ and σ are the mean and standard deviation for the IF function value of gene pair *k* in a biological state in the training set, respectively. Thus, the phenotype score 𝑃*_s,t_*_,’_ is actually the sum of absolute z-scores of IFF values of each gene pair in the test sample *t* as compared to those in the training set. The predicted biological state of sample *t* is the one with smallest phenotype score 𝑃*_s,t_*_,’_. The training-testing procedures were repeated for another 4 folds until all 5 folds had been used as testing sets. The same procedure was used for IFF2. Performance of IF1_1000_ and IF2_1000_ is presented as percentage of predictions that correctly assign samples to their actual biological states. From the five folds of training results, we selected the gene pairs that appear in all five folds (i.e., in every fold), and were unique to a given cancer type, excluding any gene pairs present in other cancer types. These are called conserved unique gene pairs for that cancer type. The hold-set performance was assessed in comparison to the training data from all five folds. We calculated the IF values for the conserved unique gene pairs in each cancer type using the gene expression data in the training set and then compared these values to the hold-out set, to predict the cancer type of each sample in the hold-out set.

### Pathway Enrichment with Over-Representation Analyses

A total of 9,273 gene sets from Gene Ontology Biological Processes (GO-BP) and REACTOME were used for the functional analysis. The significance of over-representation was calculated using Fisher’s exact test and corrected using the Benjamini–Hochberg method to produce false discovery rates (FDR). We filtered the results to only keep the gene sets that had at least one significant result (FDR<0.05) in each dataset. Heatmaps were plotted based on -log10(FDR) values of enriched pathways.

### Reconstruction of Expression Values of Invariant Molecular Feature Genes from network Hubs

We built disease specific unweighted networks, that we named invariant feature networks (IF-Nets), using IF1_1000_ and IF2_1000_ gene pairs for each phenotype, respectively. IF-Nets for IF1_1000_ and IF2_1000_ were construct independently for each phenotype and aggregated based on shared genes among identified IFGs gene pairs. In this study, genes with connectivity degree larger than the 90th percentile of all genes, i.e., top 10% most connected genes were selected and defined as hubs. Hub genes were tested for their ability to act as expression encoders. Using the gene expression of hub genes, we reasoned that we could reconstruct the expression values for the remaining genes within IF-Nets for each individual samples under a given biological phenotype. For reconstructing the gene expression for each gene (genes in IF1_1000_ and IF2_1000_ that are not a hub), the sample-wise reconstructed expression value is calculated by

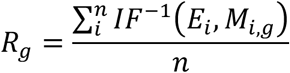

Where 𝑅*_g_* is the reconstructed expression value of gene *g* for a given sample using all hub genes that paired with gene g in IF1_1000_ and IF2_1000_ (here, hub-*i* is paired with gene *g* in this case), 𝐸*_i_* is the expression value of hub-*i*, 𝑀*_i,g_* is the mean of the invariant function value for gene *g* and hub-*i* pair, 𝐼𝐹^-1)^ is the inverse function of IFF1 or IFF2, and *n* is the number of total connecting hubs to gene *g*. Note that for the case of IFF1, its inverse function will yield two solutions thus the reconstructed expression value of gene *g* is the averaged values of these two solutions.

### Drug Information and Target Identification

The information on drug names, drug targets, and clinical phase were obtained from the Drug Repurposing Hub^8^. All genes within IF-Nets, constructed using IF1_1000_ and IF2_1000_ for each biological phenotype were matched to known drug targets, and targetable IFGs were identified. Results for drug-targeted genes within IF-Nets across 25 normal organs and 25 cancer types can be explored at life.hulilab.org.

## AUTHORS’ CONTRIBUTIONS

CZ and CC contributed equally to this work. CZ, CC, CYU and HL contributed to the conception and design of the study. CZ, CC, CYU, and HL contributed to the acquisition of data. CZ, CC, TW, SHT, ZZ, KSY, SZ, CYU and HL contributed to the analysis and interpretation of data. CZ, CC, CYU, and HL drafted the manuscript. All authors revised and accepte the final manuscript. CYU and HL supervised the study.

## ACKNOWLEDGEMENTS

This research was funded by generous support from the Mayo Clinic Comprehensive Cancer Center (P30CA015083), the Mayo Clinic Center for Cell Signaling in Gastroenterology (NIH: P30DK084567), the David F. and Margaret T. Grohne Cancer Immunology and Immunotherapy Program, the Mayo Clinic Nutrition Obesity Research Program, the Eric & Wendy Schmidt Fund for AI Research & Innovation, the Glenn Foundation for Medical Research (HL), Mayo Clinic Center for Biomedical Discovery (HL and SZ), the Mayo Clinic Center for Individualized Medicine (HL and SZ), Mayo Clinic DERIVE Office (SZ), the V Foundation for Cancer Research (SZ), and the National Institutes of Health (NIH): U19AG74879, P50CA136393, U54AG79779, R03OD038392 (HL) and R01CA240323 (SZ).

## SUPPLEMENTARY FIGURES

**Supplementary Figure 1.** Multiclass classification performance for 25 normal organ types derived with IFF1.

**Supplementary Figure 2.** Multiclass classification performance for 25 normal organ types derived with IFF2.

**Supplementary Figure 3.** Multiclass classification performance for 25 cancer types derived with IFF1.

**Supplementary Figure 4.** Multiclass classification performance for 25 cancer types derived with IFF2.

**Supplementary Figure 5.** Confusion matrix of multiclass classification for 25 normal organ types subjected to IFF1.

**Supplementary Figure 6.** Confusion matrix of multiclass classification for 25 normal organ types subjected to IFF2.

**Supplementary Figure 7.** Confusion matrix of multiclass classification for 25 cancer types subjected to IFF1.

**Supplementary Figure 8.** Confusion matrix of multiclass classification for 25 cancer types subjected to IFF2.

**Supplementary Figure 9.** Multiclass classification performance for blood-borne IF1_1000_ for Alzheimer’s disease (AD) and mild cognitive impair (MCI) and control (normal) cases.

**Supplementary Figure 10.** Multiclass classification performance for blood-borne IF2_1000_ for Alzheimer’s disease (AD) and mild cognitive impair (MCI) cases and matched controls.

**Supplementary Figure 11.** Classification performance for blood-borne IF1_1000_ in idiopathic Parkinson’s disease (IPD) and control cases.

**Supplementary Figure 12.** Classification performance for blood-borne IF2_1000_ in idiopathic Parkinson’s disease (IPD) and control cases.

**Supplementary Figure 13.** Classification performance for blood-borne IF1_1000_ in amyotrophic lateral sclerosis (ALS) and control cases.

**Supplementary Figure 14.** Classification performance for blood-borne IF2_1000_ in amyotrophic lateral sclerosis (ALS) and control cases.

**Supplementary Figure 15.** Heatmap showing commonly enriched pathways for IF1_1000_ gene-pairs across at least 4 normal organ types.

**Supplementary Figure 16.** Heatmap showing commonly enriched pathways for IF2_1000_ gene-pairs across at least 4 organ types.

**Supplementary Figure 17.** Heatmap showing commonly enriched pathways for IF1_1000_ gene-pairs across at least 4 cancer types. Number of samples correspond to each cancer type is provided along with the data label.

**Supplementary Figure 18.** Heatmap showing commonly enriched pathways for IF2_1000_ gene-pairs across at least 4 cancer types. Number of samples correspond to each cancer type is provided along with the data label.

**Supplementary Figure 19.** Venn diagram depicting of commonly IFG-enriched pathways among different normal organs and cancer types. IF1: *IFF1*; IF2: *IFF2*.

## SUPPLEMENTARY DATA

**Supplementary Data 1**. Summary of transcriptomic data used in this work. GTEx: Transcriptomic data from 25 types for normal organs; TCGA: Transcriptomic data corresponding to 25 cancer types.

**Supplementary Data 2**. IF1_1000_ for 25 normal organs (A) and 25 cancer types (B).

**Supplementary Data 3**. IF2_1000_ for 25 normal organs (A) and 25 cancer types (B).

**Supplementary Data 4**. Gene-gene expression plots for top 20 IF1_1000_ across 25 normal organs.

**Supplementary Data 5**. Gene-gene expression plots for top 20 IF2_1000_ across to 25 normal organs.

**Supplementary Data 6**. Gene-gene expression plots for top 20 IF1_1000_ across 25 cancer types. TCGA-25 is the pooled normal samples with respect to tissues of 25 cancer types.

**Supplementary Data 7**. Gene-gene expression plots for top 20 IF2_1000_ for 25 cancer types. TCGA-25 is the pooled normal sample for each tissue of 25 cancer types.

**Supplementary Data 8**. Blood-borne IF1_1000_ with respect to four neurodegenerative diseases. AD: Alzheimer’s disease; MCI: mild cognitive impair; IPD: idiopathic Parkinson’s disease; and ALS: amyotrophic lateral sclerosis.

**Supplementary Data 9**. Blood-borne IF2_1000_ with respect to four neurodegenerative diseases. AD: Alzheimer’s disease; MCI: mild cognitive impair; IPD: idiopathic Parkinson’s disease; and ALS: amyotrophic lateral sclerosis.

**Supplementary Data 10**. Enriched pathways of IF1_1000_ across 25 normal organ types.

**Supplementary Data 11**. Enriched pathways of IF2_1000_ across 25 normal organ types.

**Supplementary Data 12**. Enriched pathways of IF1_1000_ across 25 cancer types.

**Supplementary Data 13**. Enriched pathways of IF2_1000_ across 25 cancer types.

**Supplementary Data 14**. List of overlapped IFG-enriched pathways among normal organs and cancers.

