## Supplemental Data 5 for "Symmetry as a Fundamental Principle in Defining Gene Expression and Phenotypic Traits"

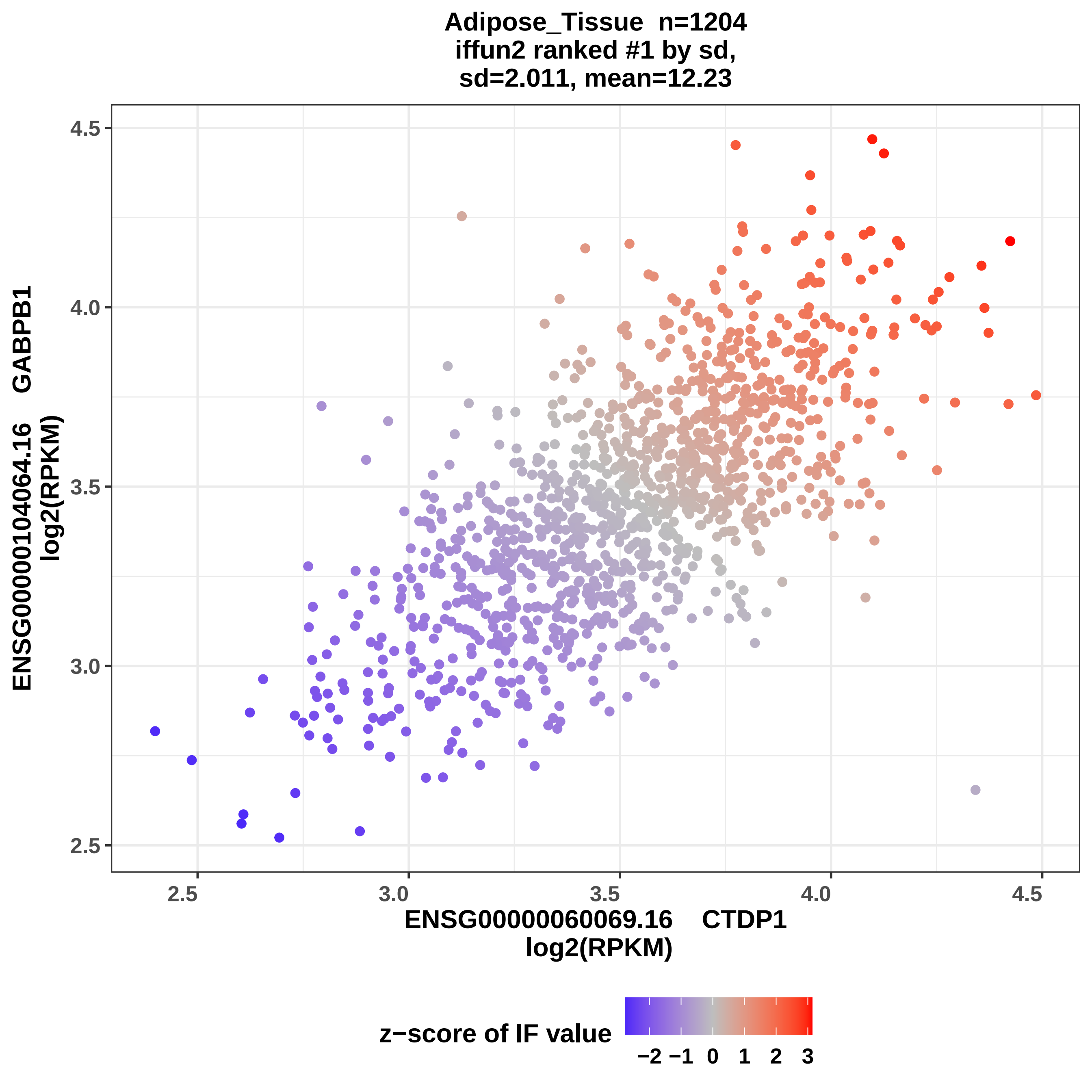
Adipose_Tissue n=1204
iffun2 ranked #1 by sd,
sd=2.011, mean=12.23
ENSG00000104064.16 GABPB1
log2(RPKM)
4.5
4.0
3.5
3.0
2.5
2.5
3.0
3.5
4.0
4.5
ENSG00000060069.16 CTDP1
log2(RPKM)
z−score of IF value
−2 −1 0 1 2 3

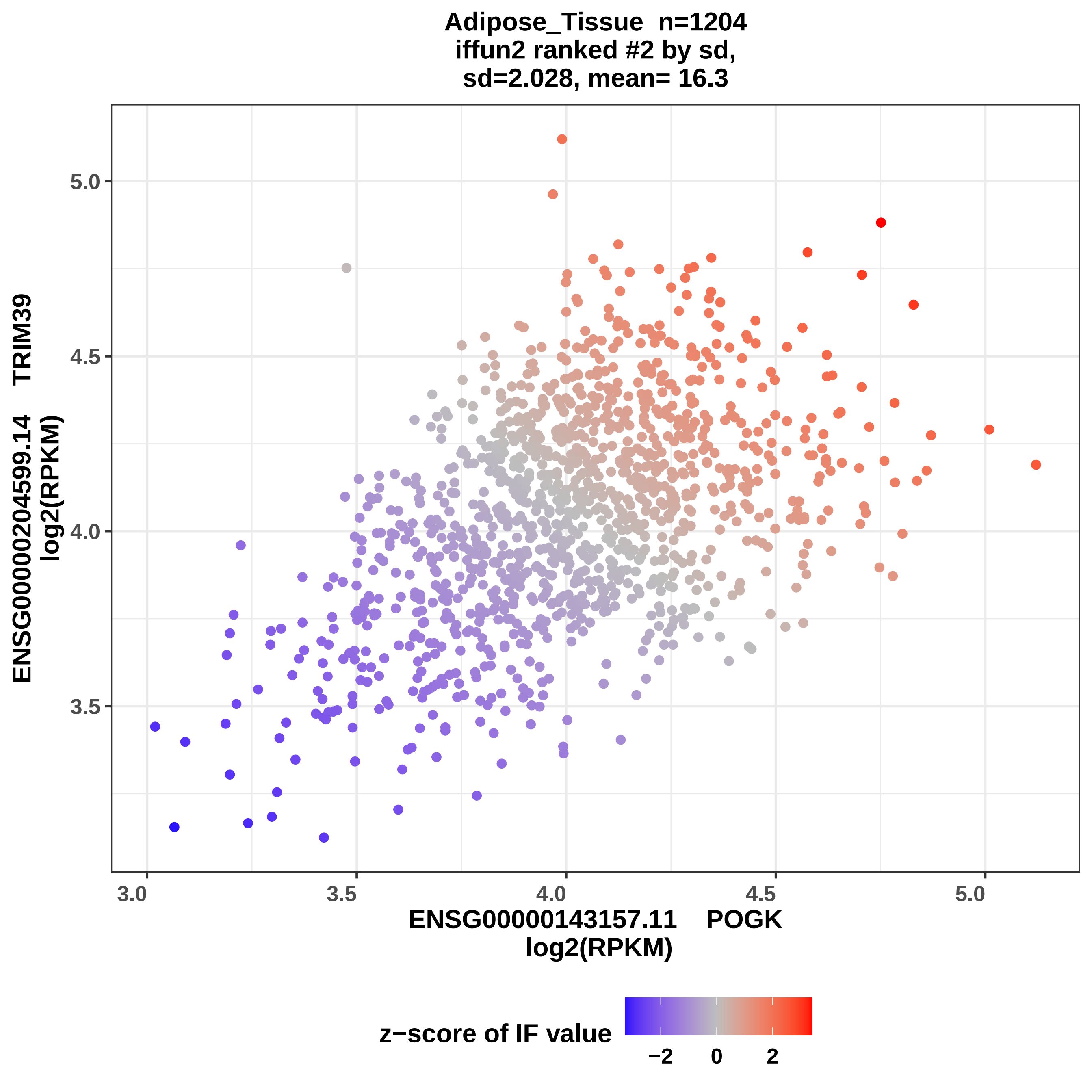

Adipose_Tissue n=1204
iffun2 ranked #2 by sd,
sd=2.028, mean= 16.3
ENSG00000204599.14 TRIM39
log2(RPKM)
5.0
4.5
4.0
3.5
3.0
3.5
4.0
4.5
5.0
ENSG00000143157.11 POGK
log2(RPKM)
z−score of IF value
−2
0
2

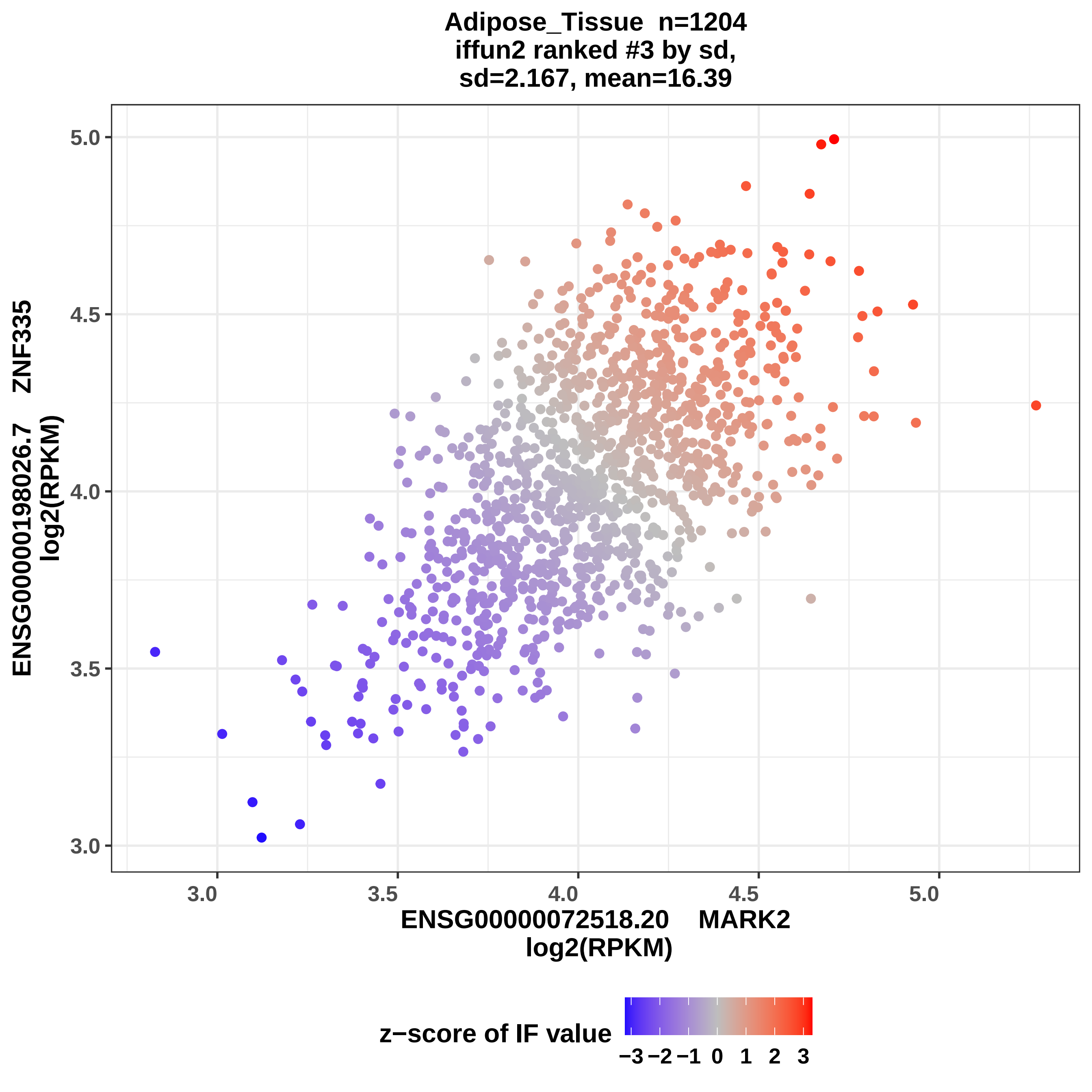
Adipose_Tissue n=1204
iffun2 ranked #3 by sd,
sd=2.167, mean=16.39
ENSG00000198026.7 ZNF335
log2(RPKM)
5.0
4.5
4.0
3.5
3.0
3.0
3.5
4.0
4.5
5.0
ENSG00000072518.20 MARK2
log2(RPKM)
z−score of IF value
−3 −2 −1 0 1 2 3

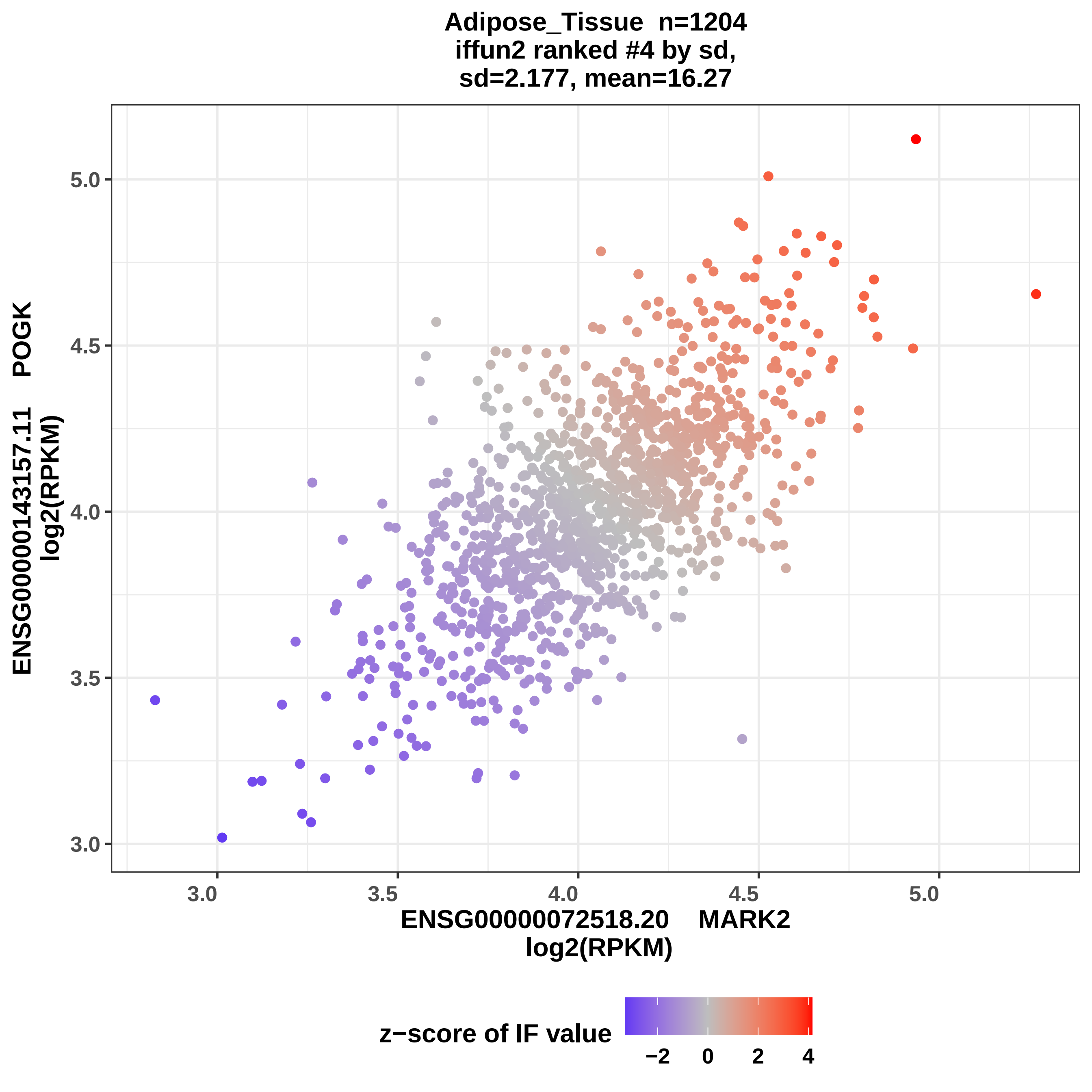
Adipose_Tissue n=1204
iffun2 ranked #4 by sd,
sd=2.177, mean=16.27
ENSG00000143157.11 POGK
log2(RPKM)
ENSG00000072518.20 MARK2
log2(RPKM)
3.0
3.5
4.0
4.5
5.0
z−score of IF value
−2
0
2
4

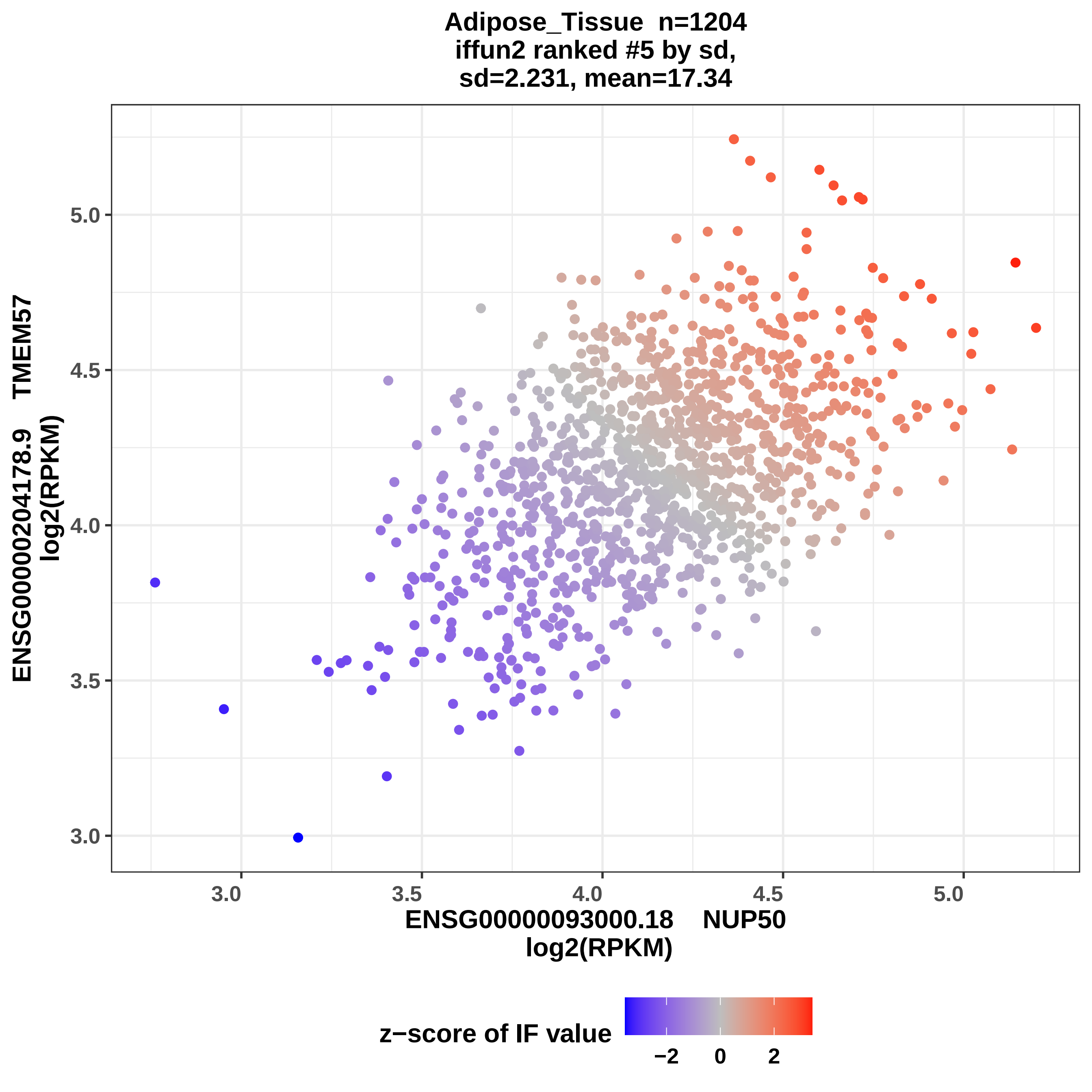

Adipose_Tissue n=1204
iffun2 ranked #5 by sd,
sd=2.231, mean=17.34
ENSG00000204178.9 TMEM57
log2(RPKM)
ENSG00000093000.18 NUP50
log2(RPKM)
5.0
4.5
4.0
3.5
3.0
3.0
3.5
4.0
4.5
5.0
z−score of IF value
−2
0
2

Adipose_Tissue n=1204
iffun2 ranked #6 by sd,
sd=2.237, mean=16.67

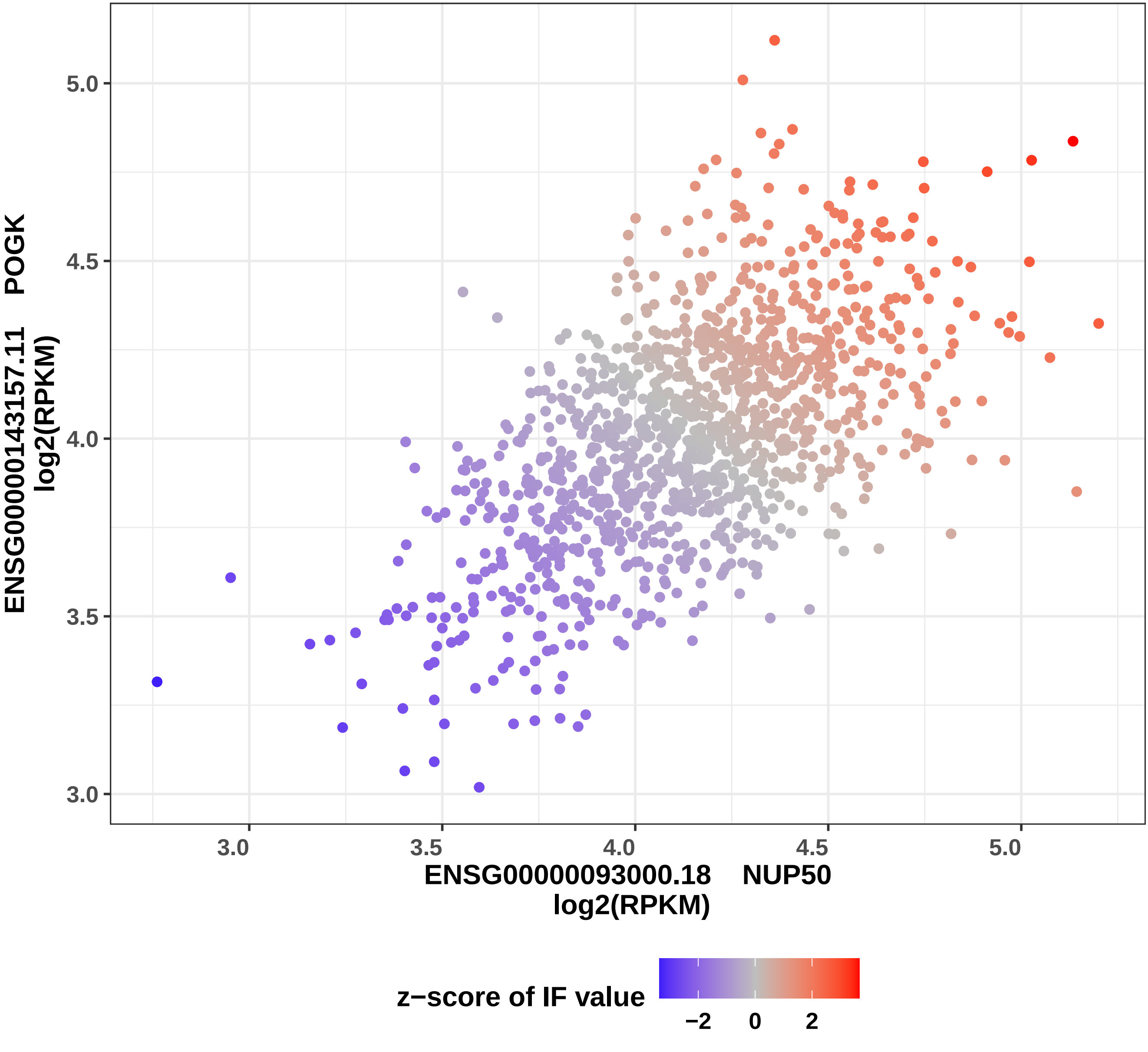

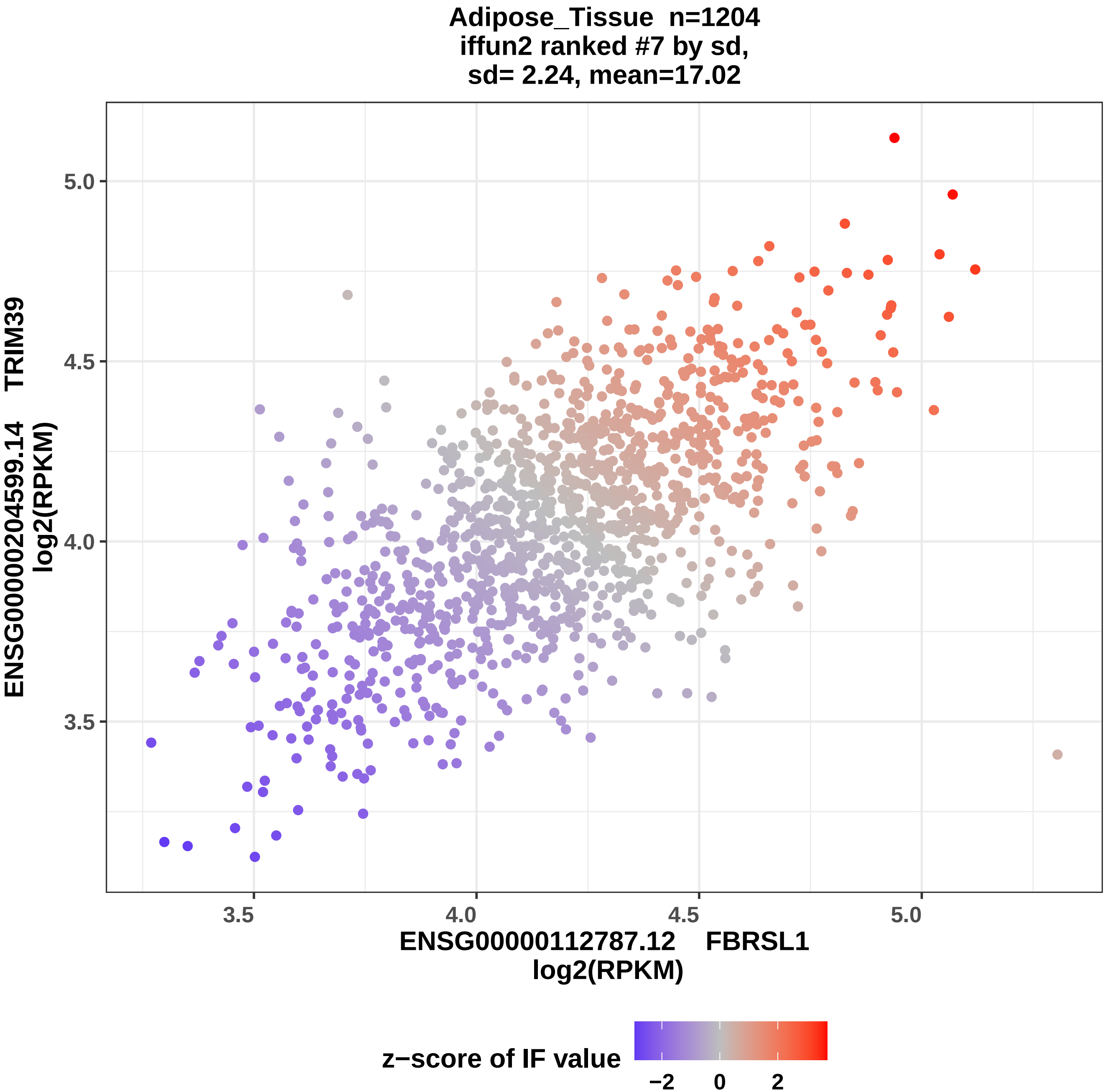
Adipose_Tissue n=1204
iffun2 ranked #7 by sd,
sd= 2.24, mean=17.02
ENSG00000204599.14 TRIM39
log2(RPKM)
5.0
4.5
4.0
3.5
3.5
4.0
4.5
5.0
ENSG00000112787.12 FBRSL1
log2(RPKM)
z−score of IF value
−2
0
2

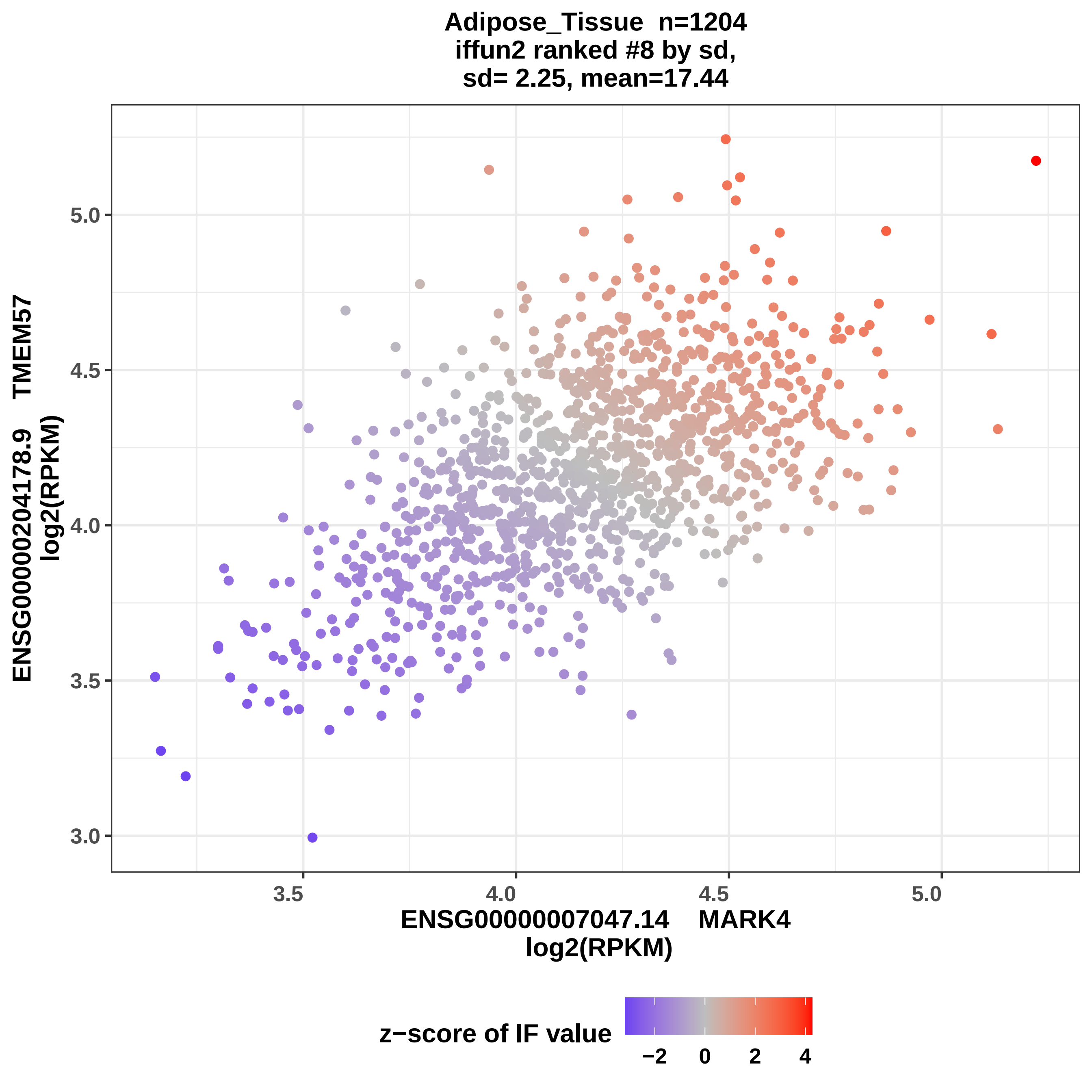
Adipose_Tissue n=1204
iffun2 ranked #8 by sd,
sd= 2.25, mean=17.44
ENSG00000204178.9 TMEM57
log2(RPKM)
5.0
4.5
4.0
3.5
3.0
3.5
4.0
4.5
5.0
ENSG00000007047.14 MARK4
log2(RPKM)
z−score of IF value
−2
0
2
4

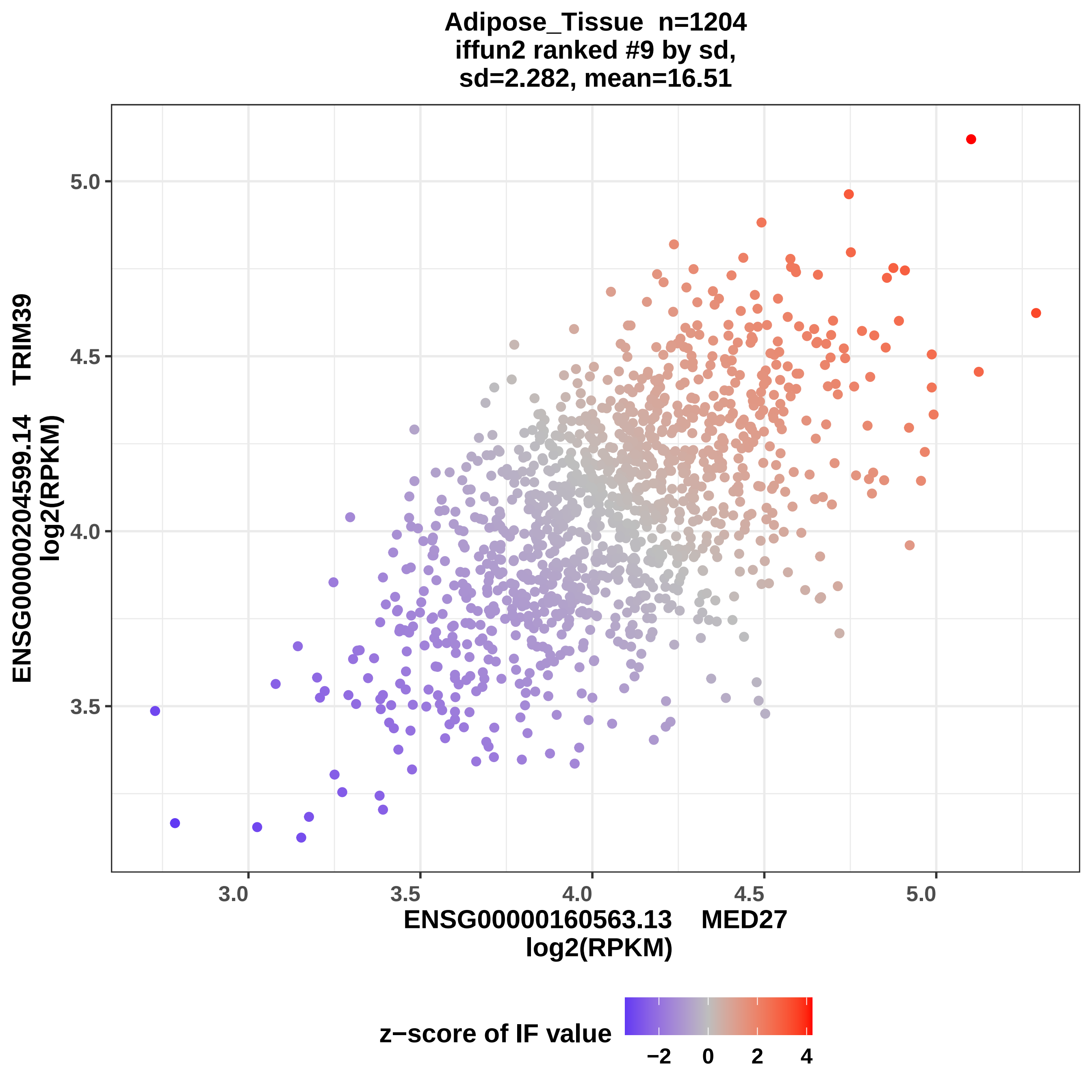

Adipose_Tissue n=1204
iffun2 ranked #9 by sd,
sd=2.282, mean=16.51
ENSG00000204599.14 TRIM39
log2(RPKM)
5.0
4.5
4.0
3.5
3.0
3.5
4.0
4.5
5.0
ENSG00000160563.13 MED27
log2(RPKM)
z−score of IF value
−2
0
2
4

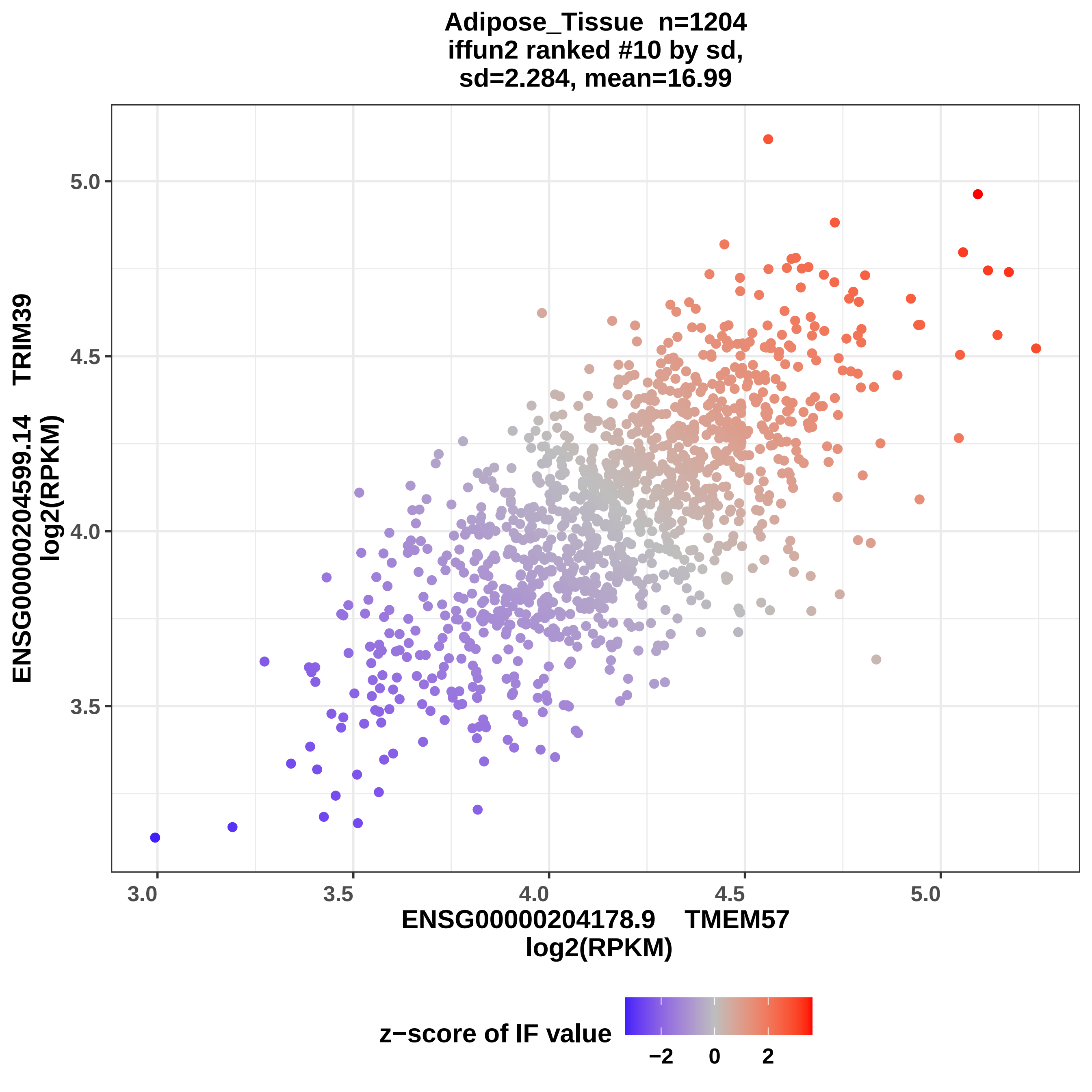

Adipose_Tissue n=1204
iffun2 ranked #10 by sd,
sd=2.284, mean=16.99
ENSG00000204599.14 TRIM39
log2(RPKM)
5.0
4.5
4.0
3.5
3.0
3.5
4.0
4.5
5.0
ENSG00000204178.9 TMEM57
log2(RPKM)
z−score of IF value
−2
0
2

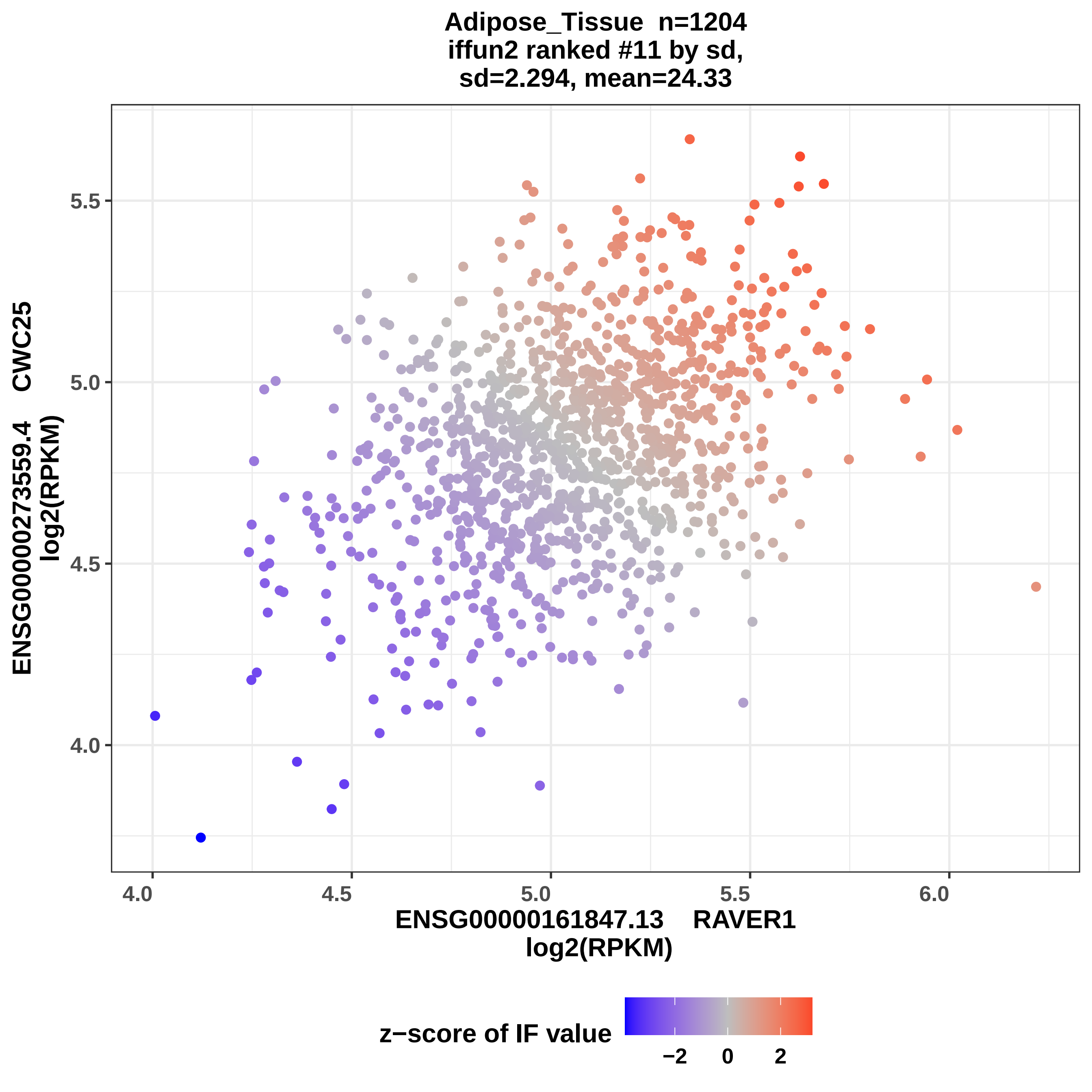

Adipose_Tissue n=1204
iffun2 ranked #11 by sd,
sd=2.294, mean=24.33
ENSG00000273559.4 CWC25
log2(RPKM)
5.5
5.0
4.5
4.0
4.0
4.5
5.0
5.5
6.0
ENSG00000161847.13 RAVER1
log2(RPKM)
z−score of IF value
−2
0
2

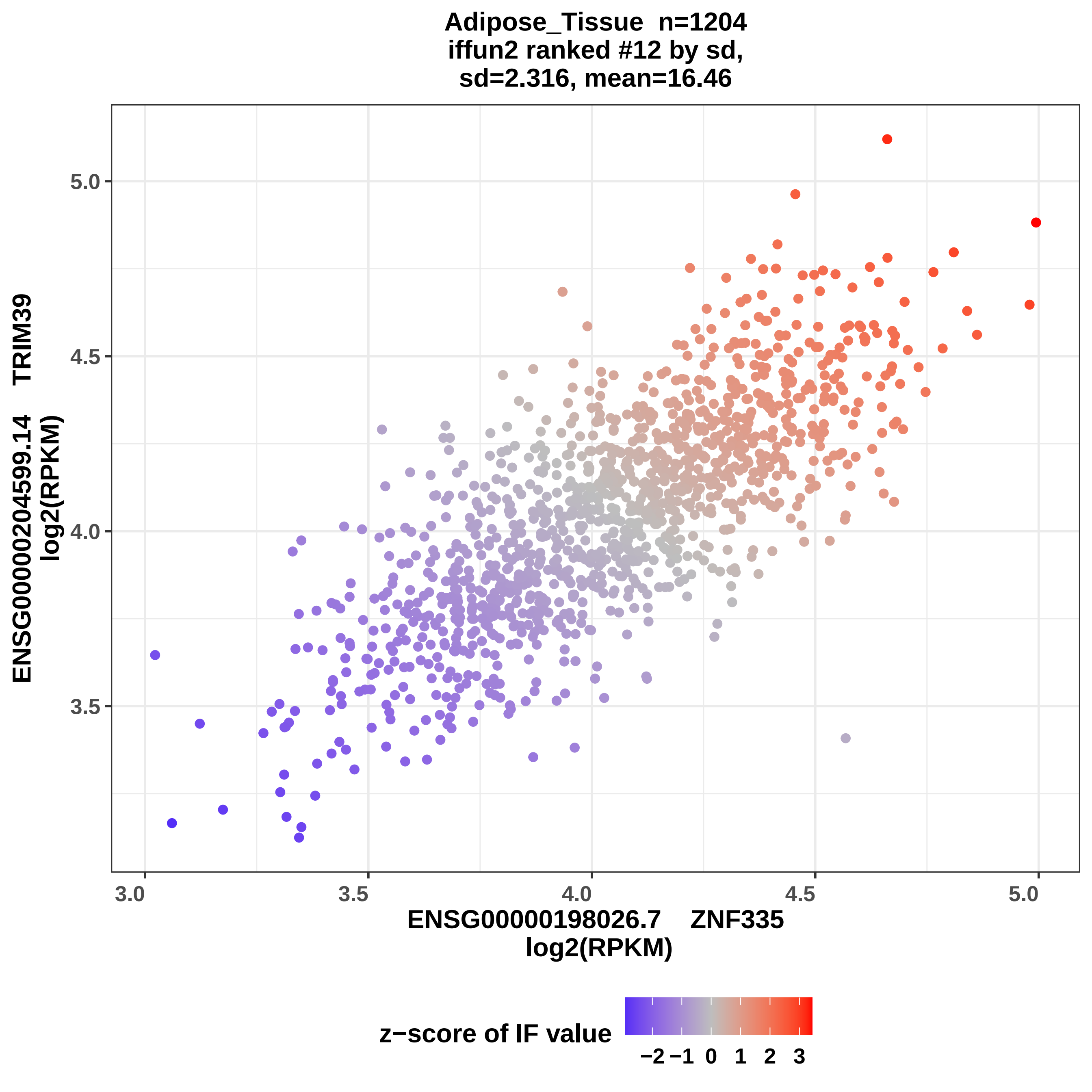
Adipose_Tissue n=1204
iffun2 ranked #12 by sd,
sd=2.316, mean=16.46
ENSG00000204599.14 TRIM39
log2(RPKM)
ENSG00000198026.7 ZNF335
log2(RPKM)
3.0
3.5
4.0
4.5
5.0
z−score of IF value
−2
−1
0
1
2
3

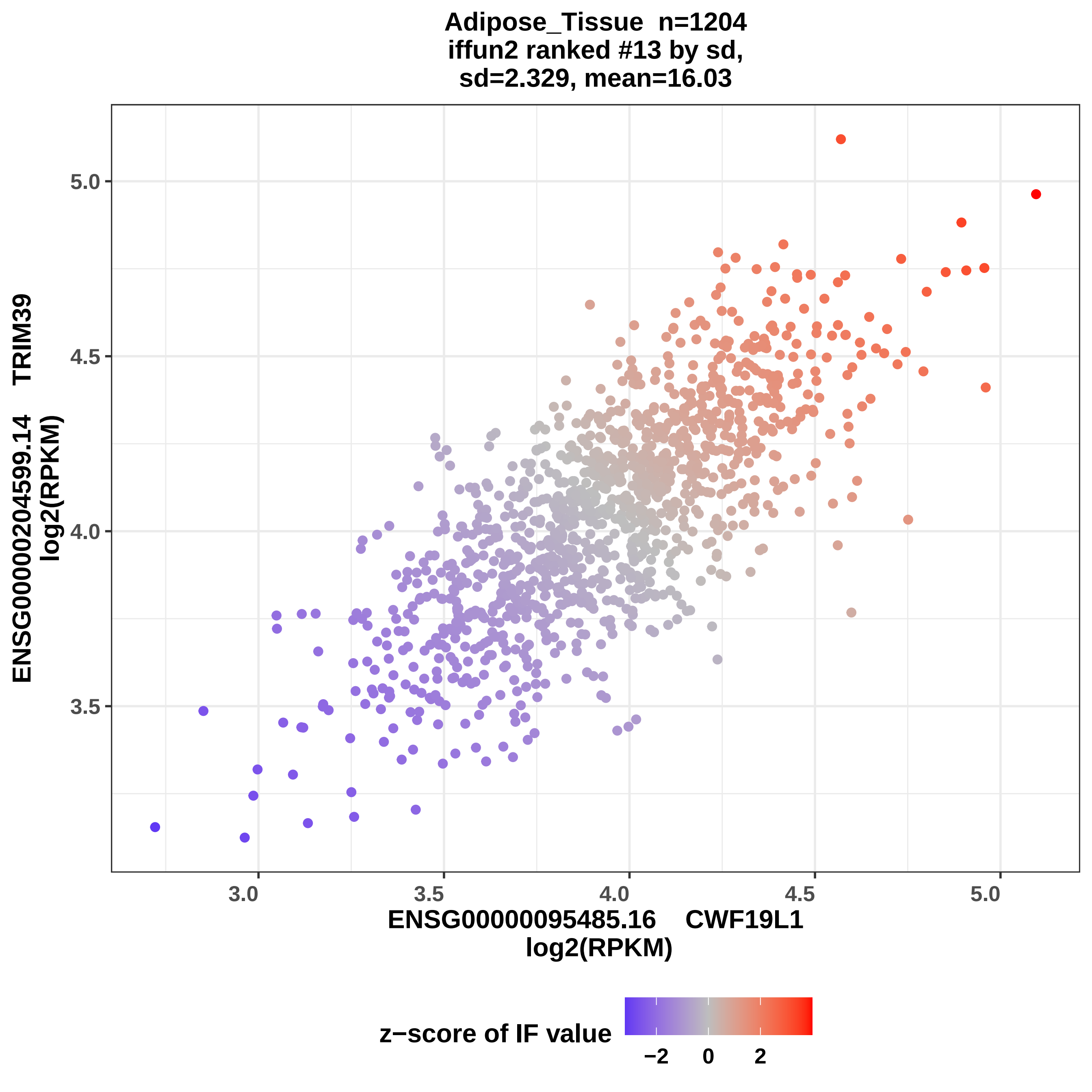
Adipose_Tissue n=1204
iffun2 ranked #13 by sd,
sd=2.329, mean=16.03
ENSG00000204599.14 TRIM39
log2(RPKM)
ENSG00000095485.16 CWF19L1
log2(RPKM)
z−score of IF value
5.0
4.5
4.0
3.5
3.0
3.5
4.0
4.5
5.0
−2
0
2

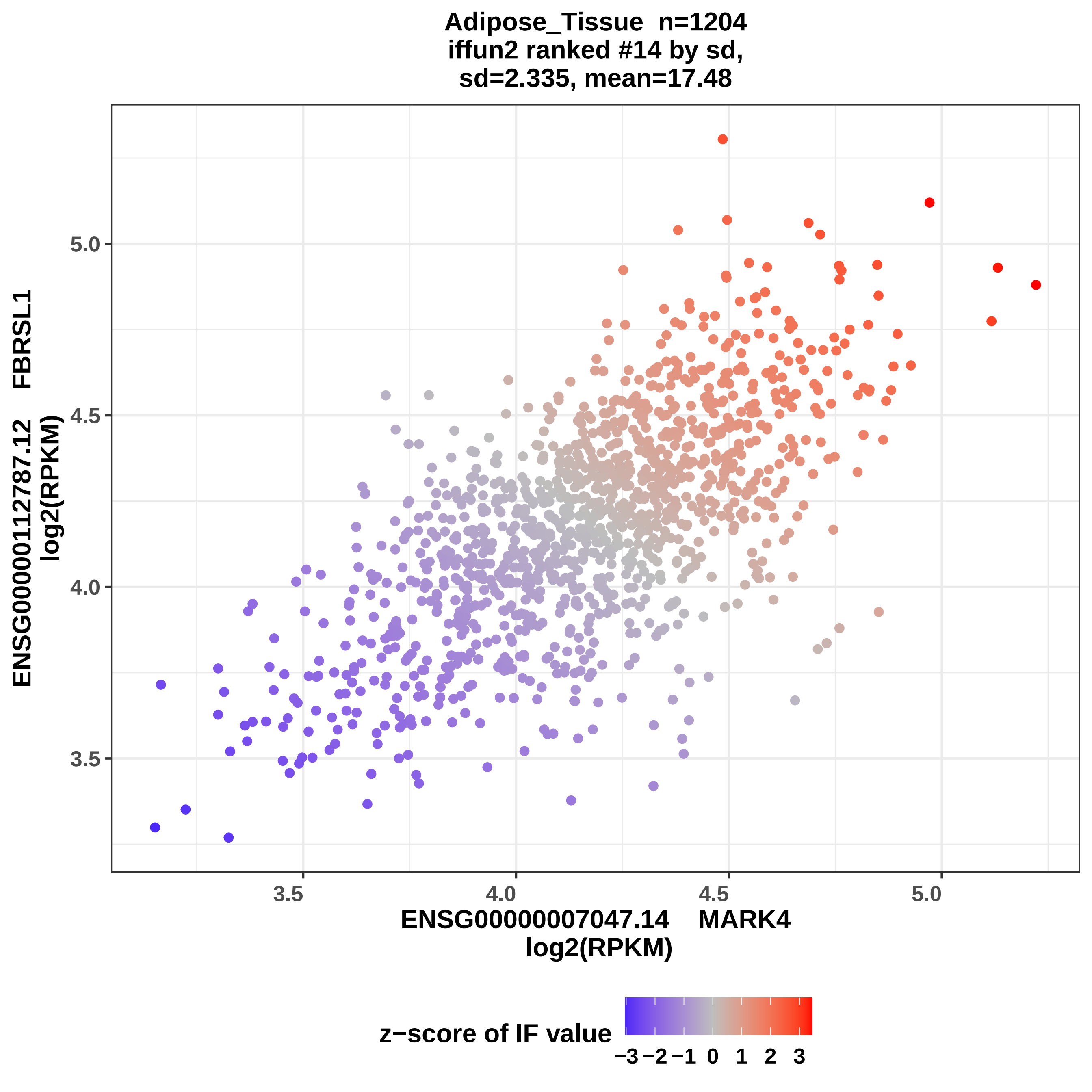

Adipose_Tissue n=1204
iffun2 ranked #14 by sd,
sd=2.335, mean=17.48
ENSG00000112787.12 FBRSL1
log2(RPKM)
5.0
4.5
4.0
3.5
3.5
4.0
4.5
5.0
ENSG00000007047.14 MARK4
log2(RPKM)
z−score of IF value
−3 −2 −1 0 1 2 3

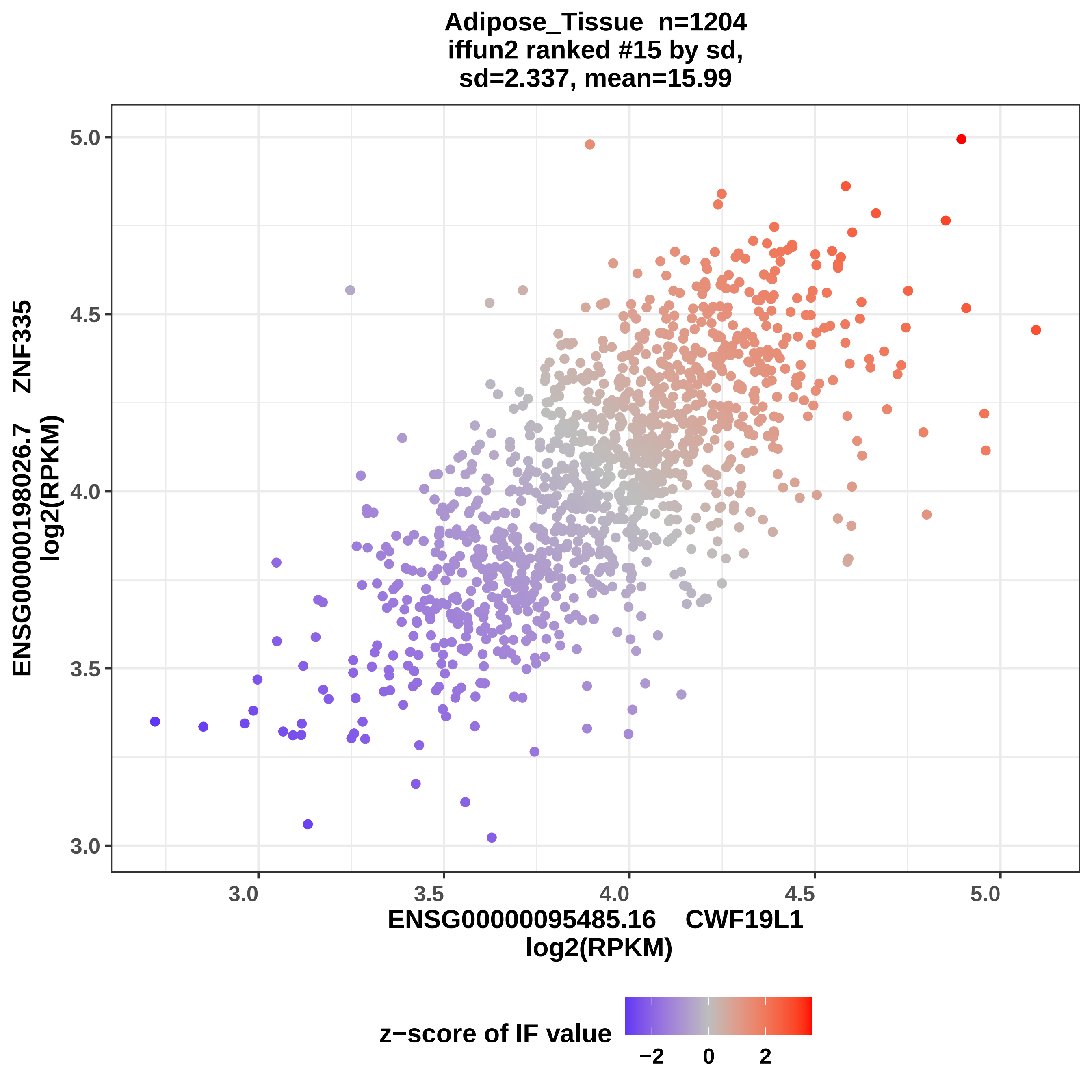
Adipose_Tissue n=1204
iffun2 ranked #15 by sd,
sd=2.337, mean=15.99
ENSG00000198026.7 ZNF335
log2(RPKM)
ENSG00000095485.16 CWF19L1
log2(RPKM)
3.0
3.5
4.0
4.5
5.0
z−score of IF value
−2
0
2

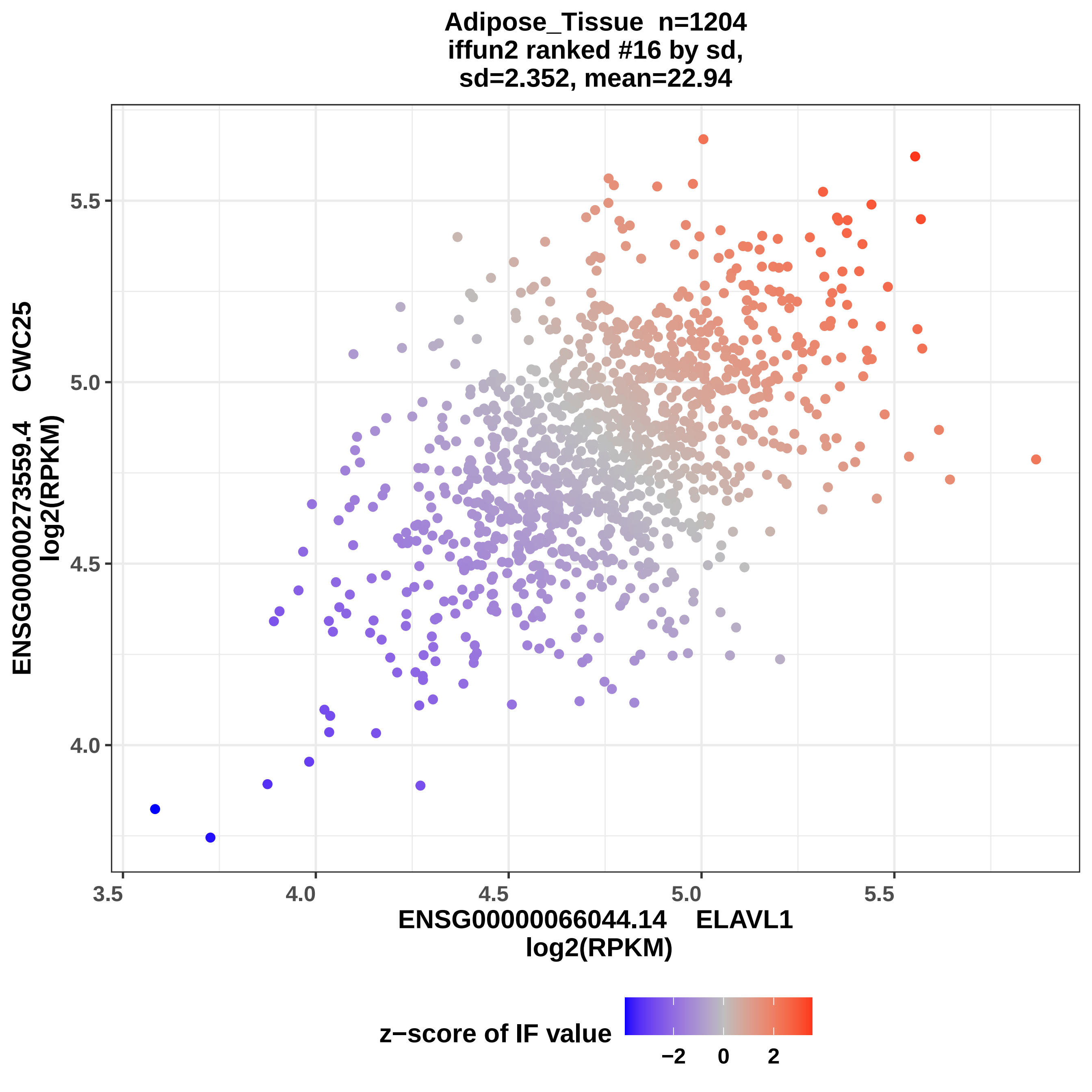

Adipose_Tissue n=1204
iffun2 ranked #16 by sd,
sd=2.352, mean=22.94
ENSG00000273559.4 CWC25
log2(RPKM)
5.5
5.0
4.5
4.0
3.5
4.0
4.5
5.0
5.5
ENSG00000066044.14 ELAVL1
log2(RPKM)
z−score of IF value
−2
0
2

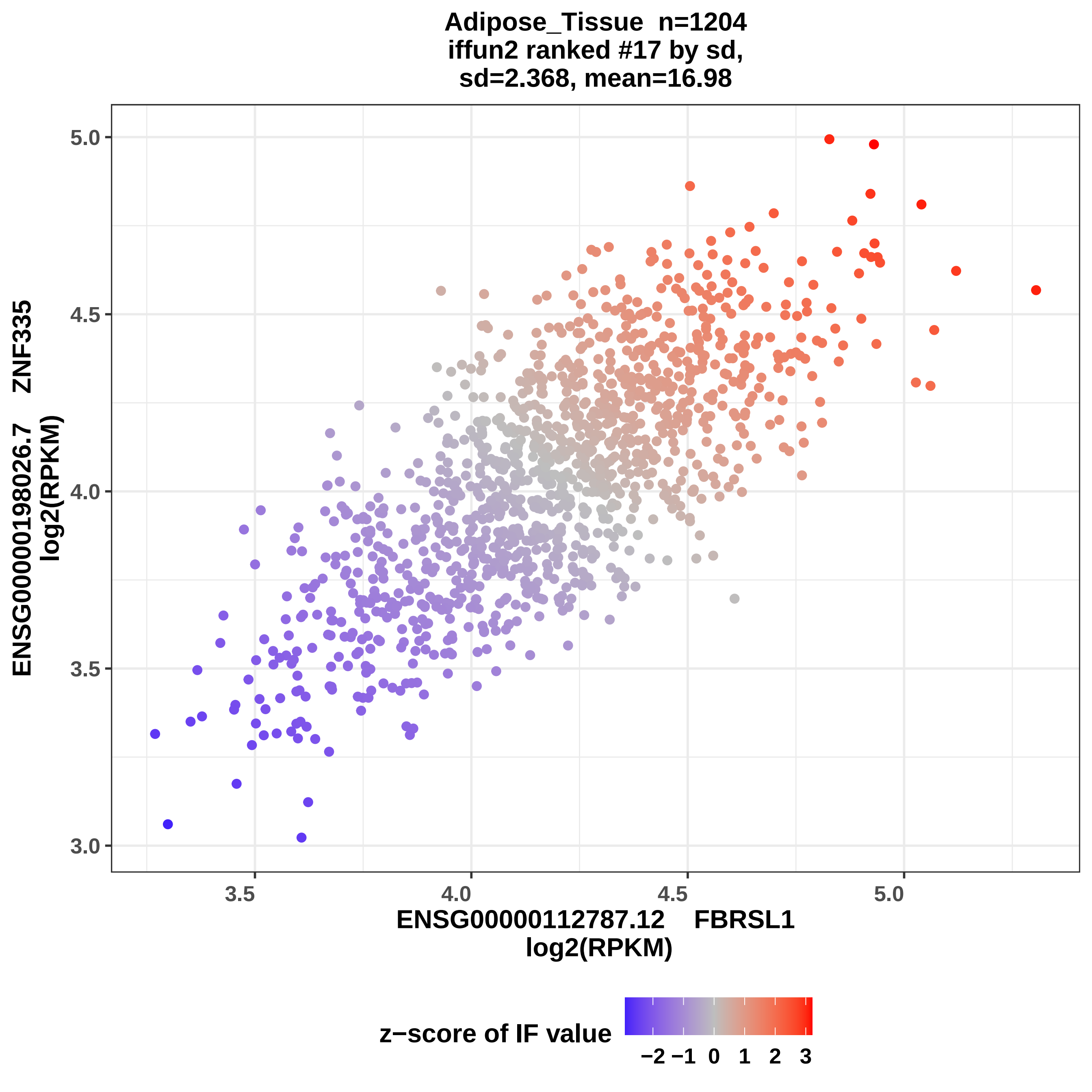

Adipose_Tissue n=1204
iffun2 ranked #17 by sd,
sd=2.368, mean=16.98
ENSG00000198026.7 ZNF335
log2(RPKM)
5.0
4.5
4.0
3.5
3.0
3.5
4.0
4.5
5.0
ENSG00000112787.12 FBRSL1
log2(RPKM)
z−score of IF value
−2 −1 0 1 2 3

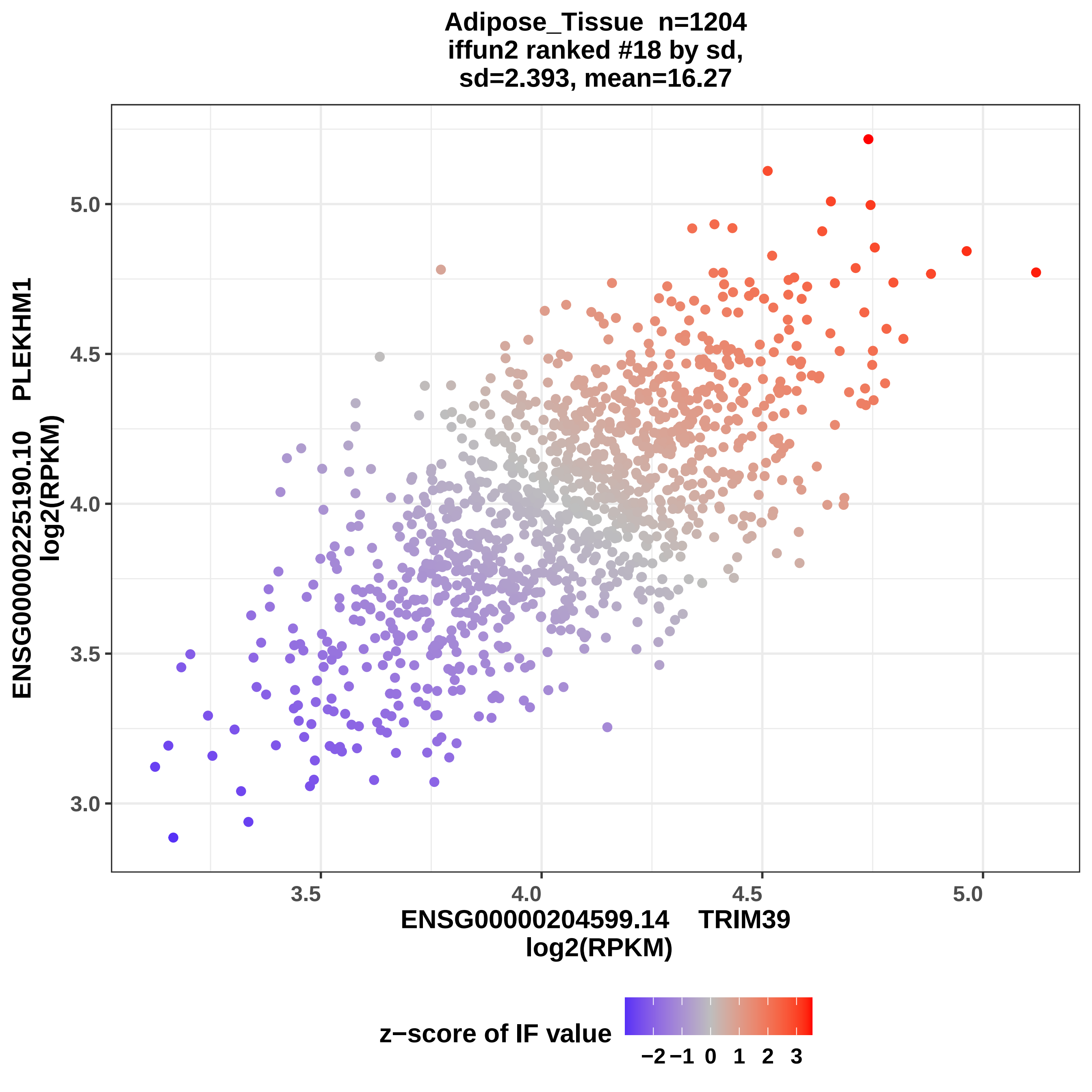
Adipose_Tissue n=1204
iffun2 ranked #18 by sd,
sd=2.393, mean=16.27
ENSG00000225190.10 PLEKHM1
log2(RPKM)
5.0
4.5
4.0
3.5
3.0
3.5
4.0
4.5
5.0
ENSG00000204599.14 TRIM39
log2(RPKM)
z−score of IF value
−2 −1 0 1 2 3

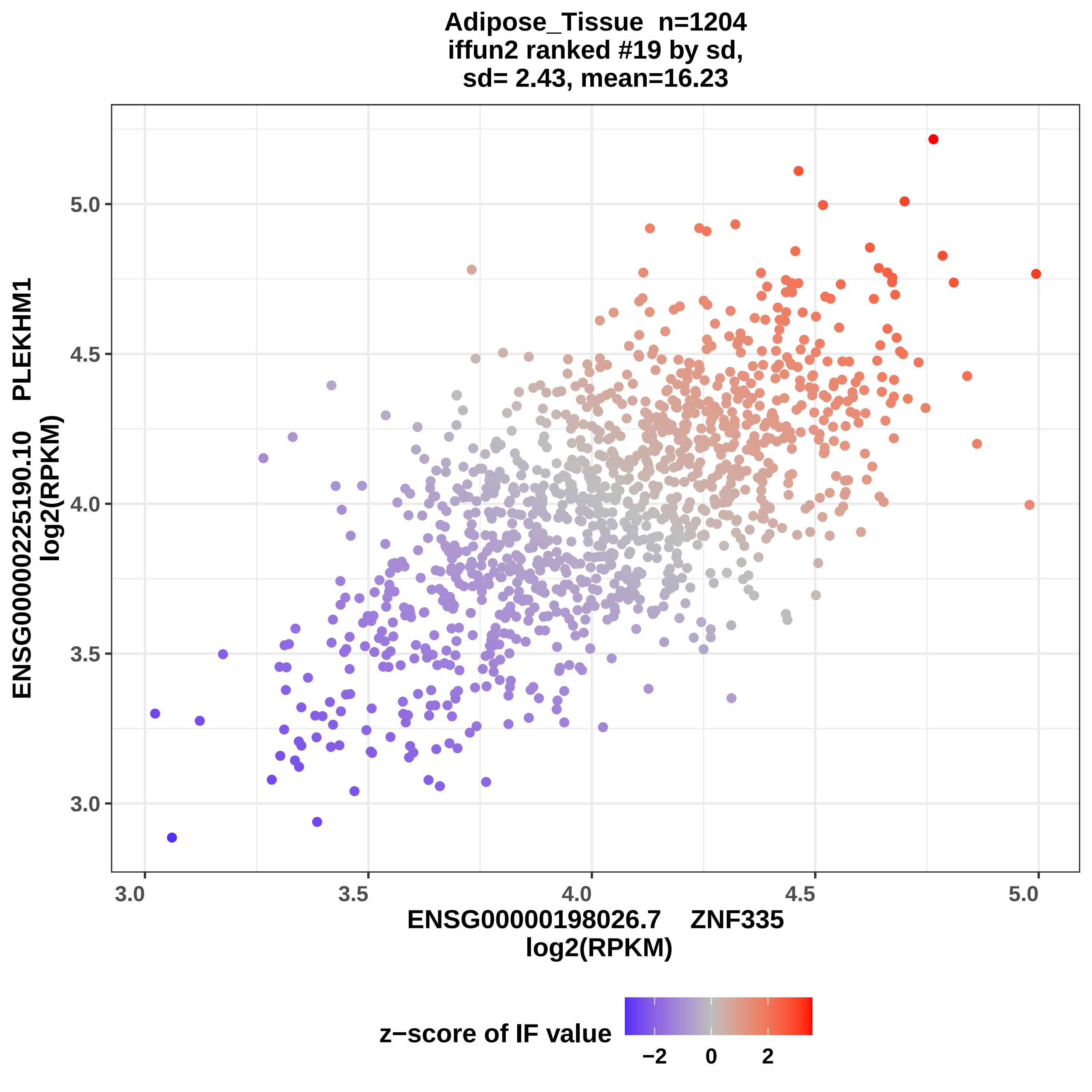
Adipose_Tissue n=1204
iffun2 ranked #19 by sd,
sd= 2.43, mean=16.23
ENSG00000225190.10 PLEKHM1
log2(RPKM)
5.0
4.5
4.0
3.5
3.0
3.0
3.5
4.0
4.5
5.0
ENSG00000198026.7 ZNF335
log2(RPKM)
z−score of IF value
−2
0
2

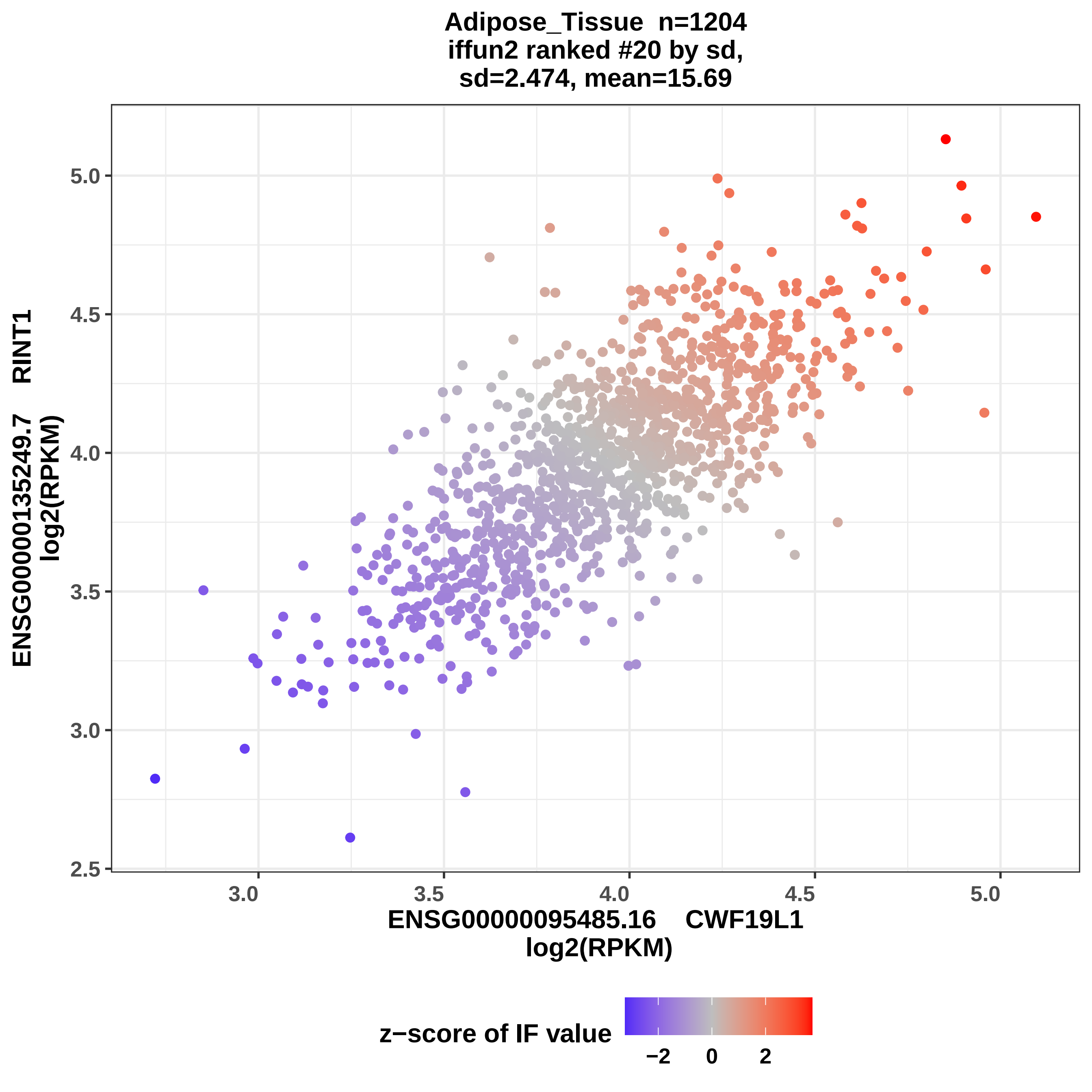
Adipose_Tissue n=1204
iffun2 ranked #20 by sd,
sd=2.474, mean=15.69
ENSG00000135249.7 RINT1
log2(RPKM)
5.0
4.5
4.0
3.5
3.0
2.5
3.0
3.5
4.0
4.5
5.0
ENSG00000095485.16 CWF19L1
log2(RPKM)
z−score of IF value
−2
0
2

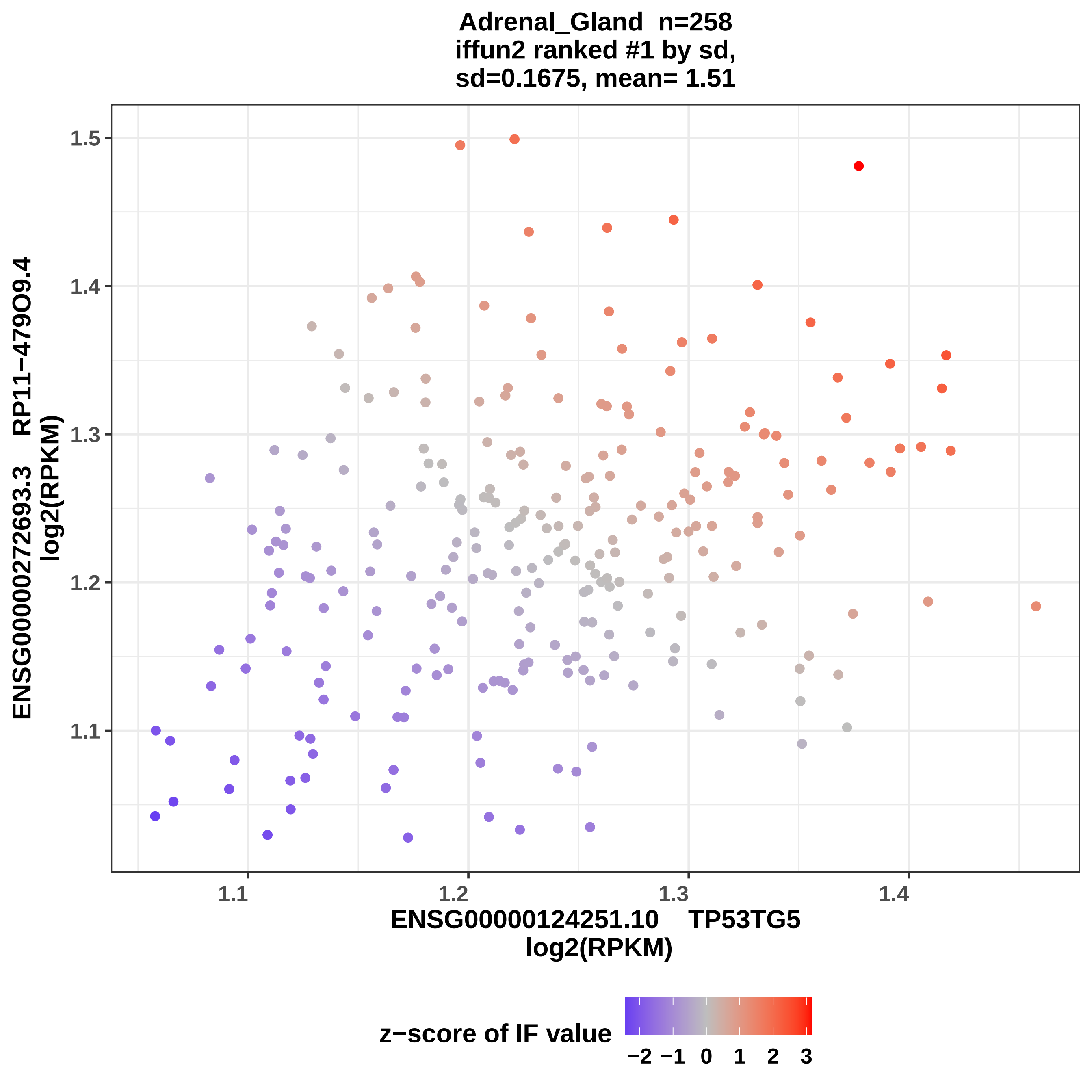

Adrenal_Gland n=258
iffun2 ranked #1 by sd,
sd=0.1675, mean= 1.51
ENSG00000272693.3 RP11−479O9.4
log2(RPKM)
1.5
1.4
1.3
1.2
1.1
1.1
1.2
1.3
1.4
ENSG00000124251.10 TP53TG5
log2(RPKM)
z−score of IF value
−2
−1
0
1
2
3

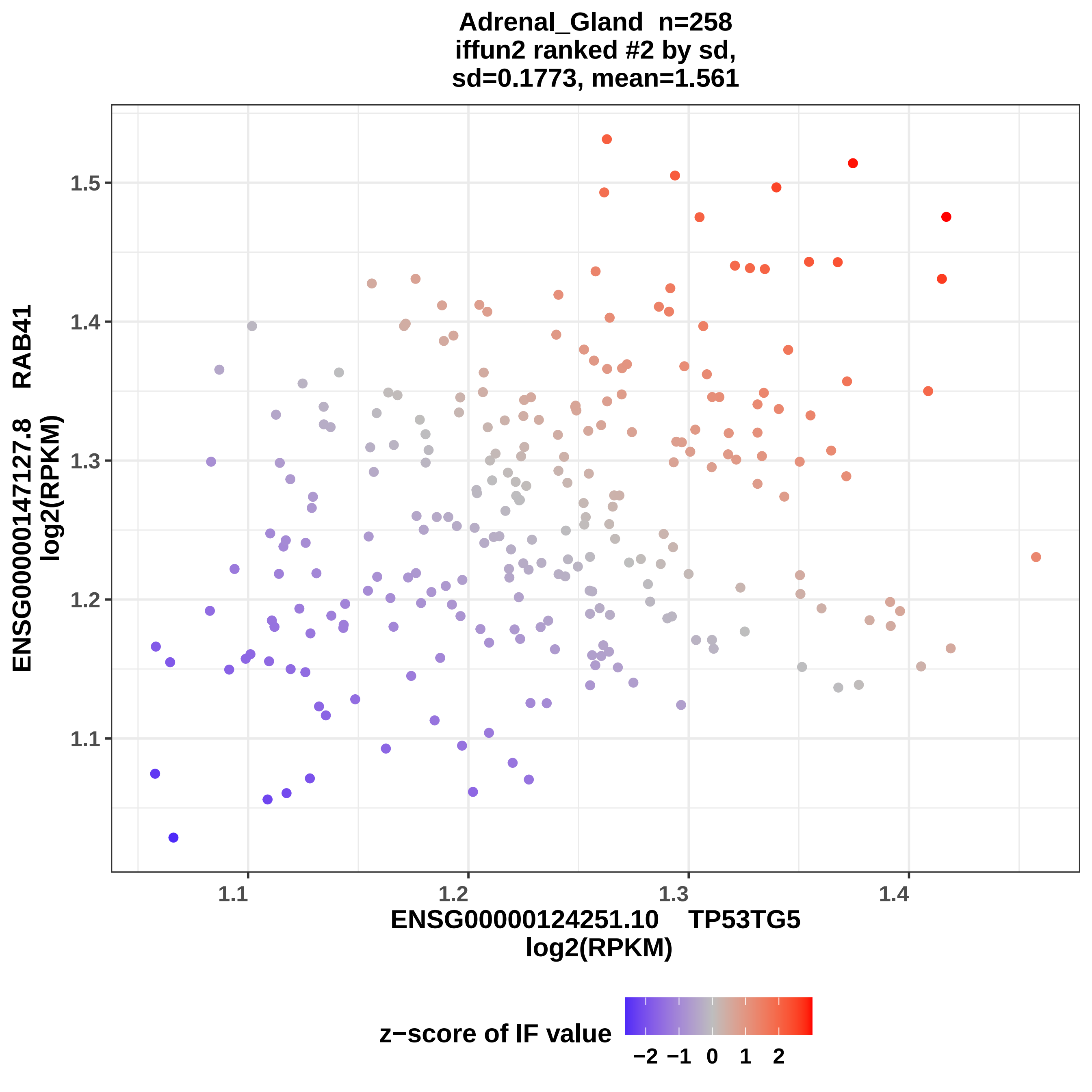

Adrenal_Gland n=258
iffun2 ranked #2 by sd,
sd=0.1773, mean=1.561
1.5
1.4
1.3
1.2
1.1
ENSG00000147127.8 RAB41
log2(RPKM)
1.1
1.2
1.3
1.4
ENSG00000124251.10 TP53TG5
log2(RPKM)
z−score of IF value
−2 −1 0 1 2

### Adrenal_Gland n=258
### iffun2 ranked #3 by sd,
### sd=0.1872, mean=1.608

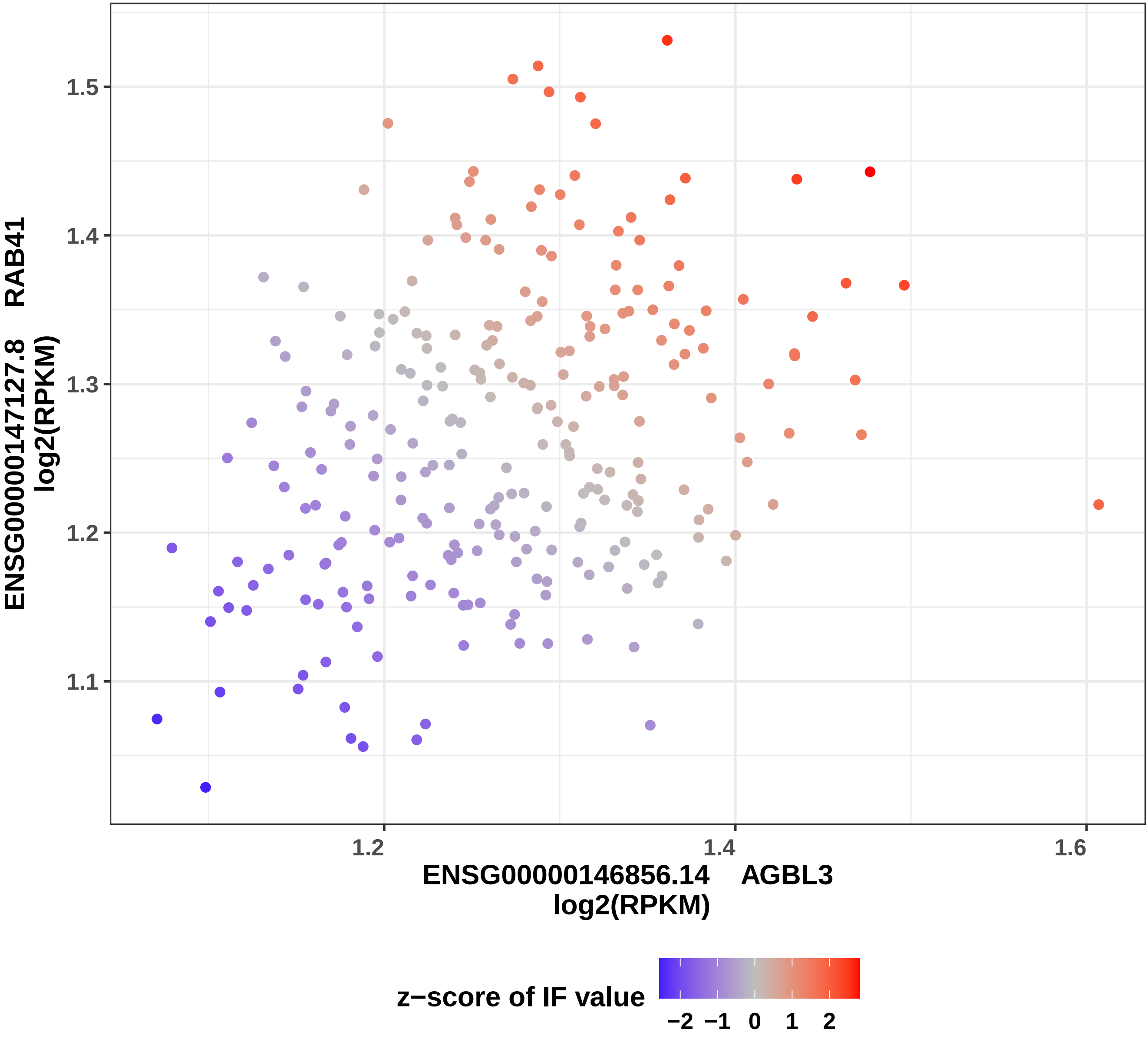

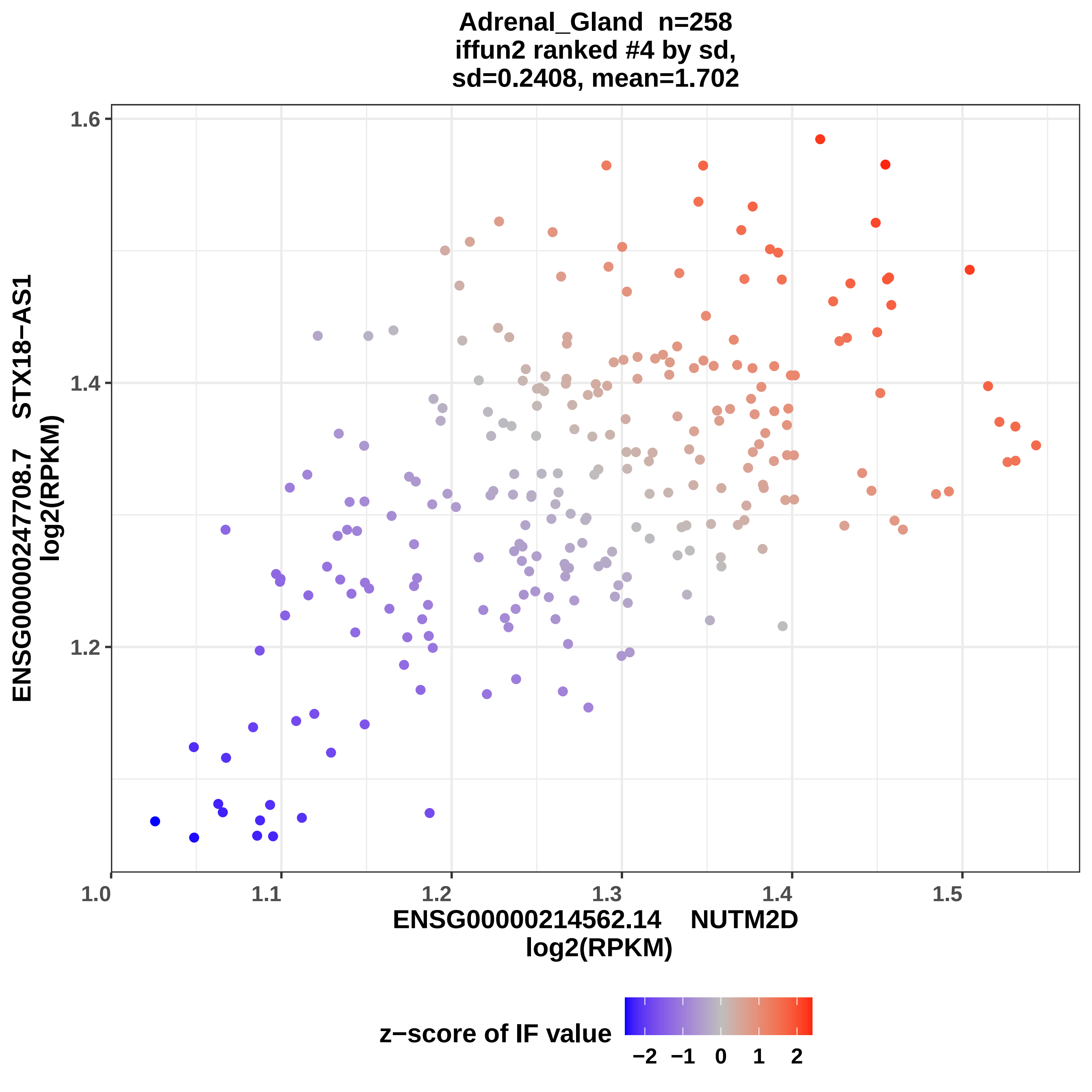

Adrenal_Gland n=258
iffun2 ranked #4 by sd,
sd=0.2408, mean=1.702
ENSG00000247708.7 STX18−AS1
log2(RPKM)
1.6
1.4
1.2
1.0
1.1
1.2
1.3
1.4
1.5
ENSG00000214562.14 NUTM2D
log2(RPKM)
z−score of IF value
−2
−1
0
1
2

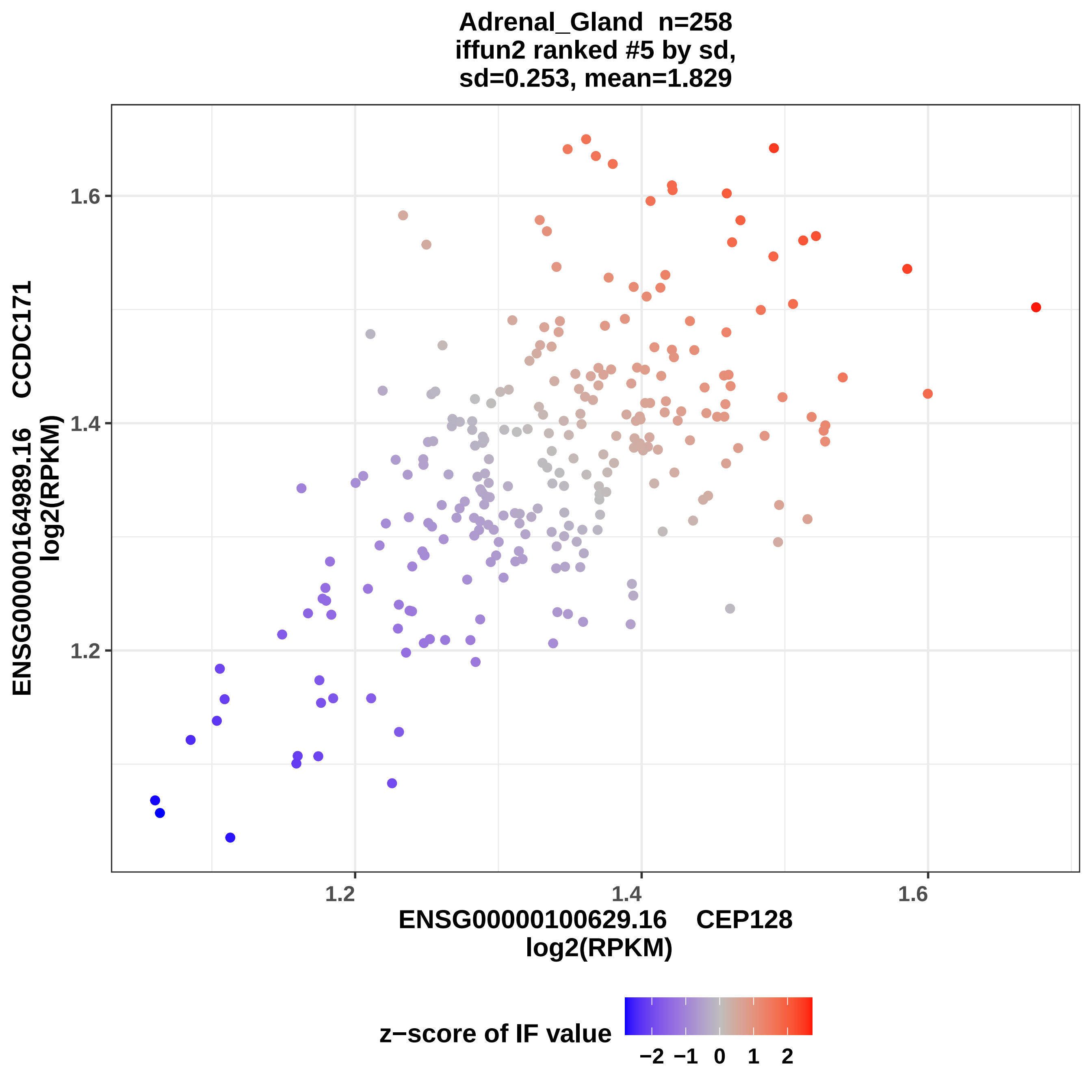
Adrenal_Gland n=258
iffun2 ranked #5 by sd,
sd=0.253, mean=1.829
ENSG00000164989.16 CCDC171
log2(RPKM)
1.6
1.4
1.2
1.2
1.4
1.6
ENSG00000100629.16 CEP128
log2(RPKM)
z−score of IF value
−2 −1 0 1 2

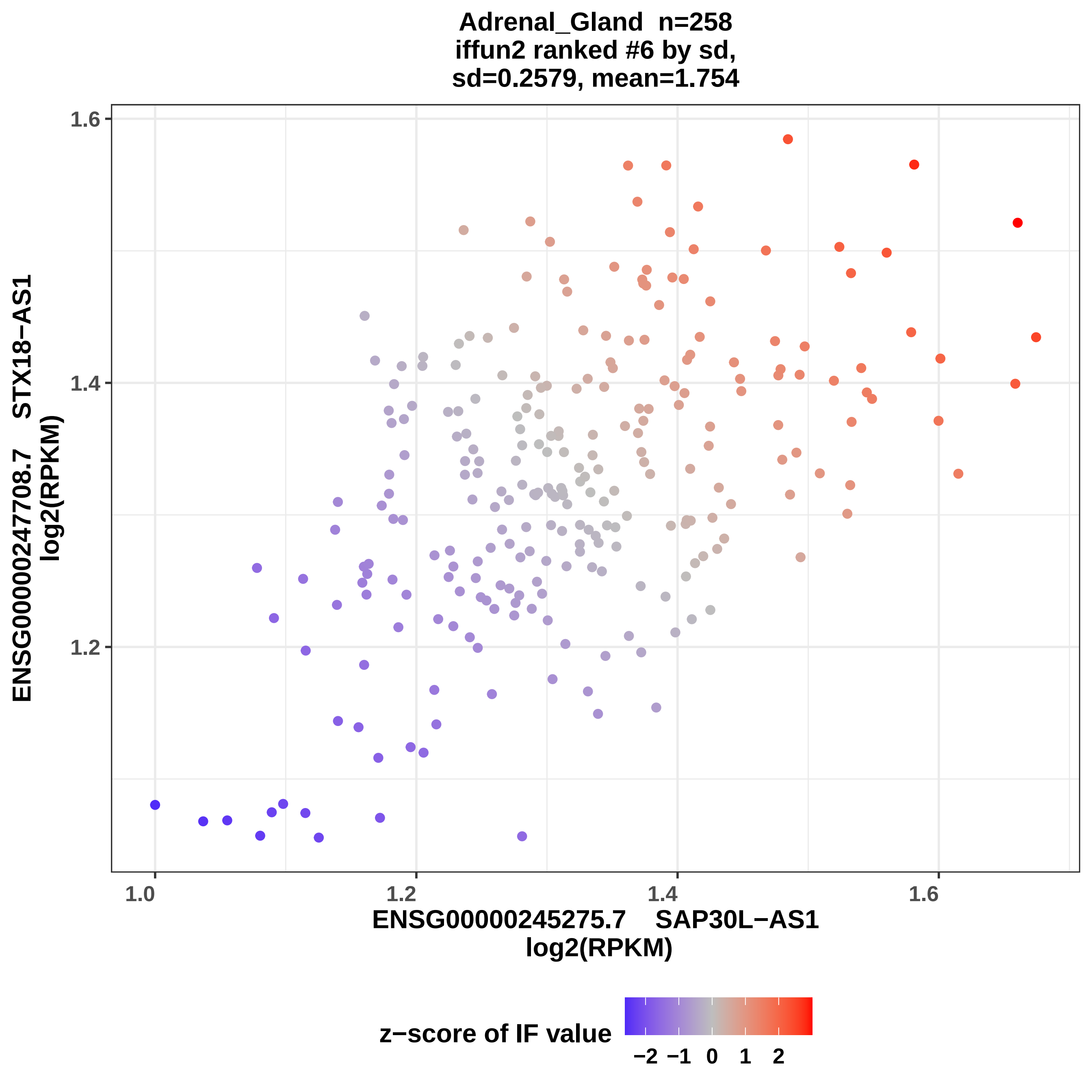

Adrenal_Gland n=258
iffun2 ranked #6 by sd,
sd=0.2579, mean=1.754
1.6
1.4
1.2
ENSG00000247708.7 STX18−AS1
log2(RPKM)
1.0
1.2
1.4
1.6
ENSG00000245275.7 SAP30L−AS1
log2(RPKM)
z−score of IF value
−2 −1 0 1 2

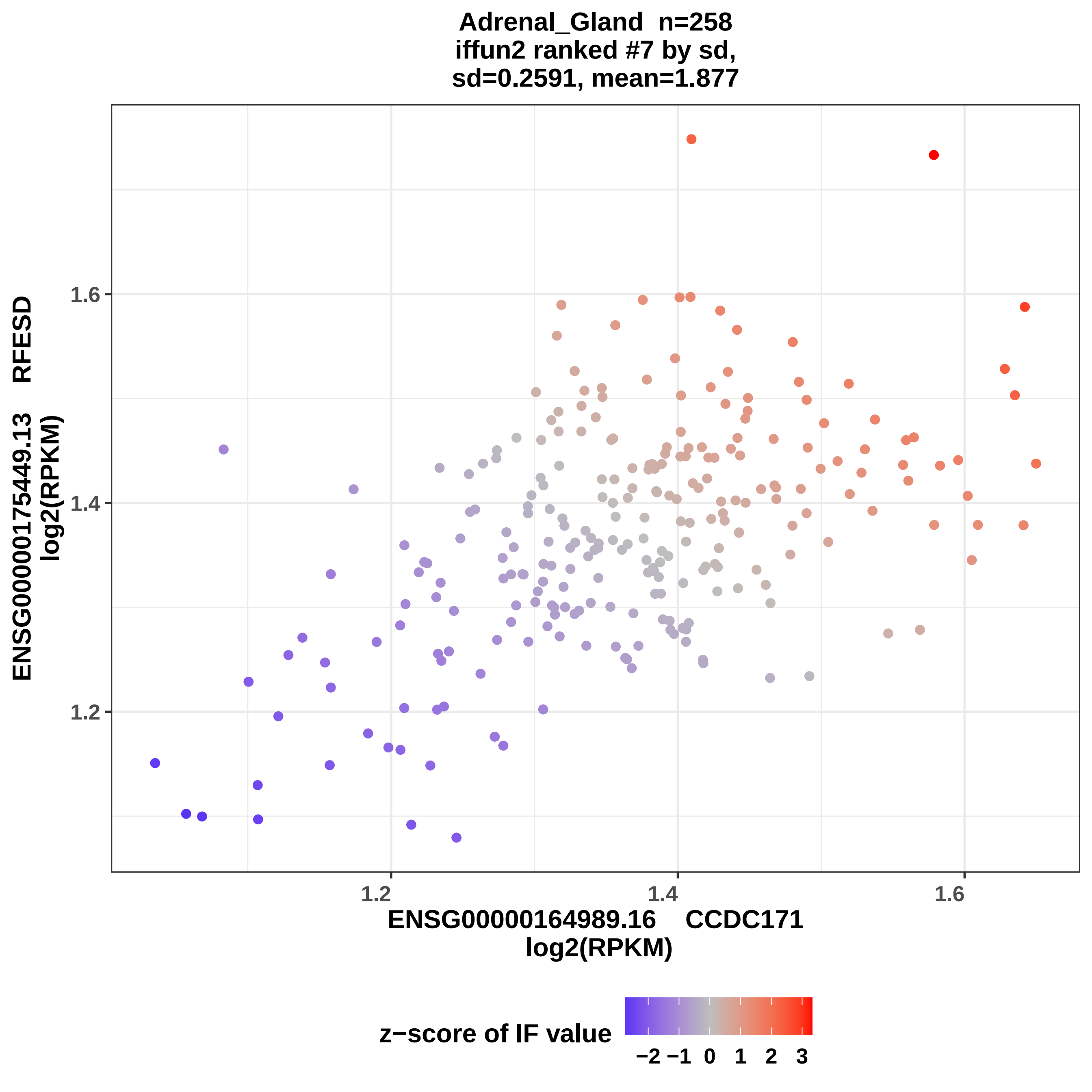
Adrenal_Gland n=258
iffun2 ranked #7 by sd,
sd=0.2591, mean=1.877
ENSG00000175449.13 RFESD
log2(RPKM)
1.6
1.4
1.2
1.2
1.4
1.6
ENSG00000164989.16 CCDC171
log2(RPKM)
z−score of IF value
−2
−1
0
1
2
3

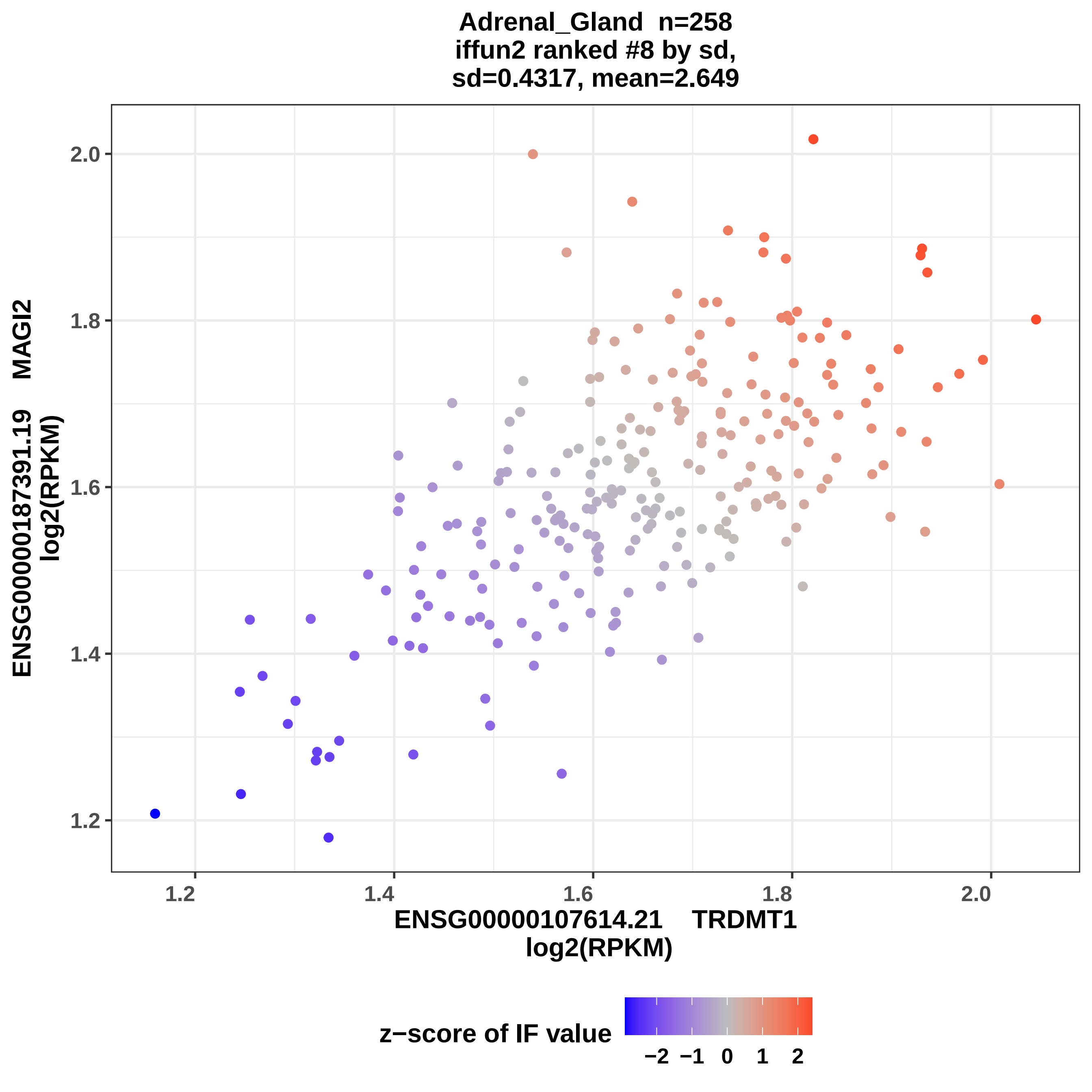

Adrenal_Gland n=258
iffun2 ranked #8 by sd,
sd=0.4317, mean=2.649
ENSG00000187391.19 MAGI2
log2(RPKM)
ENSG00000107614.21 TRDMT1
log2(RPKM)
1.2
1.4
1.6
1.8
2.0
z−score of IF value
−2
−1
0
1
2

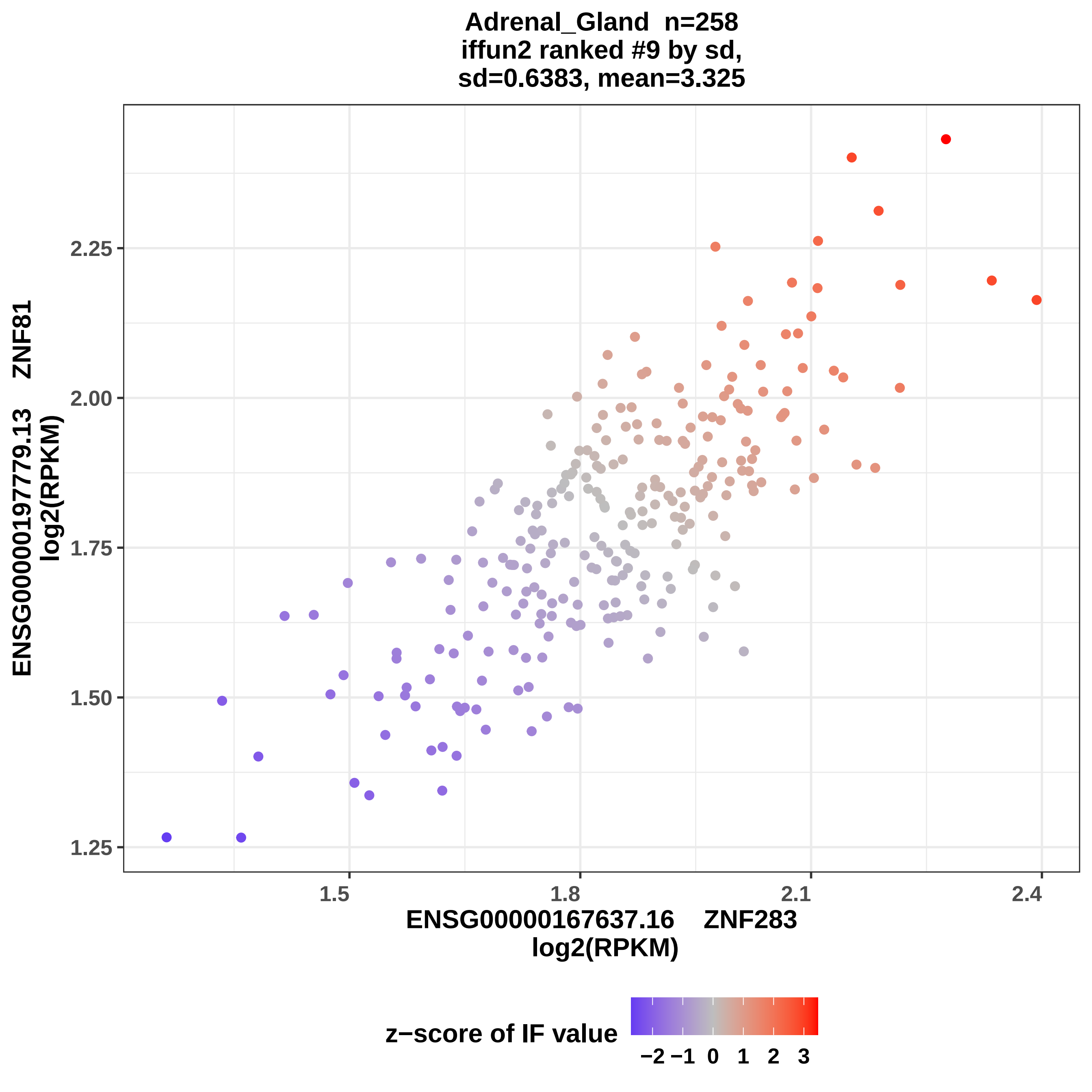
Adrenal_Gland n=258
iffun2 ranked #9 by sd,
sd=0.6383, mean=3.325
ENSG00000197779.13 ZNF81
log2(RPKM)
2.25
2.00
1.75
1.50
1.25
1.5
1.8
2.1
2.4
ENSG00000167637.16 ZNF283
log2(RPKM)
z−score of IF value
−2 −1 0 1 2 3

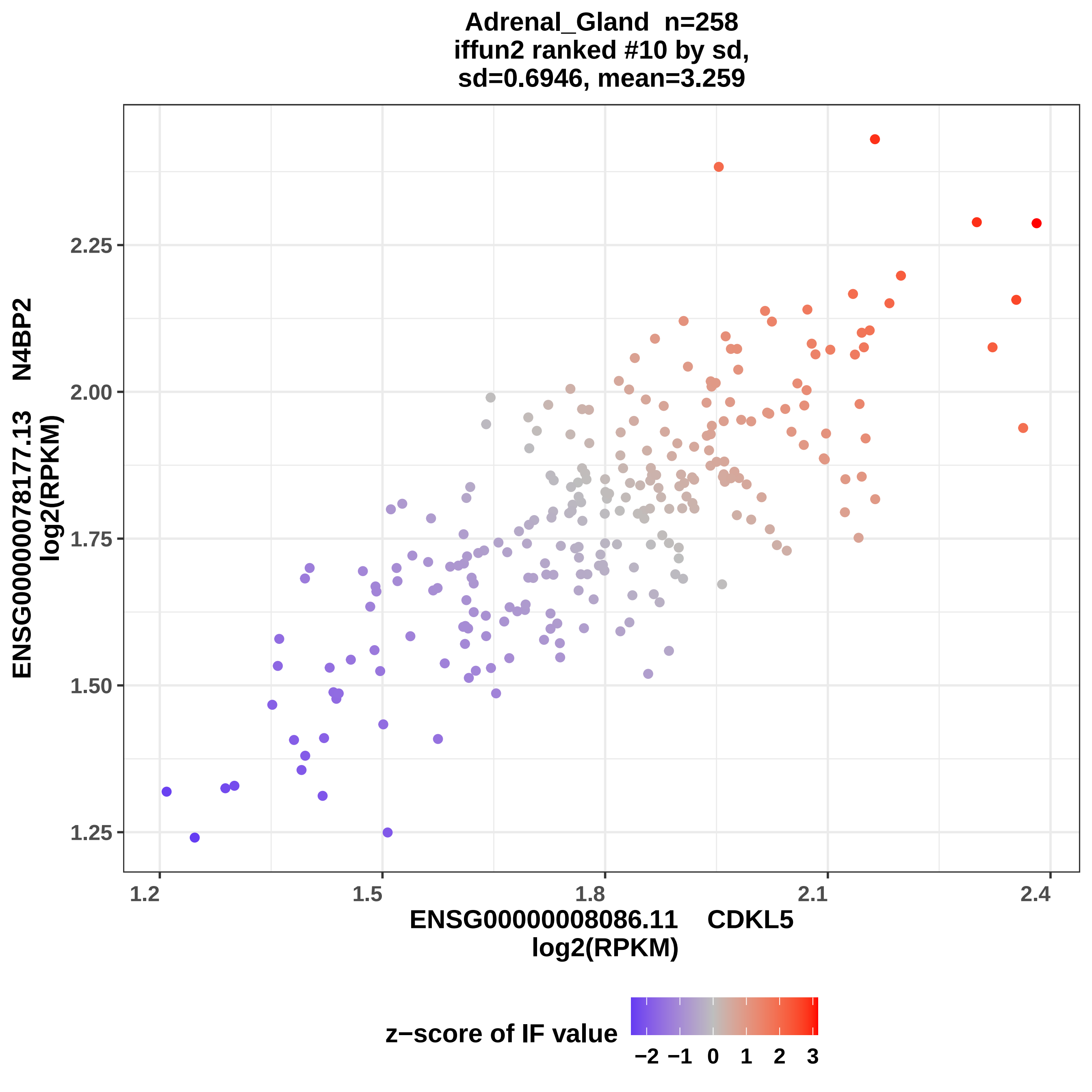

Adrenal_Gland n=258
iffun2 ranked #10 by sd,
sd=0.6946, mean=3.259
ENSG00000078177.13 N4BP2
log2(RPKM)
2.25
2.00
1.75
1.50
1.25
1.2
1.5
1.8
2.1
2.4
ENSG00000008086.11 CDKL5
log2(RPKM)
z−score of IF value
−2
−1
0
1
2
3

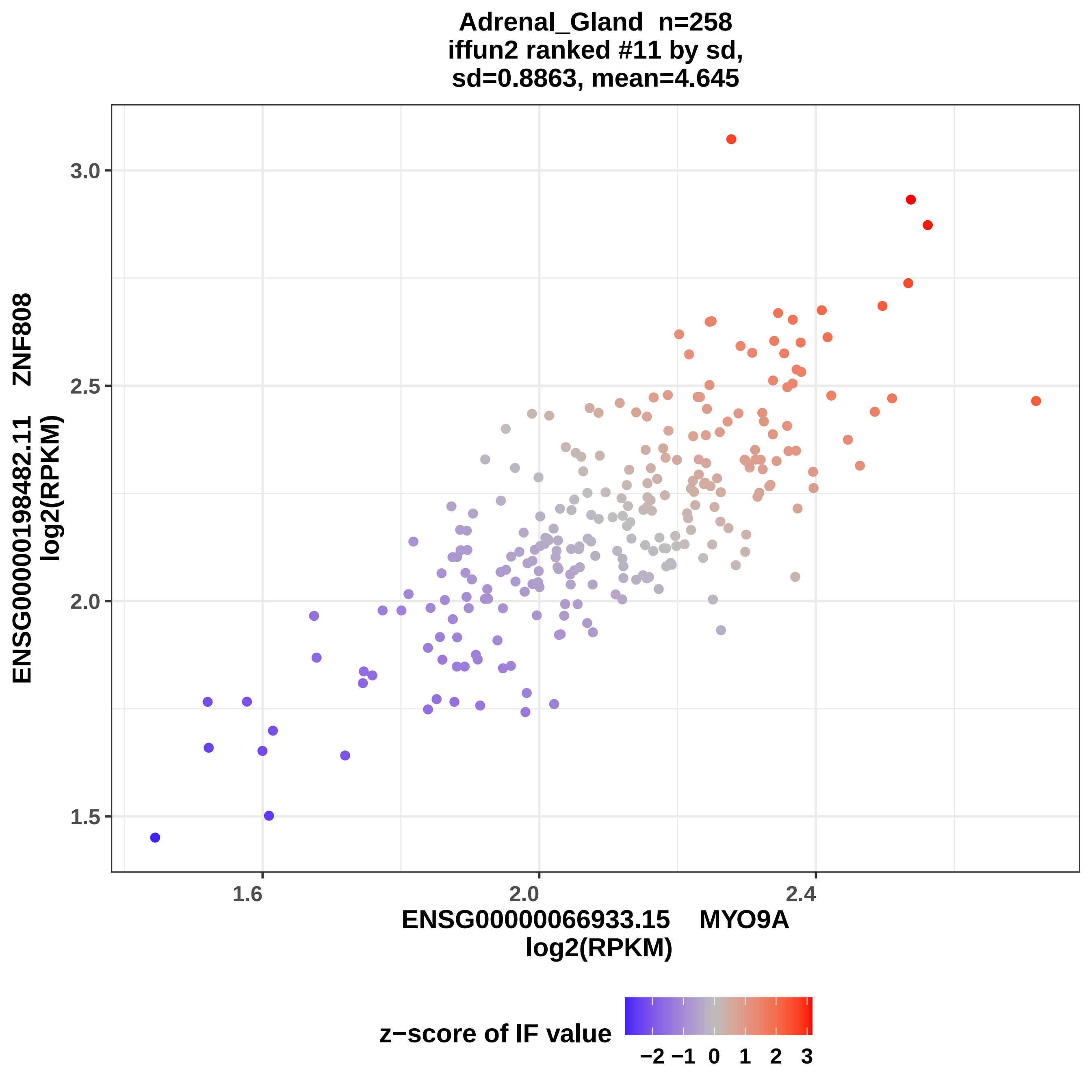

Adrenal_Gland n=258
iffun2 ranked #11 by sd,
sd=0.8863, mean=4.645
ENSG00000198482.11 ZNF808
log2(RPKM)
3.0
2.5
2.0
1.5
1.6
2.0
2.4
ENSG00000066933.15 MYO9A
log2(RPKM)
z−score of IF value
−2 −1 0 1 2 3

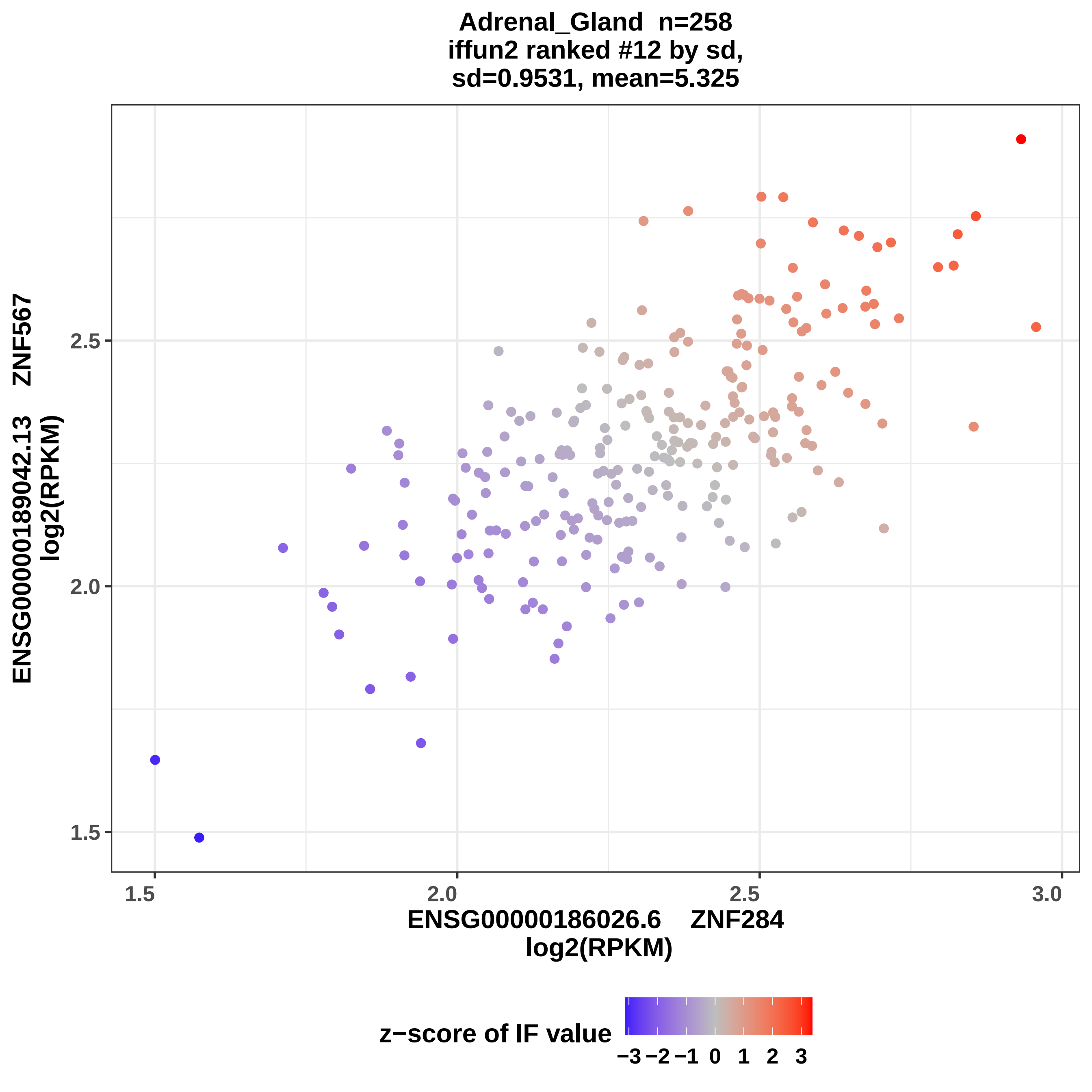
Adrenal_Gland n=258
iffun2 ranked #12 by sd,
sd=0.9531, mean=5.325
ENSG00000189042.13 ZNF567
log2(RPKM)
1.5
2.0
2.5
ENSG00000186026.6 ZNF284
log2(RPKM)
1.5
2.0
2.5
3.0
z−score of IF value
−3 −2 −1 0 1 2 3

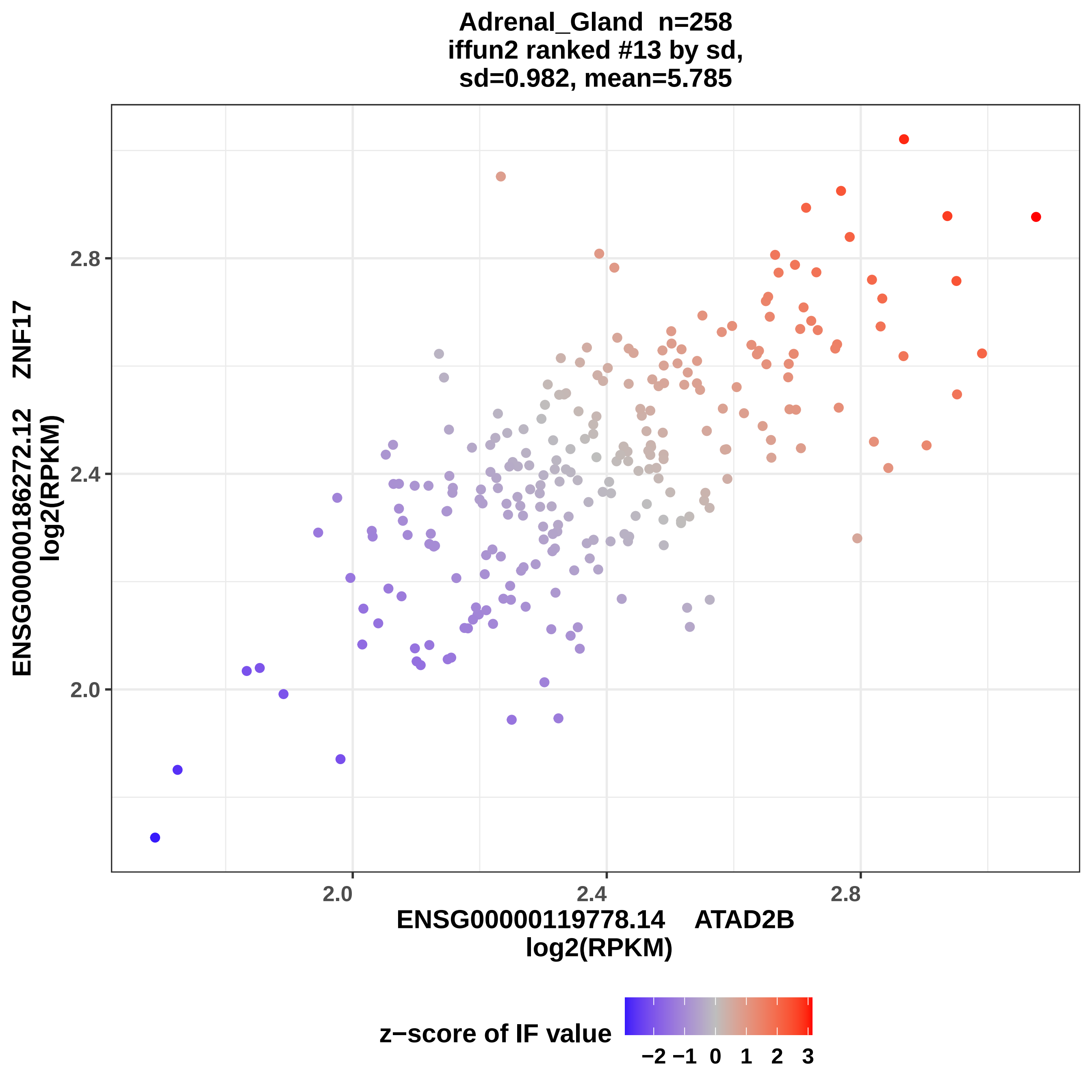

Adrenal_Gland n=258
iffun2 ranked #13 by sd,
sd=0.982, mean=5.785
ZNF17
ENSG00000186272.12
log2(RPKM)
2.8
2.4
2.0
2.0
2.4
2.8
ENSG00000119778.14 ATAD2B
log2(RPKM)
z−score of IF value
−2 −1 0 1 2 3

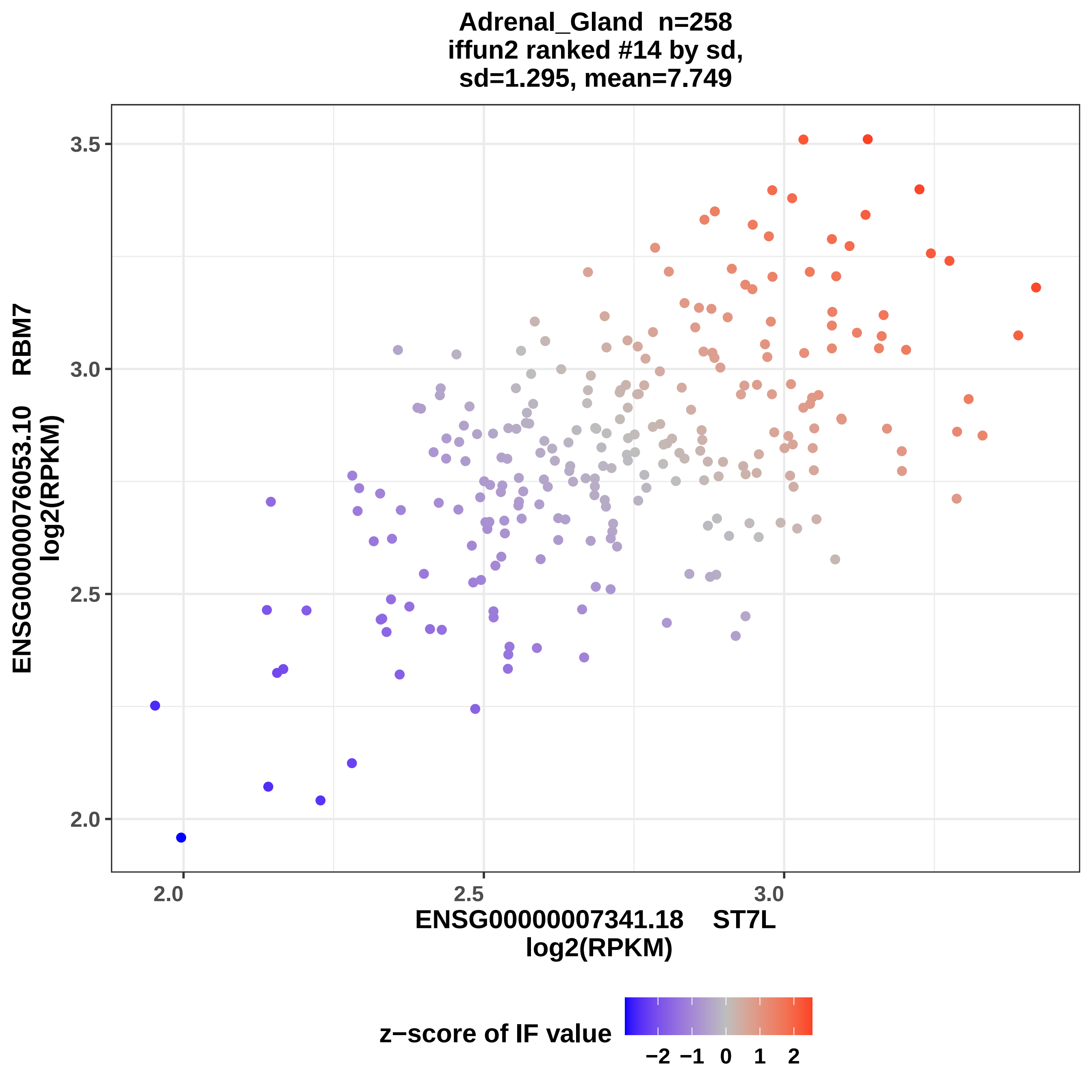
Adrenal_Gland n=258
iffun2 ranked #14 by sd,
sd=1.295, mean=7.749
ENSG00000076053.10 RBM7
log2(RPKM)
3.5
3.0
2.5
2.0
2.0
2.5
3.0
ENSG00000007341.18 ST7L
log2(RPKM)
z−score of IF value
−2 −1 0 1 2

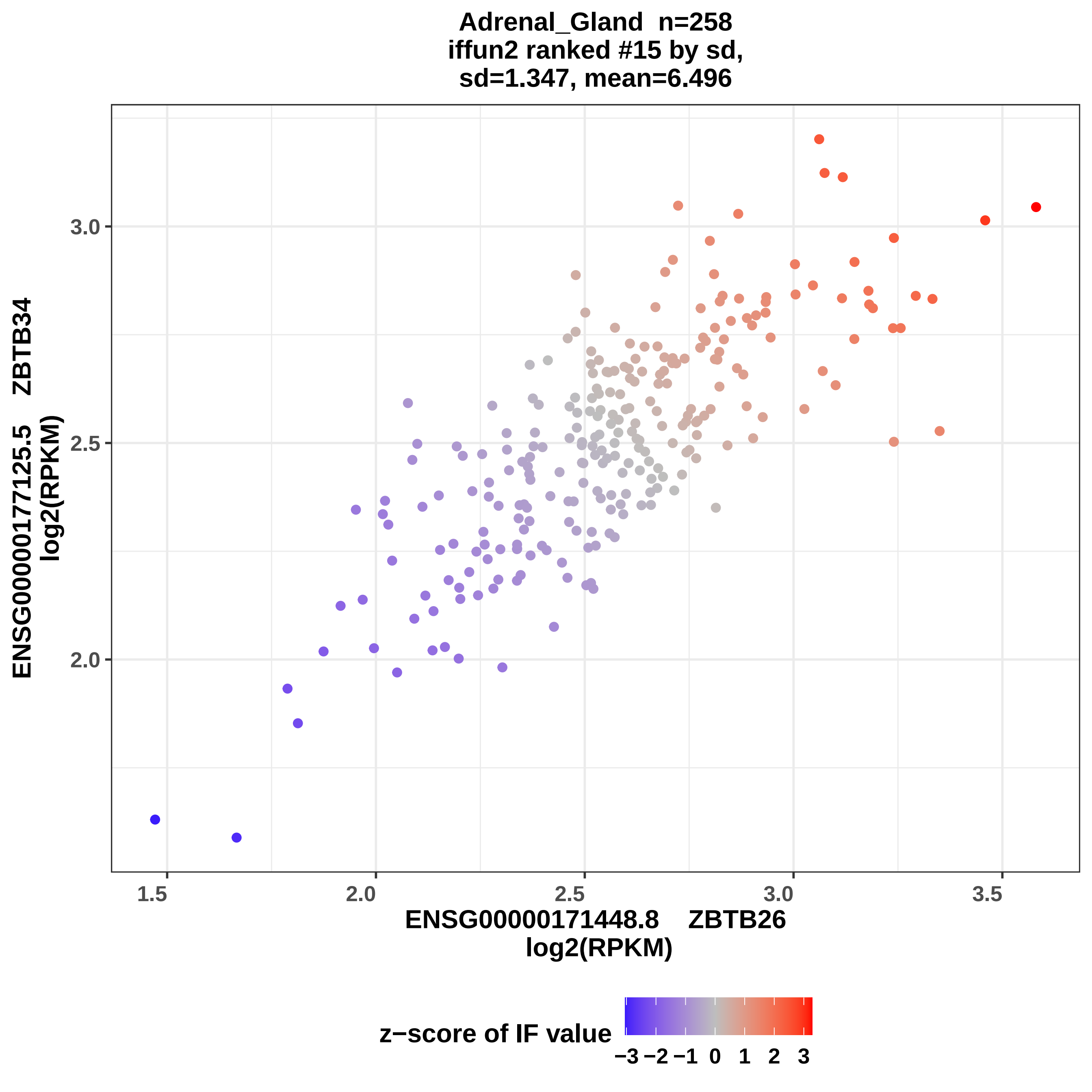
Adrenal_Gland n=258
iffun2 ranked #15 by sd,
sd=1.347, mean=6.496
ENSG00000177125.5 ZBTB34
log2(RPKM)
3.0
2.5
2.0
1.5
2.0
2.5
3.0
3.5
ENSG00000171448.8 ZBTB26
log2(RPKM)
z−score of IF value
−3 −2 −1 0 1 2 3

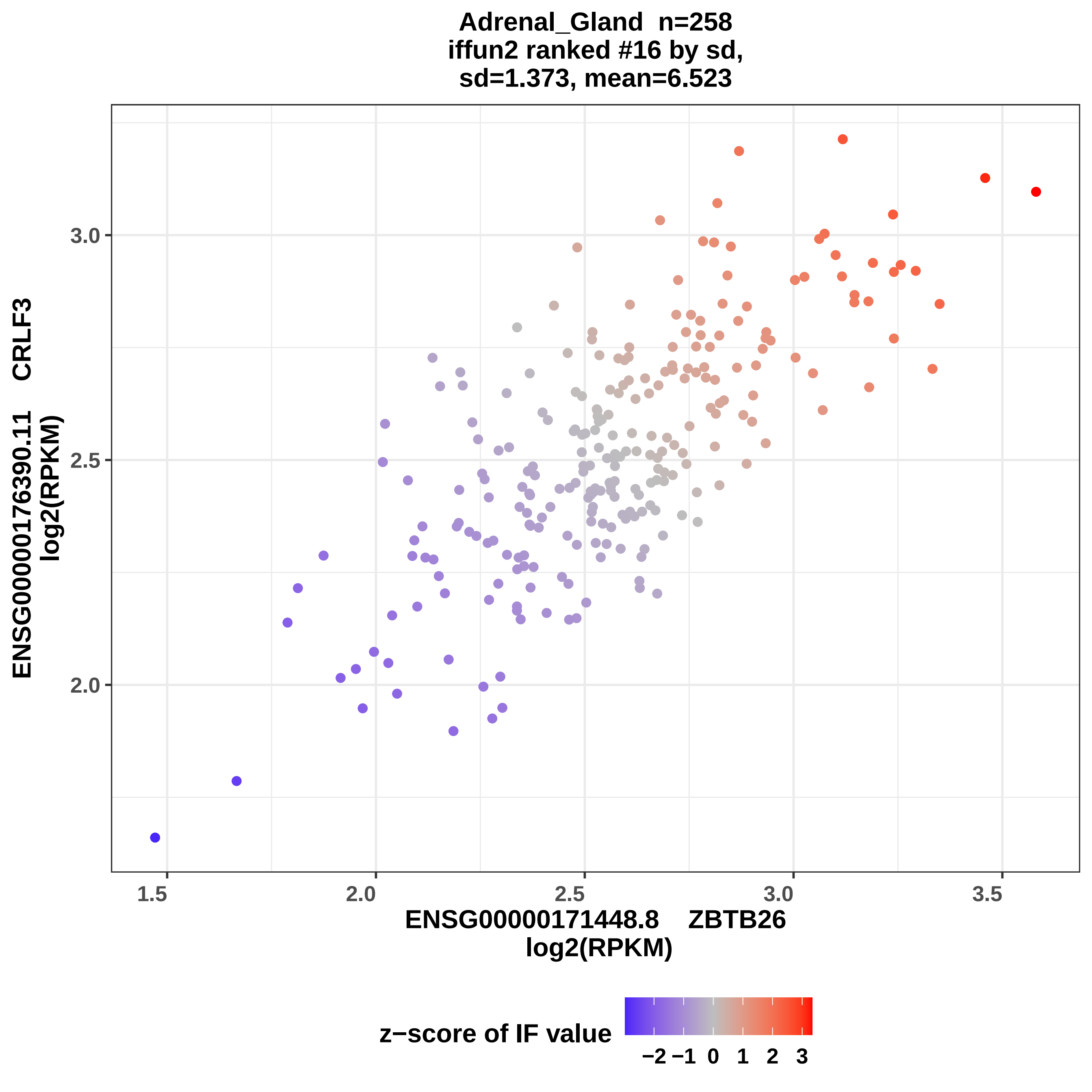
Adrenal_Gland n=258
iffun2 ranked #16 by sd,
sd=1.373, mean=6.523
ENSG00000176390.11 CRLF3
log2(RPKM)
3.0
2.5
2.0
1.5
2.0
2.5
3.0
3.5
ENSG00000171448.8 ZBTB26
log2(RPKM)
z−score of IF value
−2 −1 0 1 2 3

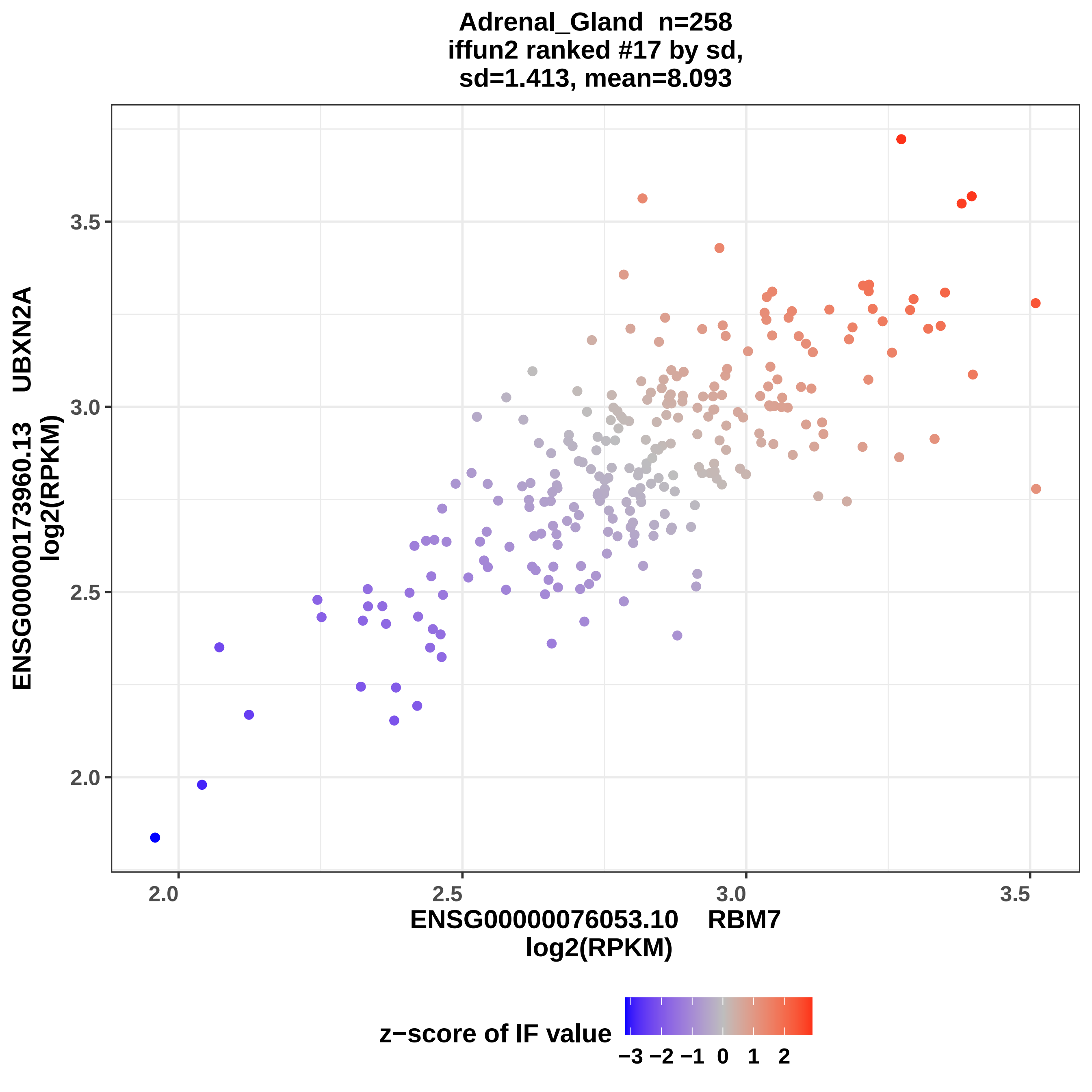

Adrenal_Gland n=258
iffun2 ranked #17 by sd,
sd=1.413, mean=8.093
ENSG00000173960.13 UBXN2A
log2(RPKM)
ENSG00000076053.10 RBM7
log2(RPKM)
2.0
2.5
3.0
3.5
z−score of IF value
−3 −2 −1 0 1 2

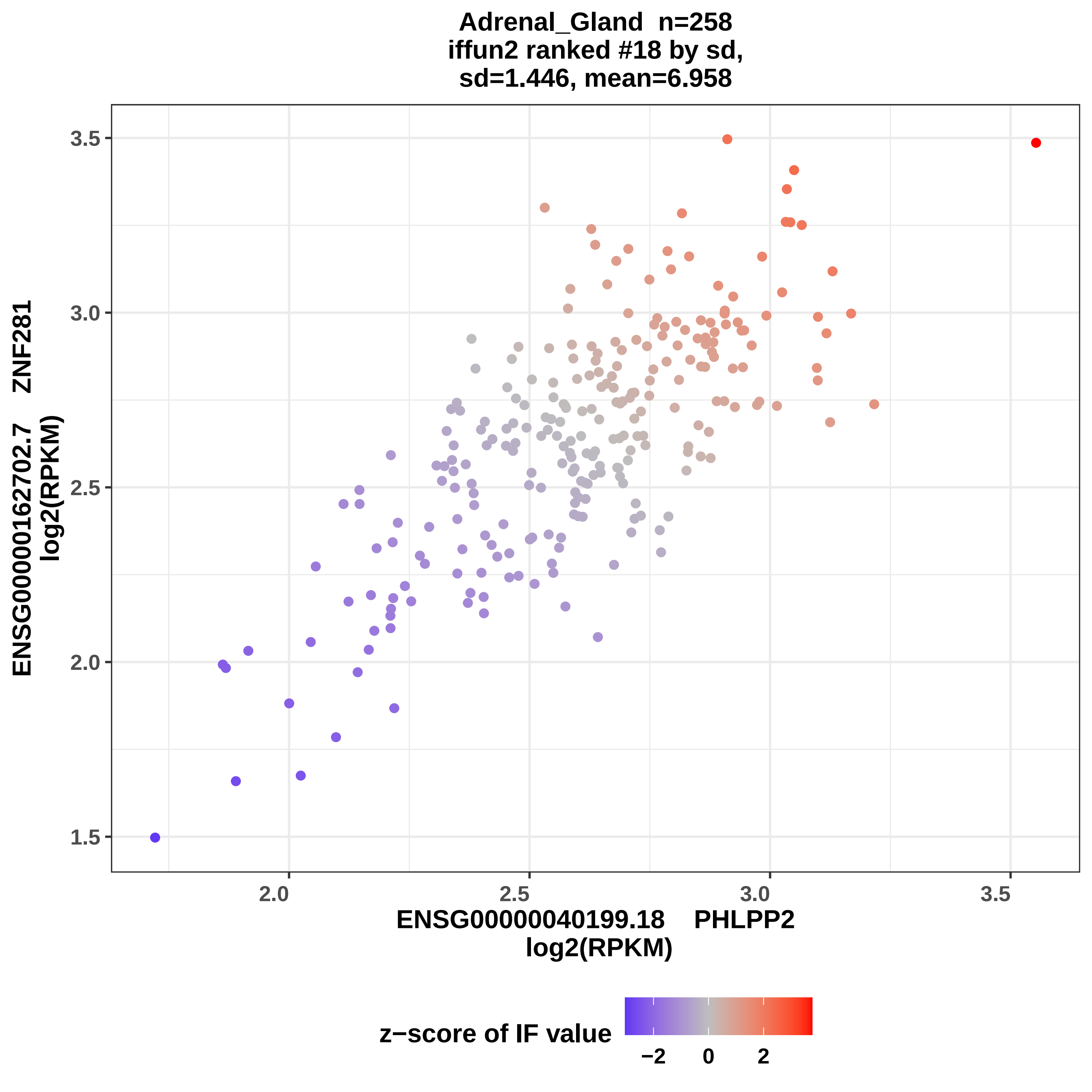

Adrenal_Gland n=258
iffun2 ranked #18 by sd,
sd=1.446, mean=6.958
ENSG00000162702.7 ZNF281
log2(RPKM)
3.5
3.0
2.5
2.0
1.5
2.0
2.5
3.0
3.5
ENSG00000040199.18 PHLPP2
log2(RPKM)
z−score of IF value
−2
0
2

Adrenal_Gland n=258
iffun2 ranked #19 by sd,
sd=1.528, mean=10.69

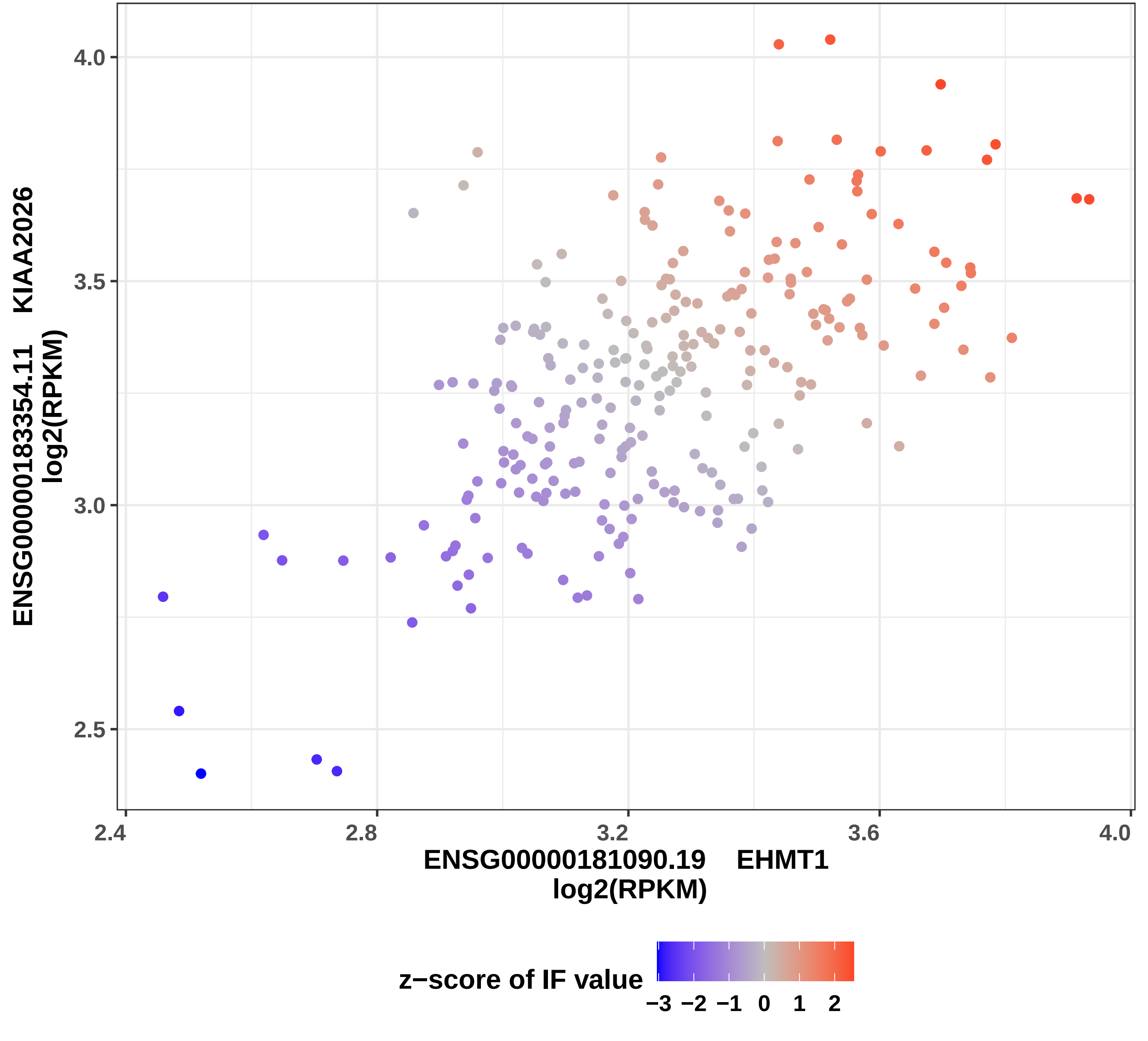

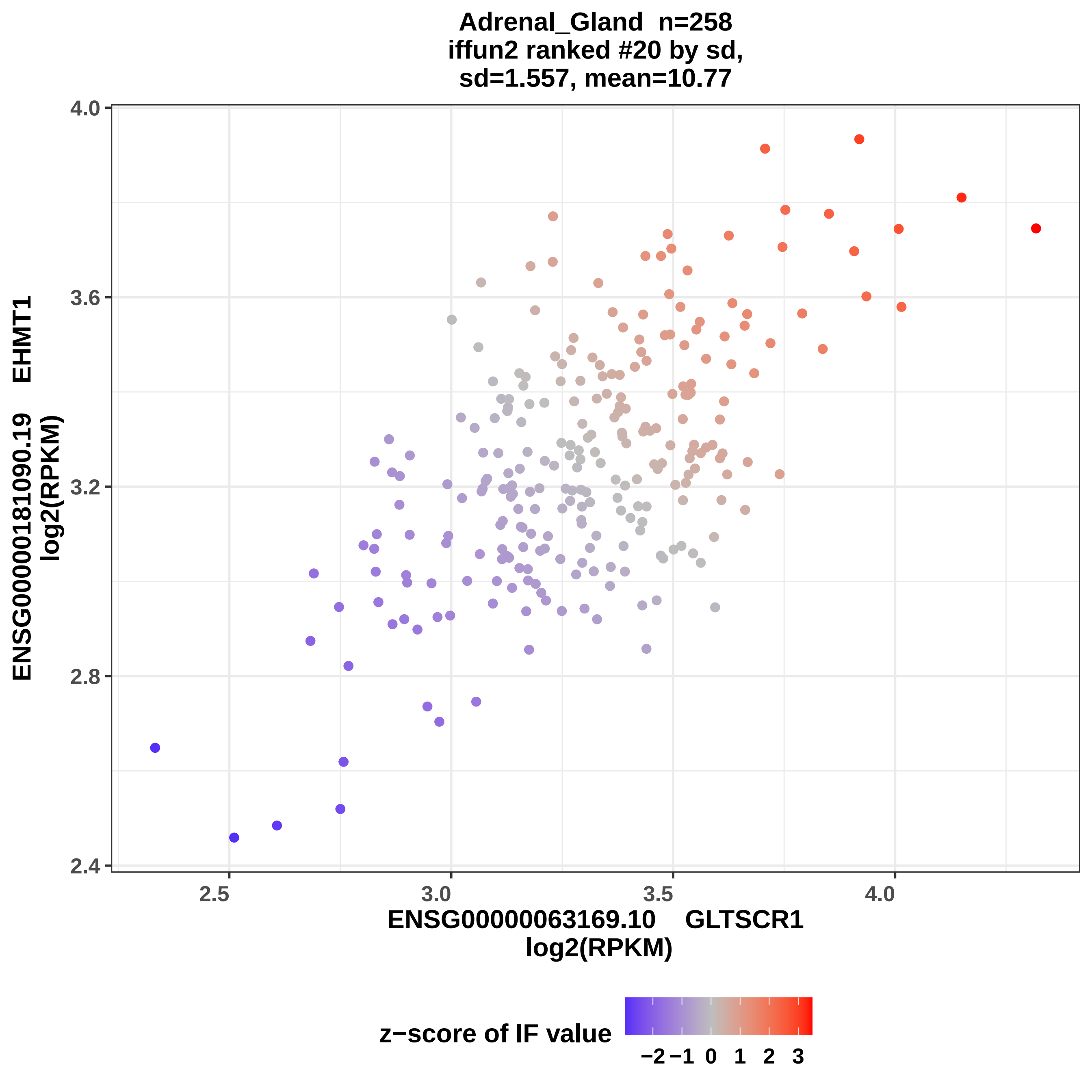

Adrenal_Gland n=258
iffun2 ranked #20 by sd,
sd=1.557, mean=10.77
ENSG00000181090.19 EHMT1
log2(RPKM)
2.4
2.8
3.2
3.6
4.0
2.5
3.0
3.5
4.0
ENSG00000063169.10 GLTSCR1
log2(RPKM)
z−score of IF value
−2
−1
0
1
2
3

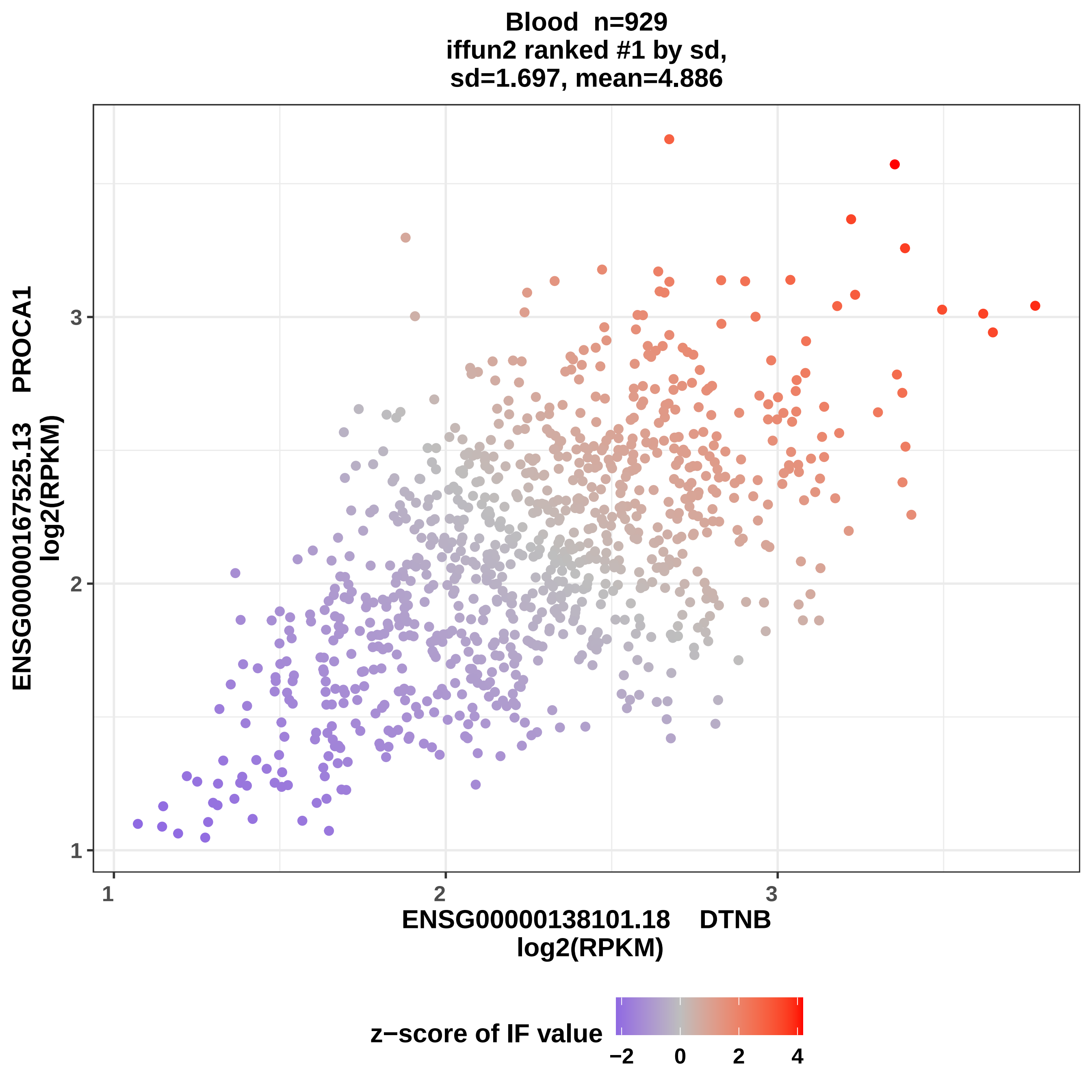
Blood n=929
iffun2 ranked #1 by sd,
sd=1.697, mean=4.886
ENSG00000167525.13 PROCA1
log2(RPKM)
ENSG00000138101.18 DTNB
log2(RPKM)
1
2
3
1
2
3
z−score of IF value
−2
0
2
4

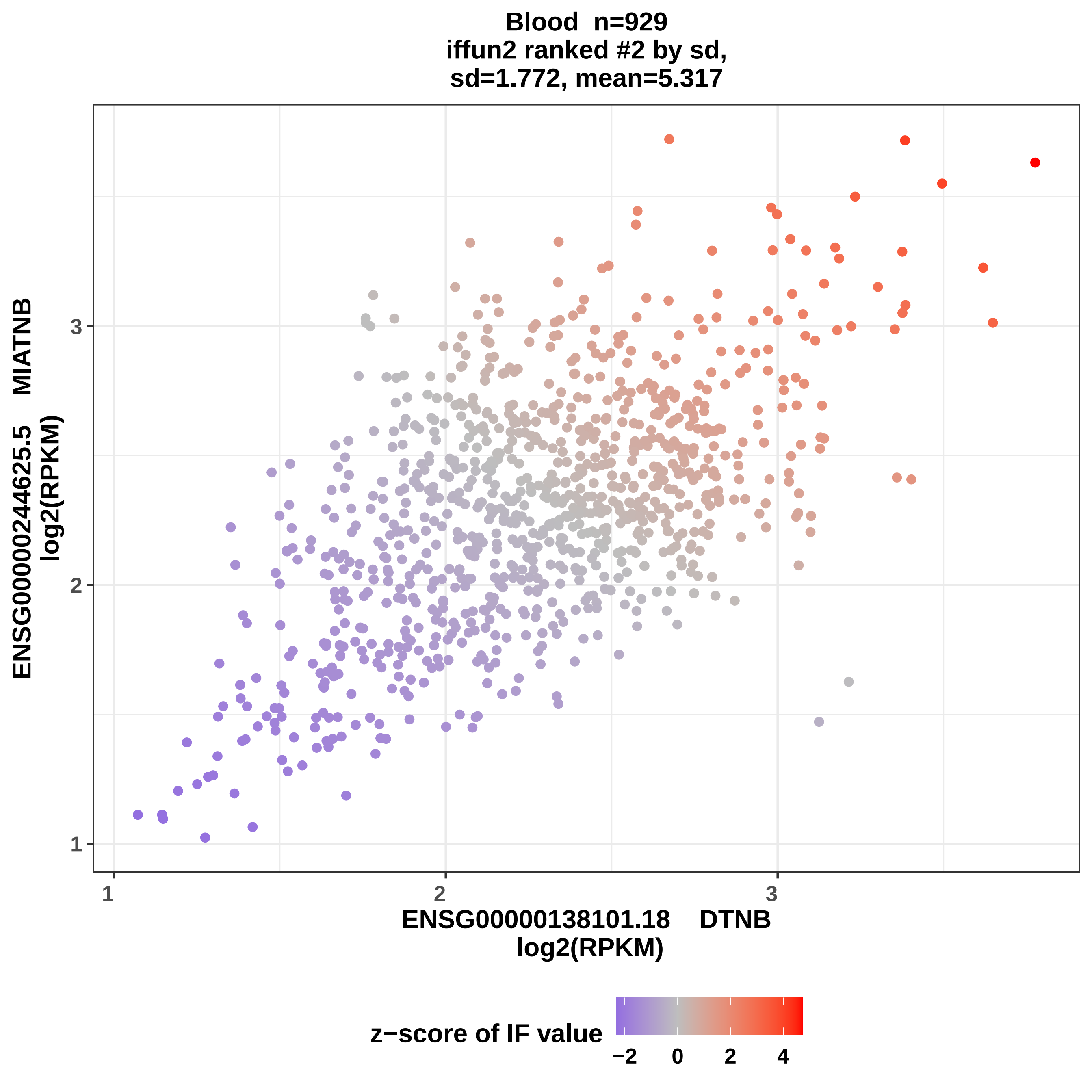
Blood n=929
iffun2 ranked #2 by sd,
sd=1.772, mean=5.317
ENSG00000244625.5 MIATNB
log2(RPKM)
3
2
1
1
2
3
ENSG00000138101.18 DTNB
log2(RPKM)
z−score of IF value
−2
0
2
4

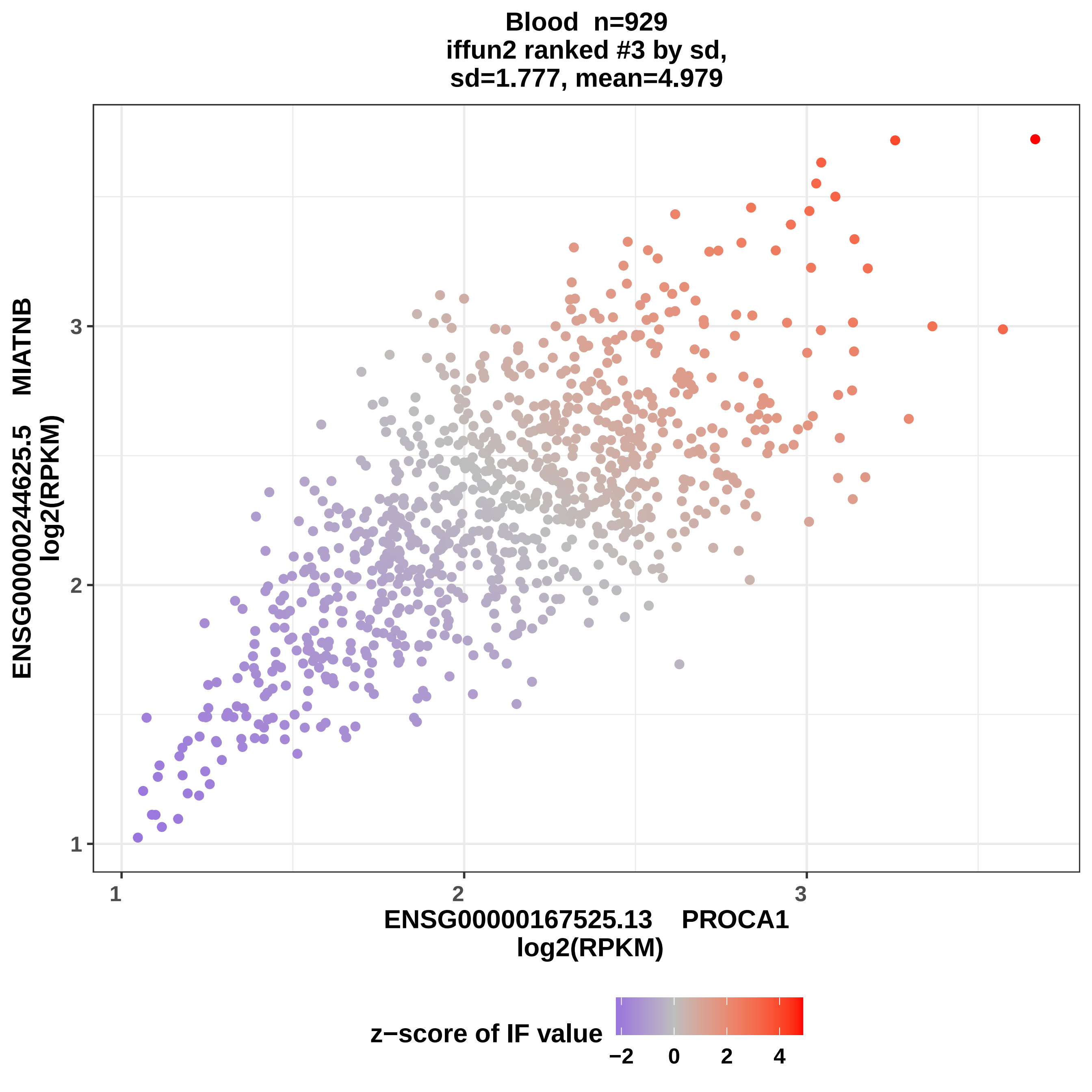
Blood n=929
iffun2 ranked #3 by sd,
sd=1.777, mean=4.979
ENSG00000244625.5 MIATNB
log2(RPKM)
ENSG00000167525.13 PROCA1
log2(RPKM)
1
2
3
z−score of IF value
−2
0
2
4

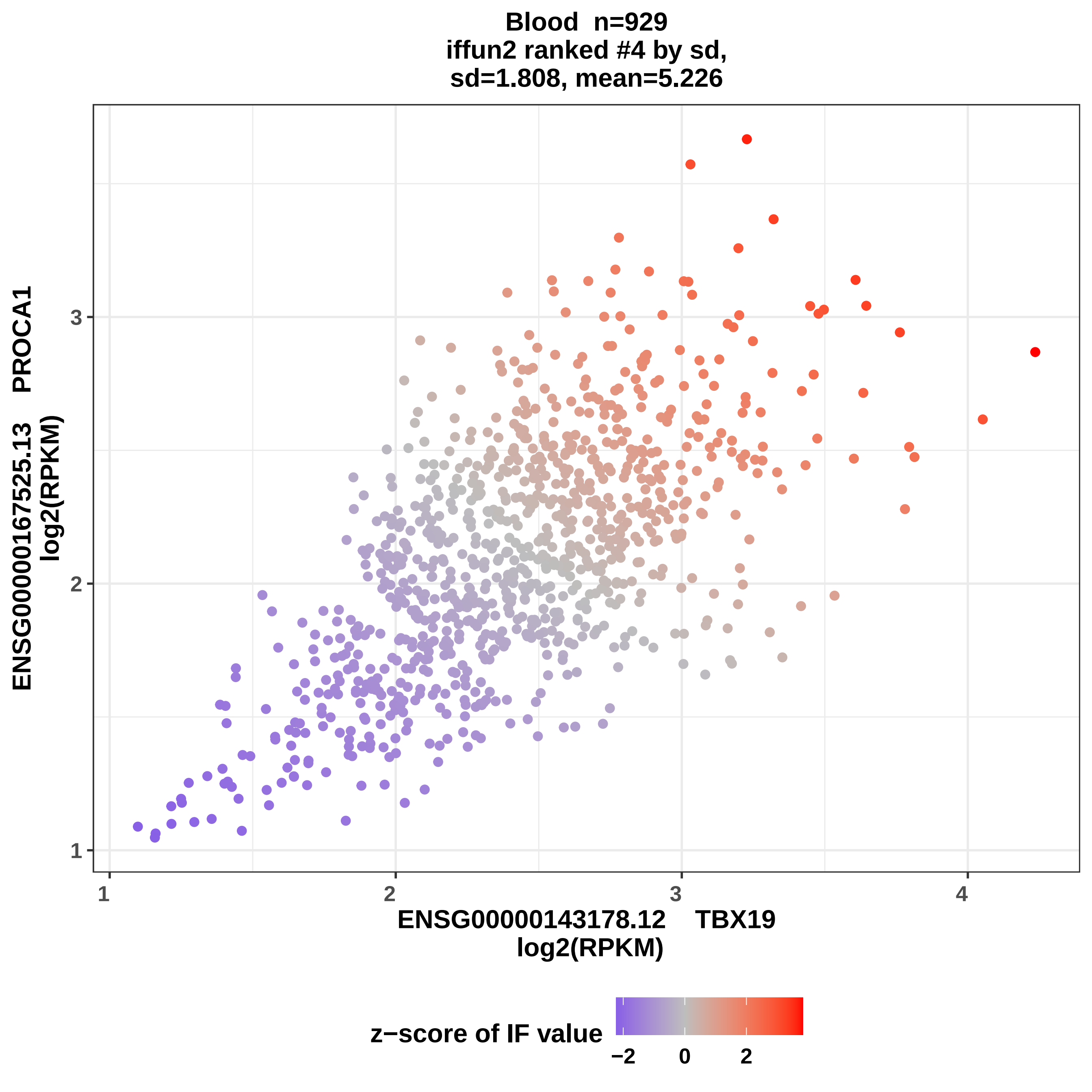

Blood n=929
iffun2 ranked #4 by sd,
sd=1.808, mean=5.226
ENSG00000167525.13 PROCA1
log2(RPKM)
1
2
3
1
2
3
4
ENSG00000143178.12 TBX19
log2(RPKM)
z−score of IF value
−2
0
2

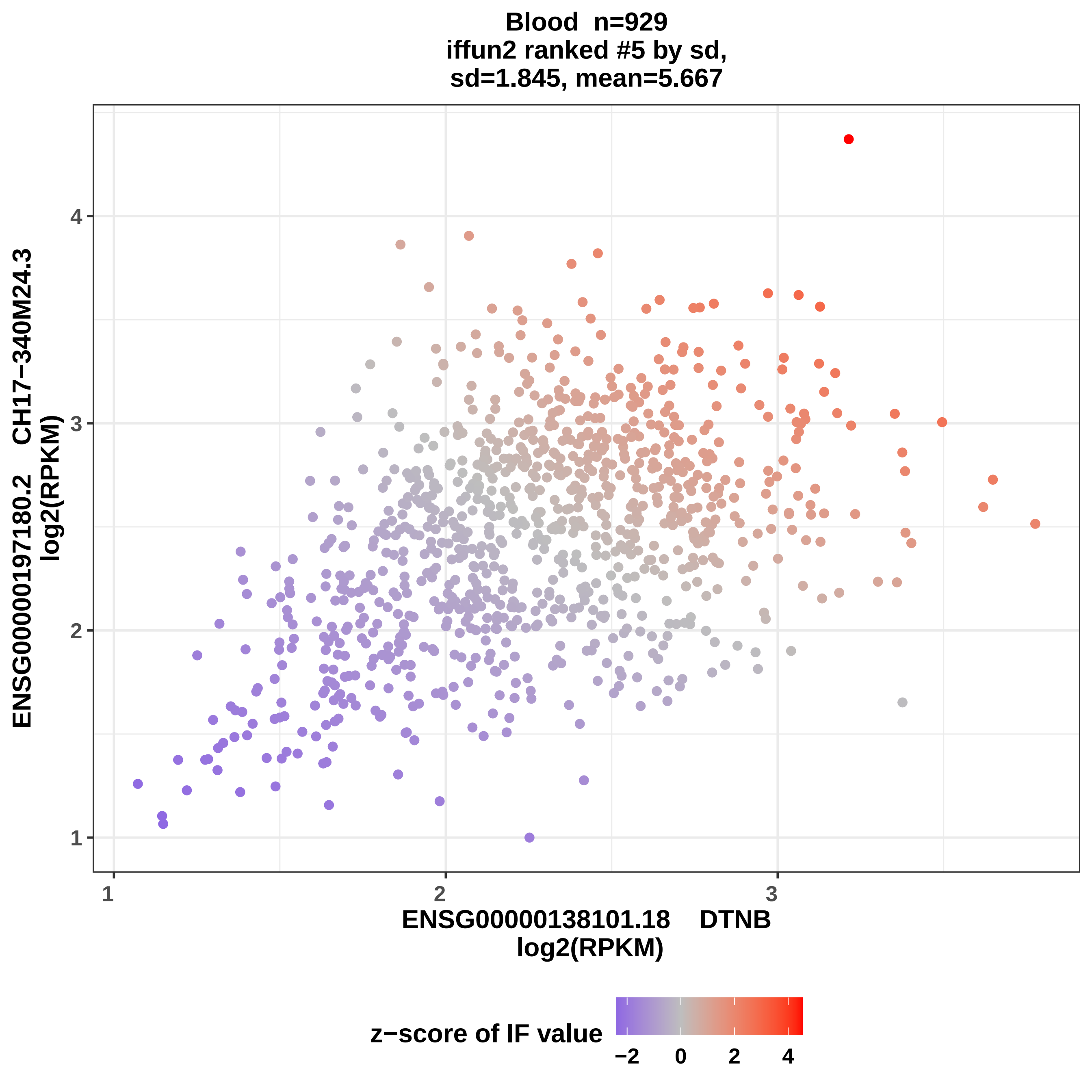
Blood n=929
iffun2 ranked #5 by sd,
sd=1.845, mean=5.667
ENSG00000197180.2 CH17−340M24.3
log2(RPKM)
ENSG00000138101.18 DTNB
log2(RPKM)
z−score of IF value
1
2
3
4
−2
0

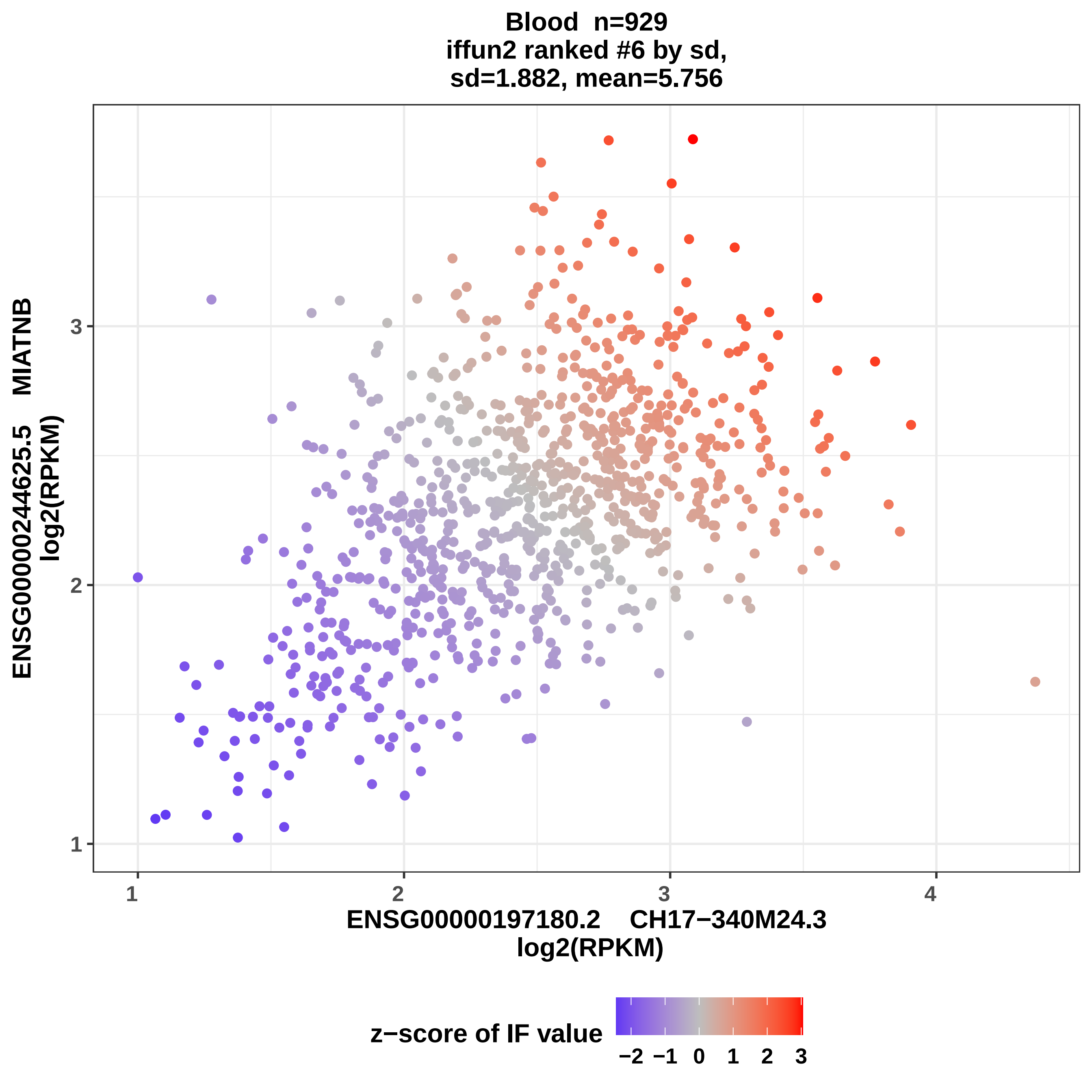

Blood n=929
iffun2 ranked #6 by sd,
sd=1.882, mean=5.756
ENSG00000244625.5 MIATNB
log2(RPKM)
3
2
1
1
2
3
4
ENSG00000197180.2 CH17−340M24.3
log2(RPKM)
z−score of IF value
−2 −1 0 1 2 3

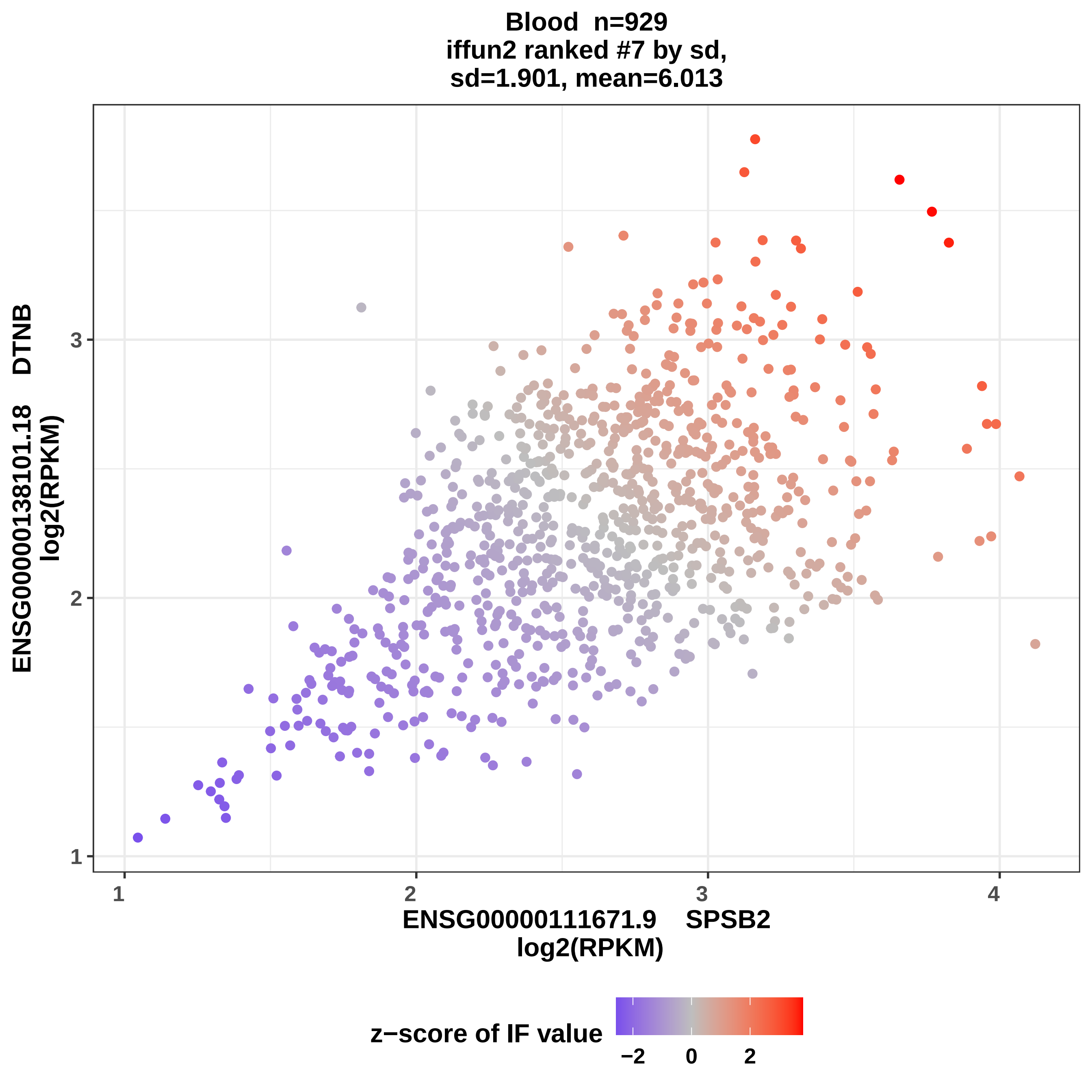

Blood n=929
iffun2 ranked #7 by sd,
sd=1.901, mean=6.013
ENSG00000138101.18 DTNB
log2(RPKM)
1
2
3
1
2
3
4
ENSG00000111671.9 SPSB2
log2(RPKM)
z−score of IF value
−2
0
2

Blood  n=929
iffun2 ranked #8 by sd,
sd=1.915, mean=5.697

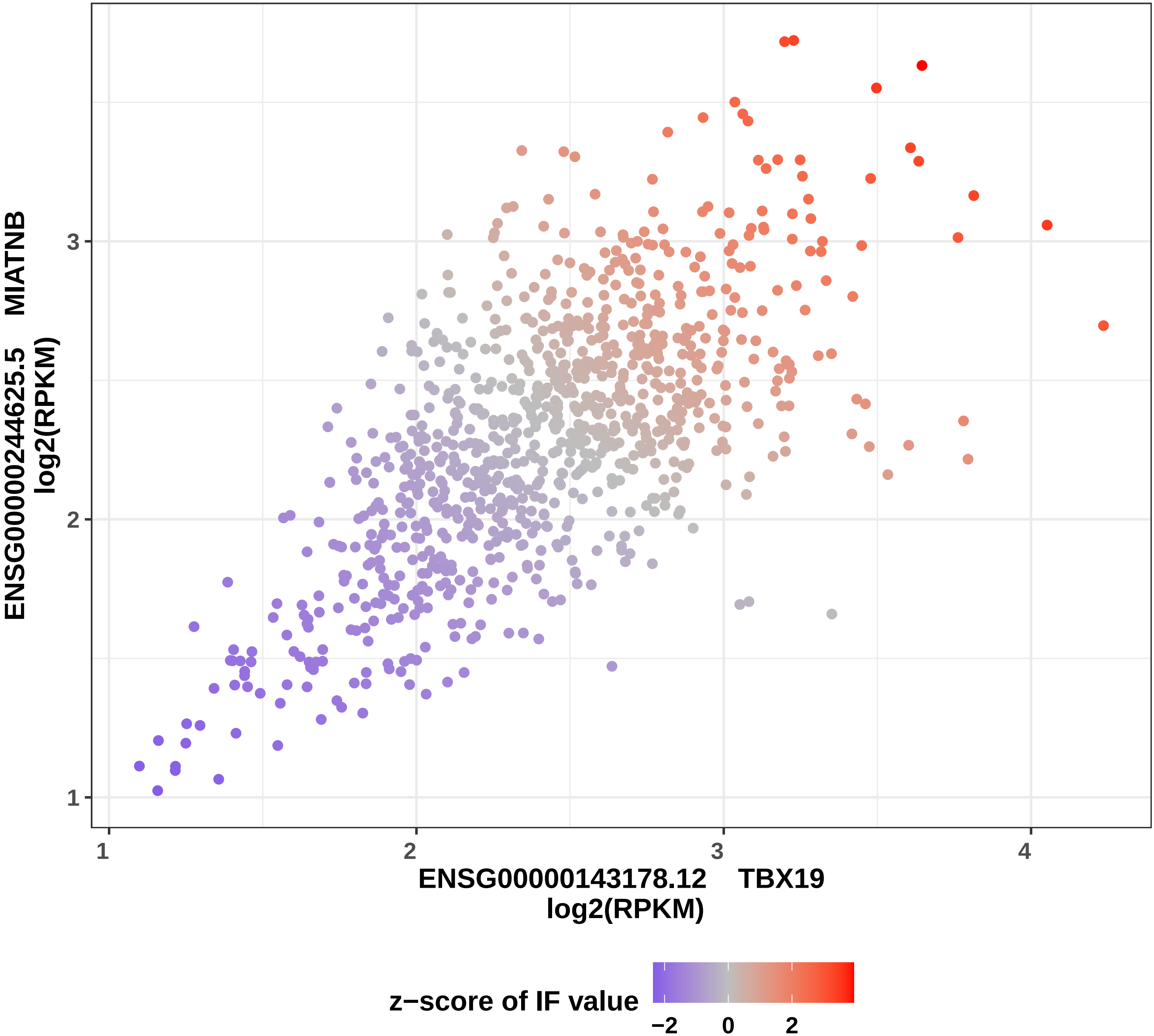

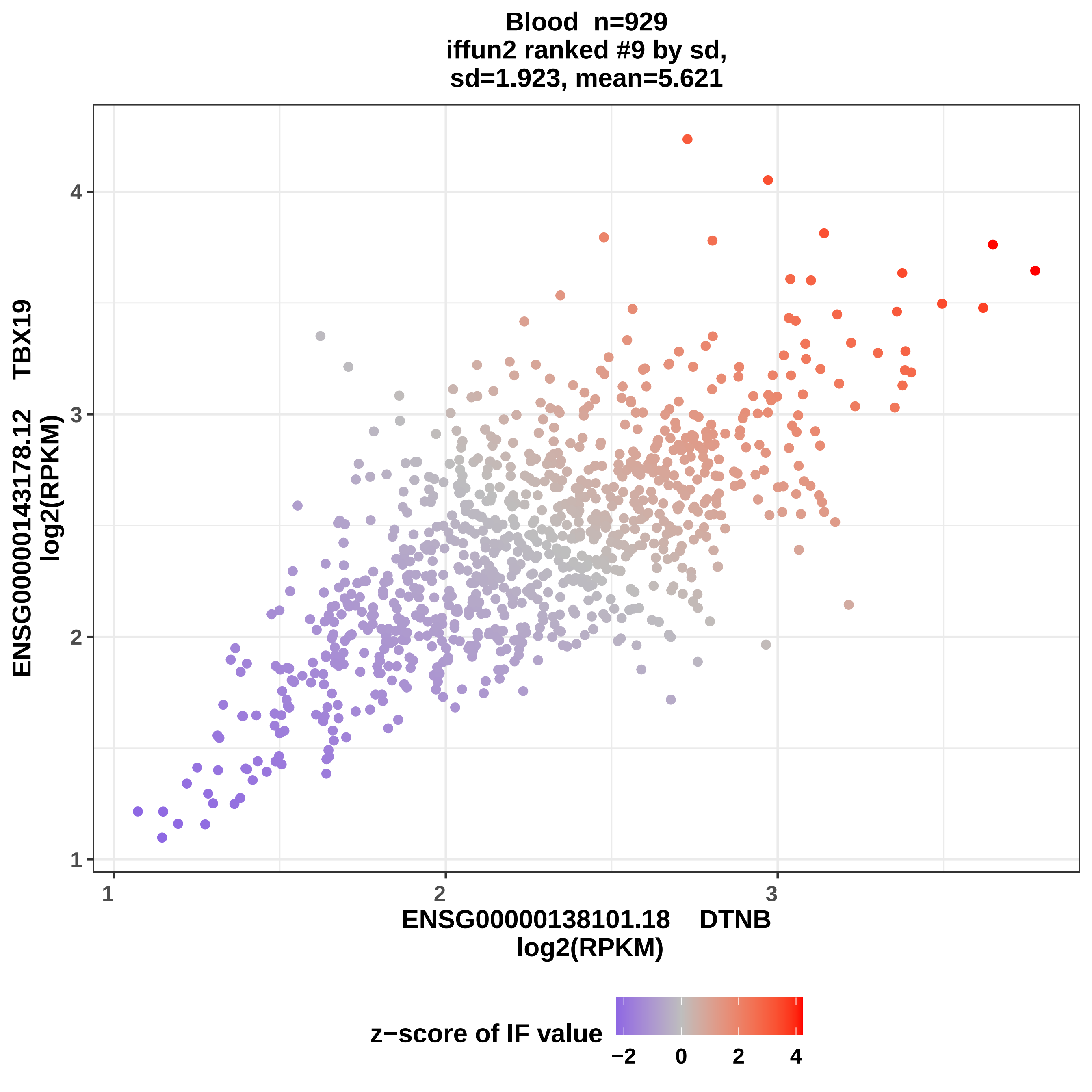

Blood n=929
iffun2 ranked #9 by sd,
sd=1.923, mean=5.621
ENSG00000143178.12 TBX19
log2(RPKM)
ENSG00000138101.18 DTNB
log2(RPKM)
1
2
3
4
z−score of IF value
−2
0
2
4

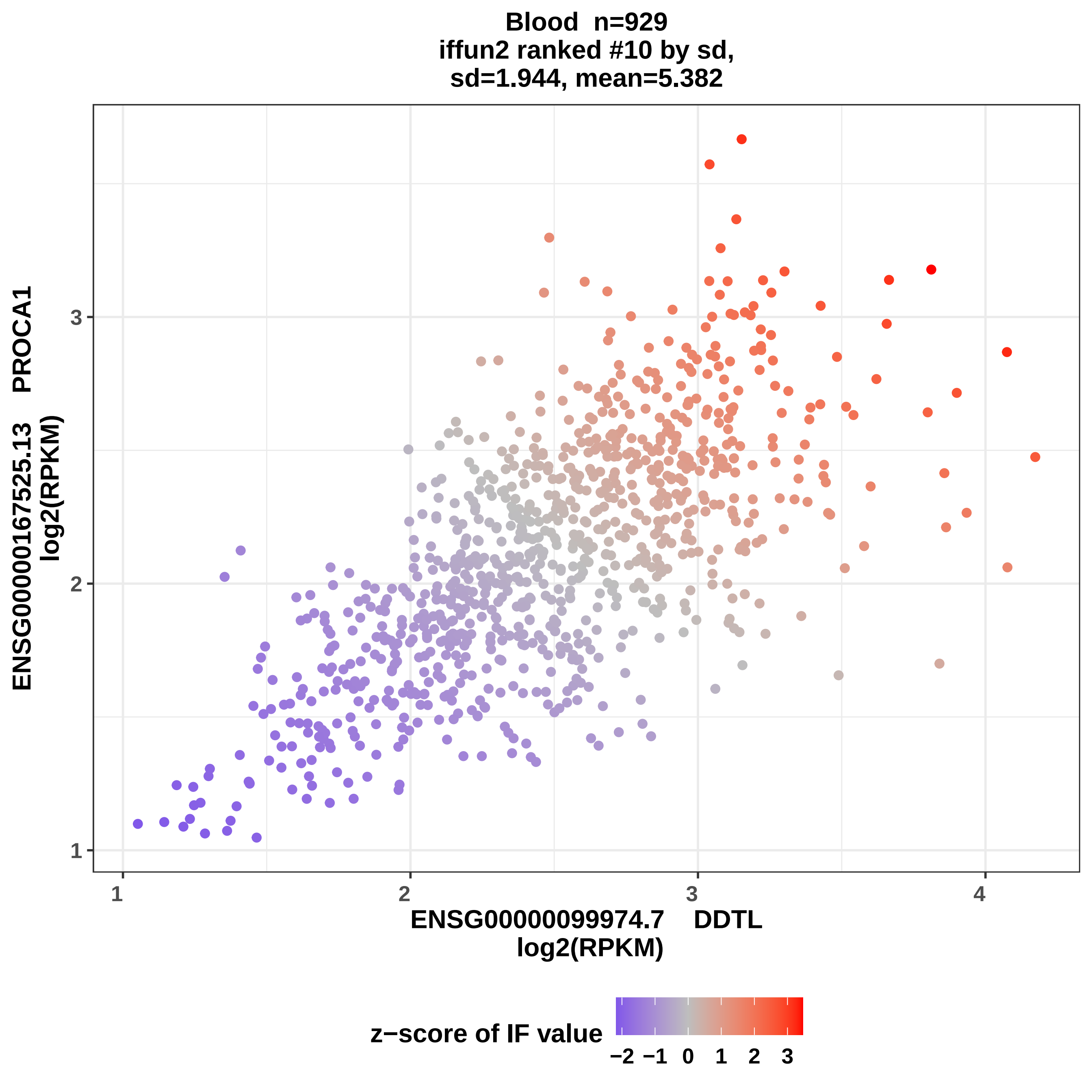

Blood n=929
iffun2 ranked #10 by sd,
sd=1.944, mean=5.382
ENSG00000167525.13 PROCA1
log2(RPKM)
1
2
3
4
ENSG00000099974.7 DDTL
log2(RPKM)
z−score of IF value
−2
−1
0
1
2
3

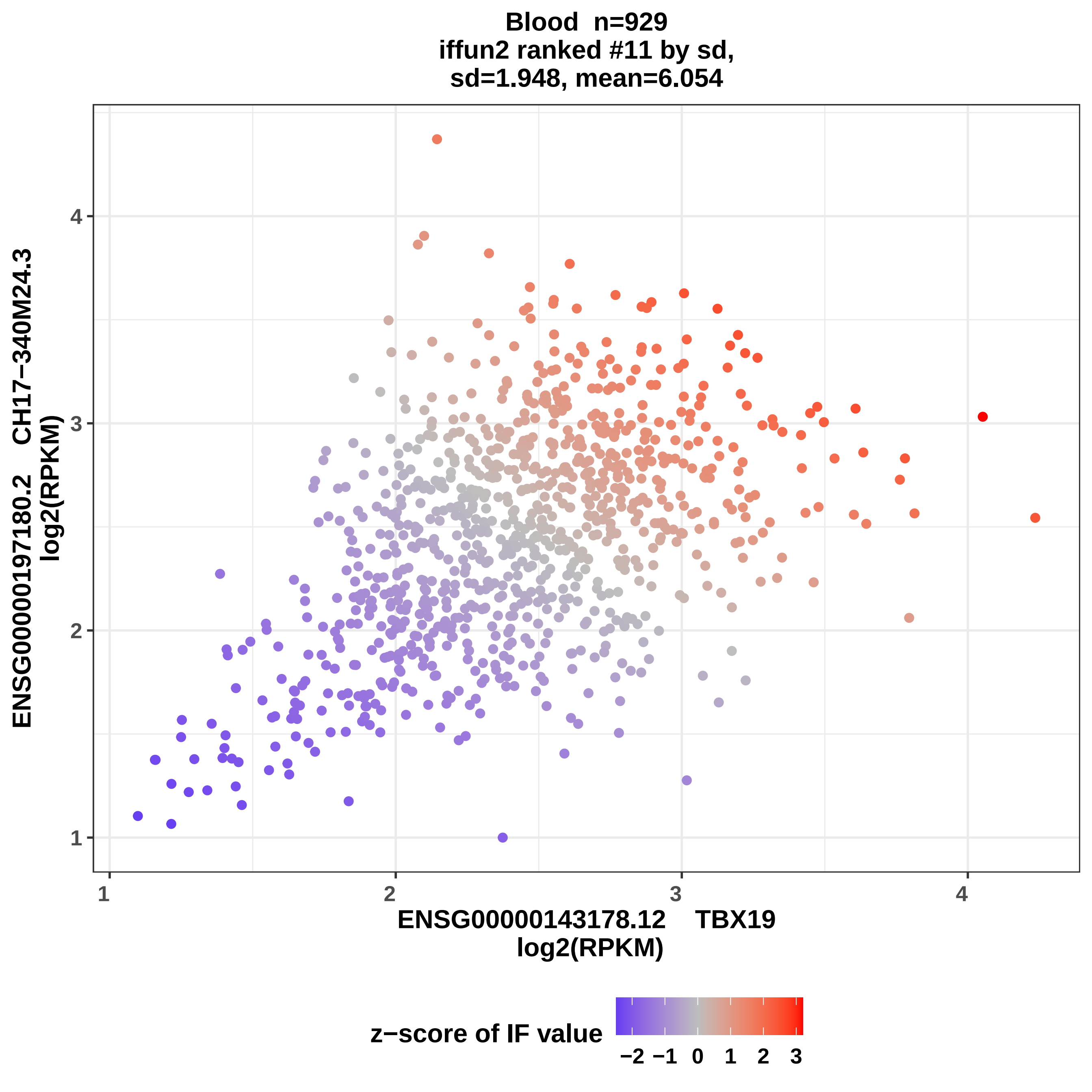
Blood n=929
iffun2 ranked #11 by sd,
sd=1.948, mean=6.054
ENSG00000197180.2 CH17−340M24.3
log2(RPKM)
ENSG00000143178.12 TBX19
log2(RPKM)
z−score of IF value
−2 −1 0 1 2 3
1
2
3
4

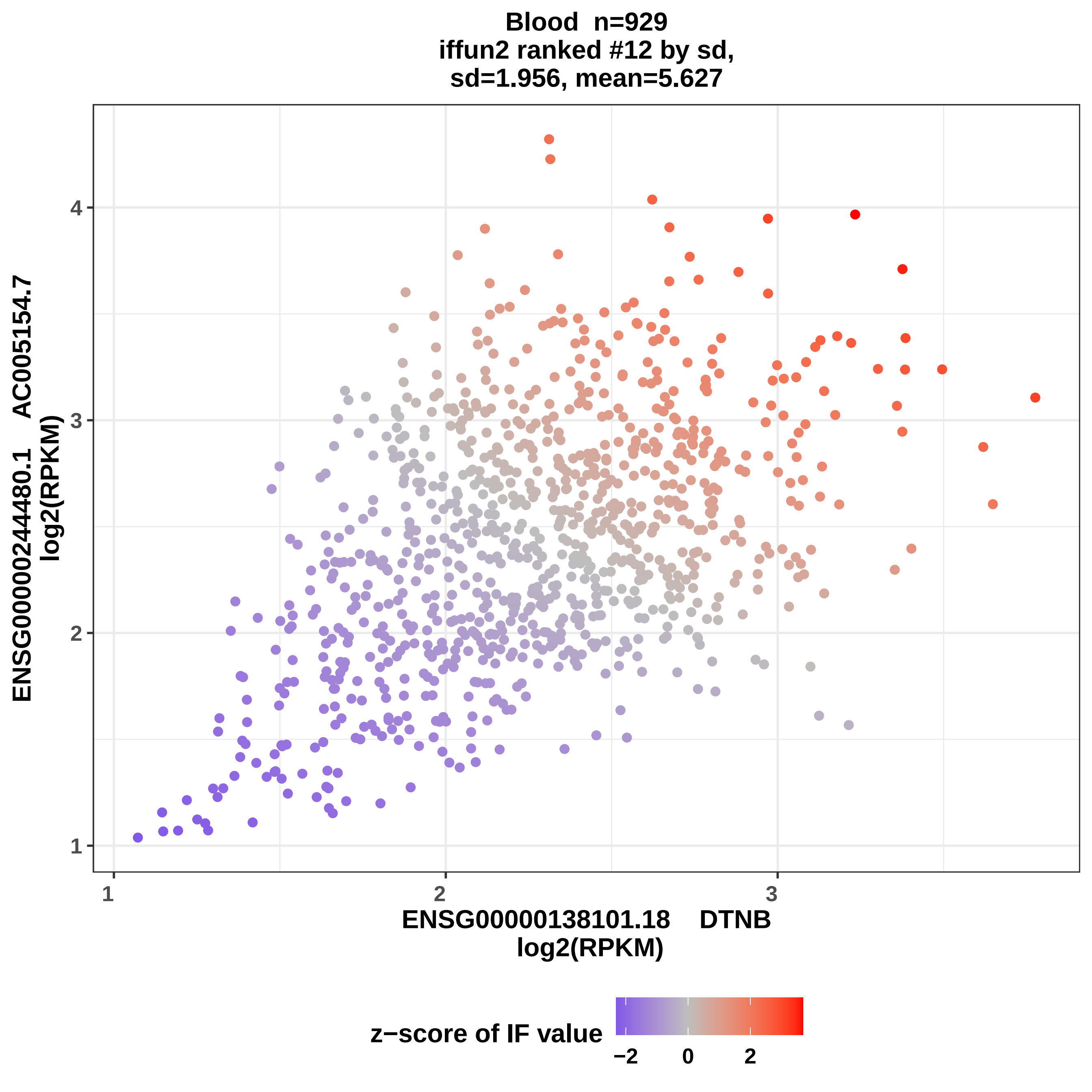
Blood n=929
iffun2 ranked #12 by sd,
sd=1.956, mean=5.627
ENSG00000244480.1 AC005154.7
log2(RPKM)
ENSG00000138101.18 DTNB
log2(RPKM)
1
2
3
4
z−score of IF value
−2
0
2

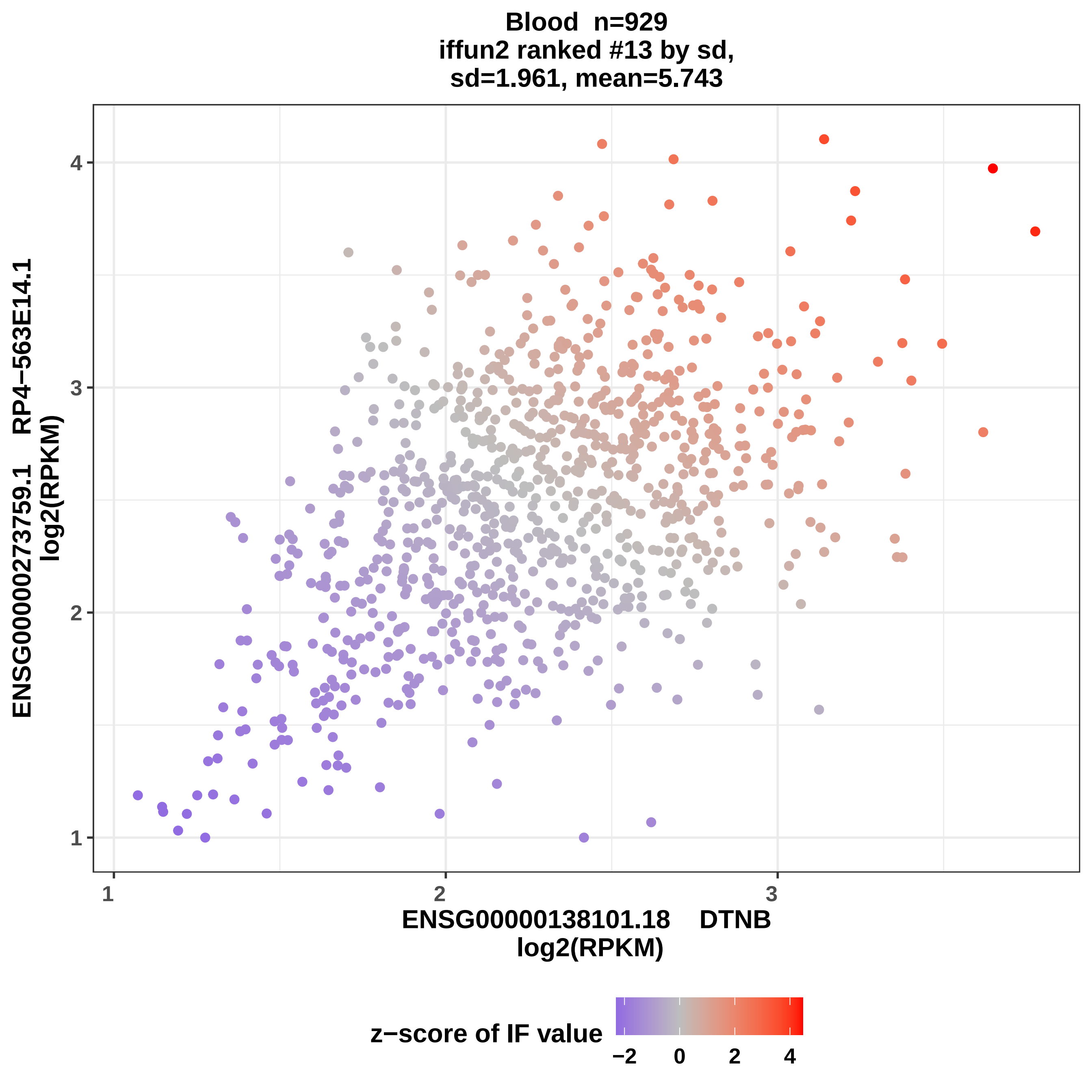
Blood n=929
iffun2 ranked #13 by sd,
sd=1.961, mean=5.743
ENSG00000273759.1 RP4−563E14.1
log2(RPKM)
ENSG00000138101.18 DTNB
log2(RPKM)
z−score of IF value
1
2
3
4
−2
0
2
4

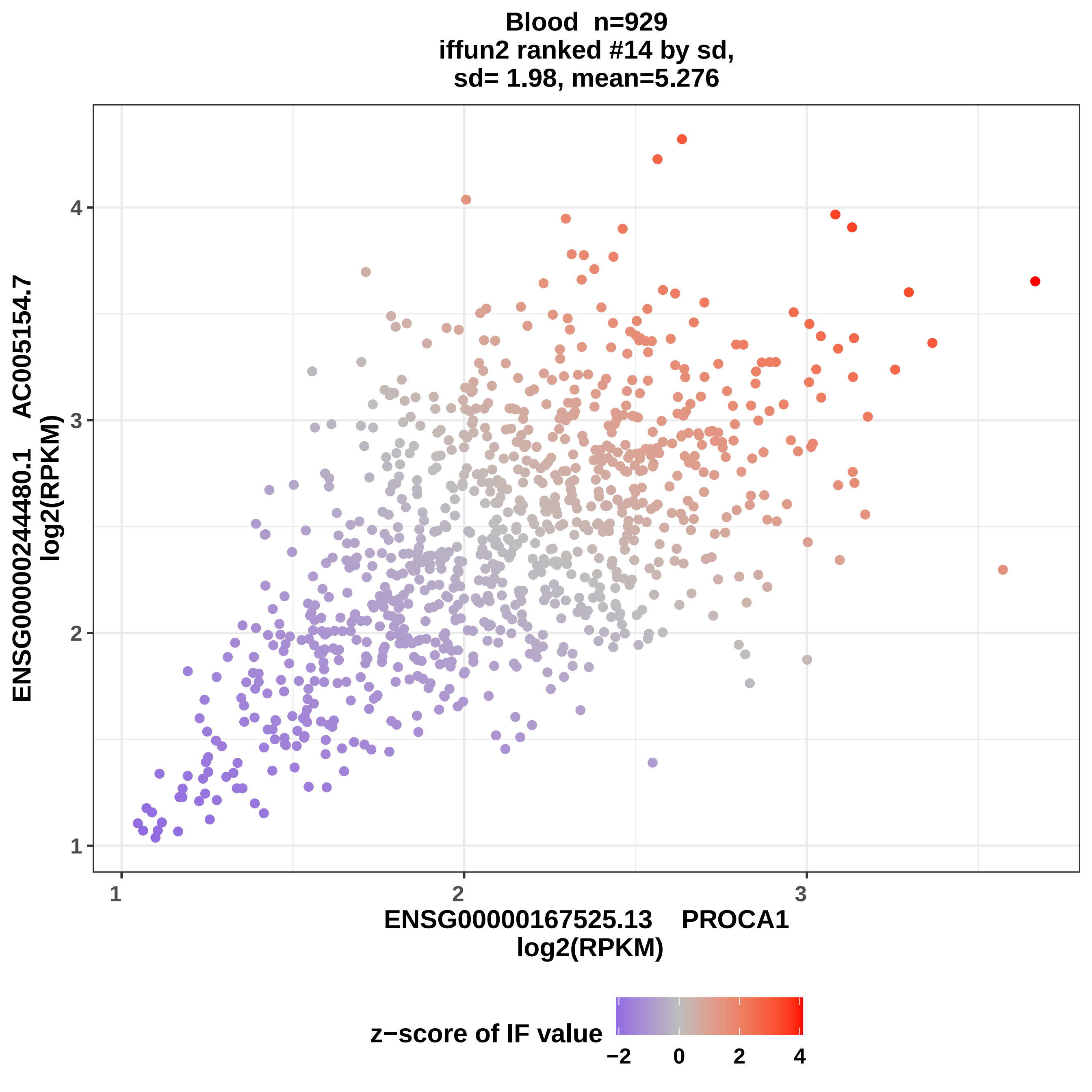
Blood n=929
iffun2 ranked #14 by sd,
sd= 1.98, mean=5.276
ENSG00000244480.1 AC005154.7
log2(RPKM)
1
2
3
4
ENSG00000167525.13 PROCA1
log2(RPKM)
z−score of IF value
−2
0
2
4

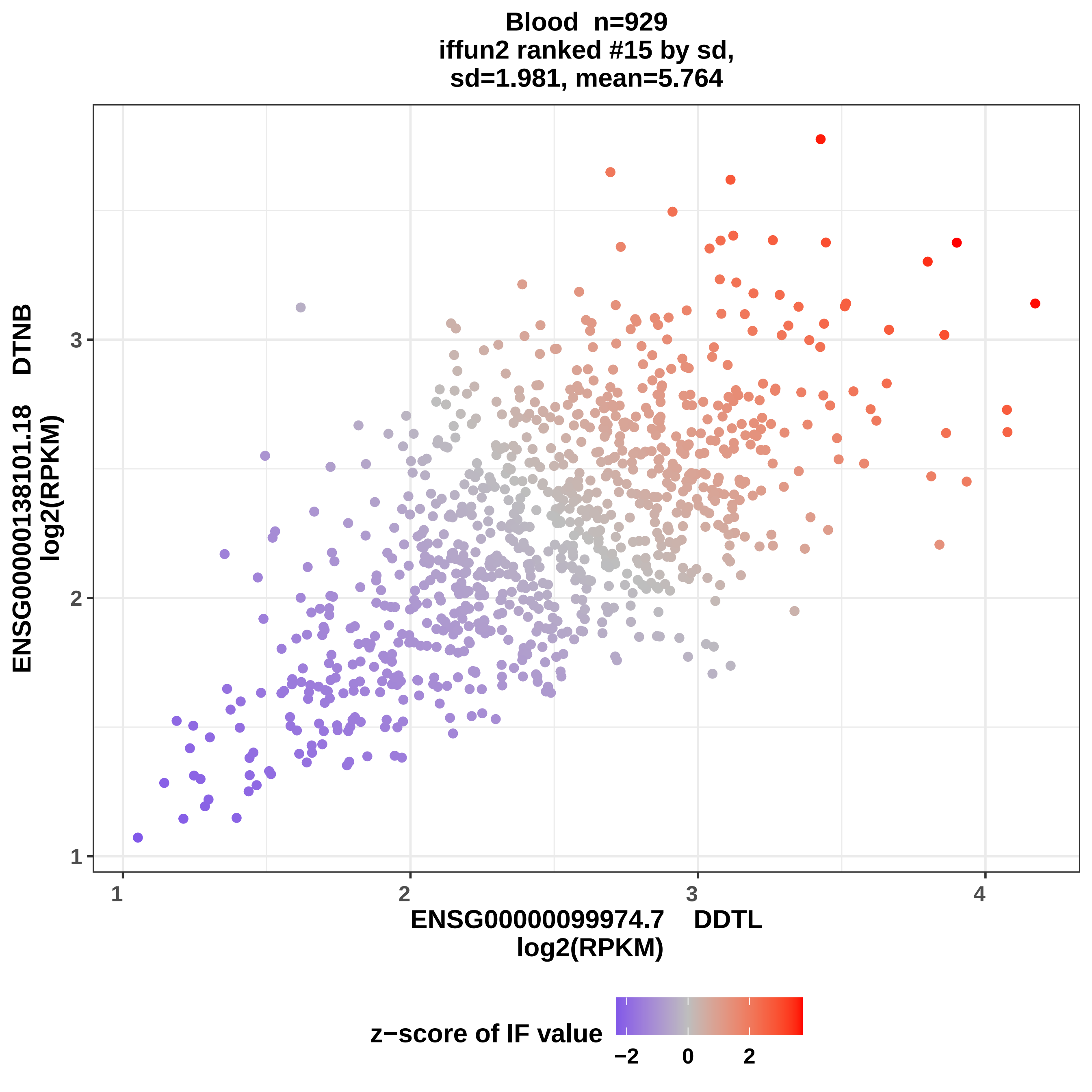
Blood n=929
iffun2 ranked #15 by sd,
sd=1.981, mean=5.764
ENSG00000138101.18 DTNB
log2(RPKM)
ENSG00000099974.7 DDTL
log2(RPKM)
1
2
3
4
z−score of IF value
−2
0
2

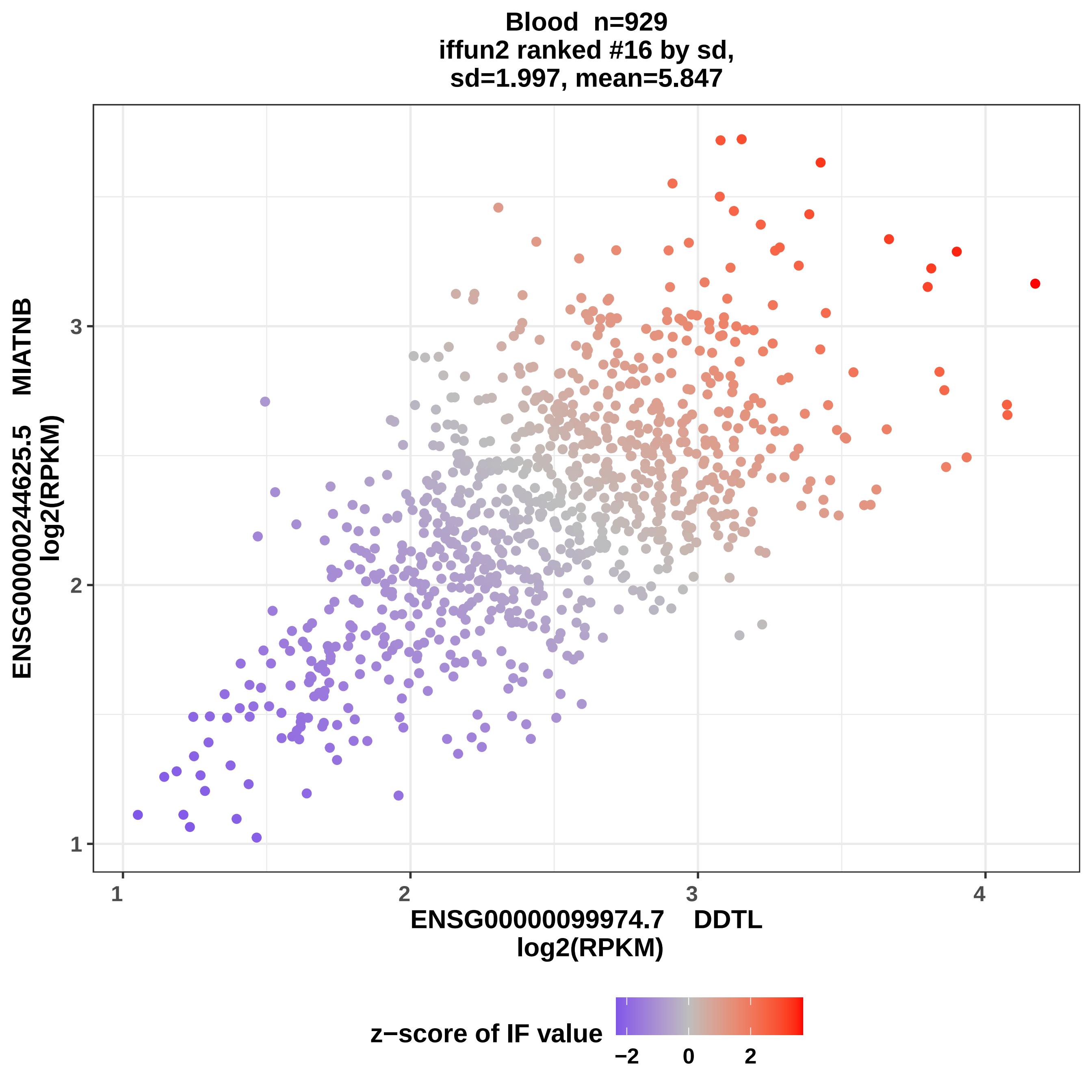
Blood n=929
iffun2 ranked #16 by sd,
sd=1.997, mean=5.847
ENSG00000244625.5 MIATNB
log2(RPKM)
ENSG00000099974.7 DDTL
log2(RPKM)
1
2
3
4
z−score of IF value
−2
0
2

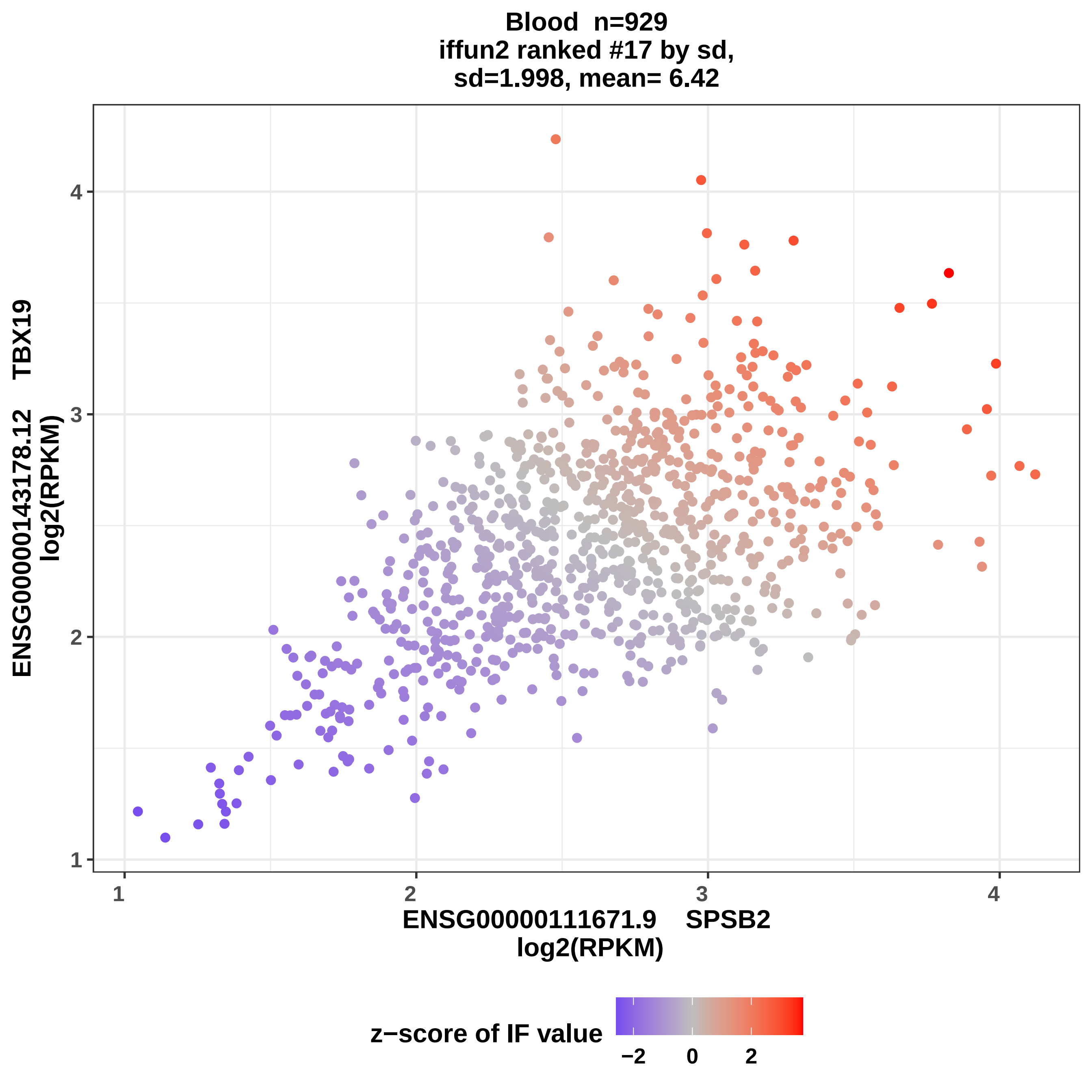
Blood n=929
iffun2 ranked #17 by sd,
sd=1.998, mean= 6.42
ENSG00000143178.12 TBX19
log2(RPKM)
ENSG00000111671.9 SPSB2
log2(RPKM)
1
2
3
4
z−score of IF value
−2
0
2

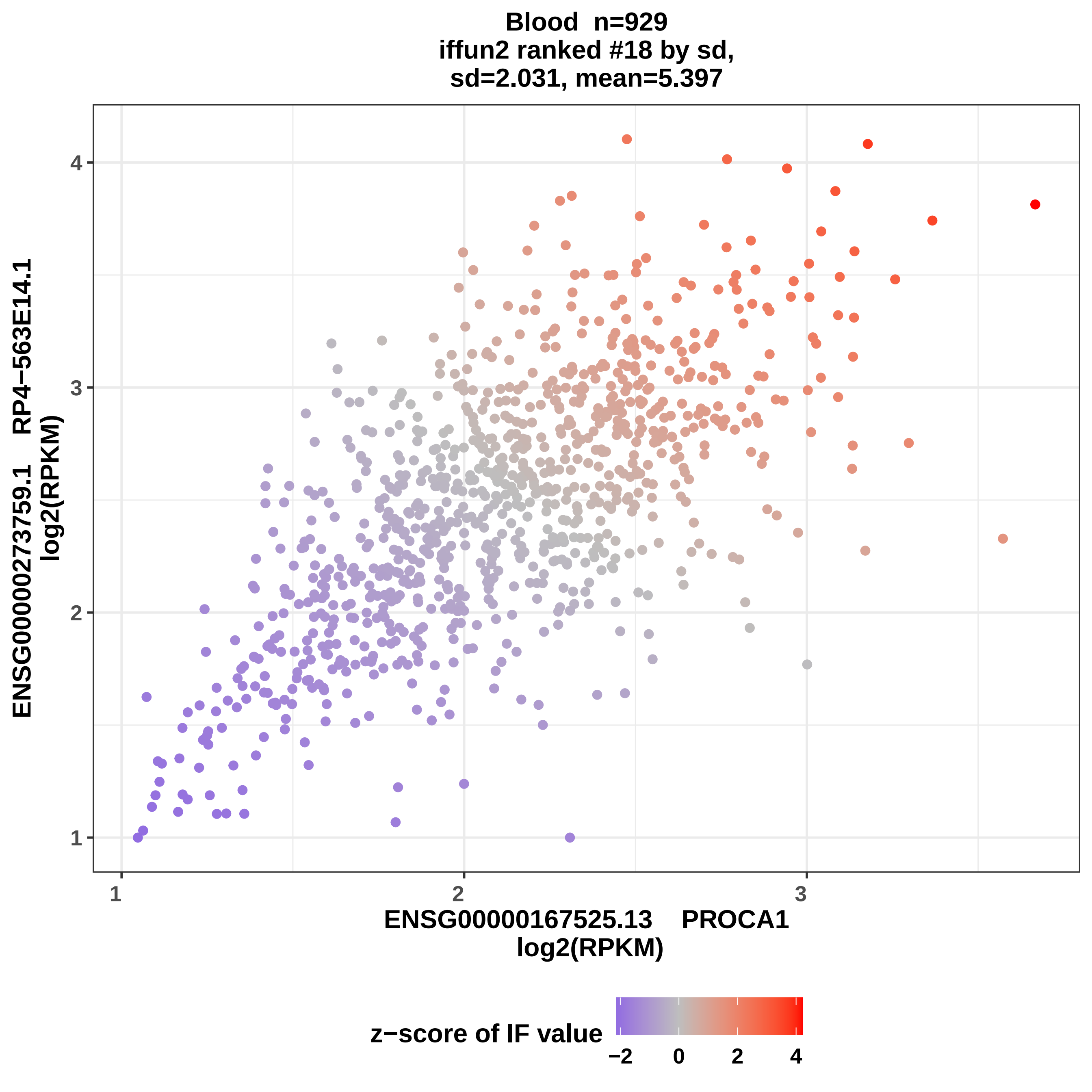

Blood n=929
iffun2 ranked #18 by sd,
sd=2.031, mean=5.397
ENSG00000273759.1 RP4−563E14.1
log2(RPKM)
ENSG00000167525.13 PROCA1
log2(RPKM)
1
2
3
4
z−score of IF value
−2
0
2
4

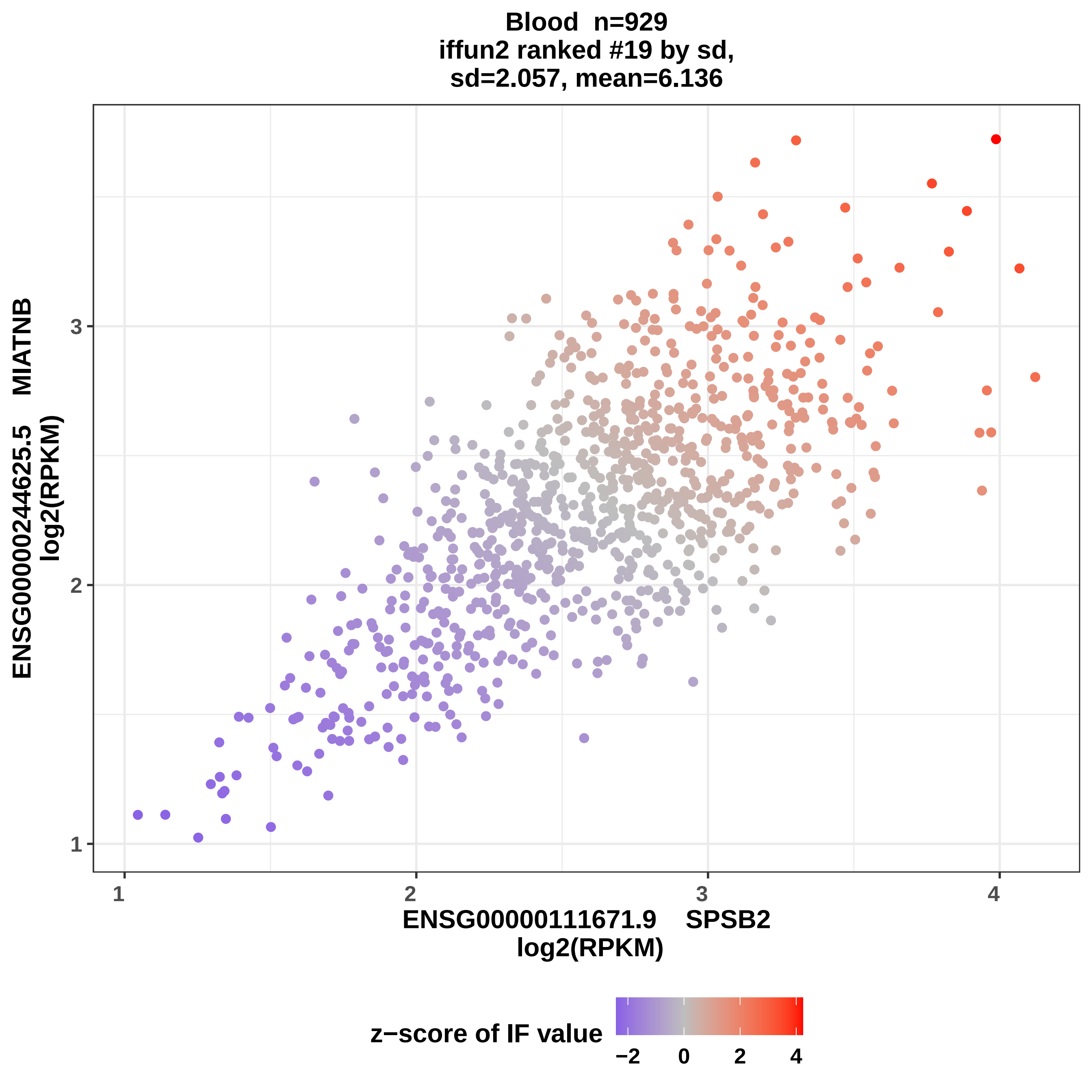

Blood n=929
iffun2 ranked #19 by sd,
sd=2.057, mean=6.136
ENSG00000244625.5 MIATNB
log2(RPKM)
ENSG00000111671.9 SPSB2
log2(RPKM)
1
2
3
4
z−score of IF value
−2
0
2
4

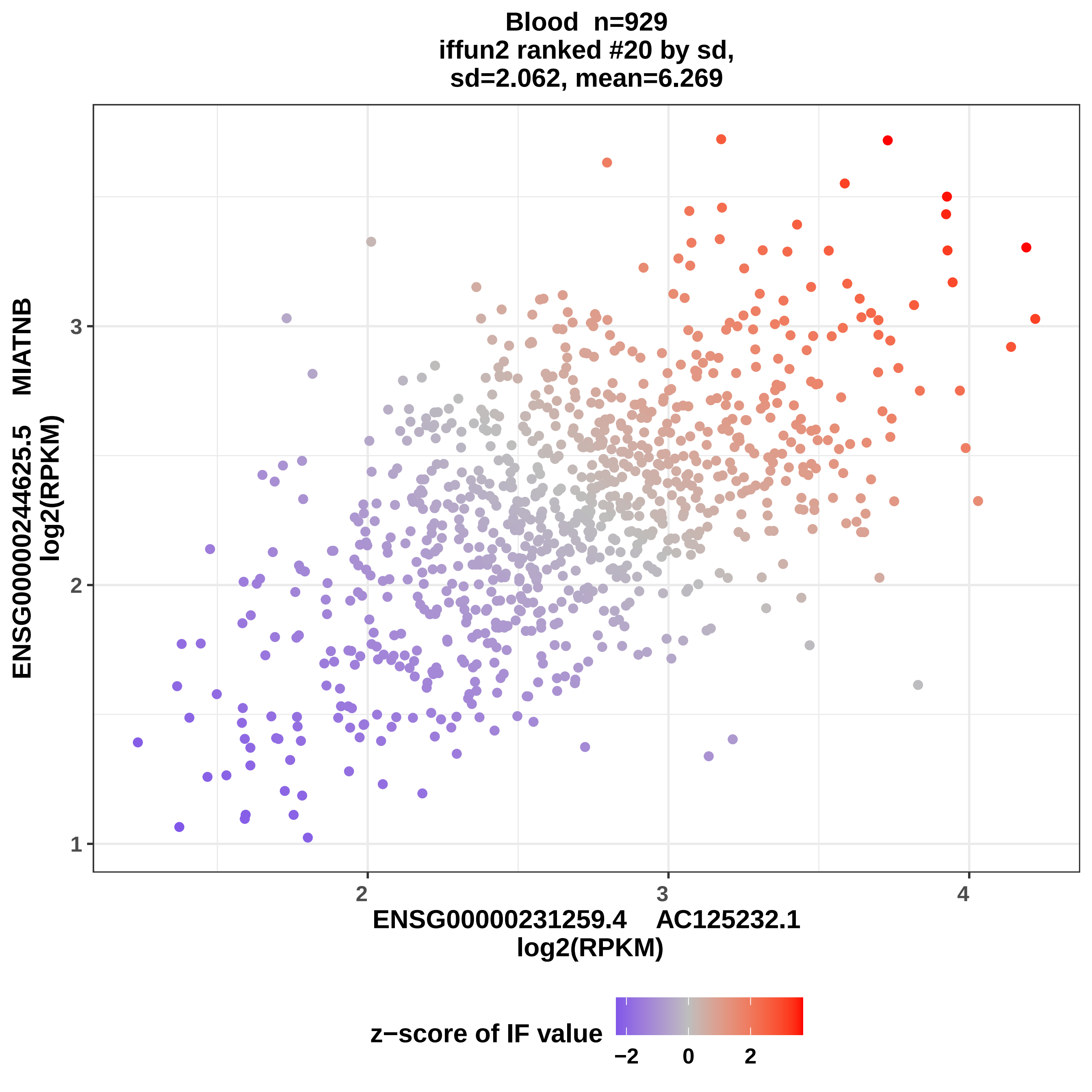

Blood n=929
iffun2 ranked #20 by sd,
sd=2.062, mean=6.269
ENSG00000244625.5 MIATNB
log2(RPKM)
3
2
1
2
3
4
ENSG00000231259.4 AC125232.1
log2(RPKM)
z−score of IF value
−2
0
2

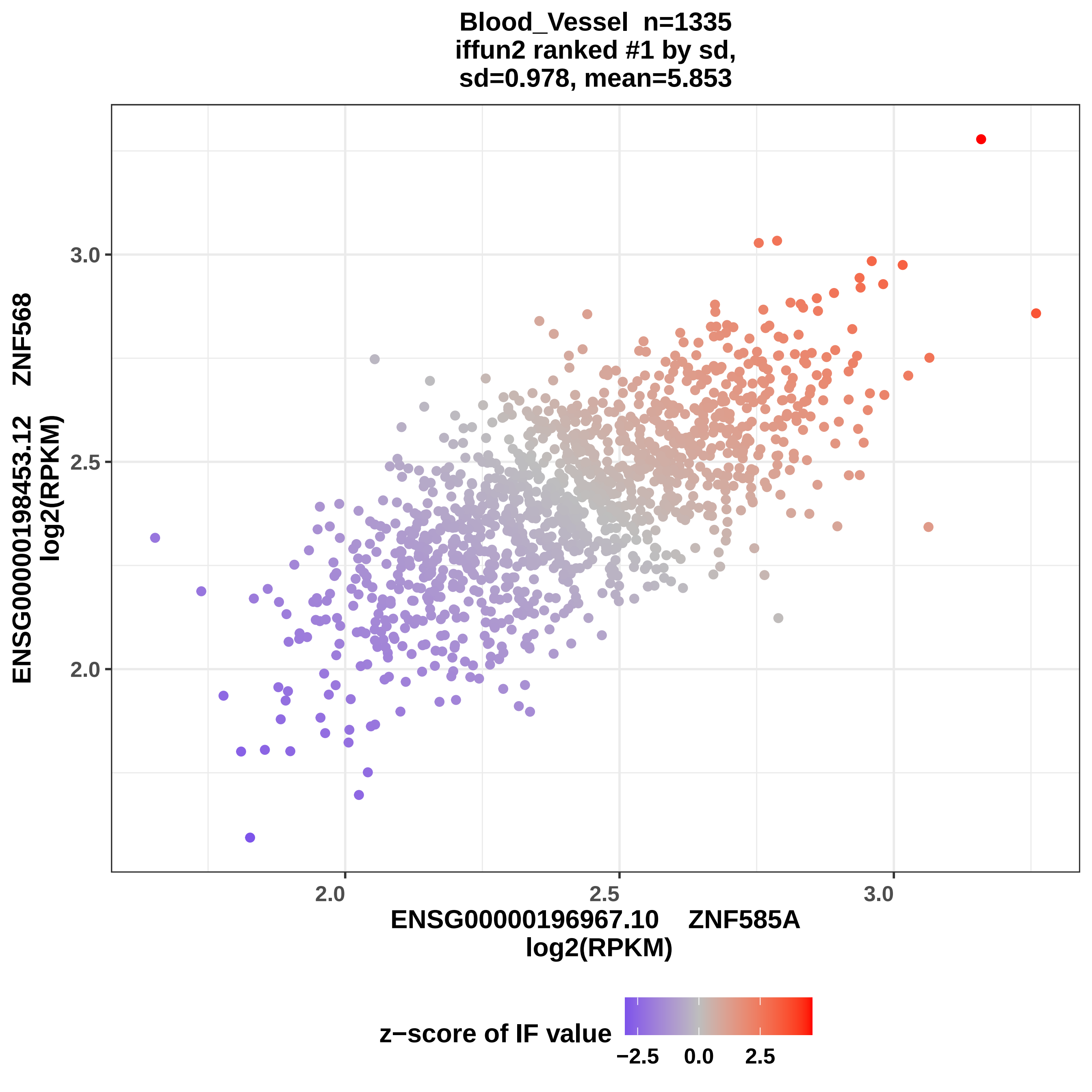

Blood_Vessel n=1335
iffun2 ranked #1 by sd,
sd=0.978, mean=5.853
ENSG00000198453.12 ZNF568
log2(RPKM)
3.0
2.5
2.0
2.0
2.5
3.0
ENSG00000196967.10 ZNF585A
log2(RPKM)
z−score of IF value
−2.5
0.0
2.5

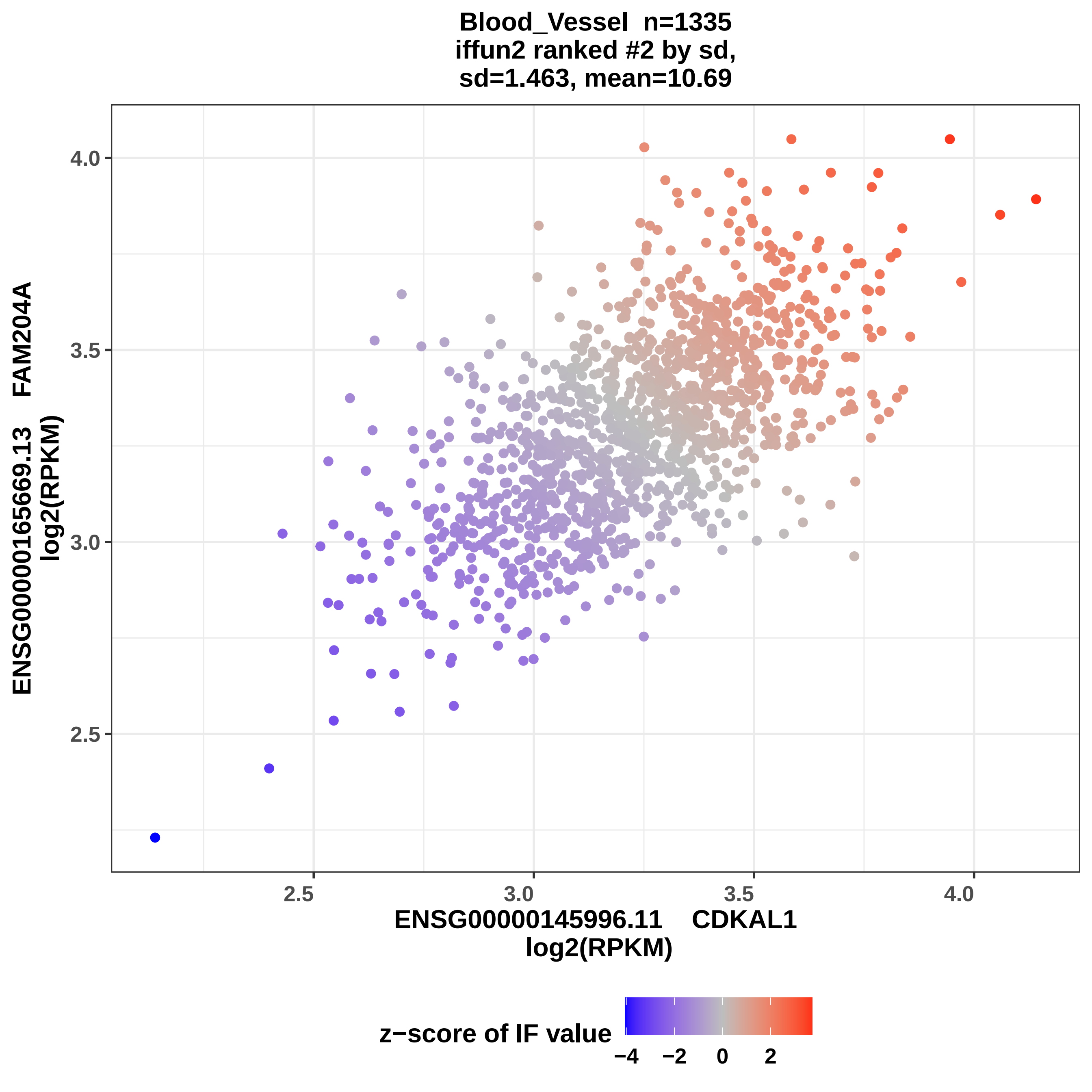
Blood_Vessel n=1335
iffun2 ranked #2 by sd,
sd=1.463, mean=10.69
ENSG00000165669.13 FAM204A
log2(RPKM)
4.0
3.5
3.0
2.5
2.5
3.0
3.5
4.0
ENSG00000145996.11 CDKAL1
log2(RPKM)
z−score of IF value
−4
−2
0
2

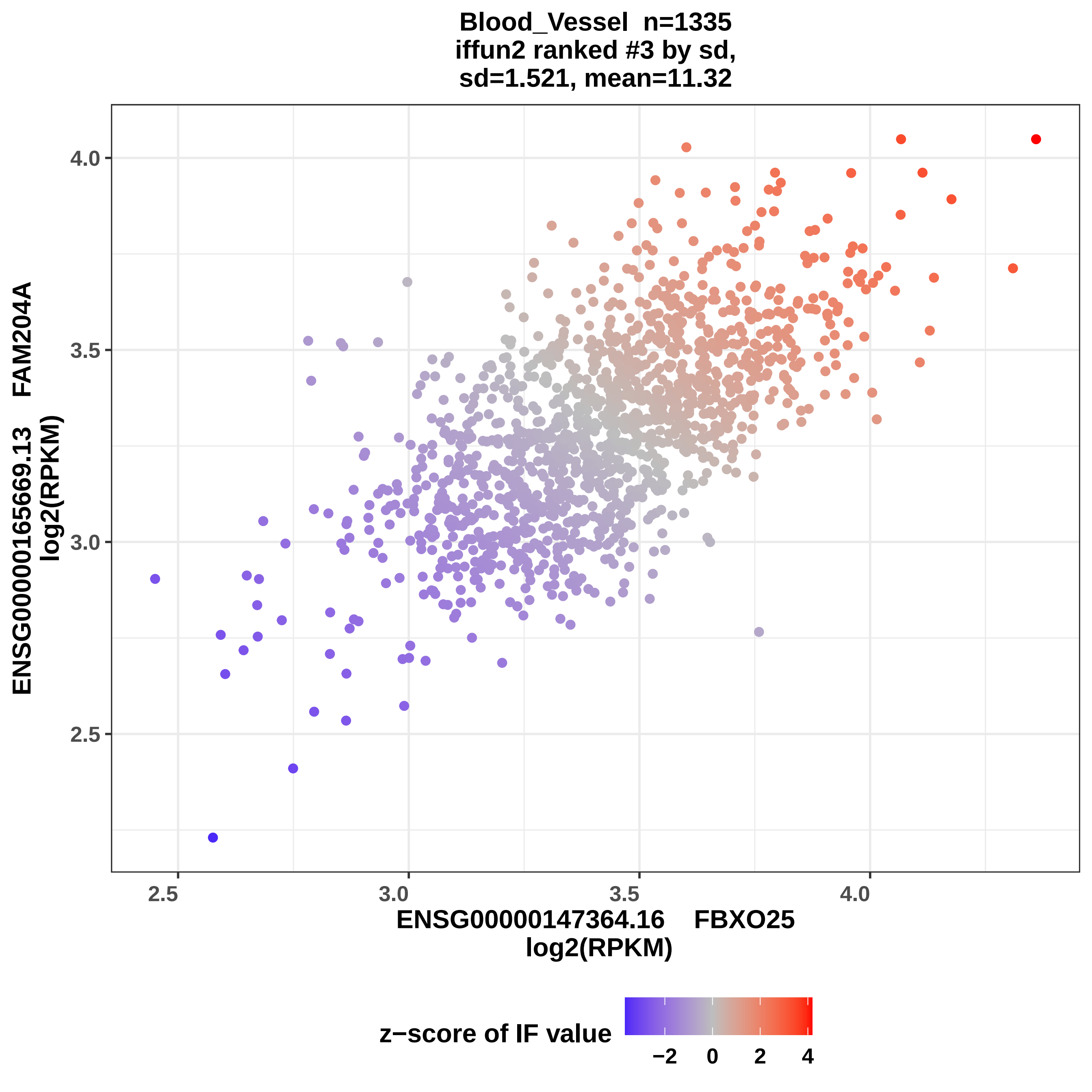
Blood_Vessel n=1335
iffun2 ranked #3 by sd,
sd=1.521, mean=11.32
ENSG00000165669.13 FAM204A
log2(RPKM)
4.0
3.5
3.0
2.5
2.5
3.0
3.5
4.0
ENSG00000147364.16 FBXO25
log2(RPKM)
z−score of IF value
−2
0
2
4

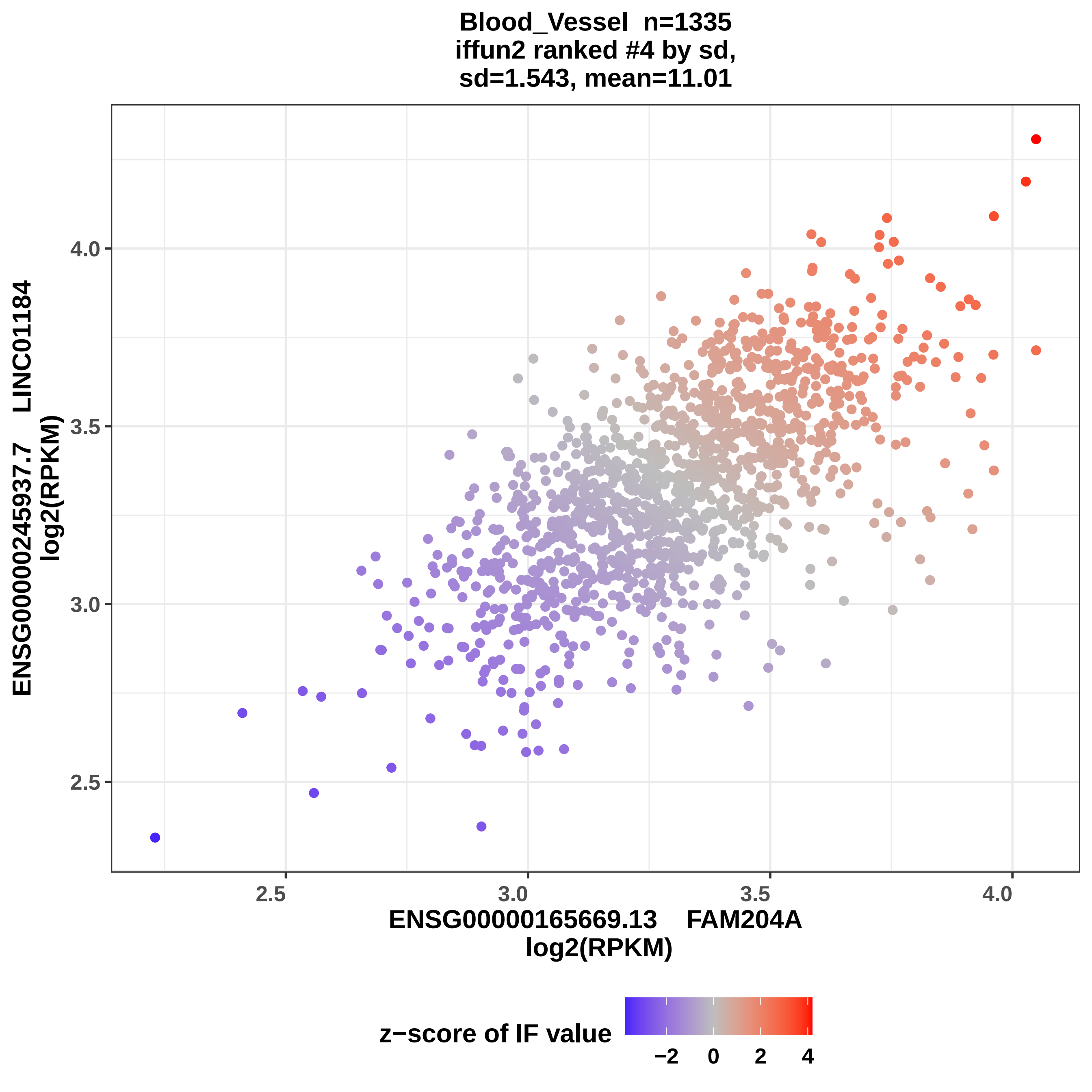

Blood_Vessel n=1335
iffun2 ranked #4 by sd,
sd=1.543, mean=11.01
ENSG00000245937.7 LINC01184
log2(RPKM)
4.0
3.5
3.0
2.5
2.5
3.0
3.5
4.0
ENSG00000165669.13 FAM204A
log2(RPKM)
z−score of IF value
−2
0
2
4

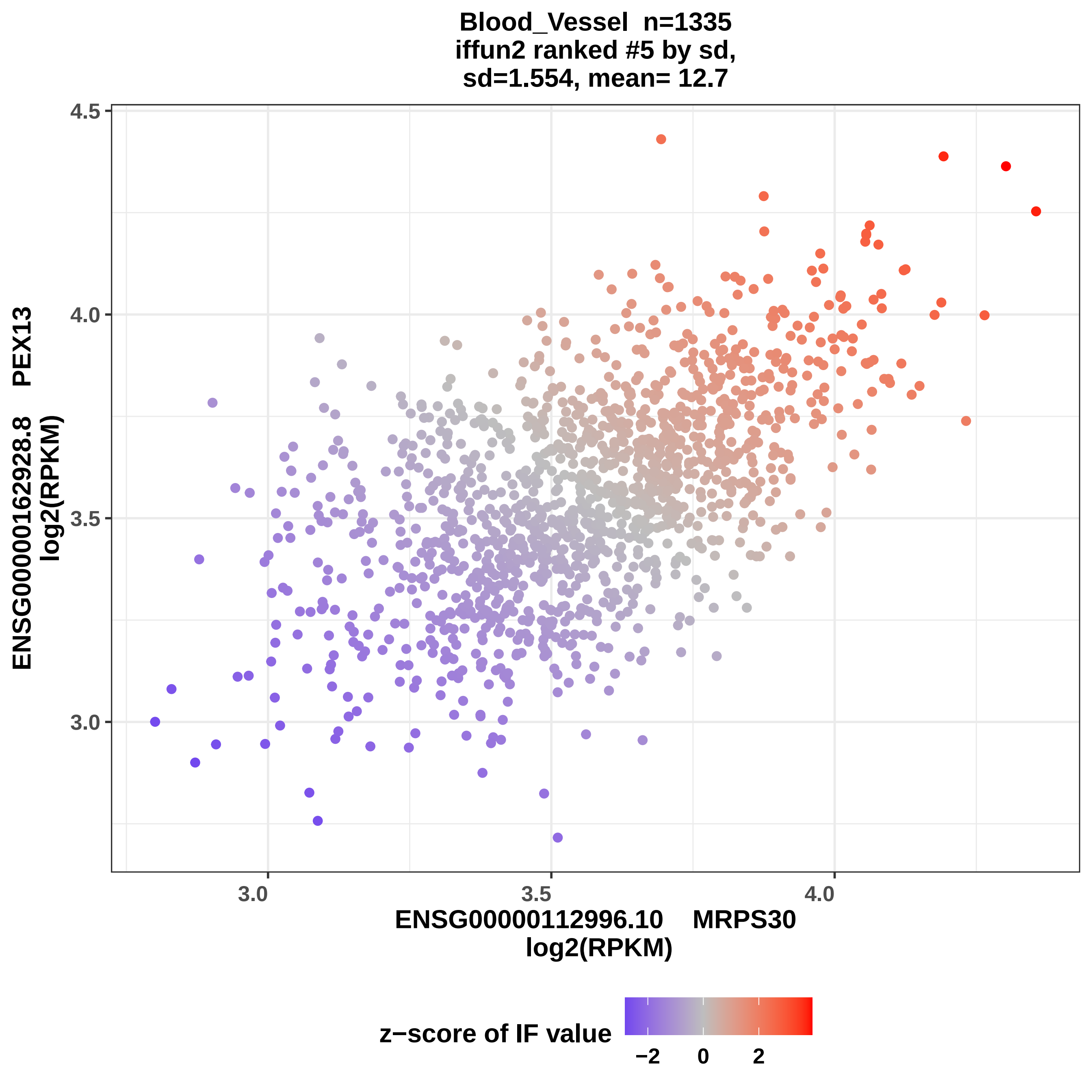

Blood_Vessel n=1335
iffun2 ranked #5 by sd,
sd=1.554, mean= 12.7
ENSG00000162928.8 PEX13
log2(RPKM)
4.5
4.0
3.5
3.0
3.0
3.5
4.0
ENSG00000112996.10 MRPS30
log2(RPKM)
z−score of IF value
−2
0
2

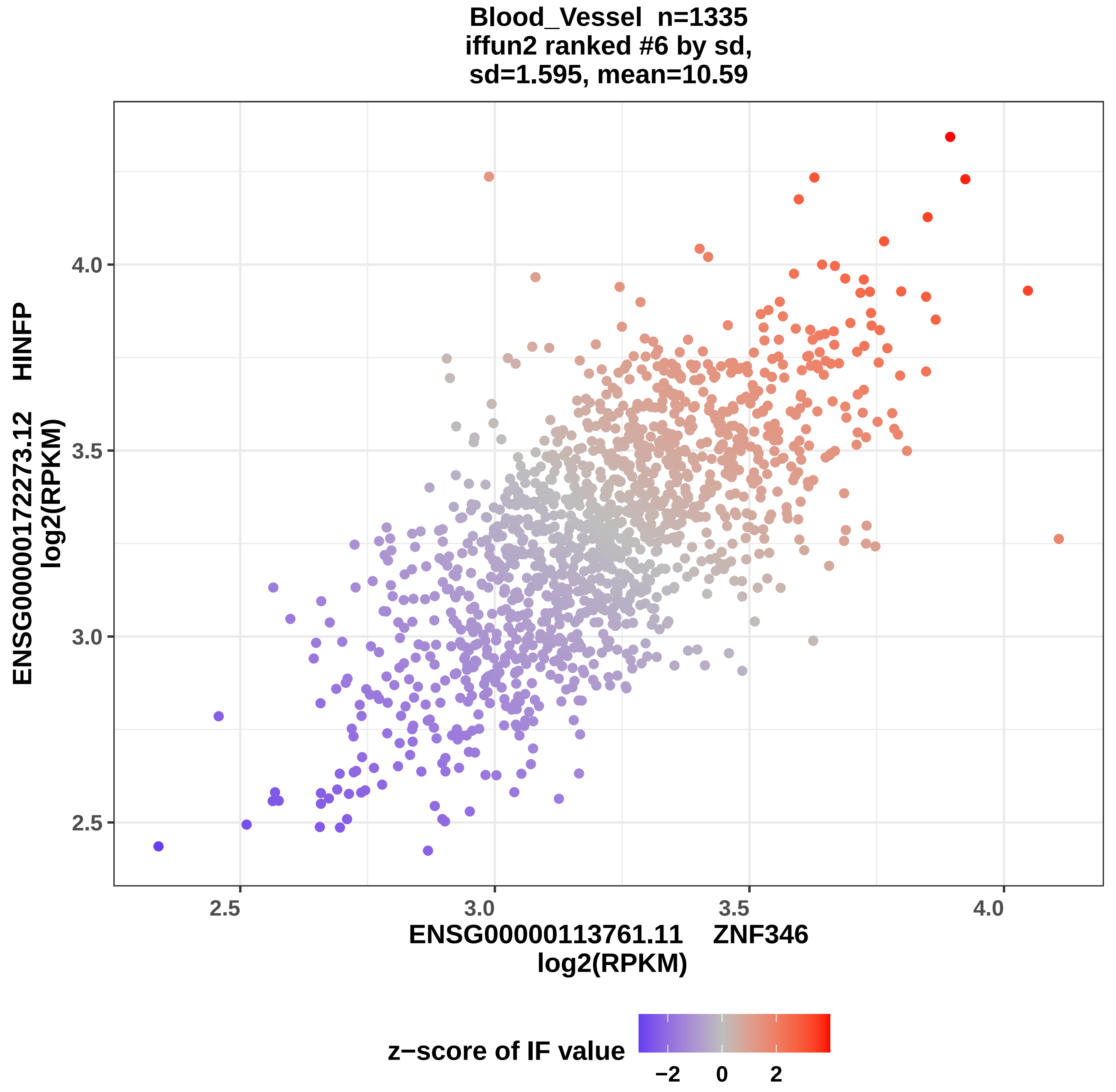
Blood_Vessel n=1335
iffun2 ranked #6 by sd,
sd=1.595, mean=10.59
ENSG00000172273.12 HINFP
log2(RPKM)
4.0
3.5
3.0
2.5
2.5
3.0
3.5
4.0
ENSG00000113761.11 ZNF346
log2(RPKM)
z−score of IF value
−2
0
2

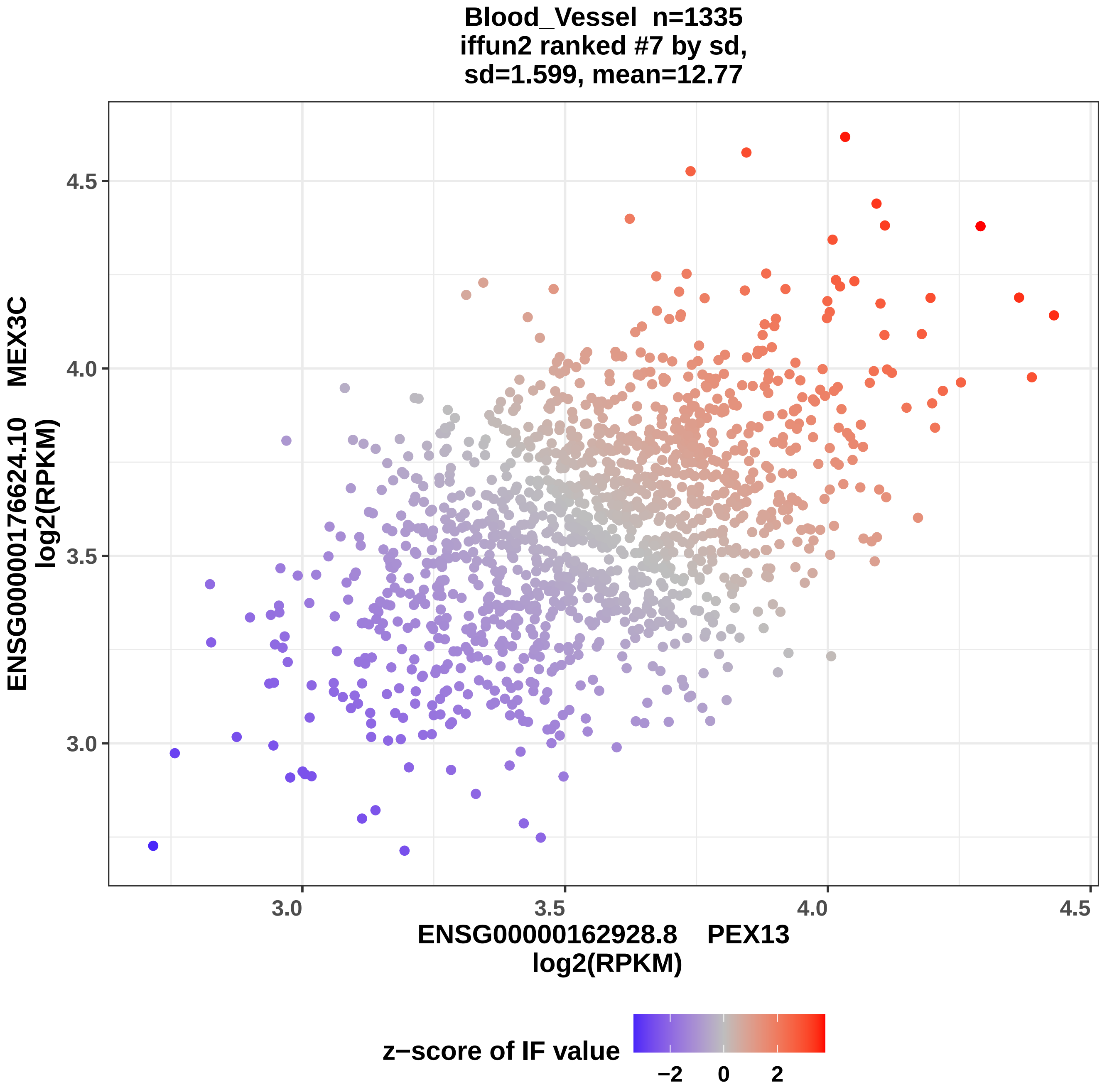
Blood_Vessel n=1335
iffun2 ranked #7 by sd,
sd=1.599, mean=12.77
ENSG00000176624.10 MEX3C
log2(RPKM)
ENSG00000162928.8 PEX13
log2(RPKM)
4.5
4.0
3.5
3.0
3.0
3.5
4.0
4.5
z−score of IF value
−2
0
2

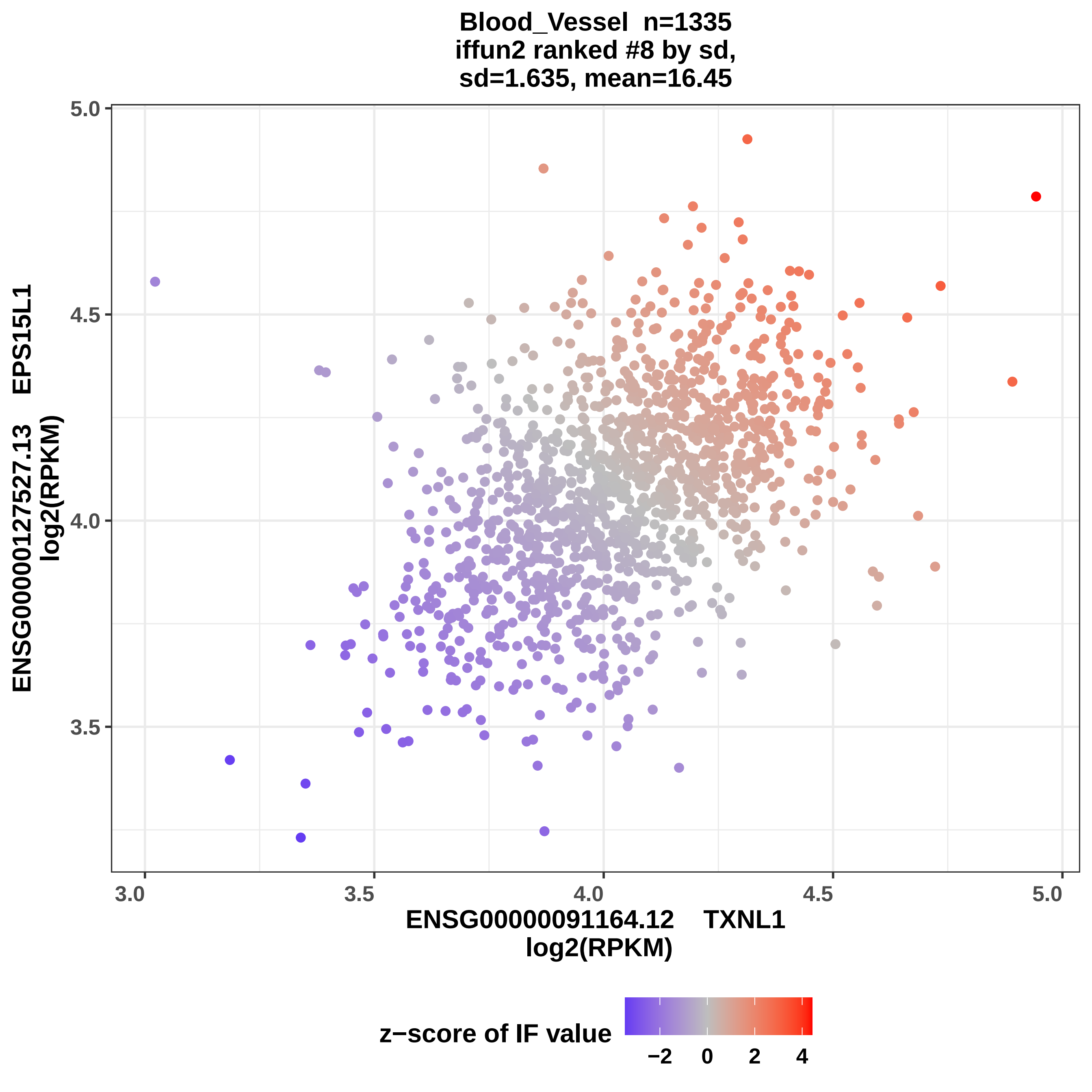

Blood_Vessel n=1335
iffun2 ranked #8 by sd,
sd=1.635, mean=16.45
ENSG00000127527.13 EPS15L1
log2(RPKM)
5.0
4.5
4.0
3.5
3.0
3.5
4.0
4.5
5.0
ENSG00000091164.12 TXNL1
log2(RPKM)
z−score of IF value
−2
0
2
4

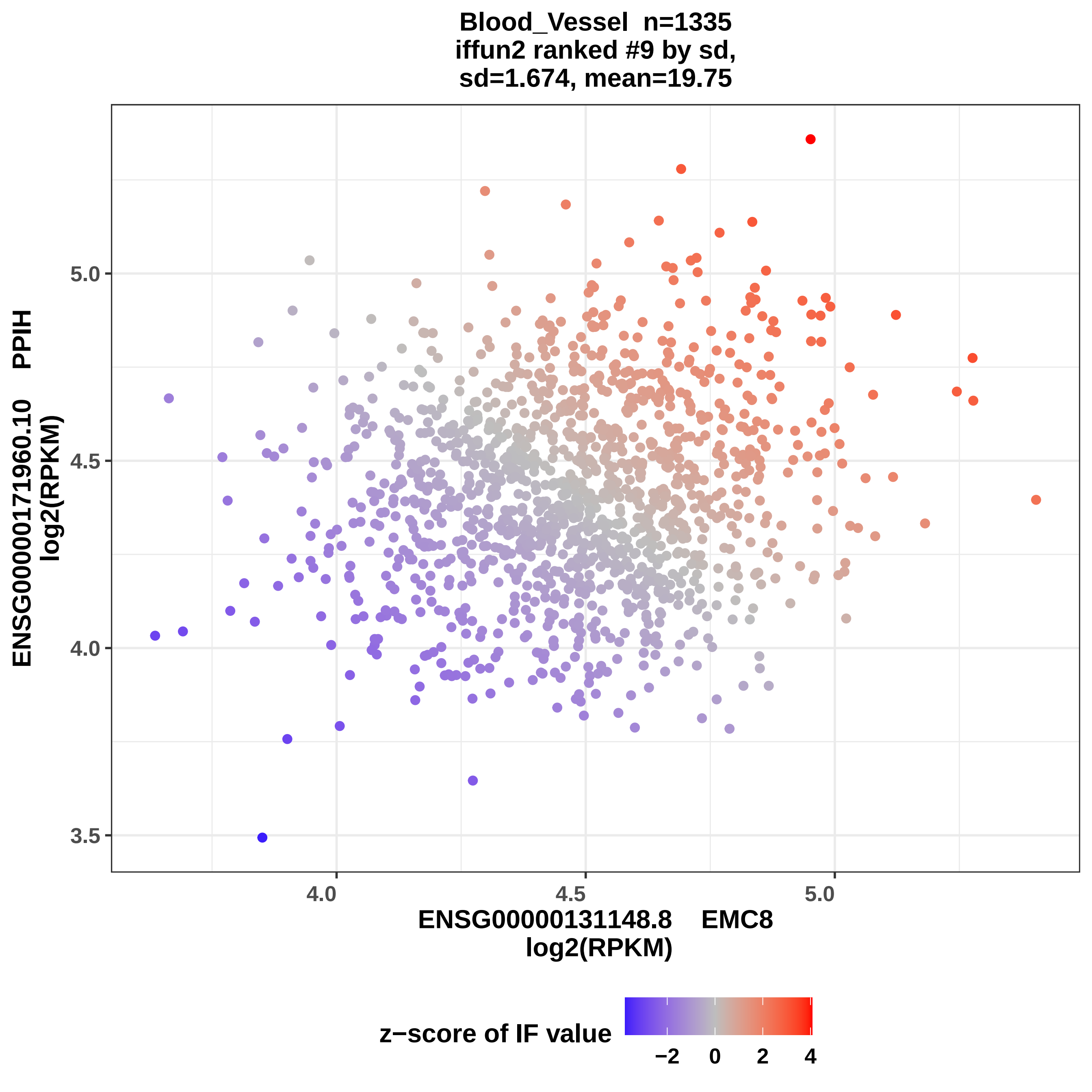
Blood_Vessel n=1335
iffun2 ranked #9 by sd,
sd=1.674, mean=19.75
ENSG00000171960.10 PPIH
log2(RPKM)
5.0
4.5
4.0
3.5
4.0
4.5
5.0
ENSG00000131148.8 EMC8
log2(RPKM)
z−score of IF value
−2
0
2
4

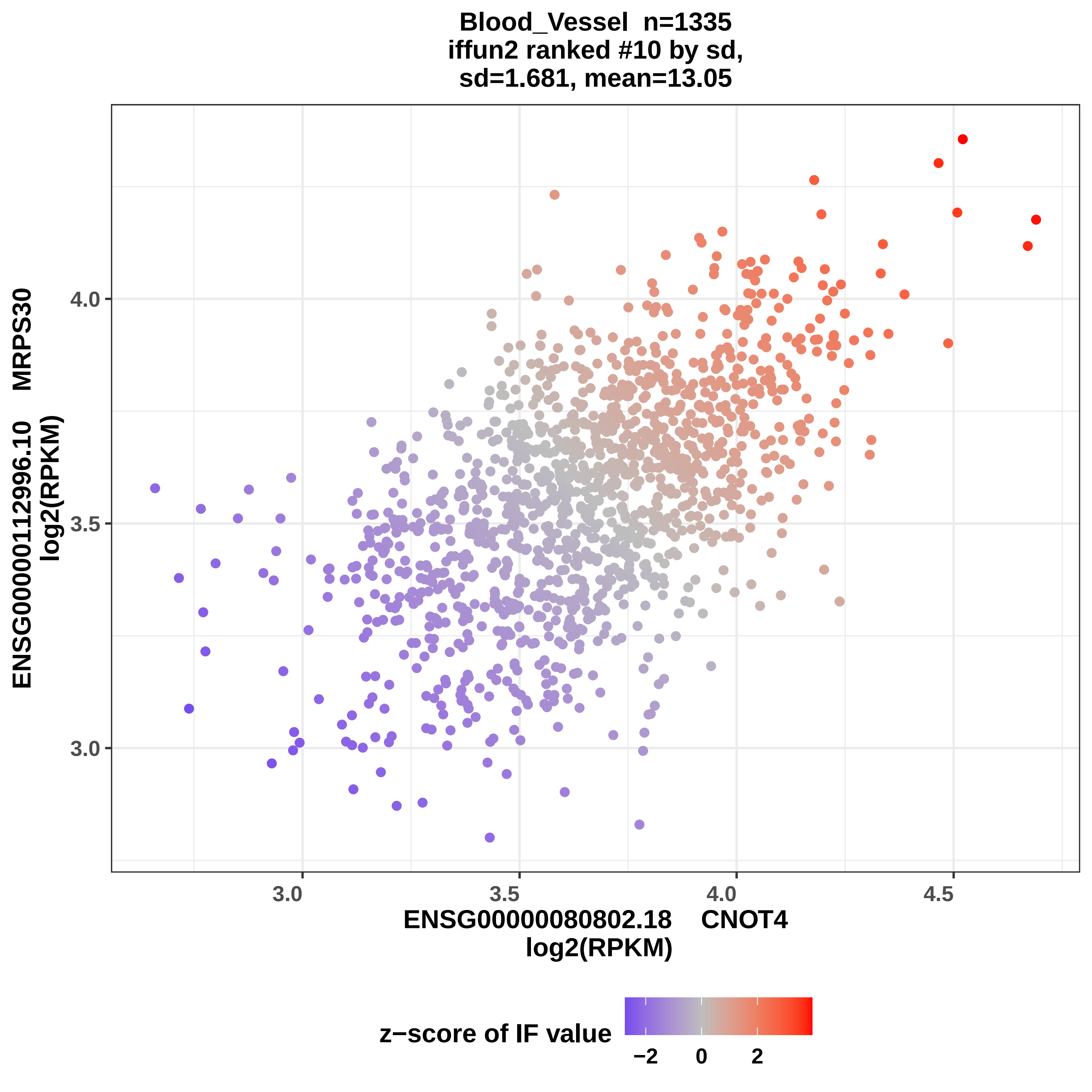
Blood_Vessel n=1335
iffun2 ranked #10 by sd,
sd=1.681, mean=13.05
ENSG00000112996.10 MRPS30
log2(RPKM)
4.0
3.5
3.0
3.0
3.5
4.0
4.5
ENSG00000080802.18 CNOT4
log2(RPKM)
z−score of IF value
−2
0
2

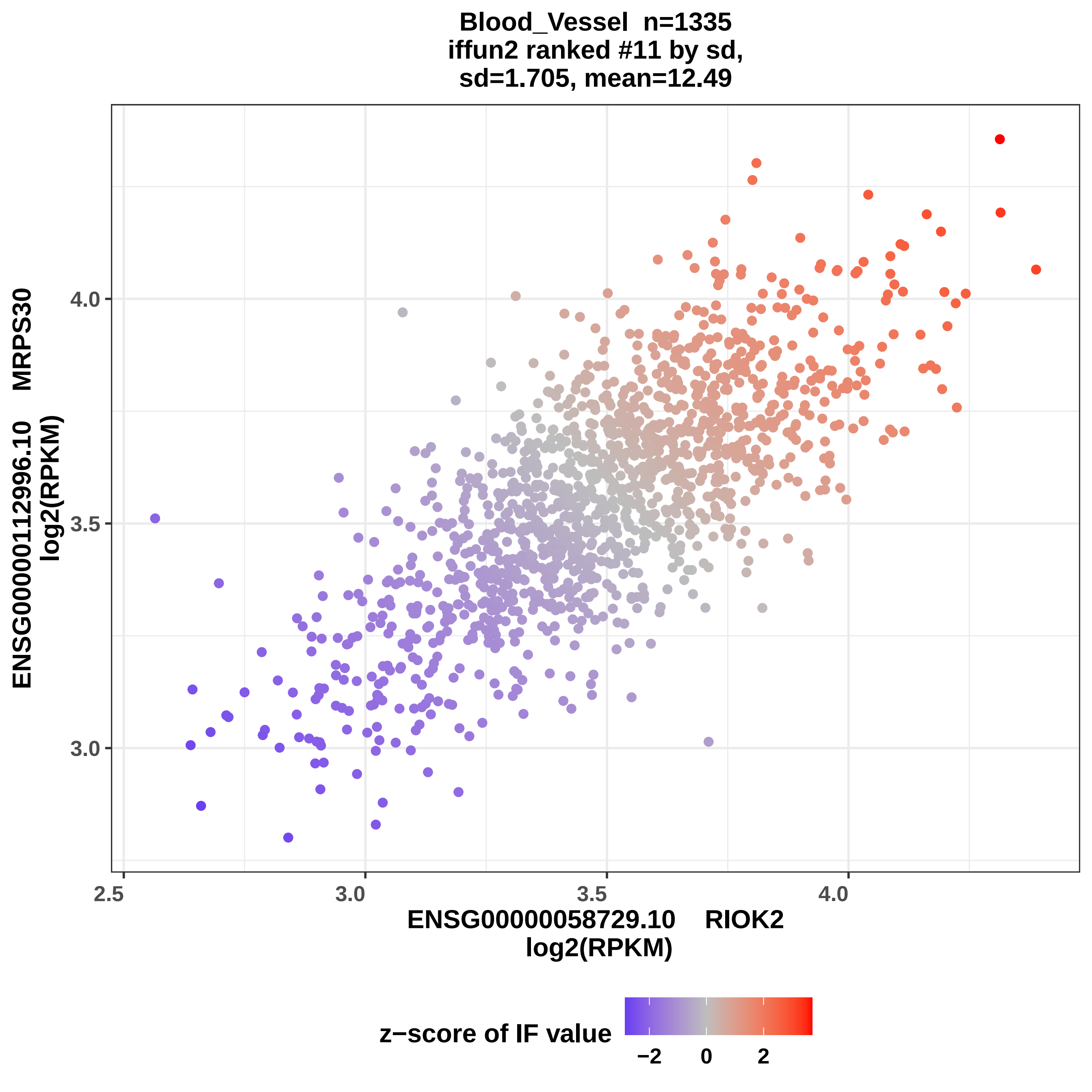
Blood_Vessel n=1335
iffun2 ranked #11 by sd,
sd=1.705, mean=12.49
ENSG00000112996.10 MRPS30
log2(RPKM)
4.0
3.5
3.0
2.5
3.0
3.5
4.0
ENSG00000058729.10 RIOK2
log2(RPKM)
z−score of IF value
−2
0
2

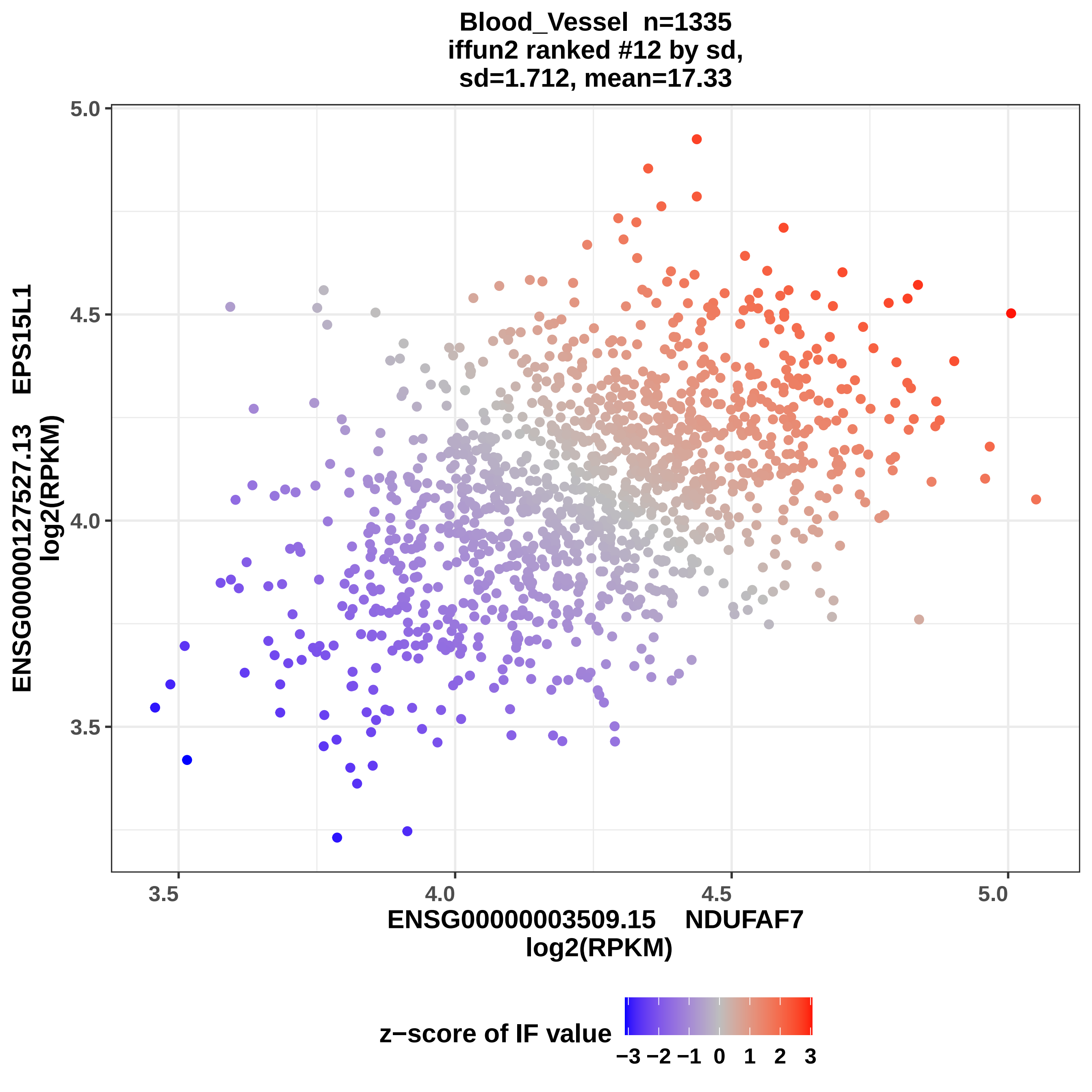
Blood_Vessel n=1335
iffun2 ranked #12 by sd,
sd=1.712, mean=17.33
ENSG00000127527.13 EPS15L1
log2(RPKM)
ENSG00000003509.15 NDUFAF7
log2(RPKM)
5.0
4.5
4.0
3.5
3.5
4.0
4.5
5.0
z−score of IF value
−3 −2 −1 0 1 2 3

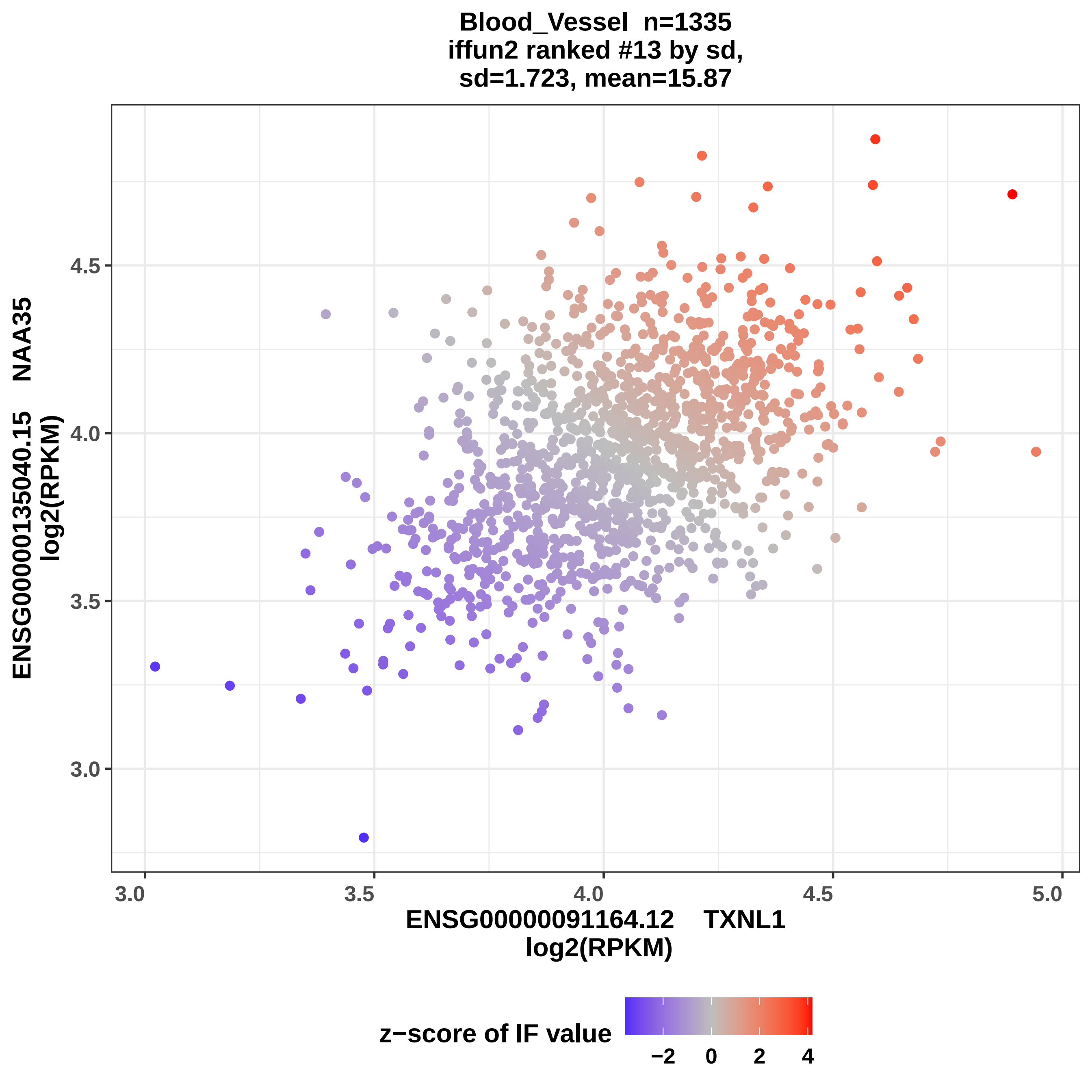

Blood_Vessel n=1335
iffun2 ranked #13 by sd,
sd=1.723, mean=15.87
ENSG00000135040.15 NAA35
log2(RPKM)
4.5
4.0
3.5
3.0
3.0
3.5
4.0
4.5
5.0
ENSG00000091164.12 TXNL1
log2(RPKM)
z−score of IF value
−2
0
2
4

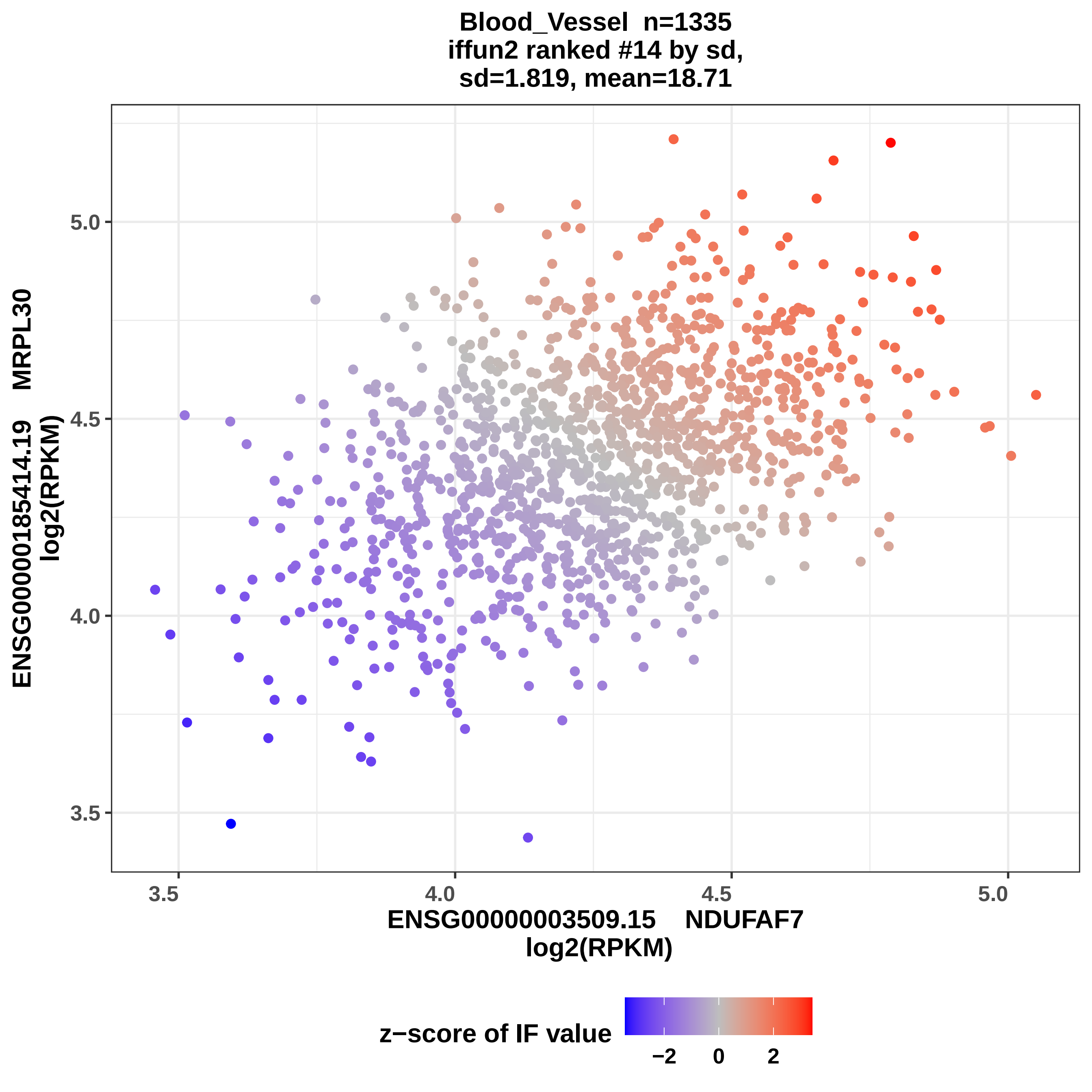

Blood_Vessel n=1335
iffun2 ranked #14 by sd,
sd=1.819, mean=18.71
ENSG00000185414.19 MRPL30
log2(RPKM)
ENSG00000003509.15 NDUFAF7
log2(RPKM)
z−score of IF value
−2
0
2
3.5
4.0
4.5
5.0

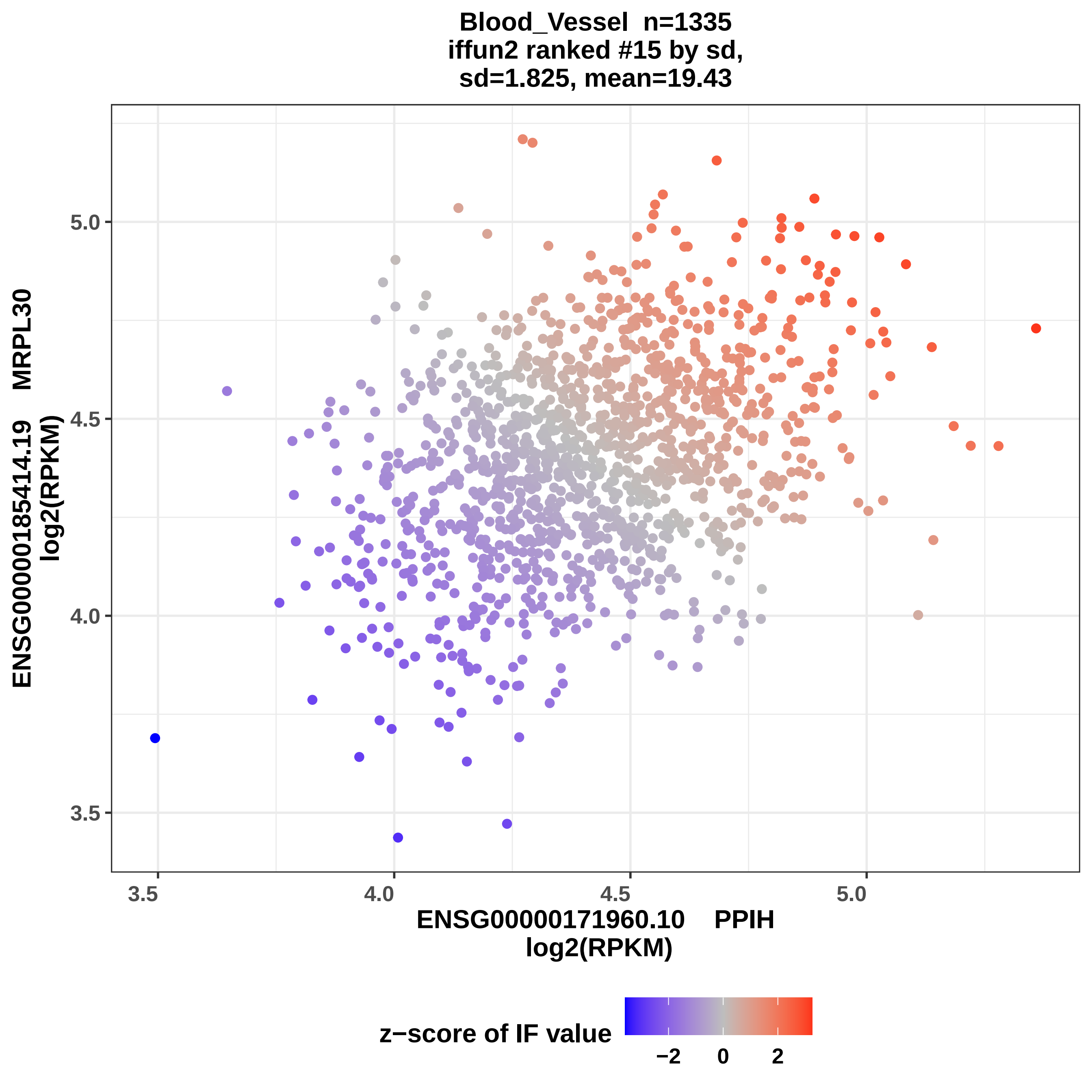

Blood_Vessel n=1335
iffun2 ranked #15 by sd,
sd=1.825, mean=19.43
ENSG00000185414.19 MRPL30
log2(RPKM)
5.0
4.5
4.0
3.5
3.5
4.0
4.5
5.0
ENSG00000171960.10 PPIH
log2(RPKM)
z−score of IF value
−2
0
2

**Blood_Vessel n=1335**
**iffun2 ranked #16 by sd,**
**sd=1.827, mean=17.05**

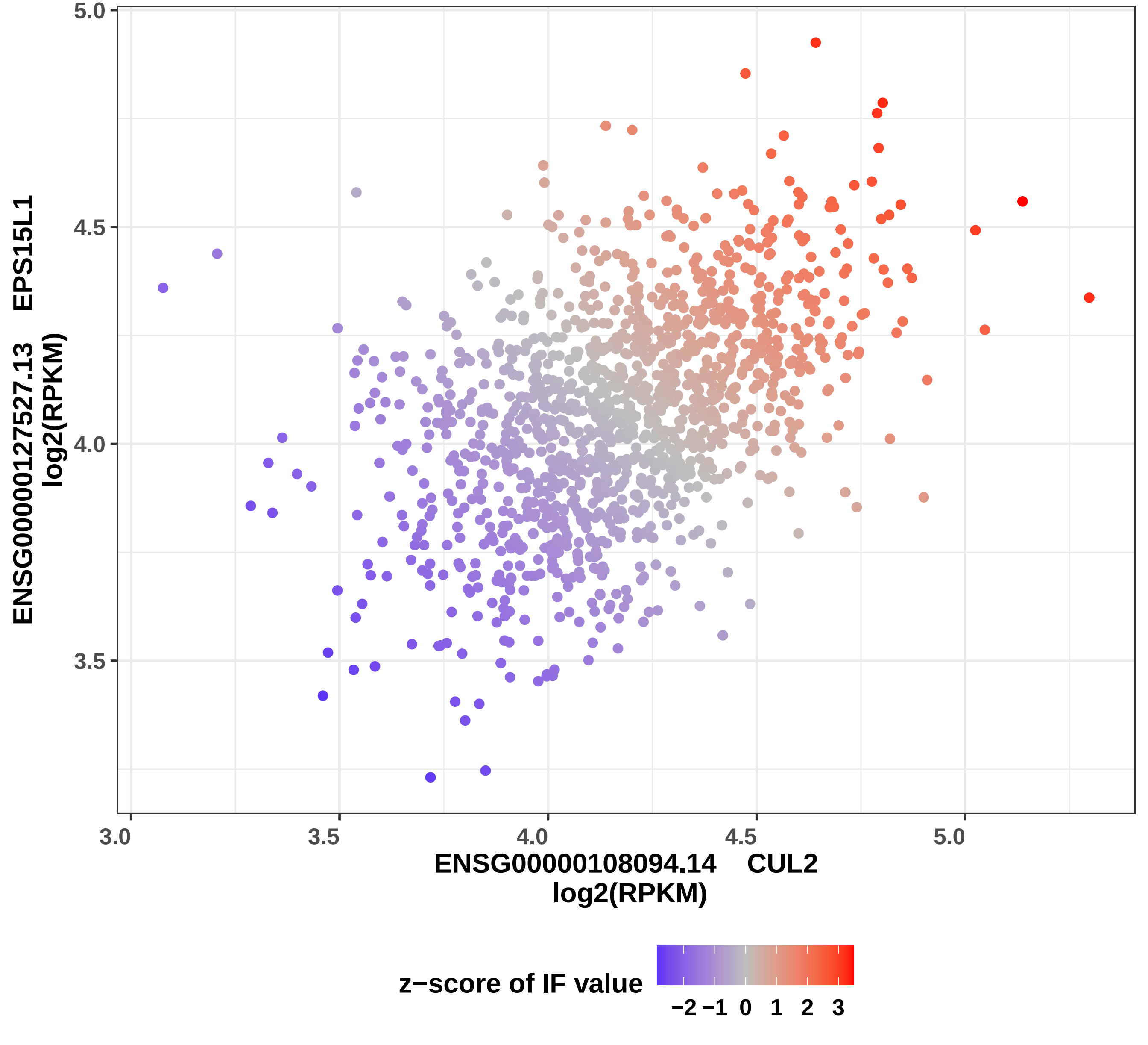

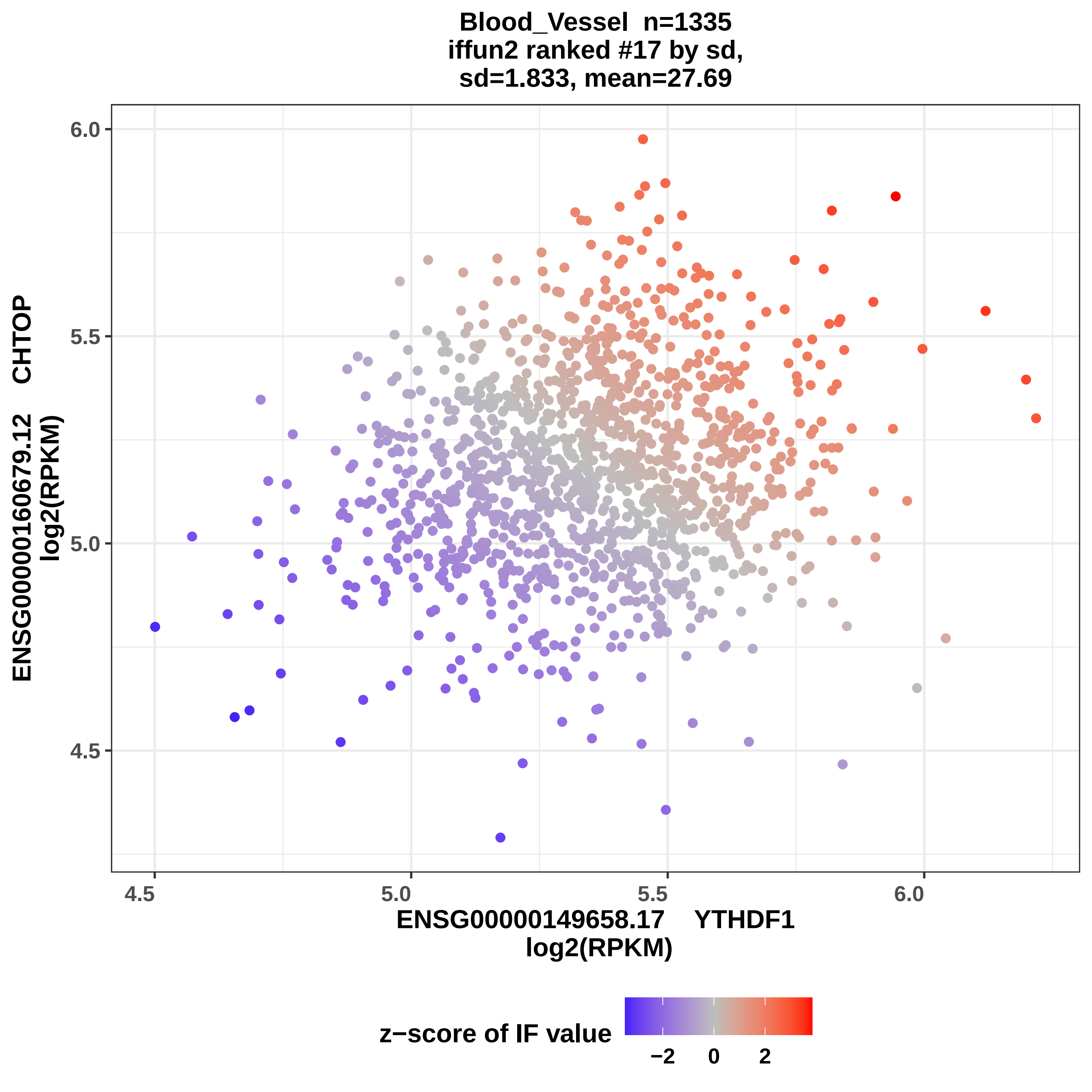
Blood_Vessel n=1335
iffun2 ranked #17 by sd,
sd=1.833, mean=27.69
ENSG00000160679.12 CHTOP
log2(RPKM)
6.0
5.5
5.0
4.5
4.5
5.0
5.5
6.0
ENSG00000149658.17 YTHDF1
log2(RPKM)
z−score of IF value
−2
0
2

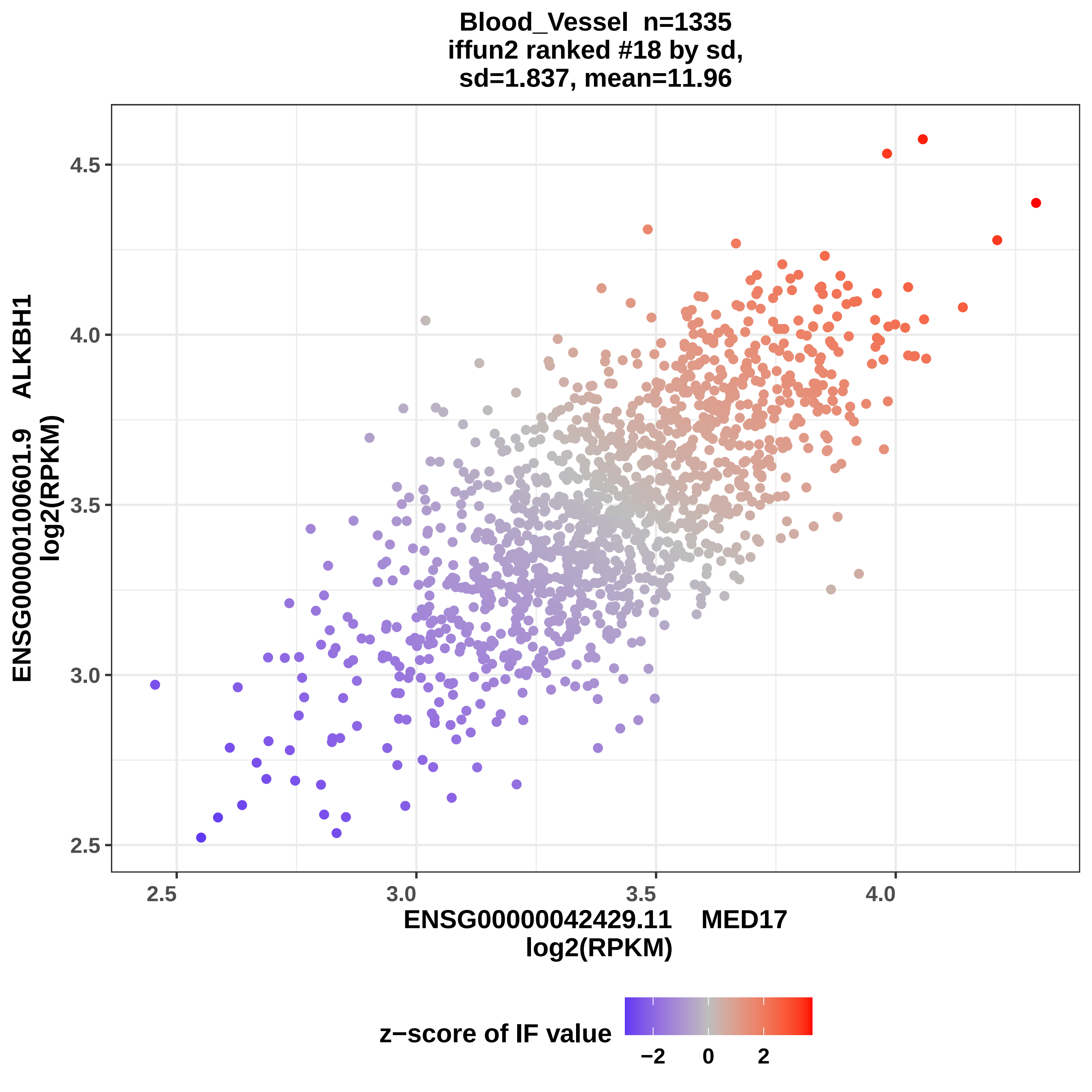
Blood_Vessel n=1335
iffun2 ranked #18 by sd,
sd=1.837, mean=11.96
ENSG00000100601.9 ALKBH1
log2(RPKM)
4.5
4.0
3.5
3.0
2.5
2.5
3.0
3.5
4.0
ENSG00000042429.11 MED17
log2(RPKM)
z−score of IF value
−2
0
2

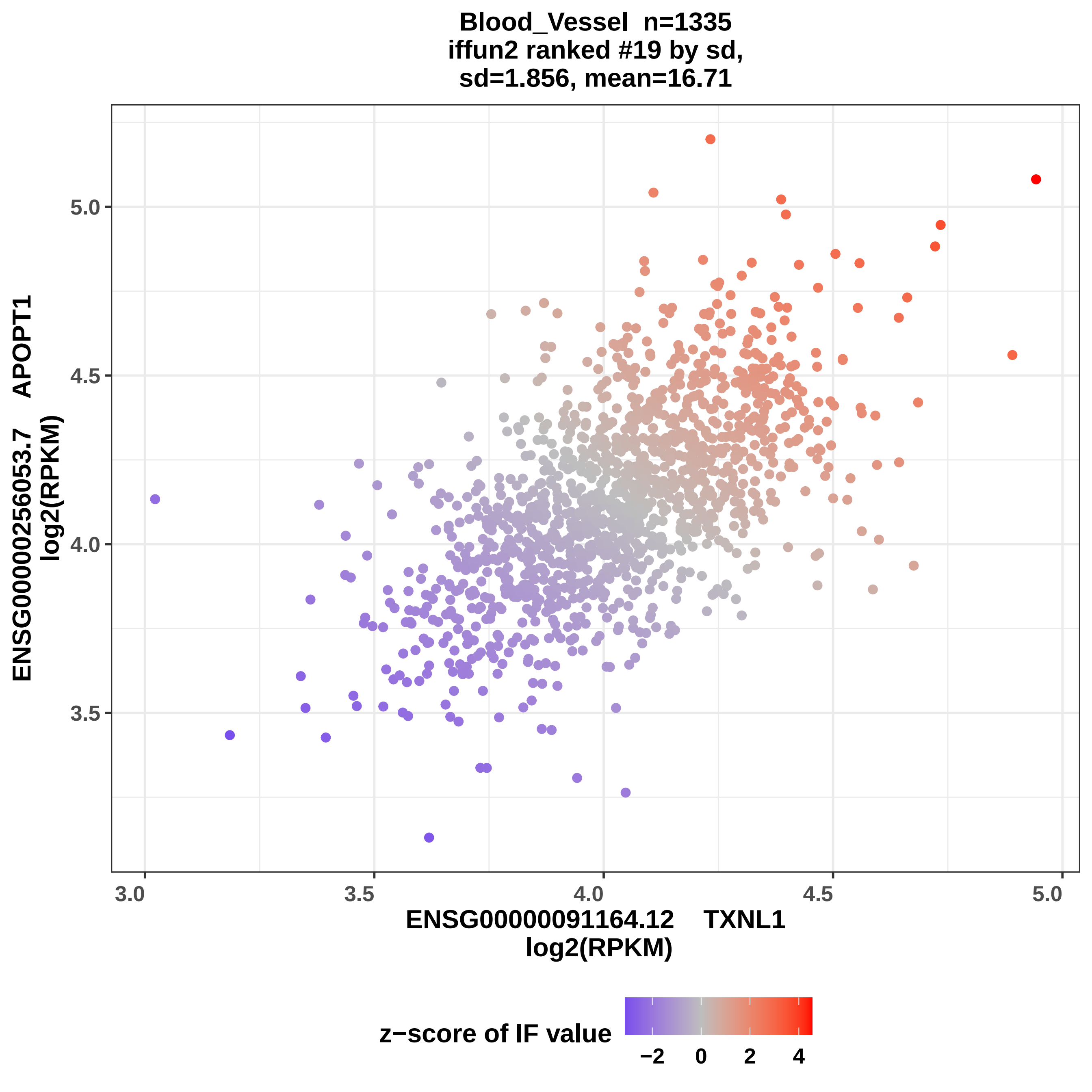
Blood_Vessel n=1335
iffun2 ranked #19 by sd,
sd=1.856, mean=16.71
ENSG00000256053.7 APOPT1
log2(RPKM)
5.0
4.5
4.0
3.5
3.0
3.5
4.0
4.5
5.0
ENSG00000091164.12 TXNL1
log2(RPKM)
z−score of IF value
−2
0
2
4

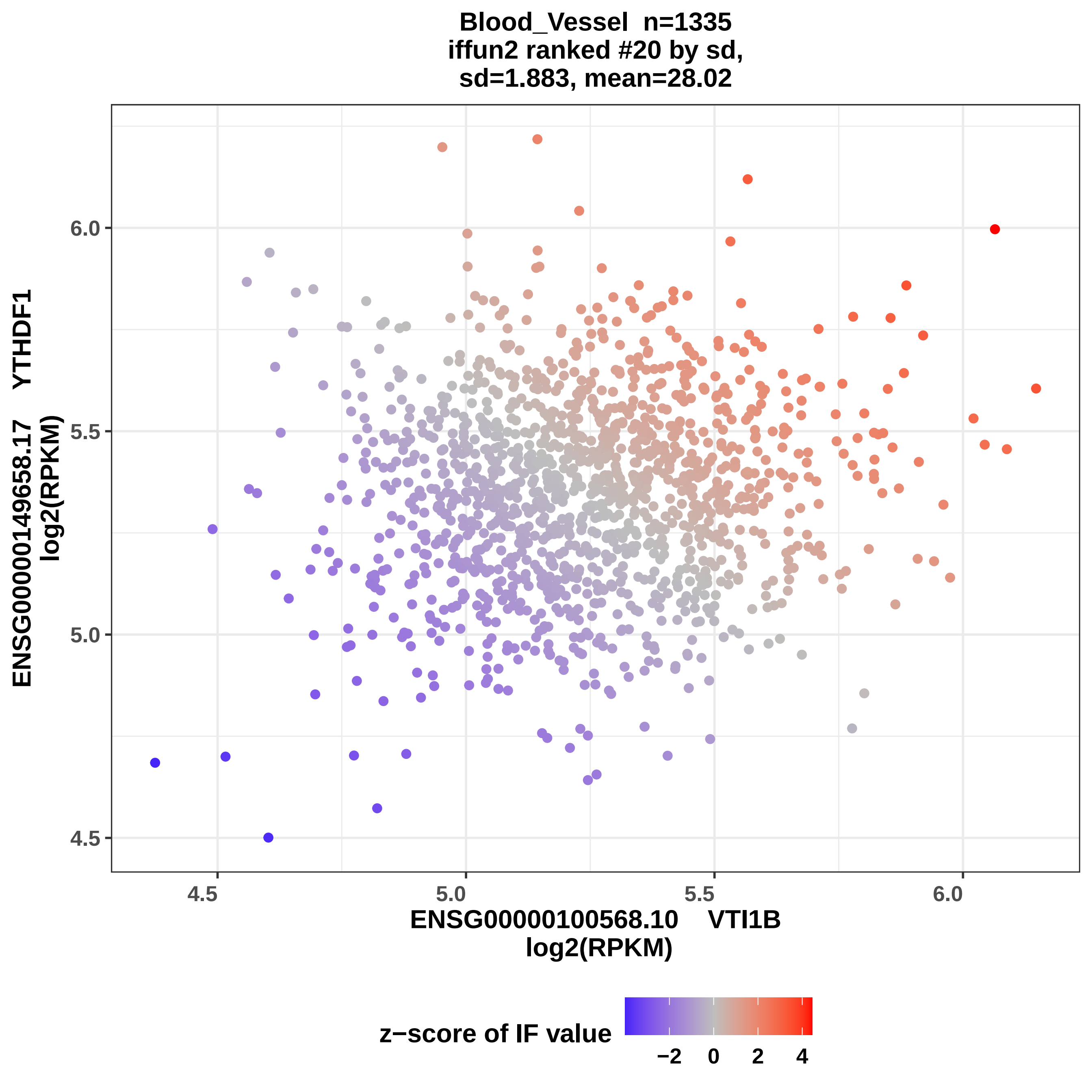
Blood_Vessel n=1335
iffun2 ranked #20 by sd,
sd=1.883, mean=28.02
ENSG00000149658.17 YTHDF1
log2(RPKM)
6.0
5.5
5.0
4.5
4.5
5.0
5.5
6.0
ENSG00000100568.10 VTI1B
log2(RPKM)
z−score of IF value
−2
0
2
4

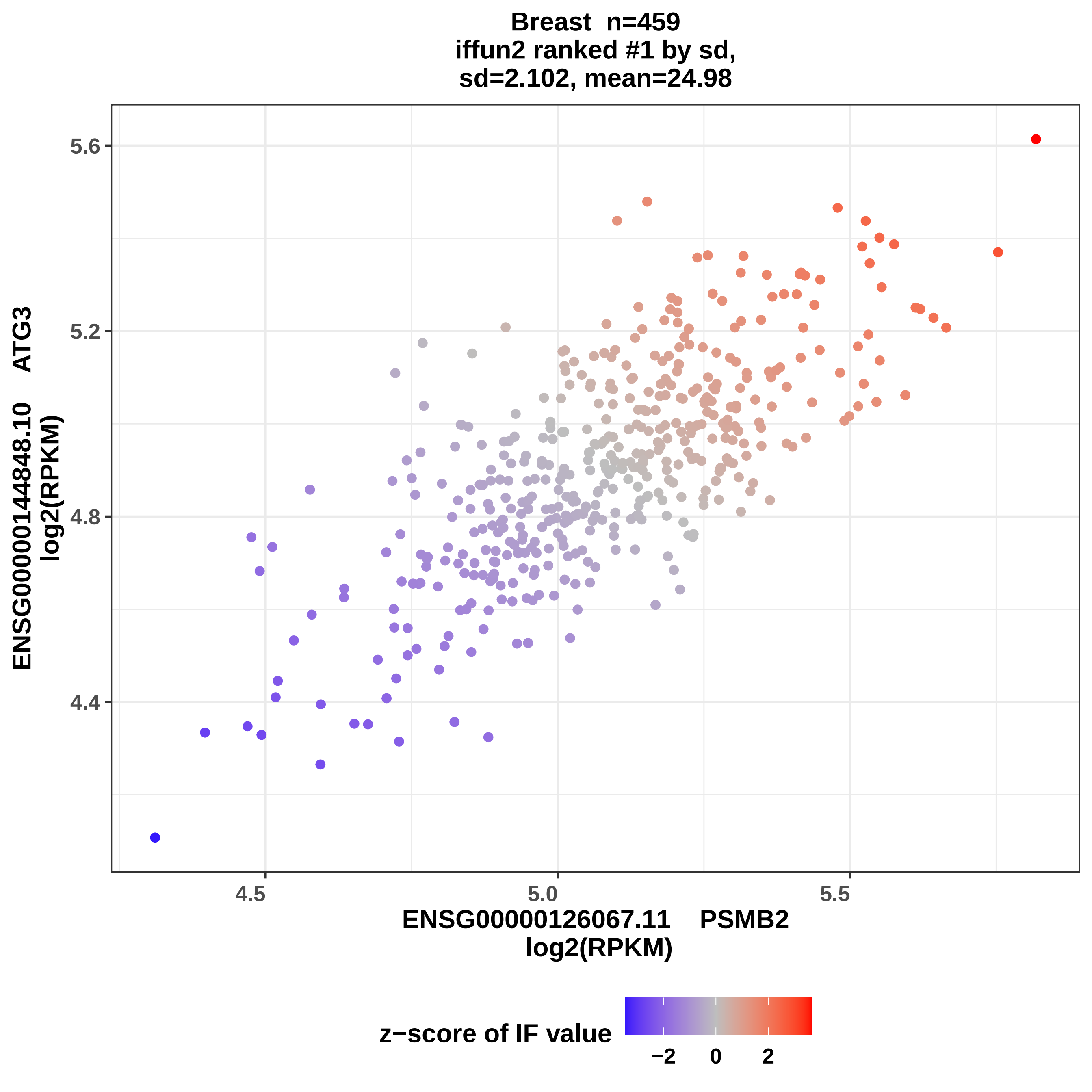

Breast n=459
iffun2 ranked #1 by sd,
sd=2.102, mean=24.98
ENSG00000144848.10 ATG3
log2(RPKM)
5.6
5.2
4.8
4.4
4.5
5.0
5.5
ENSG00000126067.11 PSMB2
log2(RPKM)
z−score of IF value
−2
0
2

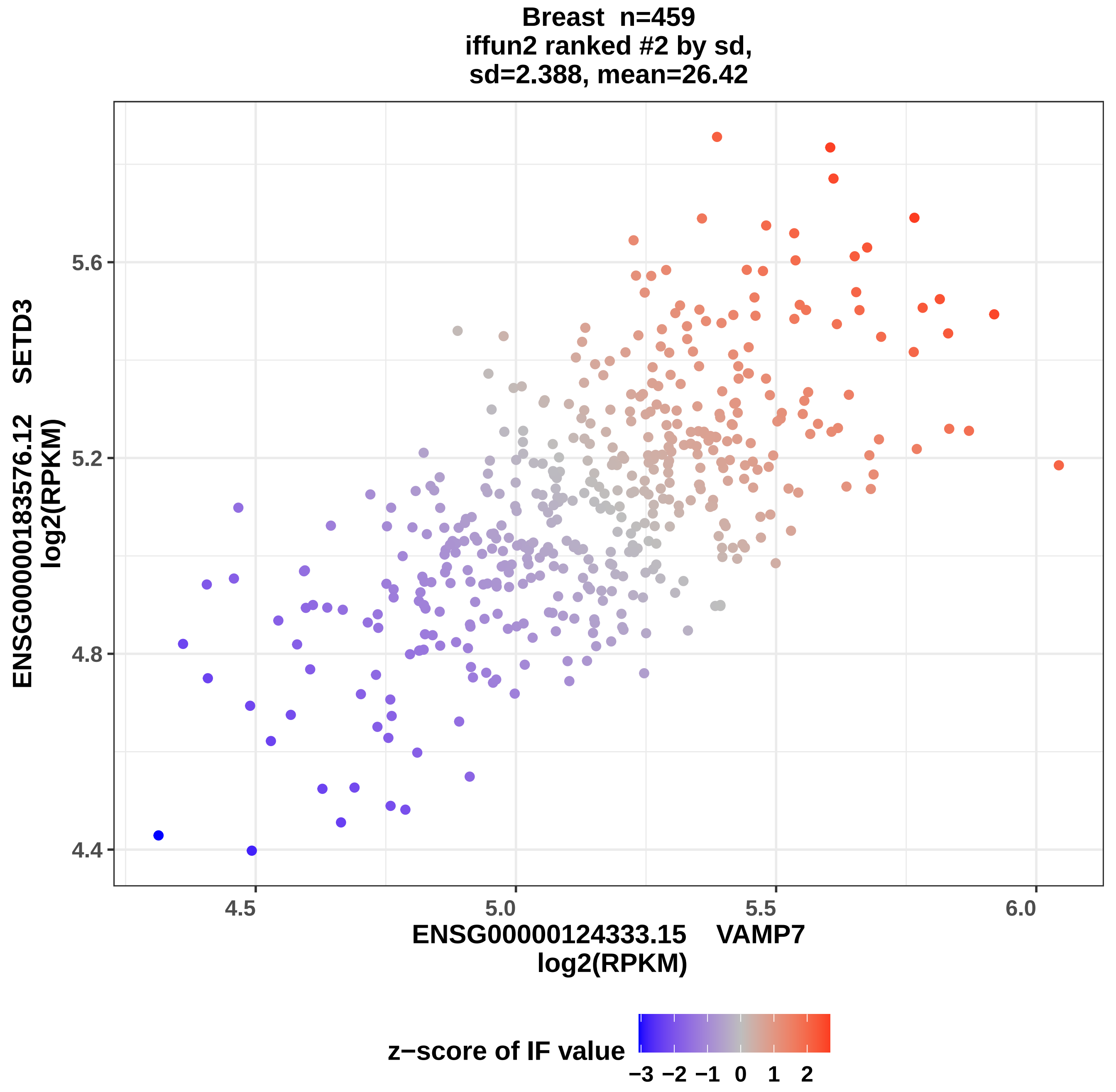

Breast n=459
iffun2 ranked #2 by sd,
sd=2.388, mean=26.42
ENSG00000183576.12 SETD3
log2(RPKM)
5.6
5.2
4.8
4.4
4.5
5.0
5.5
6.0
ENSG00000124333.15 VAMP7
log2(RPKM)
z−score of IF value
−3 −2 −1 0 1 2

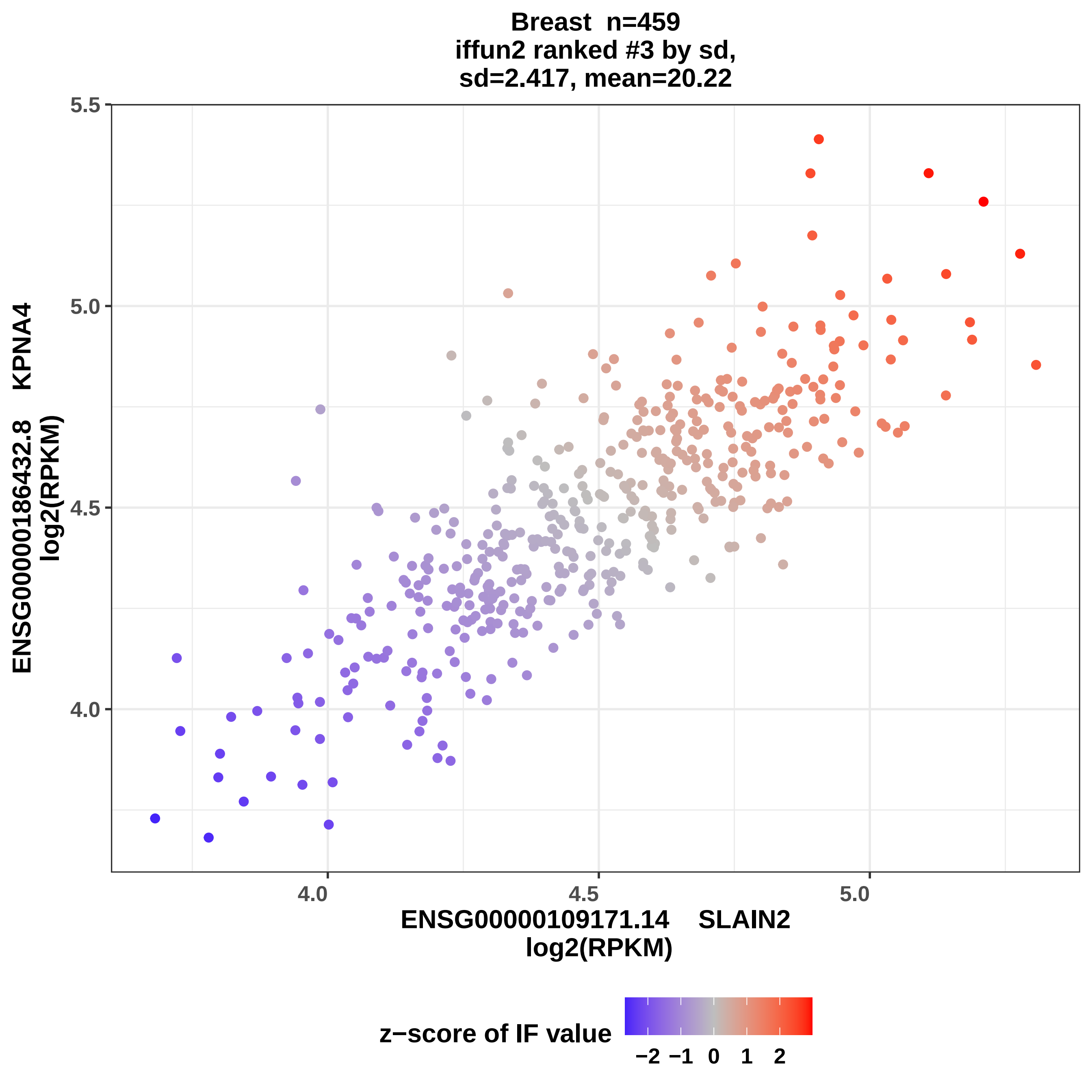
Breast n=459
iffun2 ranked #3 by sd,
sd=2.417, mean=20.22
5.5
5.0
4.5
4.0
ENSG00000186432.8 KPNA4
log2(RPKM)
4.0
4.5
5.0
ENSG00000109171.14 SLAIN2
log2(RPKM)
z−score of IF value
−2 −1 0 1 2

Breast n=459
iffun2 ranked #4 by sd,
sd=2.459, mean= 26.4

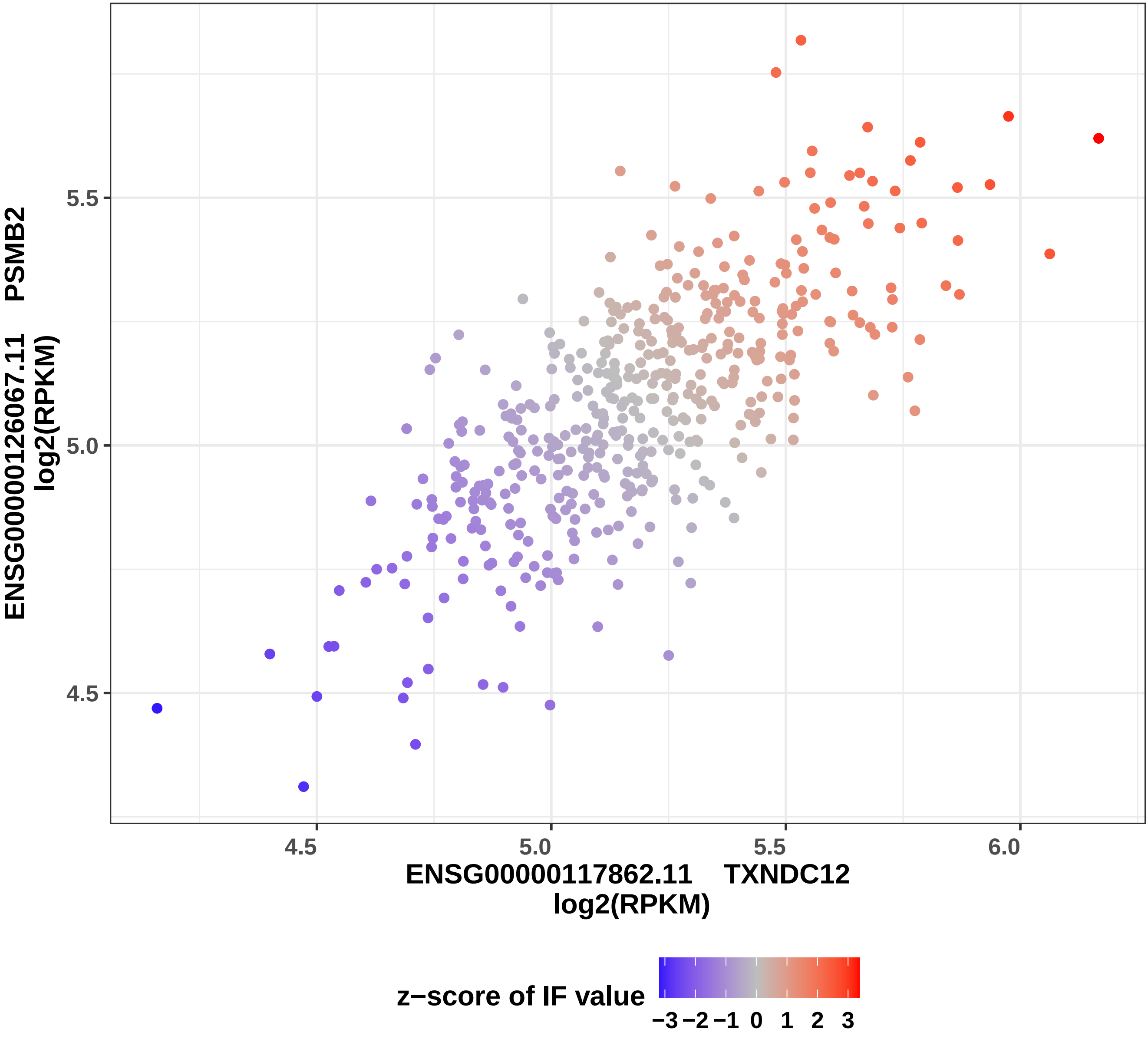

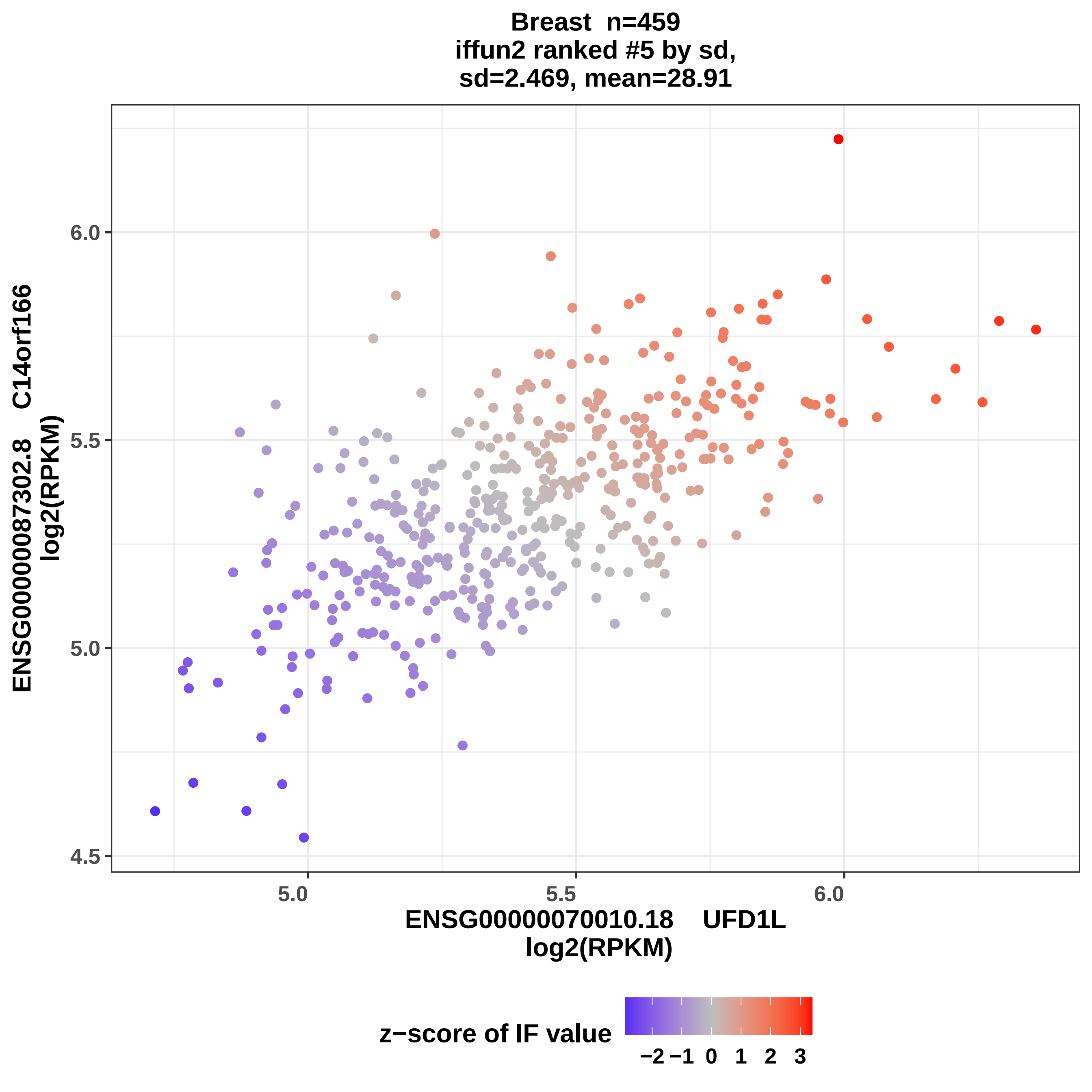
Breast n=459
iffun2 ranked #5 by sd,
sd=2.469, mean=28.91
ENSG00000087302.8 C14orf166
log2(RPKM)
6.0
5.5
5.0
4.5
5.0
5.5
6.0
ENSG00000070010.18 UFD1L
log2(RPKM)
z−score of IF value
−2 −1 0 1 2 3

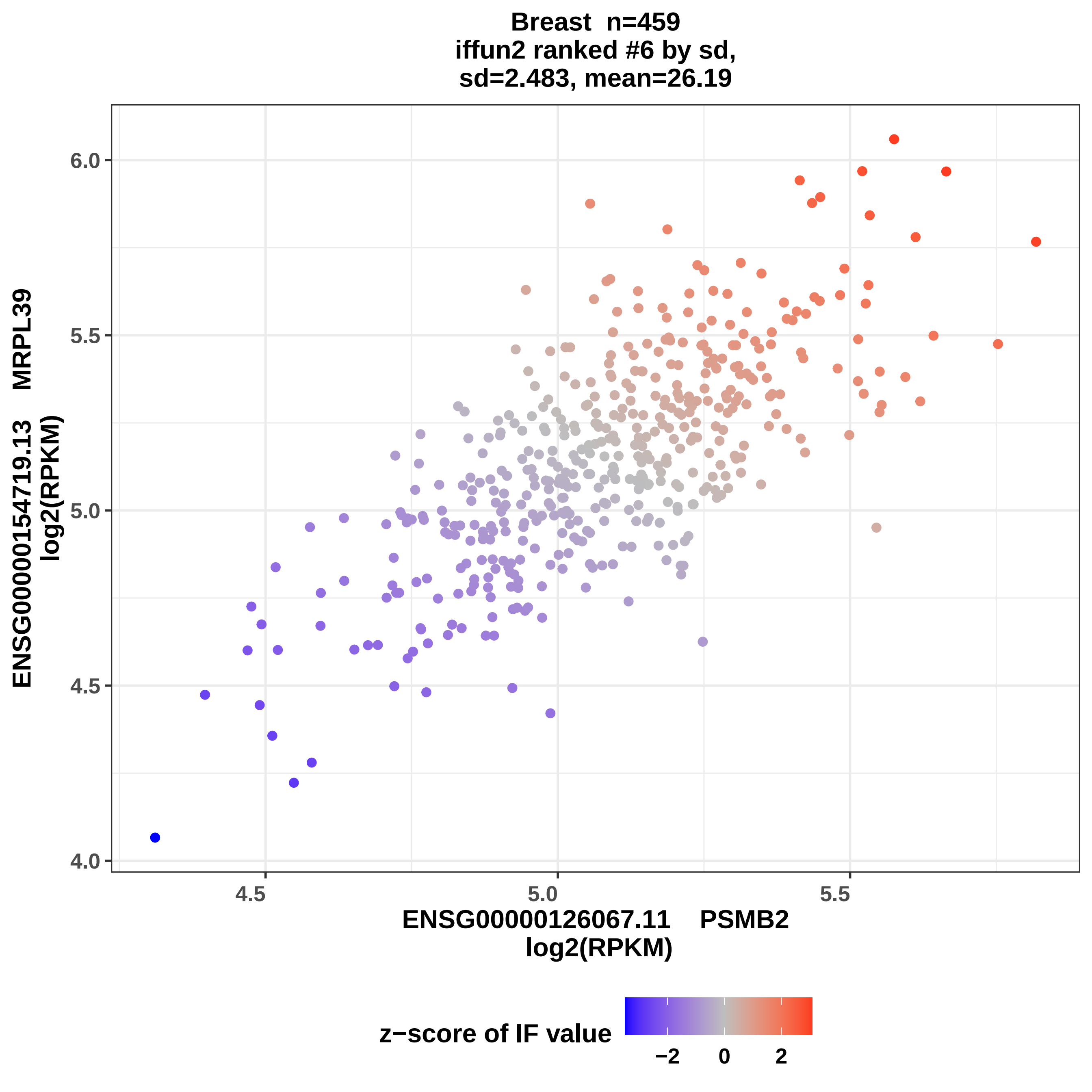

Breast n=459
iffun2 ranked #6 by sd,
sd=2.483, mean=26.19
ENSG00000154719.13 MRPL39
log2(RPKM)
6.0
5.5
5.0
4.5
4.0
4.5
5.0
5.5
ENSG00000126067.11 PSMB2
log2(RPKM)
z−score of IF value
−2
0
2

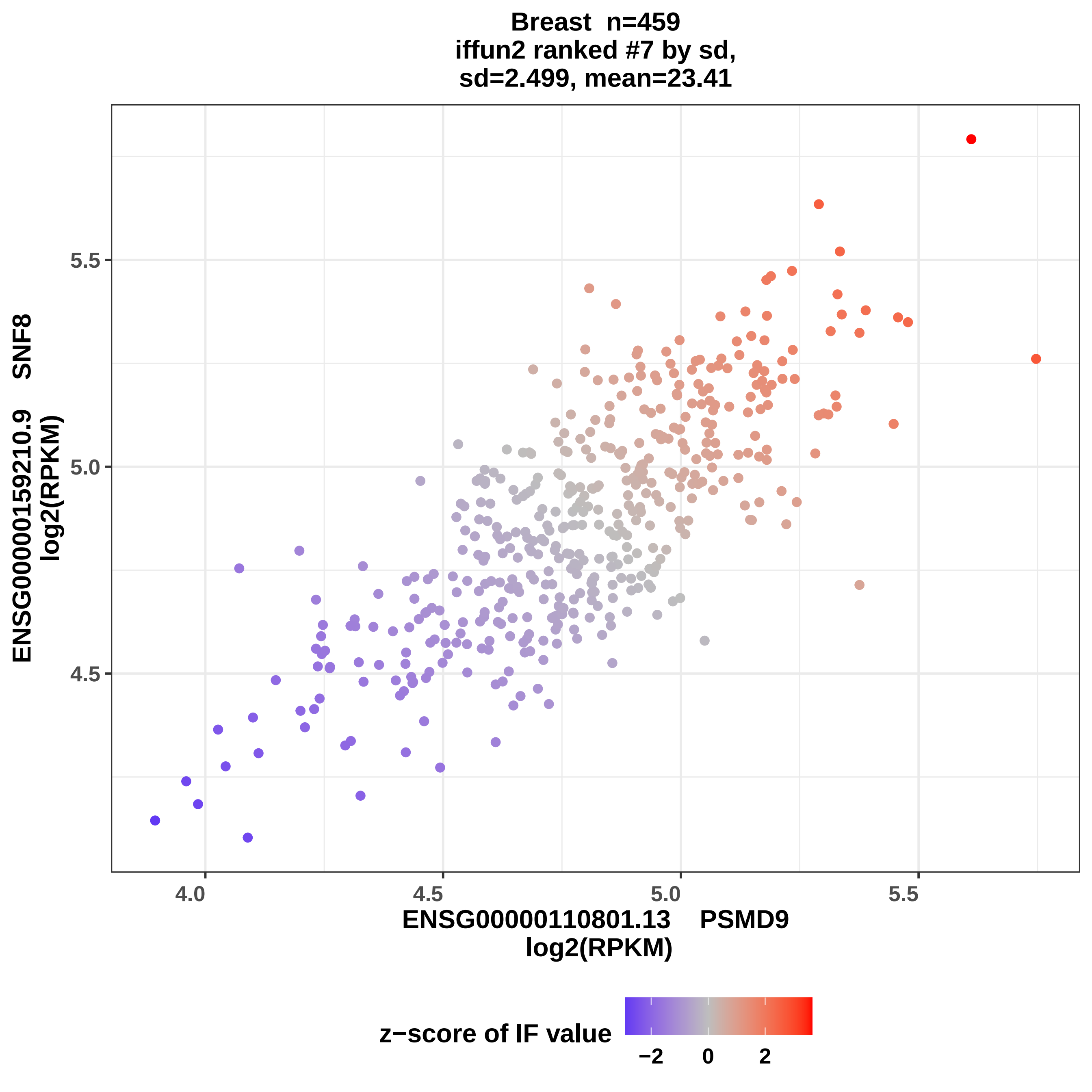
Breast n=459
iffun2 ranked #7 by sd,
sd=2.499, mean=23.41
ENSG00000159210.9 SNF8
log2(RPKM)
5.5
5.0
4.5
4.0
4.5
5.0
5.5
ENSG00000110801.13 PSMD9
log2(RPKM)
z−score of IF value
−2
0
2

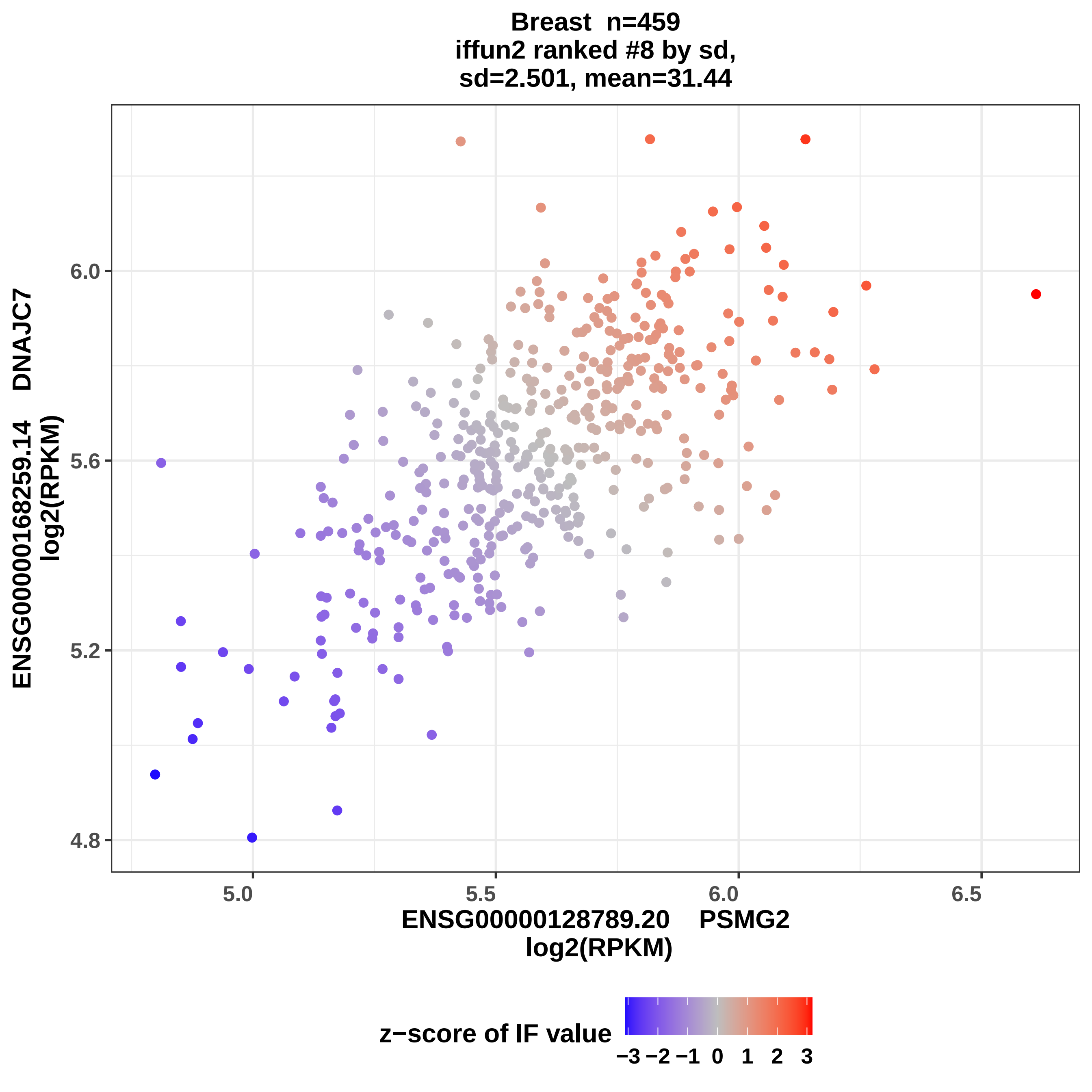
Breast n=459
iffun2 ranked #8 by sd,
sd=2.501, mean=31.44
ENSG00000168259.14 DNAJC7
log2(RPKM)
6.0
5.6
5.2
4.8
5.0
5.5
6.0
6.5
ENSG00000128789.20 PSMG2
log2(RPKM)
z−score of IF value
−3 −2 −1 0 1 2 3

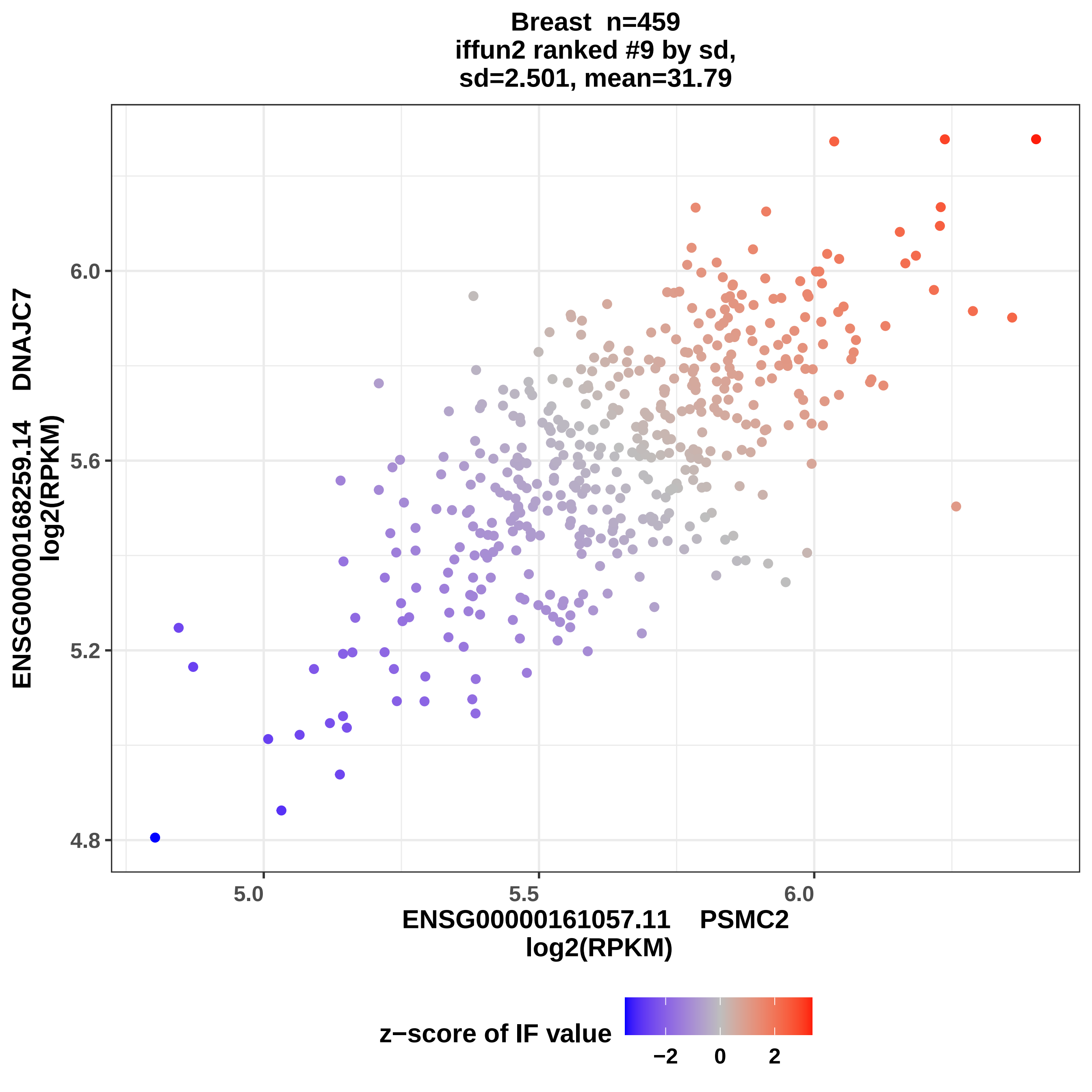
Breast n=459
iffun2 ranked #9 by sd,
sd=2.501, mean=31.79
ENSG00000168259.14 DNAJC7
log2(RPKM)
6.0
5.6
5.2
4.8
5.0
5.5
6.0
ENSG00000161057.11 PSMC2
log2(RPKM)
z−score of IF value
−2
0
2

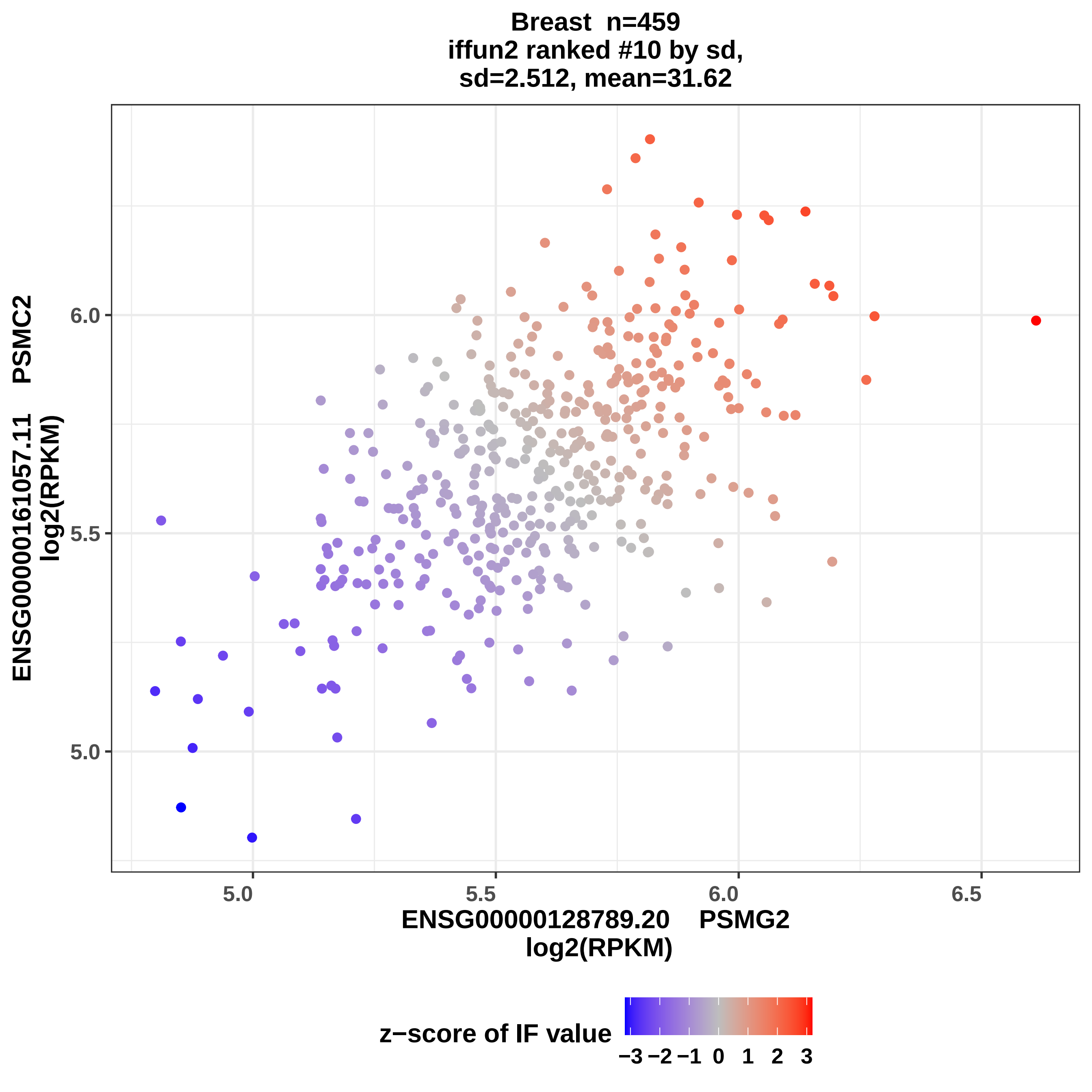

Breast n=459
iffun2 ranked #10 by sd,
sd=2.512, mean=31.62
ENSG00000161057.11 PSMC2
log2(RPKM)
6.0
5.5
5.0
5.0
5.5
6.0
6.5
ENSG00000128789.20 PSMG2
log2(RPKM)
z−score of IF value
−3 −2 −1 0 1 2 3

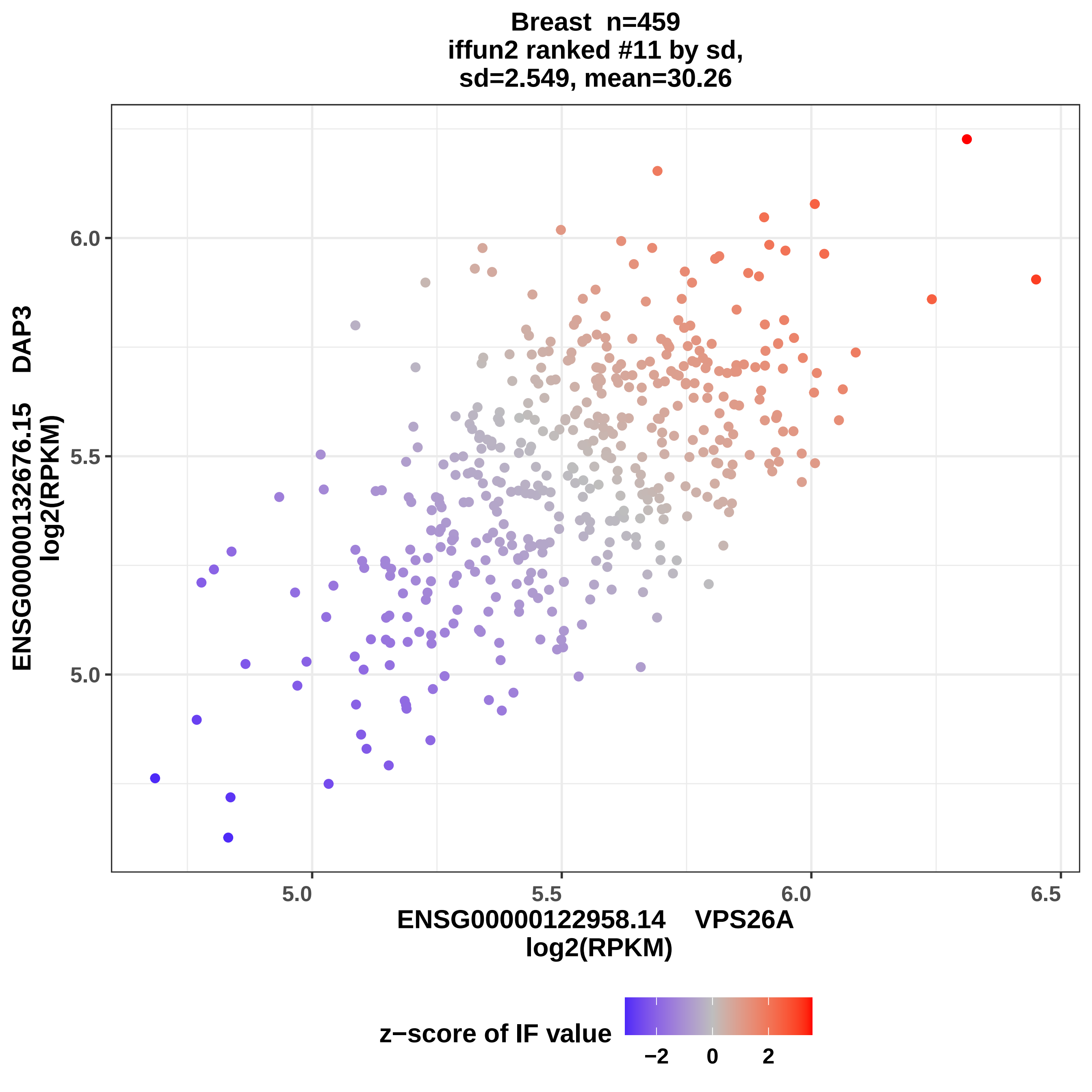

Breast n=459
iffun2 ranked #11 by sd,
sd=2.549, mean=30.26
ENSG00000132676.15 DAP3
log2(RPKM)
6.0
5.5
5.0
5.0
5.5
6.0
6.5
ENSG00000122958.14 VPS26A
log2(RPKM)
z−score of IF value
−2
0
2

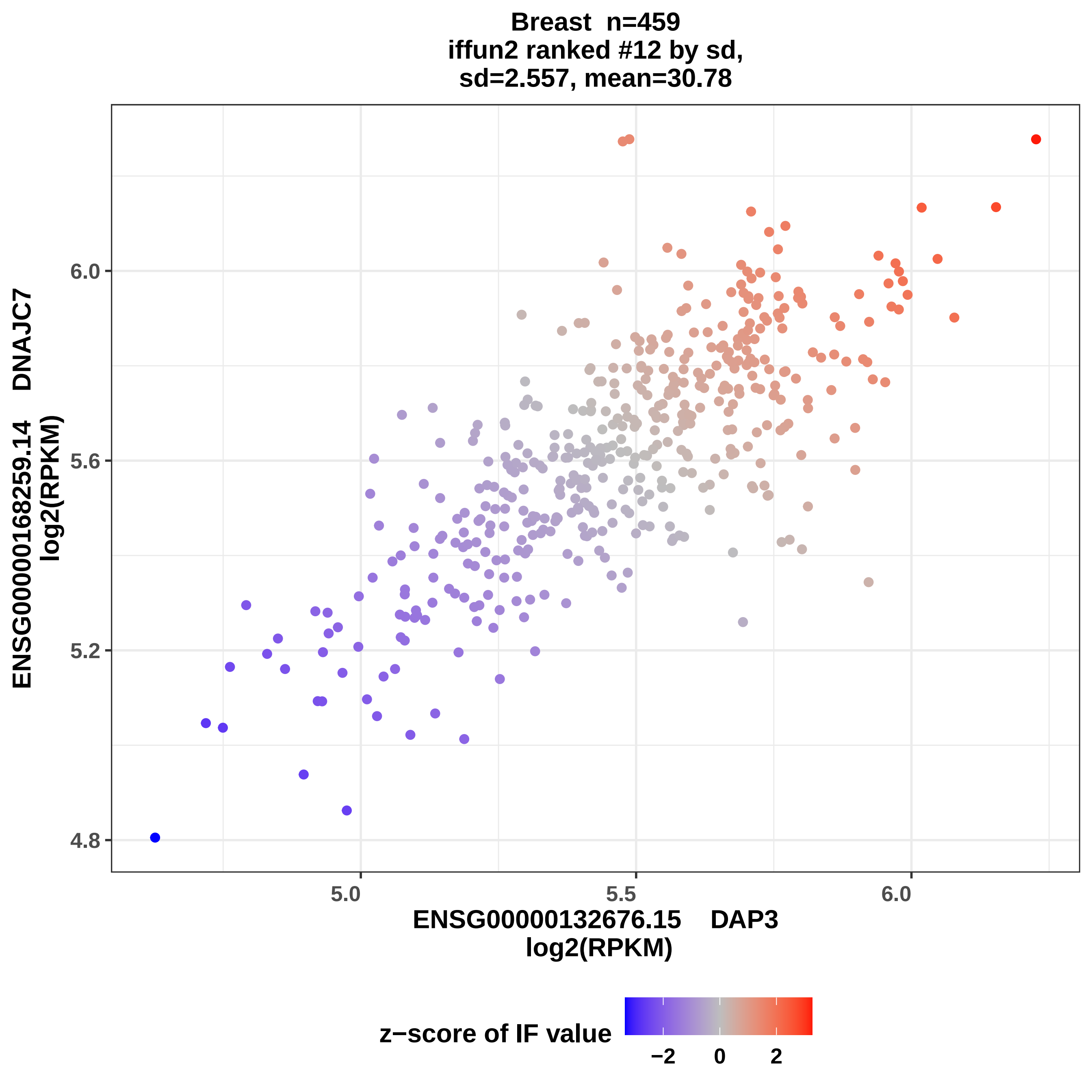

Breast n=459
iffun2 ranked #12 by sd,
sd=2.557, mean=30.78
ENSG00000168259.14 DNAJC7
log2(RPKM)
6.0
5.6
5.2
4.8
5.0
5.5
6.0
ENSG00000132676.15 DAP3
log2(RPKM)
z−score of IF value
−2
0
2

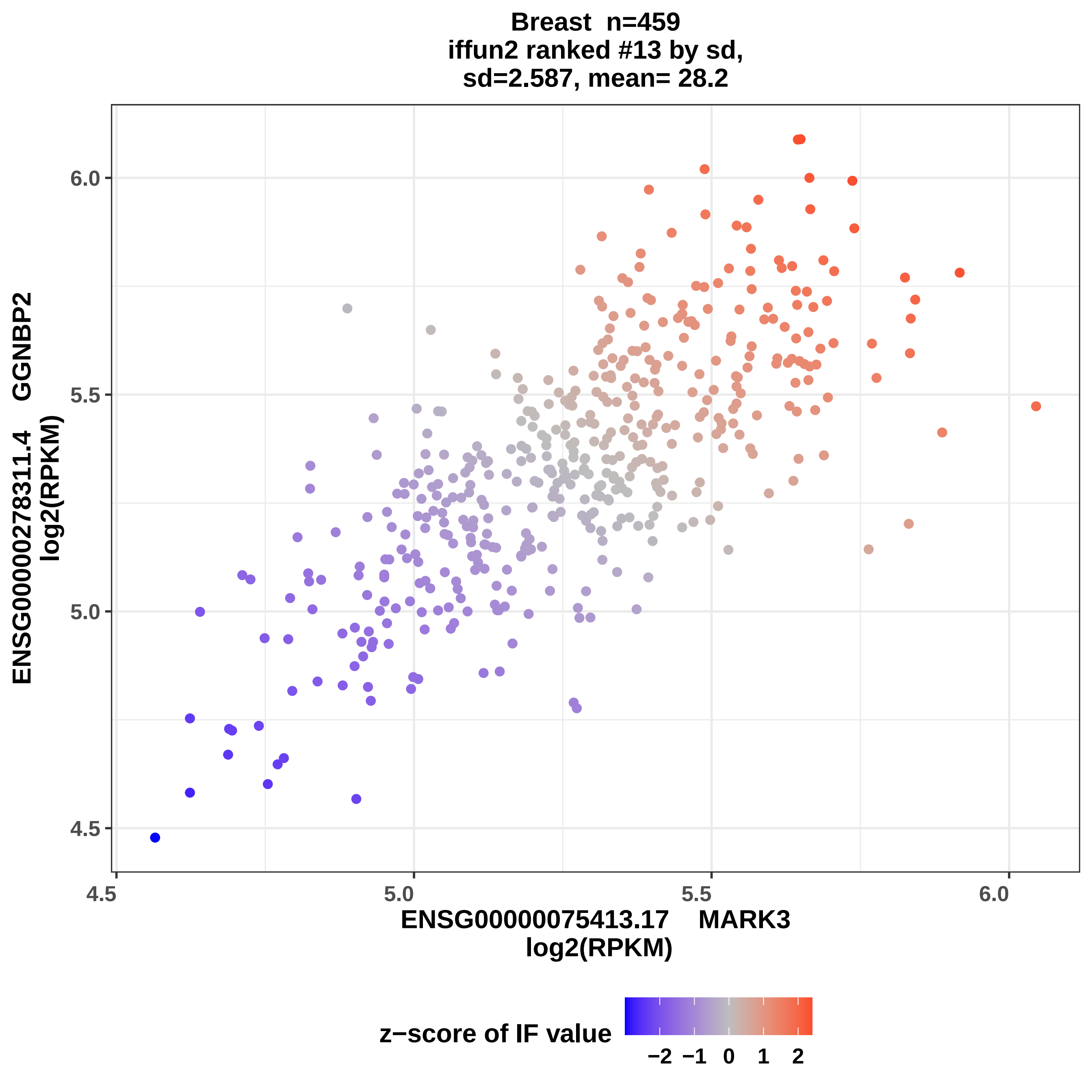
Breast n=459
iffun2 ranked #13 by sd,
sd=2.587, mean= 28.2
ENSG00000278311.4 GGNBP2
log2(RPKM)
ENSG00000075413.17 MARK3
log2(RPKM)
4.5
5.0
5.5
6.0
z−score of IF value
−2 −1 0 1 2

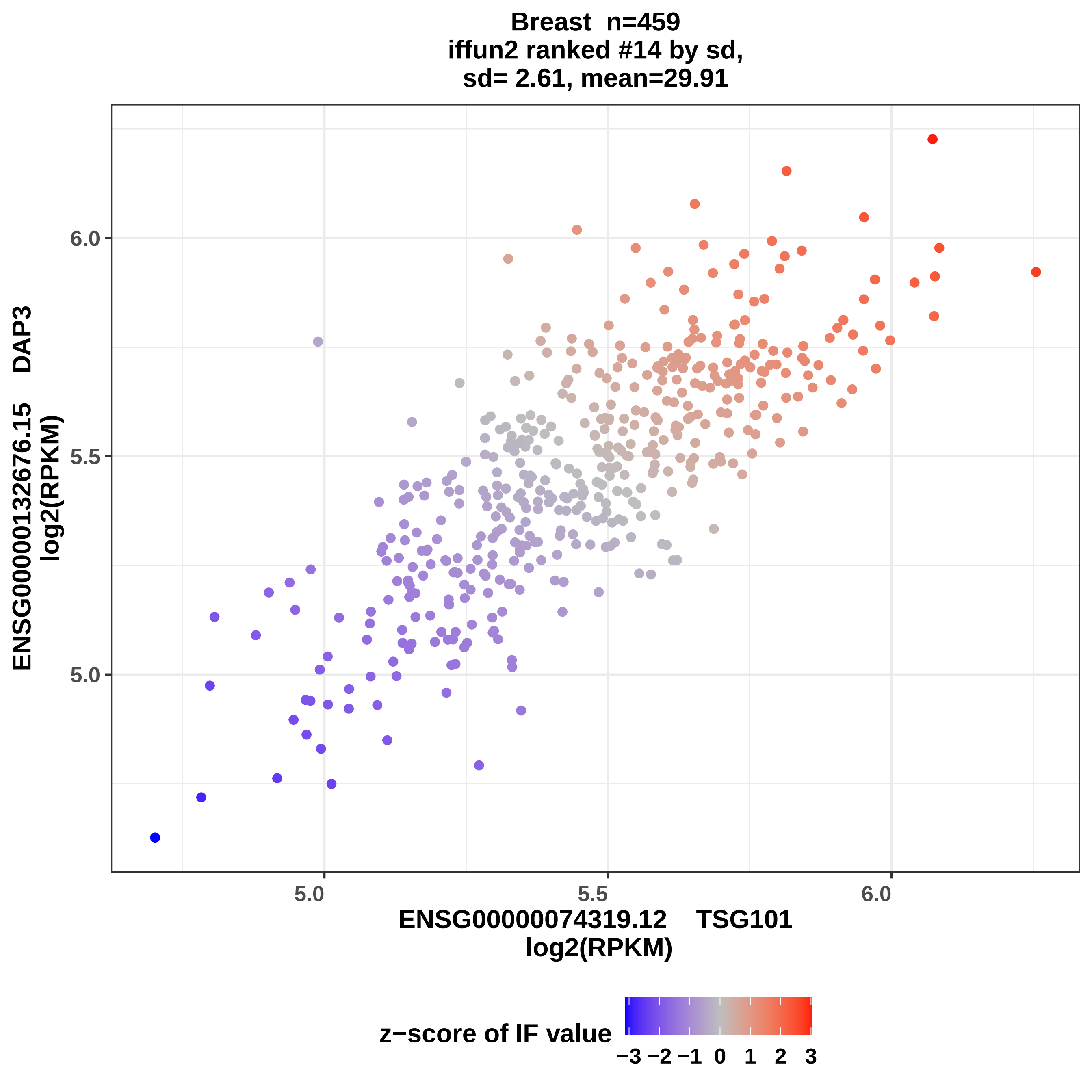

Breast n=459
iffun2 ranked #14 by sd,
sd= 2.61, mean=29.91
ENSG00000132676.15 DAP3
log2(RPKM)
6.0
5.5
5.0
5.0
5.5
6.0
ENSG00000074319.12 TSG101
log2(RPKM)
z−score of IF value
−3 −2 −1 0 1 2 3

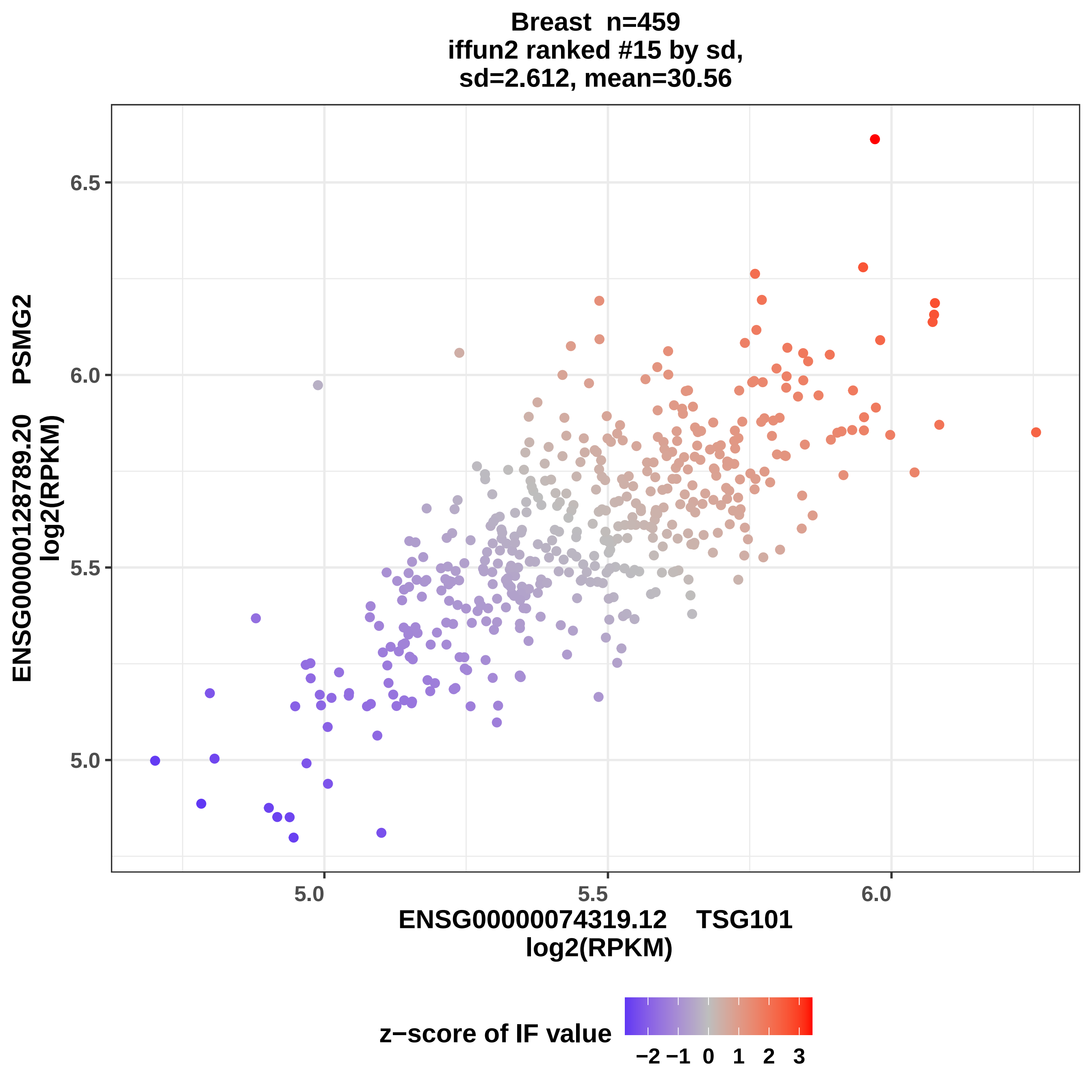
Breast n=459
iffun2 ranked #15 by sd,
sd=2.612, mean=30.56
ENSG00000128789.20 PSMG2
log2(RPKM)
6.5
6.0
5.5
5.0
5.0
5.5
6.0
ENSG00000074319.12 TSG101
log2(RPKM)
z−score of IF value
−2 −1 0 1 2 3

Breast n=459
iffun2 ranked #16 by sd,
sd=2.622, mean=30.62

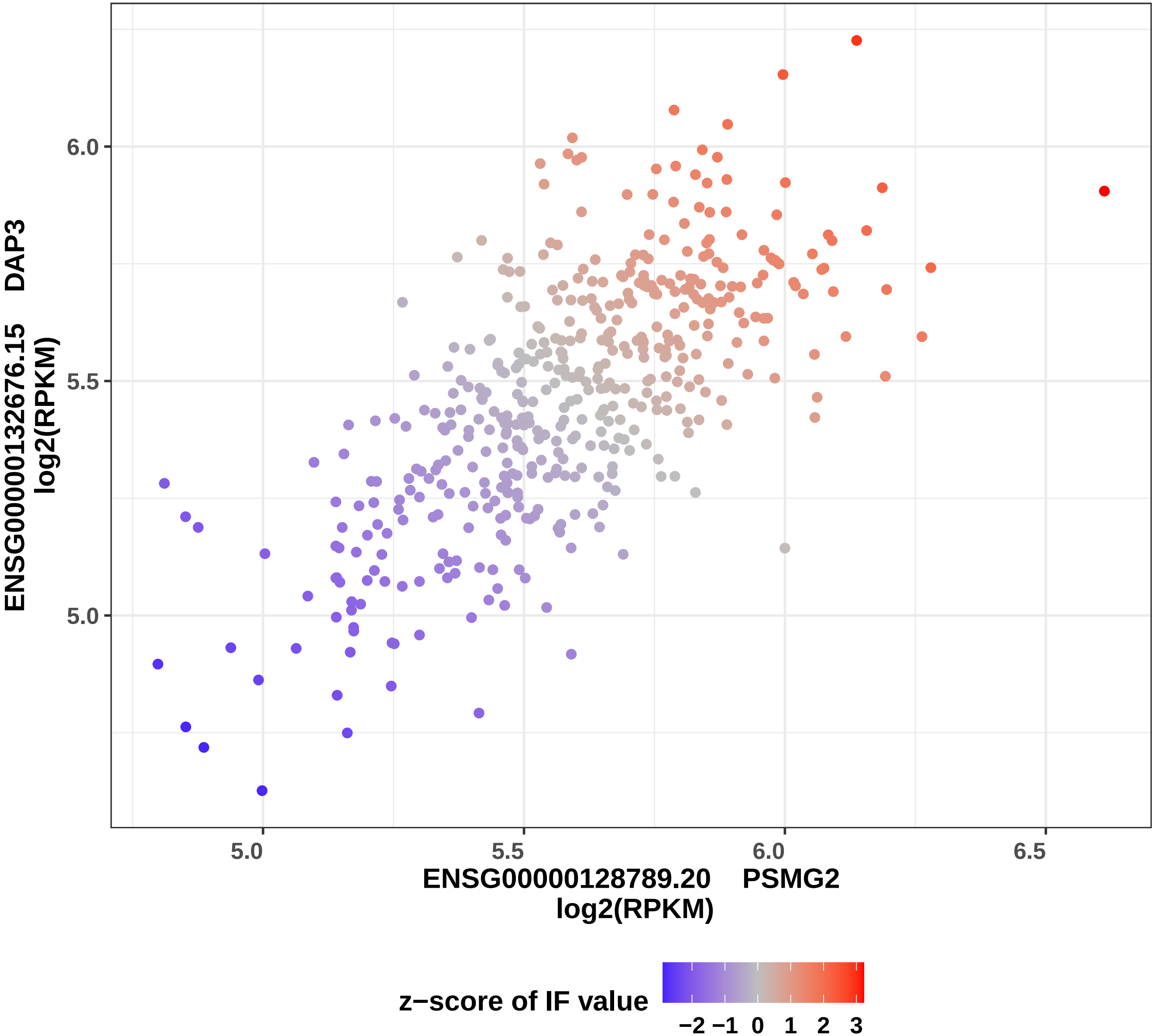

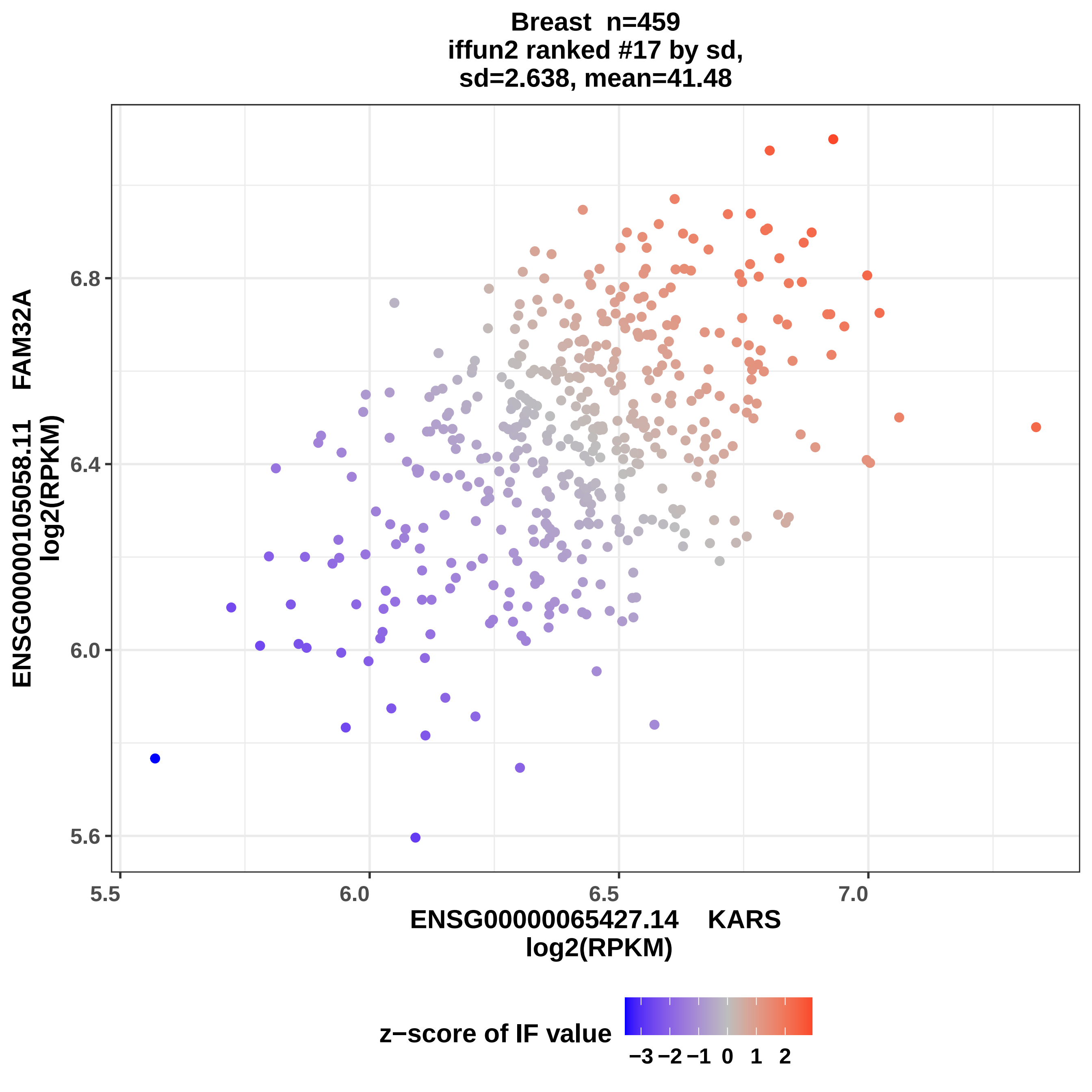
Breast n=459
iffun2 ranked #17 by sd,
sd=2.638, mean=41.48
ENSG00000105058.11 FAM32A
log2(RPKM)
6.8
6.4
6.0
5.6
5.5
6.0
6.5
7.0
ENSG00000065427.14 KARS
log2(RPKM)
z−score of IF value
−3 −2 −1 0 1 2

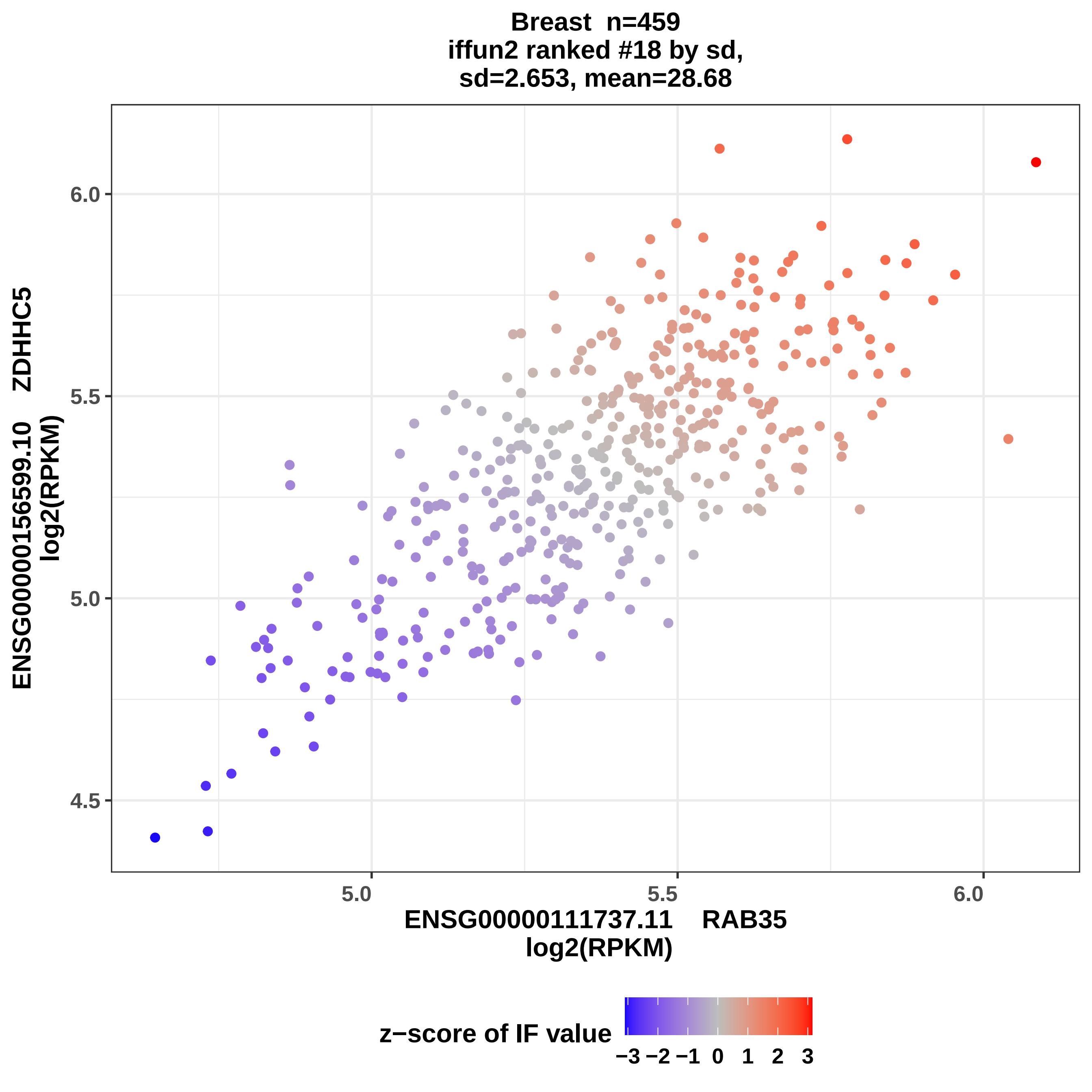
Breast n=459
iffun2 ranked #18 by sd,
sd=2.653, mean=28.68
ENSG00000156599.10 ZDHHC5
log2(RPKM)
6.0
5.5
5.0
4.5
5.0
5.5
6.0
ENSG00000111737.11 RAB35
log2(RPKM)
z−score of IF value
−3 −2 −1 0 1 2 3

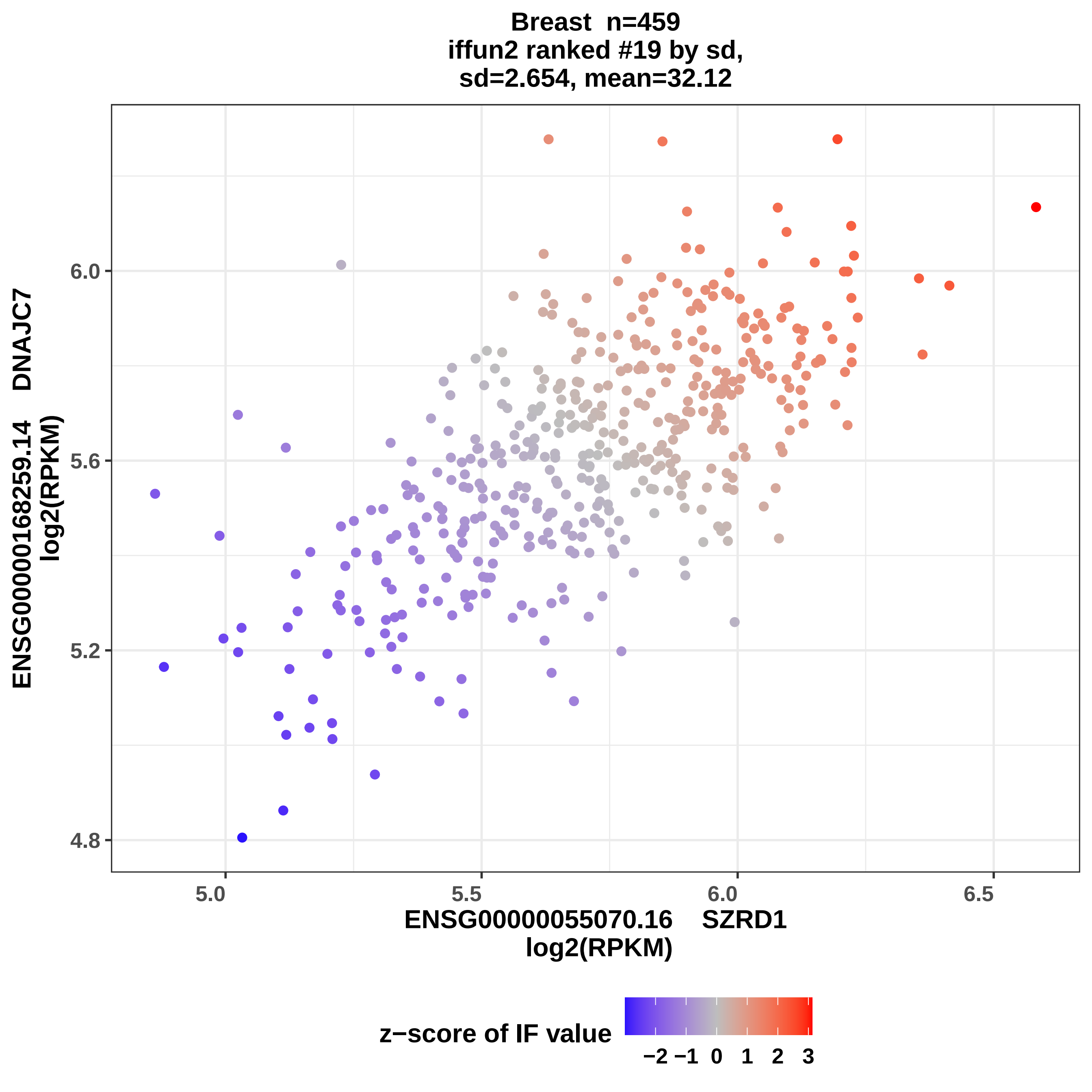
Breast n=459
iffun2 ranked #19 by sd,
sd=2.654, mean=32.12
ENSG00000168259.14 DNAJC7
log2(RPKM)
6.0
5.6
5.2
4.8
5.0
5.5
6.0
6.5
ENSG00000055070.16 SZRD1
log2(RPKM)
z−score of IF value
−2
−1
0
1
2
3

Breast n=459
iffun2 ranked #20 by sd,
sd= 2.66, mean=30.93

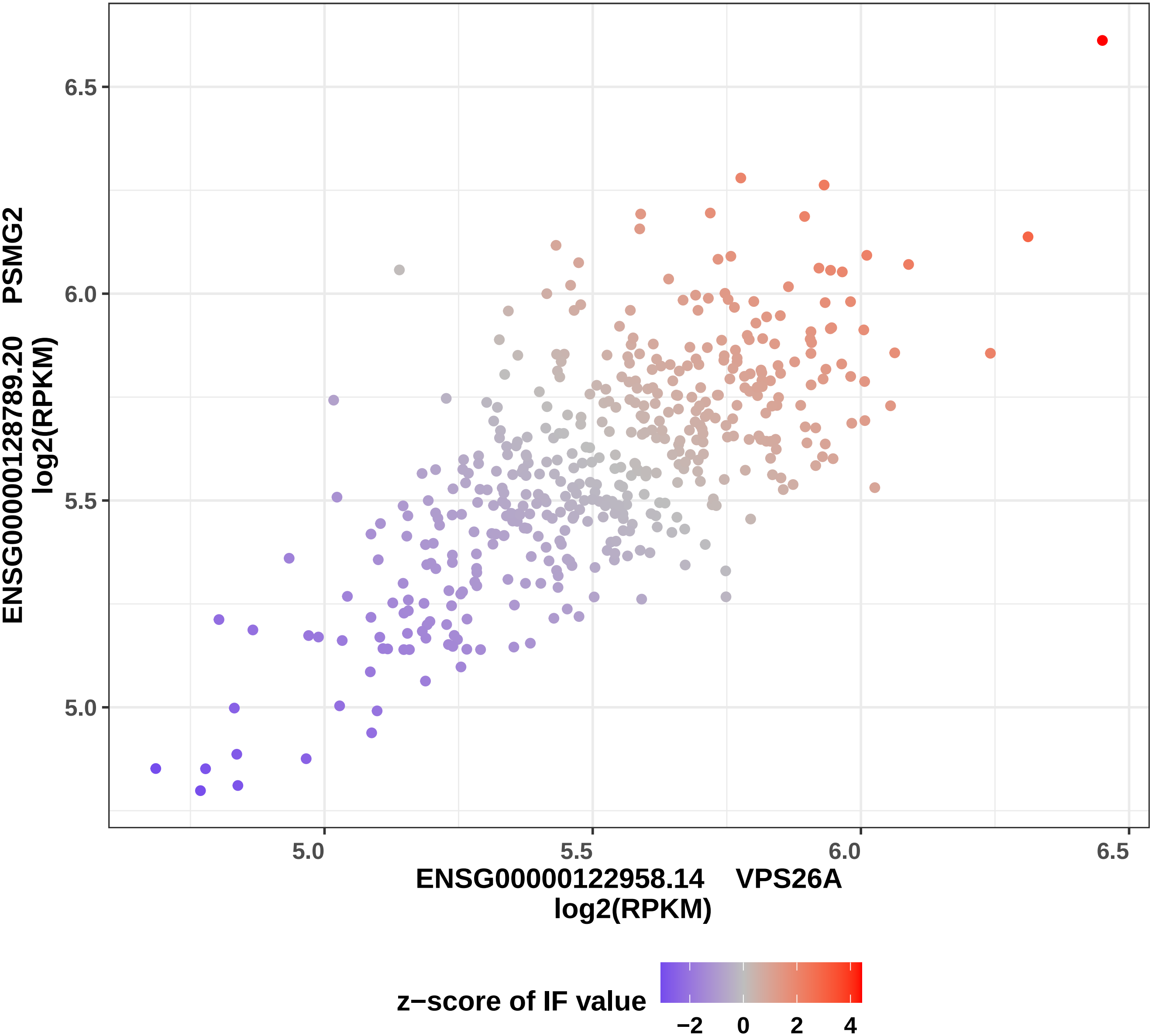

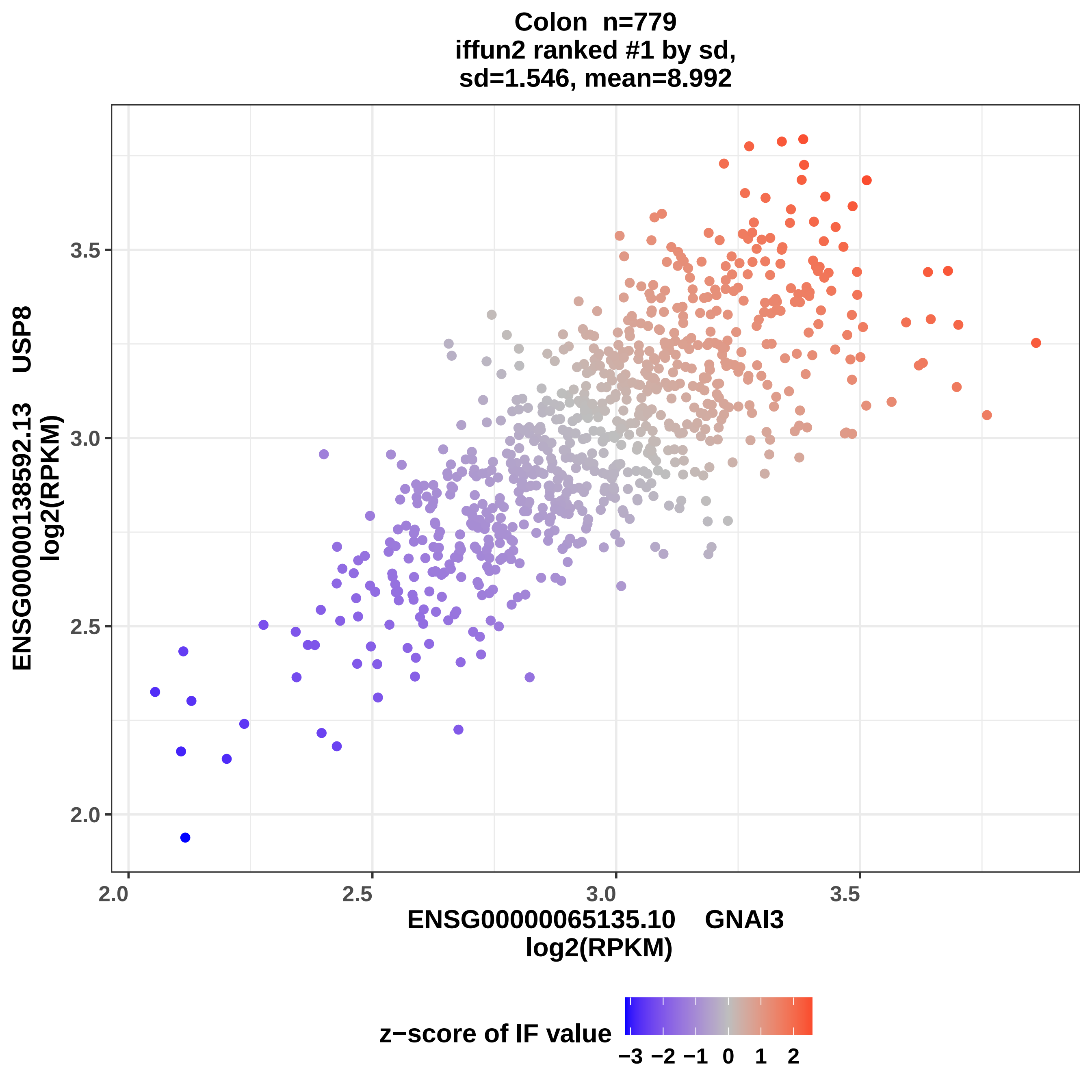

Colon n=779
iffun2 ranked #1 by sd,
sd=1.546, mean=8.992
ENSG00000138592.13 USP8
log2(RPKM)
ENSG00000065135.10 GNAI3
log2(RPKM)
2.0
2.5
3.0
3.5
z−score of IF value
−3 −2 −1 0 1 2

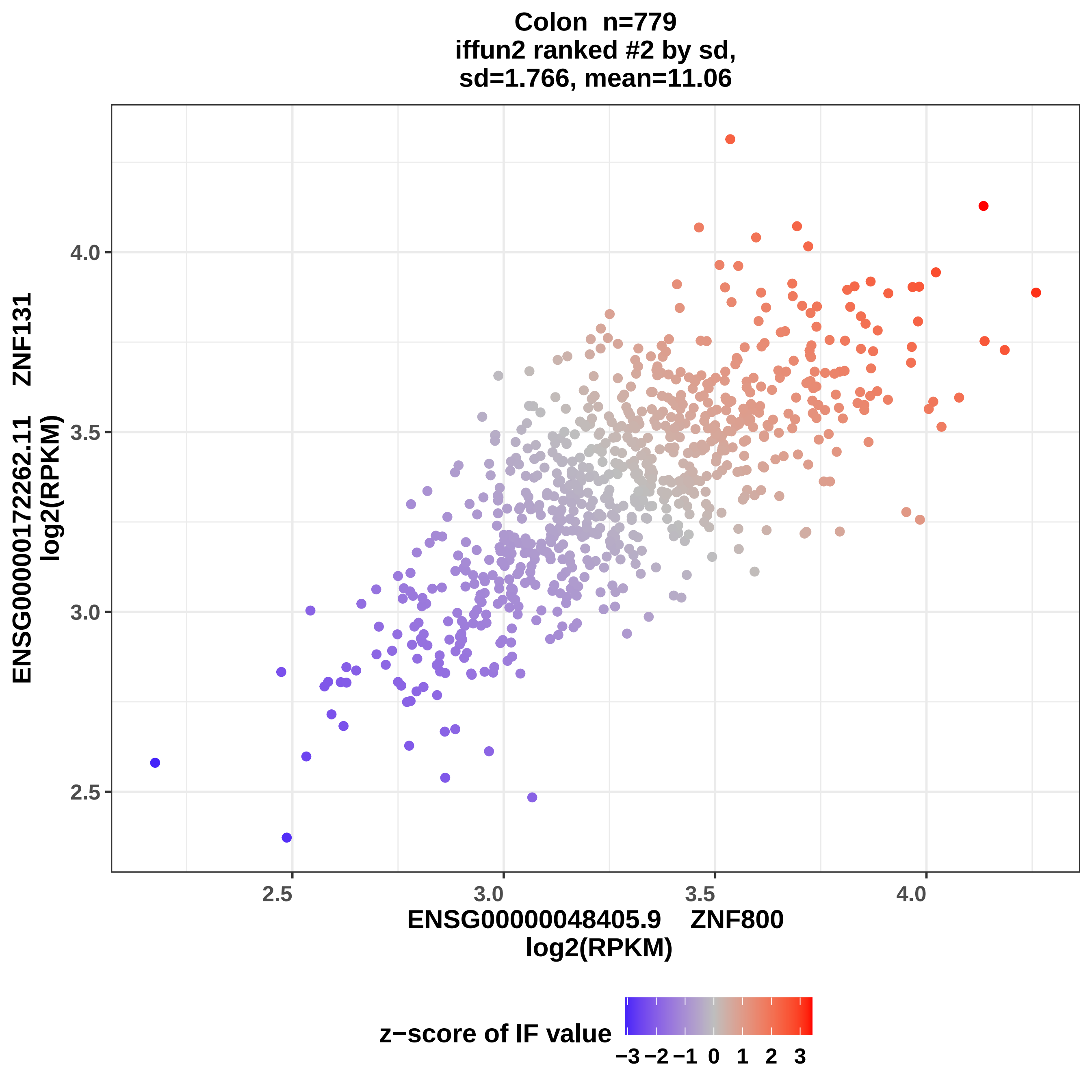

Colon n=779
iffun2 ranked #2 by sd,
sd=1.766, mean=11.06
ENSG00000172262.11 ZNF131
log2(RPKM)
4.0
3.5
3.0
2.5
2.5
3.0
3.5
4.0
ENSG00000048405.9 ZNF800
log2(RPKM)
z−score of IF value
−3 −2 −1 0 1 2 3

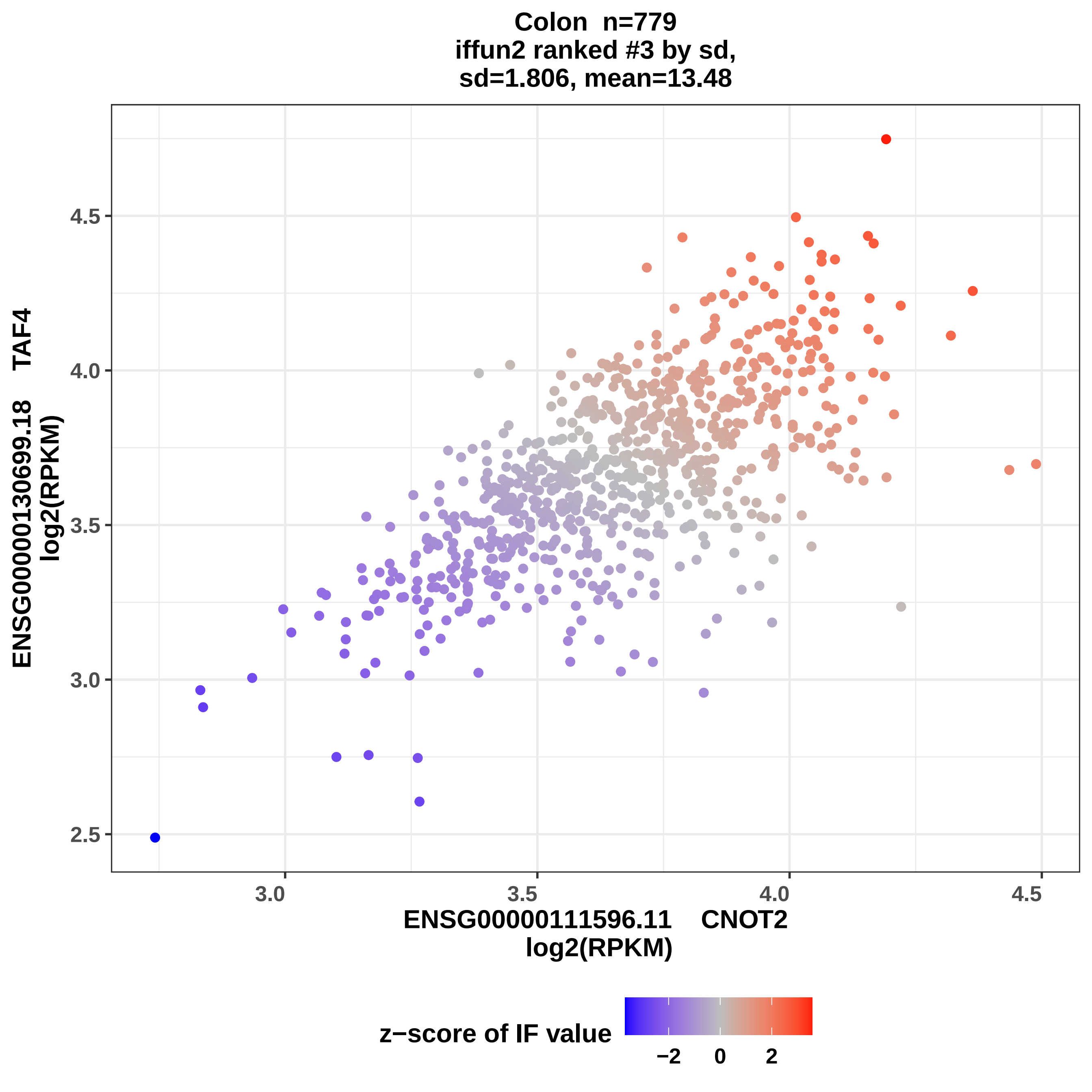

Colon n=779
iffun2 ranked #3 by sd,
sd=1.806, mean=13.48
ENSG00000130699.18 TAF4
log2(RPKM)
ENSG00000111596.11 CNOT2
log2(RPKM)
2.5
3.0
3.5
4.0
4.5
z−score of IF value
−2
0
2

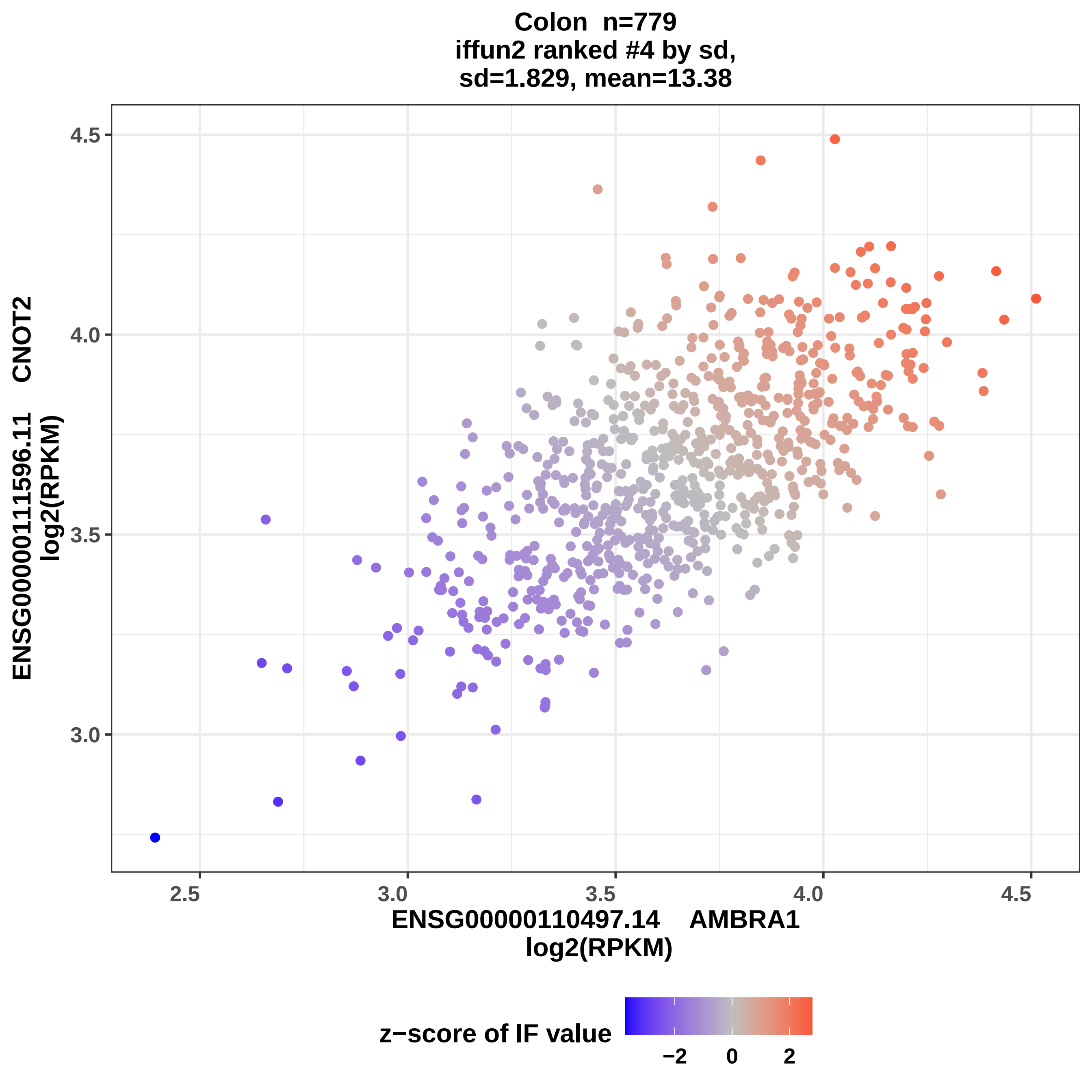

Colon n=779
iffun2 ranked #4 by sd,
sd=1.829, mean=13.38
ENSG00000111596.11 CNOT2
log2(RPKM)
4.5
4.0
3.5
3.0
2.5
3.0
3.5
4.0
4.5
ENSG00000110497.14 AMBRA1
log2(RPKM)
z−score of IF value
−2
0
2

**Colon  n=779**
**iffun2 ranked #5 by sd,**
**sd=1.831, mean=12.91**

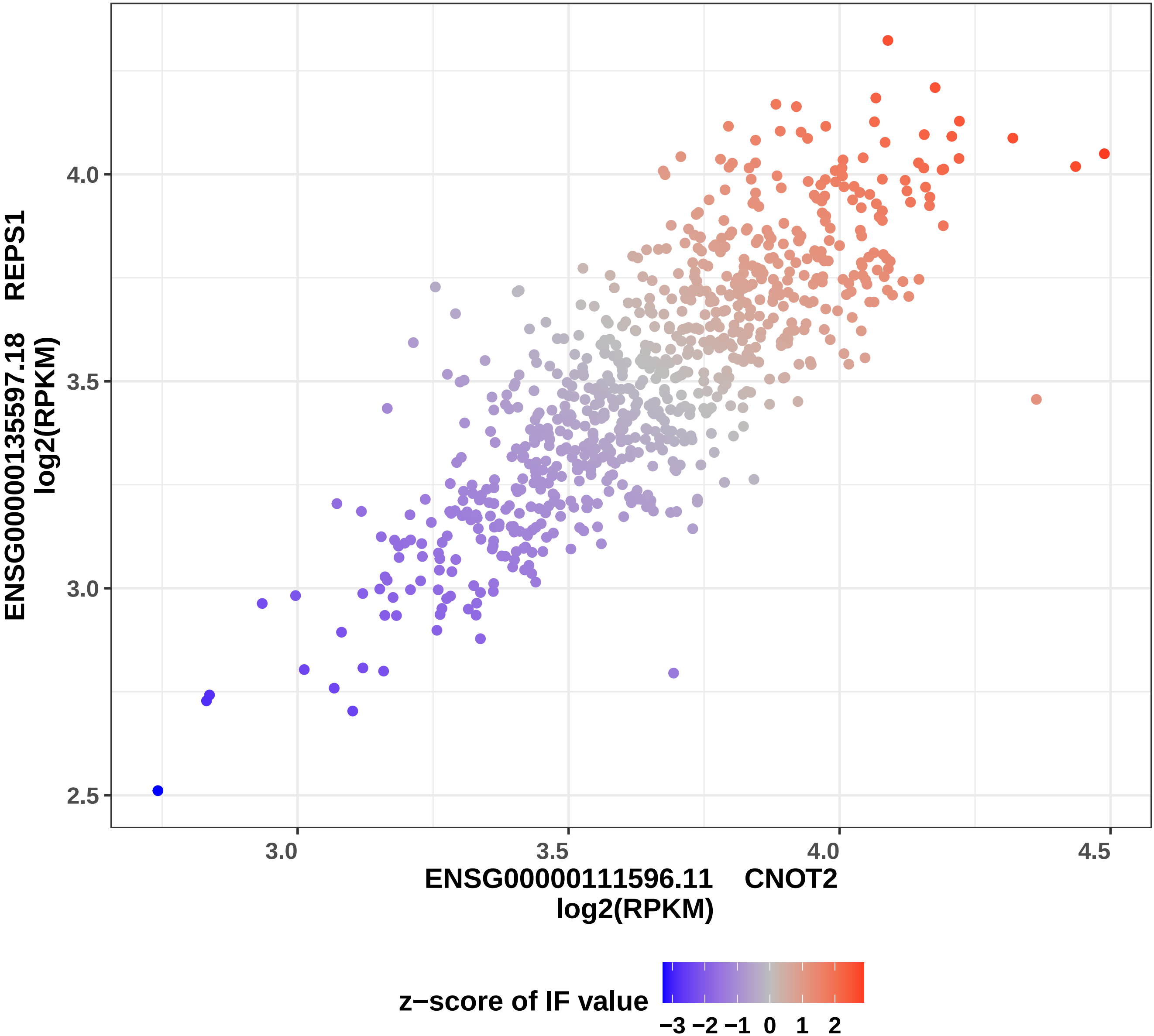

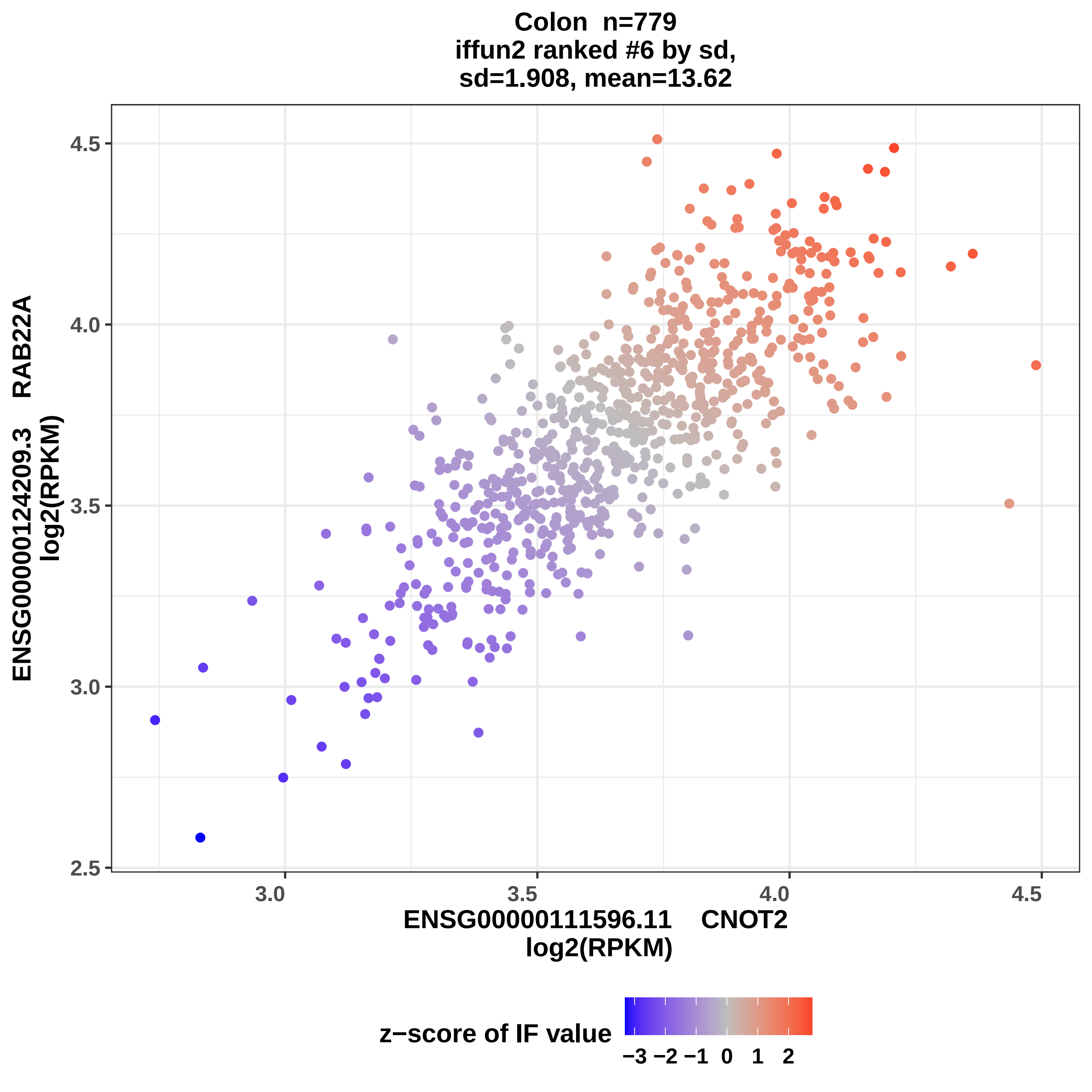

Colon n=779
iffun2 ranked #6 by sd,
sd=1.908, mean=13.62
ENSG00000124209.3 RAB22A
log2(RPKM)
4.5
4.0
3.5
3.0
2.5
3.0
3.5
4.0
4.5
ENSG00000111596.11 CNOT2
log2(RPKM)
z−score of IF value
−3 −2 −1 0 1 2

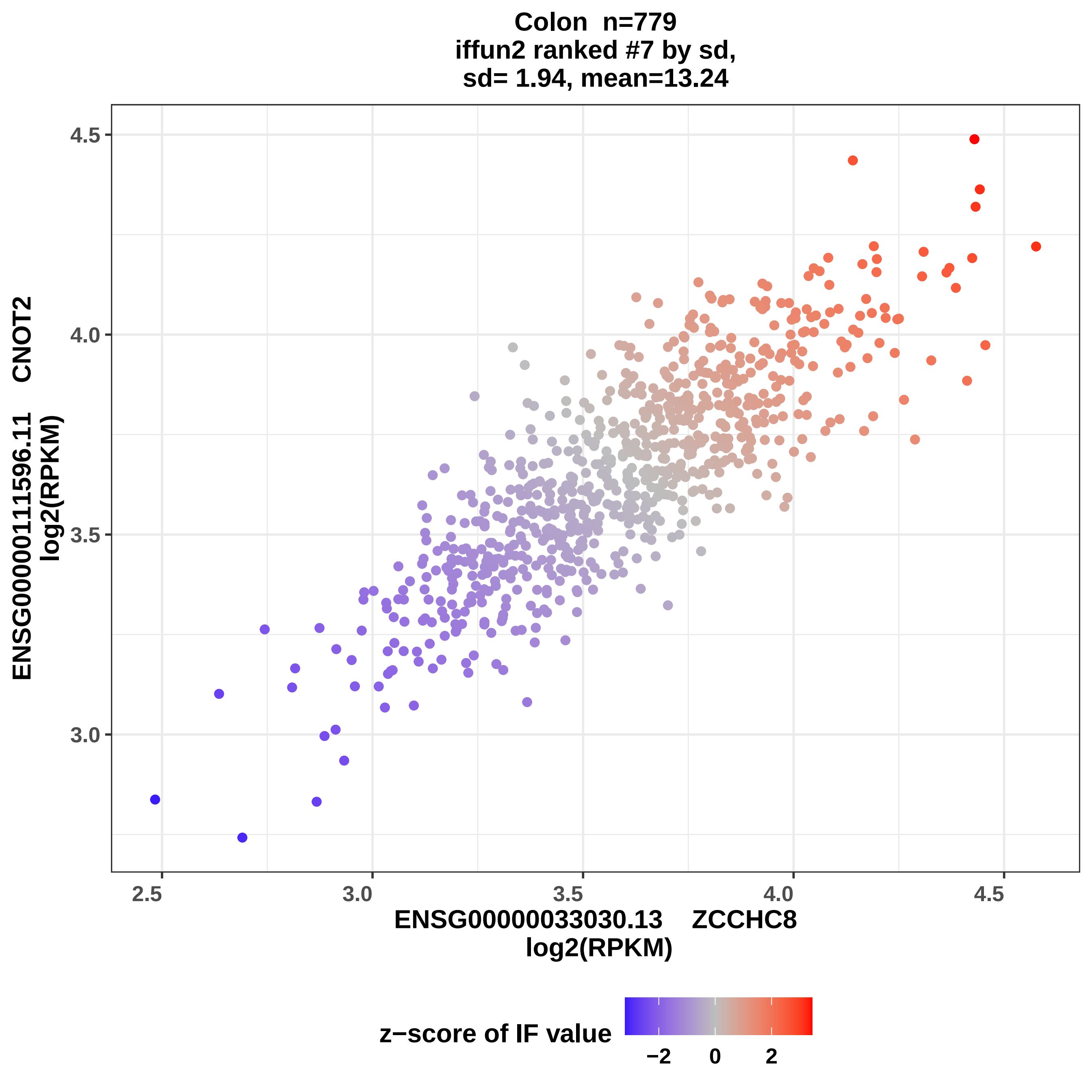
Colon n=779
iffun2 ranked #7 by sd,
sd= 1.94, mean=13.24
4.5
4.0
3.5
3.0
ENSG00000111596.11 CNOT2
log2(RPKM)
2.5
3.0
3.5
4.0
4.5
ENSG00000033030.13 ZCCHC8
log2(RPKM)
z−score of IF value
−2
0
2

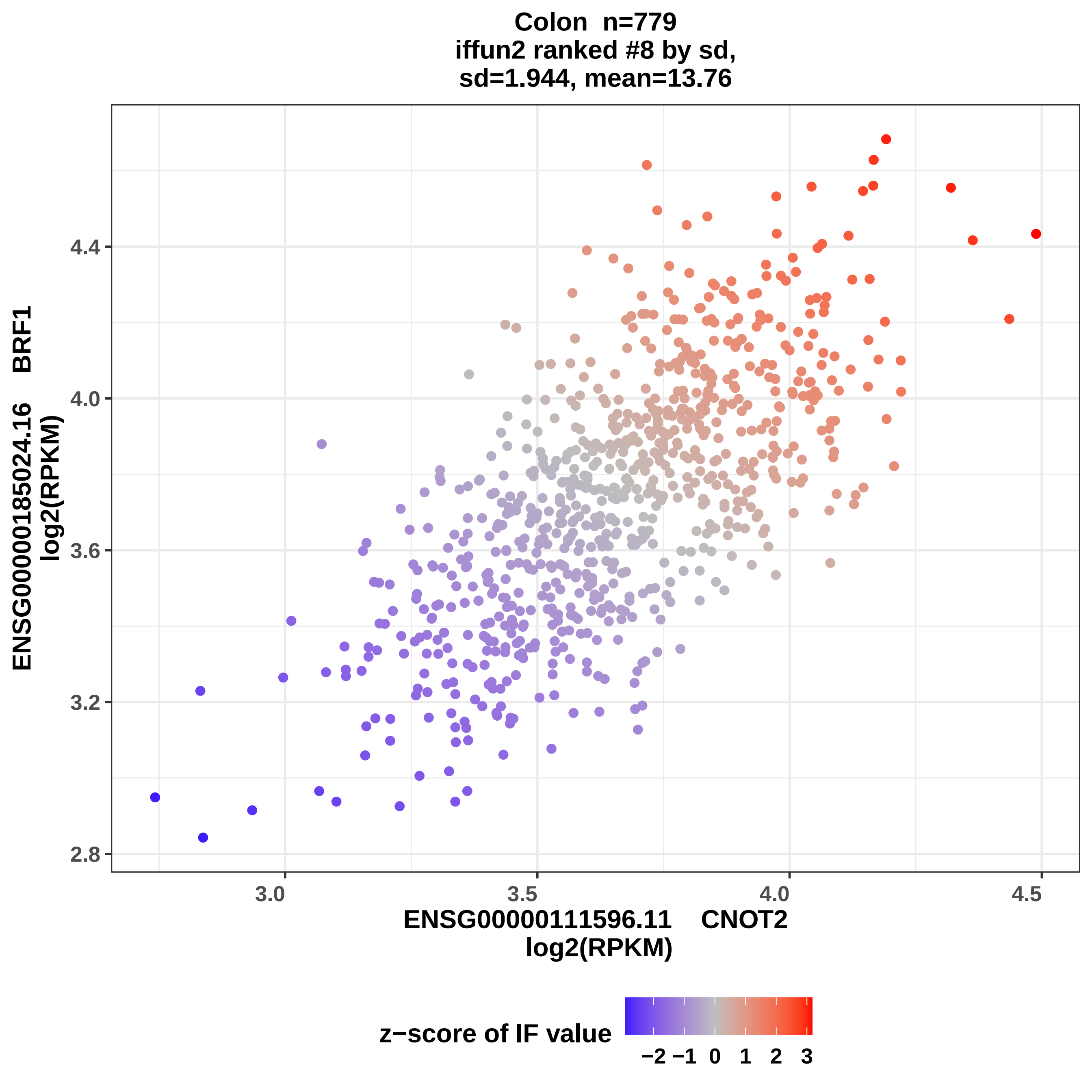
Colon n=779
iffun2 ranked #8 by sd,
sd=1.944, mean=13.76
ENSG00000185024.16 BRF1
log2(RPKM)
4.4
4.0
3.6
3.2
2.8
3.0
3.5
4.0
4.5
ENSG00000111596.11 CNOT2
log2(RPKM)
z−score of IF value
−2 −1 0 1 2 3

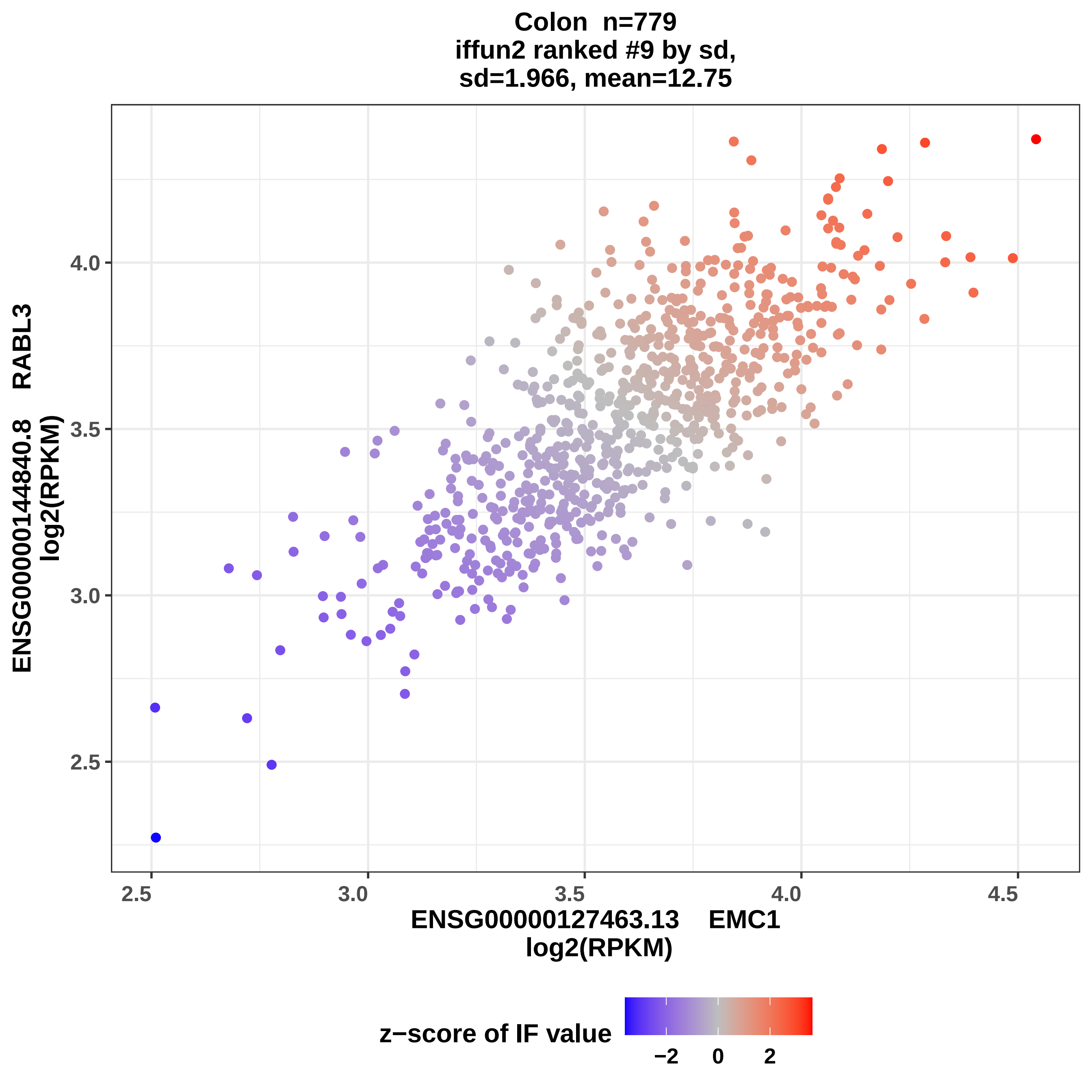
Colon n=779
iffun2 ranked #9 by sd,
sd=1.966, mean=12.75
ENSG00000144840.8 RABL3
log2(RPKM)
ENSG00000127463.13 EMC1
log2(RPKM)
2.5
3.0
3.5
4.0
4.5
z−score of IF value
−2
0
2

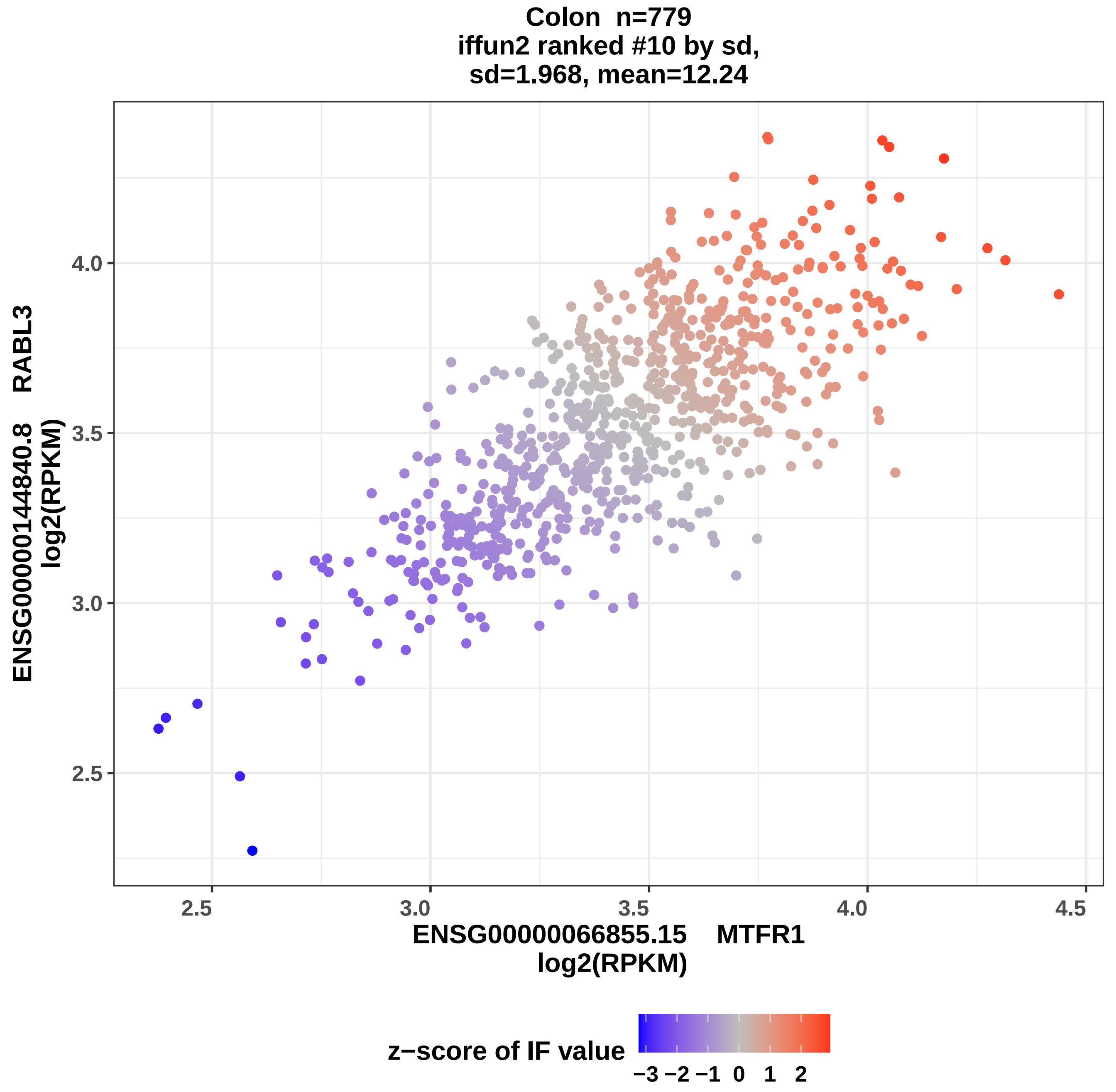

Colon n=779
iffun2 ranked #10 by sd,
sd=1.968, mean=12.24
ENSG00000144840.8 RABL3
log2(RPKM)
4.0
3.5
3.0
2.5
2.5
3.0
3.5
4.0
4.5
ENSG00000066855.15 MTFR1
log2(RPKM)
z−score of IF value
−3 −2 −1 0 1 2

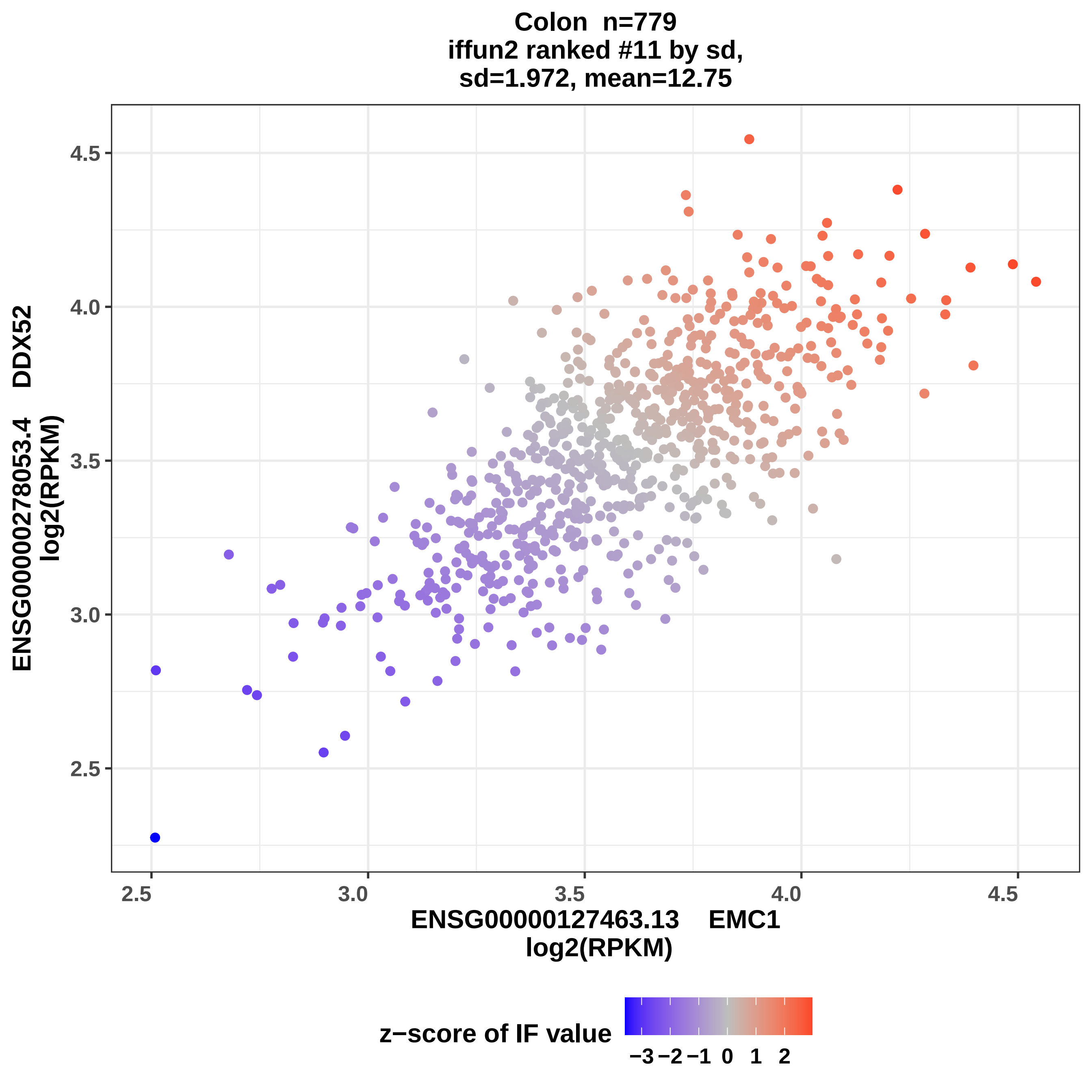
Colon n=779
iffun2 ranked #11 by sd,
sd=1.972, mean=12.75
ENSG00000278053.4 DDX52
log2(RPKM)
ENSG00000127463.13 EMC1
log2(RPKM)
4.5
4.0
3.5
3.0
2.5
2.5
3.0
3.5
4.0
4.5
z−score of IF value
−3
−2
−1
0
1
2

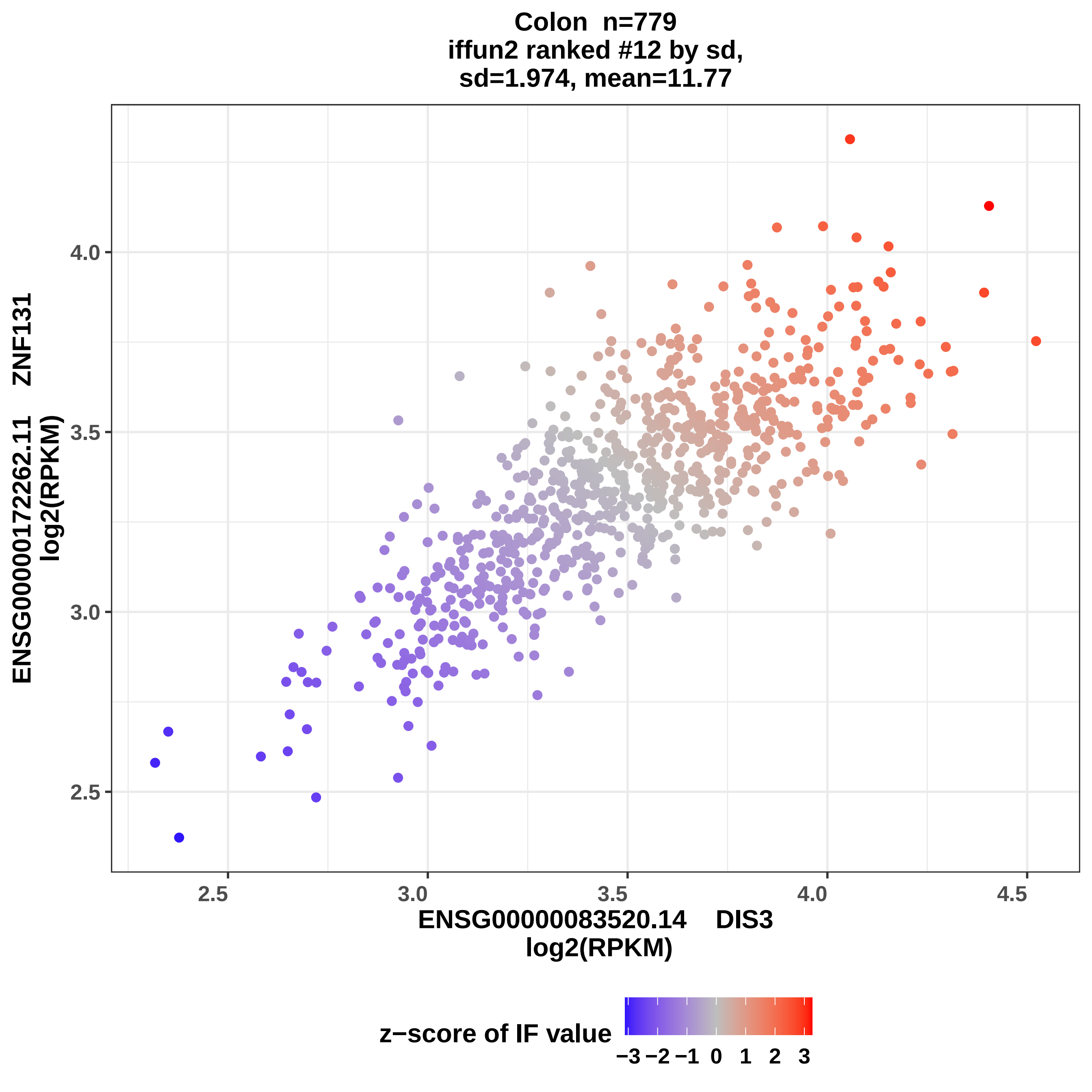
Colon n=779
iffun2 ranked #12 by sd,
sd=1.974, mean=11.77
ENSG00000172262.11 ZNF131
log2(RPKM)
4.0
3.5
3.0
2.5
2.5
3.0
3.5
4.0
4.5
ENSG00000083520.14 DIS3
log2(RPKM)
z−score of IF value
−3 −2 −1 0 1 2 3

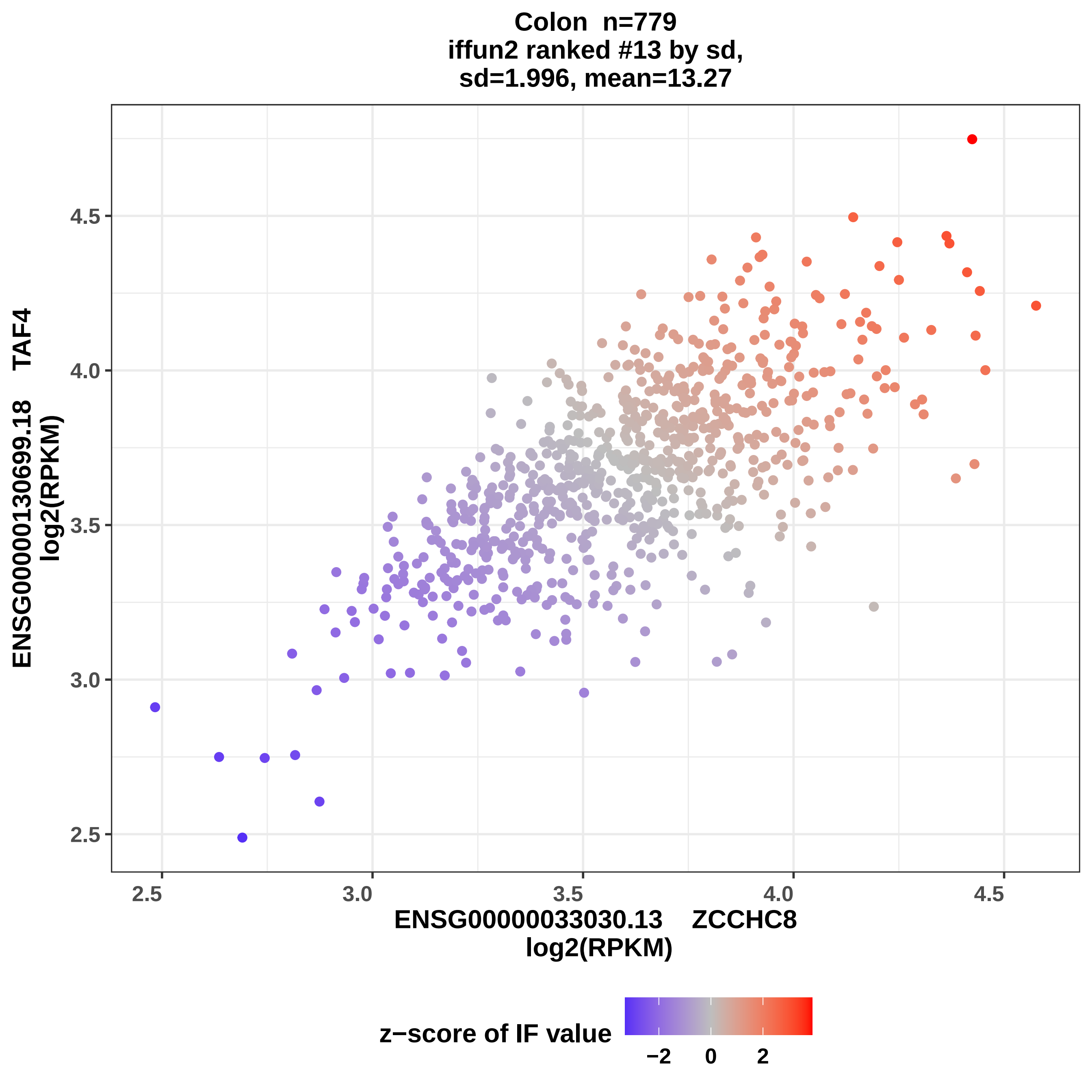

Colon n=779
iffun2 ranked #13 by sd,
sd=1.996, mean=13.27
ENSG00000130699.18 TAF4
log2(RPKM)
4.5
4.0
3.5
3.0
2.5
2.5
3.0
3.5
4.0
4.5
ENSG00000033030.13 ZCCHC8
log2(RPKM)
z−score of IF value
−2
0
2

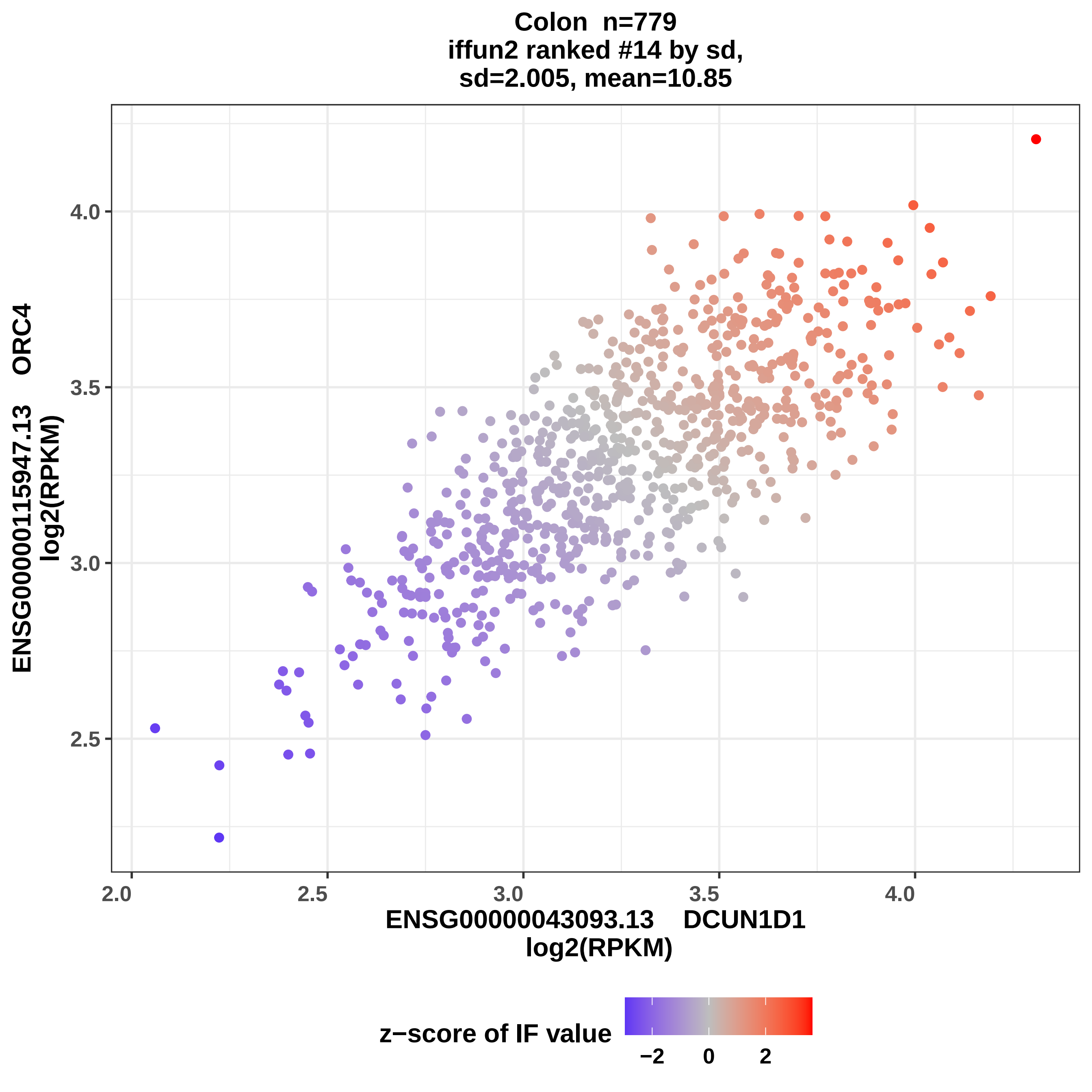

Colon n=779
iffun2 ranked #14 by sd,
sd=2.005, mean=10.85
ENSG00000115947.13 ORC4
log2(RPKM)
4.0
3.5
3.0
2.5
2.0
2.5
3.0
3.5
4.0
ENSG00000043093.13 DCUN1D1
log2(RPKM)
z−score of IF value
−2
0
2

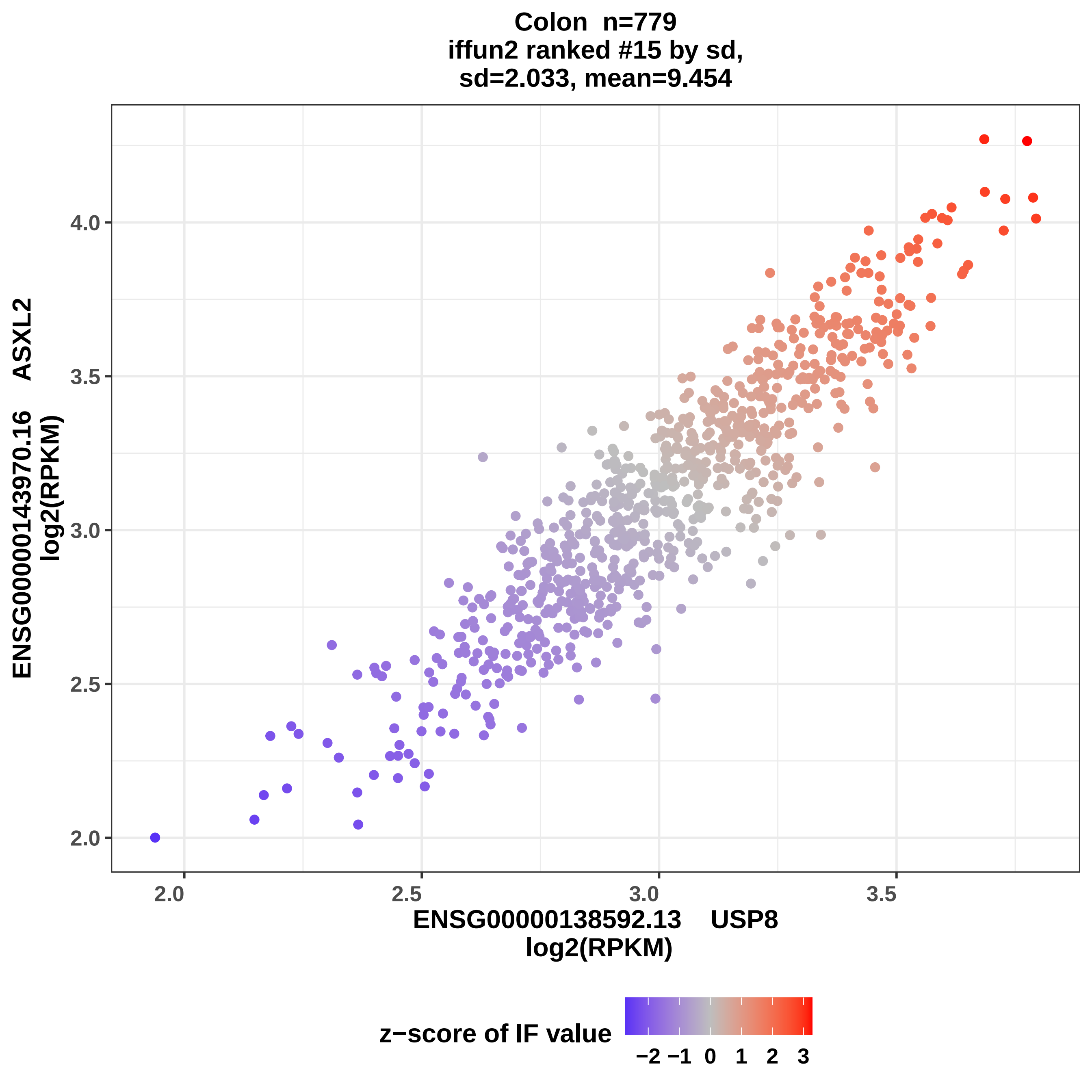
Colon n=779
iffun2 ranked #15 by sd,
sd=2.033, mean=9.454
ENSG00000143970.16 ASXL2
log2(RPKM)
2.0
2.5
3.0
3.5
4.0
2.0
2.5
3.0
3.5
ENSG00000138592.13 USP8
log2(RPKM)
z−score of IF value
−2 −1 0 1 2 3

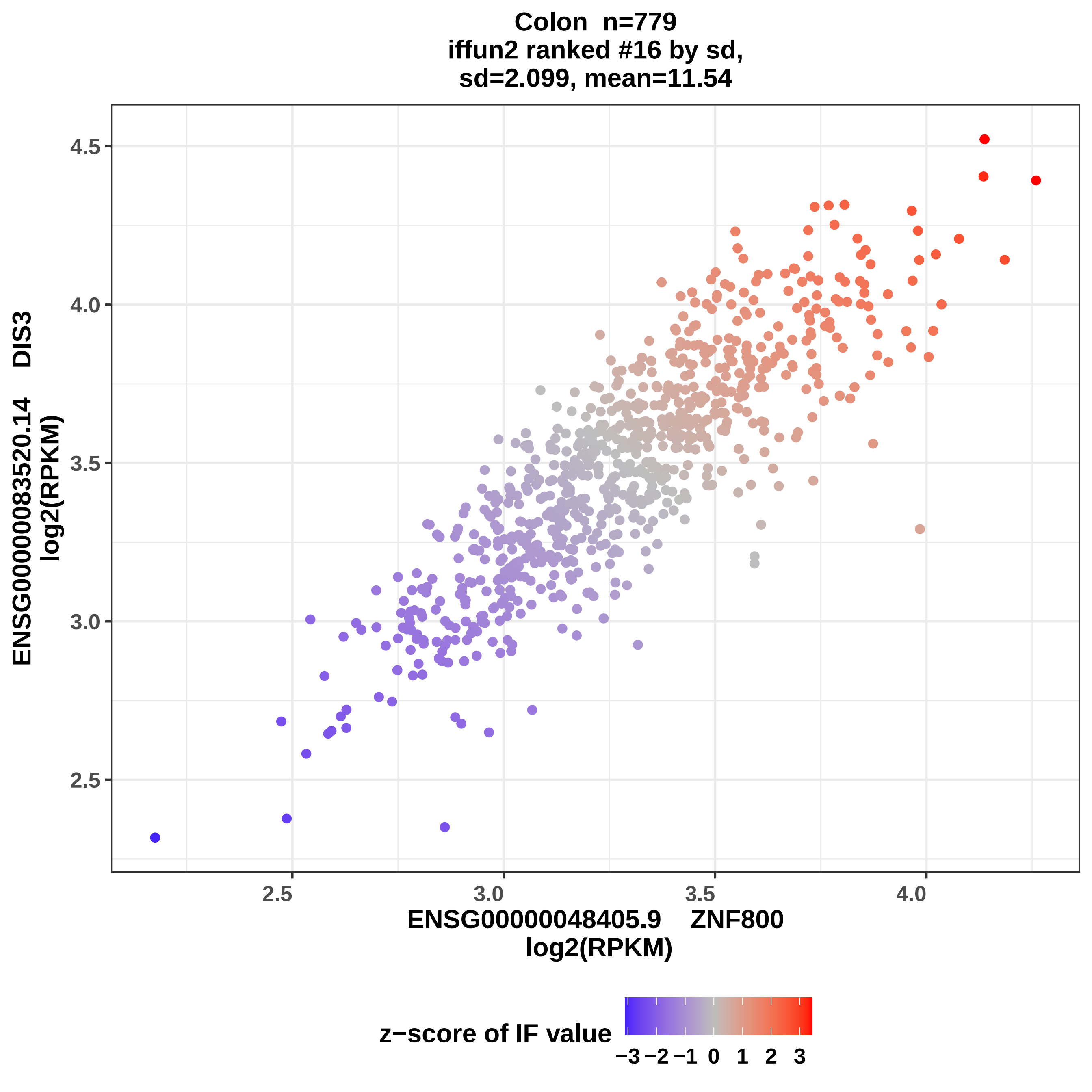

Colon n=779
iffun2 ranked #16 by sd,
sd=2.099, mean=11.54
ENSG00000083520.14 DIS3
log2(RPKM)
4.5
4.0
3.5
3.0
2.5
2.5
3.0
3.5
4.0
ENSG00000048405.9 ZNF800
log2(RPKM)
z−score of IF value
−3 −2 −1 0 1 2 3

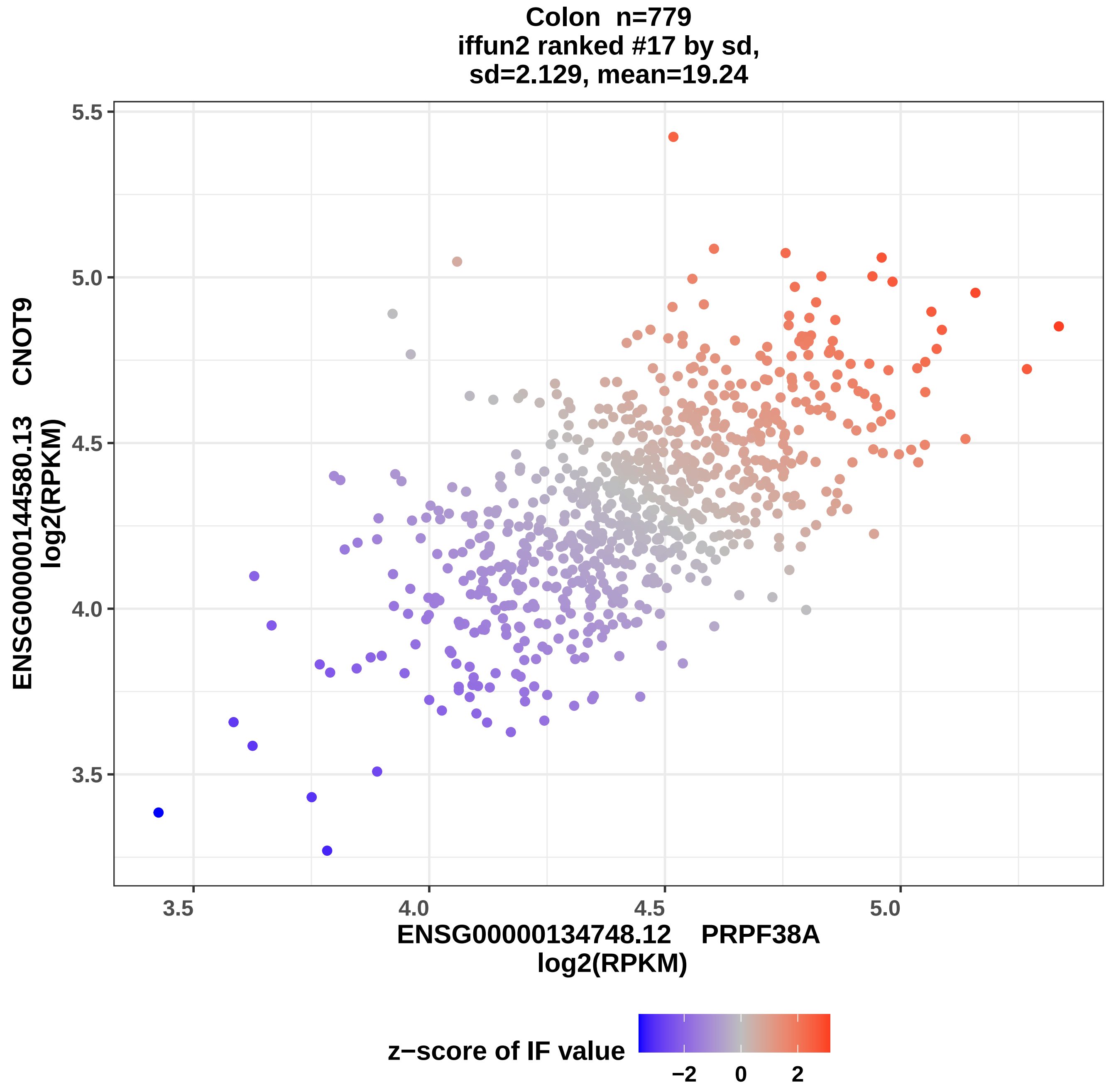

Colon n=779
iffun2 ranked #17 by sd,
sd=2.129, mean=19.24
ENSG00000144580.13 CNOT9
log2(RPKM)
ENSG00000134748.12 PRPF38A
log2(RPKM)
5.5
5.0
4.5
4.0
3.5
3.5
4.0
4.5
5.0
z−score of IF value
−2
0
2

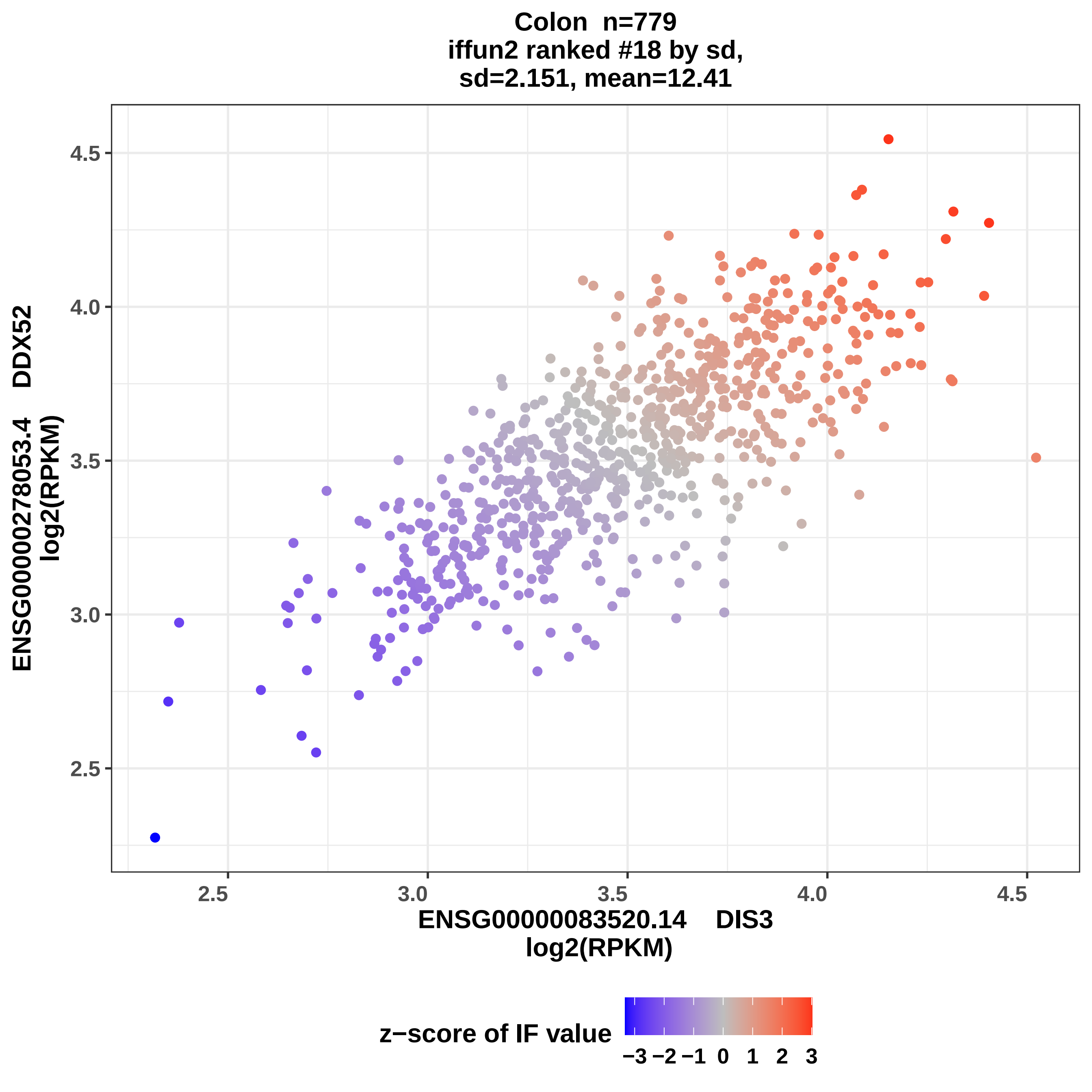
Colon n=779
iffun2 ranked #18 by sd,
sd=2.151, mean=12.41
ENSG00000278053.4 DDX52
log2(RPKM)
4.5
4.0
3.5
3.0
2.5
2.5
3.0
3.5
4.0
4.5
ENSG00000083520.14 DIS3
log2(RPKM)
z−score of IF value
−3 −2 −1 0 1 2 3

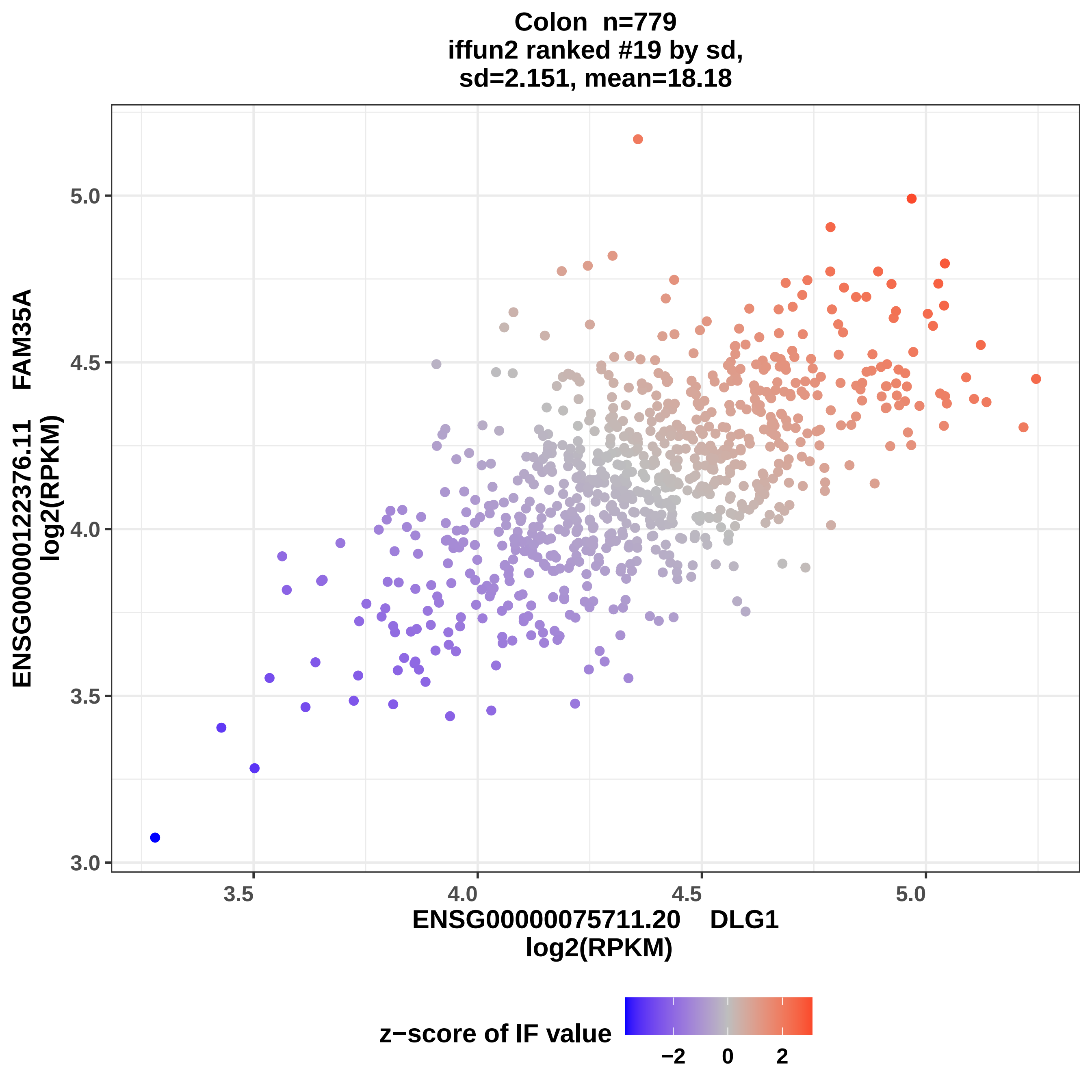

Colon n=779
iffun2 ranked #19 by sd,
sd=2.151, mean=18.18
ENSG00000122376.11 FAM35A
log2(RPKM)
ENSG00000075711.20 DLG1
log2(RPKM)
3.0
3.5
4.0
4.5
5.0
3.5
4.0
4.5
5.0
z−score of IF value
−2
0
2

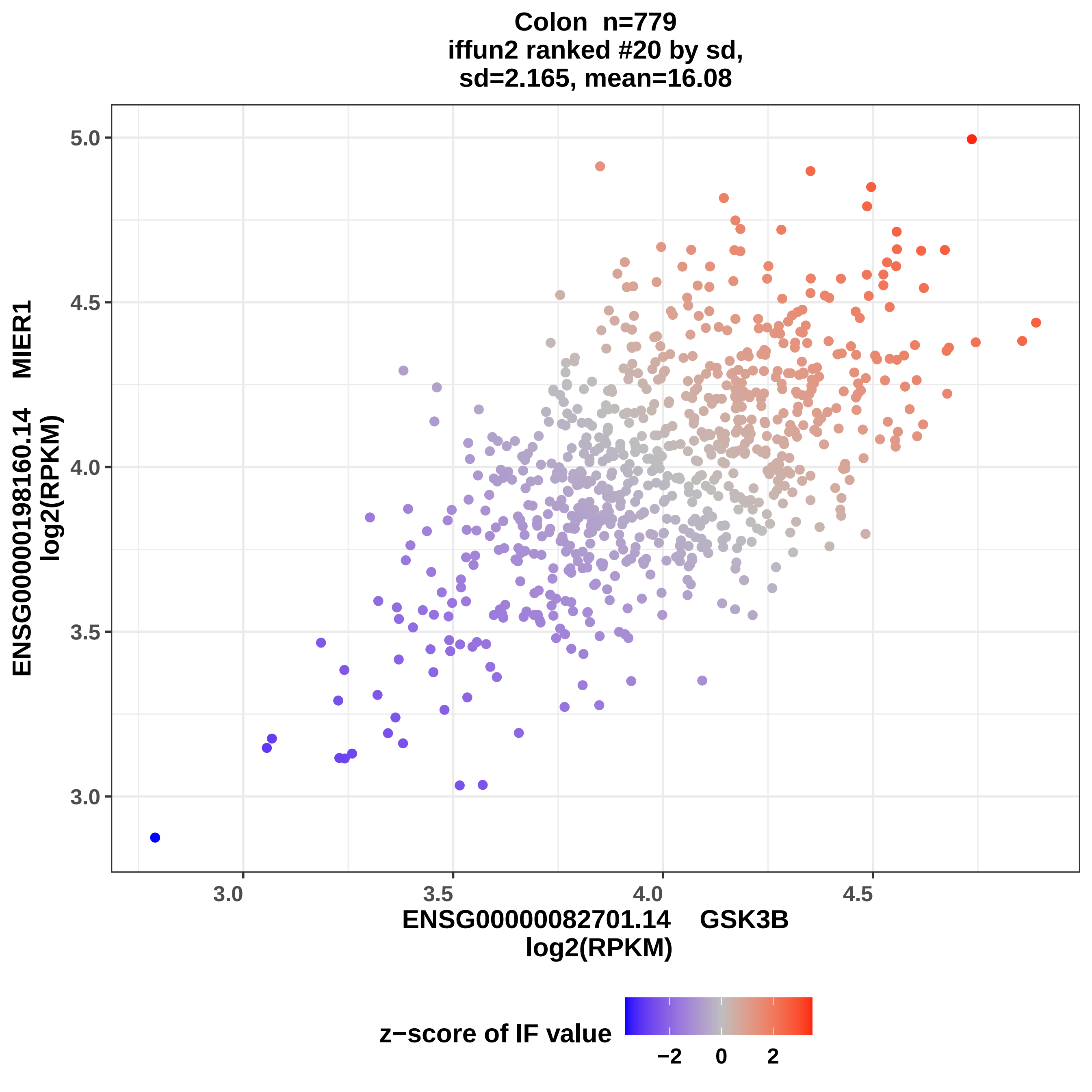
Colon n=779
iffun2 ranked #20 by sd,
sd=2.165, mean=16.08
ENSG00000198160.14 MIER1
log2(RPKM)
5.0
4.5
4.0
3.5
3.0
3.0
3.5
4.0
4.5
ENSG00000082701.14 GSK3B
log2(RPKM)
z−score of IF value
−2
0
2

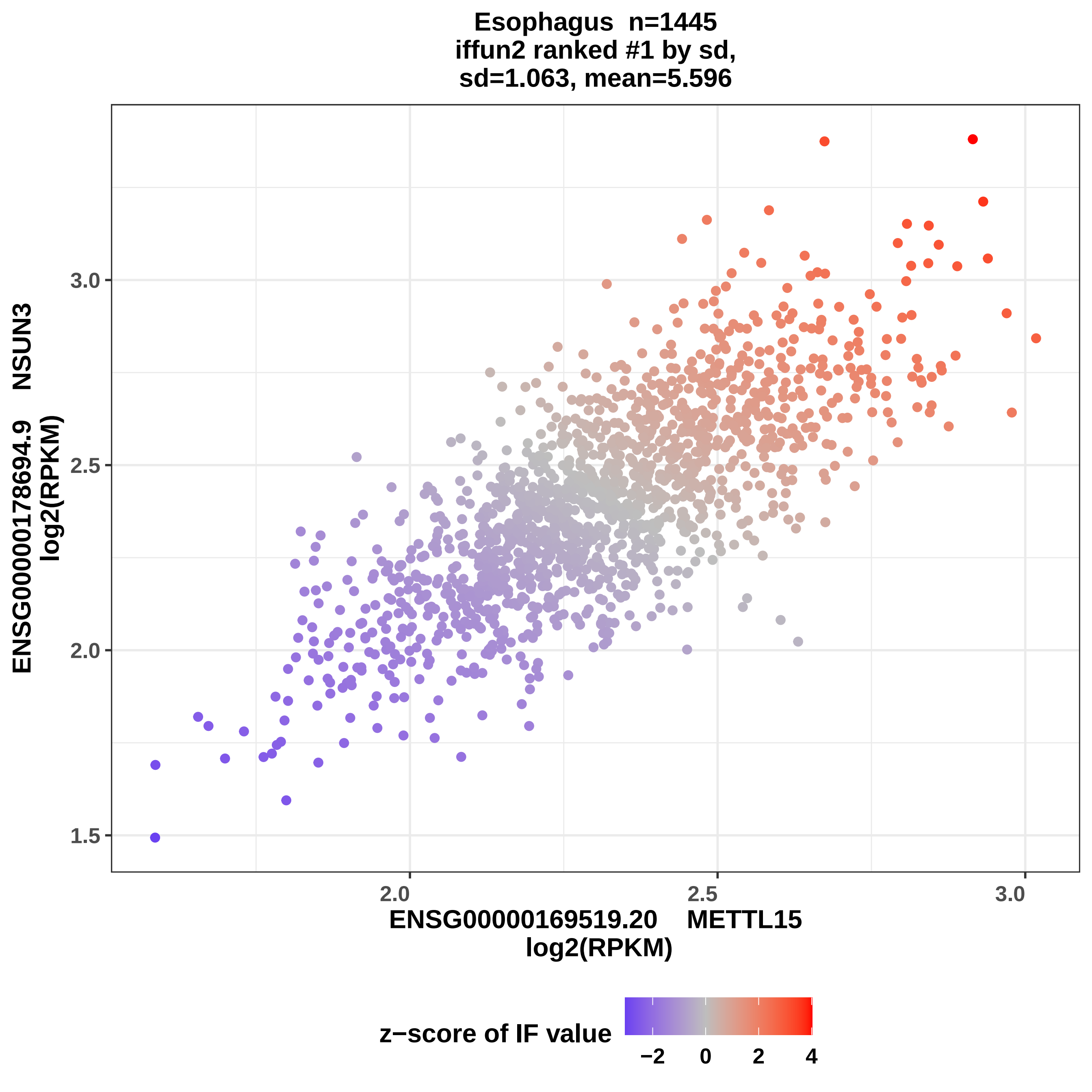
Esophagus n=1445
iffun2 ranked #1 by sd,
sd=1.063, mean=5.596
ENSG00000178694.9 NSUN3
log2(RPKM)
3.0
2.5
2.0
1.5
2.0
2.5
3.0
ENSG00000169519.20 METTL15
log2(RPKM)
z−score of IF value
−2
0
2
4

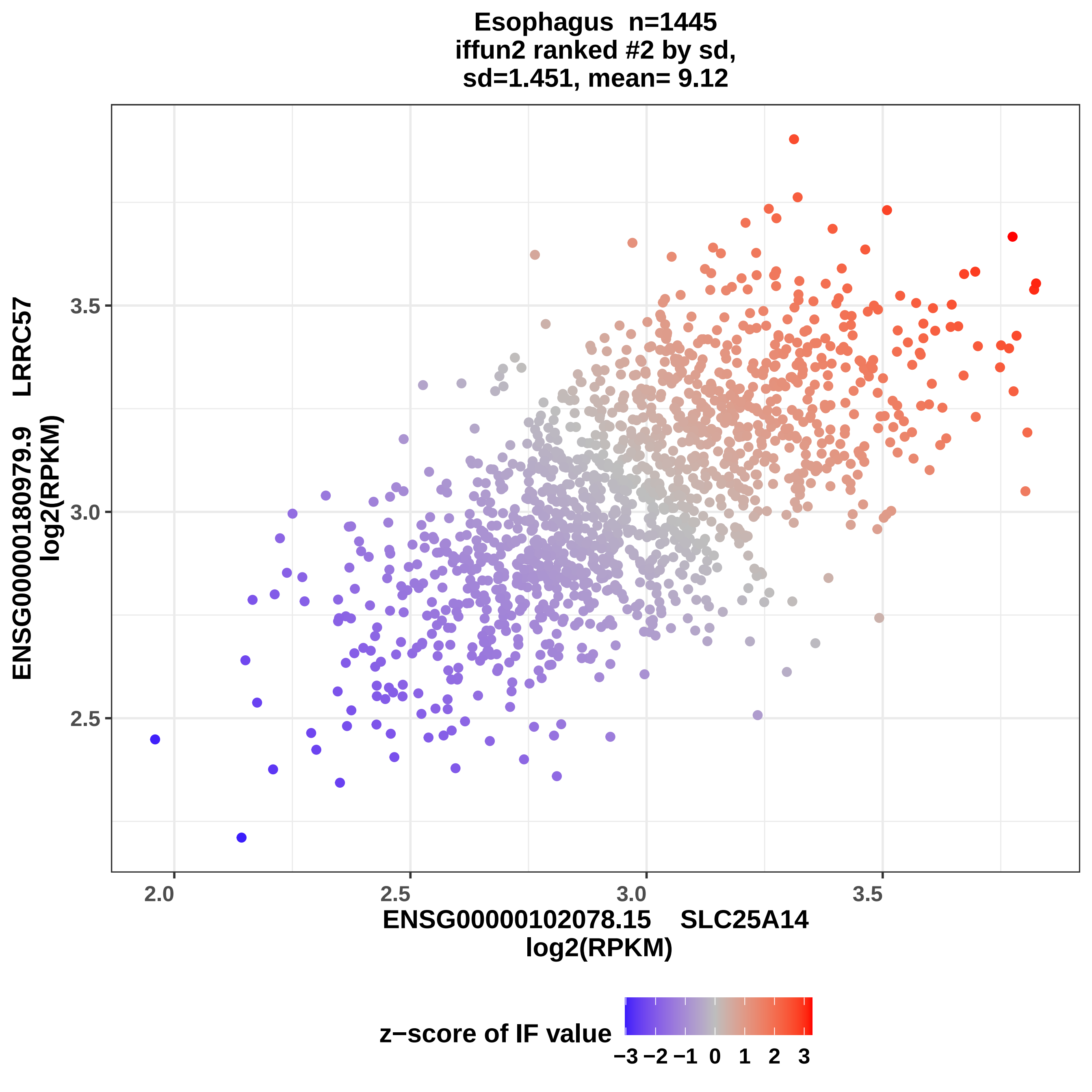
Esophagus n=1445
iffun2 ranked #2 by sd,
sd=1.451, mean= 9.12
ENSG00000180979.9 LRRC57
log2(RPKM)
3.5
3.0
2.5
2.0
2.5
3.0
3.5
ENSG00000102078.15 SLC25A14
log2(RPKM)
z−score of IF value
−3 −2 −1 0 1 2 3

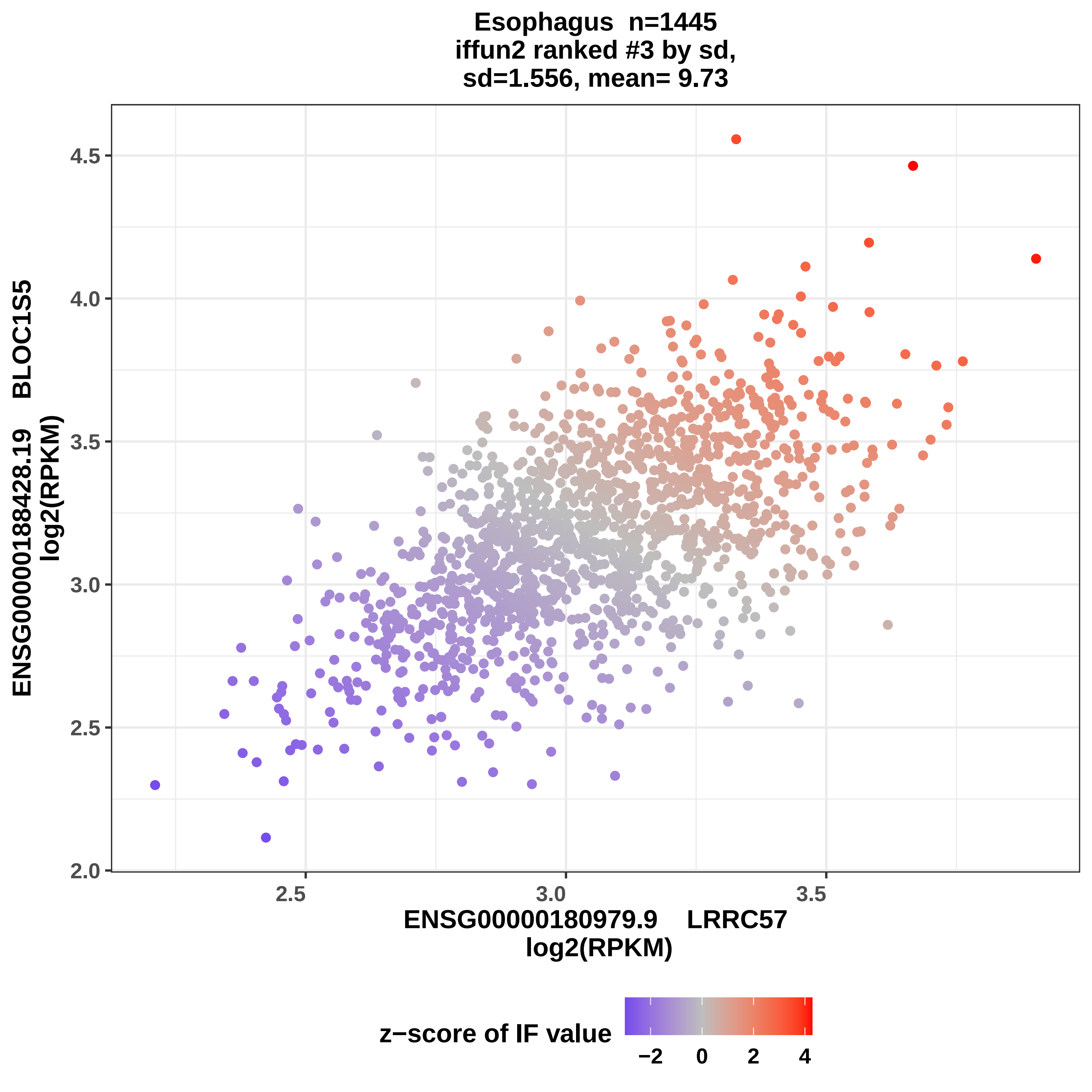
Esophagus n=1445
iffun2 ranked #3 by sd,
sd=1.556, mean= 9.73
ENSG00000188428.19 BLOC1S5
log2(RPKM)
4.5
4.0
3.5
3.0
2.5
2.0
2.5
3.0
3.5
ENSG00000180979.9 LRRC57
log2(RPKM)
z−score of IF value
−2
0
2
4

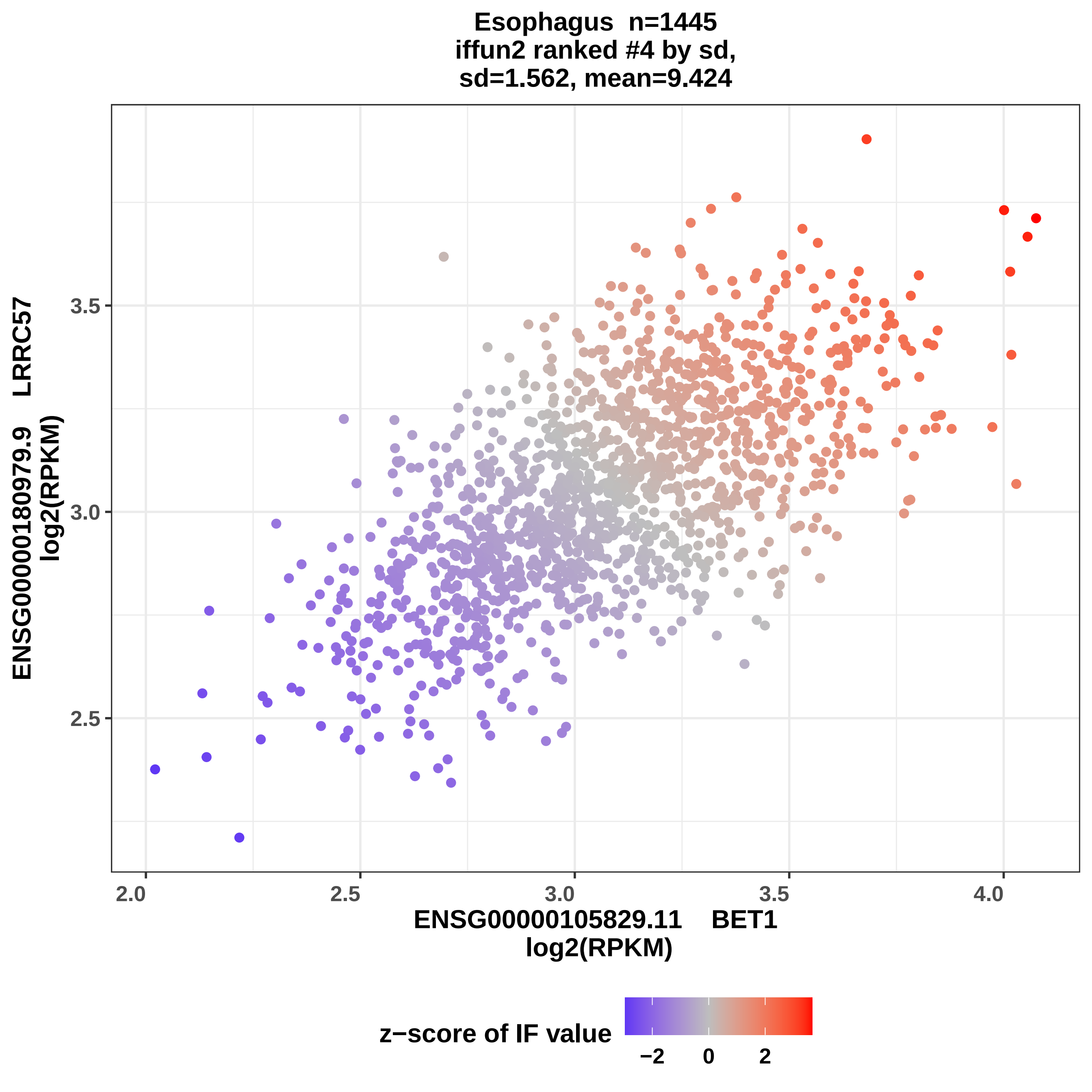
Esophagus n=1445
iffun2 ranked #4 by sd,
sd=1.562, mean=9.424
ENSG00000180979.9 LRRC57
log2(RPKM)
3.5
3.0
2.5
2.0
2.5
3.0
3.5
4.0
ENSG00000105829.11 BET1
log2(RPKM)
z−score of IF value
−2
0
2

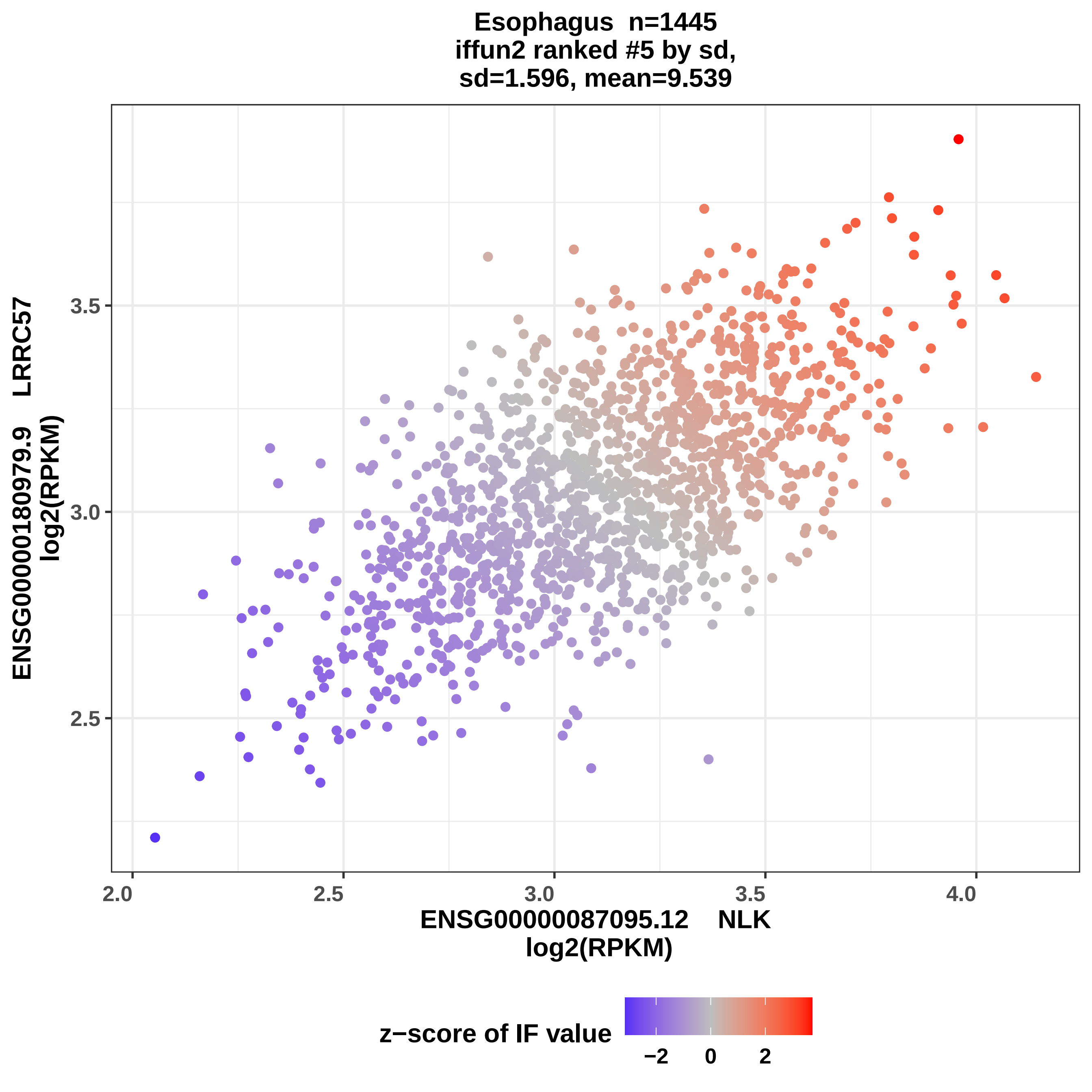
Esophagus n=1445
iffun2 ranked #5 by sd,
sd=1.596, mean=9.539
ENSG00000180979.9 LRRC57
log2(RPKM)
3.5
3.0
2.5
2.0
2.5
3.0
3.5
4.0
ENSG00000087095.12 NLK
log2(RPKM)
z−score of IF value
−2
0
2

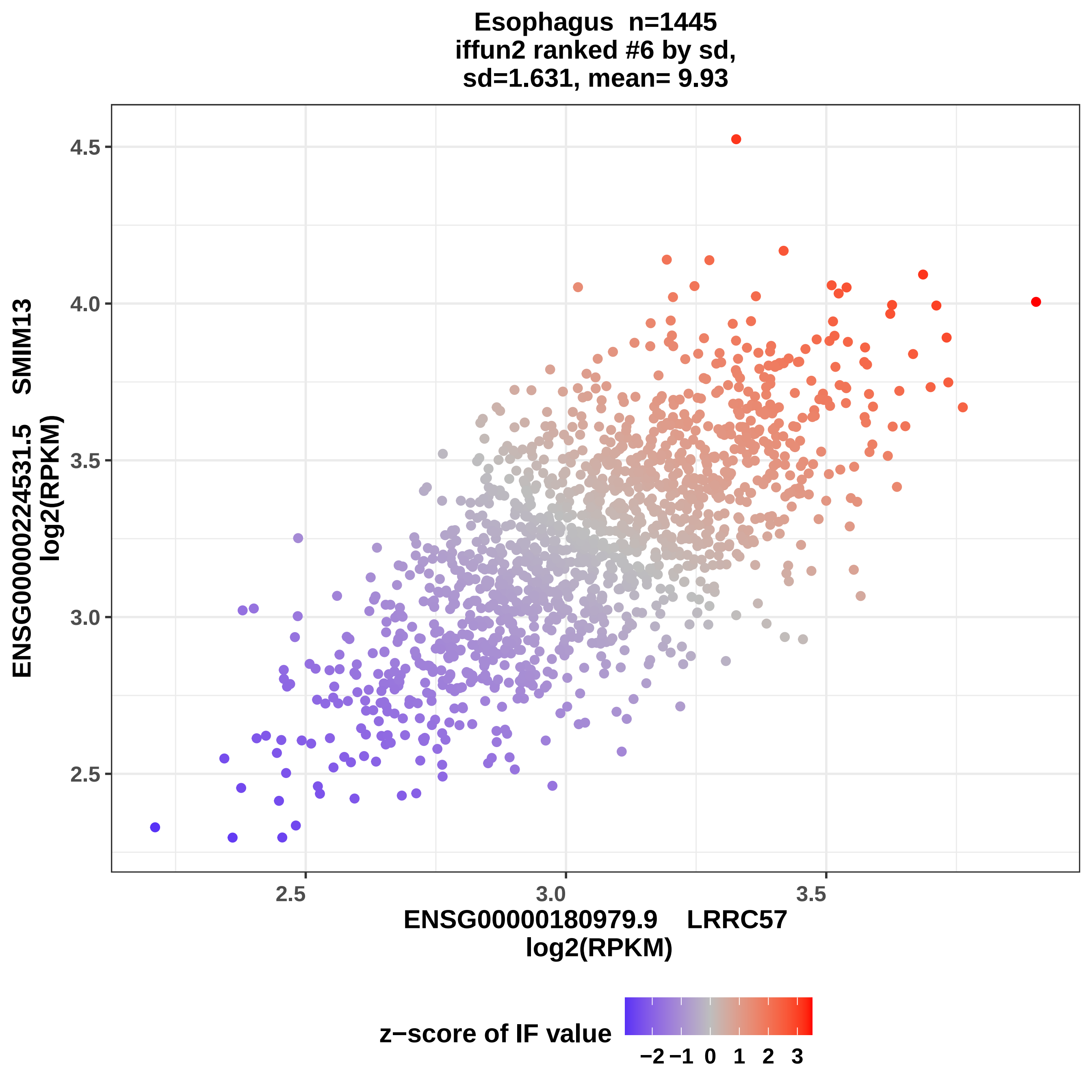
Esophagus n=1445
iffun2 ranked #6 by sd,
sd=1.631, mean= 9.93
4.5
4.0
3.5
3.0
2.5
2.5
3.0
3.5
ENSG00000224531.5 SMIM13
log2(RPKM)
ENSG00000180979.9 LRRC57
log2(RPKM)
z−score of IF value
−2 −1 0 1 2 3

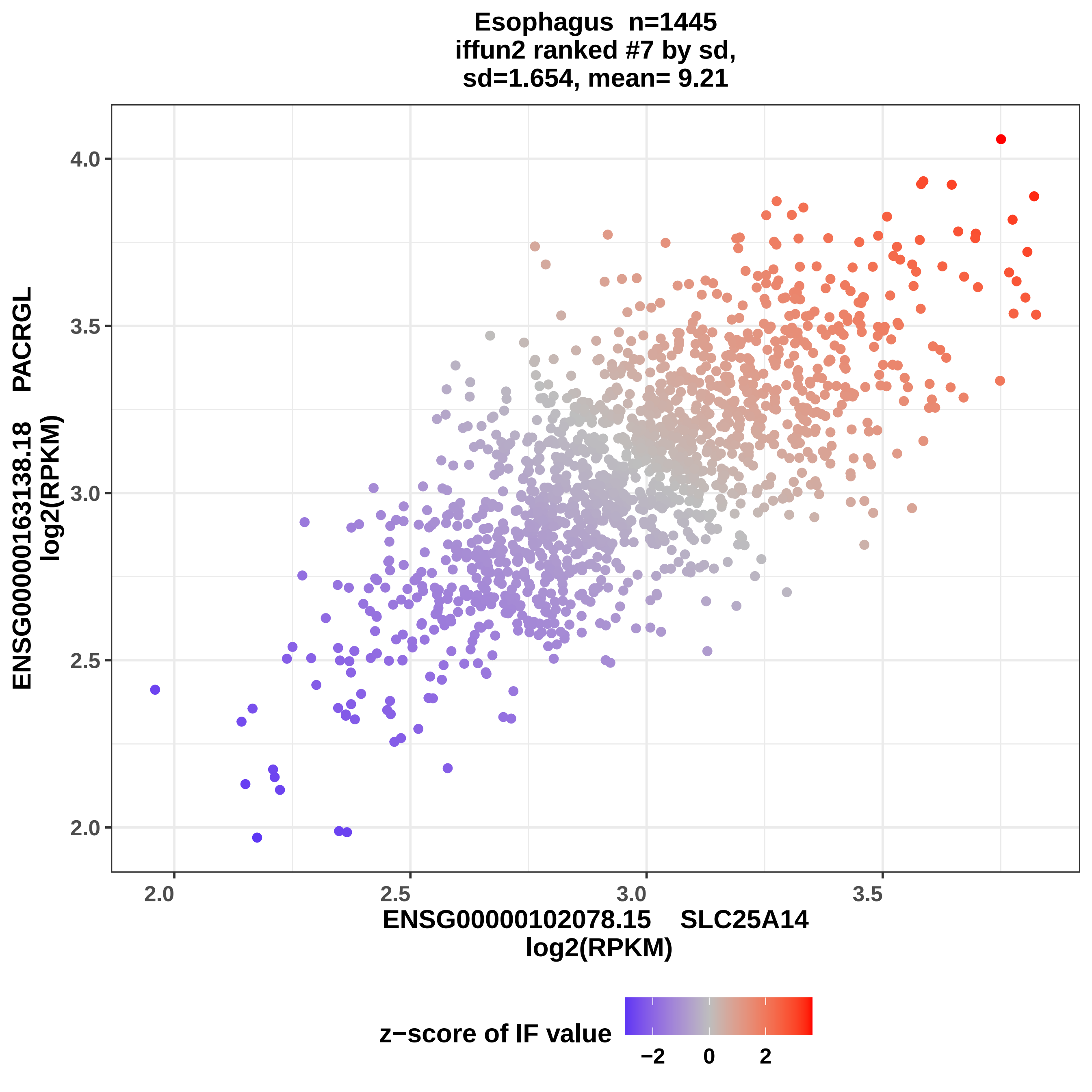
Esophagus n=1445
iffun2 ranked #7 by sd,
sd=1.654, mean= 9.21
ENSG00000163138.18 PACRGL
log2(RPKM)
4.0
3.5
3.0
2.5
2.0
2.0
2.5
3.0
3.5
ENSG00000102078.15 SLC25A14
log2(RPKM)
z−score of IF value
−2
0
2

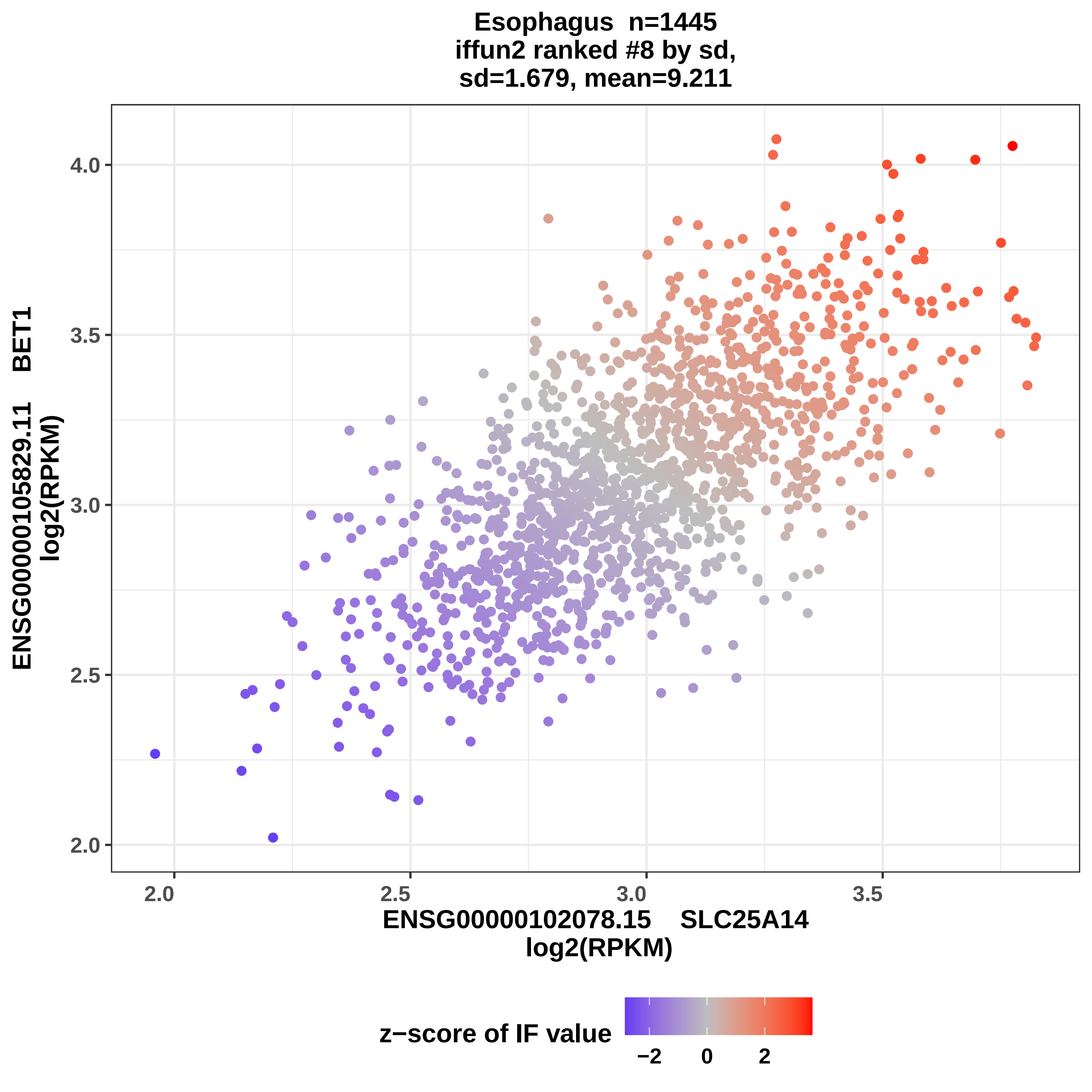

Esophagus n=1445
iffun2 ranked #8 by sd,
sd=1.679, mean=9.211
ENSG00000105829.11 BET1
log2(RPKM)
ENSG00000102078.15 SLC25A14
log2(RPKM)
4.0
3.5
3.0
2.5
2.0
2.0
2.5
3.0
3.5
z−score of IF value
−2
0
2

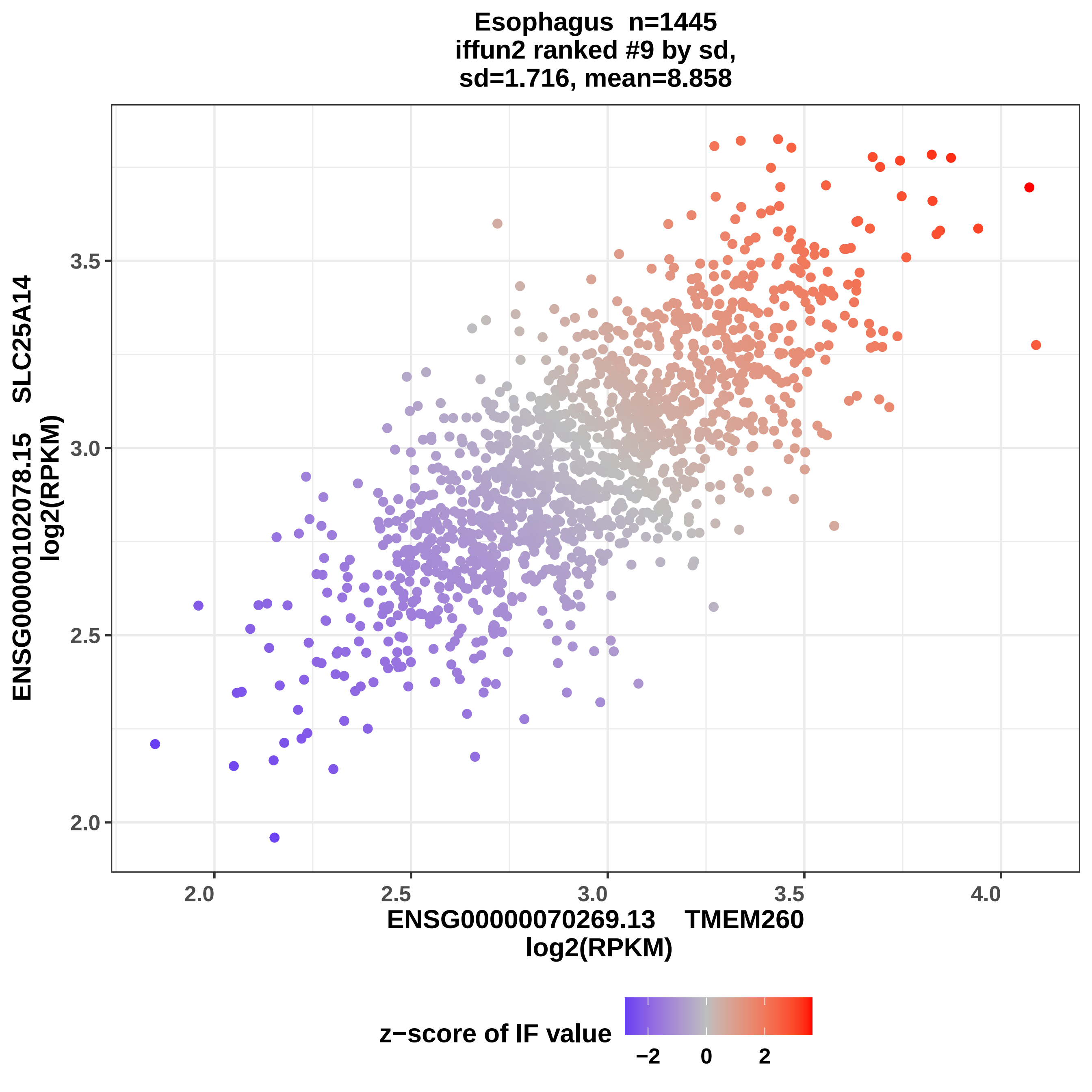
Esophagus n=1445
iffun2 ranked #9 by sd,
sd=1.716, mean=8.858
ENSG00000102078.15 SLC25A14
log2(RPKM)
ENSG00000070269.13 TMEM260
log2(RPKM)
2.0
2.5
3.0
3.5
4.0
z−score of IF value
−2
0
2

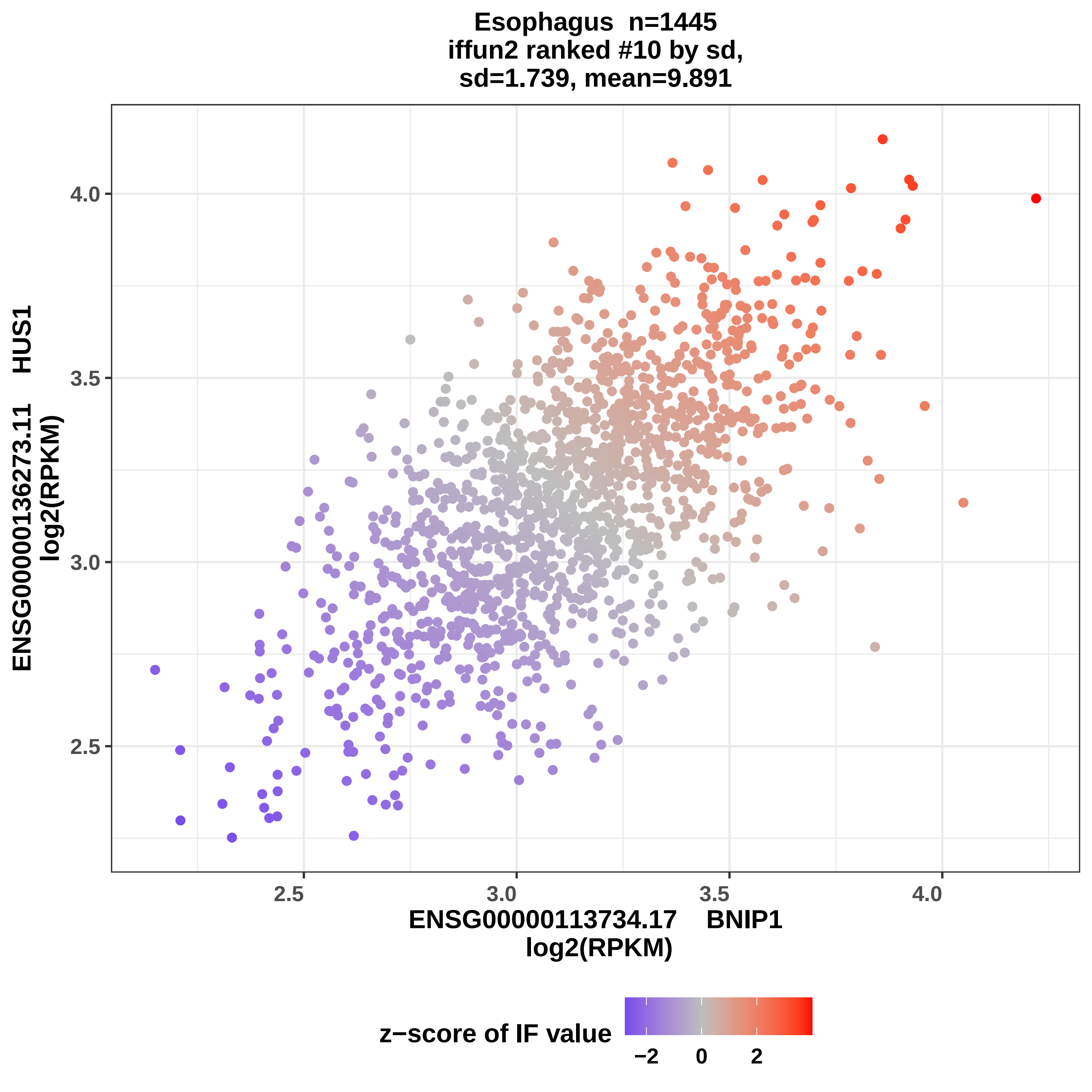

Esophagus n=1445
iffun2 ranked #10 by sd,
sd=1.739, mean=9.891
ENSG00000136273.11 HUS1
log2(RPKM)
ENSG00000113734.17 BNIP1
log2(RPKM)
4.0
3.5
3.0
2.5
2.5
3.0
3.5
4.0
z−score of IF value
−2
0
2

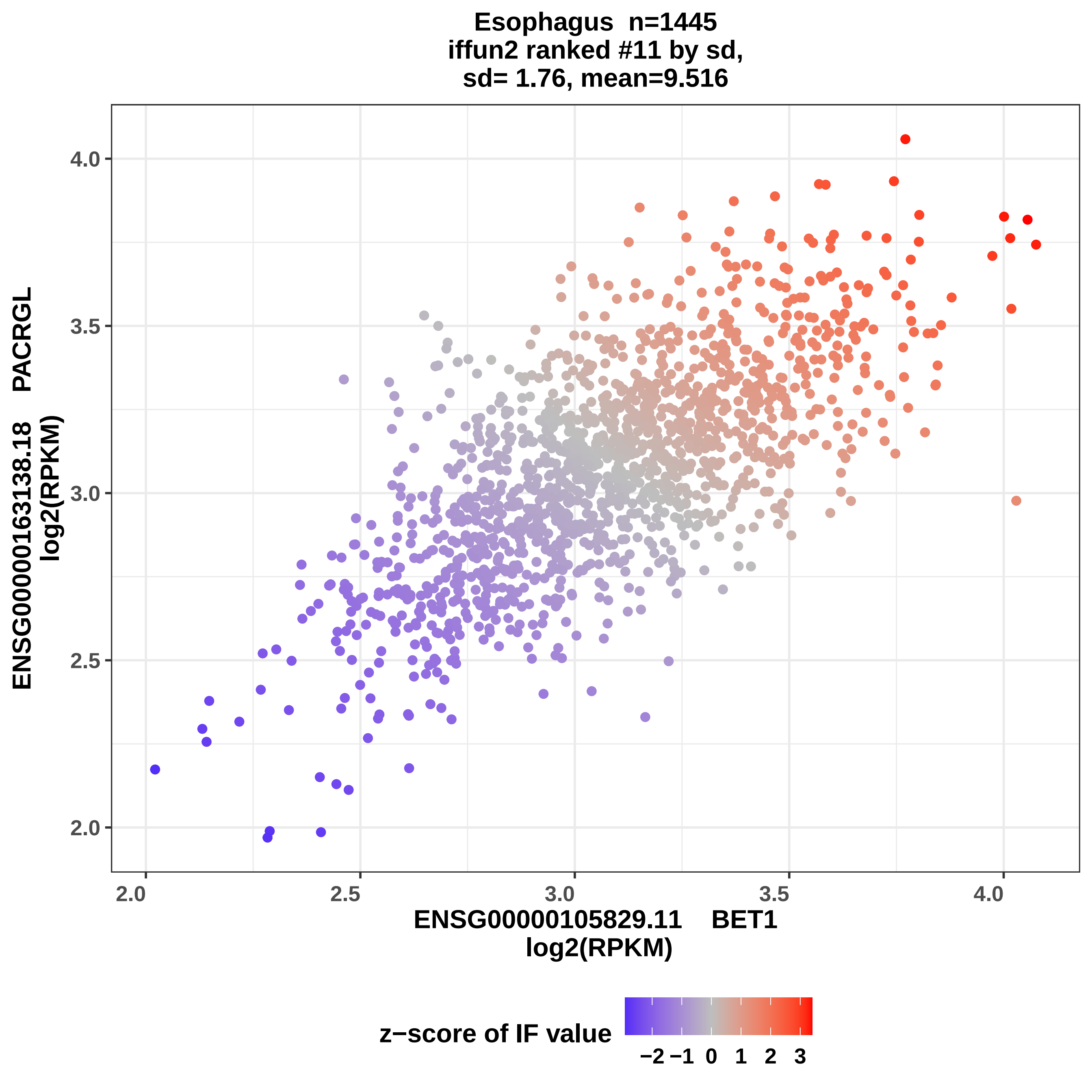

Esophagus n=1445
iffun2 ranked #11 by sd,
sd= 1.76, mean=9.516
ENSG00000163138.18 PACRGL
log2(RPKM)
4.0
3.5
3.0
2.5
2.0
2.0
2.5
3.0
3.5
4.0
ENSG00000105829.11 BET1
log2(RPKM)
z−score of IF value
−2 −1 0 1 2 3

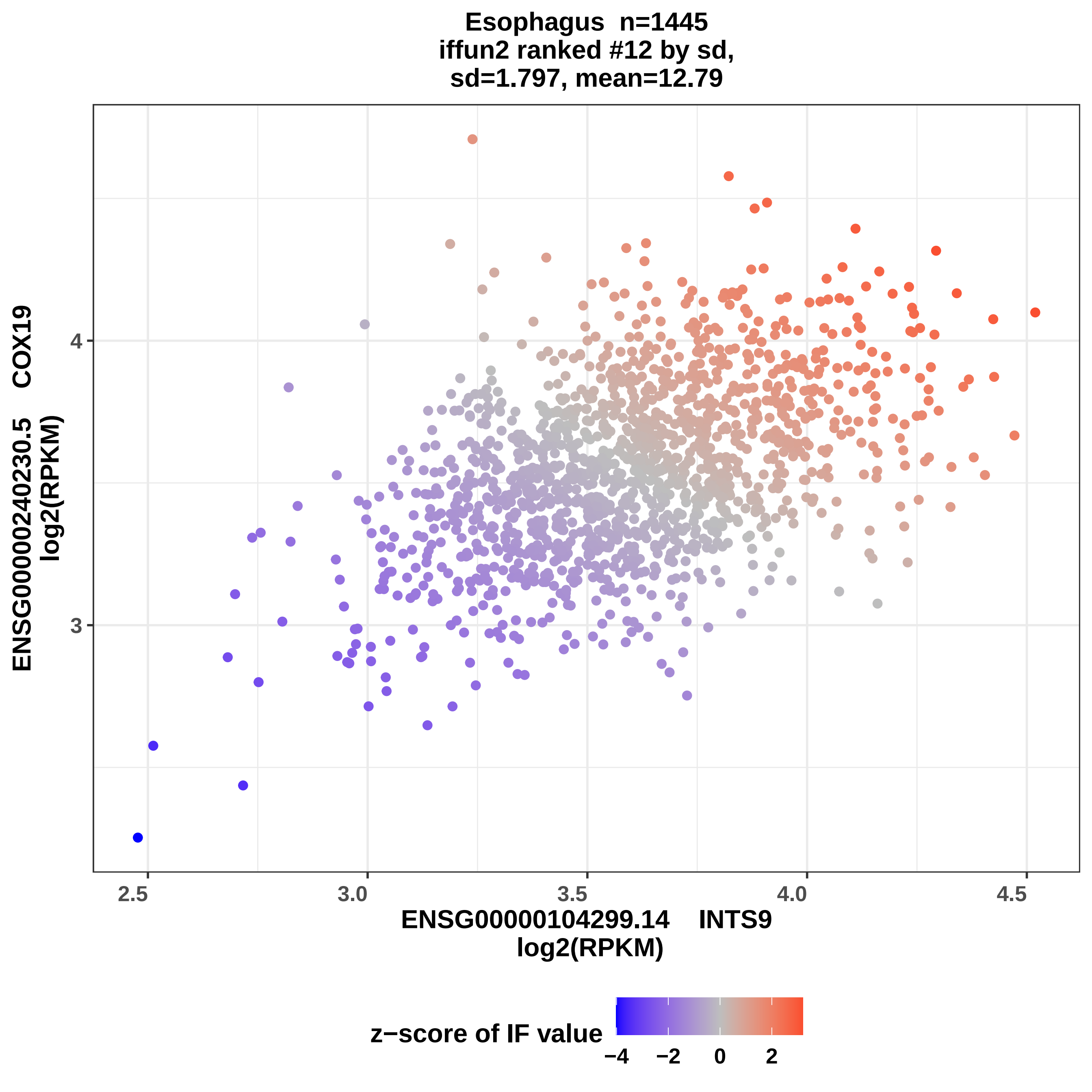

Esophagus n=1445
iffun2 ranked #12 by sd,
sd=1.797, mean=12.79
ENSG00000240230.5 COX19
log2(RPKM)
4
3
2.5
3.0
3.5
4.0
4.5
ENSG00000104299.14 INTS9
log2(RPKM)
z−score of IF value
−4
−2
0
2

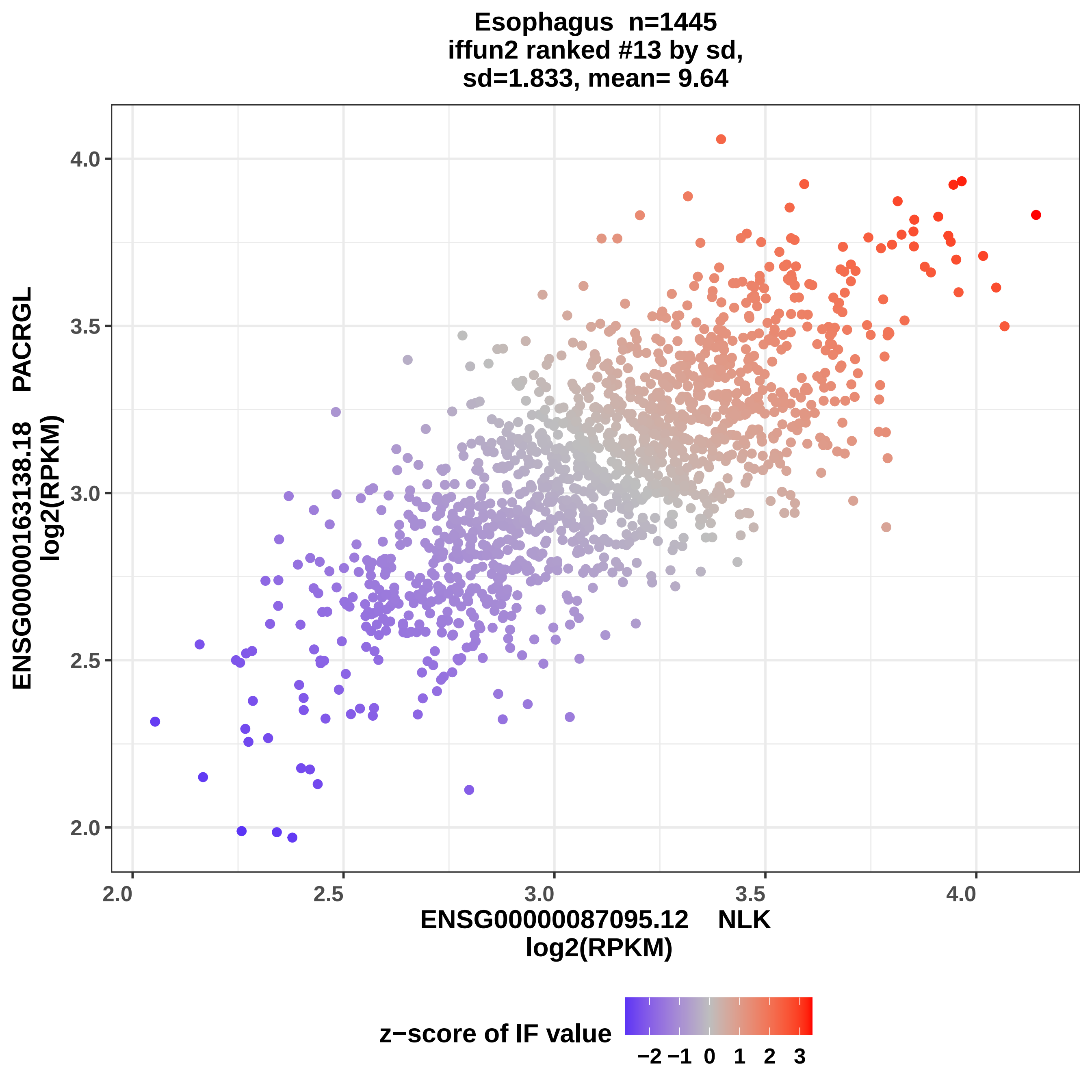

Esophagus n=1445
iffun2 ranked #13 by sd,
sd=1.833, mean= 9.64
ENSG00000163138.18 PACRGL
log2(RPKM)
4.0
3.5
3.0
2.5
2.0
2.0
2.5
3.0
3.5
4.0
ENSG00000087095.12 NLK
log2(RPKM)
z−score of IF value
−2 −1 0 1 2 3

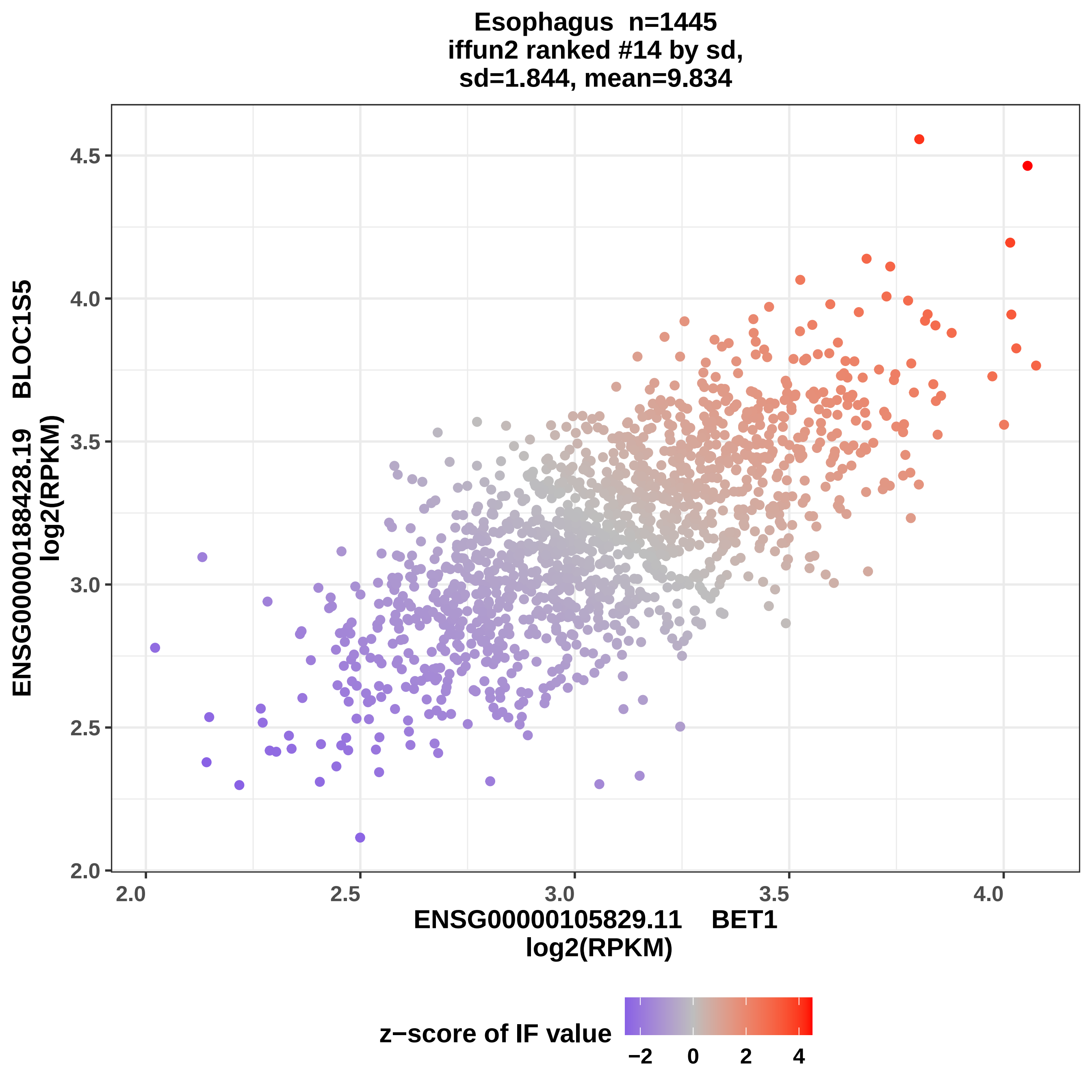

Esophagus n=1445
iffun2 ranked #14 by sd,
sd=1.844, mean=9.834
ENSG00000188428.19 BLOC1S5
log2(RPKM)
2.0
2.5
3.0
3.5
4.0
4.5
ENSG00000105829.11 BET1
log2(RPKM)
2.0
2.5
3.0
3.5
4.0
z−score of IF value
−2
0
2
4

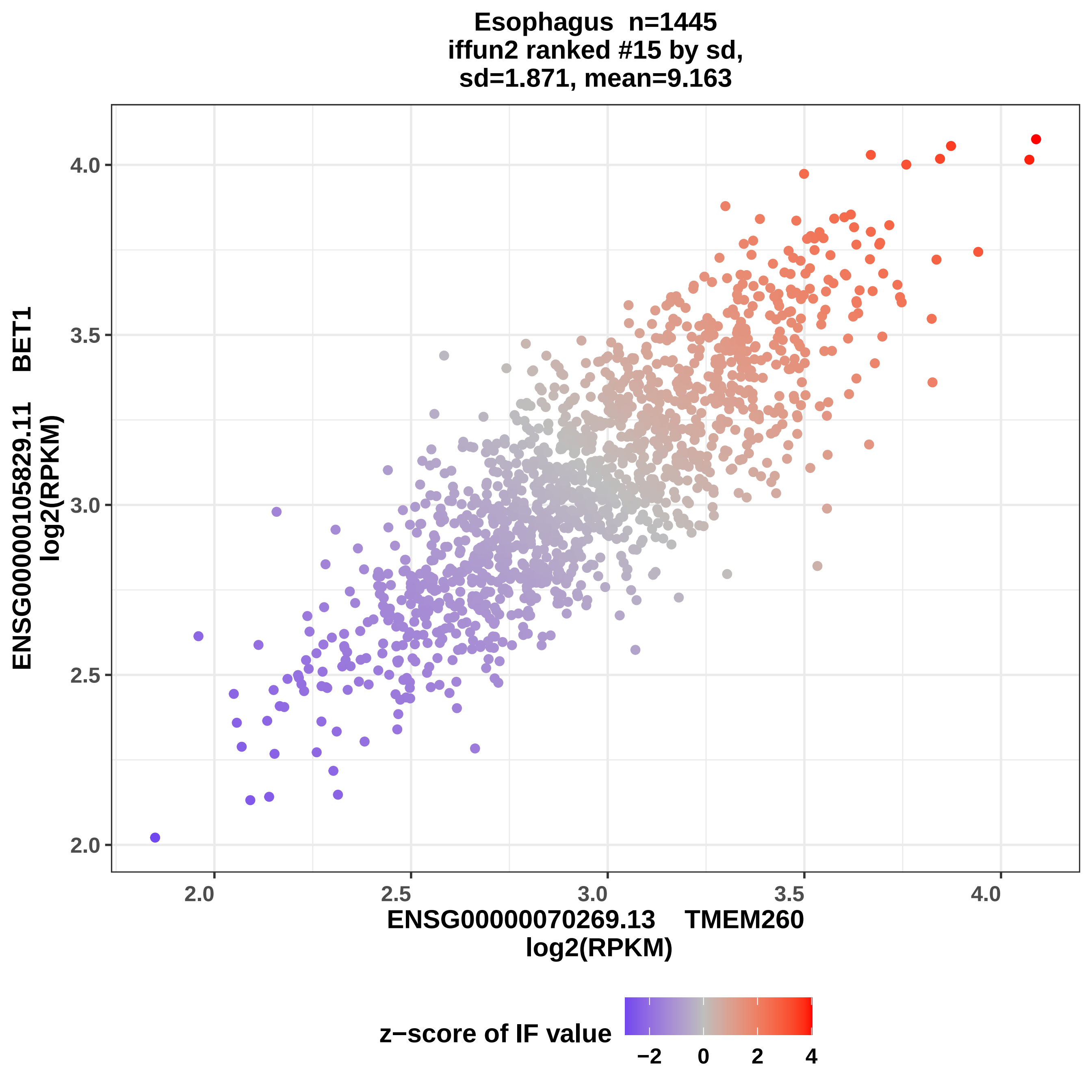

Esophagus n=1445
iffun2 ranked #15 by sd,
sd=1.871, mean=9.163
ENSG00000105829.11 BET1
log2(RPKM)
4.0
3.5
3.0
2.5
2.0
2.0
2.5
3.0
3.5
4.0
ENSG00000070269.13 TMEM260
log2(RPKM)
z−score of IF value
−2
0
2
4

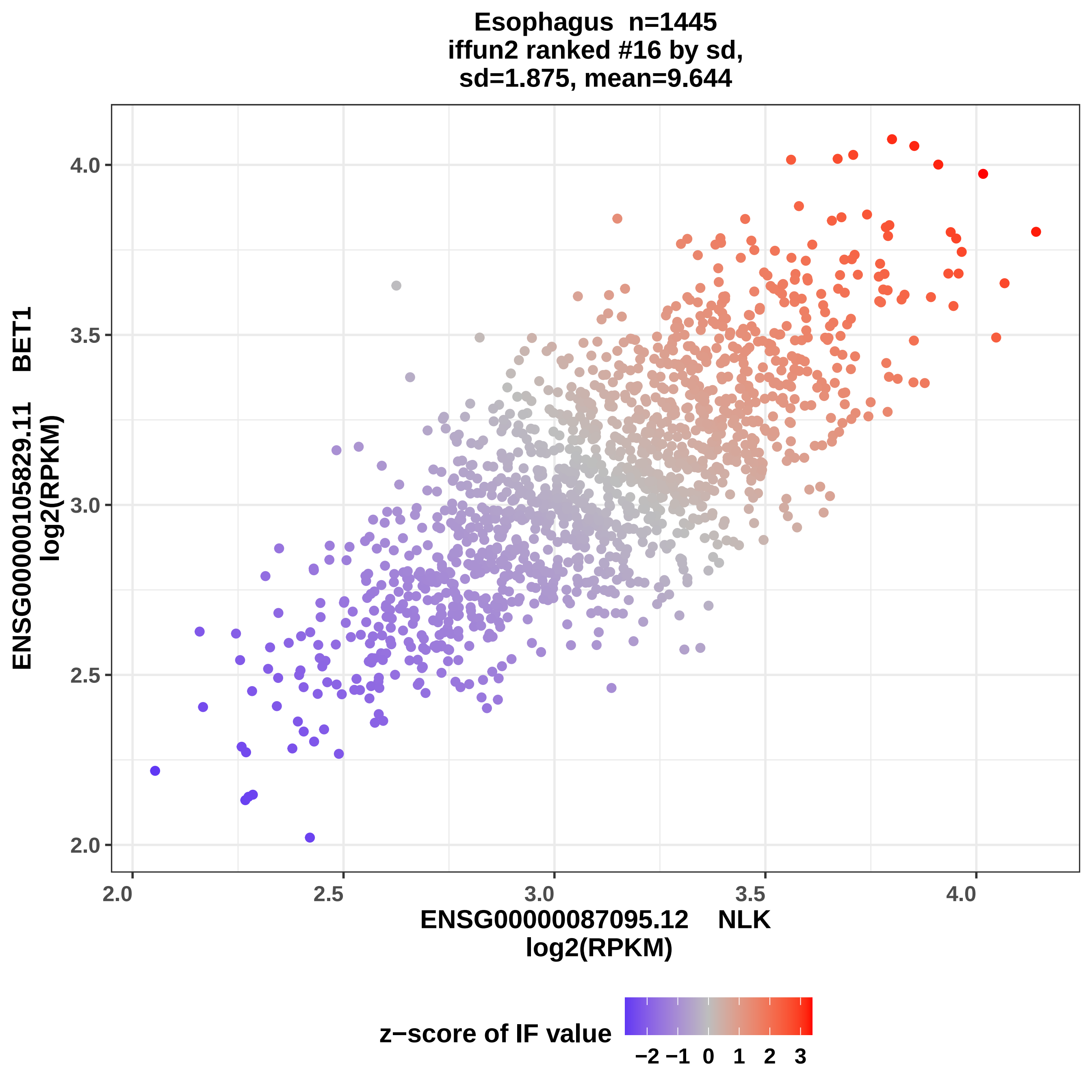
Esophagus n=1445
iffun2 ranked #16 by sd,
sd=1.875, mean=9.644
ENSG00000105829.11 BET1
log2(RPKM)
4.0
3.5
3.0
2.5
2.0
2.0
2.5
3.0
3.5
4.0
ENSG00000087095.12 NLK
log2(RPKM)
z−score of IF value
−2
−1
0
1
2
3

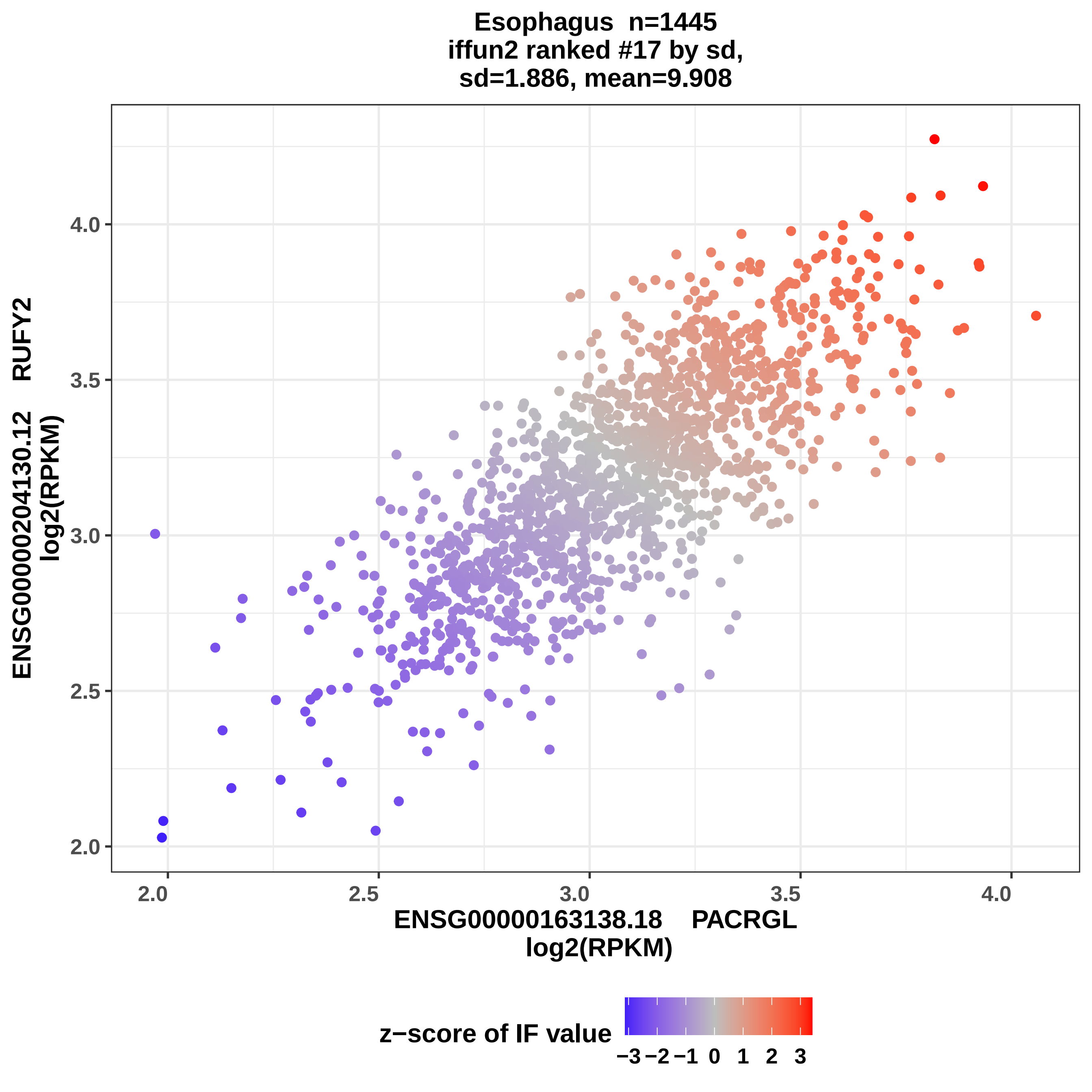
Esophagus n=1445
iffun2 ranked #17 by sd,
sd=1.886, mean=9.908
ENSG00000204130.12 RUFY2
log2(RPKM)
4.0
3.5
3.0
2.5
2.0
2.0
2.5
3.0
3.5
4.0
ENSG00000163138.18 PACRGL
log2(RPKM)
z−score of IF value
−3 −2 −1 0 1 2 3

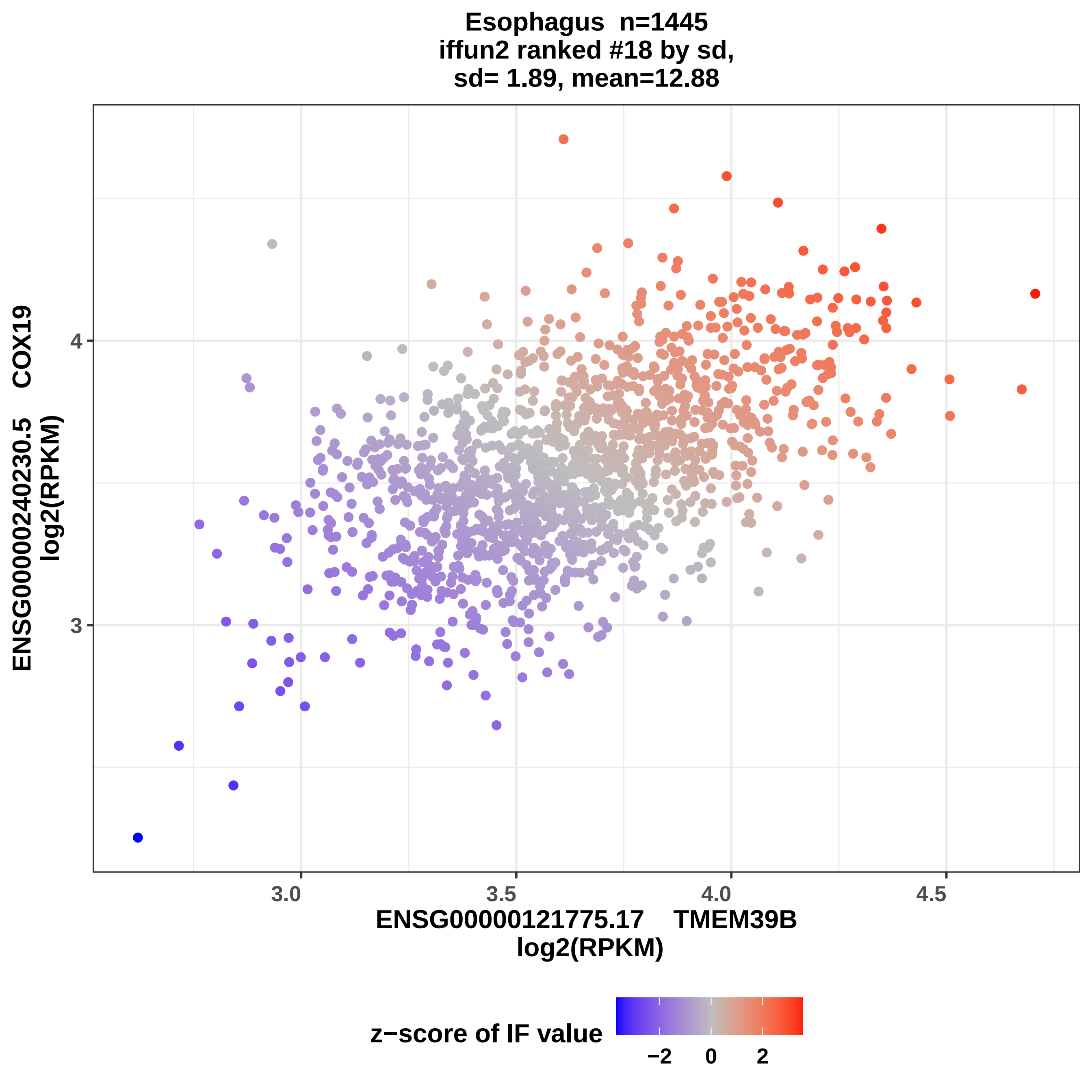
Esophagus n=1445
iffun2 ranked #18 by sd,
sd= 1.89, mean=12.88
ENSG00000240230.5 COX19
log2(RPKM)
4
3
3.0
3.5
4.0
4.5
ENSG00000121775.17 TMEM39B
log2(RPKM)
z−score of IF value
−2
0
2

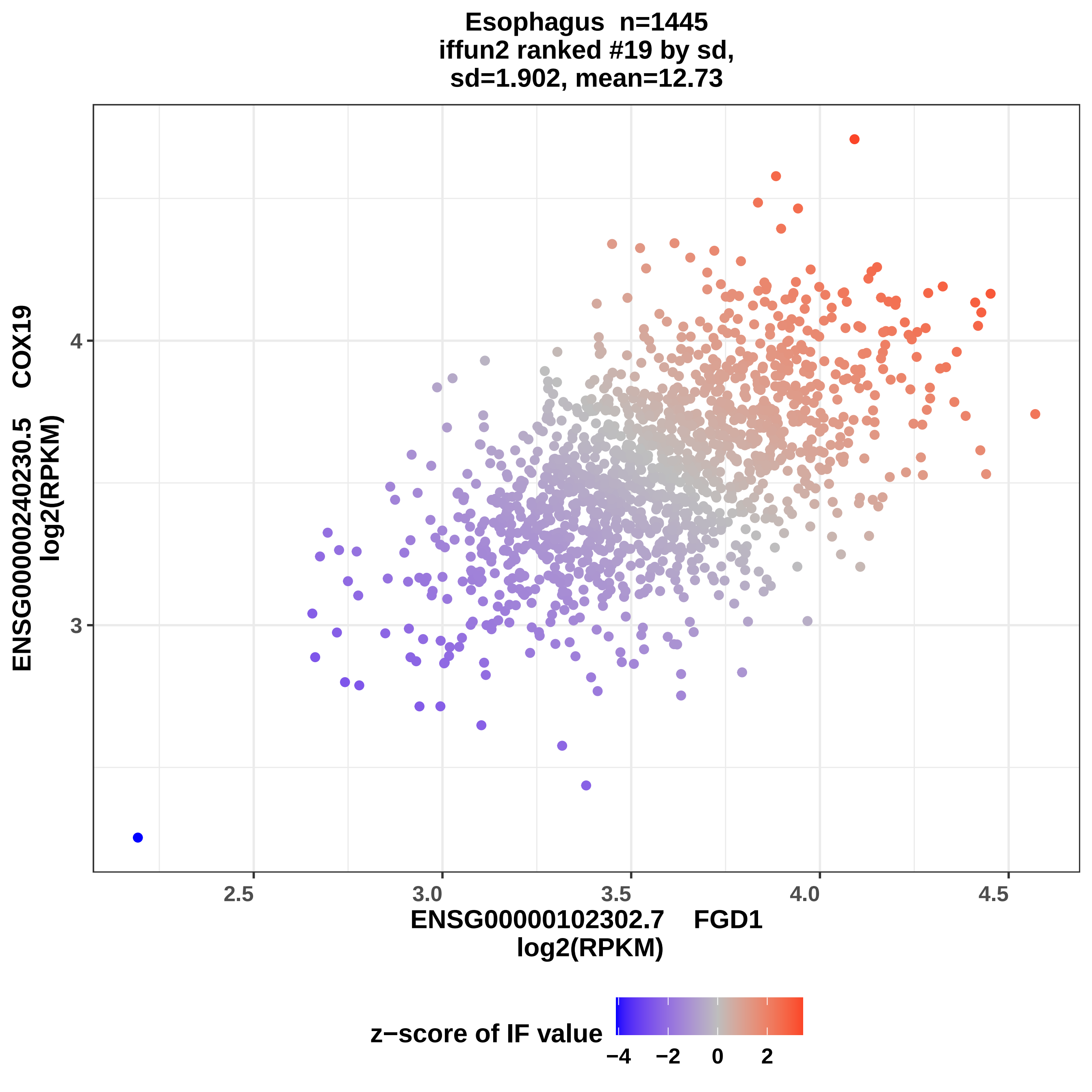
Esophagus n=1445
iffun2 ranked #19 by sd,
sd=1.902, mean=12.73
ENSG00000240230.5 COX19
log2(RPKM)
4
3
2.5
3.0
3.5
4.0
4.5
ENSG00000102302.7 FGD1
log2(RPKM)
z−score of IF value
−4
−2
0
2

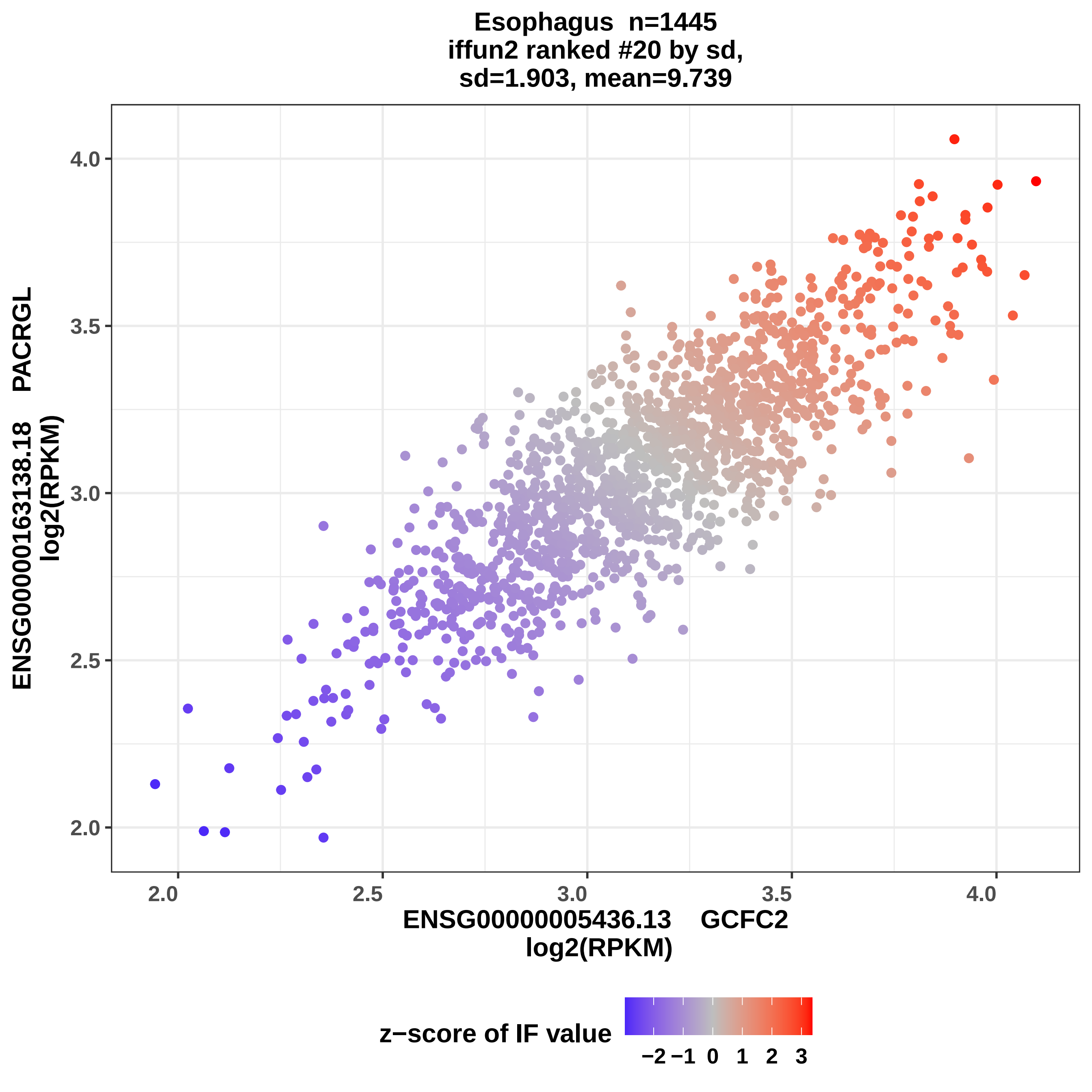

Esophagus n=1445
iffun2 ranked #20 by sd,
sd=1.903, mean=9.739
ENSG00000163138.18 PACRGL
log2(RPKM)
4.0
3.5
3.0
2.5
2.0
2.0
2.5
3.0
3.5
4.0
ENSG00000005436.13 GCFC2
log2(RPKM)
z−score of IF value
−2 −1 0 1 2 3

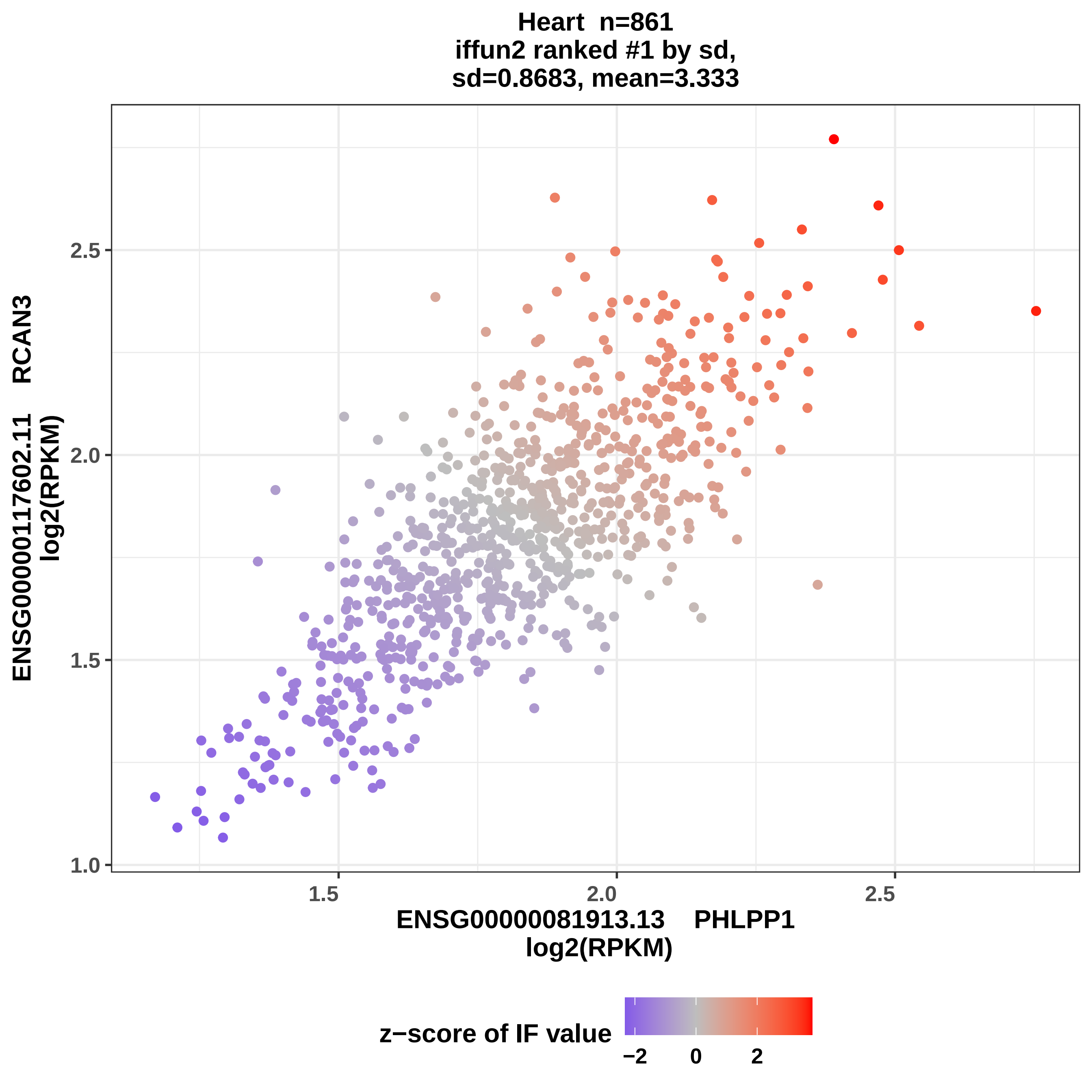

Heart n=861
iffun2 ranked #1 by sd,
sd=0.8683, mean=3.333
ENSG00000117602.11 RCAN3
log2(RPKM)
1.0
1.5
2.0
2.5
1.5
2.0
2.5
ENSG00000081913.13 PHLPP1
log2(RPKM)
z−score of IF value
−2
0
2

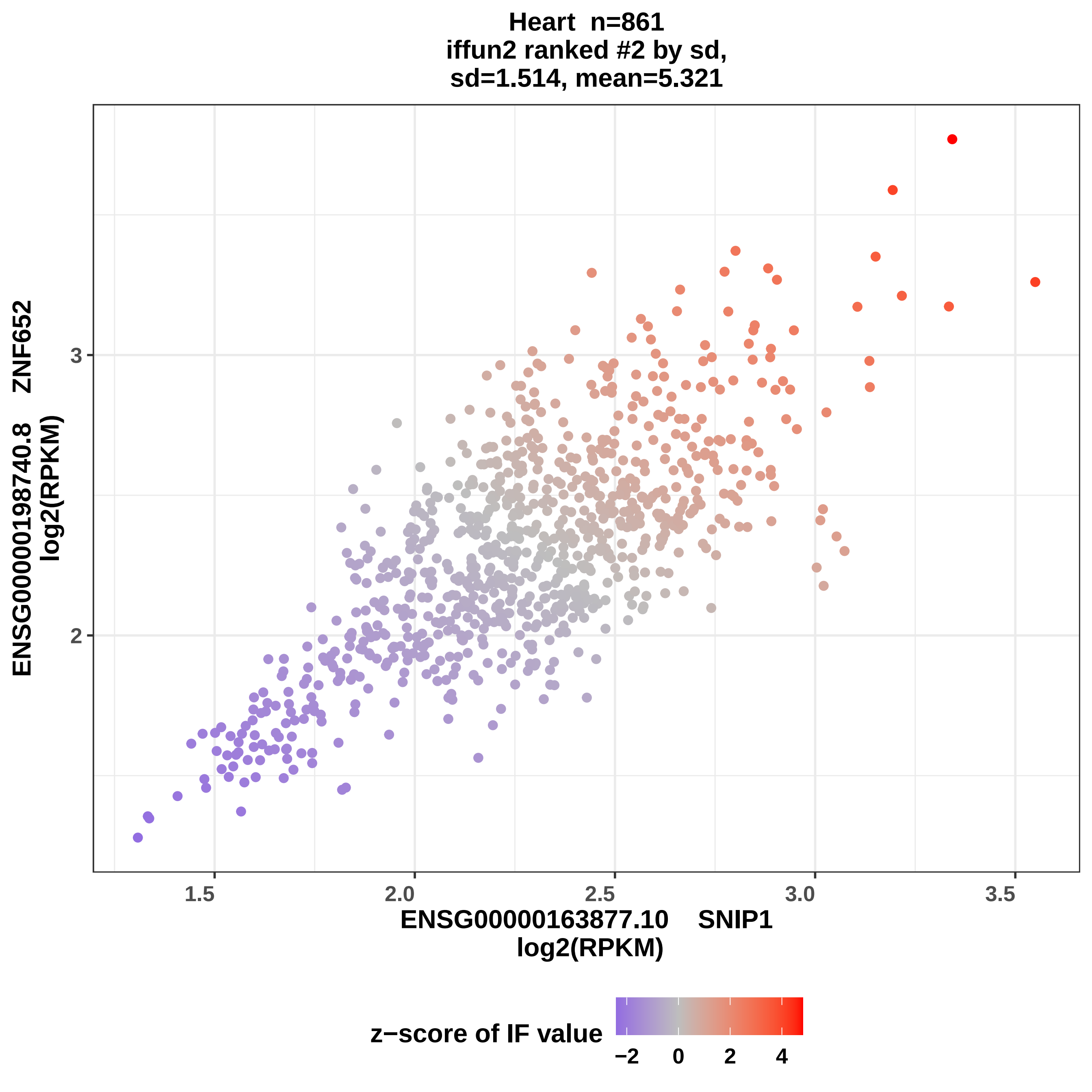

Heart n=861
iffun2 ranked #2 by sd,
sd=1.514, mean=5.321
ENSG00000198740.8 ZNF652
log2(RPKM)
3
2
1.5
2.0
2.5
3.0
3.5
ENSG00000163877.10 SNIP1
log2(RPKM)
z−score of IF value
−2
0
2
4

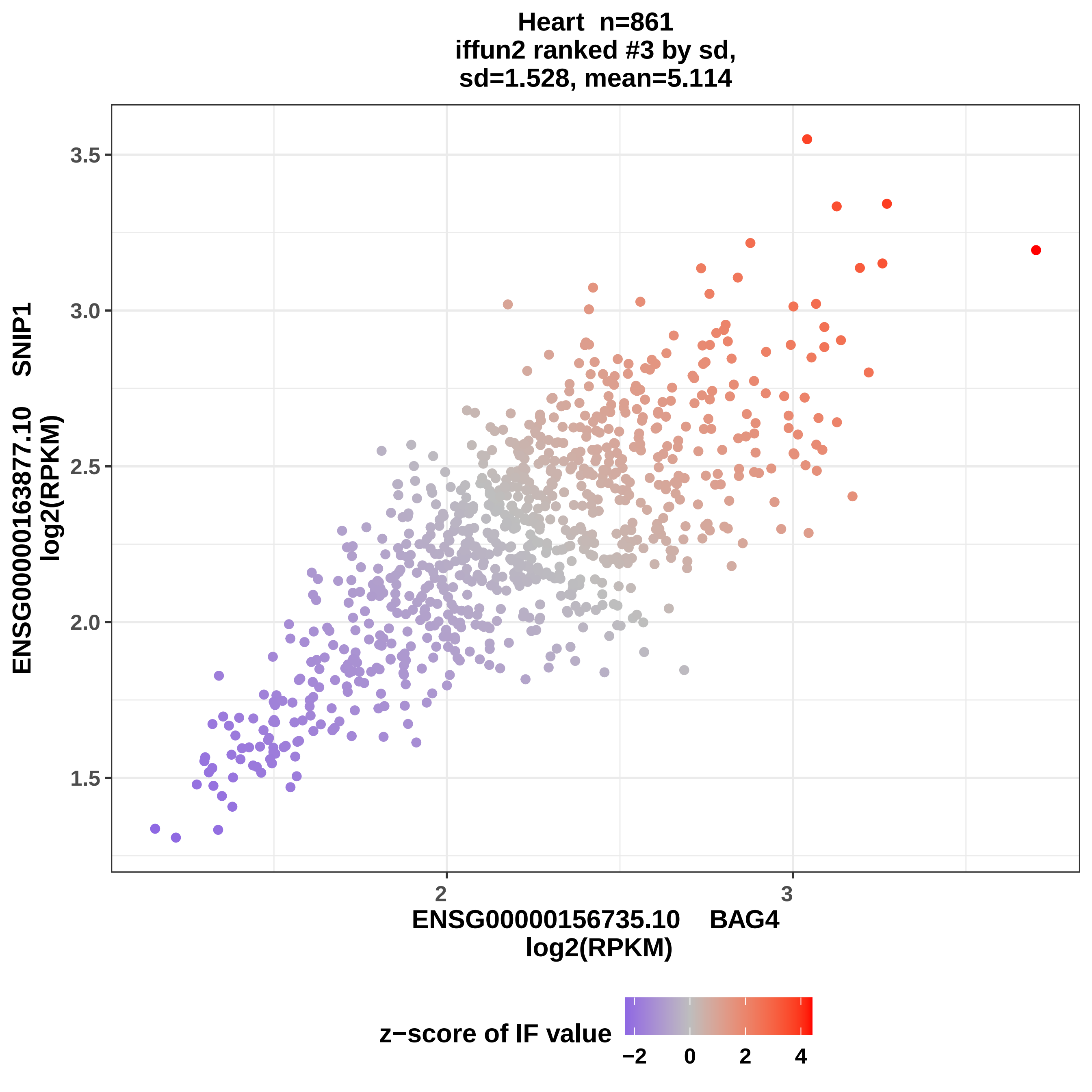

Heart n=861
iffun2 ranked #3 by sd,
sd=1.528, mean=5.114
3.5
3.0
2.5
2.0
1.5
ENSG00000163877.10 SNIP1
log2(RPKM)
2
3
ENSG0000156735.10 BAG4
log2(RPKM)
z−score of IF value
−2
0
2
4

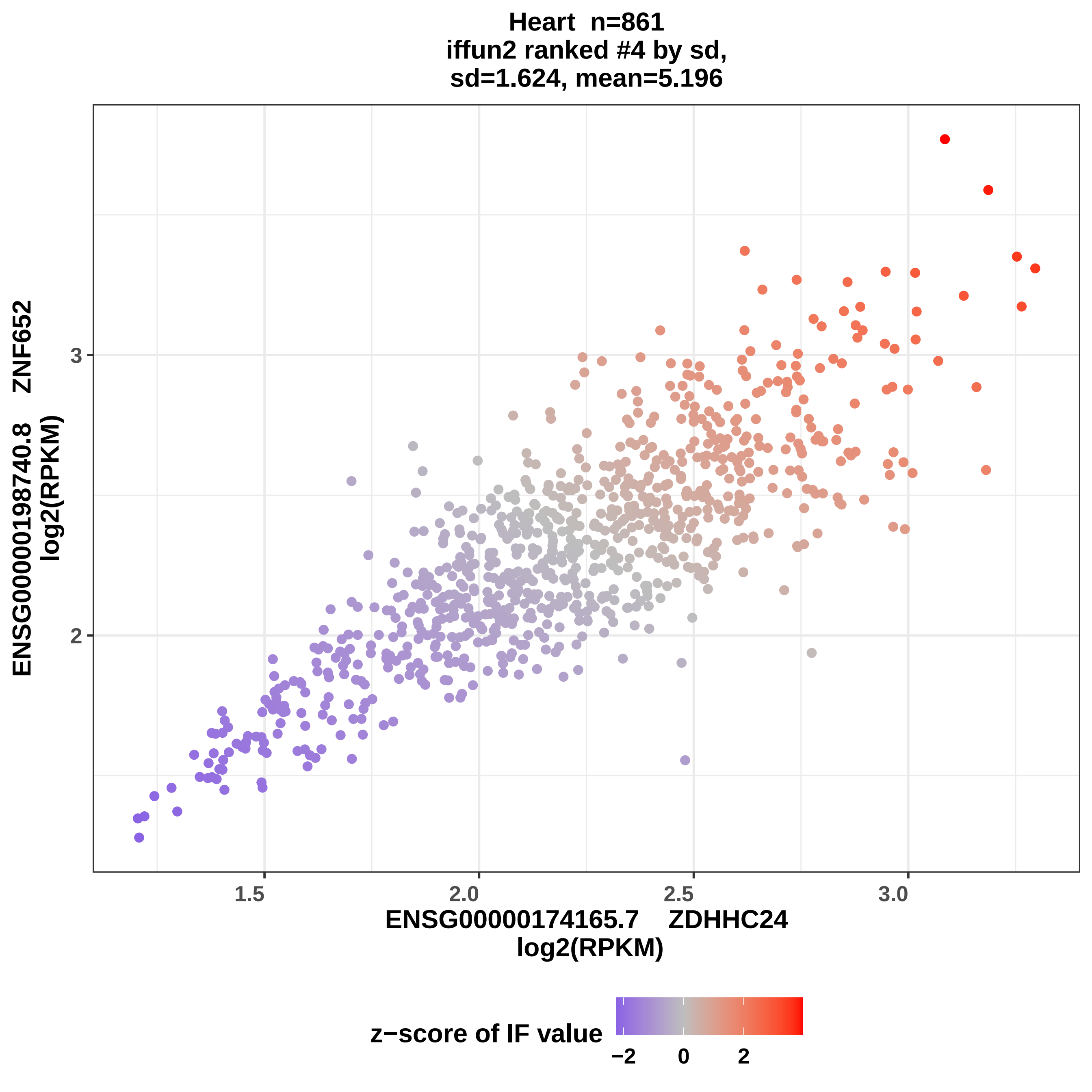

Heart n=861
iffun2 ranked #4 by sd,
sd=1.624, mean=5.196
ENSG00000198740.8 ZNF652
log2(RPKM)
3
2
1.5
2.0
2.5
3.0
ENSG00000174165.7 ZDHHC24
log2(RPKM)
z−score of IF value
−2
0
2

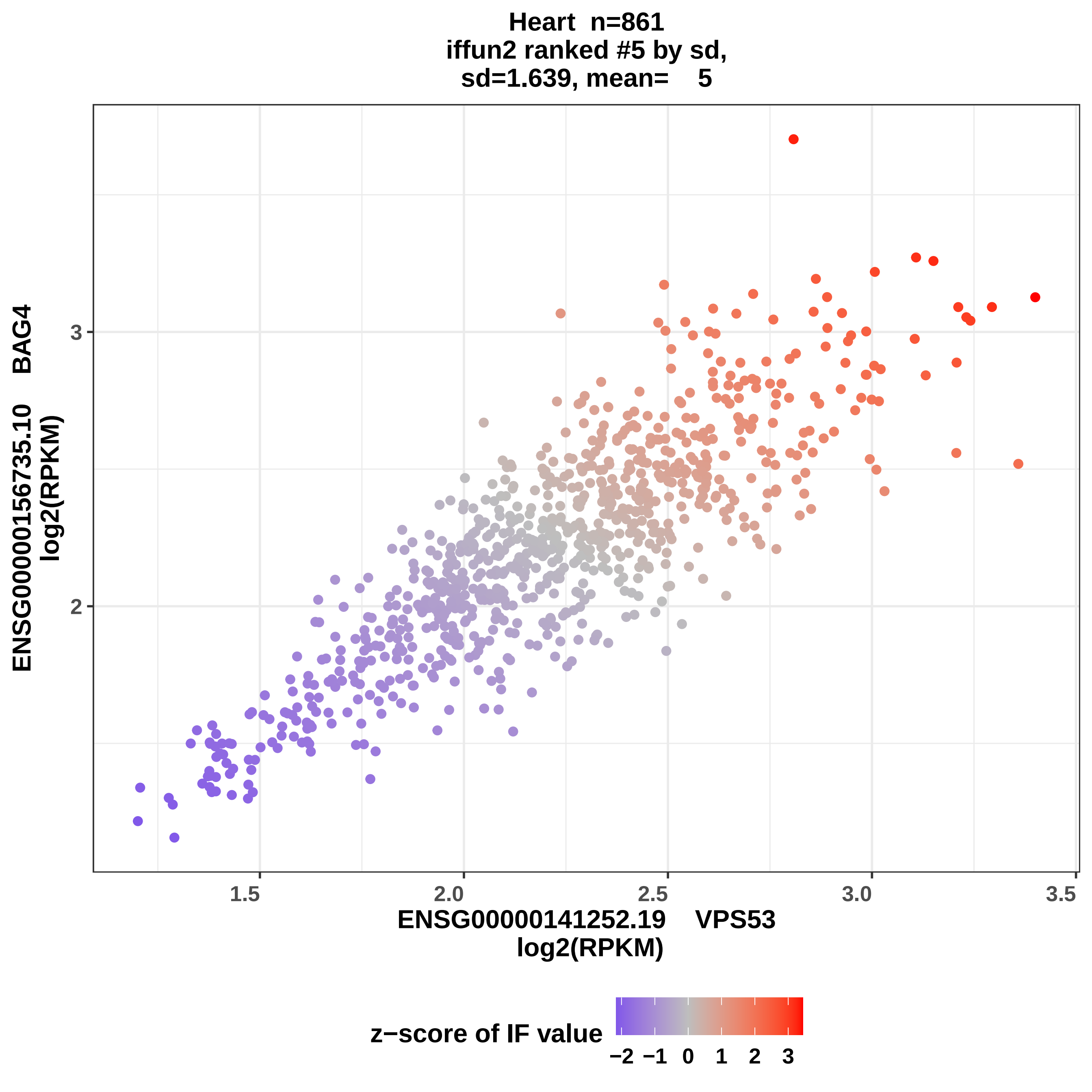
Heart n=861
iffun2 ranked #5 by sd,
sd=1.639, mean= 5
ENSG00000156735.10 BAG4
log2(RPKM)
3
2
1.5
2.0
2.5
3.0
3.5
ENSG00000141252.19 VPS53
log2(RPKM)
z−score of IF value
−2
−1
0
1
2
3

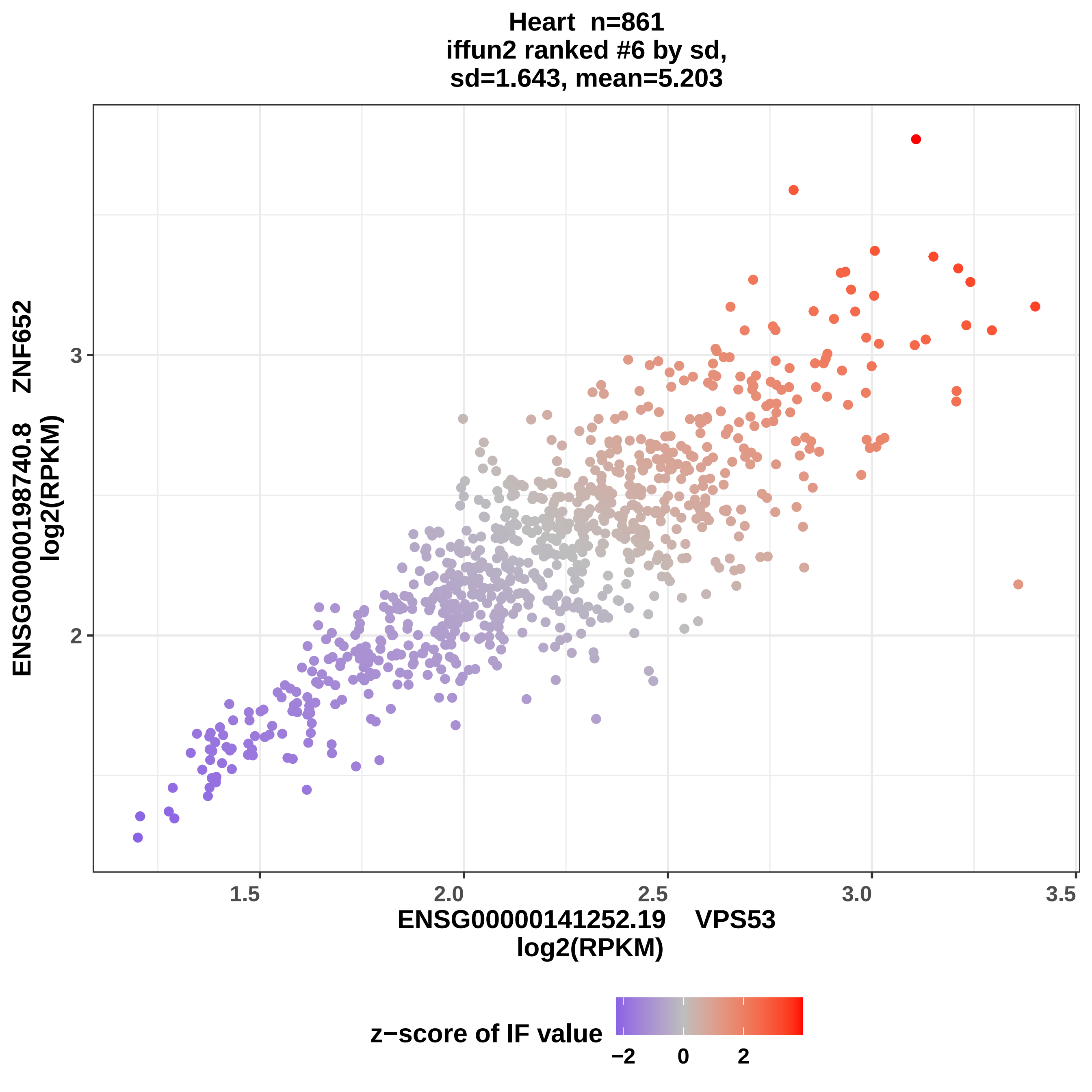
Heart n=861
iffun2 ranked #6 by sd,
sd=1.643, mean=5.203
ENSG00000198740.8 ZNF652
log2(RPKM)
3
2
1.5
2.0
2.5
3.0
3.5
ENSG00000141252.19 VPS53
log2(RPKM)
z−score of IF value
−2
0
2

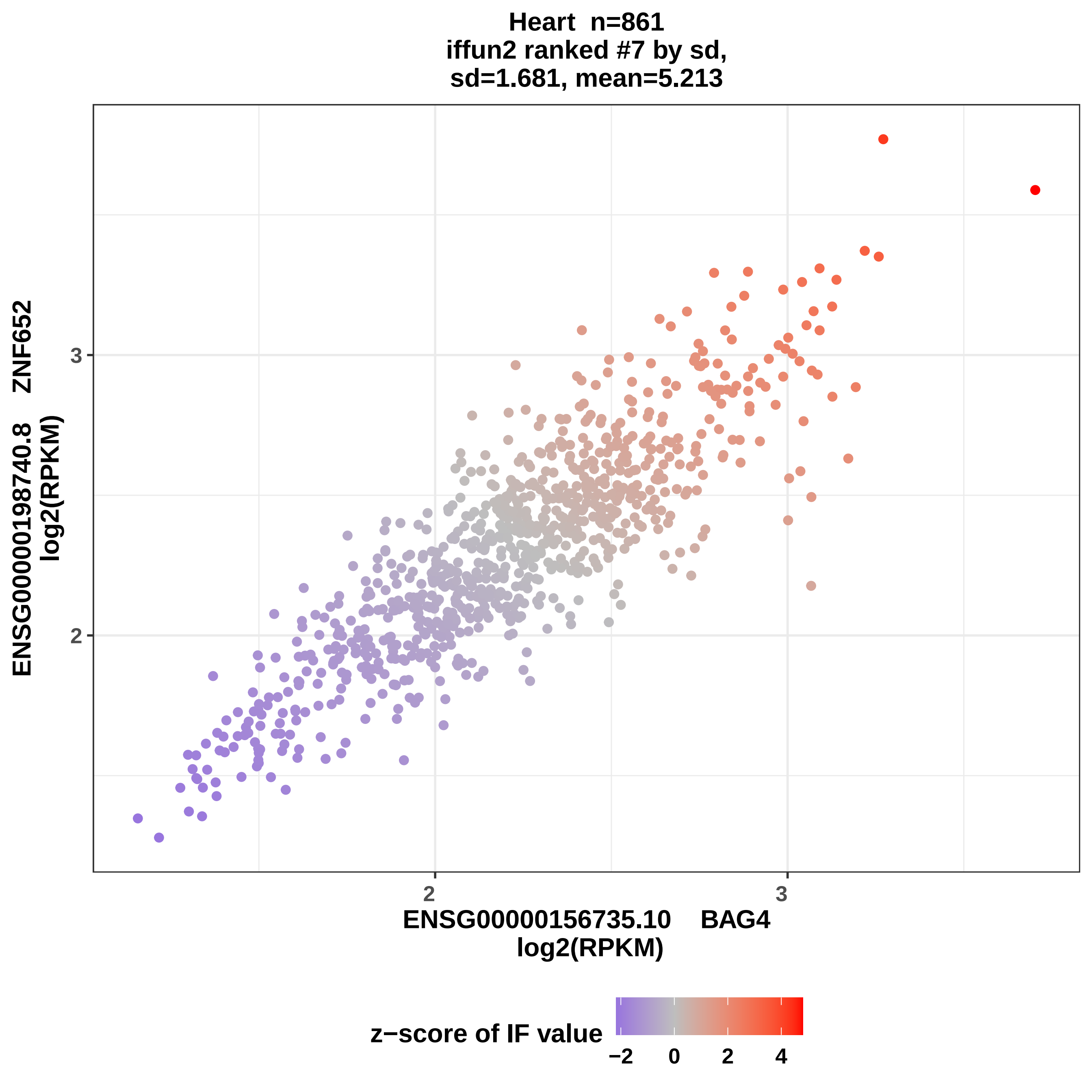
Heart n=861
iffun2 ranked #7 by sd,
sd=1.681, mean=5.213
ENSG00000198740.8 ZNF652
log2(RPKM)
3
2
2
3
ENSG00000156735.10 BAG4
log2(RPKM)
z−score of IF value
−2
0
2
4

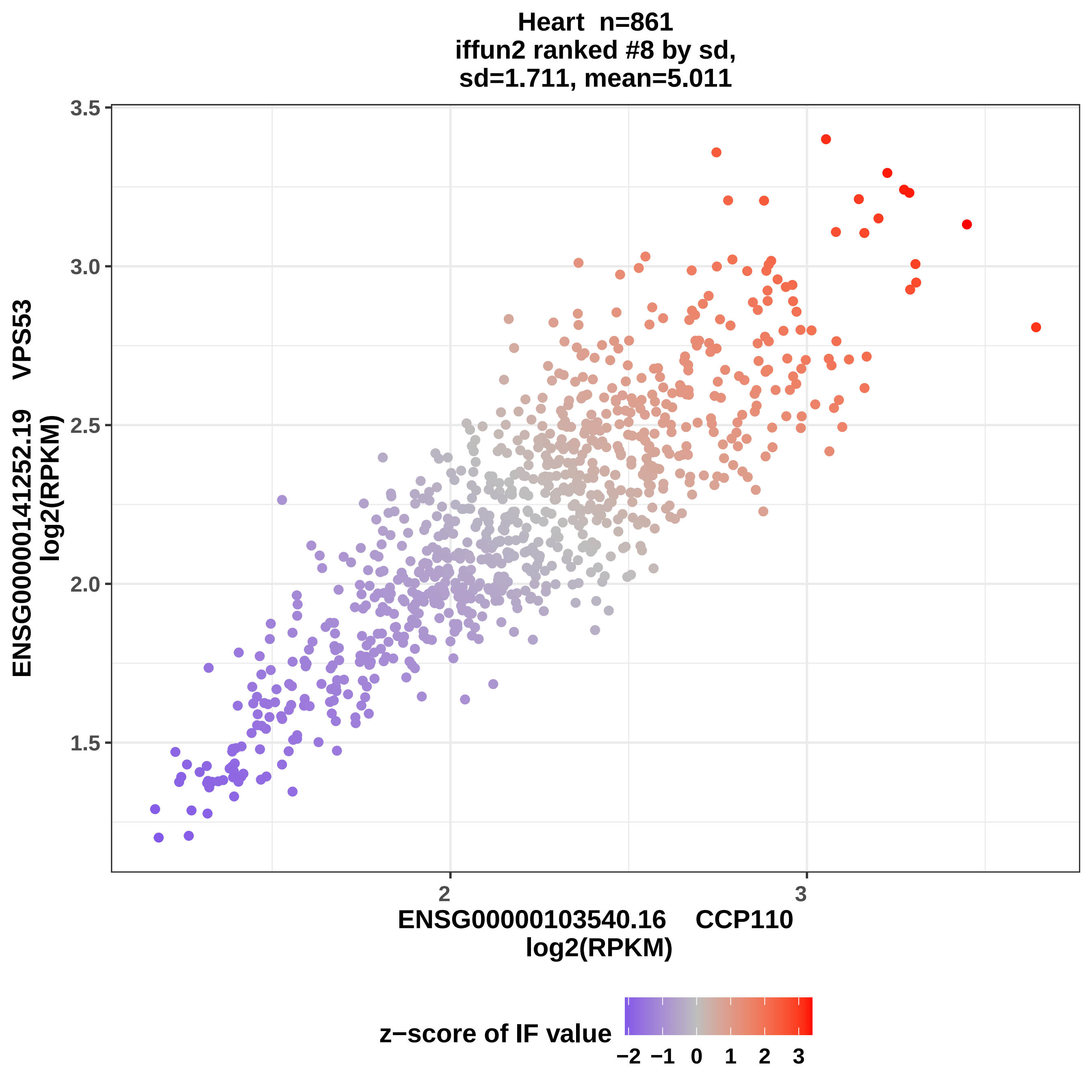

Heart n=861
iffun2 ranked #8 by sd,
sd=1.711, mean=5.011
ENSG00000141252.19 VPS53
log2(RPKM)
3.5
3.0
2.5
2.0
1.5
2
3
ENSG00000103540.16 CCP110
log2(RPKM)
z−score of IF value
−2 −1 0 1 2 3

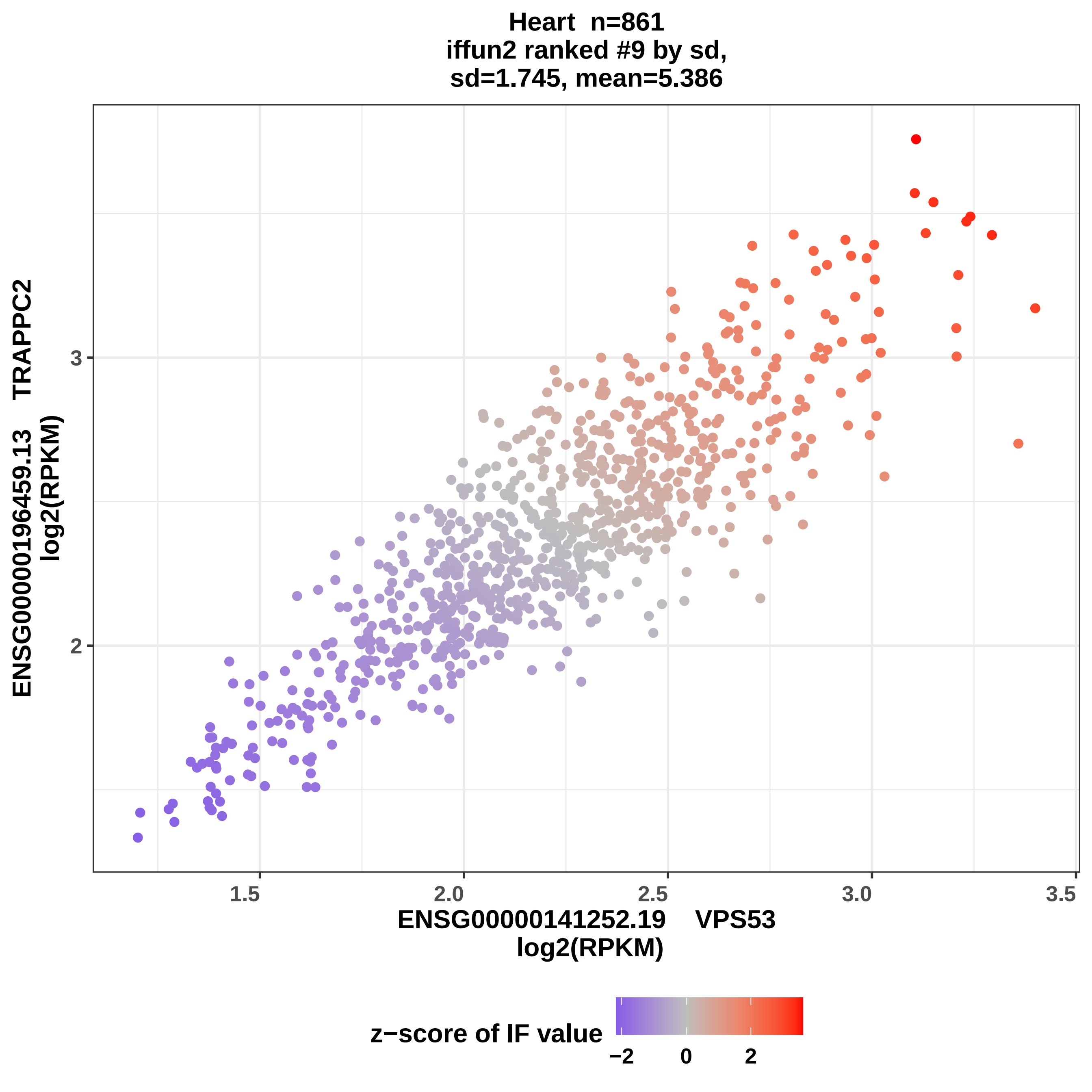

Heart n=861
iffun2 ranked #9 by sd,
sd=1.745, mean=5.386
ENSG00000196459.13 TRAPPC2
log2(RPKM)
3
2
1.5
2.0
2.5
3.0
3.5
ENSG00000141252.19 VPS53
log2(RPKM)
z−score of IF value
−2
0
2

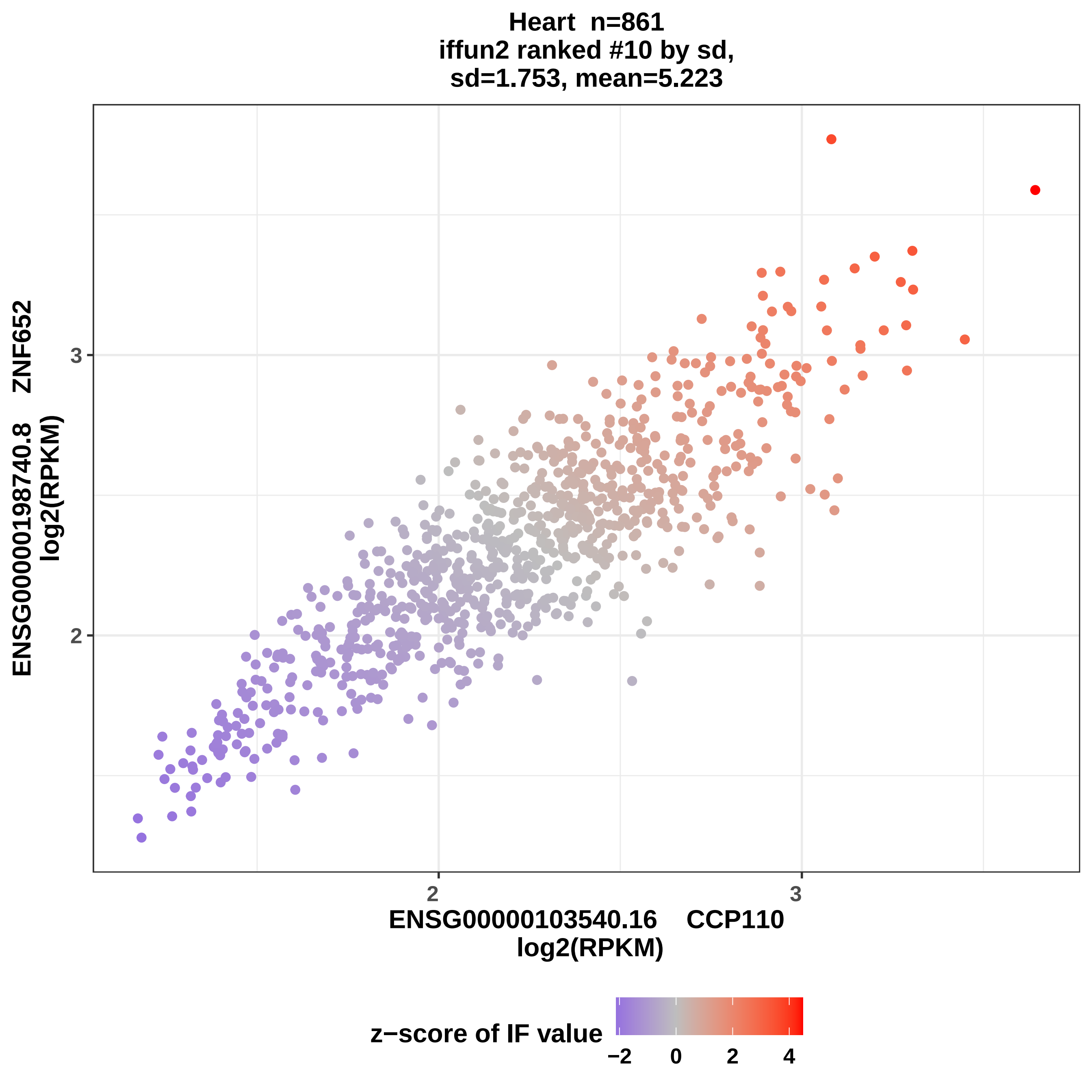

Heart n=861
iffun2 ranked #10 by sd,
sd=1.753, mean=5.223
ENSG00000198740.8 ZNF652
log2(RPKM)
3
2
ENSG00000103540.16 CCP110
log2(RPKM)
z−score of IF value
−2
0
2
4

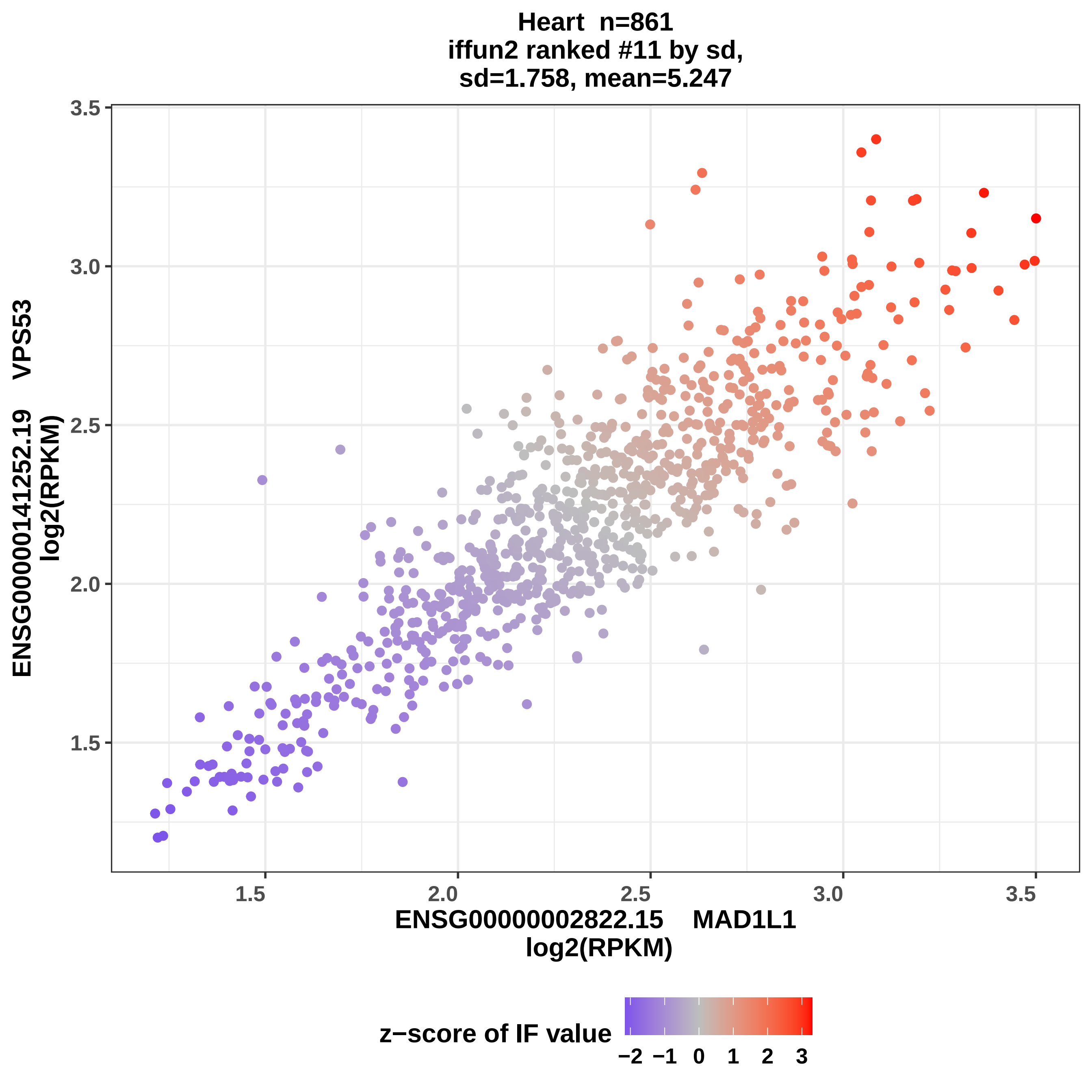
Heart n=861
iffun2 ranked #11 by sd,
sd=1.758, mean=5.247
ENSG00000141252.19 VPS53
log2(RPKM)
ENSG00000002822.15 MAD1L1
log2(RPKM)
1.5
2.0
2.5
3.0
3.5
z−score of IF value
−2
−1
0
1
2
3

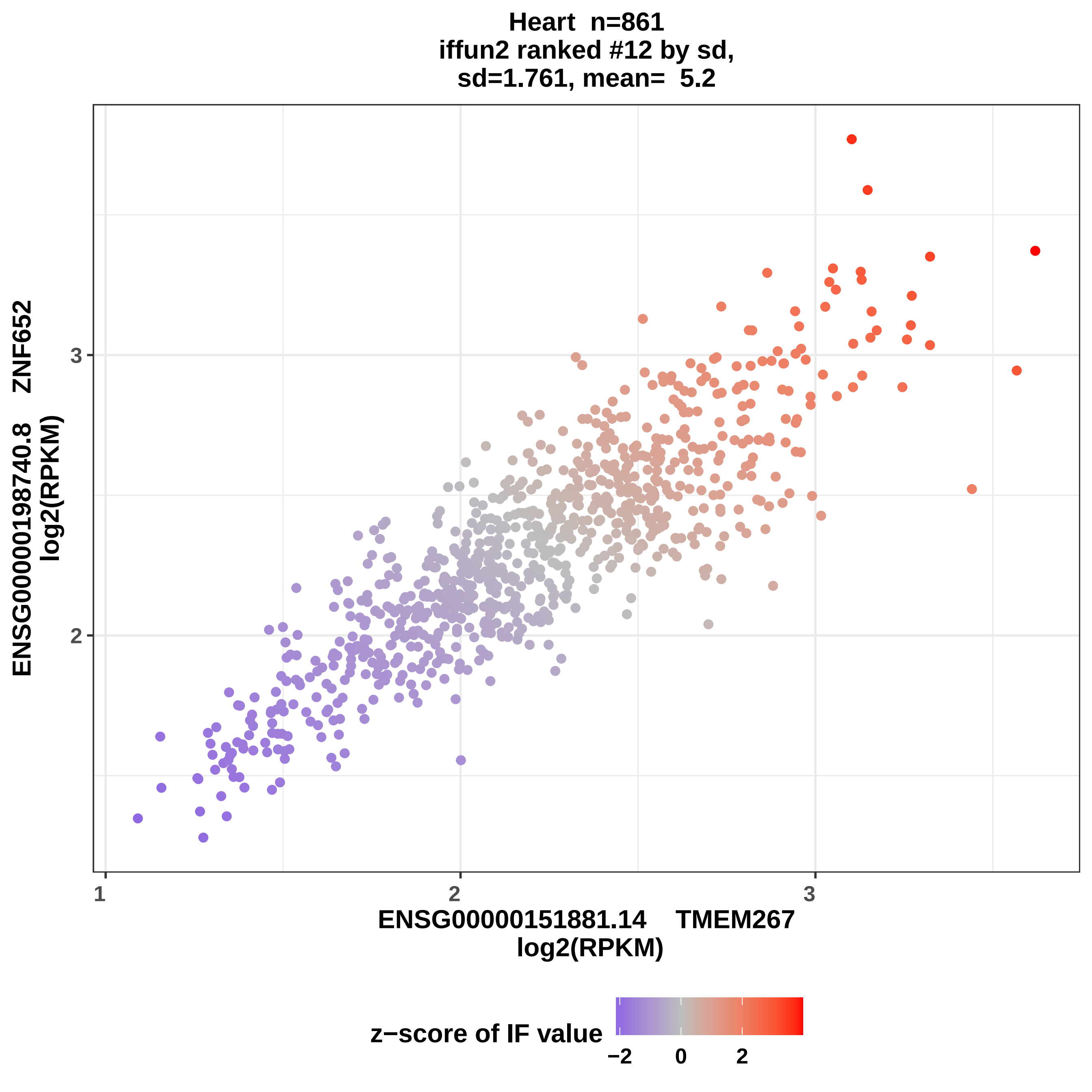

Heart n=861
iffun2 ranked #12 by sd,
sd=1.761, mean= 5.2
ENSG00000198740.8 ZNF652
log2(RPKM)
ENSG00000151881.14 TMEM267
log2(RPKM)
1
2
3
2
3
z−score of IF value
−2
0
2

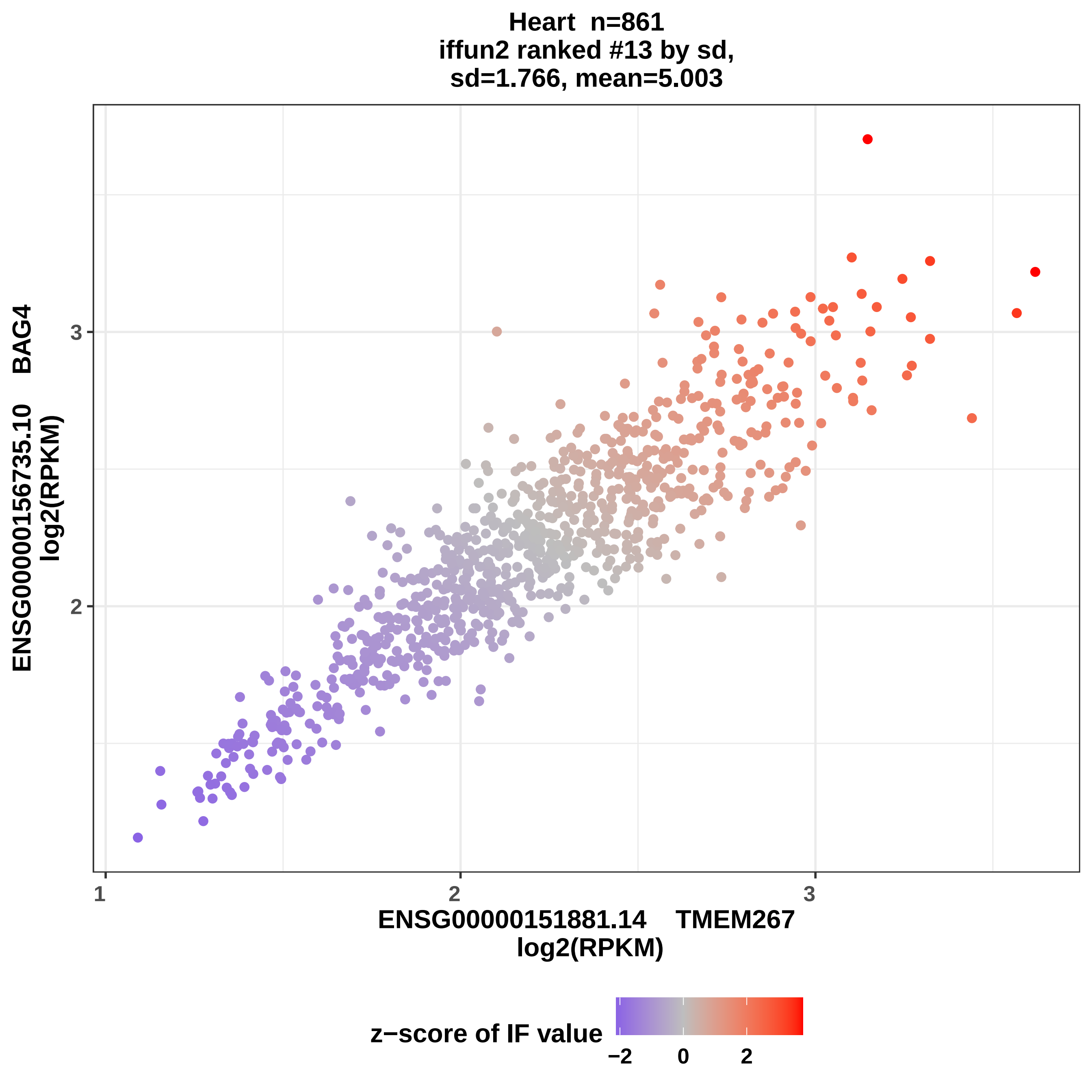
Heart n=861
iffun2 ranked #13 by sd,
sd=1.766, mean=5.003
ENSG00000156735.10 BAG4
log2(RPKM)
ENSG00000151881.14 TMEM267
log2(RPKM)
1
2
3
2
3
z−score of IF value
−2
0
2

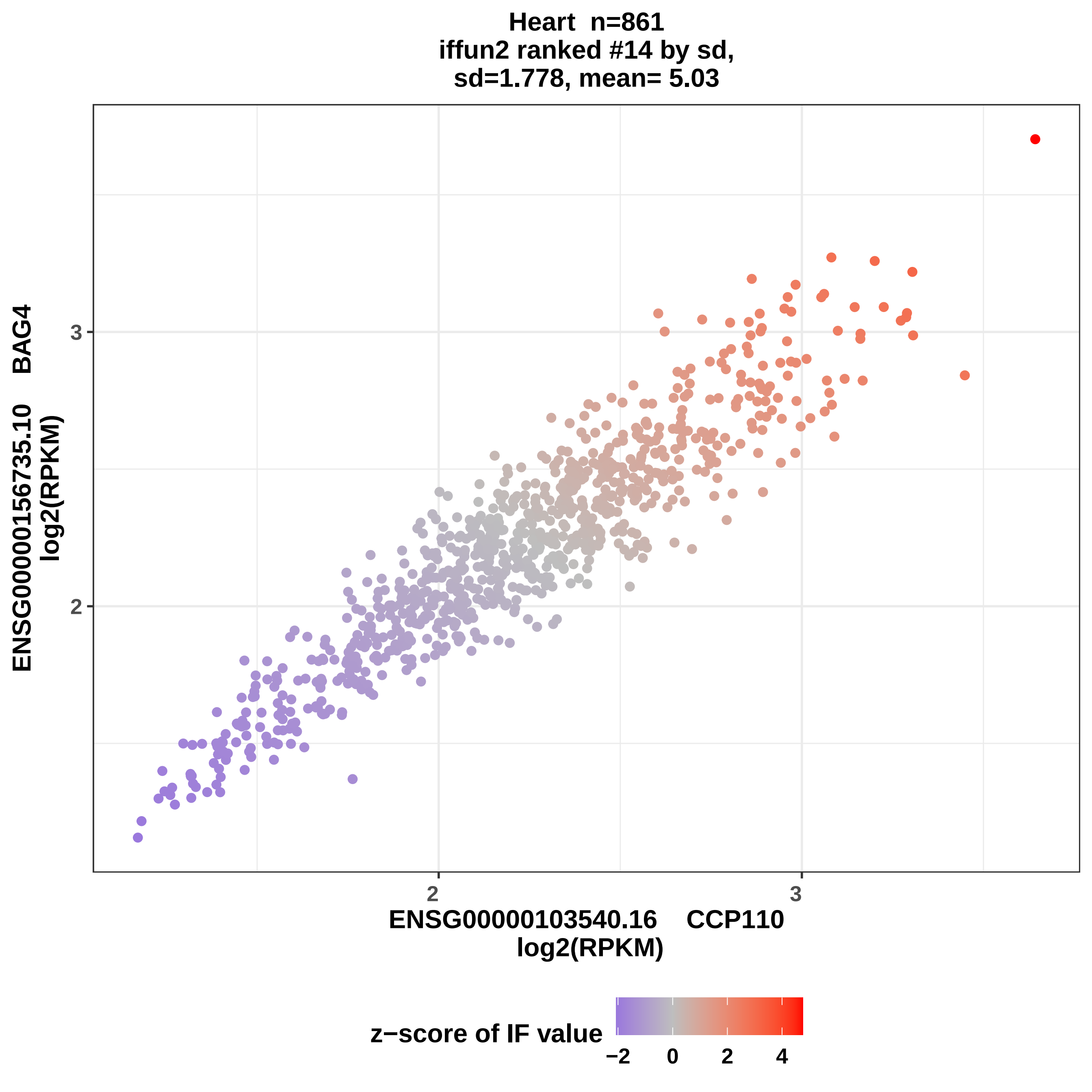

Heart n=861
iffun2 ranked #14 by sd,
sd=1.778, mean= 5.03
ENSG00000156735.10 BAG4
log2(RPKM)
3
2
2
3
ENSG00000103540.16 CCP110
log2(RPKM)
z−score of IF value
−2
0
2
4

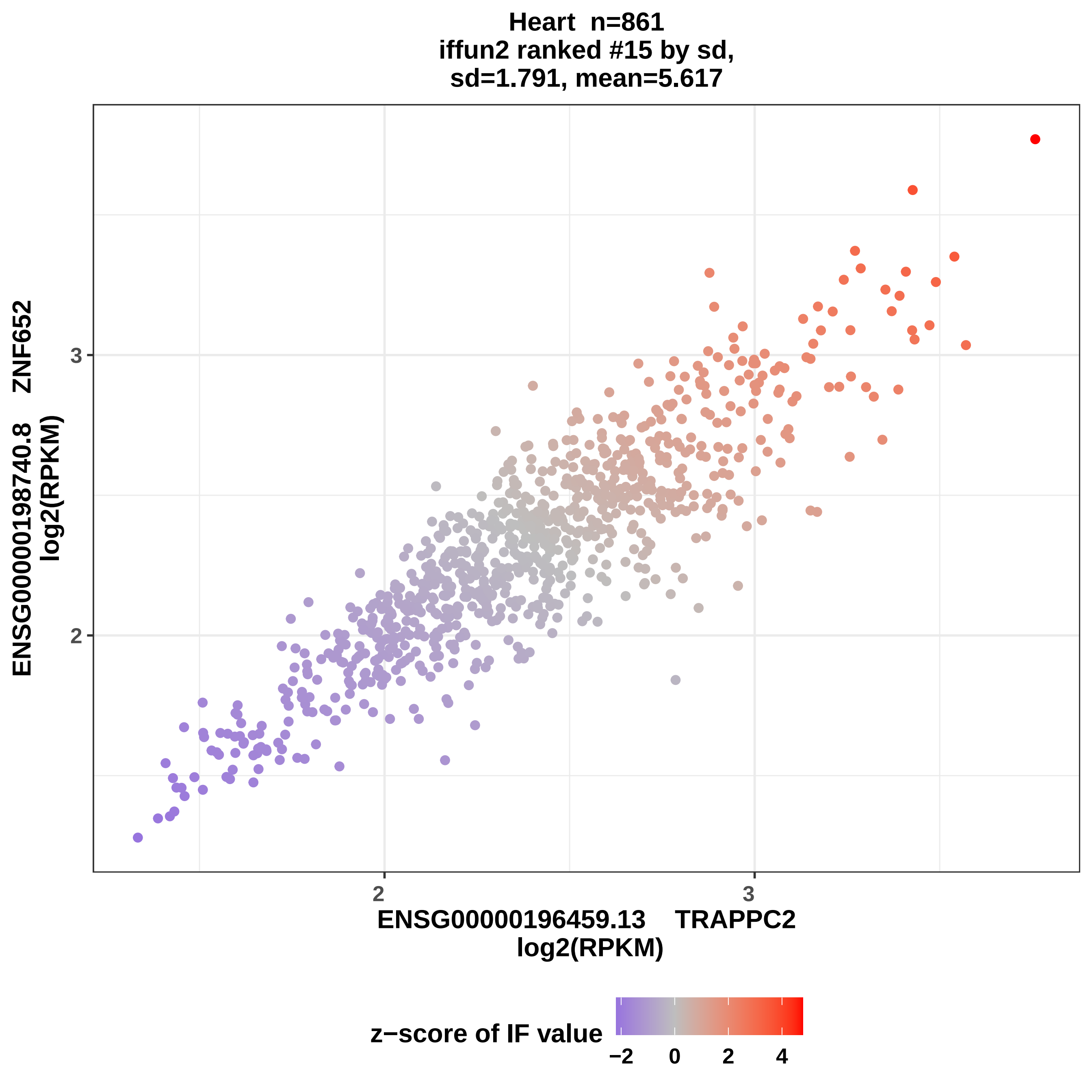

Heart n=861
iffun2 ranked #15 by sd,
sd=1.791, mean=5.617
ENSG00000198740.8 ZNF652
log2(RPKM)
3
2
ENSG00000196459.13 TRAPPC2
log2(RPKM)
z−score of IF value
−2
0
2
4

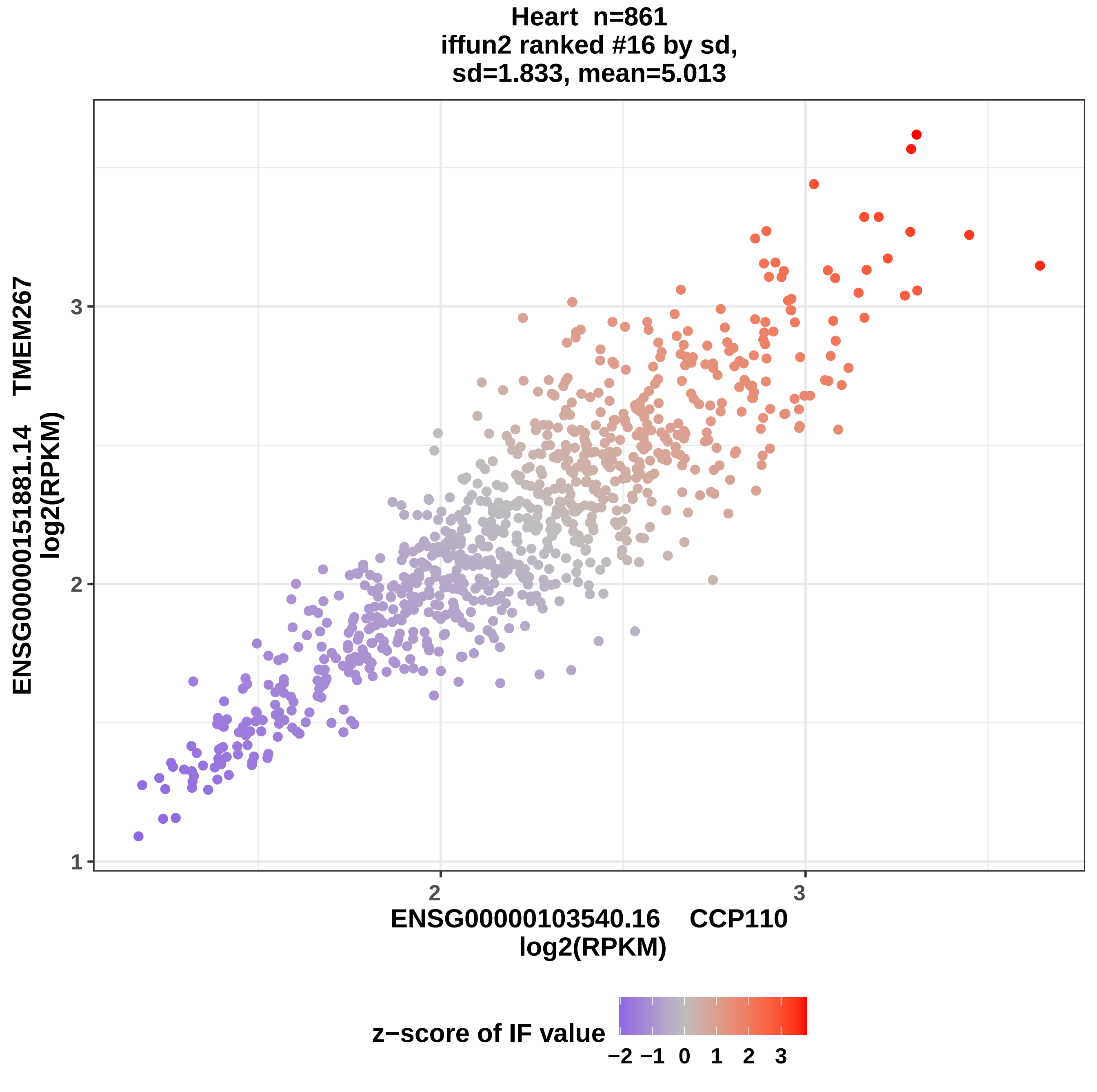
Heart n=861
iffun2 ranked #16 by sd,
sd=1.833, mean=5.013
ENSG00000151881.14 TMEM267
log2(RPKM)
3
2
1
2
3
ENSG00000103540.16 CCP110
log2(RPKM)
z−score of IF value
−2 −1 0 1 2 3

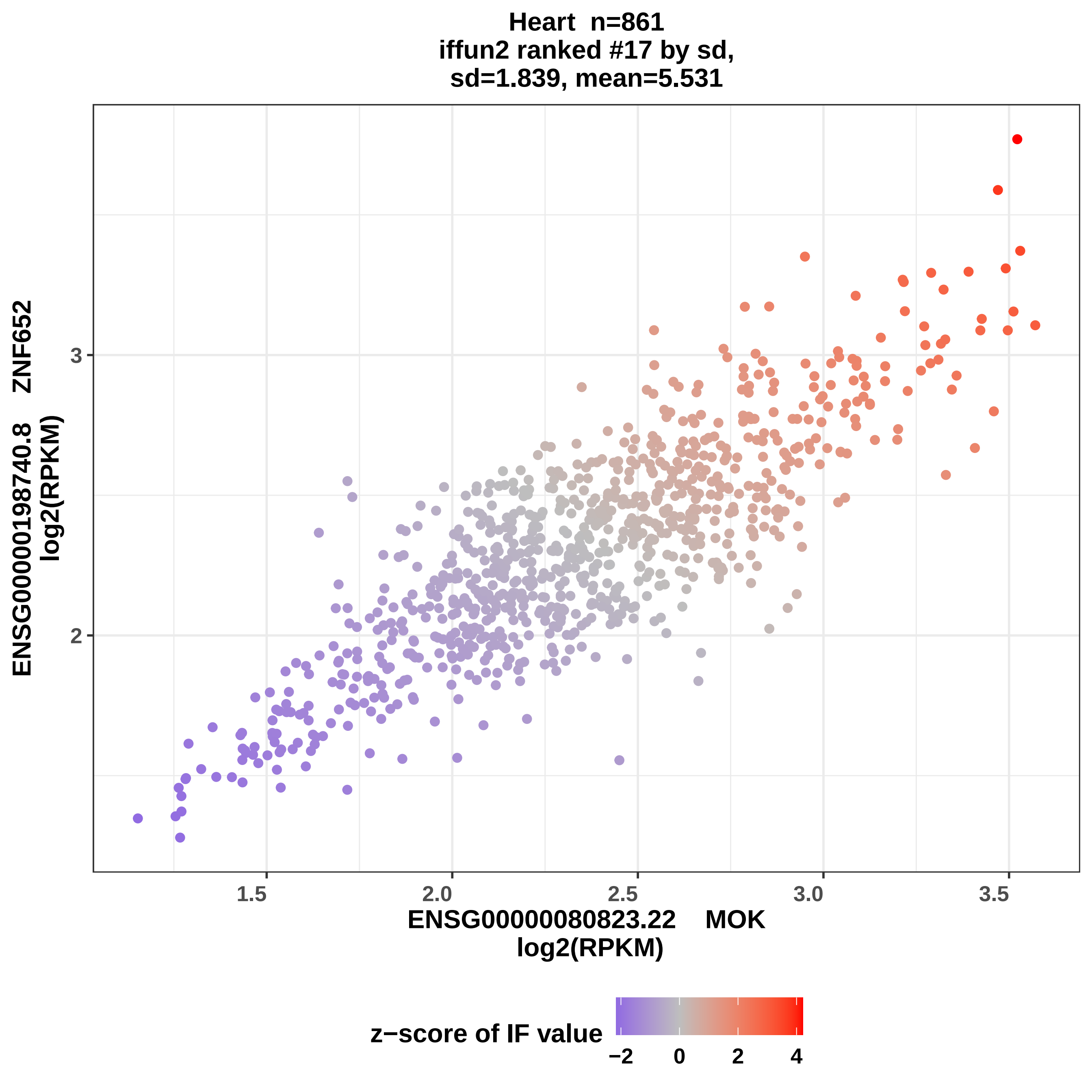

Heart n=861
iffun2 ranked #17 by sd,
sd=1.839, mean=5.531
ENSG00000198740.8 ZNF652
log2(RPKM)
3
2
1.5
2.0
2.5
3.0
3.5
ENSG00000080823.22 MOK
log2(RPKM)
z−score of IF value
−2
0
2
4

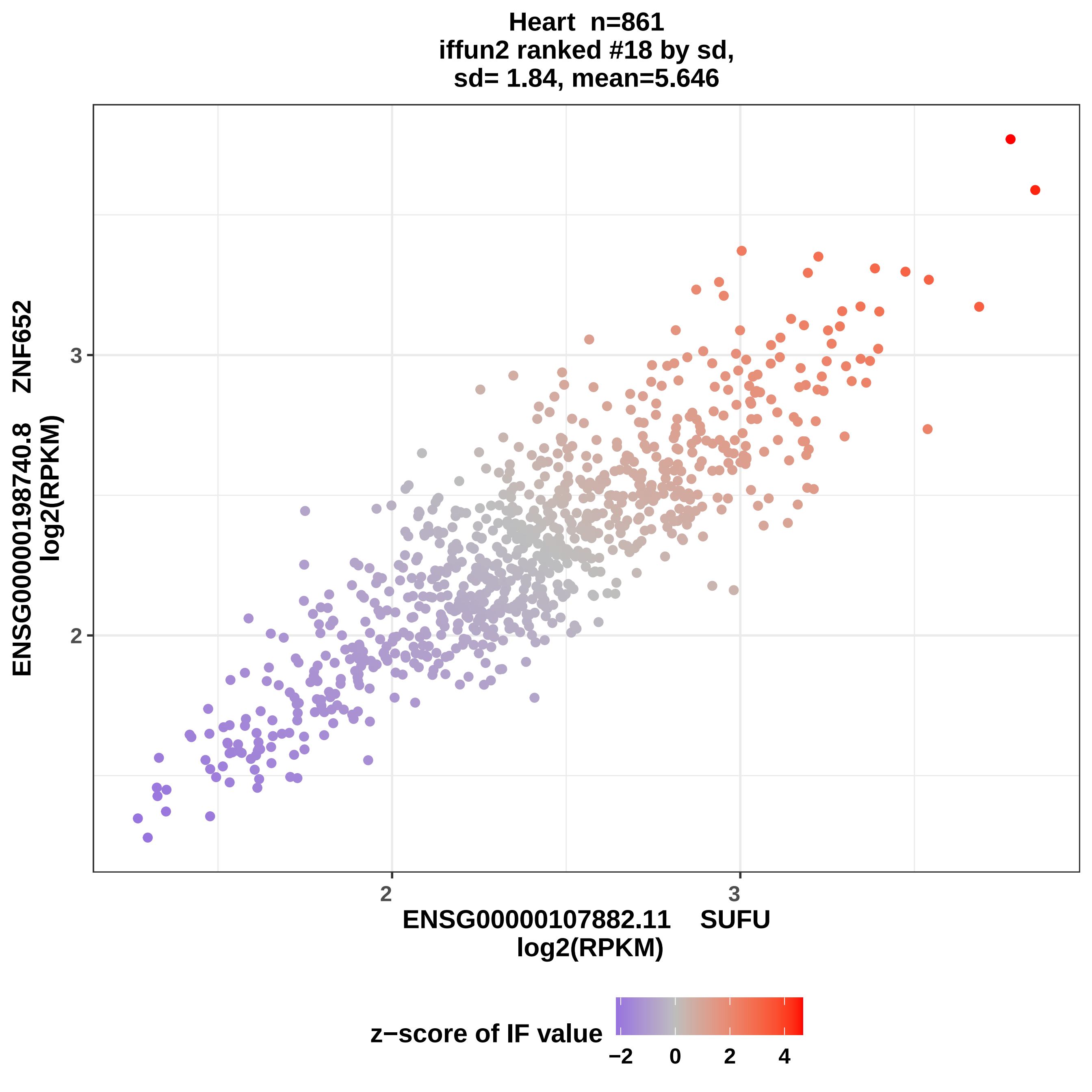
Heart n=861
iffun2 ranked #18 by sd,
sd= 1.84, mean=5.646
ENSG00000198740.8 ZNF652
log2(RPKM)
3
2
ENSG00000107882.11 SUFU
log2(RPKM)
z−score of IF value
−2
0
2
4

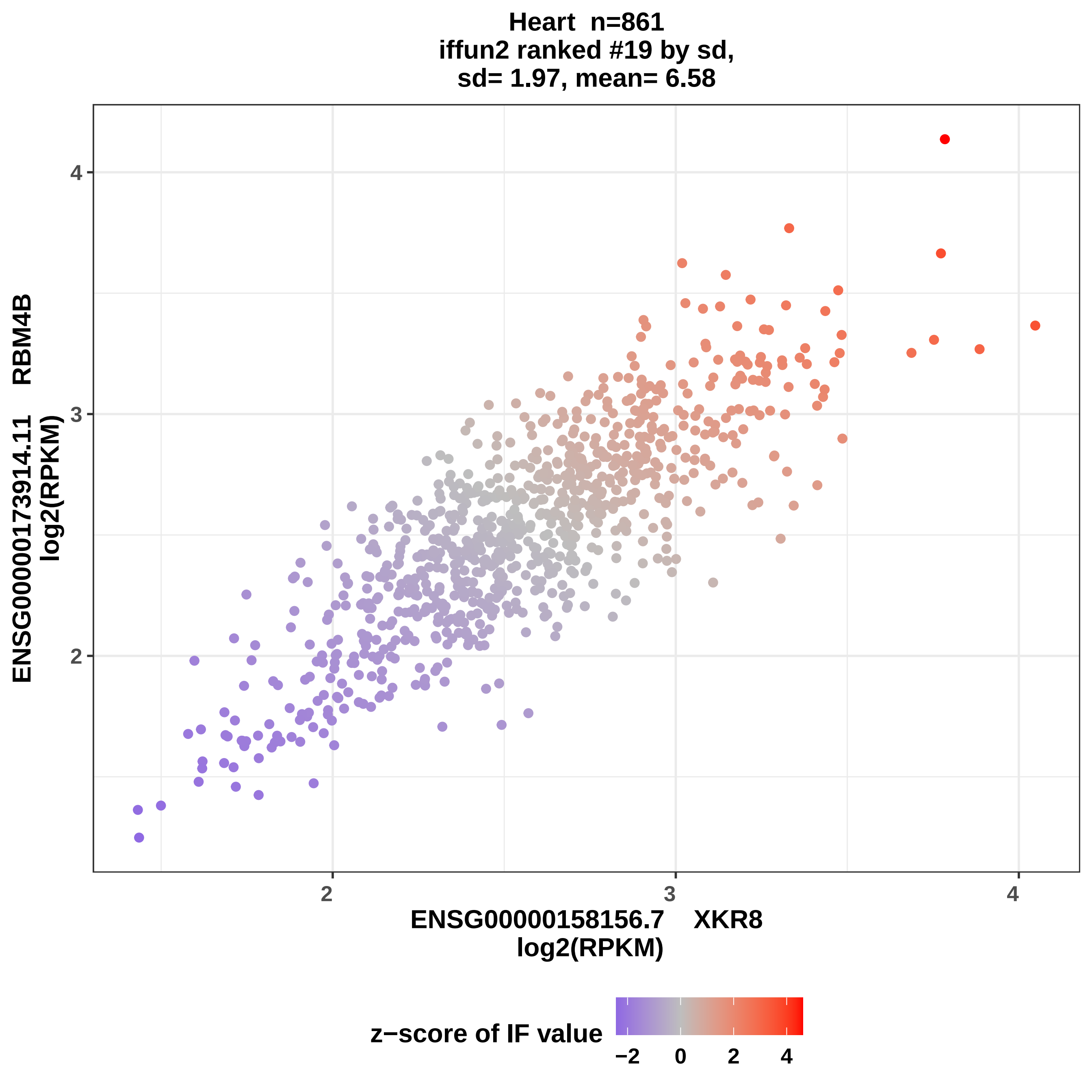
Heart n=861
iffun2 ranked #19 by sd,
sd= 1.97, mean= 6.58
ENSG00000173914.11 RBM4B
log2(RPKM)
4
3
2
2
3
4
ENSG00000158156.7 XKR8
log2(RPKM)
z−score of IF value
−2
0
2
4

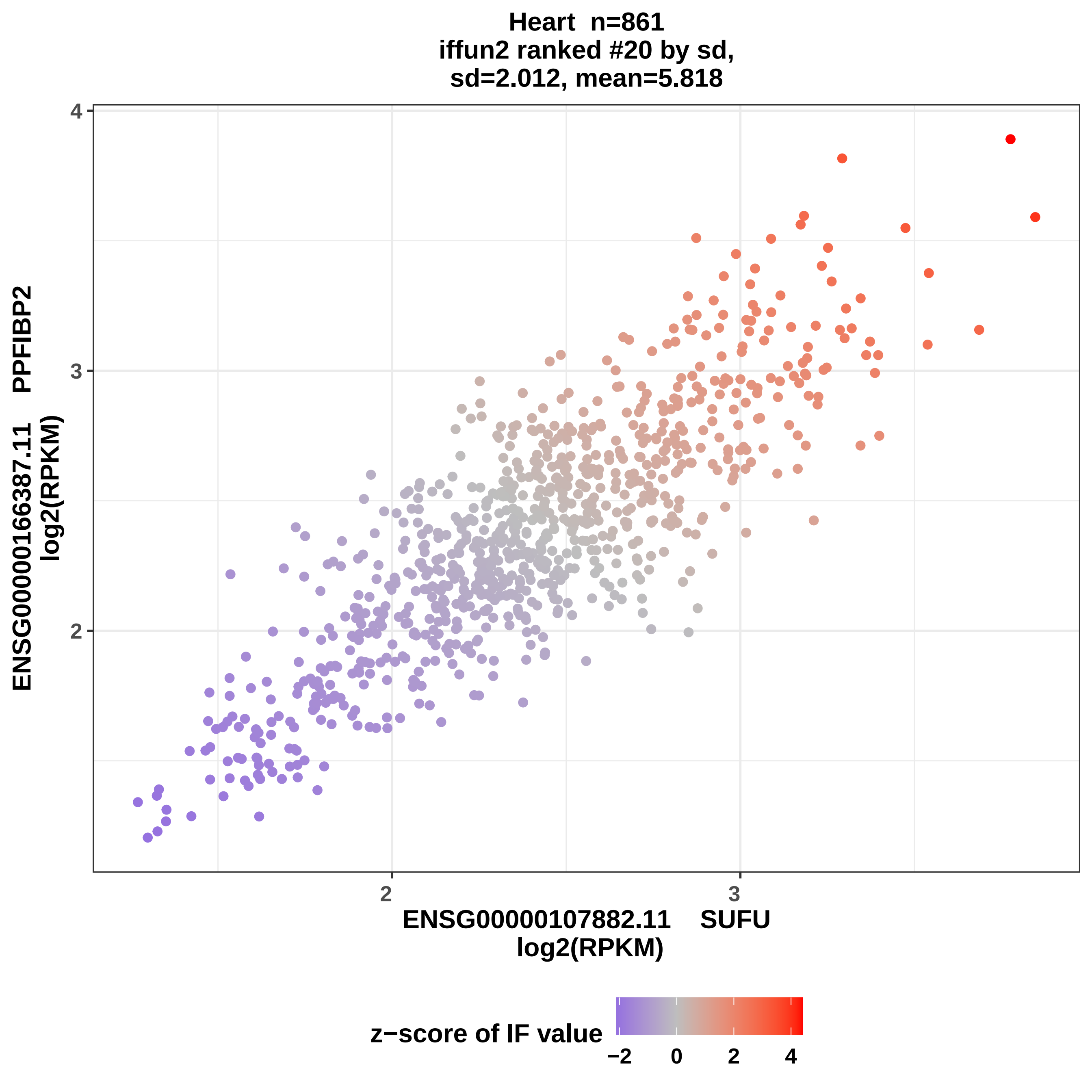
Heart n=861
iffun2 ranked #20 by sd,
sd=2.012, mean=5.818
ENSG00000166387.11 PPFIBP2
log2(RPKM)
4
3
2
2
3
ENSG00000107882.11 SUFU
log2(RPKM)
z−score of IF value
−2
0
2
4

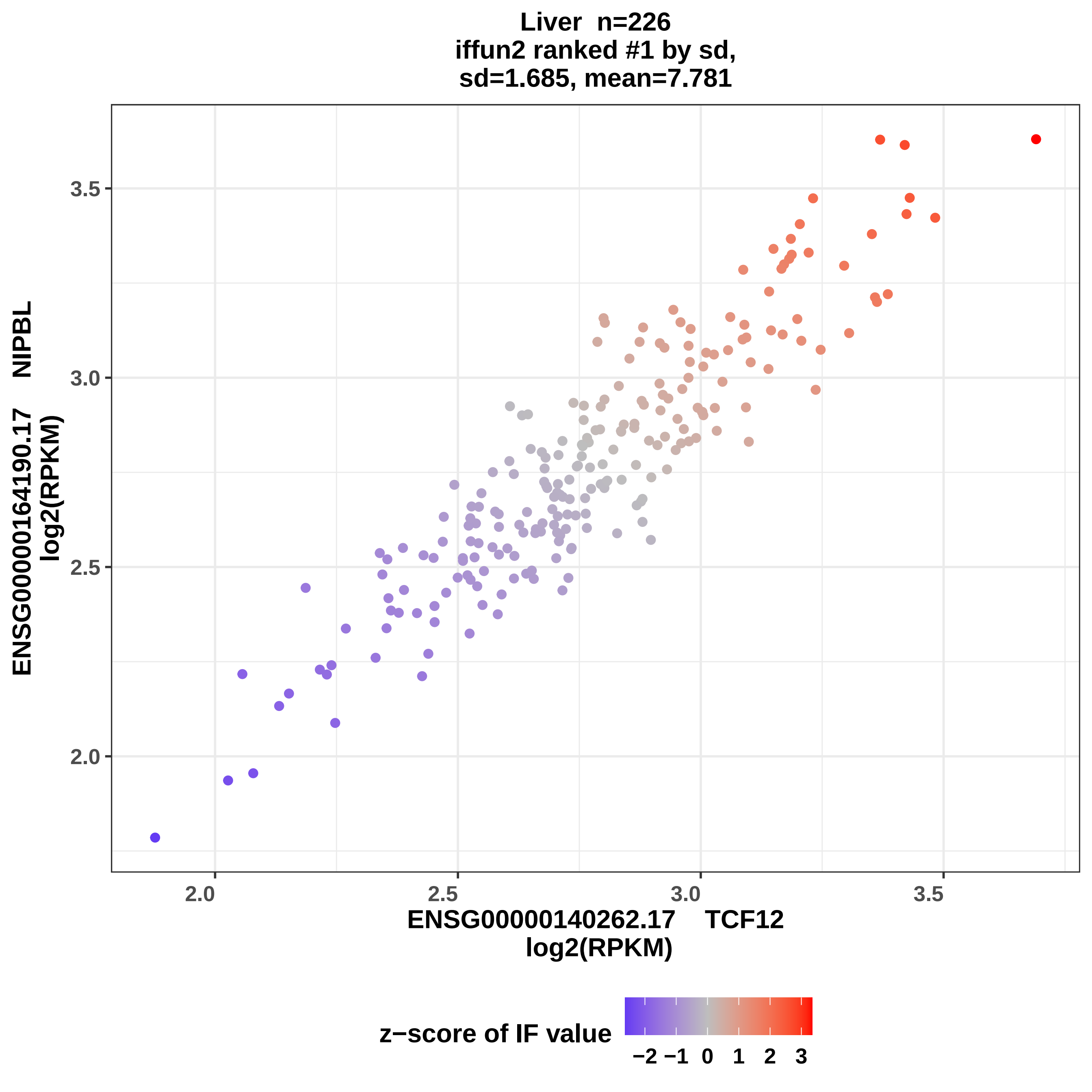
Liver n=226
iffun2 ranked #1 by sd,
sd=1.685, mean=7.781
ENSG00000164190.17 NIPBL
log2(RPKM)
3.5
3.0
2.5
2.0
2.0
2.5
3.0
3.5
ENSG00000140262.17 TCF12
log2(RPKM)
z−score of IF value
−2 −1 0 1 2 3

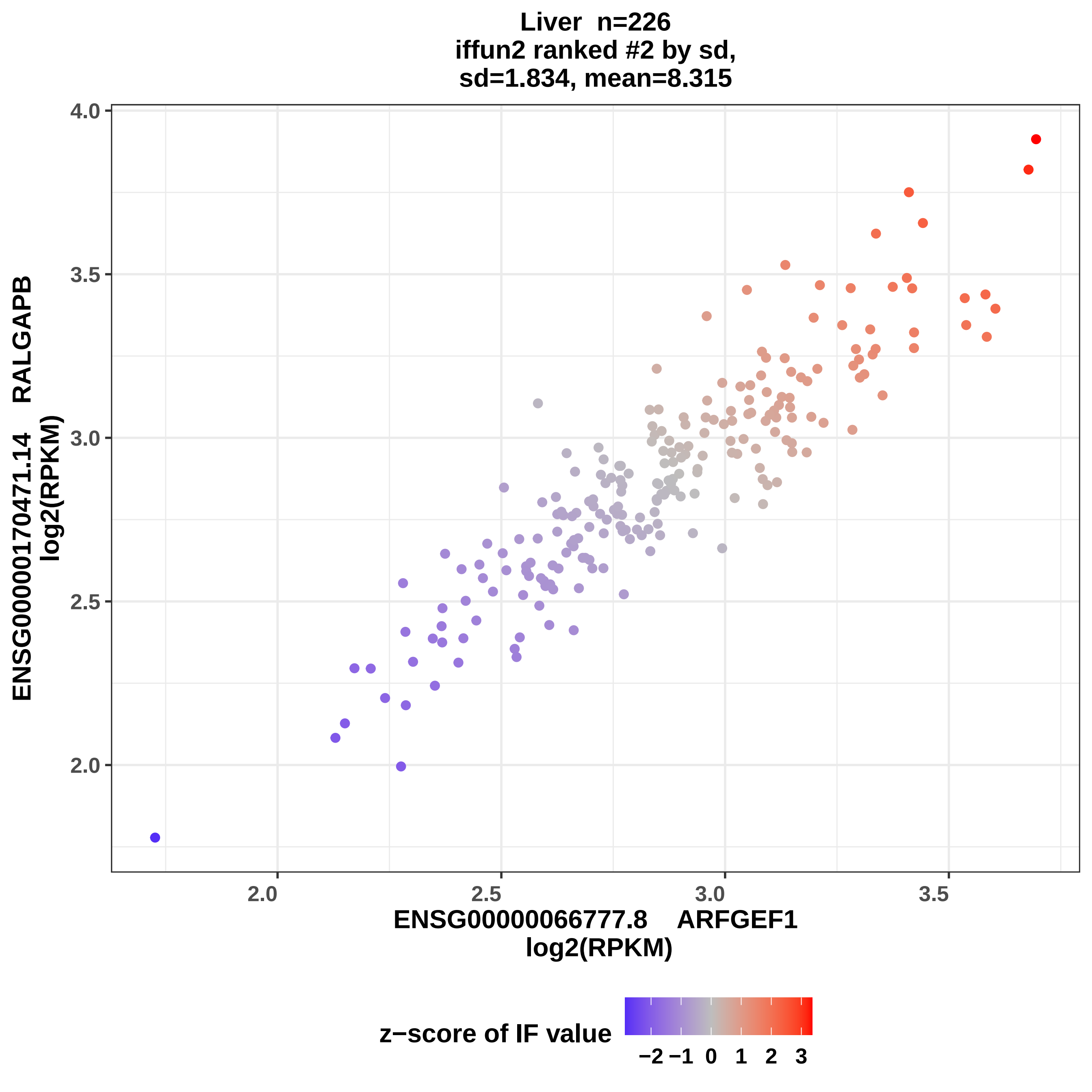
Liver n=226
iffun2 ranked #2 by sd,
sd=1.834, mean=8.315
ENSG00000170471.14 RALGAPB
log2(RPKM)
ENSG00000066777.8 ARFGEF1
log2(RPKM)
4.0
3.5
3.0
2.5
2.0
2.0
2.5
3.0
3.5
z−score of IF value
−2 −1 0 1 2 3

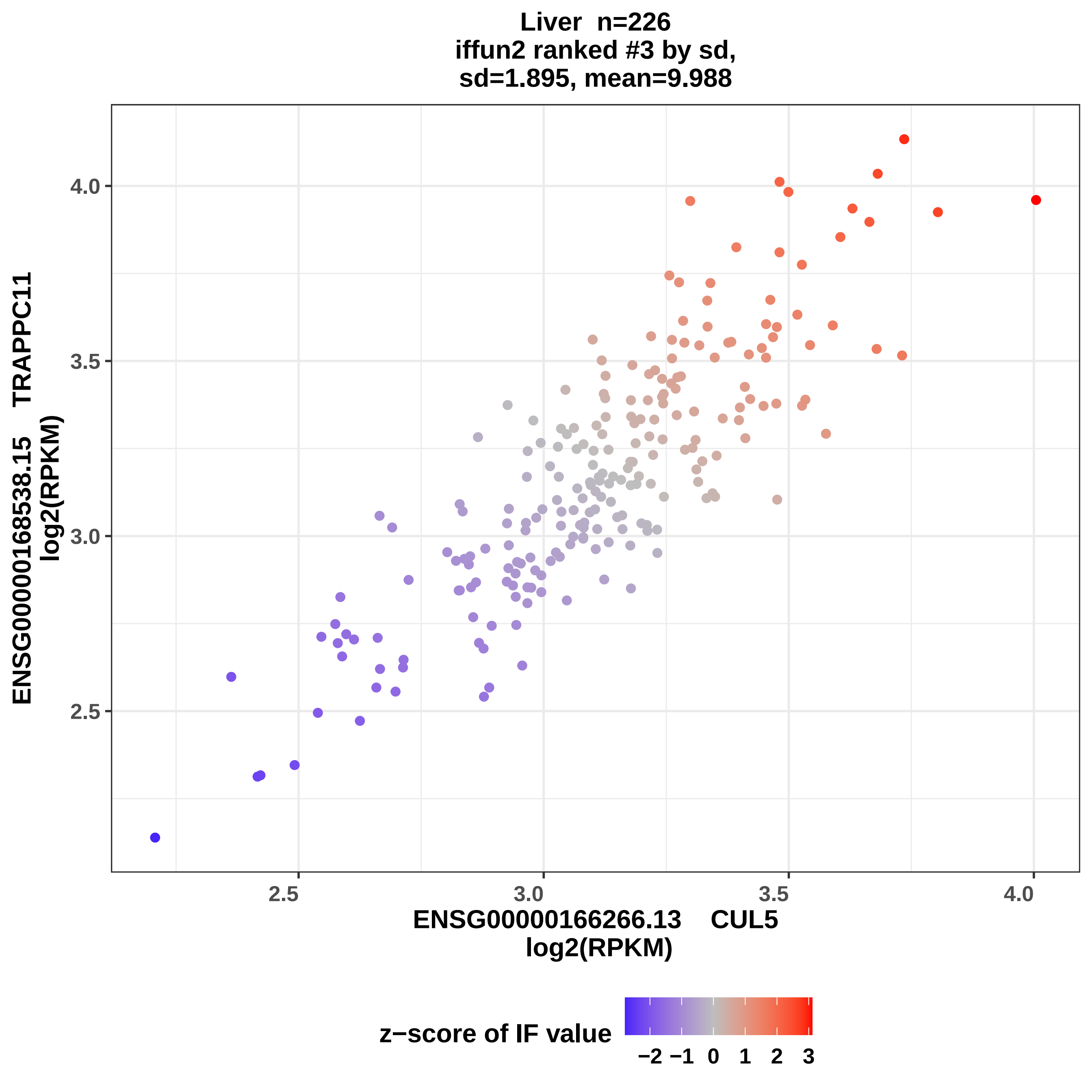

Liver n=226
iffun2 ranked #3 by sd,
sd=1.895, mean=9.988
ENSG00000168538.15 TRAPPC11
log2(RPKM)
4.0
3.5
3.0
2.5
2.5
3.0
3.5
4.0
ENSG00000166266.13 CUL5
log2(RPKM)
z−score of IF value
−2 −1 0 1 2 3

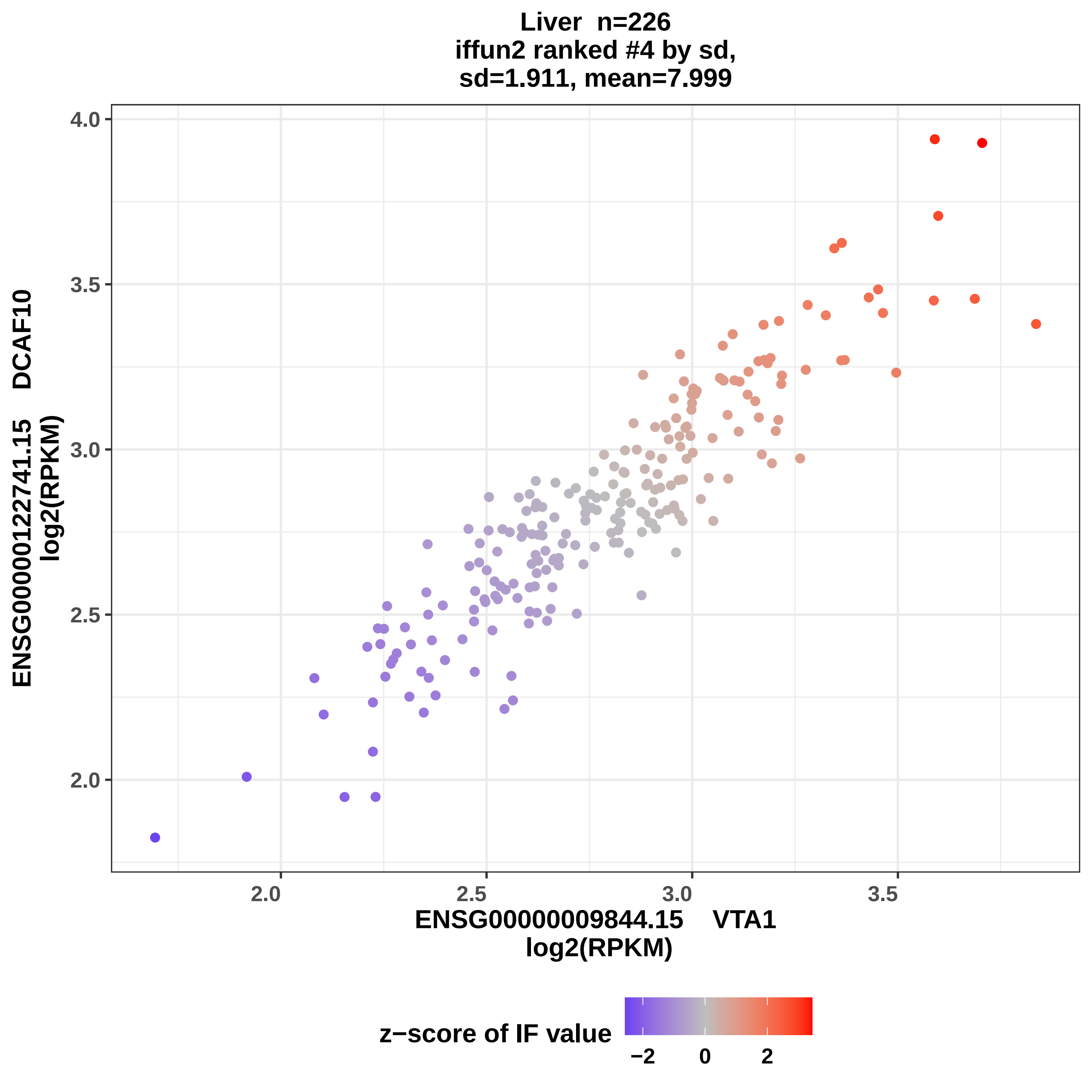
Liver n=226
iffun2 ranked #4 by sd,
sd=1.911, mean=7.999
ENSG00000122741.15 DCAF10
log2(RPKM)
ENSG00000009844.15 VTA1
log2(RPKM)
z−score of IF value
4.0
3.5
3.0
2.5
2.0
2.0
2.5
3.0
3.5
−2
0
2

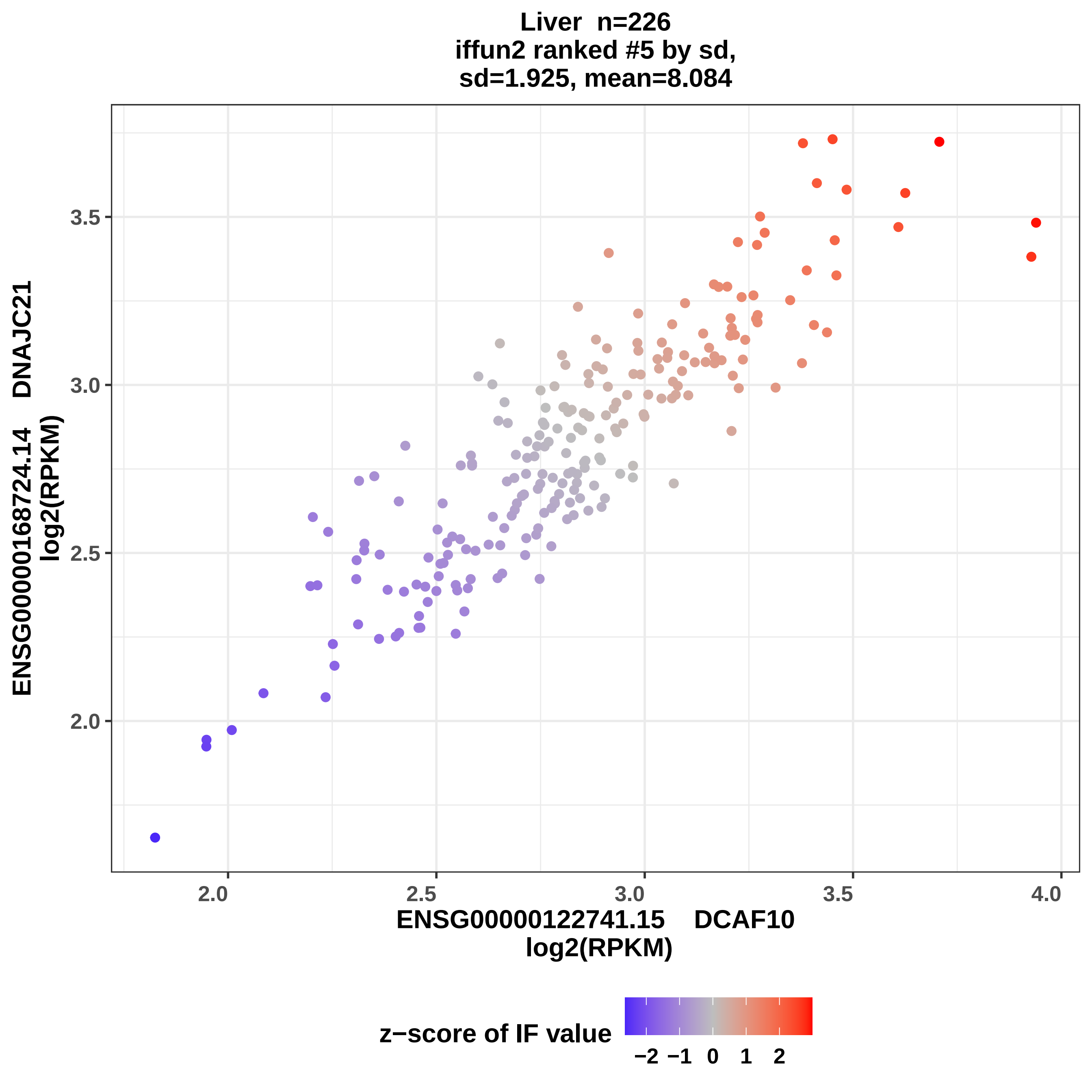
Liver n=226
iffun2 ranked #5 by sd,
sd=1.925, mean=8.084
ENSG00000168724.14 DNAJC21
log2(RPKM)
3.5
3.0
2.5
2.0
2.0
2.5
3.0
3.5
4.0
ENSG00000122741.15 DCAF10
log2(RPKM)
z−score of IF value
−2
−1
0
1
2

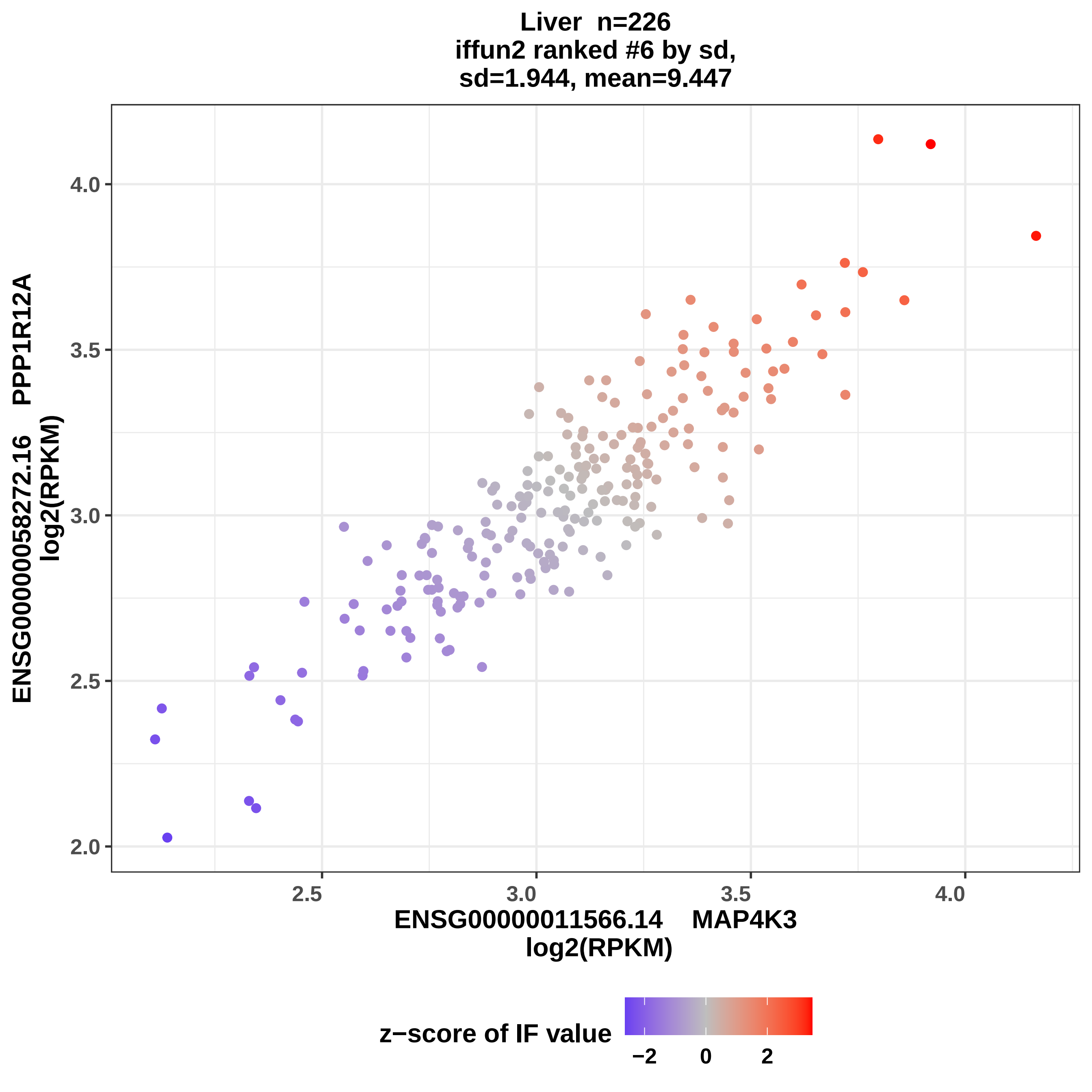

Liver n=226
iffun2 ranked #6 by sd,
sd=1.944, mean=9.447
ENSG00000058272.16 PPP1R12A
log2(RPKM)
4.0
3.5
3.0
2.5
2.0
2.5
3.0
3.5
4.0
ENSG00000011566.14 MAP4K3
log2(RPKM)
z−score of IF value
−2
0
2

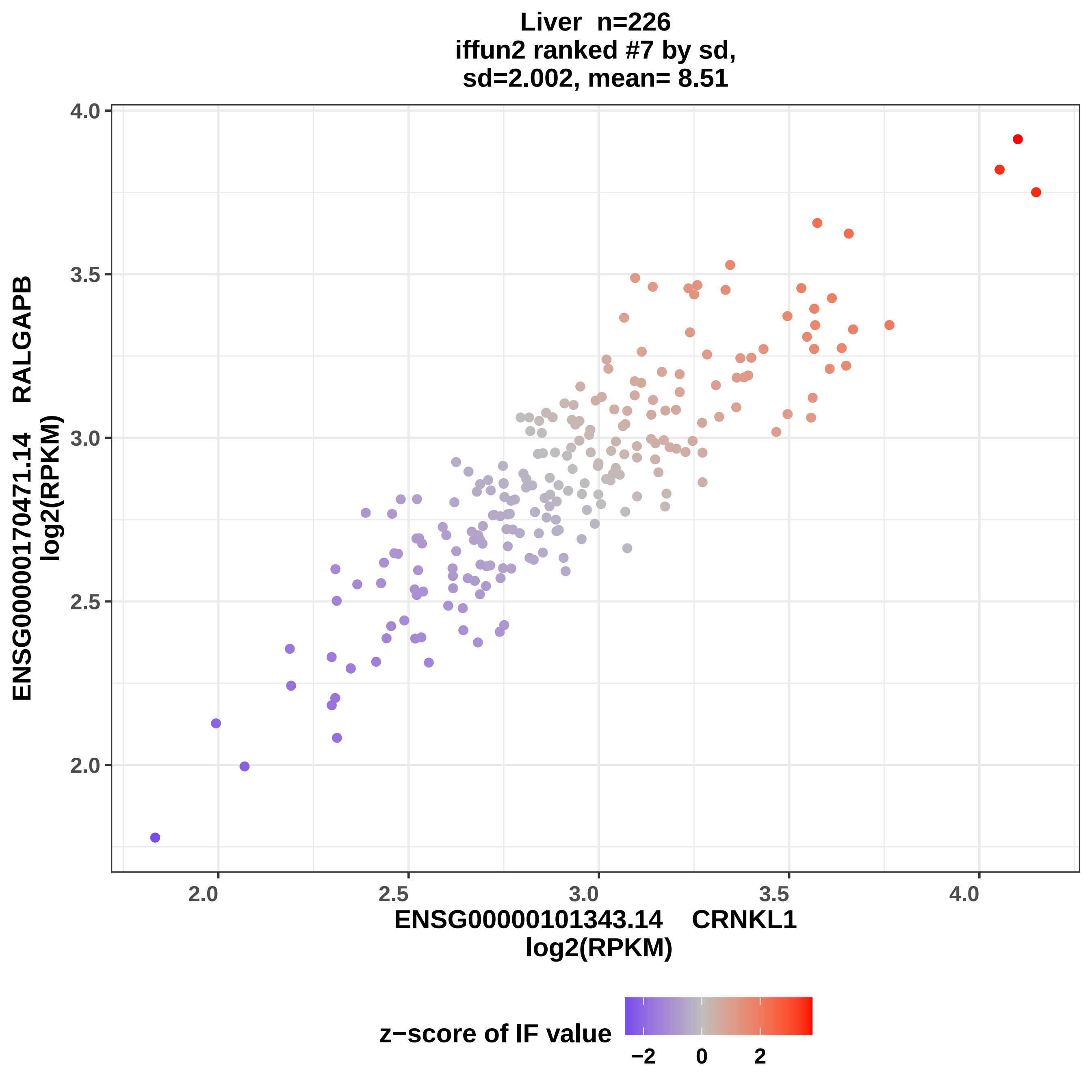

Liver n=226
iffun2 ranked #7 by sd,
sd=2.002, mean= 8.51
ENSG00000170471.14 RALGAPB
log2(RPKM)
4.0
3.5
3.0
2.5
2.0
2.0
2.5
3.0
3.5
4.0
ENSG00000101343.14 CRNKL1
log2(RPKM)
z−score of IF value
−2
0
2

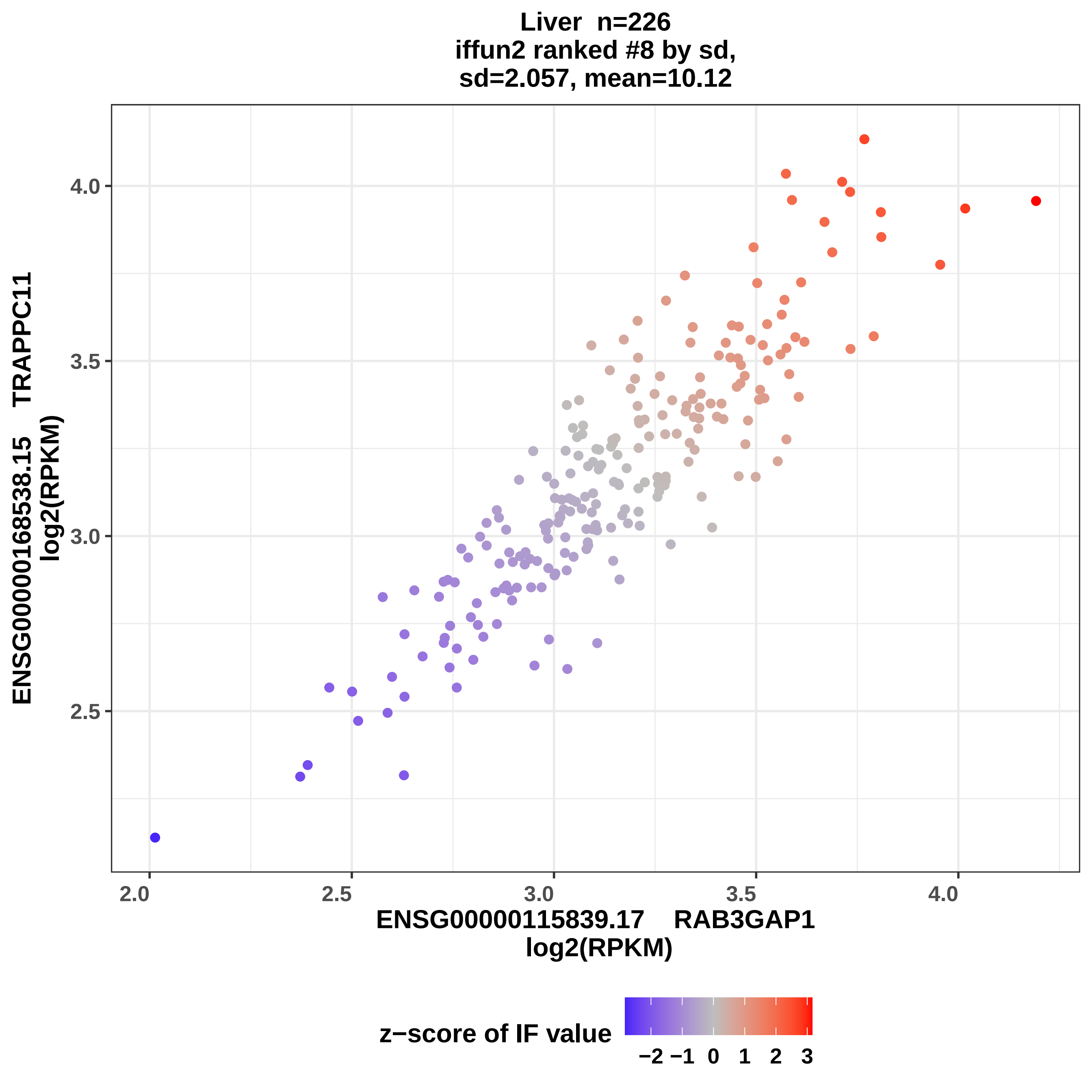
Liver n=226
iffun2 ranked #8 by sd,
sd=2.057, mean=10.12
ENSG00000168538.15 TRAPPC11
log2(RPKM)
4.0
3.5
3.0
2.5
2.0
2.5
3.0
3.5
4.0
ENSG00000115839.17 RAB3GAP1
log2(RPKM)
z−score of IF value
−2
−1
0
1
2
3

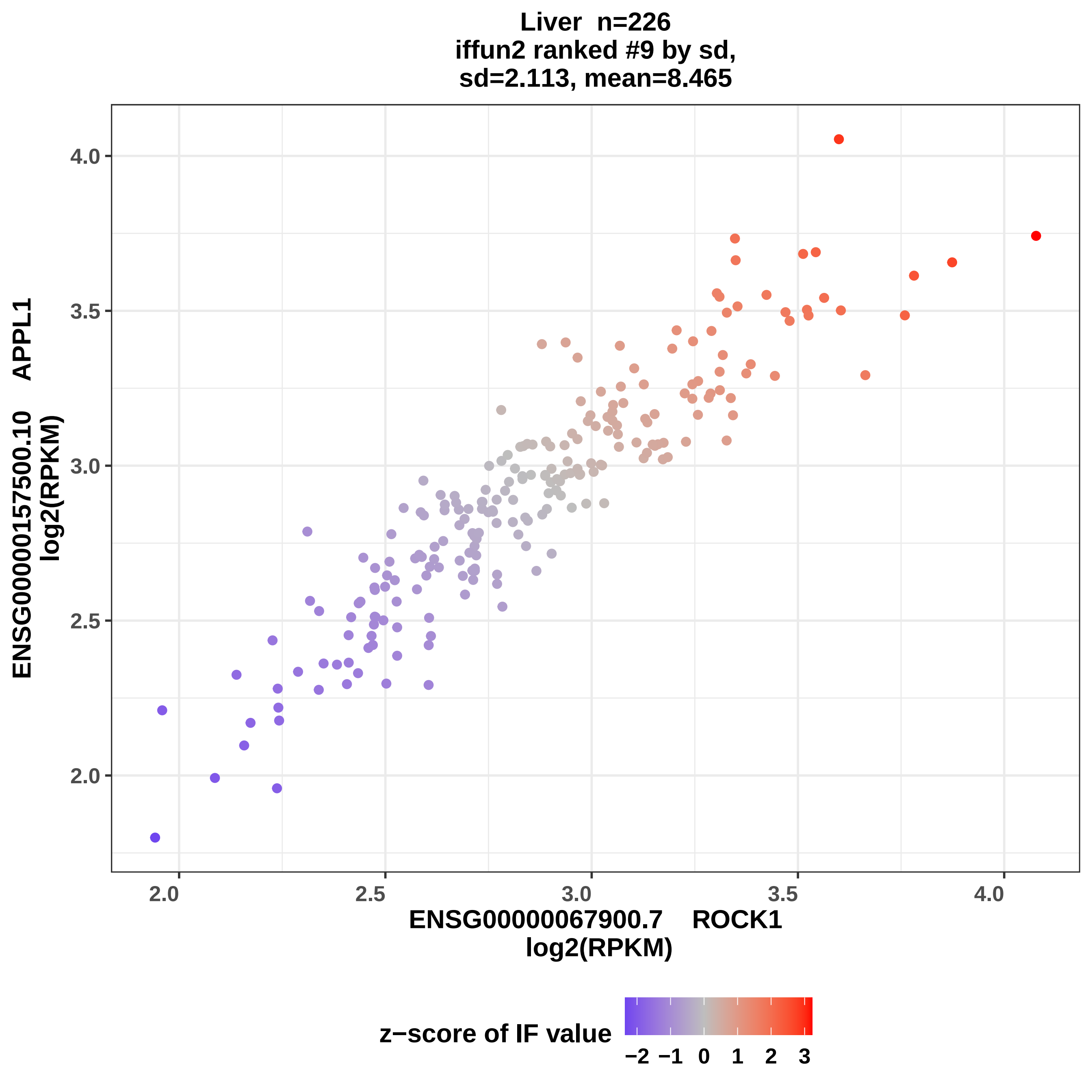
Liver n=226
iffun2 ranked #9 by sd,
sd=2.113, mean=8.465
ENSG00000157500.10 APPL1
log2(RPKM)
ENSG00000067900.7 ROCK1
log2(RPKM)
4.0
3.5
3.0
2.5
2.0
2.0
2.5
3.0
3.5
4.0
z−score of IF value
−2 −1 0 1 2 3

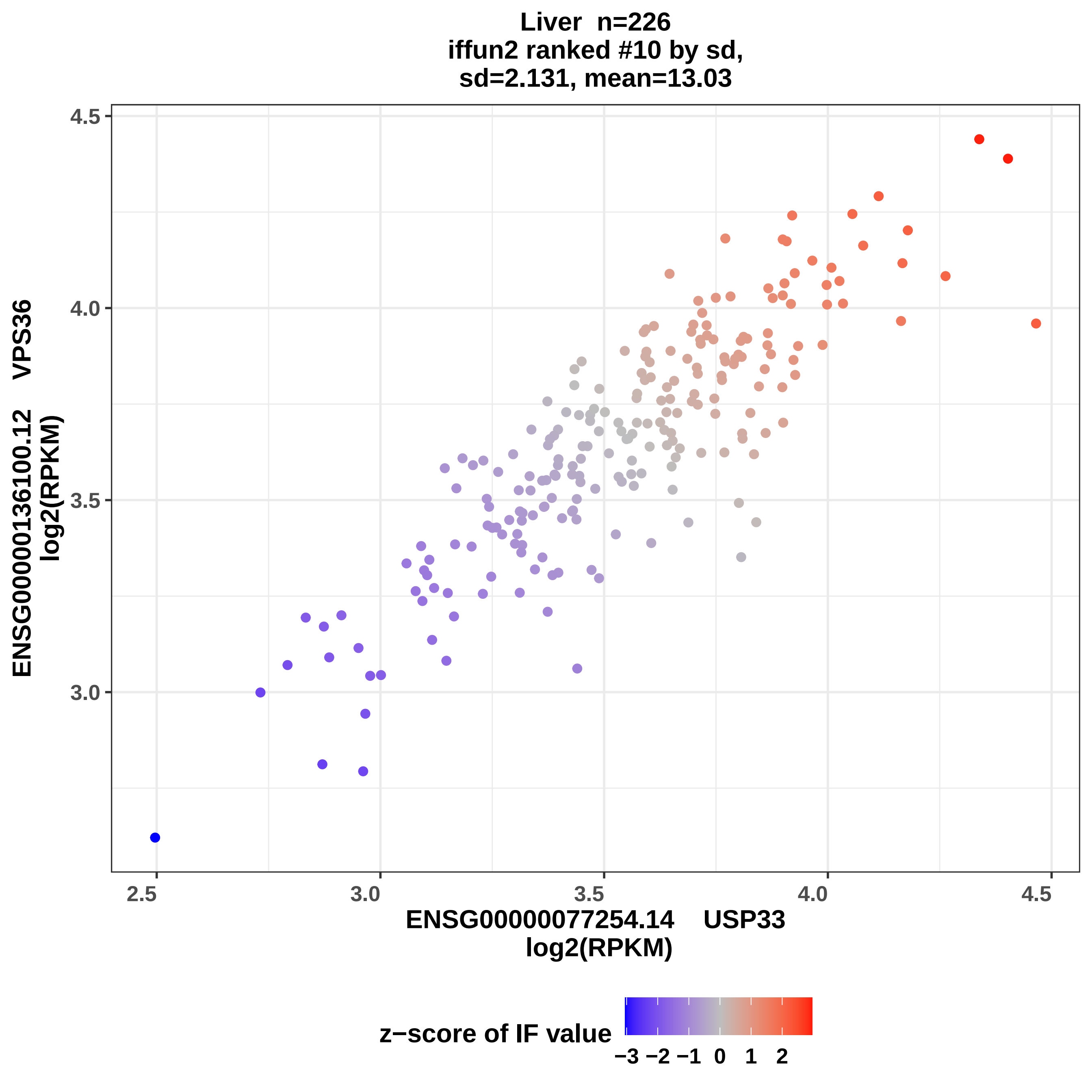

Liver n=226
iffun2 ranked #10 by sd,
sd=2.131, mean=13.03
4.5
4.0
3.5
3.0
2.5
3.0
3.5
4.0
4.5
ENSG00000136100.12 VPS36
log2(RPKM)
ENSG00000077254.14 USP33
log2(RPKM)
z−score of IF value
−3 −2 −1 0 1 2

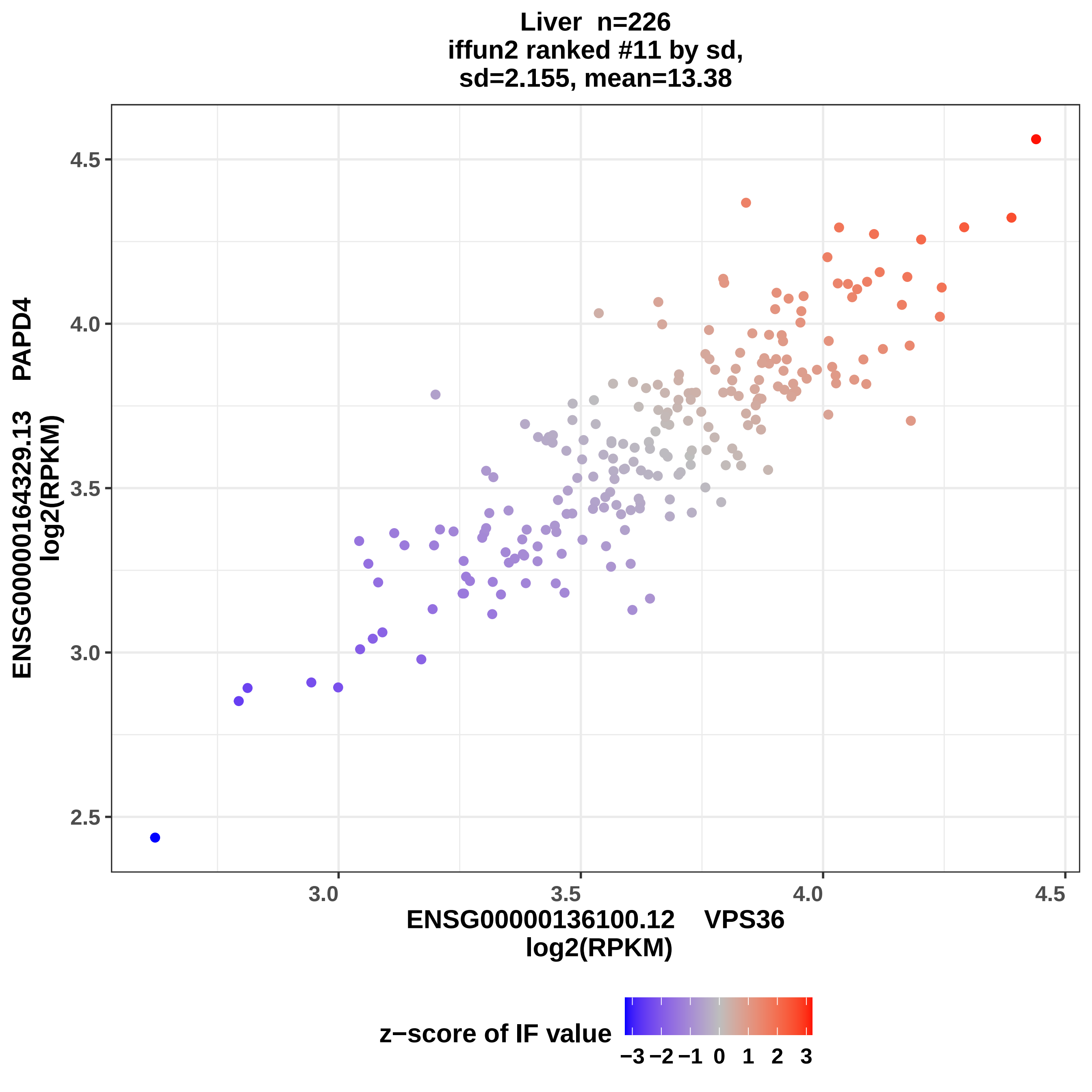

Liver n=226
iffun2 ranked #11 by sd,
sd=2.155, mean=13.38
ENSG00000164329.13 PAPD4
log2(RPKM)
4.5
4.0
3.5
3.0
2.5
3.0
3.5
4.0
4.5
ENSG00000136100.12 VPS36
log2(RPKM)
z−score of IF value
−3 −2 −1 0 1 2 3

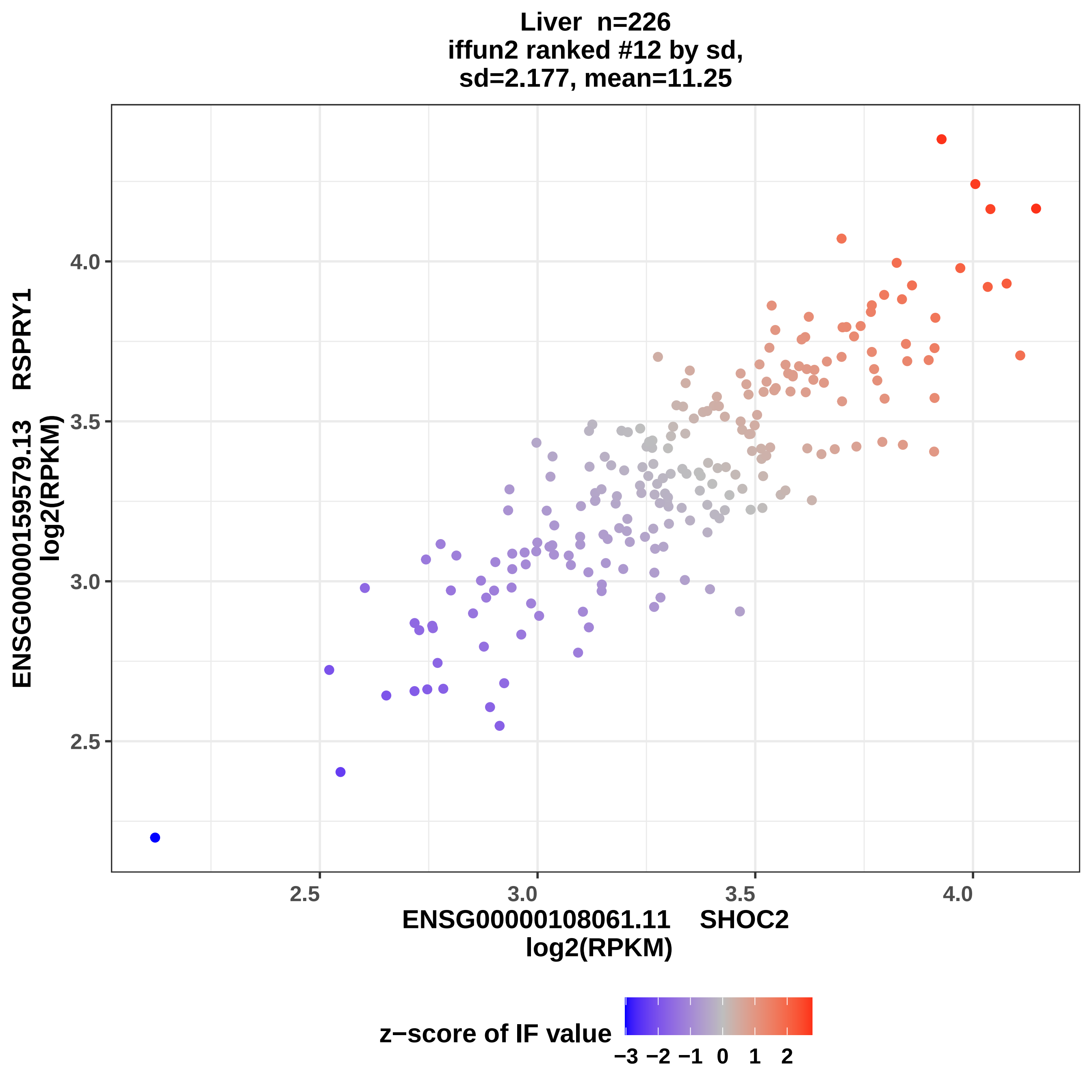

Liver n=226
iffun2 ranked #12 by sd,
sd=2.177, mean=11.25
ENSG00000159579.13 RSPRY1
log2(RPKM)
4.0
3.5
3.0
2.5
2.5
3.0
3.5
4.0
ENSG00000108061.11 SHOC2
log2(RPKM)
z−score of IF value
−3 −2 −1 0 1 2

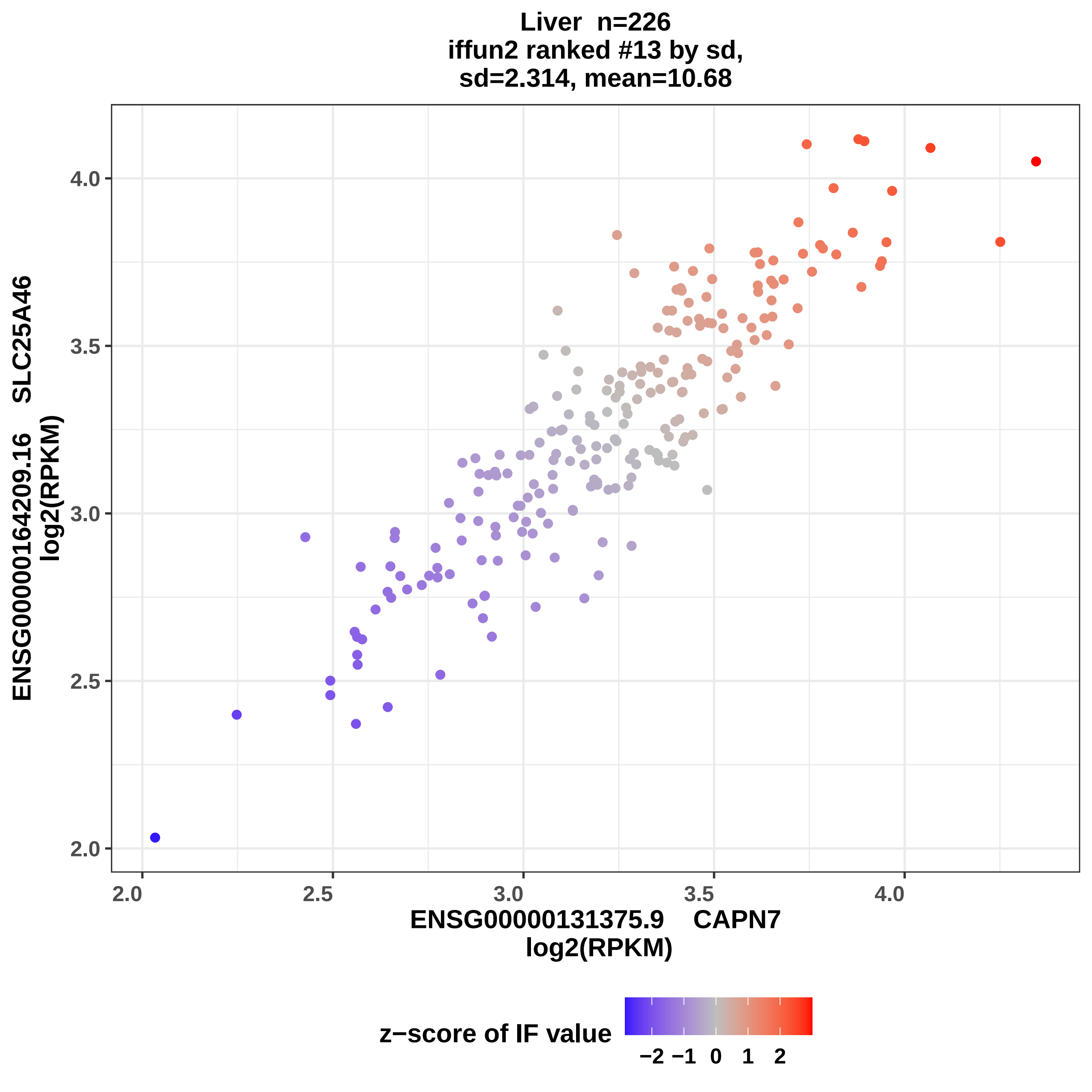

Liver n=226
iffun2 ranked #13 by sd,
sd=2.314, mean=10.68
ENSG00000164209.16 SLC25A46
log2(RPKM)
4.0
3.5
3.0
2.5
2.0
2.0
2.5
3.0
3.5
4.0
ENSG00000131375.9 CAPN7
log2(RPKM)
z−score of IF value
−2 −1 0 1 2

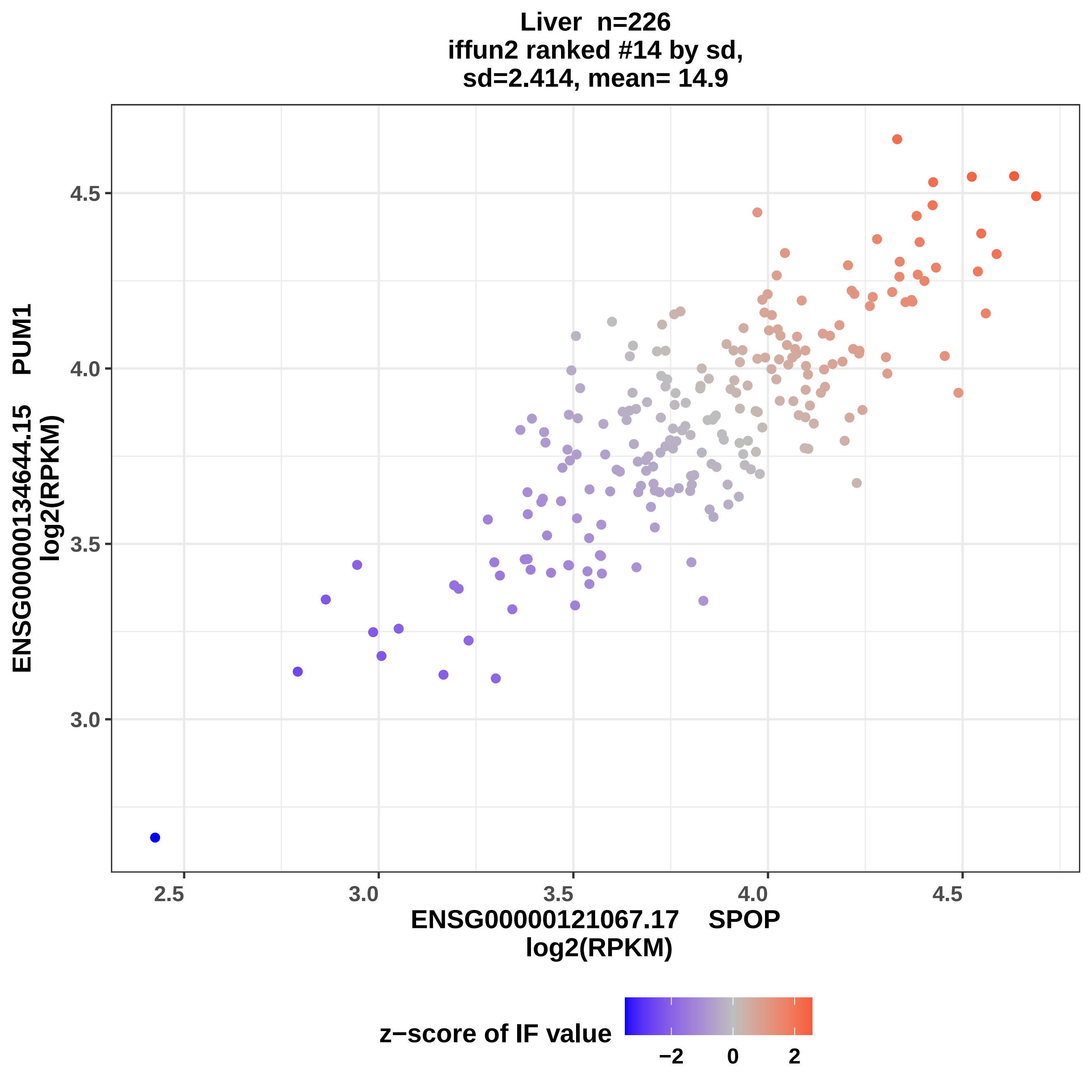

Liver n=226
iffun2 ranked #14 by sd,
sd=2.414, mean= 14.9
ENSG00000134644.15 PUM1
log2(RPKM)
4.5
4.0
3.5
3.0
2.5
3.0
3.5
4.0
4.5
ENSG00000121067.17 SPOP
log2(RPKM)
z−score of IF value
−2
0
2

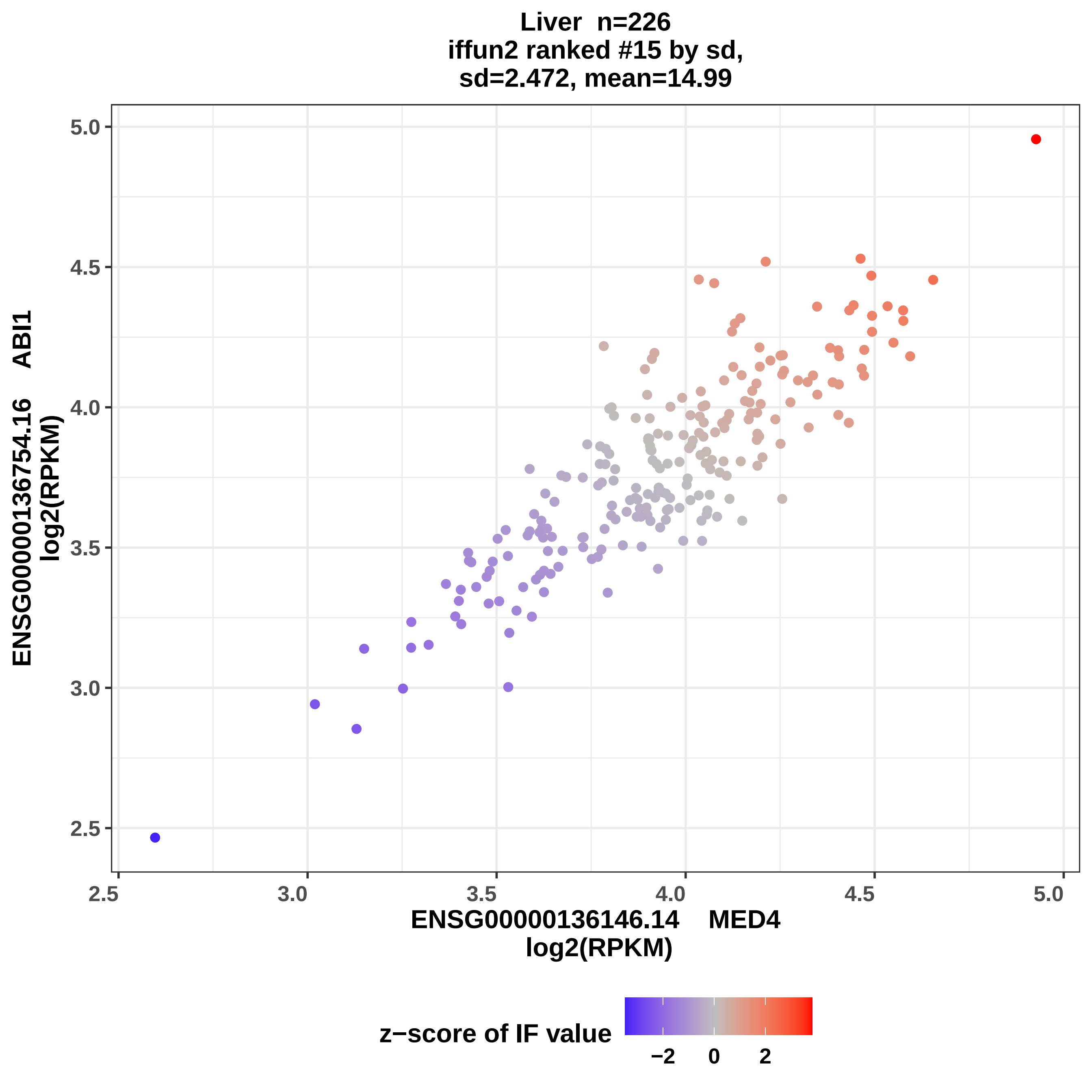

Liver n=226
iffun2 ranked #15 by sd,
sd=2.472, mean=14.99
ENSG00000136754.16 ABI1
log2(RPKM)
ENSG00000136146.14 MED4
log2(RPKM)
2.5
3.0
3.5
4.0
4.5
5.0
z−score of IF value
−2
0
2

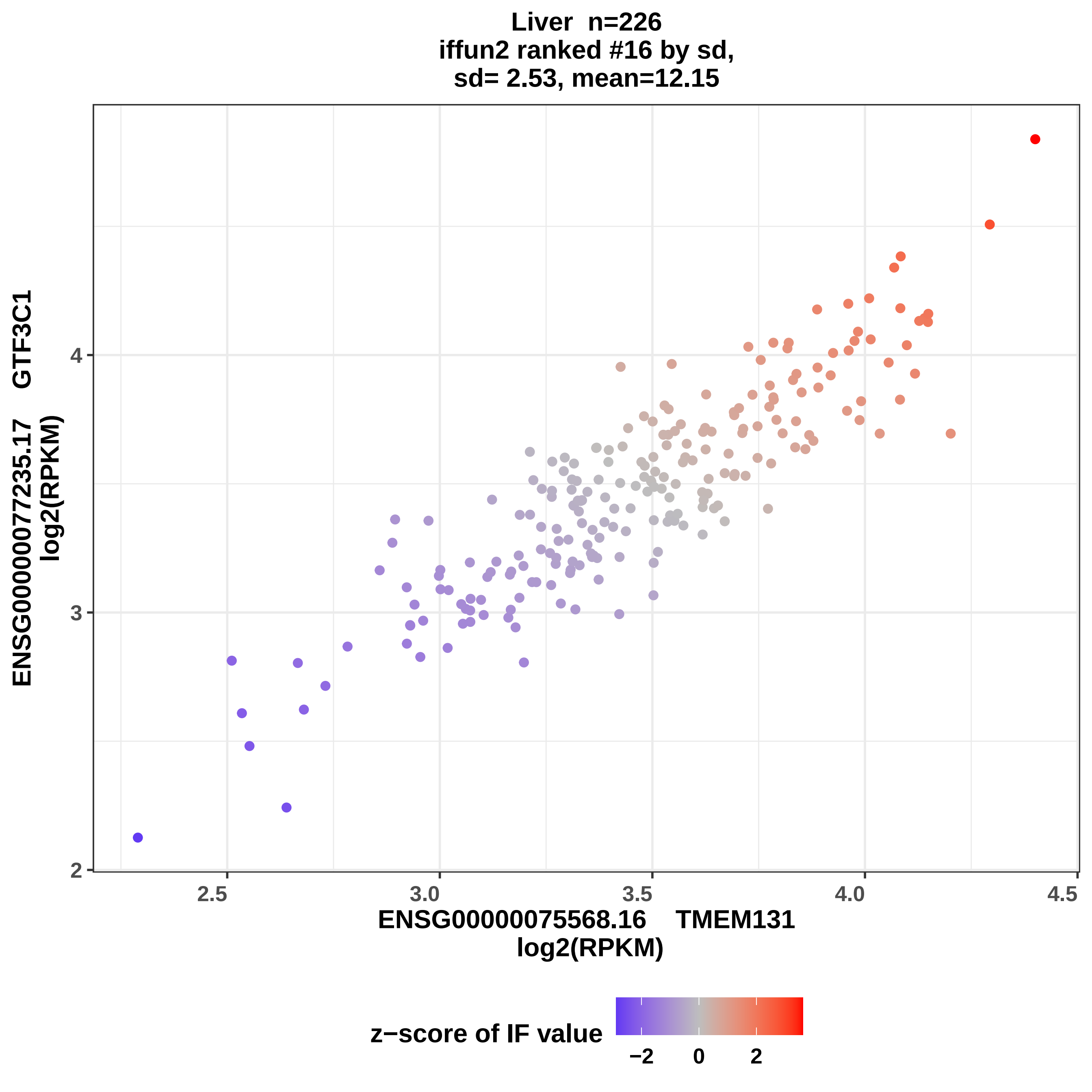

Liver n=226
iffun2 ranked #16 by sd,
sd= 2.53, mean=12.15
ENSG00000077235.17 GTF3C1
log2(RPKM)
ENSG00000075568.16 TMEM131
log2(RPKM)
2
3
4
2.5
3.0
3.5
4.0
4.5
z−score of IF value
−2
0
2

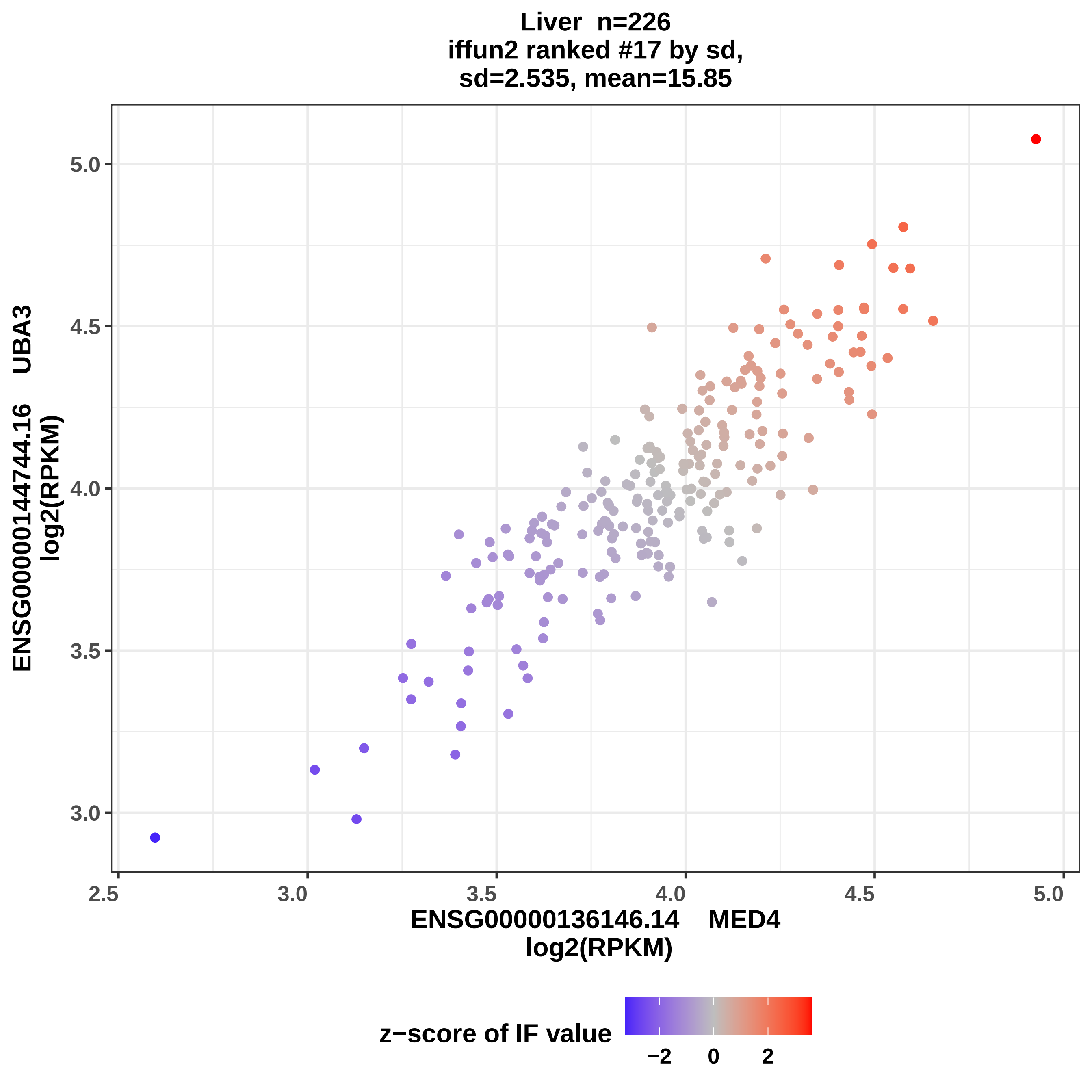

Liver n=226
iffun2 ranked #17 by sd,
sd=2.535, mean=15.85
ENSG00000144744.16 UBA3
log2(RPKM)
5.0
4.5
4.0
3.5
3.0
2.5
3.0
3.5
4.0
4.5
5.0
ENSG00000136146.14 MED4
log2(RPKM)
z−score of IF value
−2
0
2

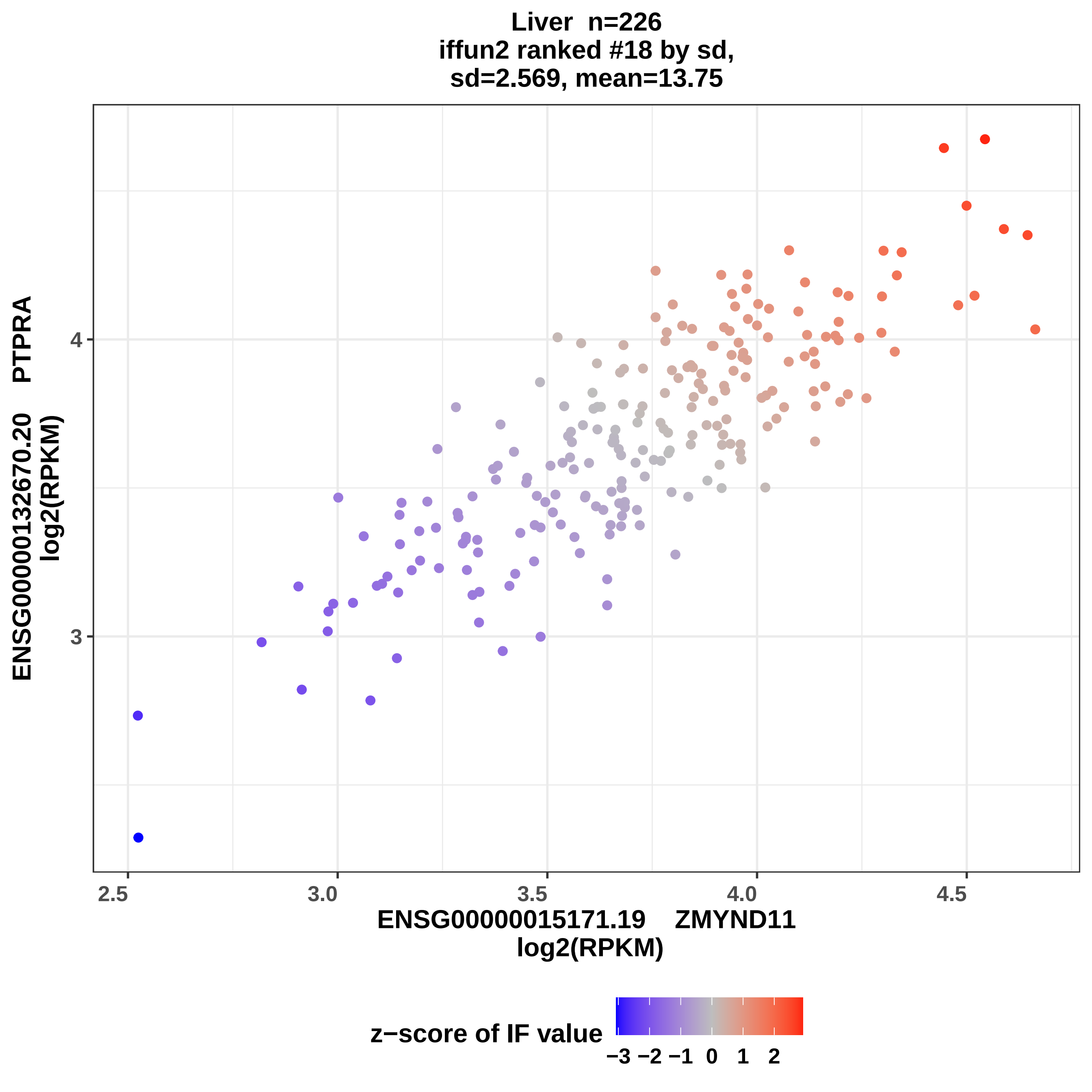

Liver n=226
iffun2 ranked #18 by sd,
sd=2.569, mean=13.75
ENSG00000132670.20 PTPRA
log2(RPKM)
4
3
2.5
3.0
3.5
4.0
4.5
ENSG00000015171.19 ZMYND11
log2(RPKM)
z−score of IF value
−3 −2 −1 0 1 2

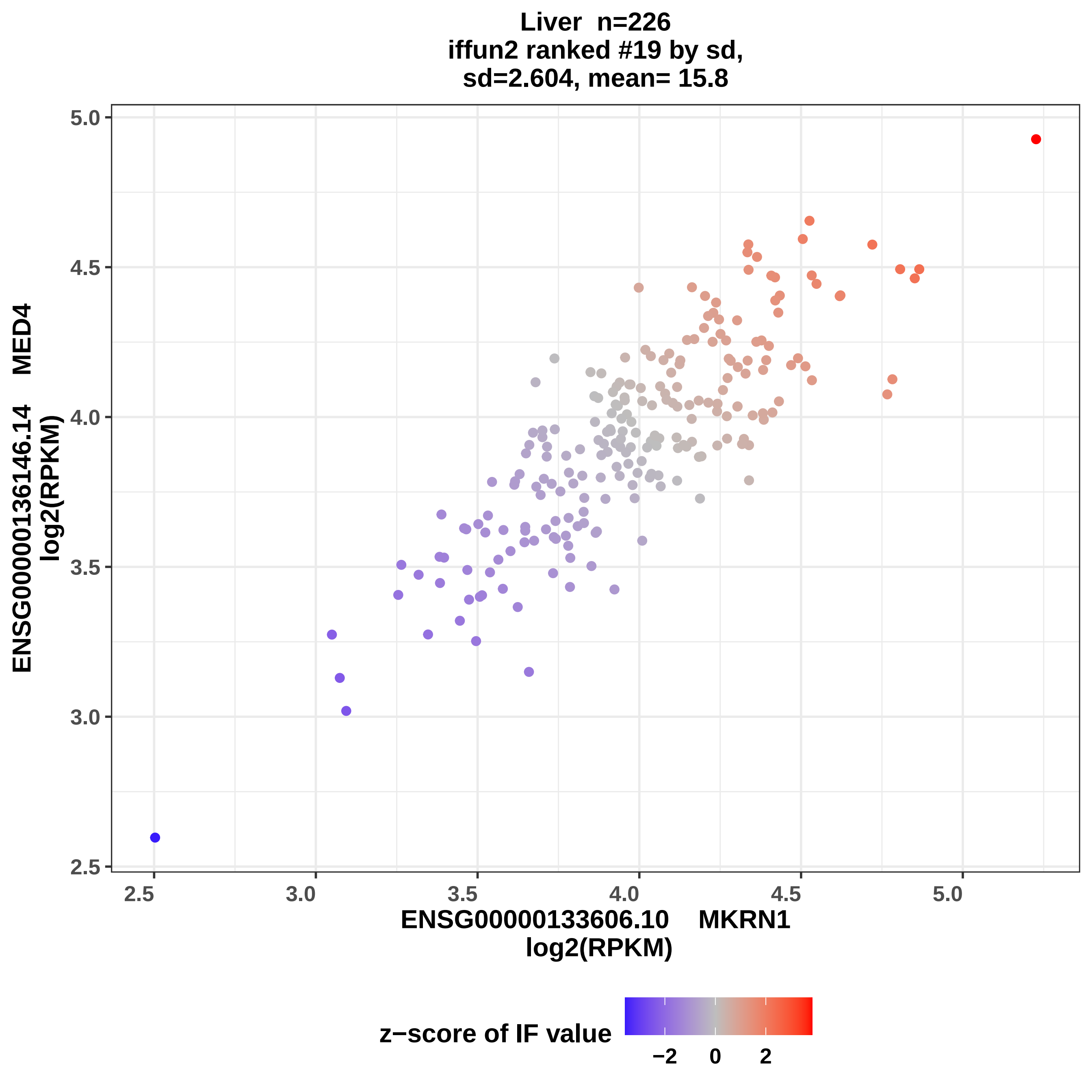

Liver n=226
iffun2 ranked #19 by sd,
sd=2.604, mean= 15.8
5.0
4.5
4.0
3.5
3.0
2.5
ENSG00000136146.14 MED4
log2(RPKM)
2.5
3.0
3.5
4.0
4.5
5.0
ENSG00000133606.10 MKRN1
log2(RPKM)
z−score of IF value
−2
0
2

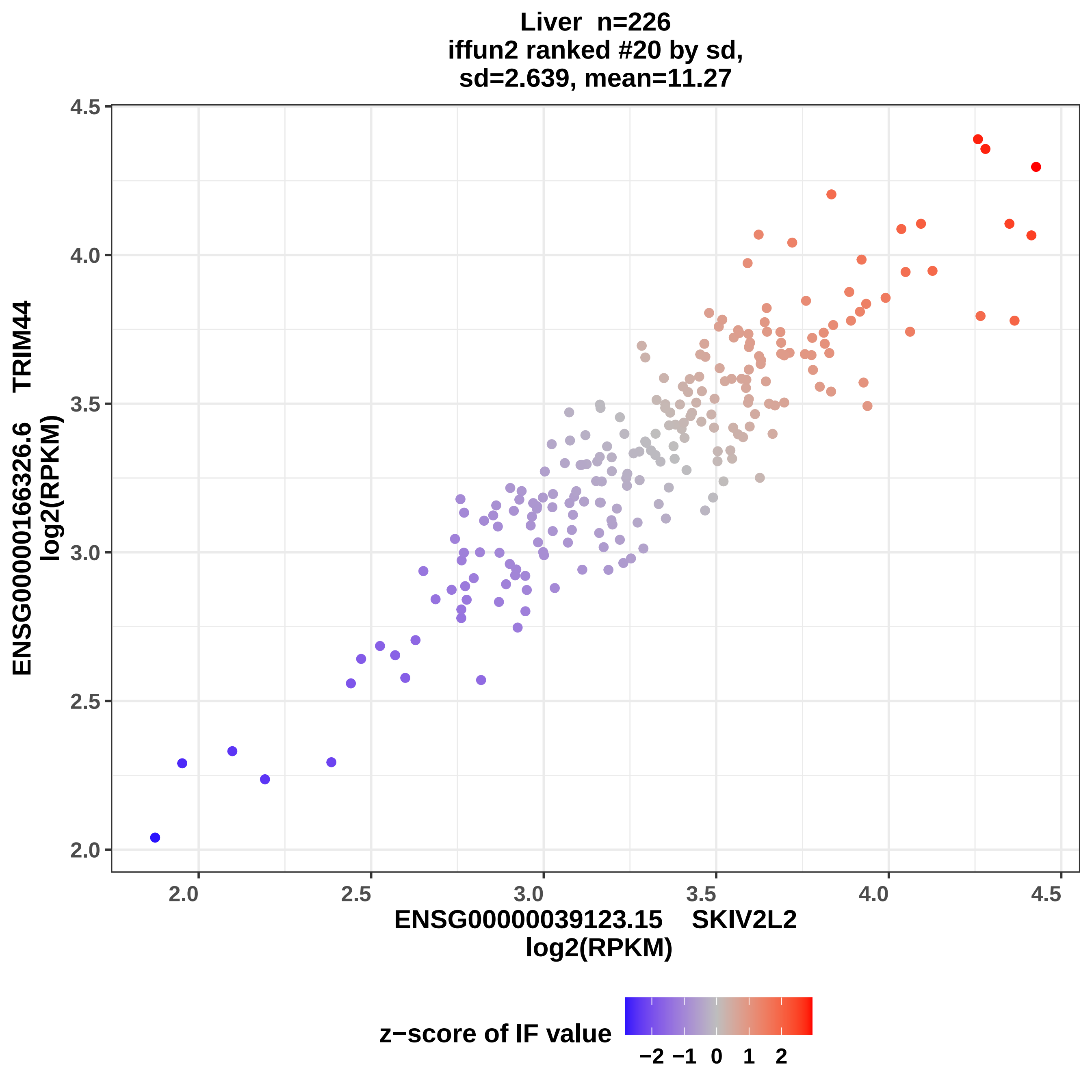

Liver n=226
iffun2 ranked #20 by sd,
sd=2.639, mean=11.27
ENSG00000166326.6 TRIM44
log2(RPKM)
4.5
4.0
3.5
3.0
2.5
2.0
2.0
2.5
3.0
3.5
4.0
4.5
ENSG00000039123.15 SKIV2L2
log2(RPKM)
z−score of IF value
−2 −1 0 1 2

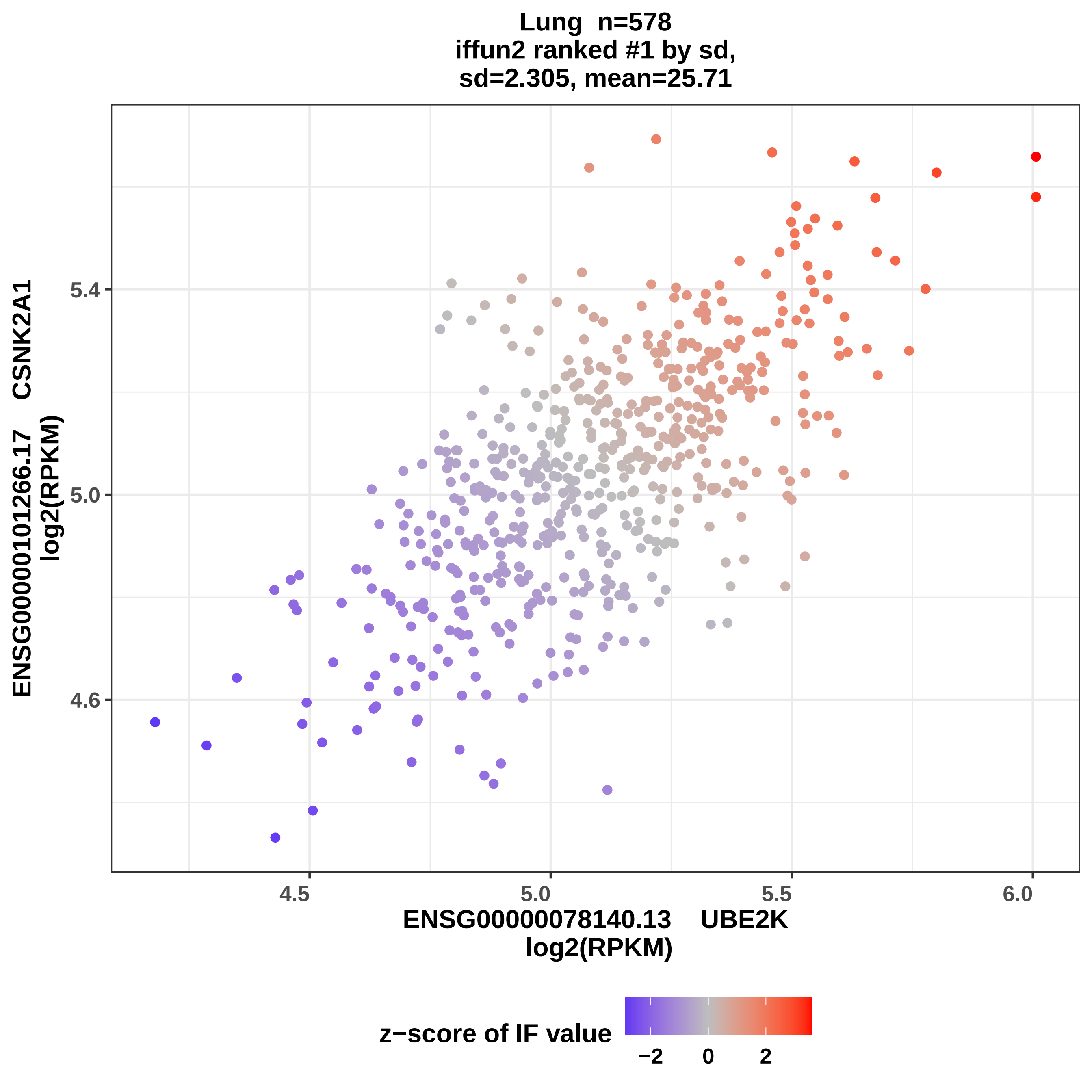

Lung n=578
iffun2 ranked #1 by sd,
sd=2.305, mean=25.71
ENSG00000101266.17 CSNK2A1
log2(RPKM)
5.4
5.0
4.6
4.5
5.0
5.5
6.0
ENSG00000078140.13 UBE2K
log2(RPKM)
z−score of IF value
−2
0
2

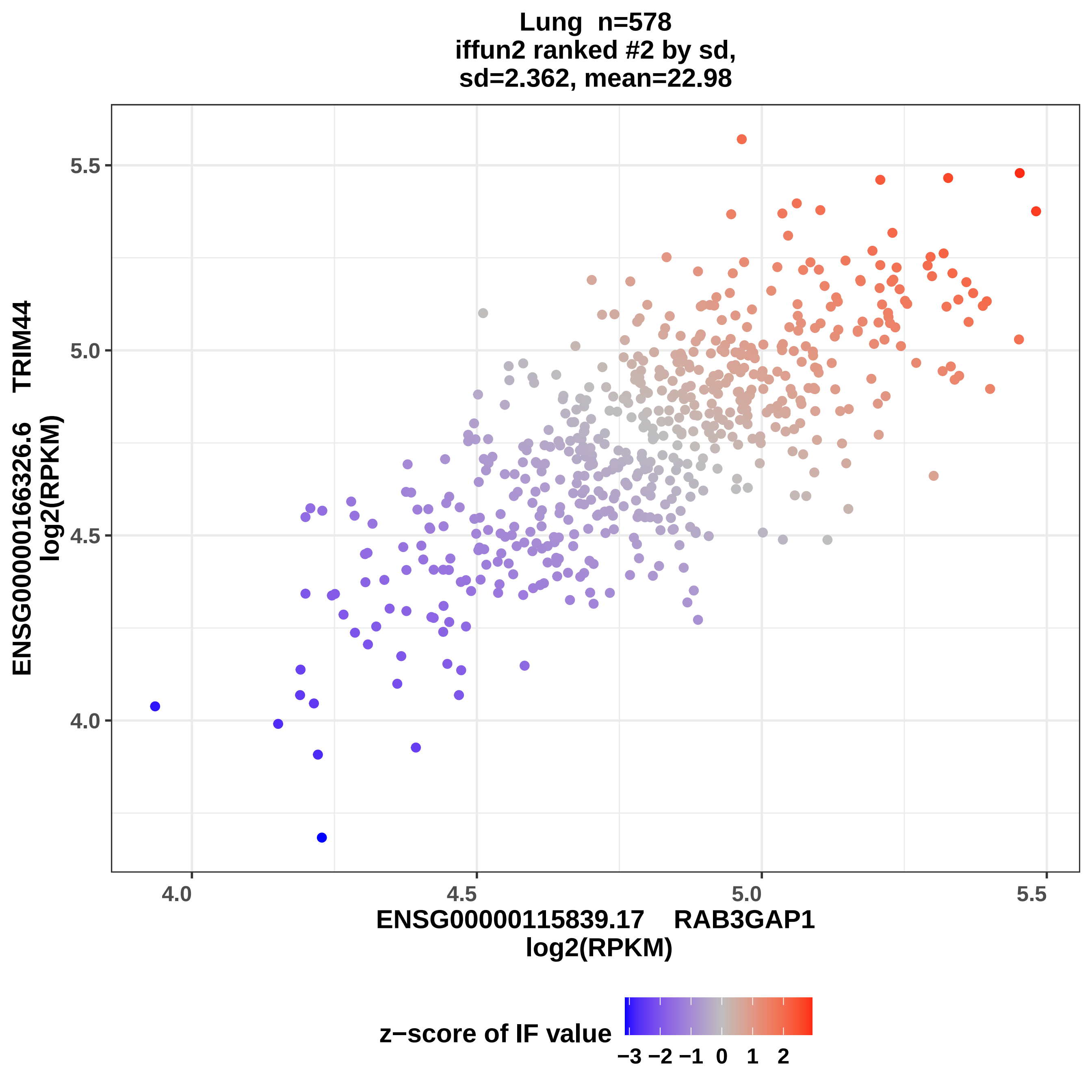

Lung n=578
iffun2 ranked #2 by sd,
sd=2.362, mean=22.98
ENSG00000166326.6 TRIM44
log2(RPKM)
ENSG00000115839.17 RAB3GAP1
log2(RPKM)
5.5
5.0
4.5
4.0
4.0
4.5
5.0
5.5
z−score of IF value
−3 −2 −1 0 1 2

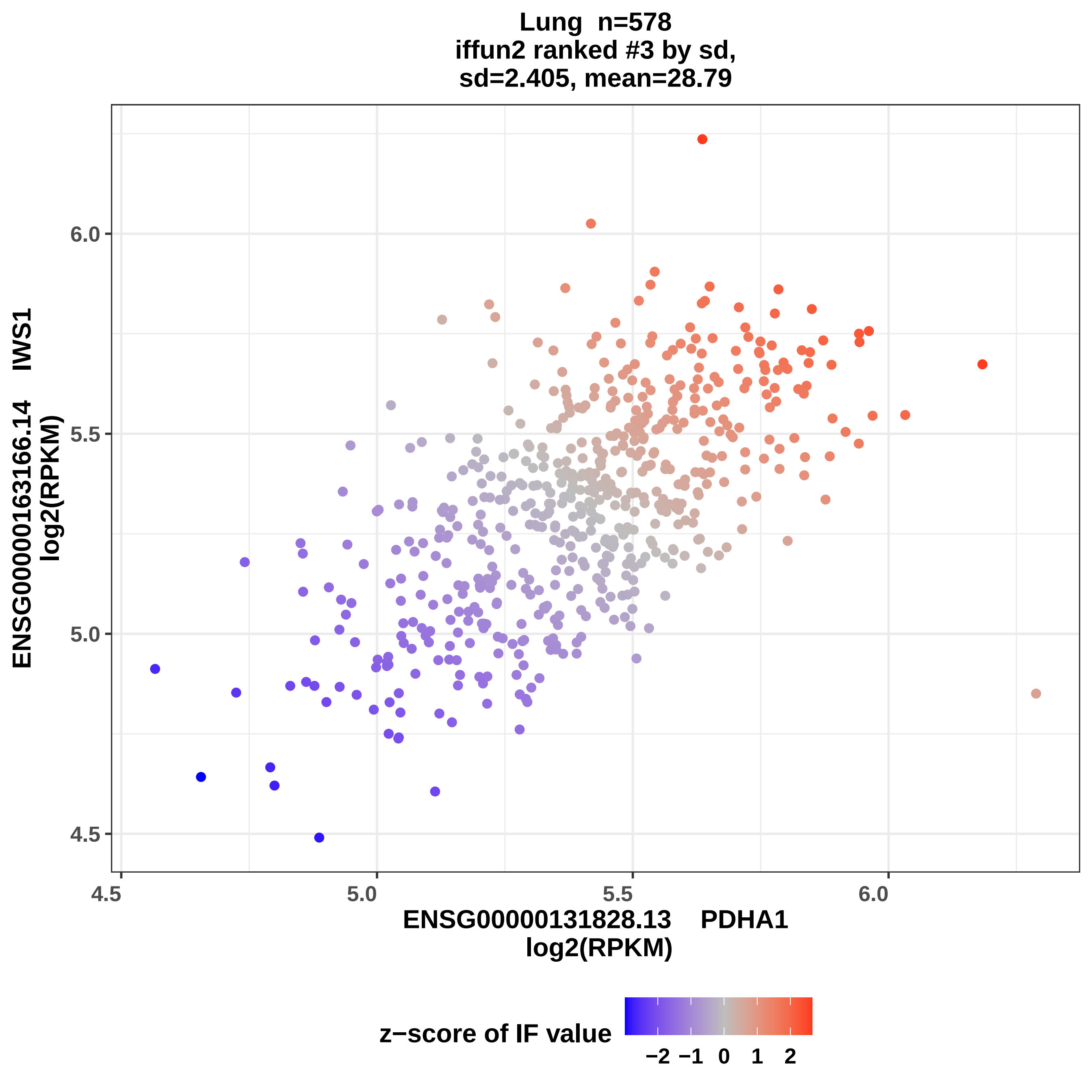

Lung n=578
iffun2 ranked #3 by sd,
sd=2.405, mean=28.79
ENSG00000163166.14 IWS1
log2(RPKM)
6.0
5.5
5.0
4.5
4.5
5.0
5.5
6.0
ENSG00000131828.13 PDHA1
log2(RPKM)
z−score of IF value
−2 −1 0 1 2

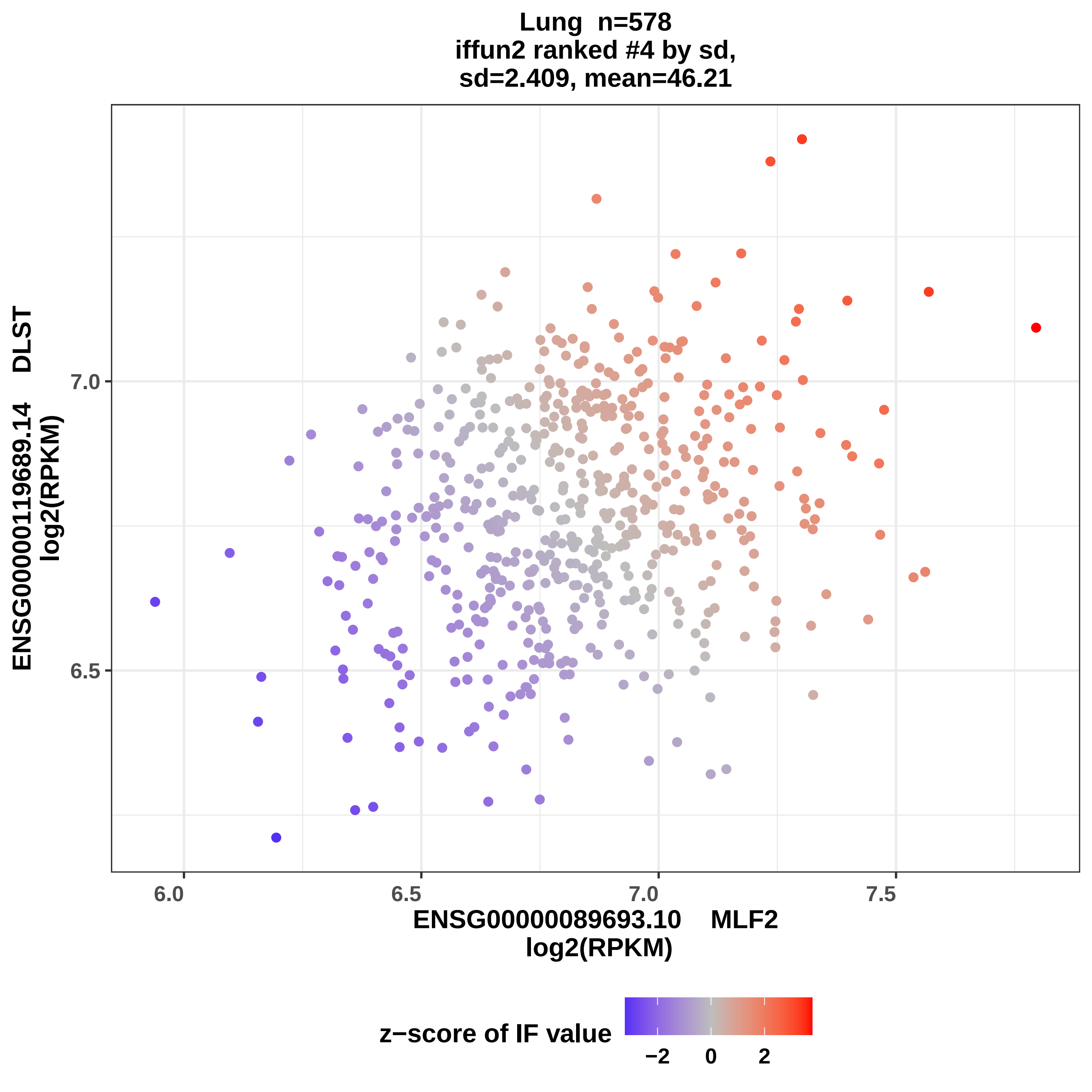

Lung n=578
iffun2 ranked #4 by sd,
sd=2.409, mean=46.21
ENSG00000119689.14 DLST
log2(RPKM)
7.0
6.5
6.0
6.5
7.0
7.5
ENSG00000089693.10 MLF2
log2(RPKM)
z−score of IF value
−2
0
2

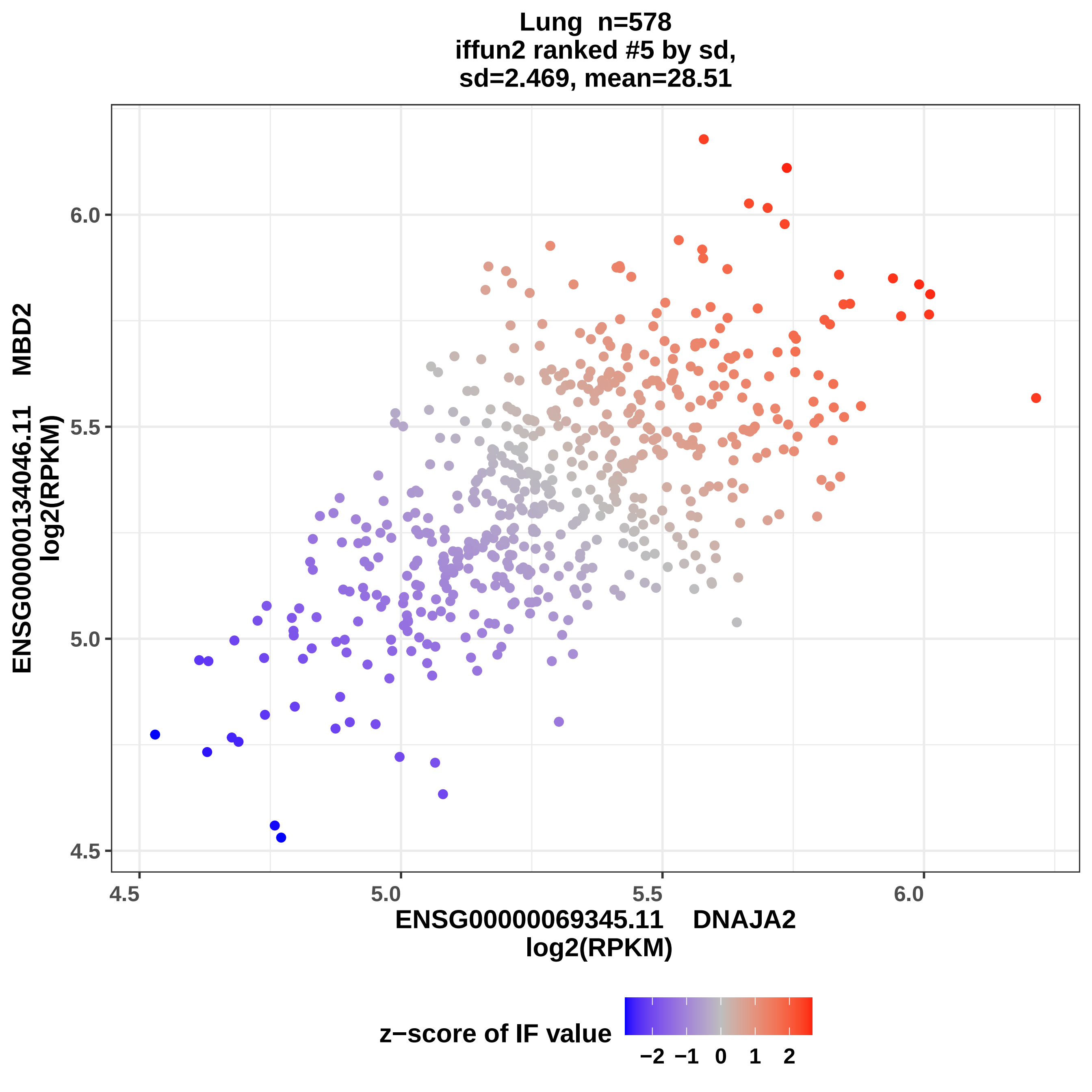
Lung n=578
iffun2 ranked #5 by sd,
sd=2.469, mean=28.51
ENSG00000134046.11 MBD2
log2(RPKM)
6.0
5.5
5.0
4.5
4.5
5.0
5.5
6.0
ENSG00000069345.11 DNAJA2
log2(RPKM)
z−score of IF value
−2 −1 0 1 2

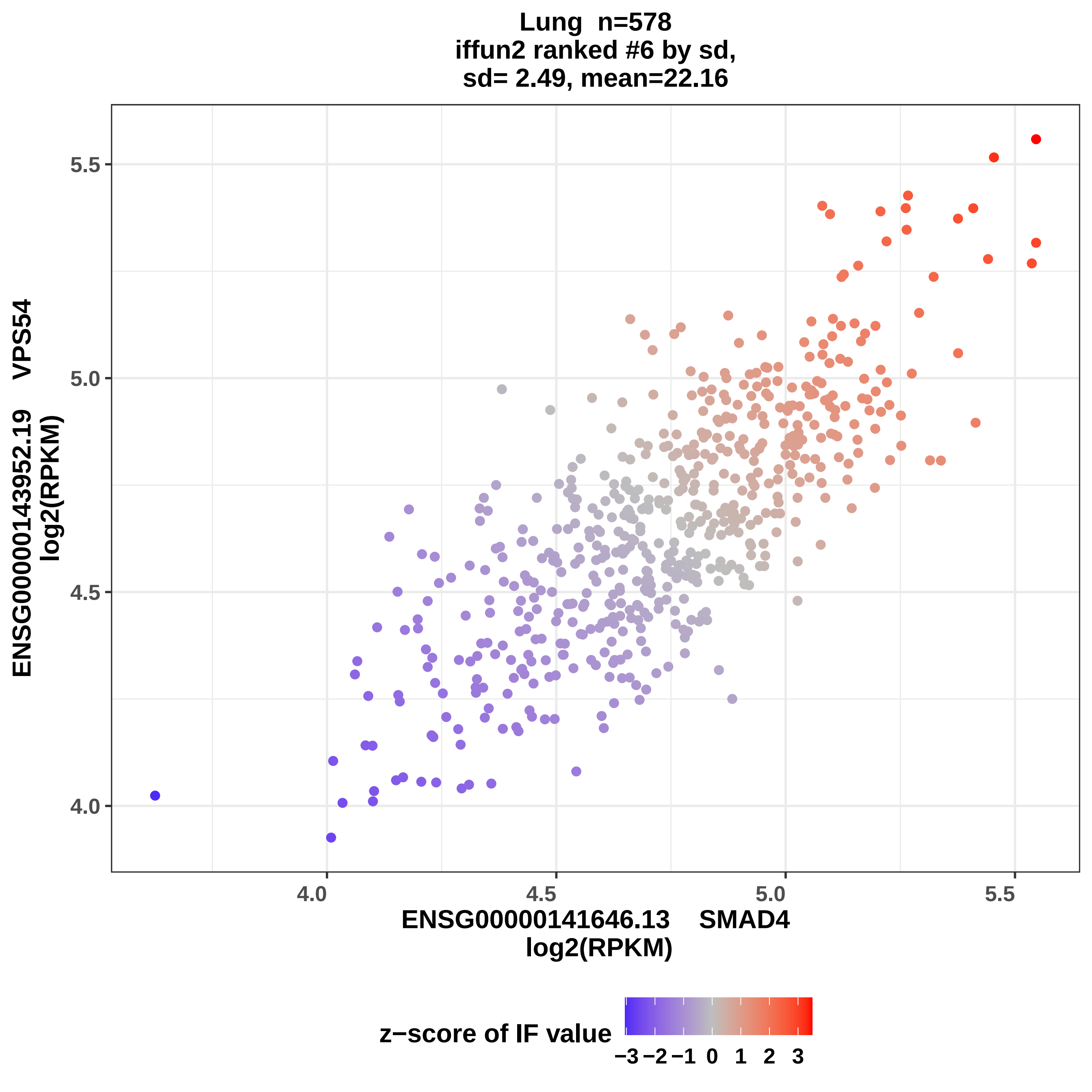

Lung n=578
iffun2 ranked #6 by sd,
sd= 2.49, mean=22.16
ENSG00000143952.19 VPS54
log2(RPKM)
5.5
5.0
4.5
4.0
4.0
4.5
5.0
5.5
ENSG00000141646.13 SMAD4
log2(RPKM)
z−score of IF value
−3 −2 −1 0 1 2 3

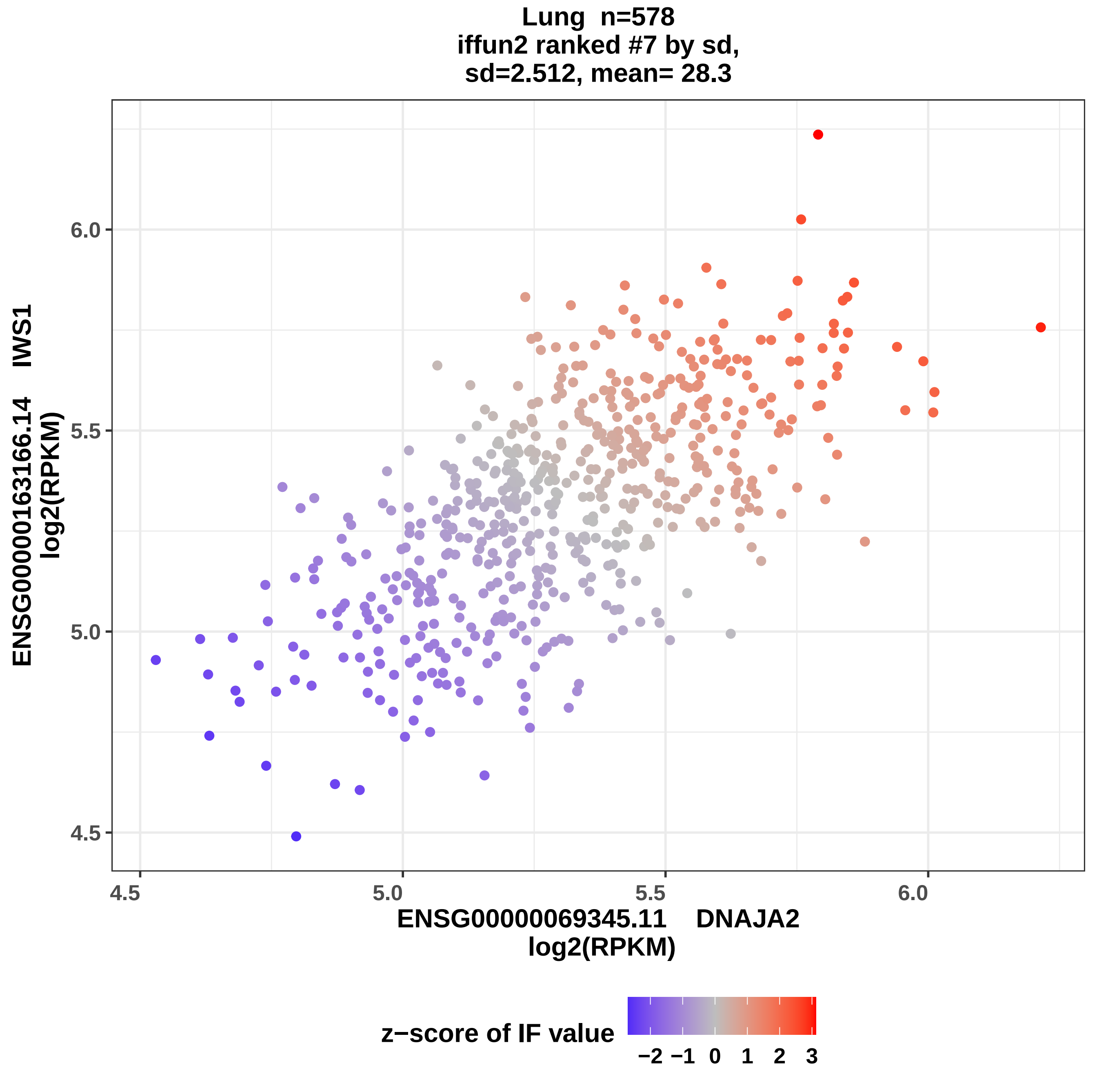

Lung n=578
iffun2 ranked #7 by sd,
sd=2.512, mean= 28.3
ENSG00000163166.14 IWS1
log2(RPKM)
6.0
5.5
5.0
4.5
4.5
5.0
5.5
6.0
ENSG00000069345.11 DNAJA2
log2(RPKM)
z−score of IF value
−2
−1
0
1
2
3

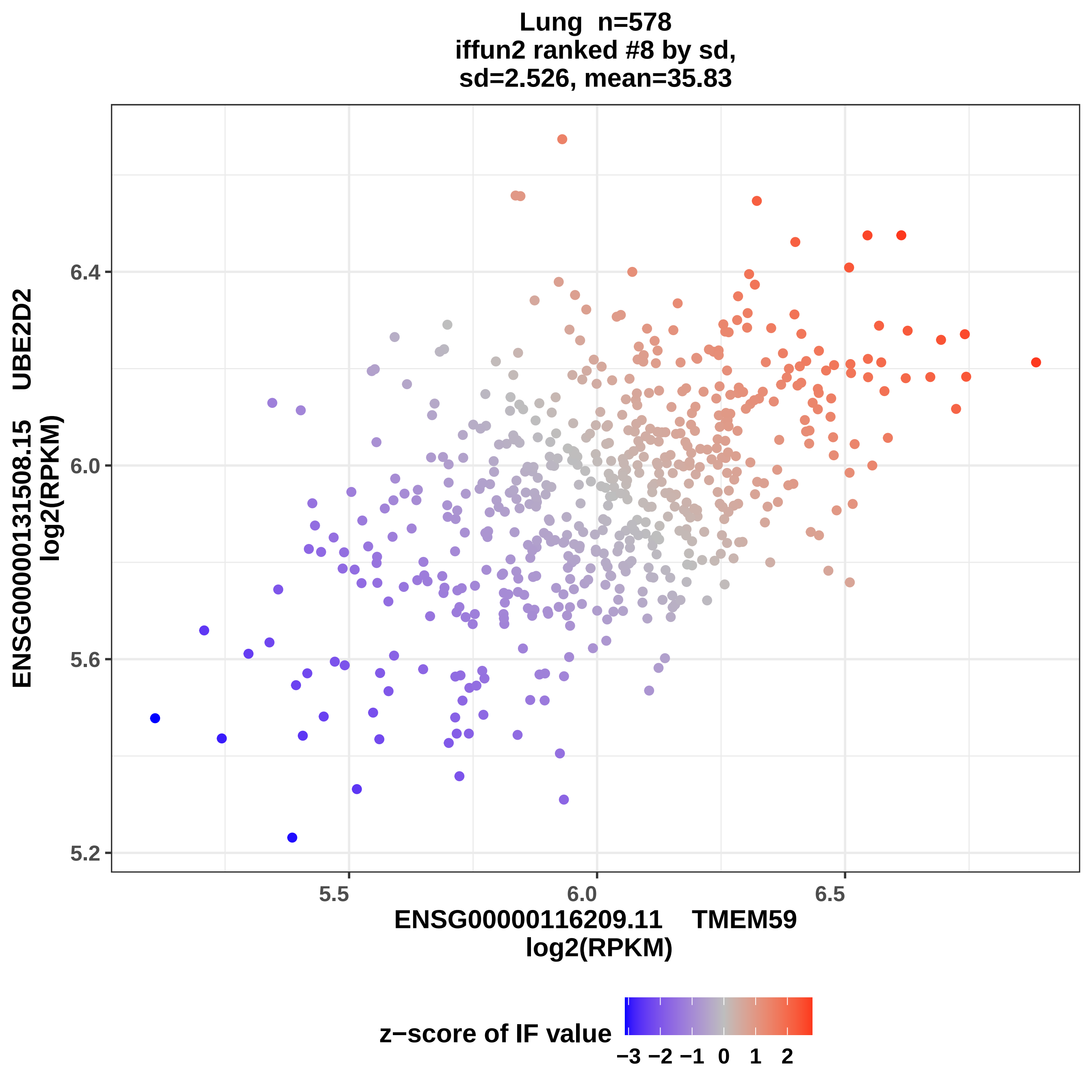
Lung n=578
iffun2 ranked #8 by sd,
sd=2.526, mean=35.83
ENSG00000131508.15 UBE2D2
log2(RPKM)
6.4
6.0
5.6
5.2
5.5
6.0
6.5
ENSG00000116209.11 TMEM59
log2(RPKM)
z−score of IF value
−3 −2 −1 0 1 2

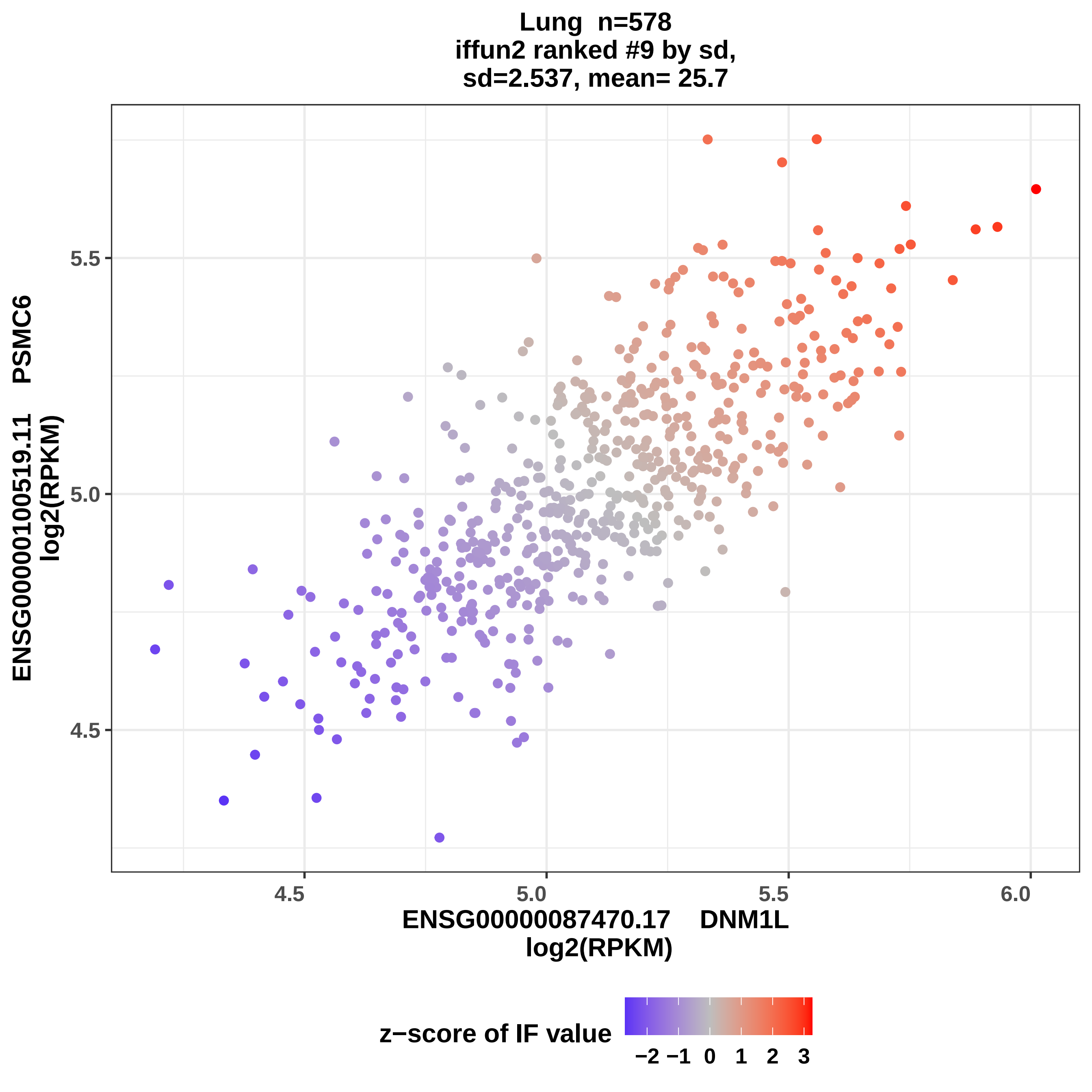
Lung n=578
iffun2 ranked #9 by sd,
sd=2.537, mean= 25.7
ENSG00000100519.11 PSMC6
log2(RPKM)
5.5
5.0
4.5
4.5
5.0
5.5
6.0
ENSG00000087470.17 DNM1L
log2(RPKM)
z−score of IF value
−2 −1 0 1 2 3

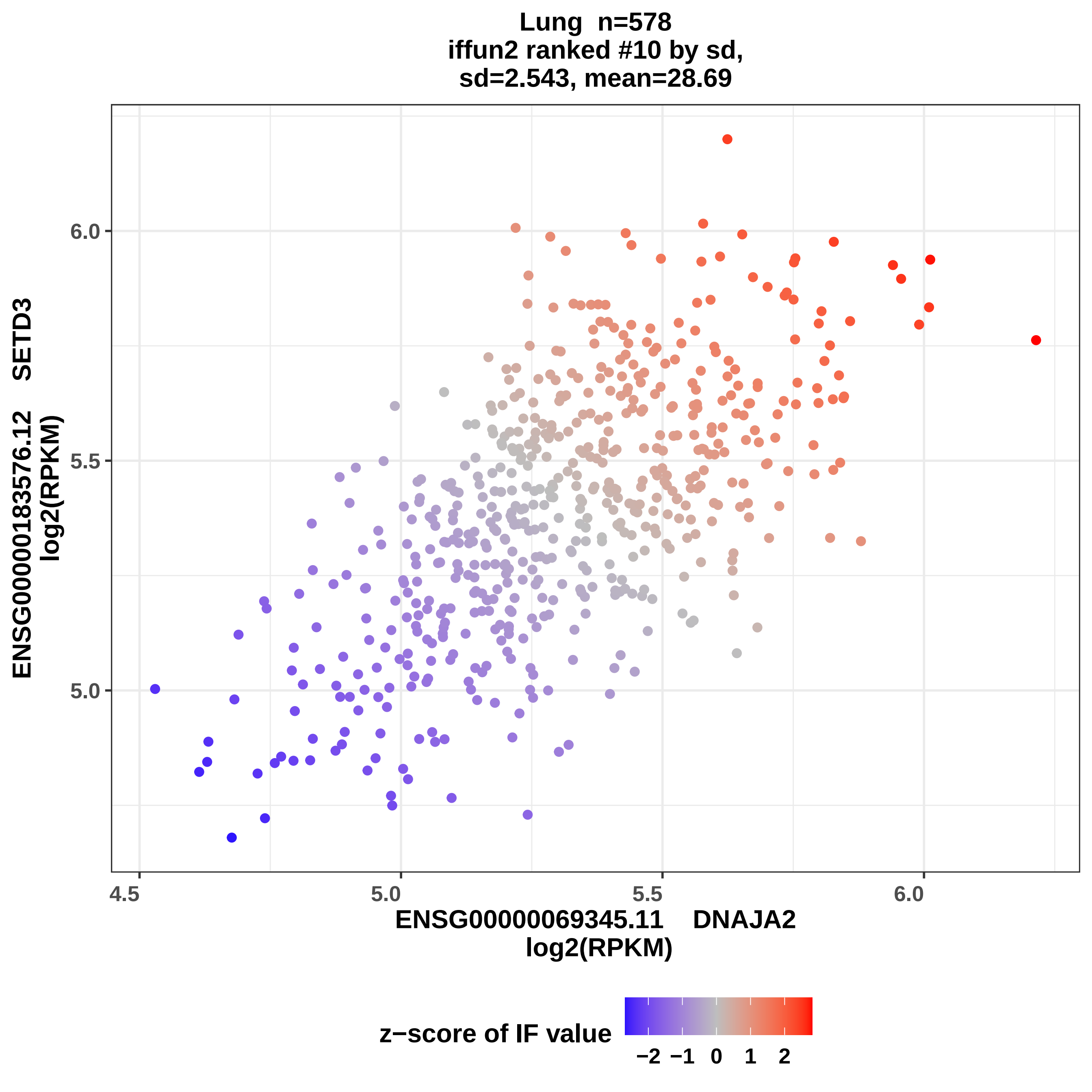
Lung n=578
iffun2 ranked #10 by sd,
sd=2.543, mean=28.69
ENSG00000183576.12 SETD3
log2(RPKM)
6.0
5.5
5.0
4.5
5.0
5.5
6.0
ENSG00000069345.11 DNAJA2
log2(RPKM)
z−score of IF value
−2 −1 0 1 2

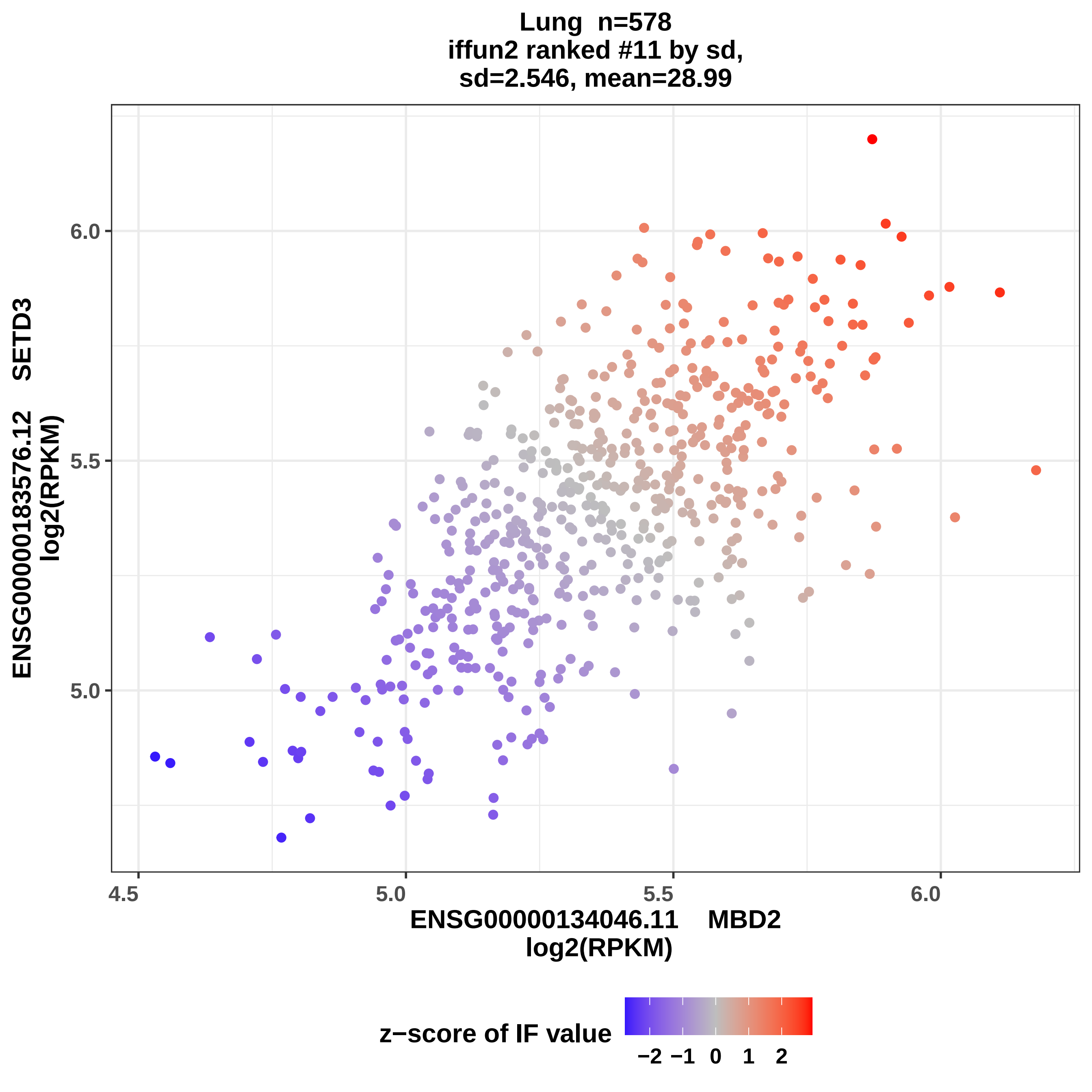

Lung n=578
iffun2 ranked #11 by sd,
sd=2.546, mean=28.99
ENSG00000183576.12 SETD3
log2(RPKM)
6.0
5.5
5.0
4.5
5.0
5.5
6.0
ENSG00000134046.11 MBD2
log2(RPKM)
z−score of IF value
−2
−1
0
1
2

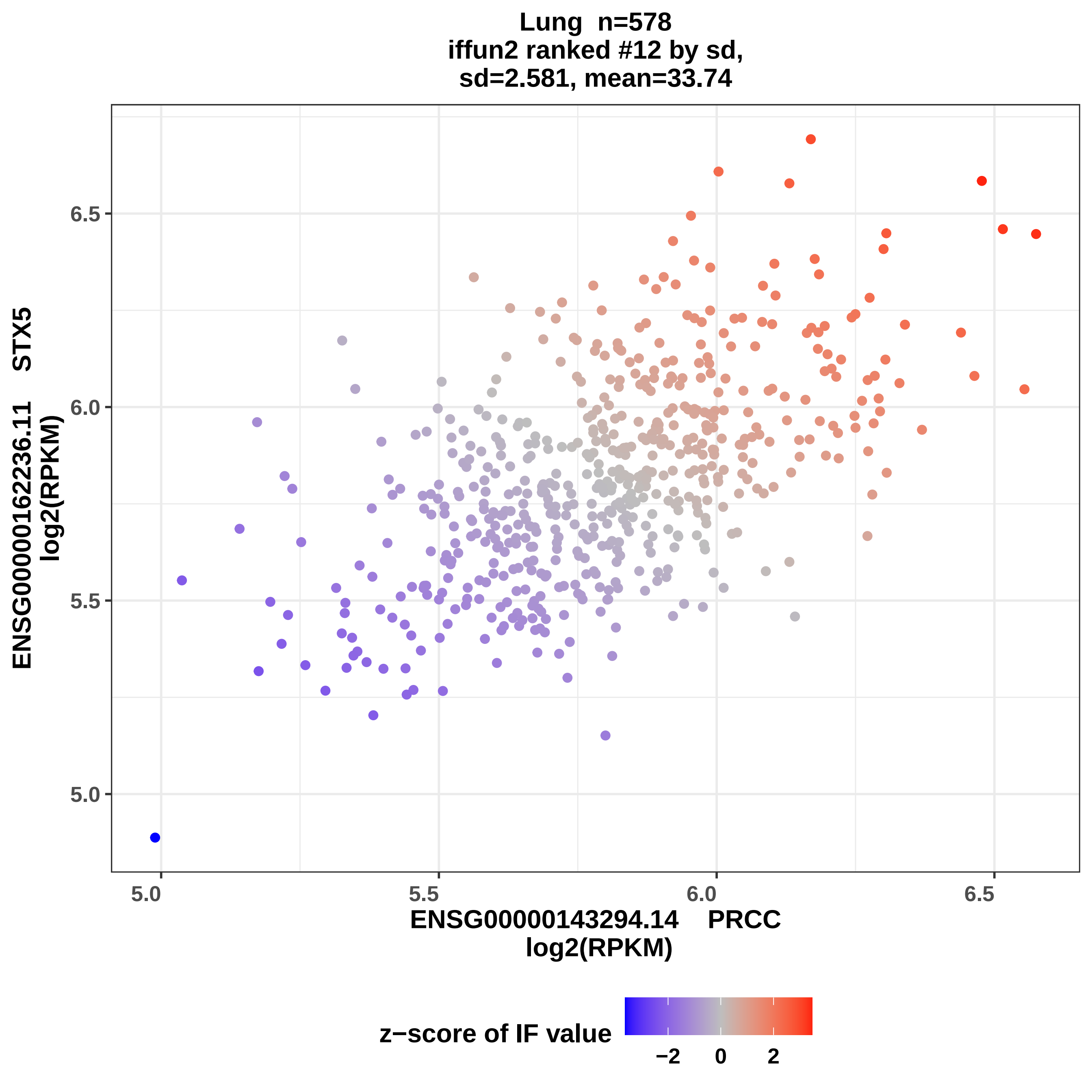

Lung n=578
iffun2 ranked #12 by sd,
sd=2.581, mean=33.74
ENSG00000162236.11 STX5
log2(RPKM)
6.5
6.0
5.5
5.0
ENSG00000143294.14 PRCC
log2(RPKM)
z−score of IF value
−2
0
2

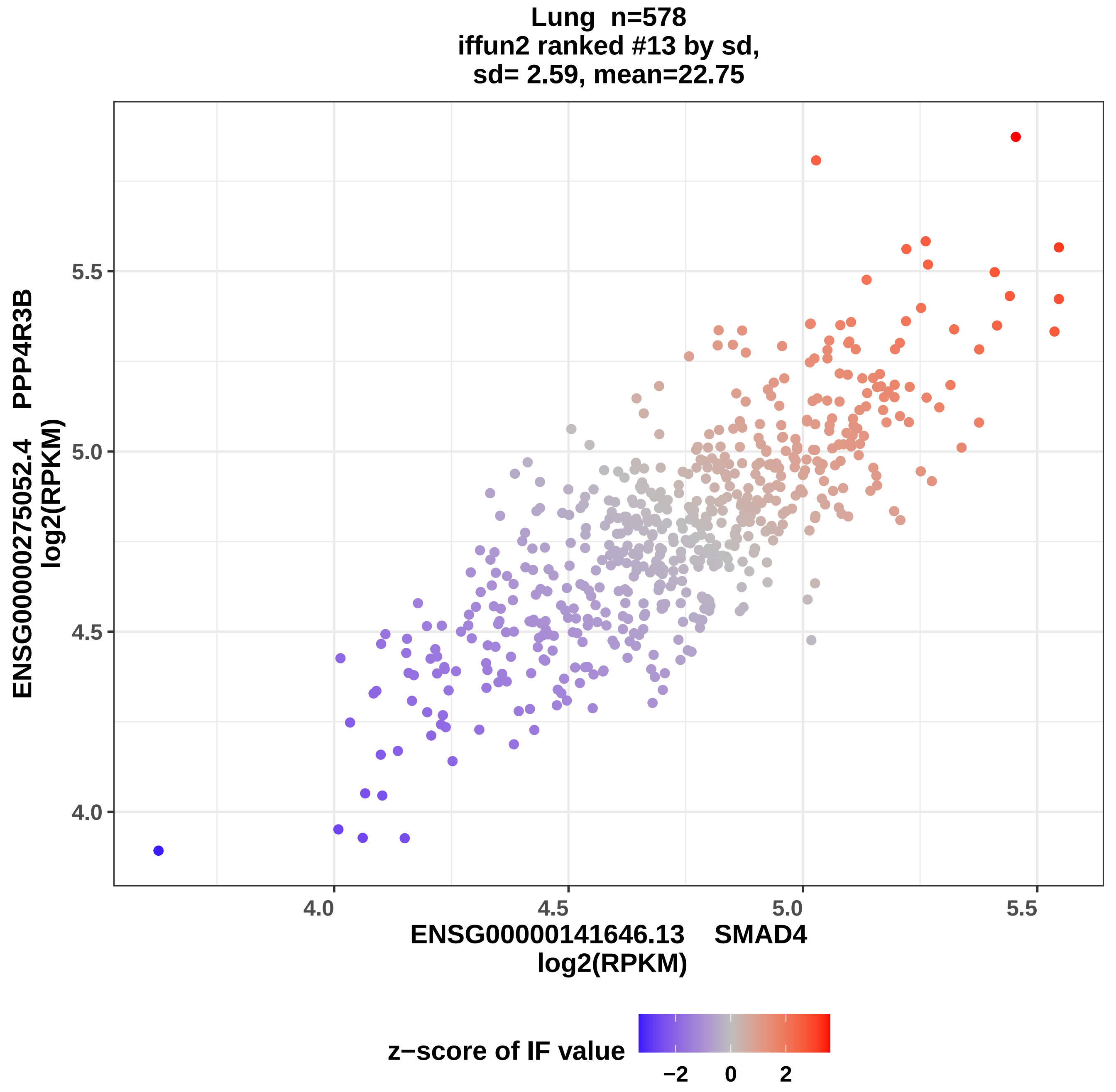

Lung n=578
iffun2 ranked #13 by sd,
sd= 2.59, mean=22.75
5.5
5.0
4.5
4.0
ENSG00000275052.4 PPP4R3B
log2(RPKM)
4.0
4.5
5.0
5.5
ENSG00000141646.13 SMAD4
log2(RPKM)
z−score of IF value
−2
0
2

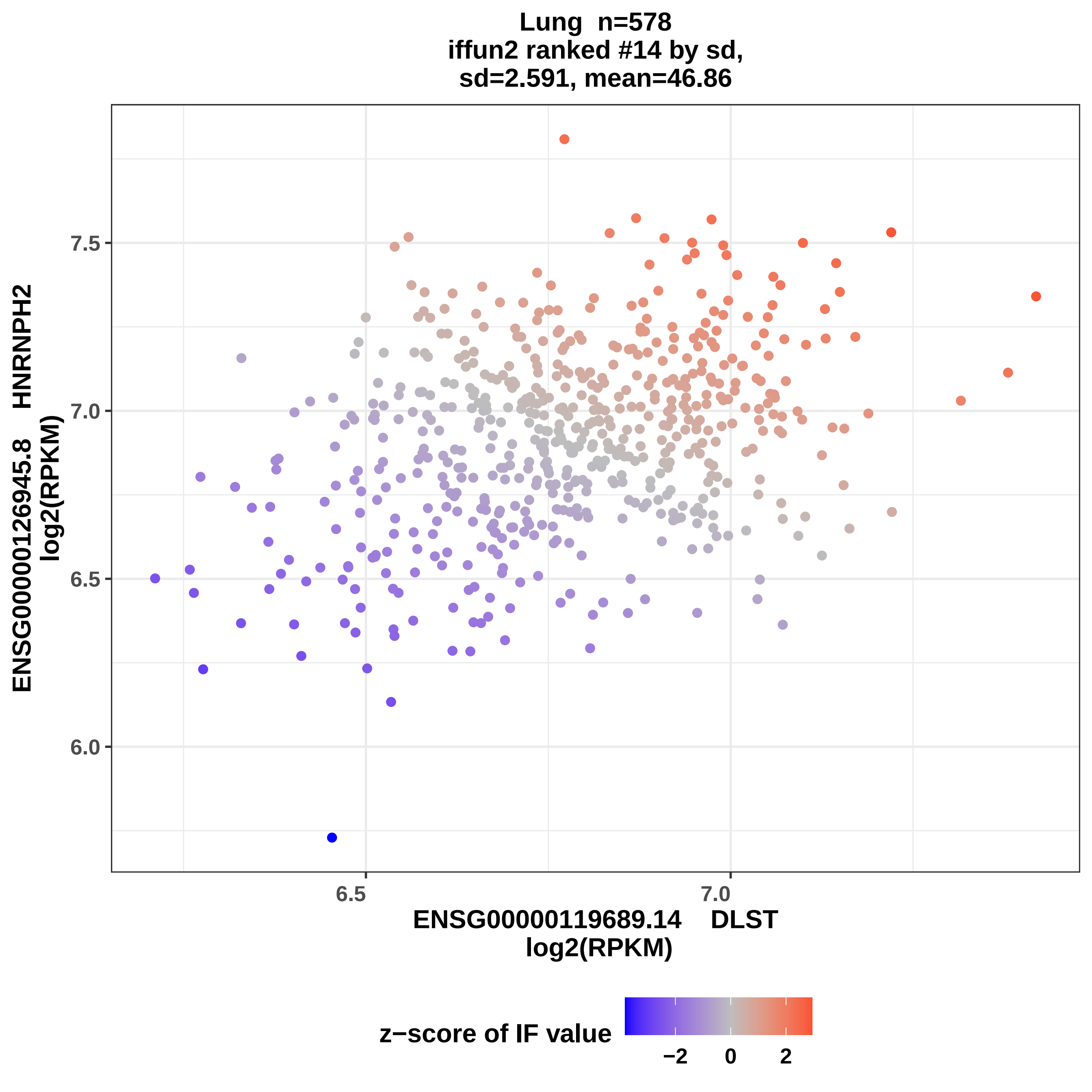
Lung n=578
iffun2 ranked #14 by sd,
sd=2.591, mean=46.86
ENSG00000126945.8 HNRNPH2
log2(RPKM)
7.5
7.0
6.5
6.0
6.5
7.0
ENSG00000119689.14 DLST
log2(RPKM)
z−score of IF value
−2
0
2

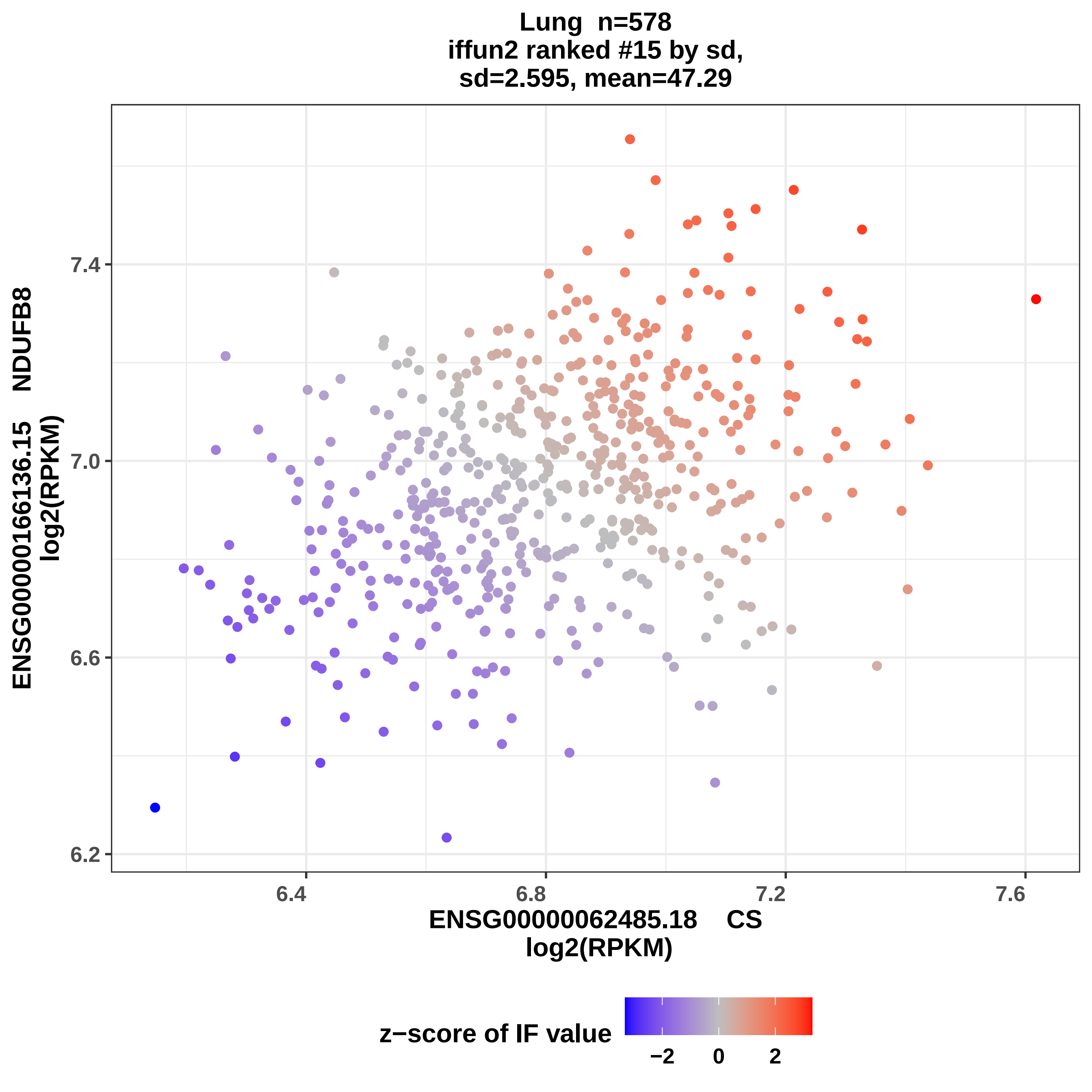
Lung n=578
iffun2 ranked #15 by sd,
sd=2.595, mean=47.29
ENSG00000166136.15 NDUFB8
log2(RPKM)
7.4
7.0
6.6
6.2
6.4
6.8
7.2
7.6
ENSG00000062485.18 CS
log2(RPKM)
z−score of IF value
−2
0
2

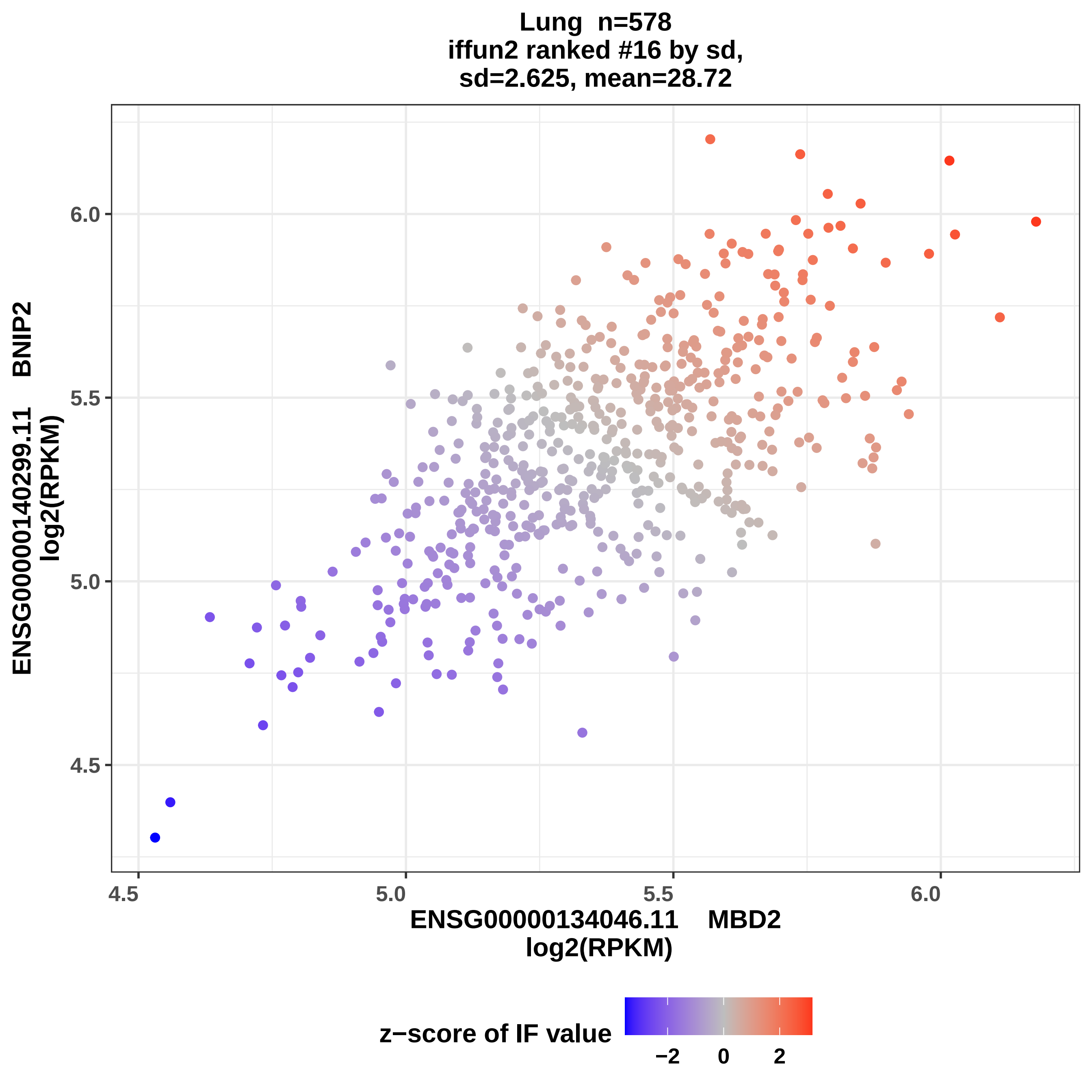

Lung n=578
iffun2 ranked #16 by sd,
sd=2.625, mean=28.72
ENSG00000140299.11 BNIP2
log2(RPKM)
6.0
5.5
5.0
4.5
4.5
5.0
5.5
6.0
ENSG00000134046.11 MBD2
log2(RPKM)
z−score of IF value
−2
0
2

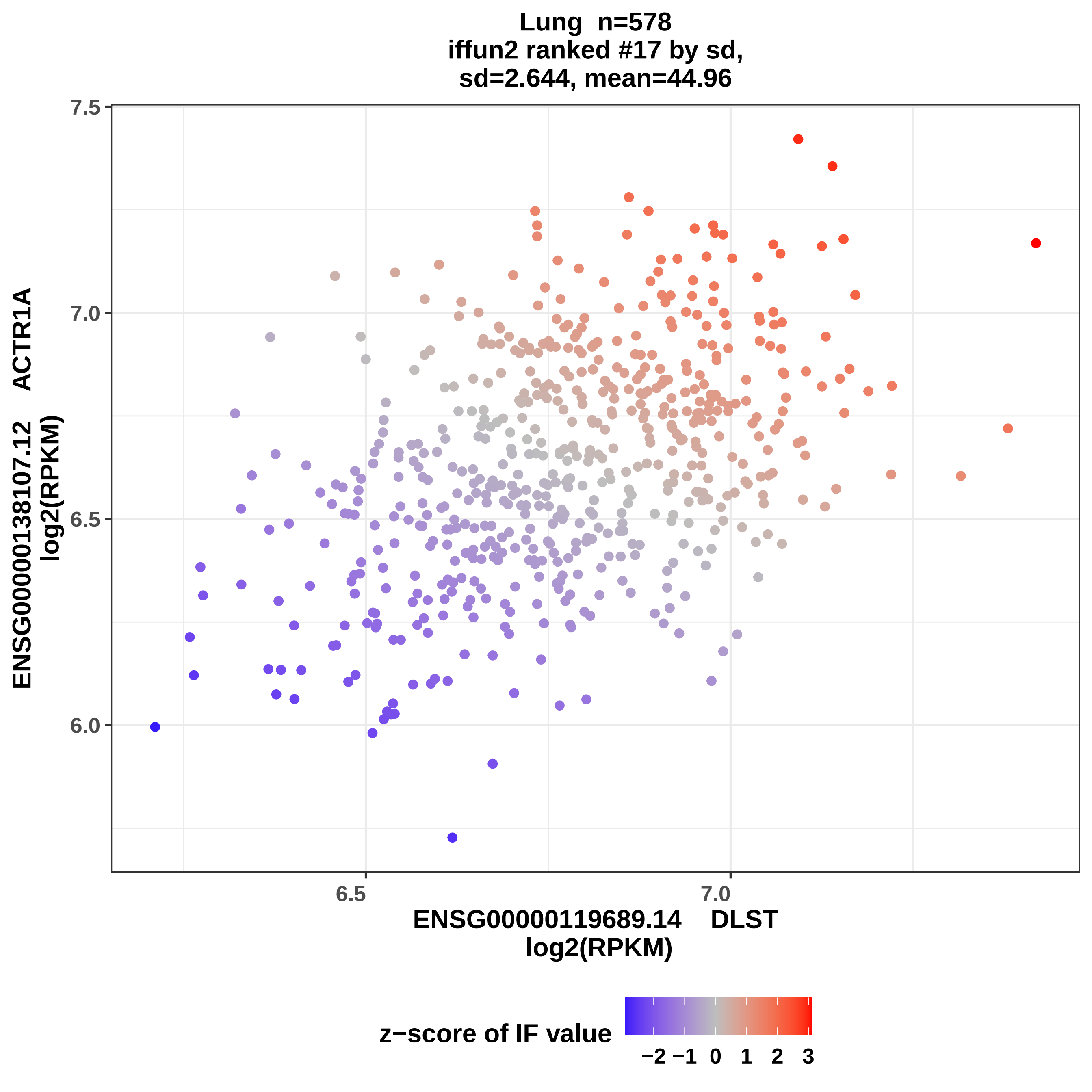
Lung n=578
iffun2 ranked #17 by sd,
sd=2.644, mean=44.96
ENSG00000138107.12 ACTR1A
log2(RPKM)
7.5
7.0
6.5
6.0
6.5
7.0
ENSG00000119689.14 DLST
log2(RPKM)
z−score of IF value
−2 −1 0 1 2 3

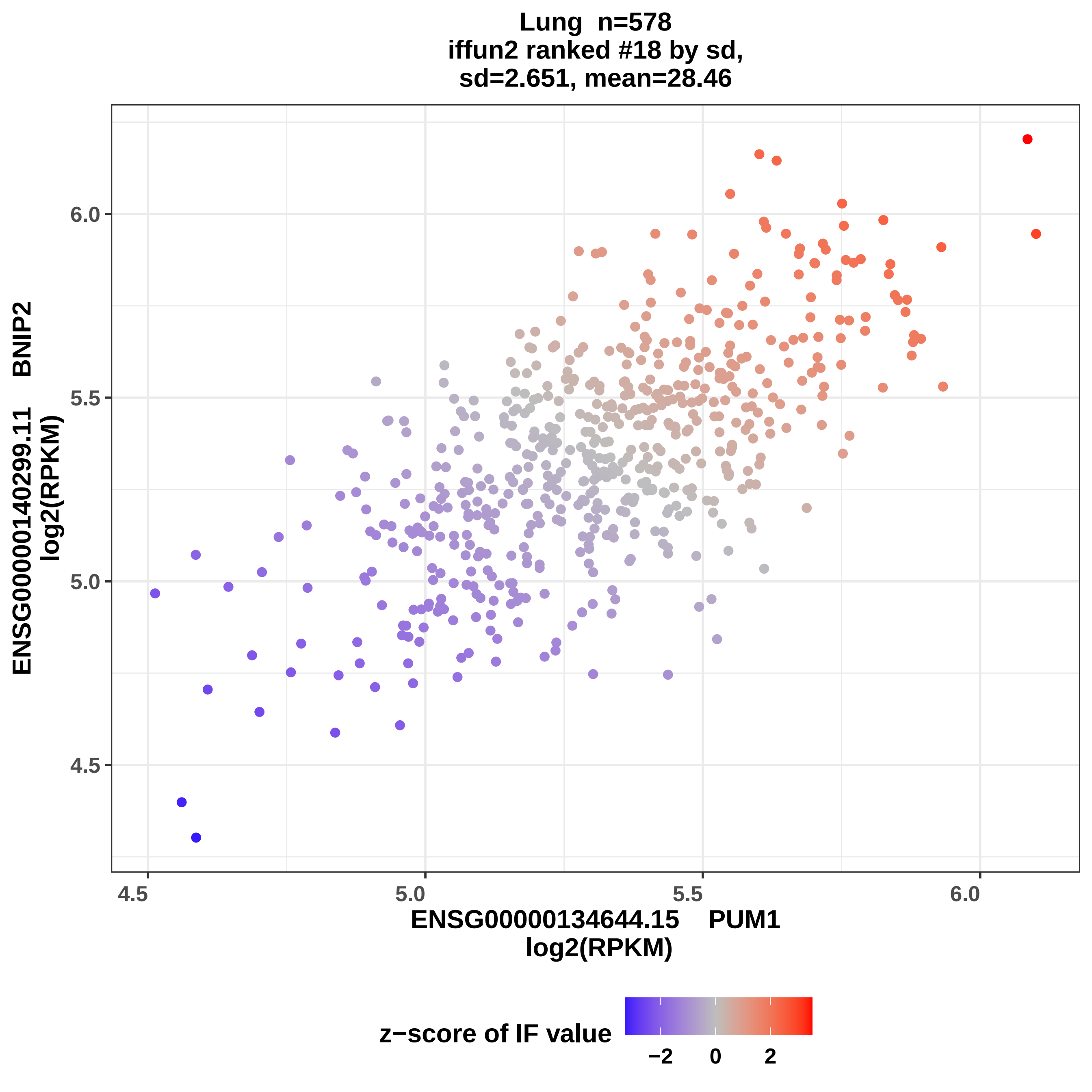
Lung n=578
iffun2 ranked #18 by sd,
sd=2.651, mean=28.46
ENSG00000140299.11 BNIP2
log2(RPKM)
ENSG00000134644.15 PUM1
log2(RPKM)
4.5
5.0
5.5
6.0
4.5
5.0
5.5
6.0
z−score of IF value
−2
0
2

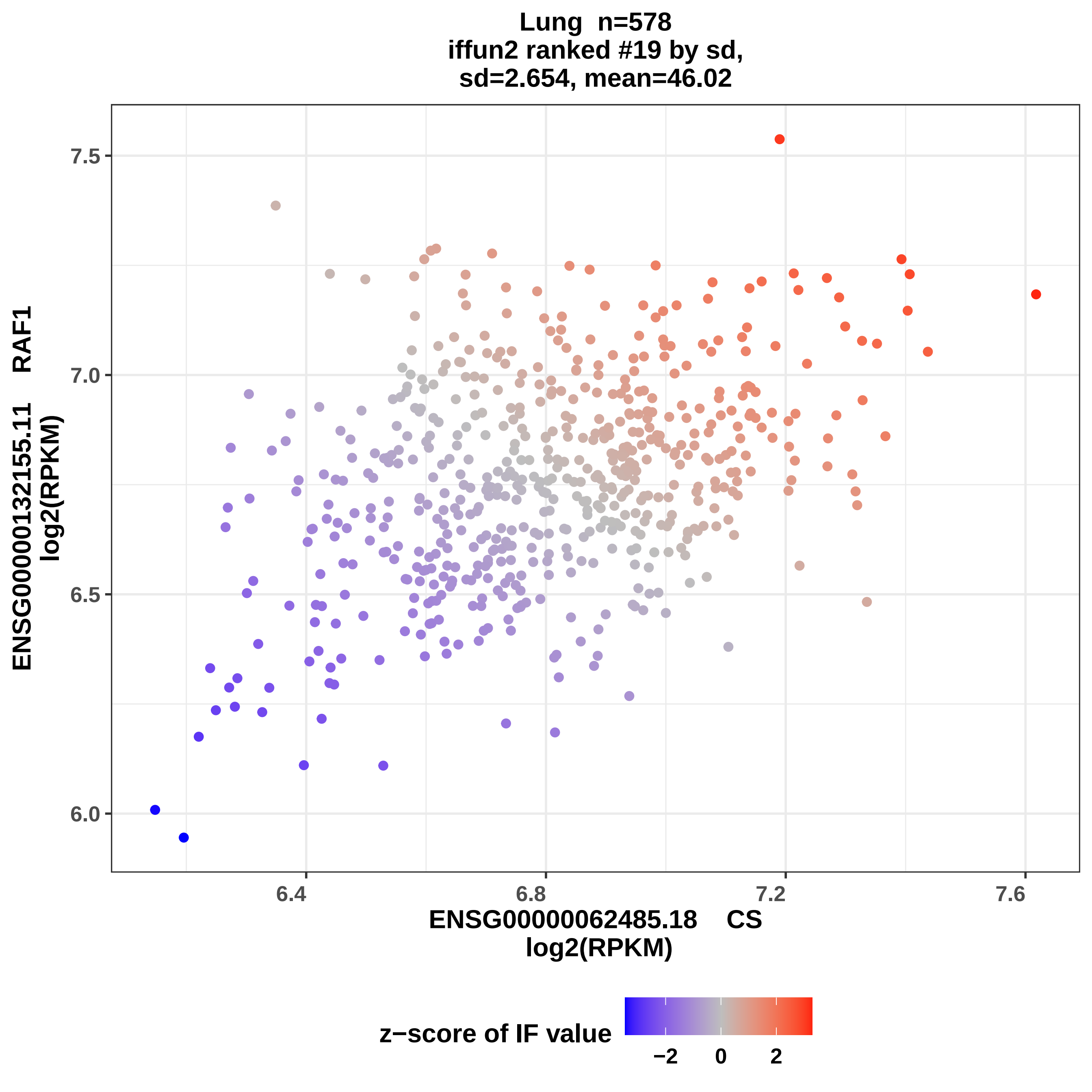

Lung n=578
iffun2 ranked #19 by sd,
sd=2.654, mean=46.02
ENSG00000132155.11 RAF1
log2(RPKM)
7.5
7.0
6.5
6.0
6.4
6.8
7.2
7.6
ENSG00000062485.18 CS
log2(RPKM)
z−score of IF value
−2
0
2

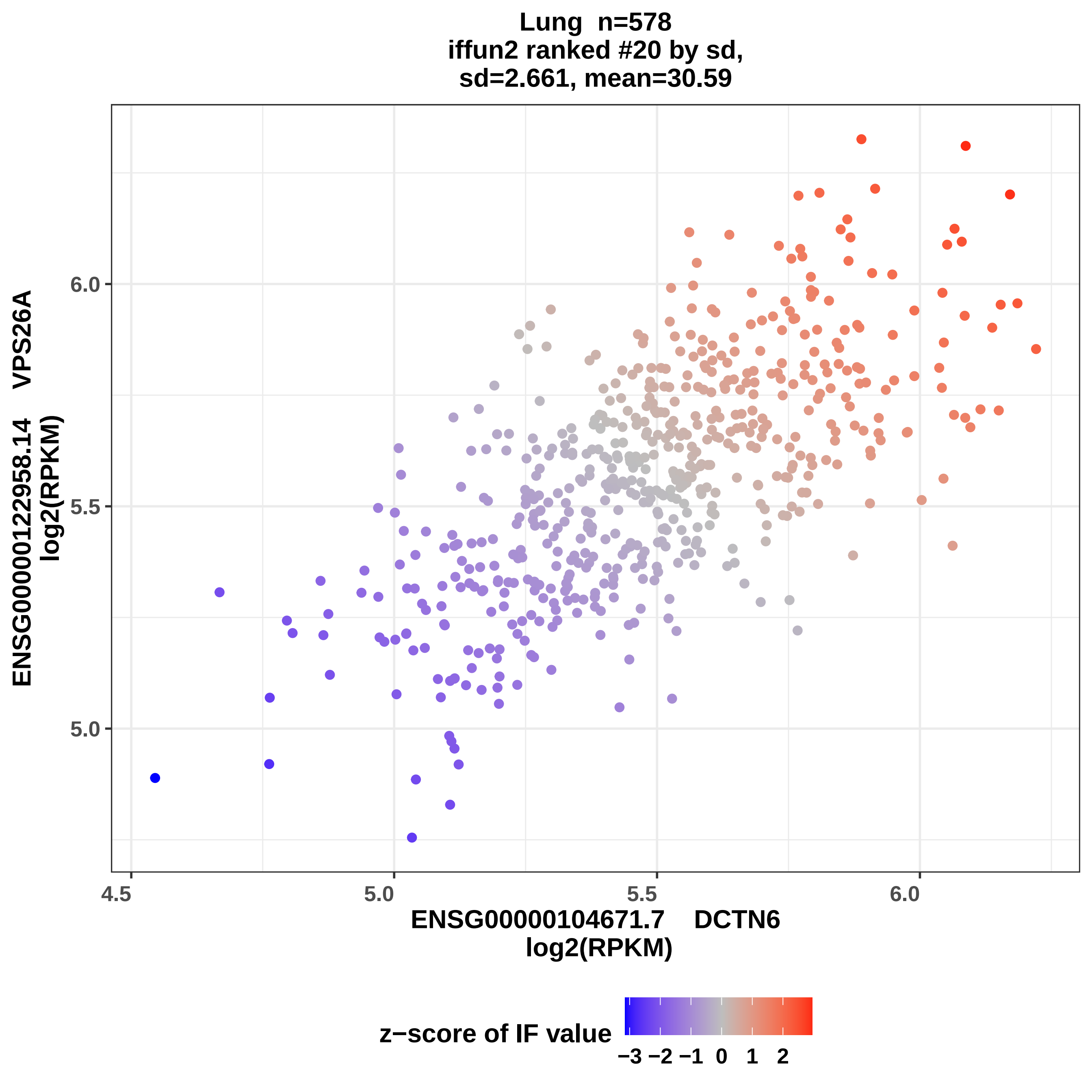

Lung n=578
iffun2 ranked #20 by sd,
sd=2.661, mean=30.59
ENSG00000122958.14 VPS26A
log2(RPKM)
6.0
5.5
5.0
4.5
5.0
5.5
6.0
ENSG00000104671.7 DCTN6
log2(RPKM)
z−score of IF value
−3 −2 −1 0 1 2

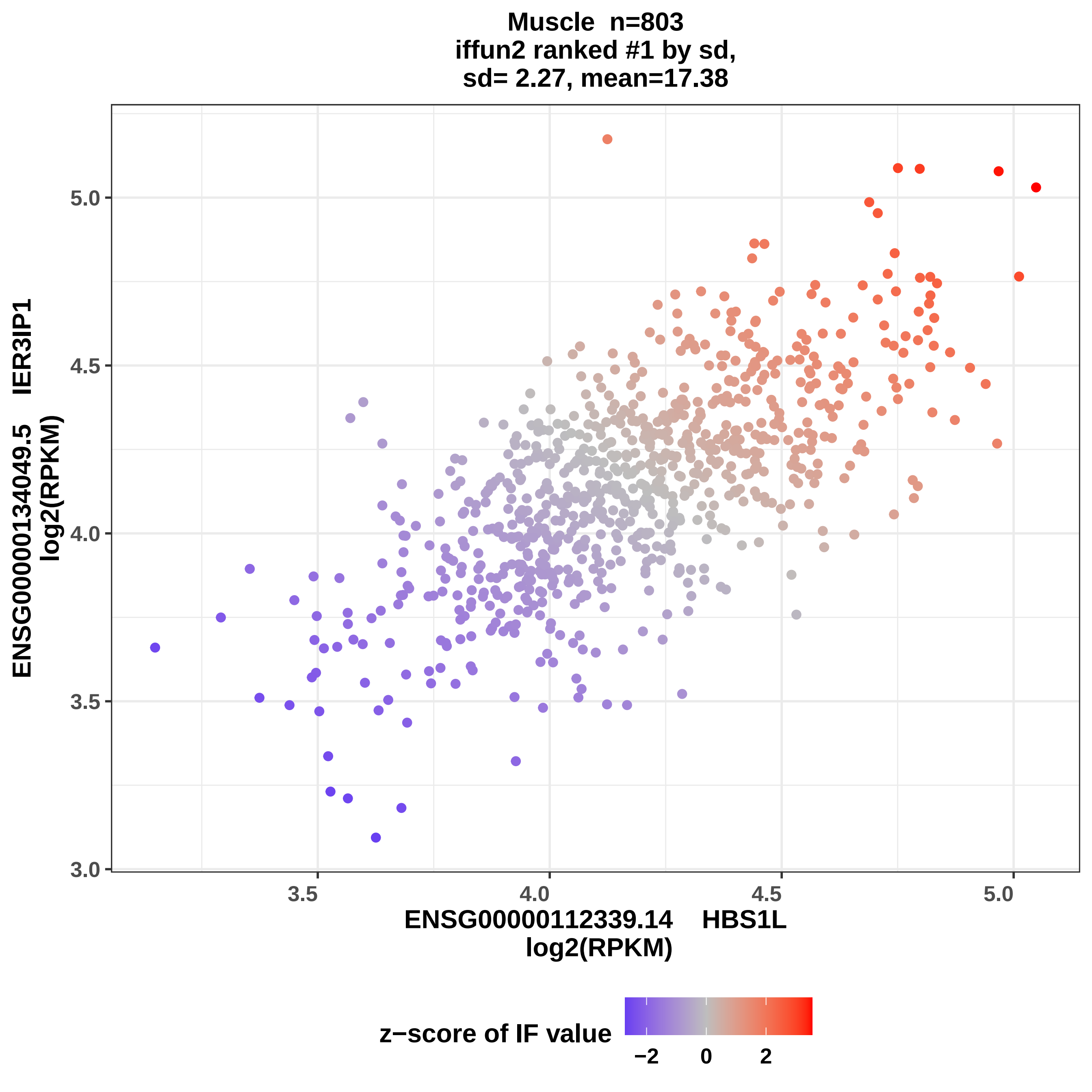

Muscle n=803
iffun2 ranked #1 by sd,
sd= 2.27, mean=17.38
ENSG00000134049.5 IER3IP1
log2(RPKM)
5.0
4.5
4.0
3.5
3.0
3.5
4.0
4.5
5.0
ENSG00000112339.14 HBS1L
log2(RPKM)
z−score of IF value
−2
0
2

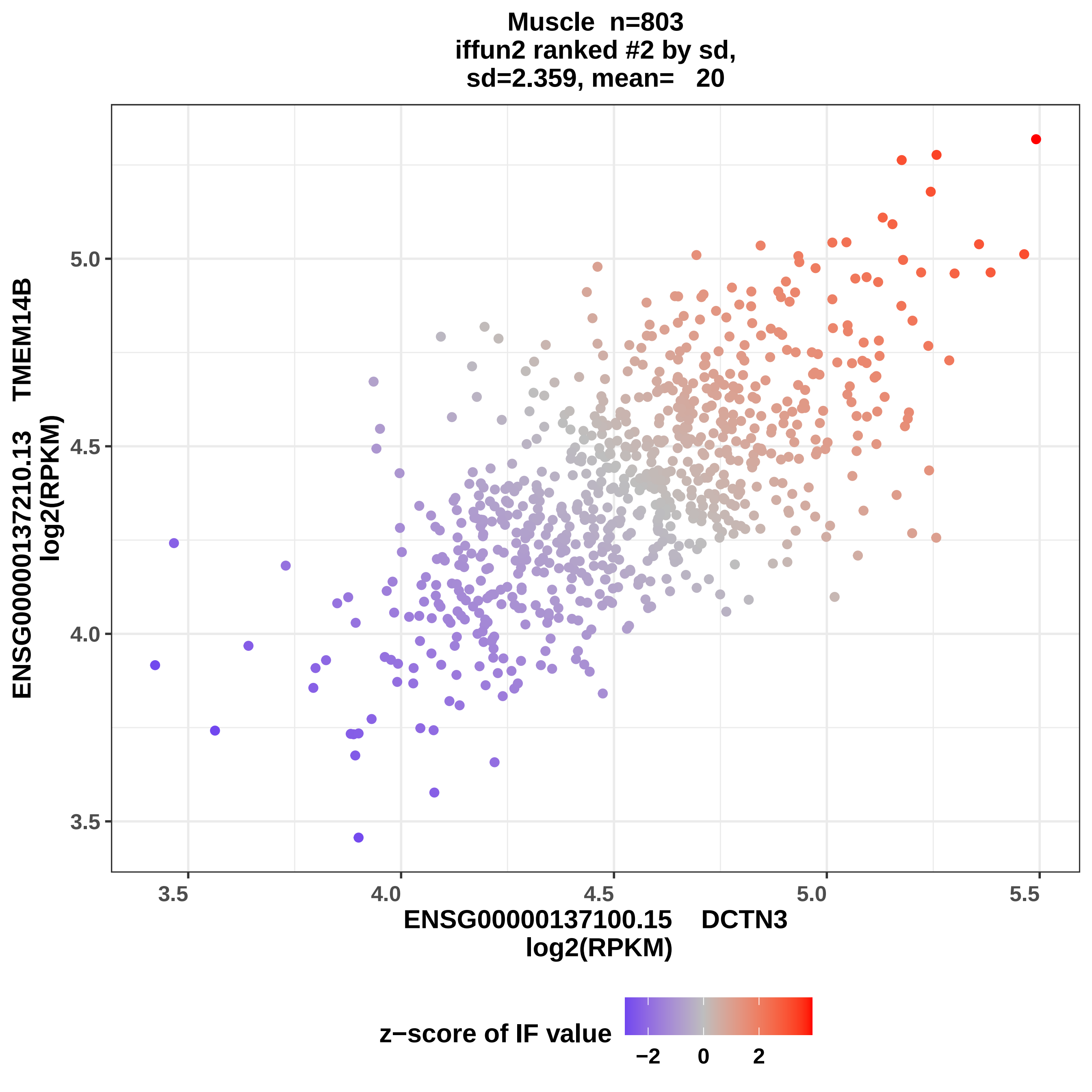

Muscle n=803
iffun2 ranked #2 by sd,
sd=2.359, mean= 20
ENSG00000137210.13 TMEM14B
log2(RPKM)
5.0
4.5
4.0
3.5
3.5
4.0
4.5
5.0
5.5
ENSG00000137100.15 DCTN3
log2(RPKM)
z−score of IF value
−2
0
2

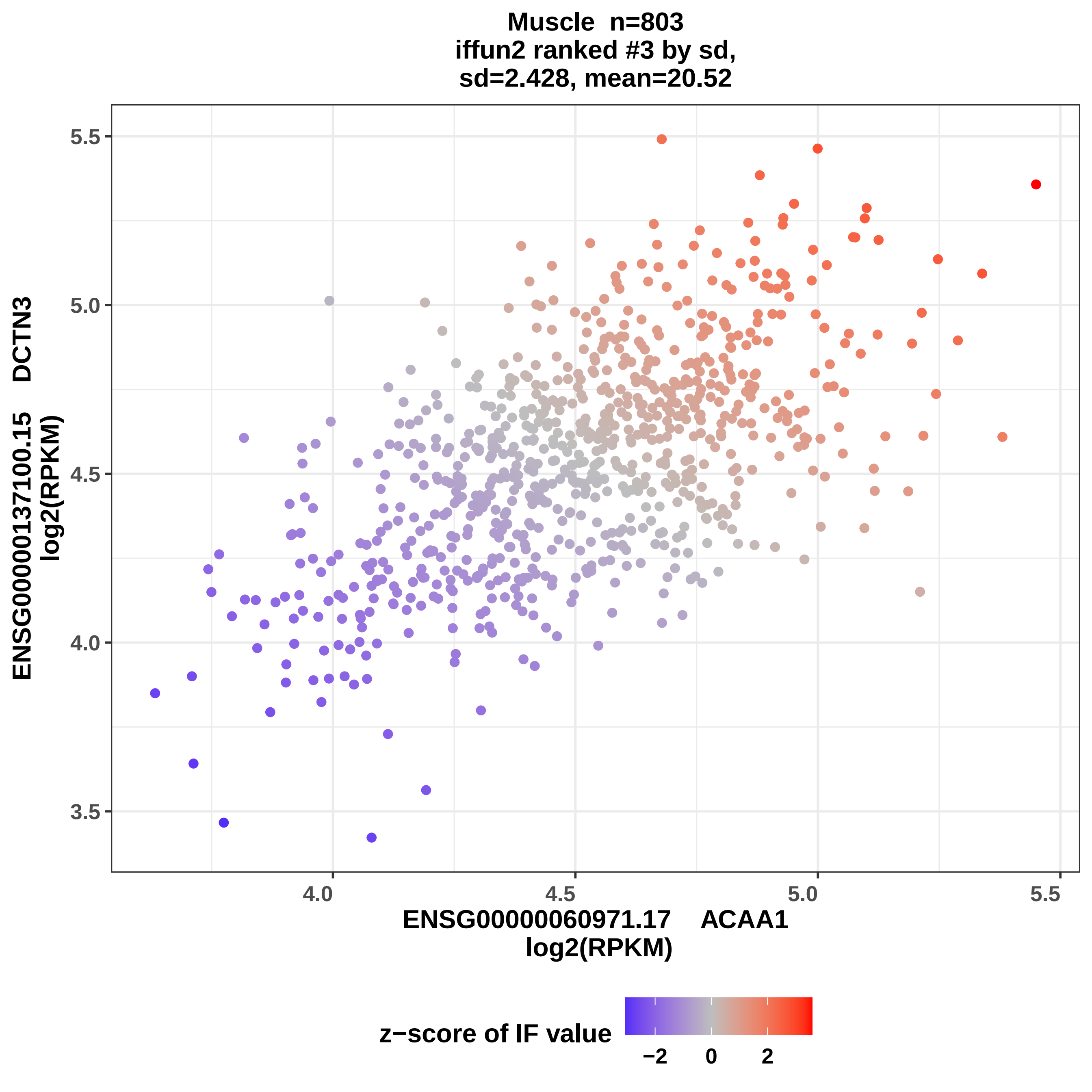
Muscle n=803
iffun2 ranked #3 by sd,
sd=2.428, mean=20.52
ENSG00000137100.15 DCTN3
log2(RPKM)
5.5
5.0
4.5
4.0
3.5
4.0
4.5
5.0
5.5
ENSG00000060971.17 ACAA1
log2(RPKM)
z−score of IF value
−2
0
2

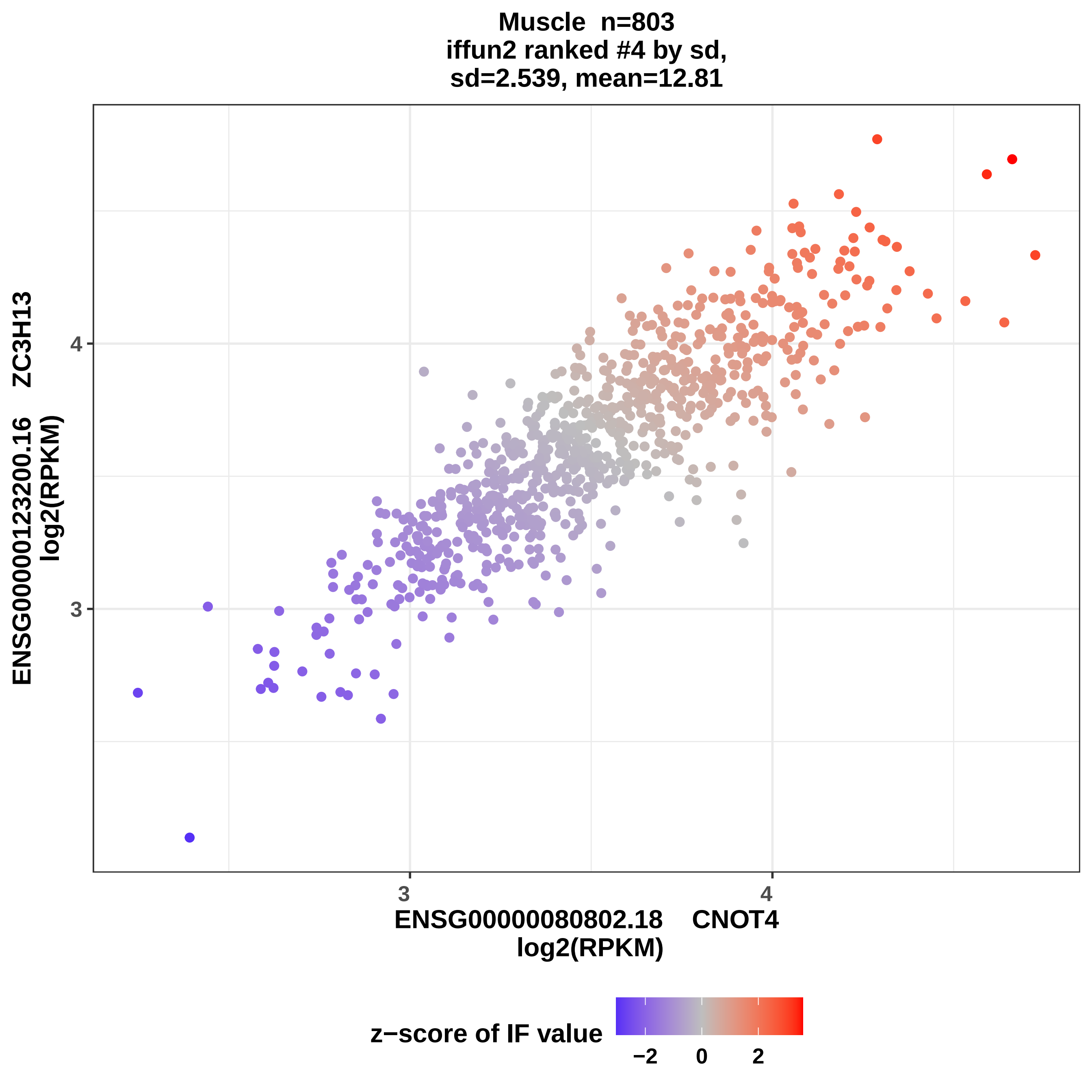
Muscle n=803
iffun2 ranked #4 by sd,
sd=2.539, mean=12.81
ENSG00000123200.16 ZC3H13
log2(RPKM)
4
3
3
4
ENSG00000080802.18 CNOT4
log2(RPKM)
z−score of IF value
−2
0
2

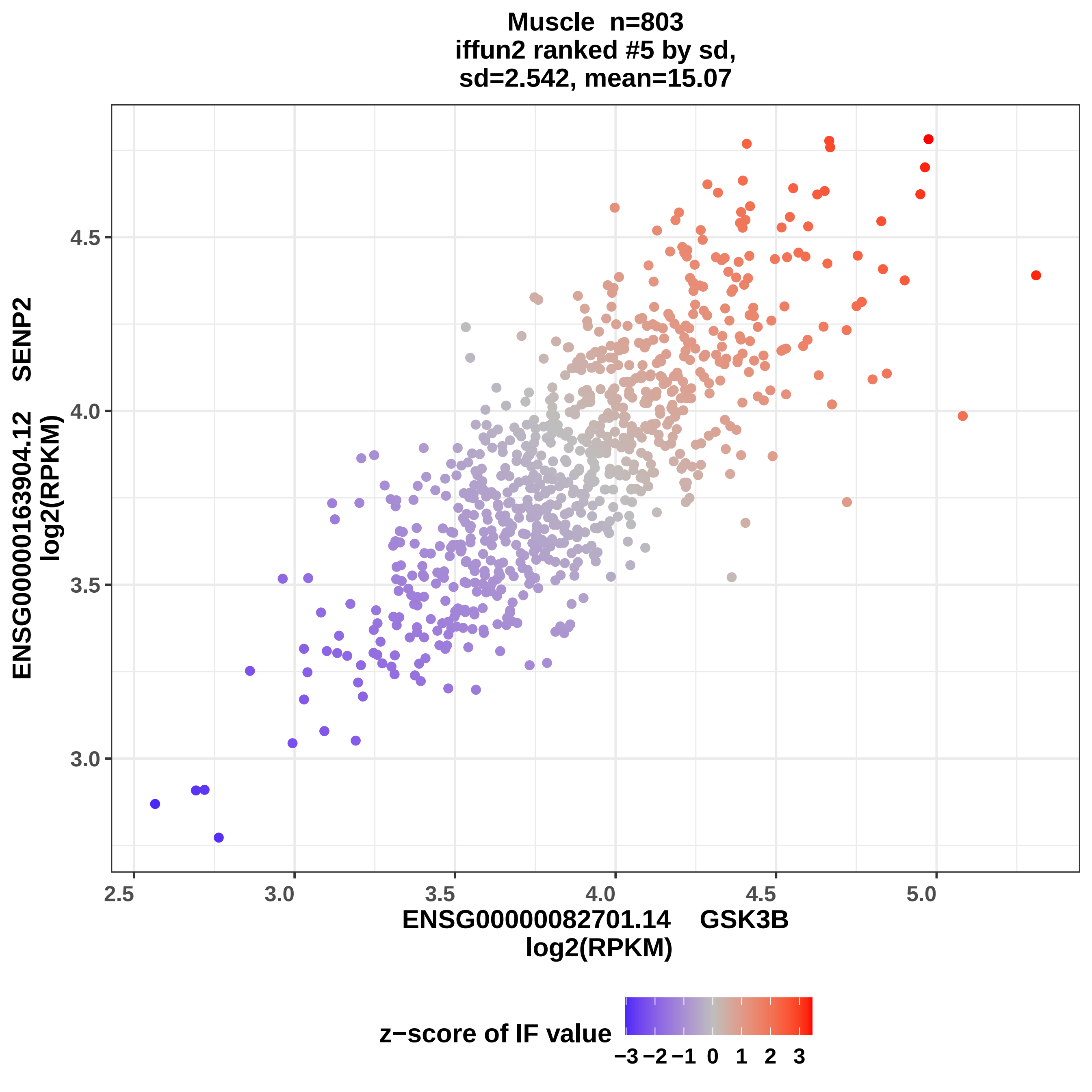

Muscle n=803
iffun2 ranked #5 by sd,
sd=2.542, mean=15.07
ENSG00000163904.12 SENP2
log2(RPKM)
4.5
4.0
3.5
3.0
2.5
3.0
3.5
4.0
4.5
5.0
ENSG00000082701.14 GSK3B
log2(RPKM)
z−score of IF value
−3 −2 −1 0 1 2 3

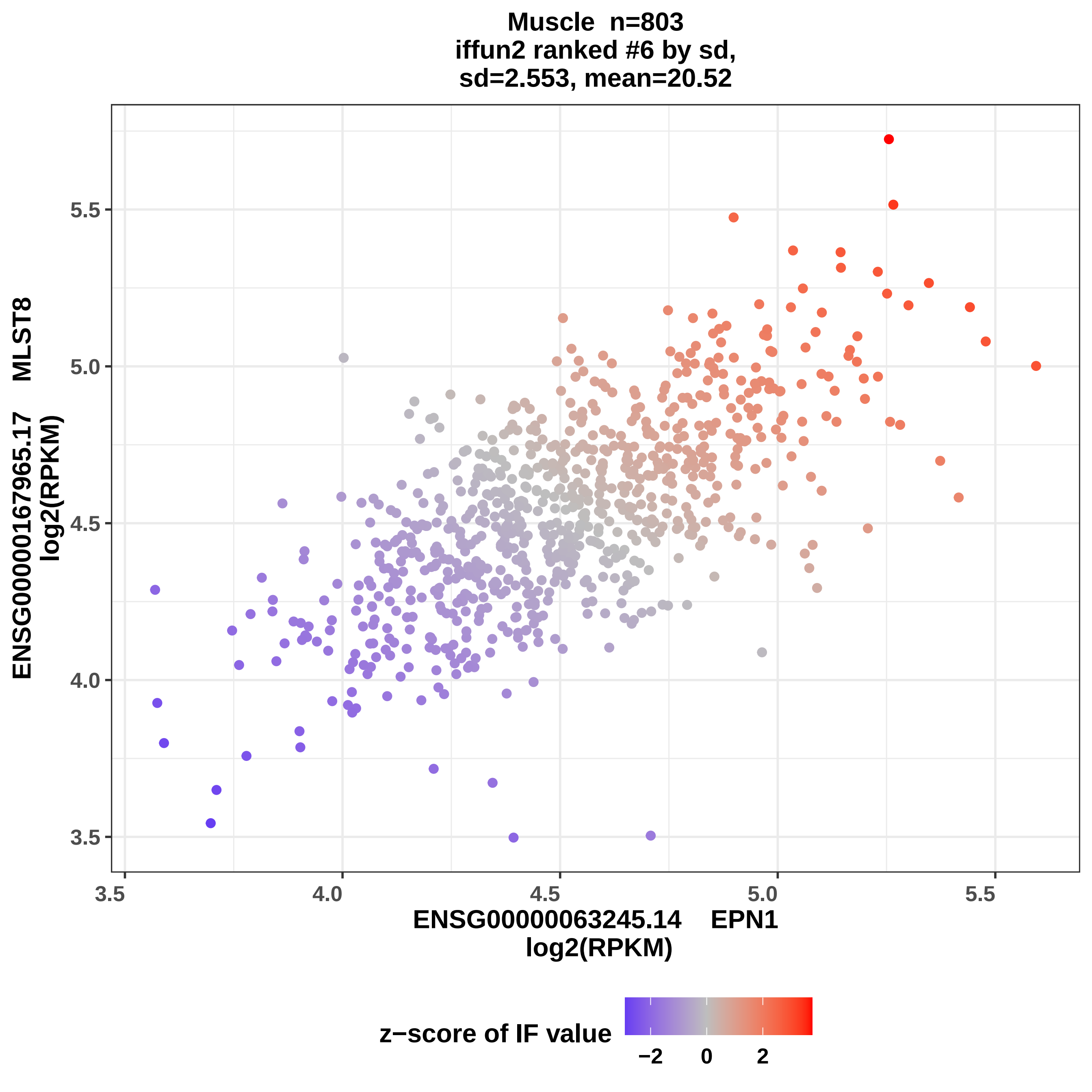
Muscle n=803
iffun2 ranked #6 by sd,
sd=2.553, mean=20.52
ENSG00000167965.17 MLST8
log2(RPKM)
5.5
5.0
4.5
4.0
3.5
3.5
4.0
4.5
5.0
5.5
ENSG00000063245.14 EPN1
log2(RPKM)
z−score of IF value
−2
0
2

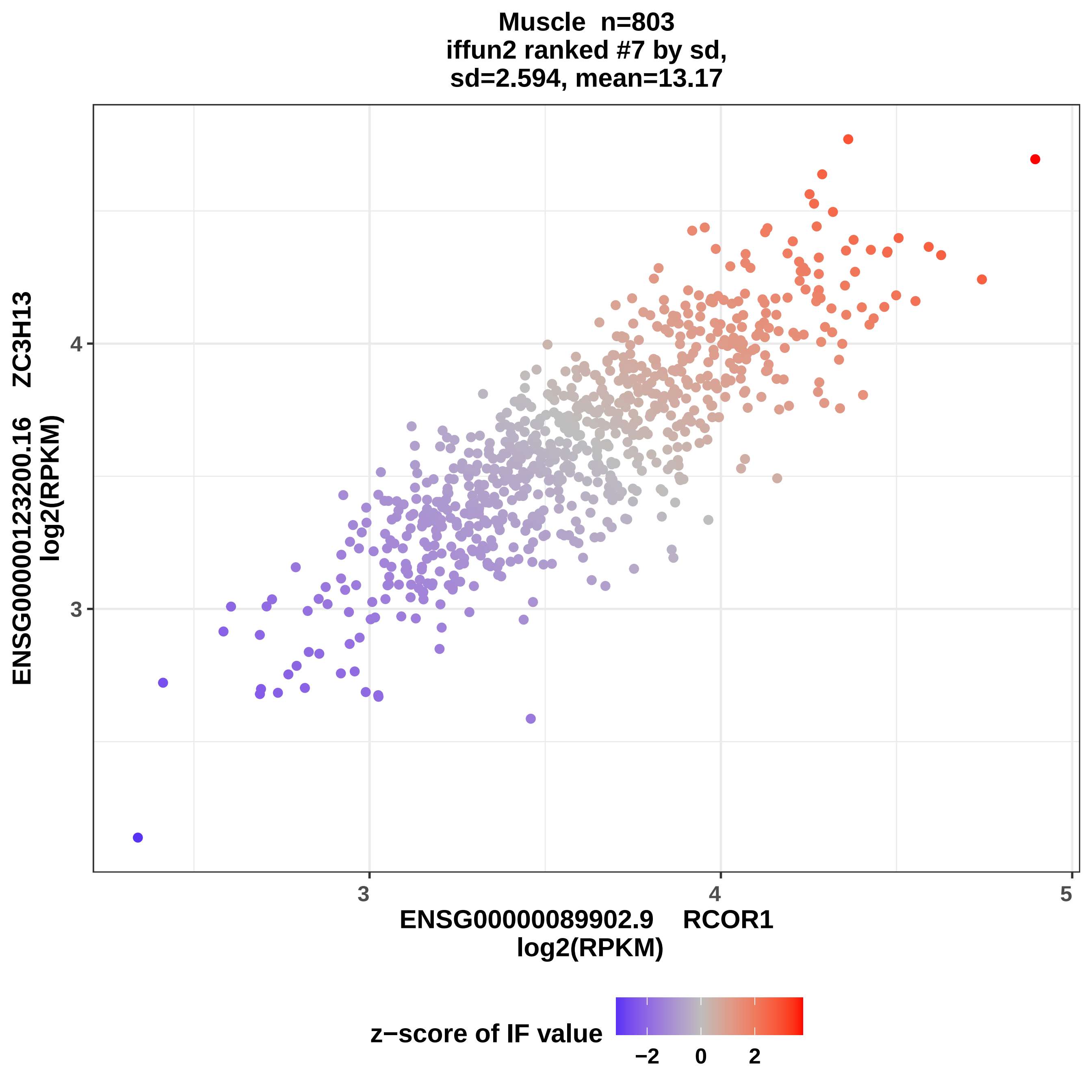
Muscle n=803
iffun2 ranked #7 by sd,
sd=2.594, mean=13.17
ENSG00000123200.16 ZC3H13
log2(RPKM)
4
3
3
4
5
ENSG00000089902.9 RCOR1
log2(RPKM)
z−score of IF value
−2
0
2

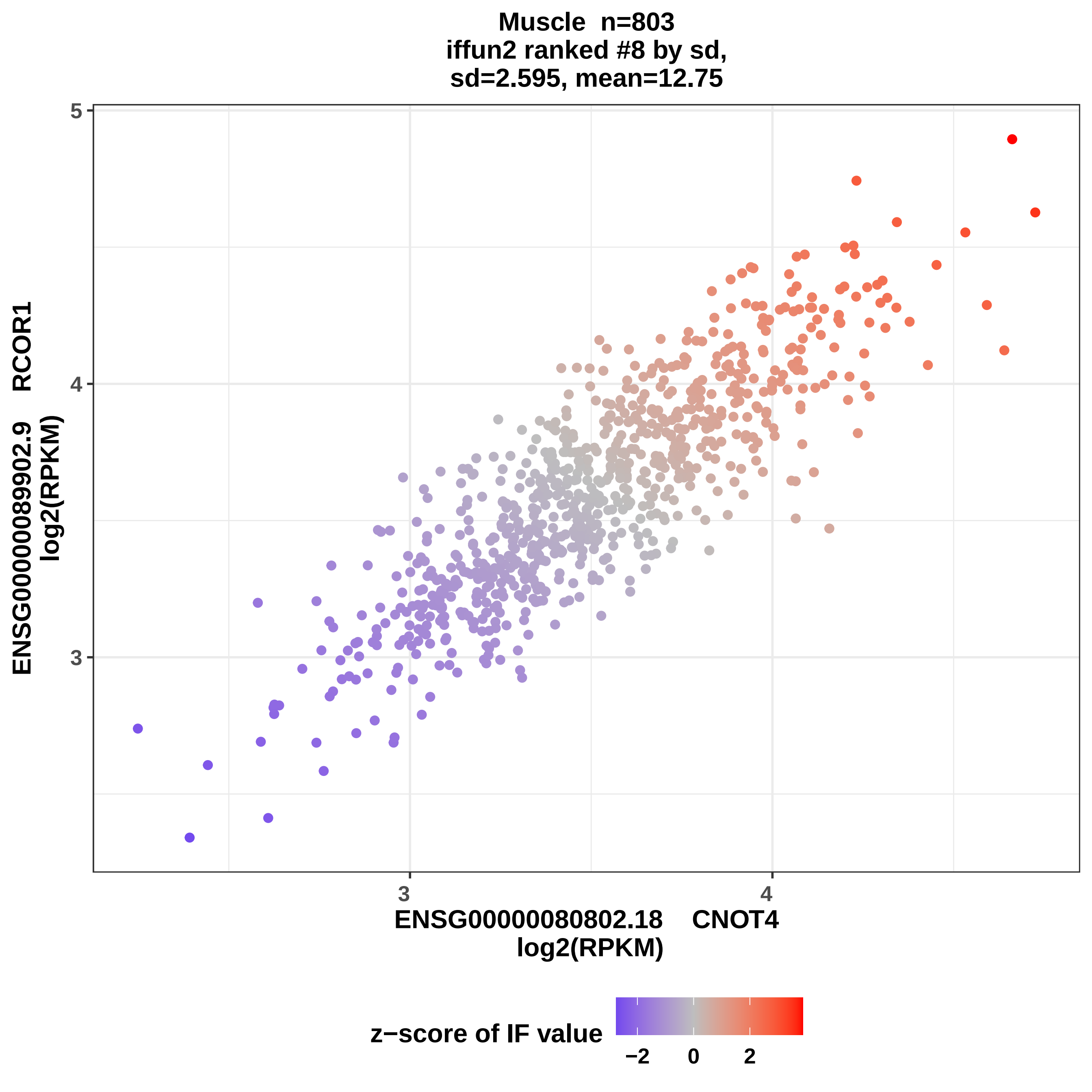

Muscle n=803
iffun2 ranked #8 by sd,
sd=2.595, mean=12.75
5
4
3
3
4
ENSG00000080802.18 CNOT4
log2(RPKM)
ENSG00000089902.9 RCOR1
log2(RPKM)
z−score of IF value
−2
0
2

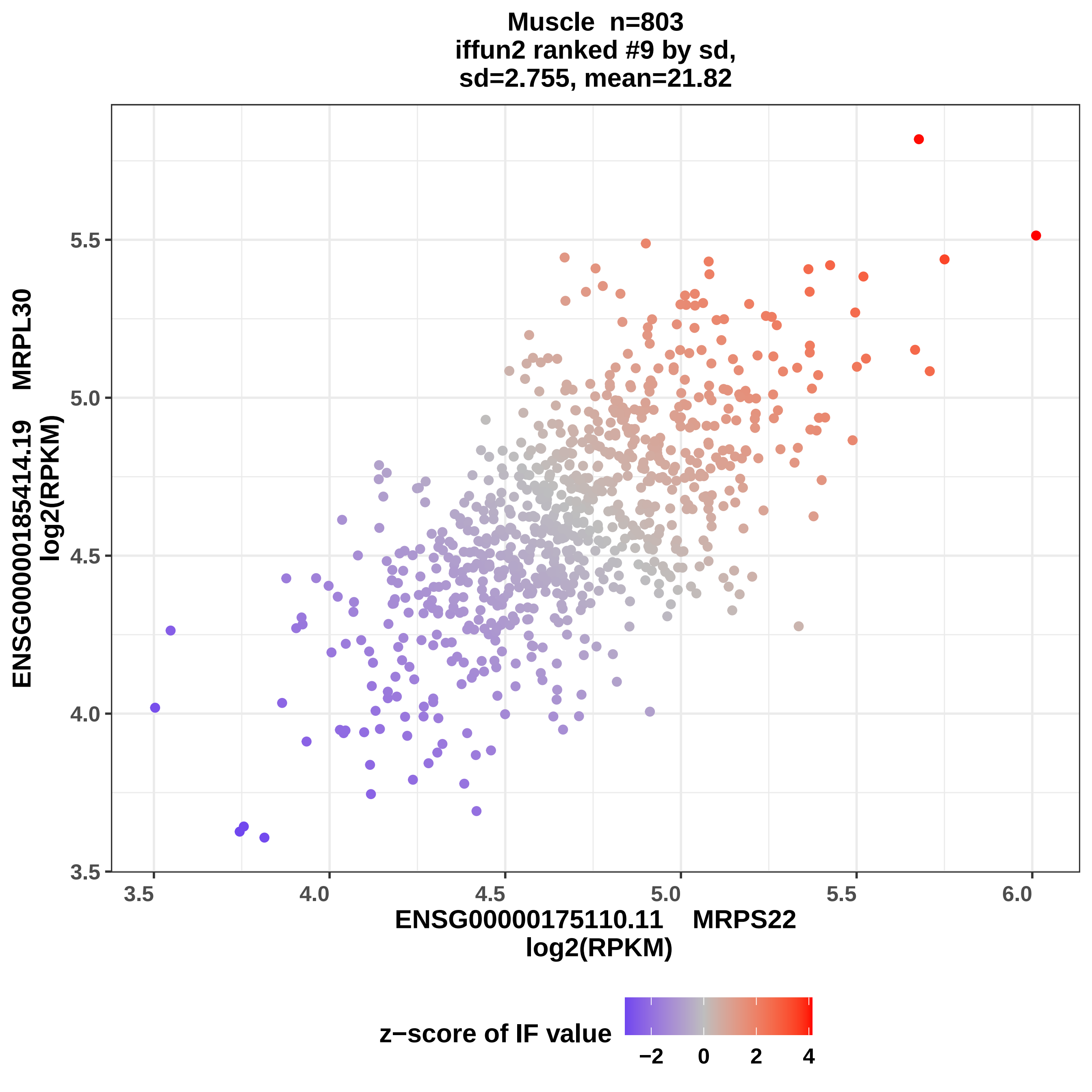
Muscle n=803
iffun2 ranked #9 by sd,
sd=2.755, mean=21.82
ENSG00000185414.19 MRPL30
log2(RPKM)
3.5
4.0
4.5
5.0
5.5
3.5
4.0
4.5
5.0
5.5
6.0
ENSG00000175110.11 MRPS22
log2(RPKM)
z−score of IF value
−2
0
2
4

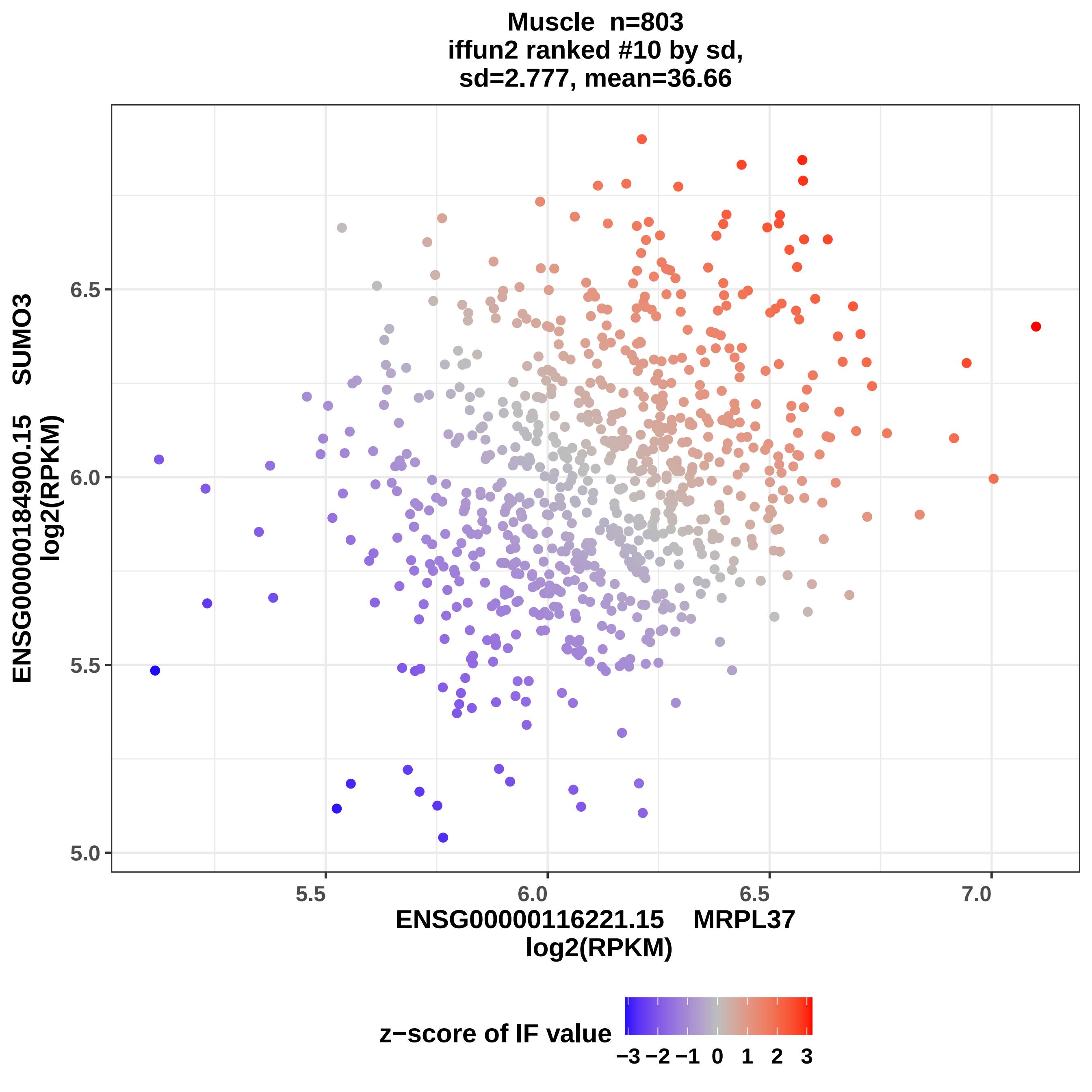
Muscle n=803
iffun2 ranked #10 by sd,
sd=2.777, mean=36.66
ENSG00000184900.15 SUMO3
log2(RPKM)
6.5
6.0
5.5
5.0
5.5
6.0
6.5
7.0
ENSG00000116221.15 MRPL37
log2(RPKM)
z−score of IF value
−3 −2 −1 0 1 2 3

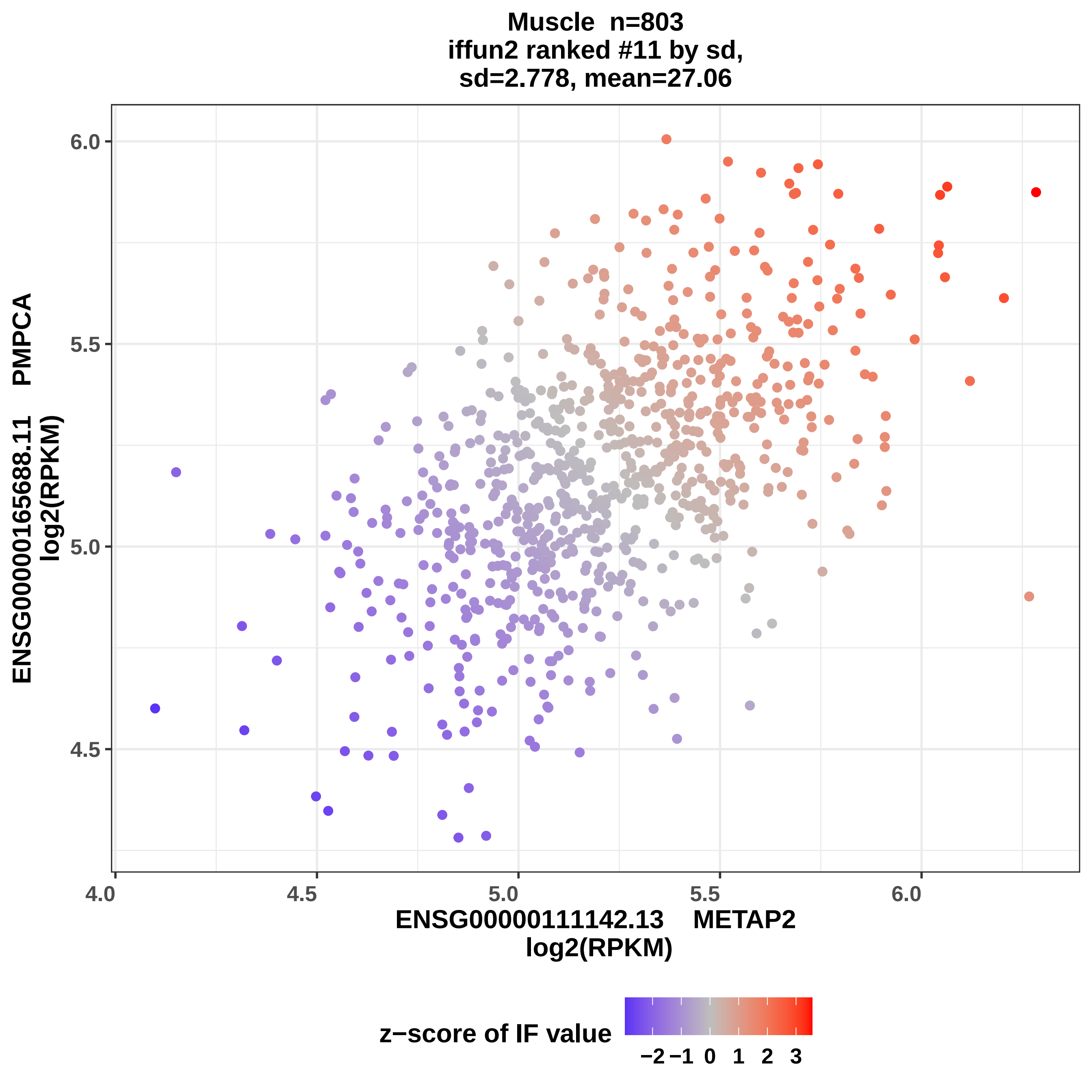

Muscle n=803
iffun2 ranked #11 by sd,
sd=2.778, mean=27.06
ENSG00000165688.11 PMPCA
log2(RPKM)
6.0
5.5
5.0
4.5
4.0
4.5
5.0
5.5
6.0
ENSG00000111142.13 METAP2
log2(RPKM)
z−score of IF value
−2
−1
0
1
2
3

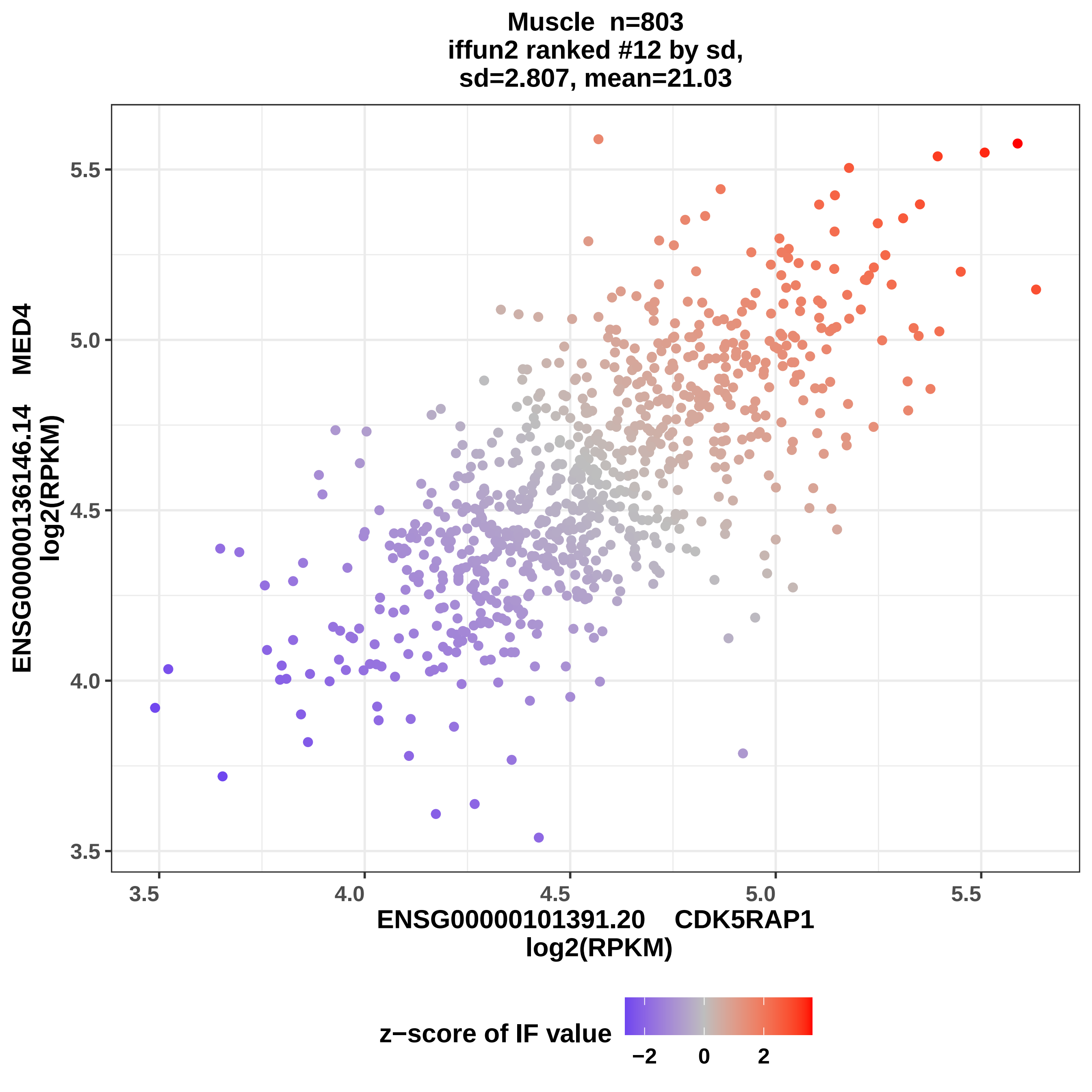

Muscle n=803
iffun2 ranked #12 by sd,
sd=2.807, mean=21.03
ENSG00000136146.14 MED4
log2(RPKM)
5.5
5.0
4.5
4.0
3.5
3.5
4.0
4.5
5.0
5.5
ENSG00000101391.20 CDK5RAP1
log2(RPKM)
z−score of IF value
−2
0
2

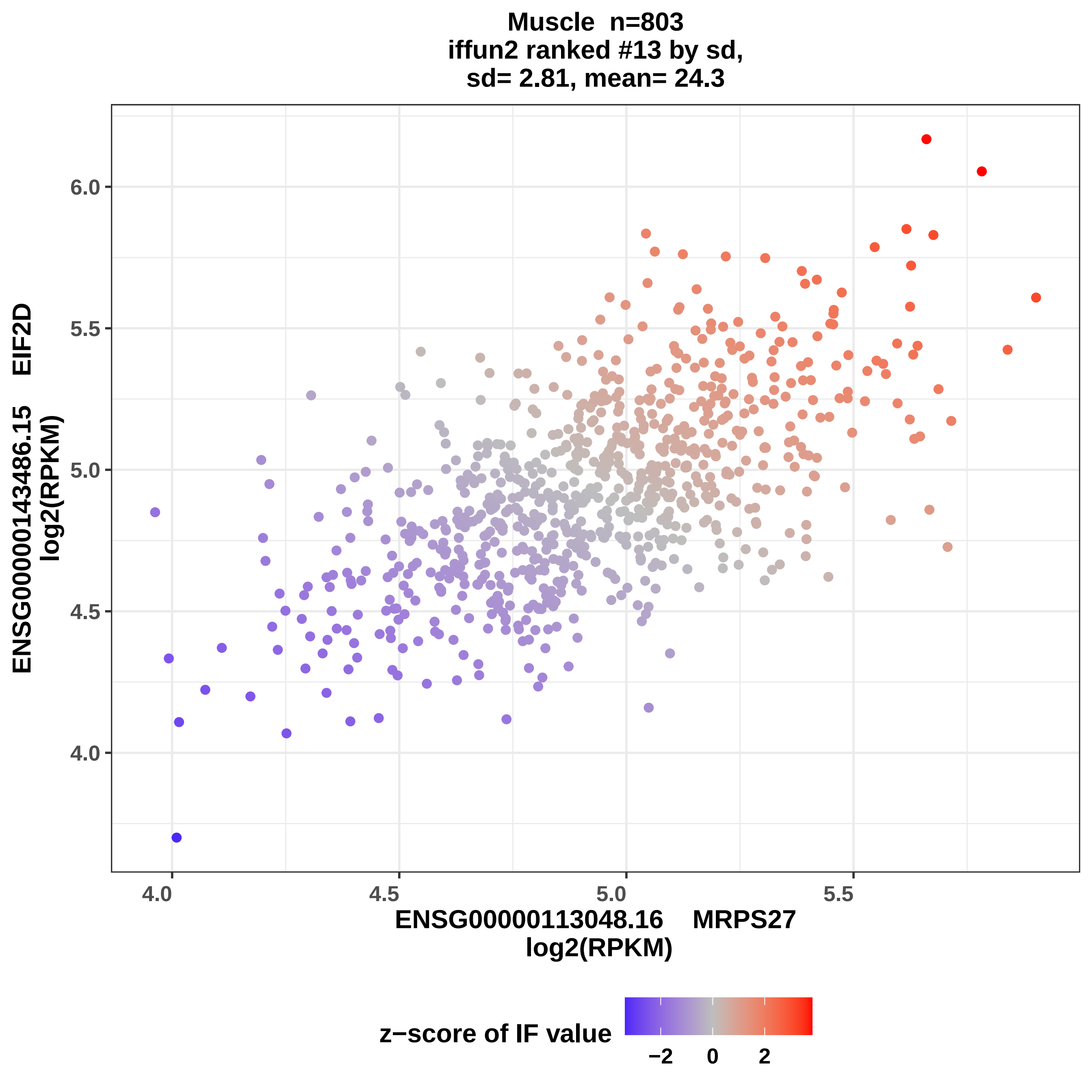
Muscle n=803
iffun2 ranked #13 by sd,
sd= 2.81, mean= 24.3
ENSG00000143486.15 EIF2D
log2(RPKM)
6.0
5.5
5.0
4.5
4.0
4.0
4.5
5.0
5.5
ENSG00000113048.16 MRPS27
log2(RPKM)
z−score of IF value
−2
0
2

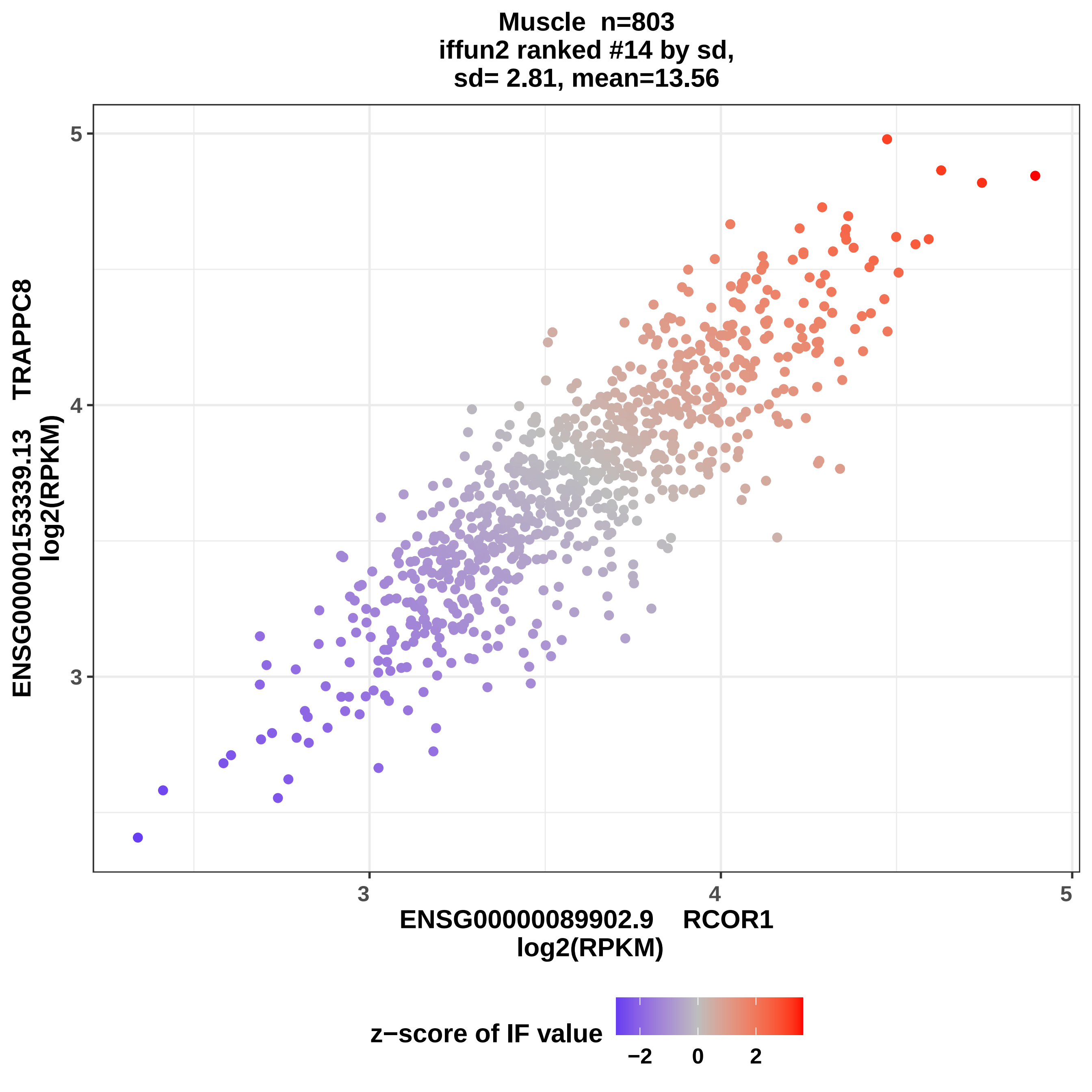
Muscle n=803
iffun2 ranked #14 by sd,
sd= 2.81, mean=13.56
ENSG00000153339.13 TRAPPC8
log2(RPKM)
5
4
3
3
4
5
ENSG00000089902.9 RCOR1
log2(RPKM)
z−score of IF value
−2
0
2

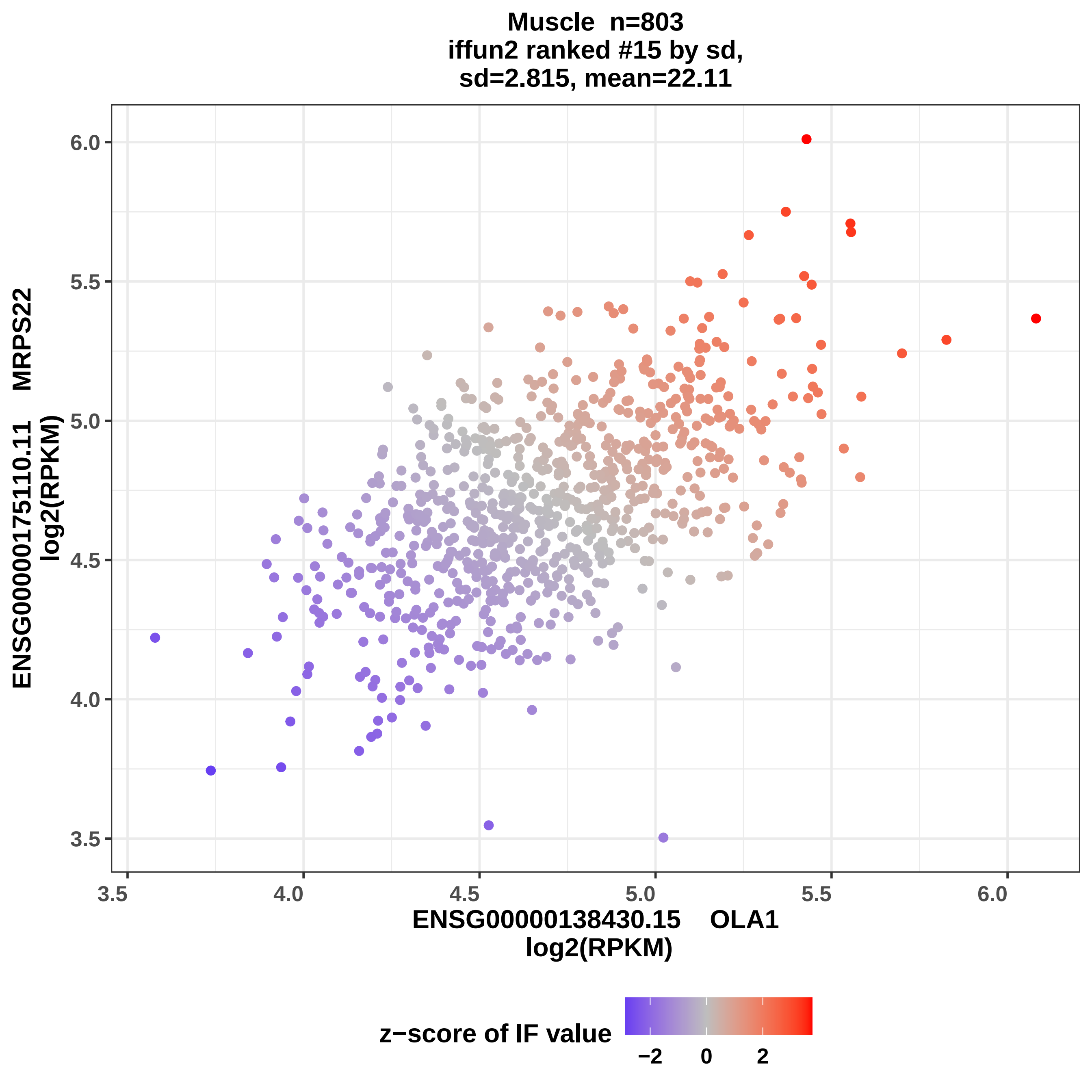
Muscle n=803
iffun2 ranked #15 by sd,
sd=2.815, mean=22.11
ENSG00000175110.11 MRPS22
log2(RPKM)
6.0
5.5
5.0
4.5
4.0
3.5
3.5
4.0
4.5
5.0
5.5
6.0
ENSG00000138430.15 OLA1
log2(RPKM)
z−score of IF value
−2
0
2

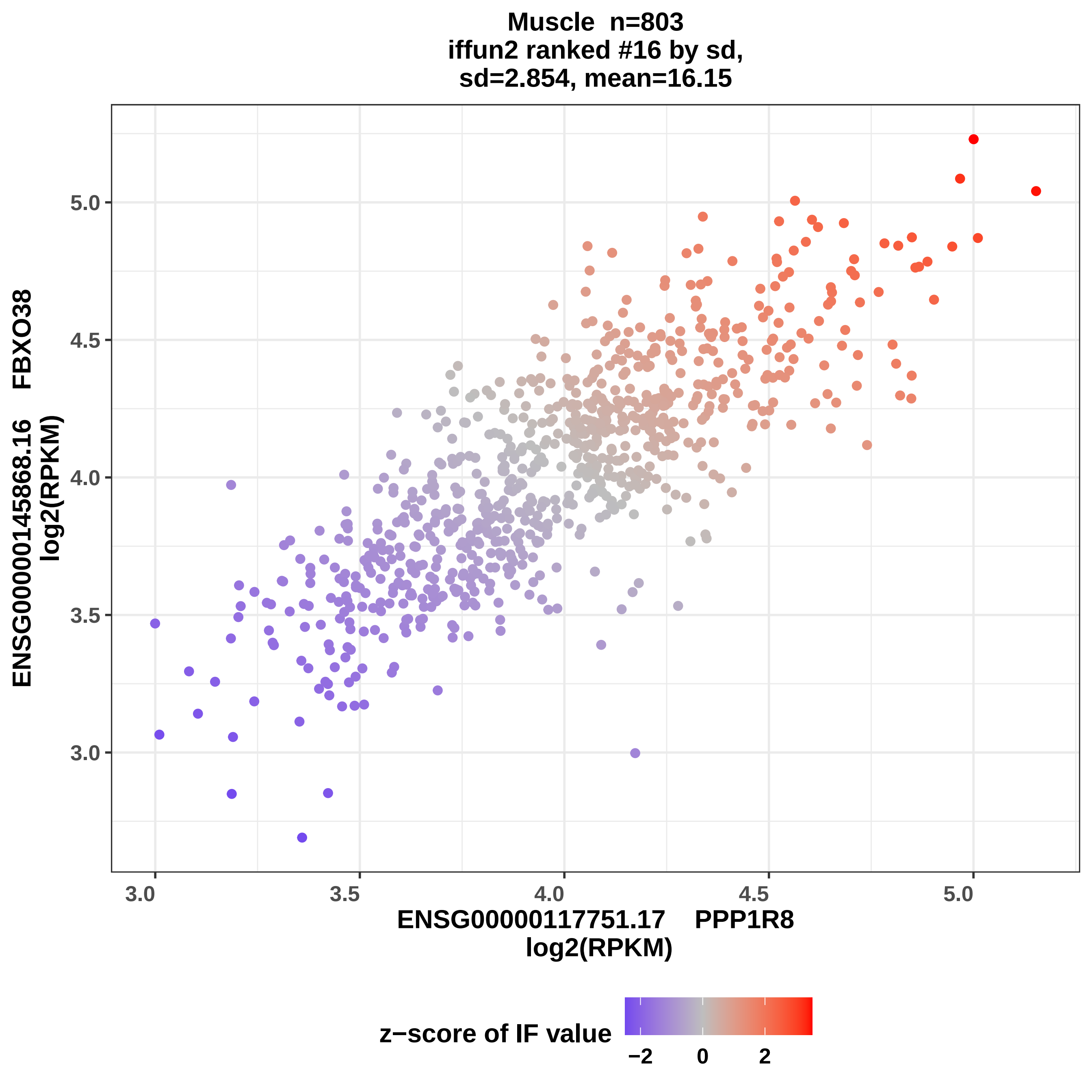

Muscle n=803
iffun2 ranked #16 by sd,
sd=2.854, mean=16.15
ENSG00000145868.16 FBXO38
log2(RPKM)
ENSG00000117751.17 PPP1R8
log2(RPKM)
3.0
3.5
4.0
4.5
5.0
z−score of IF value
−2
0
2

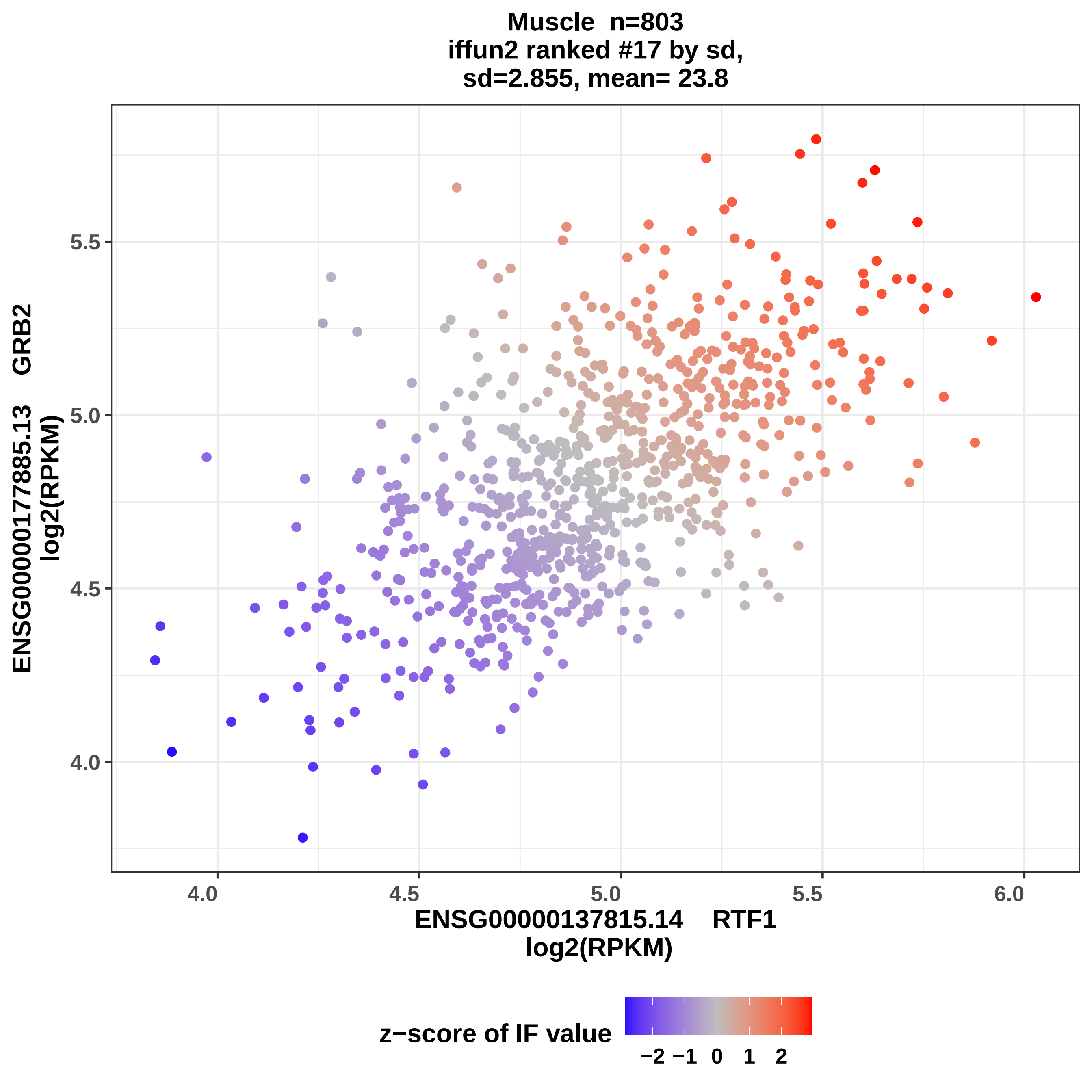
Muscle n=803
iffun2 ranked #17 by sd,
sd=2.855, mean= 23.8
ENSG00000177885.13 GRB2
log2(RPKM)
5.5
5.0
4.5
4.0
4.0
4.5
5.0
5.5
6.0
ENSG00000137815.14 RTF1
log2(RPKM)
z−score of IF value
−2 −1 0 1 2

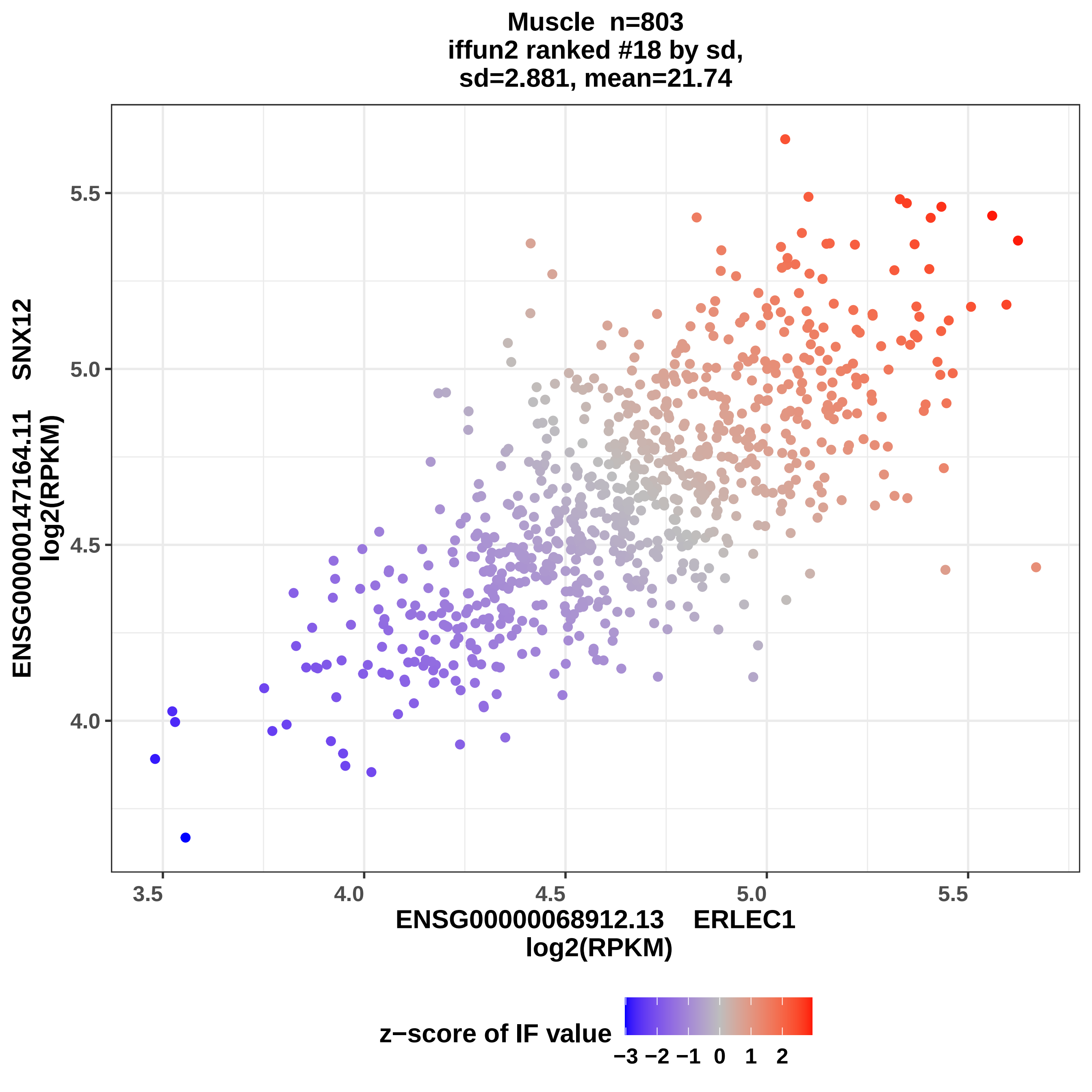

Muscle n=803
iffun2 ranked #18 by sd,
sd=2.881, mean=21.74
ENSG00000147164.11 SNX12
log2(RPKM)
5.5
5.0
4.5
4.0
3.5
4.0
4.5
5.0
5.5
ENSG00000068912.13 ERLEC1
log2(RPKM)
z−score of IF value
−3 −2 −1 0 1 2

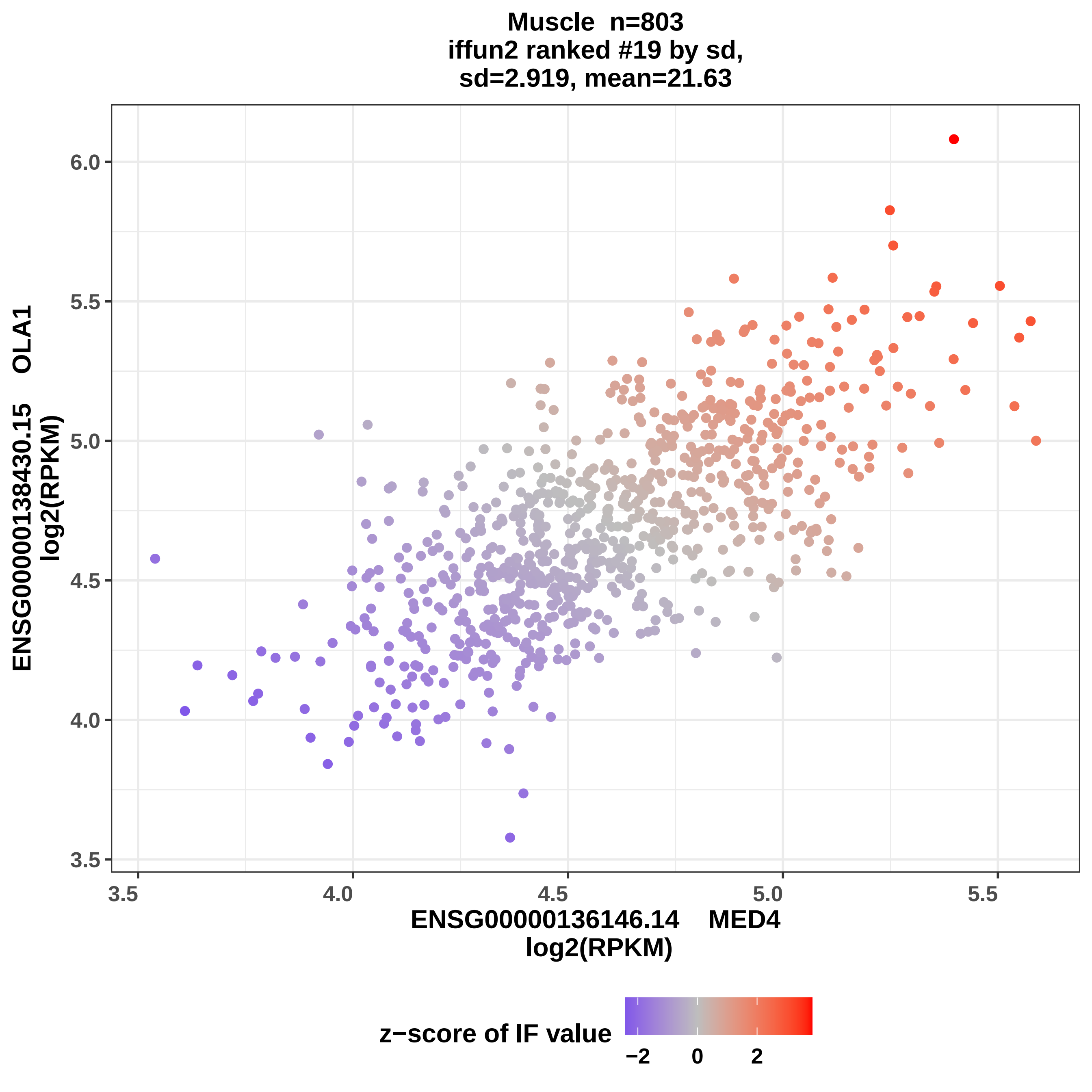
Muscle n=803
iffun2 ranked #19 by sd,
sd=2.919, mean=21.63
6.0
5.5
5.0
4.5
4.0
3.5
3.5
4.0
4.5
5.0
5.5
ENSG00000138430.15 OLA1
log2(RPKM)
ENSG00000136146.14 MED4
log2(RPKM)
z−score of IF value
−2
0
2

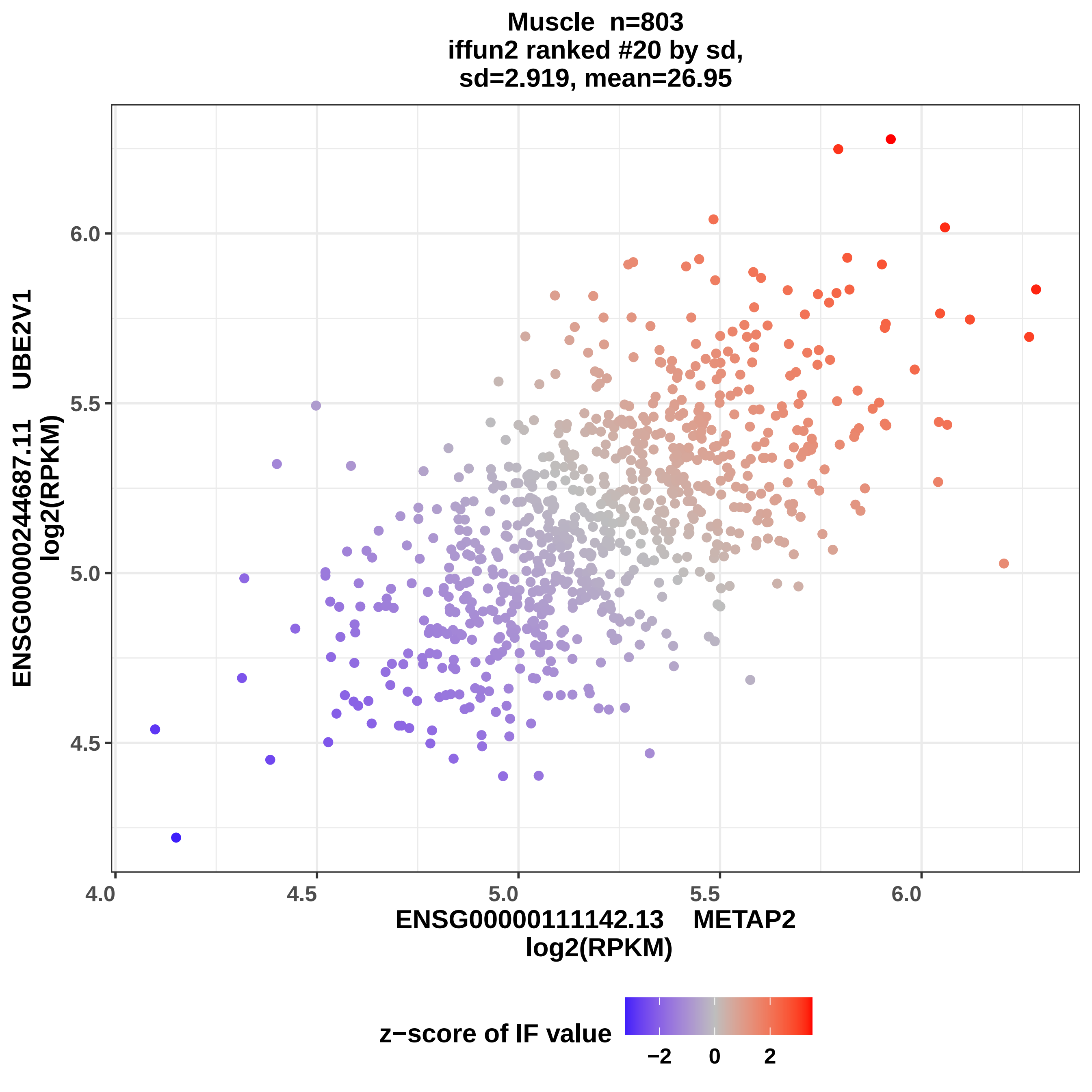
Muscle n=803
iffun2 ranked #20 by sd,
sd=2.919, mean=26.95
ENSG00000244687.11 UBE2V1
log2(RPKM)
6.0
5.5
5.0
4.5
4.0
4.5
5.0
5.5
6.0
ENSG00000111142.13 METAP2
log2(RPKM)
z−score of IF value
−2
0
2

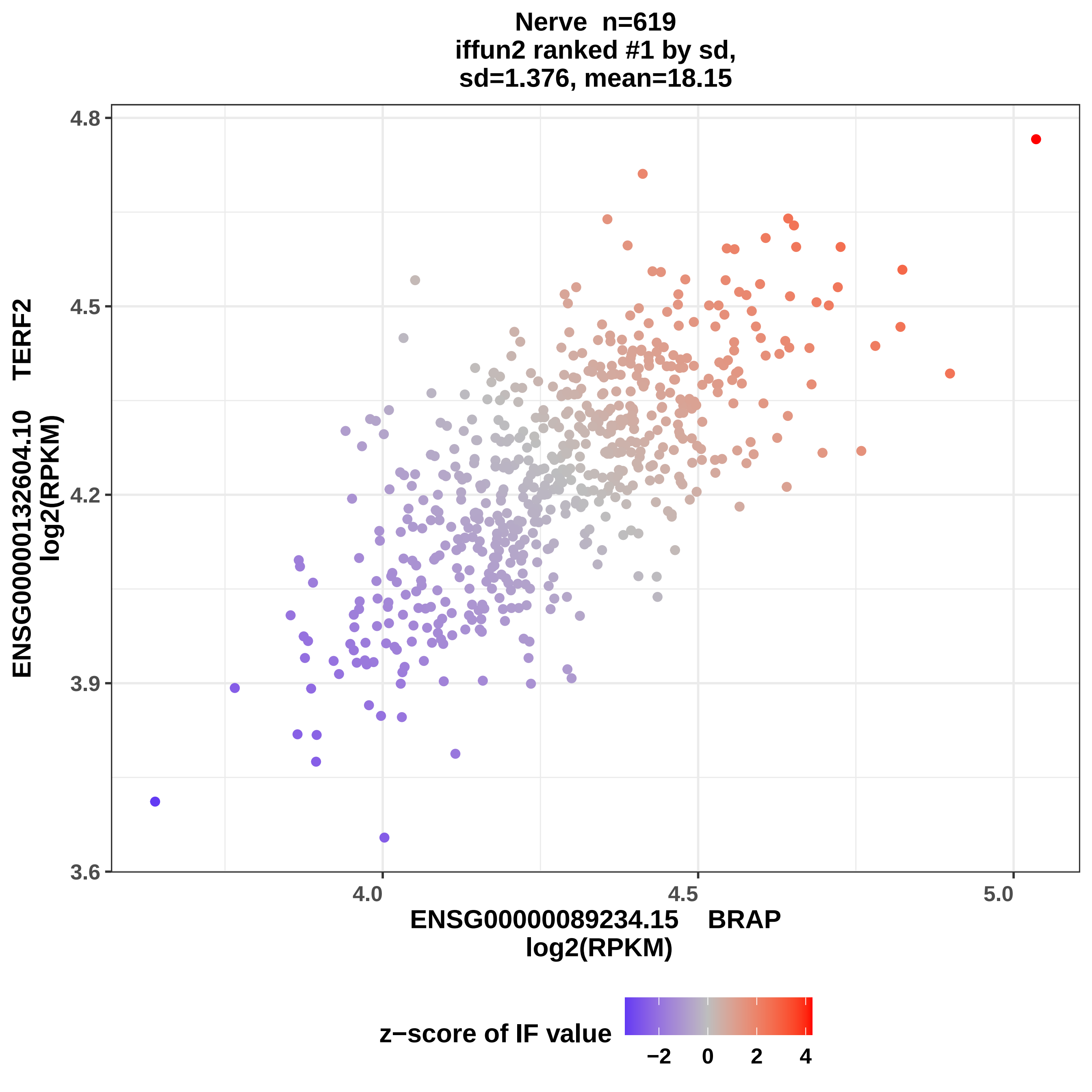
Nerve n=619
iffun2 ranked #1 by sd,
sd=1.376, mean=18.15
ENSG00000132604.10 TERF2
log2(RPKM)
4.8
4.5
4.2
3.9
3.6
4.0
4.5
5.0
ENSG00000089234.15 BRAP
log2(RPKM)
z−score of IF value
−2
0
2
4

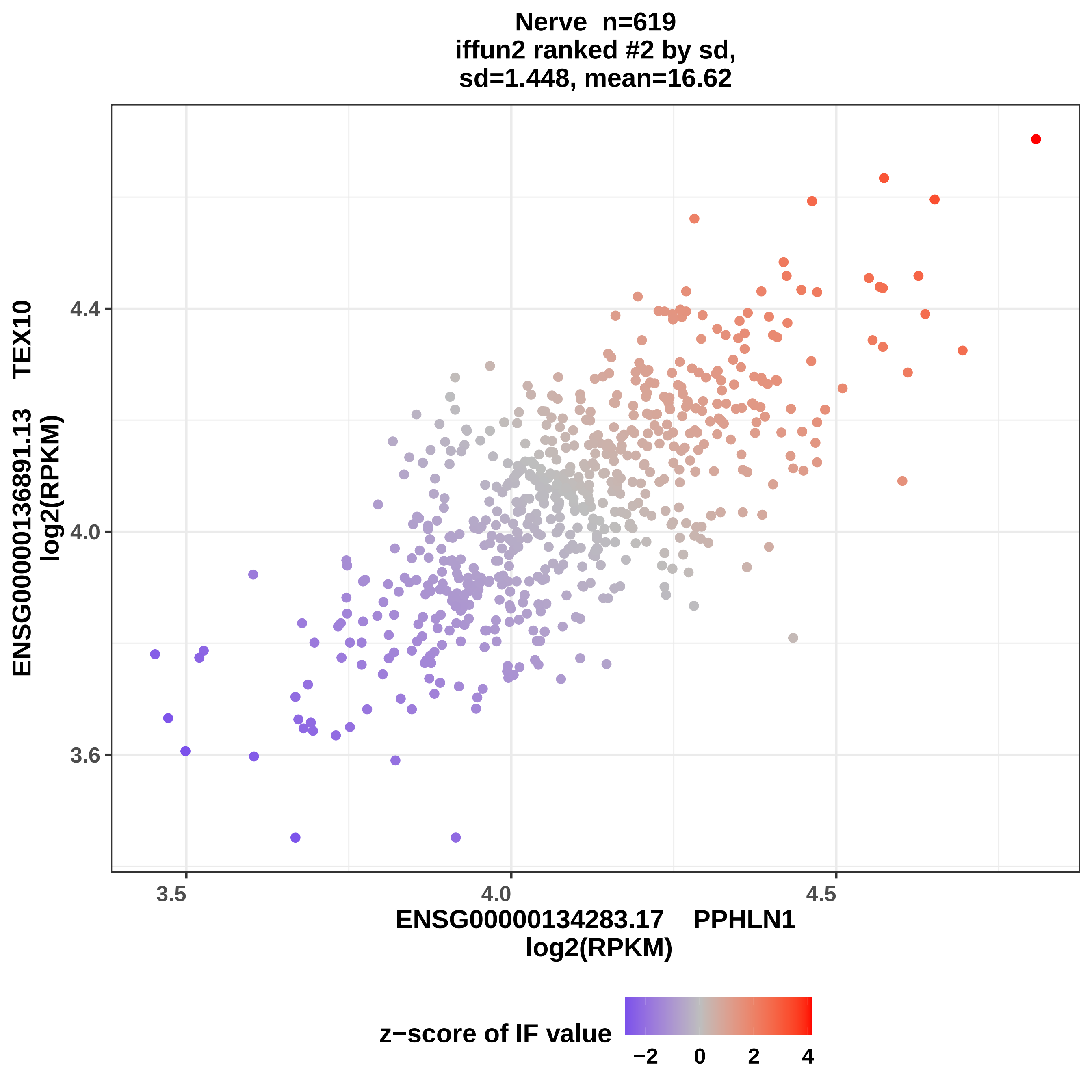

Nerve n=619
iffun2 ranked #2 by sd,
sd=1.448, mean=16.62
ENSG00000136891.13 TEX10
log2(RPKM)
4.4
4.0
3.6
3.5
4.0
4.5
ENSG00000134283.17 PPHLN1
log2(RPKM)
z−score of IF value
−2
0
2
4

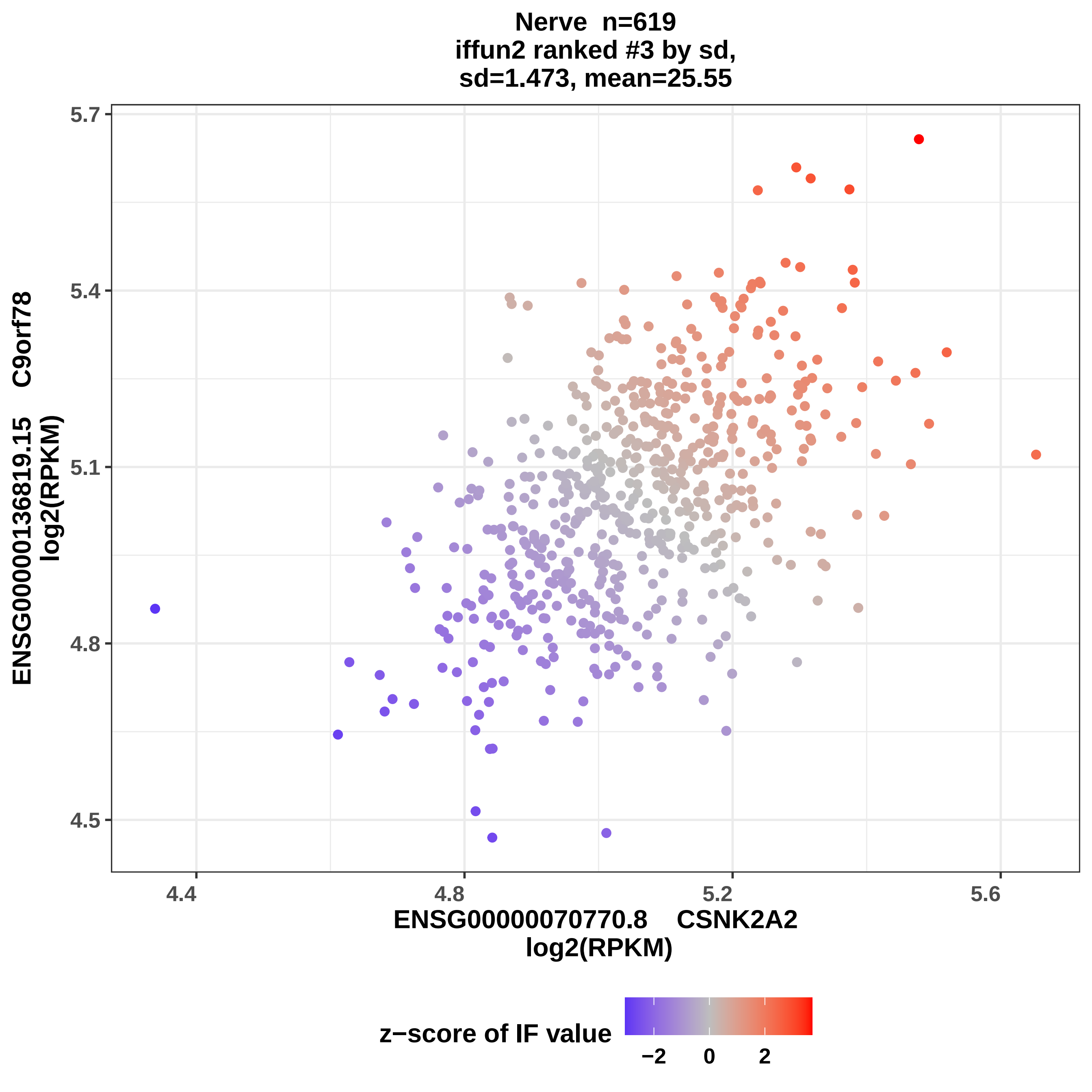

Nerve n=619
iffun2 ranked #3 by sd,
sd=1.473, mean=25.55
5.7
5.4
5.1
4.8
4.5
4.4
4.8
5.2
5.6
ENSG00000136819.15 C9orf78
log2(RPKM)
ENSG00000070770.8 CSNK2A2
log2(RPKM)
z−score of IF value
−2
0
2

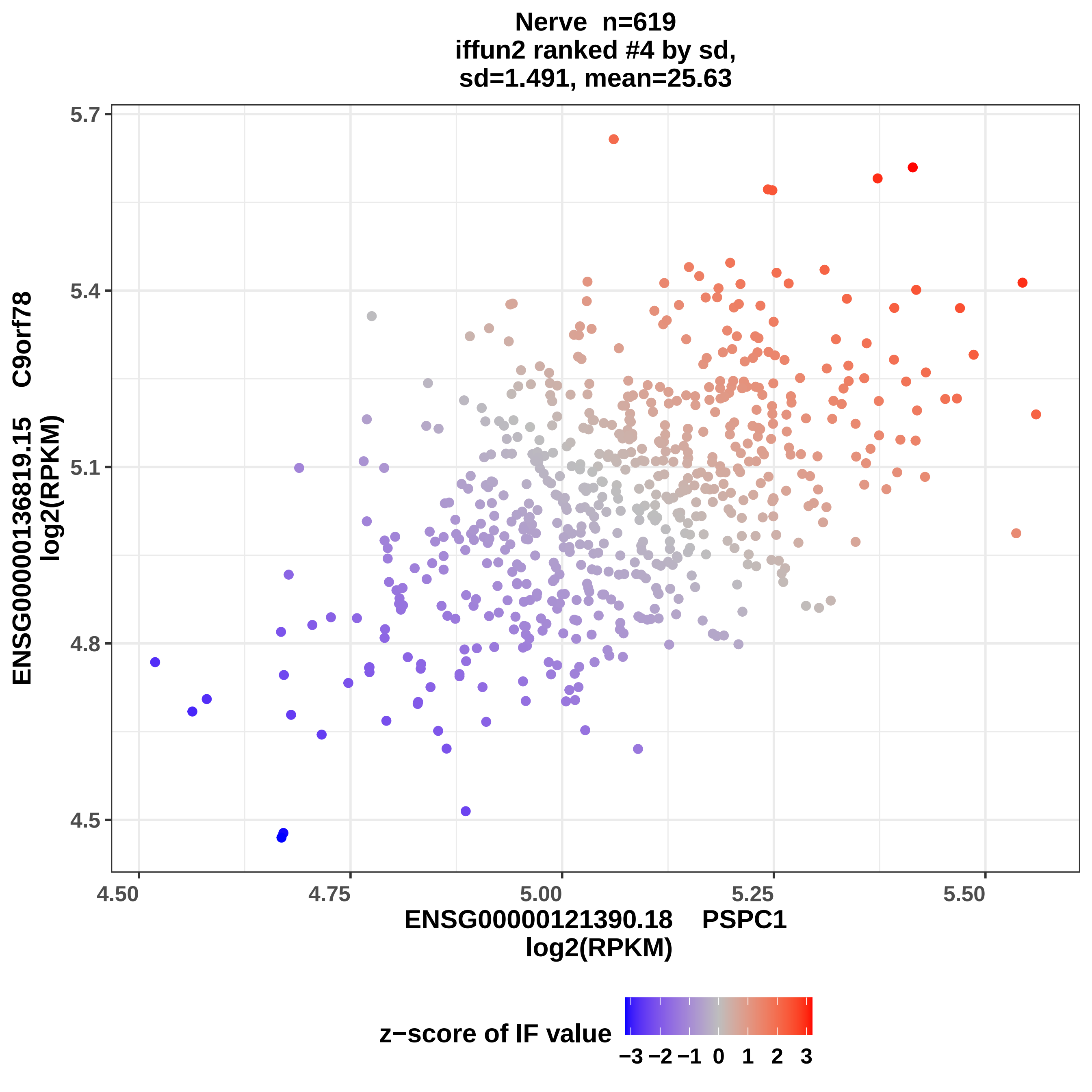

Nerve n=619
iffun2 ranked #4 by sd,
sd=1.491, mean=25.63
5.7
5.4
5.1
4.8
4.5
ENSG00000136819.15 C9orf78
log2(RPKM)
4.50
4.75
5.00
5.25
5.50
ENSG00000121390.18 PSPC1
log2(RPKM)
z−score of IF value
−3 −2 −1 0 1 2 3

Nerve n=619
iffun2 ranked #5 by sd,
sd= 1.52, mean=25.77

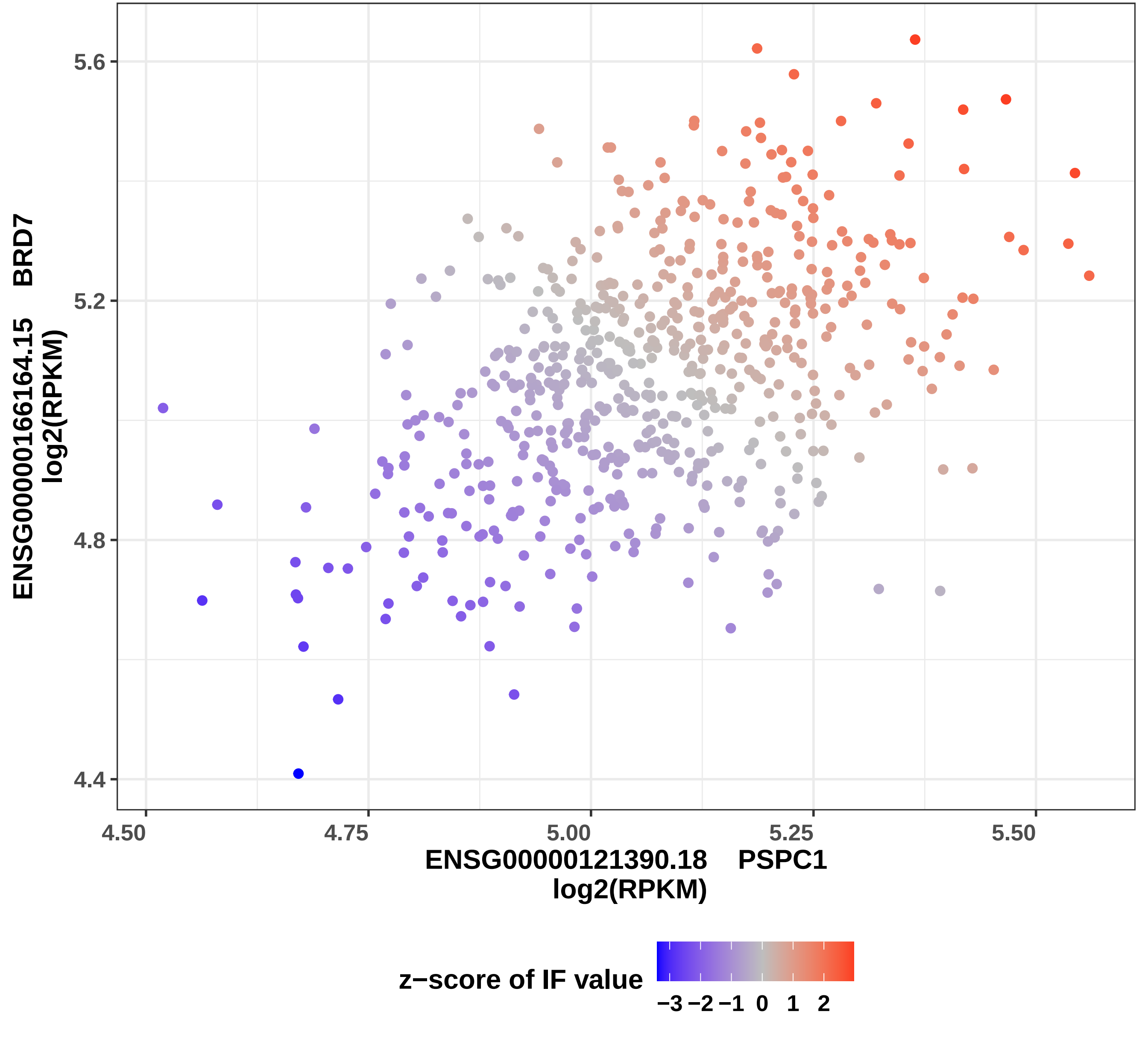

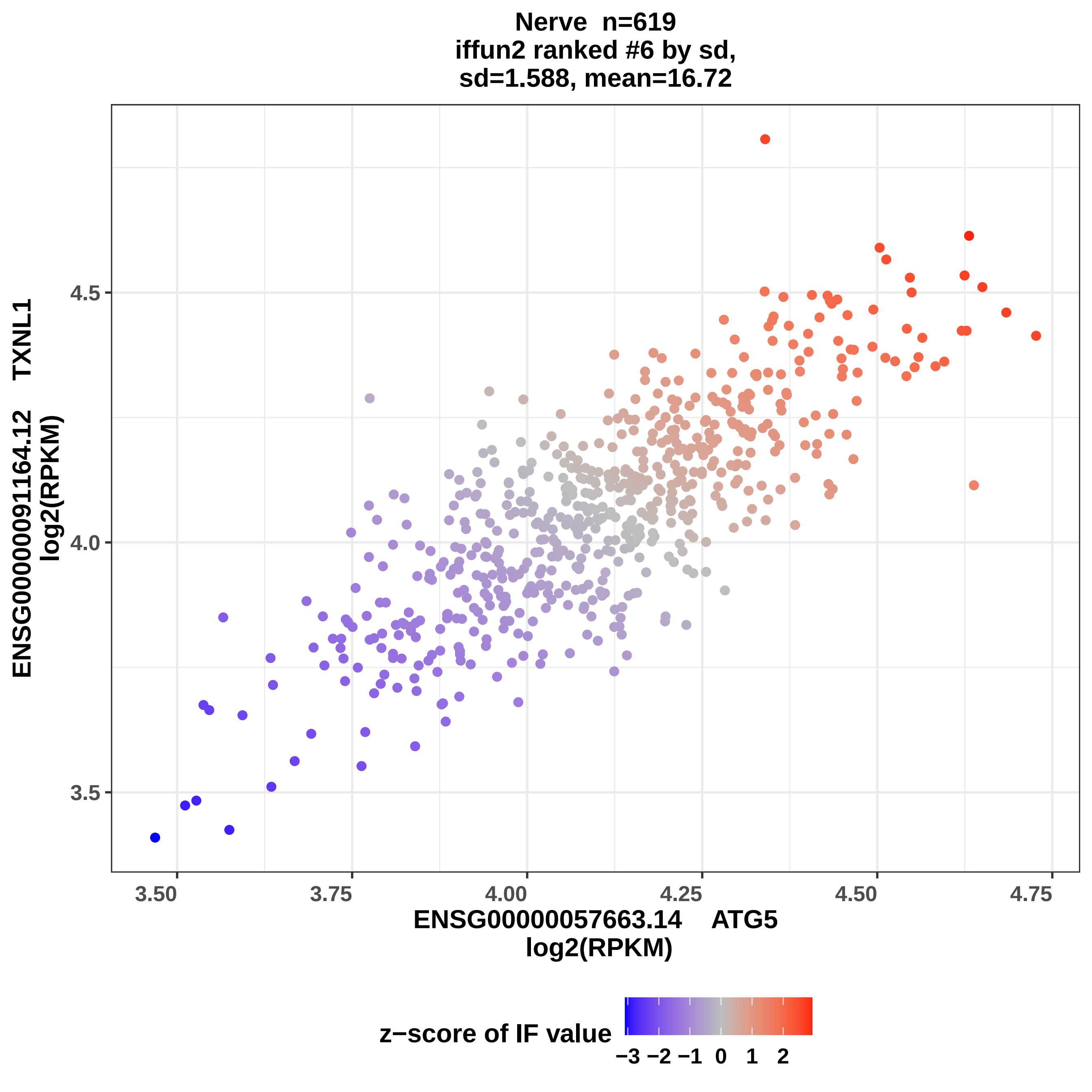

Nerve n=619
iffun2 ranked #6 by sd,
sd=1.588, mean=16.72
ENSG00000091164.12 TXNL1
log2(RPKM)
4.5
4.0
3.5
3.50
3.75
4.00
4.25
4.50
4.75
ENSG00000057663.14 ATG5
log2(RPKM)
z−score of IF value
−3 −2 −1 0 1 2

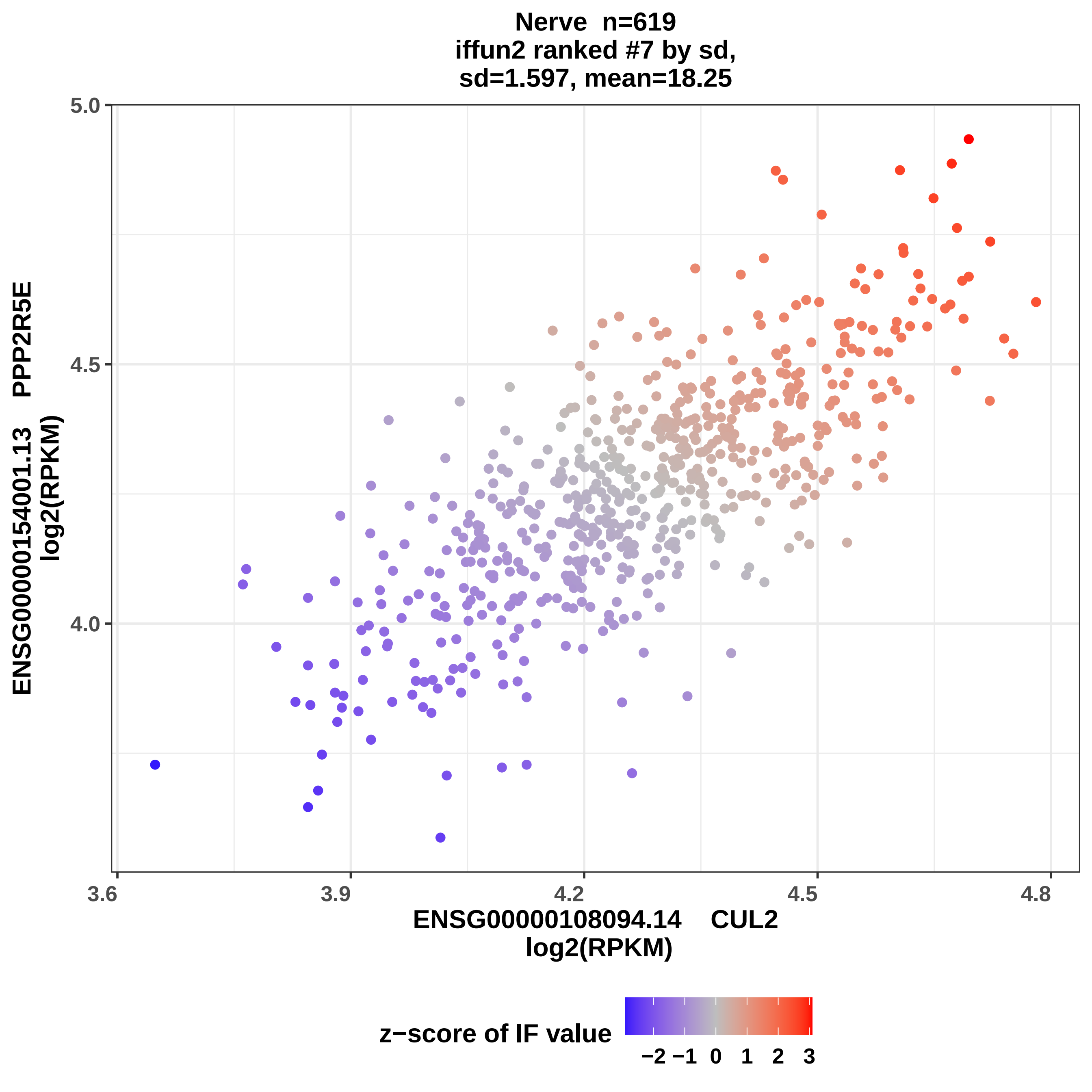
Nerve n=619
iffun2 ranked #7 by sd,
sd=1.597, mean=18.25
ENSG00000154001.13 PPP2R5E
log2(RPKM)
5.0
4.5
4.0
3.6
3.9
4.2
4.5
4.8
ENSG00000108094.14 CUL2
log2(RPKM)
z−score of IF value
−2 −1 0 1 2 3

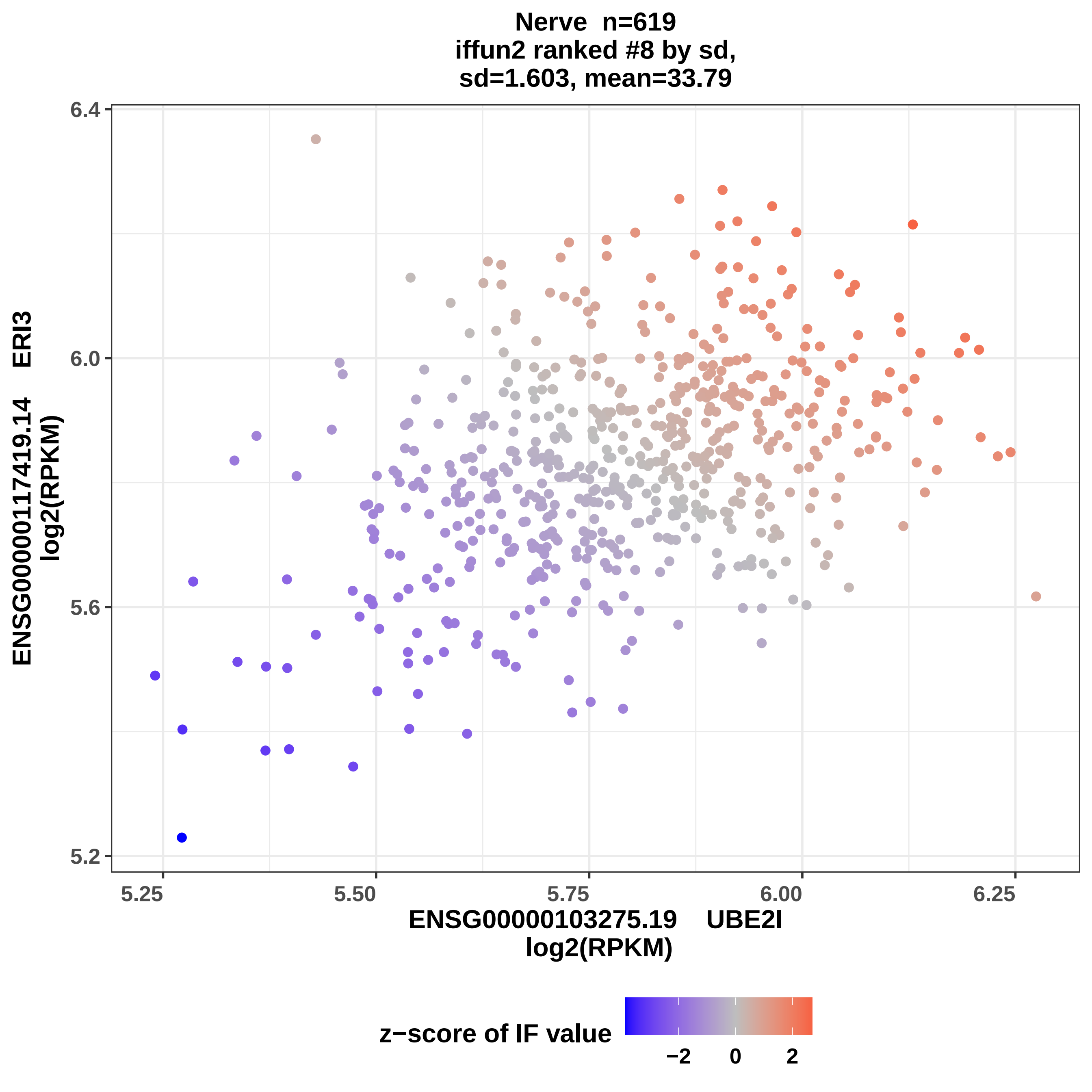
Nerve n=619
iffun2 ranked #8 by sd,
sd=1.603, mean=33.79
ENSG00000117419.14 ERI3
log2(RPKM)
6.4
6.0
5.6
5.2
5.25
5.50
5.75
6.00
6.25
ENSG00000103275.19 UBE2I
log2(RPKM)
z−score of IF value
−2
0
2

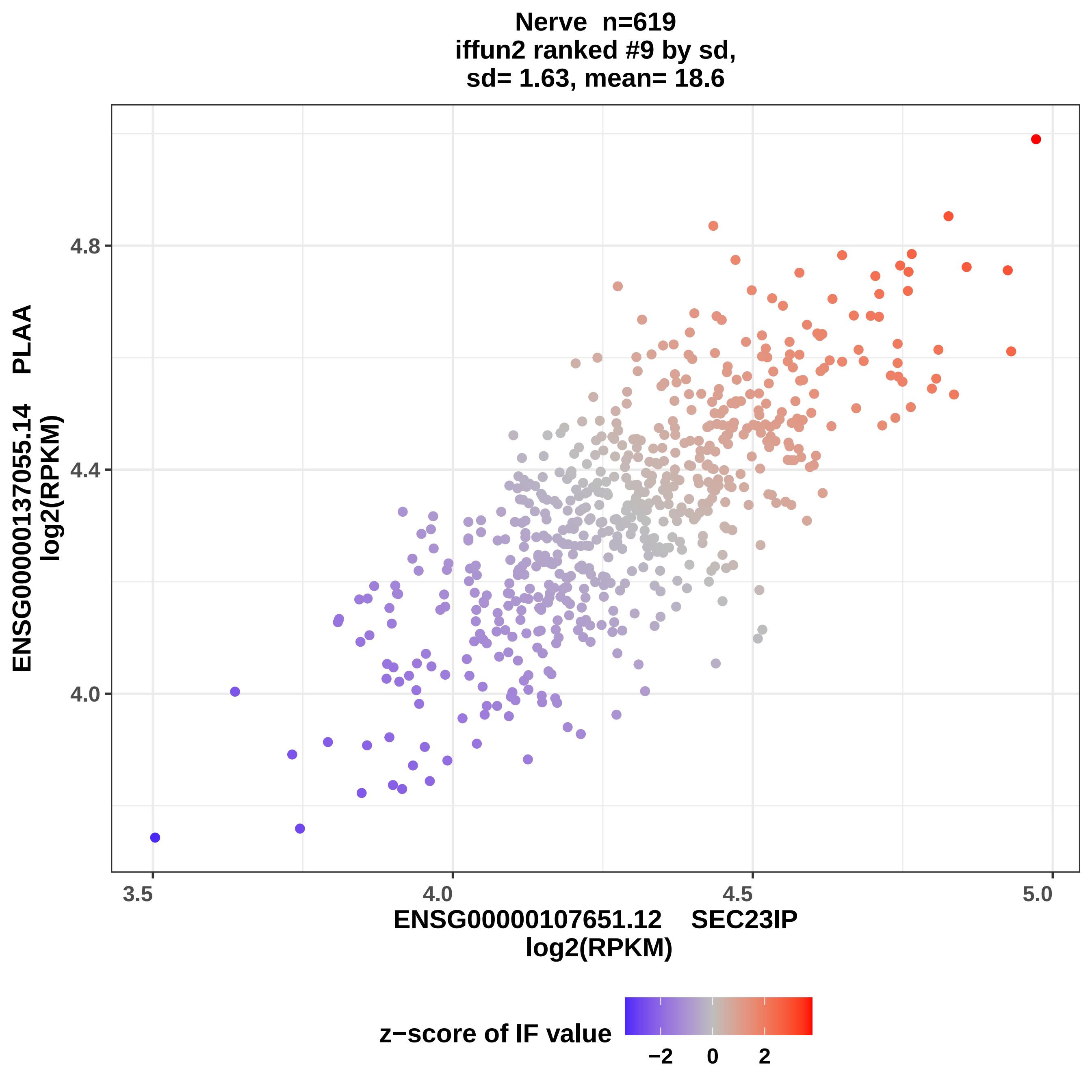
Nerve n=619
iffun2 ranked #9 by sd,
sd= 1.63, mean= 18.6
ENSG00000137055.14 PLAA
log2(RPKM)
4.8
4.4
4.0
3.5
4.0
4.5
5.0
ENSG00000107651.12 SEC23IP
log2(RPKM)
z−score of IF value
−2
0
2

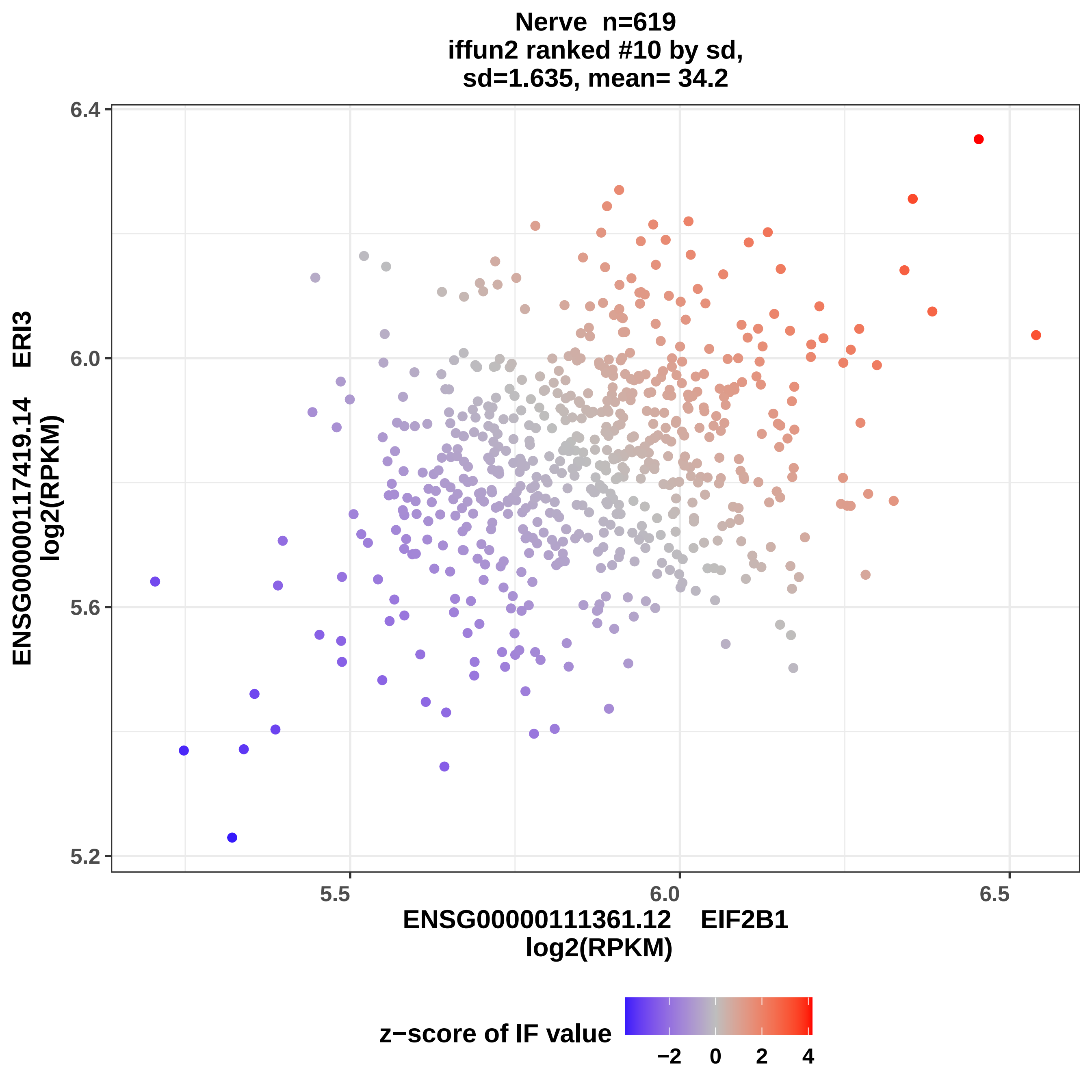

Nerve n=619
iffun2 ranked #10 by sd,
sd=1.635, mean= 34.2
ENSG00000117419.14 ERI3
log2(RPKM)
6.4
6.0
5.6
5.2
5.5
6.0
6.5
ENSG00000111361.12 EIF2B1
log2(RPKM)
z−score of IF value
−2
0
2
4

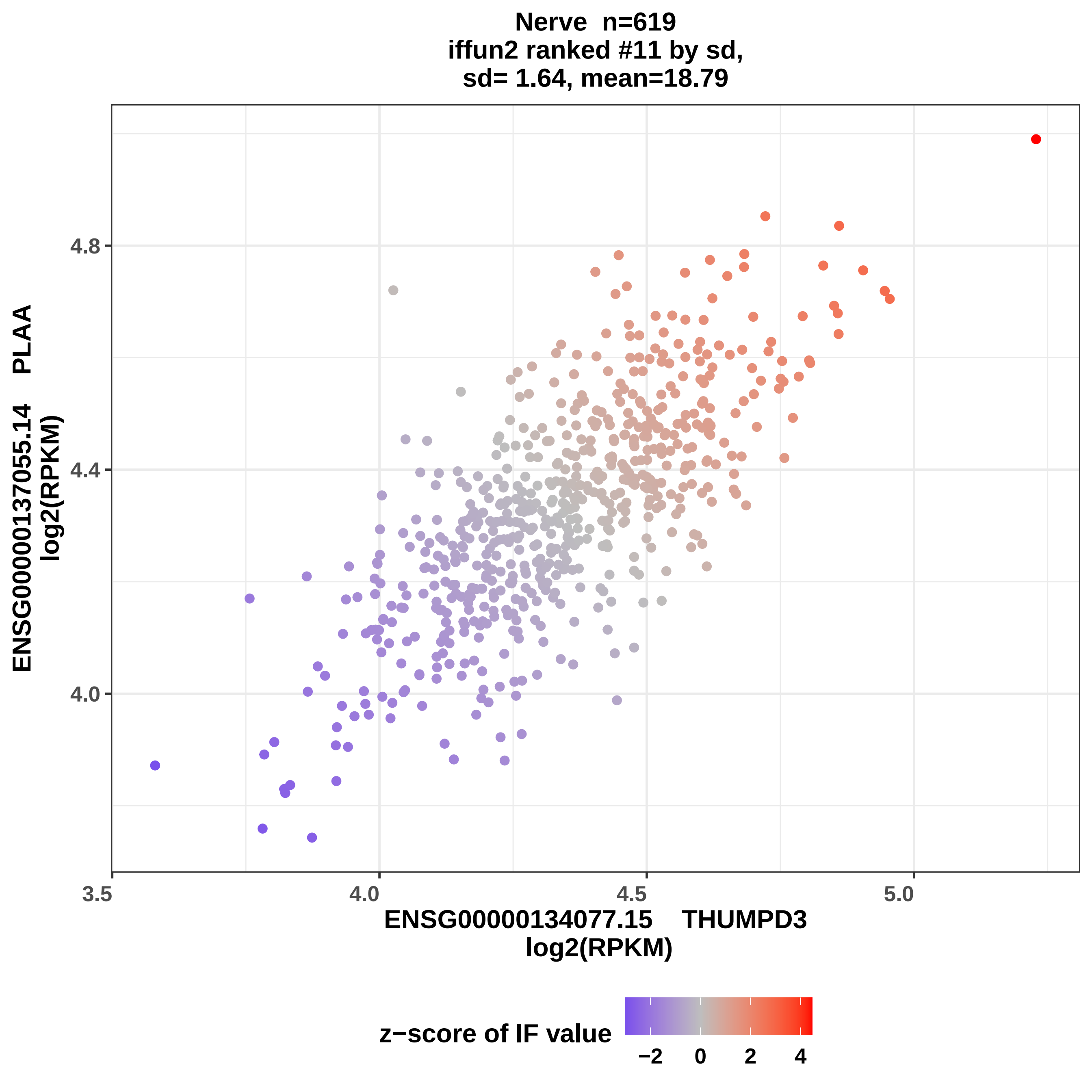
Nerve n=619
iffun2 ranked #11 by sd,
sd= 1.64, mean=18.79
ENSG00000137055.14 PLAA
log2(RPKM)
4.0
4.4
4.8
3.5
4.0
4.5
5.0
ENSG00000134077.15 THUMPD3
log2(RPKM)
z−score of IF value
−2
0
2
4

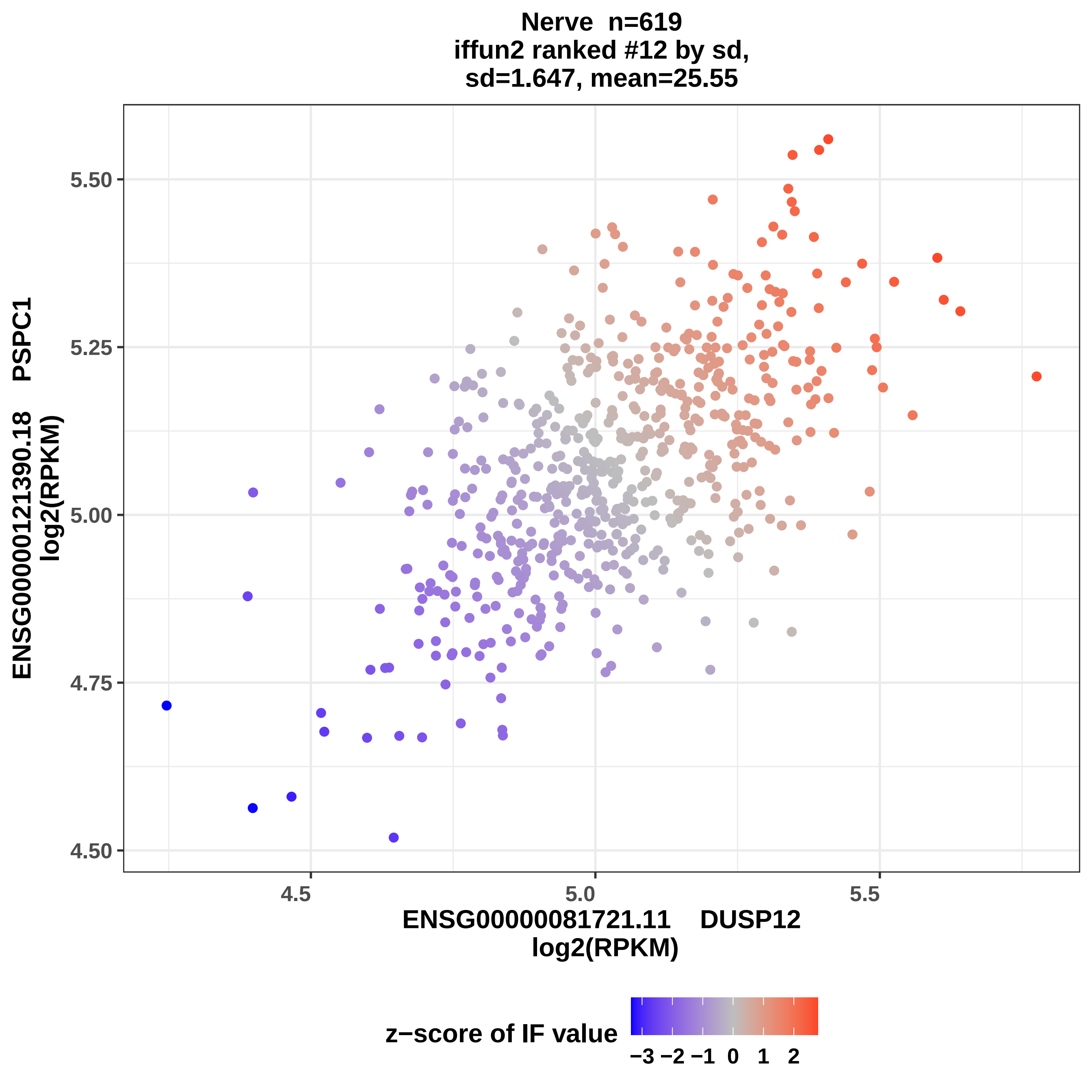

Nerve n=619
iffun2 ranked #12 by sd,
sd=1.647, mean=25.55
ENSG00000121390.18 PSPC1
log2(RPKM)
5.50
5.25
5.00
4.75
4.50
4.5
5.0
5.5
ENSG00000081721.11 DUSP12
log2(RPKM)
z−score of IF value
−3 −2 −1 0 1 2

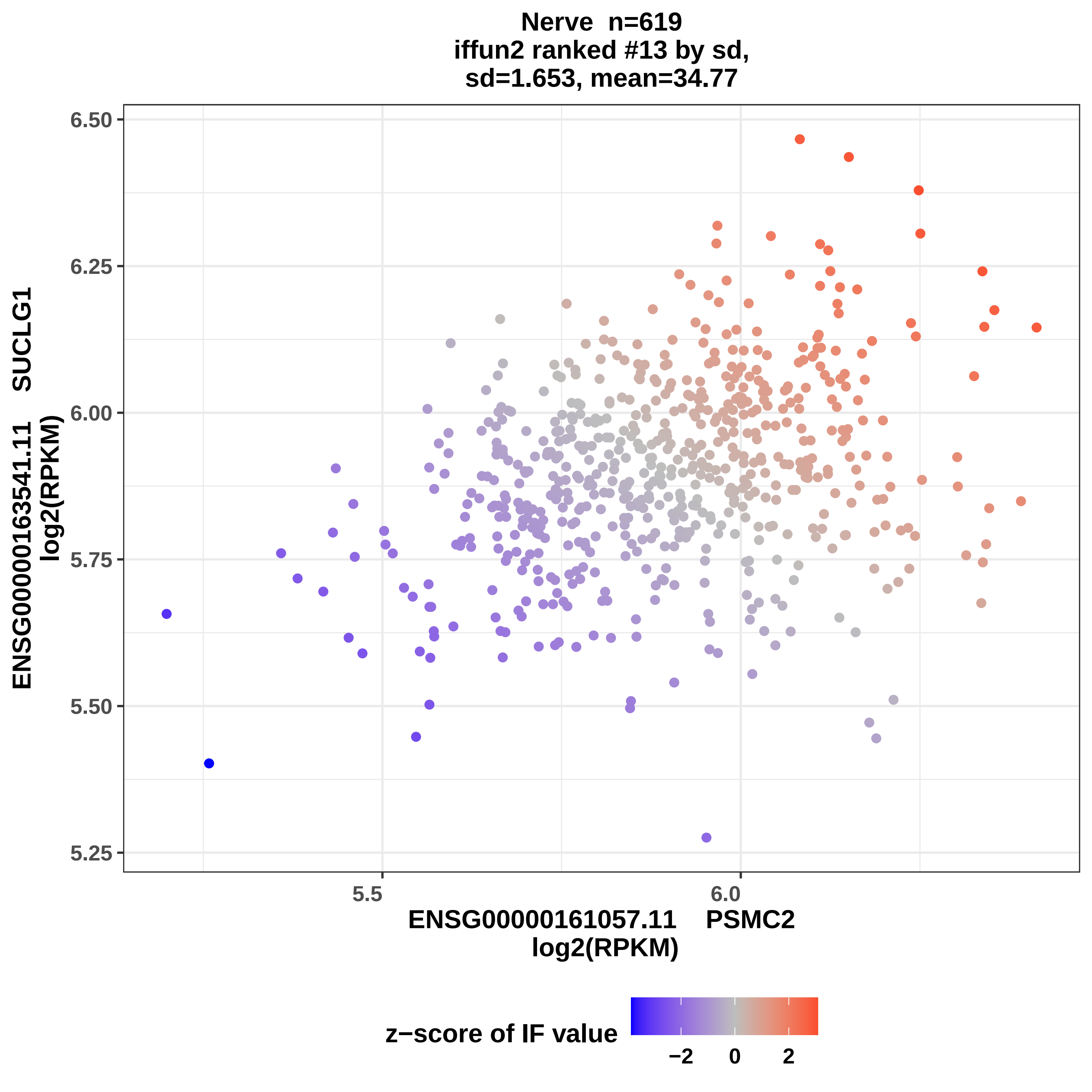
Nerve n=619
iffun2 ranked #13 by sd,
sd=1.653, mean=34.77
6.50
6.25
6.00
5.75
5.50
5.25
5.5
6.0
ENSG00000163541.11 SUCLG1
log2(RPKM)
ENSG00000161057.11 PSMC2
log2(RPKM)
z−score of IF value
−2
0
2

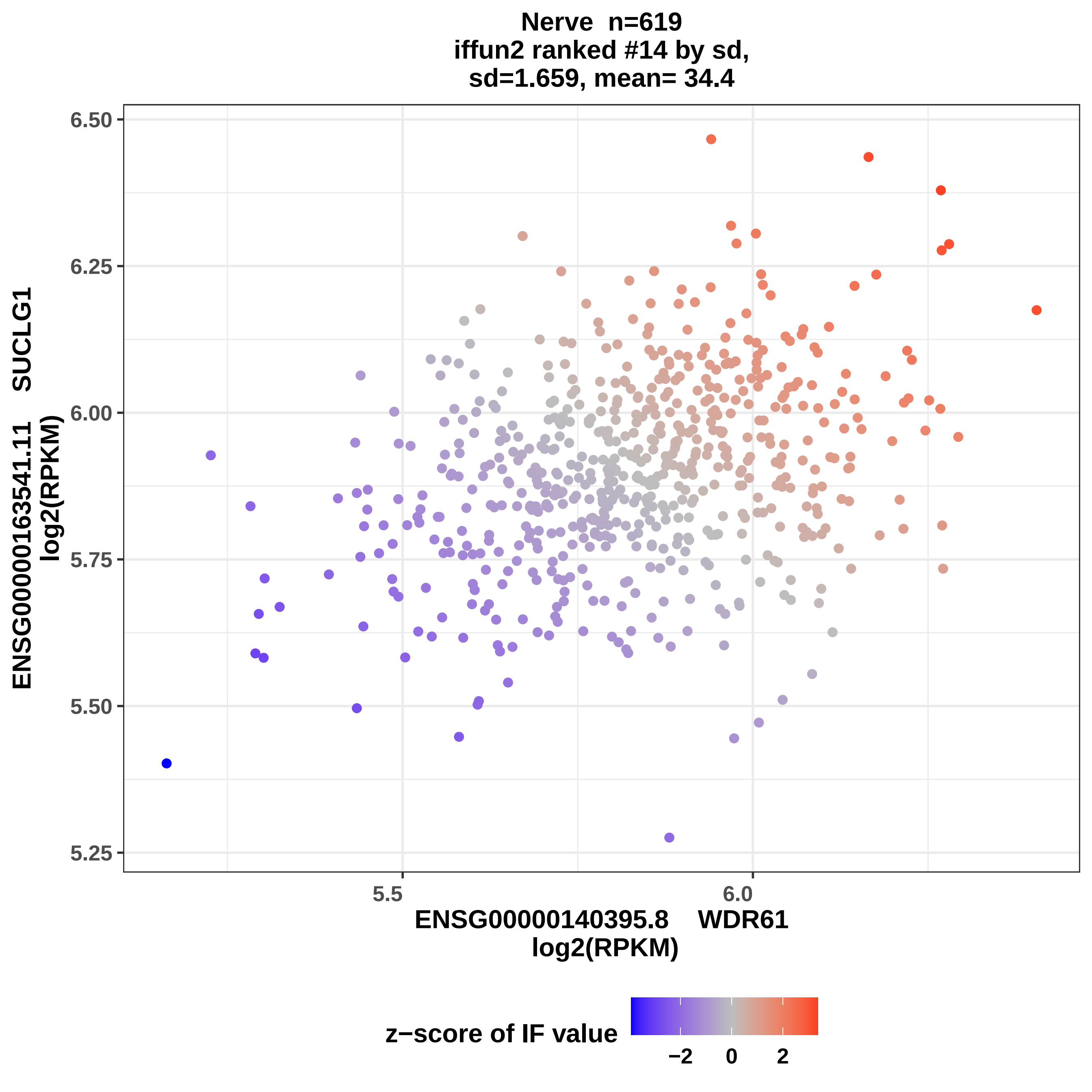

Nerve n=619
iffun2 ranked #14 by sd,
sd=1.659, mean= 34.4
6.50
6.25
6.00
5.75
5.50
5.25
ENSG00000163541.11 SUCLG1
log2(RPKM)
5.5
6.0
ENSG00000140395.8 WDR61
log2(RPKM)
z−score of IF value
−2
0
2

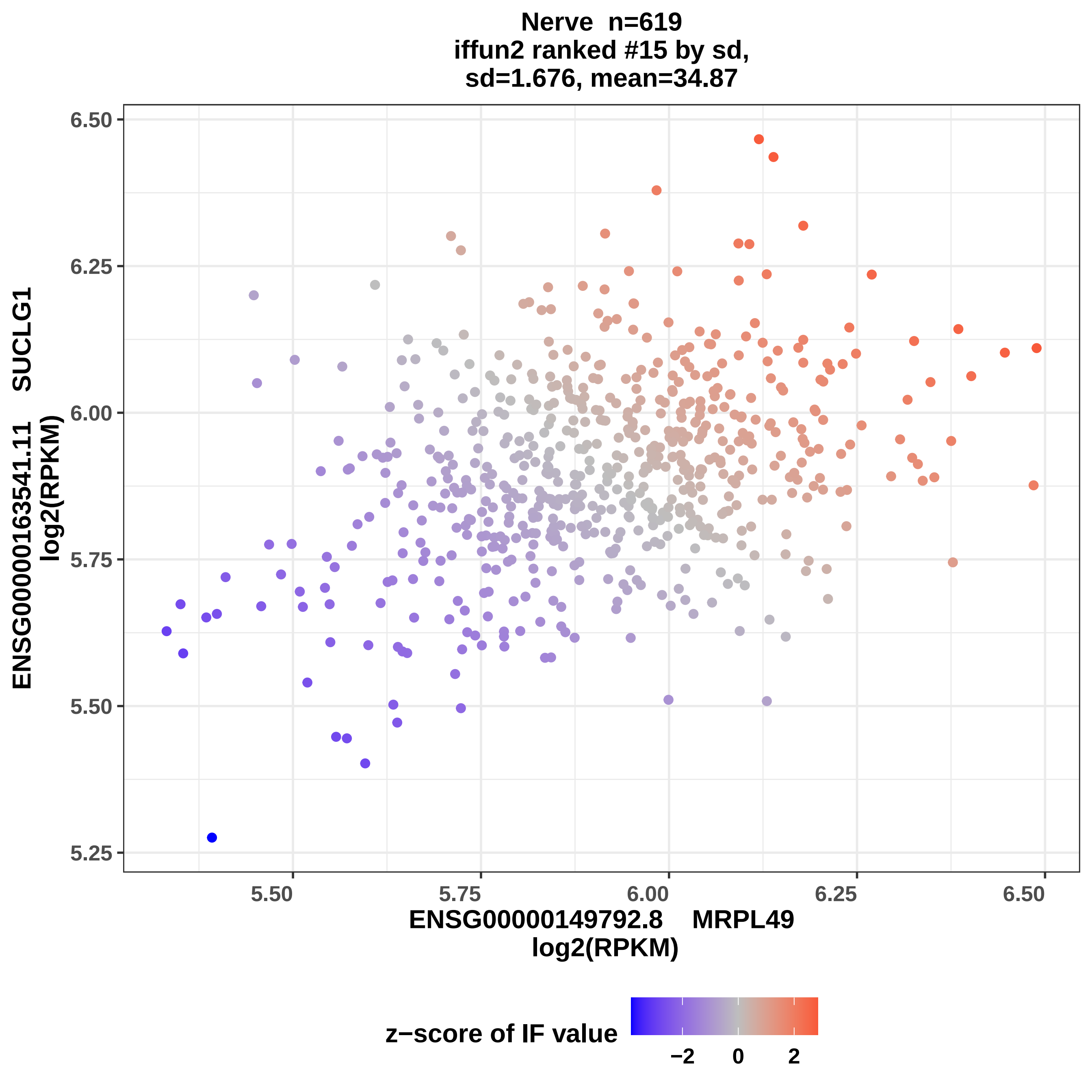

Nerve n=619
iffun2 ranked #15 by sd,
sd=1.676, mean=34.87
6.50
6.25
6.00
5.75
5.50
5.25
5.50
5.75
6.00
6.25
6.50
ENSG00000163541.11 SUCLG1
log2(RPKM)
ENSG00000149792.8 MRPL49
log2(RPKM)
z−score of IF value
−2
0
2

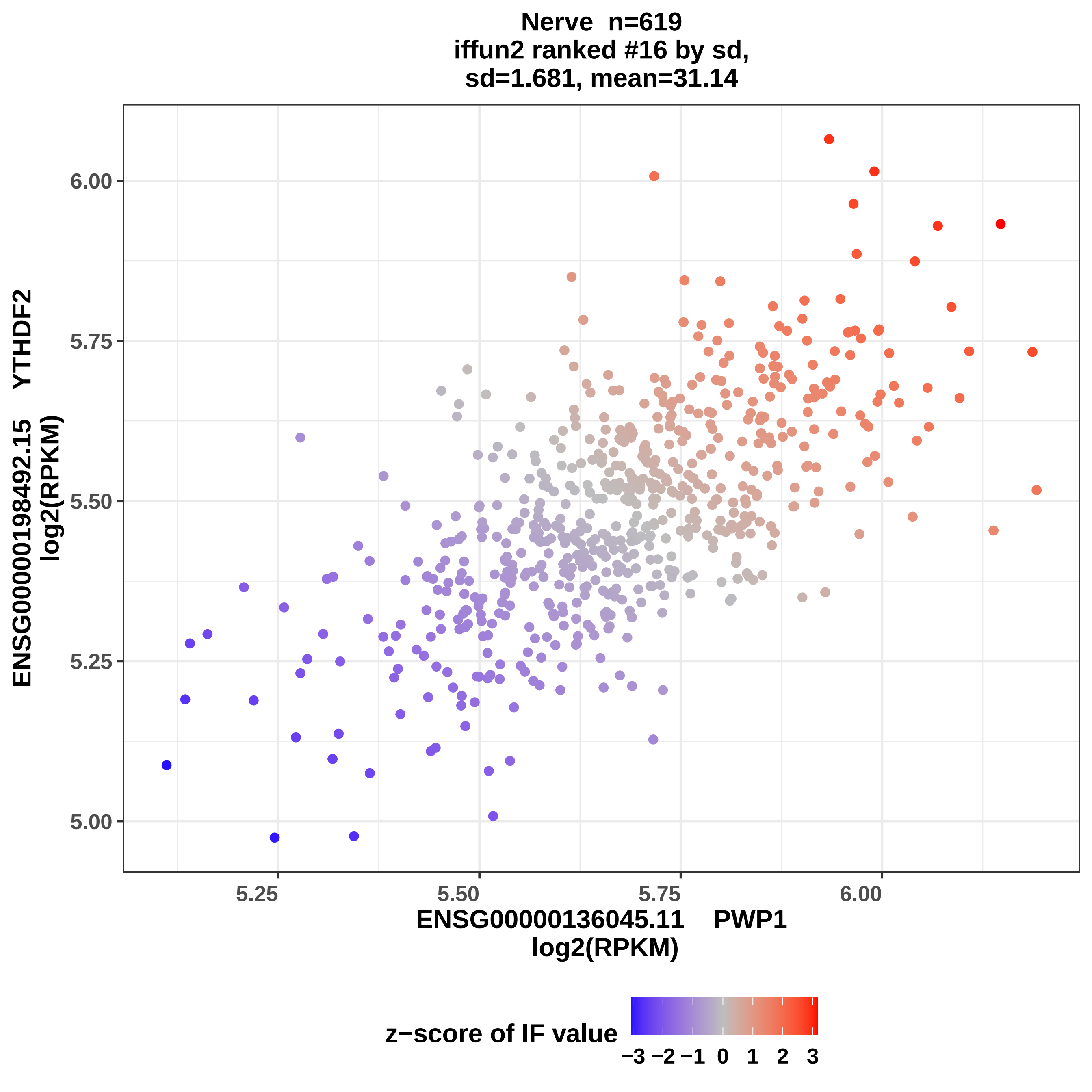
Nerve n=619
iffun2 ranked #16 by sd,
sd=1.681, mean=31.14
6.00
5.75
5.50
5.25
5.00
5.25
5.50
5.75
6.00
ENSG00000136045.11 PWP1
log2(RPKM)
ENSG00000198492.15 YTHDF2
log2(RPKM)
z−score of IF value
−3 −2 −1 0 1 2 3

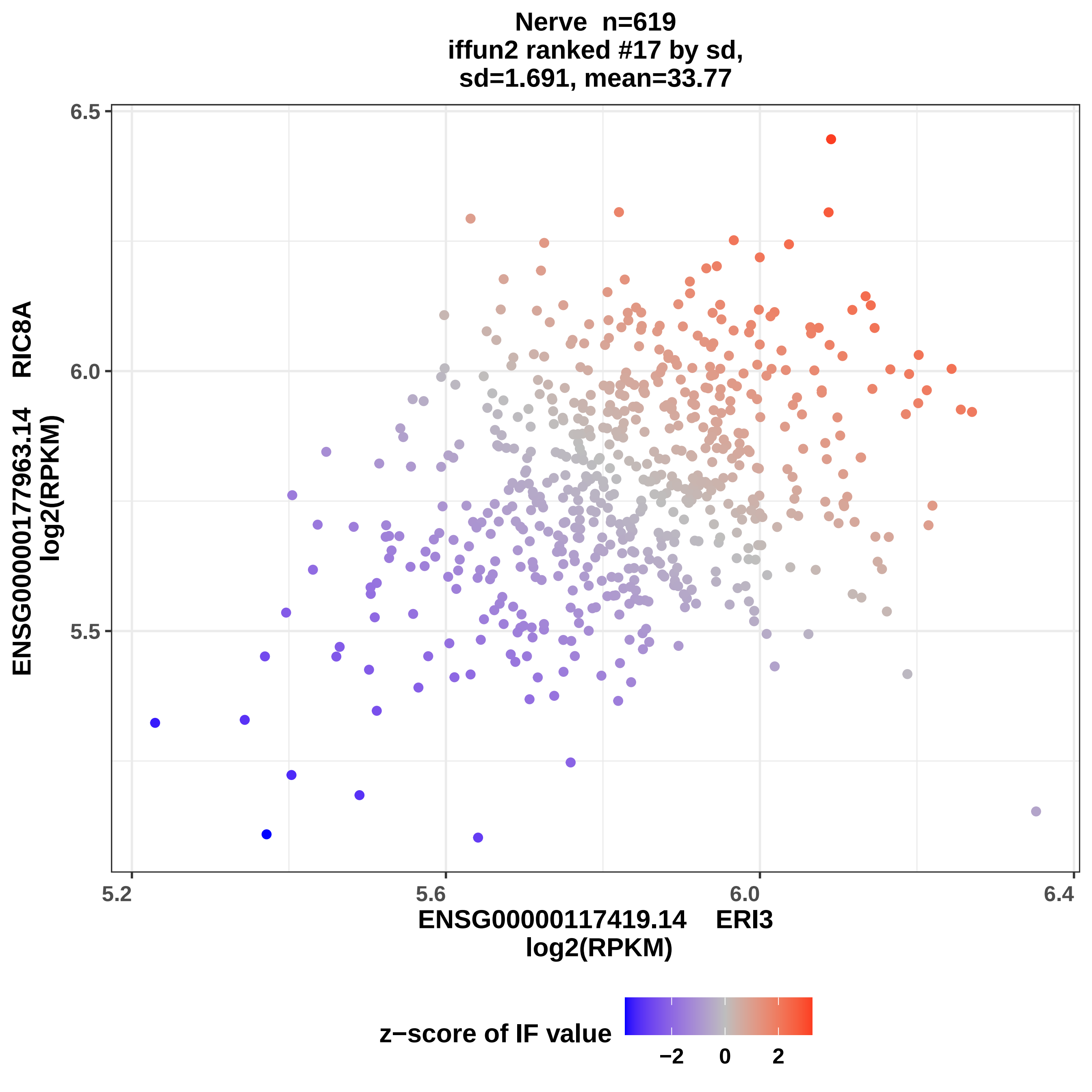
Nerve n=619
iffun2 ranked #17 by sd,
sd=1.691, mean=33.77
6.5
6.0
5.5
5.2
5.6
6.0
6.4
ENSG00000177963.14 RIC8A
log2(RPKM)
ENSG00000117419.14 ERI3
log2(RPKM)
z−score of IF value
−2
0
2

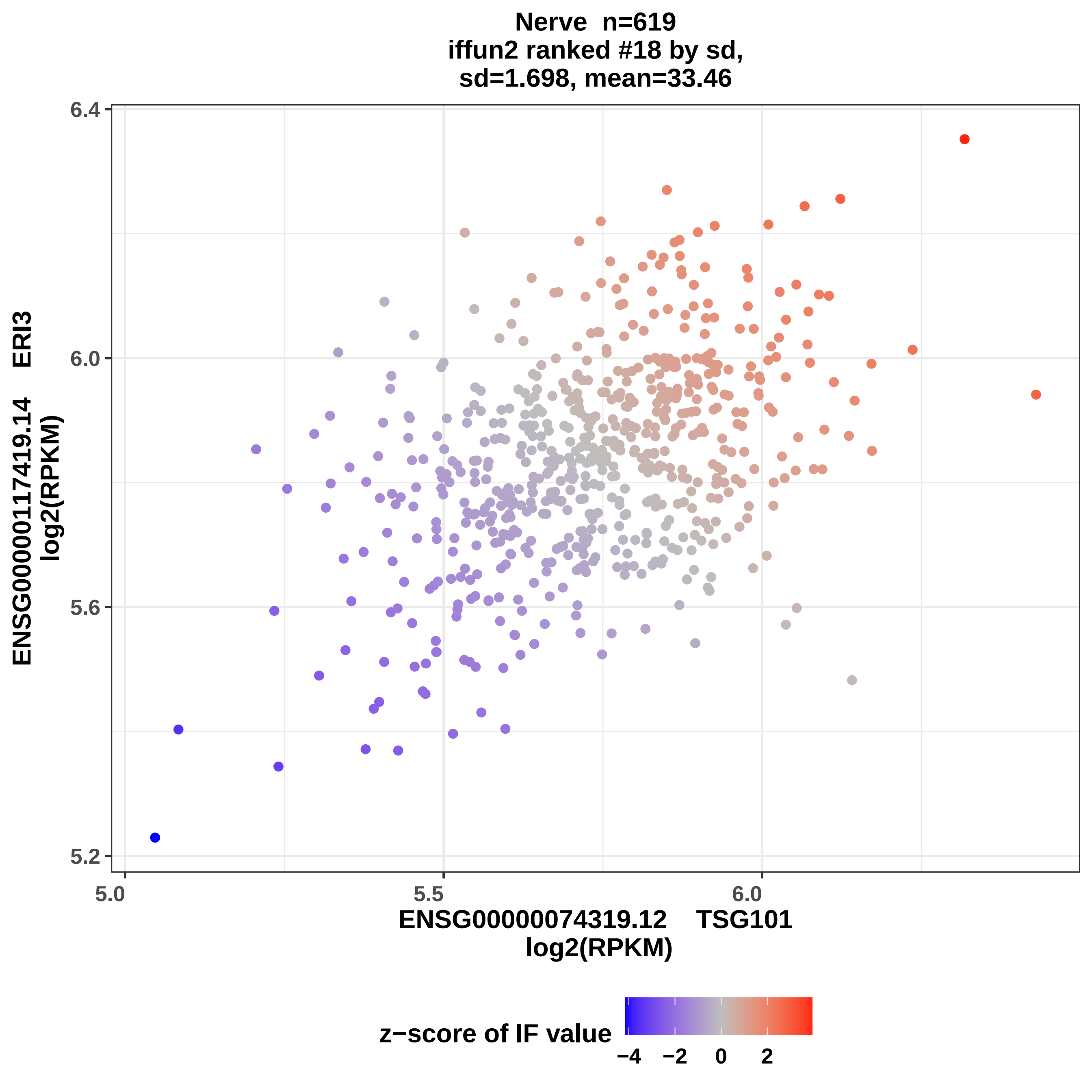
Nerve n=619
iffun2 ranked #18 by sd,
sd=1.698, mean=33.46
ENSG00000117419.14 ERI3
log2(RPKM)
6.4
6.0
5.6
5.2
5.0
5.5
6.0
ENSG00000074319.12 TSG101
log2(RPKM)
z−score of IF value
−4
−2
0
2

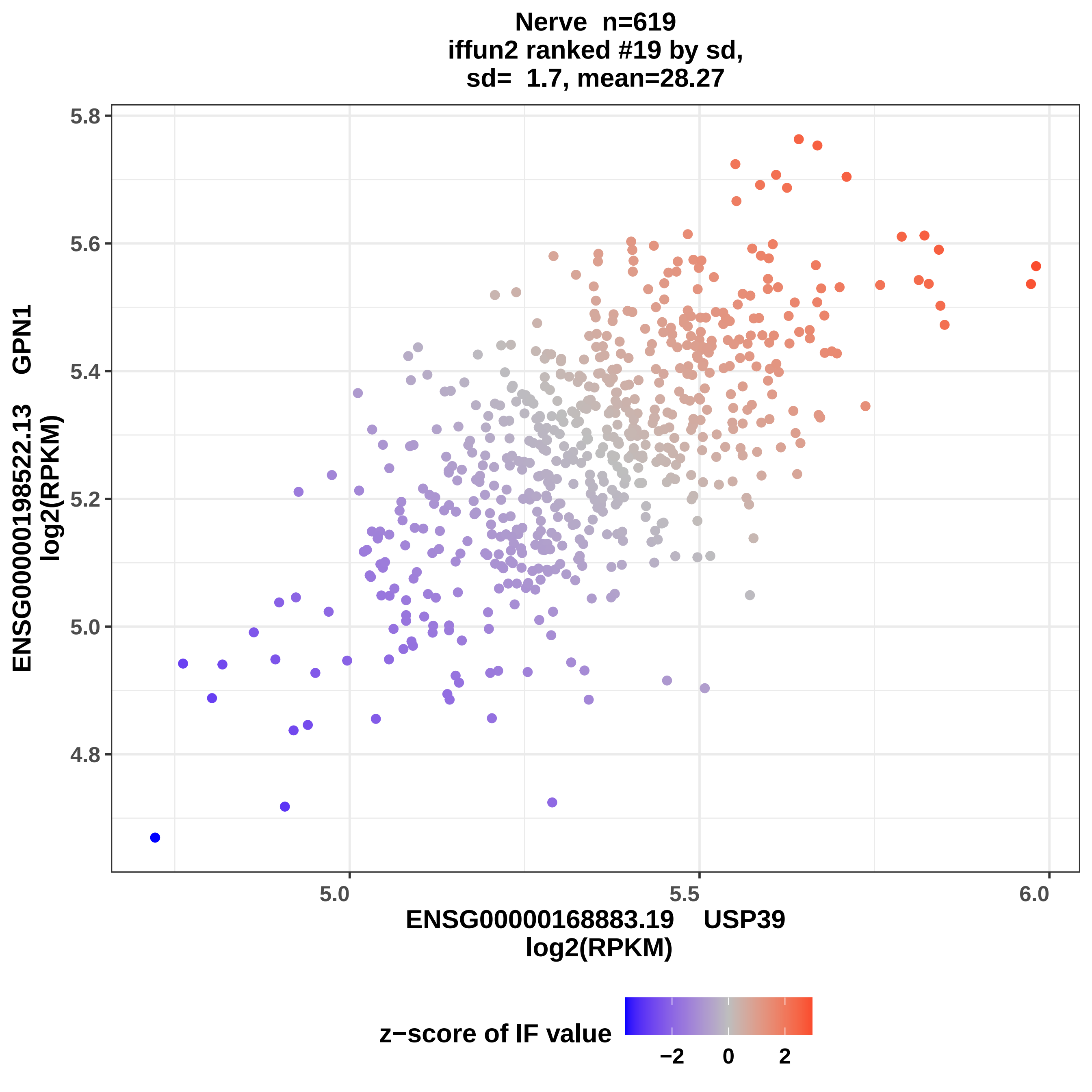

Nerve n=619
iffun2 ranked #19 by sd,
sd= 1.7, mean=28.27
ENSG00000198522.13 GPN1
log2(RPKM)
ENSG00000168883.19 USP39
log2(RPKM)
5.8
5.6
5.4
5.2
5.0
4.8
5.0
5.5
6.0
z−score of IF value
−2
0
2

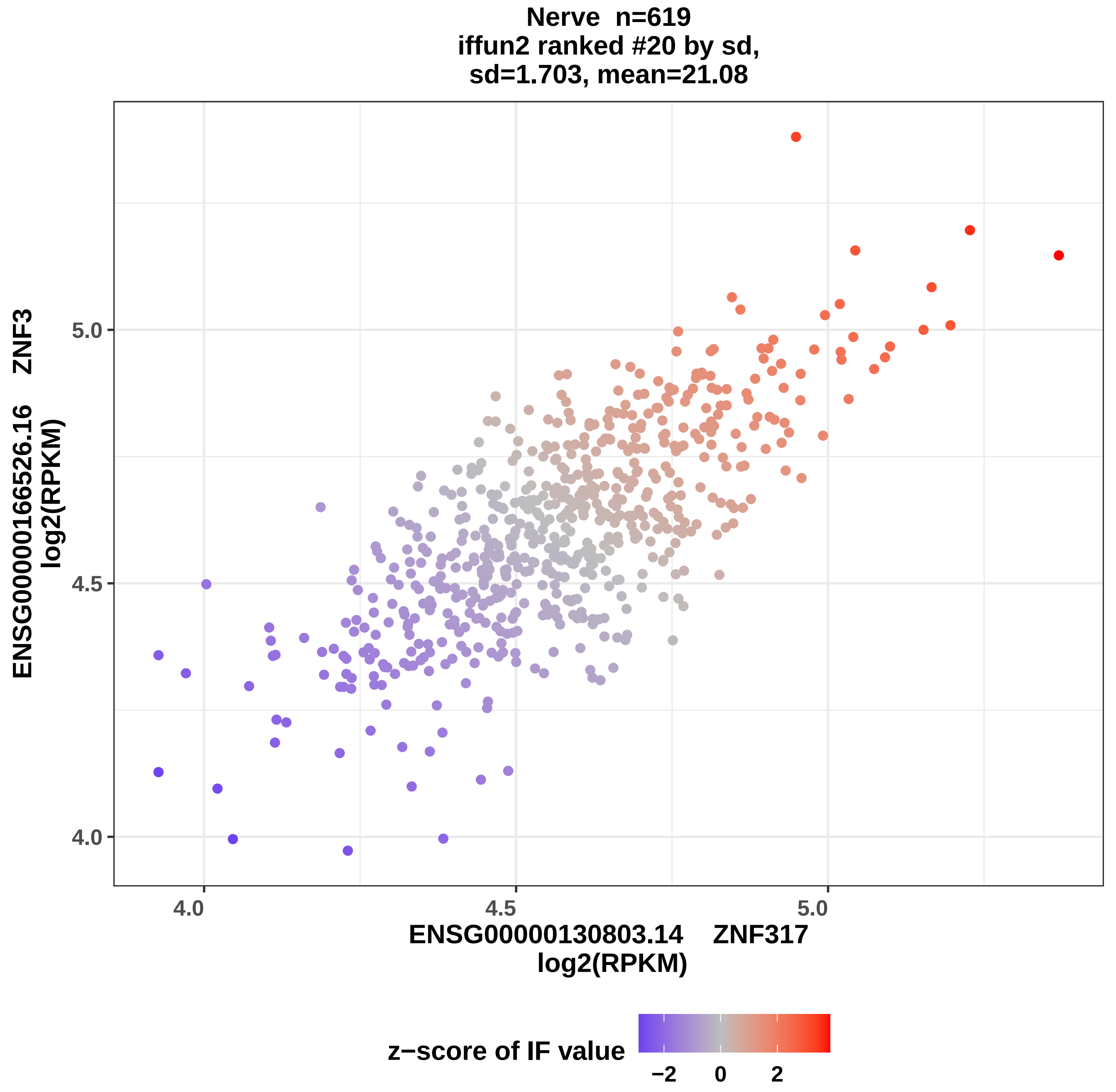
Nerve n=619
iffun2 ranked #20 by sd,
sd=1.703, mean=21.08
ENSG00000166526.16 ZNF3
log2(RPKM)
5.0
4.5
4.0
4.0
4.5
5.0
ENSG00000130803.14 ZNF317
log2(RPKM)
z−score of IF value
−2
0
2

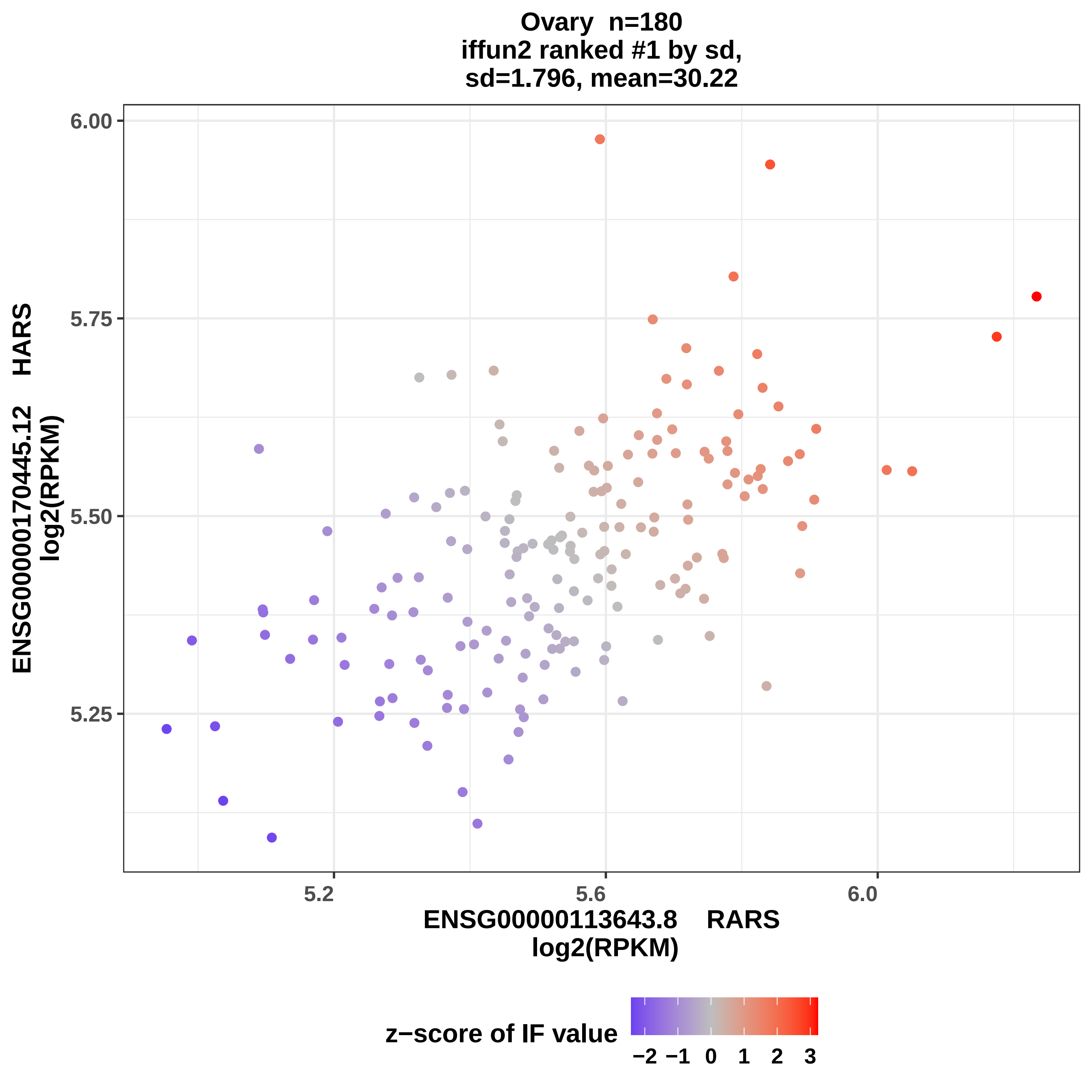
Ovary n=180
iffun2 ranked #1 by sd,
sd=1.796, mean=30.22
6.00
5.75
5.50
5.25
5.2
5.6
6.0
ENSG00000170445.12 HARS
log2(RPKM)
ENSG00000113643.8 RARS
log2(RPKM)
z−score of IF value
−2 −1 0 1 2 3

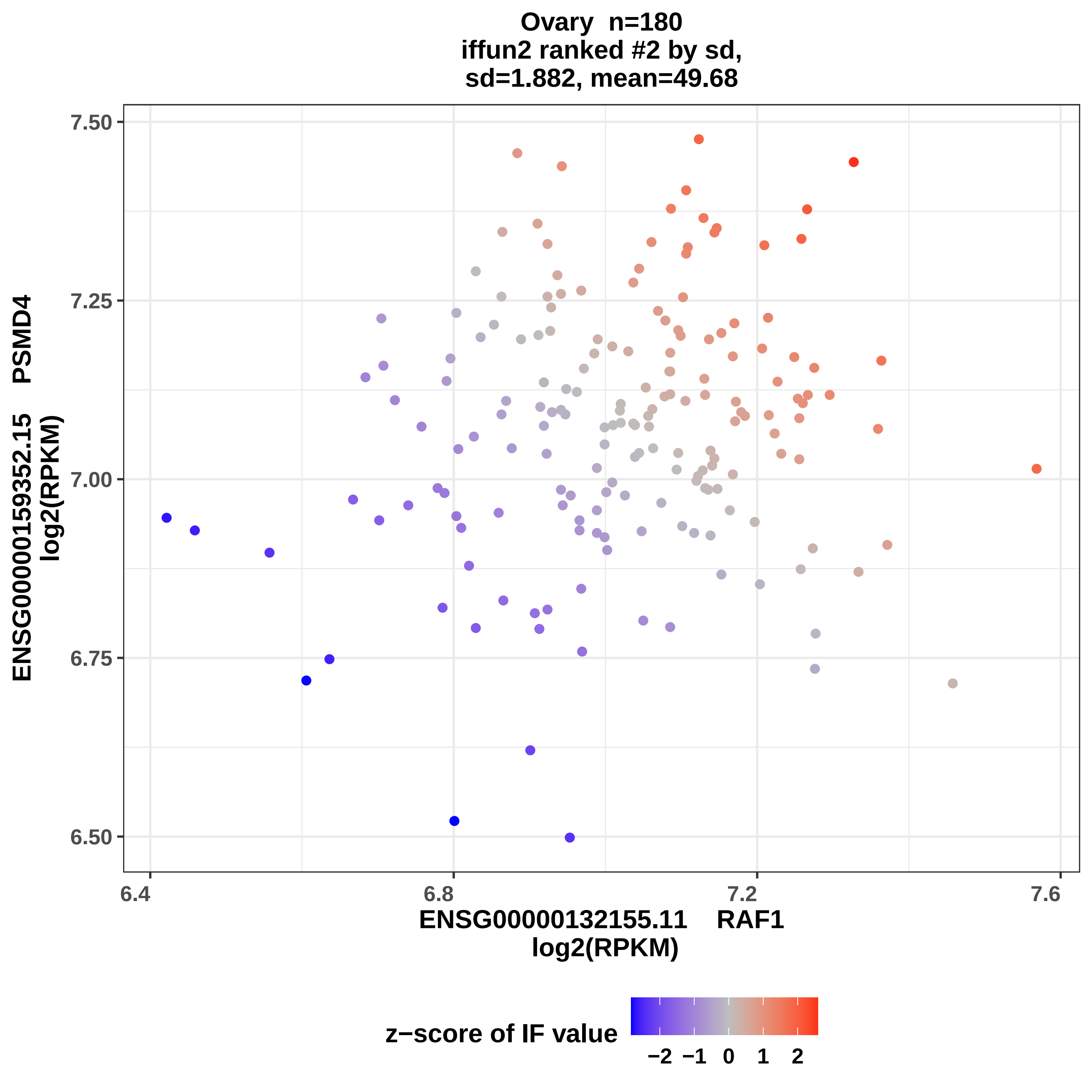
Ovary n=180
iffun2 ranked #2 by sd,
sd=1.882, mean=49.68
ENSG00000159352.15 PSMD4
log2(RPKM)
7.50
7.25
7.00
6.75
6.50
6.4
6.8
7.2
7.6
ENSG00000132155.11 RAF1
log2(RPKM)
z−score of IF value
−2 −1 0 1 2

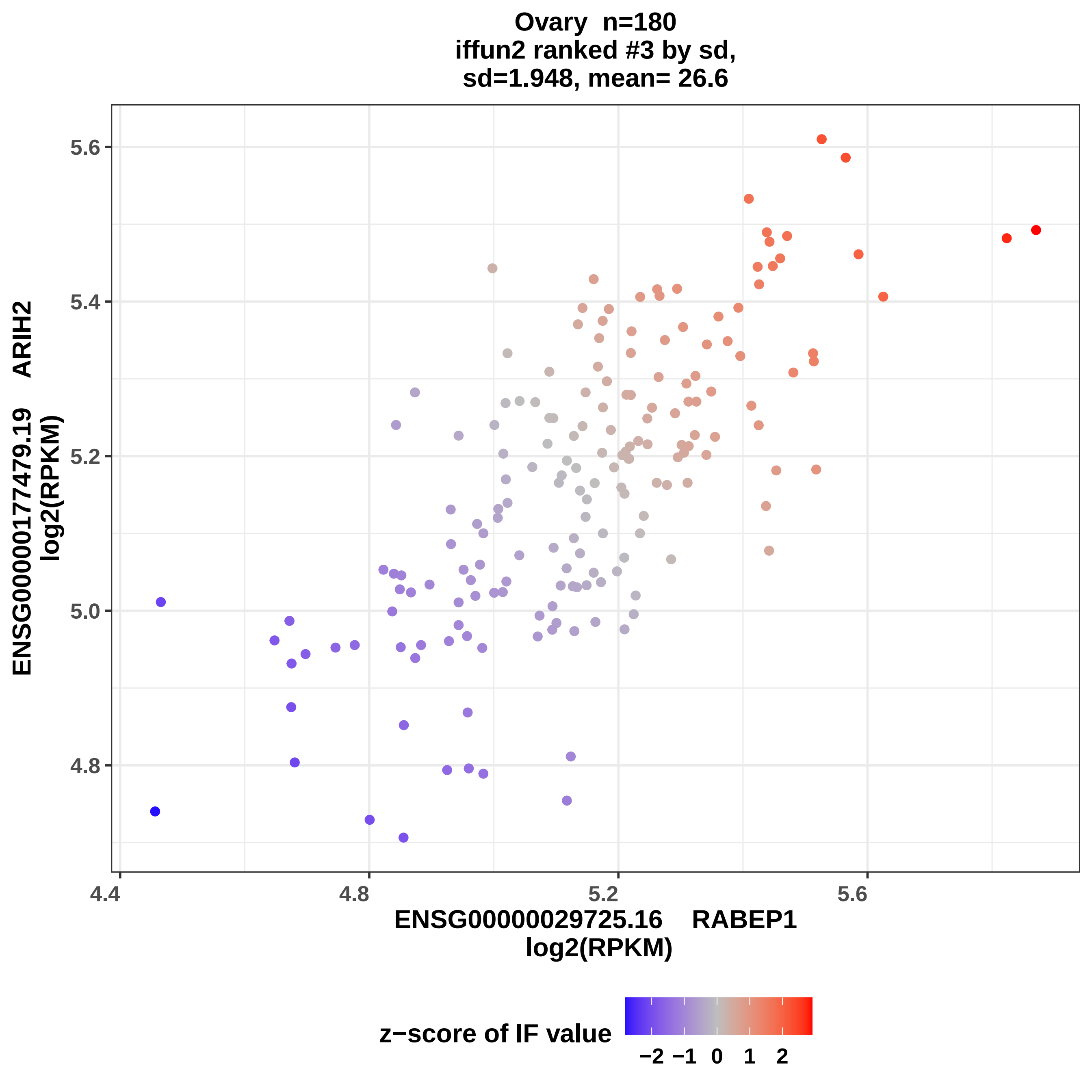

Ovary n=180
iffun2 ranked #3 by sd,
sd=1.948, mean= 26.6
5.6
5.4
5.2
5.0
4.8
ENSG00000177479.19 ARIH2
log2(RPKM)
4.4
4.8
5.2
5.6
ENSG00000029725.16 RABEP1
log2(RPKM)
z−score of IF value
−2 −1 0 1 2

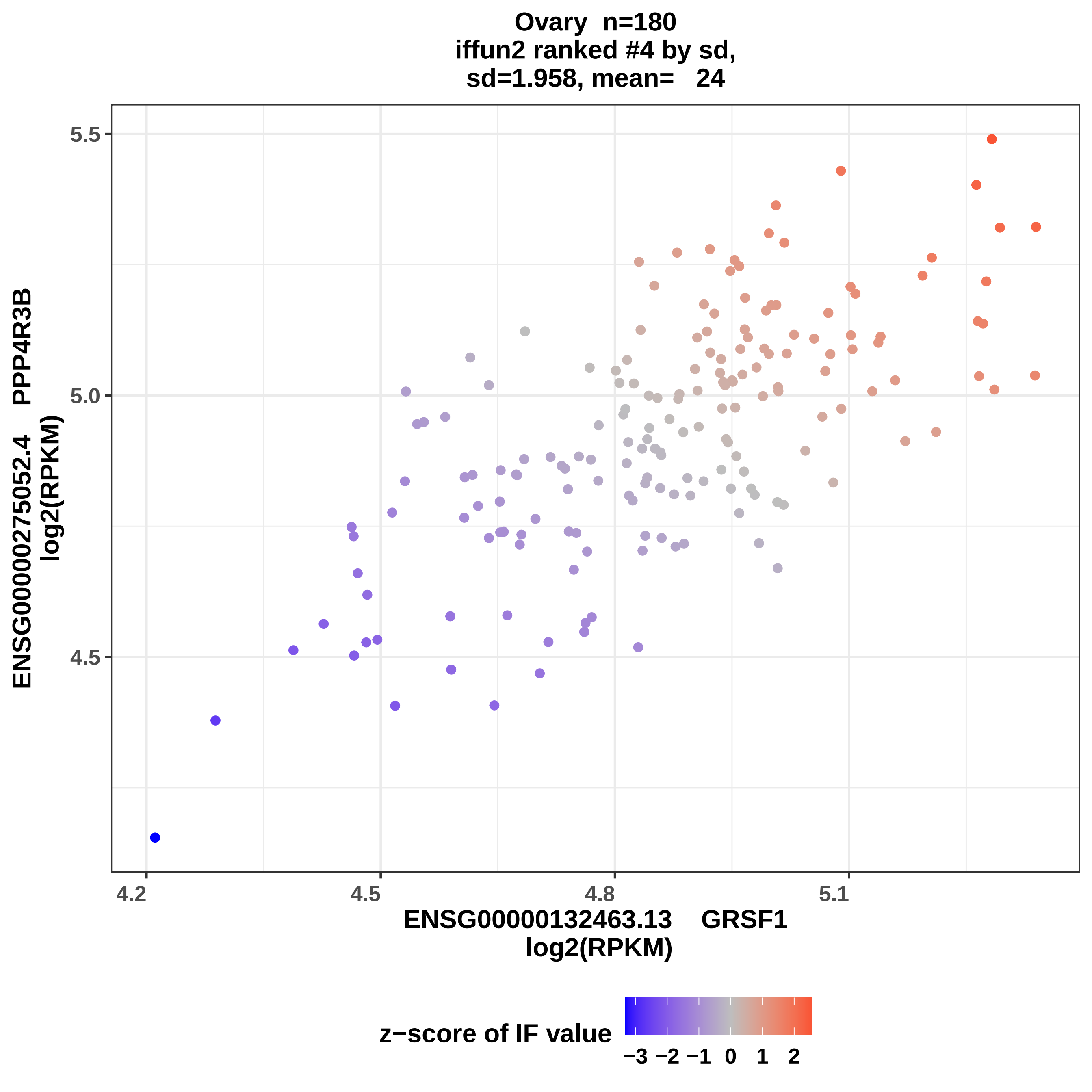
Ovary n=180
iffun2 ranked #4 by sd,
sd=1.958, mean= 24
5.5
5.0
4.5
ENSG00000275052.4 PPP4R3B
log2(RPKM)
4.2
4.5
4.8
5.1
ENSG00000132463.13 GRSF1
log2(RPKM)
z−score of IF value
−3 −2 −1 0 1 2

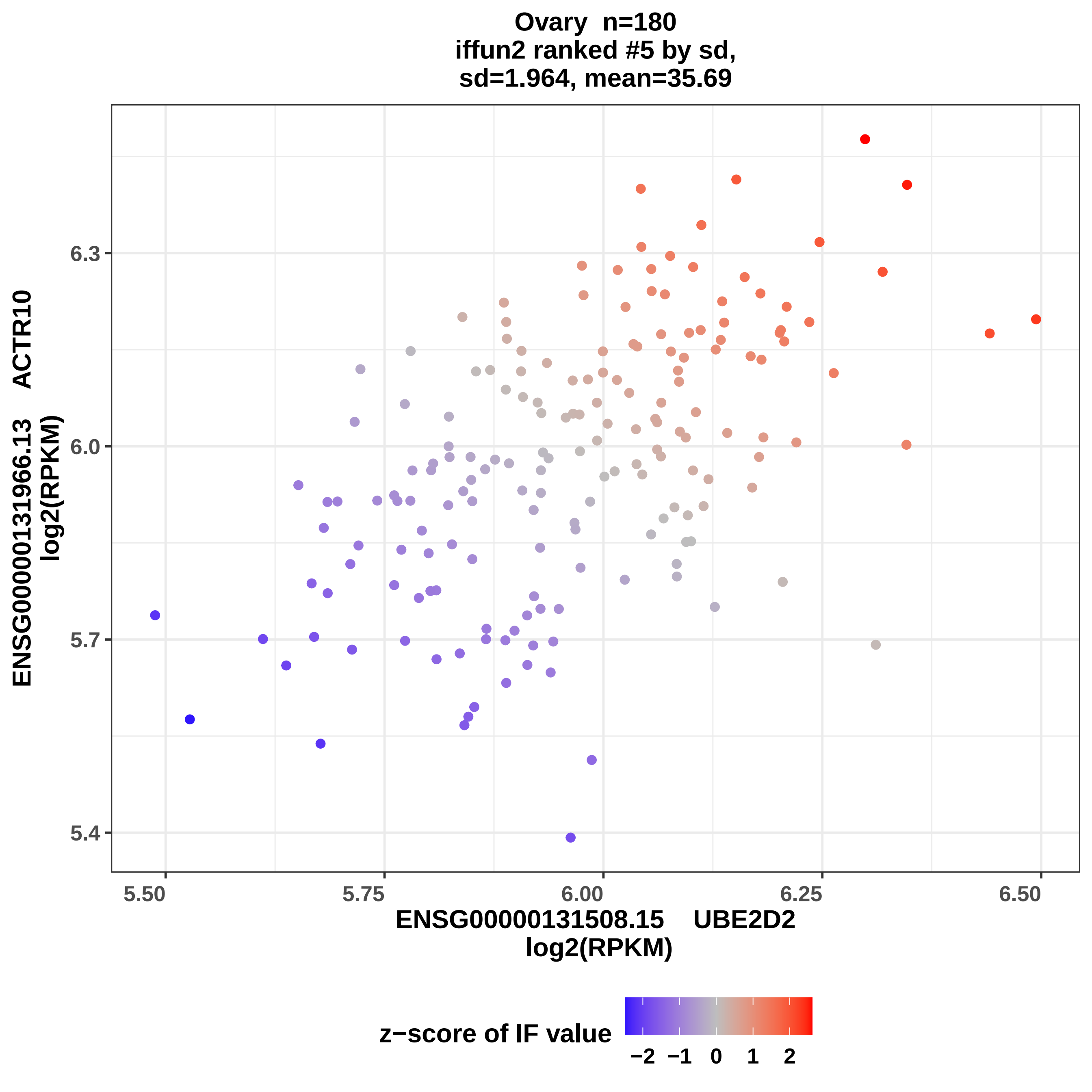

Ovary n=180
iffun2 ranked #5 by sd,
sd=1.964, mean=35.69
ENSG00000131966.13 ACTR10
log2(RPKM)
6.3
6.0
5.7
5.4
5.50
5.75
6.00
6.25
6.50
ENSG00000131508.15 UBE2D2
log2(RPKM)
z−score of IF value
−2
−1
0
1
2

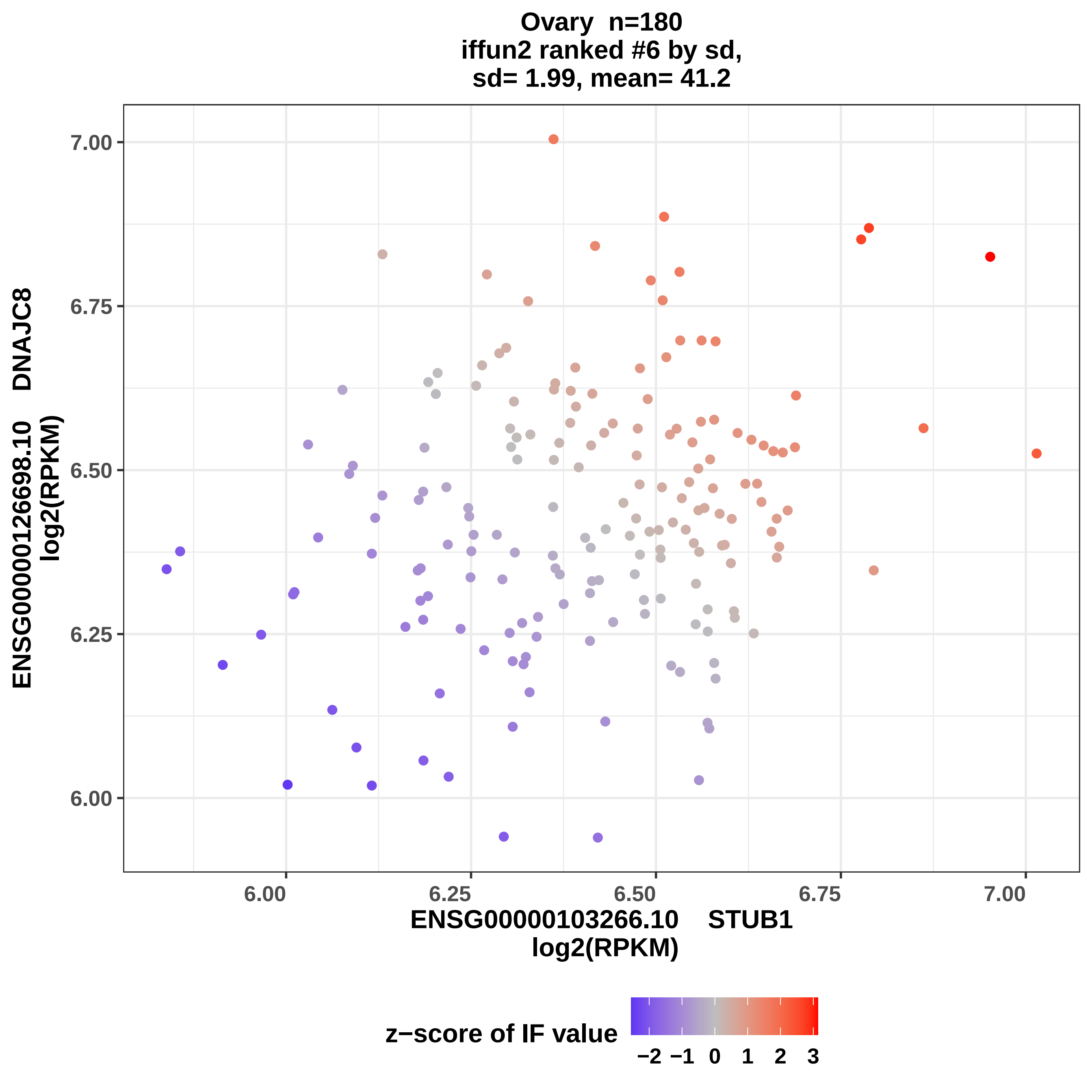

Ovary n=180
iffun2 ranked #6 by sd,
sd= 1.99, mean= 41.2
ENSG00000126698.10 DNAJC8
log2(RPKM)
7.00
6.75
6.50
6.25
6.00
6.00
6.25
6.50
6.75
7.00
ENSG00000103266.10 STUB1
log2(RPKM)
z−score of IF value
−2
−1
0
1
2
3

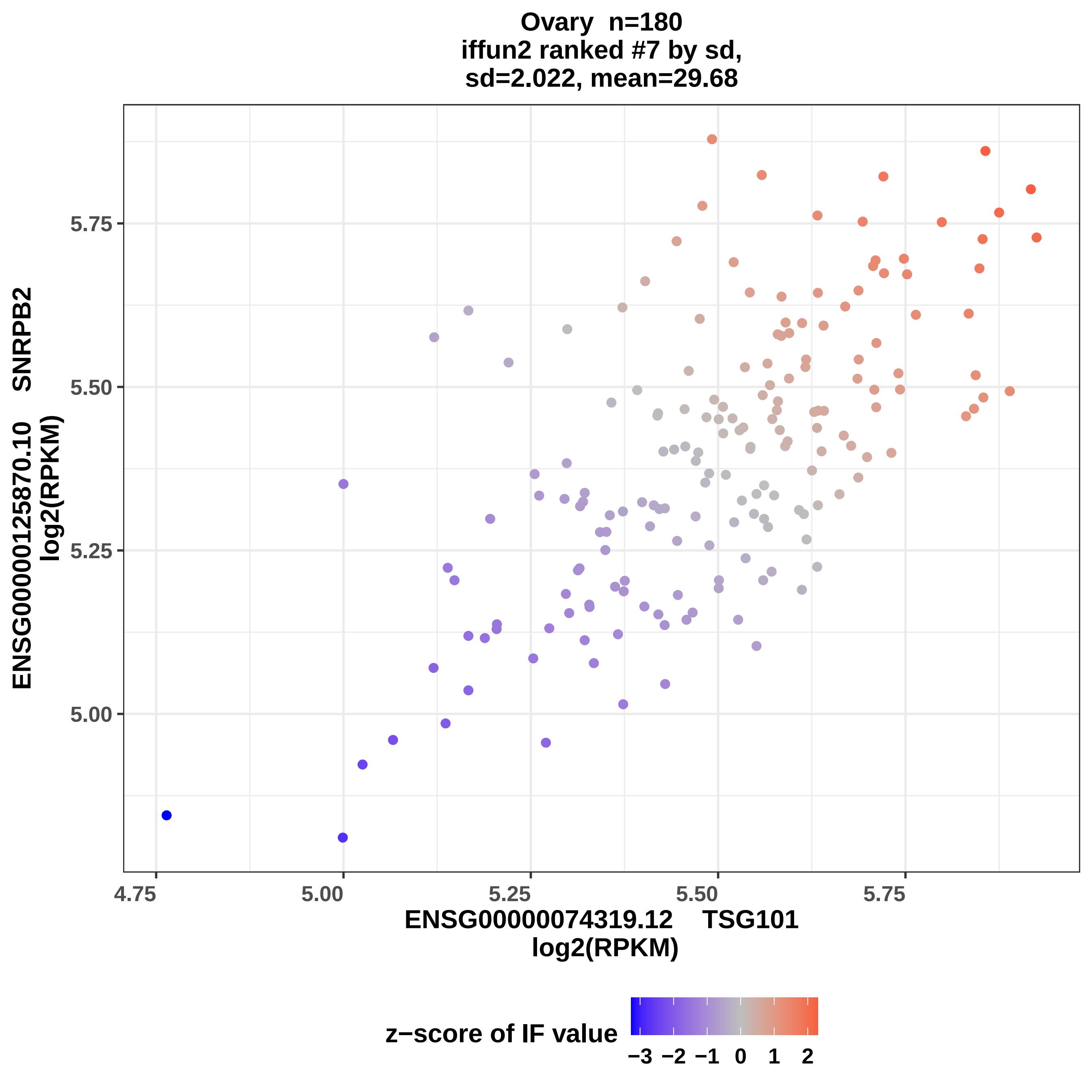
Ovary n=180
iffun2 ranked #7 by sd,
sd=2.022, mean=29.68
ENSG00000125870.10 SNRPB2
log2(RPKM)
5.75
5.50
5.25
5.00
4.75
5.00
5.25
5.50
5.75
ENSG00000074319.12 TSG101
log2(RPKM)
z−score of IF value
−3 −2 −1 0 1 2

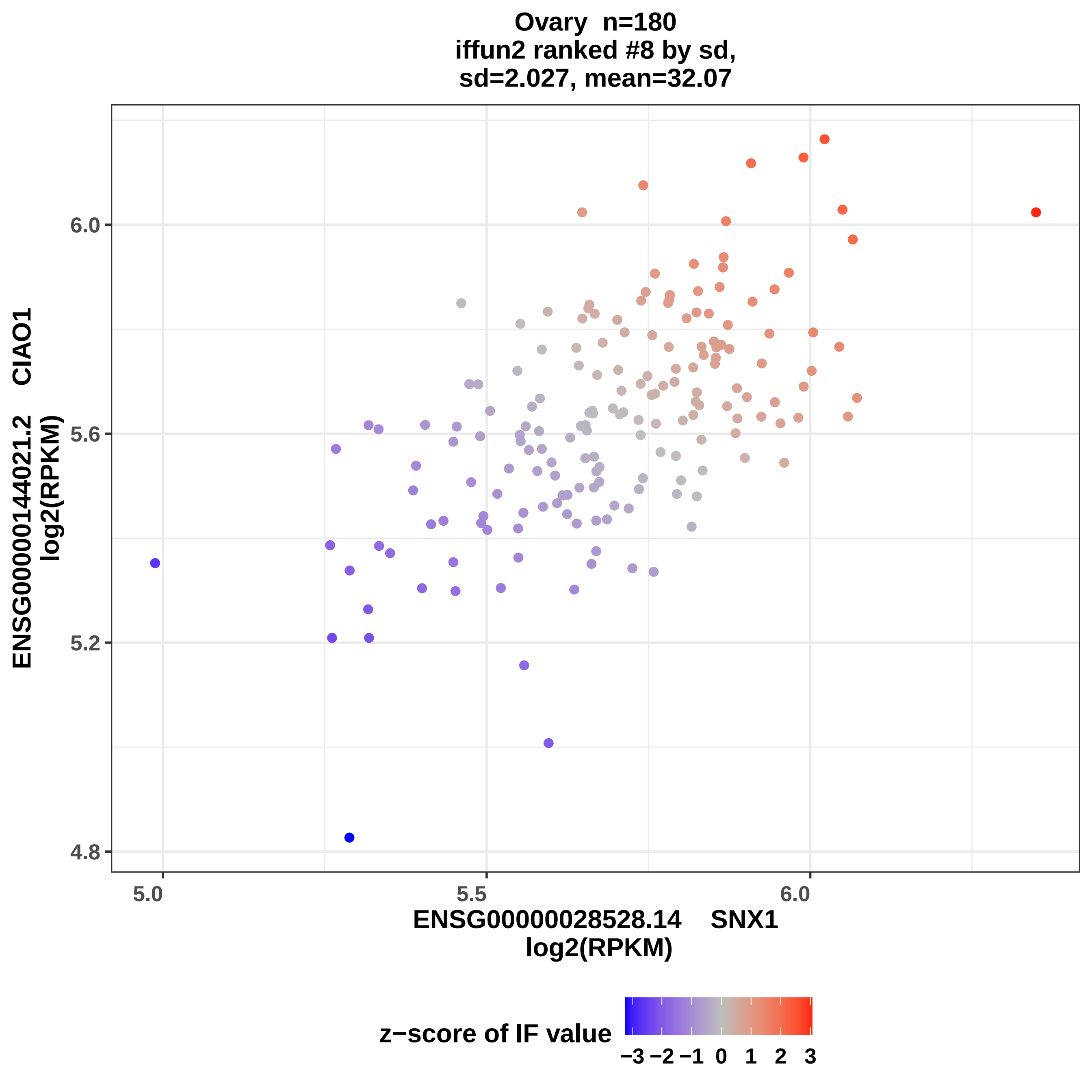

Ovary n=180
iffun2 ranked #8 by sd,
sd=2.027, mean=32.07
ENSG00000144021.2 CIAO1
log2(RPKM)
6.0
5.6
5.2
4.8
5.0
5.5
6.0
ENSG00000028528.14 SNX1
log2(RPKM)
z−score of IF value
−3 −2 −1 0 1 2 3

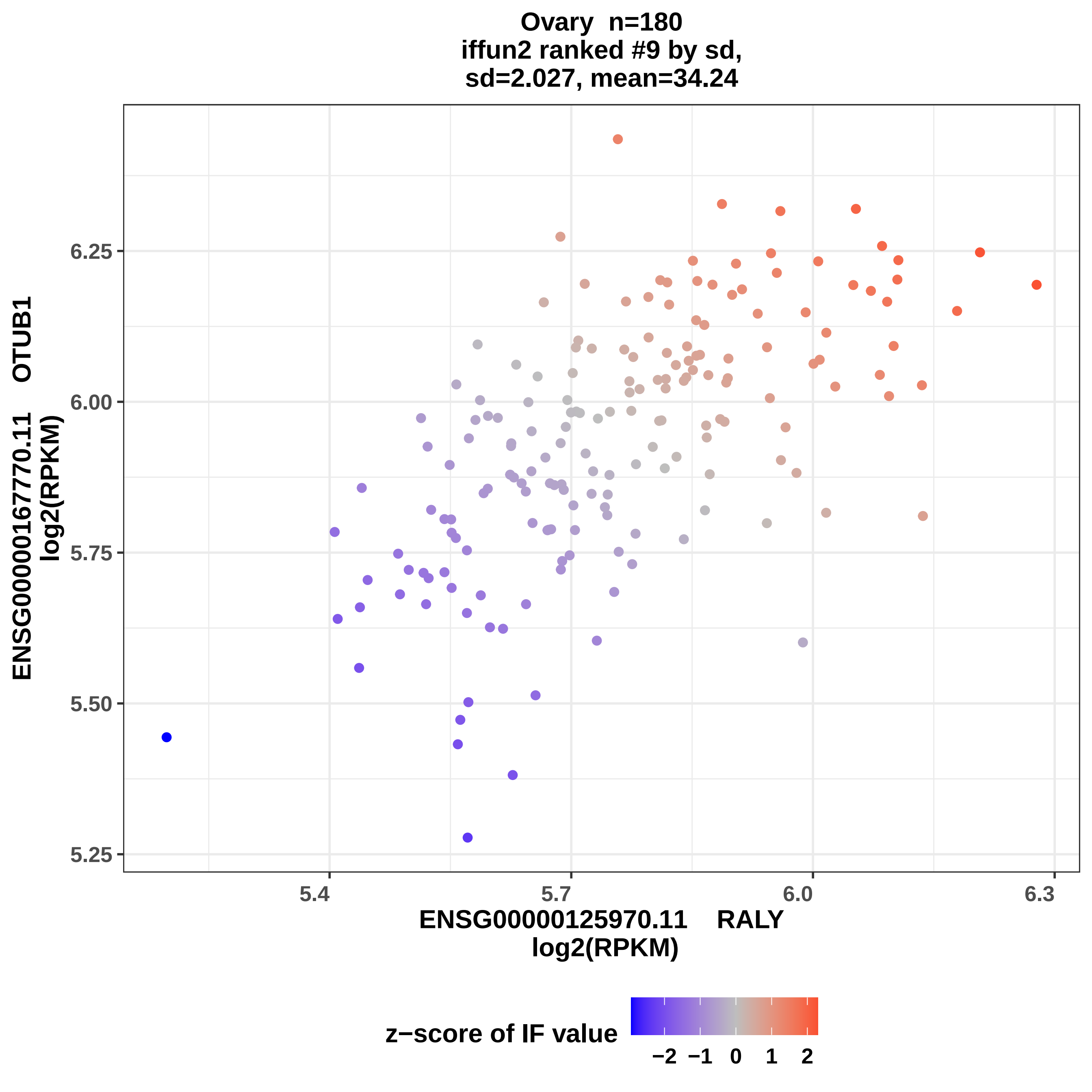

Ovary n=180
iffun2 ranked #9 by sd,
sd=2.027, mean=34.24
ENSG00000167770.11 OTUB1
log2(RPKM)
6.25
6.00
5.75
5.50
5.25
5.4
5.7
6.0
6.3
ENSG00000125970.11 RALY
log2(RPKM)
z−score of IF value
−2
−1
0
1
2

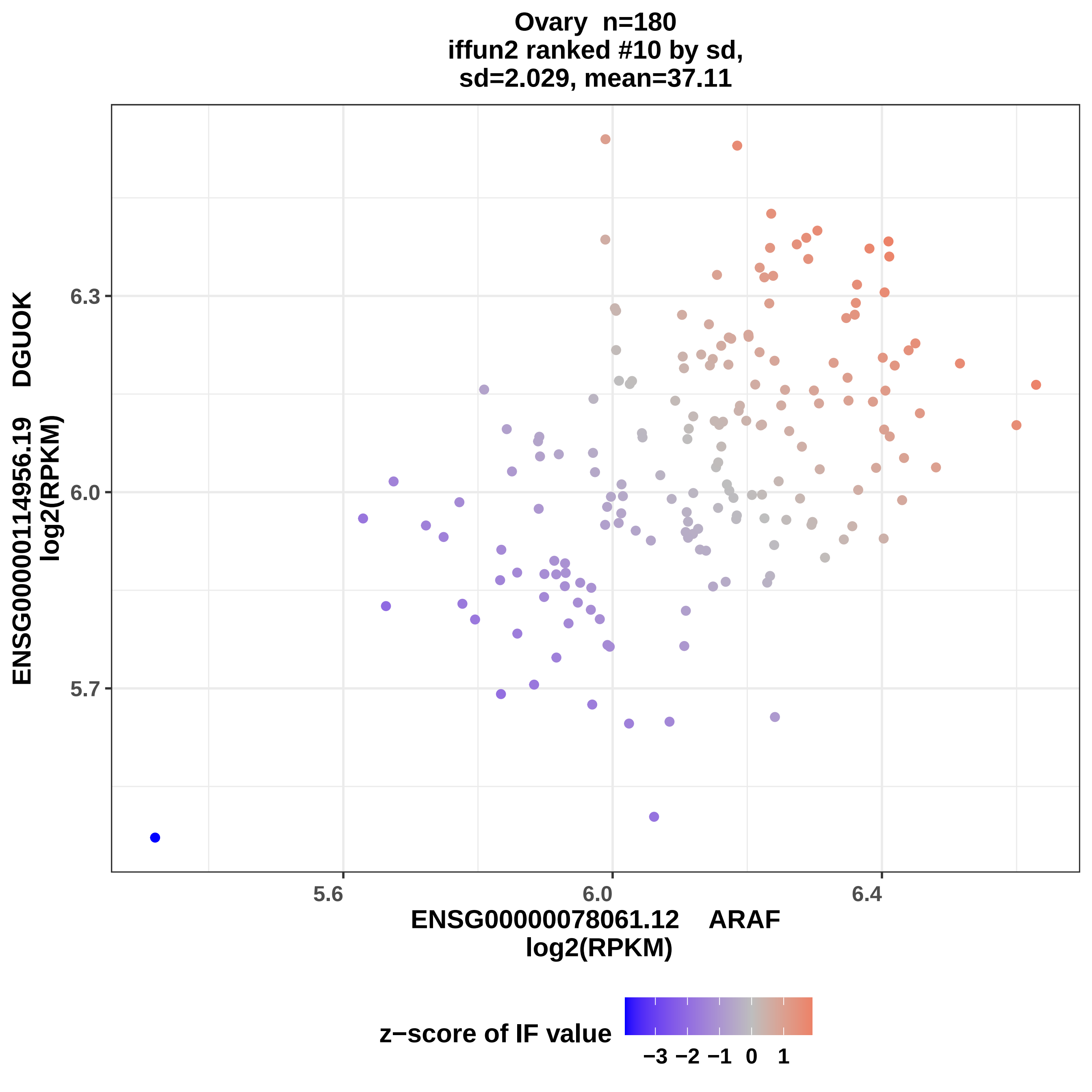
Ovary n=180
iffun2 ranked #10 by sd,
sd=2.029, mean=37.11
ENSG00000114956.19 DGUOK
log2(RPKM)
6.3
6.0
5.7
5.6
6.0
6.4
ENSG00000078061.12 ARAF
log2(RPKM)
z−score of IF value
−3 −2 −1 0 1

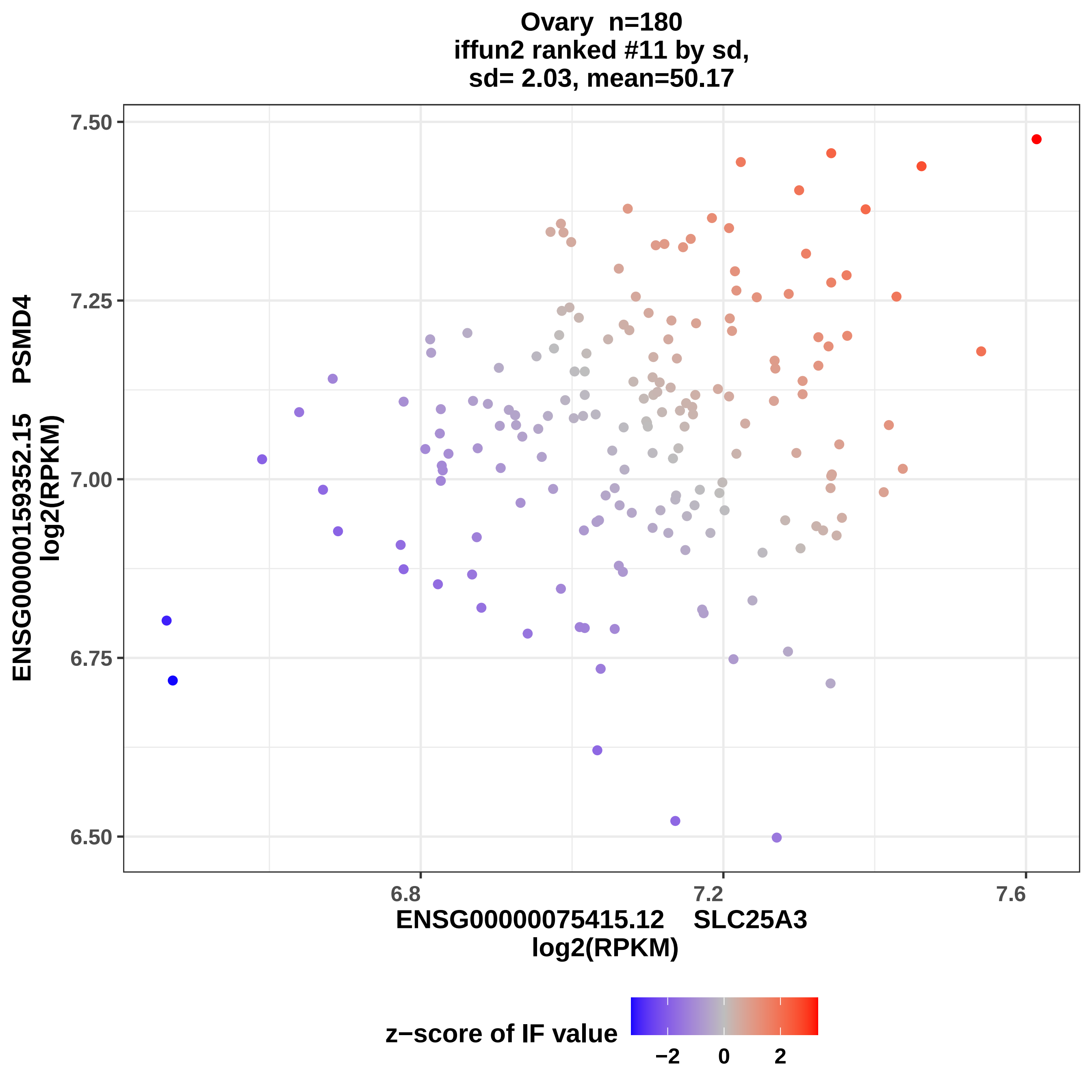
Ovary n=180
iffun2 ranked #11 by sd,
sd= 2.03, mean=50.17
7.50
7.25
7.00
6.75
6.50
6.8
7.2
7.6
ENSG00000159352.15 PSMD4
log2(RPKM)
ENSG00000075415.12 SLC25A3
log2(RPKM)
z−score of IF value
−2
0
2

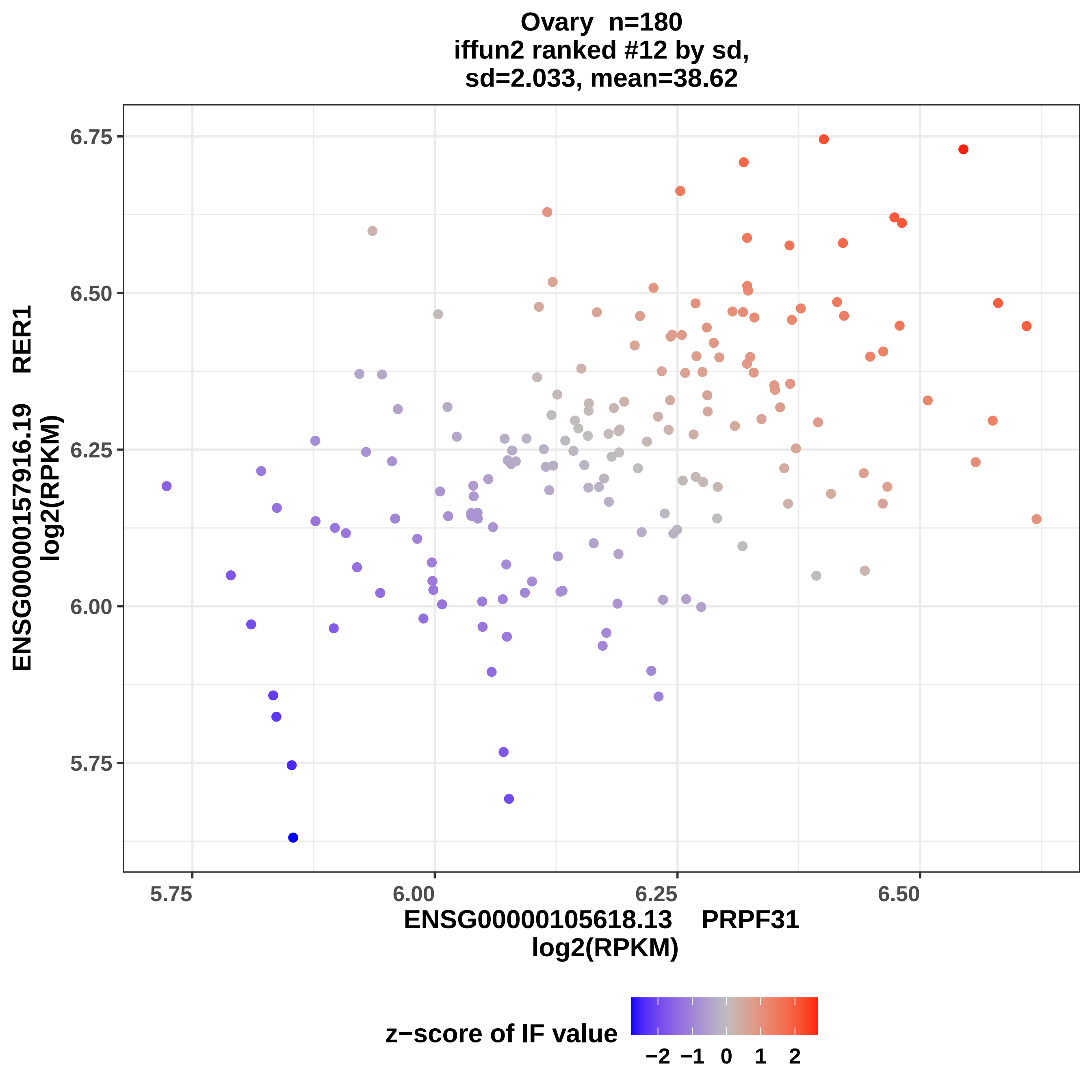

Ovary n=180
iffun2 ranked #12 by sd,
sd=2.033, mean=38.62
ENSG00000157916.19 RER1
log2(RPKM)
6.75
6.50
6.25
6.00
5.75
5.75
6.00
6.25
6.50
ENSG00000105618.13 PRPF31
log2(RPKM)
z−score of IF value
−2 −1 0 1 2

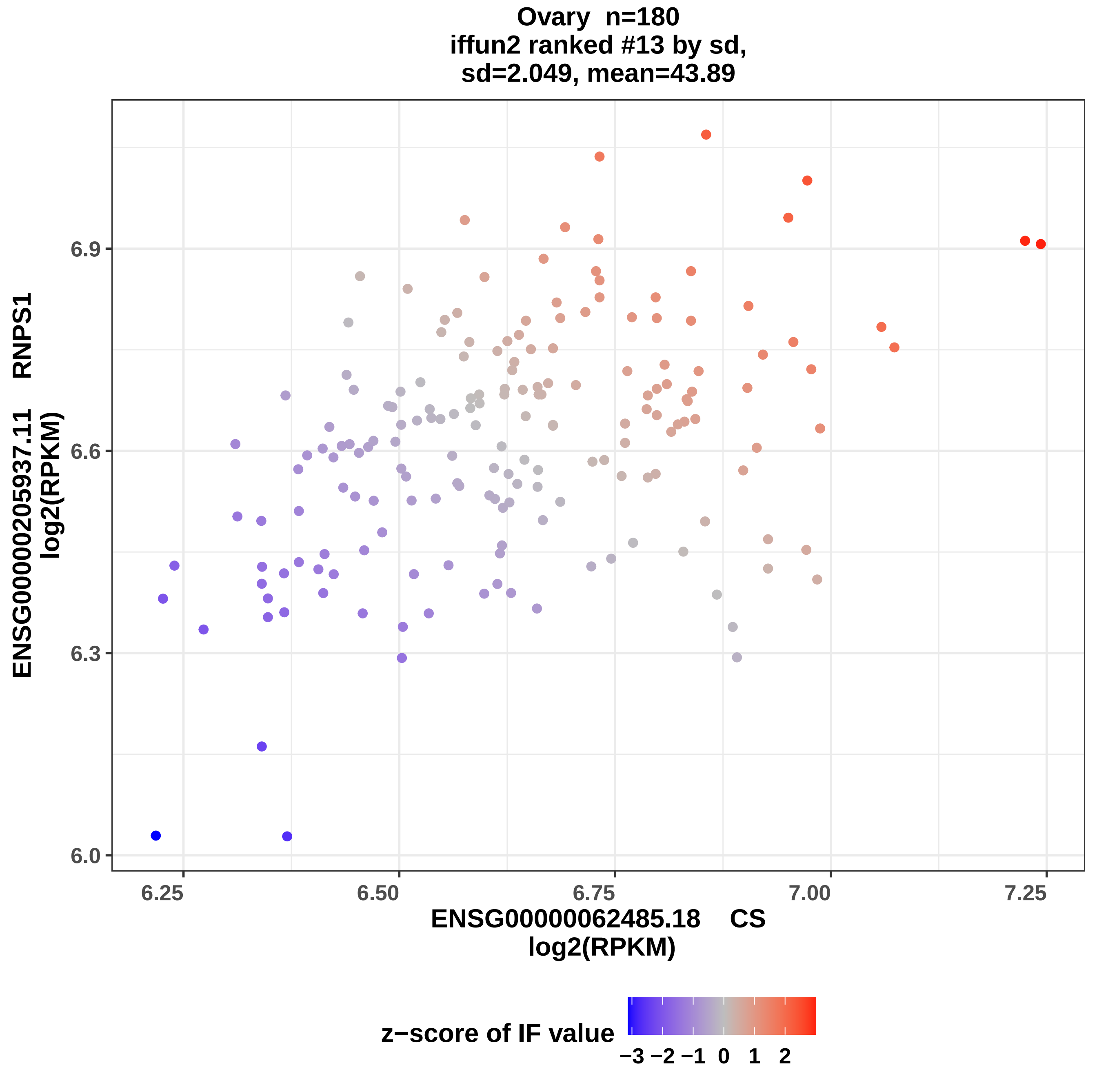

Ovary n=180
iffun2 ranked #13 by sd,
sd=2.049, mean=43.89
ENSG00000205937.11 RNPS1
log2(RPKM)
6.0
6.3
6.6
6.9
6.25
6.50
6.75
7.00
7.25
ENSG00000062485.18 CS
log2(RPKM)
z−score of IF value
−3 −2 −1 0 1 2

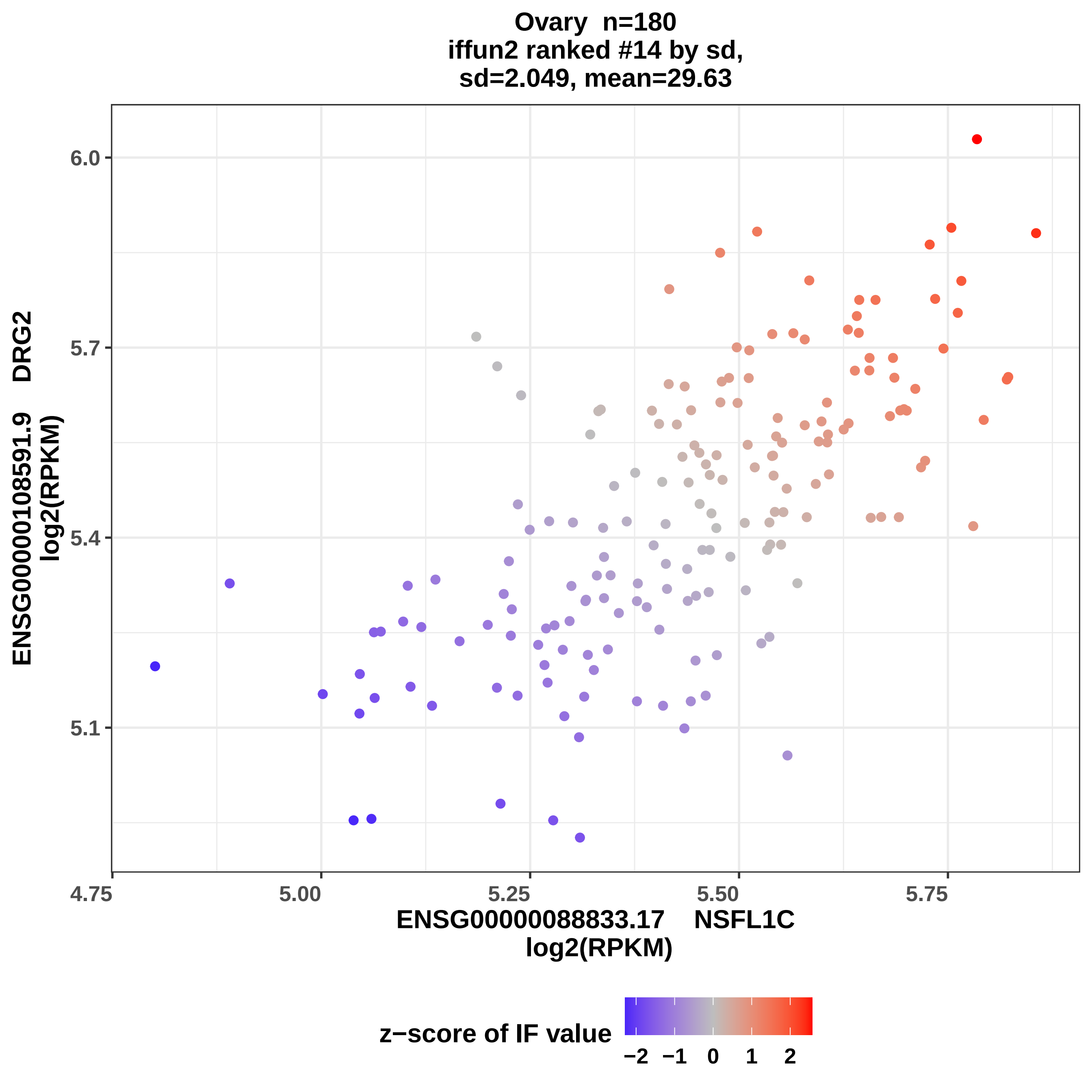
Ovary n=180
iffun2 ranked #14 by sd,
sd=2.049, mean=29.63
ENSG00000108591.9 DRG2
log2(RPKM)
6.0
5.7
5.4
5.1
4.75
5.00
5.25
5.50
5.75
ENSG00000088833.17 NSFL1C
log2(RPKM)
z−score of IF value
−2
−1
0
1
2

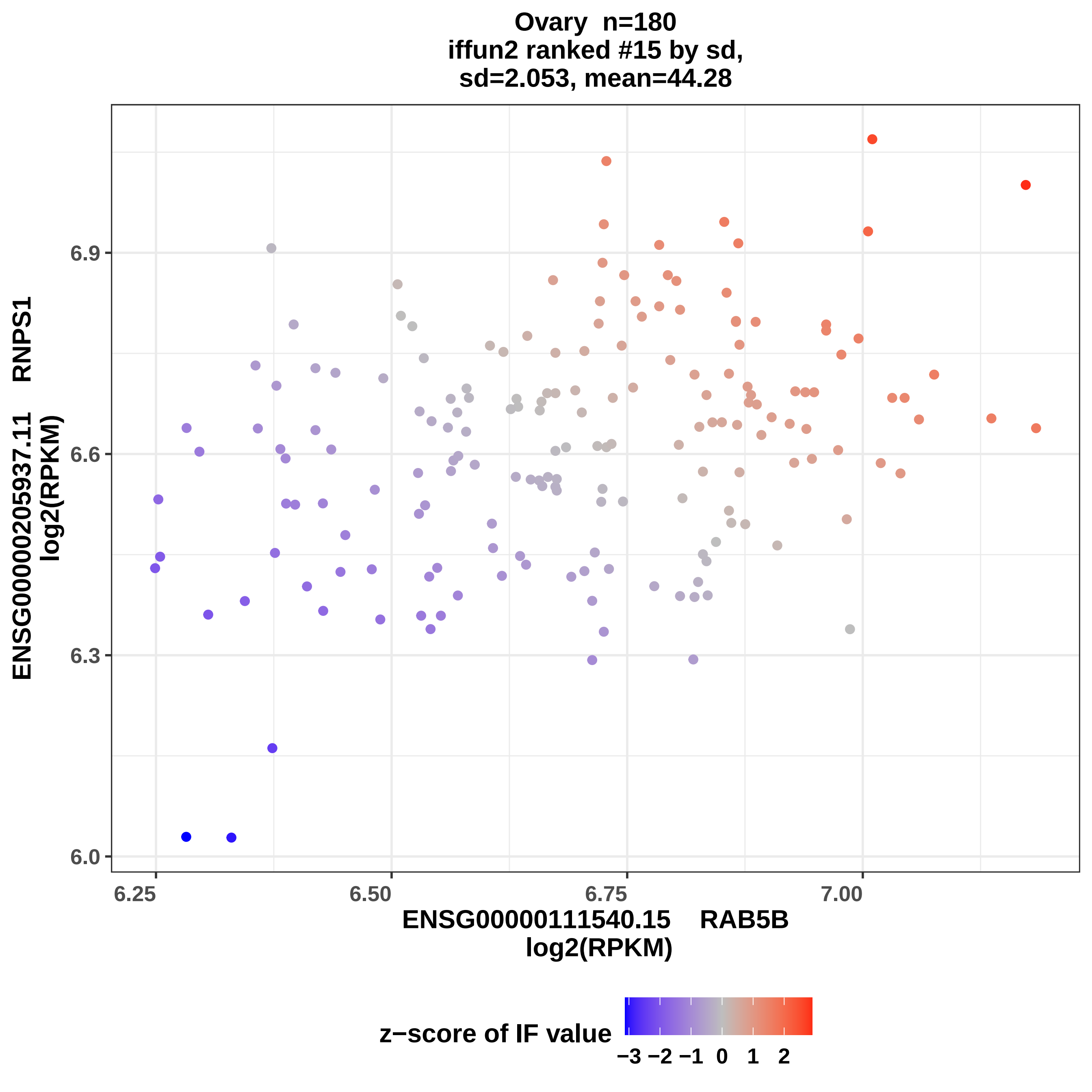

Ovary n=180
iffun2 ranked #15 by sd,
sd=2.053, mean=44.28
ENSG00000205937.11 RNPS1
log2(RPKM)
6.9
6.6
6.3
6.0
6.25
6.50
6.75
7.00
ENSG00000111540.15 RAB5B
log2(RPKM)
z−score of IF value
−3 −2 −1 0 1 2

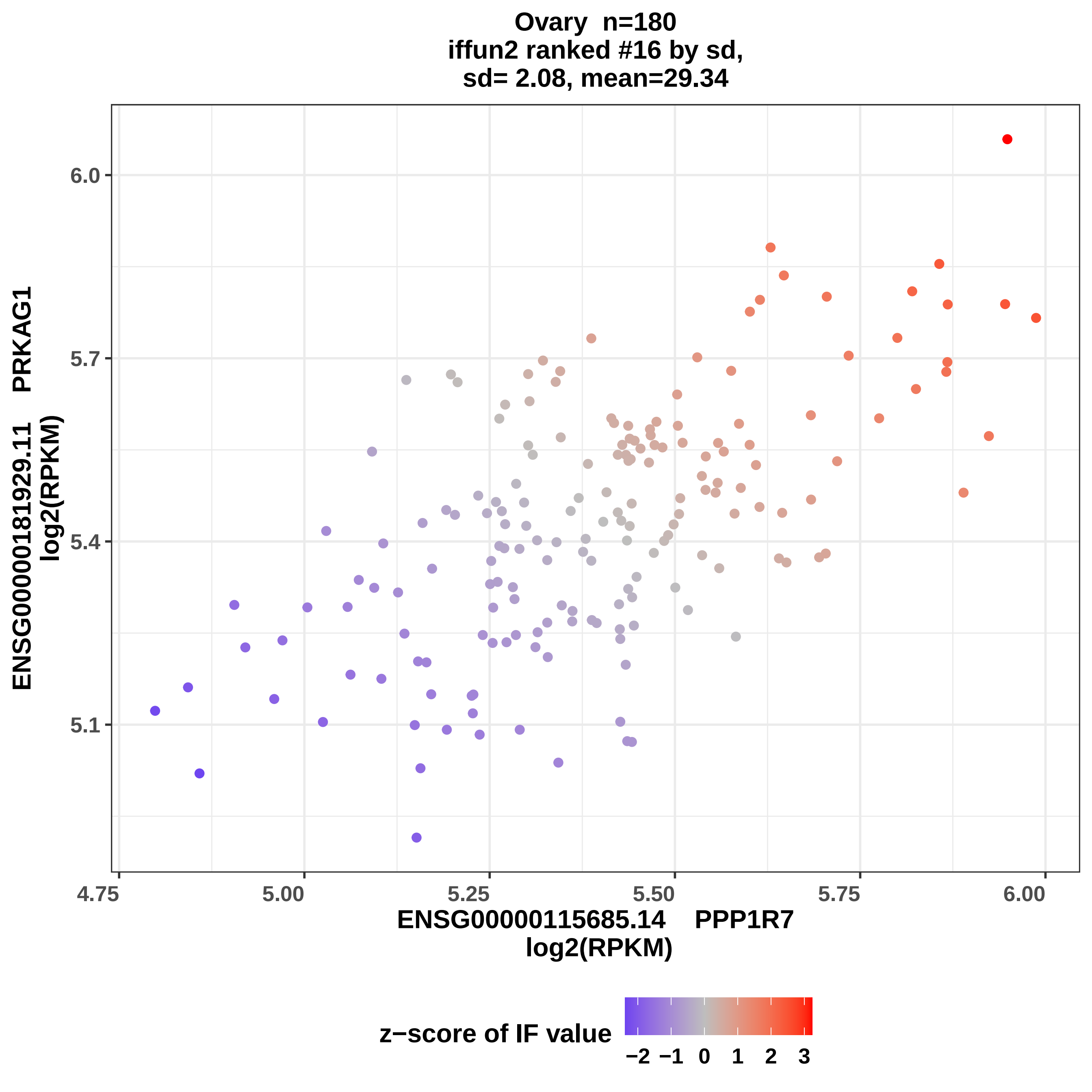

Ovary n=180
iffun2 ranked #16 by sd,
sd= 2.08, mean=29.34
ENSG00000181929.11 PRKAG1
log2(RPKM)
6.0
5.7
5.4
5.1
4.75
5.00
5.25
5.50
5.75
6.00
ENSG00000115685.14 PPP1R7
log2(RPKM)
z−score of IF value
−2 −1 0 1 2 3

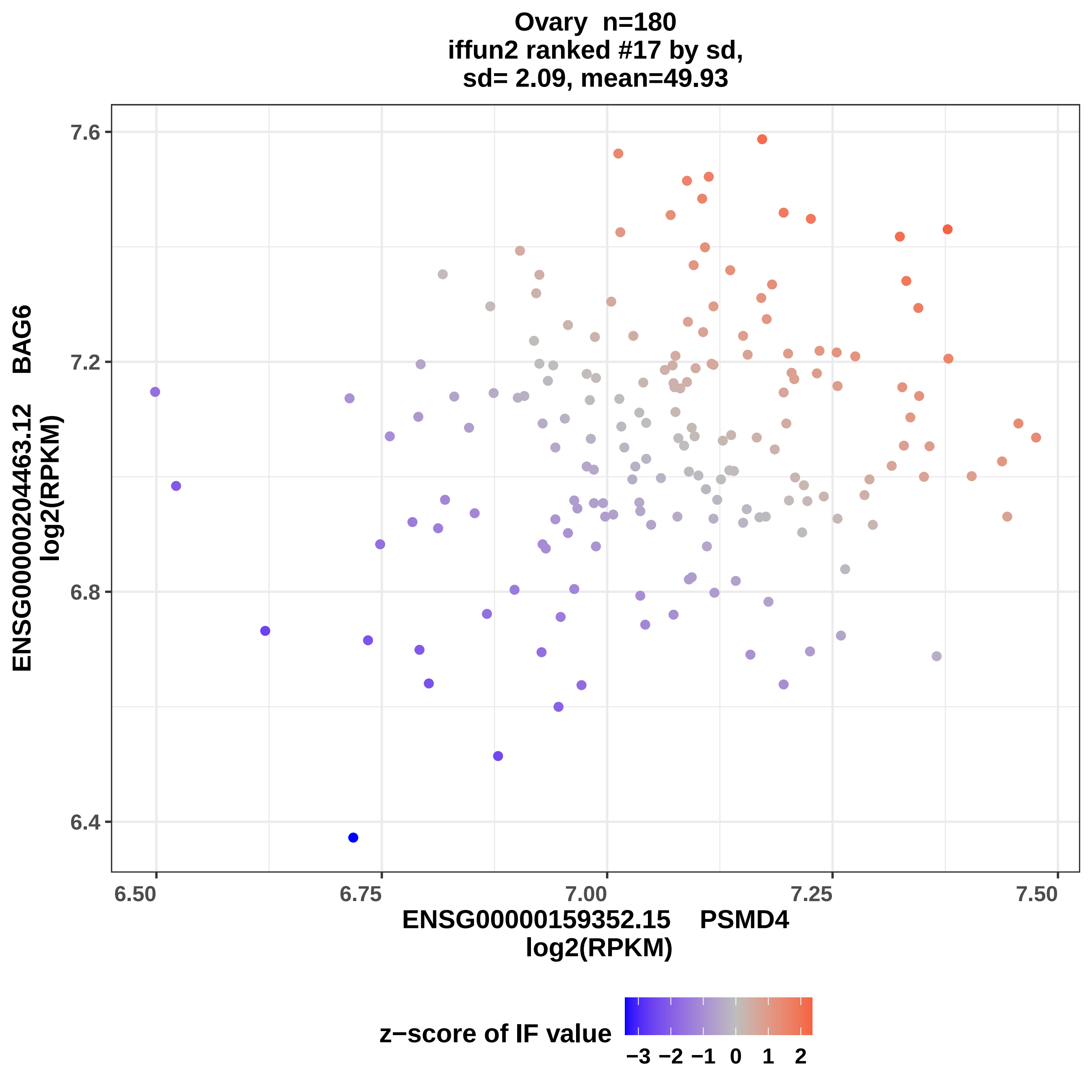

Ovary n=180
iffun2 ranked #17 by sd,
sd= 2.09, mean=49.93
ENSG00000204463.12 BAG6
log2(RPKM)
7.6
7.2
6.8
6.4
6.50
6.75
7.00
7.25
7.50
ENSG00000159352.15 PSMD4
log2(RPKM)
z−score of IF value
−3 −2 −1 0 1 2

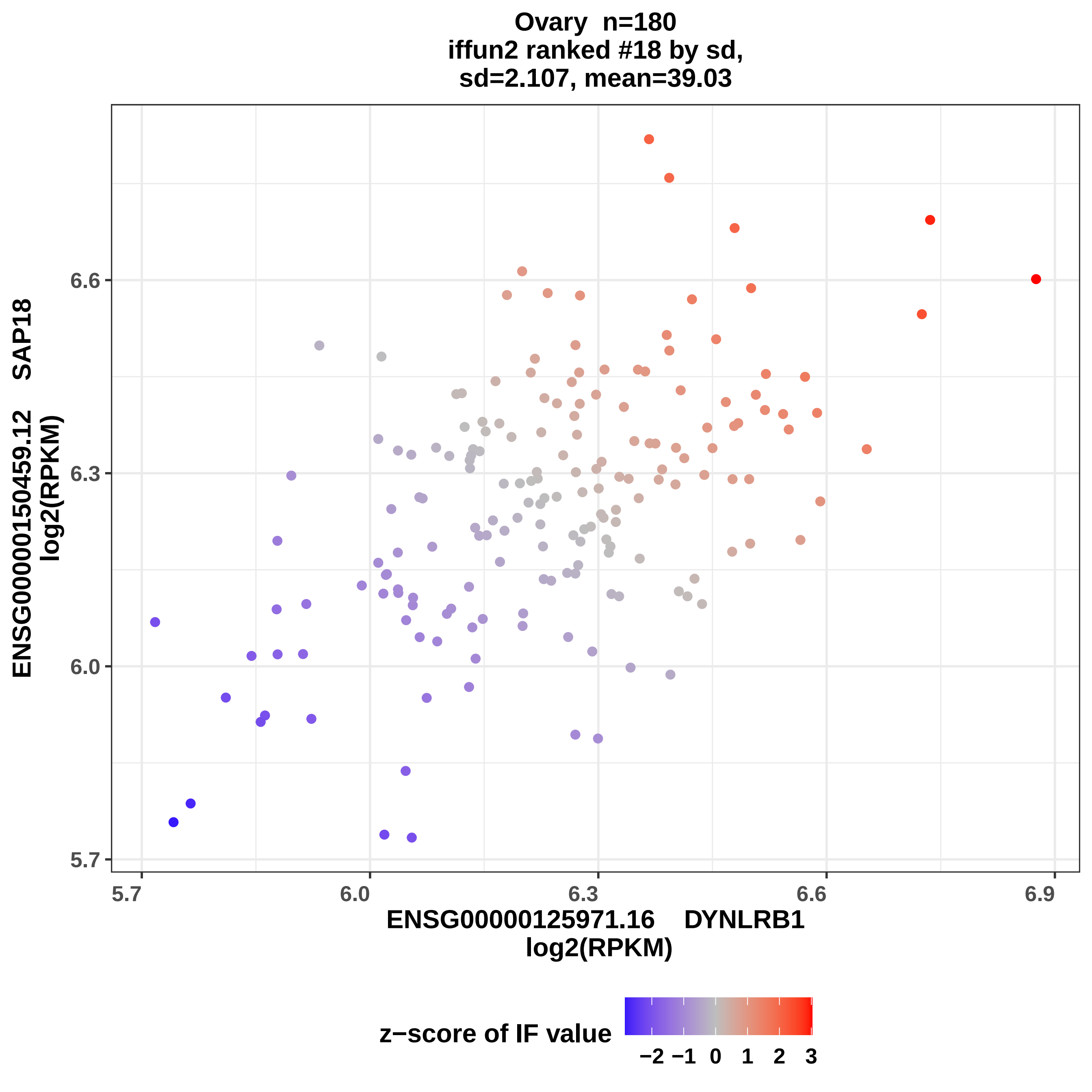

Ovary n=180
iffun2 ranked #18 by sd,
sd=2.107, mean=39.03
ENSG00000150459.12 SAP18
log2(RPKM)
6.6
6.3
6.0
5.7
5.7
6.0
6.3
6.6
6.9
ENSG00000125971.16 DYNLRB1
log2(RPKM)
z−score of IF value
−2 −1 0 1 2 3

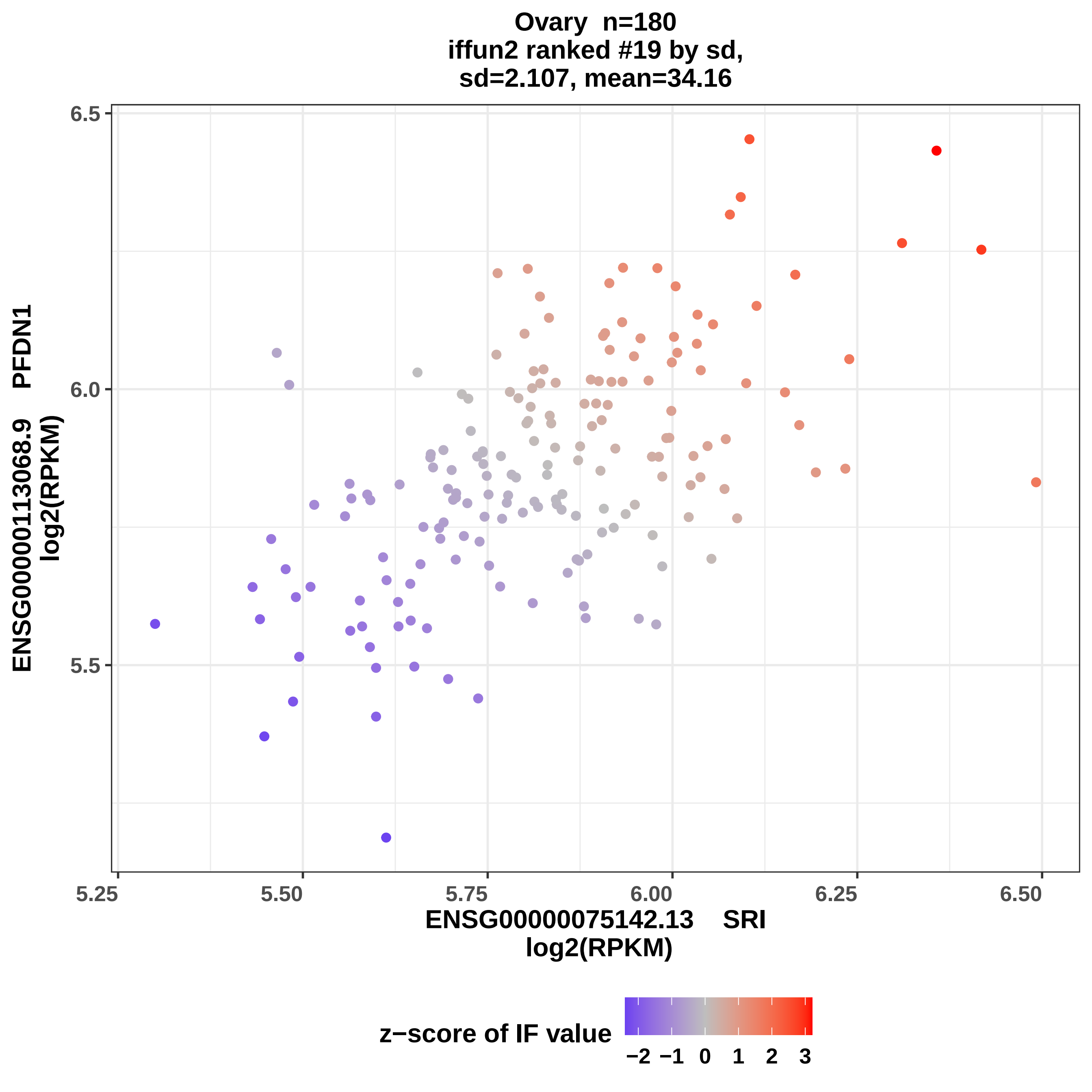

Ovary n=180
iffun2 ranked #19 by sd,
sd=2.107, mean=34.16
6.5
6.0
5.5
ENSG00000113068.9 PFDN1
log2(RPKM)
5.25
5.50
5.75
6.00
6.25
6.50
ENSG00000075142.13 SRI
log2(RPKM)
z−score of IF value
−2
−1
0
1
2
3

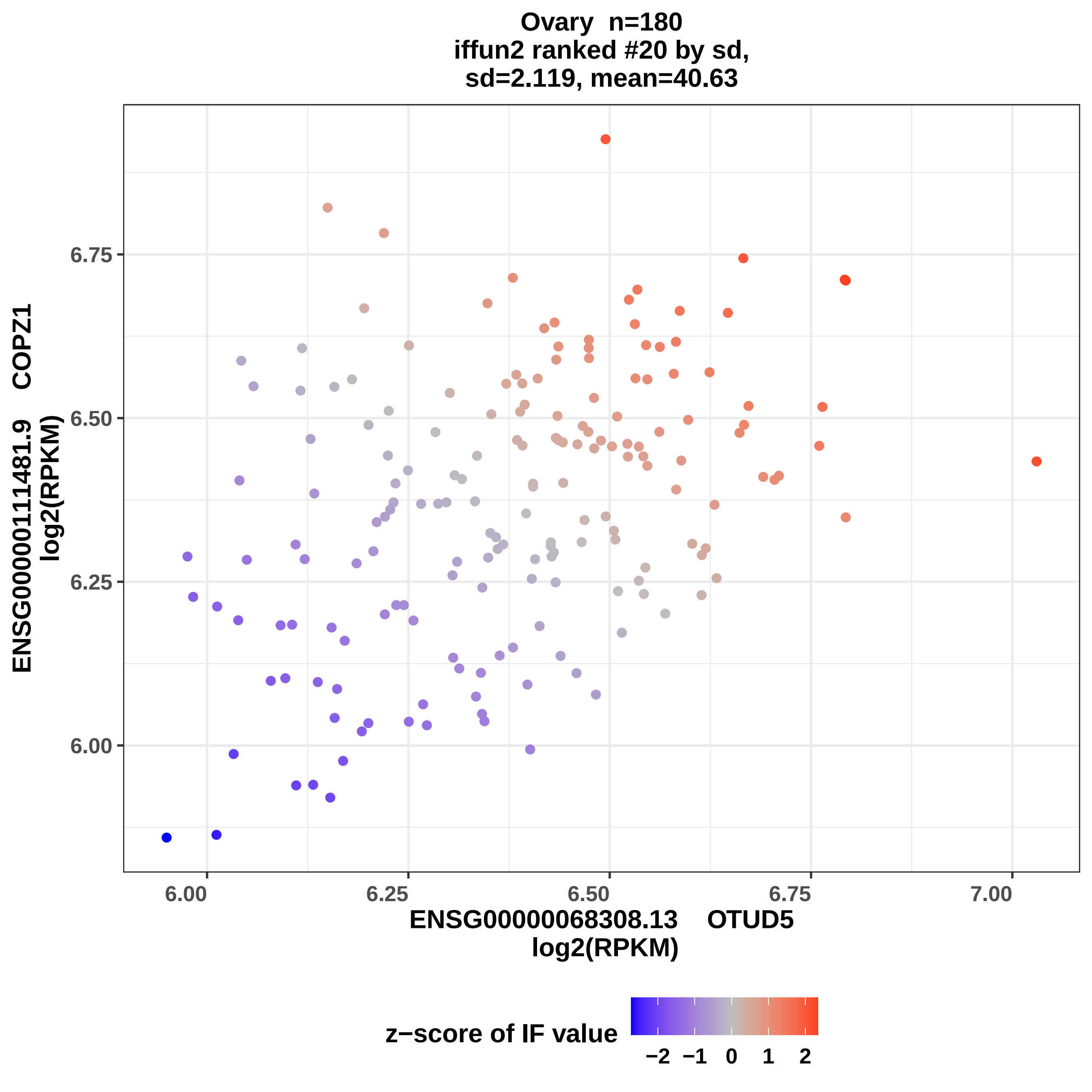

Ovary n=180
iffun2 ranked #20 by sd,
sd=2.119, mean=40.63
ENSG00000111481.9 COPZ1
log2(RPKM)
6.75
6.50
6.25
6.00
6.00
6.25
6.50
6.75
7.00
ENSG00000068308.13 OTUD5
log2(RPKM)
z−score of IF value
−2
−1
0
1
2

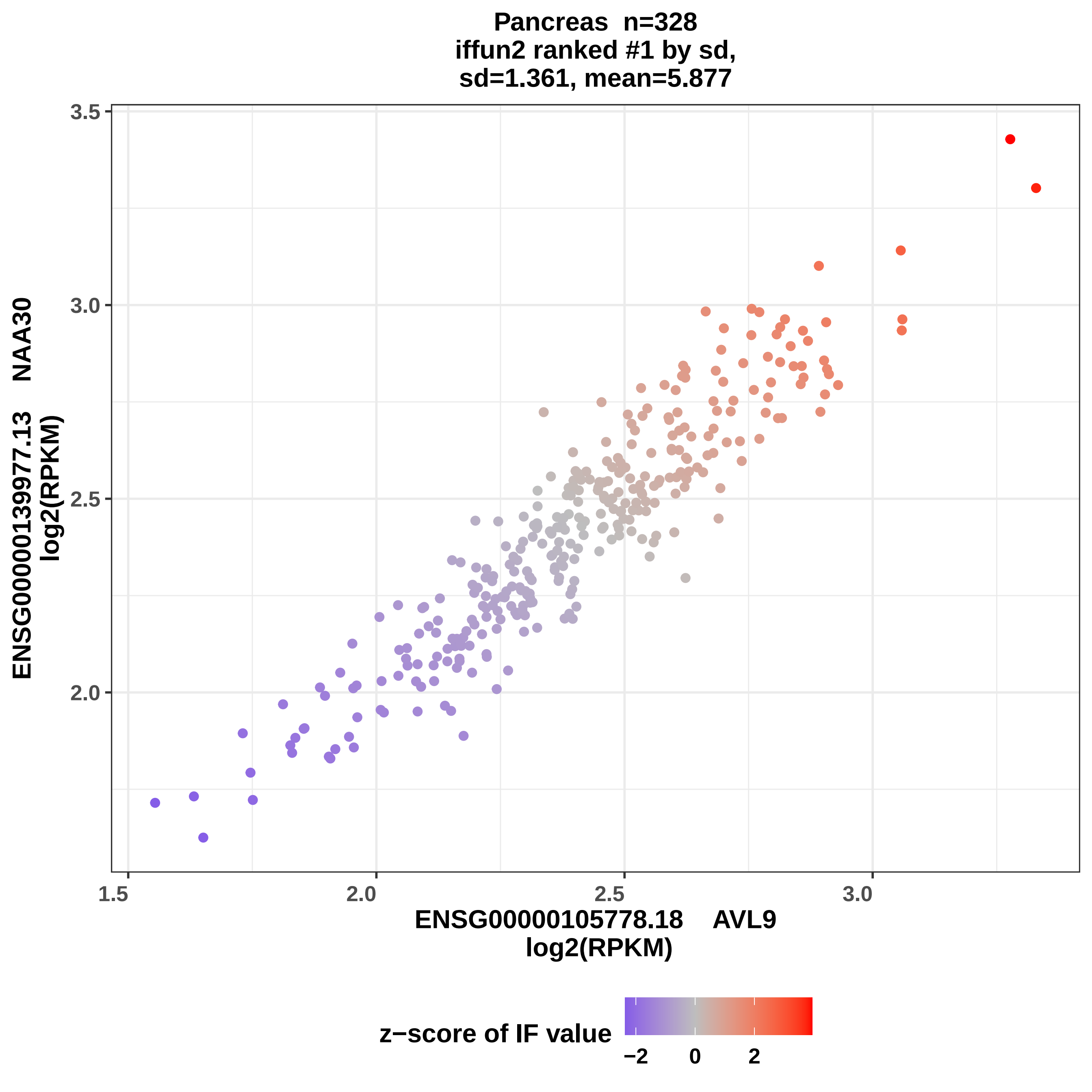

Pancreas n=328
iffun2 ranked #1 by sd,
sd=1.361, mean=5.877
ENSG00000139977.13 NAA30
log2(RPKM)
3.5
3.0
2.5
2.0
1.5
2.0
2.5
3.0
ENSG00000105778.18 AVL9
log2(RPKM)
z−score of IF value
−2
0
2

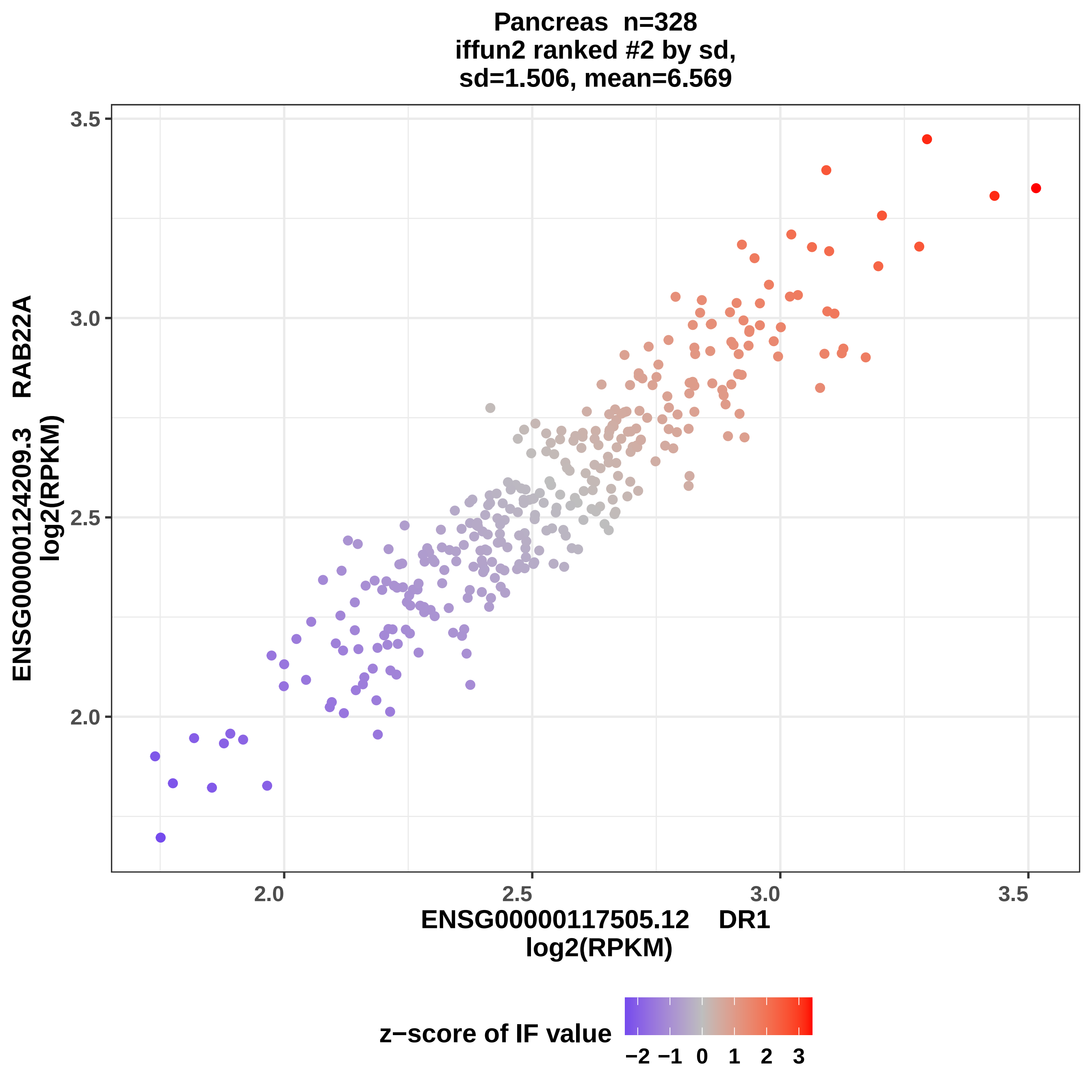

Pancreas n=328
iffun2 ranked #2 by sd,
sd=1.506, mean=6.569
ENSG00000124209.3 RAB22A
log2(RPKM)
3.5
3.0
2.5
2.0
2.0
2.5
3.0
3.5
ENSG00000117505.12 DR1
log2(RPKM)
z−score of IF value
−2 −1 0 1 2 3

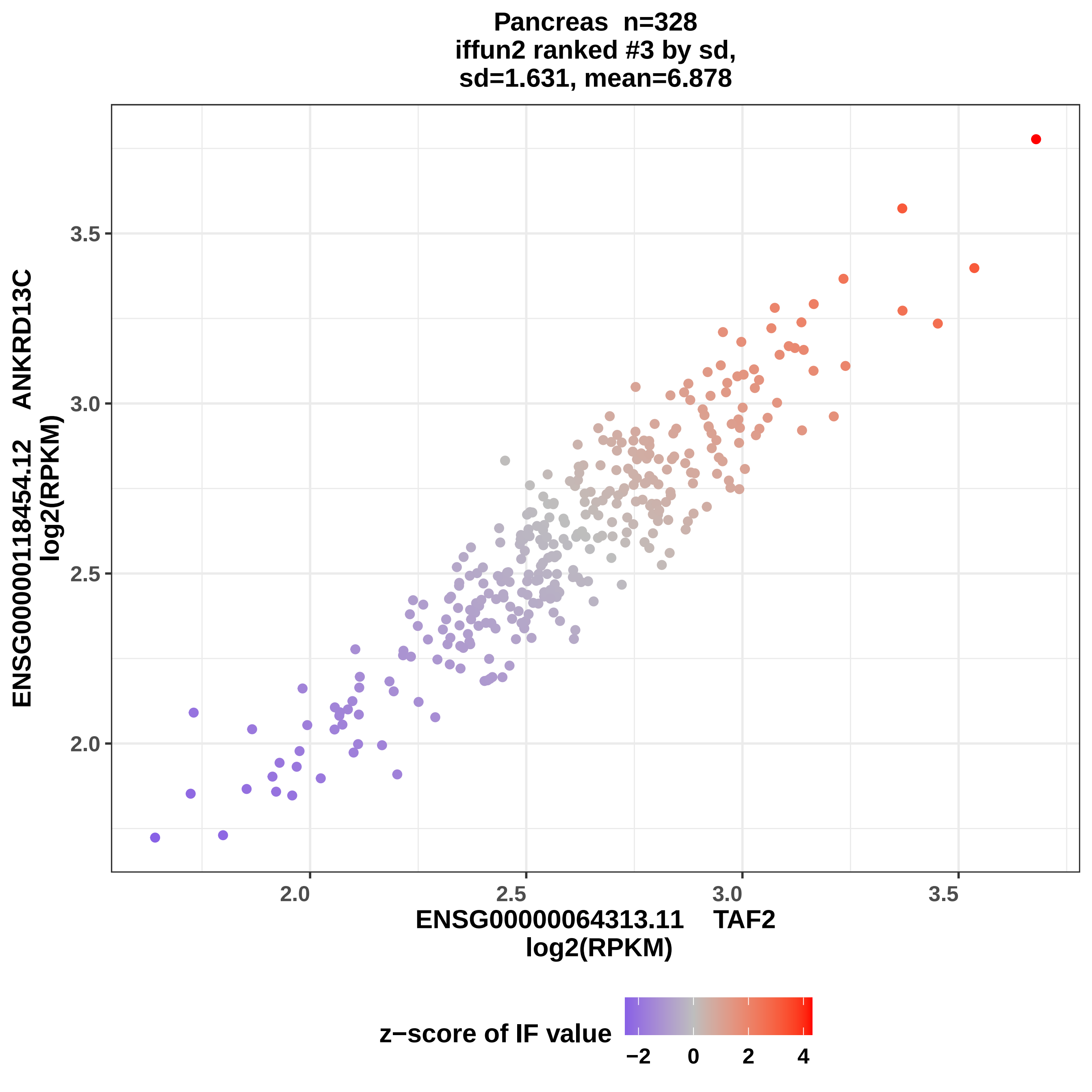
Pancreas n=328
iffun2 ranked #3 by sd,
sd=1.631, mean=6.878
ENSG00000118454.12 ANKRD13C
log2(RPKM)
3.5
3.0
2.5
2.0
2.0
2.5
3.0
3.5
ENSG00000064313.11 TAF2
log2(RPKM)
z−score of IF value
−2
0
2
4

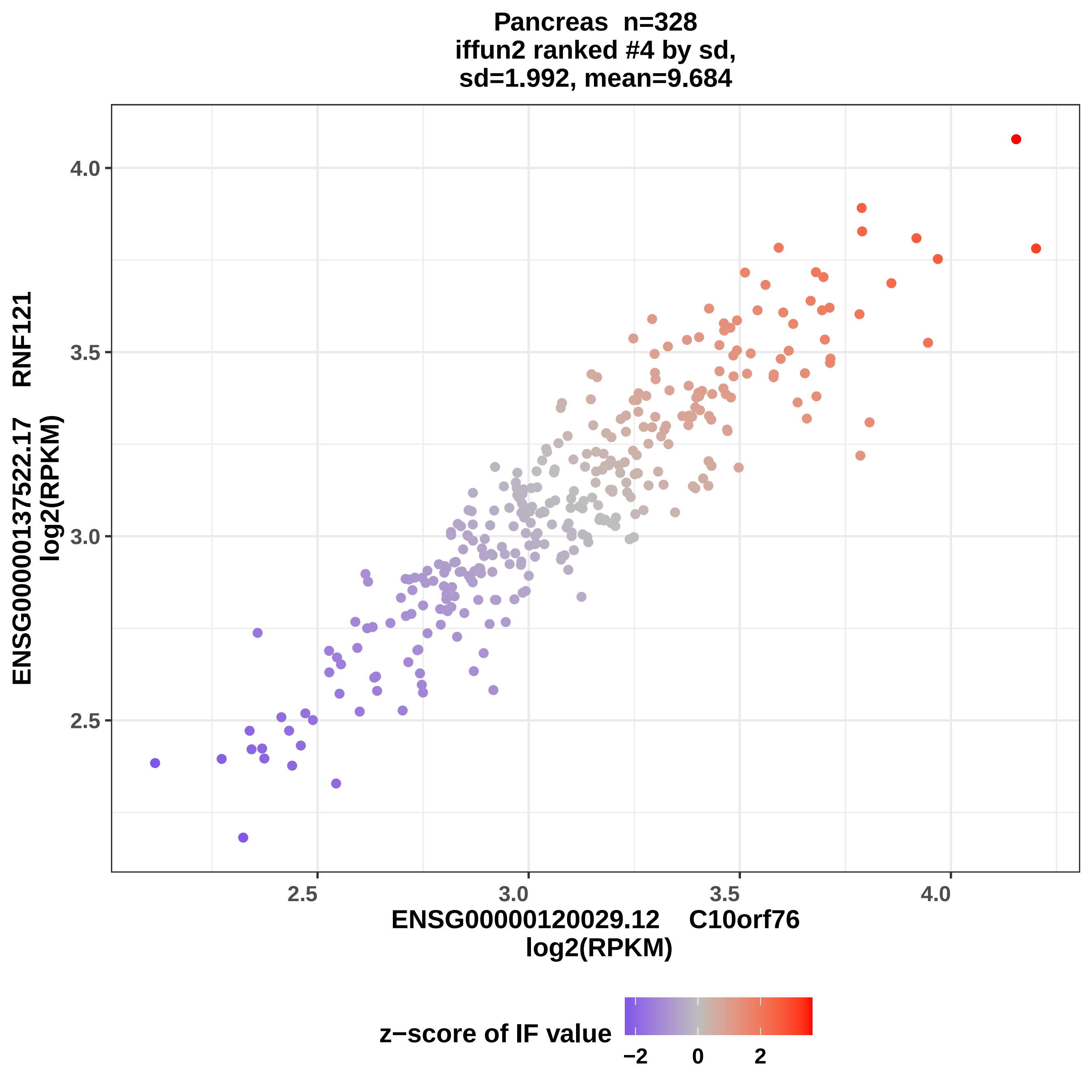

Pancreas n=328
iffun2 ranked #4 by sd,
sd=1.992, mean=9.684
ENSG00000137522.17 RNF121
log2(RPKM)
4.0
3.5
3.0
2.5
2.5
3.0
3.5
4.0
ENSG00000120029.12 C10orf76
log2(RPKM)
z−score of IF value
−2
0
2

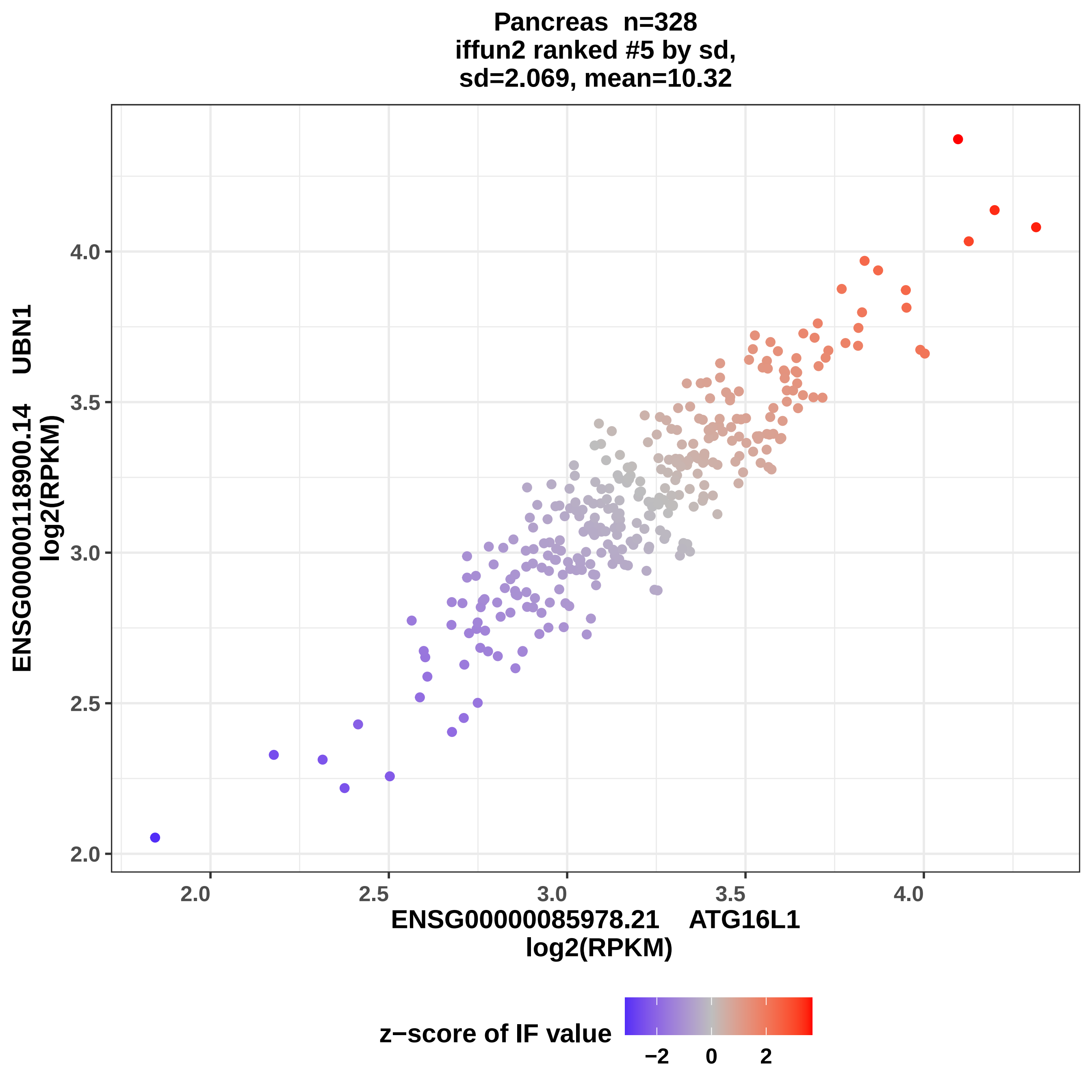
Pancreas n=328
iffun2 ranked #5 by sd,
sd=2.069, mean=10.32
ENSG00000118900.14 UBN1
log2(RPKM)
4.0
3.5
3.0
2.5
2.0
2.0
2.5
3.0
3.5
4.0
ENSG00000085978.21 ATG16L1
log2(RPKM)
z−score of IF value
−2
0
2

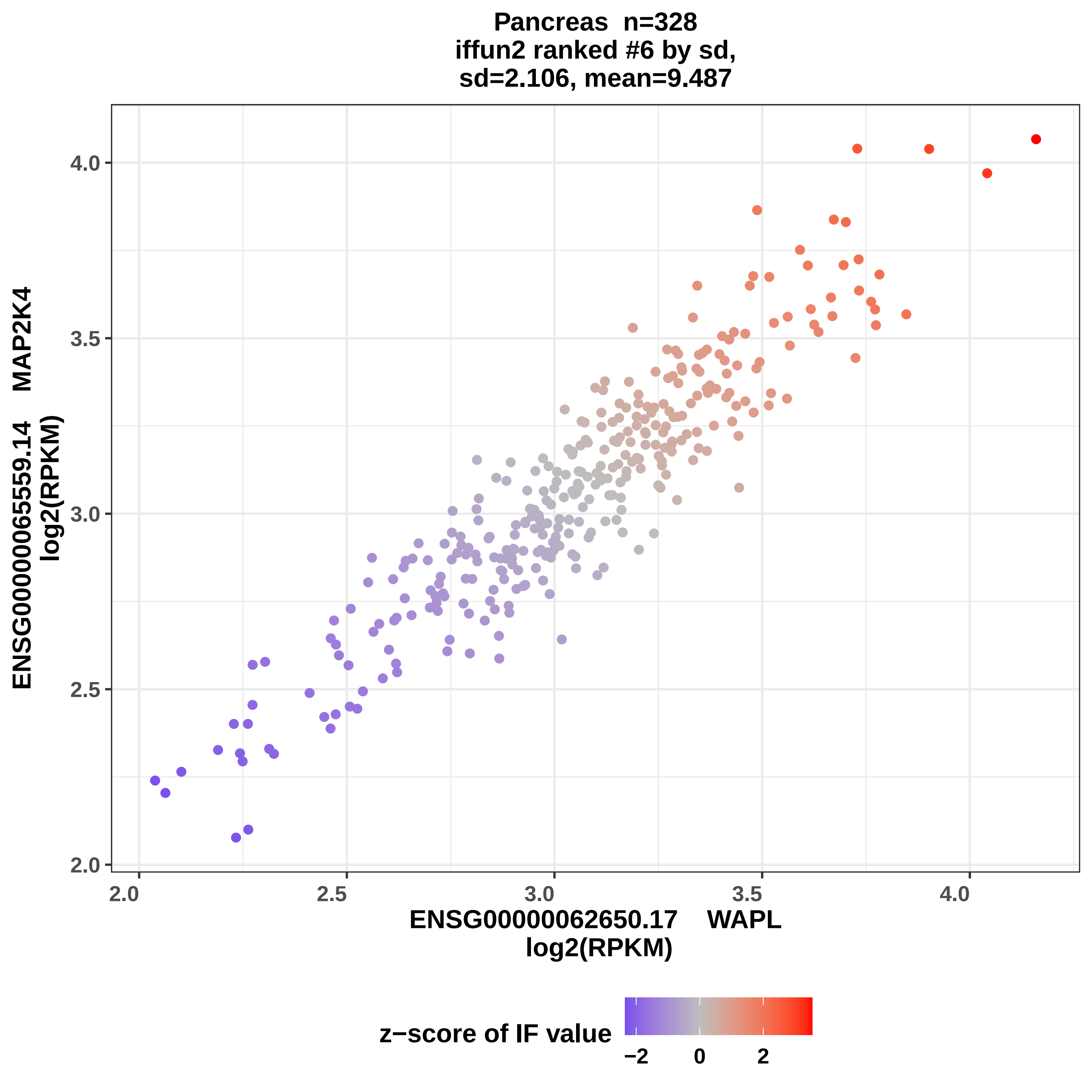
Pancreas n=328
iffun2 ranked #6 by sd,
sd=2.106, mean=9.487
ENSG00000065559.14 MAP2K4
log2(RPKM)
4.0
3.5
3.0
2.5
2.0
2.0
2.5
3.0
3.5
4.0
ENSG00000062650.17 WAPL
log2(RPKM)
z−score of IF value
−2
0
2

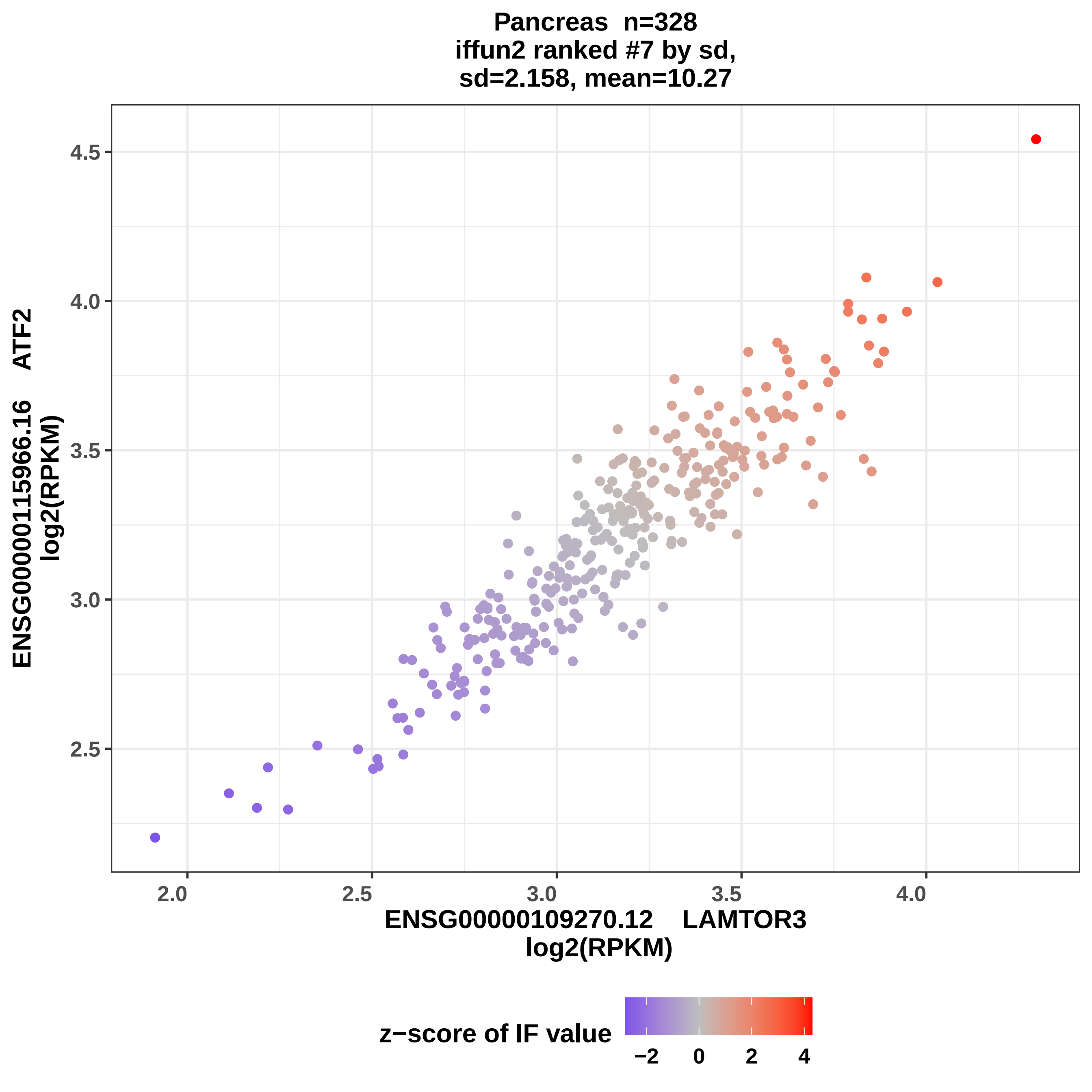

Pancreas n=328
iffun2 ranked #7 by sd,
sd=2.158, mean=10.27
ENSG00000115966.16 ATF2
log2(RPKM)
ENSG00000109270.12 LAMTOR3
log2(RPKM)
4.5
4.0
3.5
3.0
2.5
2.0
2.5
3.0
3.5
4.0
z−score of IF value
−2
0
2
4

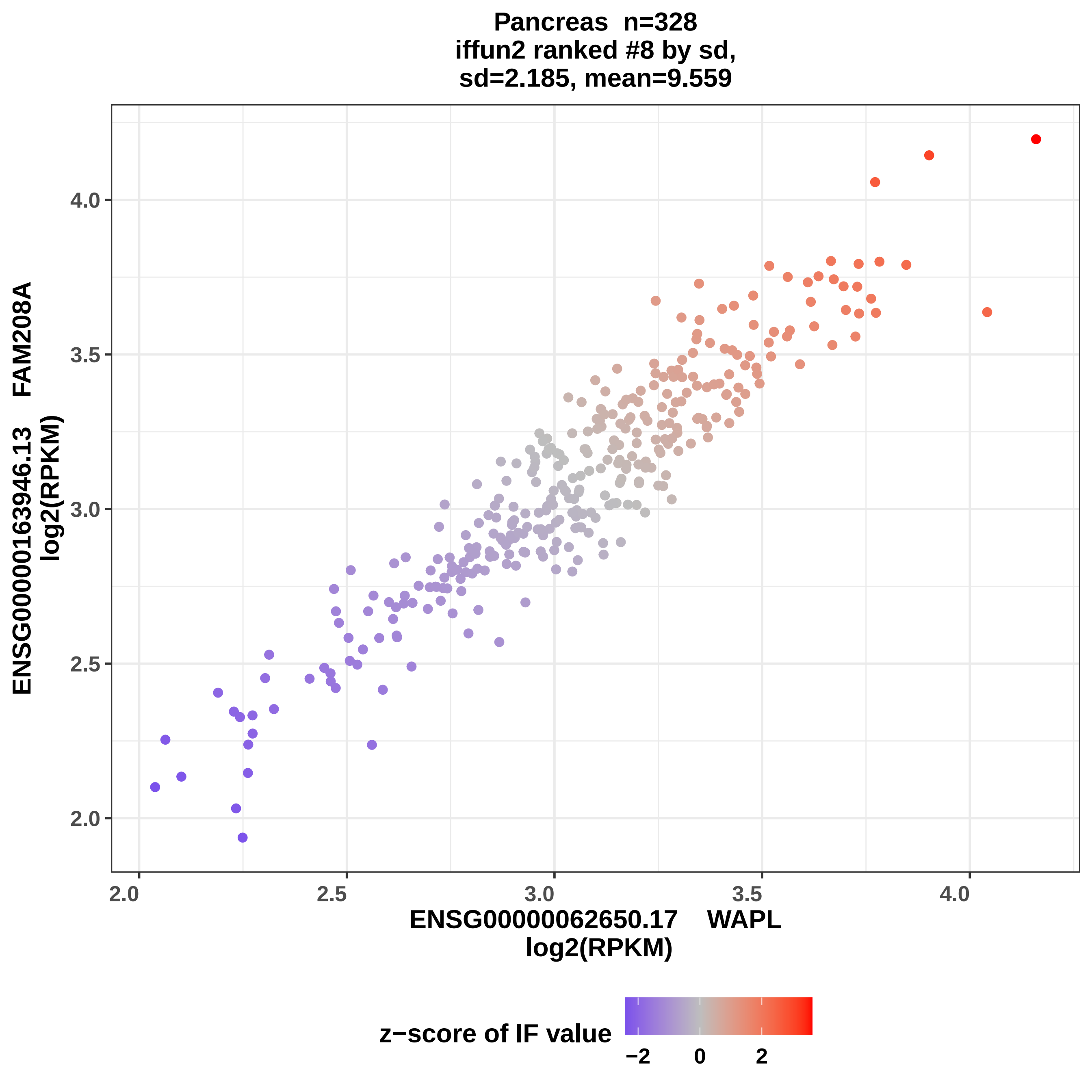
Pancreas n=328
iffun2 ranked #8 by sd,
sd=2.185, mean=9.559
ENSG00000163946.13 FAM208A
log2(RPKM)
ENSG00000062650.17 WAPL
log2(RPKM)
2.0
2.5
3.0
3.5
4.0
z−score of IF value
−2
0
2

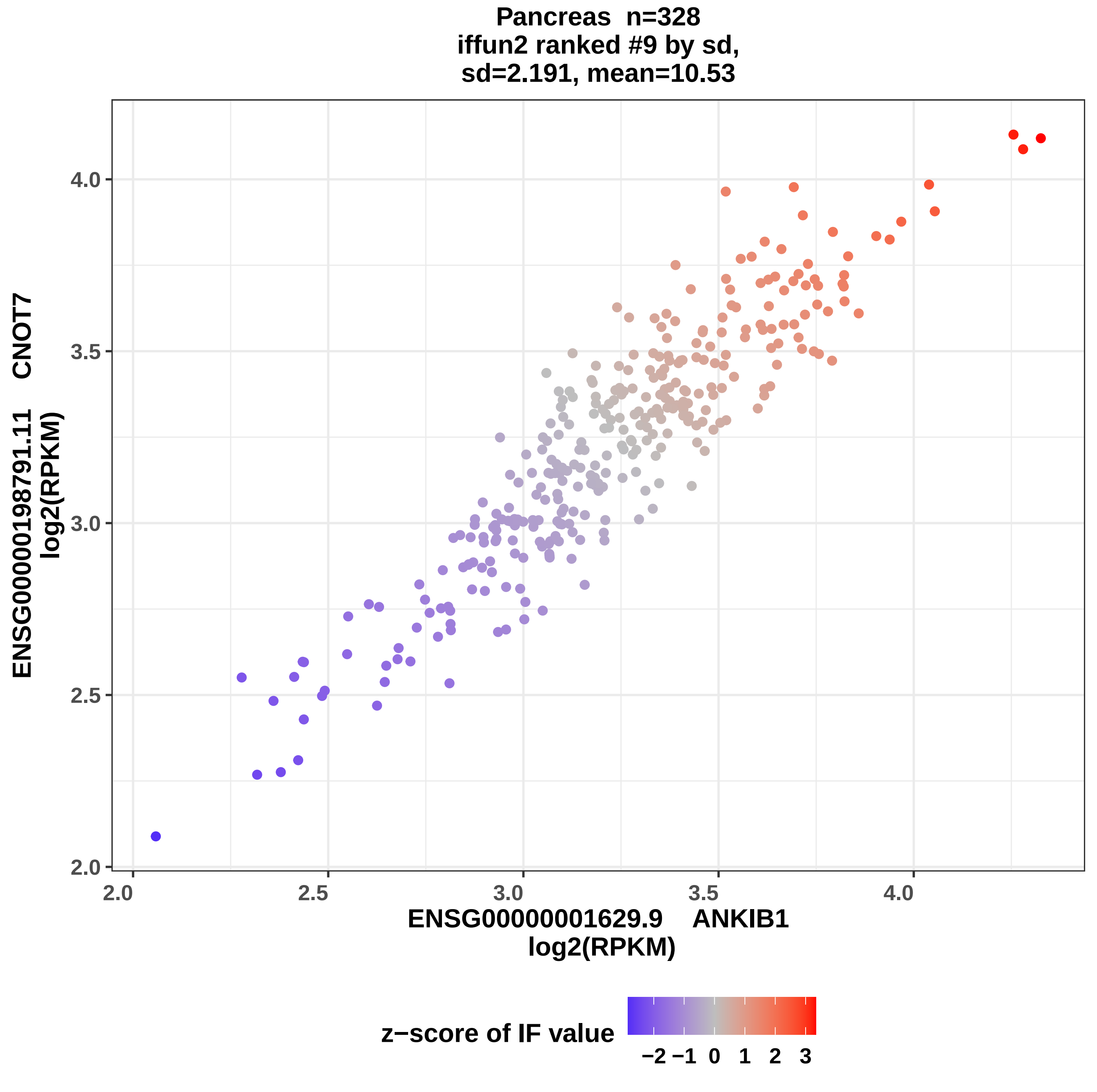
Pancreas n=328
iffun2 ranked #9 by sd,
sd=2.191, mean=10.53
ENSG00000198791.11 CNOT7
log2(RPKM)
ENSG00000001629.9 ANKIB1
log2(RPKM)
2.0
2.5
3.0
3.5
4.0
z−score of IF value
−2 −1 0 1 2 3

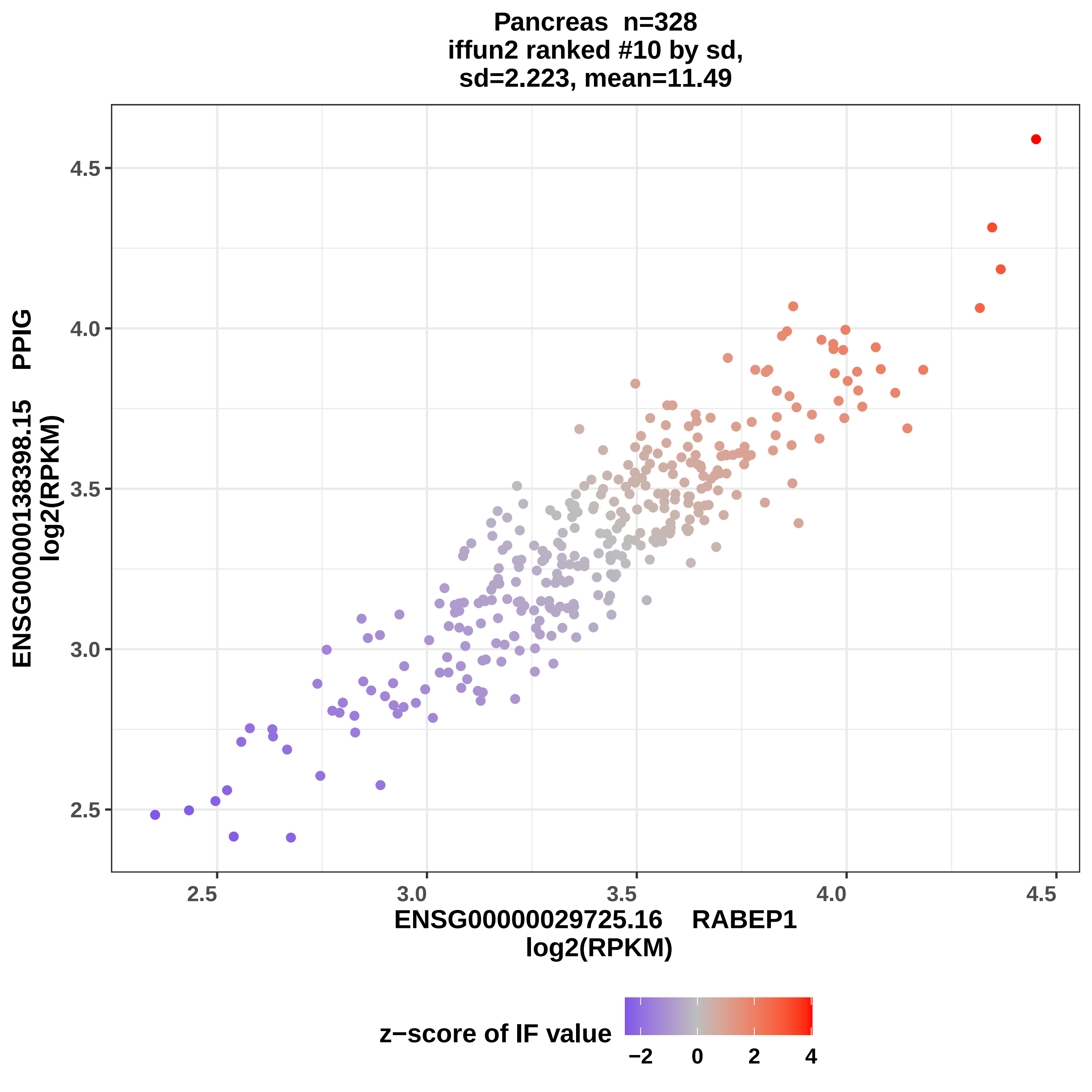

Pancreas n=328
iffun2 ranked #10 by sd,
sd=2.223, mean=11.49
ENSG00000138398.15 PPIG
log2(RPKM)
4.5
4.0
3.5
3.0
2.5
2.5
3.0
3.5
4.0
4.5
ENSG00000029725.16 RABEP1
log2(RPKM)
z−score of IF value
−2
0
2
4

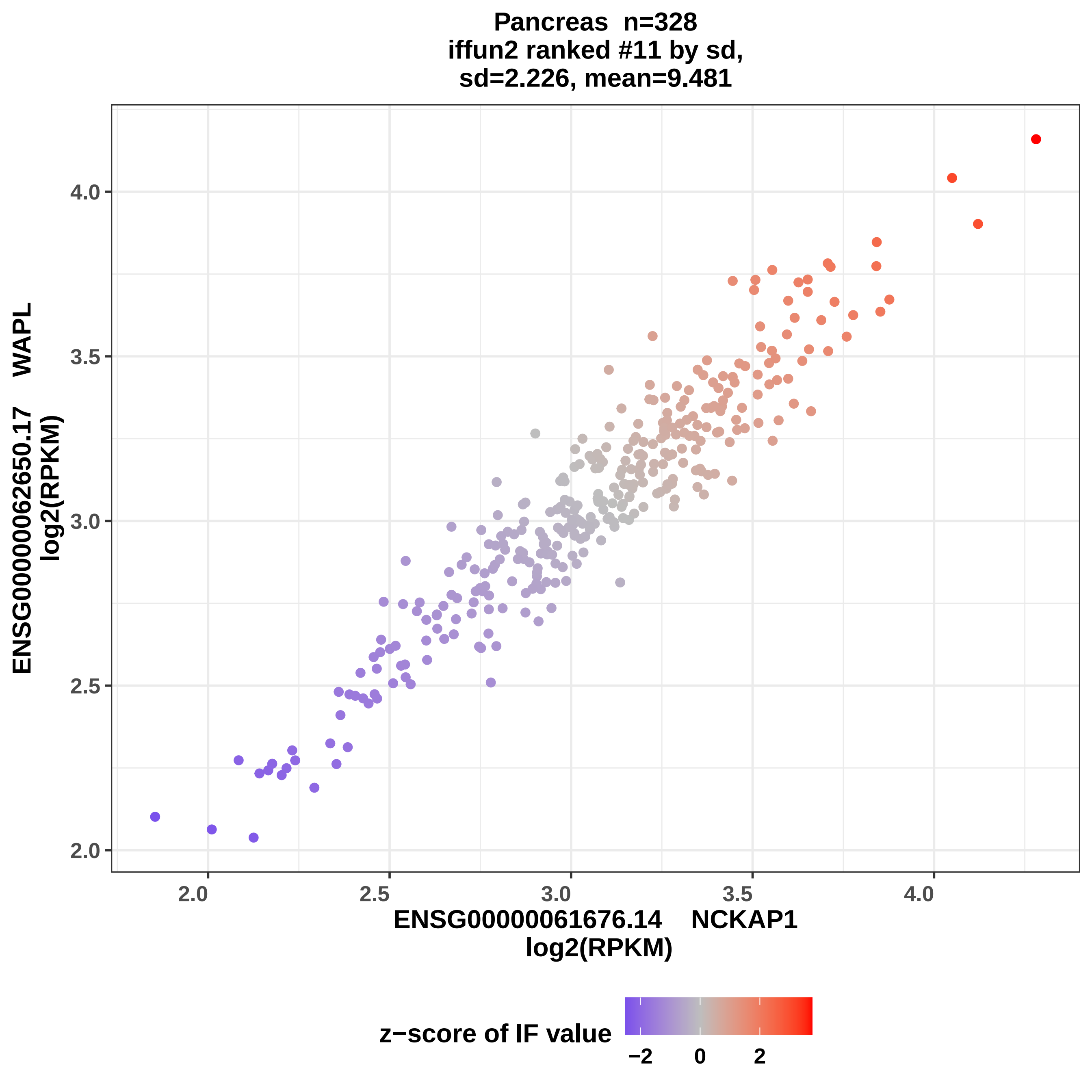

Pancreas n=328
iffun2 ranked #11 by sd,
sd=2.226, mean=9.481
ENSG00000062650.17 WAPL
log2(RPKM)
2.0
2.5
3.0
3.5
4.0
ENSG00000061676.14 NCKAP1
log2(RPKM)
z−score of IF value
−2
0
2

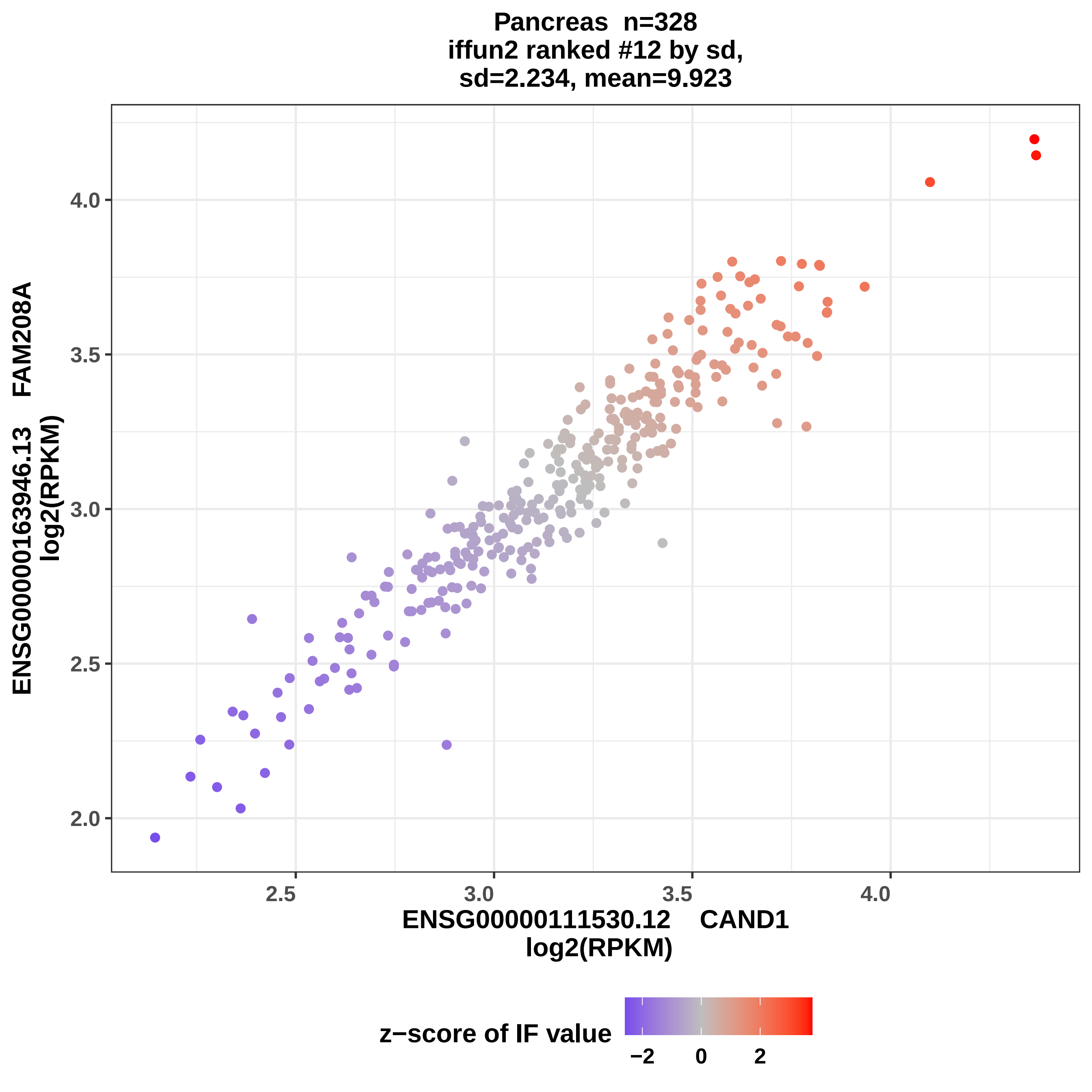
Pancreas n=328
iffun2 ranked #12 by sd,
sd=2.234, mean=9.923
ENSG00000163946.13 FAM208A
log2(RPKM)
4.0
3.5
3.0
2.5
2.0
2.5
3.0
3.5
4.0
ENSG00000111530.12 CAND1
log2(RPKM)
z−score of IF value
−2
0
2

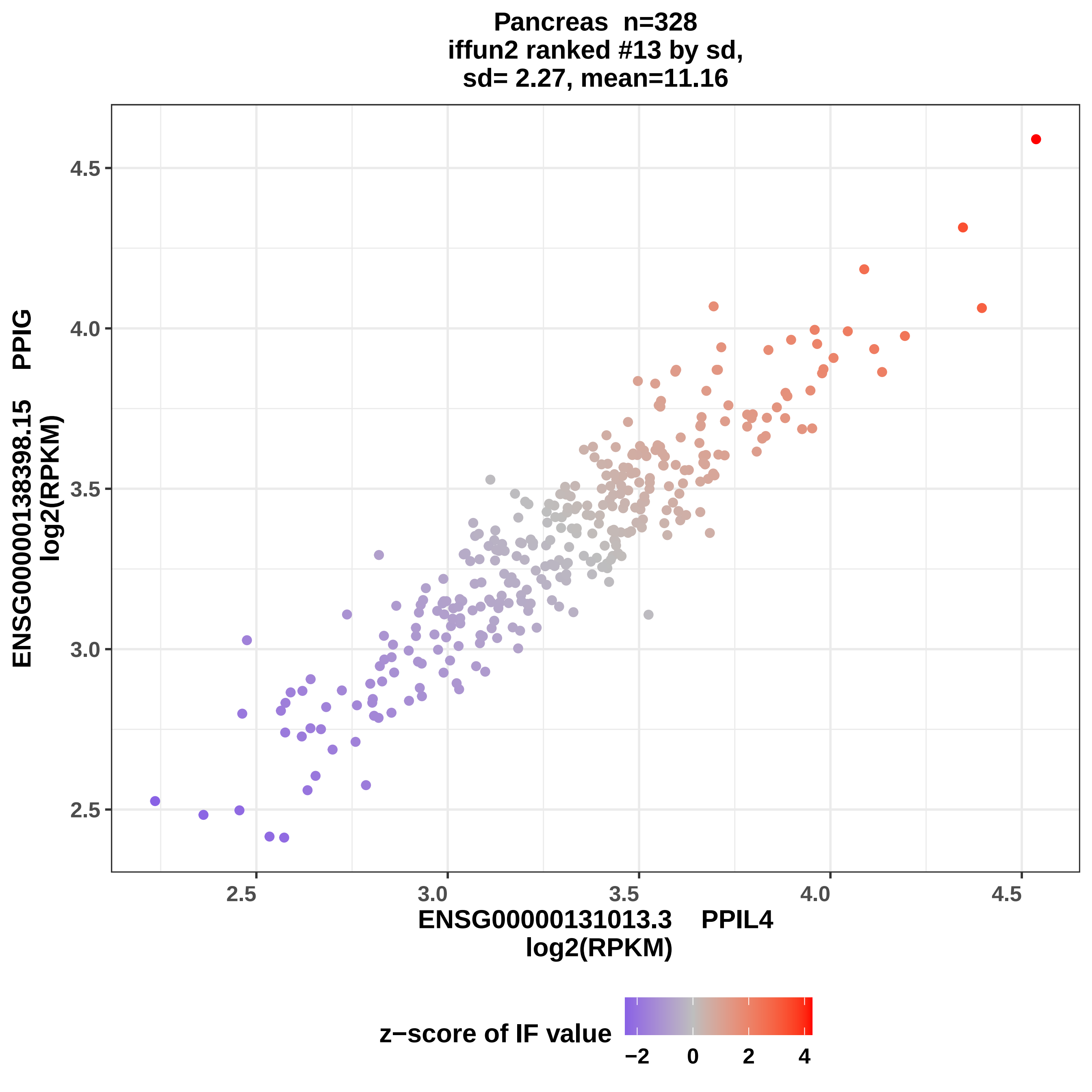

Pancreas n=328
iffun2 ranked #13 by sd,
sd= 2.27, mean=11.16
ENSG00000138398.15 PPIG
log2(RPKM)
4.5
4.0
3.5
3.0
2.5
2.5
3.0
3.5
4.0
4.5
ENSG00000131013.3 PPIL4
log2(RPKM)
z−score of IF value
−2
0
2
4

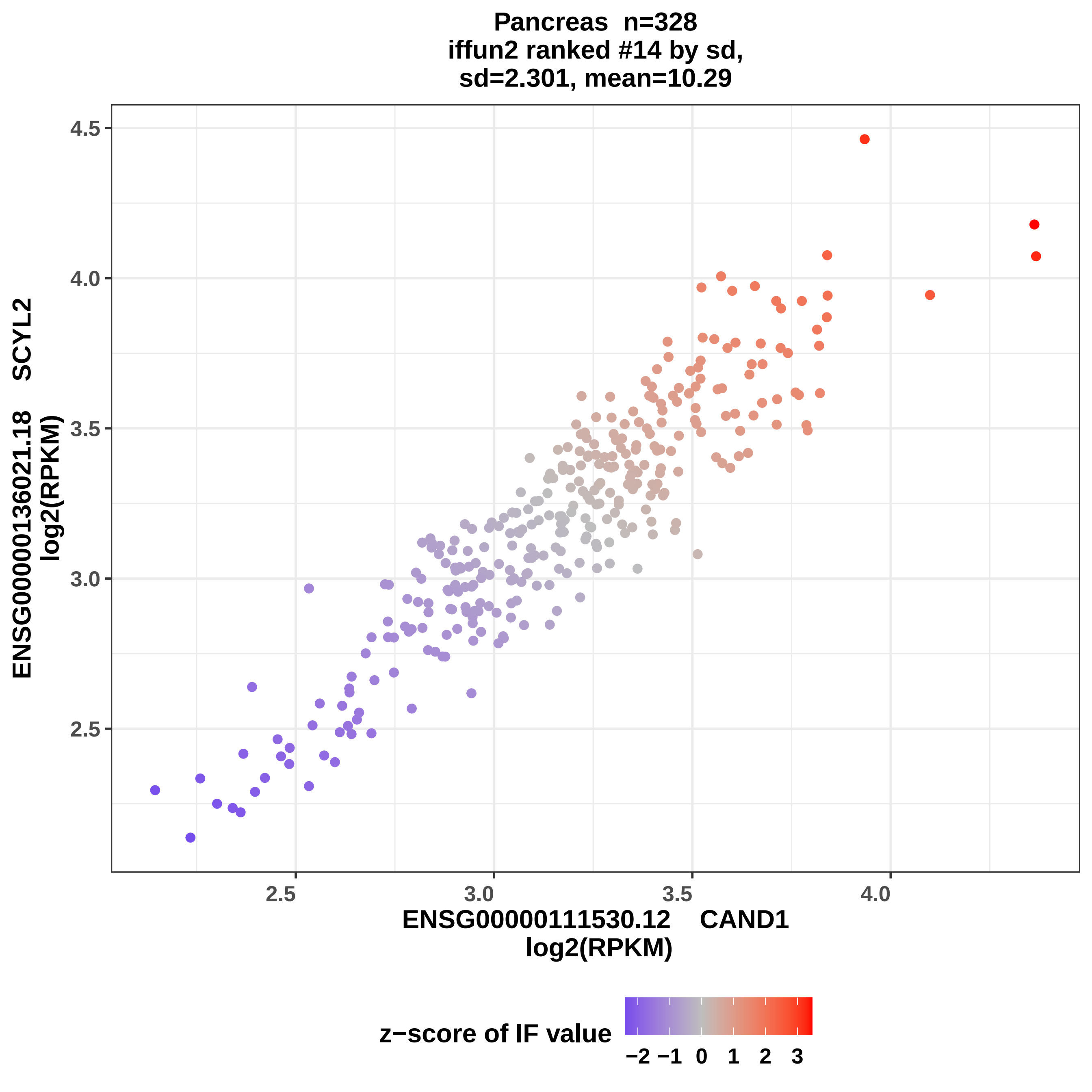
Pancreas n=328
iffun2 ranked #14 by sd,
sd=2.301, mean=10.29
ENSG00000136021.18 SCYL2
log2(RPKM)
4.5
4.0
3.5
3.0
2.5
2.5
3.0
3.5
4.0
ENSG00000111530.12 CAND1
log2(RPKM)
z−score of IF value
−2 −1 0 1 2 3

Pancreas n=328
iffun2 ranked #15 by sd,
sd=2.419, mean=11.72

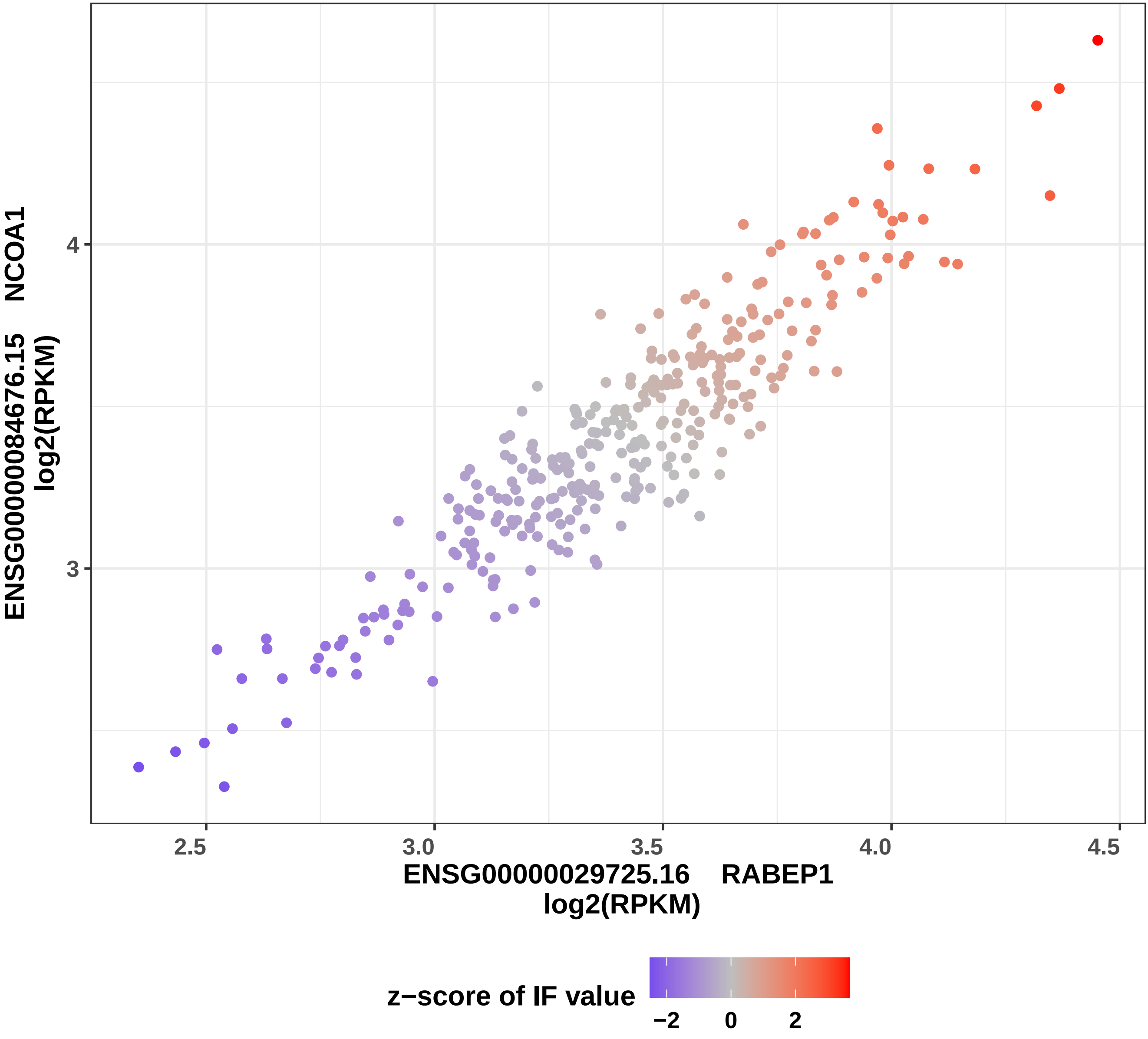

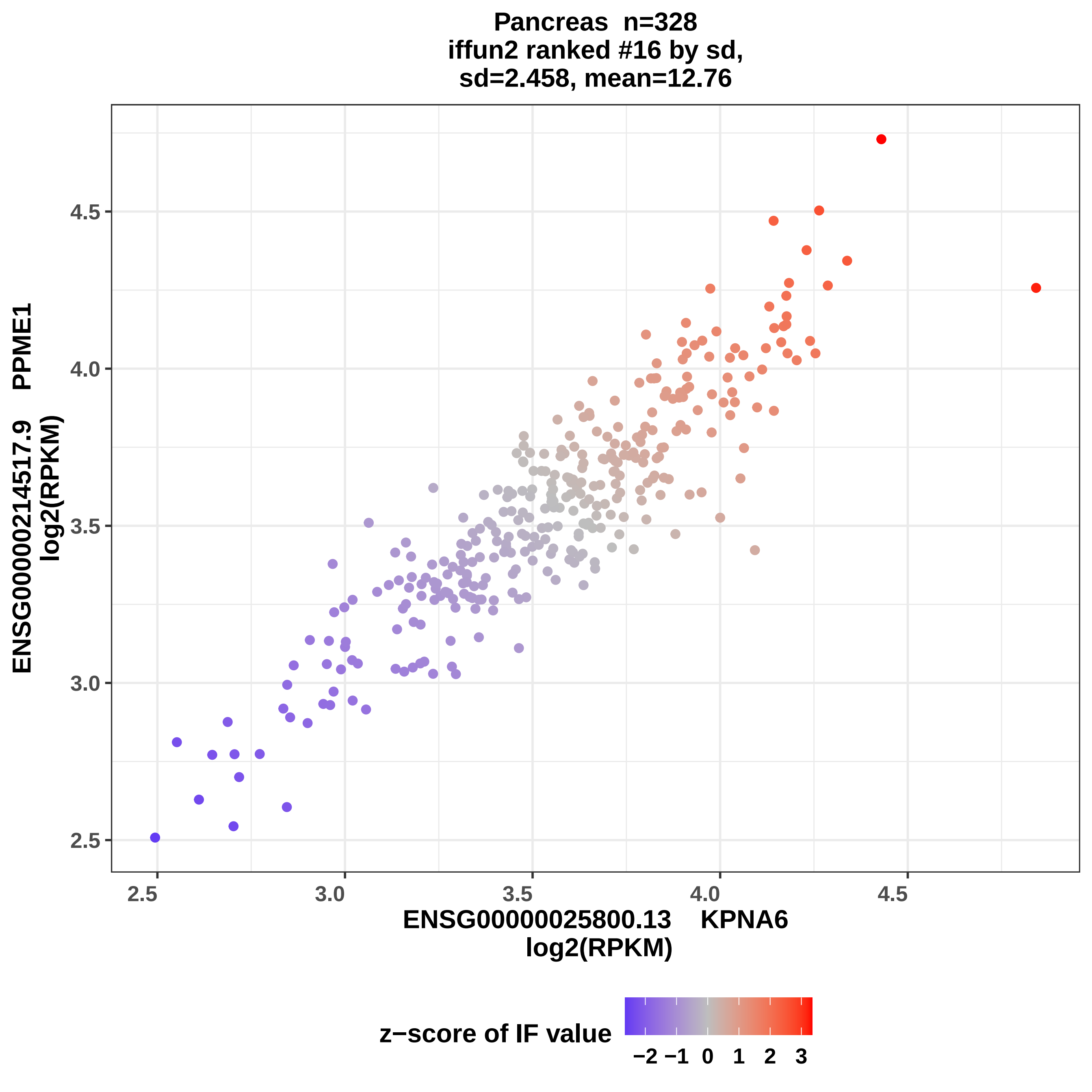
Pancreas n=328
iffun2 ranked #16 by sd,
sd=2.458, mean=12.76
ENSG00000214517.9 PPME1
log2(RPKM)
4.5
4.0
3.5
3.0
2.5
2.5
3.0
3.5
4.0
4.5
ENSG00000025800.13 KPNA6
log2(RPKM)
z−score of IF value
−2 −1 0 1 2 3

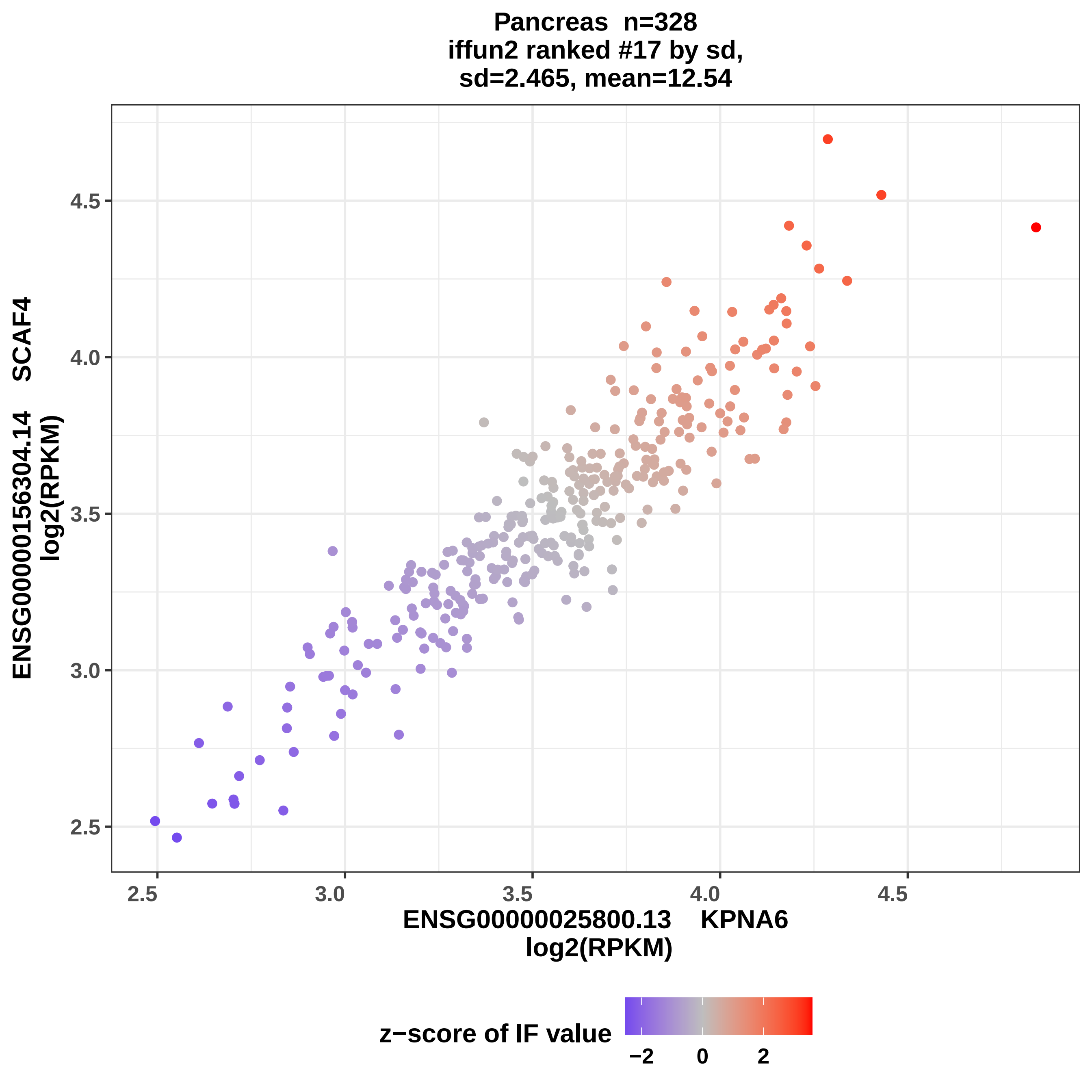

Pancreas n=328
iffun2 ranked #17 by sd,
sd=2.465, mean=12.54
ENSG00000156304.14 SCAF4
log2(RPKM)
2.5
3.0
3.5
4.0
4.5
ENSG00000025800.13 KPNA6
log2(RPKM)
z−score of IF value
−2
0
2

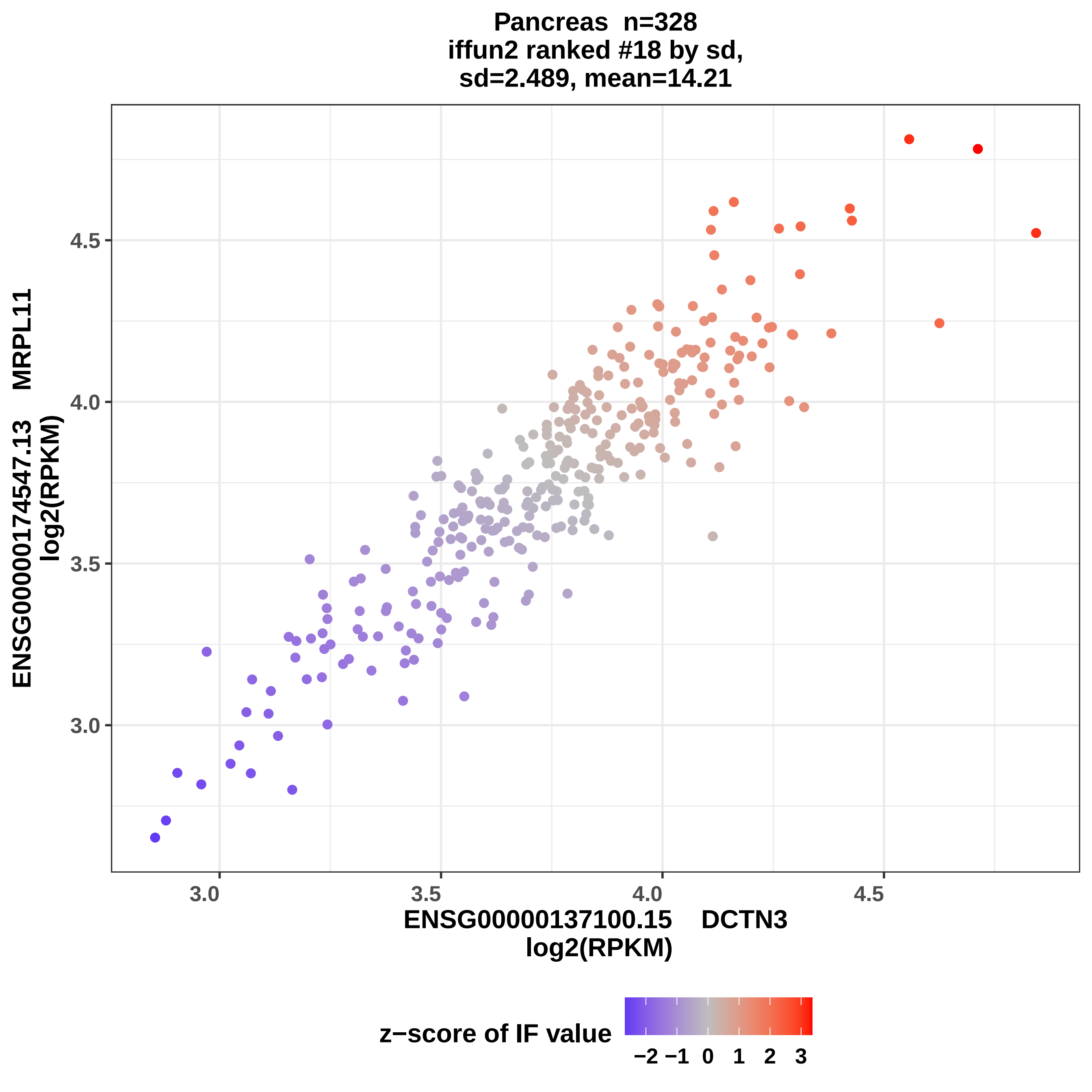
Pancreas n=328
iffun2 ranked #18 by sd,
sd=2.489, mean=14.21
ENSG00000174547.13 MRPL11
log2(RPKM)
4.5
4.0
3.5
3.0
3.0
3.5
4.0
4.5
ENSG00000137100.15 DCTN3
log2(RPKM)
z−score of IF value
−2
−1
0
1
2
3

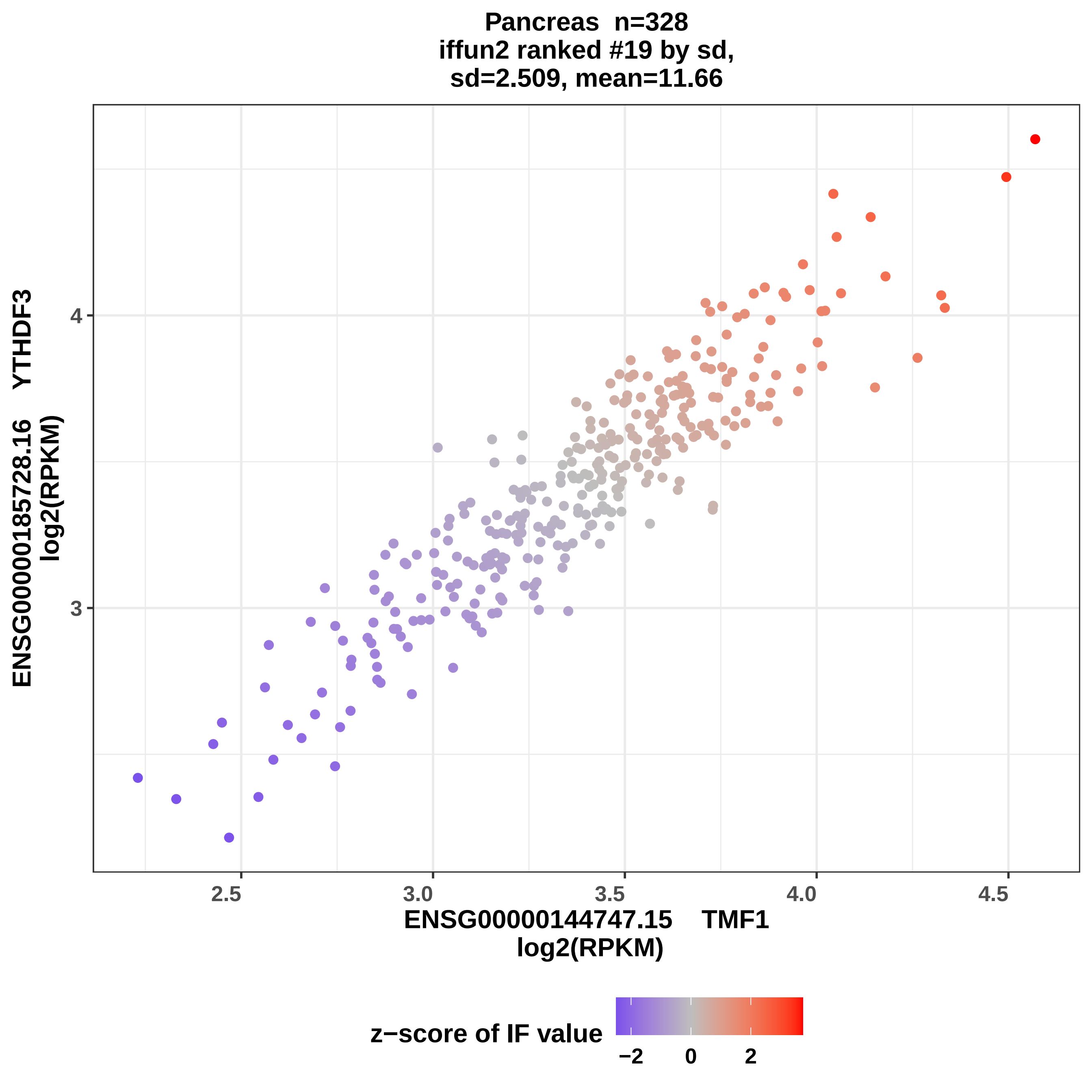

Pancreas n=328
iffun2 ranked #19 by sd,
sd=2.509, mean=11.66
ENSG00000185728.16 YTHDF3
log2(RPKM)
4
3
2.5
3.0
3.5
4.0
4.5
ENSG00000144747.15 TMF1
log2(RPKM)
z−score of IF value
−2
0
2

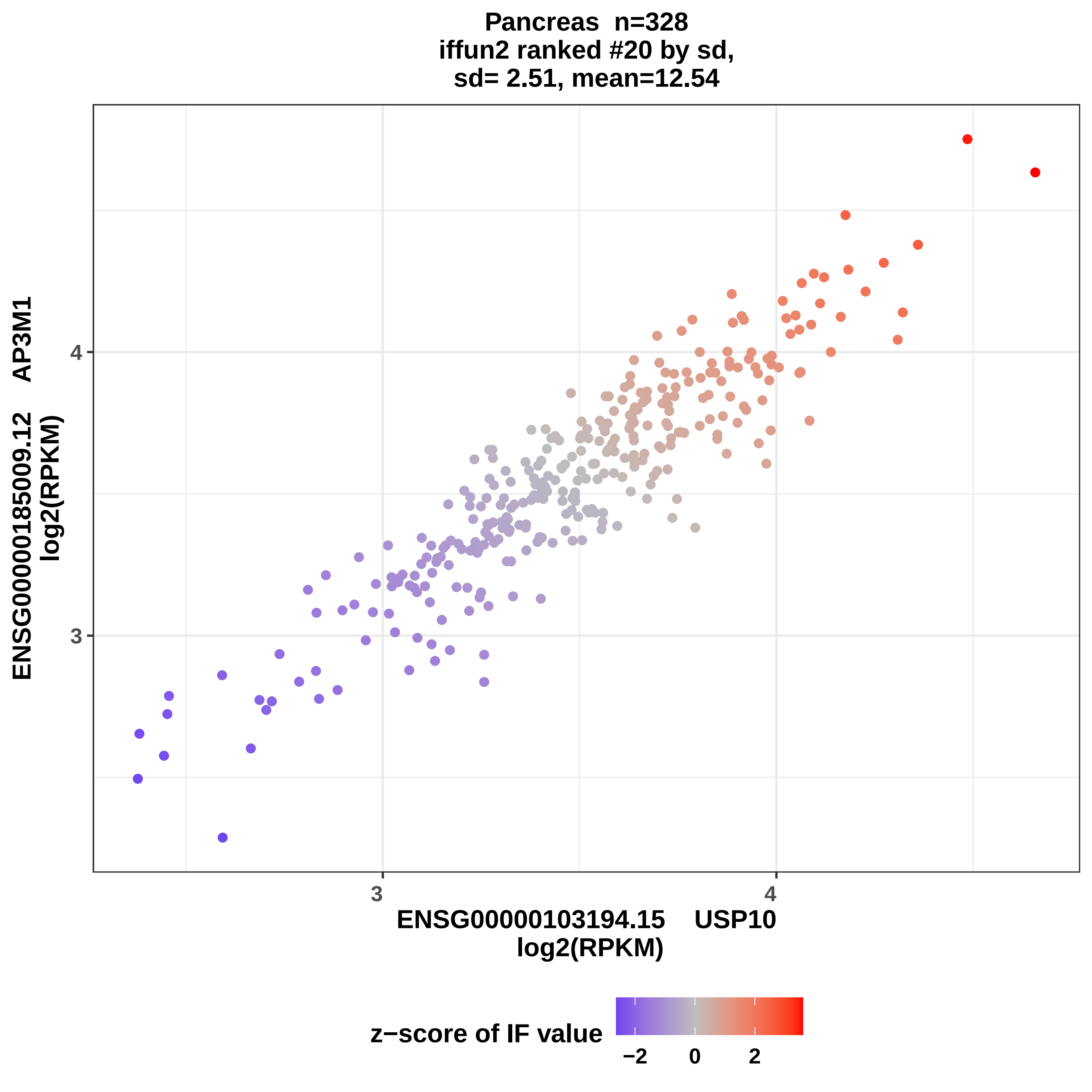
Pancreas n=328
iffun2 ranked #20 by sd,
sd= 2.51, mean=12.54
ENSG00000185009.12 AP3M1
log2(RPKM)
4
3
3
4
ENSG00000103194.15 USP10
log2(RPKM)
z−score of IF value
−2
0
2

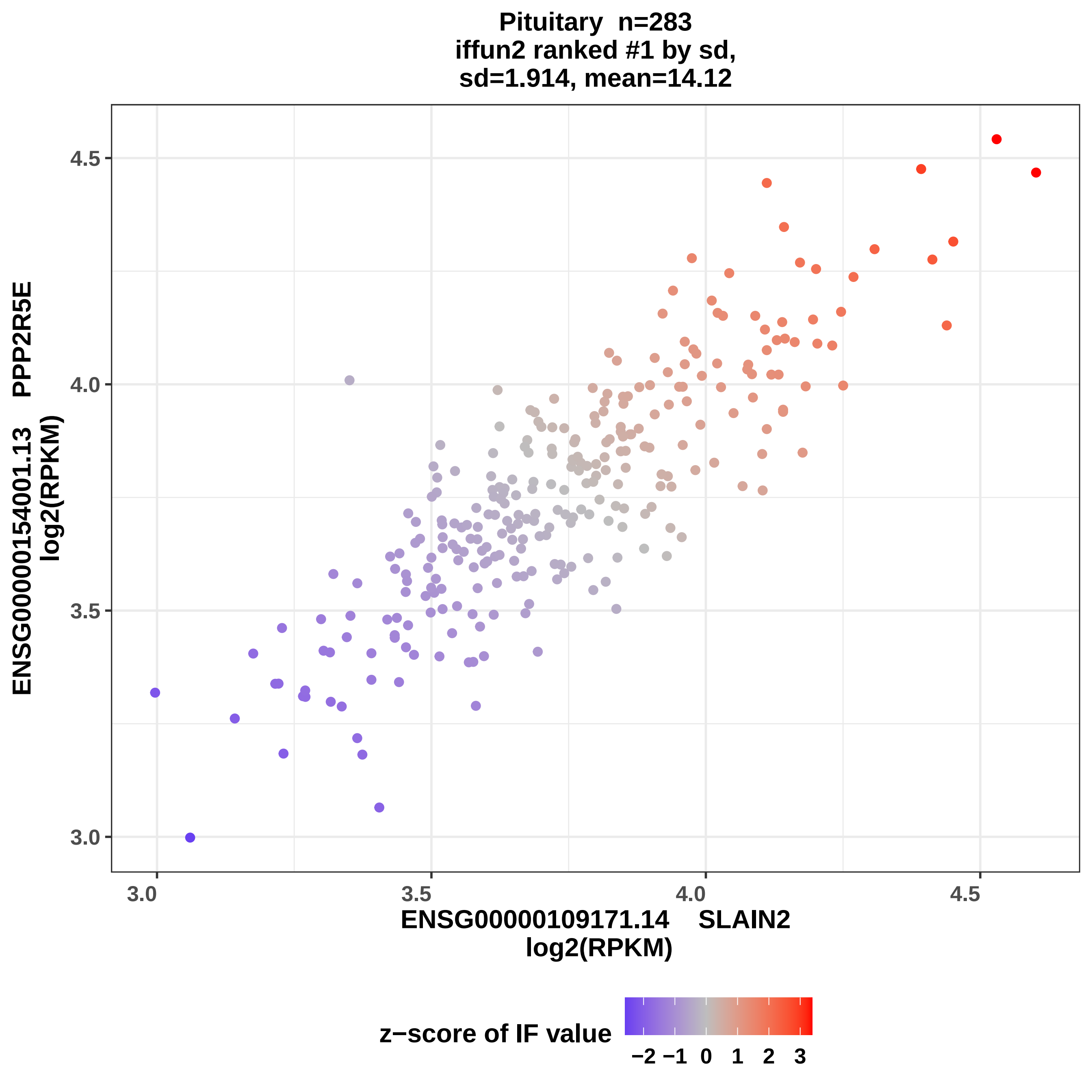

Pituitary n=283
iffun2 ranked #1 by sd,
sd=1.914, mean=14.12
ENSG00000154001.13 PPP2R5E
log2(RPKM)
4.5
4.0
3.5
3.0
3.0
3.5
4.0
4.5
ENSG00000109171.14 SLAIN2
log2(RPKM)
z−score of IF value
−2 −1 0 1 2 3

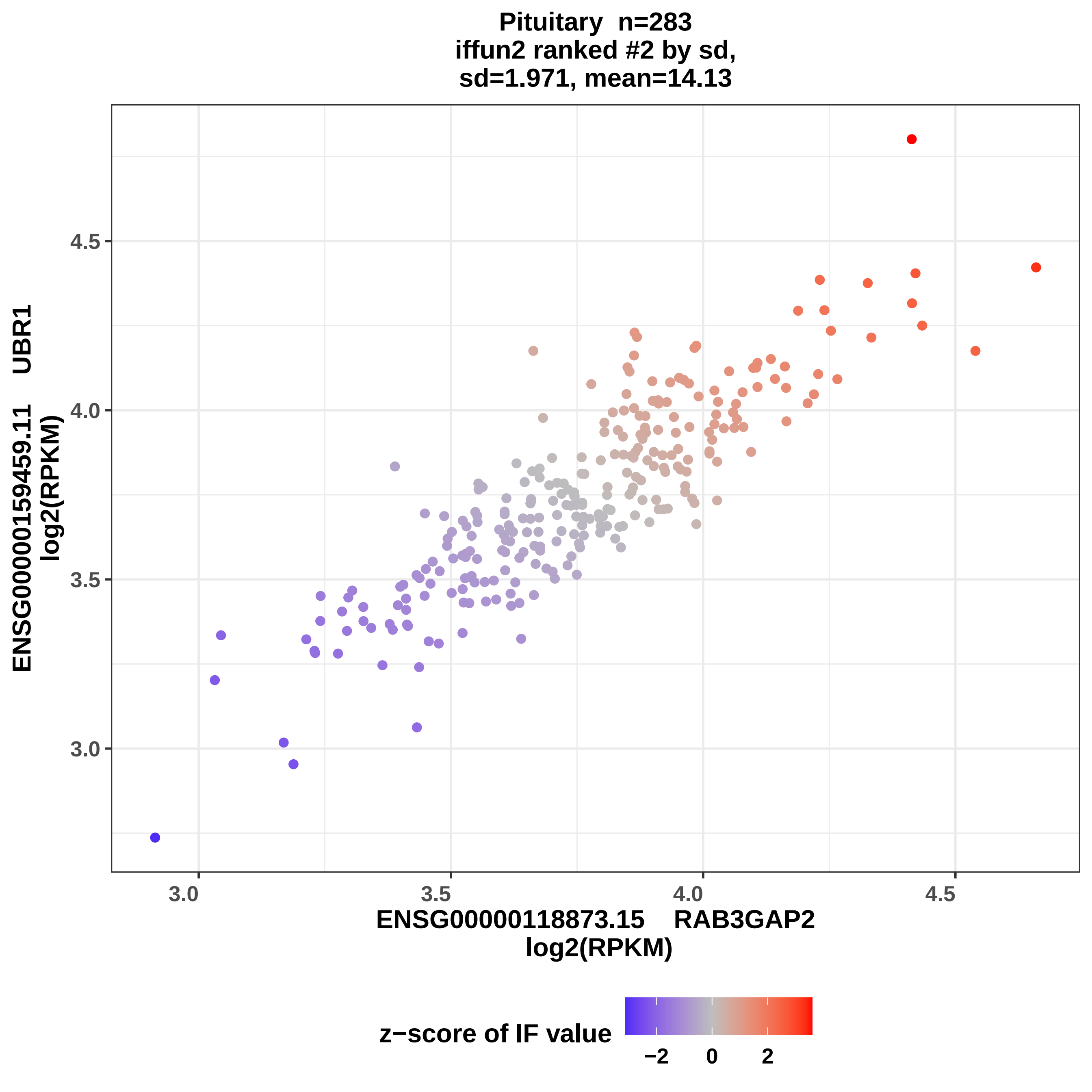
Pituitary n=283
iffun2 ranked #2 by sd,
sd=1.971, mean=14.13
ENSG00000159459.11 UBR1
log2(RPKM)
4.5
4.0
3.5
3.0
ENSG00000118873.15 RAB3GAP2
log2(RPKM)
3.0
3.5
4.0
4.5
z−score of IF value
−2
0
2

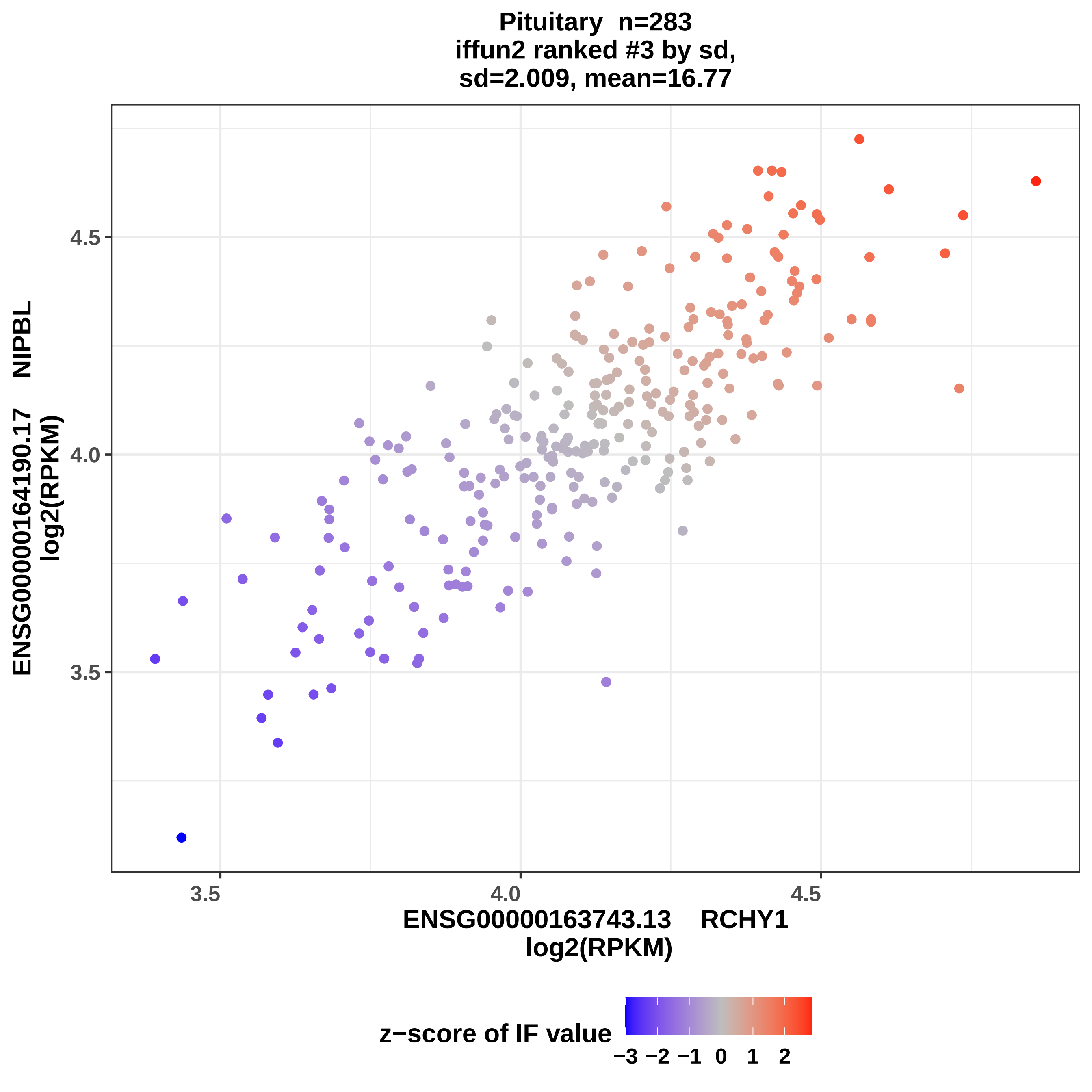

Pituitary n=283
iffun2 ranked #3 by sd,
sd=2.009, mean=16.77
ENSG00000164190.17 NIPBL
log2(RPKM)
4.5
4.0
3.5
3.5
4.0
4.5
ENSG00000163743.13 RCHY1
log2(RPKM)
z−score of IF value
−3 −2 −1 0 1 2

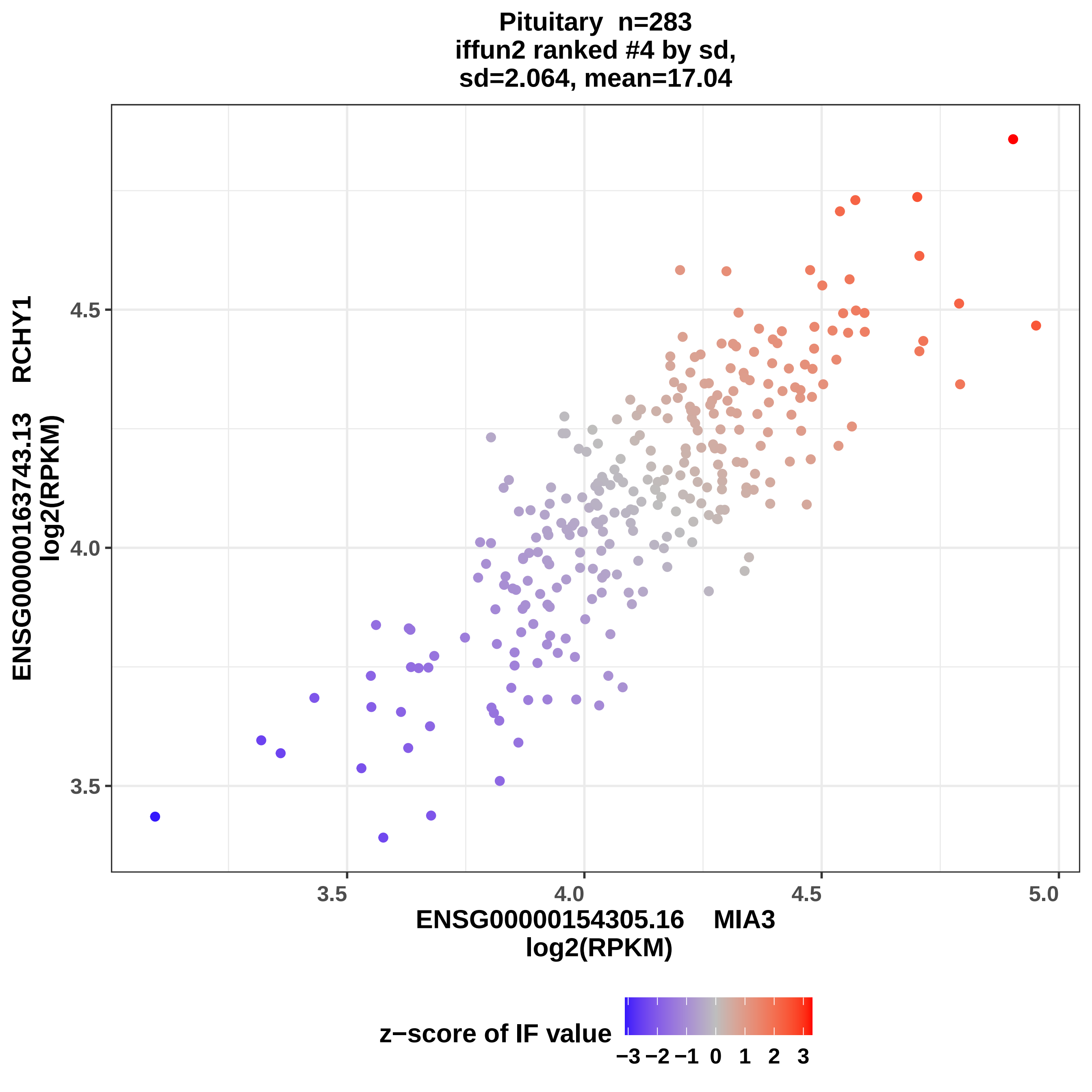

Pituitary n=283
iffun2 ranked #4 by sd,
sd=2.064, mean=17.04
ENSG00000163743.13 RCHY1
log2(RPKM)
3.5
4.0
4.5
3.5
4.0
4.5
5.0
ENSG00000154305.16 MIA3
log2(RPKM)
z−score of IF value
−3 −2 −1 0 1 2 3

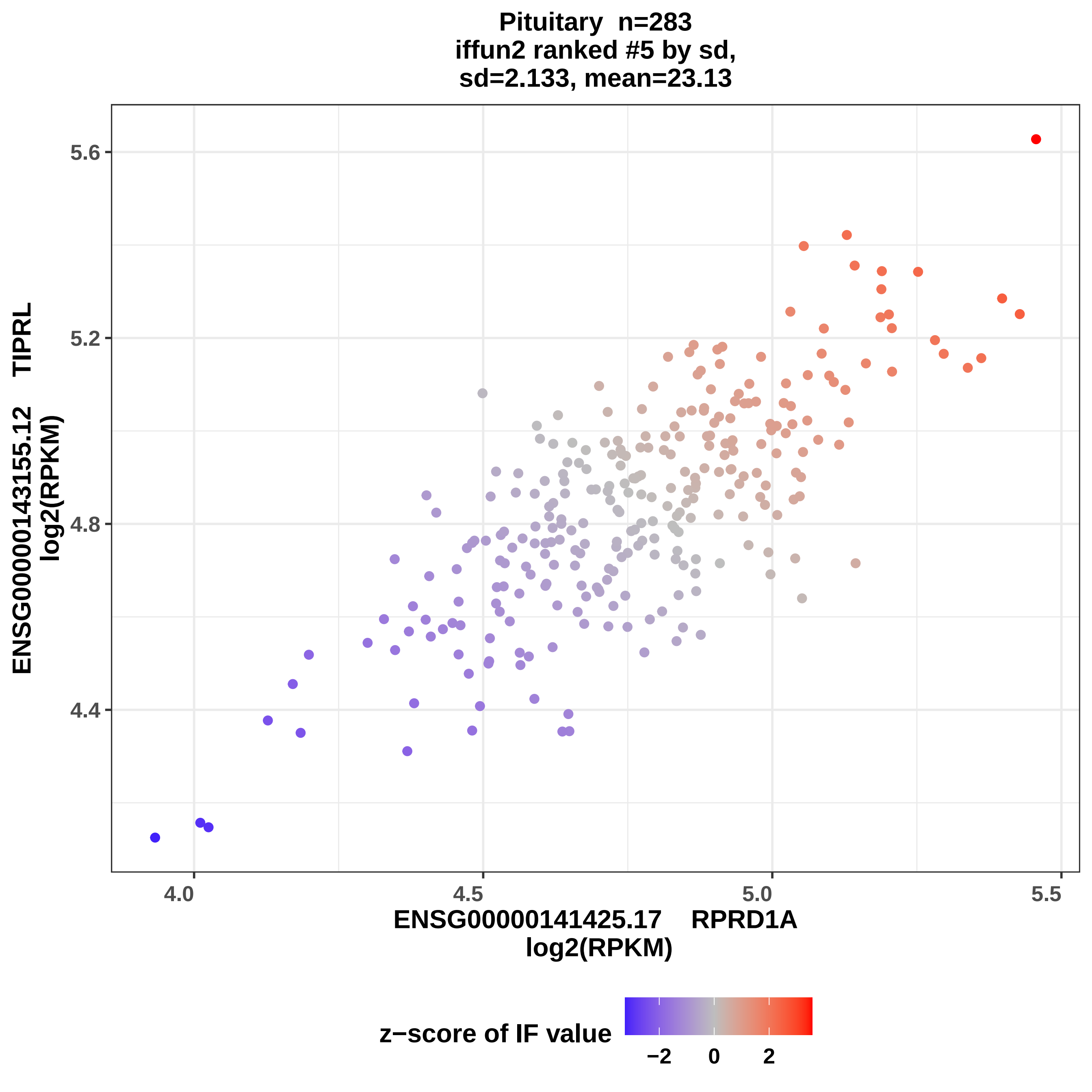

Pituitary n=283
iffun2 ranked #5 by sd,
sd=2.133, mean=23.13
ENSG00000143155.12 TIPRL
log2(RPKM)
5.6
5.2
4.8
4.4
4.0
4.5
5.0
5.5
ENSG00000141425.17 RPRD1A
log2(RPKM)
z−score of IF value
−2
0
2

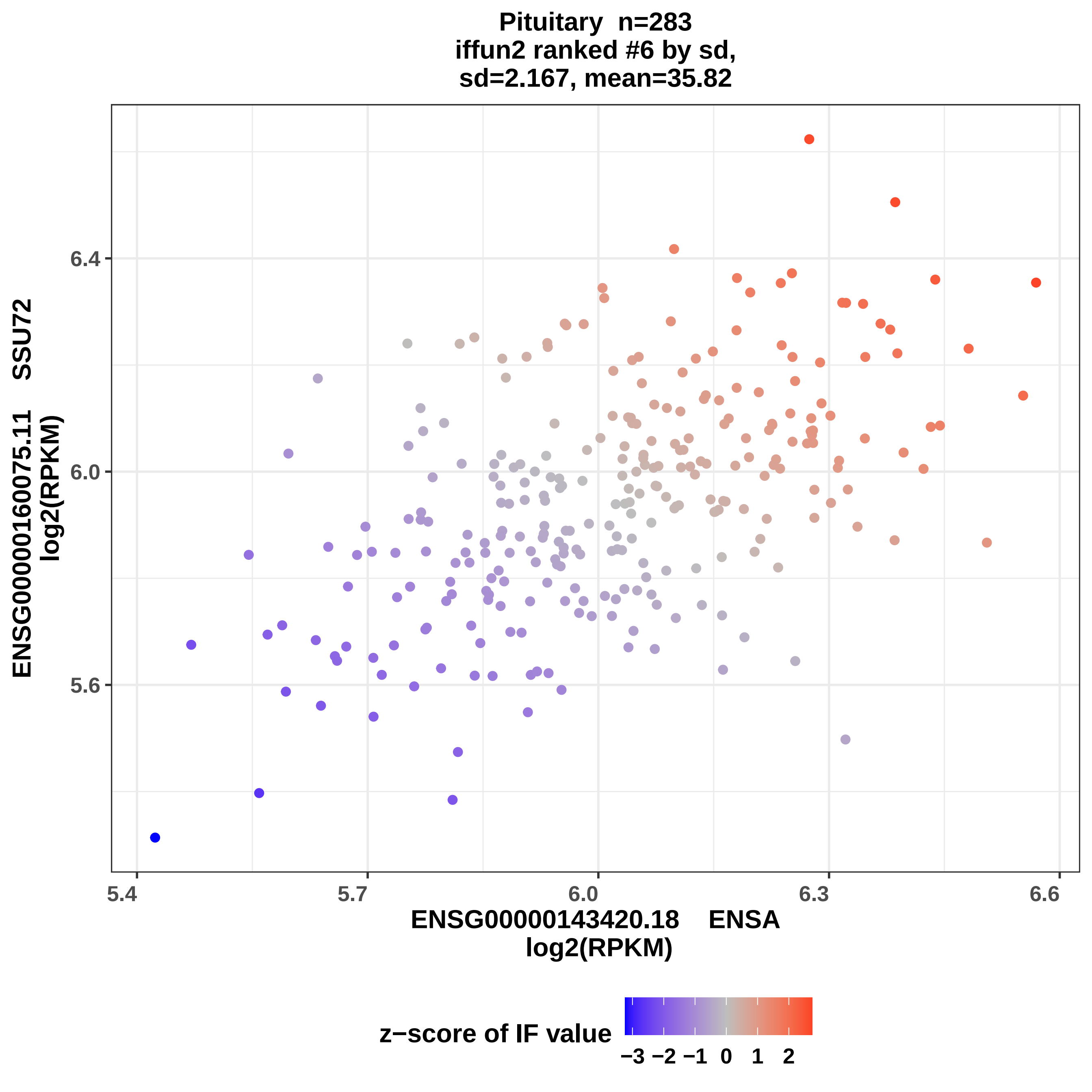
Pituitary n=283
iffun2 ranked #6 by sd,
sd=2.167, mean=35.82
ENSG00000160075.11 SSU72
log2(RPKM)
6.4
6.0
5.6
5.4
5.7
6.0
6.3
6.6
ENSG00000143420.18 ENSA
log2(RPKM)
z−score of IF value
−3 −2 −1 0 1 2

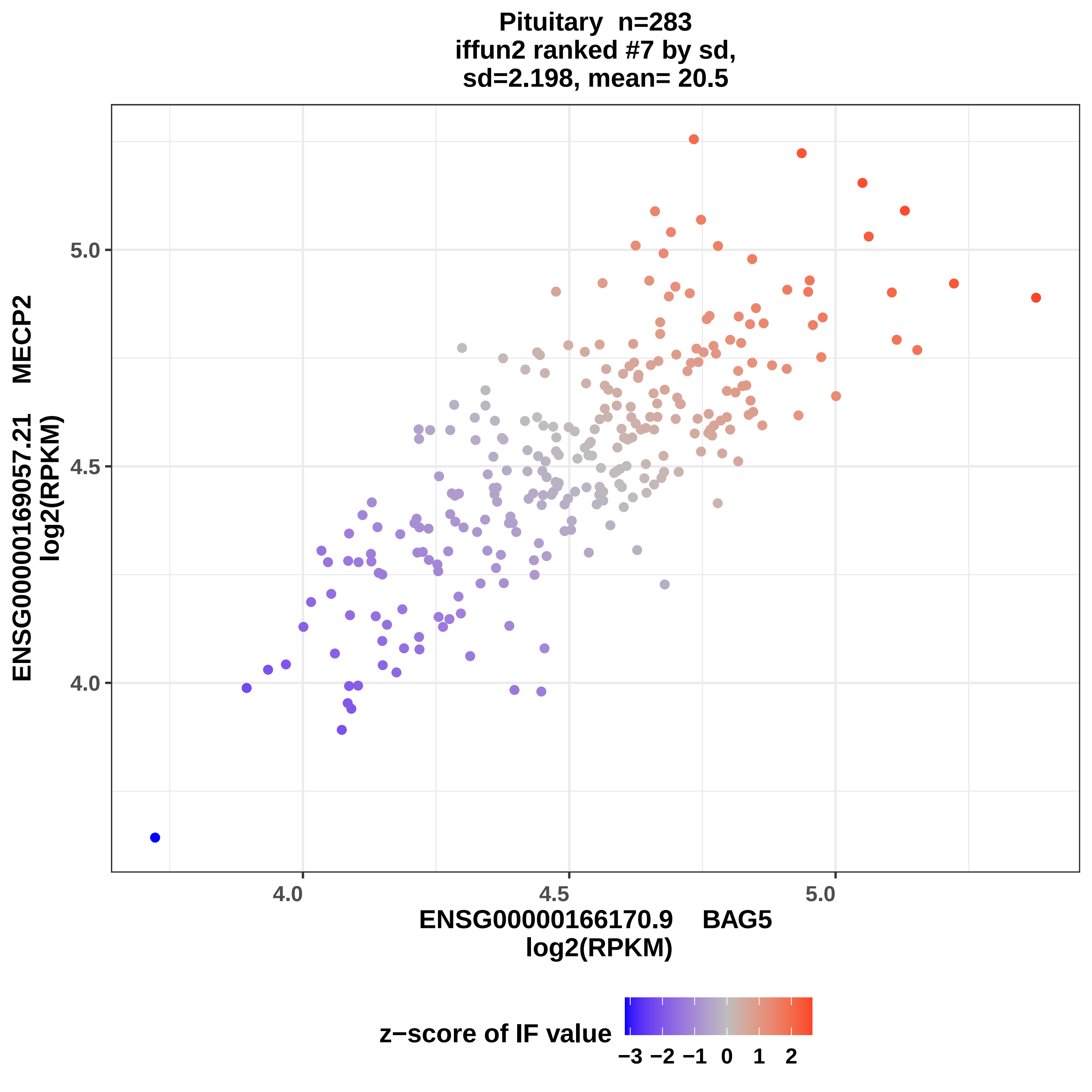

Pituitary n=283
iffun2 ranked #7 by sd,
sd=2.198, mean= 20.5
ENSG00000169057.21 MECP2
log2(RPKM)
5.0
4.5
4.0
4.0
4.5
5.0
ENSG00000166170.9 BAG5
log2(RPKM)
z−score of IF value
−3
−2
−1
0
1
2

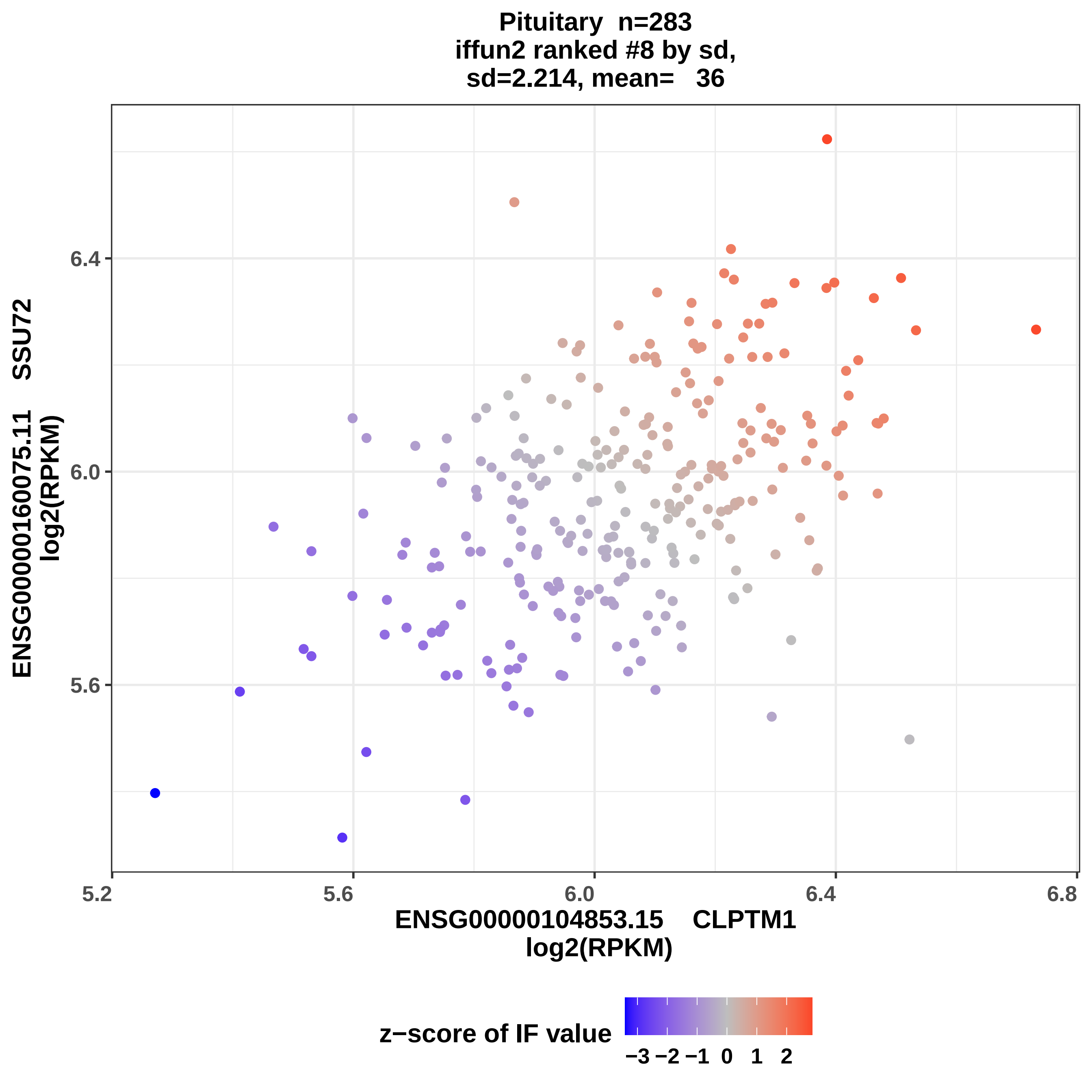

Pituitary n=283
iffun2 ranked #8 by sd,
sd=2.214, mean= 36
6.4
6.0
5.6
5.2
5.6
6.0
6.4
6.8
ENSG00000160075.11 SSU72
log2(RPKM)
ENSG00000104853.15 CLPTM1
log2(RPKM)
z−score of IF value
−3 −2 −1 0 1 2

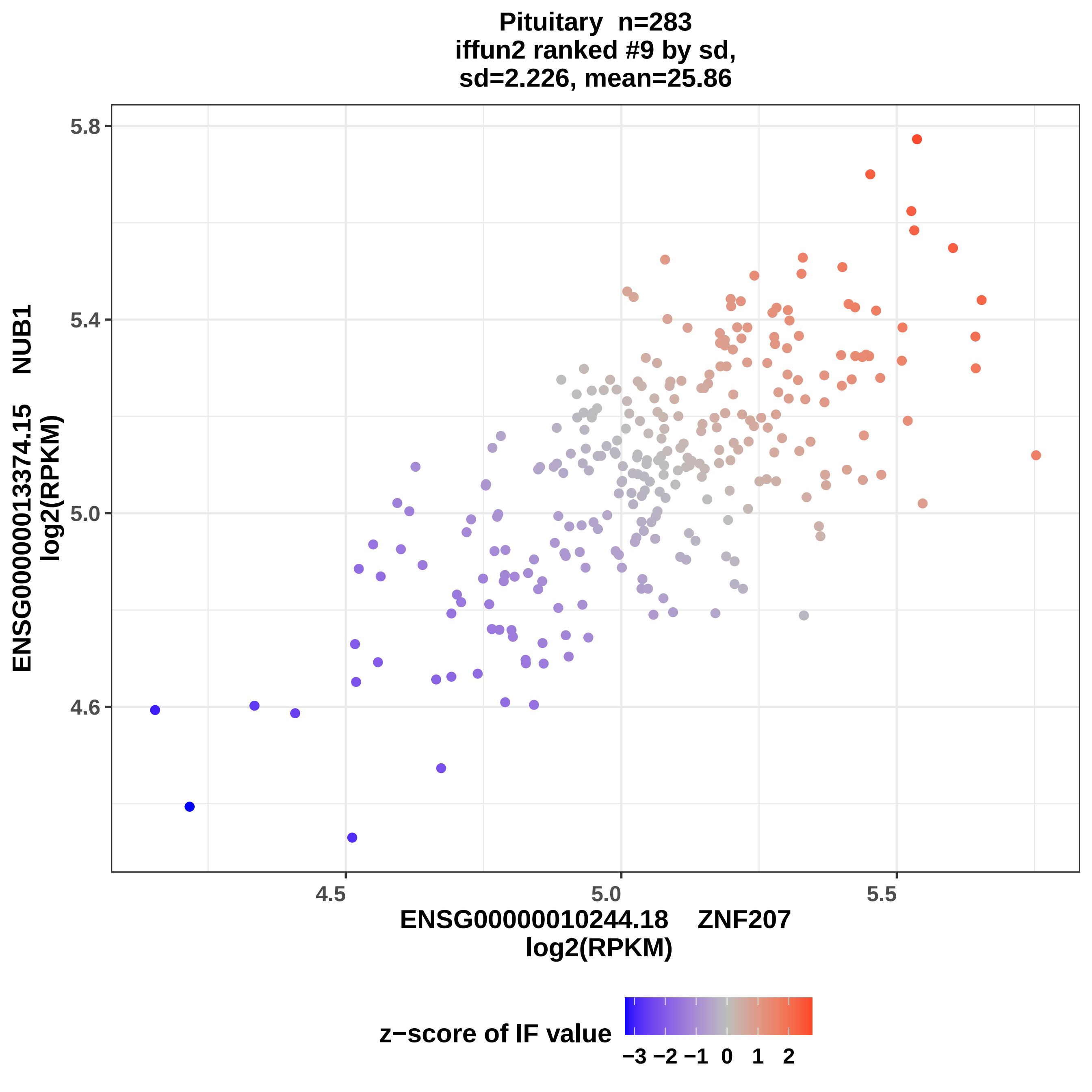

Pituitary n=283
iffun2 ranked #9 by sd,
sd=2.226, mean=25.86
5.8
5.4
5.0
4.6
4.5
5.0
5.5
ENSG00000013374.15 NUB1
log2(RPKM)
ENSG00000010244.18 ZNF207
log2(RPKM)
z−score of IF value
−3 −2 −1 0 1 2

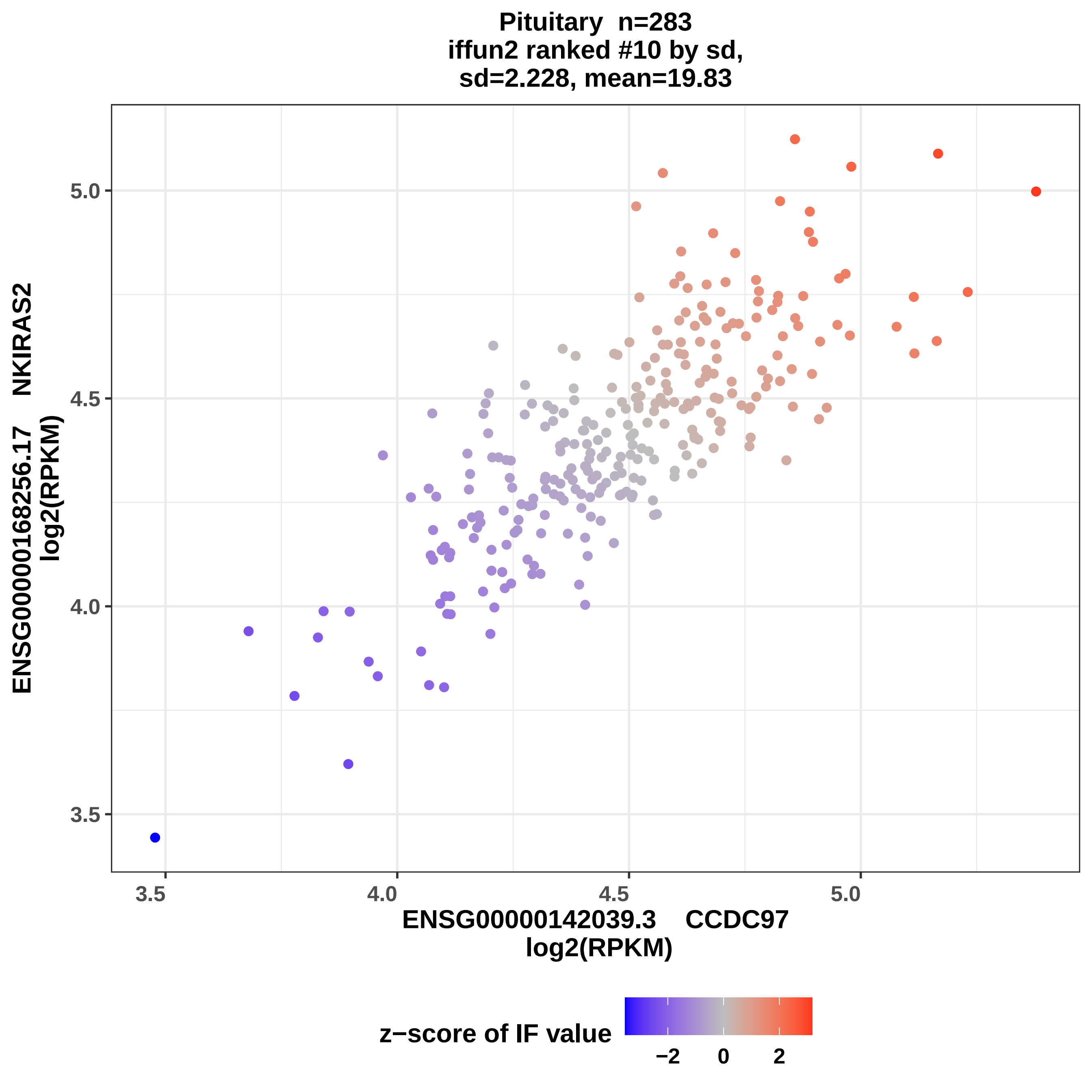
Pituitary n=283
iffun2 ranked #10 by sd,
sd=2.228, mean=19.83
ENSG00000168256.17 NKIRAS2
log2(RPKM)
5.0
4.5
4.0
3.5
3.5
4.0
4.5
5.0
ENSG00000142039.3 CCDC97
log2(RPKM)
z−score of IF value
−2
0
2

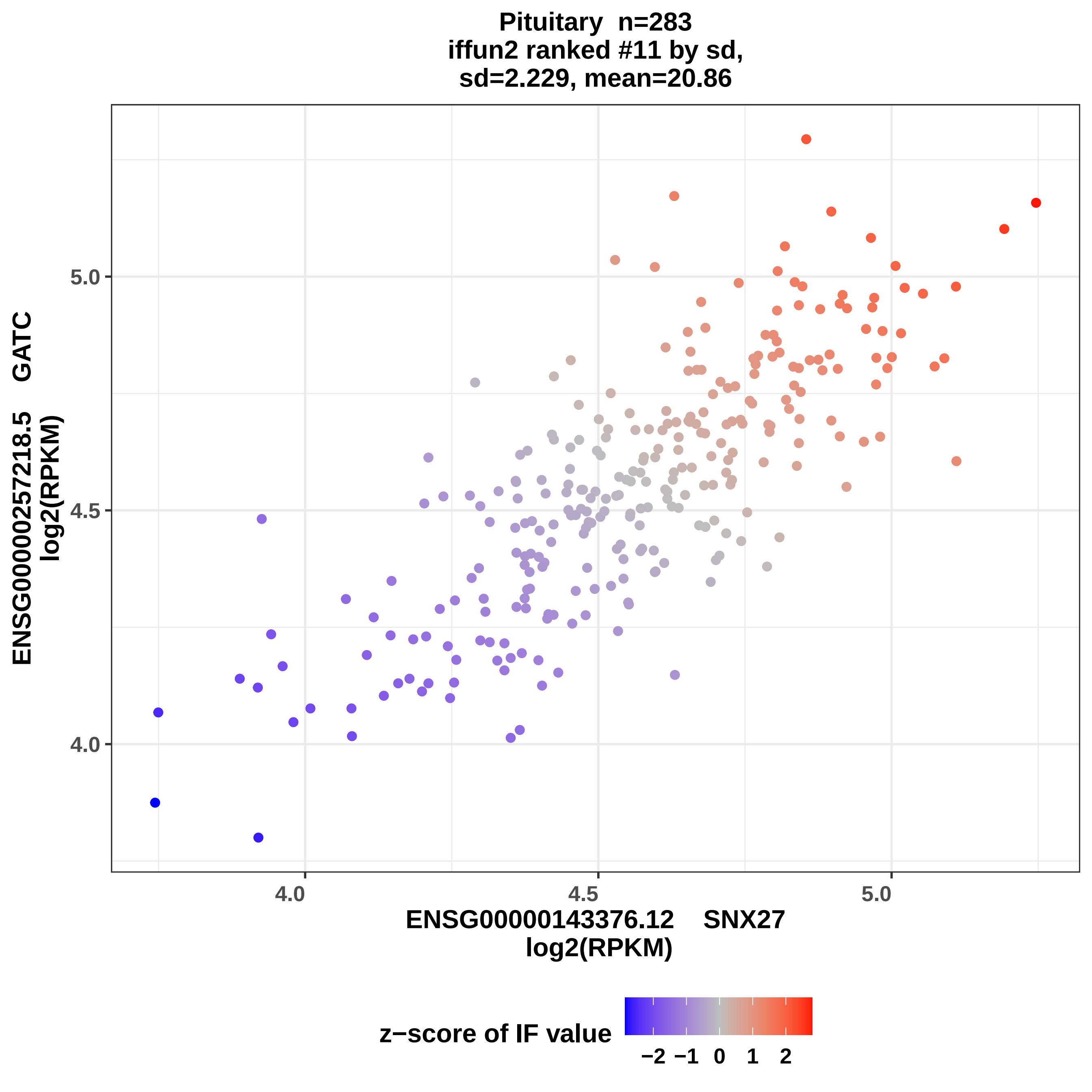

Pituitary n=283
iffun2 ranked #11 by sd,
sd=2.229, mean=20.86
ENSG00000257218.5 GATC
log2(RPKM)
5.0
4.5
4.0
ENSG00000143376.12 SNX27
log2(RPKM)
z−score of IF value
−2 −1 0 1 2

Pituitary n=283
iffun2 ranked #12 by sd,
sd=2.242, mean=25.55

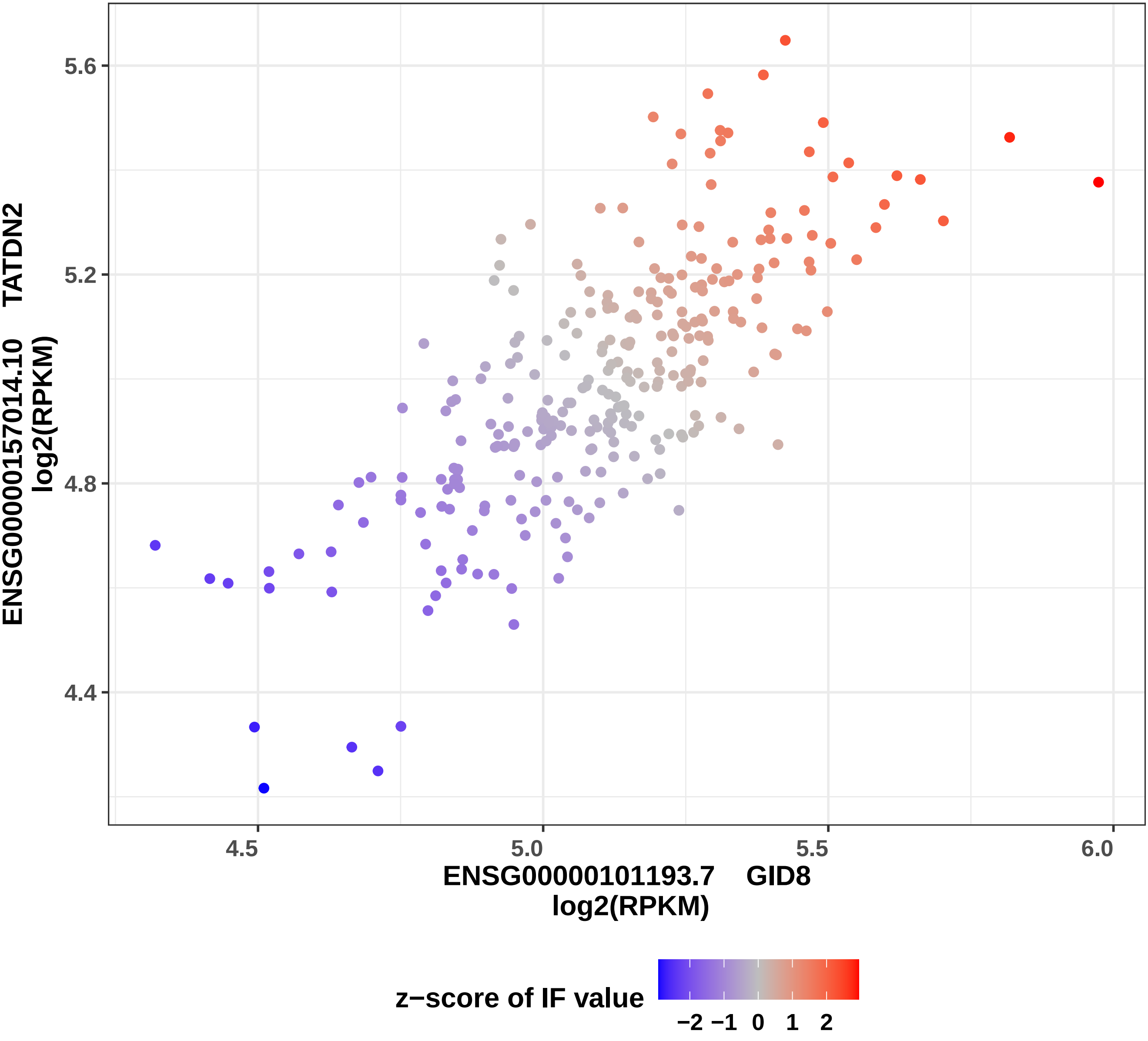

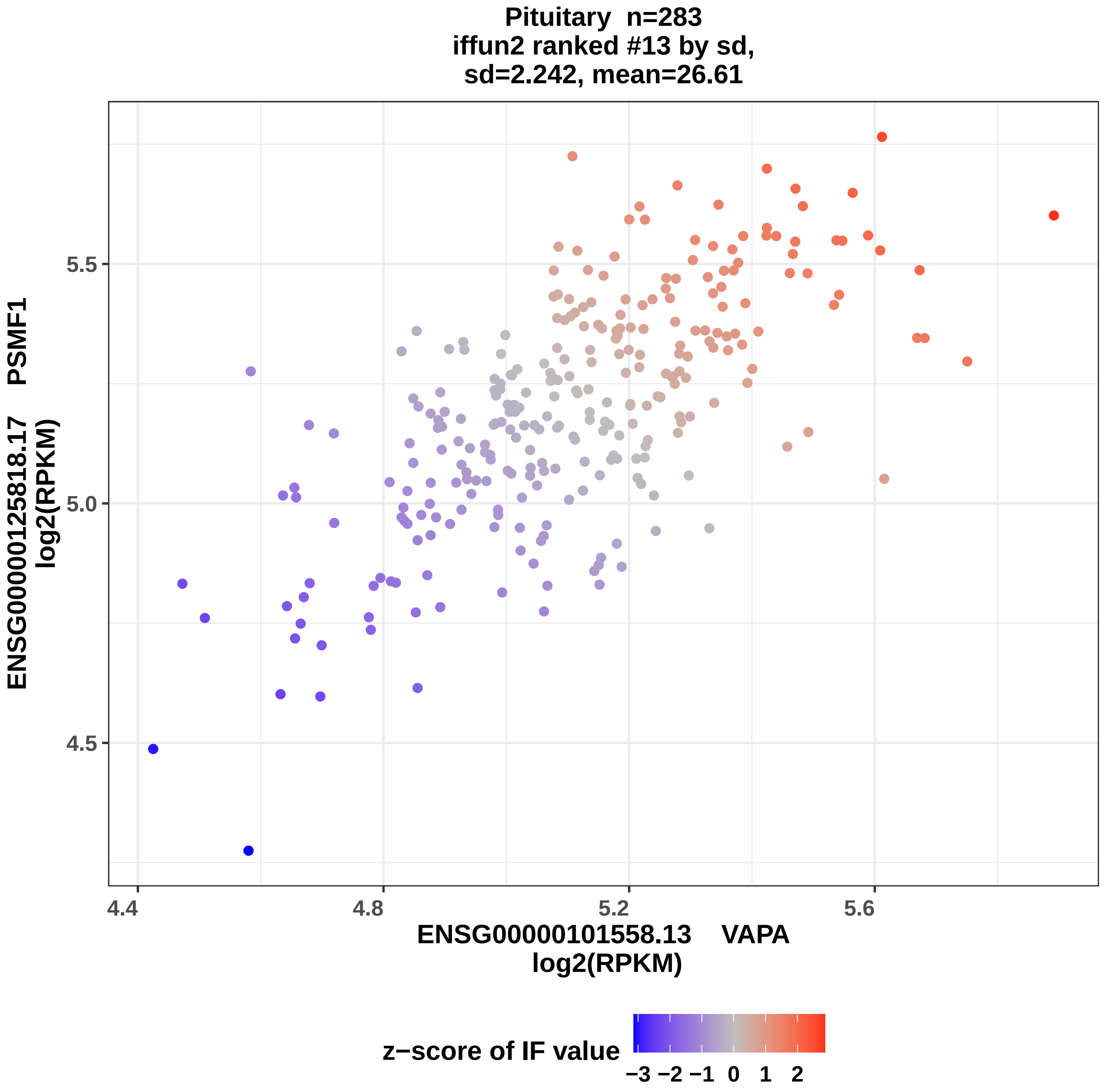
Pituitary n=283
iffun2 ranked #13 by sd,
sd=2.242, mean=26.61
ENSG00000125818.17 PSMF1
log2(RPKM)
4.5
5.0
5.5
4.4
4.8
5.2
5.6
ENSG00000101558.13 VAPA
log2(RPKM)
z−score of IF value
−3 −2 −1 0 1 2

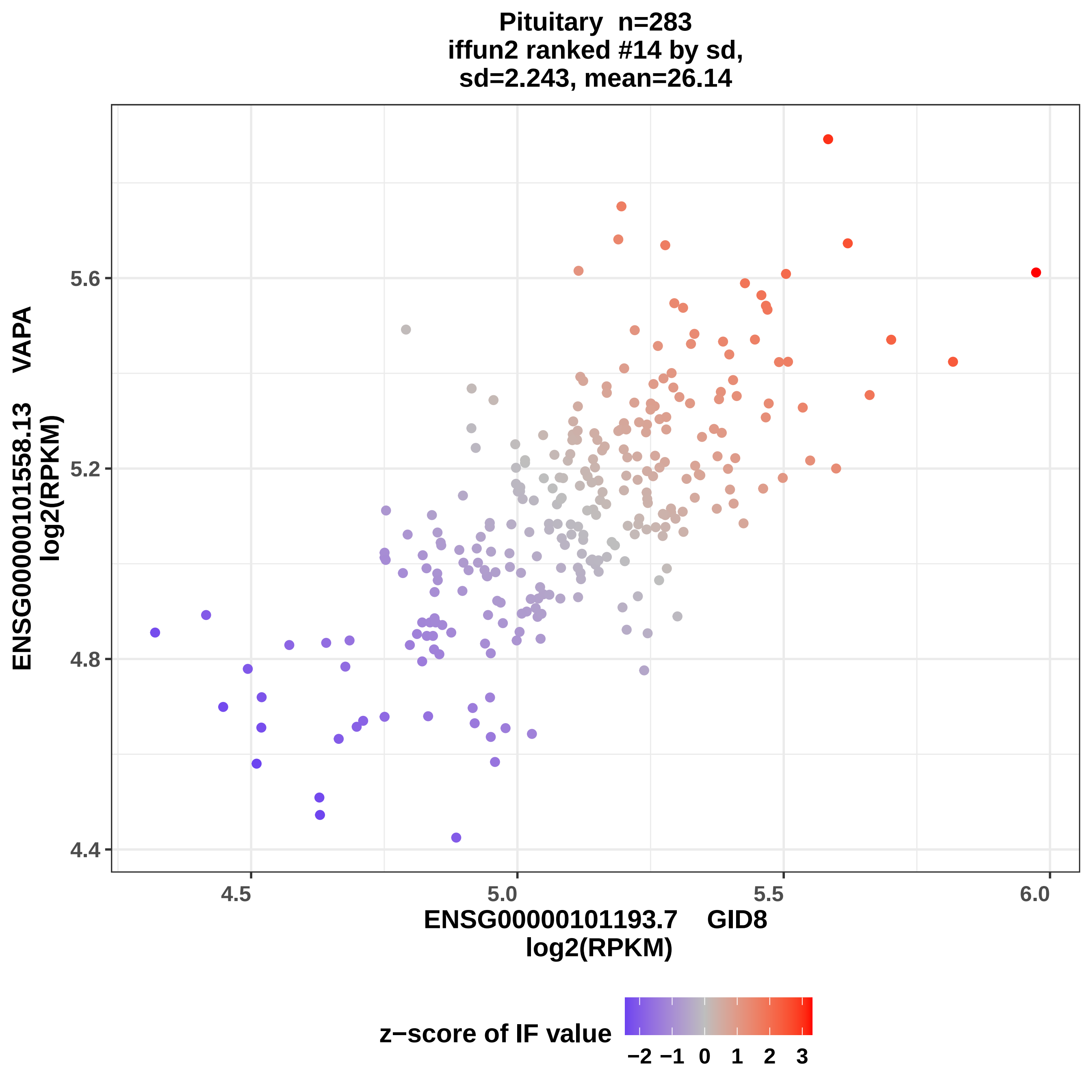

Pituitary n=283
iffun2 ranked #14 by sd,
sd=2.243, mean=26.14
ENSG00000101558.13 VAPA
log2(RPKM)
5.6
5.2
4.8
4.4
4.5
5.0
5.5
6.0
ENSG00000101193.7 GID8
log2(RPKM)
z−score of IF value
−2 −1 0 1 2 3

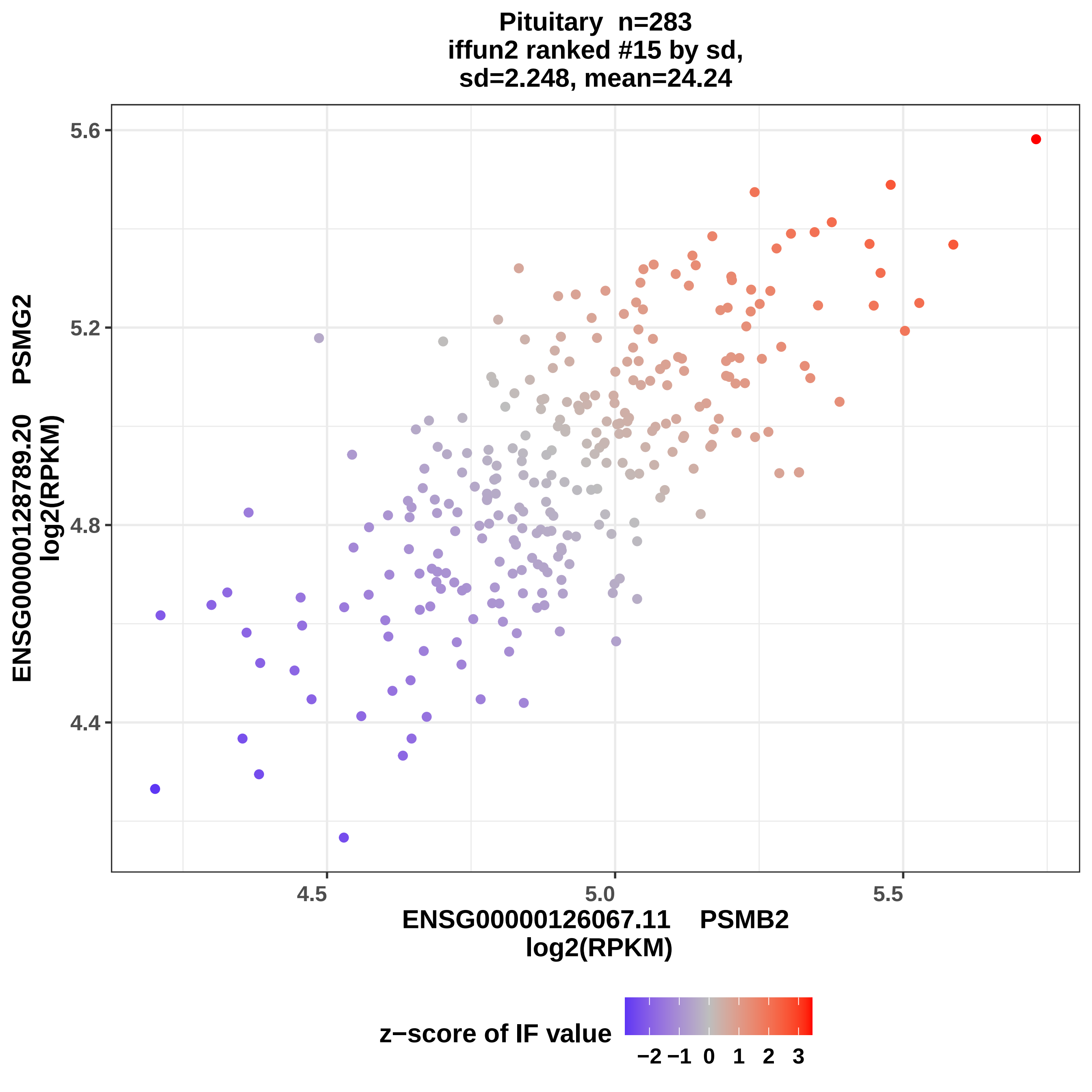

Pituitary n=283
iffun2 ranked #15 by sd,
sd=2.248, mean=24.24
5.6
5.2
4.8
4.4
4.5
5.0
5.5
ENSG00000128789.20 PSMG2
log2(RPKM)
ENSG00000126067.11 PSMB2
log2(RPKM)
z−score of IF value
−2 −1 0 1 2 3

Pituitary n=283
iffun2 ranked #16 by sd,
sd=2.268, mean=25.25

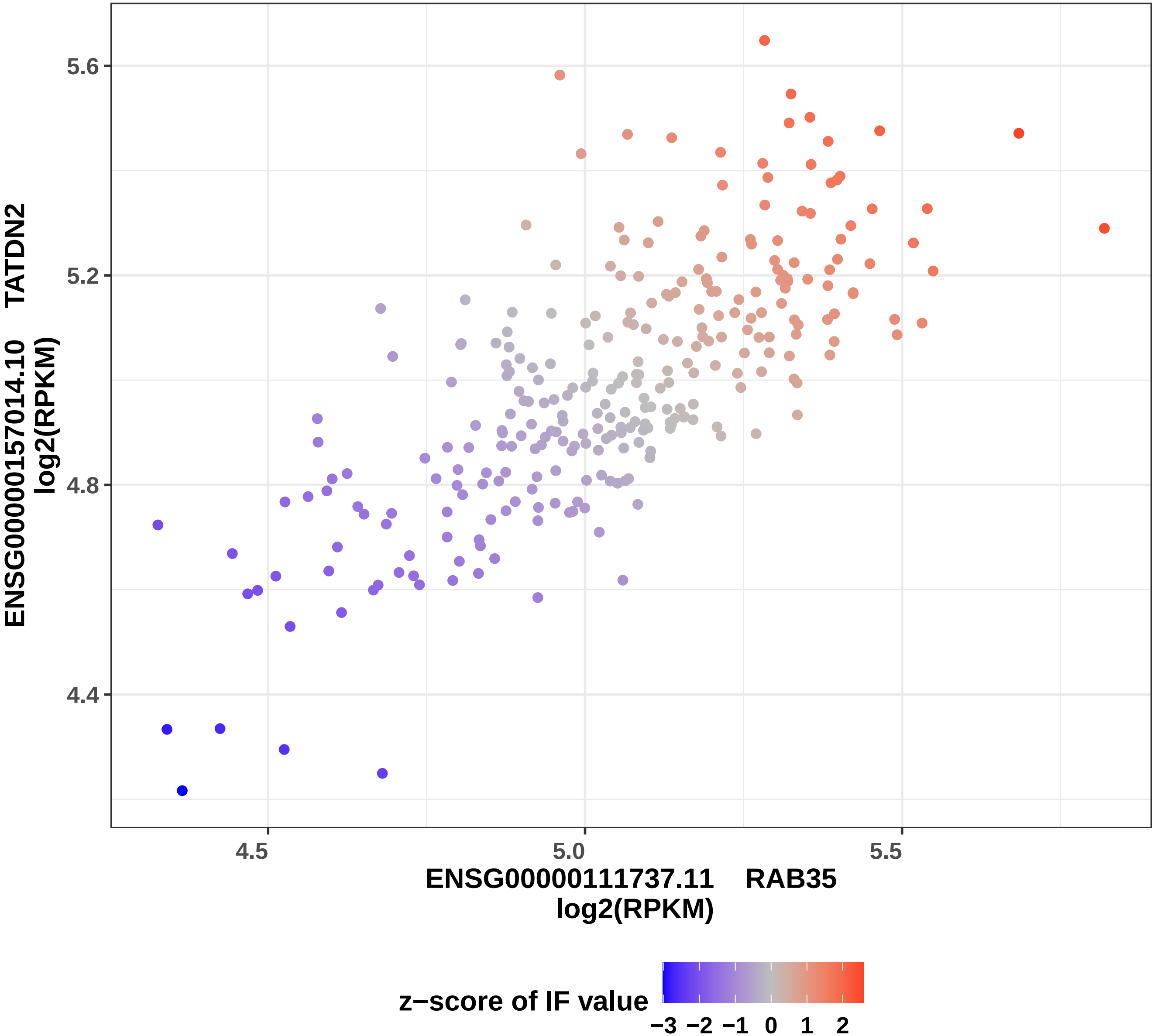

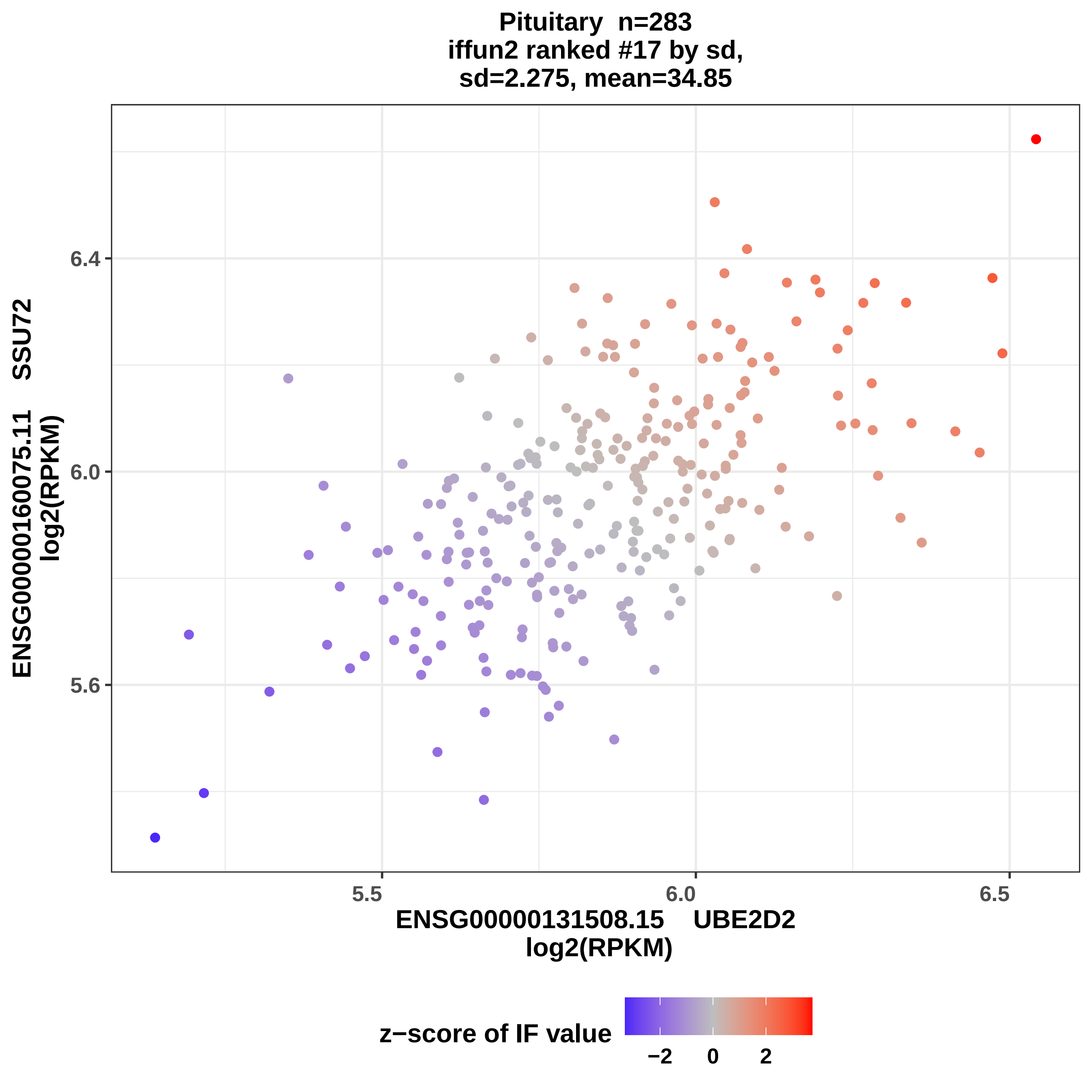

Pituitary n=283
iffun2 ranked #17 by sd,
sd=2.275, mean=34.85
6.4
6.0
5.6
5.5
6.0
6.5
ENSG00000160075.11 SSU72
log2(RPKM)
ENSG00000131508.15 UBE2D2
log2(RPKM)
z−score of IF value
−2
0
2

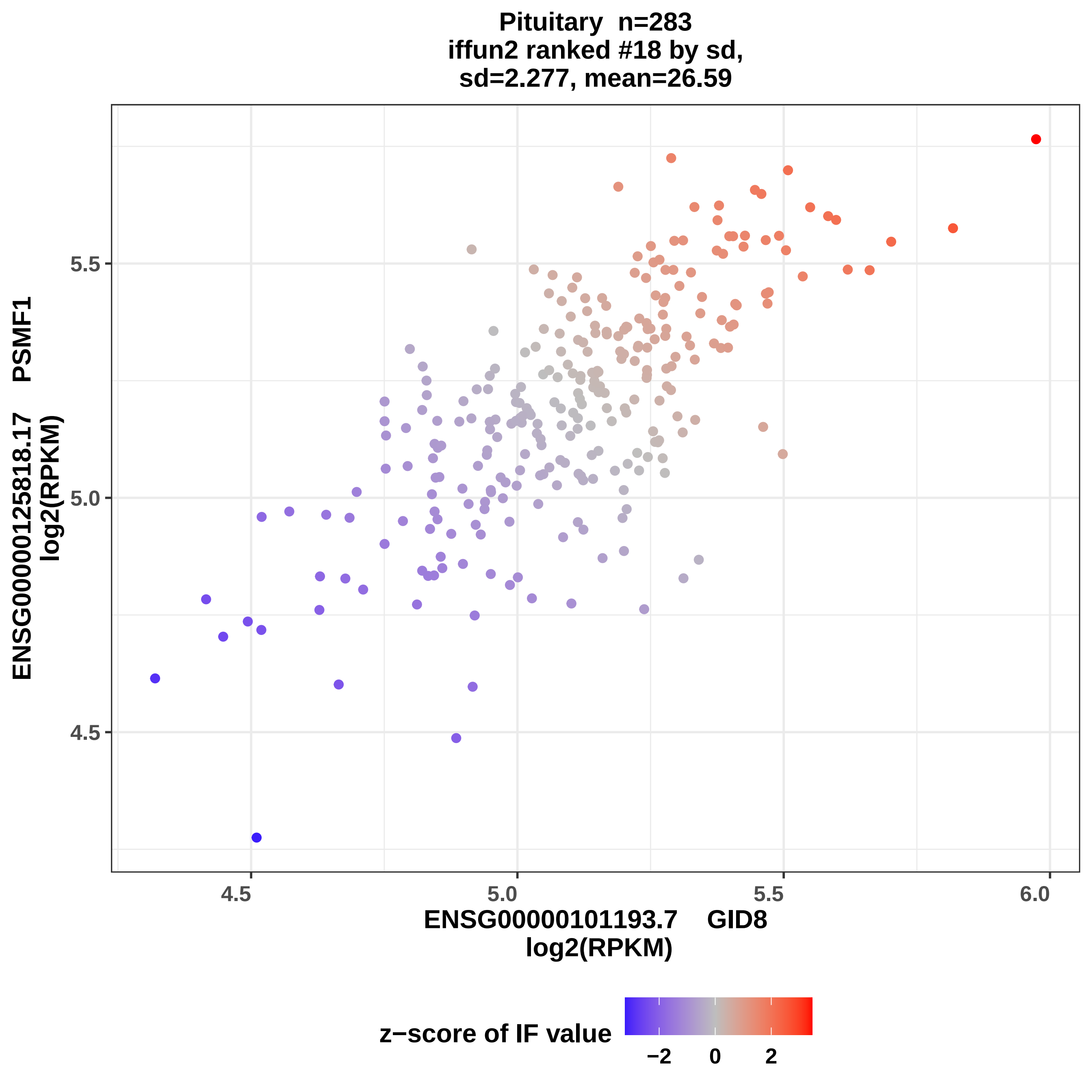
Pituitary n=283
iffun2 ranked #18 by sd,
sd=2.277, mean=26.59
ENSG00000125818.17 PSMF1
log2(RPKM)
5.5
5.0
4.5
4.5
5.0
5.5
6.0
ENSG00000101193.7 GID8
log2(RPKM)
z−score of IF value
−2
0
2

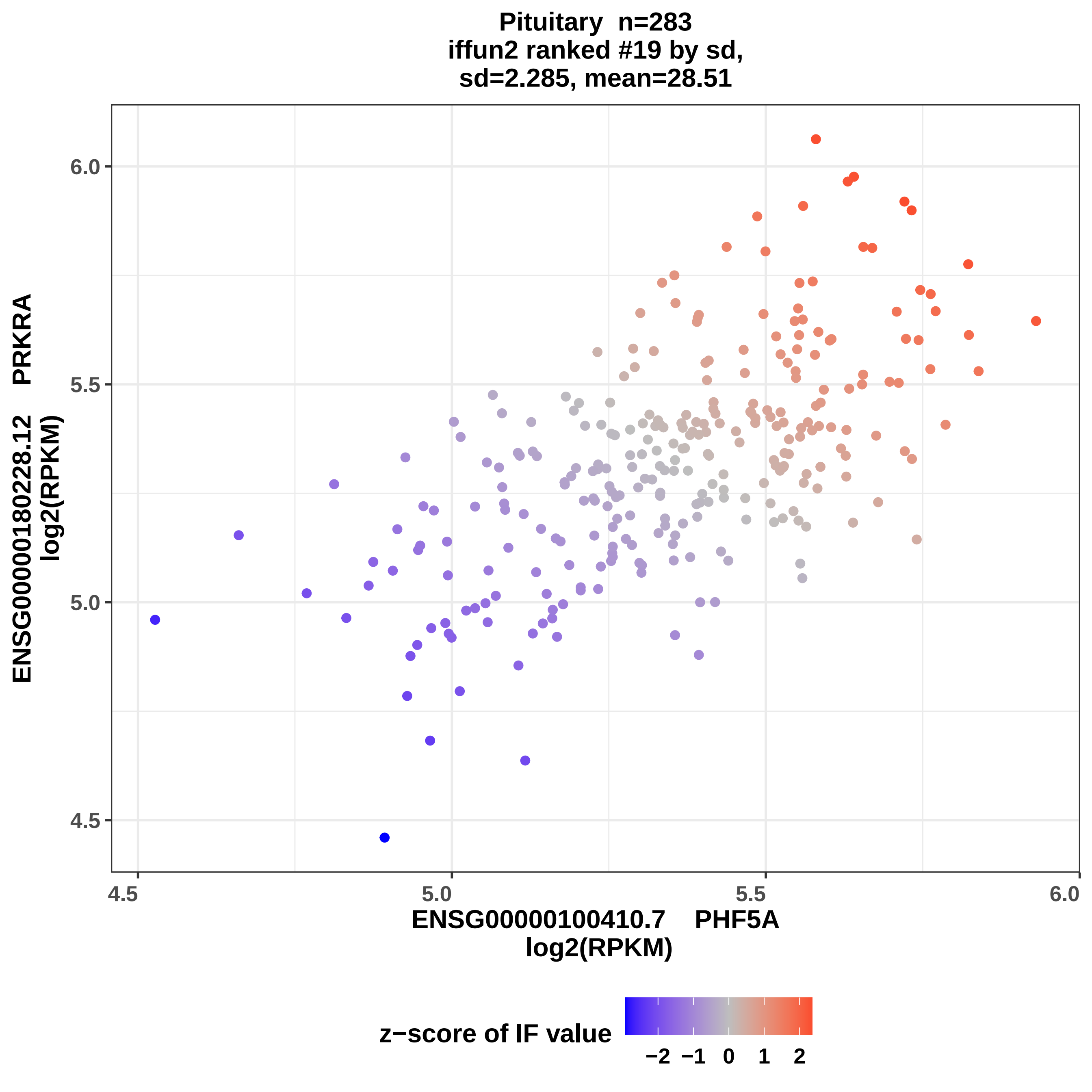
Pituitary n=283
iffun2 ranked #19 by sd,
sd=2.285, mean=28.51
ENSG00000180228.12 PRKRA
log2(RPKM)
ENSG00000100410.7 PHF5A
log2(RPKM)
4.5
5.0
5.5
6.0
z−score of IF value
−2 −1 0 1 2

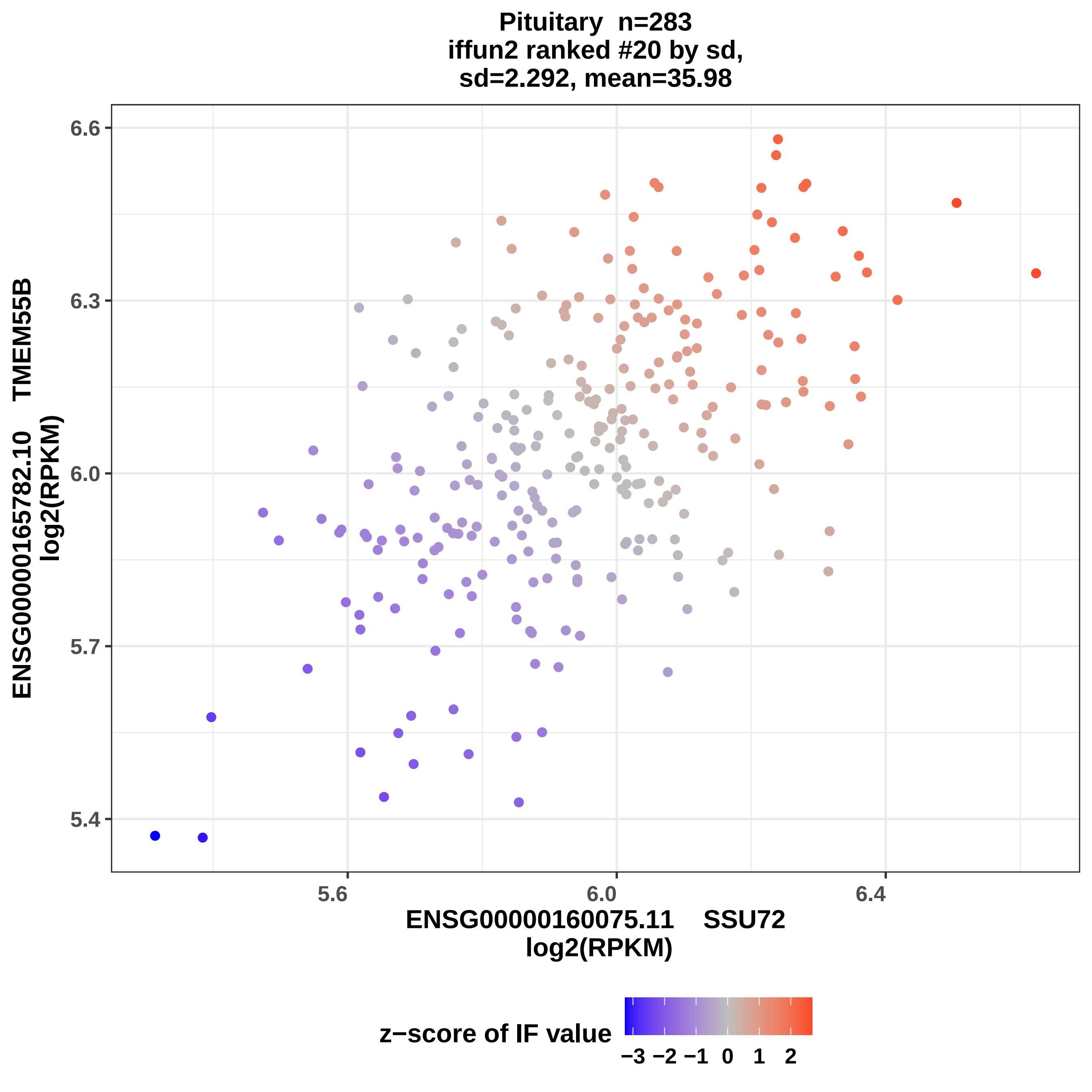

Pituitary n=283
iffun2 ranked #20 by sd,
sd=2.292, mean=35.98
ENSG00000165782.10 TMEM55B
log2(RPKM)
6.6
6.3
6.0
5.7
5.4
5.6
6.0
6.4
ENSG00000160075.11 SSU72
log2(RPKM)
z−score of IF value
−3 −2 −1 0 1 2

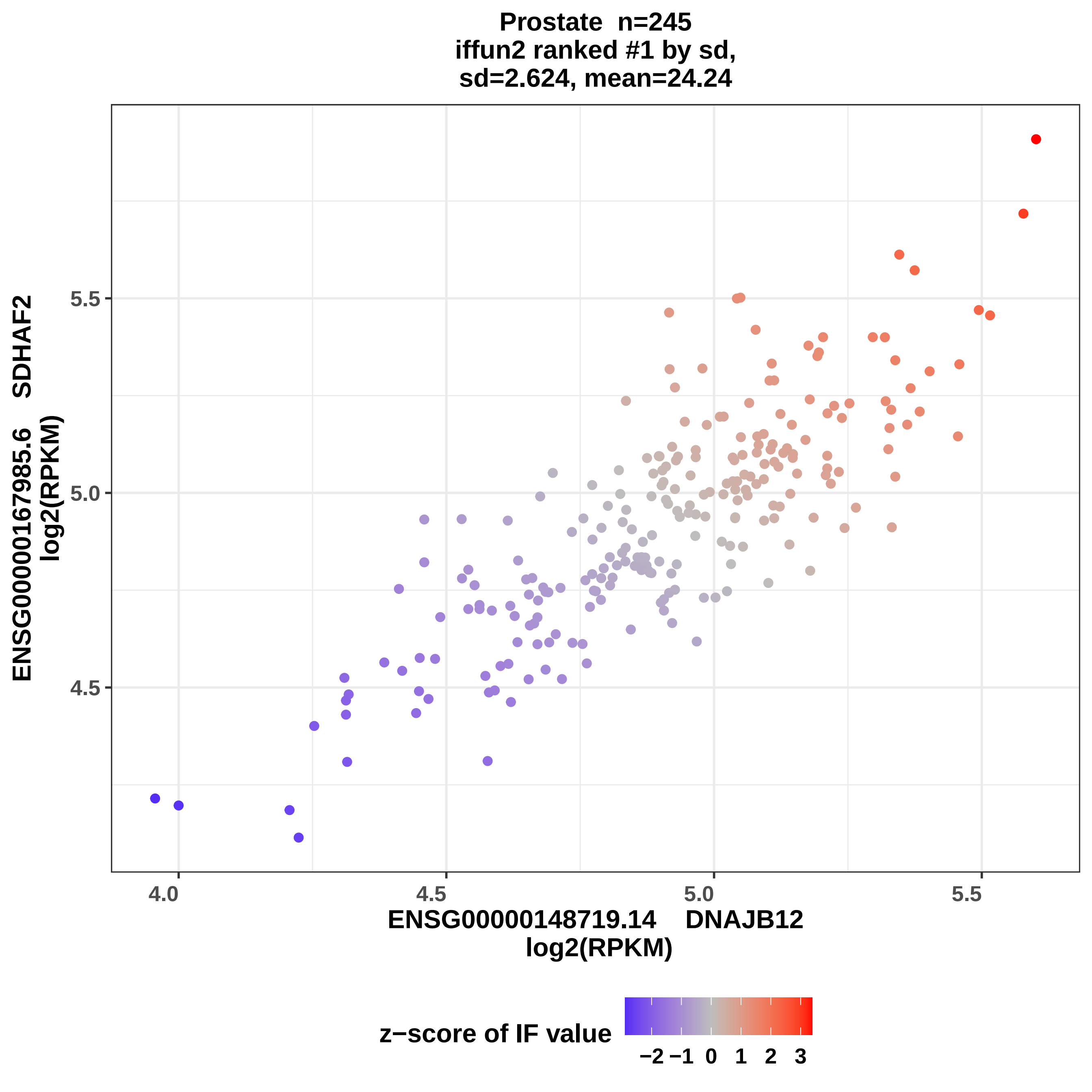

Prostate n=245
iffun2 ranked #1 by sd,
sd=2.624, mean=24.24
ENSG00000167985.6 SDHAF2
log2(RPKM)
5.5
5.0
4.5
4.0
4.5
5.0
5.5
ENSG00000148719.14 DNAJB12
log2(RPKM)
z−score of IF value
−2 −1 0 1 2 3

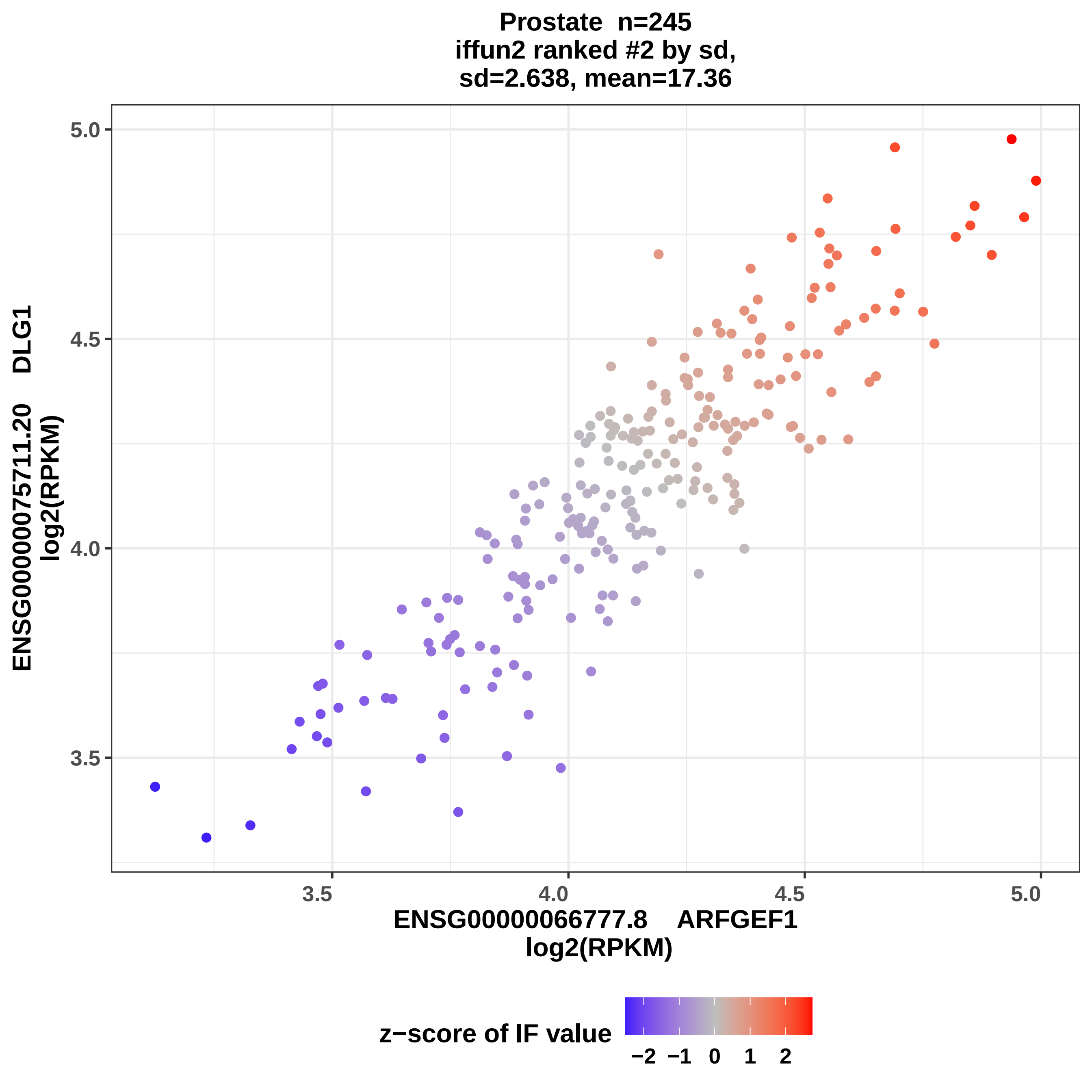
Prostate n=245
iffun2 ranked #2 by sd,
sd=2.638, mean=17.36
ENSG00000075711.20 DLG1
log2(RPKM)
5.0
4.5
4.0
3.5
3.5
4.0
4.5
5.0
ENSG00000066777.8 ARFGEF1
log2(RPKM)
z−score of IF value
−2 −1 0 1 2

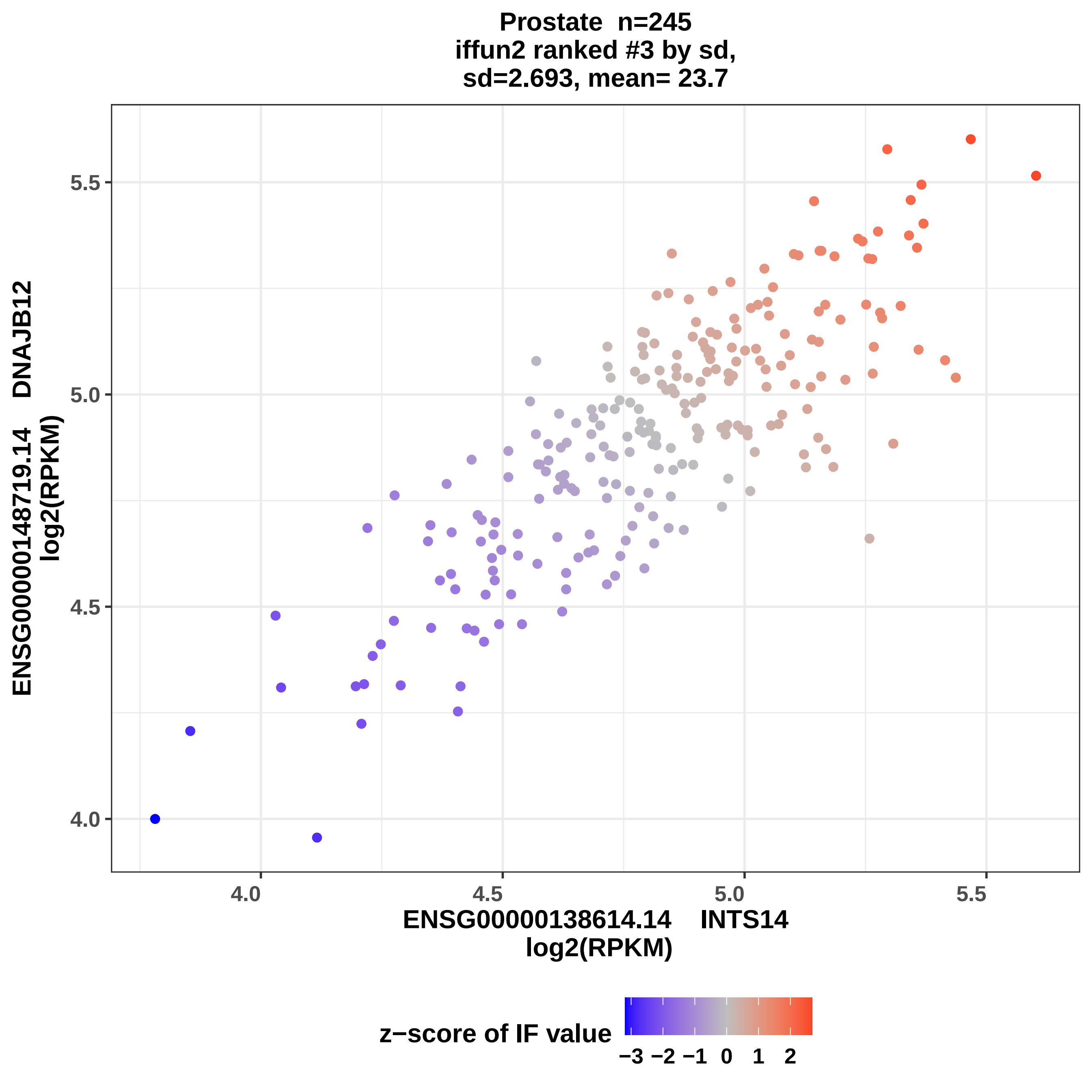

Prostate n=245
iffun2 ranked #3 by sd,
sd=2.693, mean= 23.7
5.5
5.0
4.5
4.0
4.0
4.5
5.0
5.5
ENSG00000148719.14 DNAJB12
log2(RPKM)
ENSG00000138614.14 INTS14
log2(RPKM)
z−score of IF value
−3 −2 −1 0 1 2

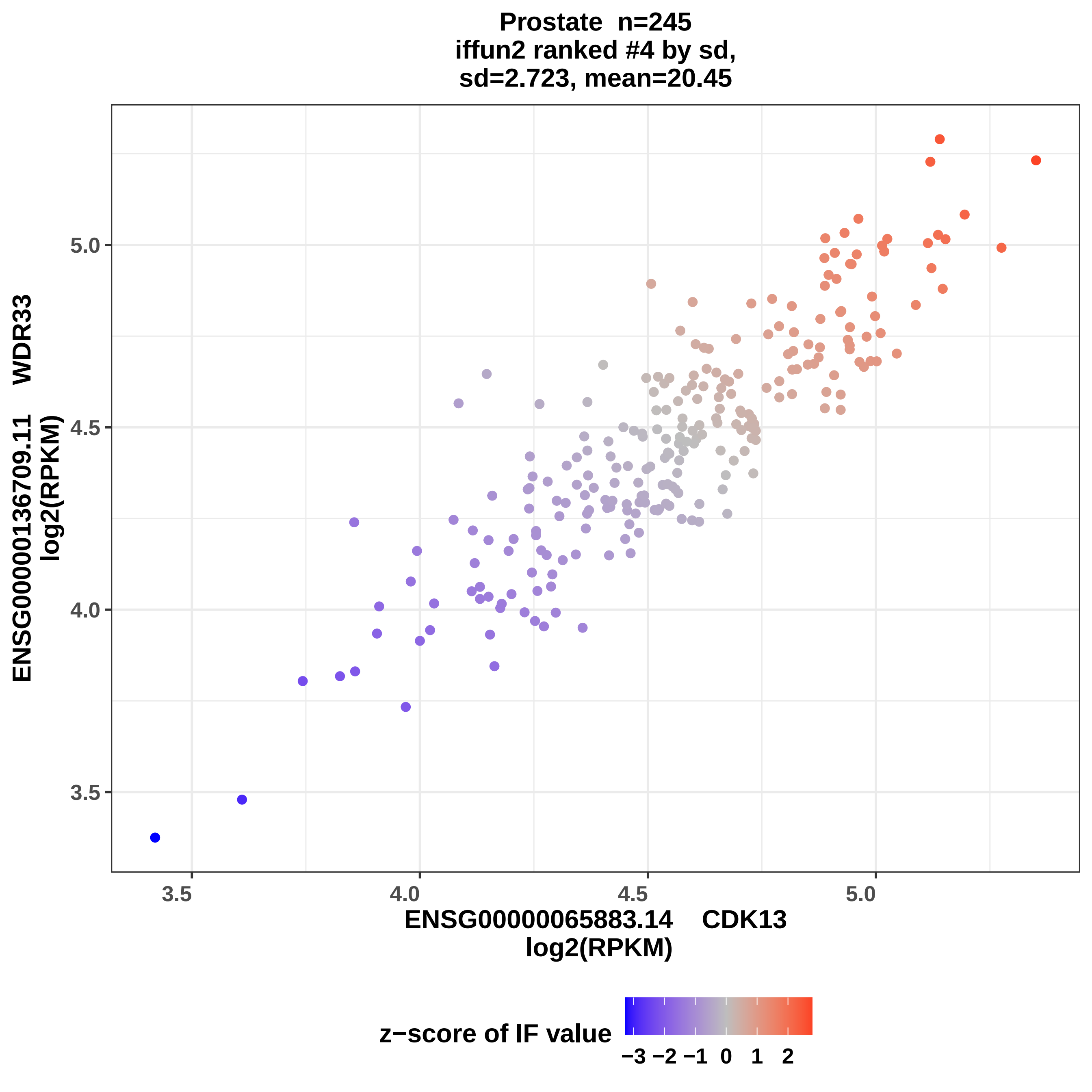

Prostate n=245
iffun2 ranked #4 by sd,
sd=2.723, mean=20.45
ENSG00000136709.11 WDR33
log2(RPKM)
3.5
4.0
4.5
5.0
ENSG00000065883.14 CDK13
log2(RPKM)
z−score of IF value
−3 −2 −1 0 1 2

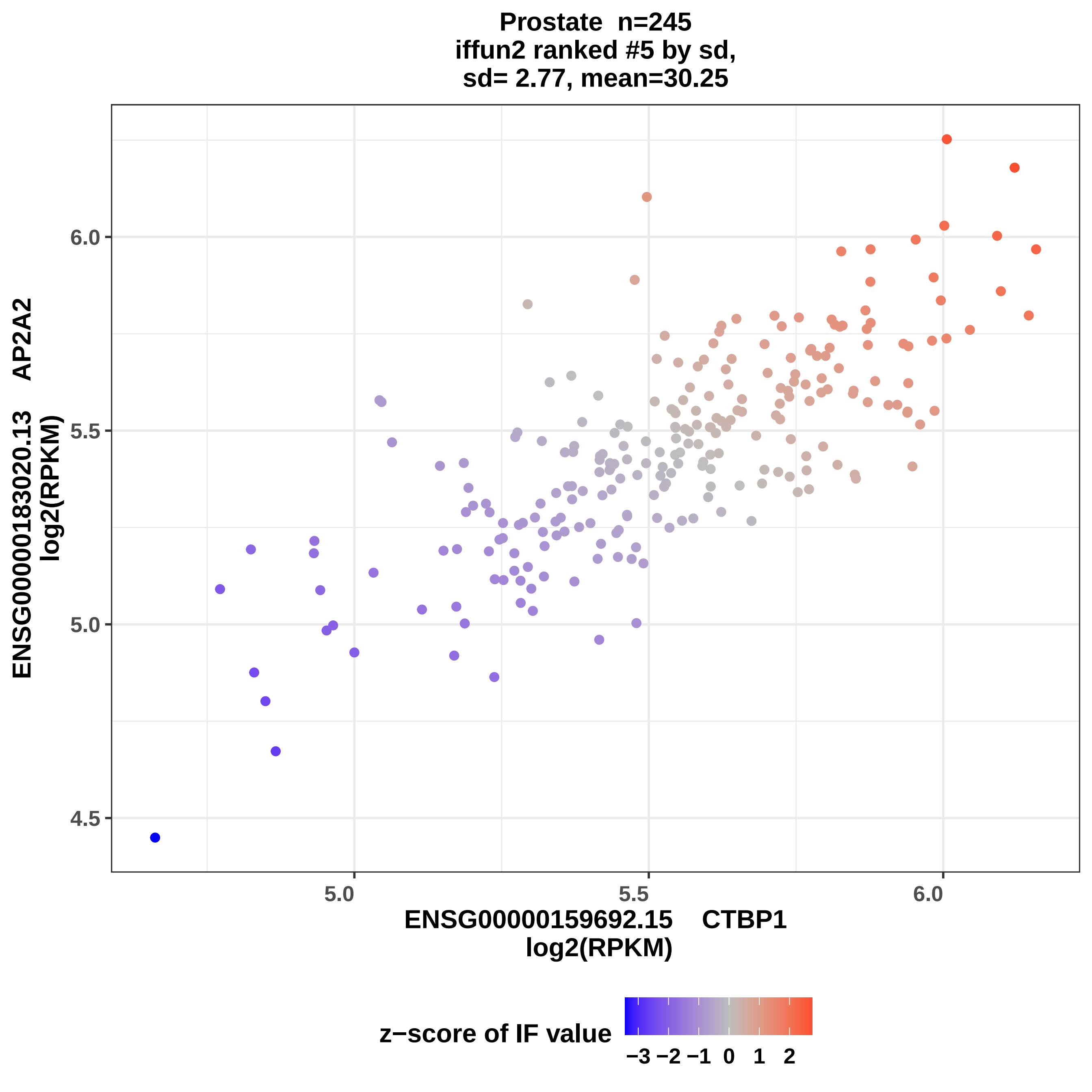
Prostate n=245
iffun2 ranked #5 by sd,
sd= 2.77, mean=30.25
ENSG00000183020.13 AP2A2
log2(RPKM)
6.0
5.5
5.0
4.5
5.0
5.5
6.0
ENSG00000159692.15 CTBP1
log2(RPKM)
z−score of IF value
−3 −2 −1 0 1 2

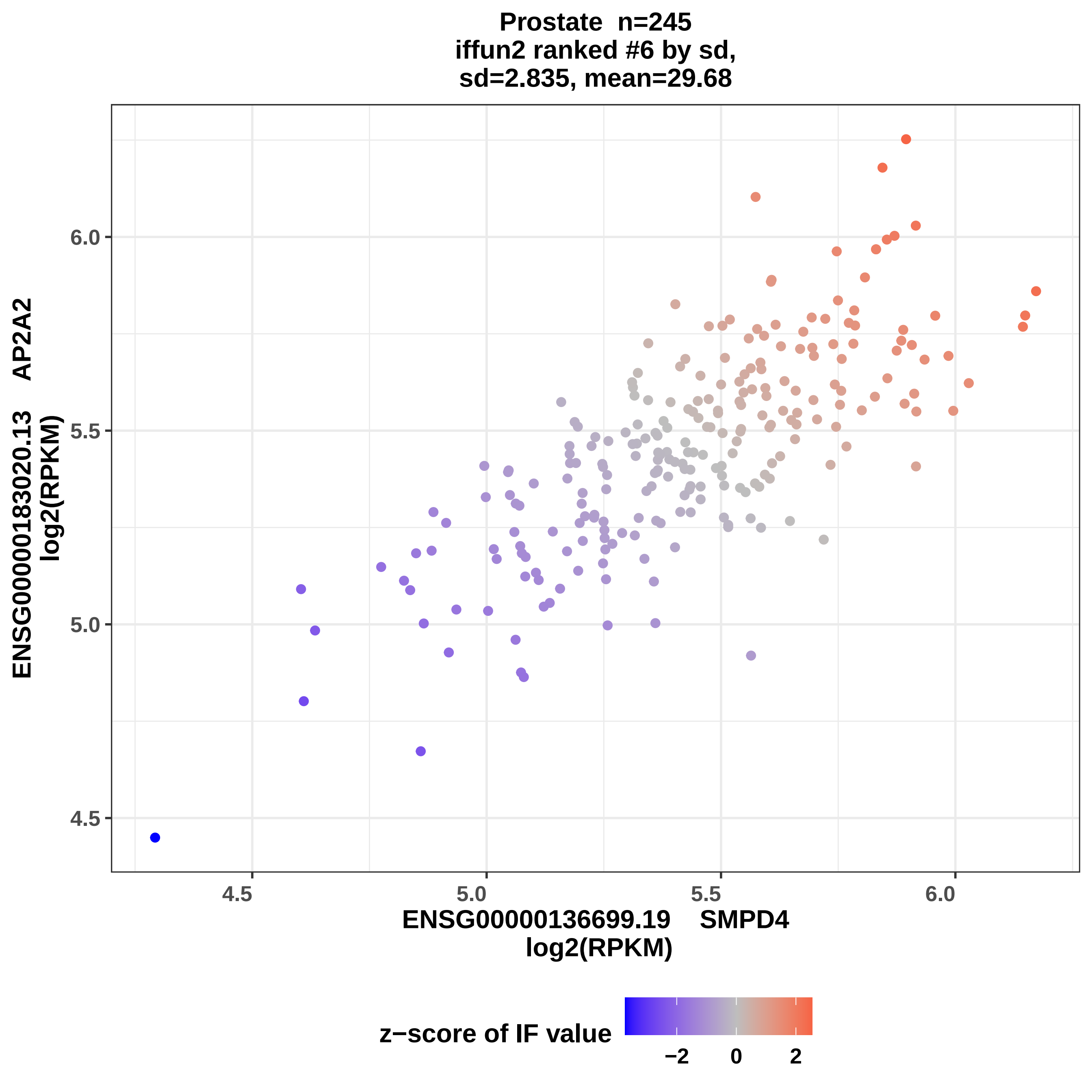
Prostate n=245
iffun2 ranked #6 by sd,
sd=2.835, mean=29.68
ENSG00000183020.13 AP2A2
log2(RPKM)
ENSG00000136699.19 SMPD4
log2(RPKM)
4.5
5.0
5.5
6.0
4.5
5.0
5.5
6.0
z−score of IF value
−2
0
2

Prostate n=245
iffun2 ranked #7 by sd,
sd=2.853, mean=29.04

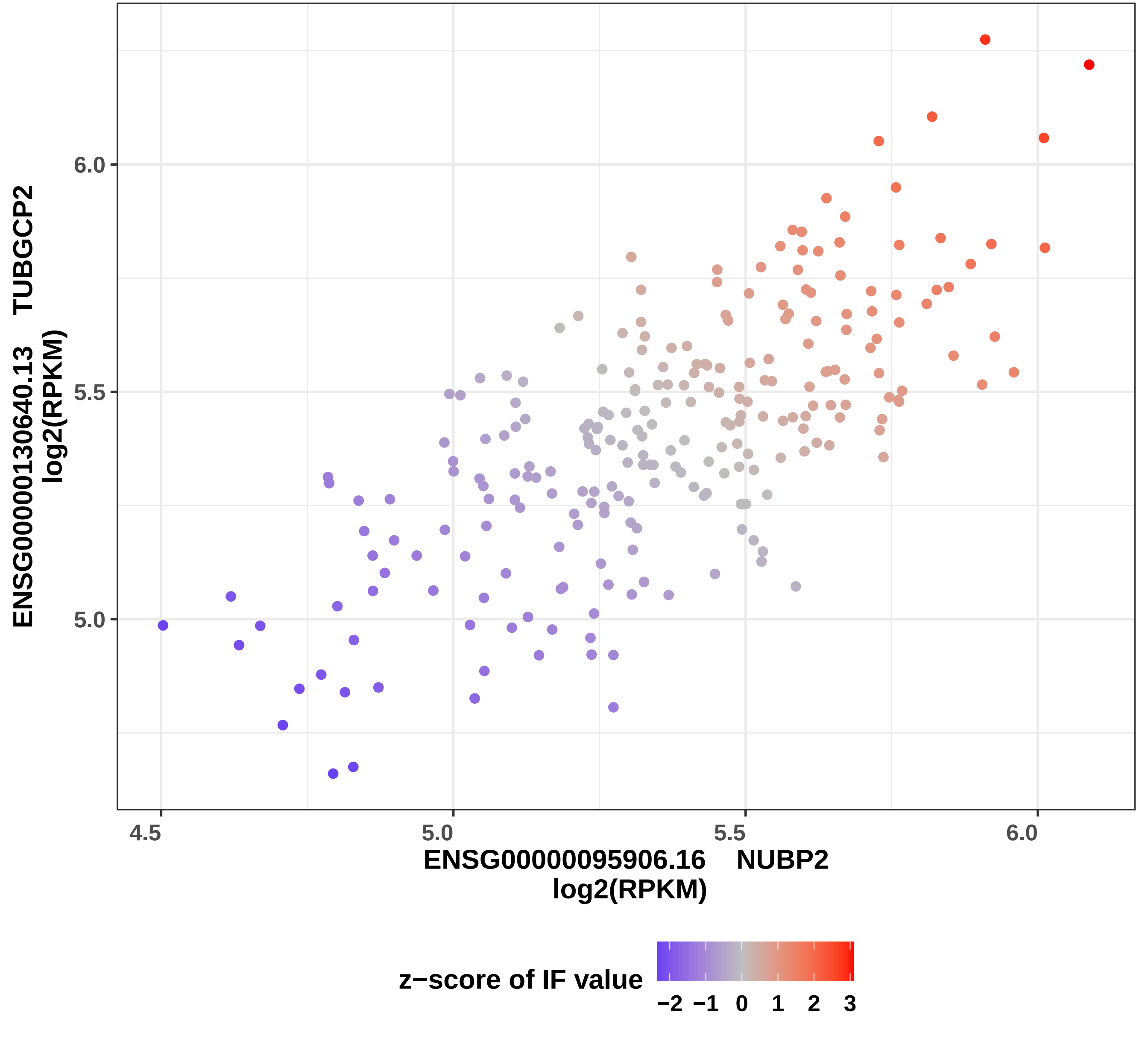

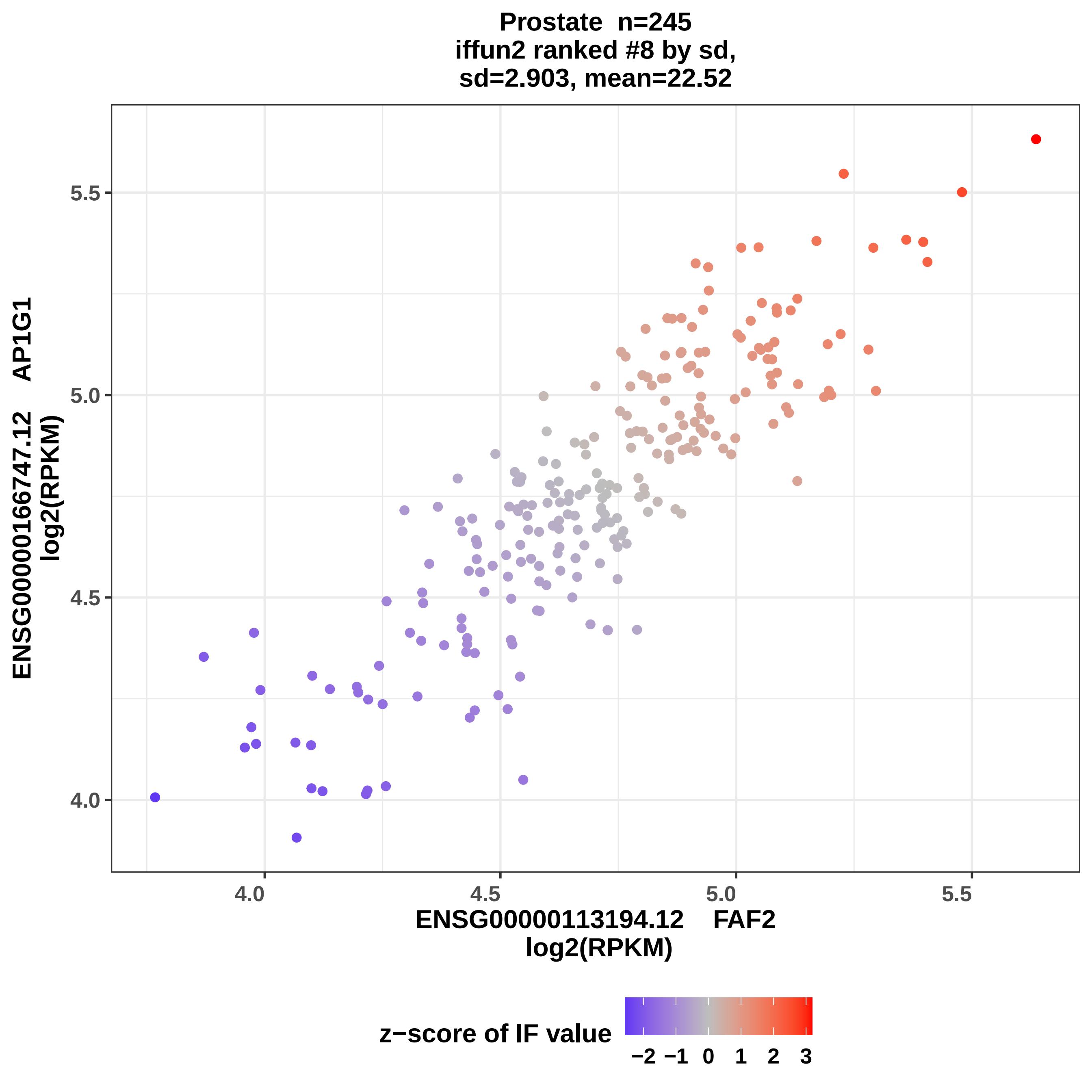
Prostate n=245
iffun2 ranked #8 by sd,
sd=2.903, mean=22.52
ENSG00000166747.12 AP1G1
log2(RPKM)
5.5
5.0
4.5
4.0
3.5
4.0
4.5
5.0
5.5
ENSG00000113194.12 FAF2
log2(RPKM)
z−score of IF value
−2 −1 0 1 2 3

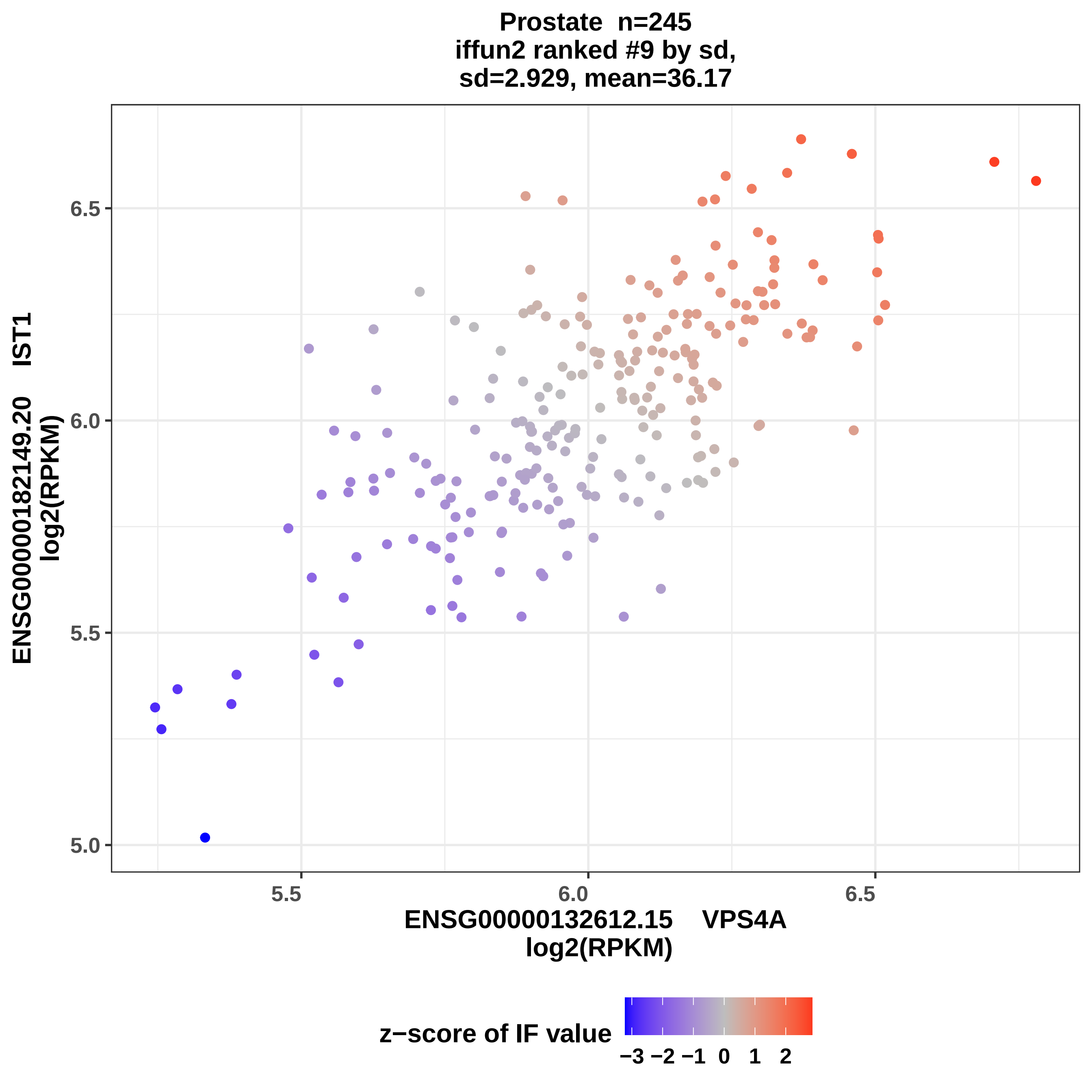
Prostate n=245
iffun2 ranked #9 by sd,
sd=2.929, mean=36.17
ENSG00000182149.20 IST1
log2(RPKM)
6.5
6.0
5.5
5.0
5.5
6.0
6.5
ENSG00000132612.15 VPS4A
log2(RPKM)
z−score of IF value
−3 −2 −1 0 1 2

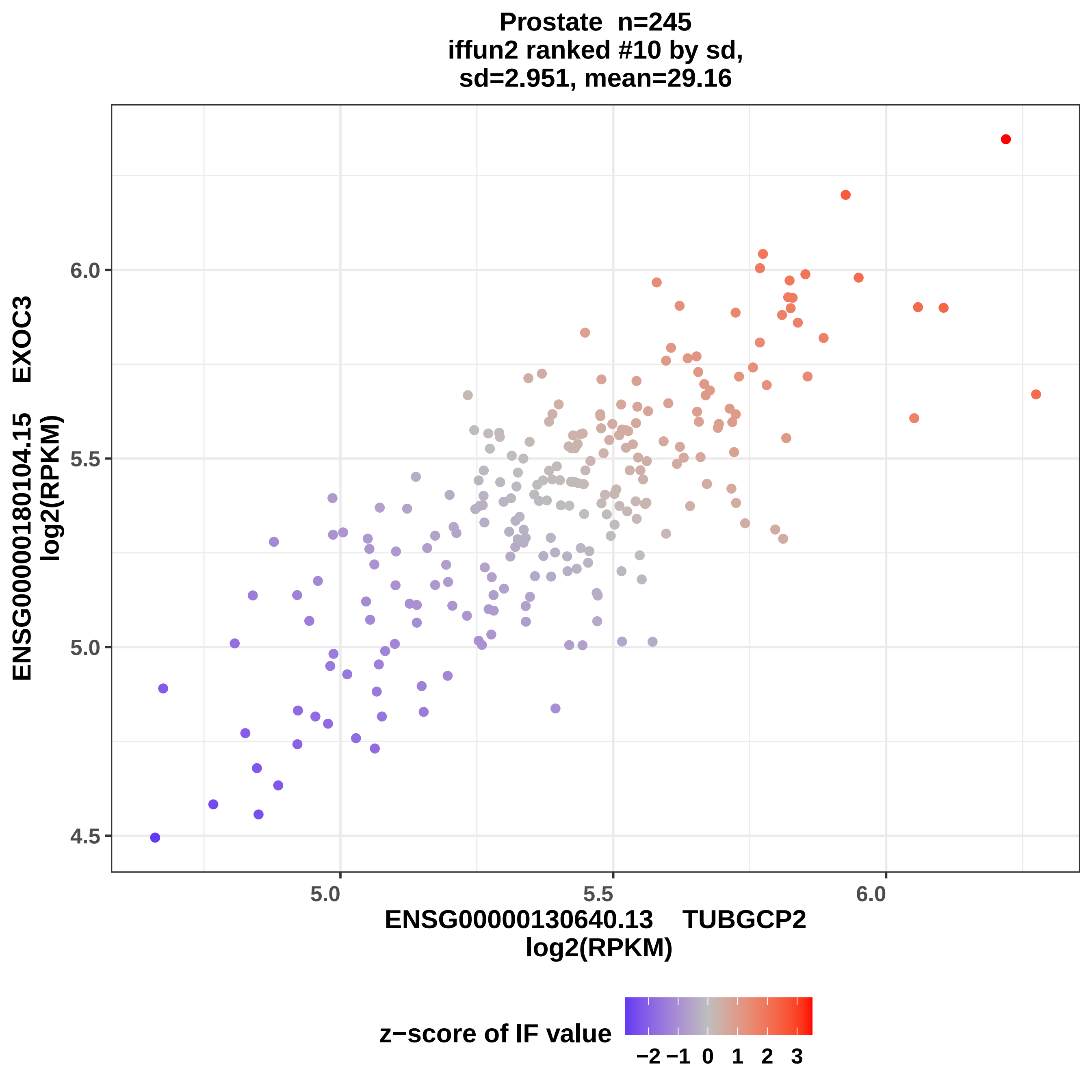

Prostate n=245
iffun2 ranked #10 by sd,
sd=2.951, mean=29.16
EXOC3
ENSG00000180104.15
log2(RPKM)
4.5
5.0
5.5
6.0
5.0
5.5
6.0
ENSG00000130640.13 TUBGCP2
log2(RPKM)
z−score of IF value
−2 −1 0 1 2 3

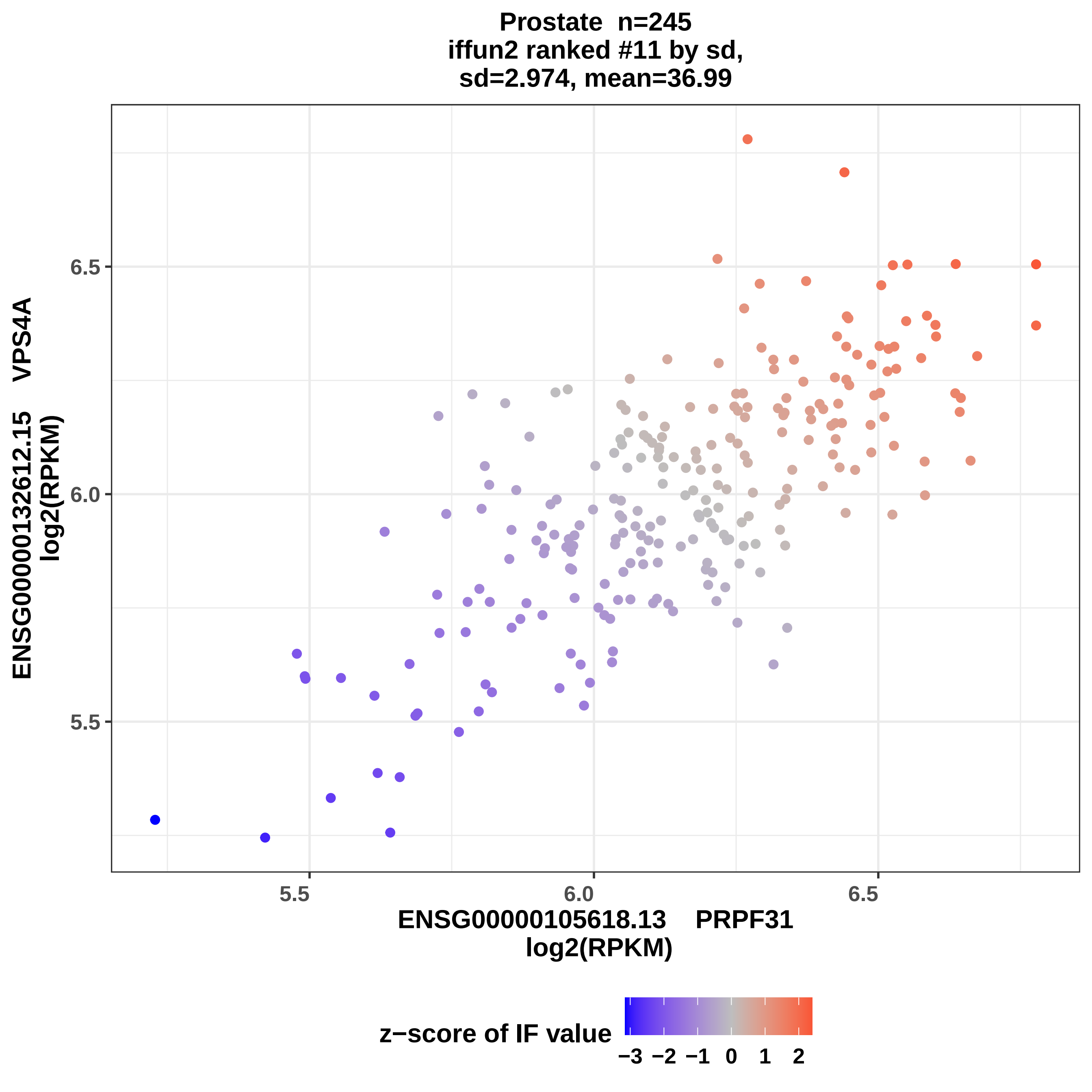

Prostate n=245
iffun2 ranked #11 by sd,
sd=2.974, mean=36.99
ENSG00000132612.15 VPS4A
log2(RPKM)
6.5
6.0
5.5
5.5
6.0
6.5
ENSG00000105618.13 PRPF31
log2(RPKM)
z−score of IF value
−3
−2
−1
0
1
2

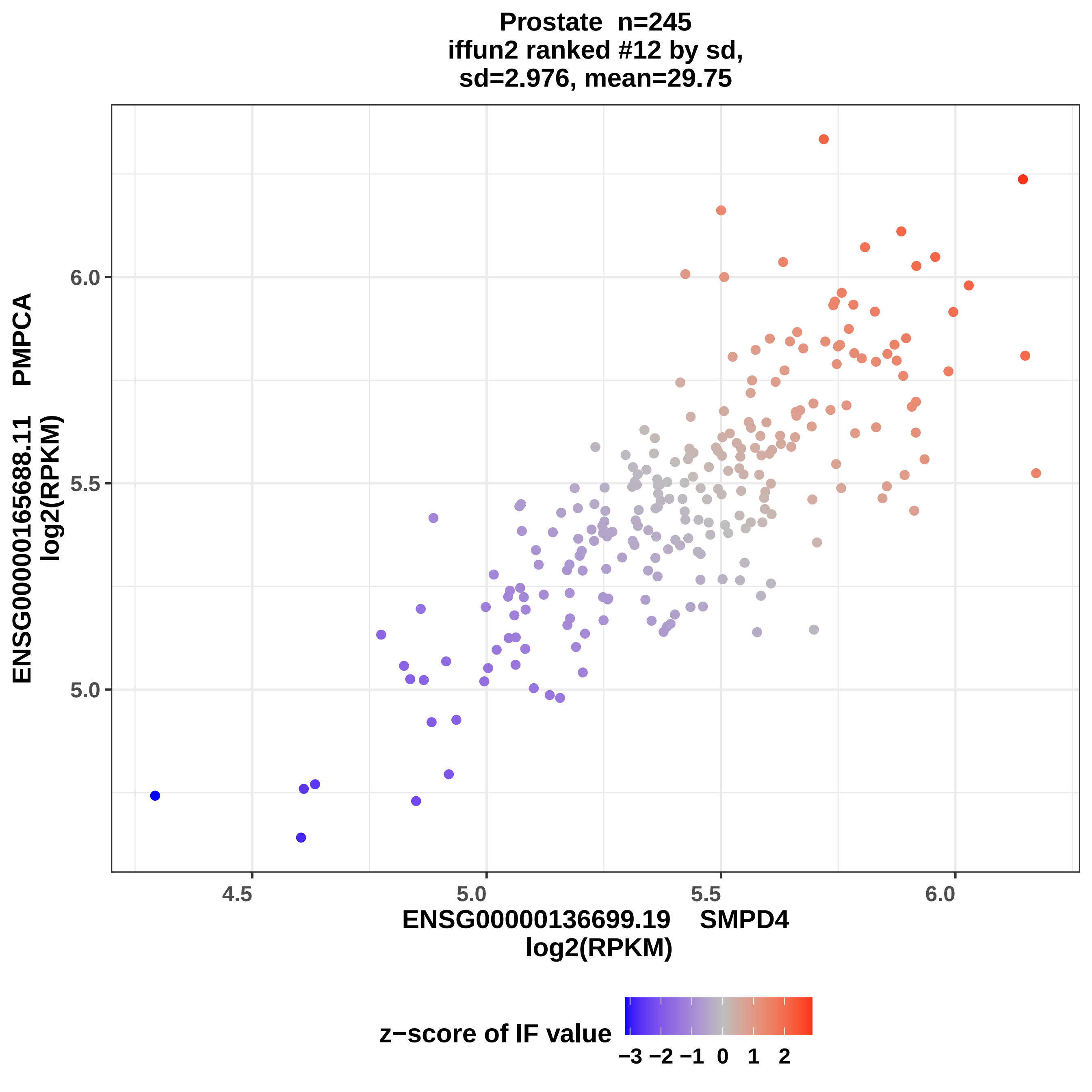
Prostate n=245
iffun2 ranked #12 by sd,
sd=2.976, mean=29.75
ENSG00000165688.11 PMPCA
log2(RPKM)
6.0
5.5
5.0
4.5
5.0
5.5
6.0
ENSG00000136699.19 SMPD4
log2(RPKM)
z−score of IF value
−3 −2 −1 0 1 2

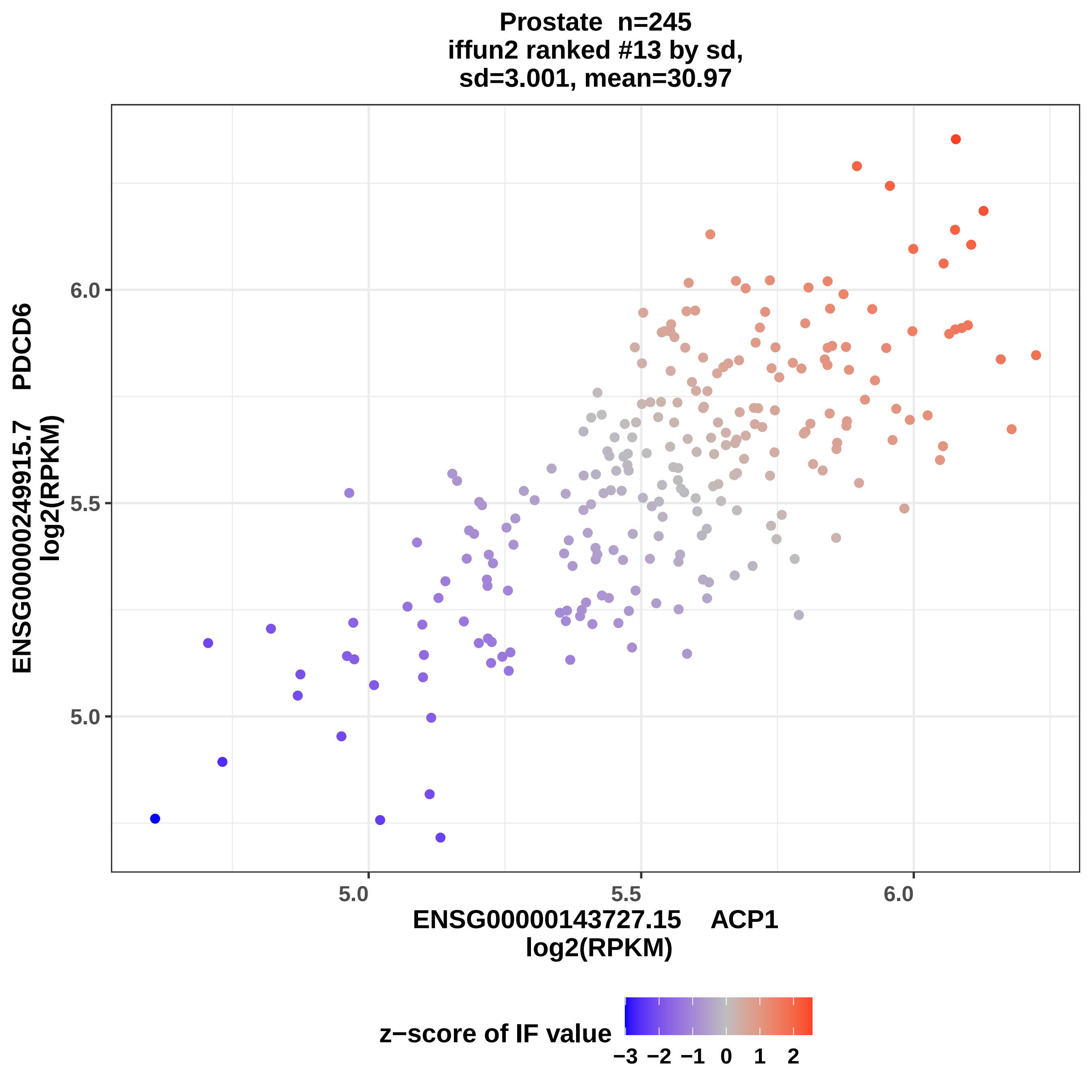

Prostate n=245
iffun2 ranked #13 by sd,
sd=3.001, mean=30.97
ENSG00000249915.7 PDCD6
log2(RPKM)
6.0
5.5
5.0
5.0
5.5
6.0
ENSG00000143727.15 ACP1
log2(RPKM)
z−score of IF value
−3
−2
−1
0
1
2

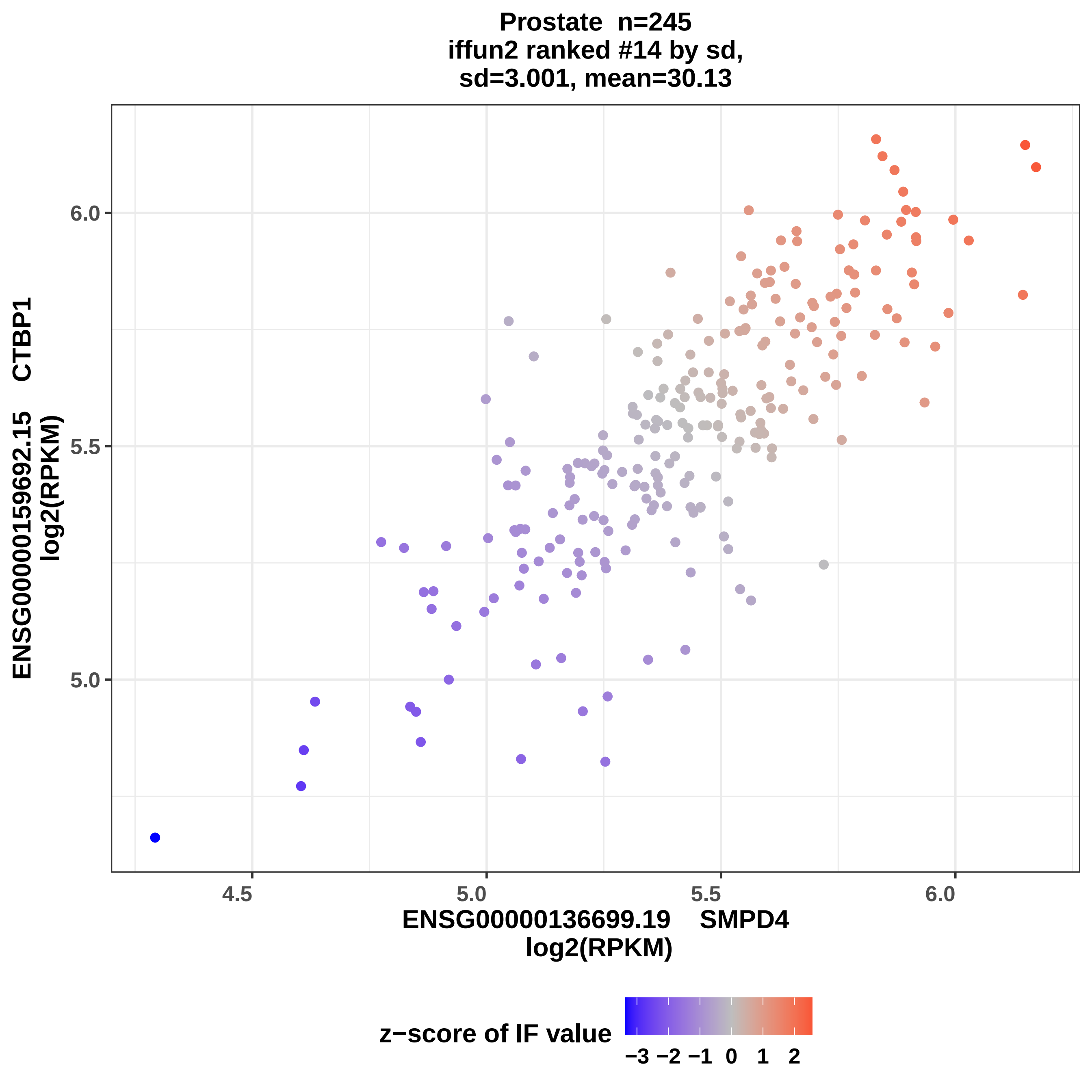
Prostate n=245
iffun2 ranked #14 by sd,
sd=3.001, mean=30.13
ENSG00000159692.15 CTBP1
log2(RPKM)
6.0
5.5
5.0
4.5
5.0
5.5
6.0
ENSG00000136699.19 SMPD4
log2(RPKM)
z−score of IF value
−3 −2 −1 0 1 2

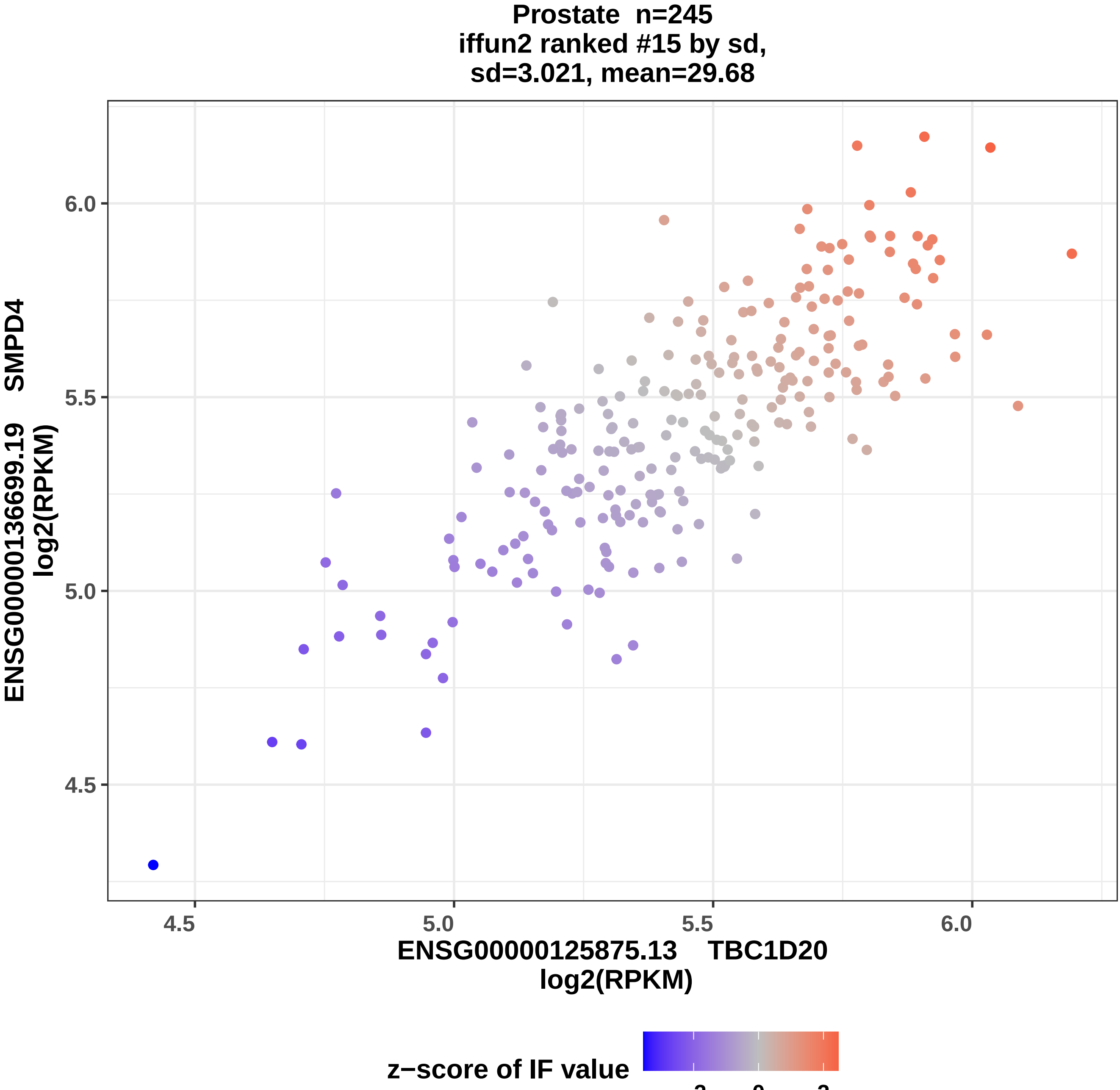

Prostate n=245
iffun2 ranked #15 by sd,
sd=3.021, mean=29.68
ENSG00000136699.19 SMPD4
log2(RPKM)
6.0
5.5
5.0
4.5
4.5
5.0
5.5
6.0
ENSG00000125875.13 TBC1D20
log2(RPKM)
z−score of IF value
−2
0
2

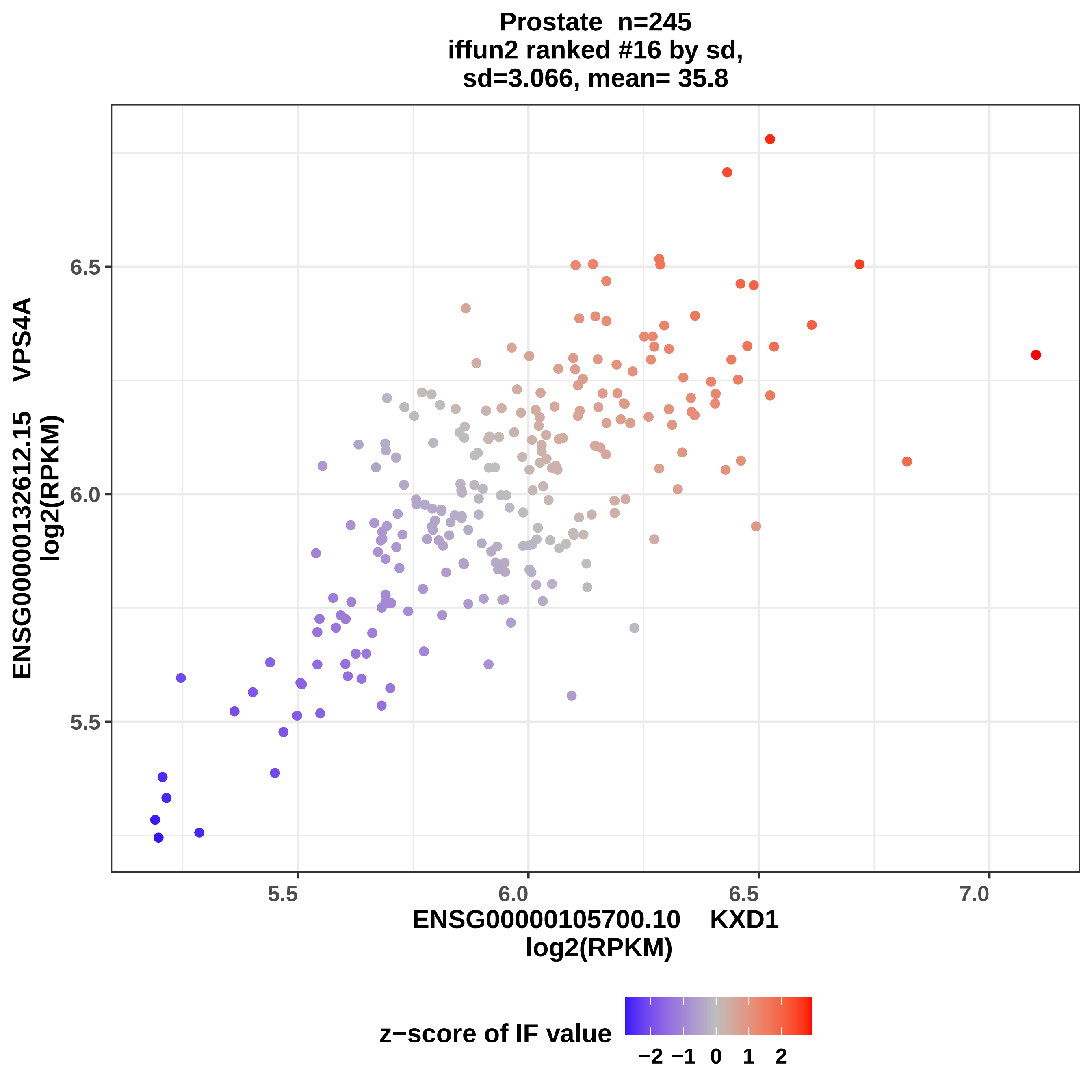
Prostate n=245
iffun2 ranked #16 by sd,
sd=3.066, mean= 35.8
ENSG00000132612.15 VPS4A
log2(RPKM)
6.5
6.0
5.5
5.5
6.0
6.5
7.0
ENSG00000105700.10 KXD1
log2(RPKM)
z−score of IF value
−2 −1 0 1 2

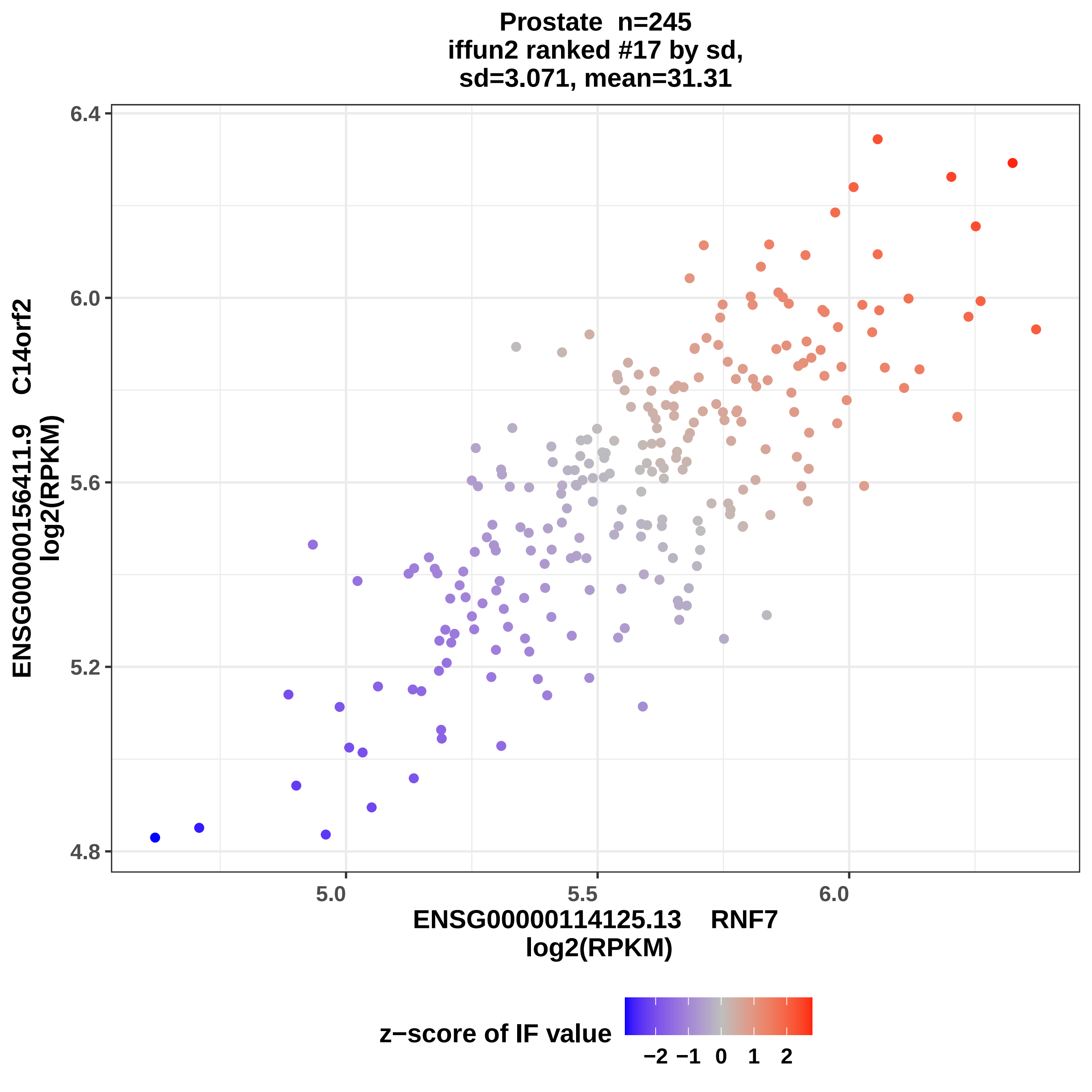
Prostate n=245
iffun2 ranked #17 by sd,
sd=3.071, mean=31.31
ENSG00000156411.9 C14orf2
log2(RPKM)
6.4
6.0
5.6
5.2
4.8
5.0
5.5
6.0
ENSG00000114125.13 RNF7
log2(RPKM)
z−score of IF value
−2 −1 0 1 2

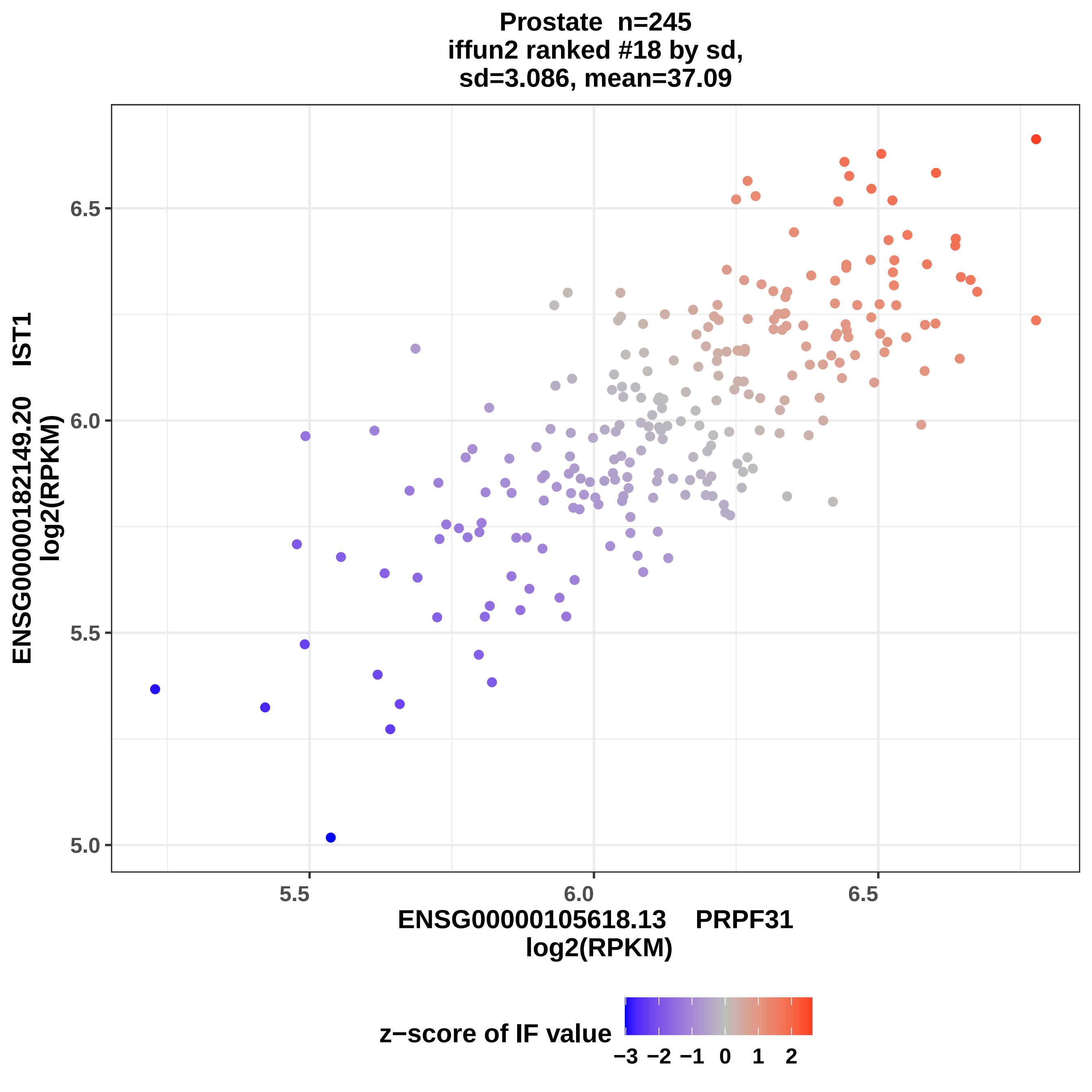

Prostate n=245
iffun2 ranked #18 by sd,
sd=3.086, mean=37.09
ENSG00000182149.20 IST1
log2(RPKM)
6.5
6.0
5.5
5.0
5.5
6.0
6.5
ENSG00000105618.13 PRPF31
log2(RPKM)
z−score of IF value
−3 −2 −1 0 1 2

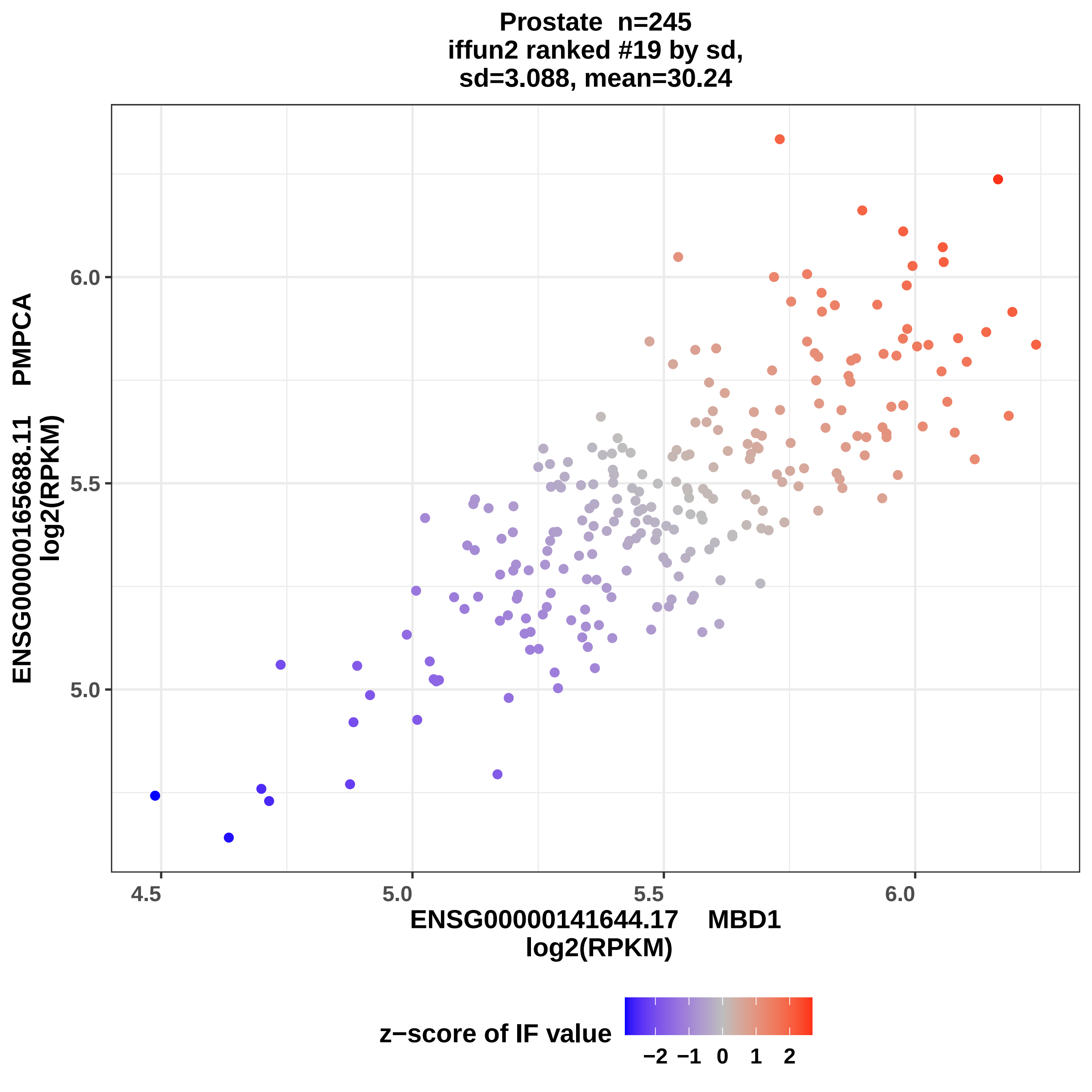

Prostate n=245
iffun2 ranked #19 by sd,
sd=3.088, mean=30.24
ENSG00000165688.11 PMPCA
log2(RPKM)
6.0
5.5
5.0
4.5
5.0
5.5
6.0
ENSG00000141644.17 MBD1
log2(RPKM)
z−score of IF value
−2
−1
0
1
2

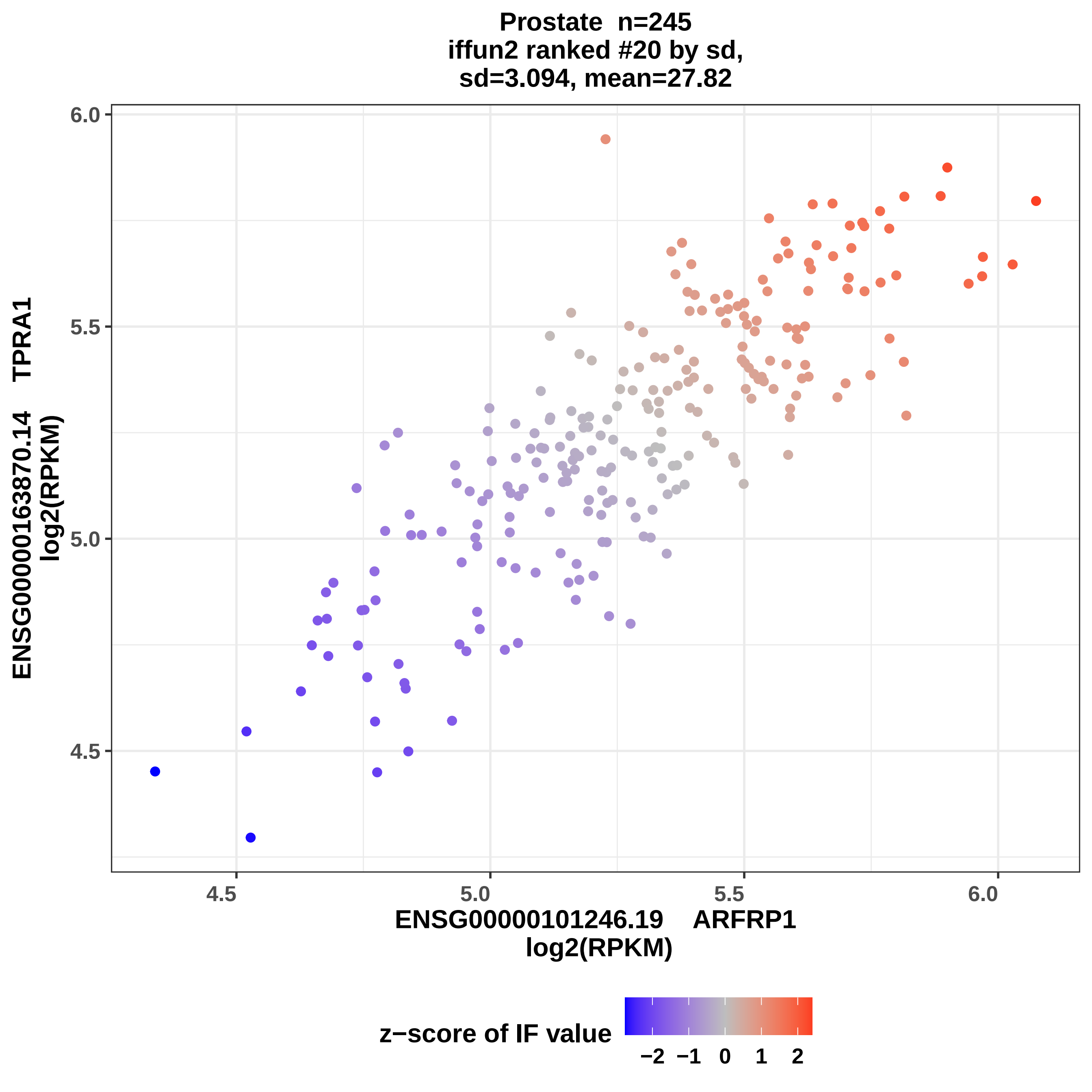

Prostate n=245
iffun2 ranked #20 by sd,
sd=3.094, mean=27.82
6.0
5.5
5.0
4.5
4.5
5.0
5.5
6.0
ENSG00000163870.14 TPRA1
log2(RPKM)
ENSG00000101246.19 ARFRP1
log2(RPKM)
z−score of IF value
−2 −1 0 1 2

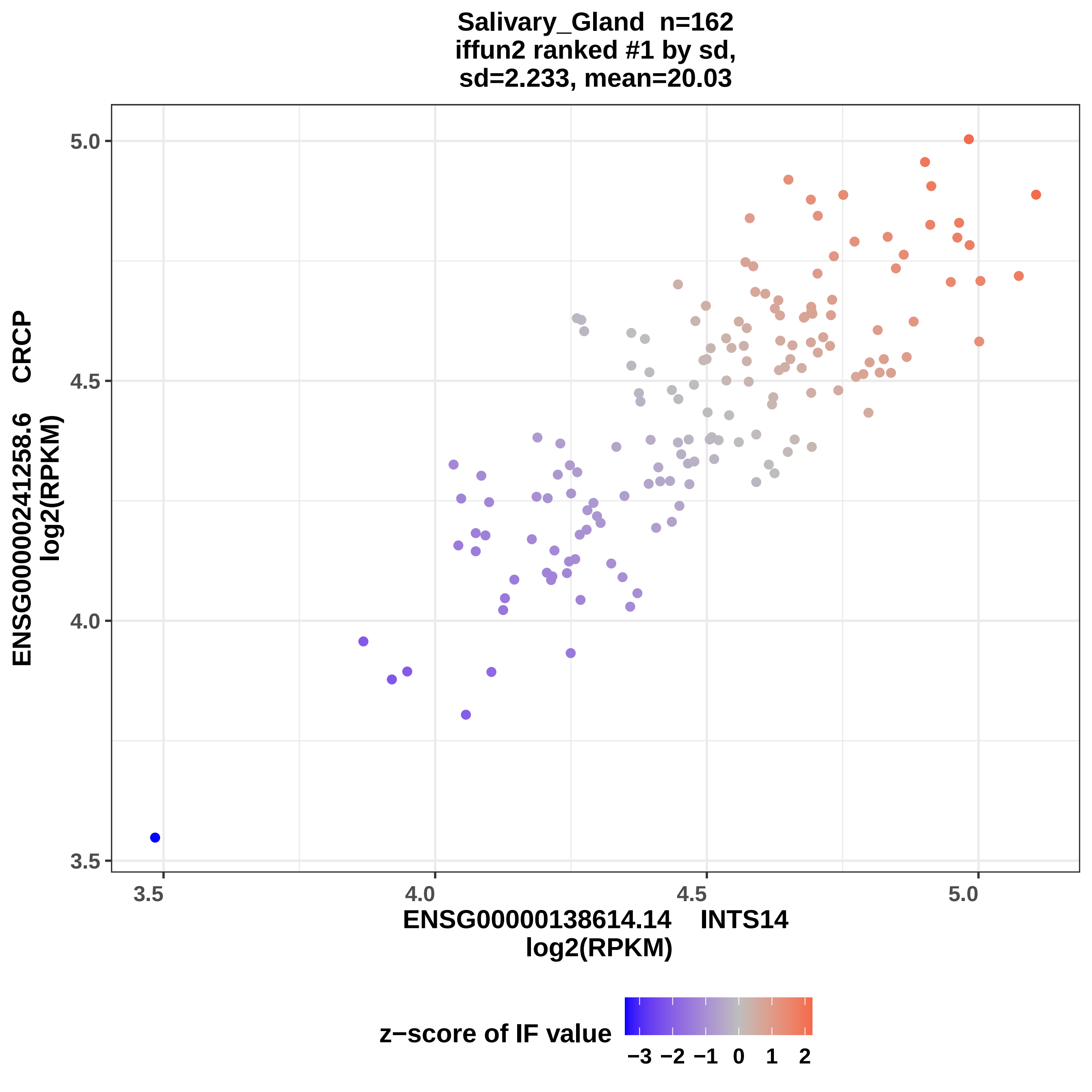

Salivary_Gland n=162
iffun2 ranked #1 by sd,
sd=2.233, mean=20.03
ENSG00000241258.6 CRCP
log2(RPKM)
ENSG00000138614.14 INTS14
log2(RPKM)
5.0
4.5
4.0
3.5
3.5
4.0
4.5
5.0
z−score of IF value
−3 −2 −1 0 1 2

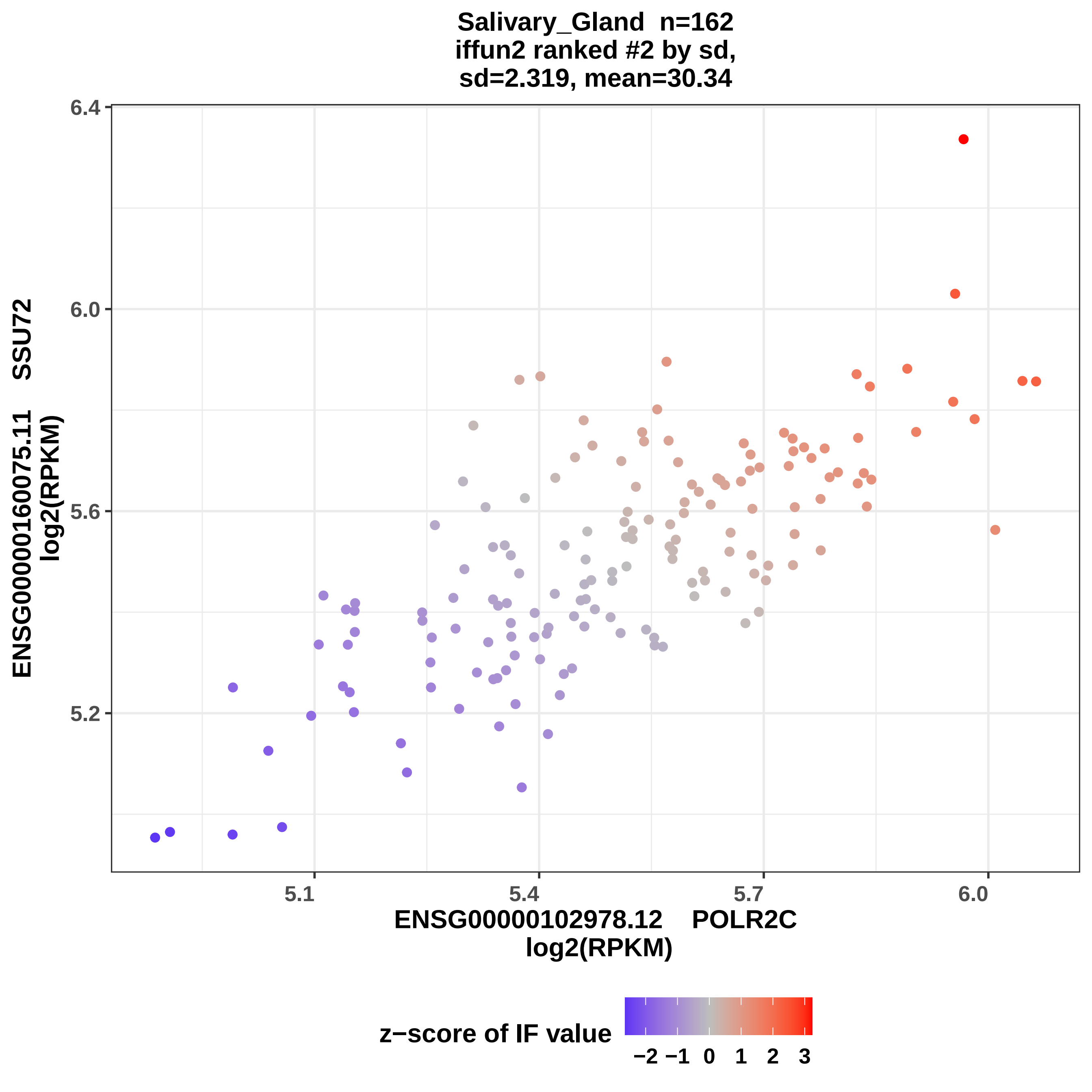

Salivary_Gland n=162
iffun2 ranked #2 by sd,
sd=2.319, mean=30.34
ENSG00000160075.11 SSU72
log2(RPKM)
6.4
6.0
5.6
5.2
5.1
5.4
5.7
6.0
ENSG00000102978.12 POLR2C
log2(RPKM)
z−score of IF value
−2 −1 0 1 2 3

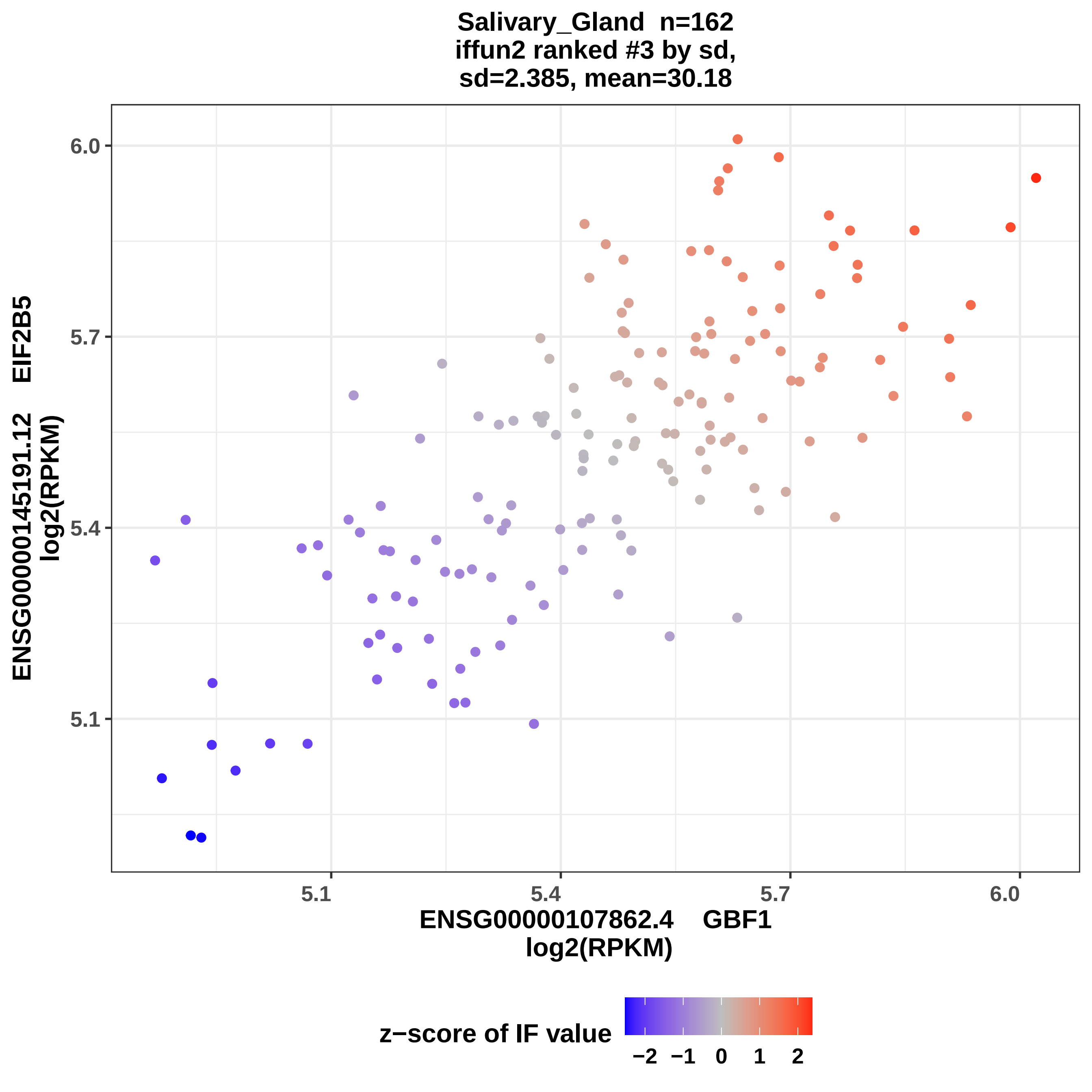
Salivary_Gland n=162
iffun2 ranked #3 by sd,
sd=2.385, mean=30.18
ENSG00000145191.12 EIF2B5
log2(RPKM)
ENSG00000107862.4 GBF1
log2(RPKM)
6.0
5.7
5.4
5.1
5.1
5.4
5.7
6.0
z−score of IF value
−2
−1
0
1
2

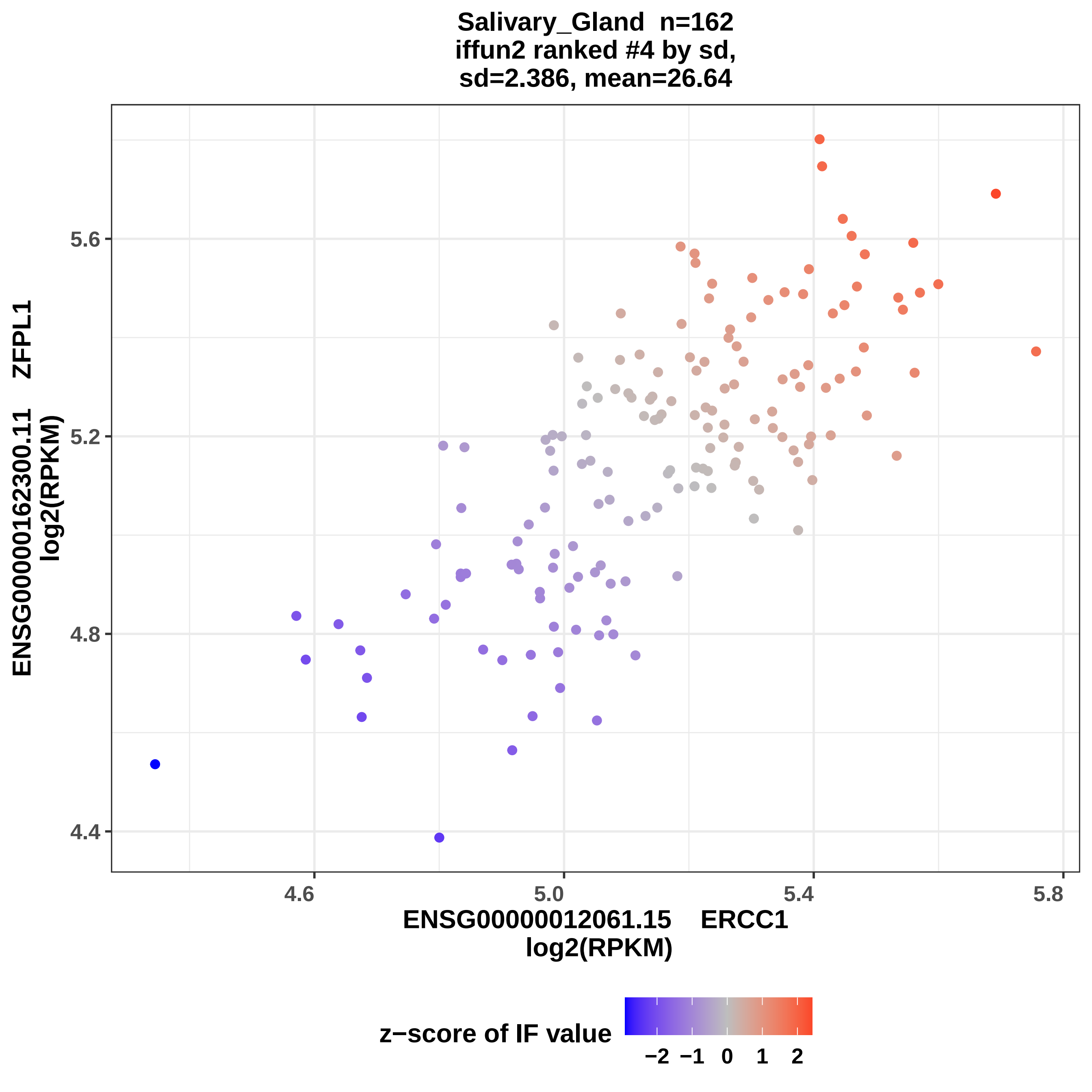

Salivary_Gland n=162
iffun2 ranked #4 by sd,
sd=2.386, mean=26.64
ENSG00000162300.11 ZFPL1
log2(RPKM)
5.6
5.2
4.8
4.4
4.6
5.0
5.4
5.8
ENSG00000012061.15 ERCC1
log2(RPKM)
z−score of IF value
−2 −1 0 1 2

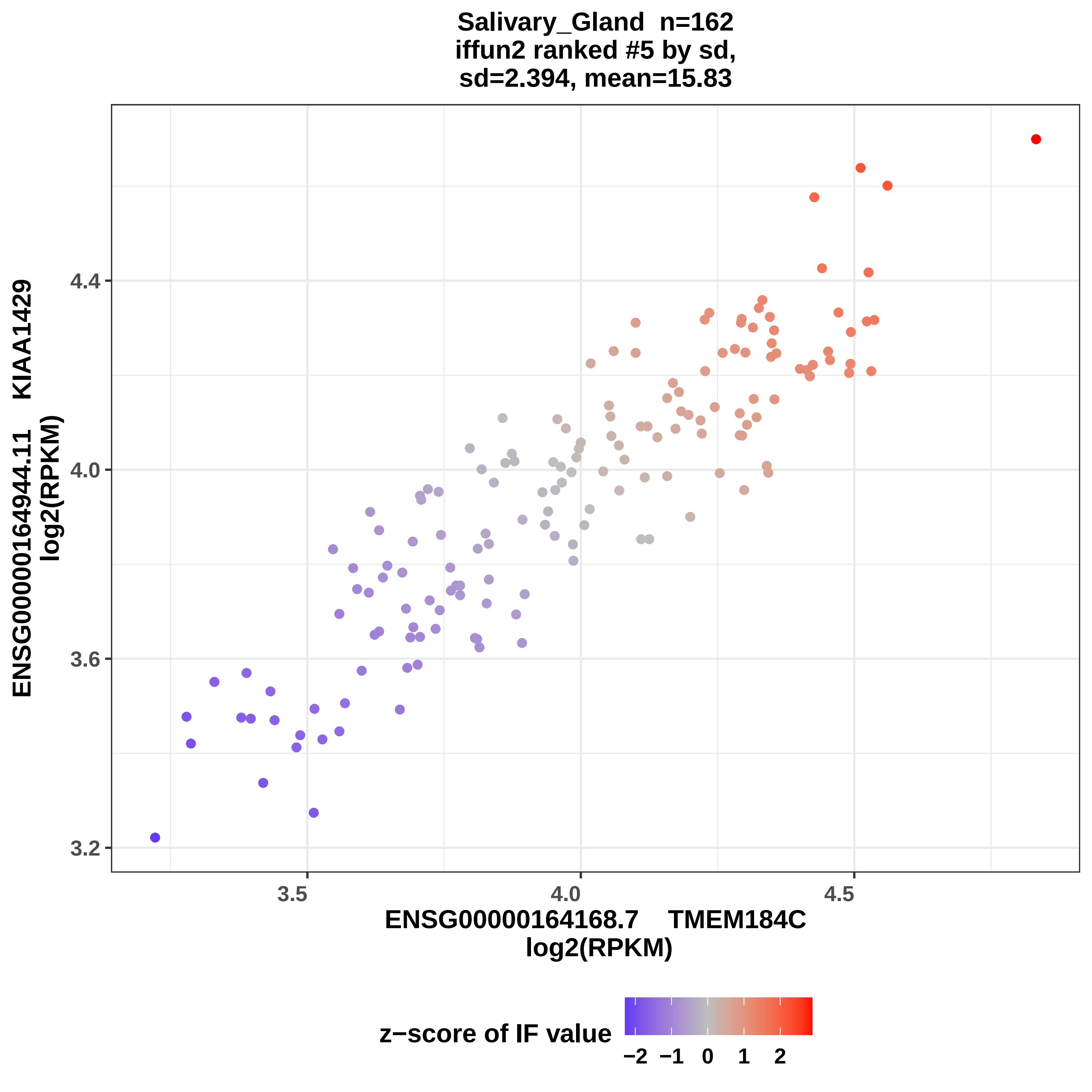
Salivary_Gland n=162
iffun2 ranked #5 by sd,
sd=2.394, mean=15.83
ENSG00000164944.11 KIAA1429
log2(RPKM)
4.4
4.0
3.6
3.2
3.5
4.0
4.5
ENSG00000164168.7 TMEM184C
log2(RPKM)
z−score of IF value
−2 −1 0 1 2

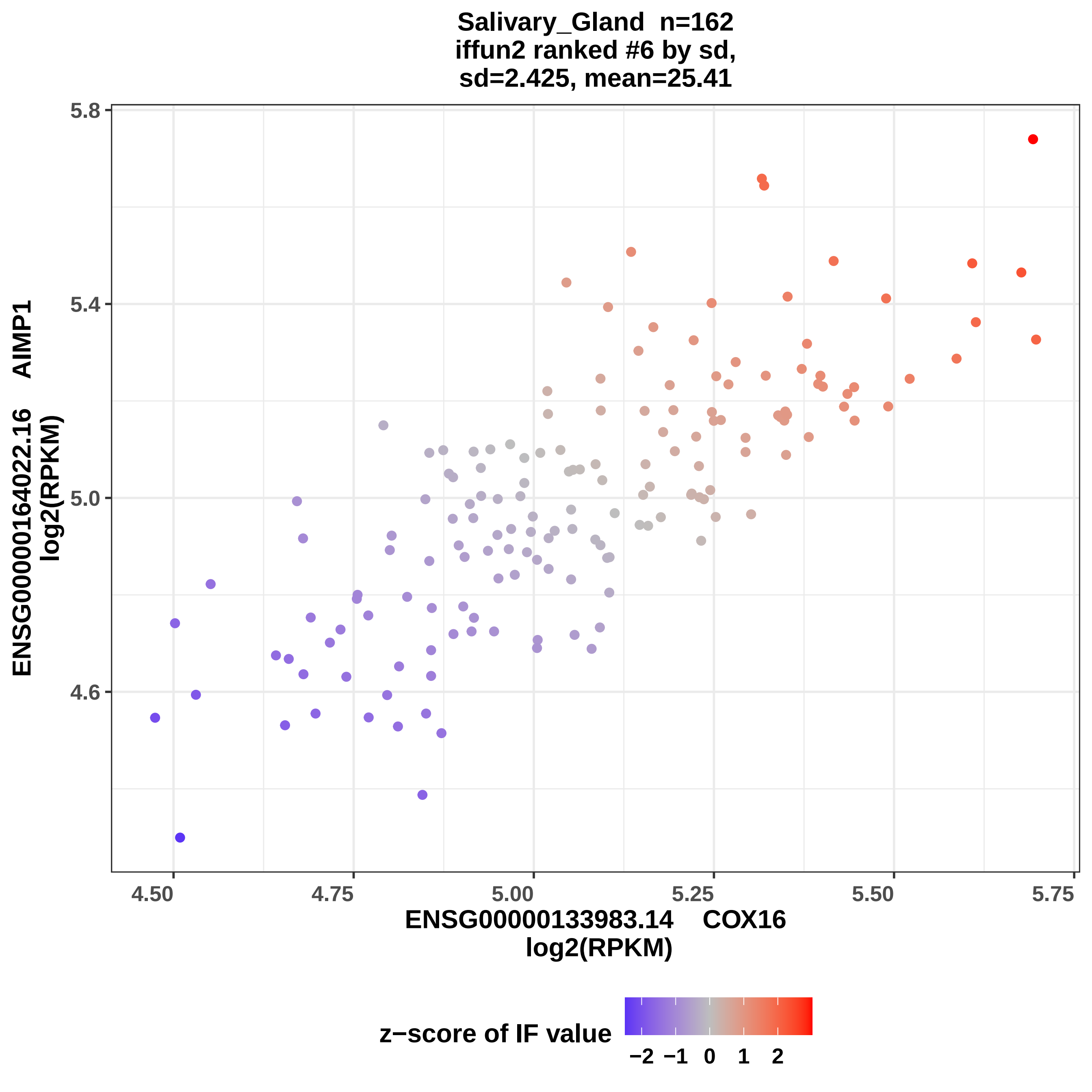
Salivary_Gland n=162
iffun2 ranked #6 by sd,
sd=2.425, mean=25.41
ENSG00000164022.16 AIMP1
log2(RPKM)
5.8
5.4
5.0
4.6
4.50
4.75
5.00
5.25
5.50
5.75
ENSG00000133983.14 COX16
log2(RPKM)
z−score of IF value
−2 −1 0 1 2

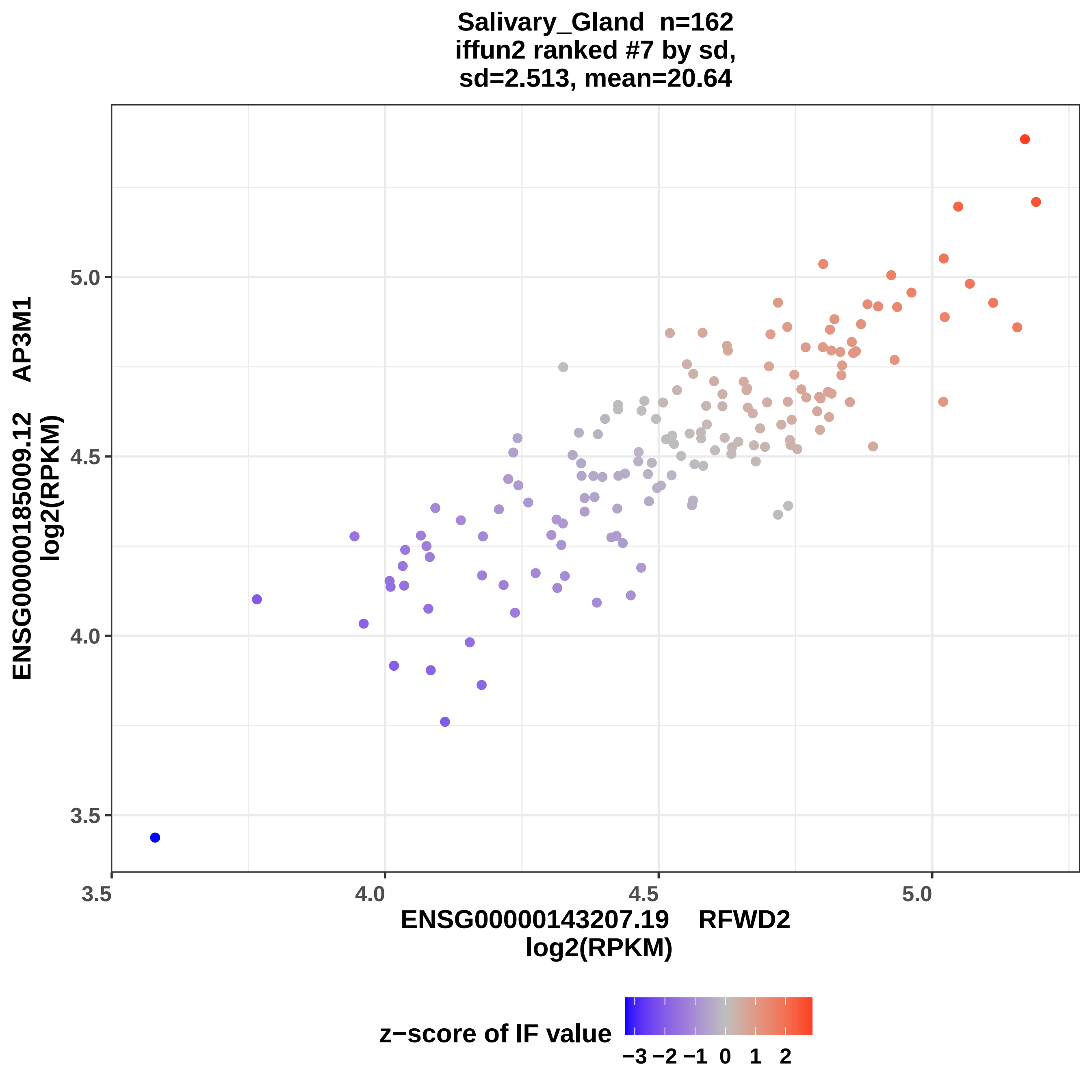
Salivary_Gland n=162
iffun2 ranked #7 by sd,
sd=2.513, mean=20.64
ENSG00000185009.12 AP3M1
log2(RPKM)
ENSG00000143207.19 RFWD2
log2(RPKM)
3.5
4.0
4.5
5.0
z−score of IF value
−3 −2 −1 0 1 2

Salivary_Gland n=162
iffun2 ranked #8 by sd,
sd=2.517, mean=35.02

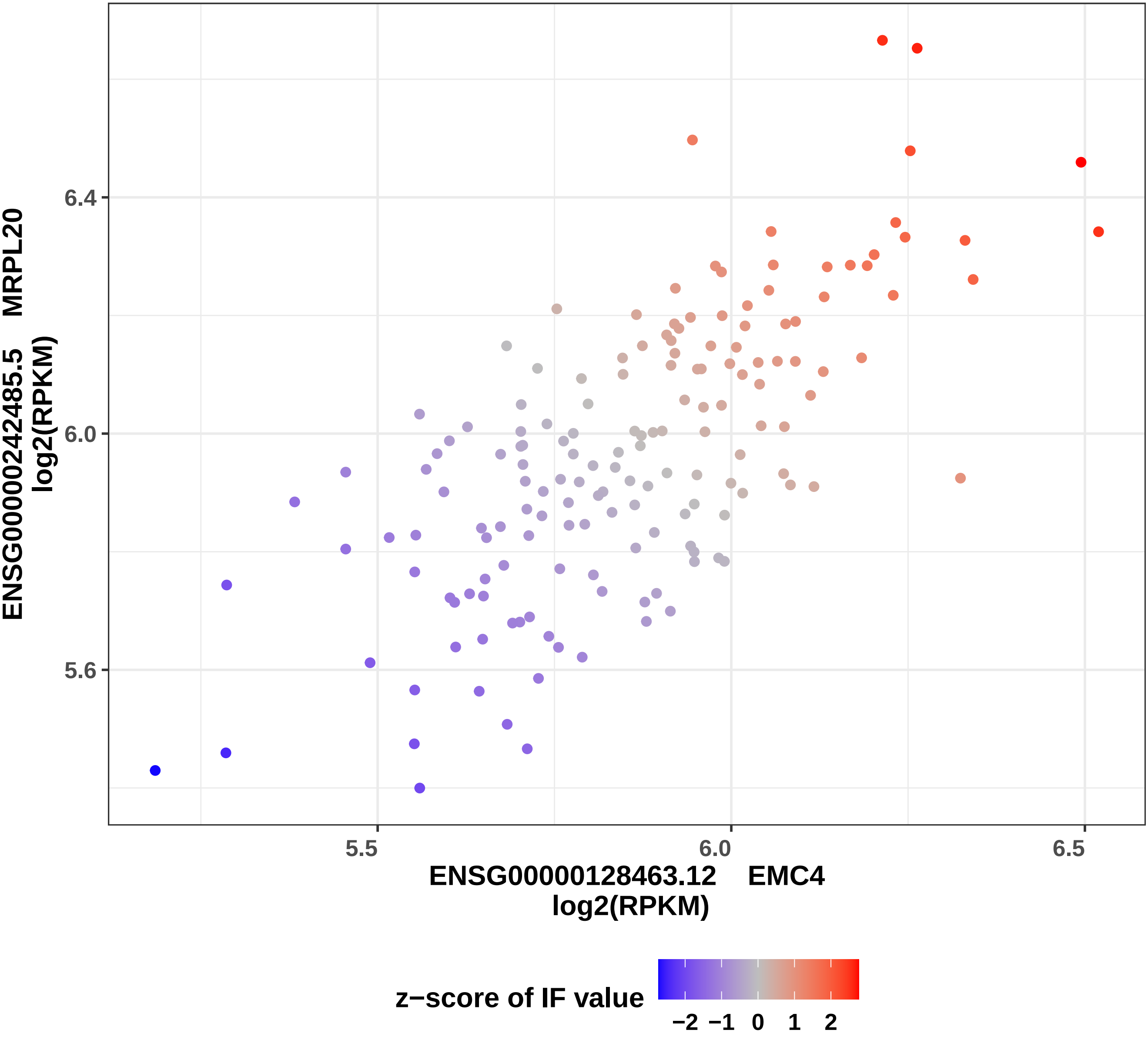

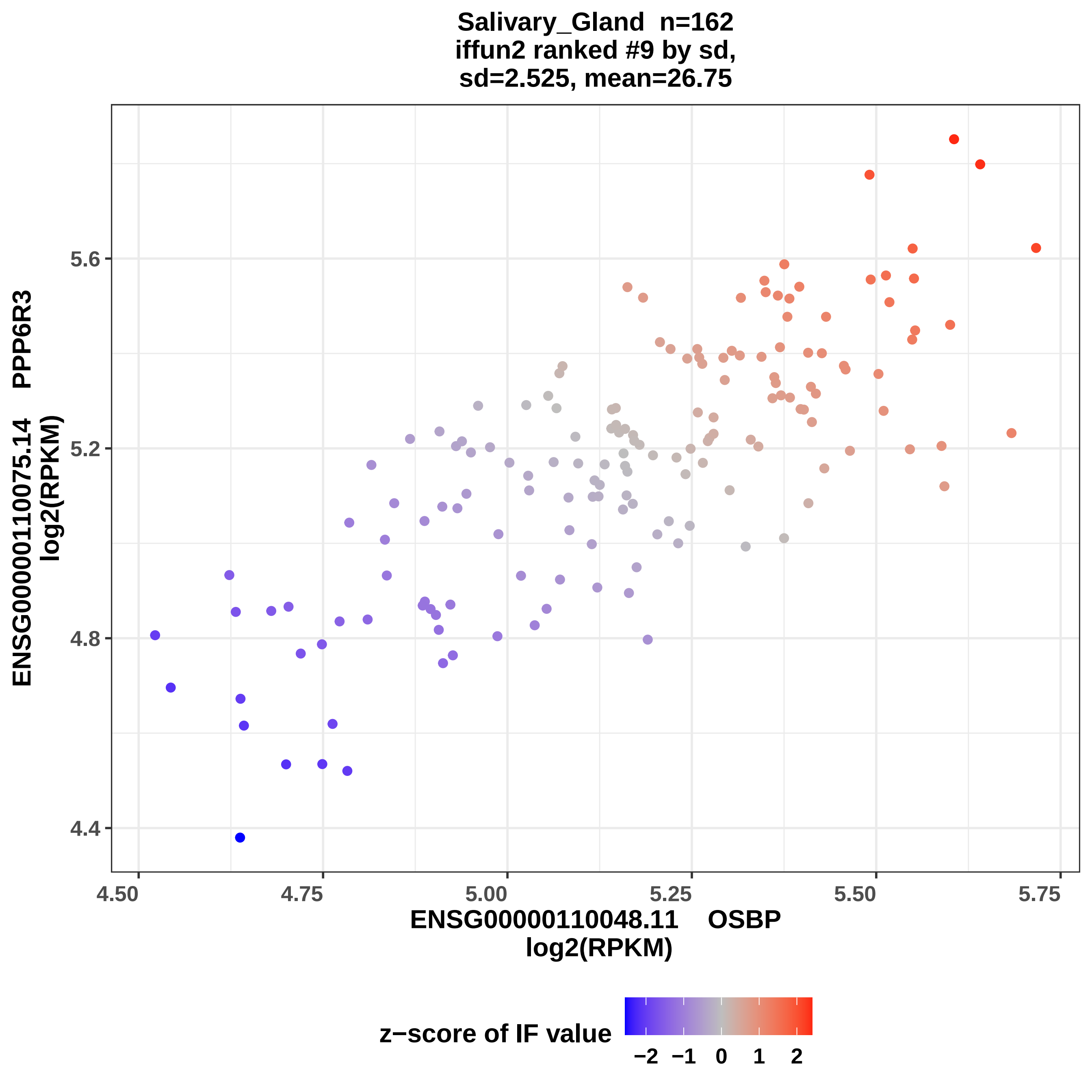

Salivary_Gland n=162
iffun2 ranked #9 by sd,
sd=2.525, mean=26.75
ENSG00000110075.14 PPP6R3
log2(RPKM)
5.6
5.2
4.8
4.4
4.50
4.75
5.00
5.25
5.50
5.75
ENSG00000110048.11 OSBP
log2(RPKM)
z−score of IF value
−2
−1
0
1
2

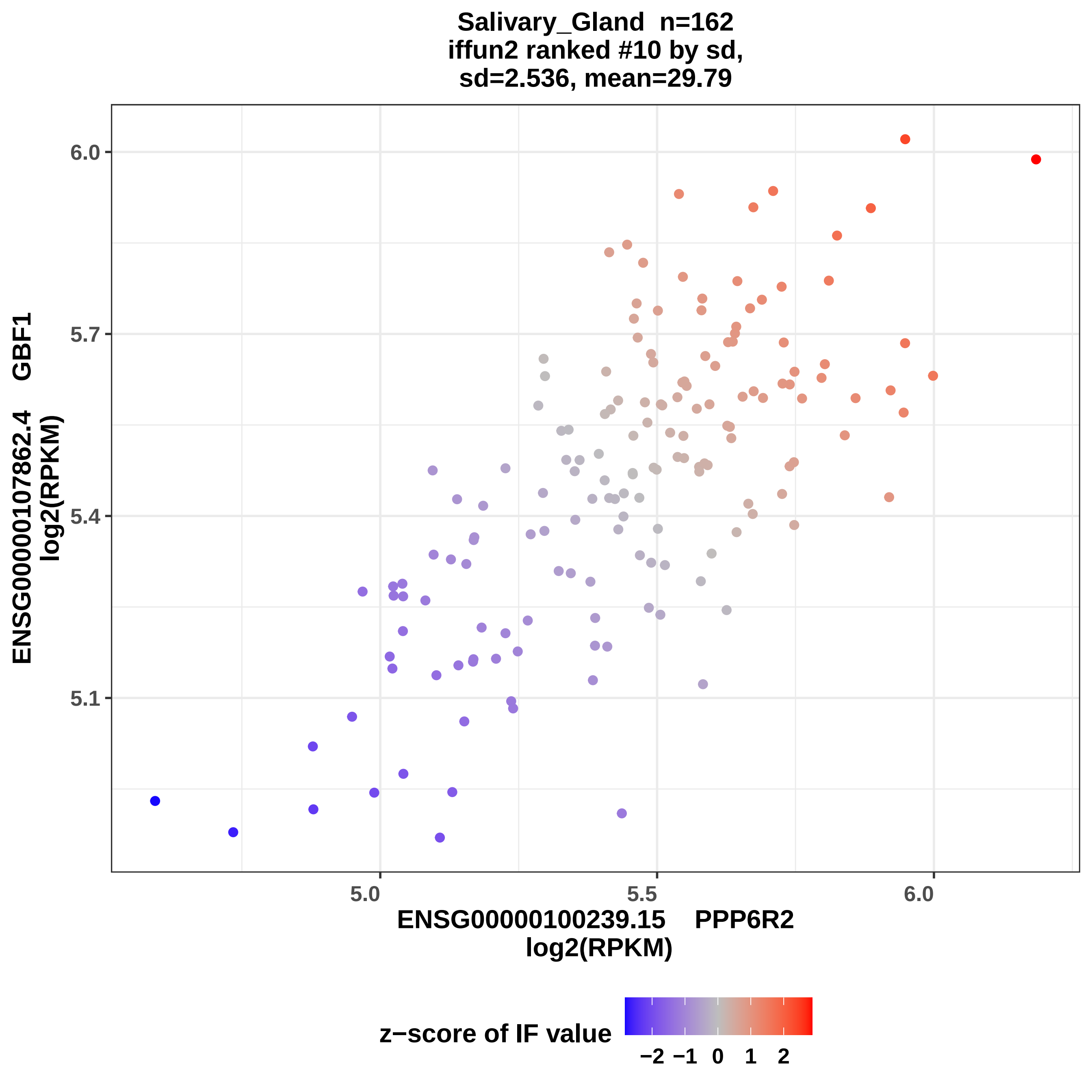

Salivary_Gland n=162
iffun2 ranked #10 by sd,
sd=2.536, mean=29.79
6.0
5.7
5.4
5.1
5.0
5.5
6.0
ENSG00000107862.4 GBF1
log2(RPKM)
ENSG00000100239.15 PPP6R2
log2(RPKM)
z−score of IF value
−2 −1 0 1 2

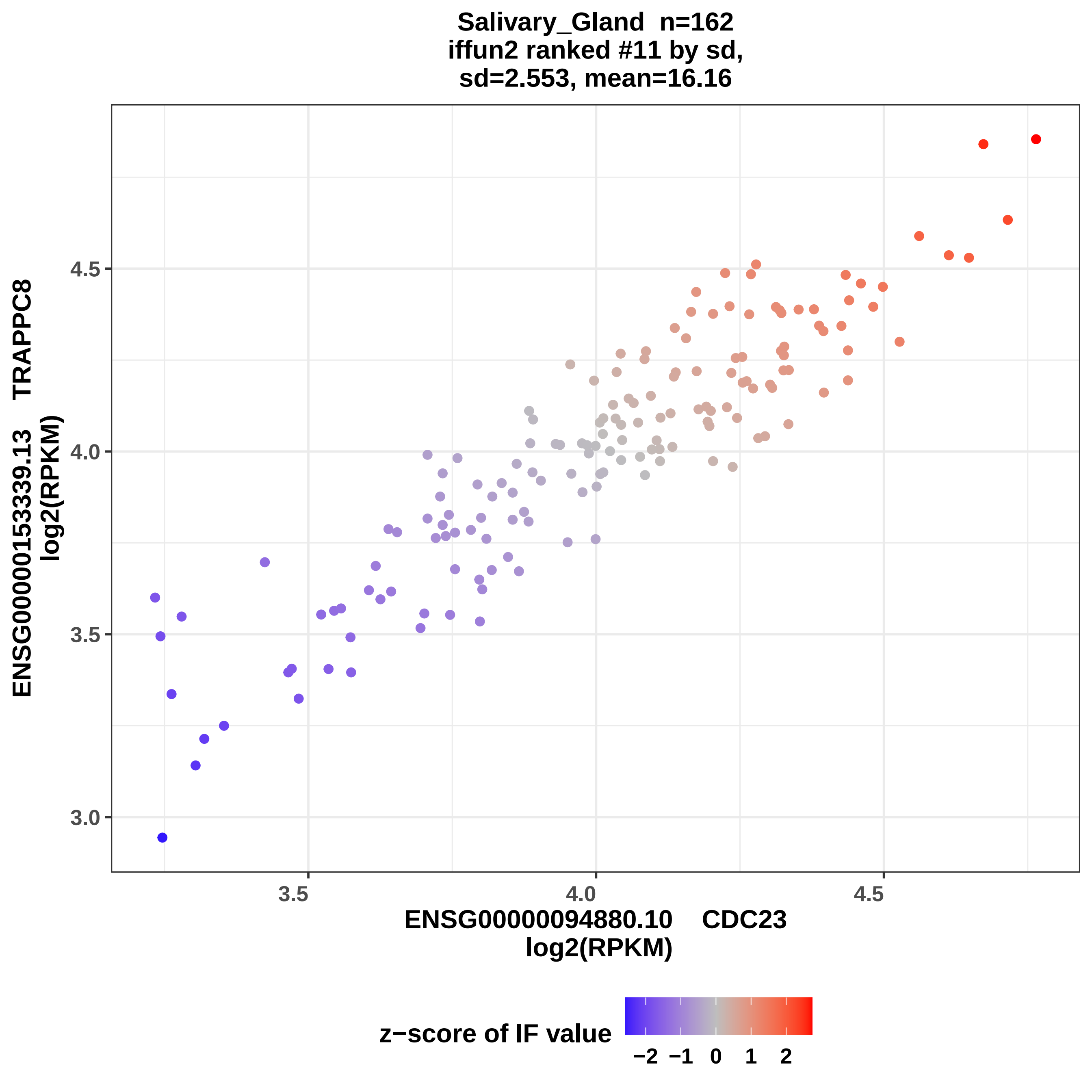

Salivary_Gland n=162
iffun2 ranked #11 by sd,
sd=2.553, mean=16.16
ENSG00000153339.13 TRAPPC8
log2(RPKM)
4.5
4.0
3.5
3.0
3.5
4.0
4.5
ENSG00000094880.10 CDC23
log2(RPKM)
z−score of IF value
−2 −1 0 1 2

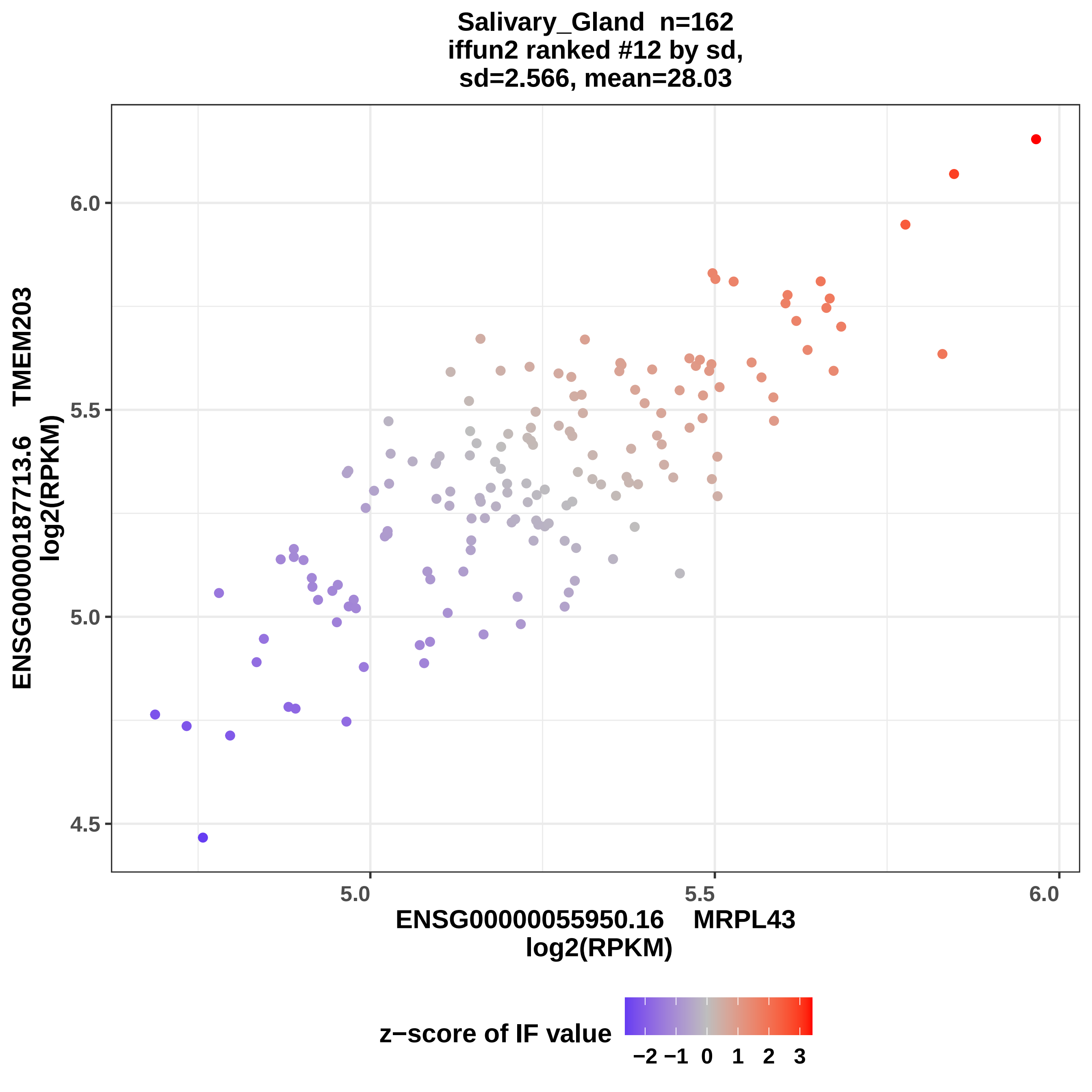
Salivary_Gland n=162
iffun2 ranked #12 by sd,
sd=2.566, mean=28.03
ENSG00000187713.6 TMEM203
log2(RPKM)
4.5
5.0
5.5
6.0
ENSG00000055950.16 MRPL43
log2(RPKM)
z−score of IF value
−2 −1 0 1 2 3

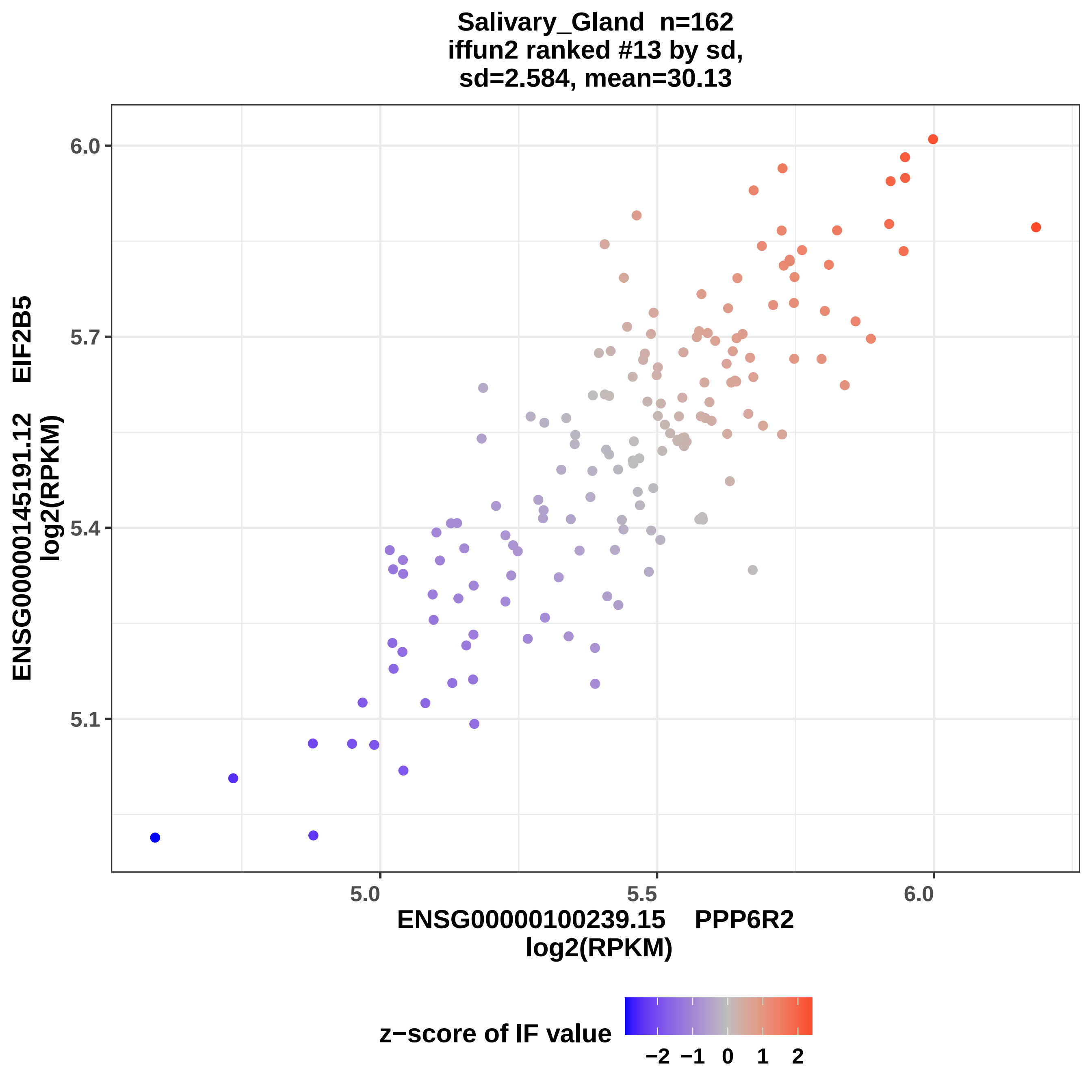

Salivary_Gland n=162
iffun2 ranked #13 by sd,
sd=2.584, mean=30.13
6.0
5.7
5.4
5.1
EIF2B5
ENSG00000145191.12
log2(RPKM)
5.0
5.5
6.0
ENSG00000100239.15 PPP6R2
log2(RPKM)
z−score of IF value
−2 −1 0 1 2

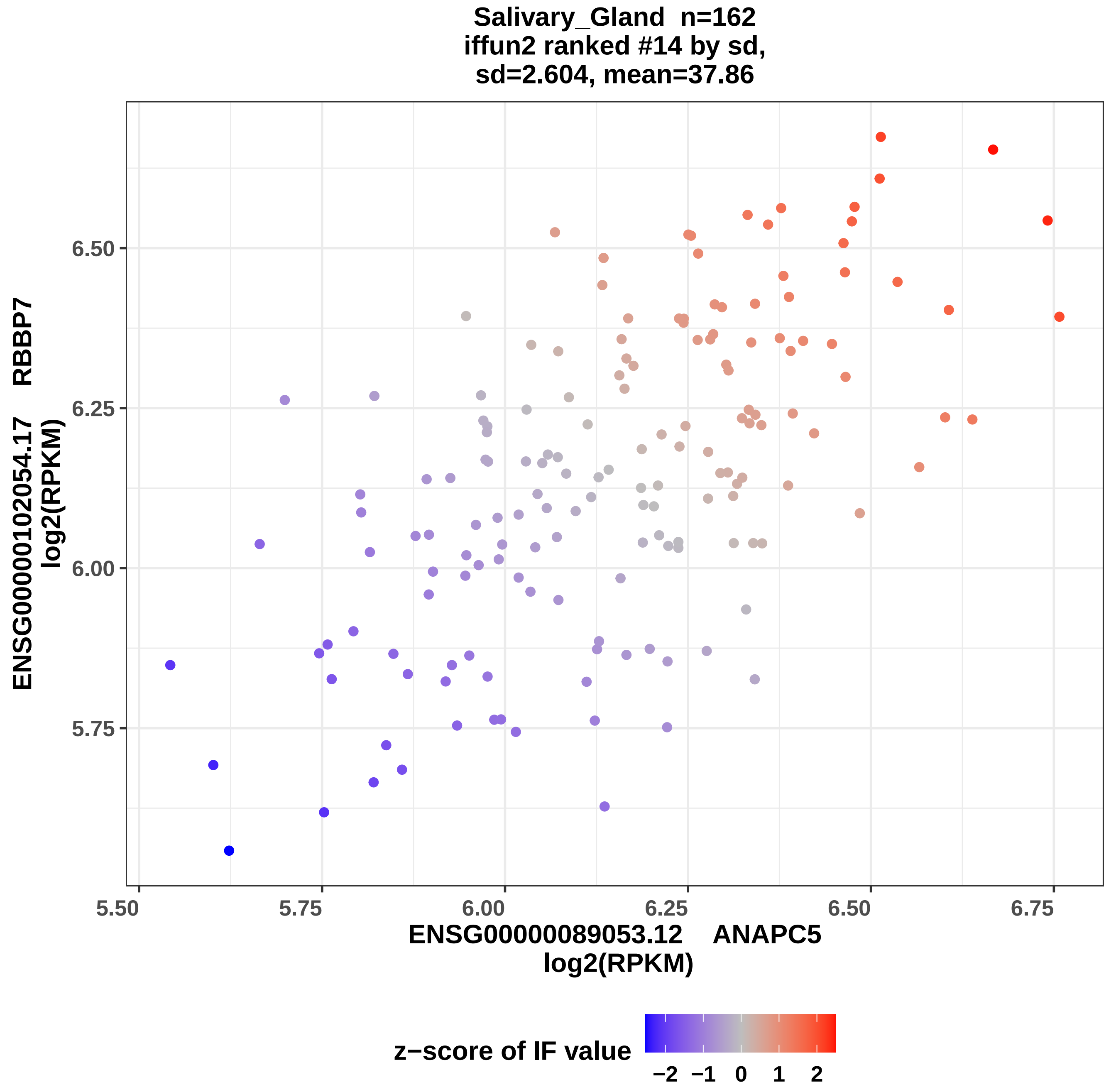
Salivary_Gland n=162
iffun2 ranked #14 by sd,
sd=2.604, mean=37.86
ENSG00000102054.17 RBBP7
log2(RPKM)
6.50
6.25
6.00
5.75
5.50
5.75
6.00
6.25
6.50
6.75
ENSG00000089053.12 ANAPC5
log2(RPKM)
z−score of IF value
−2
−1
0
1
2

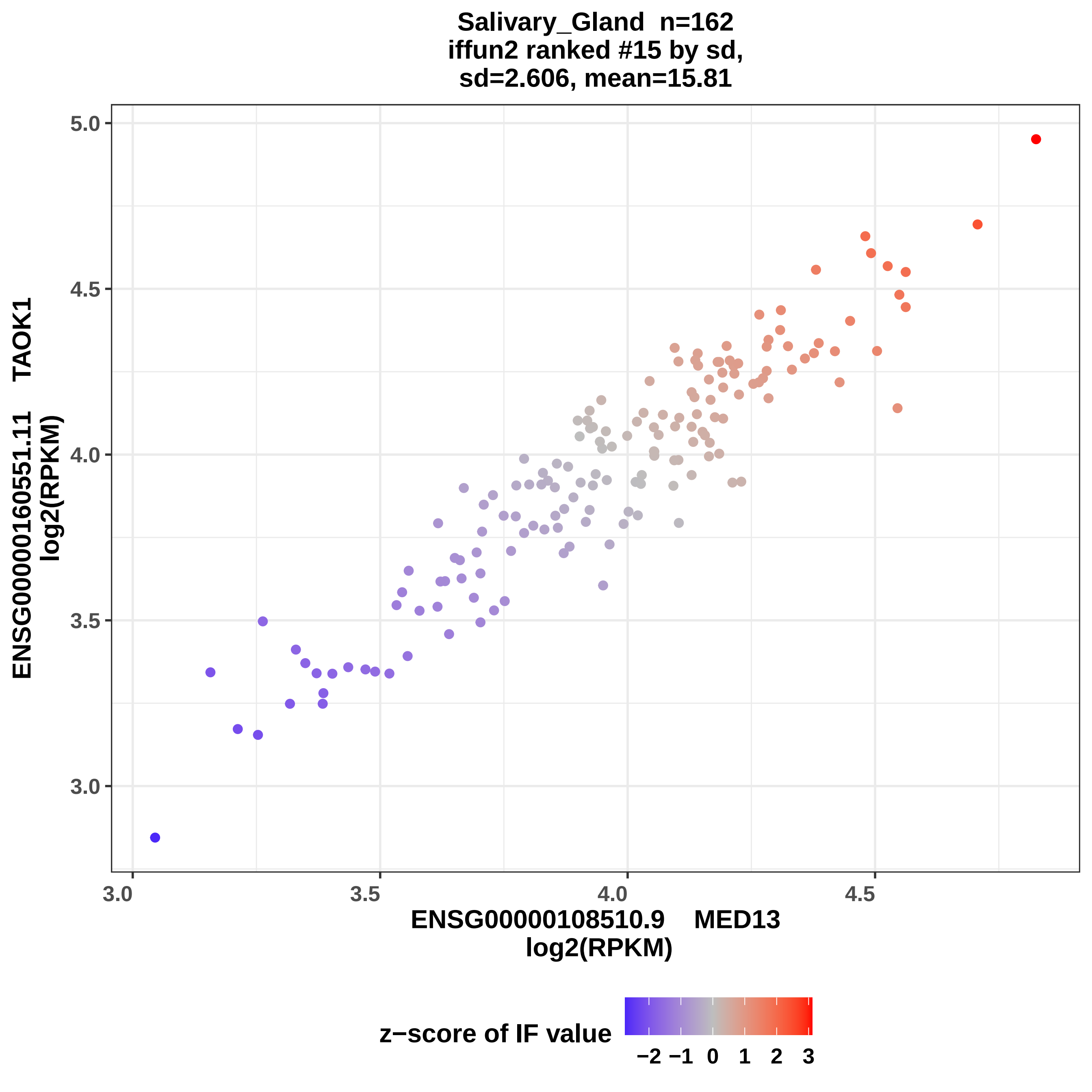

Salivary_Gland n=162
iffun2 ranked #15 by sd,
sd=2.606, mean=15.81
ENSG00000160551.11 TAOK1
log2(RPKM)
5.0
4.5
4.0
3.5
3.0
3.0
3.5
4.0
4.5
ENSG00000108510.9 MED13
log2(RPKM)
z−score of IF value
−2 −1 0 1 2 3

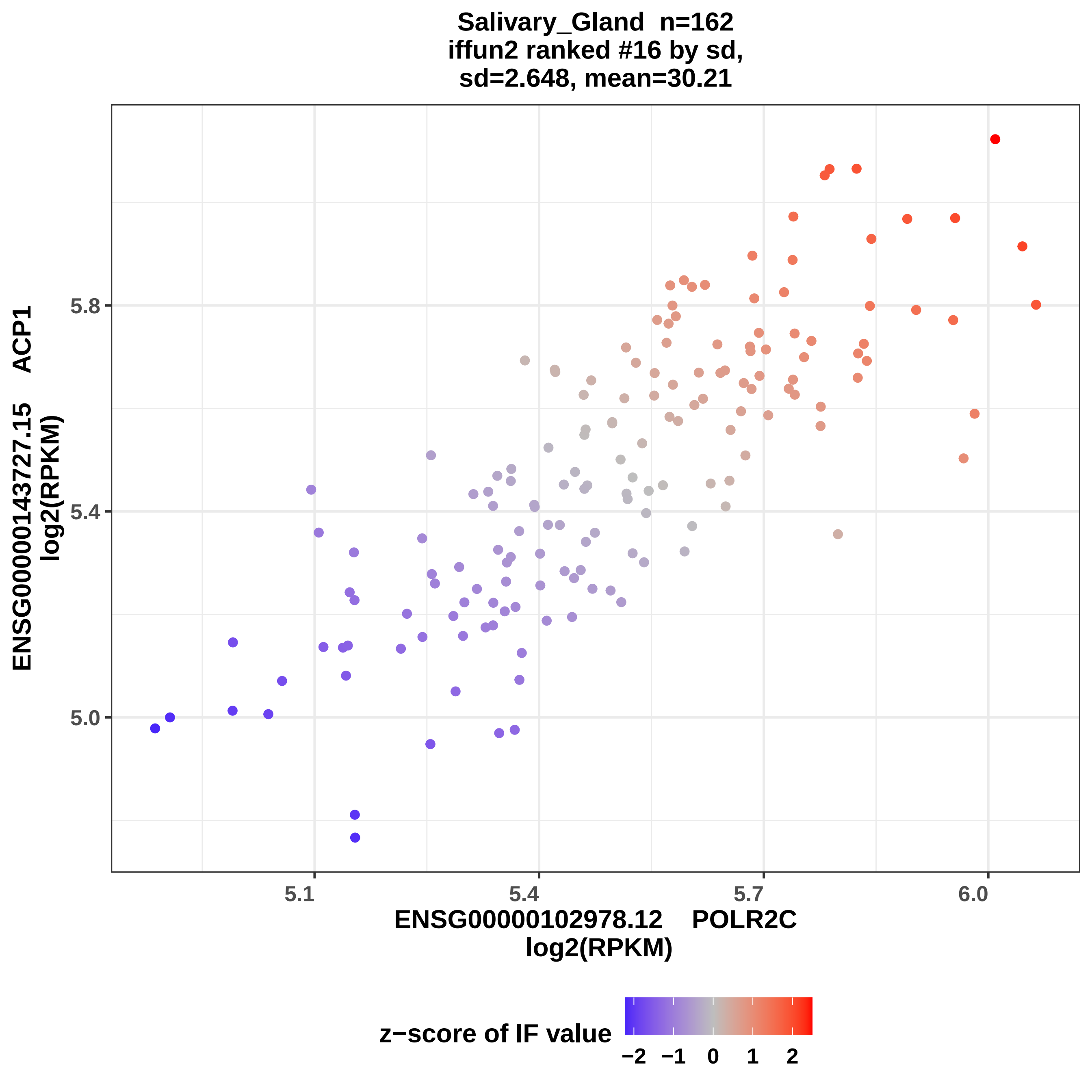

Salivary_Gland n=162
iffun2 ranked #16 by sd,
sd=2.648, mean=30.21
ENSG00000143727.15 ACP1
log2(RPKM)
5.8
5.4
5.0
5.1
5.4
5.7
6.0
ENSG00000102978.12 POLR2C
log2(RPKM)
z−score of IF value
−2
−1
0
1
2

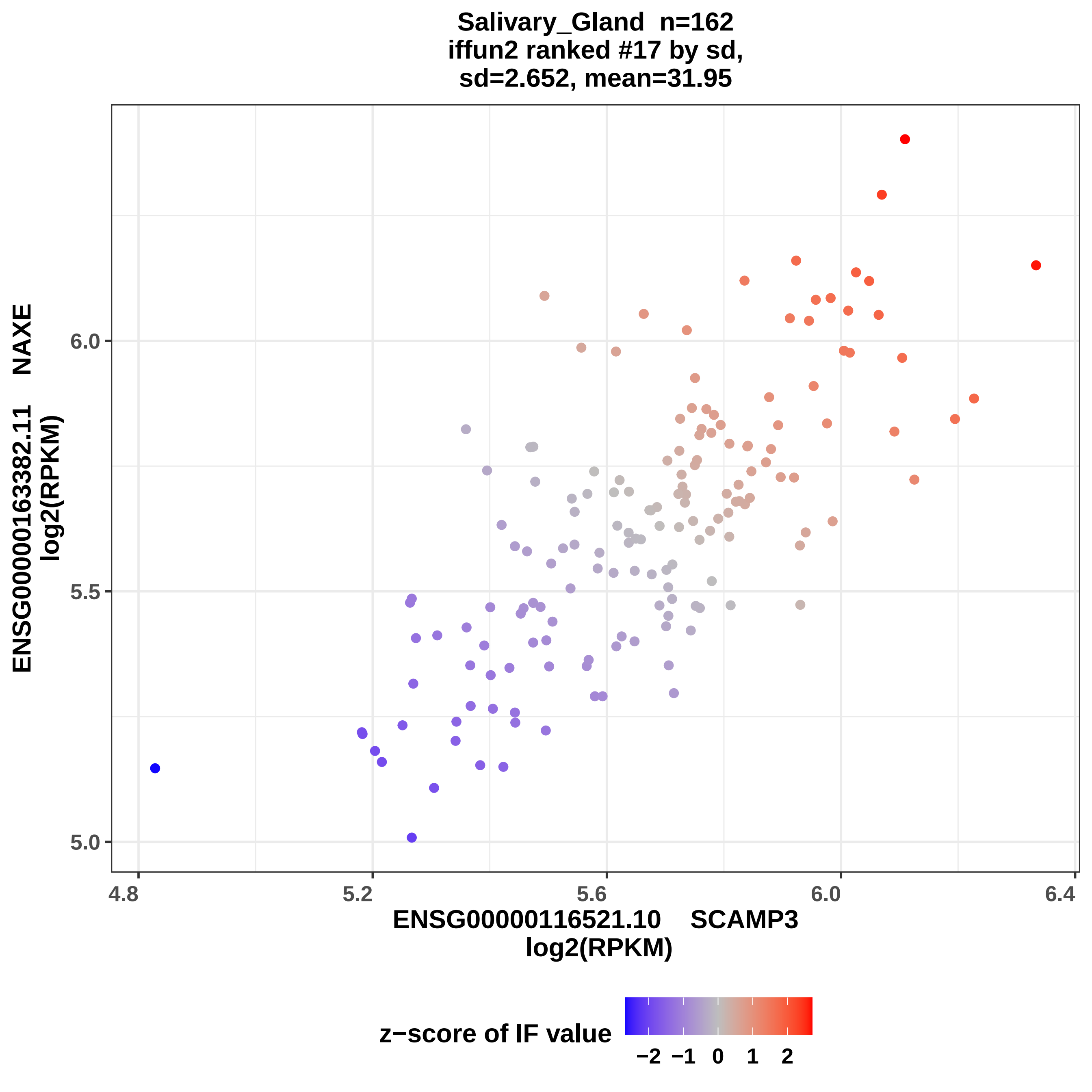

Salivary_Gland n=162
iffun2 ranked #17 by sd,
sd=2.652, mean=31.95
ENSG00000163382.11 NAXE
log2(RPKM)
6.0
5.5
5.0
4.8
5.2
5.6
6.0
6.4
ENSG00000116521.10 SCAMP3
log2(RPKM)
z−score of IF value
−2
−1
0
1
2

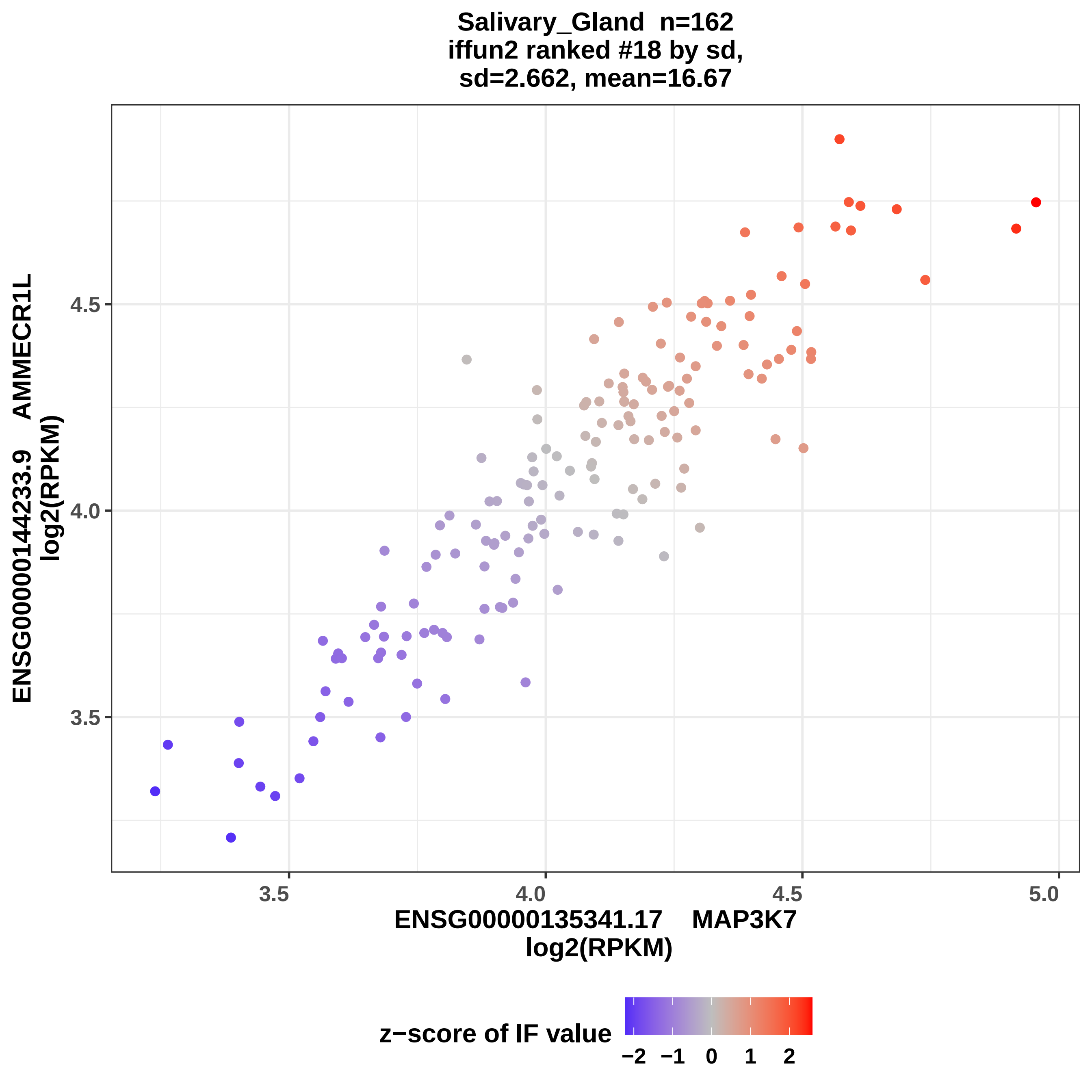

Salivary_Gland n=162
iffun2 ranked #18 by sd,
sd=2.662, mean=16.67
ENSG00000144233.9 AMMECR1L
log2(RPKM)
ENSG00000135341.17 MAP3K7
log2(RPKM)
3.5
4.0
4.5
5.0
z−score of IF value
−2
−1
0
1
2

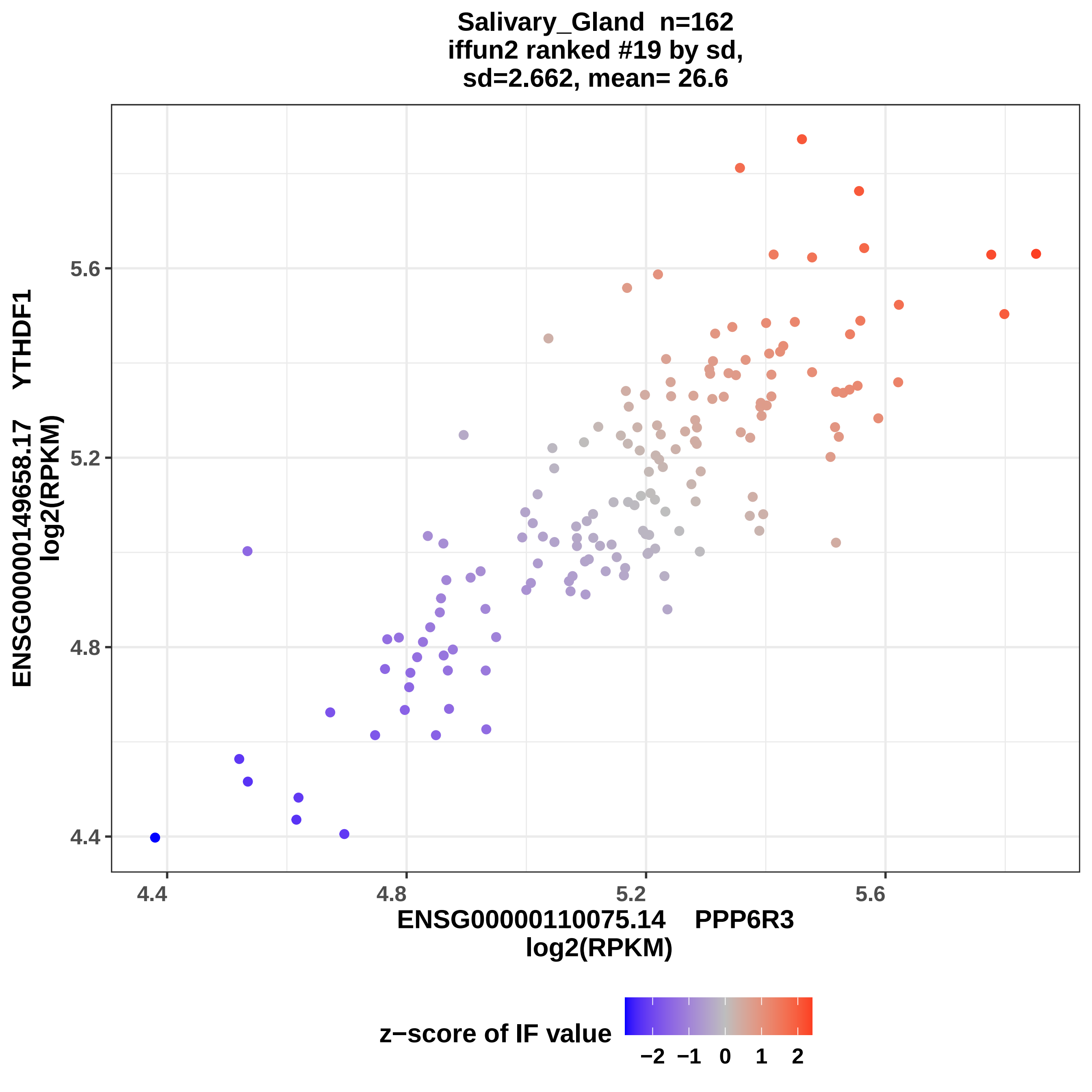
Salivary_Gland n=162
iffun2 ranked #19 by sd,
sd=2.662, mean= 26.6
ENSG00000149658.17 YTHDF1
log2(RPKM)
5.6
5.2
4.8
4.4
4.4
4.8
5.2
5.6
ENSG00000110075.14 PPP6R3
log2(RPKM)
z−score of IF value
−2
−1
0
1
2

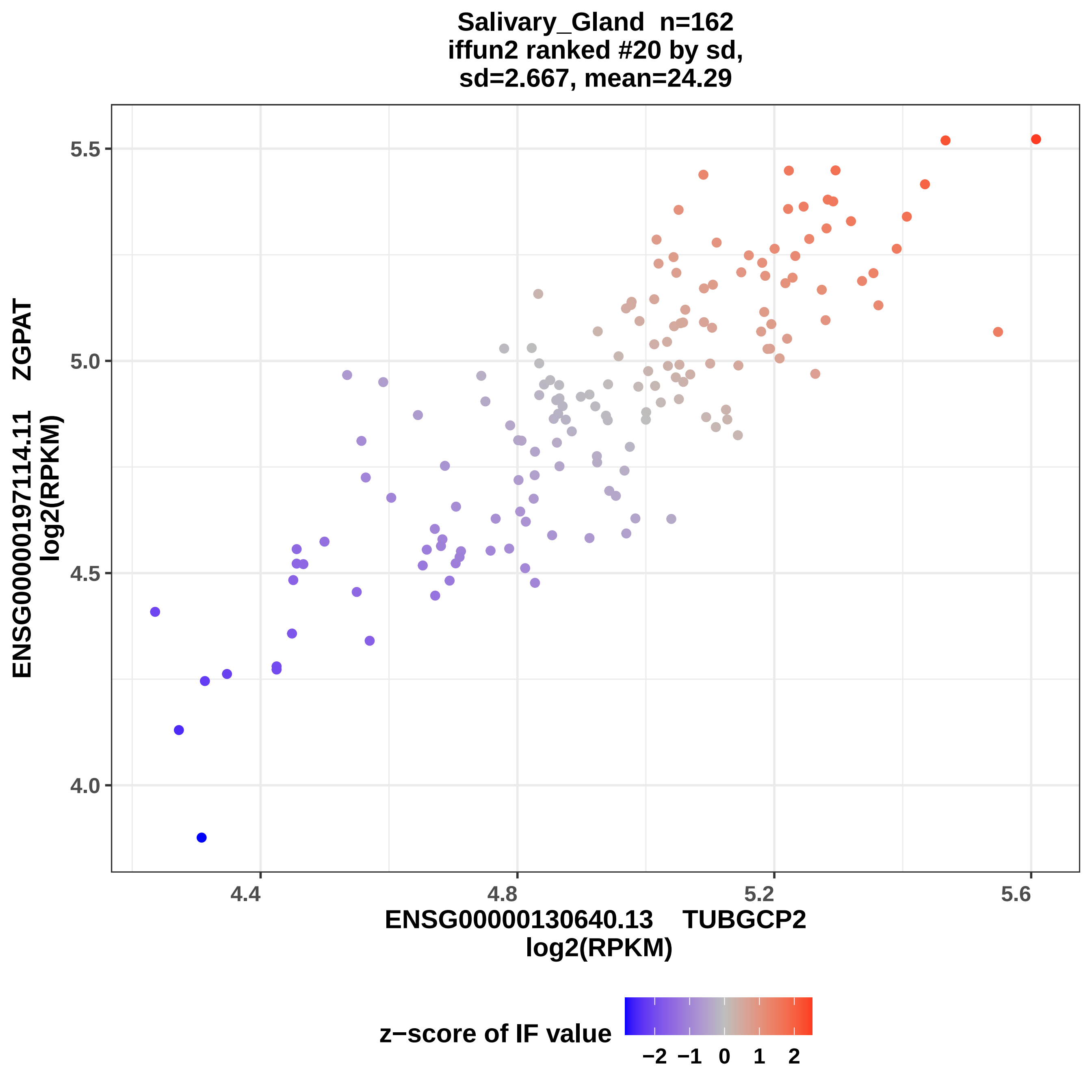

Salivary_Gland n=162
iffun2 ranked #20 by sd,
sd=2.667, mean=24.29
ENSG00000197114.11 ZGPAT
log2(RPKM)
5.5
5.0
4.5
4.0
4.4
4.8
5.2
5.6
ENSG00000130640.13 TUBGCP2
log2(RPKM)
z−score of IF value
−2
−1
0
1
2

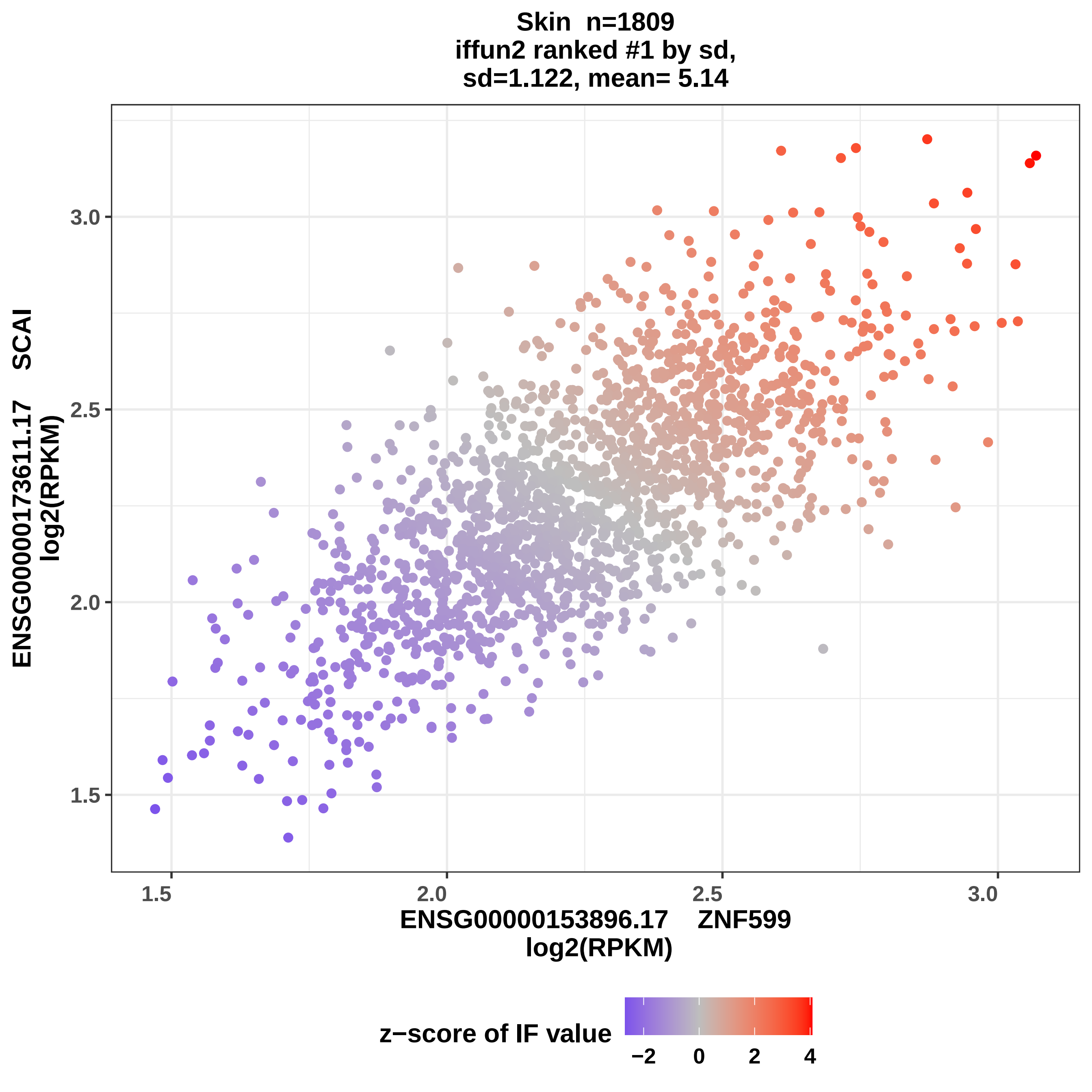
Skin n=1809
iffun2 ranked #1 by sd,
sd=1.122, mean= 5.14
ENSG00000173611.17 SCAI
log2(RPKM)
3.0
2.5
2.0
1.5
1.5
2.0
2.5
3.0
ENSG00000153896.17 ZNF599
log2(RPKM)
z−score of IF value
−2
0
2
4

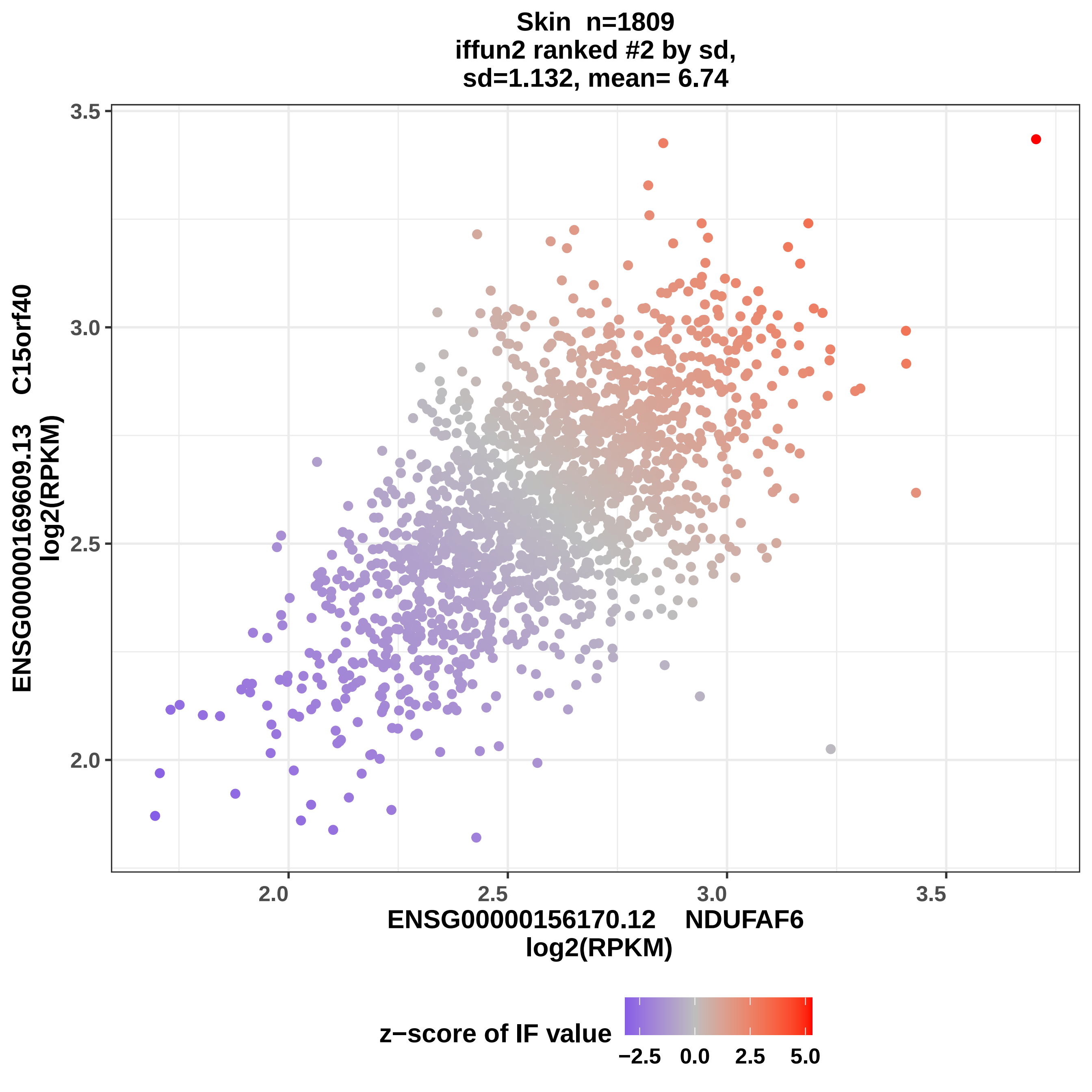

Skin n=1809
iffun2 ranked #2 by sd,
sd=1.132, mean= 6.74
ENSG00000169609.13 C15orf40
log2(RPKM)
3.5
3.0
2.5
2.0
ENSG00000156170.12 NDUFAF6
log2(RPKM)
z−score of IF value
−2.5
0.0
2.5
5.0

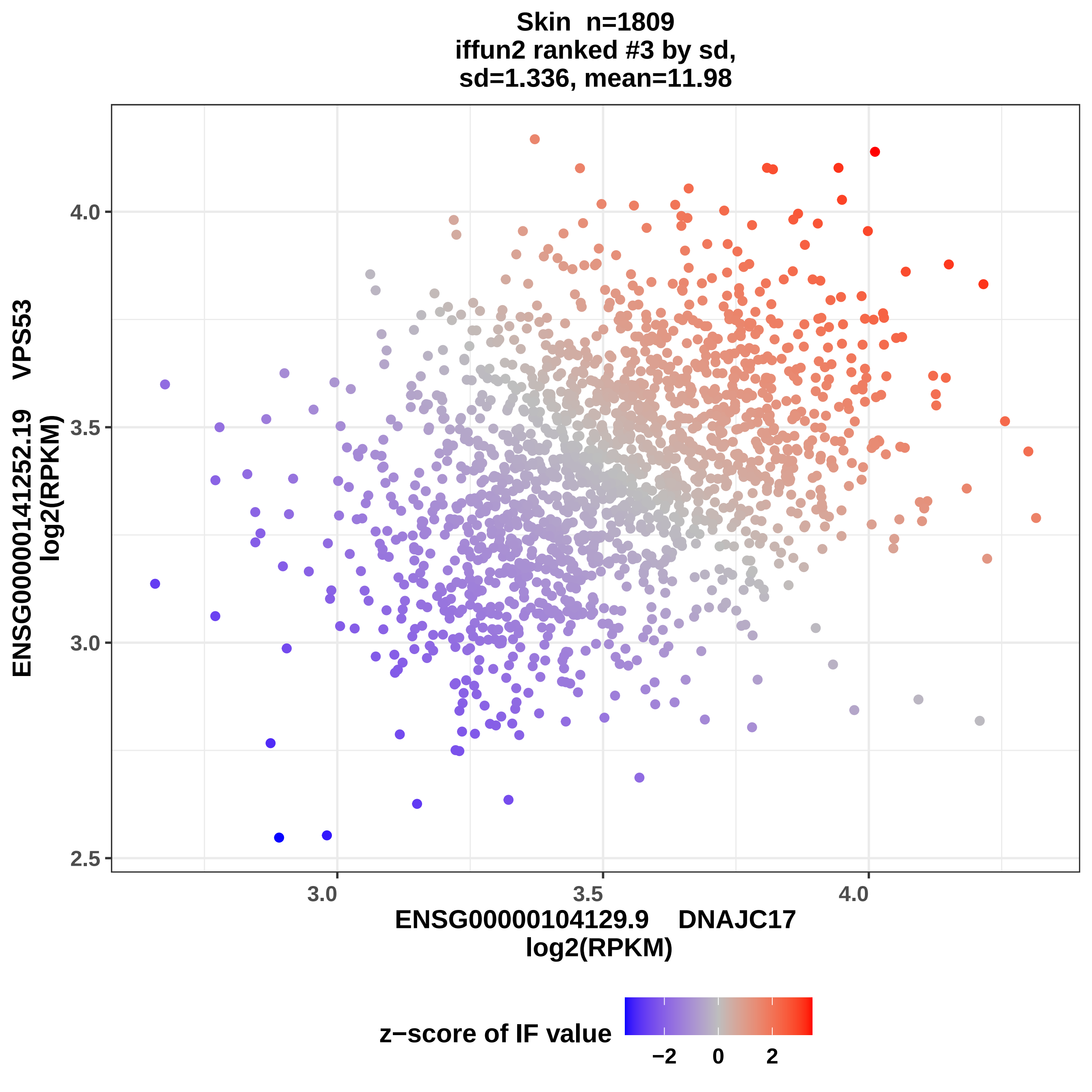
Skin n=1809
iffun2 ranked #3 by sd,
sd=1.336, mean=11.98
ENSG00000141252.19 VPS53
log2(RPKM)
4.0
3.5
3.0
2.5
3.0
3.5
4.0
ENSG00000104129.9 DNAJC17
log2(RPKM)
z−score of IF value
−2
0
2

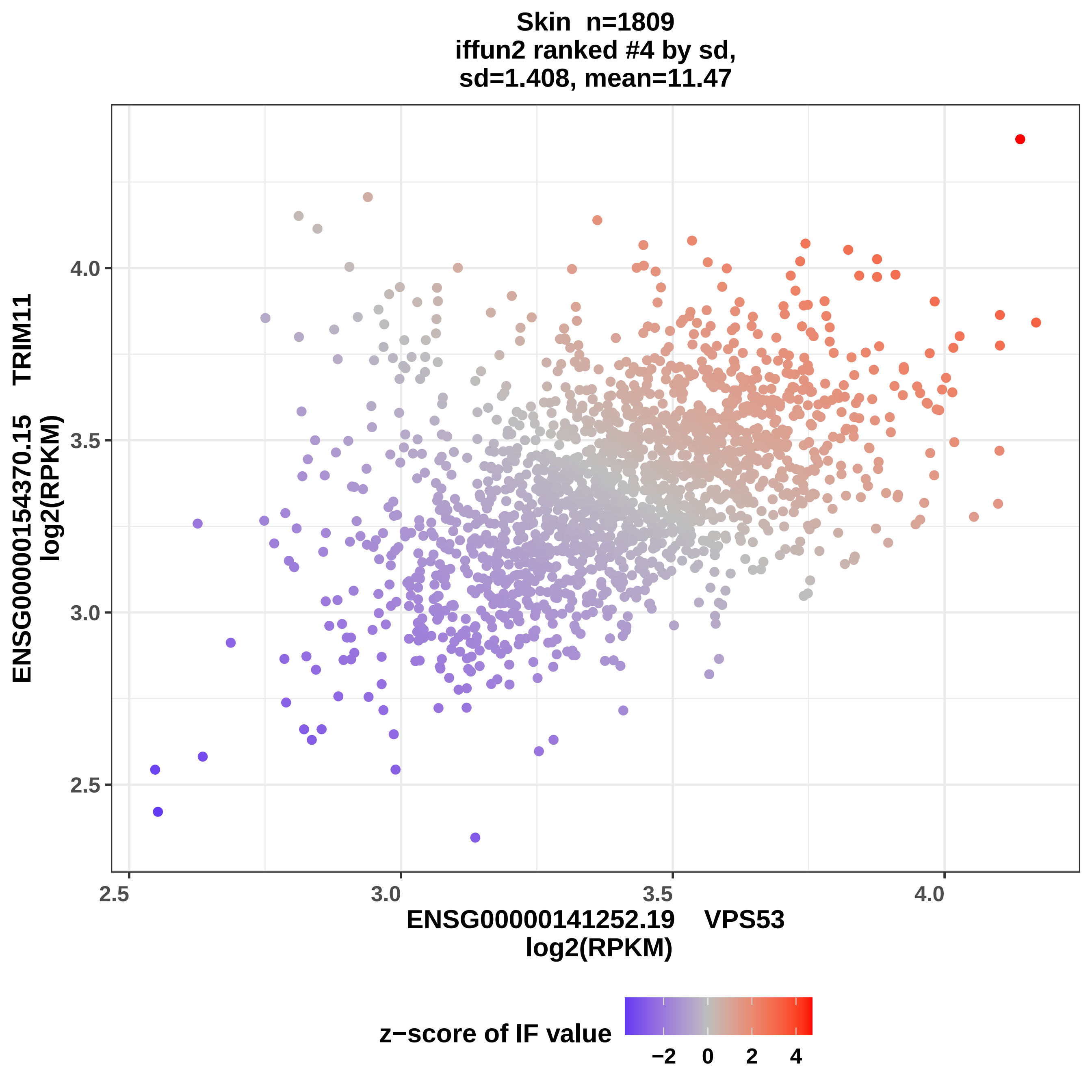
Skin n=1809
iffun2 ranked #4 by sd,
sd=1.408, mean=11.47
ENSG00000154370.15 TRIM11
log2(RPKM)
ENSG00000141252.19 VPS53
log2(RPKM)
2.5
3.0
3.5
4.0
z−score of IF value
−2
0
2
4

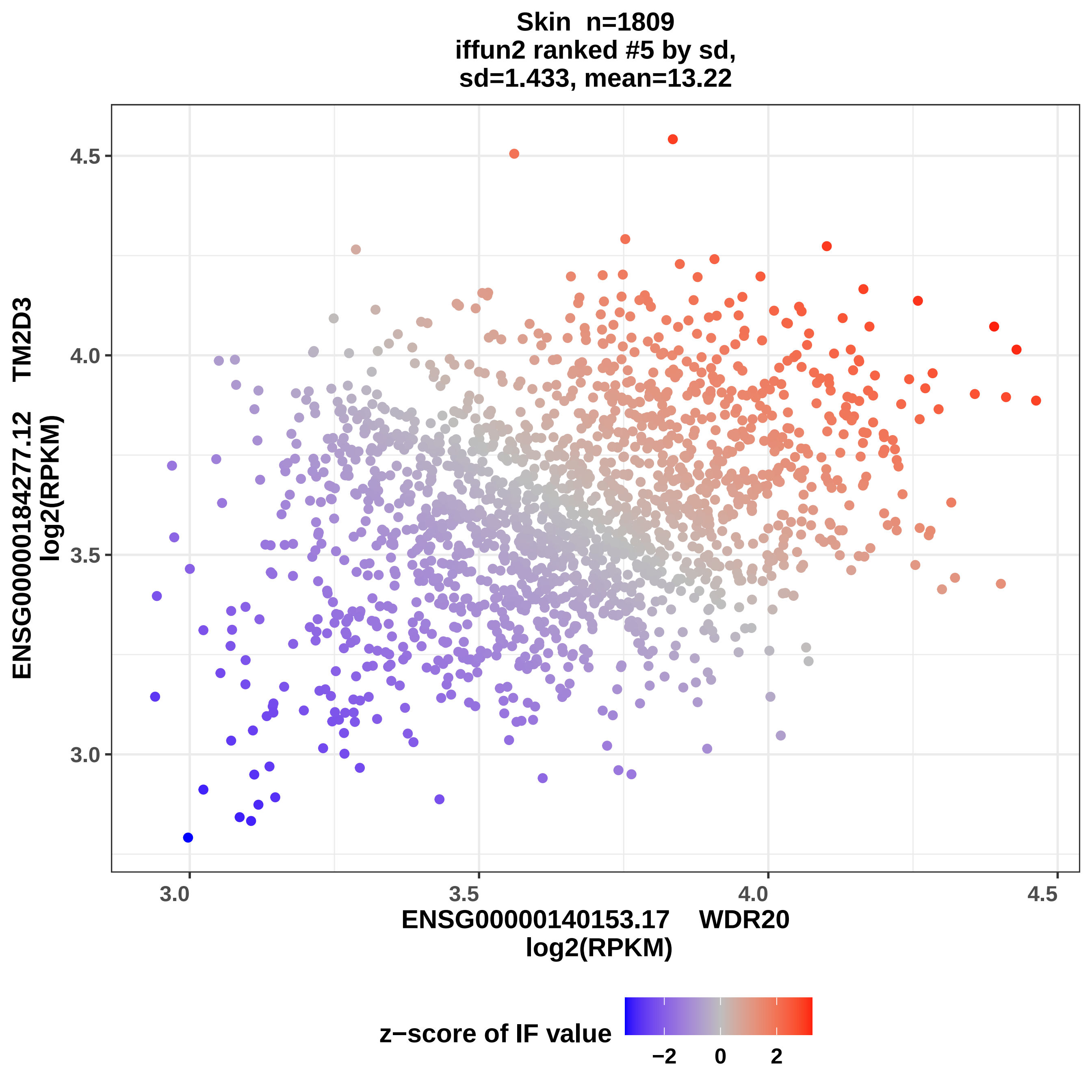

Skin n=1809
iffun2 ranked #5 by sd,
sd=1.433, mean=13.22
ENSG00000184277.12 TM2D3
log2(RPKM)
4.5
4.0
3.5
3.0
3.0
3.5
4.0
4.5
ENSG00000140153.17 WDR20
log2(RPKM)
z−score of IF value
−2
0
2

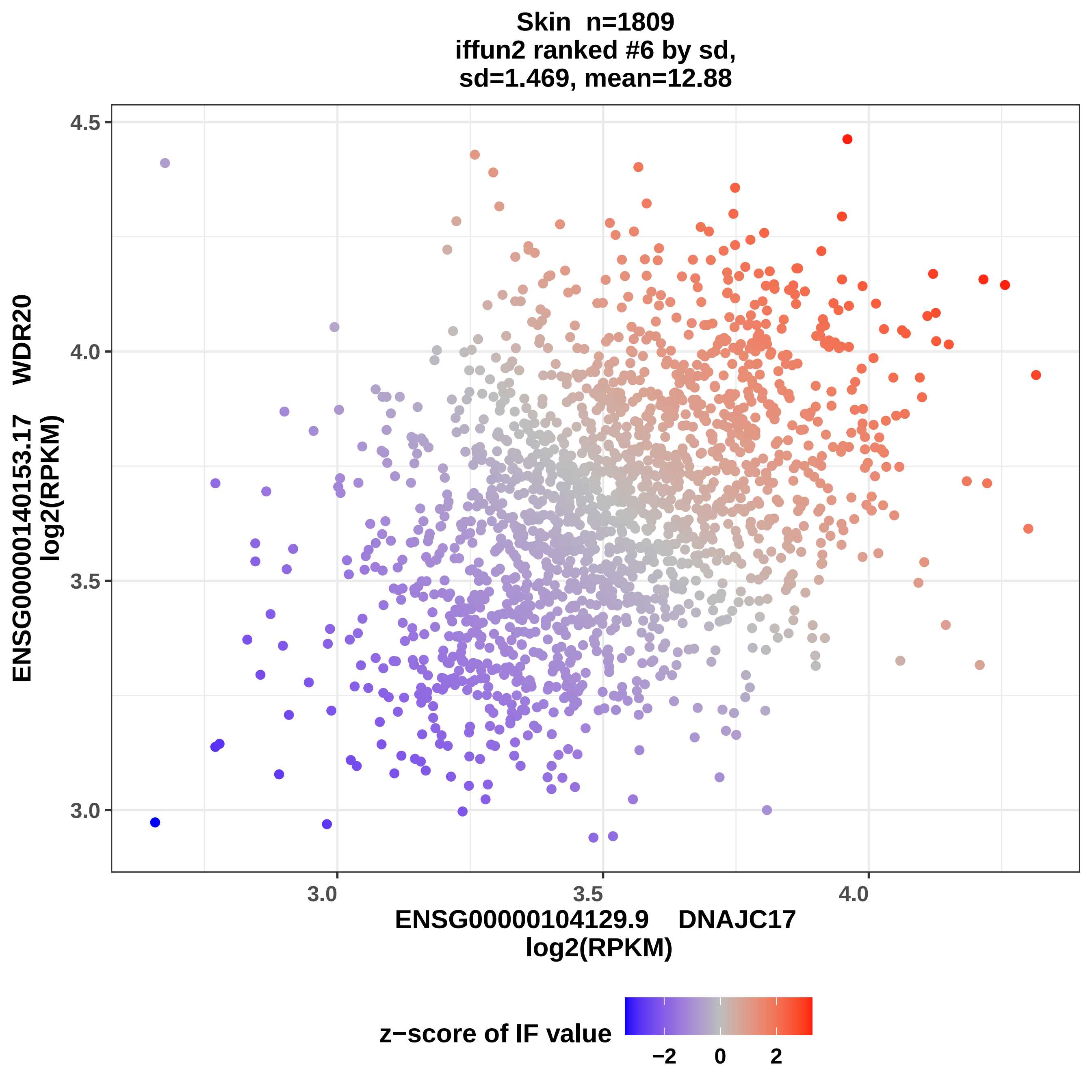

Skin n=1809
iffun2 ranked #6 by sd,
sd=1.469, mean=12.88
4.5
4.0
3.5
3.0
ENSG00000140153.17 WDR20
log2(RPKM)
3.0
3.5
4.0
ENSG00000104129.9 DNAJC17
log2(RPKM)
z−score of IF value
−2
0
2

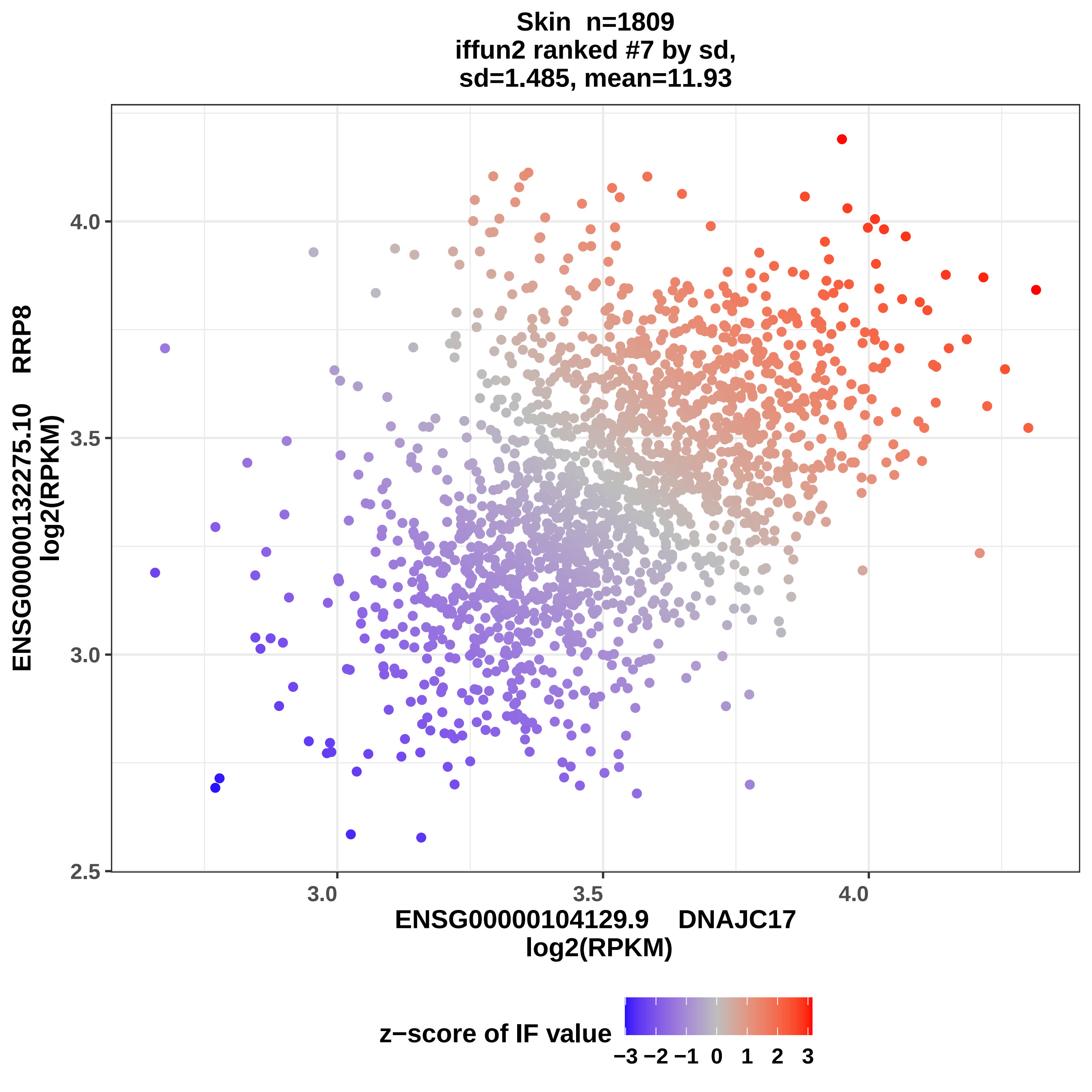

Skin n=1809
iffun2 ranked #7 by sd,
sd=1.485, mean=11.93
ENSG00000132275.10 RRP8
log2(RPKM)
4.0
3.5
3.0
2.5
3.0
3.5
4.0
ENSG00000104129.9 DNAJC17
log2(RPKM)
z−score of IF value
−3 −2 −1 0 1 2 3

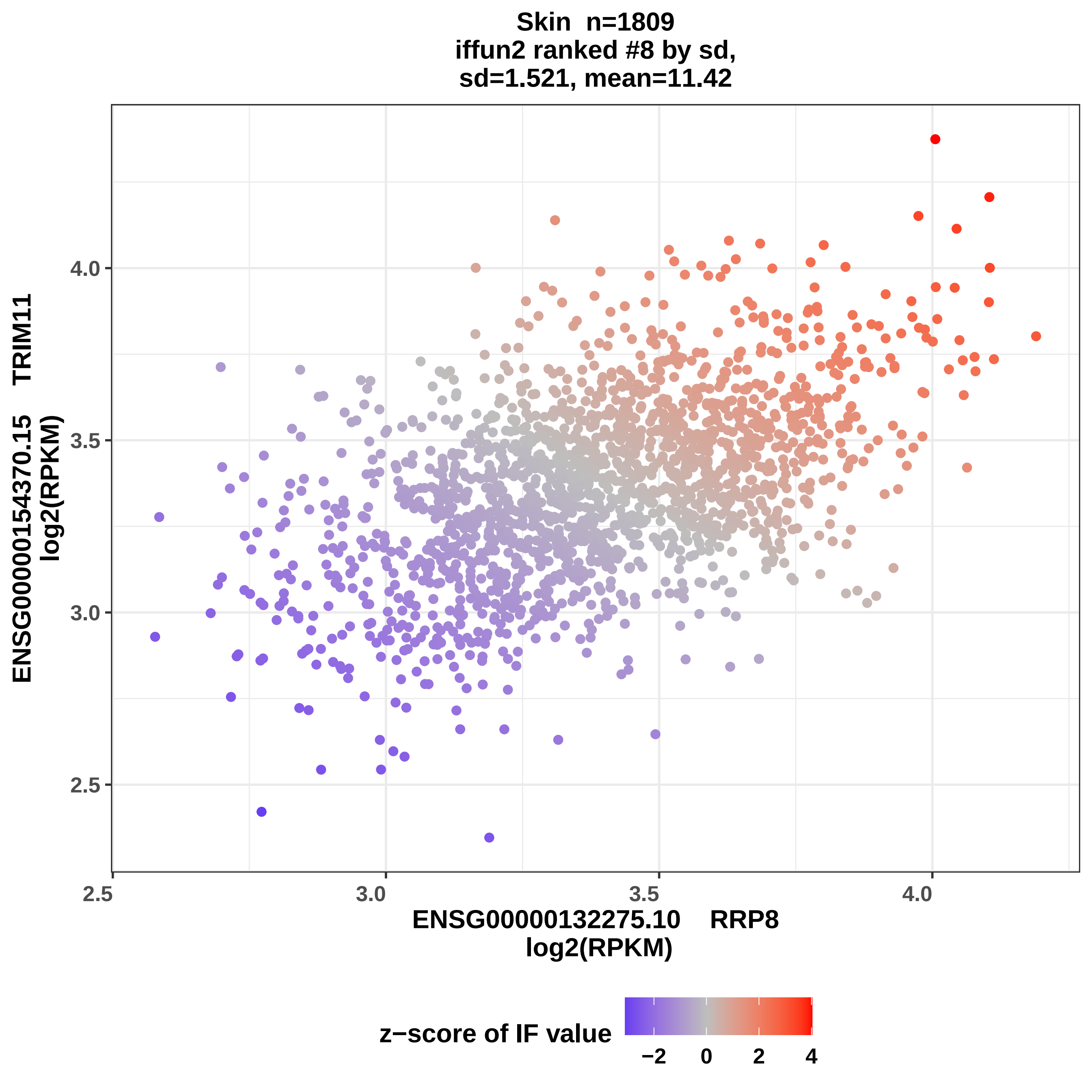
Skin n=1809
iffun2 ranked #8 by sd,
sd=1.521, mean=11.42
ENSG00000154370.15 TRIM11
log2(RPKM)
4.0
3.5
3.0
2.5
2.5
3.0
3.5
4.0
ENSG00000132275.10 RRP8
log2(RPKM)
z−score of IF value
−2
0
2
4

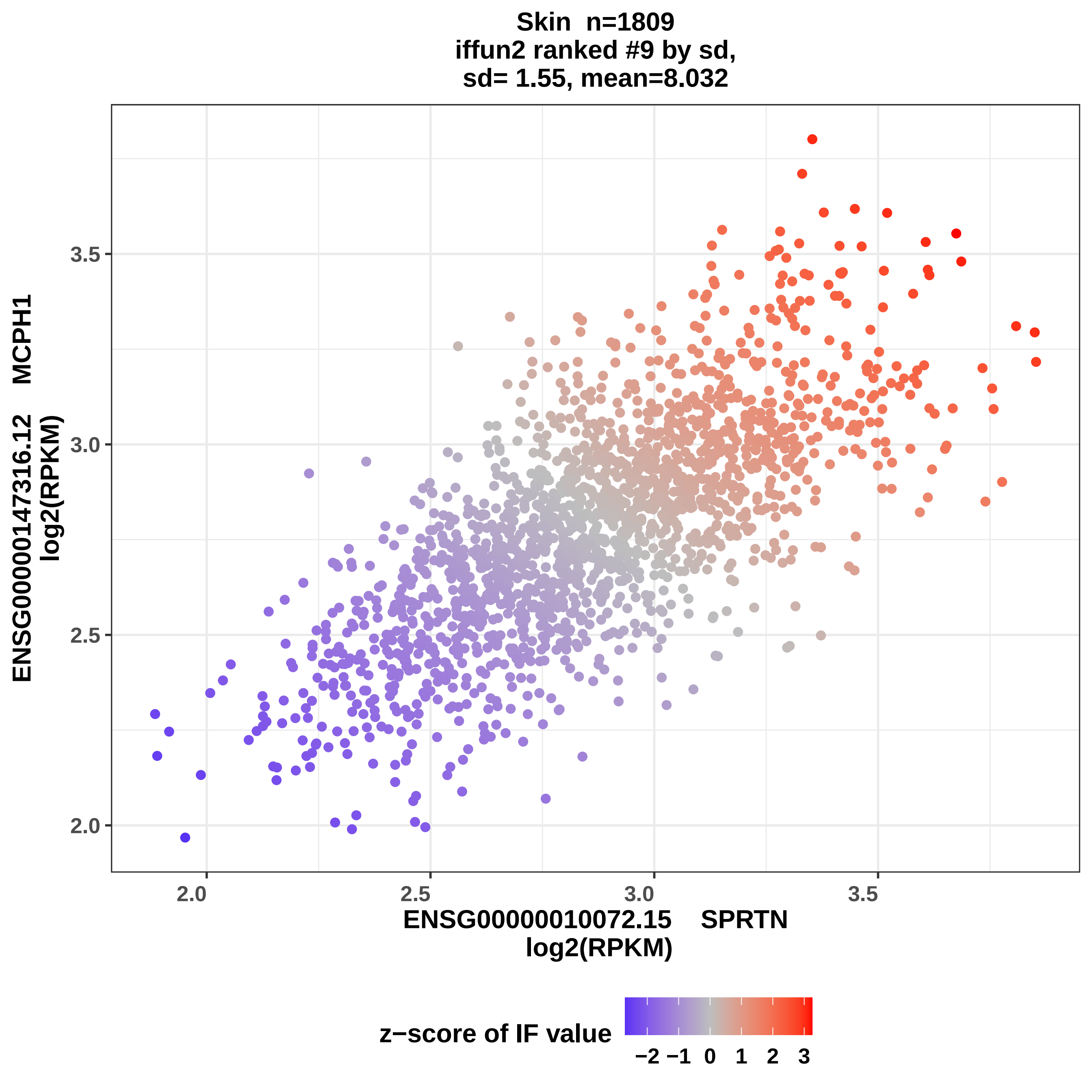

Skin n=1809
iffun2 ranked #9 by sd,
sd= 1.55, mean=8.032
ENSG00000147316.12 MCPH1
log2(RPKM)
3.5
3.0
2.5
2.0
2.0
2.5
3.0
3.5
ENSG00000010072.15 SPRTN
log2(RPKM)
z−score of IF value
−2 −1 0 1 2 3

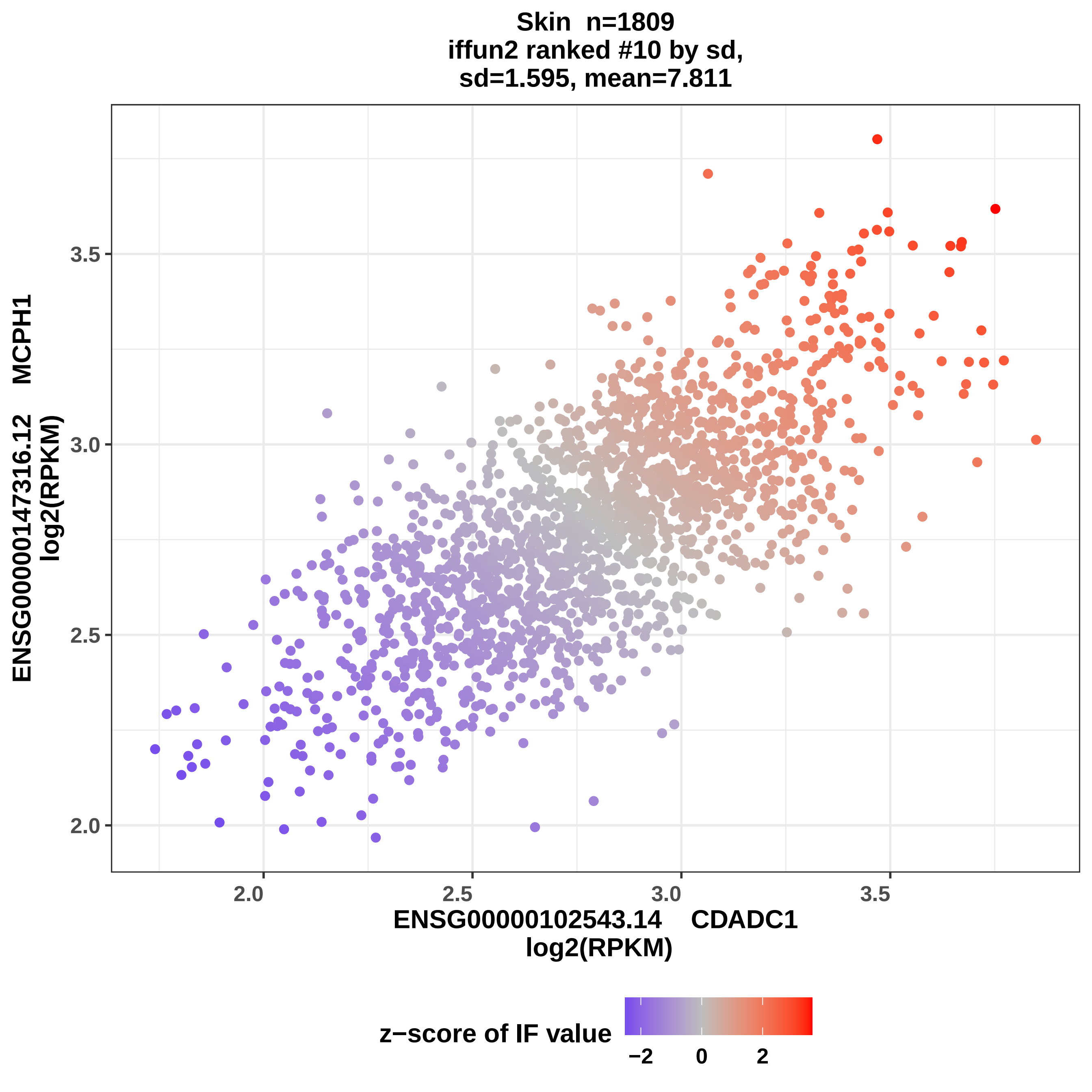

Skin n=1809
iffun2 ranked #10 by sd,
sd=1.595, mean=7.811
ENSG00000147316.12 MCPH1
log2(RPKM)
3.5
3.0
2.5
2.0
2.0
2.5
3.0
3.5
ENSG00000102543.14 CDADC1
log2(RPKM)
z−score of IF value
−2
0
2

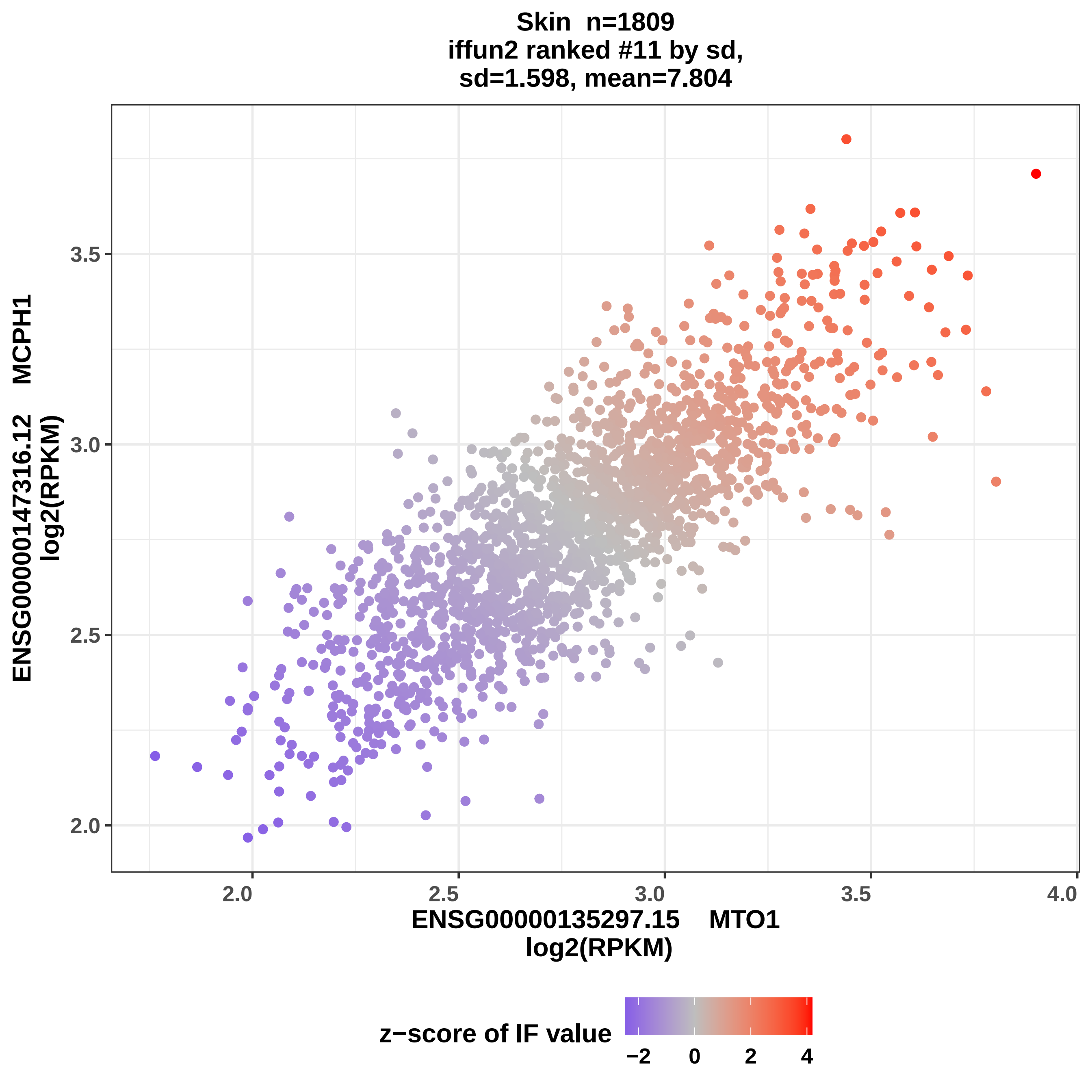

Skin n=1809
iffun2 ranked #11 by sd,
sd=1.598, mean=7.804
ENSG00000147316.12 MCPH1
log2(RPKM)
3.5
3.0
2.5
2.0
2.0
2.5
3.0
3.5
4.0
ENSG00000135297.15 MTO1
log2(RPKM)
z−score of IF value
−2
0
2
4

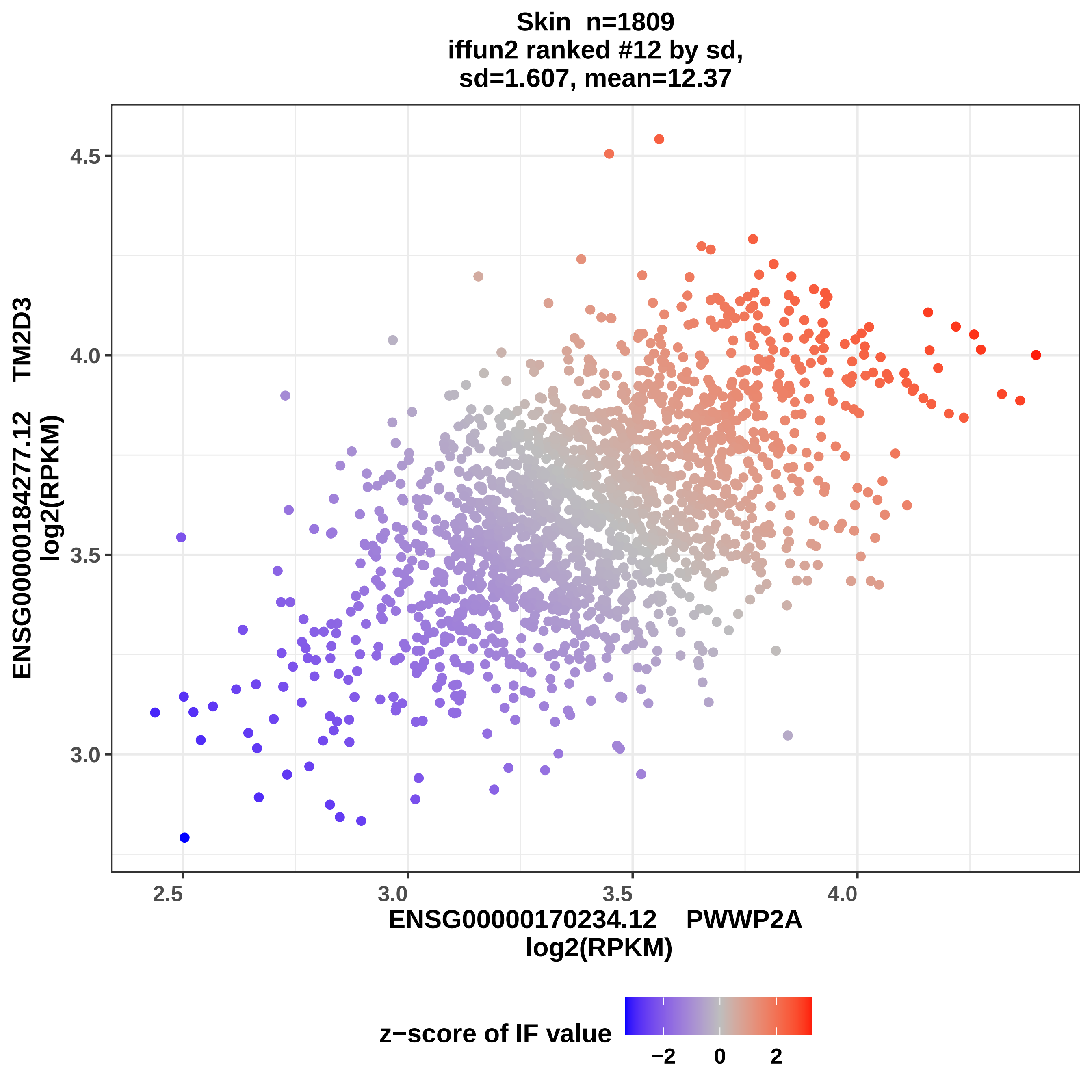

Skin n=1809
iffun2 ranked #12 by sd,
sd=1.607, mean=12.37
ENSG00000184277.12 TM2D3
log2(RPKM)
4.5
4.0
3.5
3.0
2.5
3.0
3.5
4.0
ENSG00000170234.12 PWWP2A
log2(RPKM)
z−score of IF value
−2
0
2

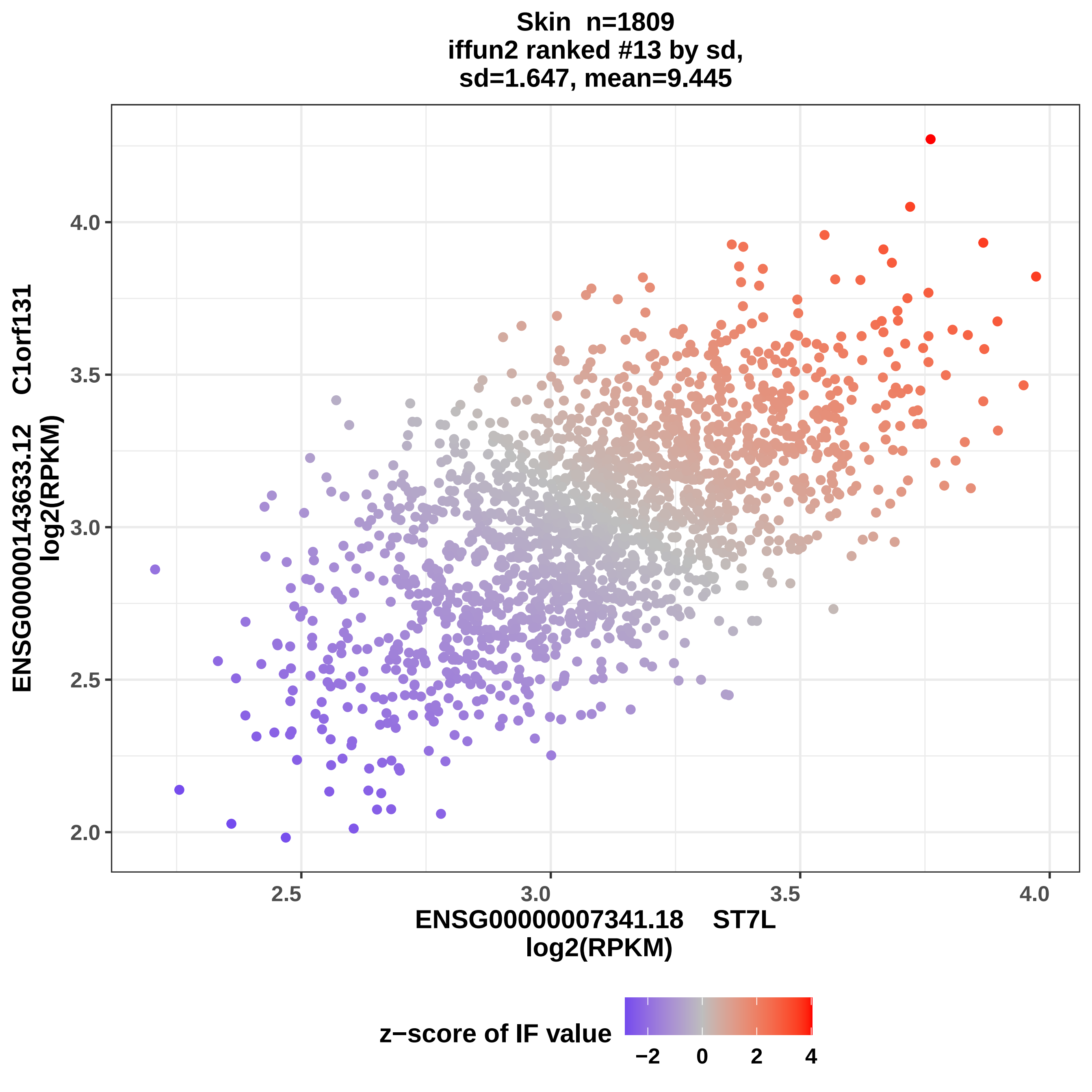

Skin n=1809
iffun2 ranked #13 by sd,
sd=1.647, mean=9.445
ENSG00000143633.12 C1orf131
log2(RPKM)
4.0
3.5
3.0
2.5
2.0
2.5
3.0
3.5
4.0
ENSG00000007341.18 ST7L
log2(RPKM)
z−score of IF value
−2
0
2
4

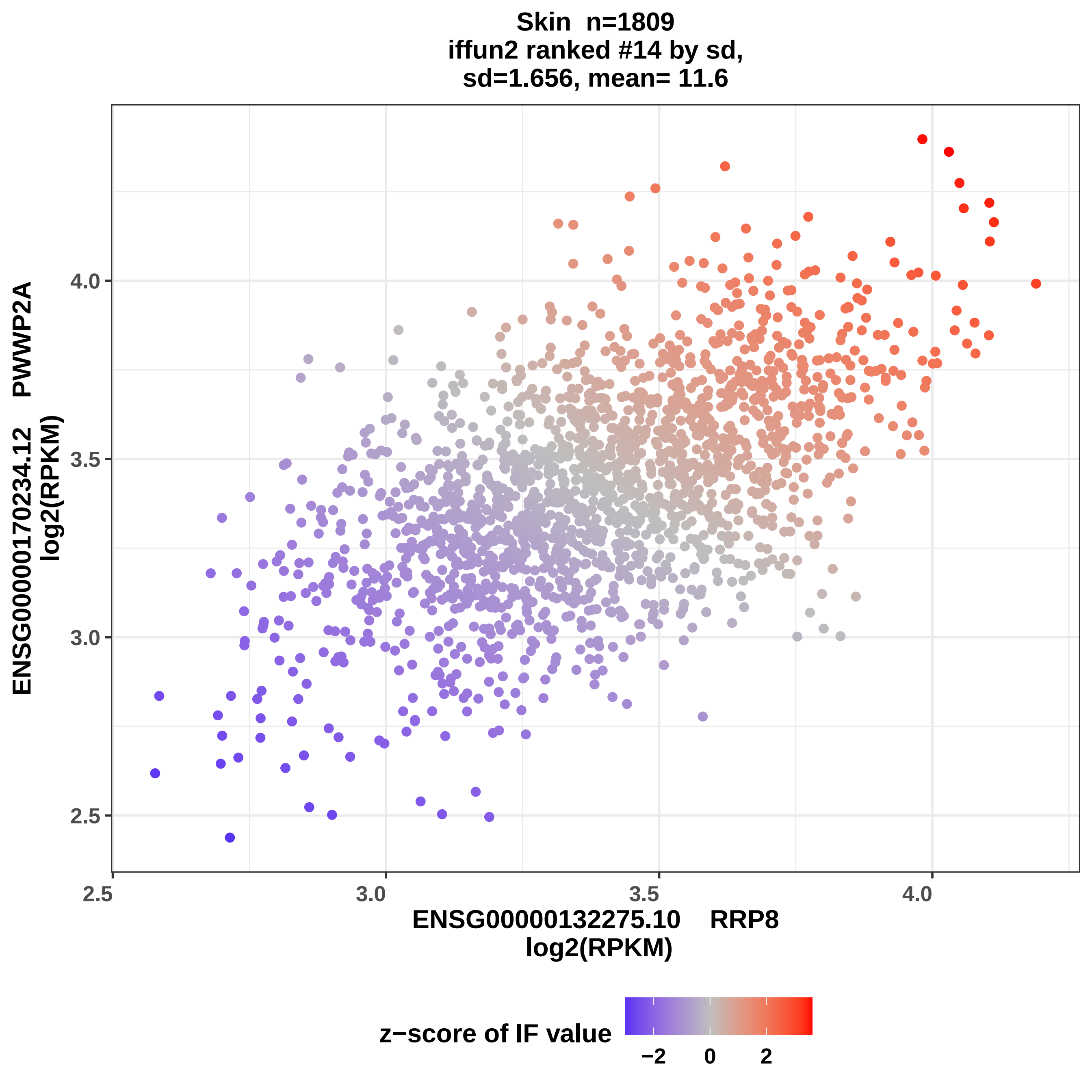
Skin n=1809
iffun2 ranked #14 by sd,
sd=1.656, mean= 11.6
ENSG00000170234.12 PWWP2A
log2(RPKM)
4.0
3.5
3.0
2.5
2.5
3.0
3.5
4.0
ENSG00000132275.10 RRP8
log2(RPKM)
z−score of IF value
−2
0
2

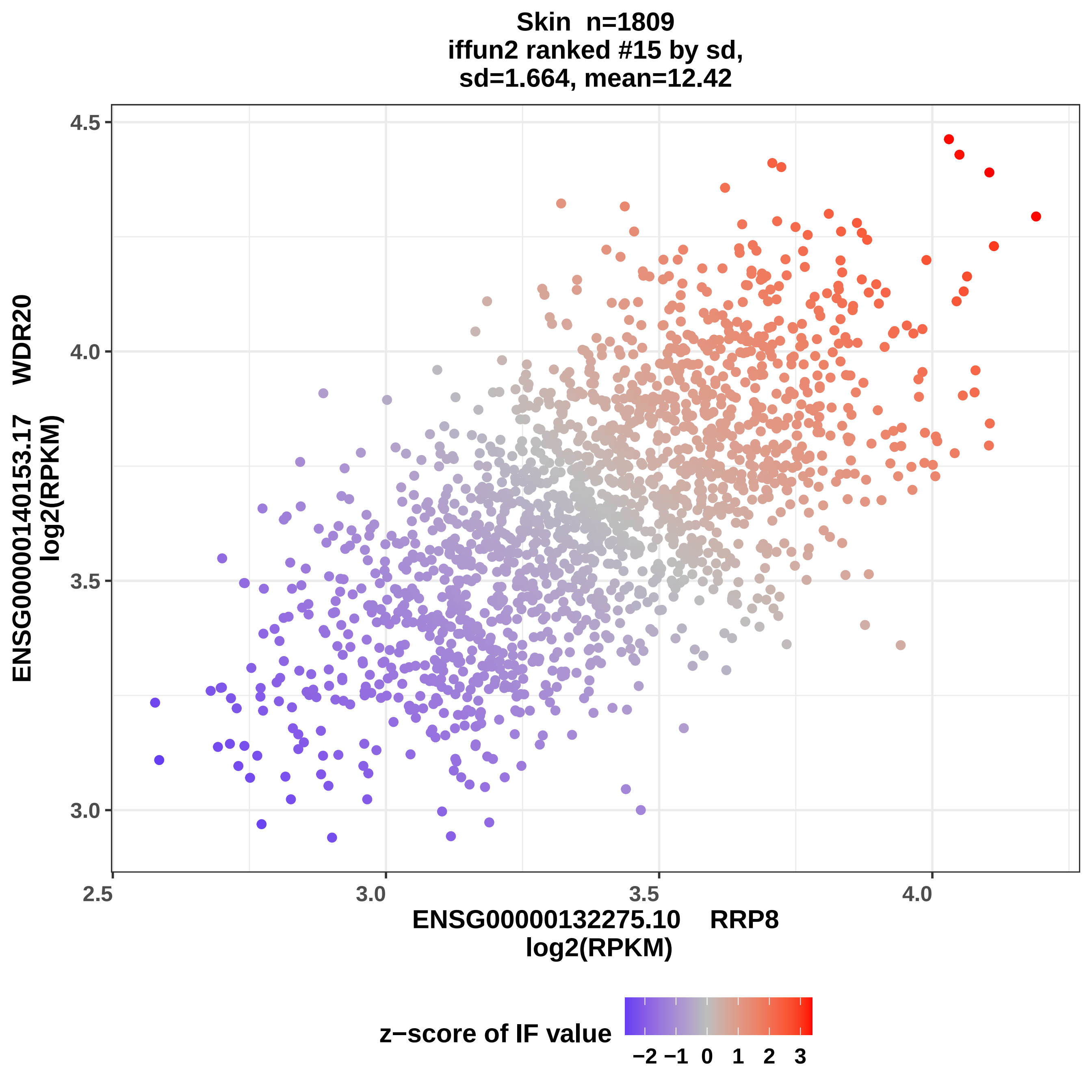
Skin n=1809
iffun2 ranked #15 by sd,
sd=1.664, mean=12.42
ENSG00000140153.17 WDR20
log2(RPKM)
ENSG00000132275.10 RRP8
log2(RPKM)
4.5
4.0
3.5
3.0
2.5
3.0
3.5
4.0
z−score of IF value
−2 −1 0 1 2 3

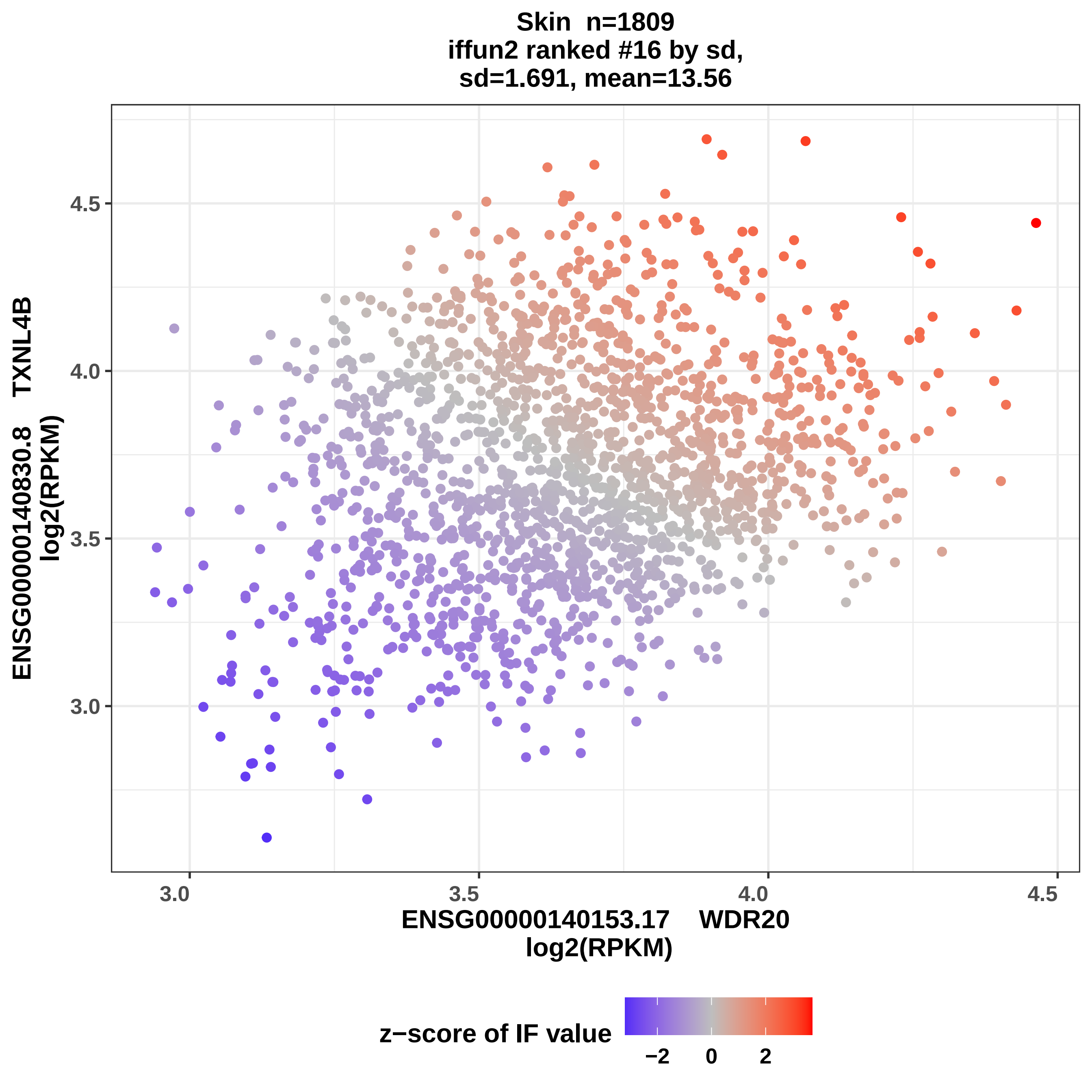
Skin n=1809
iffun2 ranked #16 by sd,
sd=1.691, mean=13.56
ENSG00000140830.8 TXNL4B
log2(RPKM)
4.5
4.0
3.5
3.0
3.0
3.5
4.0
4.5
ENSG00000140153.17 WDR20
log2(RPKM)
z−score of IF value
−2
0
2

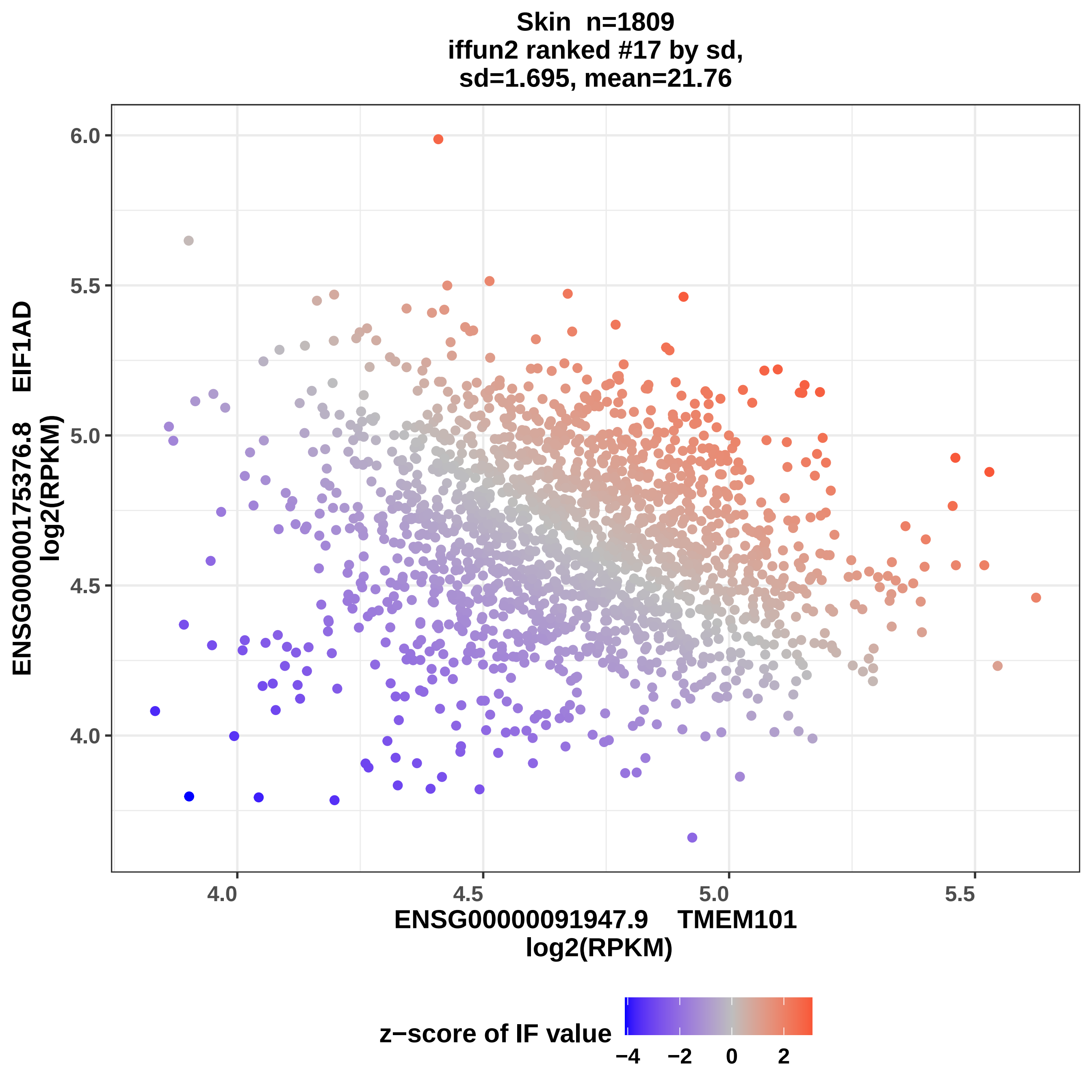

Skin n=1809
iffun2 ranked #17 by sd,
sd=1.695, mean=21.76
ENSG00000175376.8 EIF1AD
log2(RPKM)
ENSG00000091947.9 TMEM101
log2(RPKM)
6.0
5.5
5.0
4.5
4.0
4.0
4.5
5.0
5.5
z−score of IF value
−4
−2
0
2

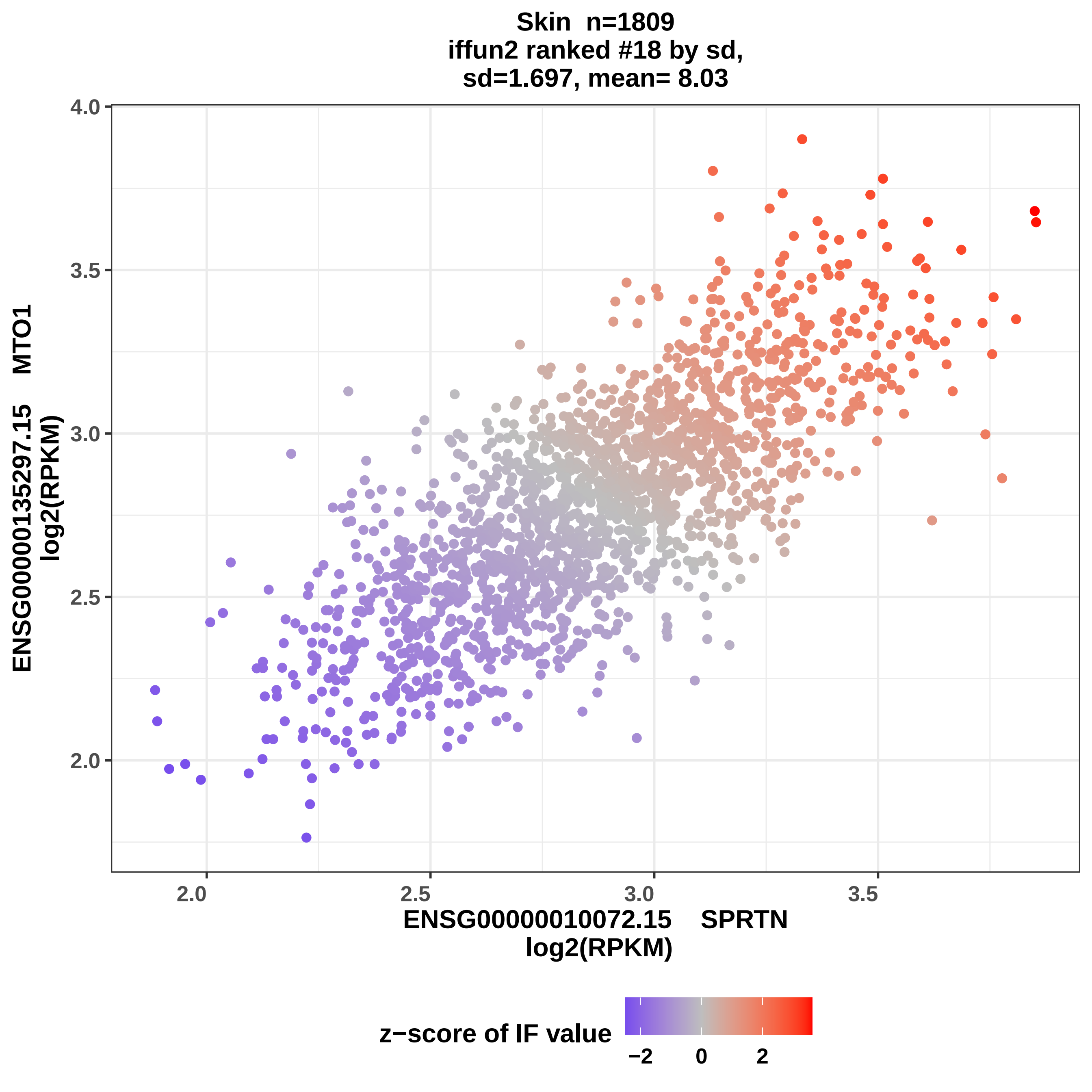
Skin n=1809
iffun2 ranked #18 by sd,
sd=1.697, mean= 8.03
ENSG00000135297.15 MTO1
log2(RPKM)
ENSG00000010072.15 SPRTN
log2(RPKM)
4.0
3.5
3.0
2.5
2.0
2.0
2.5
3.0
3.5
z−score of IF value
−2
0
2

Skin n=1809
iffun2 ranked #19 by sd,
sd=1.711, mean= 7.81

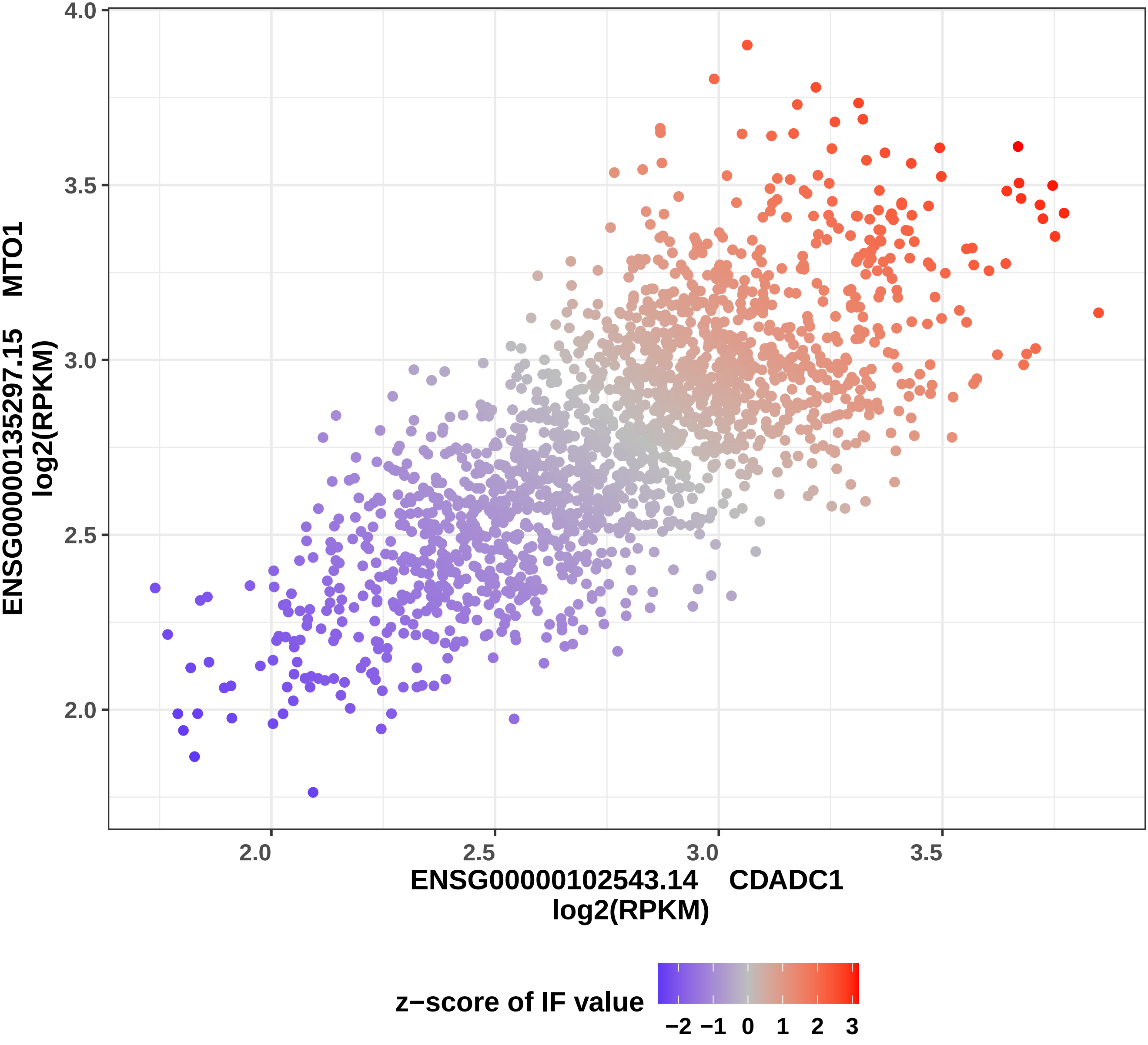

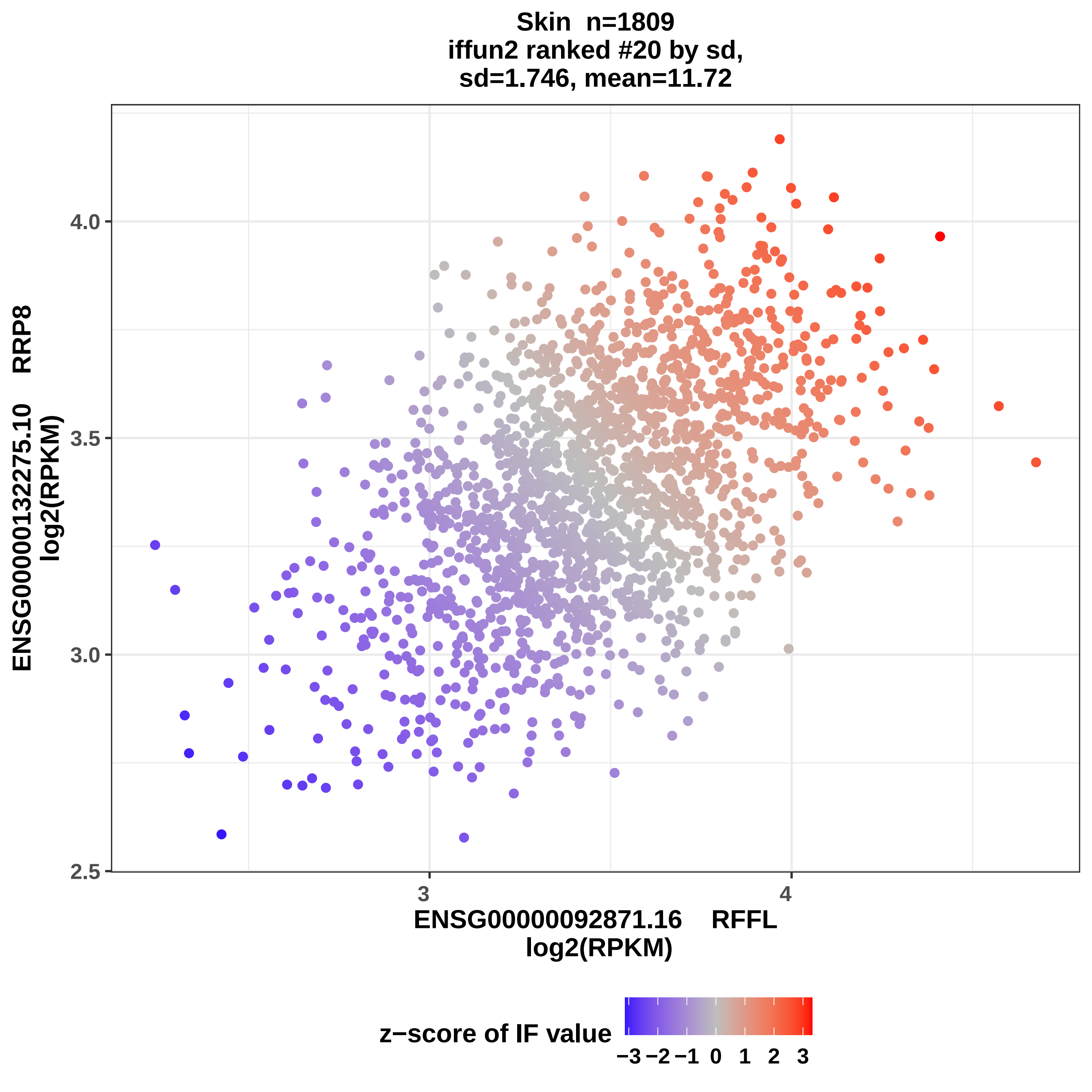

Skin n=1809
iffun2 ranked #20 by sd,
sd=1.746, mean=11.72
ENSG00000132275.10 RRP8
log2(RPKM)
4.0
3.5
3.0
2.5
3
4
ENSG00000092871.16 RFFL
log2(RPKM)
z−score of IF value
−3 −2 −1 0 1 2 3

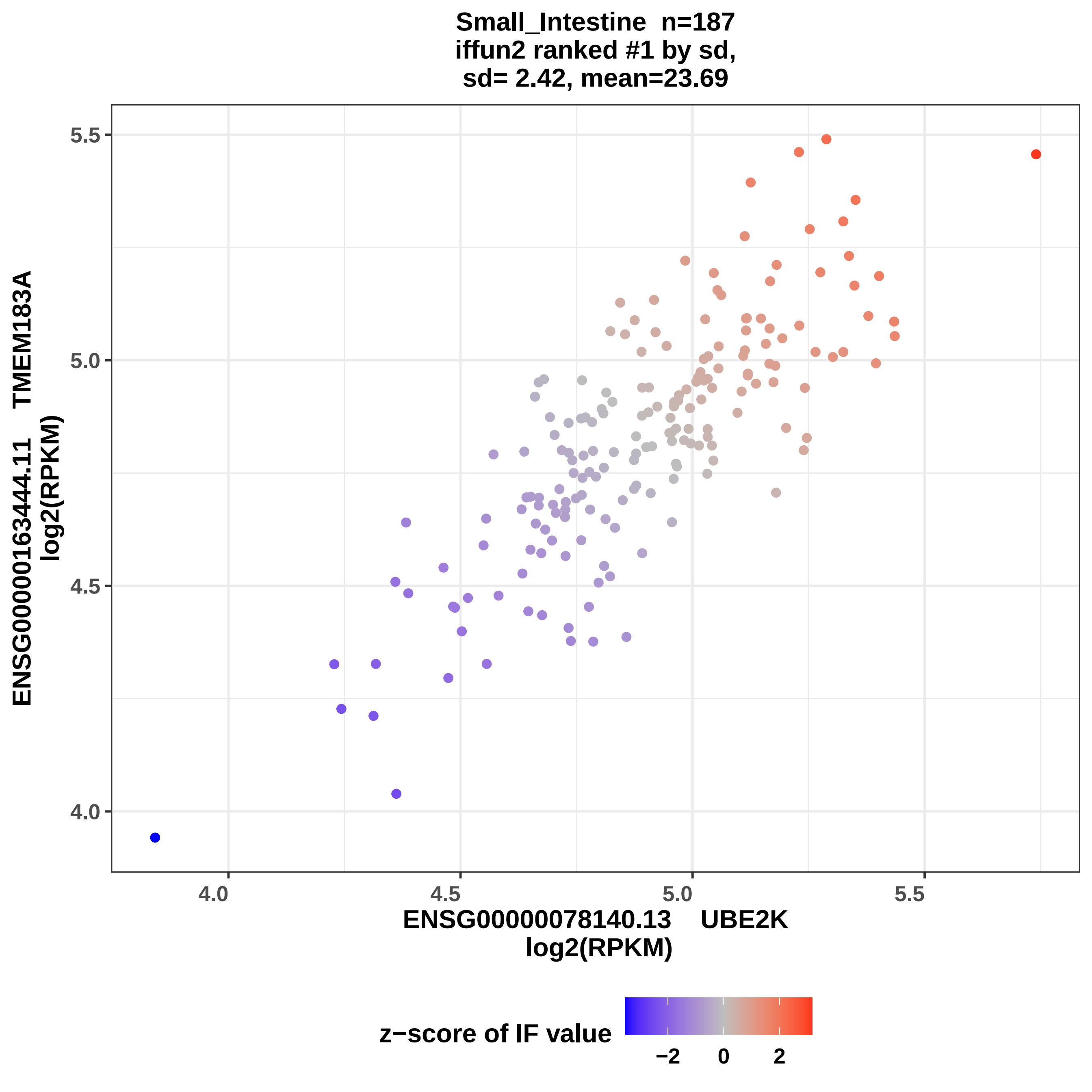
Small_Intestine n=187
iffun2 ranked #1 by sd,
sd= 2.42, mean=23.69
ENSG00000163444.11 TMEM183A
log2(RPKM)
5.5
5.0
4.5
4.0
4.0
4.5
5.0
5.5
ENSG00000078140.13 UBE2K
log2(RPKM)
z−score of IF value
−2
0
2

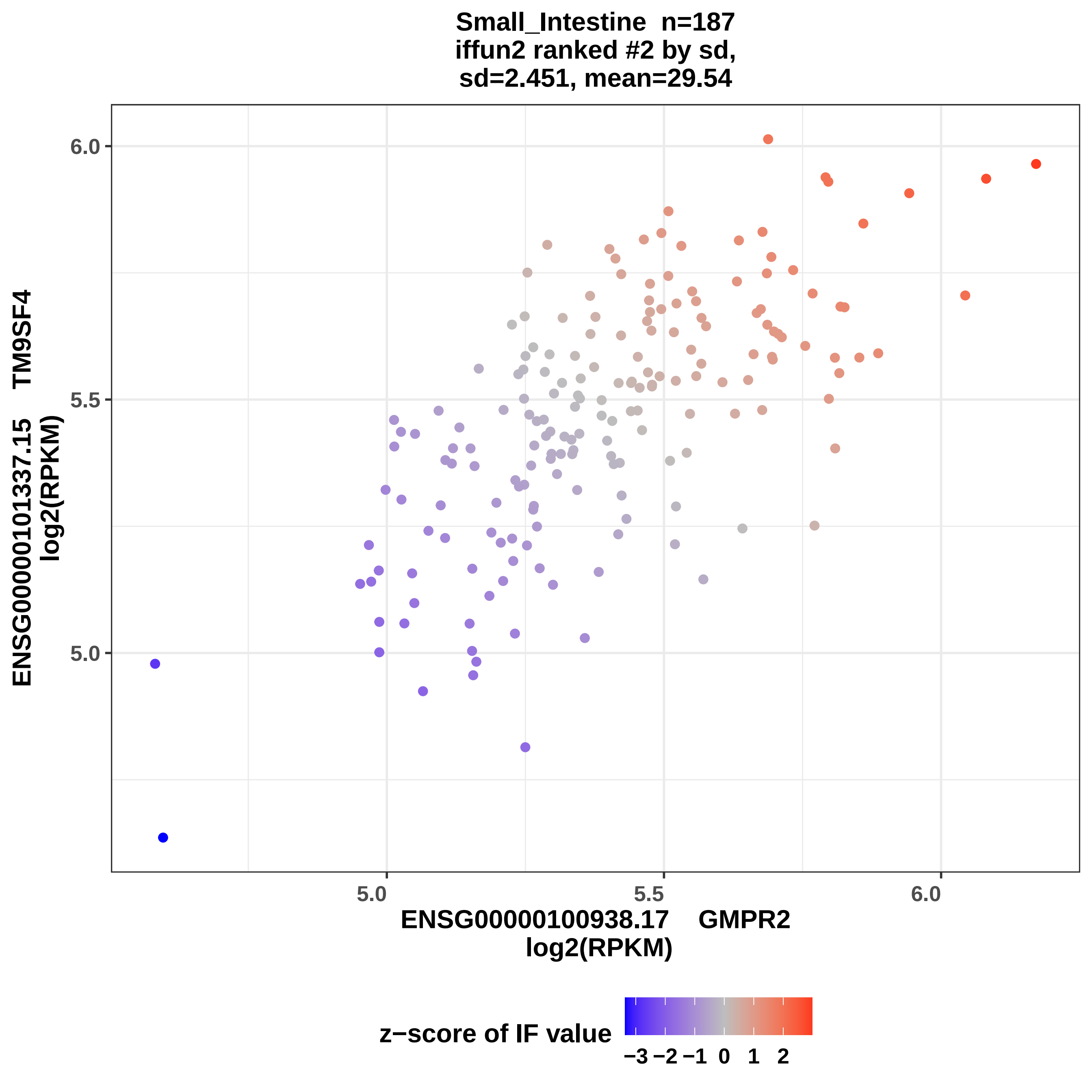
Small_Intestine n=187
iffun2 ranked #2 by sd,
sd=2.451, mean=29.54
ENSG00000101337.15 TM9SF4
log2(RPKM)
6.0
5.5
5.0
5.0
5.5
6.0
ENSG00000100938.17 GMPR2
log2(RPKM)
z−score of IF value
−3 −2 −1 0 1 2

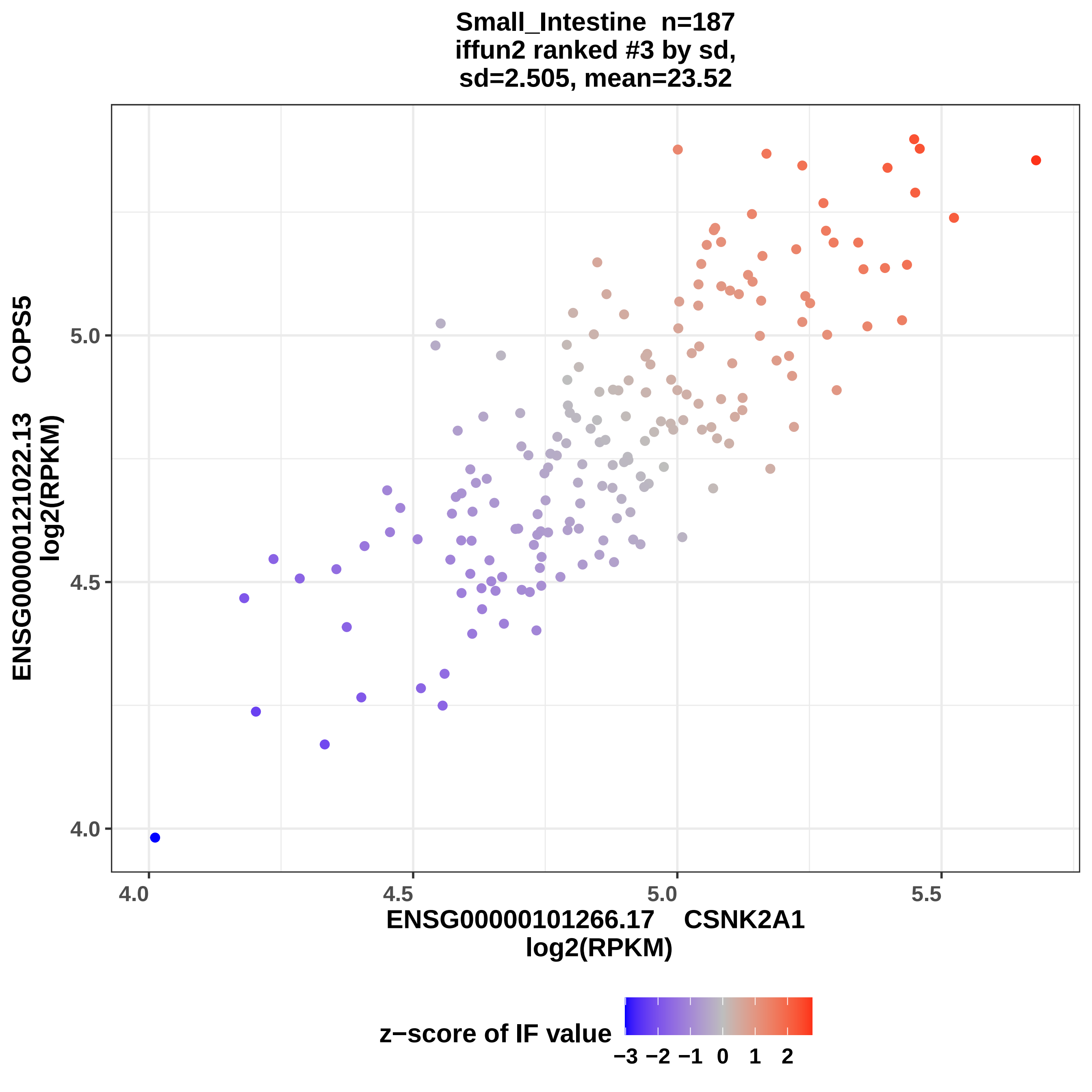

Small_Intestine n=187
iffun2 ranked #3 by sd,
sd=2.505, mean=23.52
ENSG00000121022.13 COPS5
log2(RPKM)
5.0
4.5
4.0
4.0
4.5
5.0
5.5
ENSG00000101266.17 CSNK2A1
log2(RPKM)
z−score of IF value
−3
−2
−1
0
1
2

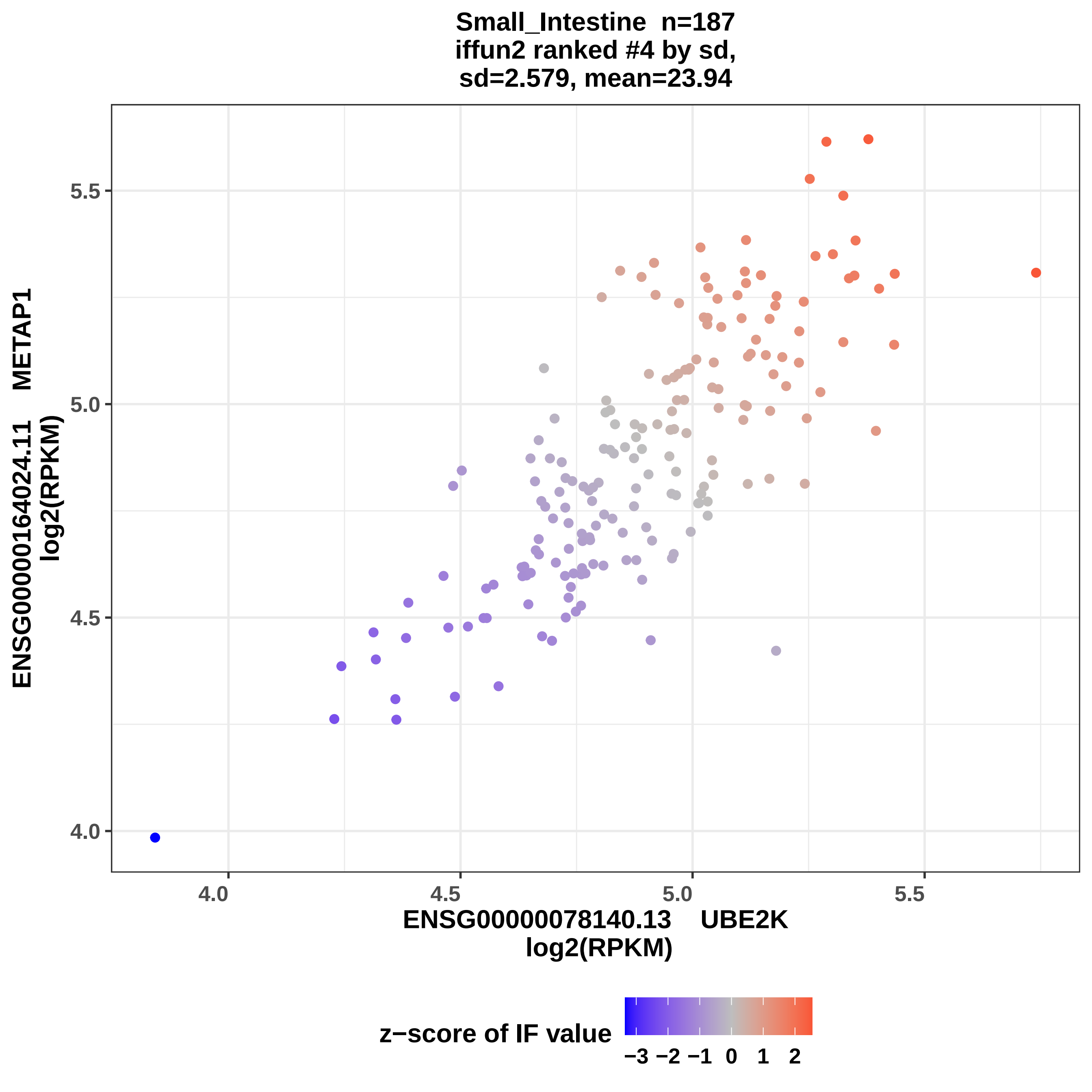

Small_Intestine n=187
iffun2 ranked #4 by sd,
sd=2.579, mean=23.94
ENSG00000164024.11 METAP1
log2(RPKM)
5.5
5.0
4.5
4.0
4.0
4.5
5.0
5.5
ENSG00000078140.13 UBE2K
log2(RPKM)
z−score of IF value
−3 −2 −1 0 1 2

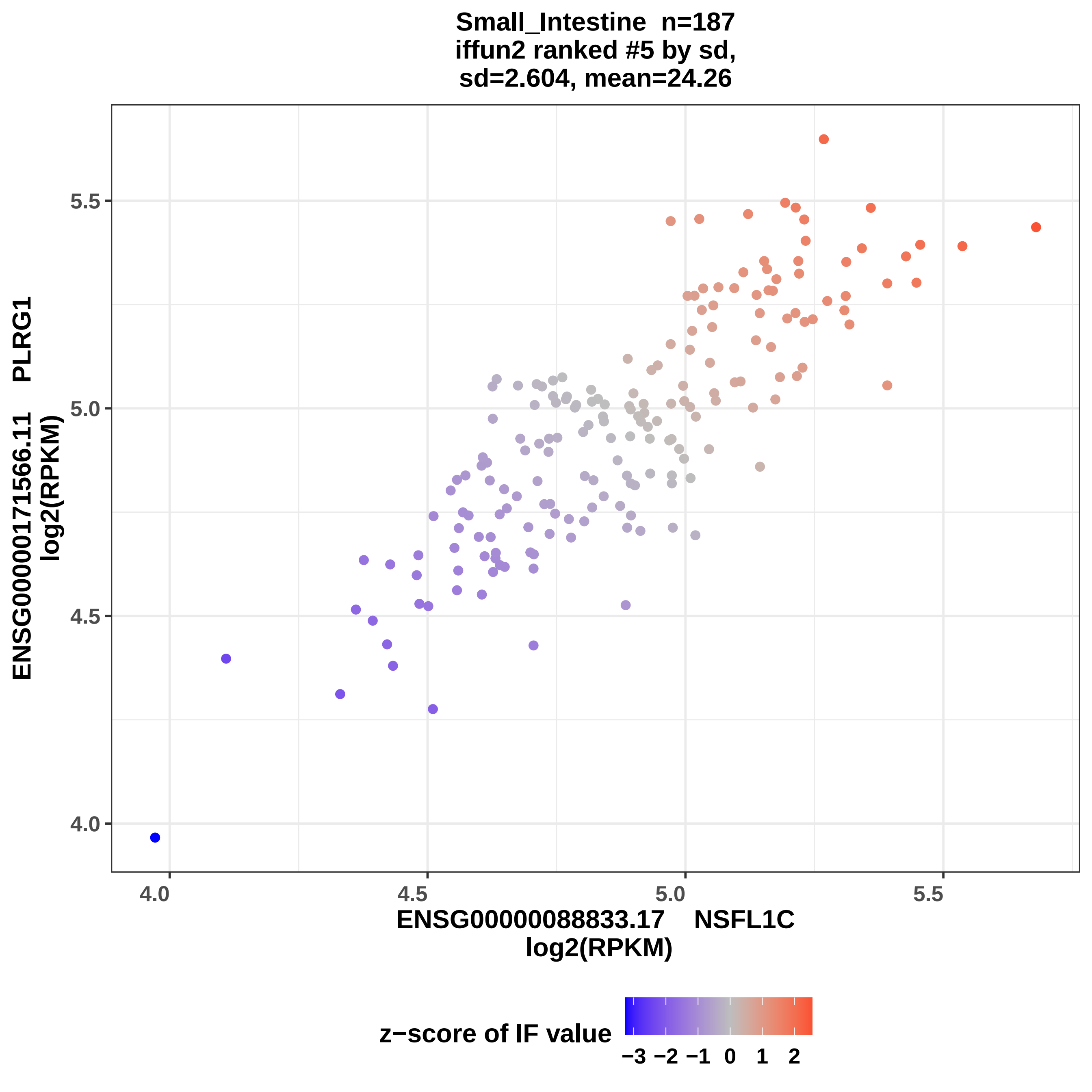

Small_Intestine n=187
iffun2 ranked #5 by sd,
sd=2.604, mean=24.26
ENSG00000171566.11 PLRG1
log2(RPKM)
5.5
5.0
4.5
4.0
4.0
4.5
5.0
5.5
ENSG00000088833.17 NSFL1C
log2(RPKM)
z−score of IF value
−3 −2 −1 0 1 2

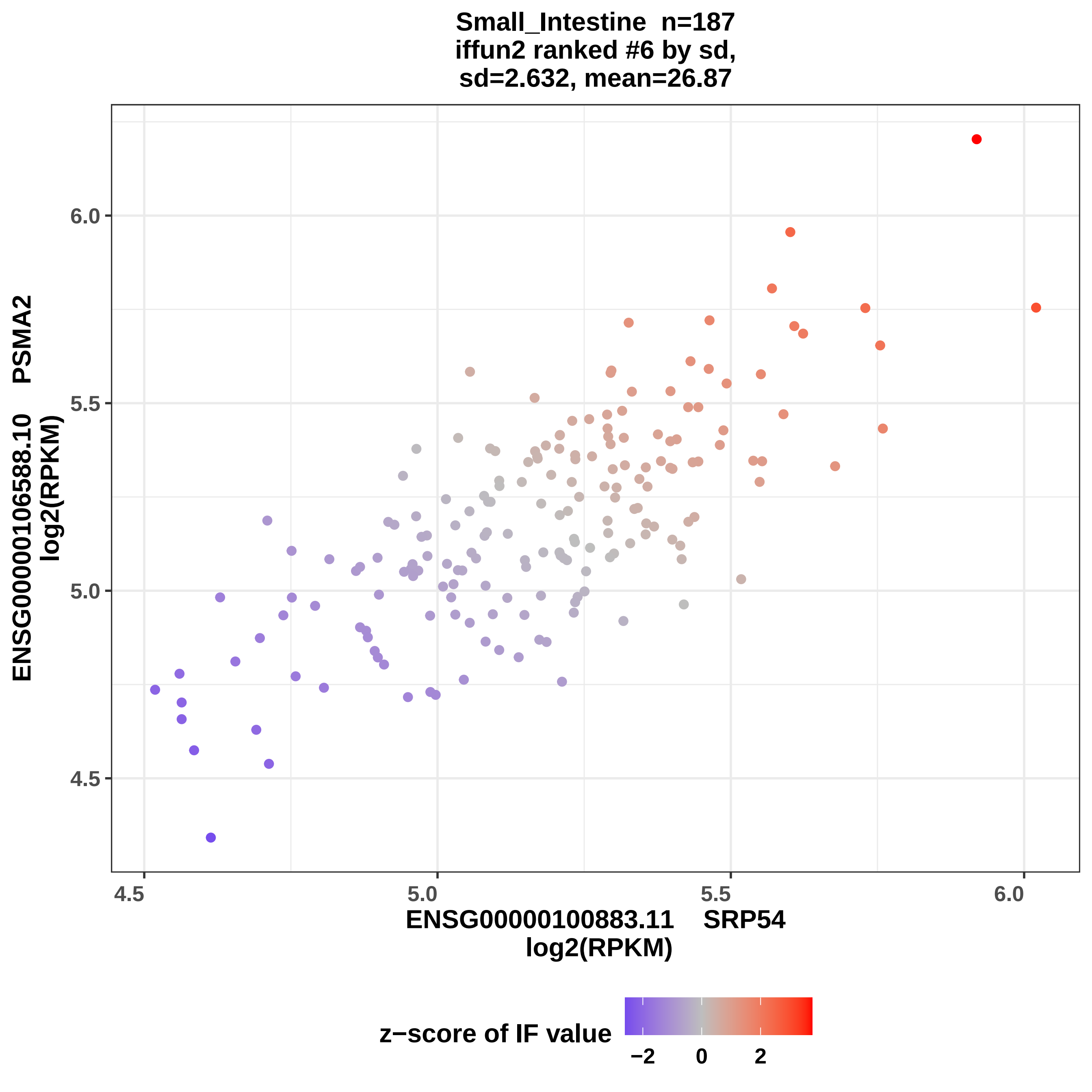
Small_Intestine n=187
iffun2 ranked #6 by sd,
sd=2.632, mean=26.87
ENSG00000106588.10 PSMA2
log2(RPKM)
6.0
5.5
5.0
4.5
4.5
5.0
5.5
6.0
ENSG00000100883.11 SRP54
log2(RPKM)
z−score of IF value
−2
0
2

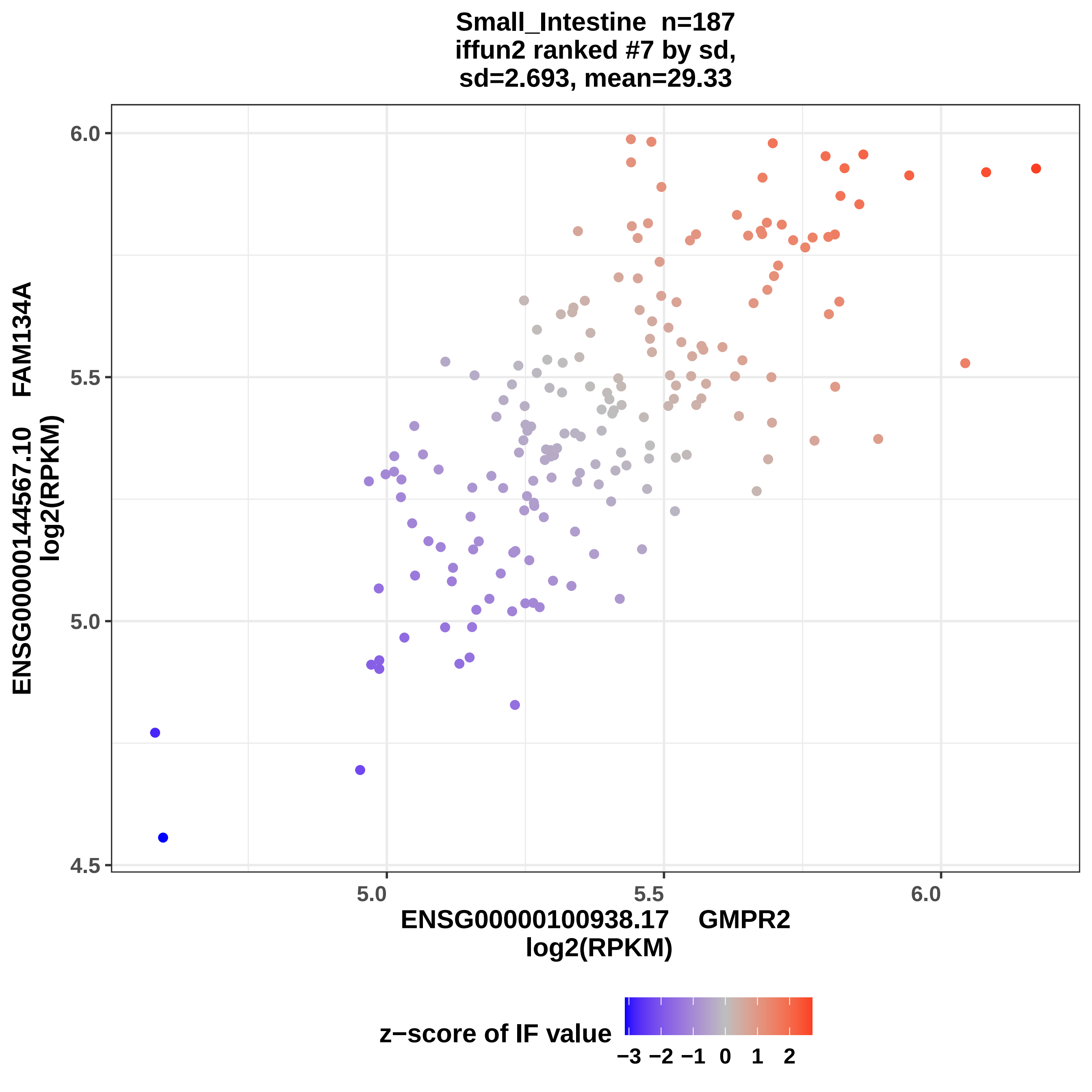
Small_Intestine n=187
iffun2 ranked #7 by sd,
sd=2.693, mean=29.33
ENSG00000144567.10 FAM134A
log2(RPKM)
6.0
5.5
5.0
4.5
5.0
5.5
6.0
ENSG00000100938.17 GMPR2
log2(RPKM)
z−score of IF value
−3 −2 −1 0 1 2

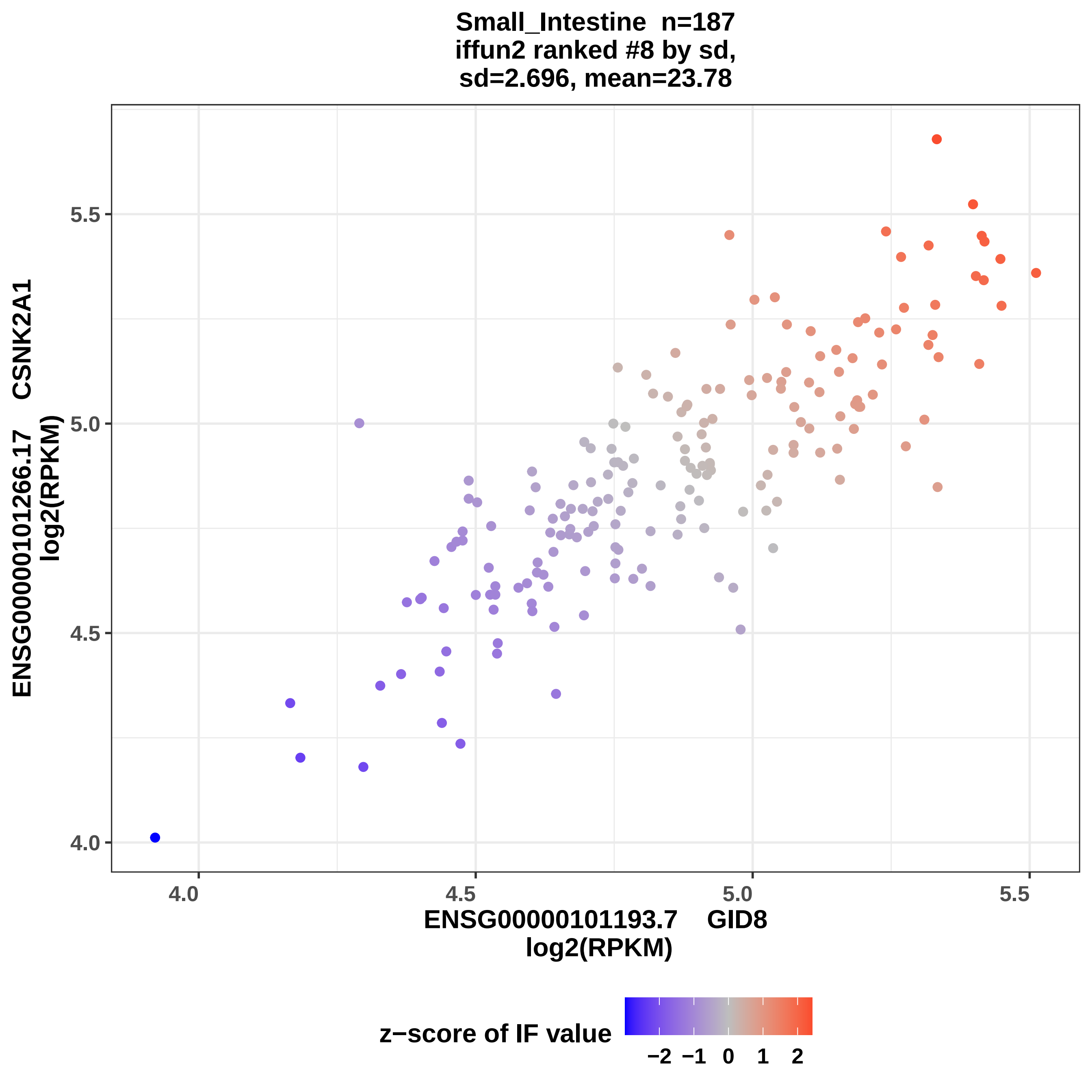
Small_Intestine n=187
iffun2 ranked #8 by sd,
sd=2.696, mean=23.78
ENSG00000101266.17 CSNK2A1
log2(RPKM)
ENSG00000101193.7 GID8
log2(RPKM)
4.0
4.5
5.0
5.5
z−score of IF value
−2
−1
0
1
2

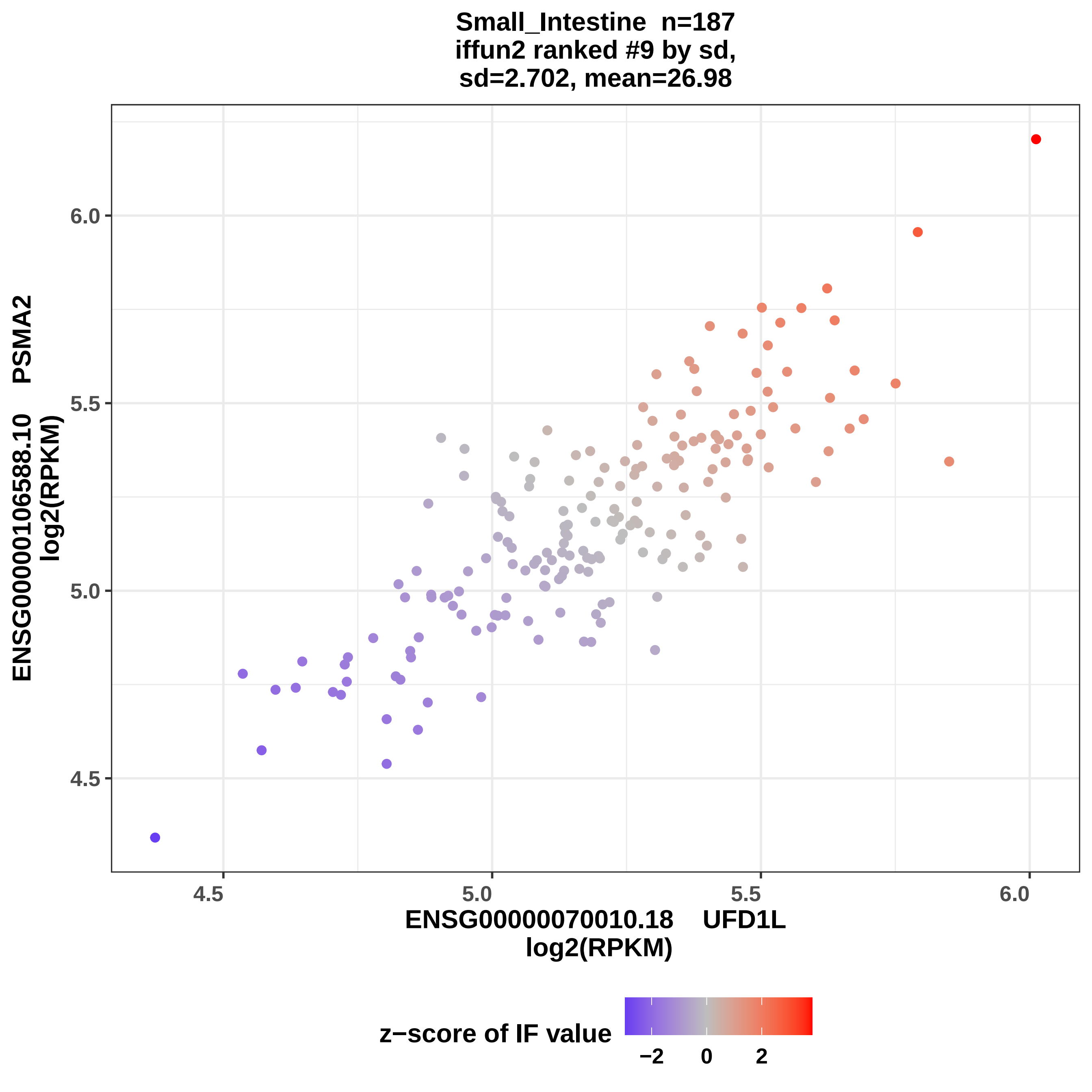
Small_Intestine n=187
iffun2 ranked #9 by sd,
sd=2.702, mean=26.98
ENSG00000106588.10 PSMA2
log2(RPKM)
6.0
5.5
5.0
4.5
4.5
5.0
5.5
6.0
ENSG00000070010.18 UFD1L
log2(RPKM)
z−score of IF value
−2
0
2

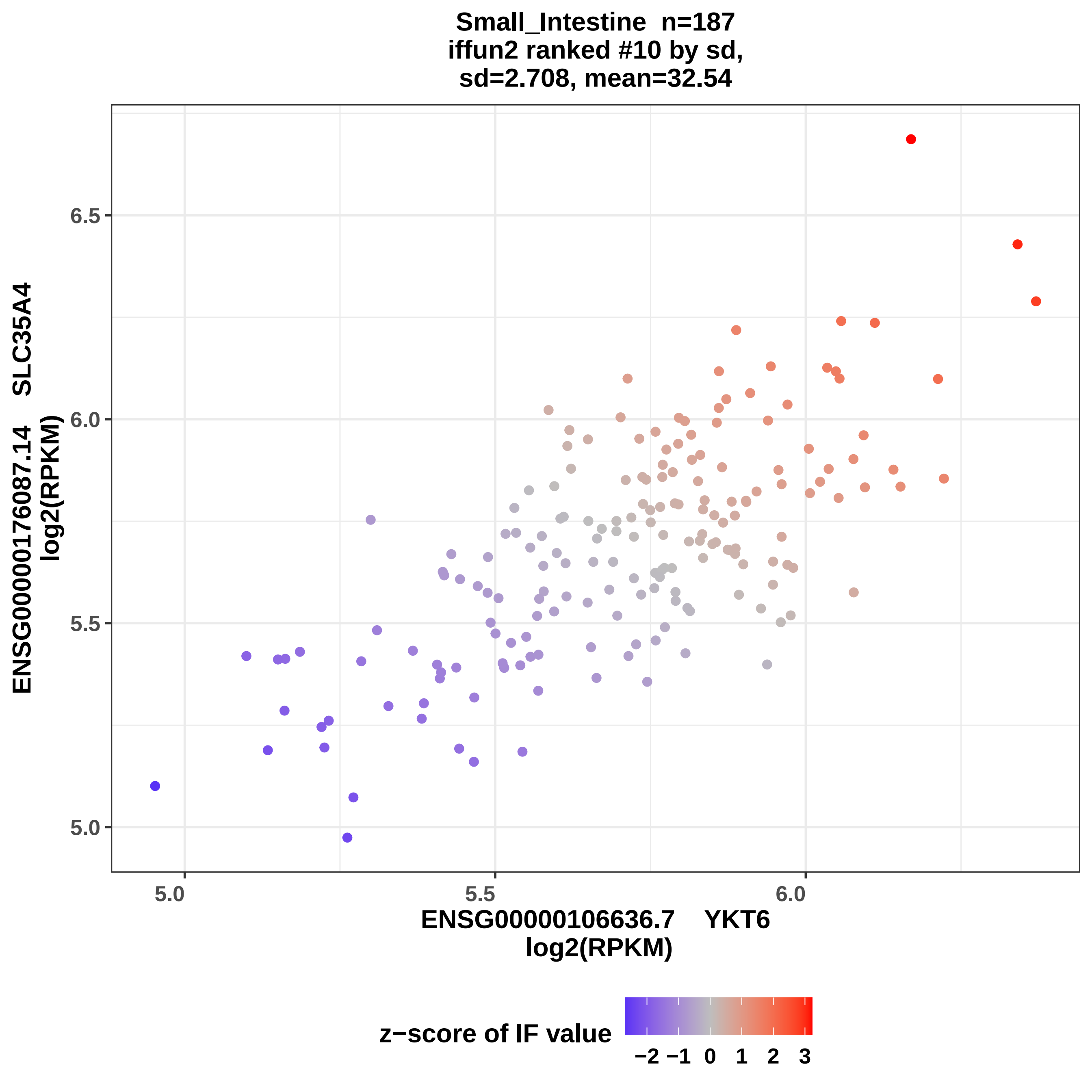
Small_Intestine n=187
iffun2 ranked #10 by sd,
sd=2.708, mean=32.54
ENSG00000176087.14 SLC35A4
log2(RPKM)
6.5
6.0
5.5
5.0
5.0
5.5
6.0
ENSG00000106636.7 YKT6
log2(RPKM)
z−score of IF value
−2 −1 0 1 2 3

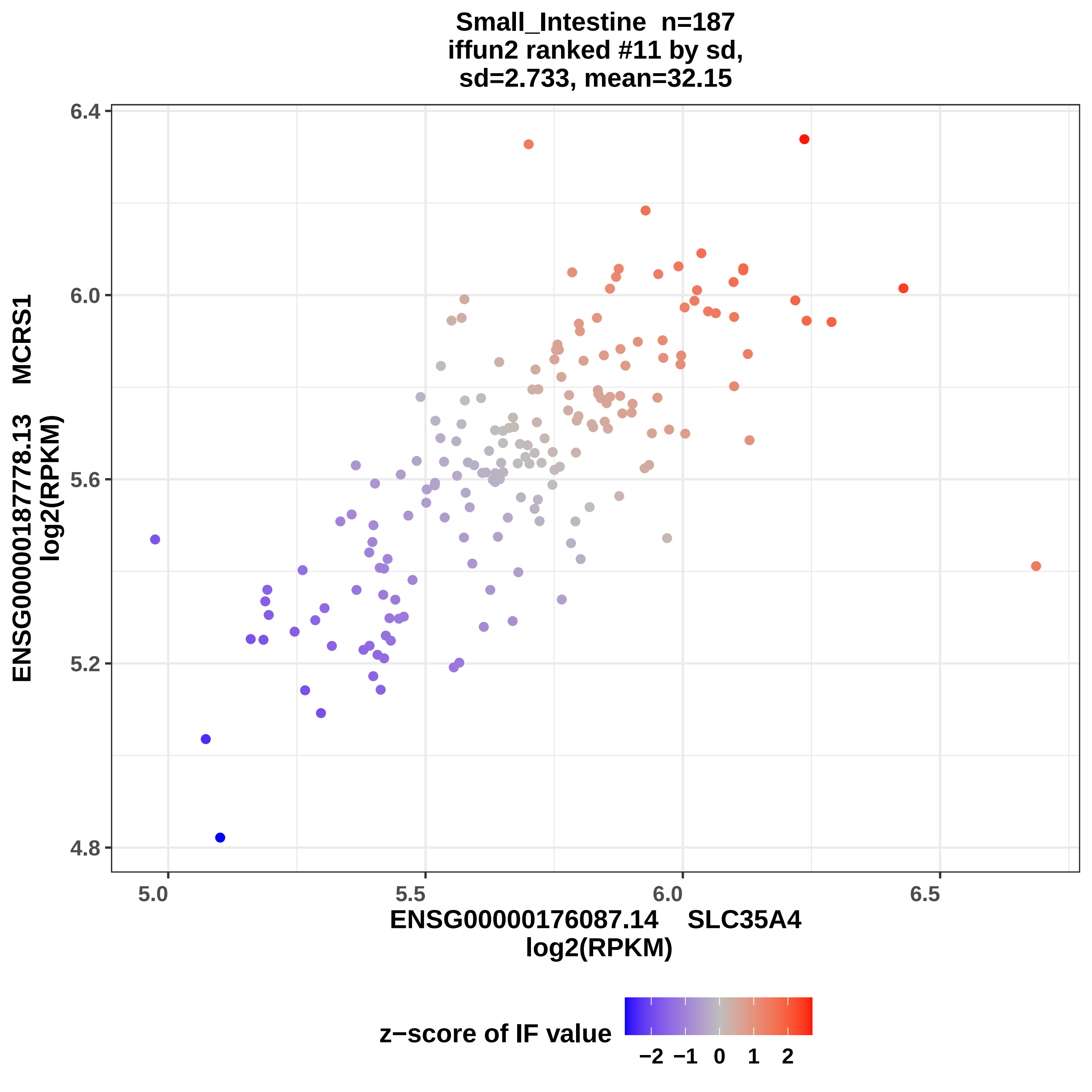
Small_Intestine n=187
iffun2 ranked #11 by sd,
sd=2.733, mean=32.15
ENSG00000187778.13 MCRS1
log2(RPKM)
ENSG00000176087.14 SLC35A4
log2(RPKM)
6.4
6.0
5.6
5.2
4.8
5.0
5.5
6.0
6.5
z−score of IF value
−2 −1 0 1 2

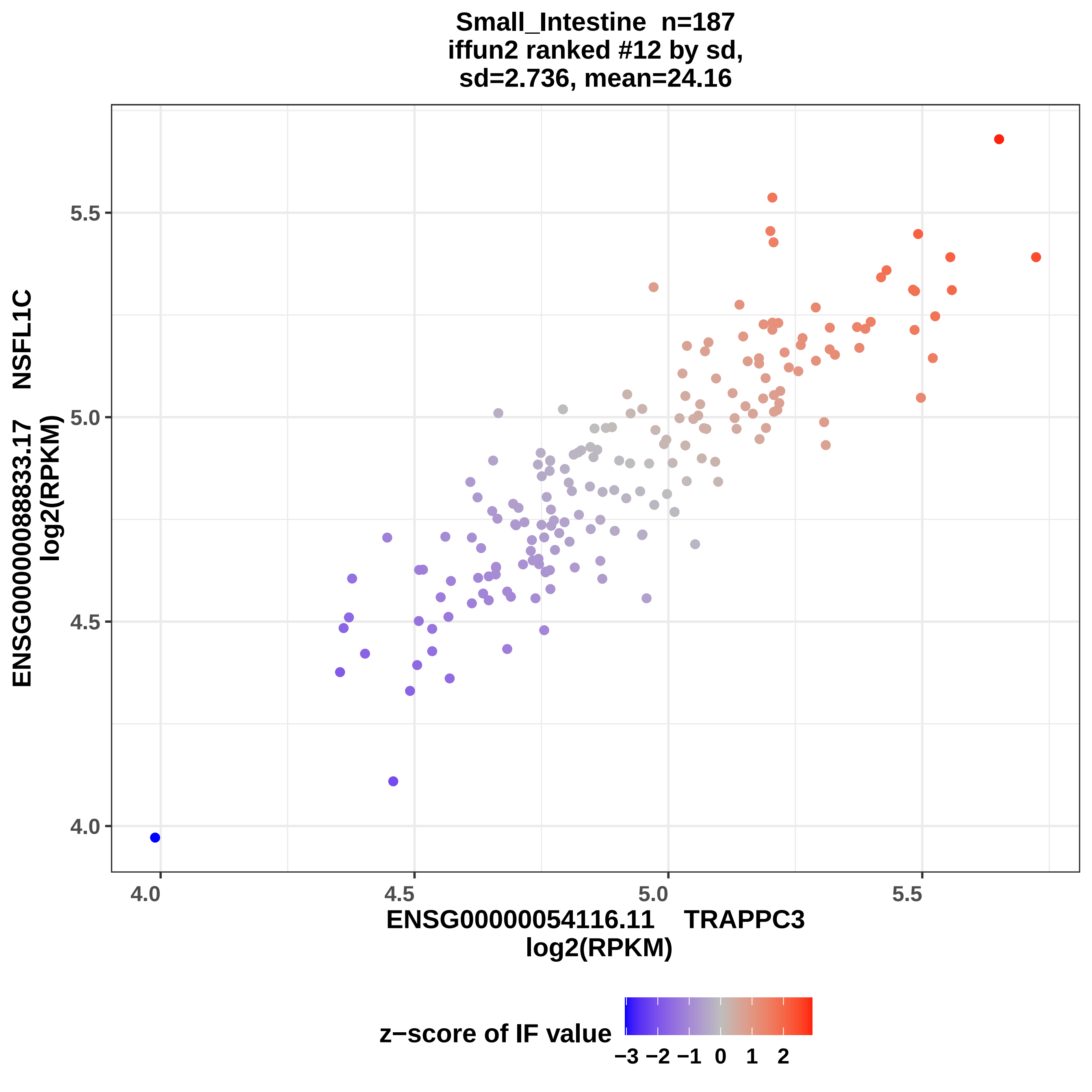

Small_Intestine n=187
iffun2 ranked #12 by sd,
sd=2.736, mean=24.16
ENSG00000088833.17 NSFL1C
log2(RPKM)
5.5
5.0
4.5
4.0
4.0
4.5
5.0
5.5
ENSG00000054116.11 TRAPPC3
log2(RPKM)
z−score of IF value
−3 −2 −1 0 1 2

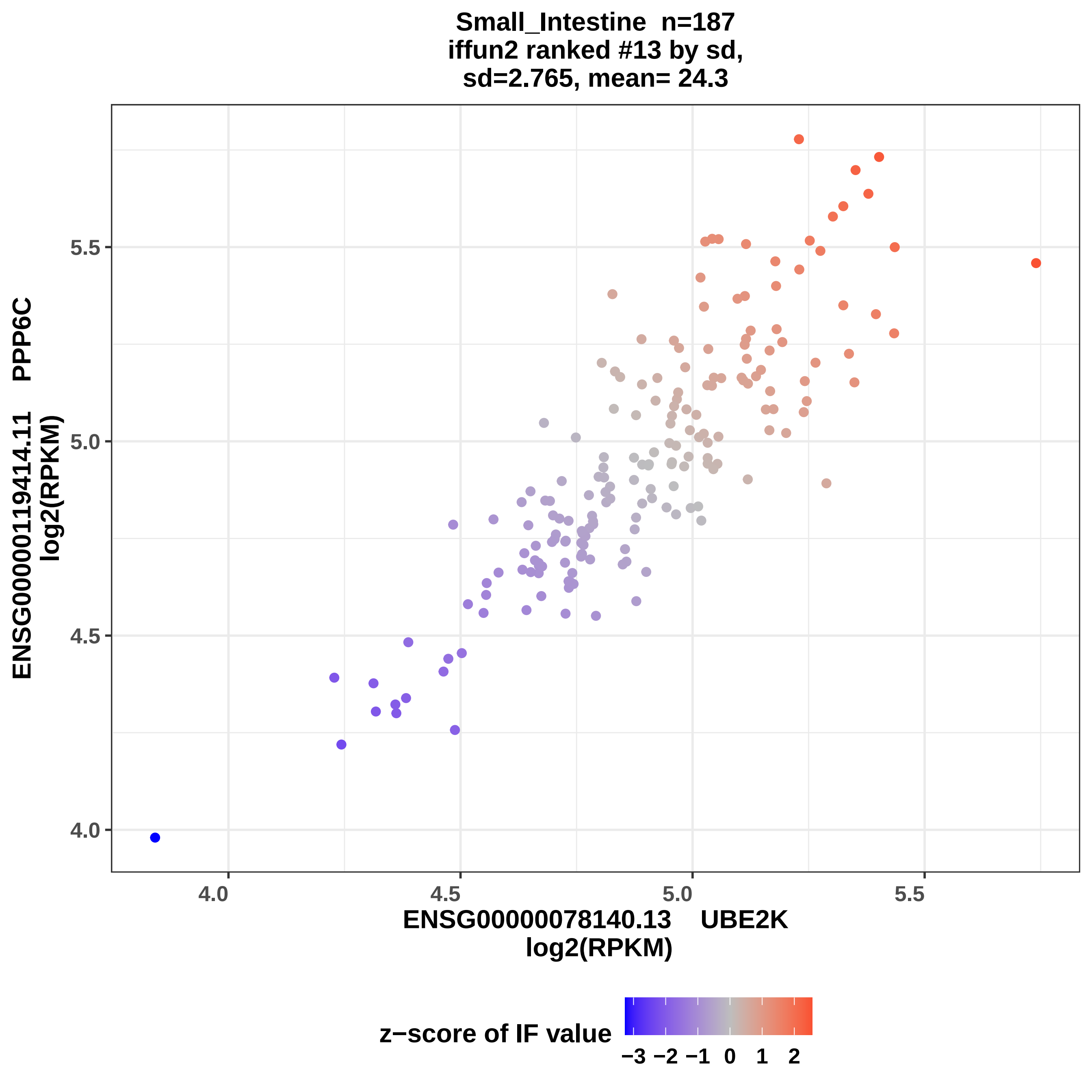
Small_Intestine n=187
iffun2 ranked #13 by sd,
sd=2.765, mean= 24.3
ENSG00000119414.11 PPP6C
log2(RPKM)
4.0
4.5
5.0
5.5
ENSG00000078140.13 UBE2K
log2(RPKM)
z−score of IF value
−3 −2 −1 0 1 2

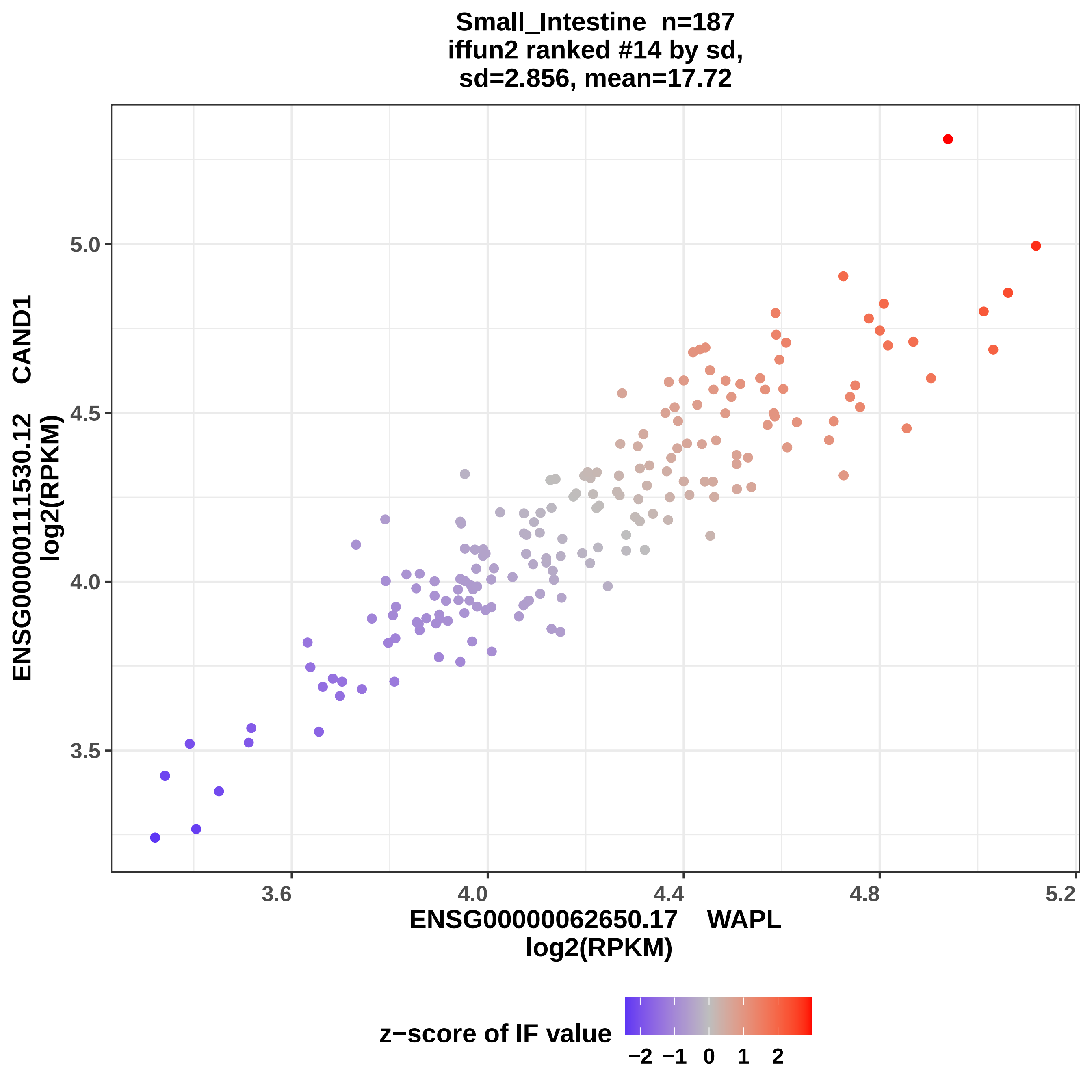

Small_Intestine n=187
iffun2 ranked #14 by sd,
sd=2.856, mean=17.72
ENSG00000111530.12 CAND1
log2(RPKM)
5.0
4.5
4.0
3.5
3.6
4.0
4.4
4.8
5.2
ENSG00000062650.17 WAPL
log2(RPKM)
z−score of IF value
−2
−1
0
1
2

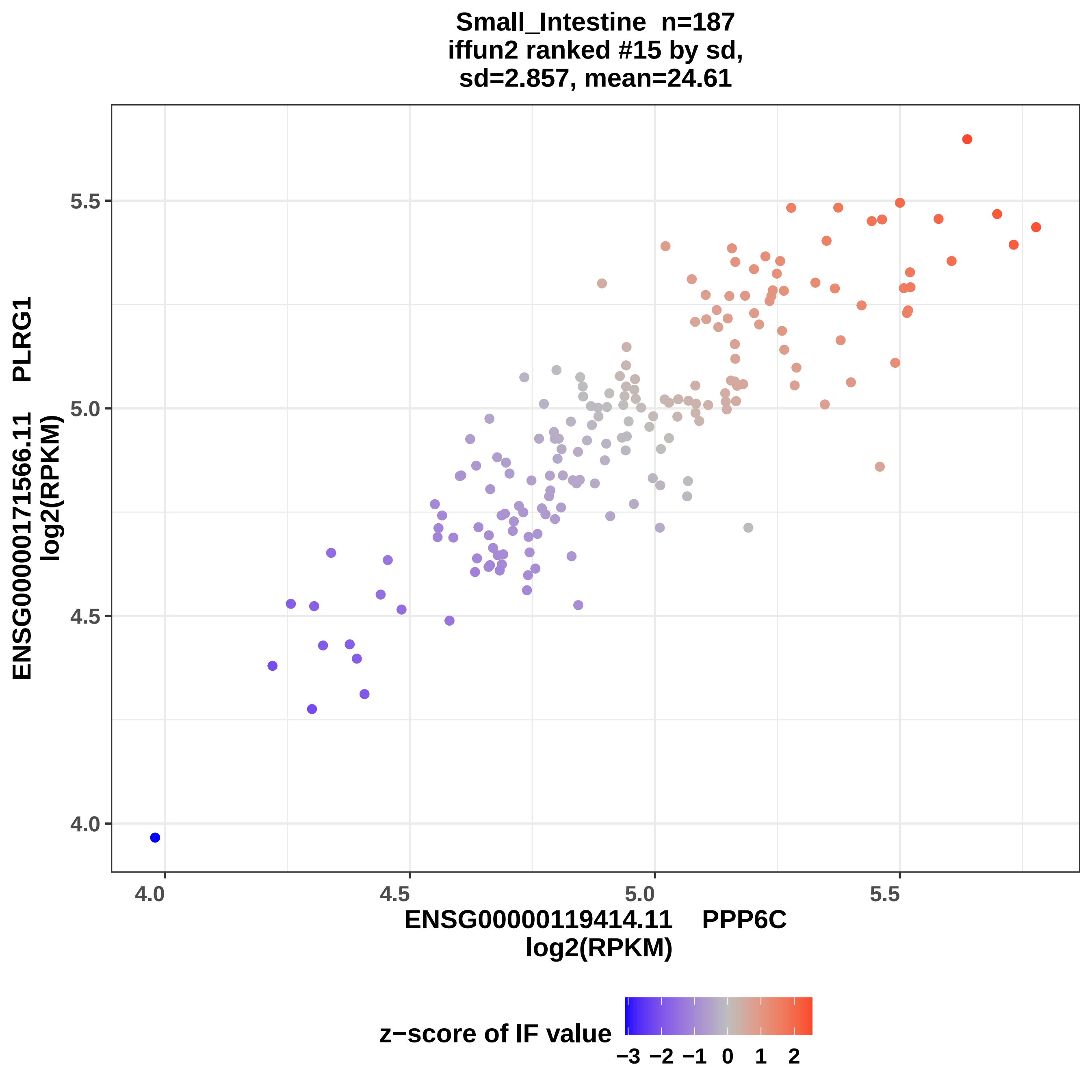

Small_Intestine n=187
iffun2 ranked #15 by sd,
sd=2.857, mean=24.61
ENSG00000171566.11 PLRG1
log2(RPKM)
5.5
5.0
4.5
4.0
4.0
4.5
5.0
5.5
ENSG00000119414.11 PPP6C
log2(RPKM)
z−score of IF value
−3 −2 −1 0 1 2

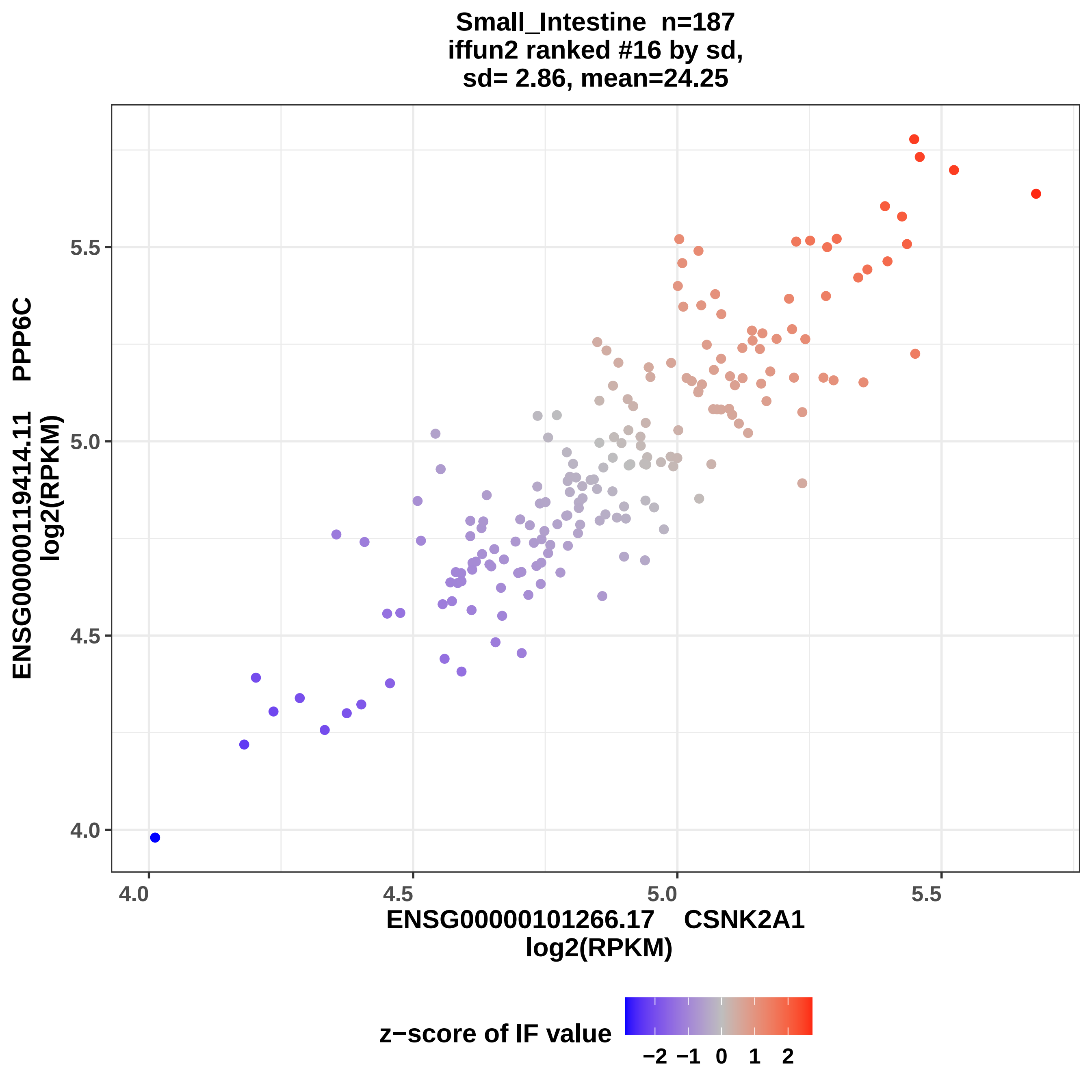

Small_Intestine n=187
iffun2 ranked #16 by sd,
sd= 2.86, mean=24.25
ENSG00000119414.11 PPP6C
log2(RPKM)
4.0
4.5
5.0
5.5
ENSG00000101266.17 CSNK2A1
log2(RPKM)
z−score of IF value
−2
−1
0
1
2

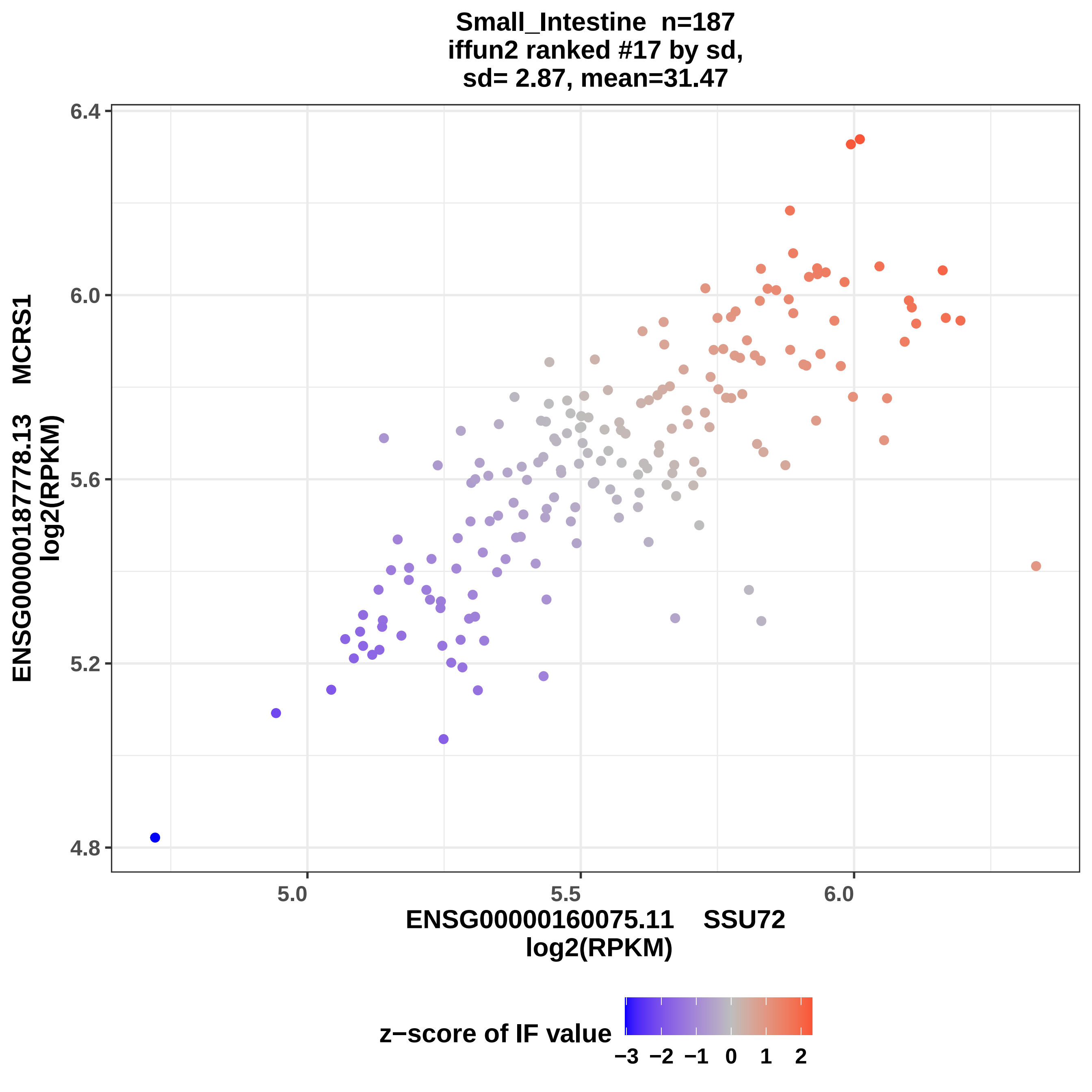

Small_Intestine n=187
iffun2 ranked #17 by sd,
sd= 2.87, mean=31.47
ENSG00000187778.13 MCRS1
log2(RPKM)
6.4
6.0
5.6
5.2
4.8
5.0
5.5
6.0
ENSG00000160075.11 SSU72
log2(RPKM)
z−score of IF value
−3 −2 −1 0 1 2

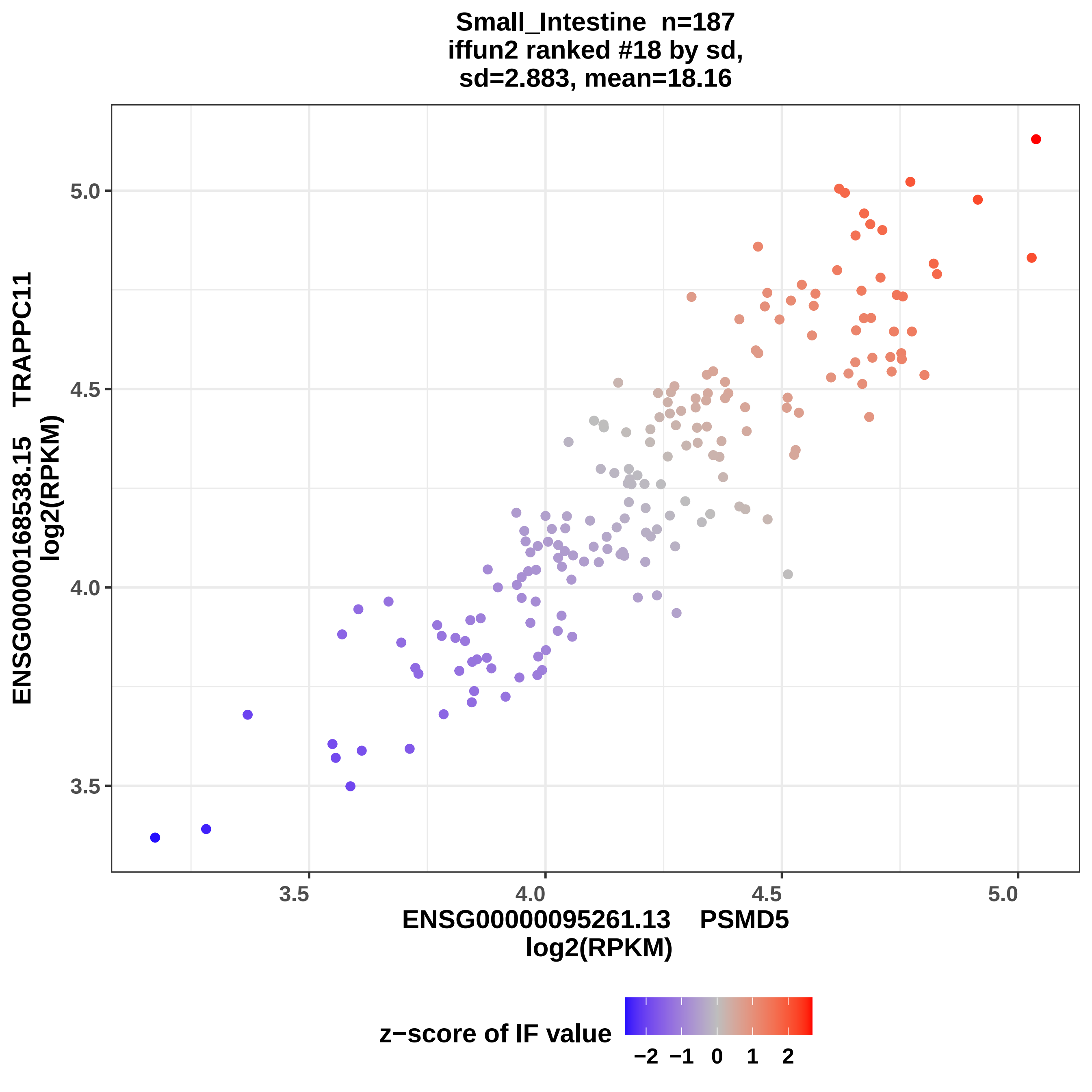
Small_Intestine n=187
iffun2 ranked #18 by sd,
sd=2.883, mean=18.16
ENSG00000168538.15 TRAPPC11
log2(RPKM)
5.0
4.5
4.0
3.5
3.5
4.0
4.5
5.0
ENSG00000095261.13 PSMD5
log2(RPKM)
z−score of IF value
−2
−1
0
1
2

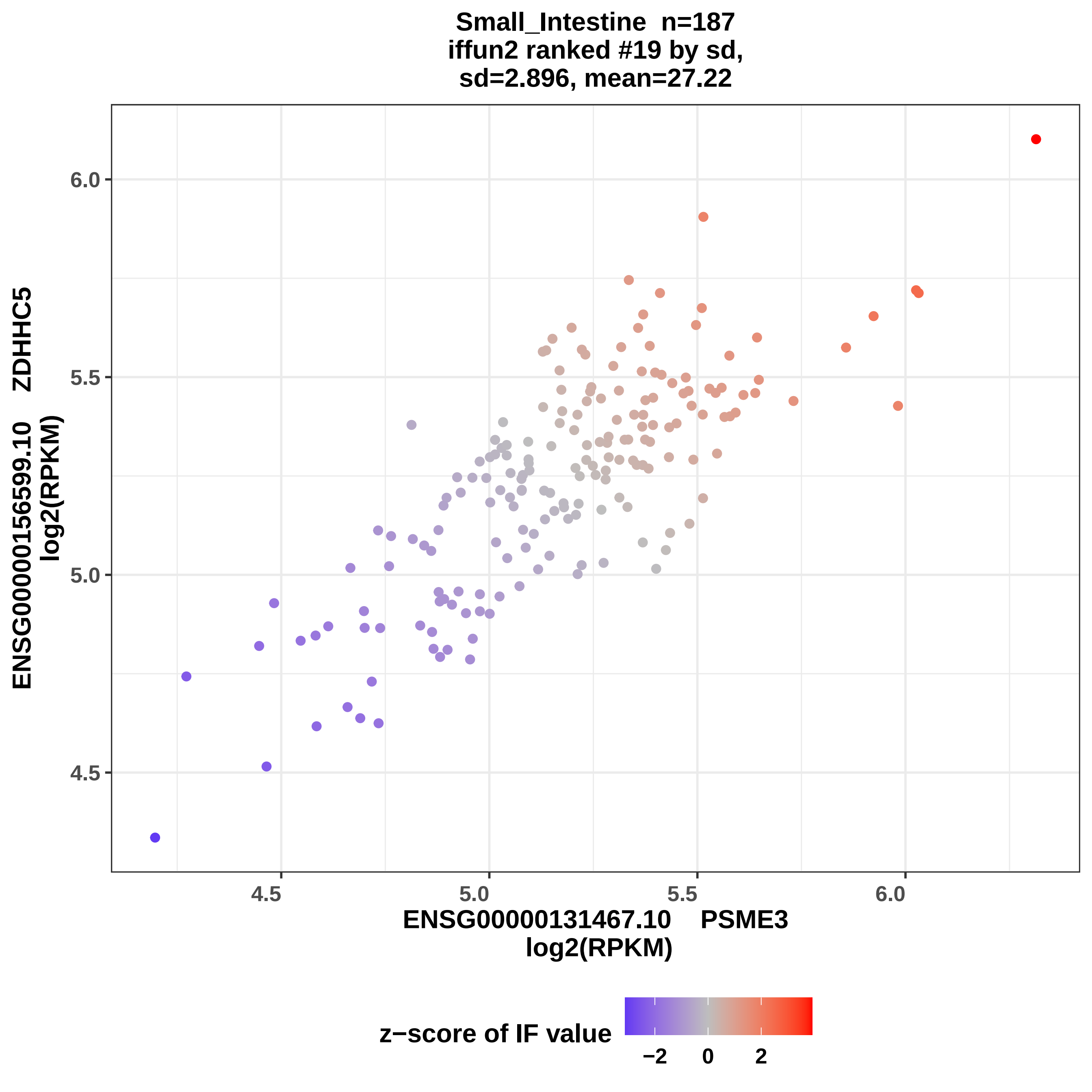
Small_Intestine n=187
iffun2 ranked #19 by sd,
sd=2.896, mean=27.22
ENSG00000156599.10 ZDHHC5
log2(RPKM)
ENSG00000131467.10 PSME3
log2(RPKM)
6.0
5.5
5.0
4.5
4.5
5.0
5.5
6.0
z−score of IF value
−2
0
2

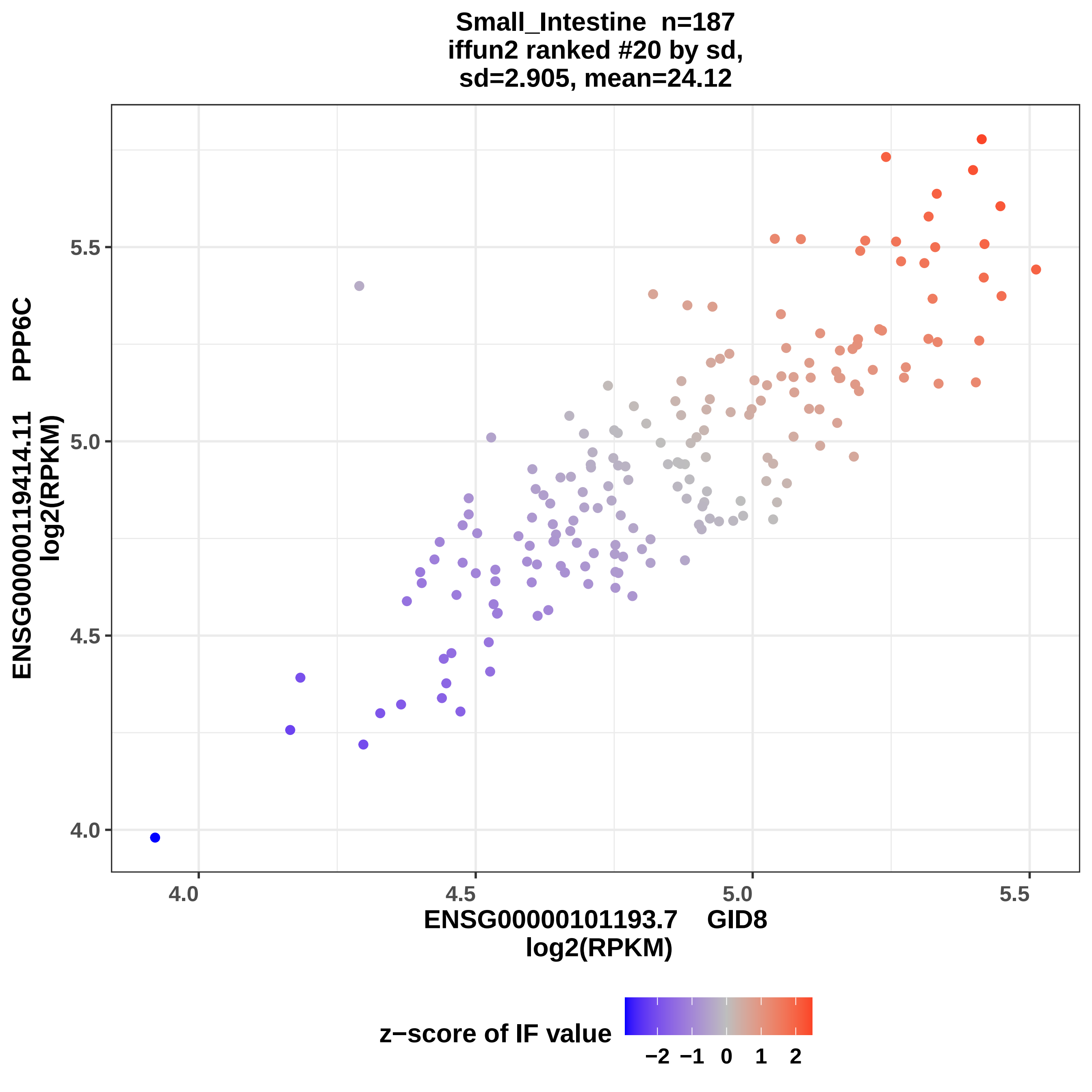
Small_Intestine n=187
iffun2 ranked #20 by sd,
sd=2.905, mean=24.12
ENSG00000119414.11 PPP6C
log2(RPKM)
5.5
5.0
4.5
4.0
4.0
4.5
5.0
5.5
ENSG00000101193.7 GID8
log2(RPKM)
z−score of IF value
−2 −1 0 1 2

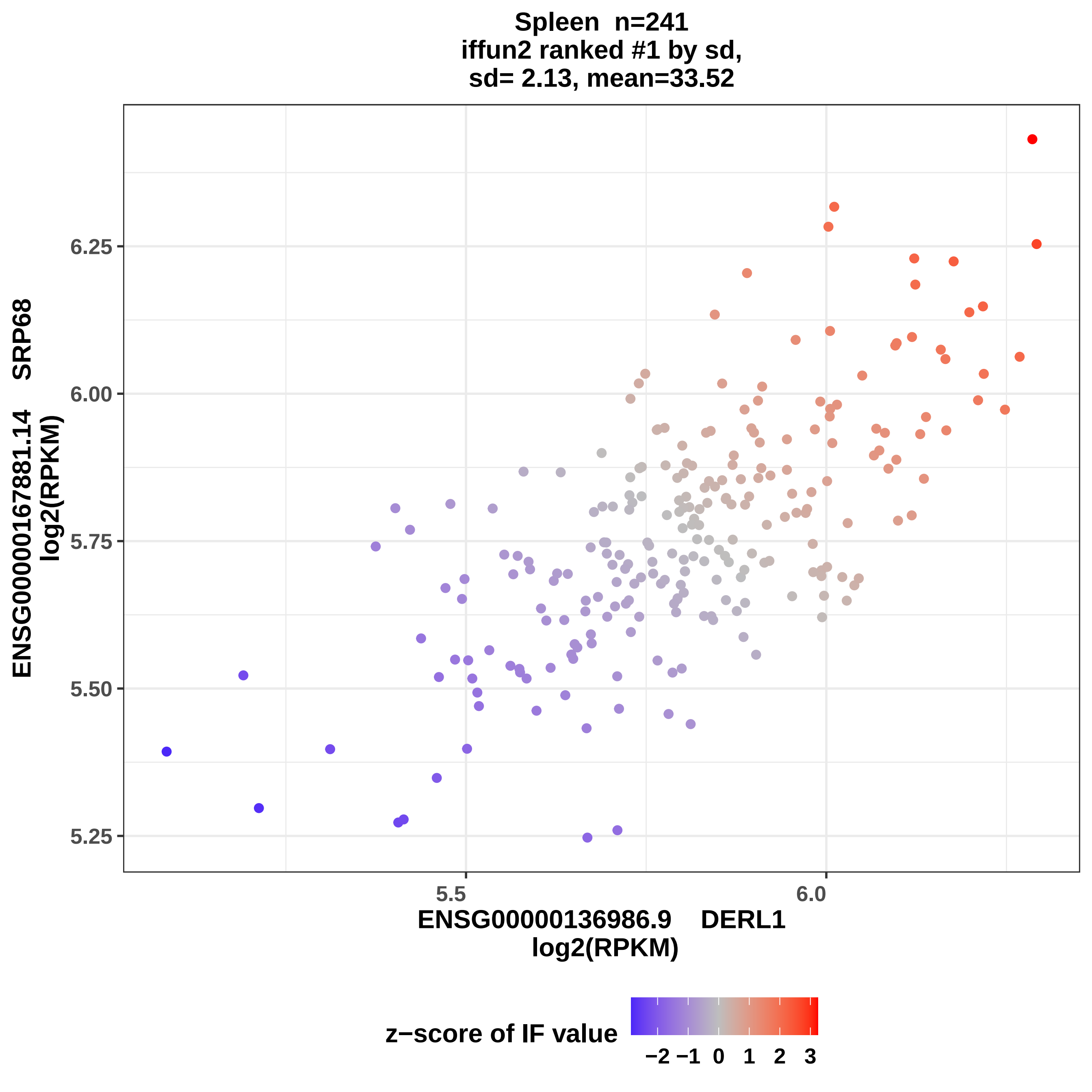
Spleen n=241
iffun2 ranked #1 by sd,
sd= 2.13, mean=33.52
ENSG00000167881.14 SRP68
log2(RPKM)
6.25
6.00
5.75
5.50
5.25
5.5
6.0
ENSG00000136986.9 DERL1
log2(RPKM)
z−score of IF value
−2 −1 0 1 2 3

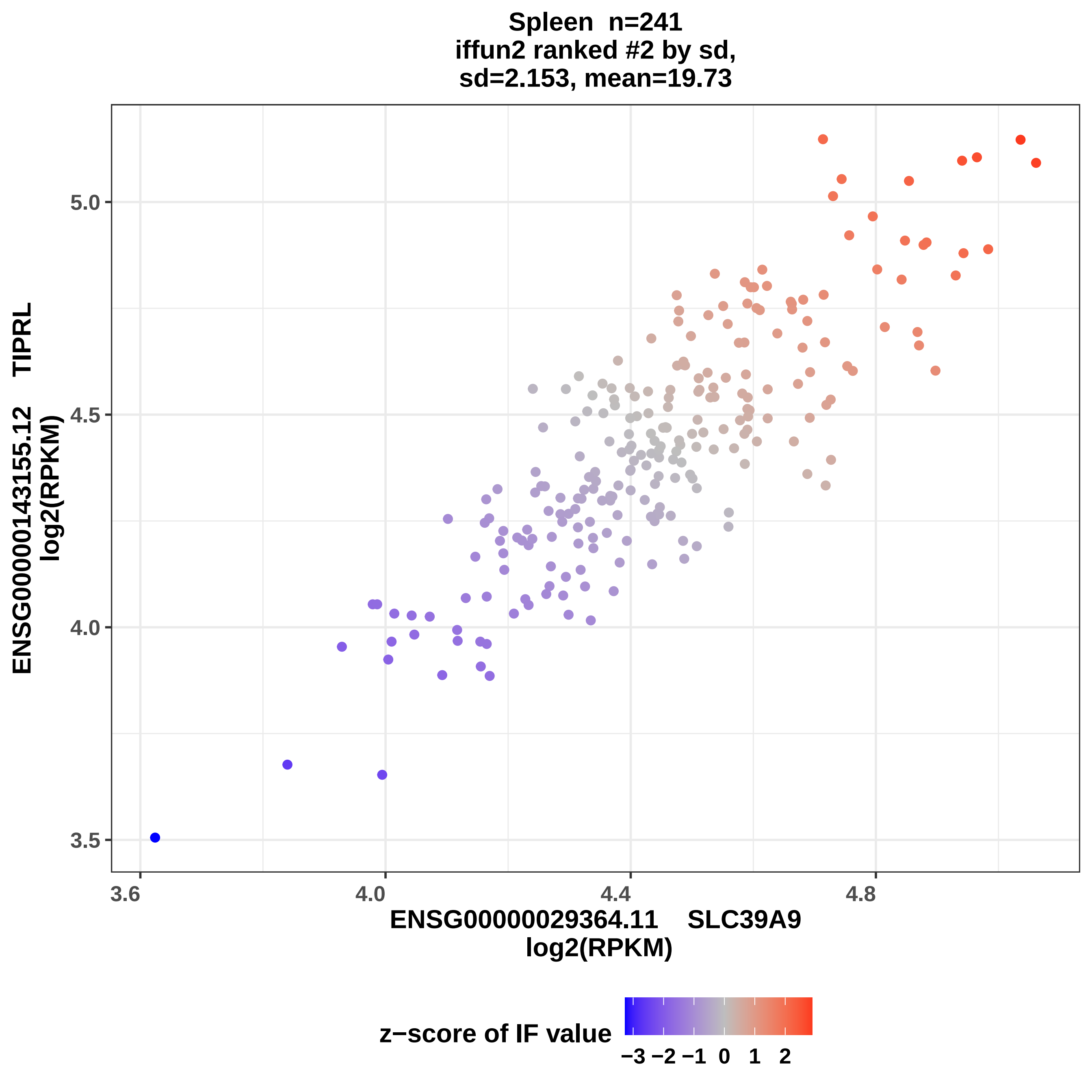

Spleen n=241
iffun2 ranked #2 by sd,
sd=2.153, mean=19.73
ENSG00000143155.12 TIPRL
log2(RPKM)
5.0
4.5
4.0
3.5
3.6
4.0
4.4
4.8
ENSG00000029364.11 SLC39A9
log2(RPKM)
z−score of IF value
−3 −2 −1 0 1 2

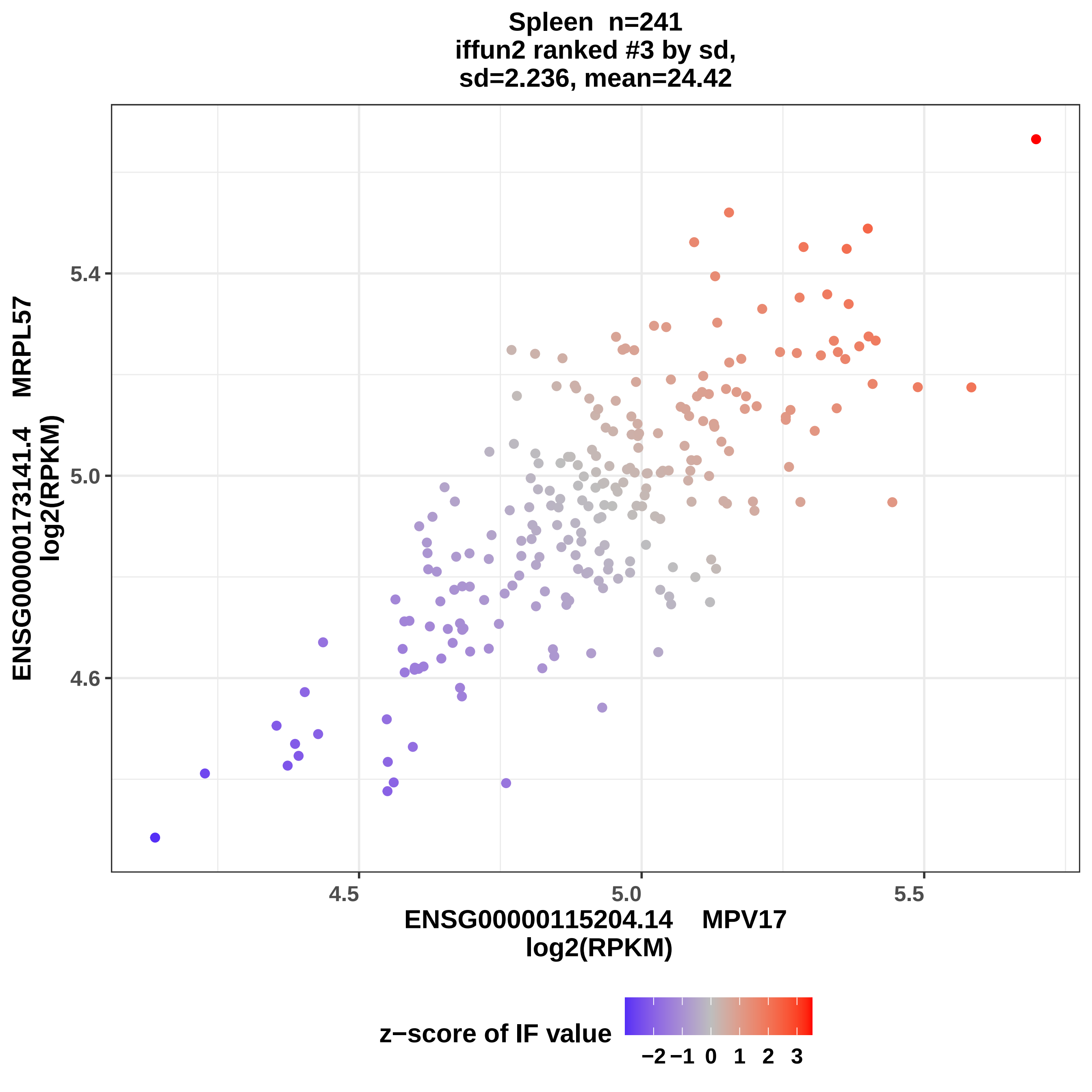

Spleen n=241
iffun2 ranked #3 by sd,
sd=2.236, mean=24.42
ENSG00000173141.4 MRPL57
log2(RPKM)
5.4
5.0
4.6
4.5
5.0
5.5
ENSG00000115204.14 MPV17
log2(RPKM)
z−score of IF value
−2 −1 0 1 2 3

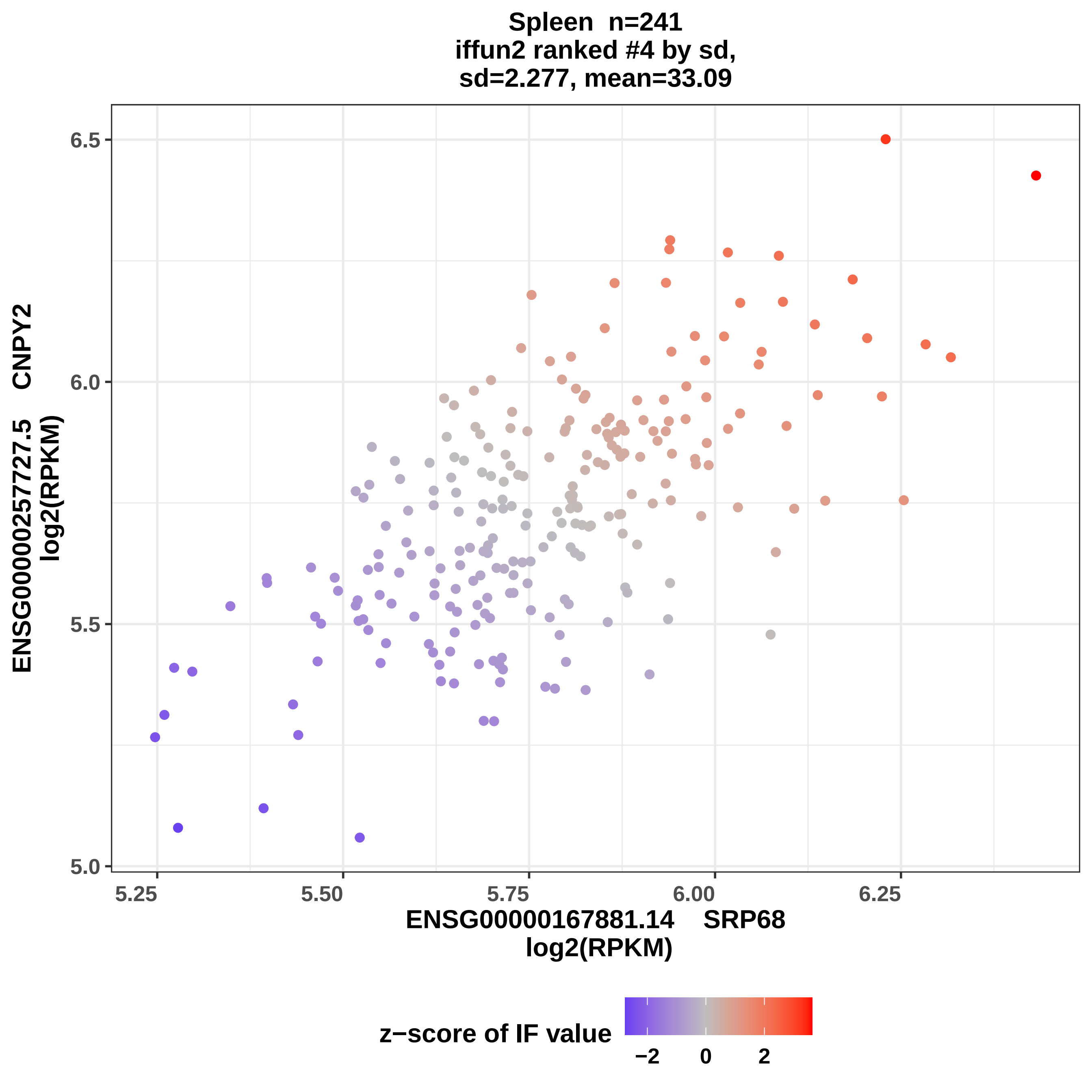
Spleen n=241
iffun2 ranked #4 by sd,
sd=2.277, mean=33.09
ENSG00000257727.5 CNPY2
log2(RPKM)
6.5
6.0
5.5
5.0
5.25
5.50
5.75
6.00
6.25
ENSG00000167881.14 SRP68
log2(RPKM)
z−score of IF value
−2
0
2

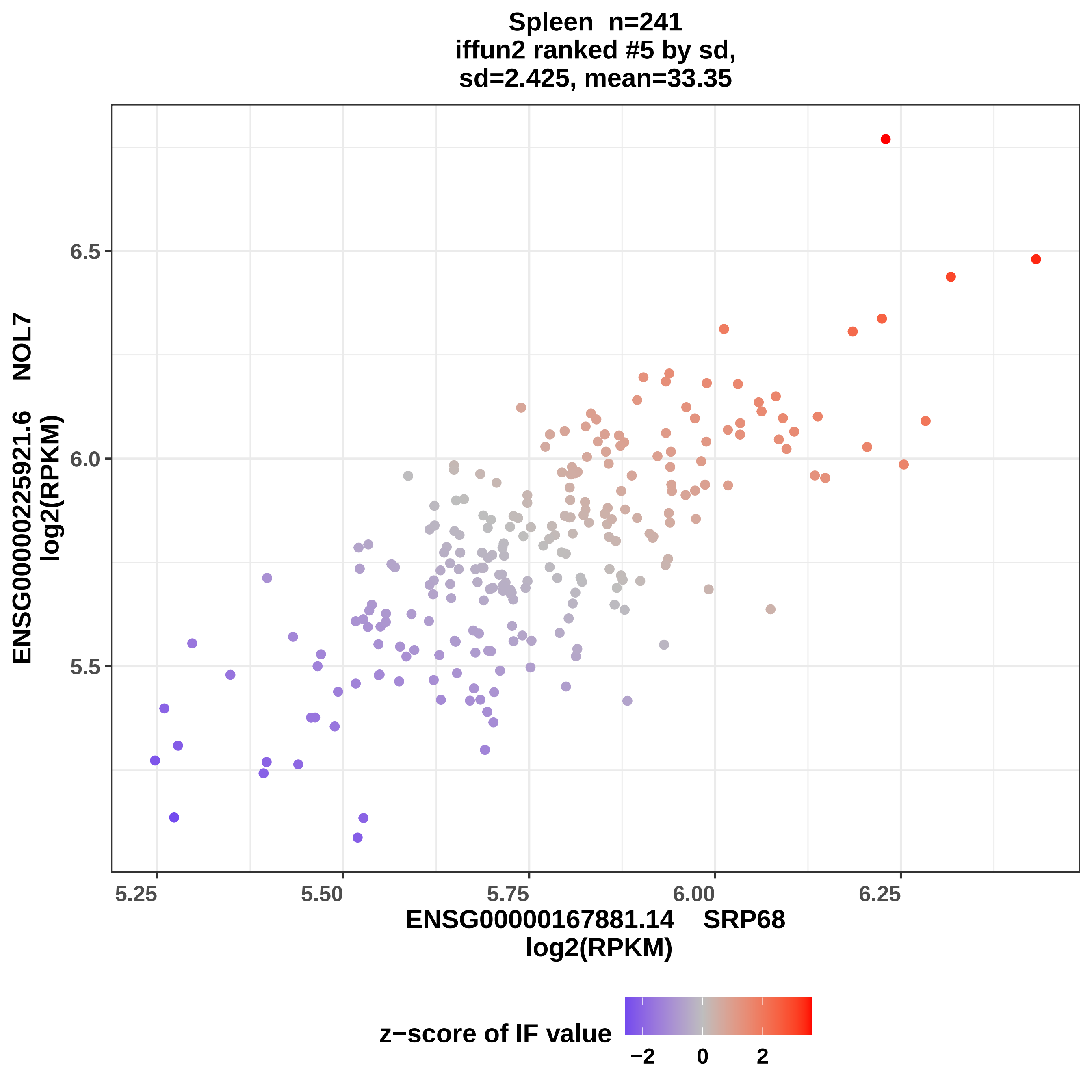

Spleen n=241
iffun2 ranked #5 by sd,
sd=2.425, mean=33.35
ENSG00000225921.6 NOL7
log2(RPKM)
6.5
6.0
5.5
5.25
5.50
5.75
6.00
6.25
ENSG00000167881.14 SRP68
log2(RPKM)
z−score of IF value
−2
0
2

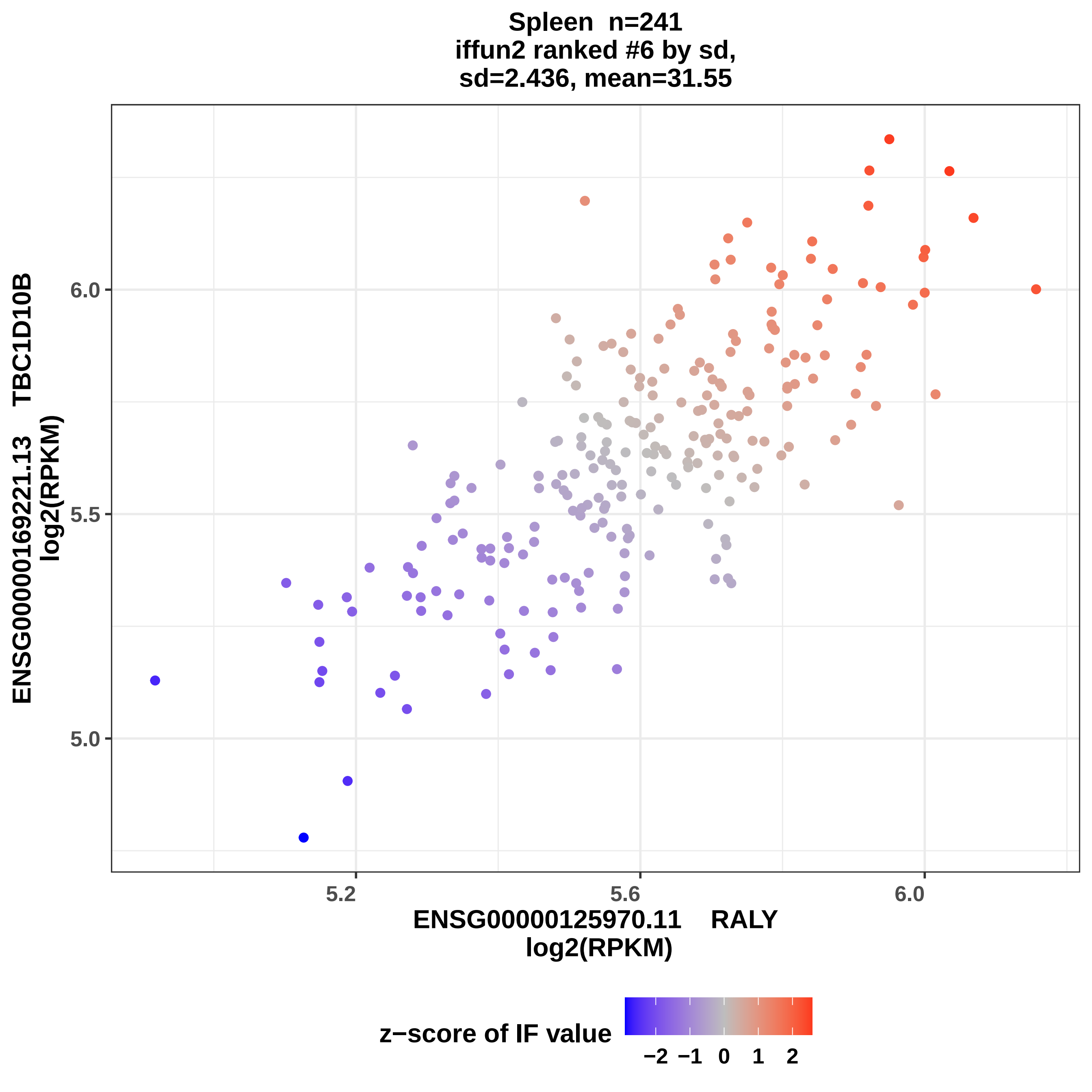

Spleen n=241
iffun2 ranked #6 by sd,
sd=2.436, mean=31.55
ENSG00000169221.13 TBC1D10B
log2(RPKM)
6.0
5.5
5.0
5.2
5.6
6.0
ENSG00000125970.11 RALY
log2(RPKM)
z−score of IF value
−2 −1 0 1 2

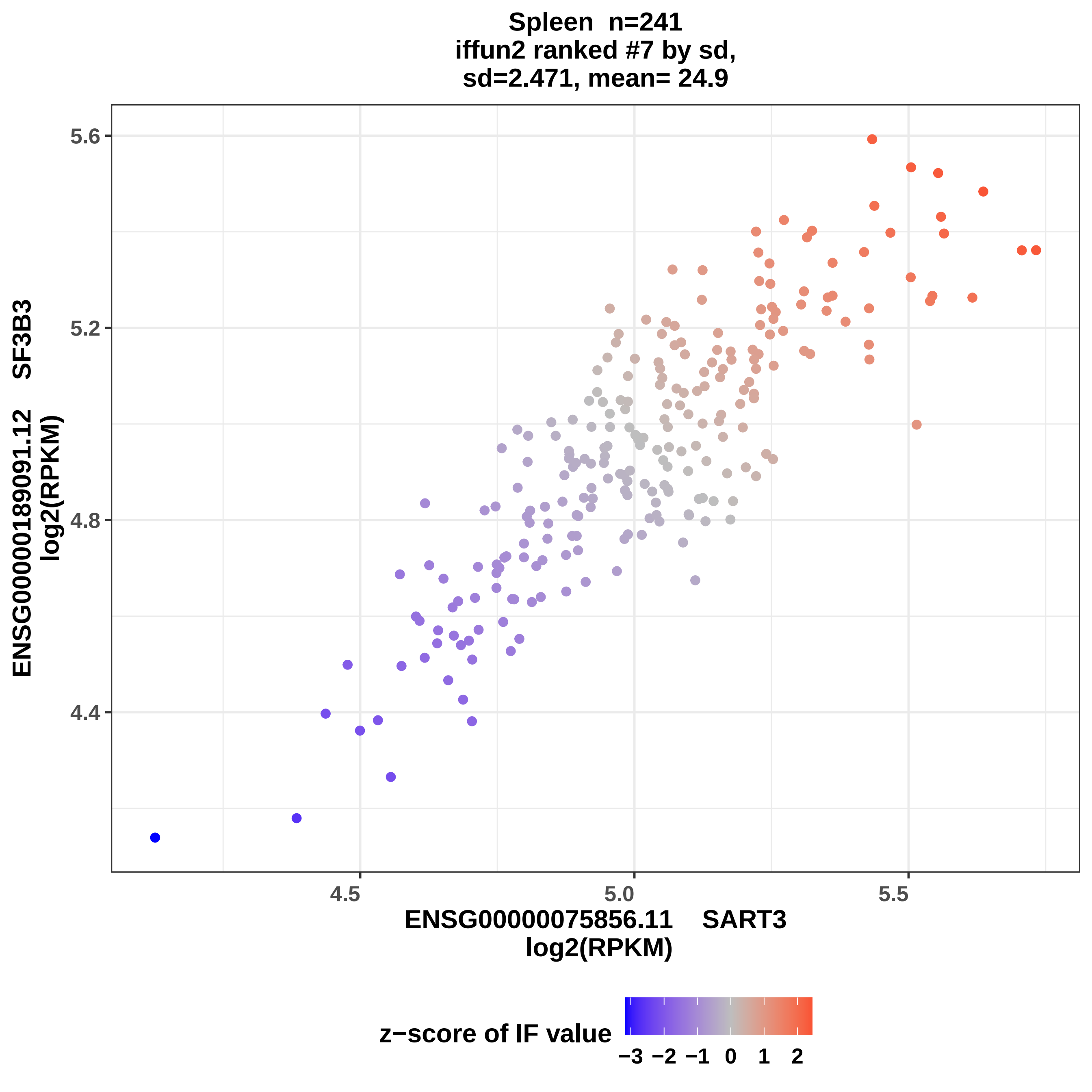
Spleen n=241
iffun2 ranked #7 by sd,
sd=2.471, mean= 24.9
ENSG00000189091.12 SF3B3
log2(RPKM)
5.6
5.2
4.8
4.4
4.5
5.0
5.5
ENSG00000075856.11 SART3
log2(RPKM)
z−score of IF value
−3
−2
−1
0
1
2

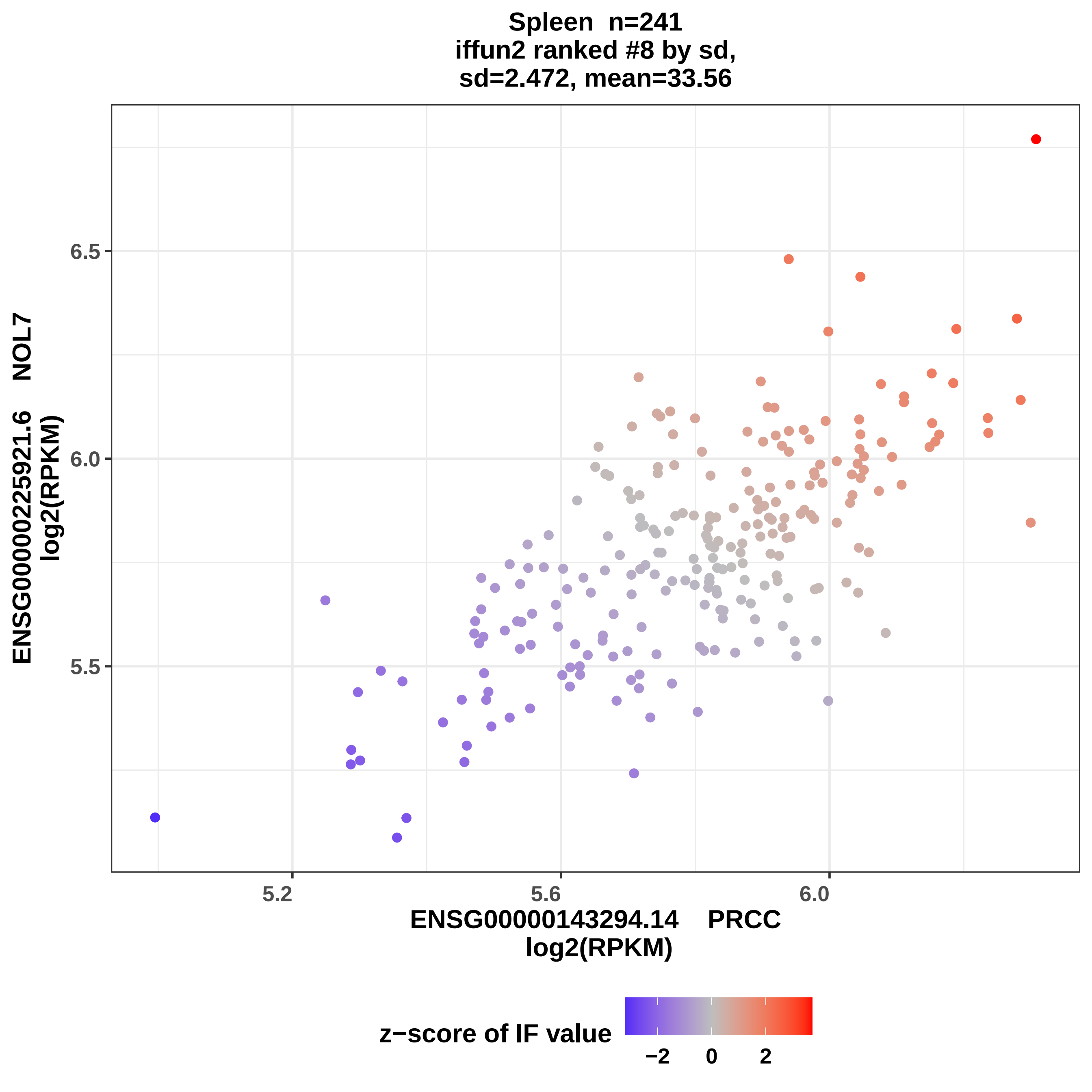

Spleen n=241
iffun2 ranked #8 by sd,
sd=2.472, mean=33.56
ENSG00000225921.6 NOL7
log2(RPKM)
6.5
6.0
5.5
5.2
5.6
6.0
ENSG00000143294.14 PRCC
log2(RPKM)
z−score of IF value
−2
0
2

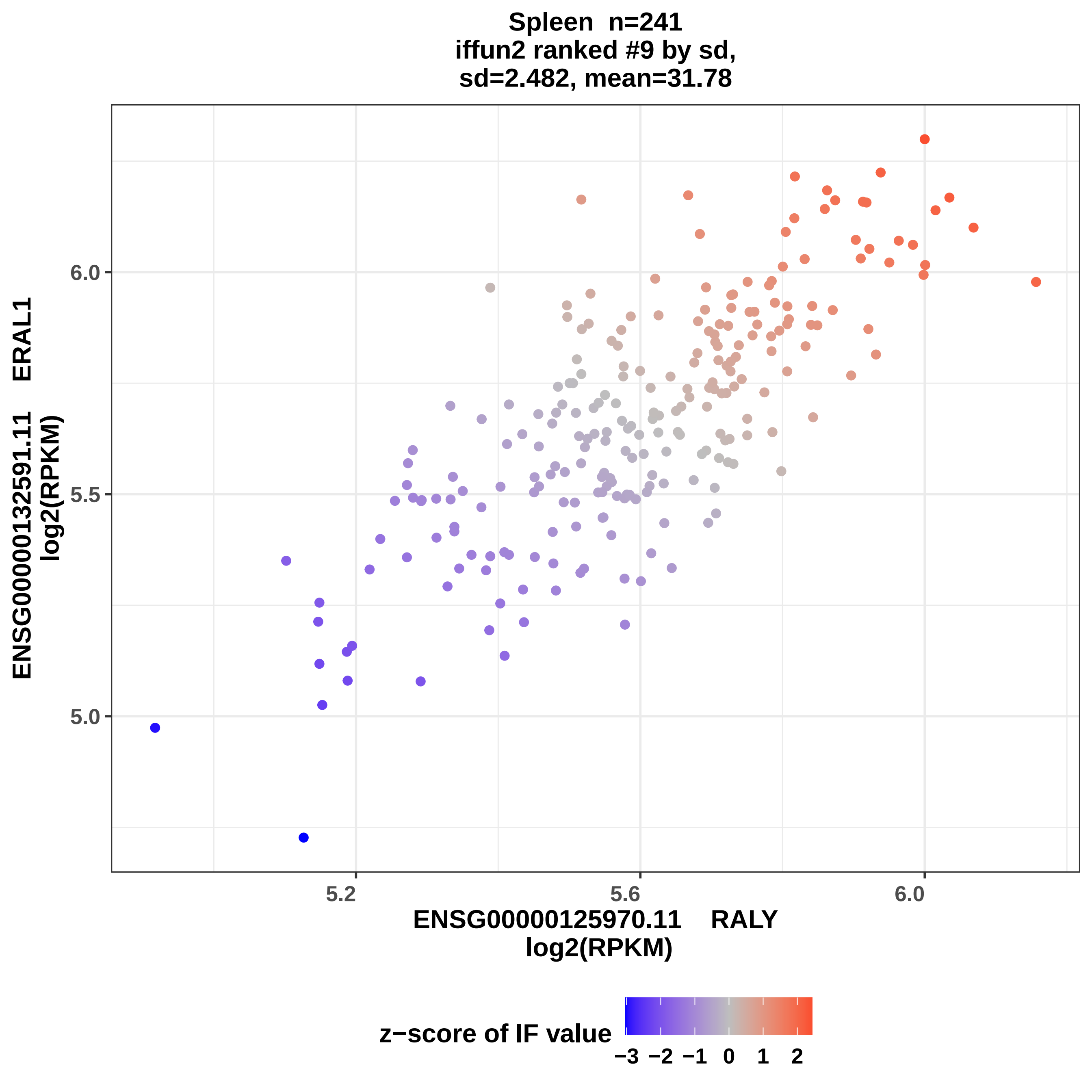
Spleen n=241
iffun2 ranked #9 by sd,
sd=2.482, mean=31.78
ENSG00000132591.11 ERAL1
log2(RPKM)
ENSG00000125970.11 RALY
log2(RPKM)
5.0
5.5
6.0
5.2
5.6
6.0
z−score of IF value
−3 −2 −1 0 1 2

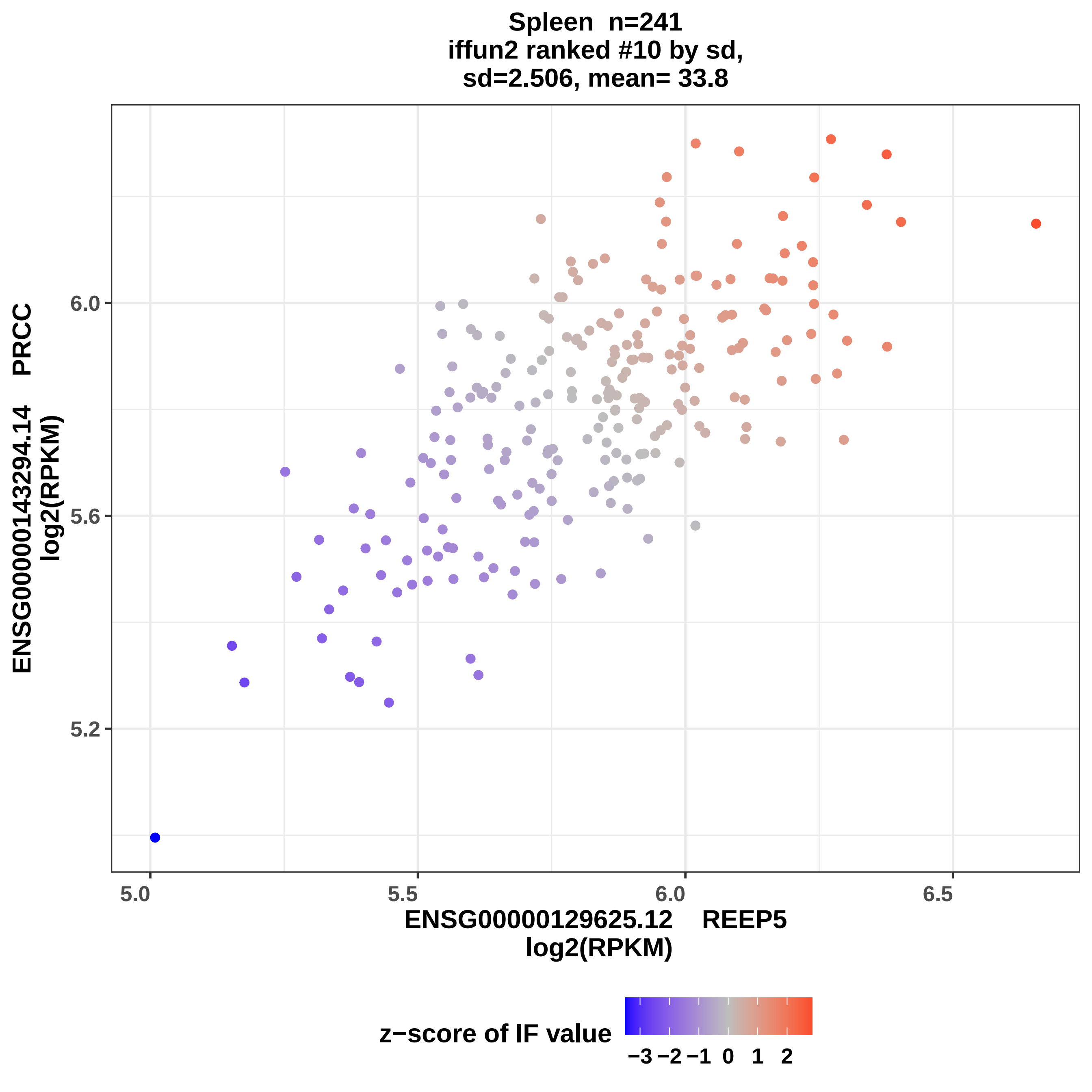

Spleen n=241
iffun2 ranked #10 by sd,
sd=2.506, mean= 33.8
ENSG00000143294.14 PRCC
log2(RPKM)
6.0
5.6
5.2
5.0
5.5
6.0
6.5
ENSG00000129625.12 REEP5
log2(RPKM)
z−score of IF value
−3 −2 −1 0 1 2

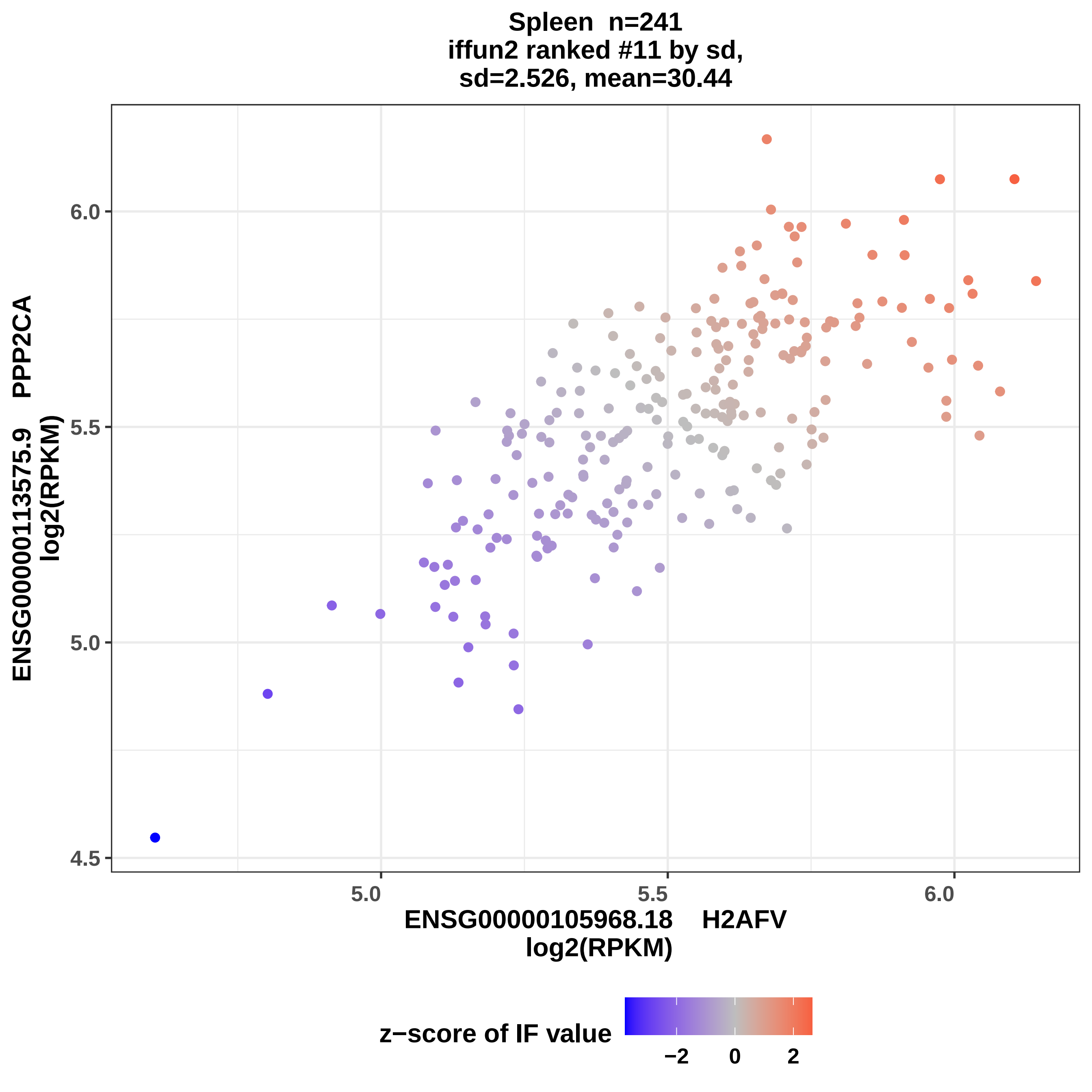
Spleen n=241
iffun2 ranked #11 by sd,
sd=2.526, mean=30.44
ENSG00000113575.9 PPP2CA
log2(RPKM)
6.0
5.5
5.0
4.5
5.0
5.5
6.0
ENSG00000105968.18 H2AFV
log2(RPKM)
z−score of IF value
−2
0
2

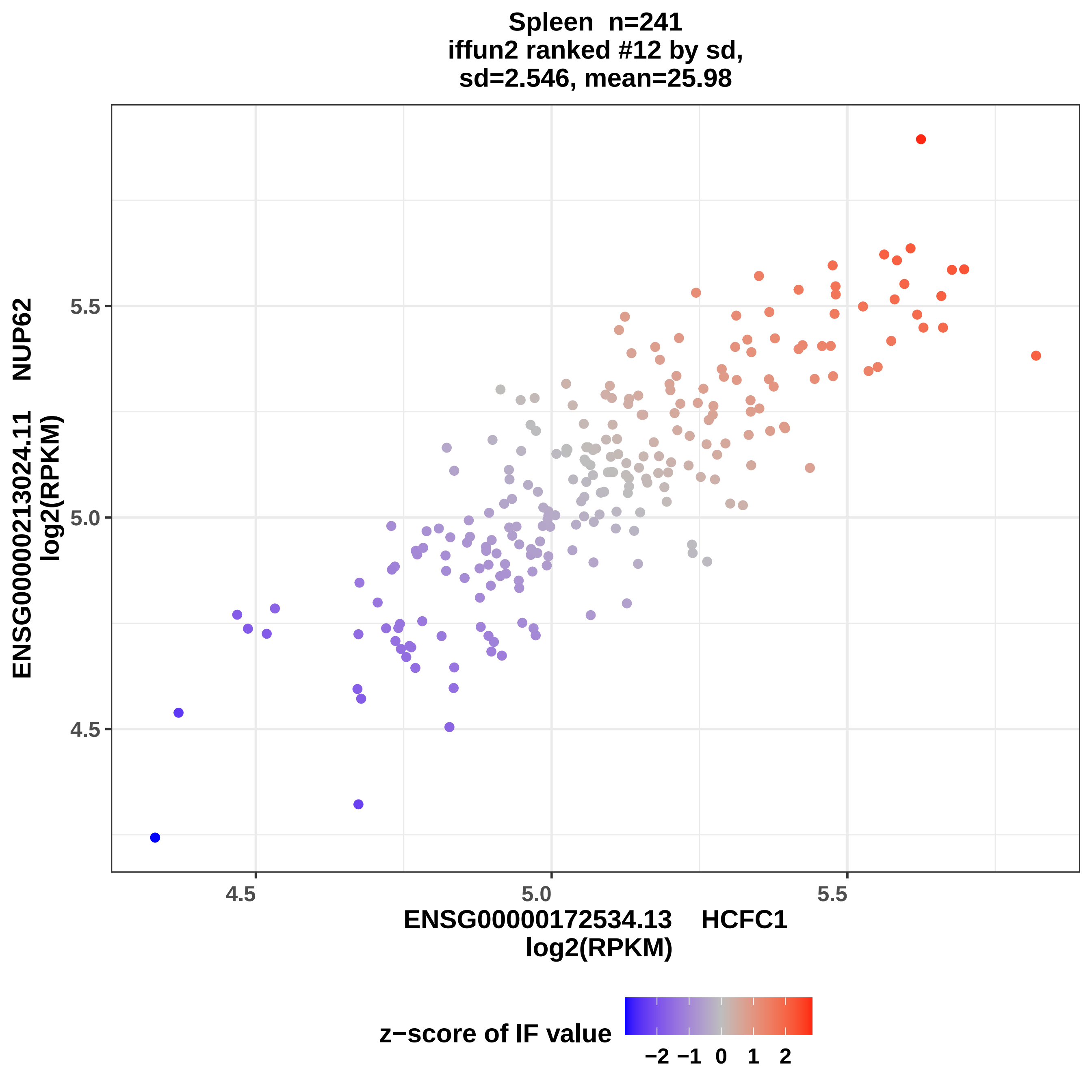
Spleen n=241
iffun2 ranked #12 by sd,
sd=2.546, mean=25.98
ENSG00000213024.11 NUP62
log2(RPKM)
ENSG00000172534.13 HCFC1
log2(RPKM)
4.5
5.0
5.5
4.5
5.0
5.5
z−score of IF value
−2 −1 0 1 2

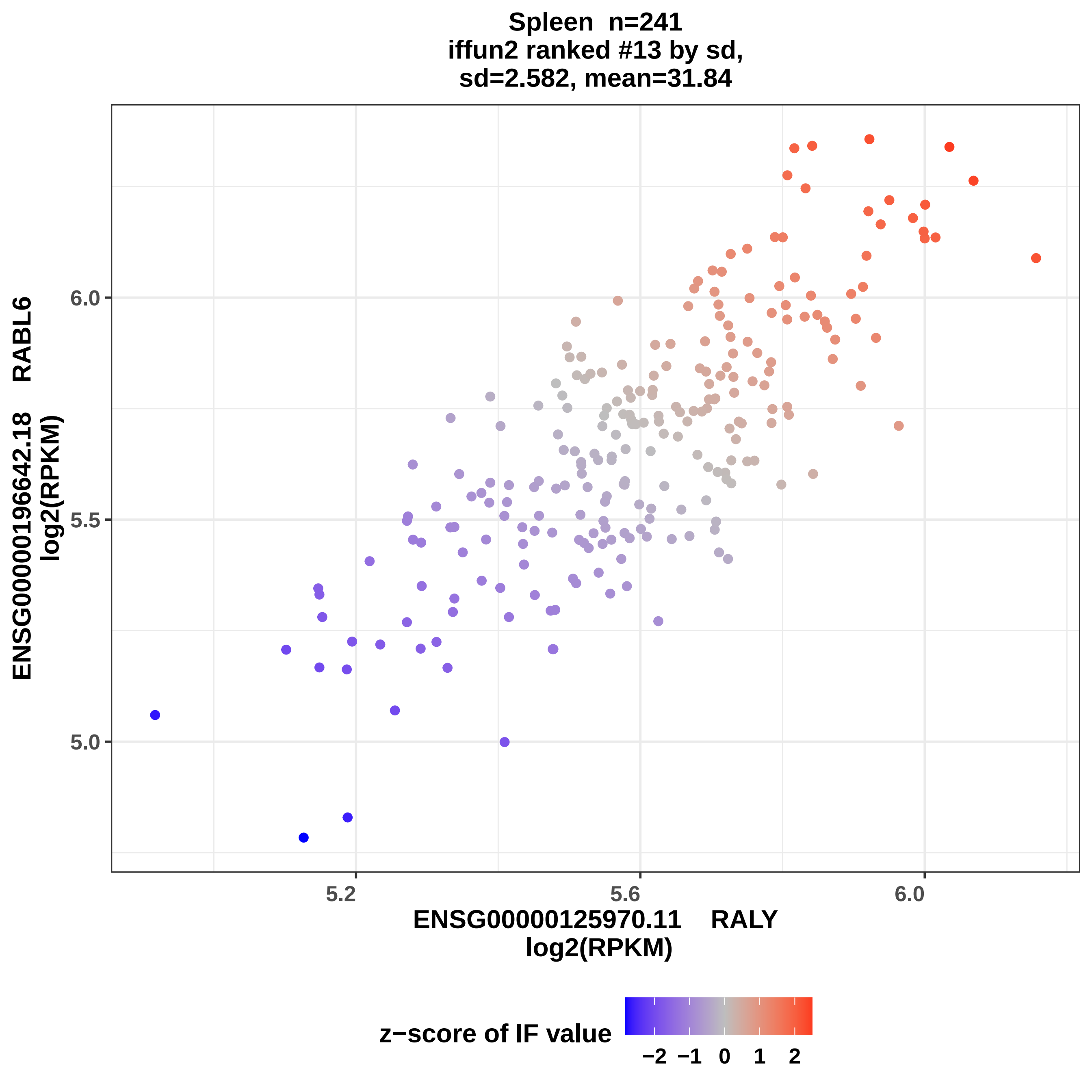

Spleen n=241
iffun2 ranked #13 by sd,
sd=2.582, mean=31.84
ENSG00000196642.18 RABL6
log2(RPKM)
6.0
5.5
5.0
5.2
5.6
6.0
ENSG00000125970.11 RALY
log2(RPKM)
z−score of IF value
−2
−1
0
1
2

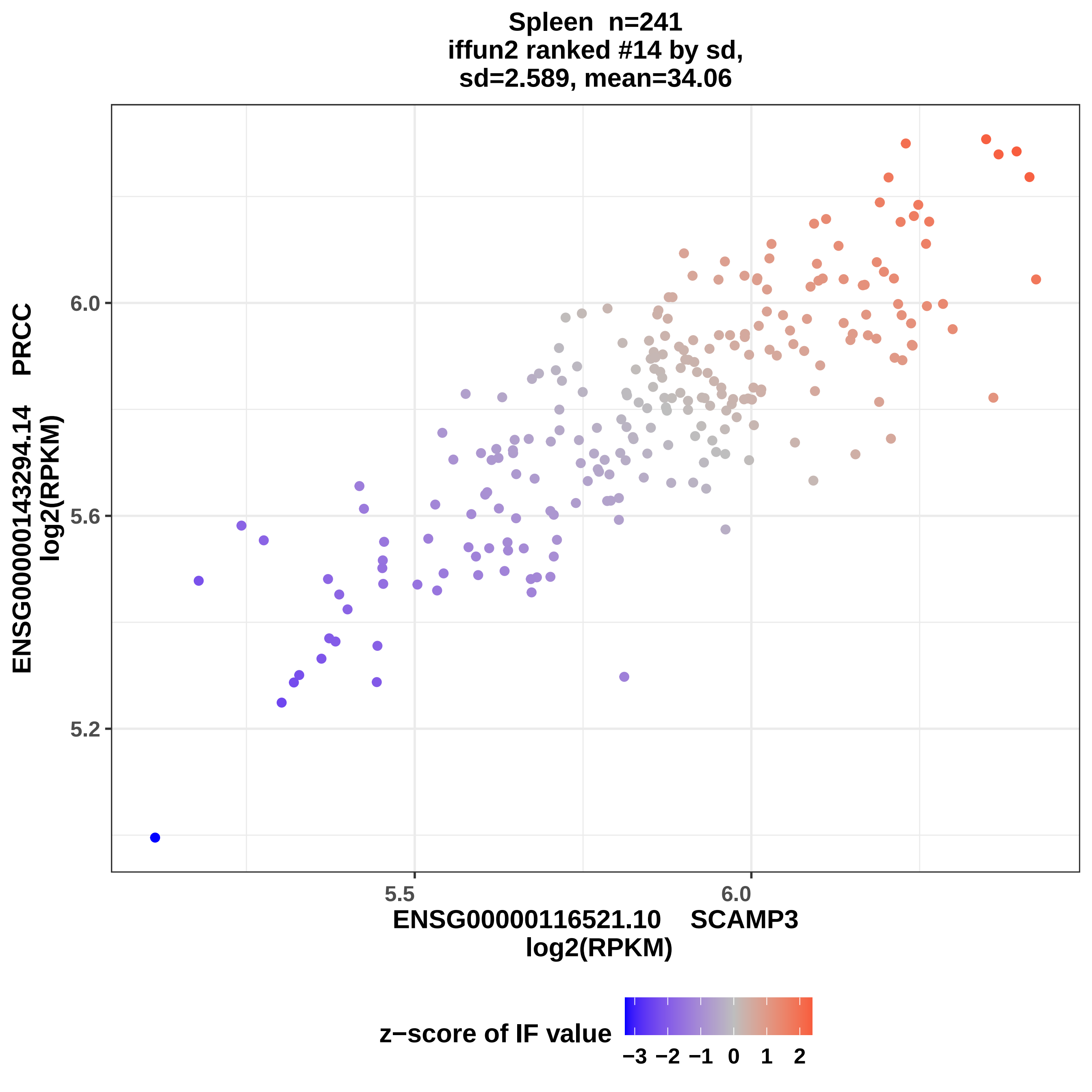
Spleen n=241
iffun2 ranked #14 by sd,
sd=2.589, mean=34.06
ENSG00000143294.14 PRCC
log2(RPKM)
6.0
5.6
5.2
5.5
6.0
ENSG00000116521.10 SCAMP3
log2(RPKM)
z−score of IF value
−3 −2 −1 0 1 2

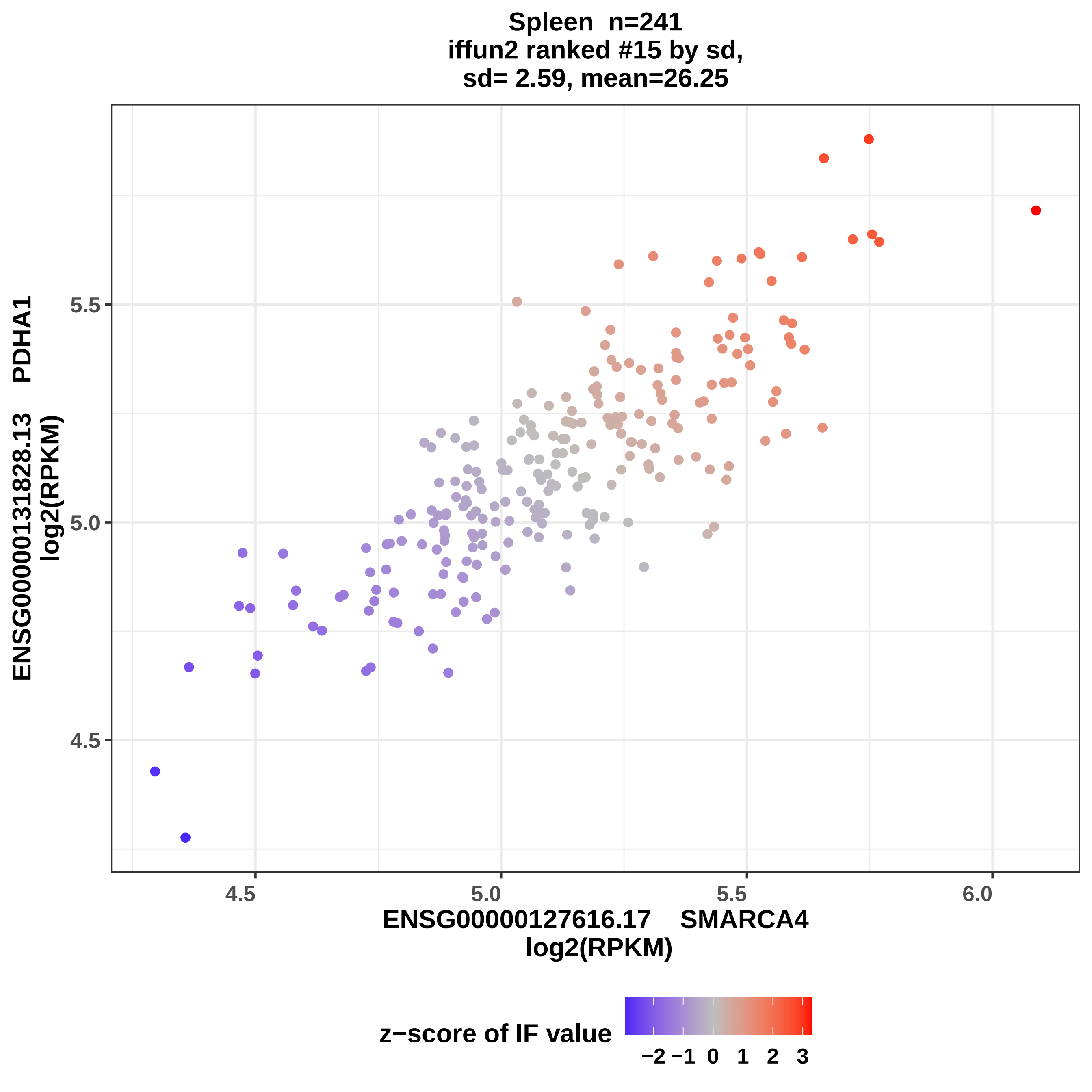
Spleen n=241
iffun2 ranked #15 by sd,
sd= 2.59, mean=26.25
ENSG00000131828.13 PDHA1
log2(RPKM)
5.5
5.0
4.5
4.5
5.0
5.5
6.0
ENSG00000127616.17 SMARCA4
log2(RPKM)
z−score of IF value
−2 −1 0 1 2 3

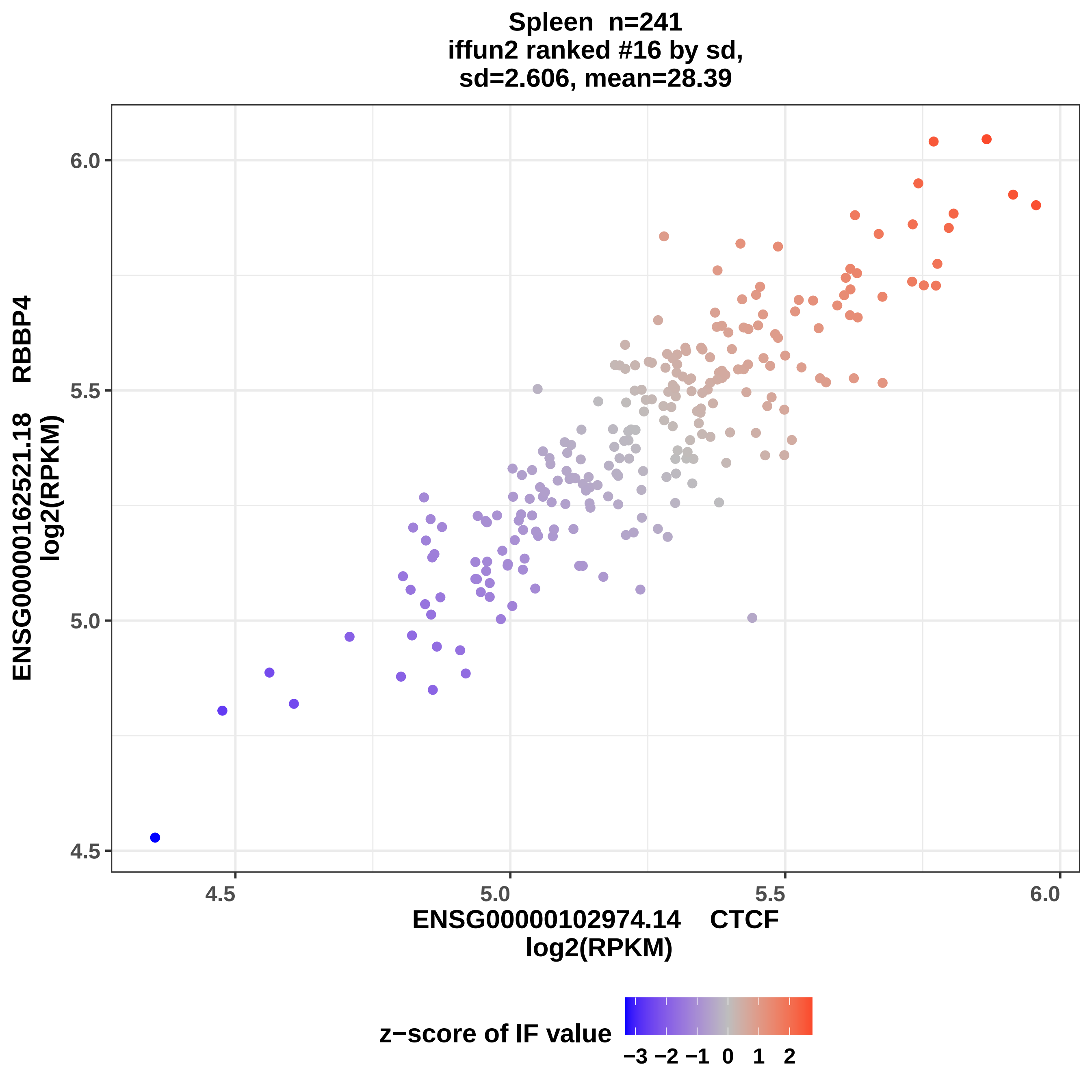
Spleen n=241
iffun2 ranked #16 by sd,
sd=2.606, mean=28.39
ENSG00000162521.18 RBBP4
log2(RPKM)
ENSG00000102974.14 CTCF
log2(RPKM)
4.5
5.0
5.5
6.0
z−score of IF value
−3 −2 −1 0 1 2

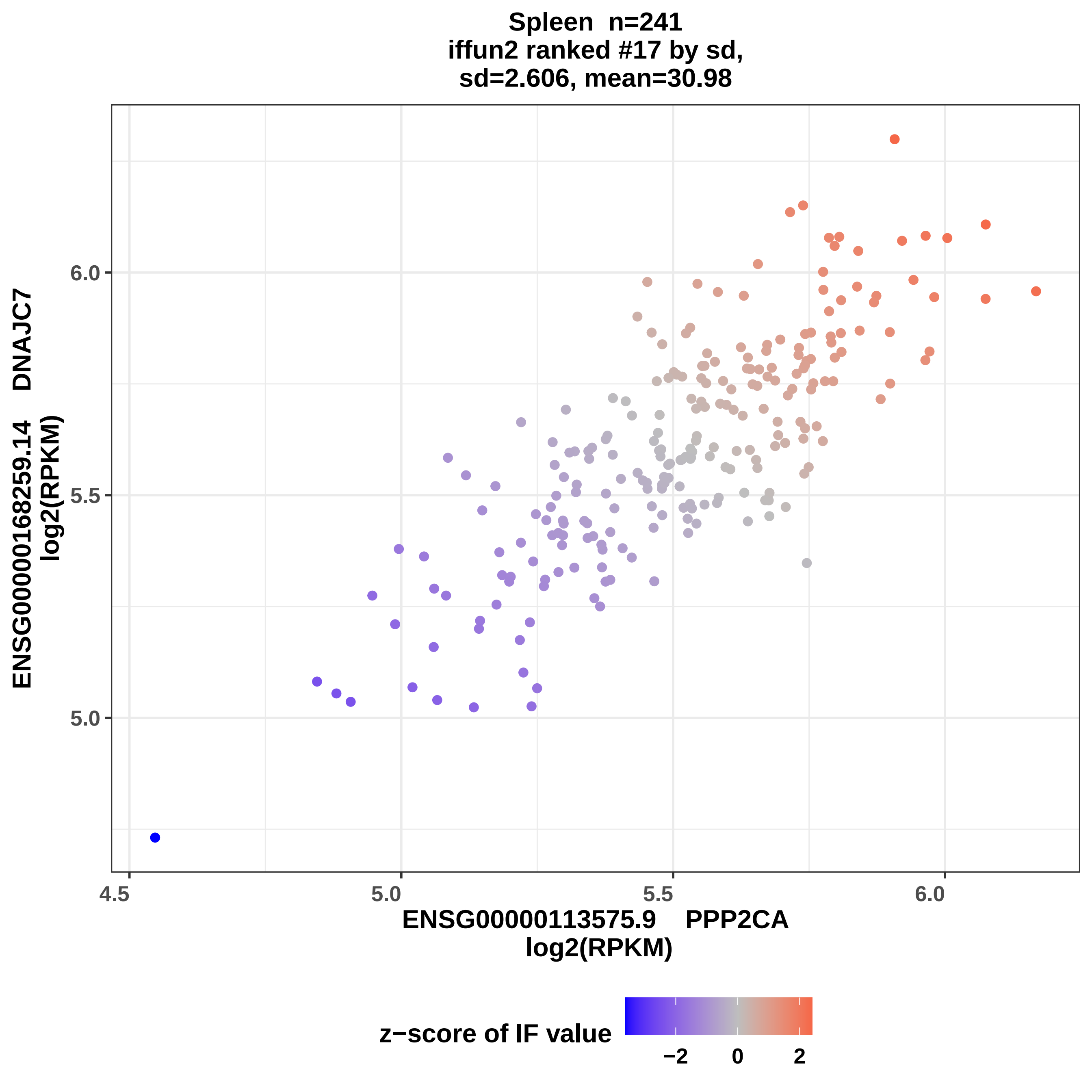
Spleen n=241
iffun2 ranked #17 by sd,
sd=2.606, mean=30.98
ENSG00000168259.14 DNAJC7
log2(RPKM)
6.0
5.5
5.0
4.5
5.0
5.5
6.0
ENSG00000113575.9 PPP2CA
log2(RPKM)
z−score of IF value
−2
0
2

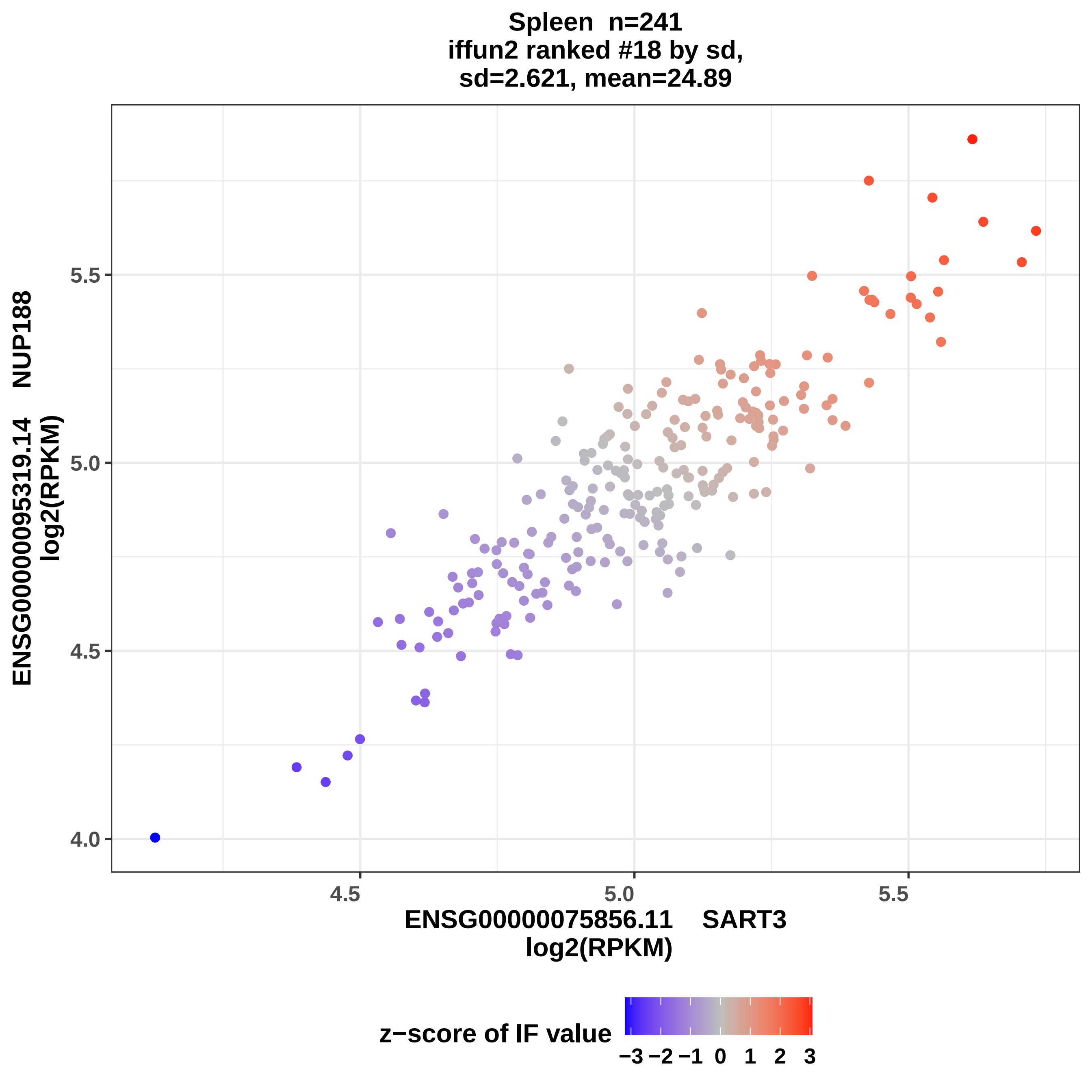
Spleen n=241
iffun2 ranked #18 by sd,
sd=2.621, mean=24.89
ENSG00000095319.14 NUP188
log2(RPKM)
5.5
5.0
4.5
4.0
4.5
5.0
5.5
ENSG00000075856.11 SART3
log2(RPKM)
z−score of IF value
−3 −2 −1 0 1 2 3

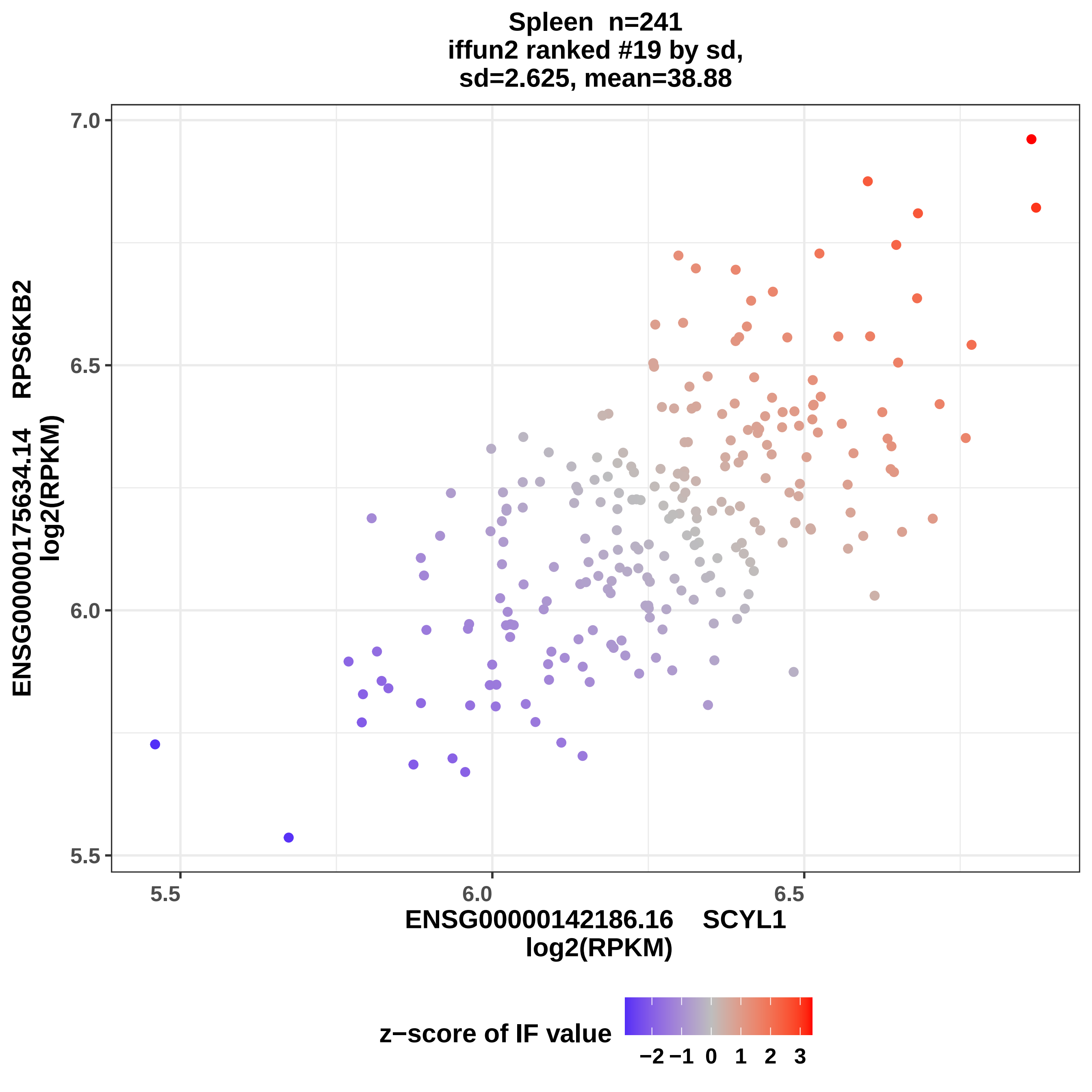
Spleen n=241
iffun2 ranked #19 by sd,
sd=2.625, mean=38.88
ENSG00000175634.14 RPS6KB2
log2(RPKM)
7.0
6.5
6.0
5.5
5.5
6.0
6.5
ENSG00000142186.16 SCYL1
log2(RPKM)
z−score of IF value
−2
−1
0
1
2
3

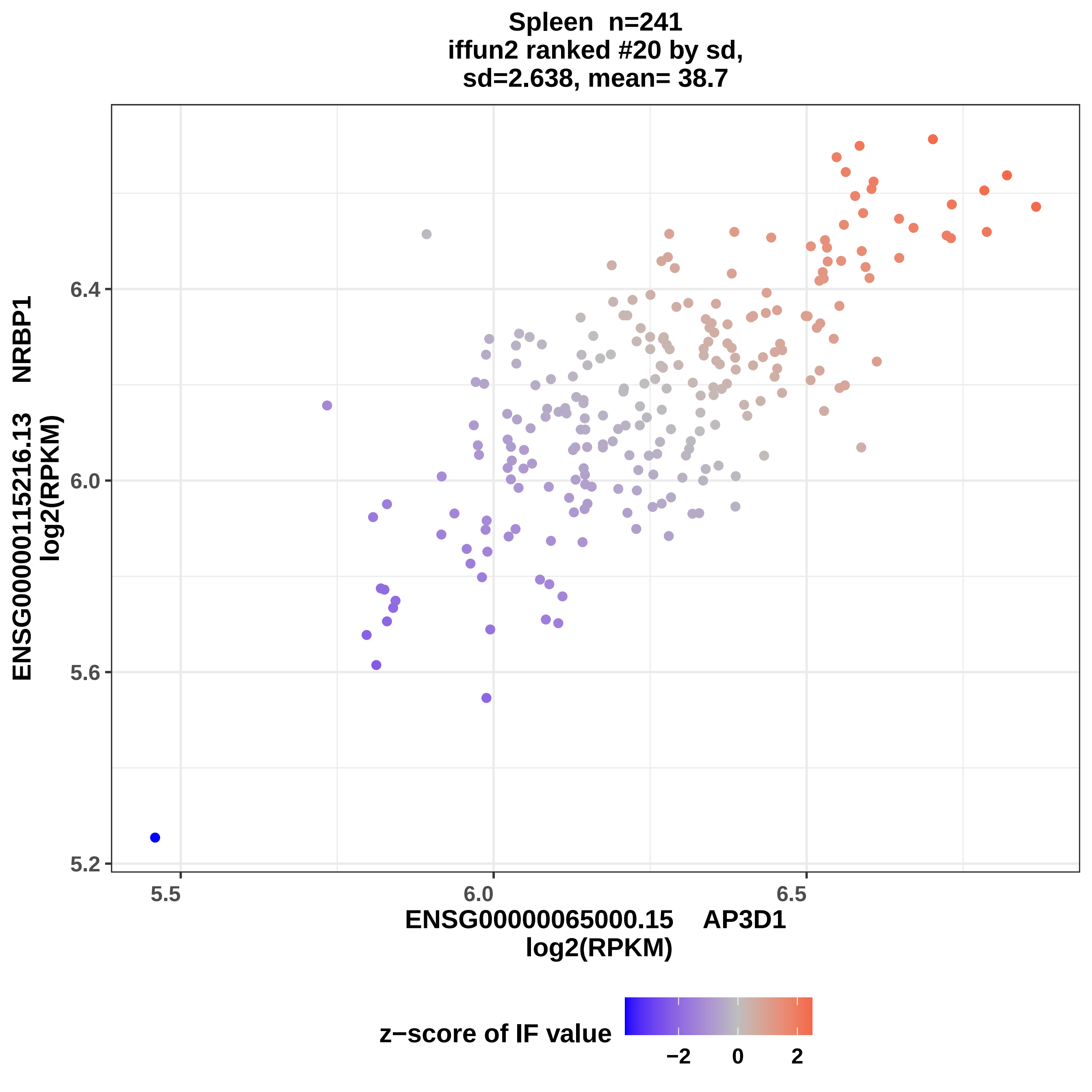
Spleen n=241
iffun2 ranked #20 by sd,
sd=2.638, mean= 38.7
ENSG00000115216.13 NRBP1
log2(RPKM)
6.4
6.0
5.6
5.2
5.5
6.0
6.5
ENSG00000065000.15 AP3D1
log2(RPKM)
z−score of IF value
−2
0
2

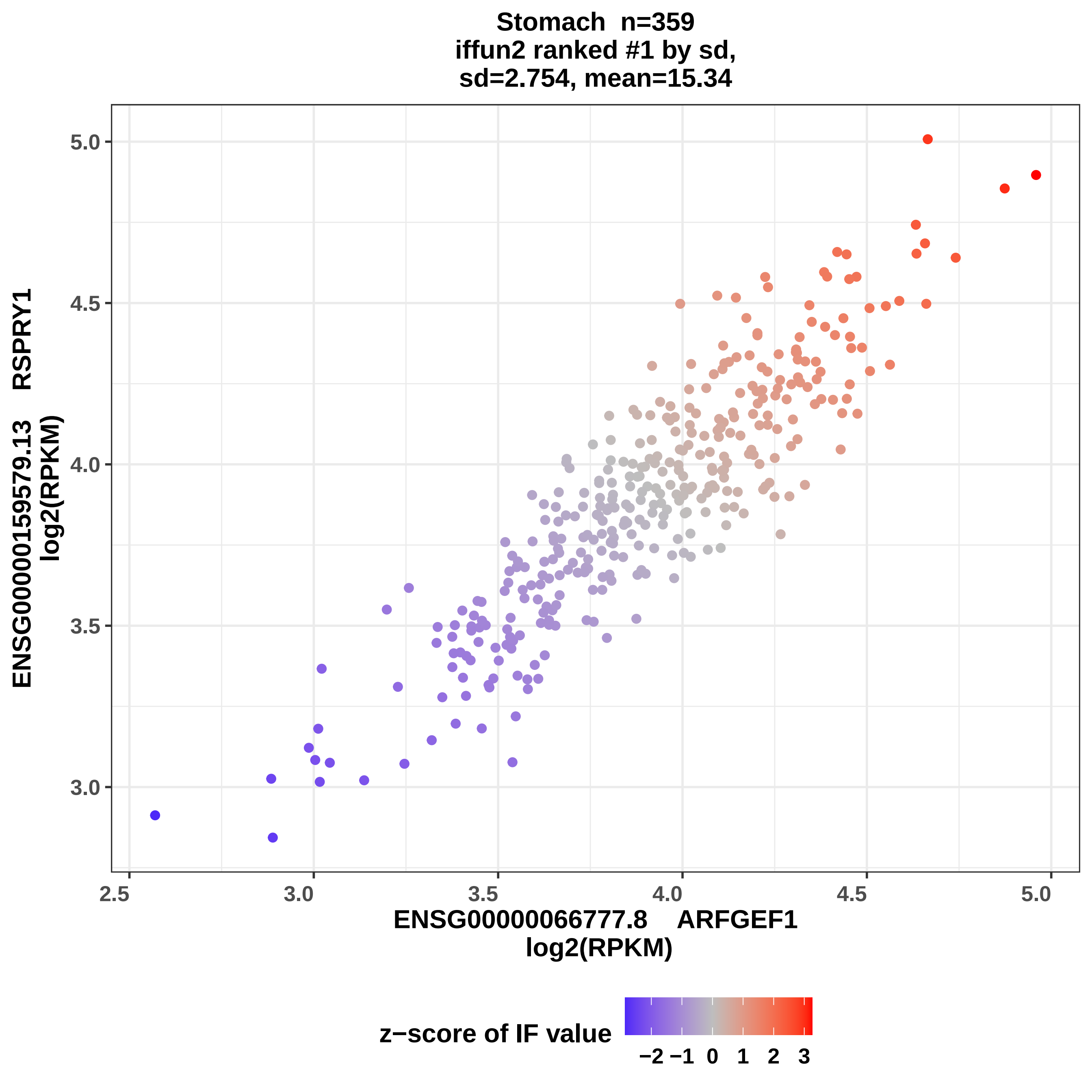

Stomach n=359
iffun2 ranked #1 by sd,
sd=2.754, mean=15.34
ENSG00000159579.13 RSPRY1
log2(RPKM)
5.0
4.5
4.0
3.5
3.0
2.5
3.0
3.5
4.0
4.5
5.0
ENSG00000066777.8 ARFGEF1
log2(RPKM)
z−score of IF value
−2 −1 0 1 2 3

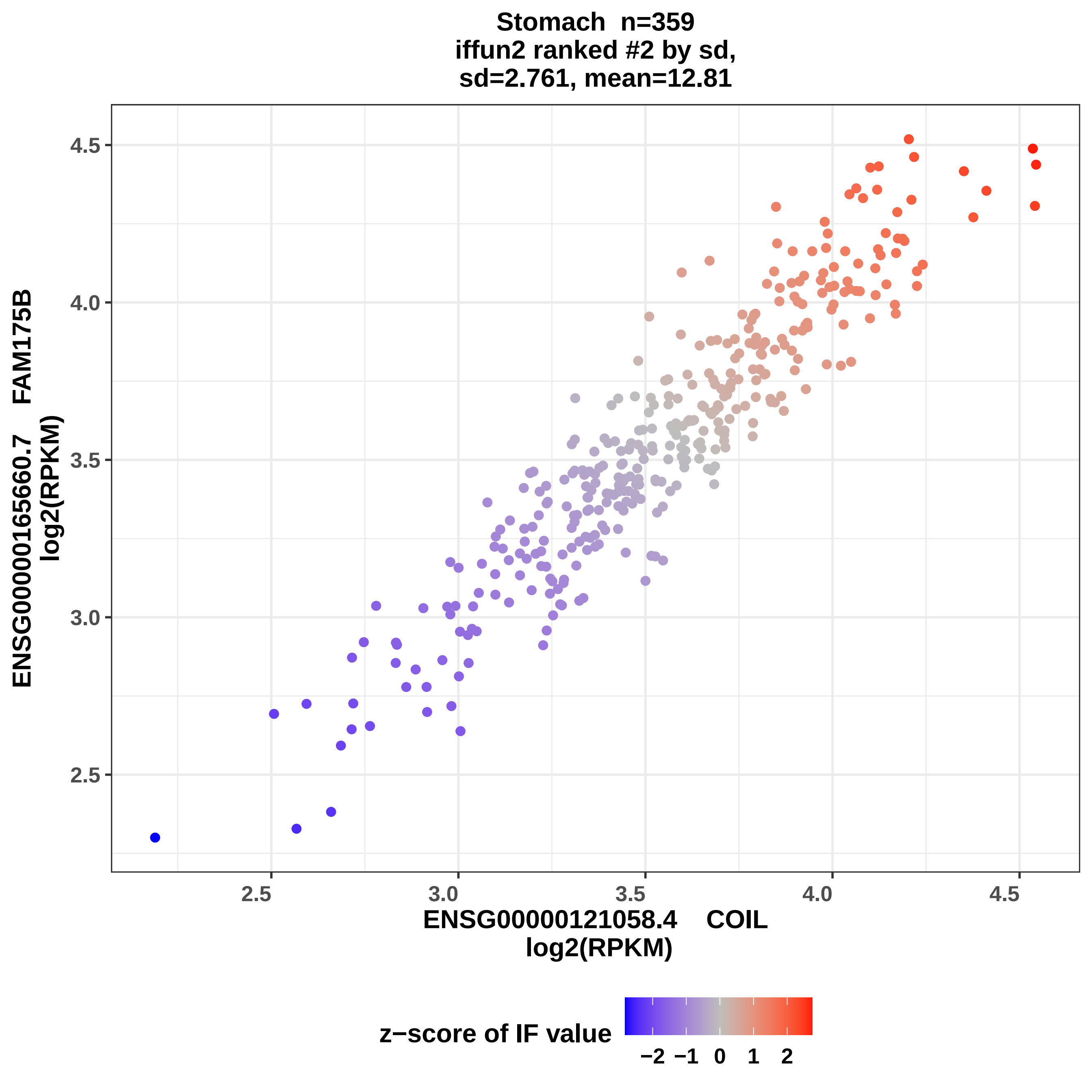

Stomach n=359
iffun2 ranked #2 by sd,
sd=2.761, mean=12.81
ENSG00000165660.7 FAM175B
log2(RPKM)
ENSG00000121058.4 COIL
log2(RPKM)
4.5
4.0
3.5
3.0
2.5
2.5
3.0
3.5
4.0
4.5
z−score of IF value
−2 −1 0 1 2

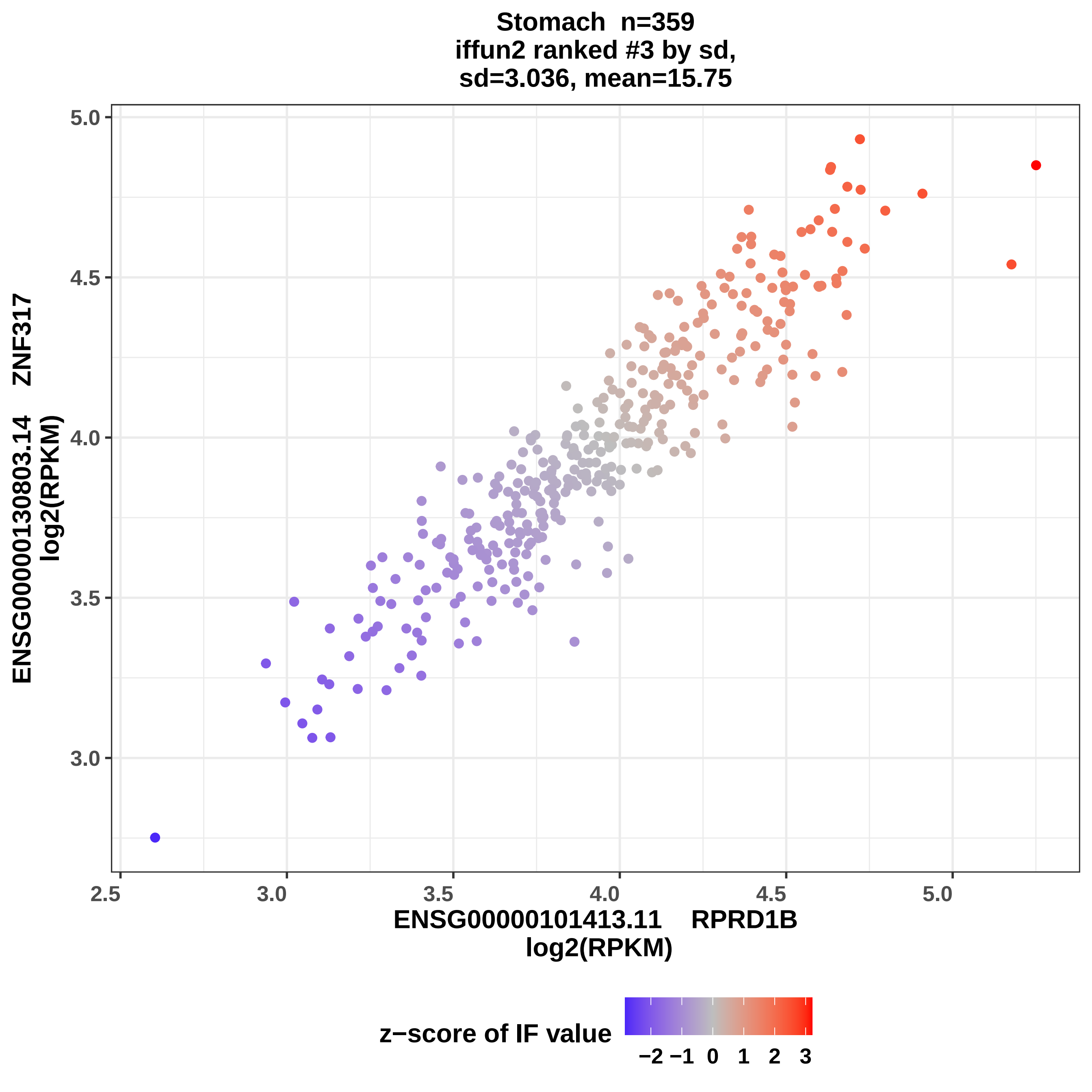
Stomach n=359
iffun2 ranked #3 by sd,
sd=3.036, mean=15.75
ENSG00000130803.14 ZNF317
log2(RPKM)
5.0
4.5
4.0
3.5
3.0
2.5
3.0
3.5
4.0
4.5
5.0
ENSG00000101413.11 RPRD1B
log2(RPKM)
z−score of IF value
−2
−1
0
1
2
3

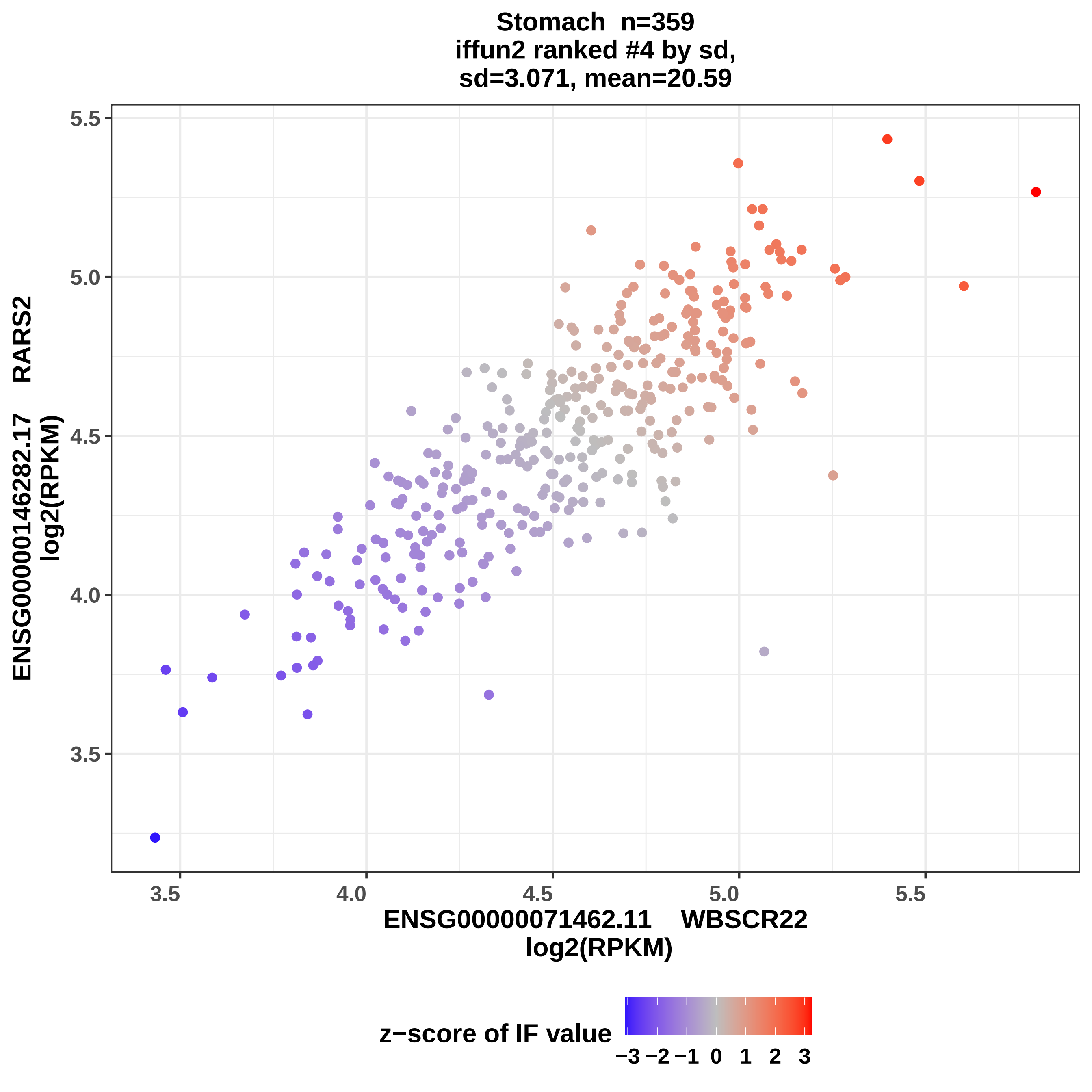

Stomach n=359
iffun2 ranked #4 by sd,
sd=3.071, mean=20.59
ENSG00000146282.17 RARS2
log2(RPKM)
5.5
5.0
4.5
4.0
3.5
3.5
4.0
4.5
5.0
5.5
ENSG00000071462.11 WBSCR22
log2(RPKM)
z−score of IF value
−3 −2 −1 0 1 2 3

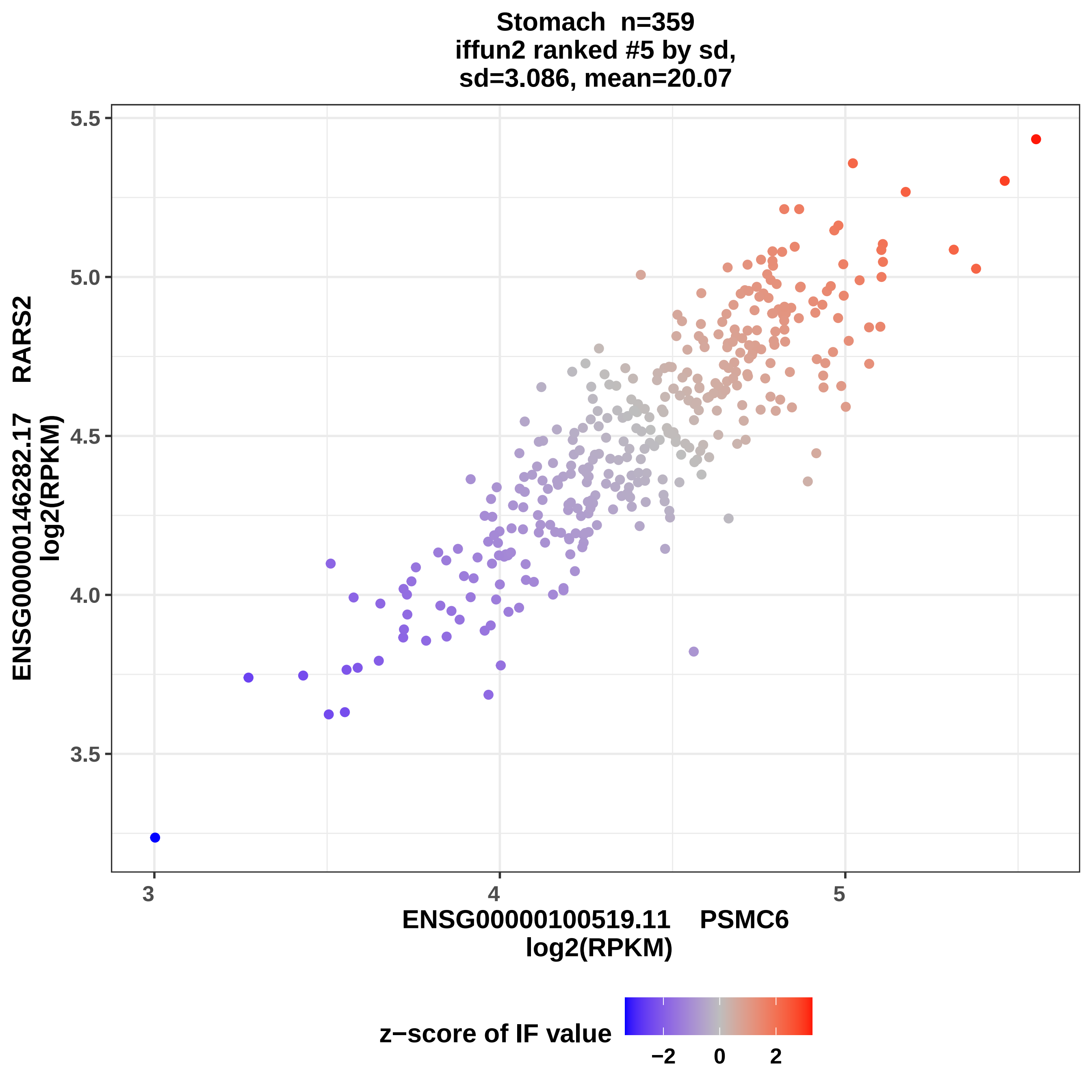
Stomach n=359
iffun2 ranked #5 by sd,
sd=3.086, mean=20.07
ENSG00000146282.17 RARS2
log2(RPKM)
5.5
5.0
4.5
4.0
3.5
3
4
5
ENSG00000100519.11 PSMC6
log2(RPKM)
z−score of IF value
−2
0
2

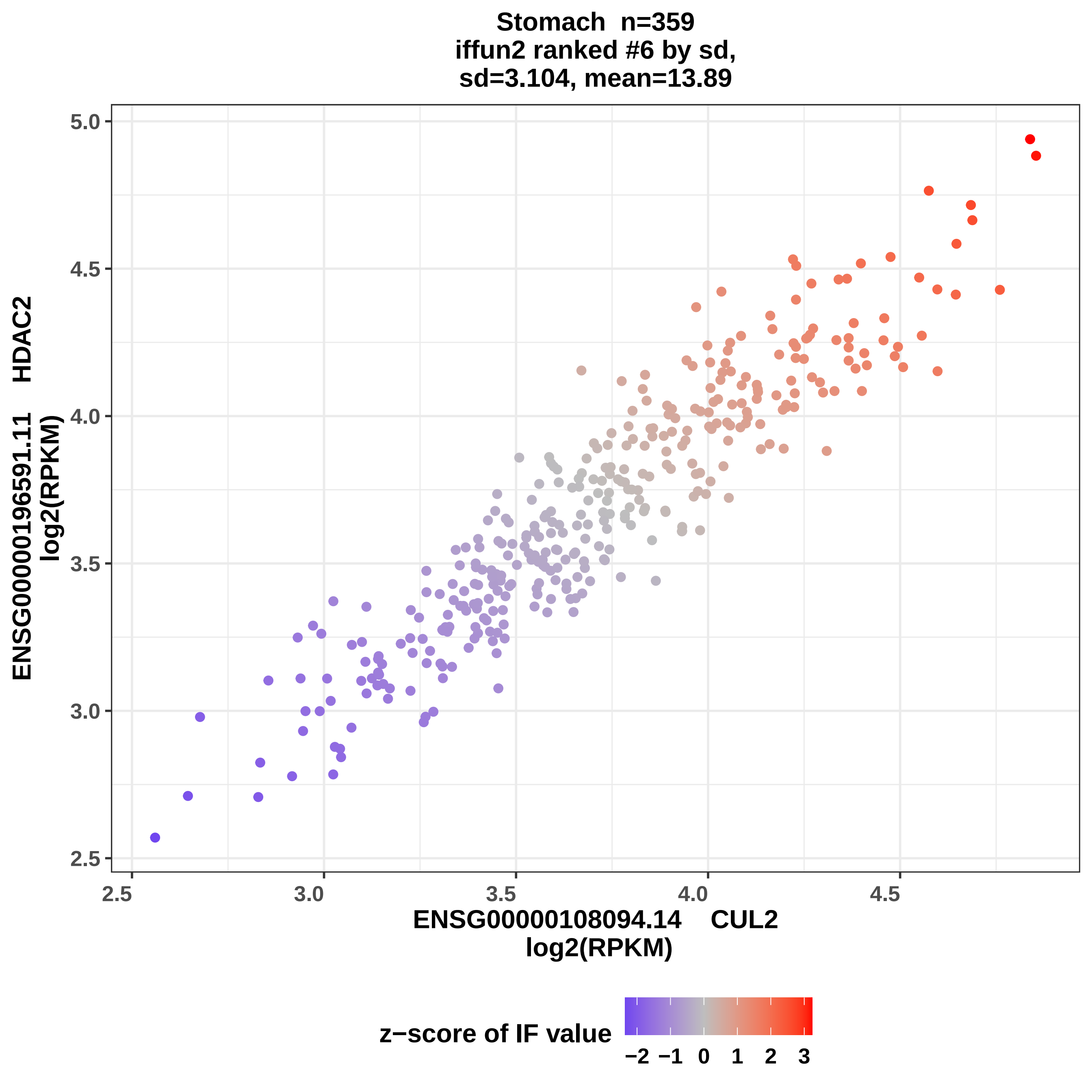
Stomach n=359
iffun2 ranked #6 by sd,
sd=3.104, mean=13.89
ENSG00000196591.11 HDAC2
log2(RPKM)
ENSG00000108094.14 CUL2
log2(RPKM)
2.5
3.0
3.5
4.0
4.5
5.0
z−score of IF value
−2
−1
0
1
2
3

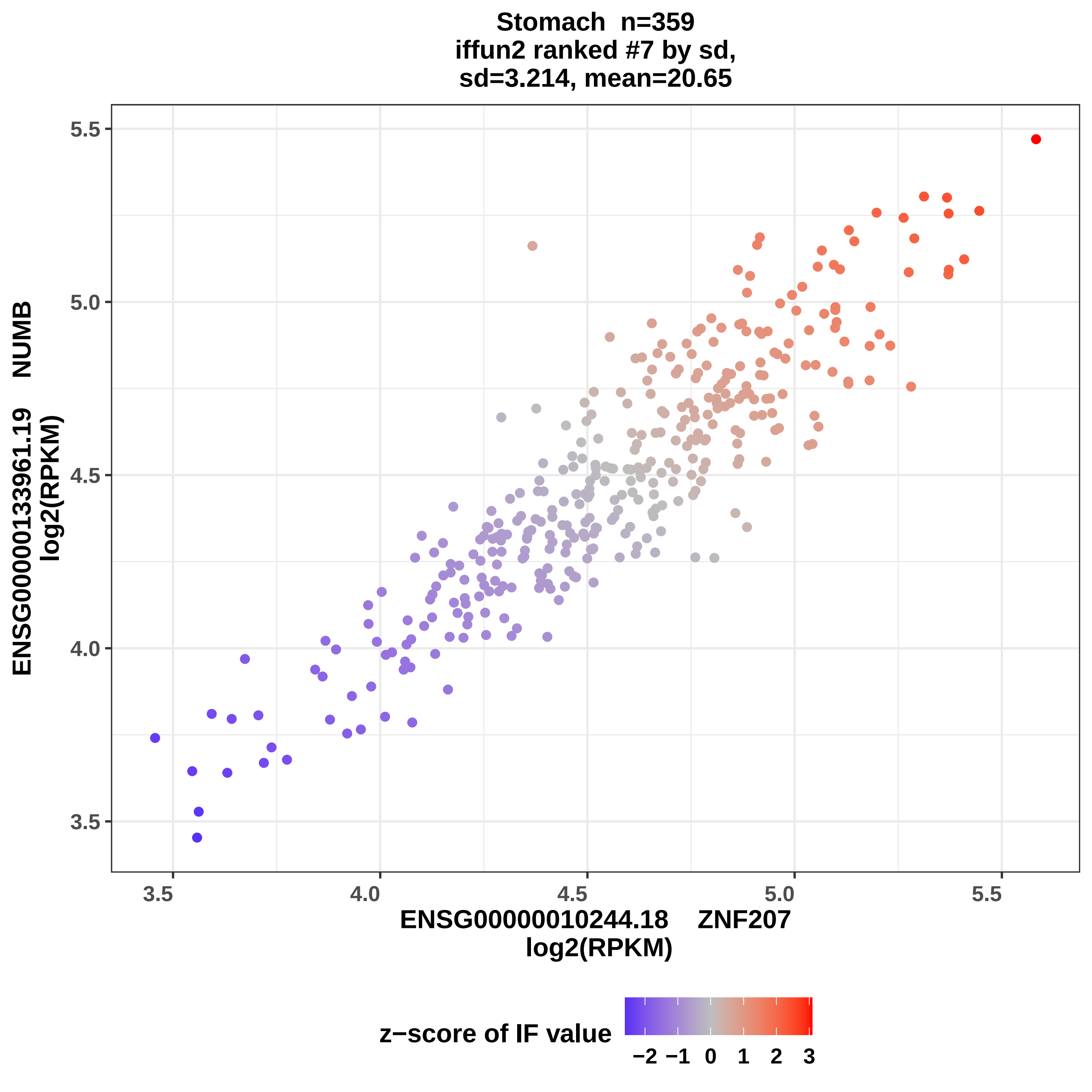
Stomach n=359
iffun2 ranked #7 by sd,
sd=3.214, mean=20.65
ENSG00000133961.19 NUMB
log2(RPKM)
ENSG00000010244.18 ZNF207
log2(RPKM)
3.5
4.0
4.5
5.0
5.5
z−score of IF value
−2 −1 0 1 2 3

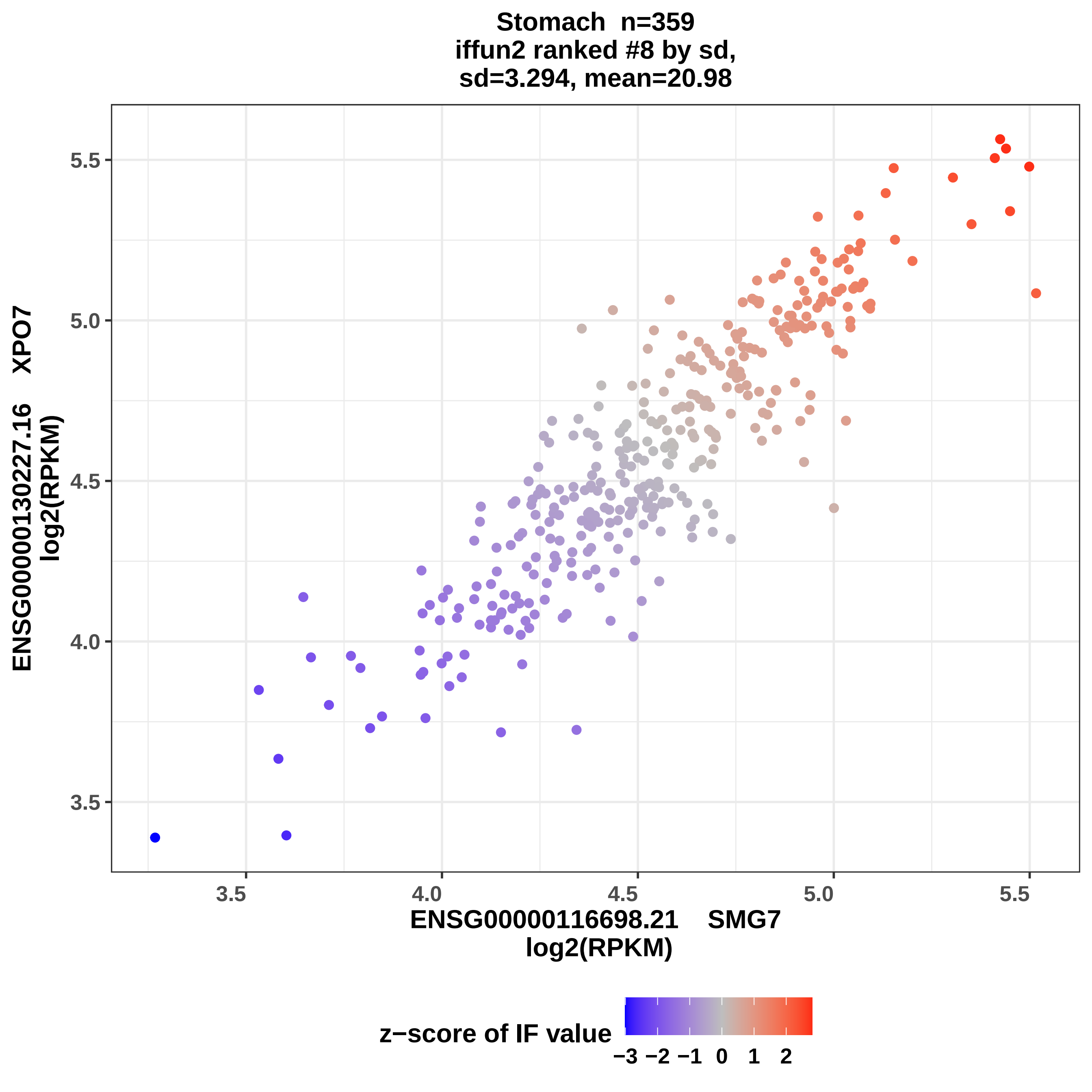

Stomach n=359
iffun2 ranked #8 by sd,
sd=3.294, mean=20.98
ENSG00000130227.16 XPO7
log2(RPKM)
5.5
5.0
4.5
4.0
3.5
3.5
4.0
4.5
5.0
5.5
ENSG00000116698.21 SMG7
log2(RPKM)
z−score of IF value
−3 −2 −1 0 1 2

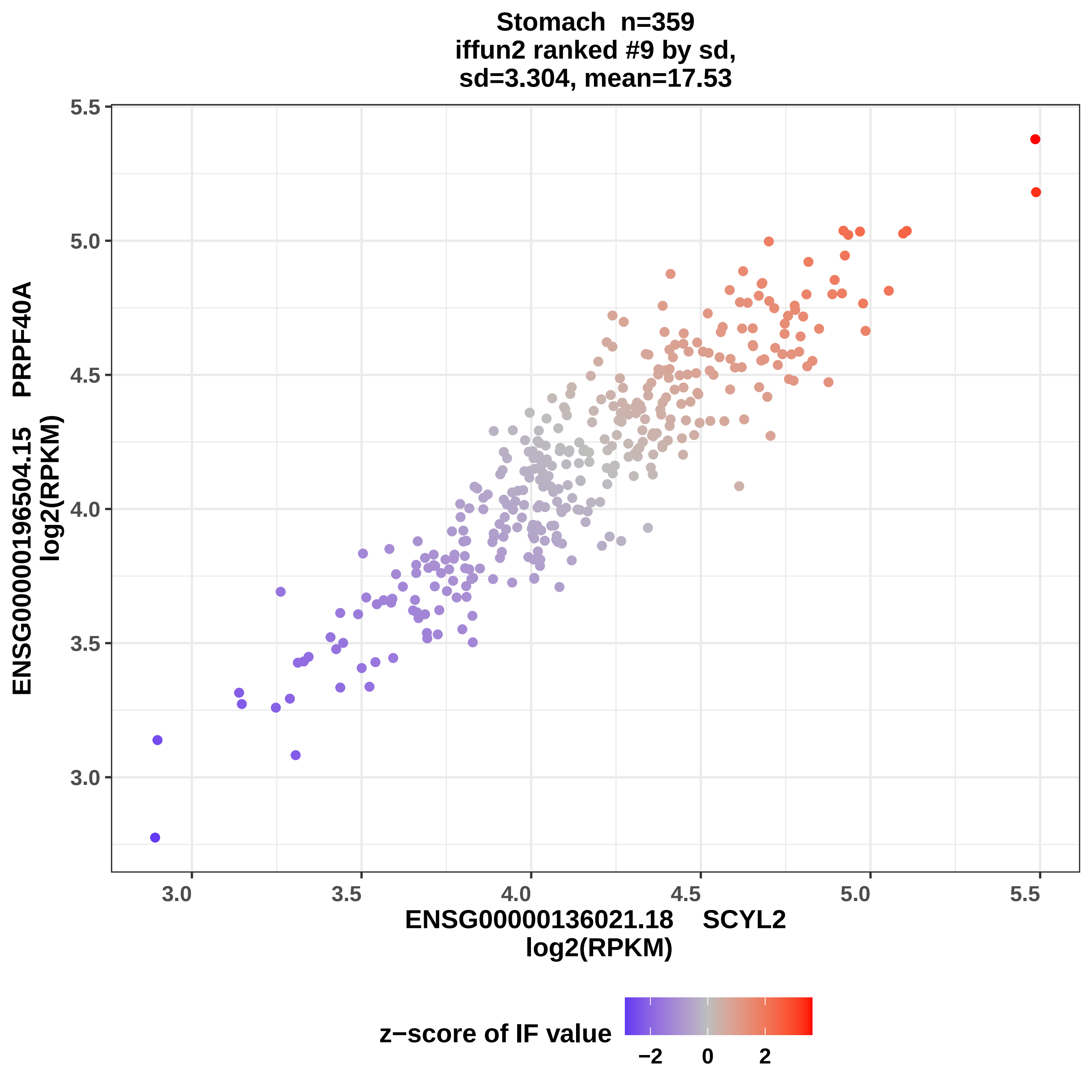
Stomach n=359
iffun2 ranked #9 by sd,
sd=3.304, mean=17.53
ENSG00000196504.15 PRPF40A
log2(RPKM)
ENSG00000136021.18 SCYL2
log2(RPKM)
5.5
5.0
4.5
4.0
3.5
3.0
z−score of IF value
−2
0
2

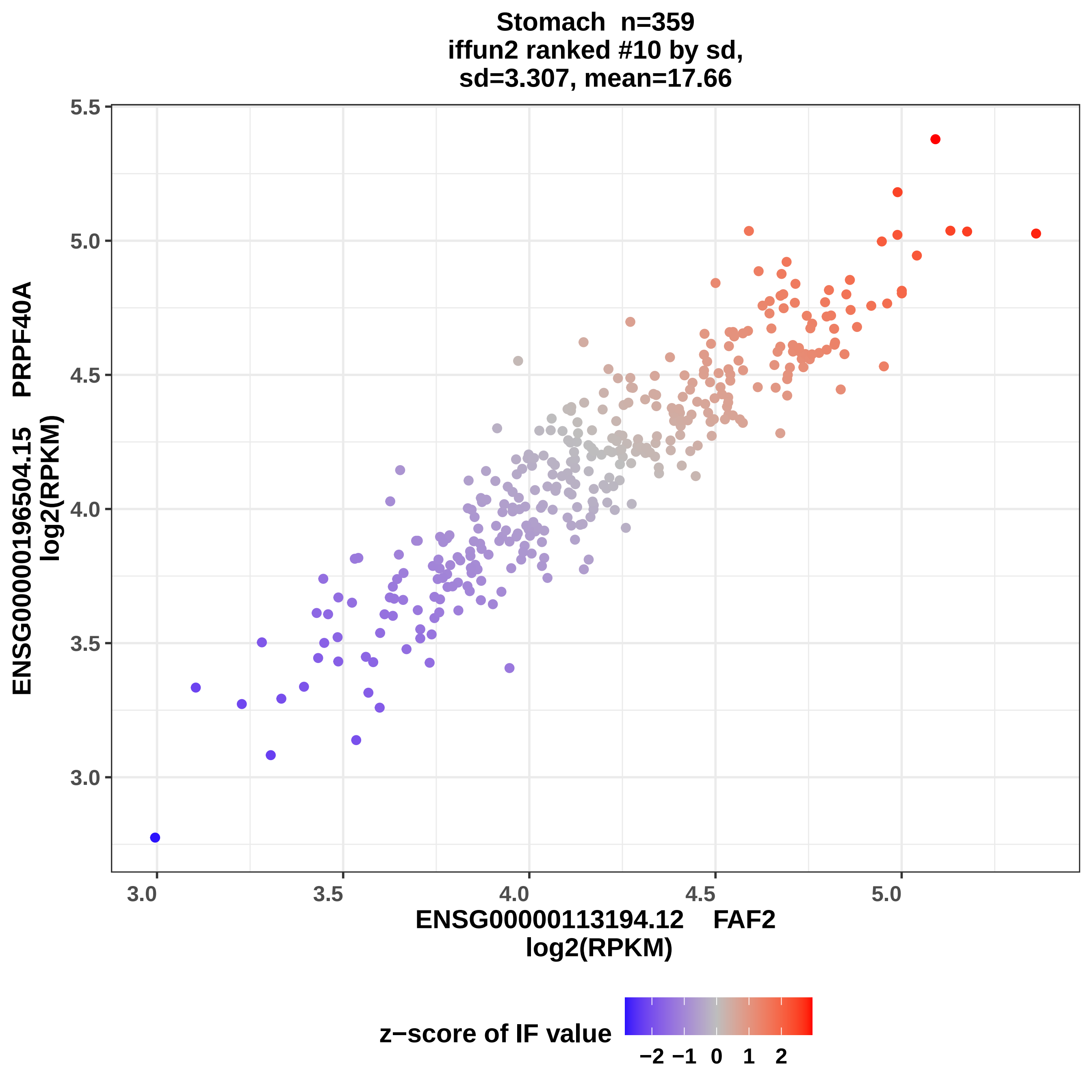
Stomach n=359
iffun2 ranked #10 by sd,
sd=3.307, mean=17.66
ENSG00000196504.15 PRPF40A
log2(RPKM)
5.5
5.0
4.5
4.0
3.5
3.0
3.0
3.5
4.0
4.5
5.0
ENSG00000113194.12 FAF2
log2(RPKM)
z−score of IF value
−2
−1
0
1
2

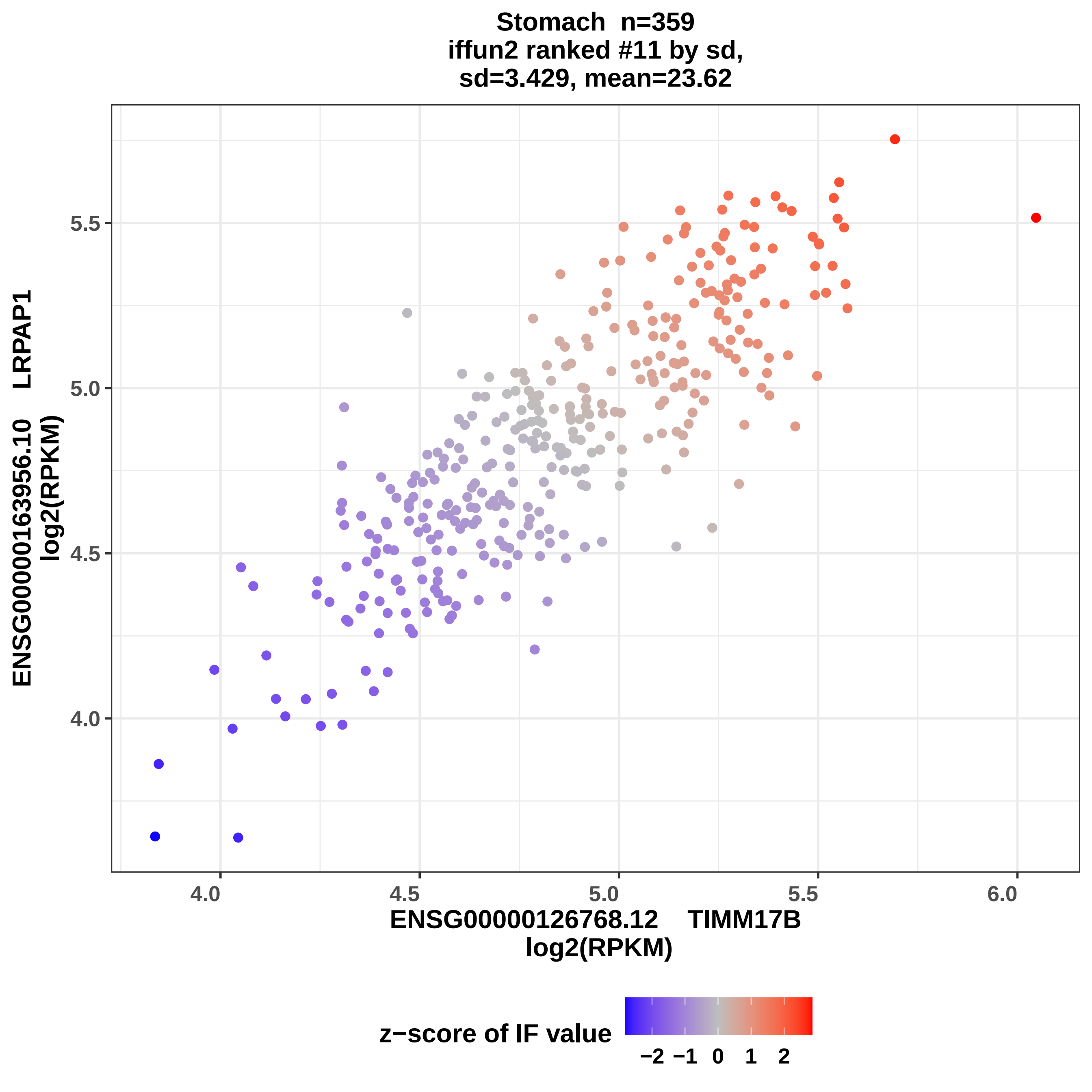

Stomach n=359
iffun2 ranked #11 by sd,
sd=3.429, mean=23.62
ENSG00000163956.10 LRPAP1
log2(RPKM)
5.5
5.0
4.5
4.0
4.0
4.5
5.0
5.5
6.0
ENSG00000126768.12 TIMM17B
log2(RPKM)
z−score of IF value
−2 −1 0 1 2

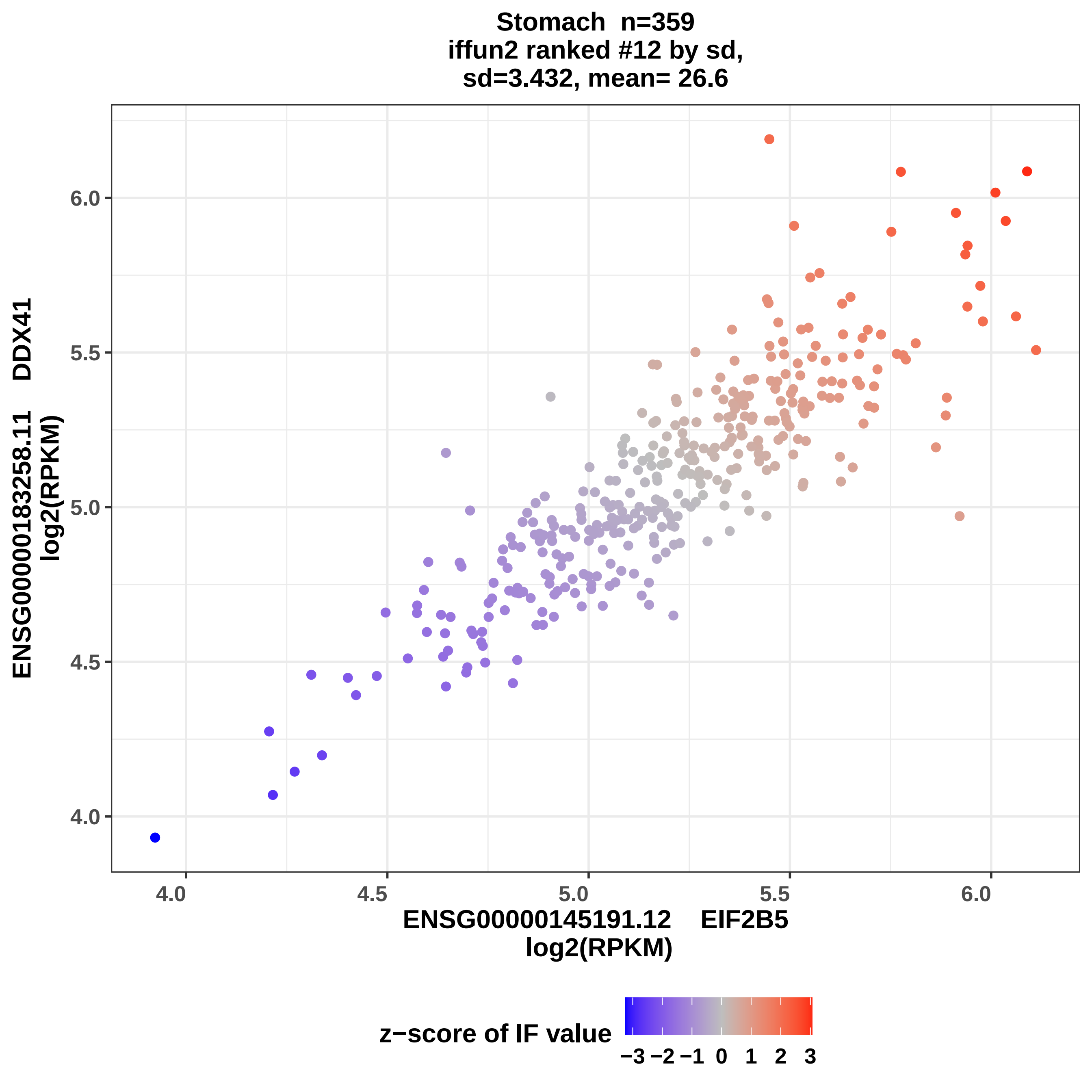
Stomach n=359
iffun2 ranked #12 by sd,
sd=3.432, mean= 26.6
ENSG00000183258.11 DDX41
log2(RPKM)
6.0
5.5
5.0
4.5
4.0
4.0
4.5
5.0
5.5
6.0
ENSG00000145191.12 EIF2B5
log2(RPKM)
z−score of IF value
−3 −2 −1 0 1 2 3

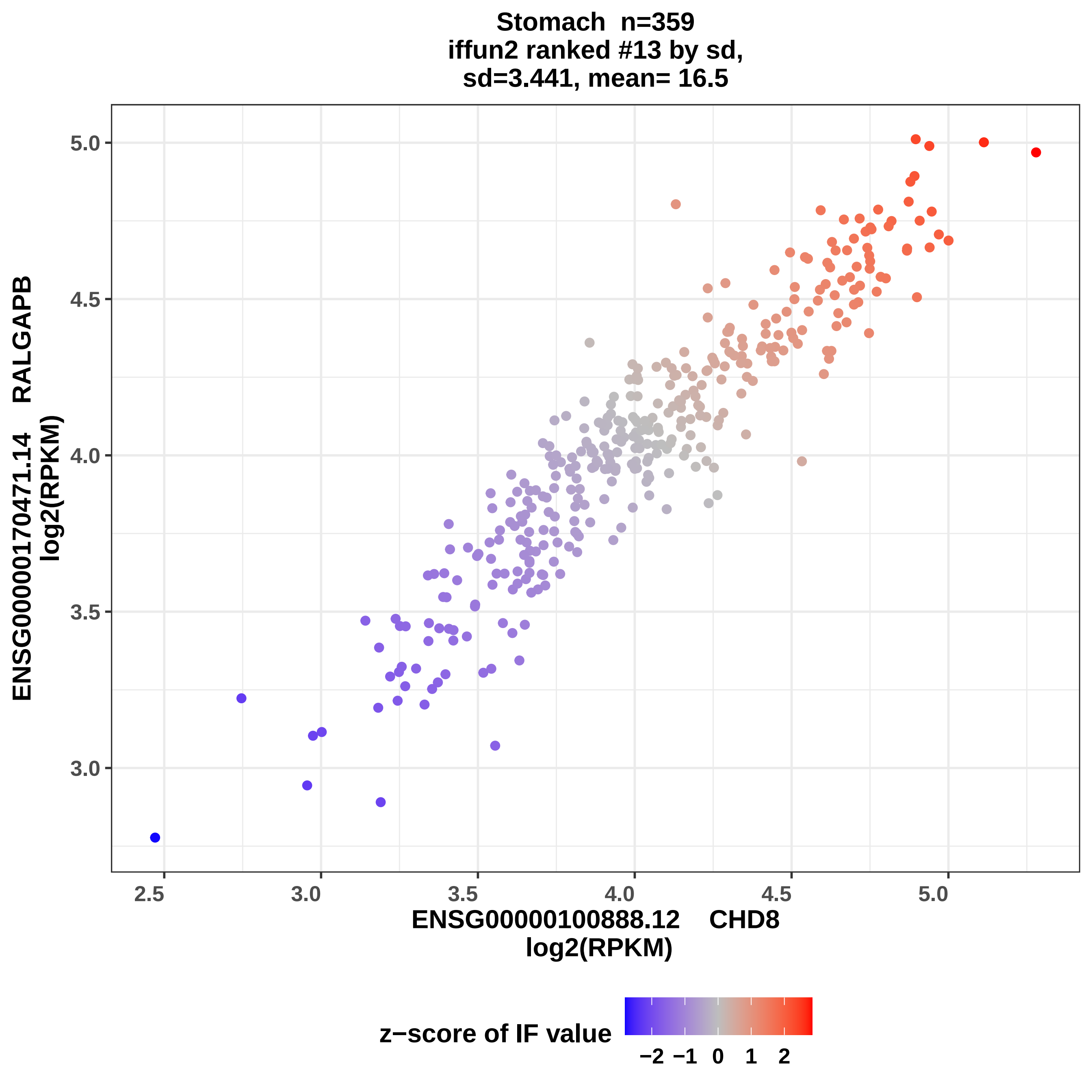
Stomach n=359
iffun2 ranked #13 by sd,
sd=3.441, mean= 16.5
ENSG00000170471.14 RALGAPB
log2(RPKM)
ENSG00000100888.12 CHD8
log2(RPKM)
z−score of IF value
−2 −1 0 1 2
5.0
4.5
4.0
3.5
3.0
2.5
3.0
3.5
4.0
4.5
5.0

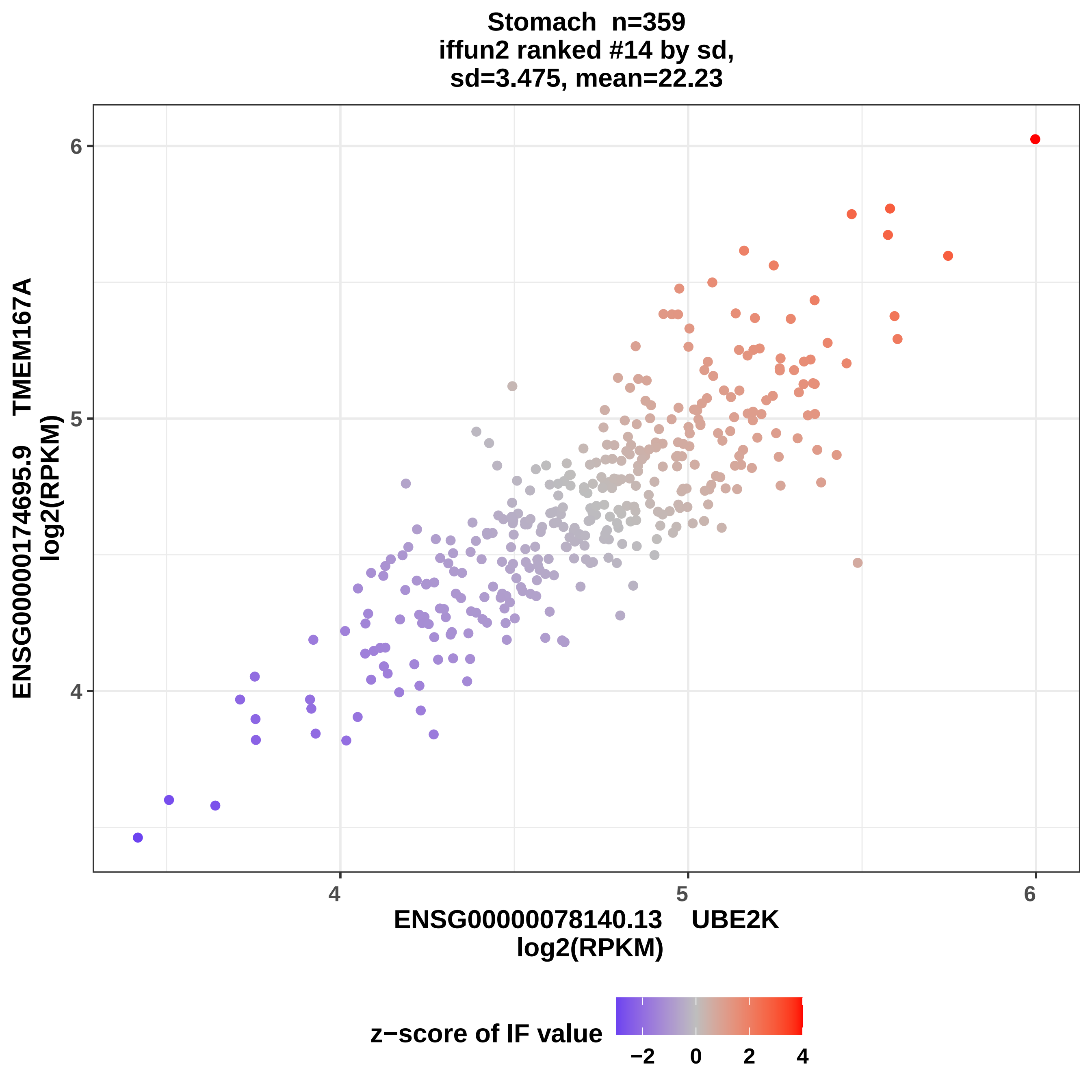

Stomach n=359
iffun2 ranked #14 by sd,
sd=3.475, mean=22.23
ENSG00000174695.9 TMEM167A
log2(RPKM)
ENSG00000078140.13 UBE2K
log2(RPKM)
6
5
4
4
5
6
z−score of IF value
−2
0
2
4

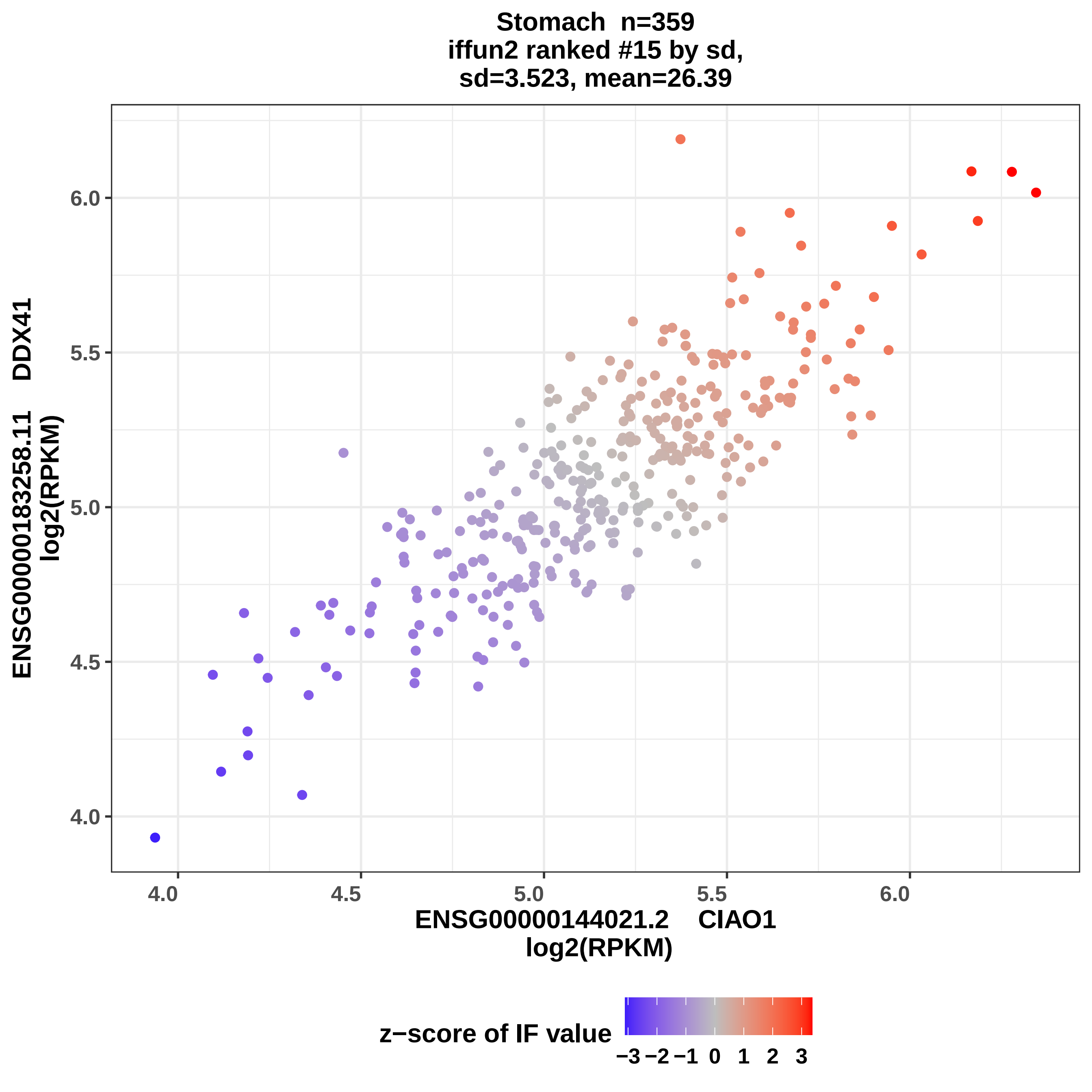

Stomach n=359
iffun2 ranked #15 by sd,
sd=3.523, mean=26.39
ENSG00000183258.11 DDX41
log2(RPKM)
6.0
5.5
5.0
4.5
4.0
4.0
4.5
5.0
5.5
6.0
ENSG00000144021.2 CIAO1
log2(RPKM)
z−score of IF value
−3 −2 −1 0 1 2 3

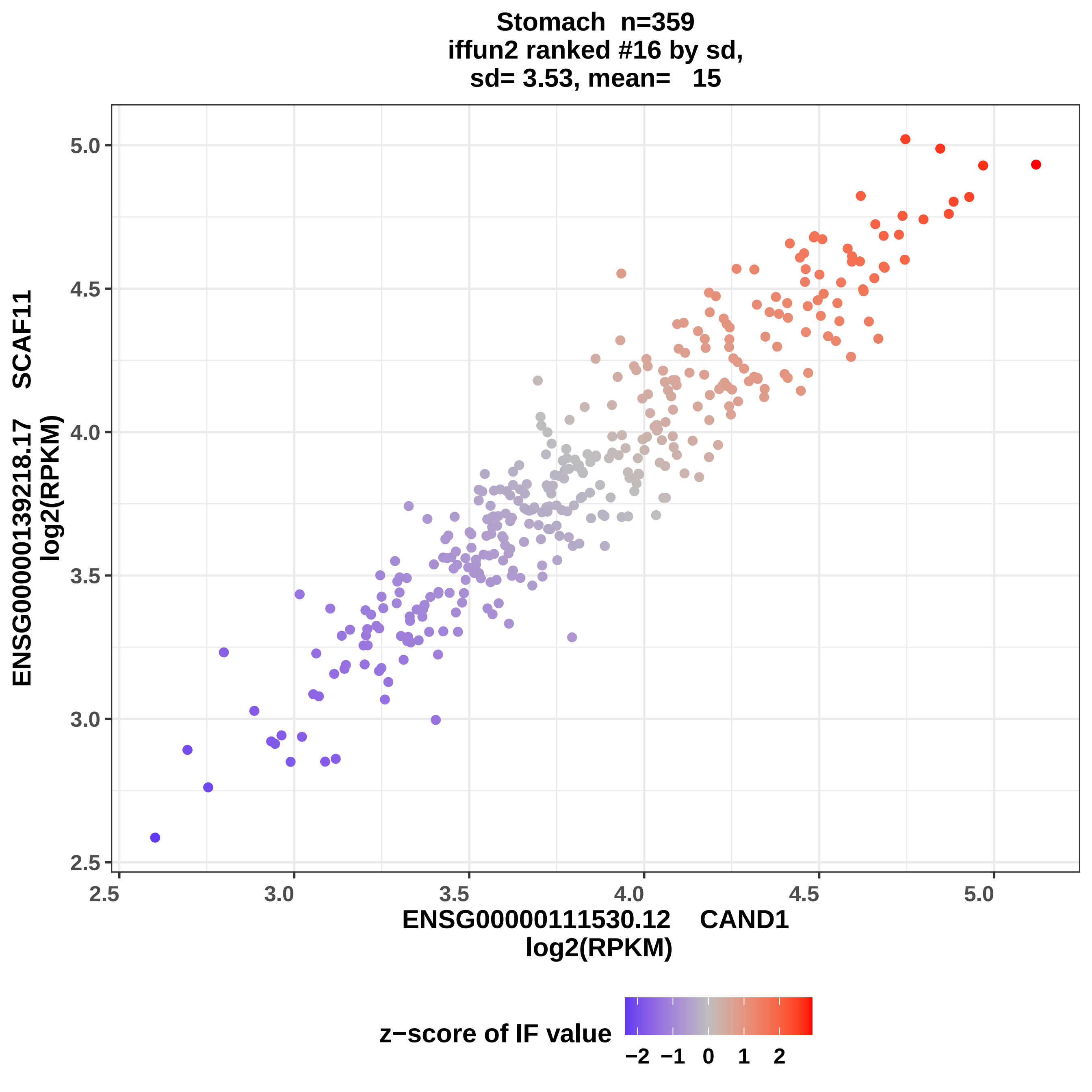
Stomach n=359
iffun2 ranked #16 by sd,
sd= 3.53, mean= 15
ENSG00000139218.17 SCAF11
log2(RPKM)
5.0
4.5
4.0
3.5
3.0
2.5
2.5
3.0
3.5
4.0
4.5
5.0
ENSG00000111530.12 CAND1
log2(RPKM)
z−score of IF value
−2
−1
0
1
2

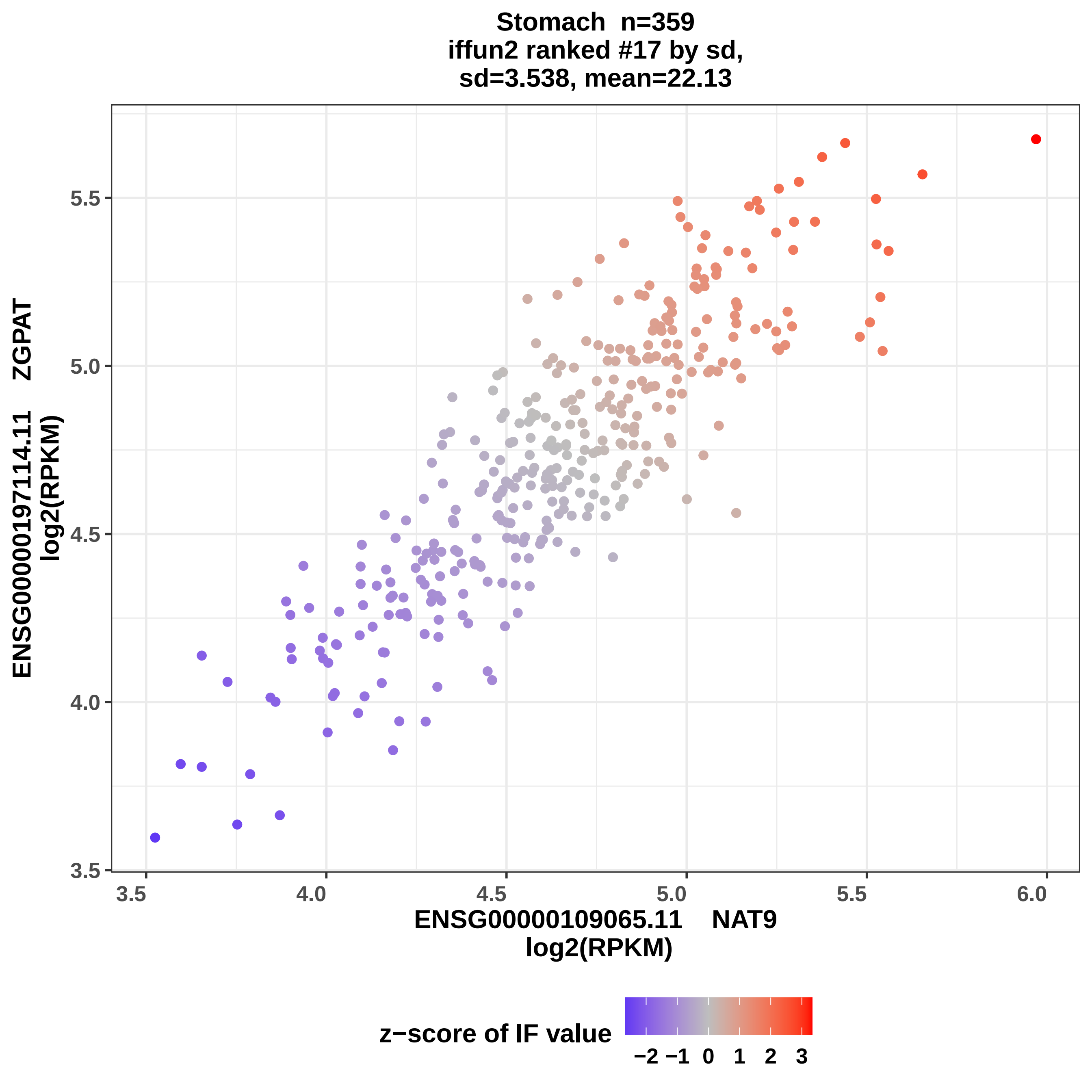
Stomach n=359
iffun2 ranked #17 by sd,
sd=3.538, mean=22.13
ENSG00000197114.11 ZGPAT
log2(RPKM)
5.5
5.0
4.5
4.0
3.5
3.5
4.0
4.5
5.0
5.5
6.0
ENSG00000109065.11 NAT9
log2(RPKM)
z−score of IF value
−2 −1 0 1 2 3

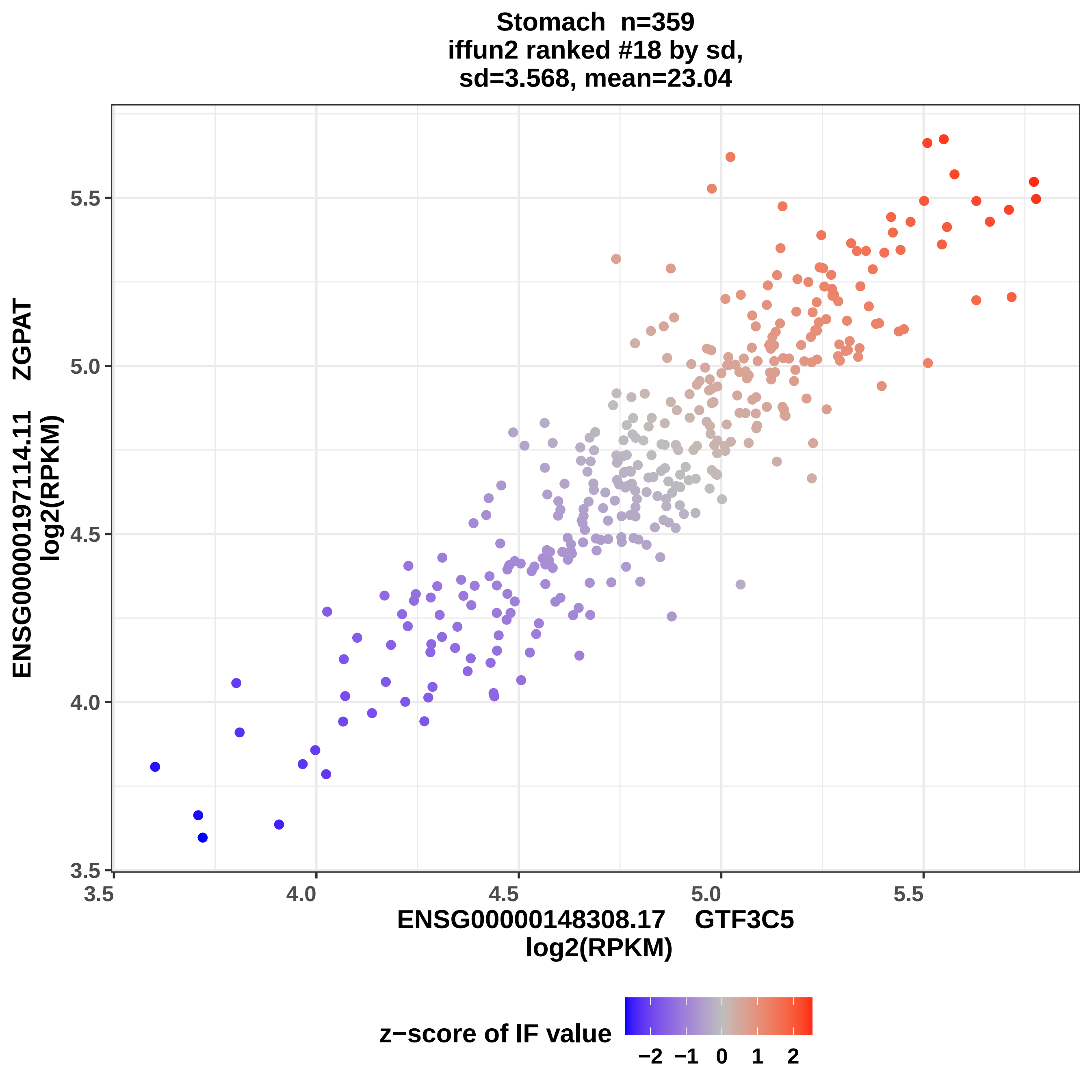
Stomach n=359
iffun2 ranked #18 by sd,
sd=3.568, mean=23.04
ENSG00000197114.11 ZGPAT
log2(RPKM)
5.5
5.0
4.5
4.0
3.5
3.5
4.0
4.5
5.0
5.5
ENSG00000148308.17 GTF3C5
log2(RPKM)
z−score of IF value
−2
−1
0
1
2

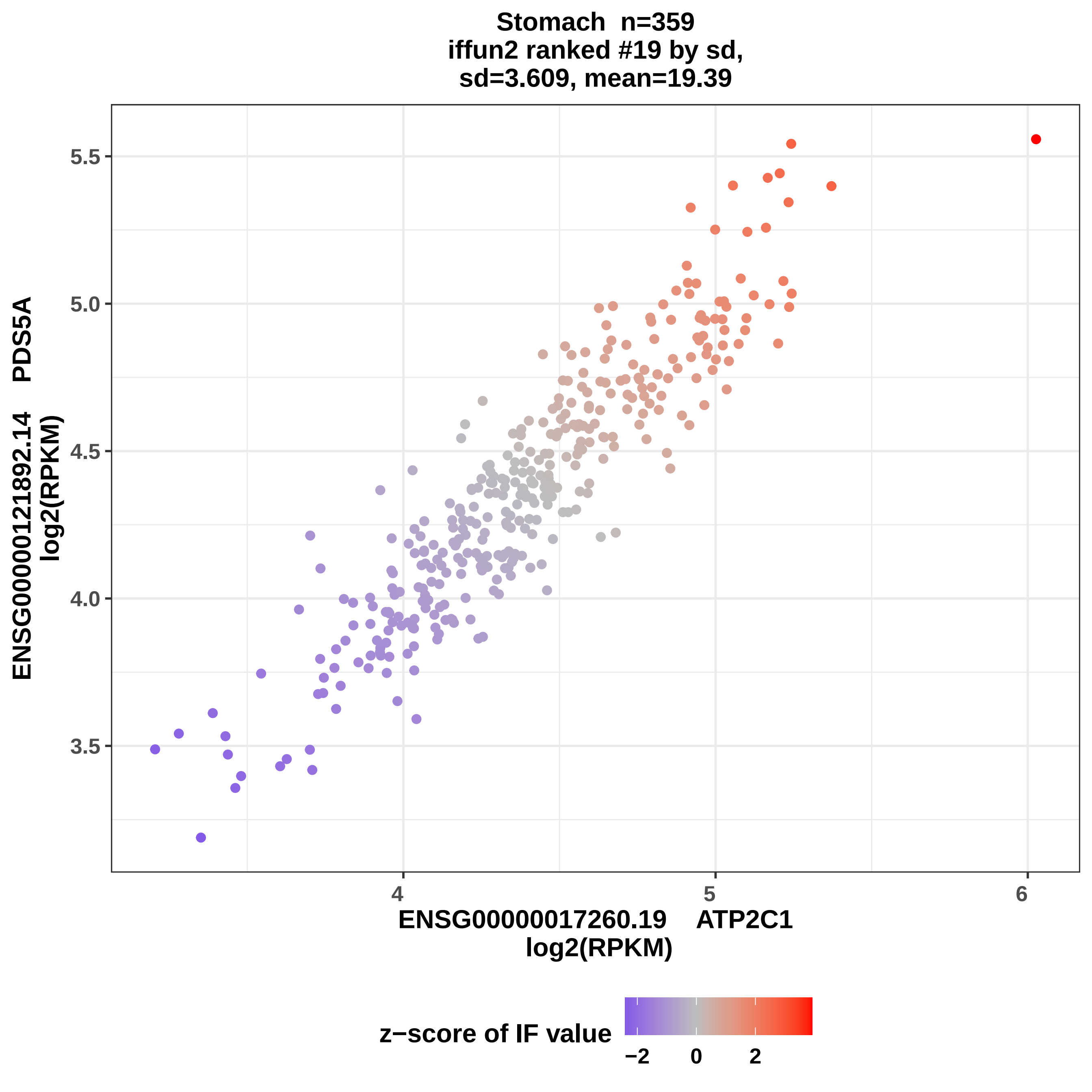

Stomach n=359
iffun2 ranked #19 by sd,
sd=3.609, mean=19.39
ENSG00000121892.14 PDS5A
log2(RPKM)
5.5
5.0
4.5
4.0
3.5
4
5
6
ENSG00000017260.19 ATP2C1
log2(RPKM)
z−score of IF value
−2
0
2

### Stomach n=359
### iffun2 ranked #20 by sd,
### sd=3.615, mean=21.15

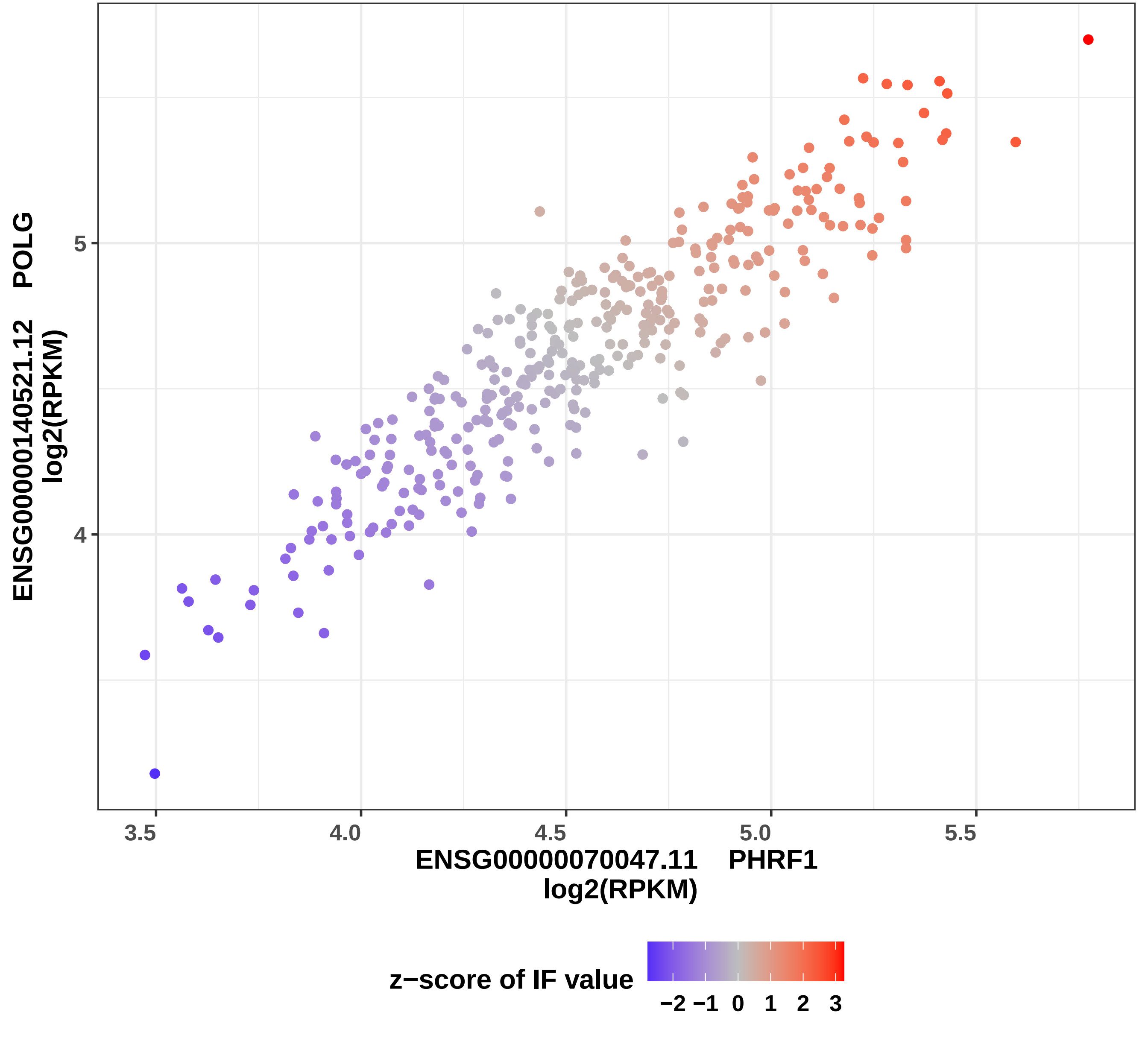

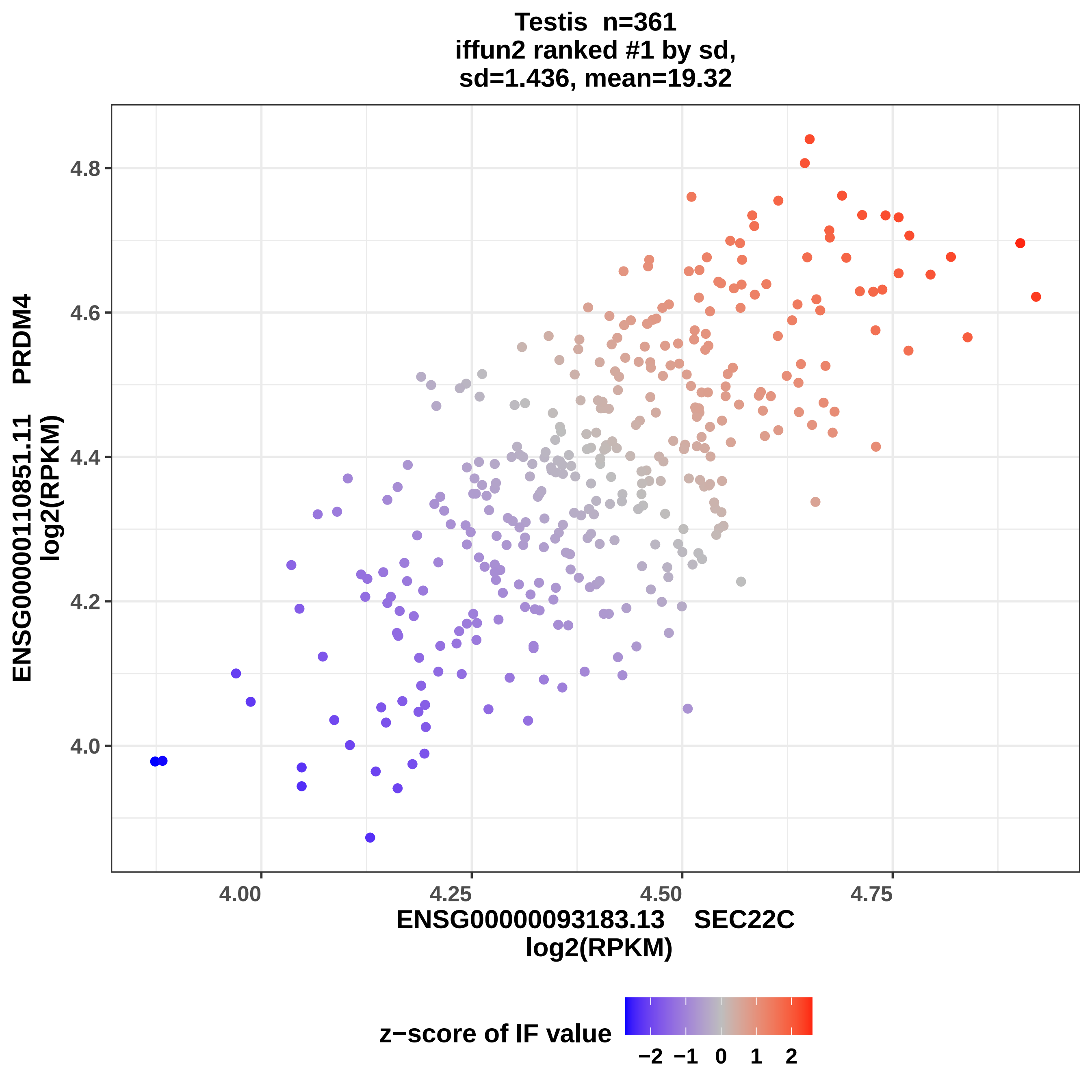
Testis n=361
iffun2 ranked #1 by sd,
sd=1.436, mean=19.32
ENSG00000110851.11 PRDM4
log2(RPKM)
4.0
4.2
4.4
4.6
4.8
4.00
4.25
4.50
4.75
ENSG00000093183.13 SEC22C
log2(RPKM)
z−score of IF value
−2 −1 0 1 2

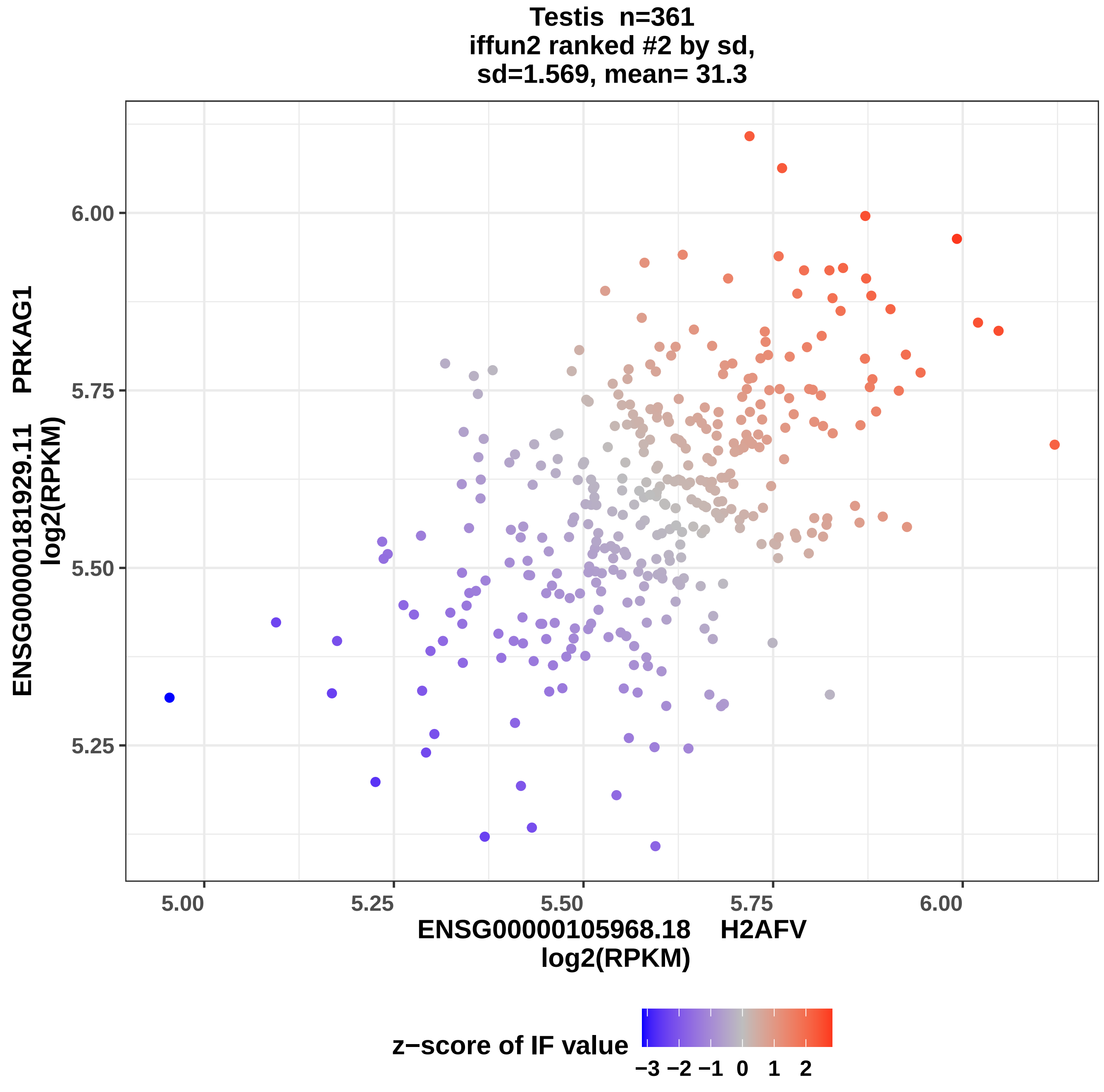

Testis n=361
iffun2 ranked #2 by sd,
sd=1.569, mean= 31.3
6.00
5.75
5.50
5.25
5.00
5.25
5.50
5.75
6.00
ENSG00000181929.11 PRKAG1
log2(RPKM)
ENSG00000105968.18 H2AFV
log2(RPKM)
z−score of IF value
−3 −2 −1 0 1 2

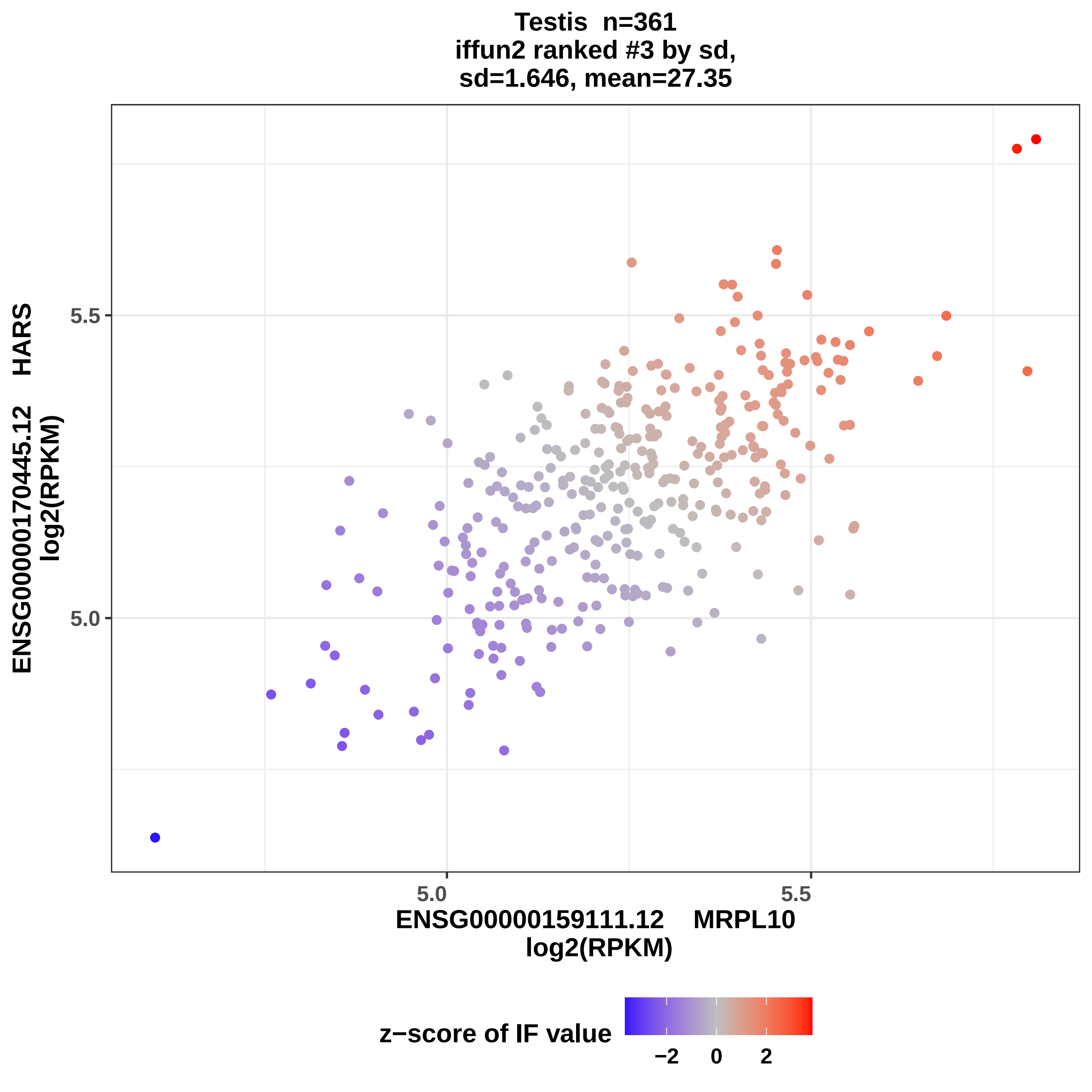
Testis n=361
iffun2 ranked #3 by sd,
sd=1.646, mean=27.35
ENSG00000170445.12 HARS
log2(RPKM)
5.5
5.0
5.0
5.5
ENSG00000159111.12 MRPL10
log2(RPKM)
z−score of IF value
−2
0
2

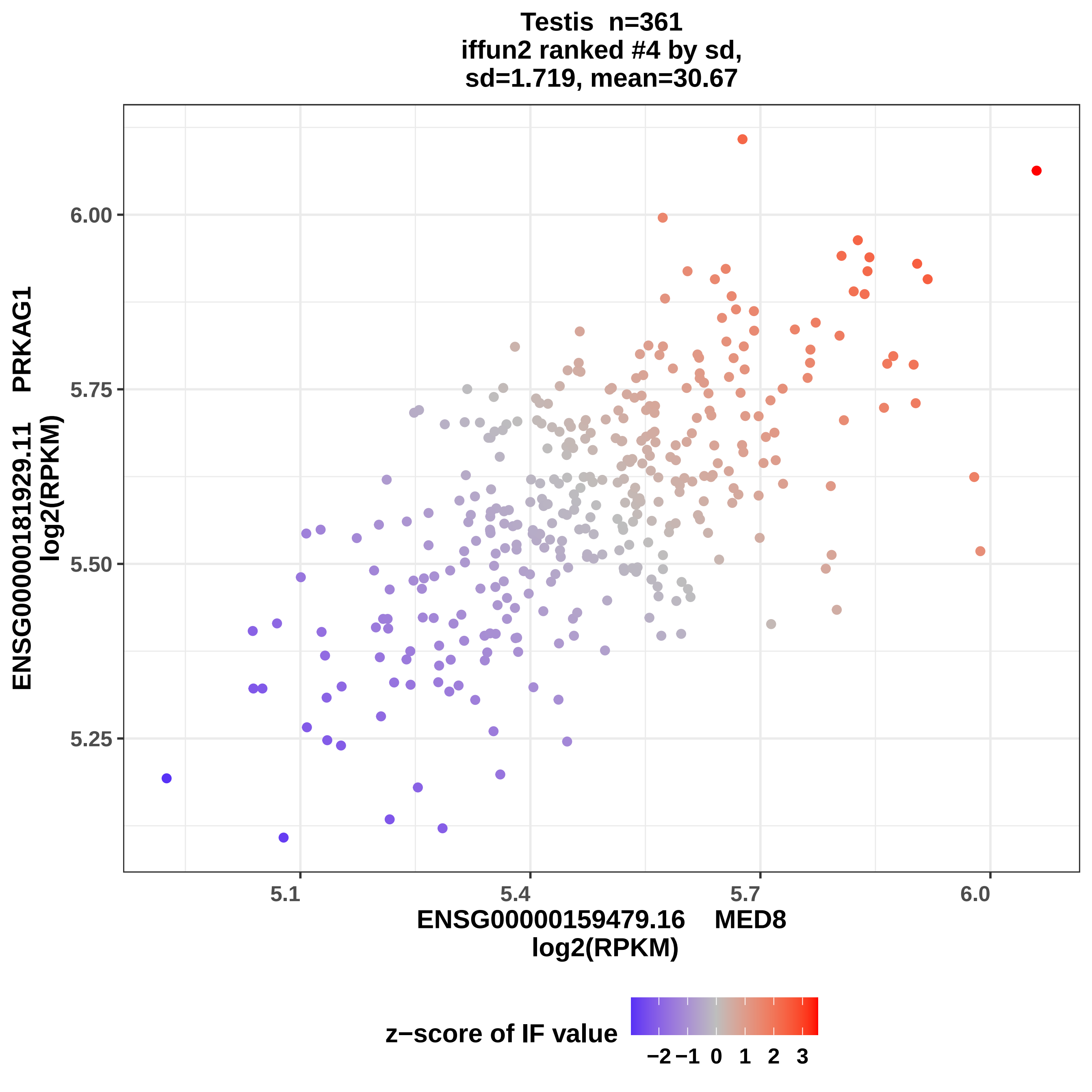
Testis n=361
iffun2 ranked #4 by sd,
sd=1.719, mean=30.67
ENSG00000181929.11 PRKAG1
log2(RPKM)
6.00
5.75
5.50
5.25
5.1
5.4
5.7
6.0
ENSG00000159479.16 MED8
log2(RPKM)
z−score of IF value
−2
−1
0
1
2
3

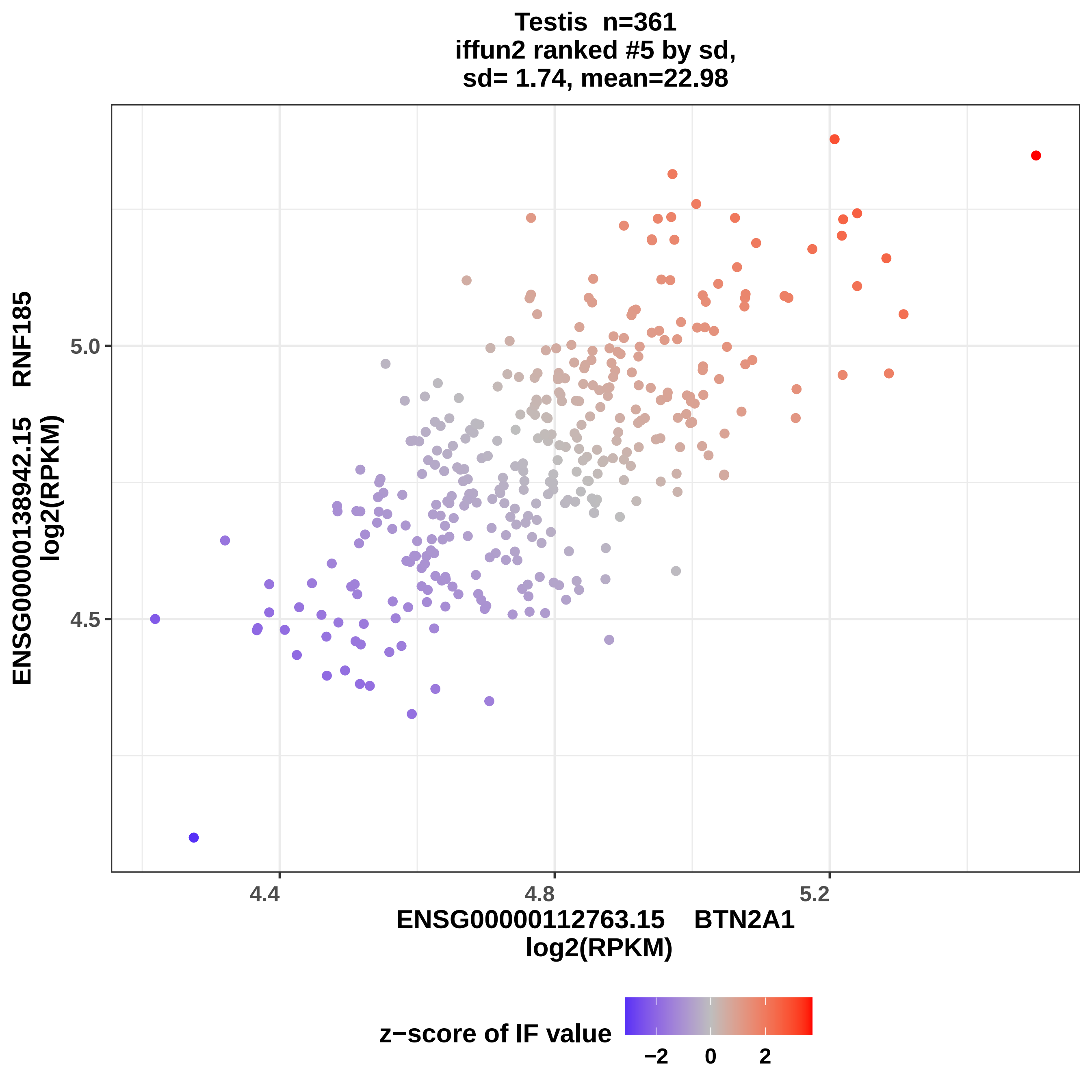
Testis n=361
iffun2 ranked #5 by sd,
sd= 1.74, mean=22.98
ENSG00000138942.15 RNF185
log2(RPKM)
5.0
4.5
4.4
4.8
5.2
ENSG00000112763.15 BTN2A1
log2(RPKM)
z−score of IF value
−2
0
2

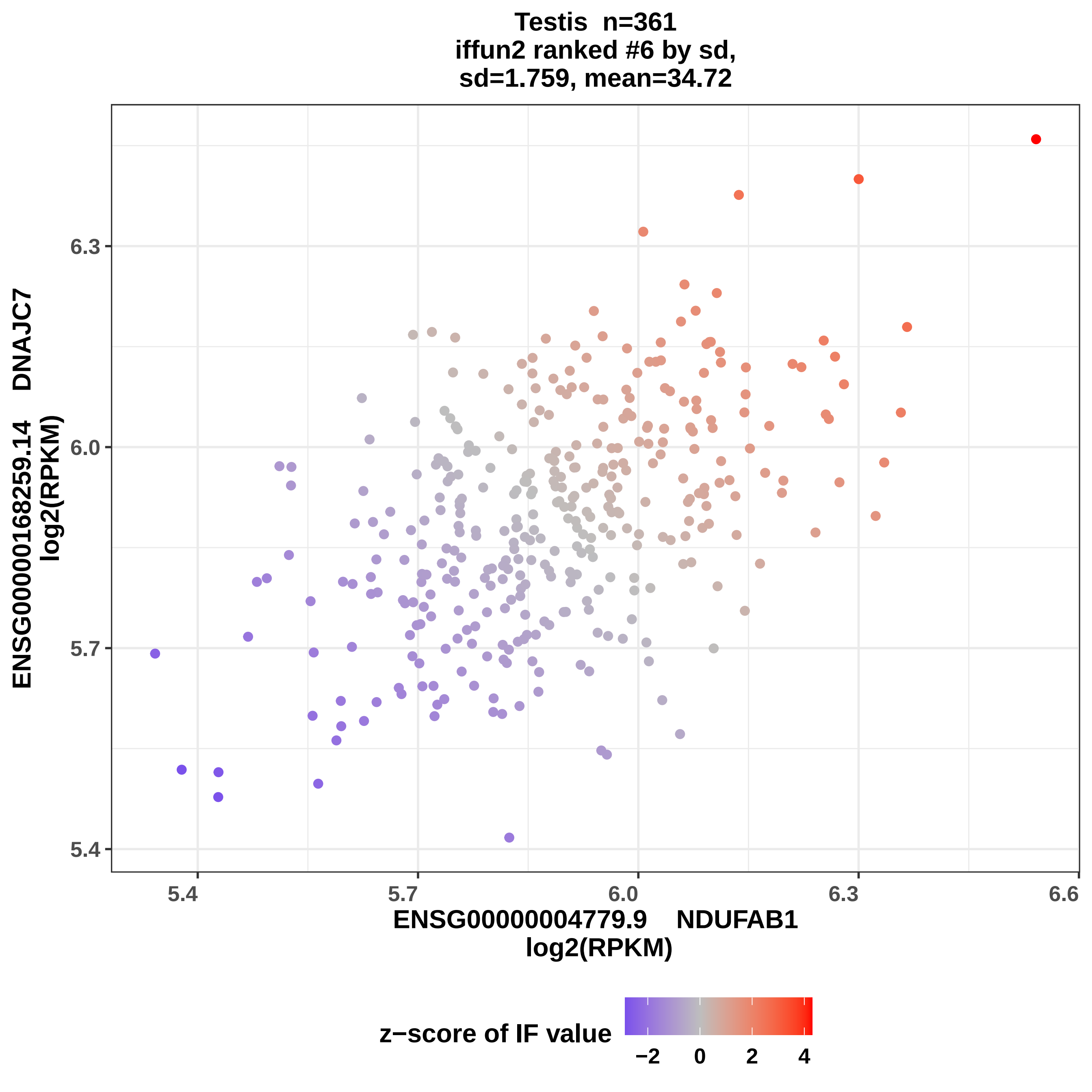
Testis n=361
iffun2 ranked #6 by sd,
sd=1.759, mean=34.72
ENSG00000168259.14 DNAJC7
log2(RPKM)
6.3
6.0
5.7
5.4
5.4
5.7
6.0
6.3
6.6
ENSG00000004779.9 NDUFAB1
log2(RPKM)
z−score of IF value
−2
0
2
4

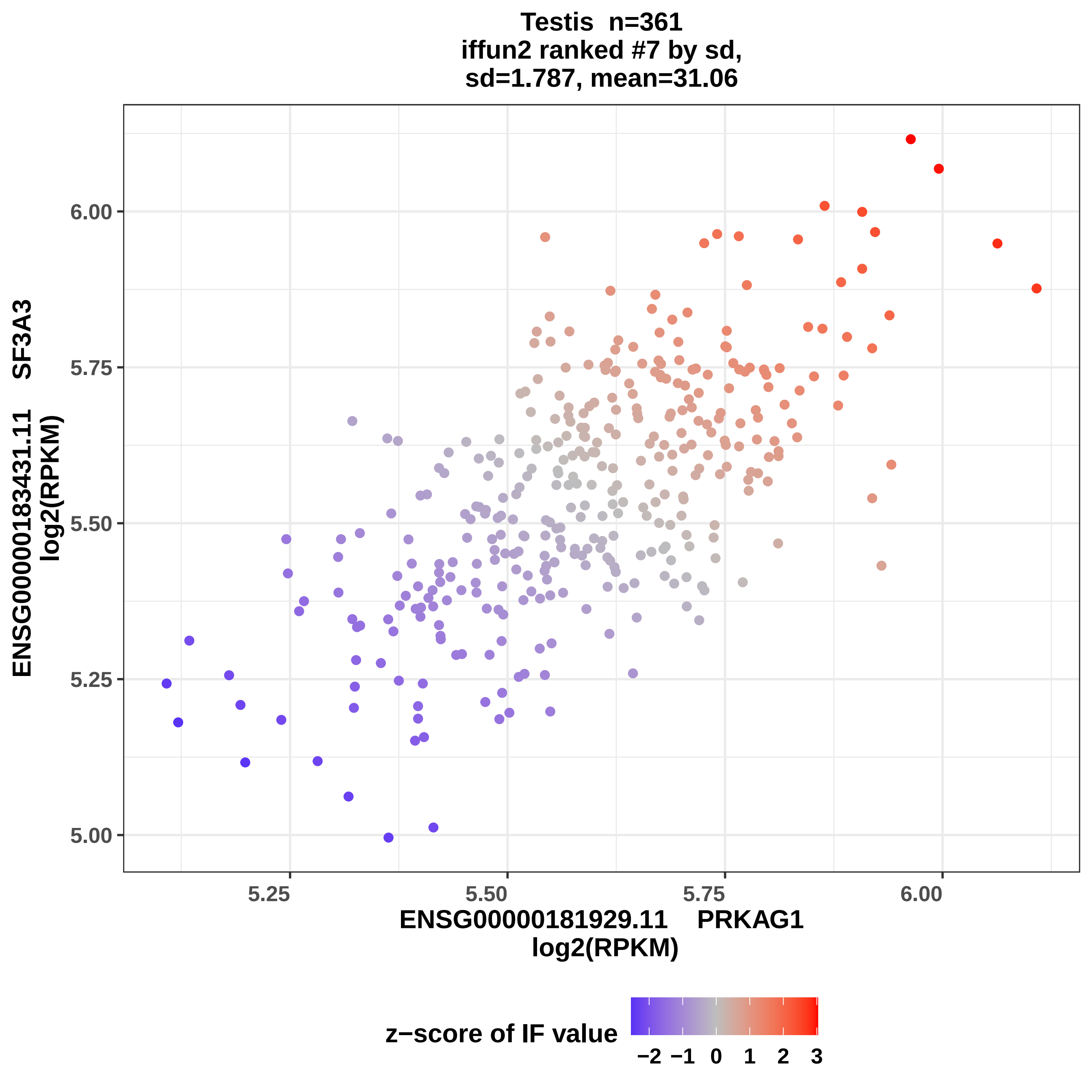
Testis n=361
iffun2 ranked #7 by sd,
sd=1.787, mean=31.06
ENSG00000183431.11 SF3A3
log2(RPKM)
6.00
5.75
5.50
5.25
5.00
5.25
5.50
5.75
6.00
ENSG00000181929.11 PRKAG1
log2(RPKM)
z−score of IF value
−2
−1
0
1
2
3

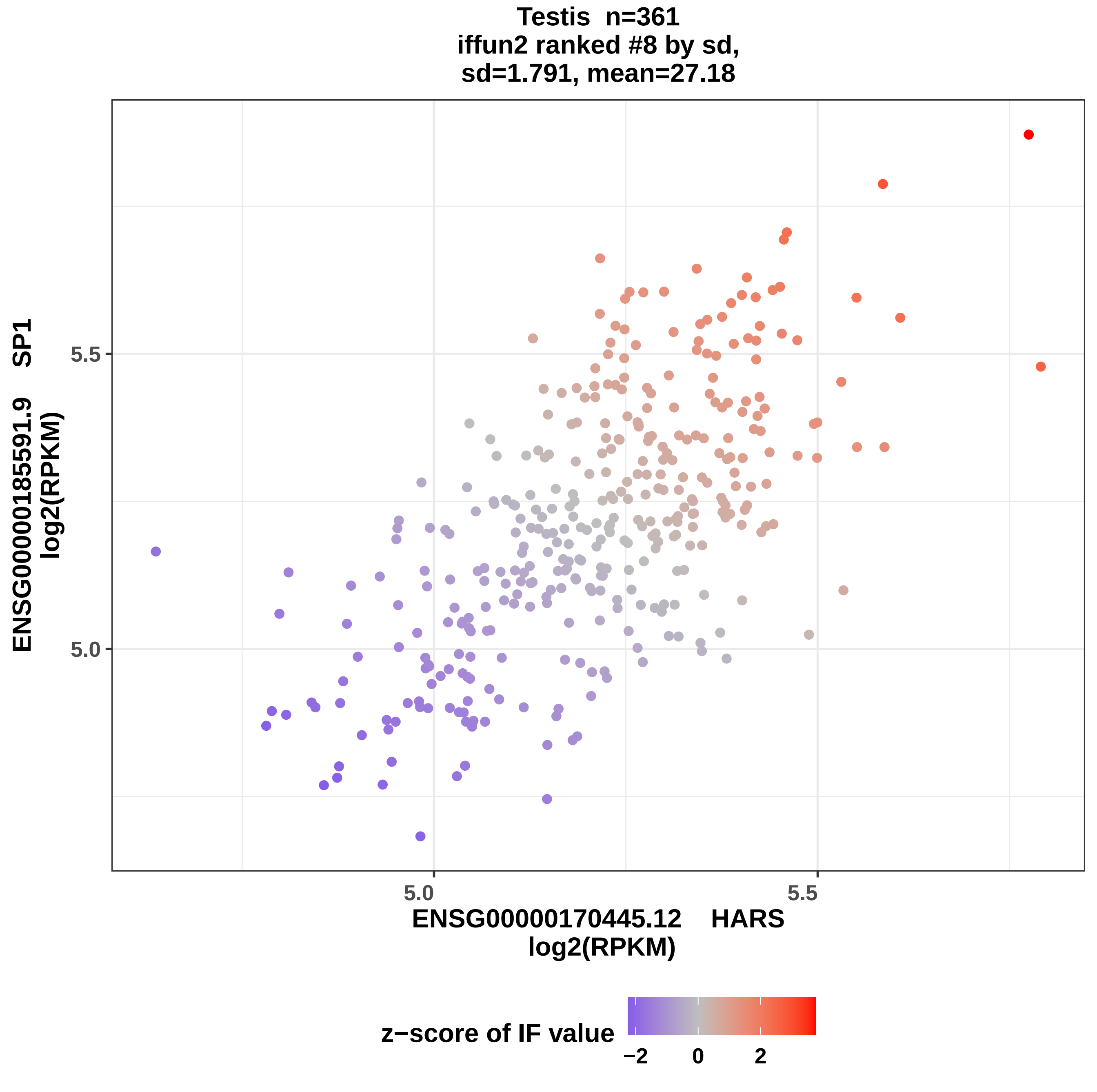
Testis n=361
iffun2 ranked #8 by sd,
sd=1.791, mean=27.18
ENSG00000185591.9 SP1
log2(RPKM)
5.5
5.0
5.0
5.5
ENSG00000170445.12 HARS
log2(RPKM)
z−score of IF value
−2
0
2

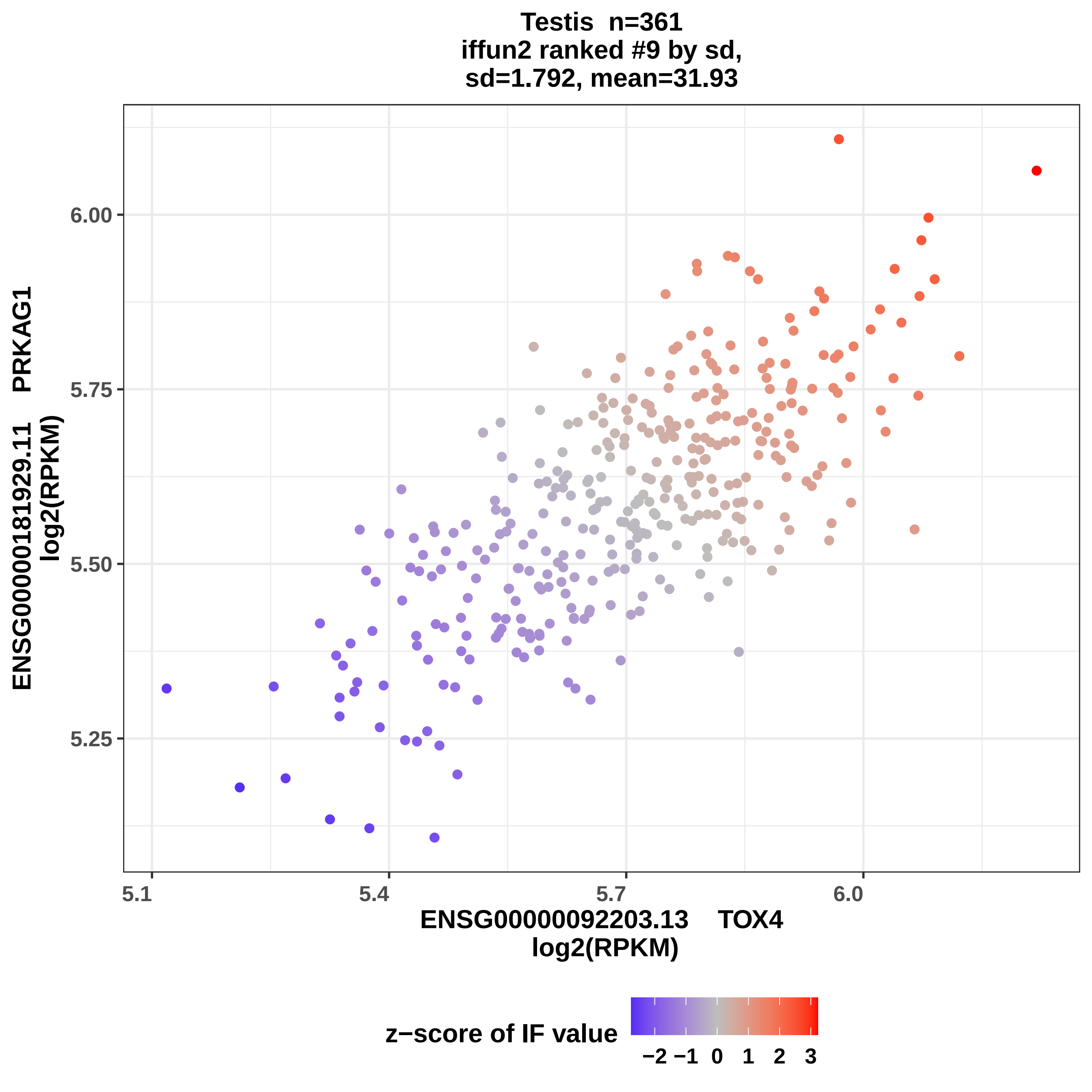

Testis n=361
iffun2 ranked #9 by sd,
sd=1.792, mean=31.93
ENSG00000181929.11 PRKAG1
log2(RPKM)
6.00
5.75
5.50
5.25
5.1
5.4
5.7
6.0
ENSG00000092203.13 TOX4
log2(RPKM)
z−score of IF value
−2 −1 0 1 2 3

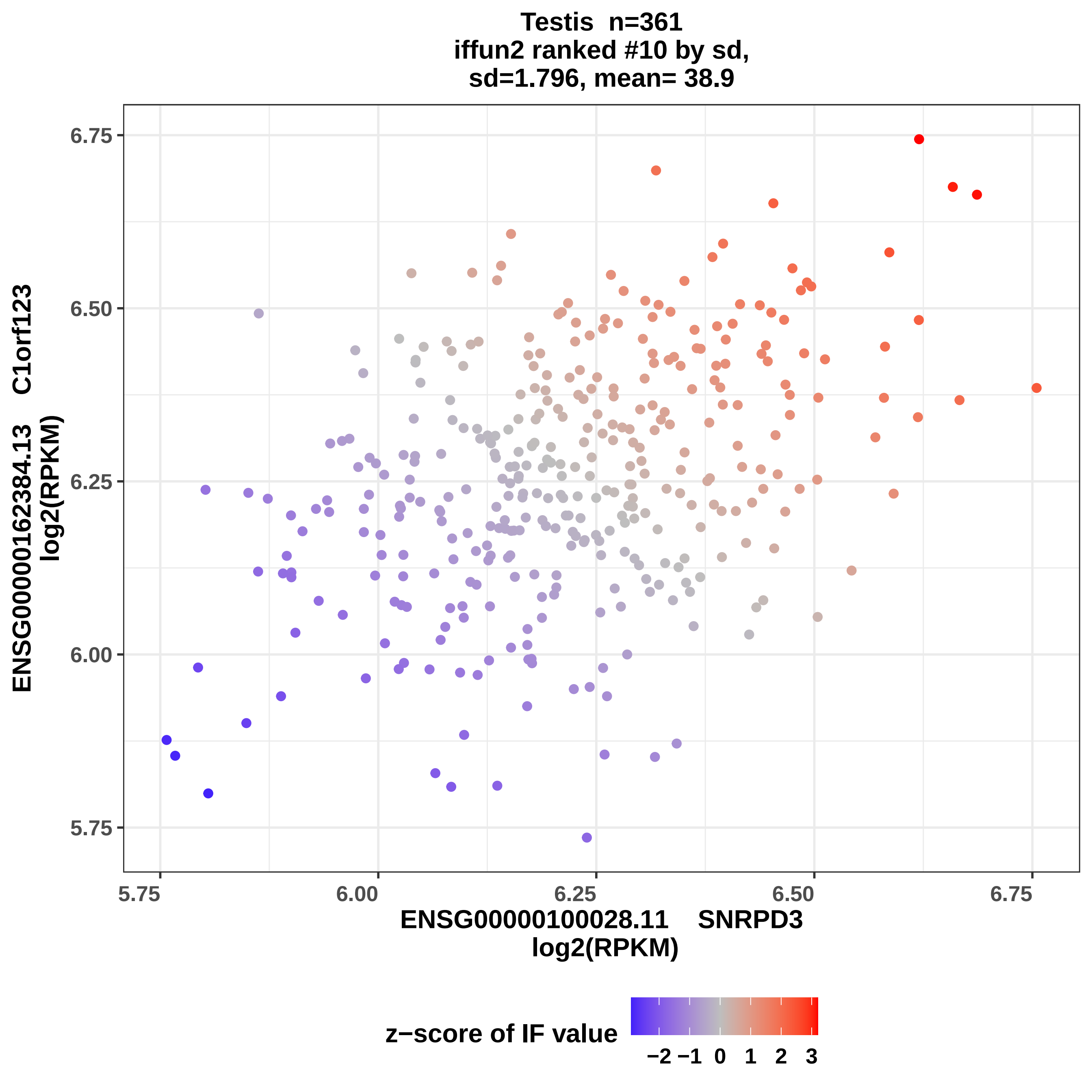
Testis n=361
iffun2 ranked #10 by sd,
sd=1.796, mean= 38.9
ENSG00000162384.13 C1orf123
log2(RPKM)
6.75
6.50
6.25
6.00
5.75
5.75
6.00
6.25
6.50
6.75
ENSG00000100028.11 SNRPD3
log2(RPKM)
z−score of IF value
−2 −1 0 1 2 3

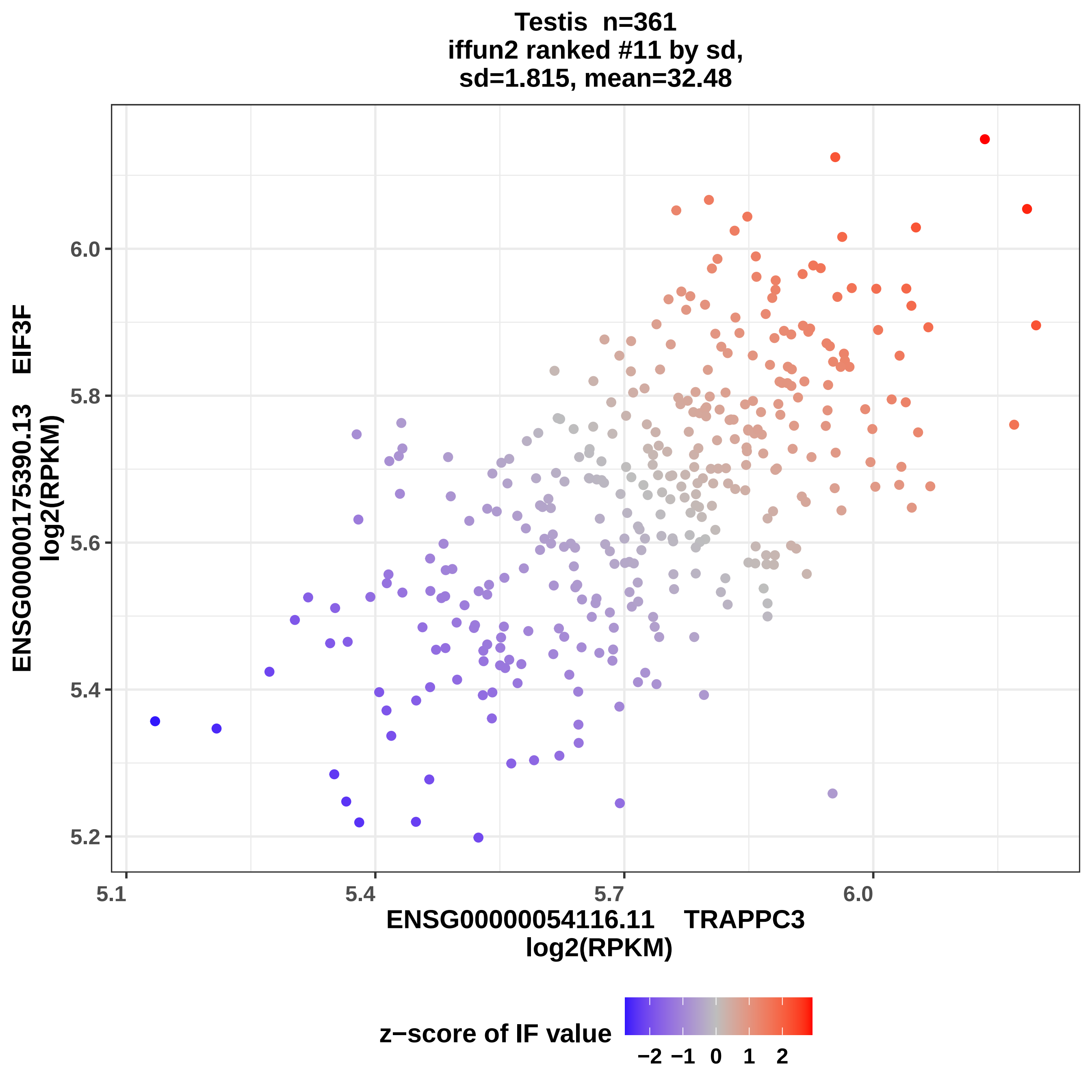
Testis n=361
iffun2 ranked #11 by sd,
sd=1.815, mean=32.48
ENSG00000175390.13 EIF3F
log2(RPKM)
6.0
5.8
5.6
5.4
5.2
5.1
5.4
5.7
6.0
ENSG00000054116.11 TRAPPC3
log2(RPKM)
z−score of IF value
−2 −1 0 1 2

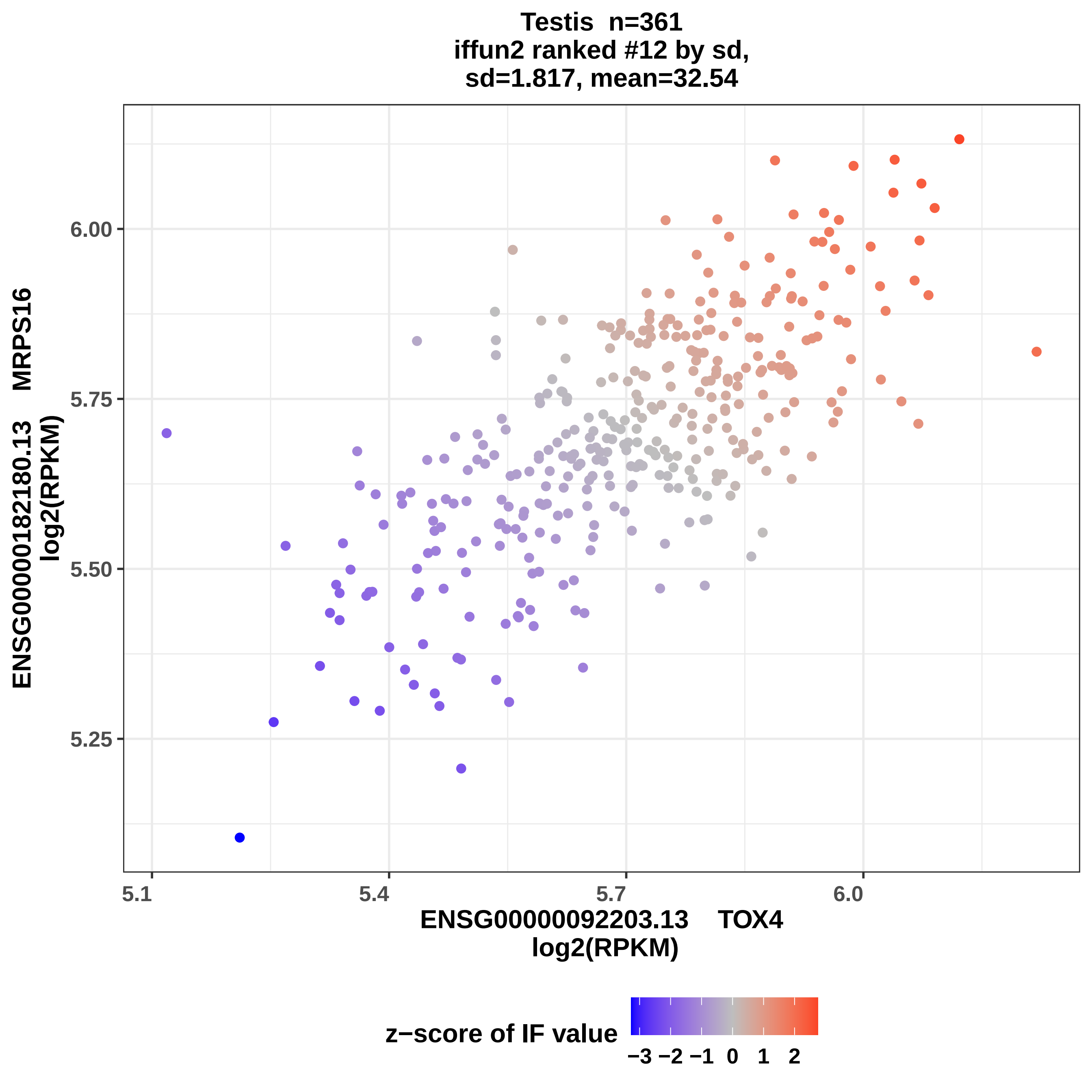

Testis n=361
iffun2 ranked #12 by sd,
sd=1.817, mean=32.54
ENSG00000182180.13 MRPS16
log2(RPKM)
6.00
5.75
5.50
5.25
5.1
5.4
5.7
6.0
ENSG00000092203.13 TOX4
log2(RPKM)
z−score of IF value
−3 −2 −1 0 1 2

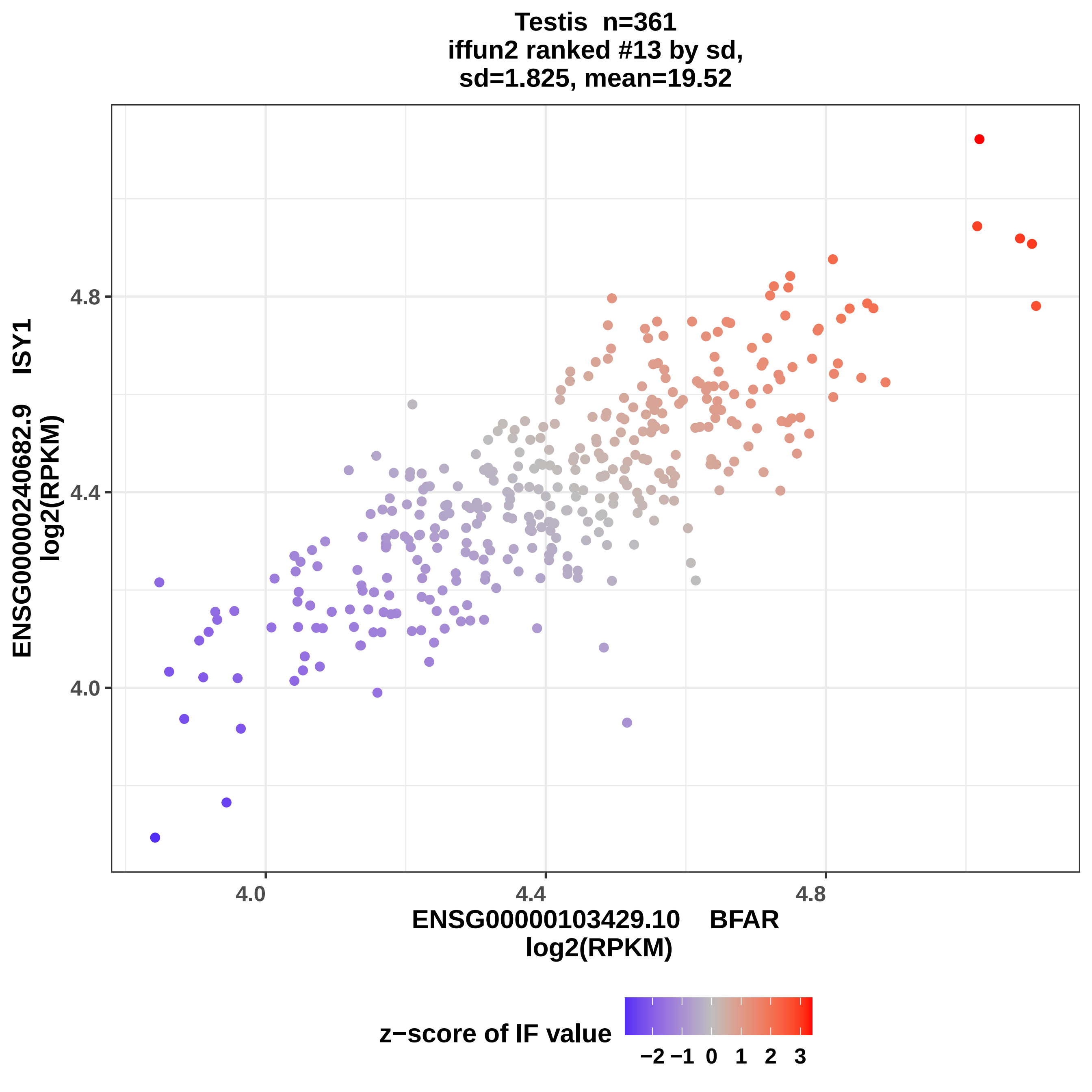
Testis n=361
iffun2 ranked #13 by sd,
sd=1.825, mean=19.52
4.8
4.4
4.0
ENSG00000240682.9 ISY1
log2(RPKM)
4.0
4.4
4.8
ENSG00000103429.10 BFAR
log2(RPKM)
z−score of IF value
−2
−1
0
1
2
3

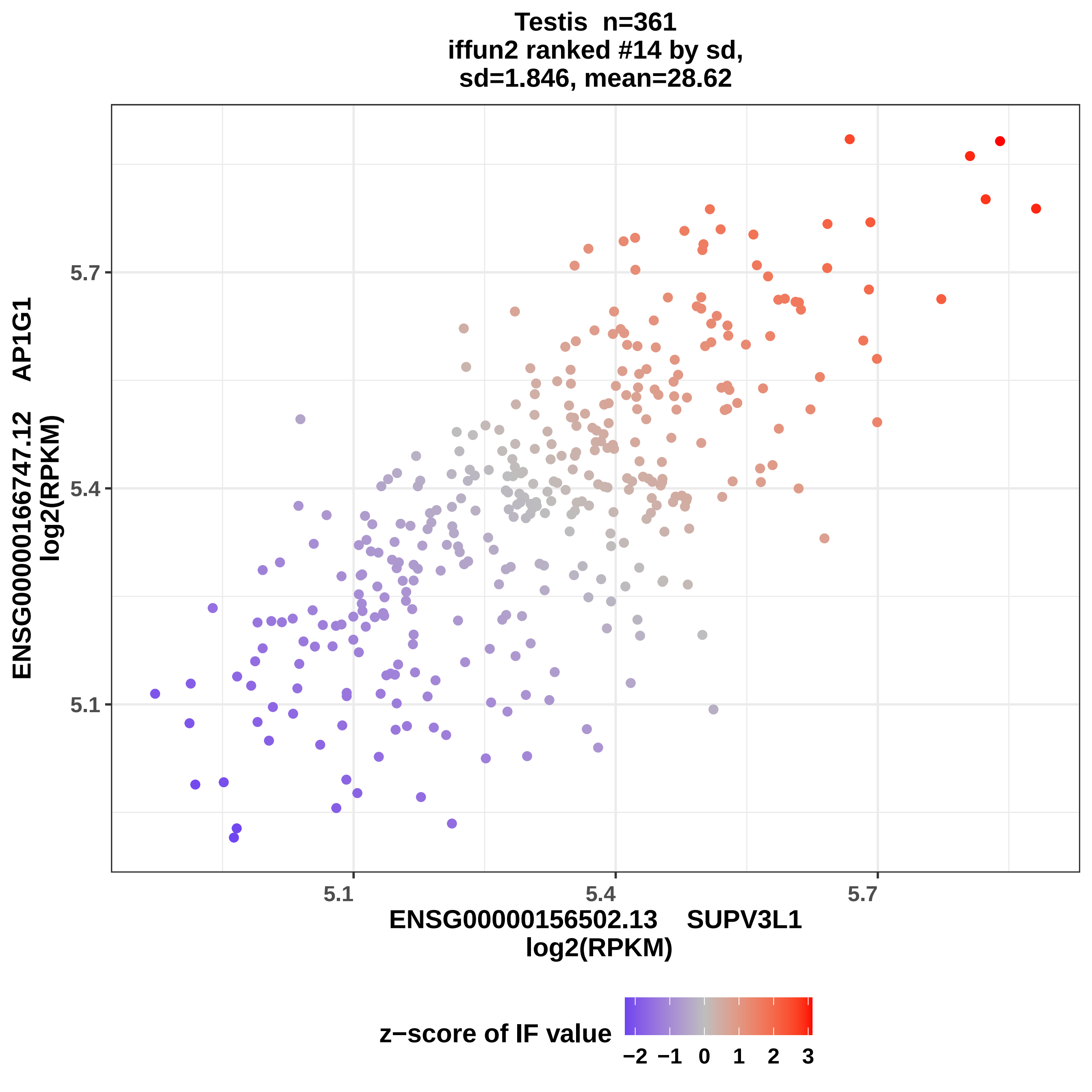

Testis n=361
iffun2 ranked #14 by sd,
sd=1.846, mean=28.62
5.7
5.4
5.1
ENSG00000166747.12 AP1G1
log2(RPKM)
5.1
5.4
5.7
ENSG00000156502.13 SUPV3L1
log2(RPKM)
z−score of IF value
−2
−1
0
1
2
3

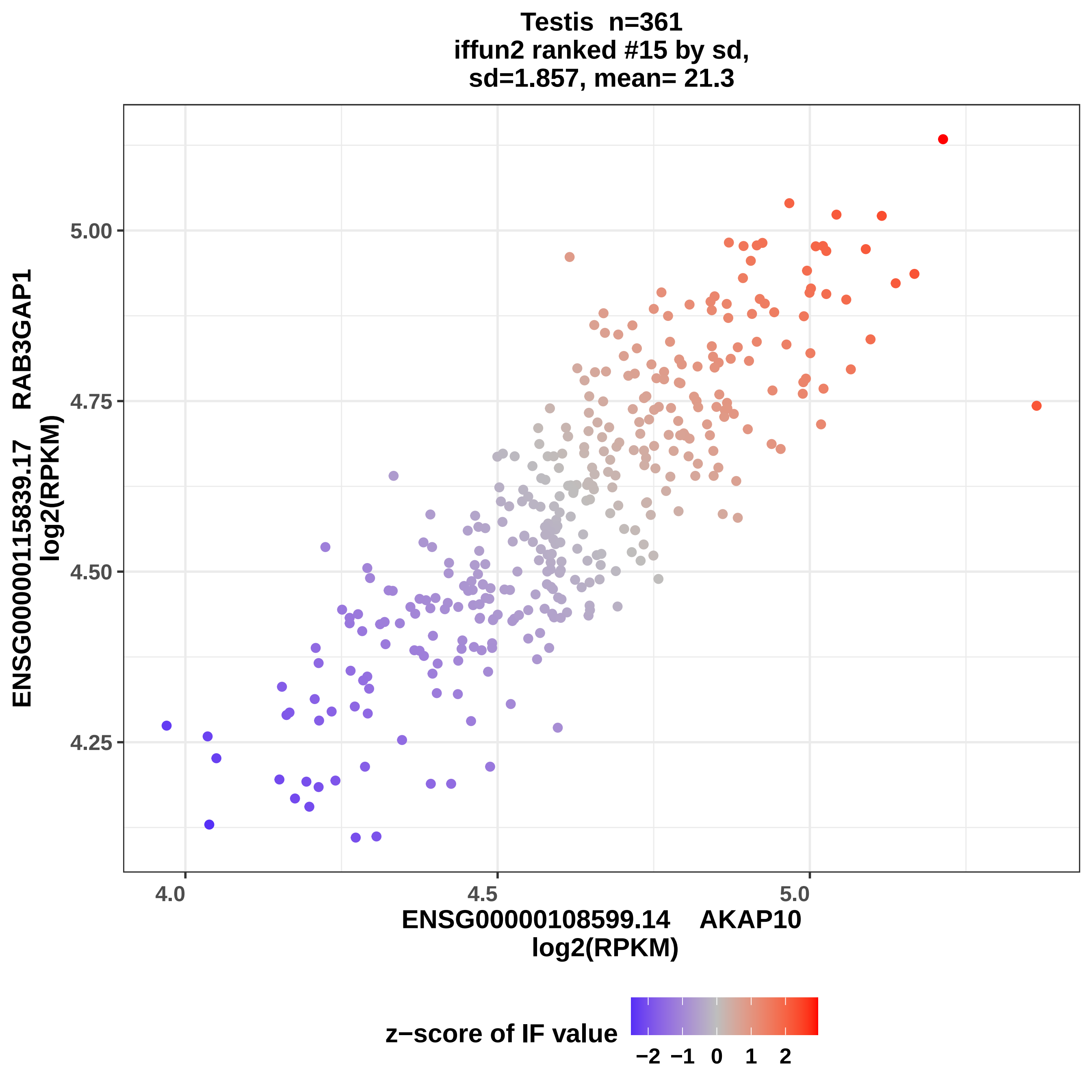

Testis n=361
iffun2 ranked #15 by sd,
sd=1.857, mean= 21.3
ENSG00000115839.17 RAB3GAP1
log2(RPKM)
5.00
4.75
4.50
4.25
4.0
4.5
5.0
ENSG00000108599.14 AKAP10
log2(RPKM)
z−score of IF value
−2
−1
0
1
2

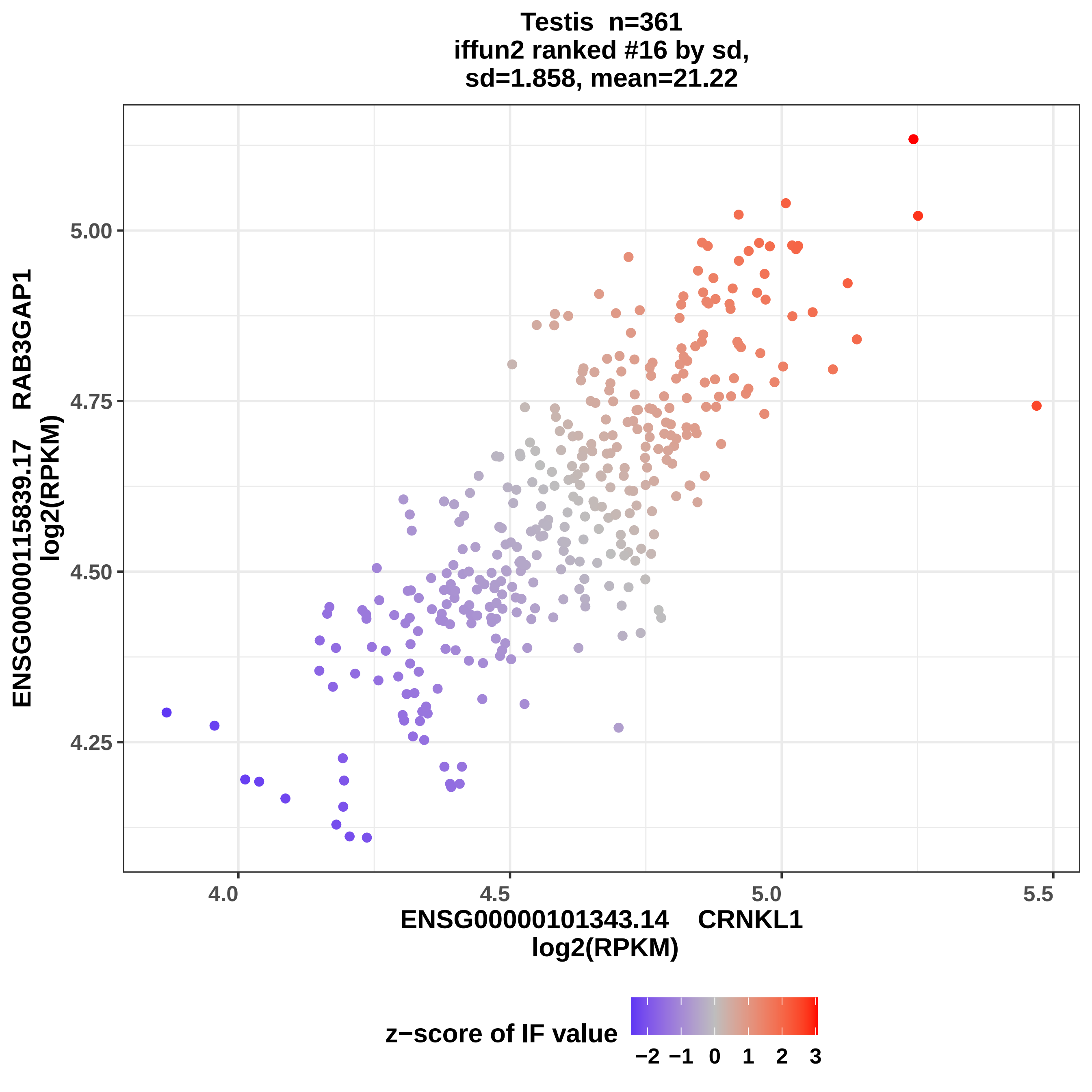

Testis n=361
iffun2 ranked #16 by sd,
sd=1.858, mean=21.22
ENSG00000115839.17 RAB3GAP1
log2(RPKM)
5.00
4.75
4.50
4.25
4.0
4.5
5.0
5.5
ENSG00000101343.14 CRNKL1
log2(RPKM)
z−score of IF value
−2
−1
0
1
2
3

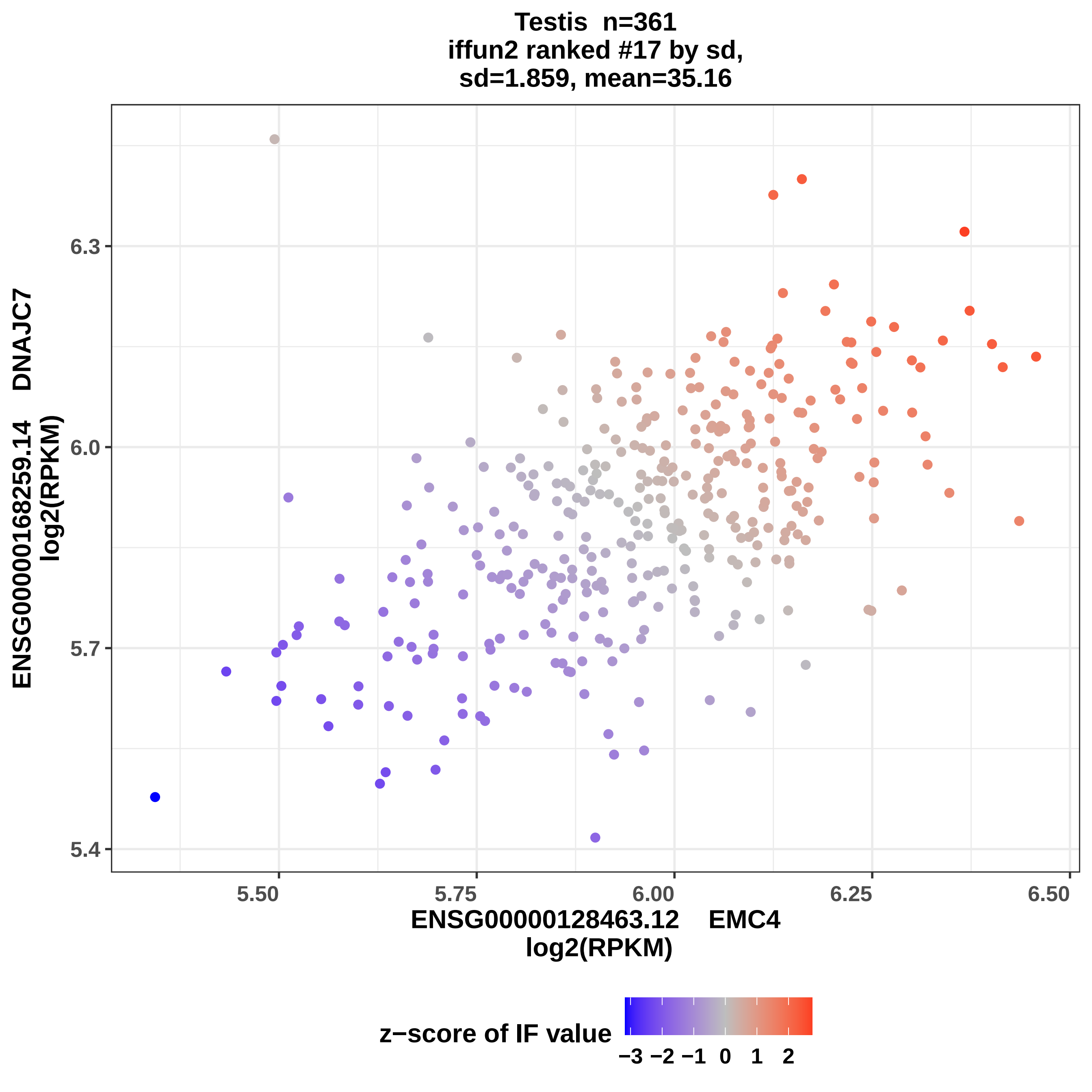

Testis n=361
iffun2 ranked #17 by sd,
sd=1.859, mean=35.16
ENSG00000168259.14 DNAJC7
log2(RPKM)
6.3
6.0
5.7
5.4
5.50
5.75
6.00
6.25
6.50
ENSG00000128463.12 EMC4
log2(RPKM)
z−score of IF value
−3 −2 −1 0 1 2

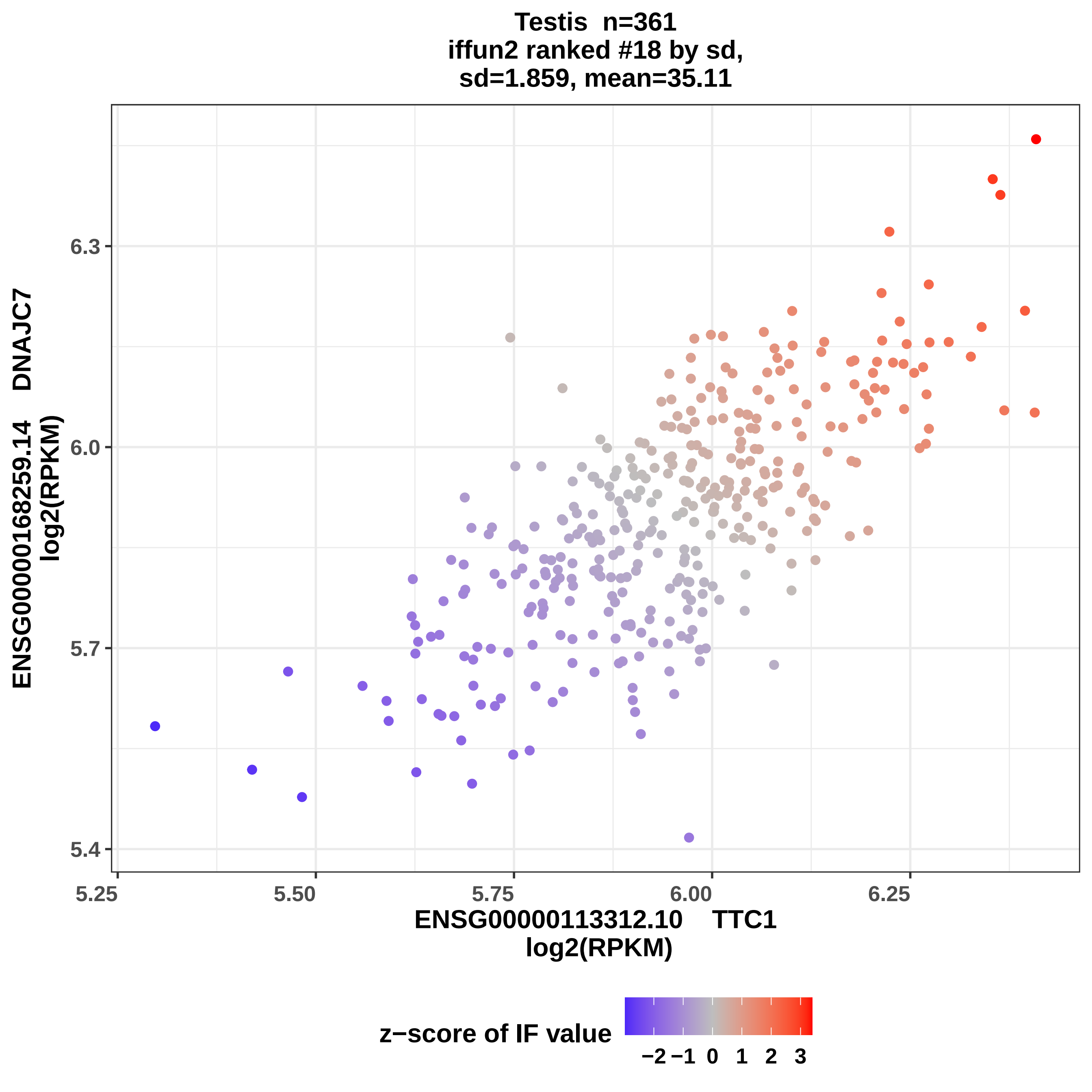
Testis n=361
iffun2 ranked #18 by sd,
sd=1.859, mean=35.11
6.3
6.0
5.7
5.4
ENSG00000168259.14 DNAJC7
log2(RPKM)
5.25
5.50
5.75
6.00
6.25
ENSG00000113312.10 TTC1
log2(RPKM)
z−score of IF value
−2
−1
0
1
2
3

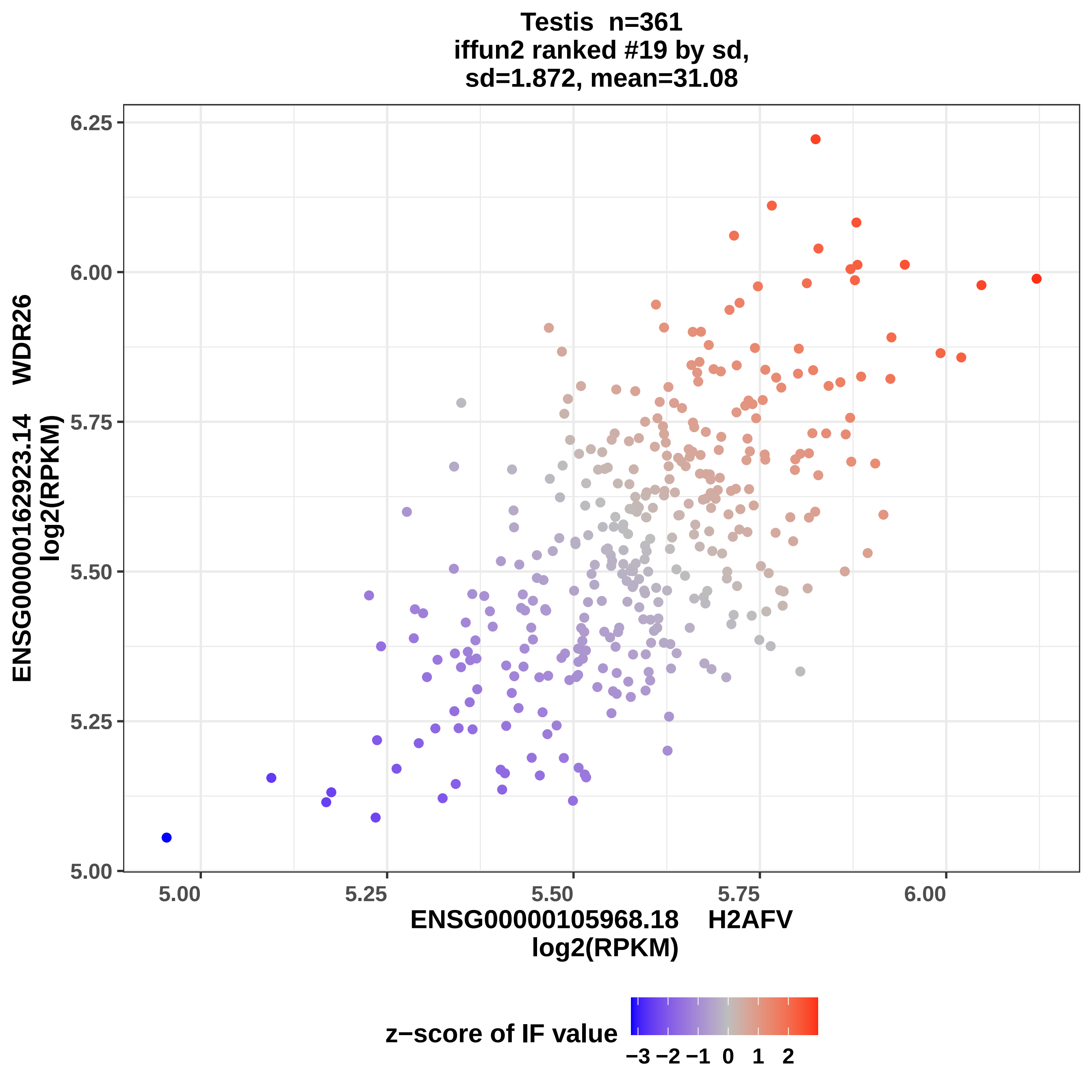
Testis n=361
iffun2 ranked #19 by sd,
sd=1.872, mean=31.08
ENSG00000162923.14 WDR26
log2(RPKM)
6.25
6.00
5.75
5.50
5.25
5.00
5.00
5.25
5.50
5.75
6.00
ENSG00000105968.18 H2AFV
log2(RPKM)
z−score of IF value
−3 −2 −1 0 1 2

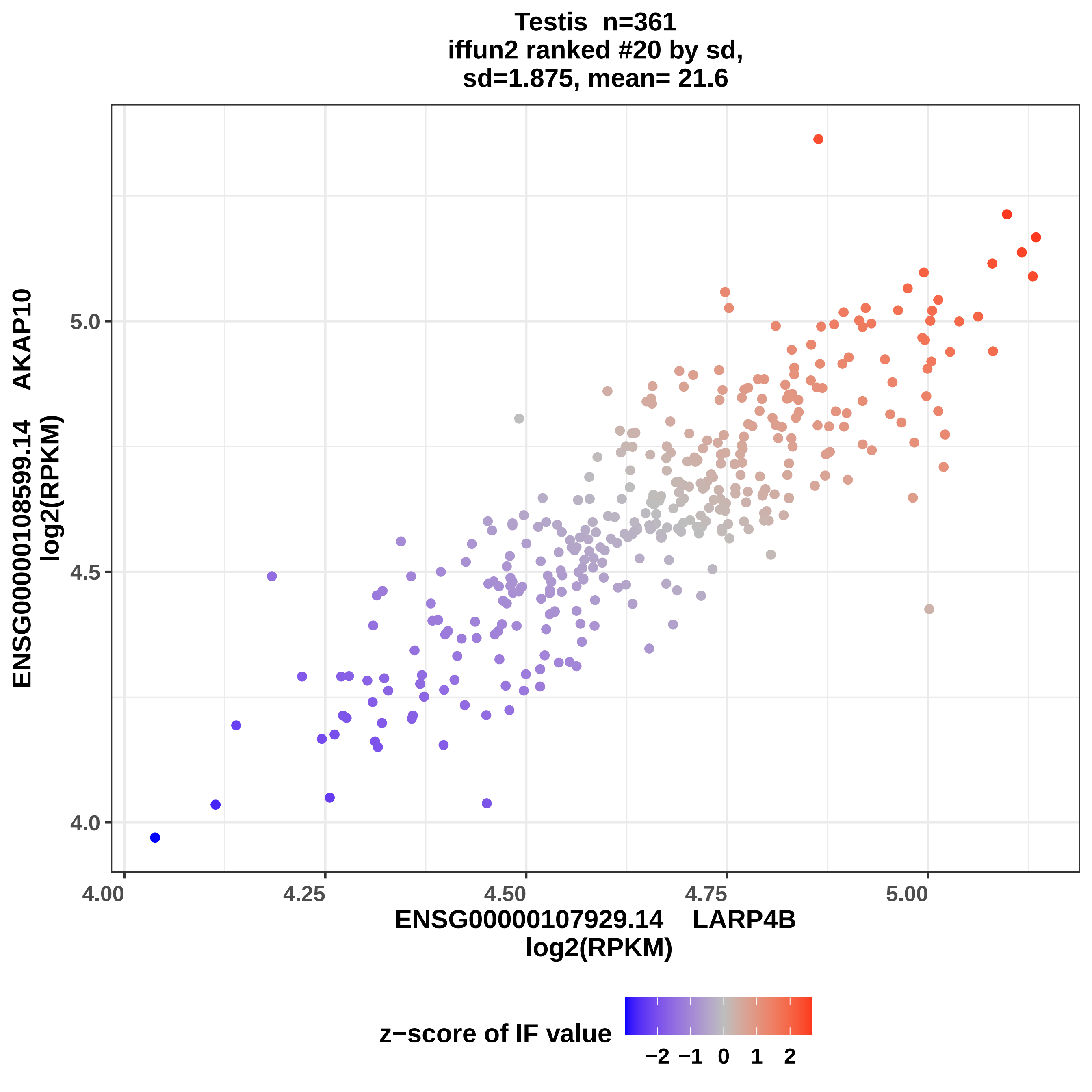
Testis n=361
iffun2 ranked #20 by sd,
sd=1.875, mean= 21.6
ENSG00000108599.14 AKAP10
log2(RPKM)
ENSG00000107929.14 LARP4B
log2(RPKM)
4.0
4.5
5.0
4.00
4.25
4.50
4.75
5.00
z−score of IF value
−2 −1 0 1 2

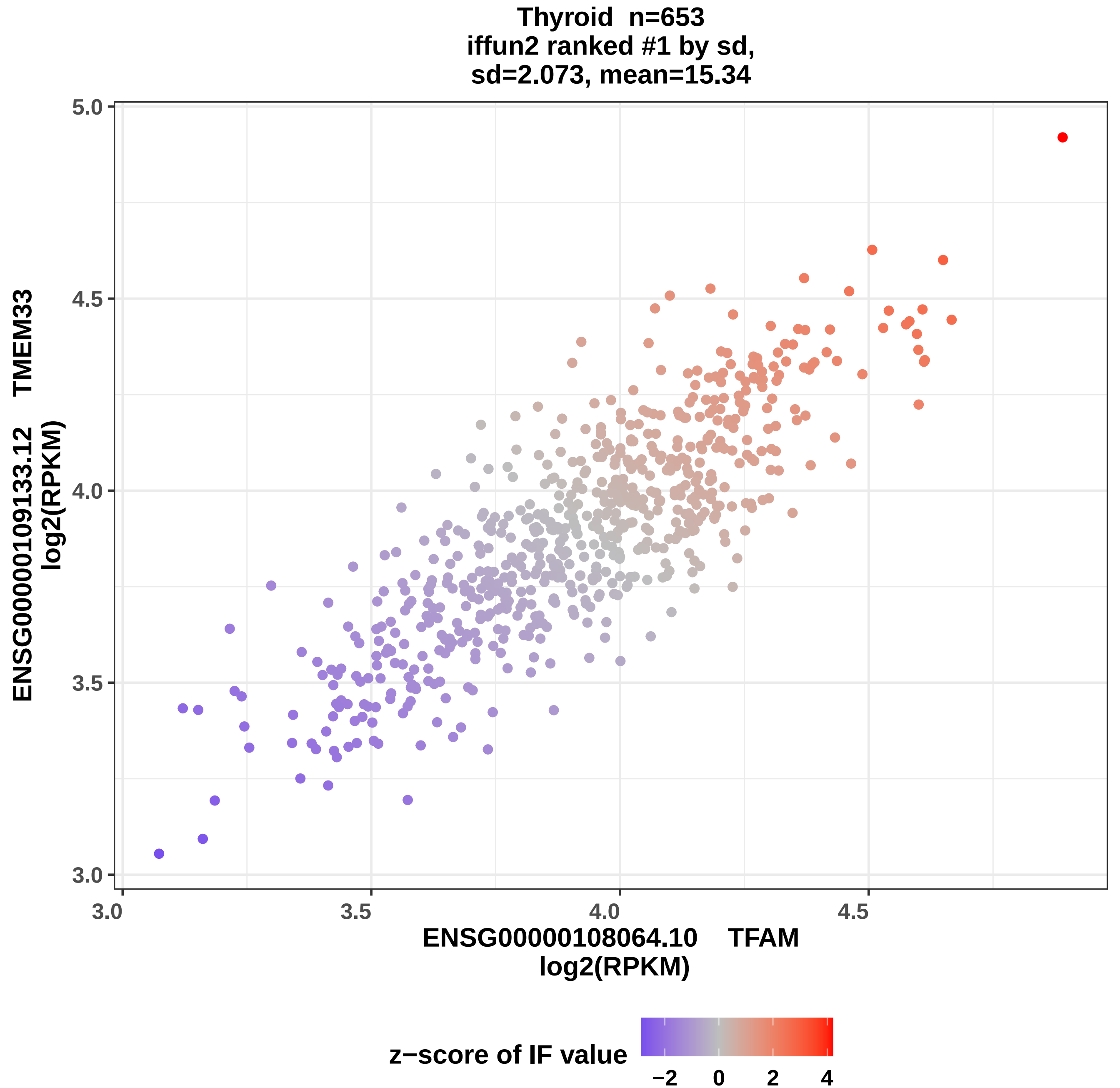

Thyroid n=653
iffun2 ranked #1 by sd,
sd=2.073, mean=15.34
ENSG00000109133.12 TMEM33
log2(RPKM)
5.0
4.5
4.0
3.5
3.0
3.0
3.5
4.0
4.5
ENSG00000108064.10 TFAM
log2(RPKM)
z−score of IF value
−2
0
2
4

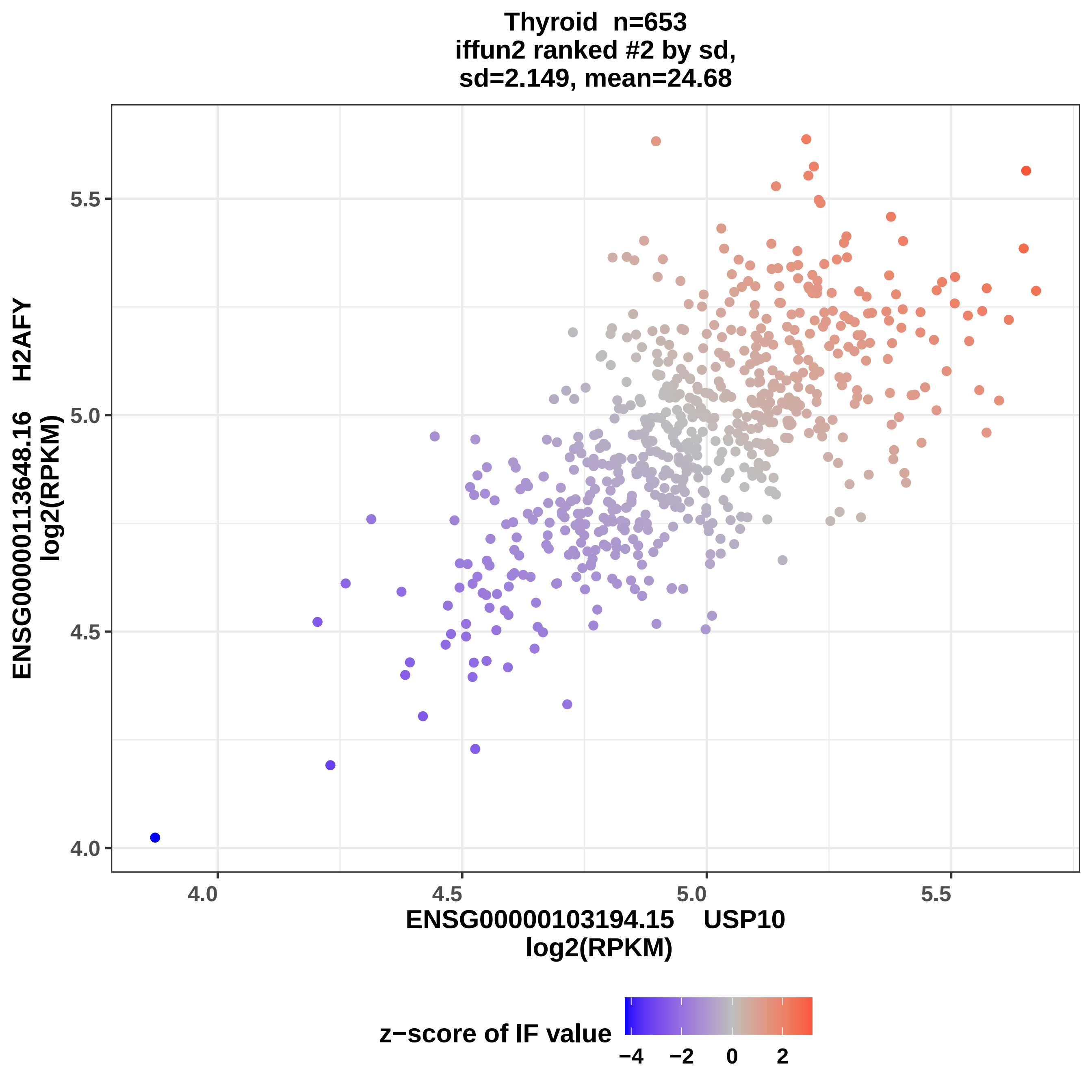
Thyroid n=653
iffun2 ranked #2 by sd,
sd=2.149, mean=24.68
5.5
5.0
4.5
4.0
4.0
4.5
5.0
5.5
ENSG00000113648.16 H2AFY
log2(RPKM)
ENSG00000103194.15 USP10
log2(RPKM)
z−score of IF value
−4
−2
0
2

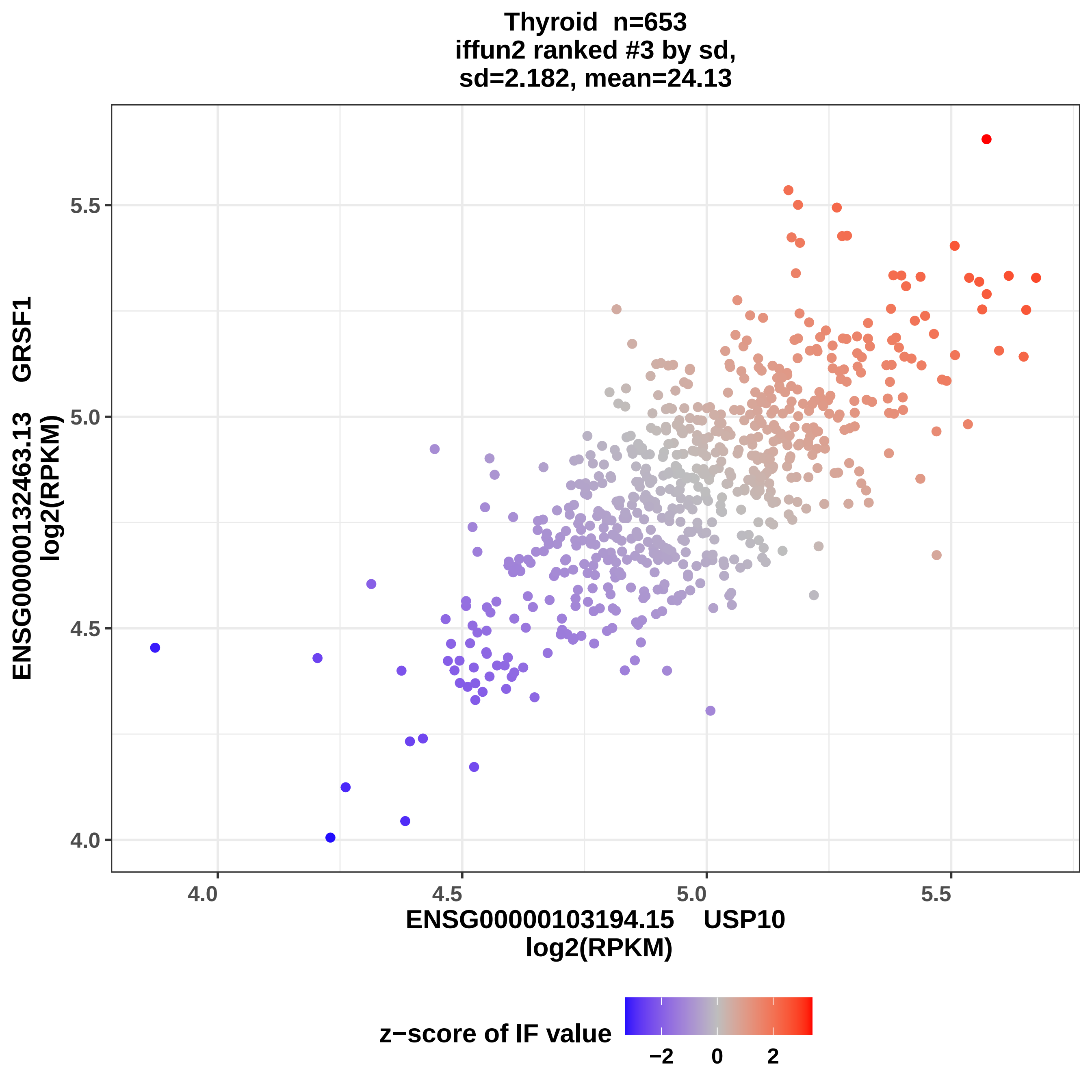

Thyroid n=653
iffun2 ranked #3 by sd,
sd=2.182, mean=24.13
ENSG00000132463.13 GRSF1
log2(RPKM)
5.5
5.0
4.5
4.0
ENSG00000103194.15 USP10
log2(RPKM)
z−score of IF value
−2
0
2

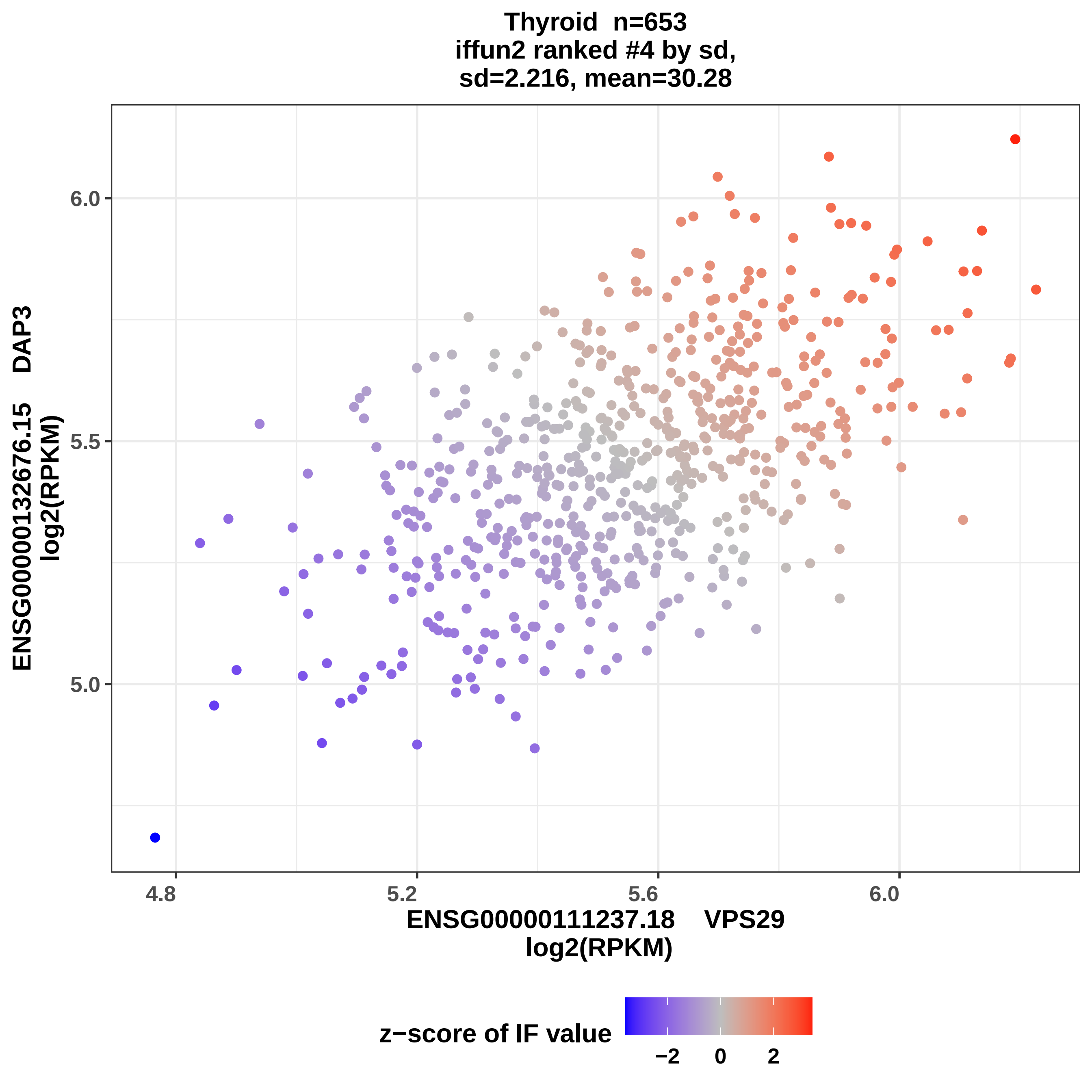

Thyroid n=653
iffun2 ranked #4 by sd,
sd=2.216, mean=30.28
ENSG00000132676.15 DAP3
log2(RPKM)
6.0
5.5
5.0
4.8
5.2
5.6
6.0
ENSG00000111237.18 VPS29
log2(RPKM)
z−score of IF value
−2
0
2

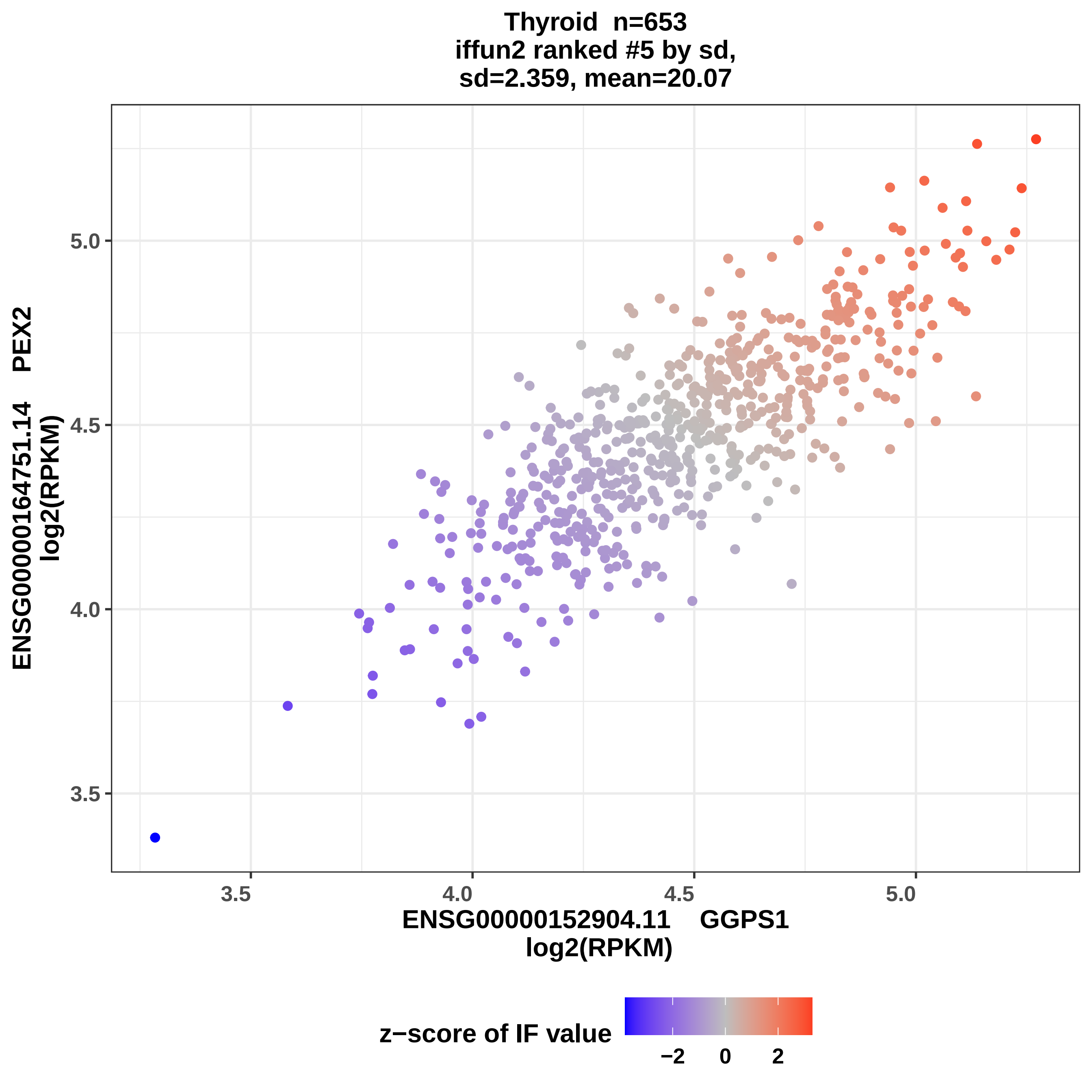
Thyroid n=653
iffun2 ranked #5 by sd,
sd=2.359, mean=20.07
ENSG00000164751.14 PEX2
log2(RPKM)
5.0
4.5
4.0
3.5
3.5
4.0
4.5
5.0
ENSG00000152904.11 GGPS1
log2(RPKM)
z−score of IF value
−2
0
2

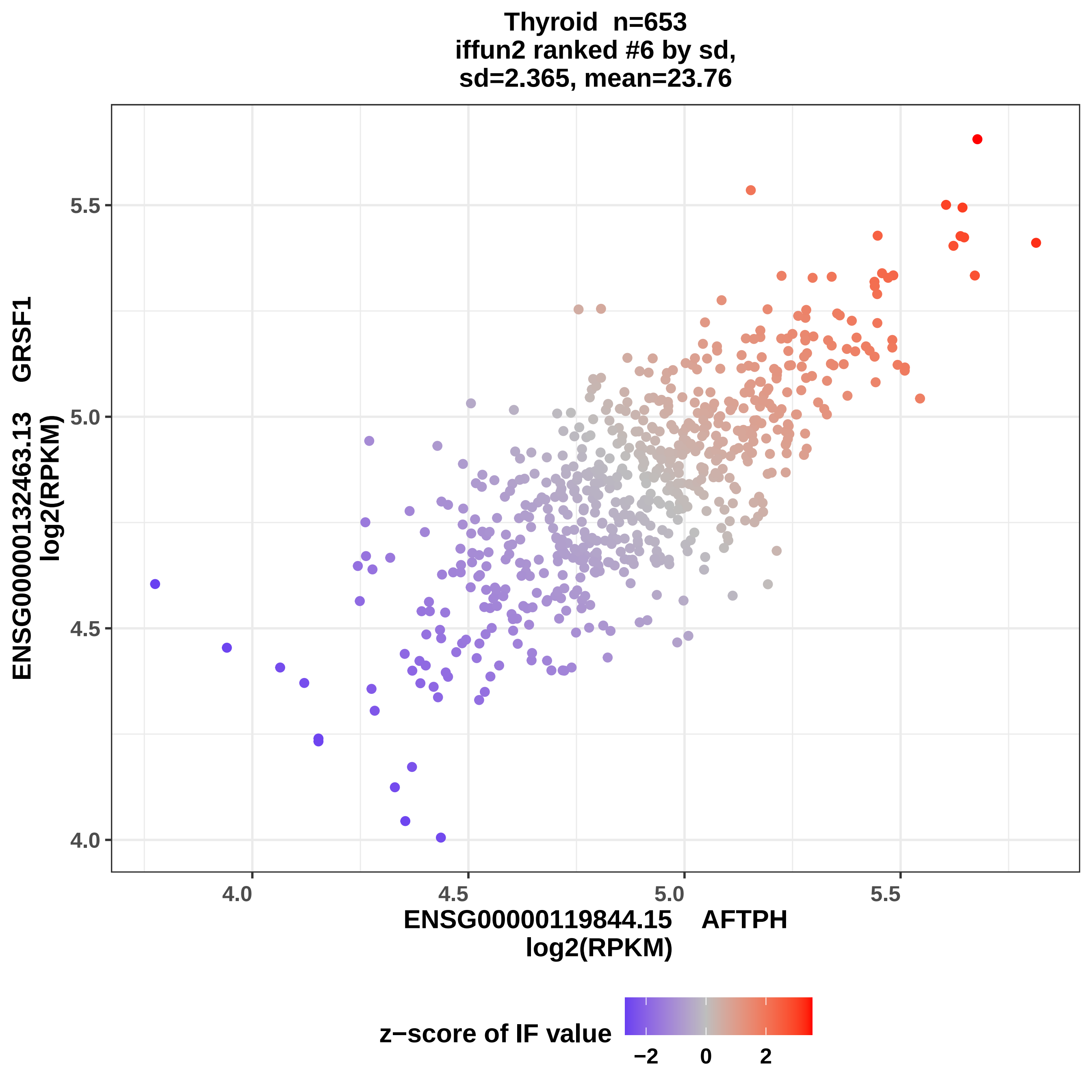

Thyroid n=653
iffun2 ranked #6 by sd,
sd=2.365, mean=23.76
ENSG00000132463.13 GRSF1
log2(RPKM)
5.5
5.0
4.5
4.0
ENSG00000119844.15 AFTPH
log2(RPKM)
z−score of IF value
−2
0
2

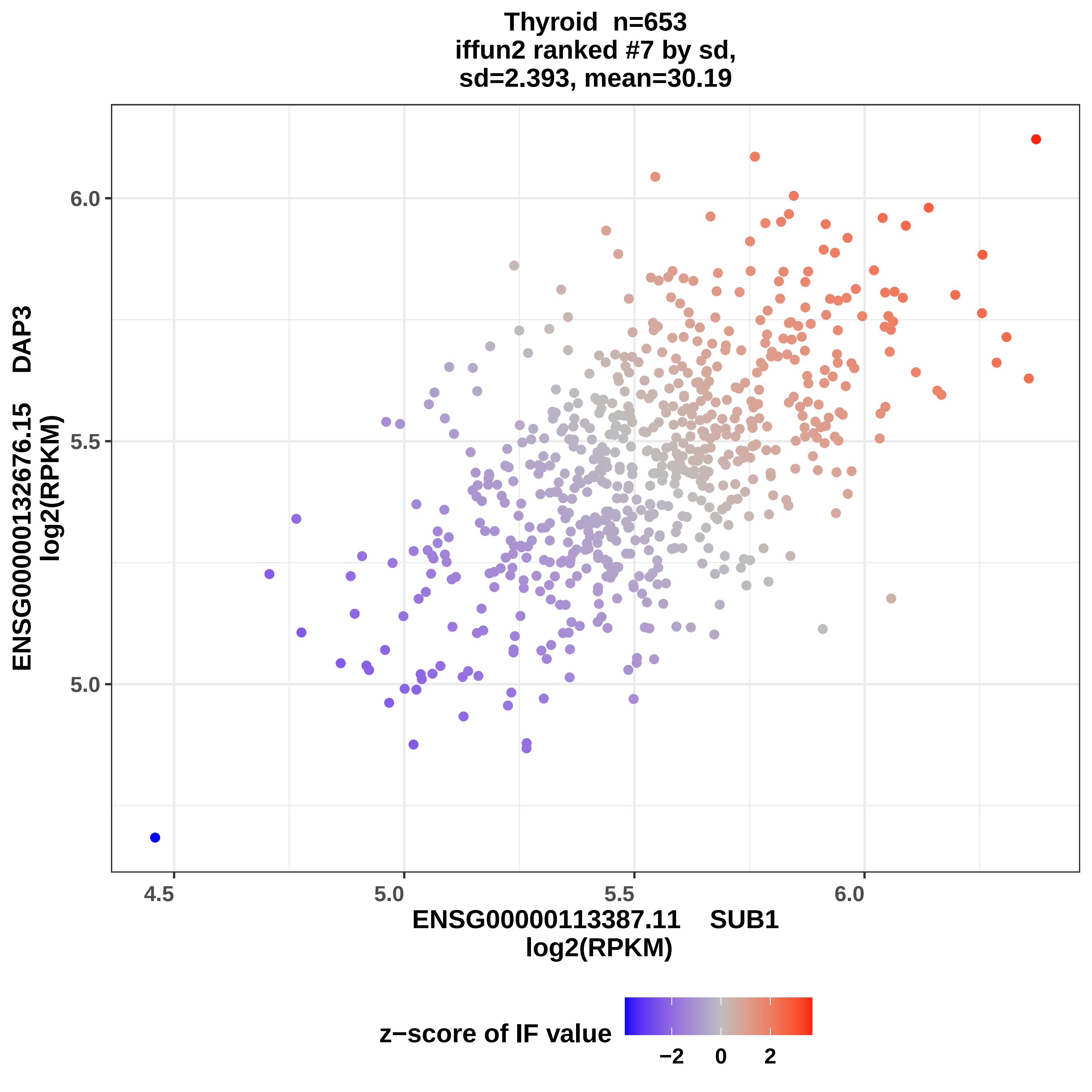

Thyroid n=653
iffun2 ranked #7 by sd,
sd=2.393, mean=30.19
ENSG00000132676.15 DAP3
log2(RPKM)
6.0
5.5
5.0
4.5
5.0
5.5
6.0
ENSG00000113387.11 SUB1
log2(RPKM)
z−score of IF value
−2
0
2

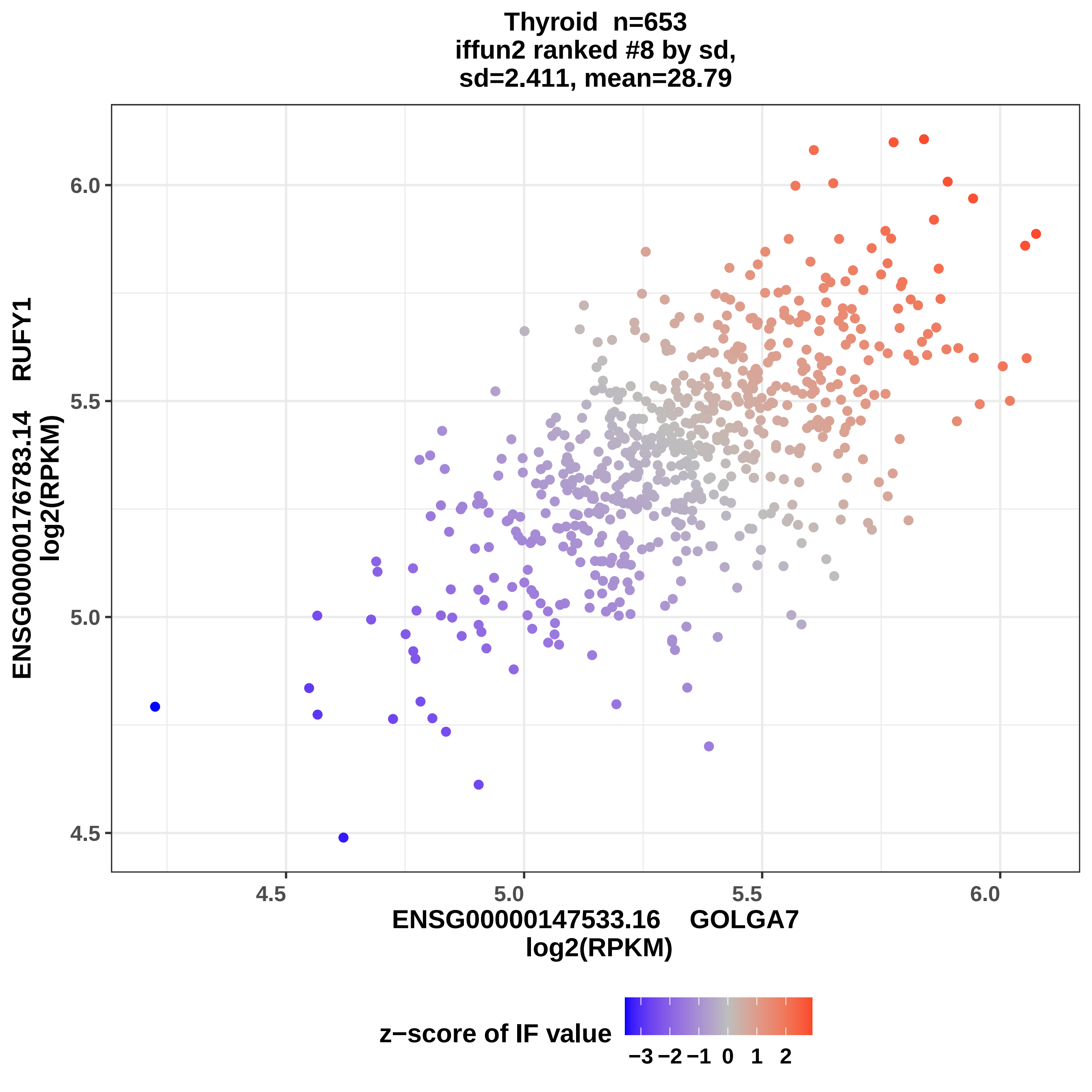
Thyroid n=653
iffun2 ranked #8 by sd,
sd=2.411, mean=28.79
ENSG00000176783.14 RUFY1
log2(RPKM)
6.0
5.5
5.0
4.5
4.5
5.0
5.5
6.0
ENSG00000147533.16 GOLGA7
log2(RPKM)
z−score of IF value
−3 −2 −1 0 1 2

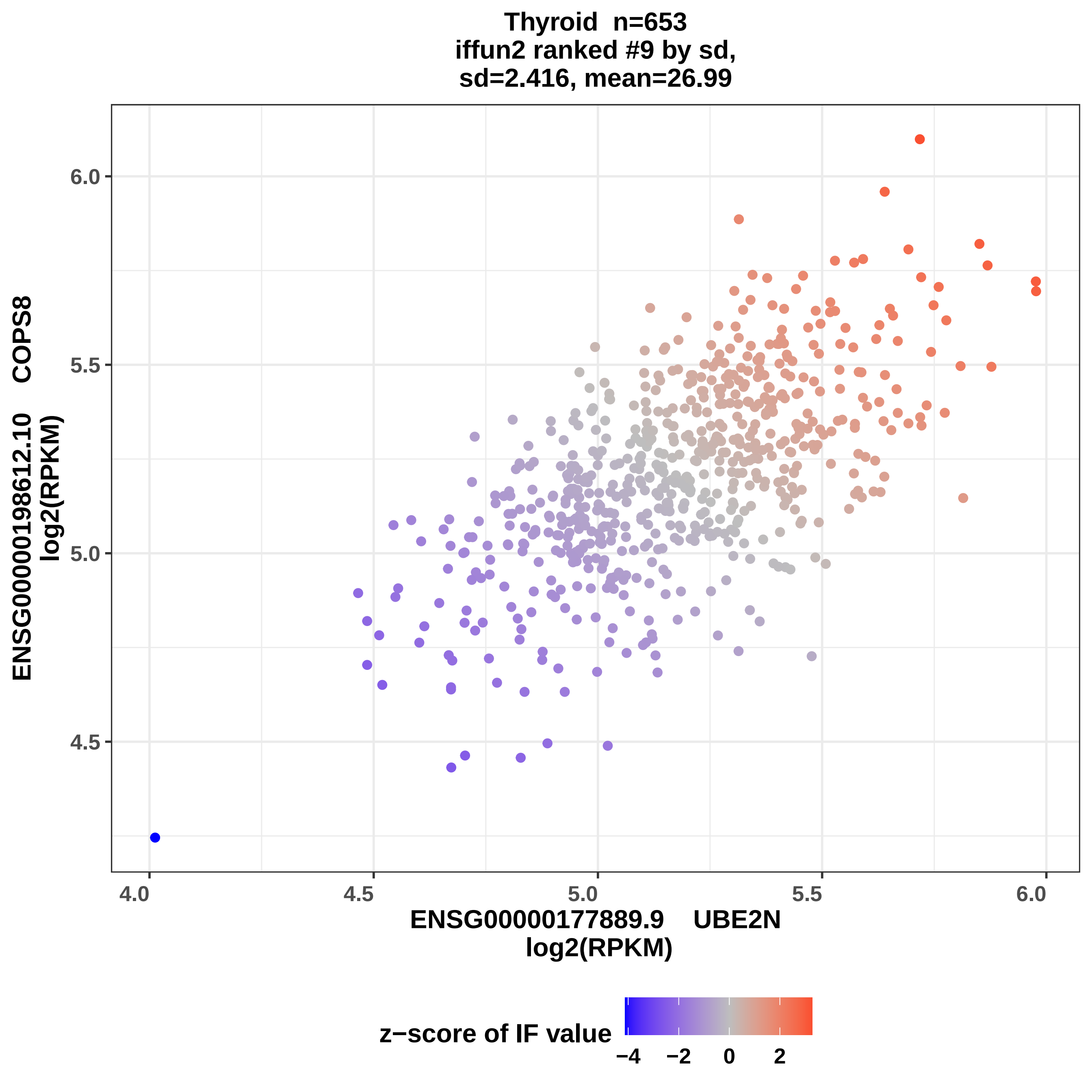
Thyroid n=653
iffun2 ranked #9 by sd,
sd=2.416, mean=26.99
ENSG00000198612.10 COPS8
log2(RPKM)
6.0
5.5
5.0
4.5
4.0
4.5
5.0
5.5
6.0
ENSG00000177889.9 UBE2N
log2(RPKM)
z−score of IF value
−4
−2
0
2

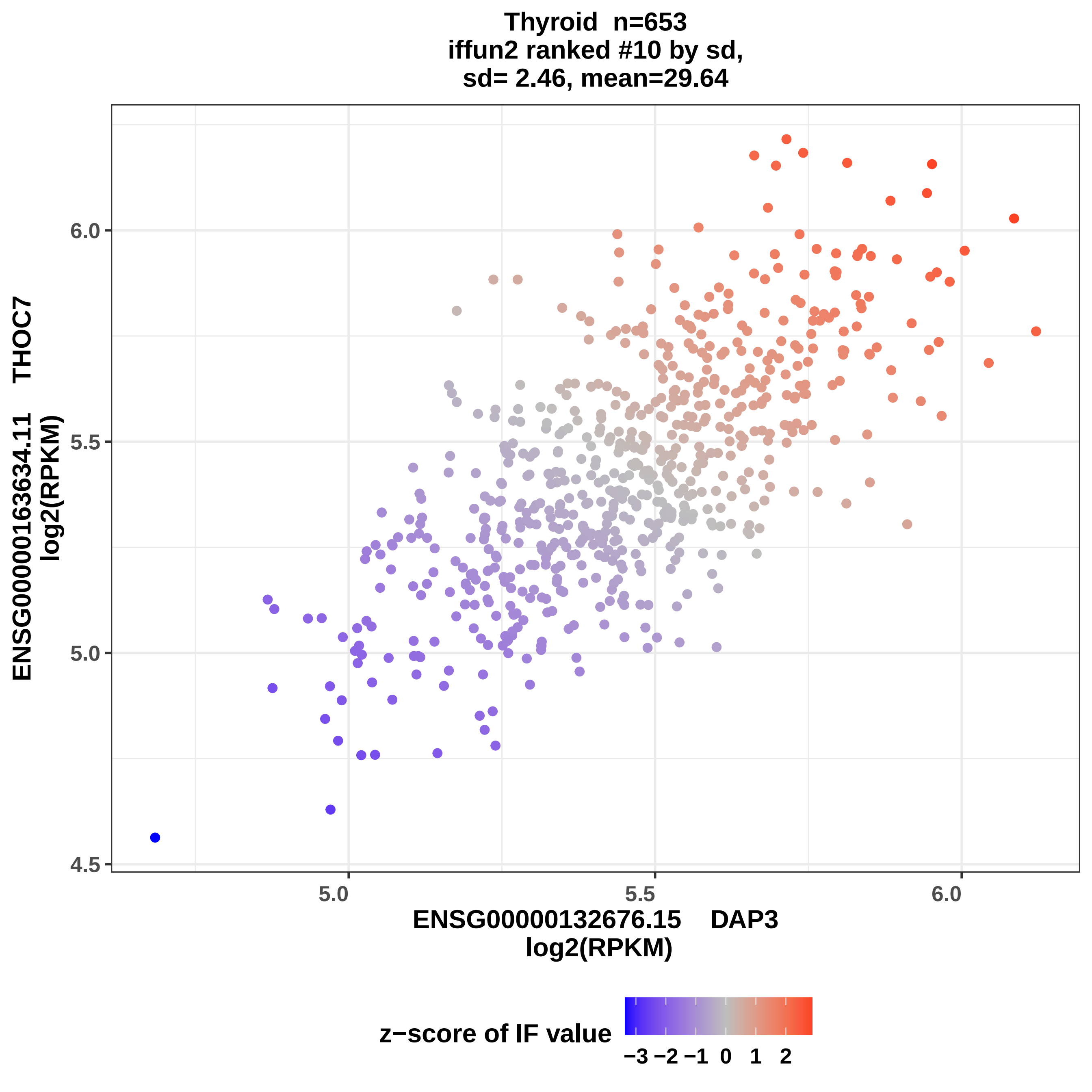
Thyroid n=653
iffun2 ranked #10 by sd,
sd= 2.46, mean=29.64
ENSG00000163634.11 THOC7
log2(RPKM)
6.0
5.5
5.0
4.5
5.0
5.5
6.0
ENSG00000132676.15 DAP3
log2(RPKM)
z−score of IF value
−3 −2 −1 0 1 2

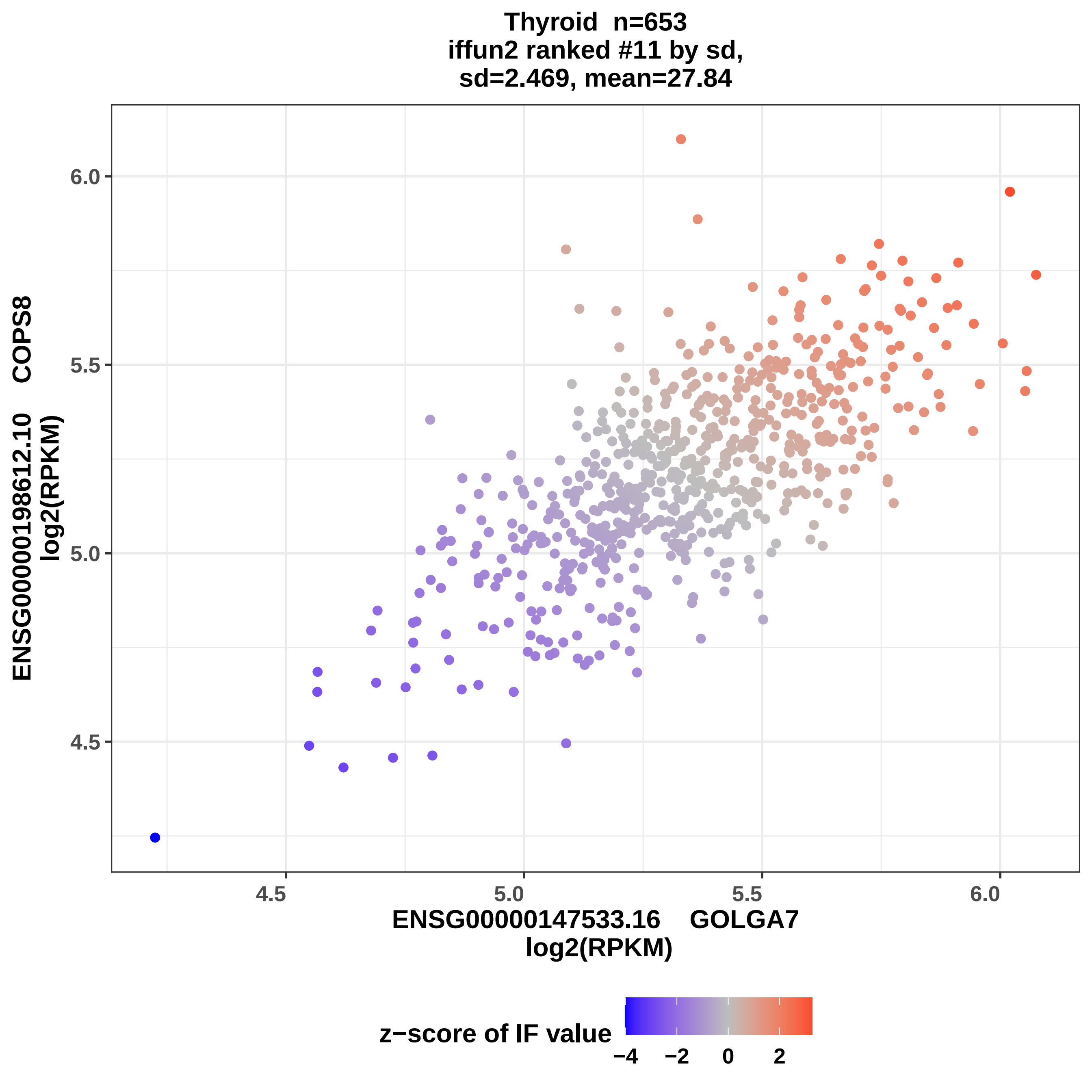
Thyroid n=653
iffun2 ranked #11 by sd,
sd=2.469, mean=27.84
ENSG00000198612.10 COPS8
log2(RPKM)
6.0
5.5
5.0
4.5
4.5
5.0
5.5
6.0
ENSG00000147533.16 GOLGA7
log2(RPKM)
z−score of IF value
−4
−2
0
2

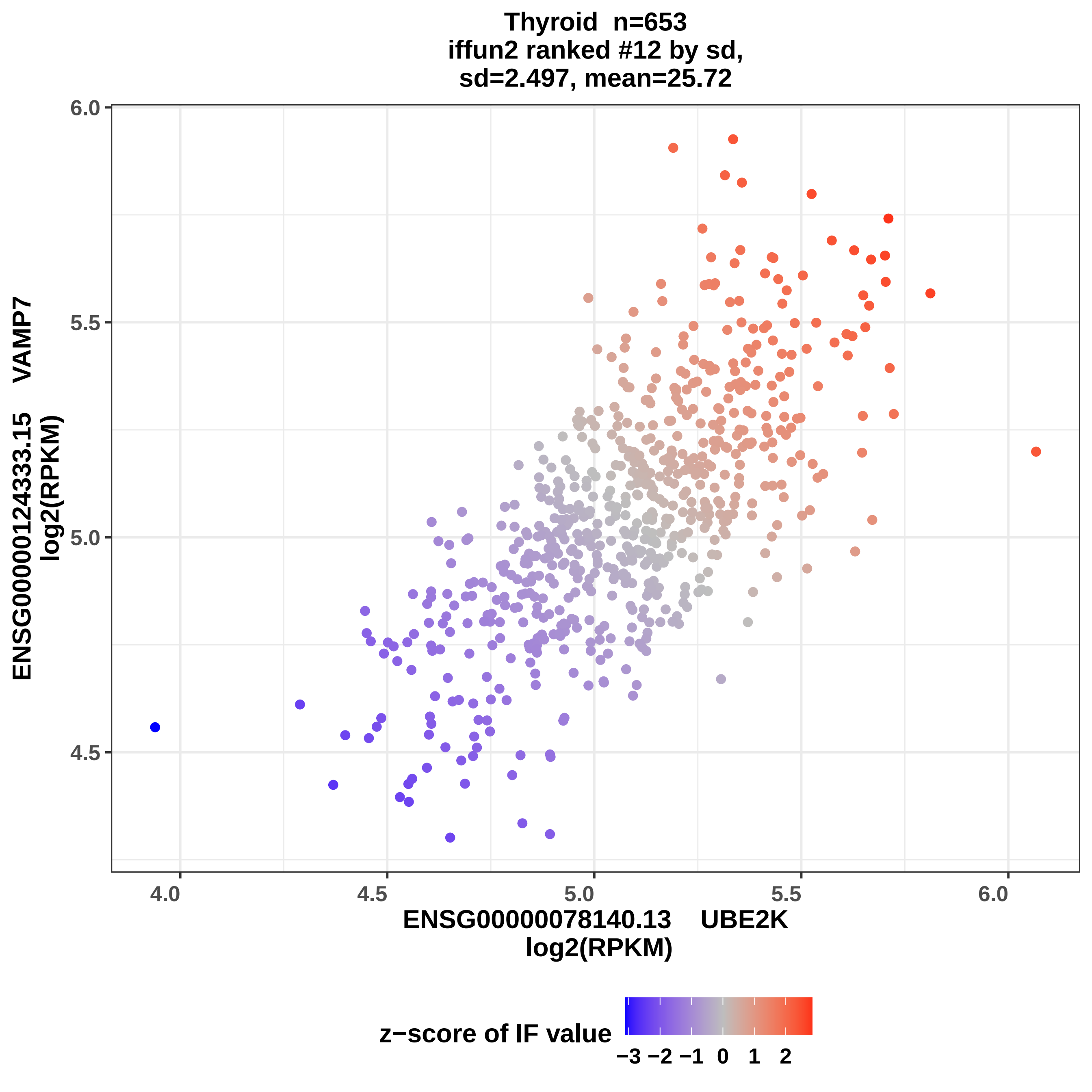
Thyroid n=653
iffun2 ranked #12 by sd,
sd=2.497, mean=25.72
ENSG00000124333.15 VAMP7
log2(RPKM)
6.0
5.5
5.0
4.5
4.0
4.5
5.0
5.5
6.0
ENSG00000078140.13 UBE2K
log2(RPKM)
z−score of IF value
−3 −2 −1 0 1 2

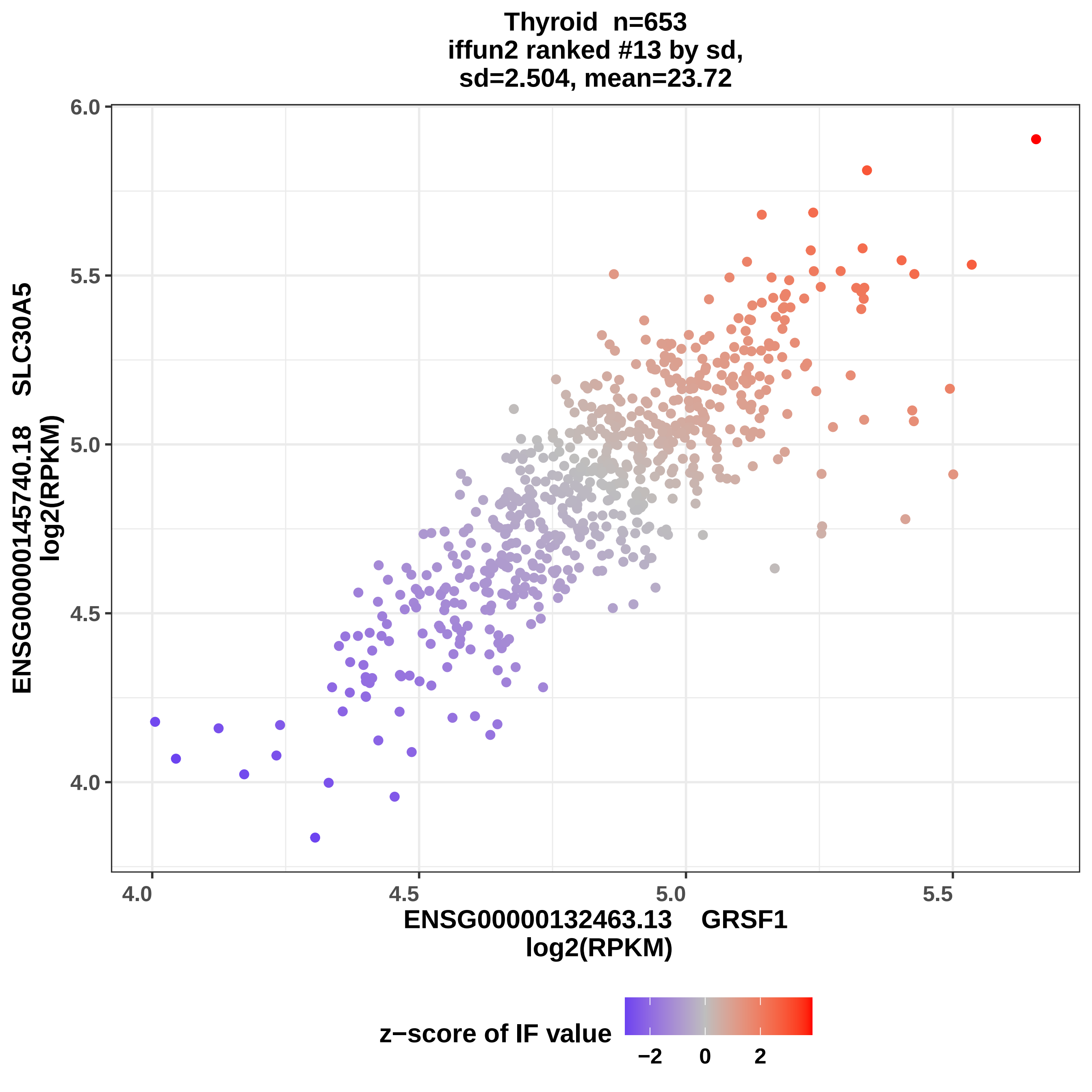

Thyroid n=653
iffun2 ranked #13 by sd,
sd=2.504, mean=23.72
ENSG00000145740.18 SLC30A5
log2(RPKM)
6.0
5.5
5.0
4.5
4.0
4.0
4.5
5.0
5.5
ENSG00000132463.13 GRSF1
log2(RPKM)
z−score of IF value
−2
0
2

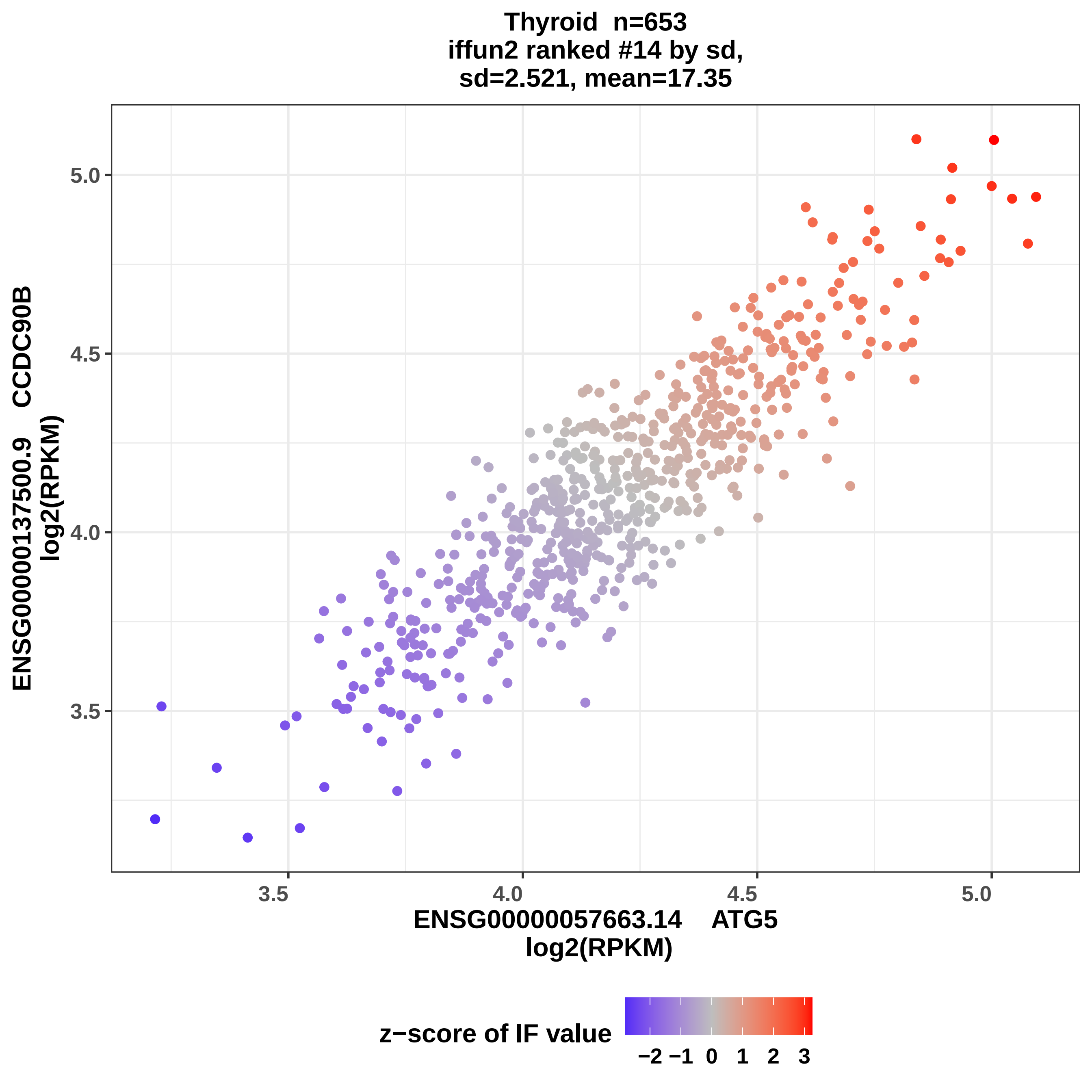
Thyroid n=653
iffun2 ranked #14 by sd,
sd=2.521, mean=17.35
ENSG00000137500.9 CCDC90B
log2(RPKM)
5.0
4.5
4.0
3.5
3.5
4.0
4.5
5.0
ENSG00000057663.14 ATG5
log2(RPKM)
z−score of IF value
−2 −1 0 1 2 3

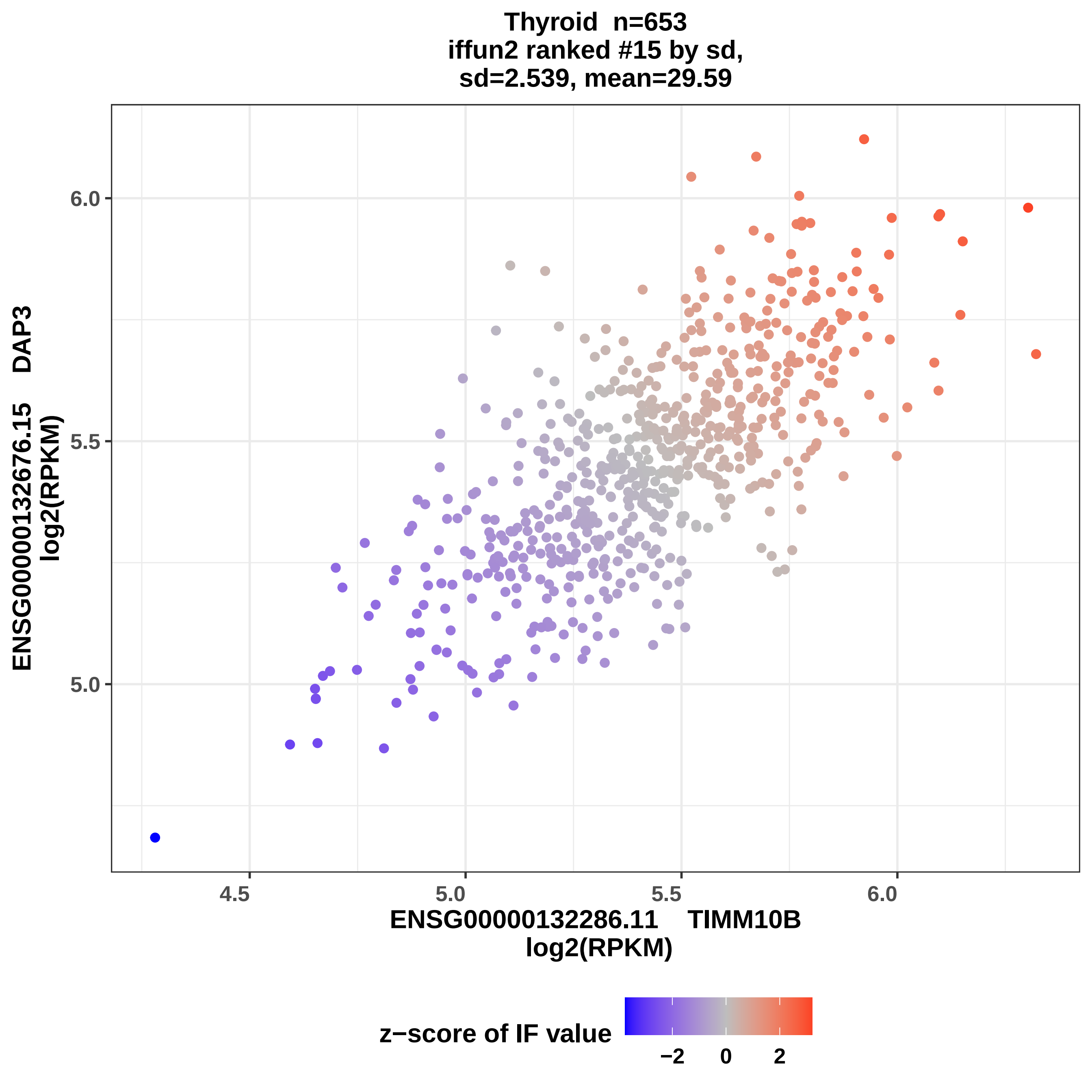

Thyroid n=653
iffun2 ranked #15 by sd,
sd=2.539, mean=29.59
ENSG00000132676.15 DAP3
log2(RPKM)
6.0
5.5
5.0
4.5
5.0
5.5
6.0
ENSG00000132286.11 TIMM10B
log2(RPKM)
z−score of IF value
−2
0
2

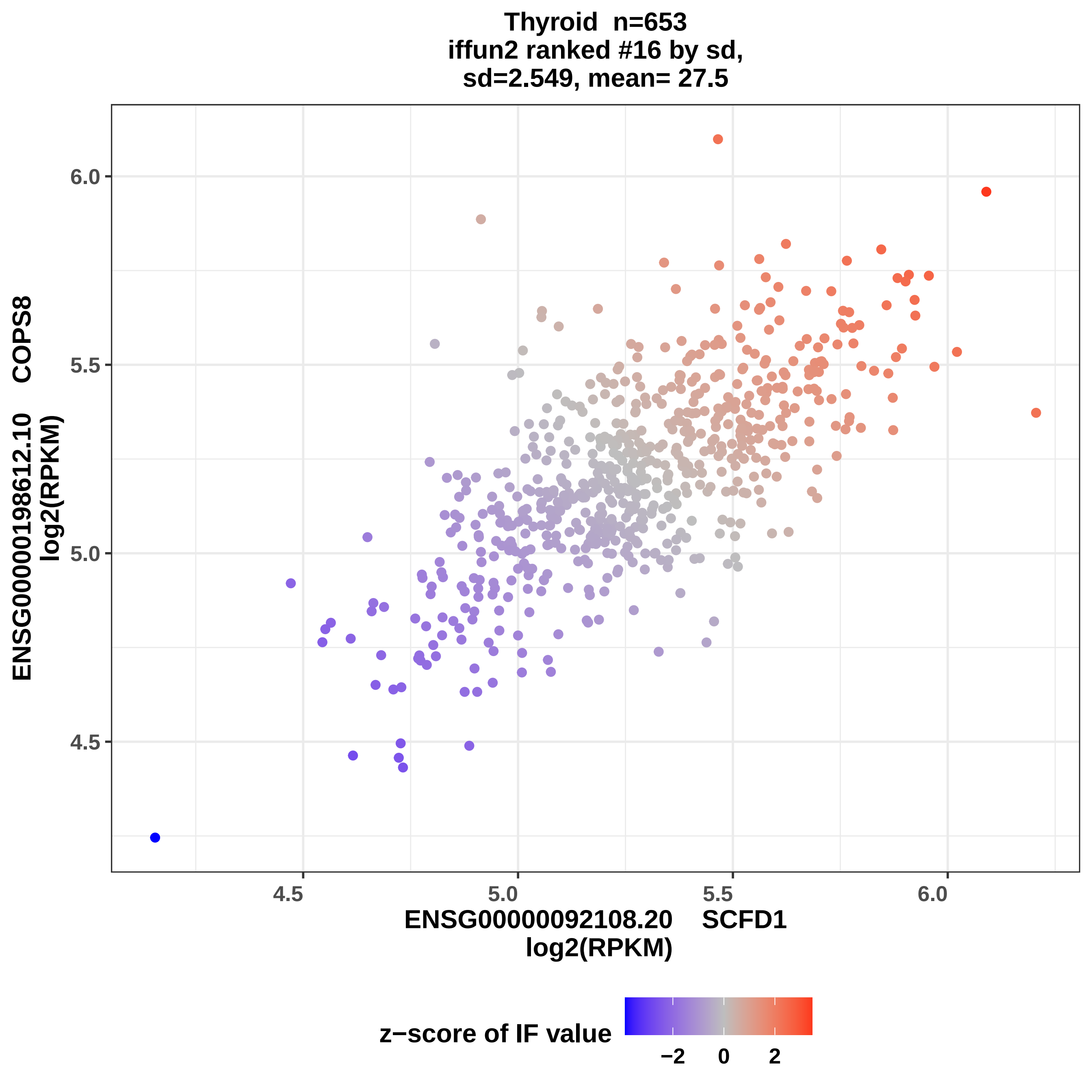
Thyroid n=653
iffun2 ranked #16 by sd,
sd=2.549, mean= 27.5
ENSG00000198612.10 COPS8
log2(RPKM)
6.0
5.5
5.0
4.5
4.5
5.0
5.5
6.0
ENSG00000092108.20 SCFD1
log2(RPKM)
z−score of IF value
−2
0
2

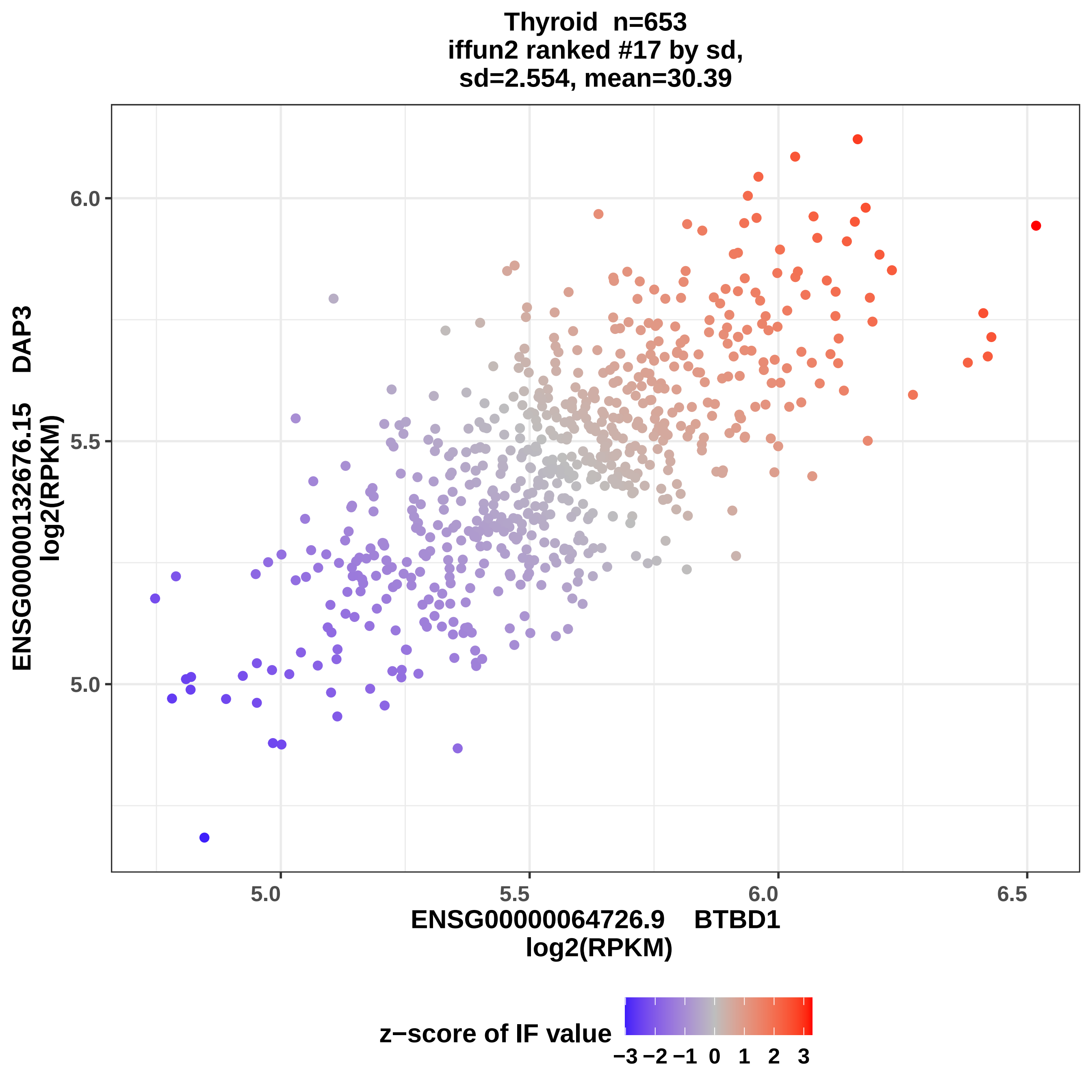

Thyroid n=653
iffun2 ranked #17 by sd,
sd=2.554, mean=30.39
ENSG00000132676.15 DAP3
log2(RPKM)
6.0
5.5
5.0
5.0
5.5
6.0
6.5
ENSG00000064726.9 BTBD1
log2(RPKM)
z−score of IF value
−3 −2 −1 0 1 2 3

Thyroid n=653
iffun2 ranked #18 by sd,
sd=2.558, mean=28.46

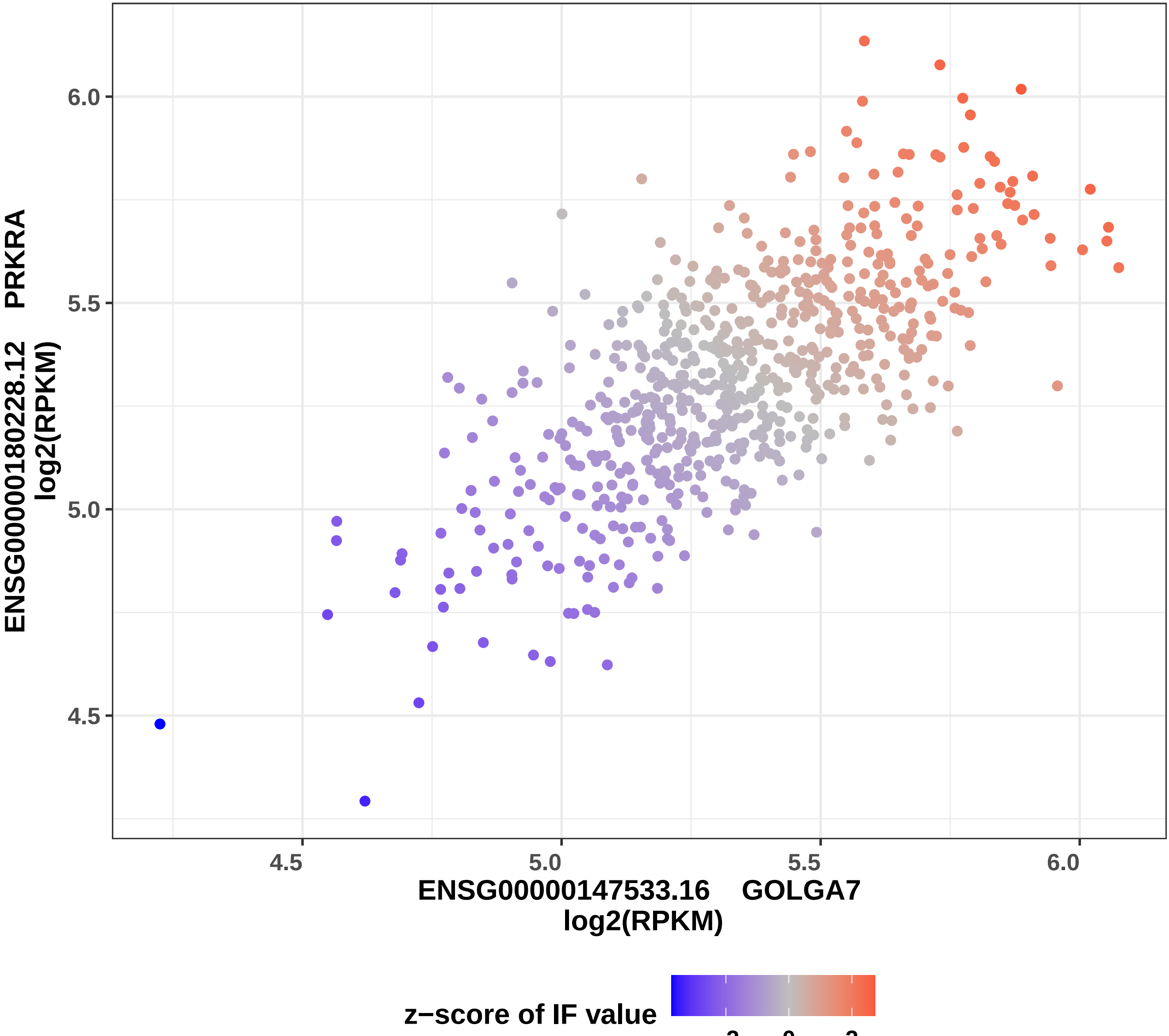

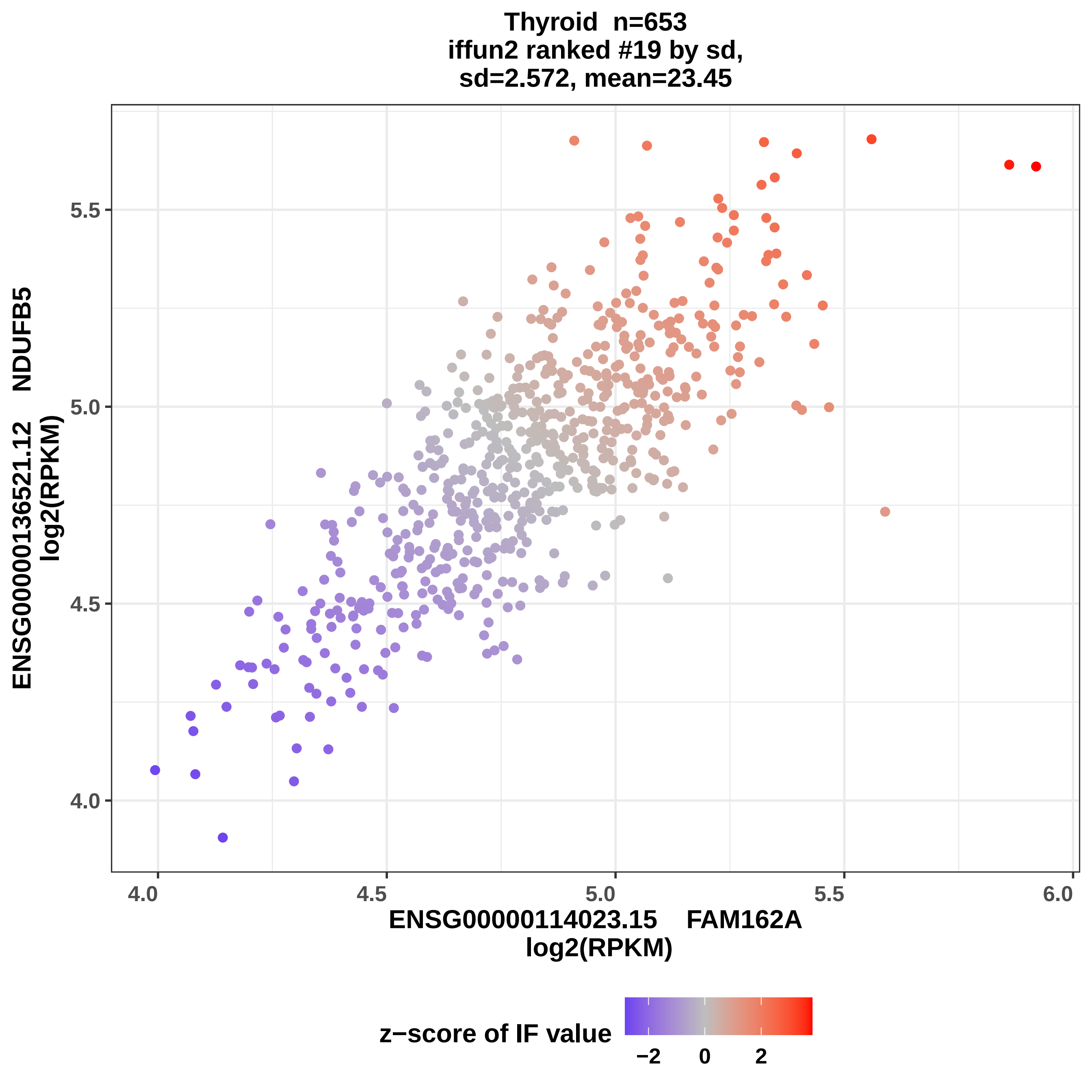

Thyroid n=653
iffun2 ranked #19 by sd,
sd=2.572, mean=23.45
ENSG00000136521.12 NDUFB5
log2(RPKM)
5.5
5.0
4.5
4.0
4.0
4.5
5.0
5.5
6.0
ENSG00000114023.15 FAM162A
log2(RPKM)
z−score of IF value
−2
0
2

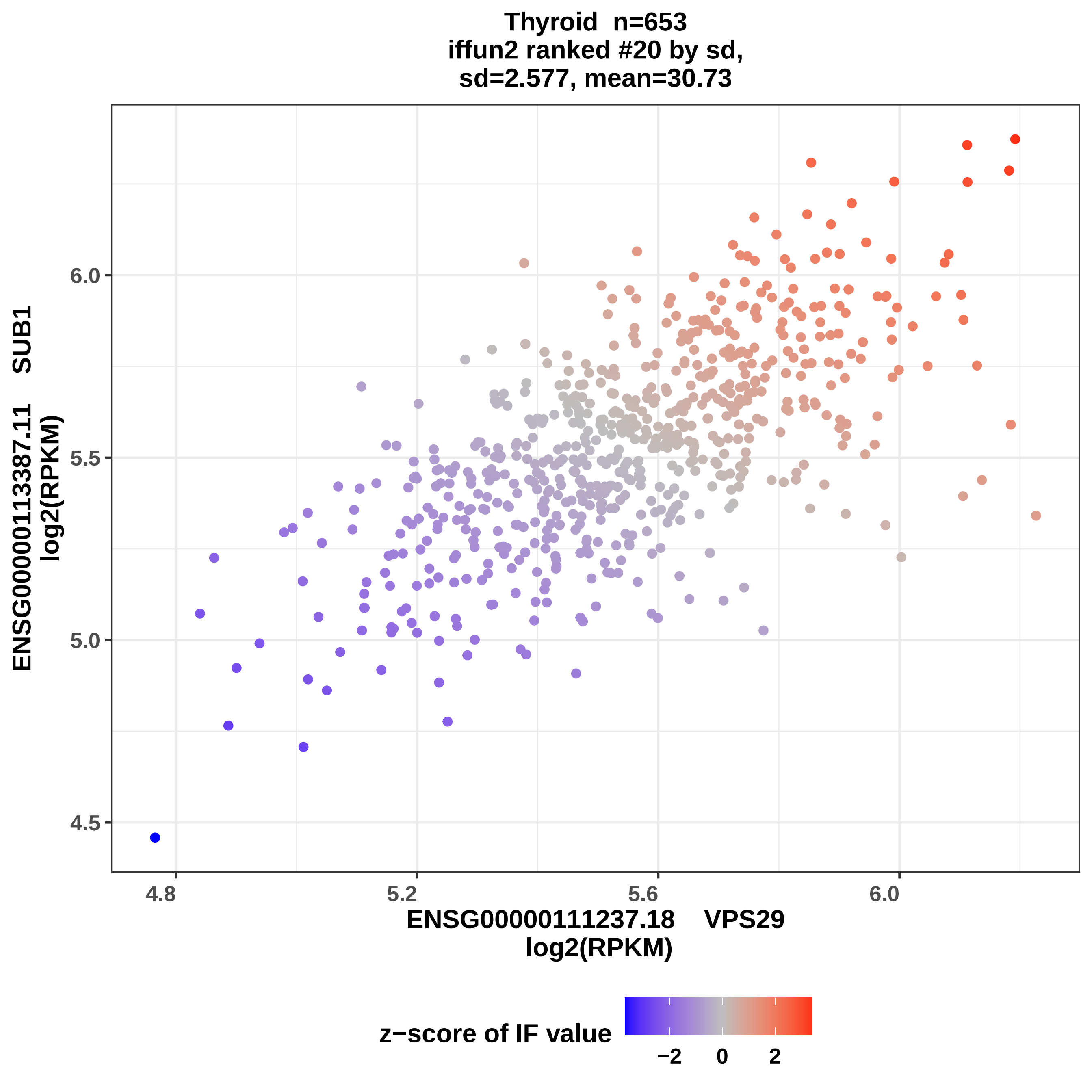

Thyroid n=653
iffun2 ranked #20 by sd,
sd=2.577, mean=30.73
ENSG00000113387.11 SUB1
log2(RPKM)
4.5
5.0
5.5
6.0
4.8
5.2
5.6
6.0
ENSG00000111237.18 VPS29
log2(RPKM)
z−score of IF value
−2
0
2

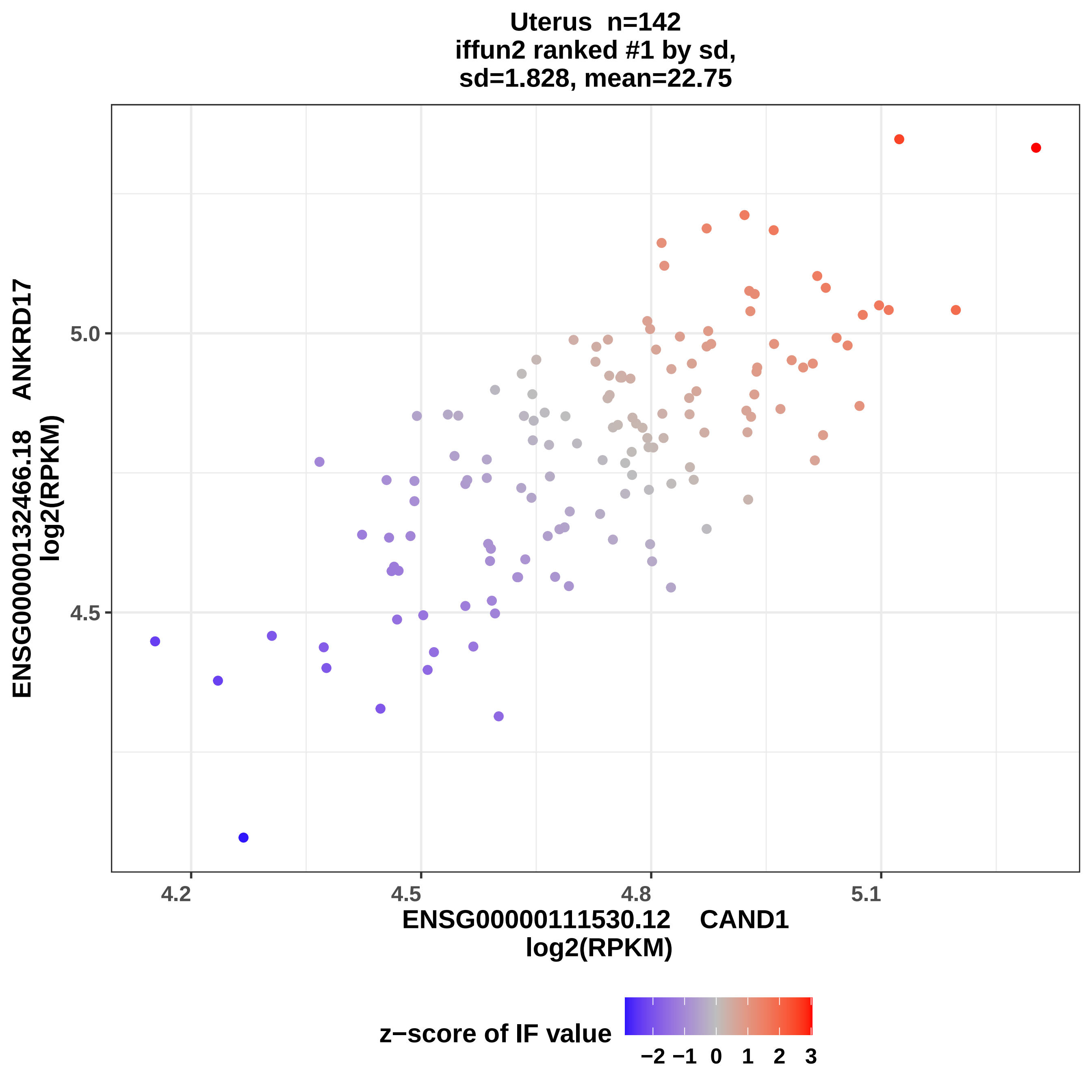

Uterus n=142
iffun2 ranked #1 by sd,
sd=1.828, mean=22.75
ENSG00000132466.18 ANKRD17
log2(RPKM)
5.0
4.5
4.2
4.5
4.8
5.1
ENSG00000111530.12 CAND1
log2(RPKM)
z−score of IF value
−2 −1 0 1 2 3

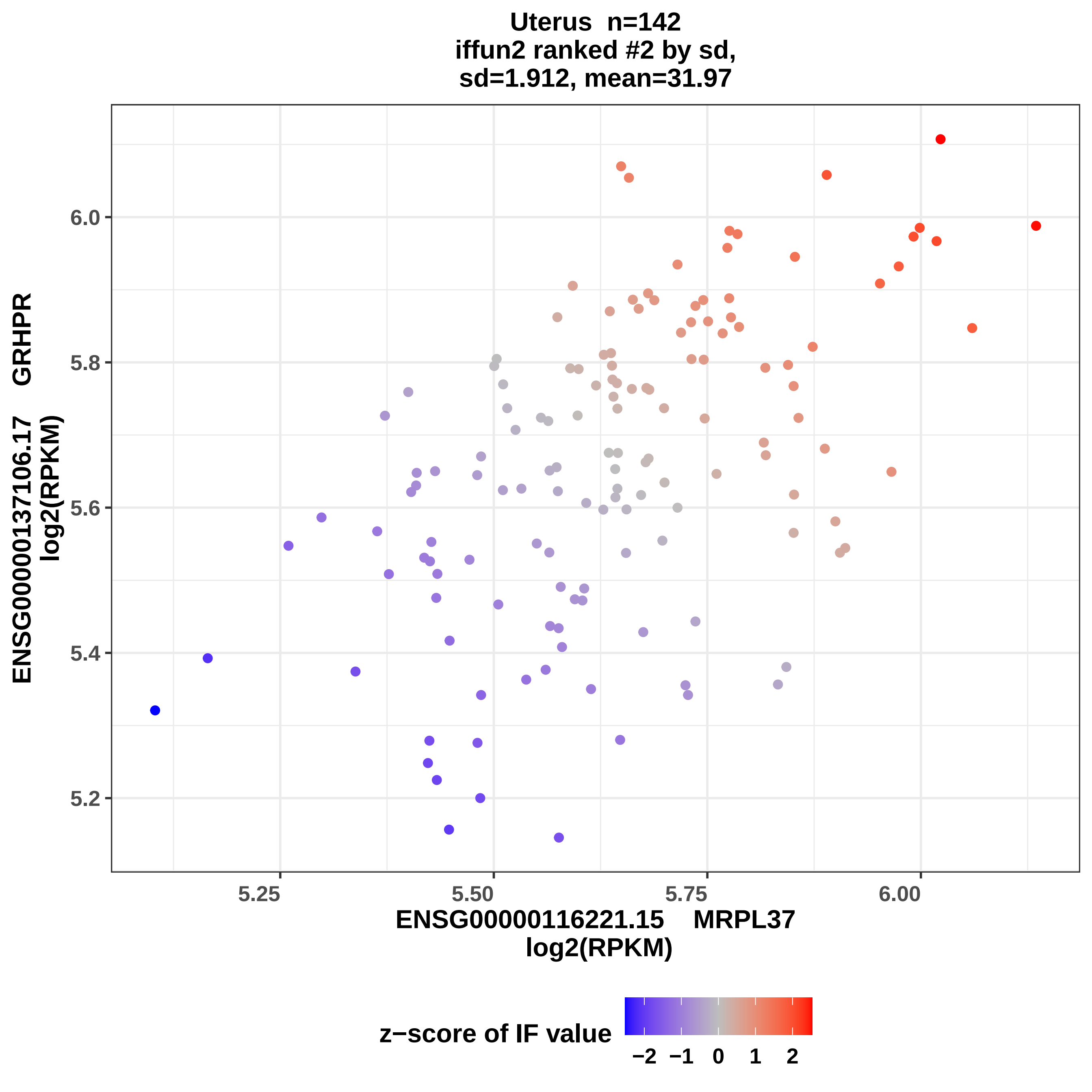
Uterus n=142
iffun2 ranked #2 by sd,
sd=1.912, mean=31.97
ENSG00000137106.17 GRHPR
log2(RPKM)
6.0
5.8
5.6
5.4
5.2
5.25
5.50
5.75
6.00
ENSG00000116221.15 MRPL37
log2(RPKM)
z−score of IF value
−2 −1 0 1 2

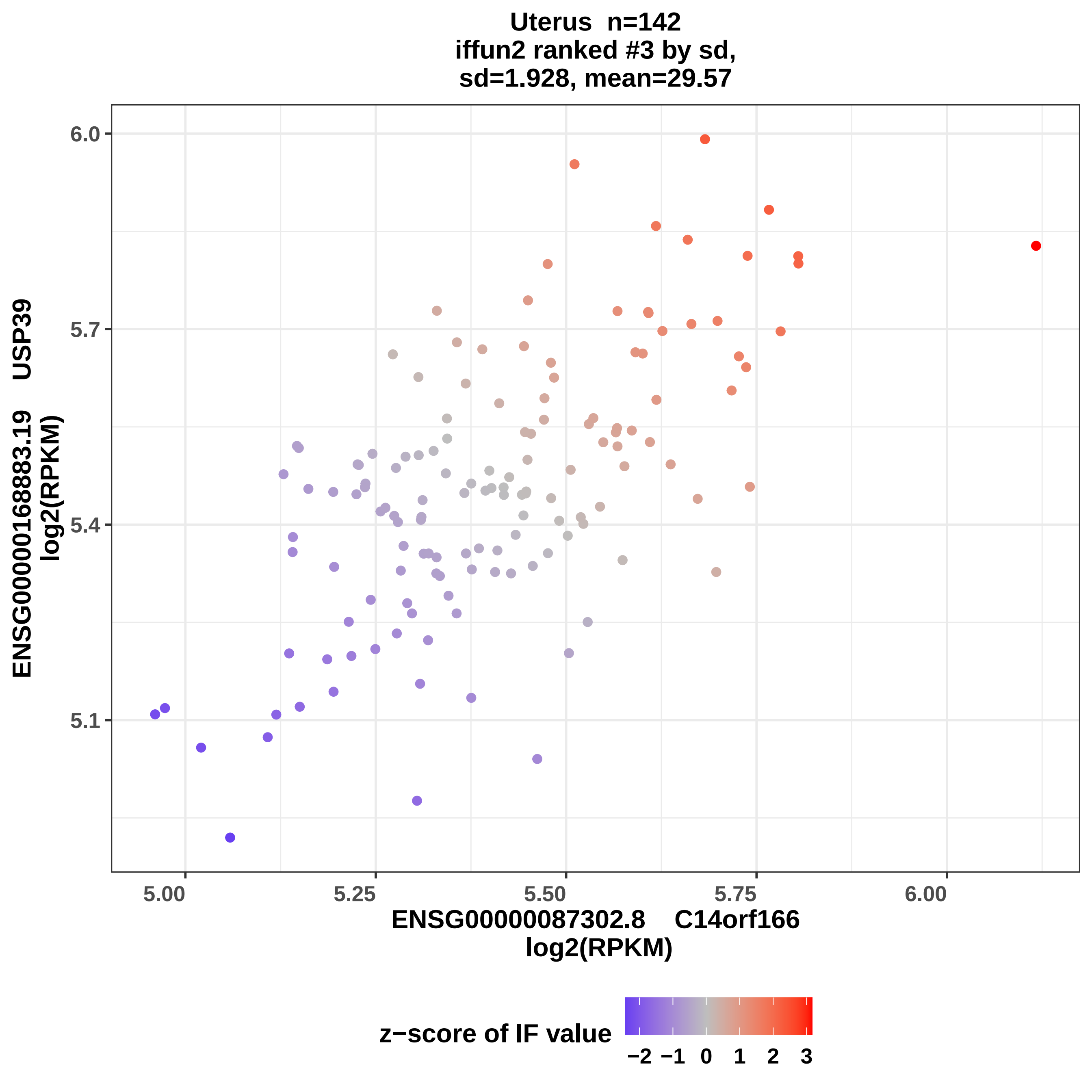
Uterus n=142
iffun2 ranked #3 by sd,
sd=1.928, mean=29.57
ENSG00000168883.19 USP39
log2(RPKM)
6.0
5.7
5.4
5.1
5.00
5.25
5.50
5.75
6.00
ENSG00000087302.8 C14orf166
log2(RPKM)
z−score of IF value
−2
−1
0
1
2
3

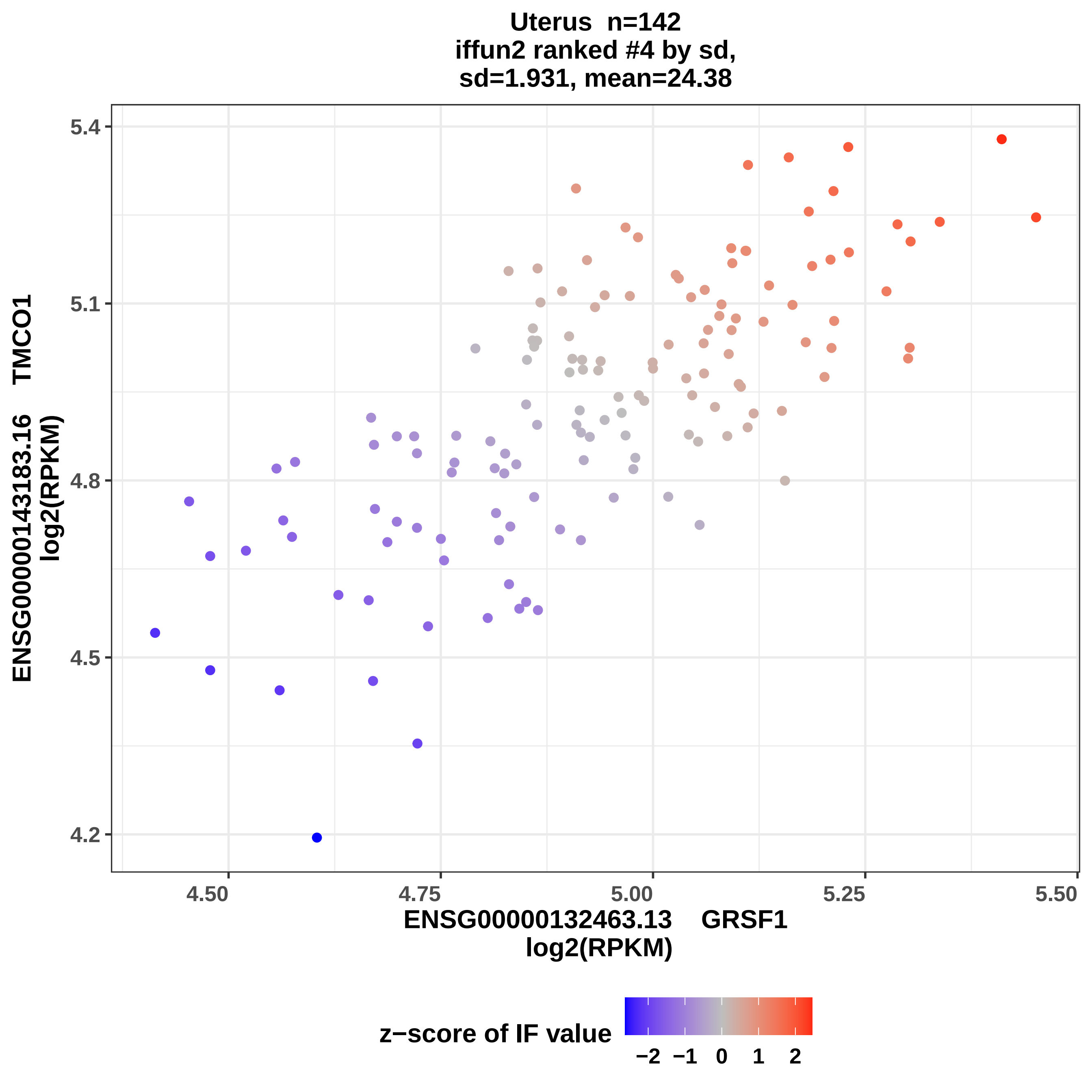
Uterus n=142
iffun2 ranked #4 by sd,
sd=1.931, mean=24.38
ENSG00000143183.16 TMCO1
log2(RPKM)
4.2
4.5
4.8
5.1
5.4
4.50
4.75
5.00
5.25
5.50
ENSG00000132463.13 GRSF1
log2(RPKM)
z−score of IF value
−2
−1
0
1
2

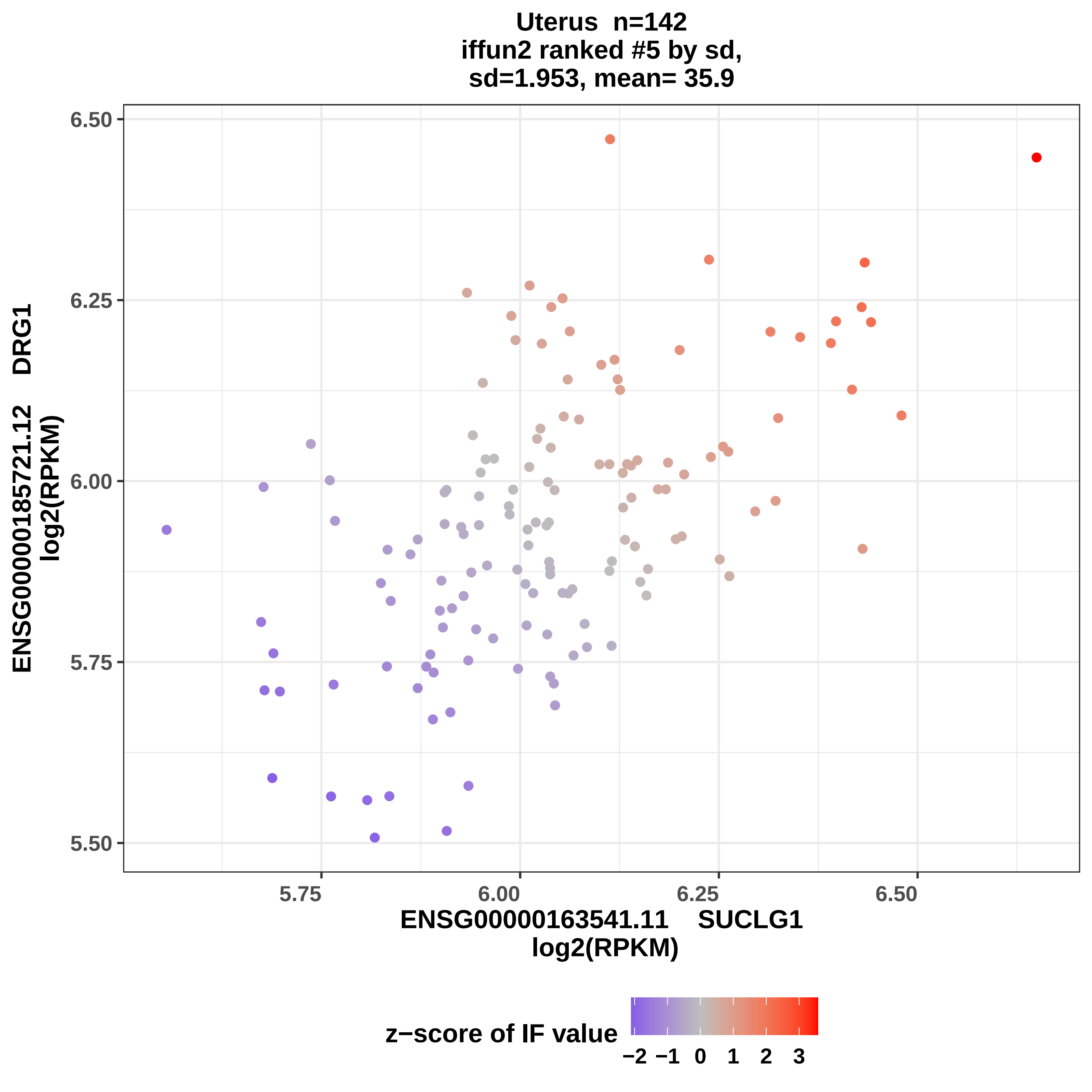

Uterus n=142
iffun2 ranked #5 by sd,
sd=1.953, mean= 35.9
ENSG00000185721.12 DRG1
log2(RPKM)
6.50
6.25
6.00
5.75
5.50
5.75
6.00
6.25
6.50
ENSG00000163541.11 SUCLG1
log2(RPKM)
z−score of IF value
−2 −1 0 1 2 3

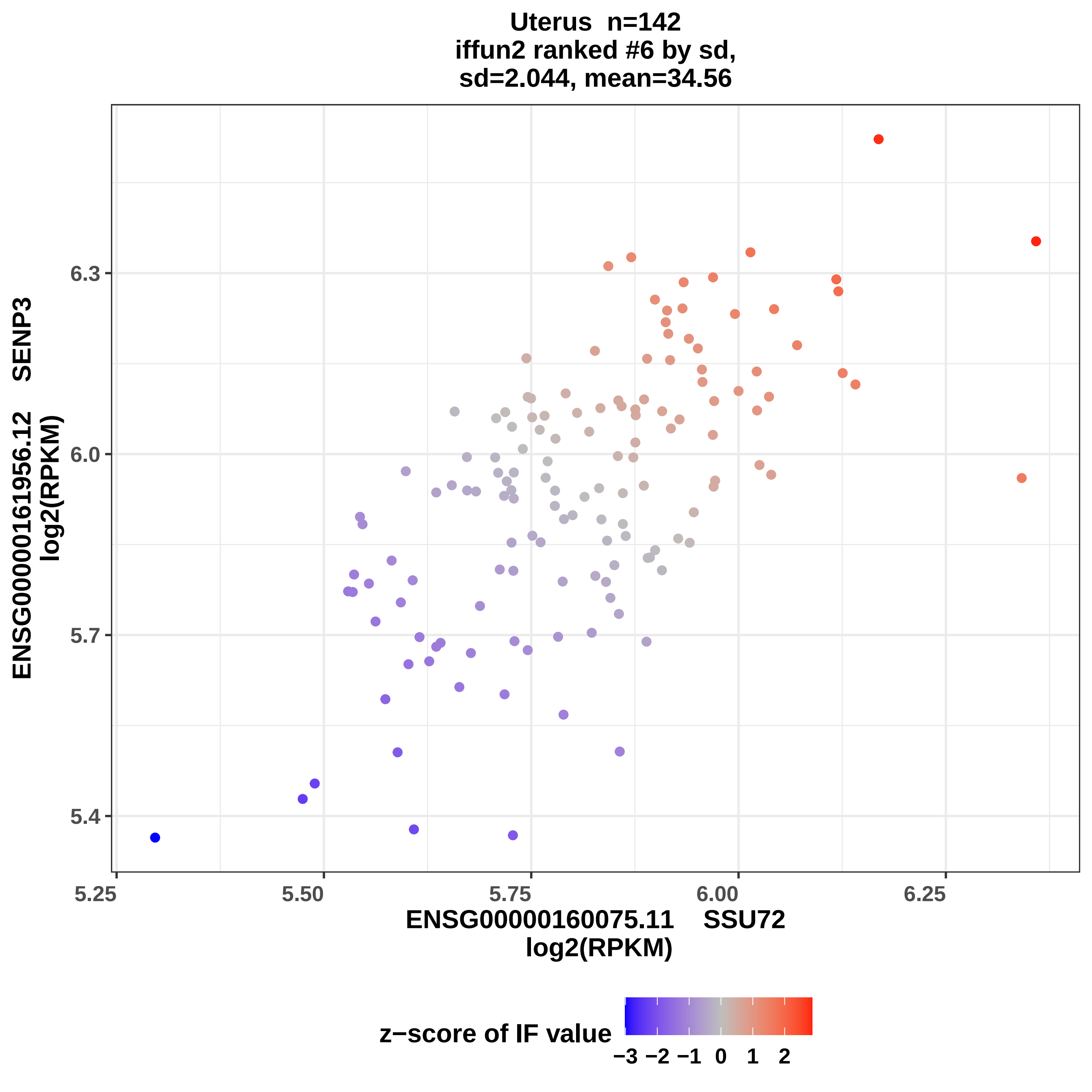
Uterus n=142
iffun2 ranked #6 by sd,
sd=2.044, mean=34.56
ENSG00000161956.12 SENP3
log2(RPKM)
6.3
6.0
5.7
5.4
5.25
5.50
5.75
6.00
6.25
ENSG00000160075.11 SSU72
log2(RPKM)
z−score of IF value
−3 −2 −1 0 1 2

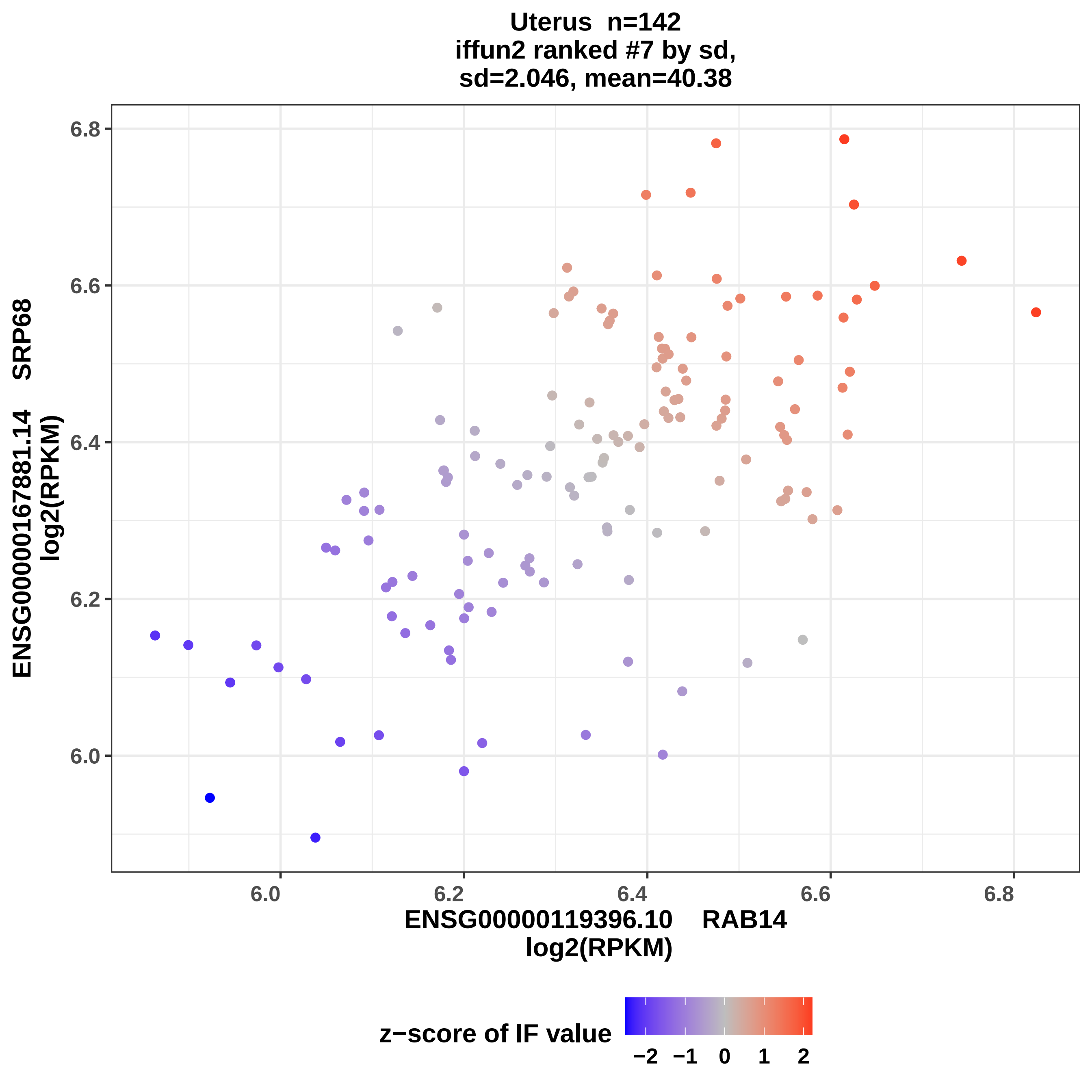
Uterus n=142
iffun2 ranked #7 by sd,
sd=2.046, mean=40.38
ENSG00000167881.14 SRP68
log2(RPKM)
ENSG00000119396.10 RAB14
log2(RPKM)
6.0
6.2
6.4
6.6
6.8
z−score of IF value
−2
−1
0
1
2

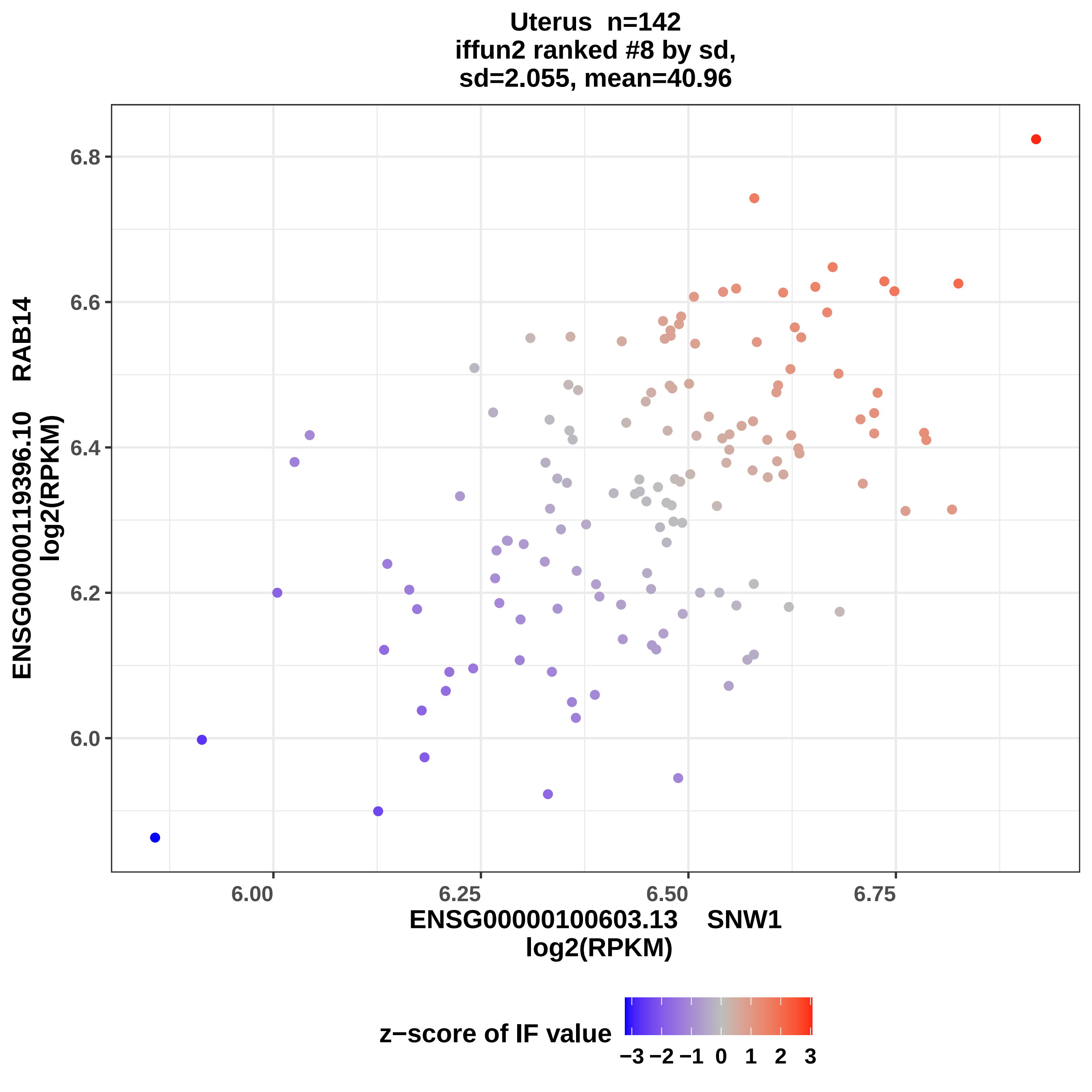
Uterus n=142
iffun2 ranked #8 by sd,
sd=2.055, mean=40.96
ENSG00000119396.10 RAB14
log2(RPKM)
6.8
6.6
6.4
6.2
6.0
6.00
6.25
6.50
6.75
ENSG00000100603.13 SNW1
log2(RPKM)
z−score of IF value
−3 −2 −1 0 1 2 3

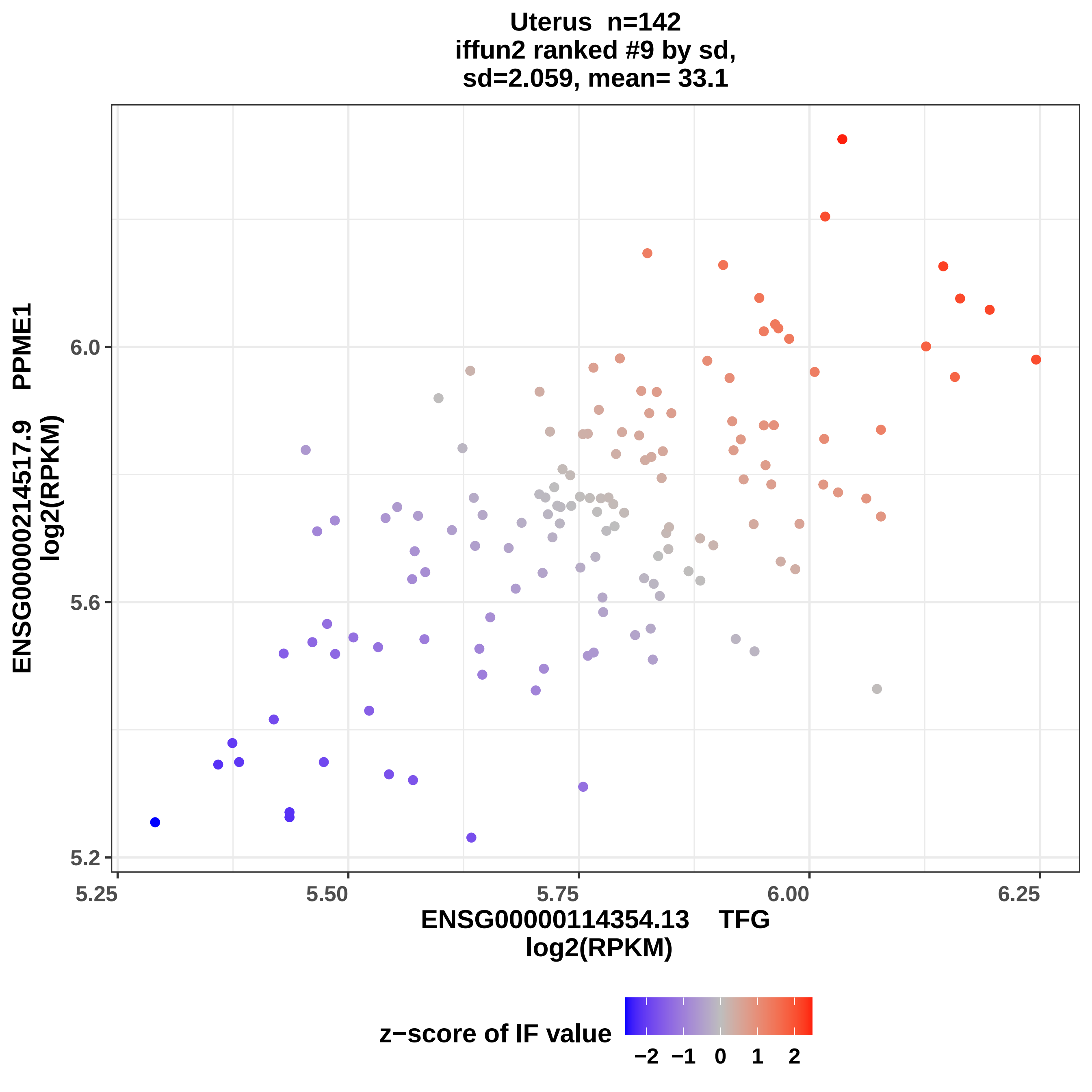
Uterus n=142
iffun2 ranked #9 by sd,
sd=2.059, mean= 33.1
ENSG00000214517.9 PPME1
log2(RPKM)
6.0
5.6
5.2
5.25
5.50
5.75
6.00
6.25
ENSG00000114354.13 TFG
log2(RPKM)
z−score of IF value
−2 −1 0 1 2

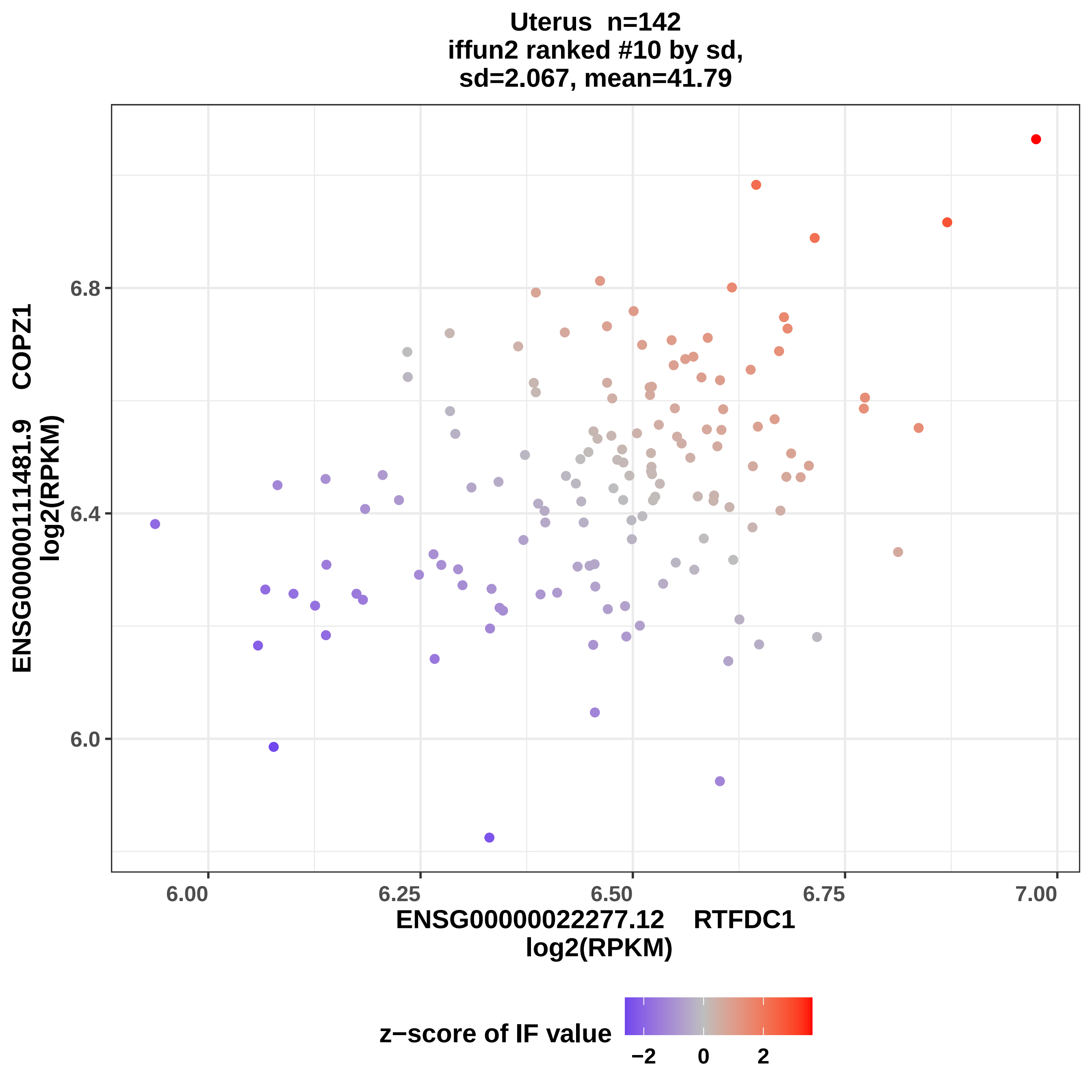

Uterus n=142
iffun2 ranked #10 by sd,
sd=2.067, mean=41.79
ENSG00000111481.9 COPZ1
log2(RPKM)
6.0
6.4
6.8
6.00
6.25
6.50
6.75
7.00
ENSG00000022277.12 RTFDC1
log2(RPKM)
z−score of IF value
−2
0
2

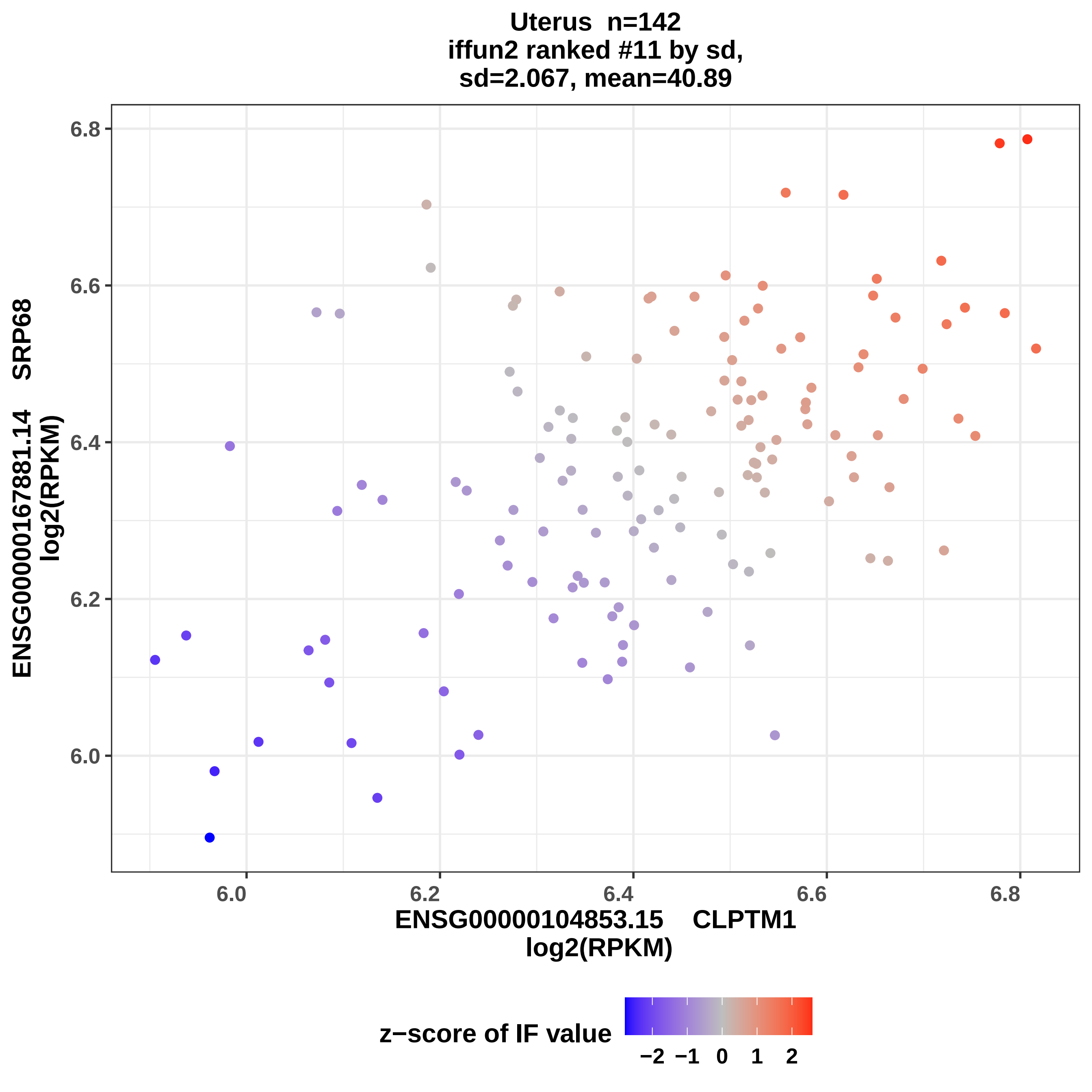

Uterus n=142
iffun2 ranked #11 by sd,
sd=2.067, mean=40.89
6.8
6.6
6.4
6.2
6.0
6.0
6.2
6.4
6.6
6.8
ENSG00000167881.14 SRP68
log2(RPKM)
ENSG00000104853.15 CLPTM1
log2(RPKM)
z−score of IF value
−2 −1 0 1 2

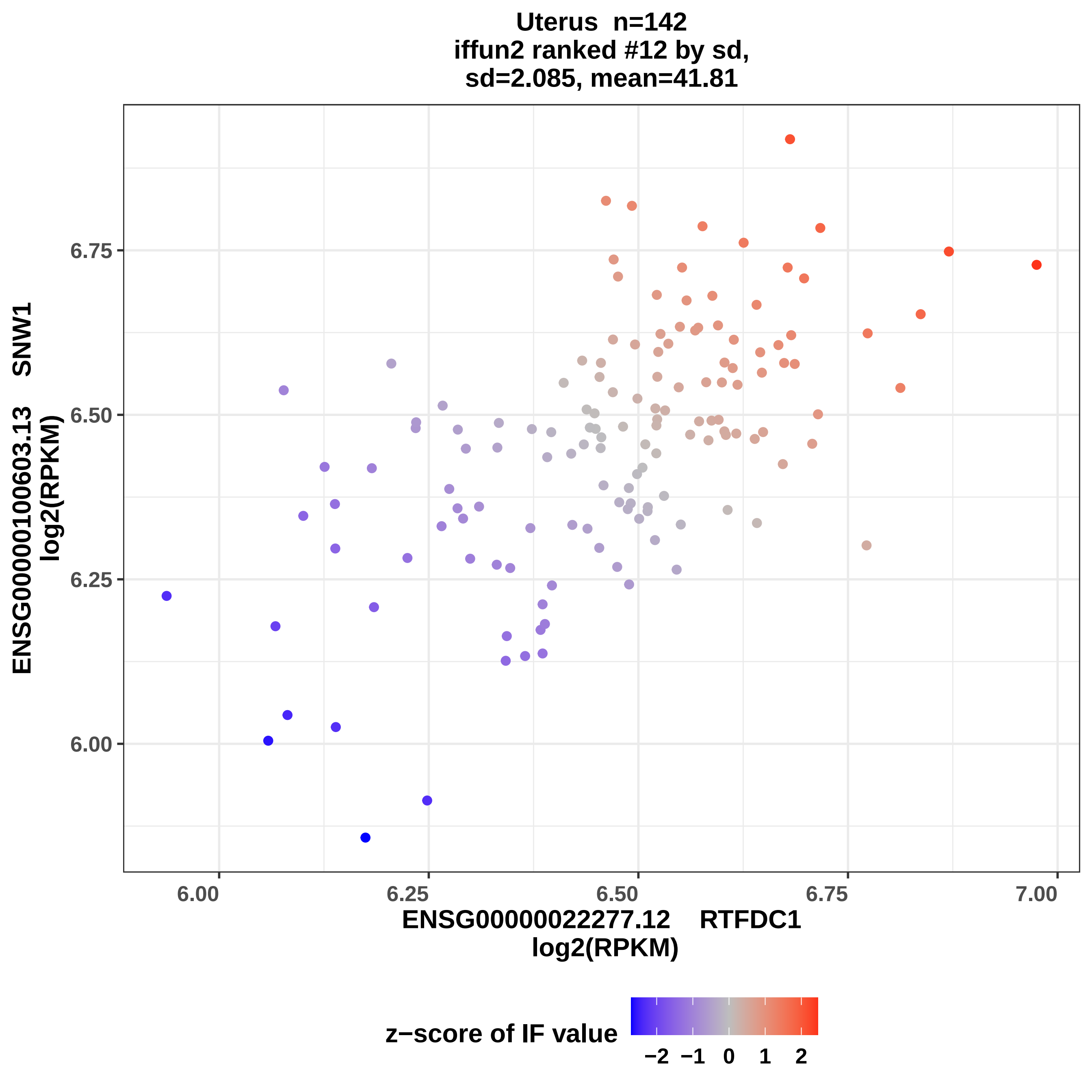
Uterus n=142
iffun2 ranked #12 by sd,
sd=2.085, mean=41.81
ENSG00000100603.13 SNW1
log2(RPKM)
6.75
6.50
6.25
6.00
6.00
6.25
6.50
6.75
7.00
ENSG00000022277.12 RTFDC1
log2(RPKM)
z−score of IF value
−2
−1
0
1
2

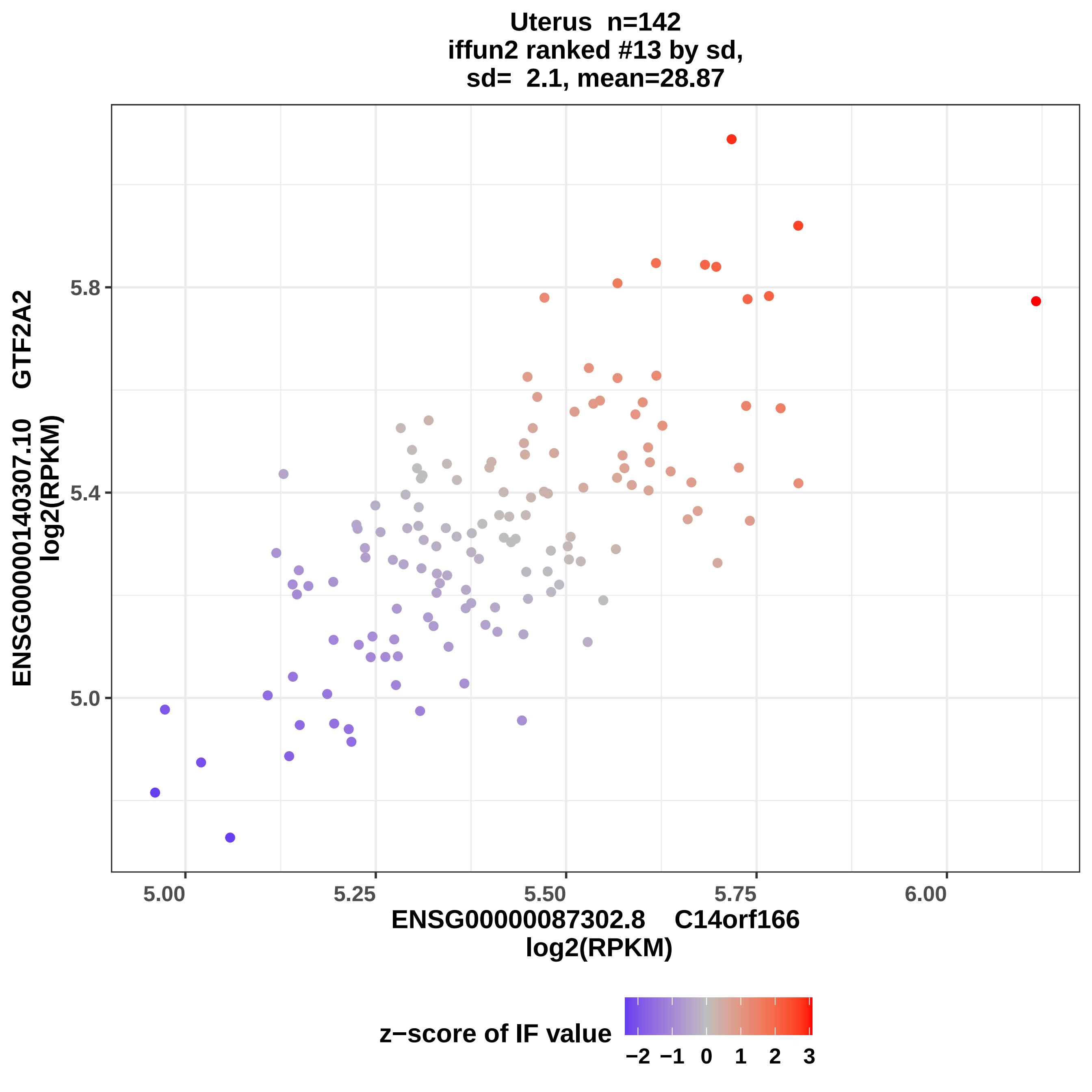

Uterus n=142
iffun2 ranked #13 by sd,
sd= 2.1, mean=28.87
ENSG00000140307.10 GTF2A2
log2(RPKM)
5.8
5.4
5.0
5.00
5.25
5.50
5.75
6.00
ENSG00000087302.8 C14orf166
log2(RPKM)
z−score of IF value
−2 −1 0 1 2 3

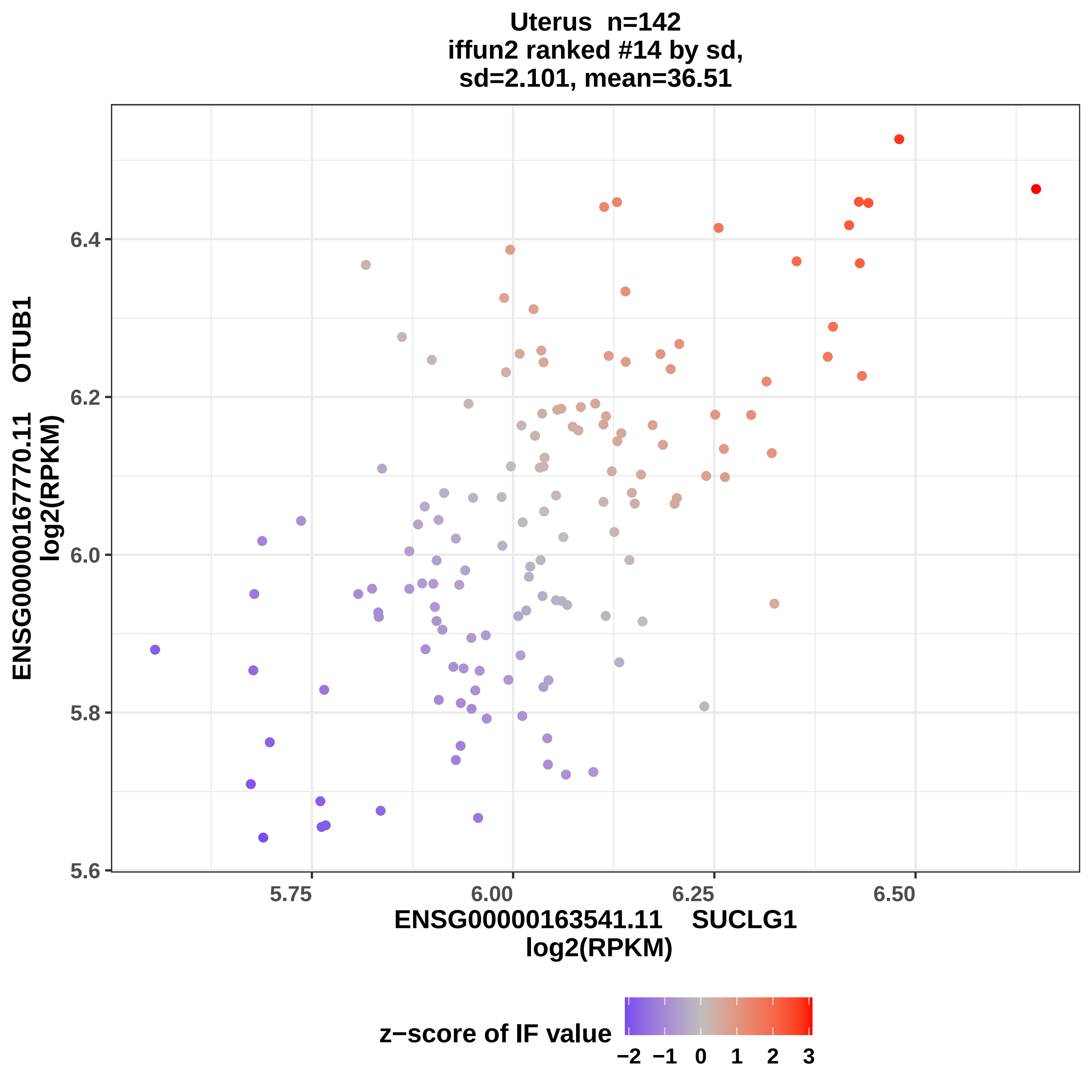
Uterus n=142
iffun2 ranked #14 by sd,
sd=2.101, mean=36.51
ENSG00000167770.11 OTUB1
log2(RPKM)
6.4
6.2
6.0
5.8
5.6
5.75
6.00
6.25
6.50
ENSG00000163541.11 SUCLG1
log2(RPKM)
z−score of IF value
−2 −1 0 1 2 3

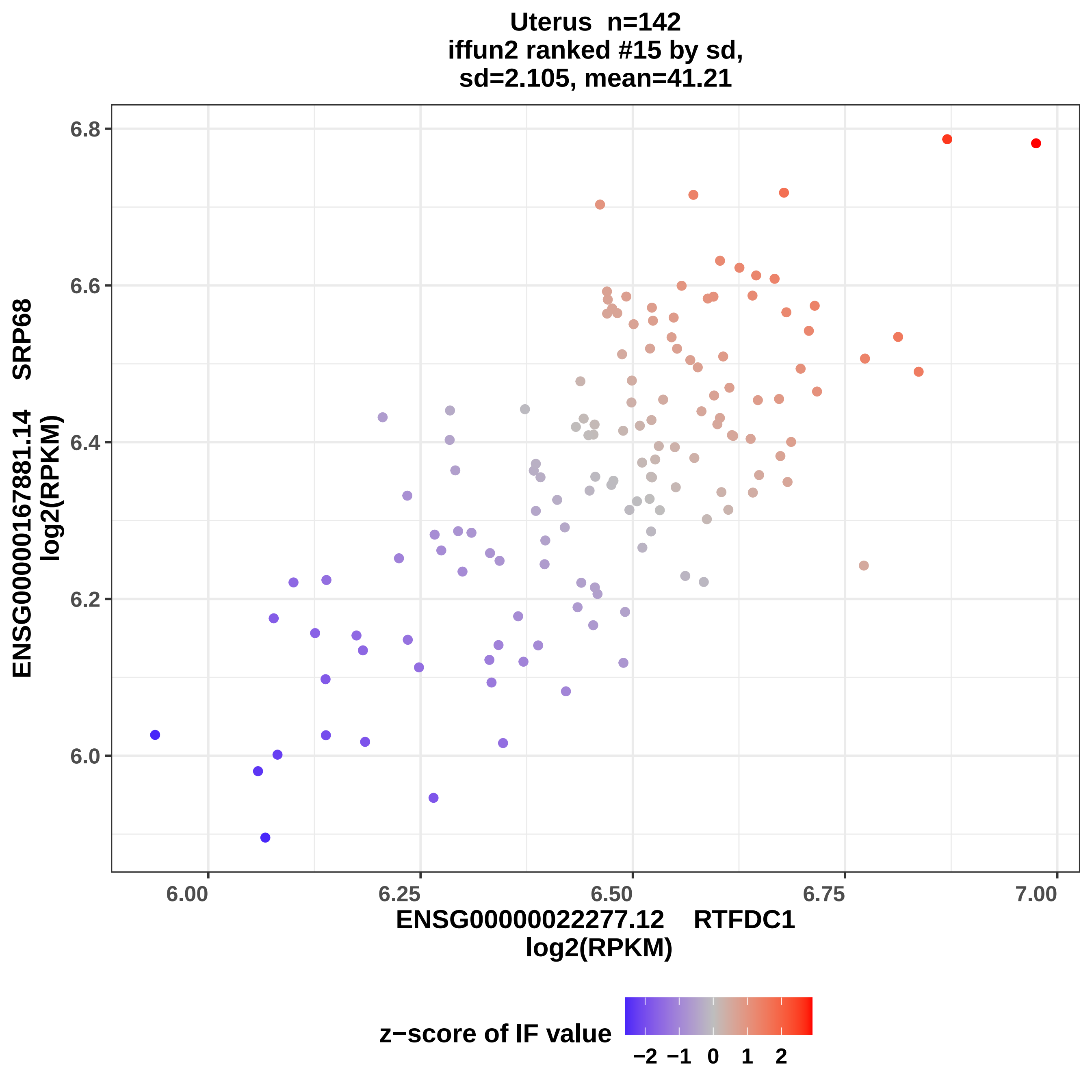
Uterus n=142
iffun2 ranked #15 by sd,
sd=2.105, mean=41.21
ENSG00000167881.14 SRP68
log2(RPKM)
6.8
6.6
6.4
6.2
6.0
6.00
6.25
6.50
6.75
7.00
ENSG00000022277.12 RTFDC1
log2(RPKM)
z−score of IF value
−2 −1 0 1 2

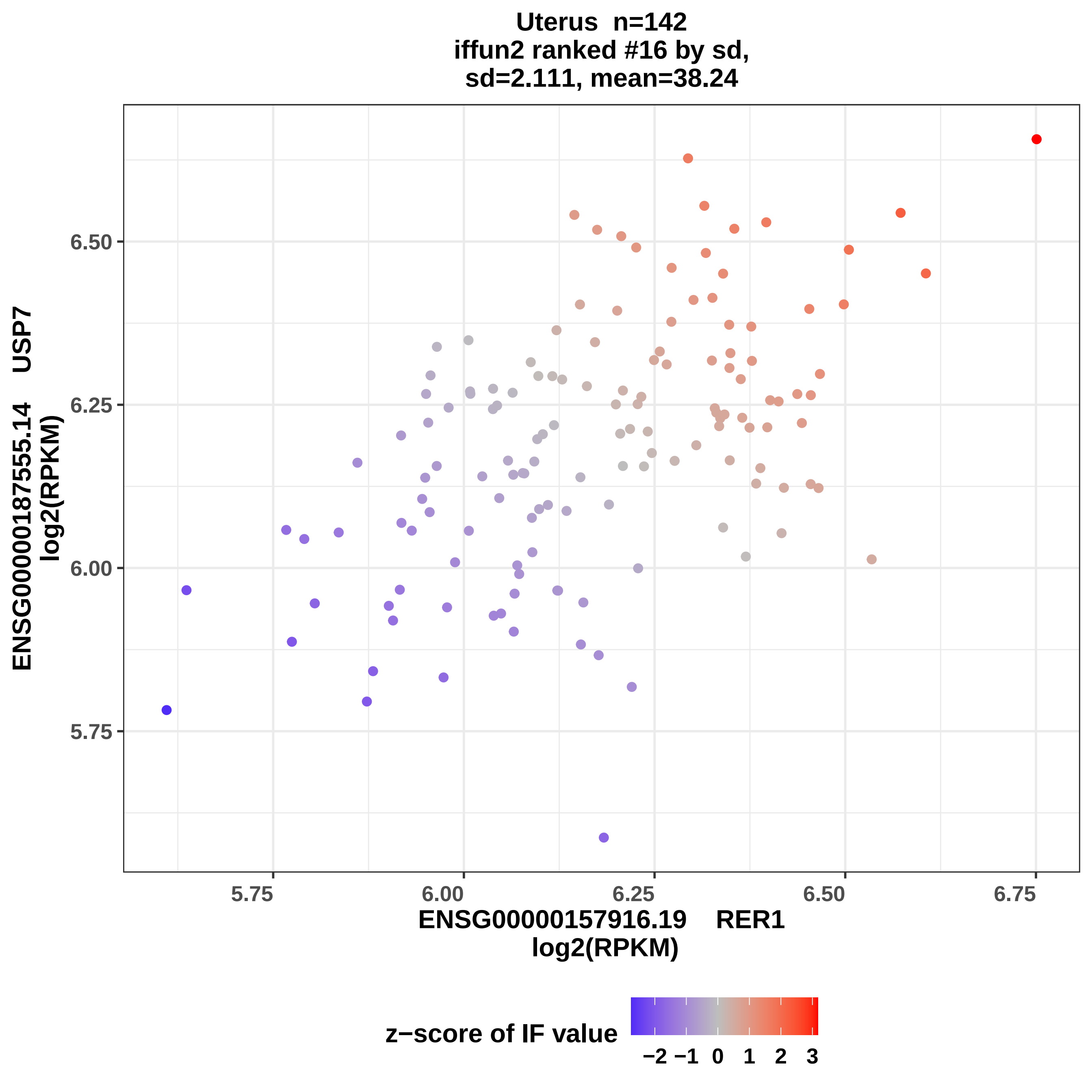

Uterus n=142
iffun2 ranked #16 by sd,
sd=2.111, mean=38.24
ENSG00000187555.14 USP7
log2(RPKM)
6.50
6.25
6.00
5.75
5.75
6.00
6.25
6.50
6.75
ENSG00000157916.19 RER1
log2(RPKM)
z−score of IF value
−2 −1 0 1 2 3

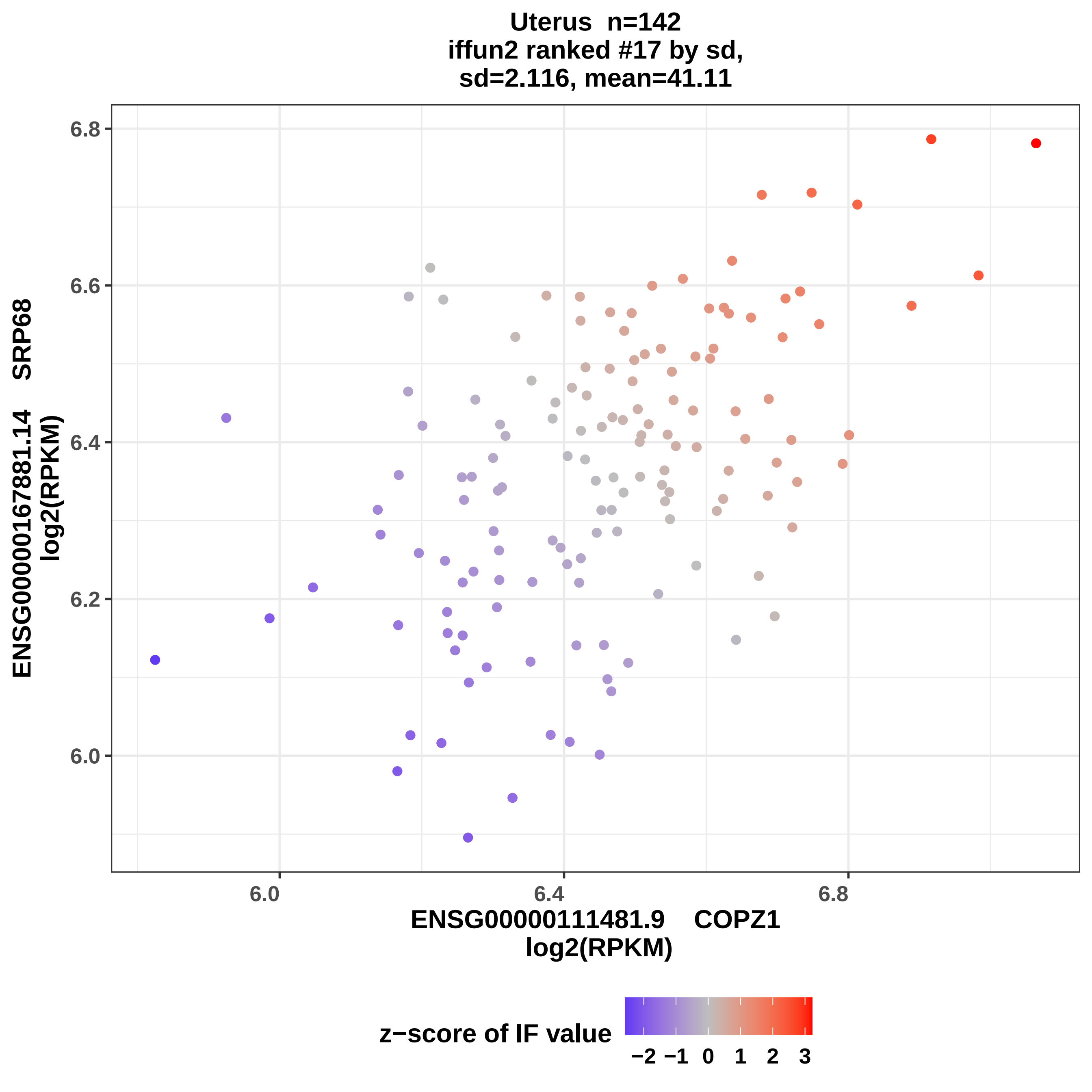
Uterus n=142
iffun2 ranked #17 by sd,
sd=2.116, mean=41.11
ENSG00000167881.14 SRP68
log2(RPKM)
6.8
6.6
6.4
6.2
6.0
6.0
6.4
6.8
ENSG00000111481.9 COPZ1
log2(RPKM)
z−score of IF value
−2 −1 0 1 2 3

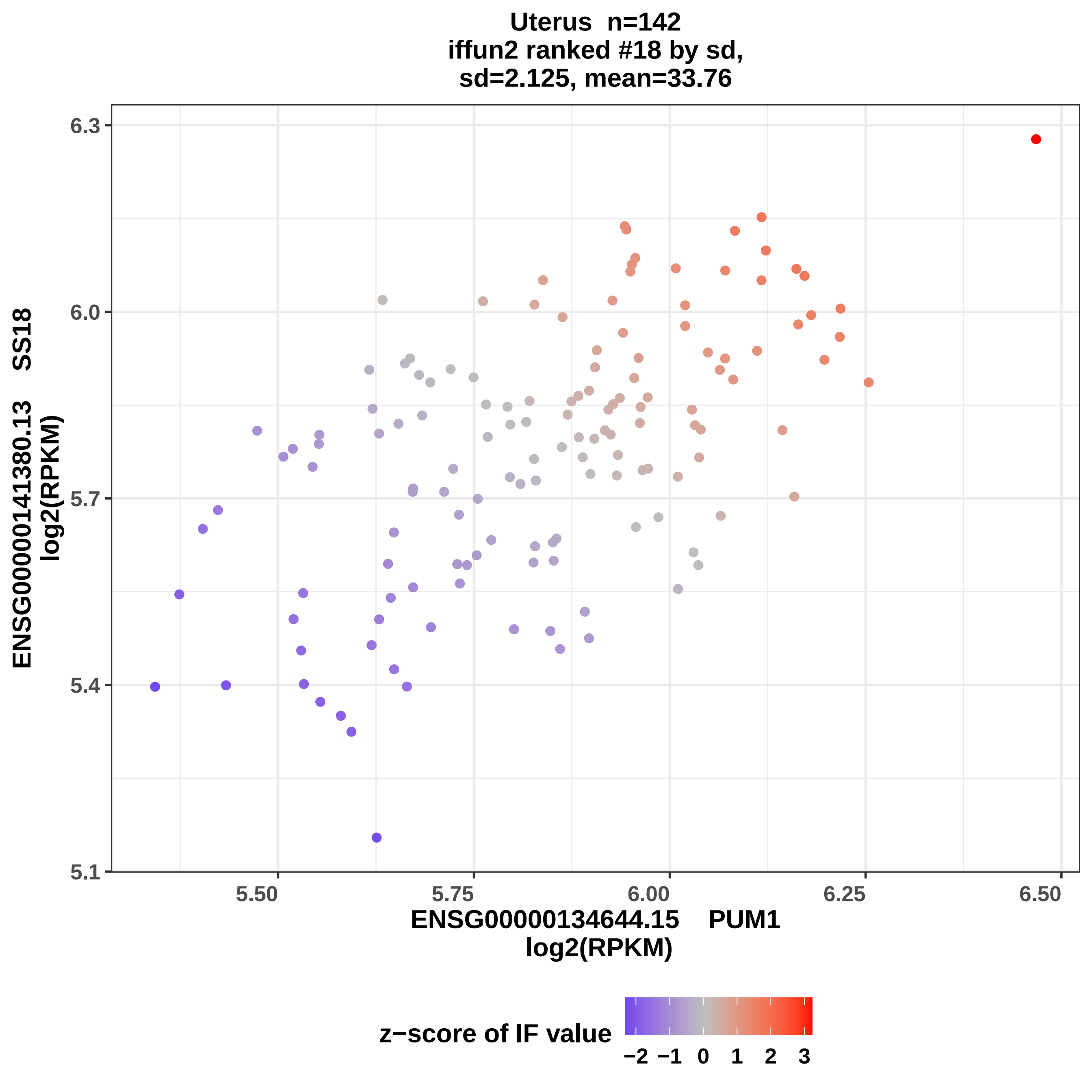
Uterus n=142
iffun2 ranked #18 by sd,
sd=2.125, mean=33.76
ENSG00000141380.13 SS18
log2(RPKM)
6.3
6.0
5.7
5.4
5.1
5.50
5.75
6.00
6.25
6.50
ENSG00000134644.15 PUM1
log2(RPKM)
z−score of IF value
−2 −1 0 1 2 3

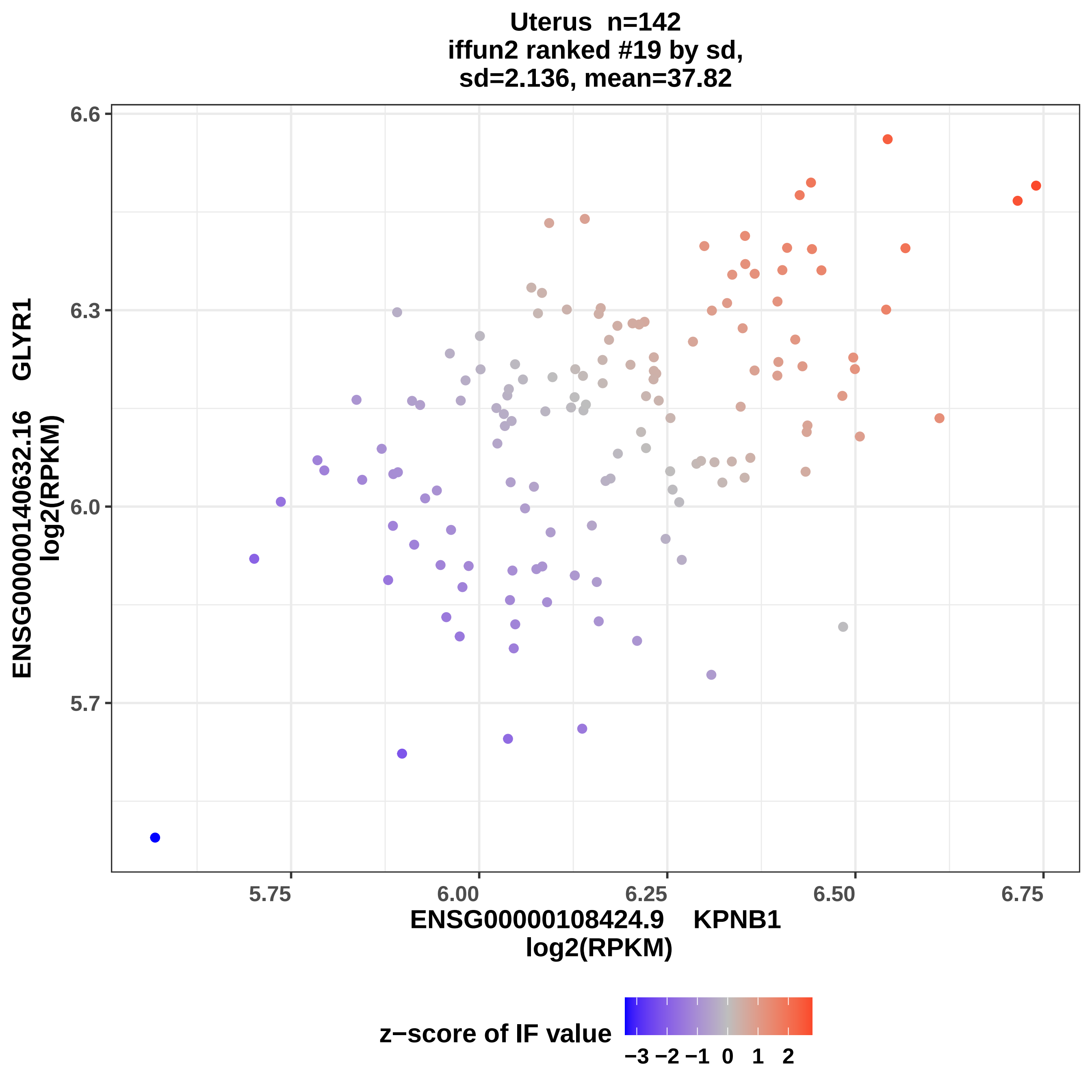
Uterus n=142
iffun2 ranked #19 by sd,
sd=2.136, mean=37.82
ENSG00000140632.16 GLYR1
log2(RPKM)
6.6
6.3
6.0
5.7
5.75
6.00
6.25
6.50
6.75
ENSG00000108424.9 KPNB1
log2(RPKM)
z−score of IF value
−3 −2 −1 0 1 2

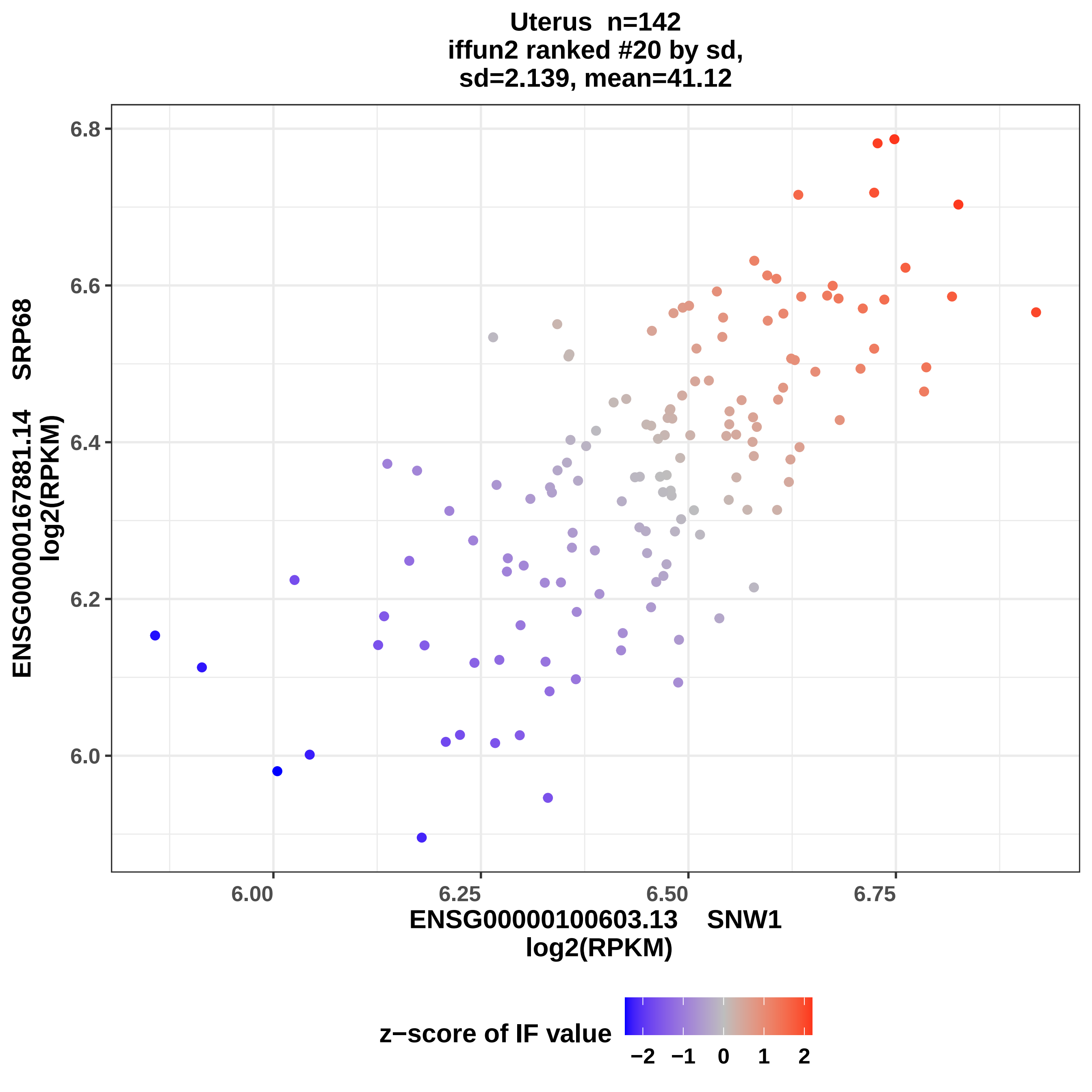
Uterus n=142
iffun2 ranked #20 by sd,
sd=2.139, mean=41.12
6.8
6.6
6.4
6.2
6.0
ENSG00000167881.14 SRP68
log2(RPKM)
6.00
6.25
6.50
6.75
ENSG00000100603.13 SNW1
log2(RPKM)
z−score of IF value
−2
−1
0
1
2

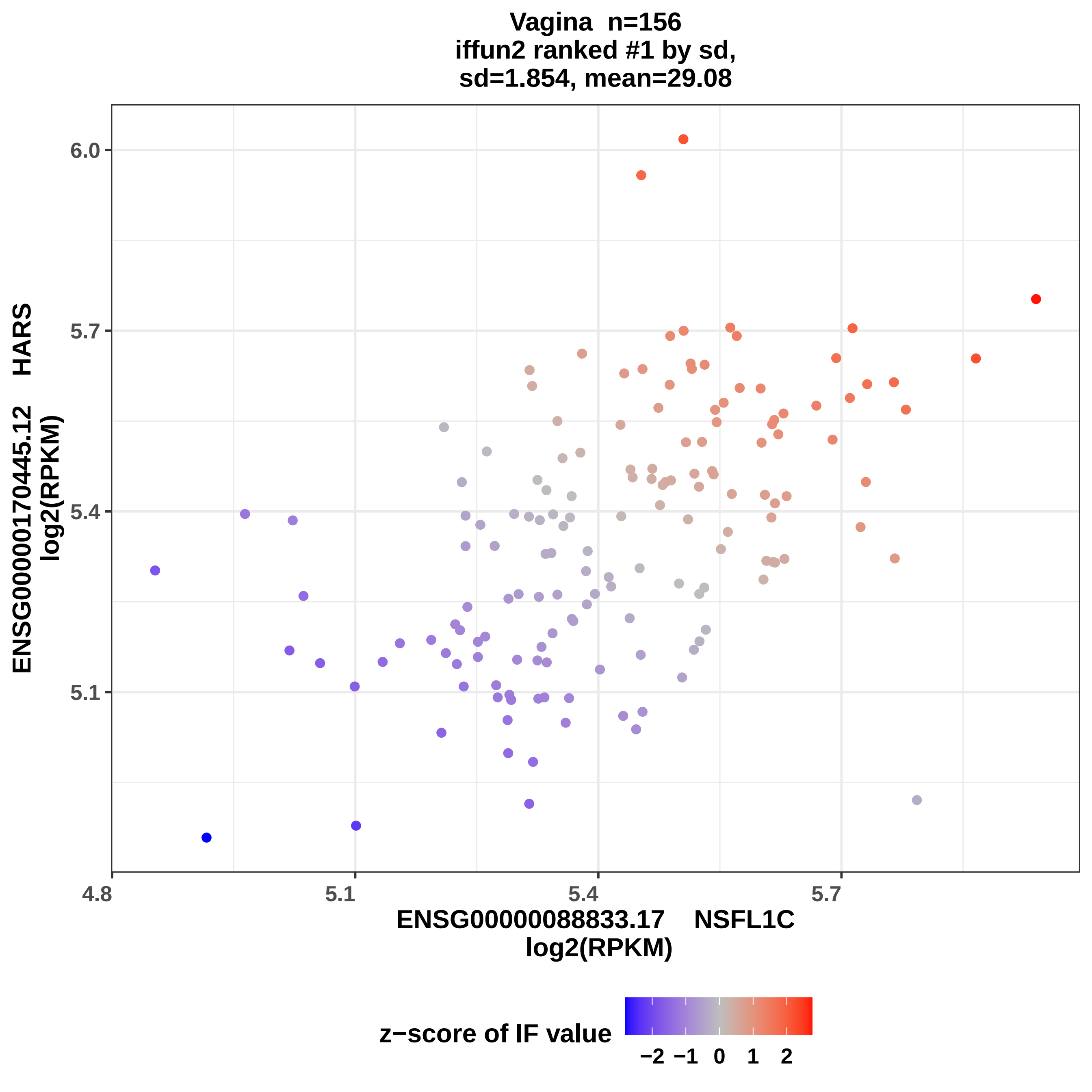

Vagina n=156
iffun2 ranked #1 by sd,
sd=1.854, mean=29.08
ENSG00000170445.12 HARS
log2(RPKM)
6.0
5.7
5.4
5.1
4.8
5.1
5.4
5.7
ENSG00000088833.17 NSFL1C
log2(RPKM)
z−score of IF value
−2 −1 0 1 2

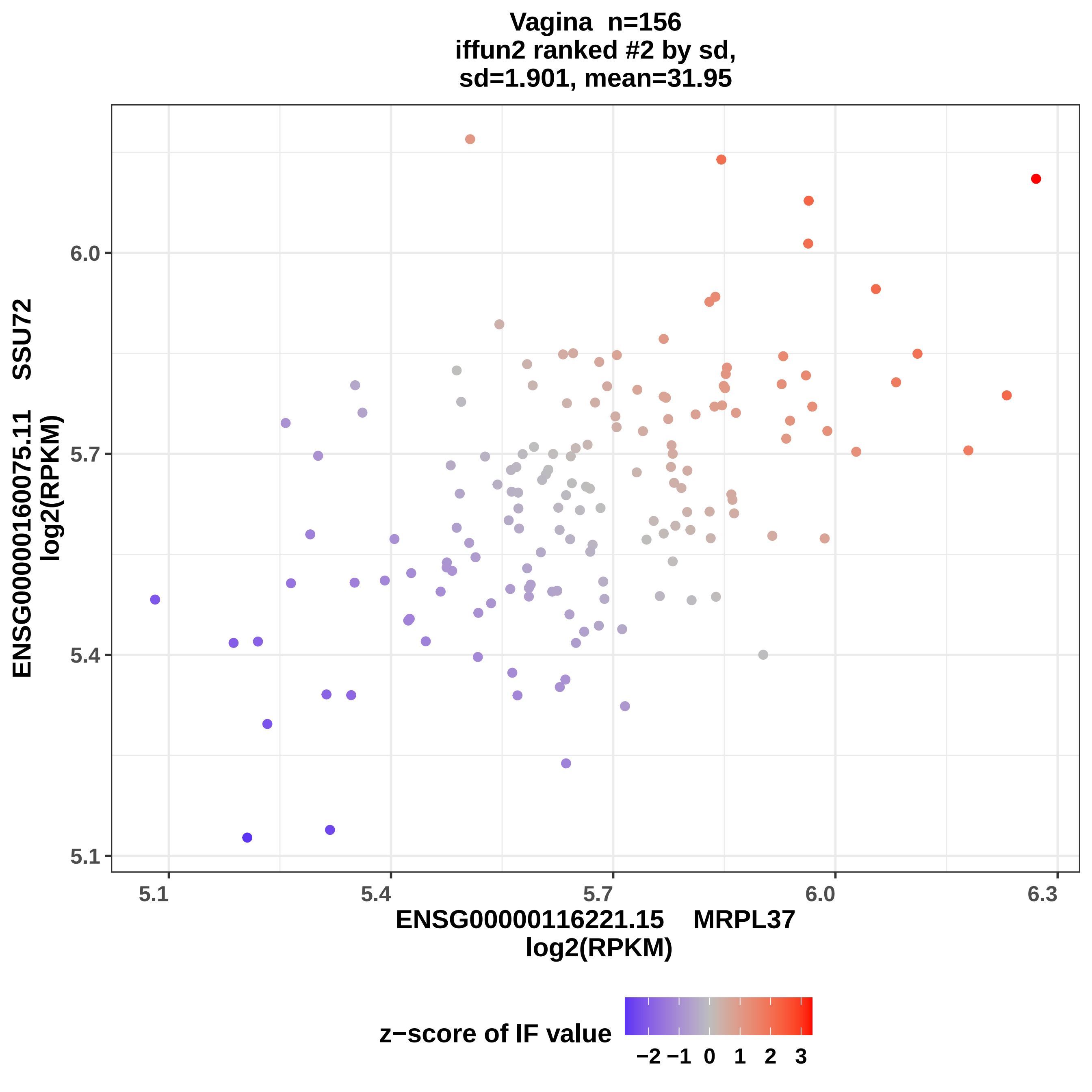
Vagina n=156
iffun2 ranked #2 by sd,
sd=1.901, mean=31.95
ENSG00000160075.11 SSU72
log2(RPKM)
6.0
5.7
5.4
5.1
5.1
5.4
5.7
6.0
6.3
ENSG00000116221.15 MRPL37
log2(RPKM)
z−score of IF value
−2 −1 0 1 2 3

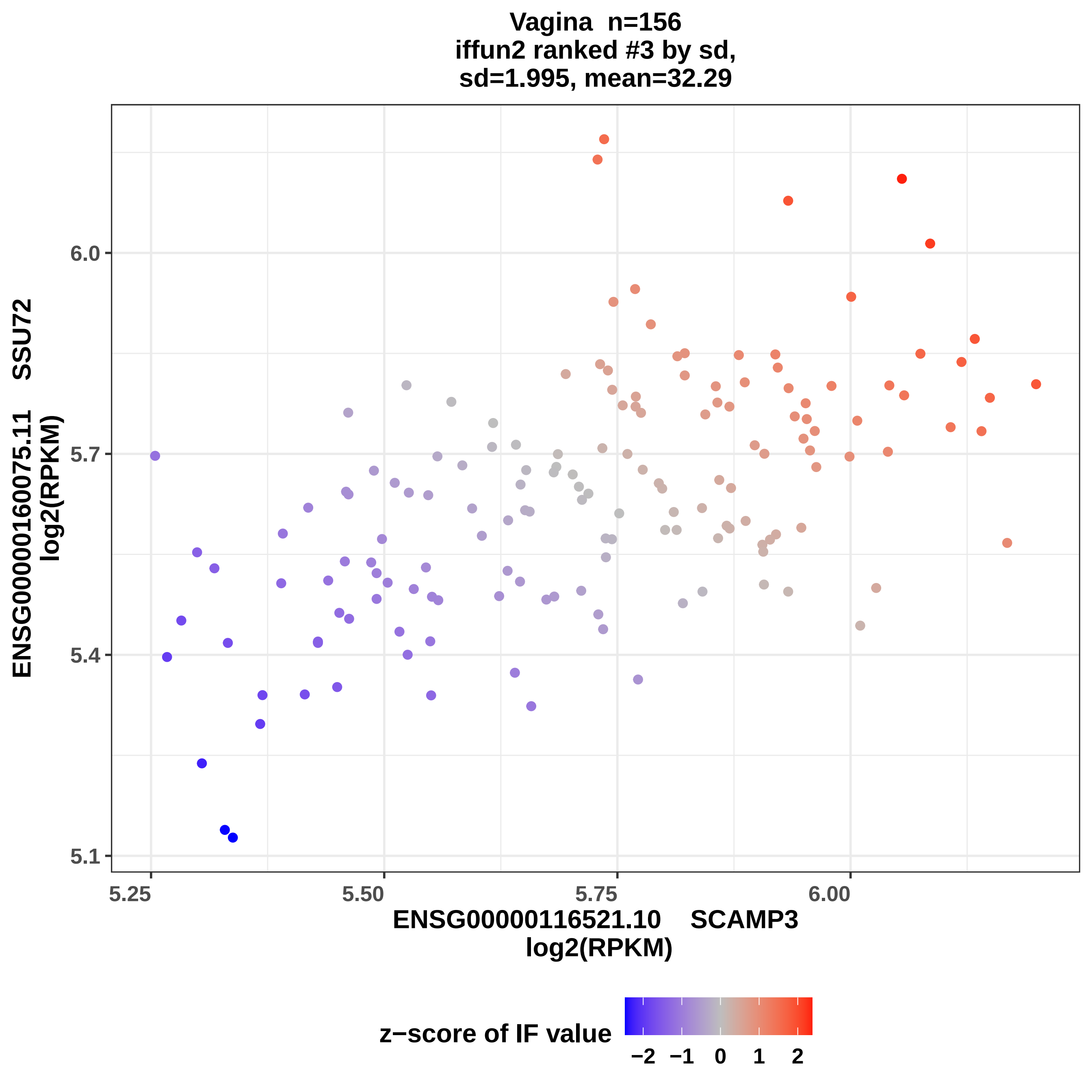
Vagina n=156
iffun2 ranked #3 by sd,
sd=1.995, mean=32.29
ENSG00000160075.11 SSU72
log2(RPKM)
6.0
5.7
5.4
5.1
5.25
5.50
5.75
6.00
ENSG00000116521.10 SCAMP3
log2(RPKM)
z−score of IF value
−2
−1
0
1
2

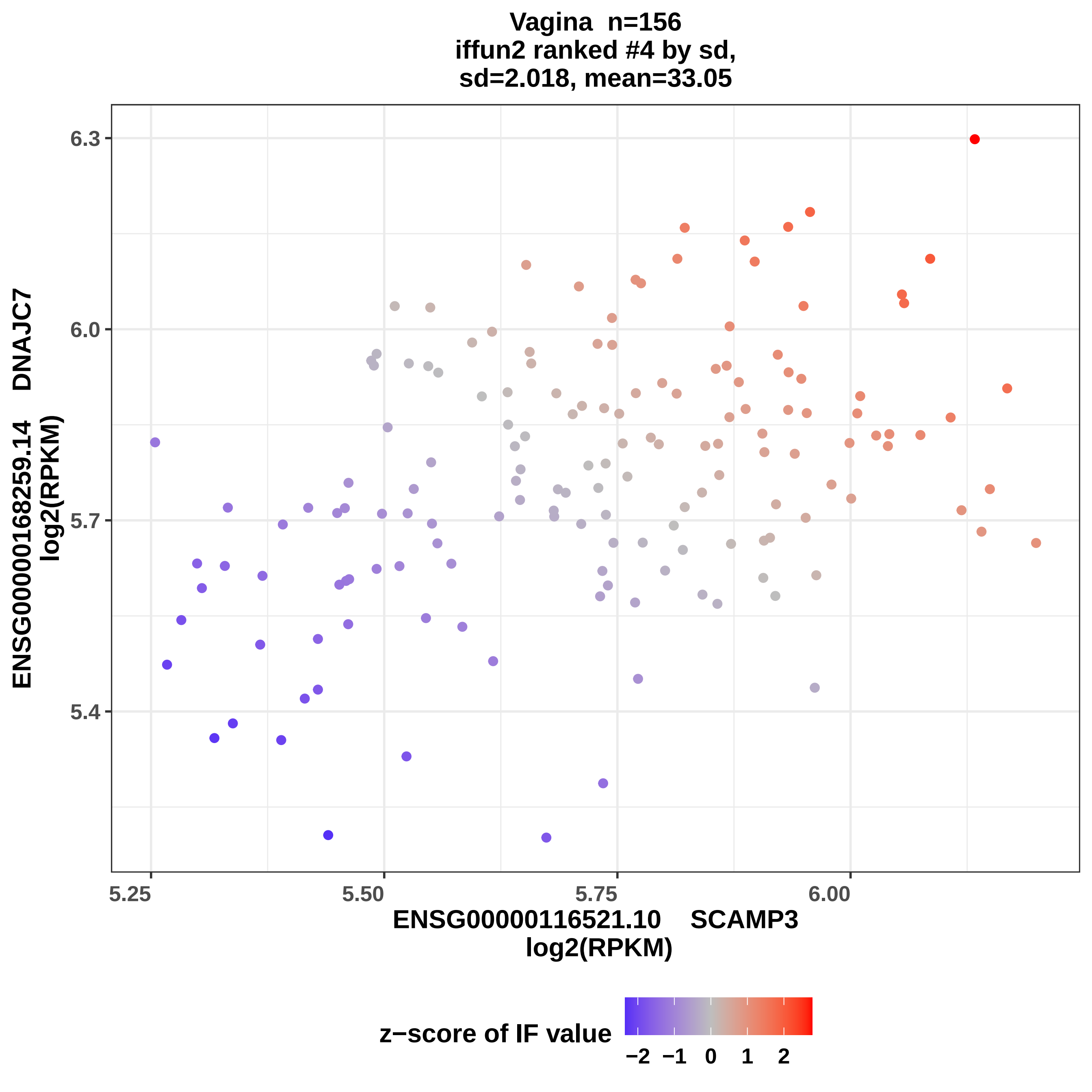

Vagina n=156
iffun2 ranked #4 by sd,
sd=2.018, mean=33.05
ENSG00000168259.14 DNAJC7
log2(RPKM)
6.3
6.0
5.7
5.4
5.25
5.50
5.75
6.00
ENSG00000116521.10 SCAMP3
log2(RPKM)
z−score of IF value
−2
−1
0
1
2

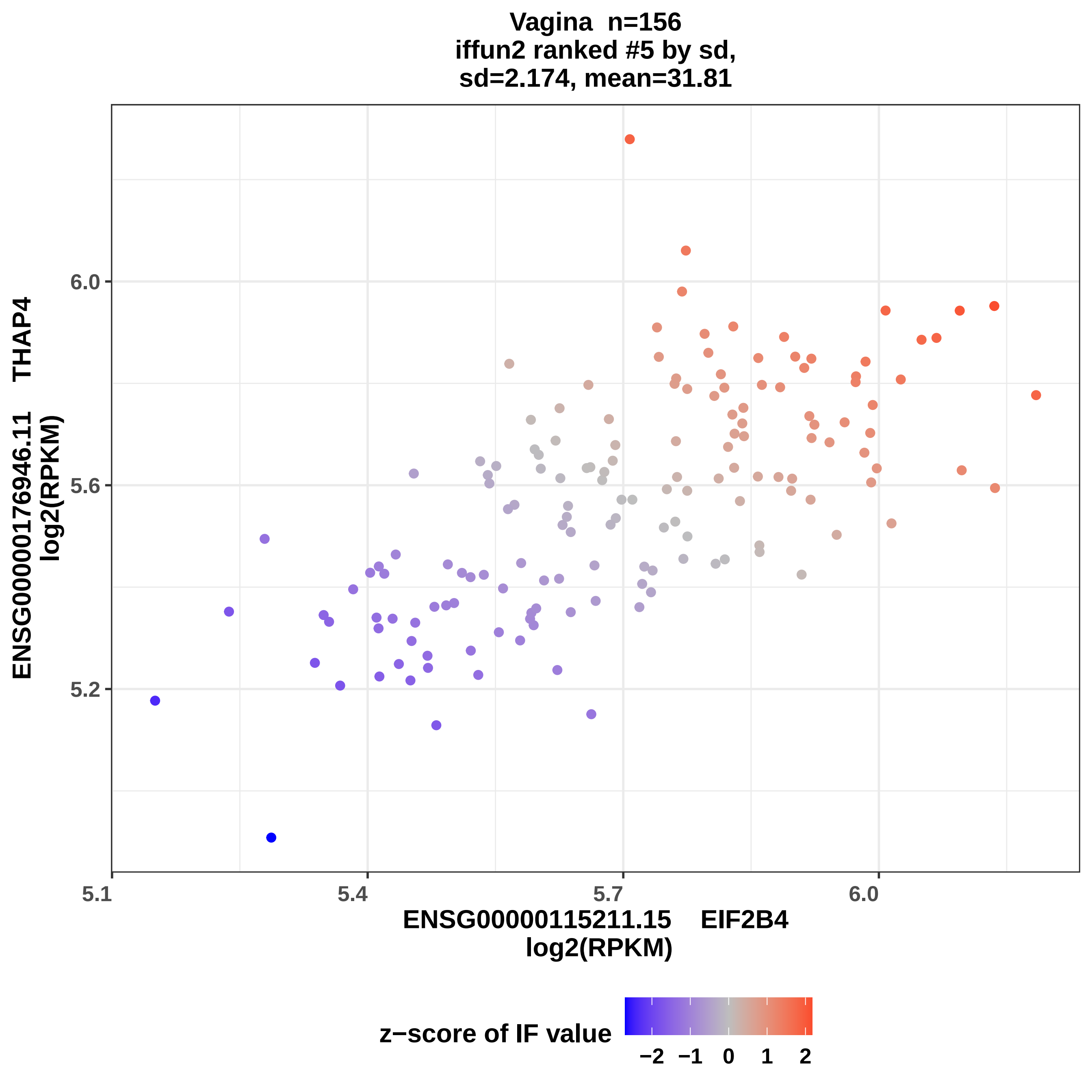
Vagina n=156
iffun2 ranked #5 by sd,
sd=2.174, mean=31.81
ENSG00000176946.11 THAP4
log2(RPKM)
6.0
5.6
5.2
5.1
5.4
5.7
6.0
ENSG00000115211.15 EIF2B4
log2(RPKM)
z−score of IF value
−2 −1 0 1 2

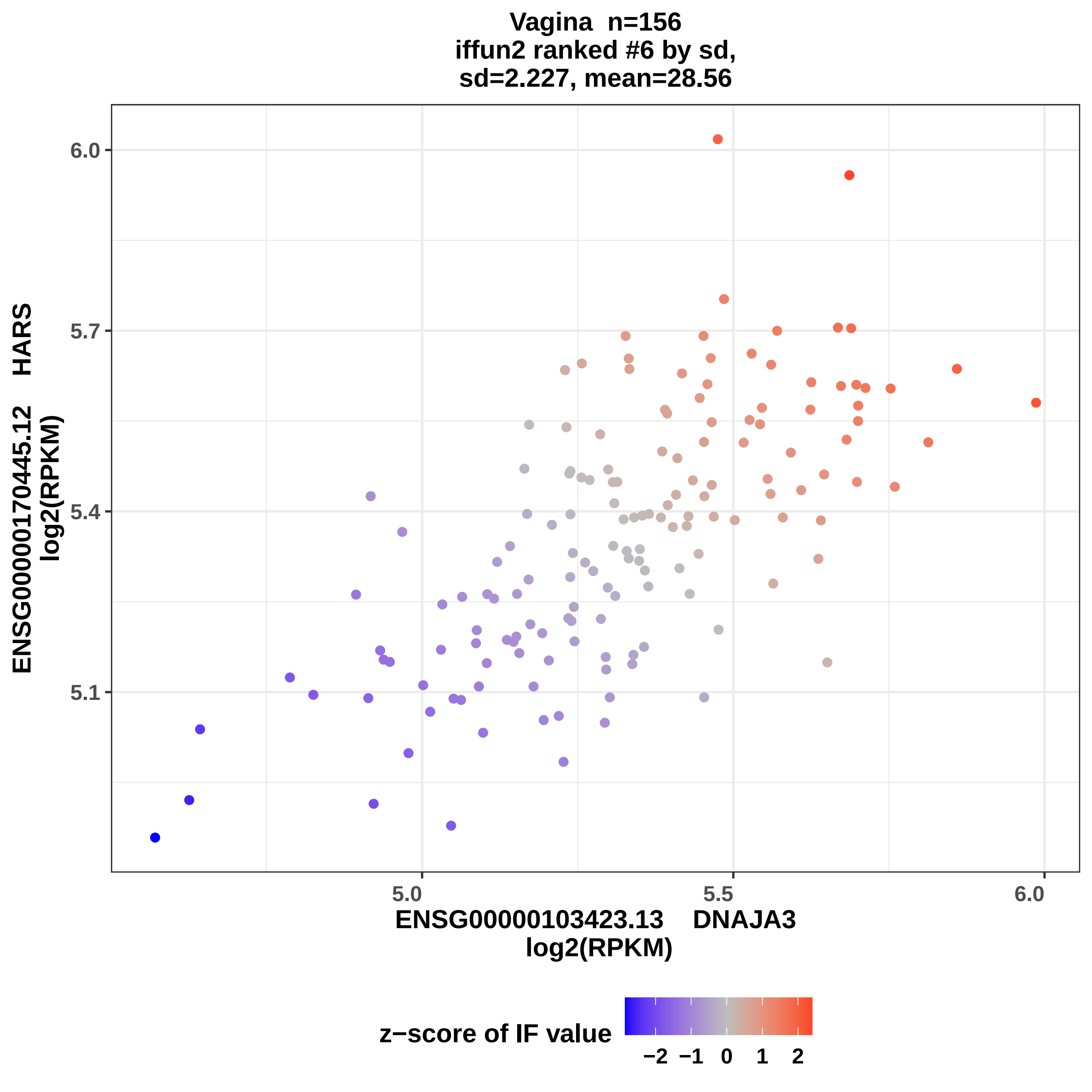

Vagina n=156
iffun2 ranked #6 by sd,
sd=2.227, mean=28.56
ENSG00000170445.12 HARS
log2(RPKM)
6.0
5.7
5.4
5.1
5.0
5.5
6.0
ENSG00000103423.13 DNAJA3
log2(RPKM)
z−score of IF value
−2 −1 0 1 2

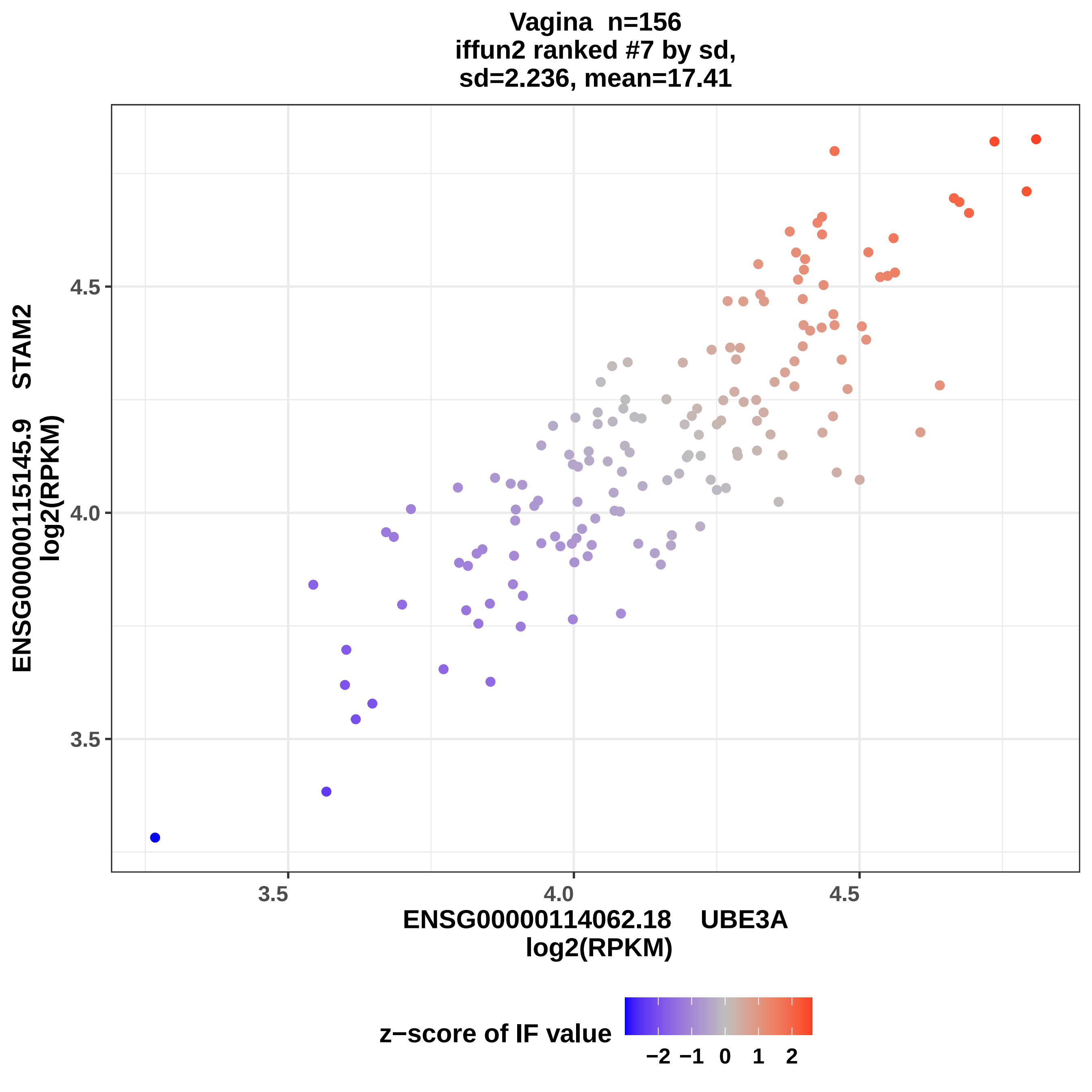
Vagina n=156
iffun2 ranked #7 by sd,
sd=2.236, mean=17.41
ENSG00000115145.9 STAM2
log2(RPKM)
4.5
4.0
3.5
3.5
4.0
4.5
ENSG00000114062.18 UBE3A
log2(RPKM)
z−score of IF value
−2 −1 0 1 2

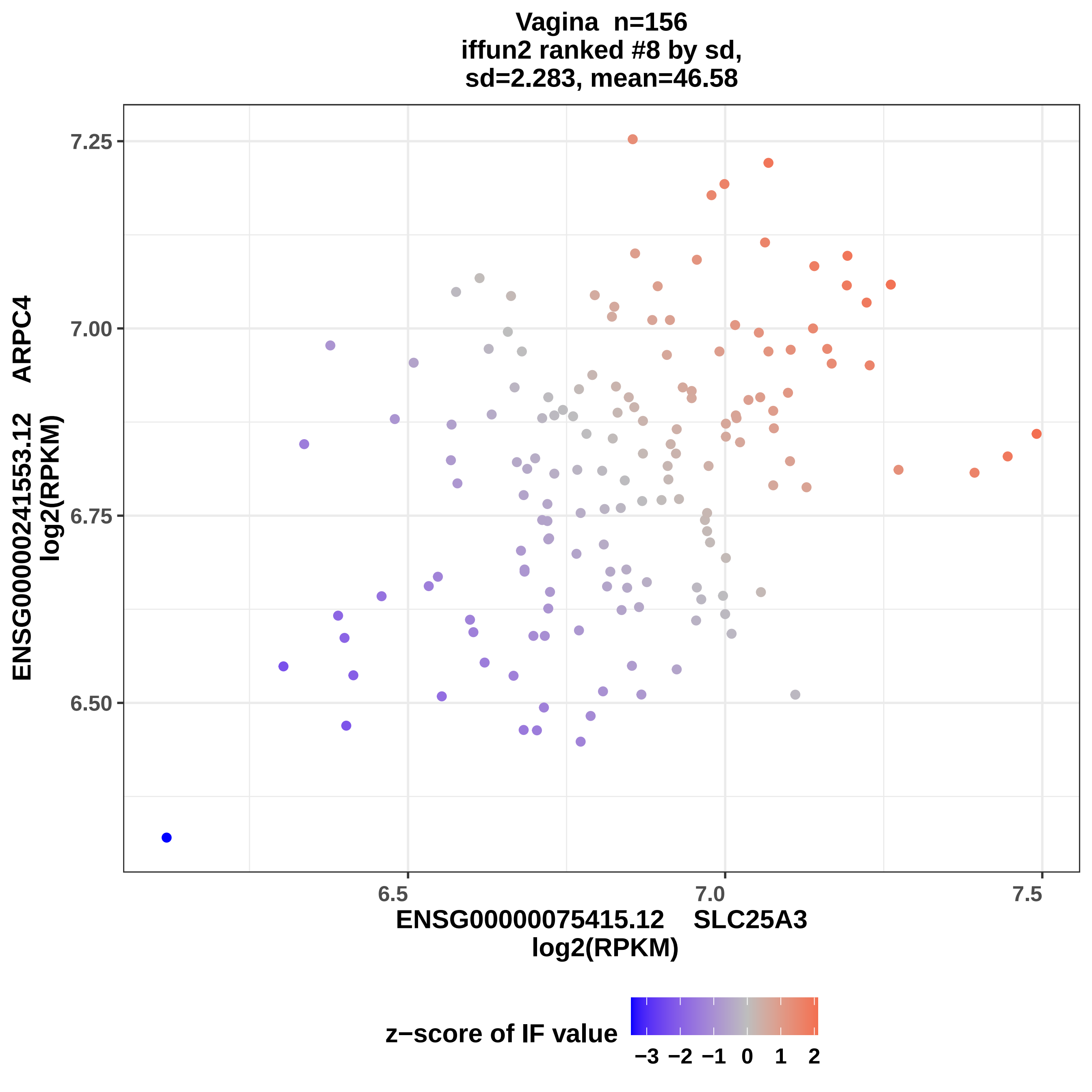

Vagina n=156
iffun2 ranked #8 by sd,
sd=2.283, mean=46.58
ENSG00000241553.12 ARPC4
log2(RPKM)
7.25
7.00
6.75
6.50
6.5
7.0
7.5
ENSG00000075415.12 SLC25A3
log2(RPKM)
z−score of IF value
−3 −2 −1 0 1 2

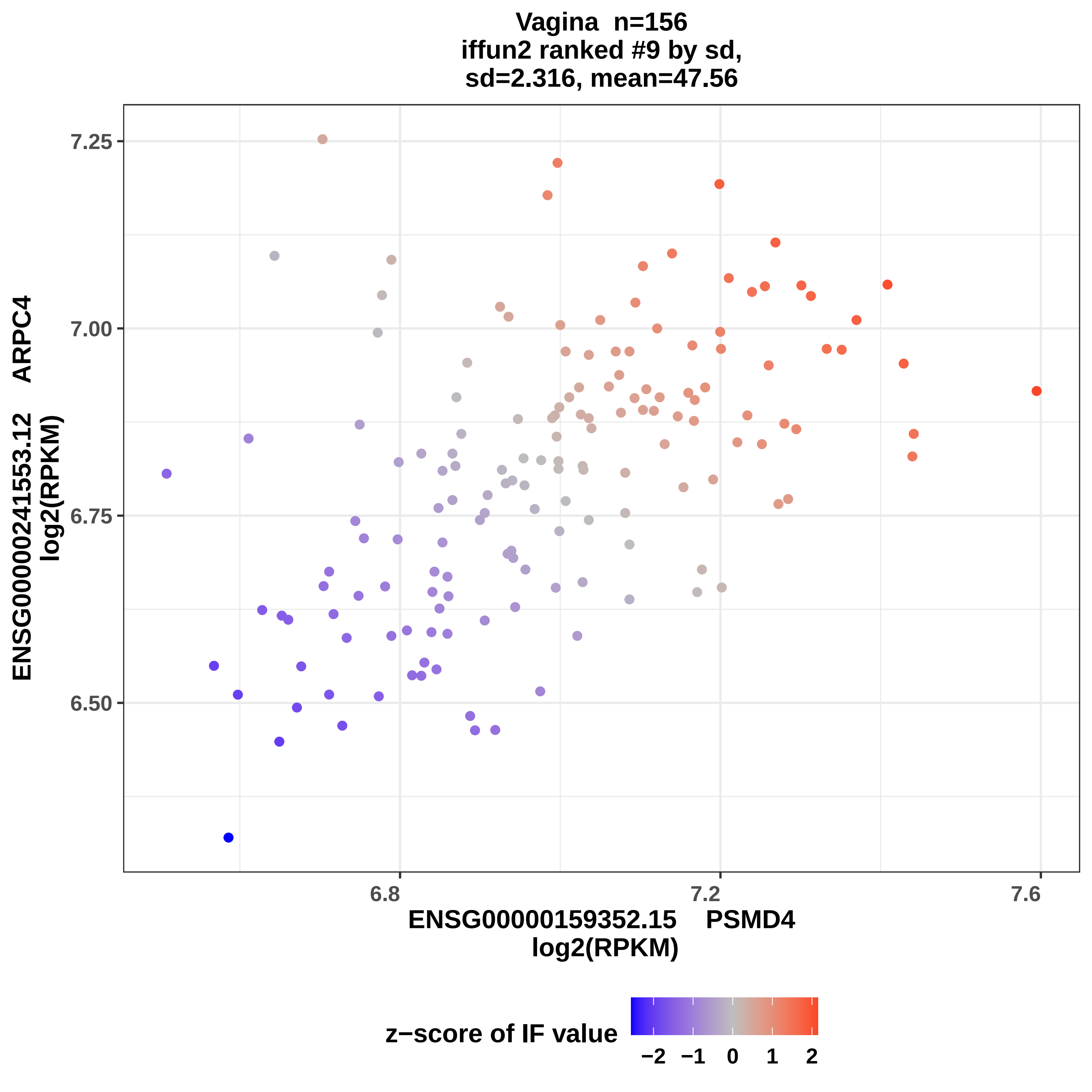

Vagina n=156
iffun2 ranked #9 by sd,
sd=2.316, mean=47.56
ENSG00000241553.12 ARPC4
log2(RPKM)
7.25
7.00
6.75
6.50
6.8
7.2
7.6
ENSG00000159352.15 PSMD4
log2(RPKM)
z−score of IF value
−2
−1
0
1
2

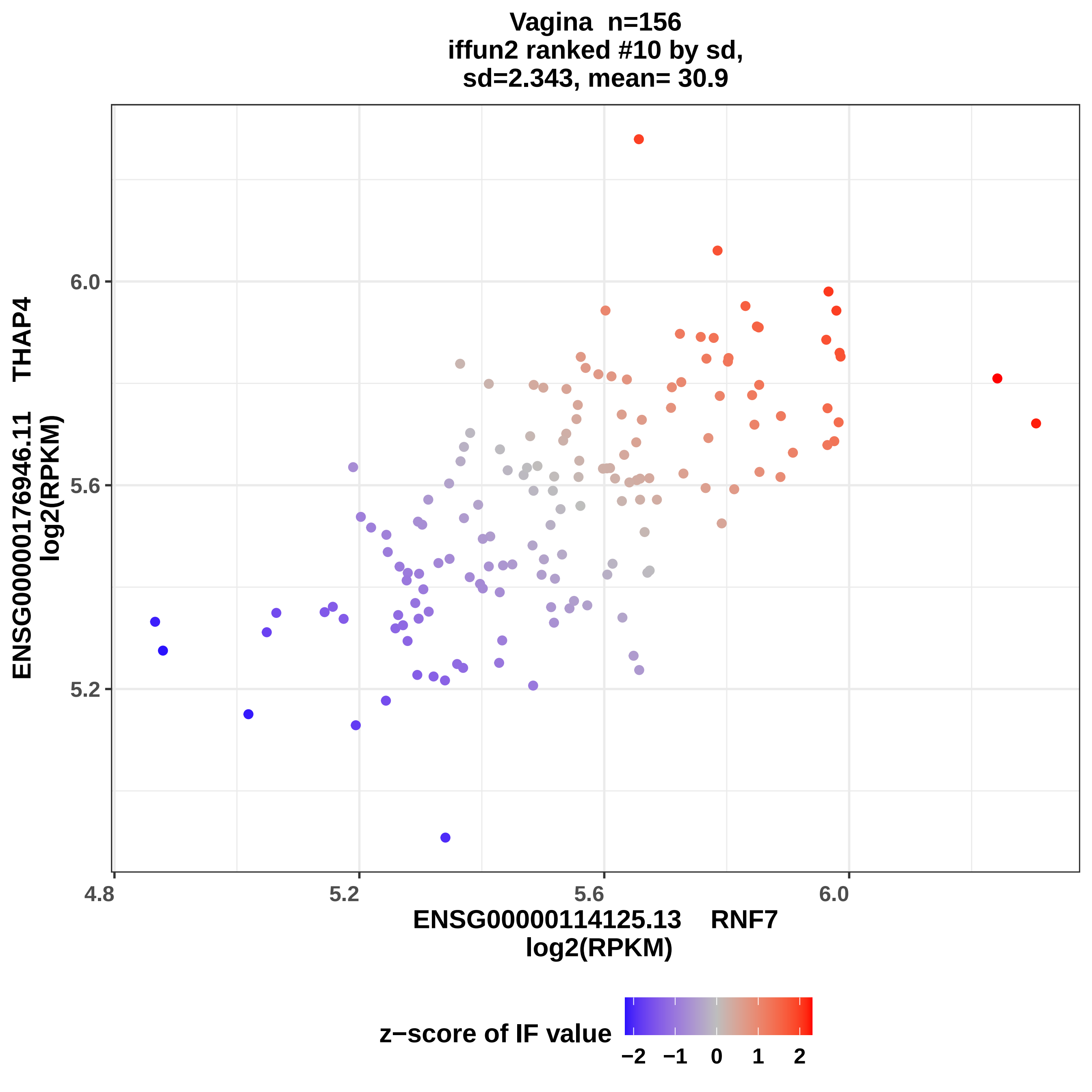

Vagina n=156
iffun2 ranked #10 by sd,
sd=2.343, mean= 30.9
ENSG00000176946.11 THAP4
log2(RPKM)
6.0
5.6
5.2
4.8
5.2
5.6
6.0
ENSG00000114125.13 RNF7
log2(RPKM)
z−score of IF value
−2
−1
0
1
2

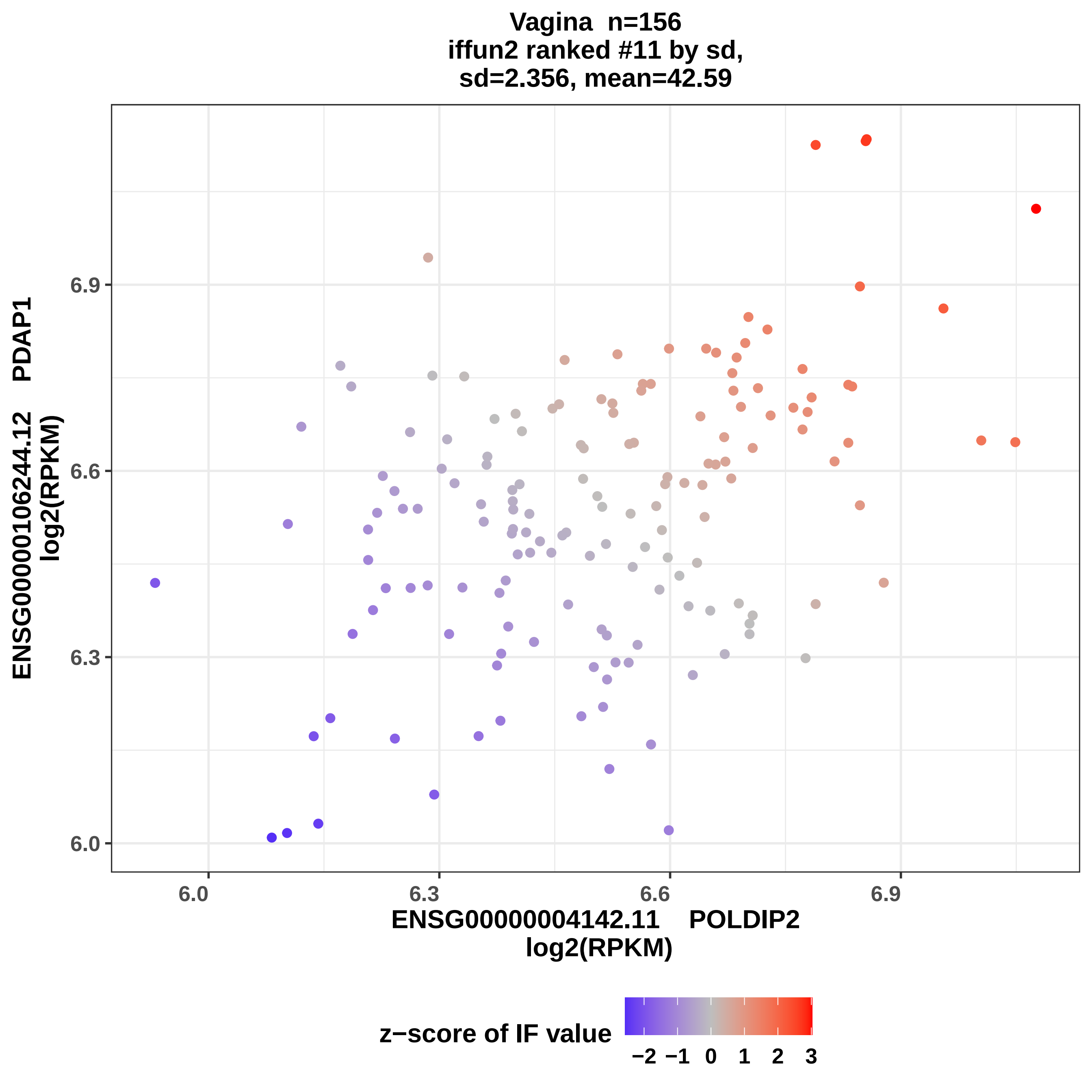

Vagina n=156
iffun2 ranked #11 by sd,
sd=2.356, mean=42.59
ENSG00000106244.12 PDAP1
log2(RPKM)
6.0
6.3
6.6
6.9
ENSG00000004142.11 POLDIP2
log2(RPKM)
z−score of IF value
−2 −1 0 1 2 3

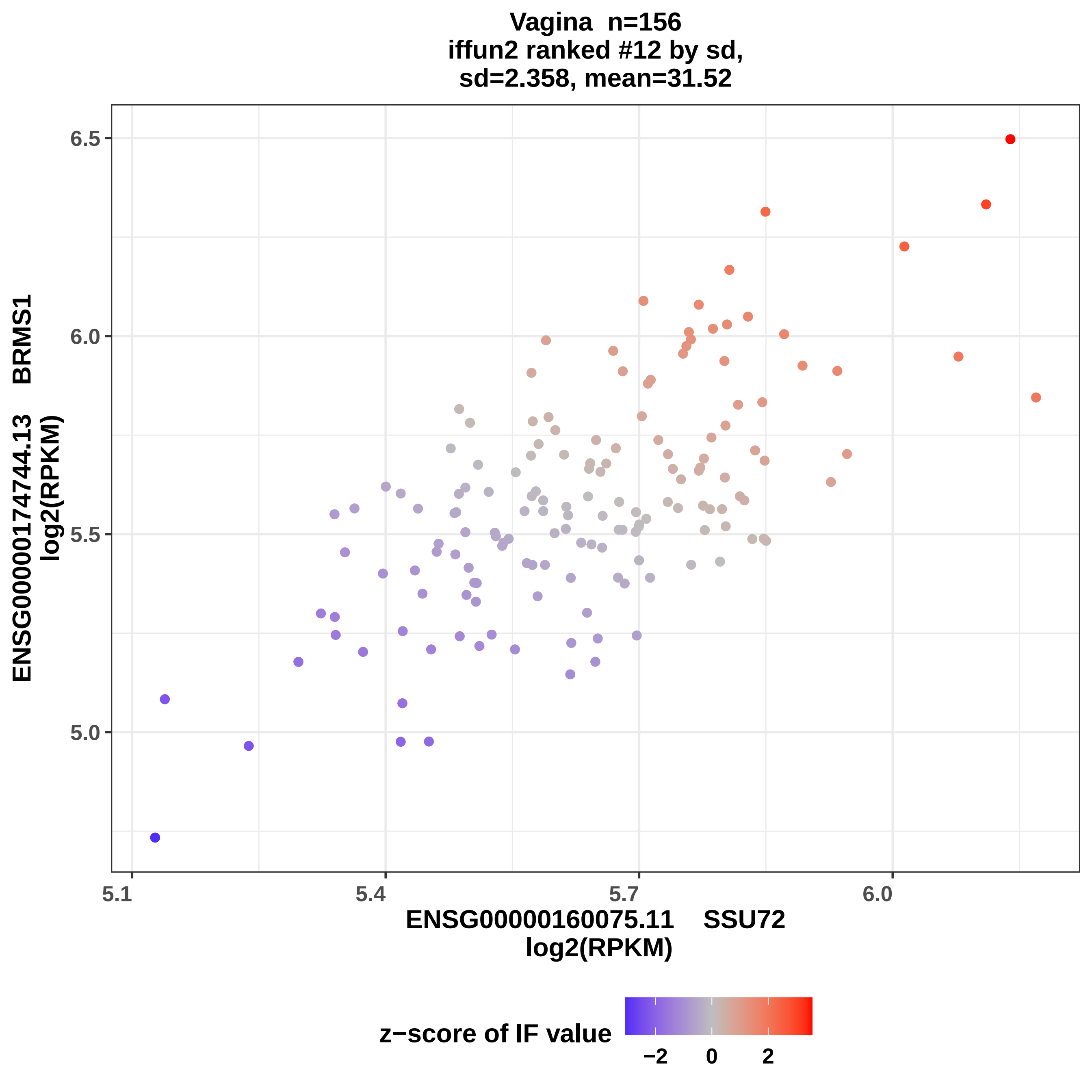

Vagina n=156
iffun2 ranked #12 by sd,
sd=2.358, mean=31.52
ENSG00000174744.13 BRMS1
log2(RPKM)
6.5
6.0
5.5
5.0
5.1
5.4
5.7
6.0
ENSG00000160075.11 SSU72
log2(RPKM)
z−score of IF value
−2
0
2

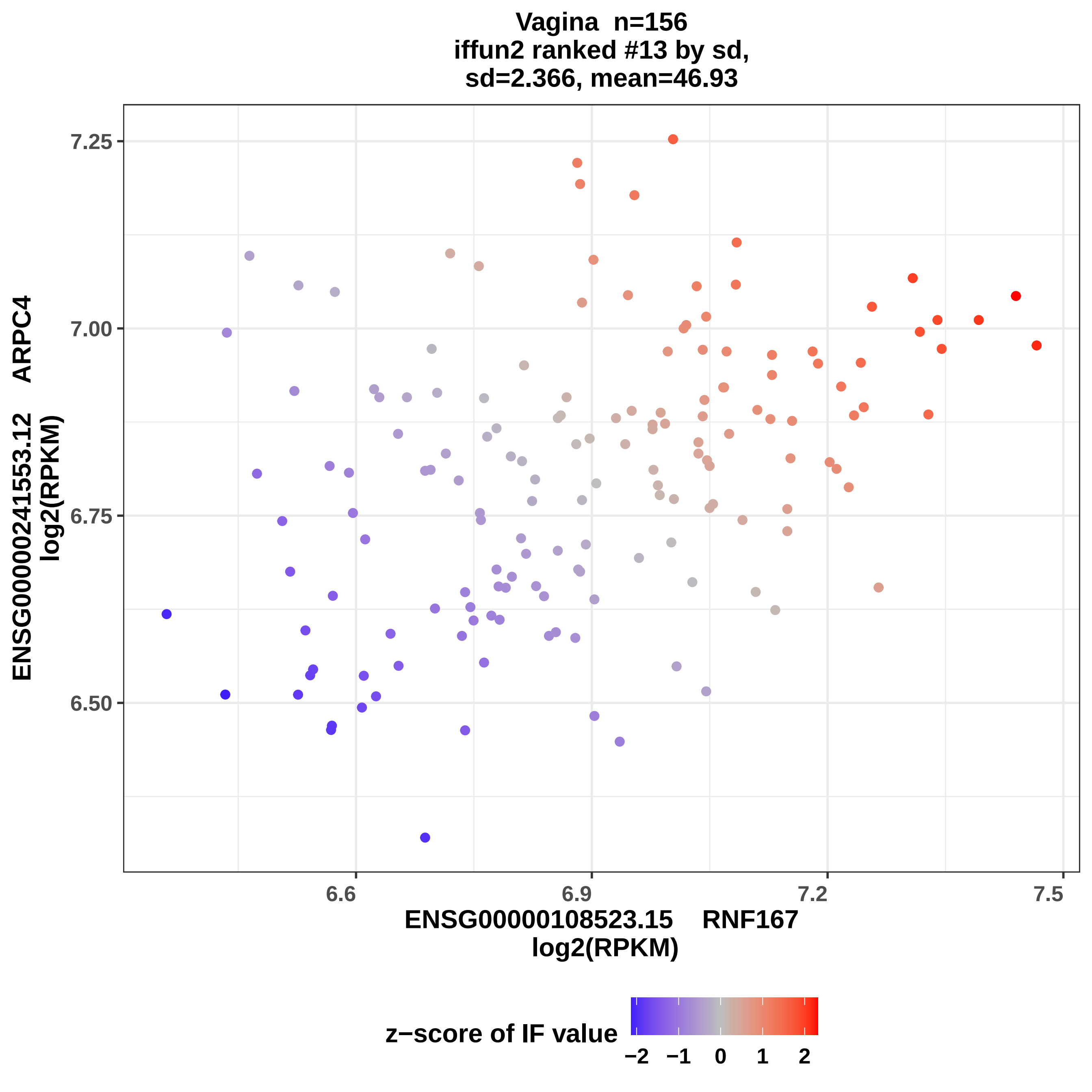

Vagina n=156
iffun2 ranked #13 by sd,
sd=2.366, mean=46.93
ENSG00000241553.12 ARPC4
log2(RPKM)
7.25
7.00
6.75
6.50
6.6
6.9
7.2
7.5
ENSG00000108523.15 RNF167
log2(RPKM)
z−score of IF value
−2
−1
0
1
2

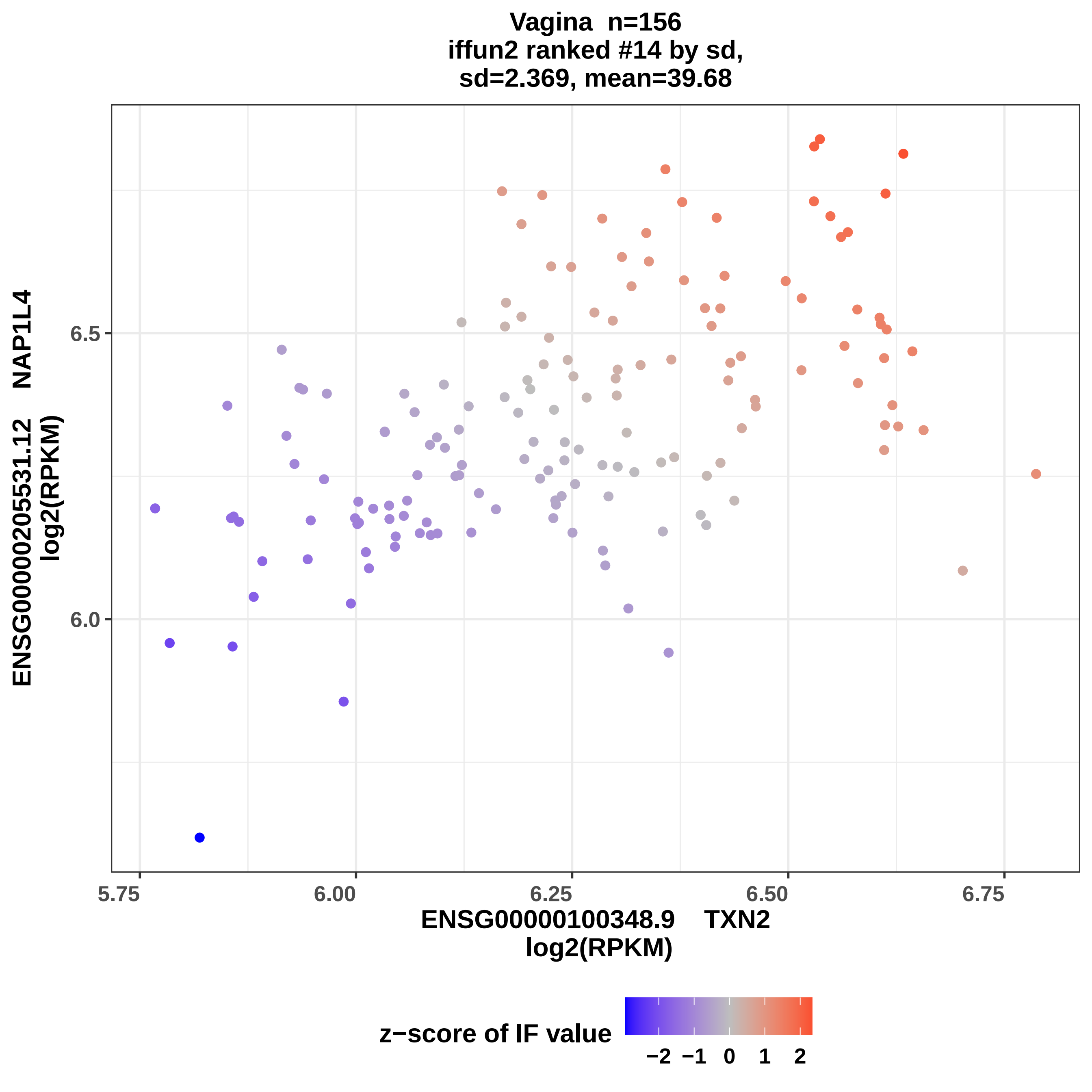

Vagina n=156
iffun2 ranked #14 by sd,
sd=2.369, mean=39.68
ENSG00000205531.12 NAP1L4
log2(RPKM)
6.5
6.0
5.75
6.00
6.25
6.50
6.75
ENSG00000100348.9 TXN2
log2(RPKM)
z−score of IF value
−2
−1
0
1
2

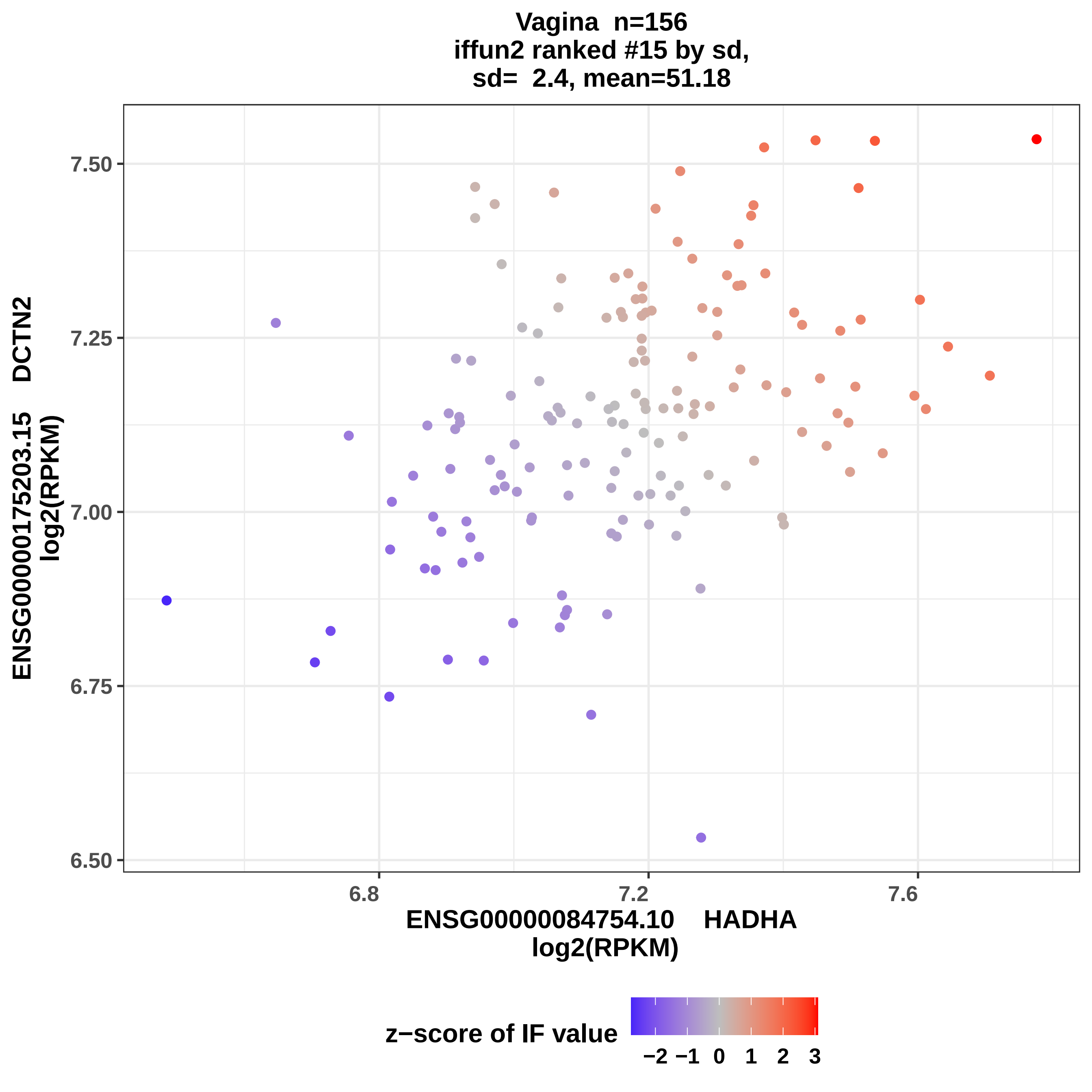

Vagina n=156
iffun2 ranked #15 by sd,
sd= 2.4, mean=51.18
ENSG00000175203.15 DCTN2
log2(RPKM)
7.50
7.25
7.00
6.75
6.50
6.8
7.2
7.6
ENSG00000084754.10 HADHA
log2(RPKM)
z−score of IF value
−2
−1
0
1
2
3

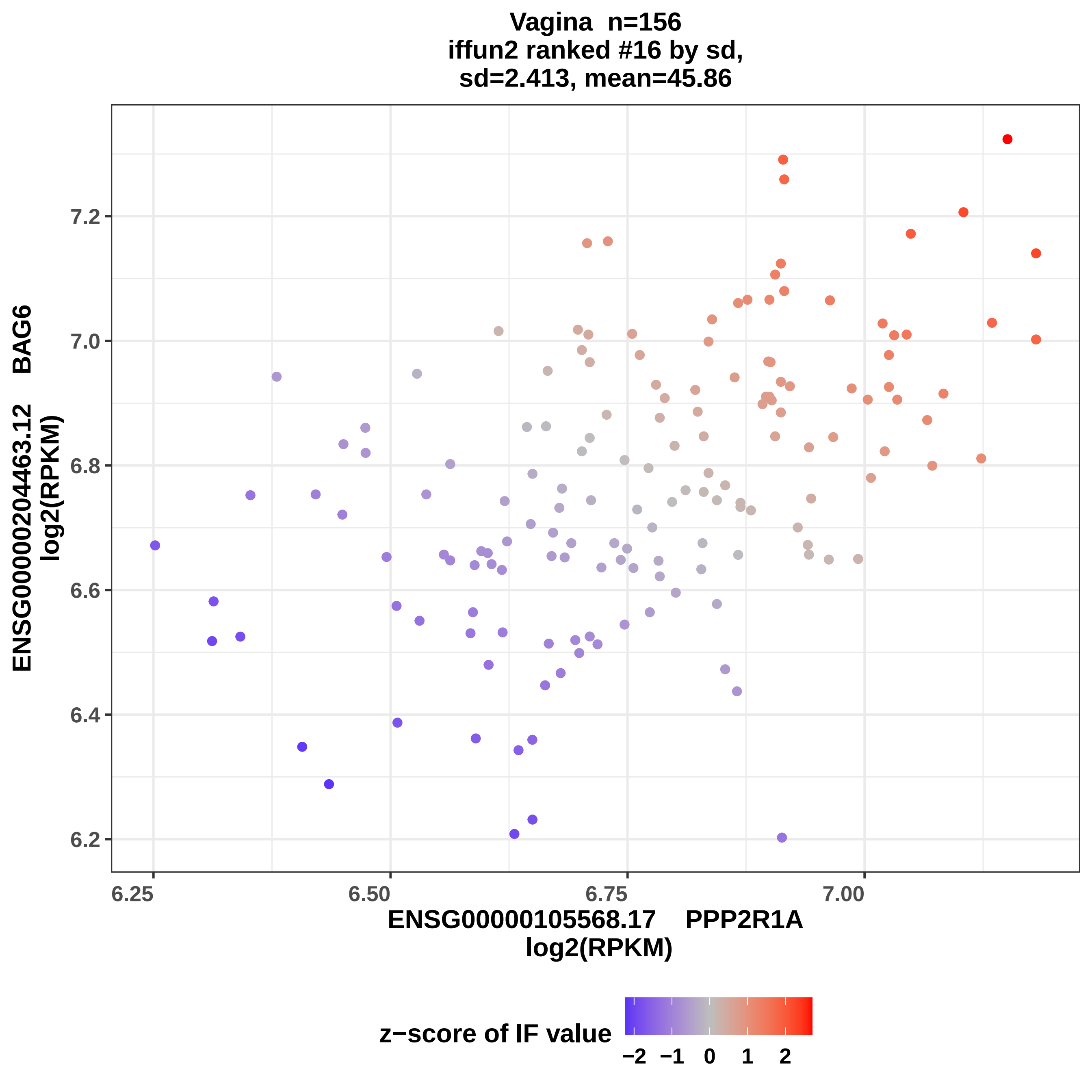

Vagina n=156
iffun2 ranked #16 by sd,
sd=2.413, mean=45.86
ENSG00000204463.12 BAG6
log2(RPKM)
7.2
7.0
6.8
6.6
6.4
6.2
6.25
6.50
6.75
7.00
ENSG00000105568.17 PPP2R1A
log2(RPKM)
z−score of IF value
−2 −1 0 1 2

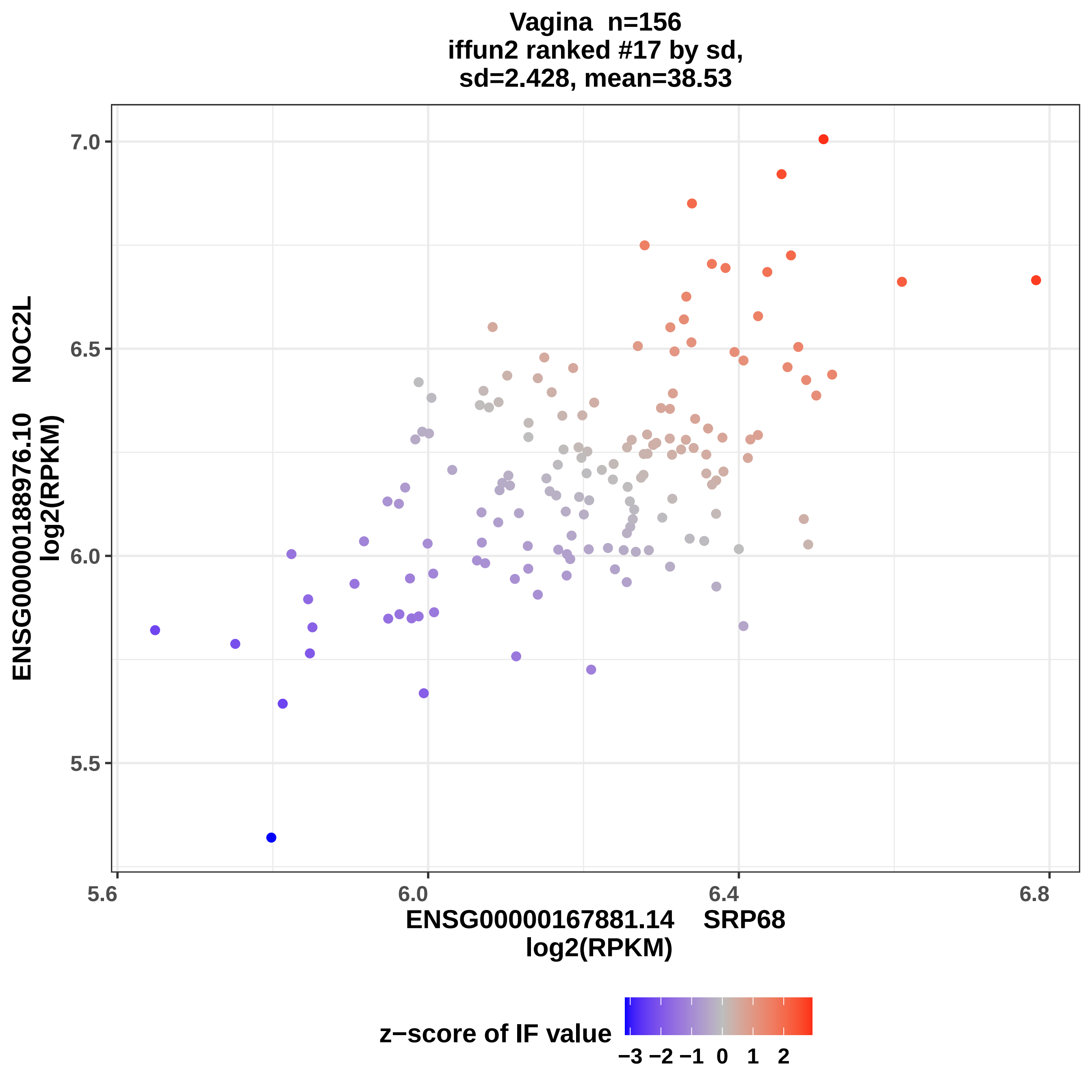

Vagina n=156
iffun2 ranked #17 by sd,
sd=2.428, mean=38.53
ENSG00000188976.10 NOC2L
log2(RPKM)
7.0
6.5
6.0
5.5
5.6
6.0
6.4
6.8
ENSG00000167881.14 SRP68
log2(RPKM)
z−score of IF value
−3 −2 −1 0 1 2

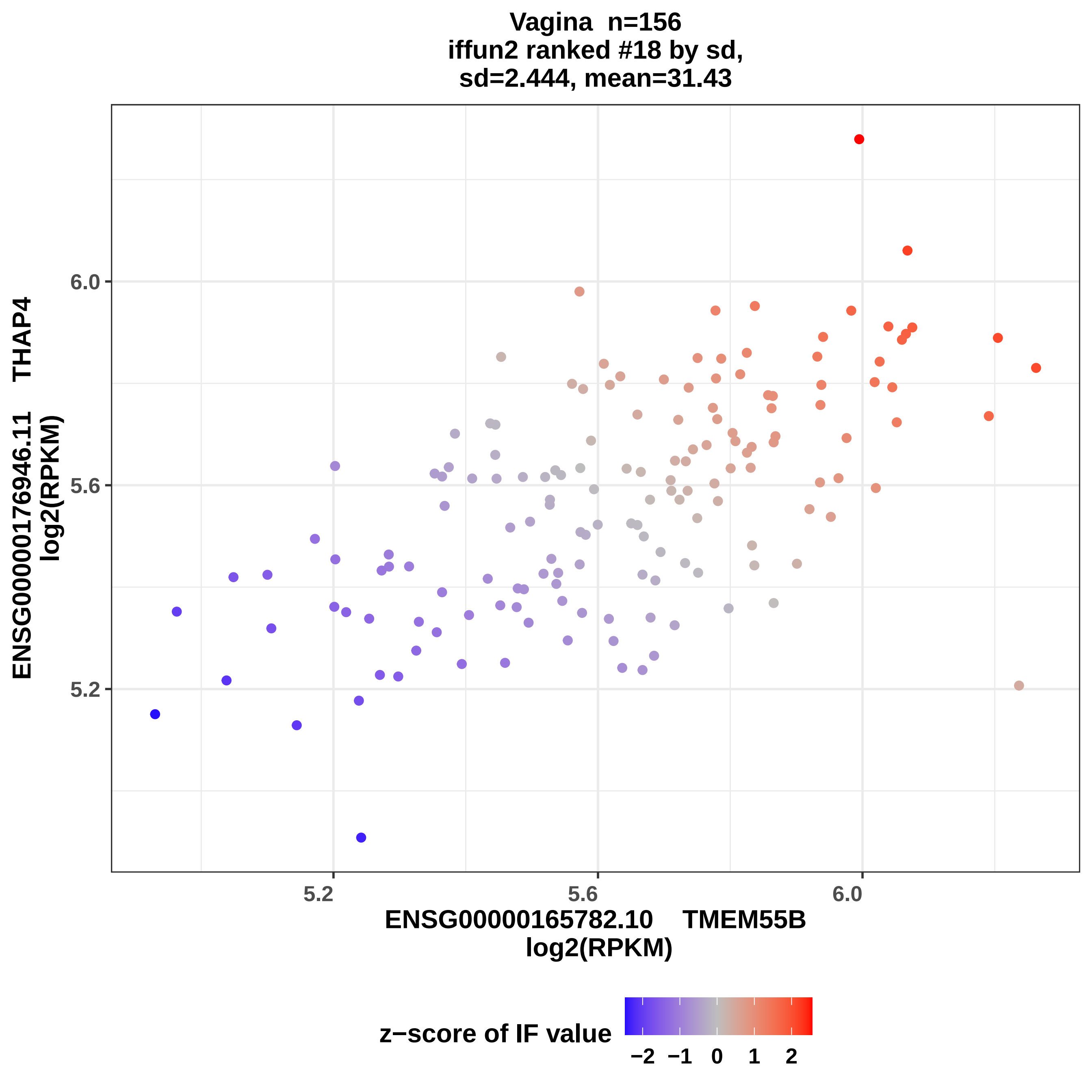

Vagina n=156
iffun2 ranked #18 by sd,
sd=2.444, mean=31.43
ENSG00000176946.11 THAP4
log2(RPKM)
6.0
5.6
5.2
5.2
5.6
6.0
ENSG00000165782.10 TMEM55B
log2(RPKM)
z−score of IF value
−2 −1 0 1 2

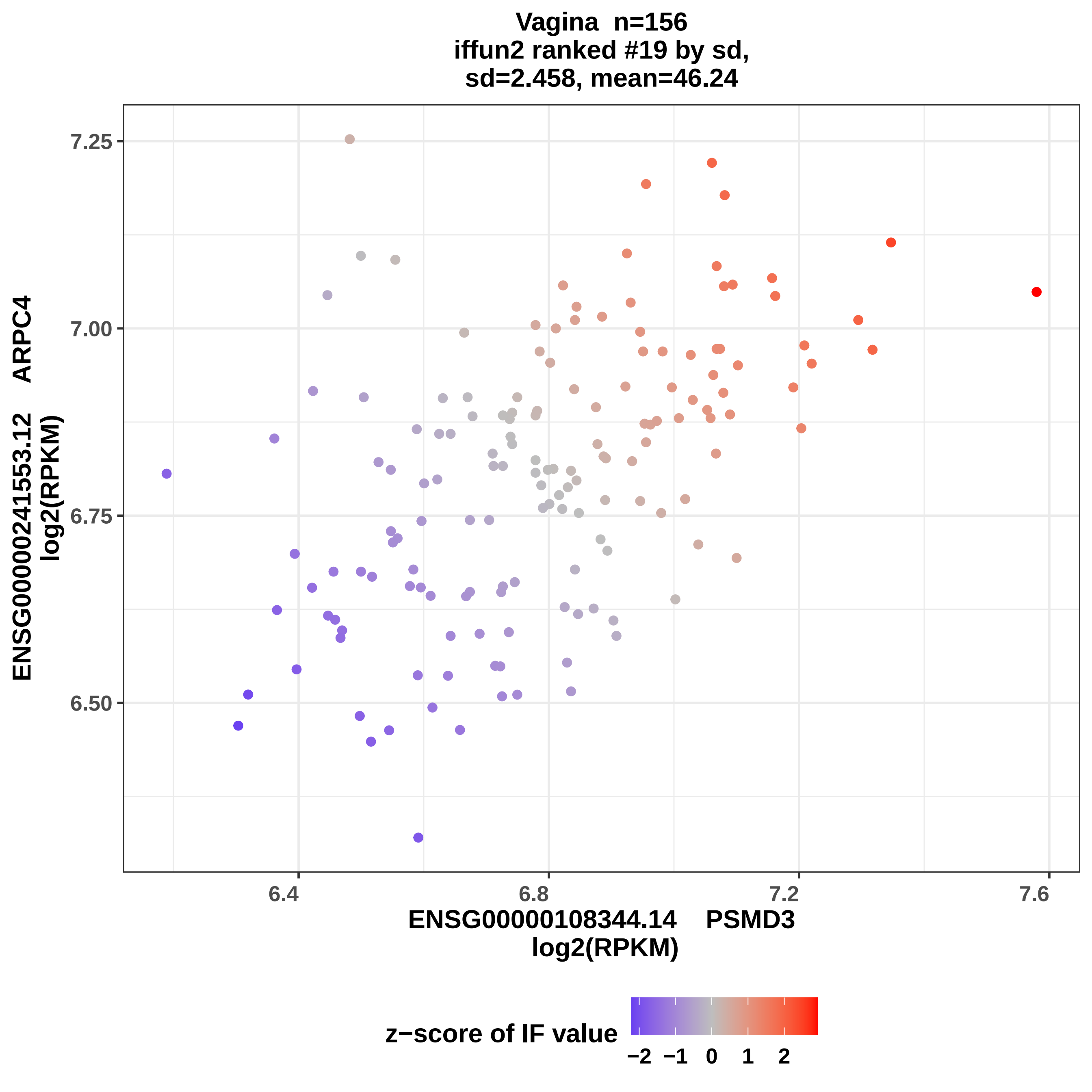

Vagina n=156
iffun2 ranked #19 by sd,
sd=2.458, mean=46.24
7.25
7.00
6.75
6.50
ENSG00000241553.12 ARPC4
log2(RPKM)
6.4
6.8
7.2
7.6
ENSG00000108344.14 PSMD3
log2(RPKM)
z−score of IF value
−2 −1 0 1 2

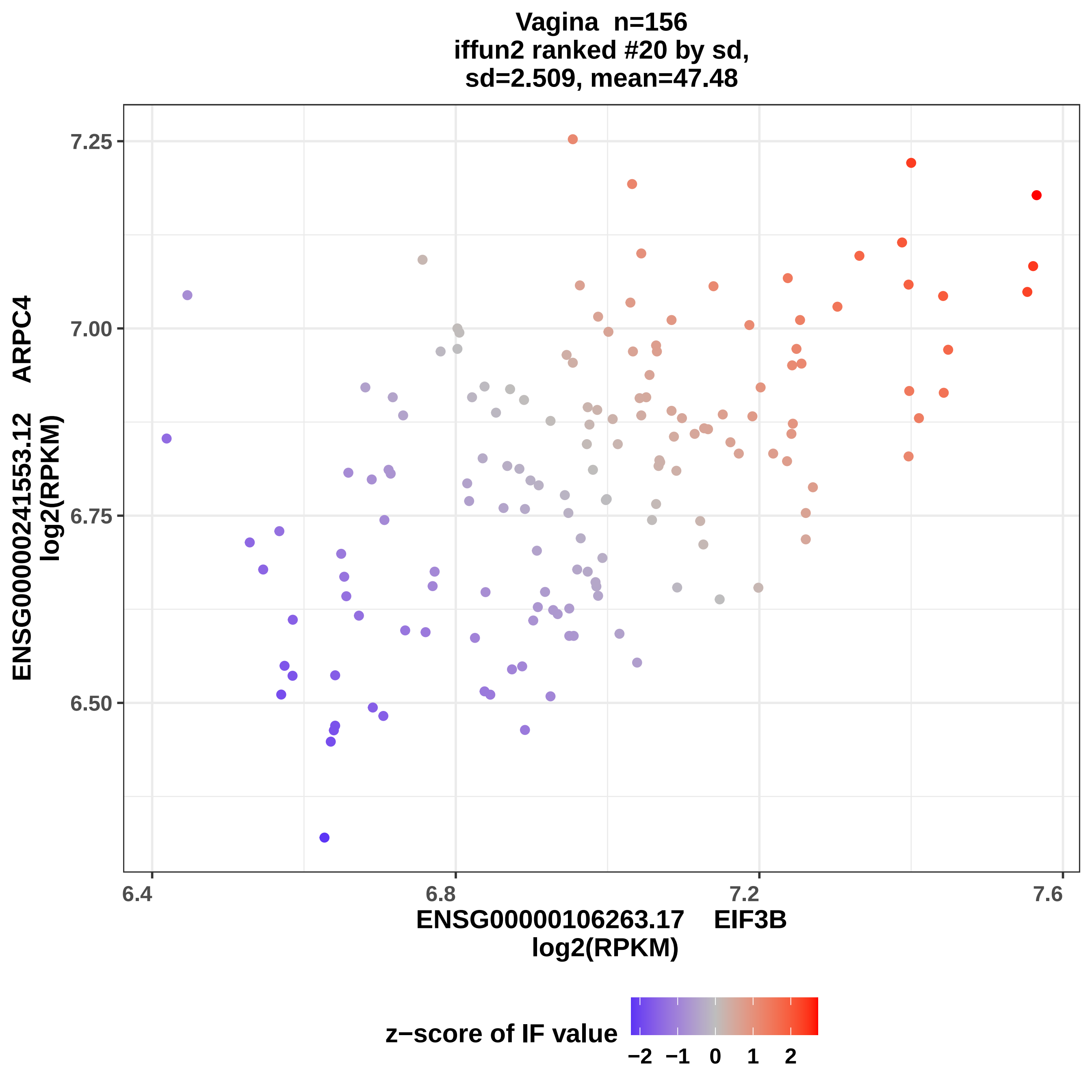

Vagina n=156
iffun2 ranked #20 by sd,
sd=2.509, mean=47.48
ENSG00000241553.12 ARPC4
log2(RPKM)
7.25
7.00
6.75
6.50
6.4
6.8
7.2
7.6
ENSG00000106263.17 EIF3B
log2(RPKM)
z−score of IF value
−2 −1 0 1 2
