## Supplemental Data 4 for "Symmetry as a Fundamental Principle in Defining Gene Expression and Phenotypic Traits"

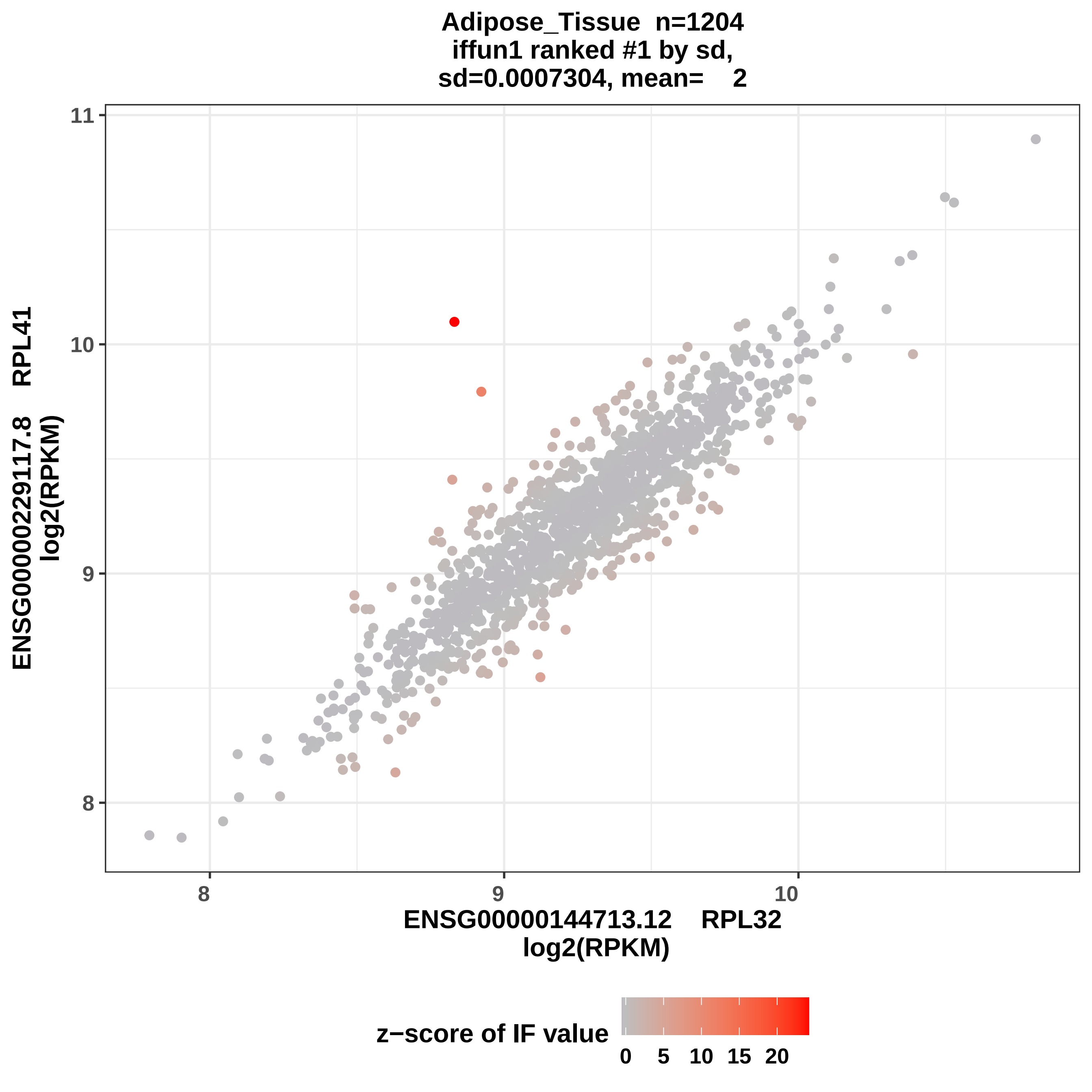

Adipose_Tissue n=1204
iffun1 ranked #1 by sd,
sd=0.0007304, mean= 2
11
10
9
8
ENSG00000229117.8 RPL41
log2(RPKM)
8
9
10
ENSG00000144713.12 RPL32
log2(RPKM)
z−score of IF value
0
5
10
15
20

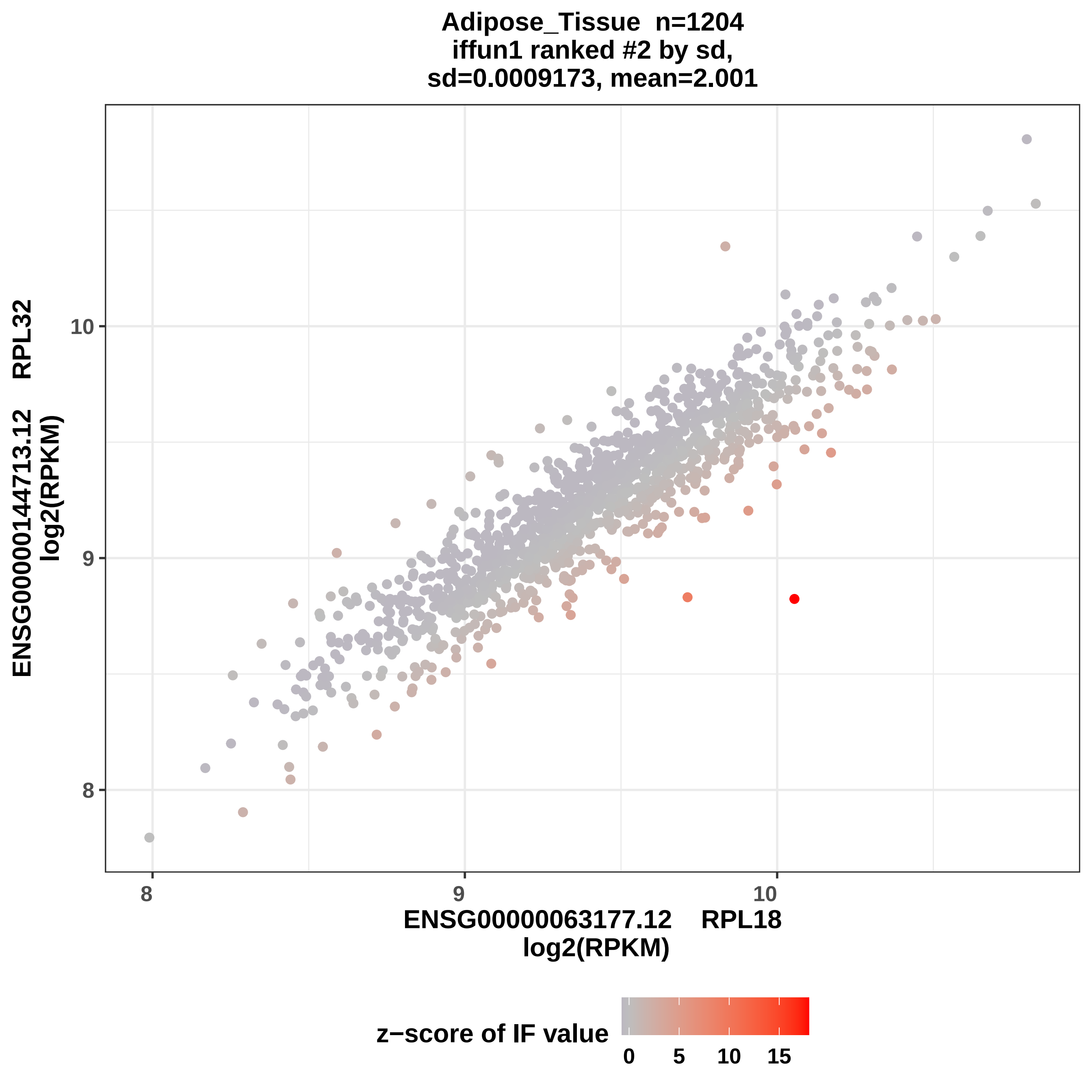

Adipose_Tissue n=1204
iffun1 ranked #2 by sd,
sd=0.0009173, mean=2.001
ENSG00000144713.12 RPL32
log2(RPKM)
10
9
8
8
9
10
ENSG00000063177.12 RPL18
log2(RPKM)
z−score of IF value
0
5
10
15

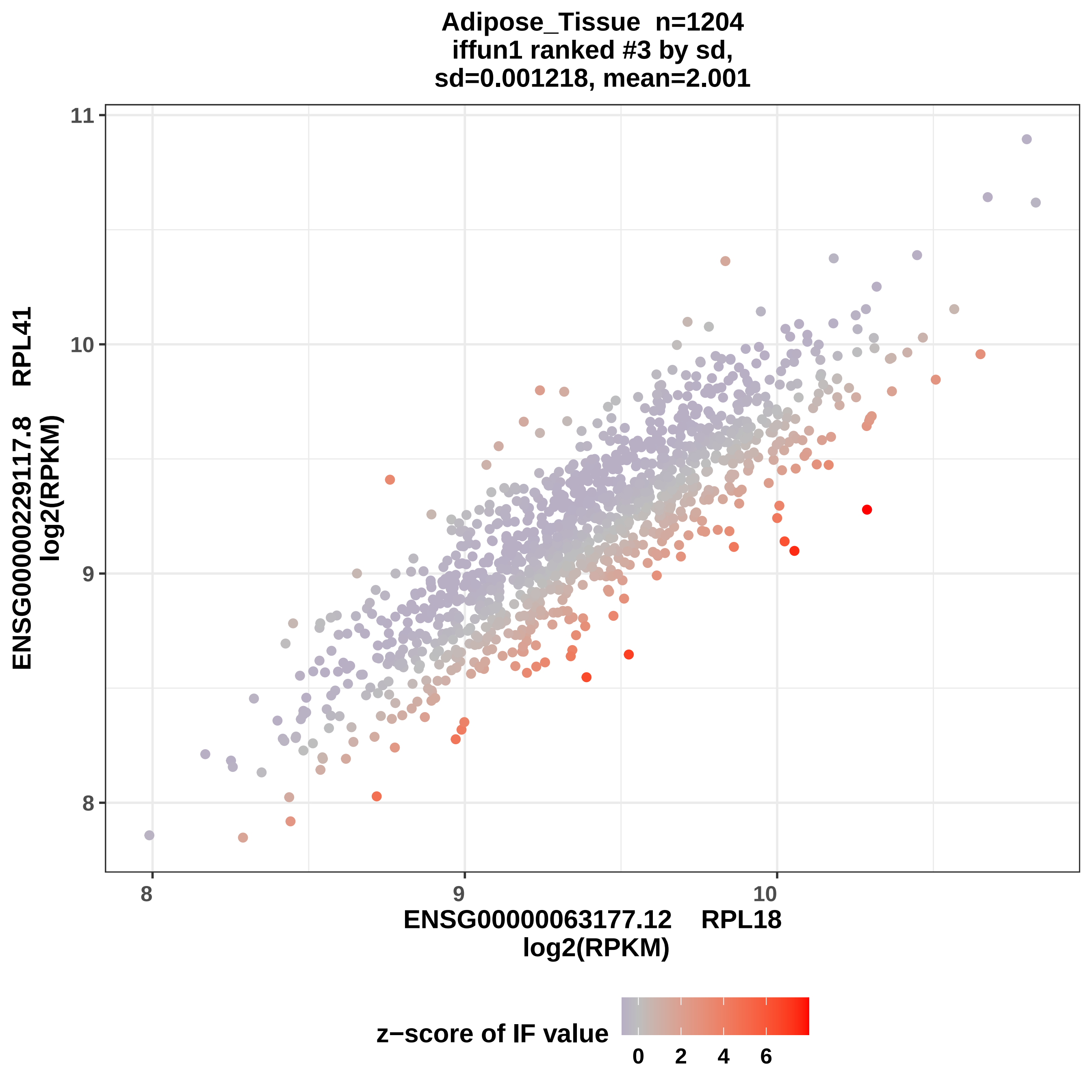
Adipose_Tissue n=1204
iffun1 ranked #3 by sd,
sd=0.001218, mean=2.001
ENSG00000229117.8 RPL41
log2(RPKM)
11
10
9
8
8
9
10
ENSG00000063177.12 RPL18
log2(RPKM)
z−score of IF value
0
2
4
6

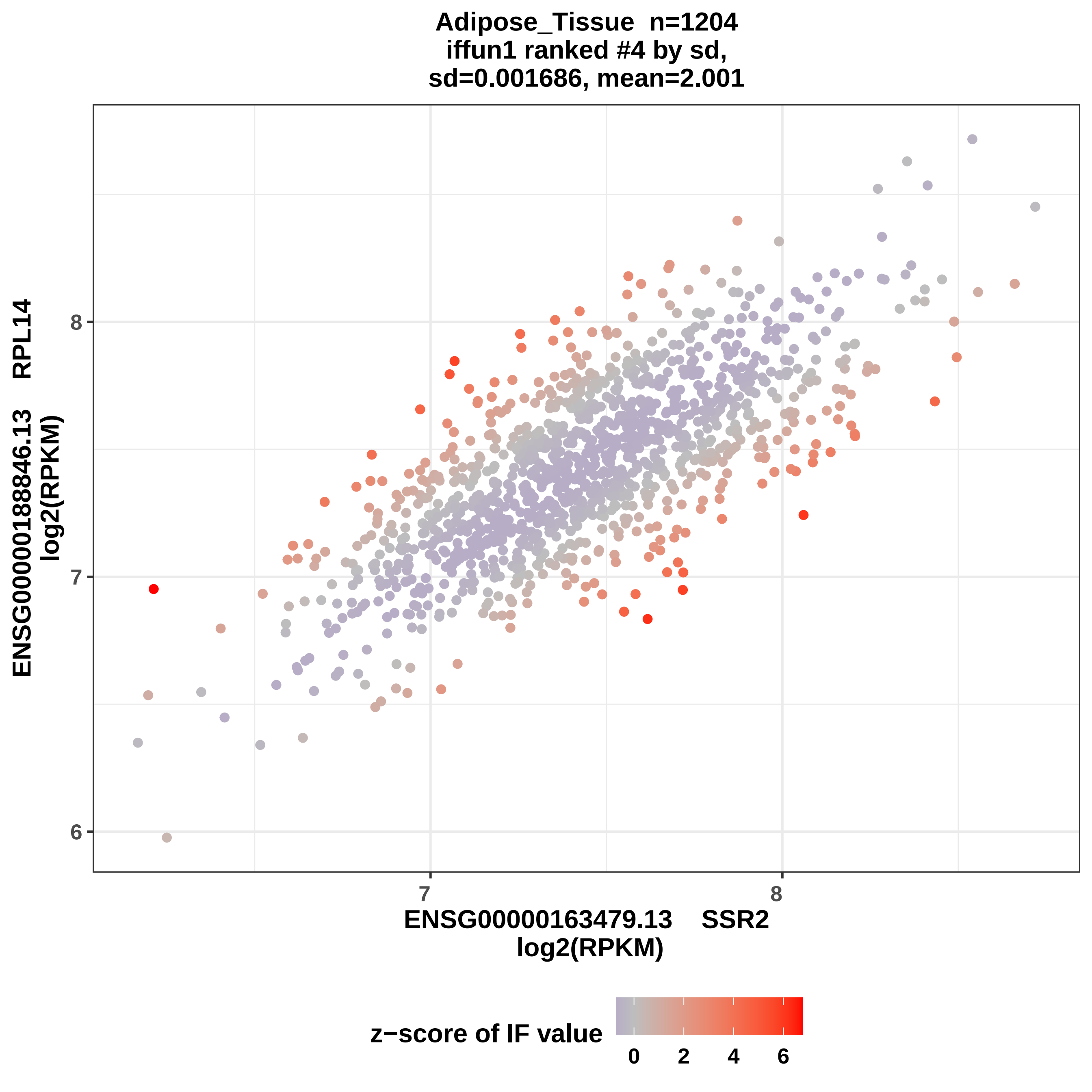

Adipose_Tissue n=1204
iffun1 ranked #4 by sd,
sd=0.001686, mean=2.001
ENSG00000188846.13 RPL14
log2(RPKM)
8
7
6
7
8
ENSG00000163479.13 SSR2
log2(RPKM)
z−score of IF value
0
2
4
6

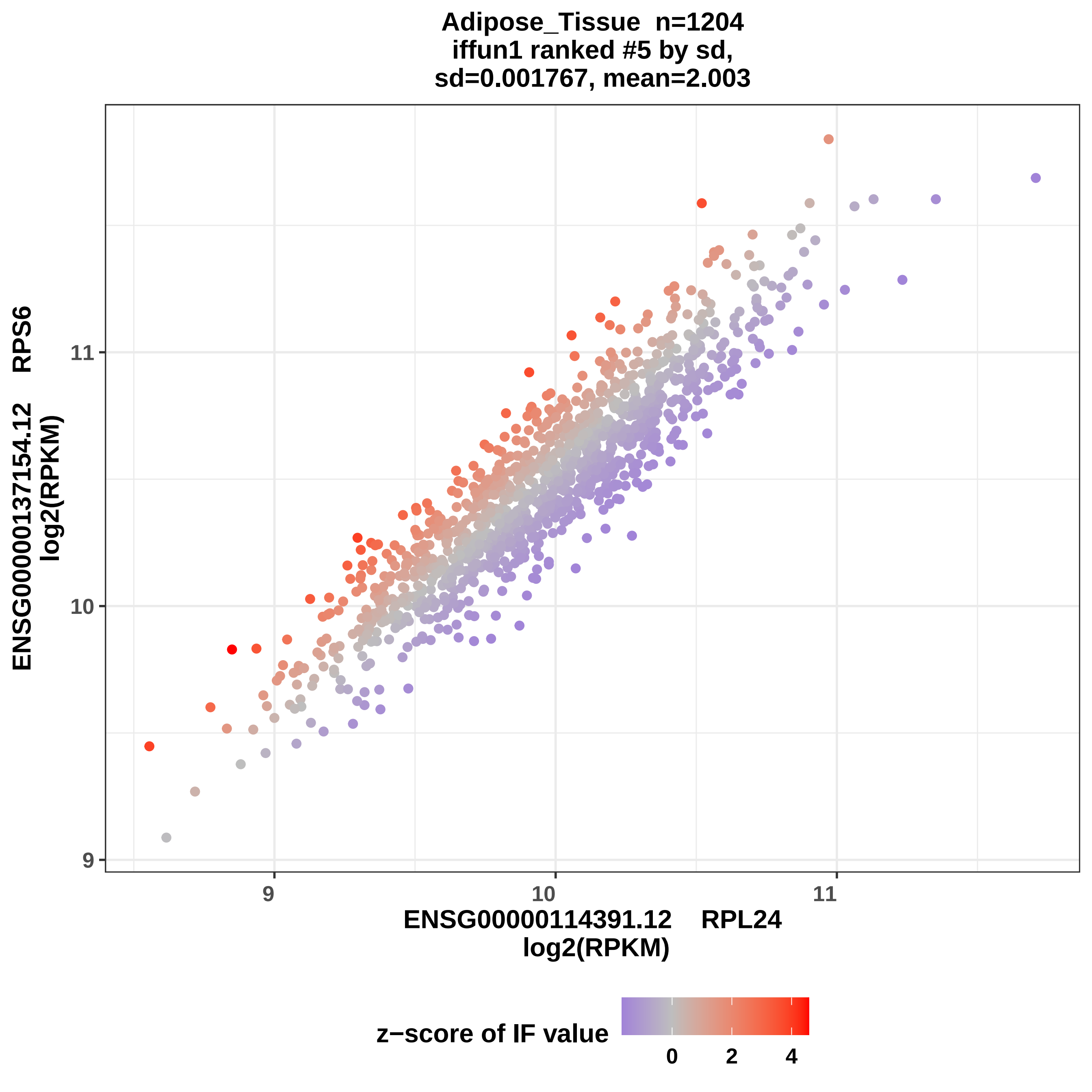
Adipose_Tissue n=1204
iffun1 ranked #5 by sd,
sd=0.001767, mean=2.003
ENSG00000137154.12 RPS6
log2(RPKM)
11
10
9
9
10
11
ENSG00000114391.12 RPL24
log2(RPKM)
z−score of IF value
0
2
4

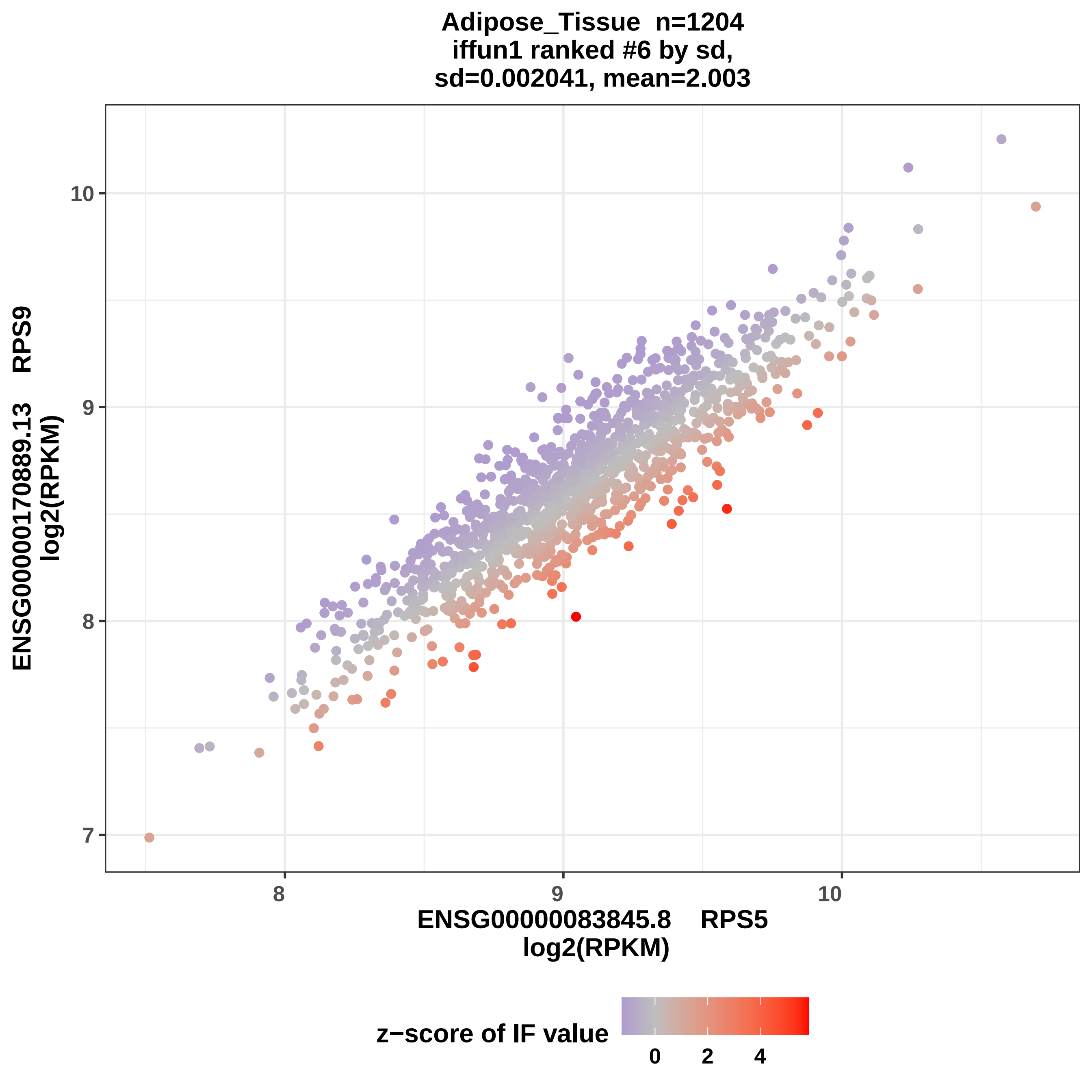

Adipose_Tissue n=1204
iffun1 ranked #6 by sd,
sd=0.002041, mean=2.003
ENSG00000170889.13 RPS9
log2(RPKM)
ENSG00000083845.8 RPS5
log2(RPKM)
7
8
9
10
8
9
10
z−score of IF value
0
2
4

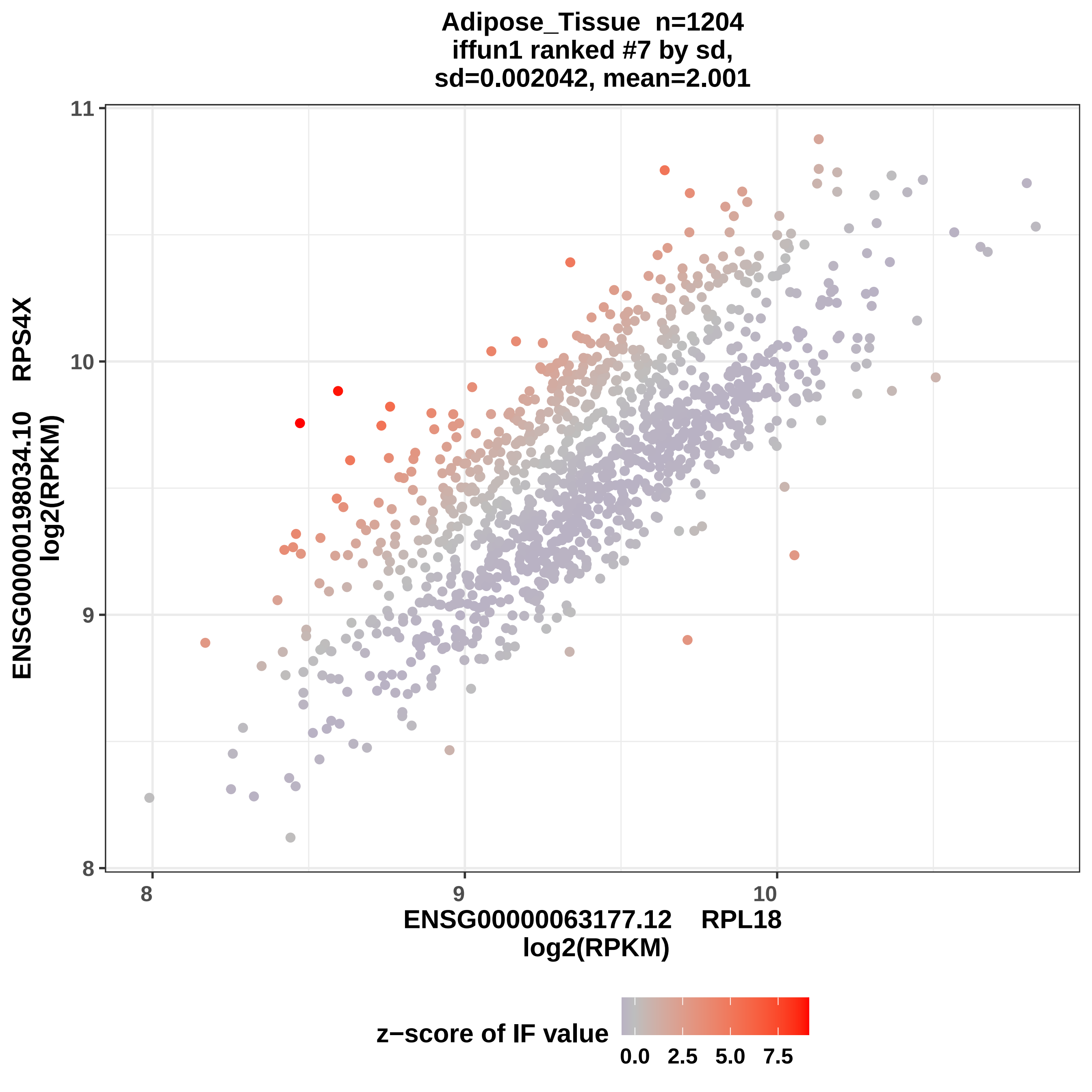
Adipose_Tissue n=1204
iffun1 ranked #7 by sd,
sd=0.002042, mean=2.001
ENSG00000198034.10 RPS4X
log2(RPKM)
ENSG00000063177.12 RPL18
log2(RPKM)
8
9
10
11
z−score of IF value
0.0
2.5
5.0
7.5

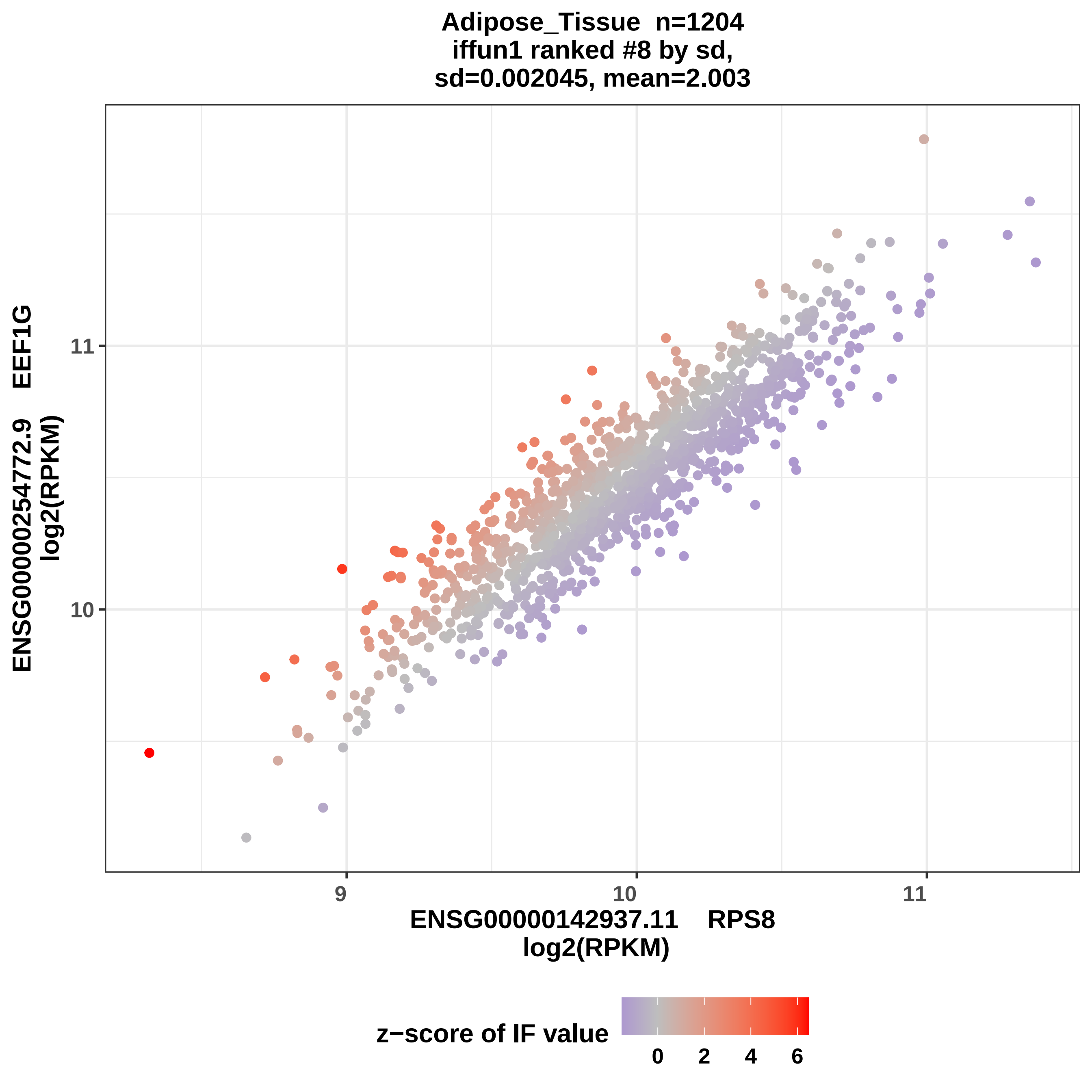
Adipose_Tissue n=1204
iffun1 ranked #8 by sd,
sd=0.002045, mean=2.003
ENSG00000254772.9 EEF1G
log2(RPKM)
11
10
9
10
11
ENSG00000142937.11 RPS8
log2(RPKM)
z−score of IF value
0
2
4
6

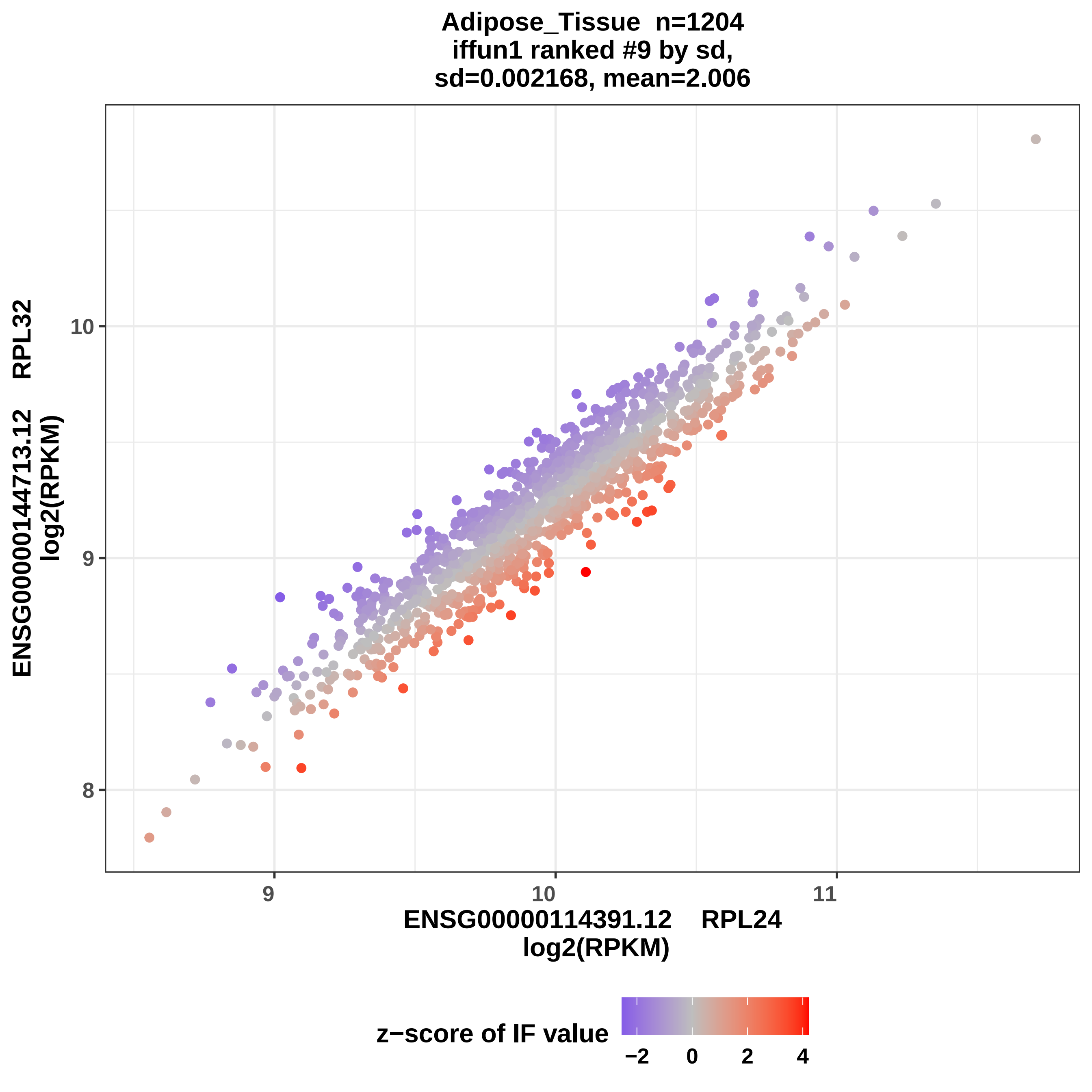
Adipose_Tissue n=1204
iffun1 ranked #9 by sd,
sd=0.002168, mean=2.006
ENSG00000144713.12 RPL32
log2(RPKM)
10
9
8
9
10
11
ENSG00000114391.12 RPL24
log2(RPKM)
z−score of IF value
−2
0
2
4

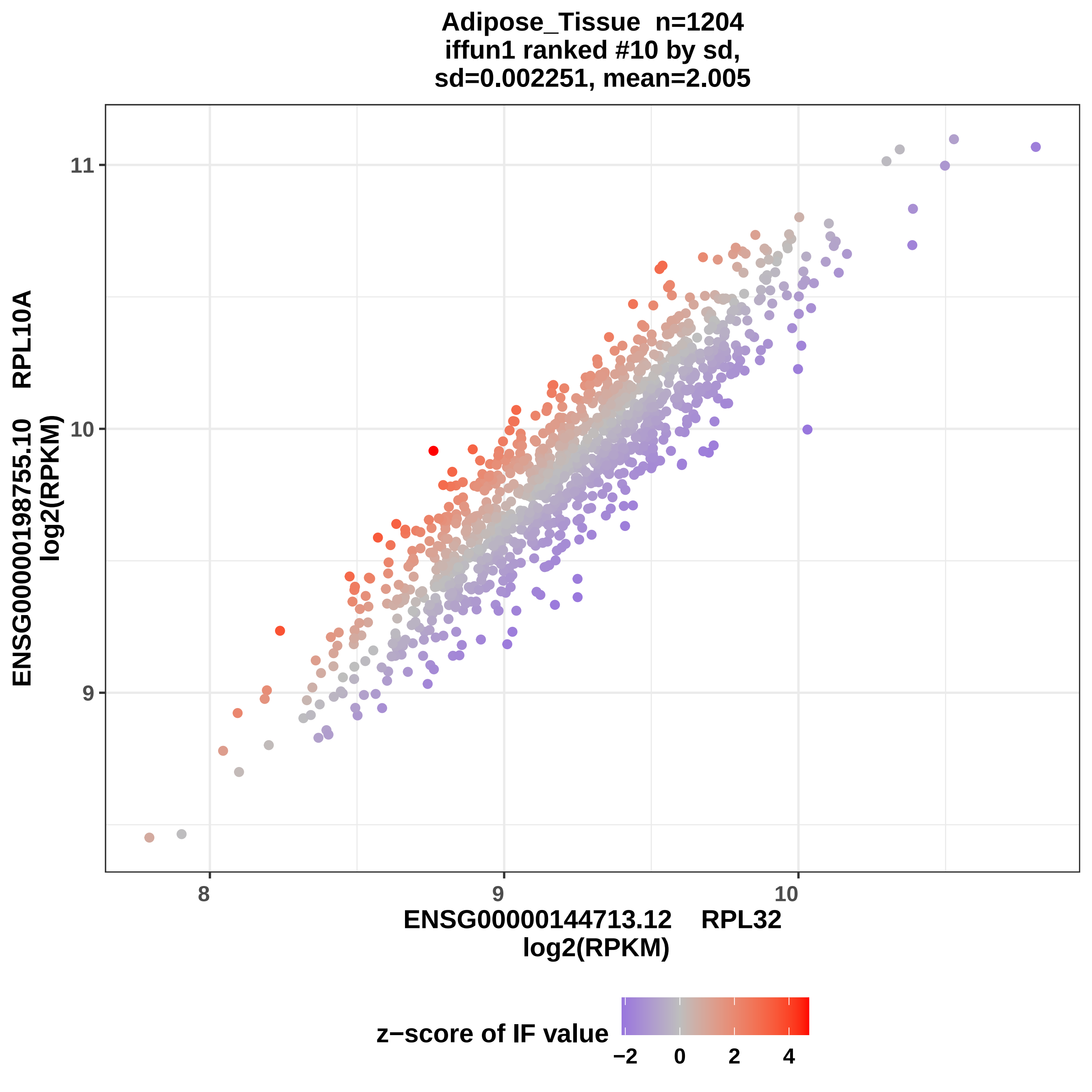
Adipose_Tissue n=1204
iffun1 ranked #10 by sd,
sd=0.002251, mean=2.005
ENSG00000198755.10 RPL10A
log2(RPKM)
11
10
9
8
9
10
ENSG00000144713.12 RPL32
log2(RPKM)
z−score of IF value
−2
0
2
4

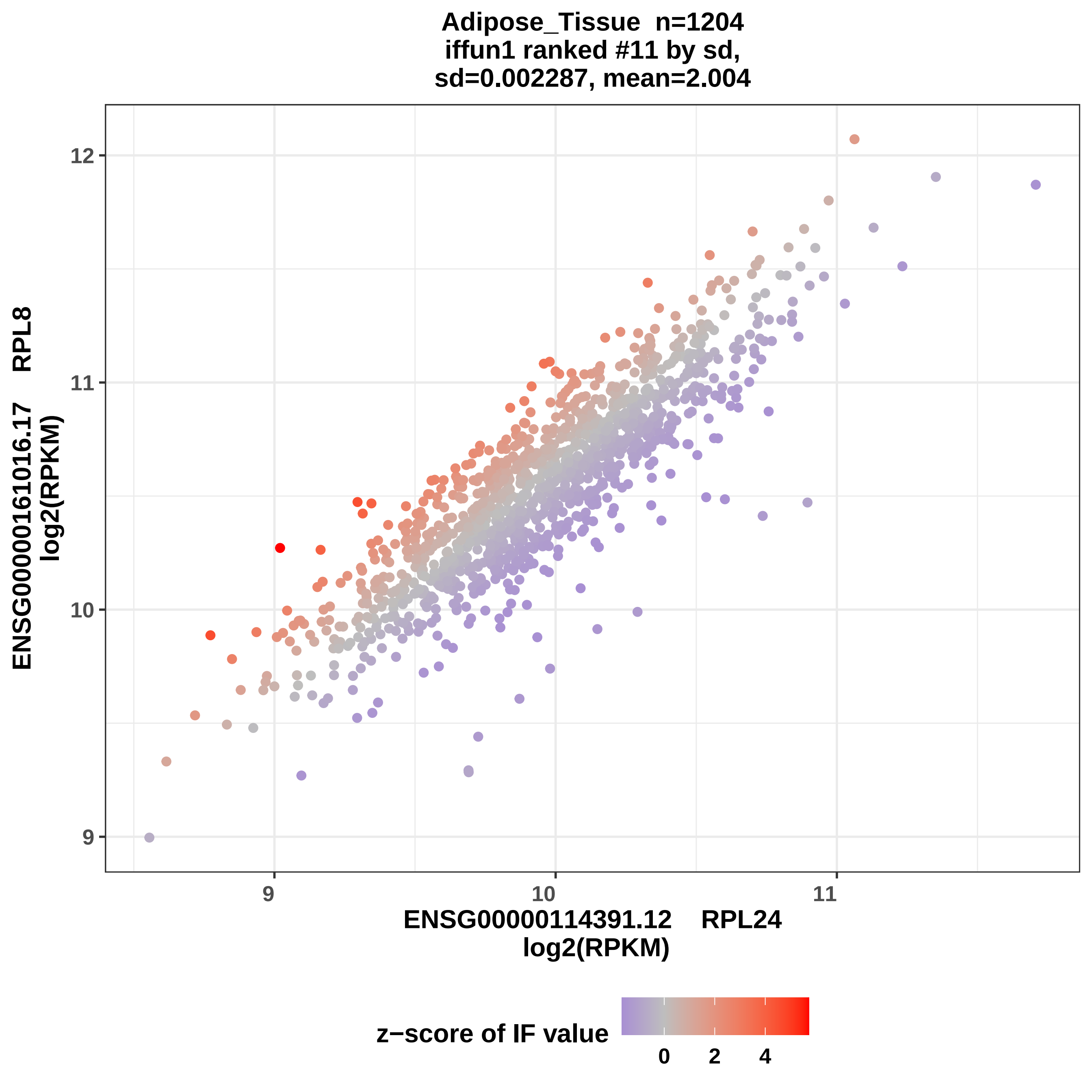
Adipose_Tissue n=1204
iffun1 ranked #11 by sd,
sd=0.002287, mean=2.004
ENSG00000161016.17 RPL8
log2(RPKM)
ENSG00000114391.12 RPL24
log2(RPKM)
12
11
10
9
9
10
11
z−score of IF value
0
2
4

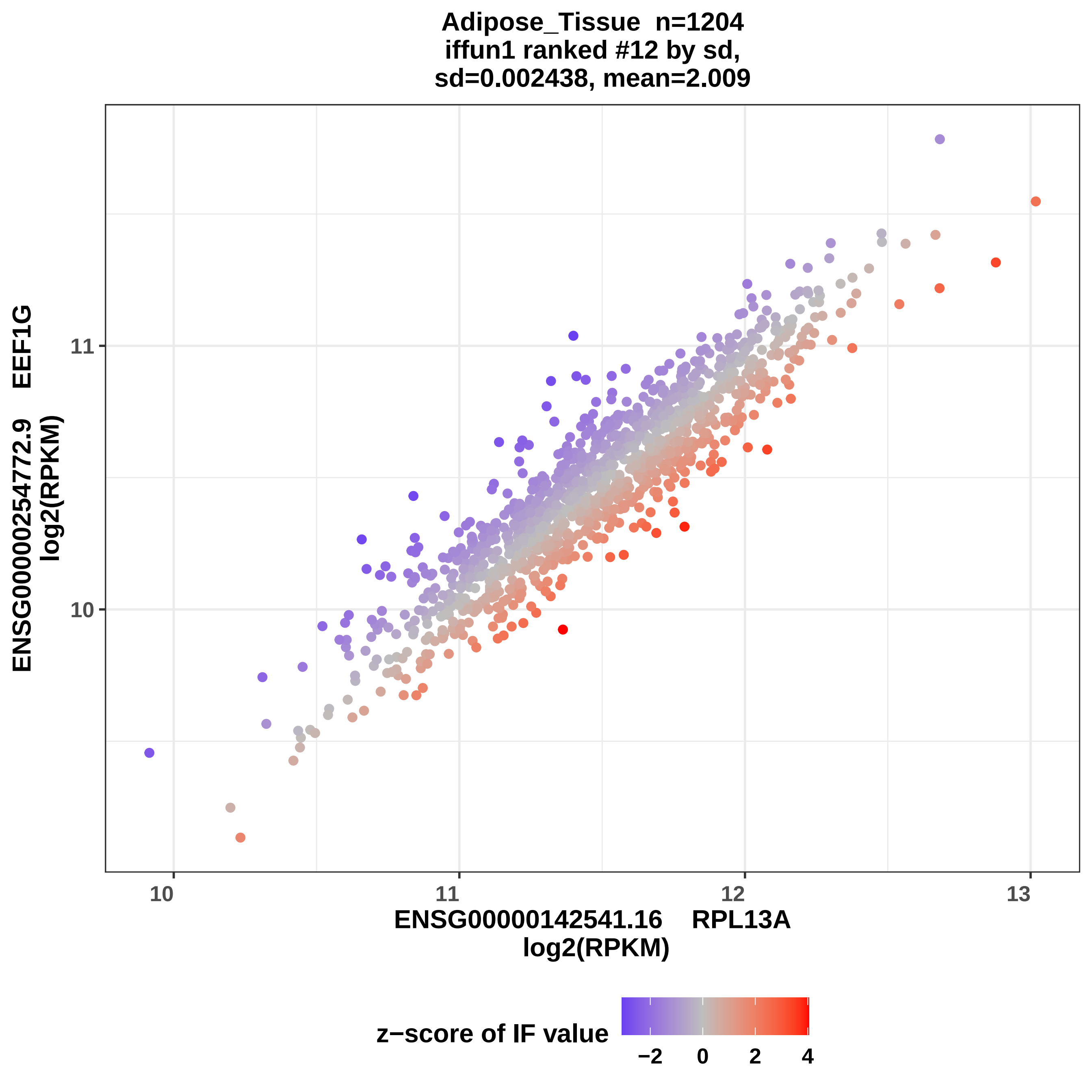

Adipose_Tissue n=1204
iffun1 ranked #12 by sd,
sd=0.002438, mean=2.009
ENSG00000254772.9 EEF1G
log2(RPKM)
11
10
10
11
12
13
ENSG00000142541.16 RPL13A
log2(RPKM)
z−score of IF value
−2
0
2
4

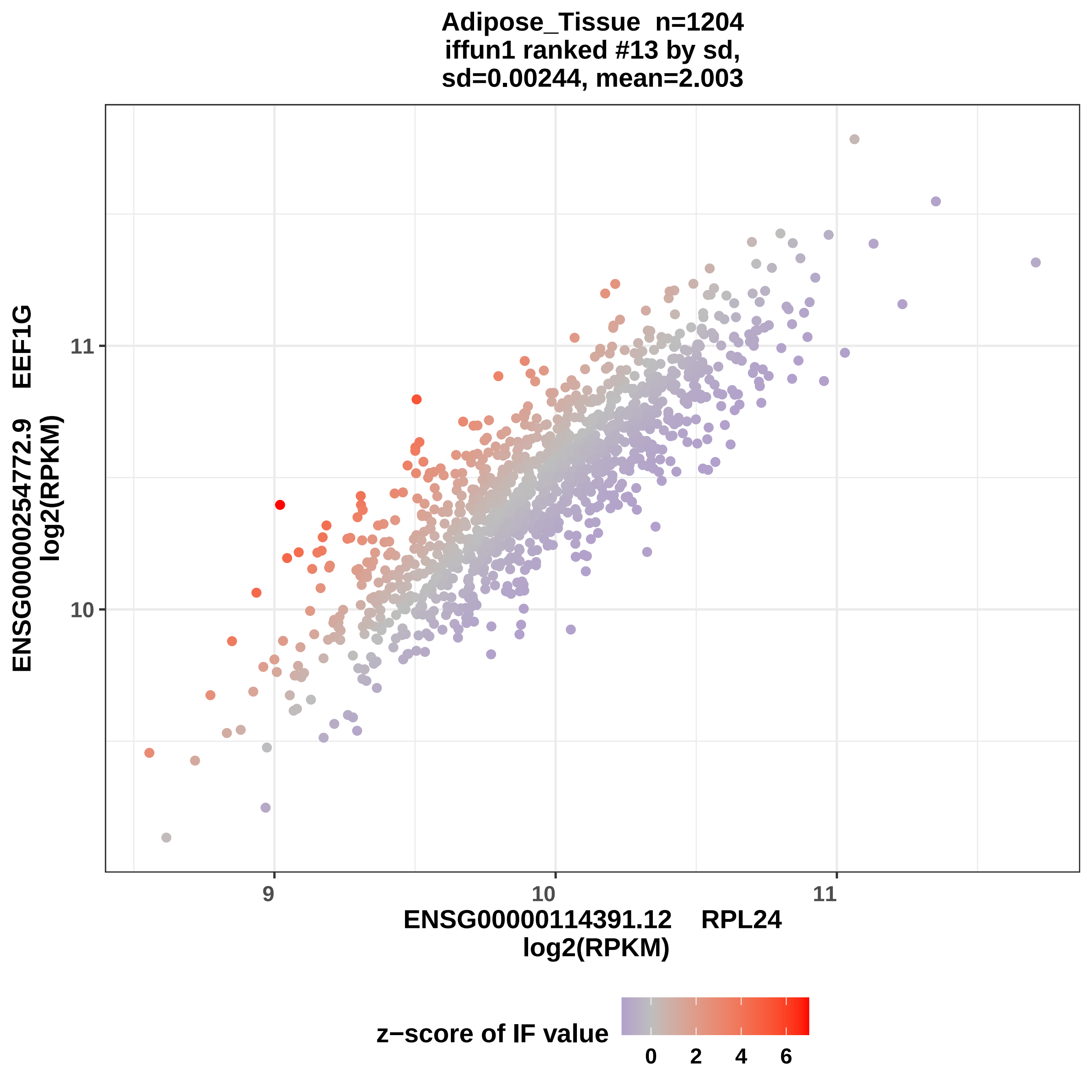

Adipose_Tissue n=1204
iffun1 ranked #13 by sd,
sd=0.00244, mean=2.003
ENSG00000254772.9 EEF1G
log2(RPKM)
11
10
9
10
11
ENSG00000114391.12 RPL24
log2(RPKM)
z−score of IF value
0
2
4
6

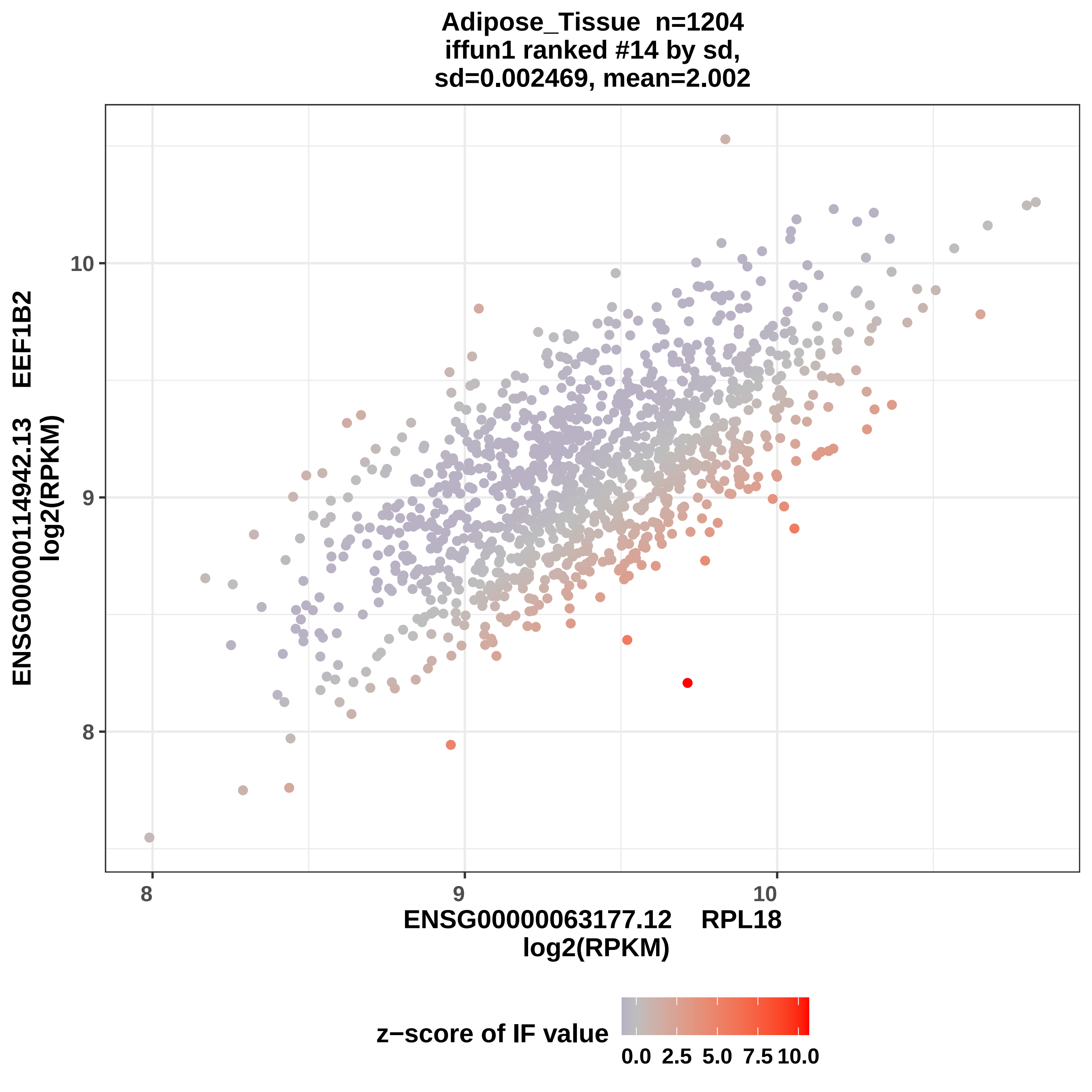

Adipose_Tissue n=1204
iffun1 ranked #14 by sd,
sd=0.002469, mean=2.002
ENSG00000114942.13 EEF1B2
log2(RPKM)
ENSG00000063177.12 RPL18
log2(RPKM)
8
9
10
8
9
10
z−score of IF value
0.0
2.5
5.0
7.5
10.0

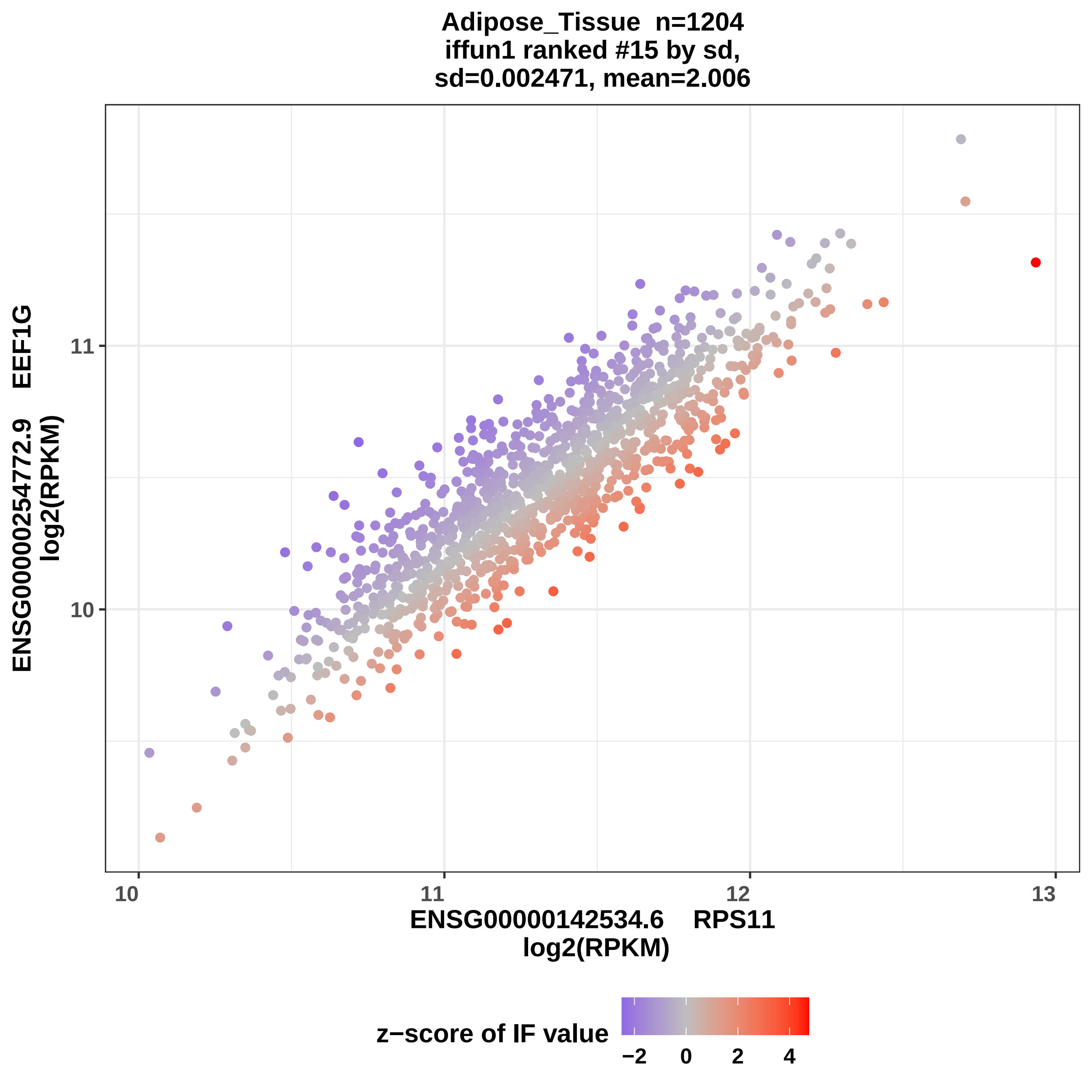

Adipose_Tissue n=1204
iffun1 ranked #15 by sd,
sd=0.002471, mean=2.006
ENSG00000254772.9 EEF1G
log2(RPKM)
11
10
10
11
12
13
ENSG00000142534.6 RPS11
log2(RPKM)
z−score of IF value
−2
0
2
4

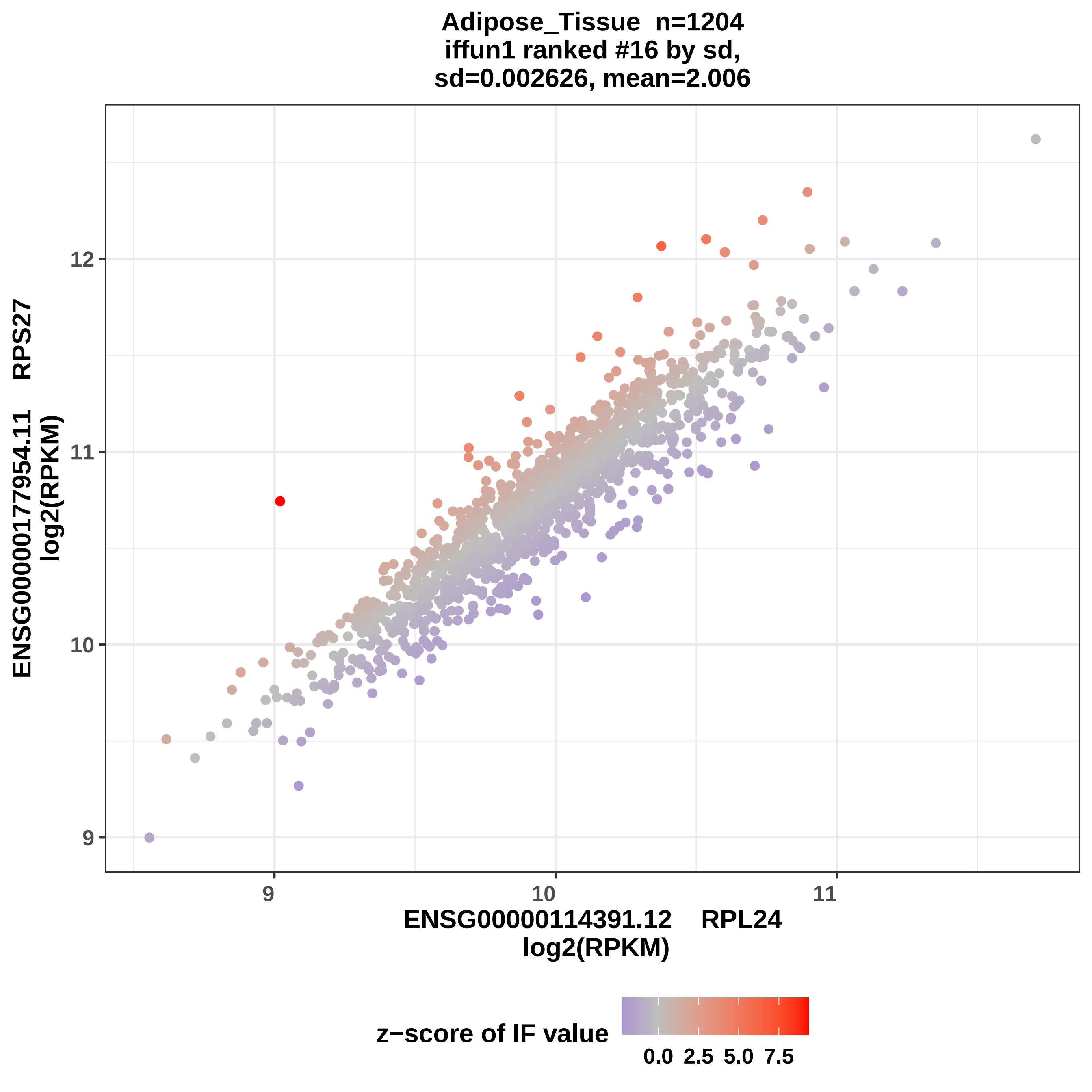
Adipose_Tissue n=1204
iffun1 ranked #16 by sd,
sd=0.002626, mean=2.006
ENSG00000177954.11 RPS27
log2(RPKM)
12
11
10
9
9
10
11
ENSG00000114391.12 RPL24
log2(RPKM)
z−score of IF value
0.0
2.5
5.0
7.5

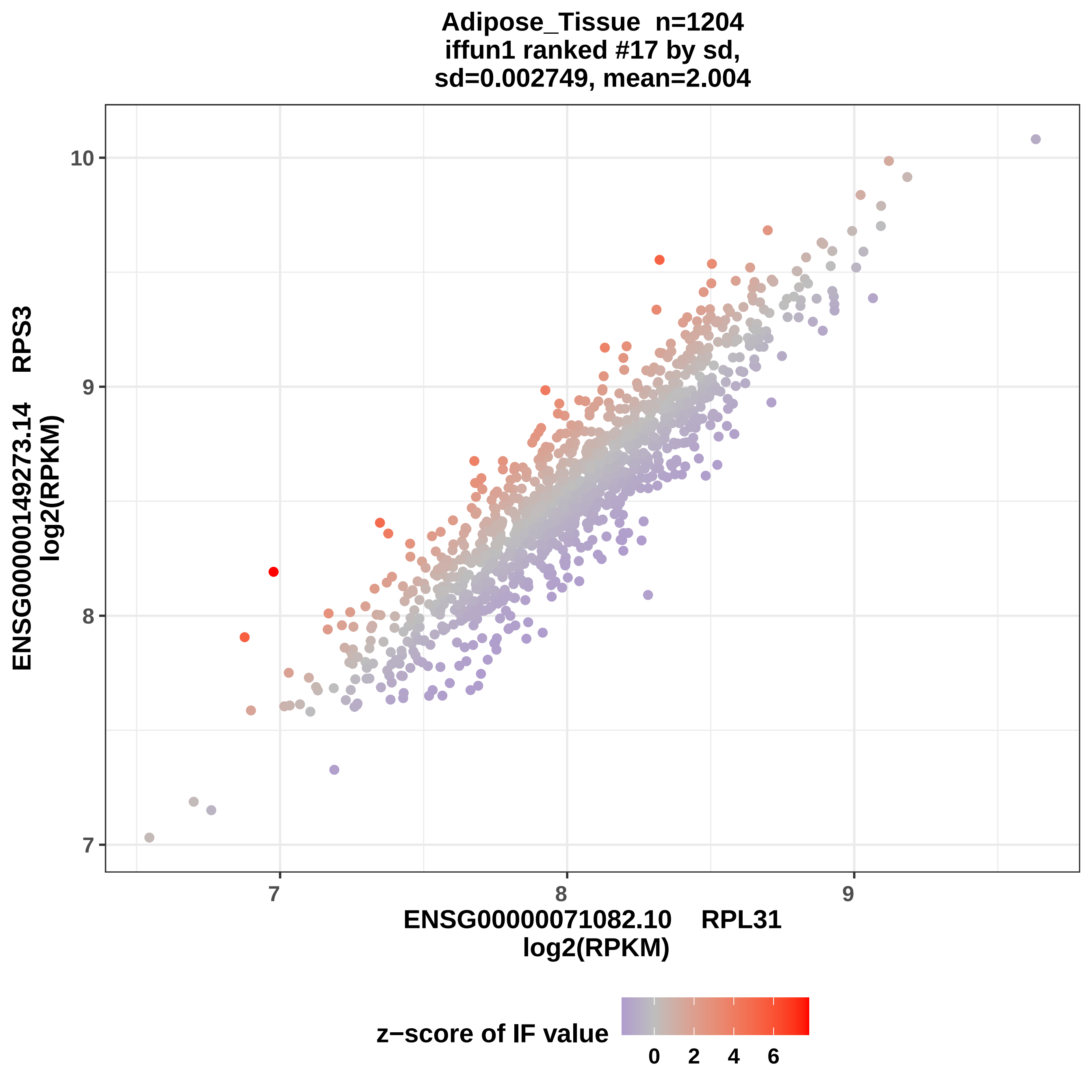

Adipose_Tissue n=1204
iffun1 ranked #17 by sd,
sd=0.002749, mean=2.004
ENSG00000149273.14 RPS3
log2(RPKM)
ENSG00000071082.10 RPL31
log2(RPKM)
10
9
8
7
7
8
9
z−score of IF value
0
2
4
6

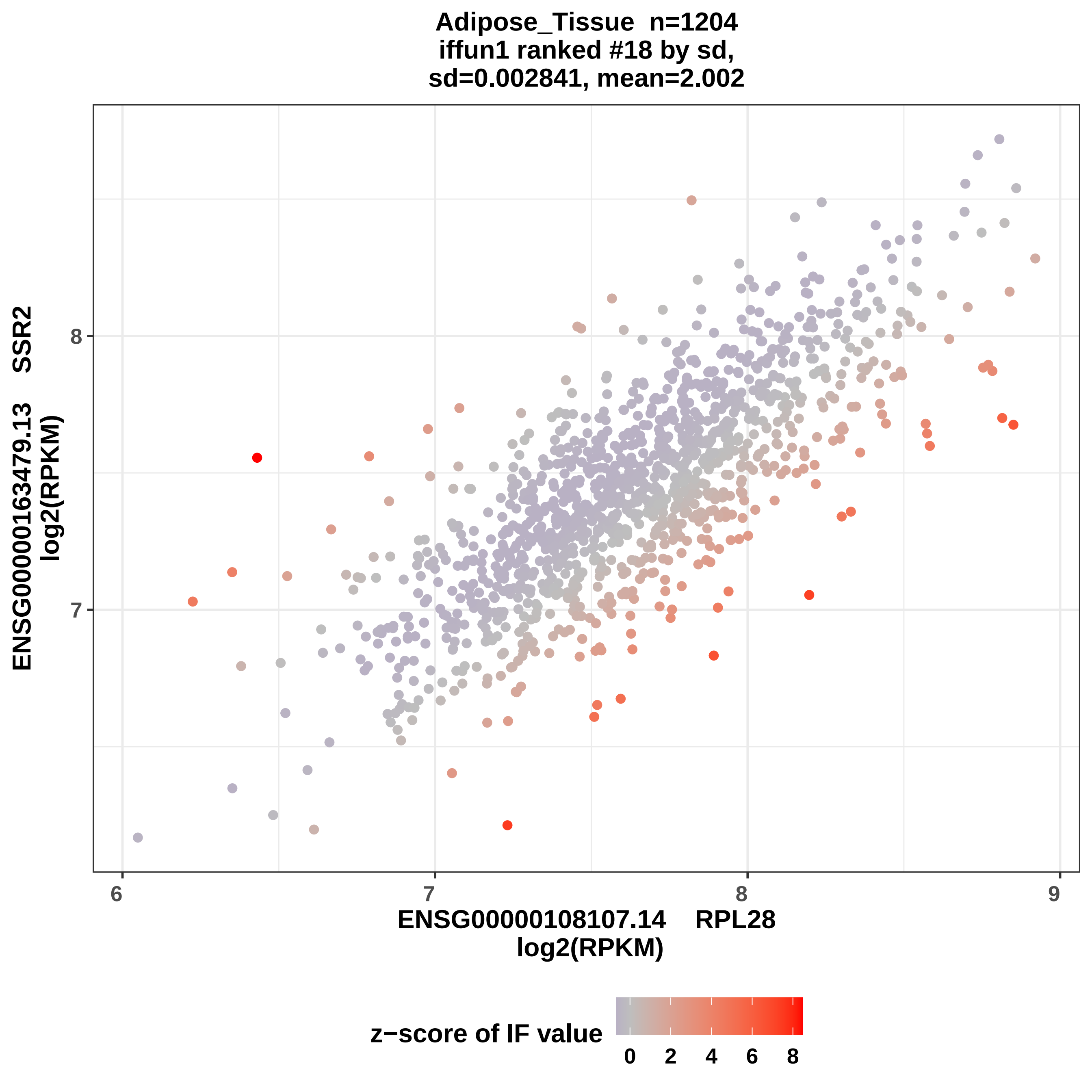
Adipose_Tissue n=1204
iffun1 ranked #18 by sd,
sd=0.002841, mean=2.002
ENSG00000163479.13 SSR2
log2(RPKM)
8
7
6
7
8
9
ENSG00000108107.14 RPL28
log2(RPKM)
z−score of IF value
0
2
4
6
8

Adipose_Tissue n=1204
iffun1 ranked #19 by sd,
sd=0.002883, mean= 2.01

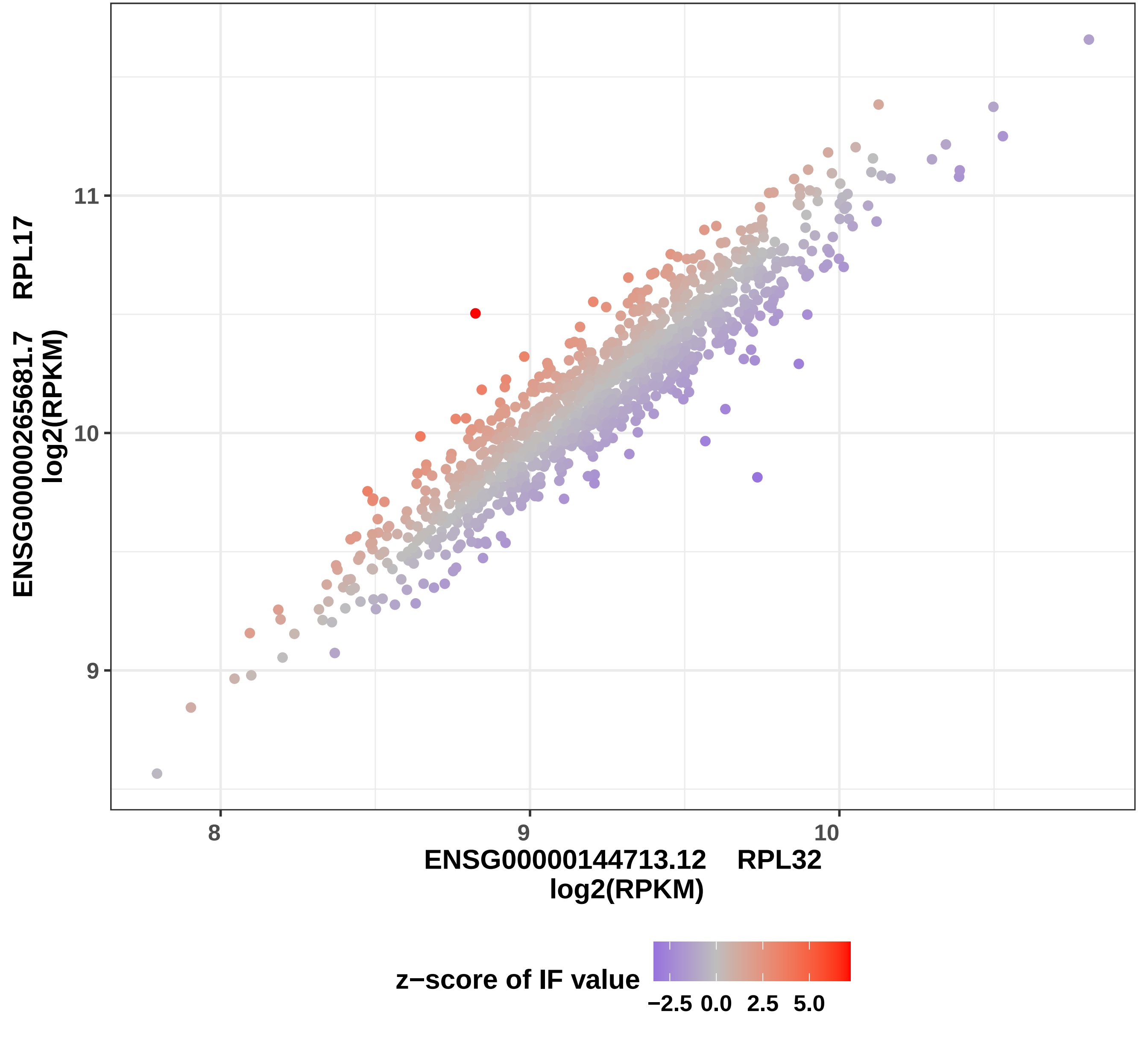

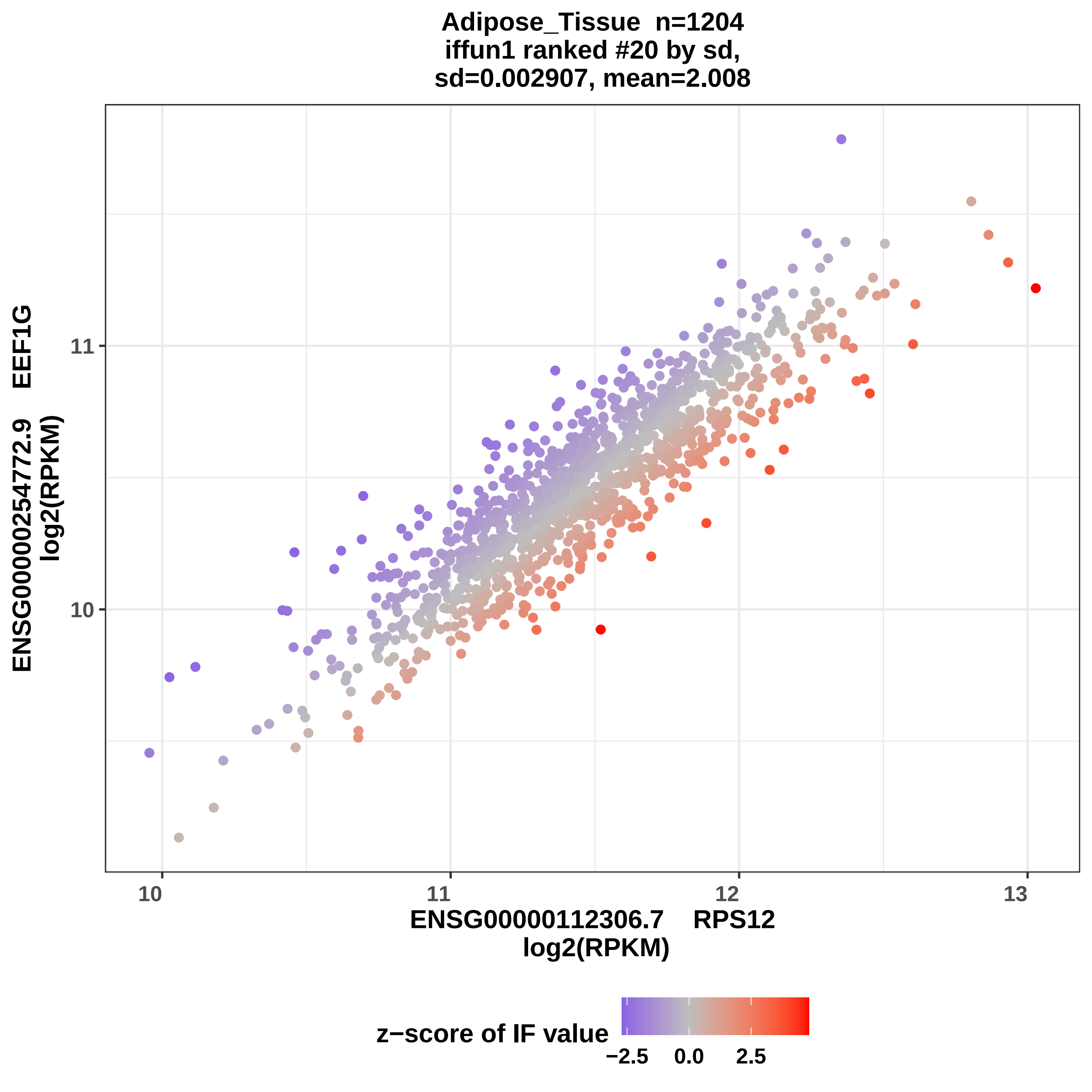
Adipose_Tissue n=1204
iffun1 ranked #20 by sd,
sd=0.002907, mean=2.008
ENSG00000254772.9 EEF1G
log2(RPKM)
11
10
10
11
12
13
ENSG00000112306.7 RPS12
log2(RPKM)
z−score of IF value
−2.5
0.0
2.5

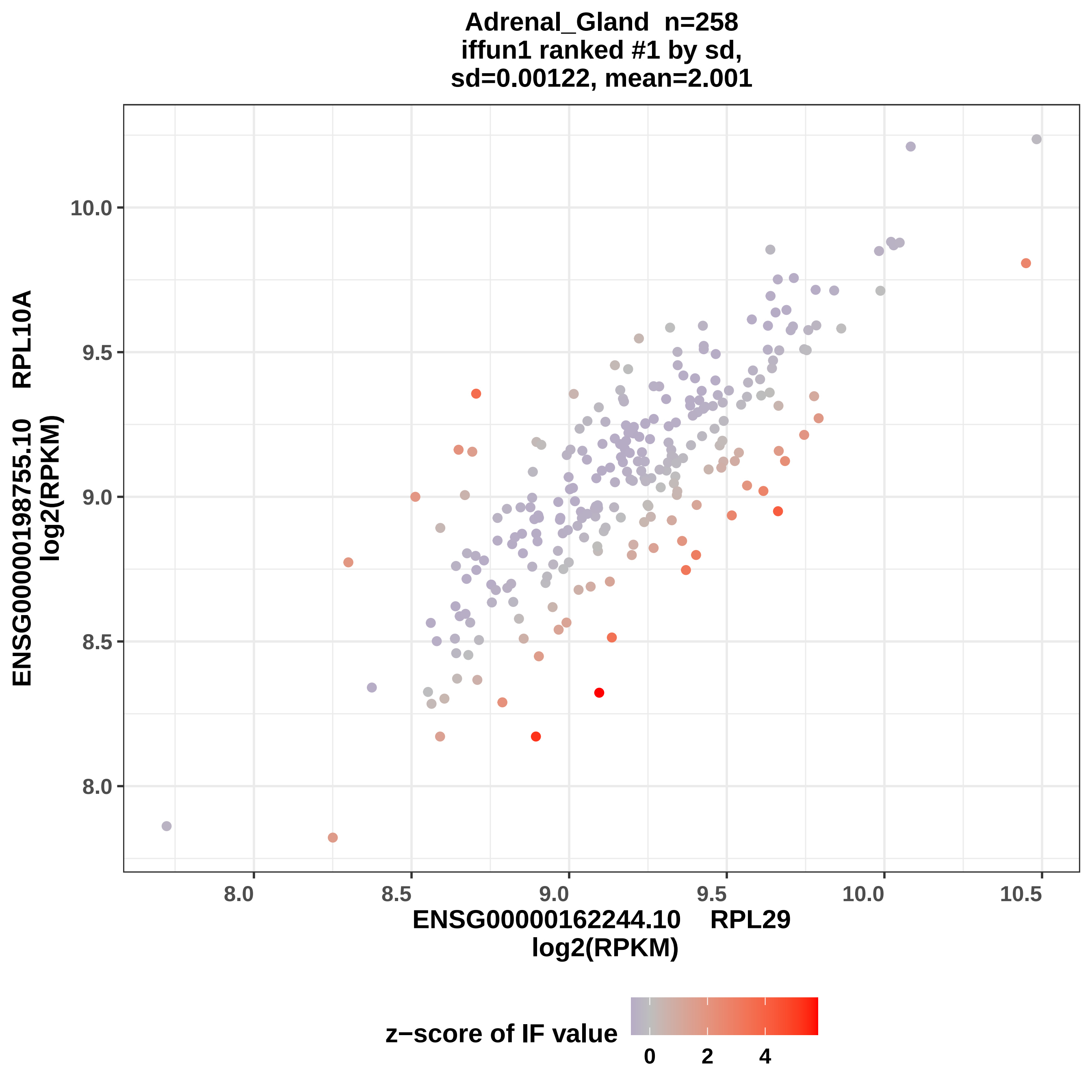

Adrenal_Gland n=258
iffun1 ranked #1 by sd,
sd=0.00122, mean=2.001
ENSG00000198755.10 RPL10A
log2(RPKM)
10.0
9.5
9.0
8.5
8.0
8.0
8.5
9.0
9.5
10.0
10.5
ENSG00000162244.10 RPL29
log2(RPKM)
z−score of IF value
0
2
4

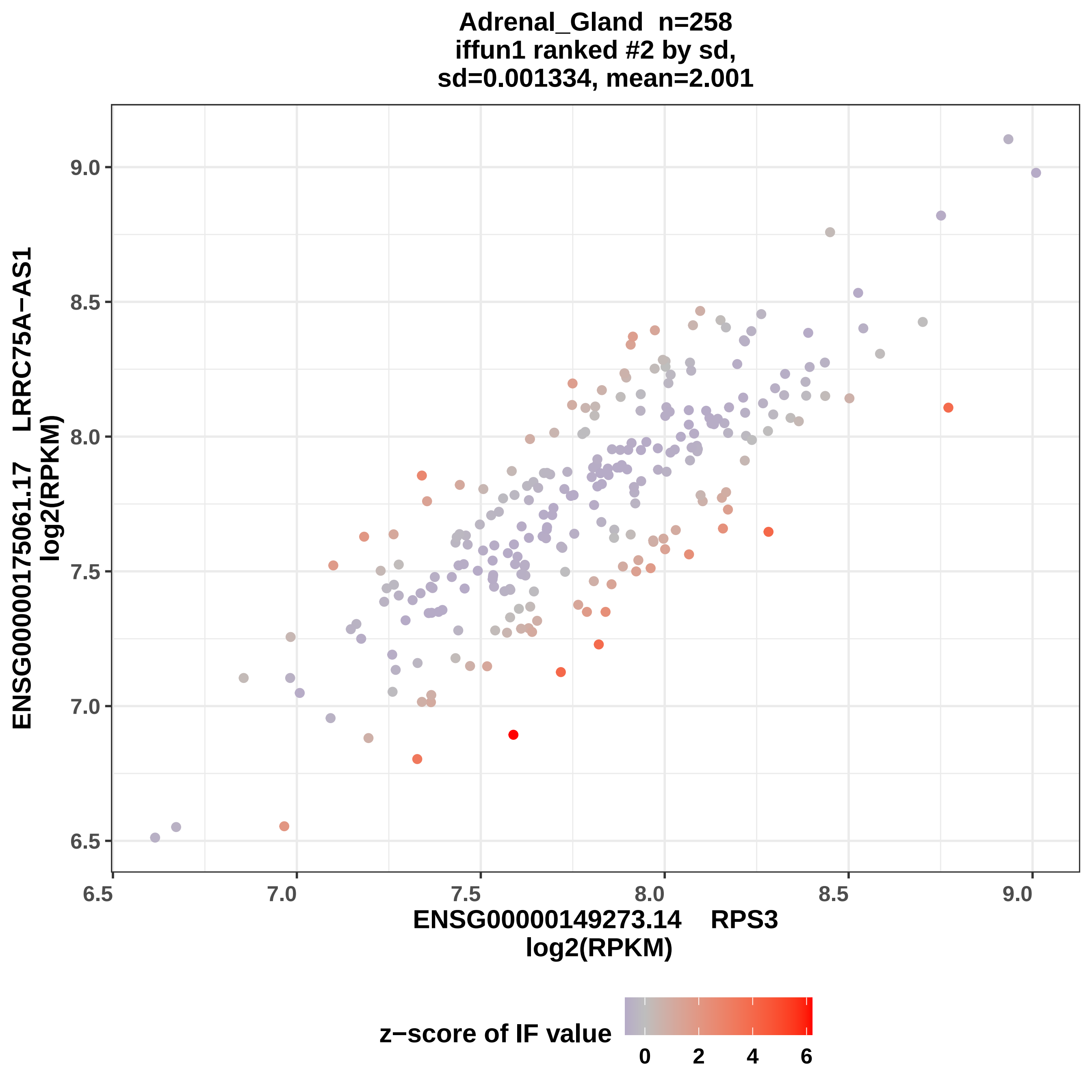
Adrenal_Gland n=258
iffun1 ranked #2 by sd,
sd=0.001334, mean=2.001
ENSG00000175061.17 LRRC75A−AS1
log2(RPKM)
ENSG00000149273.14 RPS3
log2(RPKM)
6.5
7.0
7.5
8.0
8.5
9.0
z−score of IF value
0
2
4
6

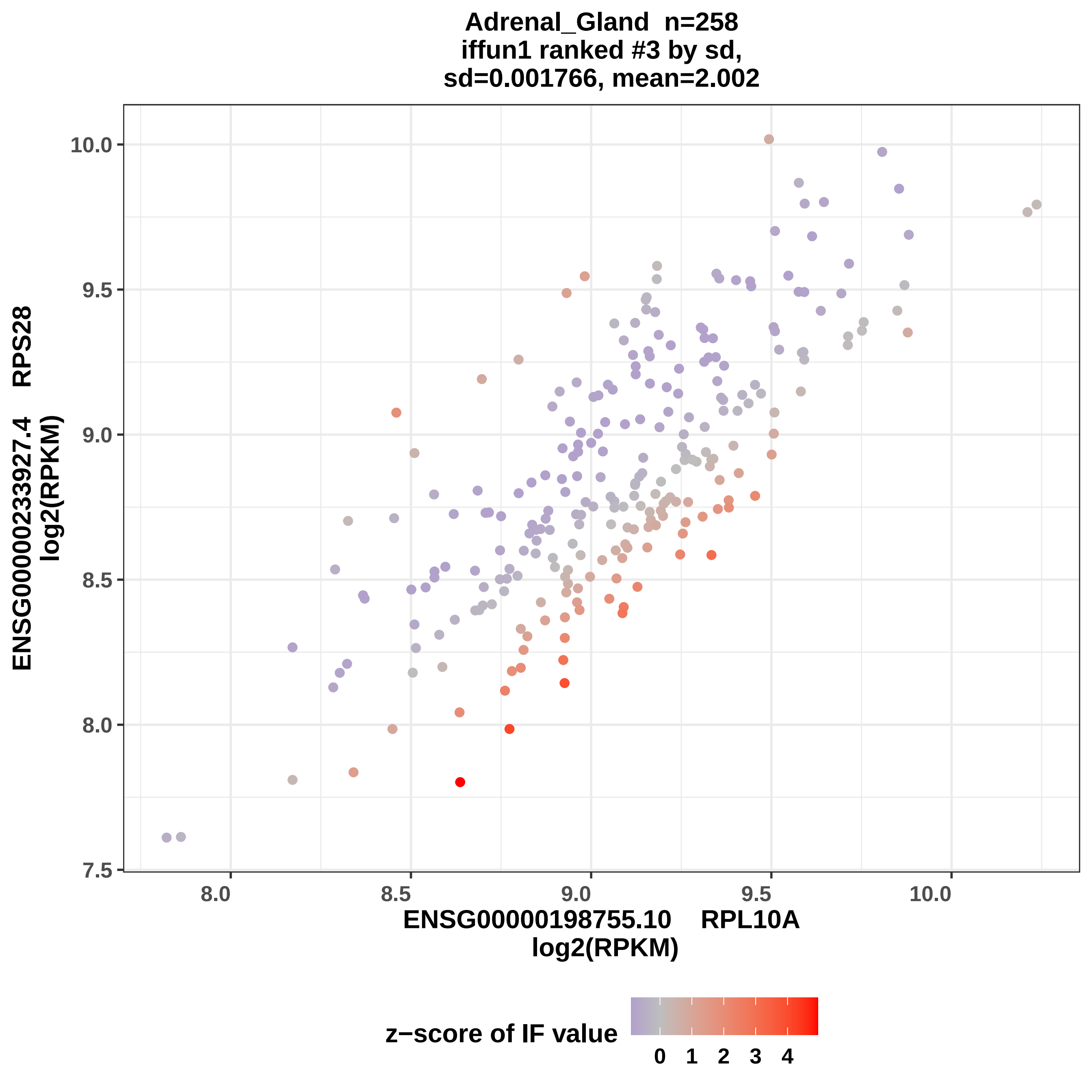

Adrenal_Gland n=258
iffun1 ranked #3 by sd,
sd=0.001766, mean=2.002
ENSG00000233927.4 RPS28
log2(RPKM)
10.0
9.5
9.0
8.5
8.0
7.5
8.0
8.5
9.0
9.5
10.0
ENSG00000198755.10 RPL10A
log2(RPKM)
z−score of IF value
0
1
2
3
4

Adrenal_Gland n=258
iffun1 ranked #4 by sd,
sd=0.001807, mean=2.001

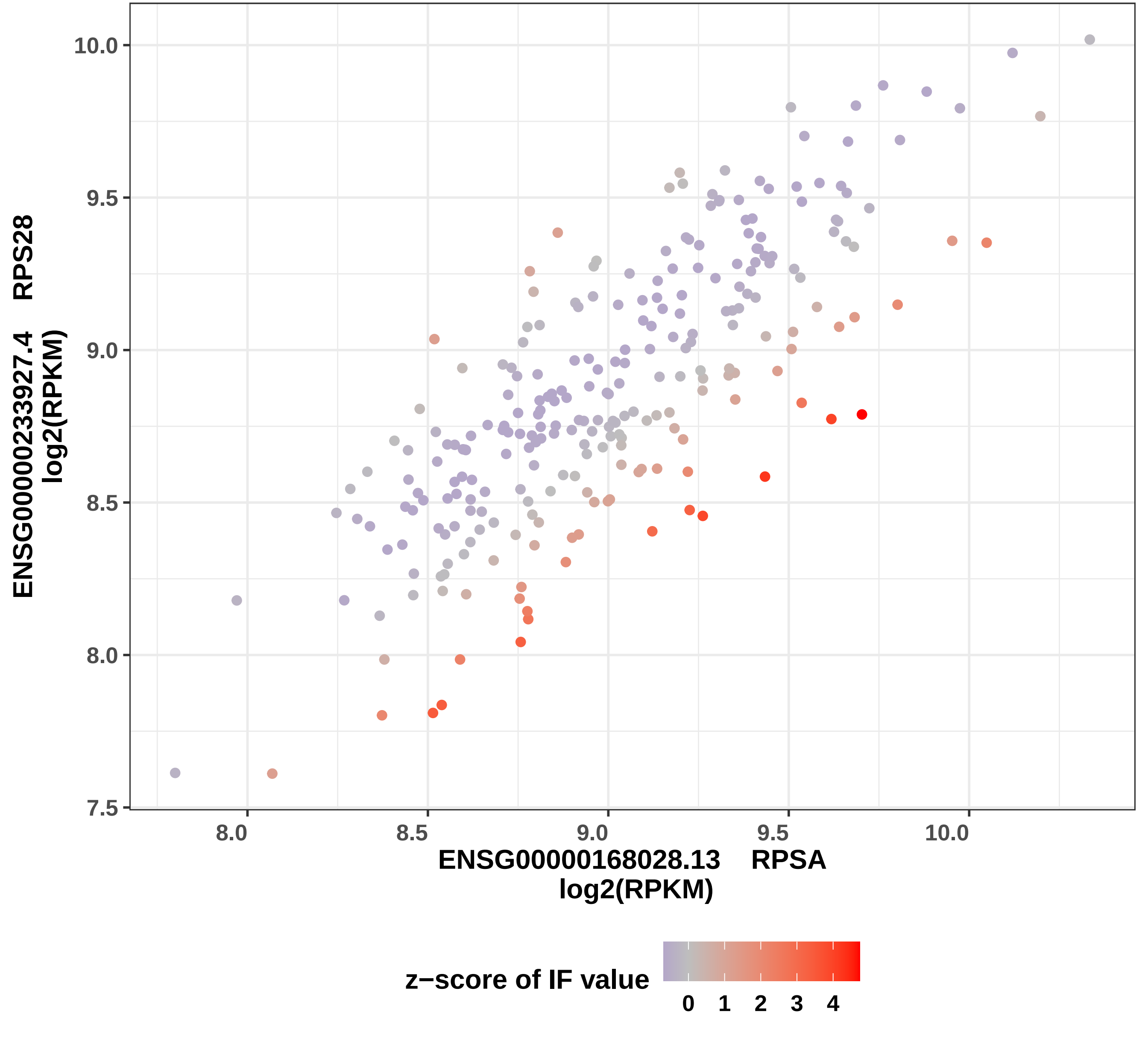

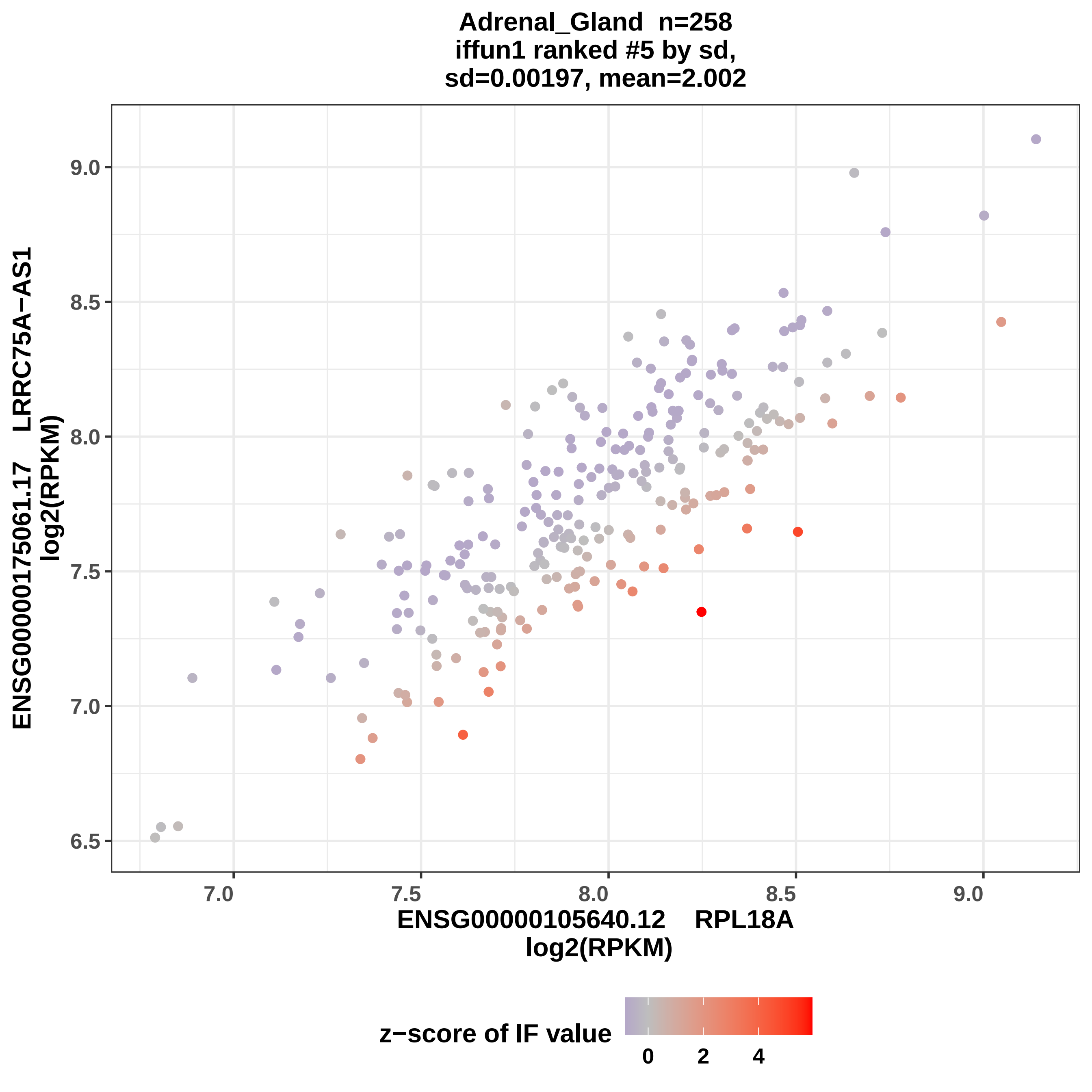
Adrenal_Gland n=258
iffun1 ranked #5 by sd,
sd=0.00197, mean=2.002
ENSG00000175061.17 LRRC75A−AS1
log2(RPKM)
ENSG00000105640.12 RPL18A
log2(RPKM)
9.0
8.5
8.0
7.5
7.0
6.5
7.0
7.5
8.0
8.5
9.0
z−score of IF value
0
2
4

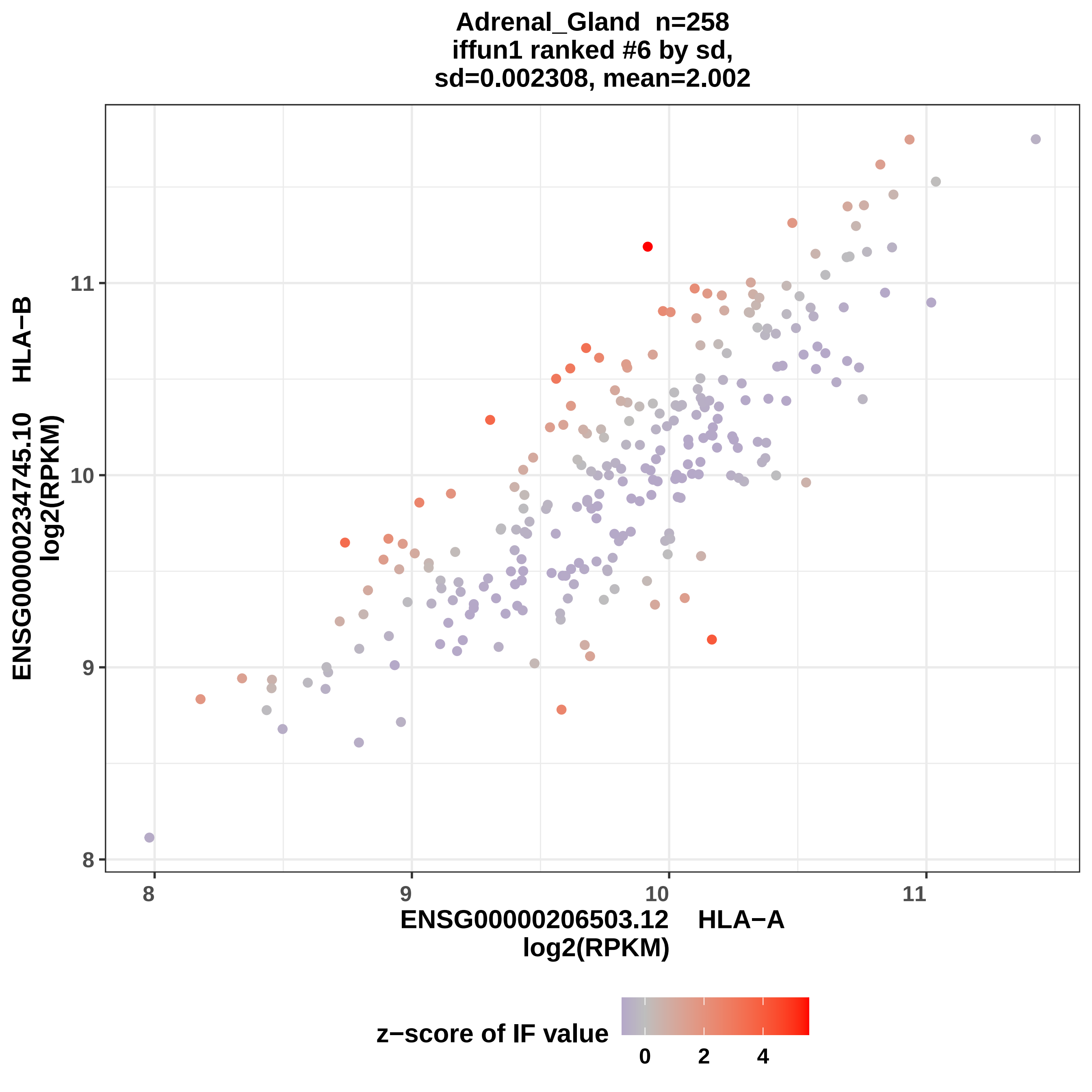
Adrenal_Gland n=258
iffun1 ranked #6 by sd,
sd=0.002308, mean=2.002
ENSG00000234745.10 HLA−B
log2(RPKM)
ENSG00000206503.12 HLA−A
log2(RPKM)
8
9
10
11
z−score of IF value
0
2
4

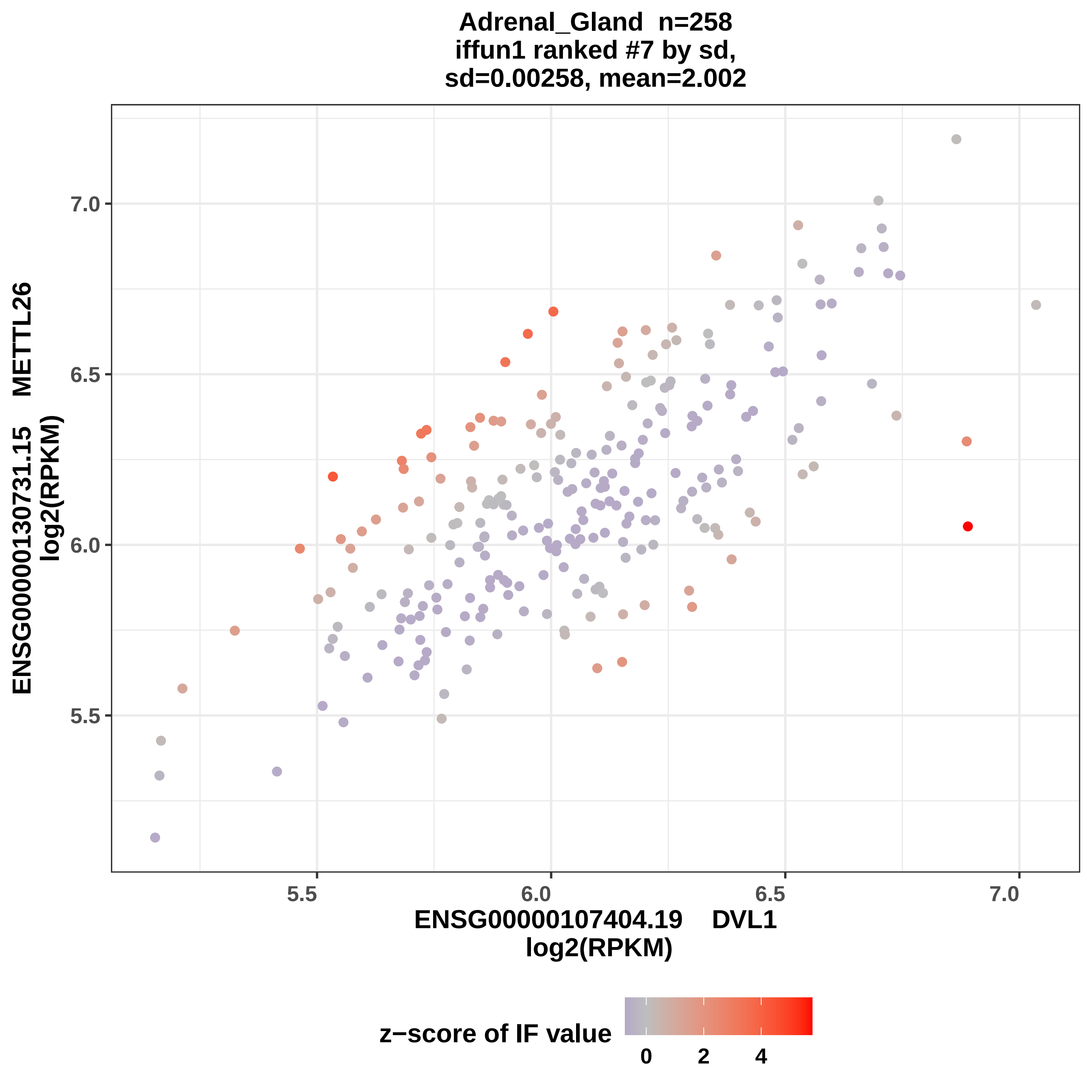
Adrenal_Gland n=258
iffun1 ranked #7 by sd,
sd=0.00258, mean=2.002
ENSG00000130731.15 METTL26
log2(RPKM)
7.0
6.5
6.0
5.5
5.5
6.0
6.5
7.0
ENSG00000107404.19 DVL1
log2(RPKM)
z−score of IF value
0
2
4

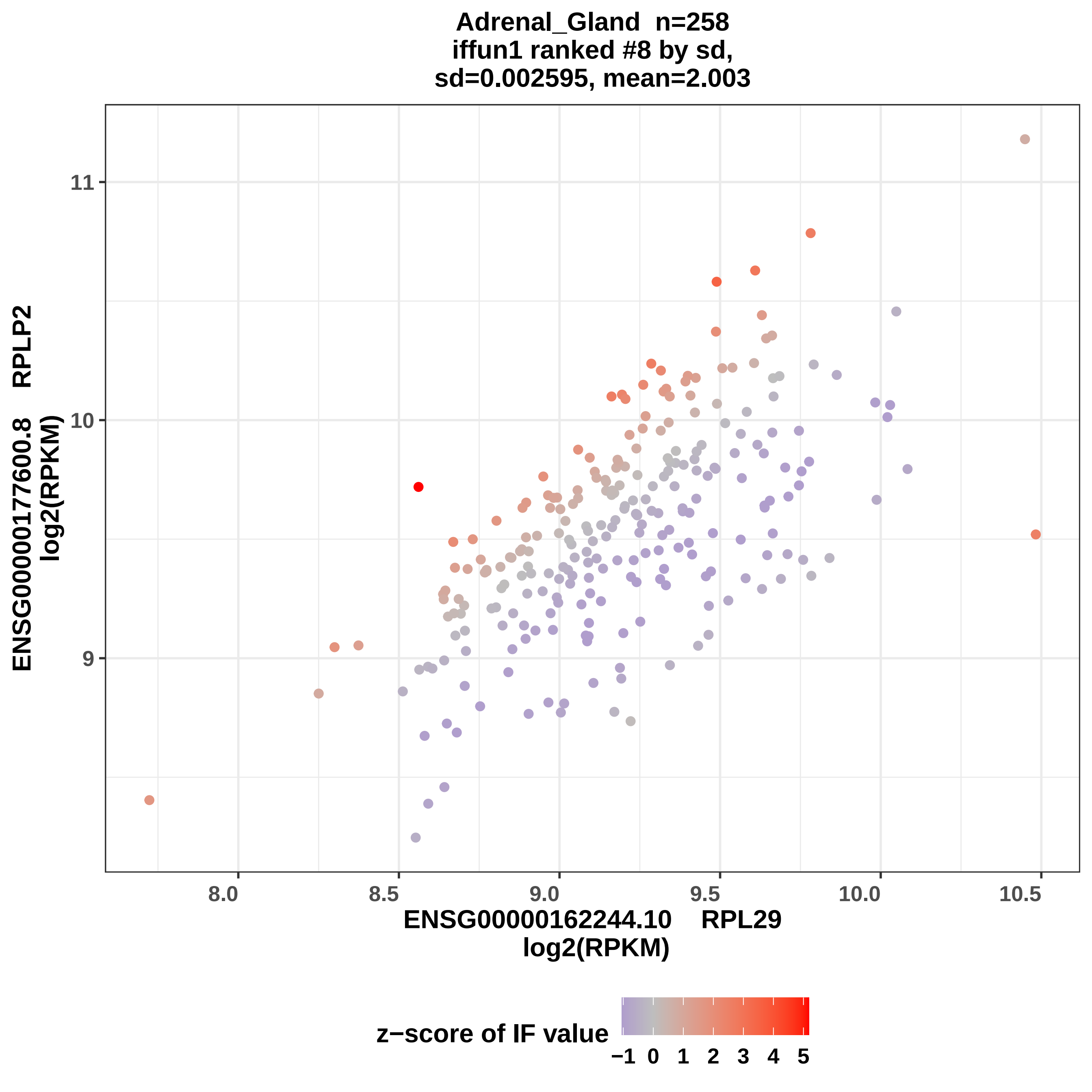
Adrenal_Gland n=258
iffun1 ranked #8 by sd,
sd=0.002595, mean=2.003
ENSG00000177600.8 RPLP2
log2(RPKM)
11
10
9
8.0
8.5
9.0
9.5
10.0
10.5
ENSG00000162244.10 RPL29
log2(RPKM)
z−score of IF value
−1
0
1
2
3
4
5

Adrenal_Gland n=258
iffun1 ranked #9 by sd,
sd=0.002597, mean=2.002

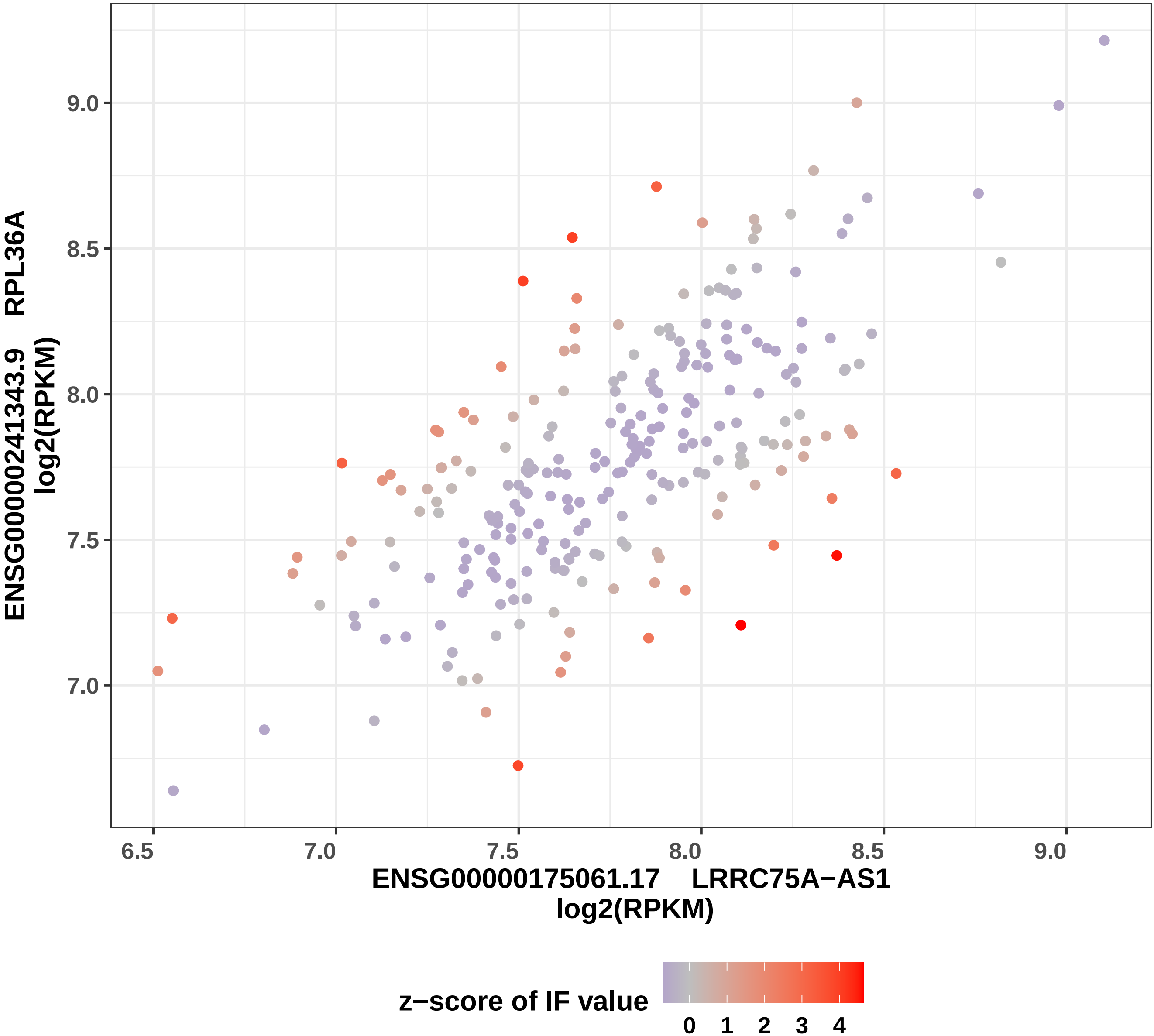

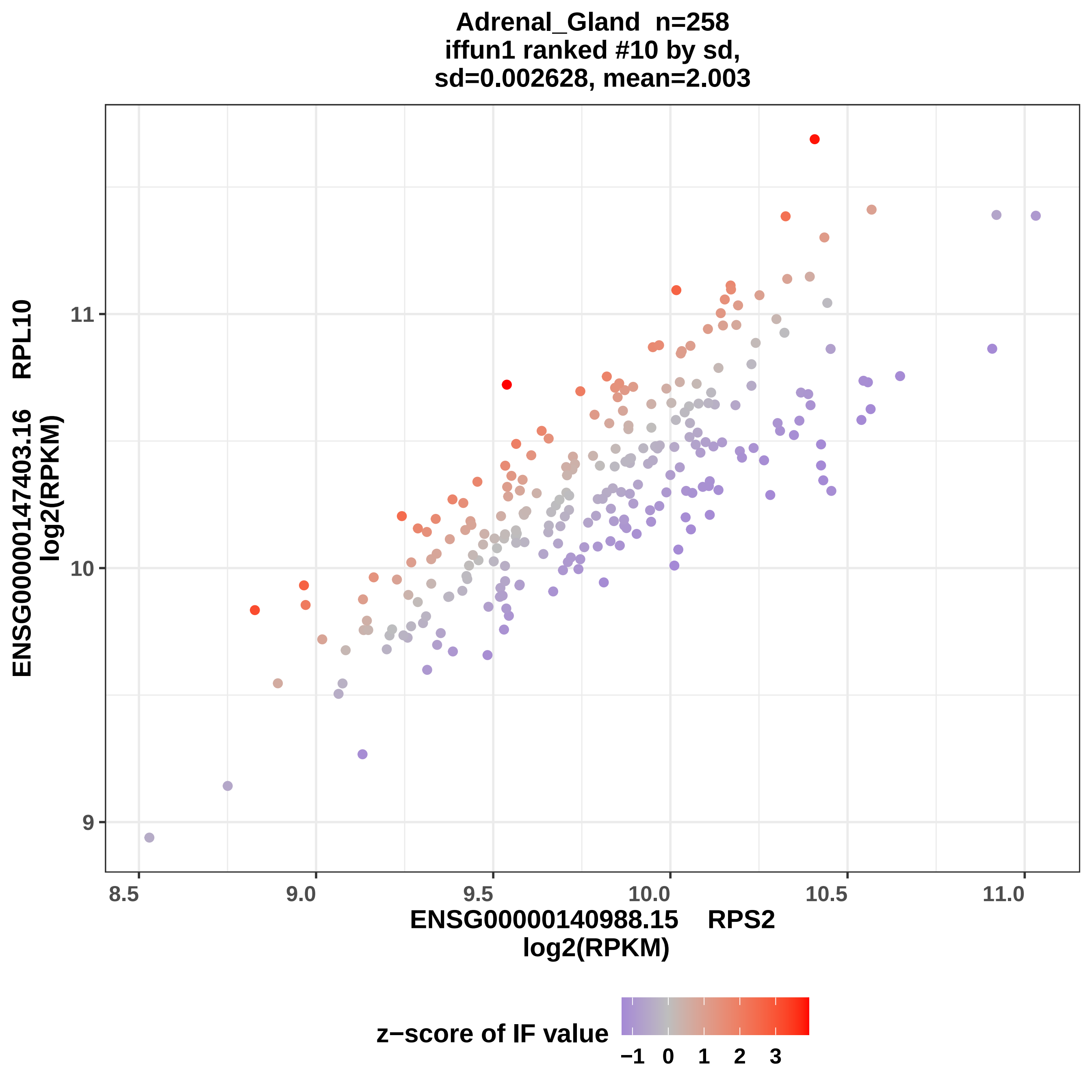
Adrenal_Gland n=258
iffun1 ranked #10 by sd,
sd=0.002628, mean=2.003
ENSG00000147403.16 RPL10
log2(RPKM)
11
10
9
8.5
9.0
9.5
10.0
10.5
11.0
ENSG00000140988.15 RPS2
log2(RPKM)
z−score of IF value
−1
0
1
2
3

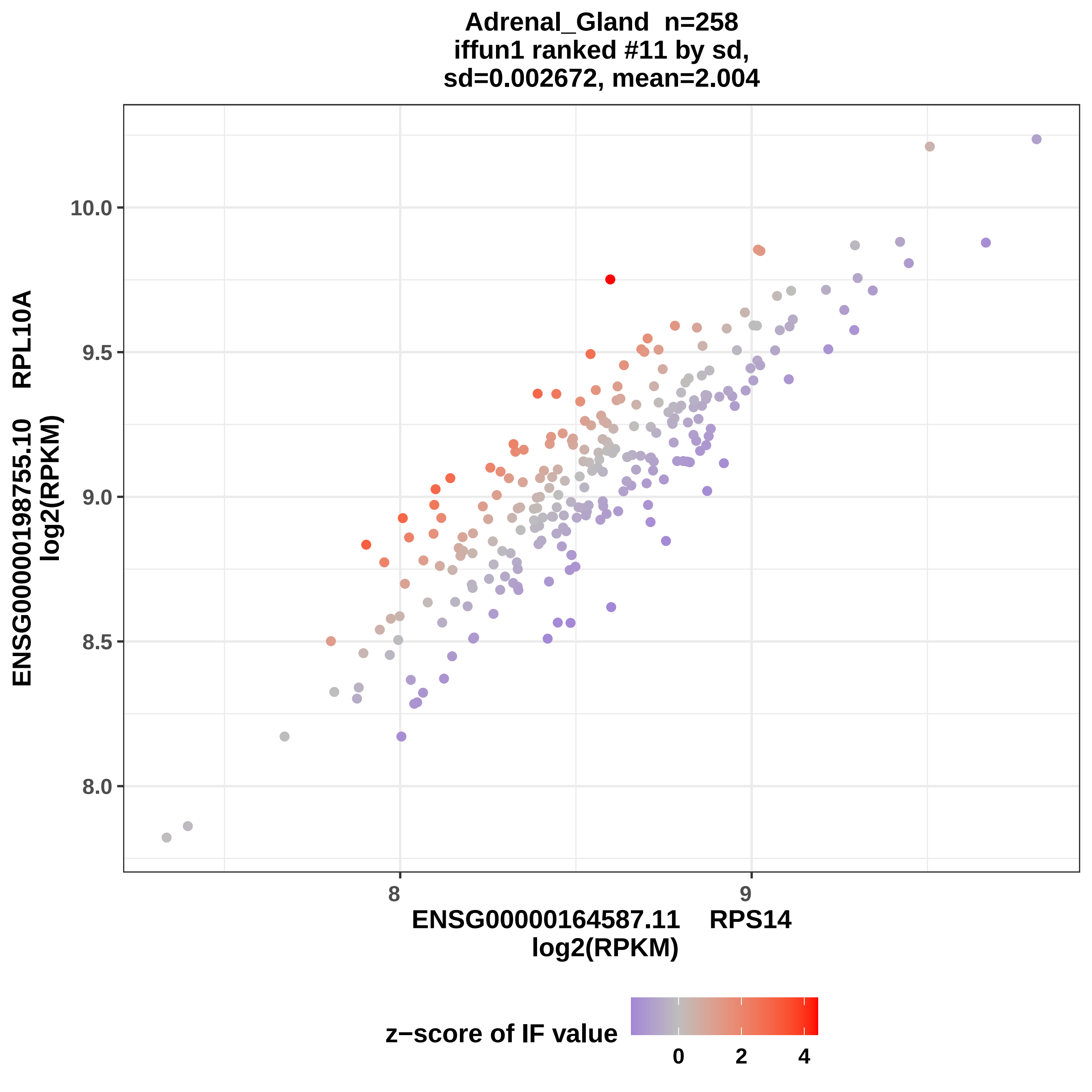

Adrenal_Gland n=258
iffun1 ranked #11 by sd,
sd=0.002672, mean=2.004
ENSG00000198755.10 RPL10A
log2(RPKM)
10.0
9.5
9.0
8.5
8.0
8
9
ENSG00000164587.11 RPS14
log2(RPKM)
z−score of IF value
0
2
4

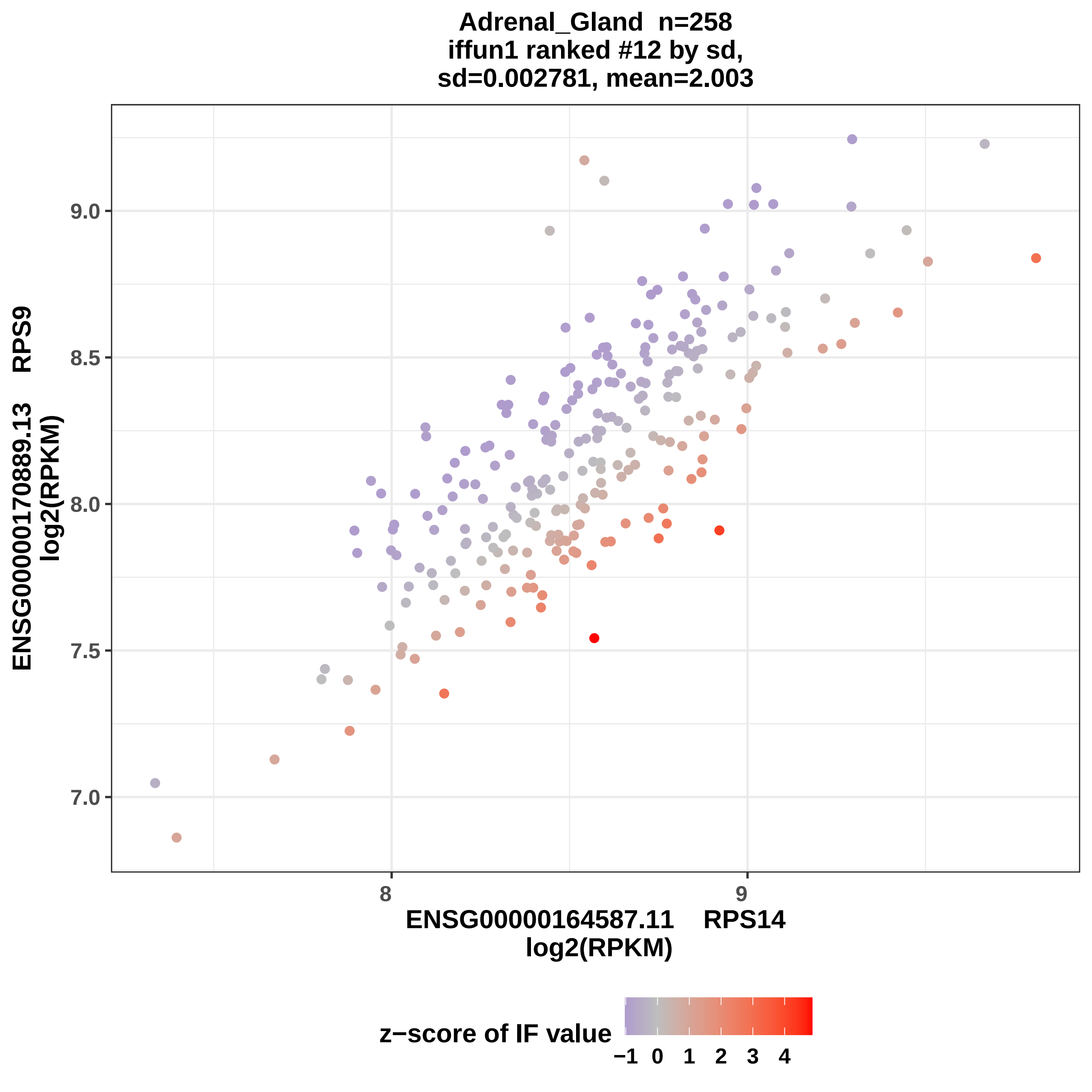

Adrenal_Gland n=258
iffun1 ranked #12 by sd,
sd=0.002781, mean=2.003
ENSG00000170889.13 RPS9
log2(RPKM)
9.0
8.5
8.0
7.5
7.0
8
9
ENSG00000164587.11 RPS14
log2(RPKM)
z−score of IF value
−1
0
1
2
3
4

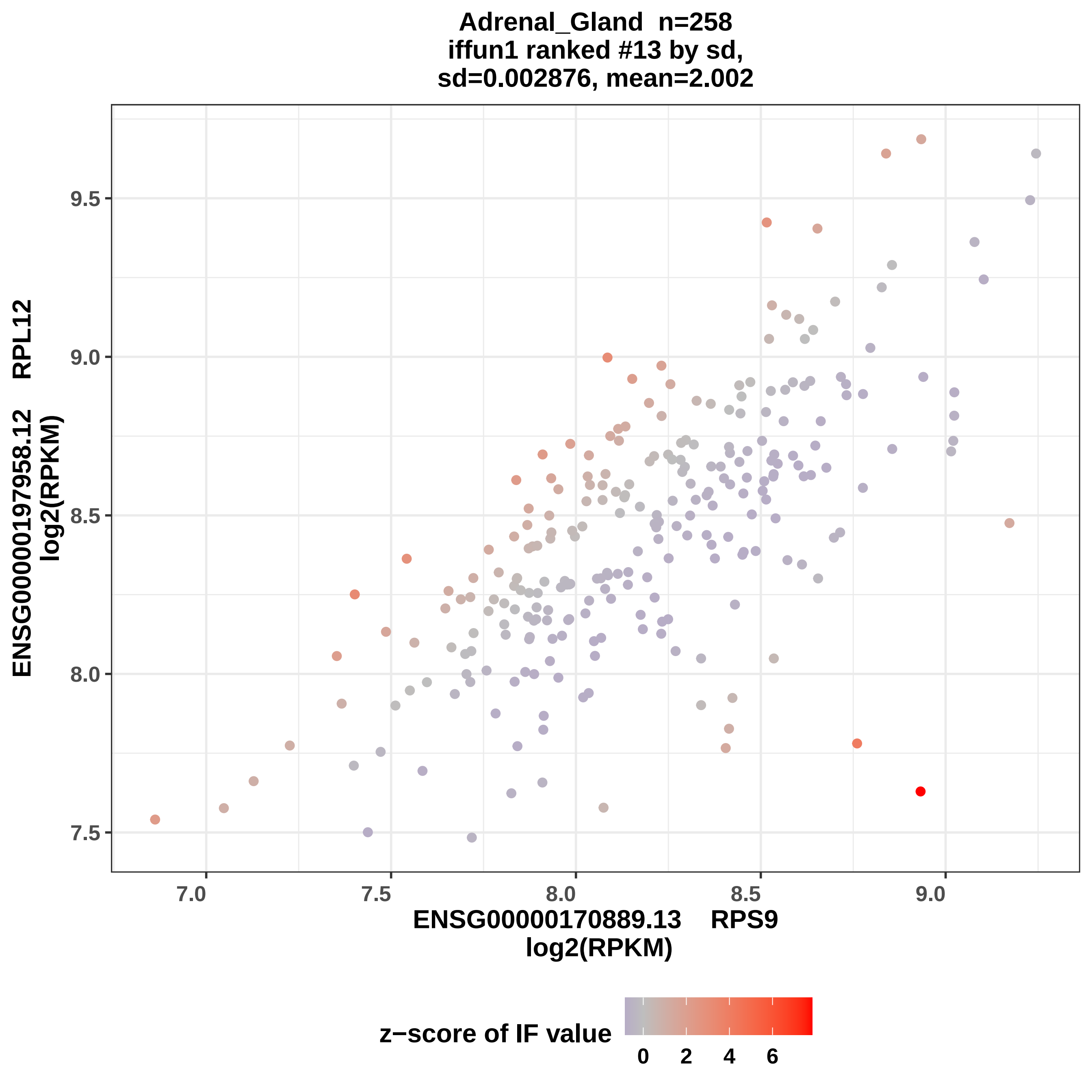
Adrenal_Gland n=258
iffun1 ranked #13 by sd,
sd=0.002876, mean=2.002
ENSG00000197958.12 RPL12
log2(RPKM)
9.5
9.0
8.5
8.0
7.5
7.0
7.5
8.0
8.5
9.0
ENSG00000170889.13 RPS9
log2(RPKM)
z−score of IF value
0
2
4
6

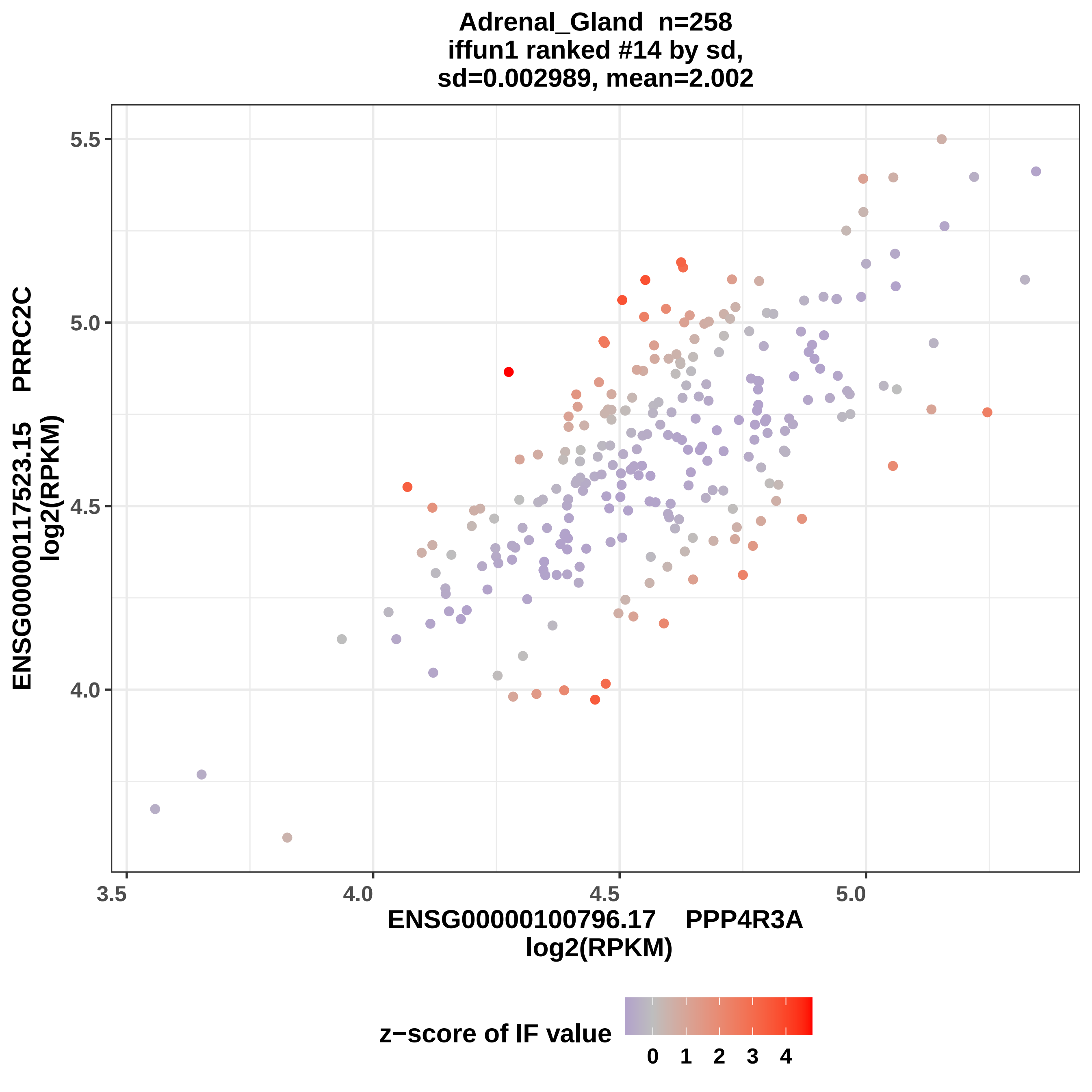
Adrenal_Gland n=258
iffun1 ranked #14 by sd,
sd=0.002989, mean=2.002
ENSG00000117523.15 PRRC2C
log2(RPKM)
5.5
5.0
4.5
4.0
3.5
4.0
4.5
5.0
ENSG00000100796.17 PPP4R3A
log2(RPKM)
z−score of IF value
0 1 2 3 4

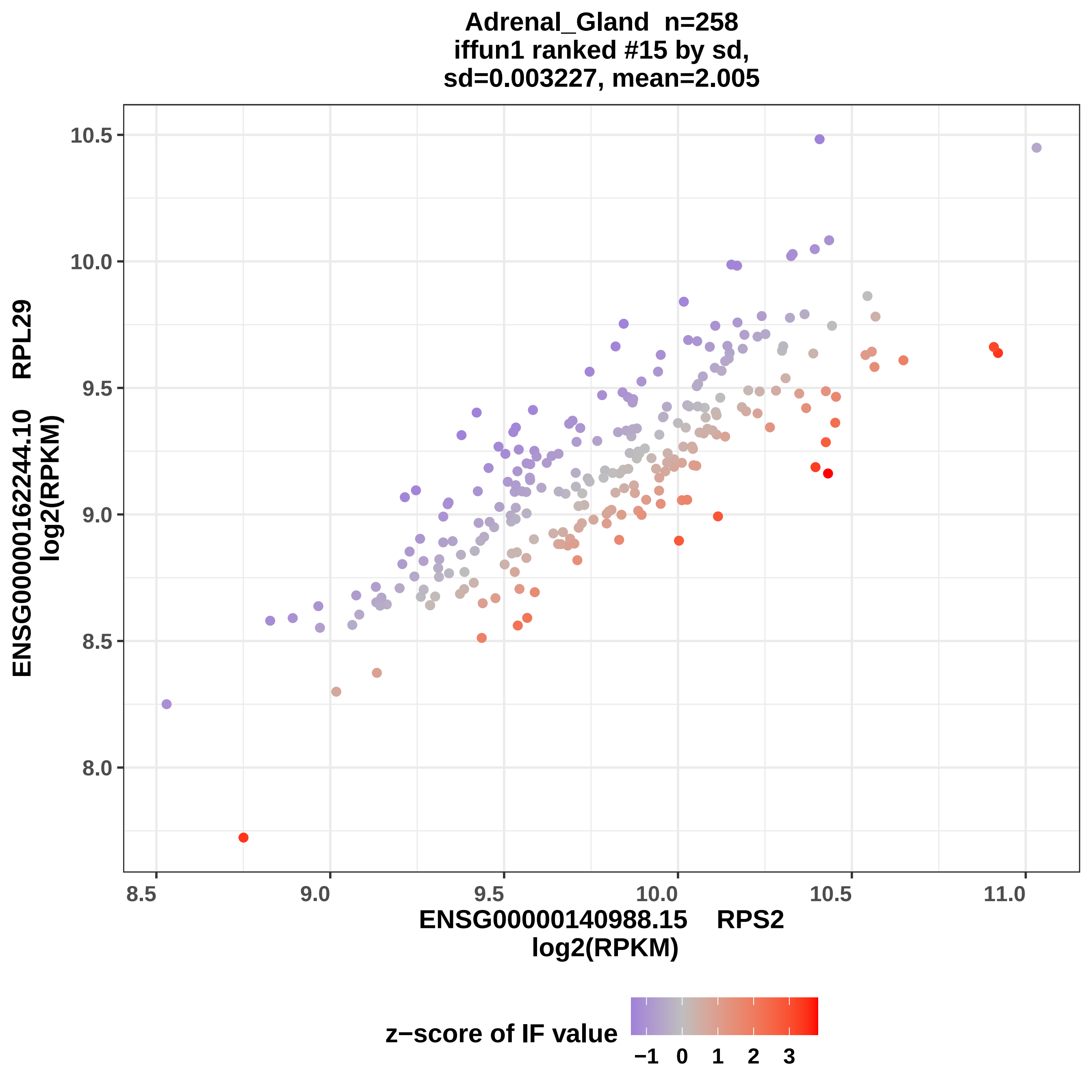
Adrenal_Gland n=258
iffun1 ranked #15 by sd,
sd=0.003227, mean=2.005
ENSG00000162244.10 RPL29
log2(RPKM)
10.5
10.0
9.5
9.0
8.5
8.0
8.5
9.0
9.5
10.0
10.5
11.0
ENSG00000140988.15 RPS2
log2(RPKM)
z−score of IF value
−1
0
1
2
3

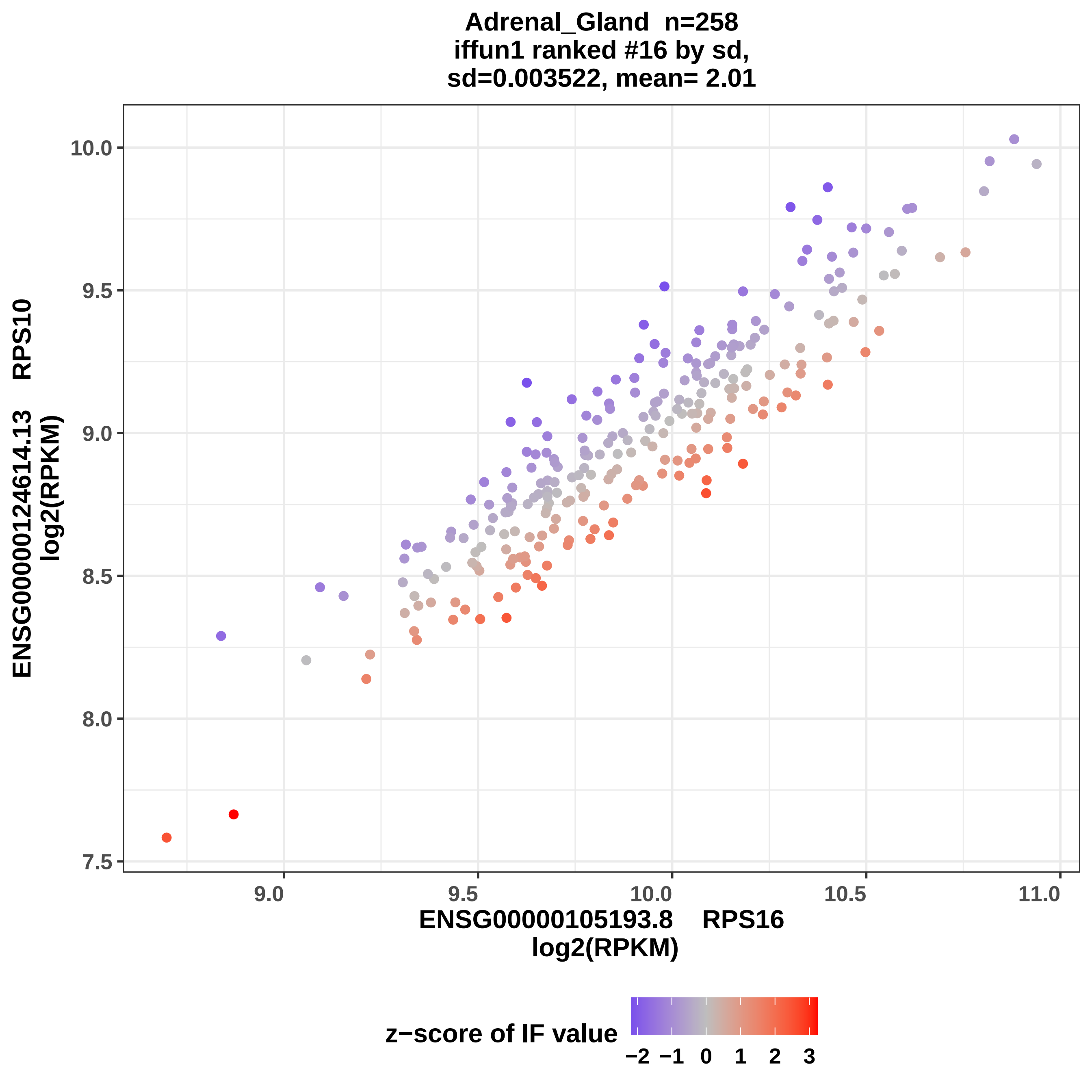
Adrenal_Gland n=258
iffun1 ranked #16 by sd,
sd=0.003522, mean= 2.01
ENSG00000124614.13 RPS10
log2(RPKM)
ENSG00000105193.8 RPS16
log2(RPKM)
z−score of IF value
10.0
9.5
9.0
8.5
8.0
7.5
9.0
9.5
10.0
10.5
11.0
−2
−1
0
1
2
3

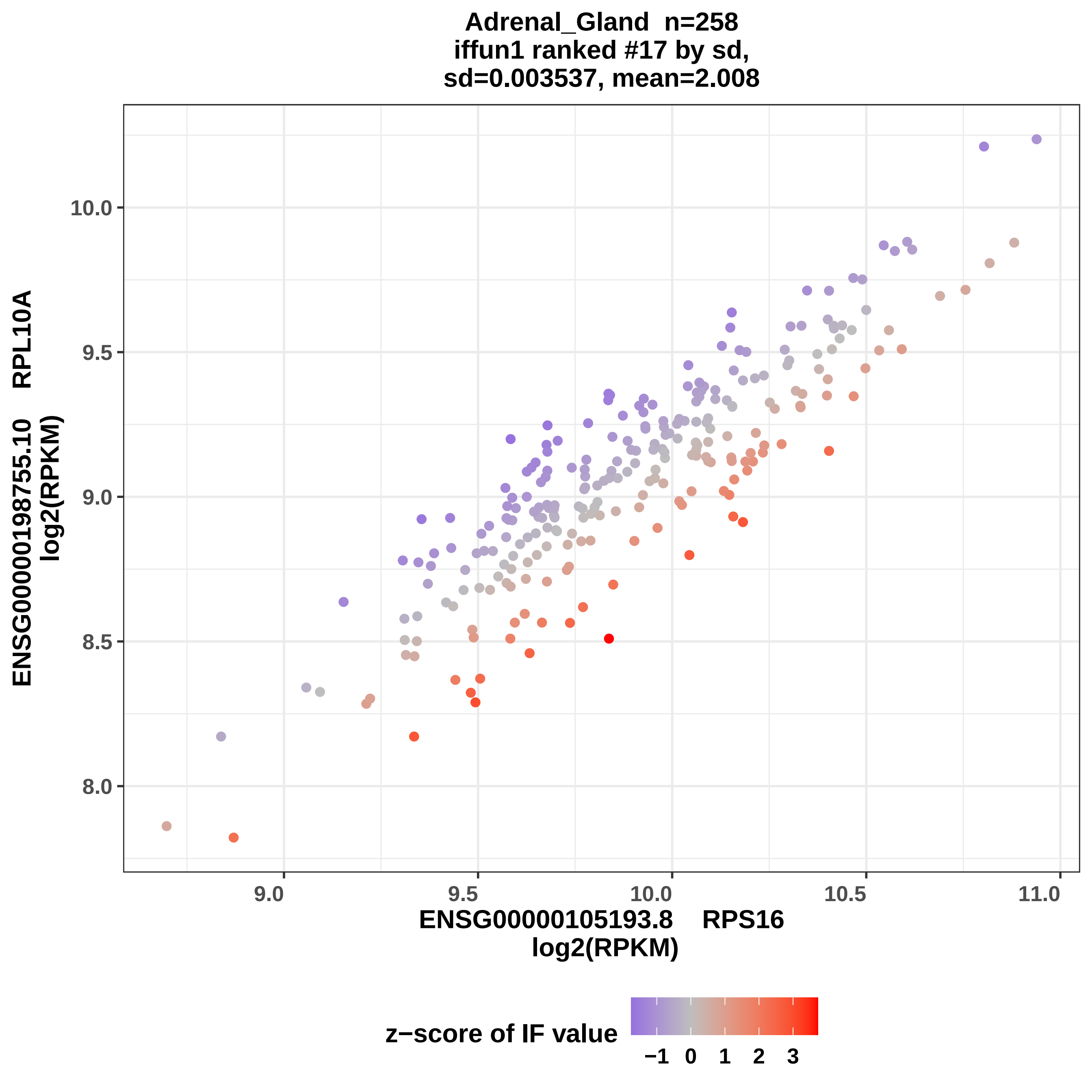
Adrenal_Gland n=258
iffun1 ranked #17 by sd,
sd=0.003537, mean=2.008
ENSG00000198755.10 RPL10A
log2(RPKM)
10.0
9.5
9.0
8.5
8.0
9.0
9.5
10.0
10.5
11.0
ENSG00000105193.8 RPS16
log2(RPKM)
z−score of IF value
−1
0
1
2
3

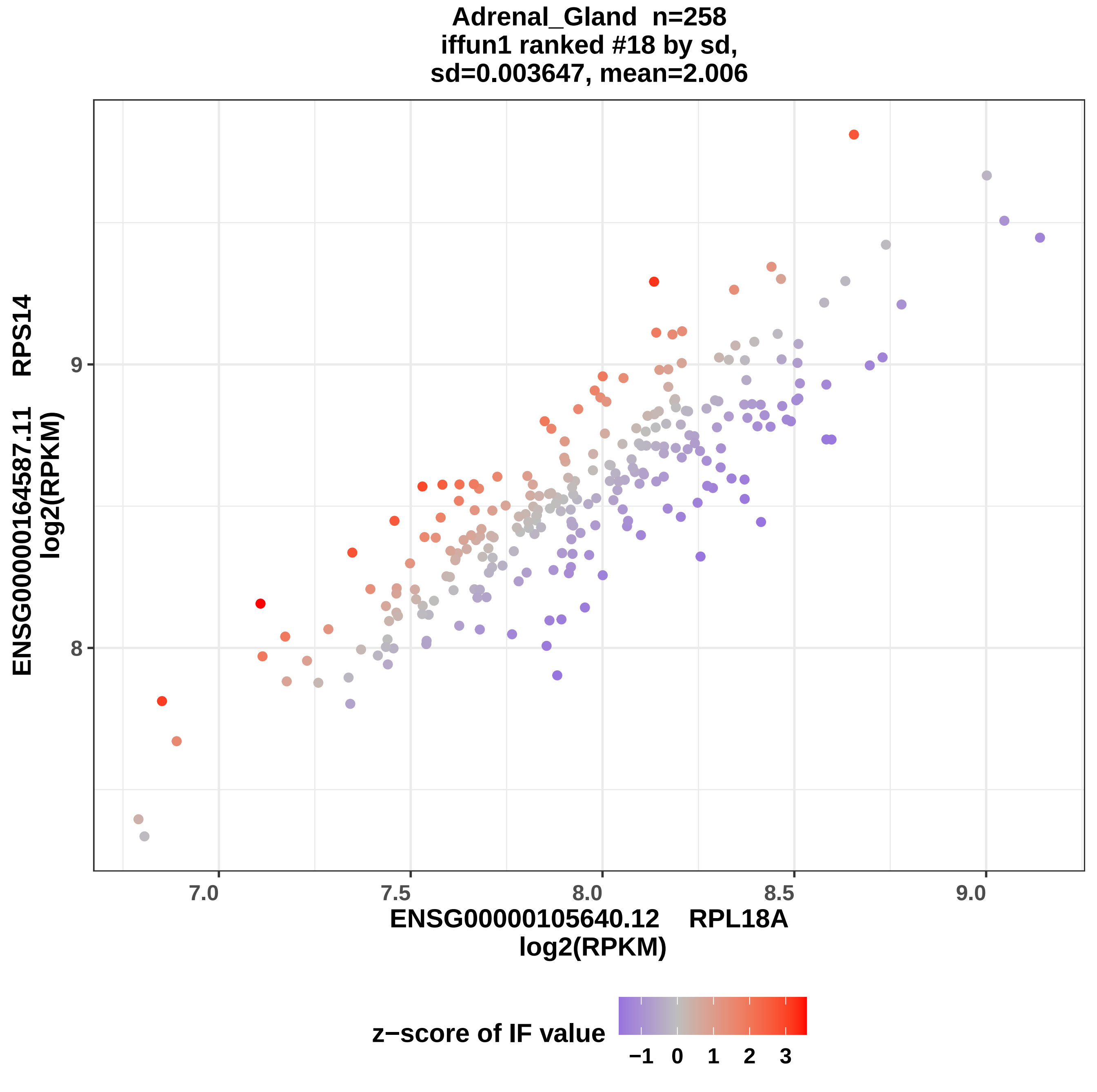

Adrenal_Gland n=258
iffun1 ranked #18 by sd,
sd=0.003647, mean=2.006
ENSG00000164587.11 RPS14
log2(RPKM)
9
8
7.0
7.5
8.0
8.5
9.0
ENSG00000105640.12 RPL18A
log2(RPKM)
z−score of IF value
−1
0
1
2
3

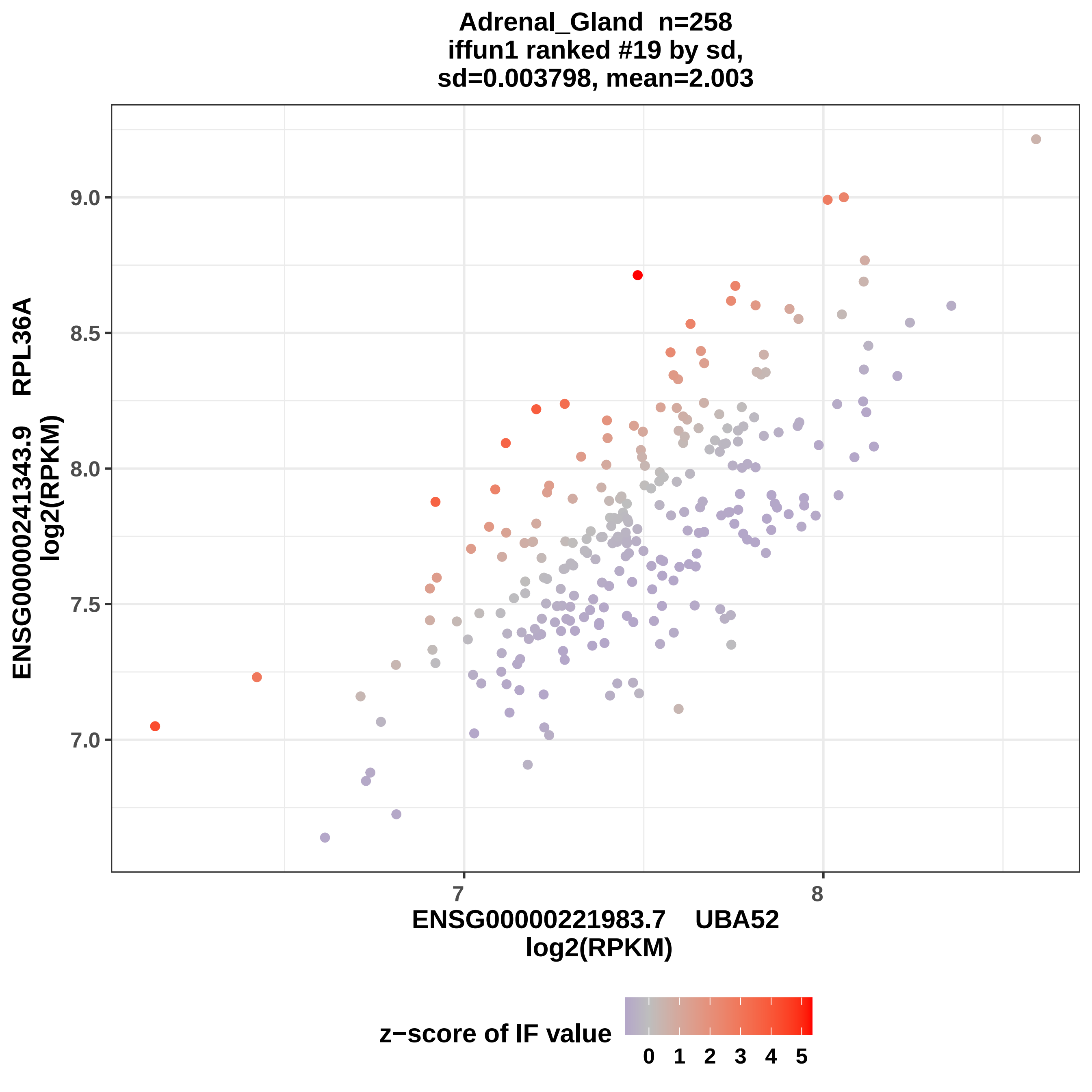

Adrenal_Gland n=258
iffun1 ranked #19 by sd,
sd=0.003798, mean=2.003
9.0
8.5
8.0
7.5
7.0
7
8
ENSG00000241343.9 RPL36A
log2(RPKM)
ENSG00000221983.7 UBA52
log2(RPKM)
z−score of IF value
0 1 2 3 4 5

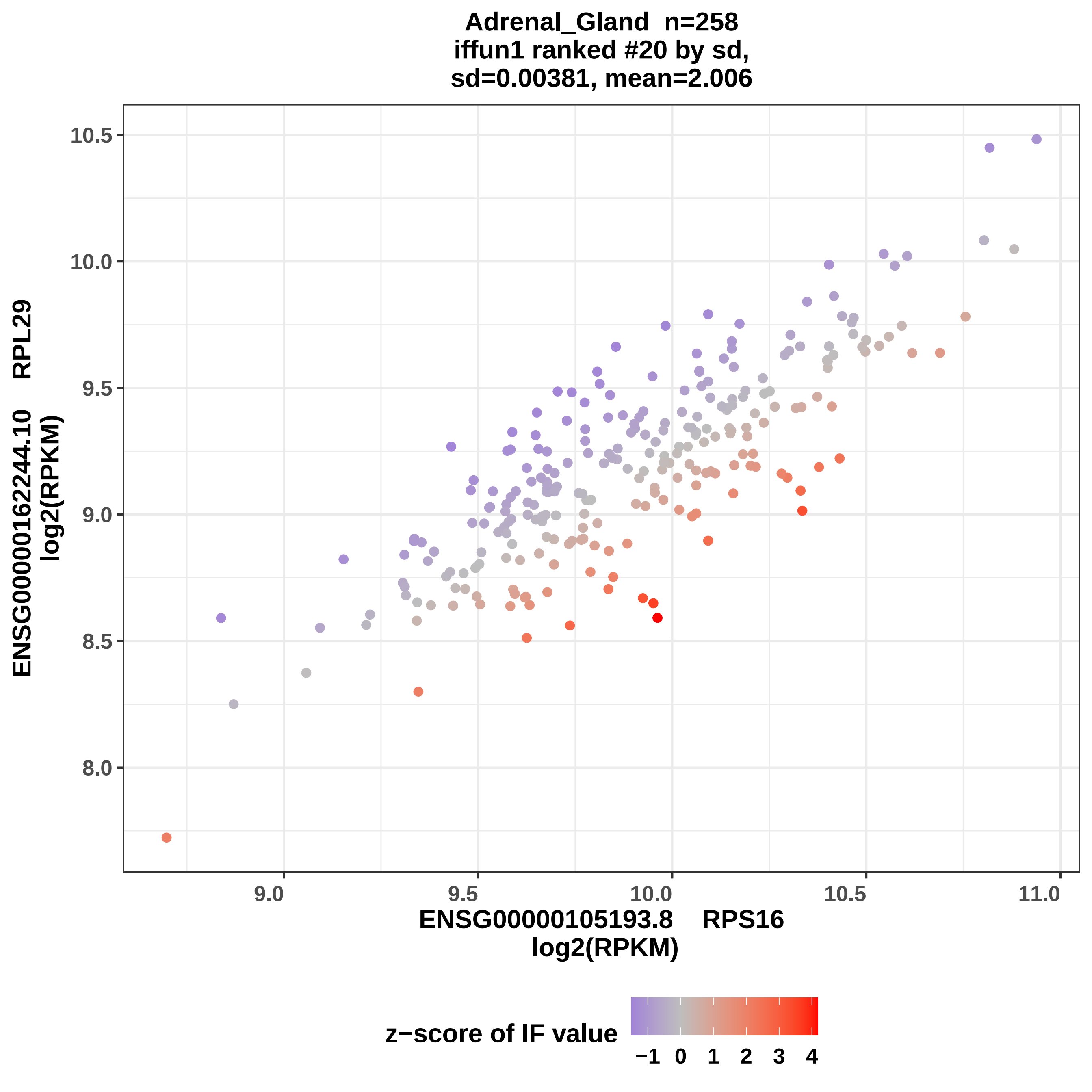
Adrenal_Gland n=258
iffun1 ranked #20 by sd,
sd=0.00381, mean=2.006
ENSG00000162244.10 RPL29
log2(RPKM)
10.5
10.0
9.5
9.0
8.5
8.0
9.0
9.5
10.0
10.5
11.0
ENSG00000105193.8 RPS16
log2(RPKM)
z−score of IF value
−1
0
1
2
3
4

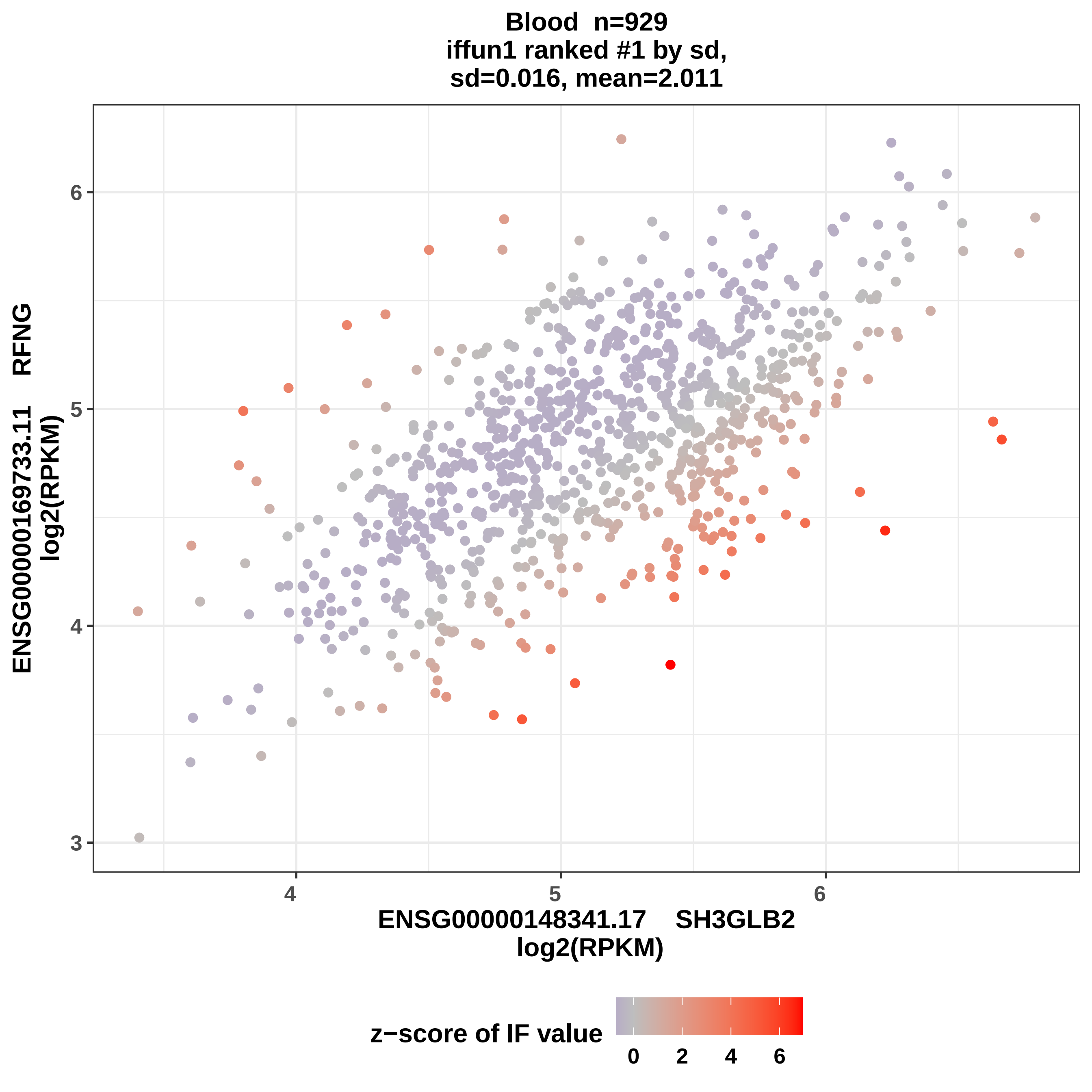

Blood n=929
iffun1 ranked #1 by sd,
sd=0.016, mean=2.011
ENSG00000169733.11 RFNG
log2(RPKM)
ENSG00000148341.17 SH3GLB2
log2(RPKM)
3
4
5
6
4
5
6
z−score of IF value
0
2
4
6

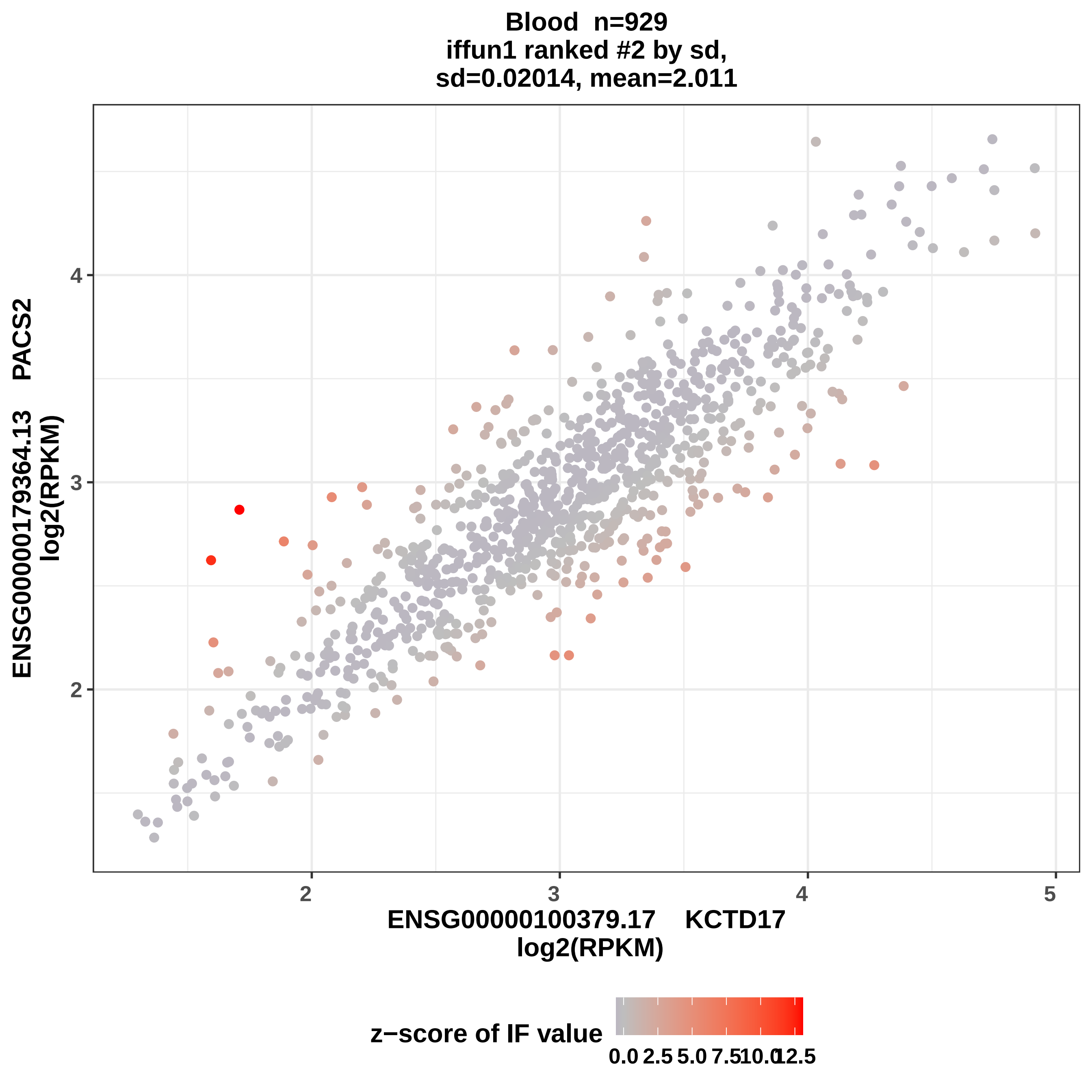
Blood n=929
iffun1 ranked #2 by sd,
sd=0.02014, mean=2.011
ENSG00000179364.13 PACS2
log2(RPKM)
ENSG00000100379.17 KCTD17
log2(RPKM)
2
3
4
5
z−score of IF value
0.0
2.5
5.0
7.5
10.0
12.5

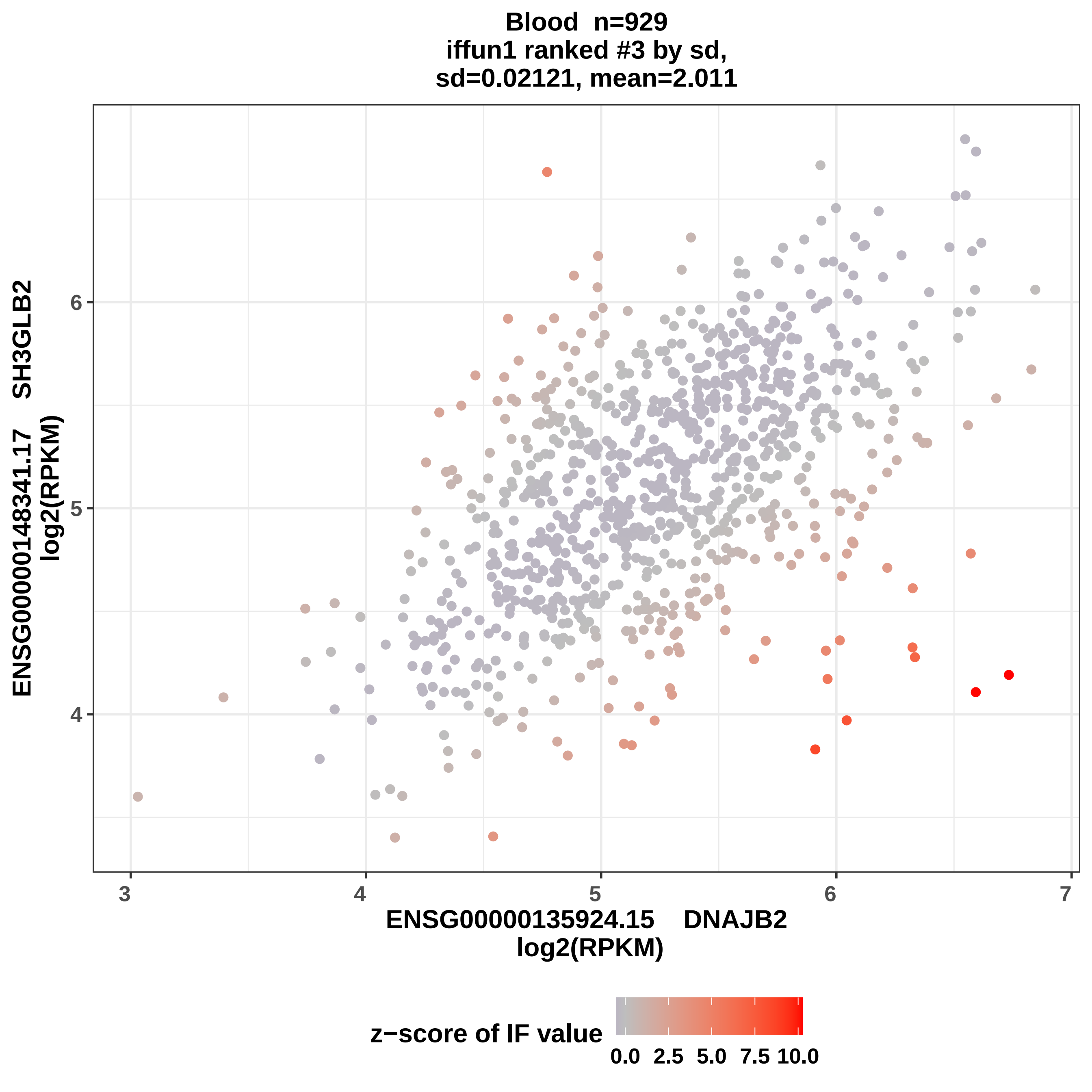

Blood n=929
iffun1 ranked #3 by sd,
sd=0.02121, mean=2.011
ENSG00000148341.17 SH3GLB2
log2(RPKM)
6
5
4
3
4
5
6
7
ENSG00000135924.15 DNAJB2
log2(RPKM)
z−score of IF value
0.0
2.5
5.0
7.5
10.0

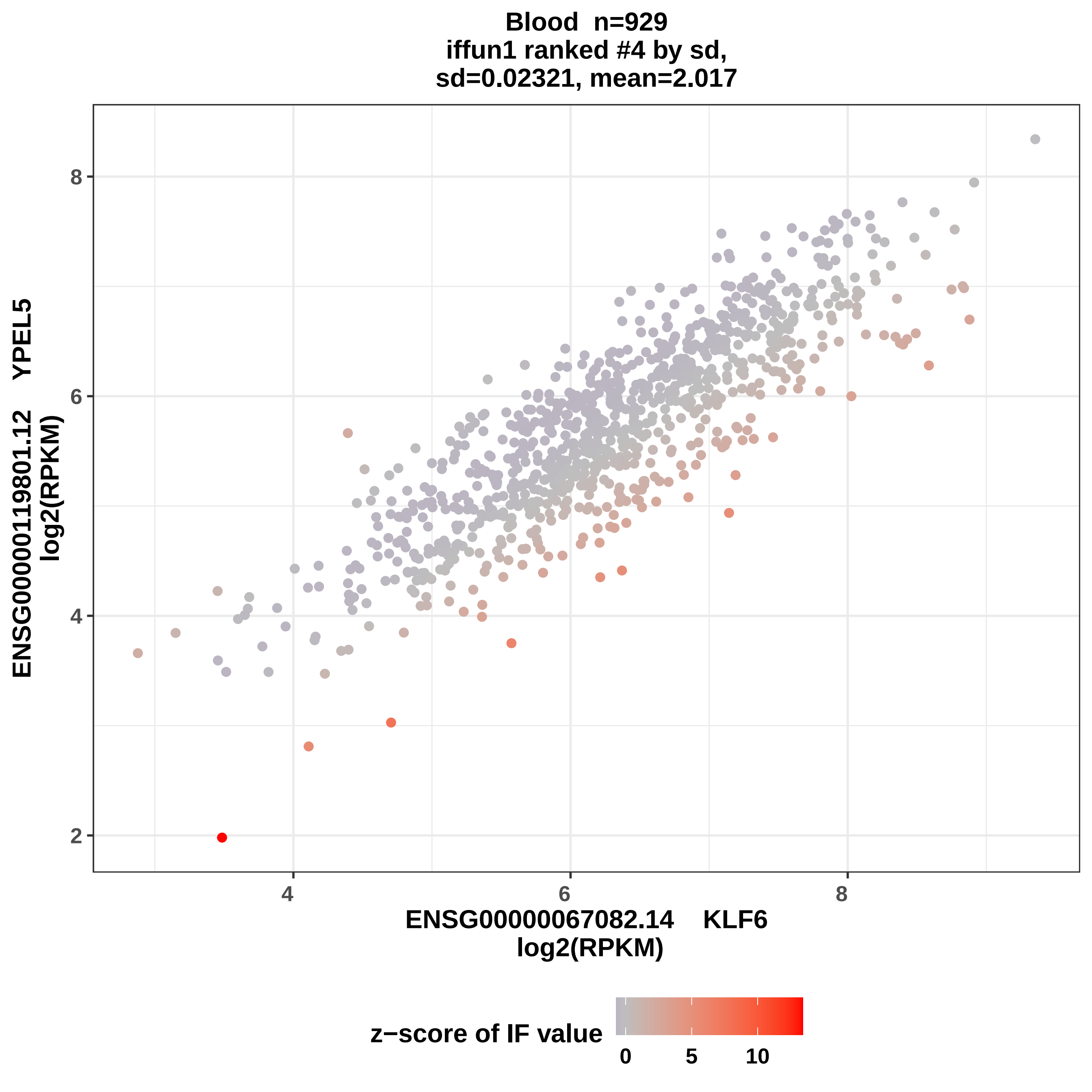
Blood n=929
iffun1 ranked #4 by sd,
sd=0.02321, mean=2.017
ENSG00000119801.12 YPEL5
log2(RPKM)
ENSG00000067082.14 KLF6
log2(RPKM)
2
4
6
8
4
6
8
z−score of IF value
0
5
10

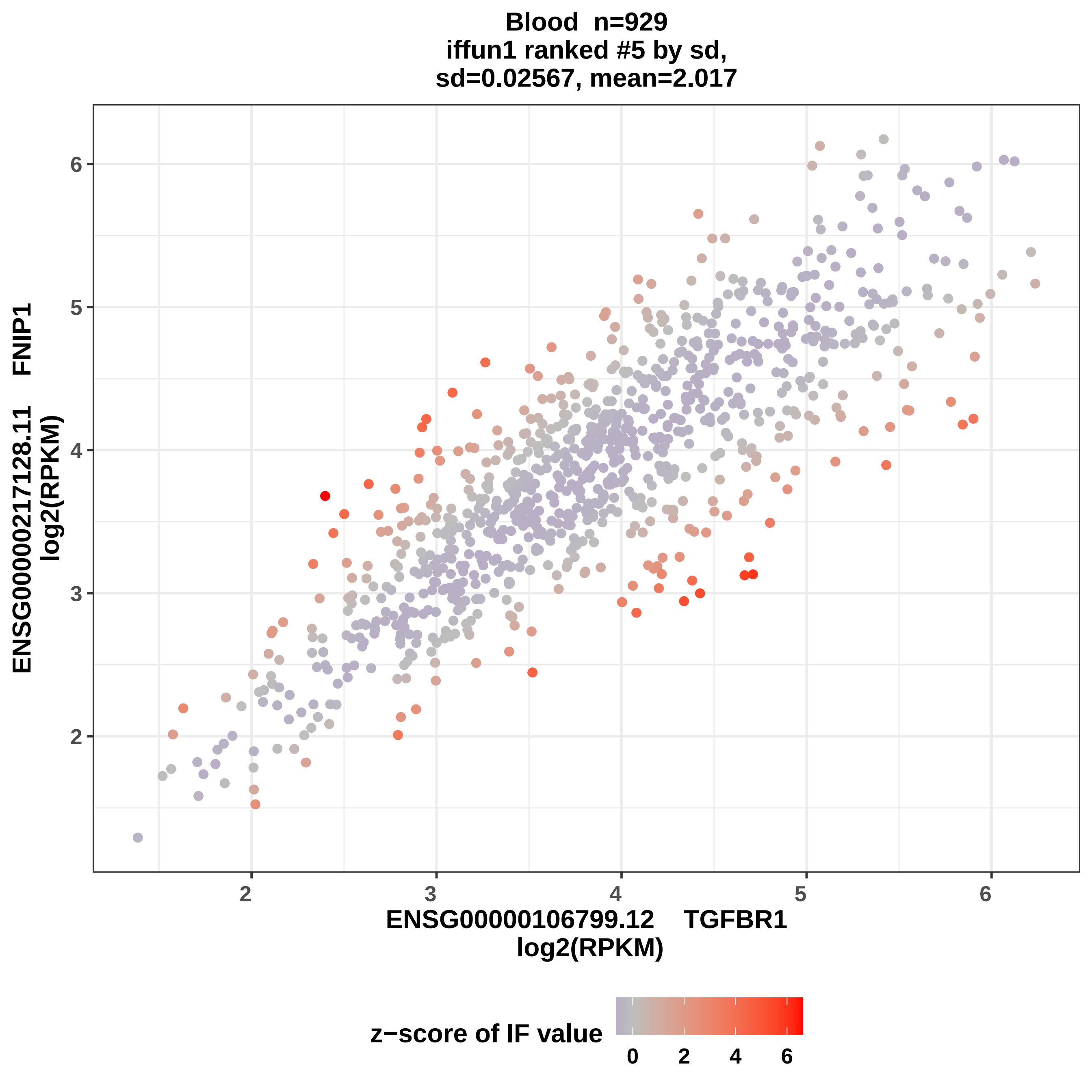

Blood n=929
iffun1 ranked #5 by sd,
sd=0.02567, mean=2.017
ENSG00000217128.11 FNIP1
log2(RPKM)
ENSG00000106799.12 TGFBR1
log2(RPKM)
2
3
4
5
6
z−score of IF value
0
2
4
6

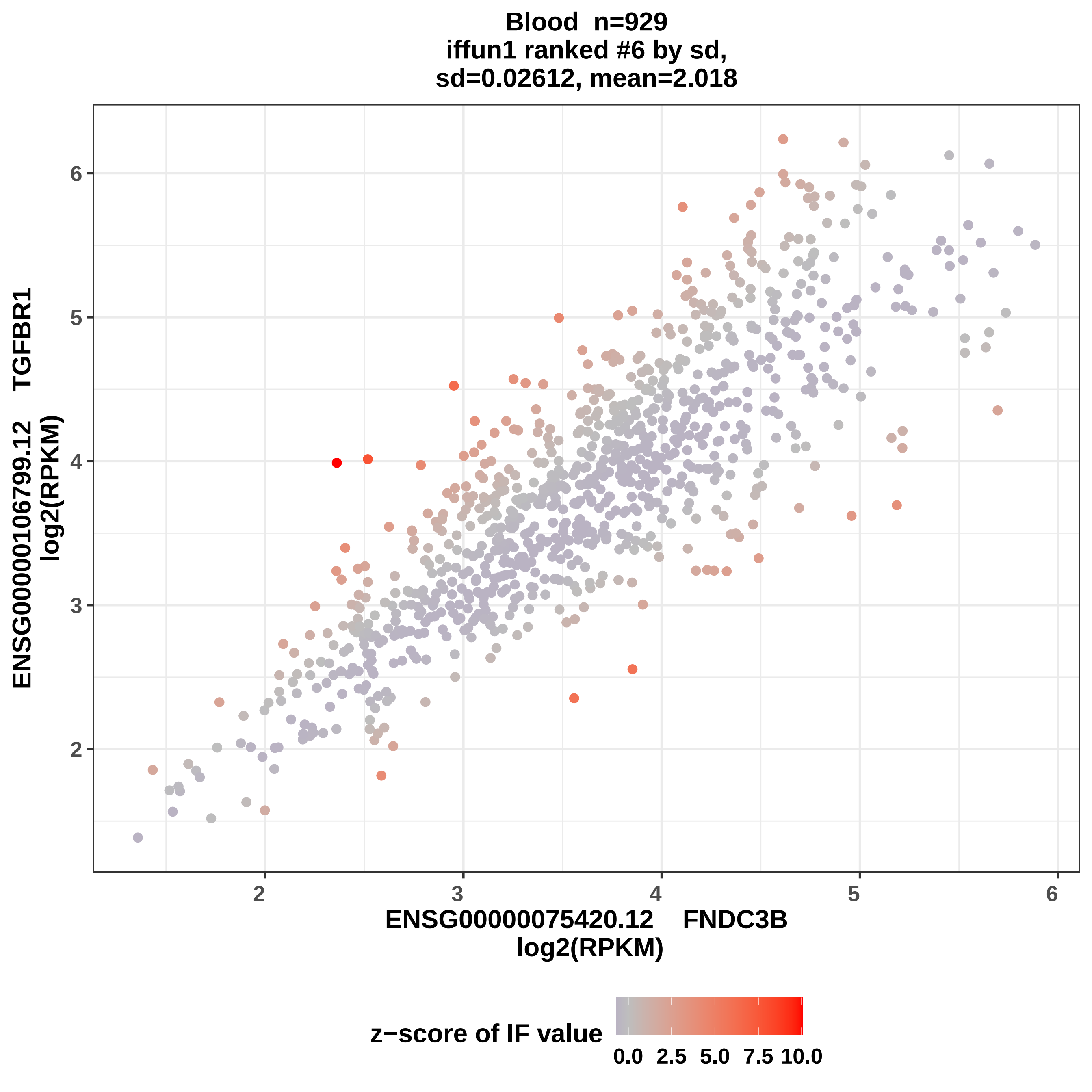
Blood n=929
iffun1 ranked #6 by sd,
sd=0.02612, mean=2.018
ENSG00000106799.12 TGFBR1
log2(RPKM)
ENSG00000075420.12 FNDC3B
log2(RPKM)
2
3
4
5
6
z−score of IF value
0.0
2.5
5.0
7.5
10.0

**Blood n=929**
**iffun1 ranked #7 by sd,**
**sd=0.02627, mean= 2.02**

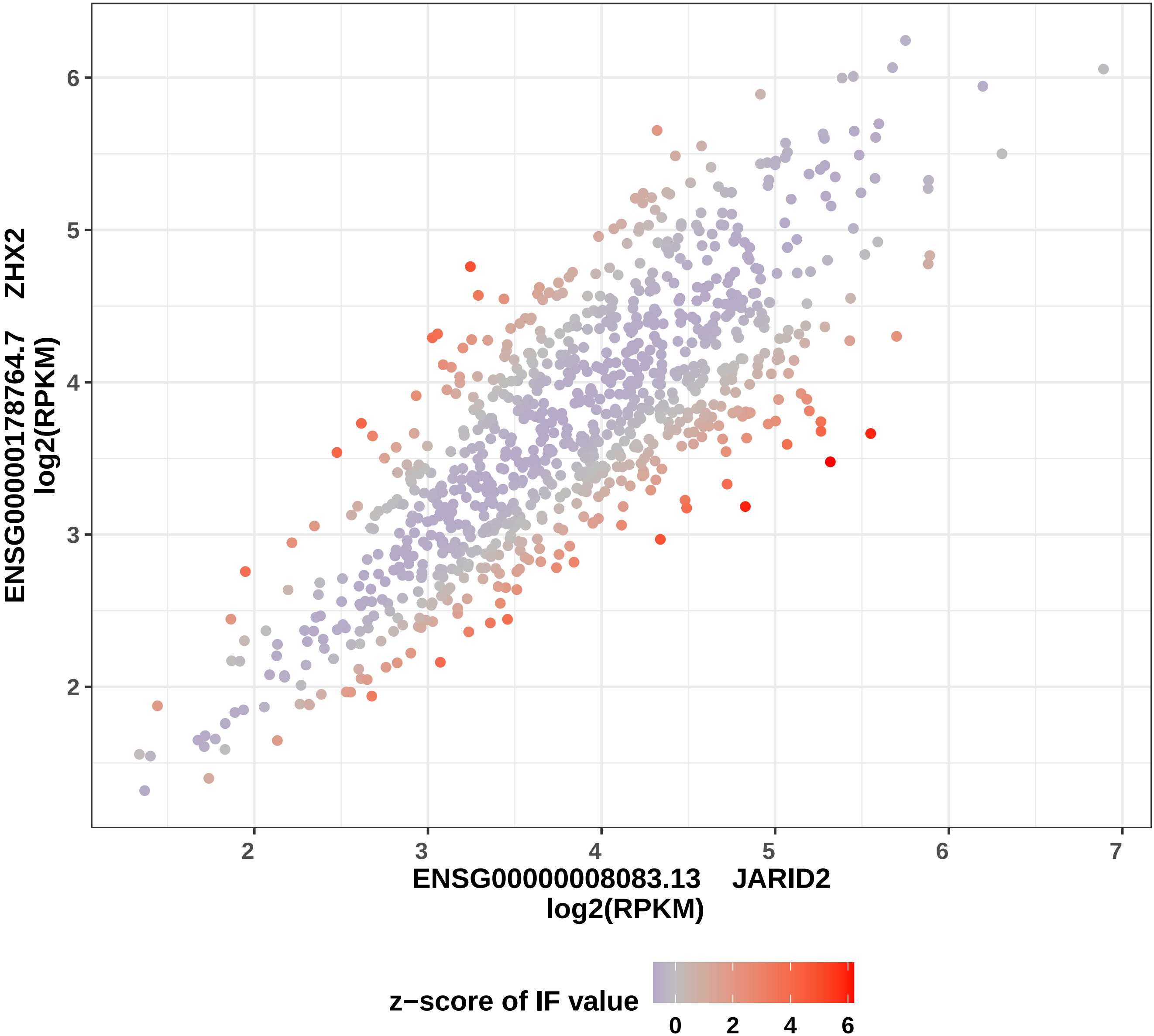

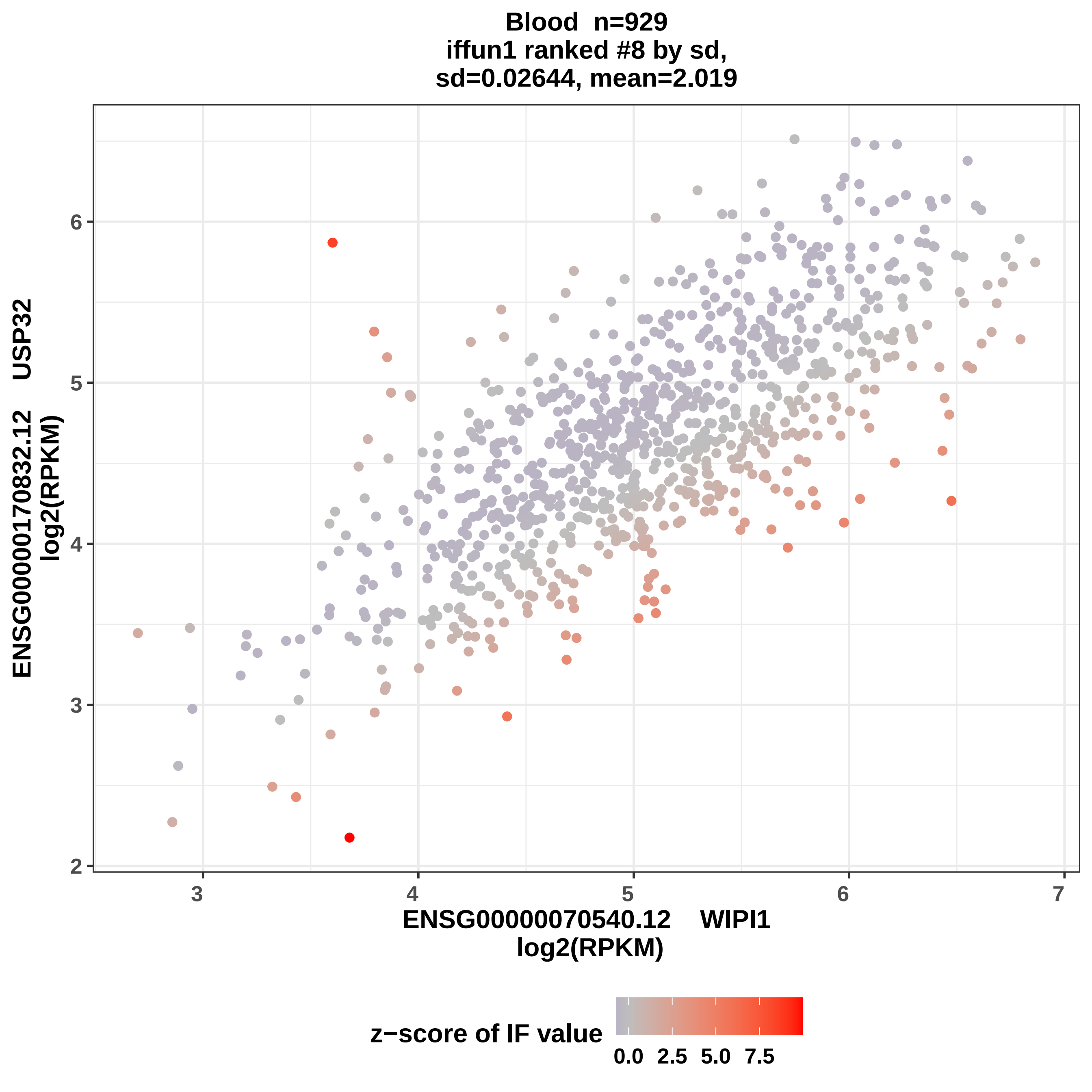
Blood n=929
iffun1 ranked #8 by sd,
sd=0.02644, mean=2.019
ENSG00000170832.12 USP32
log2(RPKM)
ENSG00000070540.12 WIPI1
log2(RPKM)
2
3
4
5
6
3
4
5
6
7
z−score of IF value
0.0
2.5
5.0
7.5

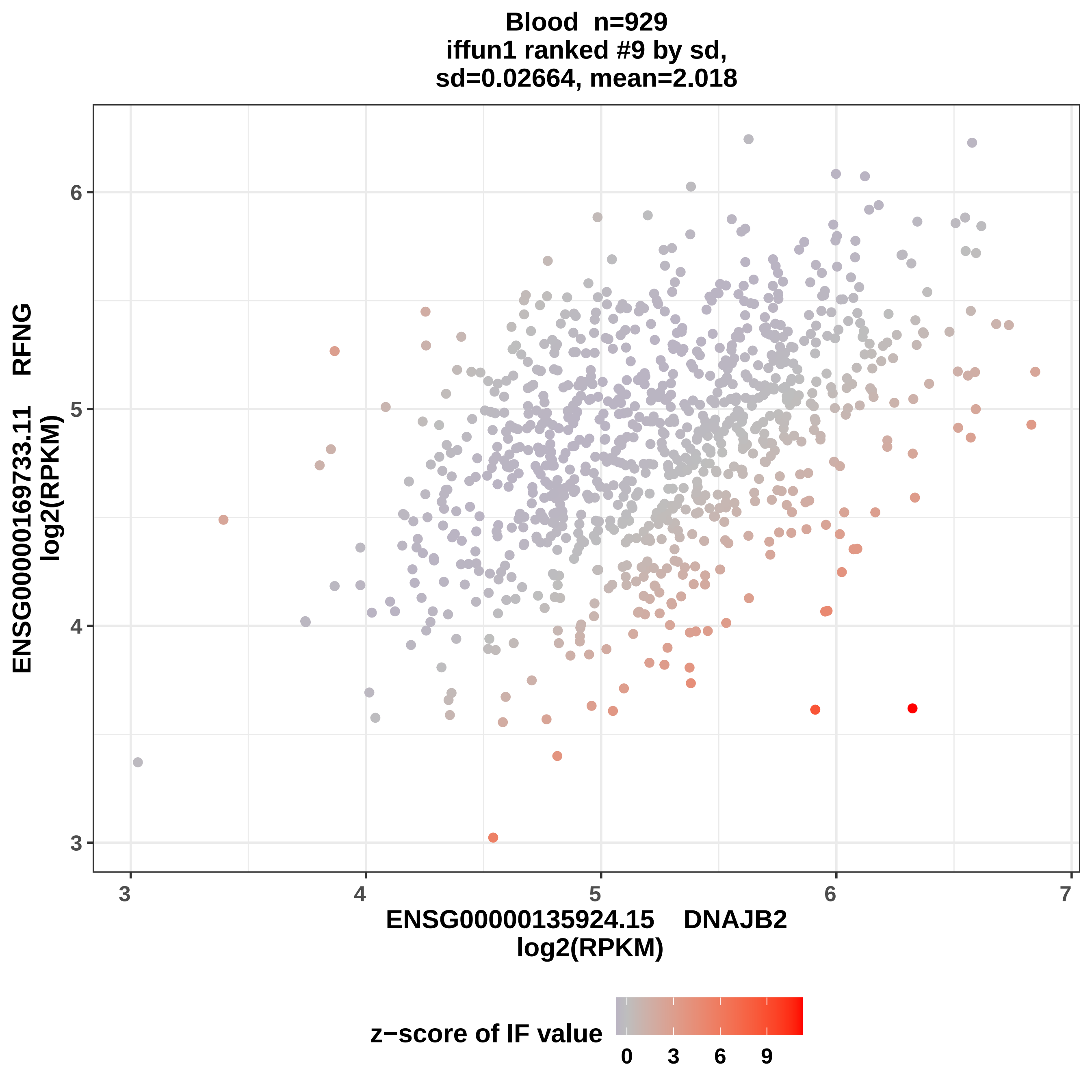
Blood n=929
iffun1 ranked #9 by sd,
sd=0.02664, mean=2.018
ENSG00000169733.11 RFNG
log2(RPKM)
ENSG00000135924.15 DNAJB2
log2(RPKM)
3
4
5
6
7
z−score of IF value
0
3
6
9

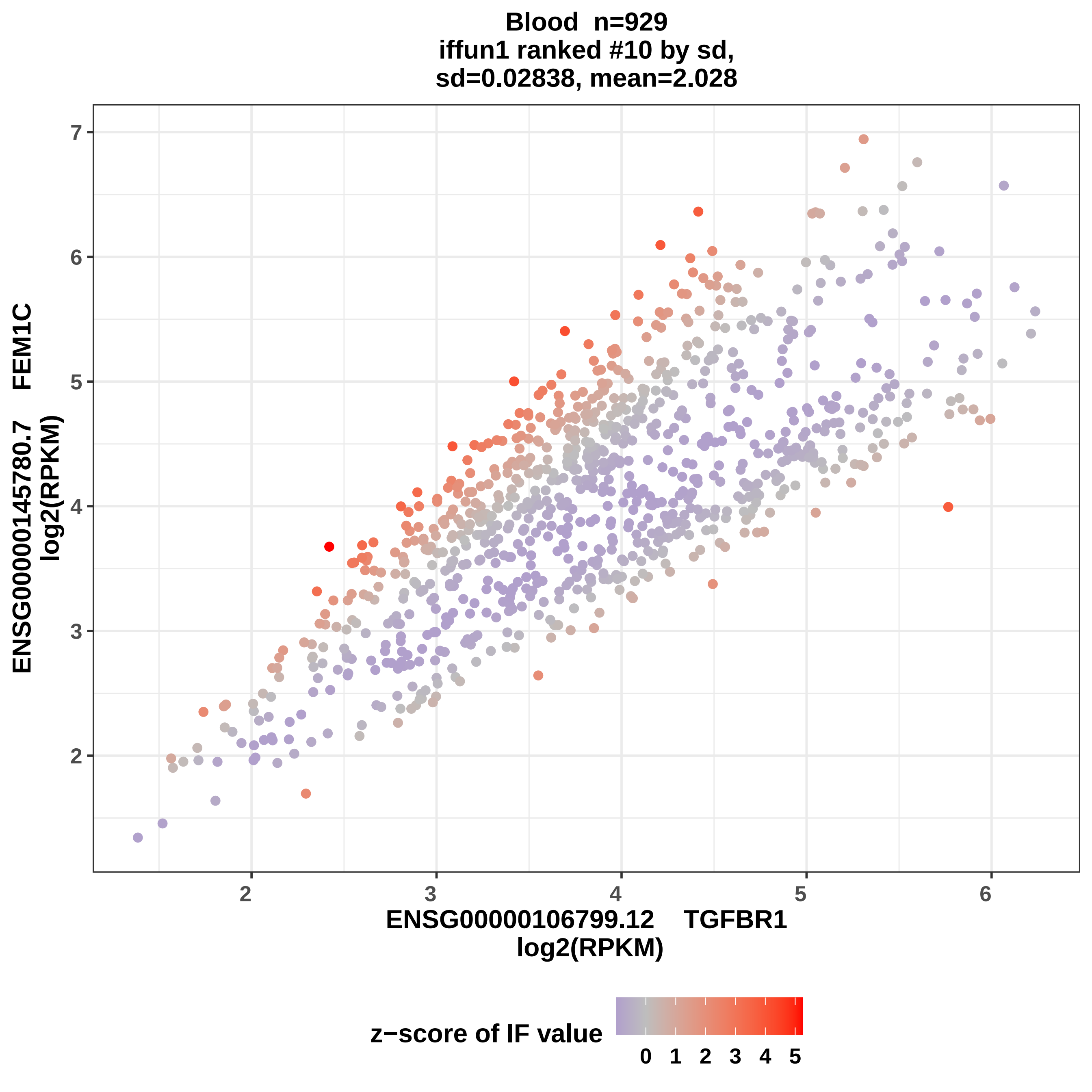

Blood n=929
iffun1 ranked #10 by sd,
sd=0.02838, mean=2.028
ENSG00000145780.7 FEM1C
log2(RPKM)
ENSG00000106799.12 TGFBR1
log2(RPKM)
2
3
4
5
6
7
z−score of IF value
0 1 2 3 4 5

**Blood n=929**
**iffun1 ranked #11 by sd,**
**sd=0.02915, mean=2.028**

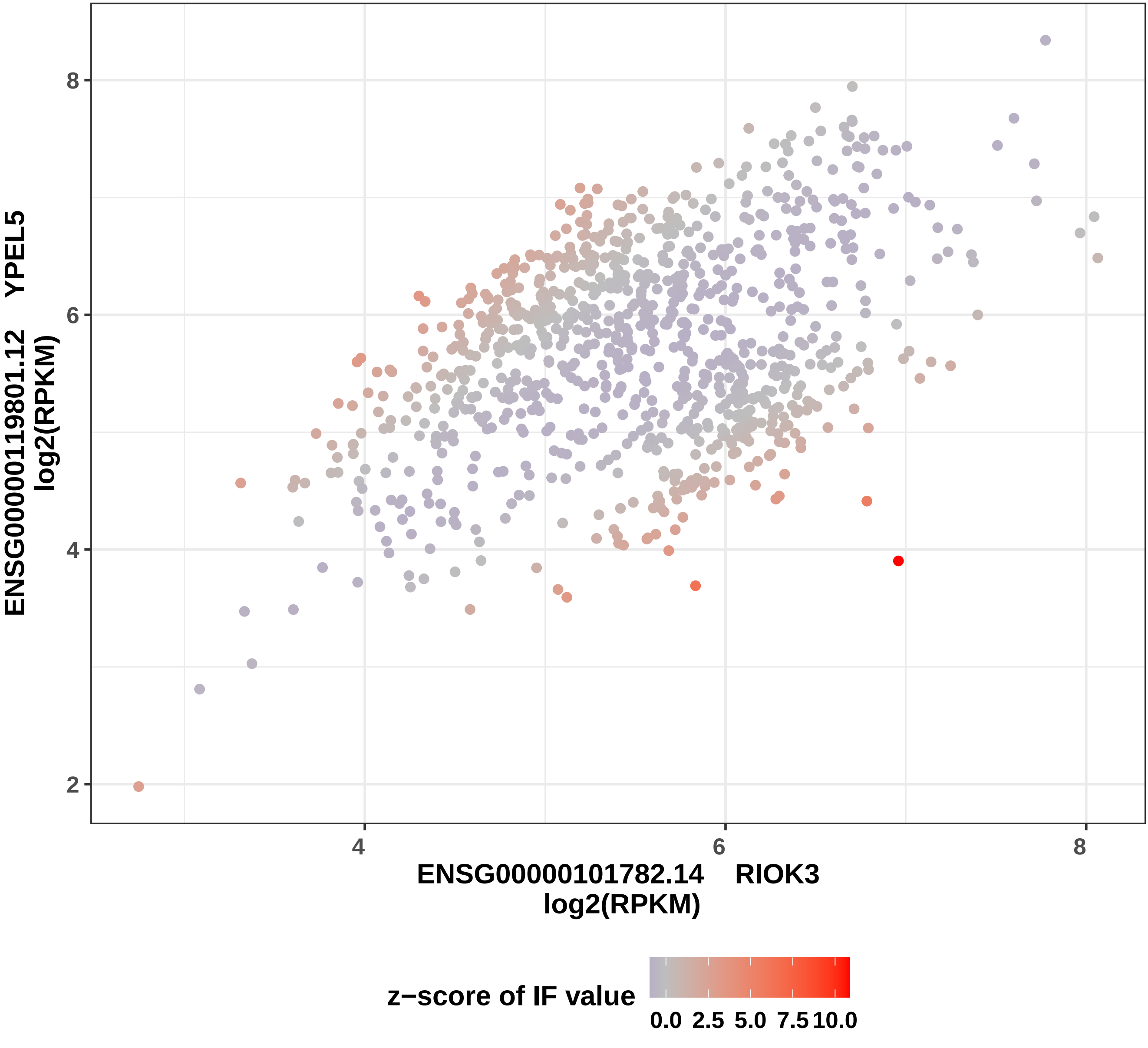

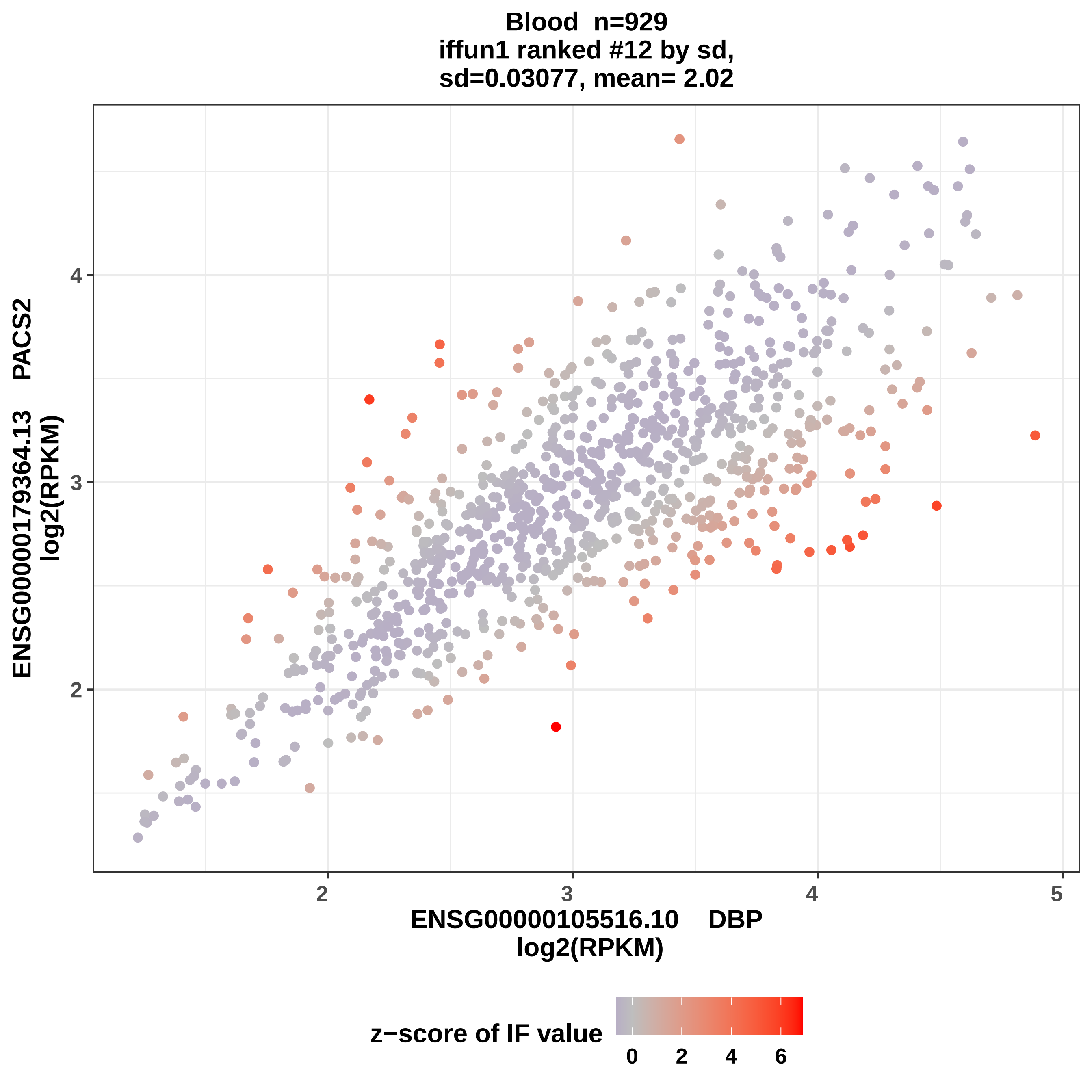
Blood n=929
iffun1 ranked #12 by sd,
sd=0.03077, mean= 2.02
ENSG00000179364.13 PACS2
log2(RPKM)
ENSG00000105516.10 DBP
log2(RPKM)
2
3
4
2
3
4
5
z−score of IF value
0
2
4
6

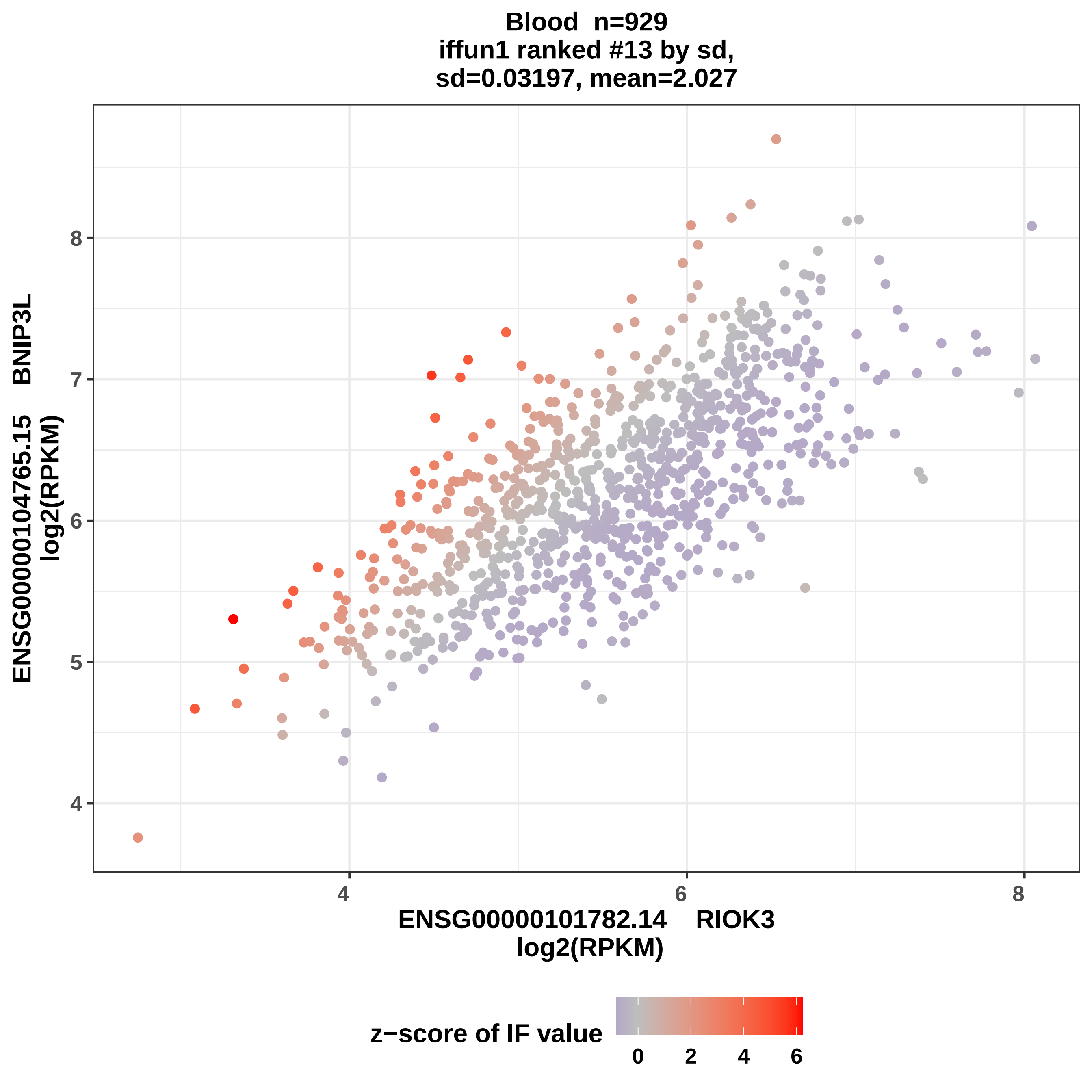

Blood n=929
iffun1 ranked #13 by sd,
sd=0.03197, mean=2.027
ENSG00000104765.15 BNIP3L
log2(RPKM)
ENSG00000101782.14 RIOK3
log2(RPKM)
4
5
6
7
8
4
6
8
z−score of IF value
0
2
4
6

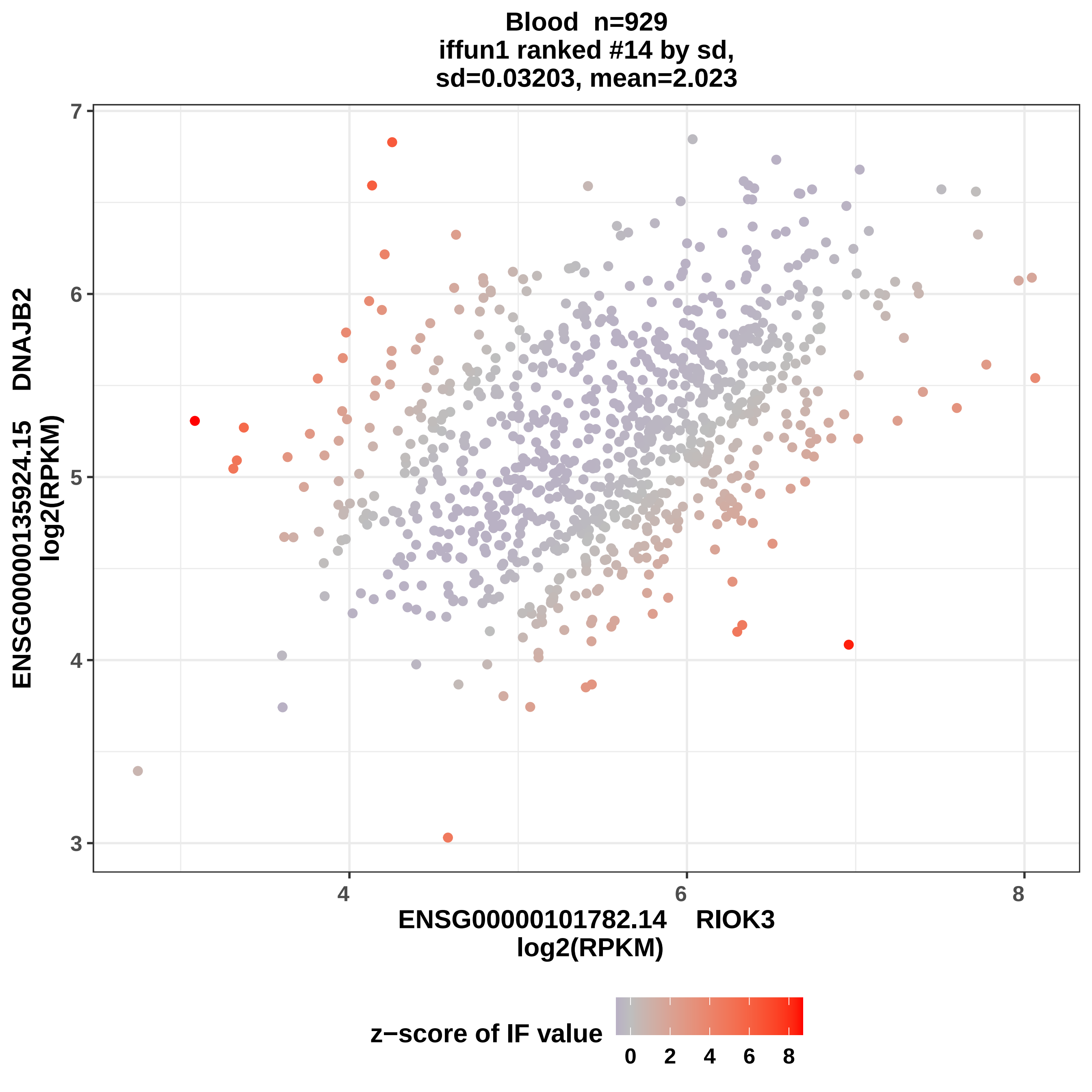
Blood n=929
iffun1 ranked #14 by sd,
sd=0.03203, mean=2.023
ENSG00000135924.15 DNAJB2
log2(RPKM)
ENSG00000101782.14 RIOK3
log2(RPKM)
z−score of IF value
3
4
5
6
7
4
6
8
0
2
4
6
8

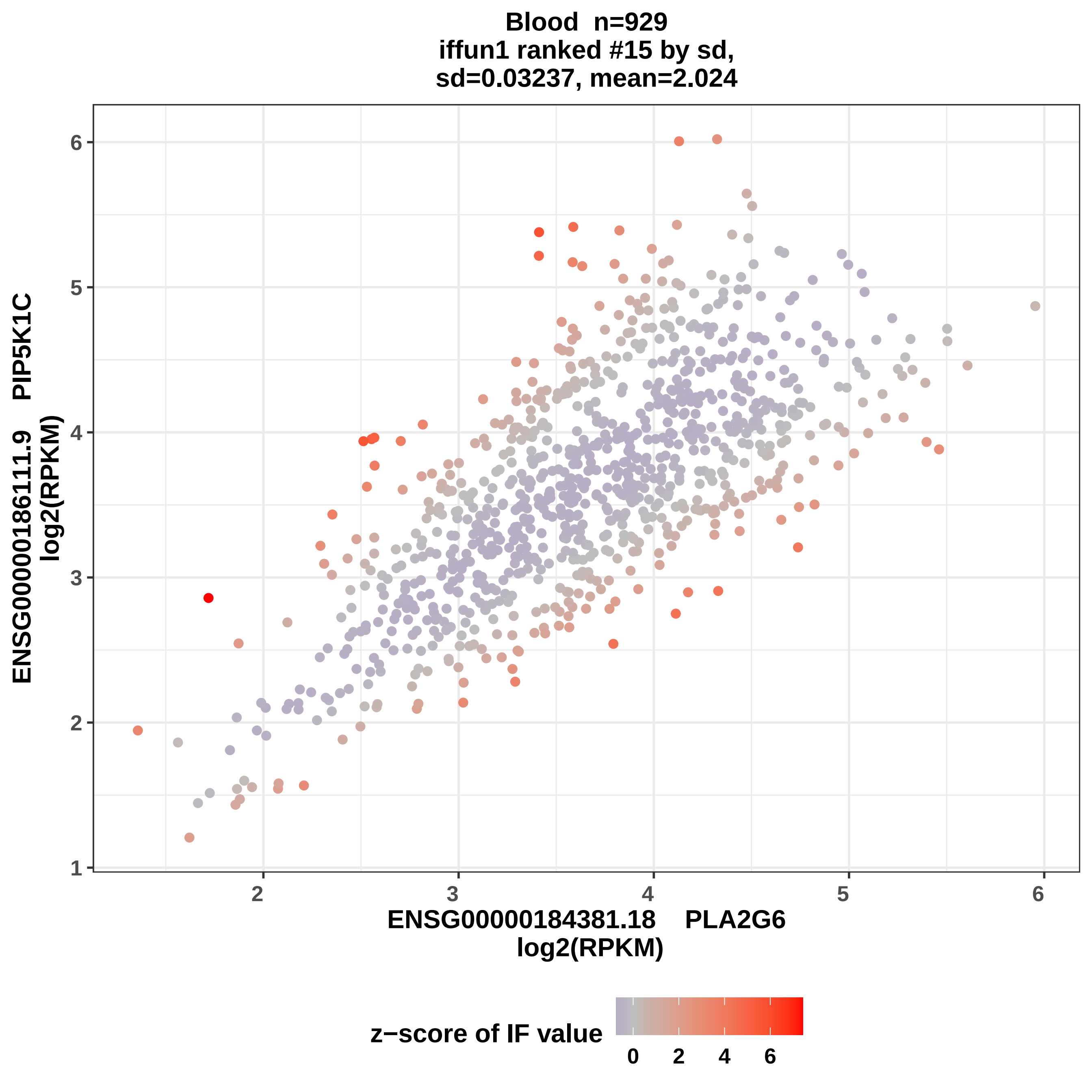
Blood n=929
iffun1 ranked #15 by sd,
sd=0.03237, mean=2.024
ENSG00000186111.9 PIP5K1C
log2(RPKM)
ENSG00000184381.18 PLA2G6
log2(RPKM)
z−score of IF value
1
2
3
4
5
6
0
2
4
6

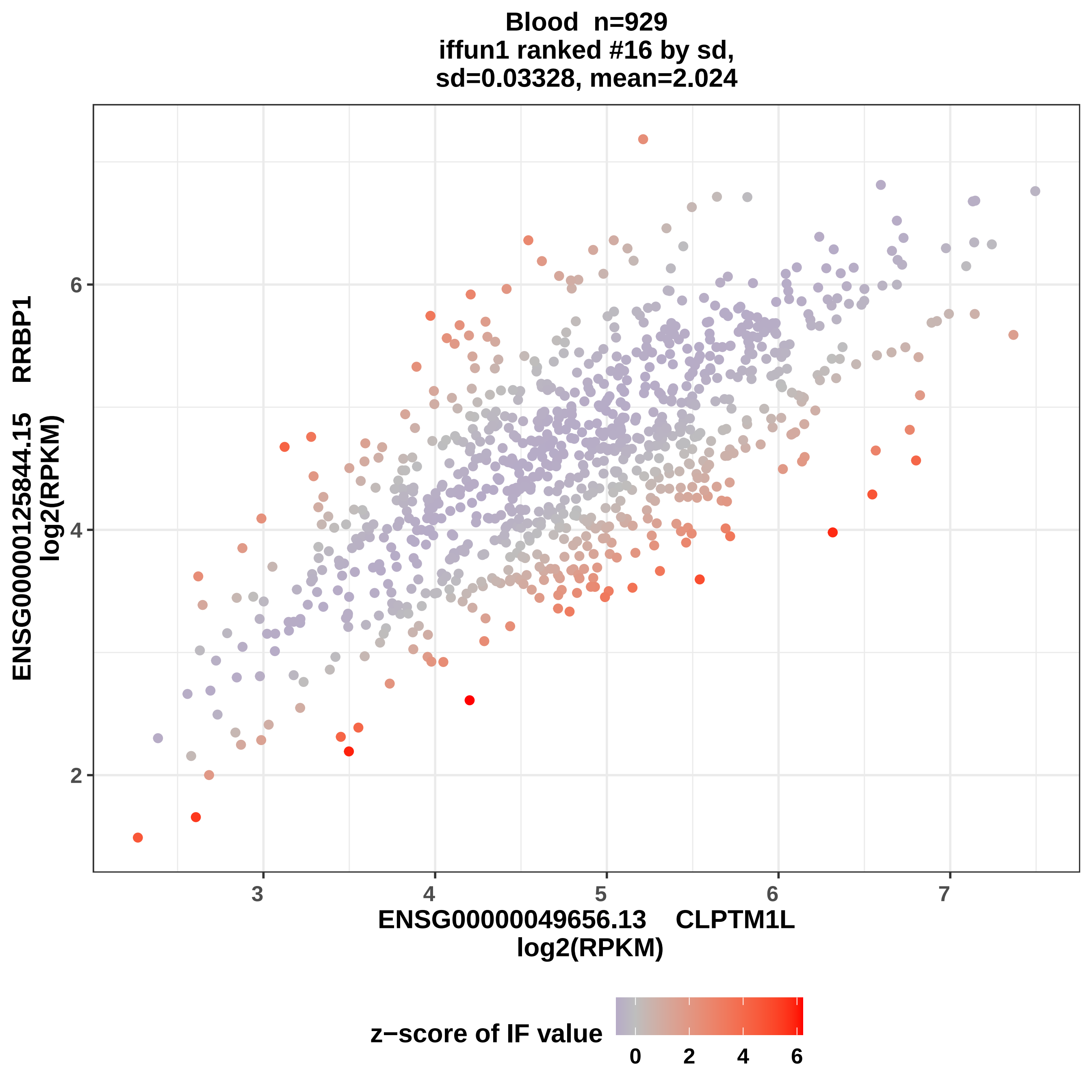

Blood n=929
iffun1 ranked #16 by sd,
sd=0.03328, mean=2.024
ENSG00000125844.15 RRBP1
log2(RPKM)
2
4
6
3
4
5
6
7
ENSG00000049656.13 CLPTM1L
log2(RPKM)
z−score of IF value
0
2
4
6

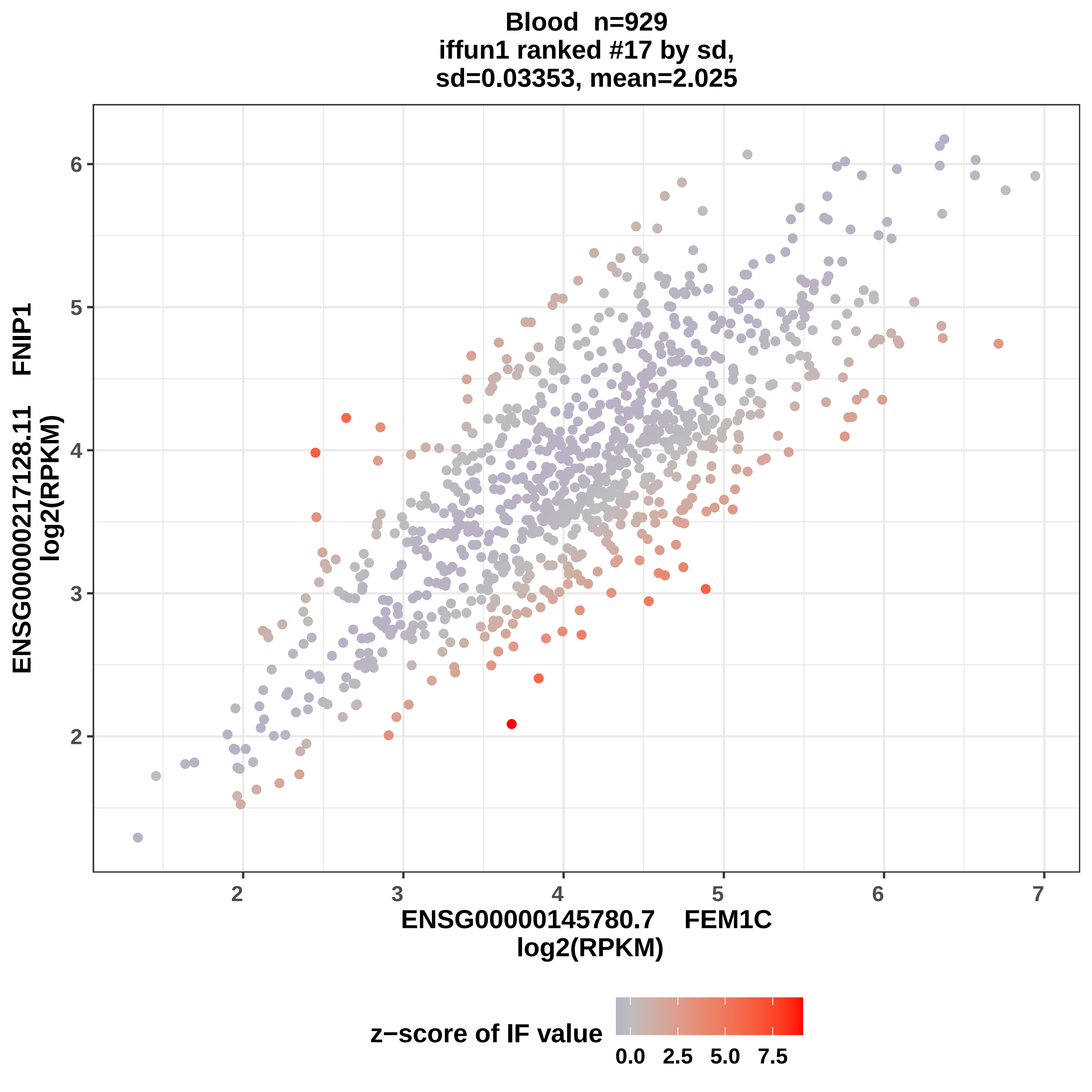

Blood n=929
iffun1 ranked #17 by sd,
sd=0.03353, mean=2.025
ENSG00000217128.11 FNIP1
log2(RPKM)
ENSG00000145780.7 FEM1C
log2(RPKM)
2
3
4
5
6
7
z−score of IF value
0.0
2.5
5.0
7.5

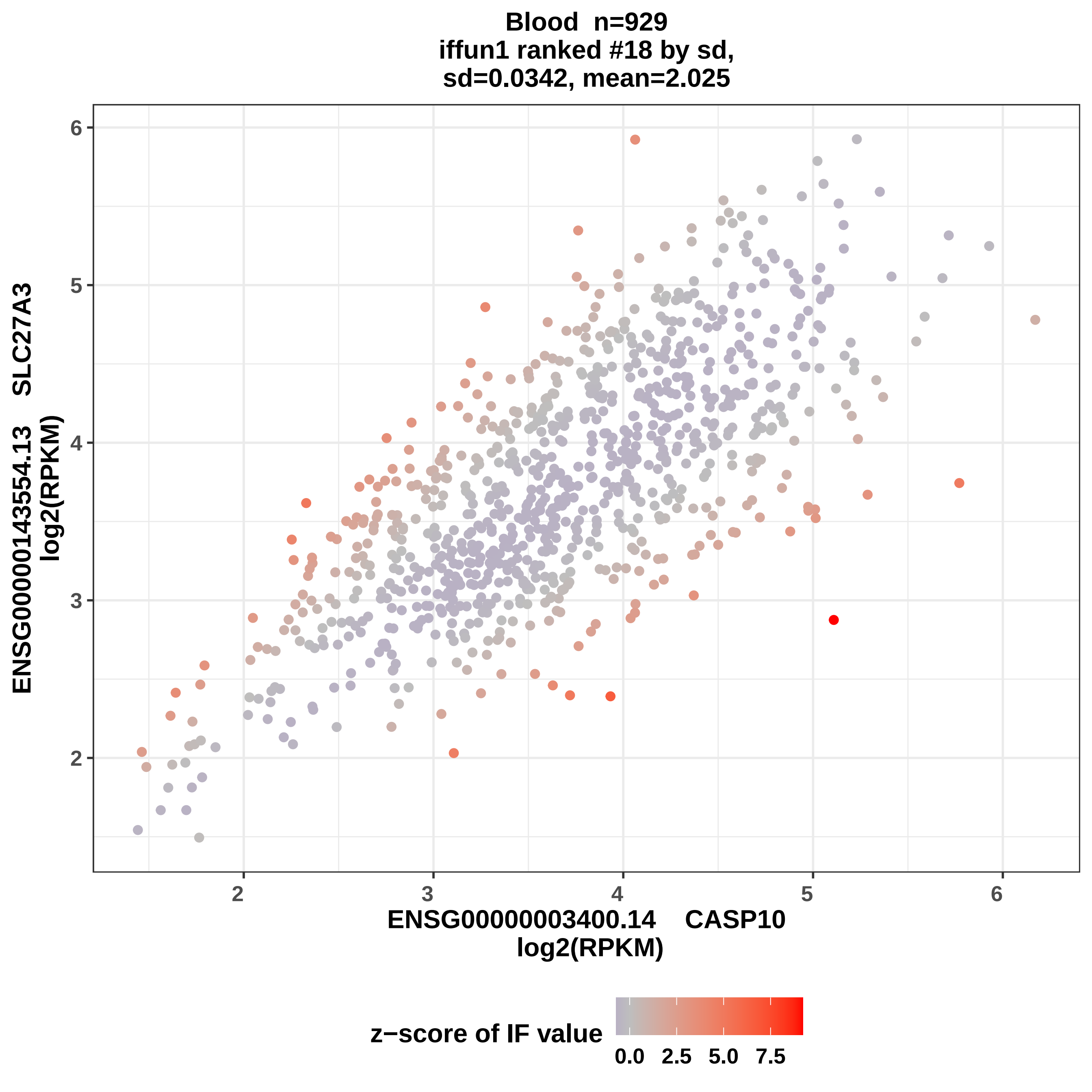

Blood n=929
iffun1 ranked #18 by sd,
sd=0.0342, mean=2.025
ENSG00000143554.13 SLC27A3
log2(RPKM)
ENSG00000003400.14 CASP10
log2(RPKM)
2
3
4
5
6
2
3
4
5
6
z−score of IF value
0.0
2.5
5.0
7.5

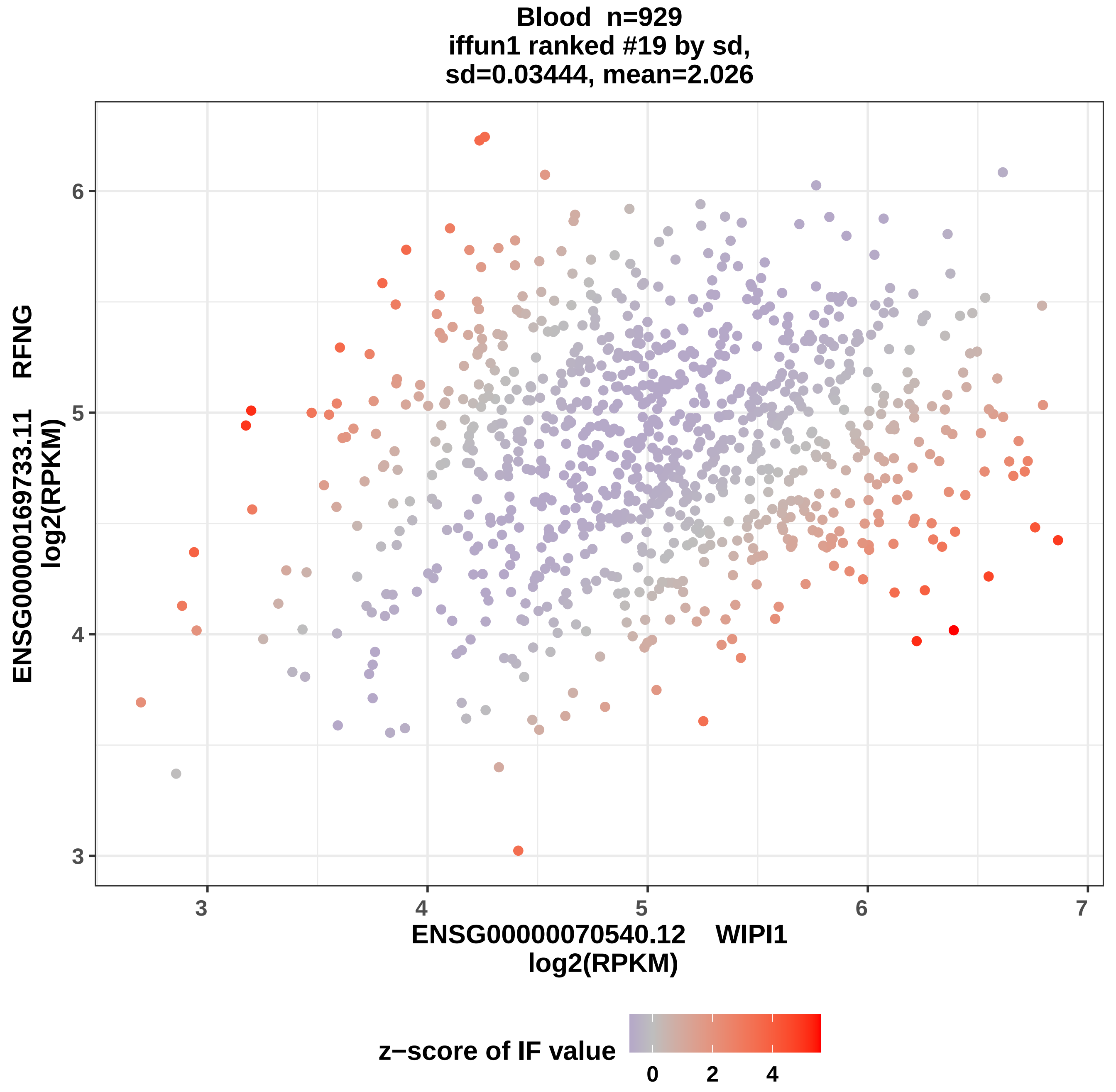

Blood n=929
iffun1 ranked #19 by sd,
sd=0.03444, mean=2.026
ENSG00000169733.11 RFNG
log2(RPKM)
ENSG00000070540.12 WIPI1
log2(RPKM)
3
4
5
6
7
z−score of IF value
0
2
4

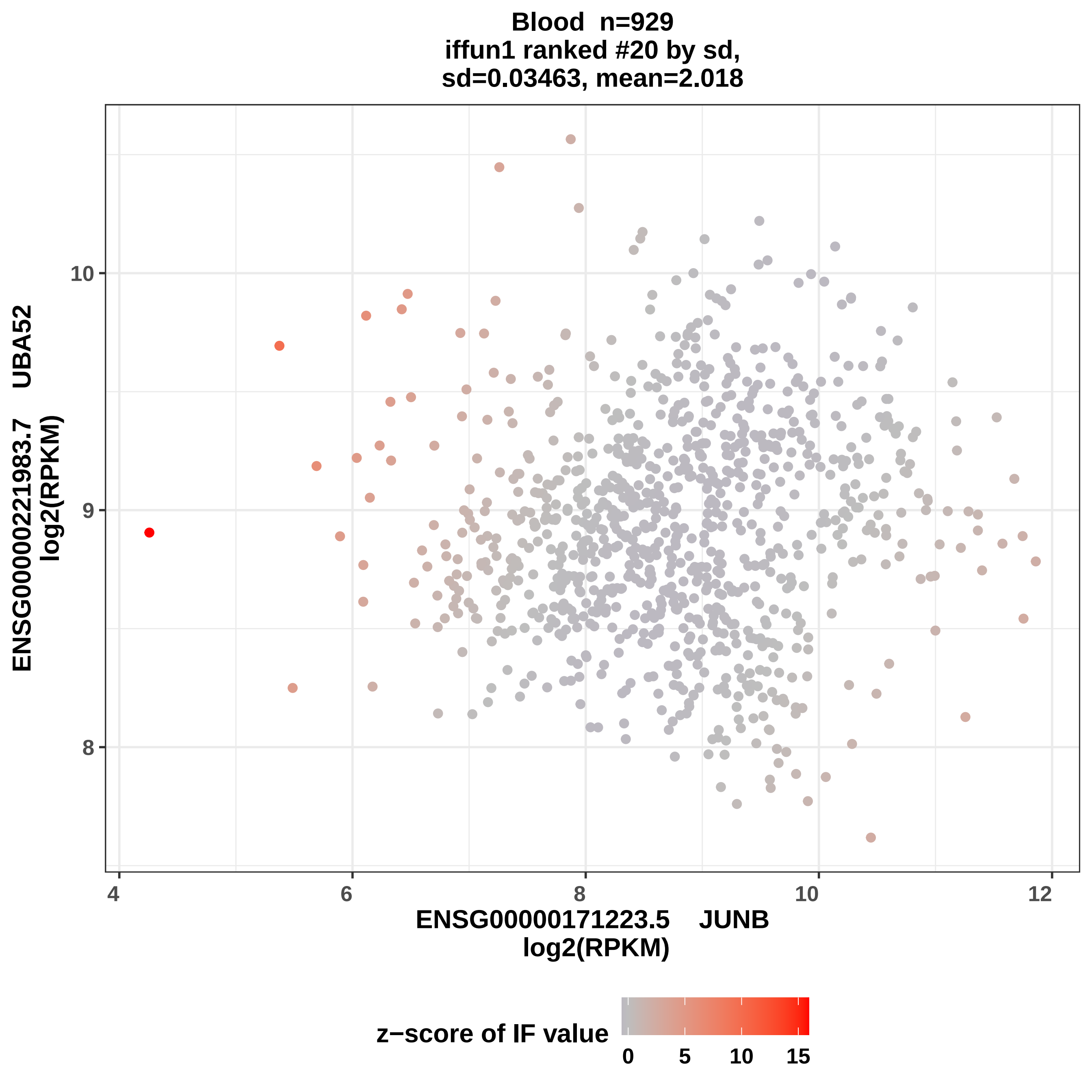
Blood n=929
iffun1 ranked #20 by sd,
sd=0.03463, mean=2.018
ENSG00000221983.7 UBA52
log2(RPKM)
ENSG00000171223.5 JUNB
log2(RPKM)
z−score of IF value
0
5
10
15

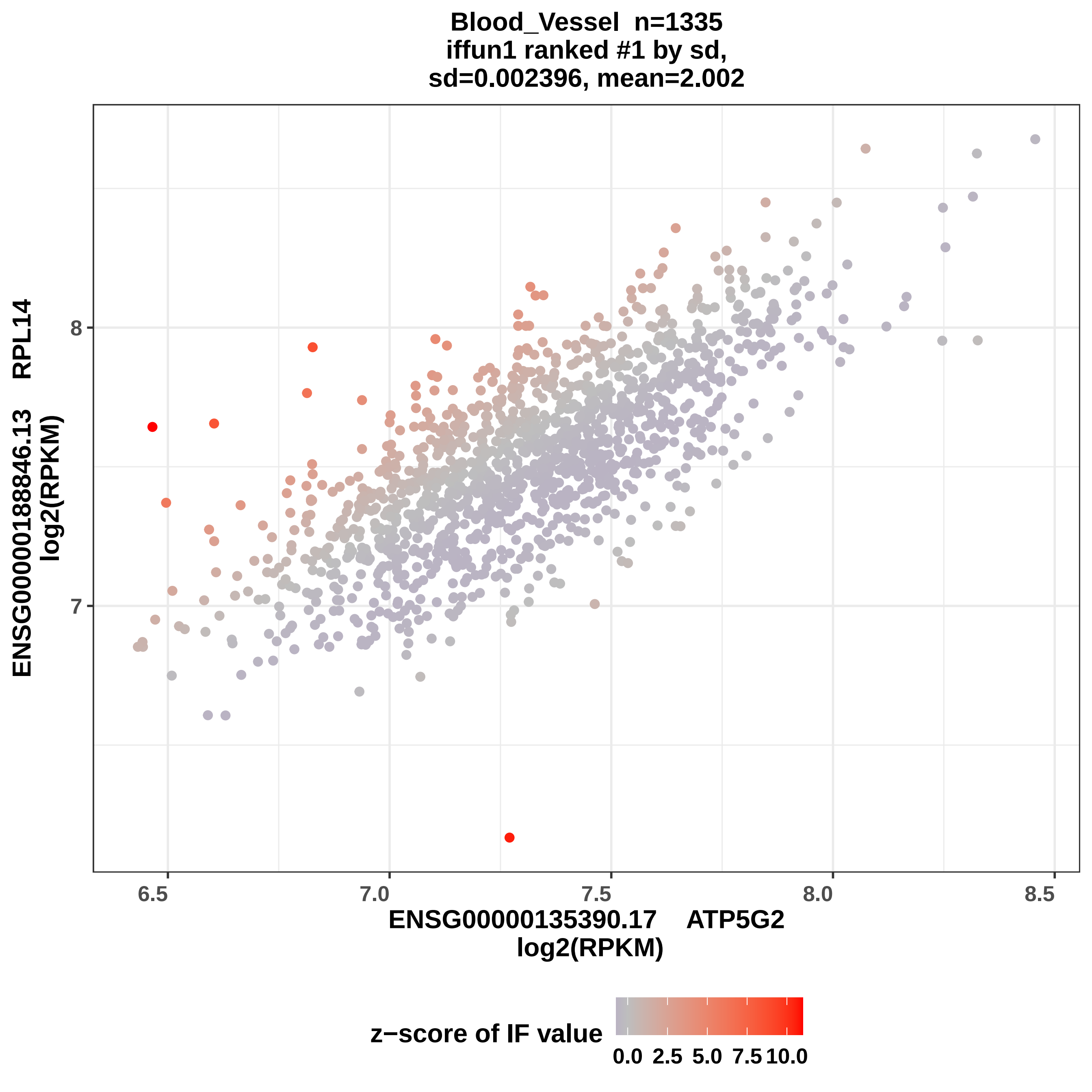
Blood_Vessel n=1335
iffun1 ranked #1 by sd,
sd=0.002396, mean=2.002
ENSG00000188846.13 RPL14
log2(RPKM)
8
7
6.5
7.0
7.5
8.0
8.5
ENSG00000135390.17 ATP5G2
log2(RPKM)
z−score of IF value
0.0
2.5
5.0
7.5
10.0

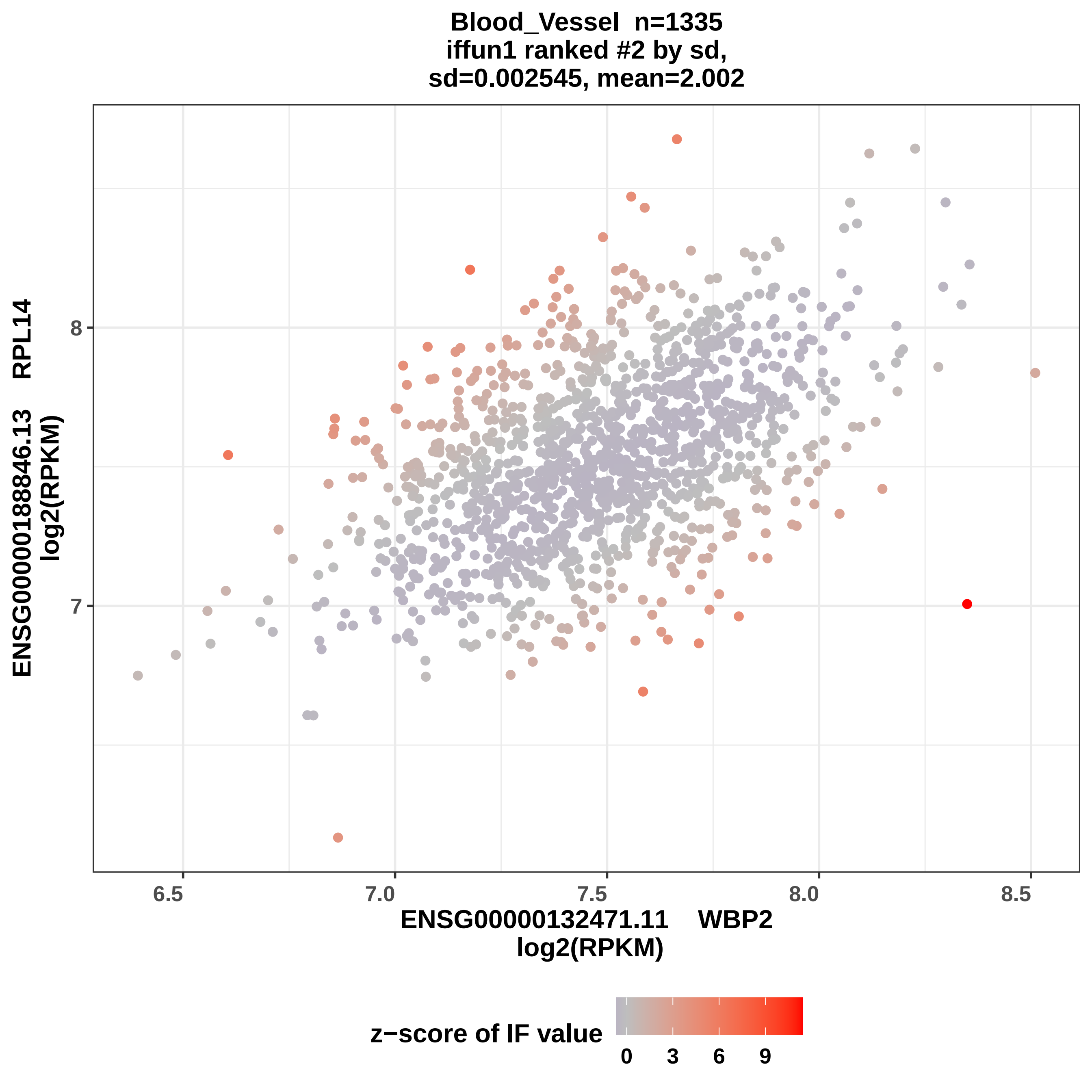

Blood_Vessel n=1335
iffun1 ranked #2 by sd,
sd=0.002545, mean=2.002
ENSG00000188846.13 RPL14
log2(RPKM)
8
7
6.5
7.0
7.5
8.0
8.5
ENSG00000132471.11 WBP2
log2(RPKM)
z−score of IF value
0
3
6
9

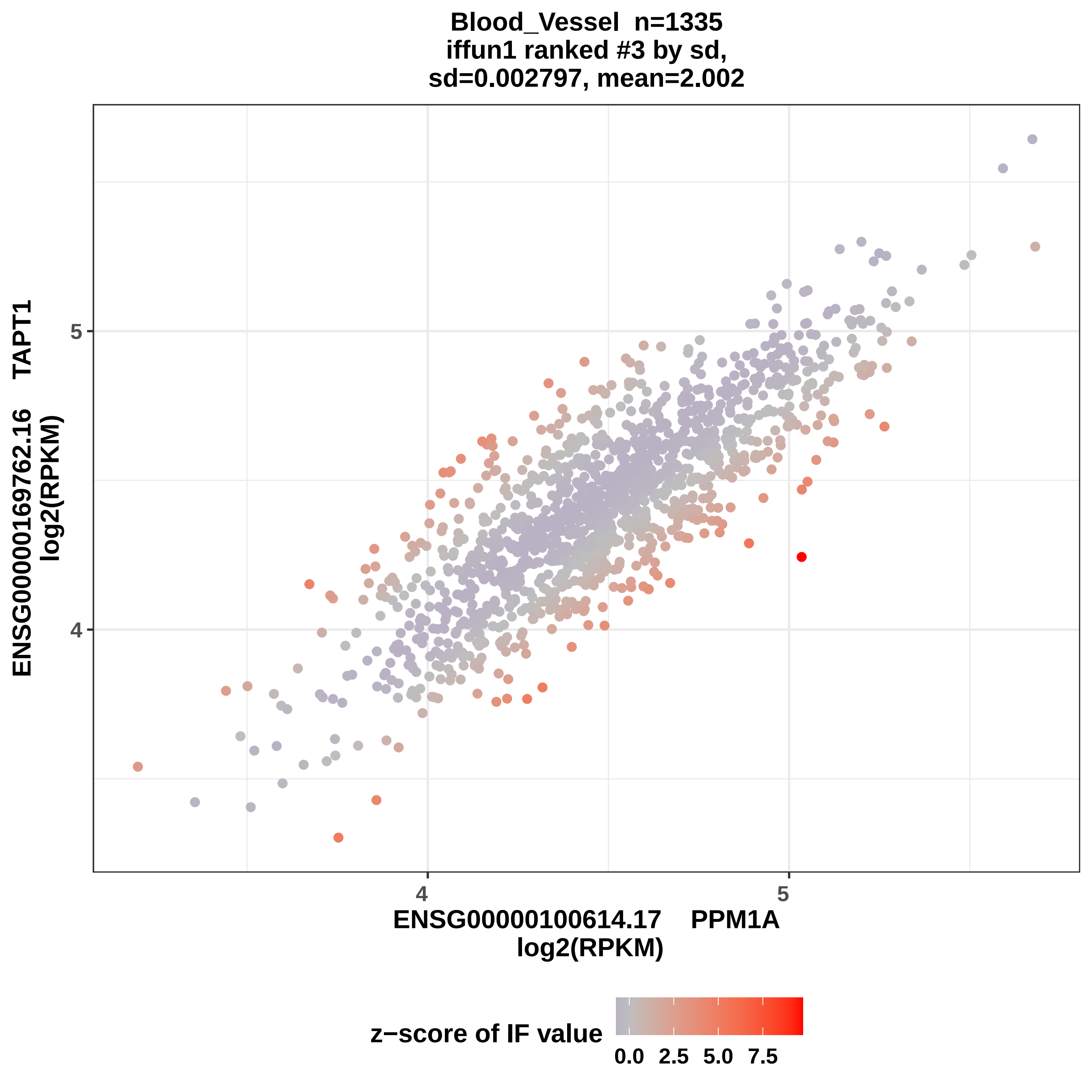

Blood_Vessel n=1335
iffun1 ranked #3 by sd,
sd=0.002797, mean=2.002
ENSG00000169762.16 TAPT1
log2(RPKM)
5
4
4
5
ENSG00000100614.17 PPM1A
log2(RPKM)
z−score of IF value
0.0
2.5
5.0
7.5

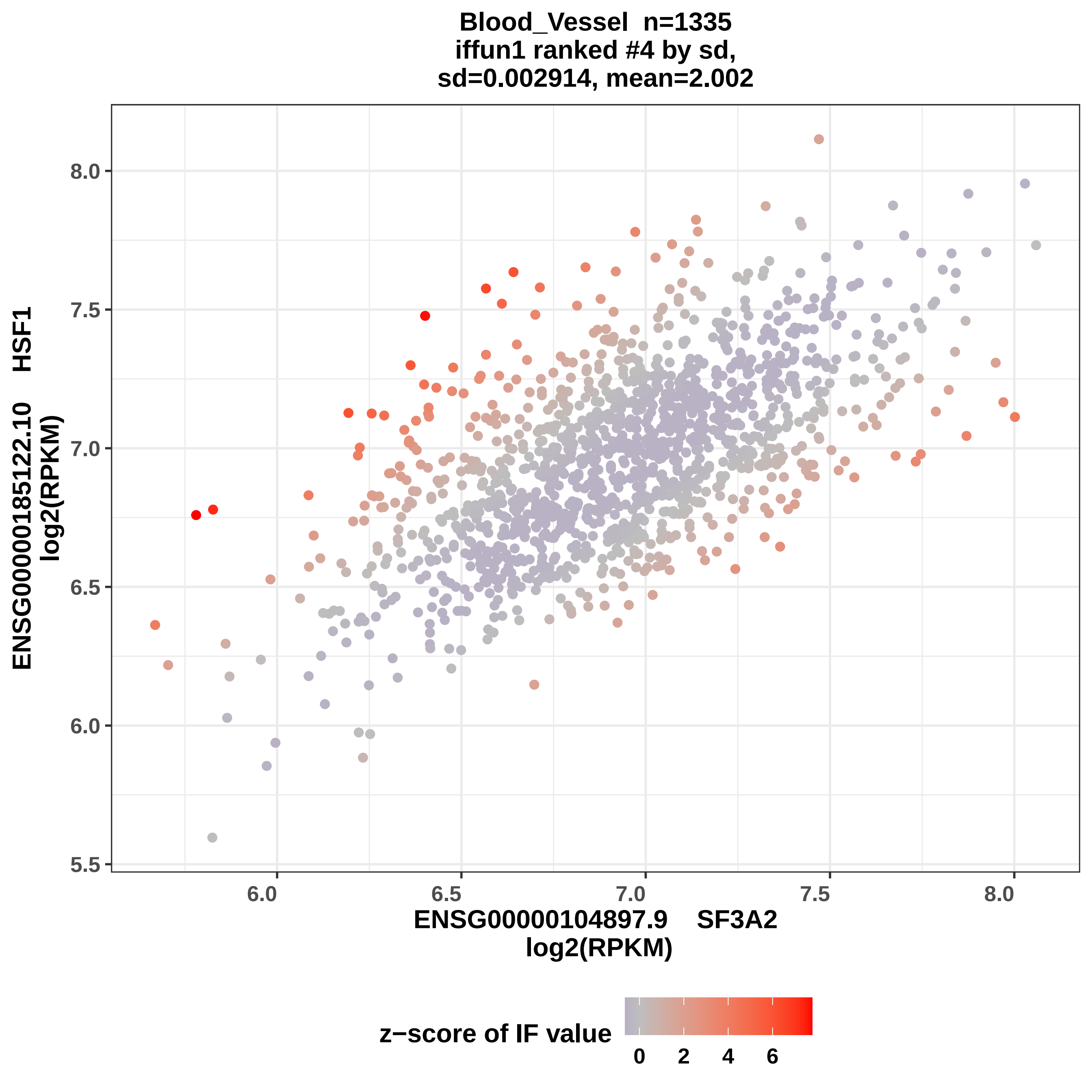

Blood_Vessel n=1335
iffun1 ranked #4 by sd,
sd=0.002914, mean=2.002
ENSG00000185122.10 HSF1
log2(RPKM)
ENSG00000104897.9 SF3A2
log2(RPKM)
8.0
7.5
7.0
6.5
6.0
5.5
6.0
6.5
7.0
7.5
8.0
z−score of IF value
0
2
4
6

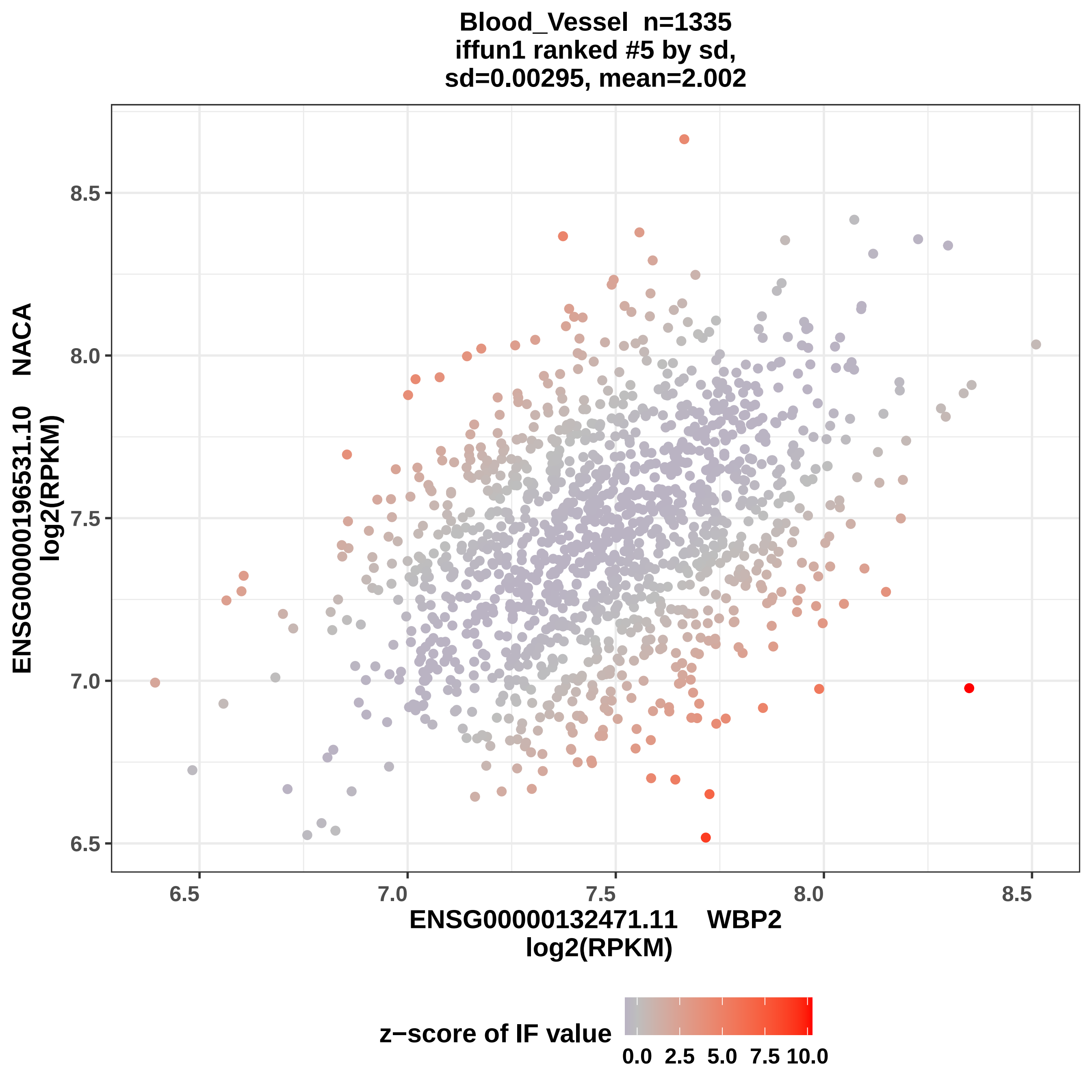
Blood_Vessel n=1335
iffun1 ranked #5 by sd,
sd=0.00295, mean=2.002
ENSG00000196531.10 NACA
log2(RPKM)
8.5
8.0
7.5
7.0
6.5
6.5
7.0
7.5
8.0
8.5
ENSG00000132471.11 WBP2
log2(RPKM)
z−score of IF value
0.0
2.5
5.0
7.5
10.0

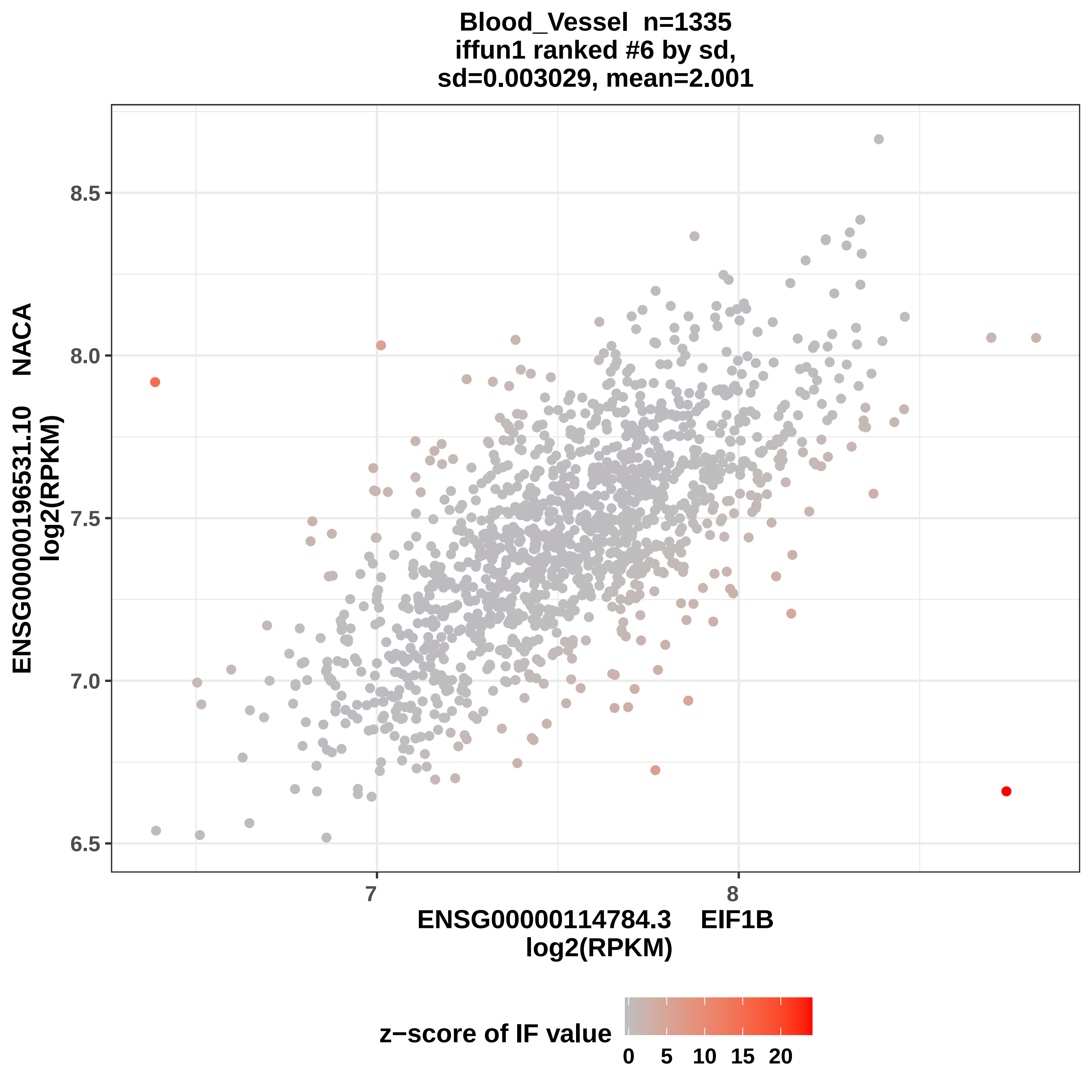
Blood_Vessel n=1335
iffun1 ranked #6 by sd,
sd=0.003029, mean=2.001
ENSG00000196531.10 NACA
log2(RPKM)
8.5
8.0
7.5
7.0
6.5
7
8
ENSG00000114784.3 EIF1B
log2(RPKM)
z−score of IF value
0
5
10
15
20

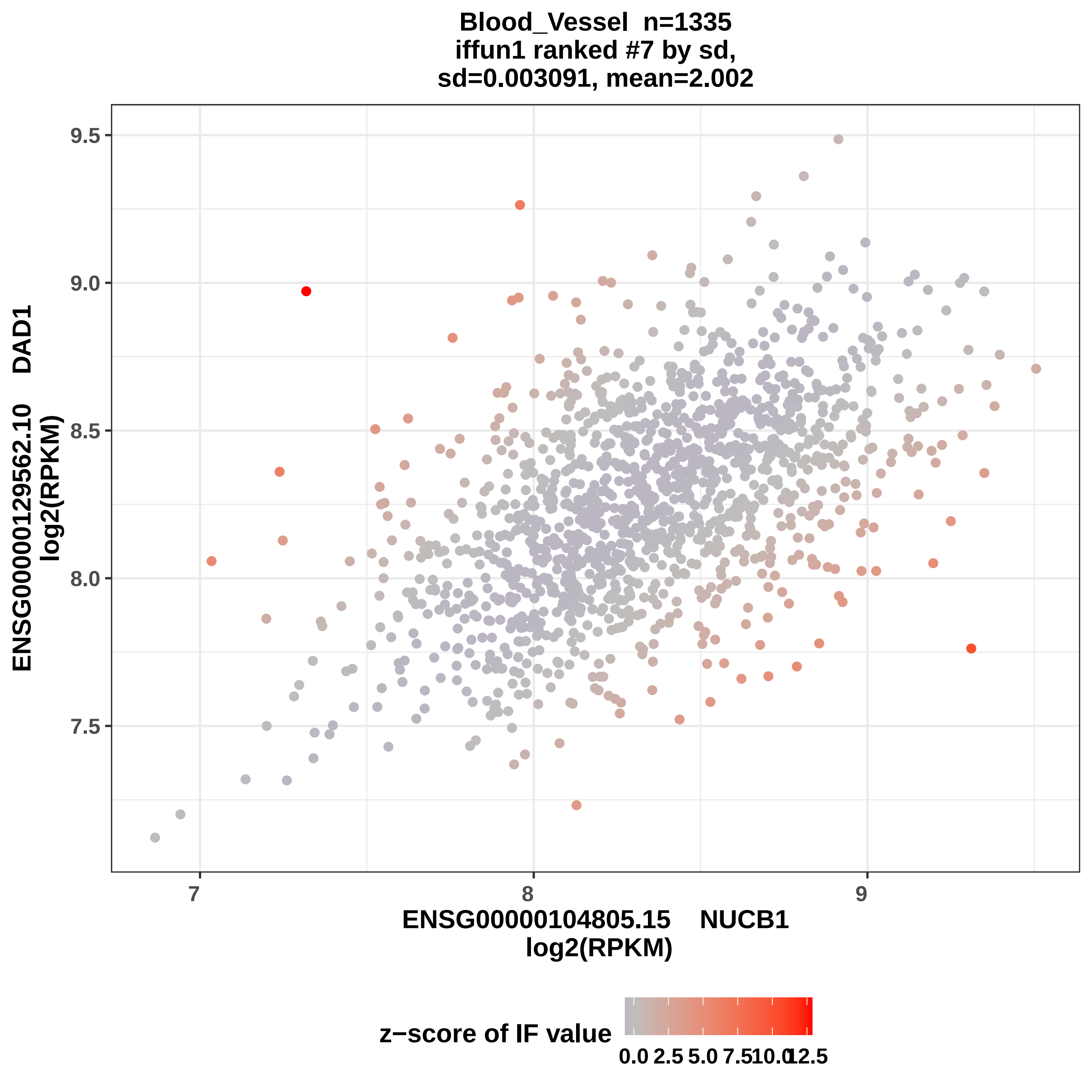
Blood_Vessel n=1335
iffun1 ranked #7 by sd,
sd=0.003091, mean=2.002
ENSG00000129562.10 DAD1
log2(RPKM)
9.5
9.0
8.5
8.0
7.5
7
8
9
ENSG00000104805.15 NUCB1
log2(RPKM)
z−score of IF value
0.0
2.5
5.0
7.5
10.0
12.5

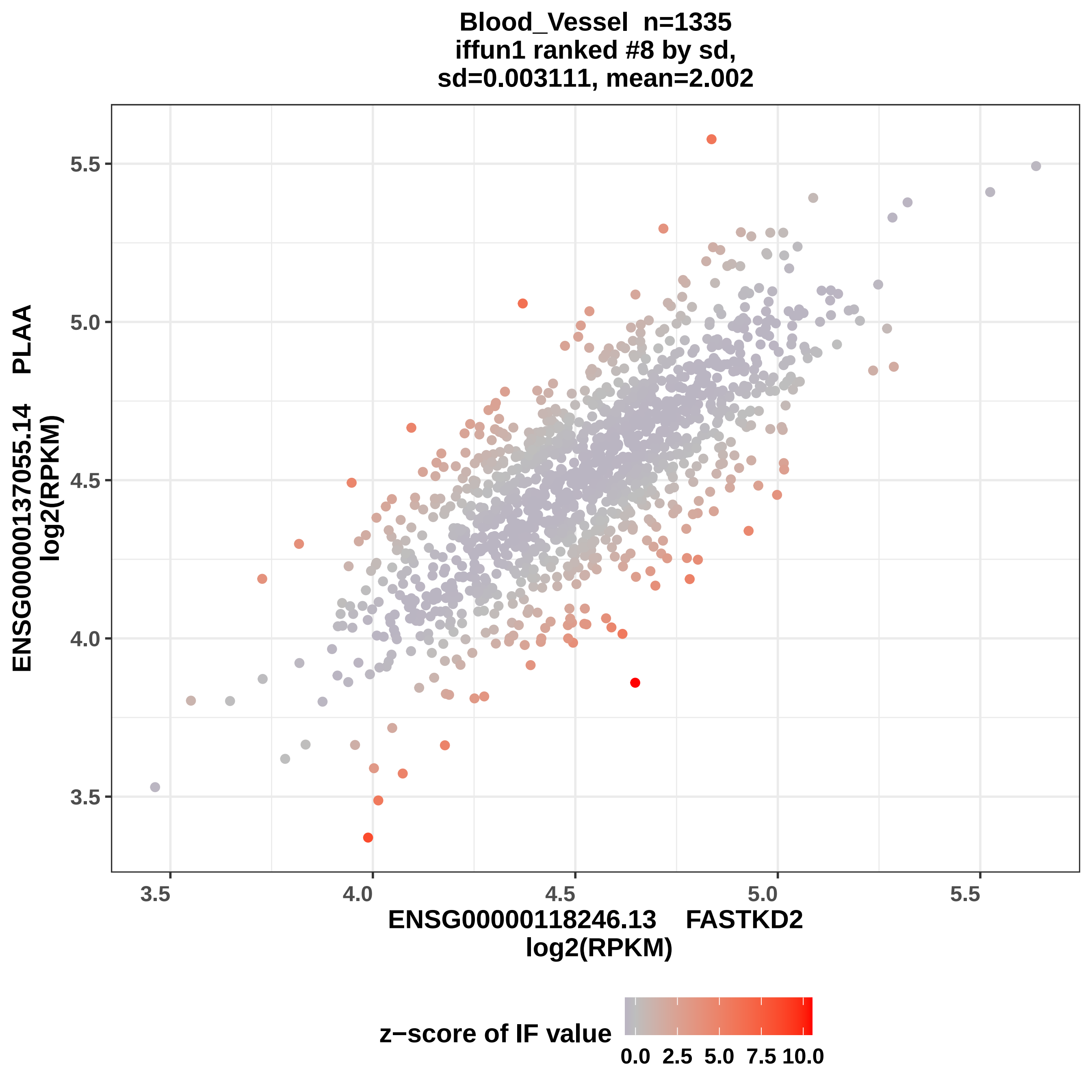
Blood_Vessel n=1335
iffun1 ranked #8 by sd,
sd=0.003111, mean=2.002
ENSG00000137055.14 PLAA
log2(RPKM)
ENSG00000118246.13 FASTKD2
log2(RPKM)
3.5
4.0
4.5
5.0
5.5
z−score of IF value
0.0 2.5 5.0 7.5 10.0

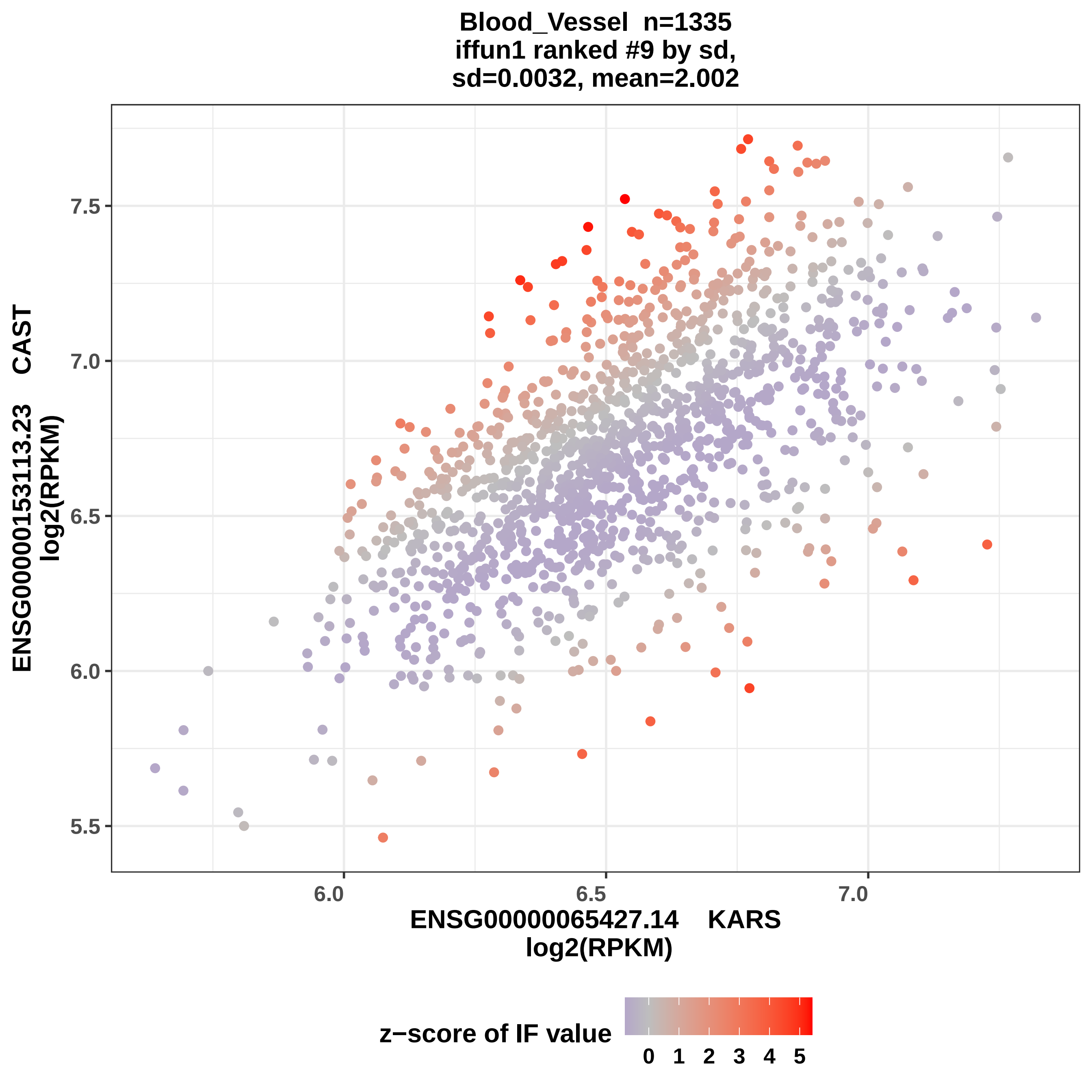

Blood_Vessel n=1335
iffun1 ranked #9 by sd,
sd=0.0032, mean=2.002
ENSG00000153113.23 CAST
log2(RPKM)
7.5
7.0
6.5
6.0
5.5
6.0
6.5
7.0
ENSG00000065427.14 KARS
log2(RPKM)
z−score of IF value
0 1 2 3 4 5

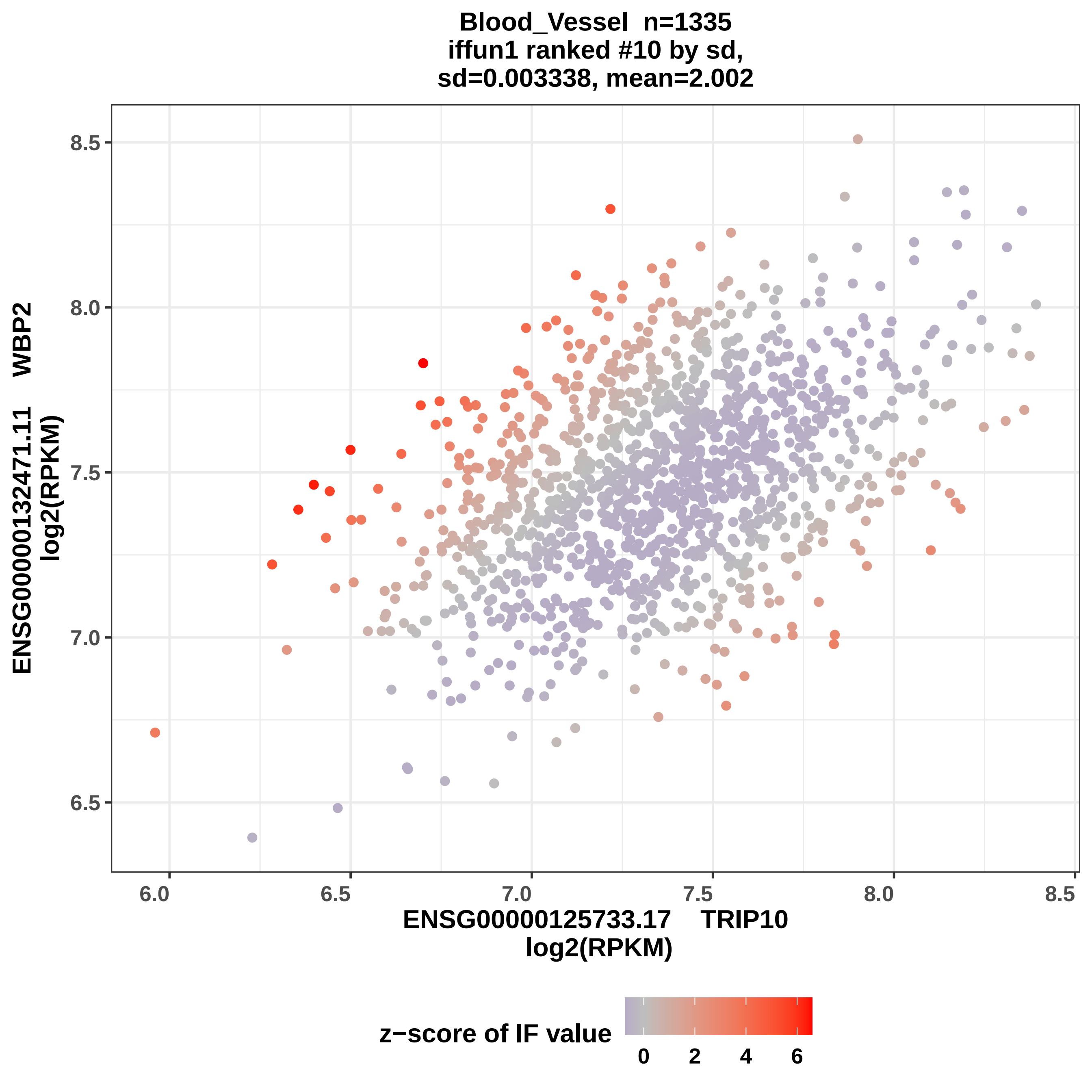

Blood_Vessel n=1335
iffun1 ranked #10 by sd,
sd=0.003338, mean=2.002
ENSG00000132471.11 WBP2
log2(RPKM)
ENSG00000125733.17 TRIP10
log2(RPKM)
z−score of IF value
8.5
8.0
7.5
7.0
6.5
6.0
6.5
7.0
7.5
8.0
8.5
0
2
4
6

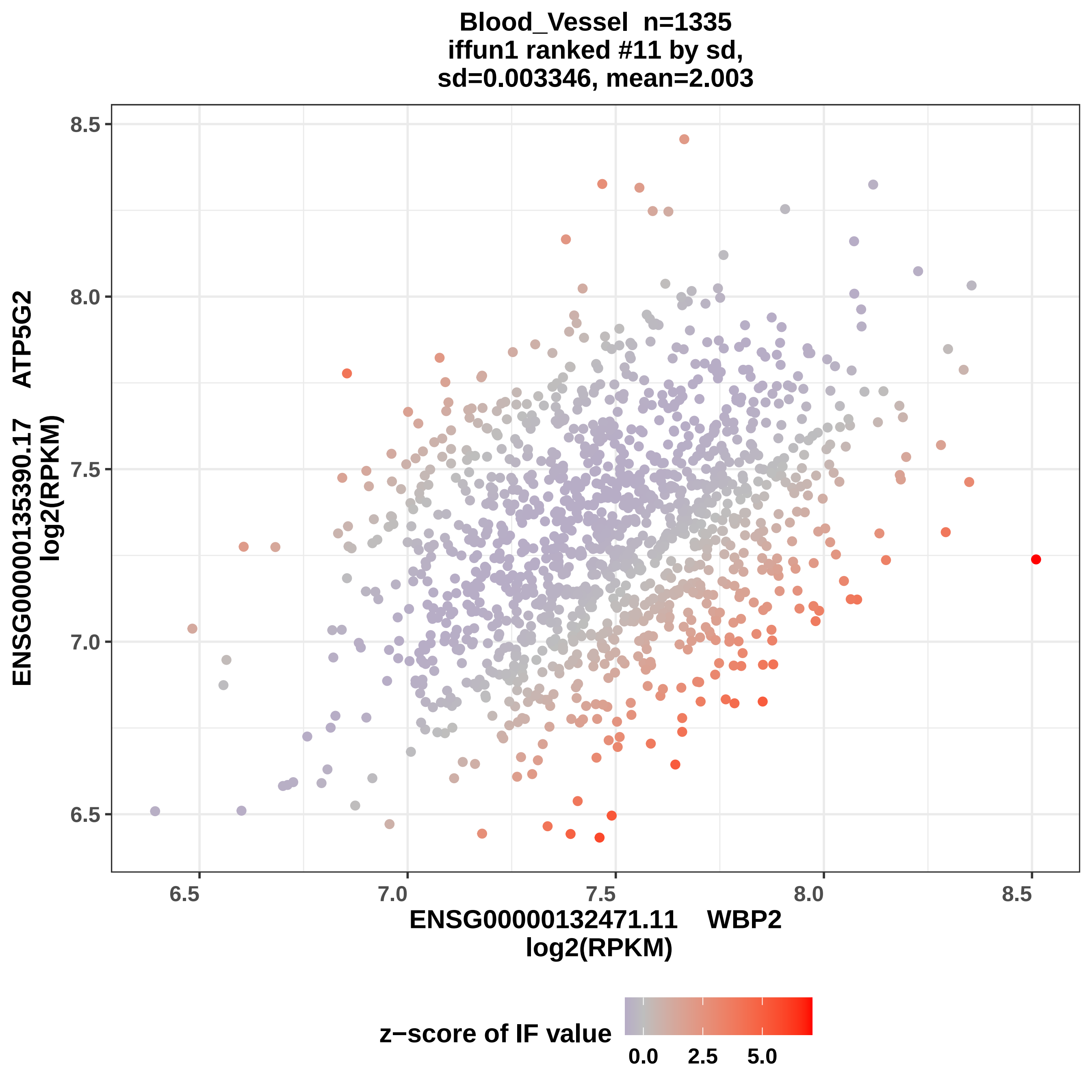
Blood_Vessel n=1335
iffun1 ranked #11 by sd,
sd=0.003346, mean=2.003
ENSG00000135390.17 ATP5G2
log2(RPKM)
8.5
8.0
7.5
7.0
6.5
6.5
7.0
7.5
8.0
8.5
ENSG00000132471.11 WBP2
log2(RPKM)
z−score of IF value
0.0
2.5
5.0

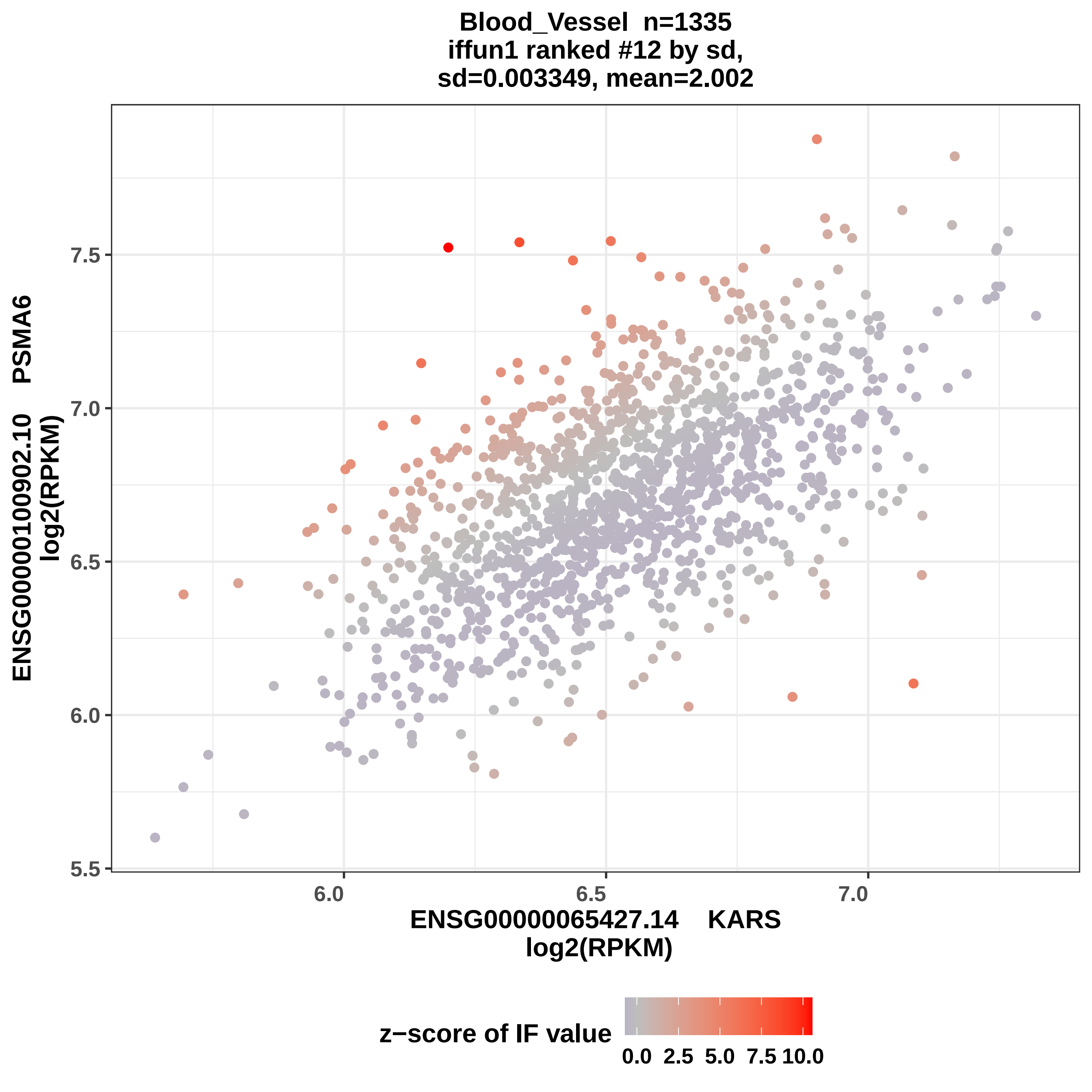

Blood_Vessel n=1335
iffun1 ranked #12 by sd,
sd=0.003349, mean=2.002
ENSG00000100902.10 PSMA6
log2(RPKM)
ENSG00000065427.14 KARS
log2(RPKM)
7.5
7.0
6.5
6.0
5.5
6.0
6.5
7.0
z−score of IF value
0.0
2.5
5.0
7.5
10.0

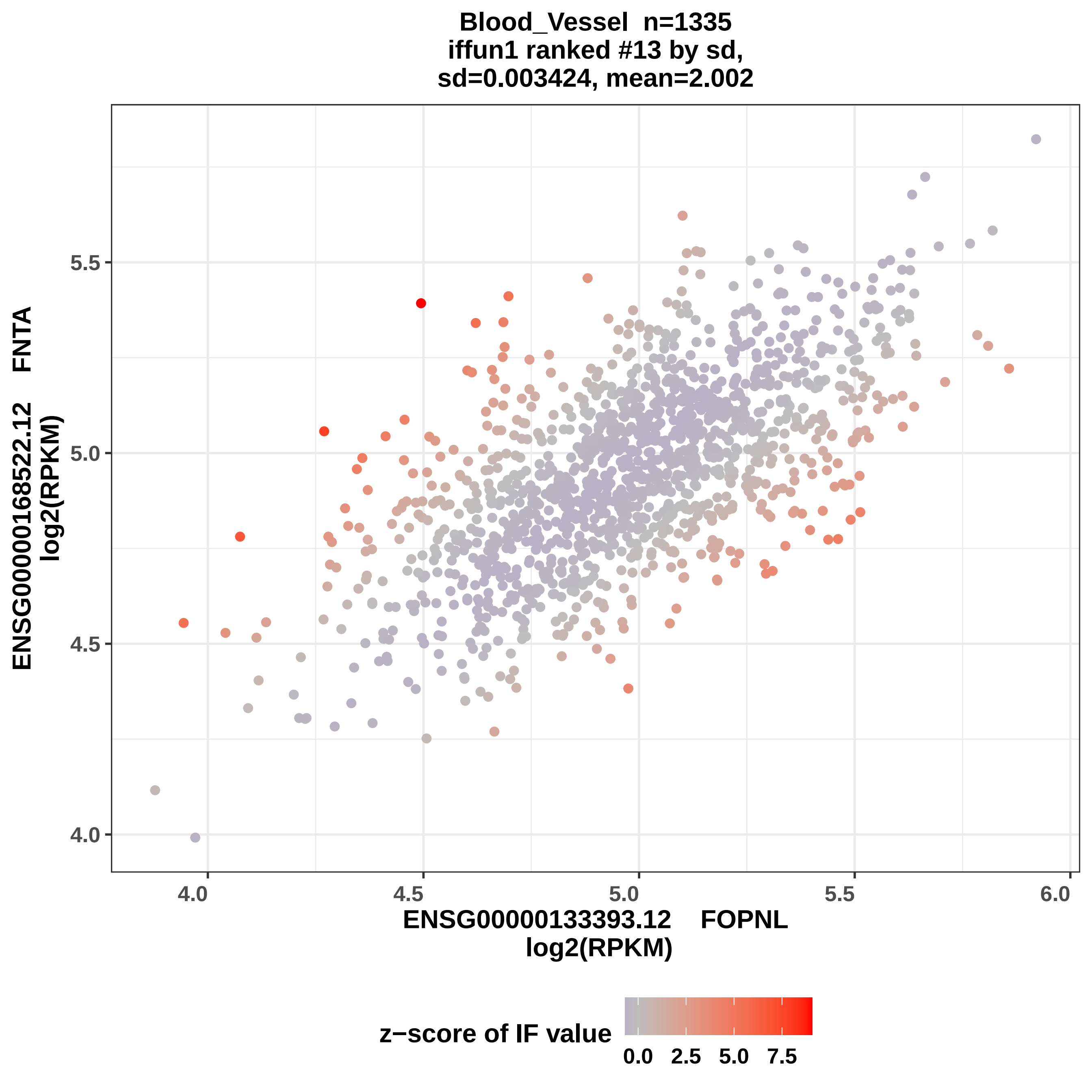

Blood_Vessel n=1335
iffun1 ranked #13 by sd,
sd=0.003424, mean=2.002
ENSG00000168522.12 FNTA
log2(RPKM)
5.5
5.0
4.5
4.0
4.0
4.5
5.0
5.5
6.0
ENSG00000133393.12 FOPNL
log2(RPKM)
z−score of IF value
0.0
2.5
5.0
7.5

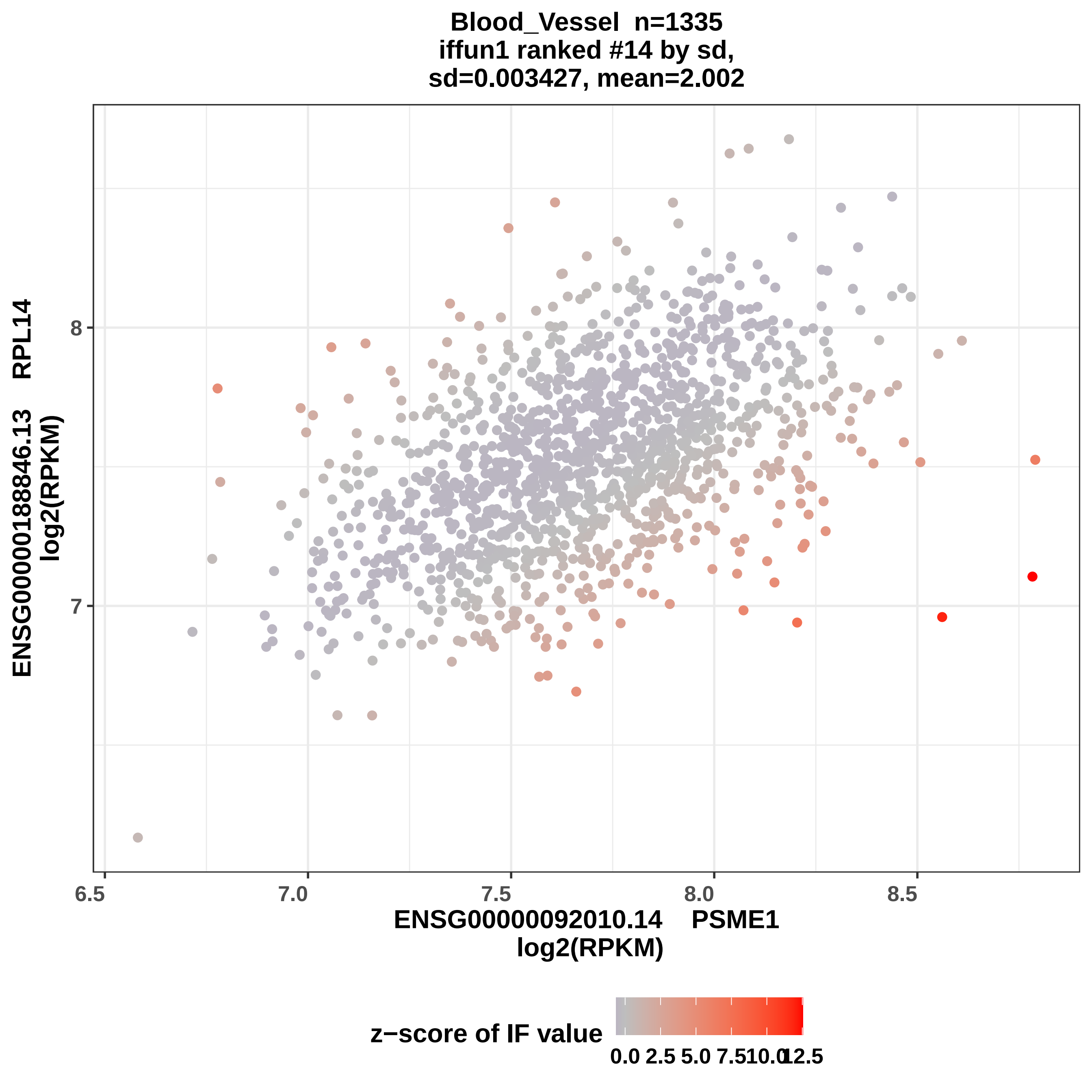

Blood_Vessel n=1335
iffun1 ranked #14 by sd,
sd=0.003427, mean=2.002
ENSG00000188846.13 RPL14
log2(RPKM)
8
7
6.5
7.0
7.5
8.0
8.5
ENSG00000092010.14 PSME1
log2(RPKM)
z−score of IF value
0.0
2.5
5.0
7.5
10.0
12.5

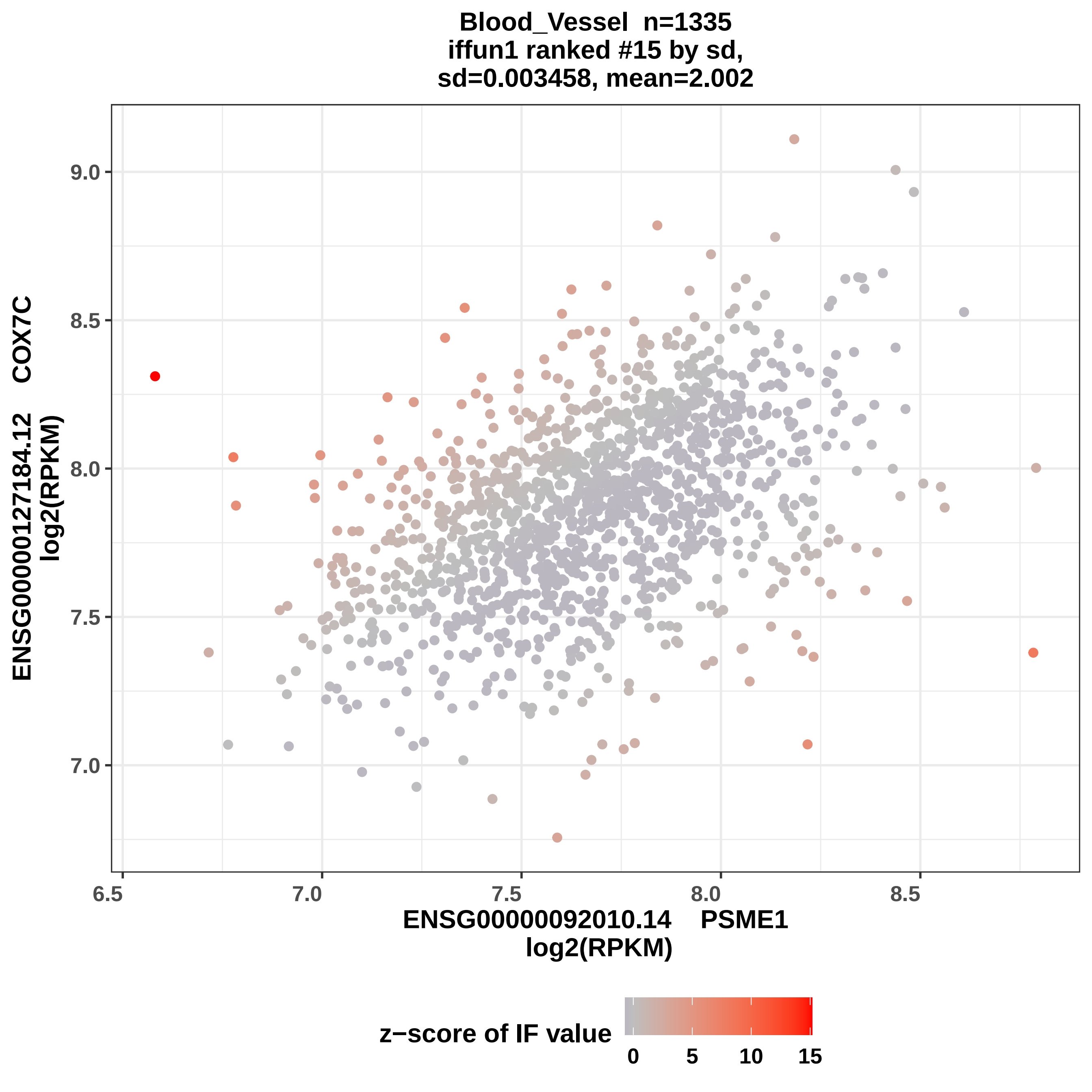

Blood_Vessel n=1335
iffun1 ranked #15 by sd,
sd=0.003458, mean=2.002
ENSG00000127184.12 COX7C
log2(RPKM)
9.0
8.5
8.0
7.5
7.0
6.5
7.0
7.5
8.0
8.5
ENSG00000092010.14 PSME1
log2(RPKM)
z−score of IF value
0
5
10
15

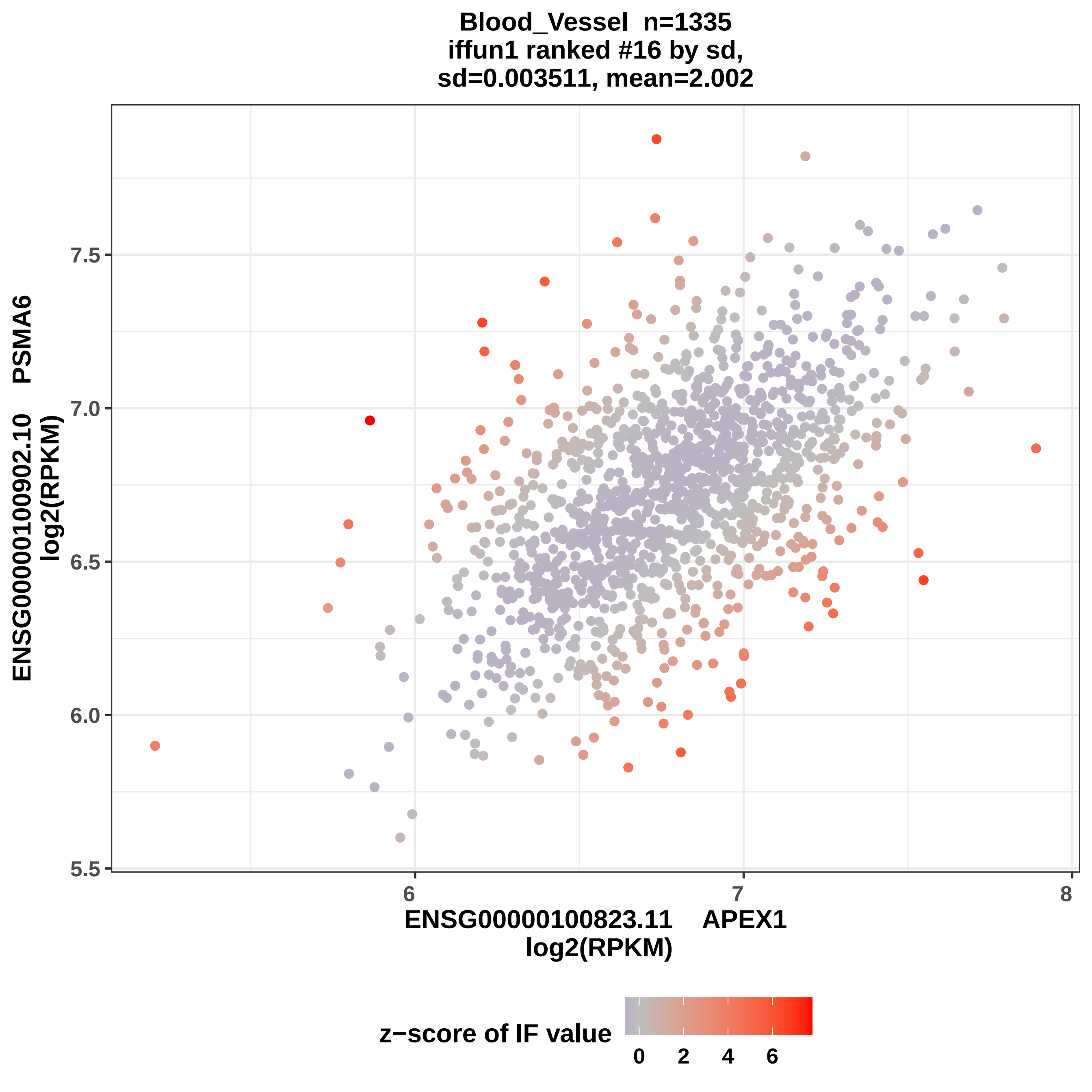

Blood_Vessel n=1335
iffun1 ranked #16 by sd,
sd=0.003511, mean=2.002
ENSG00000100902.10 PSMA6
log2(RPKM)
7.5
7.0
6.5
6.0
5.5
6
7
8
ENSG00000100823.11 APEX1
log2(RPKM)
z−score of IF value
0
2
4
6

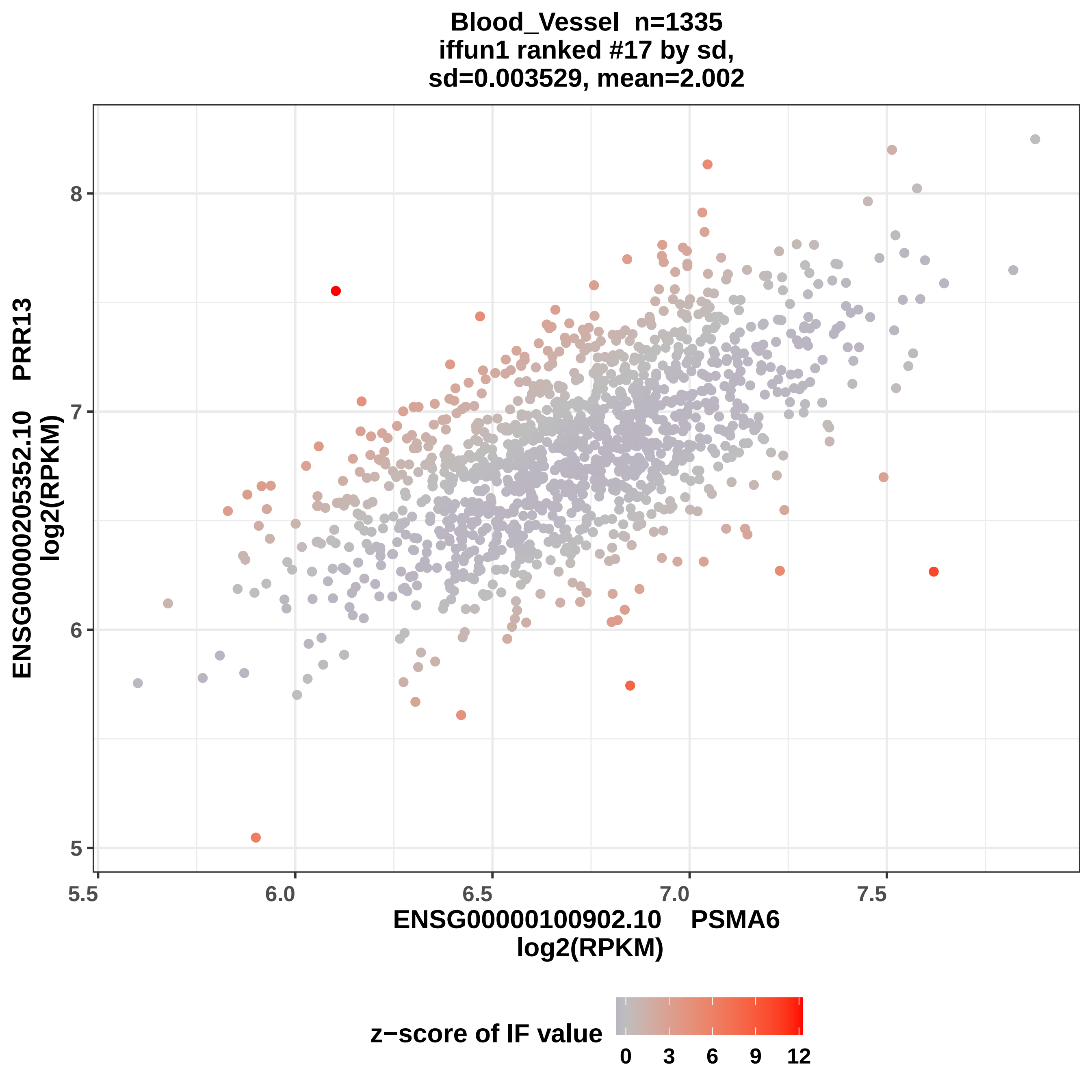

Blood_Vessel n=1335
iffun1 ranked #17 by sd,
sd=0.003529, mean=2.002
ENSG00000205352.10 PRR13
log2(RPKM)
8
7
6
5
5.5
6.0
6.5
7.0
7.5
ENSG00000100902.10 PSMA6
log2(RPKM)
z−score of IF value
0
3
6
9
12

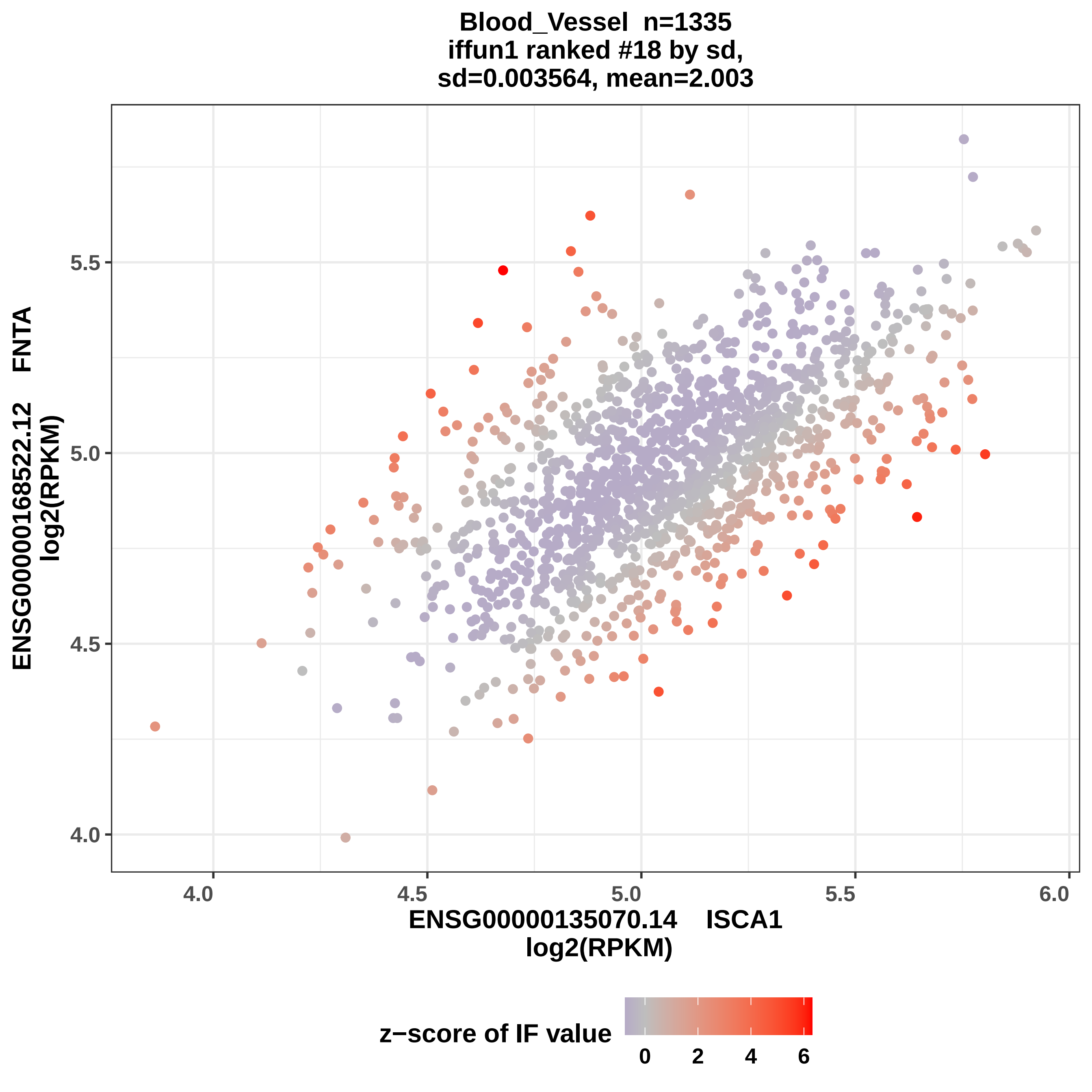
Blood_Vessel n=1335
iffun1 ranked #18 by sd,
sd=0.003564, mean=2.003
FNTA
ENSG00000168522.12
log2(RPKM)
5.5
5.0
4.5
4.0
4.0
4.5
5.0
5.5
6.0
ENSG00000135070.14 ISCA1
log2(RPKM)
z−score of IF value
0
2
4
6

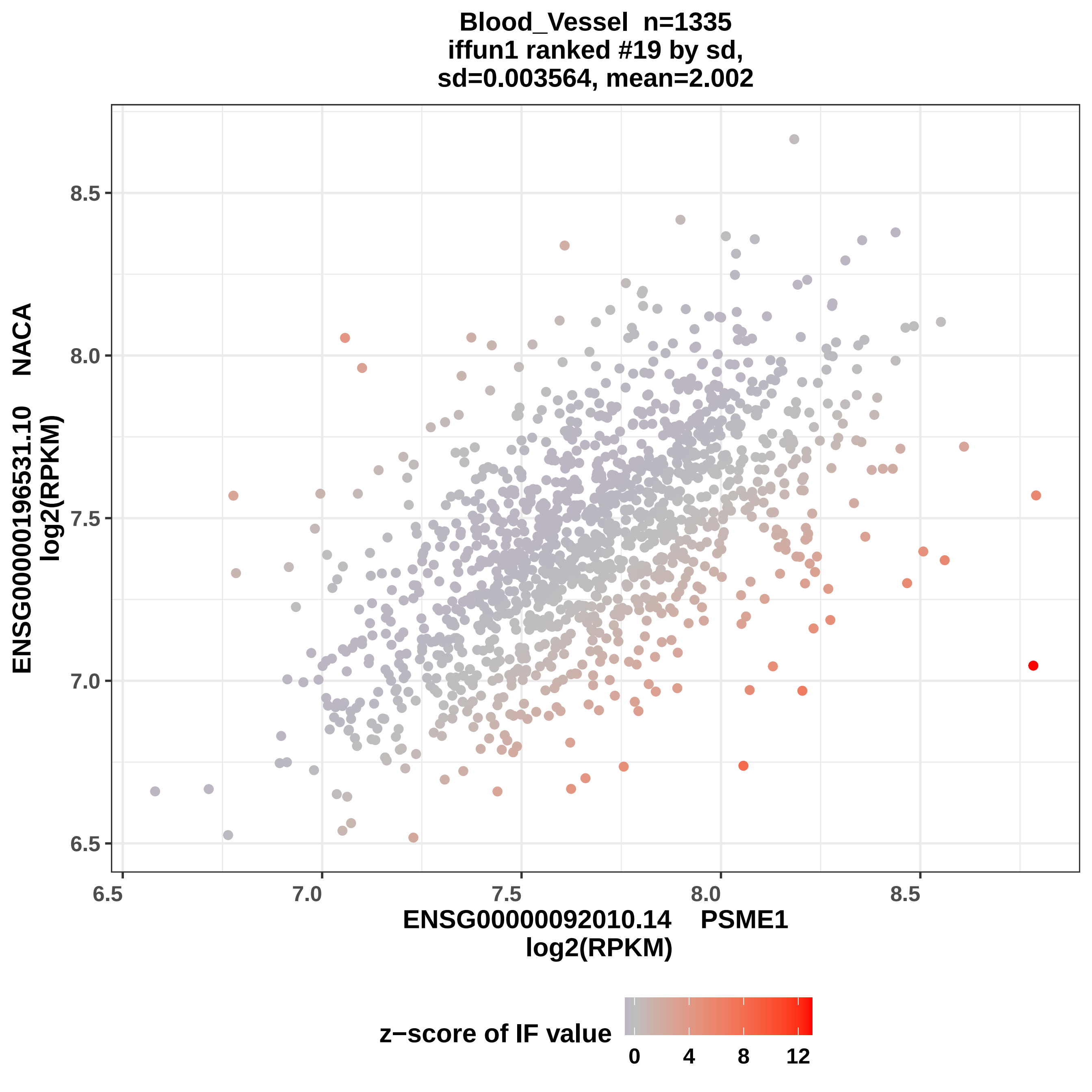

Blood_Vessel n=1335
iffun1 ranked #19 by sd,
sd=0.003564, mean=2.002
ENSG00000196531.10 NACA
log2(RPKM)
8.5
8.0
7.5
7.0
6.5
6.5
7.0
7.5
8.0
8.5
ENSG00000092010.14 PSME1
log2(RPKM)
z−score of IF value
0
4
8
12

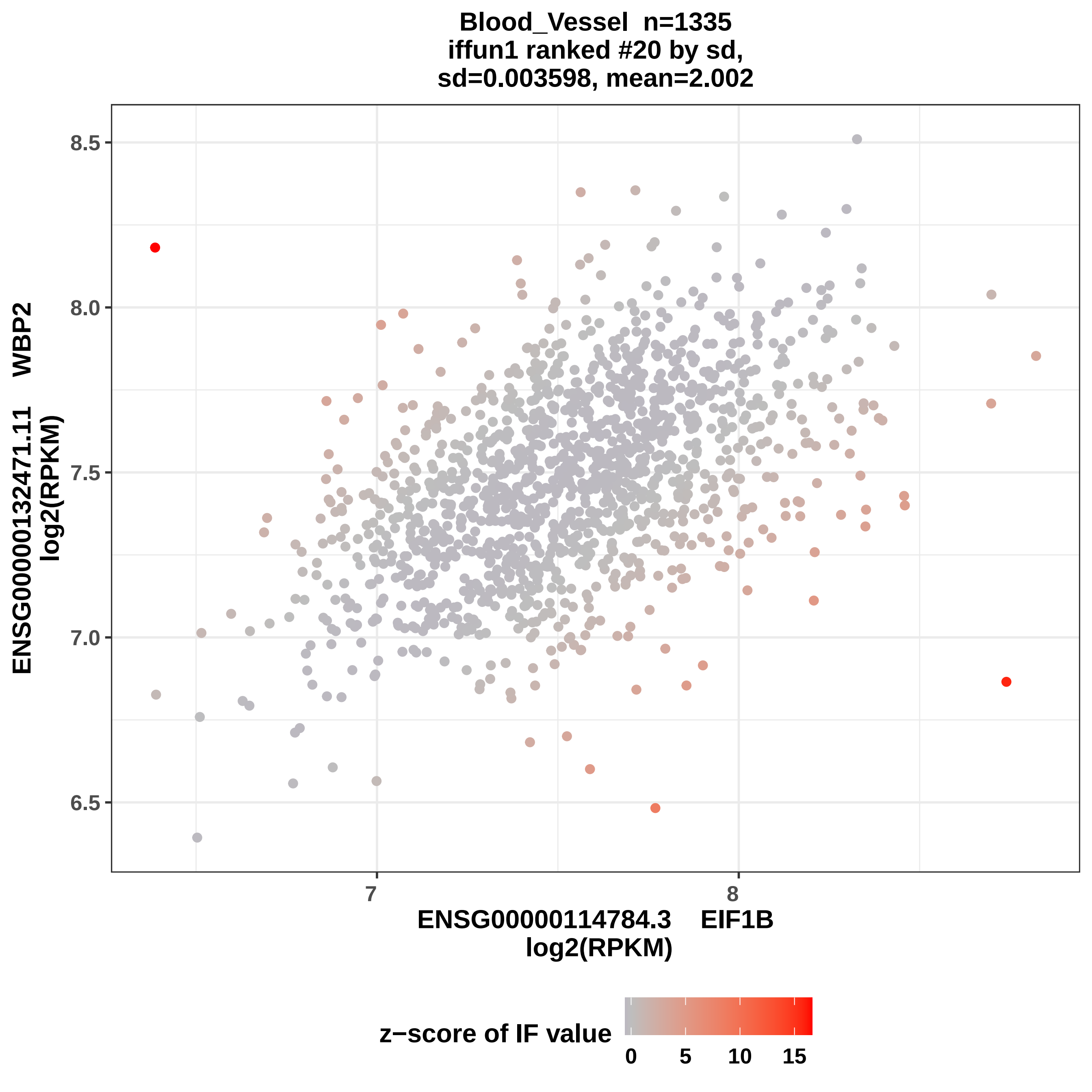

Blood_Vessel n=1335
iffun1 ranked #20 by sd,
sd=0.003598, mean=2.002
8.5
8.0
7.5
7.0
6.5
ENSG00000132471.11 WBP2
log2(RPKM)
7
8
ENSG00000114784.3 EIF1B
log2(RPKM)
z−score of IF value
0
5
10
15

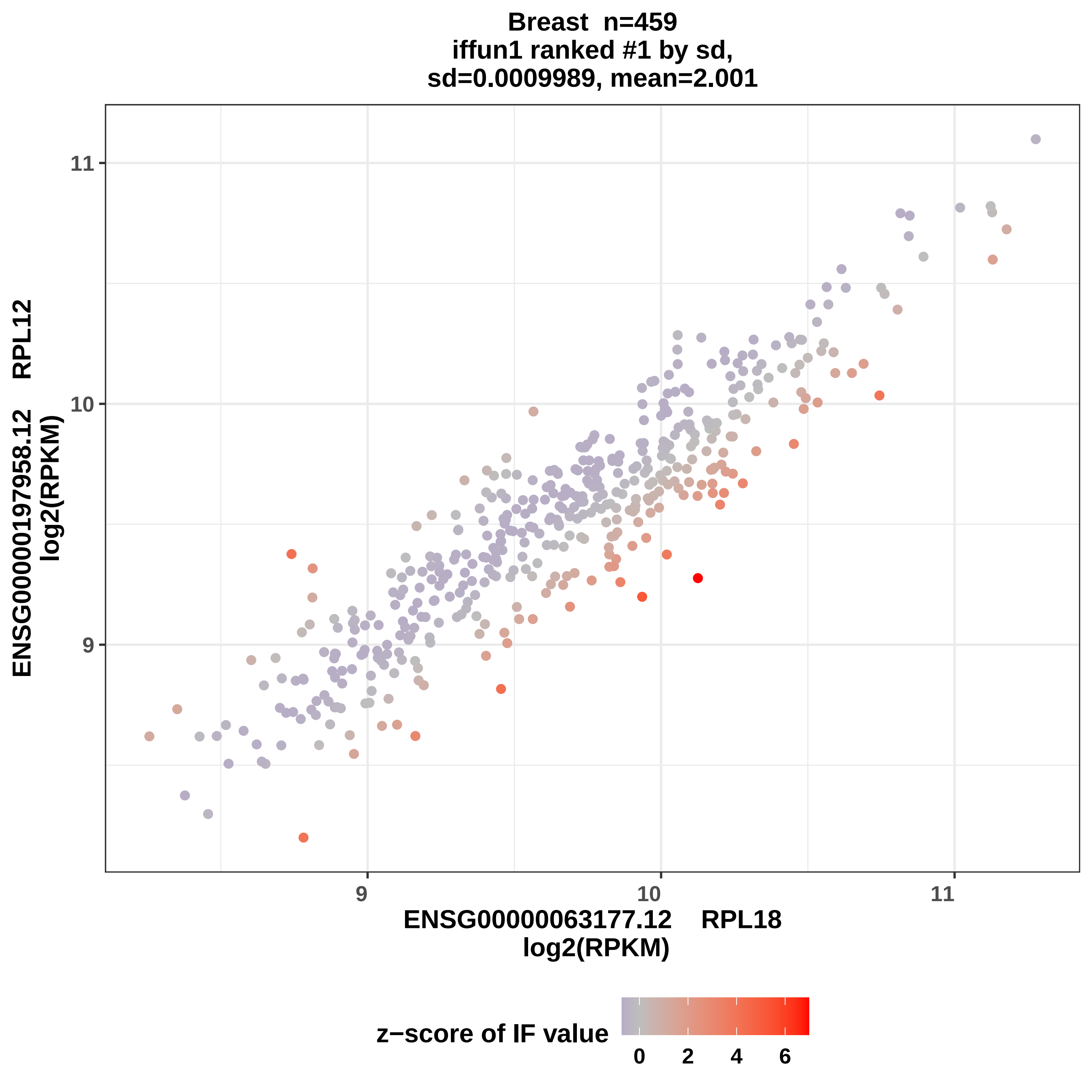

Breast n=459
iffun1 ranked #1 by sd,
sd=0.0009989, mean=2.001
ENSG00000197958.12 RPL12
log2(RPKM)
11
10
9
9
10
11
ENSG00000063177.12 RPL18
log2(RPKM)
z−score of IF value
0
2
4
6

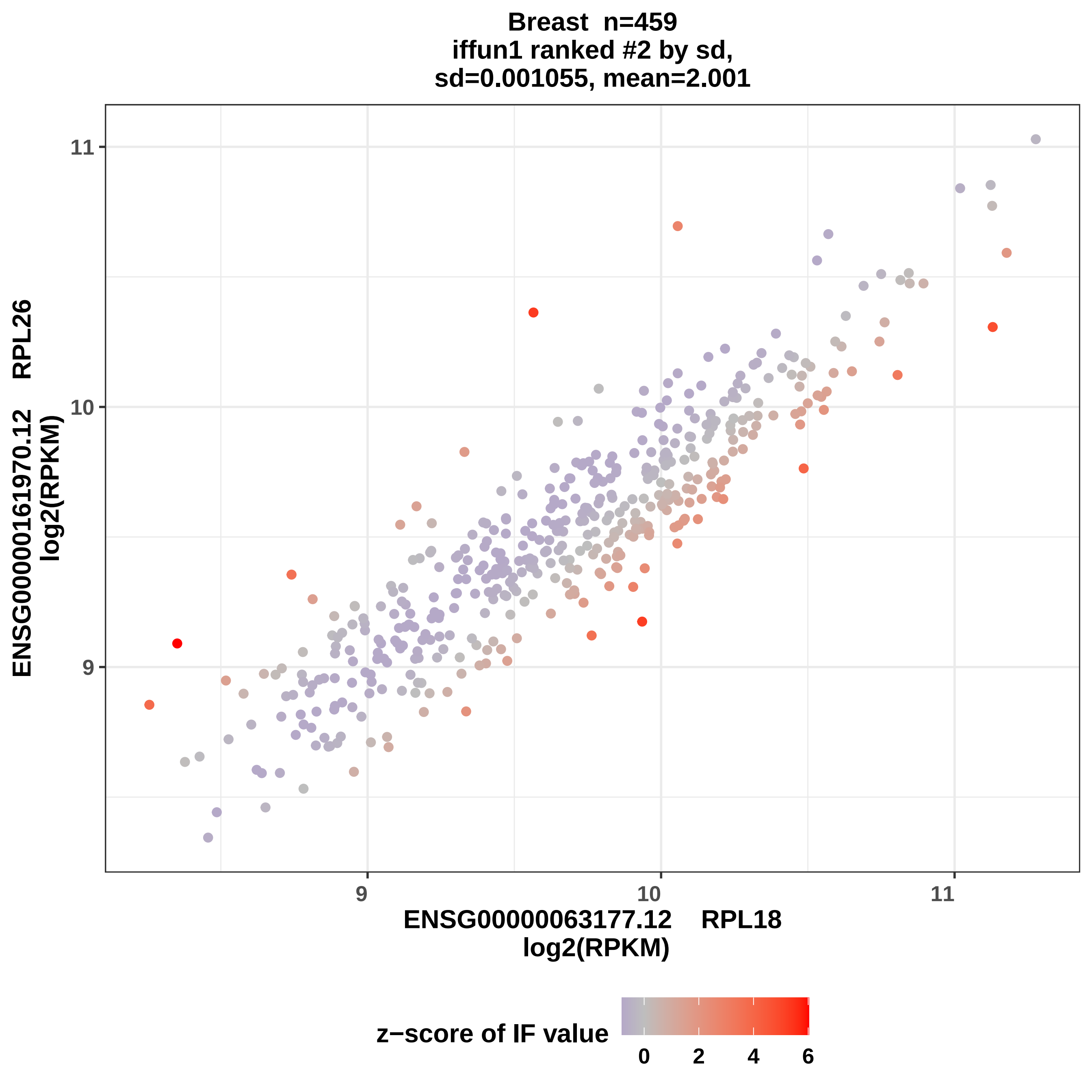

Breast n=459
iffun1 ranked #2 by sd,
sd=0.001055, mean=2.001
ENSG00000161970.12 RPL26
log2(RPKM)
ENSG00000063177.12 RPL18
log2(RPKM)
z−score of IF value
9
10
11
0
2
4
6

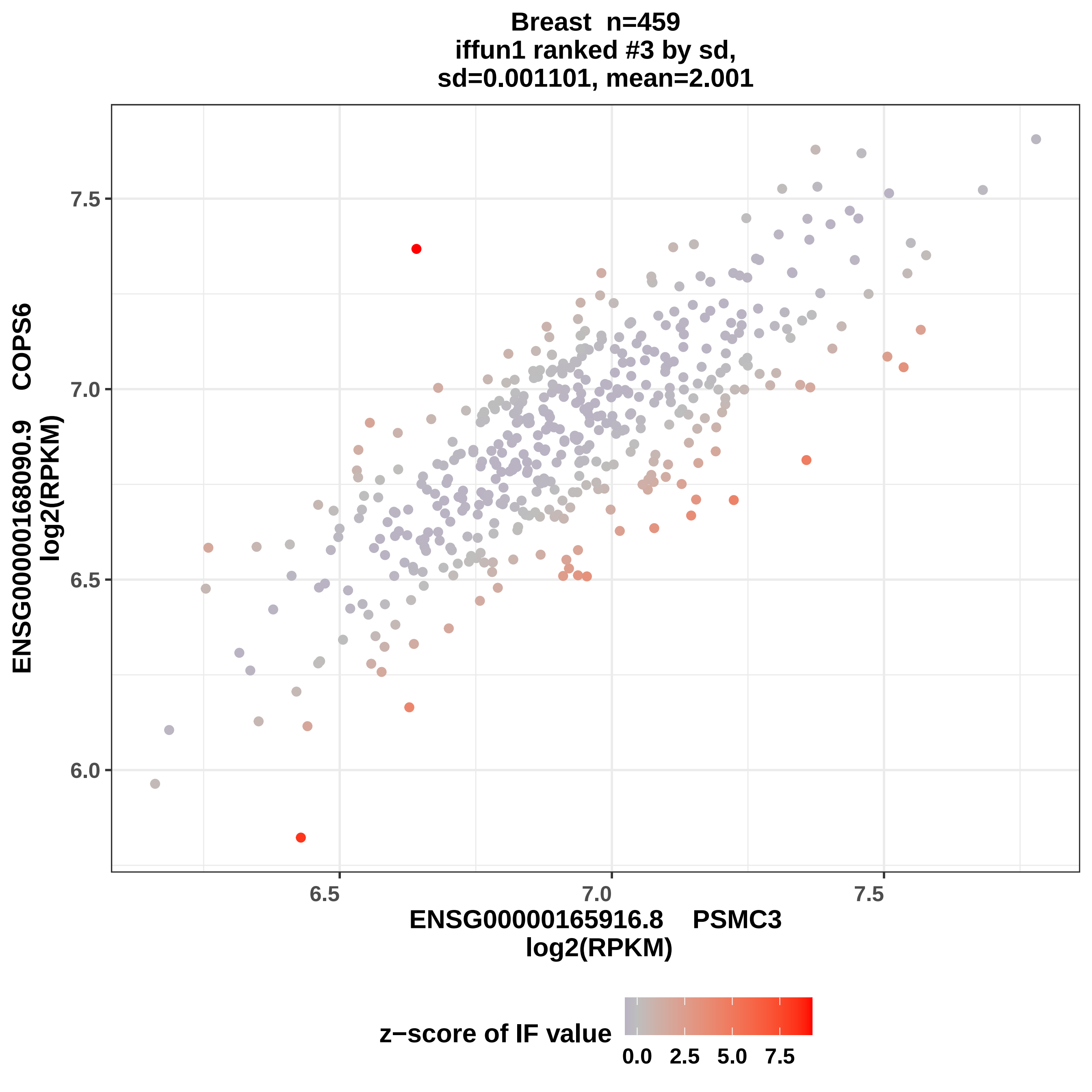
Breast n=459
iffun1 ranked #3 by sd,
sd=0.001101, mean=2.001
ENSG00000168090.9 COPS6
log2(RPKM)
7.5
7.0
6.5
6.0
6.5
7.0
7.5
ENSG00000165916.8 PSMC3
log2(RPKM)
z−score of IF value
0.0
2.5
5.0
7.5

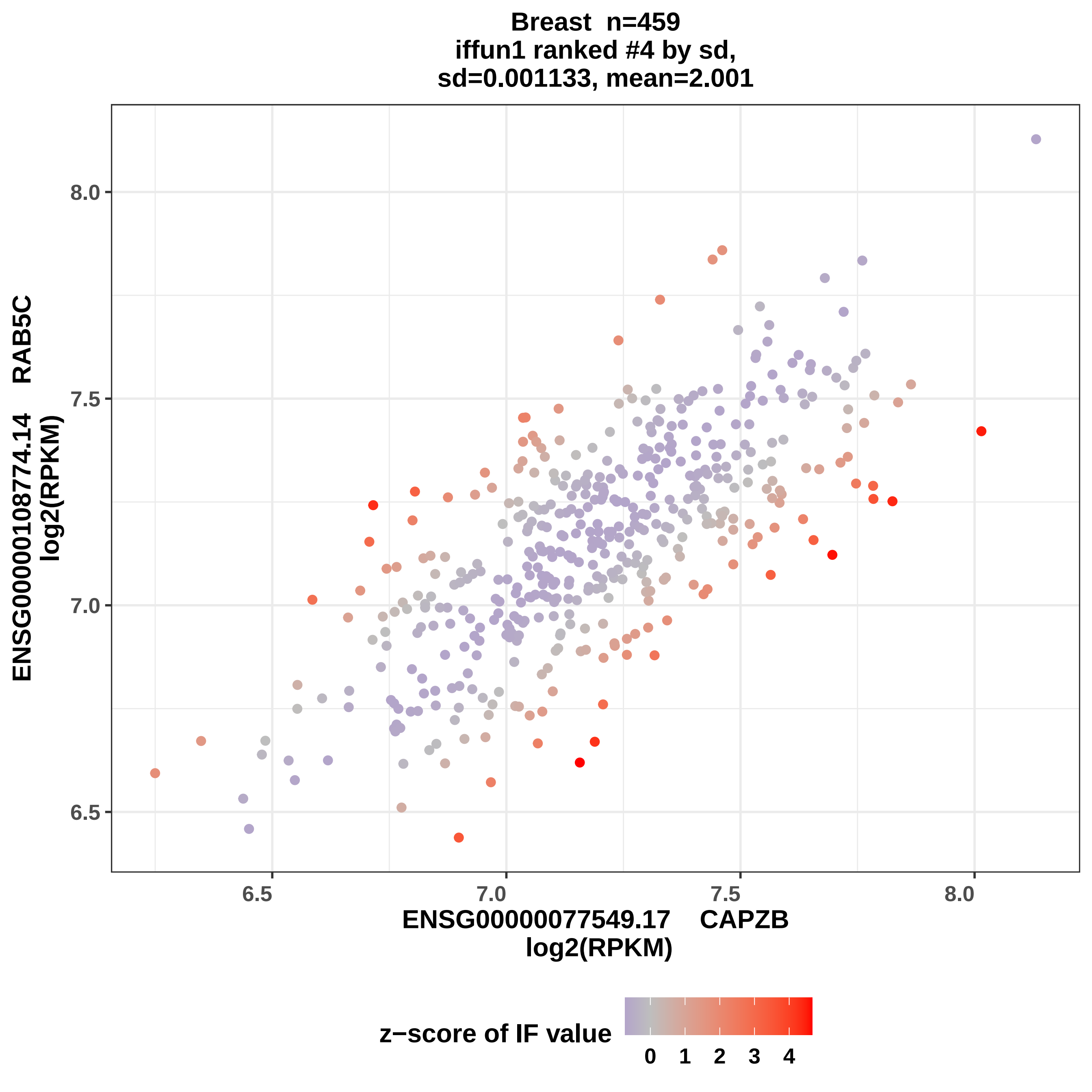
Breast n=459
iffun1 ranked #4 by sd,
sd=0.001133, mean=2.001
ENSG00000108774.14 RAB5C
log2(RPKM)
8.0
7.5
7.0
6.5
6.5
7.0
7.5
8.0
ENSG00000077549.17 CAPZB
log2(RPKM)
z−score of IF value
0
1
2
3
4

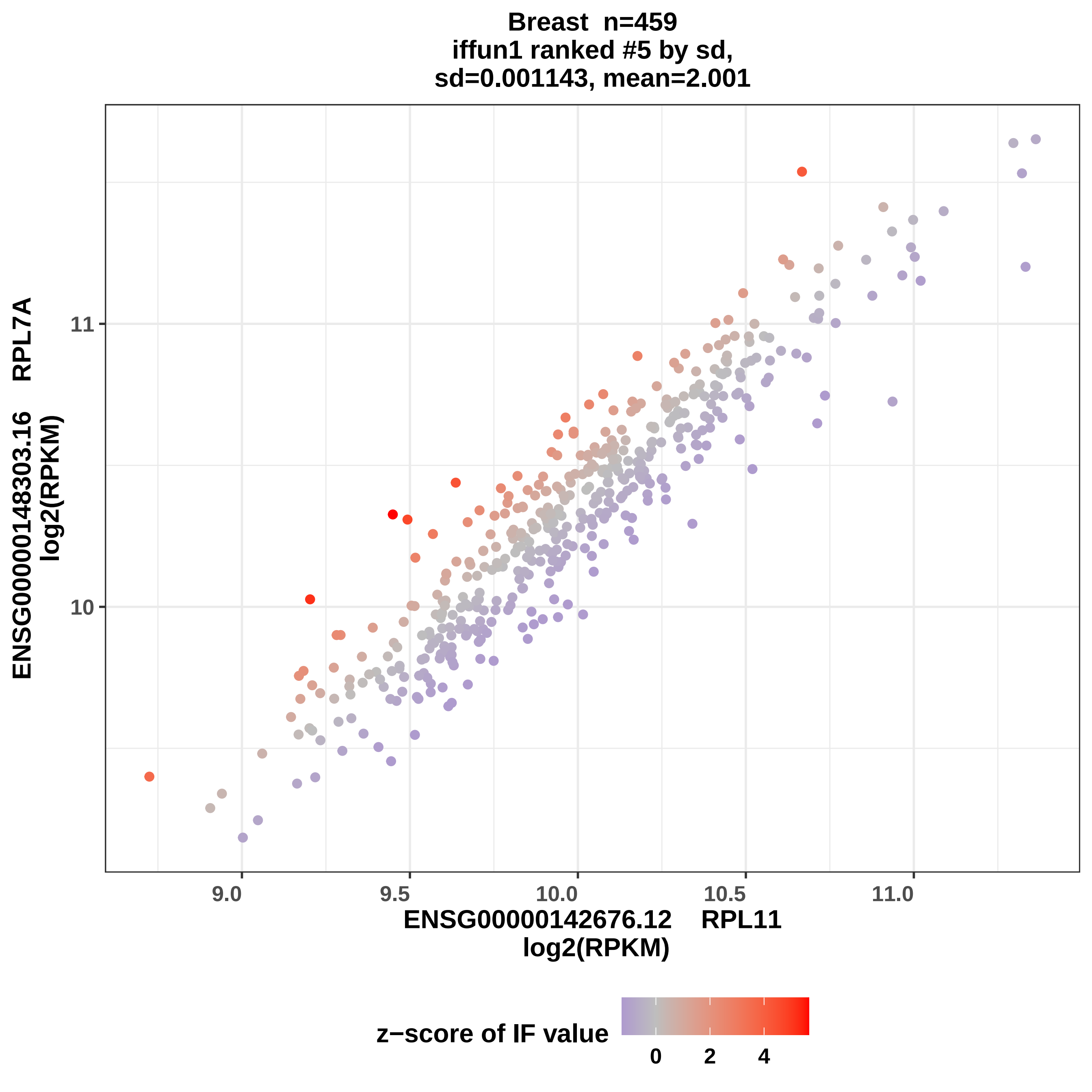

Breast n=459
iffun1 ranked #5 by sd,
sd=0.001143, mean=2.001
ENSG00000148303.16 RPL7A
log2(RPKM)
11
10
9.0
9.5
10.0
10.5
11.0
ENSG00000142676.12 RPL11
log2(RPKM)
z−score of IF value
0
2
4

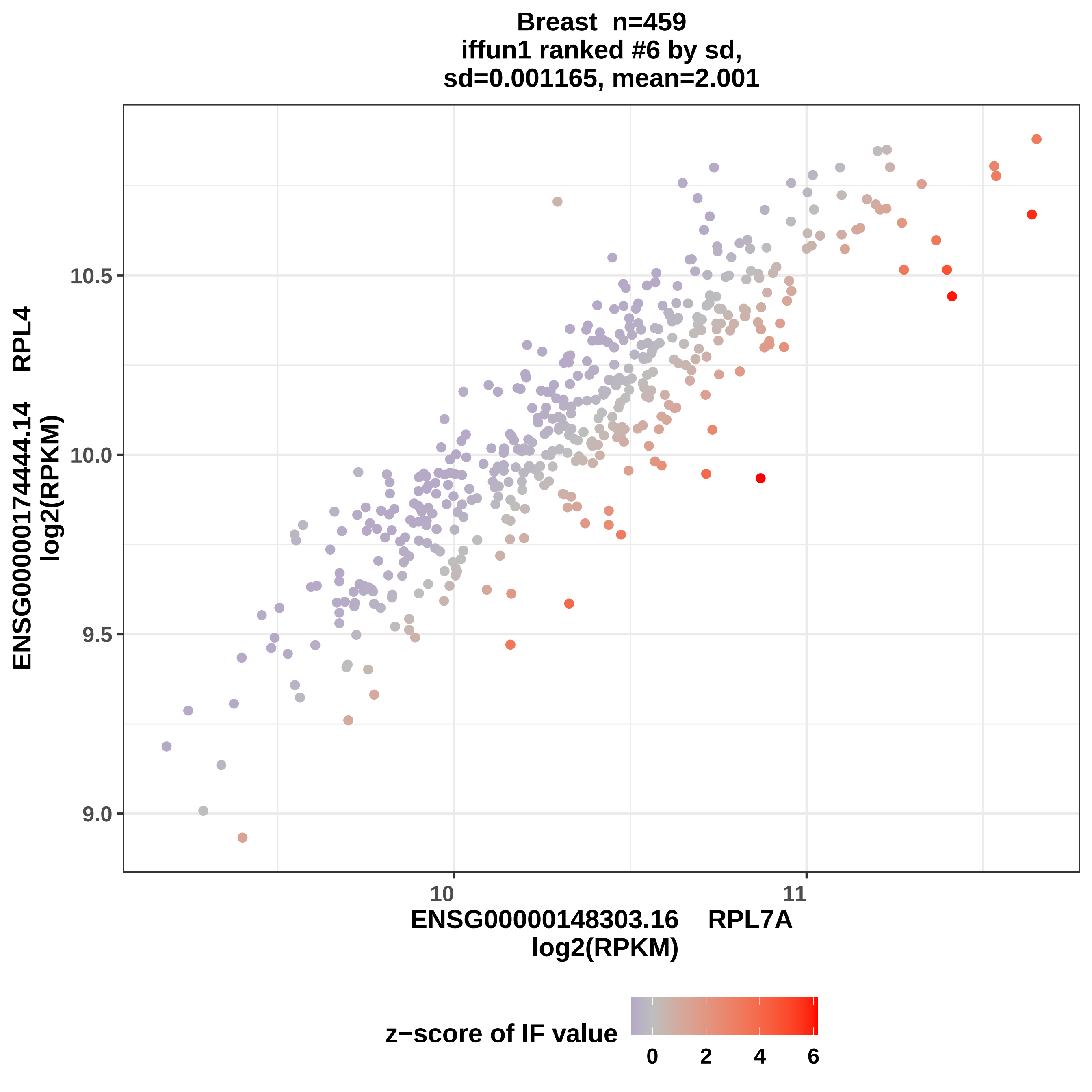
Breast n=459
iffun1 ranked #6 by sd,
sd=0.001165, mean=2.001
ENSG00000174444.14 RPL4
log2(RPKM)
10.5
10.0
9.5
9.0
10
11
ENSG00000148303.16 RPL7A
log2(RPKM)
z−score of IF value
0
2
4
6

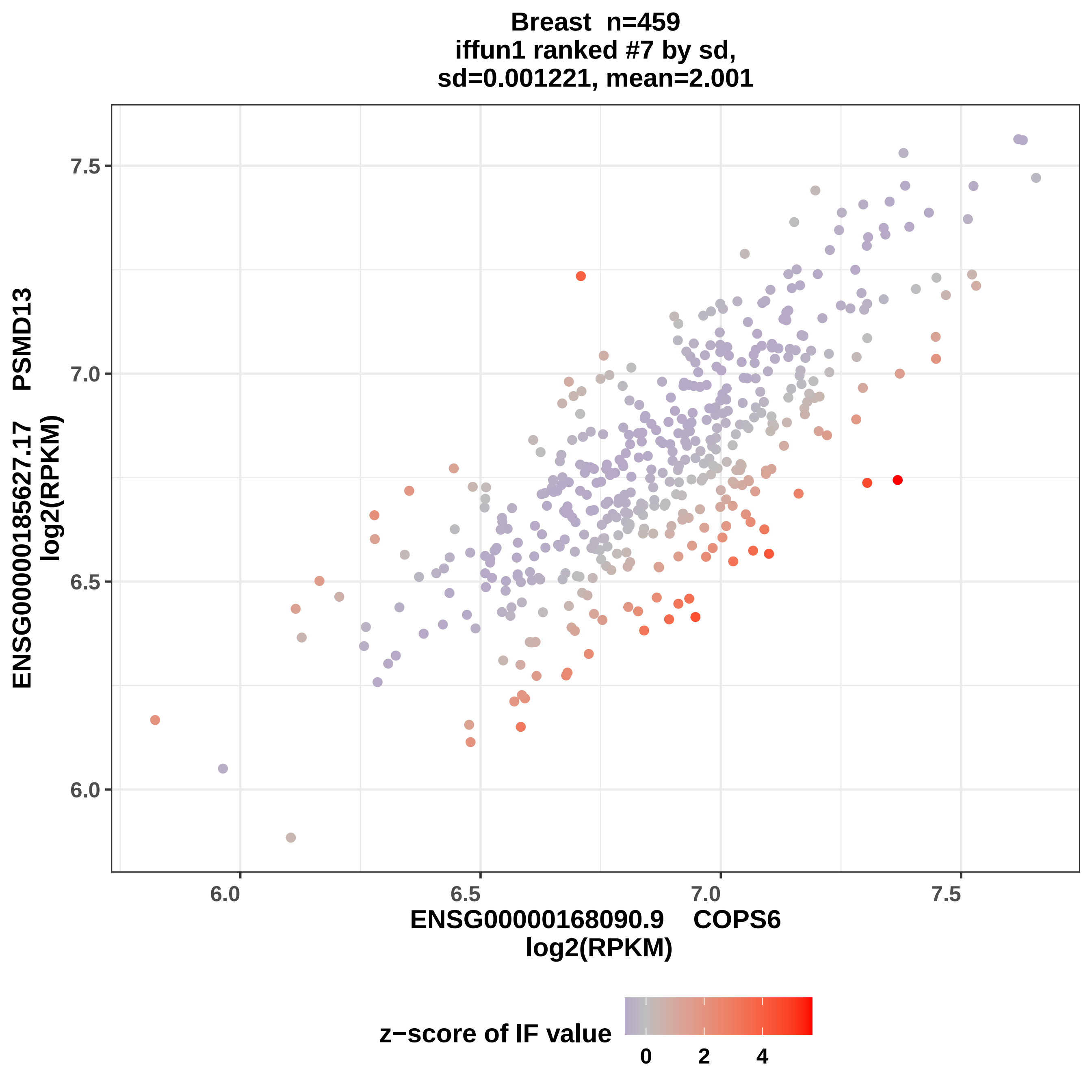

Breast n=459
iffun1 ranked #7 by sd,
sd=0.001221, mean=2.001
ENSG00000185627.17 PSMD13
log2(RPKM)
ENSG00000168090.9 COPS6
log2(RPKM)
7.5
7.0
6.5
6.0
6.0
6.5
7.0
7.5
z−score of IF value
0
2
4

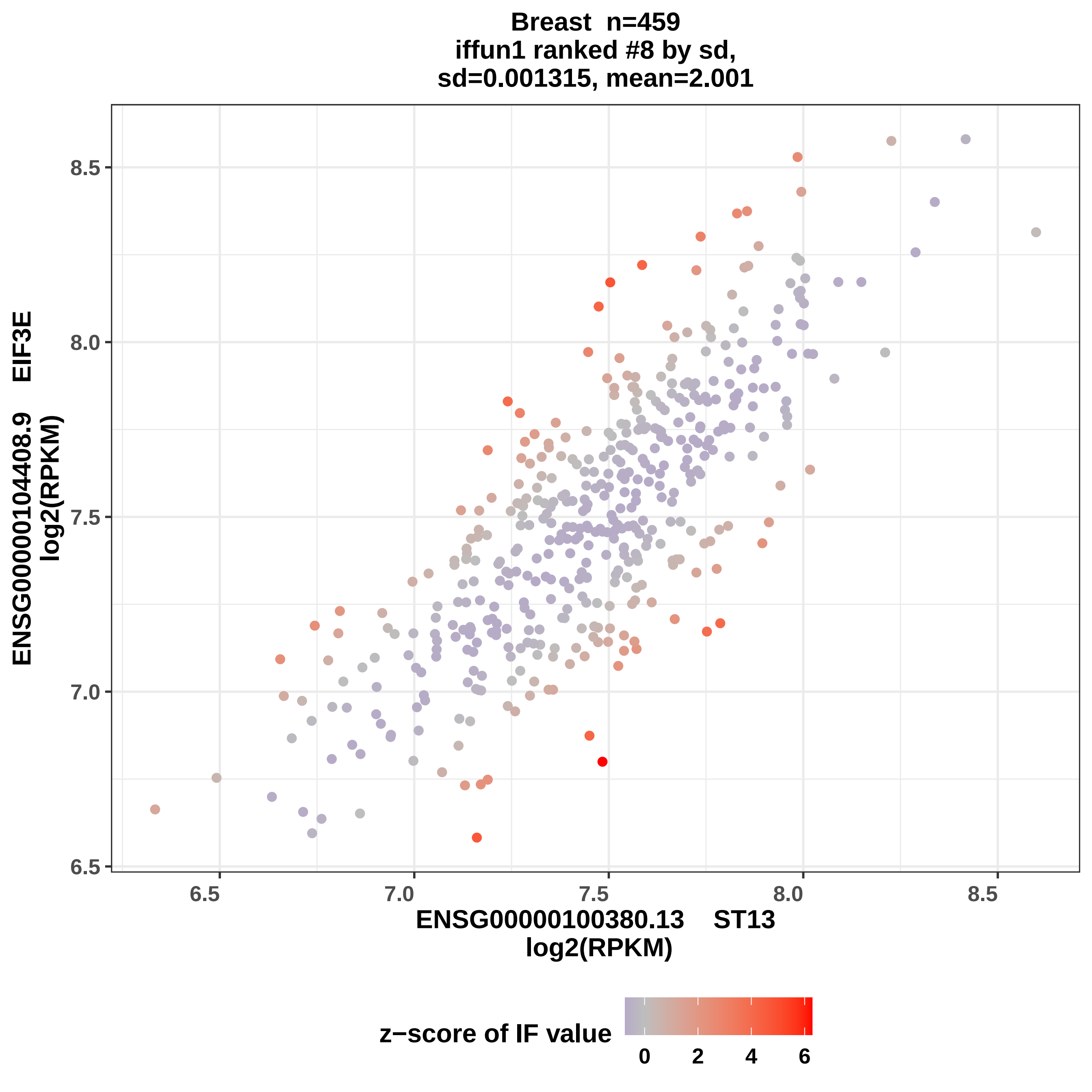

Breast n=459
iffun1 ranked #8 by sd,
sd=0.001315, mean=2.001
ENSG00000104408.9 EIF3E
log2(RPKM)
ENSG00000100380.13 ST13
log2(RPKM)
8.5
8.0
7.5
7.0
6.5
6.5
7.0
7.5
8.0
8.5
z−score of IF value
0
2
4
6

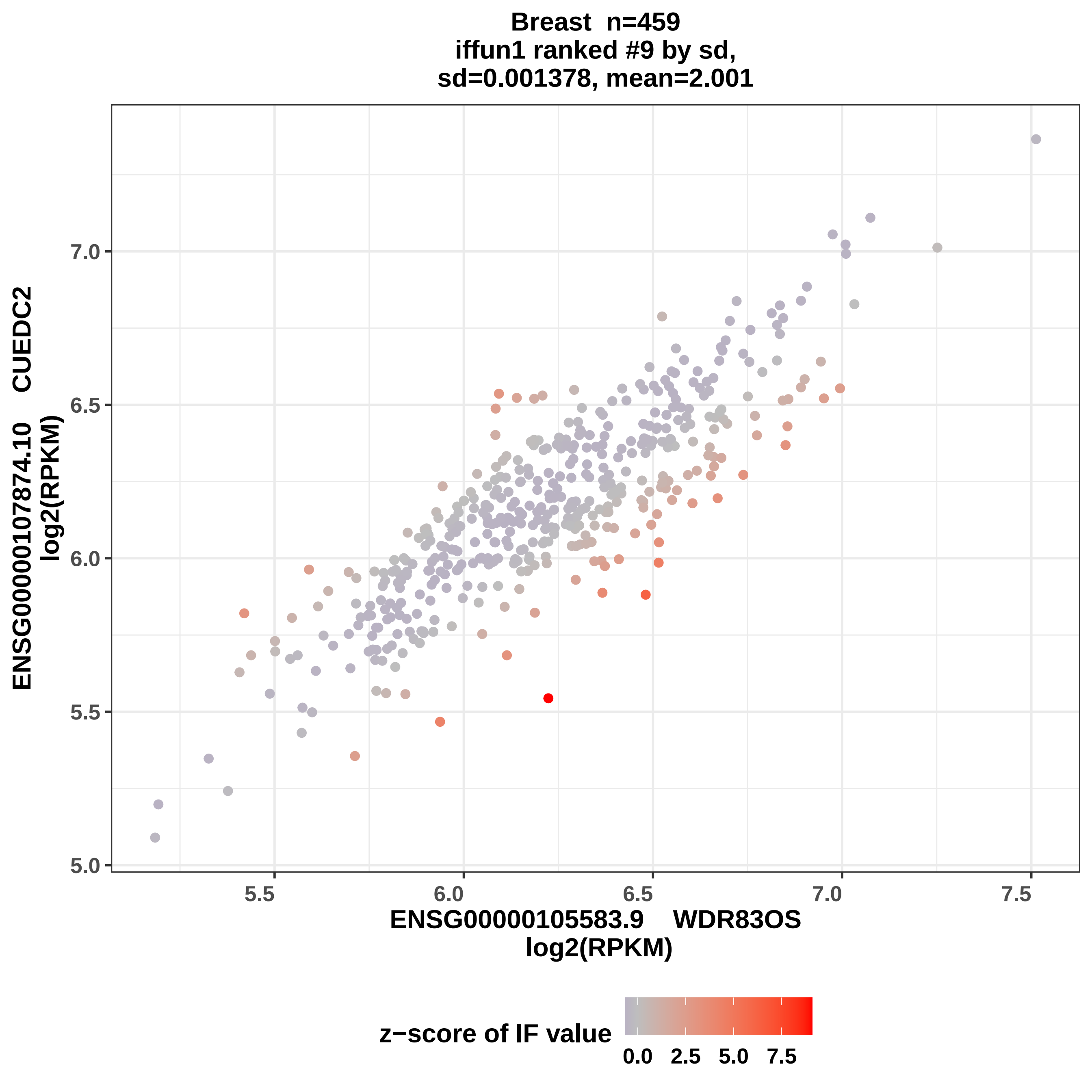

Breast n=459
iffun1 ranked #9 by sd,
sd=0.001378, mean=2.001
ENSG00000107874.10 CUEDC2
log2(RPKM)
7.0
6.5
6.0
5.5
5.0
5.5
6.0
6.5
7.0
7.5
ENSG00000105583.9 WDR83OS
log2(RPKM)
z−score of IF value
0.0
2.5
5.0
7.5

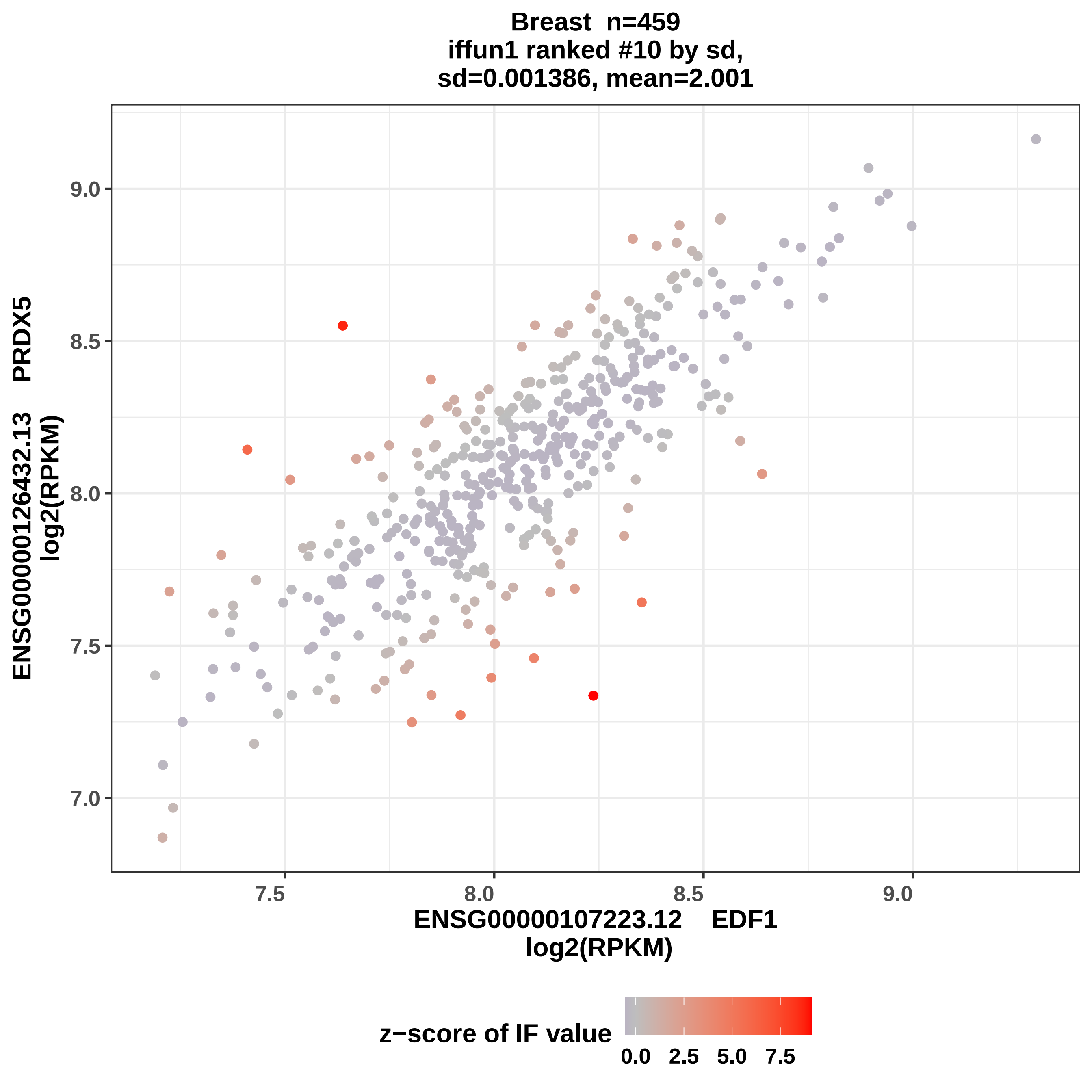
Breast n=459
iffun1 ranked #10 by sd,
sd=0.001386, mean=2.001
ENSG00000126432.13 PRDX5
log2(RPKM)
ENSG00000107223.12 EDF1
log2(RPKM)
7.0
7.5
8.0
8.5
9.0
z−score of IF value
0.0
2.5
5.0
7.5

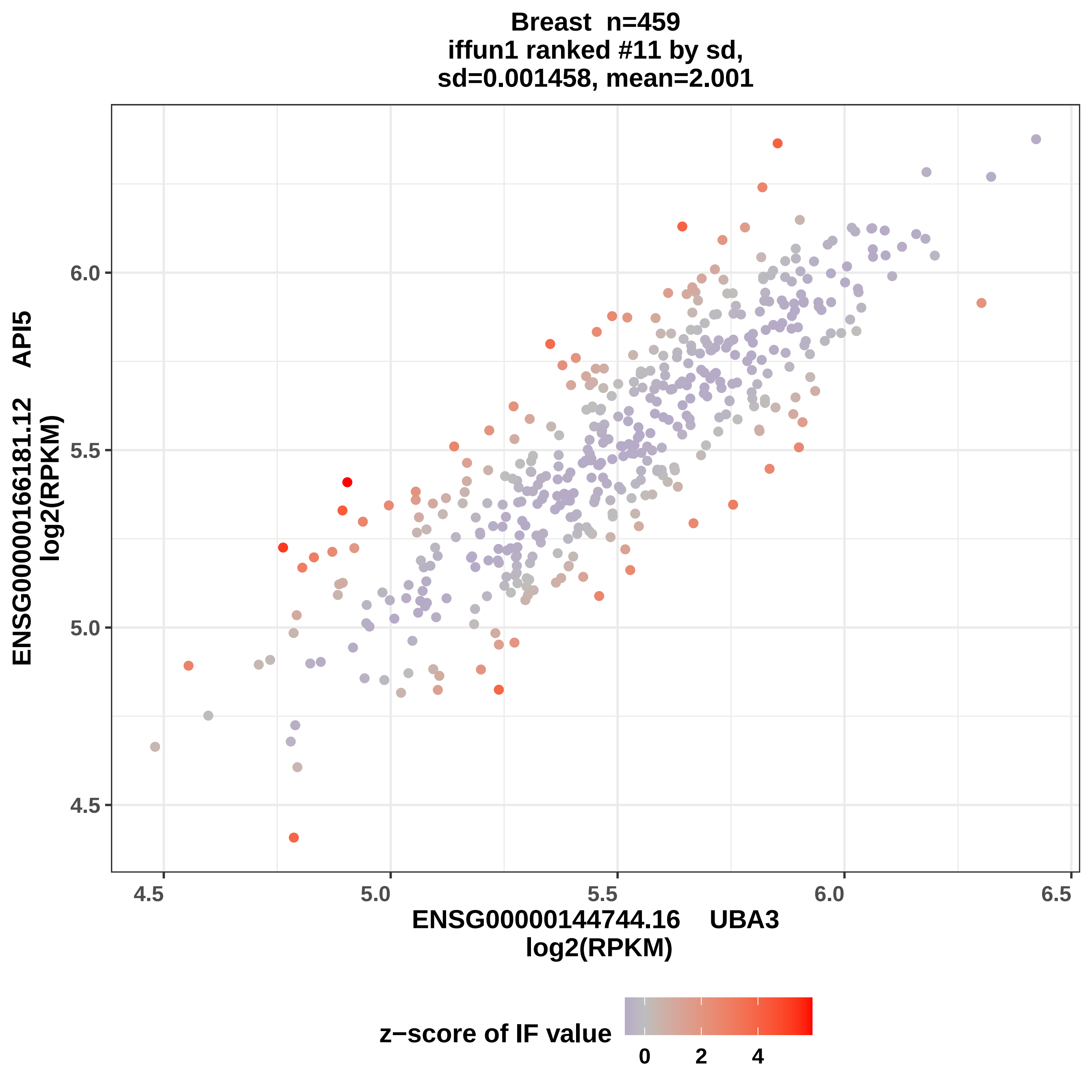
Breast n=459
iffun1 ranked #11 by sd,
sd=0.001458, mean=2.001
ENSG00000166181.12 API5
log2(RPKM)
6.0
5.5
5.0
4.5
4.5
5.0
5.5
6.0
6.5
ENSG00000144744.16 UBA3
log2(RPKM)
z−score of IF value
0
2
4

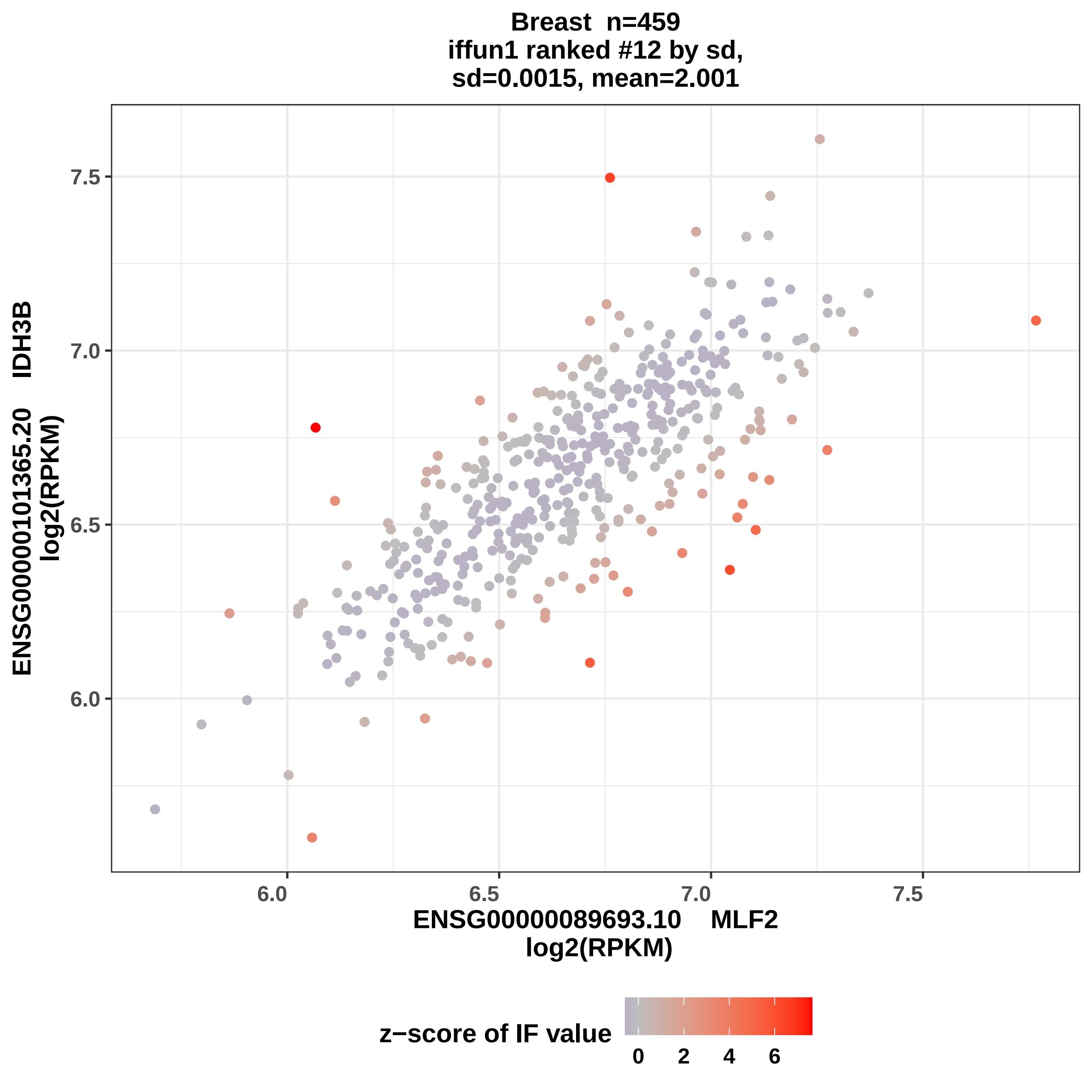

Breast n=459
iffun1 ranked #12 by sd,
sd=0.0015, mean=2.001
ENSG00000101365.20 IDH3B
log2(RPKM)
ENSG00000089693.10 MLF2
log2(RPKM)
7.5
7.0
6.5
6.0
6.0
6.5
7.0
7.5
z−score of IF value
0
2
4
6

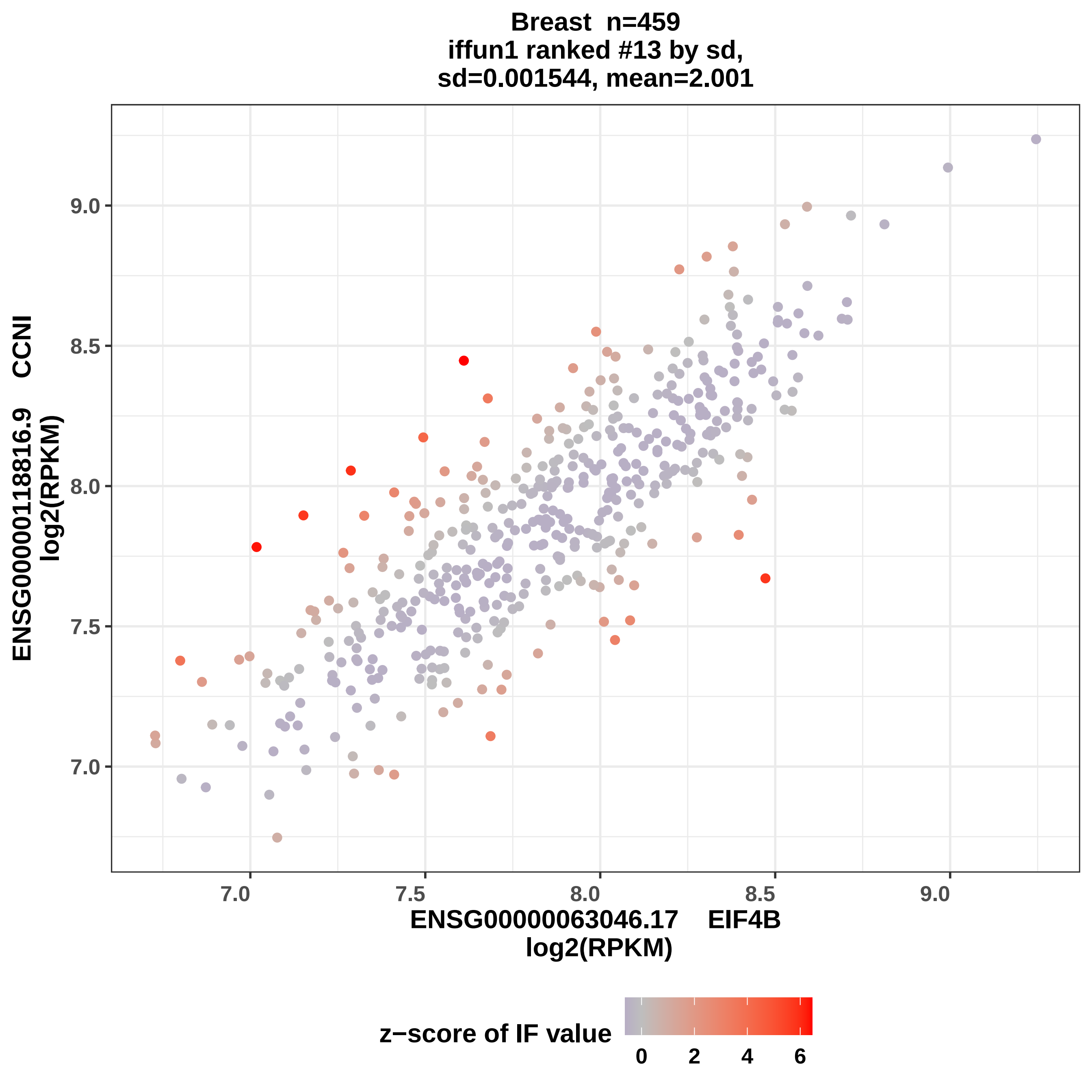
Breast n=459
iffun1 ranked #13 by sd,
sd=0.001544, mean=2.001
ENSG00000118816.9 CCNI
log2(RPKM)
9.0
8.5
8.0
7.5
7.0
ENSG00000063046.17 EIF4B
log2(RPKM)
z−score of IF value
0
2
4
6

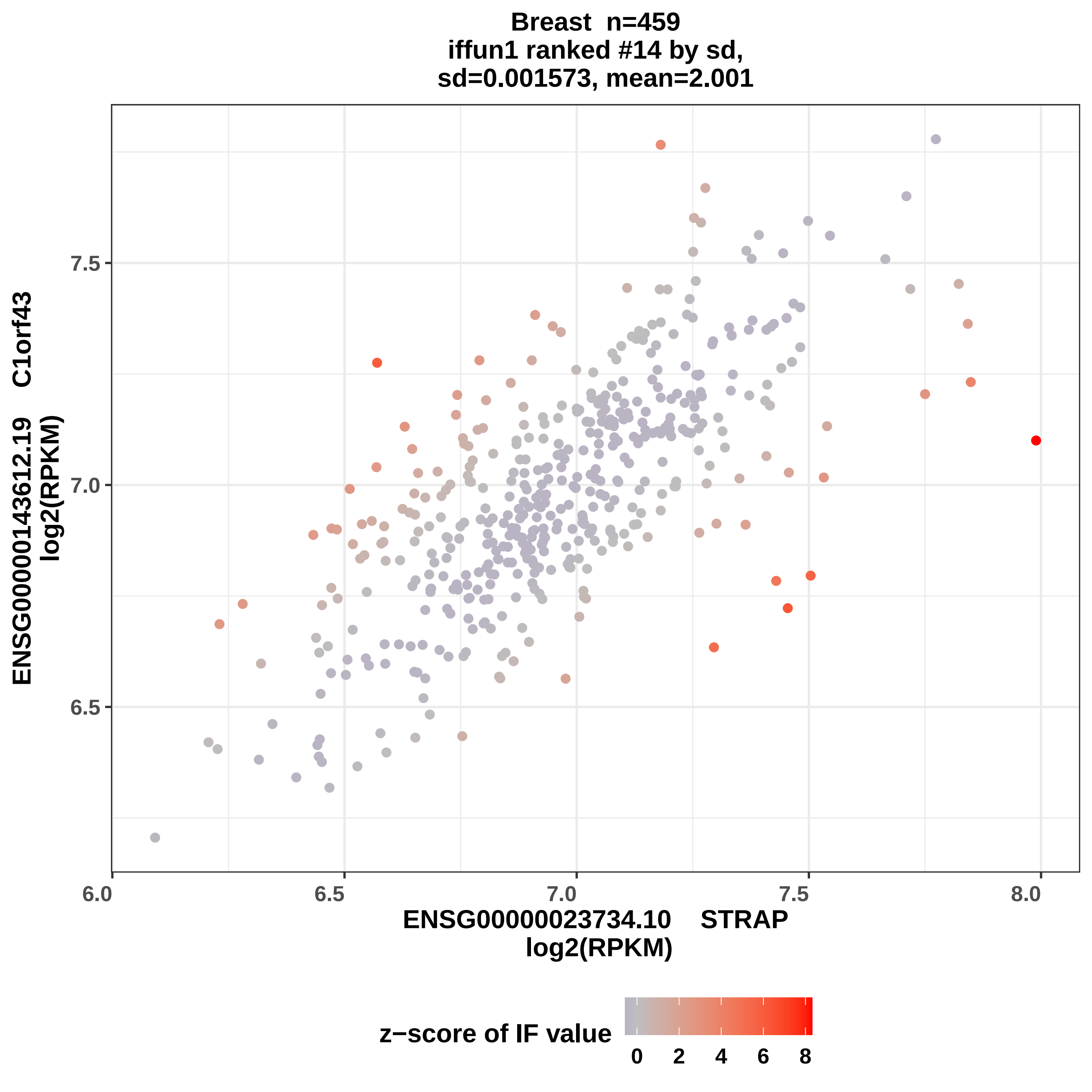
Breast n=459
iffun1 ranked #14 by sd,
sd=0.001573, mean=2.001
ENSG00000143612.19 C1orf43
log2(RPKM)
7.5
7.0
6.5
6.0
6.5
7.0
7.5
8.0
ENSG00000023734.10 STRAP
log2(RPKM)
z−score of IF value
0
2
4
6
8

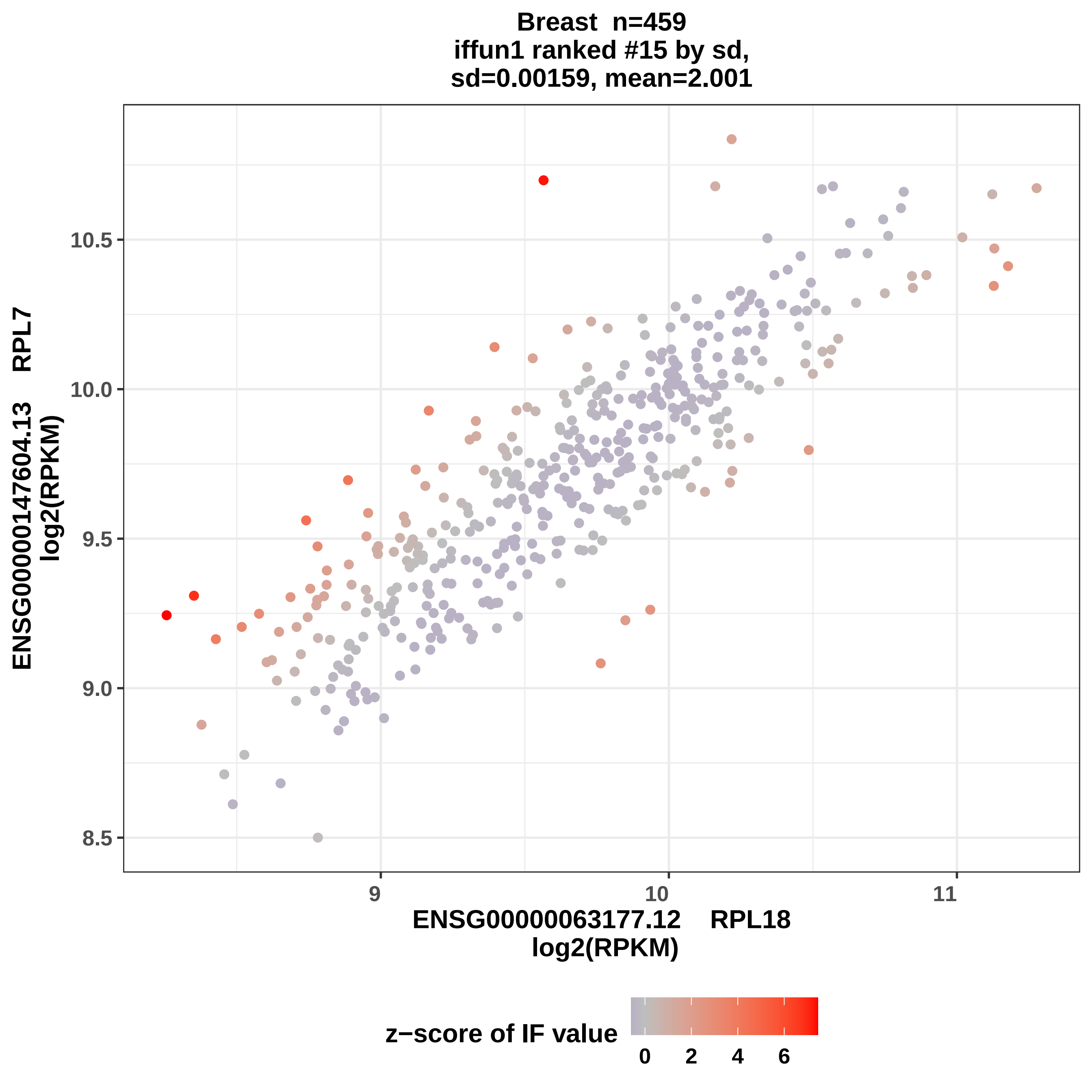

Breast n=459
iffun1 ranked #15 by sd,
sd=0.00159, mean=2.001
ENSG00000147604.13 RPL7
log2(RPKM)
10.5
10.0
9.5
9.0
8.5
9
10
11
ENSG00000063177.12 RPL18
log2(RPKM)
z−score of IF value
0
2
4
6

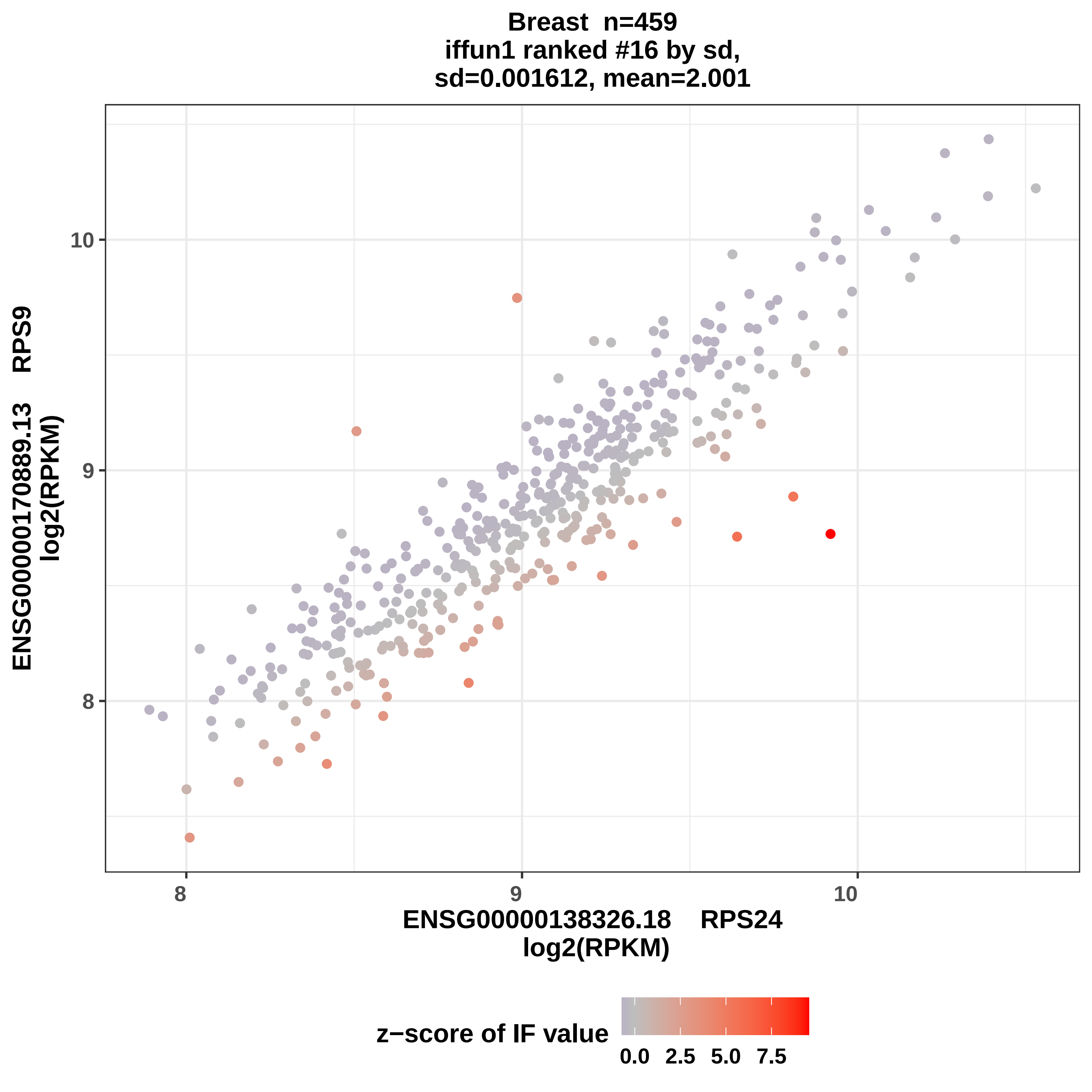
Breast n=459
iffun1 ranked #16 by sd,
sd=0.001612, mean=2.001
ENSG00000170889.13 RPS9
log2(RPKM)
10
9
8
8
9
10
ENSG00000138326.18 RPS24
log2(RPKM)
z−score of IF value
0.0
2.5
5.0
7.5

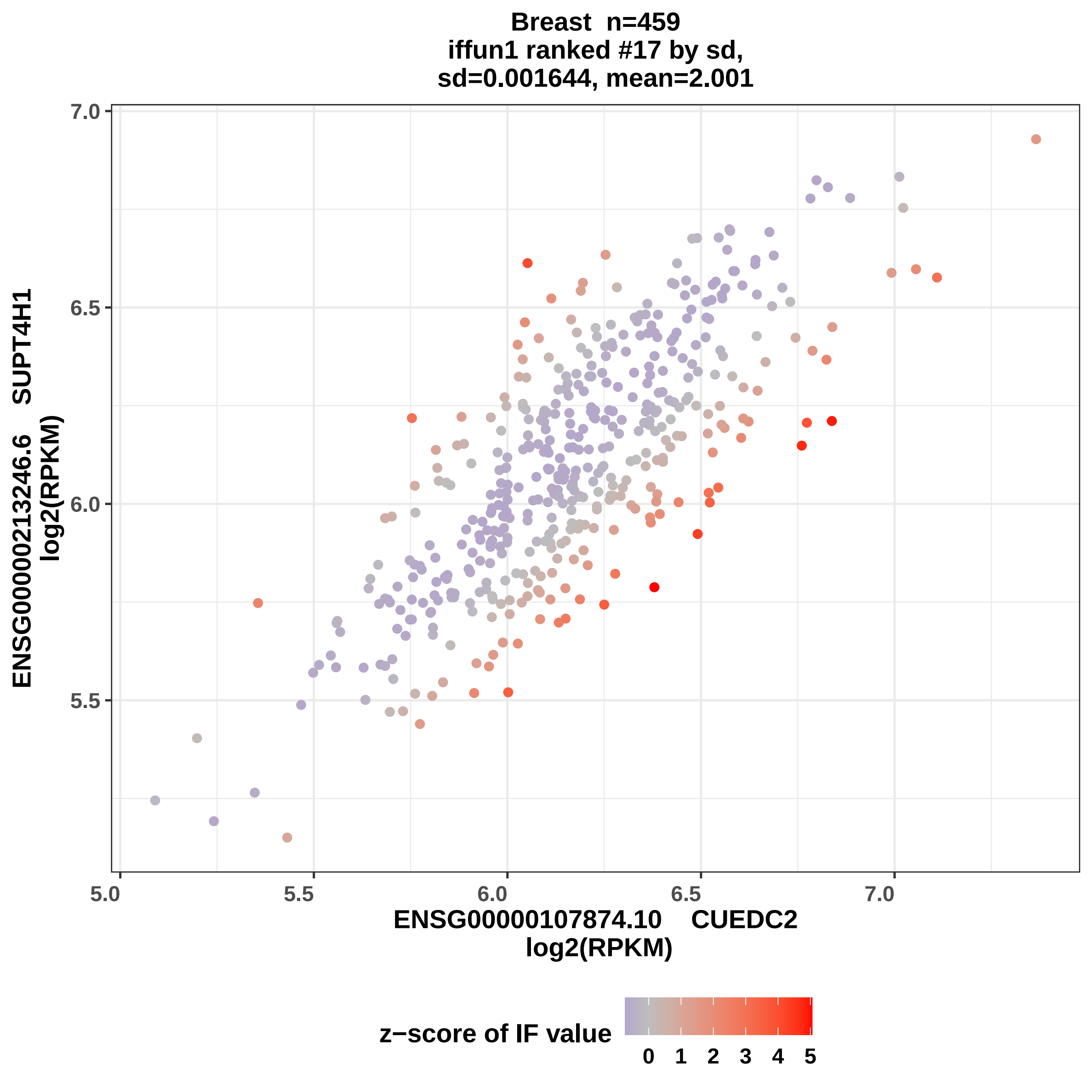

Breast n=459
iffun1 ranked #17 by sd,
sd=0.001644, mean=2.001
ENSG00000213246.6 SUPT4H1
log2(RPKM)
7.0
6.5
6.0
5.5
5.0
5.5
6.0
6.5
7.0
ENSG00000107874.10 CUEDC2
log2(RPKM)
z−score of IF value
0
1
2
3
4
5

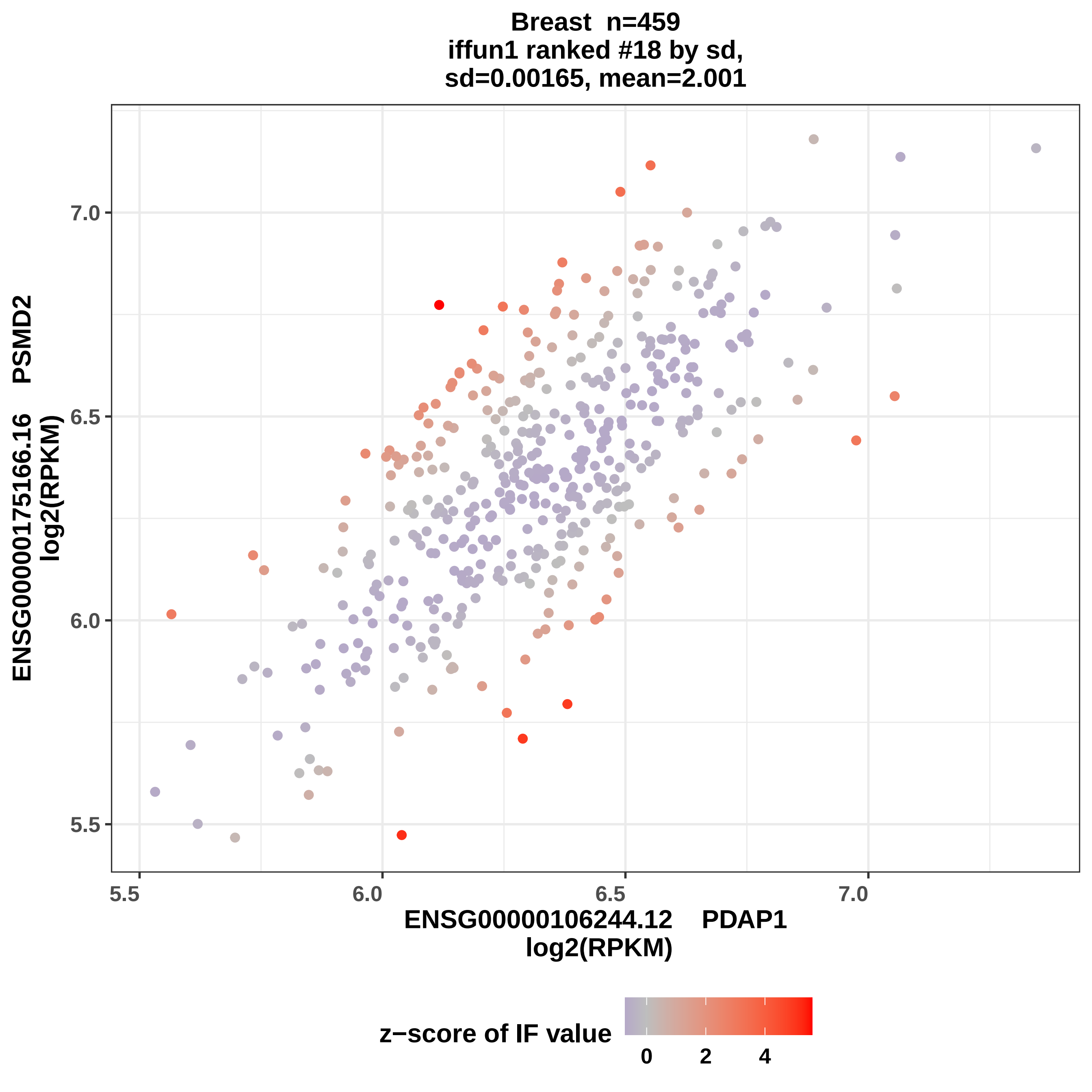

Breast n=459
iffun1 ranked #18 by sd,
sd=0.00165, mean=2.001
ENSG00000175166.16 PSMD2
log2(RPKM)
7.0
6.5
6.0
5.5
5.5
6.0
6.5
7.0
ENSG00000106244.12 PDAP1
log2(RPKM)
z−score of IF value
0
2
4

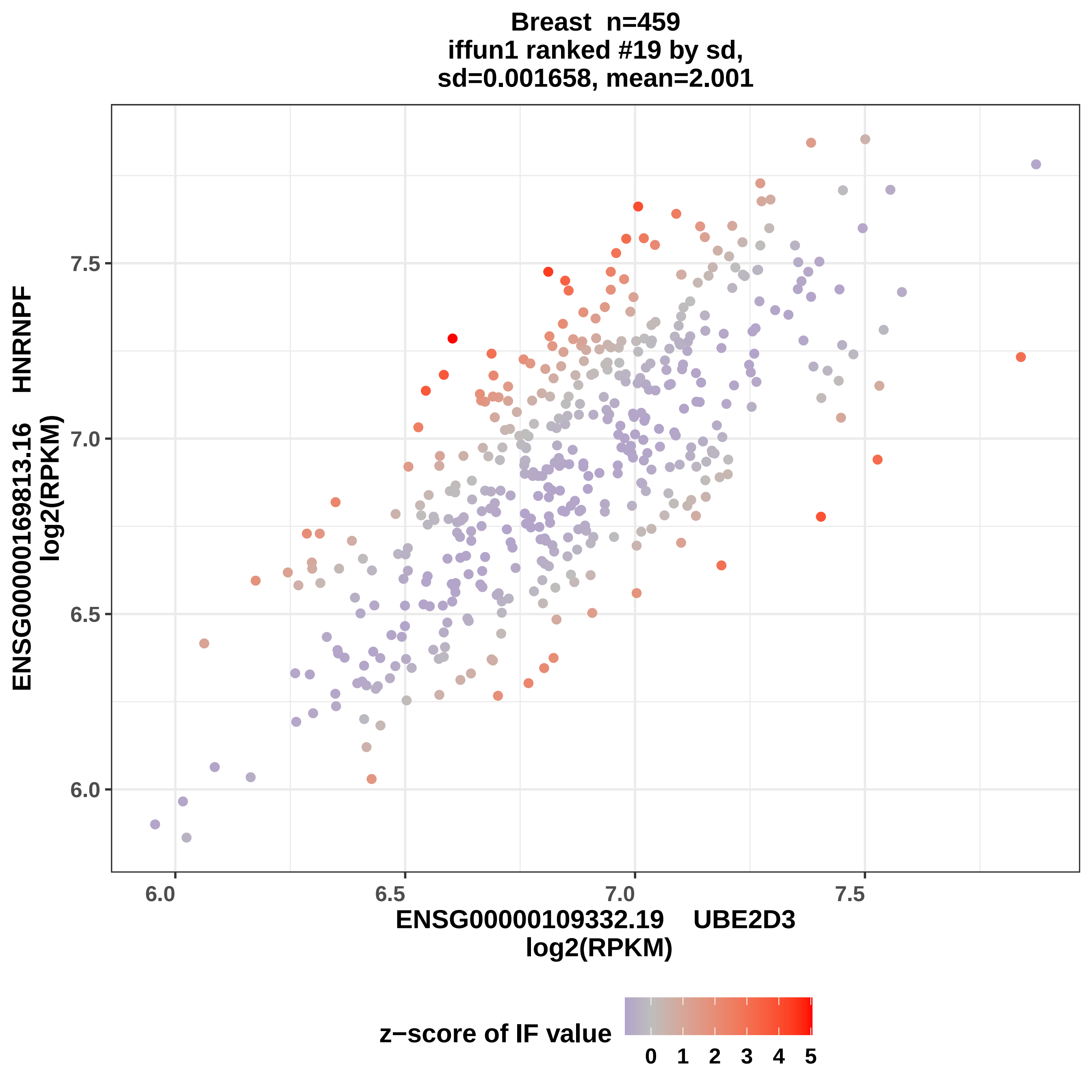
Breast n=459
iffun1 ranked #19 by sd,
sd=0.001658, mean=2.001
ENSG00000169813.16 HNRNPF
log2(RPKM)
7.5
7.0
6.5
6.0
6.0
6.5
7.0
7.5
ENSG00000109332.19 UBE2D3
log2(RPKM)
z−score of IF value
0
1
2
3
4
5

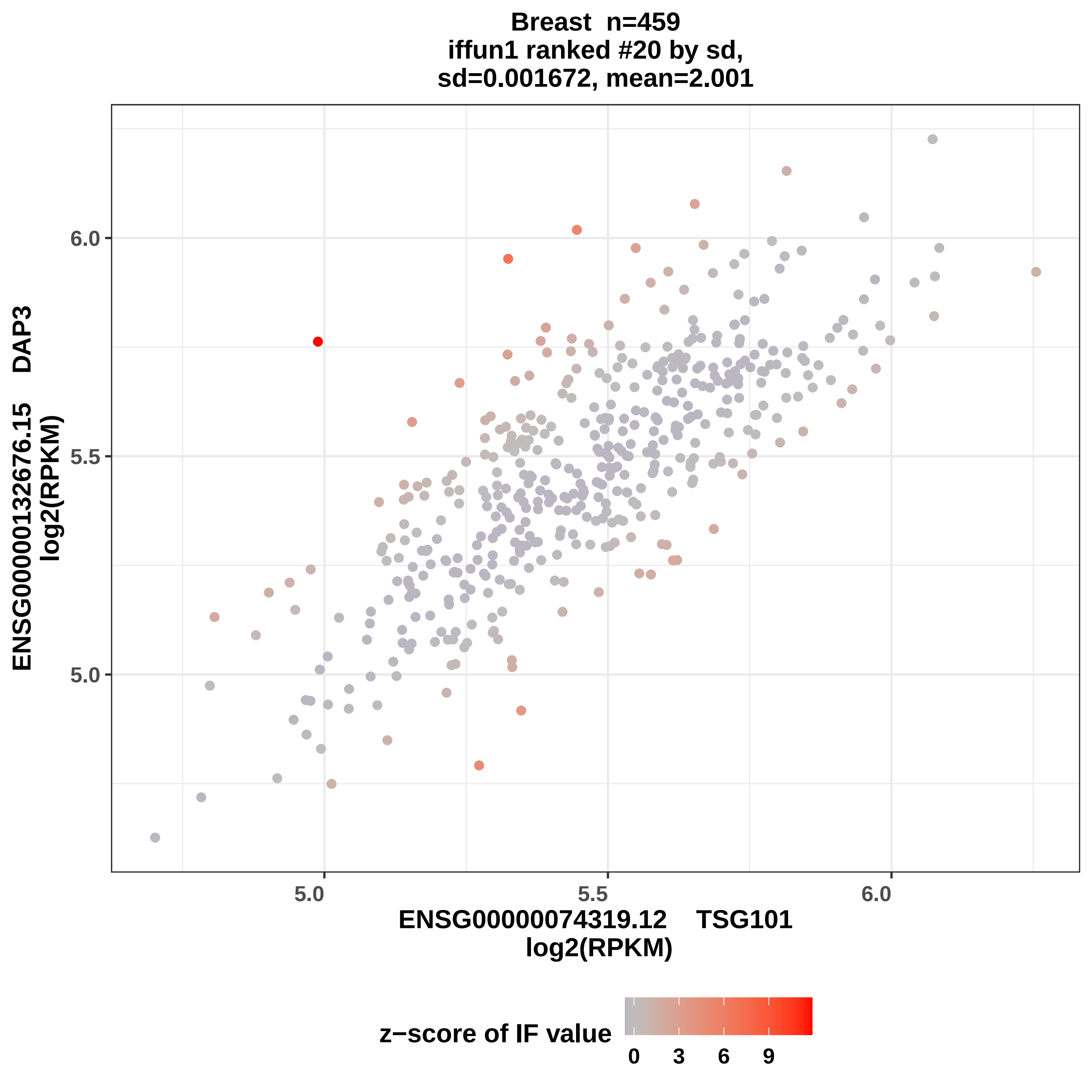

Breast n=459
iffun1 ranked #20 by sd,
sd=0.001672, mean=2.001
ENSG00000132676.15 DAP3
log2(RPKM)
6.0
5.5
5.0
5.0
5.5
6.0
ENSG00000074319.12 TSG101
log2(RPKM)
z−score of IF value
0
3
6
9

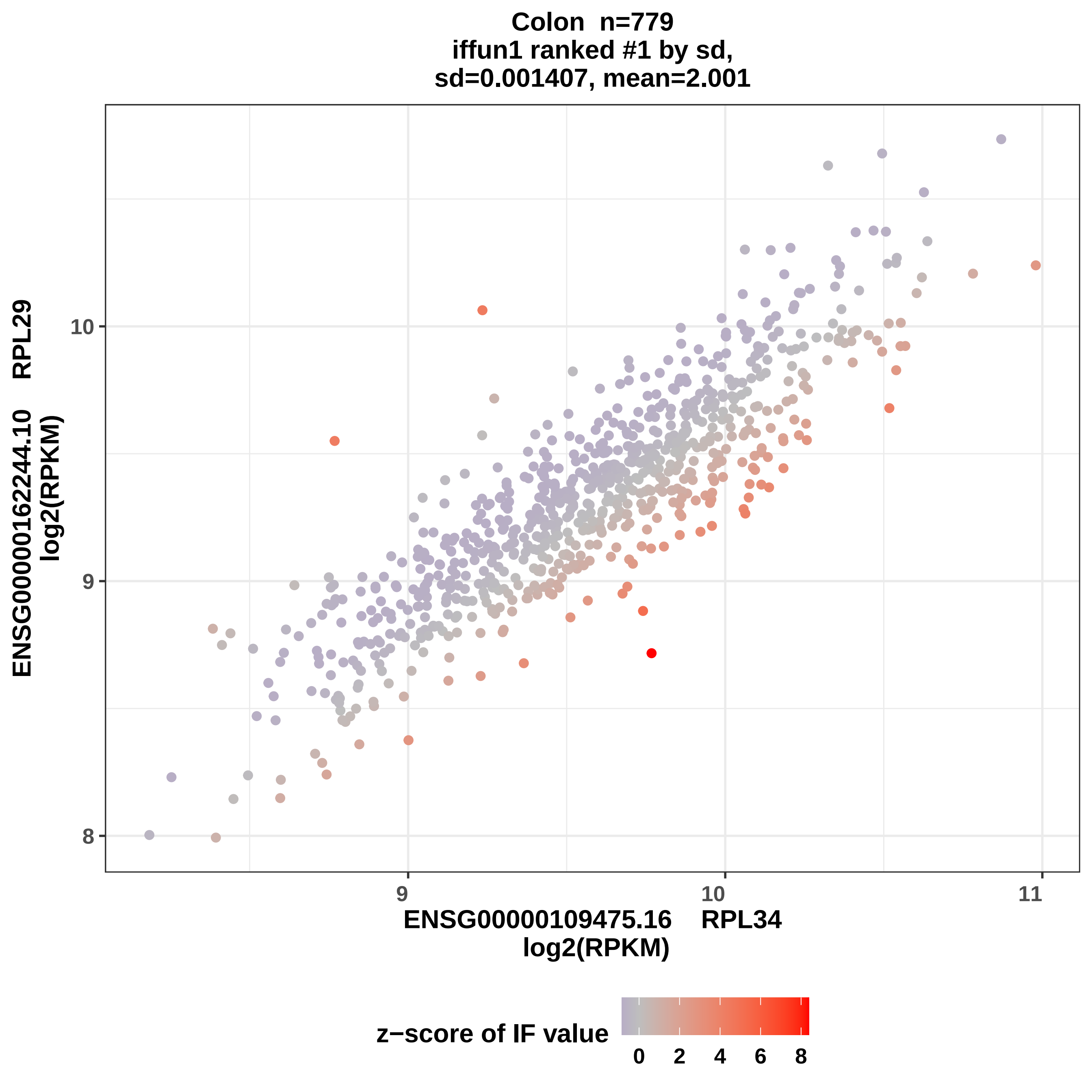
Colon n=779
iffun1 ranked #1 by sd,
sd=0.001407, mean=2.001
ENSG00000162244.10 RPL29
log2(RPKM)
ENSG00000109475.16 RPL34
log2(RPKM)
8
9
10
9
10
11
z−score of IF value
0
2
4
6
8

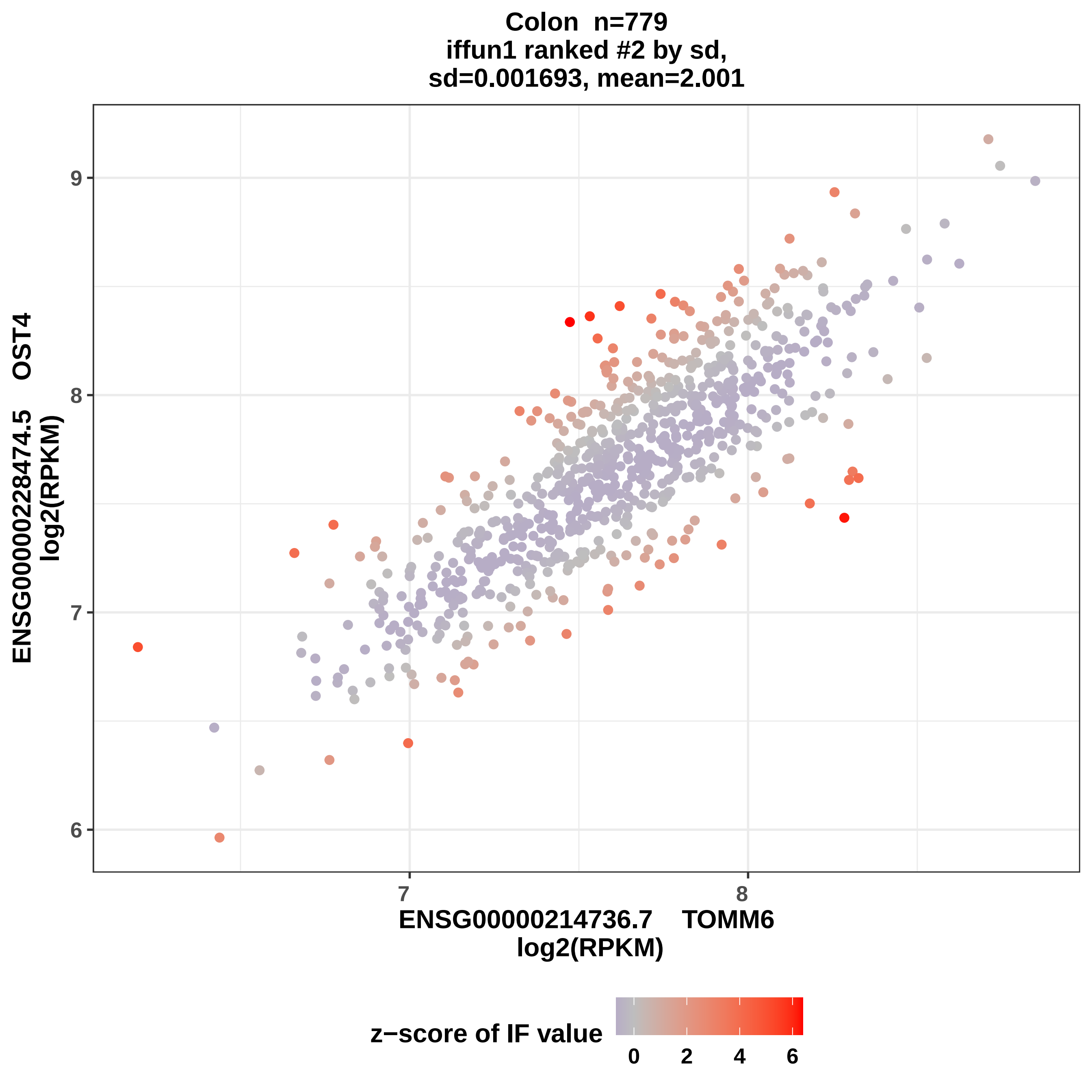

Colon n=779
iffun1 ranked #2 by sd,
sd=0.001693, mean=2.001
ENSG00000228474.5 OST4
log2(RPKM)
9
8
7
6
7
8
ENSG00000214736.7 TOMM6
log2(RPKM)
z−score of IF value
0
2
4
6

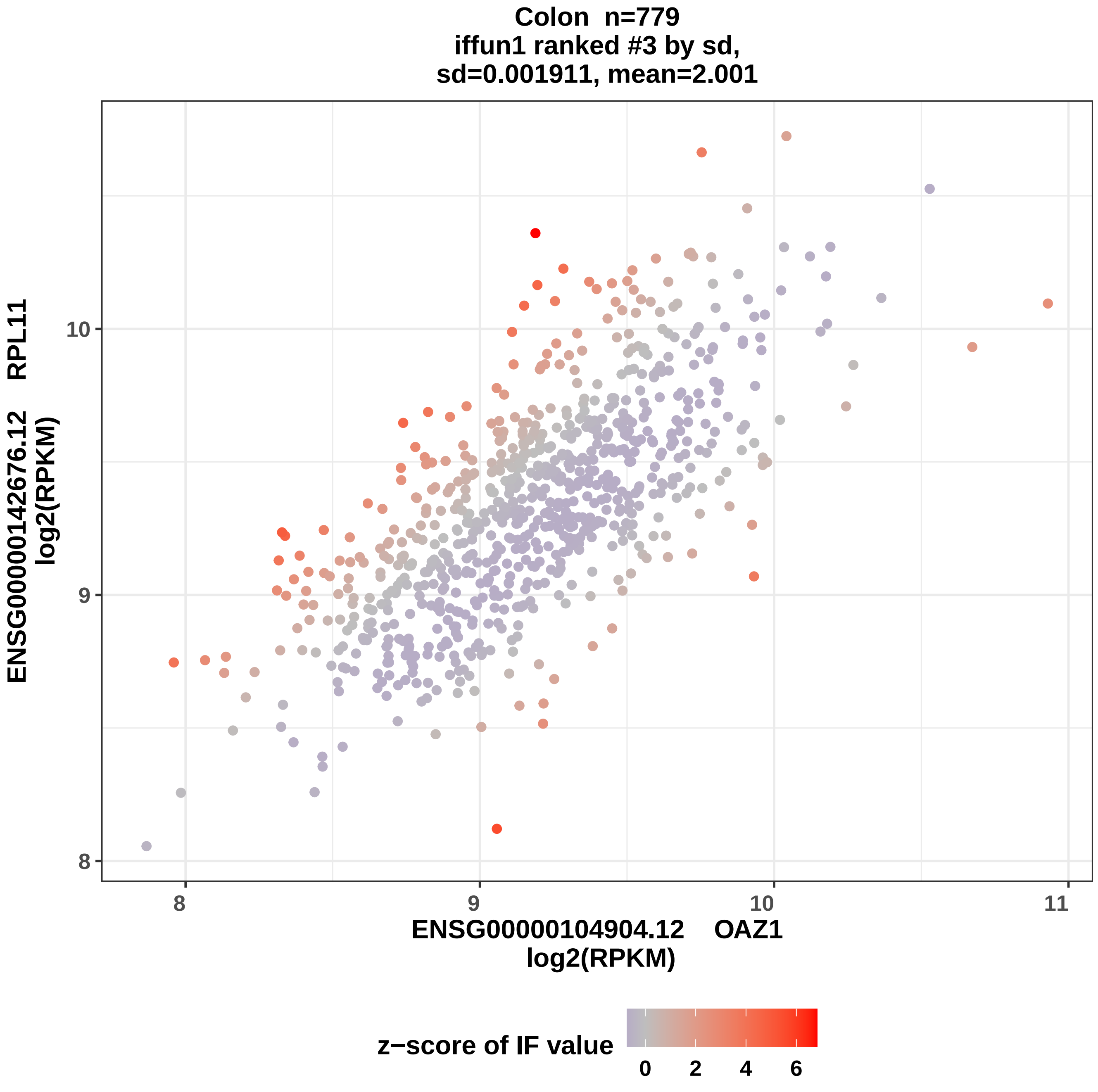

Colon n=779
iffun1 ranked #3 by sd,
sd=0.001911, mean=2.001
ENSG00000142676.12 RPL11
log2(RPKM)
10
9
8
8
9
10
11
ENSG00000104904.12 OAZ1
log2(RPKM)
z−score of IF value
0
2
4
6

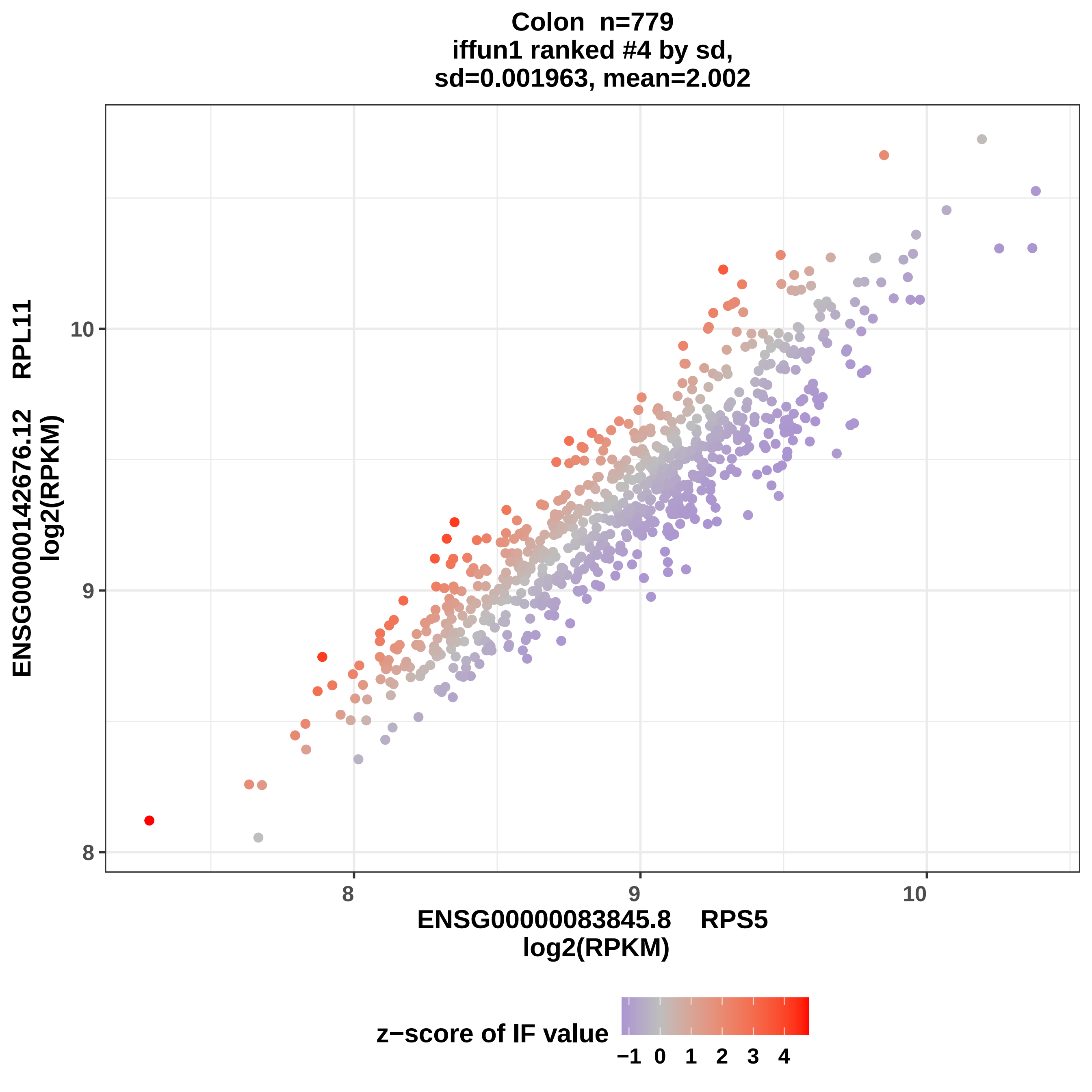
Colon n=779
iffun1 ranked #4 by sd,
sd=0.001963, mean=2.002
ENSG00000142676.12 RPL11
log2(RPKM)
10
9
8
8
9
10
ENSG00000083845.8 RPS5
log2(RPKM)
z−score of IF value
−1 0 1 2 3 4

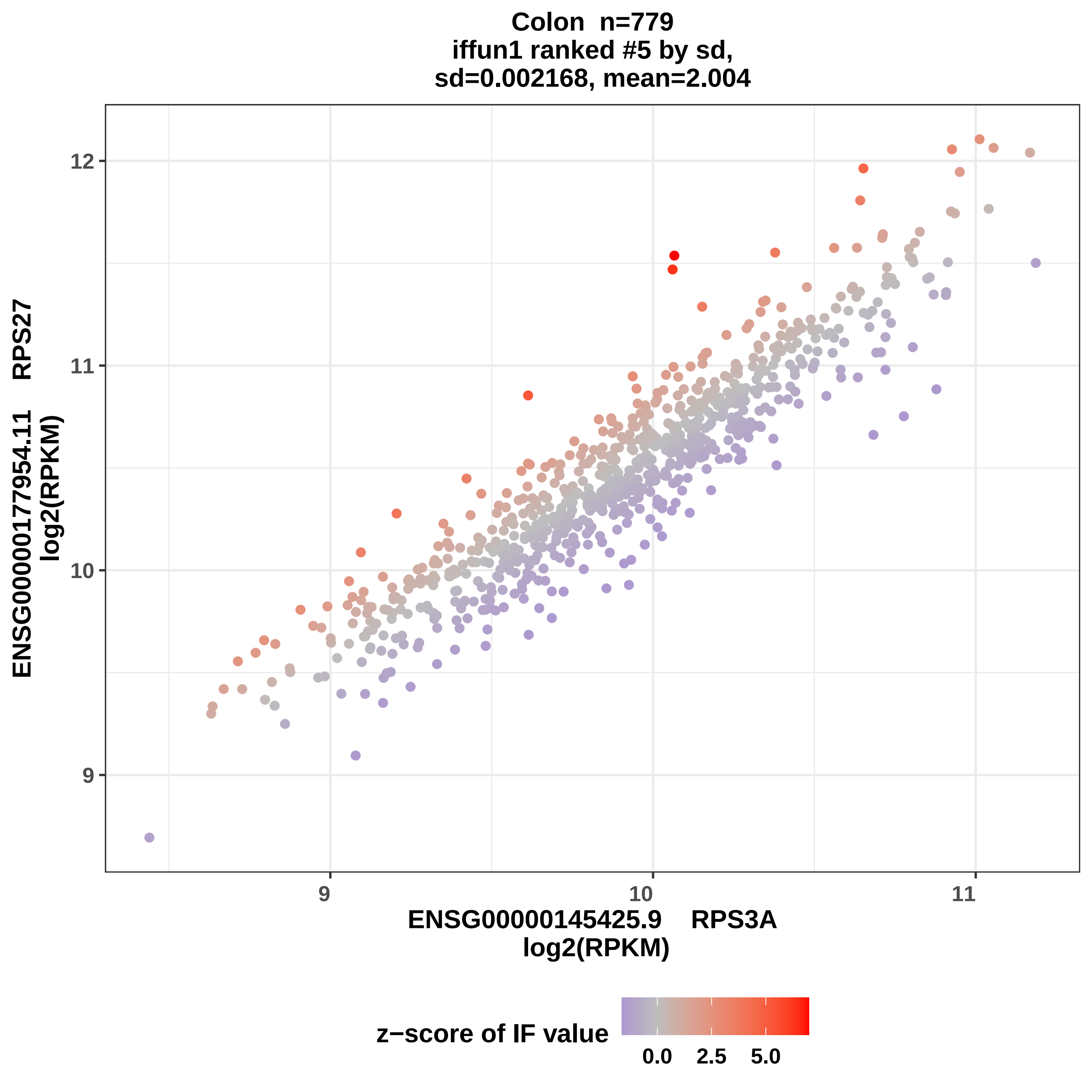

Colon n=779
iffun1 ranked #5 by sd,
sd=0.002168, mean=2.004
ENSG00000177954.11 RPS27
log2(RPKM)
12
11
10
9
9
10
11
ENSG00000145425.9 RPS3A
log2(RPKM)
z−score of IF value
0.0
2.5
5.0

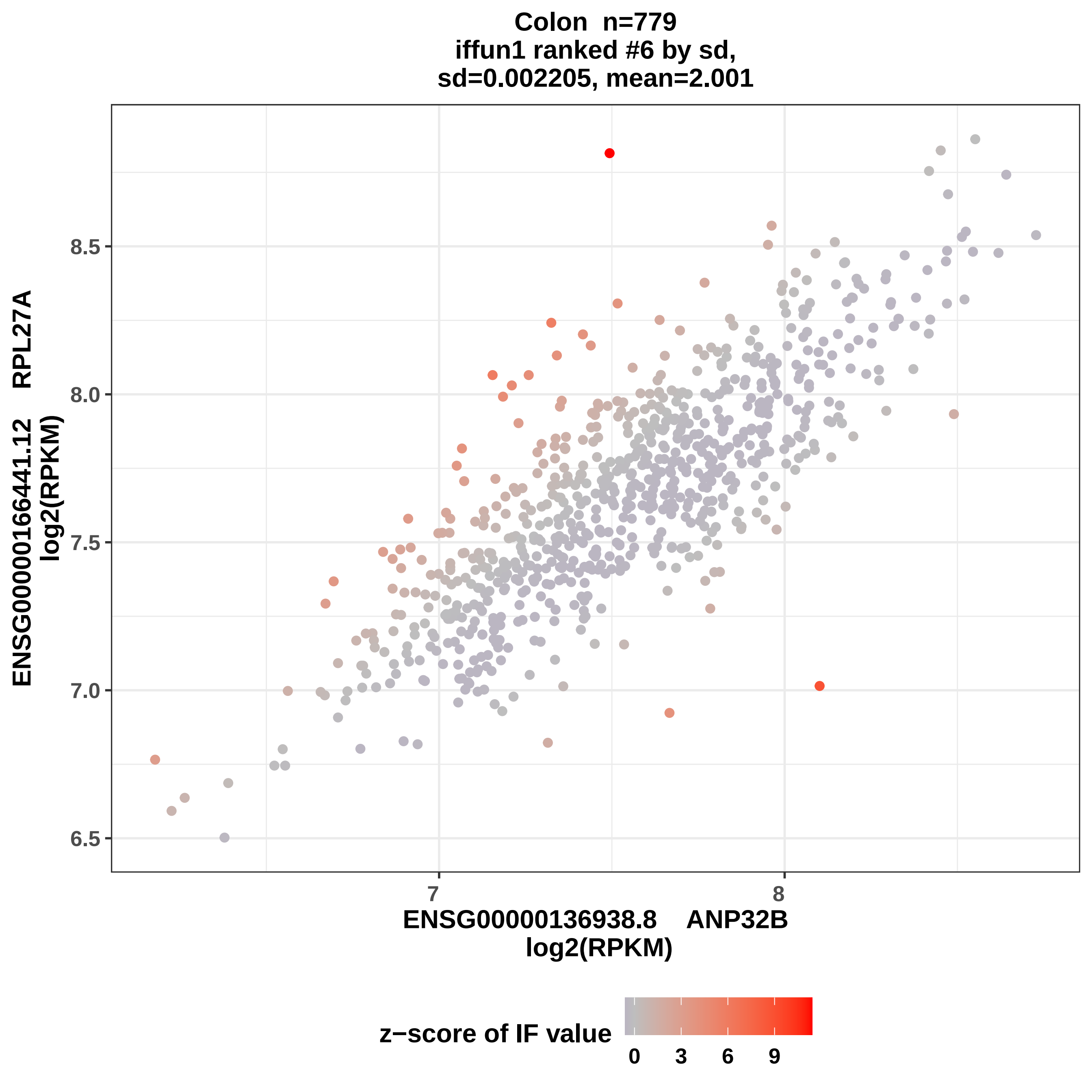
Colon n=779
iffun1 ranked #6 by sd,
sd=0.002205, mean=2.001
ENSG00000166441.12 RPL27A
log2(RPKM)
8.5
8.0
7.5
7.0
6.5
7
8
ENSG00000136938.8 ANP32B
log2(RPKM)
z−score of IF value
0
3
6
9

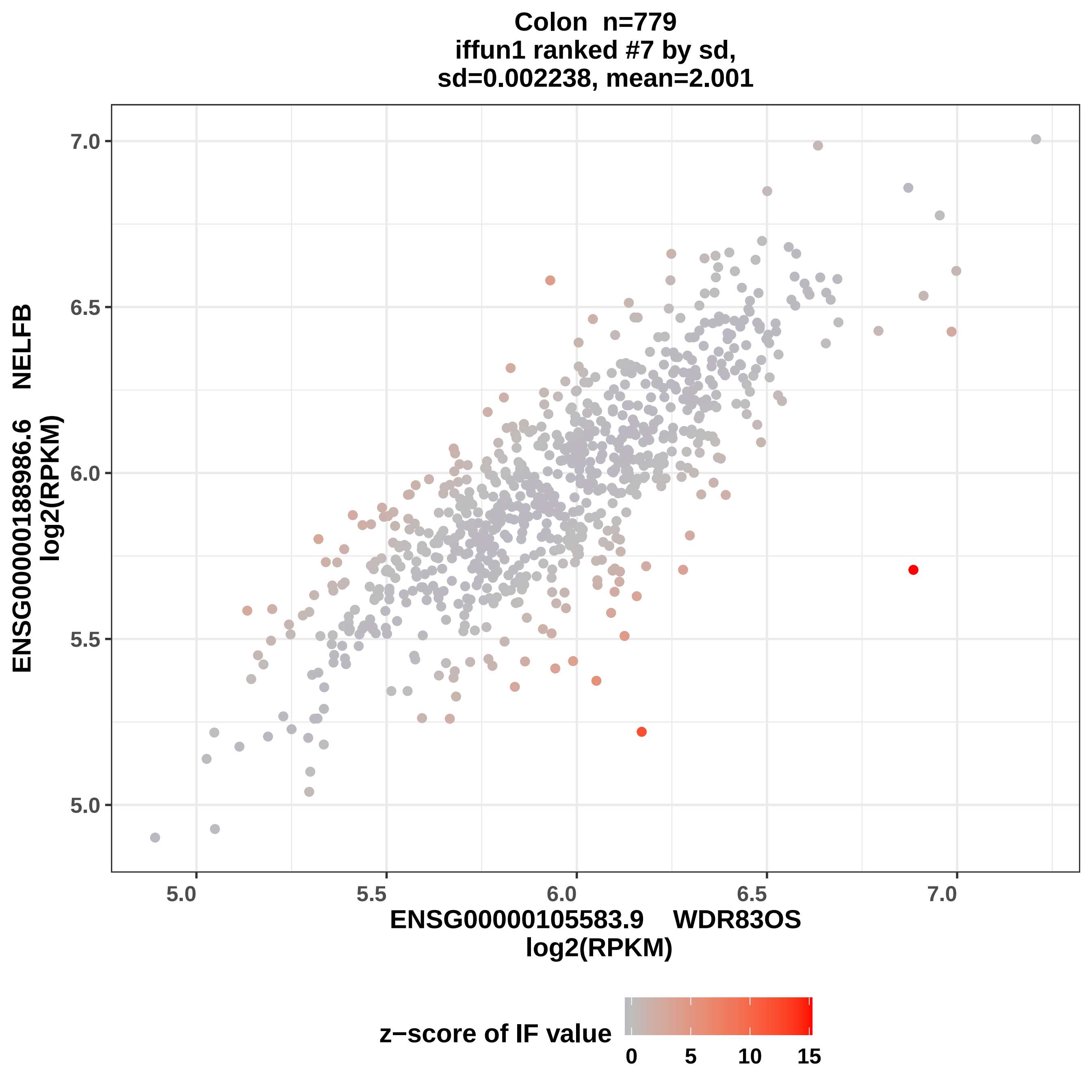
Colon n=779
iffun1 ranked #7 by sd,
sd=0.002238, mean=2.001
ENSG00000188986.6 NELFB
log2(RPKM)
ENSG00000105583.9 WDR83OS
log2(RPKM)
7.0
6.5
6.0
5.5
5.0
5.0
5.5
6.0
6.5
7.0
z−score of IF value
0
5
10
15

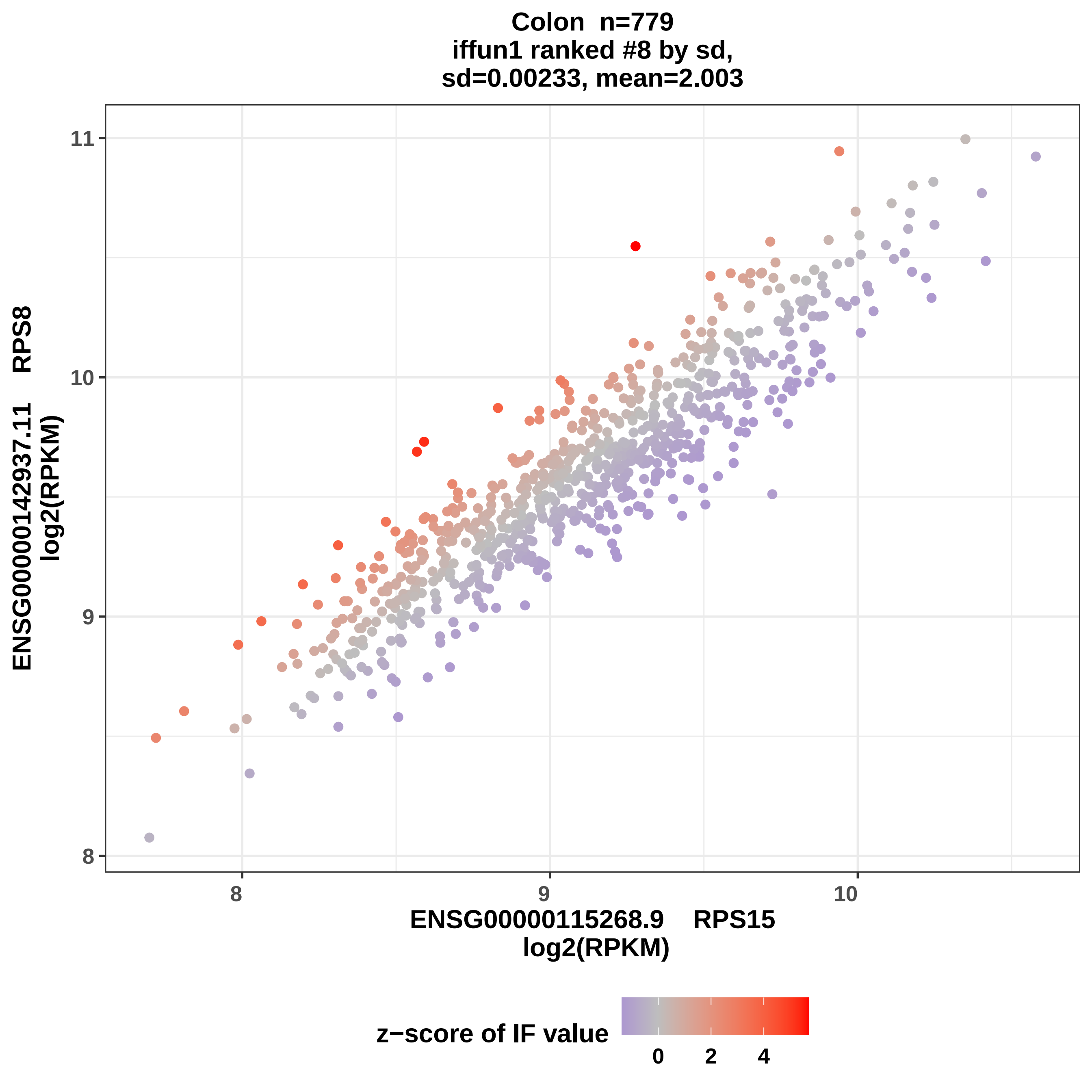

Colon n=779
iffun1 ranked #8 by sd,
sd=0.00233, mean=2.003
ENSG00000142937.11 RPS8
log2(RPKM)
ENSG00000115268.9 RPS15
log2(RPKM)
8
9
10
11
z−score of IF value
0
2
4

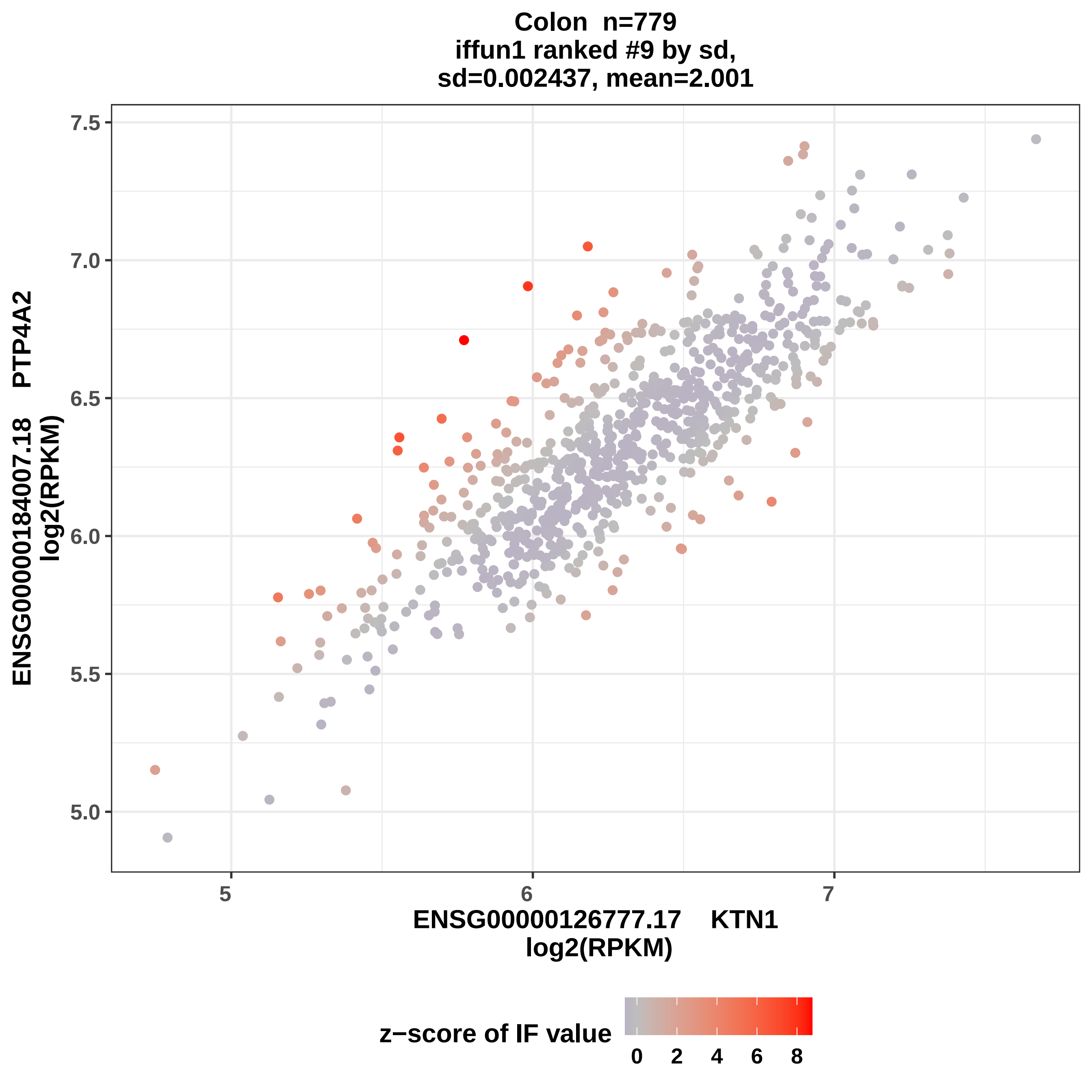

Colon n=779
iffun1 ranked #9 by sd,
sd=0.002437, mean=2.001
ENSG00000184007.18 PTP4A2
log2(RPKM)
7.5
7.0
6.5
6.0
5.5
5.0
5
6
7
ENSG00000126777.17 KTN1
log2(RPKM)
z−score of IF value
0
2
4
6
8

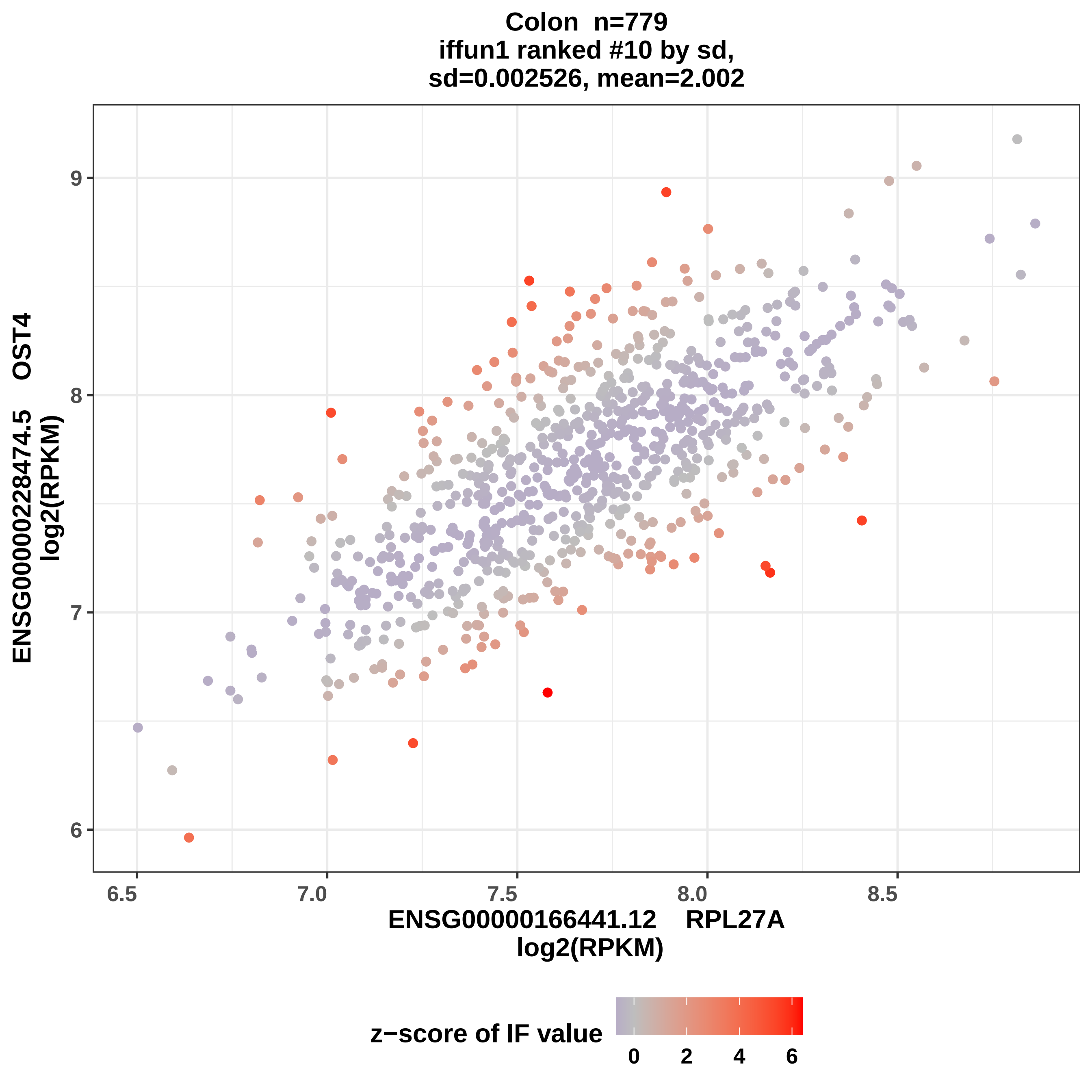

Colon n=779
iffun1 ranked #10 by sd,
sd=0.002526, mean=2.002
ENSG00000228474.5 OST4
log2(RPKM)
9
8
7
6
6.5
7.0
7.5
8.0
8.5
ENSG00000166441.12 RPL27A
log2(RPKM)
z−score of IF value
0
2
4
6

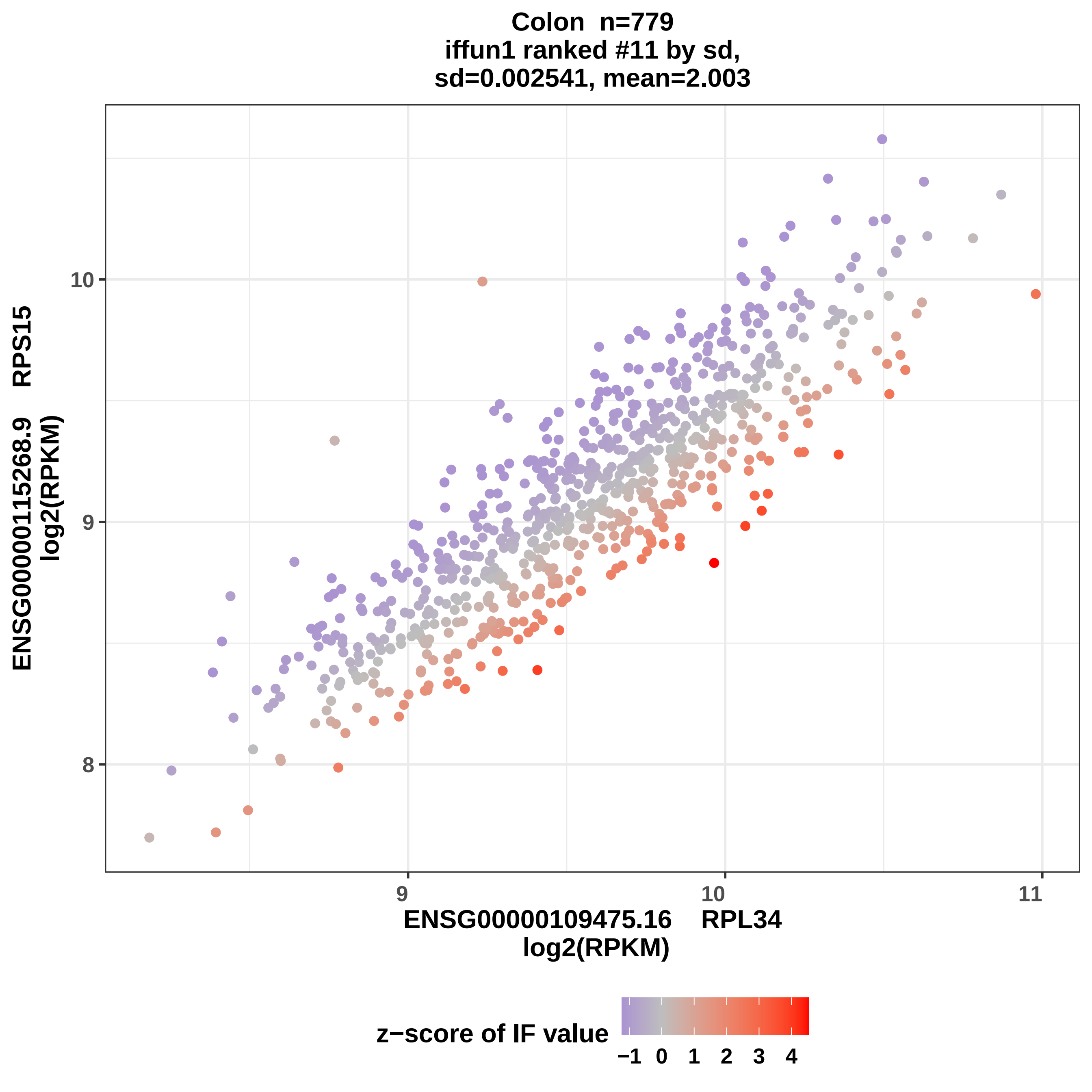
Colon n=779
iffun1 ranked #11 by sd,
sd=0.002541, mean=2.003
ENSG00000115268.9 RPS15
log2(RPKM)
10
9
8
9
10
11
ENSG00000109475.16 RPL34
log2(RPKM)
z−score of IF value
−1 0 1 2 3 4

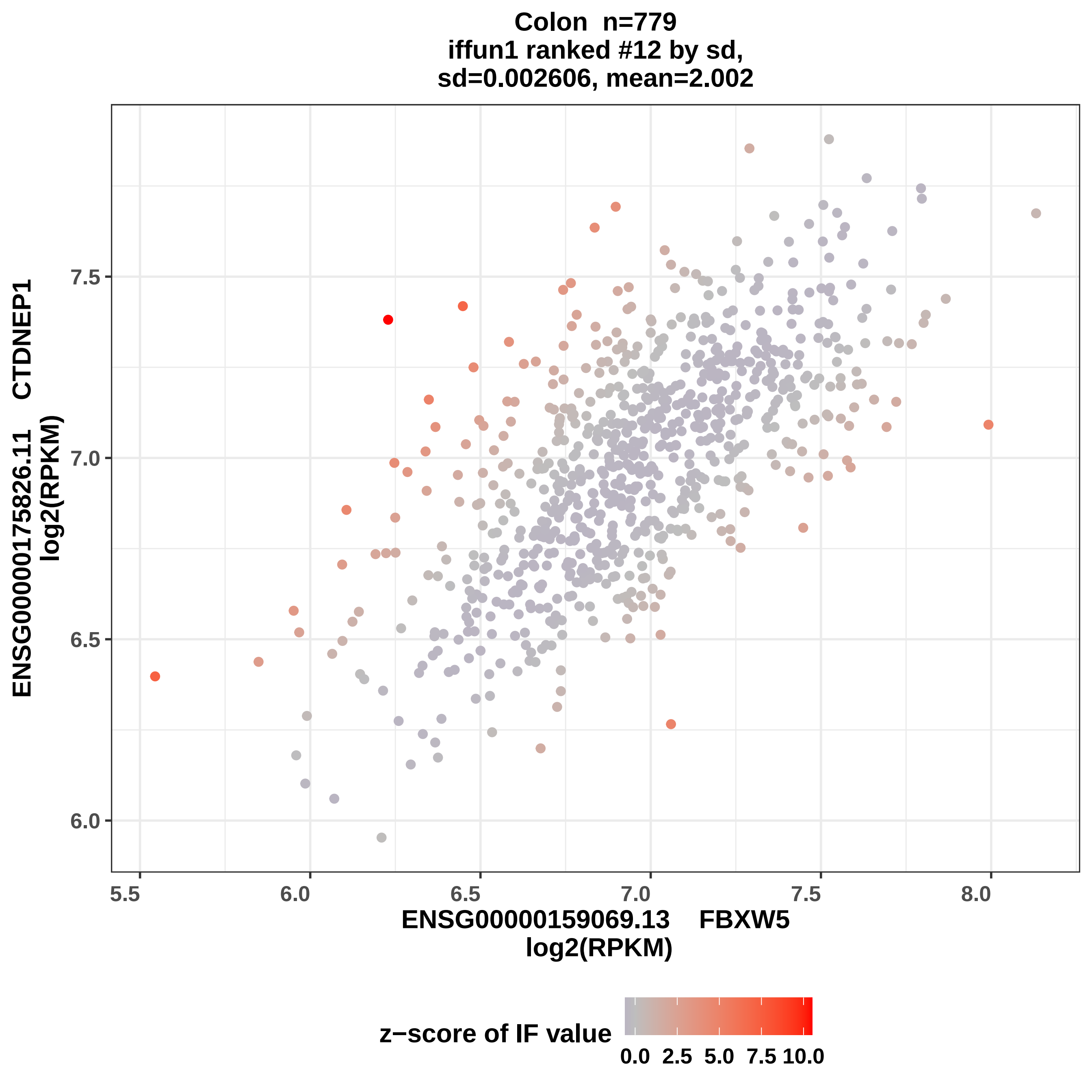
Colon n=779
iffun1 ranked #12 by sd,
sd=0.002606, mean=2.002
ENSG00000175826.11 CTDNEP1
log2(RPKM)
7.5
7.0
6.5
6.0
5.5
6.0
6.5
7.0
7.5
8.0
ENSG00000159069.13 FBXW5
log2(RPKM)
z−score of IF value
0.0
2.5
5.0
7.5
10.0

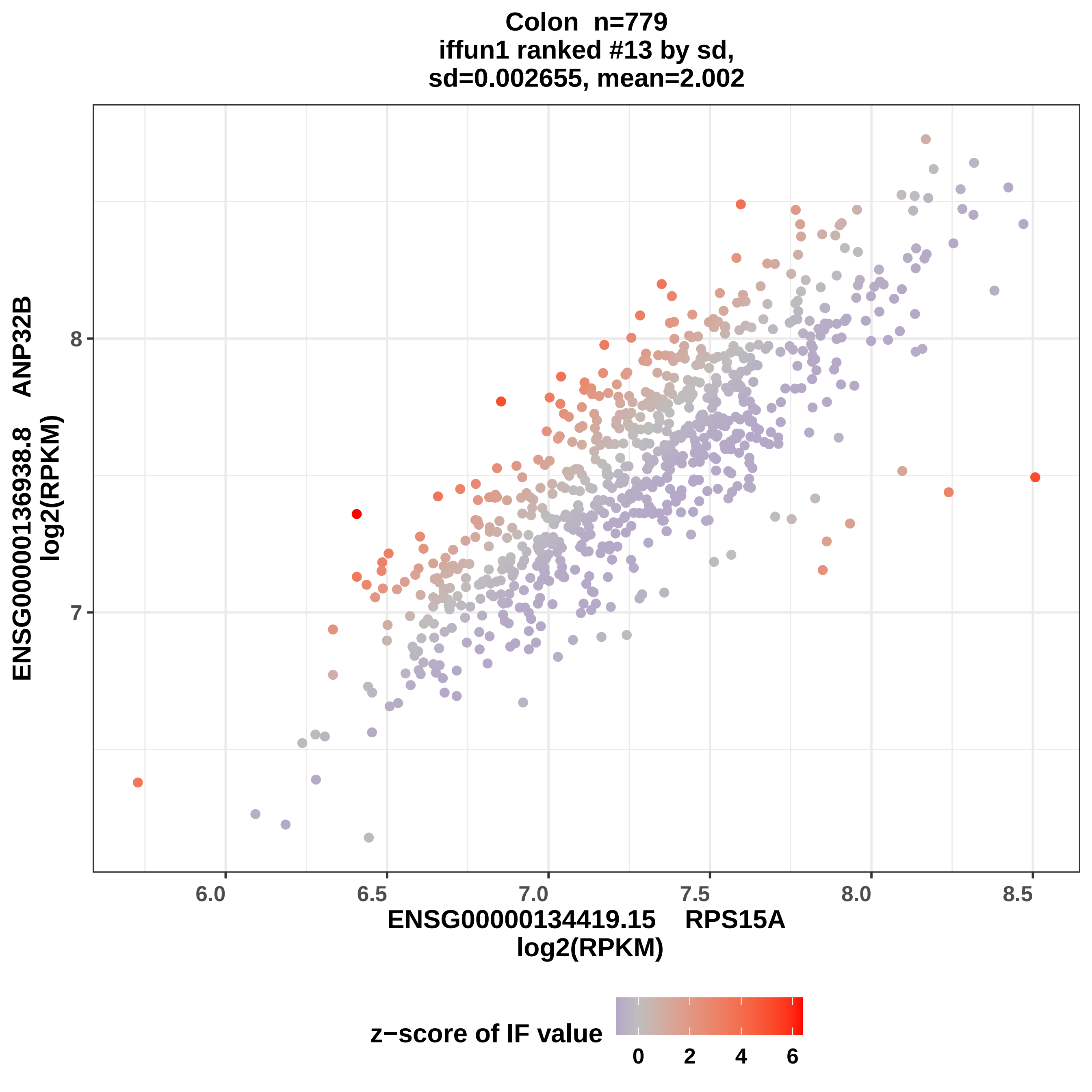
Colon n=779
iffun1 ranked #13 by sd,
sd=0.002655, mean=2.002
ENSG00000136938.8 ANP32B
log2(RPKM)
8
7
6.0
6.5
7.0
7.5
8.0
8.5
ENSG00000134419.15 RPS15A
log2(RPKM)
z−score of IF value
0
2
4
6

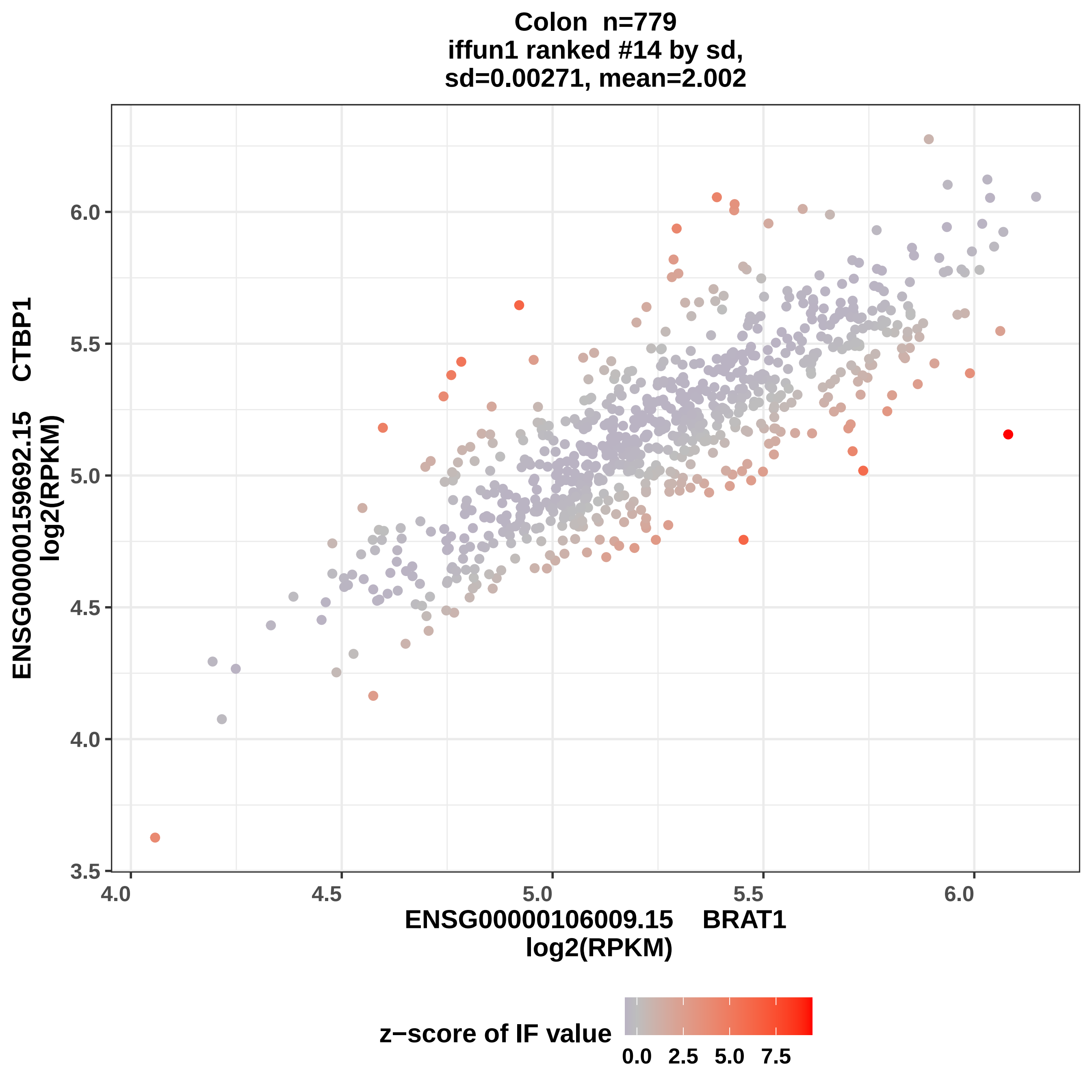
Colon n=779
iffun1 ranked #14 by sd,
sd=0.00271, mean=2.002
ENSG00000159692.15 CTBP1
log2(RPKM)
6.0
5.5
5.0
4.5
4.0
3.5
4.0
4.5
5.0
5.5
6.0
ENSG00000106009.15 BRAT1
log2(RPKM)
z−score of IF value
0.0
2.5
5.0
7.5

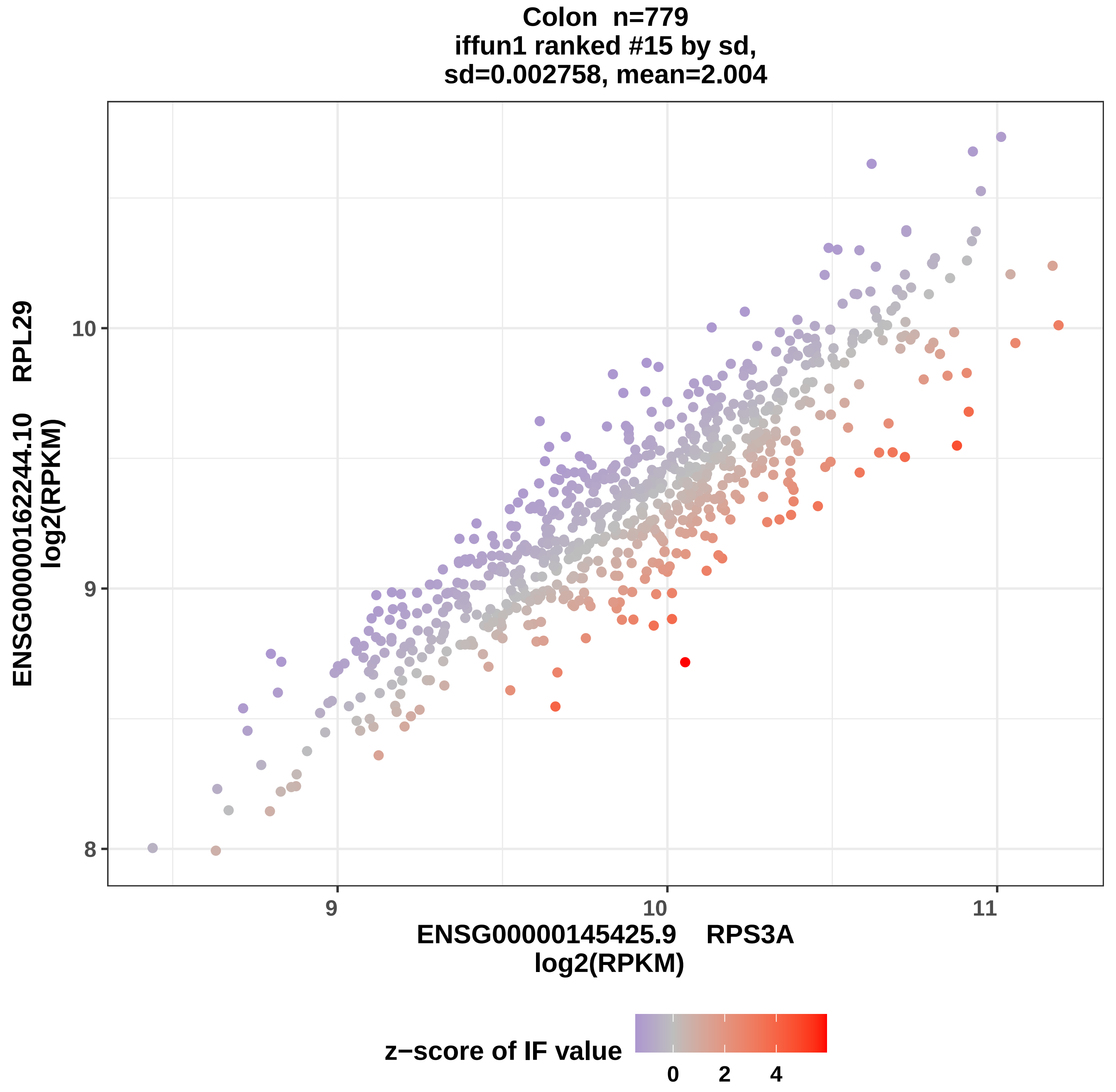

Colon n=779
iffun1 ranked #15 by sd,
sd=0.002758, mean=2.004
ENSG00000162244.10 RPL29
log2(RPKM)
10
9
8
9
10
11
ENSG00000145425.9 RPS3A
log2(RPKM)
z−score of IF value
0
2
4

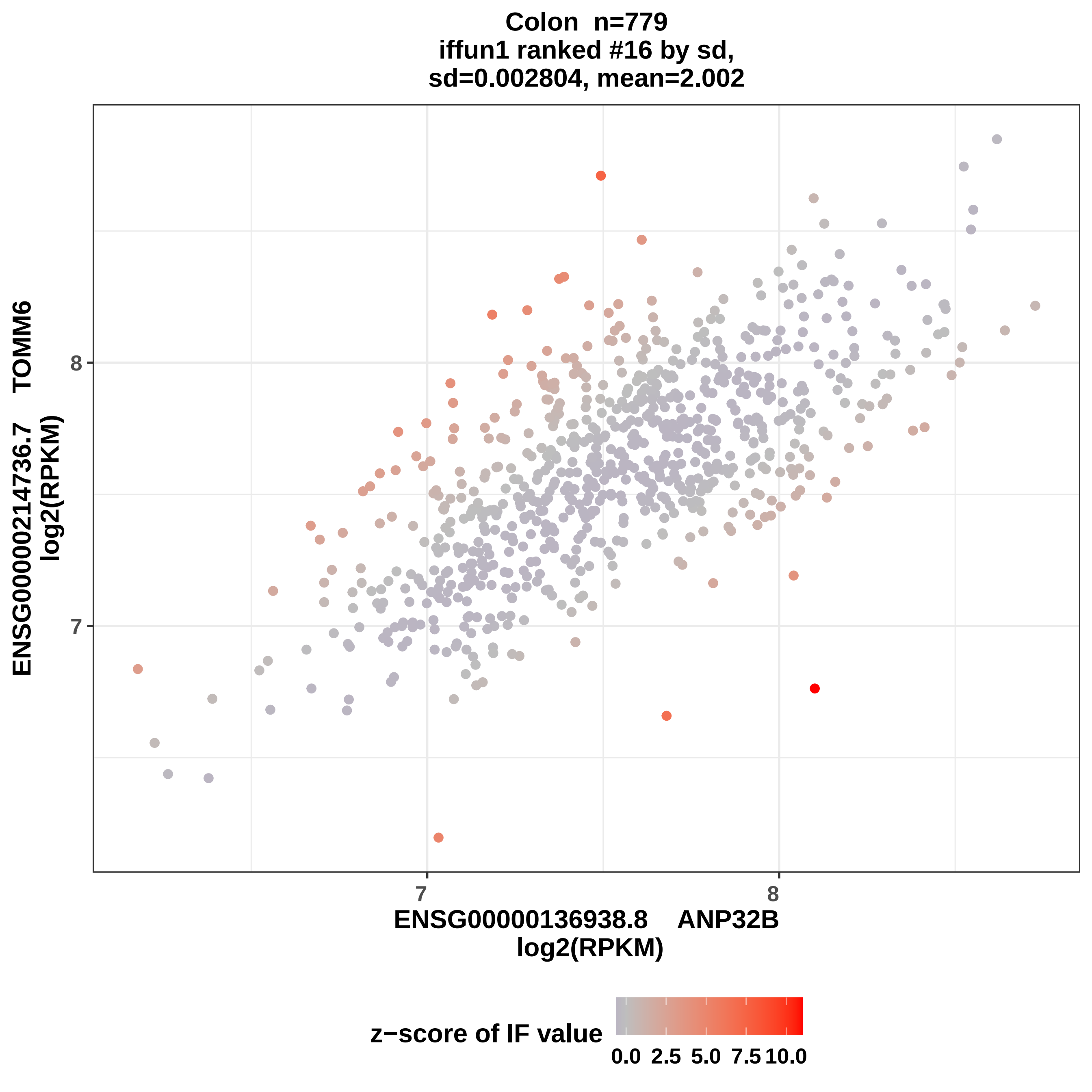
Colon n=779
iffun1 ranked #16 by sd,
sd=0.002804, mean=2.002
ENSG00000214736.7 TOMM6
log2(RPKM)
8
7
7
8
ENSG00000136938.8 ANP32B
log2(RPKM)
z−score of IF value
0.0
2.5
5.0
7.5
10.0

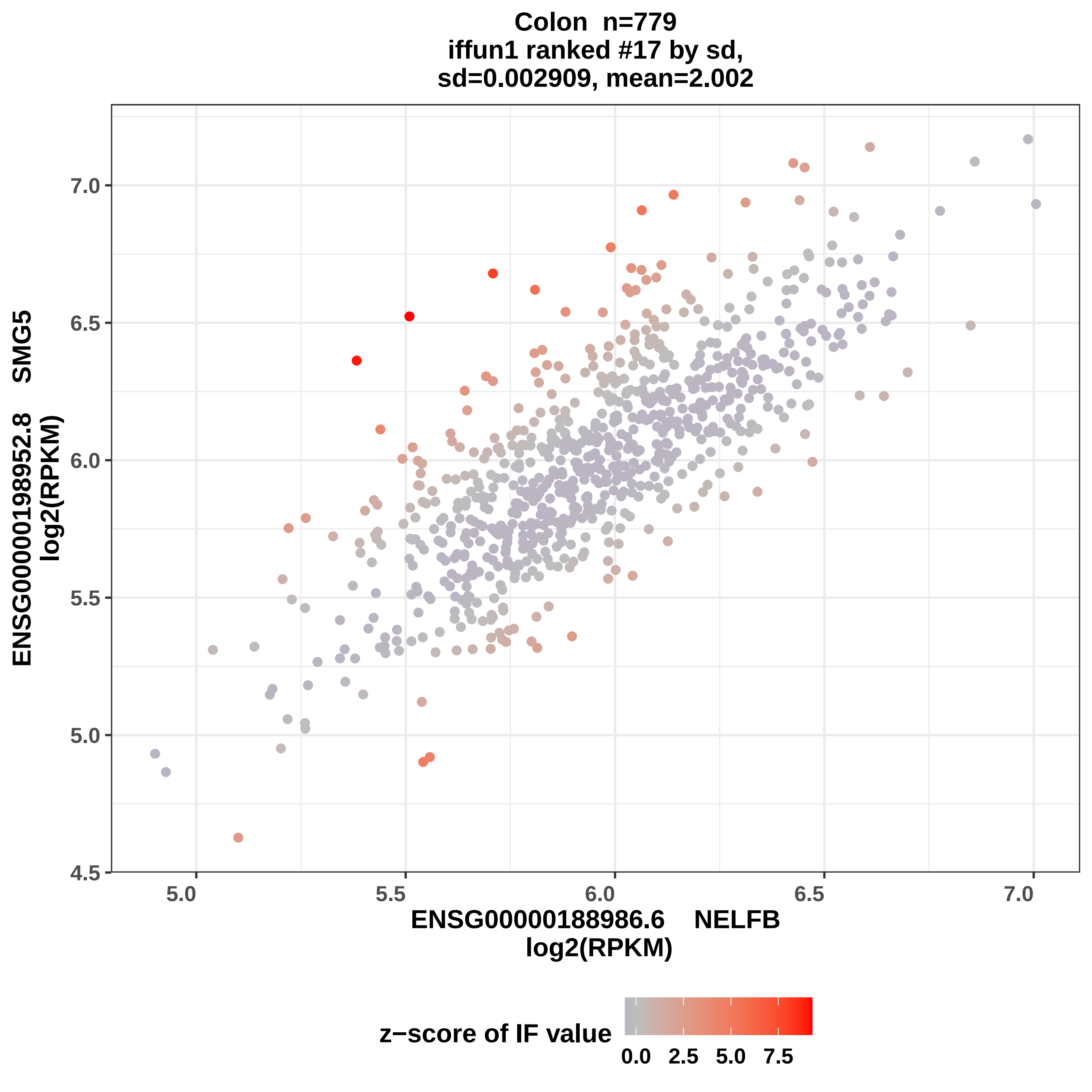

Colon n=779
iffun1 ranked #17 by sd,
sd=0.002909, mean=2.002
ENSG00000198952.8 SMG5
log2(RPKM)
ENSG00000188986.6 NELFB
log2(RPKM)
4.5
5.0
5.5
6.0
6.5
7.0
5.0
5.5
6.0
6.5
7.0
z−score of IF value
0.0
2.5
5.0
7.5

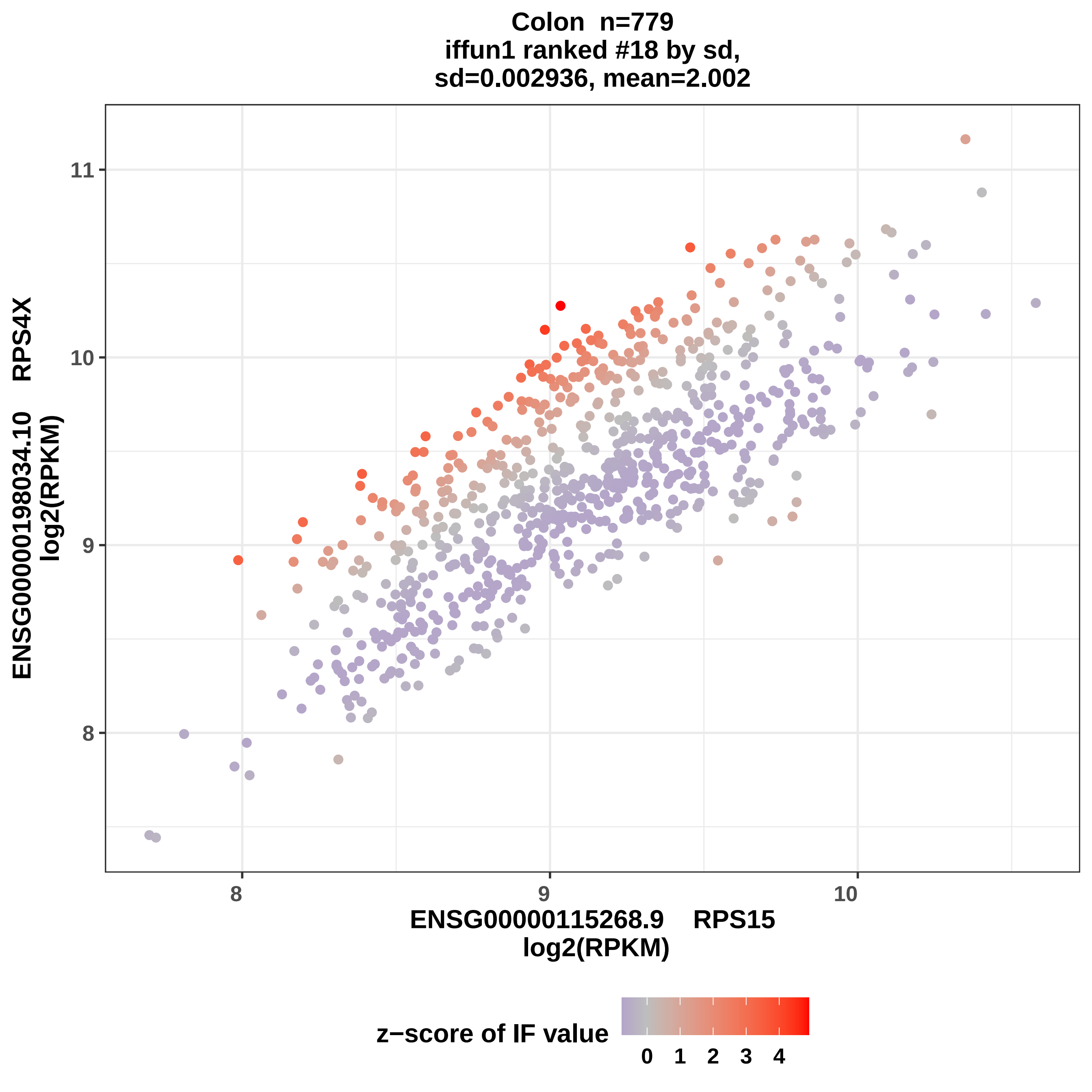
Colon n=779
iffun1 ranked #18 by sd,
sd=0.002936, mean=2.002
ENSG00000198034.10 RPS4X
log2(RPKM)
ENSG00000115268.9 RPS15
log2(RPKM)
8
9
10
11
z−score of IF value
0 1 2 3 4

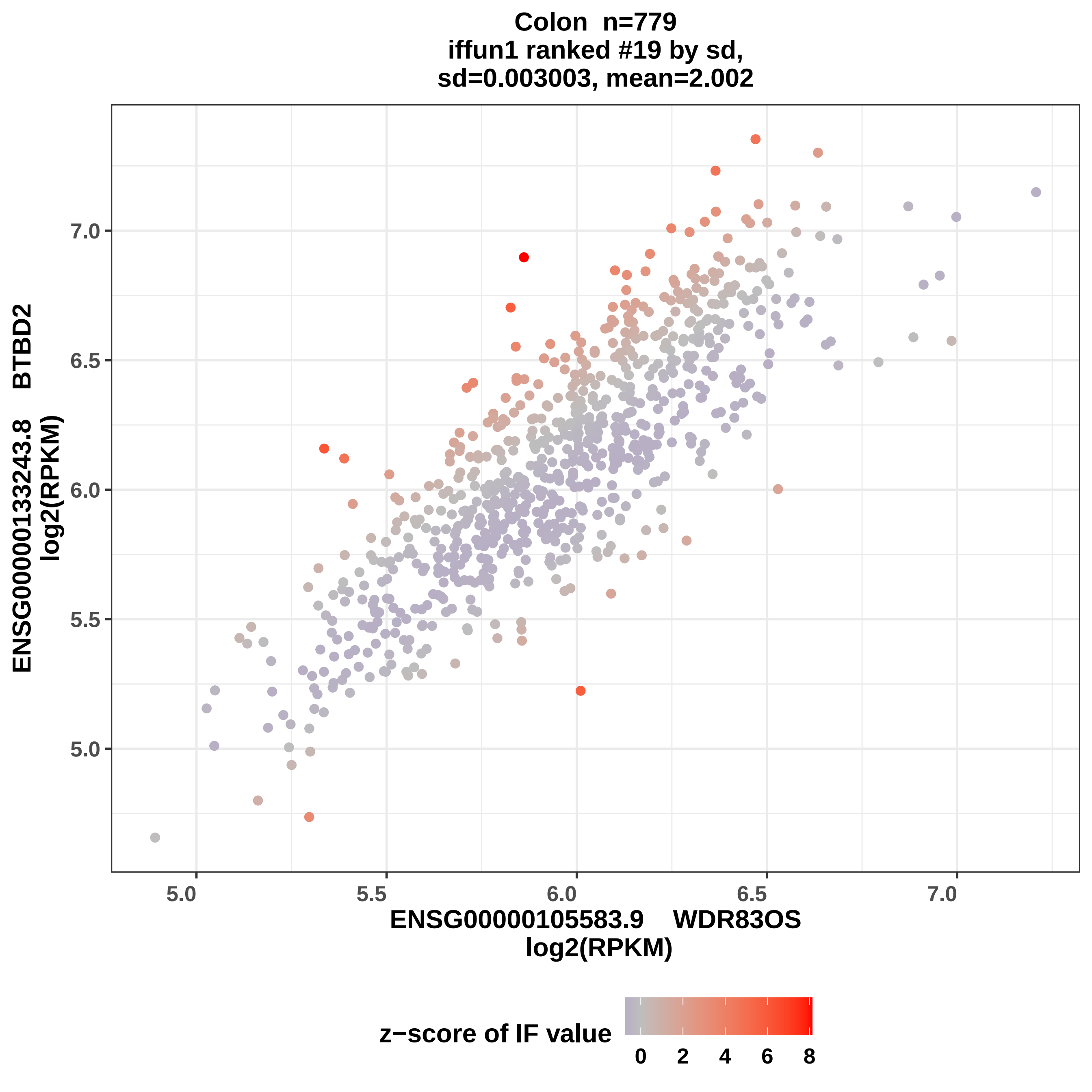

Colon n=779
iffun1 ranked #19 by sd,
sd=0.003003, mean=2.002
ENSG00000133243.8 BTBD2
log2(RPKM)
7.0
6.5
6.0
5.5
5.0
5.0
5.5
6.0
6.5
7.0
ENSG00000105583.9 WDR83OS
log2(RPKM)
z−score of IF value
0
2
4
6
8

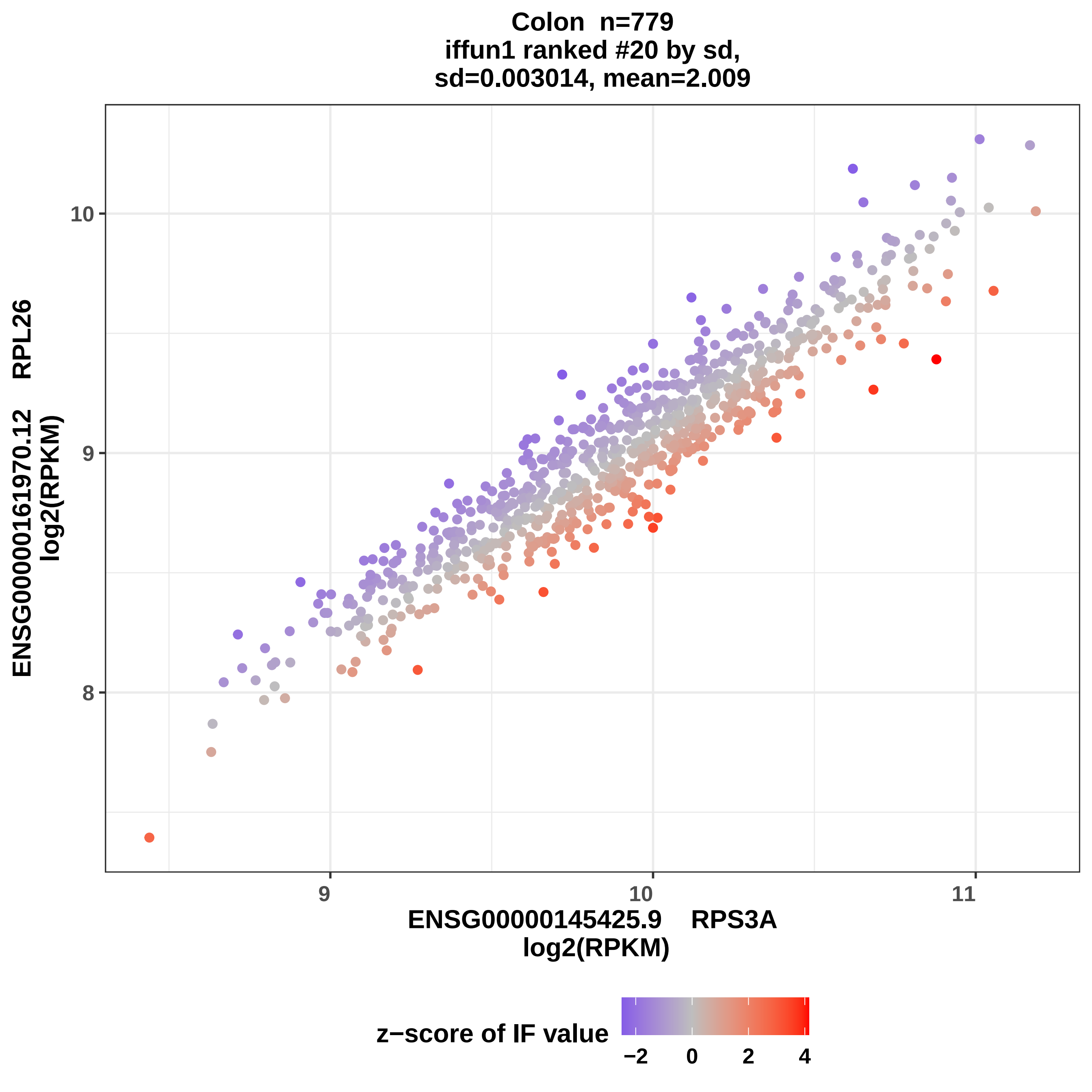

Colon n=779
iffun1 ranked #20 by sd,
sd=0.003014, mean=2.009
10
9
8
9
10
11
ENSG00000161970.12 RPL26
log2(RPKM)
ENSG00000145425.9 RPS3A
log2(RPKM)
z−score of IF value
−2
0
2
4

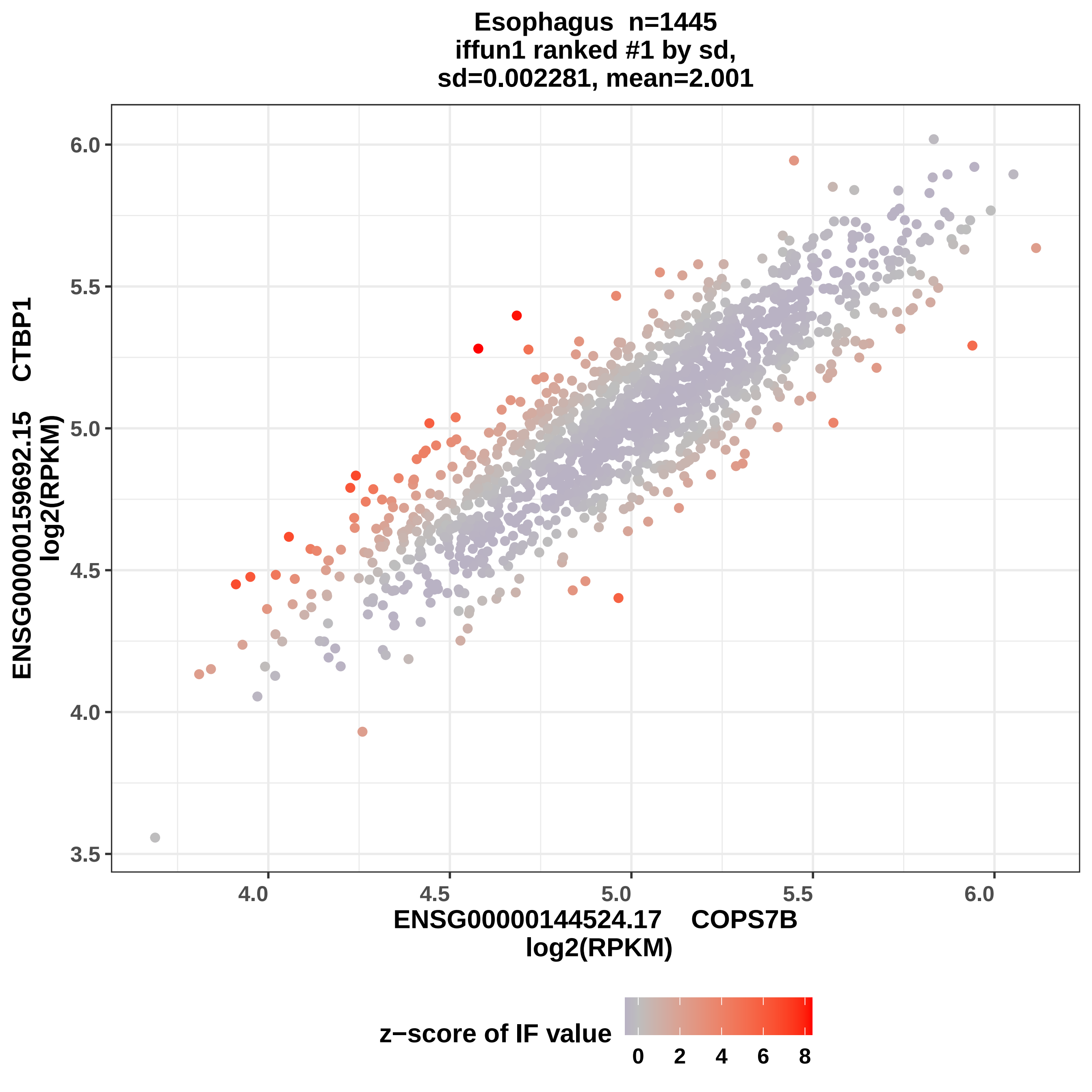
Esophagus n=1445
iffun1 ranked #1 by sd,
sd=0.002281, mean=2.001
ENSG00000159692.15 CTBP1
log2(RPKM)
ENSG00000144524.17 COPS7B
log2(RPKM)
z−score of IF value
0
2
4
6
8
6.0
5.5
5.0
4.5
4.0
3.5
4.0
4.5
5.0
5.5
6.0

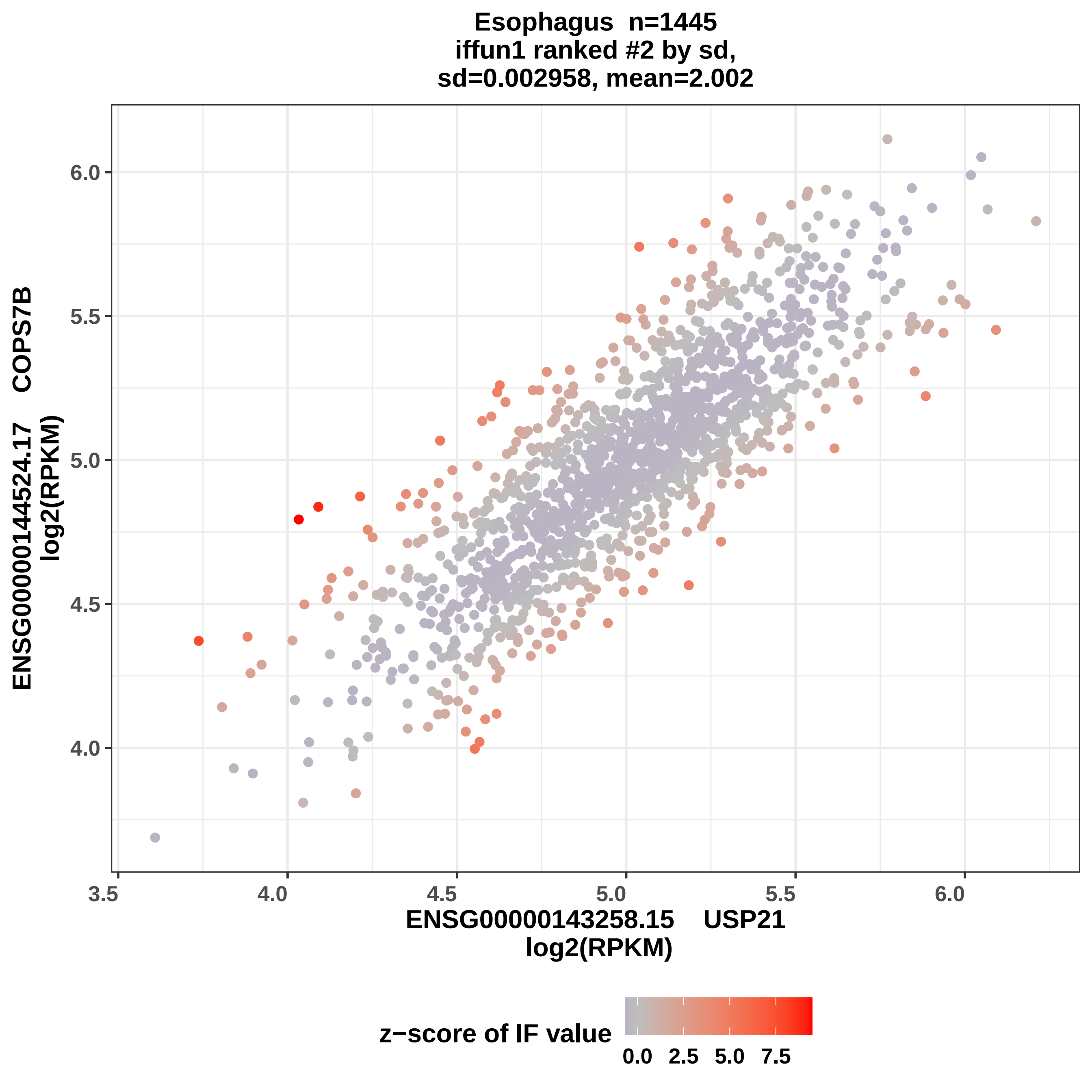

Esophagus n=1445
iffun1 ranked #2 by sd,
sd=0.002958, mean=2.002
ENSG00000144524.17 COPS7B
log2(RPKM)
ENSG00000143258.15 USP21
log2(RPKM)
6.0
5.5
5.0
4.5
4.0
3.5
z−score of IF value
0.0
2.5
5.0
7.5

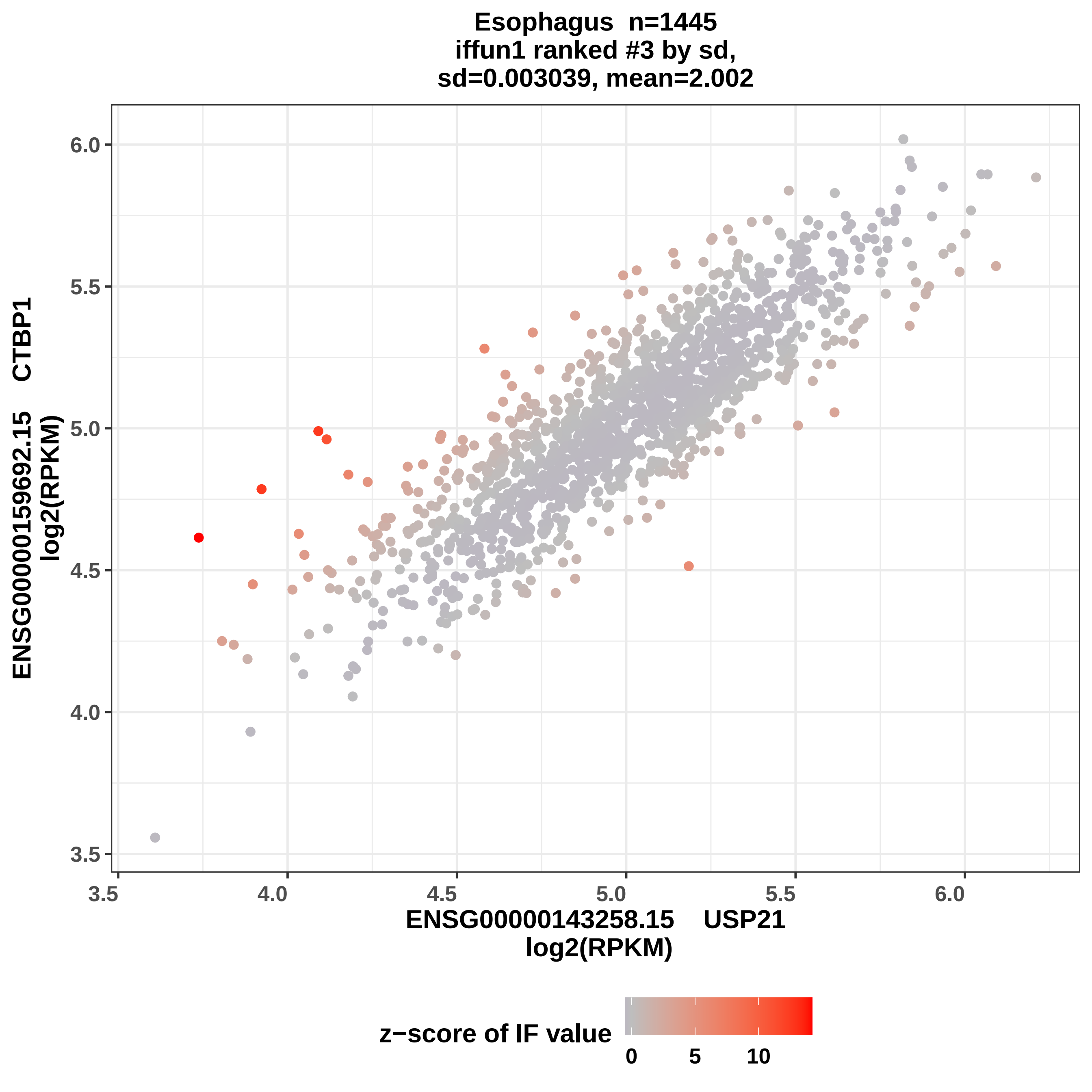
Esophagus n=1445
iffun1 ranked #3 by sd,
sd=0.003039, mean=2.002
ENSG00000159692.15 CTBP1
log2(RPKM)
ENSG00000143258.15 USP21
log2(RPKM)
z−score of IF value
0
5
10
3.5
4.0
4.5
5.0
5.5
6.0

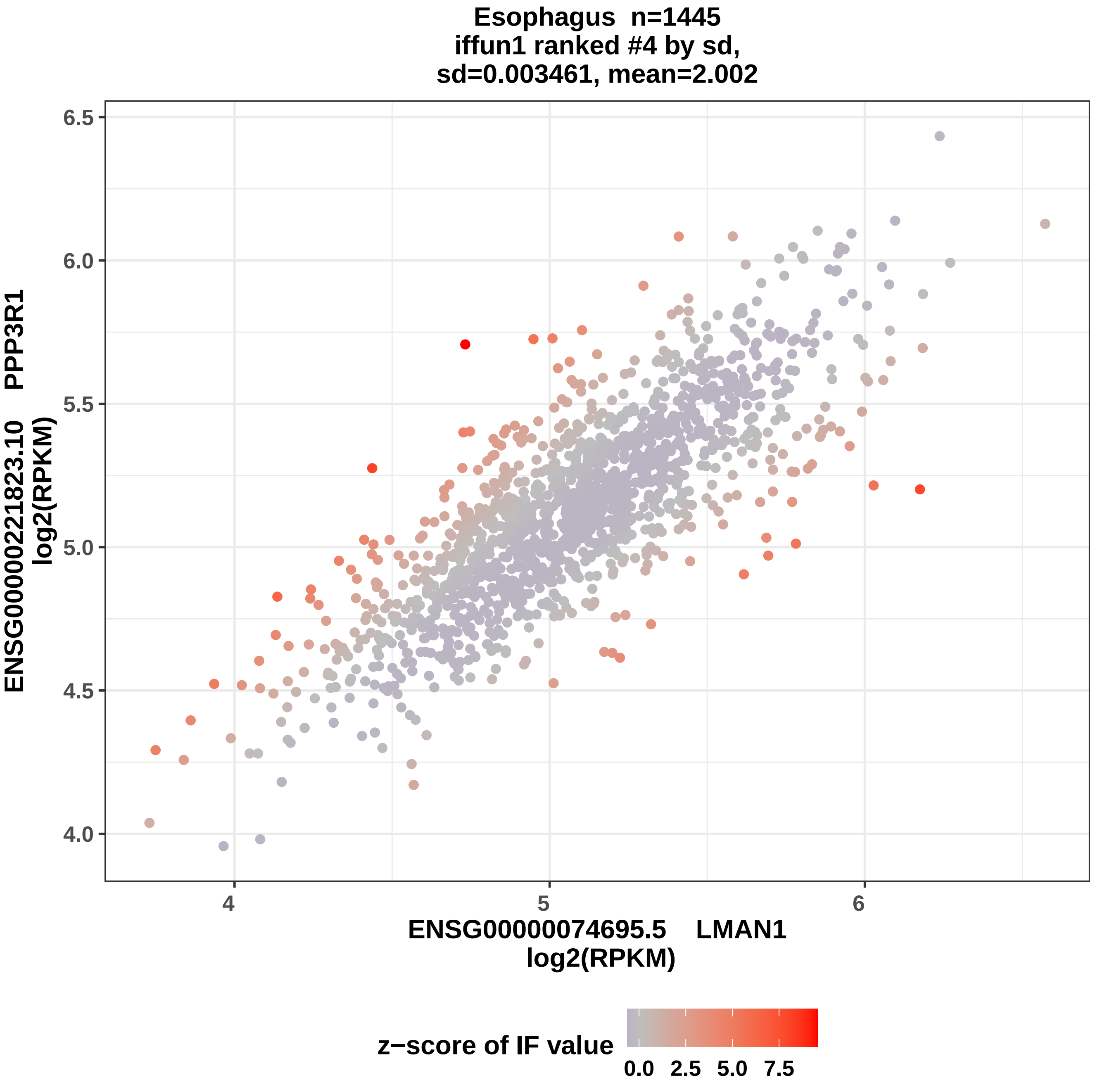
Esophagus n=1445
iffun1 ranked #4 by sd,
sd=0.003461, mean=2.002
ENSG00000221823.10 PPP3R1
log2(RPKM)
6.5
6.0
5.5
5.0
4.5
4.0
4
5
6
ENSG00000074695.5 LMAN1
log2(RPKM)
z−score of IF value
0.0
2.5
5.0
7.5

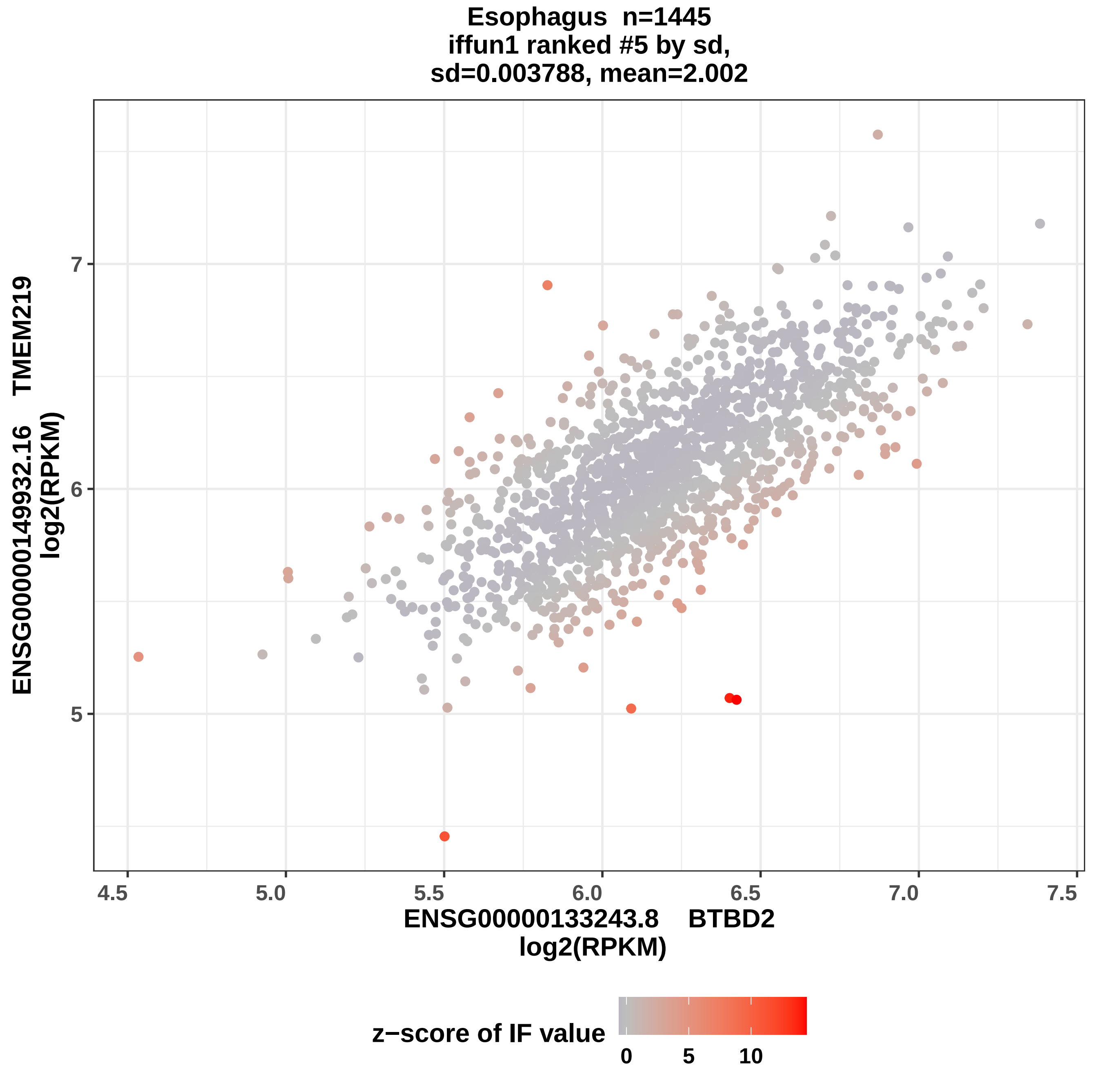

Esophagus n=1445
iffun1 ranked #5 by sd,
sd=0.003788, mean=2.002
ENSG00000149932.16 TMEM219
log2(RPKM)
7
6
5
4.5
5.0
5.5
6.0
6.5
7.0
7.5
ENSG00000133243.8 BTBD2
log2(RPKM)
z−score of IF value
0
5
10

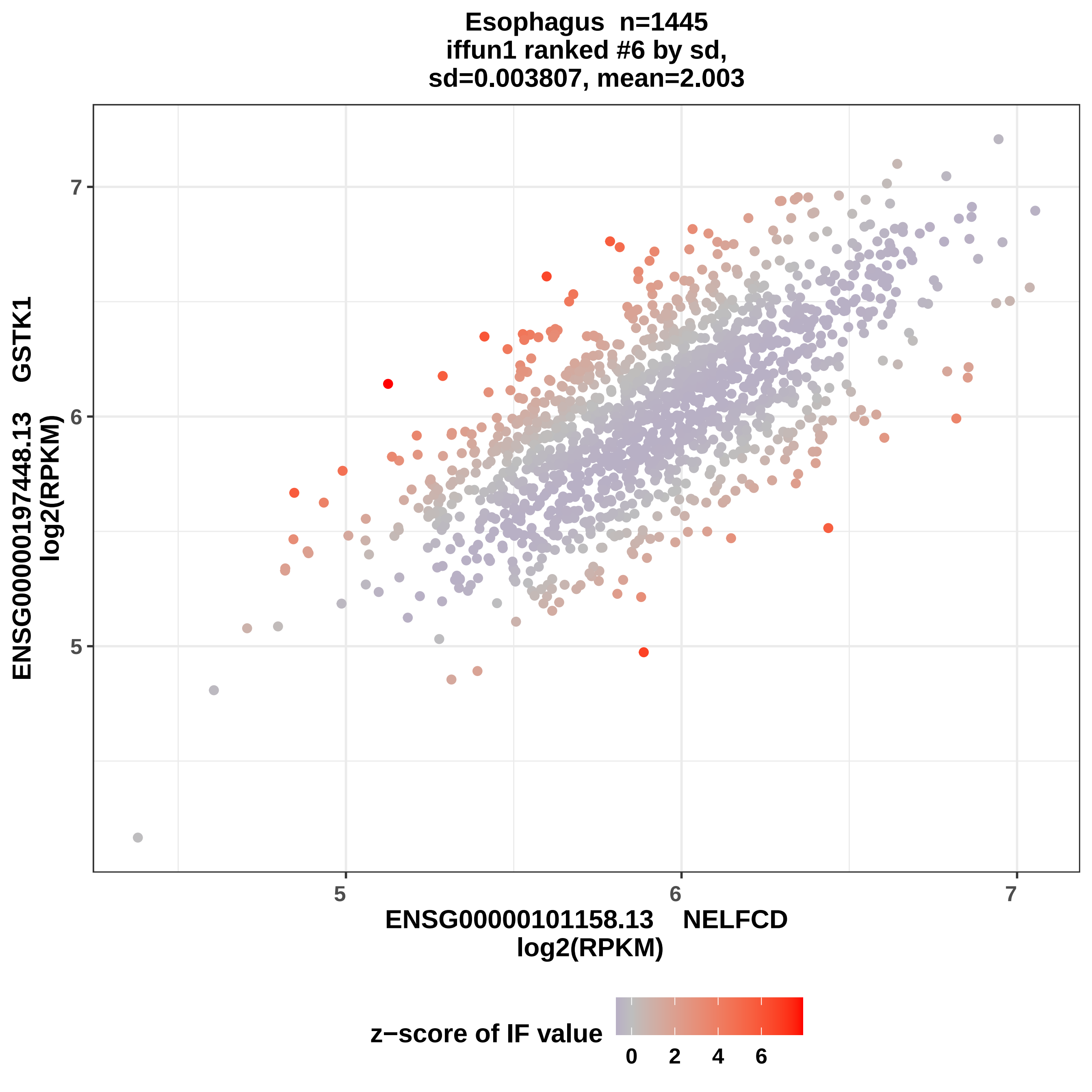
Esophagus n=1445
iffun1 ranked #6 by sd,
sd=0.003807, mean=2.003
ENSG00000197448.13 GSTK1
log2(RPKM)
ENSG00000101158.13 NELFCD
log2(RPKM)
5
6
7
5
6
7
z−score of IF value
0
2
4
6

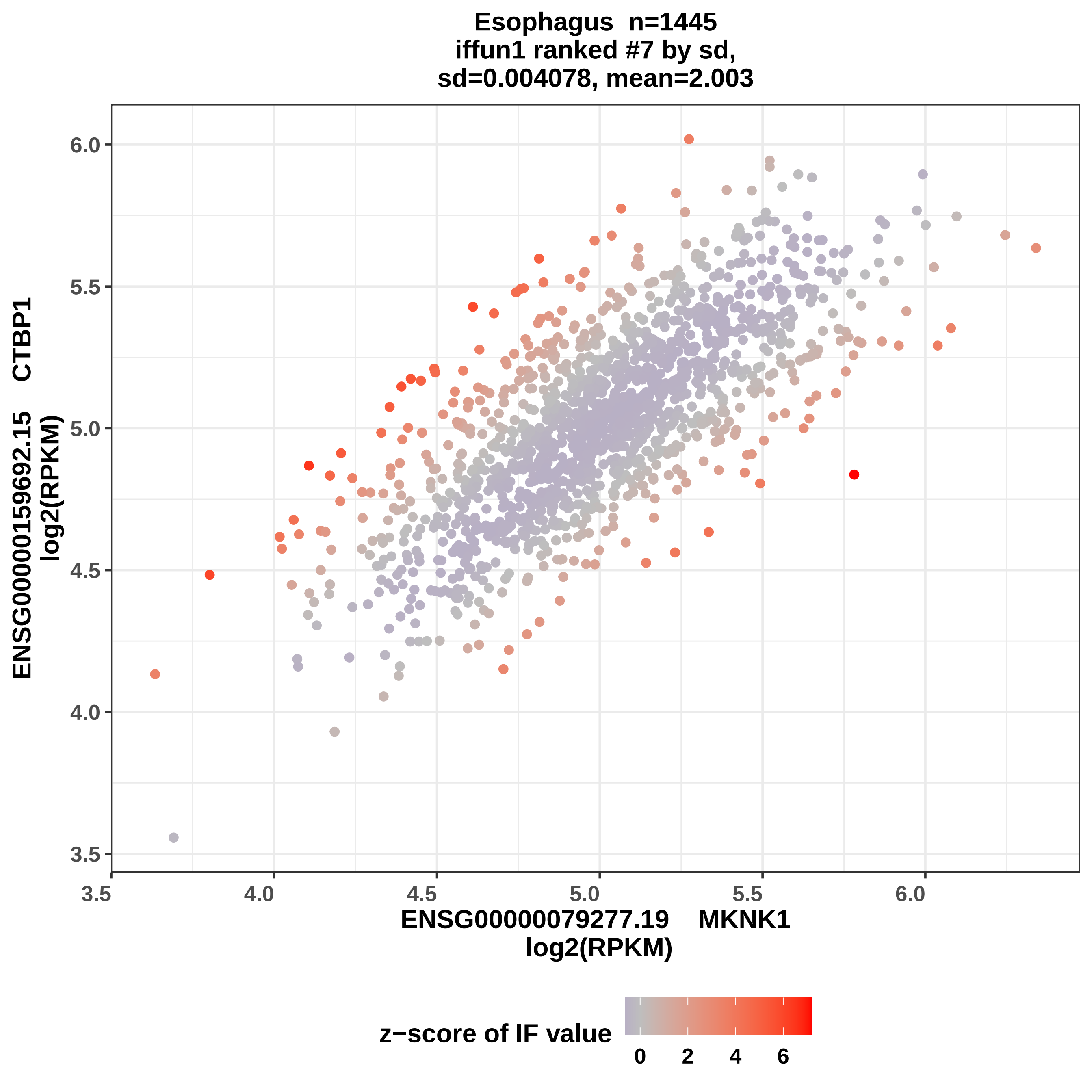

Esophagus n=1445
iffun1 ranked #7 by sd,
sd=0.004078, mean=2.003
ENSG00000159692.15 CTBP1
log2(RPKM)
ENSG00000079277.19 MKNK1
log2(RPKM)
z−score of IF value
3.5
4.0
4.5
5.0
5.5
6.0
0
2
4
6

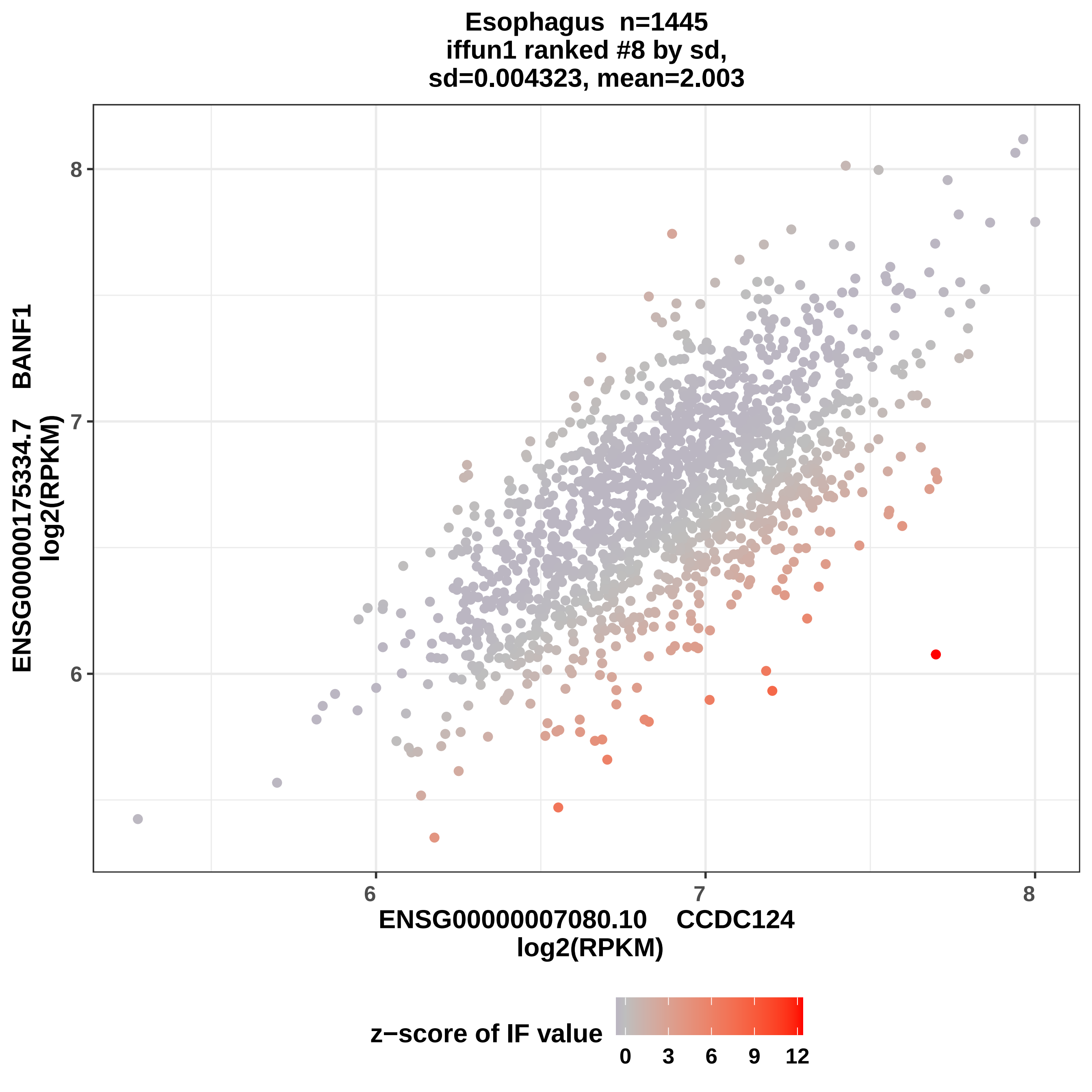

Esophagus n=1445
iffun1 ranked #8 by sd,
sd=0.004323, mean=2.003
ENSG00000175334.7 BANF1
log2(RPKM)
ENSG00000007080.10 CCDC124
log2(RPKM)
z−score of IF value
6
7
8
6
7
8
0
3
6
9
12

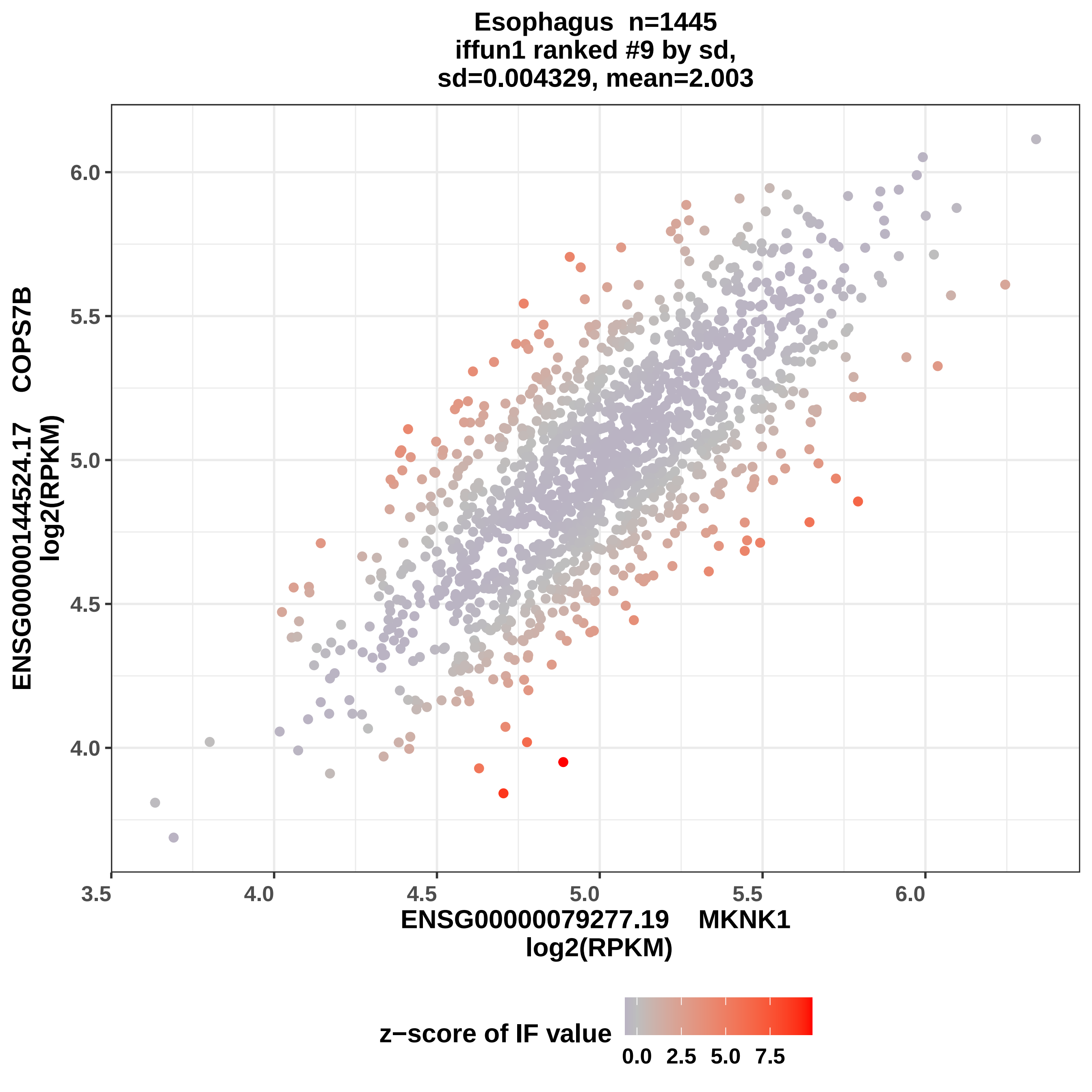

Esophagus n=1445
iffun1 ranked #9 by sd,
sd=0.004329, mean=2.003
ENSG00000144524.17 COPS7B
log2(RPKM)
ENSG00000079277.19 MKNK1
log2(RPKM)
6.0
5.5
5.0
4.5
4.0
3.5
4.0
4.5
5.0
5.5
6.0
z−score of IF value
0.0
2.5
5.0
7.5

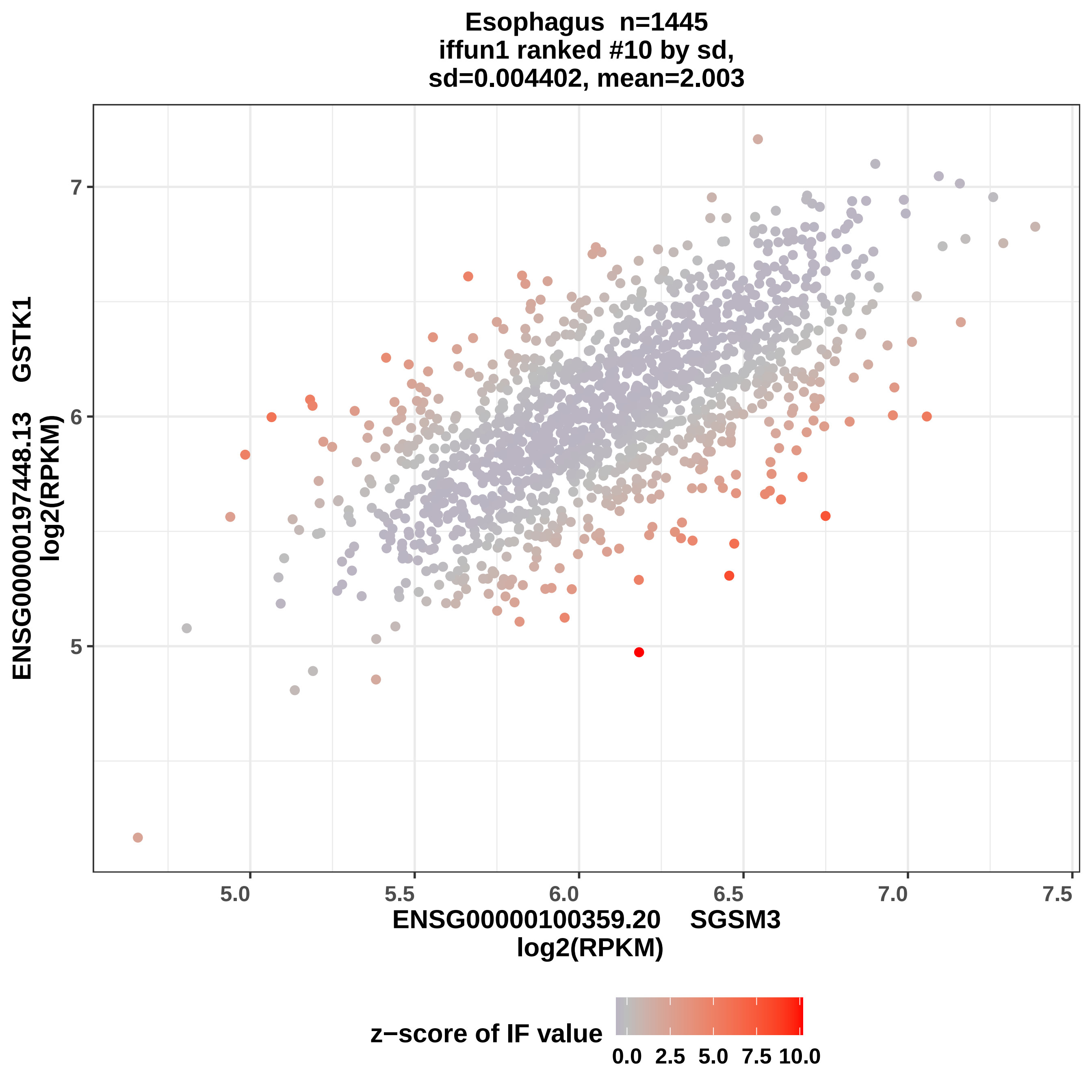

Esophagus n=1445
iffun1 ranked #10 by sd,
sd=0.004402, mean=2.003
ENSG00000197448.13 GSTK1
log2(RPKM)
7
6
5
5.0
5.5
6.0
6.5
7.0
7.5
ENSG00000100359.20 SGSM3
log2(RPKM)
z−score of IF value
0.0
2.5
5.0
7.5
10.0

Esophagus n=1445
iffun1 ranked #11 by sd,
sd=0.00466, mean=2.004

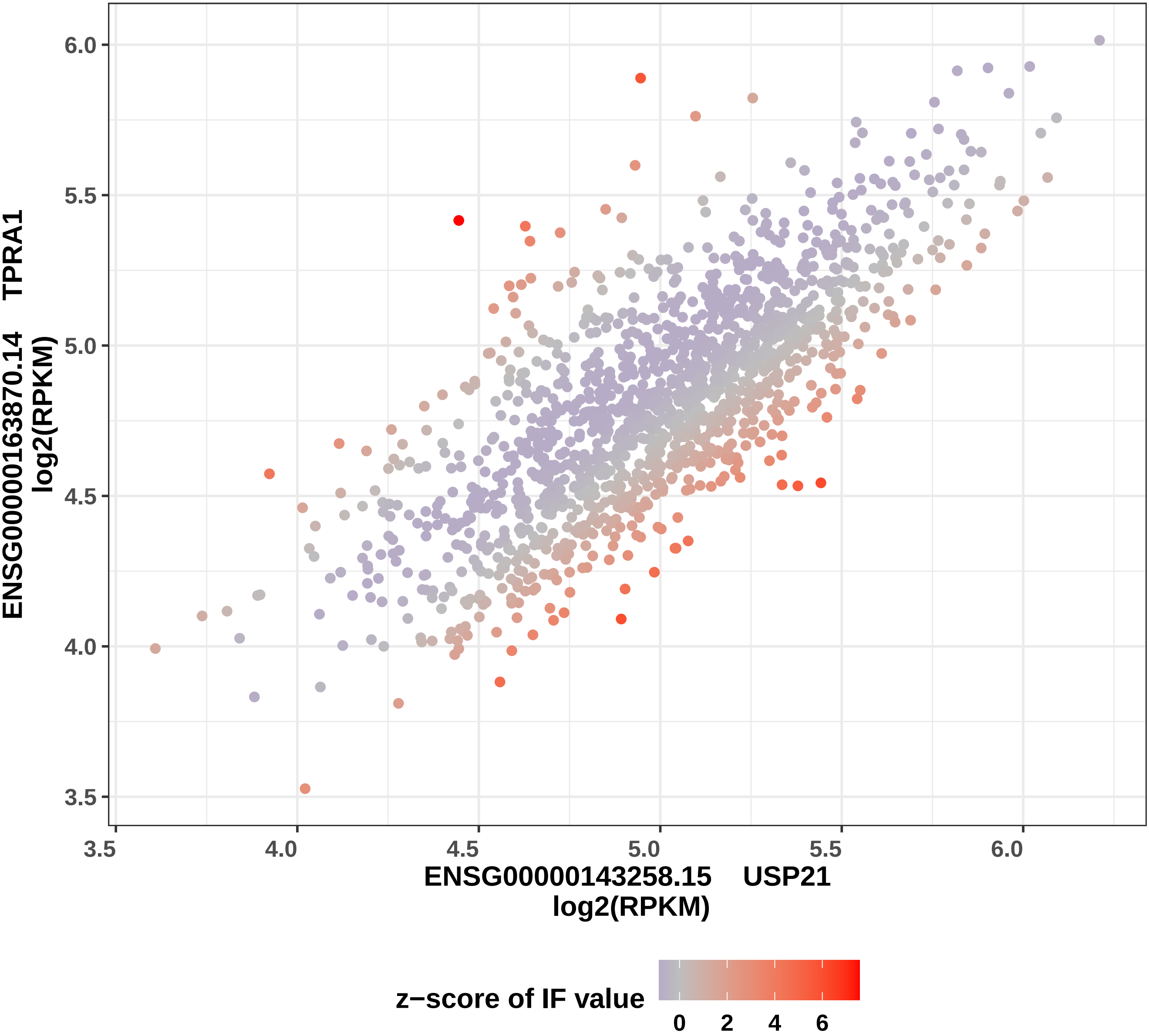

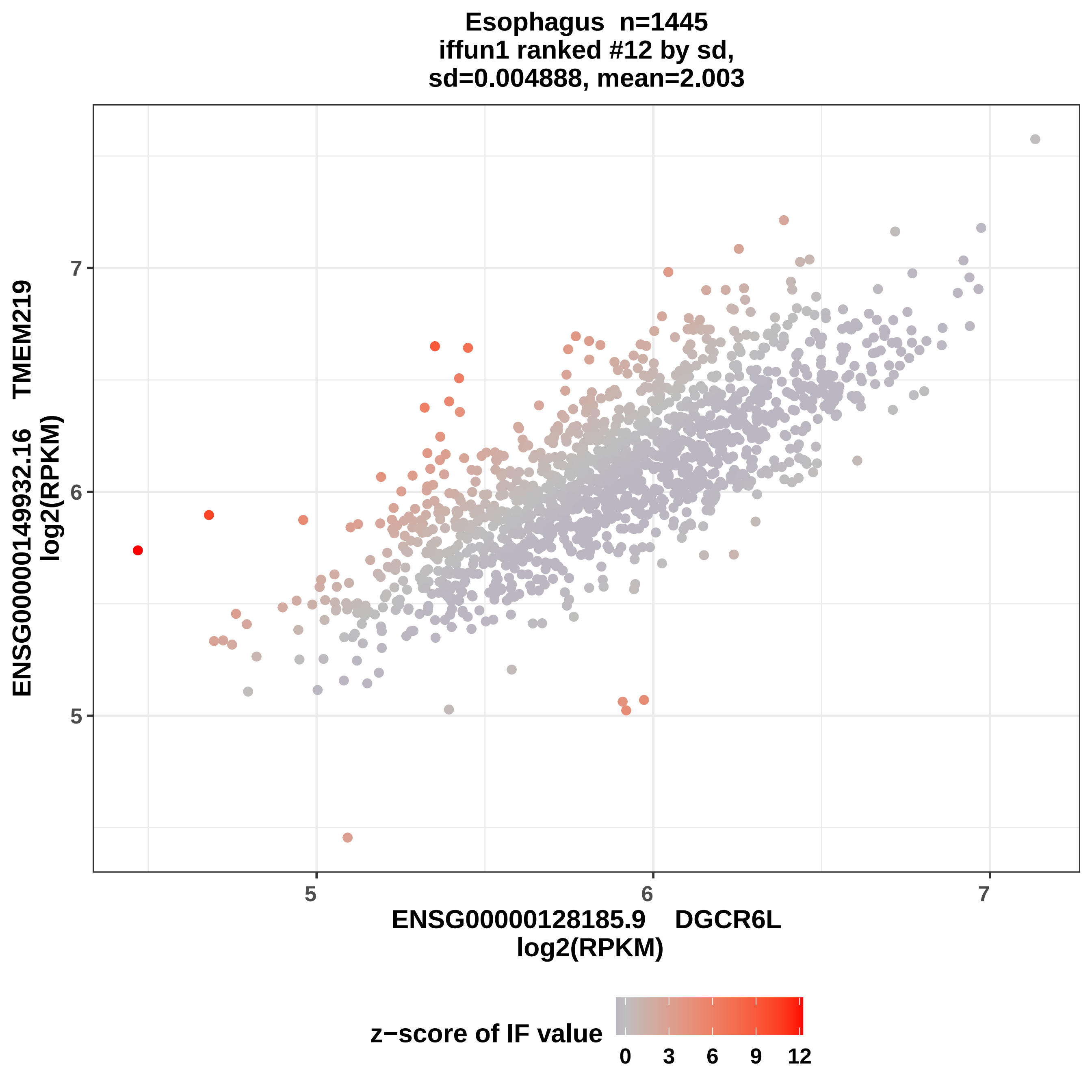
Esophagus n=1445
iffun1 ranked #12 by sd,
sd=0.004888, mean=2.003
ENSG00000149932.16 TMEM219
log2(RPKM)
ENSG00000128185.9 DGCR6L
log2(RPKM)
5
6
7
5
6
7
z−score of IF value
0
3
6
9
12

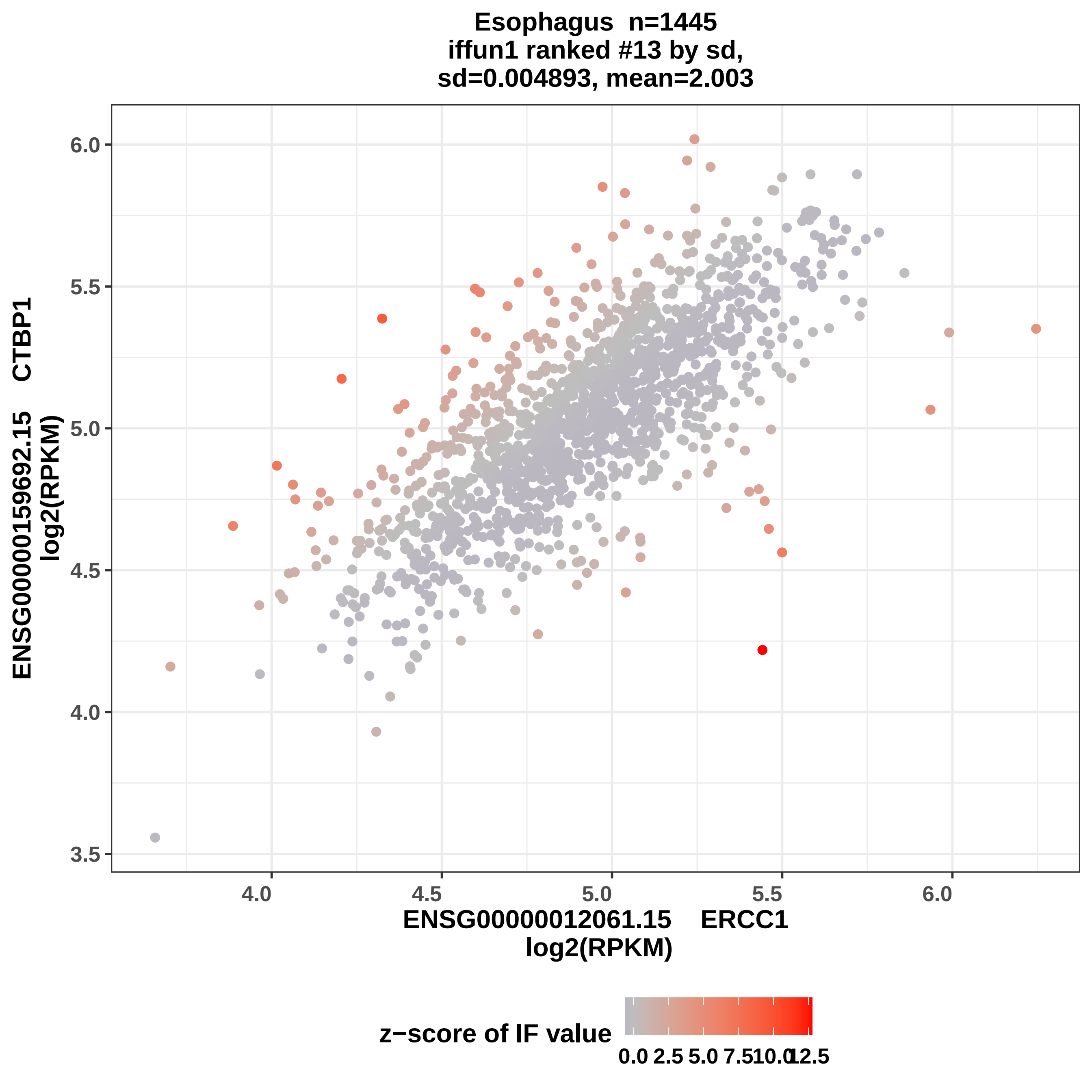

Esophagus n=1445
iffun1 ranked #13 by sd,
sd=0.004893, mean=2.003
ENSG00000159692.15 CTBP1
log2(RPKM)
ENSG00000012061.15 ERCC1
log2(RPKM)
6.0
5.5
5.0
4.5
4.0
3.5
4.0
4.5
5.0
5.5
6.0
z−score of IF value
0.0
2.5
5.0
7.5
10.0
12.5

Esophagus  n=1445
iffun1 ranked #14 by sd,
sd=0.0049, mean=2.003

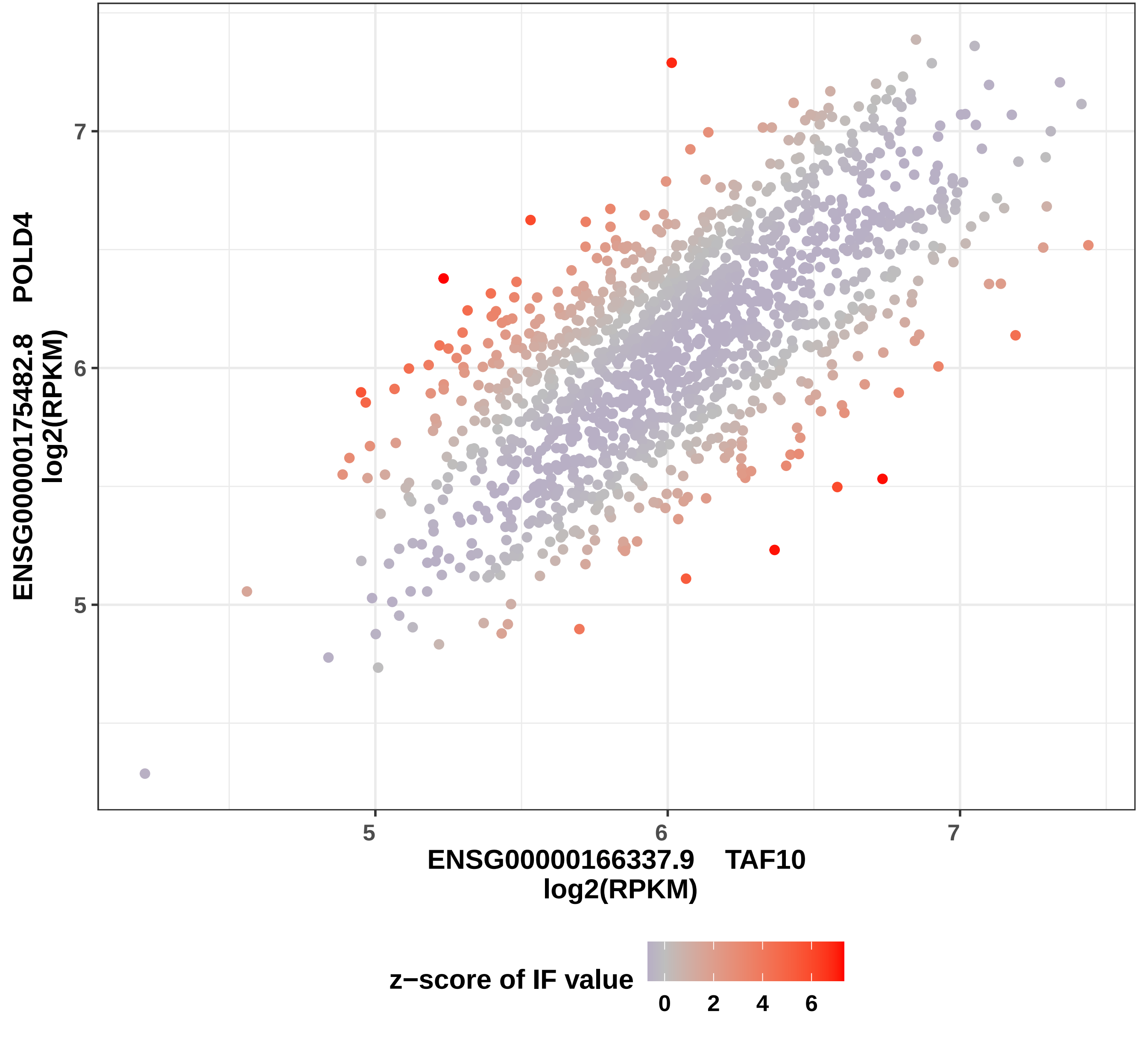

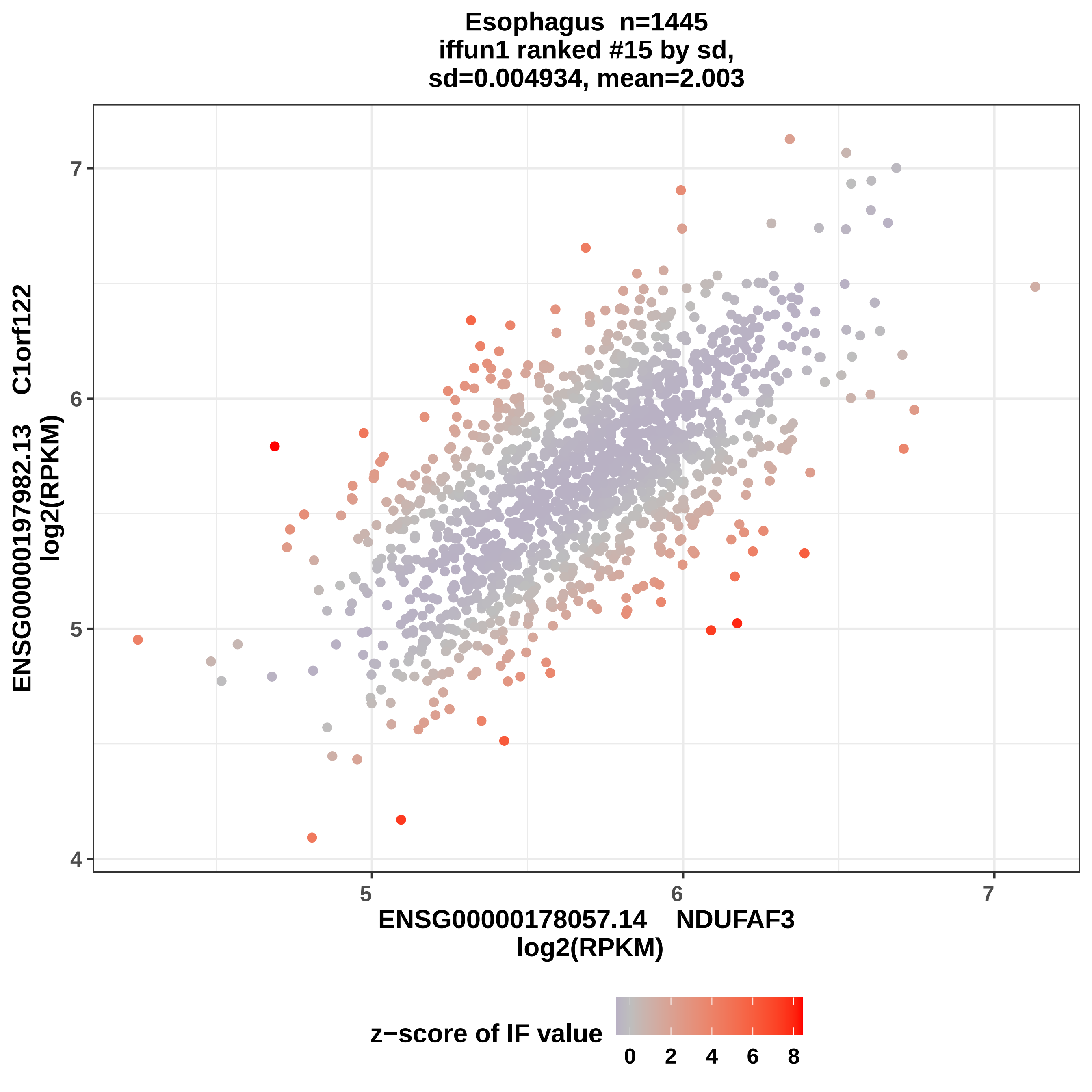

Esophagus n=1445
iffun1 ranked #15 by sd,
sd=0.004934, mean=2.003
ENSG00000197982.13 C1orf122
log2(RPKM)
ENSG00000178057.14 NDUFAF3
log2(RPKM)
4
5
6
7
z−score of IF value
0
2
4
6
8

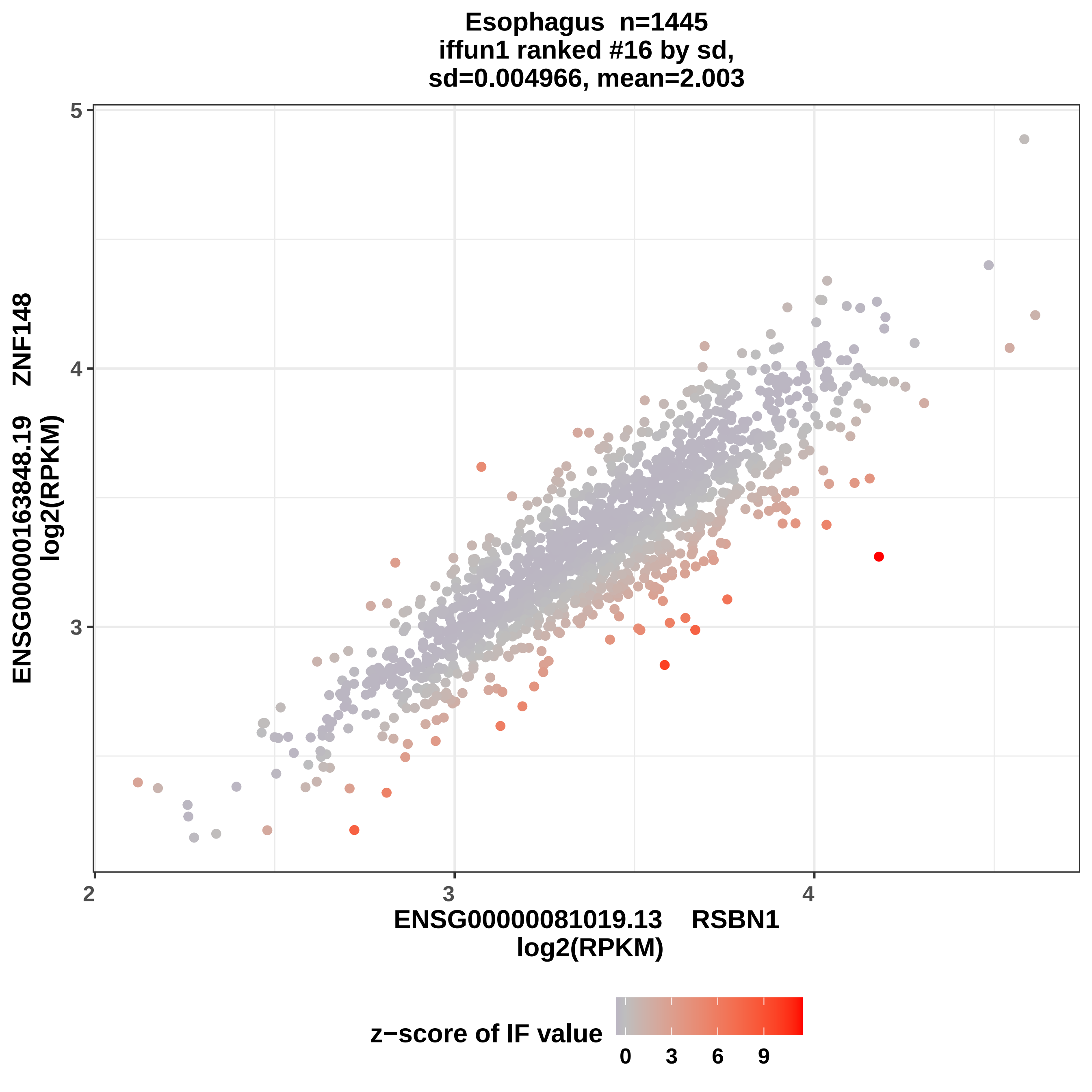
Esophagus n=1445
iffun1 ranked #16 by sd,
sd=0.004966, mean=2.003
ENSG00000163848.19 ZNF148
log2(RPKM)
ENSG00000081019.13 RSBN1
log2(RPKM)
5
4
3
2
3
4
z−score of IF value
0
3
6
9

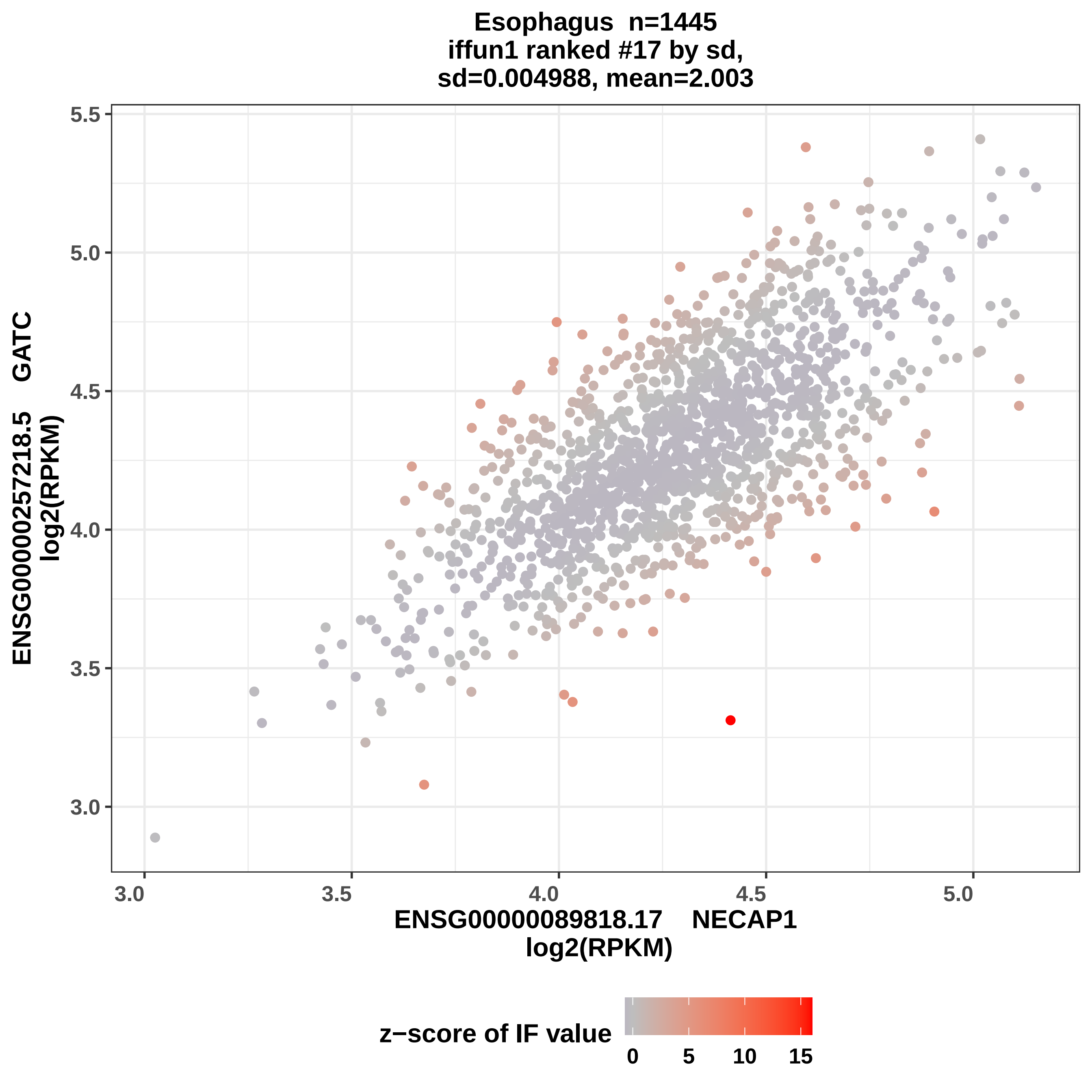
Esophagus n=1445
iffun1 ranked #17 by sd,
sd=0.004988, mean=2.003
ENSG00000257218.5 GATC
log2(RPKM)
ENSG00000089818.17 NECAP1
log2(RPKM)
3.0
3.5
4.0
4.5
5.0
5.5
z−score of IF value
0
5
10
15

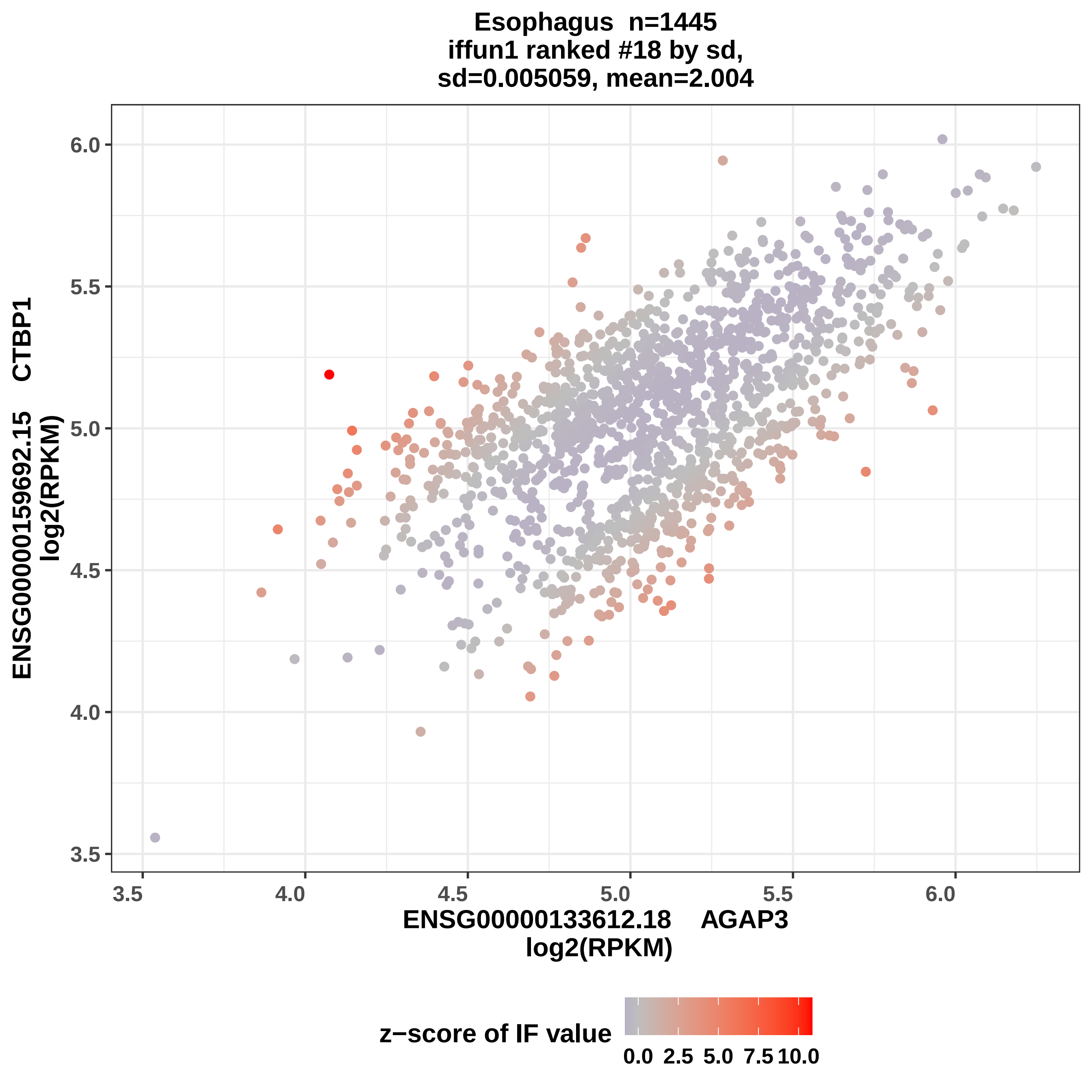

Esophagus n=1445
iffun1 ranked #18 by sd,
sd=0.005059, mean=2.004
ENSG00000159692.15 CTBP1
log2(RPKM)
6.0
5.5
5.0
4.5
4.0
3.5
3.5
4.0
4.5
5.0
5.5
6.0
ENSG00000133612.18 AGAP3
log2(RPKM)
z−score of IF value
0.0
2.5
5.0
7.5
10.0

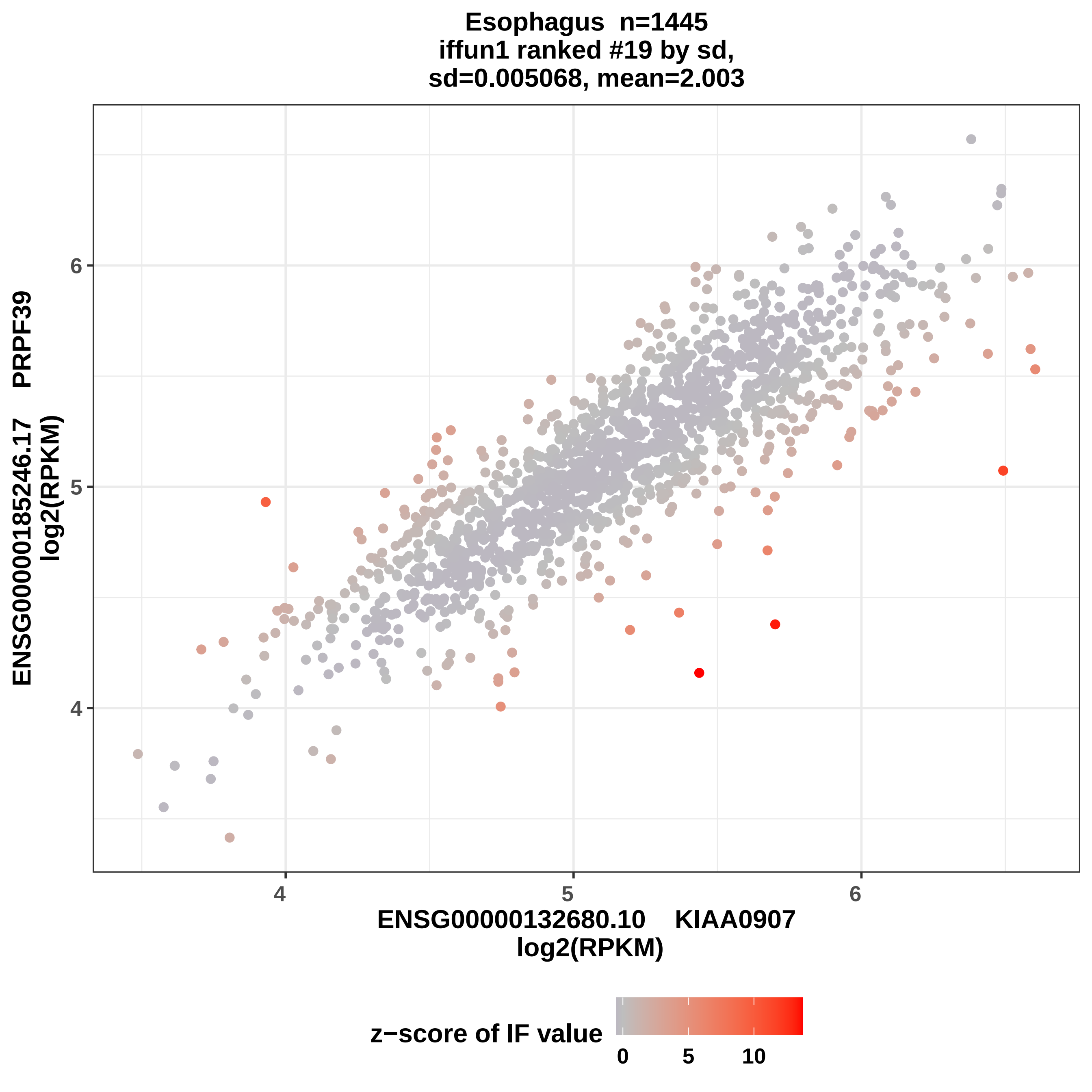

Esophagus n=1445
iffun1 ranked #19 by sd,
sd=0.005068, mean=2.003
ENSG00000185246.17 PRPF39
log2(RPKM)
6
5
4
4
5
6
ENSG00000132680.10 KIAA0907
log2(RPKM)
z−score of IF value
0
5
10

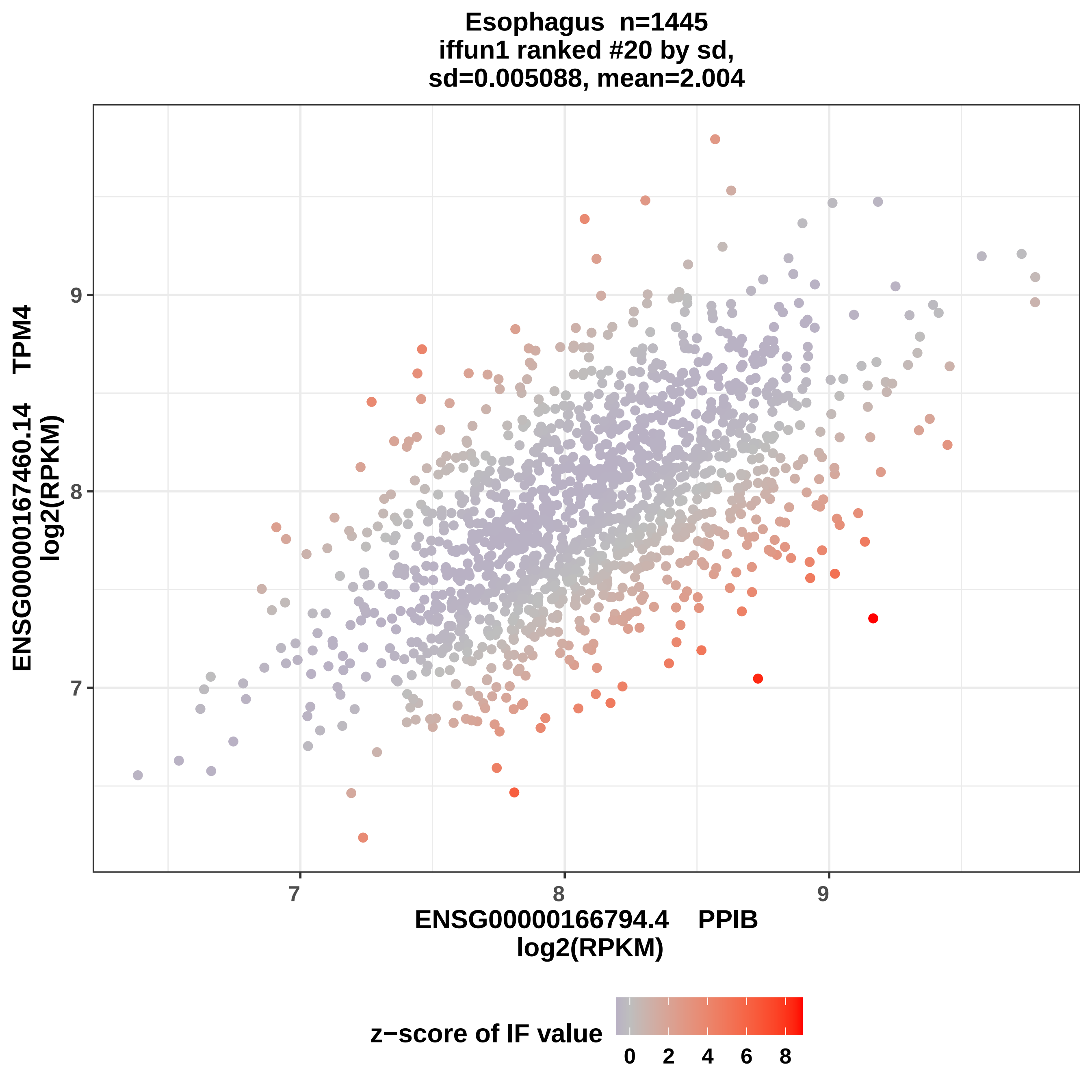
Esophagus n=1445
iffun1 ranked #20 by sd,
sd=0.005088, mean=2.004
ENSG00000167460.14 TPM4
log2(RPKM)
9
8
7
ENSG00000166794.4 PPIB
log2(RPKM)
z−score of IF value
0
2
4
6
8

### Heart n=861
### iffun1 ranked #1 by sd,
### sd=0.005338, mean=2.003

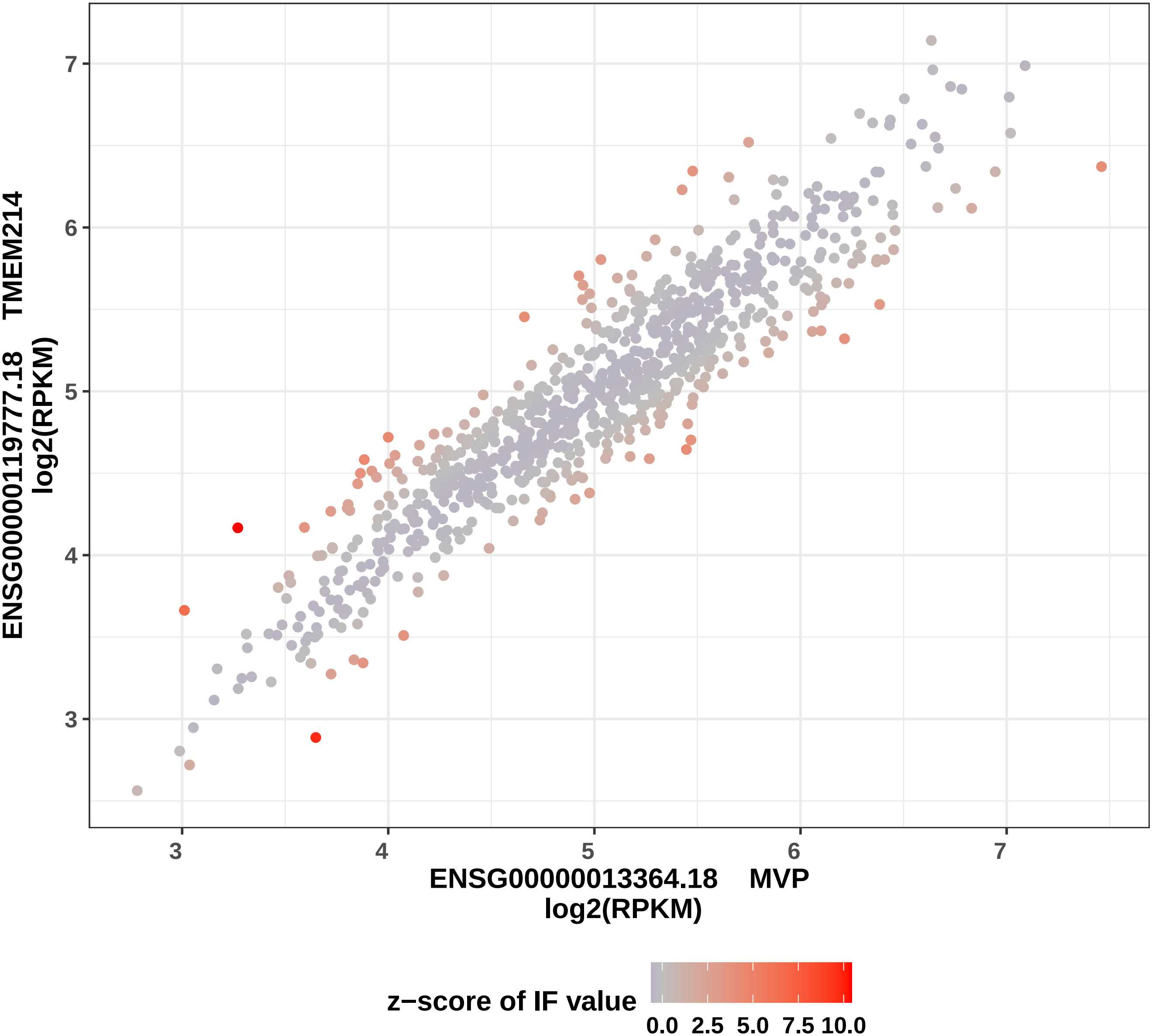

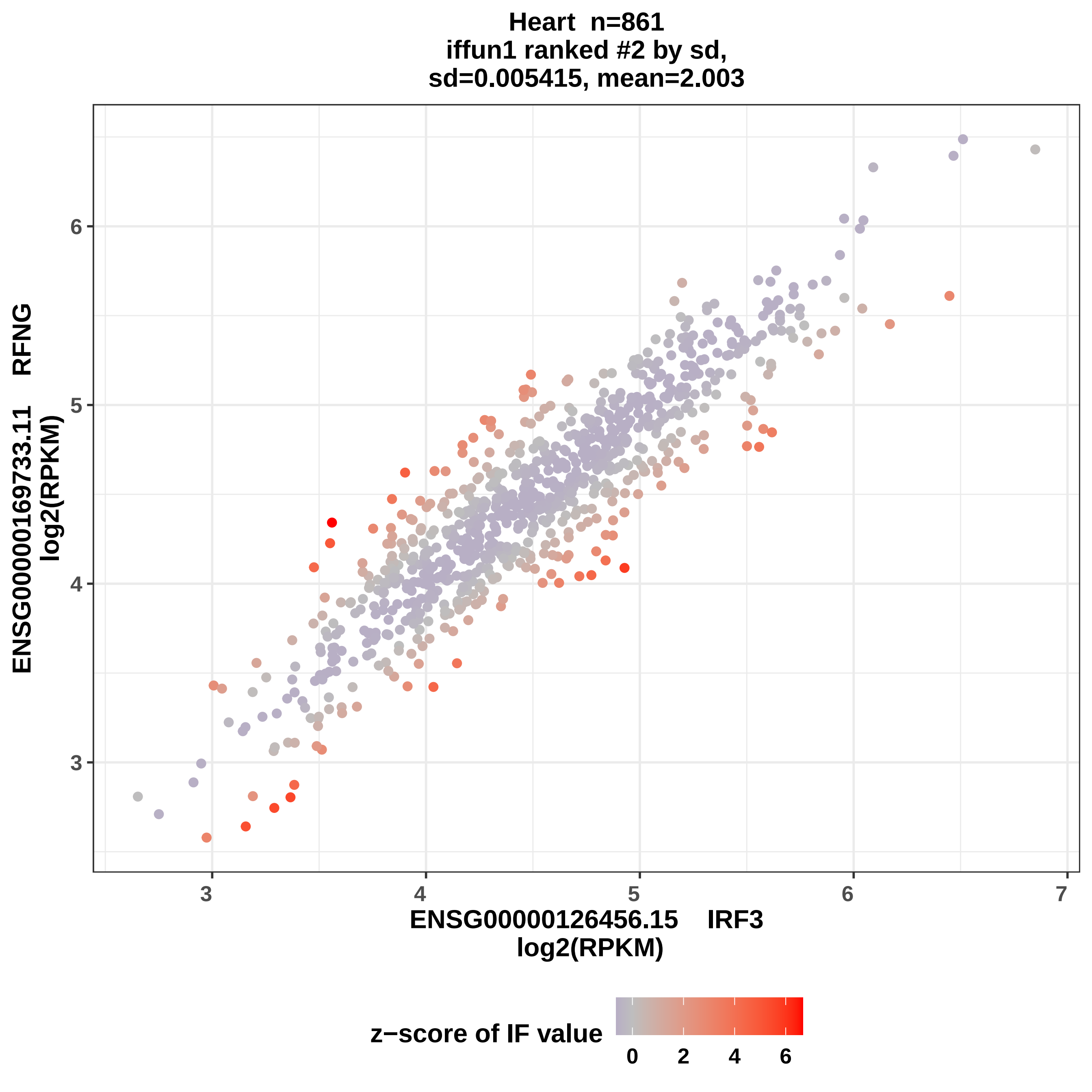

Heart n=861
iffun1 ranked #2 by sd,
sd=0.005415, mean=2.003
ENSG00000169733.11 RFNG
log2(RPKM)
ENSG00000126456.15 IRF3
log2(RPKM)
3
4
5
6
7
z−score of IF value
0
2
4
6

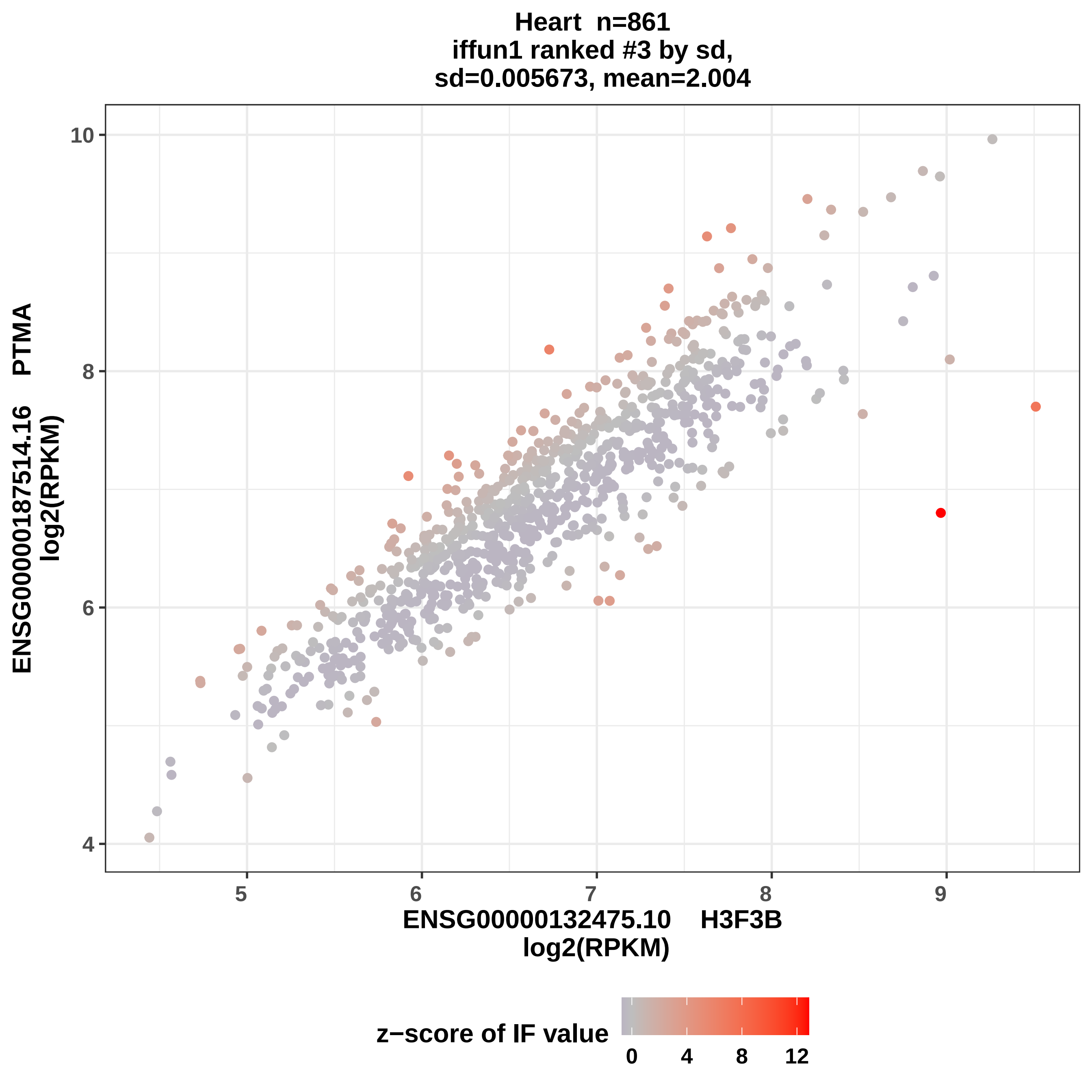
Heart n=861
iffun1 ranked #3 by sd,
sd=0.005673, mean=2.004
ENSG00000187514.16 PTMA
log2(RPKM)
10
8
6
4
5
6
7
8
9
ENSG00000132475.10 H3F3B
log2(RPKM)
z−score of IF value
0
4
8
12

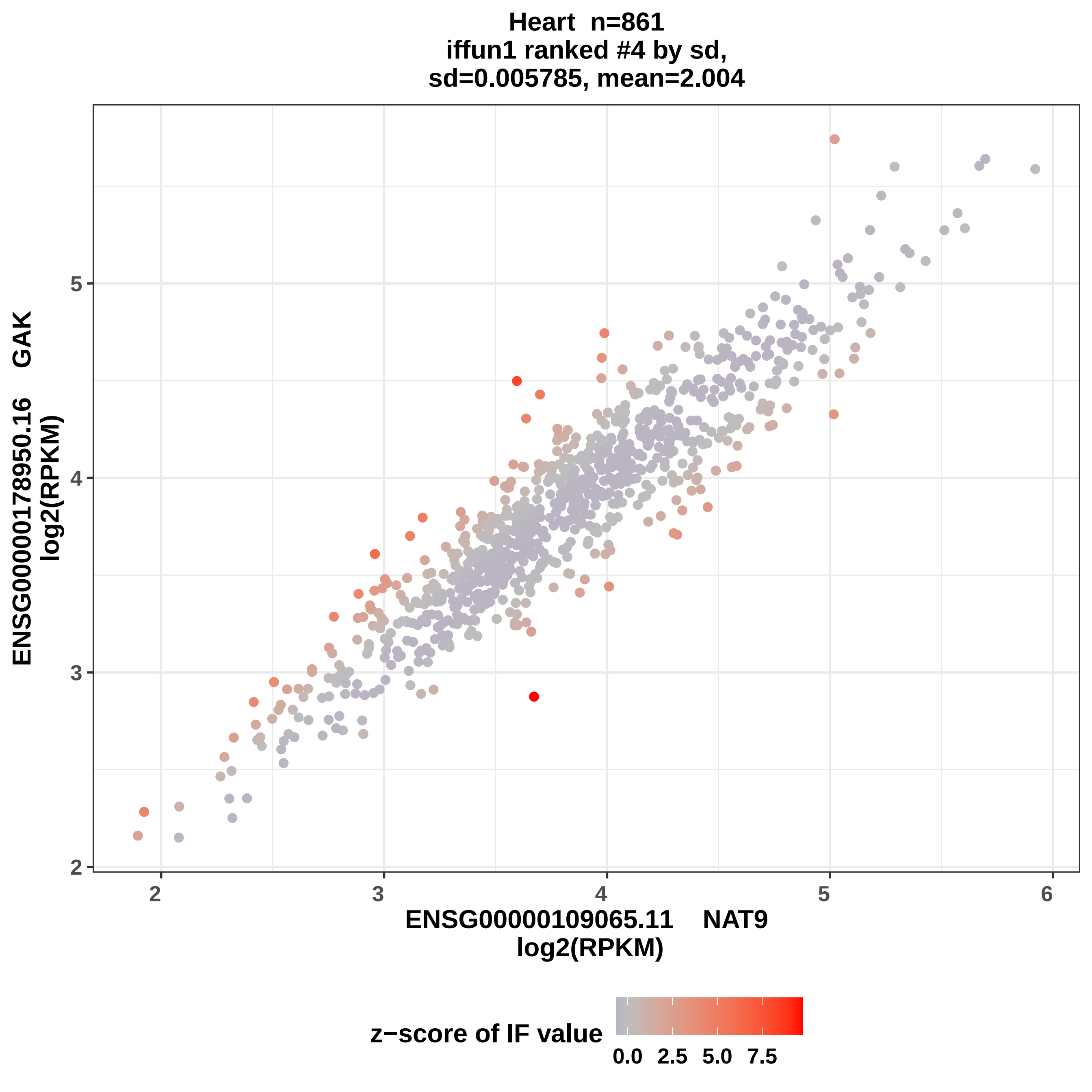
Heart n=861
iffun1 ranked #4 by sd,
sd=0.005785, mean=2.004
ENSG00000178950.16 GAK
log2(RPKM)
2
3
4
5
ENSG00000109065.11 NAT9
log2(RPKM)
2
3
4
5
6
z−score of IF value
0.0
2.5
5.0
7.5

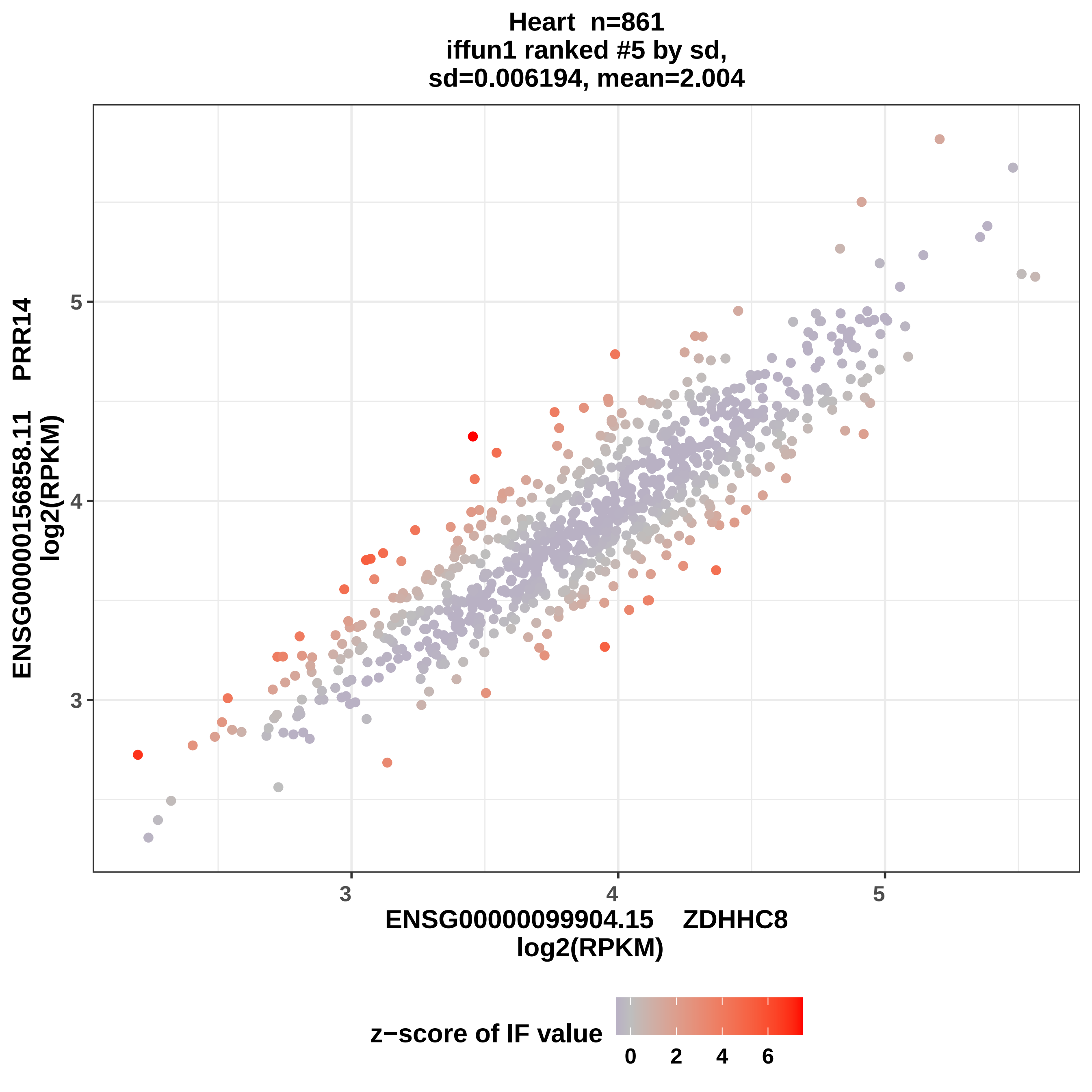
Heart n=861
iffun1 ranked #5 by sd,
sd=0.006194, mean=2.004
ENSG00000156858.11 PRR14
log2(RPKM)
ENSG00000099904.15 ZDHHC8
log2(RPKM)
3
4
5
3
4
5
z−score of IF value
0
2
4
6

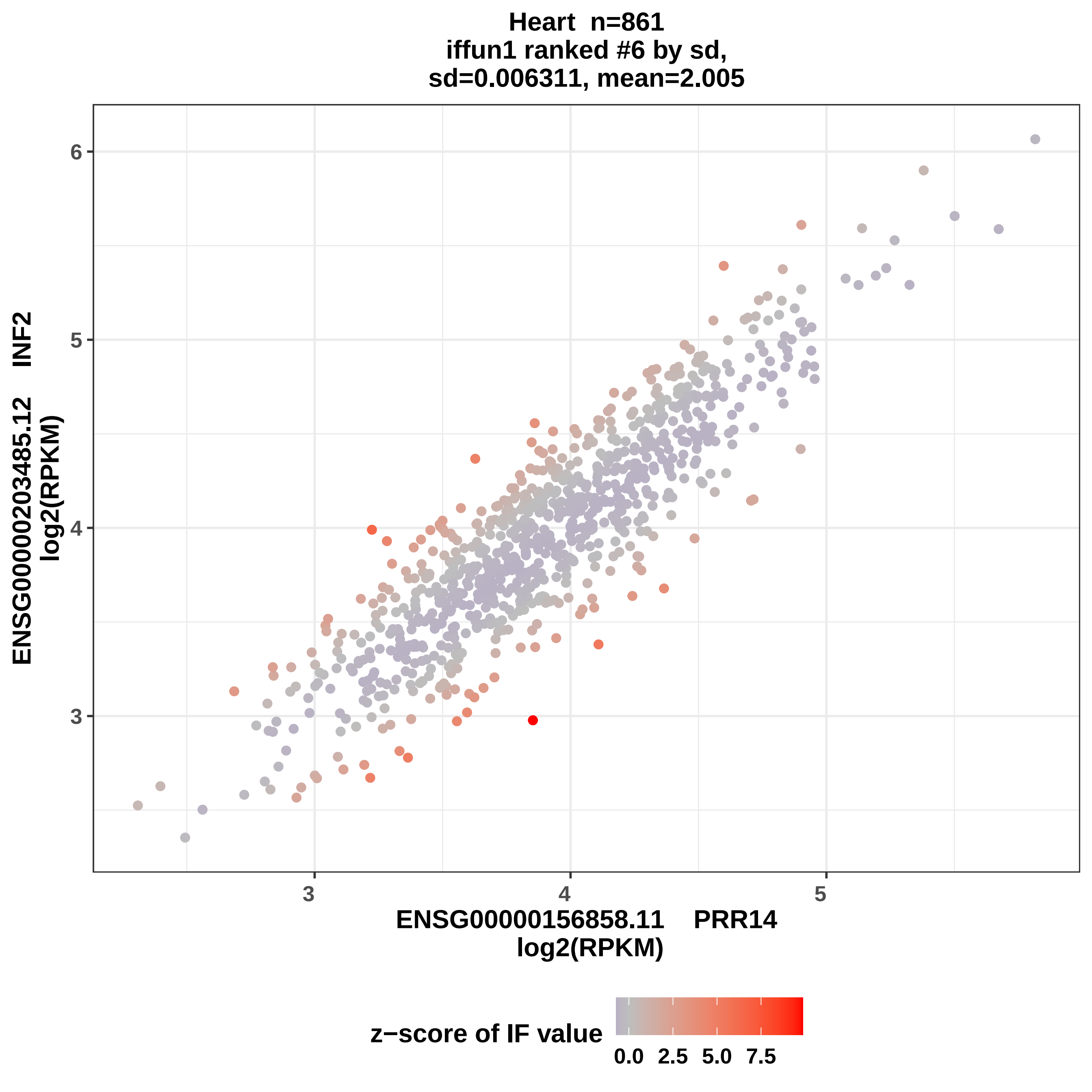

Heart n=861
iffun1 ranked #6 by sd,
sd=0.006311, mean=2.005
ENSG00000203485.12 INF2
log2(RPKM)
6
5
4
3
3
4
5
ENSG00000156858.11 PRR14
log2(RPKM)
z−score of IF value
0.0
2.5
5.0
7.5

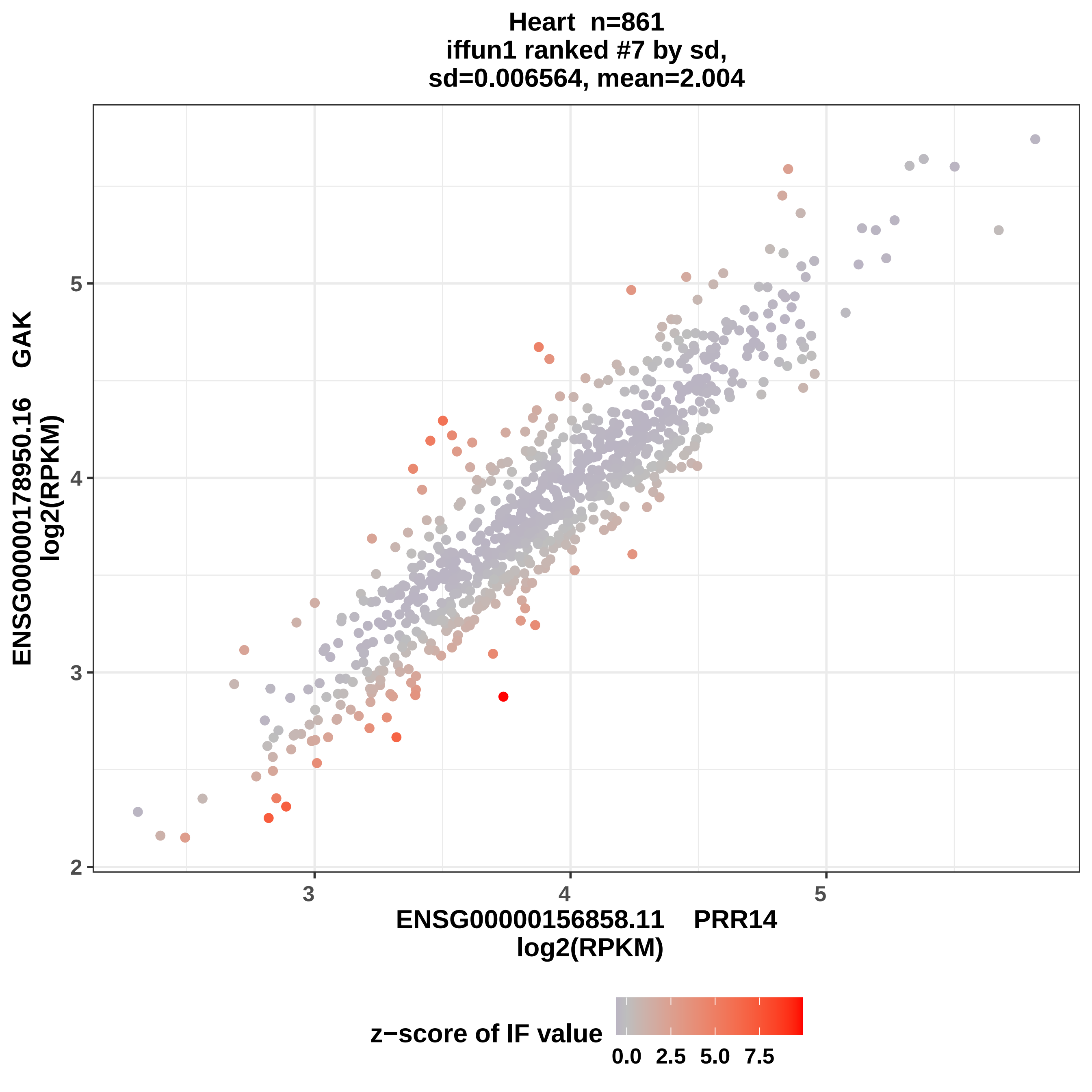

Heart n=861
iffun1 ranked #7 by sd,
sd=0.006564, mean=2.004
ENSG00000178950.16 GAK
log2(RPKM)
ENSG00000156858.11 PRR14
log2(RPKM)
2
3
4
5
3
4
5
z−score of IF value
0.0
2.5
5.0
7.5

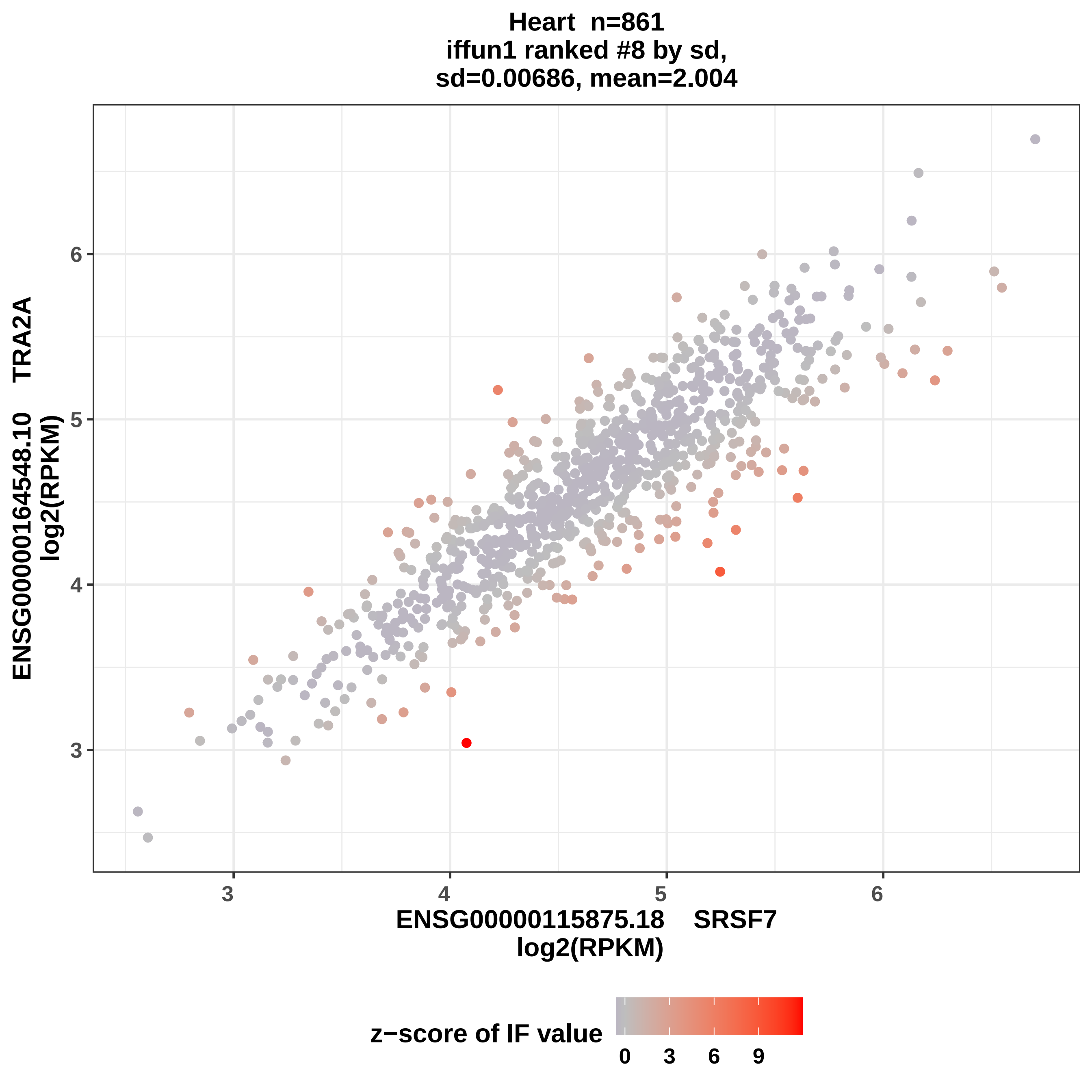
Heart n=861
iffun1 ranked #8 by sd,
sd=0.00686, mean=2.004
ENSG00000164548.10 TRA2A
log2(RPKM)
ENSG00000115875.18 SRSF7
log2(RPKM)
z−score of IF value
0 3 6 9
3 4 5 6
3 4 5 6

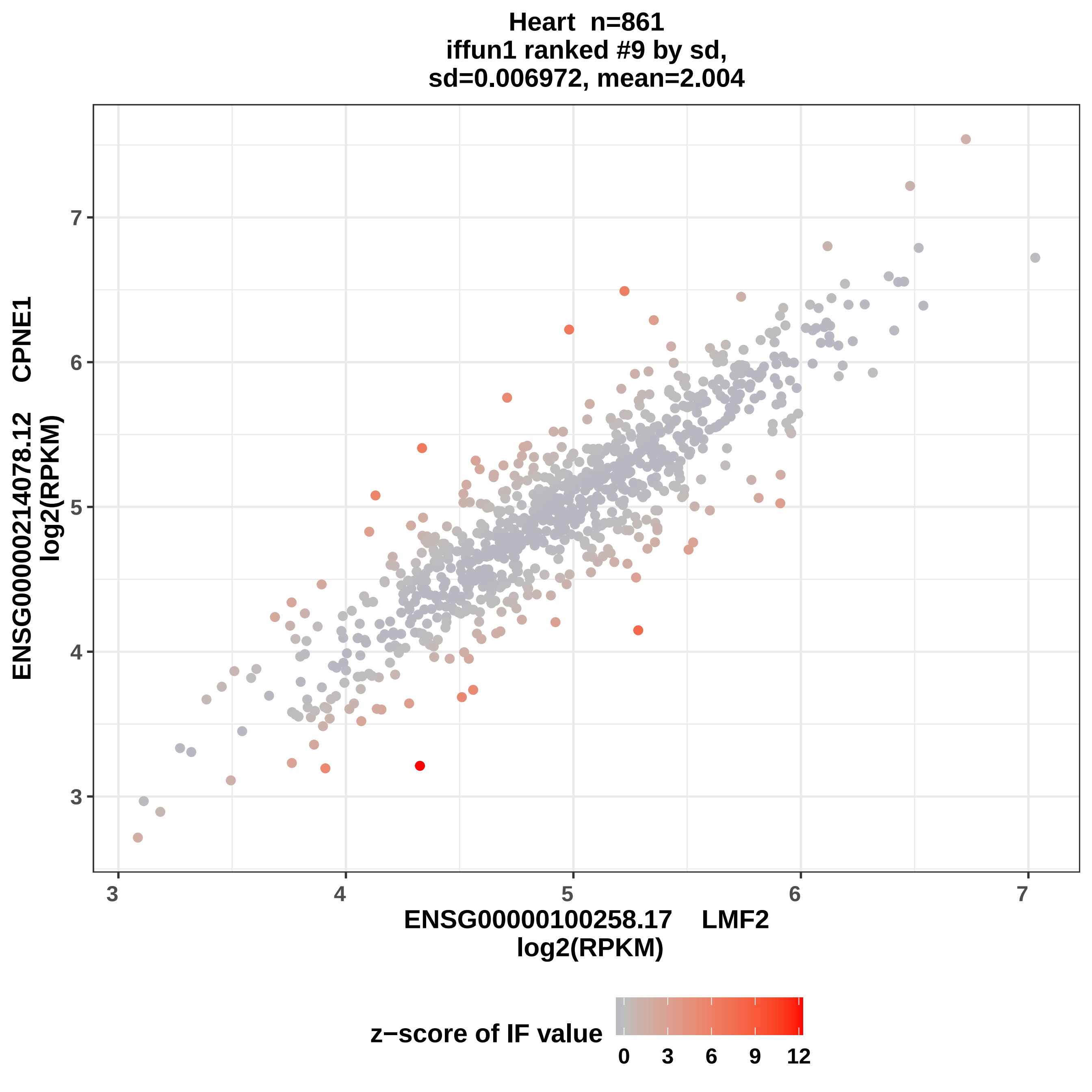
Heart n=861
iffun1 ranked #9 by sd,
sd=0.006972, mean=2.004
ENSG00000214078.12 CPNE1
log2(RPKM)
ENSG00000100258.17 LMF2
log2(RPKM)
3
4
5
6
7
z−score of IF value
0
3
6
9
12

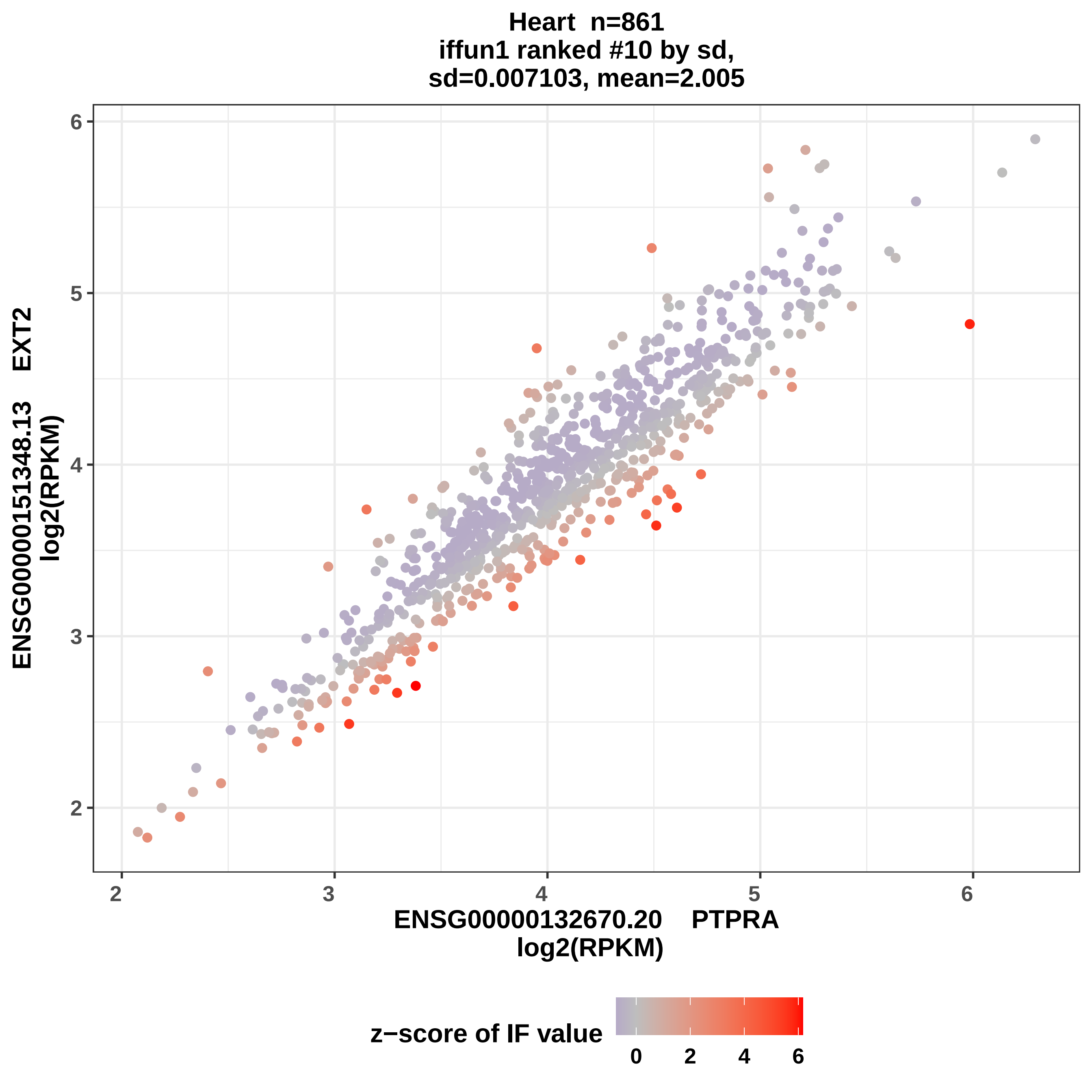

Heart n=861
iffun1 ranked #10 by sd,
sd=0.007103, mean=2.005
ENSG00000151348.13 EXT2
log2(RPKM)
ENSG00000132670.20 PTPRA
log2(RPKM)
2
3
4
5
6
z−score of IF value
0
2
4
6

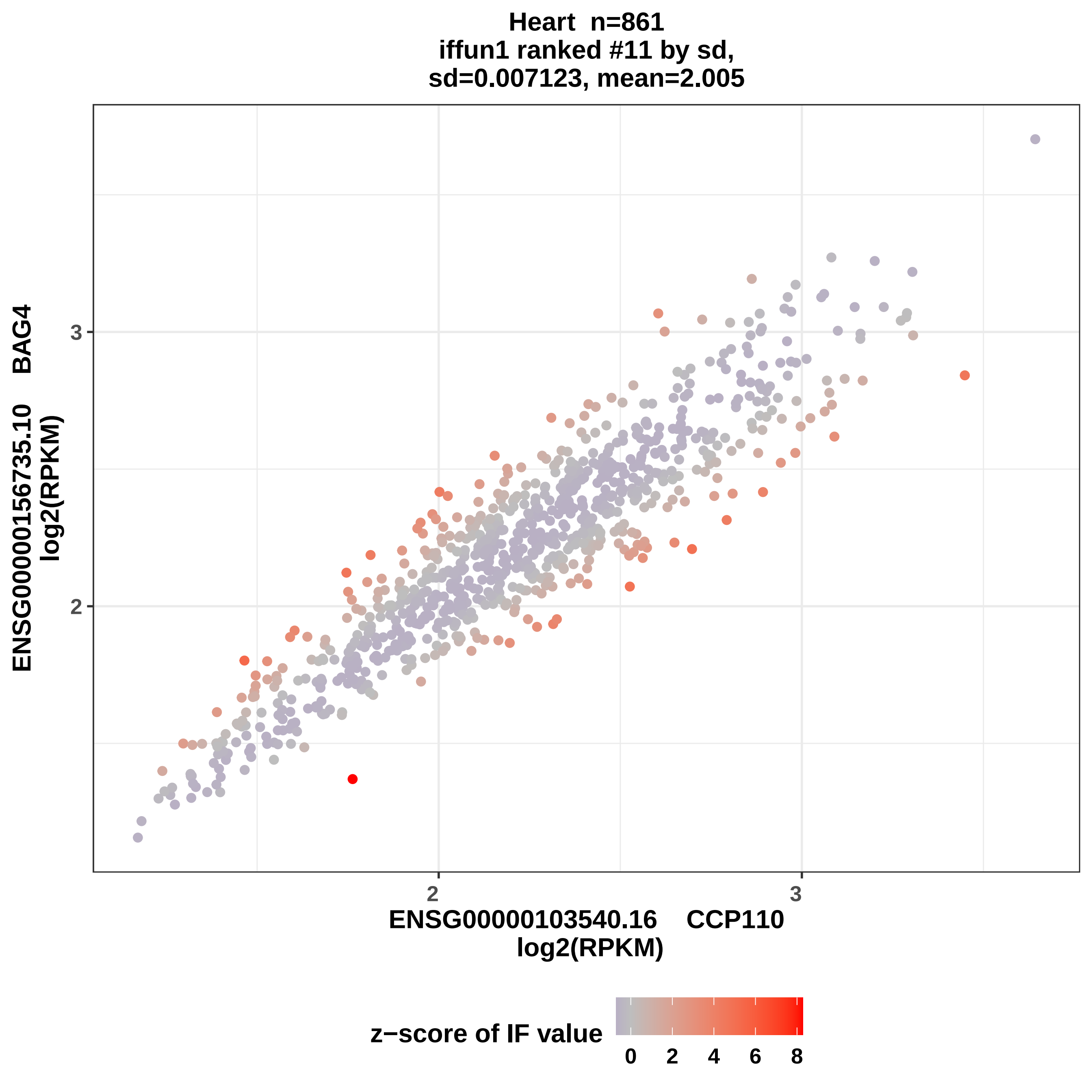

Heart n=861
iffun1 ranked #11 by sd,
sd=0.007123, mean=2.005
ENSG00000156735.10 BAG4
log2(RPKM)
3
2
2
3
ENSG00000103540.16 CCP110
log2(RPKM)
z−score of IF value
0
2
4
6
8

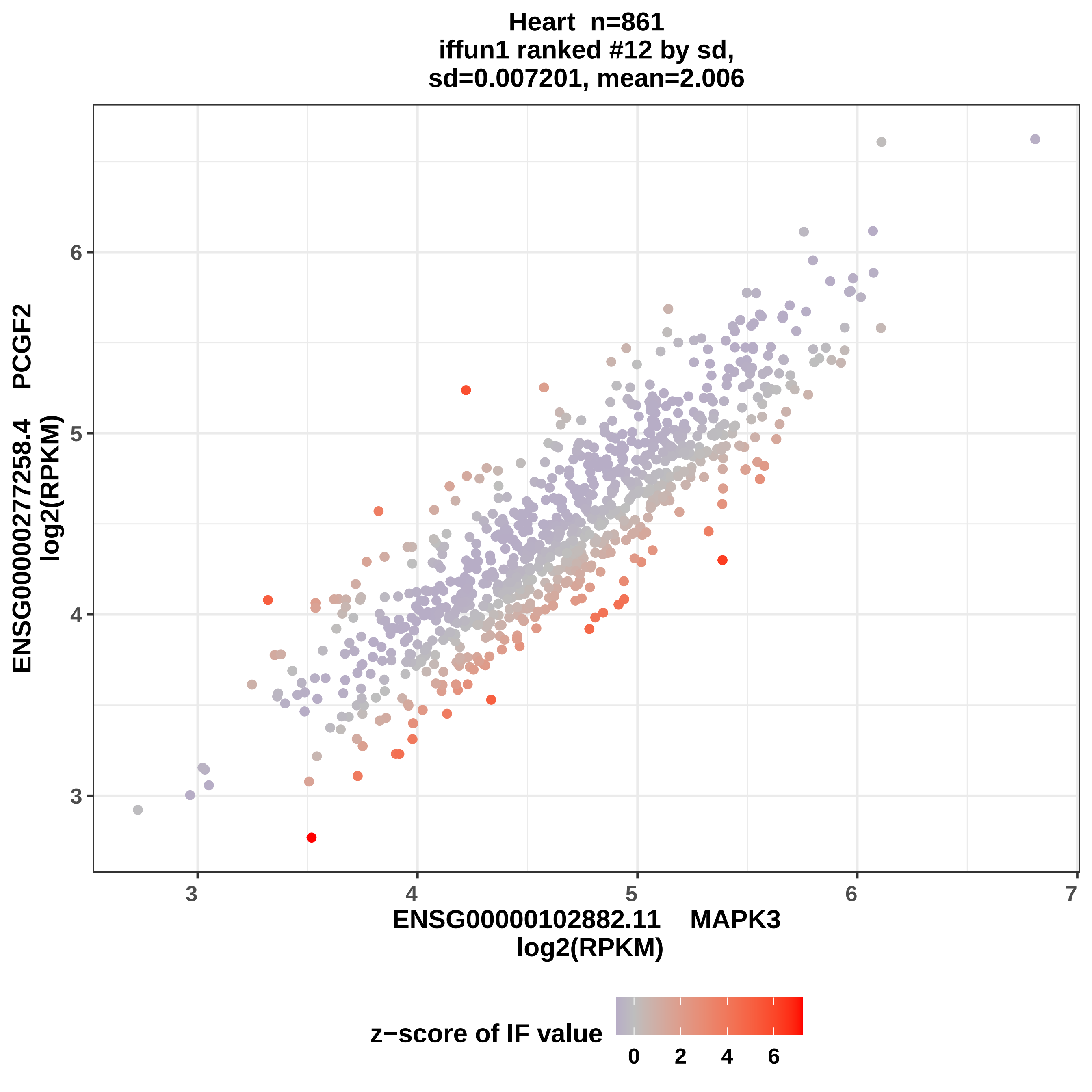
Heart n=861
iffun1 ranked #12 by sd,
sd=0.007201, mean=2.006
ENSG00000277258.4 PCGF2
log2(RPKM)
ENSG00000102882.11 MAPK3
log2(RPKM)
z−score of IF value
3
4
5
6
7
0
2
4
6

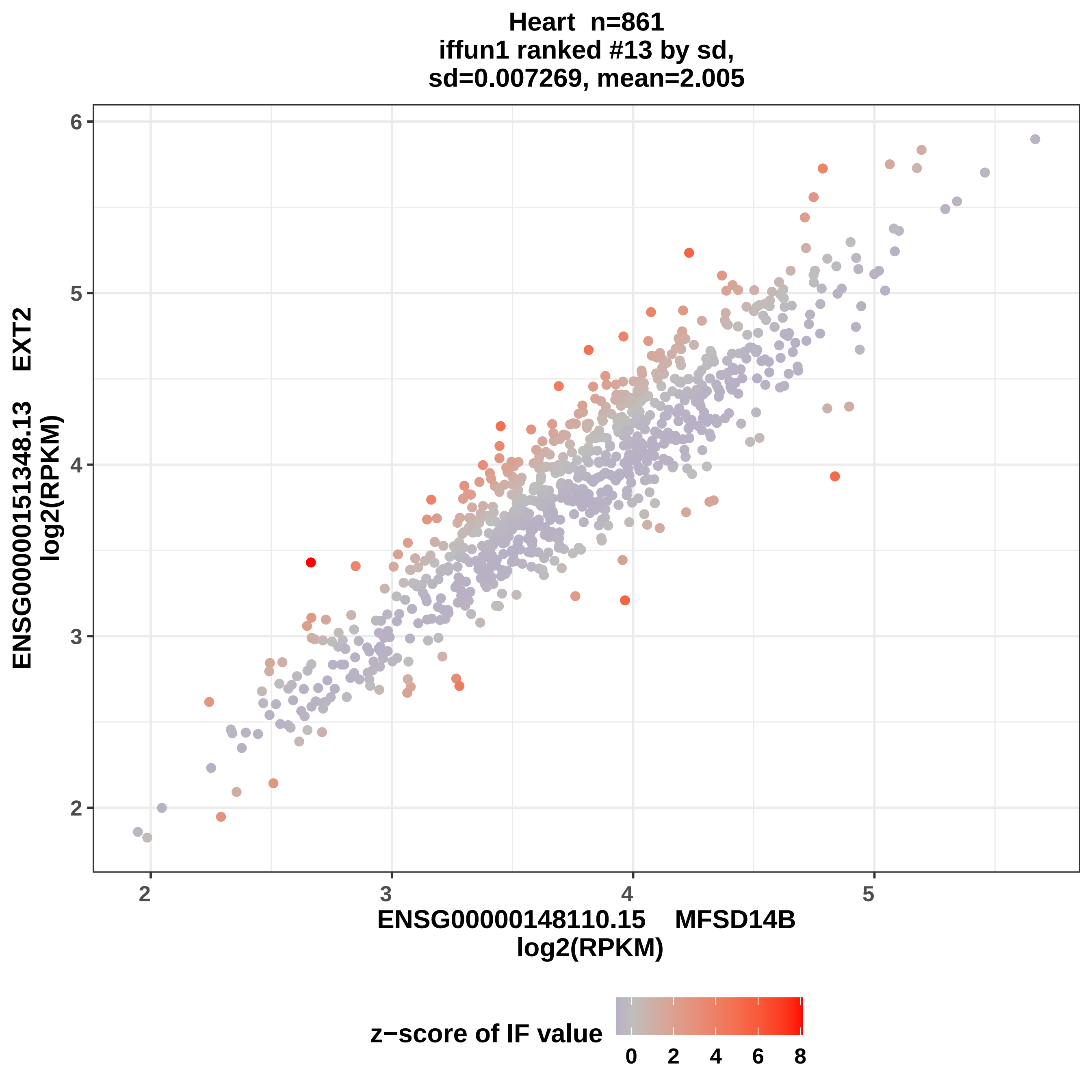

Heart n=861
iffun1 ranked #13 by sd,
sd=0.007269, mean=2.005
ENSG00000151348.13 EXT2
log2(RPKM)
ENSG00000148110.15 MFSD14B
log2(RPKM)
2
3
4
5
6
z−score of IF value
0
2
4
6
8

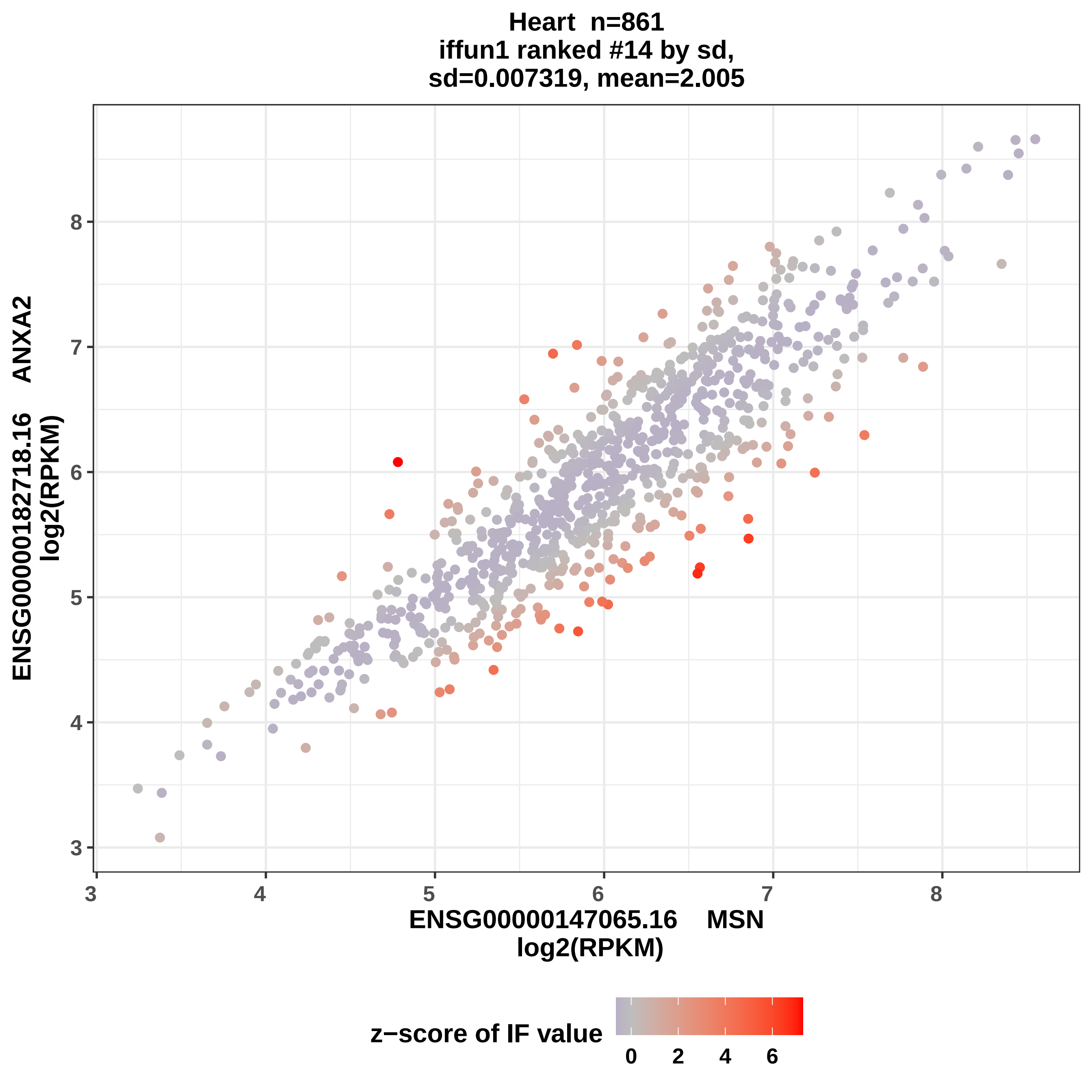
Heart n=861
iffun1 ranked #14 by sd,
sd=0.007319, mean=2.005
ENSG00000182718.16 ANXA2
log2(RPKM)
ENSG00000147065.16 MSN
log2(RPKM)
3
4
5
6
7
8
z−score of IF value
0
2
4
6

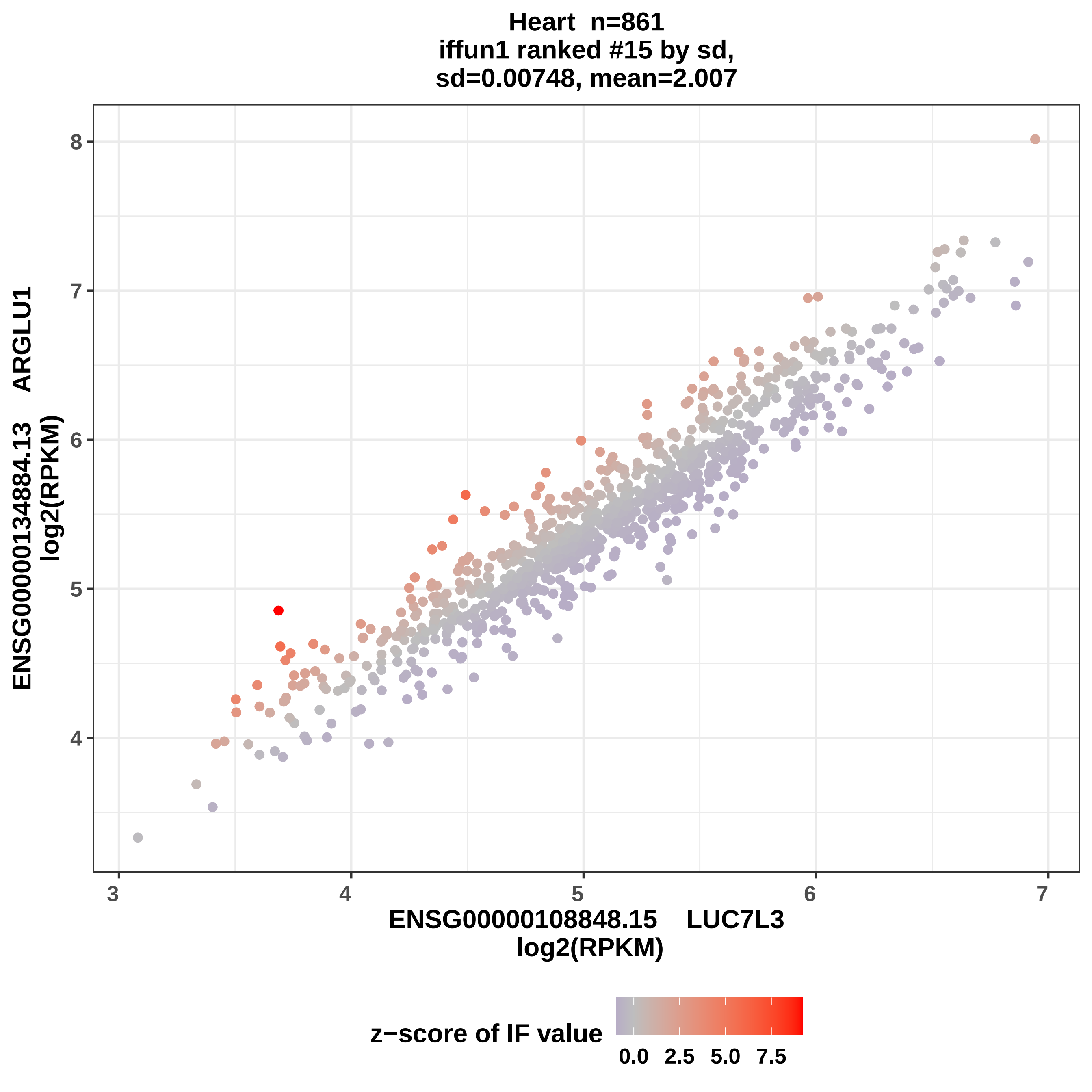
Heart n=861
iffun1 ranked #15 by sd,
sd=0.00748, mean=2.007
ENSG00000134884.13 ARGLU1
log2(RPKM)
ENSG00000108848.15 LUC7L3
log2(RPKM)
z−score of IF value
0.0 2.5 5.0 7.5
3 4 5 6 7
4 5 6 7 8

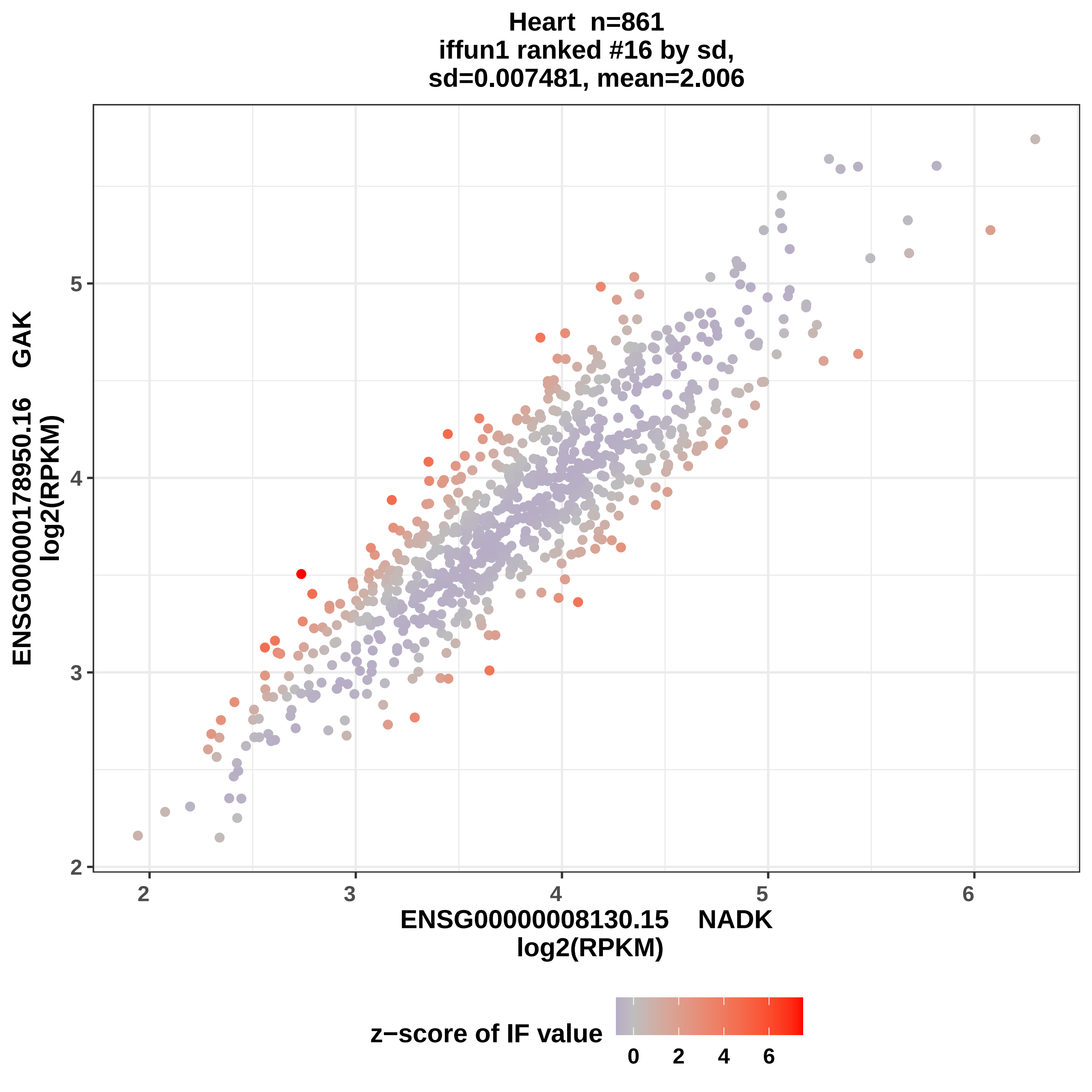

Heart n=861
iffun1 ranked #16 by sd,
sd=0.007481, mean=2.006
ENSG00000178950.16 GAK
log2(RPKM)
2
3
4
5
2
3
4
5
6
ENSG00000008130.15 NADK
log2(RPKM)
z−score of IF value
0
2
4
6

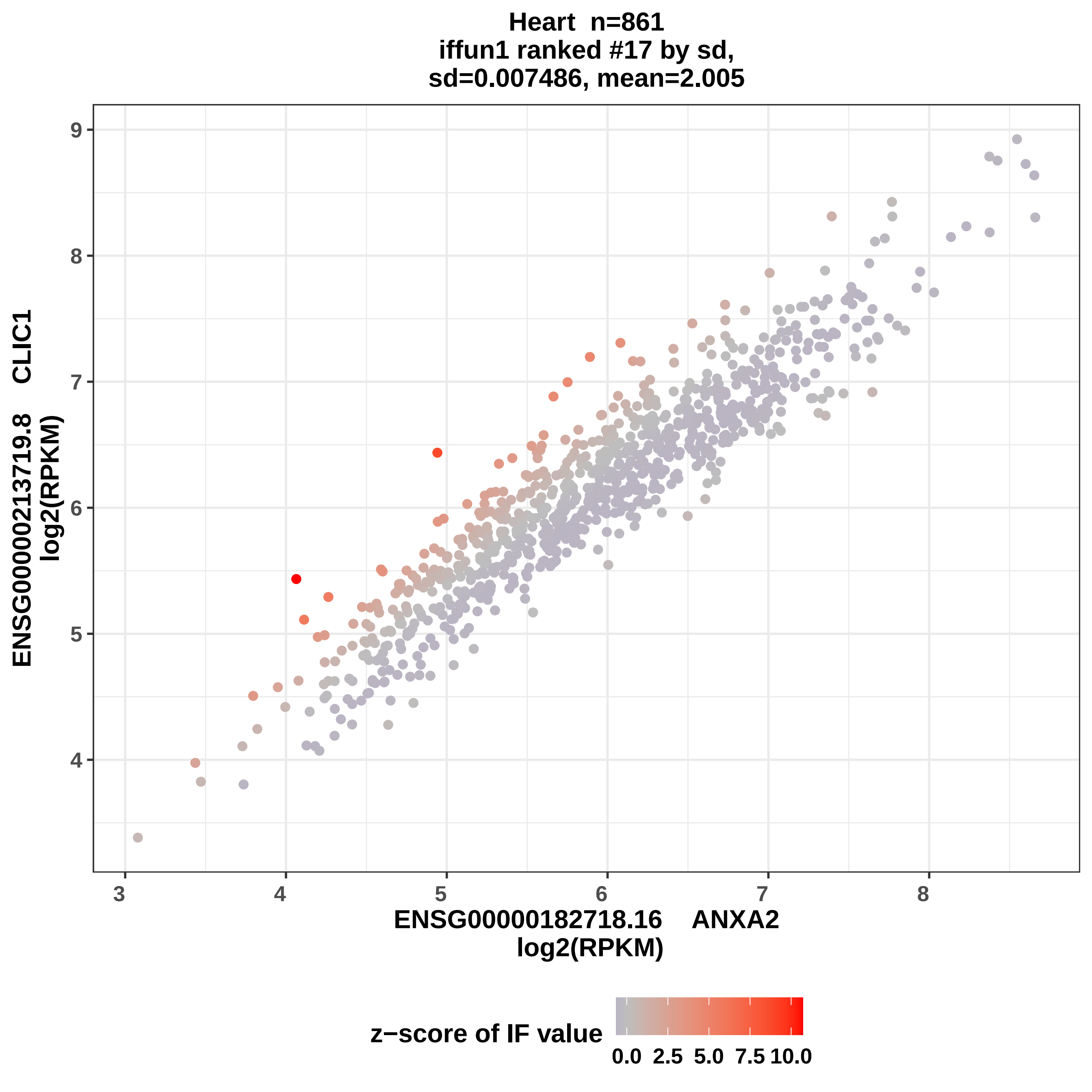
Heart n=861
iffun1 ranked #17 by sd,
sd=0.007486, mean=2.005
ENSG00000213719.8 CLIC1
log2(RPKM)
ENSG00000182718.16 ANXA2
log2(RPKM)
z−score of IF value
0.0 2.5 5.0 7.5 10.0
3 4 5 6 7 8
4 5 6 7 8 9

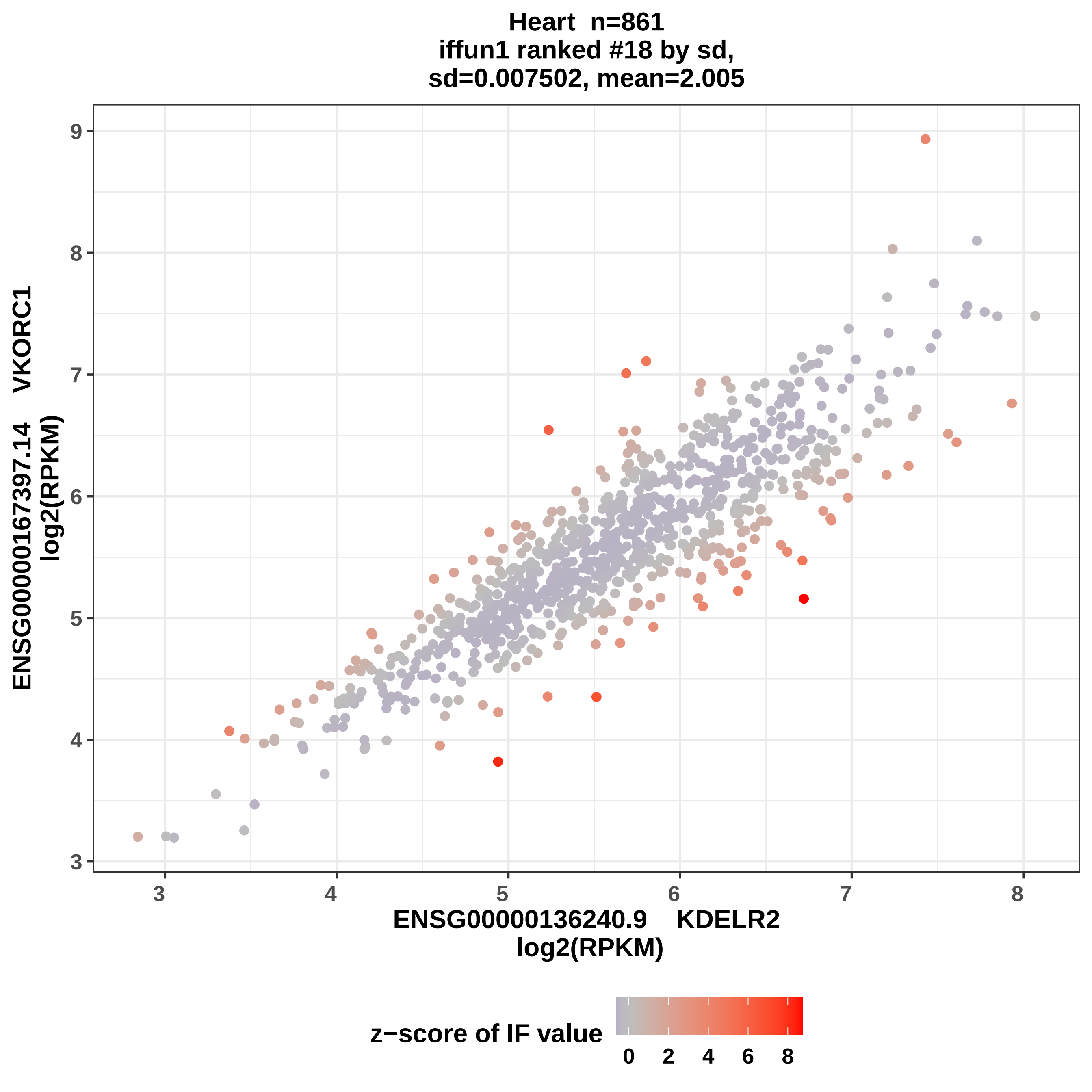

Heart n=861
iffun1 ranked #18 by sd,
sd=0.007502, mean=2.005
ENSG00000167397.14 VKORC1
log2(RPKM)
ENSG00000136240.9 KDELR2
log2(RPKM)
z−score of IF value
0 2 4 6 8
3 4 5 6 7 8
3 4 5 6 7 8 9

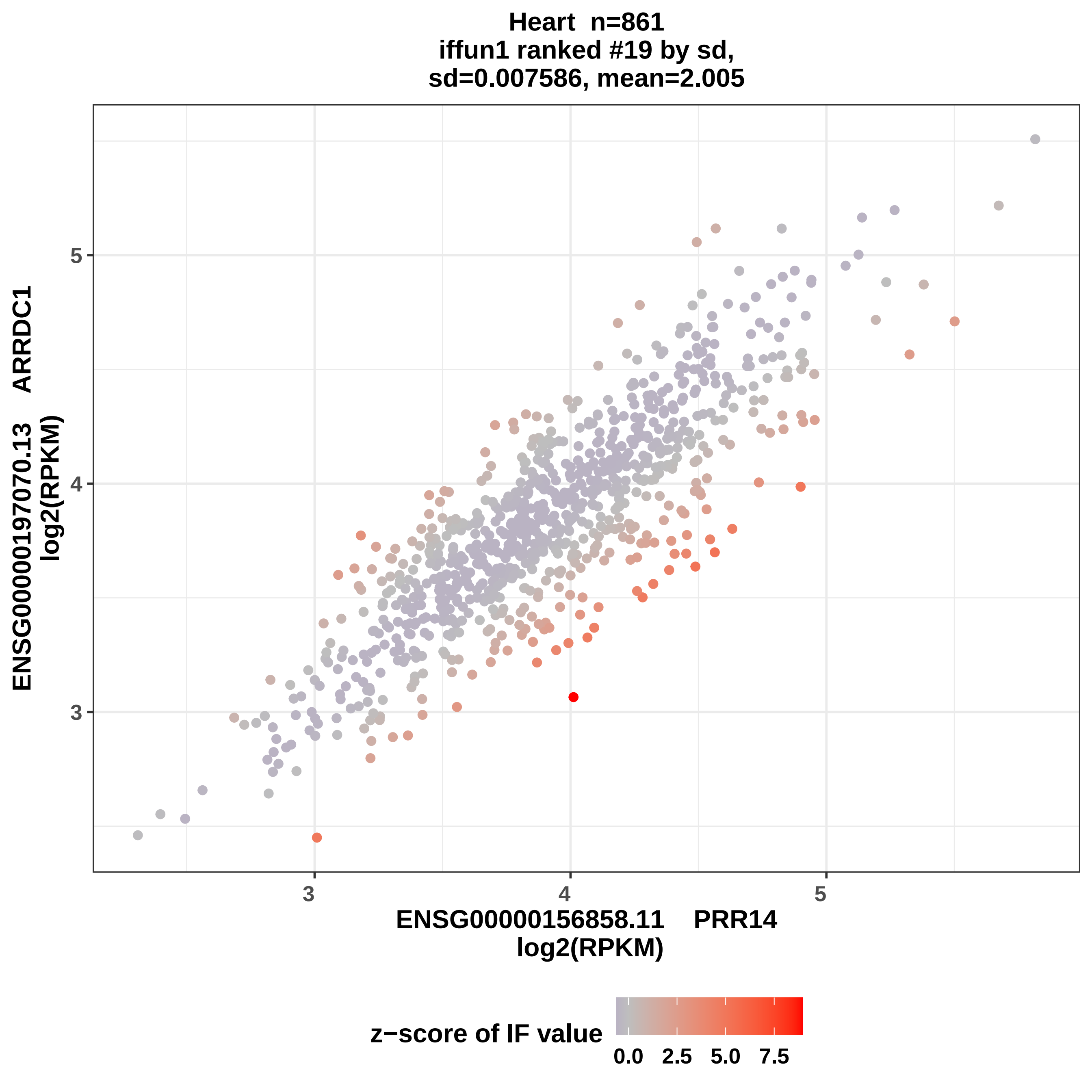

Heart n=861
iffun1 ranked #19 by sd,
sd=0.007586, mean=2.005
ENSG00000197070.13 ARRDC1
log2(RPKM)
ENSG00000156858.11 PRR14
log2(RPKM)
3
4
5
z−score of IF value
0.0
2.5
5.0
7.5

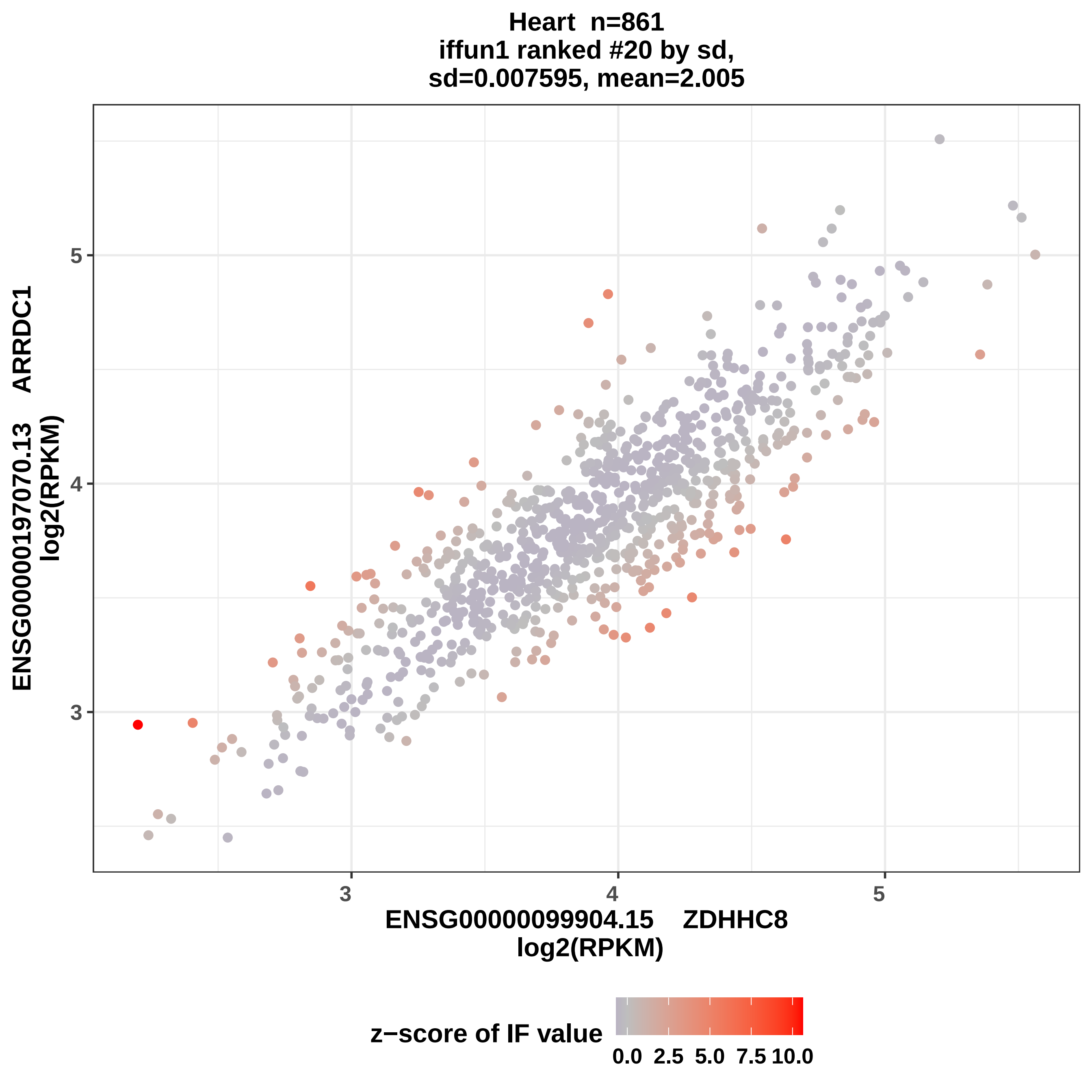
Heart n=861
iffun1 ranked #20 by sd,
sd=0.007595, mean=2.005
ENSG00000197070.13 ARRDC1
log2(RPKM)
ENSG00000099904.15 ZDHHC8
log2(RPKM)
3
4
5
3
4
5
z−score of IF value
0.0
2.5
5.0
7.5
10.0

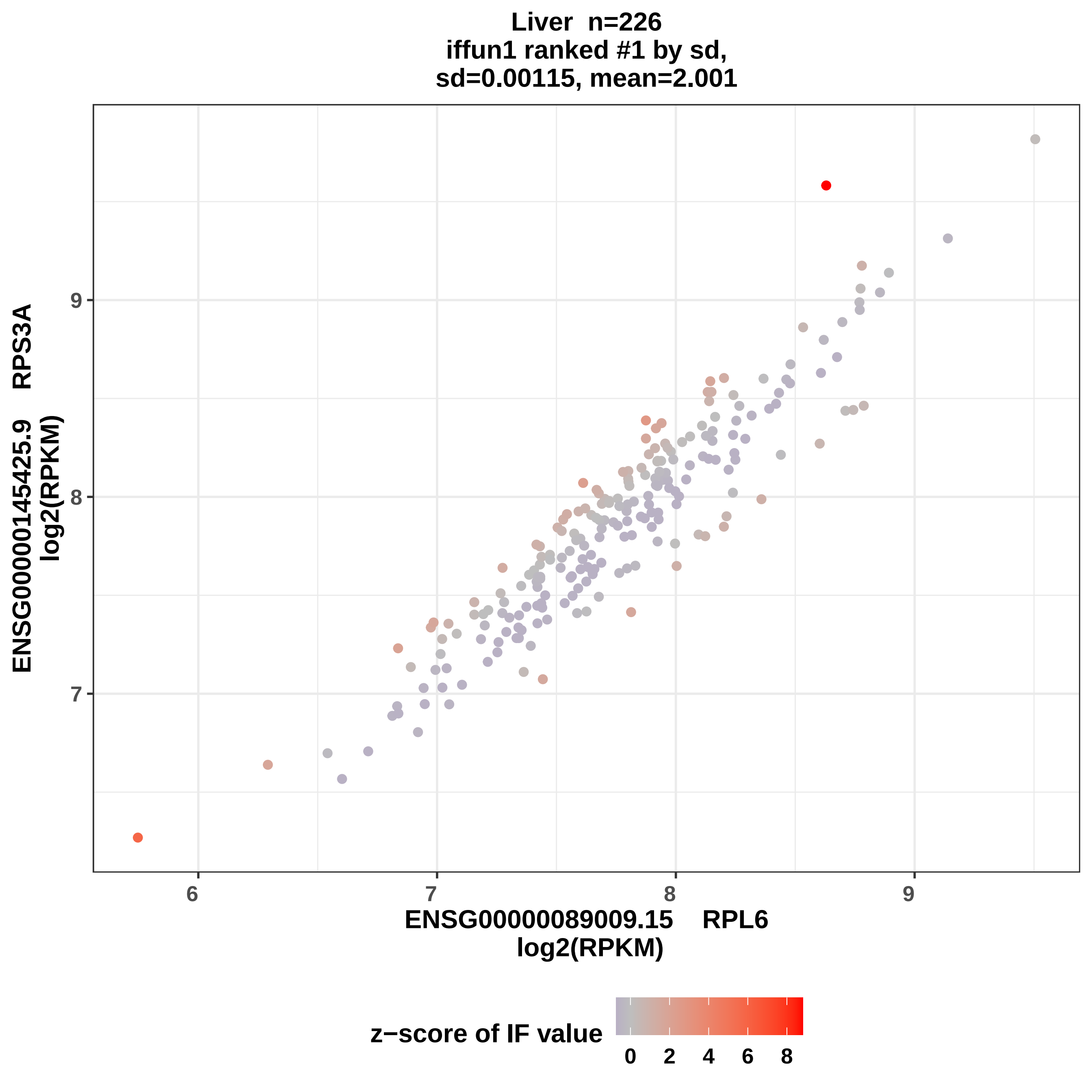

Liver n=226
iffun1 ranked #1 by sd,
sd=0.00115, mean=2.001
ENSG00000145425.9 RPS3A
log2(RPKM)
ENSG00000089009.15 RPL6
log2(RPKM)
z−score of IF value
0 2 4 6 8

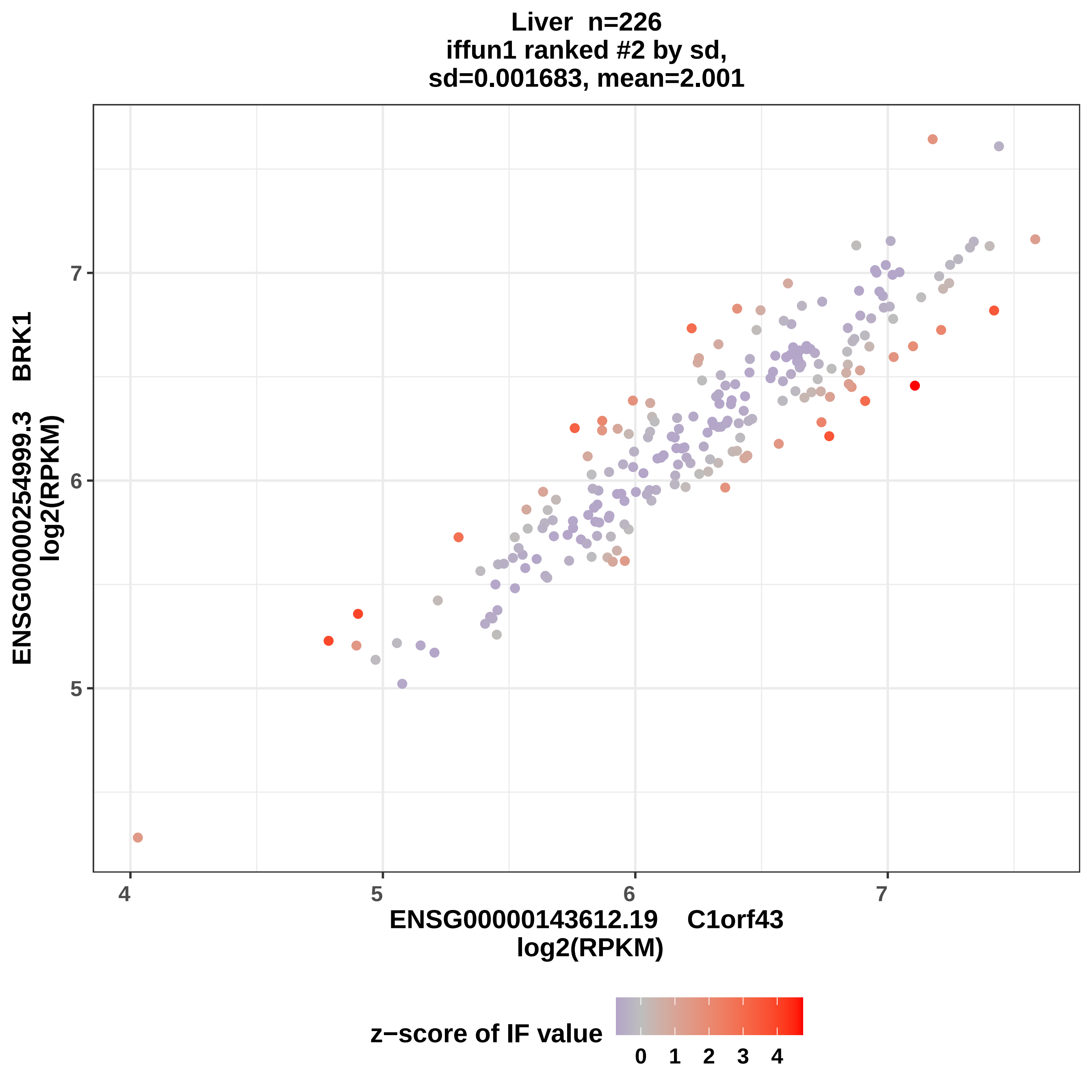

Liver n=226
iffun1 ranked #2 by sd,
sd=0.001683, mean=2.001
ENSG00000254999.3 BRK1
log2(RPKM)
7
6
5
4
5
6
7
ENSG00000143612.19 C1orf43
log2(RPKM)
z−score of IF value
0
1
2
3
4

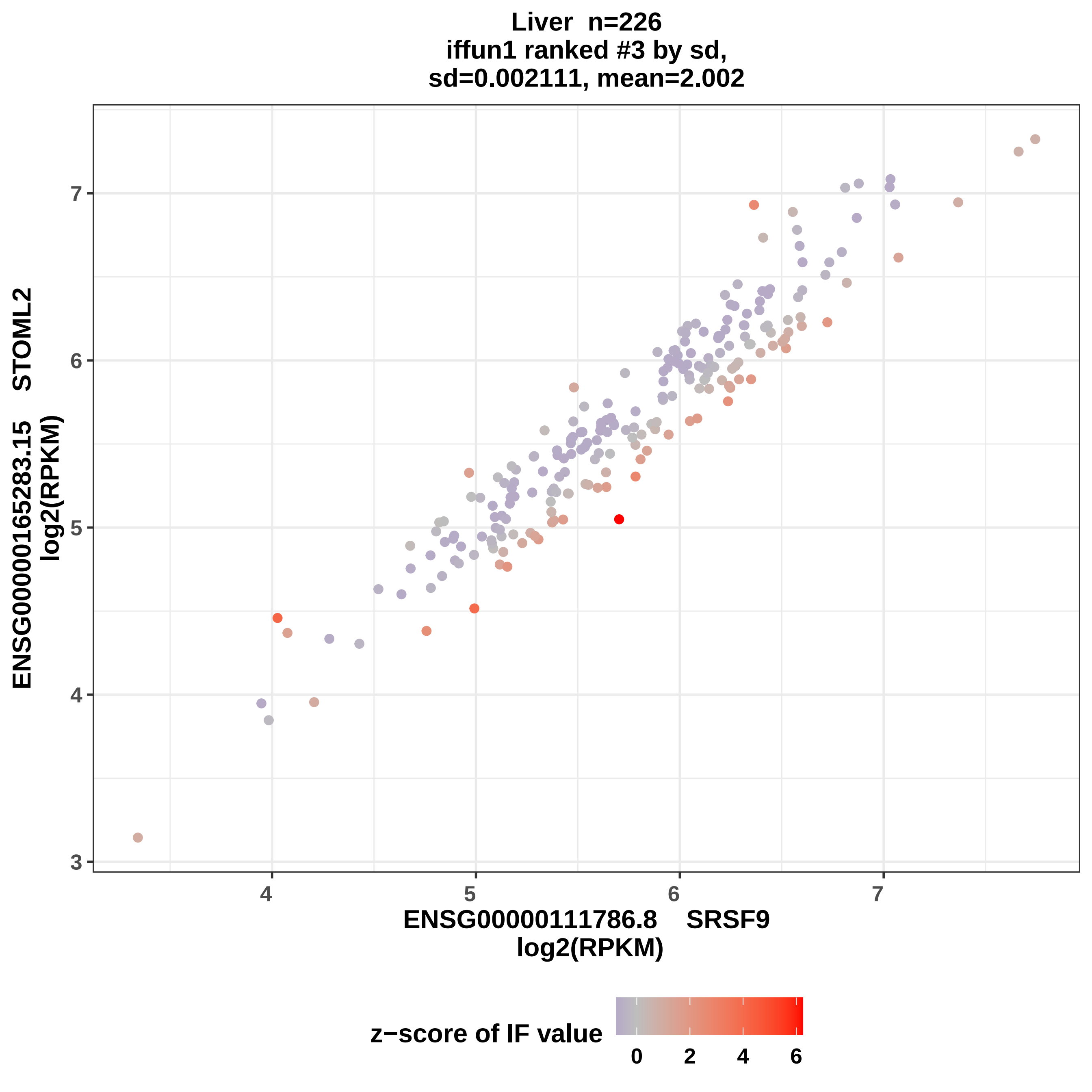

Liver n=226
iffun1 ranked #3 by sd,
sd=0.002111, mean=2.002
ENSG00000165283.15 STOML2
log2(RPKM)
ENSG00000111786.8 SRSF9
log2(RPKM)
3
4
5
6
7
4
5
6
7
z−score of IF value
0
2
4
6

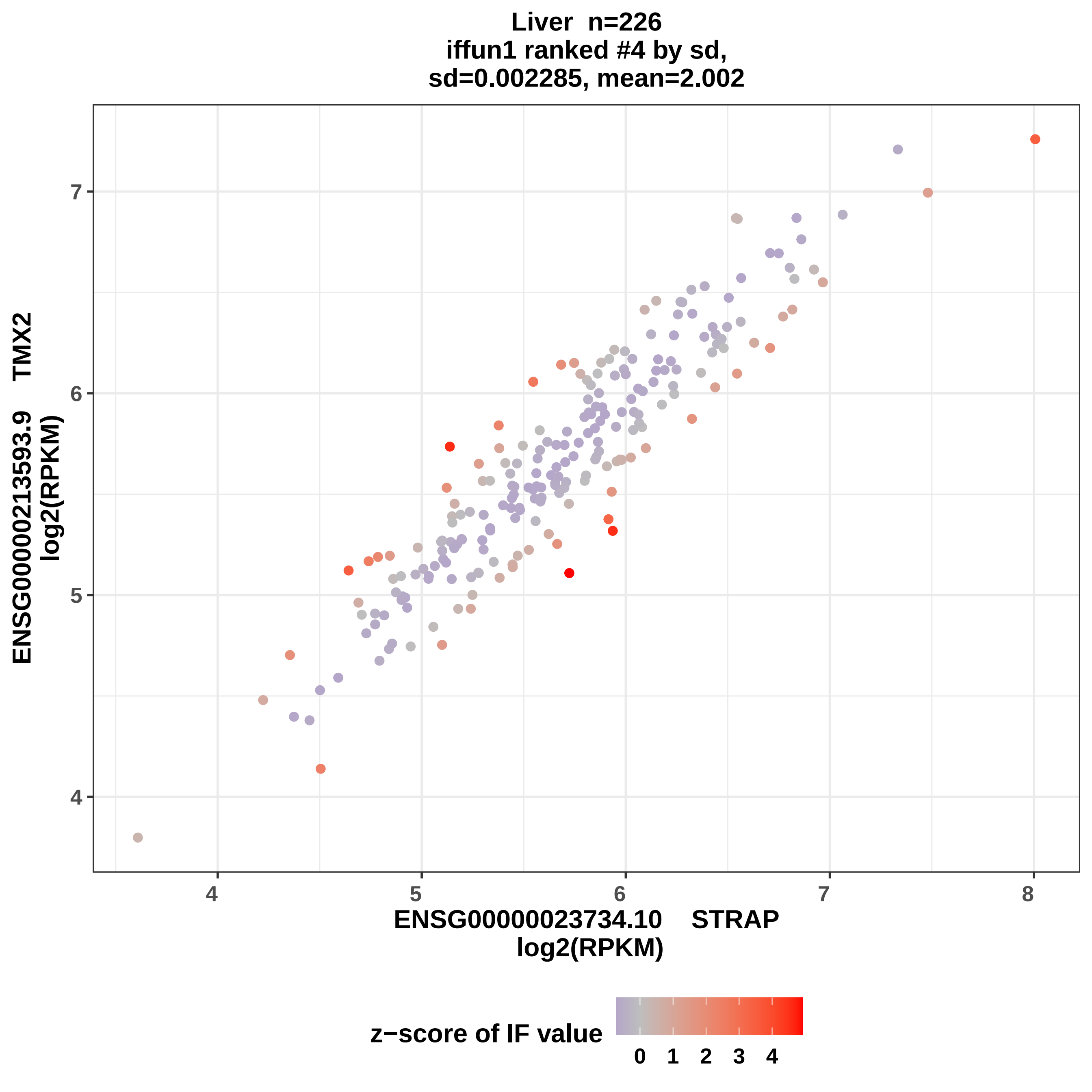
Liver n=226
iffun1 ranked #4 by sd,
sd=0.002285, mean=2.002
ENSG00000213593.9 TMX2
log2(RPKM)
ENSG00000023734.10 STRAP
log2(RPKM)
4
5
6
7
8
z−score of IF value
0
1
2
3
4

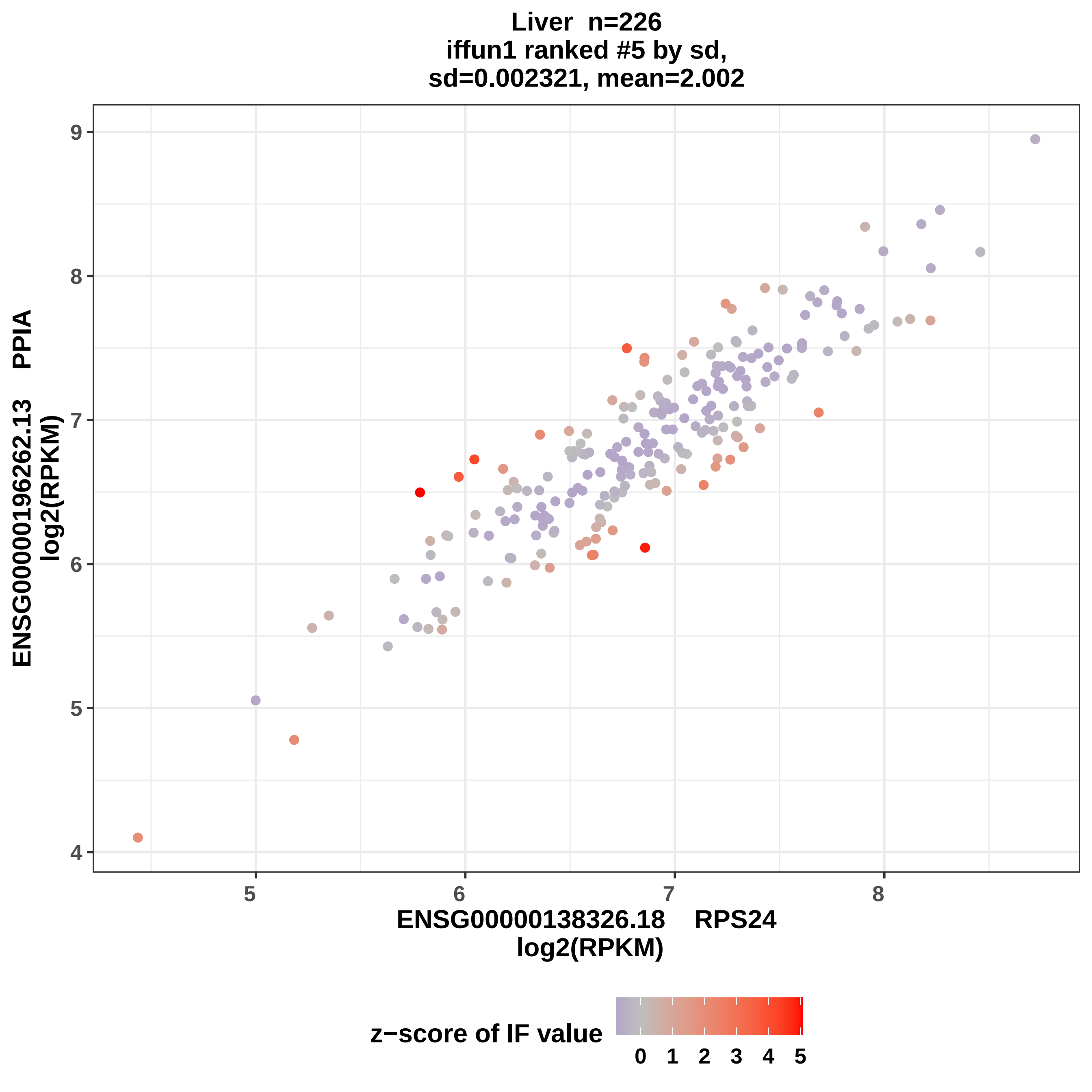
Liver n=226
iffun1 ranked #5 by sd,
sd=0.002321, mean=2.002
ENSG00000196262.13 PPIA
log2(RPKM)
ENSG00000138326.18 RPS24
log2(RPKM)
z−score of IF value
0 1 2 3 4 5

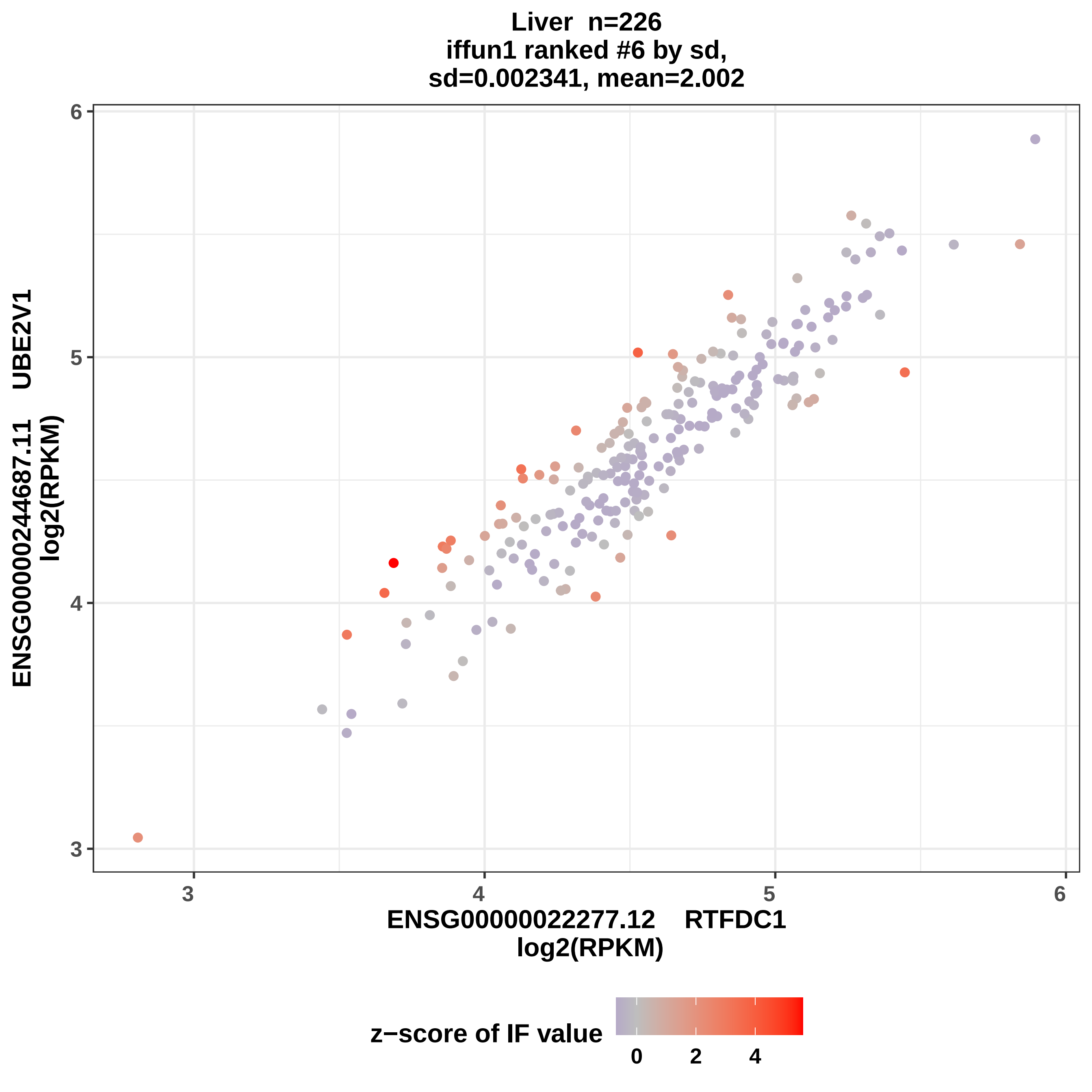

Liver n=226
iffun1 ranked #6 by sd,
sd=0.002341, mean=2.002
ENSG00000244687.11 UBE2V1
log2(RPKM)
ENSG00000022277.12 RTFDC1
log2(RPKM)
z−score of IF value
3
4
5
6
0
2
4

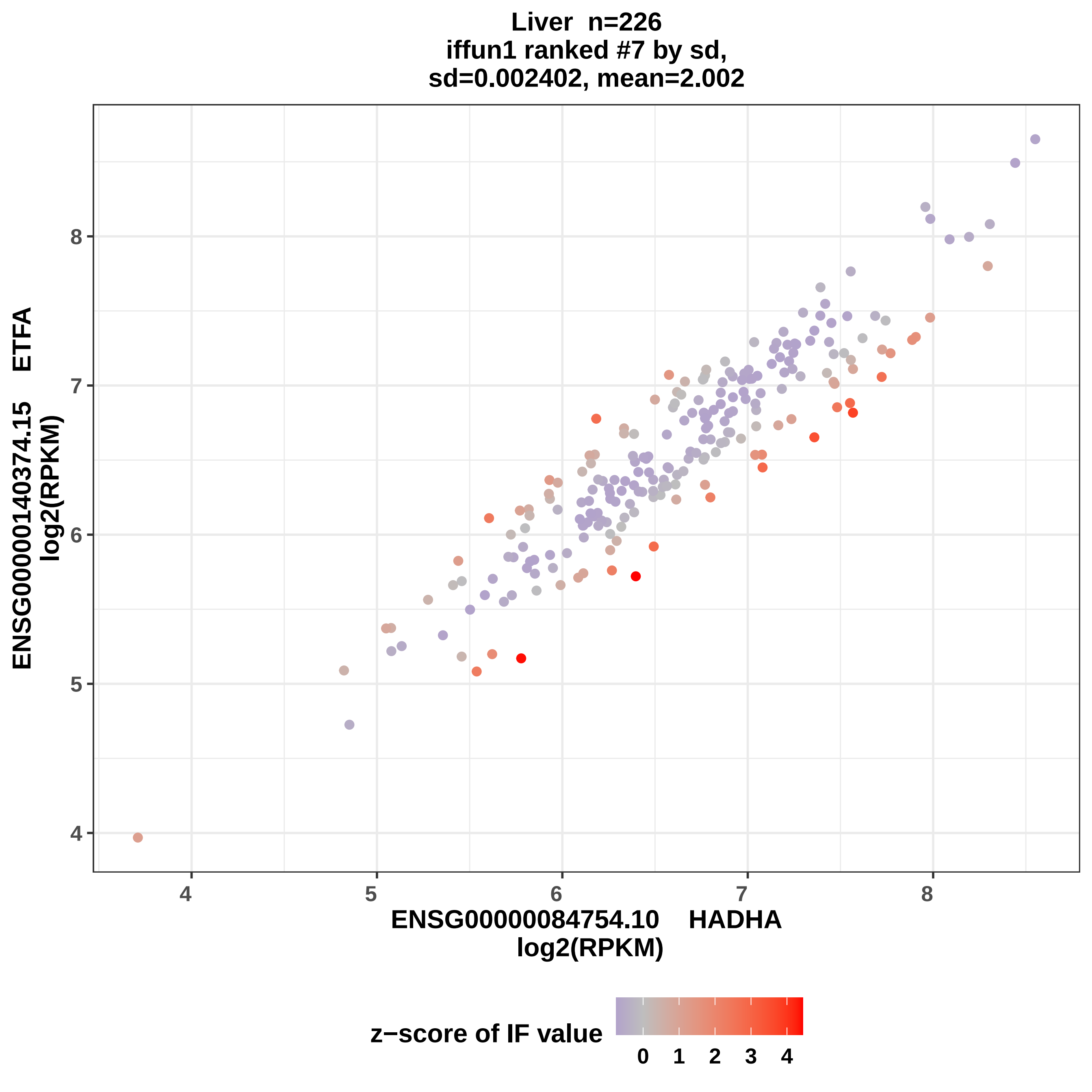

Liver n=226
iffun1 ranked #7 by sd,
sd=0.002402, mean=2.002
ENSG00000140374.15 ETFA
log2(RPKM)
8
7
6
5
4
4
5
6
7
8
ENSG00000084754.10 HADHA
log2(RPKM)
z−score of IF value
0
1
2
3
4

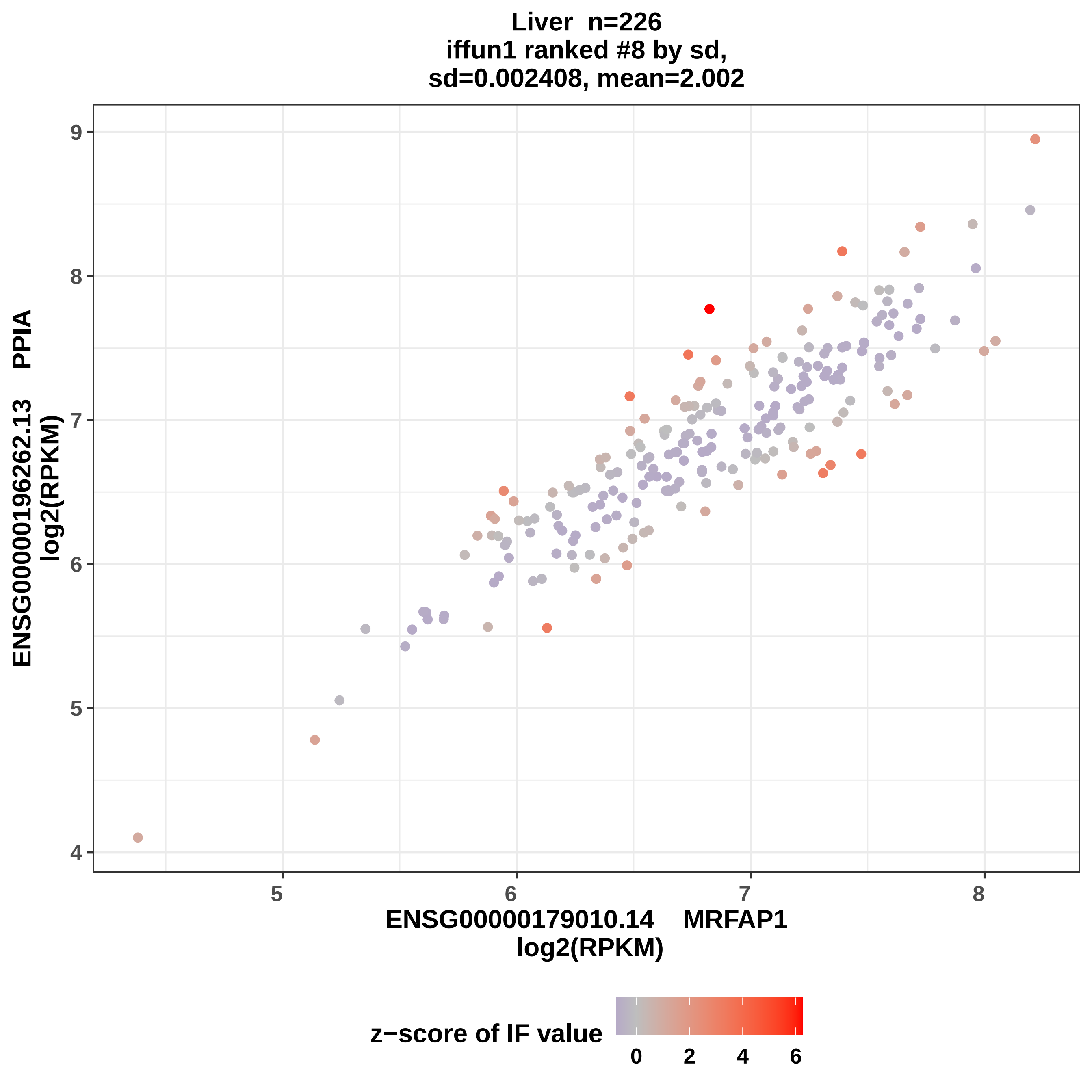

Liver n=226
iffun1 ranked #8 by sd,
sd=0.002408, mean=2.002
ENSG00000196262.13 PPIA
log2(RPKM)
ENSG00000179010.14 MRFAP1
log2(RPKM)
4
5
6
7
8
9
5
6
7
8
z−score of IF value
0
2
4
6

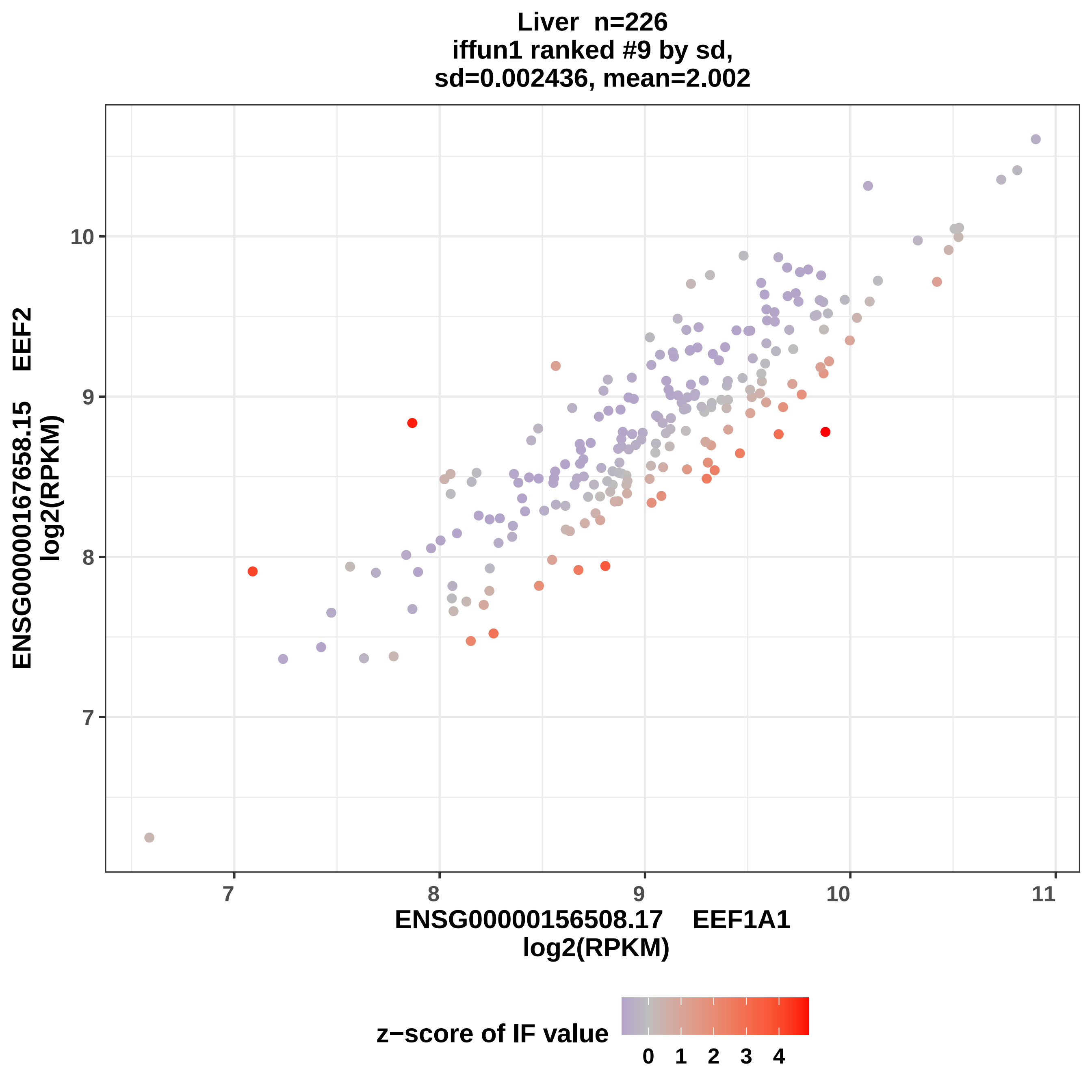

Liver n=226
iffun1 ranked #9 by sd,
sd=0.002436, mean=2.002
ENSG00000167658.15 EEF2
log2(RPKM)
10
9
8
7
7
8
9
10
11
ENSG00000156508.17 EEF1A1
log2(RPKM)
z−score of IF value
0 1 2 3 4

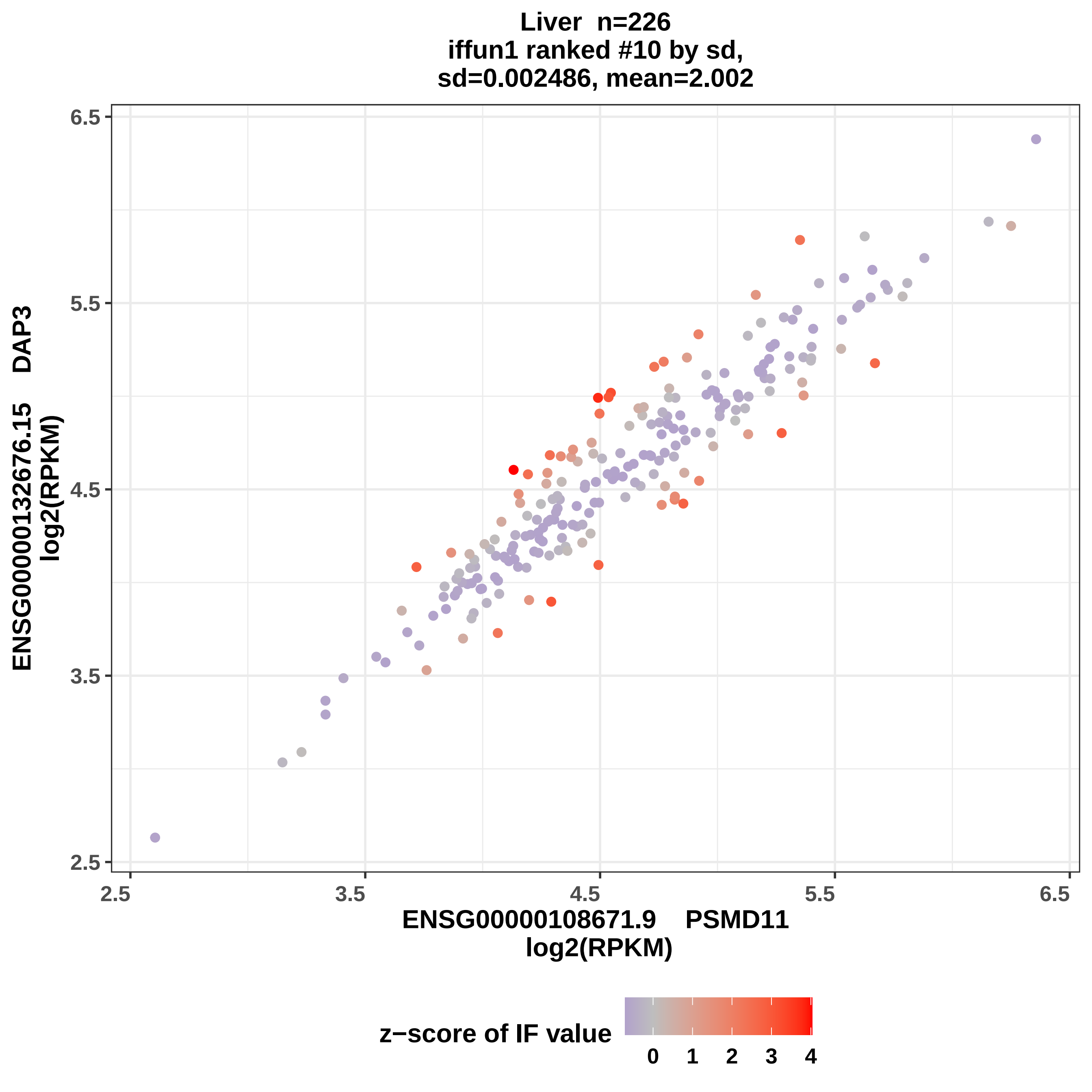

Liver n=226
iffun1 ranked #10 by sd,
sd=0.002486, mean=2.002
ENSG00000132676.15 DAP3
log2(RPKM)
ENSG00000108671.9 PSMD11
log2(RPKM)
2.5
3.5
4.5
5.5
6.5
z−score of IF value
0
1
2
3
4

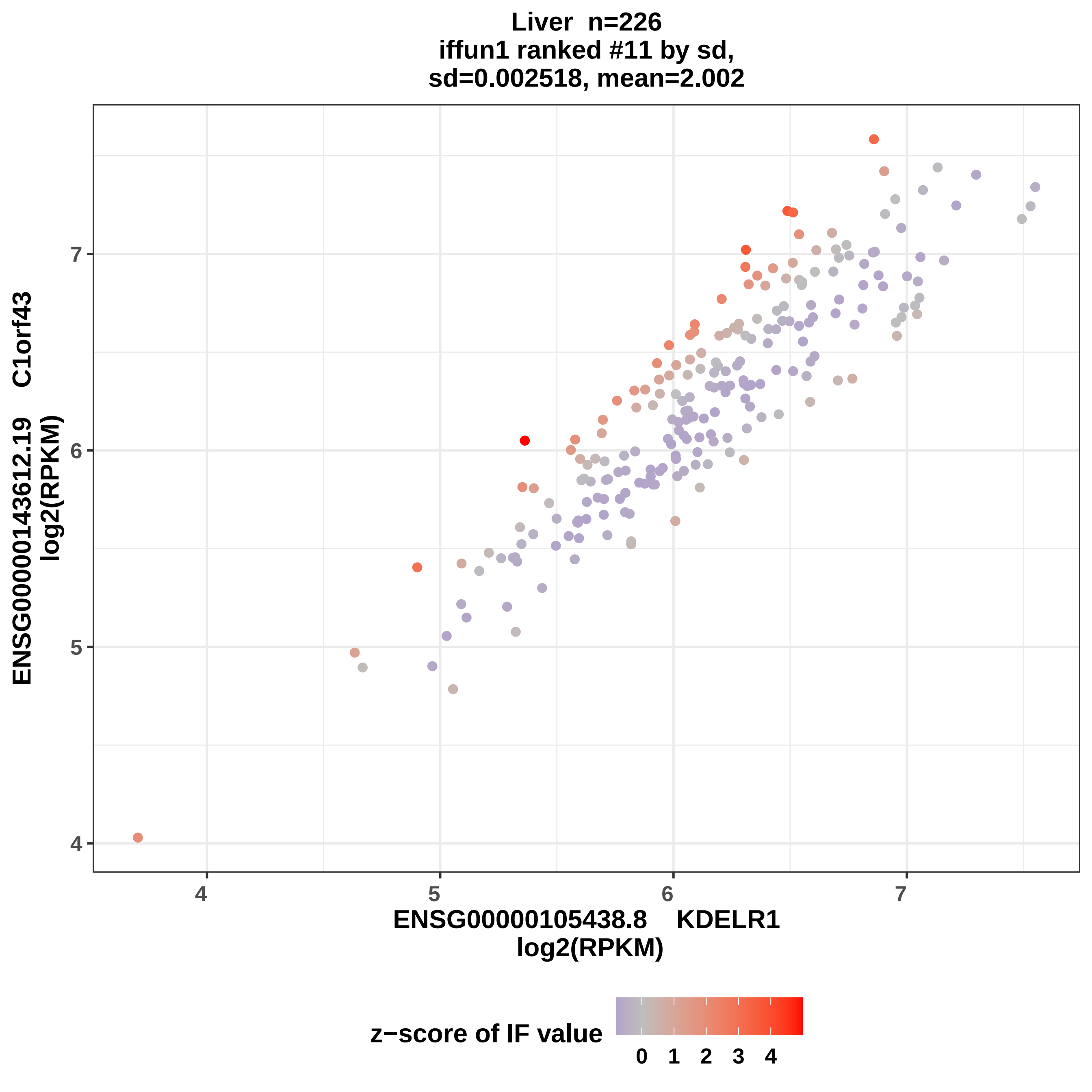
Liver n=226
iffun1 ranked #11 by sd,
sd=0.002518, mean=2.002
ENSG00000143612.19 C1orf43
log2(RPKM)
ENSG00000105438.8 KDELR1
log2(RPKM)
4
5
6
7
z−score of IF value
0 1 2 3 4

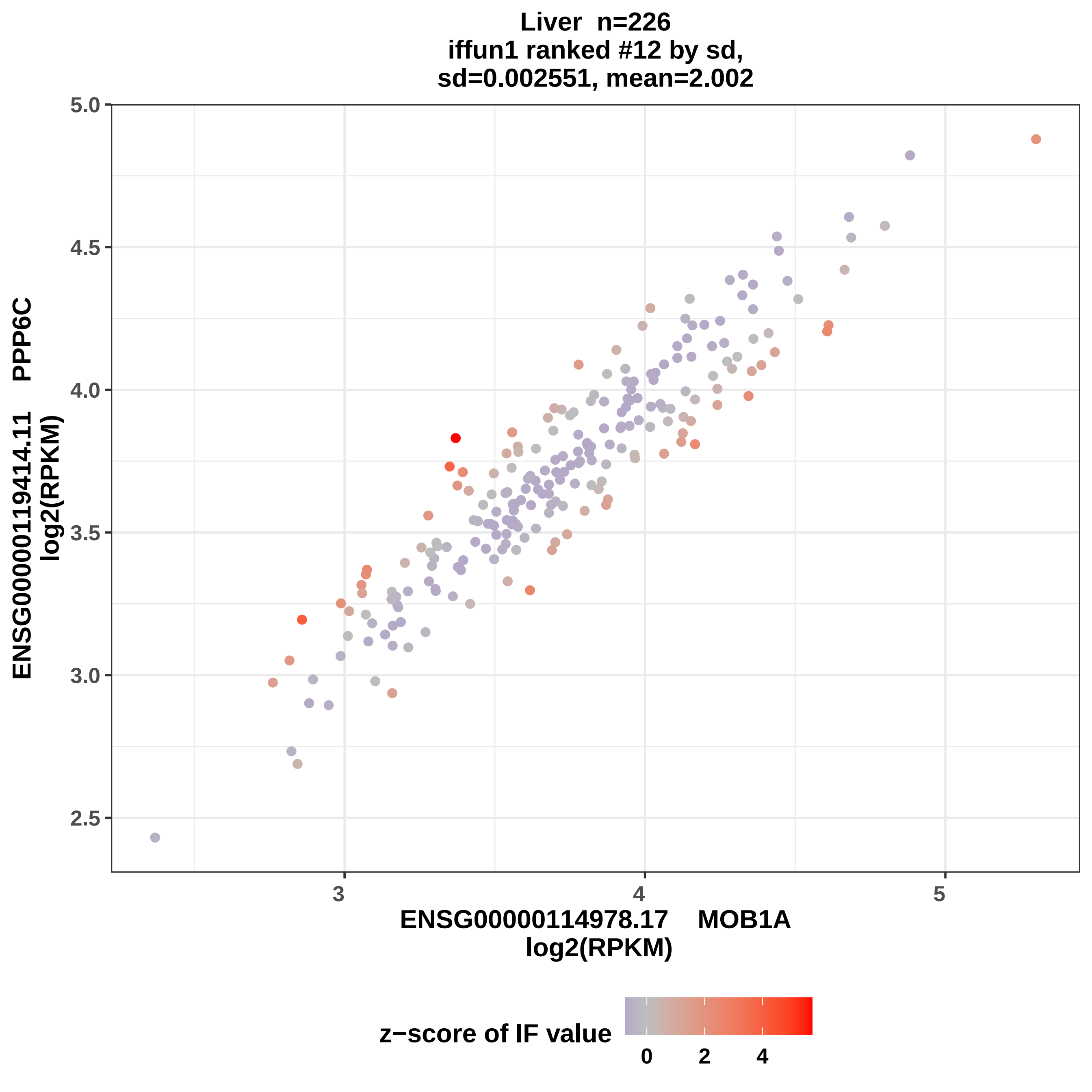

Liver n=226
iffun1 ranked #12 by sd,
sd=0.002551, mean=2.002
5.0
4.5
4.0
3.5
3.0
2.5
3
4
5
ENSG00000114978.17 MOB1A
log2(RPKM)
ENSG00000119414.11 PPP6C
log2(RPKM)
z−score of IF value
0
2
4

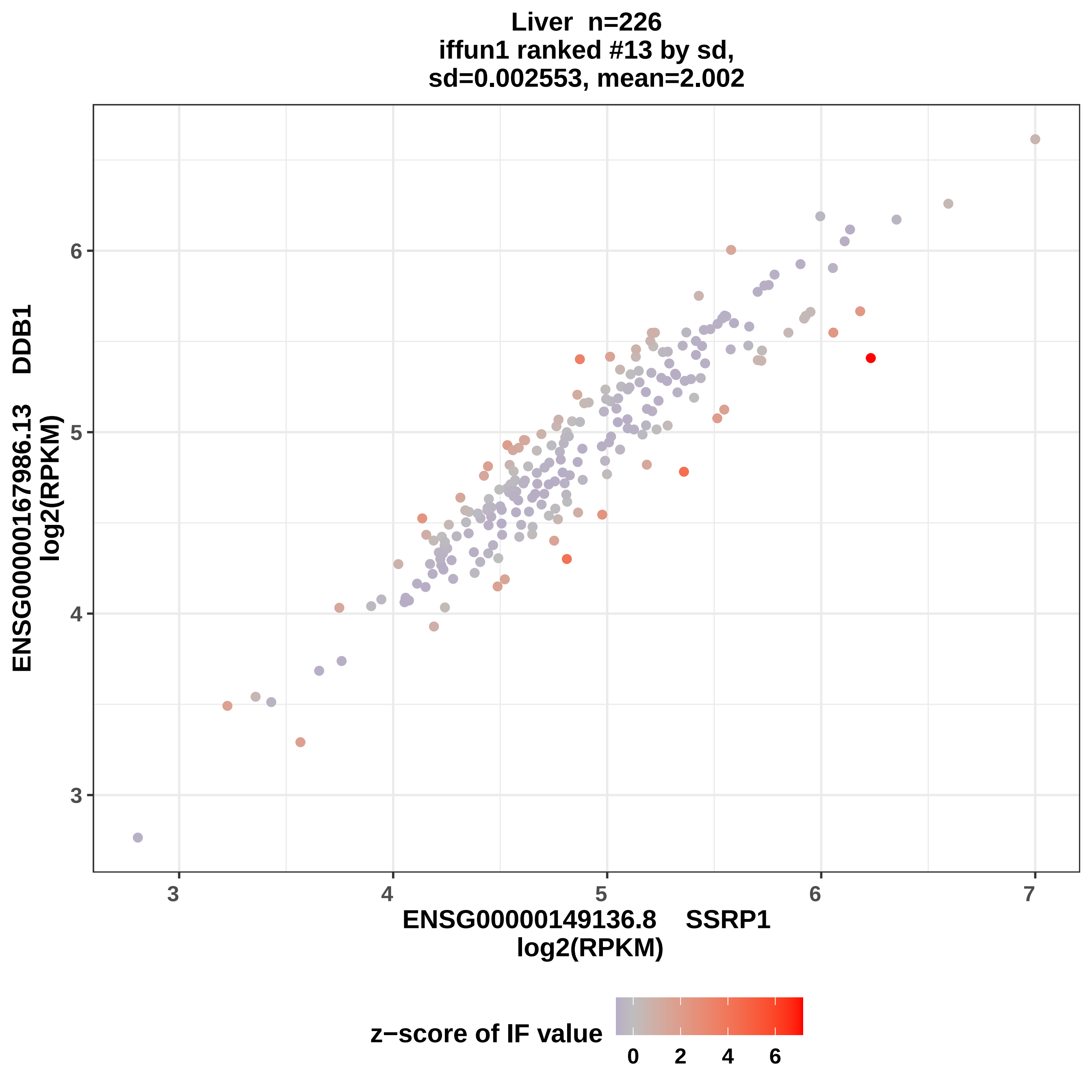
Liver n=226
iffun1 ranked #13 by sd,
sd=0.002553, mean=2.002
ENSG00000167986.13 DDB1
log2(RPKM)
ENSG00000149136.8 SSRP1
log2(RPKM)
3
4
5
6
7
z−score of IF value
0
2
4
6

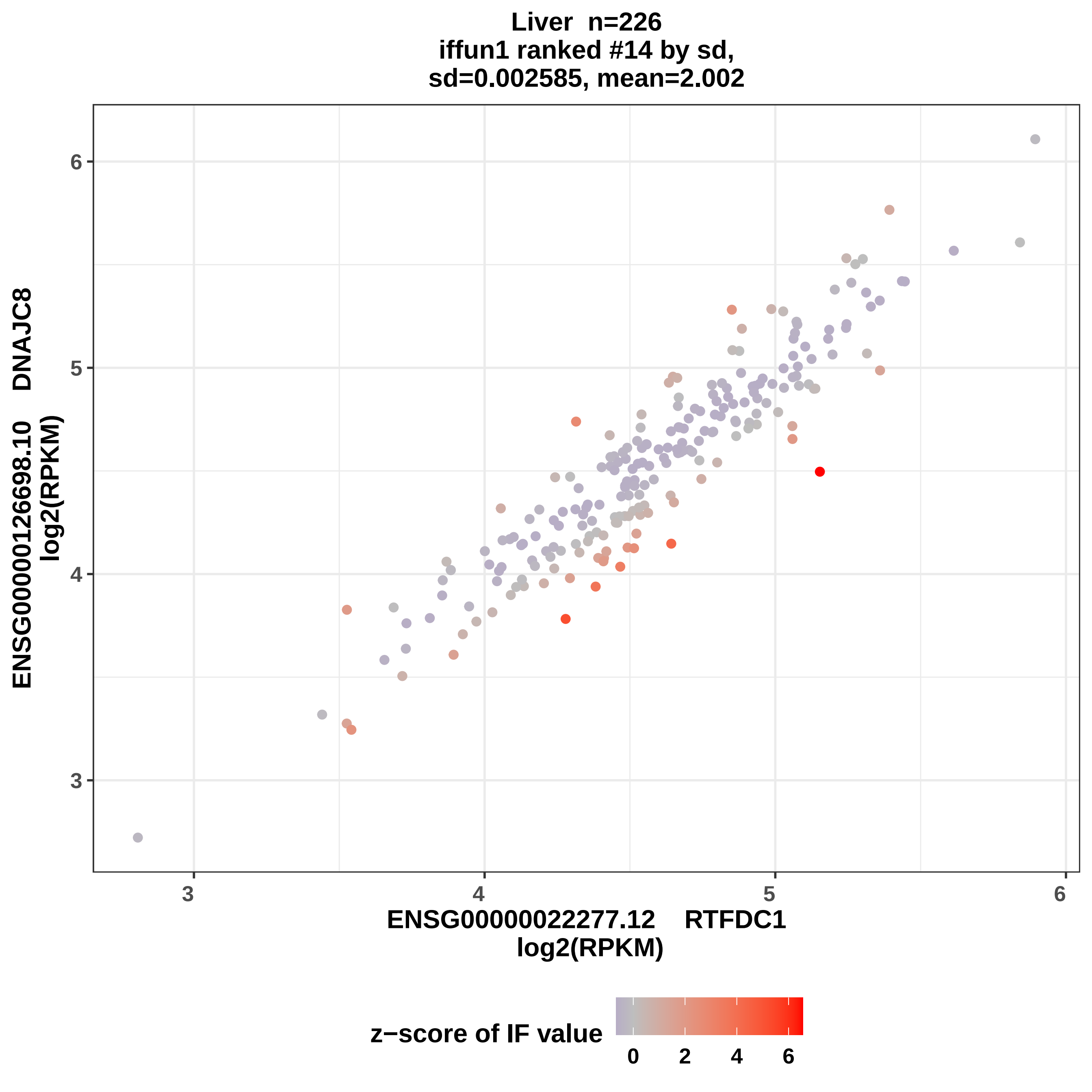

Liver n=226
iffun1 ranked #14 by sd,
sd=0.002585, mean=2.002
ENSG00000126698.10 DNAJC8
log2(RPKM)
3
4
5
6
ENSG00000022277.12 RTFDC1
log2(RPKM)
3
4
5
6
z−score of IF value
0
2
4
6

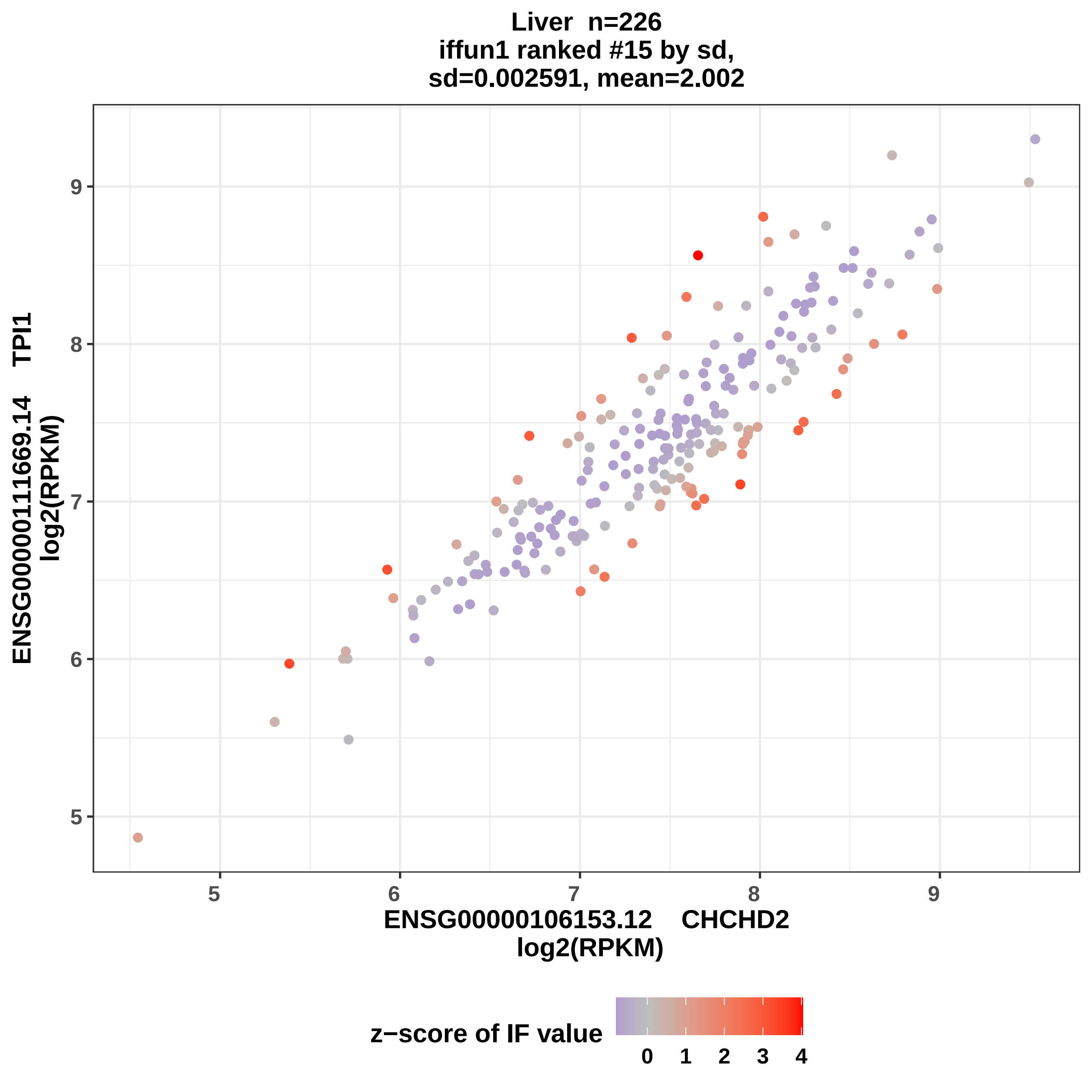

Liver n=226
iffun1 ranked #15 by sd,
sd=0.002591, mean=2.002
ENSG00000111669.14 TPI1
log2(RPKM)
ENSG00000106153.12 CHCHD2
log2(RPKM)
5
6
7
8
9
z−score of IF value
0
1
2
3
4

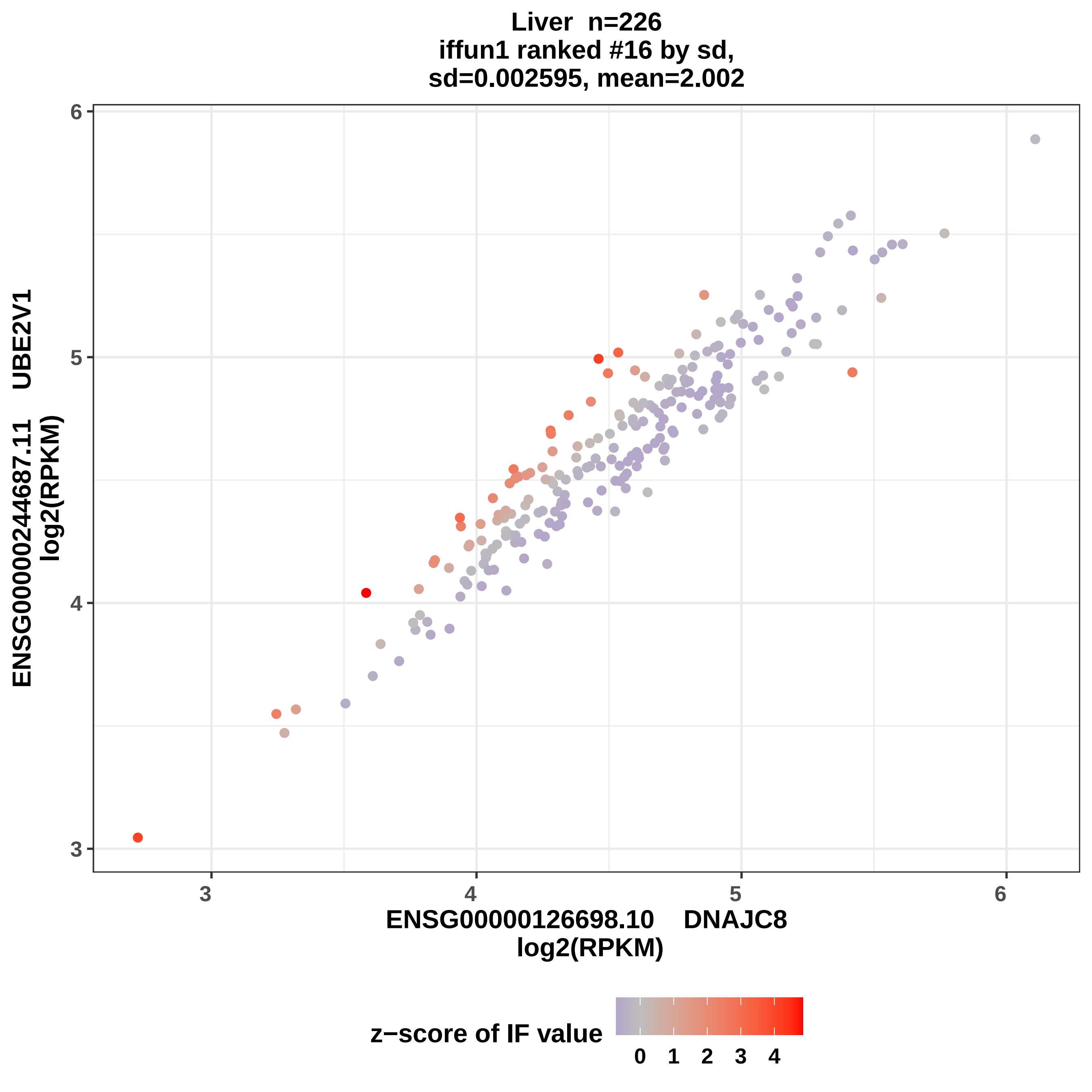

Liver n=226
iffun1 ranked #16 by sd,
sd=0.002595, mean=2.002
ENSG00000244687.11 UBE2V1
log2(RPKM)
ENSG00000126698.10 DNAJC8
log2(RPKM)
3
4
5
6
z−score of IF value
0
1
2
3
4

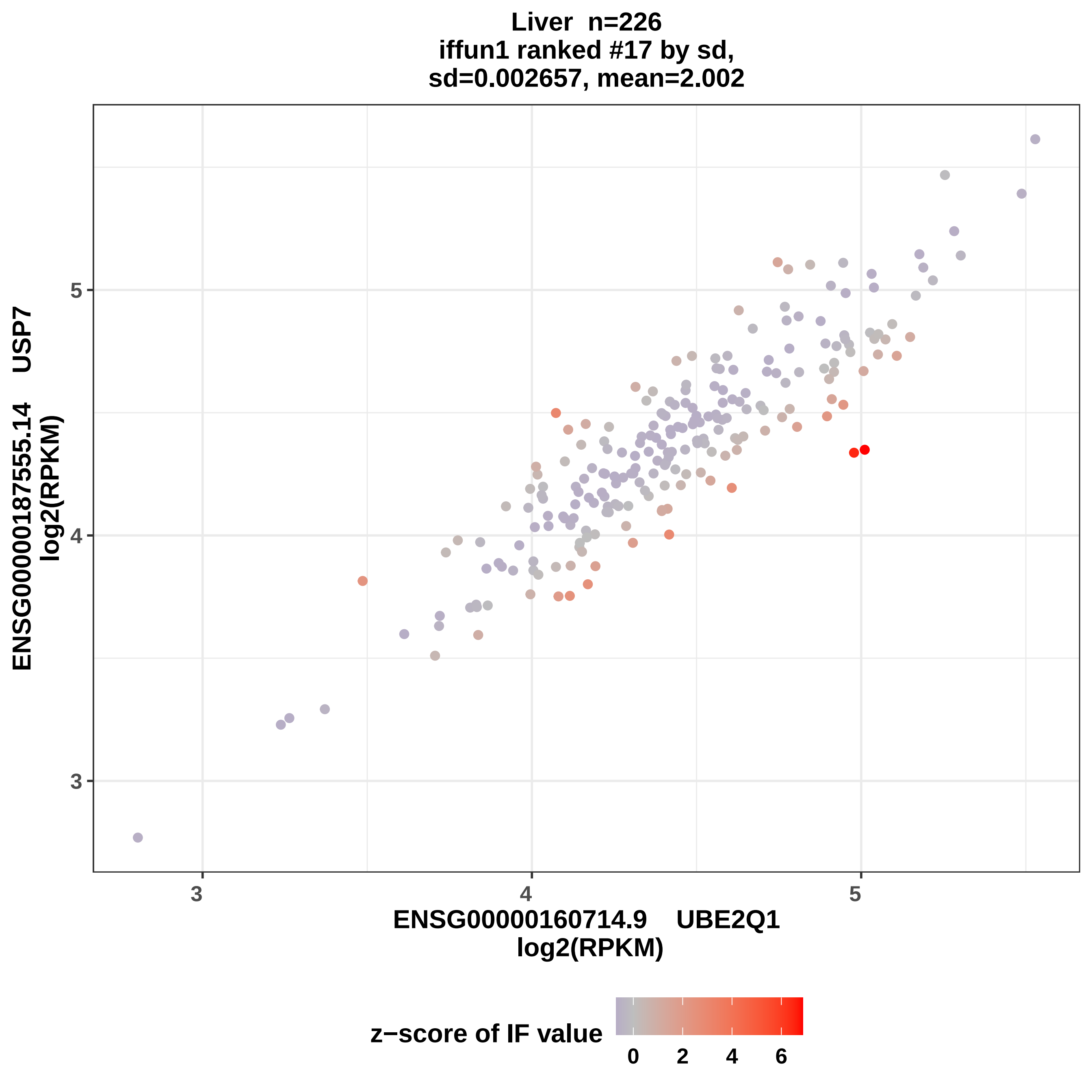
Liver n=226
iffun1 ranked #17 by sd,
sd=0.002657, mean=2.002
ENSG00000187555.14 USP7
log2(RPKM)
5
4
3
3
4
5
ENSG00000160714.9 UBE2Q1
log2(RPKM)
z−score of IF value
0
2
4
6

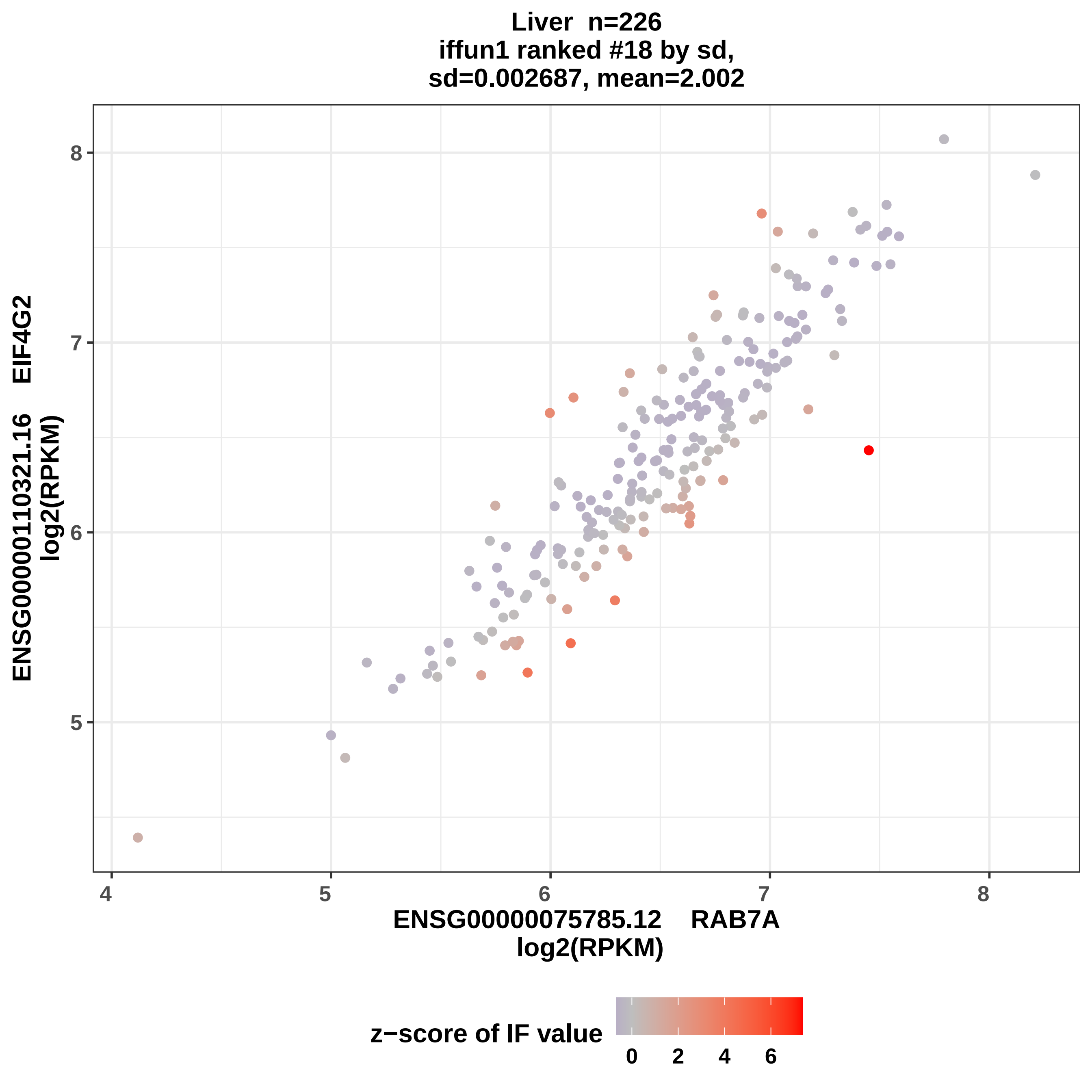
Liver n=226
iffun1 ranked #18 by sd,
sd=0.002687, mean=2.002
ENSG00000110321.16 EIF4G2
log2(RPKM)
ENSG00000075785.12 RAB7A
log2(RPKM)
4
5
6
7
8
z−score of IF value
0
2
4
6

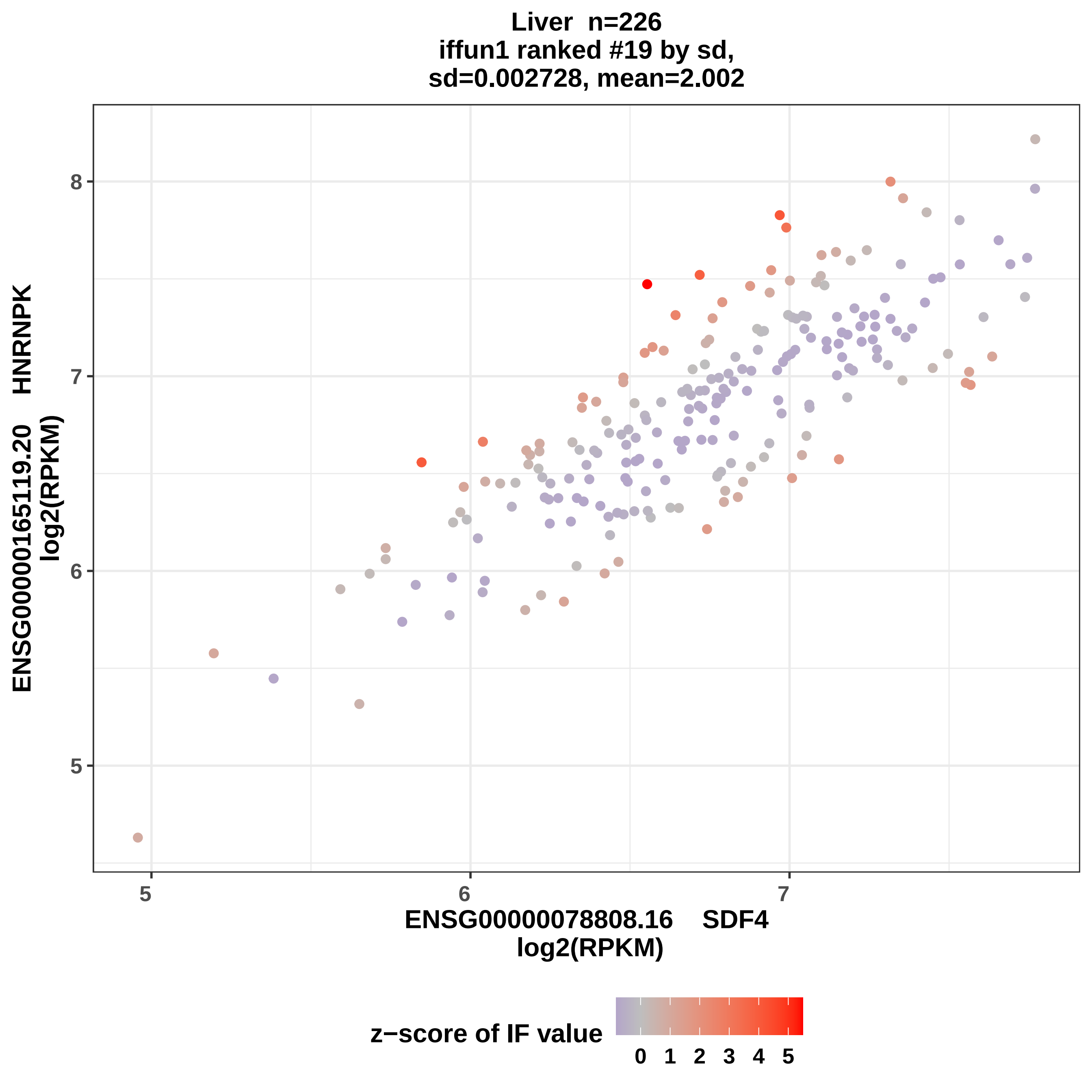
Liver n=226
iffun1 ranked #19 by sd,
sd=0.002728, mean=2.002
ENSG00000165119.20 HNRNPK
log2(RPKM)
ENSG00000078808.16 SDF4
log2(RPKM)
5
6
7
8
z−score of IF value
0 1 2 3 4 5

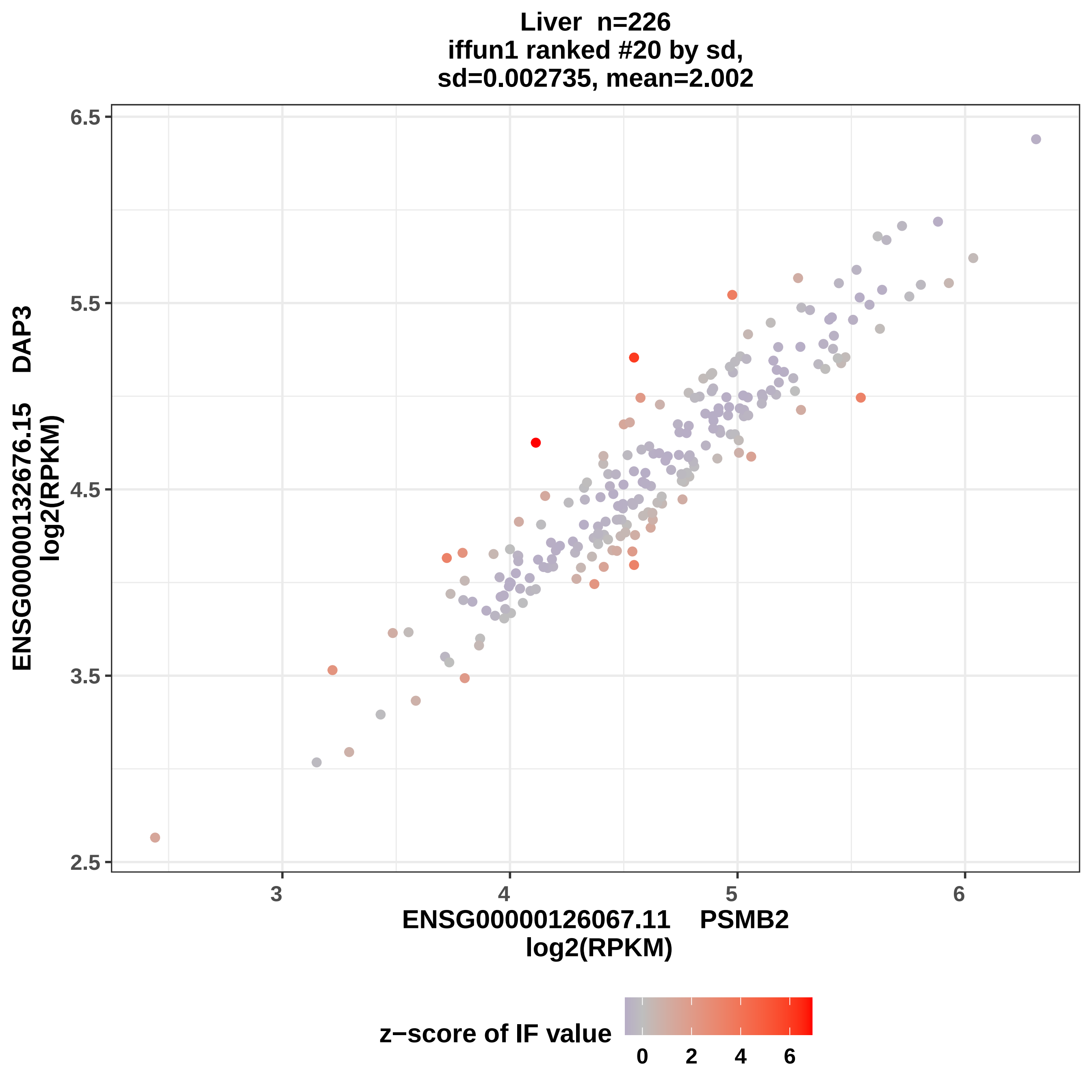
Liver n=226
iffun1 ranked #20 by sd,
sd=0.002735, mean=2.002
6.5
5.5
4.5
3.5
2.5
3
4
5
6
ENSG00000132676.15 DAP3
log2(RPKM)
ENSG00000126067.11 PSMB2
log2(RPKM)
z−score of IF value
0
2
4
6

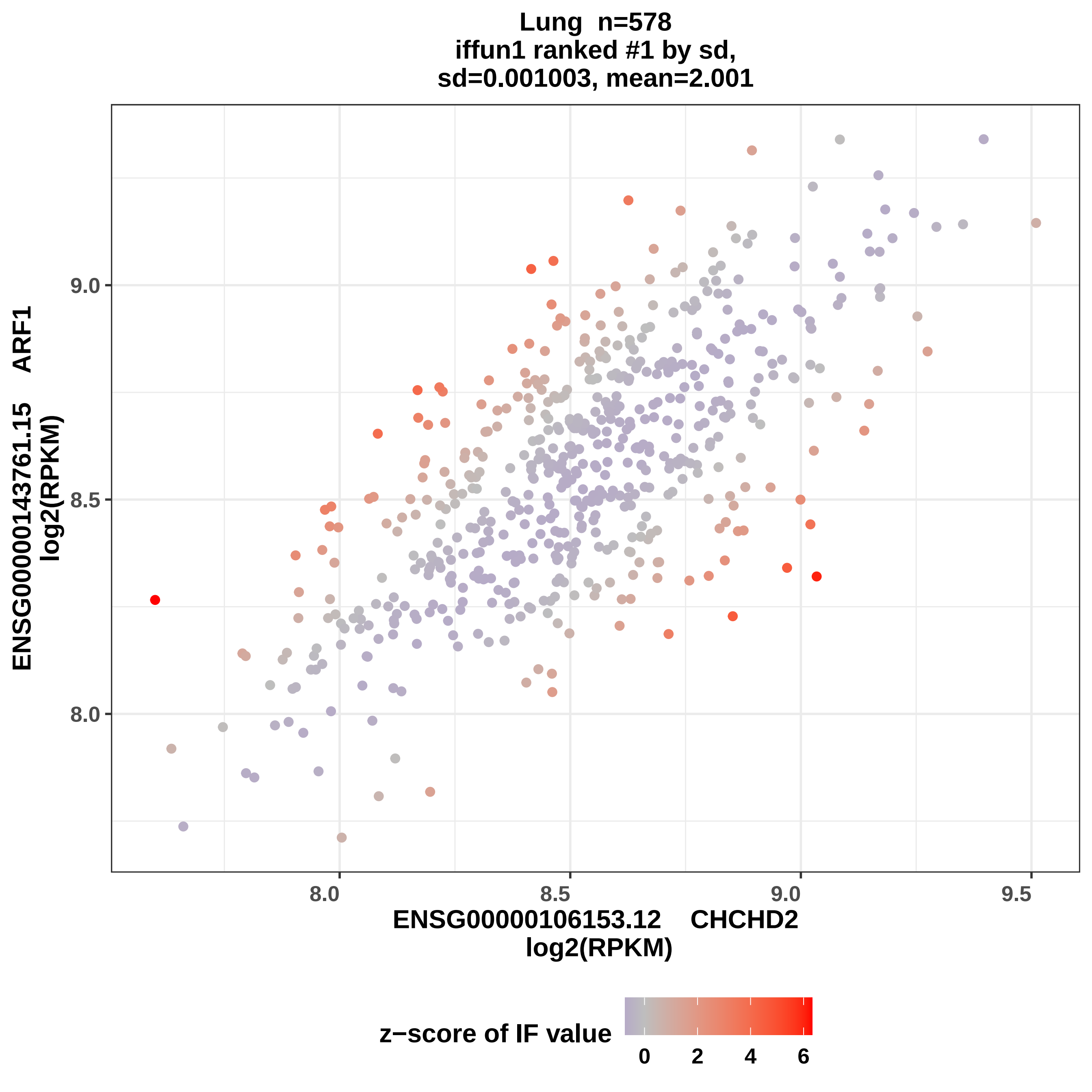

Lung n=578
iffun1 ranked #1 by sd,
sd=0.001003, mean=2.001
ENSG00000143761.15 ARF1
log2(RPKM)
9.0
8.5
8.0
8.0
8.5
9.0
9.5
ENSG00000106153.12 CHCHD2
log2(RPKM)
z−score of IF value
0
2
4
6

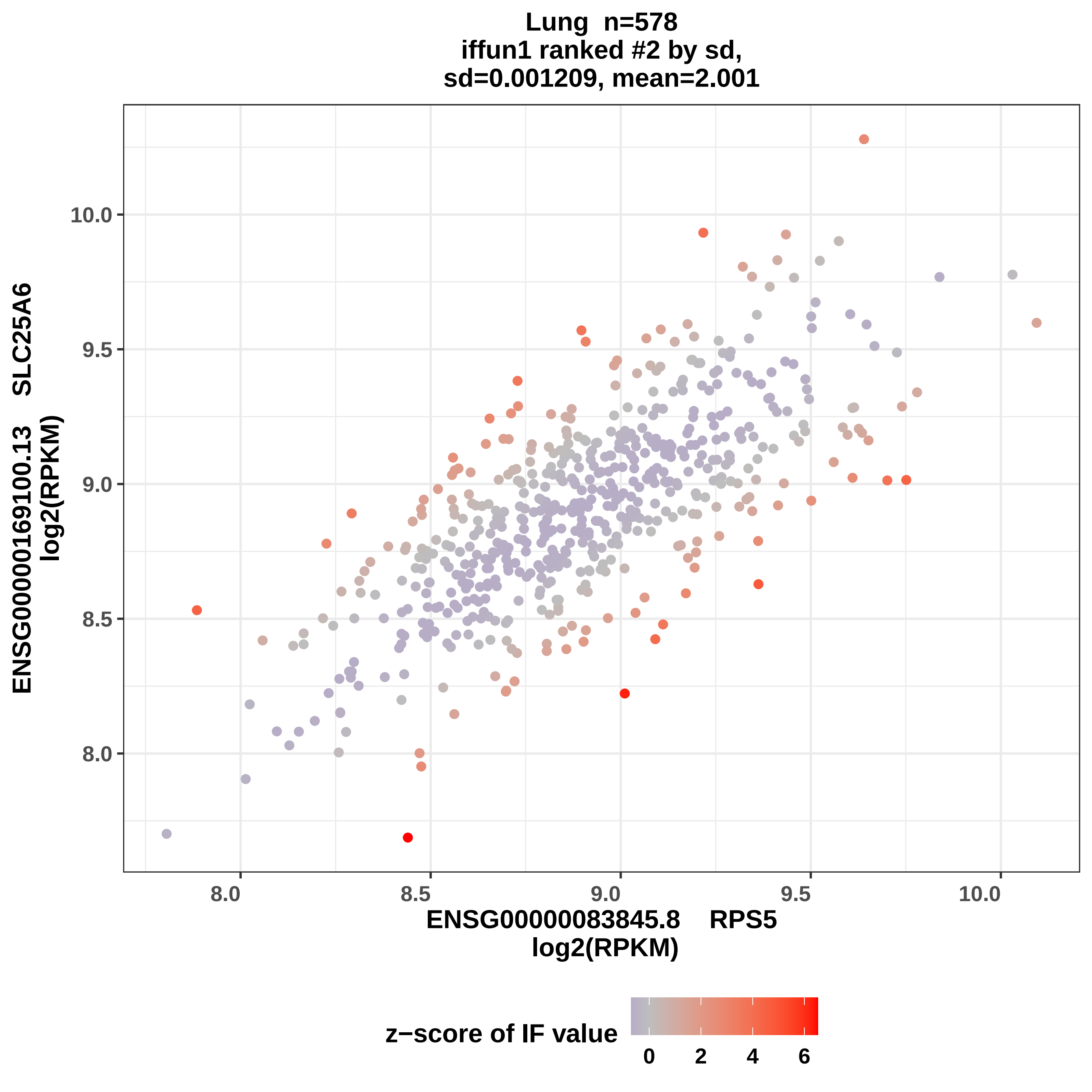
Lung n=578
iffun1 ranked #2 by sd,
sd=0.001209, mean=2.001
ENSG00000169100.13 SLC25A6
log2(RPKM)
10.0
9.5
9.0
8.5
8.0
8.0
8.5
9.0
9.5
10.0
ENSG00000083845.8 RPS5
log2(RPKM)
z−score of IF value
0
2
4
6

Lung  n=578
iffun1 ranked #3 by sd,
sd=0.001214, mean=2.001

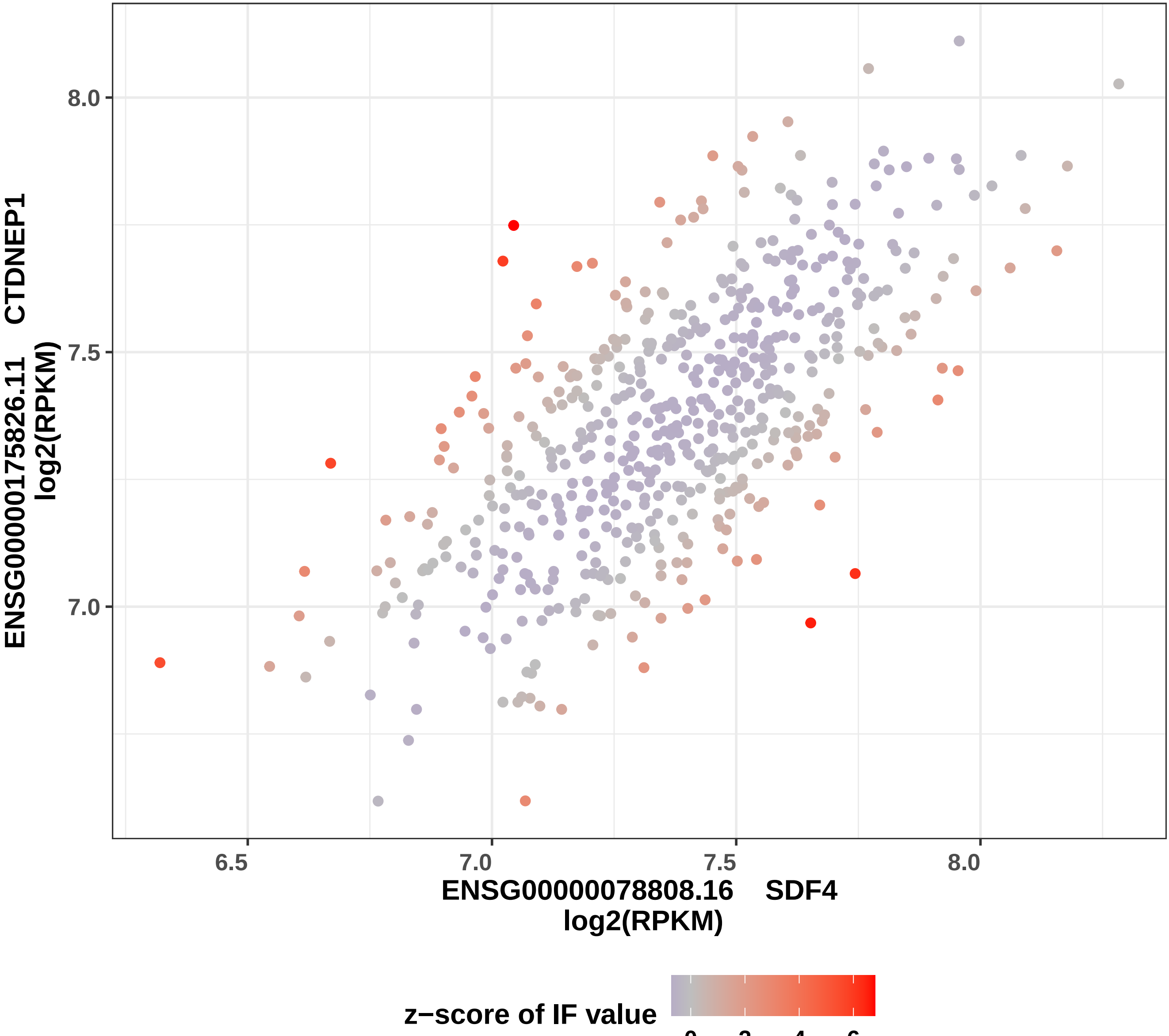

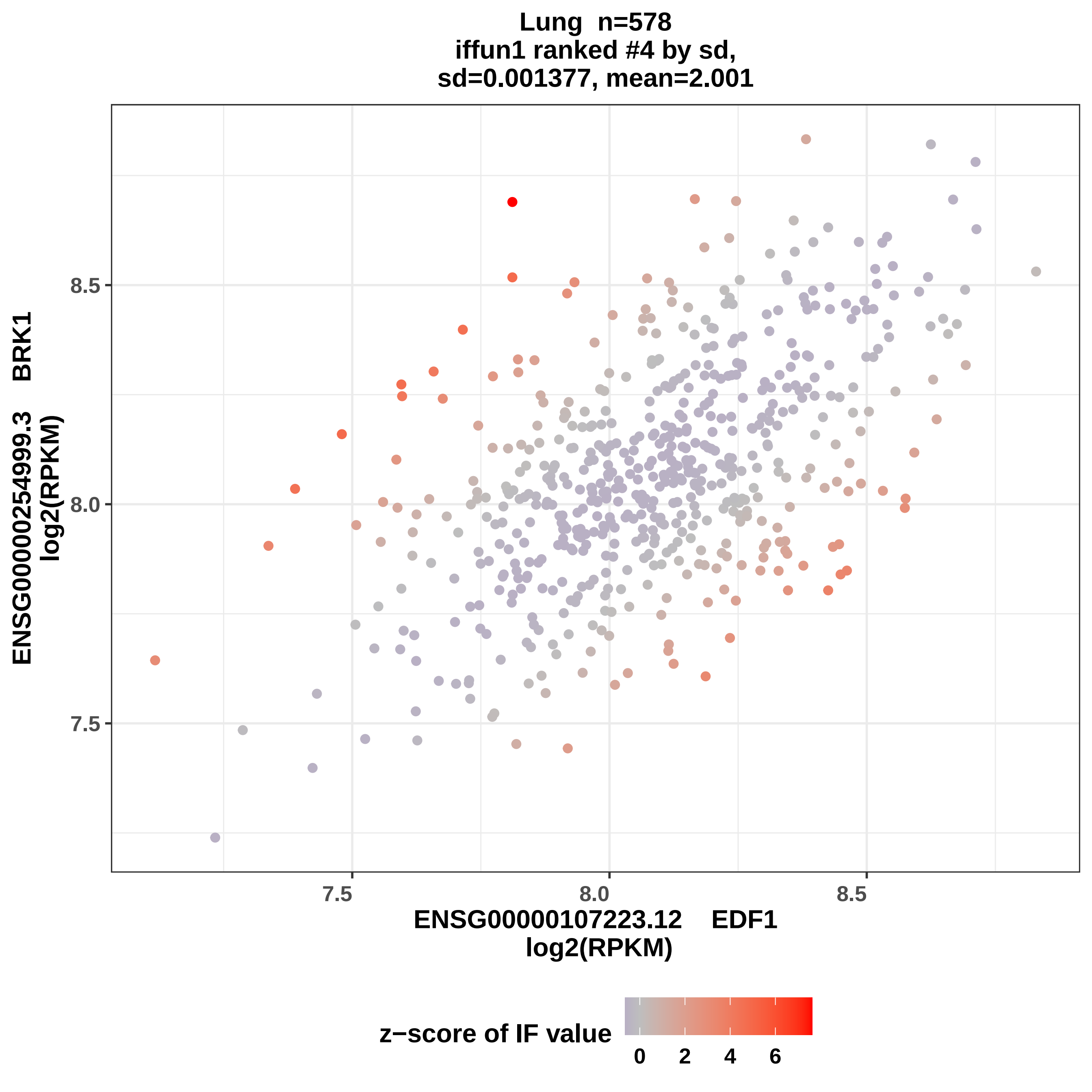

Lung n=578
iffun1 ranked #4 by sd,
sd=0.001377, mean=2.001
ENSG00000254999.3 BRK1
log2(RPKM)
8.5
8.0
7.5
7.5
8.0
8.5
ENSG00000107223.12 EDF1
log2(RPKM)
z−score of IF value
0
2
4
6

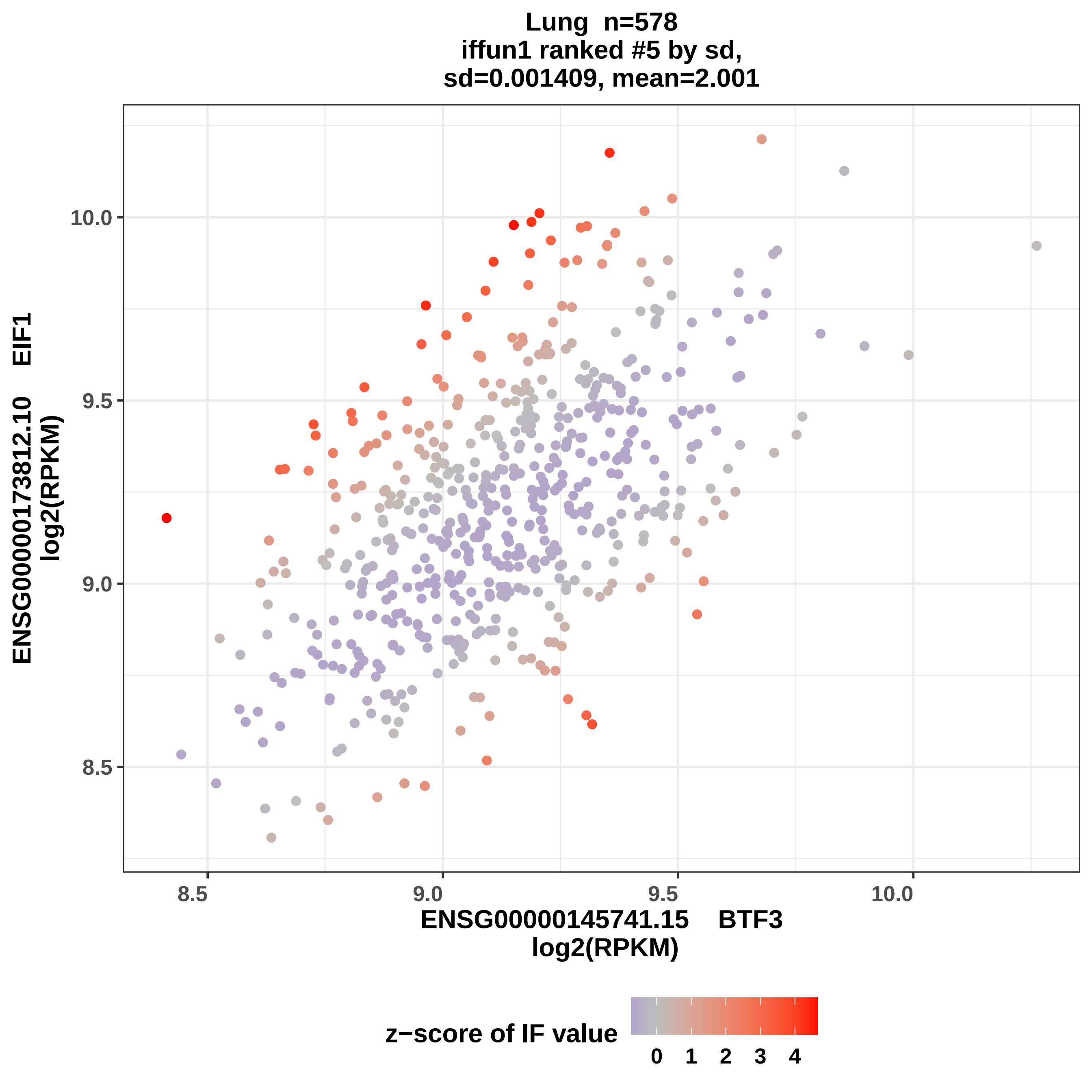
Lung n=578
iffun1 ranked #5 by sd,
sd=0.001409, mean=2.001
ENSG00000173812.10 EIF1
log2(RPKM)
10.0
9.5
9.0
8.5
8.5
9.0
9.5
10.0
ENSG00000145741.15 BTF3
log2(RPKM)
z−score of IF value
0
1
2
3
4

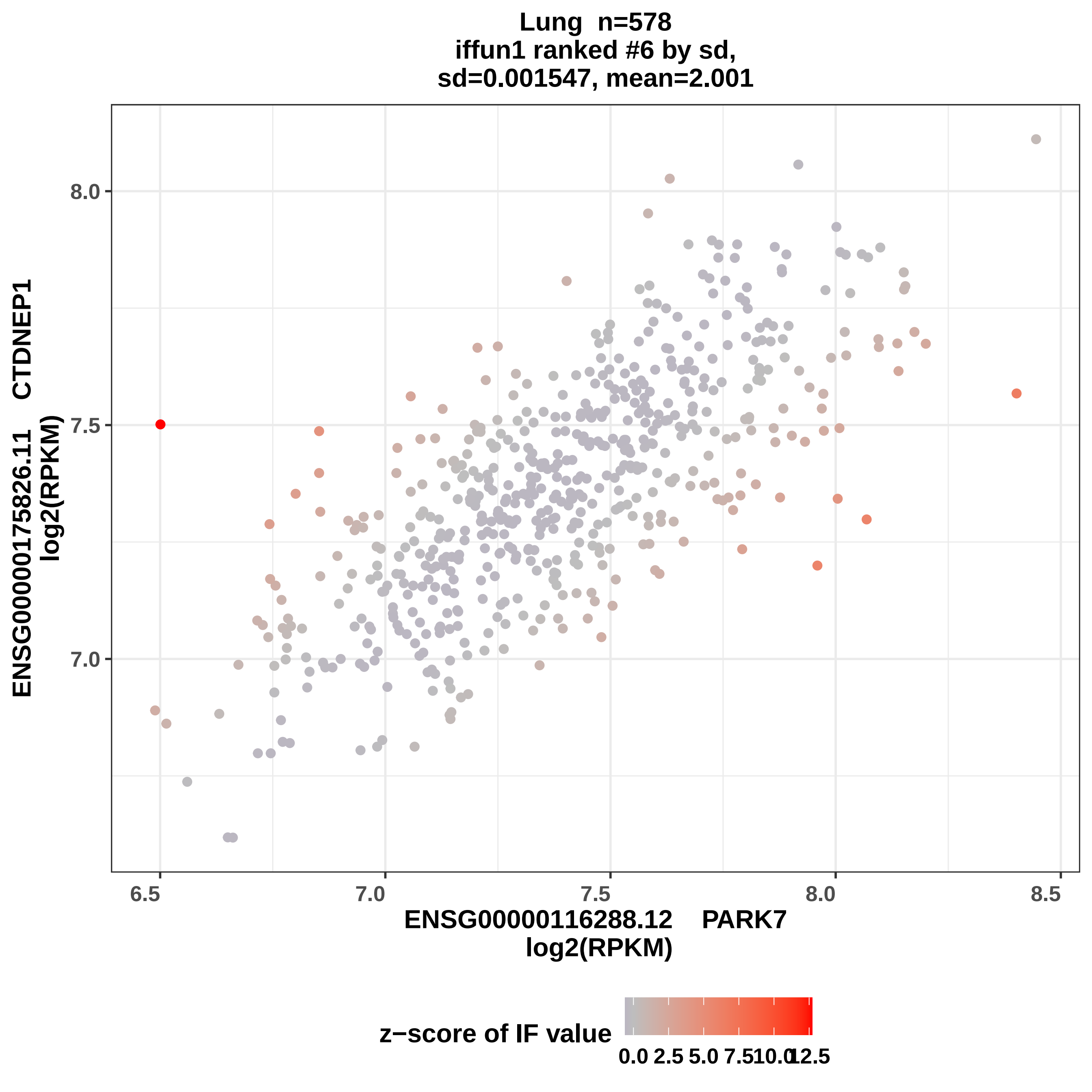
Lung n=578
iffun1 ranked #6 by sd,
sd=0.001547, mean=2.001
ENSG00000175826.11 CTDNEP1
log2(RPKM)
8.0
7.5
7.0
6.5
7.0
7.5
8.0
8.5
ENSG00000116288.12 PARK7
log2(RPKM)
z−score of IF value
0.0
2.5
5.0
7.5
10.0
12.5

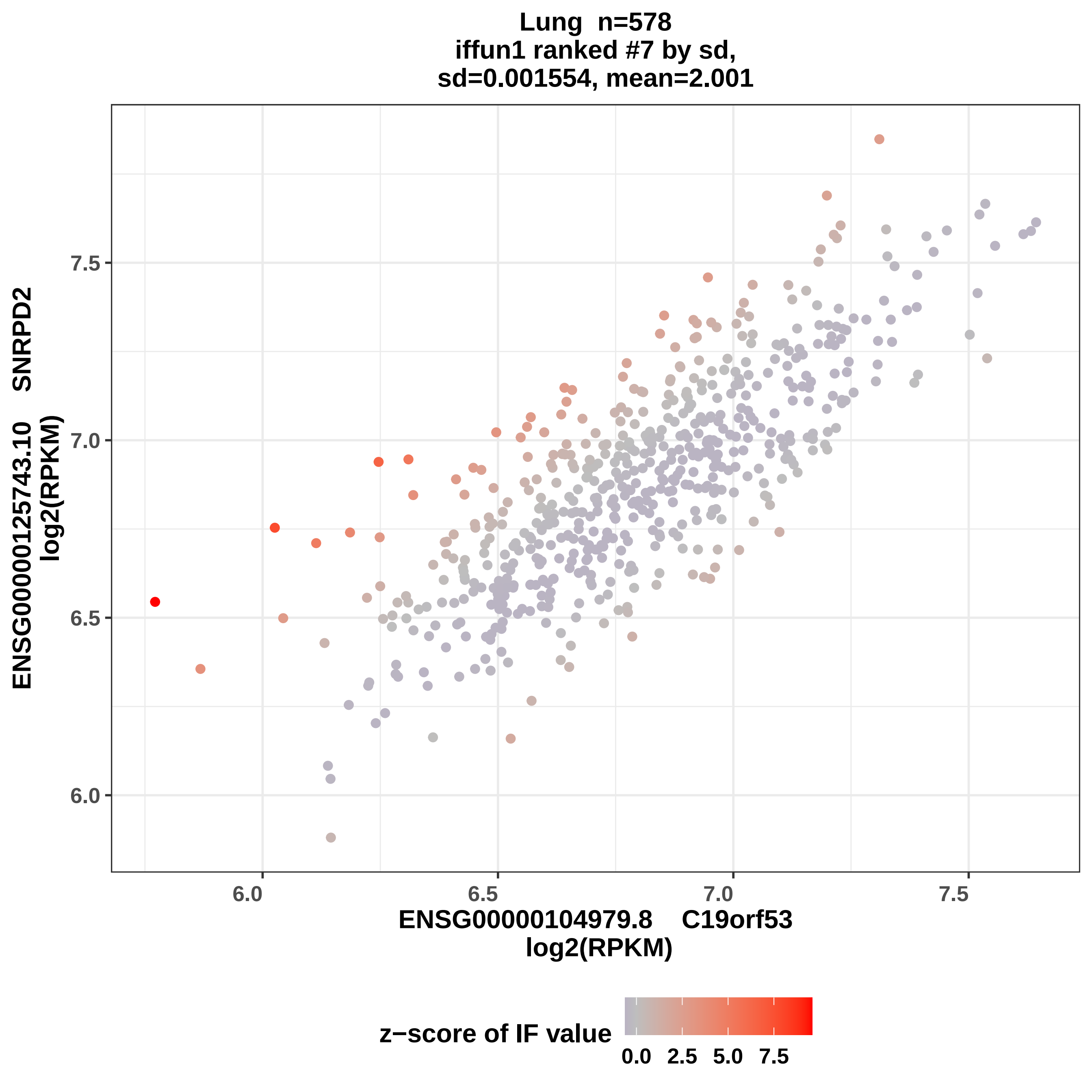

Lung n=578
iffun1 ranked #7 by sd,
sd=0.001554, mean=2.001
ENSG00000125743.10 SNRPD2
log2(RPKM)
7.5
7.0
6.5
6.0
6.0
6.5
7.0
7.5
ENSG00000104979.8 C19orf53
log2(RPKM)
z−score of IF value
0.0
2.5
5.0
7.5

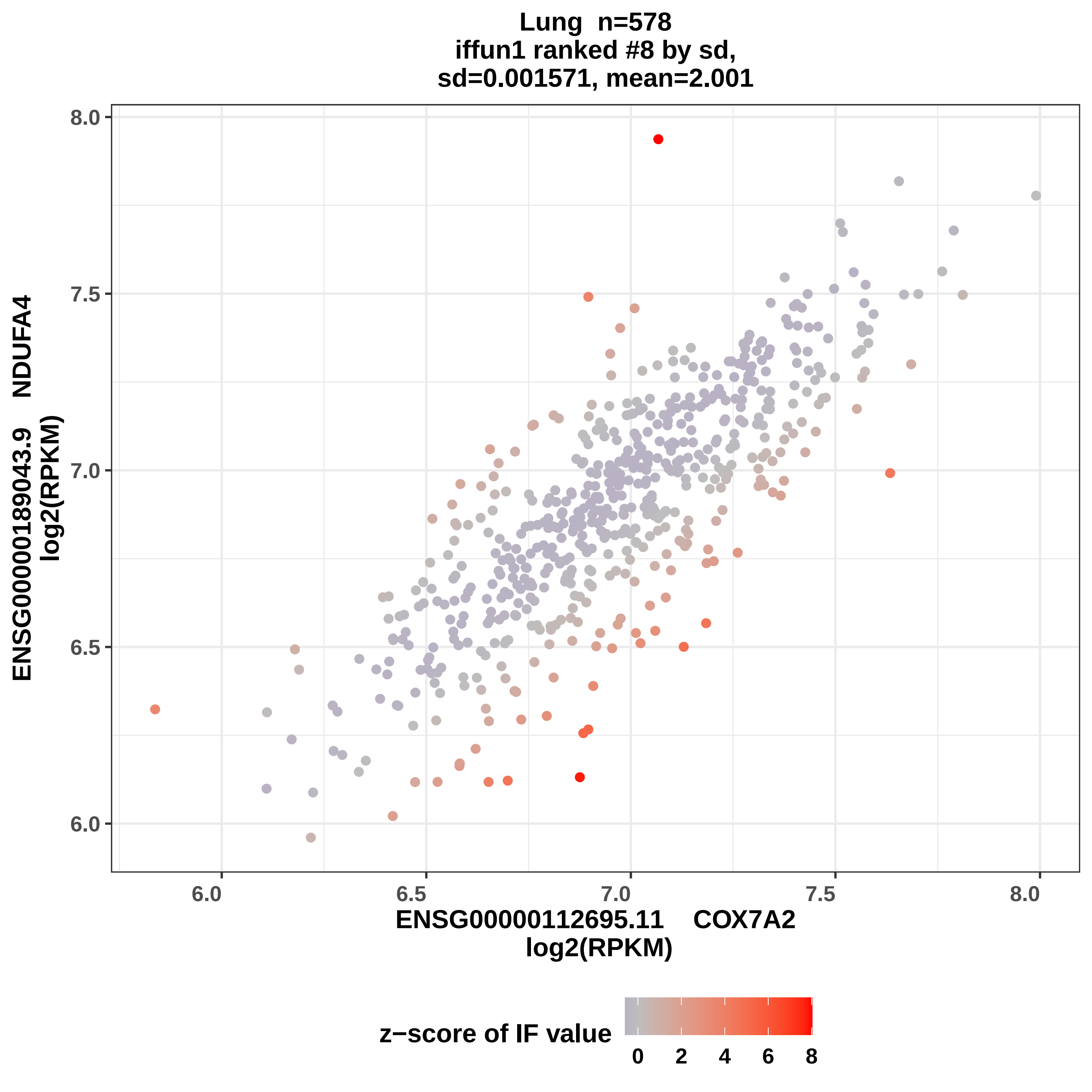
Lung n=578
iffun1 ranked #8 by sd,
sd=0.001571, mean=2.001
ENSG00000189043.9 NDUFA4
log2(RPKM)
8.0
7.5
7.0
6.5
6.0
6.0
6.5
7.0
7.5
8.0
ENSG00000112695.11 COX7A2
log2(RPKM)
z−score of IF value
0
2
4
6
8

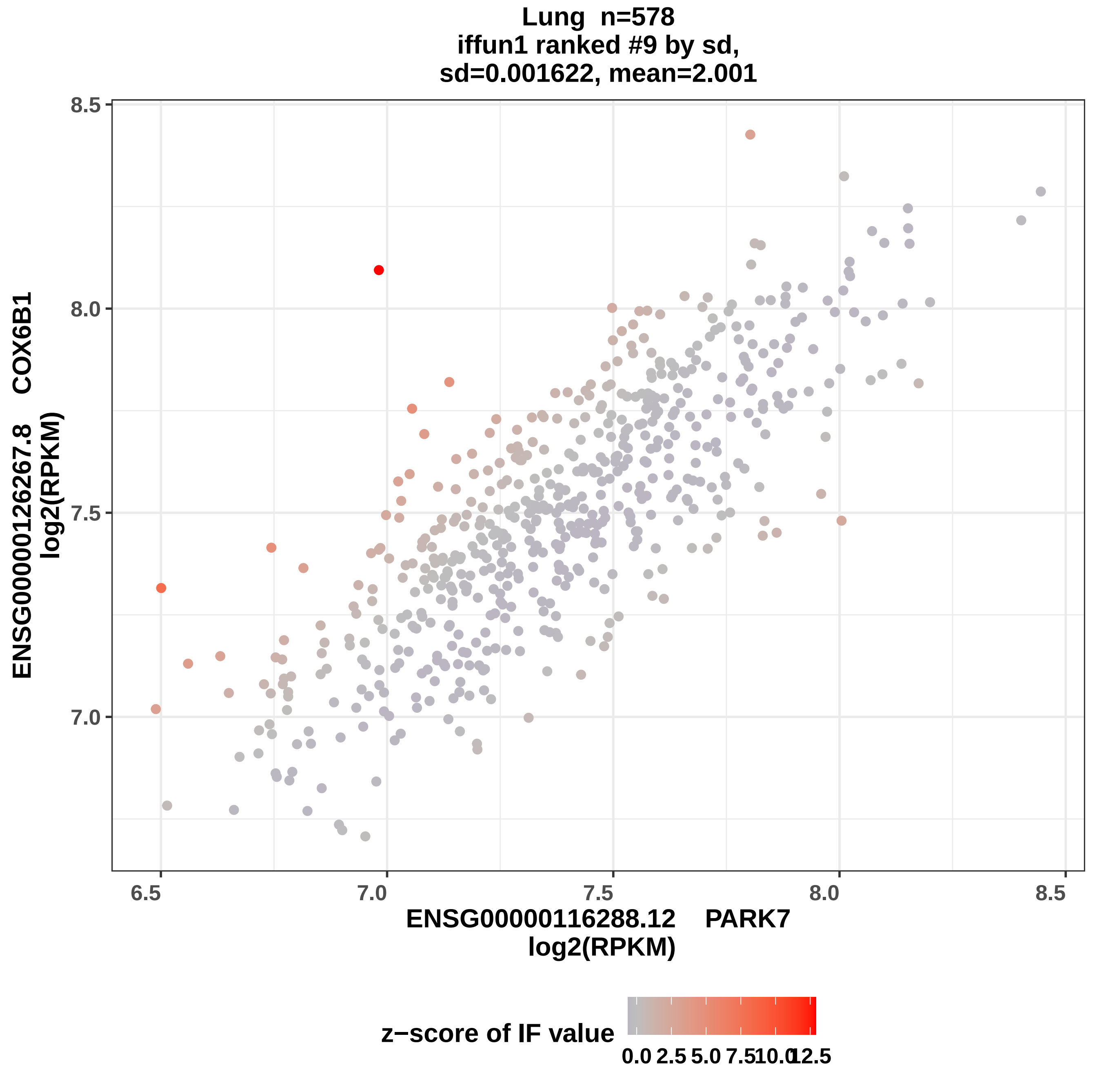
Lung n=578
iffun1 ranked #9 by sd,
sd=0.001622, mean=2.001
ENSG00000126267.8 COX6B1
log2(RPKM)
8.5
8.0
7.5
7.0
6.5
7.0
7.5
8.0
8.5
ENSG00000116288.12 PARK7
log2(RPKM)
z−score of IF value
0.0
2.5
5.0
7.5
10.0
12.5

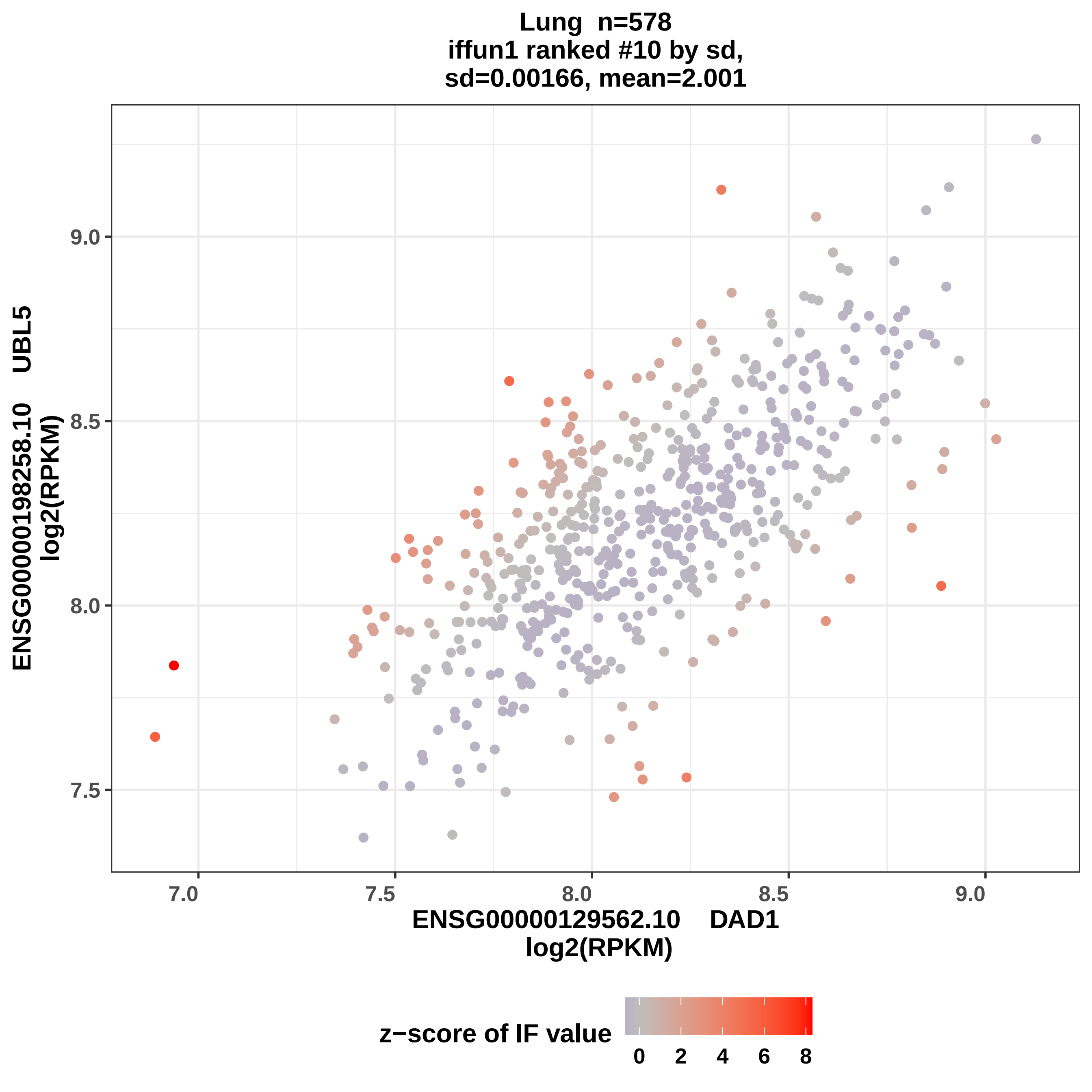

Lung n=578
iffun1 ranked #10 by sd,
sd=0.00166, mean=2.001
ENSG00000198258.10 UBL5
log2(RPKM)
ENSG00000129562.10 DAD1
log2(RPKM)
7.5
8.0
8.5
9.0
7.0
z−score of IF value
0
2
4
6
8

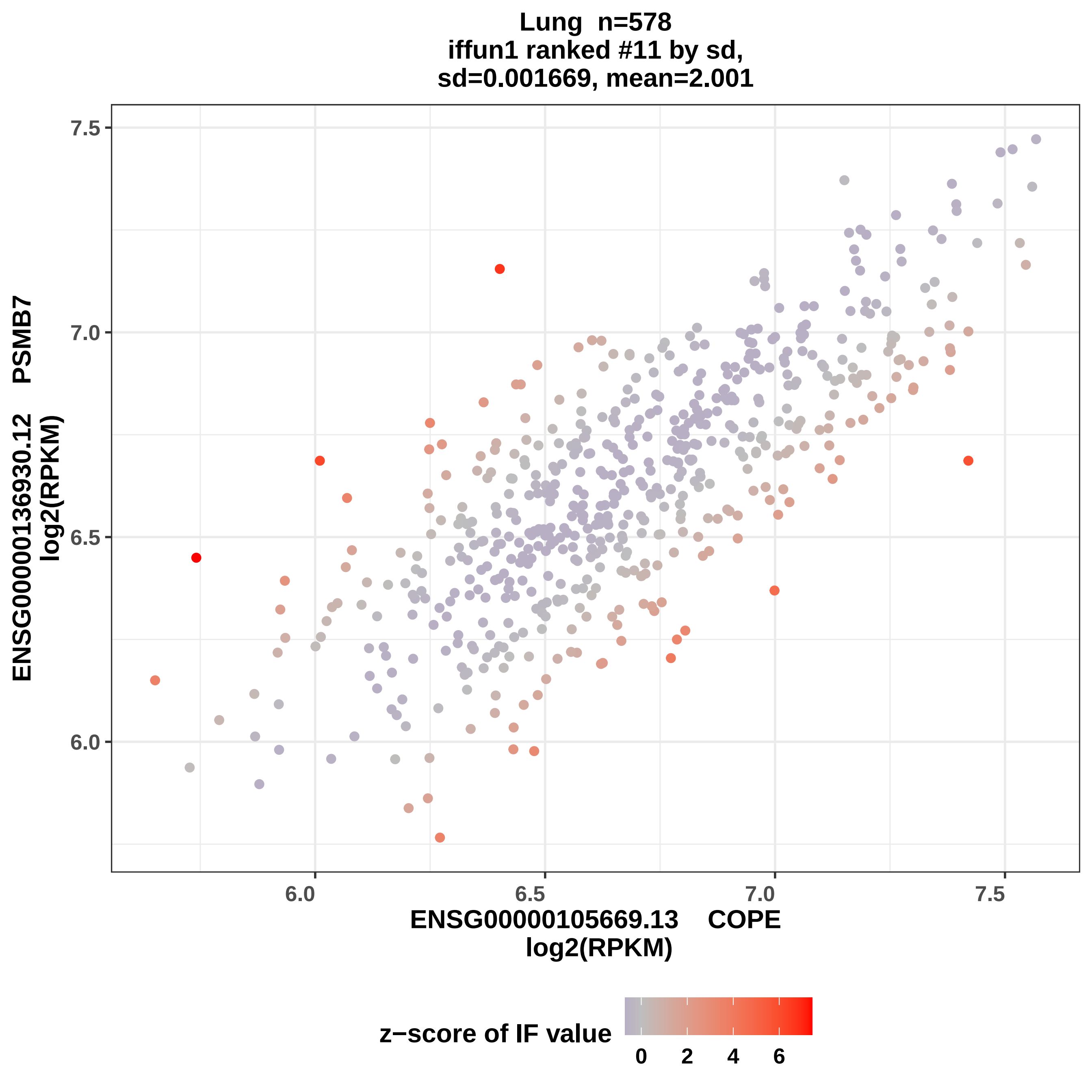

Lung n=578
iffun1 ranked #11 by sd,
sd=0.001669, mean=2.001
ENSG00000136930.12 PSMB7
log2(RPKM)
7.5
7.0
6.5
6.0
6.0
6.5
7.0
7.5
ENSG00000105669.13 COPE
log2(RPKM)
z−score of IF value
0
2
4
6

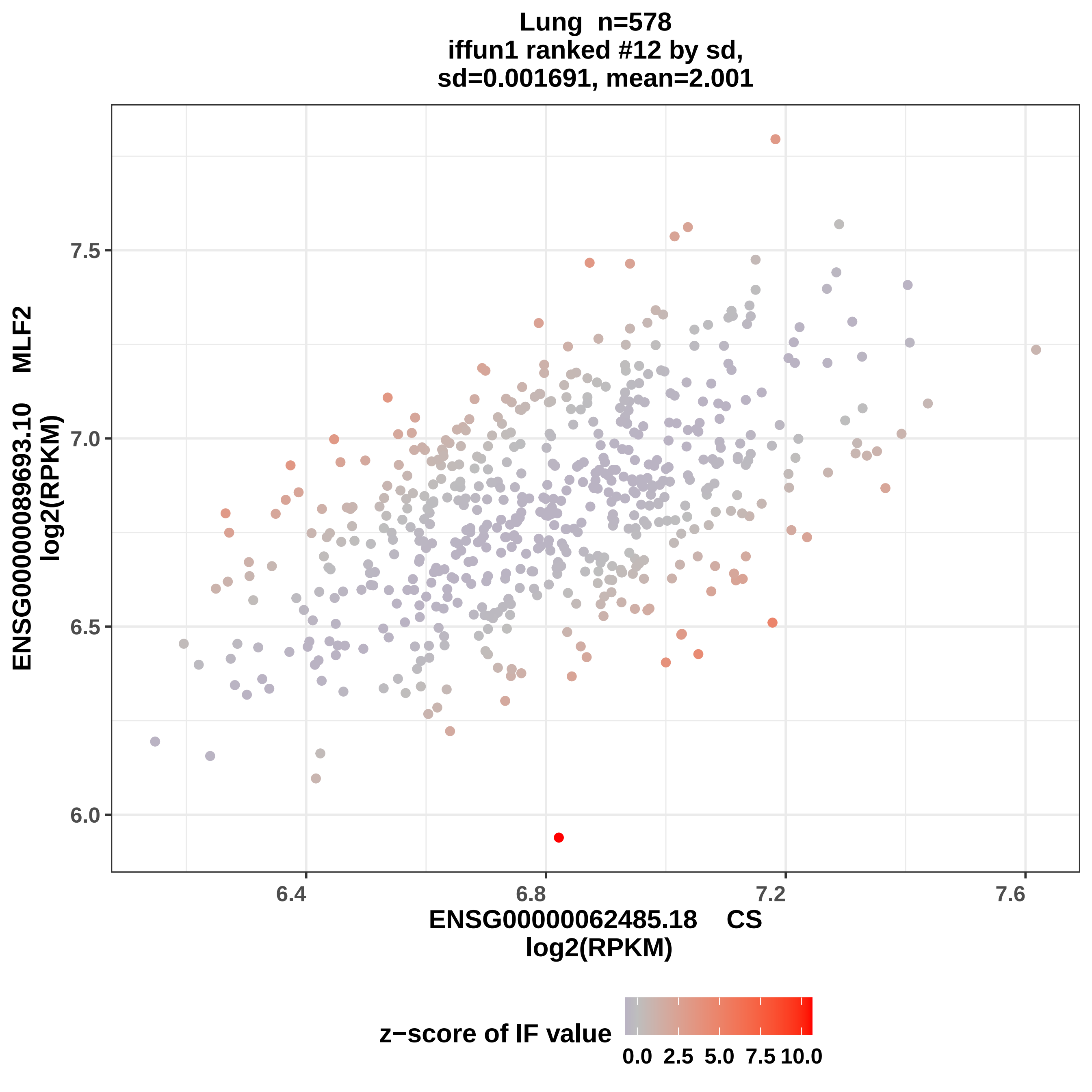
Lung n=578
iffun1 ranked #12 by sd,
sd=0.001691, mean=2.001
ENSG00000089693.10 MLF2
log2(RPKM)
7.5
7.0
6.5
6.0
6.4
6.8
7.2
7.6
ENSG00000062485.18 CS
log2(RPKM)
z−score of IF value
0.0
2.5
5.0
7.5
10.0

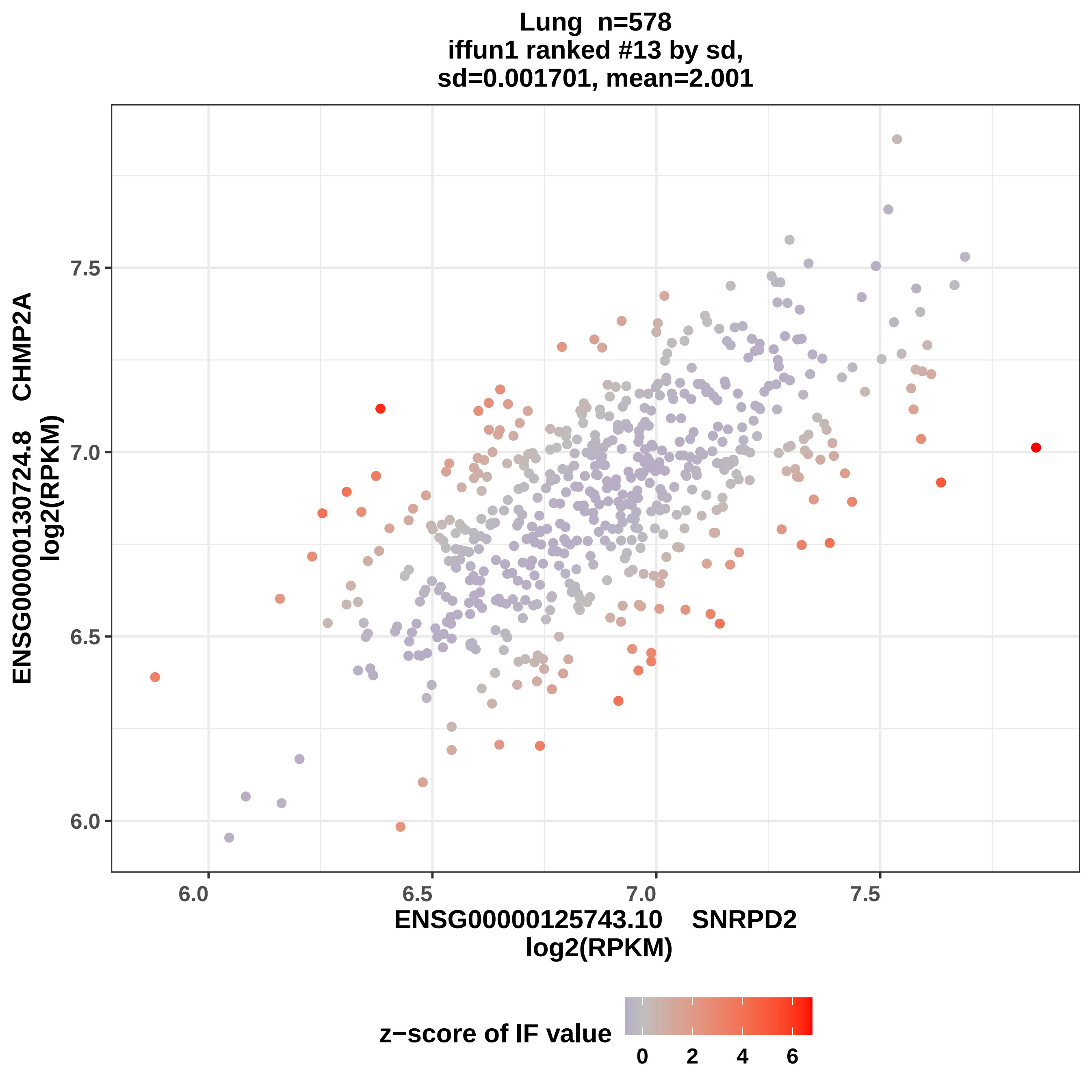
Lung n=578
iffun1 ranked #13 by sd,
sd=0.001701, mean=2.001
ENSG00000130724.8 CHMP2A
log2(RPKM)
7.5
7.0
6.5
6.0
6.0
6.5
7.0
7.5
ENSG00000125743.10 SNRPD2
log2(RPKM)
z−score of IF value
0
2
4
6

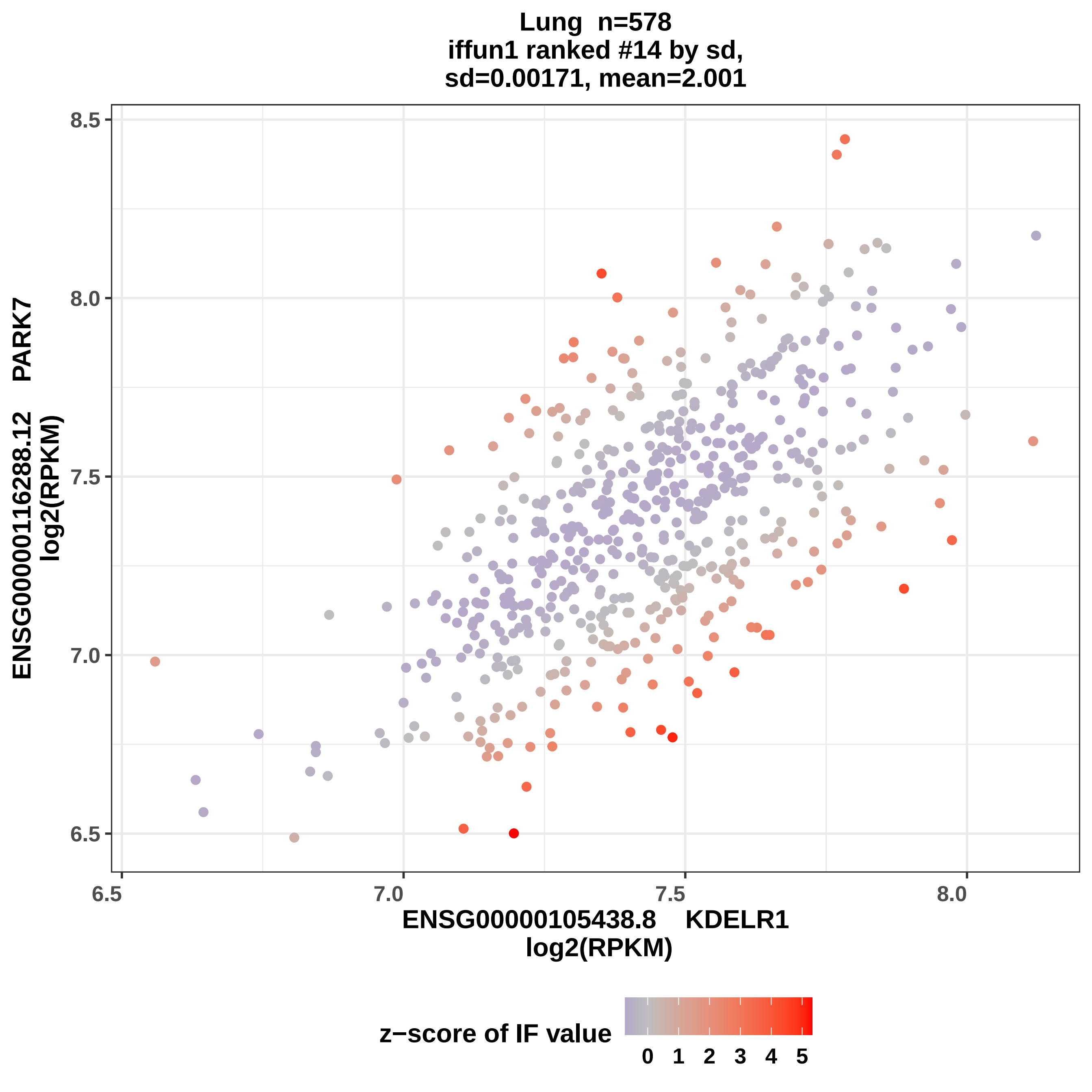

Lung n=578
iffun1 ranked #14 by sd,
sd=0.00171, mean=2.001
ENSG00000116288.12 PARK7
log2(RPKM)
8.5
8.0
7.5
7.0
6.5
6.5
7.0
7.5
8.0
ENSG00000105438.8 KDELR1
log2(RPKM)
z−score of IF value
0 1 2 3 4 5

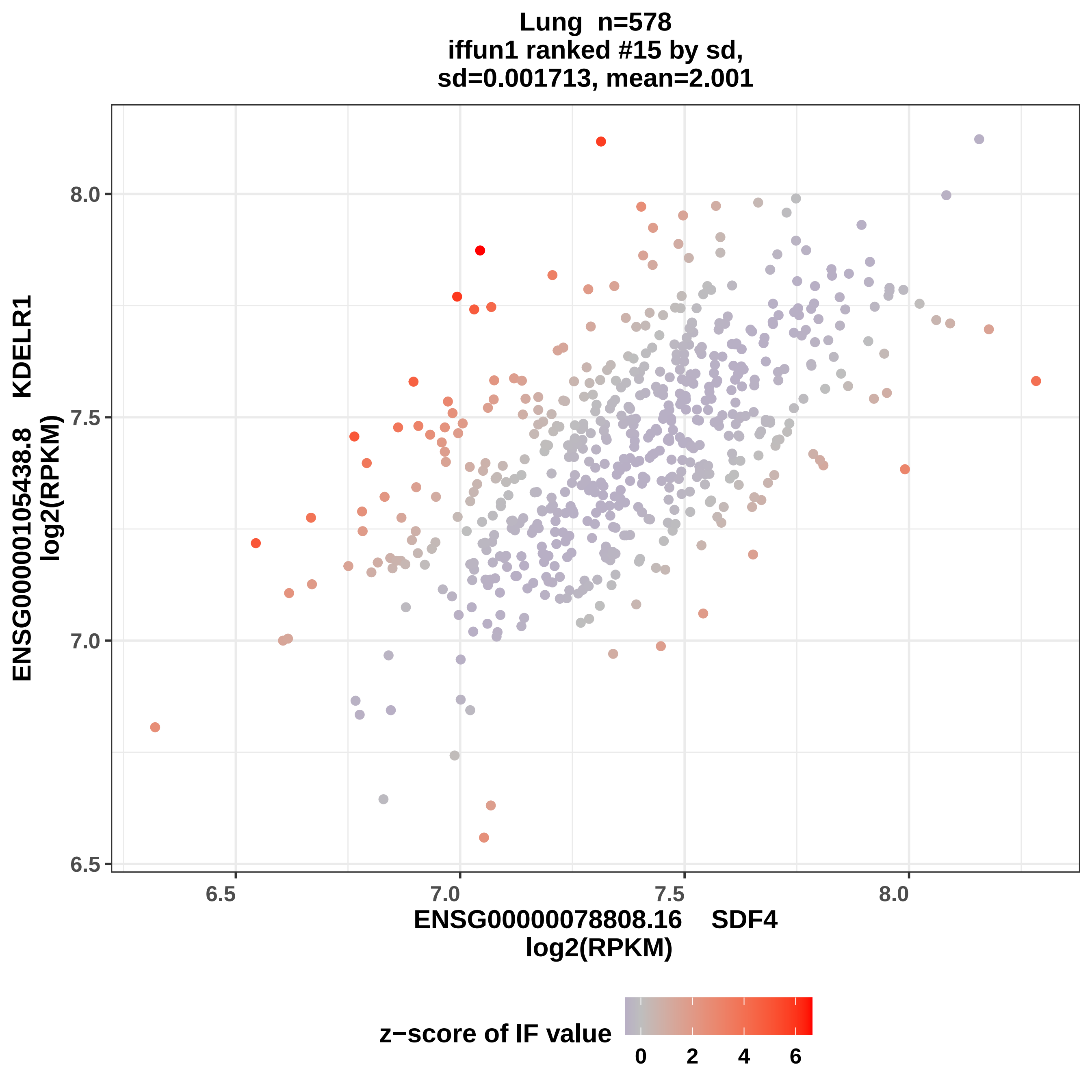
Lung n=578
iffun1 ranked #15 by sd,
sd=0.001713, mean=2.001
ENSG00000105438.8 KDELR1
log2(RPKM)
8.0
7.5
7.0
6.5
6.5
7.0
7.5
8.0
ENSG00000078808.16 SDF4
log2(RPKM)
z−score of IF value
0
2
4
6

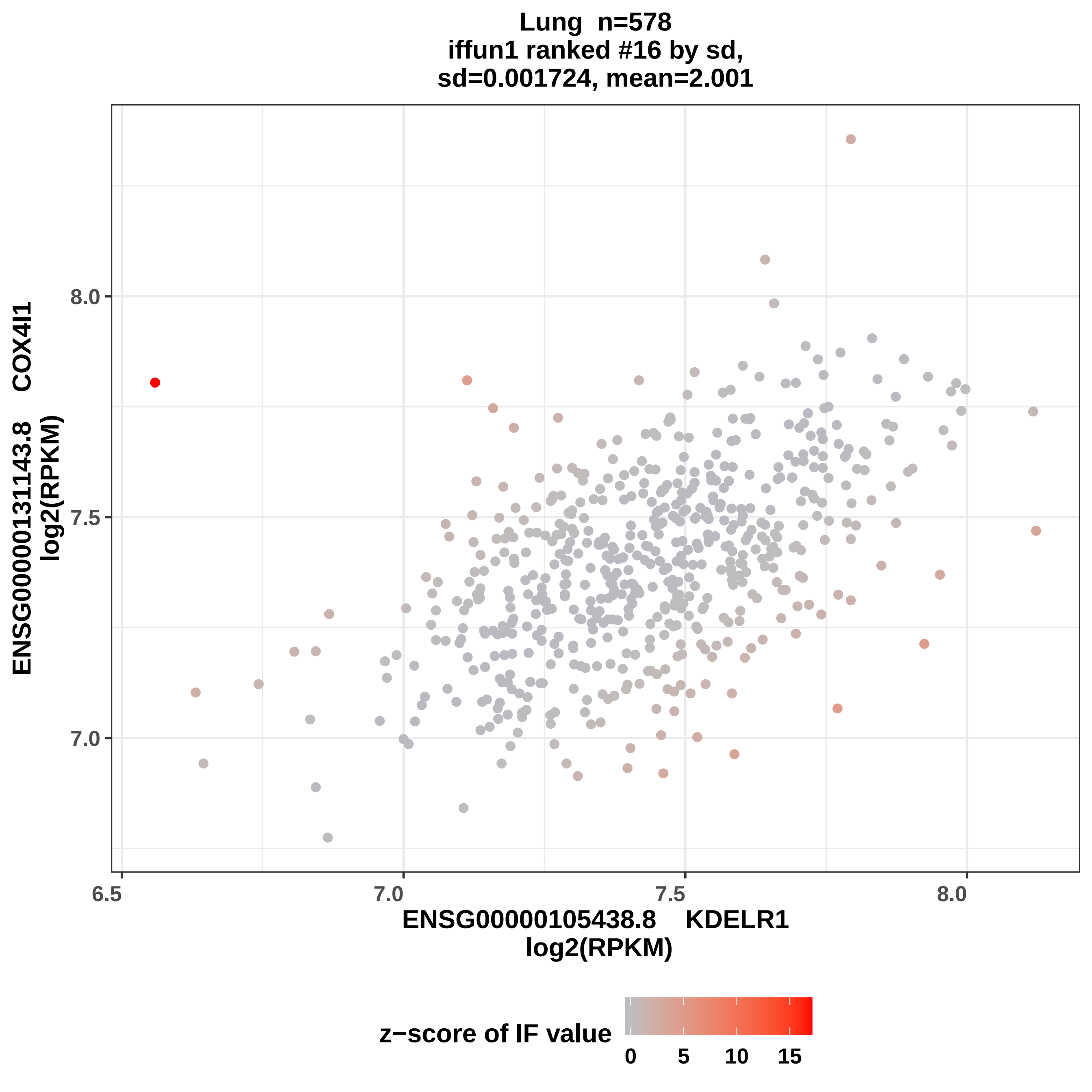

Lung n=578
iffun1 ranked #16 by sd,
sd=0.001724, mean=2.001
ENSG00000131143.8 COX4I1
log2(RPKM)
8.0
7.5
7.0
6.5
7.0
7.5
8.0
ENSG00000105438.8 KDELR1
log2(RPKM)
z−score of IF value
0
5
10
15

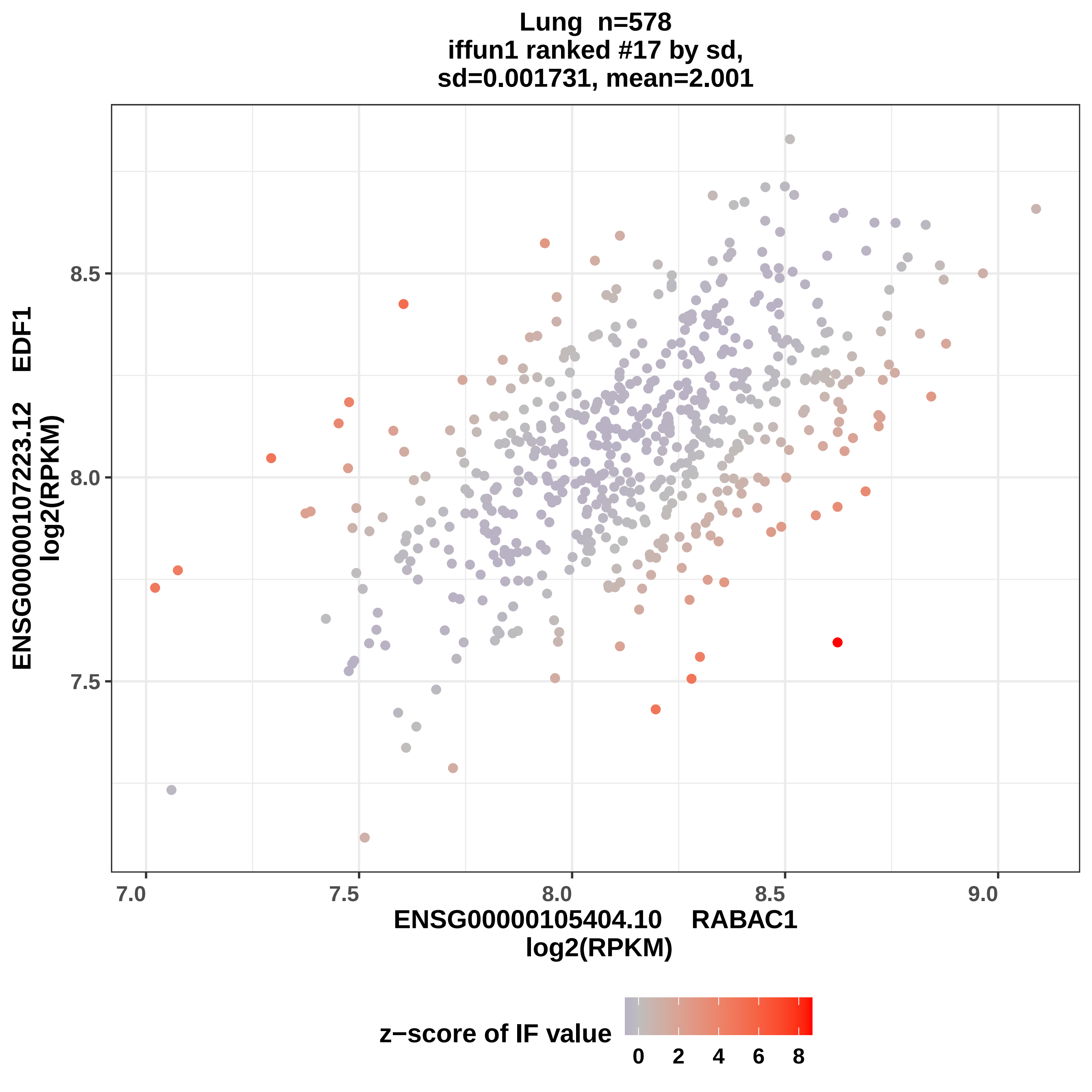
Lung n=578
iffun1 ranked #17 by sd,
sd=0.001731, mean=2.001
ENSG00000107223.12 EDF1
log2(RPKM)
8.5
8.0
7.5
7.0
7.5
8.0
8.5
9.0
ENSG00000105404.10 RABAC1
log2(RPKM)
z−score of IF value
0
2
4
6
8

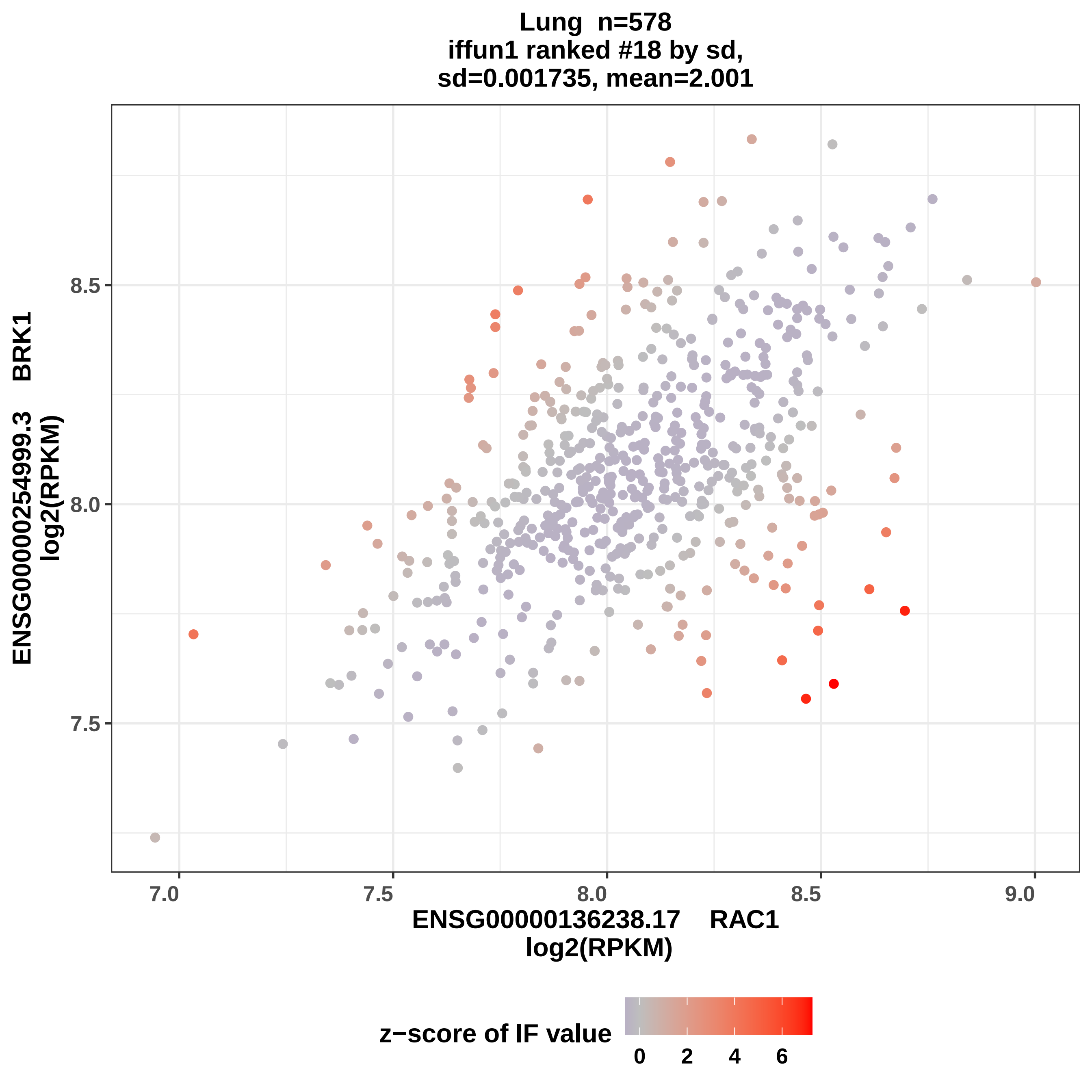
Lung n=578
iffun1 ranked #18 by sd,
sd=0.001735, mean=2.001
ENSG00000254999.3 BRK1
log2(RPKM)
8.5
8.0
7.5
7.0
7.5
8.0
8.5
9.0
ENSG00000136238.17 RAC1
log2(RPKM)
z−score of IF value
0
2
4
6

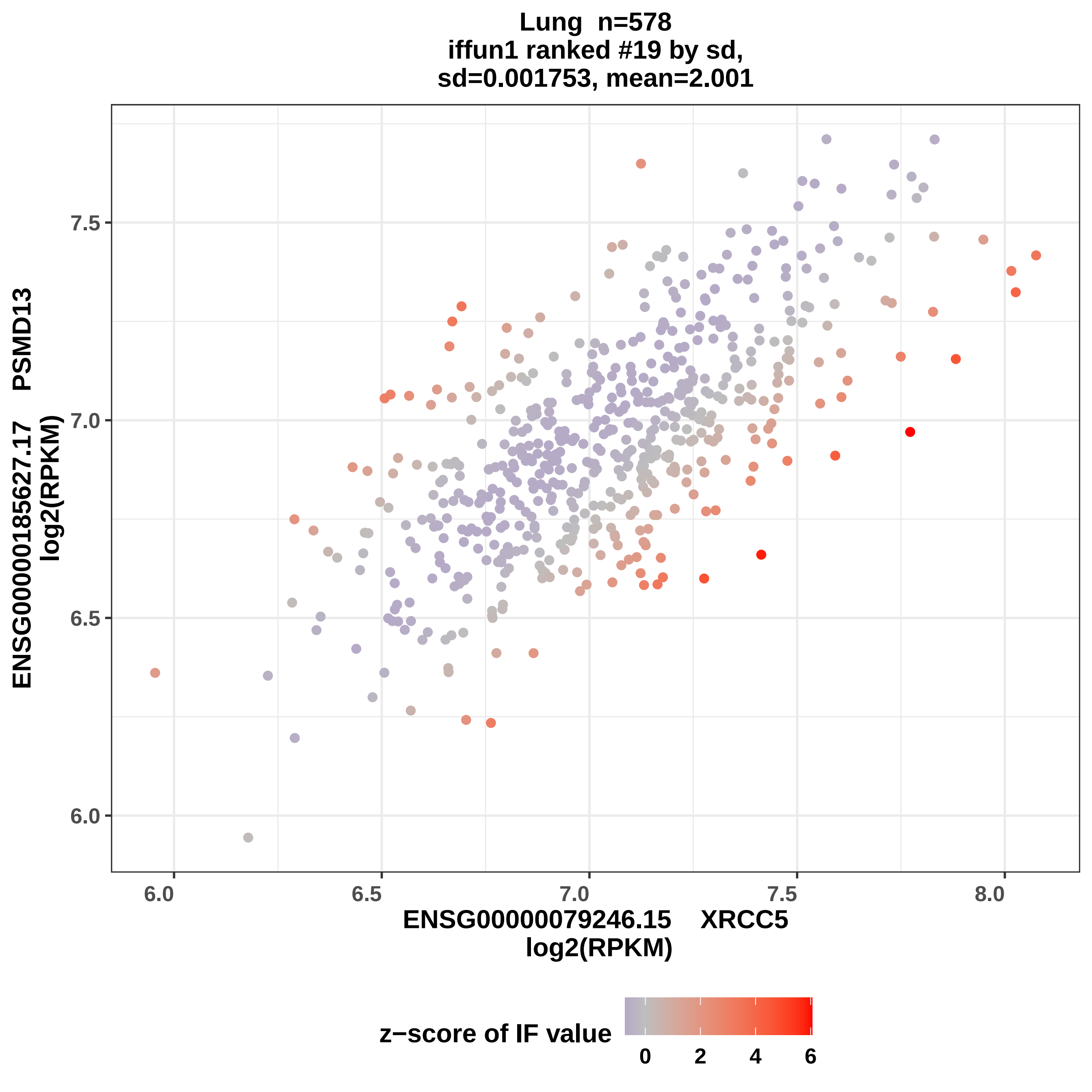

Lung n=578
iffun1 ranked #19 by sd,
sd=0.001753, mean=2.001
ENSG00000185627.17 PSMD13
log2(RPKM)
7.5
7.0
6.5
6.0
6.0
6.5
7.0
7.5
8.0
ENSG00000079246.15 XRCC5
log2(RPKM)
z−score of IF value
0
2
4
6

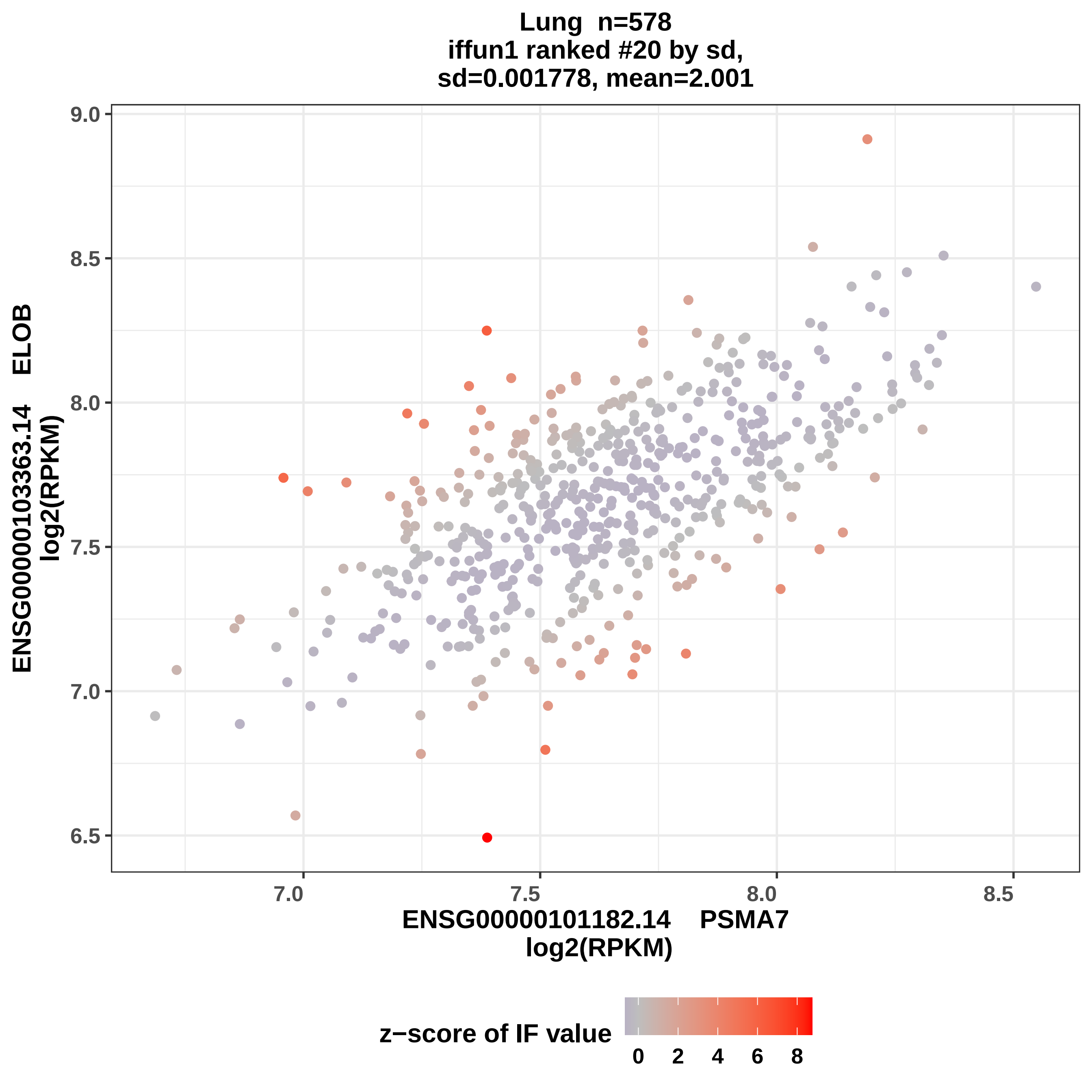
Lung n=578
iffun1 ranked #20 by sd,
sd=0.001778, mean=2.001
ENSG00000103363.14 ELOB
log2(RPKM)
ENSG00000101182.14 PSMA7
log2(RPKM)
9.0
8.5
8.0
7.5
7.0
6.5
7.0
7.5
8.0
8.5
z−score of IF value
0
2
4
6
8

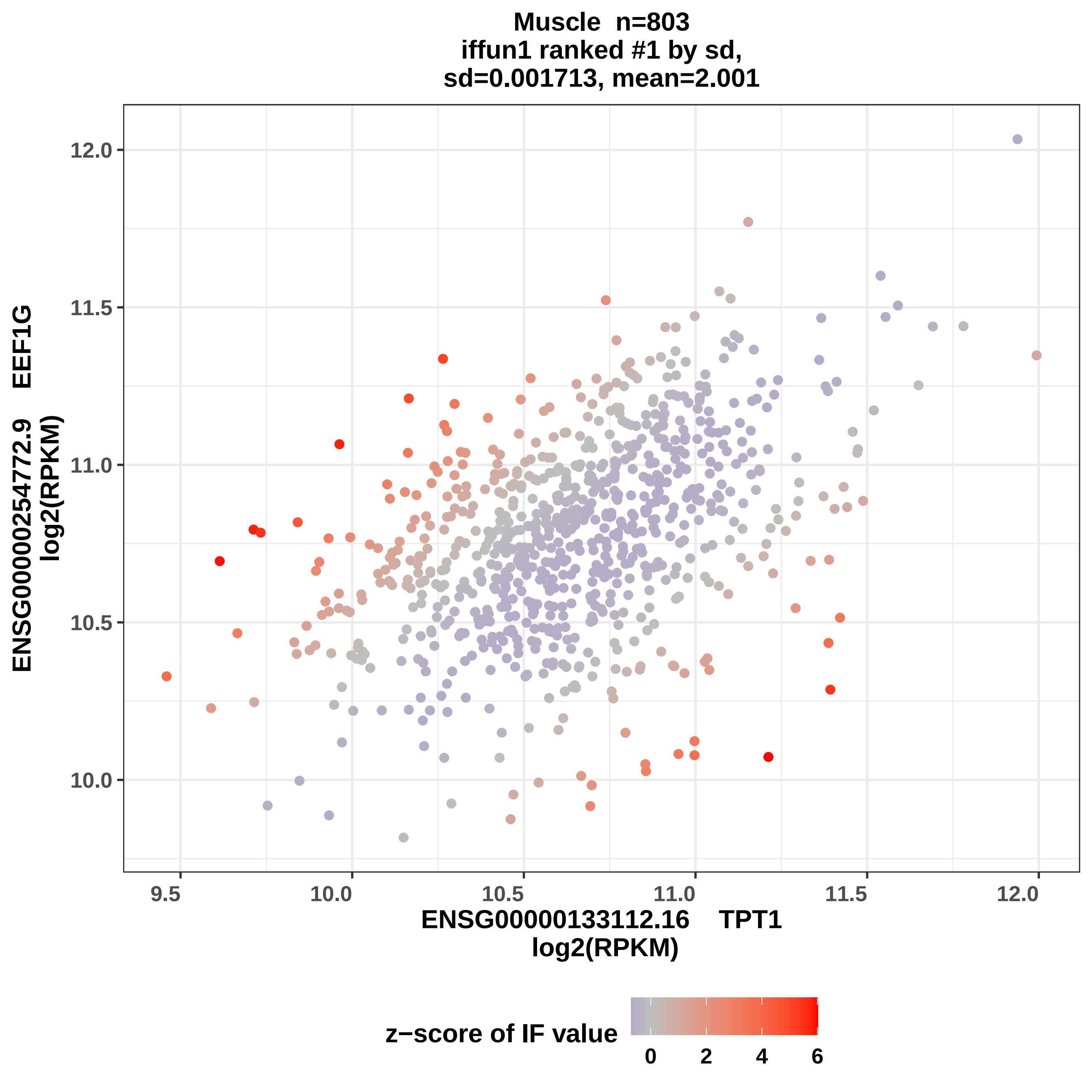
Muscle n=803
iffun1 ranked #1 by sd,
sd=0.001713, mean=2.001
ENSG00000254772.9 EEF1G
log2(RPKM)
ENSG00000133112.16 TPT1
log2(RPKM)
12.0
11.5
11.0
10.5
10.0
9.5
10.0
10.5
11.0
11.5
12.0
z−score of IF value
0
2
4
6

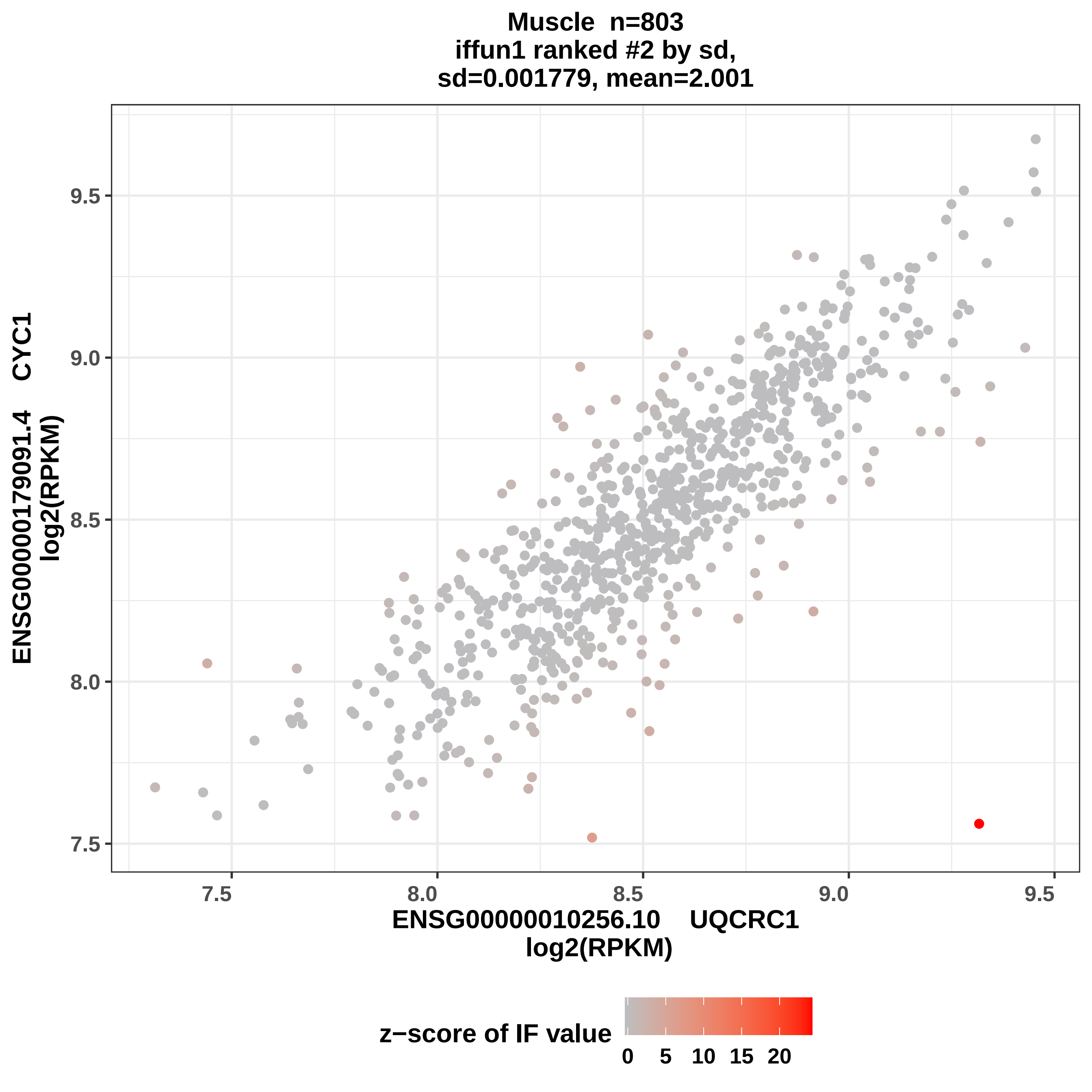
Muscle n=803
iffun1 ranked #2 by sd,
sd=0.001779, mean=2.001
9.5
9.0
8.5
8.0
7.5
ENSG00000179091.4 CYC1
log2(RPKM)
7.5
8.0
8.5
9.0
9.5
ENSG00000010256.10 UQCRC1
log2(RPKM)
z−score of IF value
0
5
10
15
20

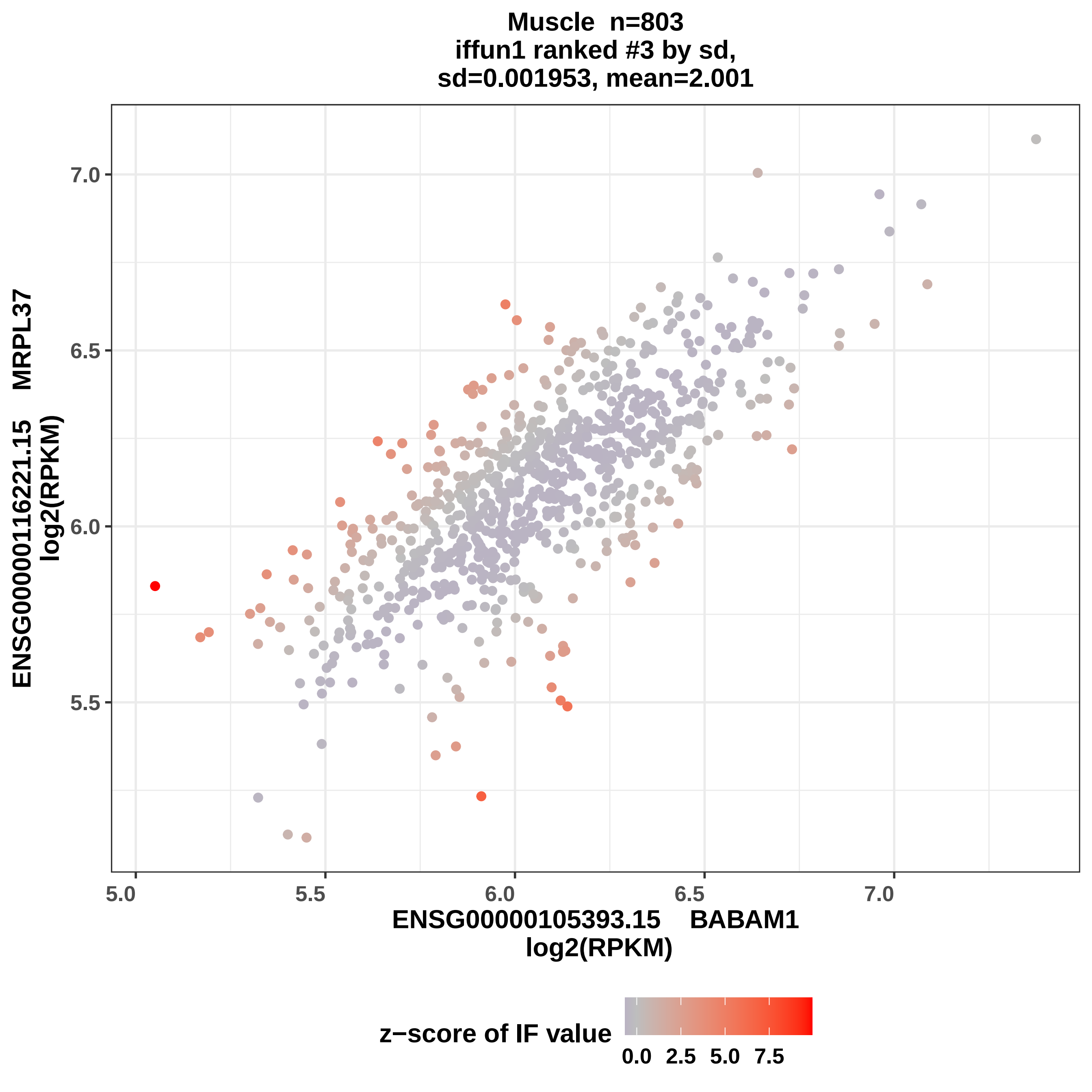
Muscle n=803
iffun1 ranked #3 by sd,
sd=0.001953, mean=2.001
ENSG00000116221.15 MRPL37
log2(RPKM)
7.0
6.5
6.0
5.5
5.0
5.5
6.0
6.5
7.0
ENSG00000105393.15 BABAM1
log2(RPKM)
z−score of IF value
0.0
2.5
5.0
7.5

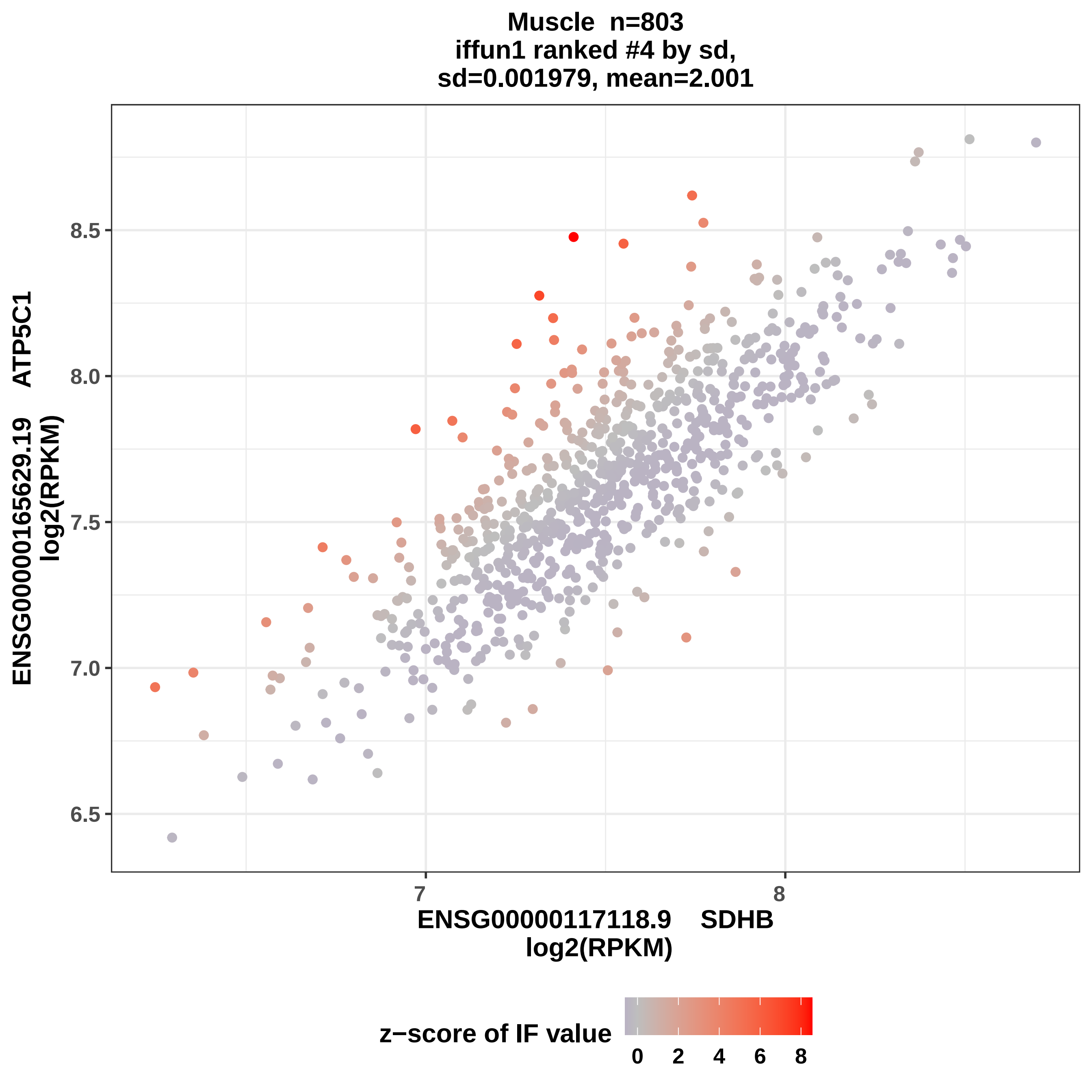
Muscle n=803
iffun1 ranked #4 by sd,
sd=0.001979, mean=2.001
ENSG00000165629.19 ATP5C1
log2(RPKM)
8.5
8.0
7.5
7.0
6.5
7
8
ENSG00000117118.9 SDHB
log2(RPKM)
z−score of IF value
0
2
4
6
8

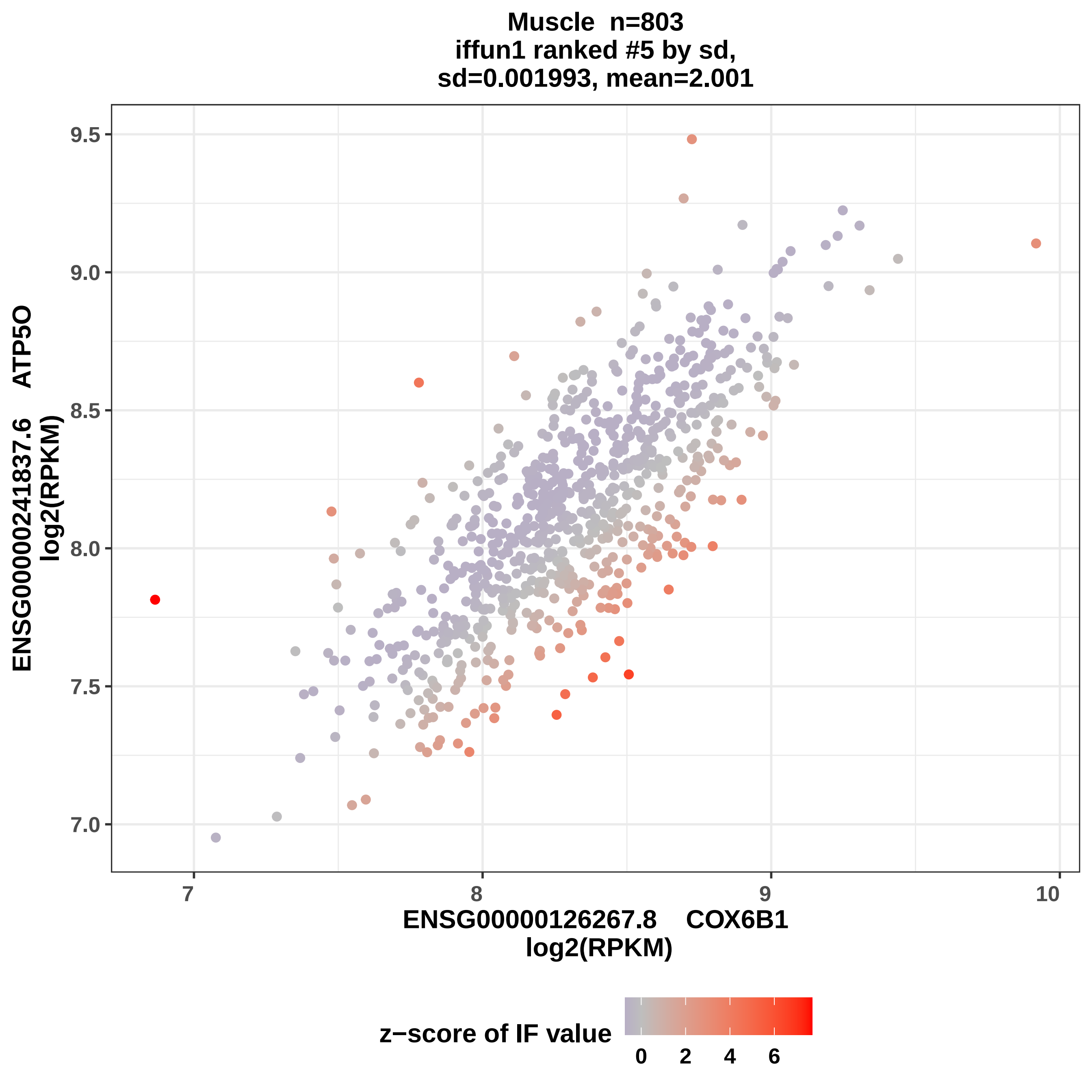

Muscle n=803
iffun1 ranked #5 by sd,
sd=0.001993, mean=2.001
ENSG00000241837.6 ATP5O
log2(RPKM)
9.5
9.0
8.5
8.0
7.5
7.0
7
8
9
10
ENSG00000126267.8 COX6B1
log2(RPKM)
z−score of IF value
0
2
4
6

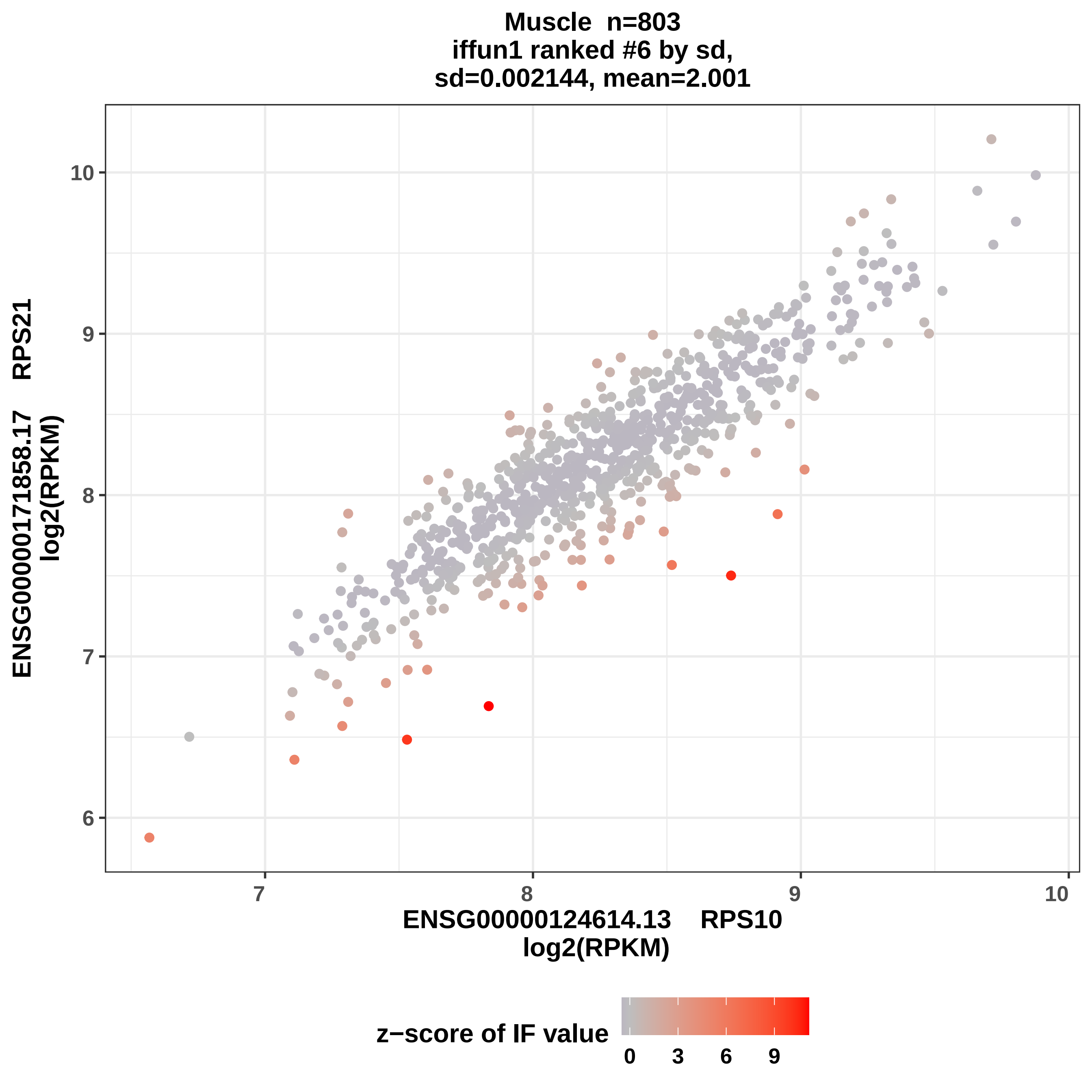
Muscle n=803
iffun1 ranked #6 by sd,
sd=0.002144, mean=2.001
ENSG00000171858.17 RPS21
log2(RPKM)
10
9
8
7
6
7
8
9
10
ENSG00000124614.13 RPS10
log2(RPKM)
z−score of IF value
0
3
6
9

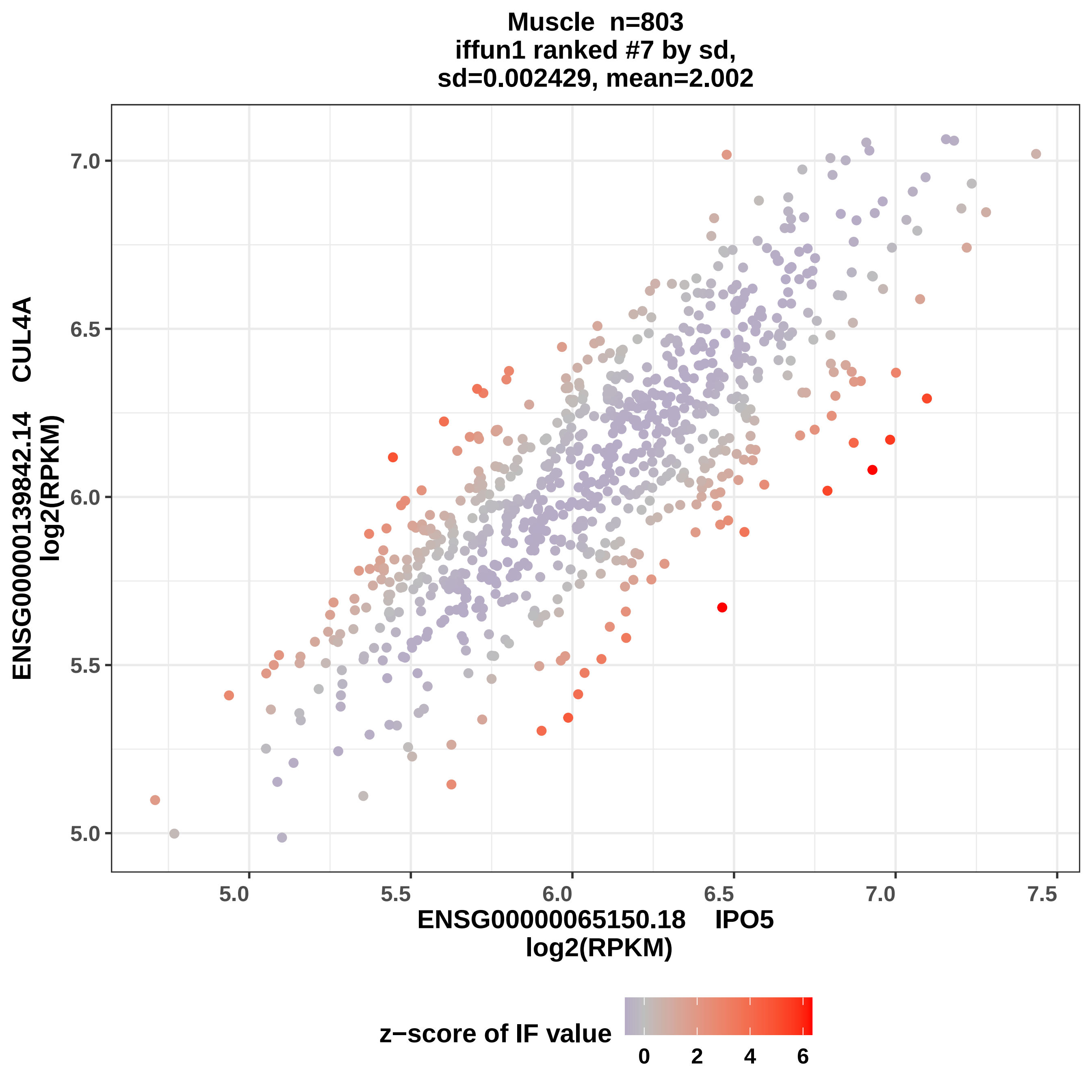

Muscle n=803
iffun1 ranked #7 by sd,
sd=0.002429, mean=2.002
ENSG00000139842.14 CUL4A
log2(RPKM)
7.0
6.5
6.0
5.5
5.0
5.0
5.5
6.0
6.5
7.0
7.5
ENSG00000065150.18 IPO5
log2(RPKM)
z−score of IF value
0
2
4
6

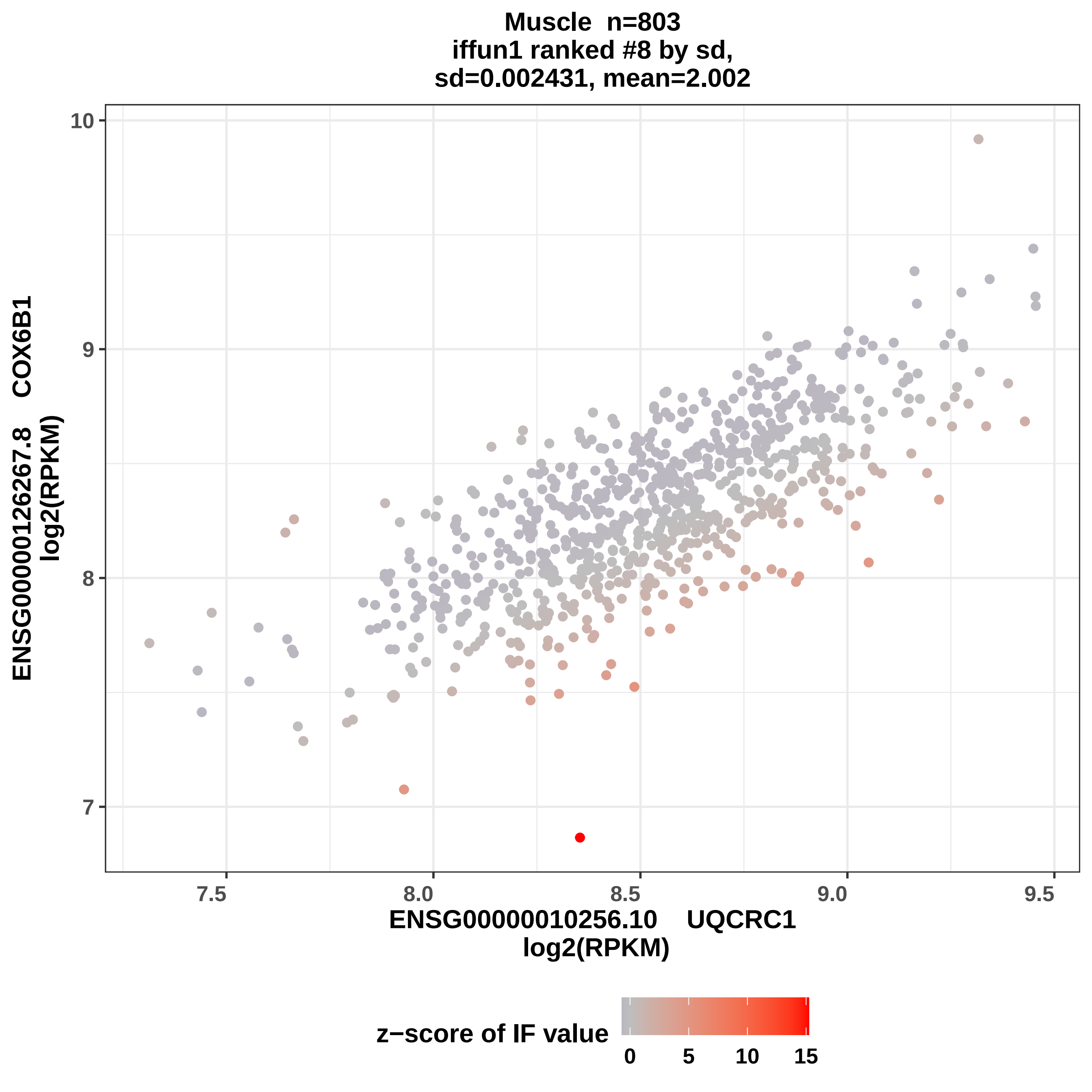

Muscle n=803
iffun1 ranked #8 by sd,
sd=0.002431, mean=2.002
10
9
8
7
7.5
8.0
8.5
9.0
9.5
ENSG00000126267.8 COX6B1
log2(RPKM)
ENSG00000010256.10 UQCRC1
log2(RPKM)
z−score of IF value
0
5
10
15

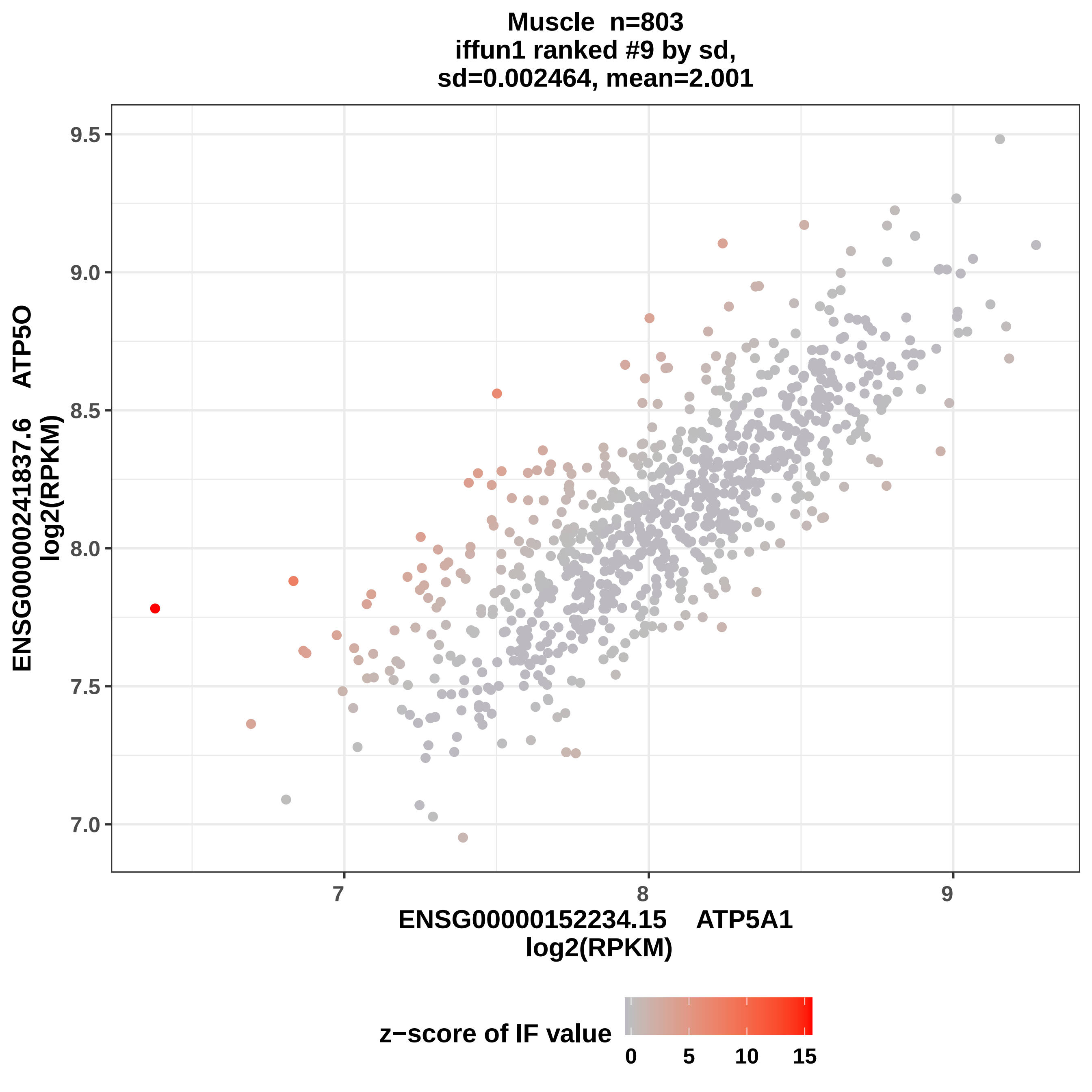

Muscle n=803
iffun1 ranked #9 by sd,
sd=0.002464, mean=2.001
ENSG00000241837.6 ATP5O
log2(RPKM)
9.5
9.0
8.5
8.0
7.5
7.0
7
8
9
ENSG00000152234.15 ATP5A1
log2(RPKM)
z−score of IF value
0
5
10
15

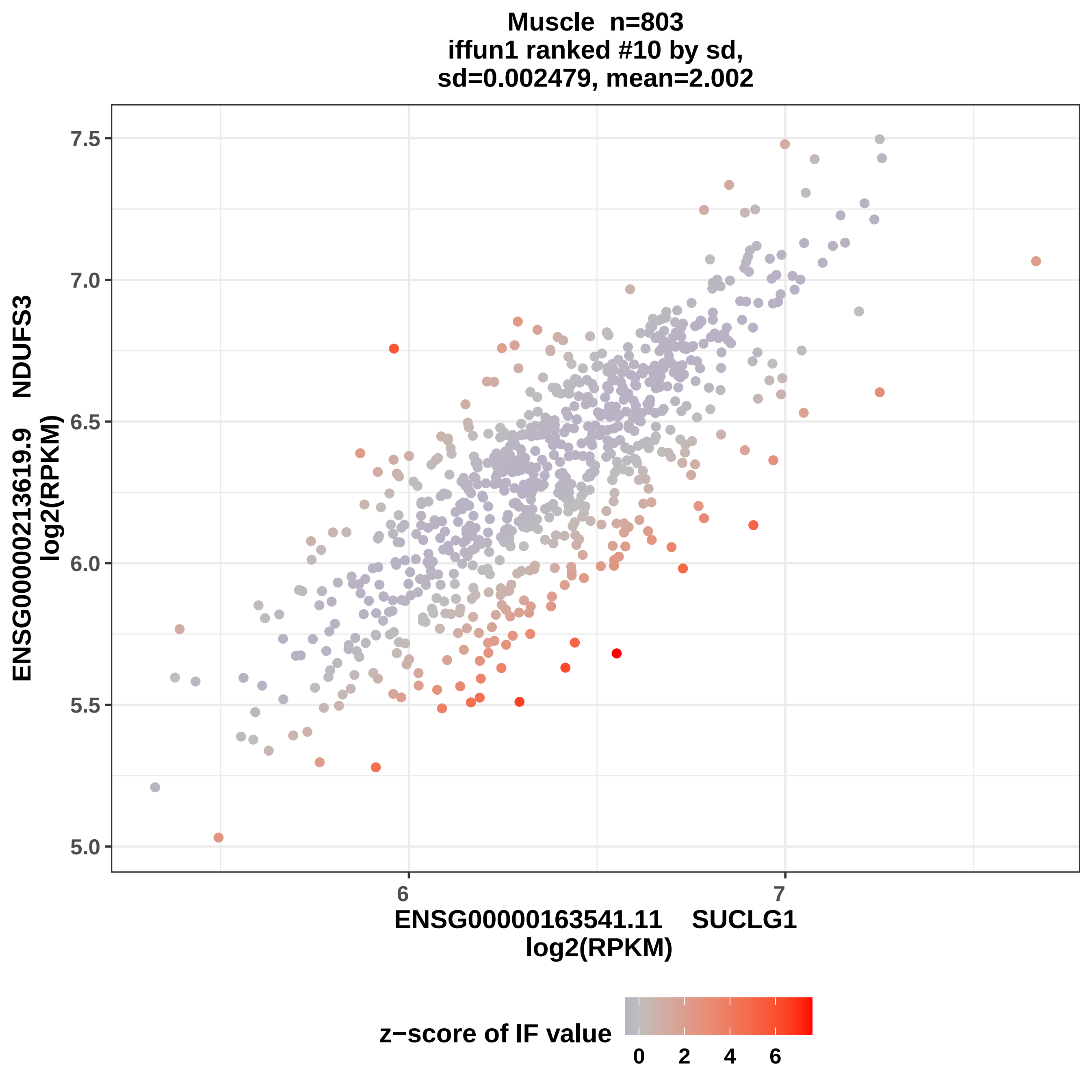
Muscle n=803
iffun1 ranked #10 by sd,
sd=0.002479, mean=2.002
ENSG00000213619.9 NDUFS3
log2(RPKM)
7.5
7.0
6.5
6.0
5.5
5.0
6
7
ENSG00000163541.11 SUCLG1
log2(RPKM)
z−score of IF value
0
2
4
6

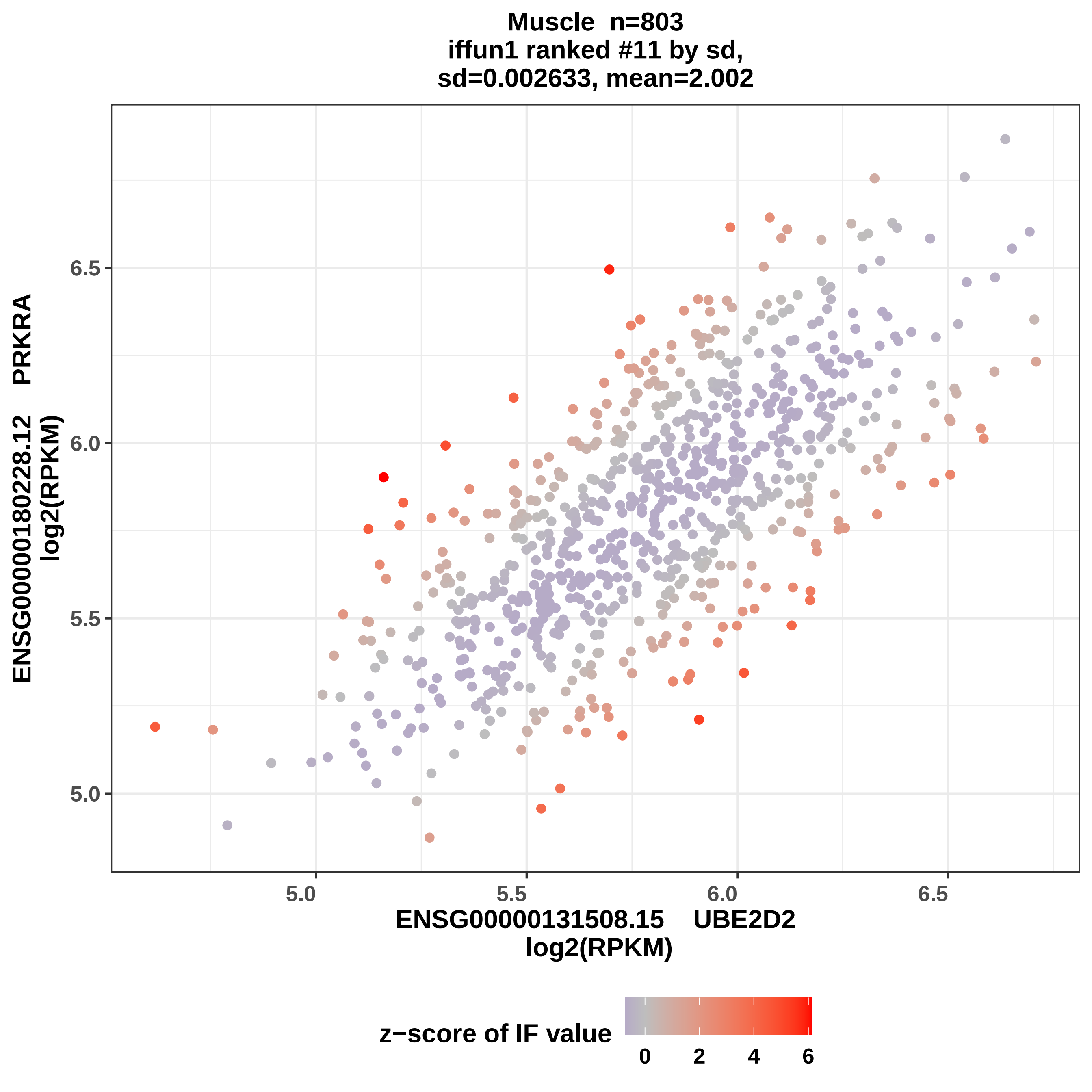
Muscle n=803
iffun1 ranked #11 by sd,
sd=0.002633, mean=2.002
ENSG00000180228.12 PRKRA
log2(RPKM)
6.5
6.0
5.5
5.0
5.0
5.5
6.0
6.5
ENSG00000131508.15 UBE2D2
log2(RPKM)
z−score of IF value
0
2
4
6

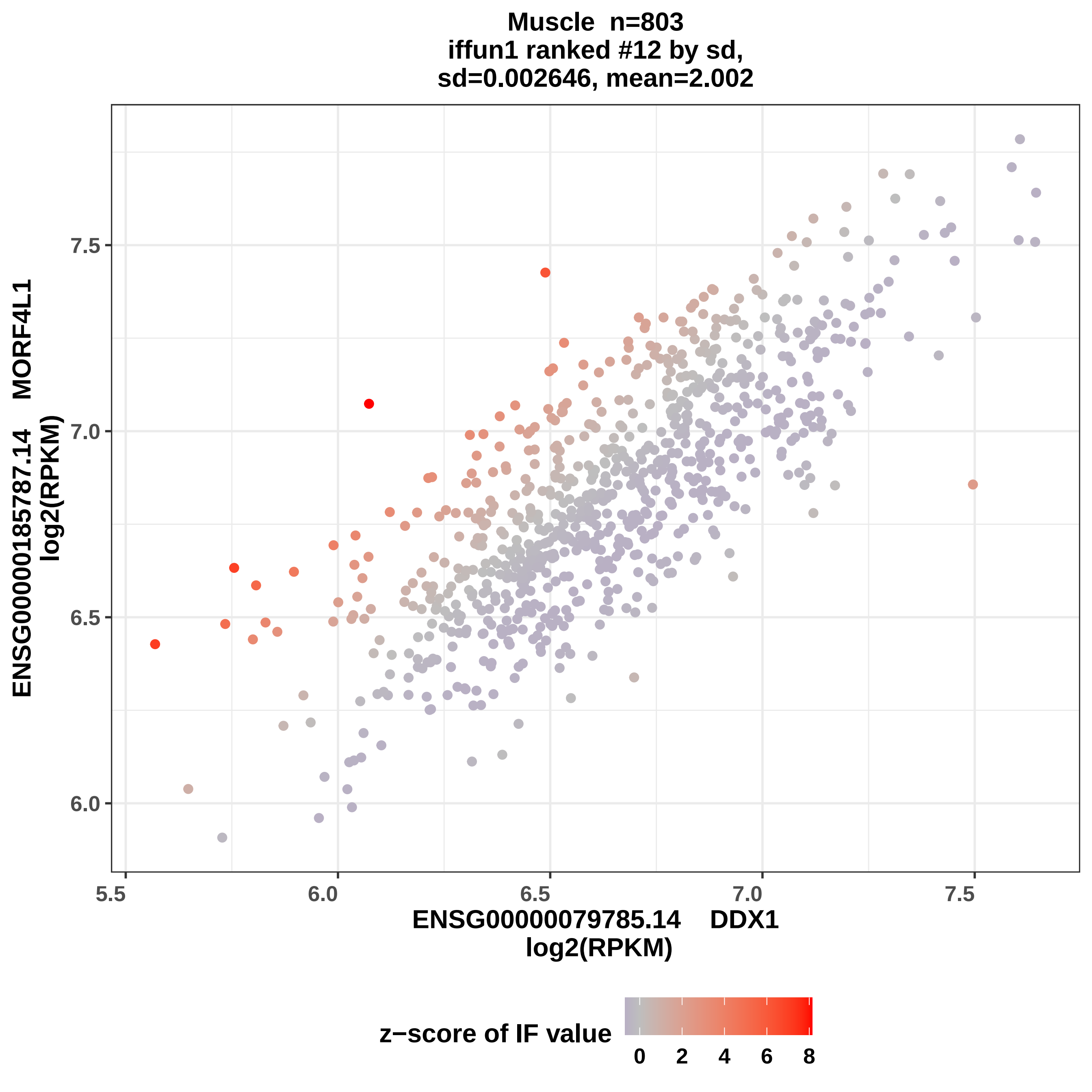

Muscle n=803
iffun1 ranked #12 by sd,
sd=0.002646, mean=2.002
ENSG00000185787.14 MORF4L1
log2(RPKM)
ENSG00000079785.14 DDX1
log2(RPKM)
6.0
6.5
7.0
7.5
5.5
z−score of IF value
0
2
4
6
8

**Muscle n=803**
**iffun1 ranked #13 by sd,**
**sd=0.002655, mean=2.002**

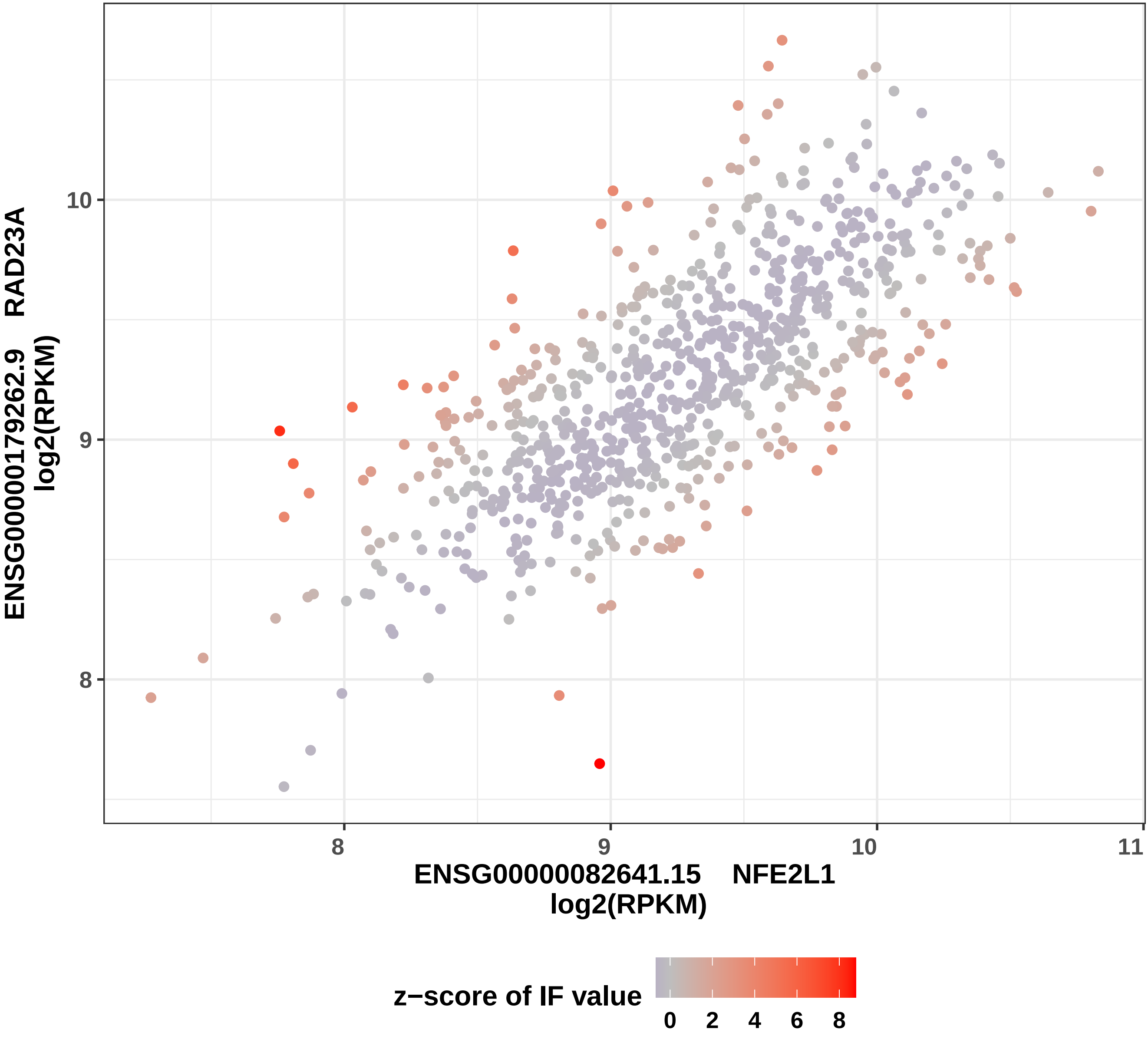

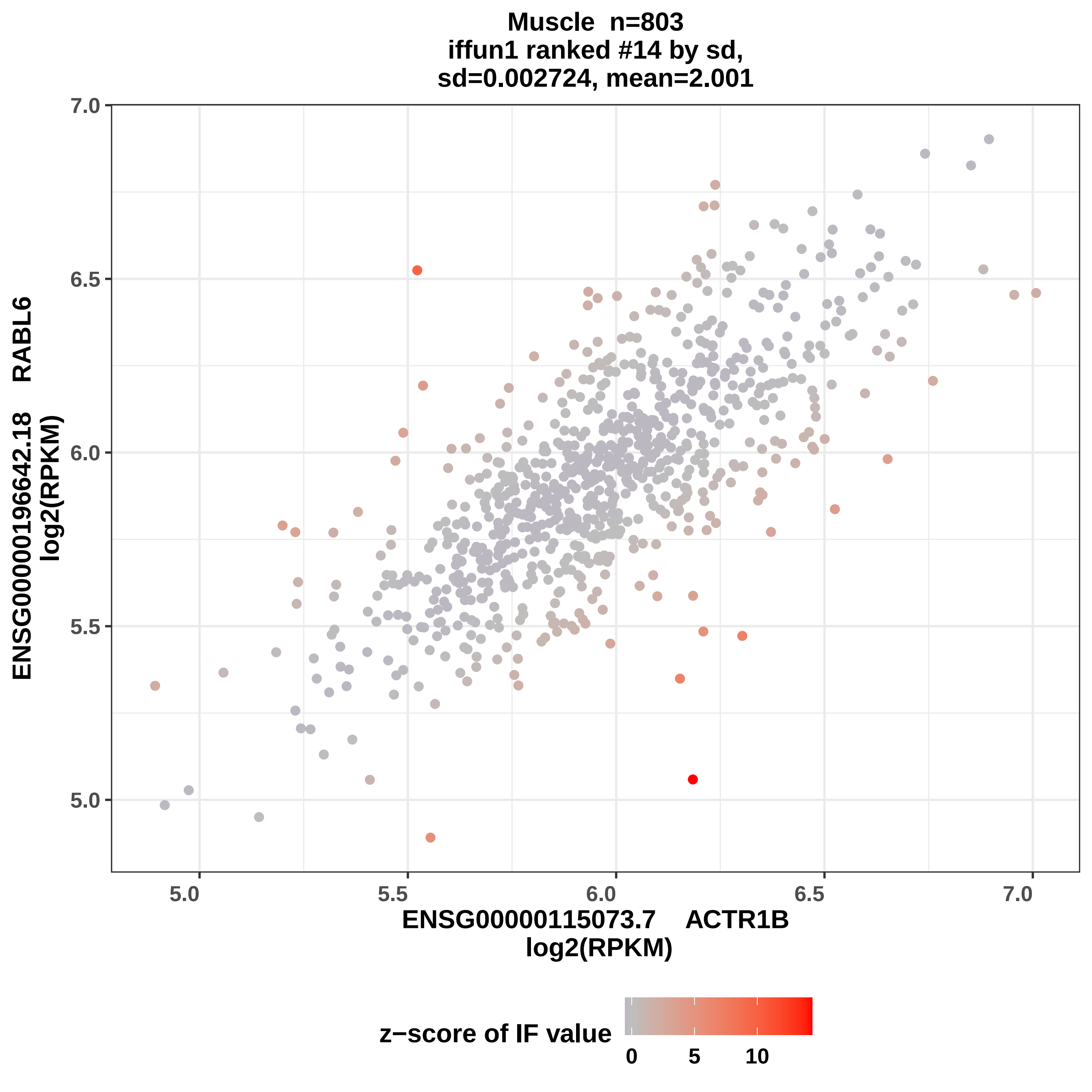
Muscle n=803
iffun1 ranked #14 by sd,
sd=0.002724, mean=2.001
7.0
6.5
6.0
5.5
5.0
ENSG00000196642.18 RABL6
log2(RPKM)
5.0
5.5
6.0
6.5
7.0
ENSG00000115073.7 ACTR1B
log2(RPKM)
z−score of IF value
0
5
10

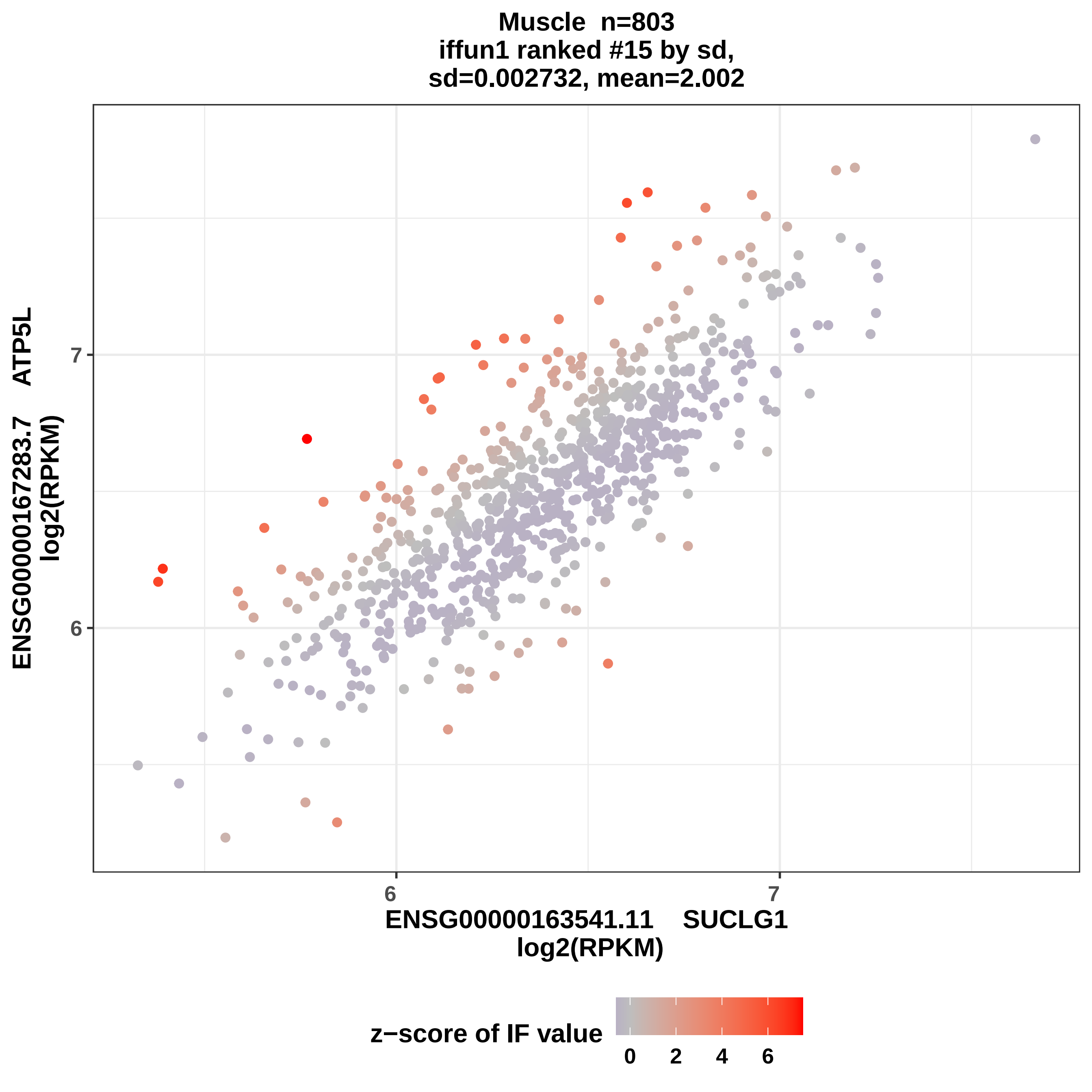
Muscle n=803
iffun1 ranked #15 by sd,
sd=0.002732, mean=2.002
ENSG00000167283.7 ATP5L
log2(RPKM)
7
6
6
7
ENSG00000163541.11 SUCLG1
log2(RPKM)
z−score of IF value
0
2
4
6

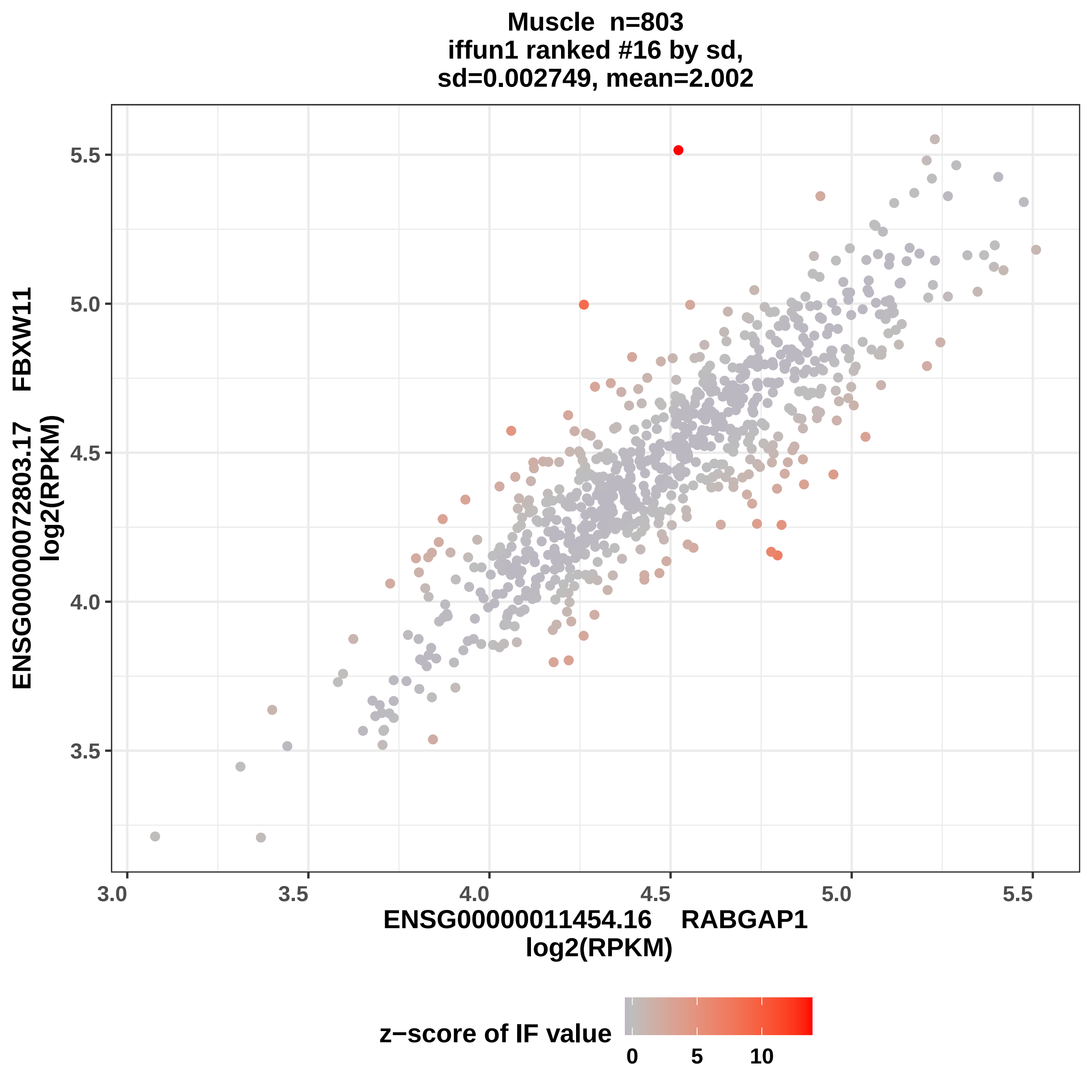

Muscle n=803
iffun1 ranked #16 by sd,
sd=0.002749, mean=2.002
ENSG00000072803.17 FBXW11
log2(RPKM)
ENSG00000011454.16 RABGAP1
log2(RPKM)
3.0
3.5
4.0
4.5
5.0
5.5
z−score of IF value
0
5
10

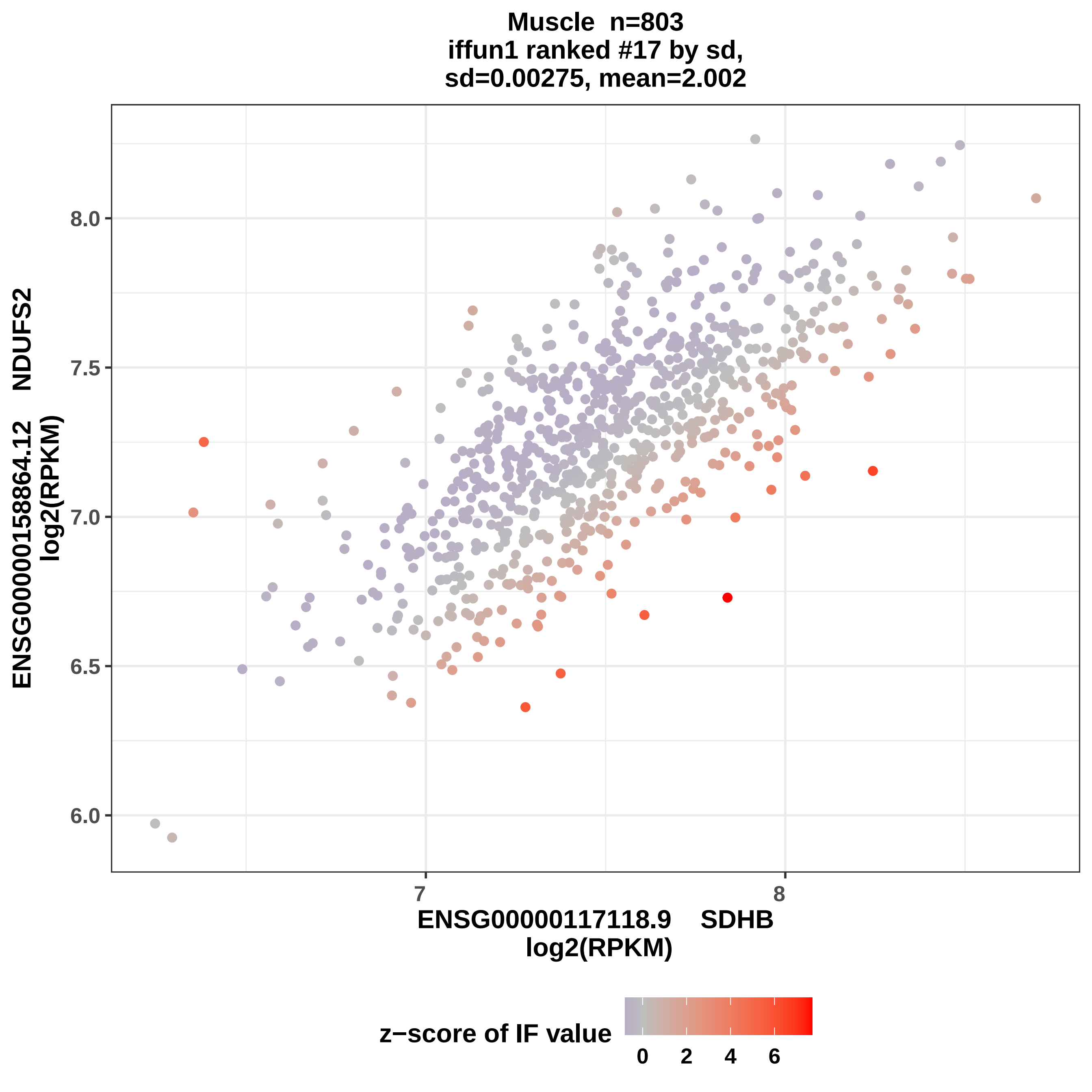

Muscle n=803
iffun1 ranked #17 by sd,
sd=0.00275, mean=2.002
ENSG00000158864.12 NDUFS2
log2(RPKM)
8.0
7.5
7.0
6.5
6.0
7
8
ENSG00000117118.9 SDHB
log2(RPKM)
z−score of IF value
0
2
4
6

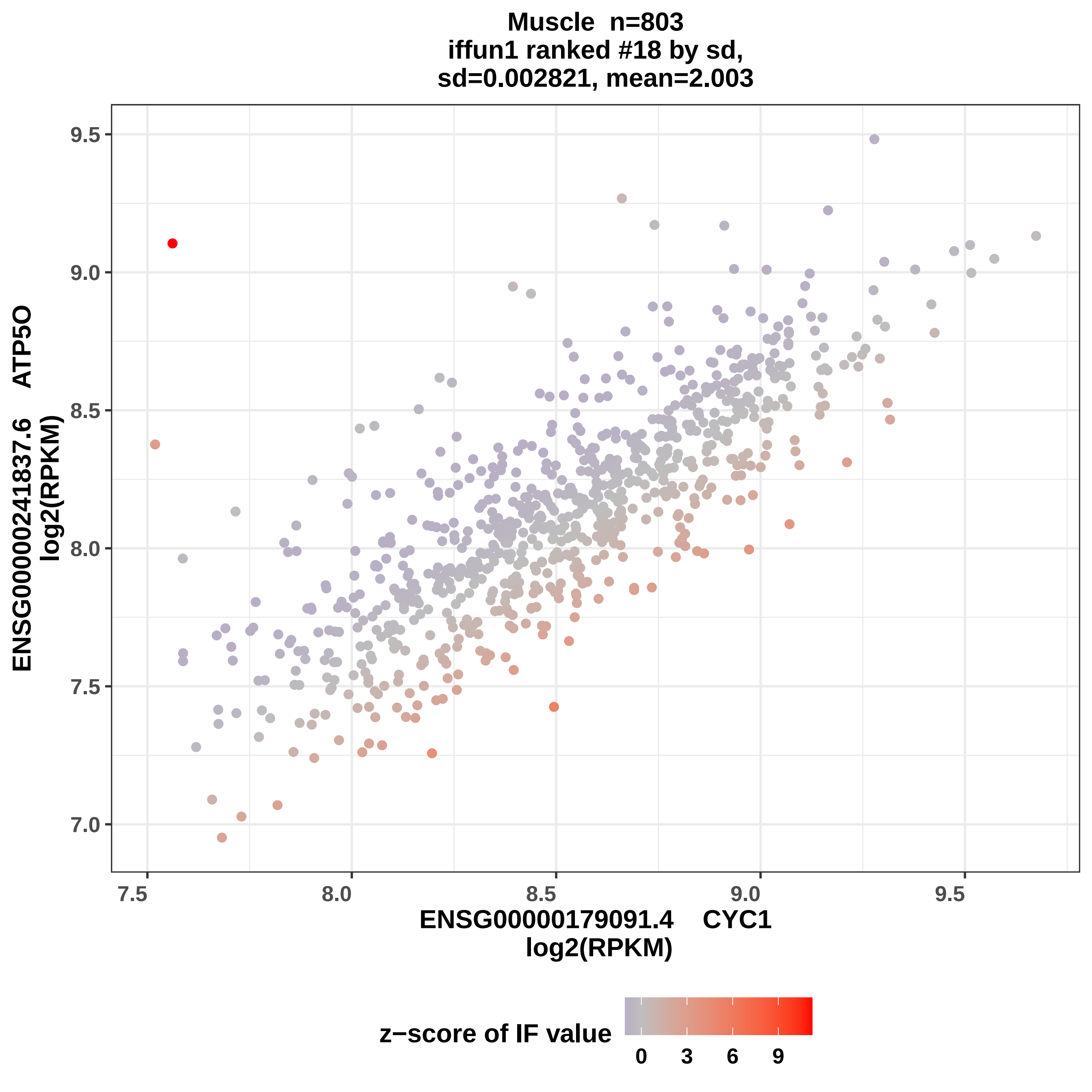

Muscle n=803
iffun1 ranked #18 by sd,
sd=0.002821, mean=2.003
ENSG00000241837.6 ATP5O
log2(RPKM)
9.5
9.0
8.5
8.0
7.5
7.0
7.5
8.0
8.5
9.0
9.5
ENSG00000179091.4 CYC1
log2(RPKM)
z−score of IF value
0
3
6
9

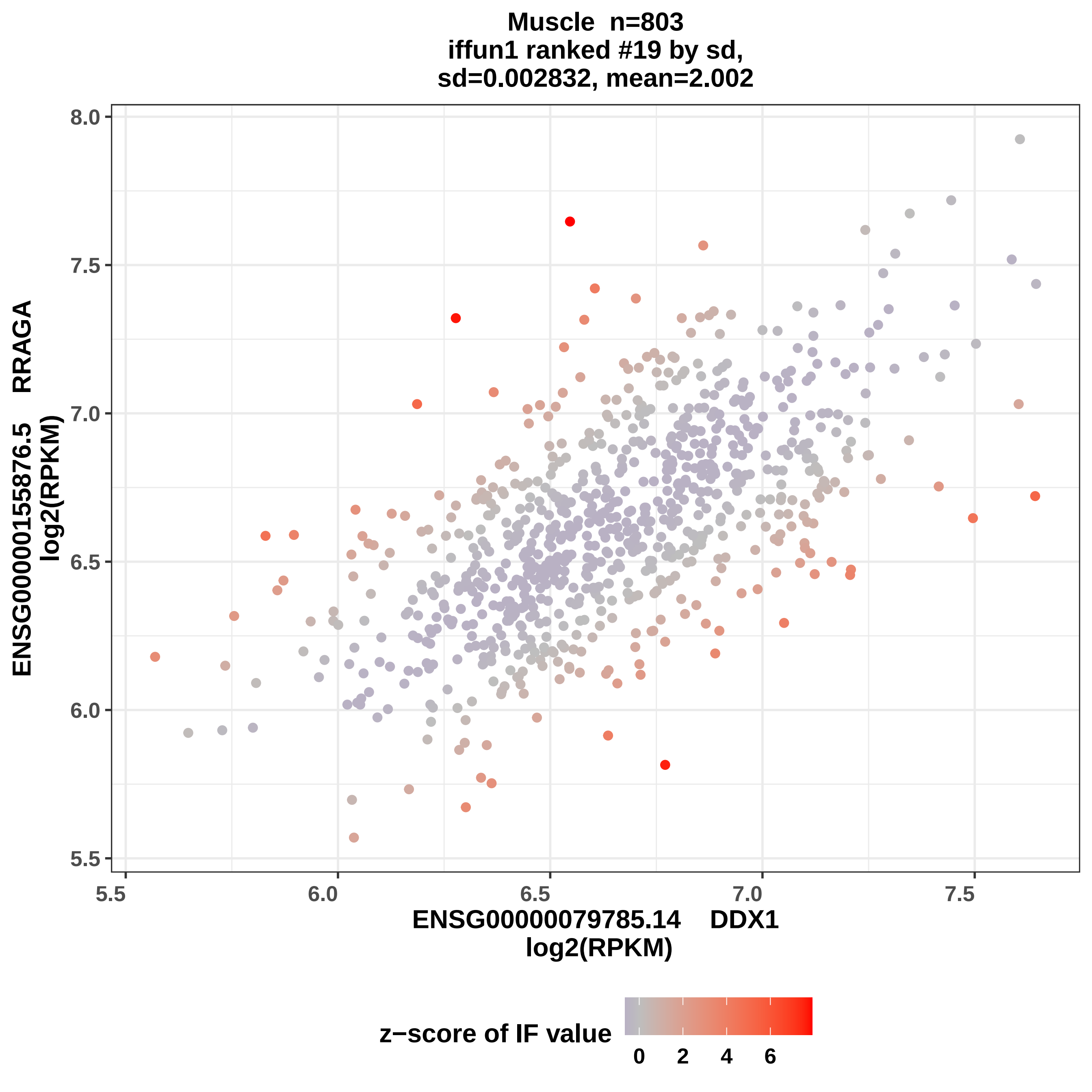

Muscle n=803
iffun1 ranked #19 by sd,
sd=0.002832, mean=2.002
ENSG00000155876.5 RRAGA
log2(RPKM)
8.0
7.5
7.0
6.5
6.0
5.5
5.5
6.0
6.5
7.0
7.5
ENSG00000079785.14 DDX1
log2(RPKM)
z−score of IF value
0
2
4
6

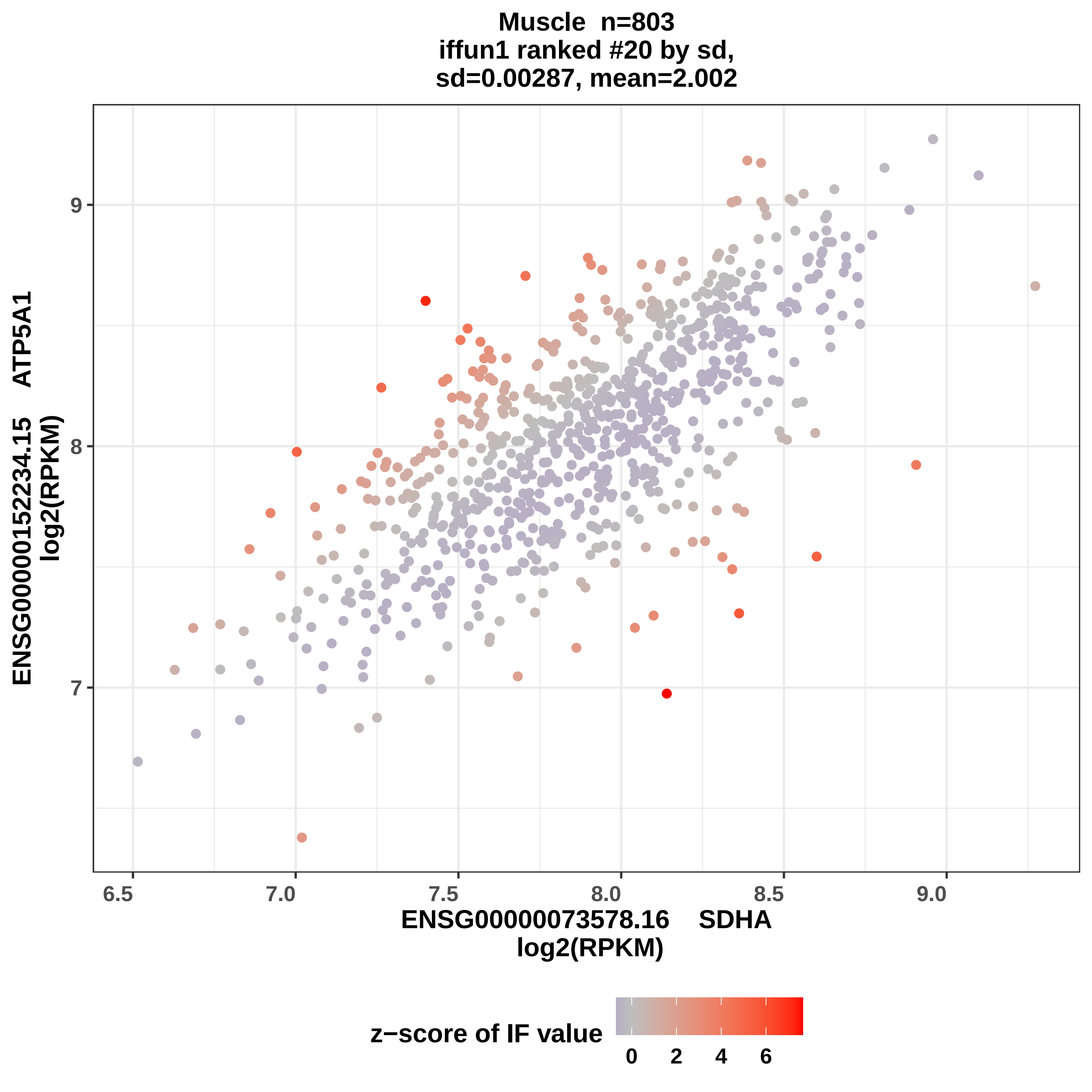

Muscle n=803
iffun1 ranked #20 by sd,
sd=0.00287, mean=2.002
ENSG00000152234.15 ATP5A1
log2(RPKM)
9
8
7
6.5
7.0
7.5
8.0
8.5
9.0
ENSG00000073578.16 SDHA
log2(RPKM)
z−score of IF value
0
2
4
6

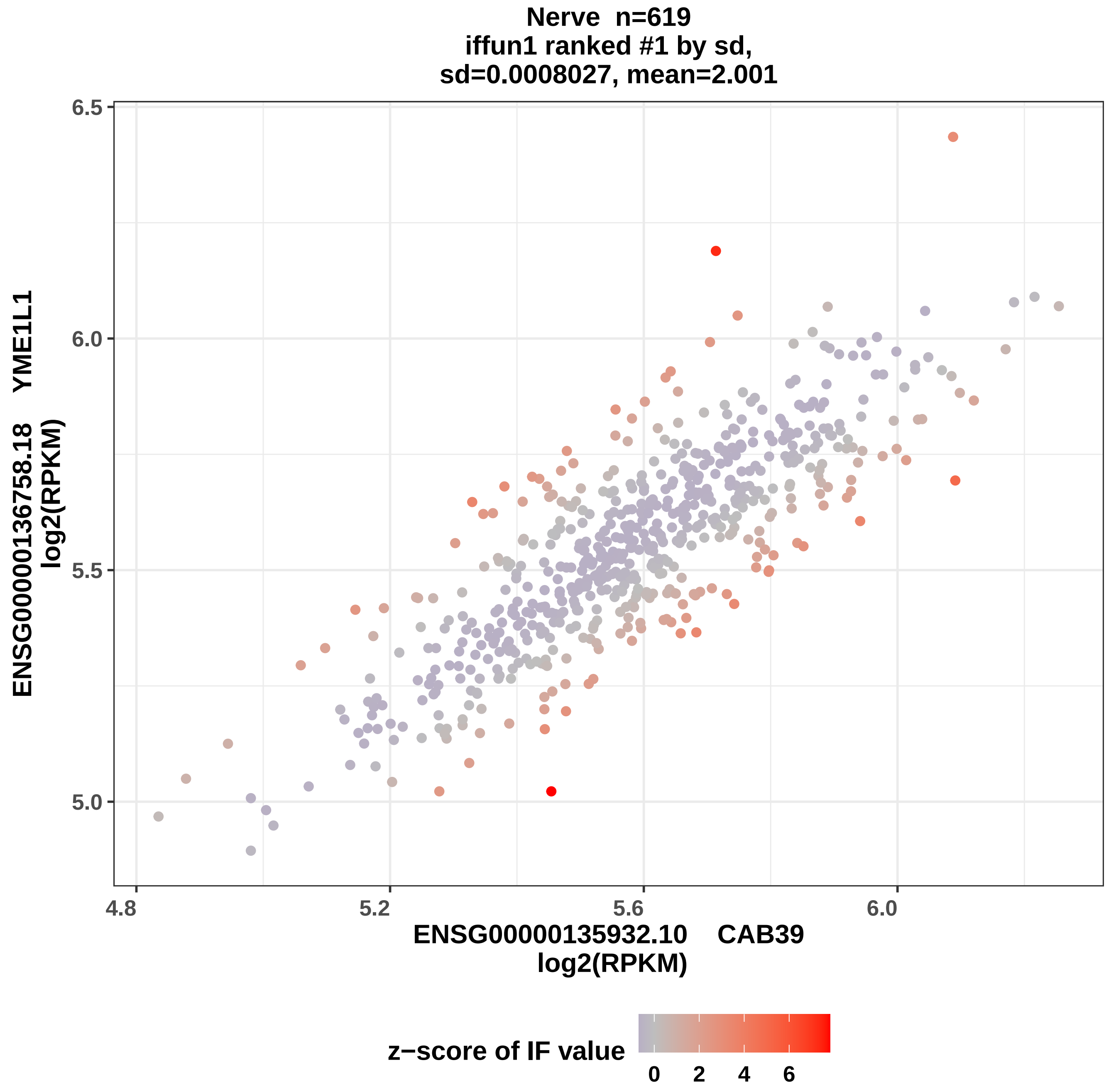

Nerve n=619
iffun1 ranked #1 by sd,
sd=0.0008027, mean=2.001
6.5
6.0
5.5
5.0
4.8
5.2
5.6
6.0
ENSG00000136758.18 YME1L1
log2(RPKM)
ENSG00000135932.10 CAB39
log2(RPKM)
z−score of IF value
0
2
4
6

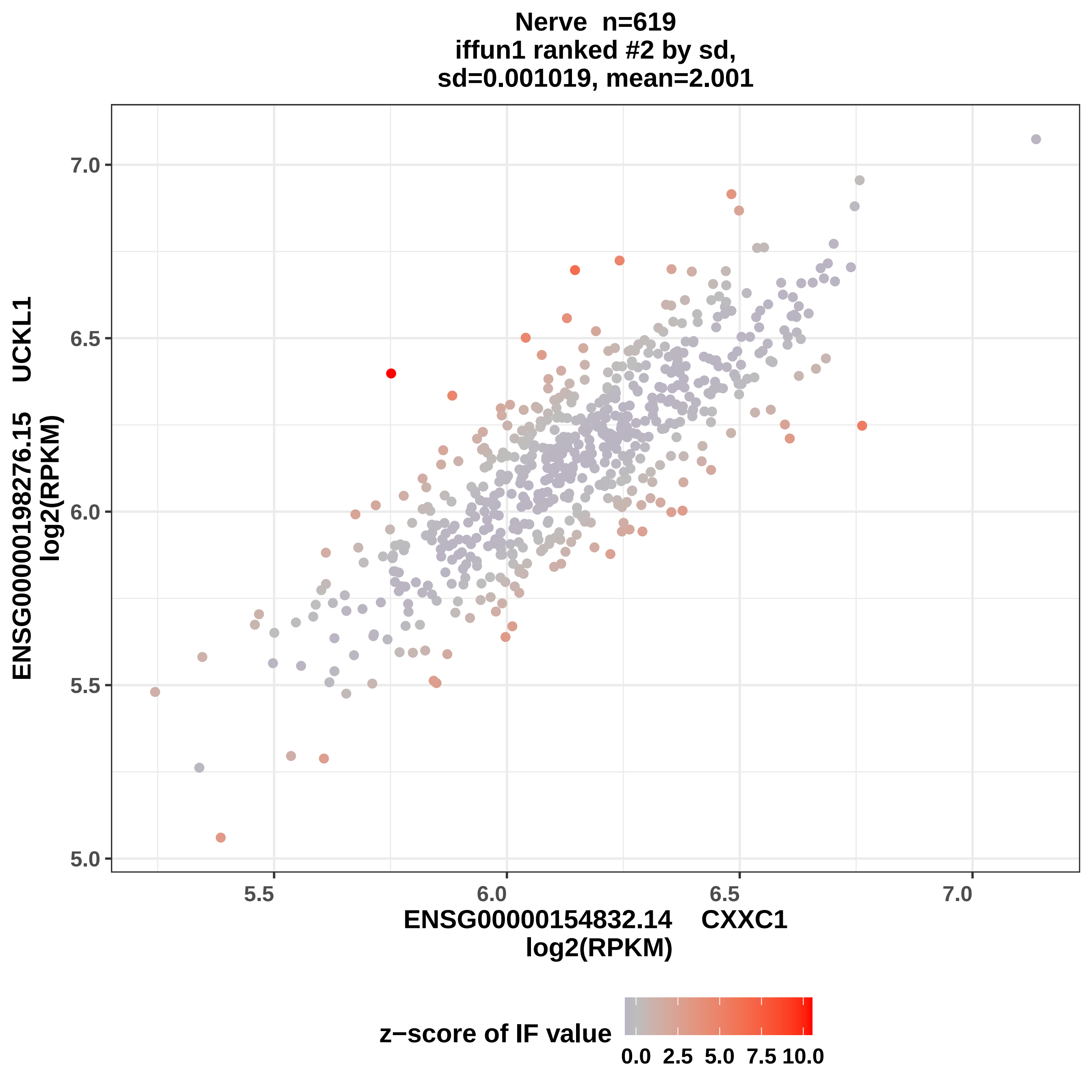
Nerve n=619
iffun1 ranked #2 by sd,
sd=0.001019, mean=2.001
ENSG00000198276.15 UCKL1
log2(RPKM)
7.0
6.5
6.0
5.5
5.0
5.5
6.0
6.5
7.0
ENSG00000154832.14 CXXC1
log2(RPKM)
z−score of IF value
0.0
2.5
5.0
7.5
10.0

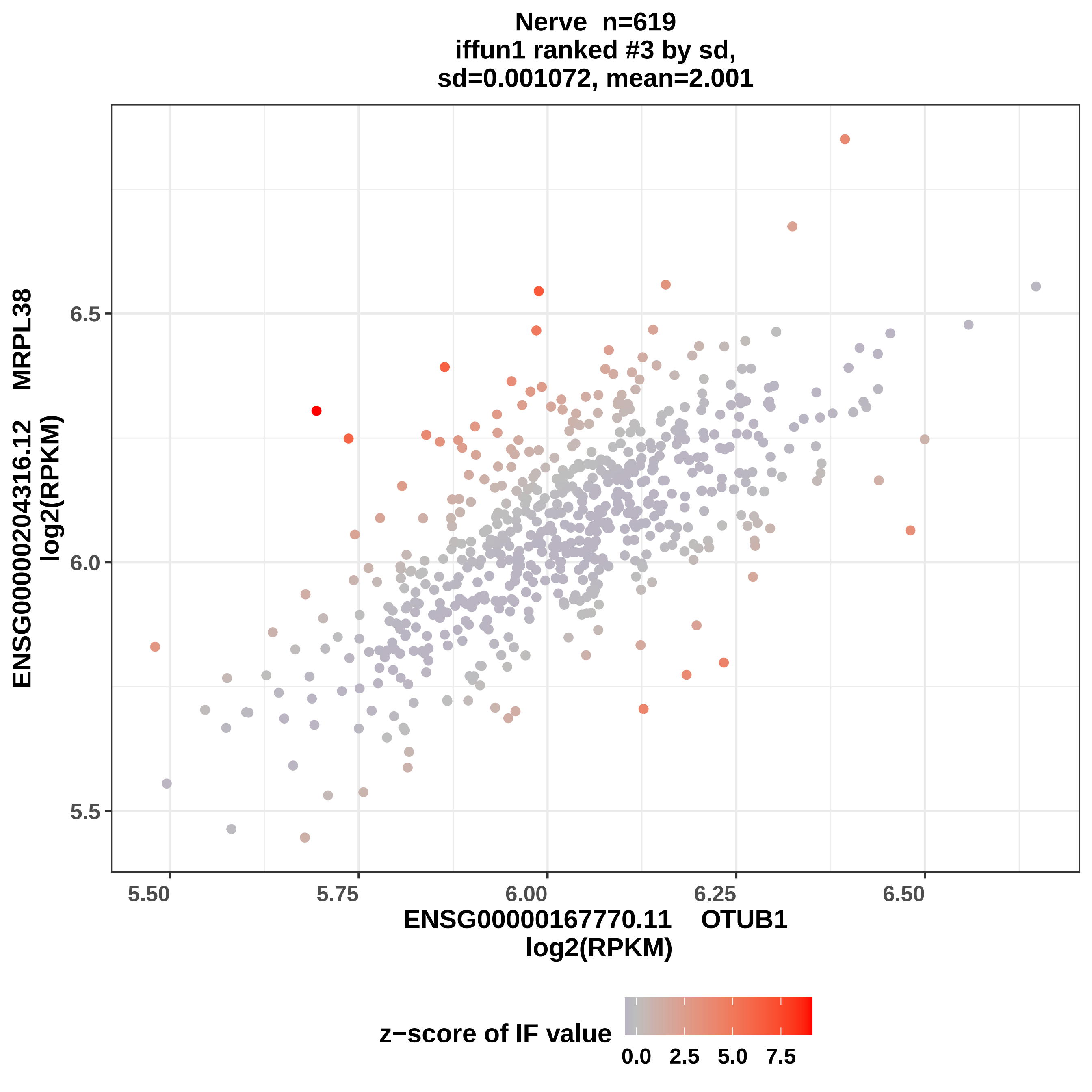

Nerve n=619
iffun1 ranked #3 by sd,
sd=0.001072, mean=2.001
ENSG00000204316.12 MRPL38
log2(RPKM)
6.5
6.0
5.5
5.50
5.75
6.00
6.25
6.50
ENSG00000167770.11 OTUB1
log2(RPKM)
z−score of IF value
0.0
2.5
5.0
7.5

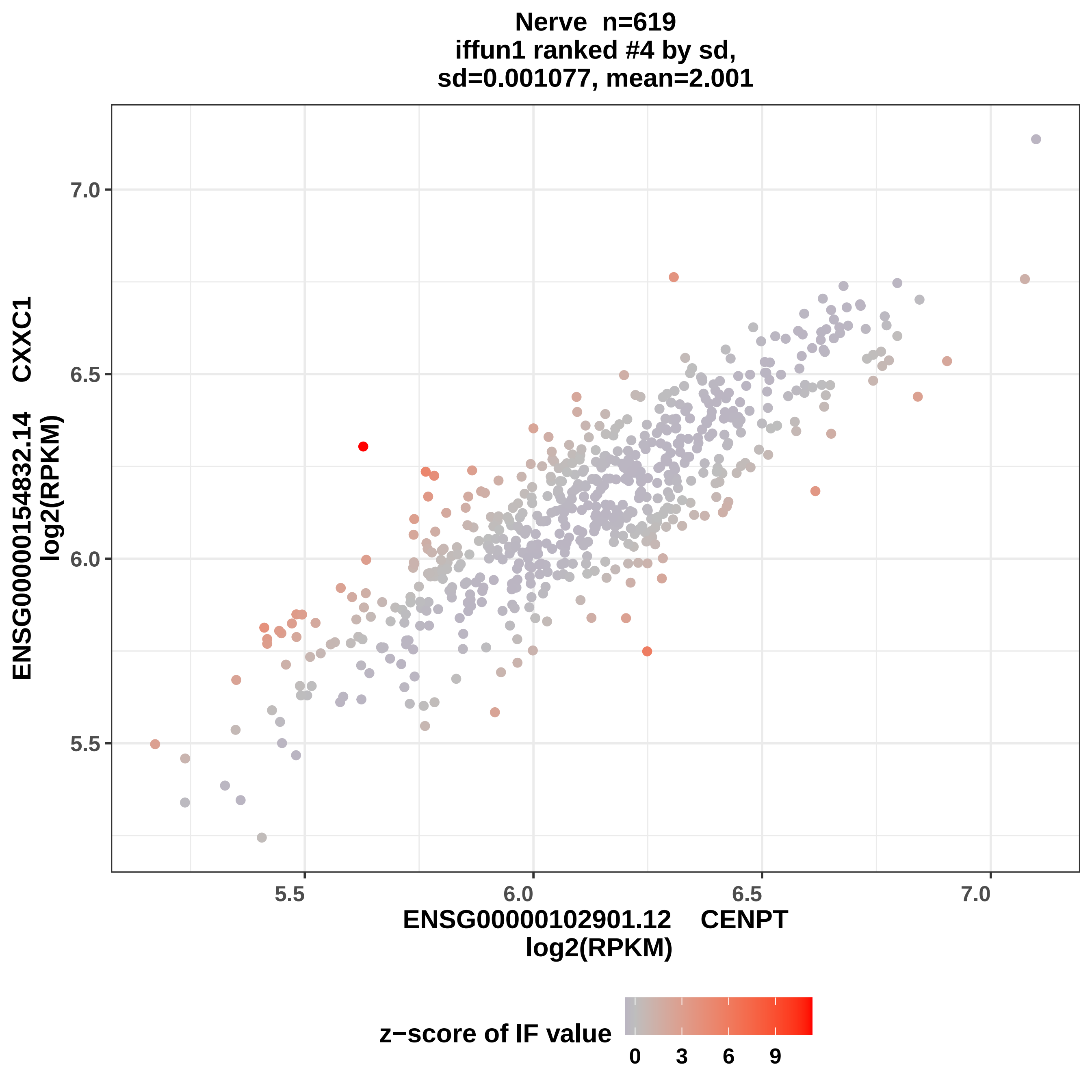

Nerve n=619
iffun1 ranked #4 by sd,
sd=0.001077, mean=2.001
ENSG00000154832.14 CXXC1
log2(RPKM)
7.0
6.5
6.0
5.5
5.5
6.0
6.5
7.0
ENSG00000102901.12 CENPT
log2(RPKM)
z−score of IF value
0
3
6
9

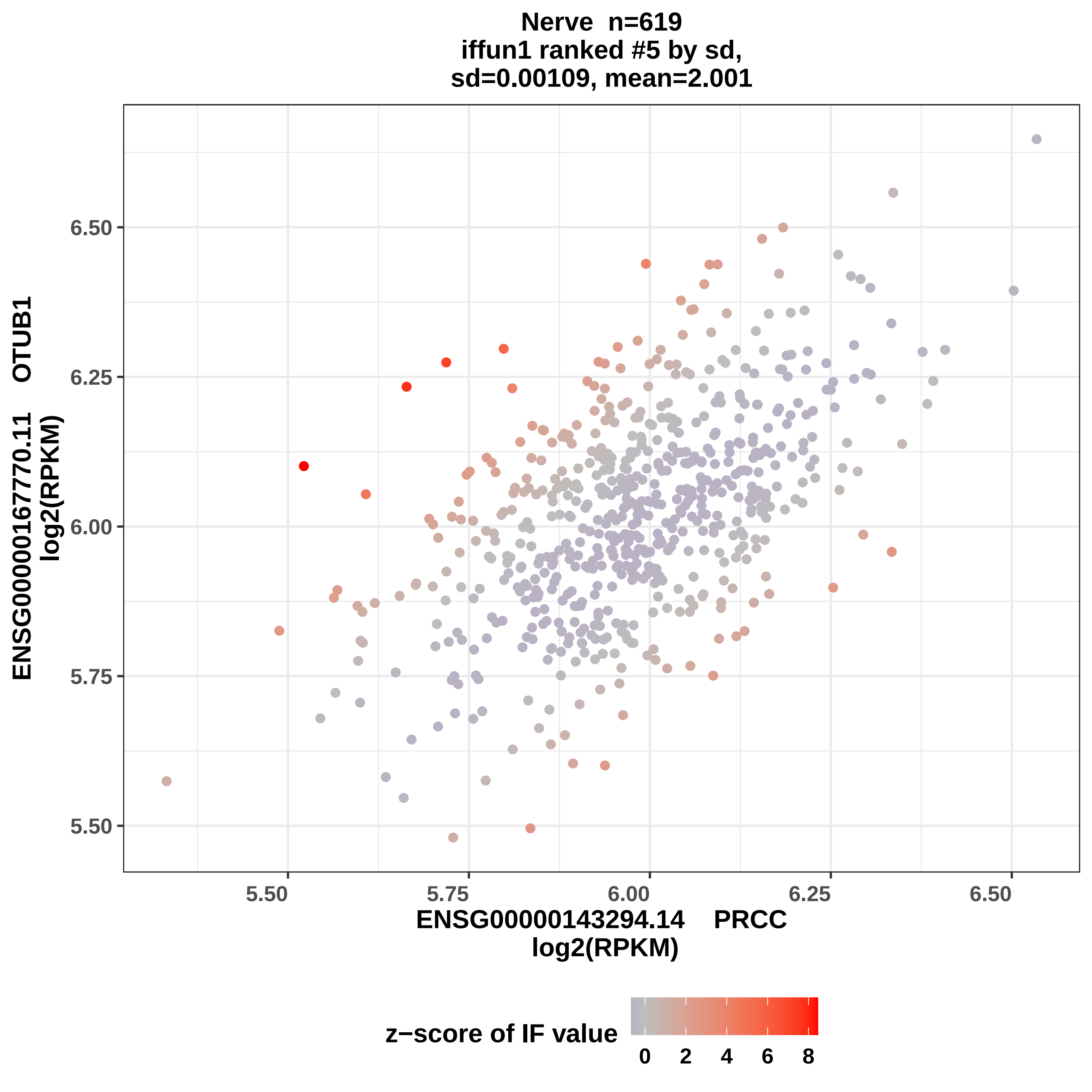

Nerve n=619
iffun1 ranked #5 by sd,
sd=0.00109, mean=2.001
ENSG00000167770.11 OTUB1
log2(RPKM)
6.50
6.25
6.00
5.75
5.50
5.50
5.75
6.00
6.25
6.50
ENSG00000143294.14 PRCC
log2(RPKM)
z−score of IF value
0
2
4
6
8

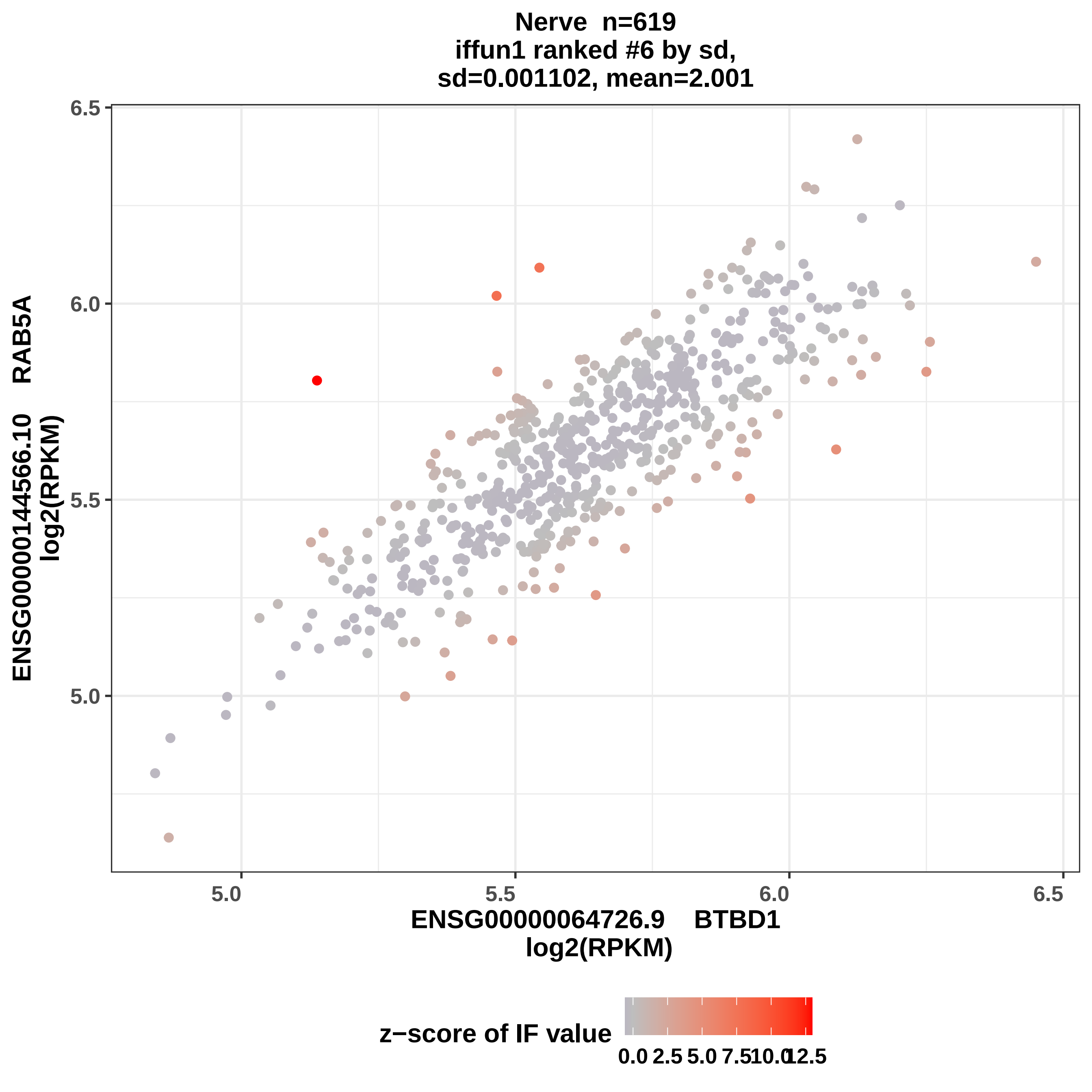
Nerve n=619
iffun1 ranked #6 by sd,
sd=0.001102, mean=2.001
ENSG00000144566.10 RAB5A
log2(RPKM)
6.5
6.0
5.5
5.0
ENSG00000064726.9 BTBD1
log2(RPKM)
5.0
5.5
6.0
6.5
z−score of IF value
0.0
2.5
5.0
7.5
10.0
12.5

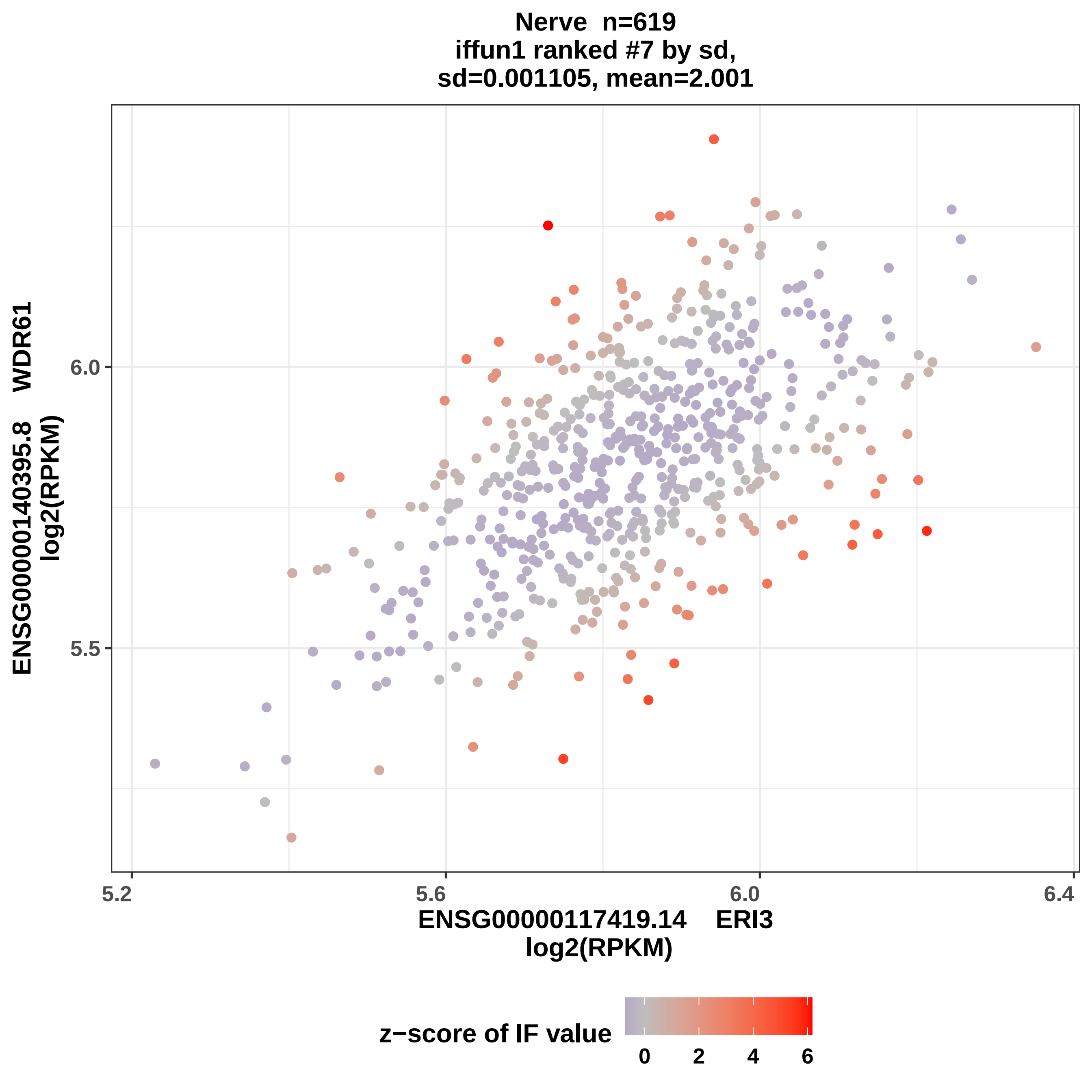

Nerve n=619
iffun1 ranked #7 by sd,
sd=0.001105, mean=2.001
ENSG00000140395.8 WDR61
log2(RPKM)
6.0
5.5
5.2
5.6
6.0
6.4
ENSG00000117419.14 ERI3
log2(RPKM)
z−score of IF value
0
2
4
6

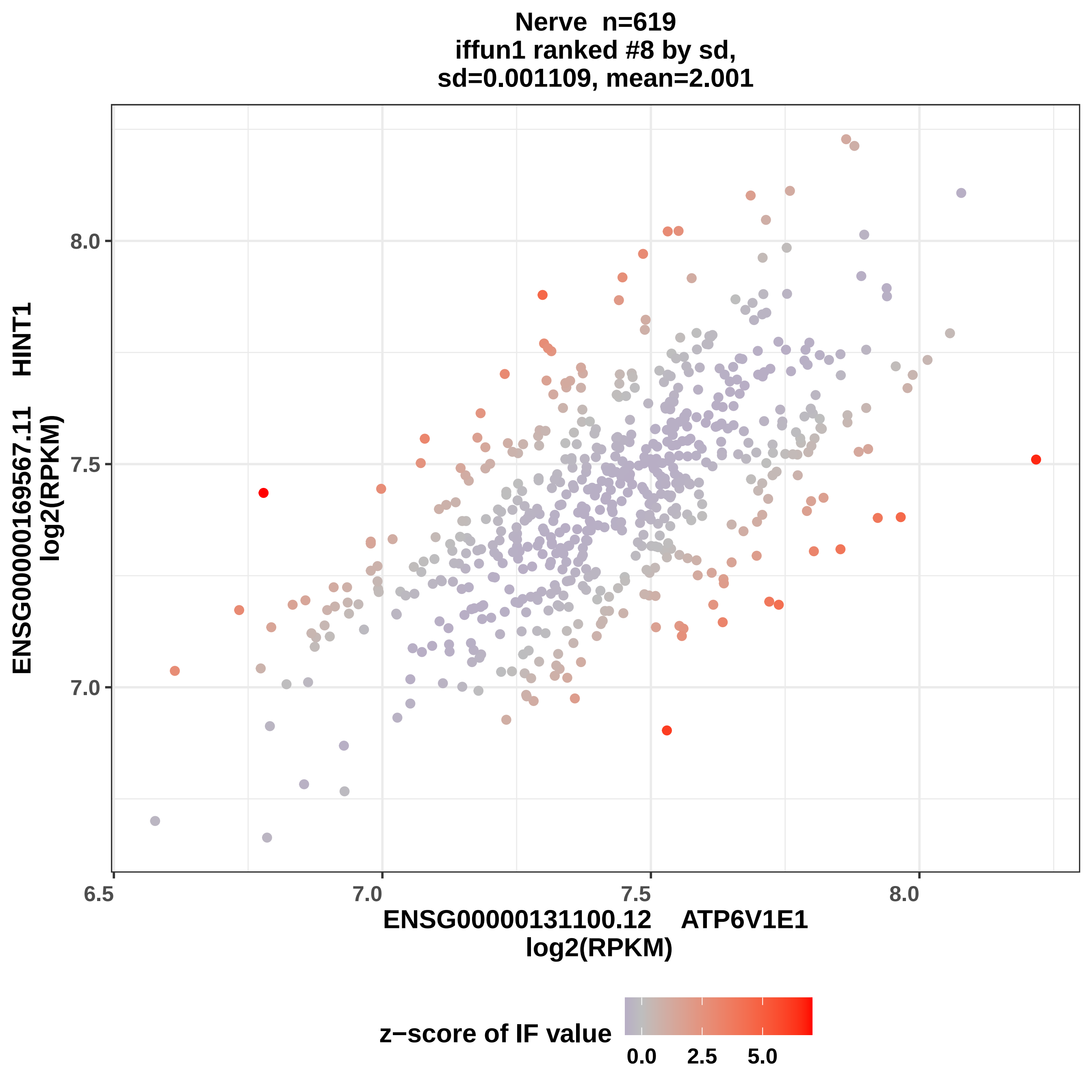
Nerve n=619
iffun1 ranked #8 by sd,
sd=0.001109, mean=2.001
ENSG00000169567.11 HINT1
log2(RPKM)
8.0
7.5
7.0
6.5
7.0
7.5
8.0
ENSG00000131100.12 ATP6V1E1
log2(RPKM)
z−score of IF value
0.0
2.5
5.0

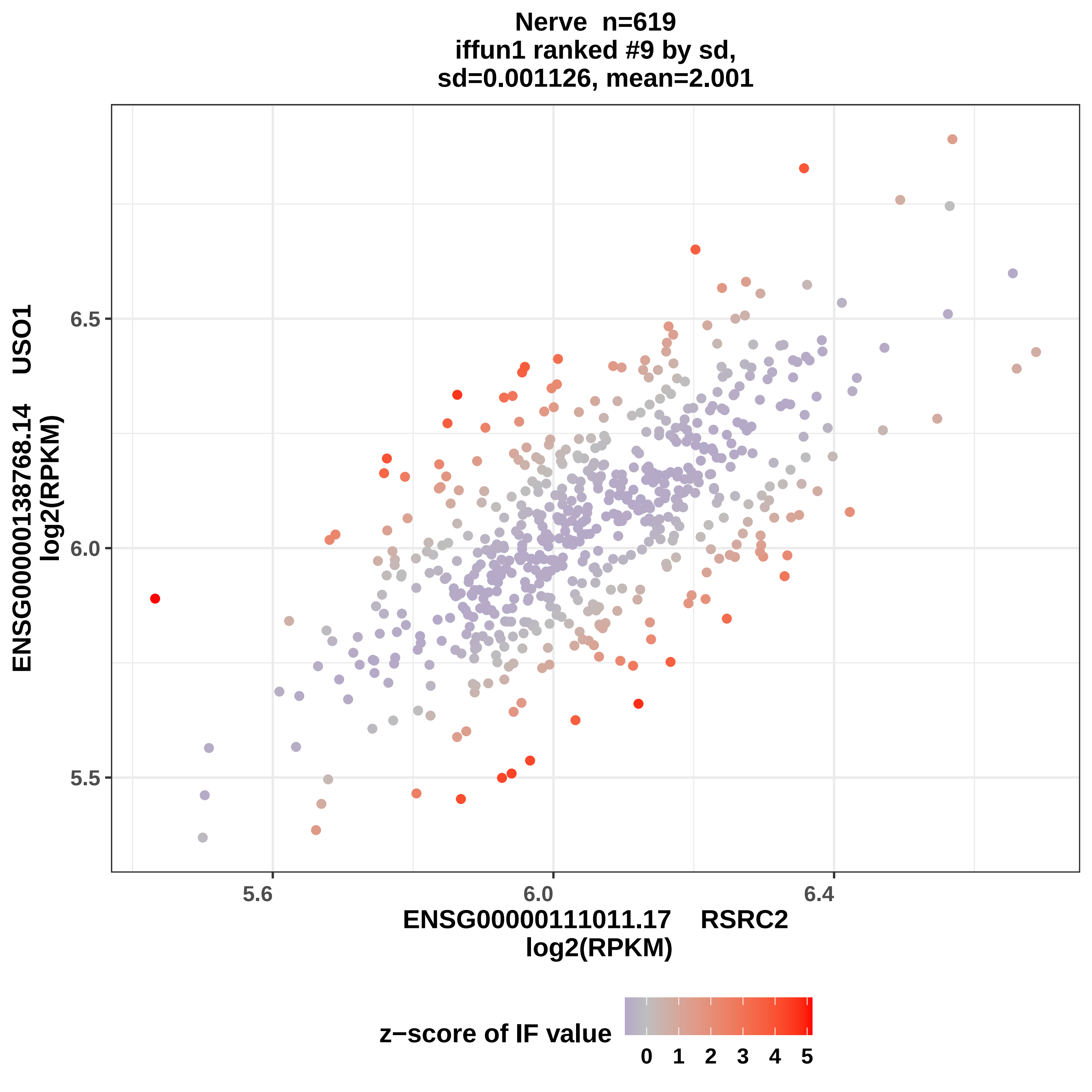

Nerve n=619
iffun1 ranked #9 by sd,
sd=0.001126, mean=2.001
ENSG00000138768.14 USO1
log2(RPKM)
6.5
6.0
5.5
5.6
6.0
6.4
ENSG00000111011.17 RSRC2
log2(RPKM)
z−score of IF value
0
1
2
3
4
5

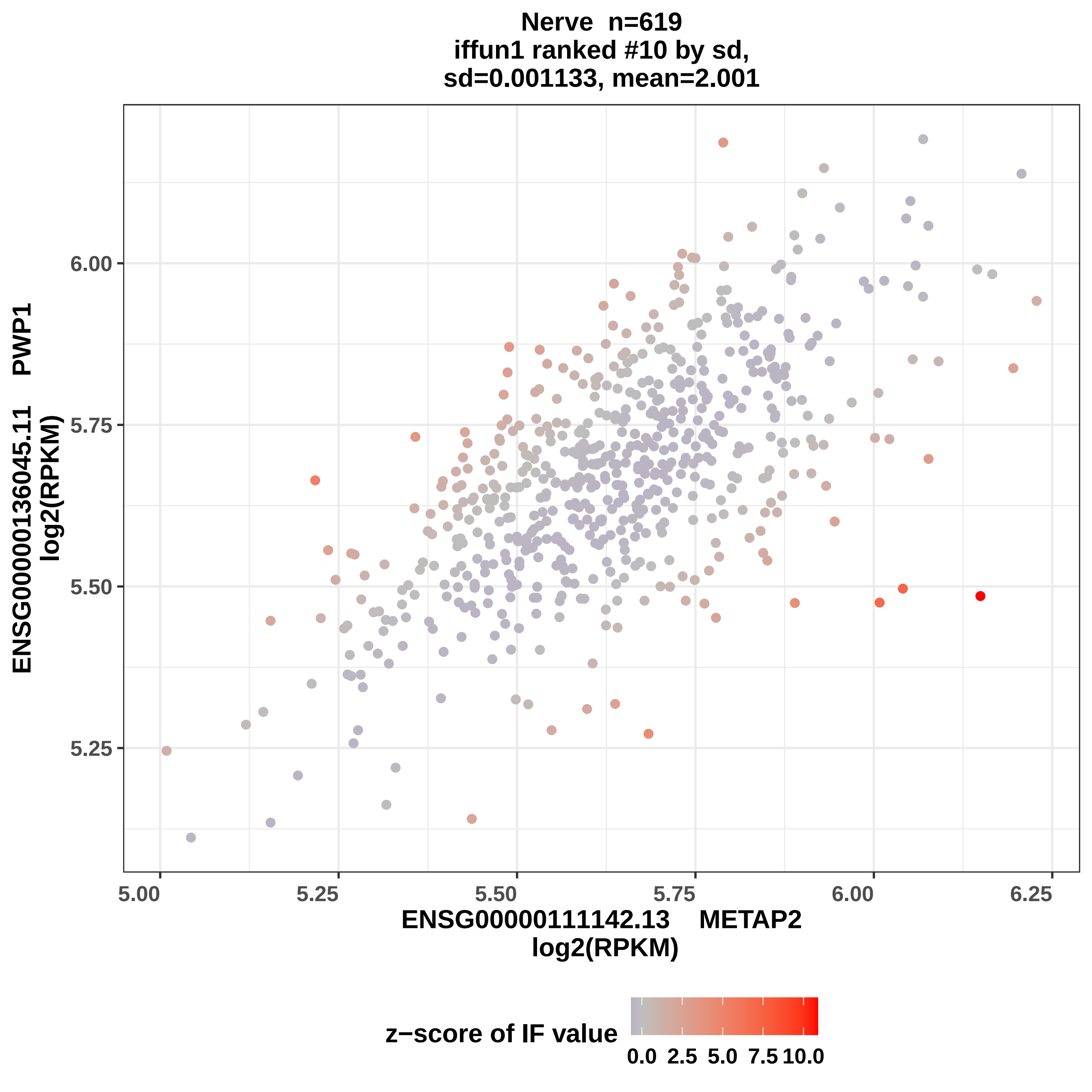

Nerve n=619
iffun1 ranked #10 by sd,
sd=0.001133, mean=2.001
ENSG00000136045.11 PWP1
log2(RPKM)
6.00
5.75
5.50
5.25
5.00
5.25
5.50
5.75
6.00
6.25
ENSG00000111142.13 METAP2
log2(RPKM)
z−score of IF value
0.0
2.5
5.0
7.5
10.0

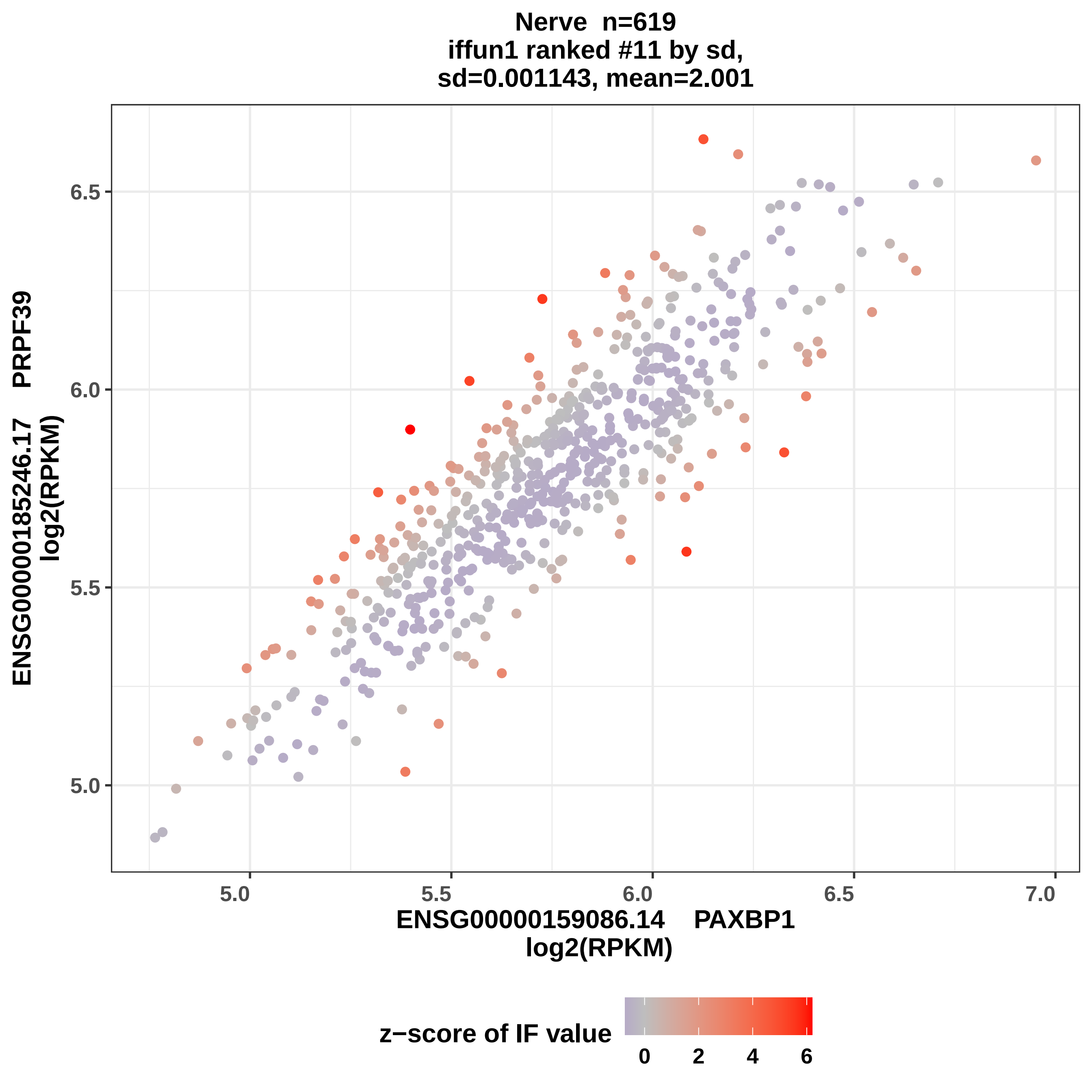
Nerve n=619
iffun1 ranked #11 by sd,
sd=0.001143, mean=2.001
ENSG00000185246.17 PRPF39
log2(RPKM)
6.5
6.0
5.5
5.0
5.0
5.5
6.0
6.5
7.0
ENSG00000159086.14 PAXBP1
log2(RPKM)
z−score of IF value
0
2
4
6

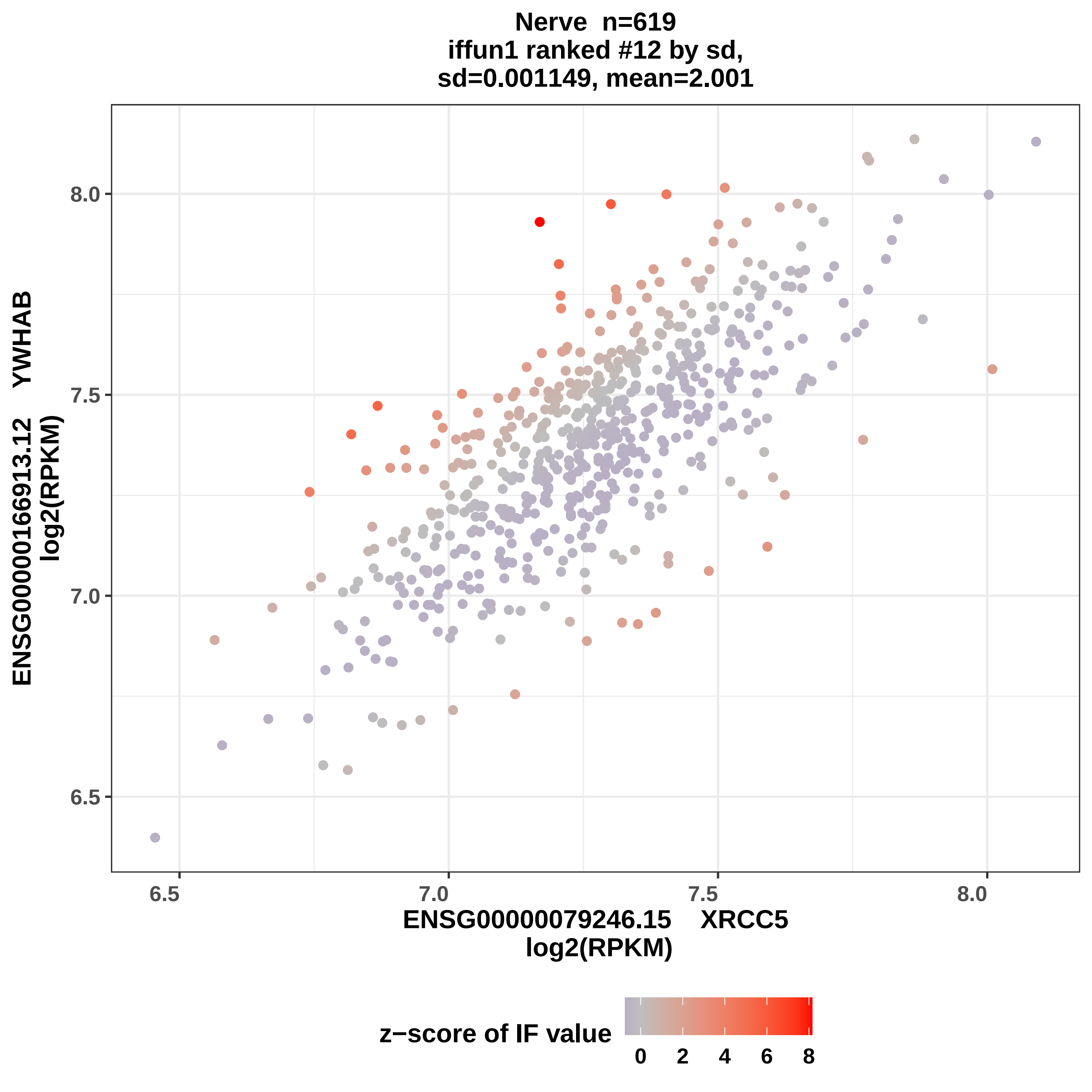

Nerve n=619
iffun1 ranked #12 by sd,
sd=0.001149, mean=2.001
ENSG00000166913.12 YWHAB
log2(RPKM)
ENSG00000079246.15 XRCC5
log2(RPKM)
6.5
7.0
7.5
8.0
z−score of IF value
0
2
4
6
8

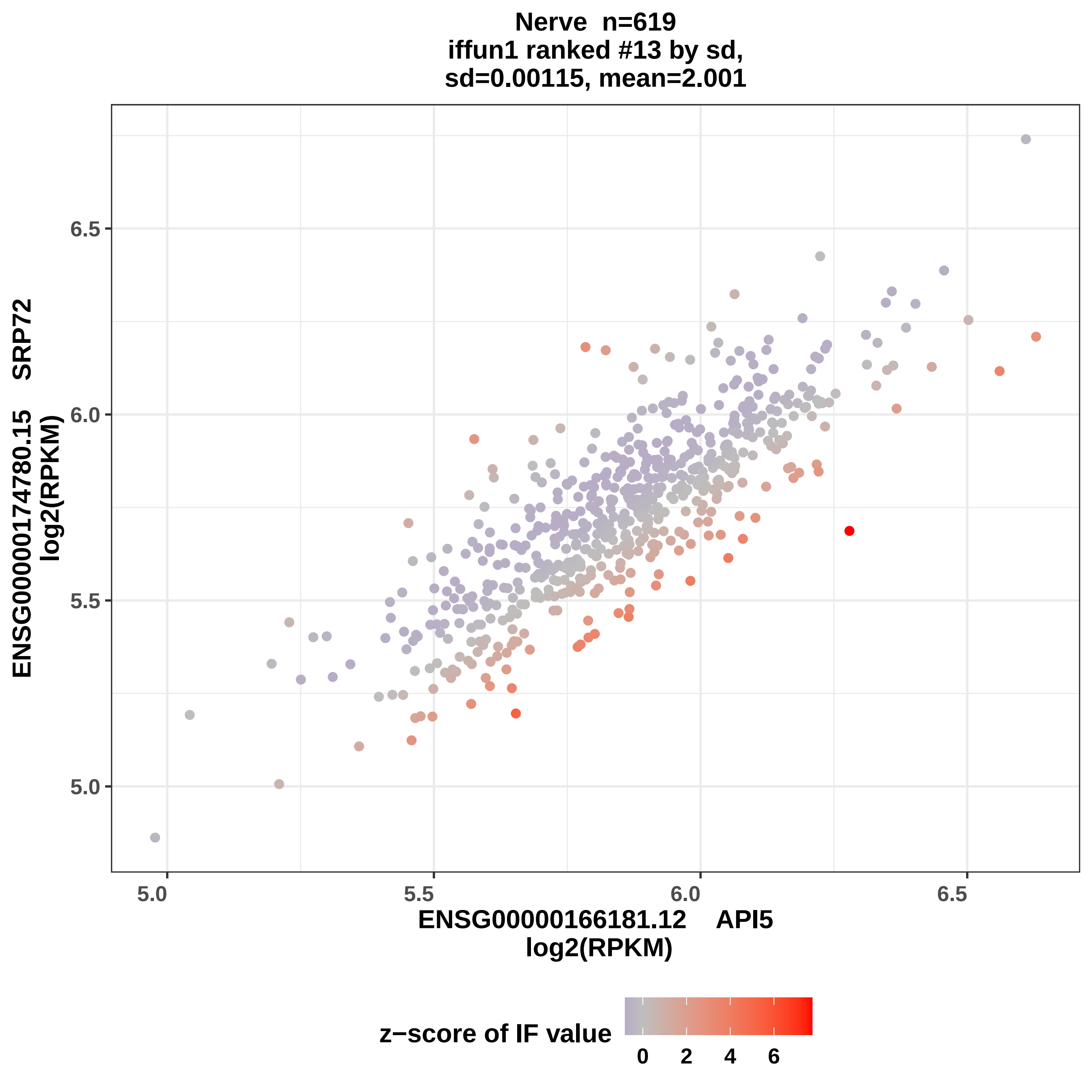

Nerve n=619
iffun1 ranked #13 by sd,
sd=0.00115, mean=2.001
ENSG00000174780.15 SRP72
log2(RPKM)
6.5
6.0
5.5
5.0
5.0
5.5
6.0
6.5
ENSG00000166181.12 API5
log2(RPKM)
z−score of IF value
0
2
4
6

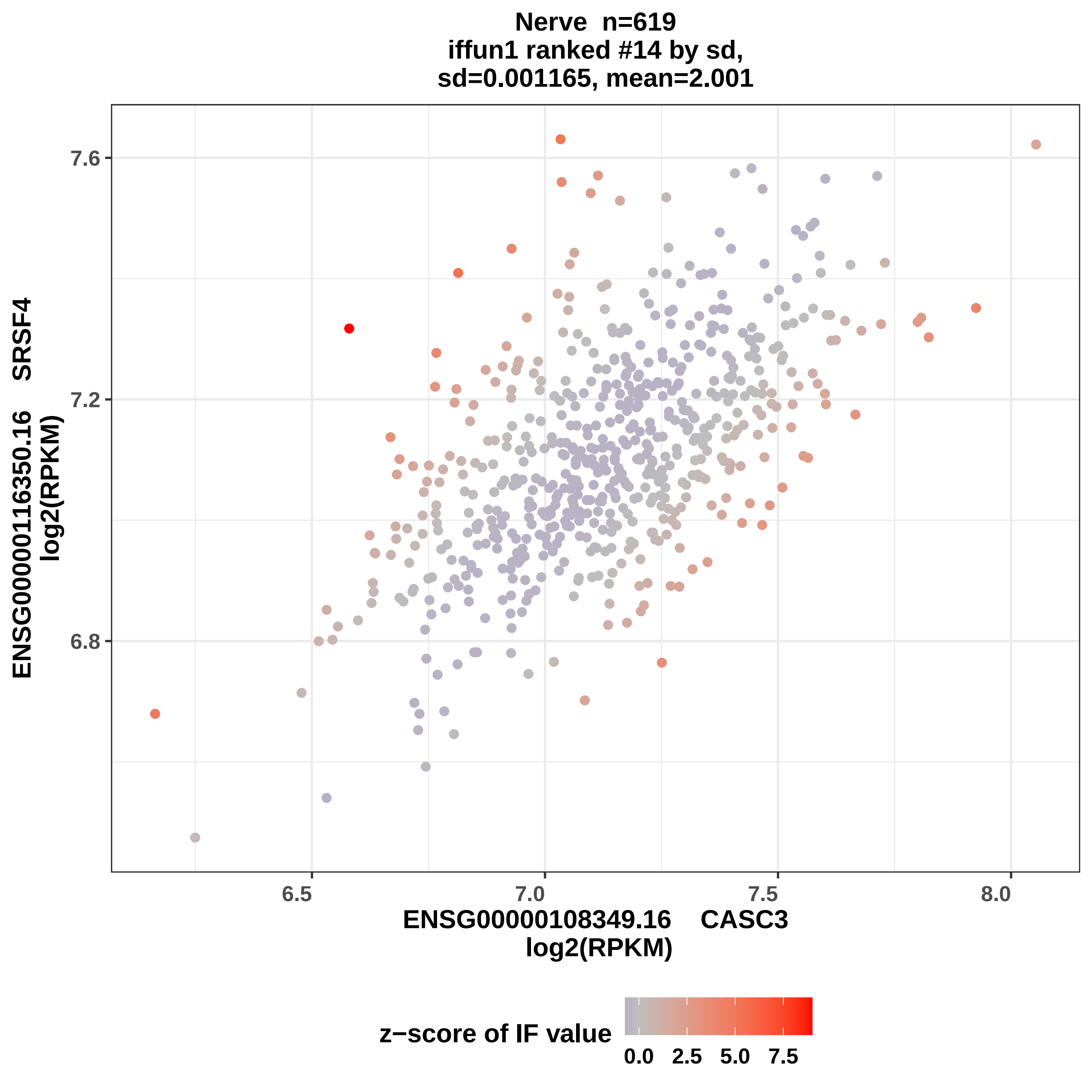

Nerve n=619
iffun1 ranked #14 by sd,
sd=0.001165, mean=2.001
7.6
7.2
6.8
6.5
7.0
7.5
8.0
ENSG00000116350.16 SRSF4
log2(RPKM)
ENSG00000108349.16 CASC3
log2(RPKM)
z−score of IF value
0.0
2.5
5.0
7.5

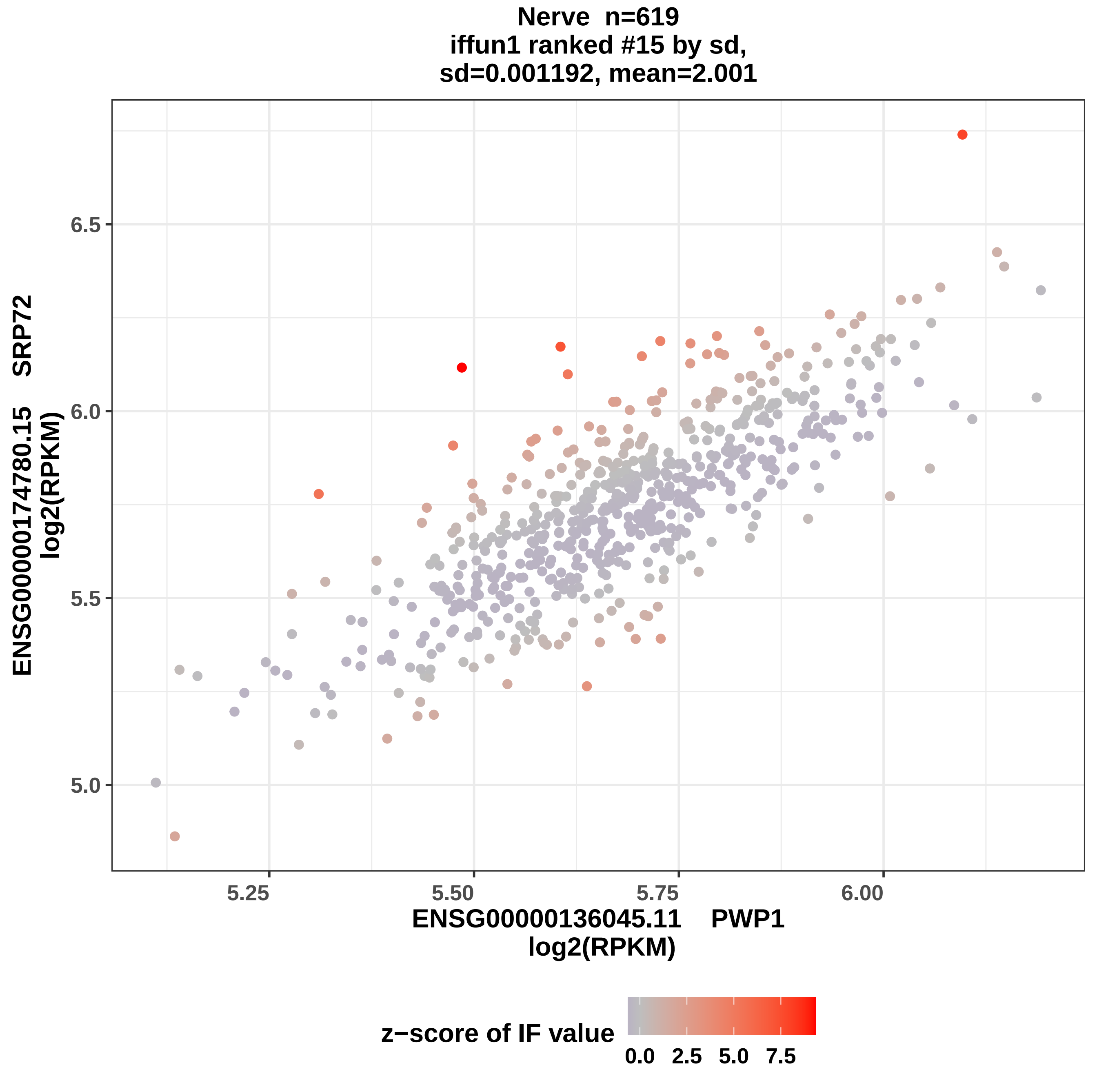
Nerve n=619
iffun1 ranked #15 by sd,
sd=0.001192, mean=2.001
ENSG00000174780.15 SRP72
log2(RPKM)
6.5
6.0
5.5
5.0
5.25
5.50
5.75
6.00
ENSG00000136045.11 PWP1
log2(RPKM)
z−score of IF value
0.0
2.5
5.0
7.5

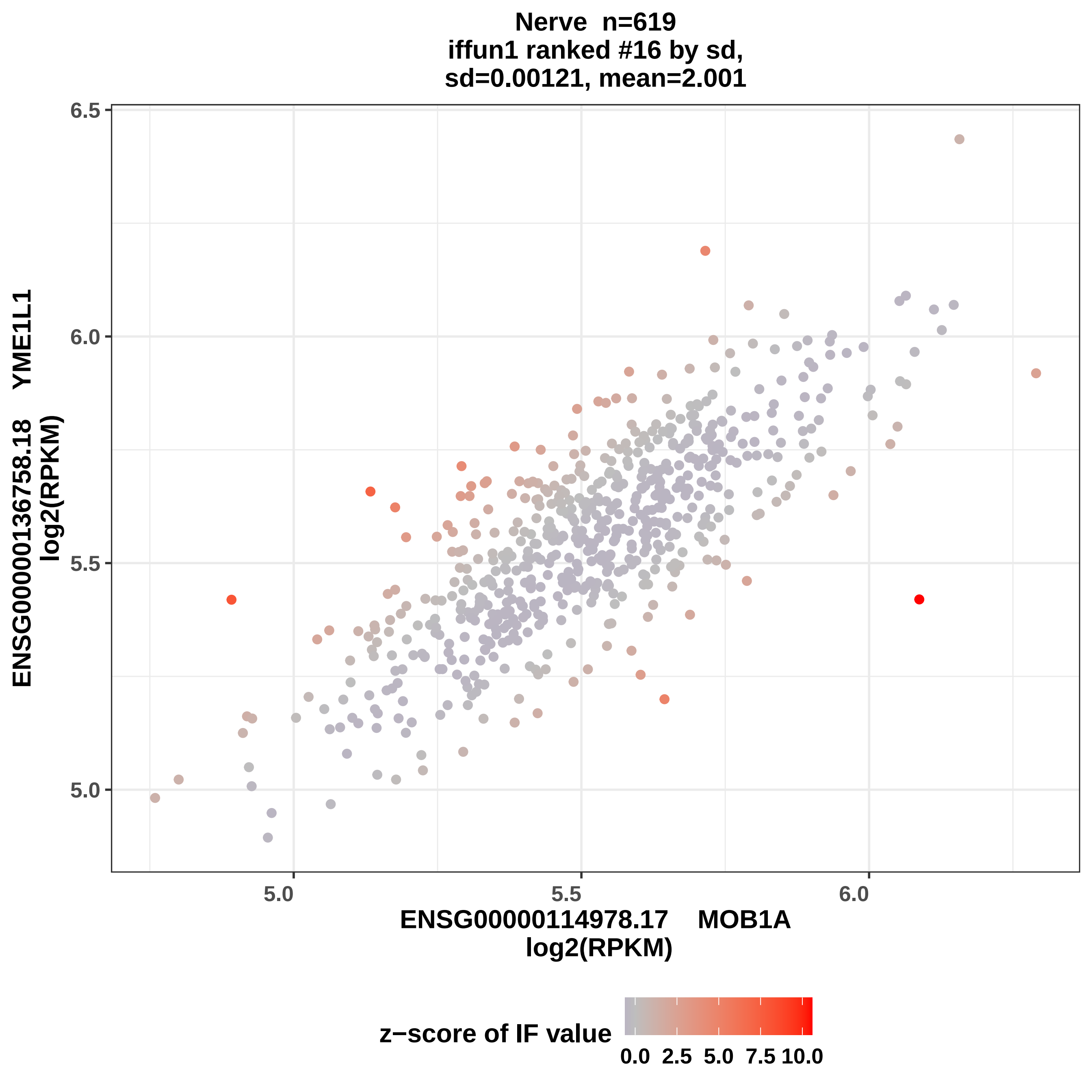
Nerve n=619
iffun1 ranked #16 by sd,
sd=0.00121, mean=2.001
ENSG00000136758.18 YME1L1
log2(RPKM)
6.5
6.0
5.5
5.0
5.0
5.5
6.0
ENSG00000114978.17 MOB1A
log2(RPKM)
z−score of IF value
0.0 2.5 5.0 7.5 10.0

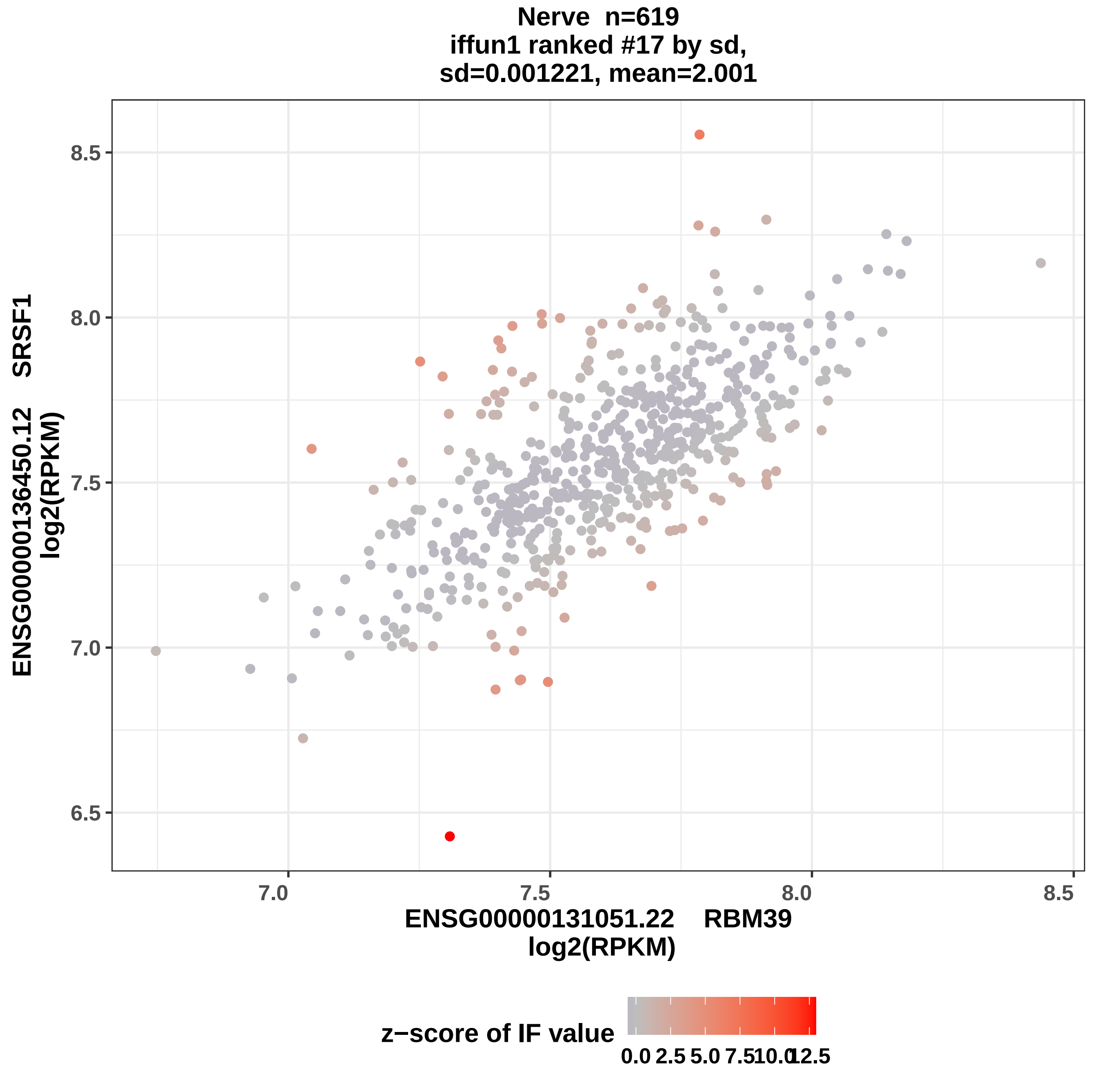
Nerve n=619
iffun1 ranked #17 by sd,
sd=0.001221, mean=2.001
ENSG00000136450.12 SRSF1
log2(RPKM)
8.5
8.0
7.5
7.0
6.5
7.0
7.5
8.0
8.5
ENSG00000131051.22 RBM39
log2(RPKM)
z−score of IF value
0.0
2.5
5.0
7.5
10.0
12.5

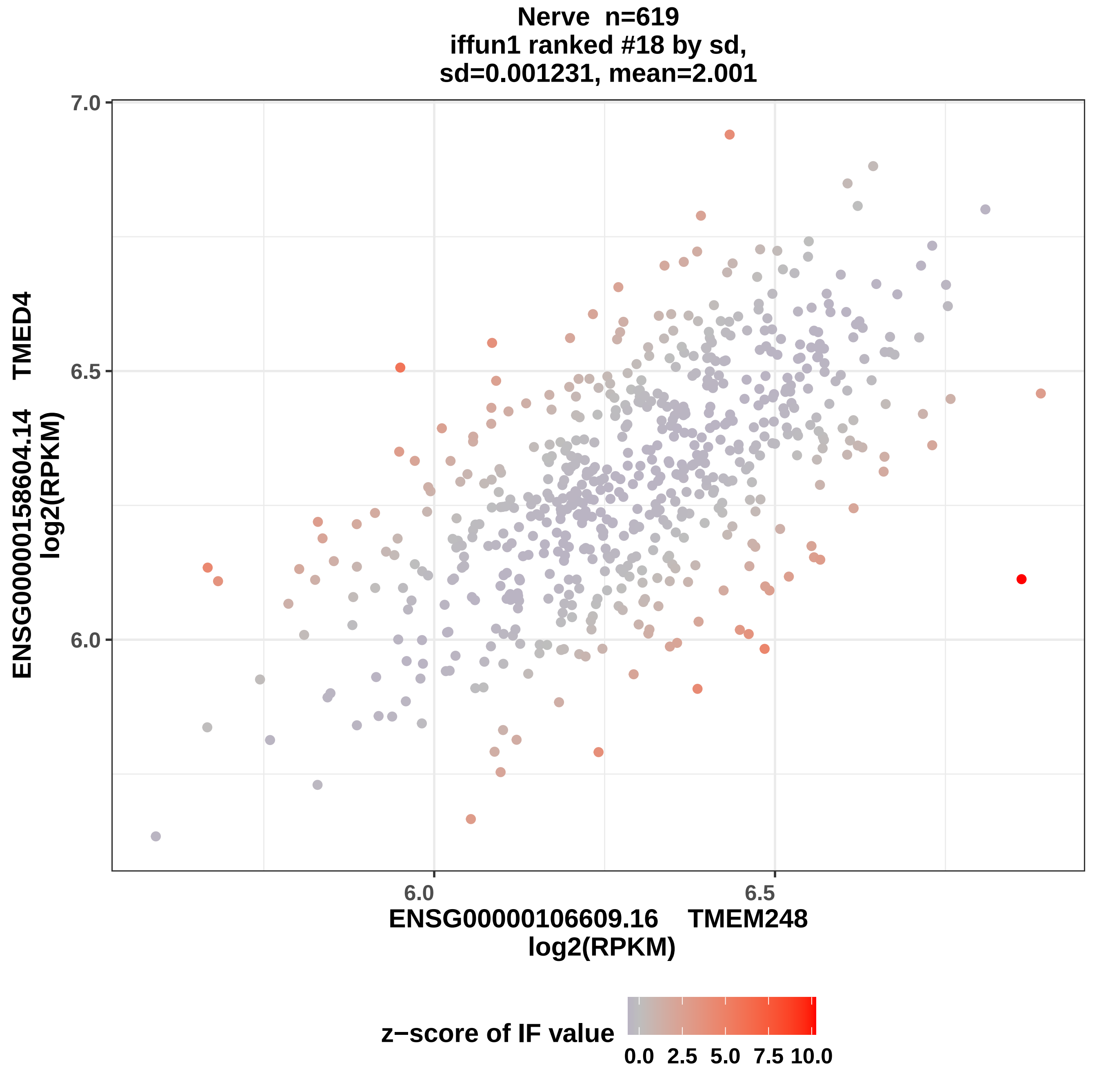
Nerve n=619
iffun1 ranked #18 by sd,
sd=0.001231, mean=2.001
7.0
6.5
6.0
6.0
6.5
ENSG00000158604.14 TMED4
log2(RPKM)
ENSG00000106609.16 TMEM248
log2(RPKM)
z−score of IF value
0.0
2.5
5.0
7.5
10.0

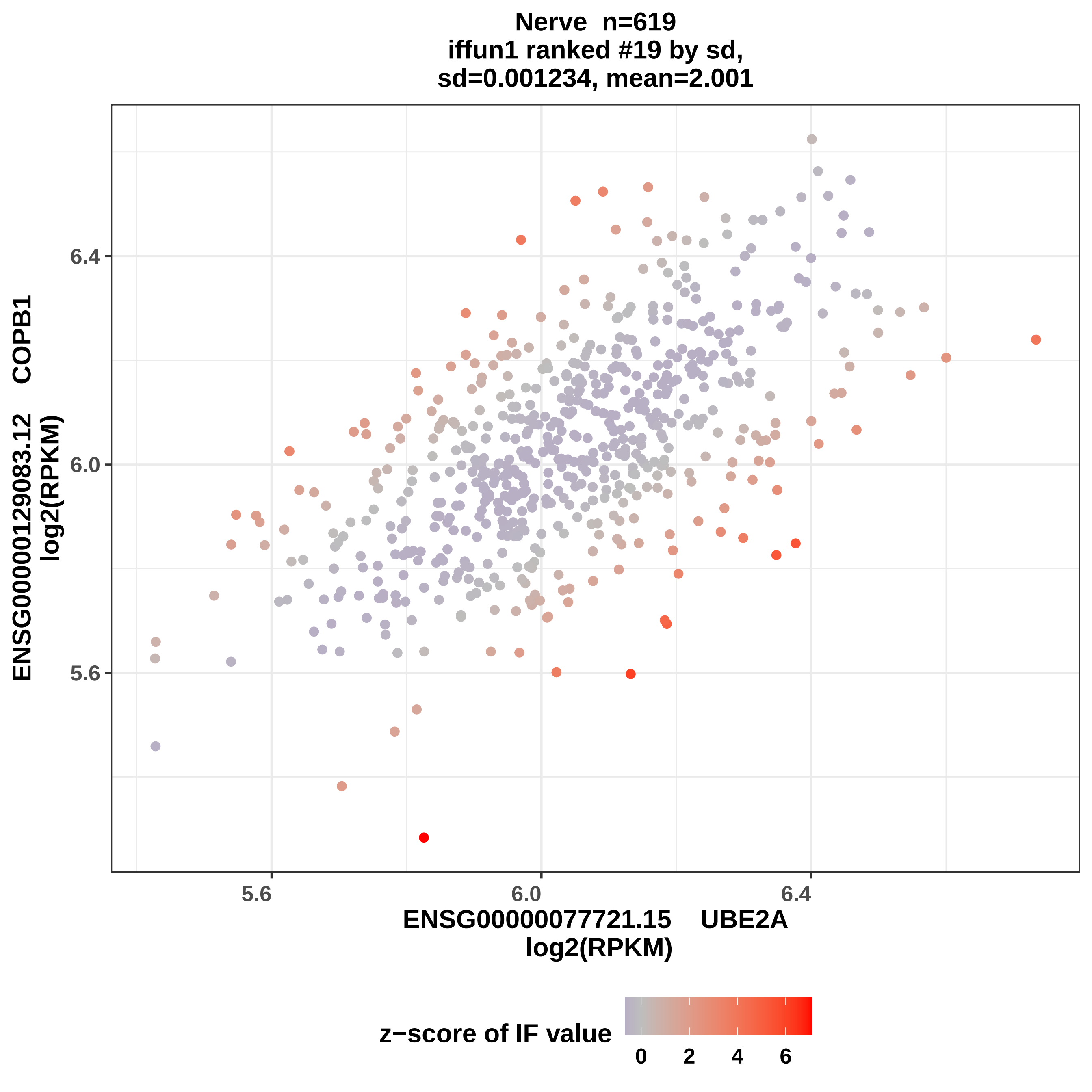

Nerve n=619
iffun1 ranked #19 by sd,
sd=0.001234, mean=2.001
ENSG00000129083.12 COPB1
log2(RPKM)
6.4
6.0
5.6
5.6
6.0
6.4
ENSG00000077721.15 UBE2A
log2(RPKM)
z−score of IF value
0
2
4
6

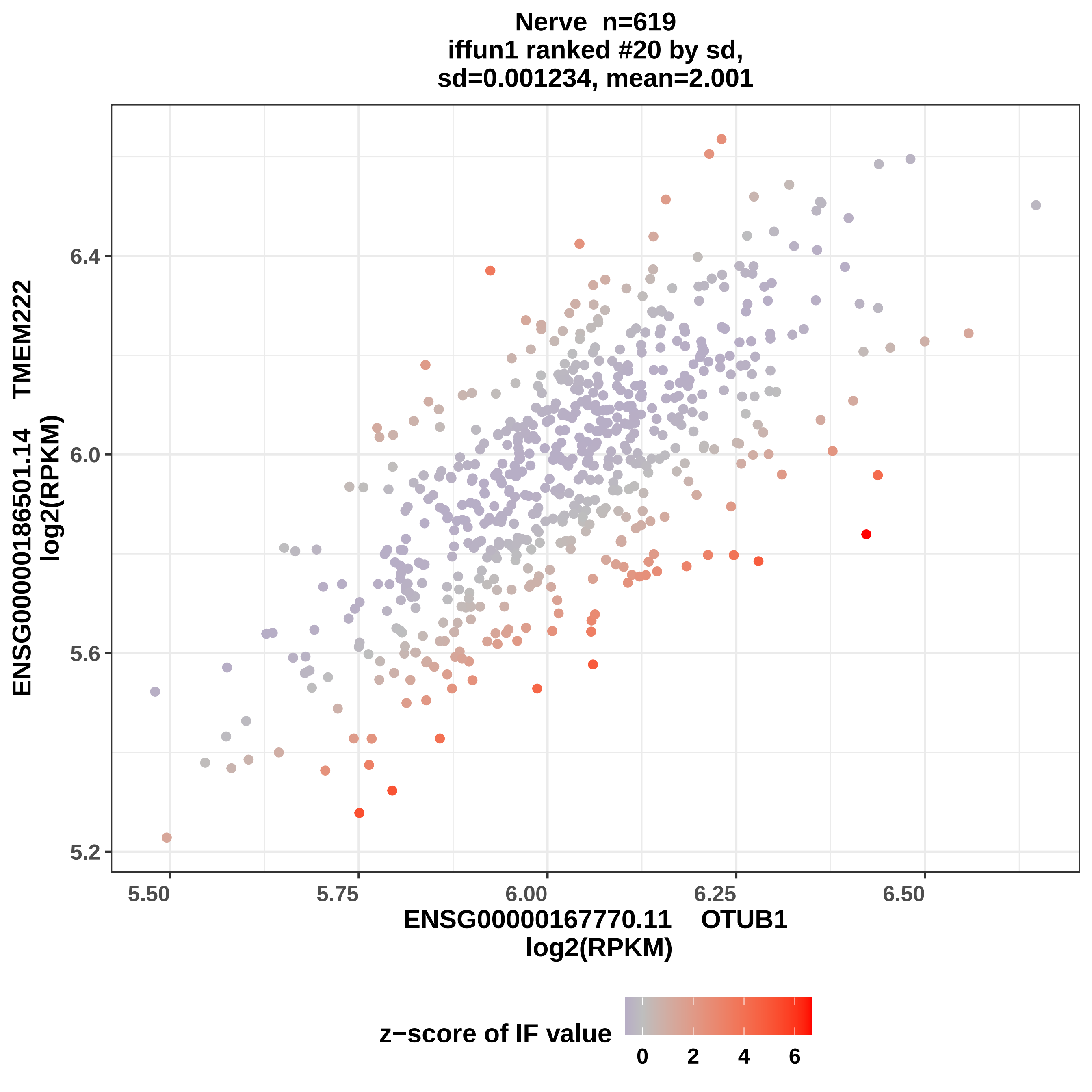

Nerve n=619
iffun1 ranked #20 by sd,
sd=0.001234, mean=2.001
ENSG00000186501.14 TMEM222
log2(RPKM)
6.4
6.0
5.6
5.2
5.50
5.75
6.00
6.25
6.50
ENSG00000167770.11 OTUB1
log2(RPKM)
z−score of IF value
0
2
4
6

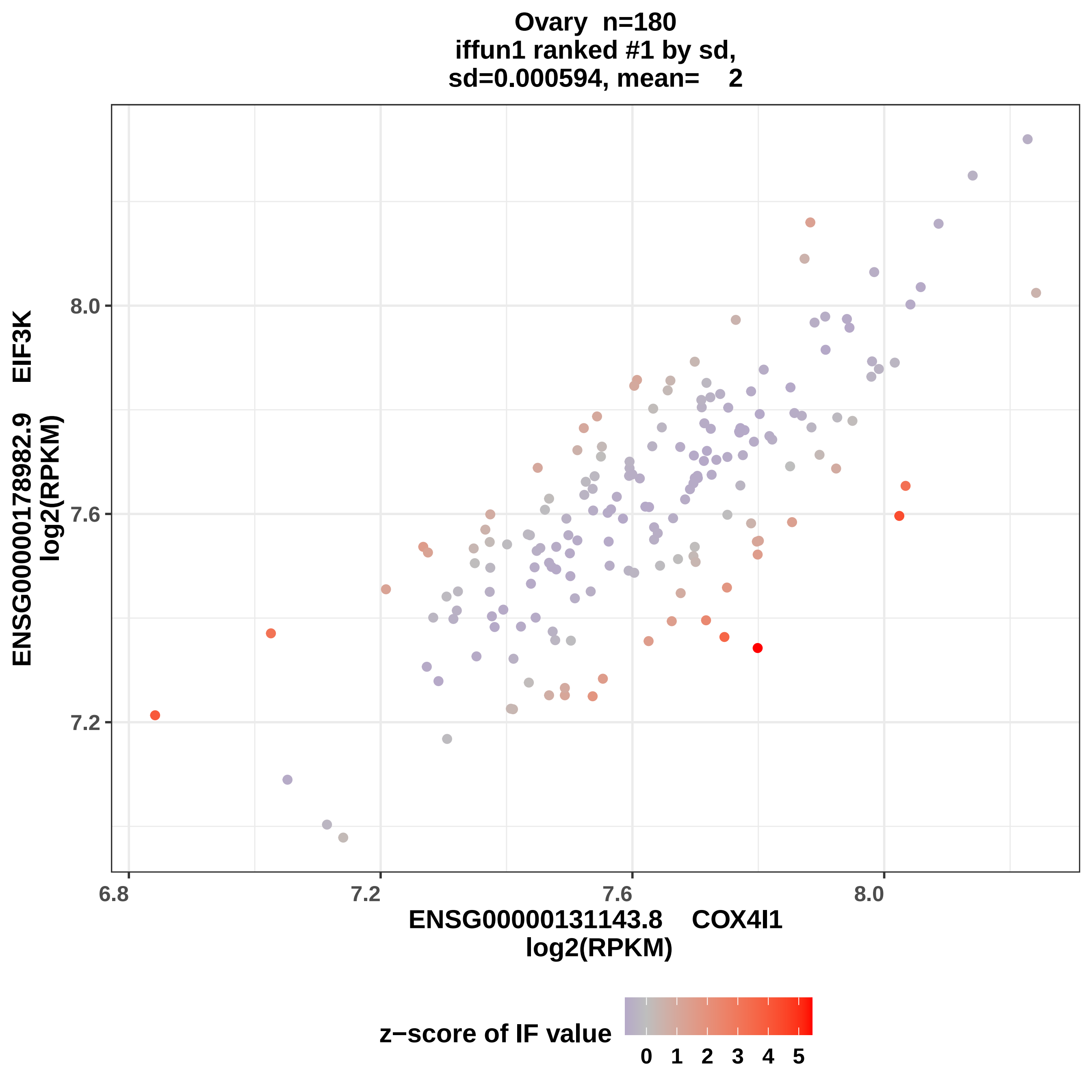

Ovary n=180
iffun1 ranked #1 by sd,
sd=0.000594, mean= 2
ENSG00000178982.9 EIF3K
log2(RPKM)
8.0
7.6
7.2
6.8
7.2
7.6
8.0
ENSG00000131143.8 COX4I1
log2(RPKM)
z−score of IF value
0 1 2 3 4 5

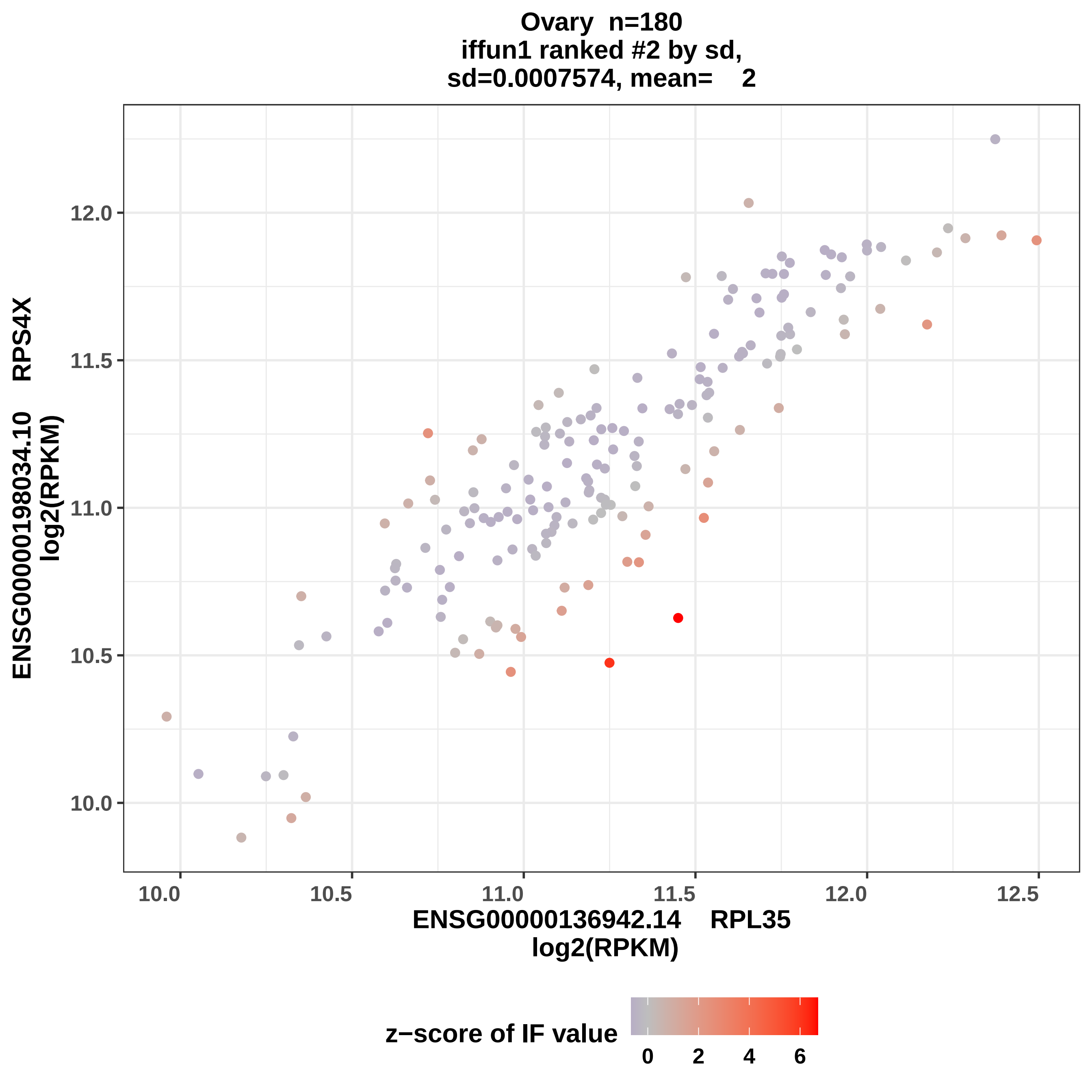

Ovary n=180
iffun1 ranked #2 by sd,
sd=0.0007574, mean= 2
ENSG00000198034.10 RPS4X
log2(RPKM)
12.0
11.5
11.0
10.5
10.0
10.0
10.5
11.0
11.5
12.0
12.5
ENSG00000136942.14 RPL35
log2(RPKM)
z−score of IF value
0
2
4
6

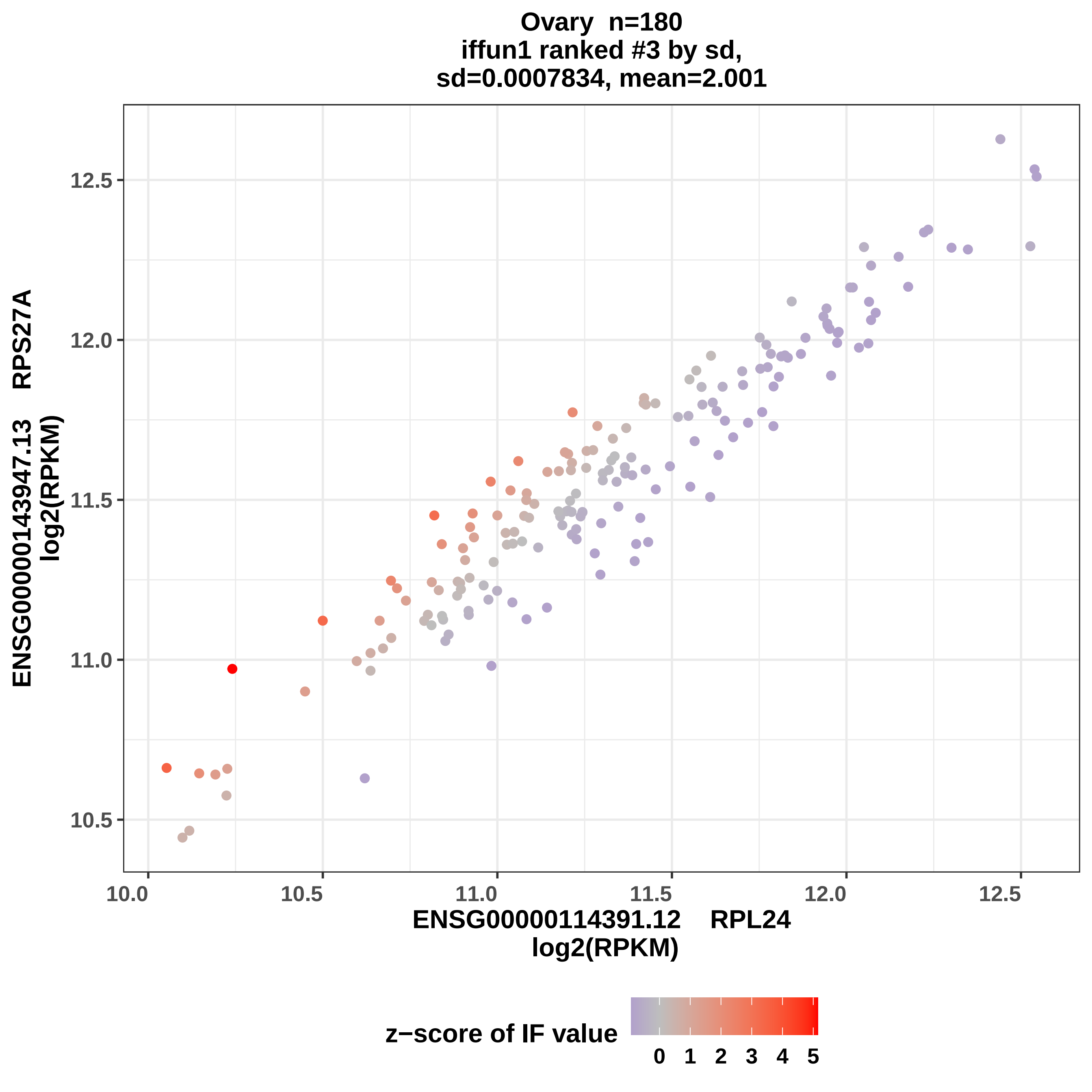

Ovary n=180
iffun1 ranked #3 by sd,
sd=0.0007834, mean=2.001
ENSG00000143947.13 RPS27A
log2(RPKM)
12.5
12.0
11.5
11.0
10.5
10.0
10.5
11.0
11.5
12.0
12.5
ENSG00000114391.12 RPL24
log2(RPKM)
z−score of IF value
0
1
2
3
4
5

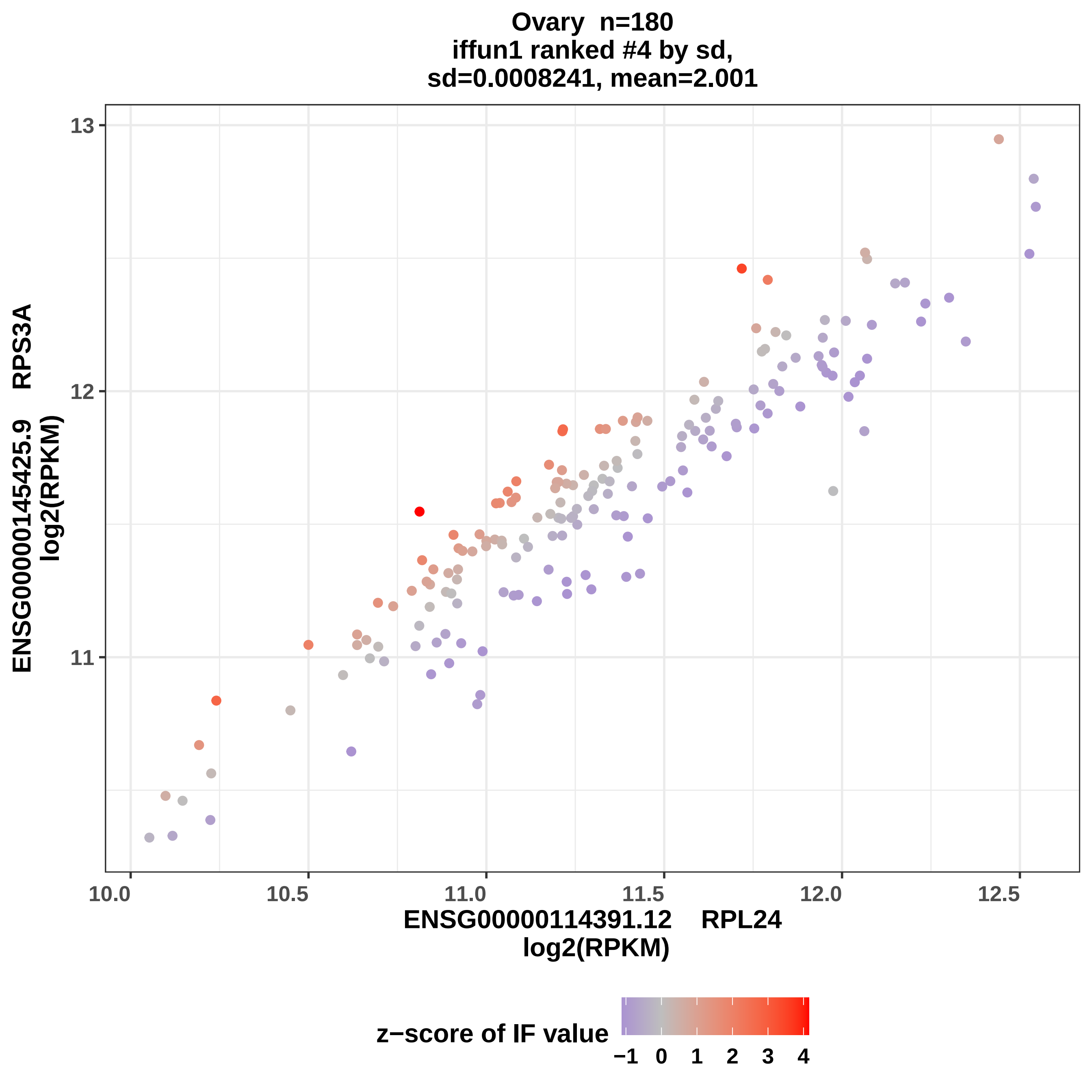

Ovary n=180
iffun1 ranked #4 by sd,
sd=0.0008241, mean=2.001
ENSG00000145425.9 RPS3A
log2(RPKM)
13
12
11
10.0
10.5
11.0
11.5
12.0
12.5
ENSG00000114391.12 RPL24
log2(RPKM)
z−score of IF value
−1
0
1
2
3
4

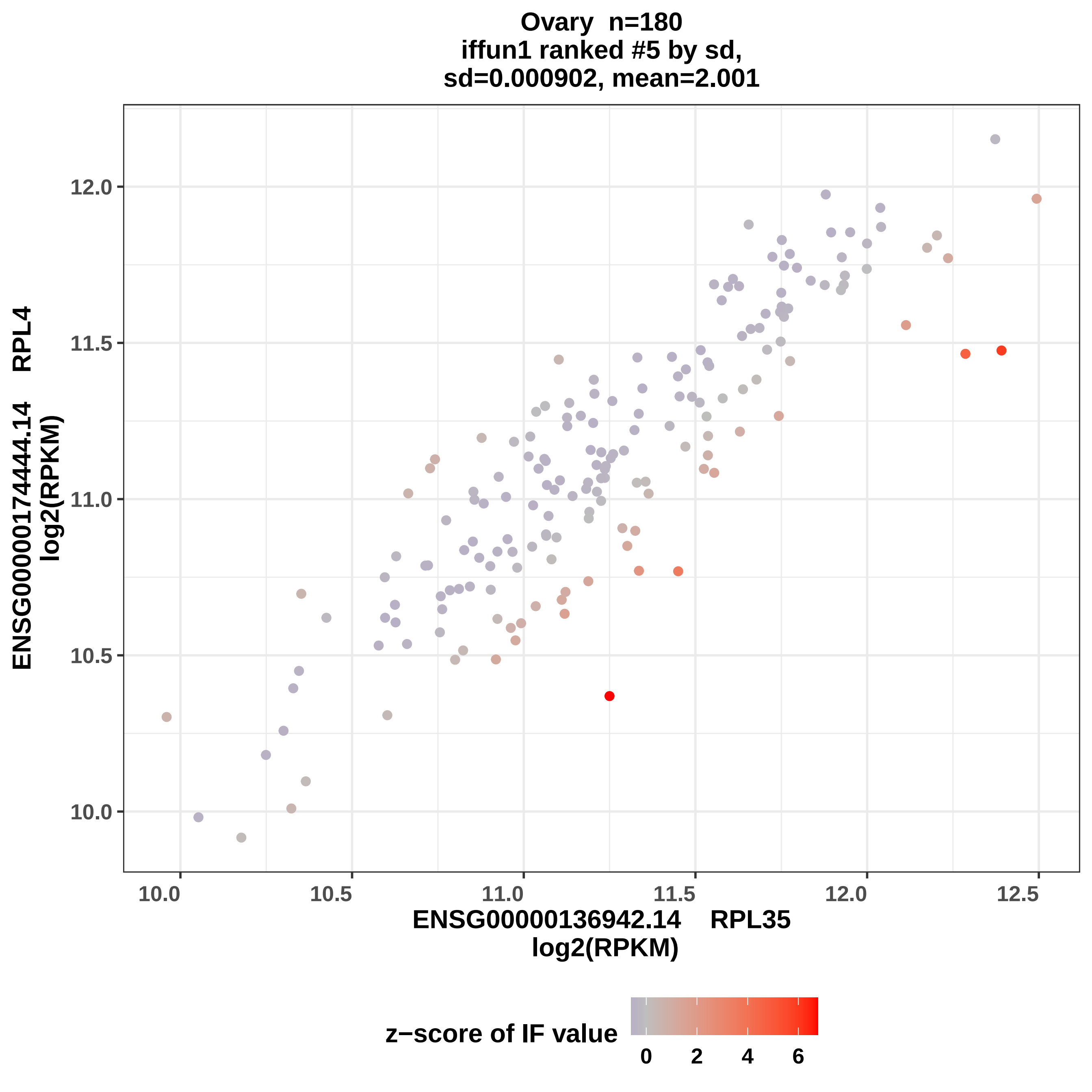
Ovary n=180
iffun1 ranked #5 by sd,
sd=0.000902, mean=2.001
ENSG00000174444.14 RPL4
log2(RPKM)
12.0
11.5
11.0
10.5
10.0
10.0
10.5
11.0
11.5
12.0
12.5
ENSG00000136942.14 RPL35
log2(RPKM)
z−score of IF value
0
2
4
6

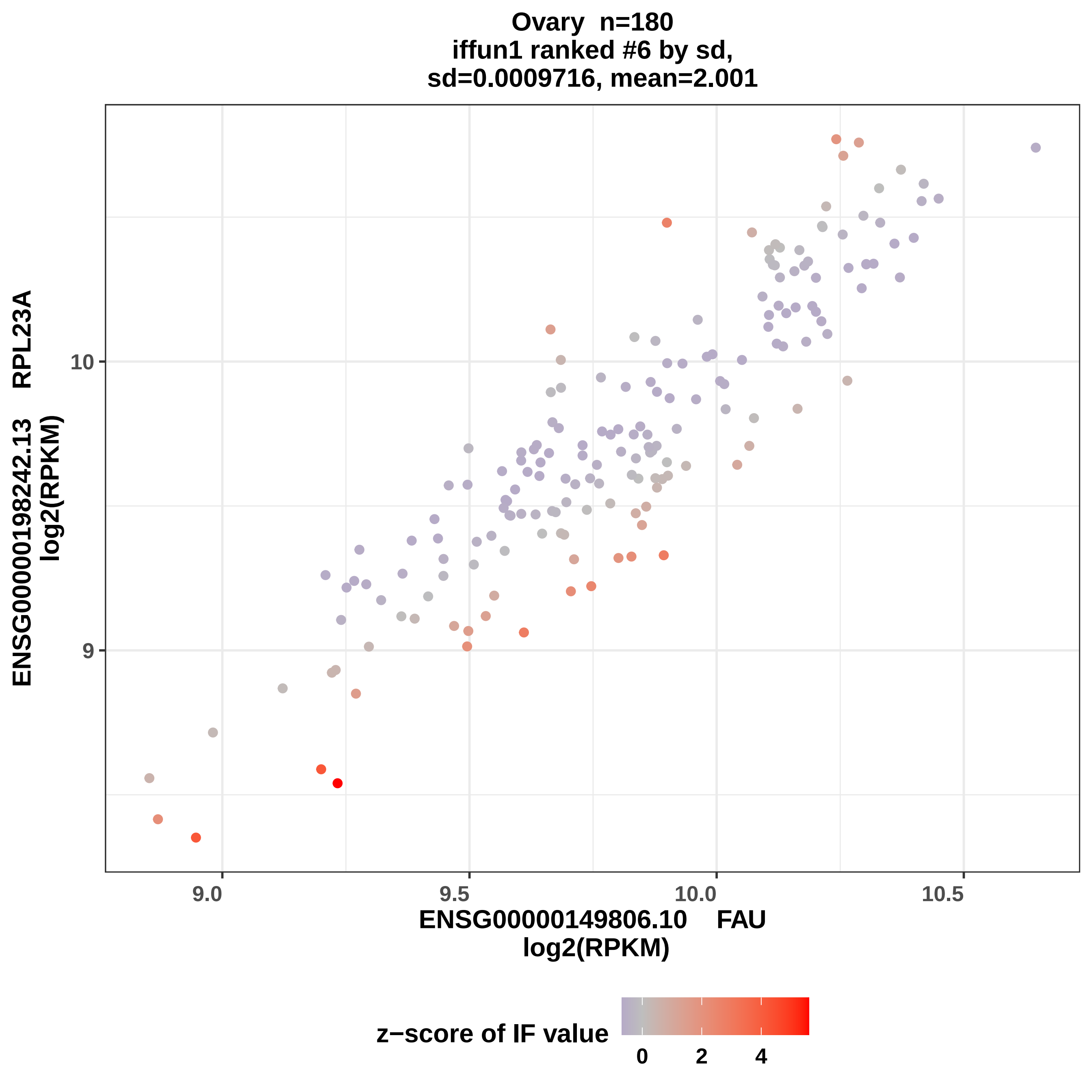
Ovary n=180
iffun1 ranked #6 by sd,
sd=0.0009716, mean=2.001
ENSG00000198242.13 RPL23A
log2(RPKM)
10
9
9.0
9.5
10.0
10.5
ENSG00000149806.10 FAU
log2(RPKM)
z−score of IF value
0
2
4

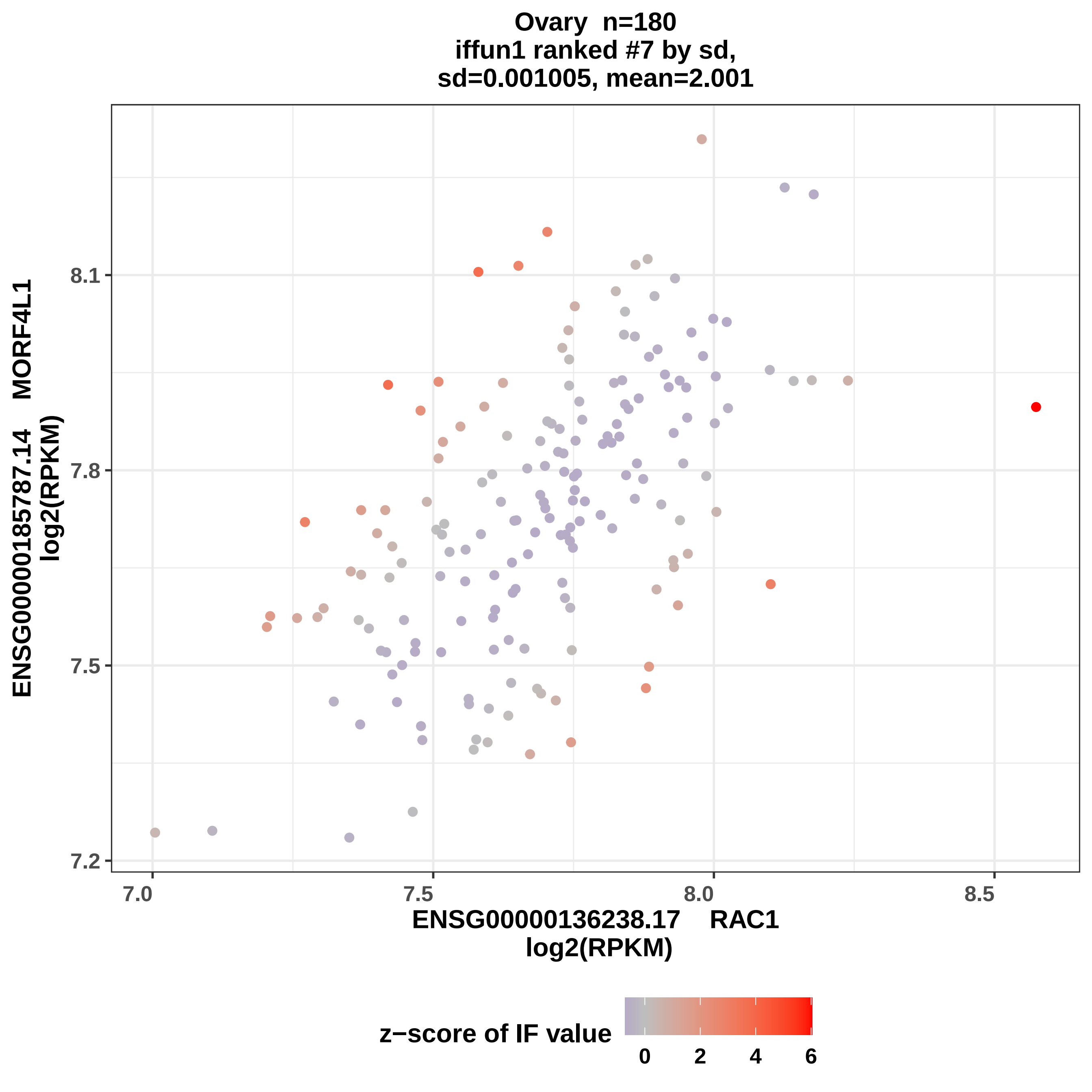
Ovary n=180
iffun1 ranked #7 by sd,
sd=0.001005, mean=2.001
ENSG00000185787.14 MORF4L1
log2(RPKM)
8.1
7.8
7.5
7.2
7.0
7.5
8.0
8.5
ENSG00000136238.17 RAC1
log2(RPKM)
z−score of IF value
0
2
4
6

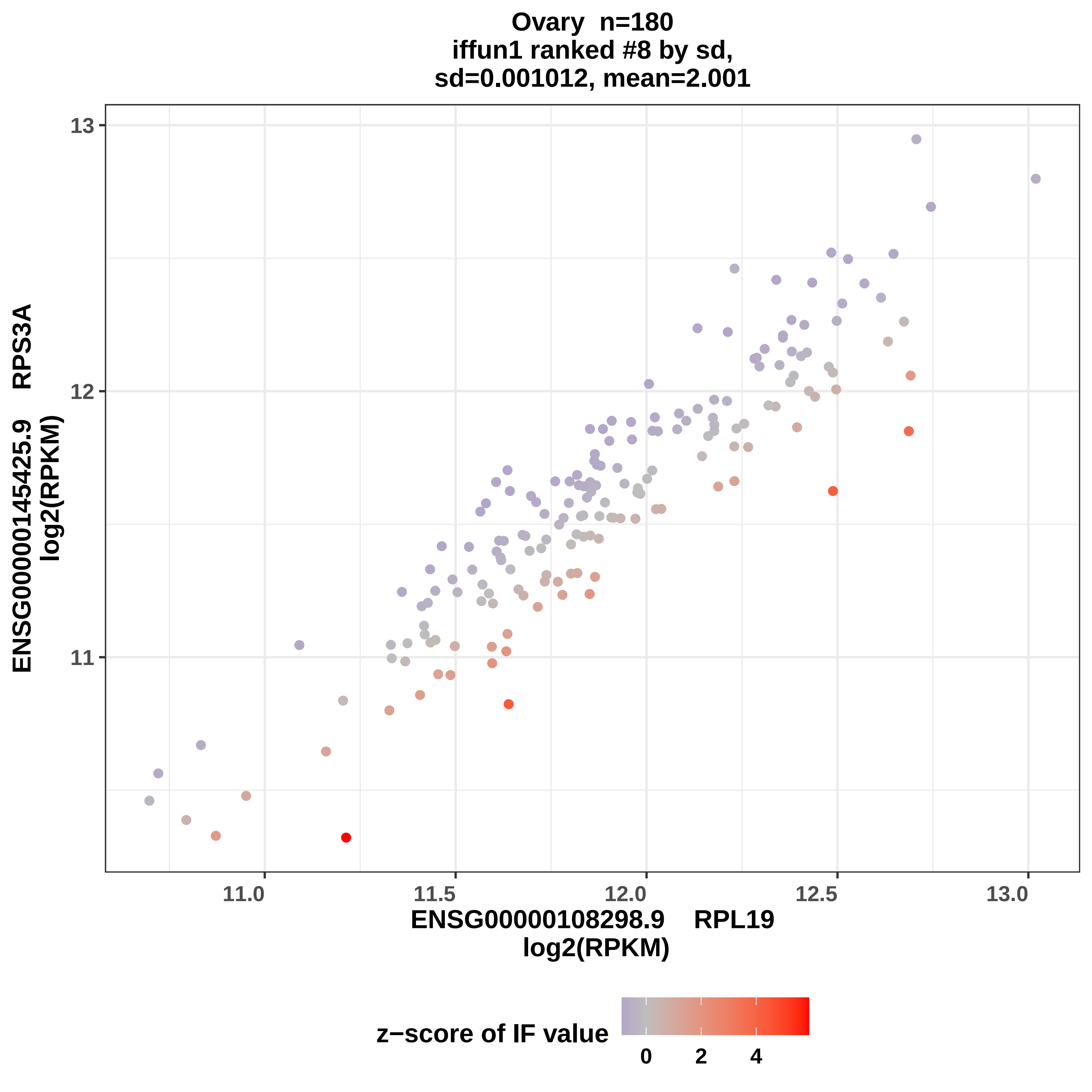

Ovary n=180
iffun1 ranked #8 by sd,
sd=0.001012, mean=2.001
13
12
11
11.0
11.5
12.0
12.5
13.0
ENSG00000145425.9 RPS3A
log2(RPKM)
ENSG00000108298.9 RPL19
log2(RPKM)
z−score of IF value
0
2
4

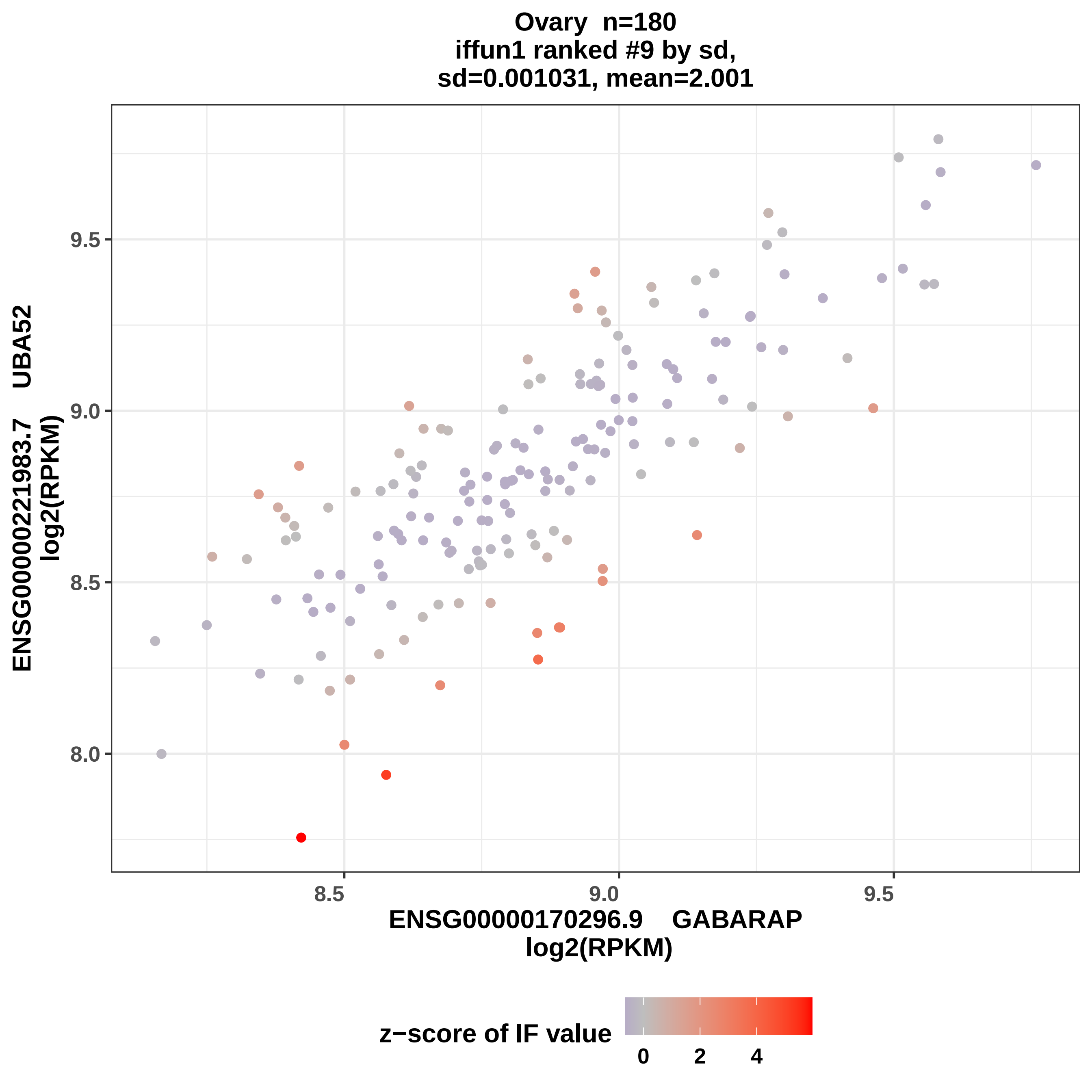

Ovary n=180
iffun1 ranked #9 by sd,
sd=0.001031, mean=2.001
ENSG00000221983.7 UBA52
log2(RPKM)
9.5
9.0
8.5
8.0
8.5
9.0
9.5
ENSG00000170296.9 GABARAP
log2(RPKM)
z−score of IF value
0
2
4

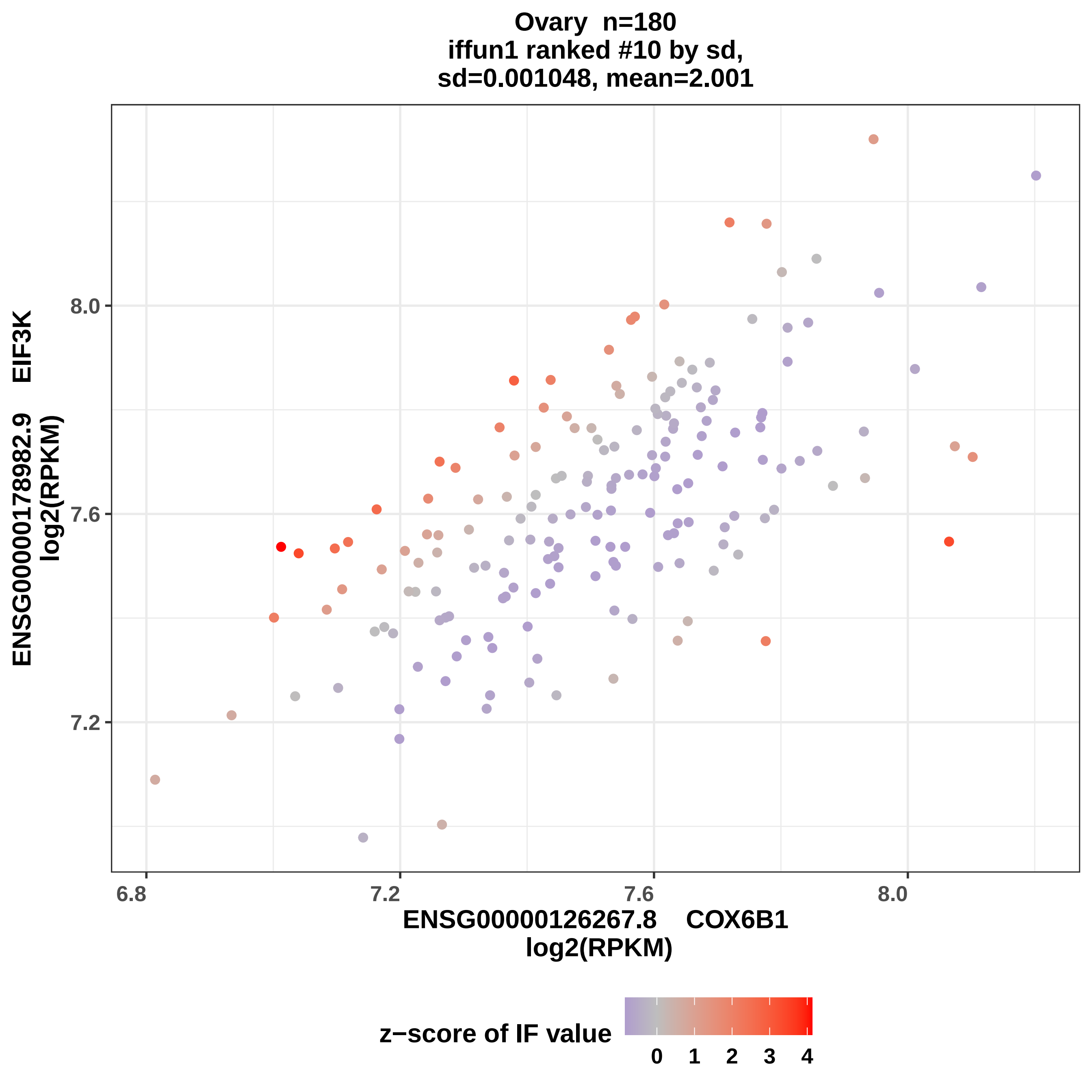

Ovary n=180
iffun1 ranked #10 by sd,
sd=0.001048, mean=2.001
ENSG00000178982.9 EIF3K
log2(RPKM)
8.0
7.6
7.2
6.8
7.2
7.6
8.0
ENSG00000126267.8 COX6B1
log2(RPKM)
z−score of IF value
0
1
2
3
4

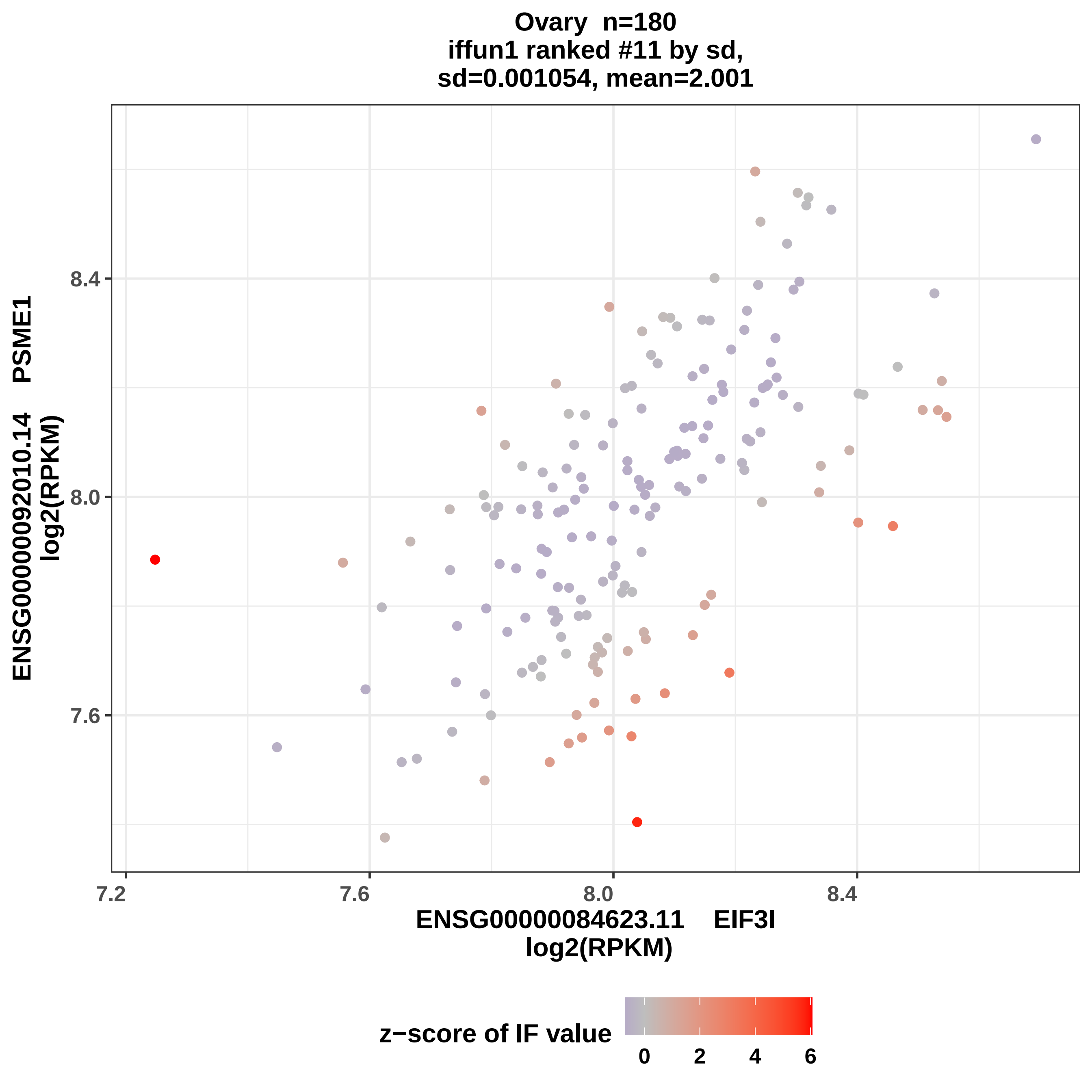
Ovary n=180
iffun1 ranked #11 by sd,
sd=0.001054, mean=2.001
ENSG00000092010.14 PSME1
log2(RPKM)
8.4
8.0
7.6
7.2
7.6
8.0
8.4
ENSG00000084623.11 EIF3I
log2(RPKM)
z−score of IF value
0
2
4
6

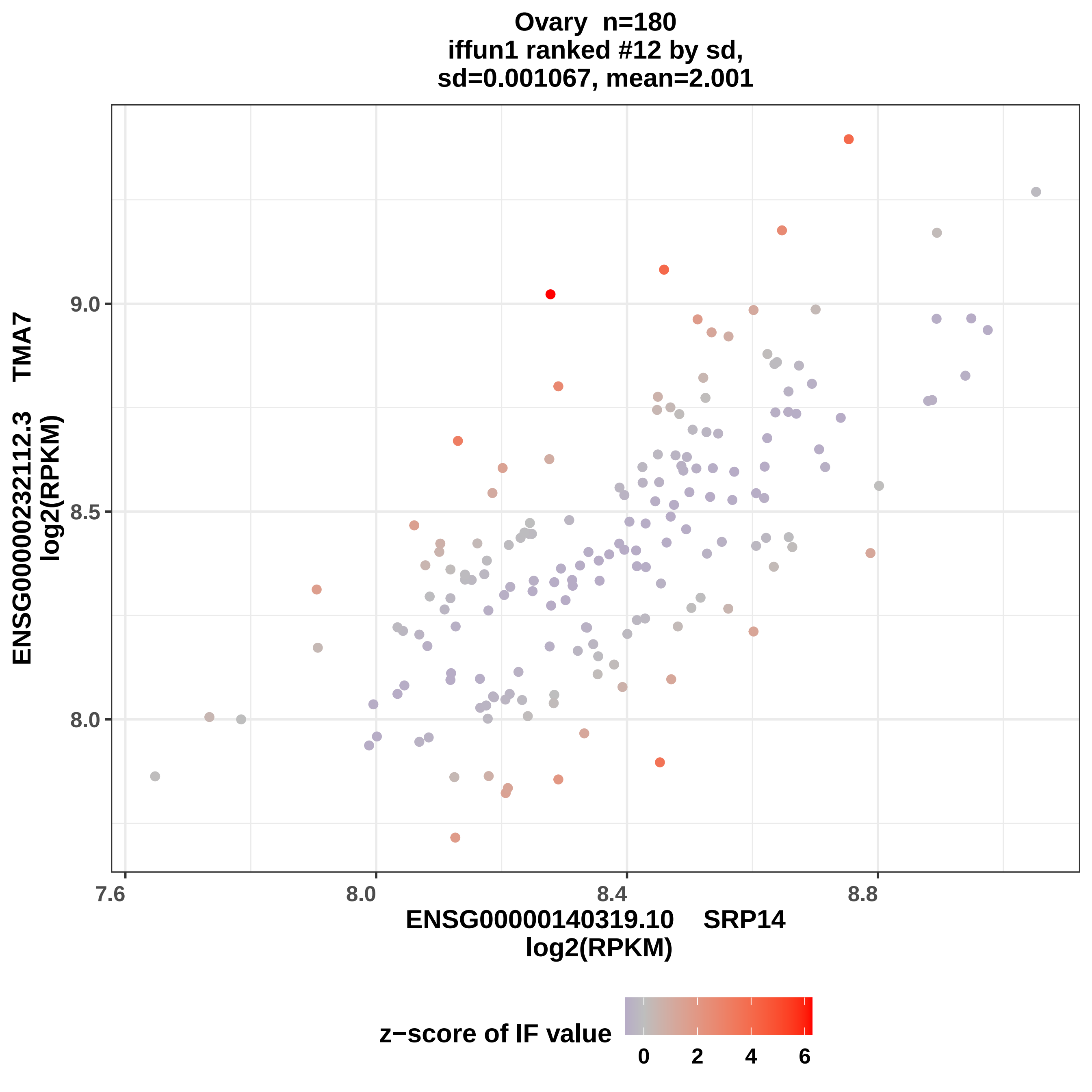

Ovary n=180
iffun1 ranked #12 by sd,
sd=0.001067, mean=2.001
ENSG00000232112.3 TMA7
log2(RPKM)
9.0
8.5
8.0
7.6
8.0
8.4
8.8
ENSG00000140319.10 SRP14
log2(RPKM)
z−score of IF value
0
2
4
6

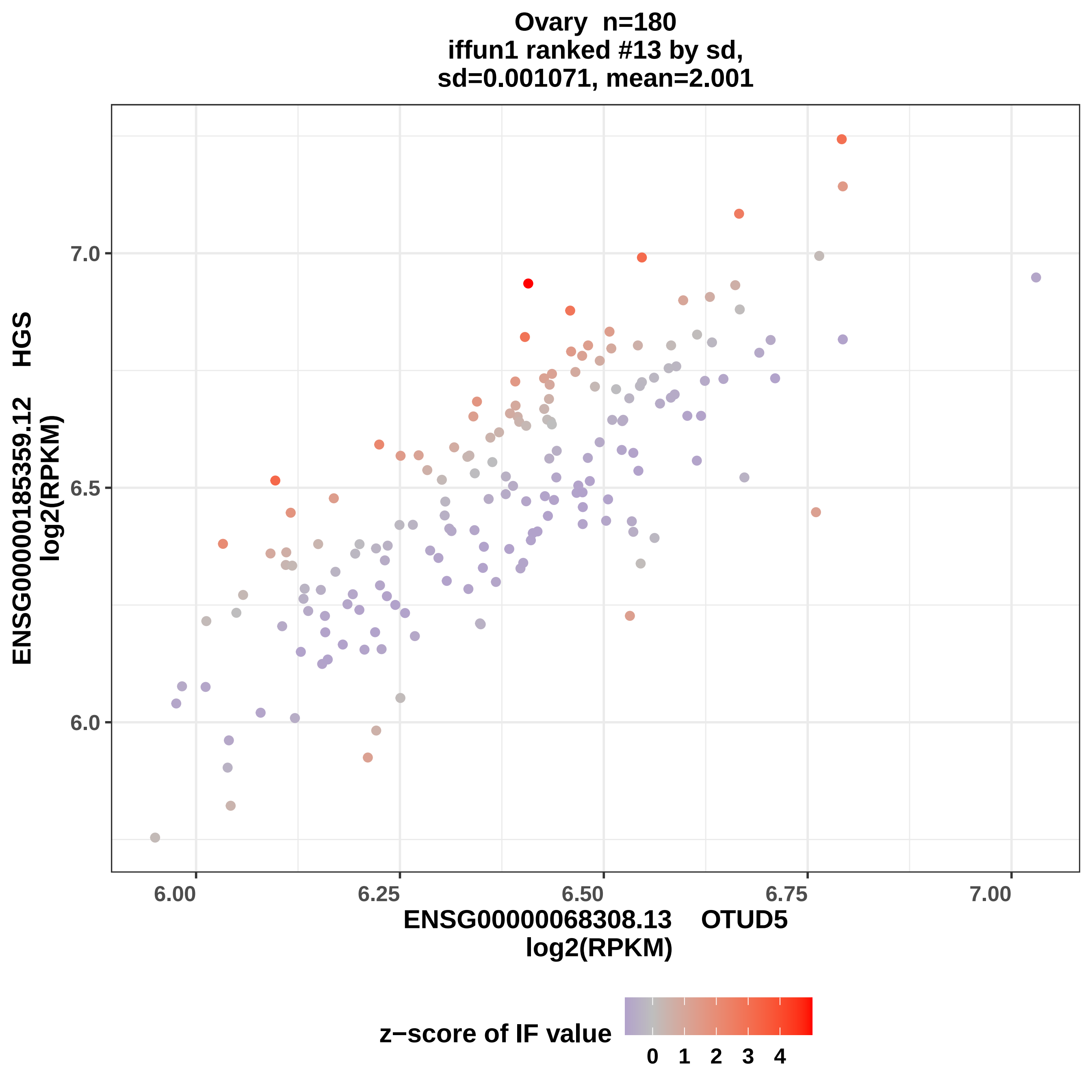
Ovary n=180
iffun1 ranked #13 by sd,
sd=0.001071, mean=2.001
ENSG00000185359.12 HGS
log2(RPKM)
7.0
6.5
6.0
6.00
6.25
6.50
6.75
7.00
ENSG00000068308.13 OTUD5
log2(RPKM)
z−score of IF value
0
1
2
3
4

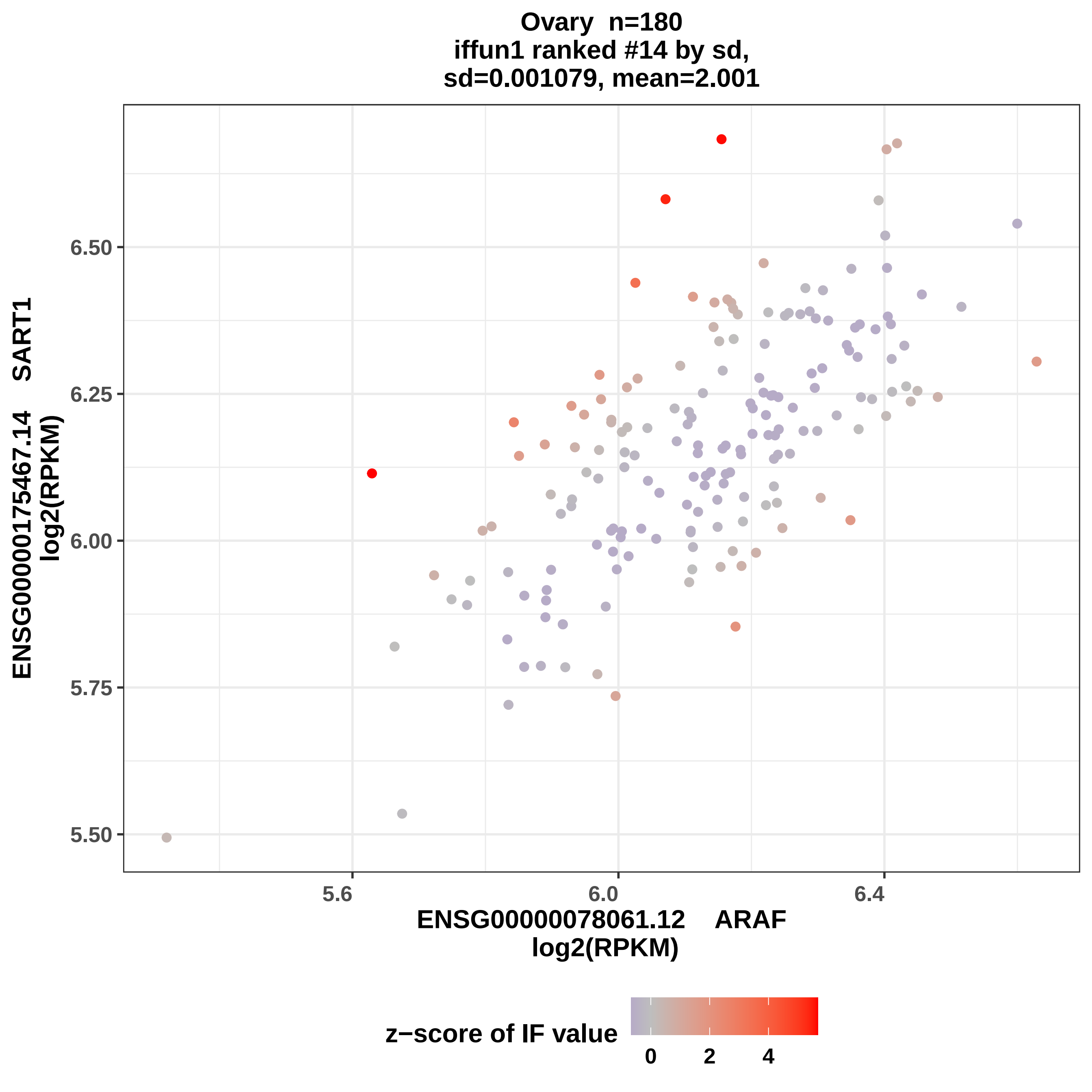

Ovary n=180
iffun1 ranked #14 by sd,
sd=0.001079, mean=2.001
ENSG00000175467.14 SART1
log2(RPKM)
6.50
6.25
6.00
5.75
5.50
5.6
6.0
6.4
ENSG00000078061.12 ARAF
log2(RPKM)
z−score of IF value
0
2
4

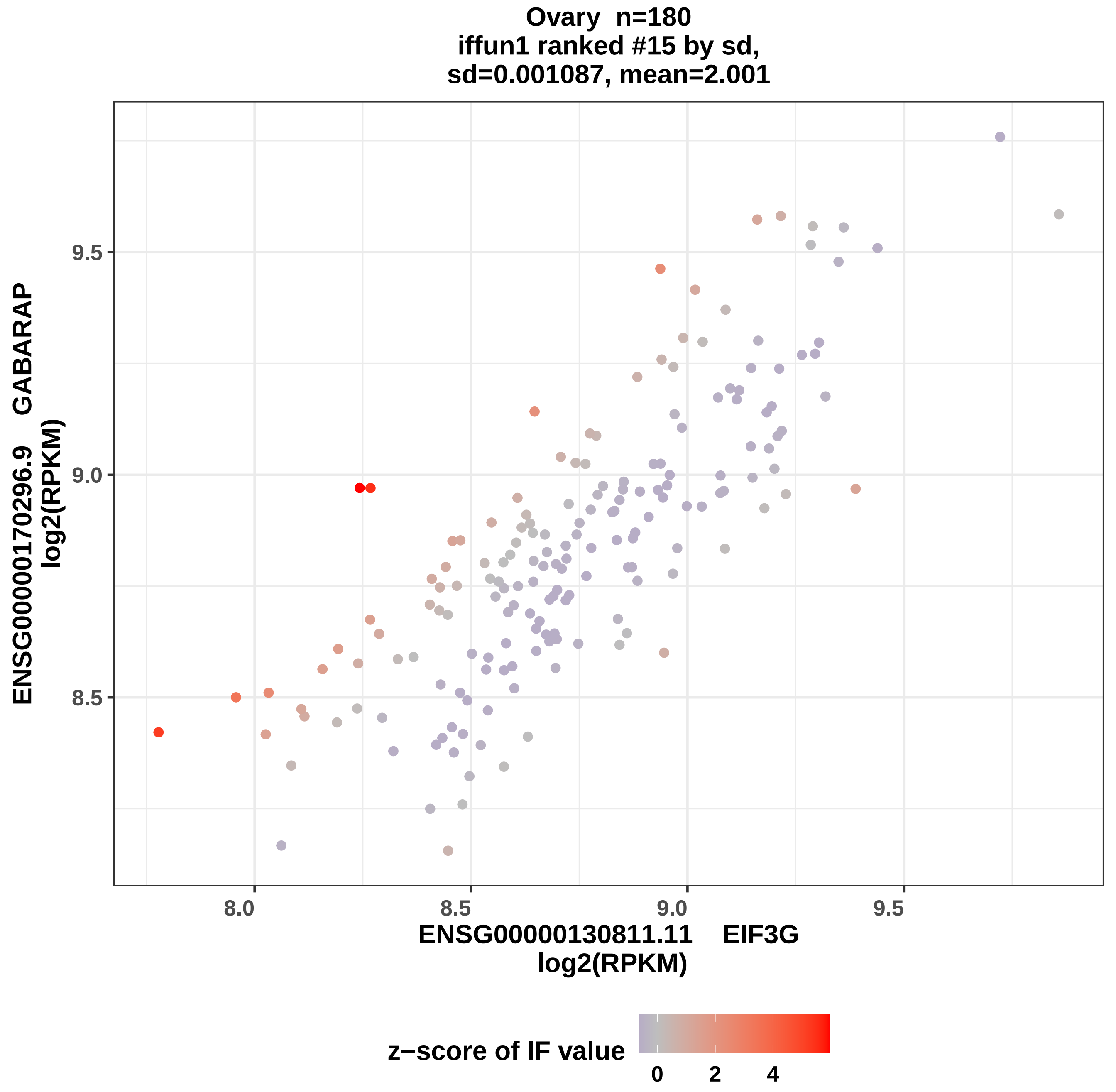
Ovary n=180
iffun1 ranked #15 by sd,
sd=0.001087, mean=2.001
ENSG00000170296.9 GABARAP
log2(RPKM)
9.5
9.0
8.5
8.0
8.5
9.0
9.5
ENSG00000130811.11 EIF3G
log2(RPKM)
z−score of IF value
0
2
4

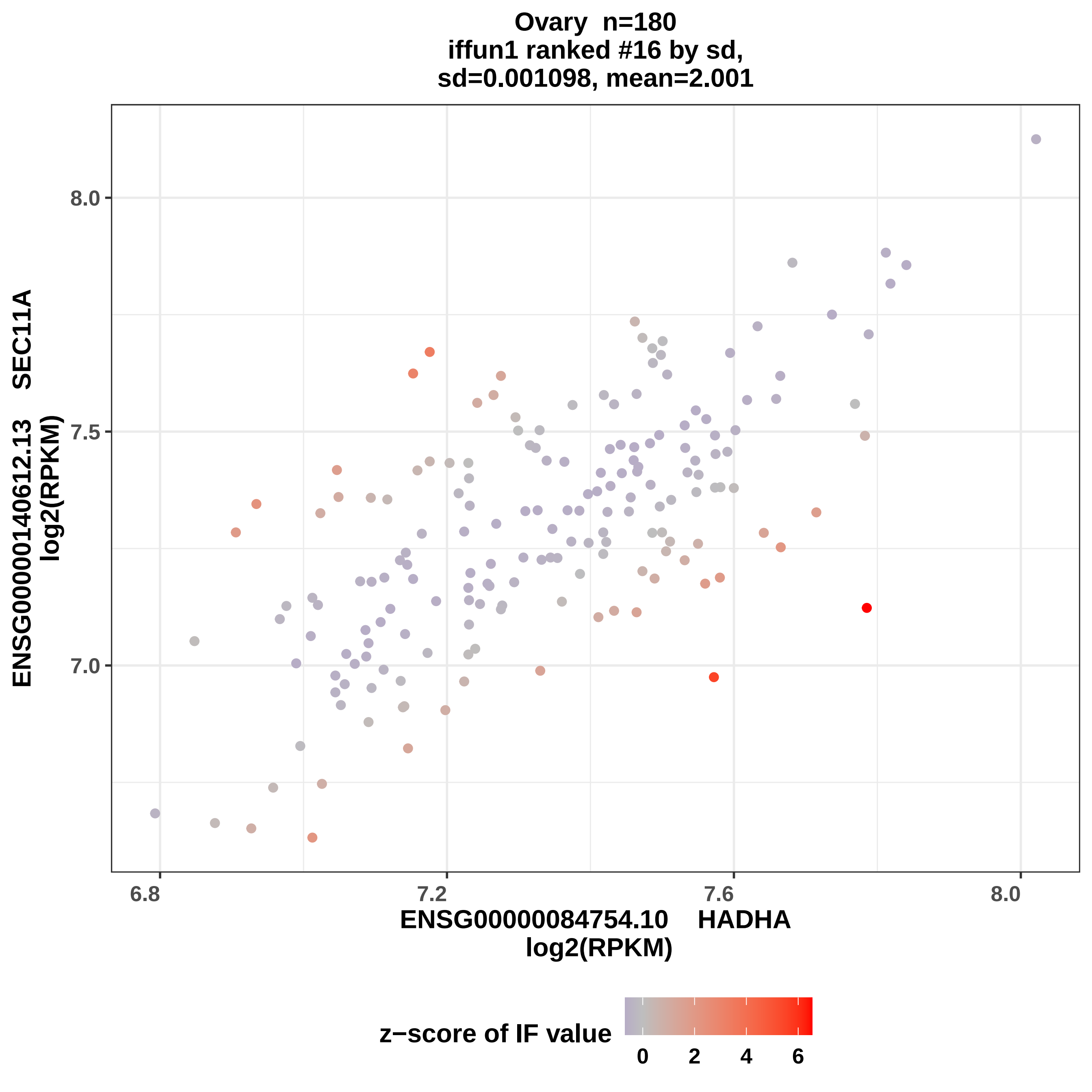

Ovary n=180
iffun1 ranked #16 by sd,
sd=0.001098, mean=2.001
ENSG00000140612.13 SEC11A
log2(RPKM)
8.0
7.5
7.0
6.8
7.2
7.6
8.0
ENSG00000084754.10 HADHA
log2(RPKM)
z−score of IF value
0
2
4
6

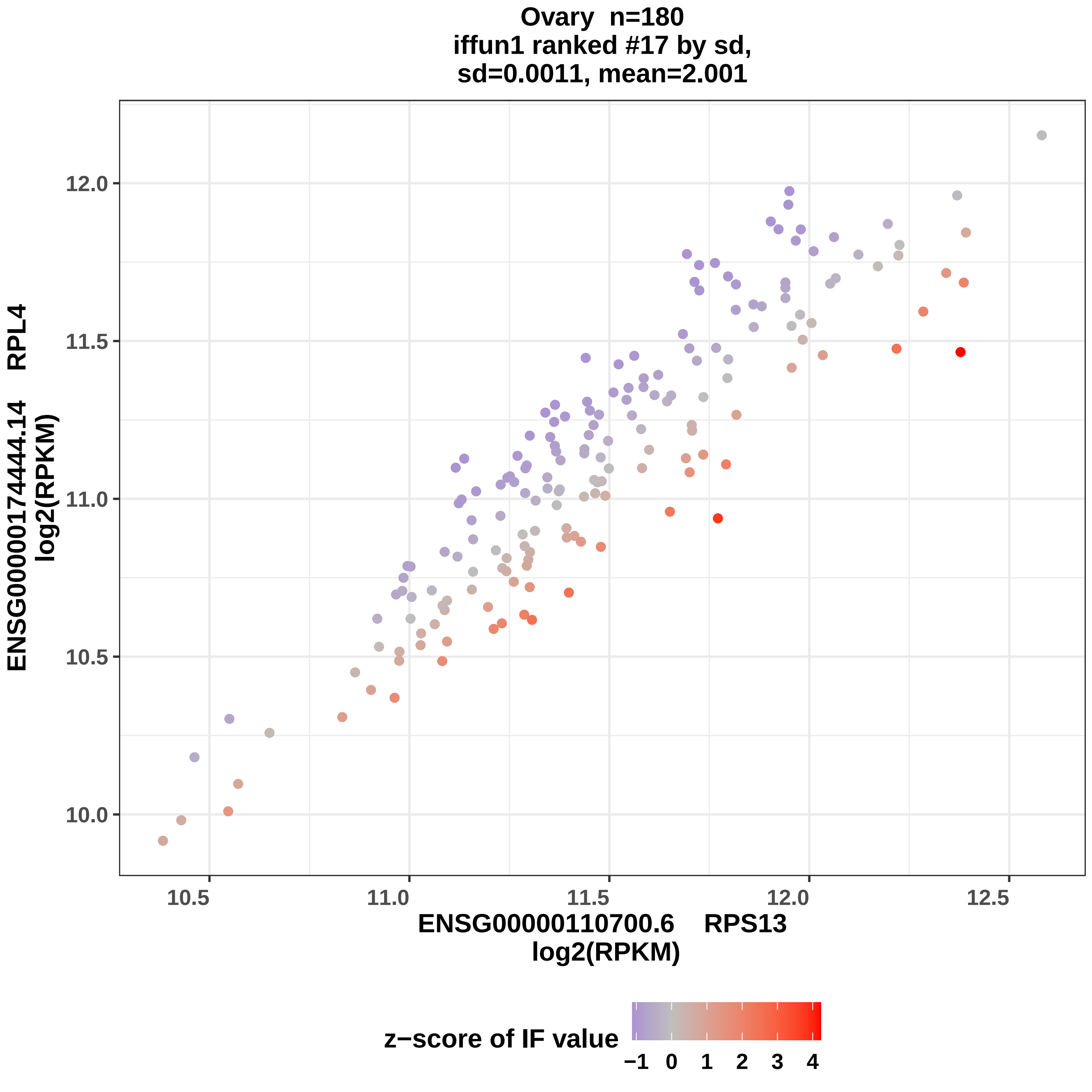

Ovary n=180
iffun1 ranked #17 by sd,
sd=0.0011, mean=2.001
ENSG00000174444.14 RPL4
log2(RPKM)
ENSG00000110700.6 RPS13
log2(RPKM)
10.0
10.5
11.0
11.5
12.0
10.5
11.0
11.5
12.0
12.5
z−score of IF value
−1
0
1
2
3
4

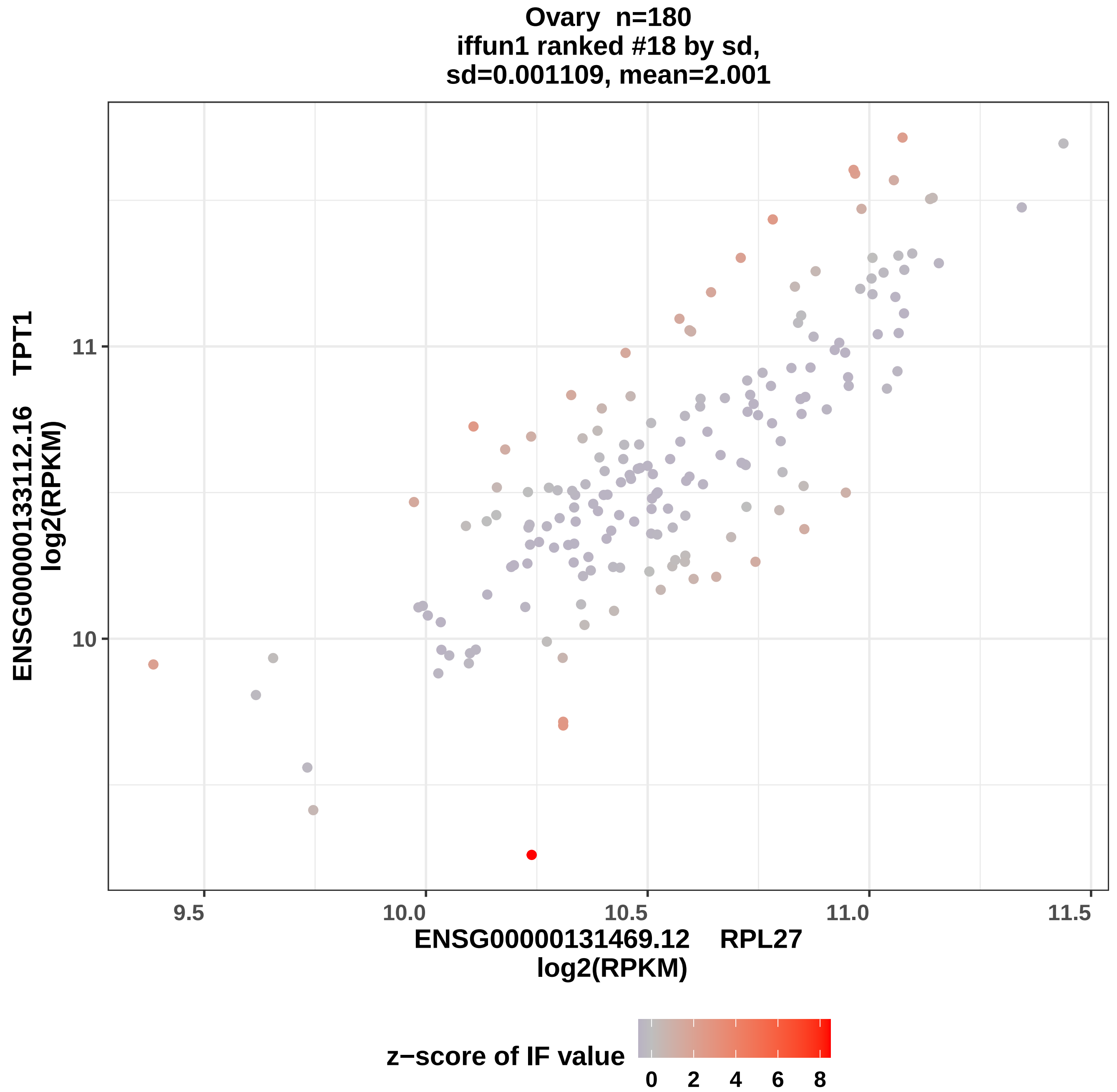
Ovary n=180
iffun1 ranked #18 by sd,
sd=0.001109, mean=2.001
ENSG00000133112.16 TPT1
log2(RPKM)
11
10
9.5
10.0
10.5
11.0
11.5
ENSG00000131469.12 RPL27
log2(RPKM)
z−score of IF value
0
2
4
6
8

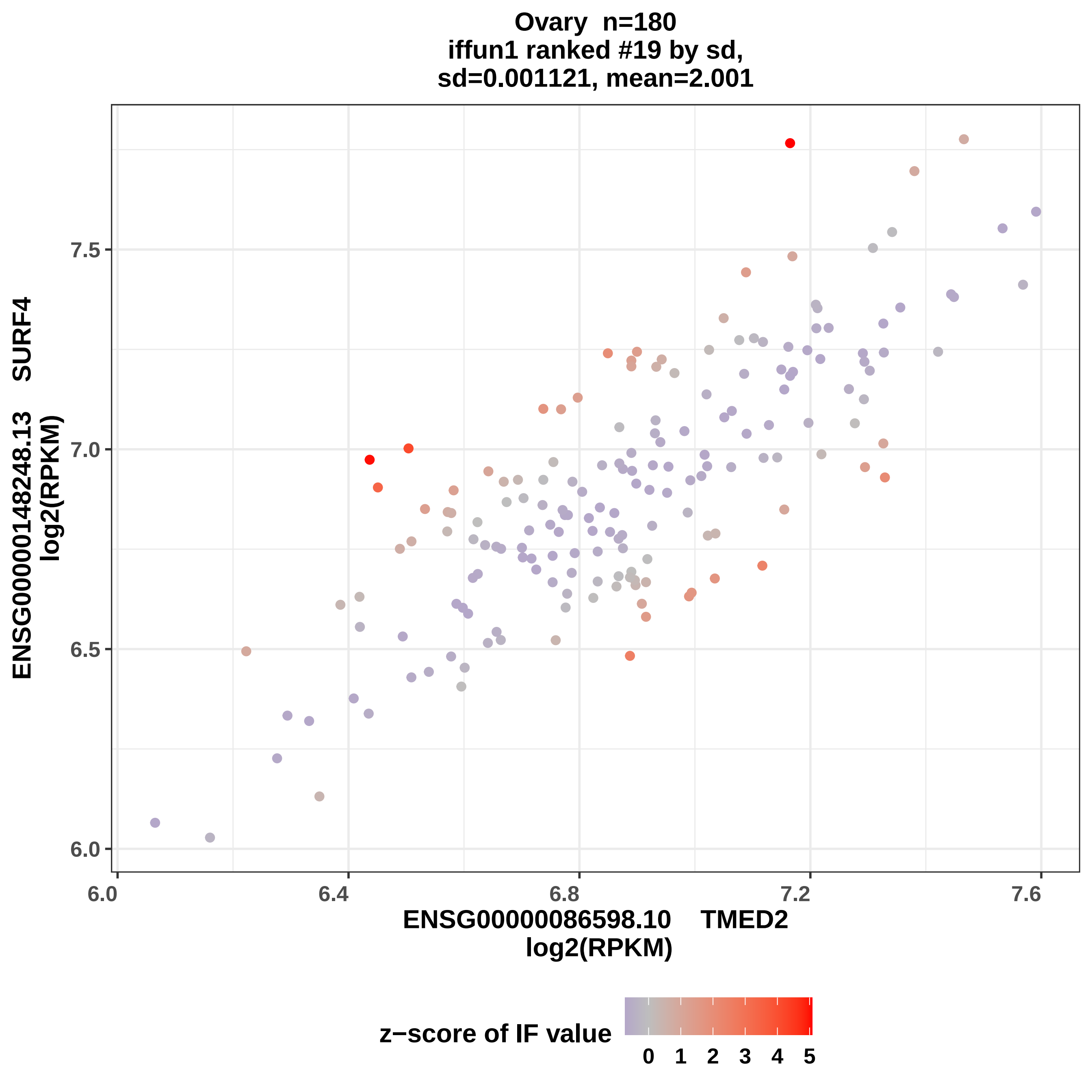
Ovary n=180
iffun1 ranked #19 by sd,
sd=0.001121, mean=2.001
ENSG00000148248.13 SURF4
log2(RPKM)
7.5
7.0
6.5
6.0
6.0
6.4
6.8
7.2
7.6
ENSG00000086598.10 TMED2
log2(RPKM)
z−score of IF value
0 1 2 3 4 5

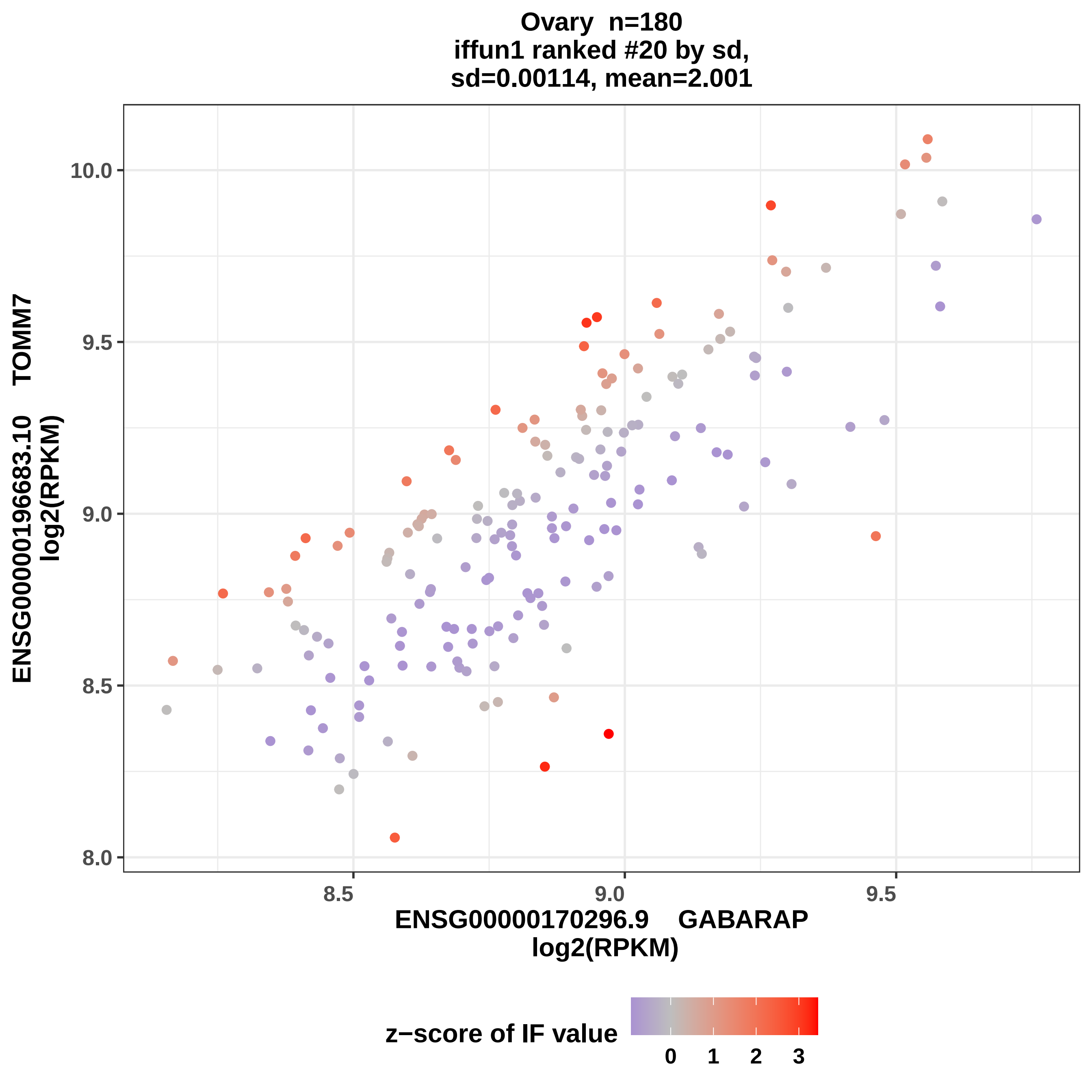

Ovary n=180
iffun1 ranked #20 by sd,
sd=0.00114, mean=2.001
ENSG00000196683.10 TOMM7
log2(RPKM)
10.0
9.5
9.0
8.5
8.0
8.5
9.0
9.5
ENSG00000170296.9 GABARAP
log2(RPKM)
z−score of IF value
0
1
2
3

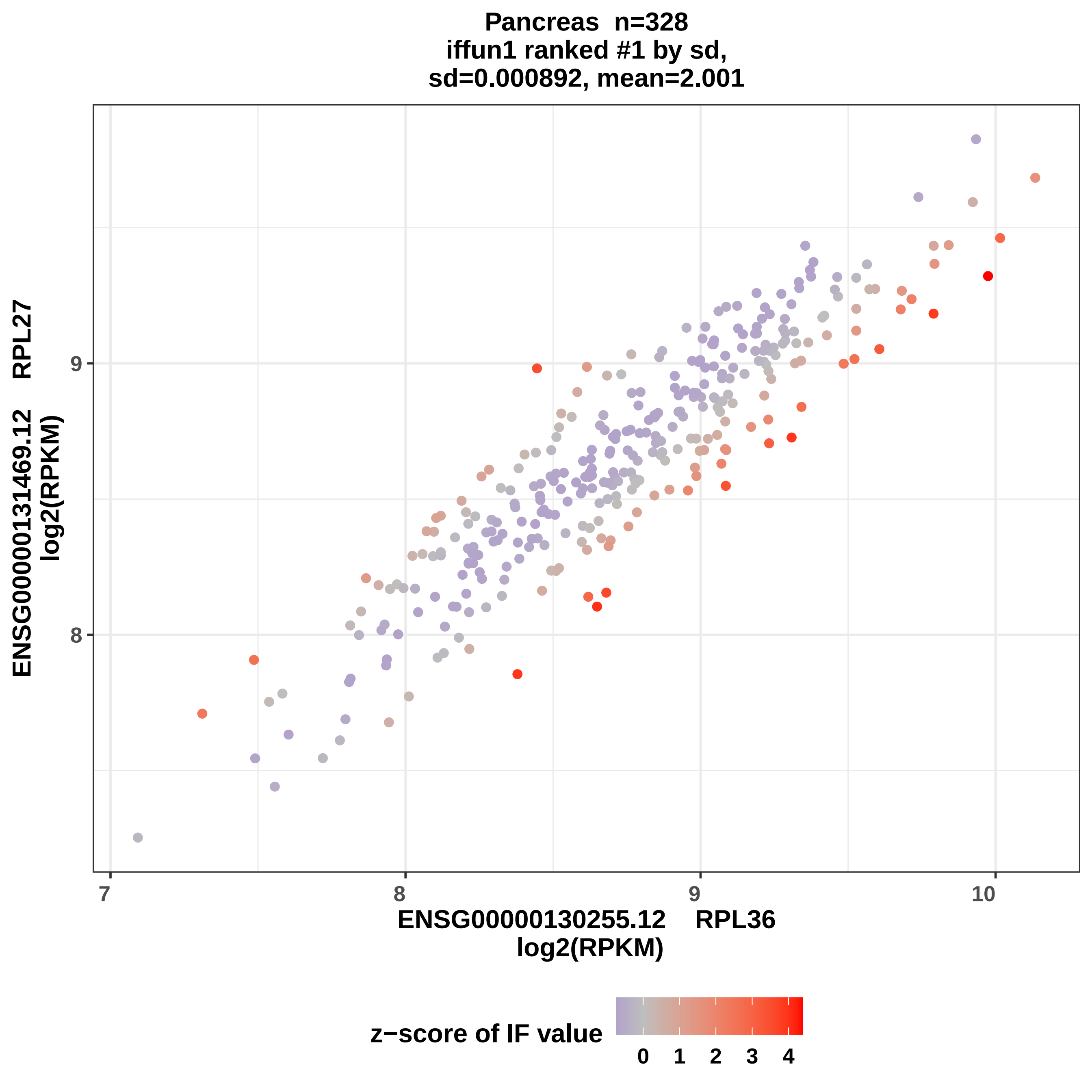

Pancreas n=328
iffun1 ranked #1 by sd,
sd=0.000892, mean=2.001
ENSG00000131469.12 RPL27
log2(RPKM)
9
8
7
8
9
10
ENSG00000130255.12 RPL36
log2(RPKM)
z−score of IF value
0
1
2
3
4

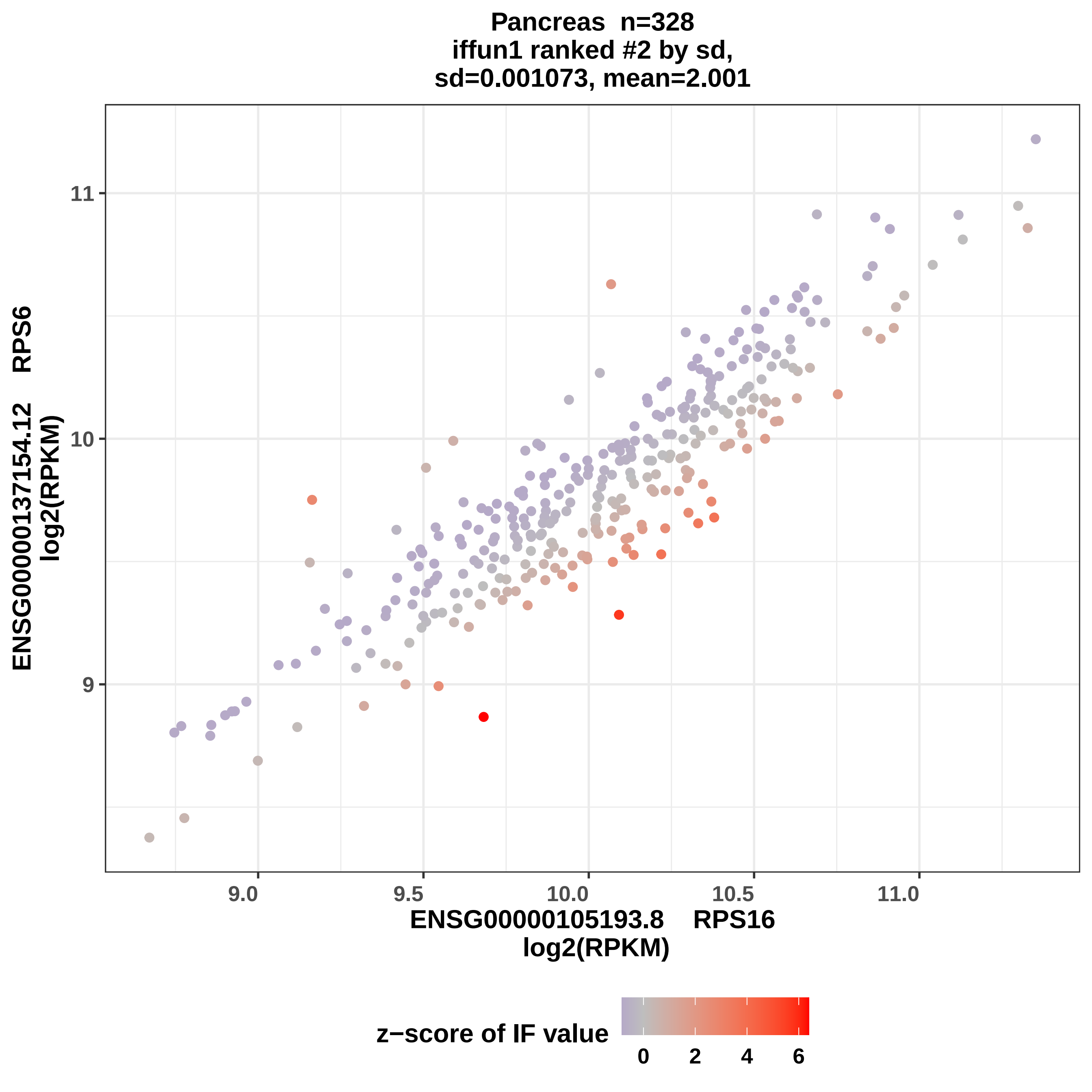

Pancreas n=328
iffun1 ranked #2 by sd,
sd=0.001073, mean=2.001
ENSG00000137154.12 RPS6
log2(RPKM)
11
10
9
9.0
9.5
10.0
10.5
11.0
ENSG00000105193.8 RPS16
log2(RPKM)
z−score of IF value
0
2
4
6

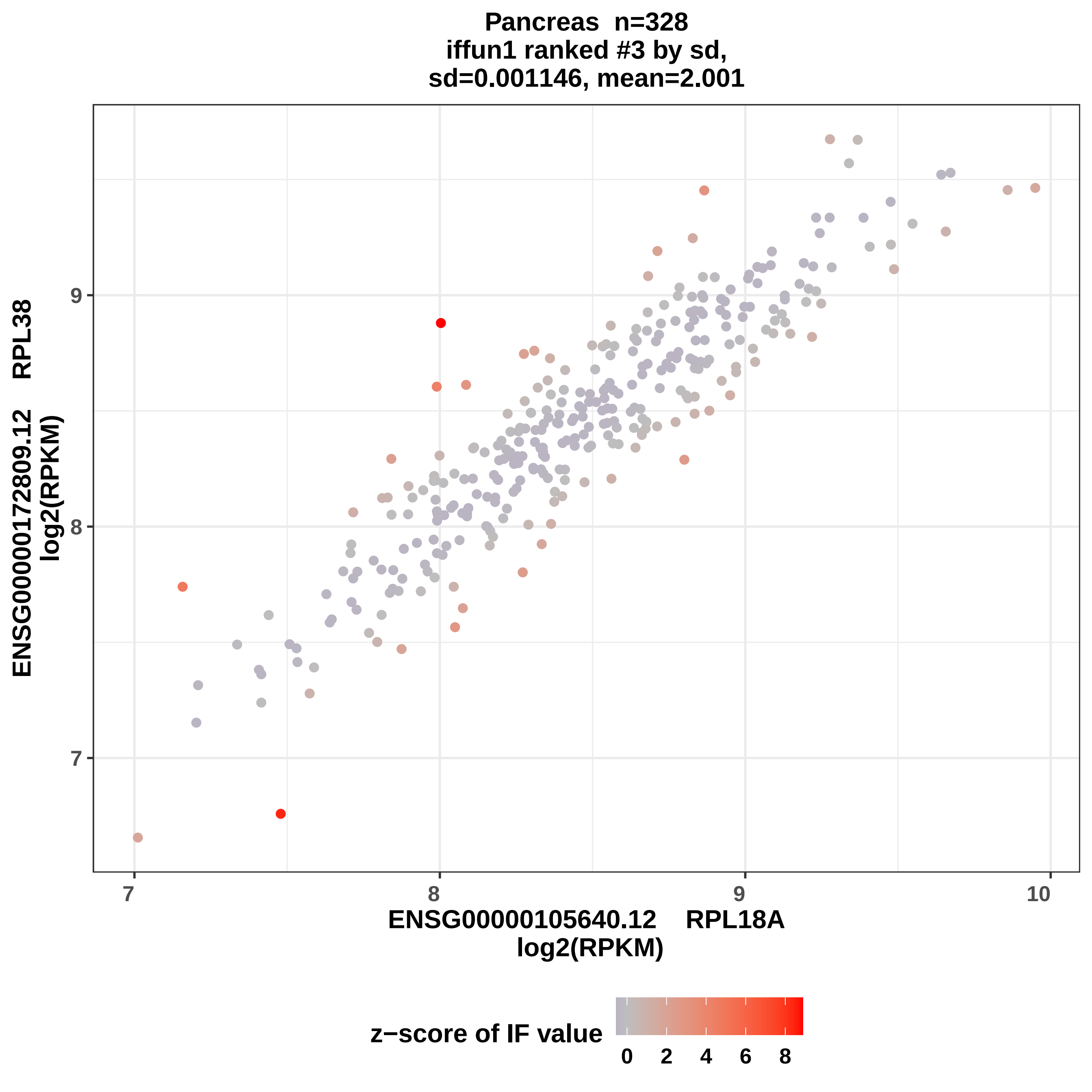

Pancreas n=328
iffun1 ranked #3 by sd,
sd=0.001146, mean=2.001
ENSG00000172809.12 RPL38
log2(RPKM)
9
8
7
7
8
9
10
ENSG00000105640.12 RPL18A
log2(RPKM)
z−score of IF value
0
2
4
6
8

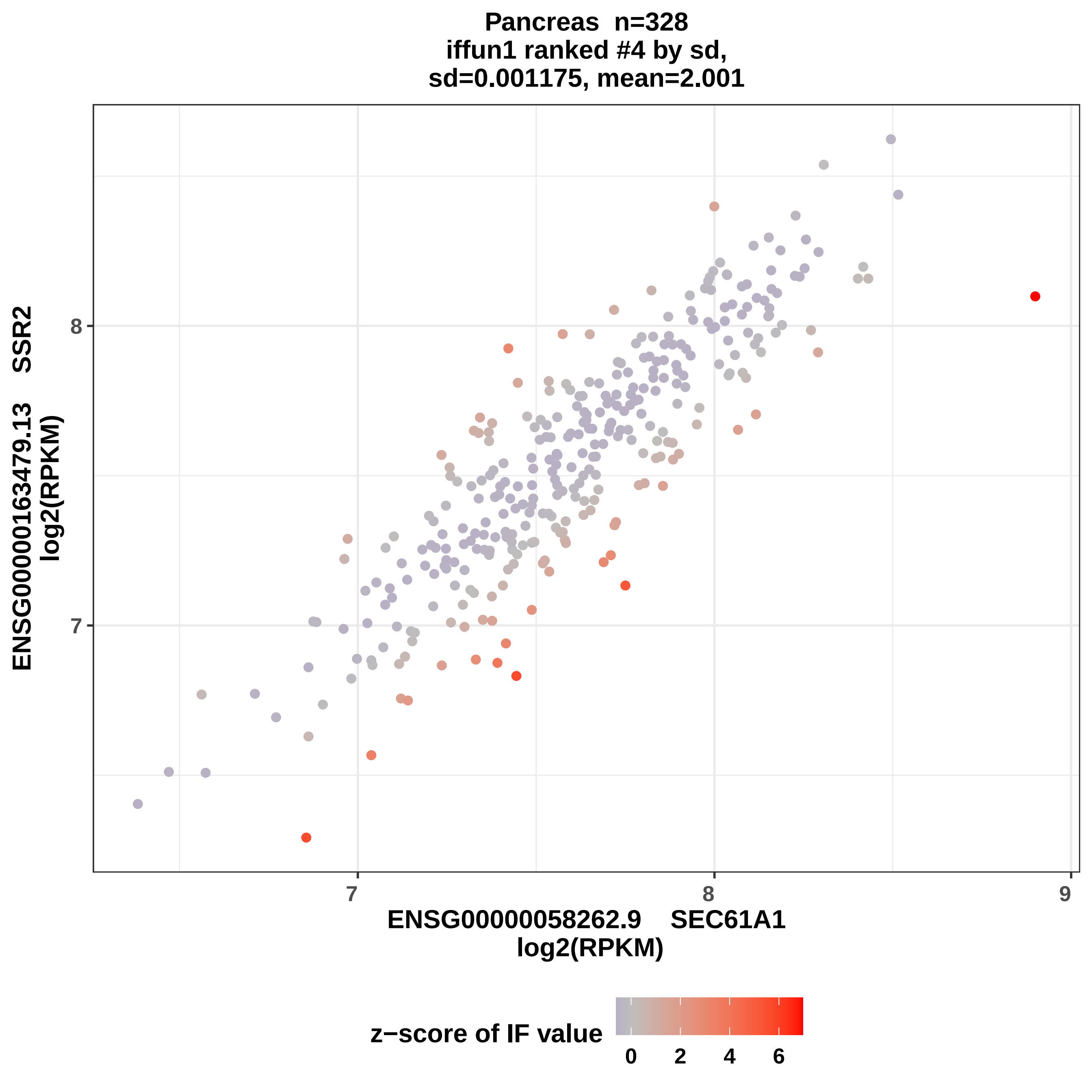
Pancreas n=328
iffun1 ranked #4 by sd,
sd=0.001175, mean=2.001
ENSG00000163479.13 SSR2
log2(RPKM)
8
7
7
8
9
ENSG00000058262.9 SEC61A1
log2(RPKM)
z−score of IF value
0
2
4
6

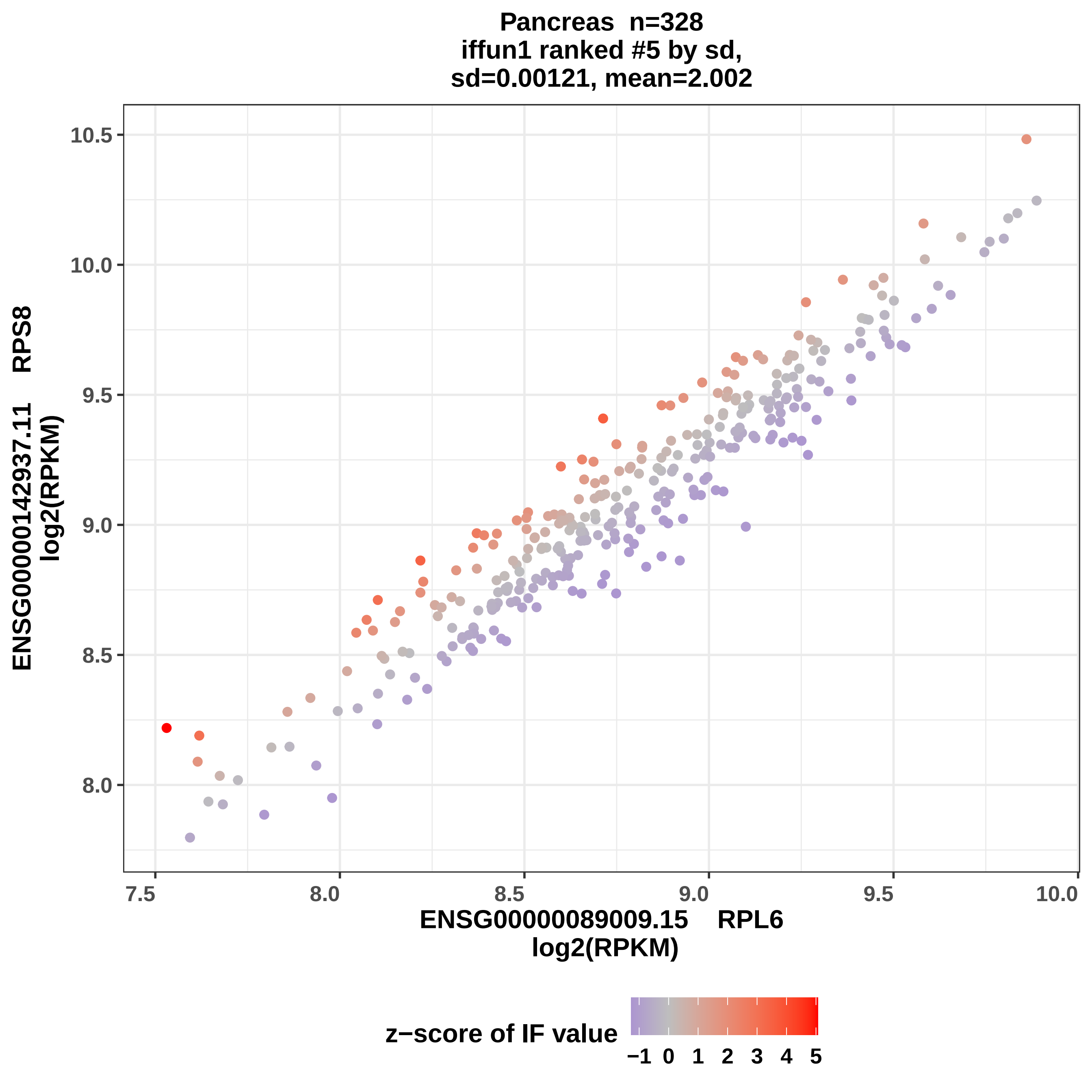

Pancreas n=328
iffun1 ranked #5 by sd,
sd=0.00121, mean=2.002
ENSG00000142937.11 RPS8
log2(RPKM)
10.5
10.0
9.5
9.0
8.5
8.0
7.5
8.0
8.5
9.0
9.5
10.0
ENSG00000089009.15 RPL6
log2(RPKM)
z−score of IF value
−1
0
1
2
3
4
5

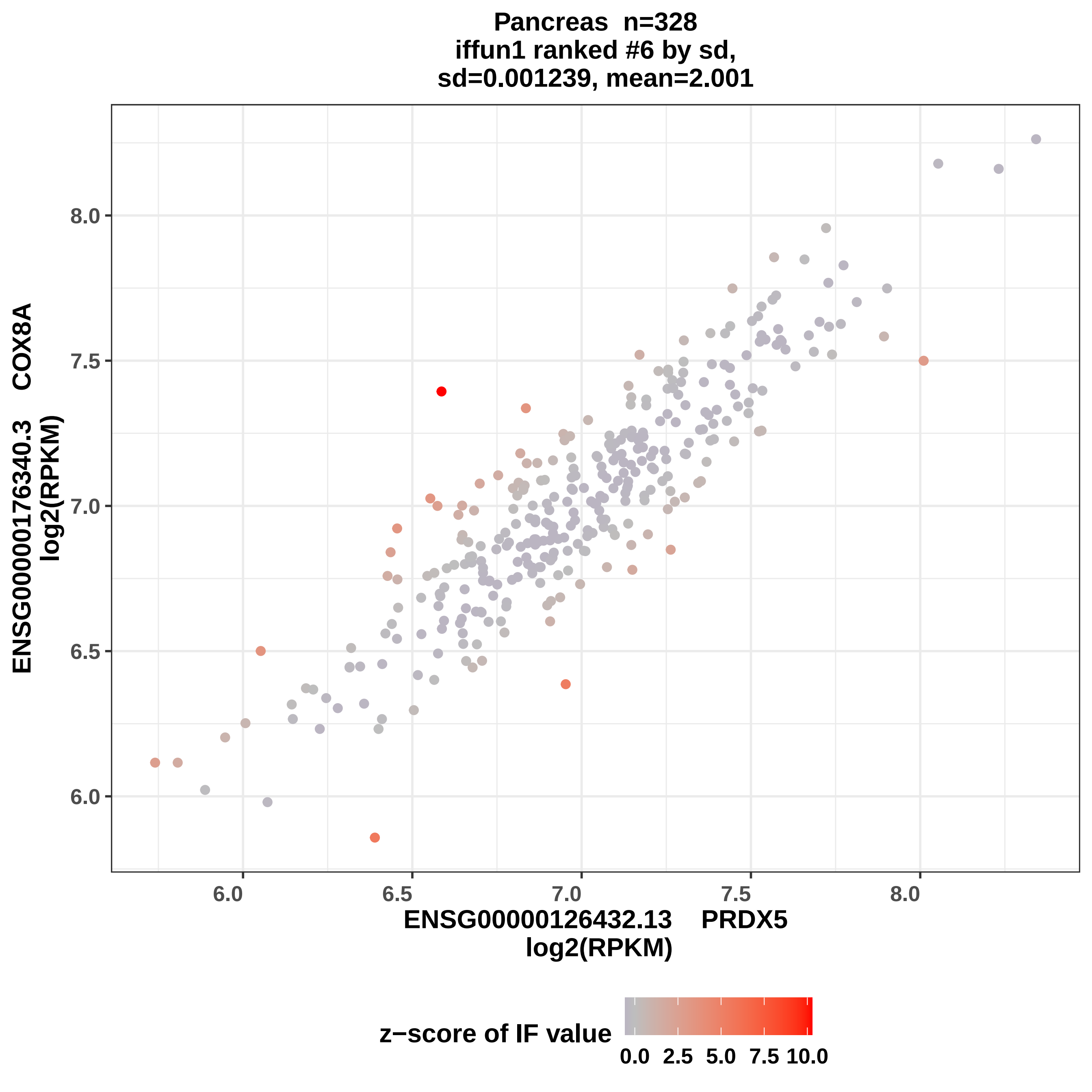

Pancreas n=328
iffun1 ranked #6 by sd,
sd=0.001239, mean=2.001
ENSG00000176340.3 COX8A
log2(RPKM)
ENSG00000126432.13 PRDX5
log2(RPKM)
z−score of IF value
0.0 2.5 5.0 7.5 10.0

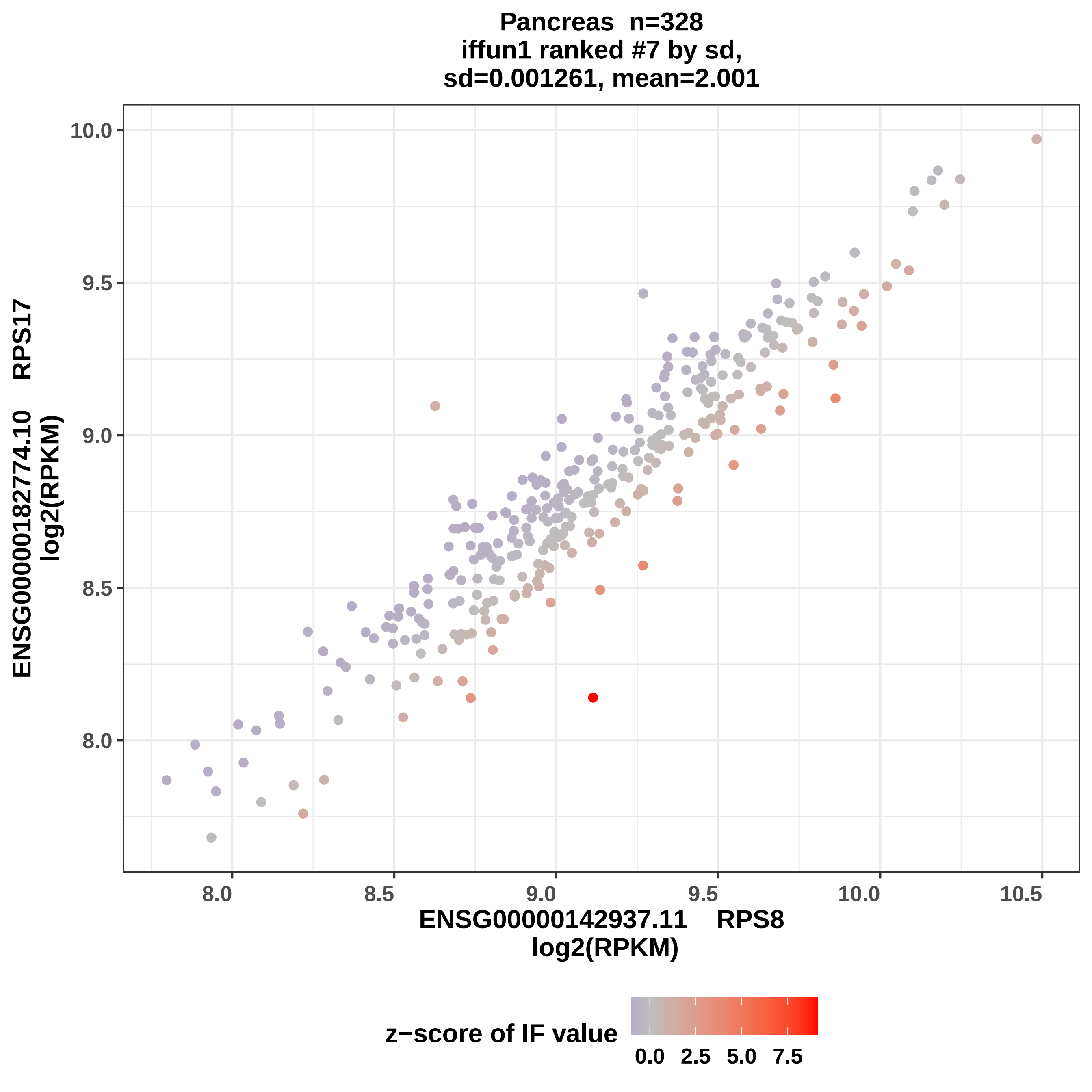

Pancreas n=328
iffun1 ranked #7 by sd,
sd=0.001261, mean=2.001
ENSG00000182774.10 RPS17
log2(RPKM)
10.0
9.5
9.0
8.5
8.0
8.0
8.5
9.0
9.5
10.0
10.5
ENSG00000142937.11 RPS8
log2(RPKM)
z−score of IF value
0.0
2.5
5.0
7.5

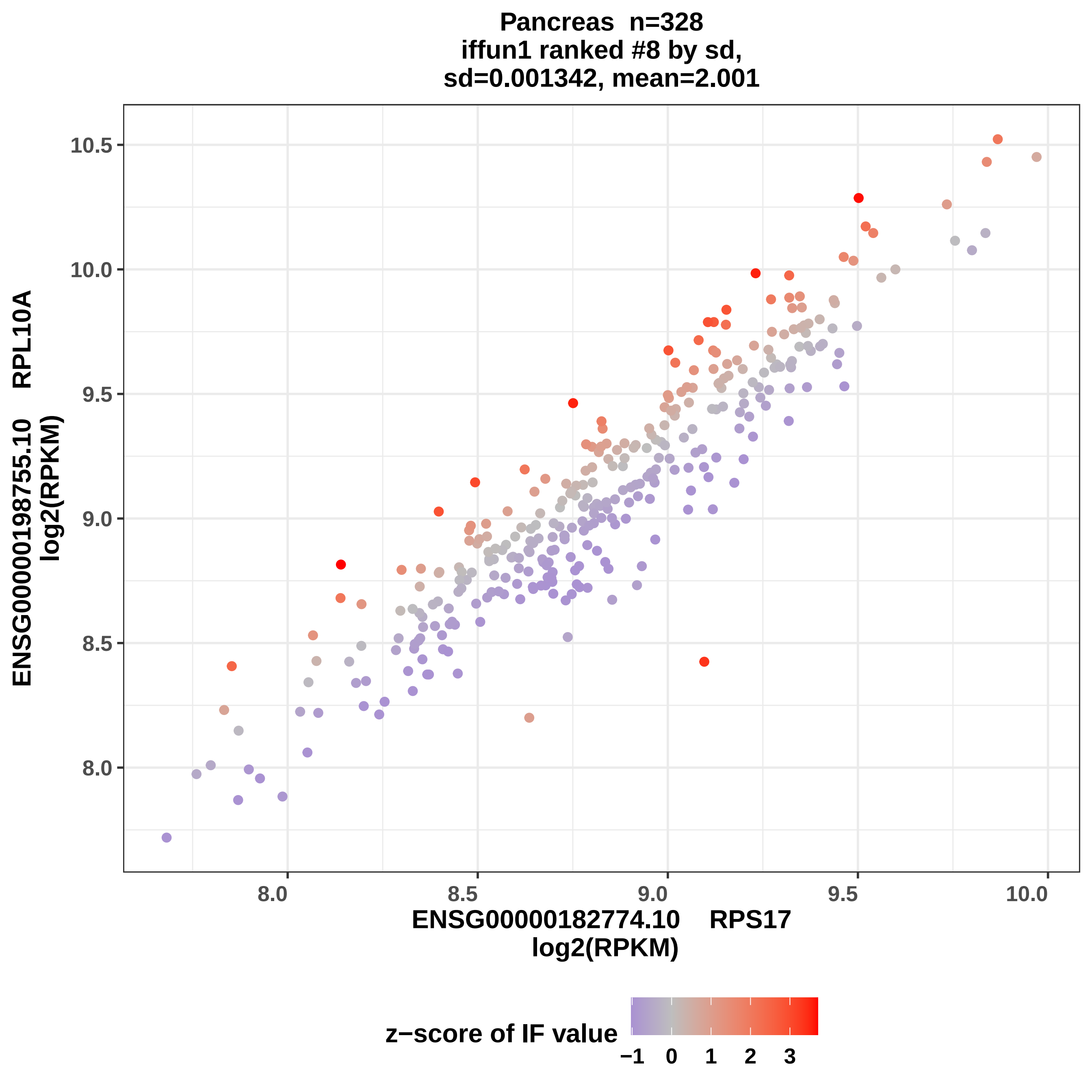
Pancreas n=328
iffun1 ranked #8 by sd,
sd=0.001342, mean=2.001
ENSG00000198755.10 RPL10A
log2(RPKM)
ENSG00000182774.10 RPS17
log2(RPKM)
10.5
10.0
9.5
9.0
8.5
8.0
8.0
8.5
9.0
9.5
10.0
z−score of IF value
−1
0
1
2
3

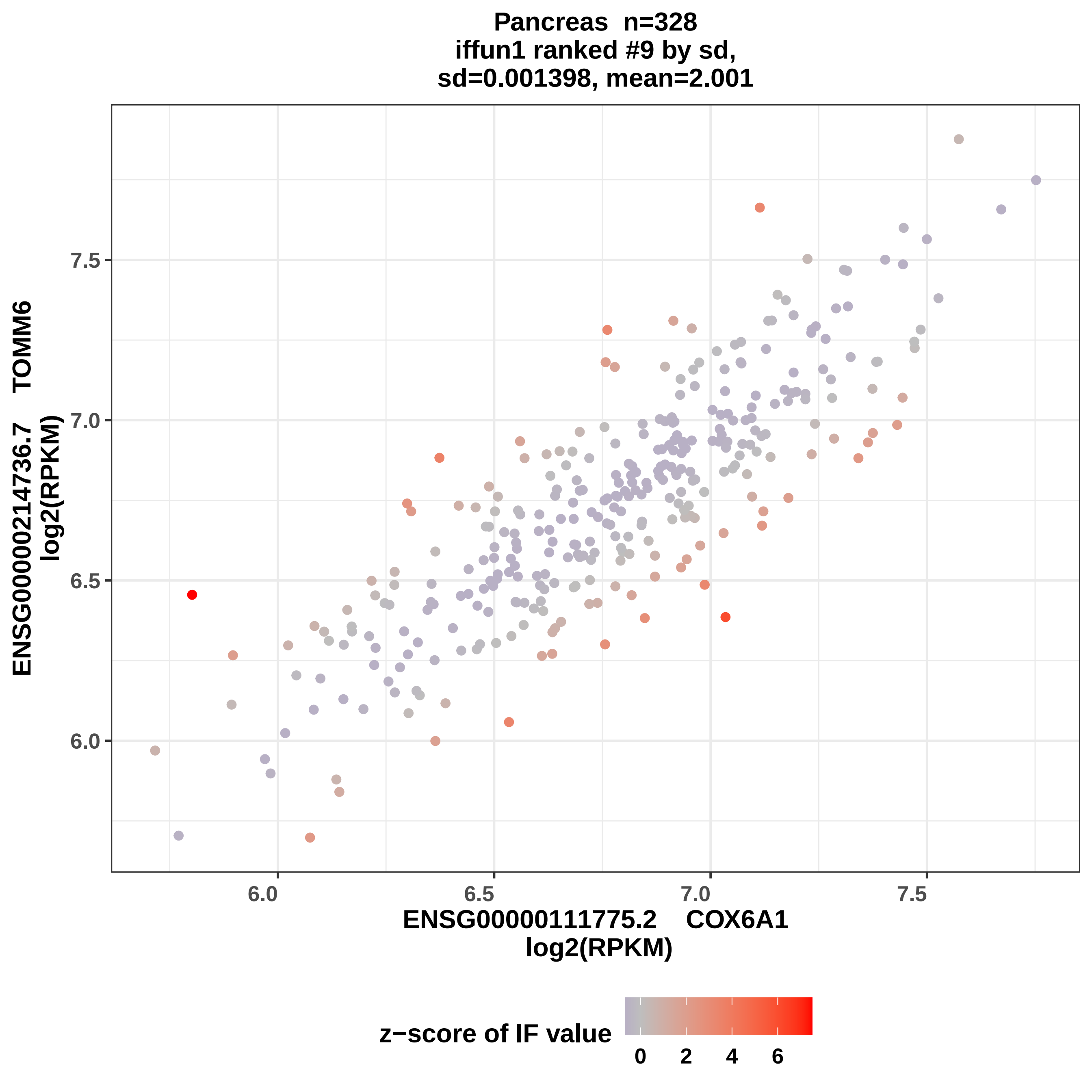

Pancreas n=328
iffun1 ranked #9 by sd,
sd=0.001398, mean=2.001
ENSG00000214736.7 TOMM6
log2(RPKM)
7.5
7.0
6.5
6.0
6.0
6.5
7.0
7.5
ENSG00000111775.2 COX6A1
log2(RPKM)
z−score of IF value
0
2
4
6

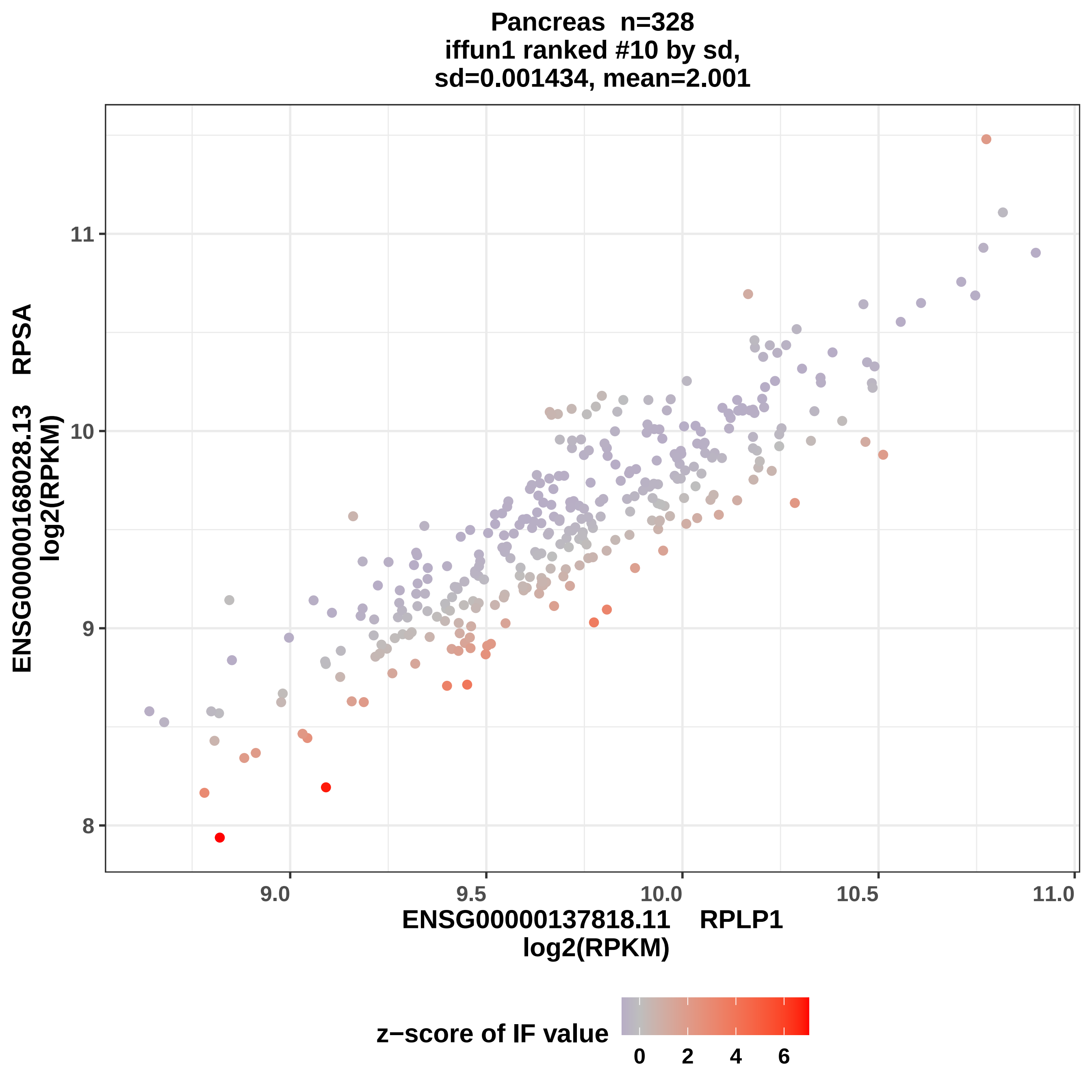

Pancreas n=328
iffun1 ranked #10 by sd,
sd=0.001434, mean=2.001
ENSG00000168028.13 RPSA
log2(RPKM)
11
10
9
8
9.0
9.5
10.0
10.5
11.0
ENSG00000137818.11 RPLP1
log2(RPKM)
z−score of IF value
0
2
4
6

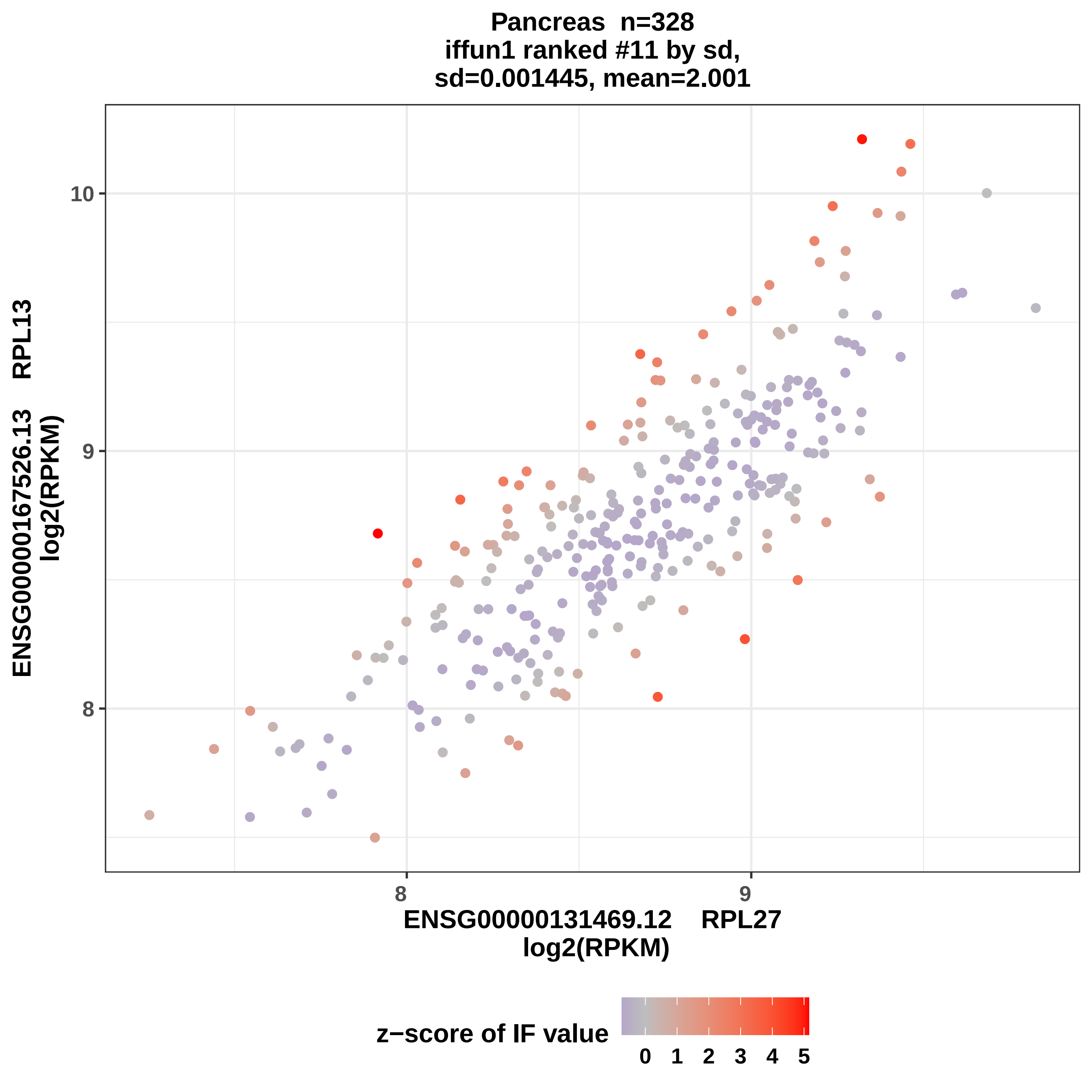

Pancreas n=328
iffun1 ranked #11 by sd,
sd=0.001445, mean=2.001
10
9
8
8
9
ENSG00000131469.12 RPL27
log2(RPKM)
ENSG00000167526.13 RPL13
log2(RPKM)
z−score of IF value
0 1 2 3 4 5

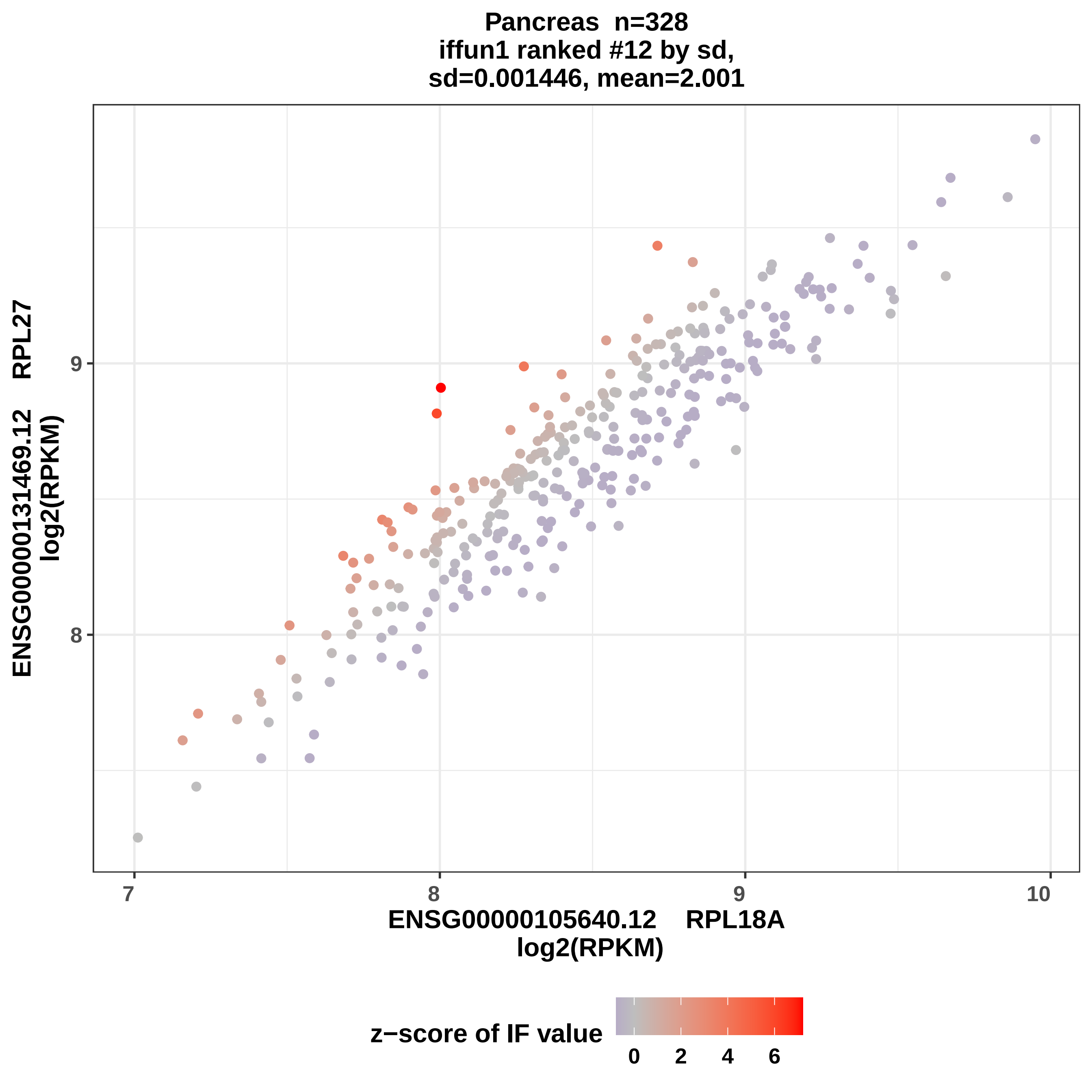
Pancreas n=328
iffun1 ranked #12 by sd,
sd=0.001446, mean=2.001
ENSG00000131469.12 RPL27
log2(RPKM)
9
8
7
8
9
10
ENSG00000105640.12 RPL18A
log2(RPKM)
z−score of IF value
0
2
4
6

Pancreas n=328
iffun1 ranked #13 by sd,
sd=0.00147, mean=2.001

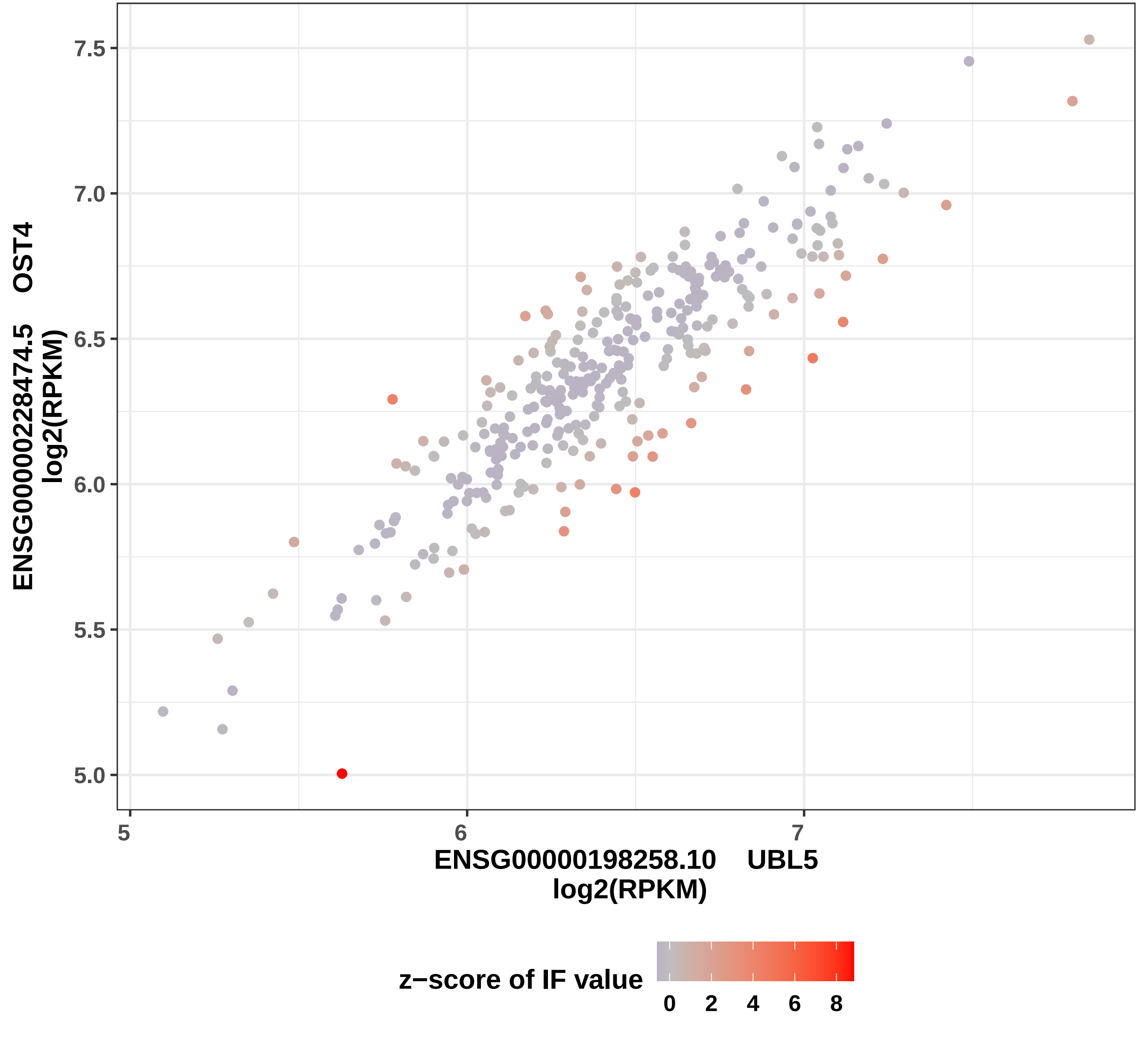

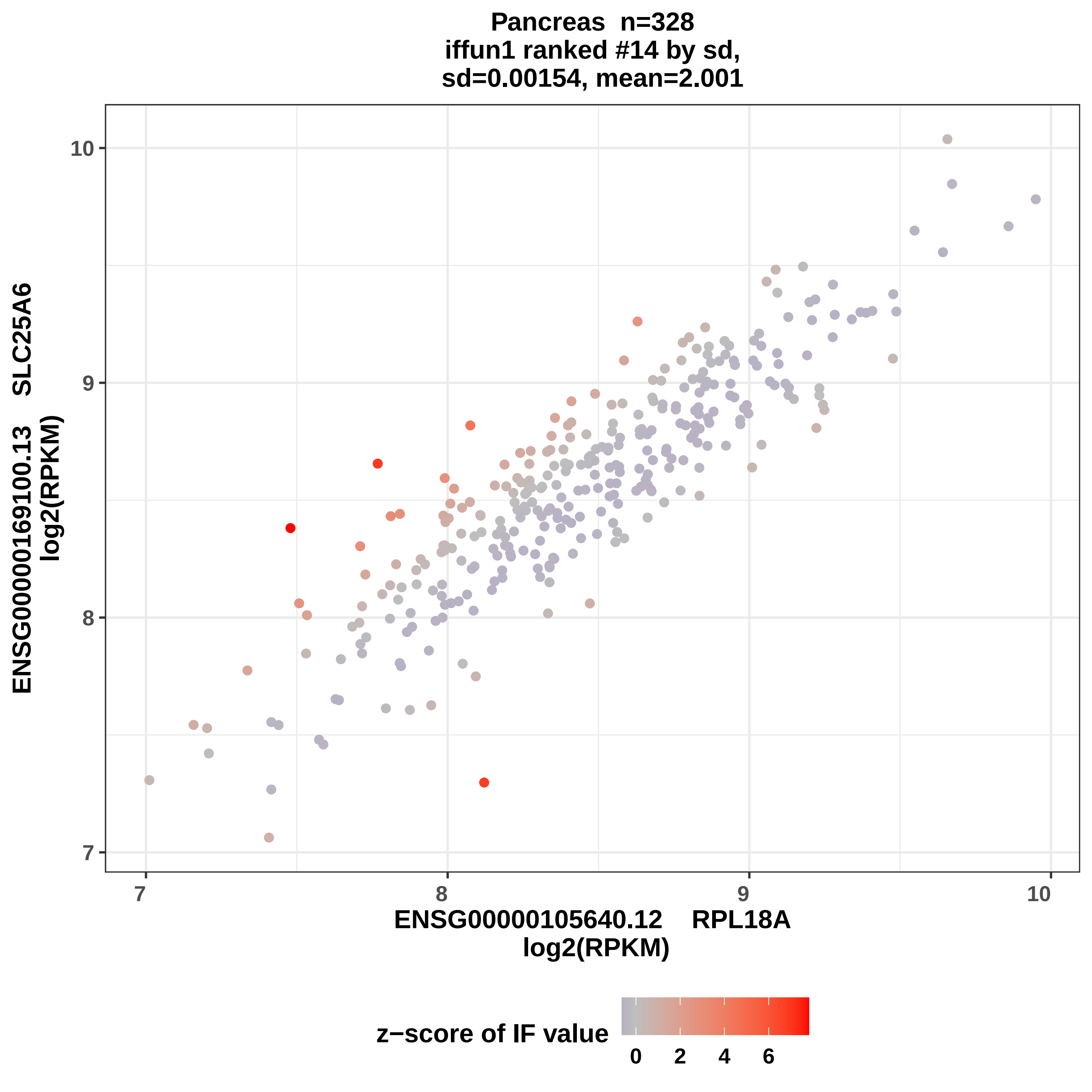

Pancreas n=328
iffun1 ranked #14 by sd,
sd=0.00154, mean=2.001
ENSG00000169100.13 SLC25A6
log2(RPKM)
ENSG00000105640.12 RPL18A
log2(RPKM)
z−score of IF value
7
8
9
10
0
2
4
6

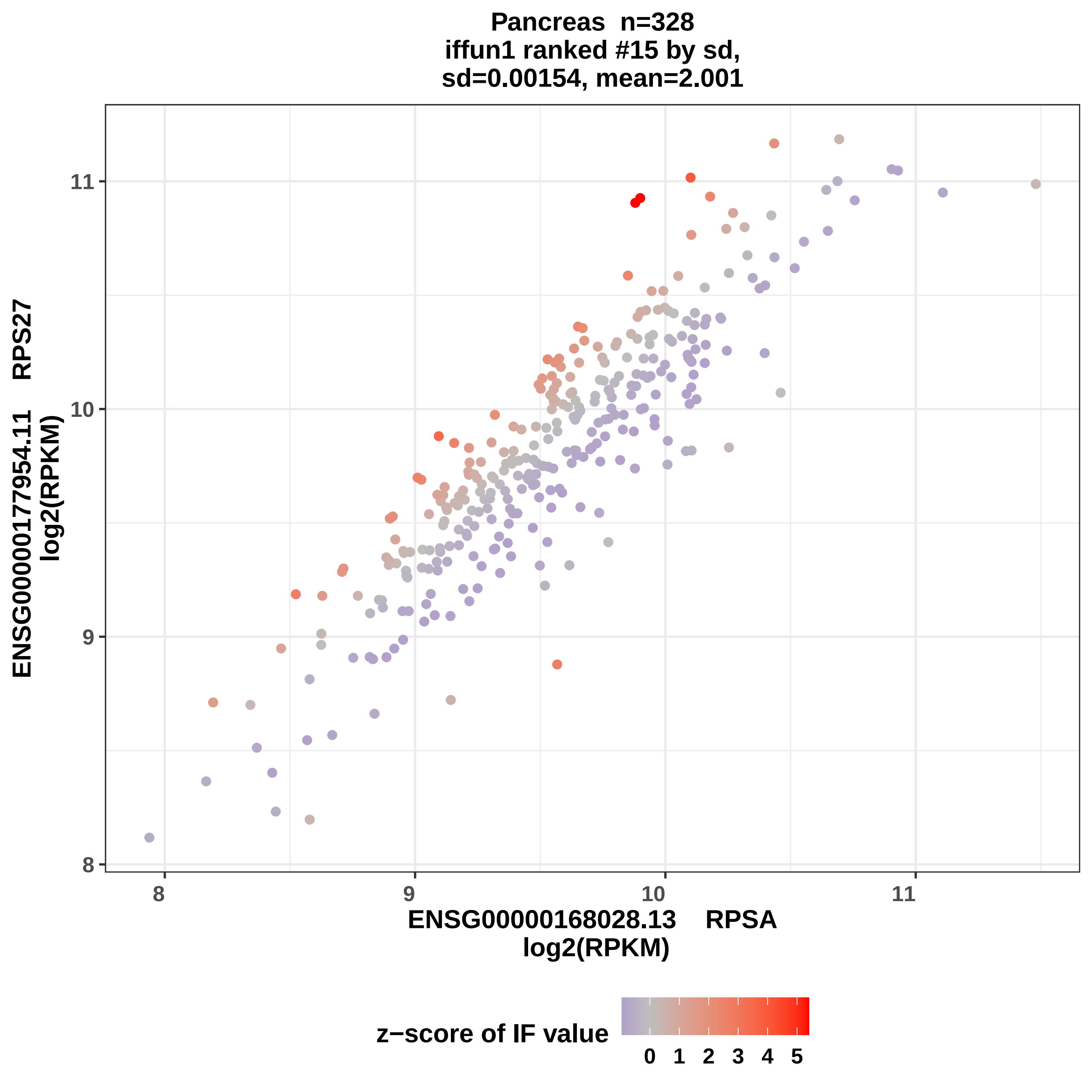
Pancreas n=328
iffun1 ranked #15 by sd,
sd=0.00154, mean=2.001
ENSG00000177954.11 RPS27
log2(RPKM)
8
9
10
11
ENSG00000168028.13 RPSA
log2(RPKM)
z−score of IF value
0 1 2 3 4 5

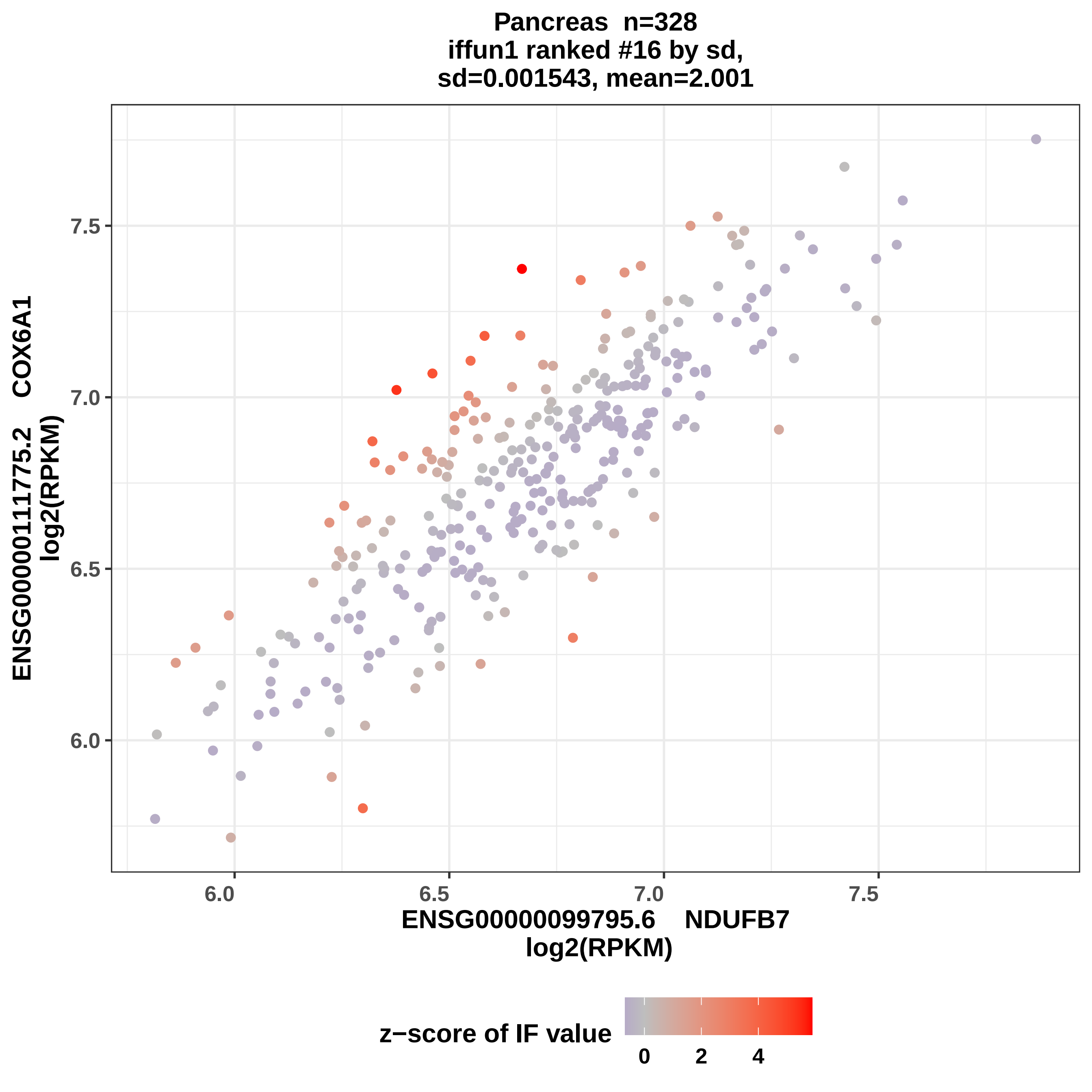

Pancreas n=328
iffun1 ranked #16 by sd,
sd=0.001543, mean=2.001
ENSG00000111775.2 COX6A1
log2(RPKM)
ENSG00000099795.6 NDUFB7
log2(RPKM)
6.0
6.5
7.0
7.5
z−score of IF value
0
2
4

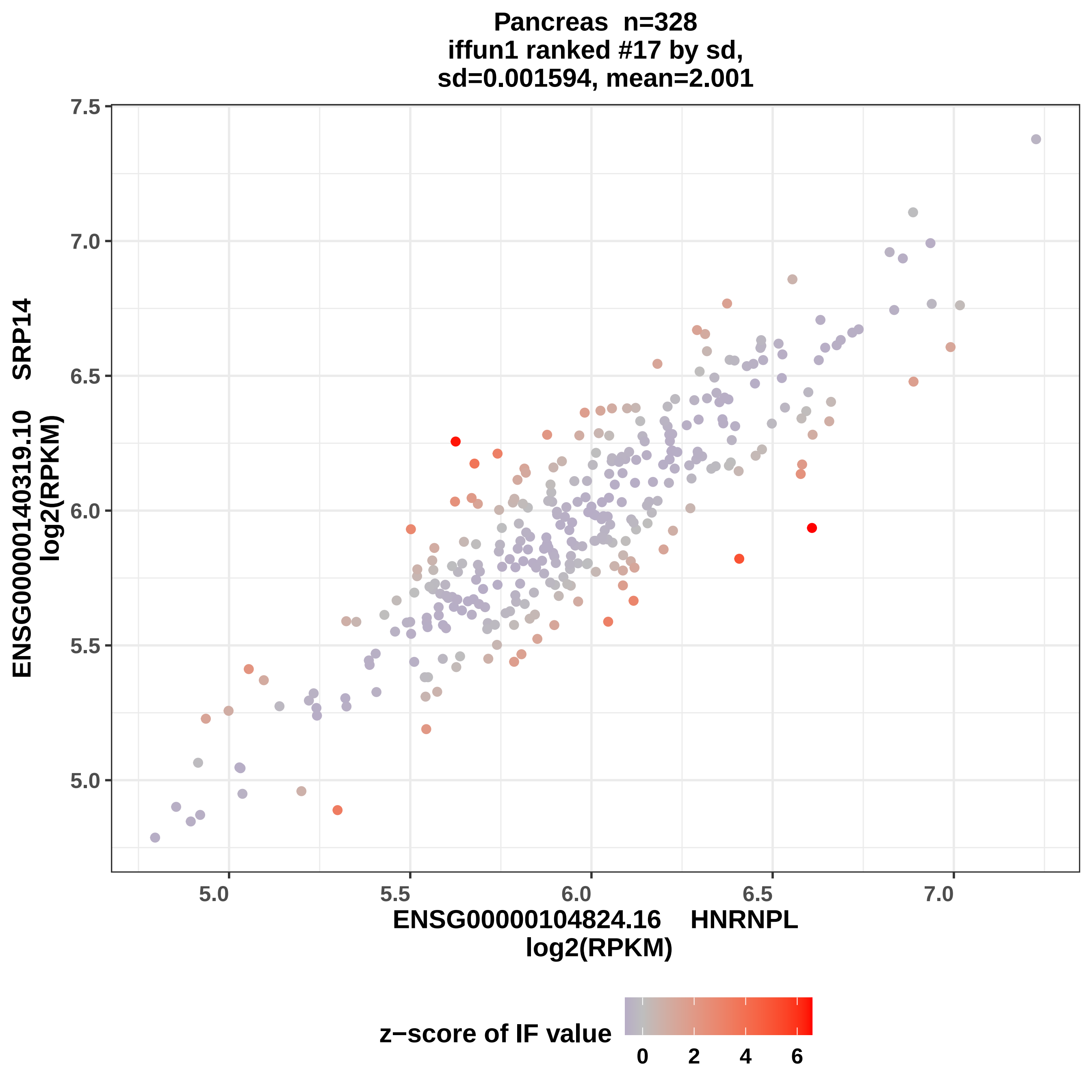

Pancreas n=328
iffun1 ranked #17 by sd,
sd=0.001594, mean=2.001
7.5
7.0
6.5
6.0
5.5
5.0
5.0
5.5
6.0
6.5
7.0
ENSG00000140319.10 SRP14
log2(RPKM)
ENSG00000104824.16 HNRNPL
log2(RPKM)
z−score of IF value
0
2
4
6

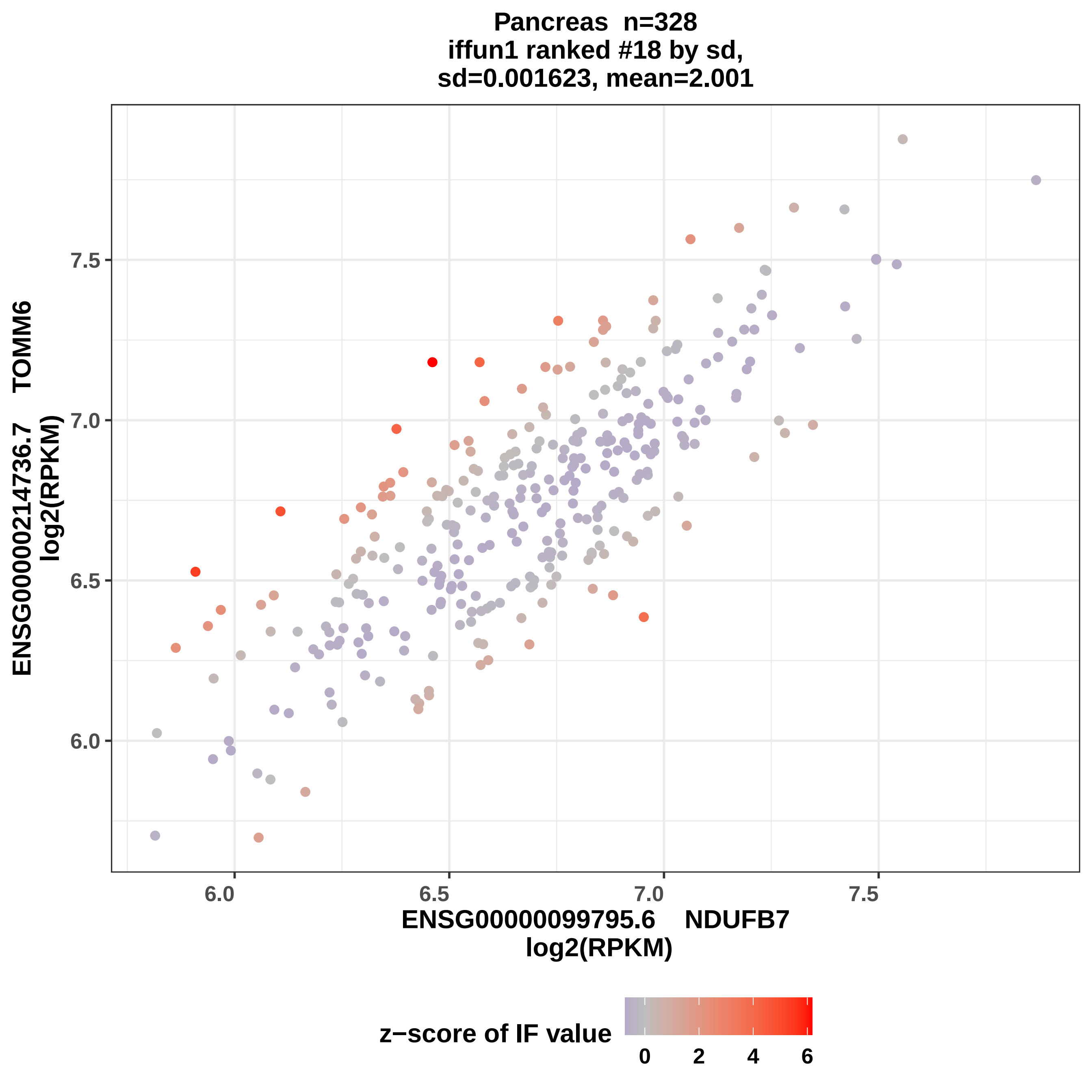

Pancreas n=328
iffun1 ranked #18 by sd,
sd=0.001623, mean=2.001
ENSG00000214736.7 TOMM6
log2(RPKM)
7.5
7.0
6.5
6.0
6.0
6.5
7.0
7.5
ENSG00000099795.6 NDUFB7
log2(RPKM)
z−score of IF value
0
2
4
6

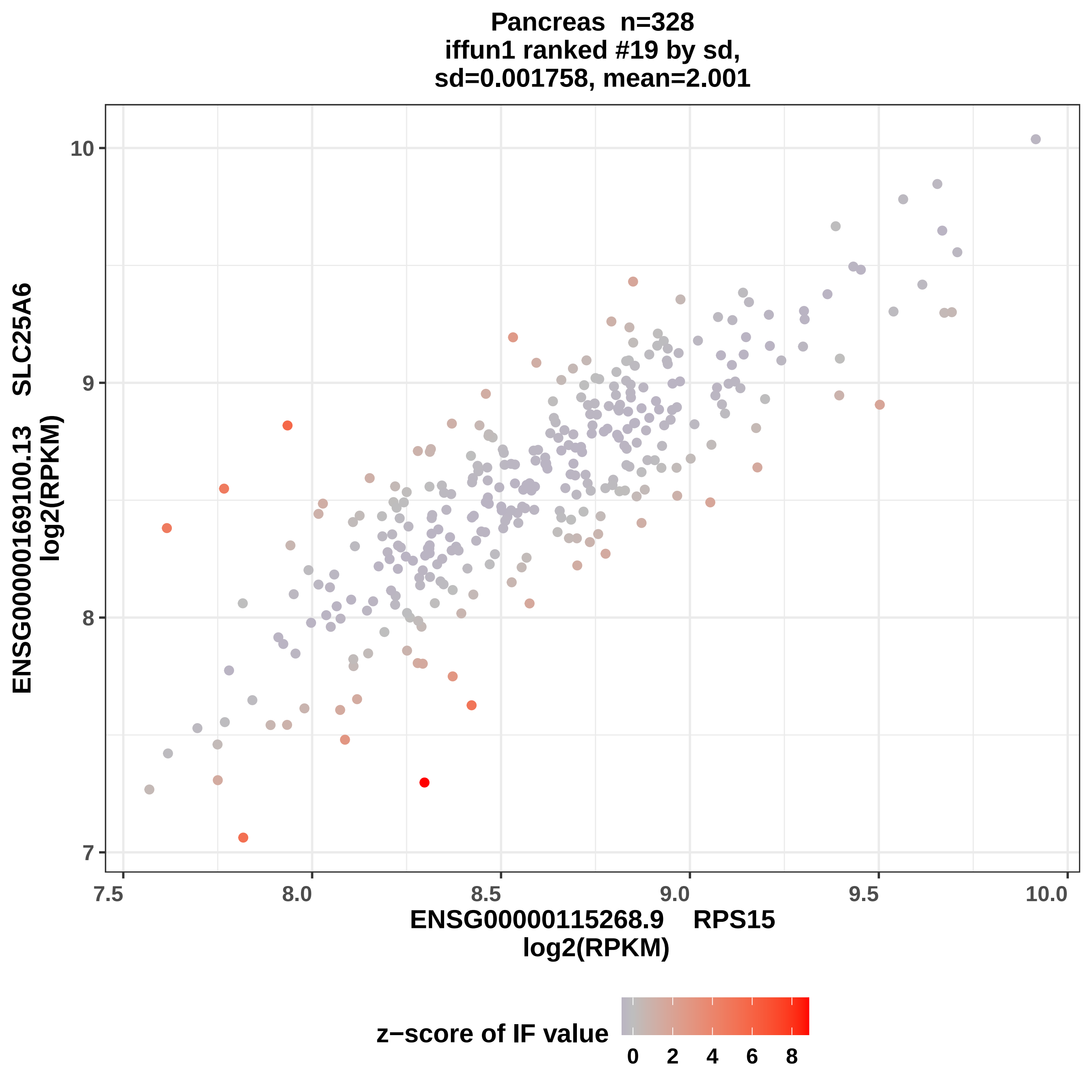
Pancreas n=328
iffun1 ranked #19 by sd,
sd=0.001758, mean=2.001
ENSG00000169100.13 SLC25A6
log2(RPKM)
ENSG00000115268.9 RPS15
log2(RPKM)
z−score of IF value
7
8
9
10
7.5
8.0
8.5
9.0
9.5
10.0
0
2
4
6
8

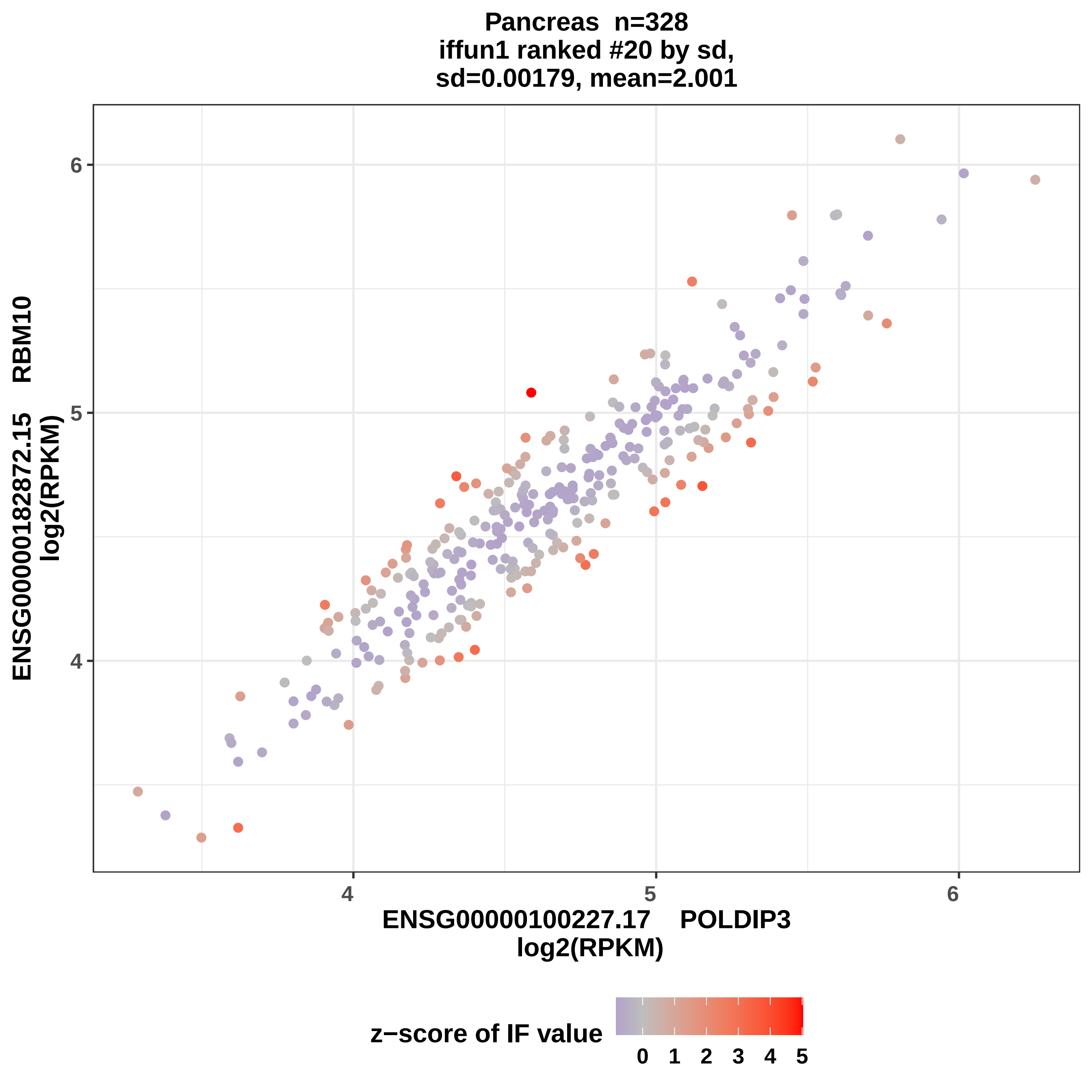
Pancreas n=328
iffun1 ranked #20 by sd,
sd=0.00179, mean=2.001
ENSG00000182872.15 RBM10
log2(RPKM)
ENSG00000100227.17 POLDIP3
log2(RPKM)
4
5
6
z−score of IF value
0 1 2 3 4 5

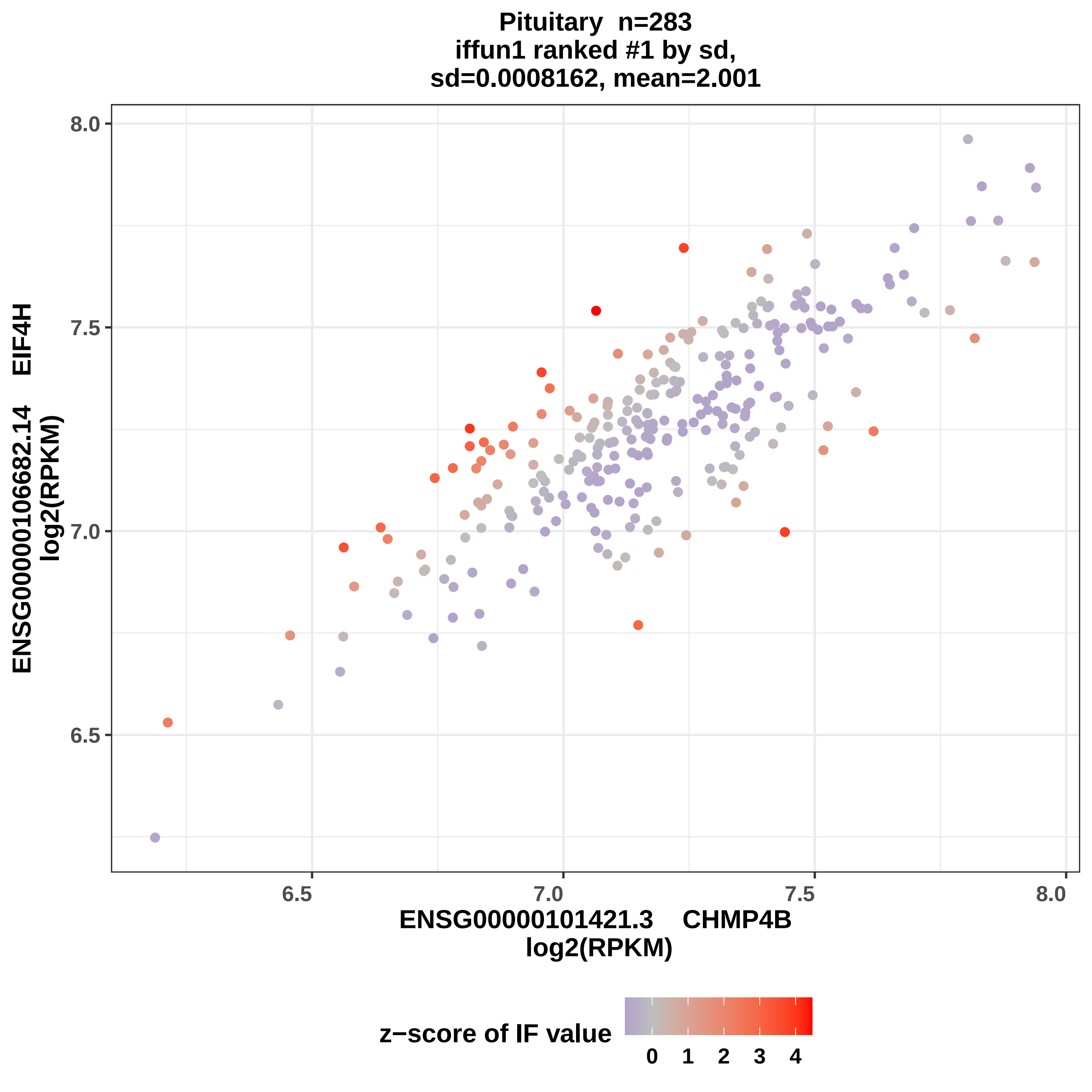
Pituitary n=283
iffun1 ranked #1 by sd,
sd=0.0008162, mean=2.001
ENSG00000106682.14 EIF4H
log2(RPKM)
ENSG00000101421.3 CHMP4B
log2(RPKM)
8.0
7.5
7.0
6.5
6.5
7.0
7.5
8.0
z−score of IF value
0
1
2
3
4

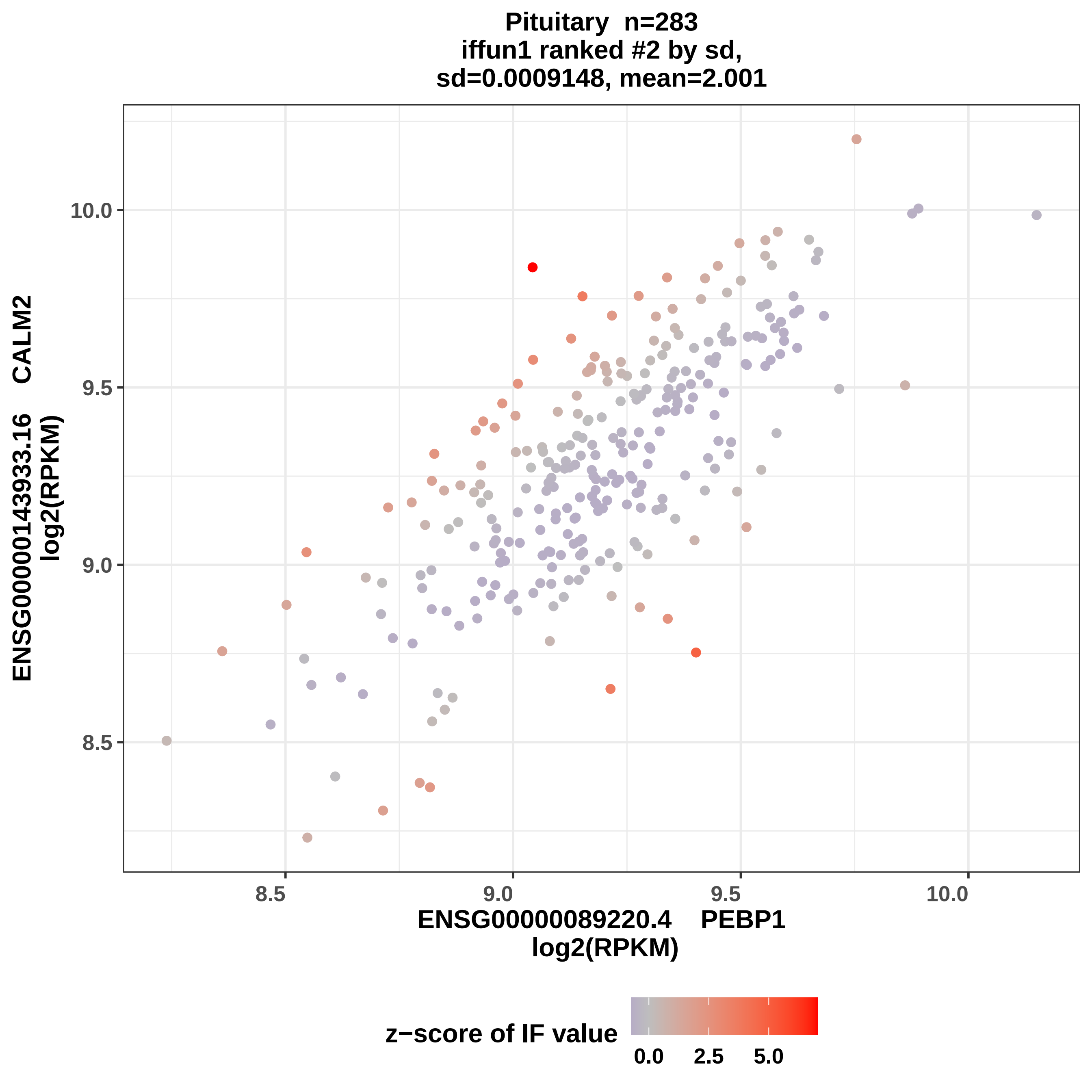

Pituitary n=283
iffun1 ranked #2 by sd,
sd=0.0009148, mean=2.001
ENSG00000143933.16 CALM2
log2(RPKM)
ENSG00000089220.4 PEBP1
log2(RPKM)
8.5
9.0
9.5
10.0
z−score of IF value
0.0
2.5
5.0

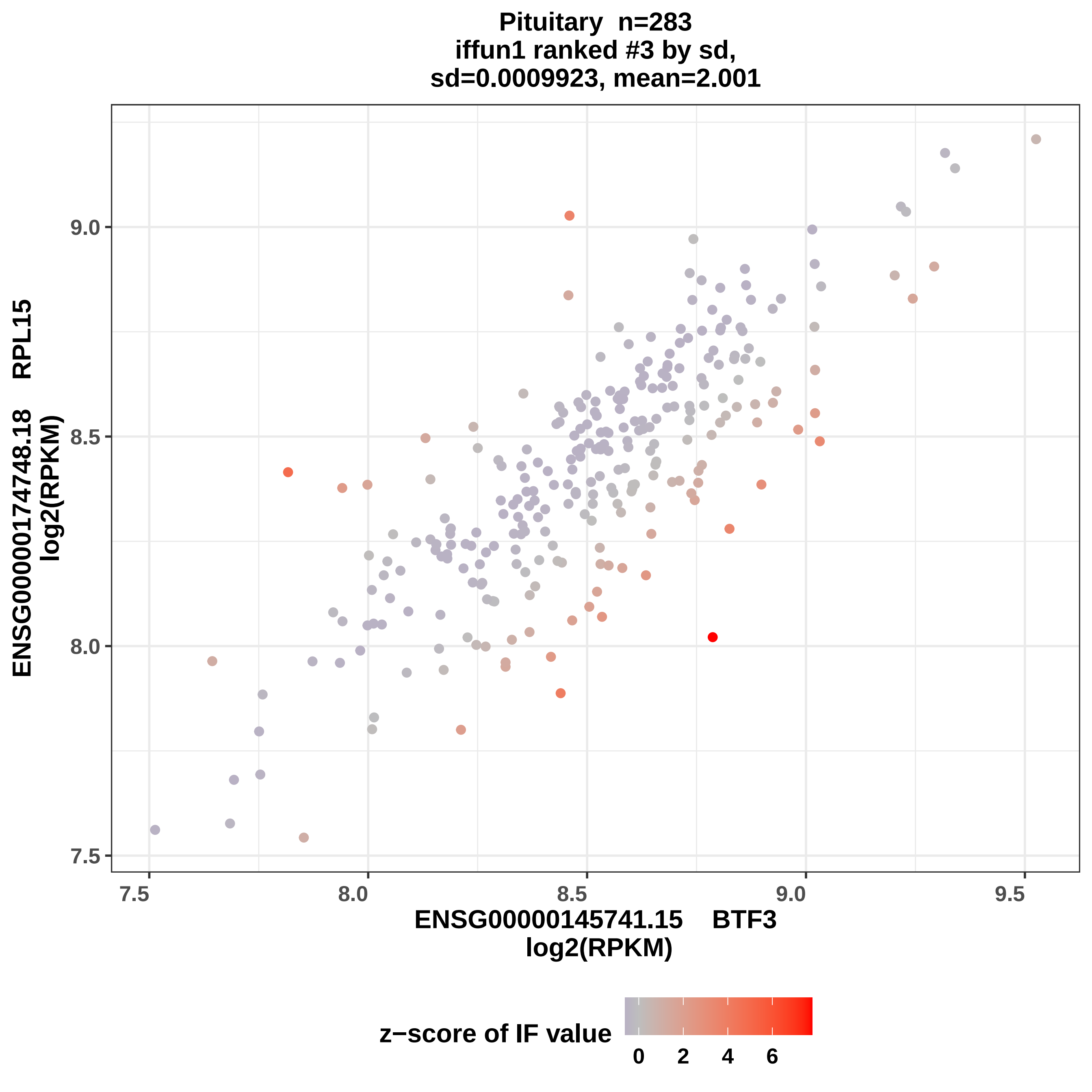

Pituitary n=283
iffun1 ranked #3 by sd,
sd=0.0009923, mean=2.001
ENSG00000174748.18 RPL15
log2(RPKM)
9.0
8.5
8.0
7.5
7.5
8.0
8.5
9.0
9.5
ENSG00000145741.15 BTF3
log2(RPKM)
z−score of IF value
0
2
4
6

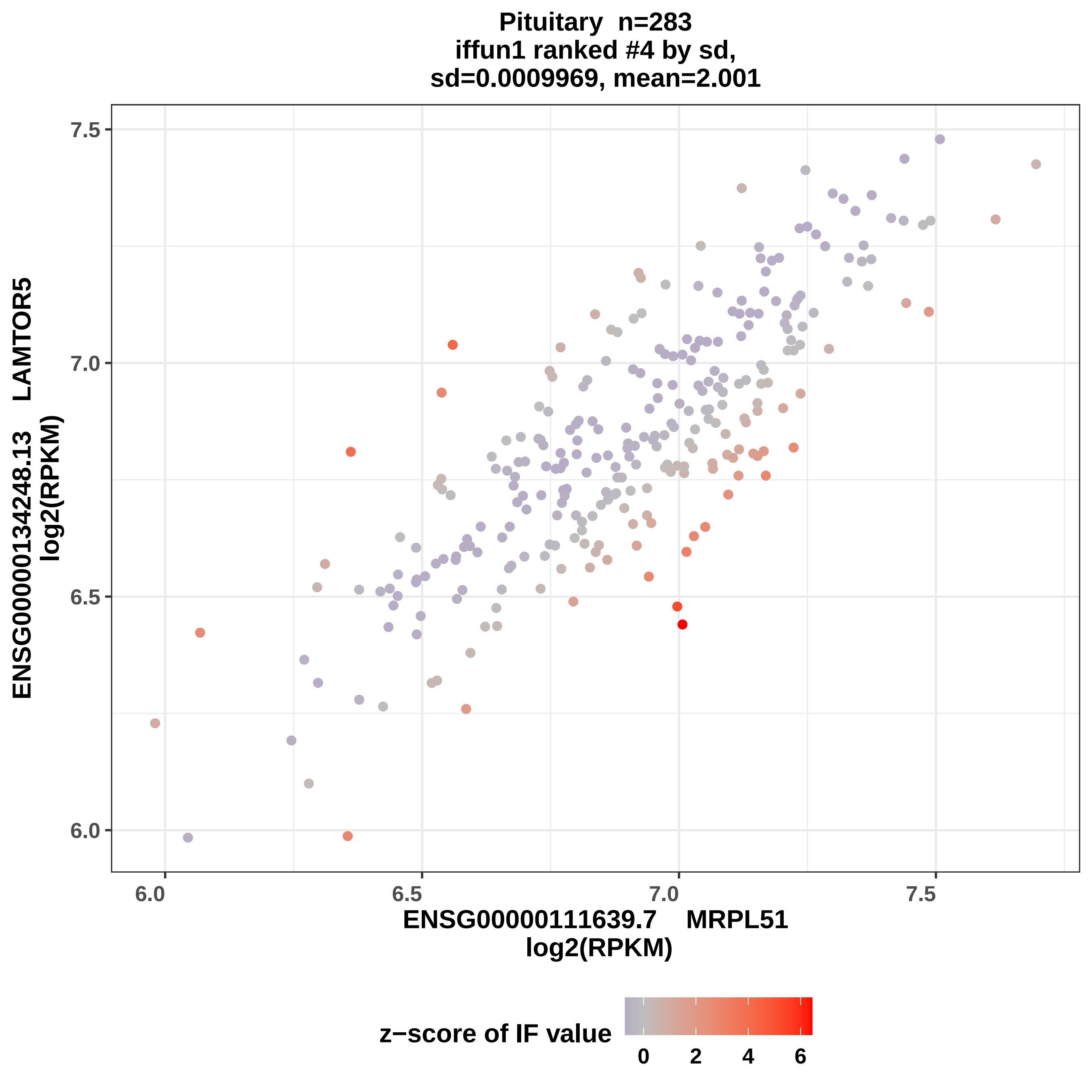
Pituitary n=283
iffun1 ranked #4 by sd,
sd=0.0009969, mean=2.001
ENSG00000134248.13 LAMTOR5
log2(RPKM)
ENSG00000111639.7 MRPL51
log2(RPKM)
7.5
7.0
6.5
6.0
6.0
6.5
7.0
7.5
z−score of IF value
0
2
4
6

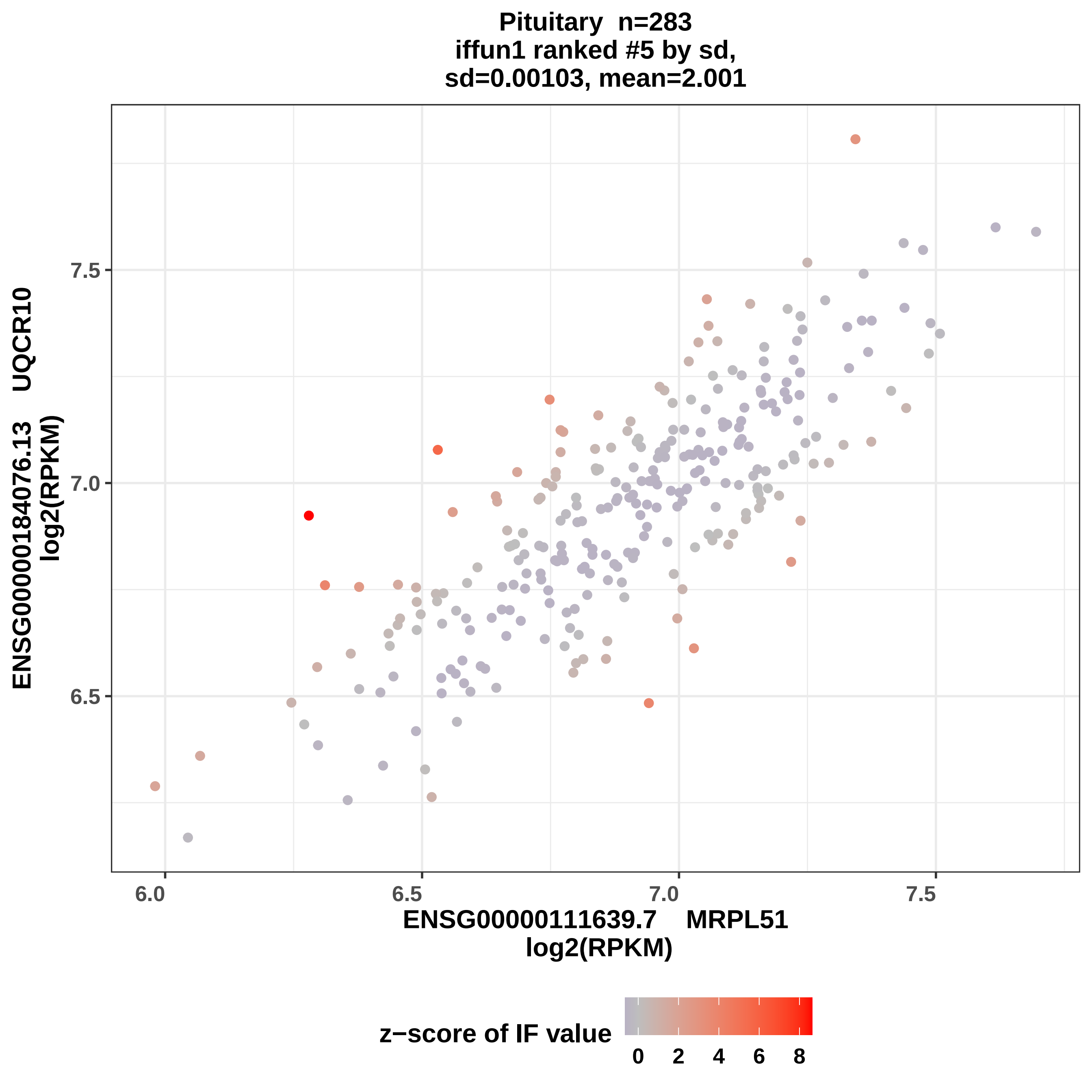

Pituitary n=283
iffun1 ranked #5 by sd,
sd=0.00103, mean=2.001
ENSG00000184076.13 UQCR10
log2(RPKM)
7.5
7.0
6.5
6.0
6.5
7.0
7.5
ENSG00000111639.7 MRPL51
log2(RPKM)
z−score of IF value
0
2
4
6
8

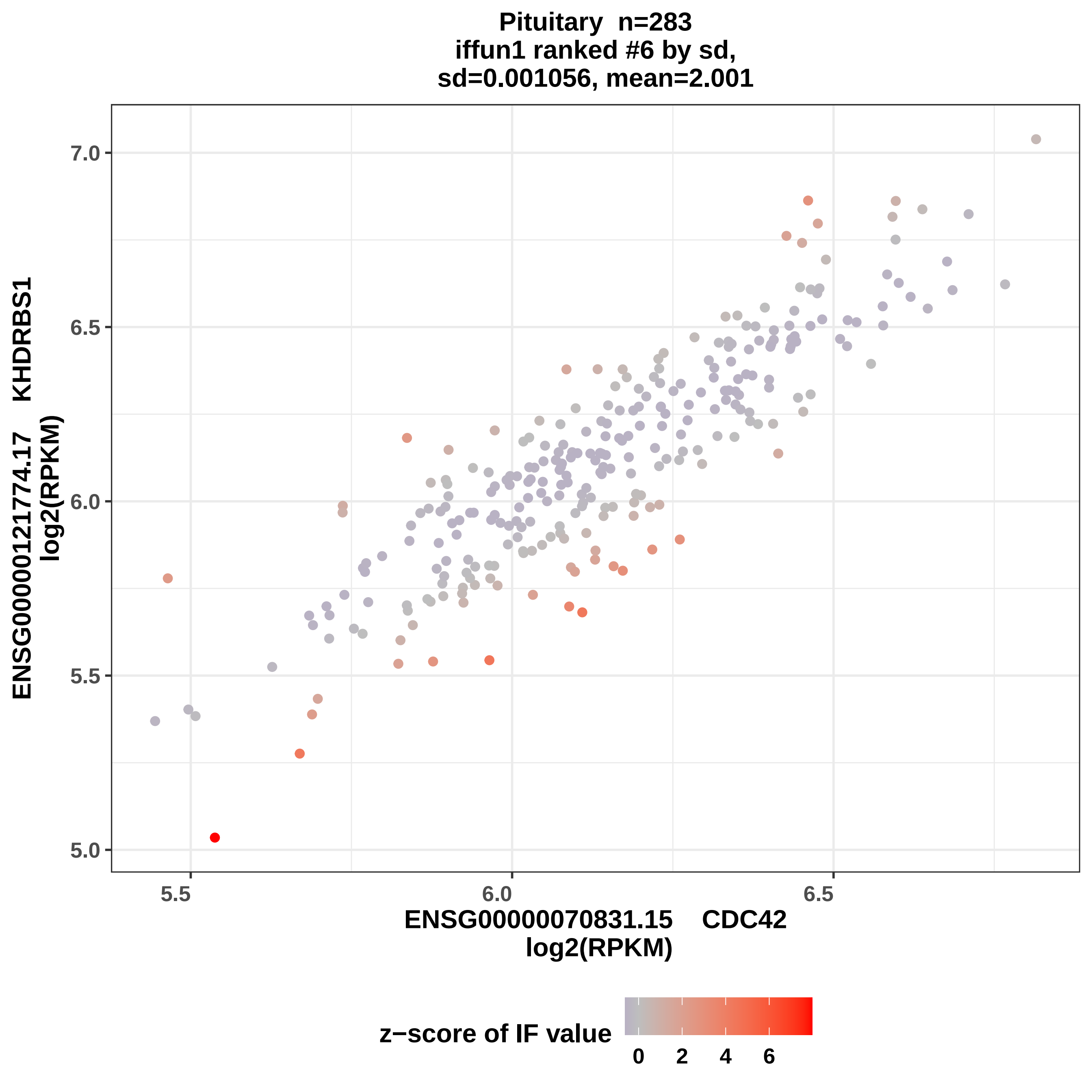
Pituitary n=283
iffun1 ranked #6 by sd,
sd=0.001056, mean=2.001
7.0
6.5
6.0
5.5
5.0
5.5
6.0
6.5
ENSG00000121774.17 KHDRBS1
log2(RPKM)
ENSG00000070831.15 CDC42
log2(RPKM)
z−score of IF value
0
2
4
6

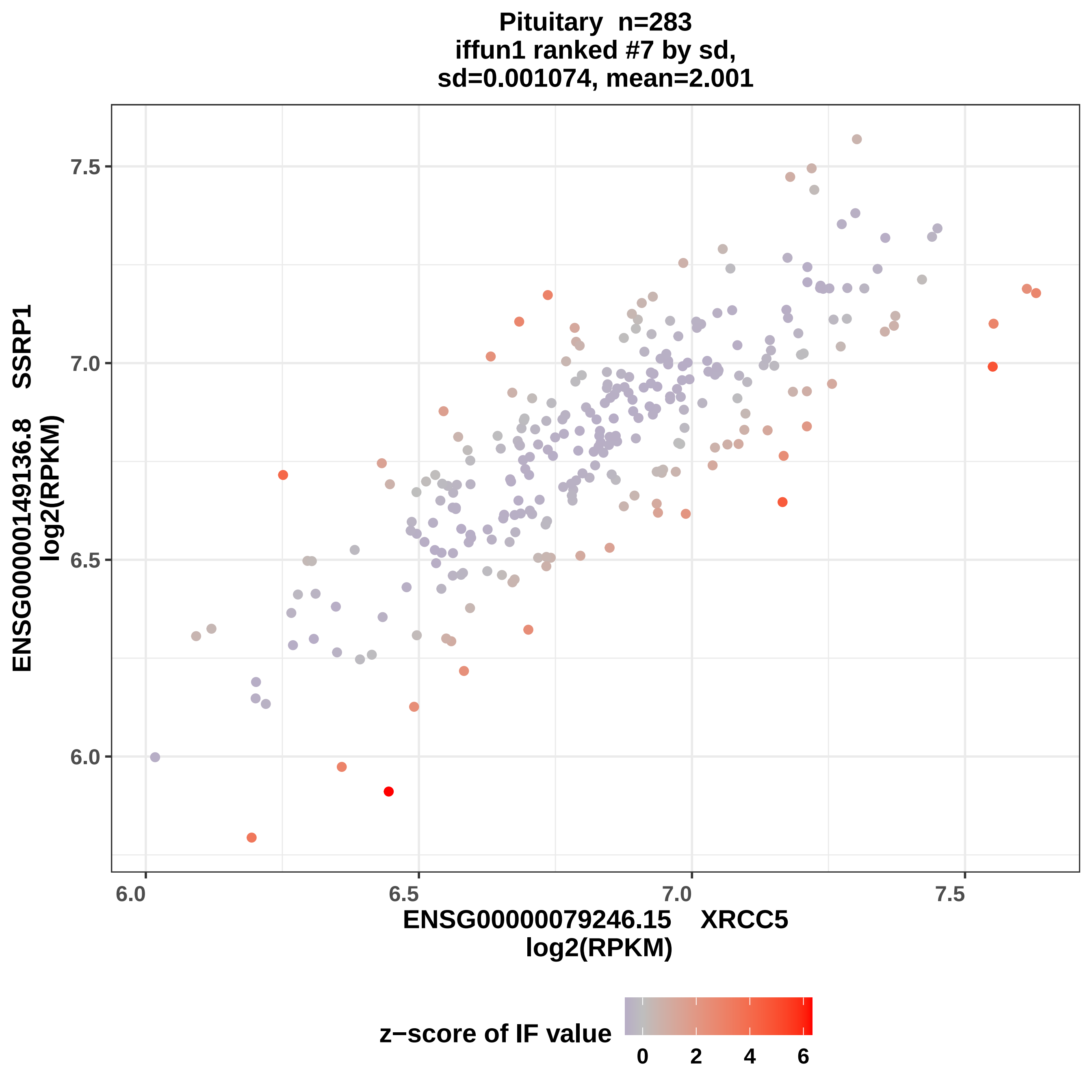

Pituitary n=283
iffun1 ranked #7 by sd,
sd=0.001074, mean=2.001
ENSG00000149136.8 SSRP1
log2(RPKM)
7.5
7.0
6.5
6.0
6.0
6.5
7.0
7.5
ENSG00000079246.15 XRCC5
log2(RPKM)
z−score of IF value
0
2
4
6

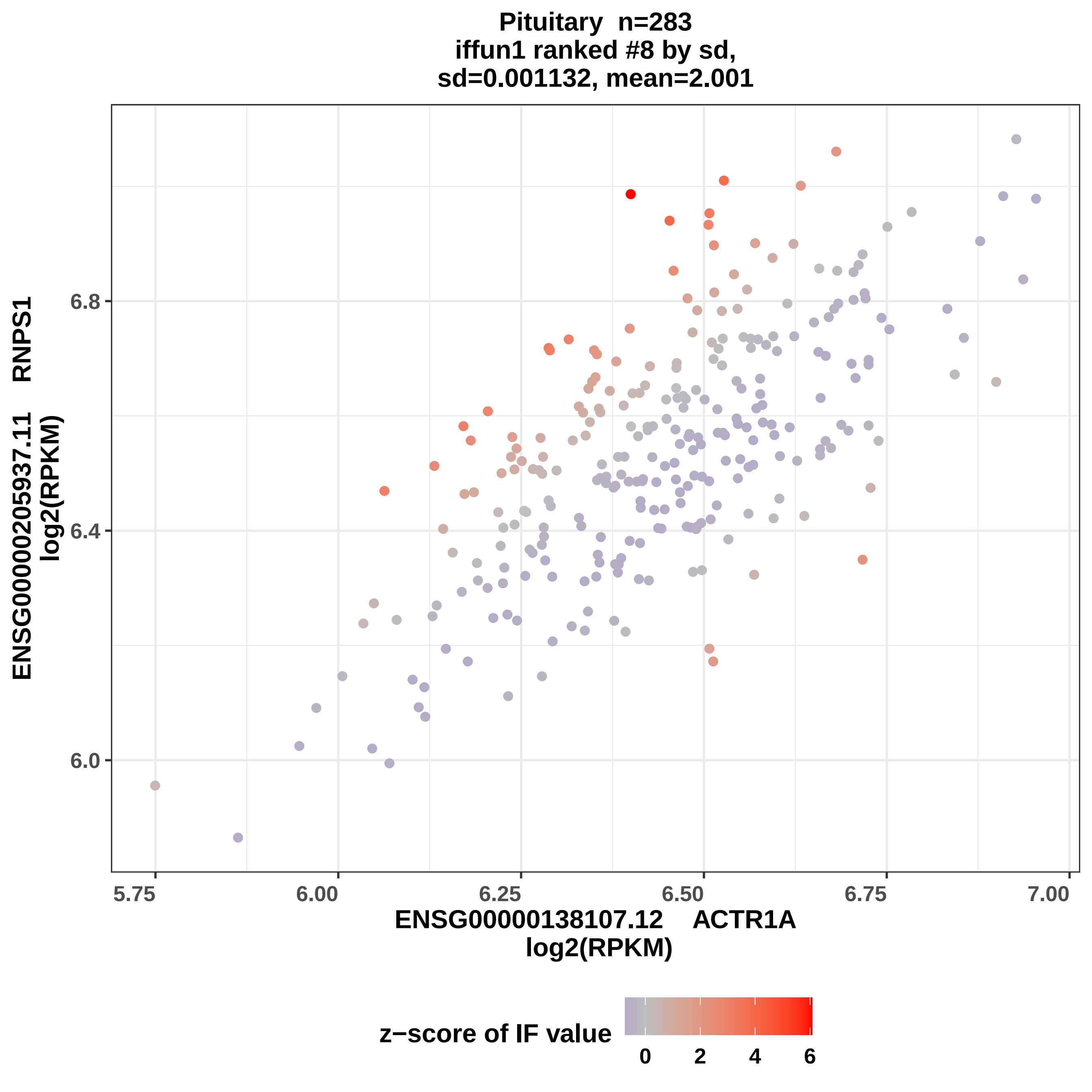

Pituitary n=283
iffun1 ranked #8 by sd,
sd=0.001132, mean=2.001
ENSG00000205937.11 RNPS1
log2(RPKM)
6.0
6.4
6.8
5.75
6.00
6.25
6.50
6.75
7.00
ENSG00000138107.12 ACTR1A
log2(RPKM)
z−score of IF value
0
2
4
6

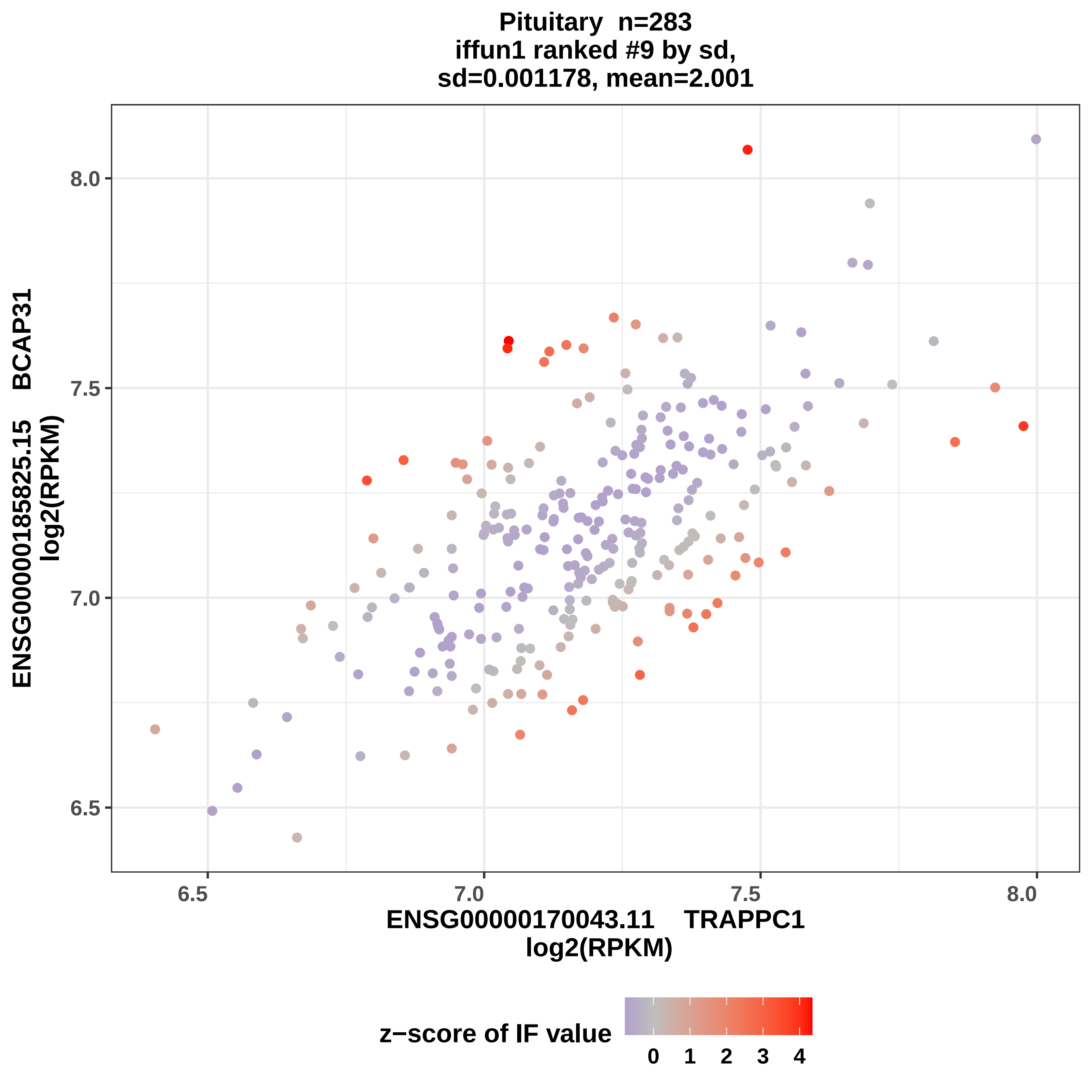

Pituitary n=283
iffun1 ranked #9 by sd,
sd=0.001178, mean=2.001
ENSG00000185825.15 BCAP31
log2(RPKM)
ENSG00000170043.11 TRAPPC1
log2(RPKM)
6.5
7.0
7.5
8.0
z−score of IF value
0
1
2
3
4

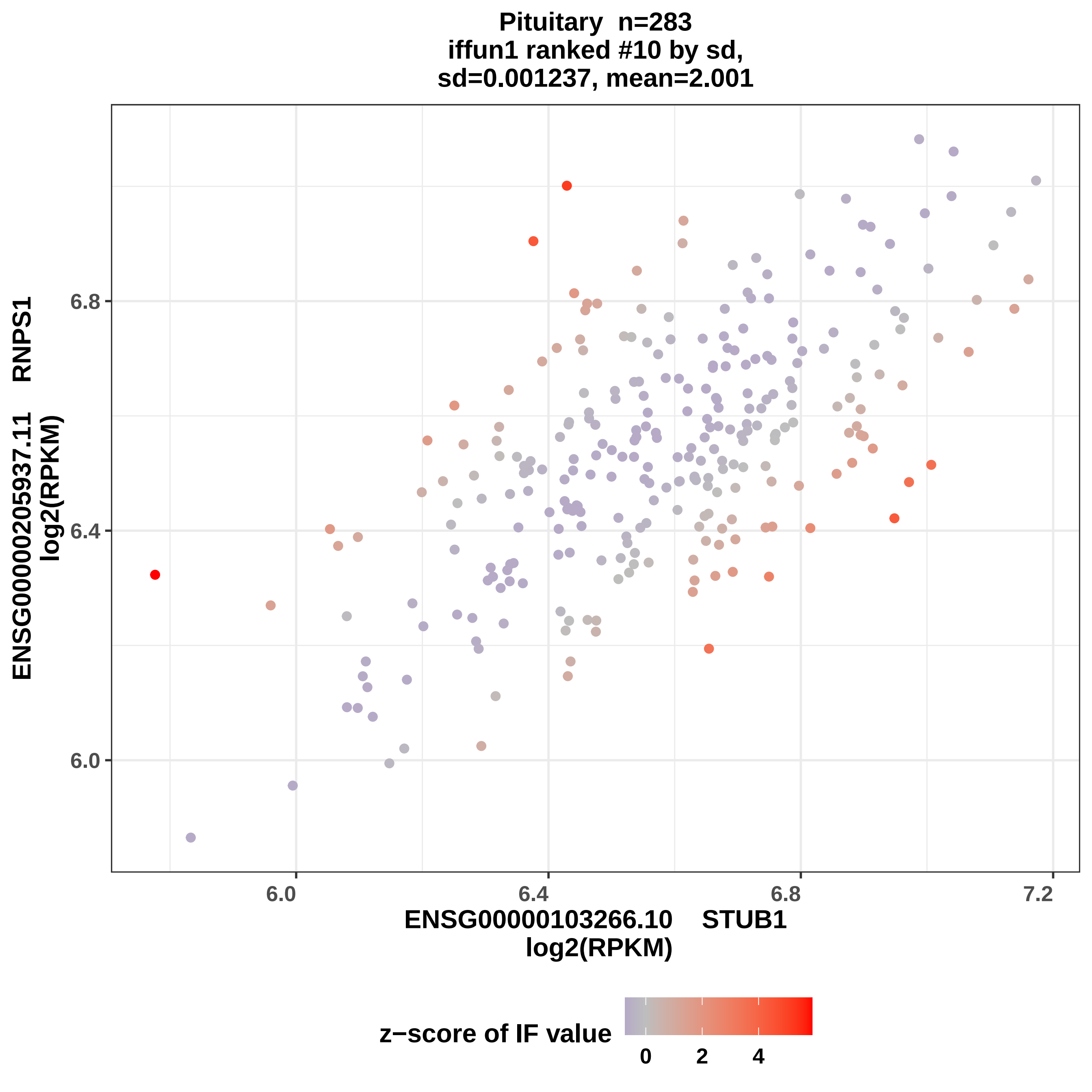
Pituitary n=283
iffun1 ranked #10 by sd,
sd=0.001237, mean=2.001
ENSG00000205937.11 RNPS1
log2(RPKM)
6.0
6.4
6.8
ENSG00000103266.10 STUB1
log2(RPKM)
6.0
6.4
6.8
7.2
z−score of IF value
0
2
4

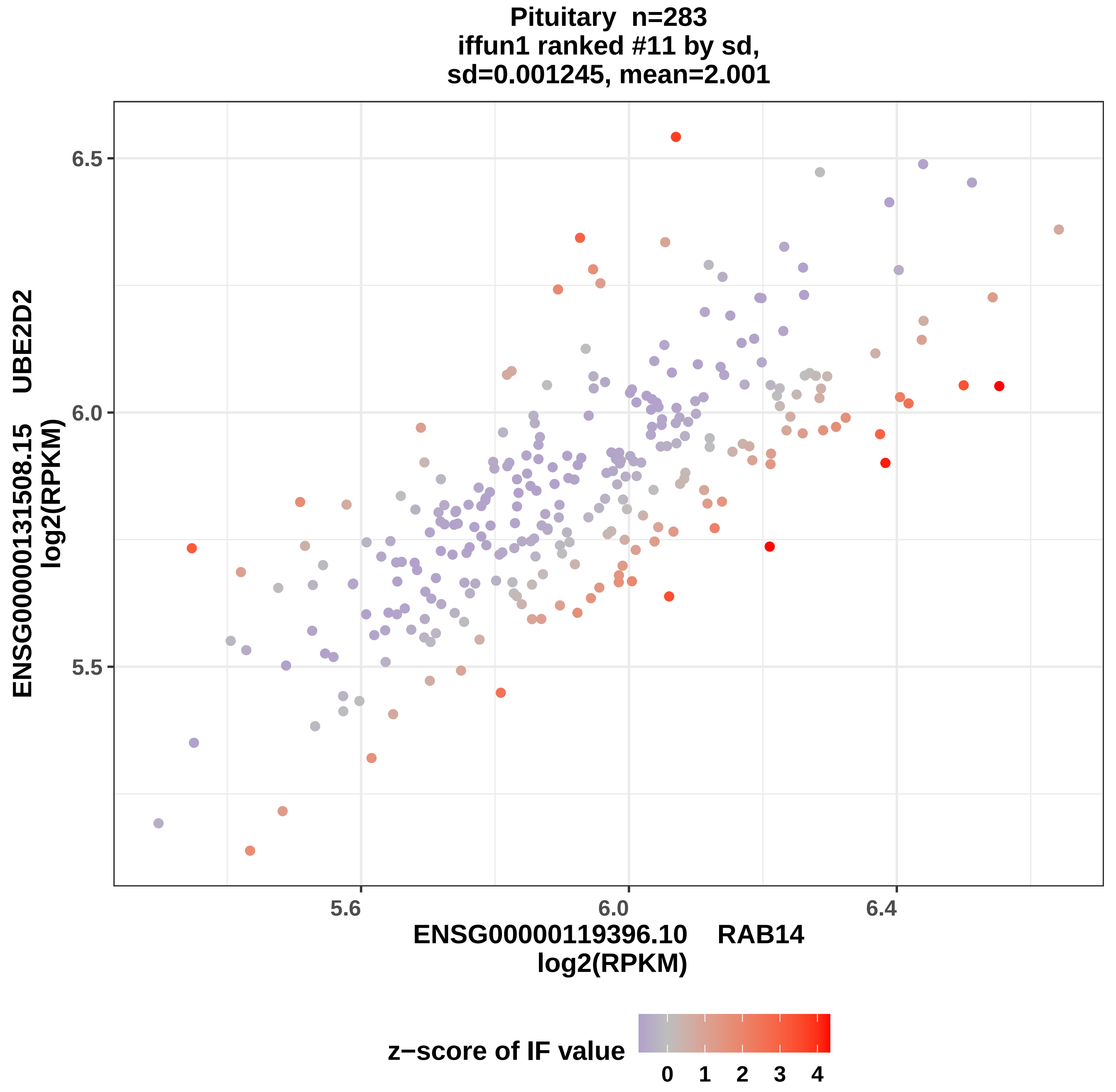
Pituitary n=283
iffun1 ranked #11 by sd,
sd=0.001245, mean=2.001
ENSG00000131508.15 UBE2D2
log2(RPKM)
6.5
6.0
5.5
5.6
6.0
6.4
ENSG00000119396.10 RAB14
log2(RPKM)
z−score of IF value
0
1
2
3
4

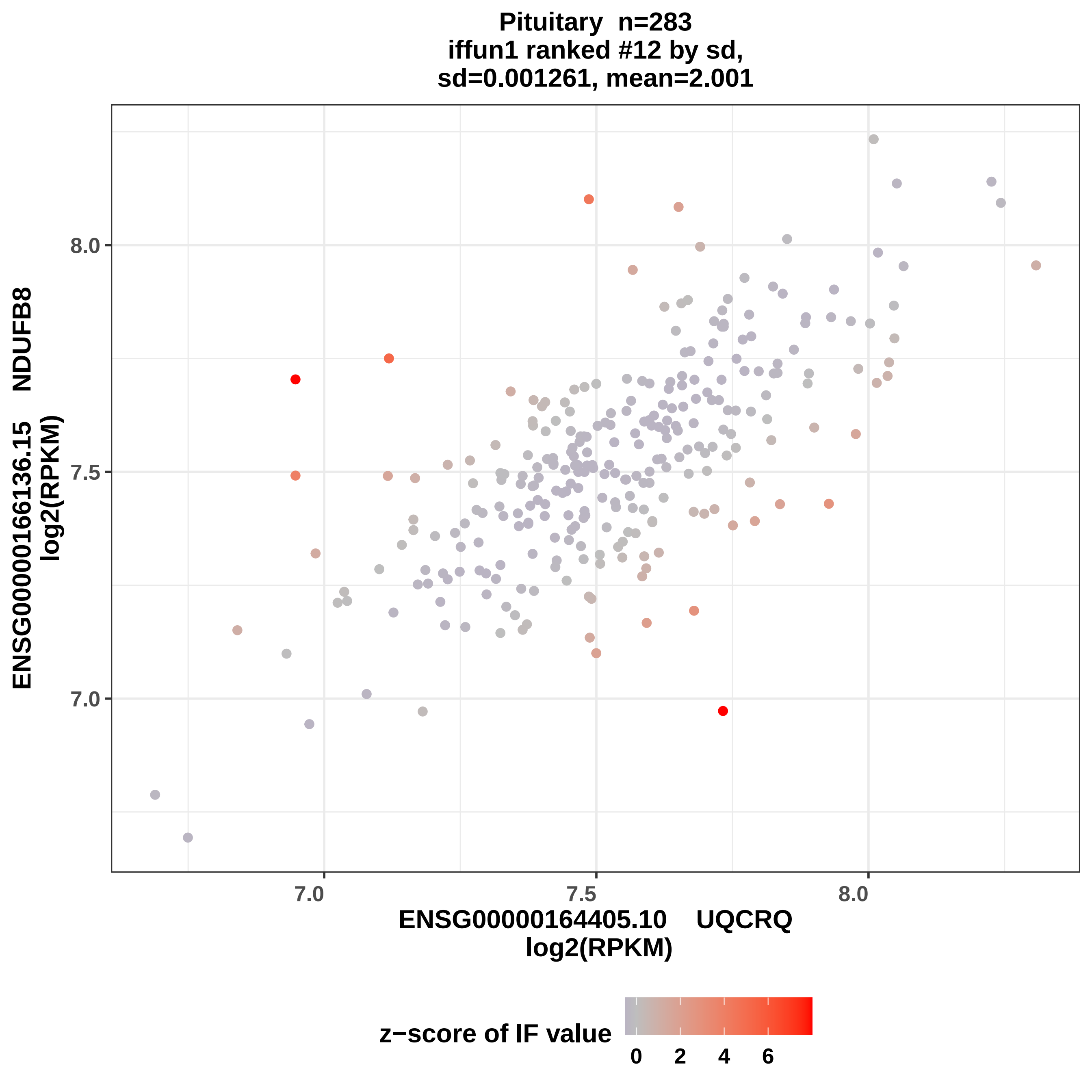
Pituitary n=283
iffun1 ranked #12 by sd,
sd=0.001261, mean=2.001
ENSG00000166136.15 NDUFB8
log2(RPKM)
ENSG00000164405.10 UQCRQ
log2(RPKM)
7.0
7.5
8.0
z−score of IF value
0
2
4
6

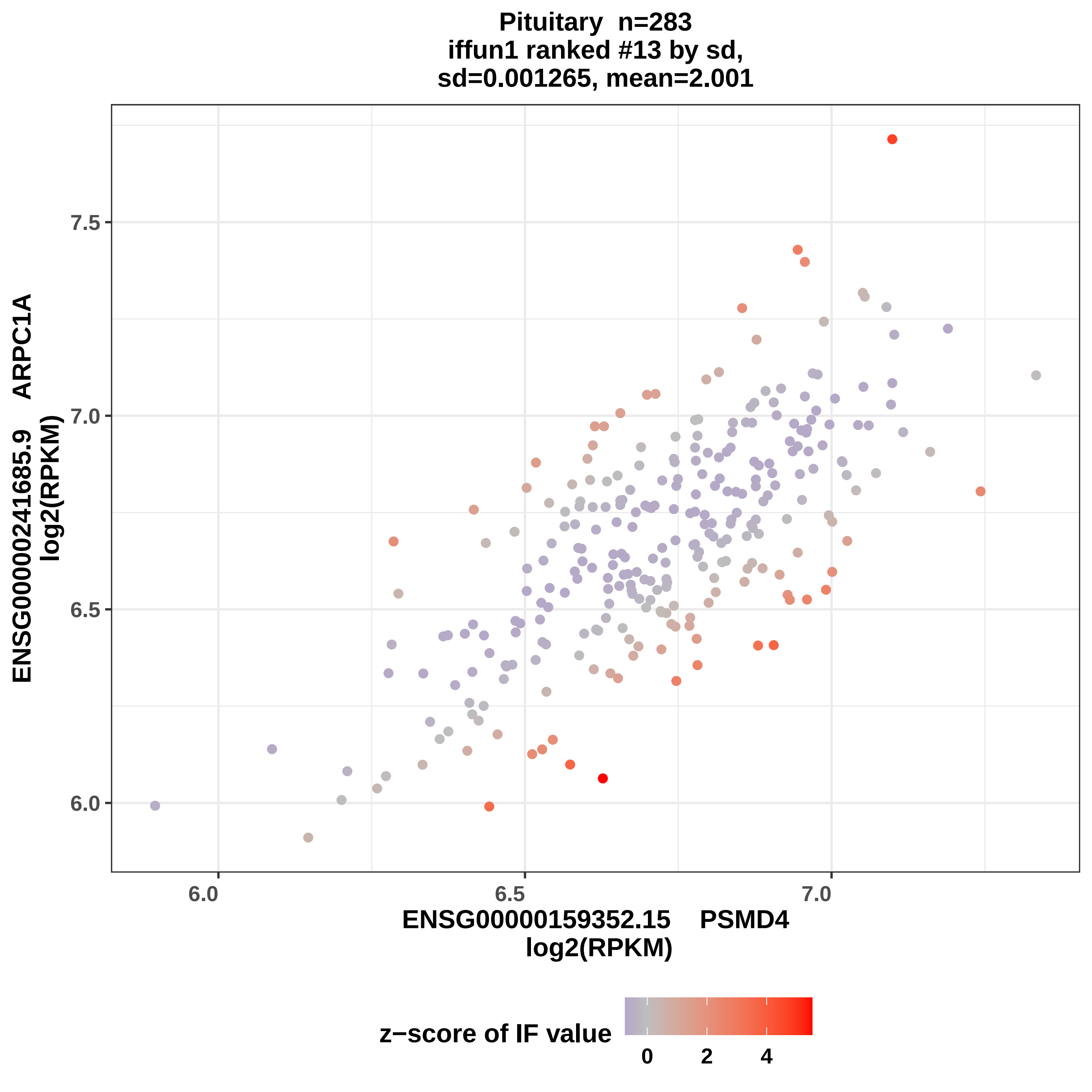
Pituitary n=283
iffun1 ranked #13 by sd,
sd=0.001265, mean=2.001
ENSG00000241685.9 ARPC1A
log2(RPKM)
7.5
7.0
6.5
6.0
6.0
6.5
7.0
ENSG00000159352.15 PSMD4
log2(RPKM)
z−score of IF value
0
2
4

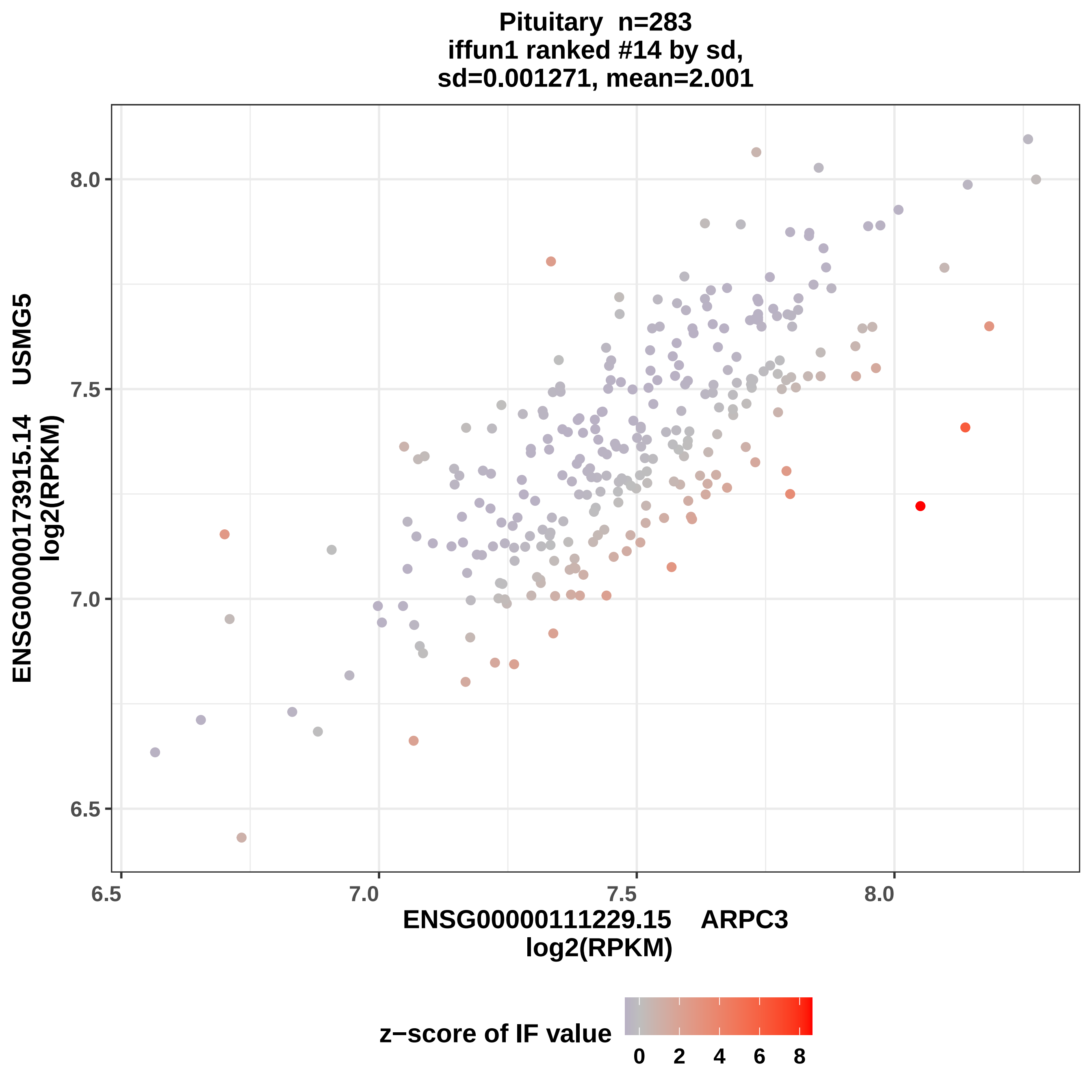
Pituitary n=283
iffun1 ranked #14 by sd,
sd=0.001271, mean=2.001
ENSG00000173915.14 USMG5
log2(RPKM)
8.0
7.5
7.0
6.5
6.5
7.0
7.5
8.0
ENSG00000111229.15 ARPC3
log2(RPKM)
z−score of IF value
0
2
4
6
8

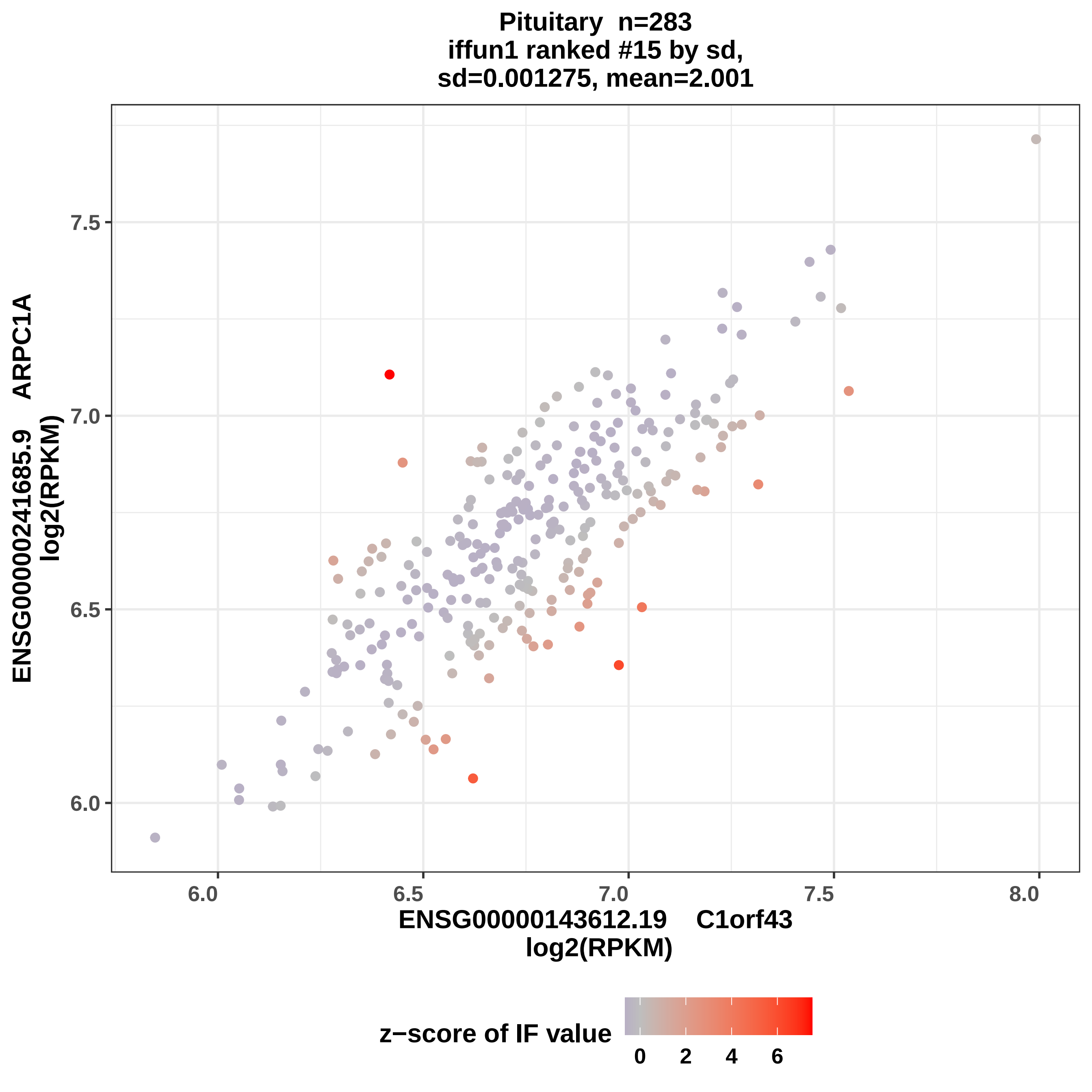
Pituitary n=283
iffun1 ranked #15 by sd,
sd=0.001275, mean=2.001
ENSG00000241685.9 ARPC1A
log2(RPKM)
7.5
7.0
6.5
6.0
6.0
6.5
7.0
7.5
8.0
ENSG00000143612.19 C1orf43
log2(RPKM)
z−score of IF value
0
2
4
6

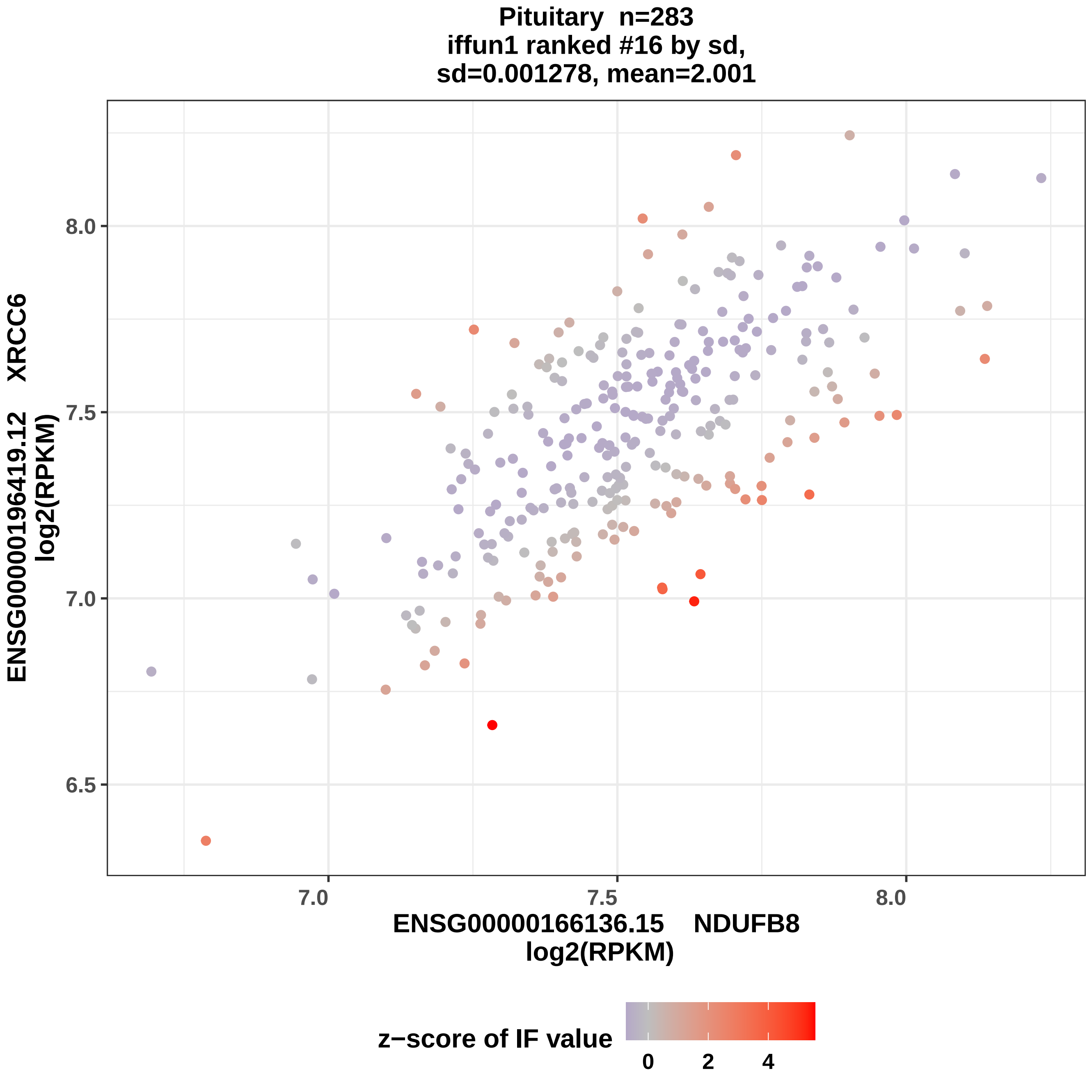

Pituitary n=283
iffun1 ranked #16 by sd,
sd=0.001278, mean=2.001
ENSG00000196419.12 XRCC6
log2(RPKM)
ENSG00000166136.15 NDUFB8
log2(RPKM)
6.5
7.0
7.5
8.0
7.0
7.5
8.0
z−score of IF value
0
2
4

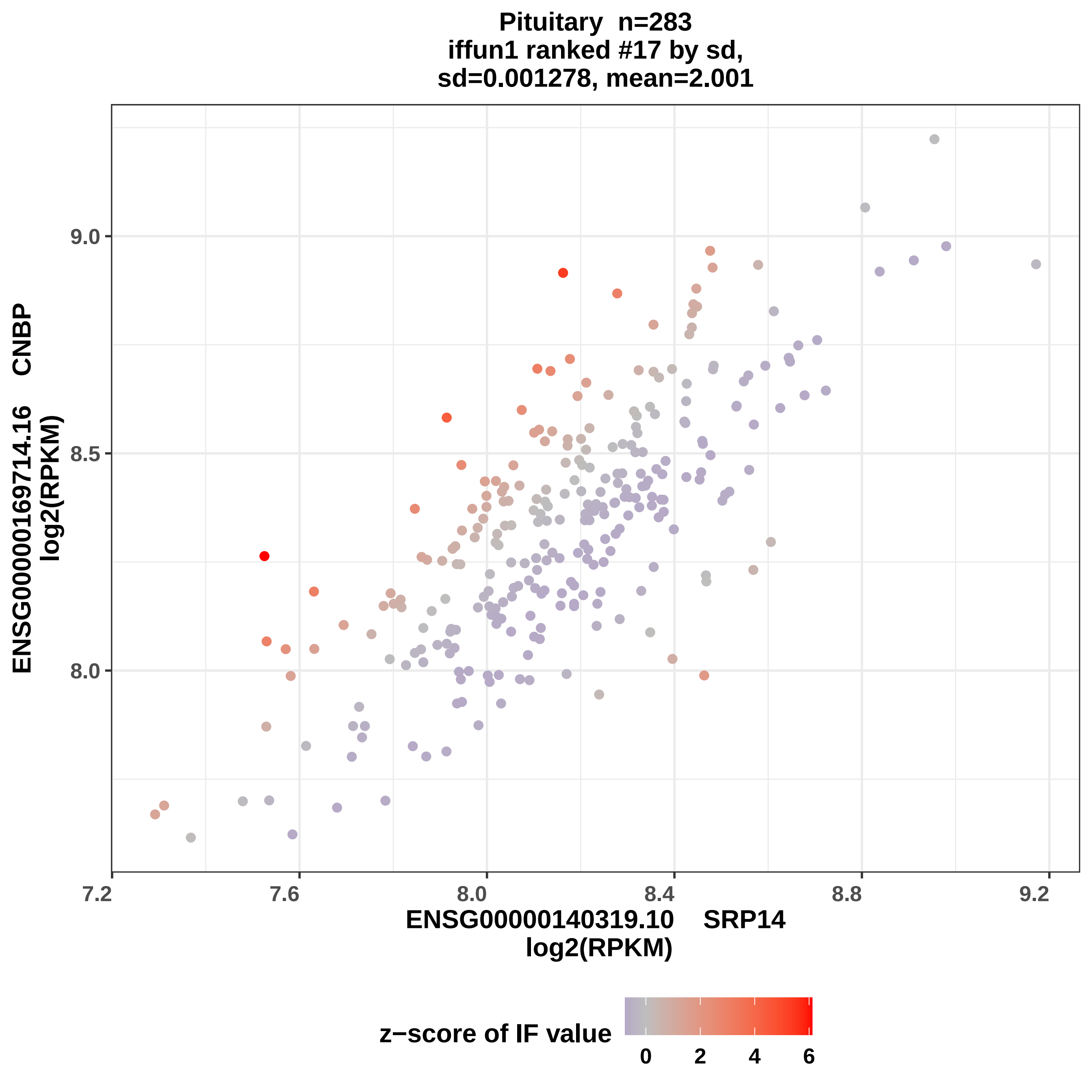
Pituitary n=283
iffun1 ranked #17 by sd,
sd=0.001278, mean=2.001
ENSG00000169714.16 CNBP
log2(RPKM)
9.0
8.5
8.0
7.2
7.6
8.0
8.4
8.8
9.2
ENSG00000140319.10 SRP14
log2(RPKM)
z−score of IF value
0
2
4
6

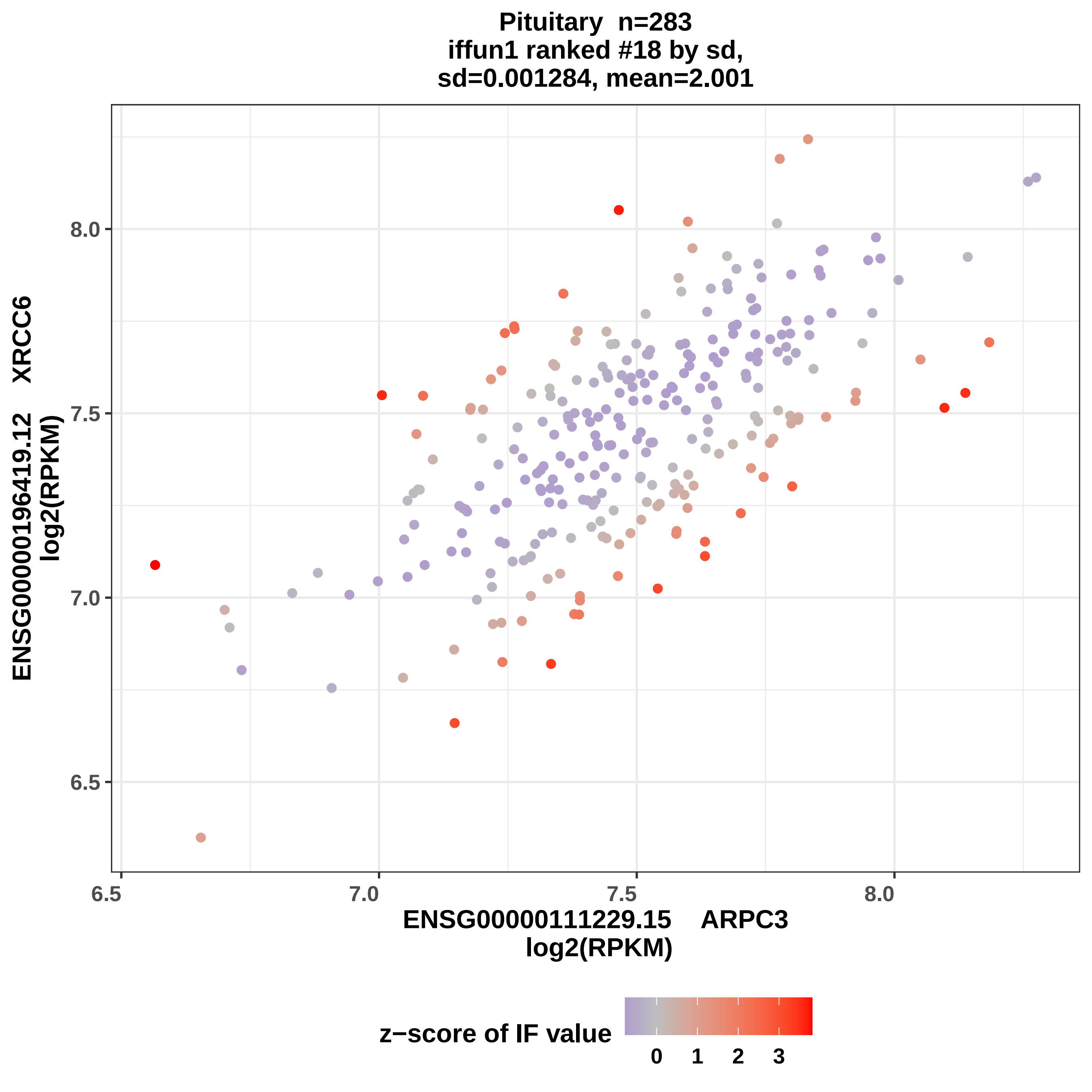

Pituitary n=283
iffun1 ranked #18 by sd,
sd=0.001284, mean=2.001
ENSG00000196419.12 XRCC6
log2(RPKM)
8.0
7.5
7.0
6.5
6.5
7.0
7.5
8.0
ENSG00000111229.15 ARPC3
log2(RPKM)
z−score of IF value
0
1
2
3

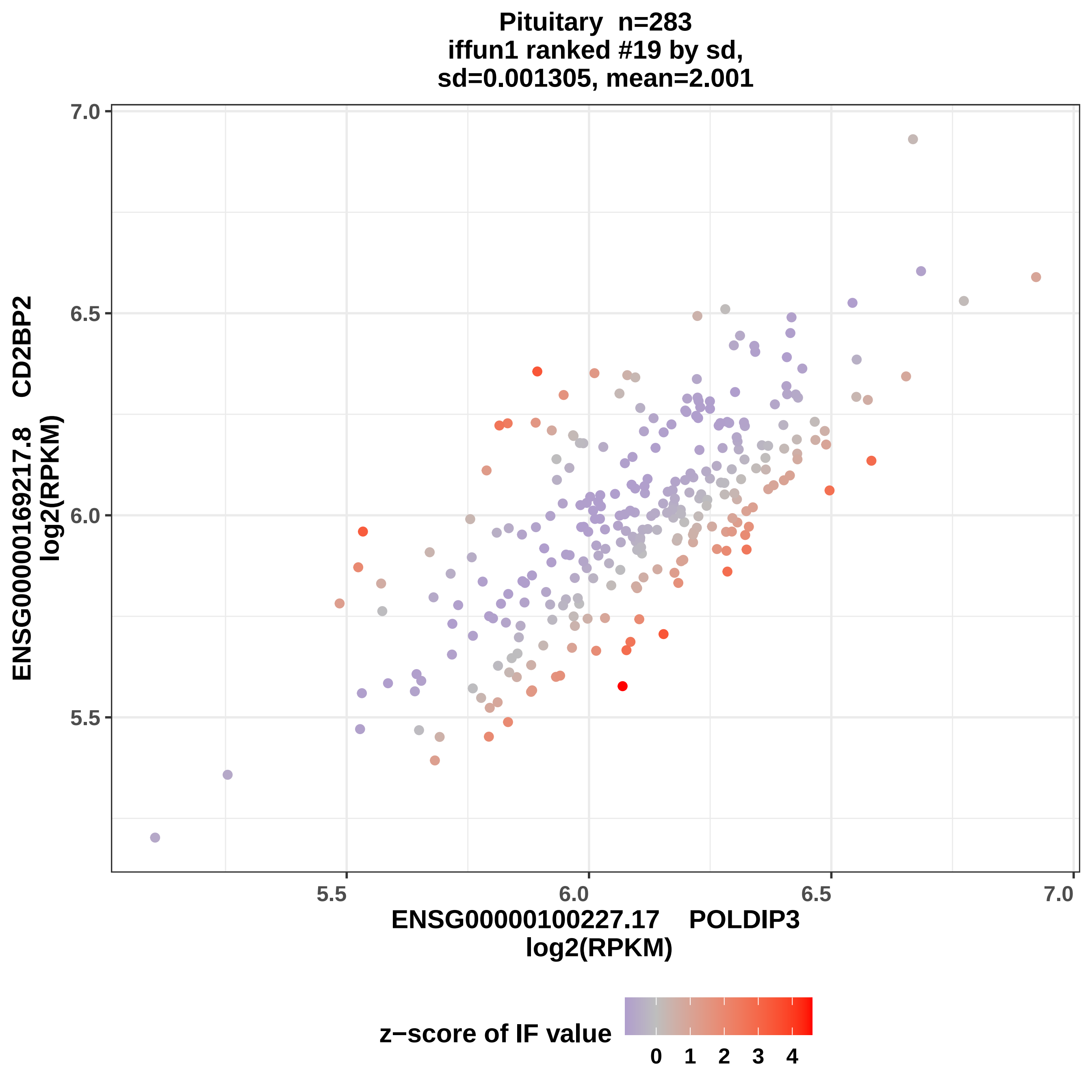
Pituitary n=283
iffun1 ranked #19 by sd,
sd=0.001305, mean=2.001
ENSG00000169217.8 CD2BP2
log2(RPKM)
7.0
6.5
6.0
5.5
5.5
6.0
6.5
7.0
ENSG00000100227.17 POLDIP3
log2(RPKM)
z−score of IF value
0 1 2 3 4

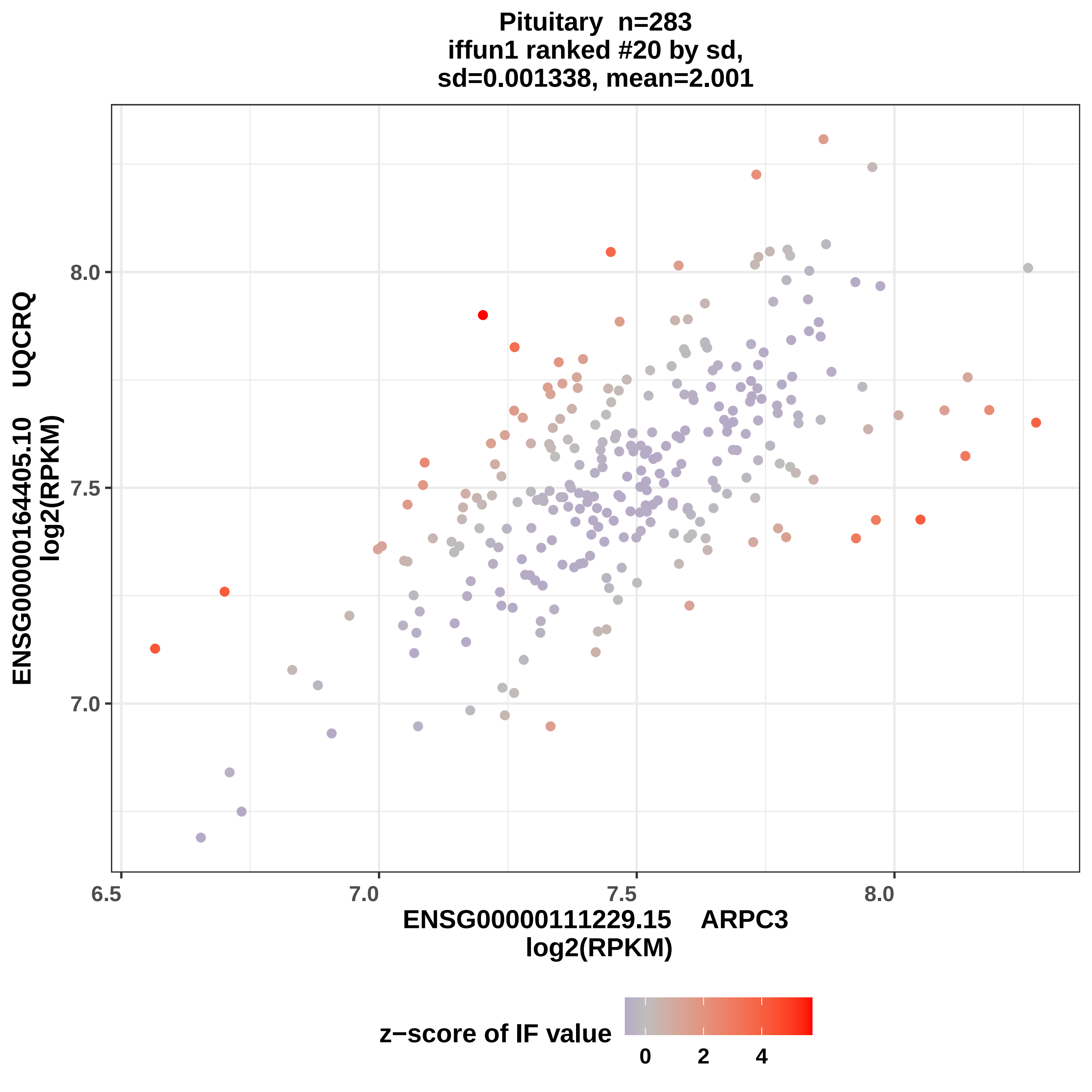

Pituitary n=283
iffun1 ranked #20 by sd,
sd=0.001338, mean=2.001
ENSG00000164405.10 UQCRQ
log2(RPKM)
8.0
7.5
7.0
6.5
7.0
7.5
8.0
ENSG00000111229.15 ARPC3
log2(RPKM)
z−score of IF value
0
2
4

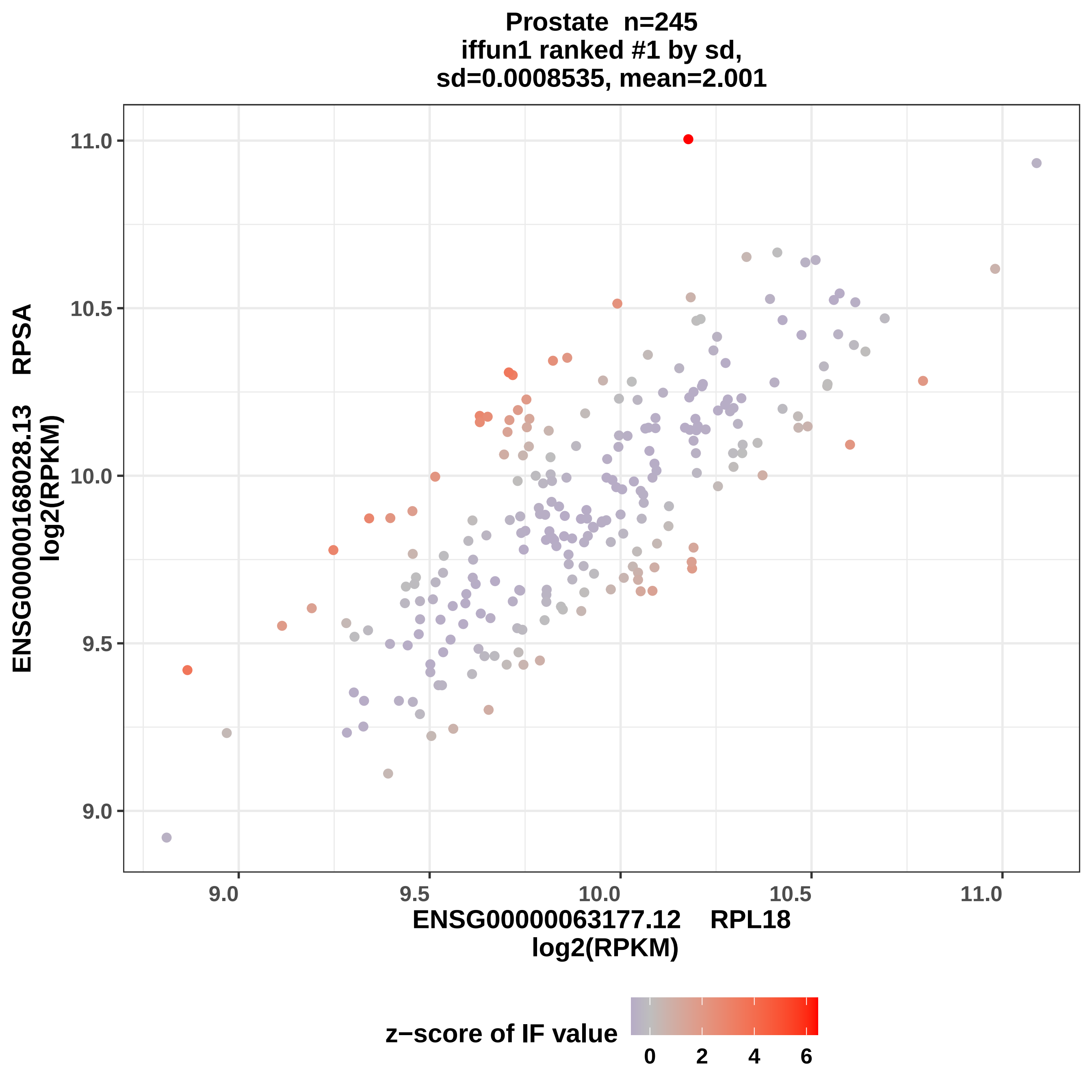

Prostate n=245
iffun1 ranked #1 by sd,
sd=0.0008535, mean=2.001
ENSG00000168028.13 RPSA
log2(RPKM)
11.0
10.5
10.0
9.5
9.0
ENSG00000063177.12 RPL18
log2(RPKM)
z−score of IF value
0
2
4
6

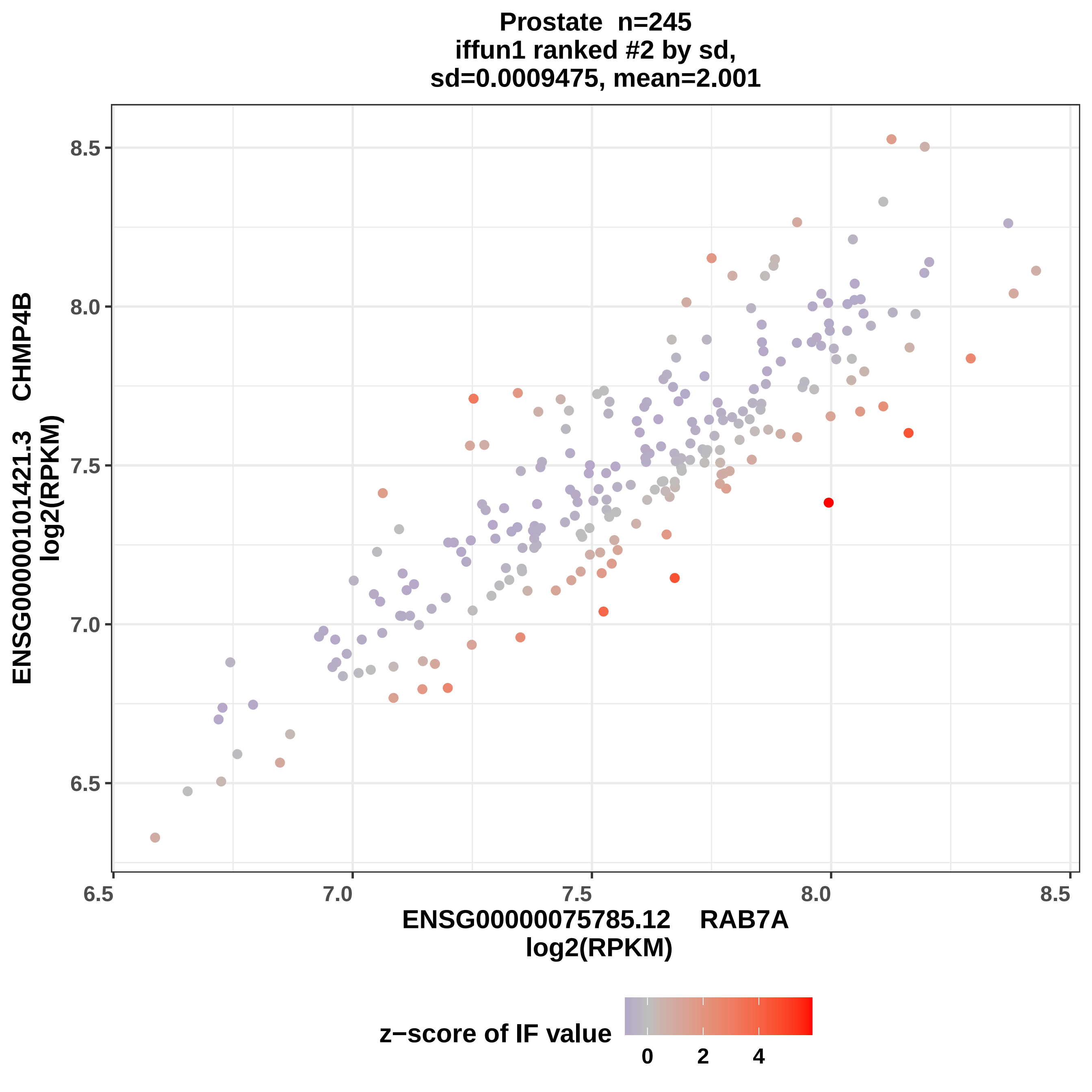
Prostate n=245
iffun1 ranked #2 by sd,
sd=0.0009475, mean=2.001
ENSG00000101421.3 CHMP4B
log2(RPKM)
8.5
8.0
7.5
7.0
6.5
6.5
7.0
7.5
8.0
8.5
ENSG00000075785.12 RAB7A
log2(RPKM)
z−score of IF value
0
2
4

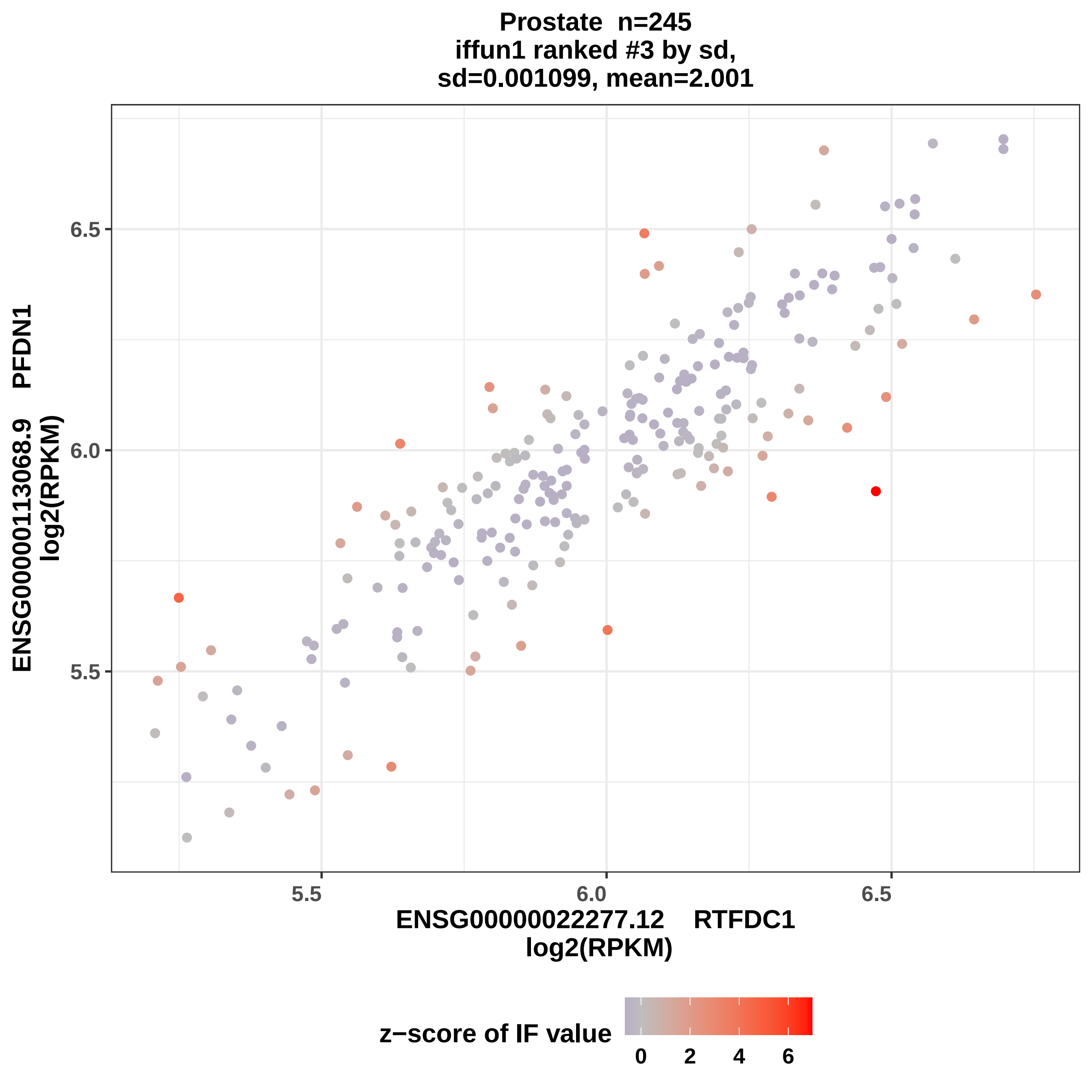

Prostate n=245
iffun1 ranked #3 by sd,
sd=0.001099, mean=2.001
ENSG00000113068.9 PFDN1
log2(RPKM)
6.5
6.0
5.5
5.5
6.0
6.5
ENSG00000022277.12 RTFDC1
log2(RPKM)
z−score of IF value
0
2
4
6

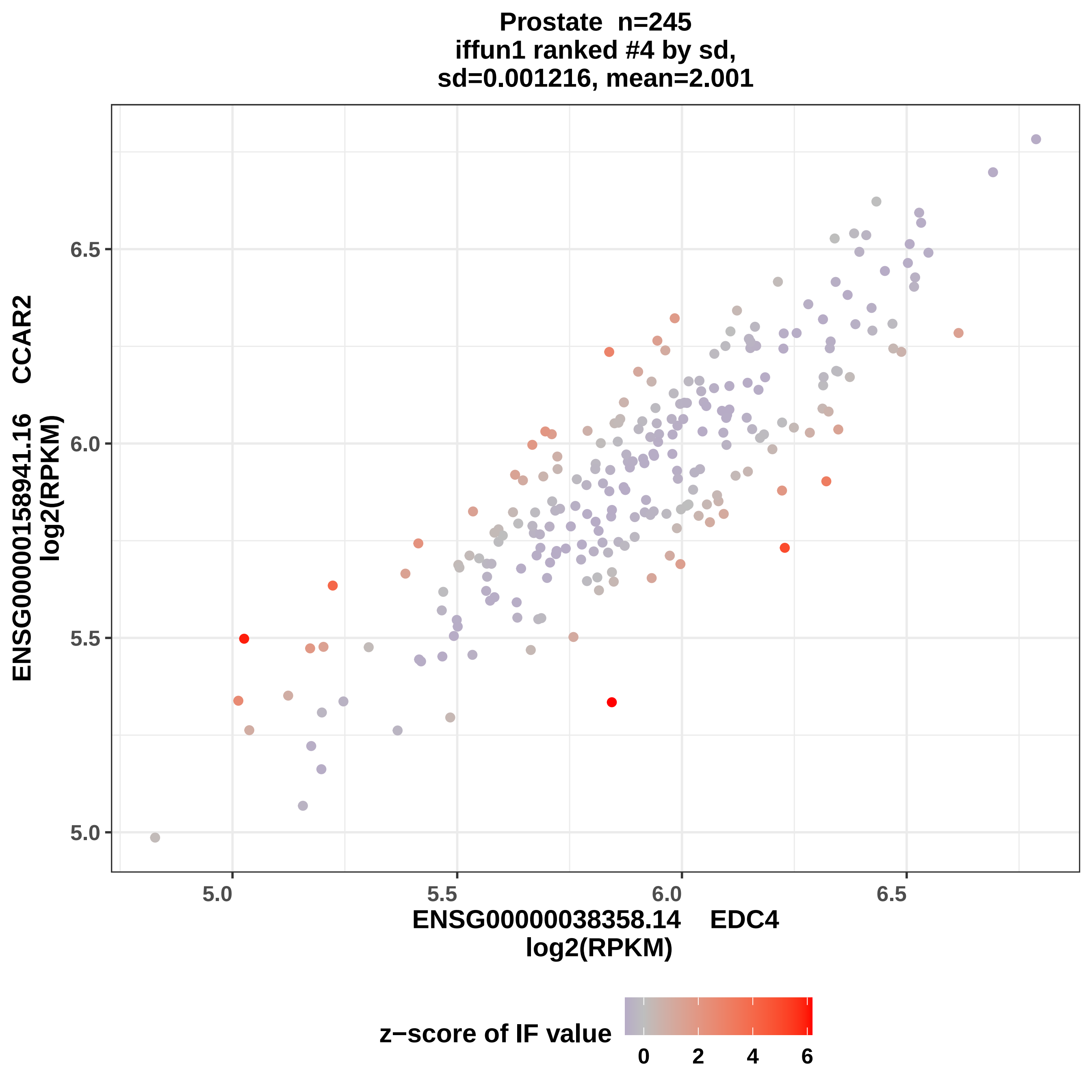
Prostate n=245
iffun1 ranked #4 by sd,
sd=0.001216, mean=2.001
ENSG00000158941.16 CCAR2
log2(RPKM)
ENSG00000038358.14 EDC4
log2(RPKM)
5.0
5.5
6.0
6.5
z−score of IF value
0
2
4
6

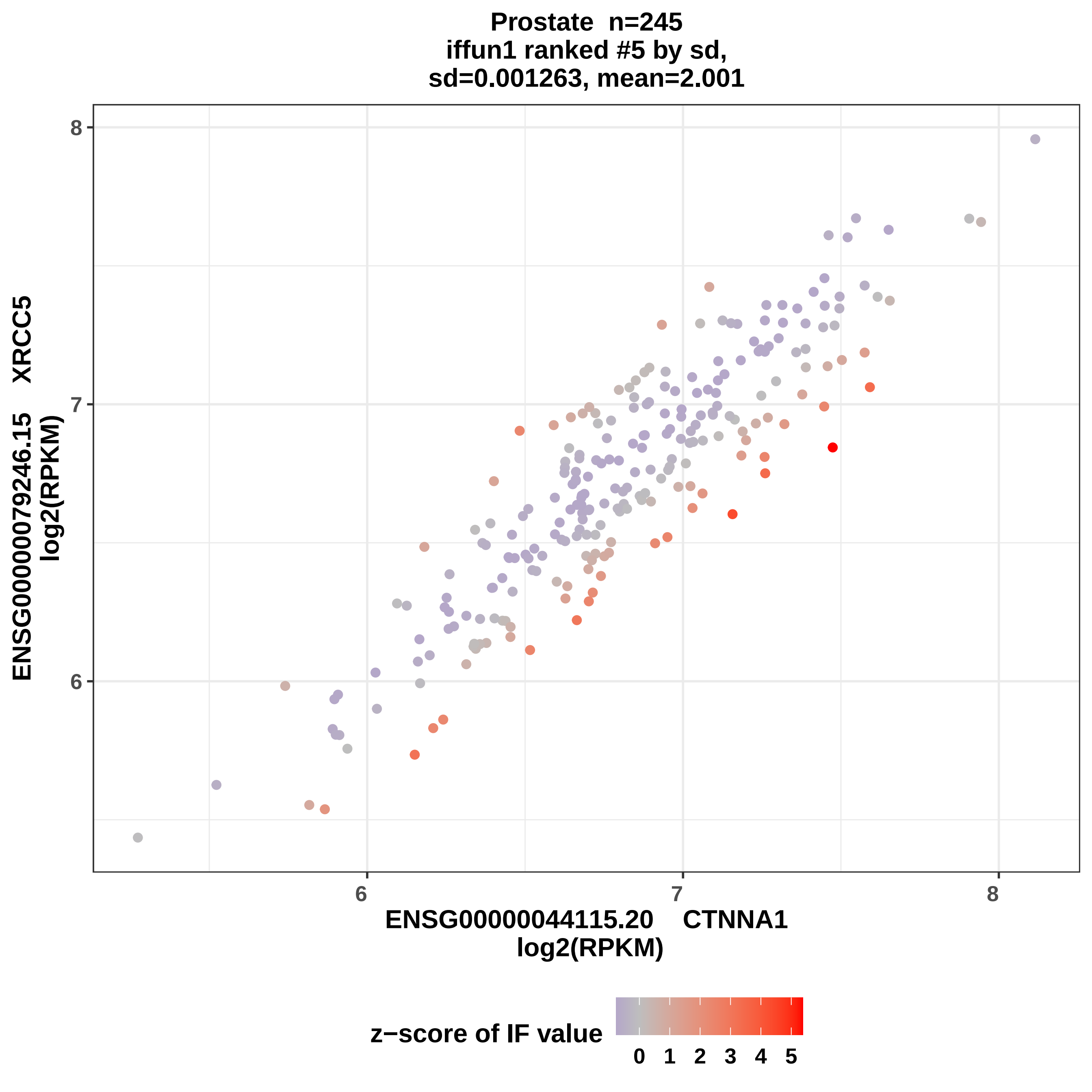

Prostate n=245
iffun1 ranked #5 by sd,
sd=0.001263, mean=2.001
ENSG00000079246.15 XRCC5
log2(RPKM)
ENSG00000044115.20 CTNNA1
log2(RPKM)
6
7
8
6
7
8
z−score of IF value
0
1
2
3
4
5

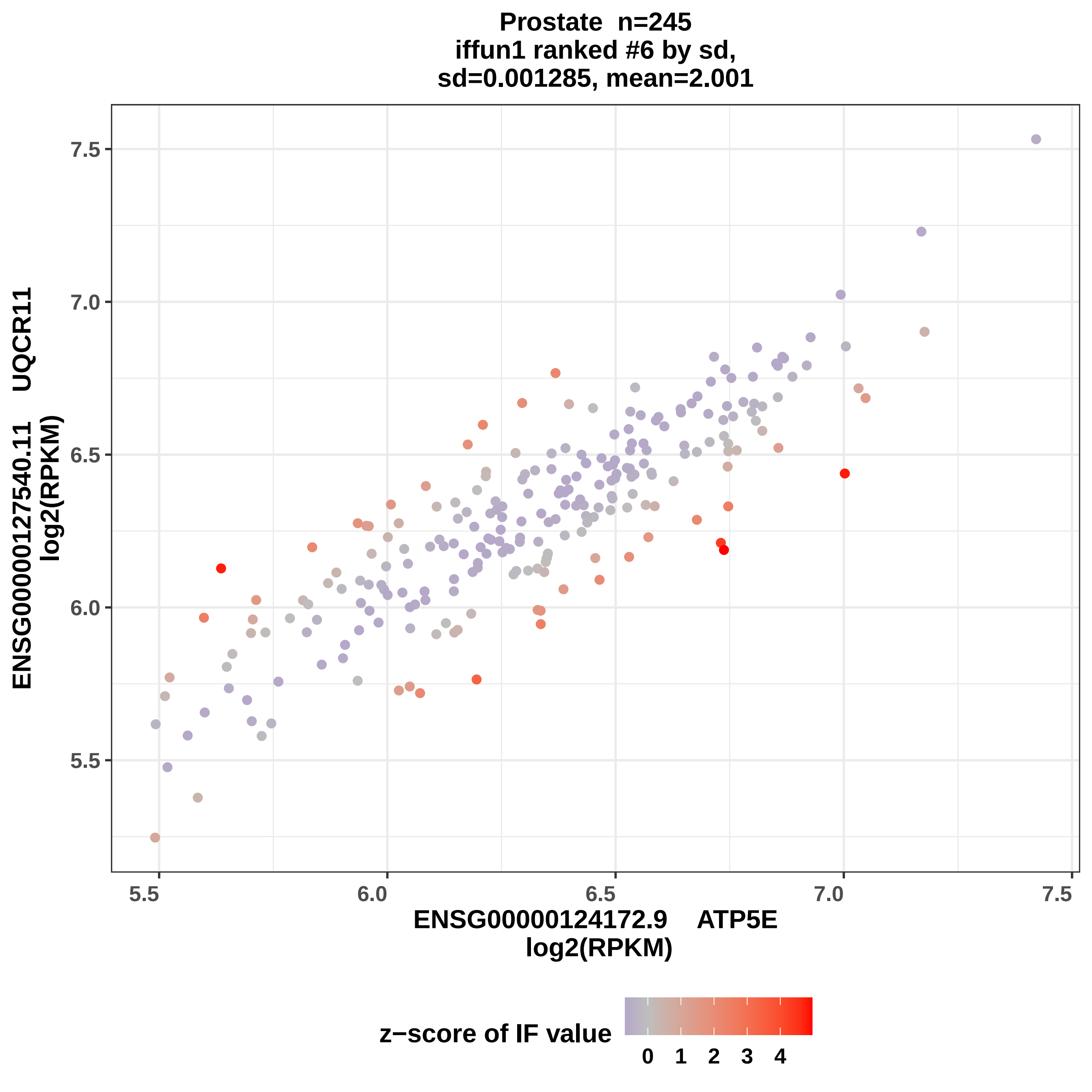

Prostate n=245
iffun1 ranked #6 by sd,
sd=0.001285, mean=2.001
ENSG00000127540.11 UQCR11
log2(RPKM)
7.5
7.0
6.5
6.0
5.5
5.5
6.0
6.5
7.0
7.5
ENSG00000124172.9 ATP5E
log2(RPKM)
z−score of IF value
0 1 2 3 4

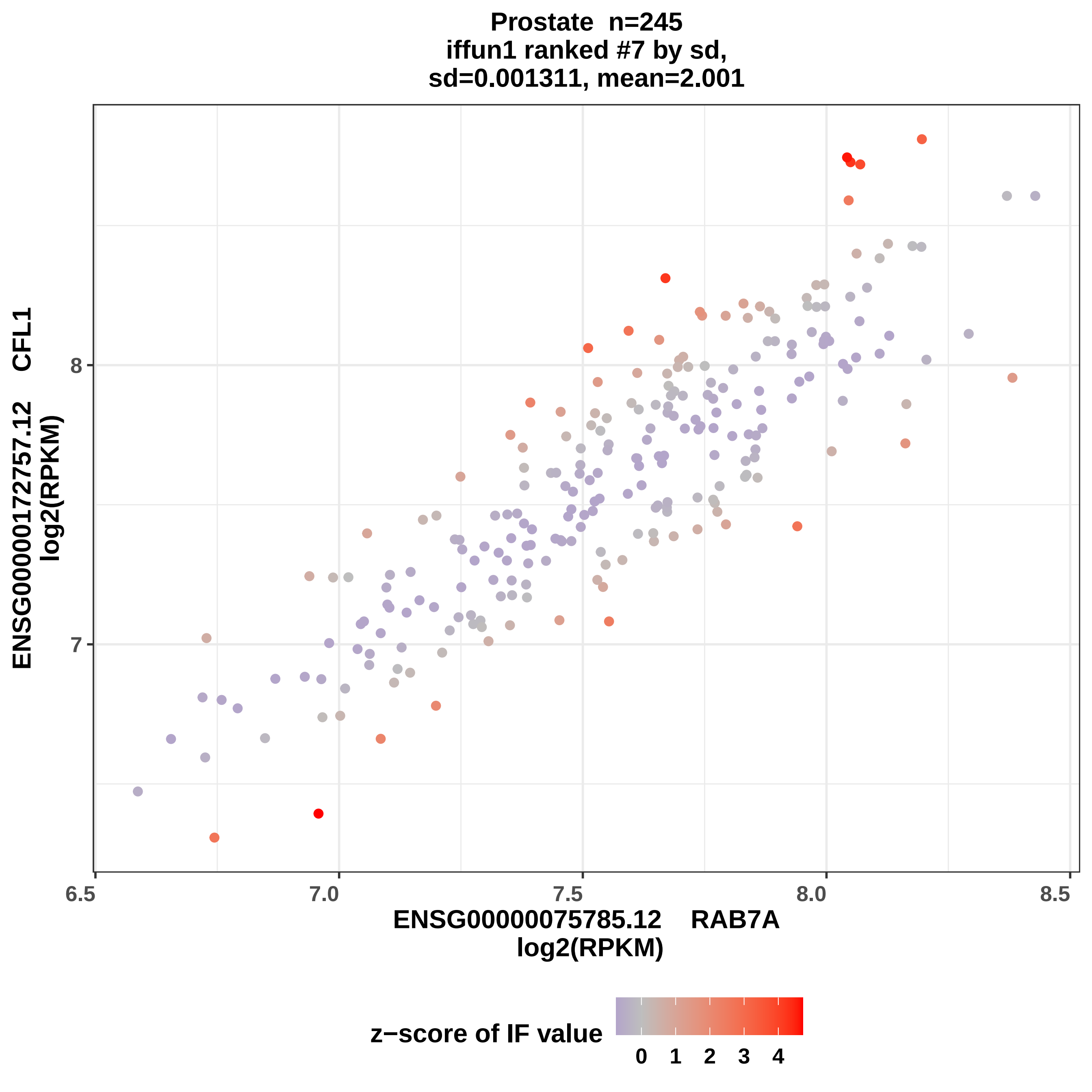

Prostate n=245
iffun1 ranked #7 by sd,
sd=0.001311, mean=2.001
ENSG00000172757.12 CFL1
log2(RPKM)
8
7
6.5
7.0
7.5
8.0
8.5
ENSG00000075785.12 RAB7A
log2(RPKM)
z−score of IF value
0
1
2
3
4

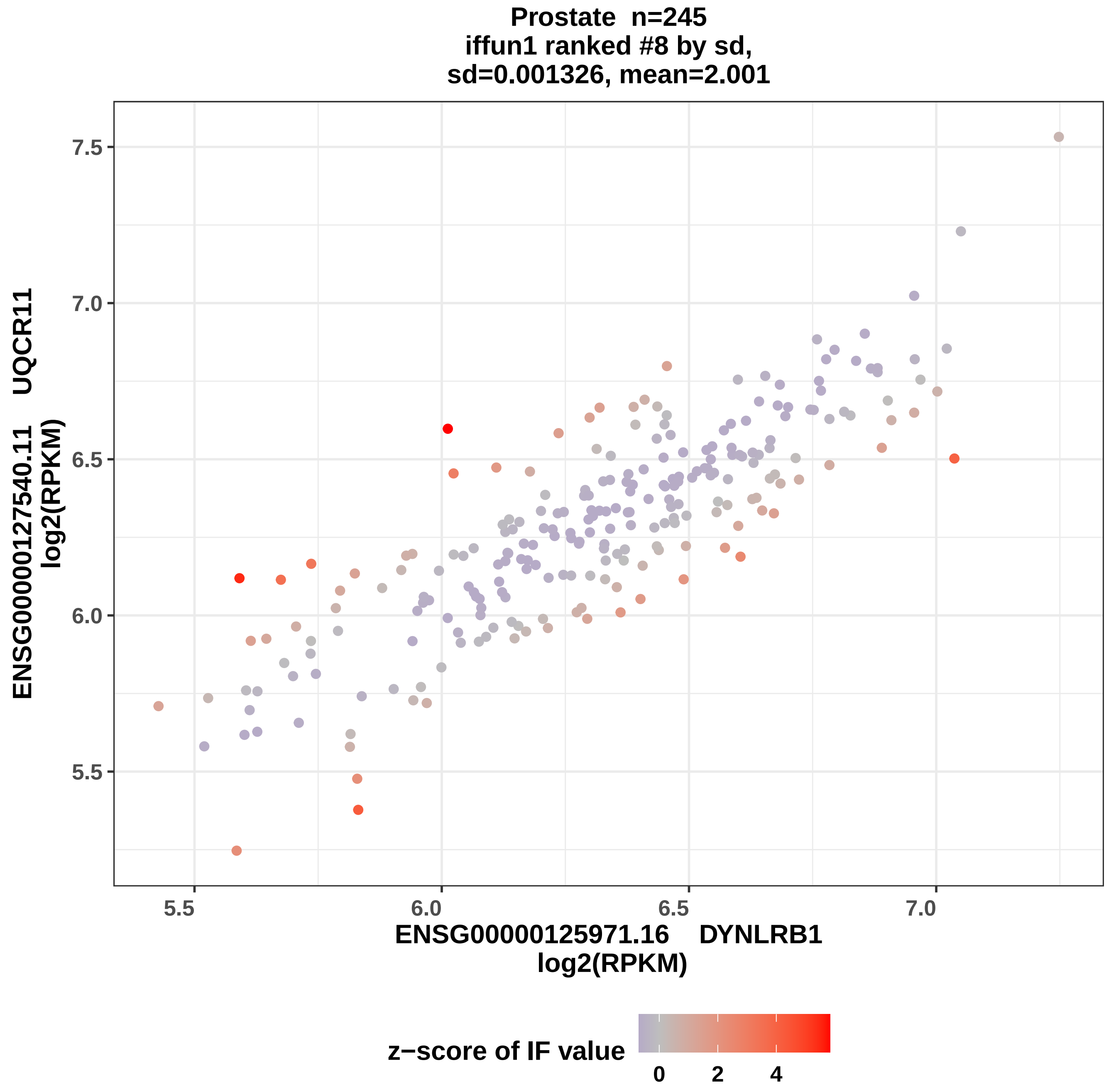
Prostate n=245
iffun1 ranked #8 by sd,
sd=0.001326, mean=2.001
ENSG00000127540.11 UQCR11
log2(RPKM)
7.5
7.0
6.5
6.0
5.5
5.5
6.0
6.5
7.0
ENSG00000125971.16 DYNLRB1
log2(RPKM)
z−score of IF value
0
2
4

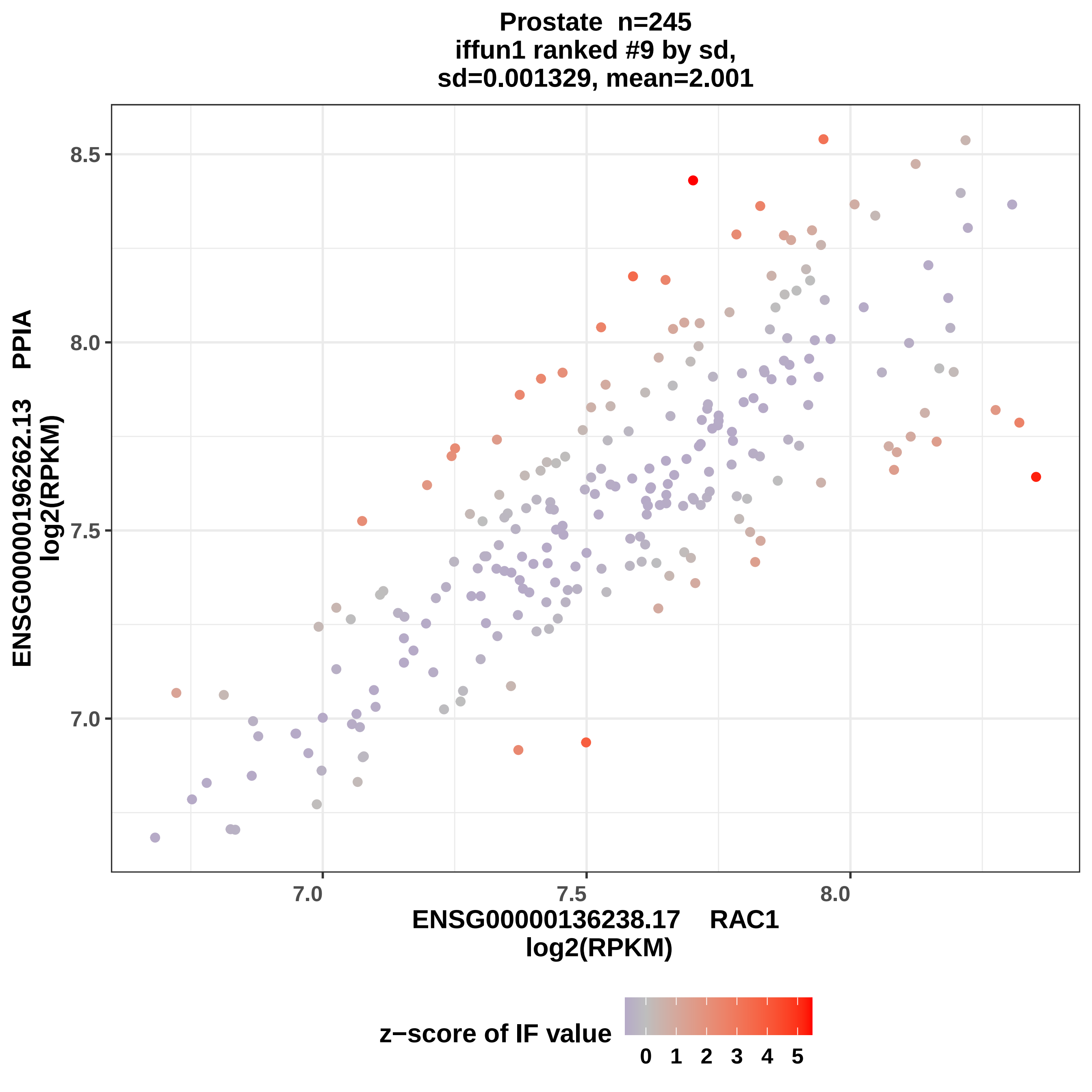
Prostate n=245
iffun1 ranked #9 by sd,
sd=0.001329, mean=2.001
ENSG00000196262.13 PPIA
log2(RPKM)
8.5
8.0
7.5
7.0
7.0
7.5
8.0
ENSG00000136238.17 RAC1
log2(RPKM)
z−score of IF value
0
1
2
3
4
5

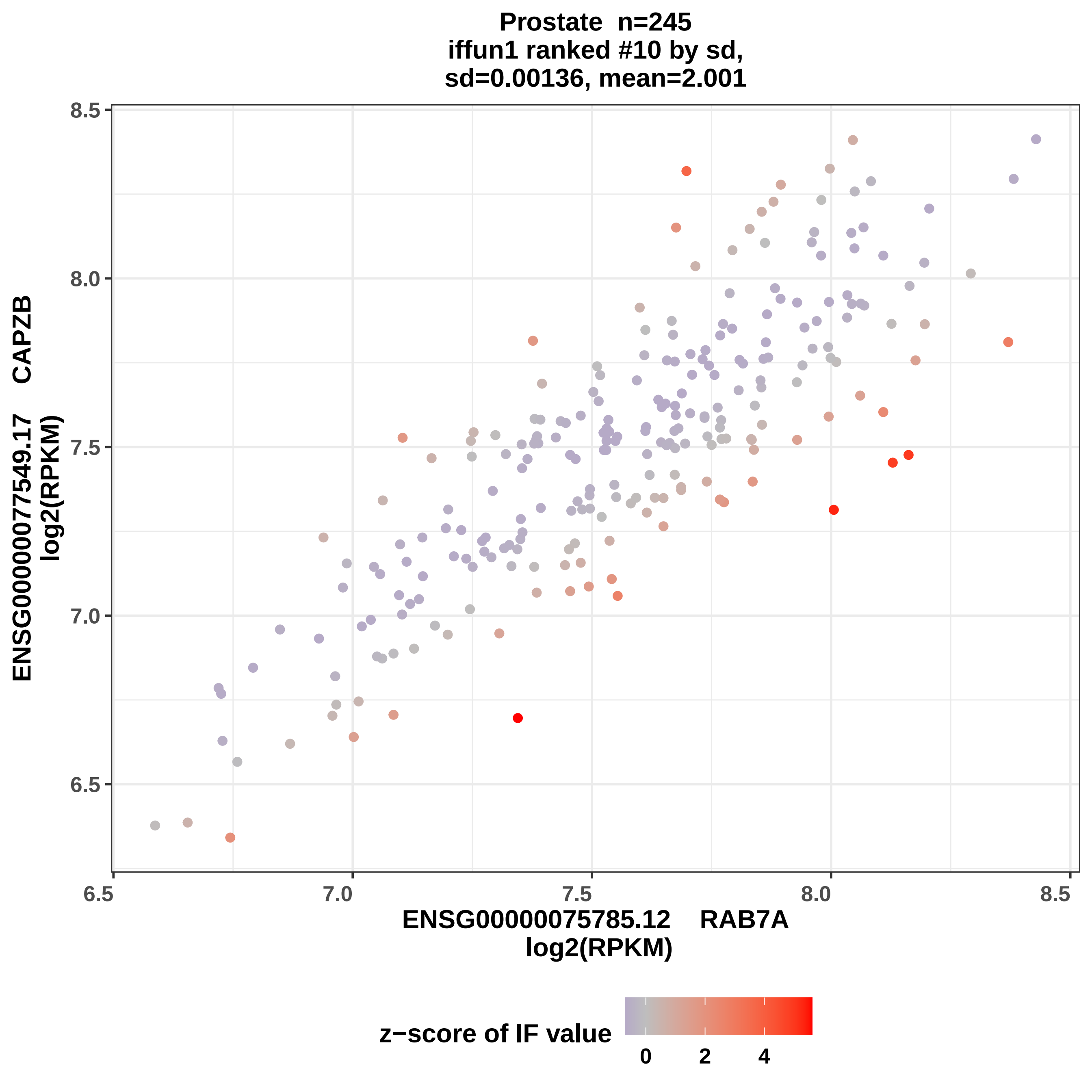
Prostate n=245
iffun1 ranked #10 by sd,
sd=0.00136, mean=2.001
ENSG00000077549.17 CAPZB
log2(RPKM)
ENSG00000075785.12 RAB7A
log2(RPKM)
6.5
7.0
7.5
8.0
8.5
z−score of IF value
0
2
4

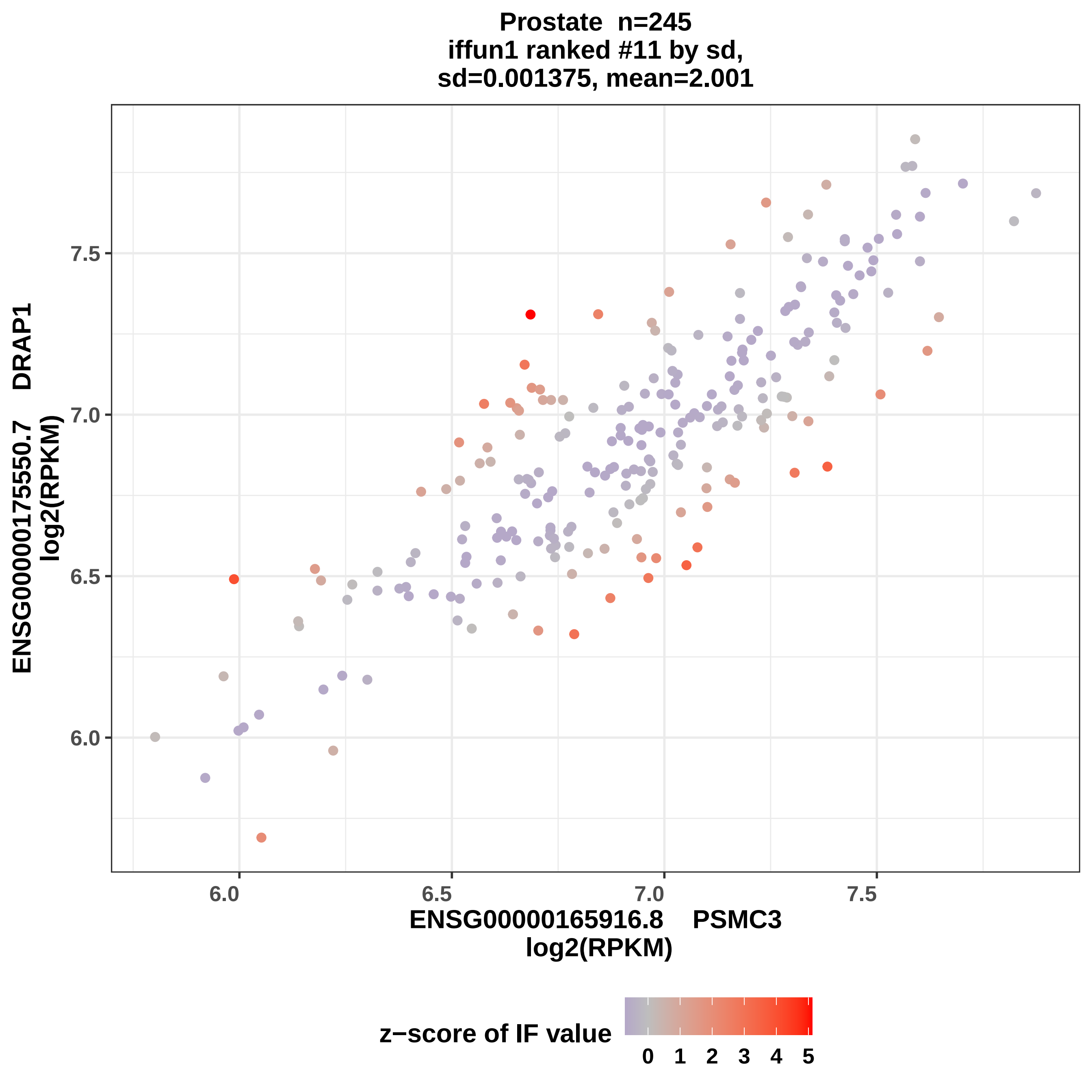
Prostate n=245
iffun1 ranked #11 by sd,
sd=0.001375, mean=2.001
ENSG00000175550.7 DRAP1
log2(RPKM)
ENSG00000165916.8 PSMC3
log2(RPKM)
6.0
6.5
7.0
7.5
z−score of IF value
0
1
2
3
4
5

Prostate n=245
iffun1 ranked #12 by sd,
sd=0.001375, mean=2.001

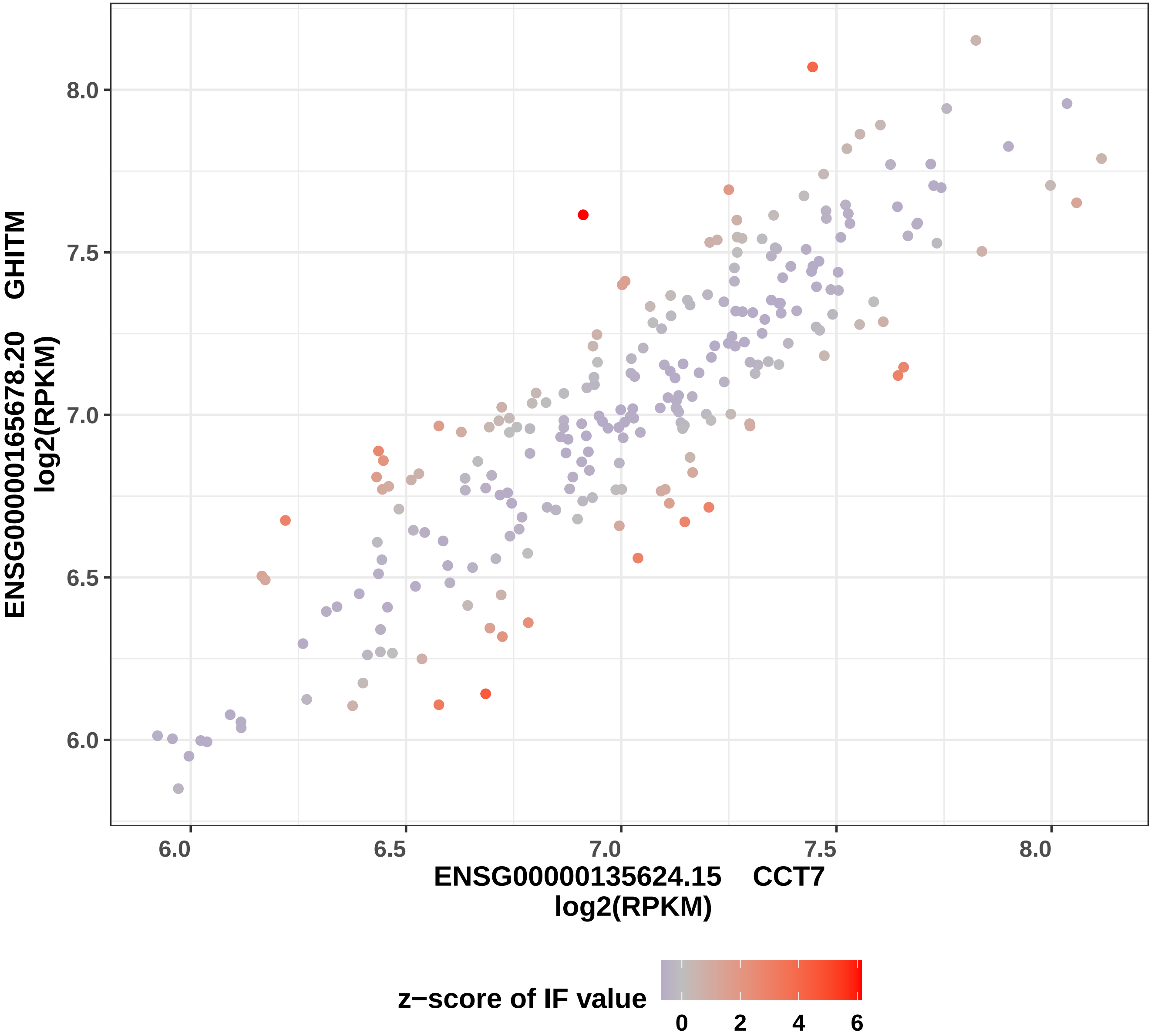

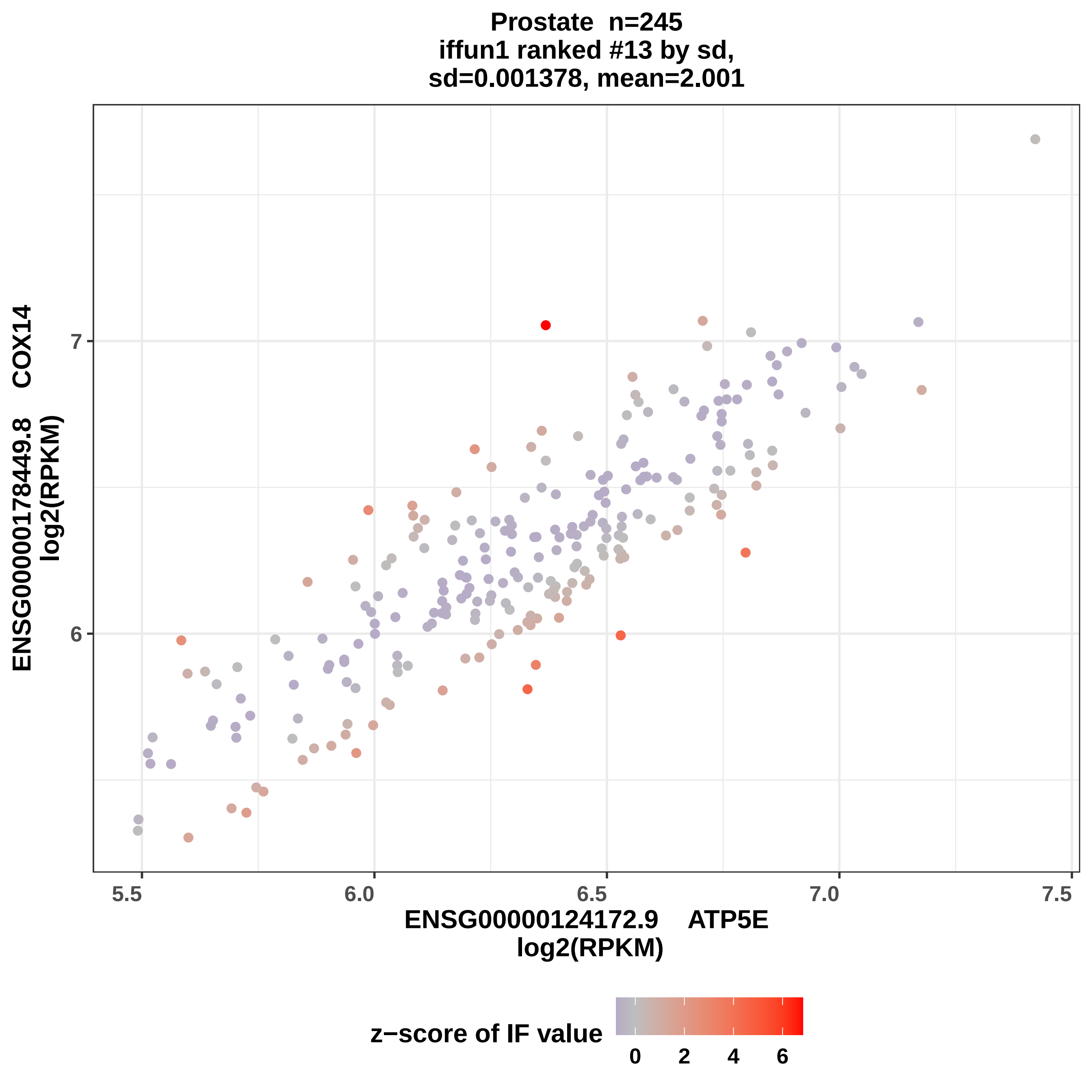
Prostate n=245
iffun1 ranked #13 by sd,
sd=0.001378, mean=2.001
ENSG00000178449.8 COX14
log2(RPKM)
7
6
5.5
6.0
6.5
7.0
7.5
ENSG00000124172.9 ATP5E
log2(RPKM)
z−score of IF value
0
2
4
6

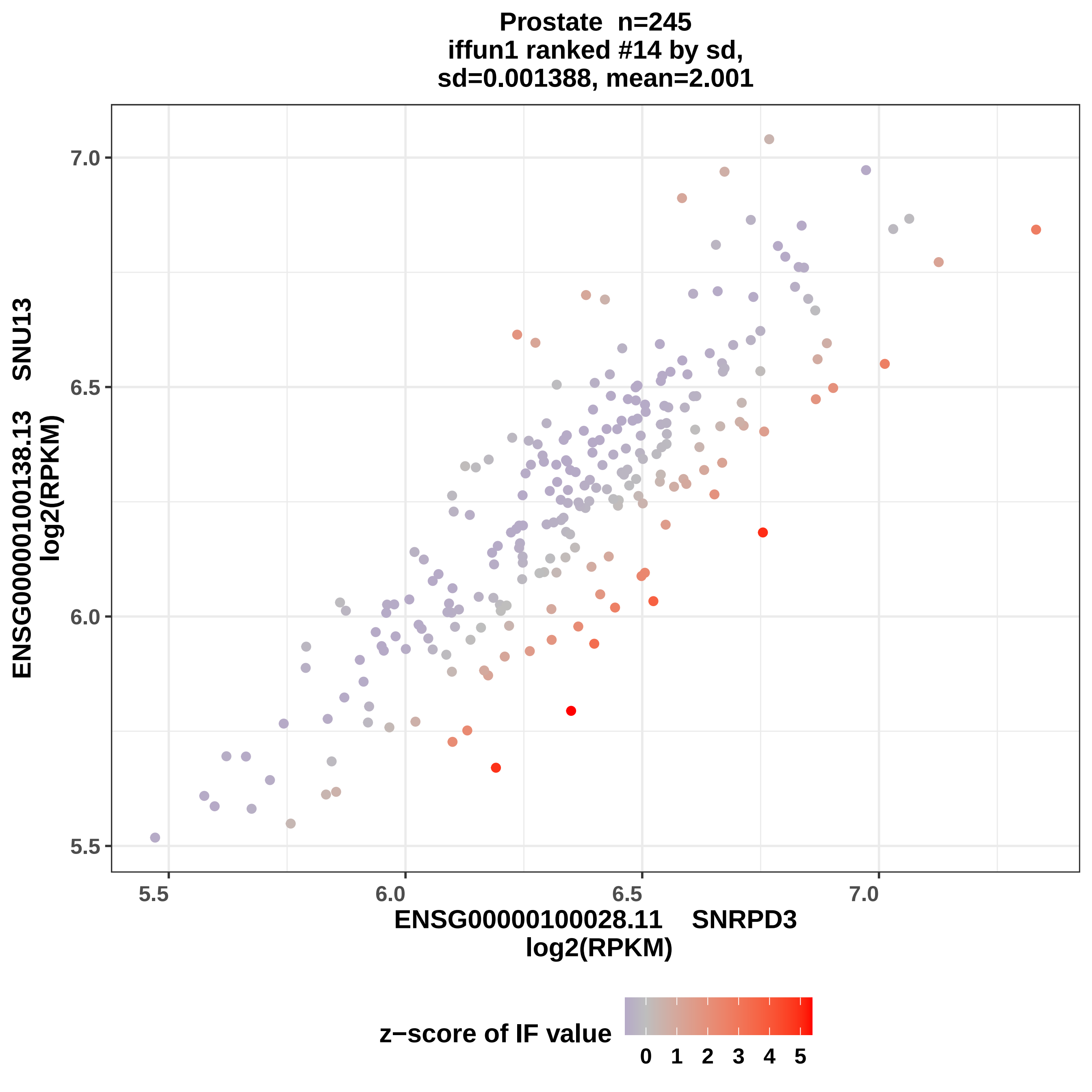
Prostate n=245
iffun1 ranked #14 by sd,
sd=0.001388, mean=2.001
ENSG00000100138.13 SNU13
log2(RPKM)
ENSG00000100028.11 SNRPD3
log2(RPKM)
5.5
6.0
6.5
7.0
z−score of IF value
0 1 2 3 4 5

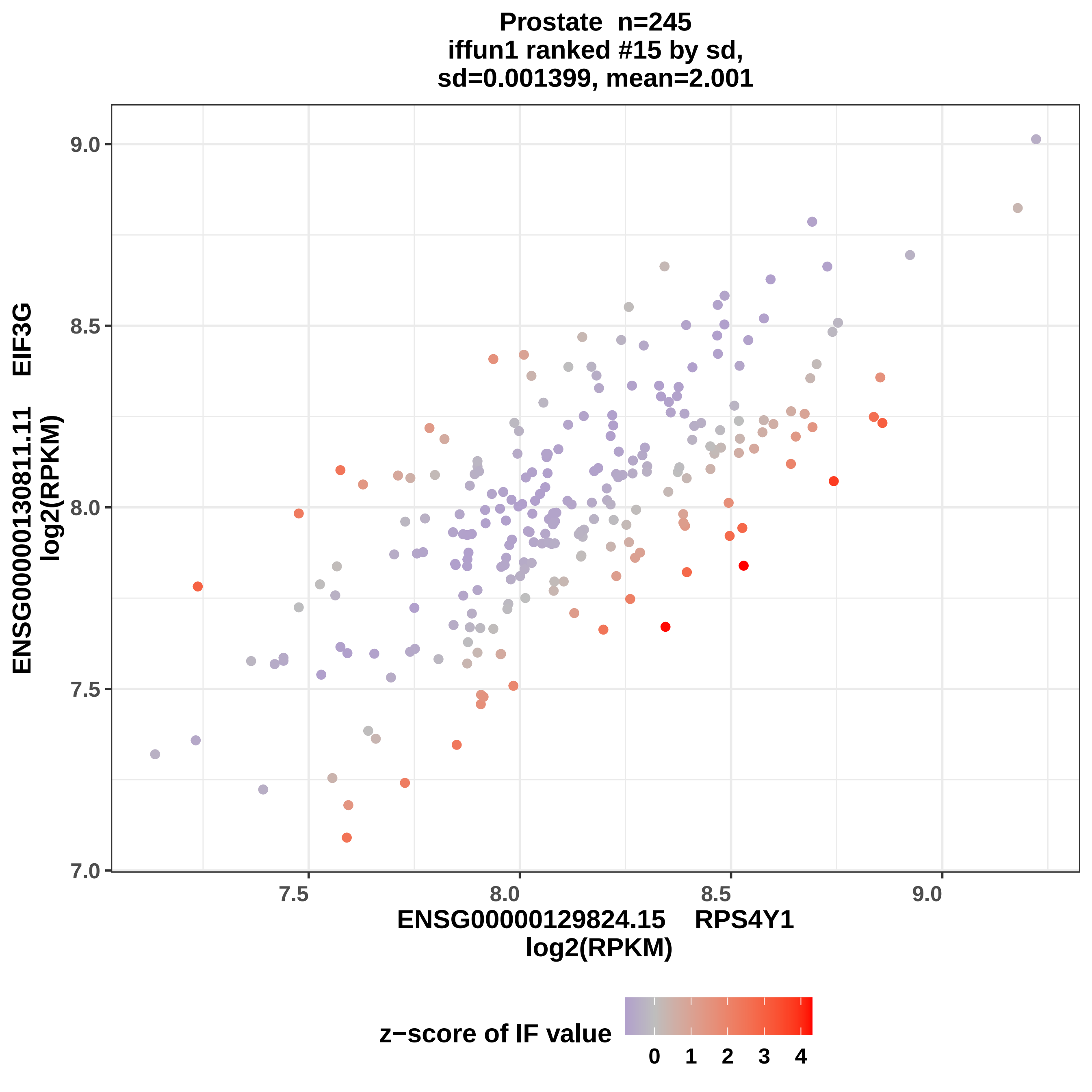

Prostate n=245
iffun1 ranked #15 by sd,
sd=0.001399, mean=2.001
ENSG00000130811.11 EIF3G
log2(RPKM)
9.0
8.5
8.0
7.5
7.0
7.5
8.0
8.5
9.0
ENSG00000129824.15 RPS4Y1
log2(RPKM)
z−score of IF value
0
1
2
3
4

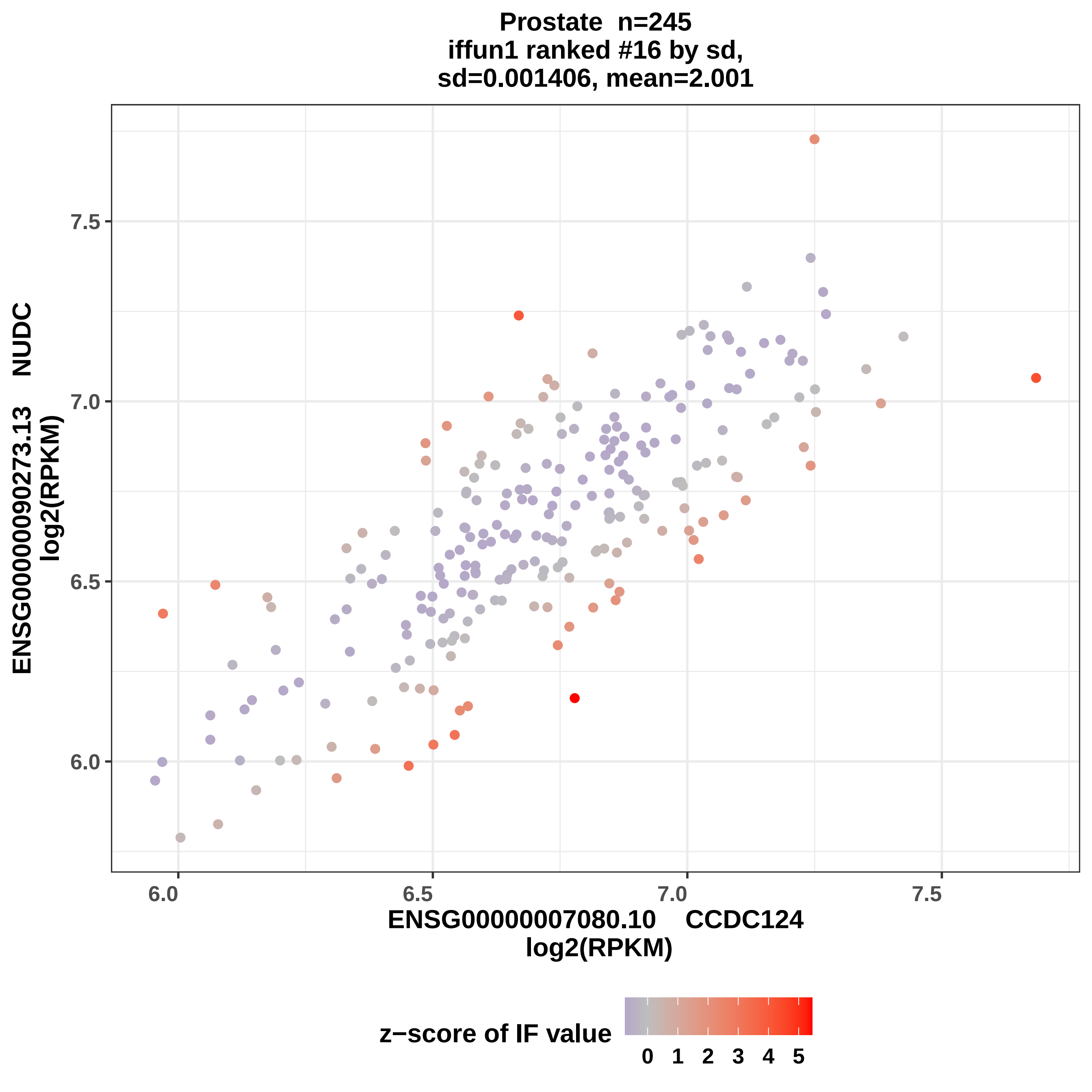

Prostate n=245
iffun1 ranked #16 by sd,
sd=0.001406, mean=2.001
ENSG00000090273.13 NUDC
log2(RPKM)
7.5
7.0
6.5
6.0
6.0
6.5
7.0
7.5
ENSG00000007080.10 CCDC124
log2(RPKM)
z−score of IF value
0 1 2 3 4 5

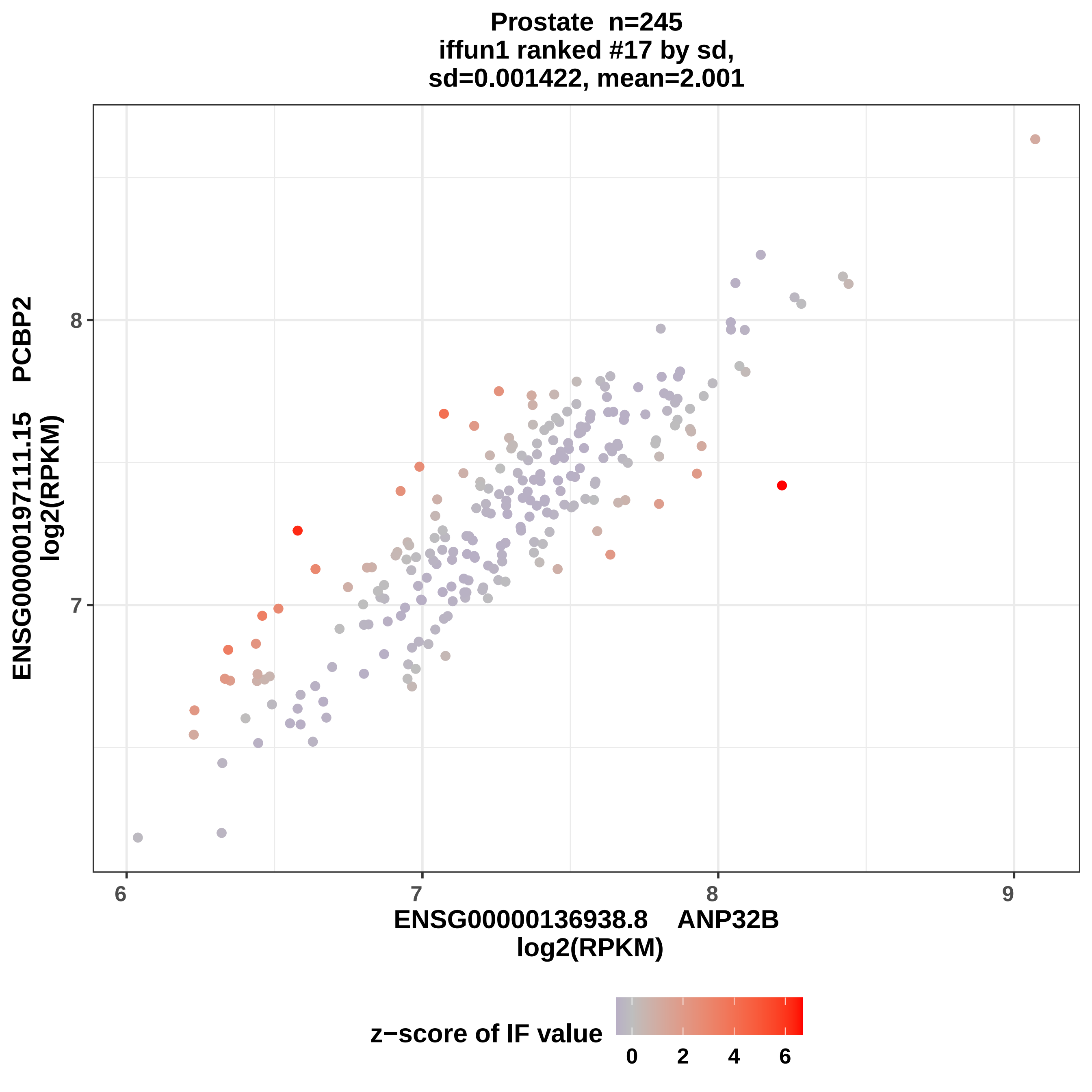

Prostate n=245
iffun1 ranked #17 by sd,
sd=0.001422, mean=2.001
ENSG00000197111.15 PCBP2
log2(RPKM)
8
7
6
7
8
9
ENSG00000136938.8 ANP32B
log2(RPKM)
z−score of IF value
0
2
4
6

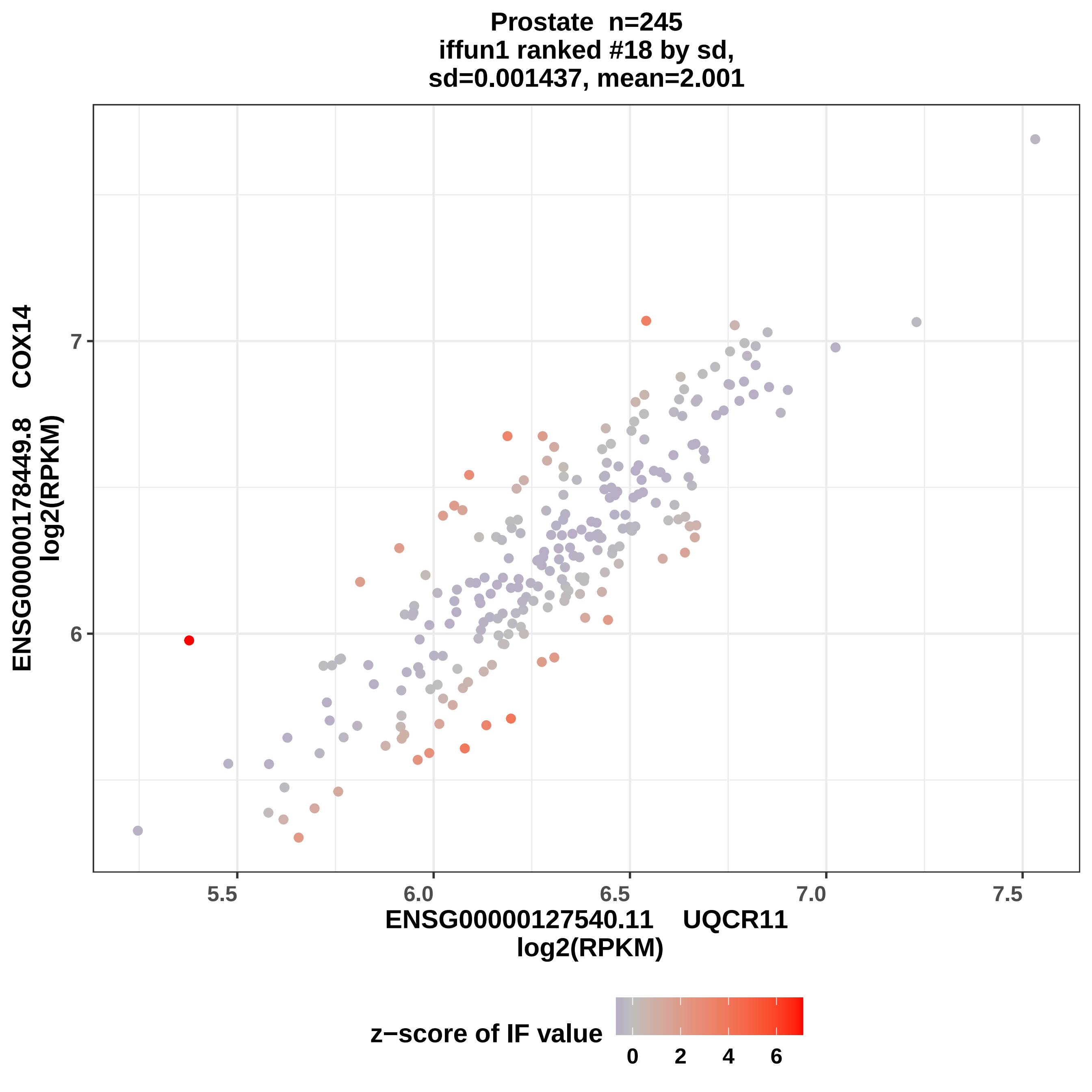

Prostate n=245
iffun1 ranked #18 by sd,
sd=0.001437, mean=2.001
ENSG00000178449.8 COX14
log2(RPKM)
7
6
5.5
6.0
6.5
7.0
7.5
ENSG00000127540.11 UQCR11
log2(RPKM)
z−score of IF value
0
2
4
6

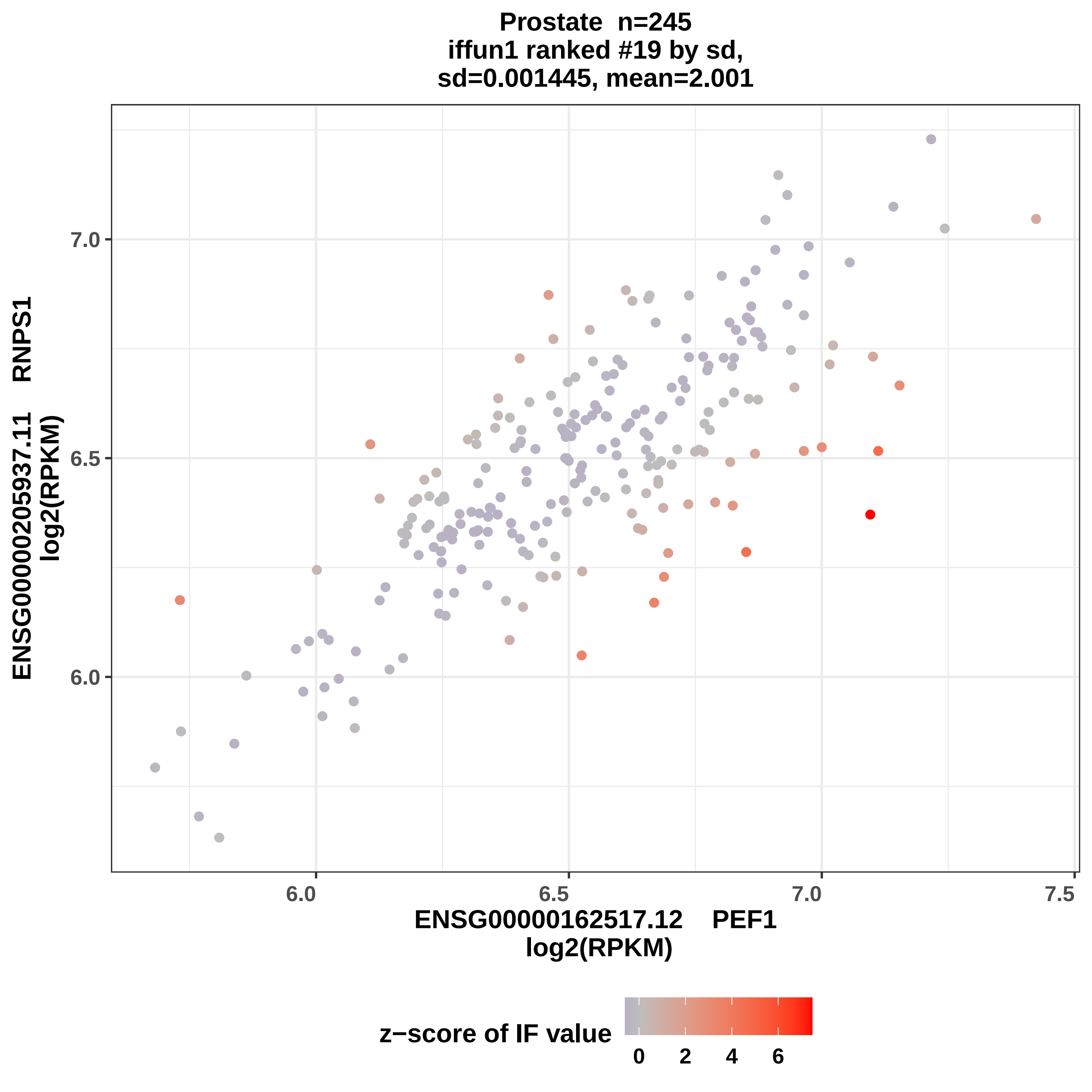

Prostate n=245
iffun1 ranked #19 by sd,
sd=0.001445, mean=2.001
ENSG00000205937.11 RNPS1
log2(RPKM)
7.0
6.5
6.0
6.0
6.5
7.0
7.5
ENSG00000162517.12 PEF1
log2(RPKM)
z−score of IF value
0
2
4
6

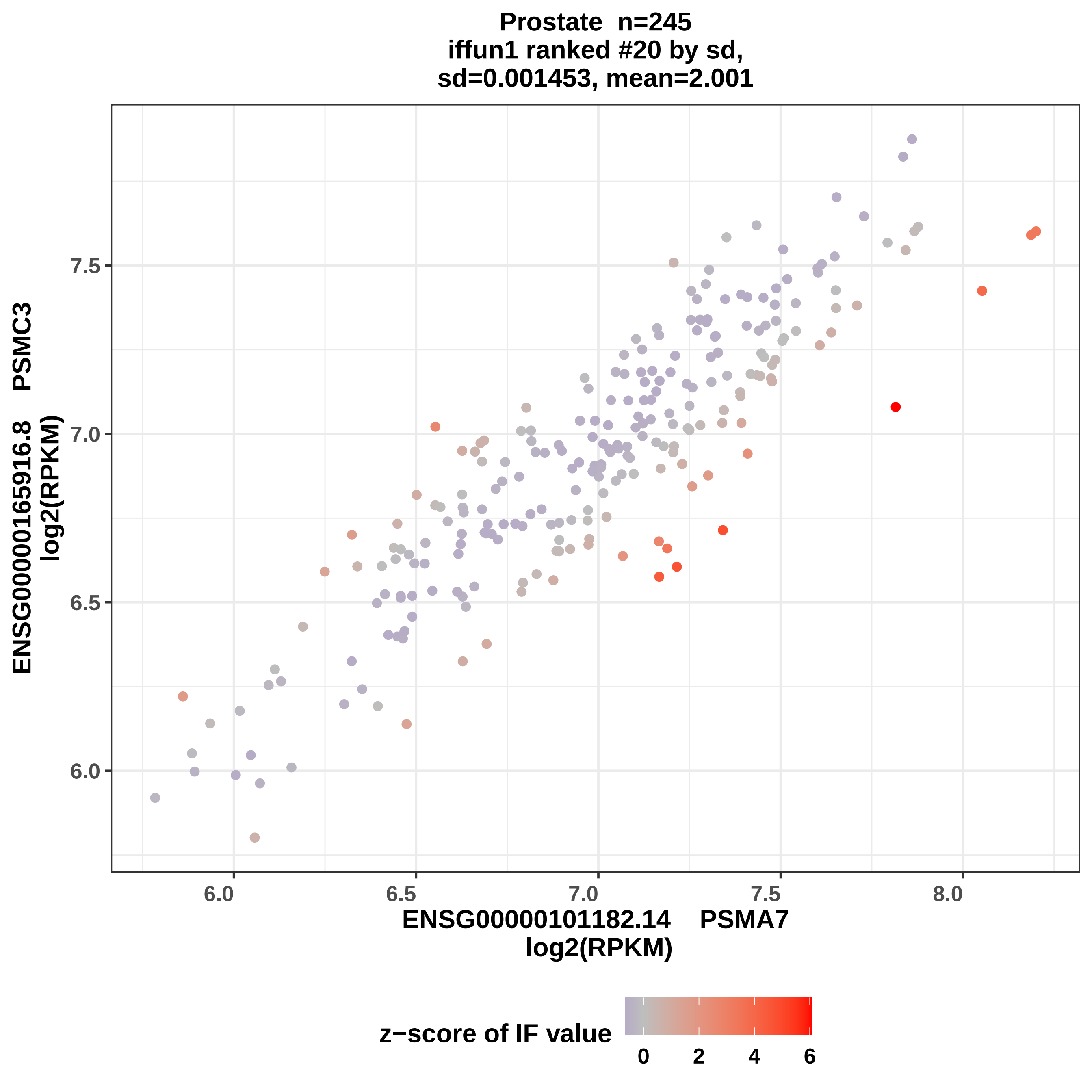
Prostate n=245
iffun1 ranked #20 by sd,
sd=0.001453, mean=2.001
ENSG00000165916.8 PSMC3
log2(RPKM)
7.5
7.0
6.5
6.0
6.0
6.5
7.0
7.5
8.0
ENSG00000101182.14 PSMA7
log2(RPKM)
z−score of IF value
0
2
4
6

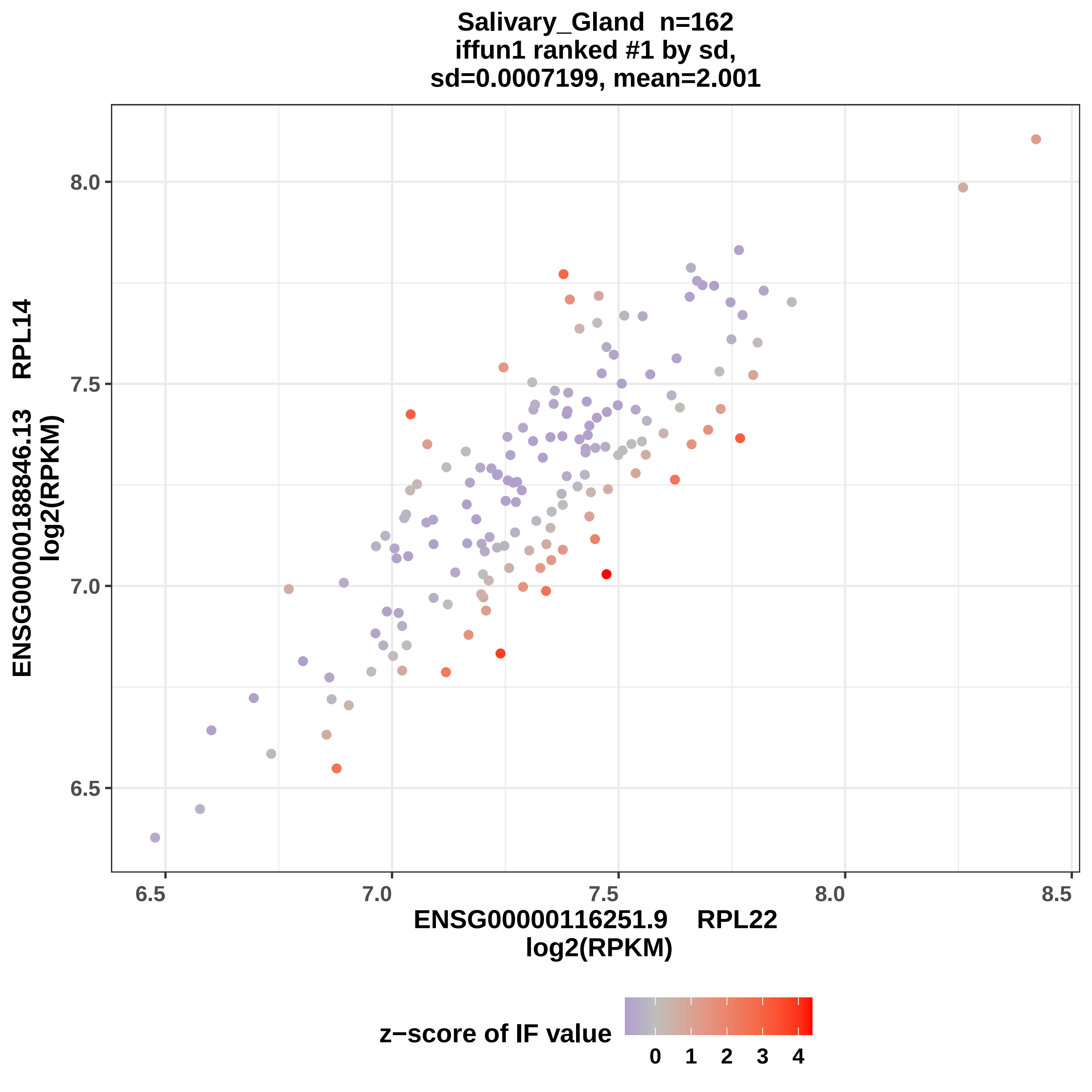

Salivary_Gland n=162
iffun1 ranked #1 by sd,
sd=0.0007199, mean=2.001
ENSG00000188846.13 RPL14
log2(RPKM)
8.0
7.5
7.0
6.5
6.5
7.0
7.5
8.0
8.5
ENSG00000116251.9 RPL22
log2(RPKM)
z−score of IF value
0
1
2
3
4

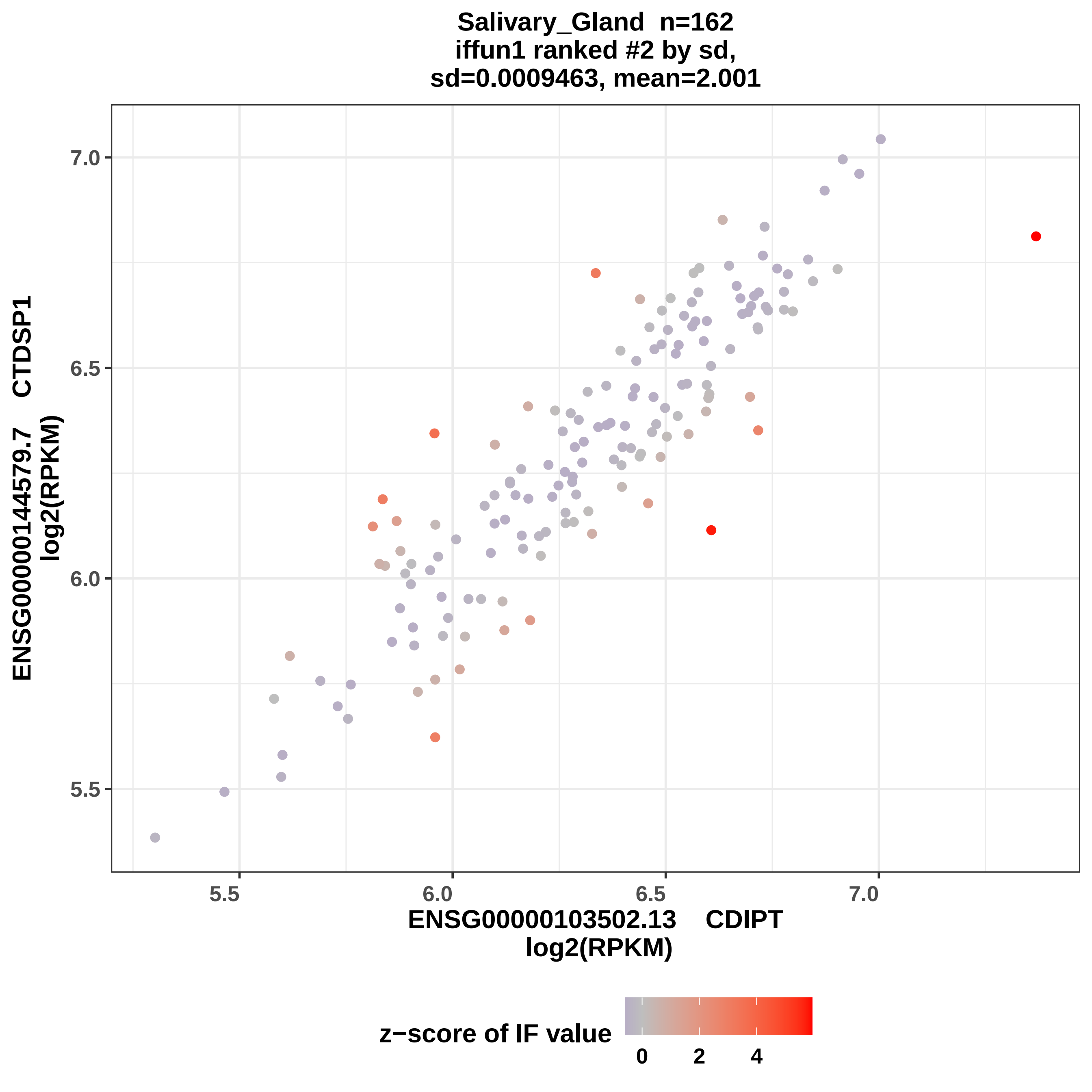
Salivary_Gland n=162
iffun1 ranked #2 by sd,
sd=0.0009463, mean=2.001
ENSG00000144579.7 CTDSP1
log2(RPKM)
ENSG00000103502.13 CDIPT
log2(RPKM)
7.0
6.5
6.0
5.5
5.5
6.0
6.5
7.0
z−score of IF value
0
2
4

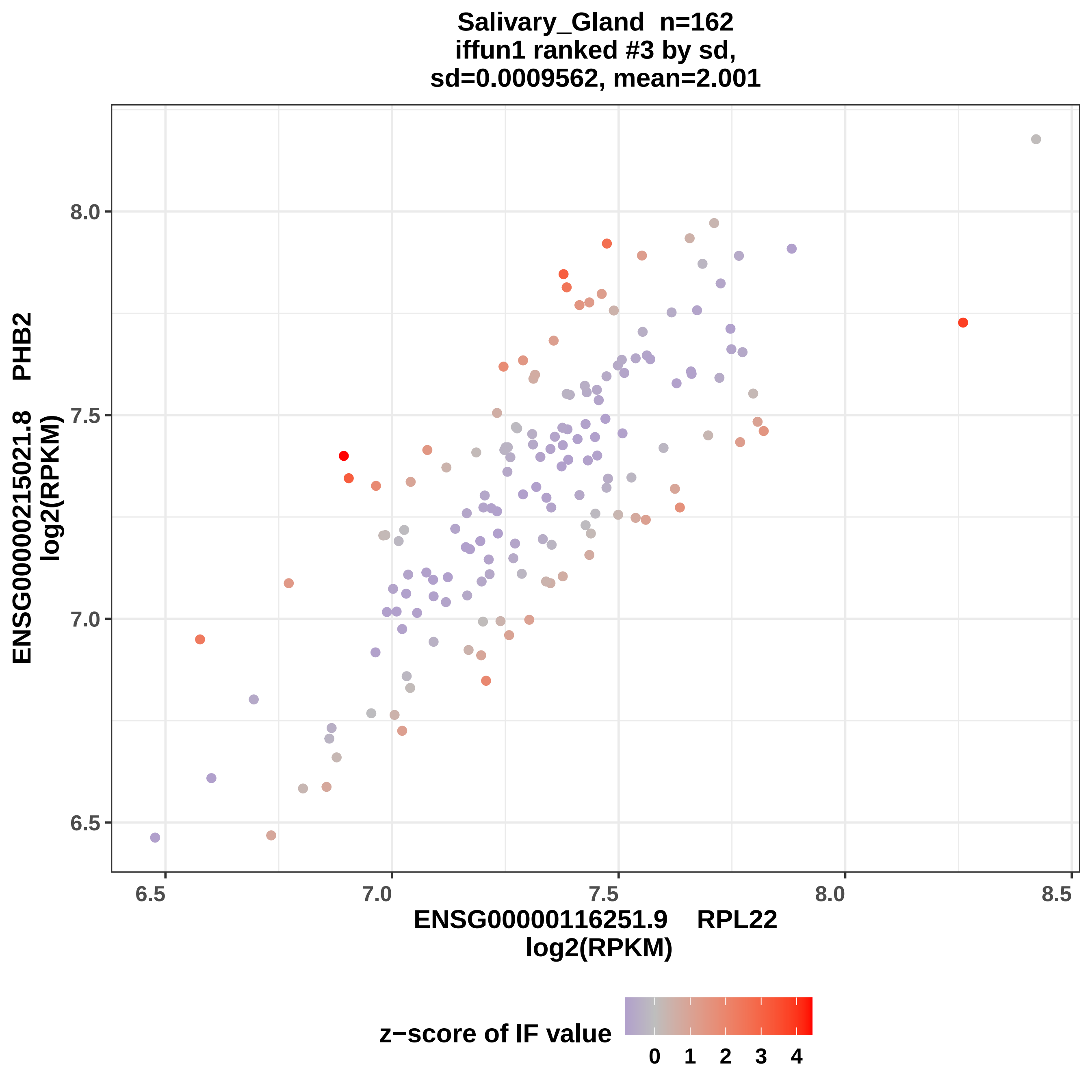
Salivary_Gland n=162
iffun1 ranked #3 by sd,
sd=0.0009562, mean=2.001
ENSG00000215021.8 PHB2
log2(RPKM)
8.0
7.5
7.0
6.5
6.5
7.0
7.5
8.0
8.5
ENSG00000116251.9 RPL22
log2(RPKM)
z−score of IF value
0
1
2
3
4

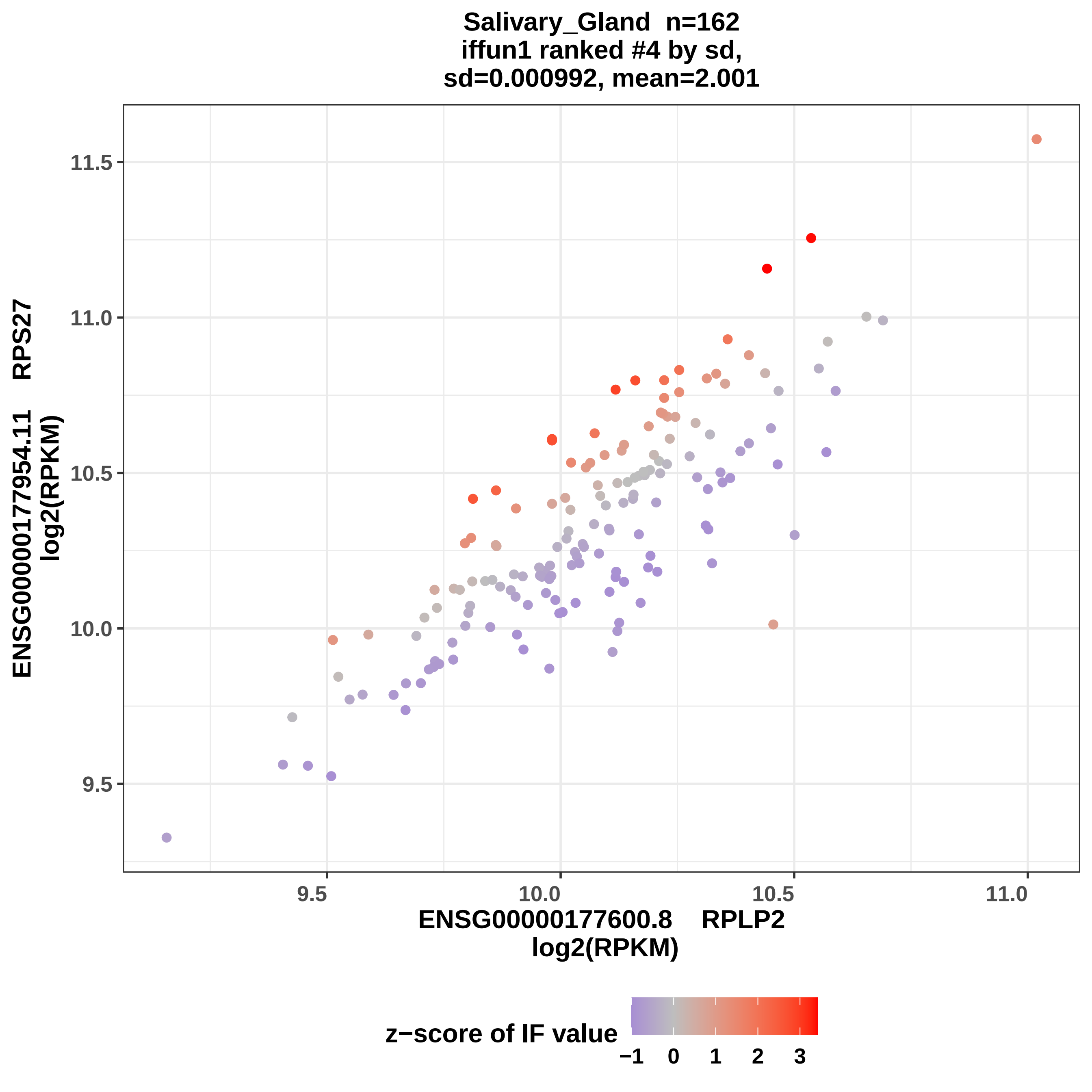
Salivary_Gland n=162
iffun1 ranked #4 by sd,
sd=0.000992, mean=2.001
ENSG00000177954.11 RPS27
log2(RPKM)
11.5
11.0
10.5
10.0
9.5
9.5
10.0
10.5
11.0
ENSG00000177600.8 RPLP2
log2(RPKM)
z−score of IF value
−1
0
1
2
3

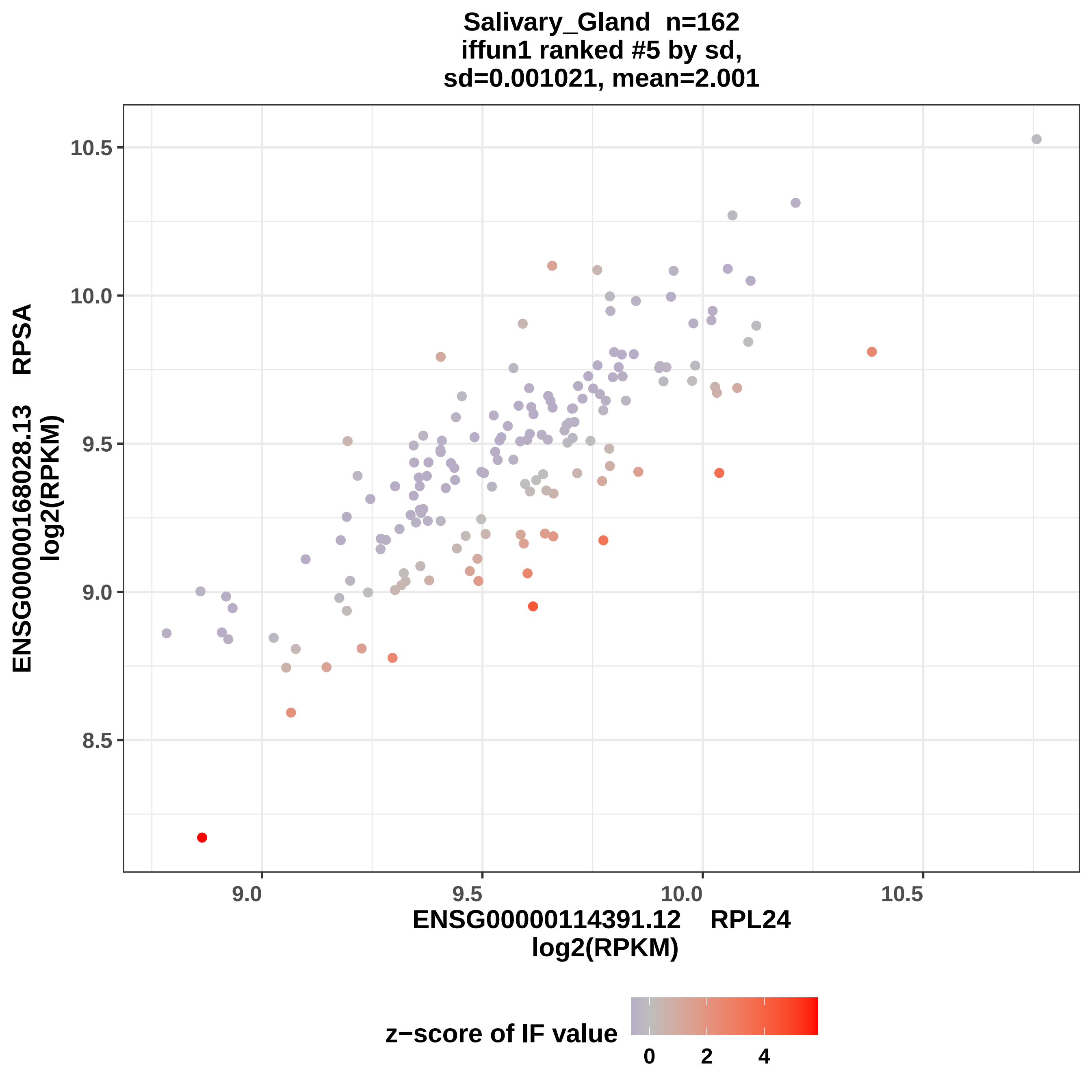
Salivary_Gland n=162
iffun1 ranked #5 by sd,
sd=0.001021, mean=2.001
ENSG00000168028.13 RPSA
log2(RPKM)
10.5
10.0
9.5
9.0
8.5
9.0
9.5
10.0
10.5
ENSG00000114391.12 RPL24
log2(RPKM)
z−score of IF value
0
2
4

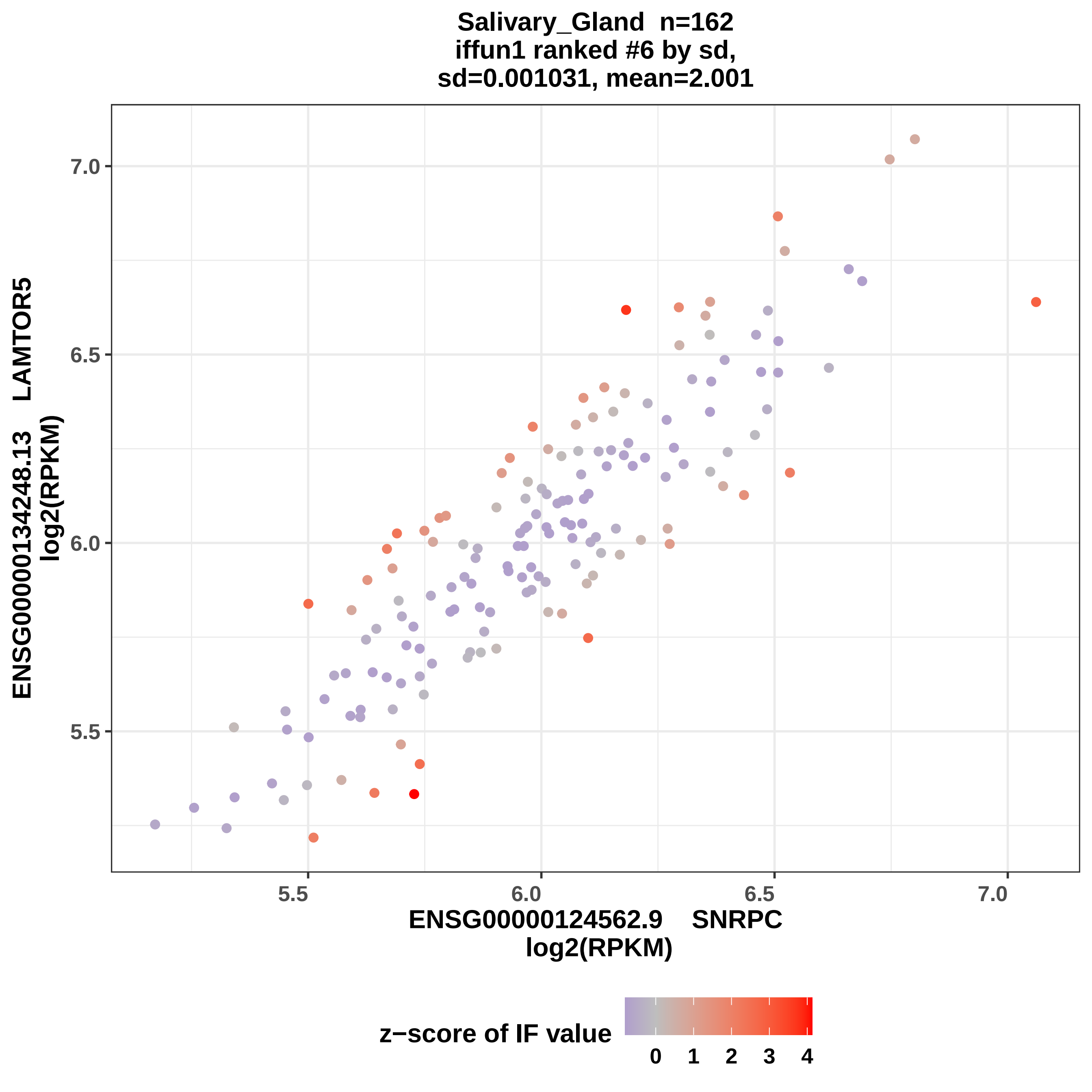
Salivary_Gland n=162
iffun1 ranked #6 by sd,
sd=0.001031, mean=2.001
ENSG00000134248.13 LAMTOR5
log2(RPKM)
7.0
6.5
6.0
5.5
5.5
6.0
6.5
7.0
ENSG00000124562.9 SNRPC
log2(RPKM)
z−score of IF value
0
1
2
3
4

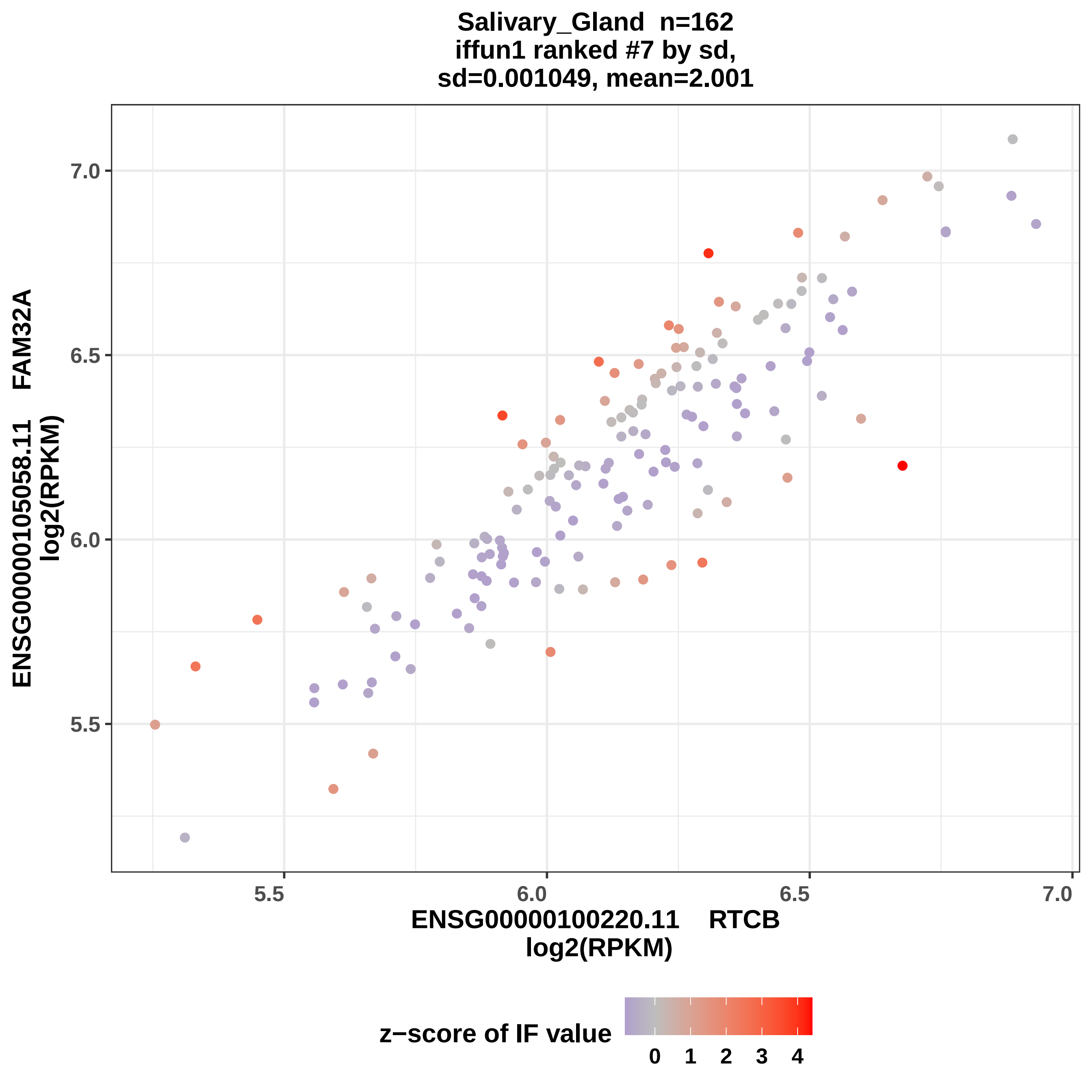
Salivary_Gland n=162
iffun1 ranked #7 by sd,
sd=0.001049, mean=2.001
ENSG00000105058.11 FAM32A
log2(RPKM)
7.0
6.5
6.0
5.5
5.5
6.0
6.5
7.0
ENSG00000100220.11 RTCB
log2(RPKM)
z−score of IF value
0
1
2
3
4

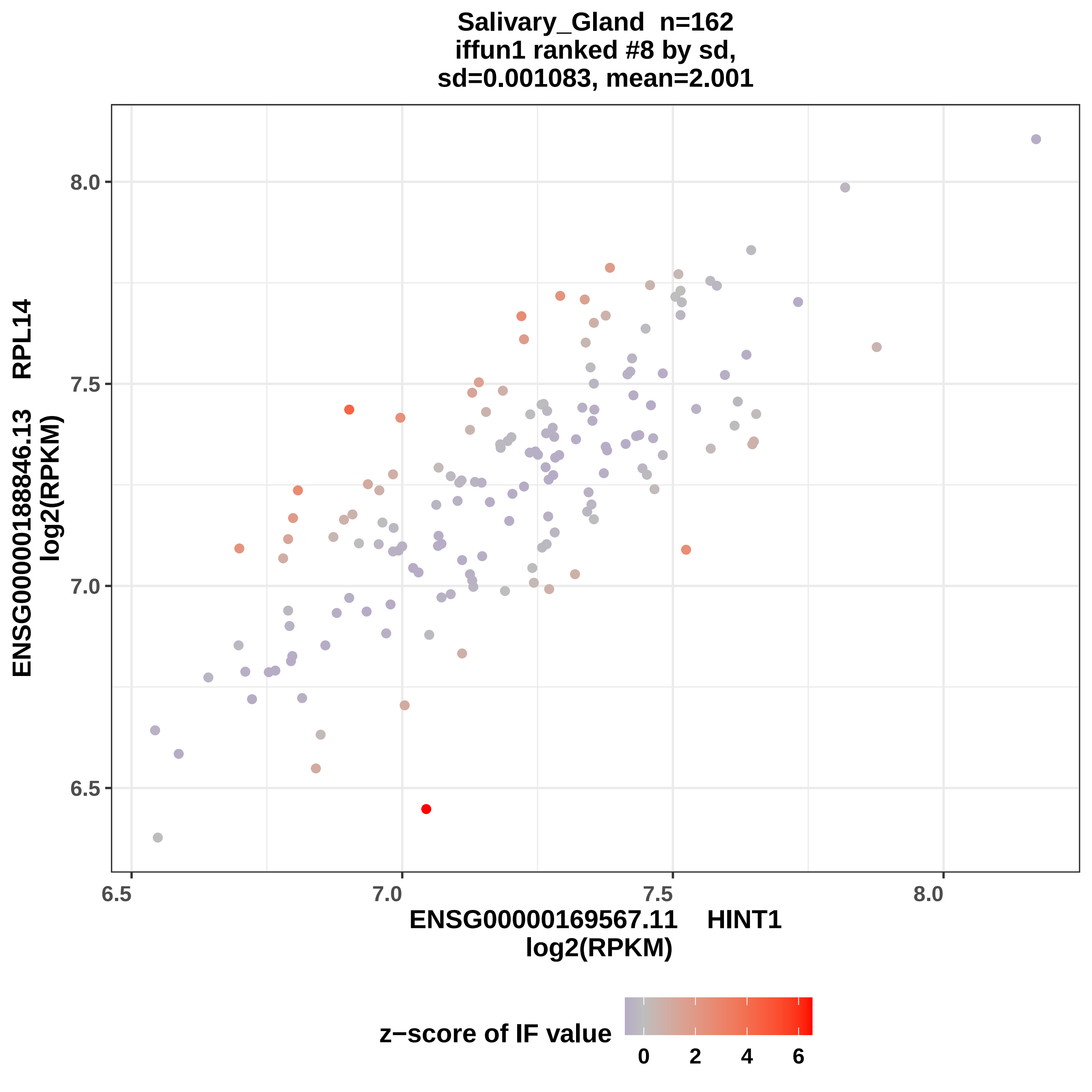
Salivary_Gland n=162
iffun1 ranked #8 by sd,
sd=0.001083, mean=2.001
ENSG00000188846.13 RPL14
log2(RPKM)
ENSG00000169567.11 HINT1
log2(RPKM)
8.0
7.5
7.0
6.5
6.5
7.0
7.5
8.0
z−score of IF value
0
2
4
6

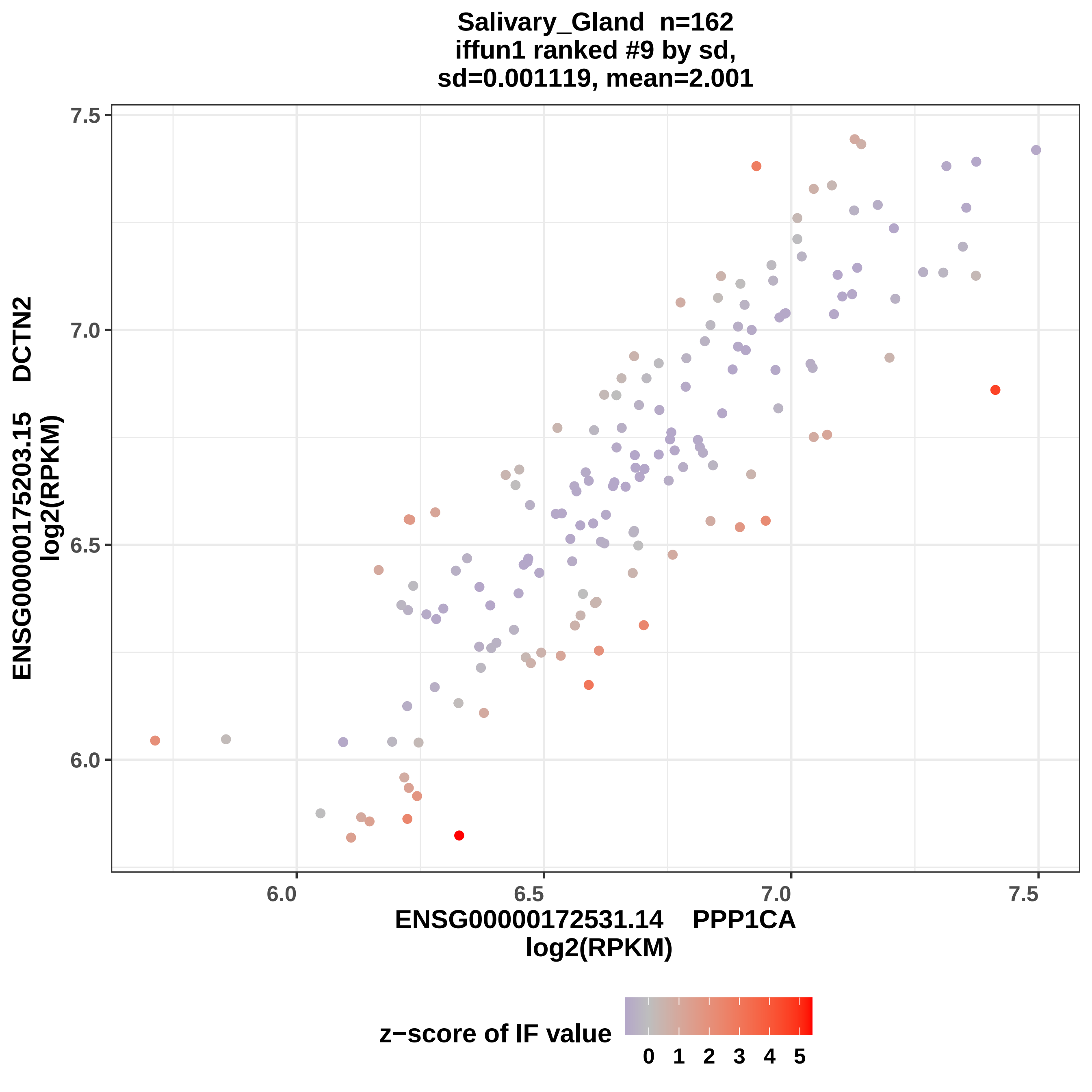
Salivary_Gland n=162
iffun1 ranked #9 by sd,
sd=0.001119, mean=2.001
7.5
7.0
6.5
6.0
ENSG00000175203.15 DCTN2
log2(RPKM)
6.0
6.5
7.0
7.5
ENSG00000172531.14 PPP1CA
log2(RPKM)
z−score of IF value
0 1 2 3 4 5

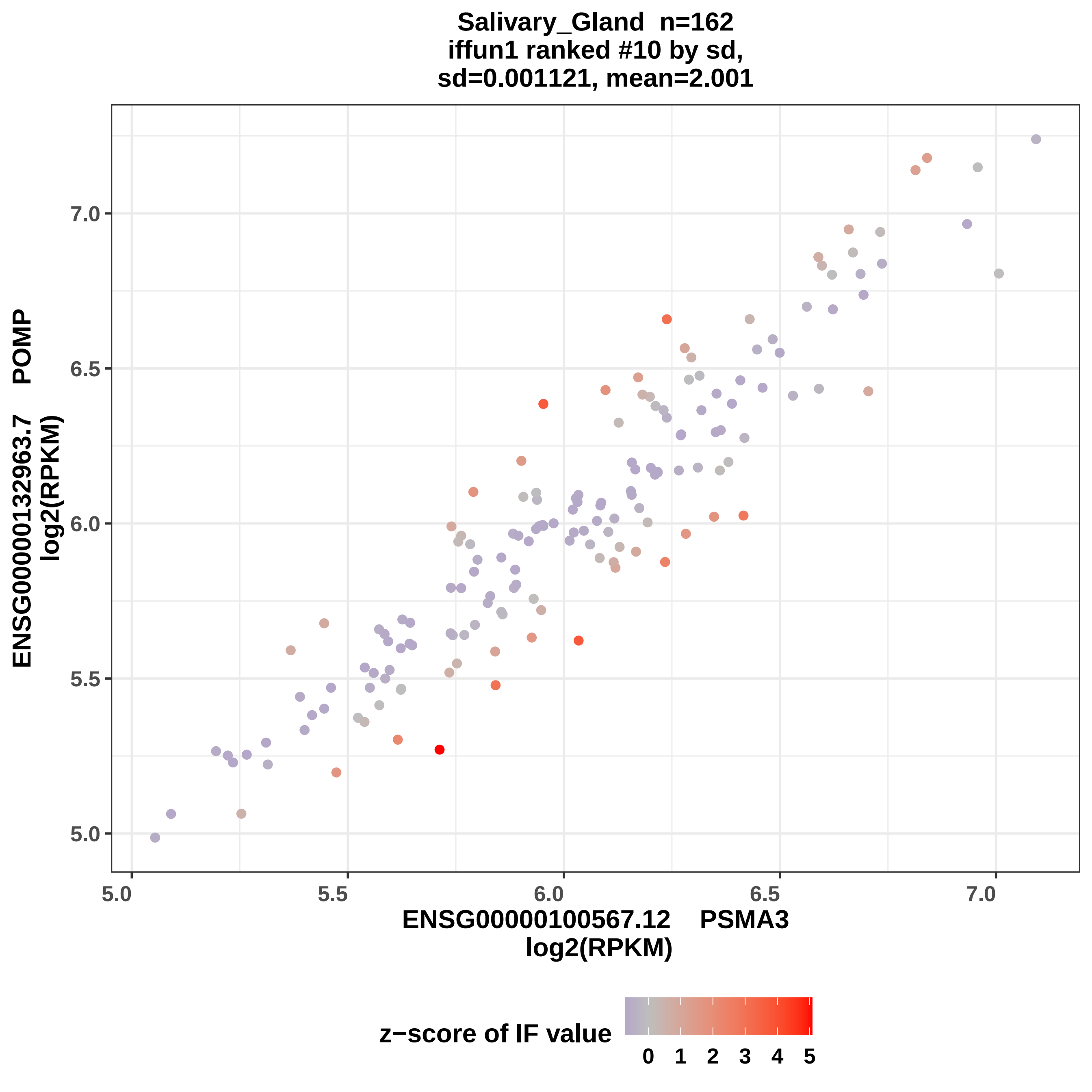
Salivary_Gland n=162
iffun1 ranked #10 by sd,
sd=0.001121, mean=2.001
ENSG00000132963.7 POMP
log2(RPKM)
ENSG00000100567.12 PSMA3
log2(RPKM)
5.0
5.5
6.0
6.5
7.0
z−score of IF value
0 1 2 3 4 5

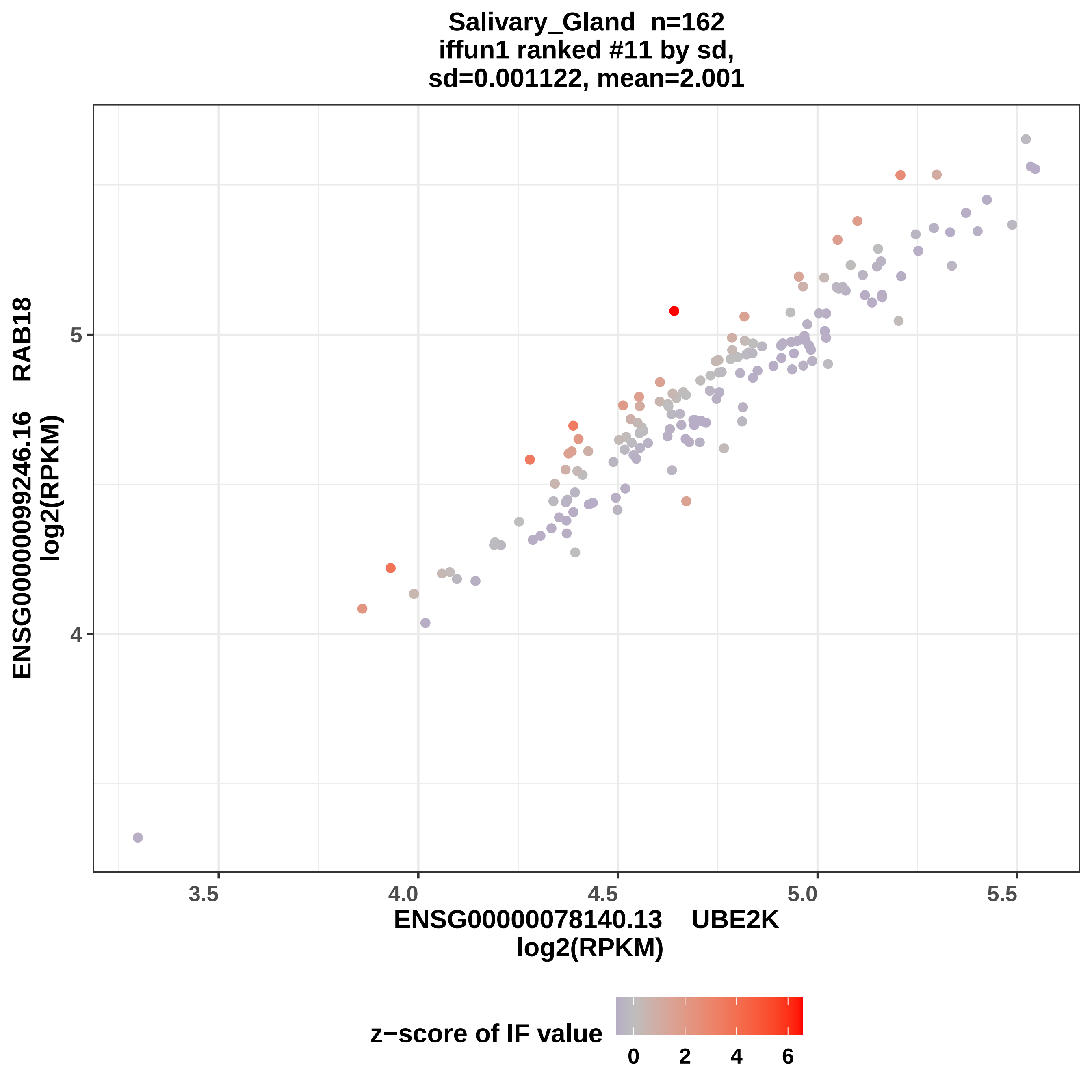

Salivary_Gland n=162
iffun1 ranked #11 by sd,
sd=0.001122, mean=2.001
ENSG00000099246.16 RAB18
log2(RPKM)
5
4
3.5
4.0
4.5
5.0
5.5
ENSG00000078140.13 UBE2K
log2(RPKM)
z−score of IF value
0
2
4
6

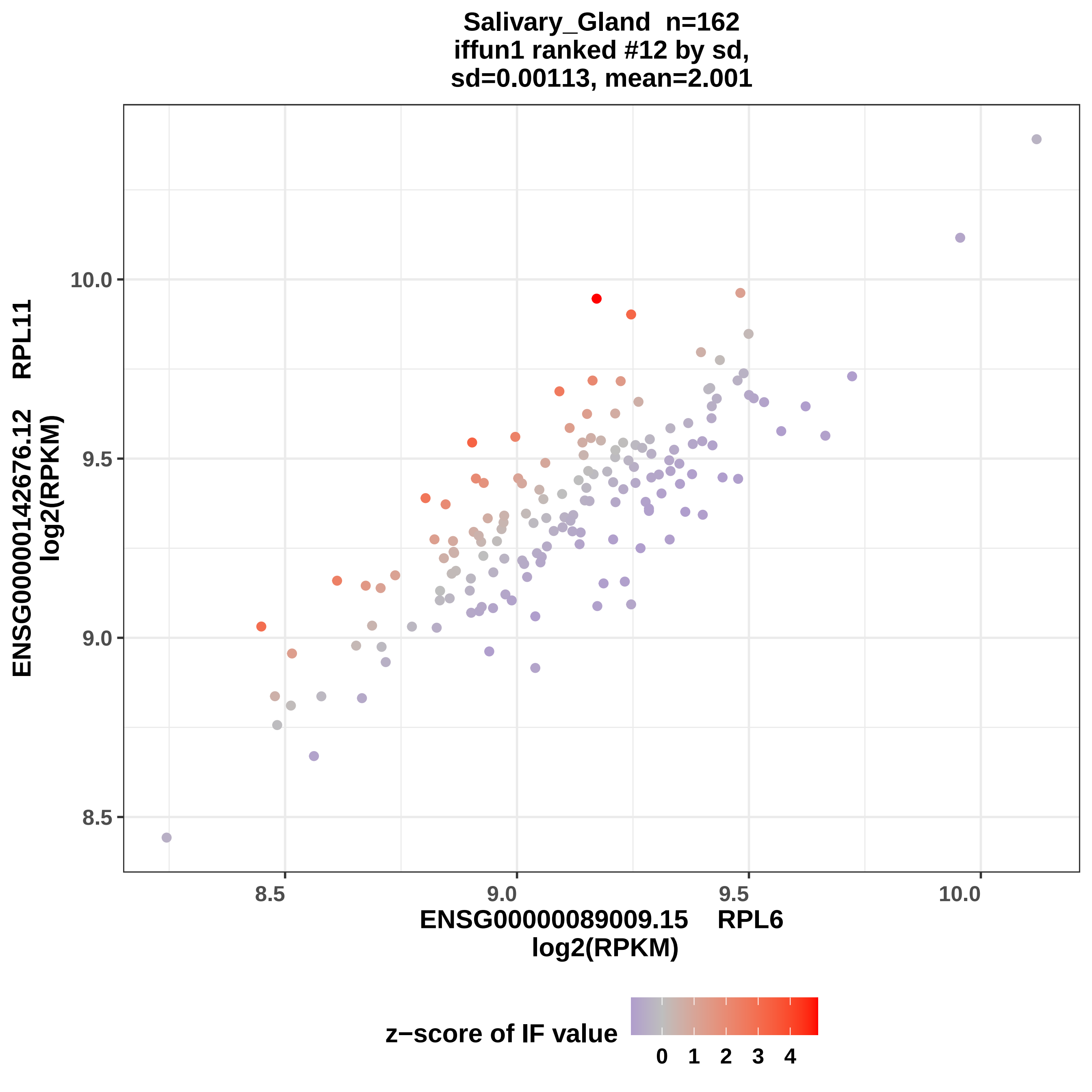
Salivary_Gland n=162
iffun1 ranked #12 by sd,
sd=0.00113, mean=2.001
ENSG00000142676.12 RPL11
log2(RPKM)
ENSG00000089009.15 RPL6
log2(RPKM)
8.5
9.0
9.5
10.0
z−score of IF value
0
1
2
3
4

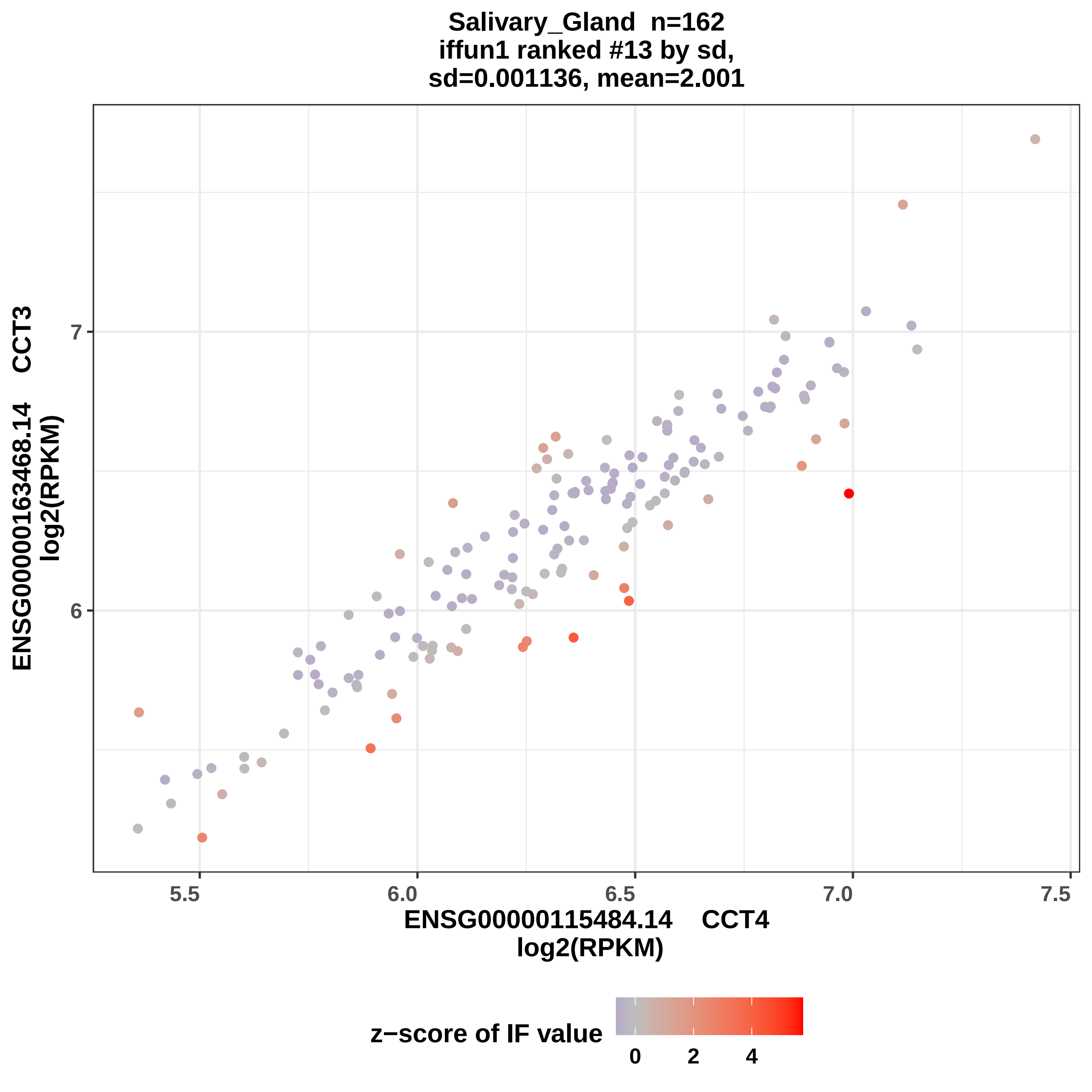
Salivary_Gland n=162
iffun1 ranked #13 by sd,
sd=0.001136, mean=2.001
ENSG00000163468.14 CCT3
log2(RPKM)
7
6
5.5
6.0
6.5
7.0
7.5
ENSG00000115484.14 CCT4
log2(RPKM)
z−score of IF value
0
2
4

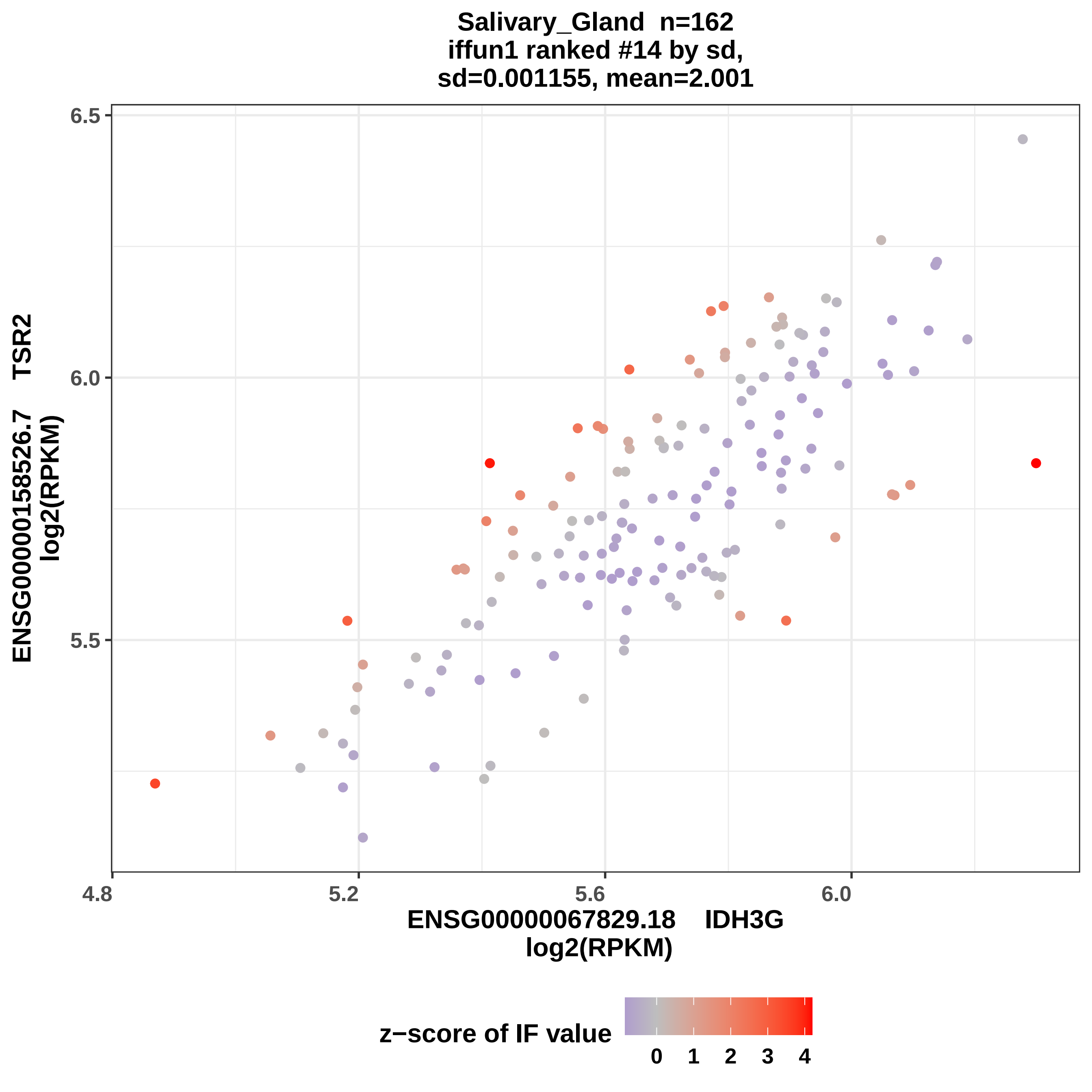
Salivary_Gland n=162
iffun1 ranked #14 by sd,
sd=0.001155, mean=2.001
6.5
6.0
5.5
4.8
5.2
5.6
6.0
ENSG00000158526.7 TSR2
log2(RPKM)
ENSG00000067829.18 IDH3G
log2(RPKM)
z−score of IF value
0
1
2
3
4

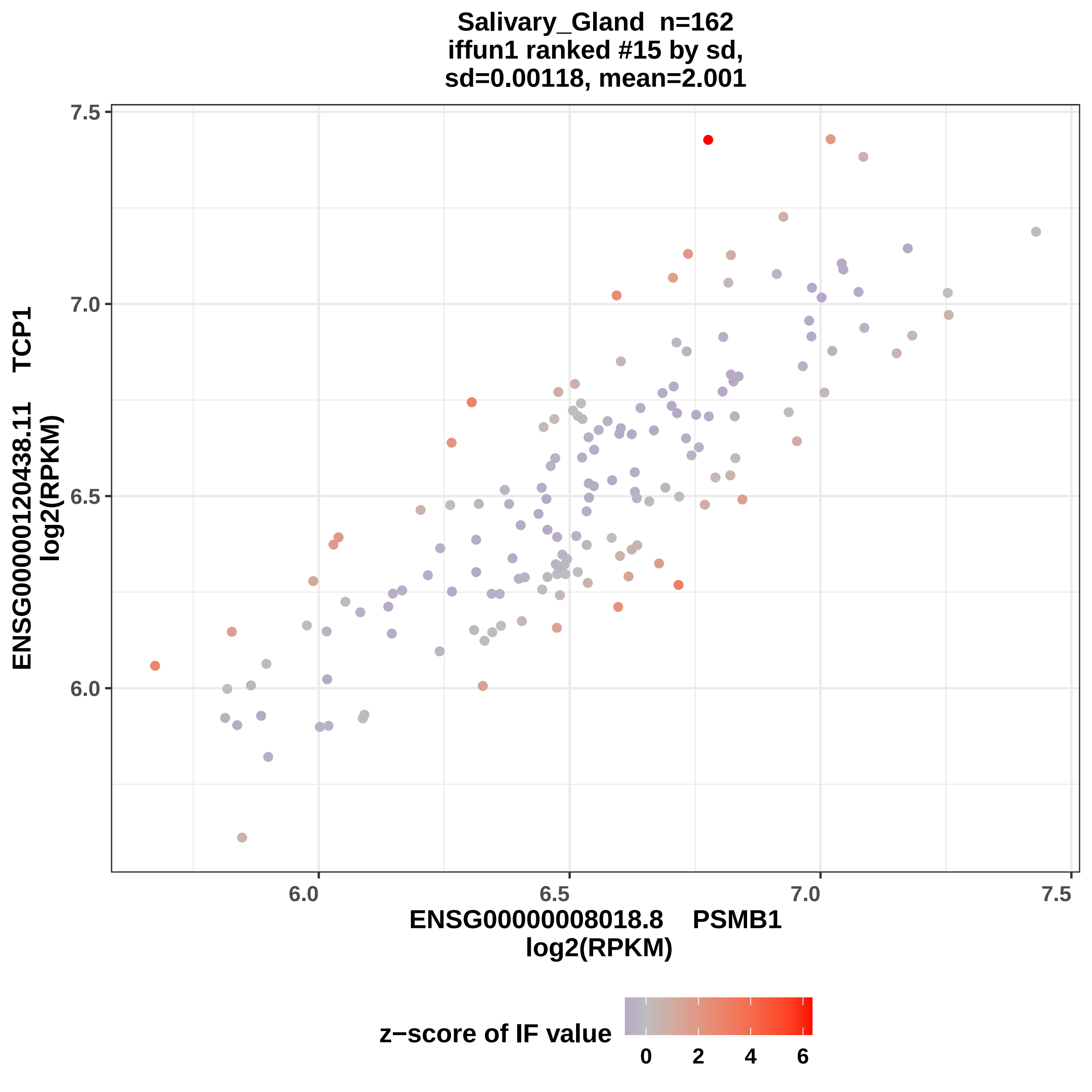
Salivary_Gland n=162
iffun1 ranked #15 by sd,
sd=0.00118, mean=2.001
ENSG00000120438.11 TCP1
log2(RPKM)
7.5
7.0
6.5
6.0
6.0
6.5
7.0
7.5
ENSG00000008018.8 PSMB1
log2(RPKM)
z−score of IF value
0
2
4
6

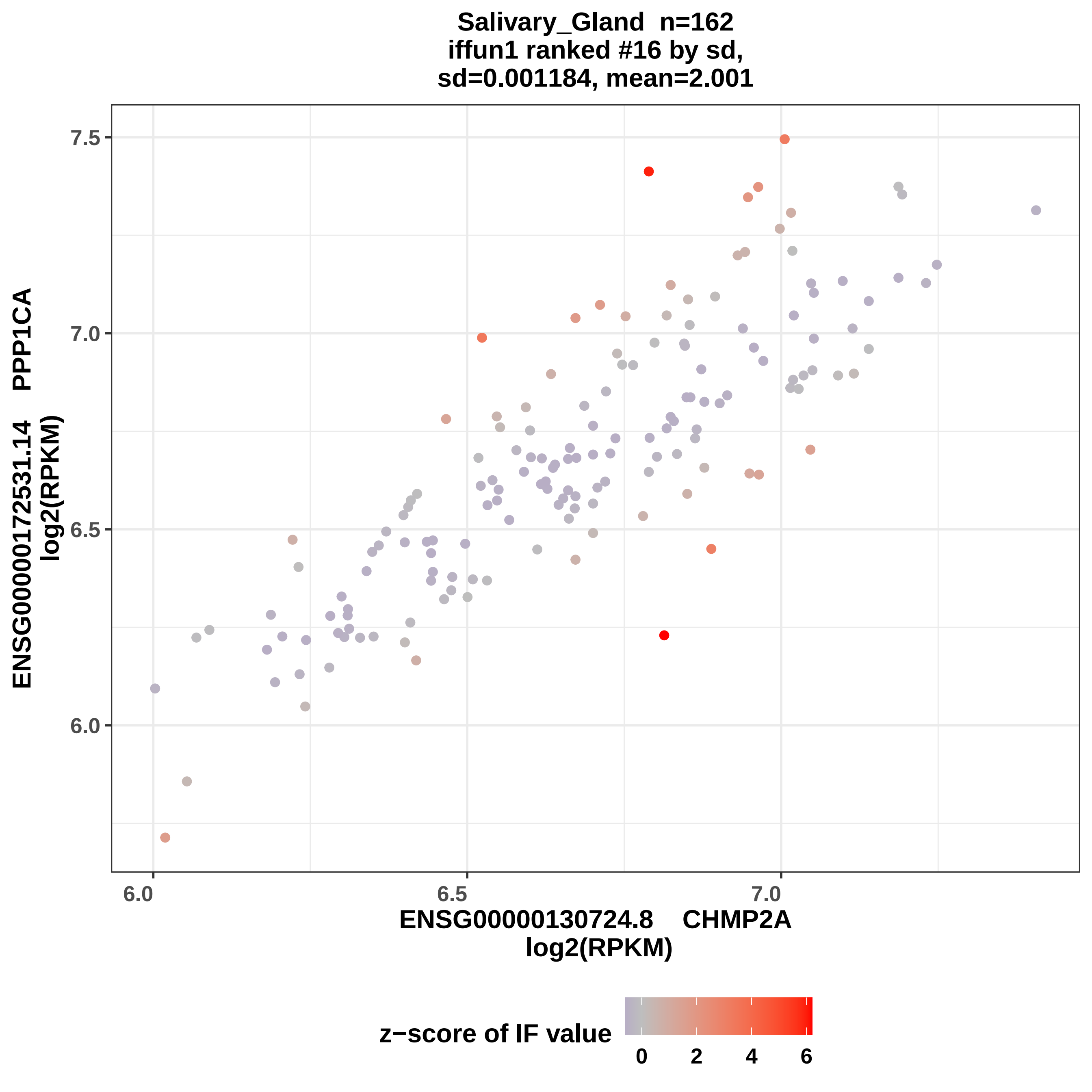

Salivary_Gland n=162
iffun1 ranked #16 by sd,
sd=0.001184, mean=2.001
ENSG00000172531.14 PPP1CA
log2(RPKM)
7.5
7.0
6.5
6.0
6.0
6.5
7.0
ENSG00000130724.8 CHMP2A
log2(RPKM)
z−score of IF value
0
2
4
6

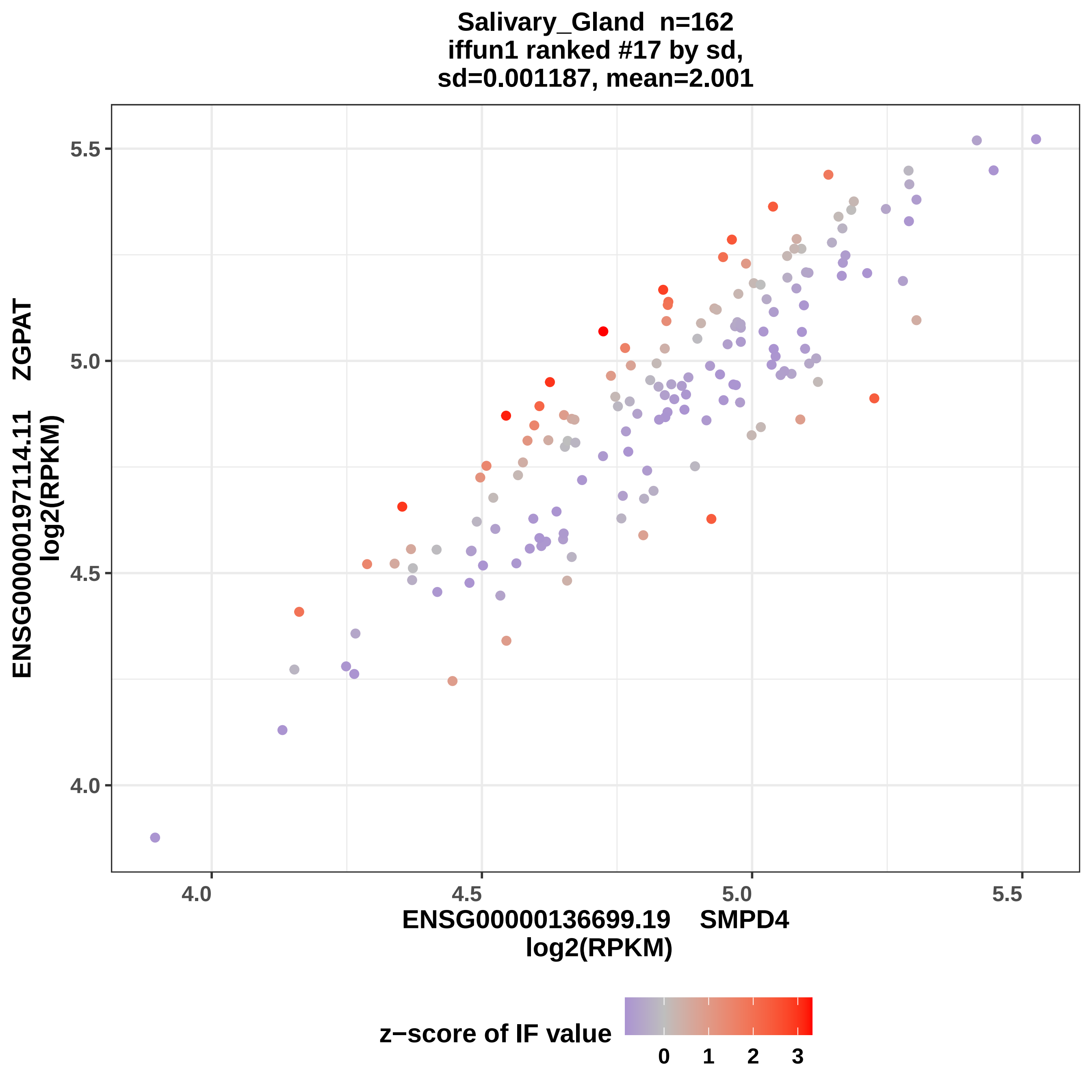
Salivary_Gland n=162
iffun1 ranked #17 by sd,
sd=0.001187, mean=2.001
ENSG00000197114.11 ZGPAT
log2(RPKM)
ENSG00000136699.19 SMPD4
log2(RPKM)
4.0
4.5
5.0
5.5
z−score of IF value
0
1
2
3

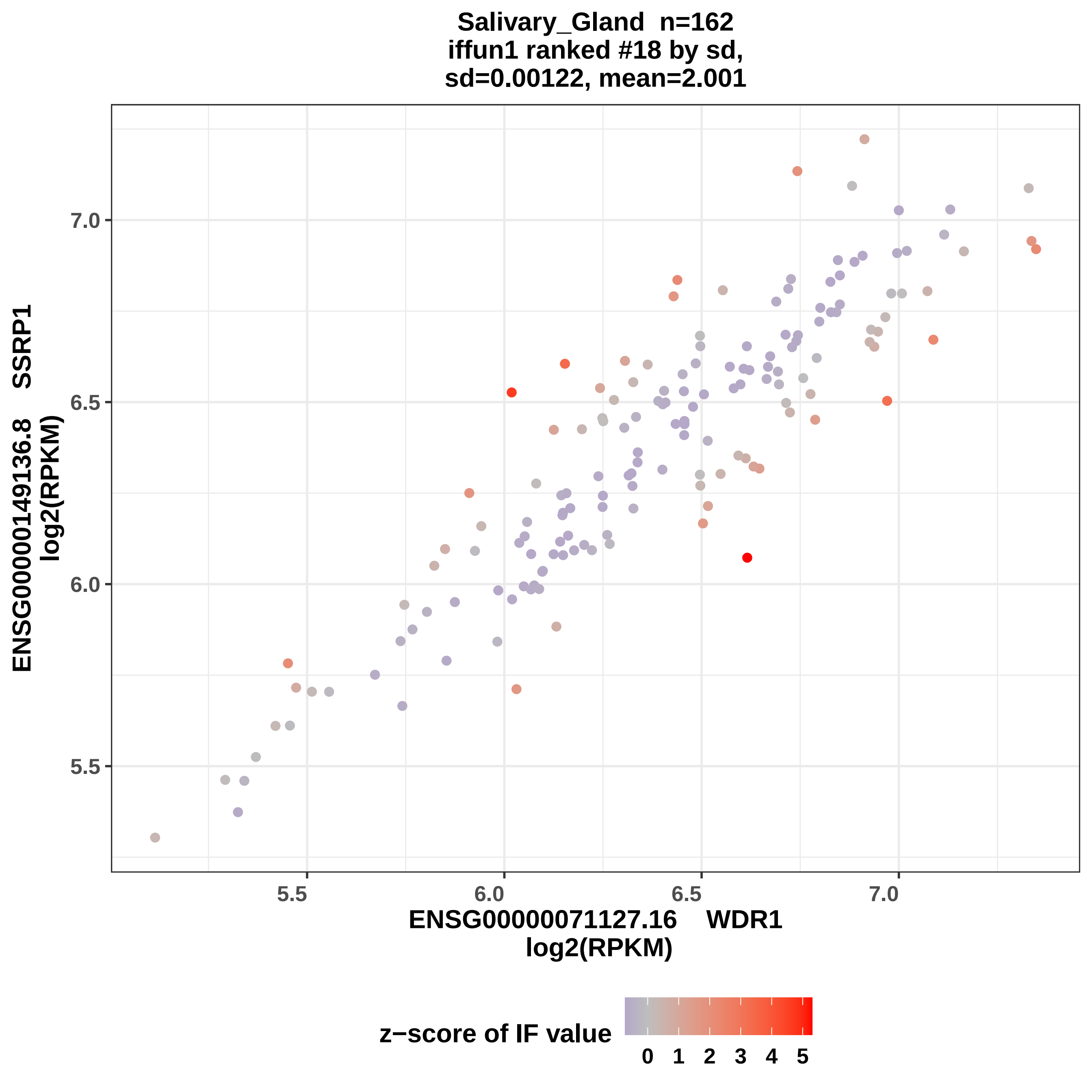

Salivary_Gland n=162
iffun1 ranked #18 by sd,
sd=0.00122, mean=2.001
ENSG00000149136.8 SSRP1
log2(RPKM)
7.0
6.5
6.0
5.5
5.5
6.0
6.5
7.0
ENSG00000071127.16 WDR1
log2(RPKM)
z−score of IF value
0 1 2 3 4 5

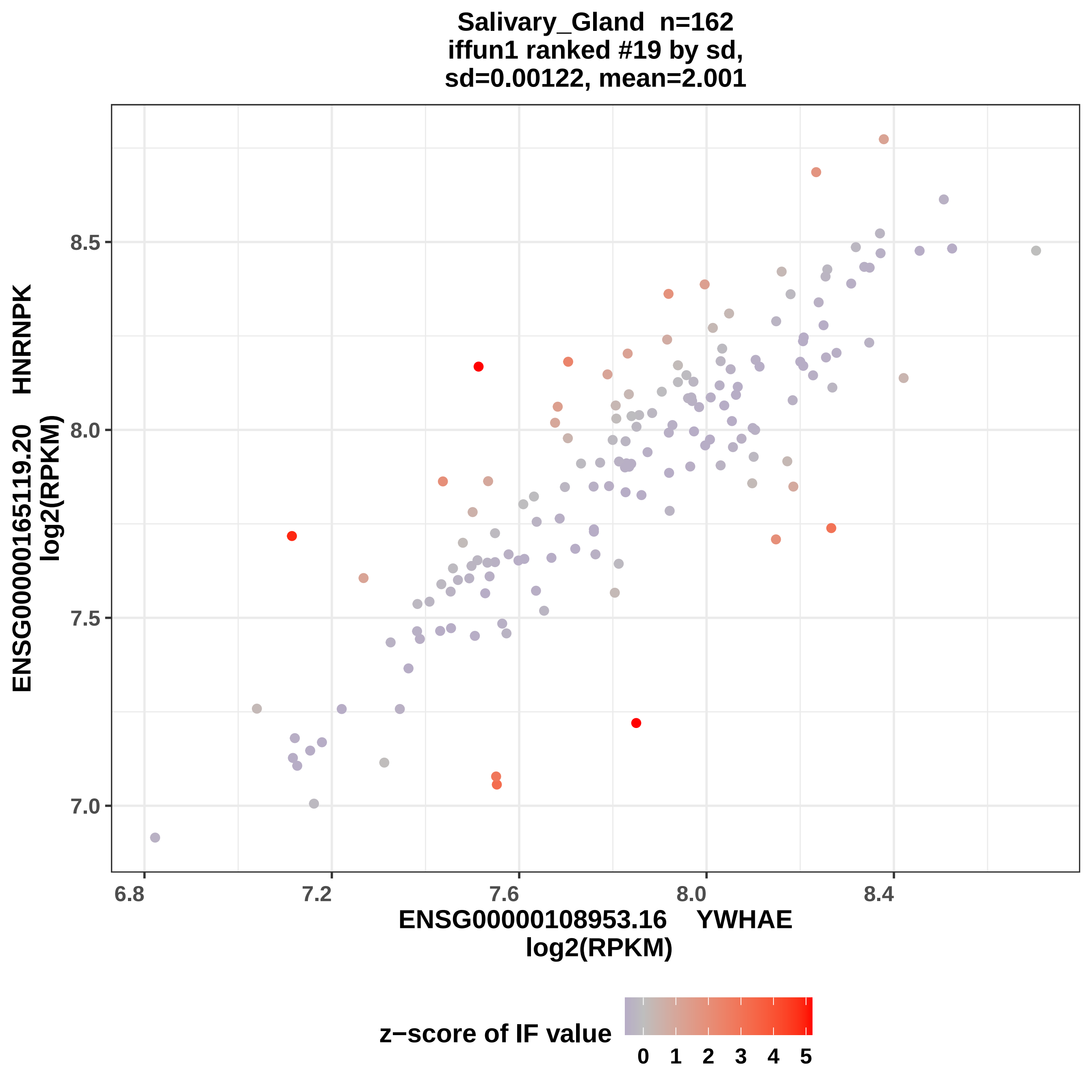
Salivary_Gland n=162
iffun1 ranked #19 by sd,
sd=0.00122, mean=2.001
ENSG00000165119.20 HNRNPK
log2(RPKM)
8.5
8.0
7.5
7.0
6.8
7.2
7.6
8.0
8.4
ENSG00000108953.16 YWHAE
log2(RPKM)
z−score of IF value
0
1
2
3
4
5

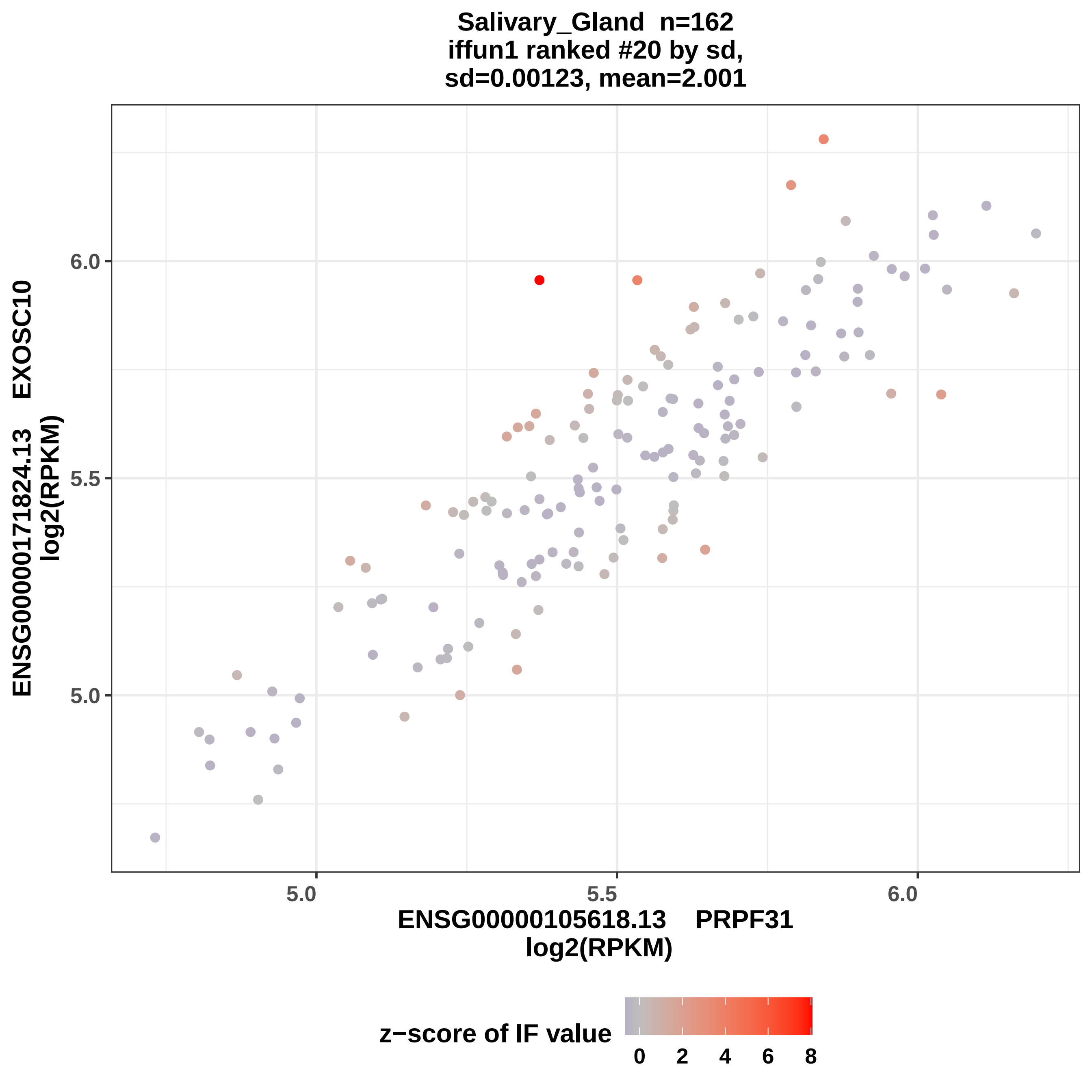

Salivary_Gland n=162
iffun1 ranked #20 by sd,
sd=0.00123, mean=2.001
ENSG00000171824.13 EXOSC10
log2(RPKM)
6.0
5.5
5.0
5.0
5.5
6.0
ENSG00000105618.13 PRPF31
log2(RPKM)
z−score of IF value
0
2
4
6
8

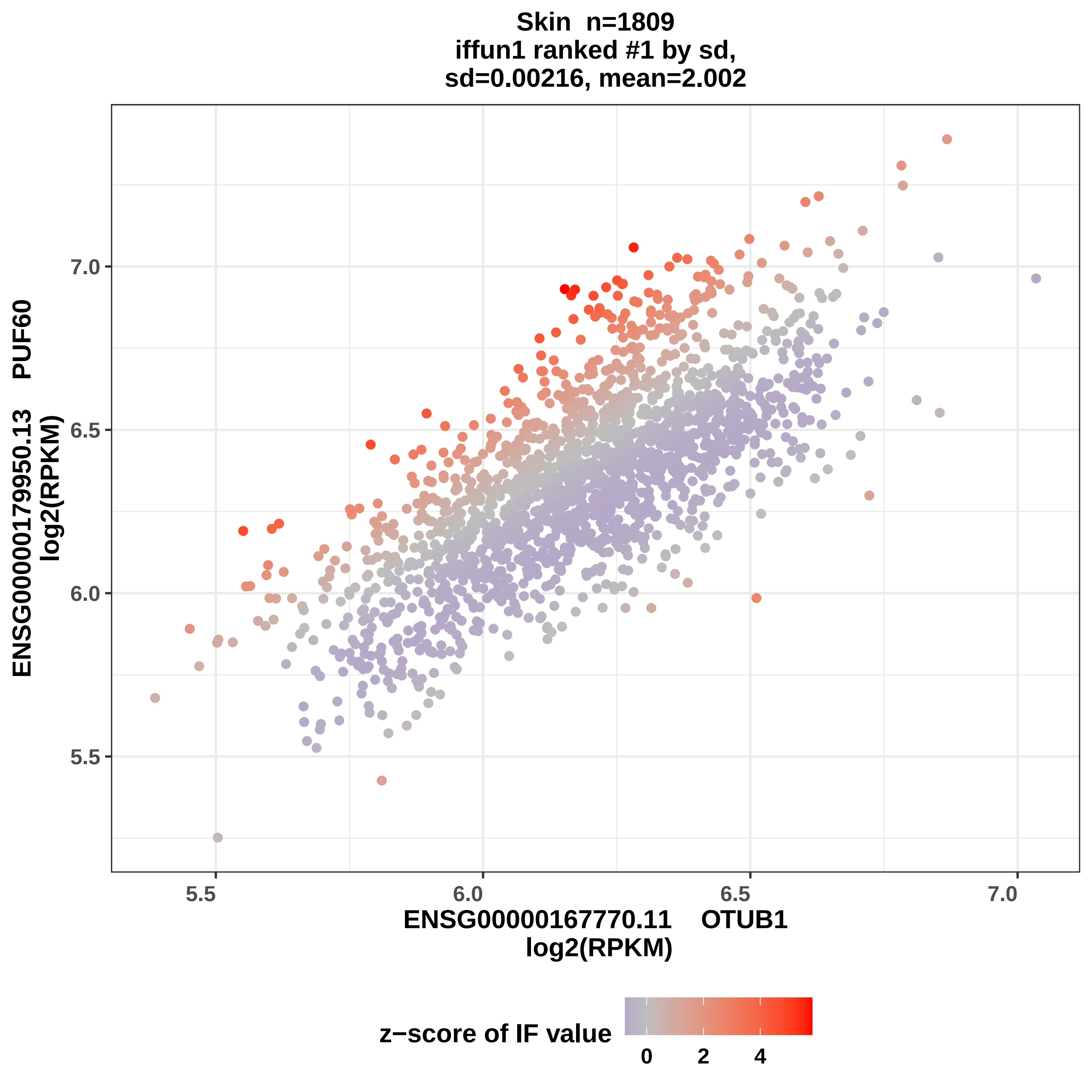

Skin n=1809
iffun1 ranked #1 by sd,
sd=0.00216, mean=2.002
ENSG00000179950.13 PUF60
log2(RPKM)
7.0
6.5
6.0
5.5
5.5
6.0
6.5
7.0
ENSG00000167770.11 OTUB1
log2(RPKM)
z−score of IF value
0
2
4

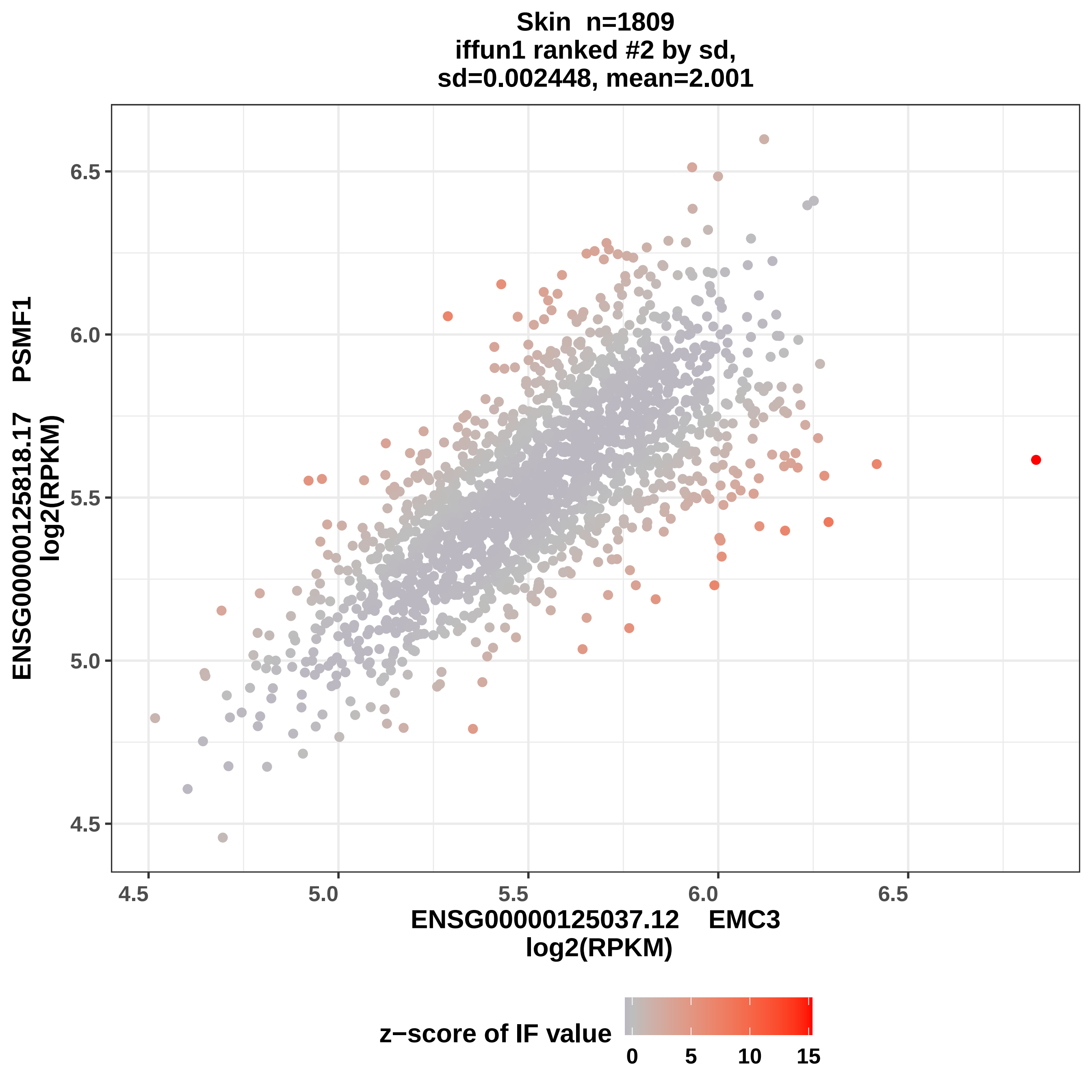

Skin n=1809
iffun1 ranked #2 by sd,
sd=0.002448, mean=2.001
ENSG00000125818.17 PSMF1
log2(RPKM)
6.5
6.0
5.5
5.0
4.5
4.5
5.0
5.5
6.0
6.5
ENSG00000125037.12 EMC3
log2(RPKM)
z−score of IF value
0
5
10
15

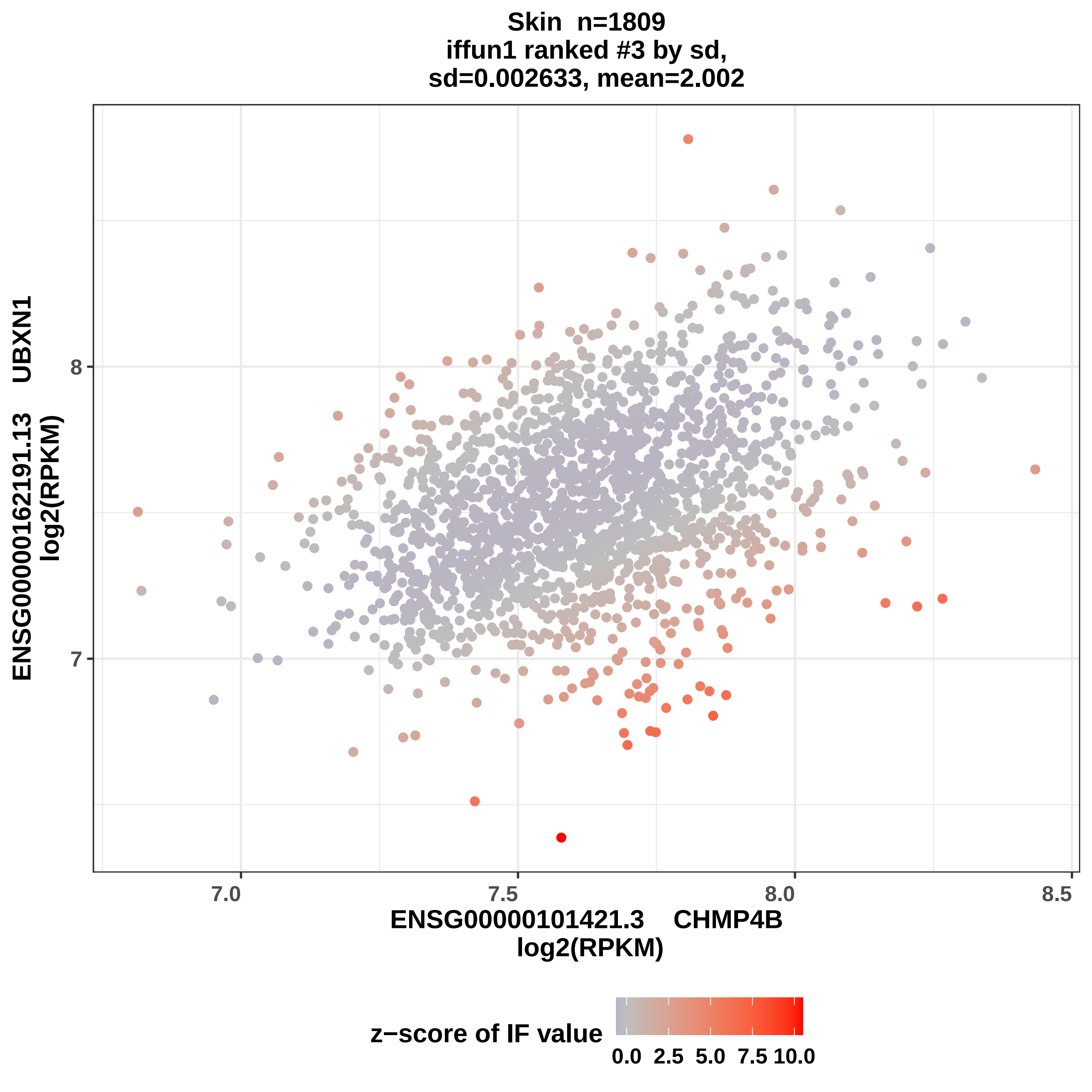
Skin n=1809
iffun1 ranked #3 by sd,
sd=0.002633, mean=2.002
ENSG00000162191.13 UBXN1
log2(RPKM)
8
7
7.0
7.5
8.0
8.5
ENSG00000101421.3 CHMP4B
log2(RPKM)
z−score of IF value
0.0
2.5
5.0
7.5
10.0

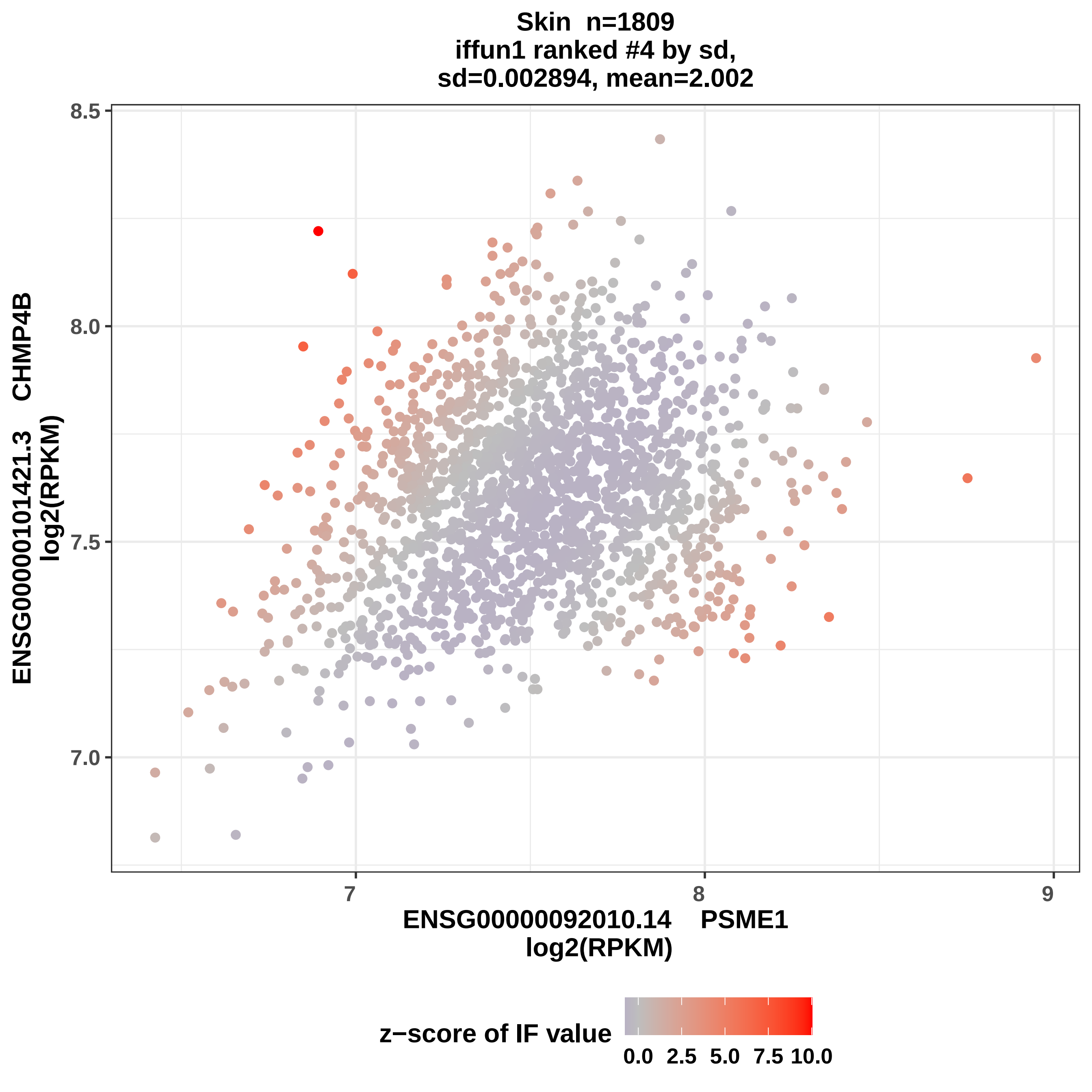
Skin n=1809
iffun1 ranked #4 by sd,
sd=0.002894, mean=2.002
8.5
8.0
7.5
7.0
ENSG00000101421.3 CHMP4B
log2(RPKM)
7
8
9
ENSG00000092010.14 PSME1
log2(RPKM)
z−score of IF value
0.0
2.5
5.0
7.5
10.0

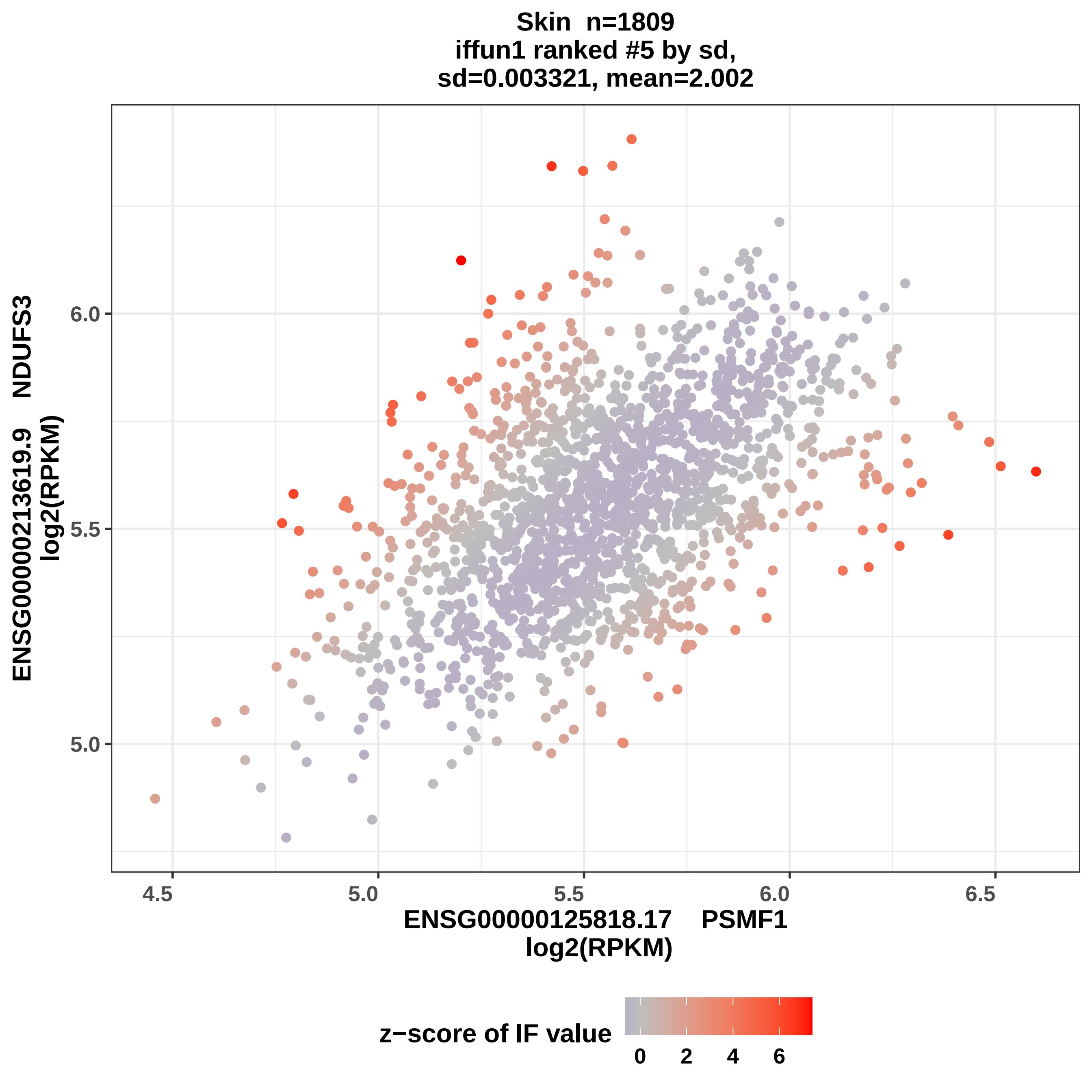
Skin n=1809
iffun1 ranked #5 by sd,
sd=0.003321, mean=2.002
ENSG00000213619.9 NDUFS3
log2(RPKM)
6.0
5.5
5.0
4.5
5.0
5.5
6.0
6.5
ENSG00000125818.17 PSMF1
log2(RPKM)
z−score of IF value
0
2
4
6

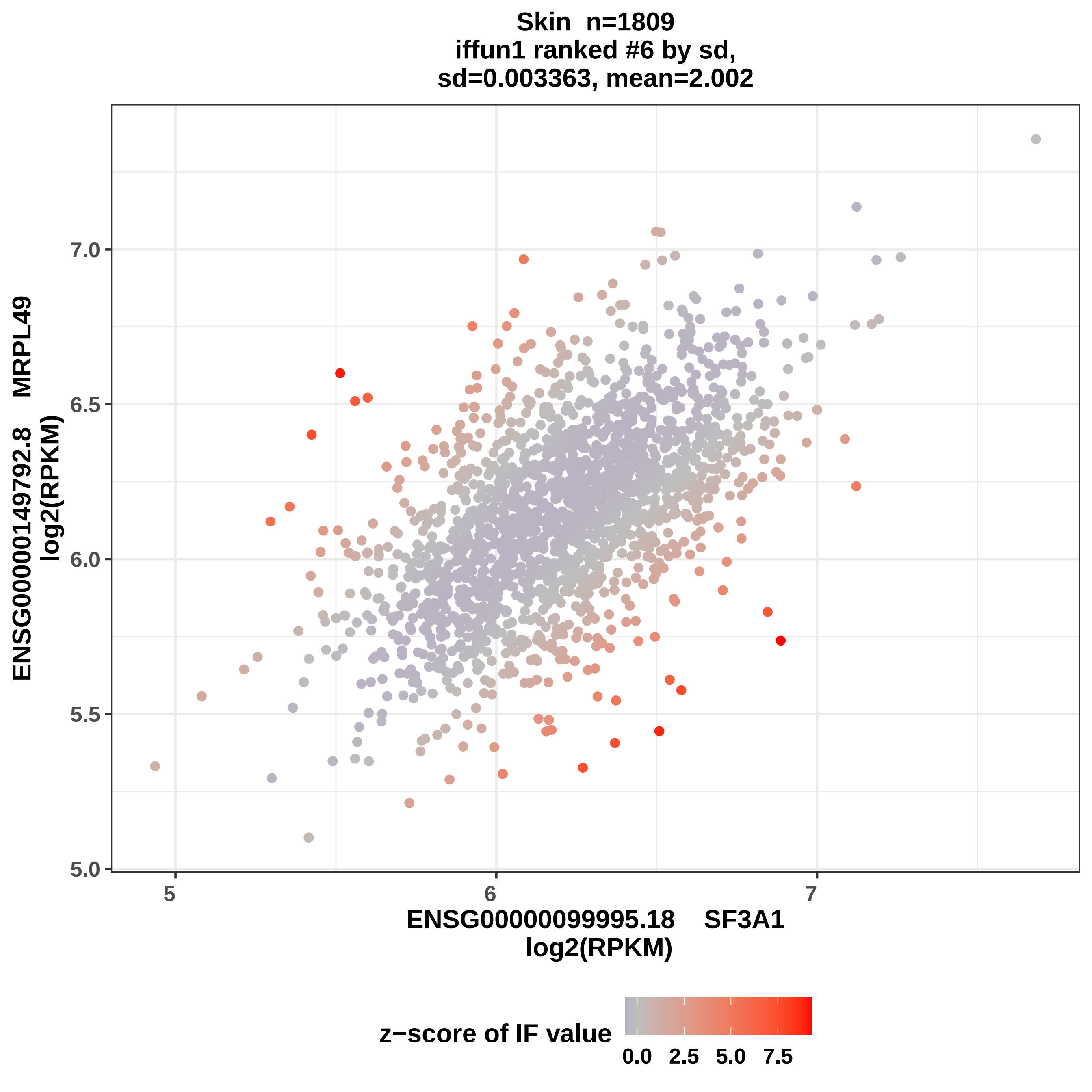
Skin n=1809
iffun1 ranked #6 by sd,
sd=0.003363, mean=2.002
ENSG00000149792.8 MRPL49
log2(RPKM)
7.0
6.5
6.0
5.5
5.0
5
6
7
ENSG00000099995.18 SF3A1
log2(RPKM)
z−score of IF value
0.0
2.5
5.0
7.5

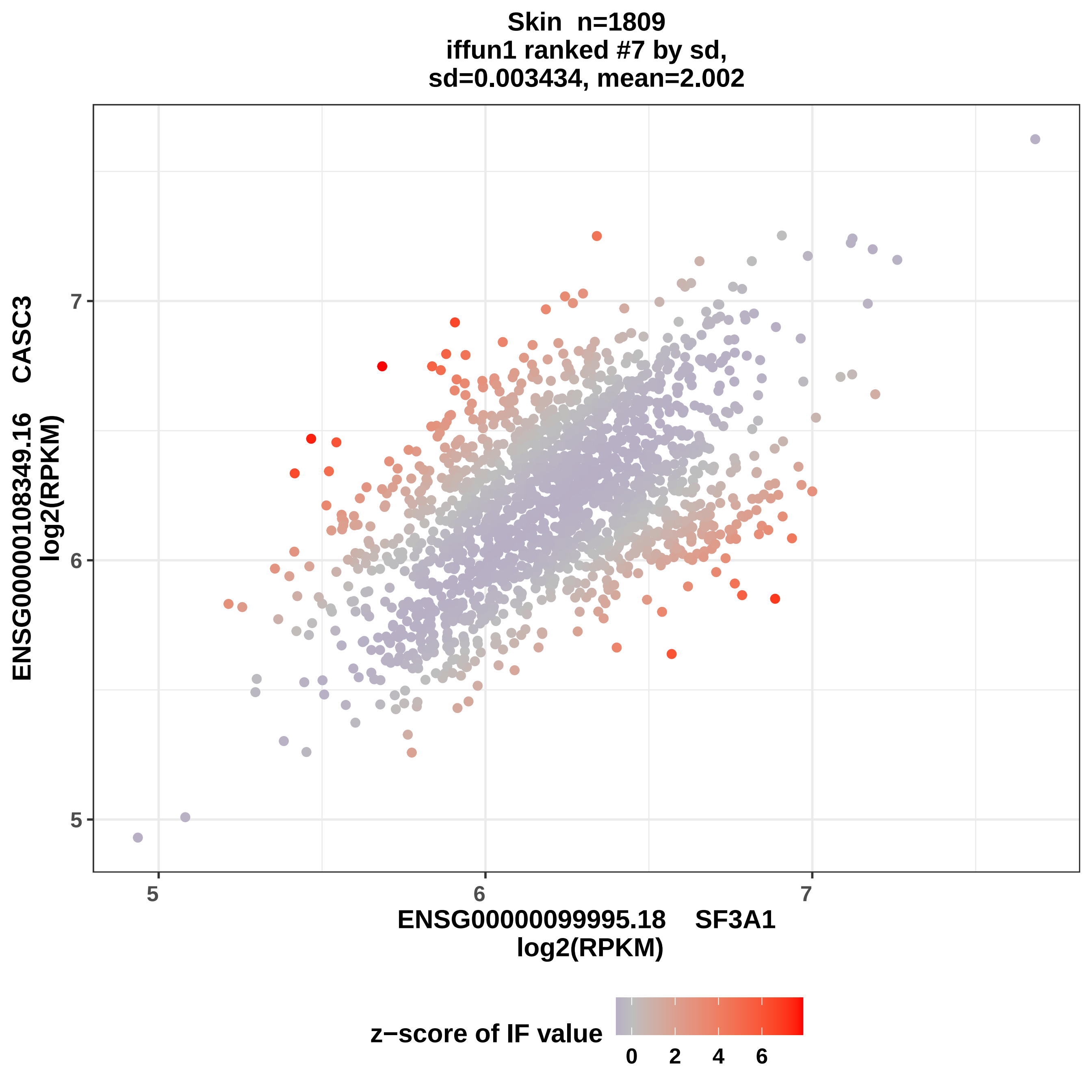
Skin n=1809
iffun1 ranked #7 by sd,
sd=0.003434, mean=2.002
ENSG00000108349.16 CASC3
log2(RPKM)
ENSG00000099995.18 SF3A1
log2(RPKM)
5
6
7
z−score of IF value
0
2
4
6

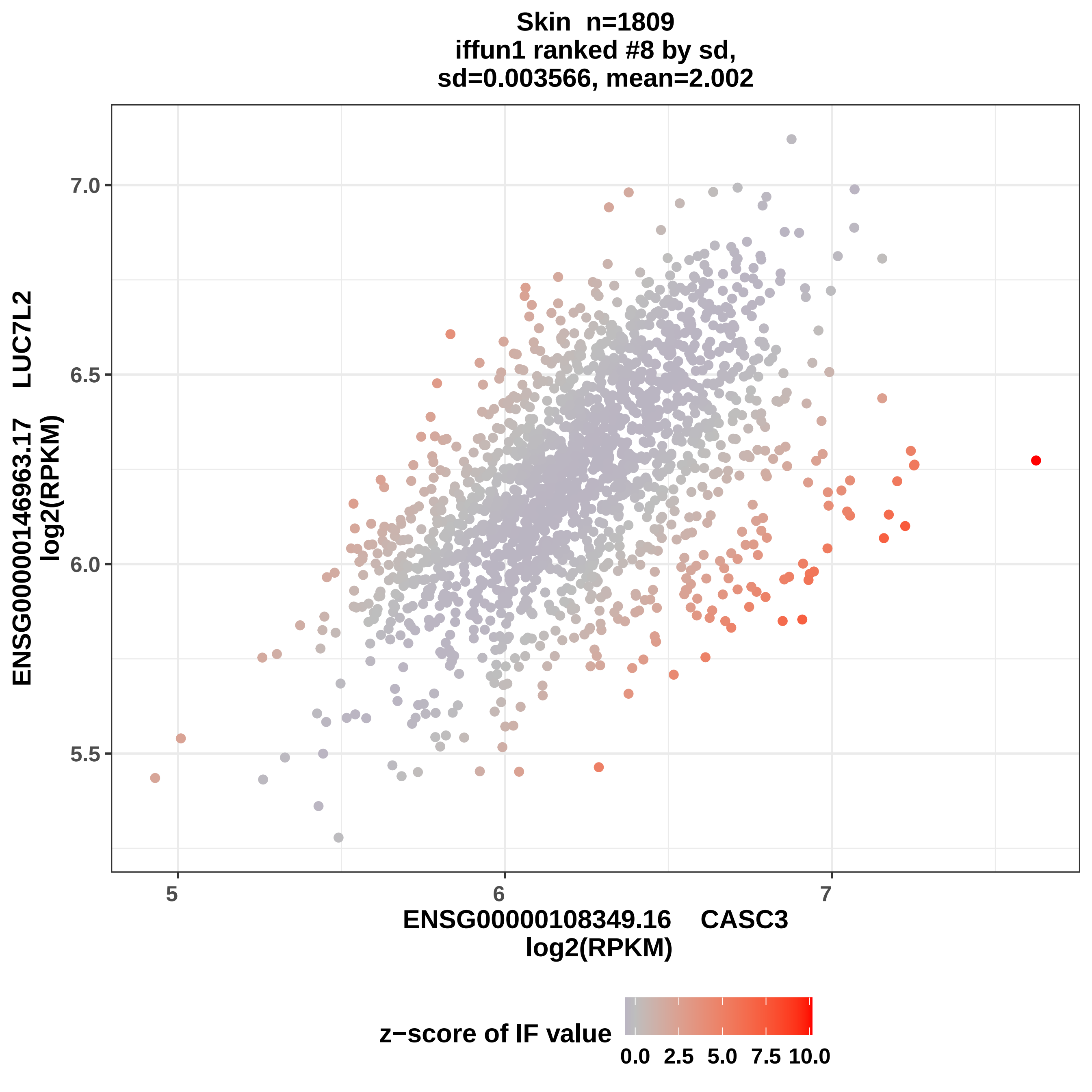

Skin n=1809
iffun1 ranked #8 by sd,
sd=0.003566, mean=2.002
7.0
6.5
6.0
5.5
5
6
7
ENSG00000146963.17 LUC7L2
log2(RPKM)
ENSG00000108349.16 CASC3
log2(RPKM)
z−score of IF value
0.0 2.5 5.0 7.5 10.0

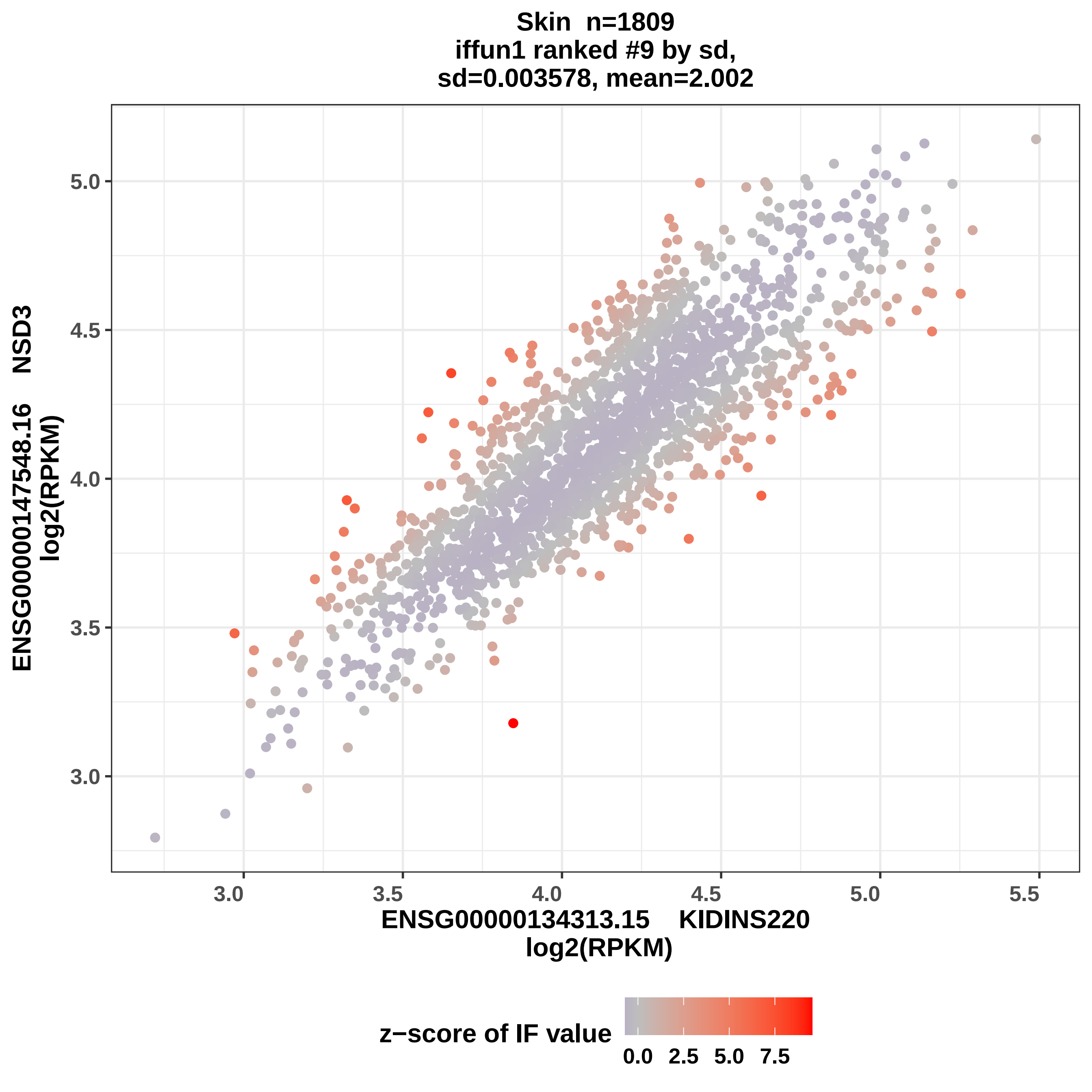

Skin n=1809
iffun1 ranked #9 by sd,
sd=0.003578, mean=2.002
ENSG00000147548.16 NSD3
log2(RPKM)
5.0
4.5
4.0
3.5
3.0
3.0
3.5
4.0
4.5
5.0
5.5
ENSG00000134313.15 KIDINS220
log2(RPKM)
z−score of IF value
0.0
2.5
5.0
7.5

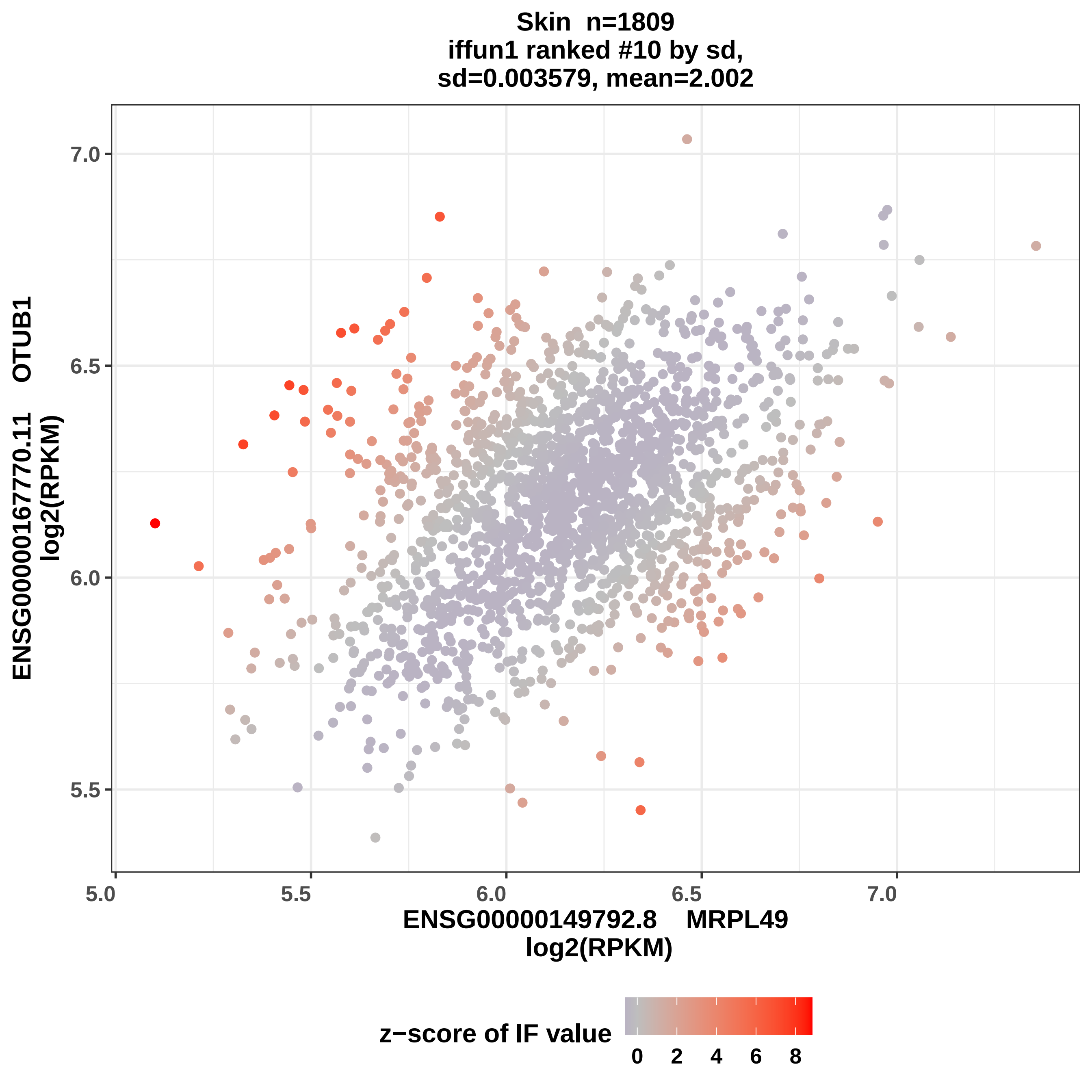

Skin n=1809
iffun1 ranked #10 by sd,
sd=0.003579, mean=2.002
ENSG00000167770.11 OTUB1
log2(RPKM)
7.0
6.5
6.0
5.5
5.0
5.5
6.0
6.5
7.0
ENSG00000149792.8 MRPL49
log2(RPKM)
z−score of IF value
0
2
4
6
8

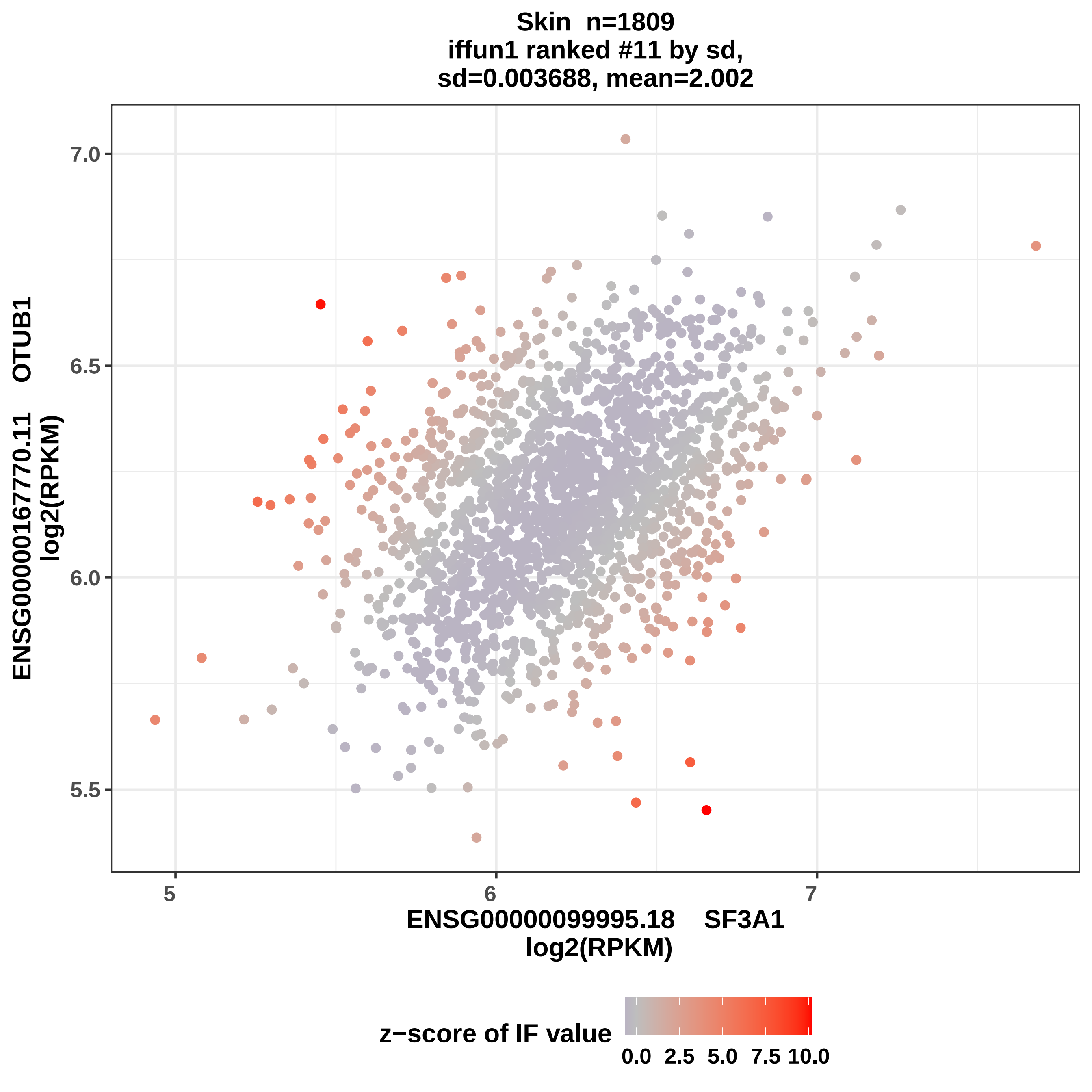

Skin n=1809
iffun1 ranked #11 by sd,
sd=0.003688, mean=2.002
7.0
6.5
6.0
5.5
5
6
7
ENSG00000167770.11 OTUB1
log2(RPKM)
ENSG00000099995.18 SF3A1
log2(RPKM)
z−score of IF value
0.0 2.5 5.0 7.5 10.0

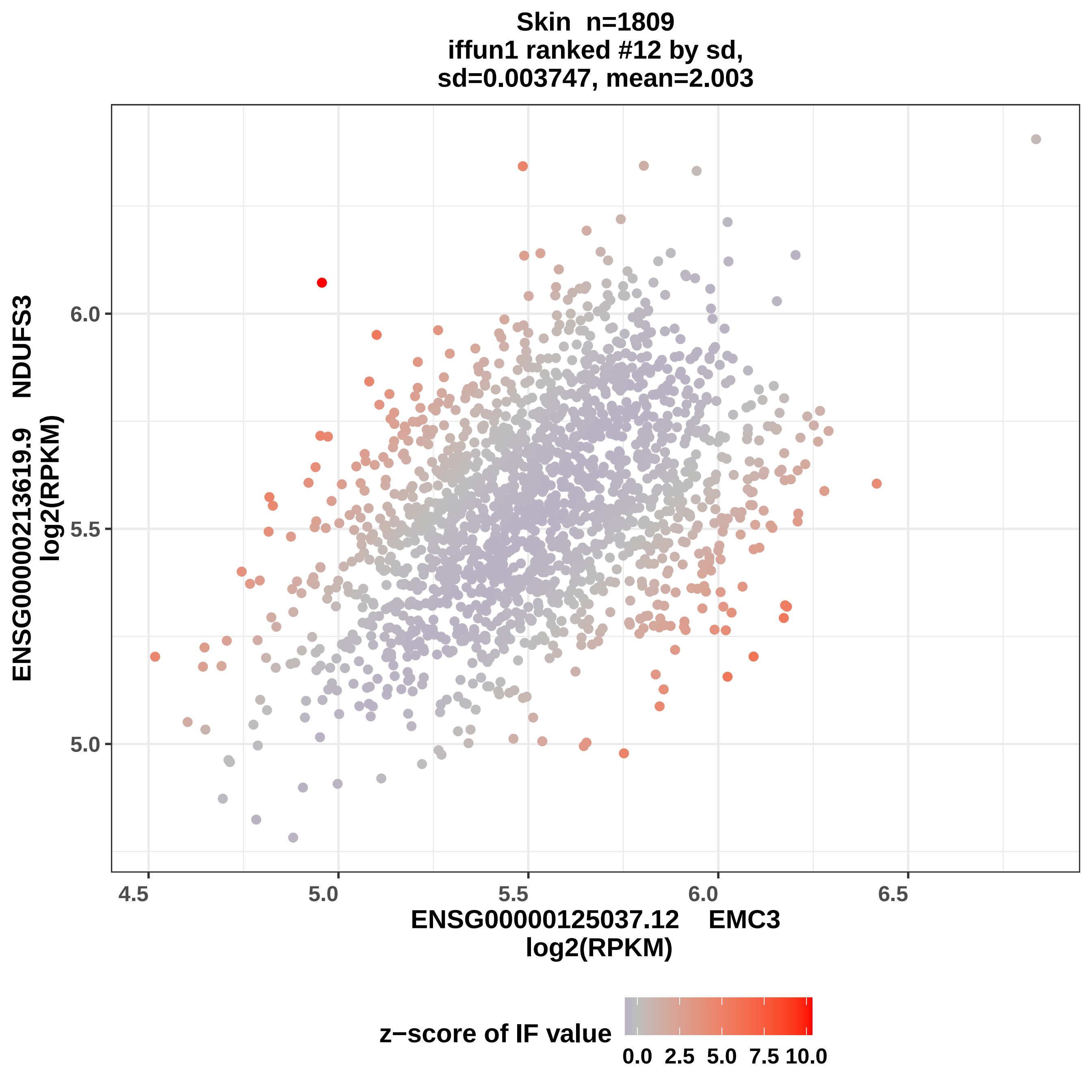
Skin n=1809
iffun1 ranked #12 by sd,
sd=0.003747, mean=2.003
ENSG00000213619.9 NDUFS3
log2(RPKM)
6.0
5.5
5.0
4.5
5.0
5.5
6.0
6.5
ENSG00000125037.12 EMC3
log2(RPKM)
z−score of IF value
0.0
2.5
5.0
7.5
10.0

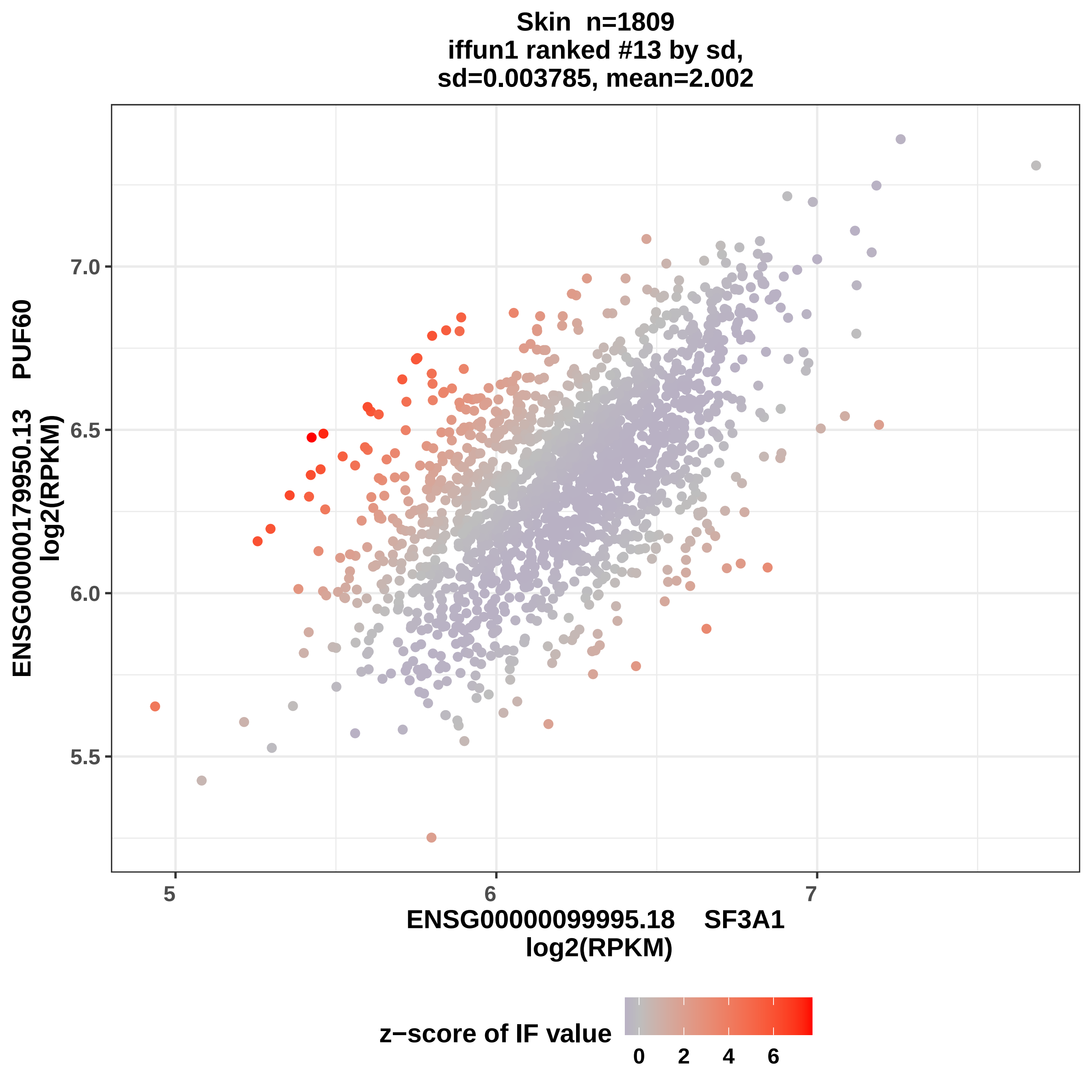

Skin n=1809
iffun1 ranked #13 by sd,
sd=0.003785, mean=2.002
ENSG00000179950.13 PUF60
log2(RPKM)
7.0
6.5
6.0
5.5
5
6
7
ENSG00000099995.18 SF3A1
log2(RPKM)
z−score of IF value
0
2
4
6

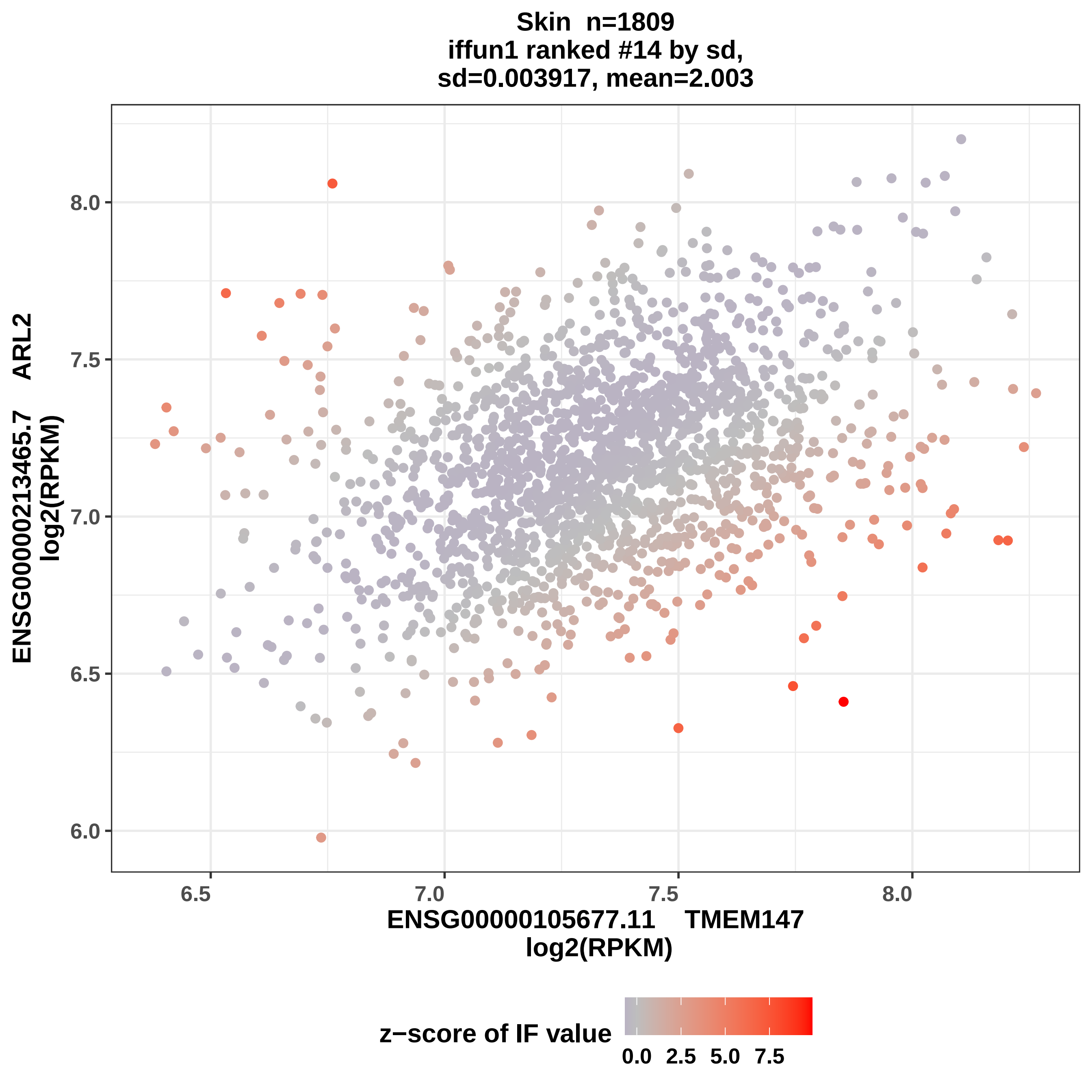

Skin n=1809
iffun1 ranked #14 by sd,
sd=0.003917, mean=2.003
8.0
7.5
7.0
6.5
6.0
6.5
7.0
7.5
8.0
ENSG00000213465.7 ARL2
log2(RPKM)
ENSG00000105677.11 TMEM147
log2(RPKM)
z−score of IF value
0.0
2.5
5.0
7.5

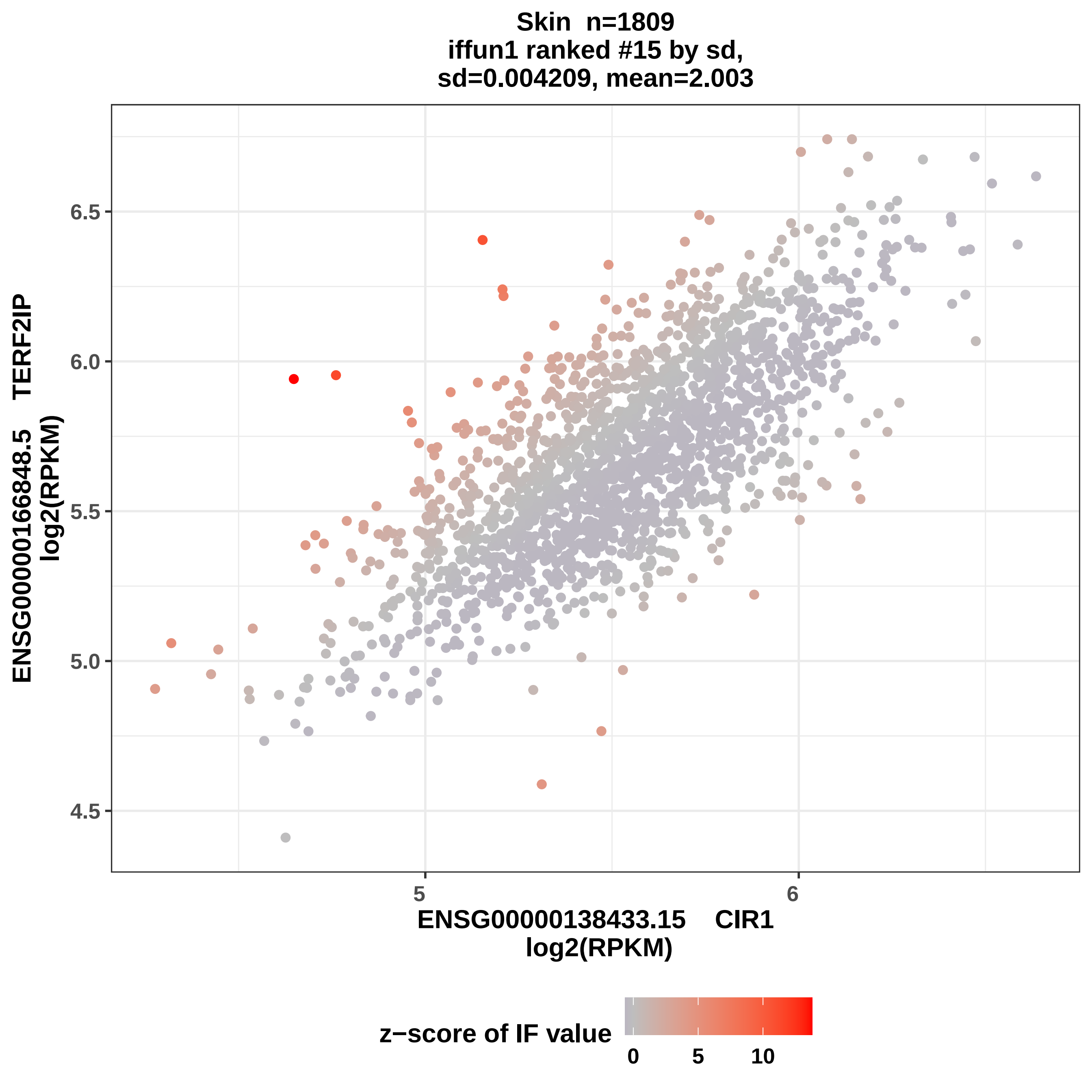
Skin n=1809
iffun1 ranked #15 by sd,
sd=0.004209, mean=2.003
ENSG00000166848.5 TERF2IP
log2(RPKM)
6.5
6.0
5.5
5.0
4.5
5
6
ENSG00000138433.15 CIR1
log2(RPKM)
z−score of IF value
0
5
10

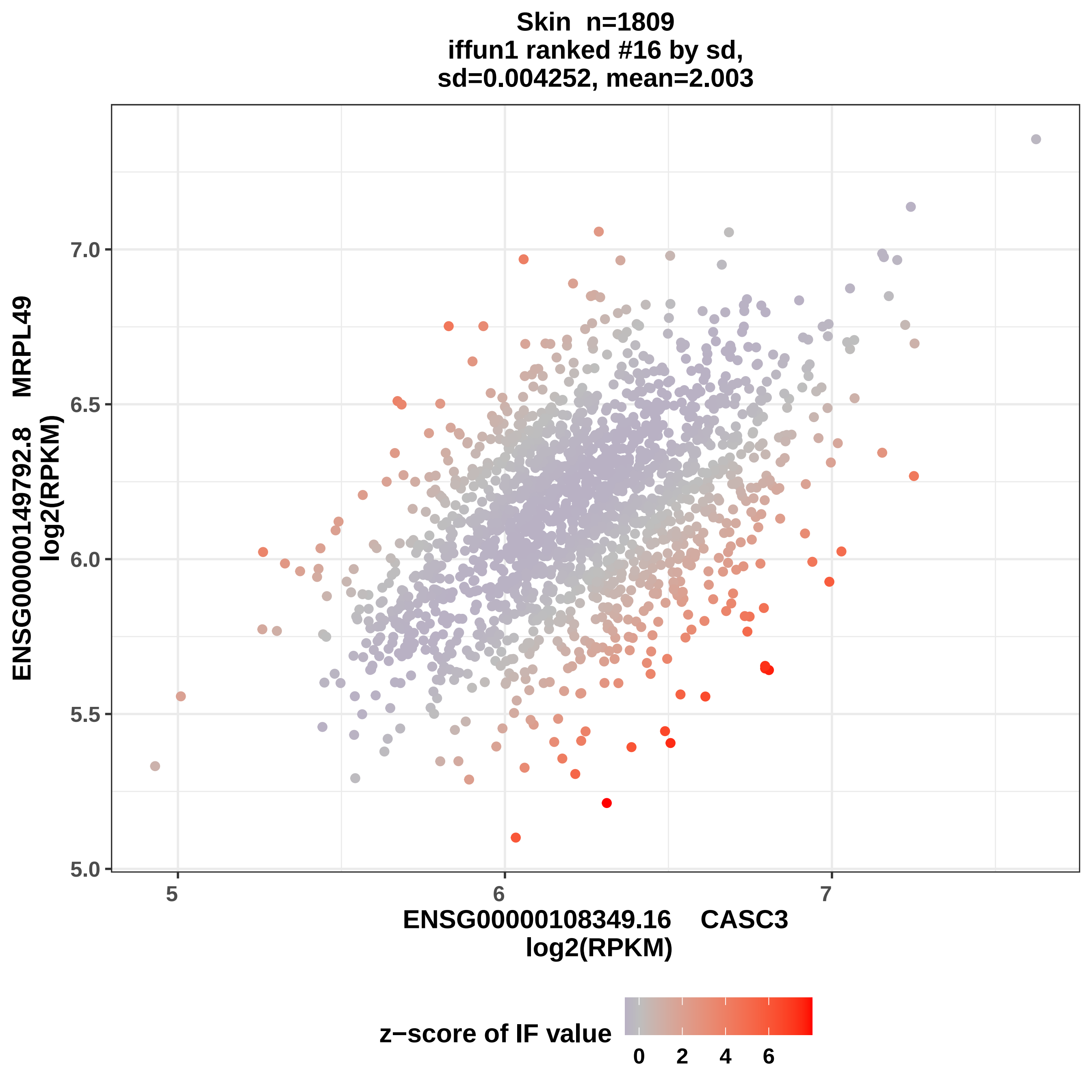
Skin n=1809
iffun1 ranked #16 by sd,
sd=0.004252, mean=2.003
ENSG00000149792.8 MRPL49
log2(RPKM)
7.0
6.5
6.0
5.5
5.0
5
6
7
ENSG00000108349.16 CASC3
log2(RPKM)
z−score of IF value
0
2
4
6

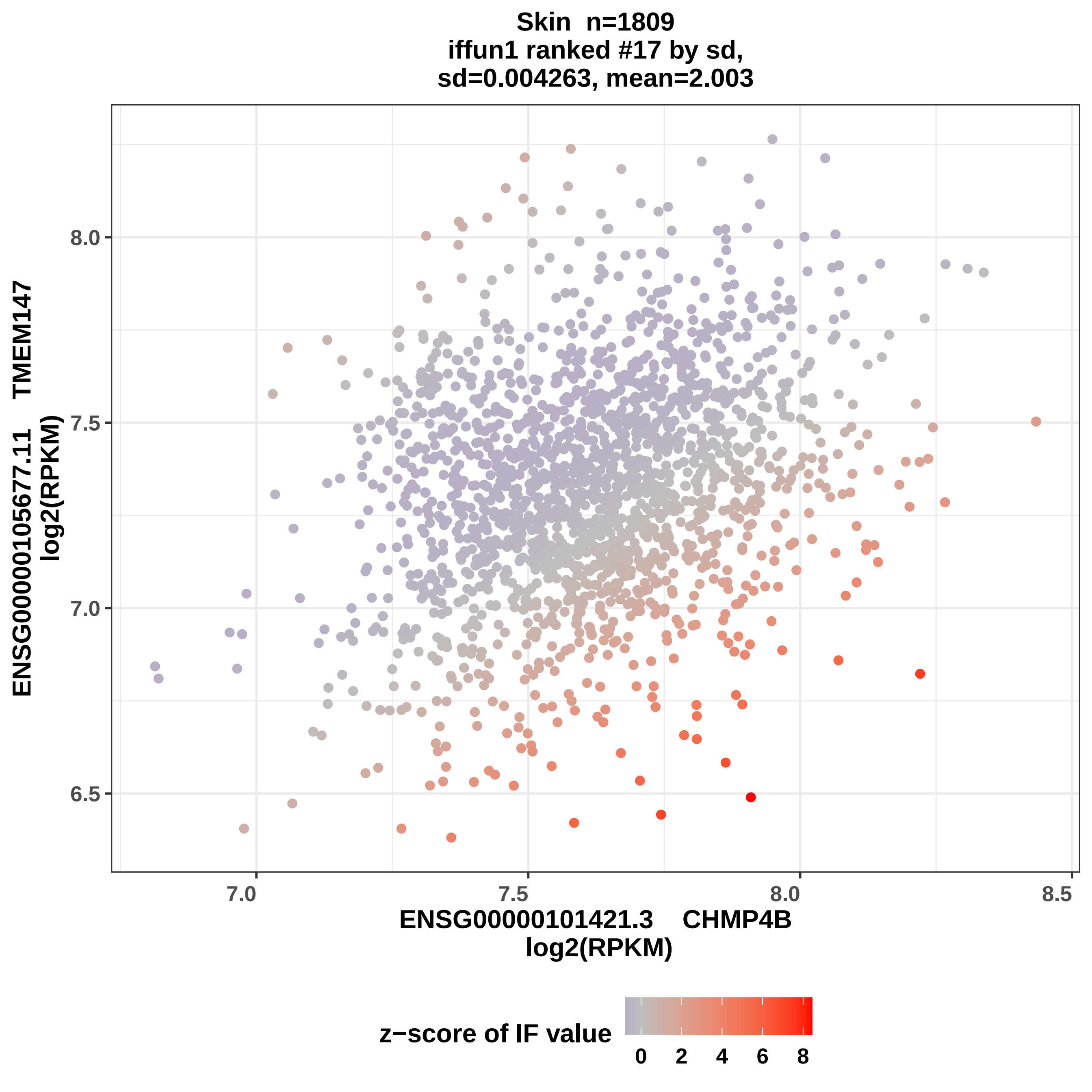

Skin n=1809
iffun1 ranked #17 by sd,
sd=0.004263, mean=2.003
ENSG00000105677.11 TMEM147
log2(RPKM)
8.0
7.5
7.0
6.5
7.0
7.5
8.0
8.5
ENSG00000101421.3 CHMP4B
log2(RPKM)
z−score of IF value
0
2
4
6
8

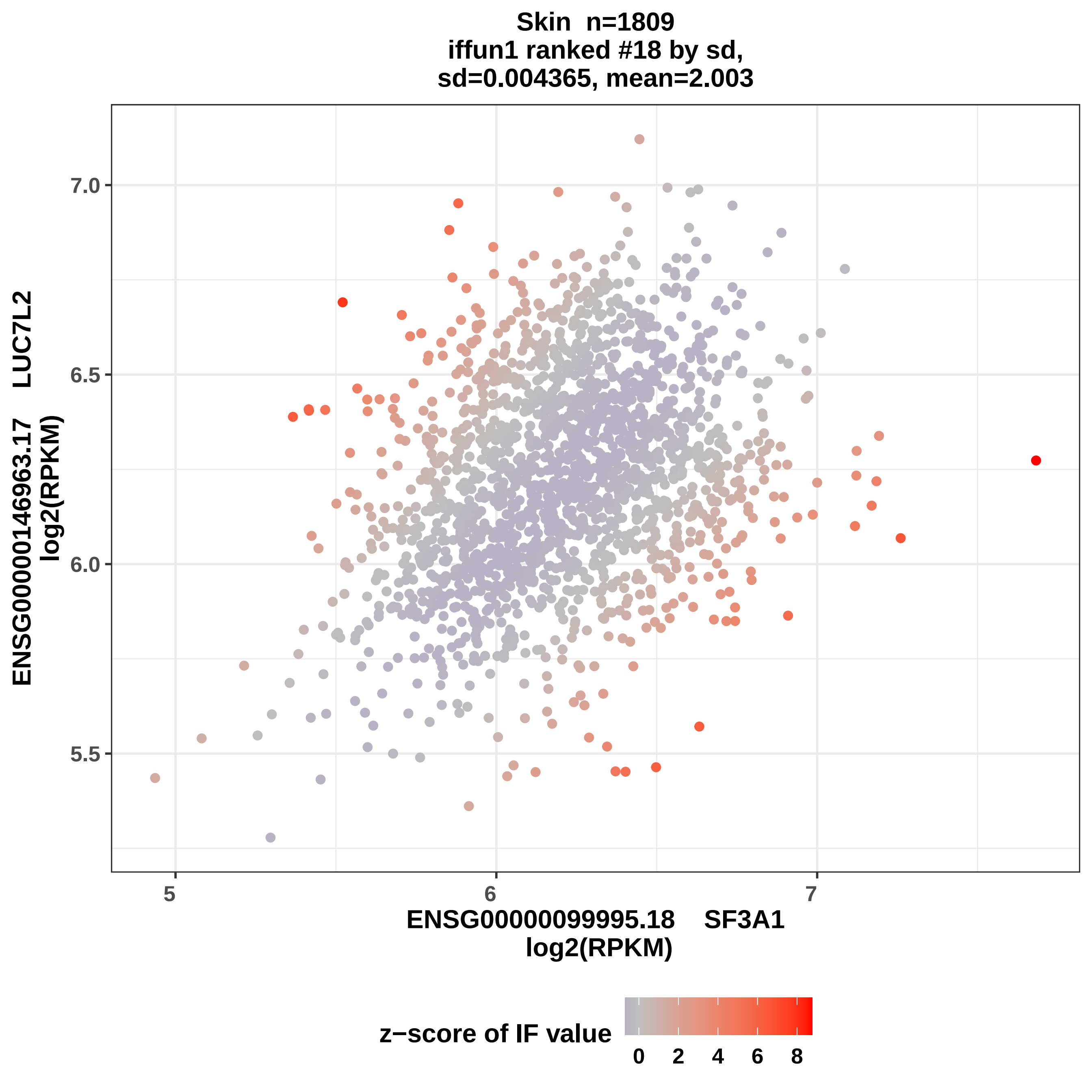
Skin n=1809
iffun1 ranked #18 by sd,
sd=0.004365, mean=2.003
7.0
6.5
6.0
5.5
5
6
7
ENSG00000146963.17 LUC7L2
log2(RPKM)
ENSG00000099995.18 SF3A1
log2(RPKM)
z−score of IF value
0
2
4
6
8

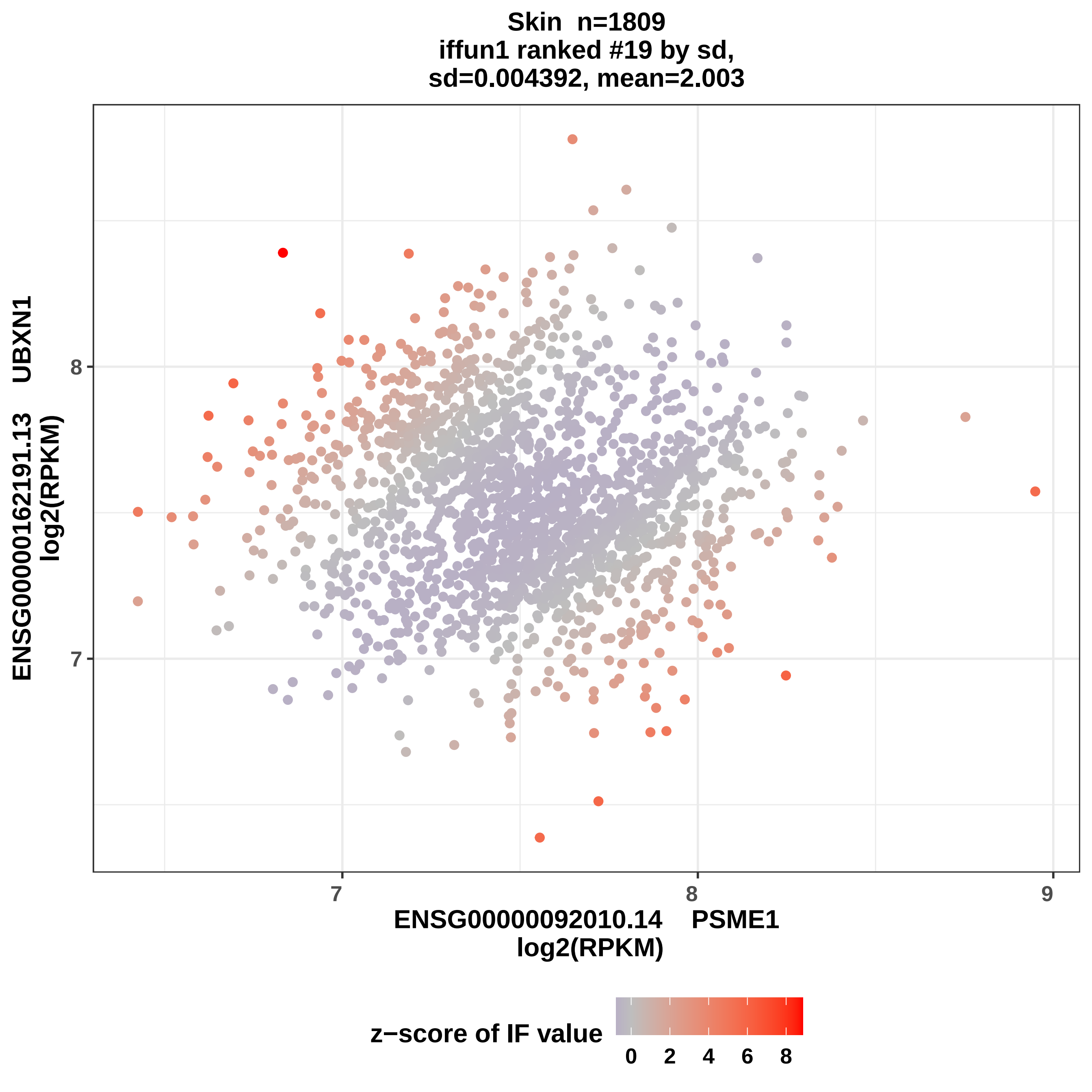

Skin n=1809
iffun1 ranked #19 by sd,
sd=0.004392, mean=2.003
ENSG00000162191.13 UBXN1
log2(RPKM)
8
7
7
8
9
ENSG00000092010.14 PSME1
log2(RPKM)
z−score of IF value
0
2
4
6
8

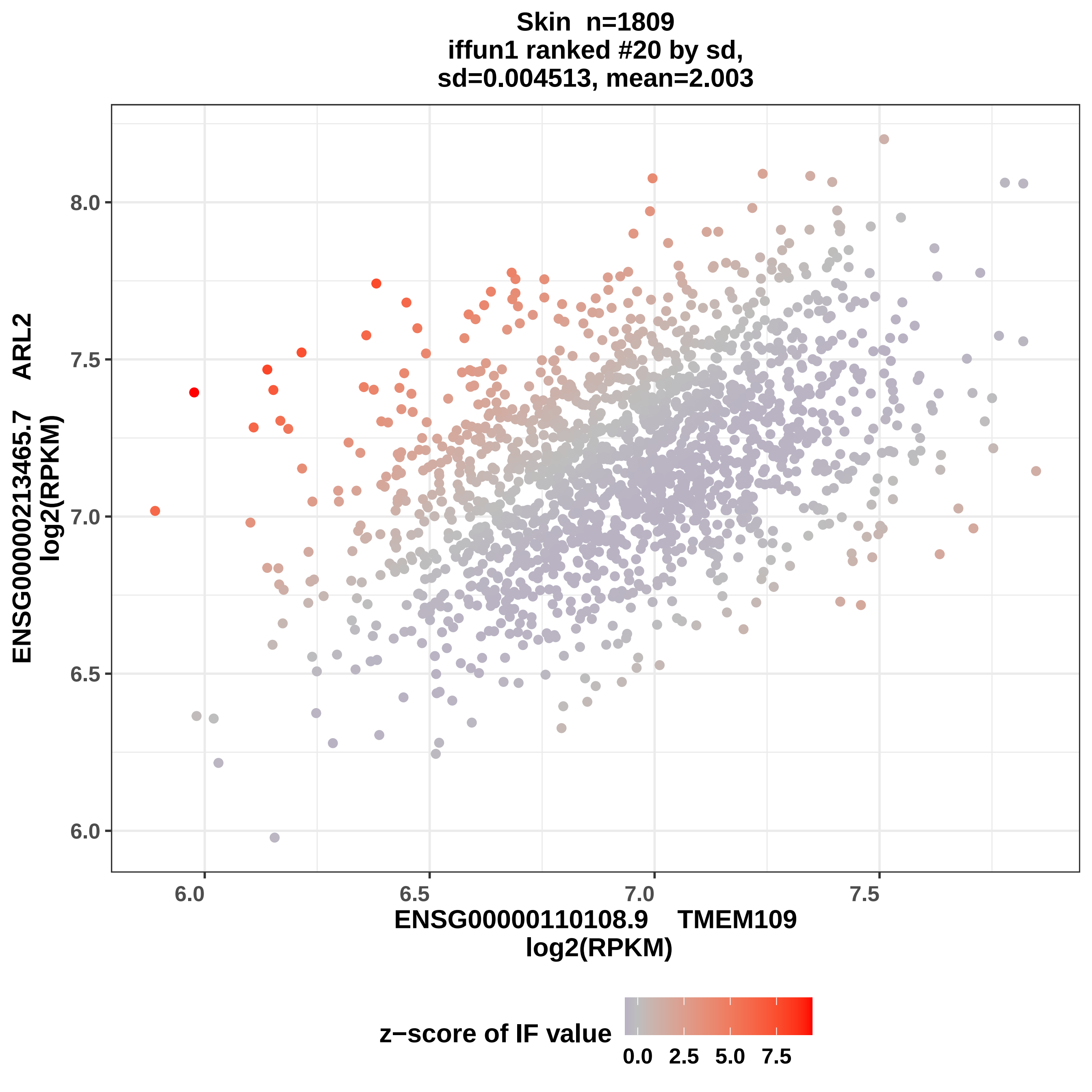
Skin n=1809
iffun1 ranked #20 by sd,
sd=0.004513, mean=2.003
ENSG00000213465.7 ARL2
log2(RPKM)
8.0
7.5
7.0
6.5
6.0
6.0
6.5
7.0
7.5
ENSG00000110108.9 TMEM109
log2(RPKM)
z−score of IF value
0.0
2.5
5.0
7.5

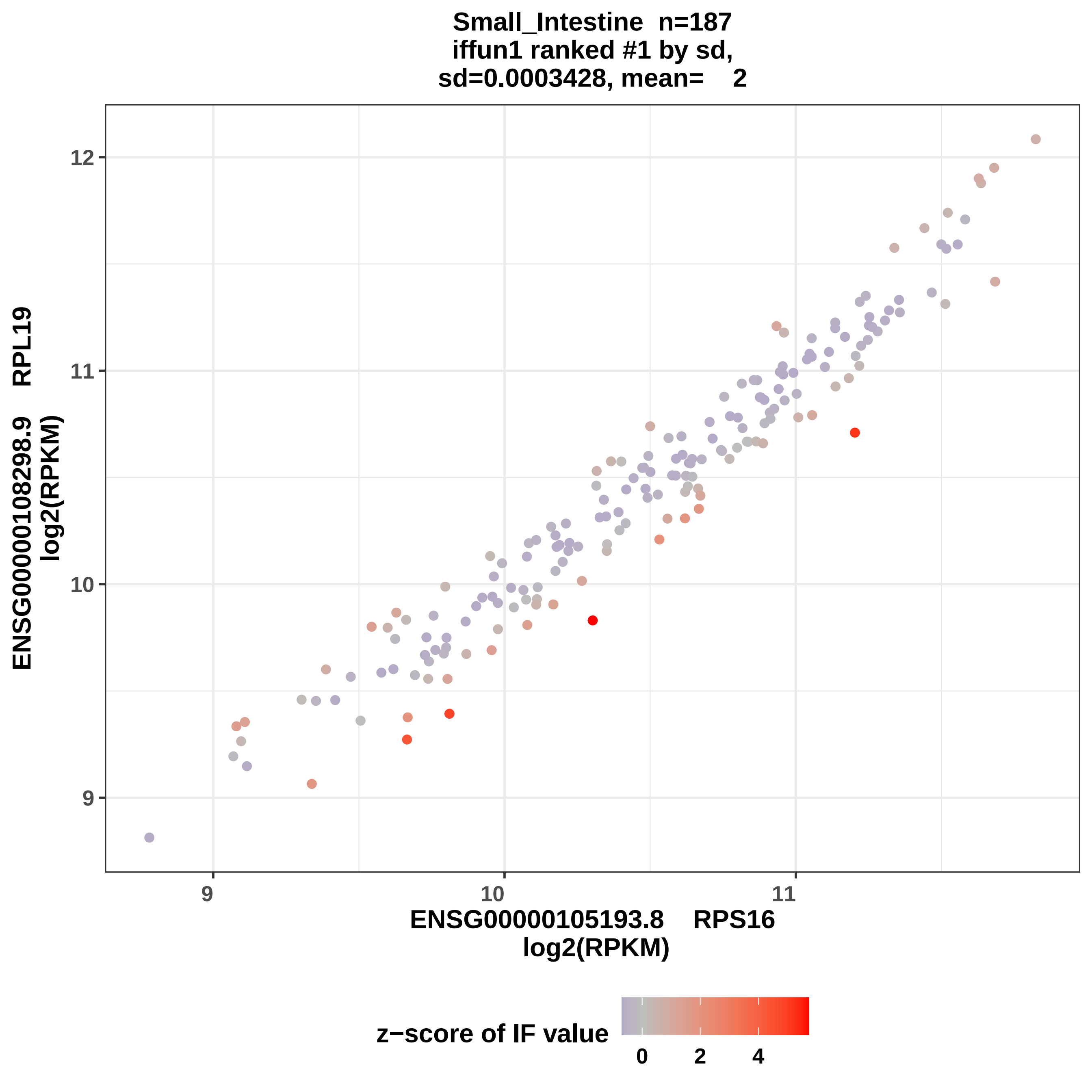
Small_Intestine n=187
iffun1 ranked #1 by sd,
sd=0.0003428, mean= 2
ENSG00000108298.9 RPL19
log2(RPKM)
12
11
10
9
9
10
11
ENSG00000105193.8 RPS16
log2(RPKM)
z−score of IF value
0
2
4

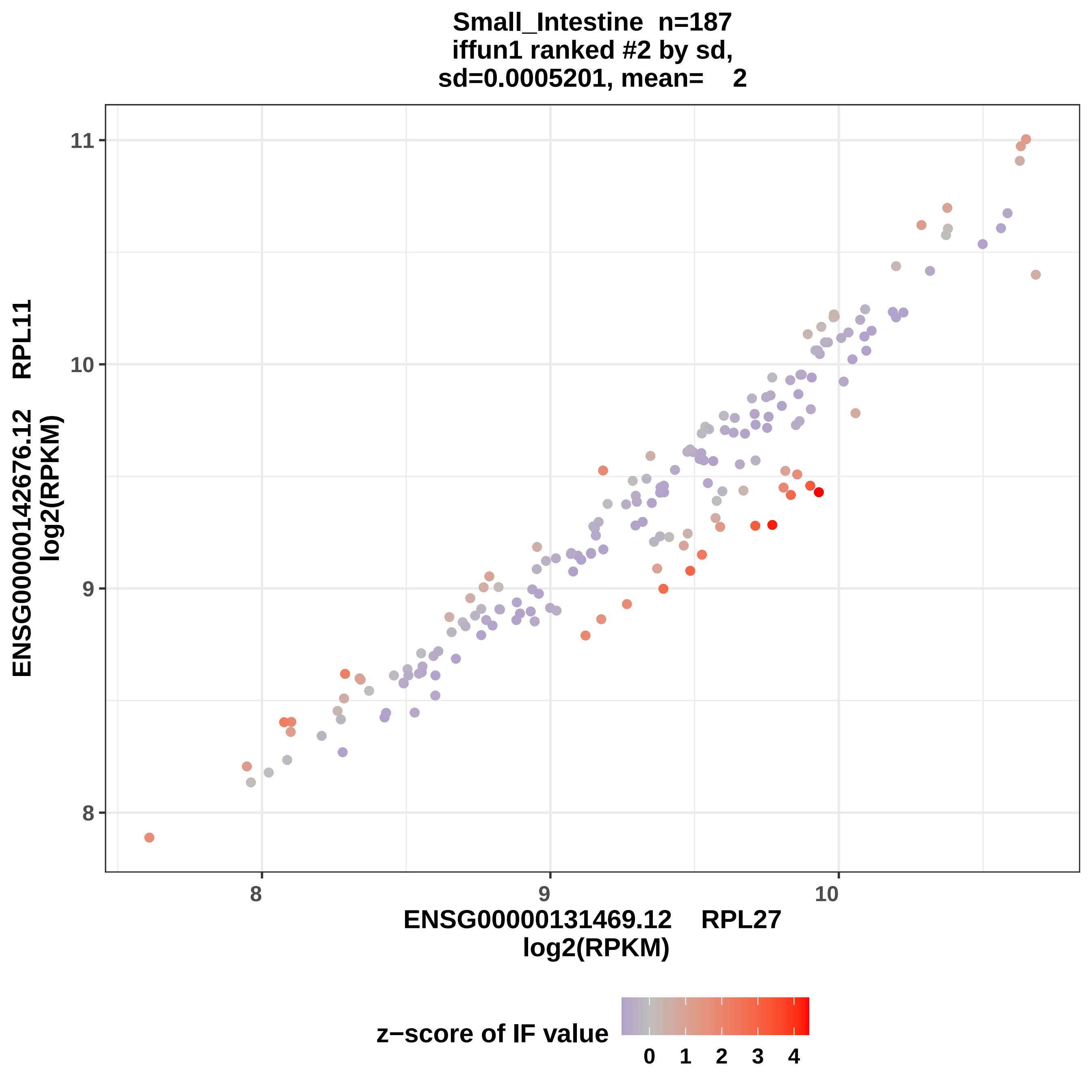
Small_Intestine n=187
iffun1 ranked #2 by sd,
sd=0.0005201, mean= 2
ENSG00000142676.12 RPL11
log2(RPKM)
ENSG00000131469.12 RPL27
log2(RPKM)
8
9
10
11
z−score of IF value
0 1 2 3 4

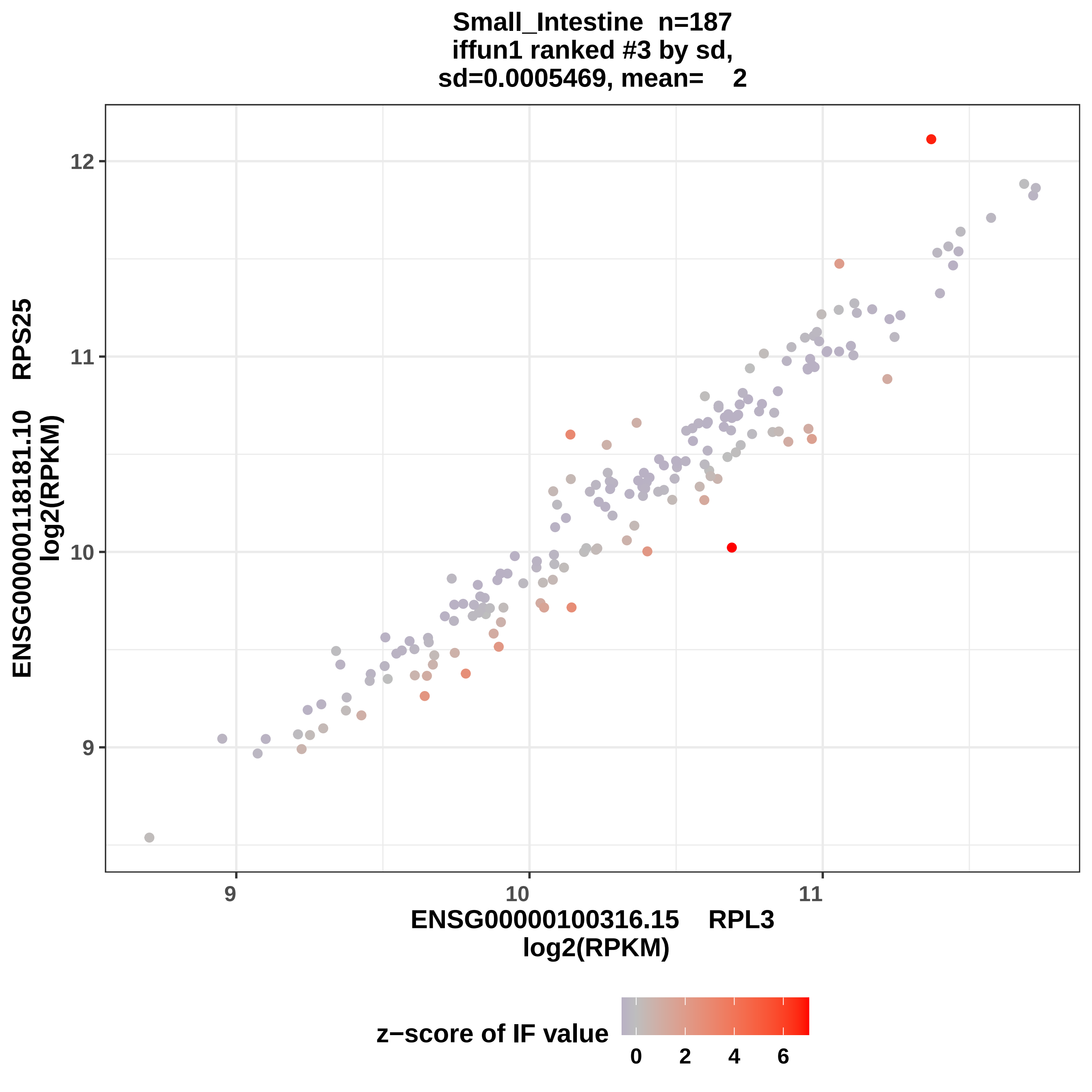
Small_Intestine n=187
iffun1 ranked #3 by sd,
sd=0.0005469, mean= 2
ENSG00000118181.10 RPS25
log2(RPKM)
12
11
10
9
9
10
11
ENSG00000100316.15 RPL3
log2(RPKM)
z−score of IF value
0
2
4
6

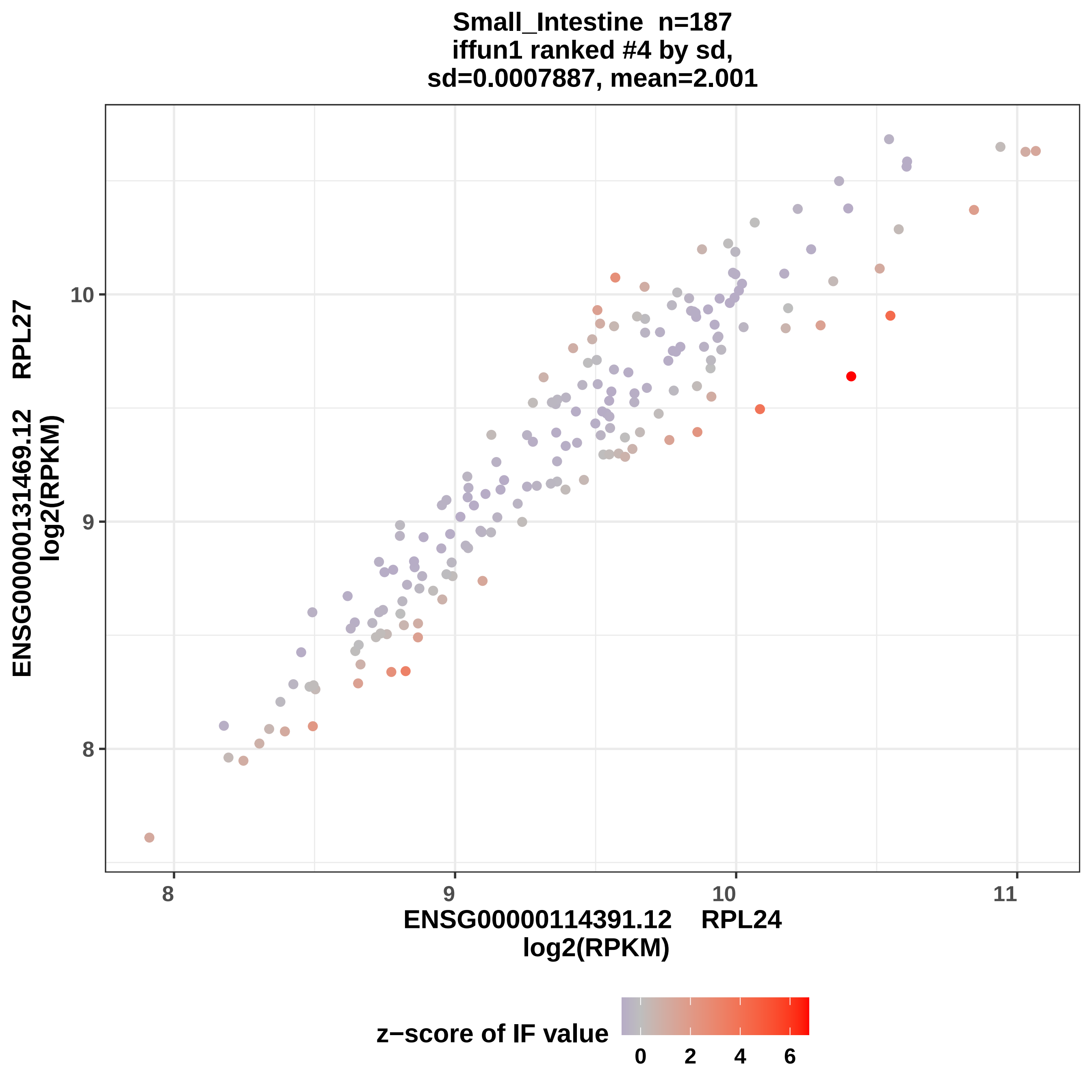
Small_Intestine n=187
iffun1 ranked #4 by sd,
sd=0.0007887, mean=2.001
ENSG00000131469.12 RPL27
log2(RPKM)
ENSG00000114391.12 RPL24
log2(RPKM)
8
9
10
11
z−score of IF value
0
2
4
6

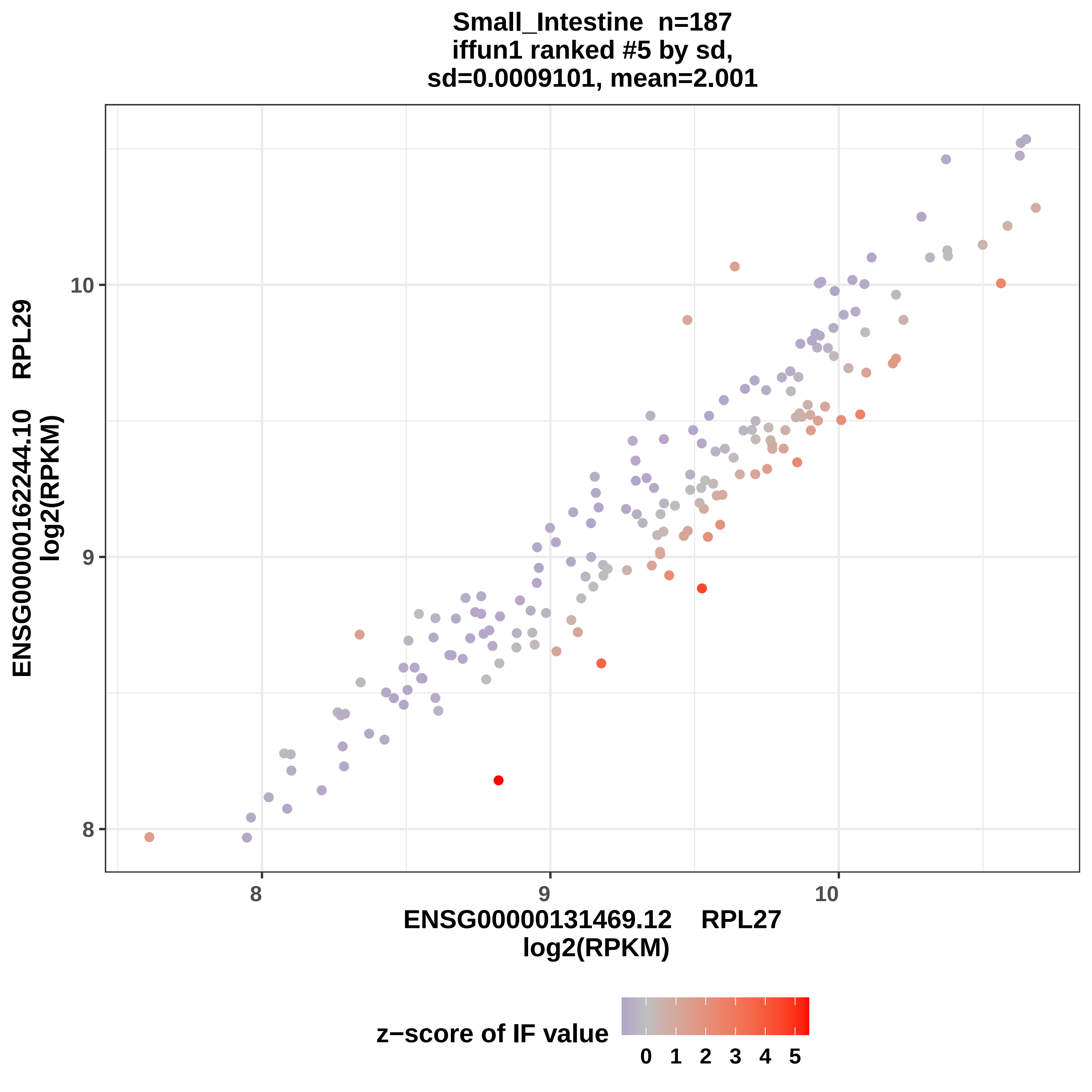

Small_Intestine n=187
iffun1 ranked #5 by sd,
sd=0.0009101, mean=2.001
ENSG00000162244.10 RPL29
log2(RPKM)
10
9
8
8
9
10
ENSG00000131469.12 RPL27
log2(RPKM)
z−score of IF value
0 1 2 3 4 5

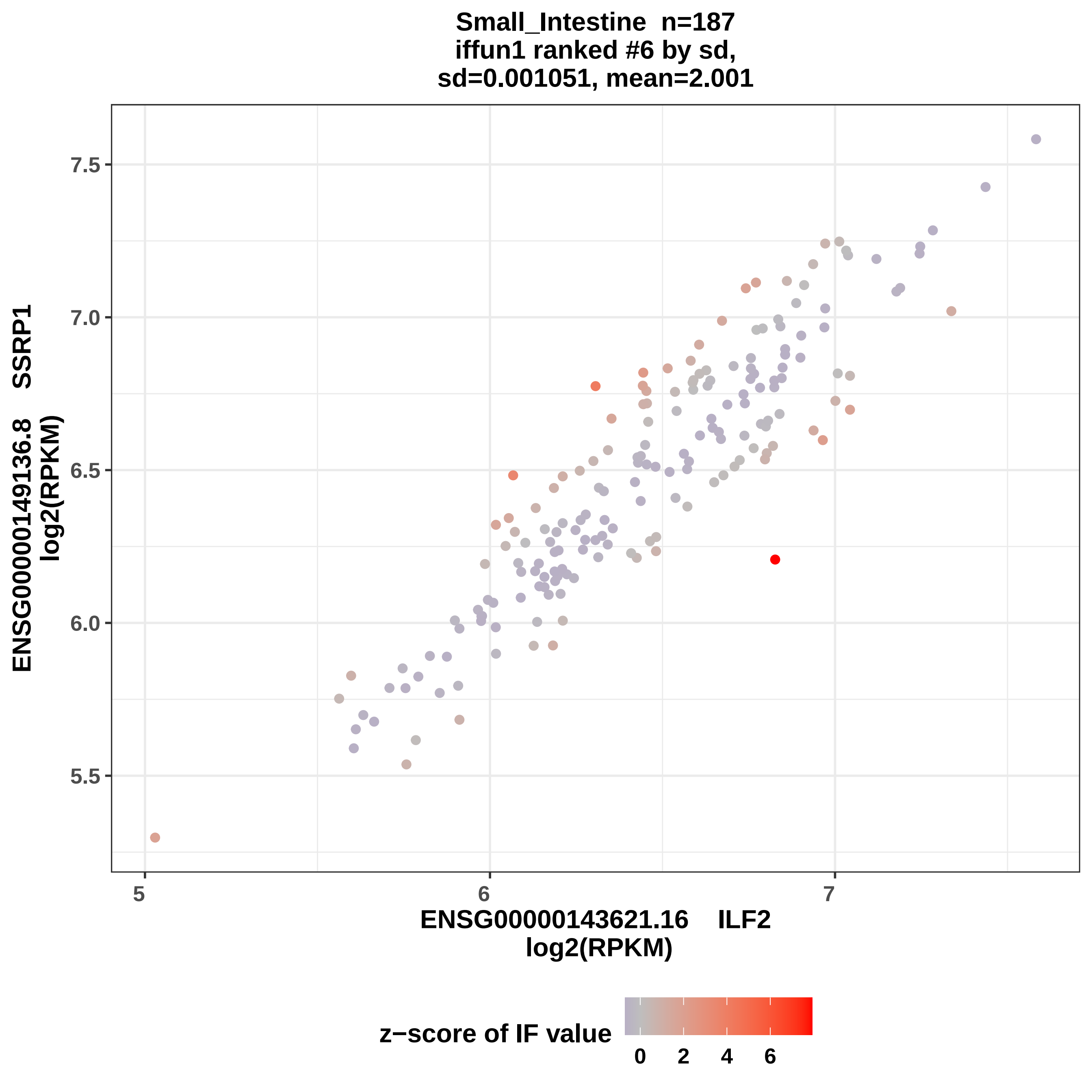

Small_Intestine n=187
iffun1 ranked #6 by sd,
sd=0.001051, mean=2.001
7.5
7.0
6.5
6.0
5.5
5
6
7
ENSG00000149136.8 SSRP1
log2(RPKM)
ENSG00000143621.16 ILF2
log2(RPKM)
z−score of IF value
0
2
4
6

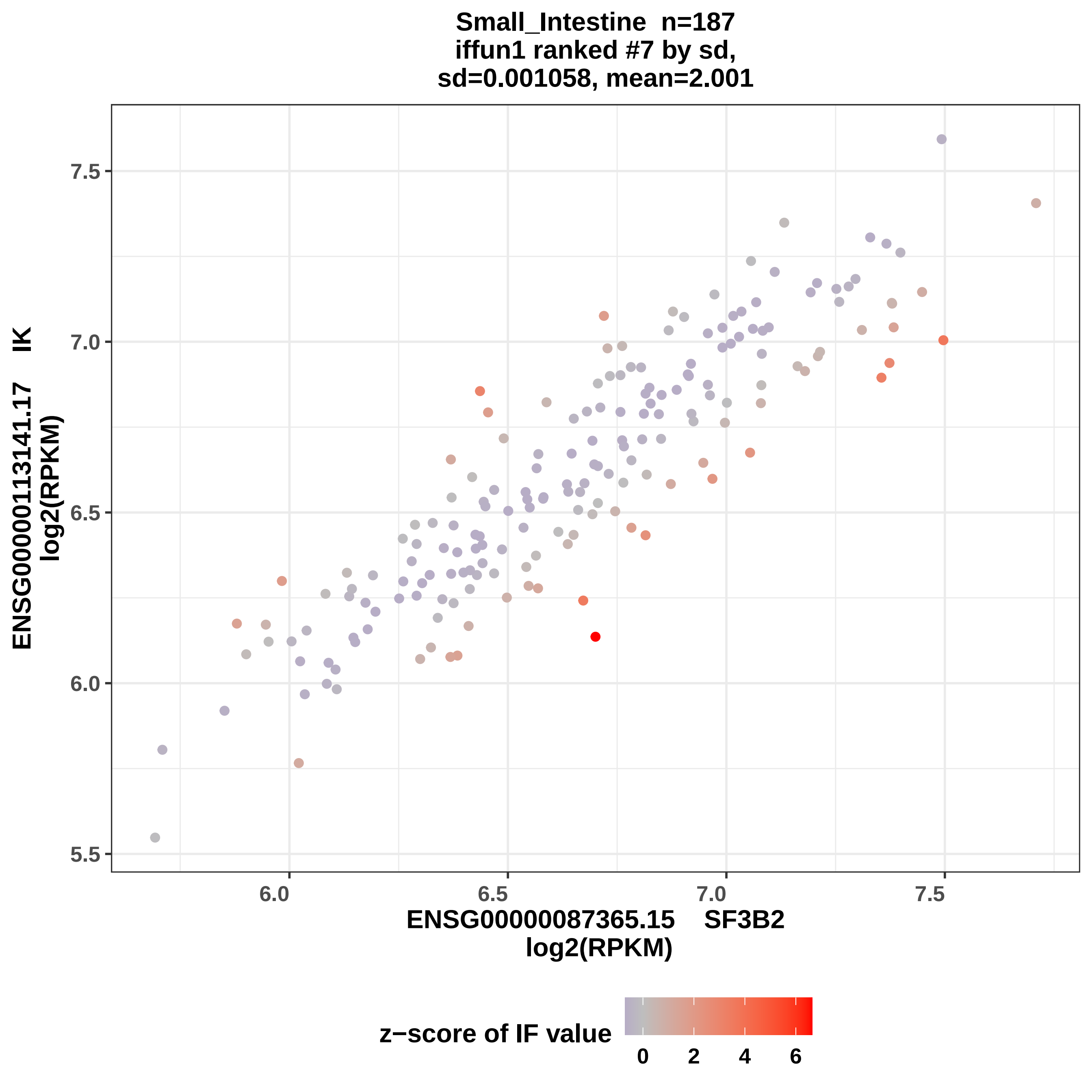

Small_Intestine n=187
iffun1 ranked #7 by sd,
sd=0.001058, mean=2.001
ENSG00000113141.17 IK
log2(RPKM)
7.5
7.0
6.5
6.0
5.5
6.0
6.5
7.0
7.5
ENSG00000087365.15 SF3B2
log2(RPKM)
z−score of IF value
0
2
4
6

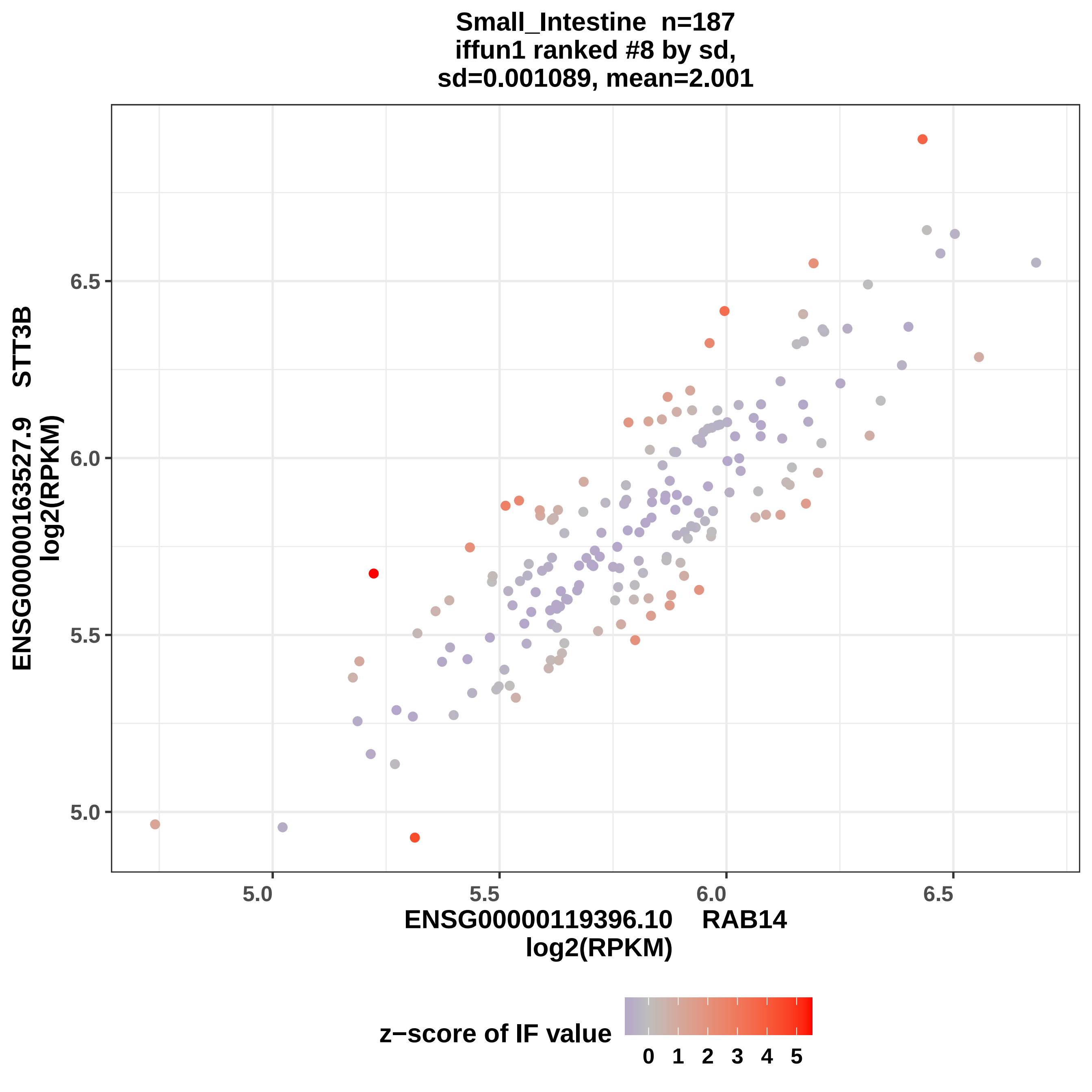

Small_Intestine n=187
iffun1 ranked #8 by sd,
sd=0.001089, mean=2.001
ENSG00000163527.9 STT3B
log2(RPKM)
6.5
6.0
5.5
5.0
5.0
5.5
6.0
6.5
ENSG00000119396.10 RAB14
log2(RPKM)
z−score of IF value
0 1 2 3 4 5

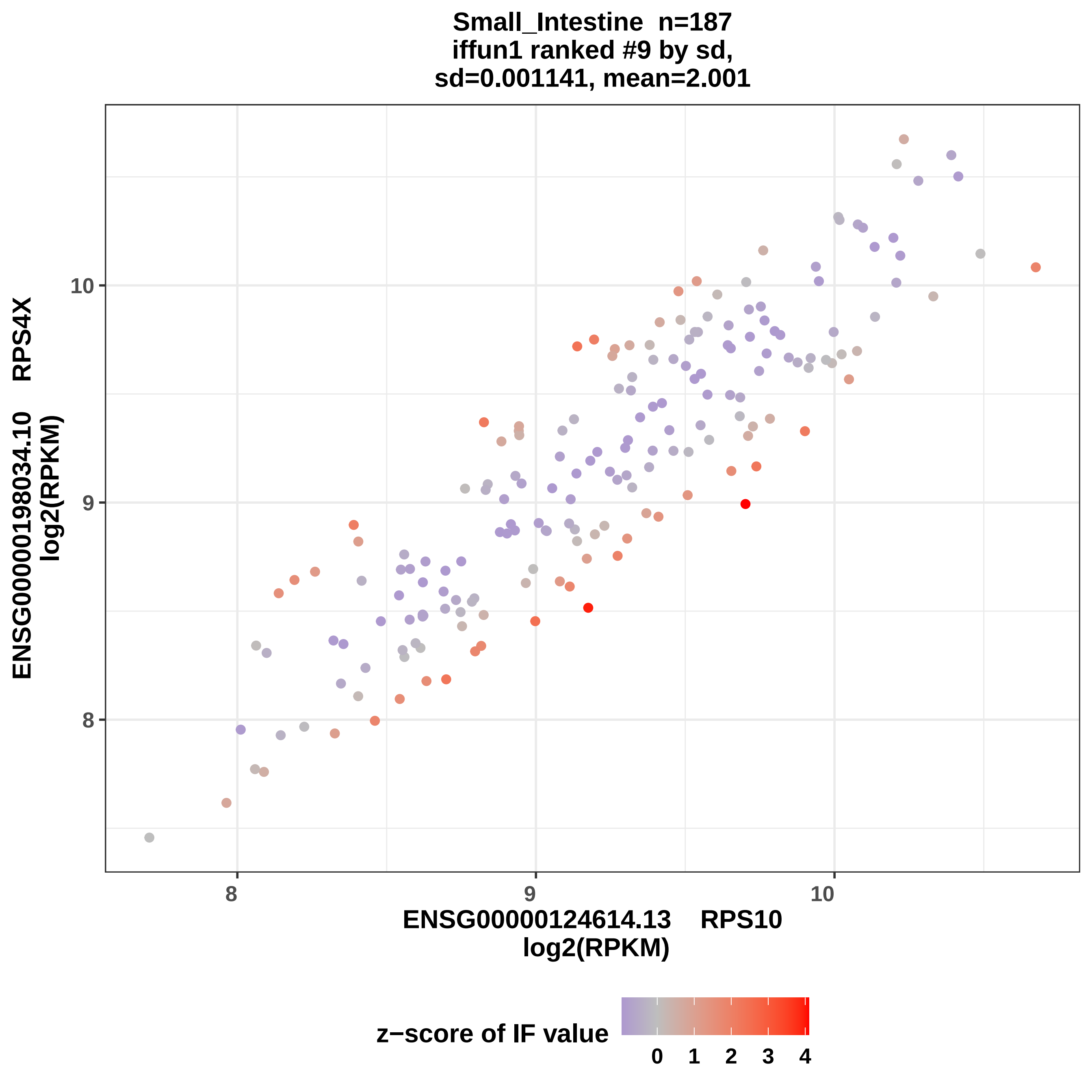

Small_Intestine n=187
iffun1 ranked #9 by sd,
sd=0.001141, mean=2.001
ENSG00000198034.10 RPS4X
log2(RPKM)
10
9
8
8
9
10
ENSG00000124614.13 RPS10
log2(RPKM)
z−score of IF value
0
1
2
3
4

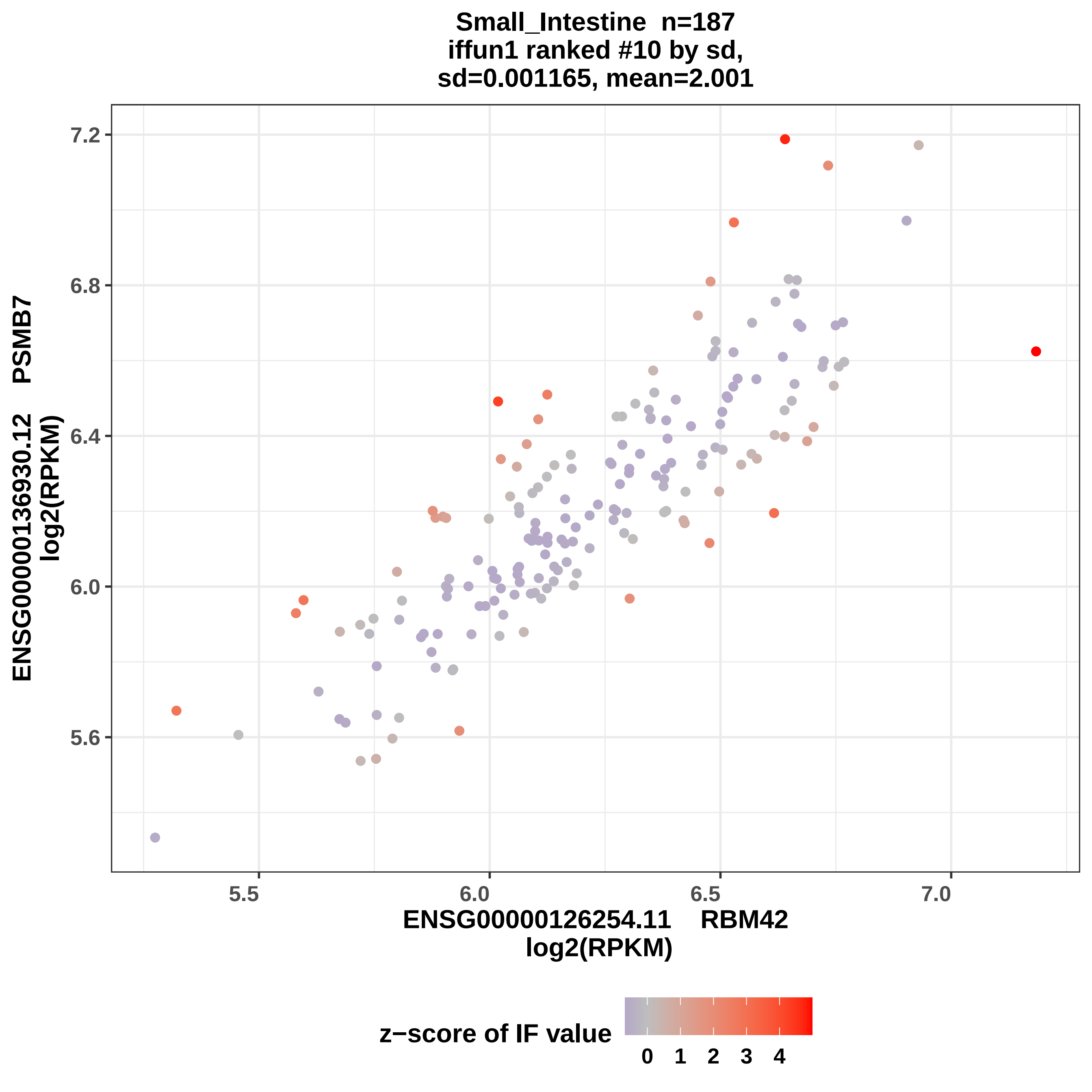
Small_Intestine n=187
iffun1 ranked #10 by sd,
sd=0.001165, mean=2.001
ENSG00000136930.12 PSMB7
log2(RPKM)
ENSG00000126254.11 RBM42
log2(RPKM)
7.2
6.8
6.4
6.0
5.6
5.5
6.0
6.5
7.0
z−score of IF value
0
1
2
3
4

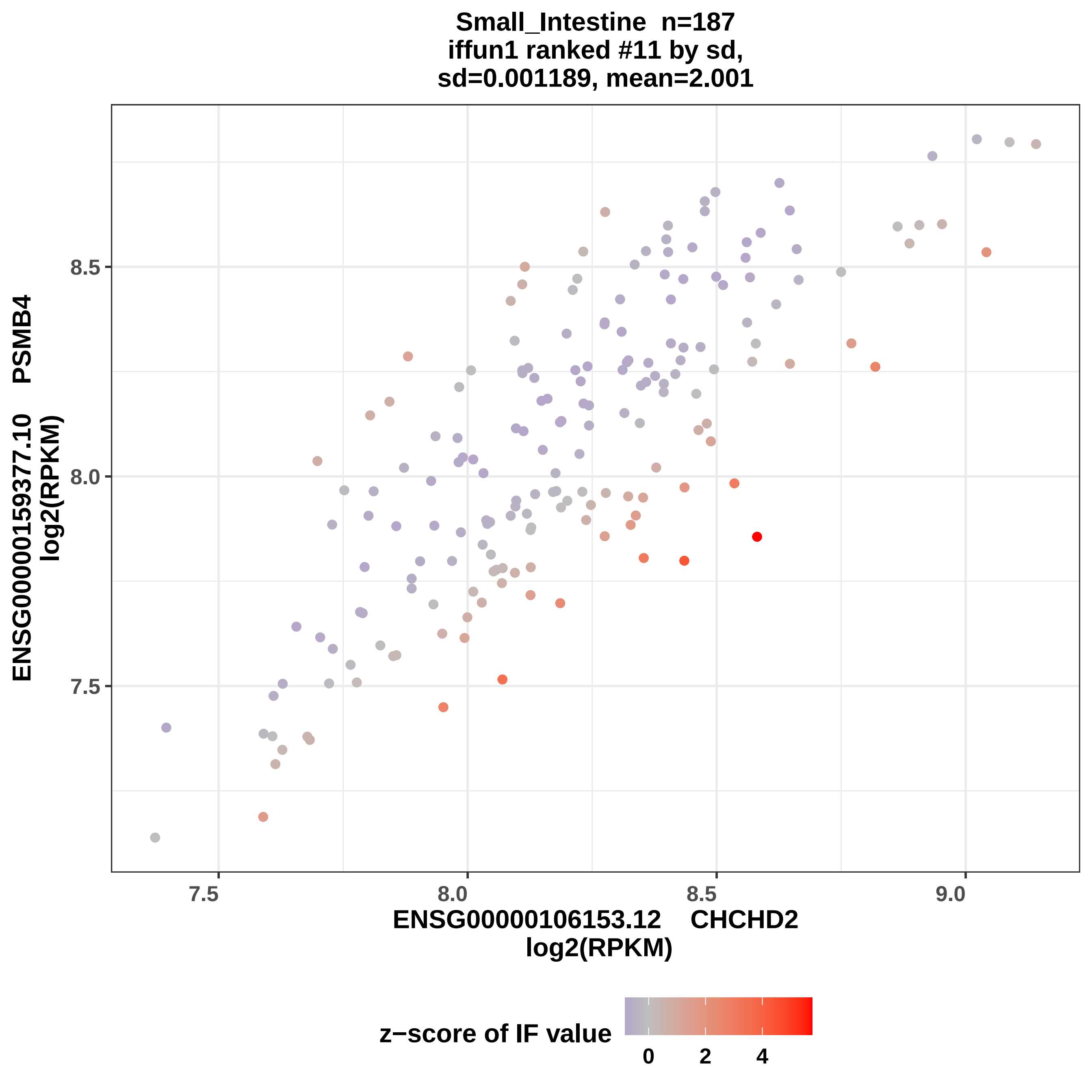

Small_Intestine n=187
iffun1 ranked #11 by sd,
sd=0.001189, mean=2.001
ENSG00000159377.10 PSMB4
log2(RPKM)
ENSG00000106153.12 CHCHD2
log2(RPKM)
z−score of IF value
7.5
8.0
8.5
9.0
0
2
4

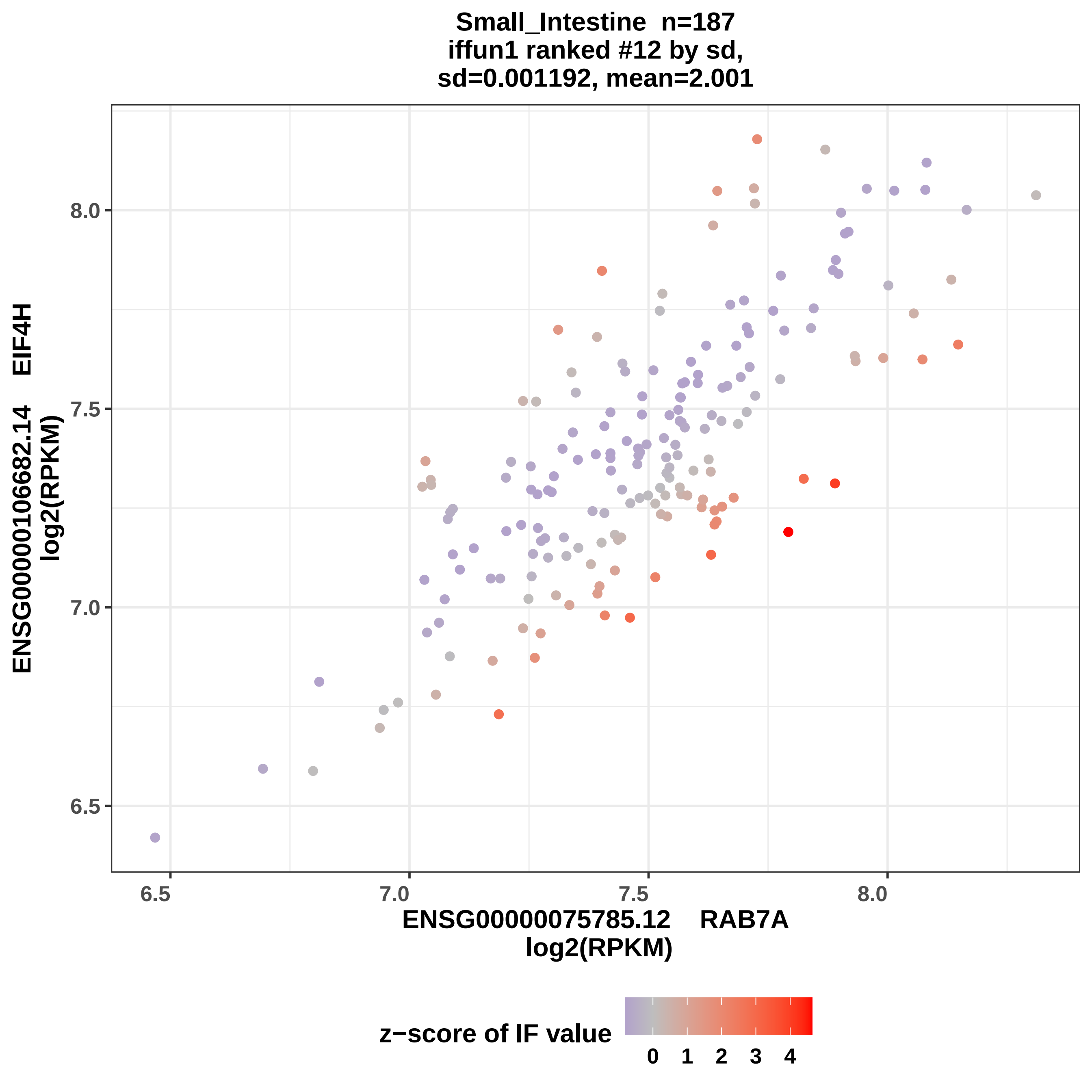
Small_Intestine n=187
iffun1 ranked #12 by sd,
sd=0.001192, mean=2.001
ENSG00000106682.14 EIF4H
log2(RPKM)
8.0
7.5
7.0
6.5
6.5
7.0
7.5
8.0
ENSG00000075785.12 RAB7A
log2(RPKM)
z−score of IF value
0 1 2 3 4

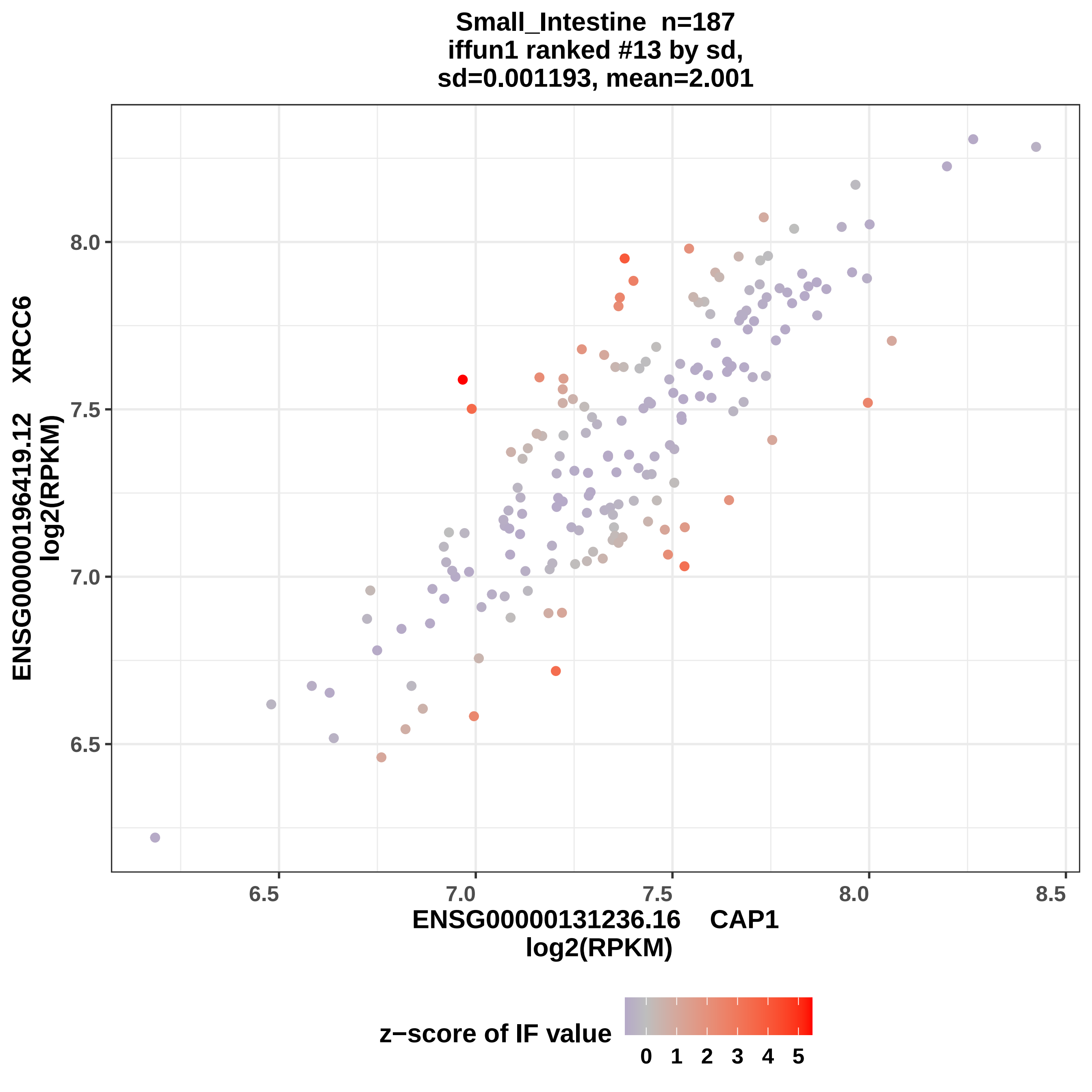

Small_Intestine n=187
iffun1 ranked #13 by sd,
sd=0.001193, mean=2.001
ENSG00000196419.12 XRCC6
log2(RPKM)
ENSG00000131236.16 CAP1
log2(RPKM)
8.0
7.5
7.0
6.5
6.5
7.0
7.5
8.0
8.5
z−score of IF value
0
1
2
3
4
5

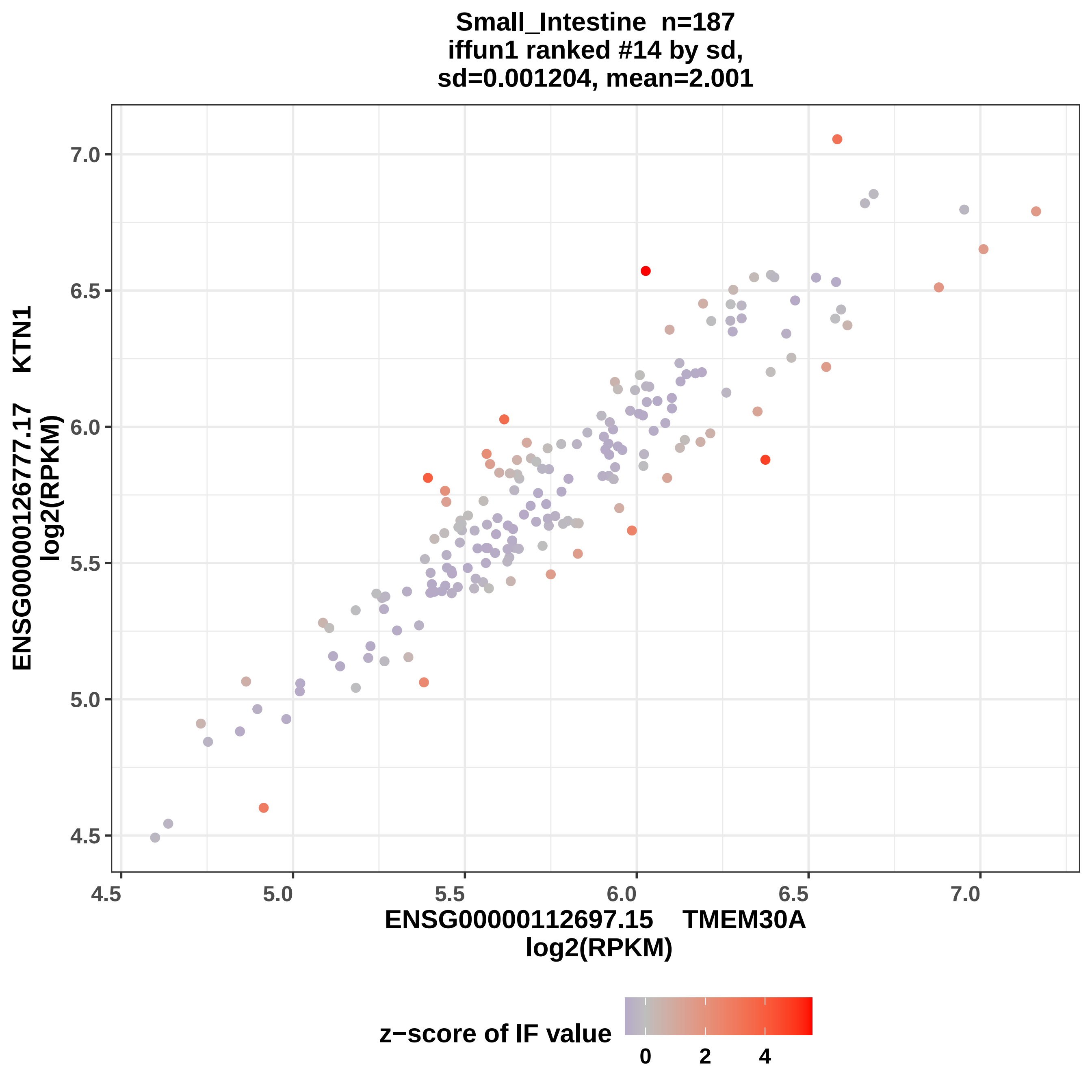

Small_Intestine n=187
iffun1 ranked #14 by sd,
sd=0.001204, mean=2.001
ENSG00000126777.17 KTN1
log2(RPKM)
7.0
6.5
6.0
5.5
5.0
4.5
4.5
5.0
5.5
6.0
6.5
7.0
ENSG00000112697.15 TMEM30A
log2(RPKM)
z−score of IF value
0
2
4

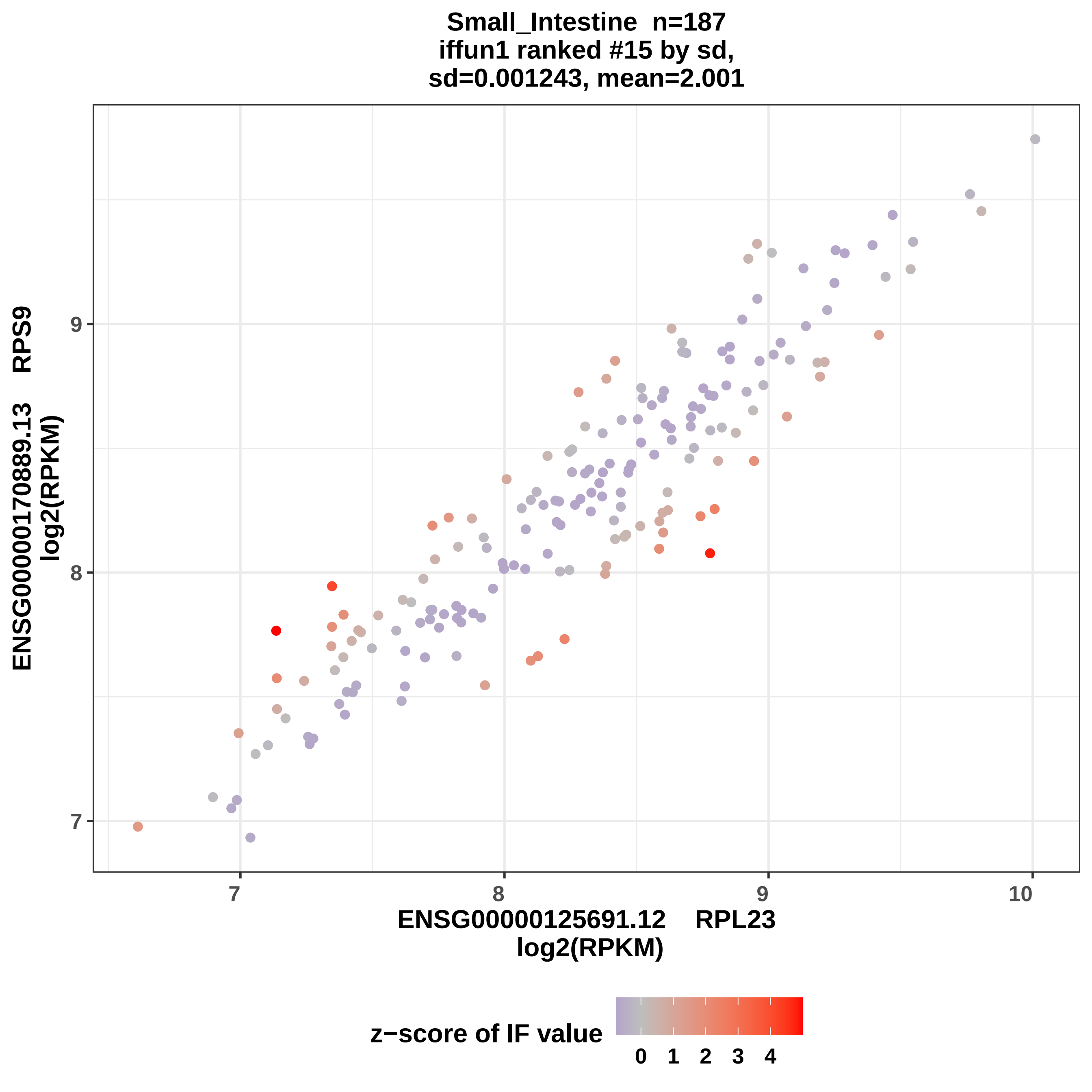
Small_Intestine n=187
iffun1 ranked #15 by sd,
sd=0.001243, mean=2.001
ENSG00000170889.13 RPS9
log2(RPKM)
ENSG00000125691.12 RPL23
log2(RPKM)
7
8
9
10
z−score of IF value
0 1 2 3 4

Small_Intestine n=187
iffun1 ranked #16 by sd,
sd=0.001256, mean=2.001

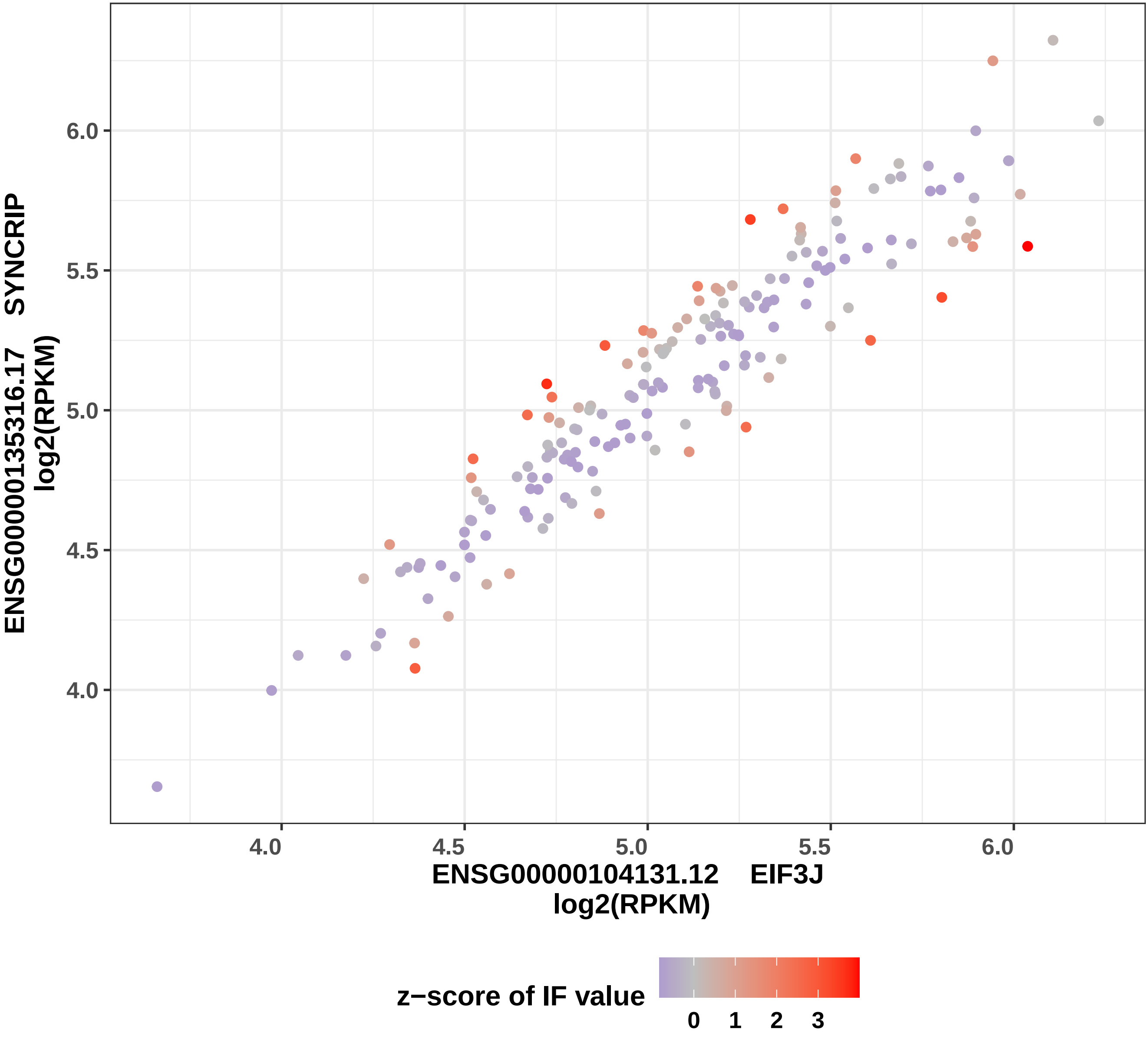

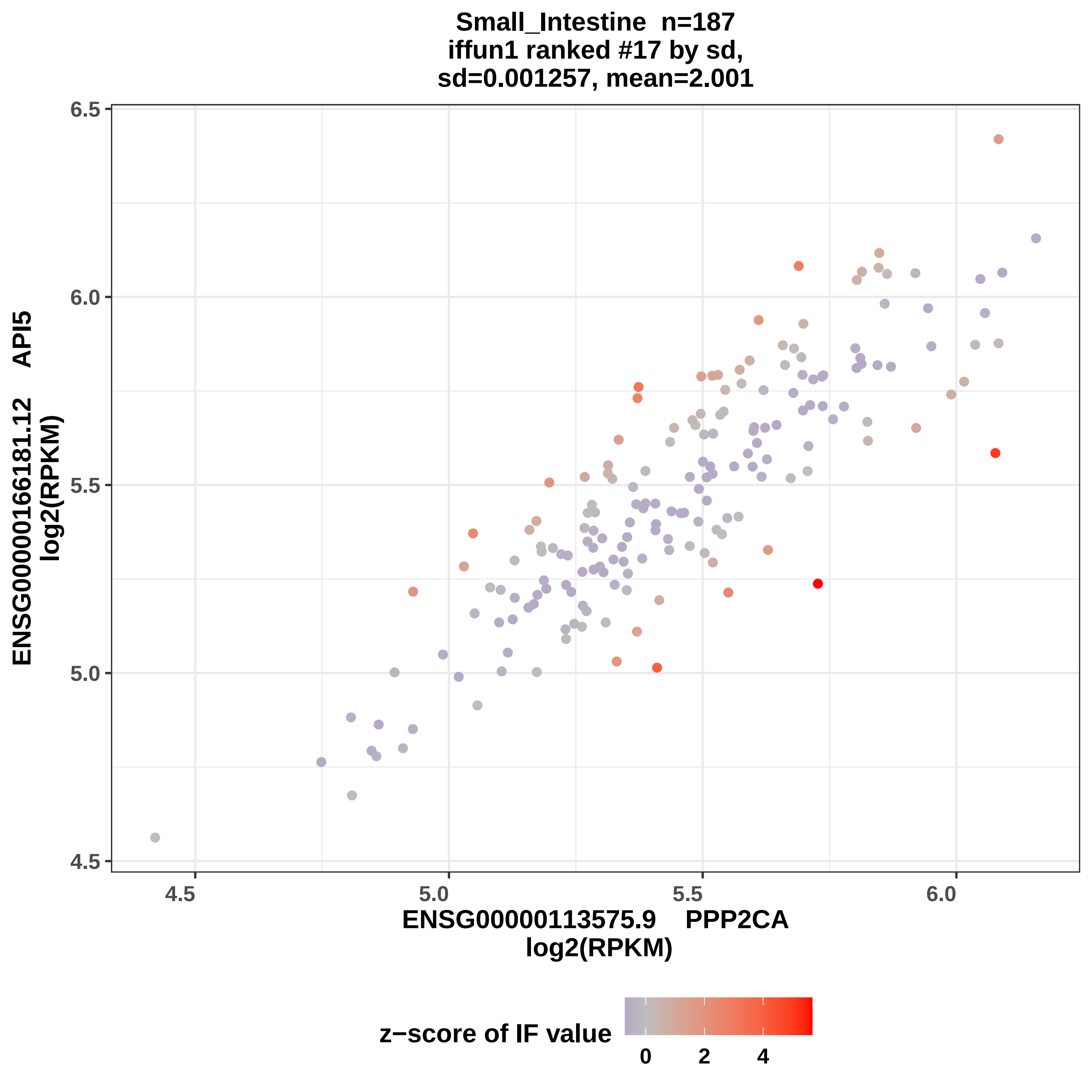

Small_Intestine n=187
iffun1 ranked #17 by sd,
sd=0.001257, mean=2.001
ENSG00000166181.12 API5
log2(RPKM)
ENSG00000113575.9 PPP2CA
log2(RPKM)
6.5
6.0
5.5
5.0
4.5
4.5
5.0
5.5
6.0
z−score of IF value
0
2
4

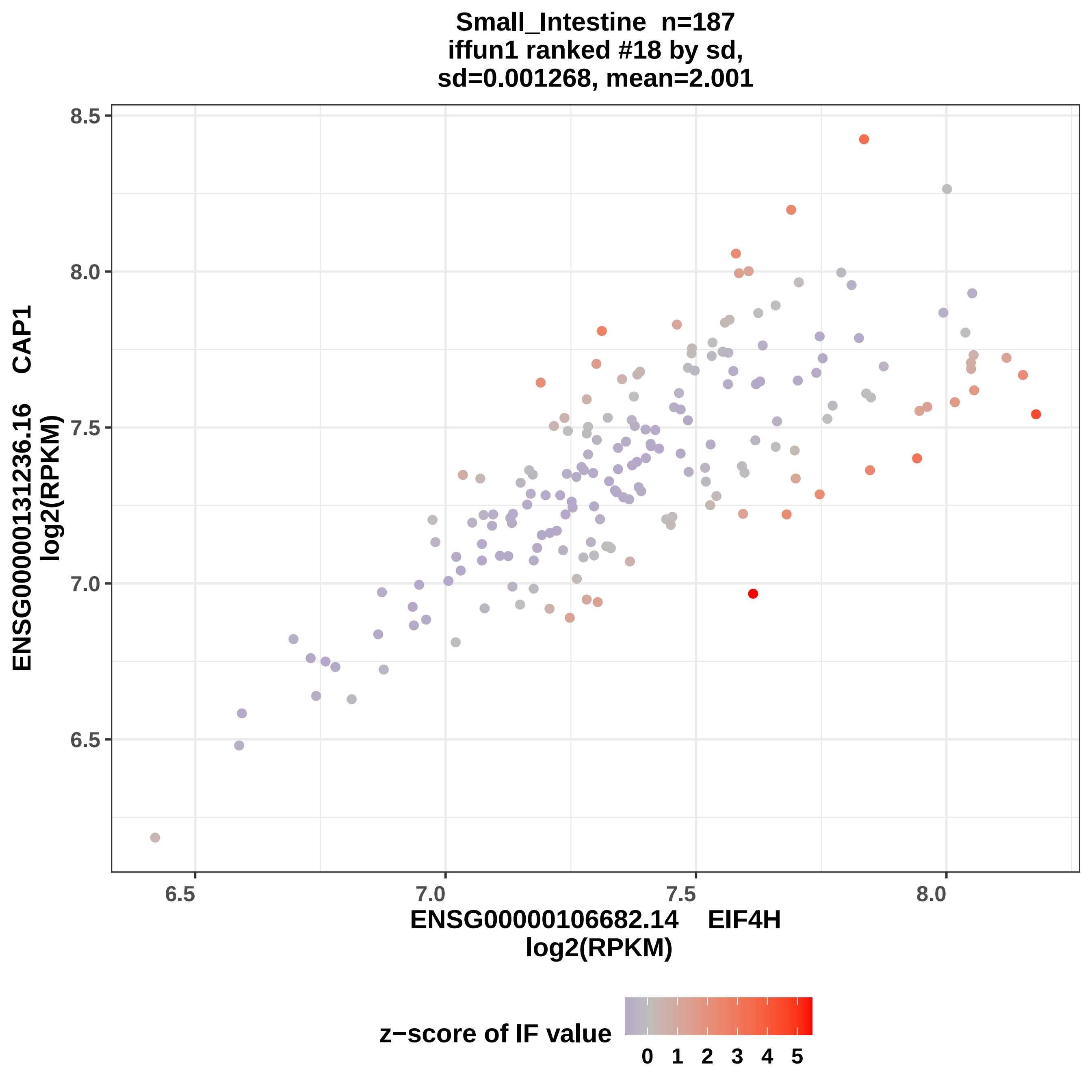
Small_Intestine n=187
iffun1 ranked #18 by sd,
sd=0.001268, mean=2.001
ENSG00000131236.16 CAP1
log2(RPKM)
ENSG00000106682.14 EIF4H
log2(RPKM)
8.5
8.0
7.5
7.0
6.5
6.5
7.0
7.5
8.0
z−score of IF value
0 1 2 3 4 5

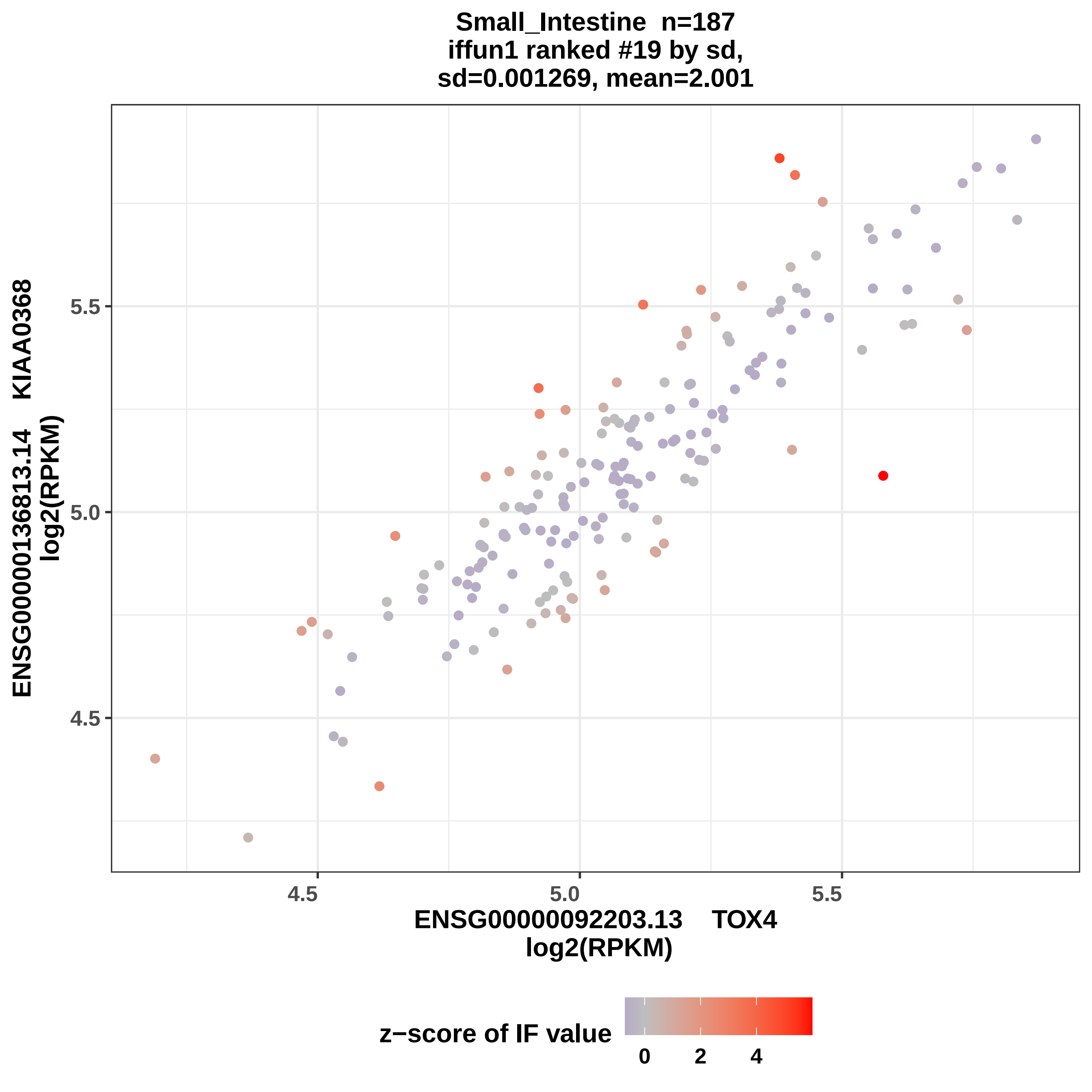
Small_Intestine n=187
iffun1 ranked #19 by sd,
sd=0.001269, mean=2.001
ENSG00000136813.14 KIAA0368
log2(RPKM)
5.5
5.0
4.5
4.5
5.0
5.5
ENSG00000092203.13 TOX4
log2(RPKM)
z−score of IF value
0
2
4

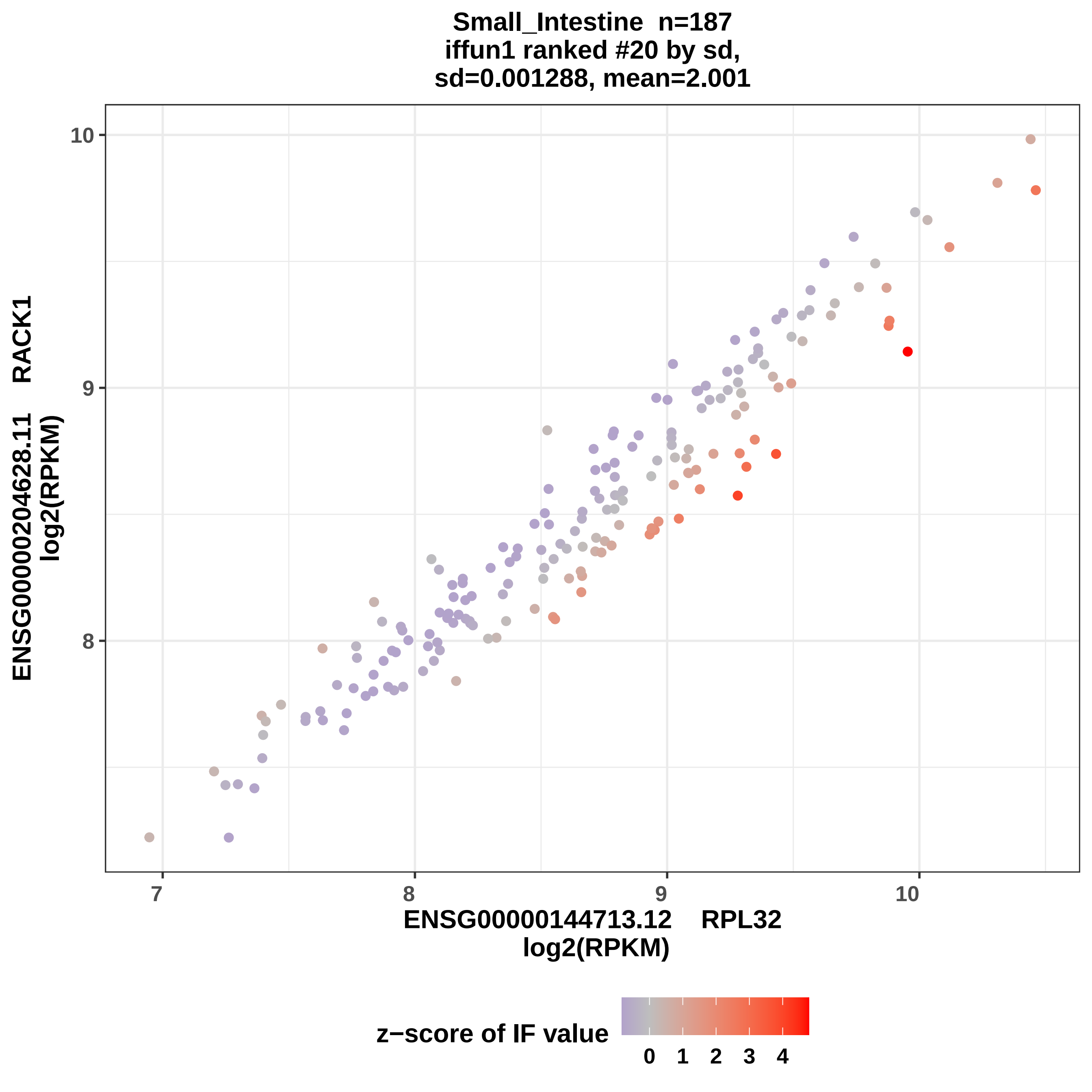
Small_Intestine n=187
iffun1 ranked #20 by sd,
sd=0.001288, mean=2.001
ENSG00000204628.11 RACK1
log2(RPKM)
ENSG00000144713.12 RPL32
log2(RPKM)
10
9
8
7
8
9
10
z−score of IF value
0
1
2
3
4

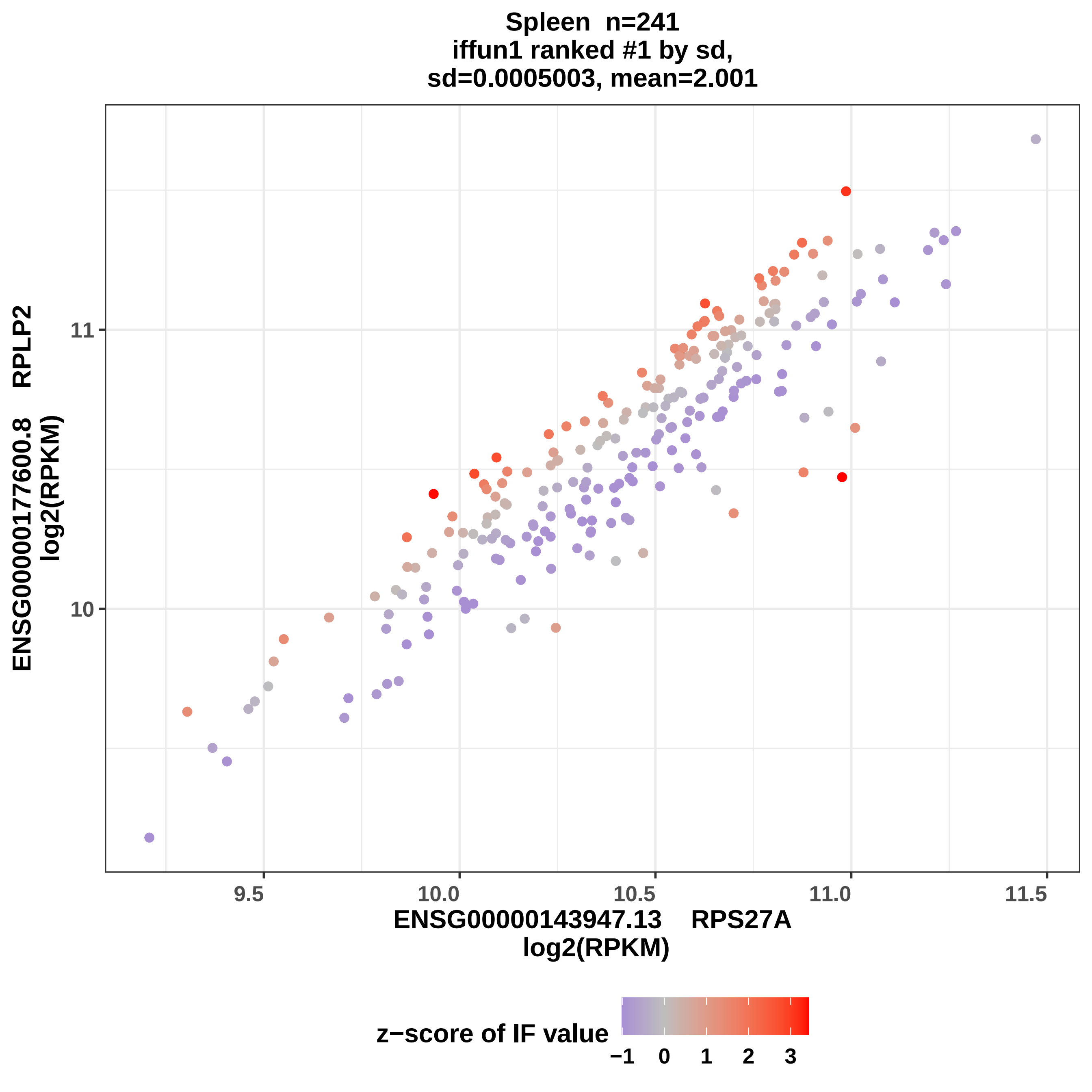
Spleen n=241
iffun1 ranked #1 by sd,
sd=0.0005003, mean=2.001
ENSG00000177600.8 RPLP2
log2(RPKM)
11
10
9.5
10.0
10.5
11.0
11.5
ENSG00000143947.13 RPS27A
log2(RPKM)
z−score of IF value
−1
0
1
2
3

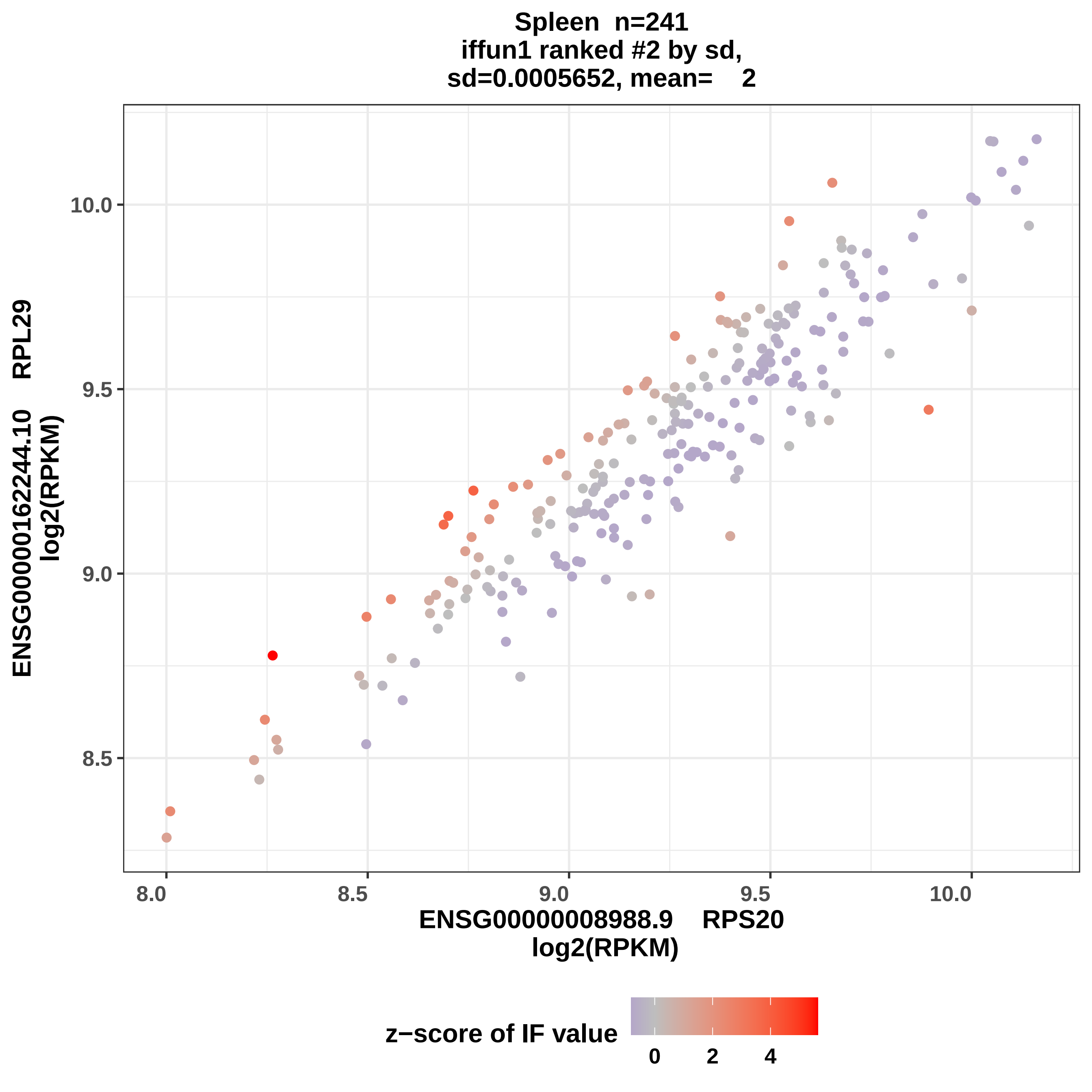
Spleen n=241
iffun1 ranked #2 by sd,
sd=0.0005652, mean= 2
ENSG00000162244.10 RPL29
log2(RPKM)
10.0
9.5
9.0
8.5
8.0
8.5
9.0
9.5
10.0
ENSG00000008988.9 RPS20
log2(RPKM)
z−score of IF value
0
2
4

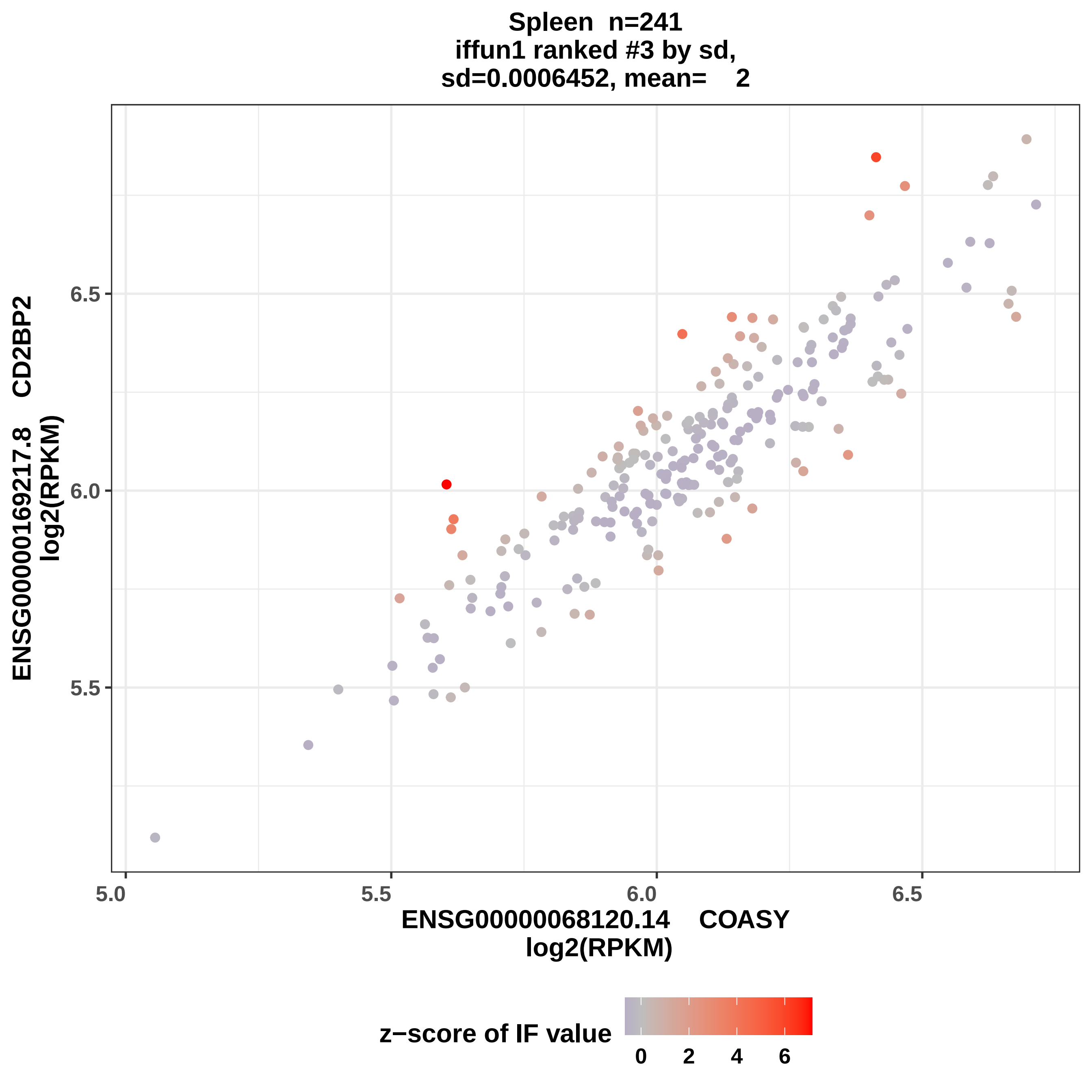
Spleen n=241
iffun1 ranked #3 by sd,
sd=0.0006452, mean= 2
ENSG00000169217.8 CD2BP2
log2(RPKM)
6.5
6.0
5.5
5.0
5.5
6.0
6.5
ENSG00000068120.14 COASY
log2(RPKM)
z−score of IF value
0
2
4
6

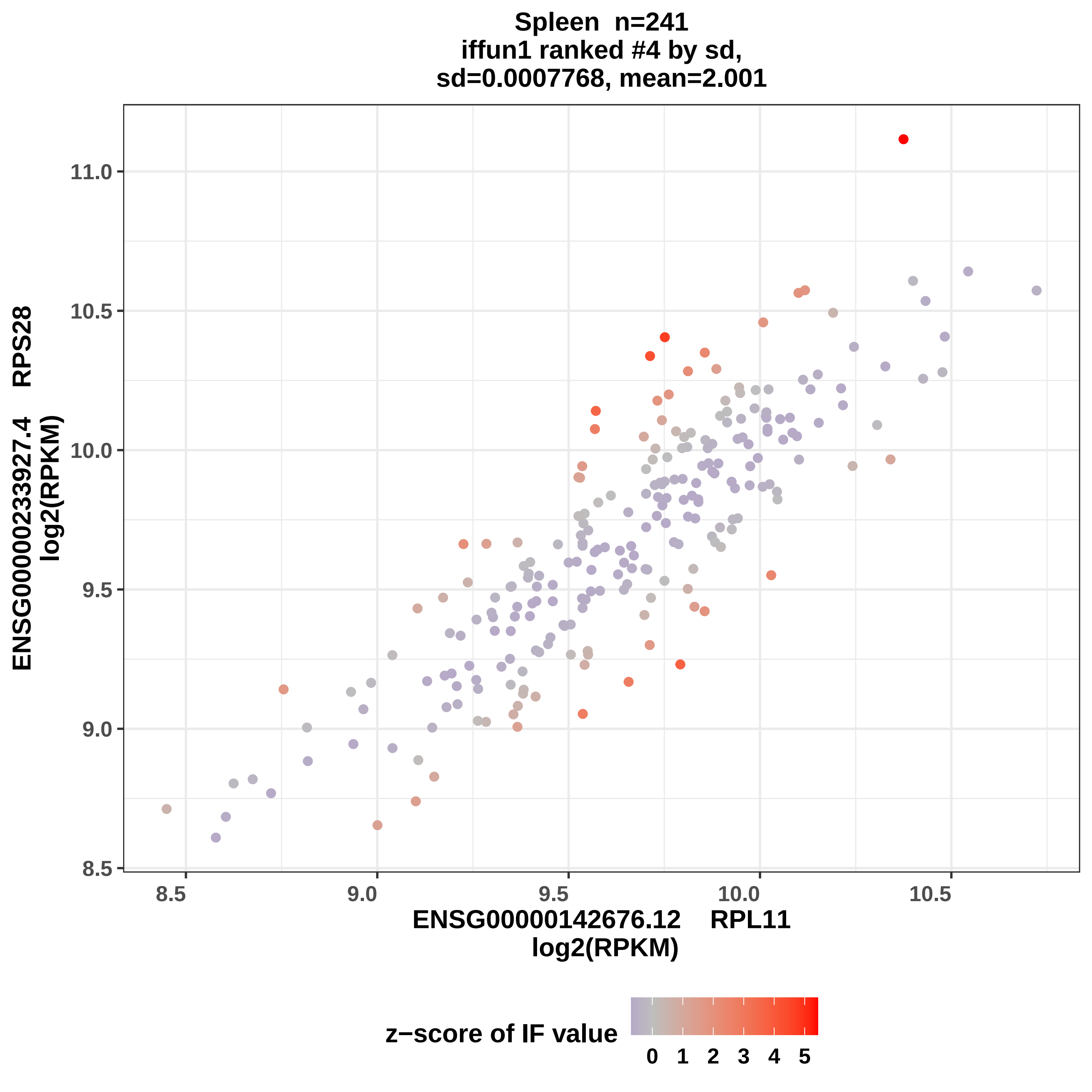
Spleen n=241
iffun1 ranked #4 by sd,
sd=0.0007768, mean=2.001
ENSG00000233927.4 RPS28
log2(RPKM)
11.0
10.5
10.0
9.5
9.0
8.5
8.5
9.0
9.5
10.0
10.5
ENSG00000142676.12 RPL11
log2(RPKM)
z−score of IF value
0
1
2
3
4
5

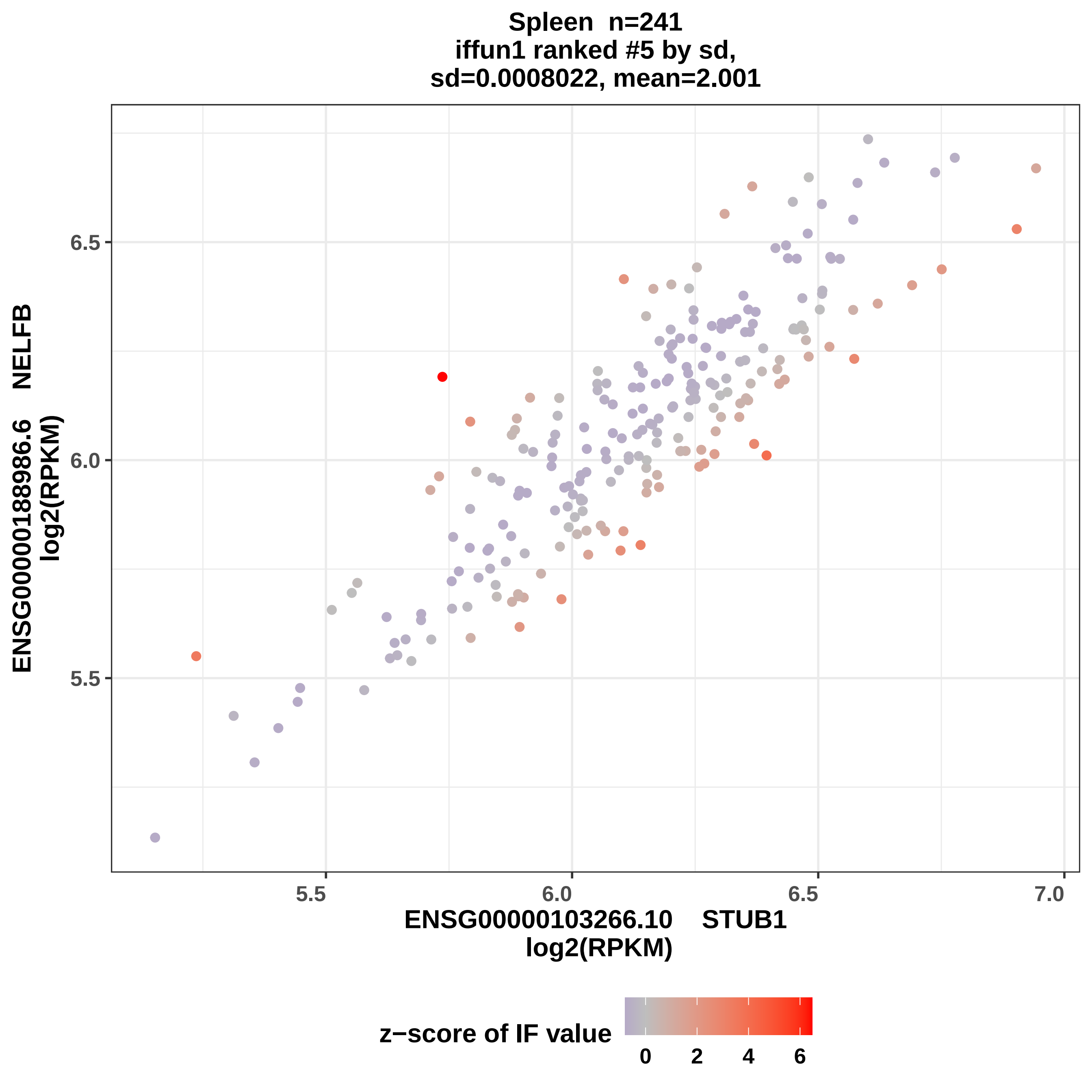
Spleen n=241
iffun1 ranked #5 by sd,
sd=0.0008022, mean=2.001
ENSG00000188986.6 NELFB
log2(RPKM)
6.5
6.0
5.5
5.5
6.0
6.5
7.0
ENSG00000103266.10 STUB1
log2(RPKM)
z−score of IF value
0
2
4
6

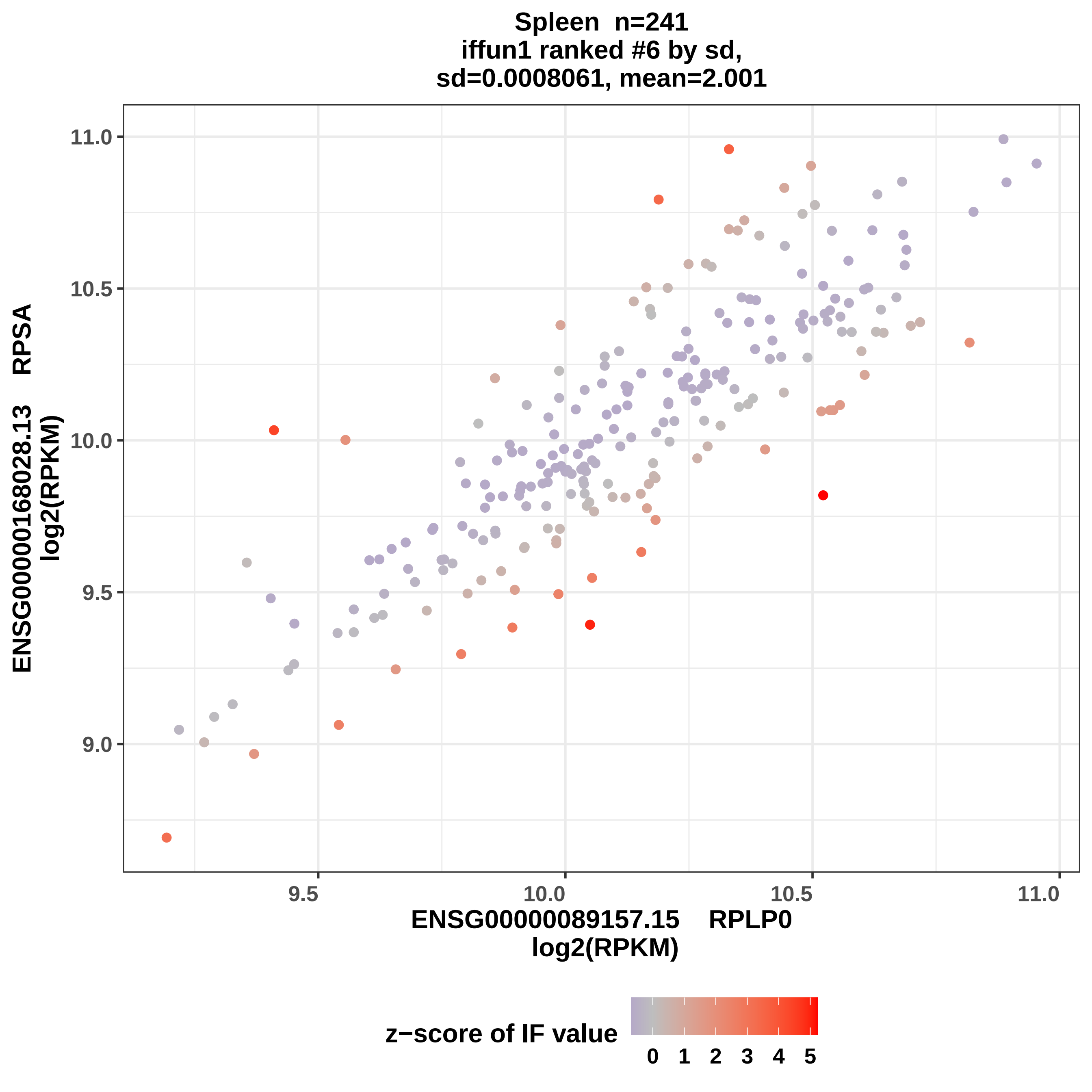

Spleen n=241
iffun1 ranked #6 by sd,
sd=0.0008061, mean=2.001
ENSG00000168028.13 RPSA
log2(RPKM)
11.0
10.5
10.0
9.5
9.0
9.5
10.0
10.5
11.0
ENSG00000089157.15 RPLP0
log2(RPKM)
z−score of IF value
0 1 2 3 4 5

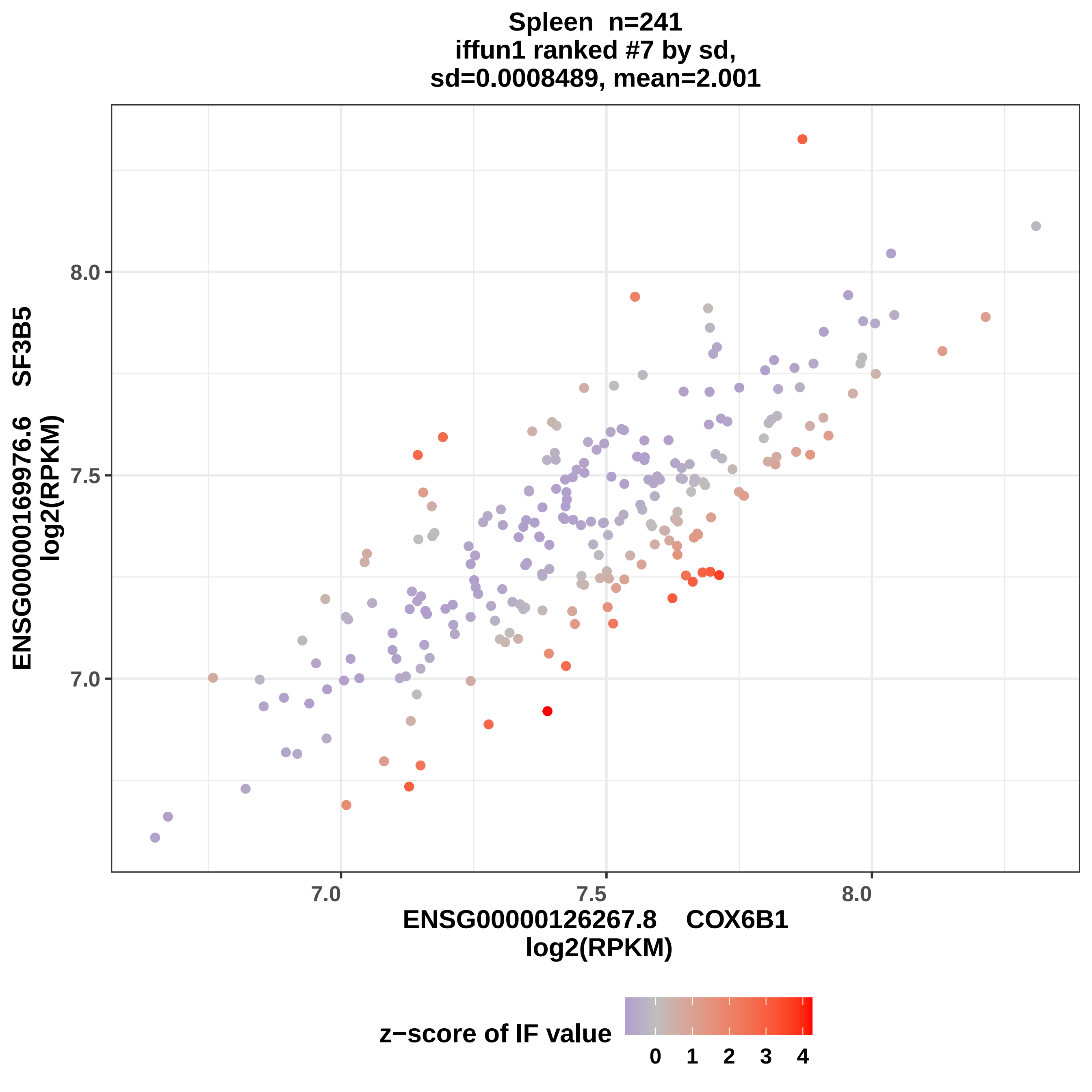

Spleen n=241
iffun1 ranked #7 by sd,
sd=0.0008489, mean=2.001
ENSG00000169976.6 SF3B5
log2(RPKM)
8.0
7.5
7.0
7.0
7.5
8.0
ENSG00000126267.8 COX6B1
log2(RPKM)
z−score of IF value
0
1
2
3
4

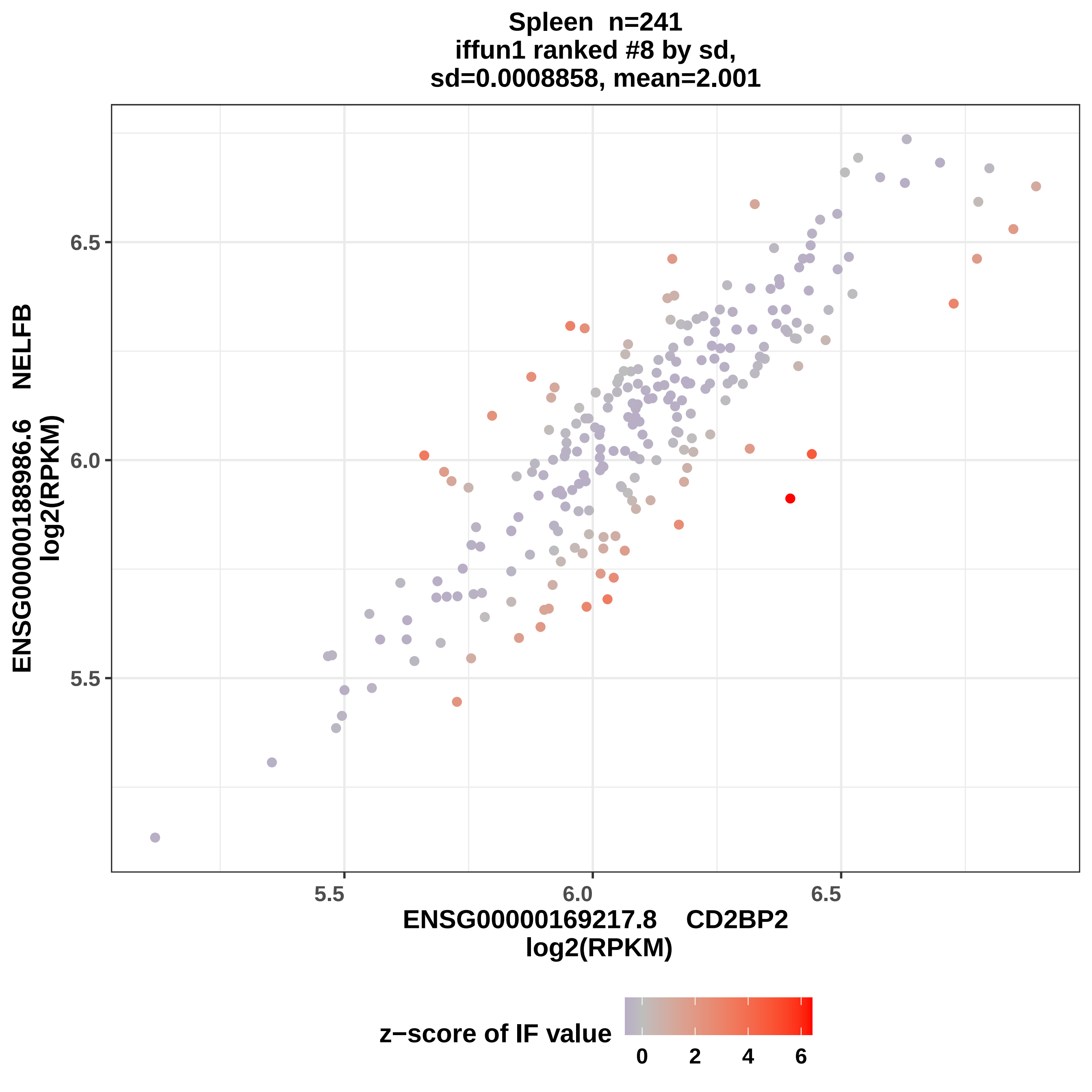
Spleen n=241
iffun1 ranked #8 by sd,
sd=0.0008858, mean=2.001
ENSG00000188986.6 NELFB
log2(RPKM)
6.5
6.0
5.5
5.5
6.0
6.5
ENSG00000169217.8 CD2BP2
log2(RPKM)
z−score of IF value
0
2
4
6

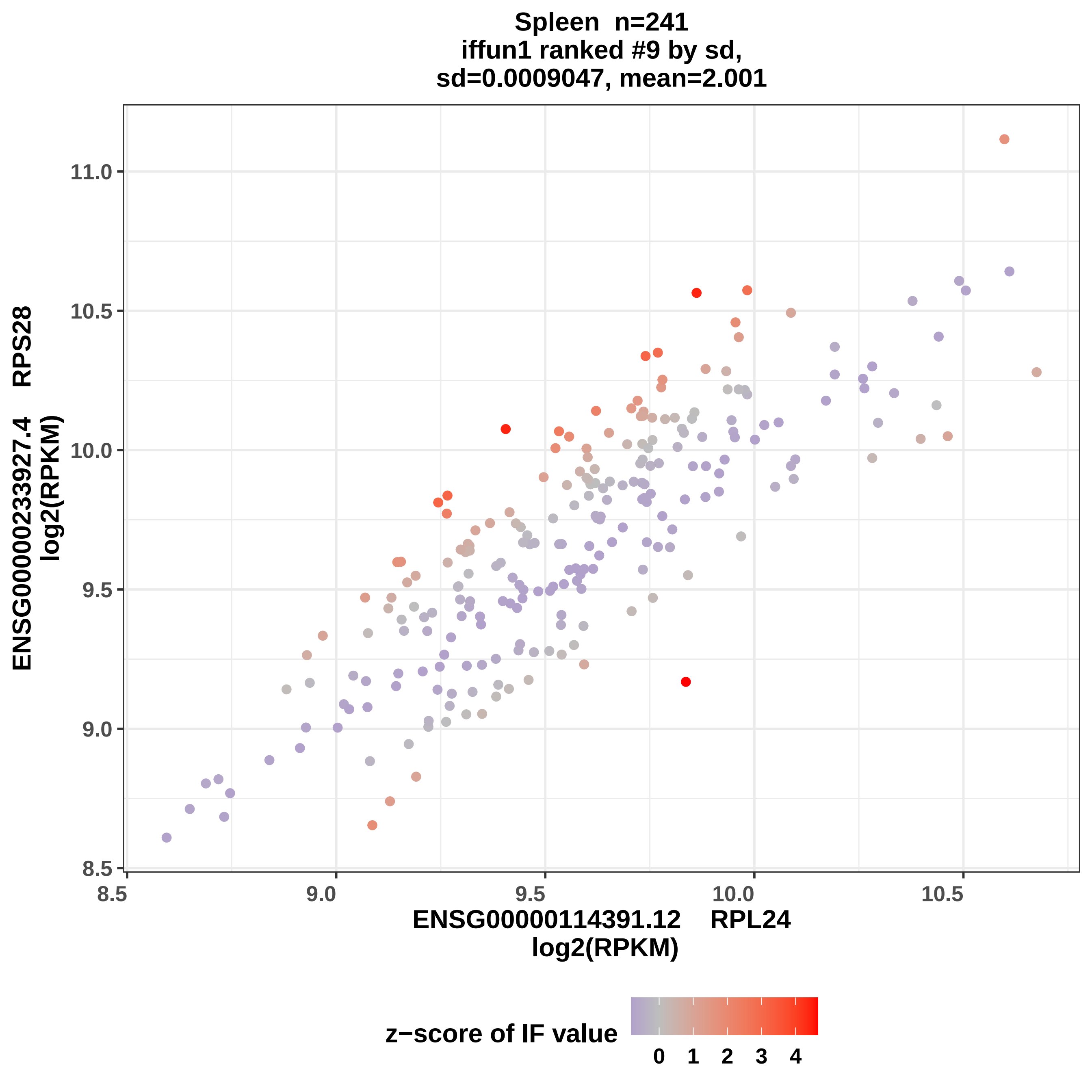
Spleen n=241
iffun1 ranked #9 by sd,
sd=0.0009047, mean=2.001
ENSG00000233927.4 RPS28
log2(RPKM)
ENSG00000114391.12 RPL24
log2(RPKM)
11.0
10.5
10.0
9.5
9.0
8.5
8.5
9.0
9.5
10.0
10.5
z−score of IF value
0
1
2
3
4

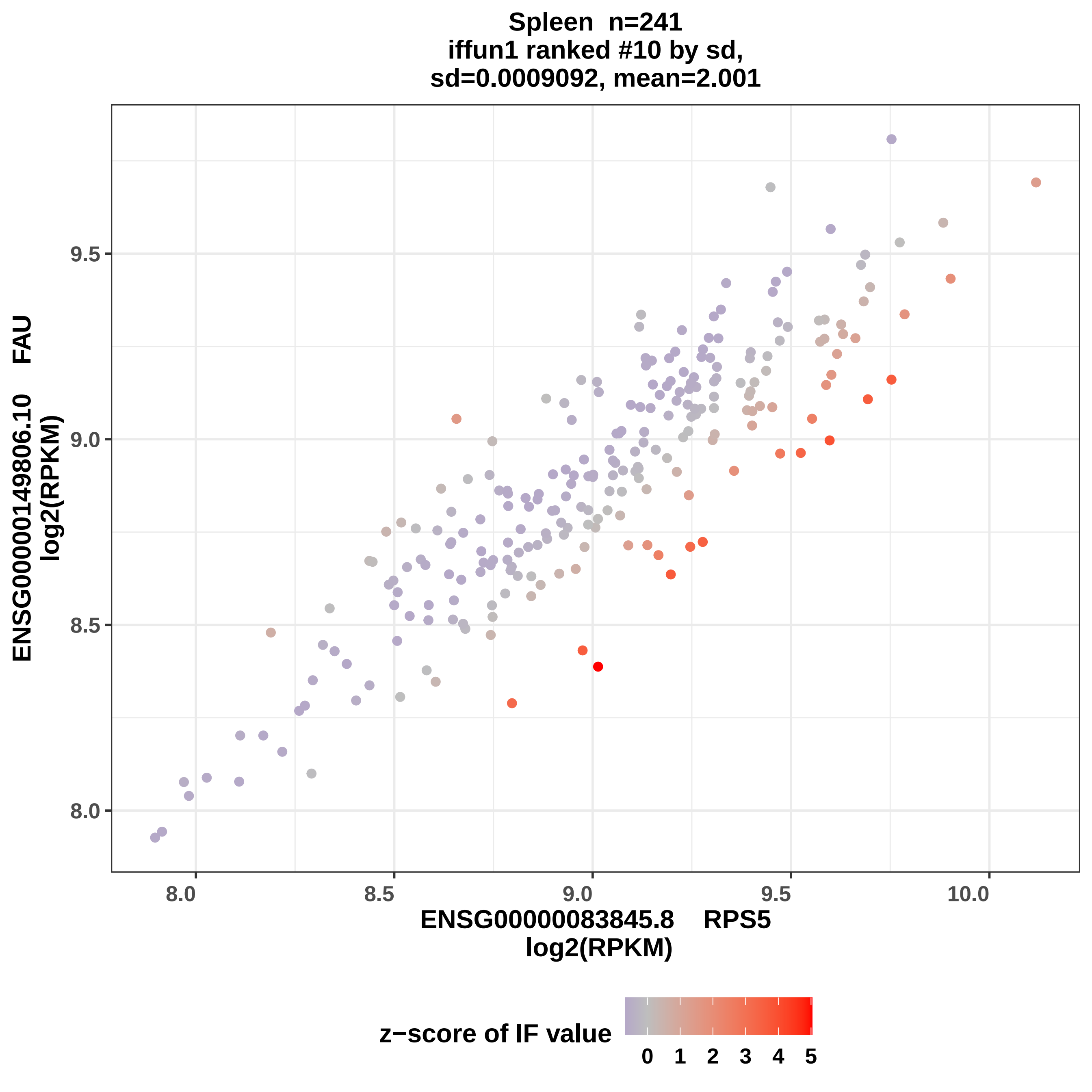
Spleen n=241
iffun1 ranked #10 by sd,
sd=0.0009092, mean=2.001
ENSG00000149806.10 FAU
log2(RPKM)
9.5
9.0
8.5
8.0
8.0
8.5
9.0
9.5
10.0
ENSG00000083845.8 RPS5
log2(RPKM)
z−score of IF value
0 1 2 3 4 5

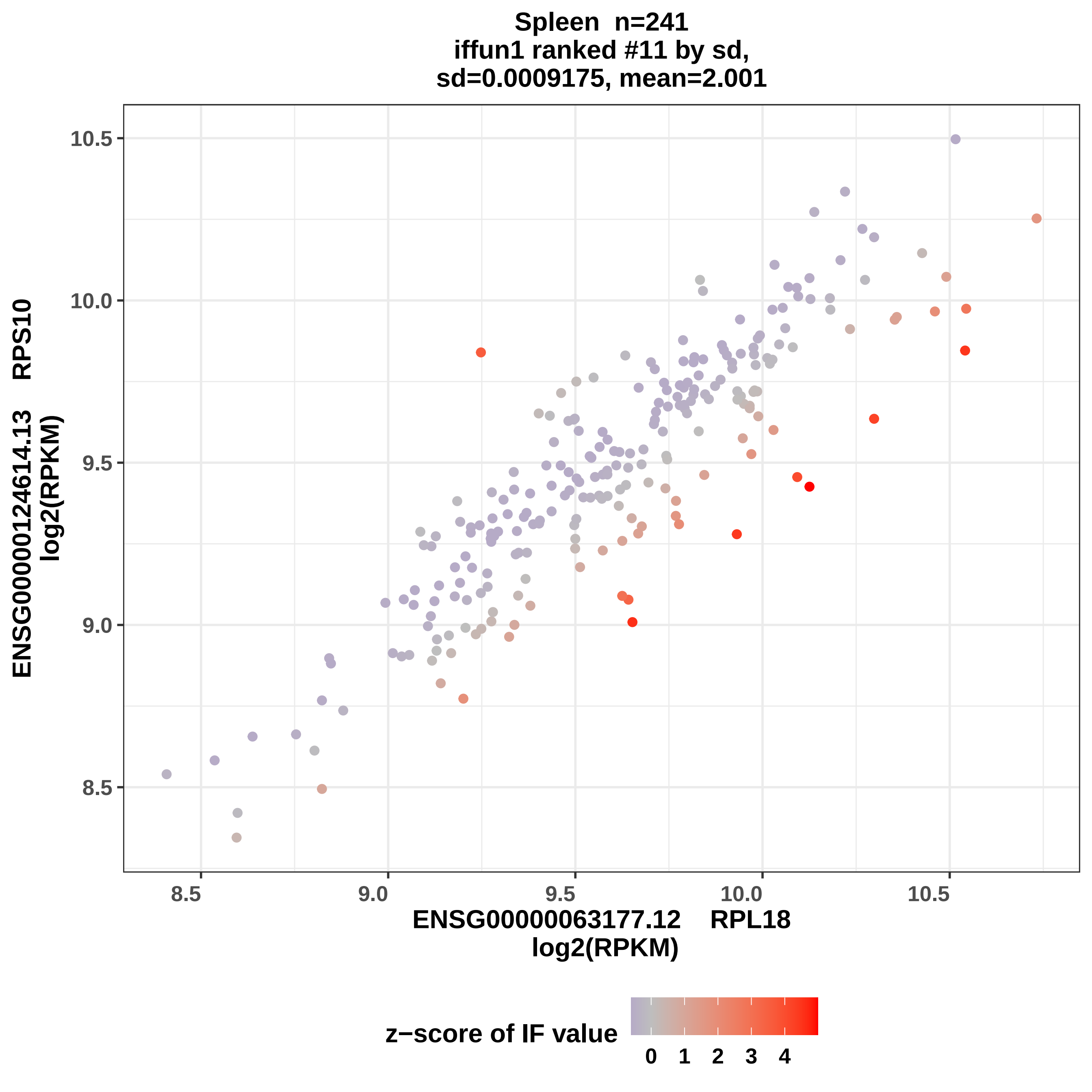

Spleen n=241
iffun1 ranked #11 by sd,
sd=0.0009175, mean=2.001
ENSG00000124614.13 RPS10
log2(RPKM)
10.5
10.0
9.5
9.0
8.5
8.5
9.0
9.5
10.0
10.5
ENSG00000063177.12 RPL18
log2(RPKM)
z−score of IF value
0
1
2
3
4

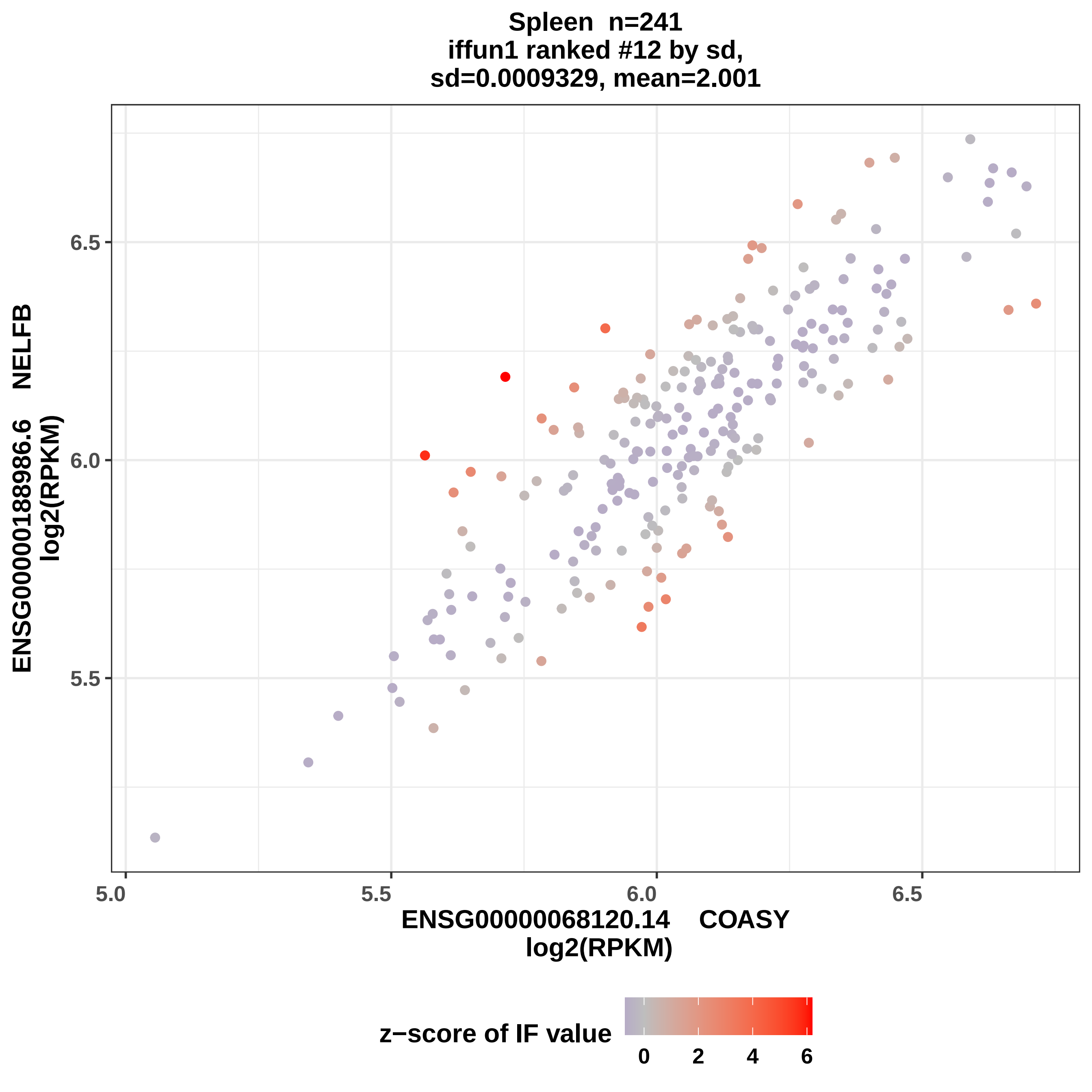

Spleen n=241
iffun1 ranked #12 by sd,
sd=0.0009329, mean=2.001
ENSG00000188986.6 NELFB
log2(RPKM)
6.5
6.0
5.5
5.0
5.5
6.0
6.5
ENSG00000068120.14 COASY
log2(RPKM)
z−score of IF value
0
2
4
6

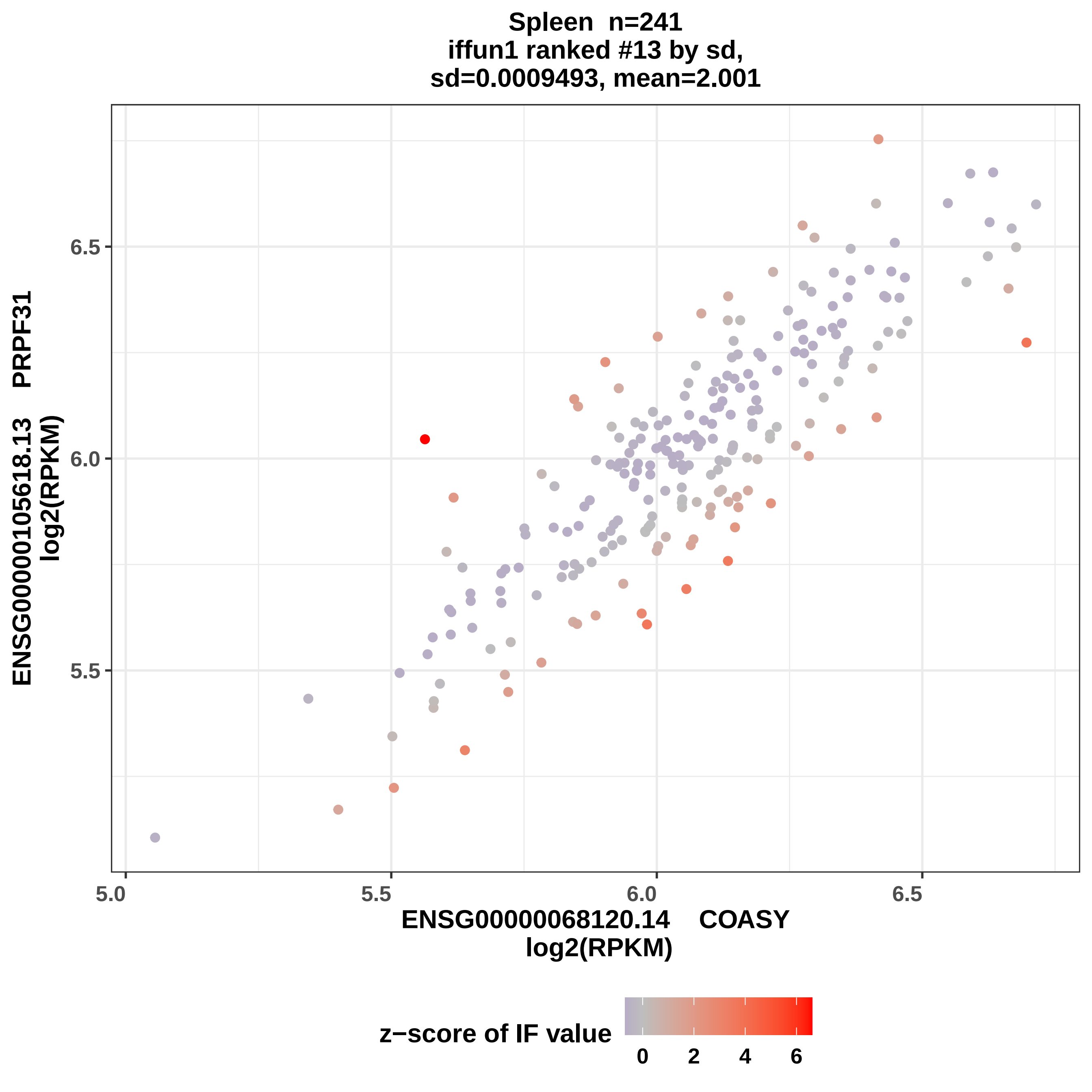

Spleen n=241
iffun1 ranked #13 by sd,
sd=0.0009493, mean=2.001
ENSG00000105618.13 PRPF31
log2(RPKM)
6.5
6.0
5.5
5.0
5.5
6.0
6.5
ENSG00000068120.14 COASY
log2(RPKM)
z−score of IF value
0
2
4
6

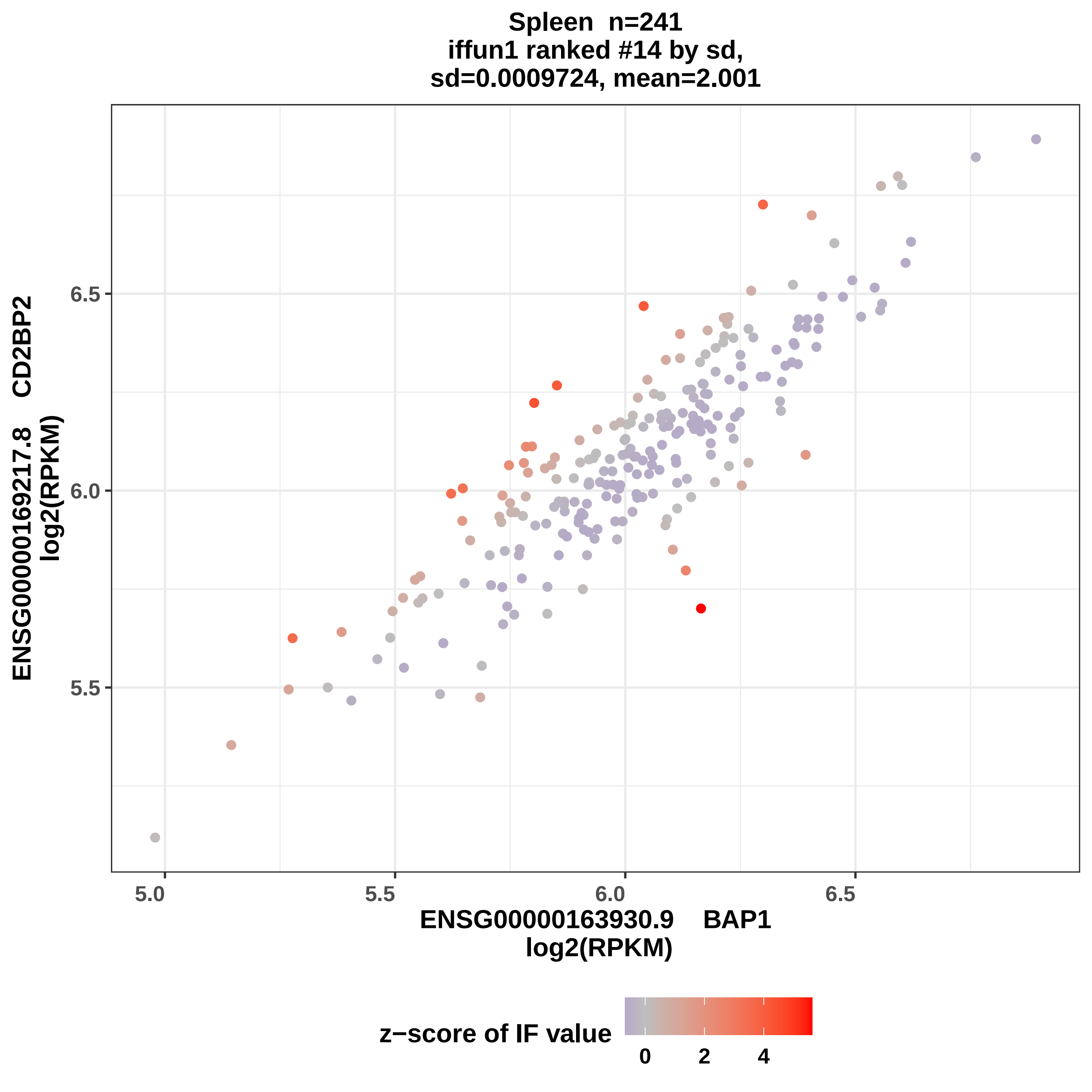

Spleen n=241
iffun1 ranked #14 by sd,
sd=0.0009724, mean=2.001
ENSG00000169217.8 CD2BP2
log2(RPKM)
6.5
6.0
5.5
5.0
5.5
6.0
6.5
ENSG00000163930.9 BAP1
log2(RPKM)
z−score of IF value
0
2
4

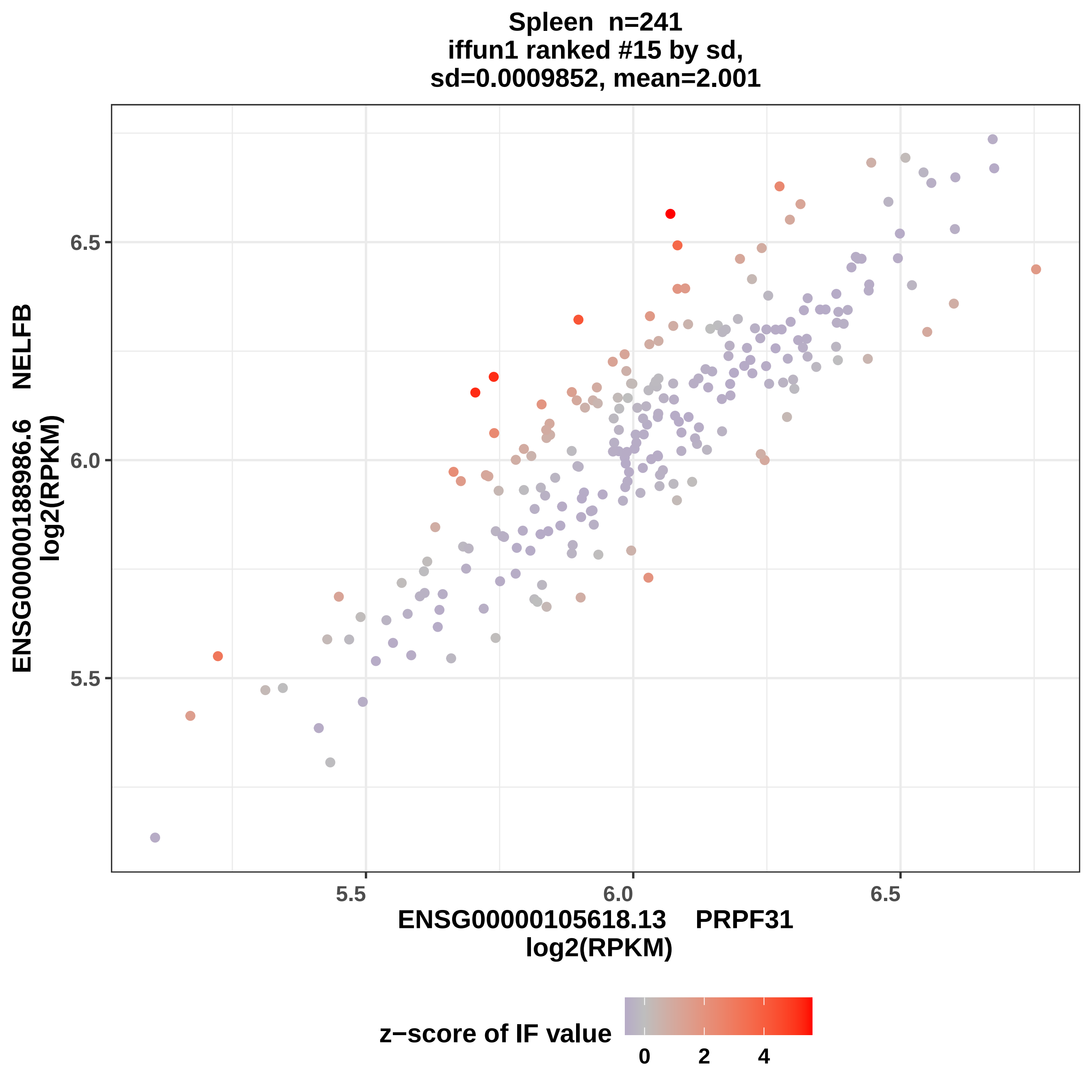

Spleen n=241
iffun1 ranked #15 by sd,
sd=0.0009852, mean=2.001
ENSG00000188986.6 NELFB
log2(RPKM)
6.5
6.0
5.5
5.5
6.0
6.5
ENSG00000105618.13 PRPF31
log2(RPKM)
z−score of IF value
0
2
4

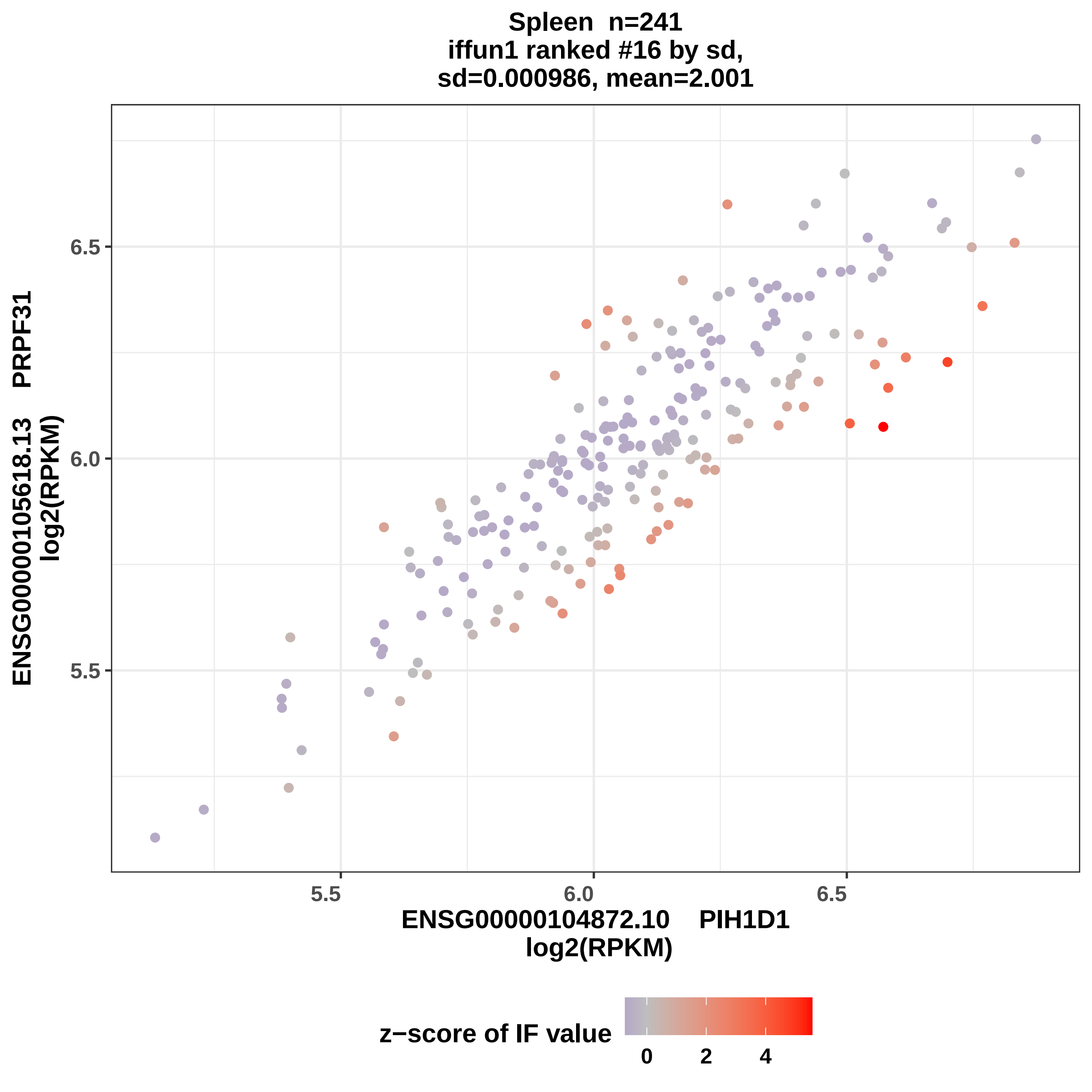
Spleen n=241
iffun1 ranked #16 by sd,
sd=0.000986, mean=2.001
ENSG00000105618.13 PRPF31
log2(RPKM)
6.5
6.0
5.5
5.5
6.0
6.5
ENSG00000104872.10 PIH1D1
log2(RPKM)
z−score of IF value
0
2
4

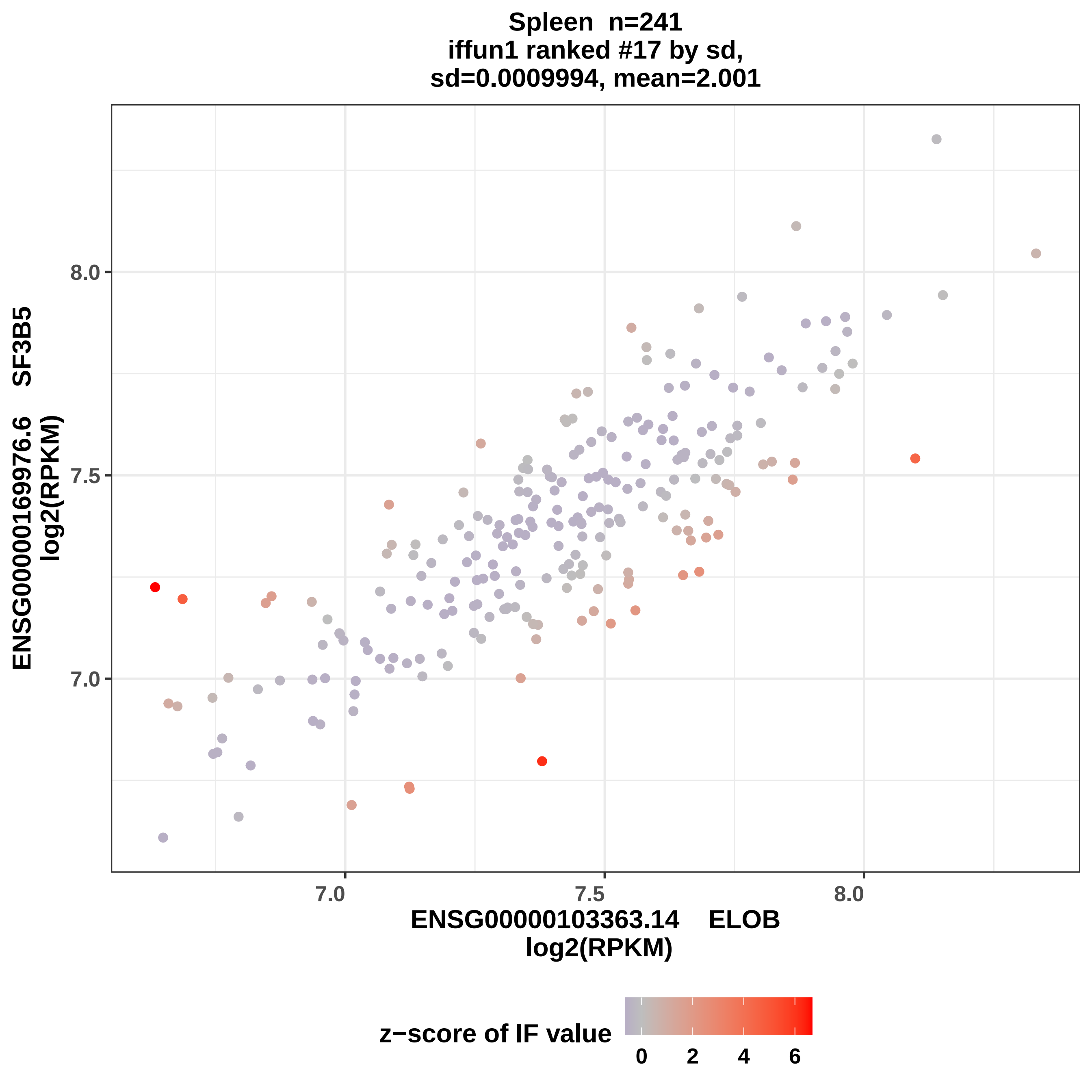

Spleen n=241
iffun1 ranked #17 by sd,
sd=0.0009994, mean=2.001
ENSG00000169976.6 SF3B5
log2(RPKM)
8.0
7.5
7.0
7.0
7.5
8.0
ENSG00000103363.14 ELOB
log2(RPKM)
z−score of IF value
0
2
4
6

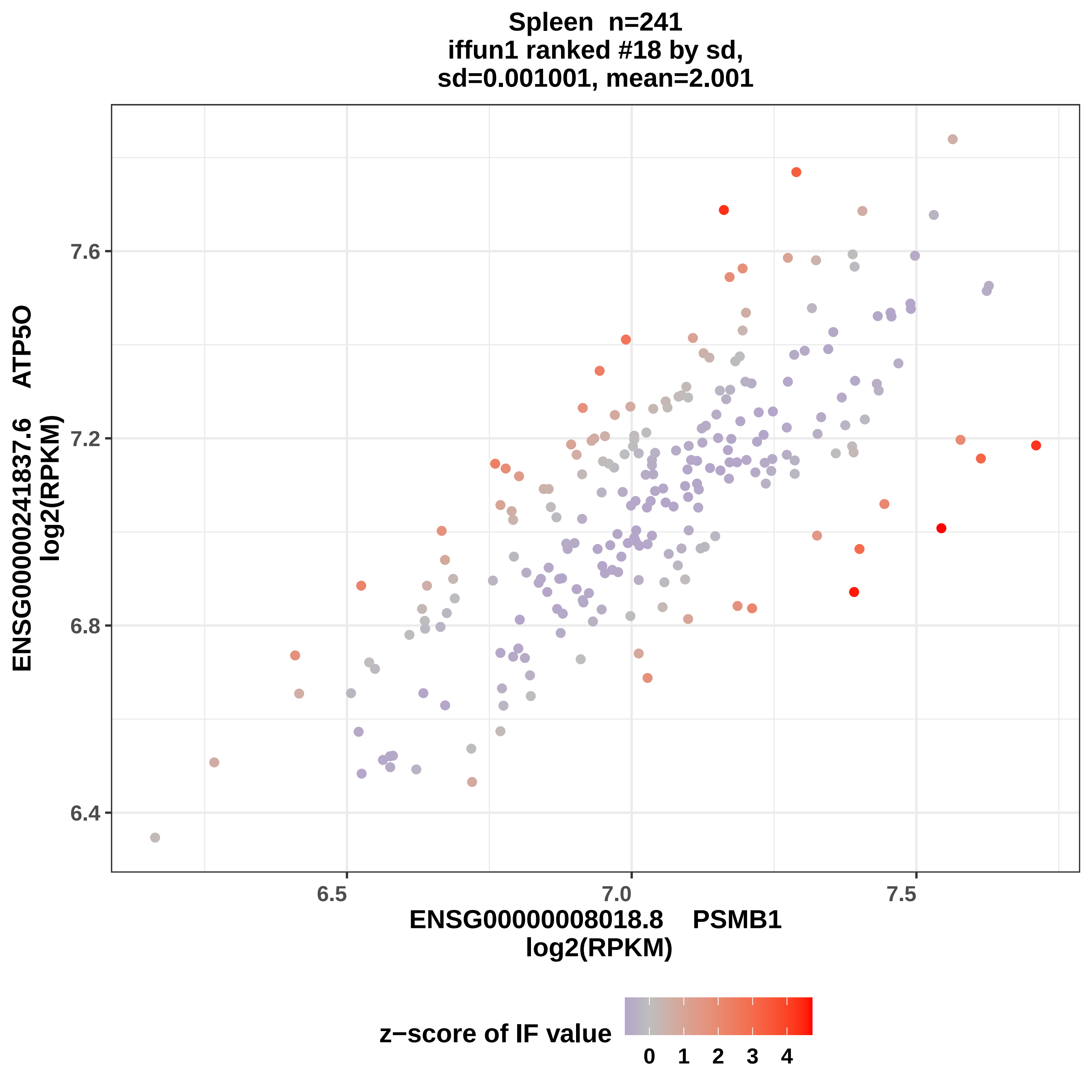

Spleen n=241
iffun1 ranked #18 by sd,
sd=0.001001, mean=2.001
ENSG00000241837.6 ATP5O
log2(RPKM)
7.6
7.2
6.8
6.4
6.5
7.0
7.5
ENSG00000008018.8 PSMB1
log2(RPKM)
z−score of IF value
0
1
2
3
4

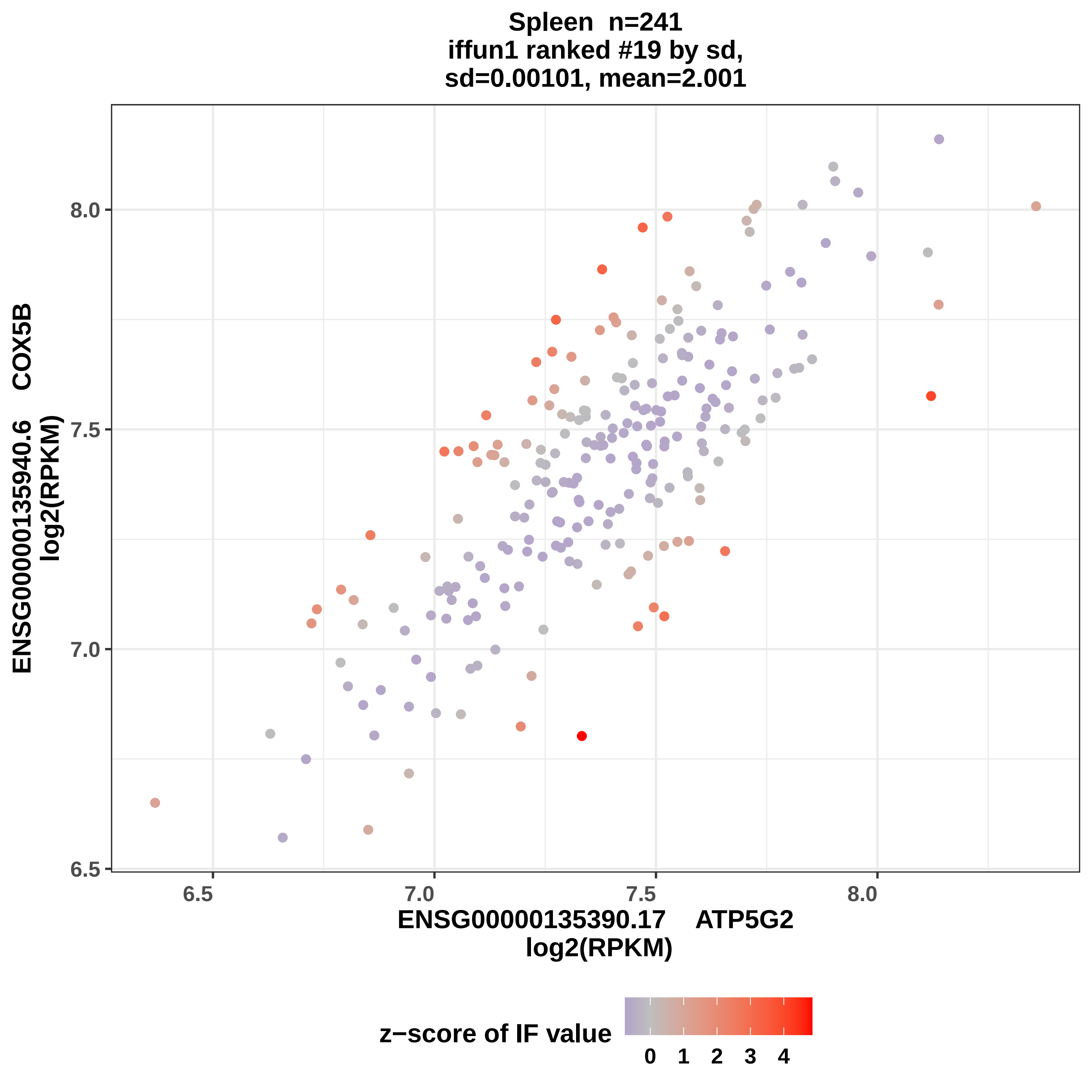

Spleen n=241
iffun1 ranked #19 by sd,
sd=0.00101, mean=2.001
ENSG00000135940.6 COX5B
log2(RPKM)
8.0
7.5
7.0
6.5
6.5
7.0
7.5
8.0
ENSG00000135390.17 ATP5G2
log2(RPKM)
z−score of IF value
0 1 2 3 4

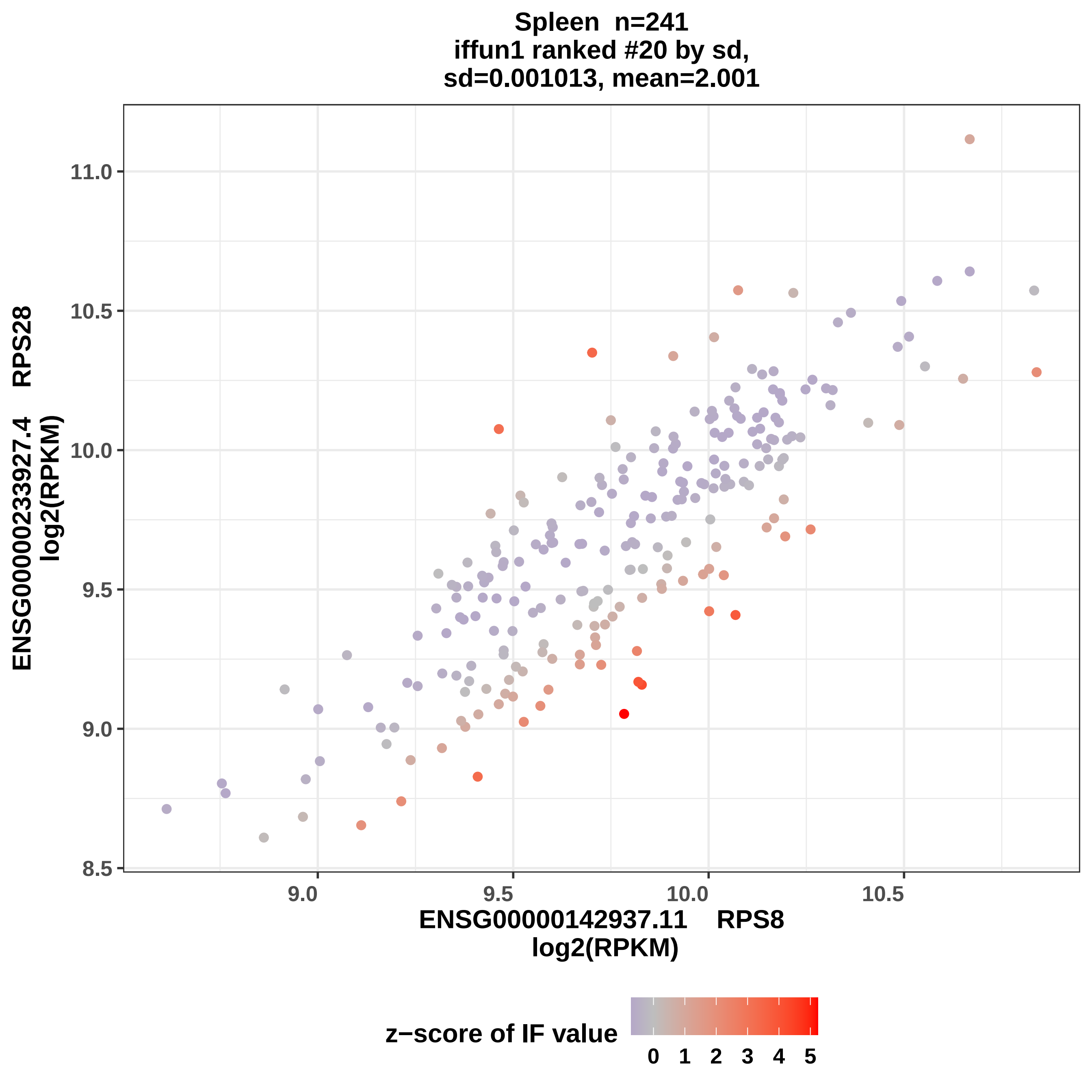
Spleen n=241
iffun1 ranked #20 by sd,
sd=0.001013, mean=2.001
ENSG00000233927.4 RPS28
log2(RPKM)
11.0
10.5
10.0
9.5
9.0
8.5
9.0
9.5
10.0
10.5
ENSG00000142937.11 RPS8
log2(RPKM)
z−score of IF value
0
1
2
3
4
5

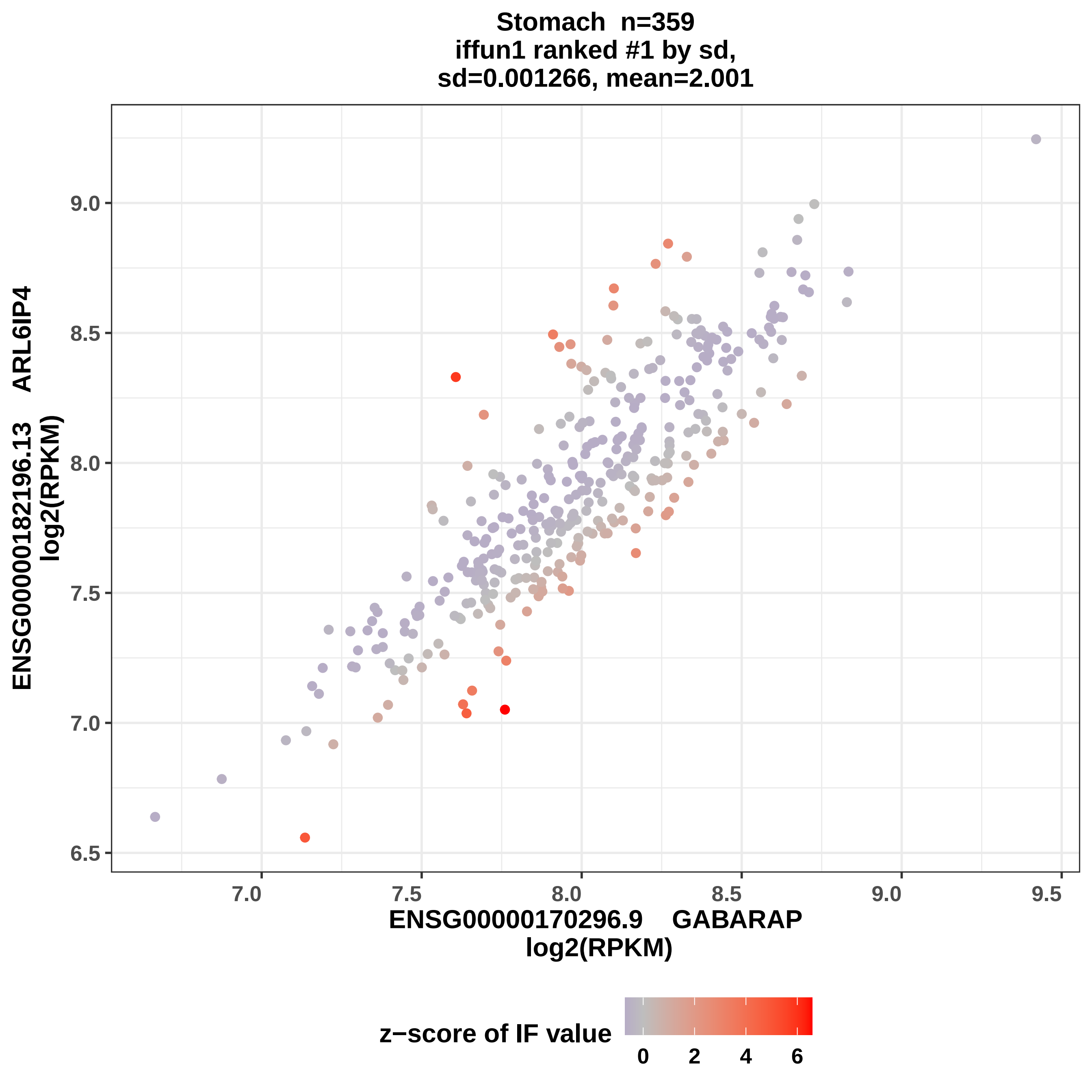

Stomach n=359
iffun1 ranked #1 by sd,
sd=0.001266, mean=2.001
ENSG00000182196.13 ARL6IP4
log2(RPKM)
9.0
8.5
8.0
7.5
7.0
6.5
7.0
7.5
8.0
8.5
9.0
9.5
ENSG00000170296.9 GABARAP
log2(RPKM)
z−score of IF value
0
2
4
6

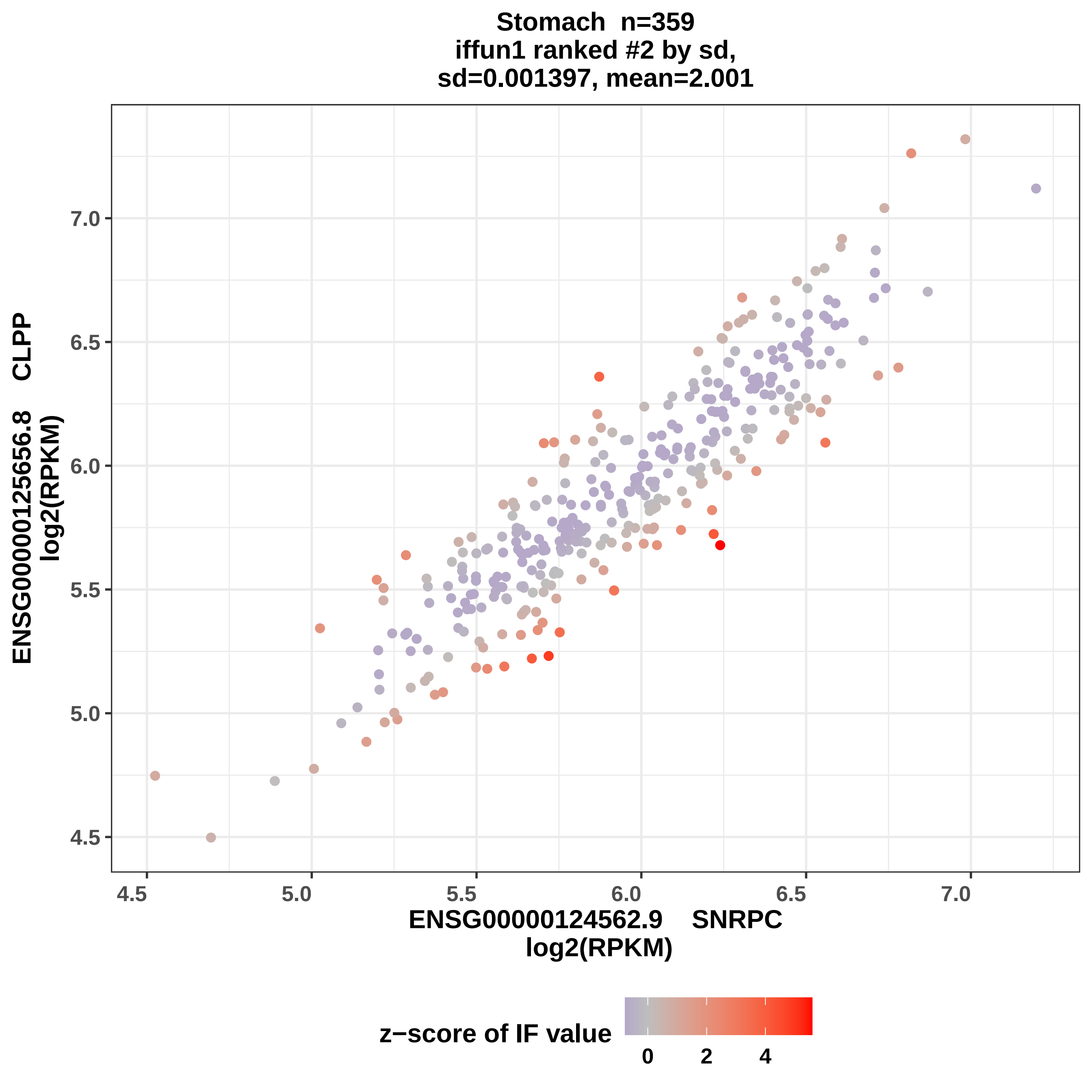
Stomach n=359
iffun1 ranked #2 by sd,
sd=0.001397, mean=2.001
ENSG00000125656.8 CLPP
log2(RPKM)
ENSG00000124562.9 SNRPC
log2(RPKM)
z−score of IF value
4.5
5.0
5.5
6.0
6.5
7.0
0
2
4

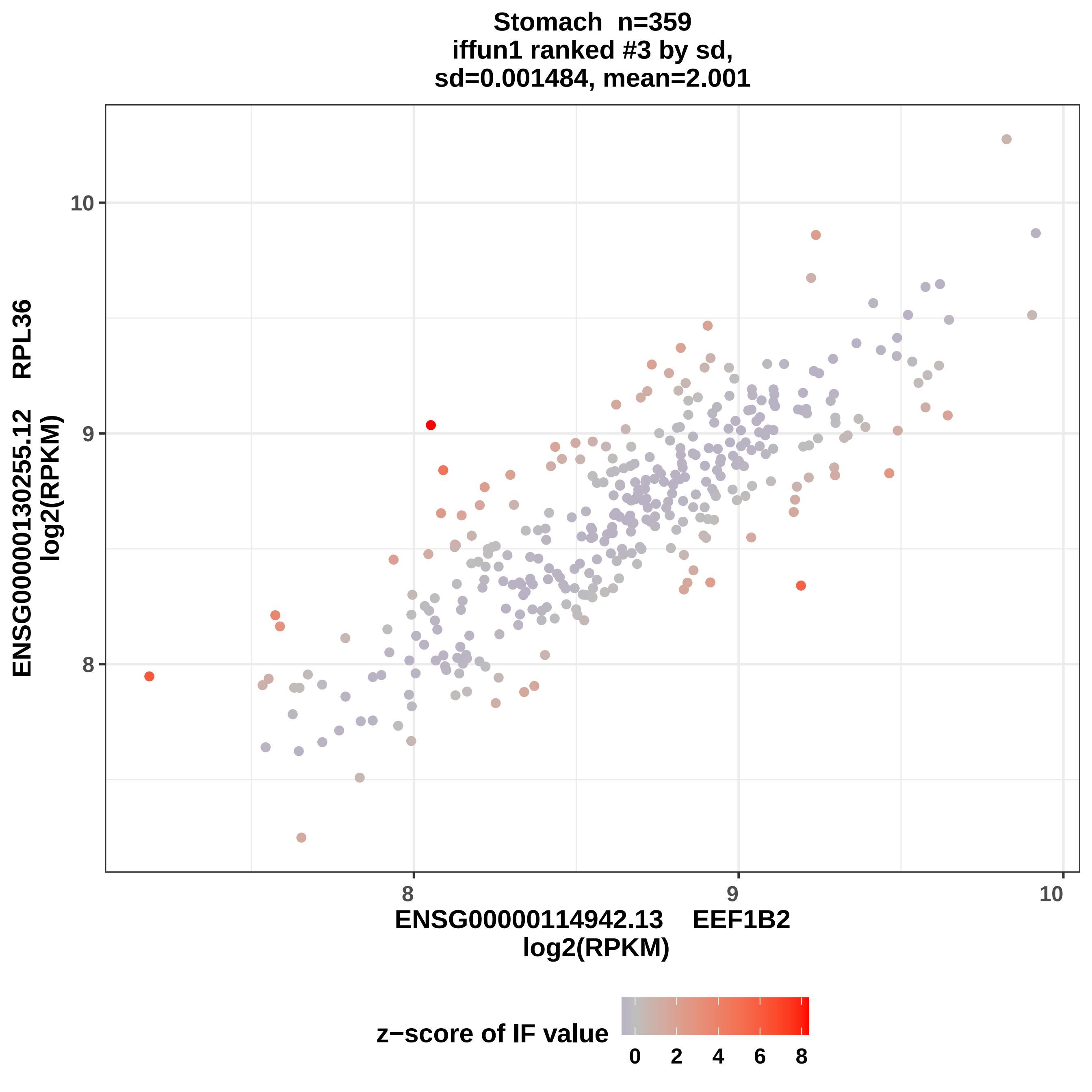

Stomach n=359
iffun1 ranked #3 by sd,
sd=0.001484, mean=2.001
ENSG00000130255.12 RPL36
log2(RPKM)
ENSG00000114942.13 EEF1B2
log2(RPKM)
10
9
8
8
9
10
z−score of IF value
0
2
4
6
8

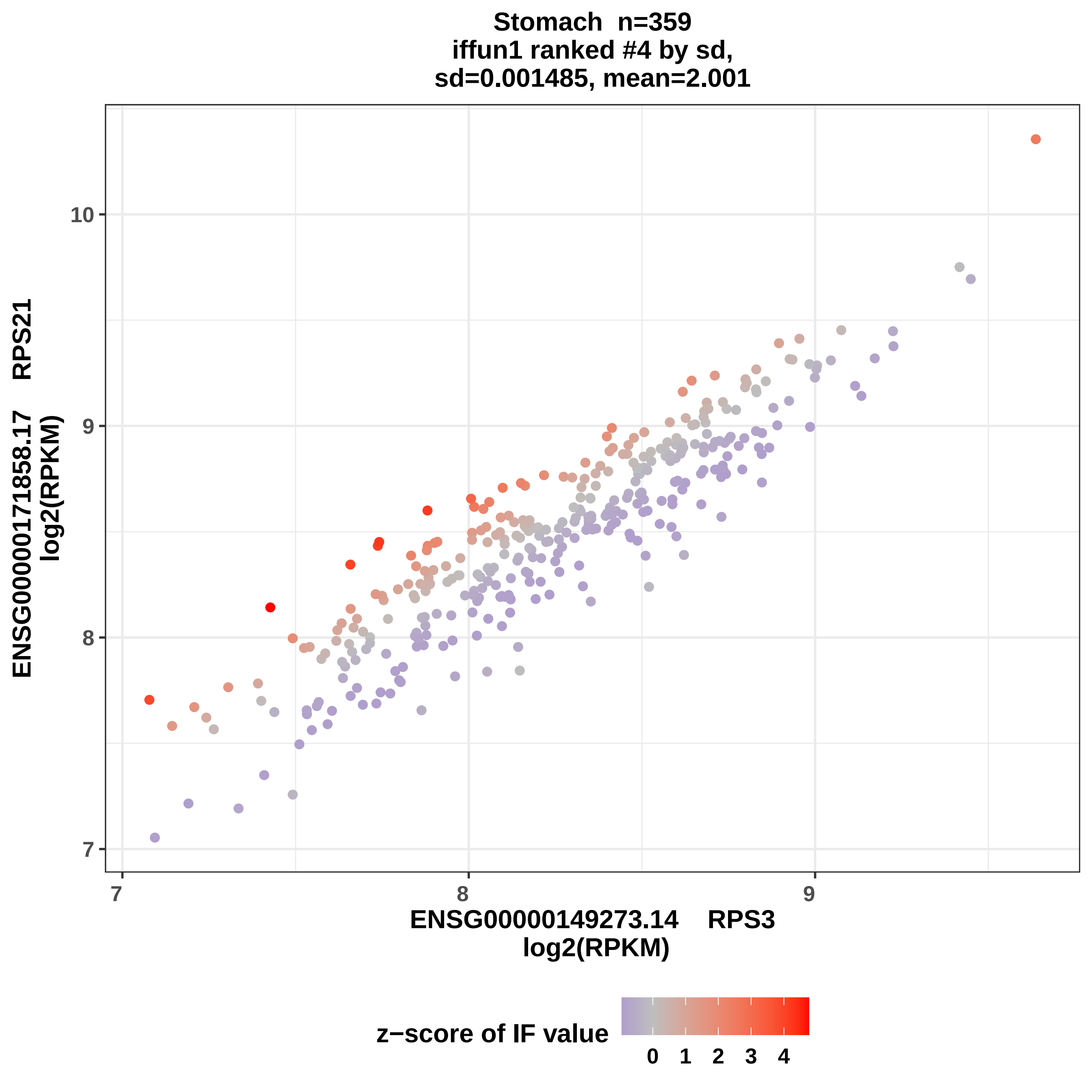

Stomach n=359
iffun1 ranked #4 by sd,
sd=0.001485, mean=2.001
ENSG00000171858.17 RPS21
log2(RPKM)
10
9
8
7
ENSG00000149273.14 RPS3
log2(RPKM)
z−score of IF value
0 1 2 3 4

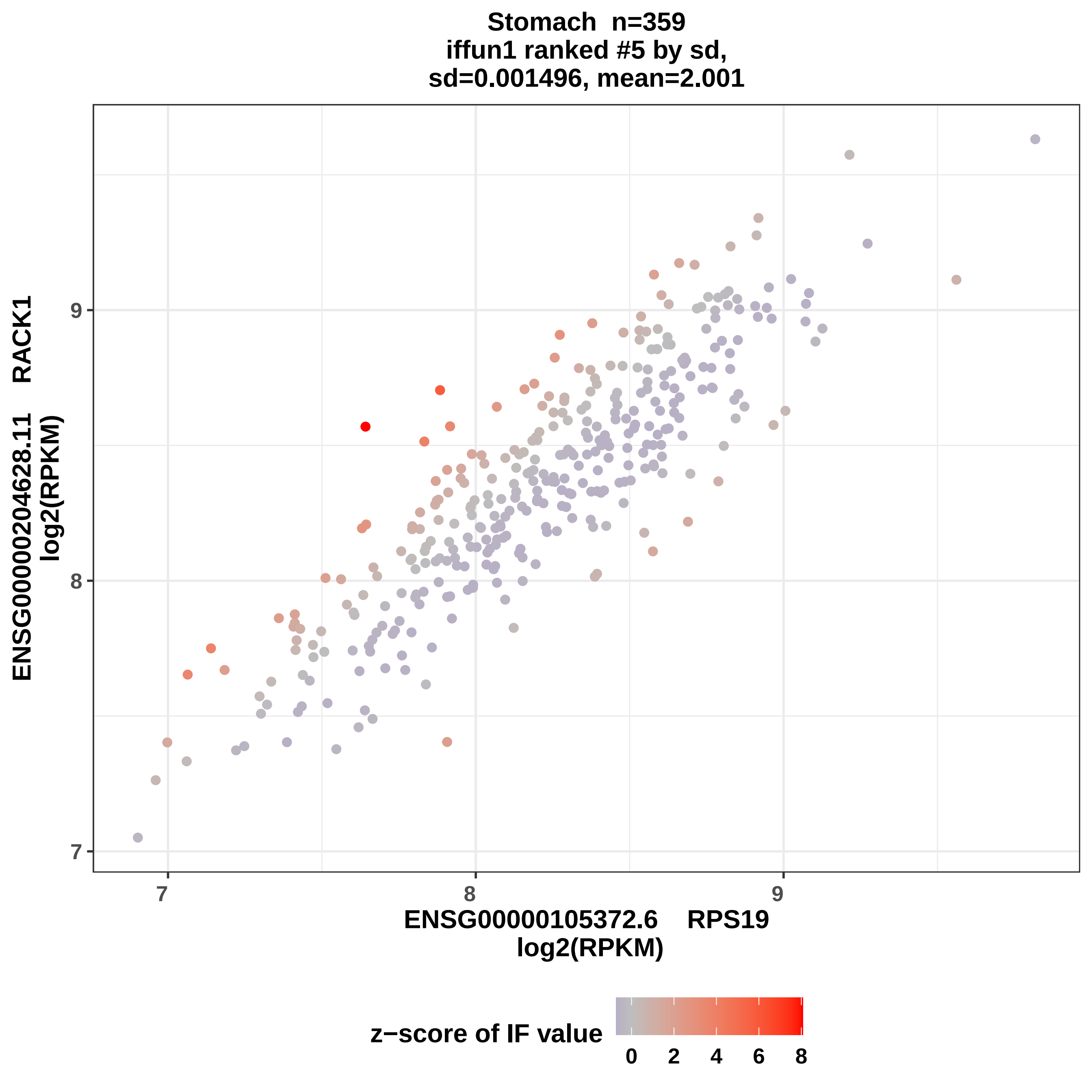

Stomach n=359
iffun1 ranked #5 by sd,
sd=0.001496, mean=2.001
ENSG00000204628.11 RACK1
log2(RPKM)
ENSG00000105372.6 RPS19
log2(RPKM)
7
8
9
z−score of IF value
0
2
4
6
8

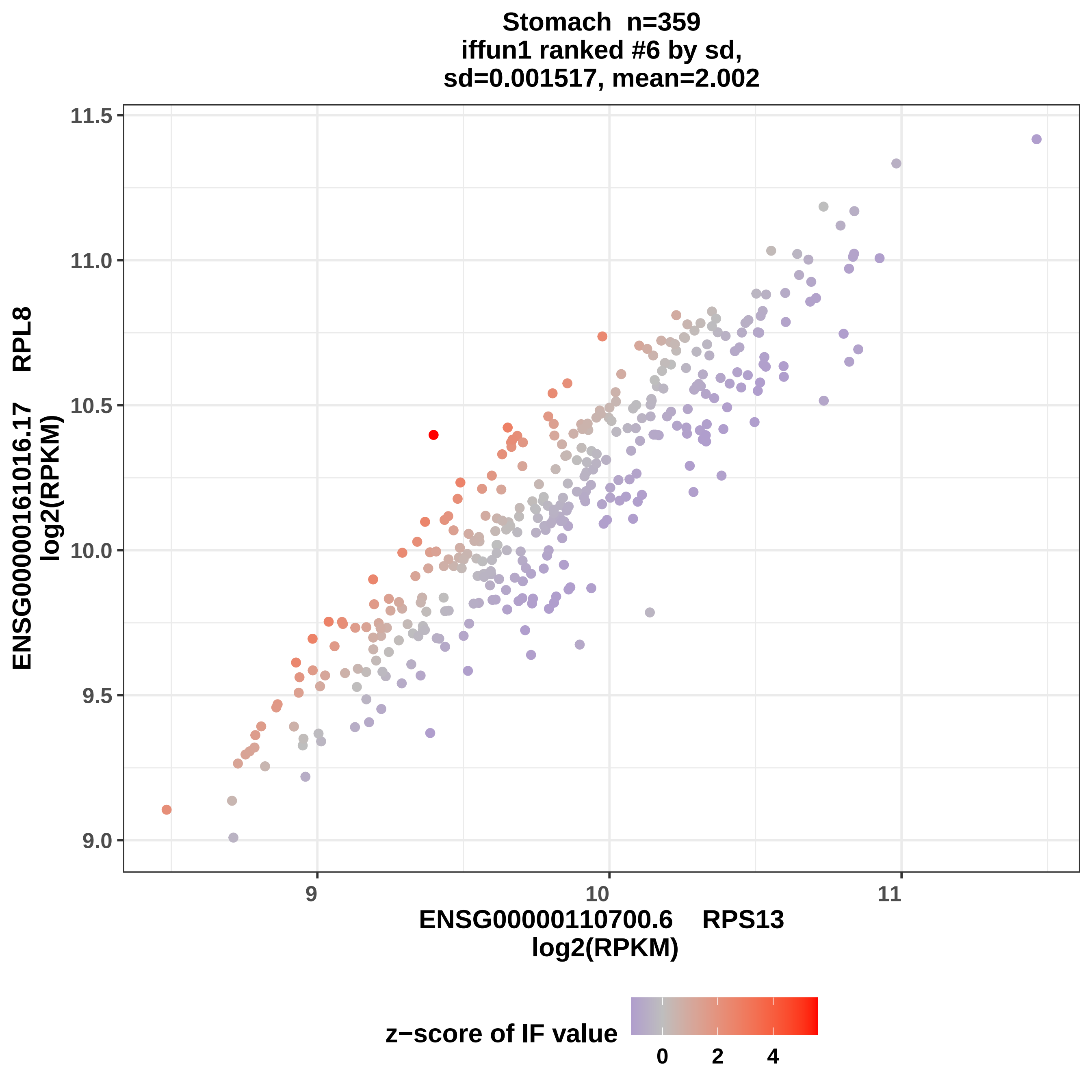

Stomach n=359
iffun1 ranked #6 by sd,
sd=0.001517, mean=2.002
11.5
11.0
10.5
10.0
9.5
9.0
ENSG00000161016.17 RPL8
log2(RPKM)
9
10
11
ENSG00000110700.6 RPS13
log2(RPKM)
z−score of IF value
0
2
4

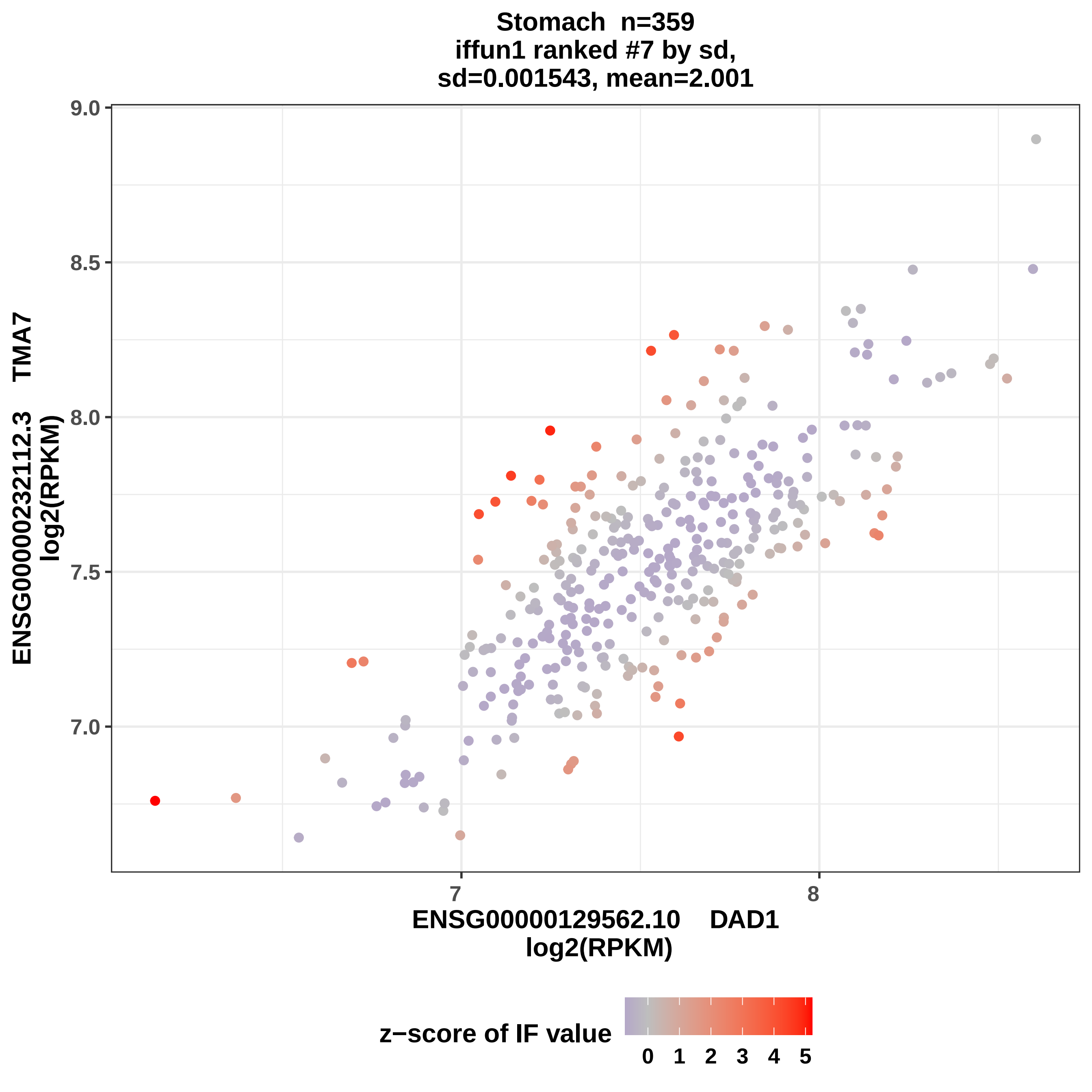
Stomach n=359
iffun1 ranked #7 by sd,
sd=0.001543, mean=2.001
9.0
8.5
8.0
7.5
7.0
ENSG00000232112.3 TMA7
log2(RPKM)
7
8
ENSG00000129562.10 DAD1
log2(RPKM)
z−score of IF value
0 1 2 3 4 5

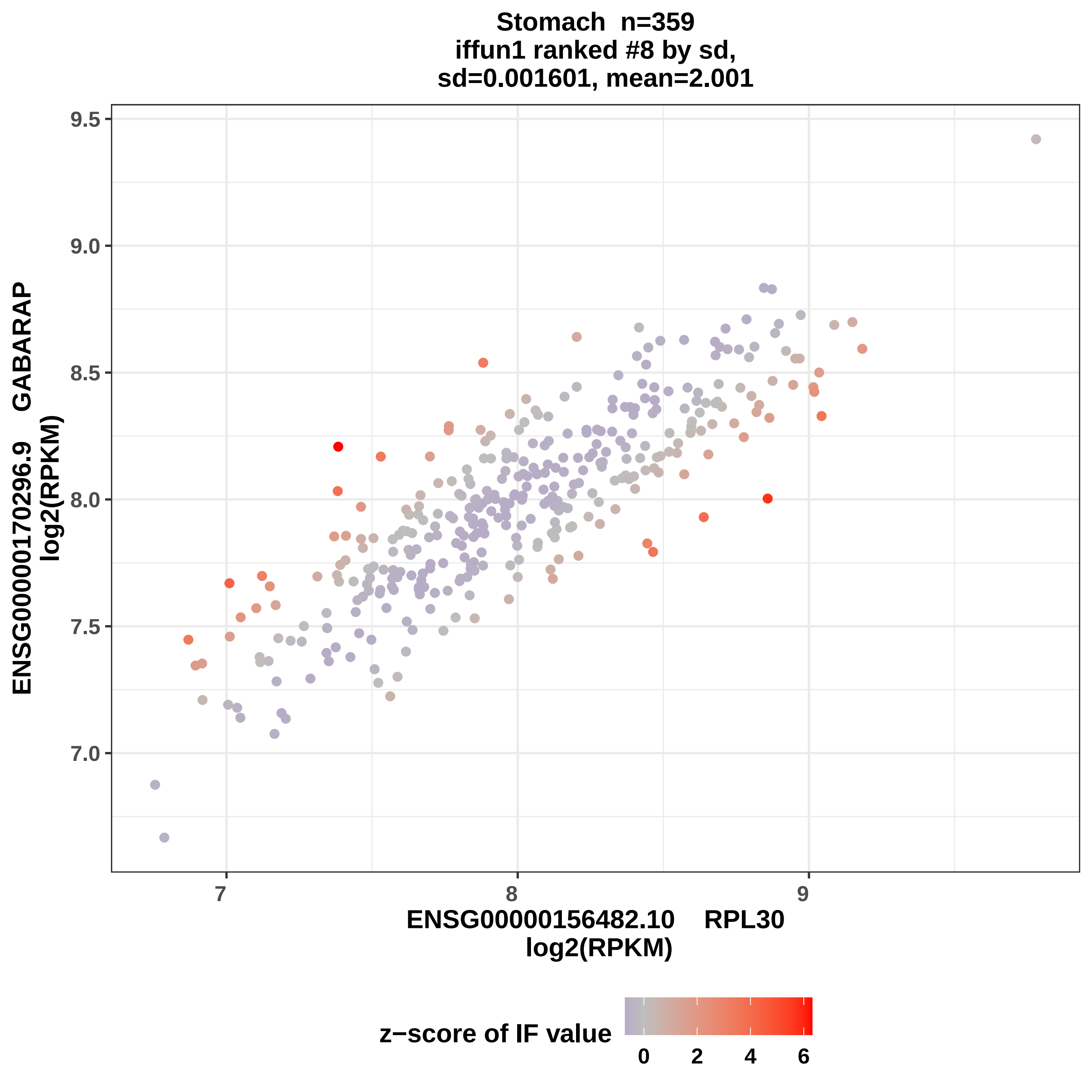
Stomach n=359
iffun1 ranked #8 by sd,
sd=0.001601, mean=2.001
ENSG00000170296.9 GABARAP
log2(RPKM)
9.5
9.0
8.5
8.0
7.5
7.0
7
8
9
ENSG00000156482.10 RPL30
log2(RPKM)
z−score of IF value
0
2
4
6

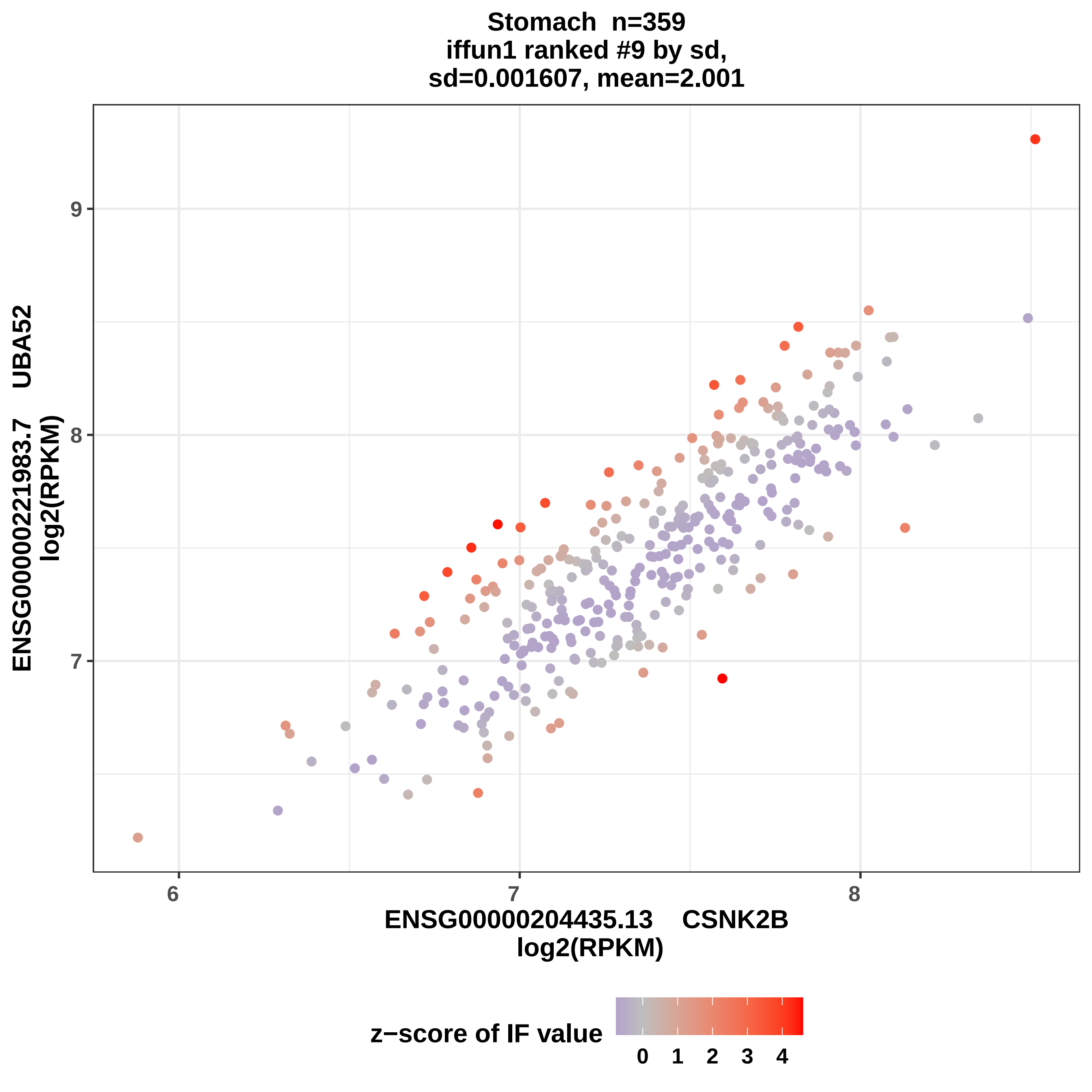
Stomach n=359
iffun1 ranked #9 by sd,
sd=0.001607, mean=2.001
9
8
7
6
7
8
ENSG00000221983.7 UBA52
log2(RPKM)
ENSG00000204435.13 CSNK2B
log2(RPKM)
z−score of IF value
0
1
2
3
4

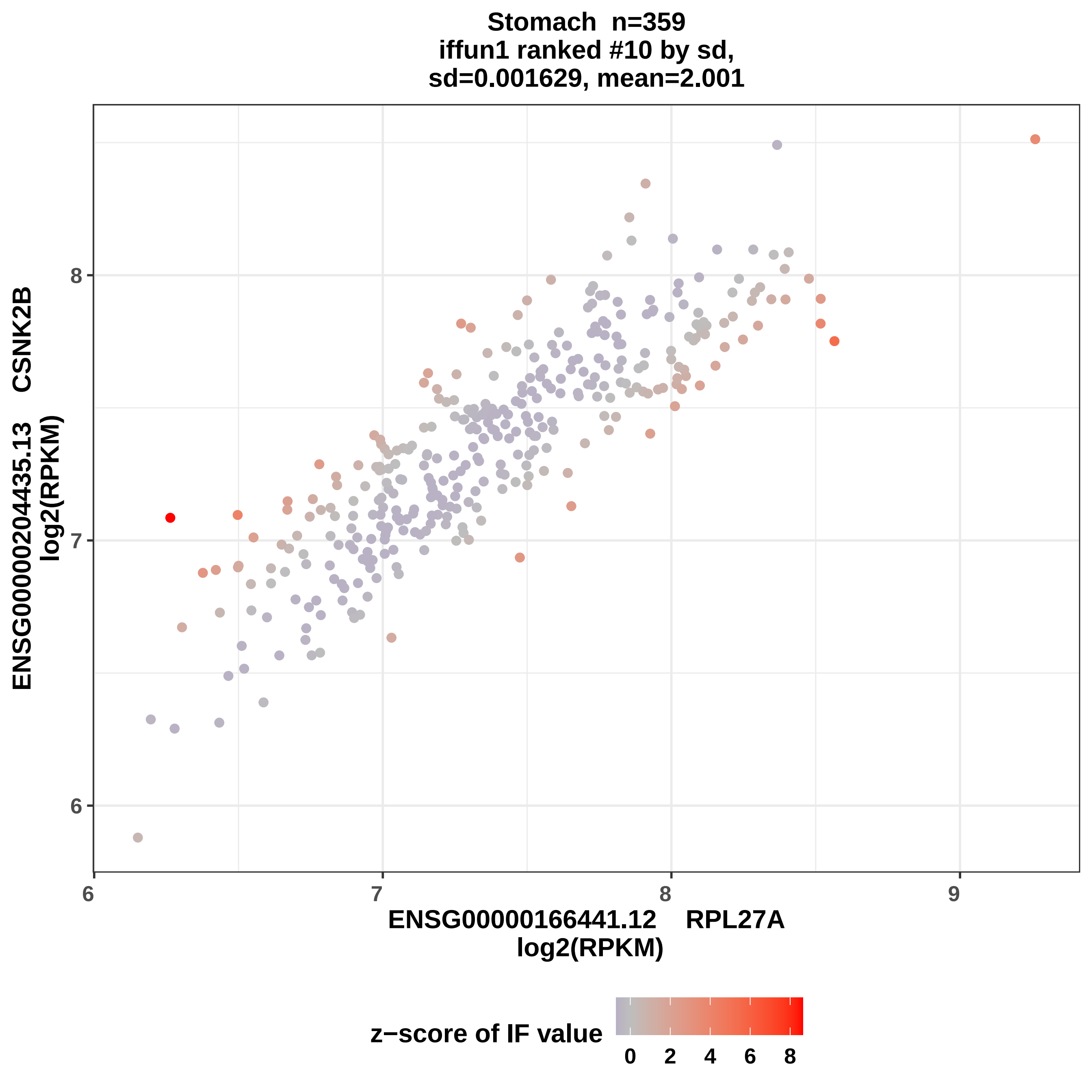

Stomach n=359
iffun1 ranked #10 by sd,
sd=0.001629, mean=2.001
ENSG00000204435.13 CSNK2B
log2(RPKM)
ENSG00000166441.12 RPL27A
log2(RPKM)
6
7
8
9
6
7
8
z−score of IF value
0
2
4
6
8

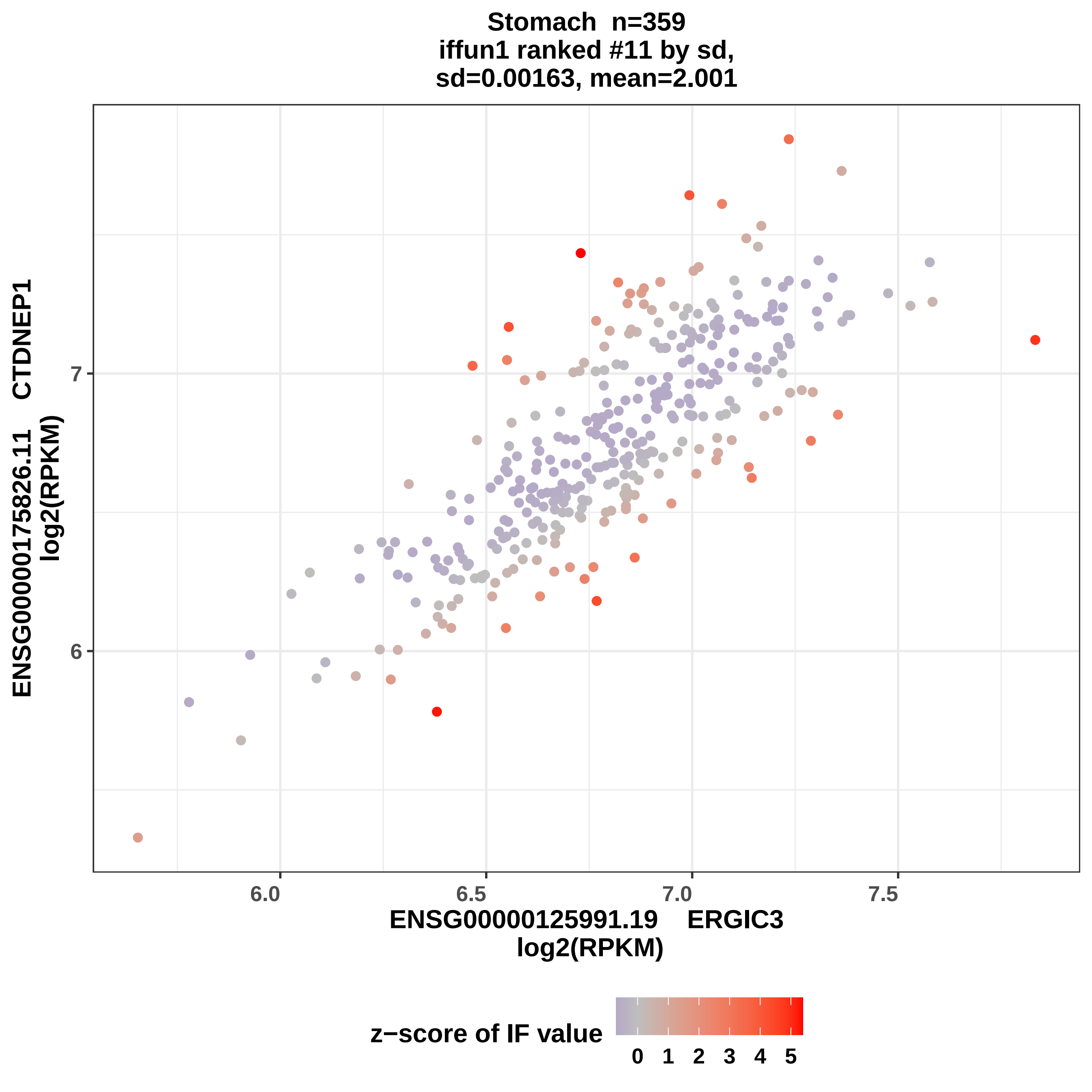
Stomach n=359
iffun1 ranked #11 by sd,
sd=0.00163, mean=2.001
ENSG00000175826.11 CTDNEP1
log2(RPKM)
ENSG00000125991.19 ERGIC3
log2(RPKM)
6.0
6.5
7.0
7.5
6
7
z−score of IF value
0
1
2
3
4
5

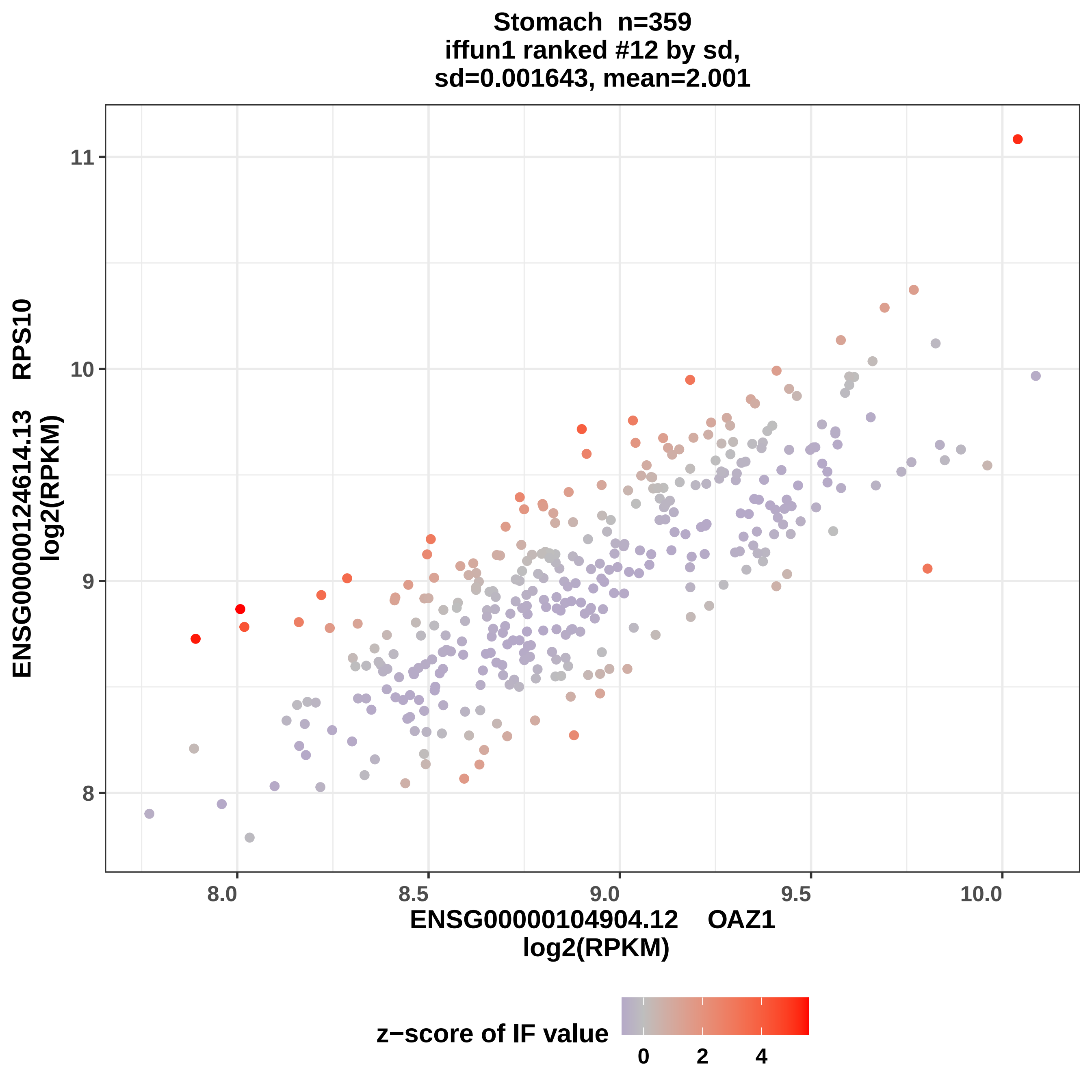

Stomach n=359
iffun1 ranked #12 by sd,
sd=0.001643, mean=2.001
ENSG00000124614.13 RPS10
log2(RPKM)
ENSG00000104904.12 OAZ1
log2(RPKM)
z−score of IF value
8
9
10
11
8.0
8.5
9.0
9.5
10.0
0
2
4

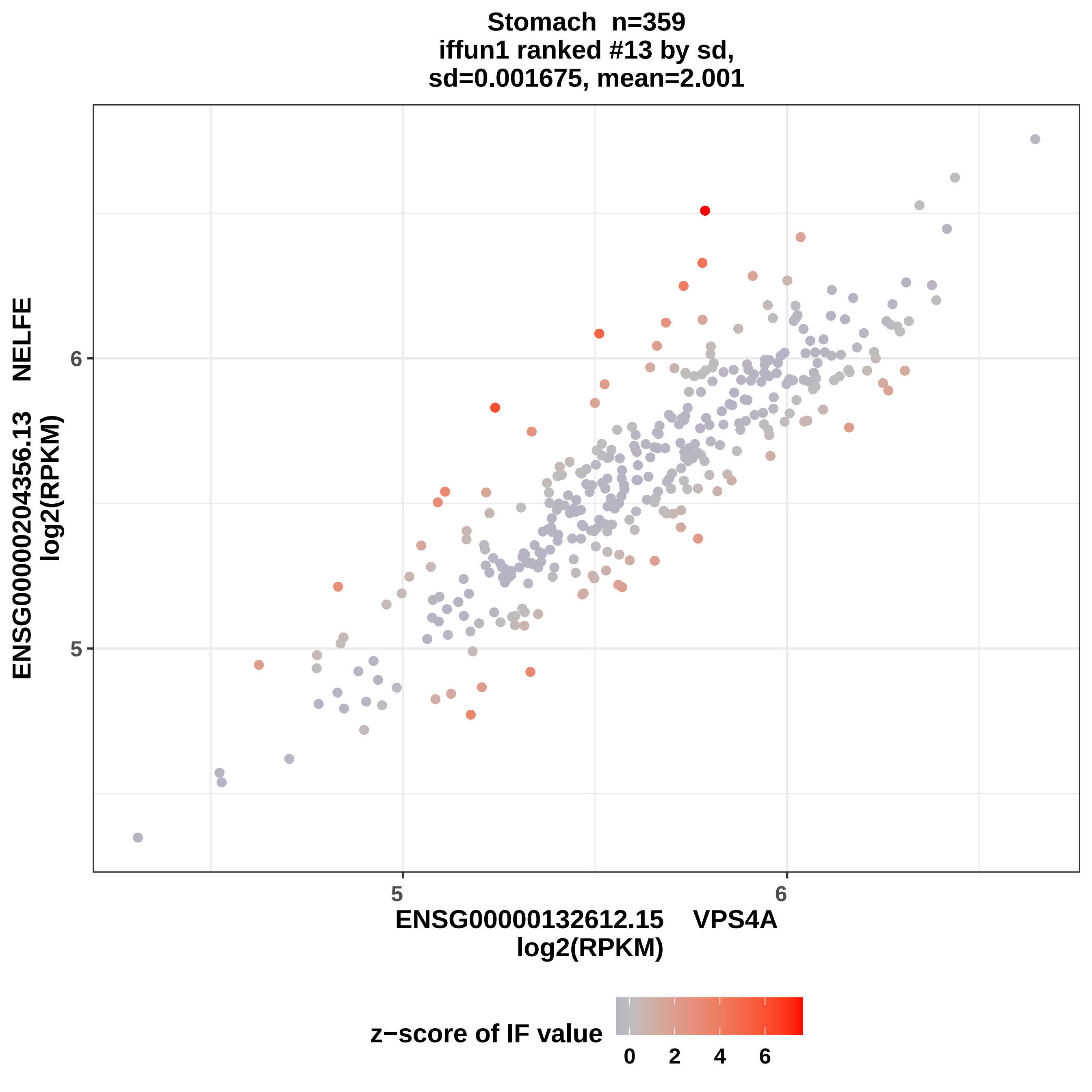

Stomach n=359
iffun1 ranked #13 by sd,
sd=0.001675, mean=2.001
6
5
ENSG00000204356.13 NELFE
log2(RPKM)
5
6
ENSG00000132612.15 VPS4A
log2(RPKM)
z−score of IF value
0
2
4
6

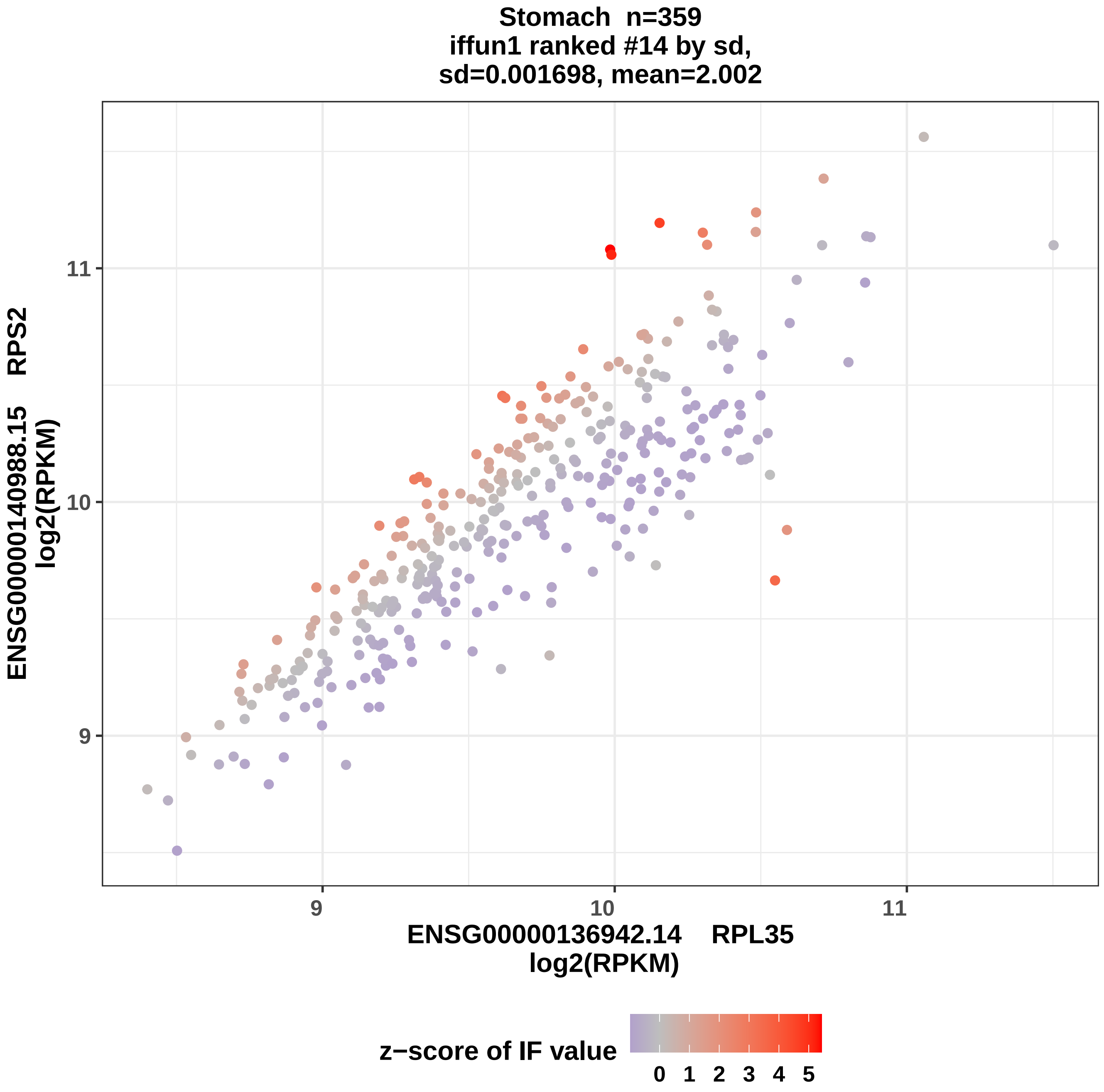

Stomach n=359
iffun1 ranked #14 by sd,
sd=0.001698, mean=2.002
ENSG00000140988.15 RPS2
log2(RPKM)
11
10
9
9
10
11
ENSG00000136942.14 RPL35
log2(RPKM)
z−score of IF value
0 1 2 3 4 5

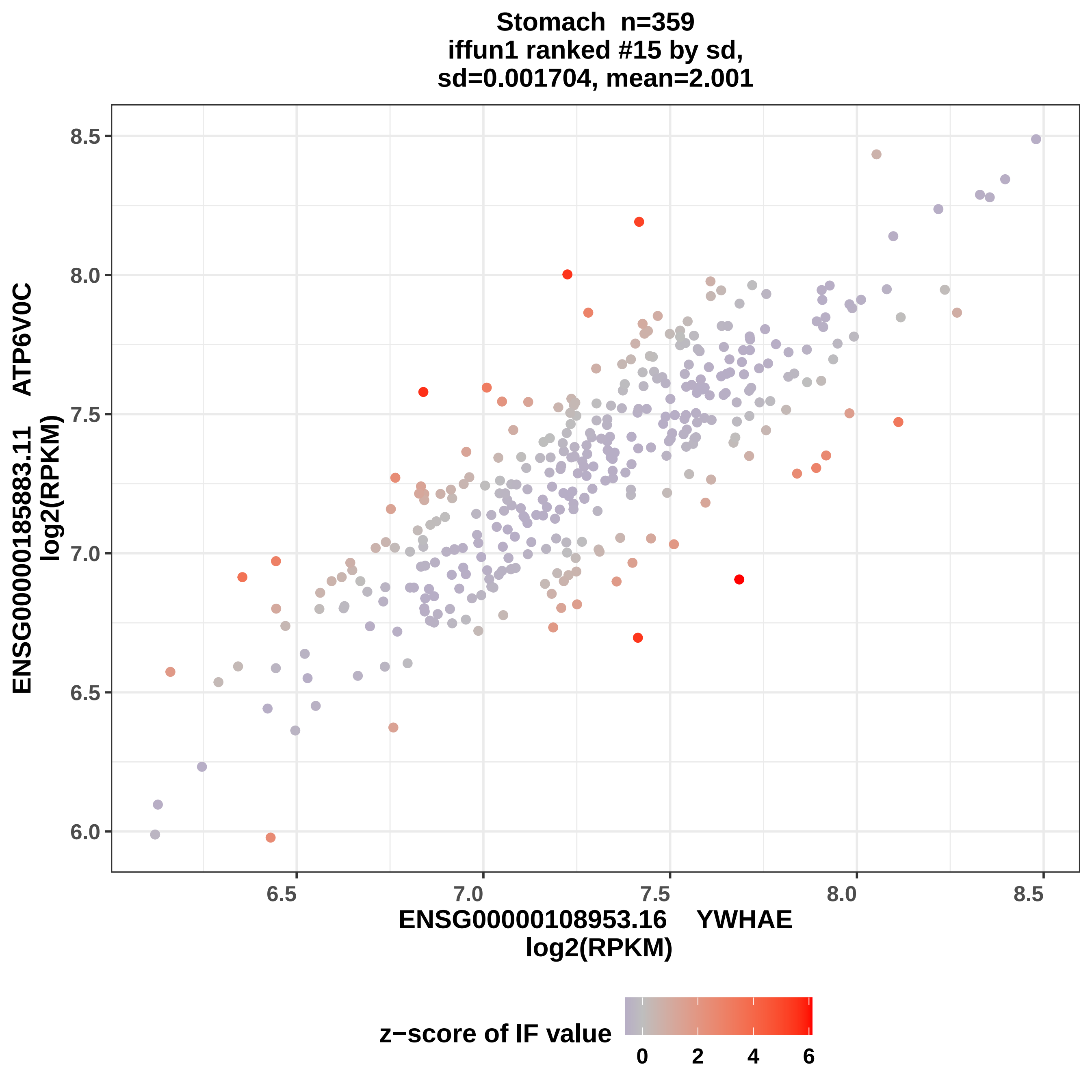
Stomach n=359
iffun1 ranked #15 by sd,
sd=0.001704, mean=2.001
ENSG00000185883.11 ATP6V0C
log2(RPKM)
6.0
6.5
7.0
7.5
8.0
8.5
ENSG00000108953.16 YWHAE
log2(RPKM)
z−score of IF value
0
2
4
6

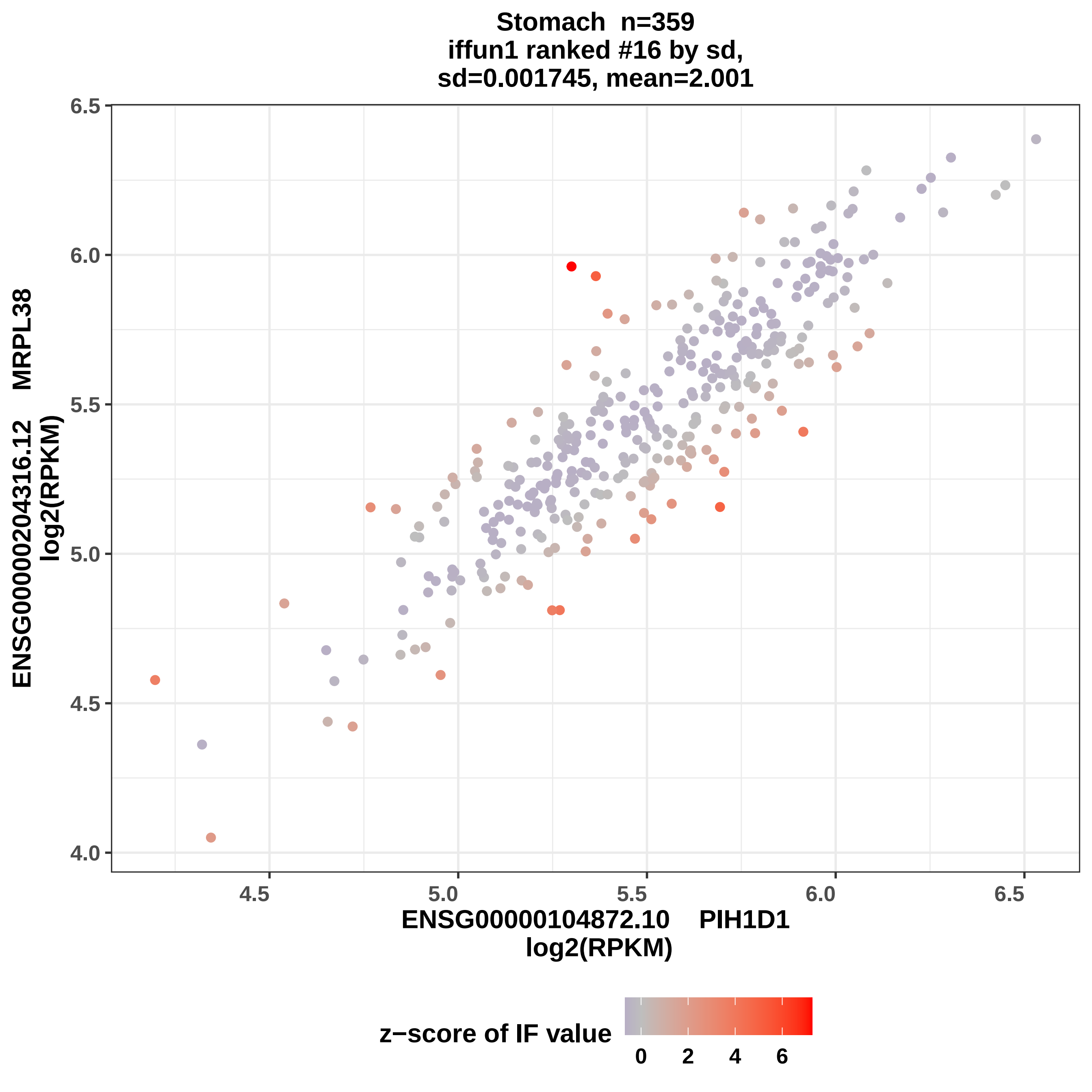

Stomach n=359
iffun1 ranked #16 by sd,
sd=0.001745, mean=2.001
ENSG00000204316.12 MRPL38
log2(RPKM)
6.5
6.0
5.5
5.0
4.5
4.0
4.5
5.0
5.5
6.0
6.5
ENSG00000104872.10 PIH1D1
log2(RPKM)
z−score of IF value
0
2
4
6

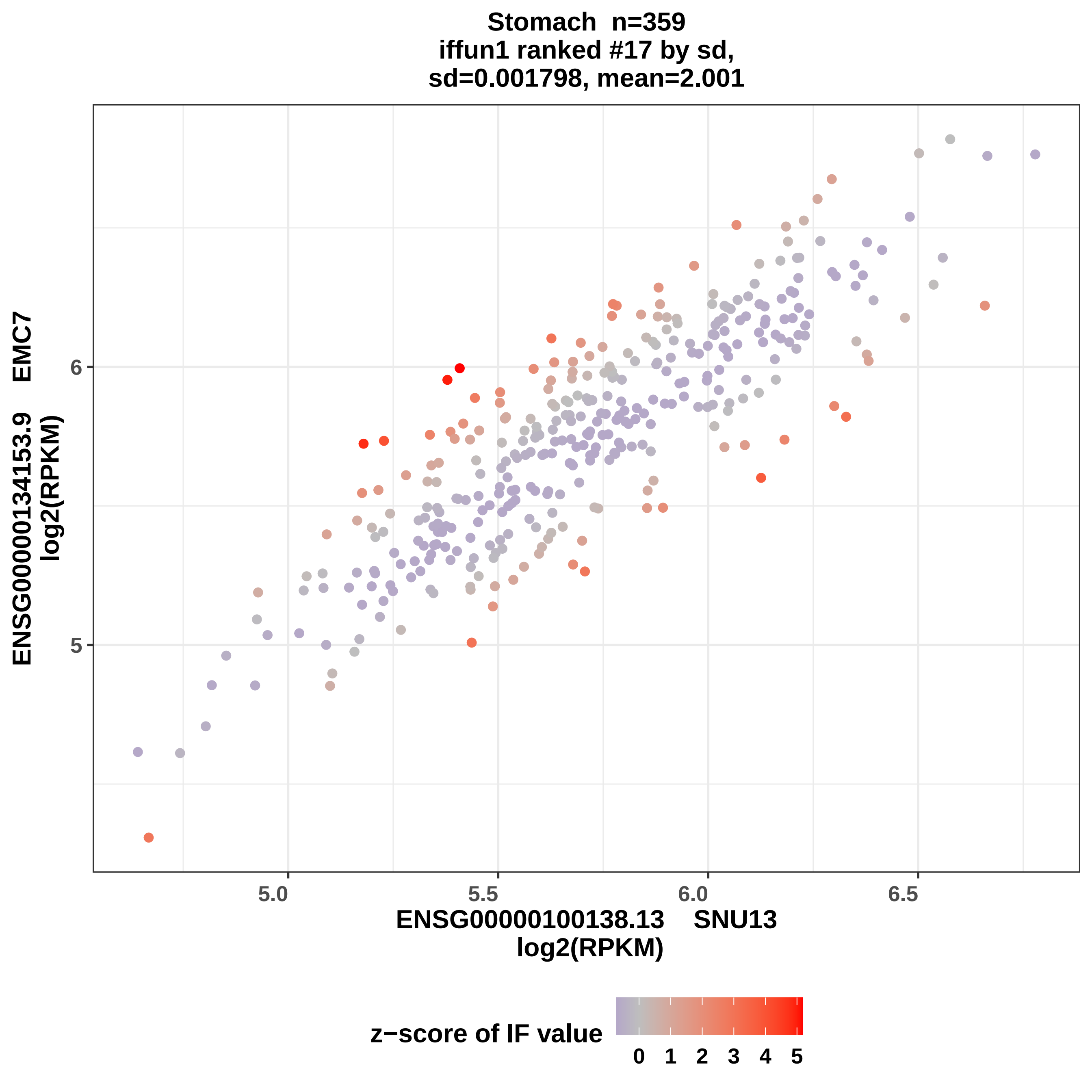
Stomach n=359
iffun1 ranked #17 by sd,
sd=0.001798, mean=2.001
ENSG00000134153.9 EMC7
log2(RPKM)
6
5
5.0
5.5
6.0
6.5
ENSG00000100138.13 SNU13
log2(RPKM)
z−score of IF value
0 1 2 3 4 5

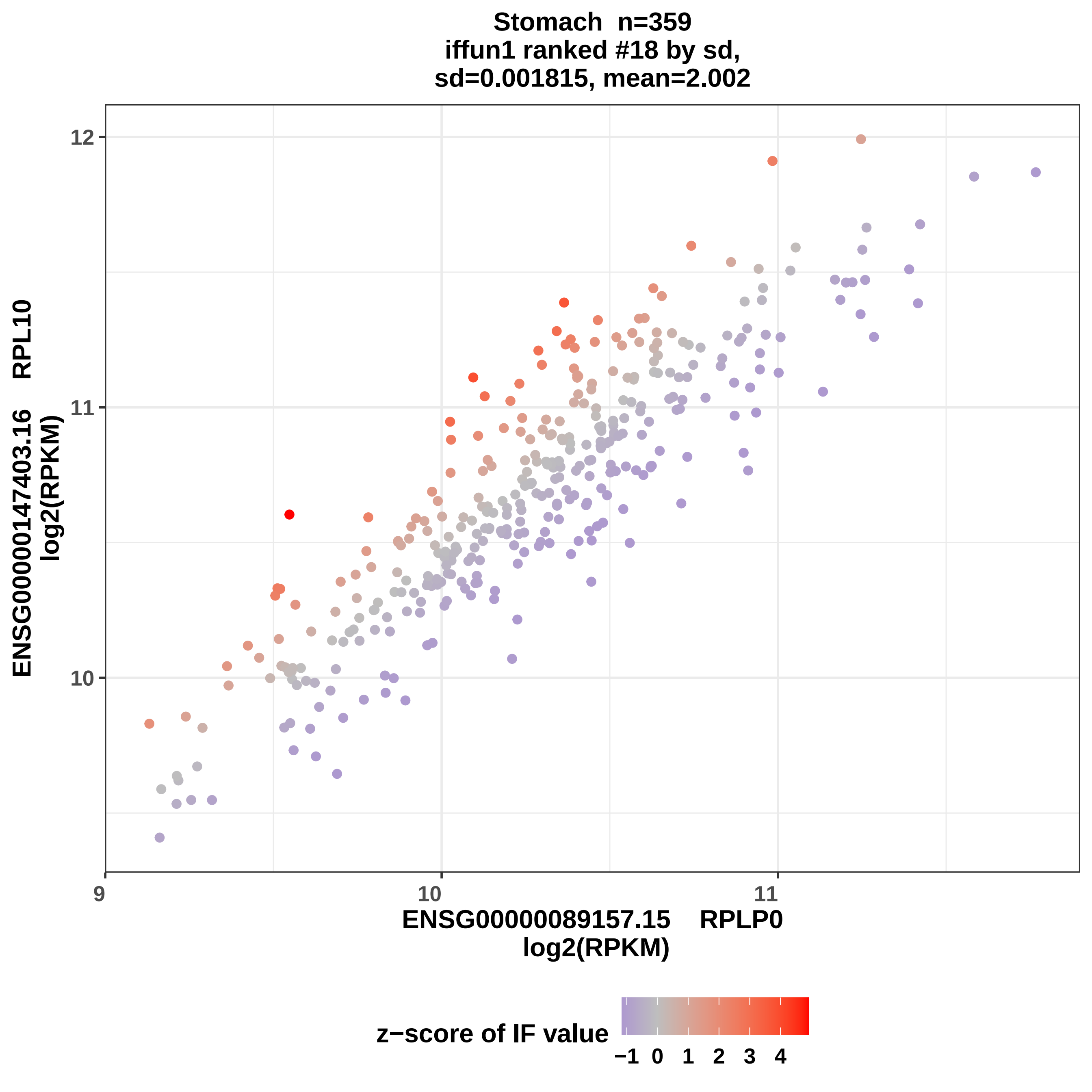
Stomach n=359
iffun1 ranked #18 by sd,
sd=0.001815, mean=2.002
ENSG00000147403.16 RPL10
log2(RPKM)
12
11
10
9
10
11
ENSG00000089157.15 RPLP0
log2(RPKM)
z−score of IF value
−1 0 1 2 3 4

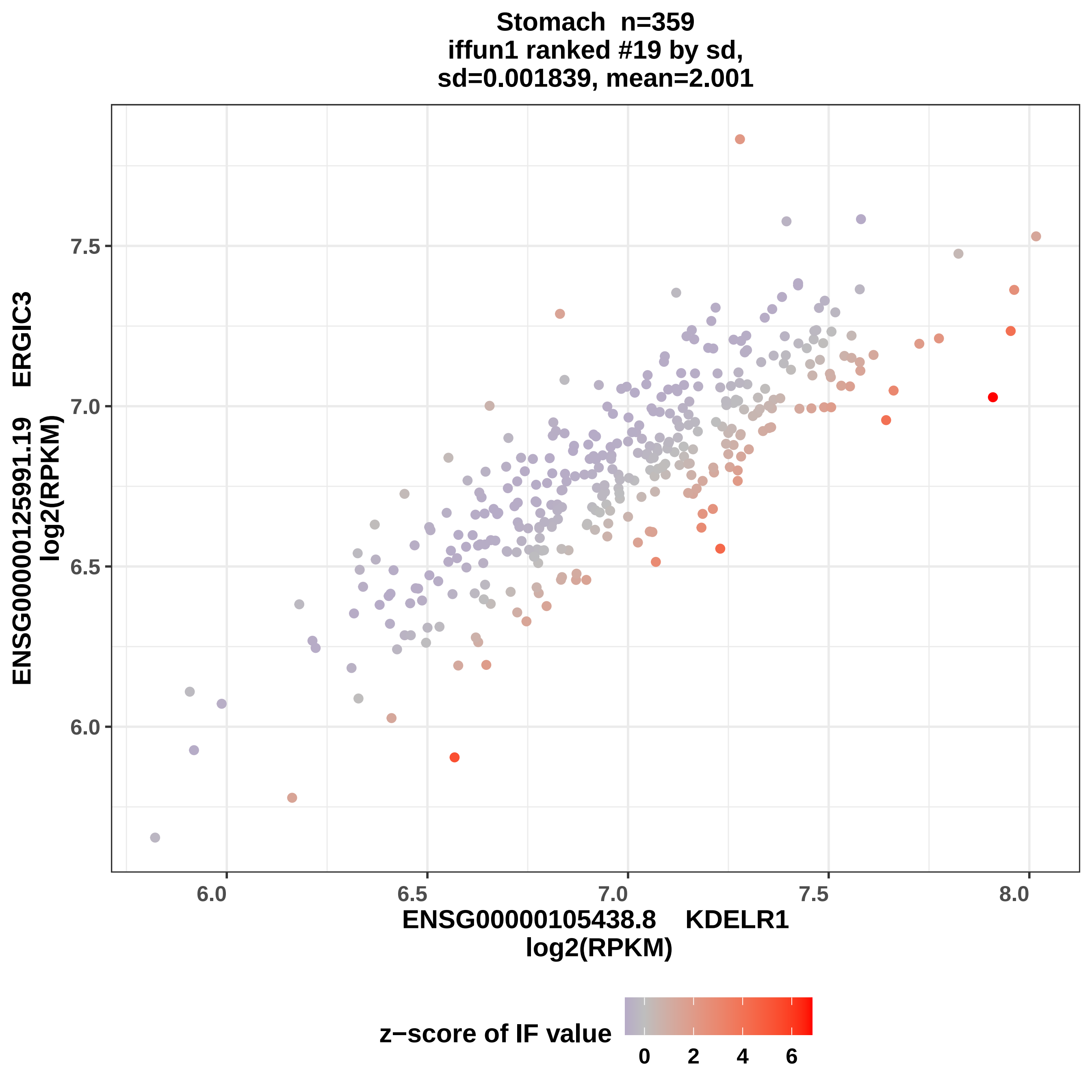
Stomach n=359
iffun1 ranked #19 by sd,
sd=0.001839, mean=2.001
ENSG00000125991.19 ERGIC3
log2(RPKM)
7.5
7.0
6.5
6.0
6.0
6.5
7.0
7.5
8.0
ENSG00000105438.8 KDELR1
log2(RPKM)
z−score of IF value
0
2
4
6

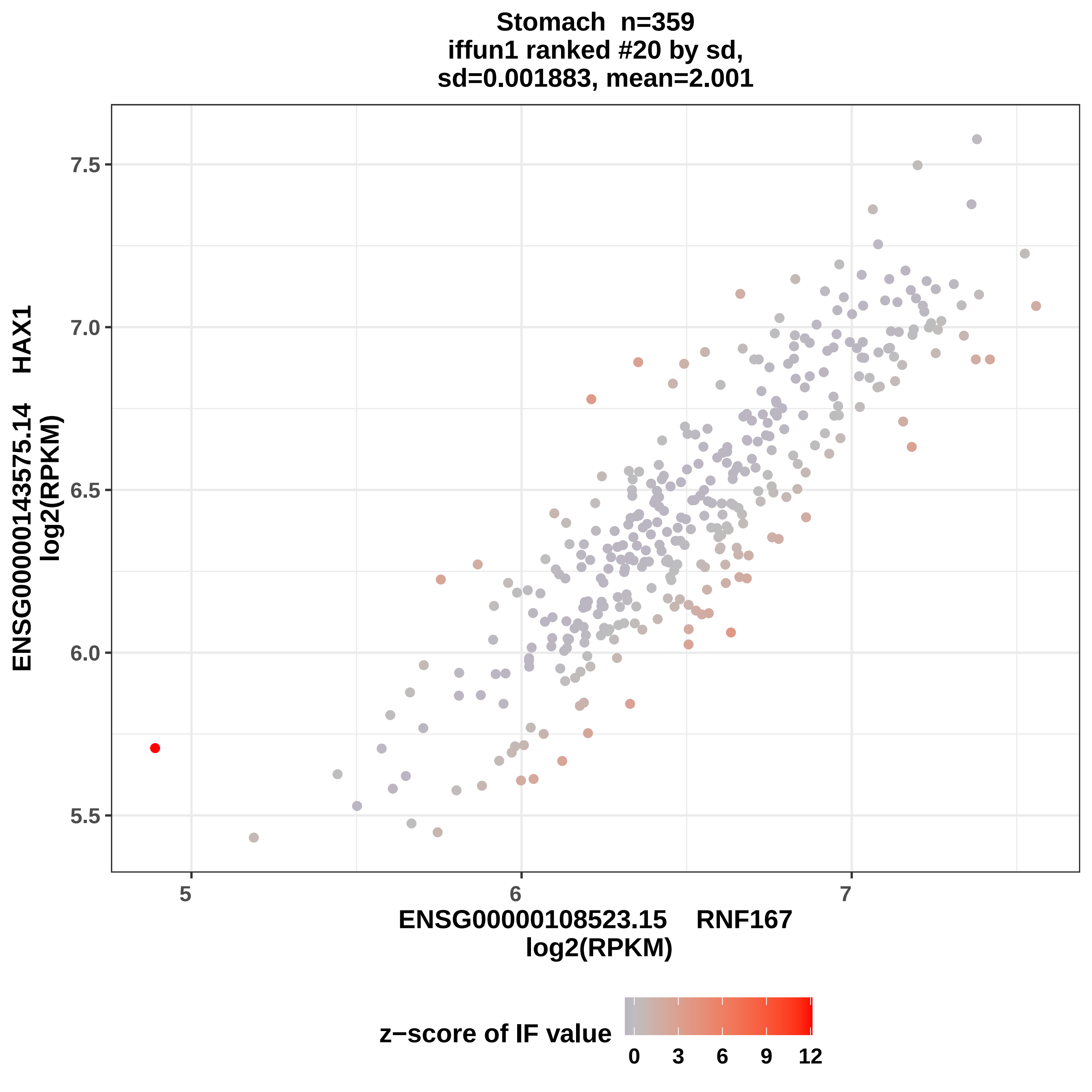
Stomach n=359
iffun1 ranked #20 by sd,
sd=0.001883, mean=2.001
ENSG00000143575.14 HAX1
log2(RPKM)
7.5
7.0
6.5
6.0
5.5
5
6
7
ENSG00000108523.15 RNF167
log2(RPKM)
z−score of IF value
0
3
6
9
12

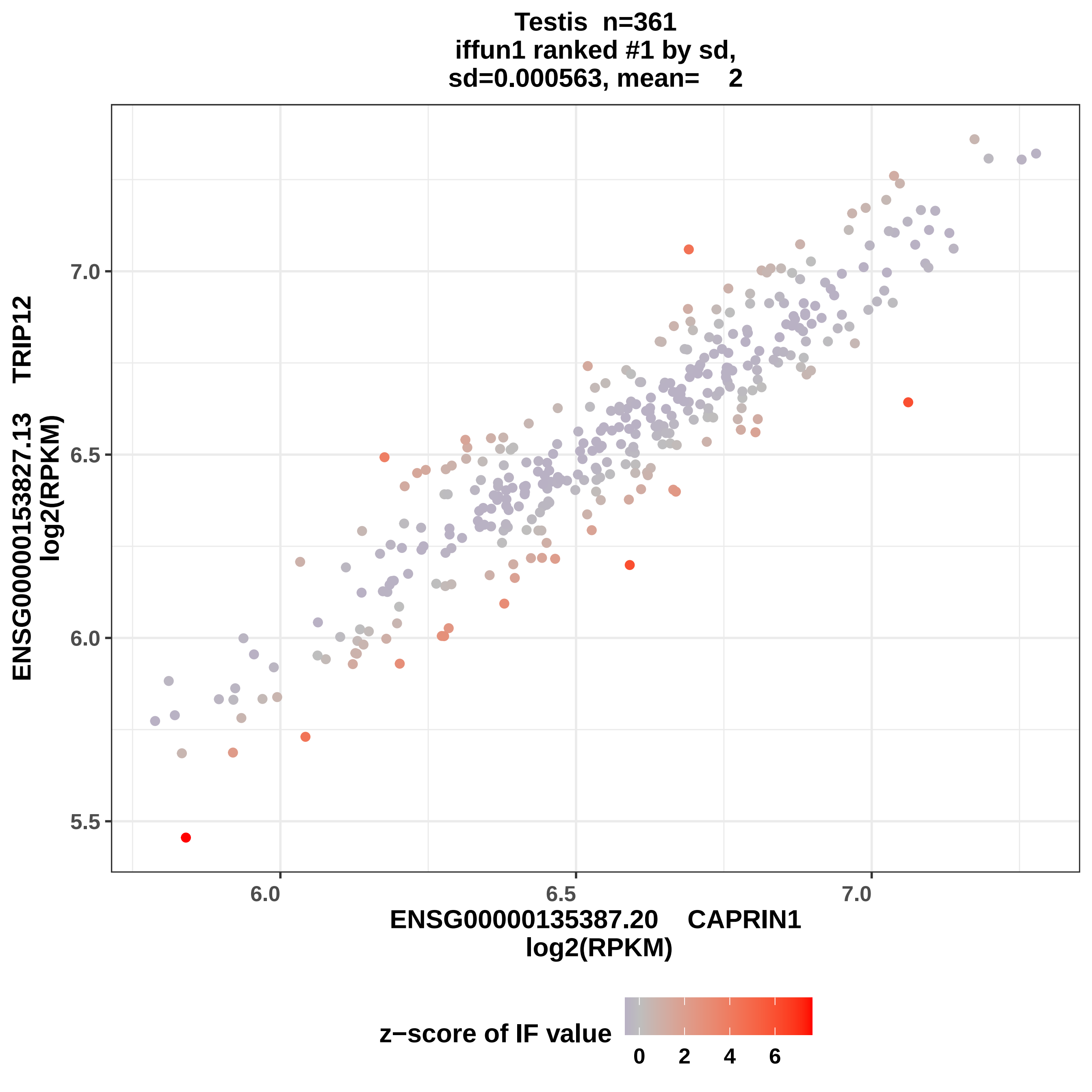

Testis n=361
iffun1 ranked #1 by sd,
sd=0.000563, mean= 2
ENSG00000153827.13 TRIP12
log2(RPKM)
7.0
6.5
6.0
5.5
6.0
6.5
7.0
ENSG00000135387.20 CAPRIN1
log2(RPKM)
z−score of IF value
0
2
4
6

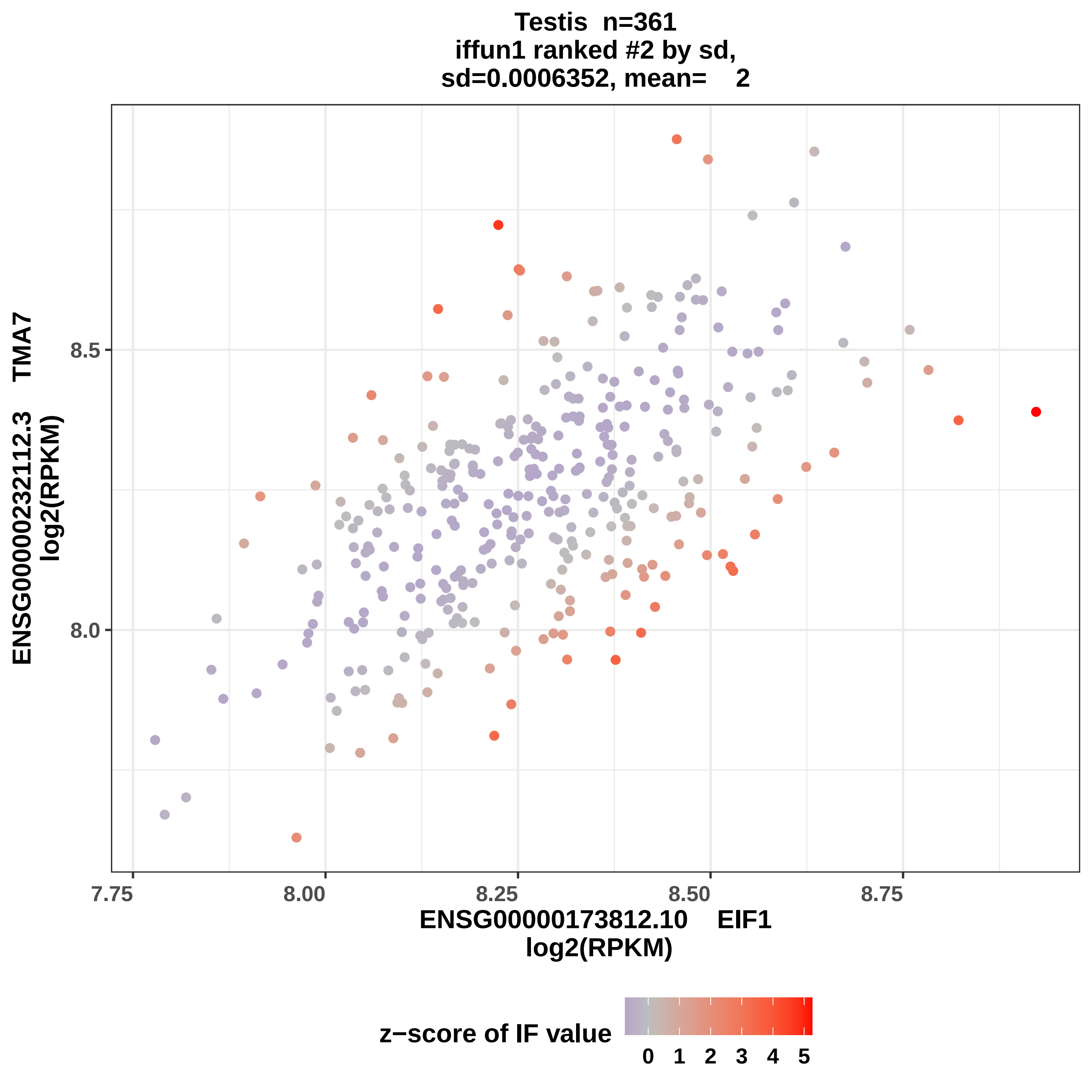

Testis n=361
iffun1 ranked #2 by sd,
sd=0.0006352, mean= 2
ENSG00000232112.3 TMA7
log2(RPKM)
8.5
8.0
7.75
8.00
8.25
8.50
8.75
ENSG00000173812.10 EIF1
log2(RPKM)
z−score of IF value
0 1 2 3 4 5

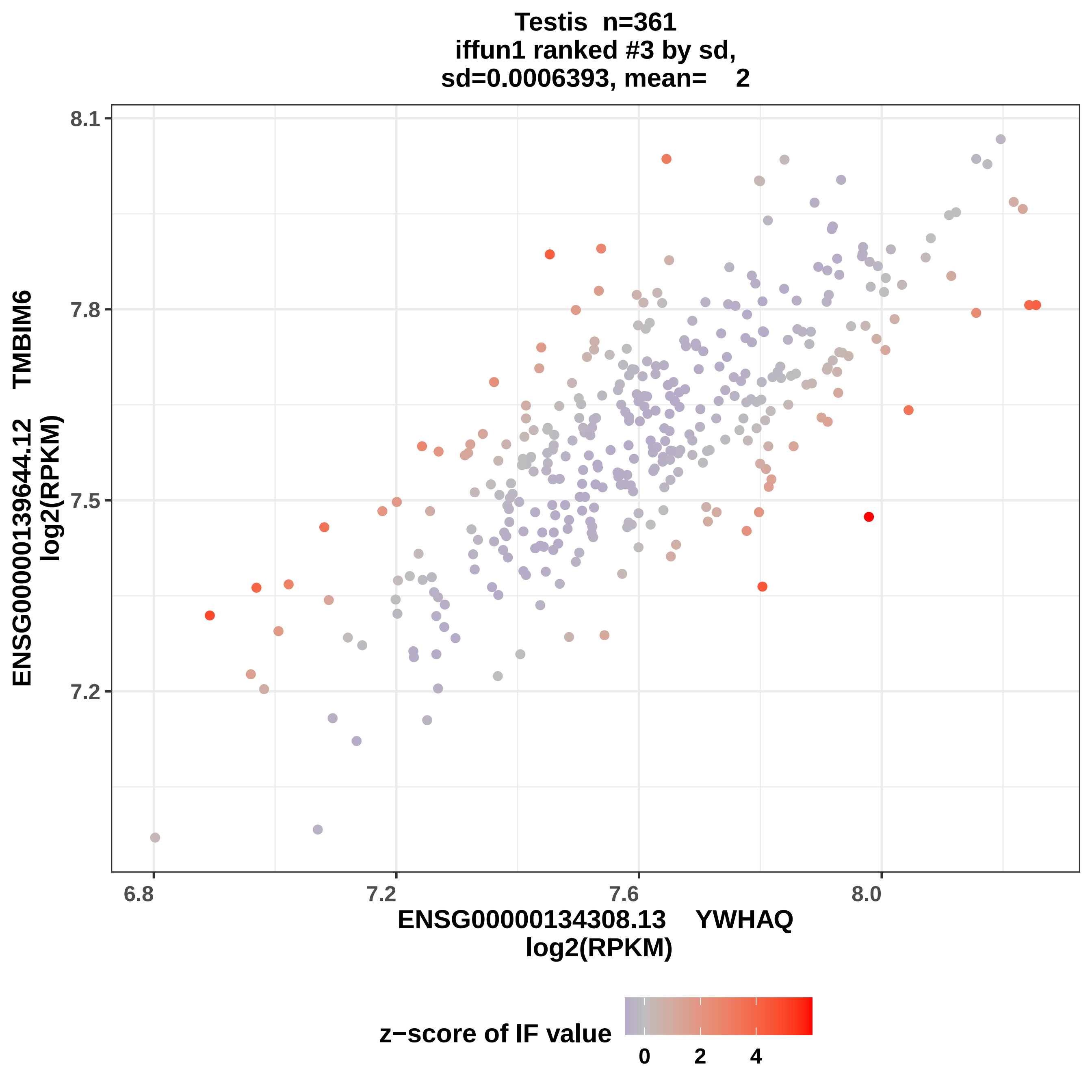
Testis n=361
iffun1 ranked #3 by sd,
sd=0.0006393, mean= 2
ENSG00000139644.12 TMBIM6
log2(RPKM)
8.1
7.8
7.5
7.2
6.8
7.2
7.6
8.0
ENSG00000134308.13 YWHAQ
log2(RPKM)
z−score of IF value
0
2
4

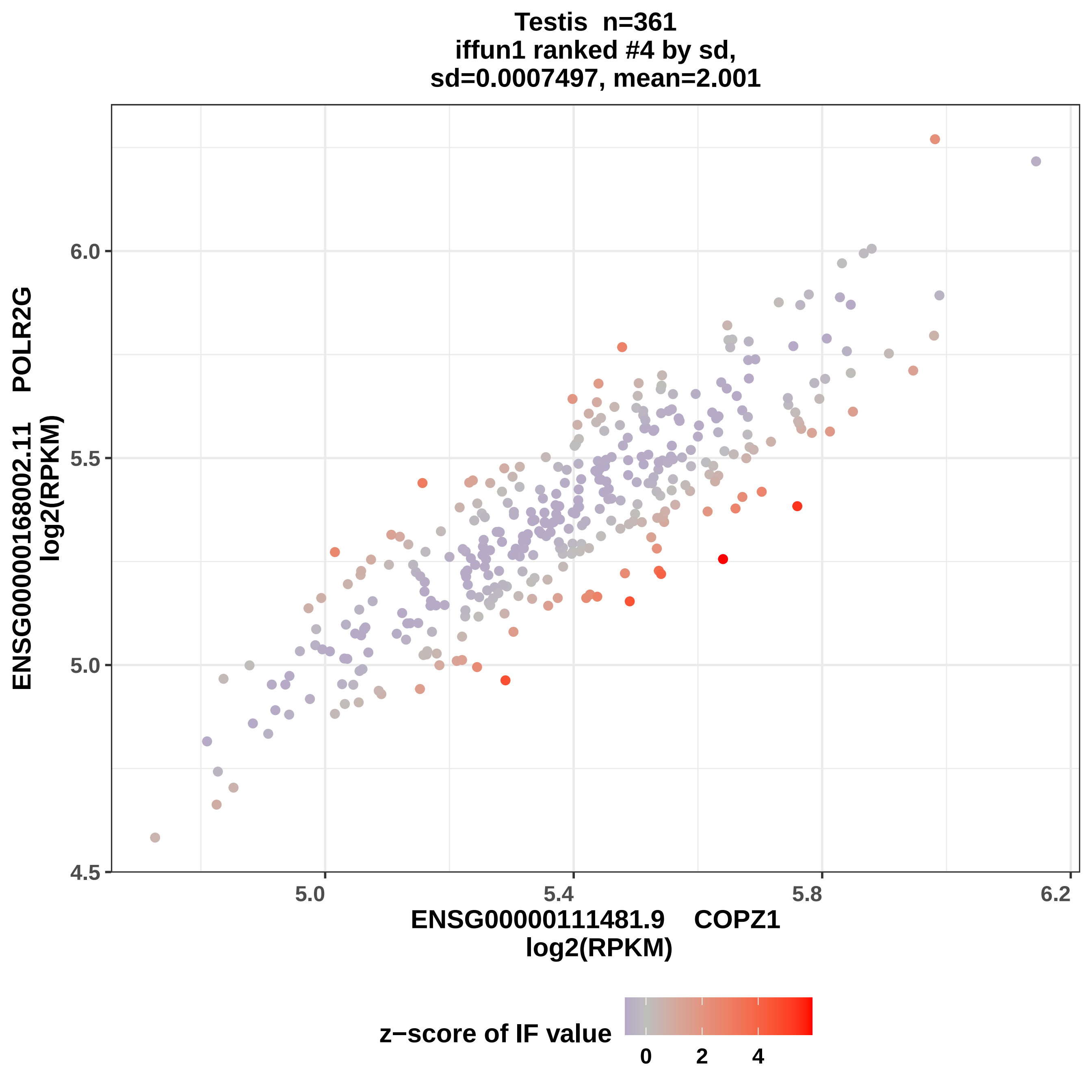

Testis n=361
iffun1 ranked #4 by sd,
sd=0.0007497, mean=2.001
ENSG00000168002.11 POLR2G
log2(RPKM)
6.0
5.5
5.0
4.5
5.0
5.4
5.8
6.2
ENSG00000111481.9 COPZ1
log2(RPKM)
z−score of IF value
0
2
4

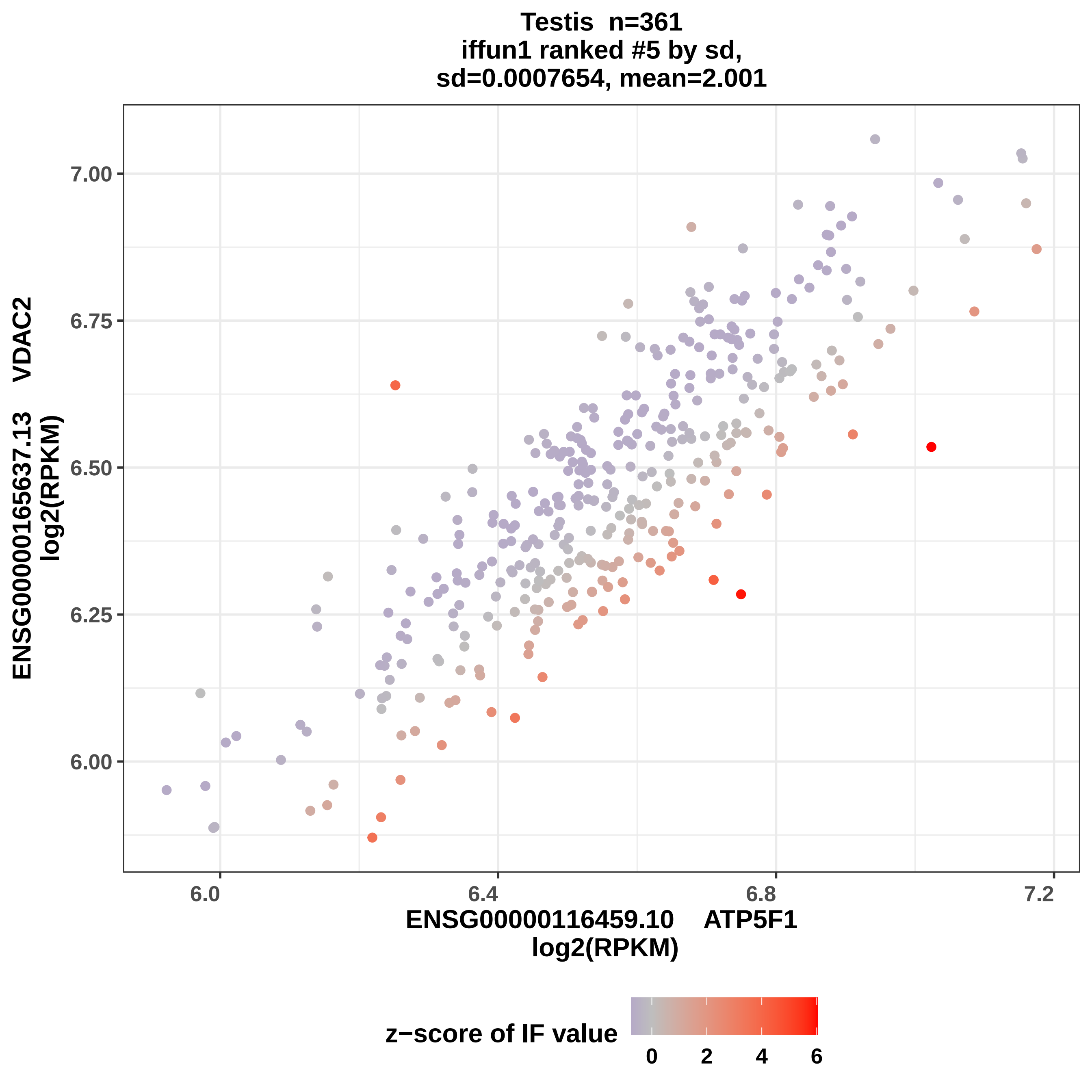
Testis n=361
iffun1 ranked #5 by sd,
sd=0.0007654, mean=2.001
ENSG00000165637.13 VDAC2
log2(RPKM)
7.00
6.75
6.50
6.25
6.00
6.0
6.4
6.8
7.2
ENSG00000116459.10 ATP5F1
log2(RPKM)
z−score of IF value
0
2
4
6

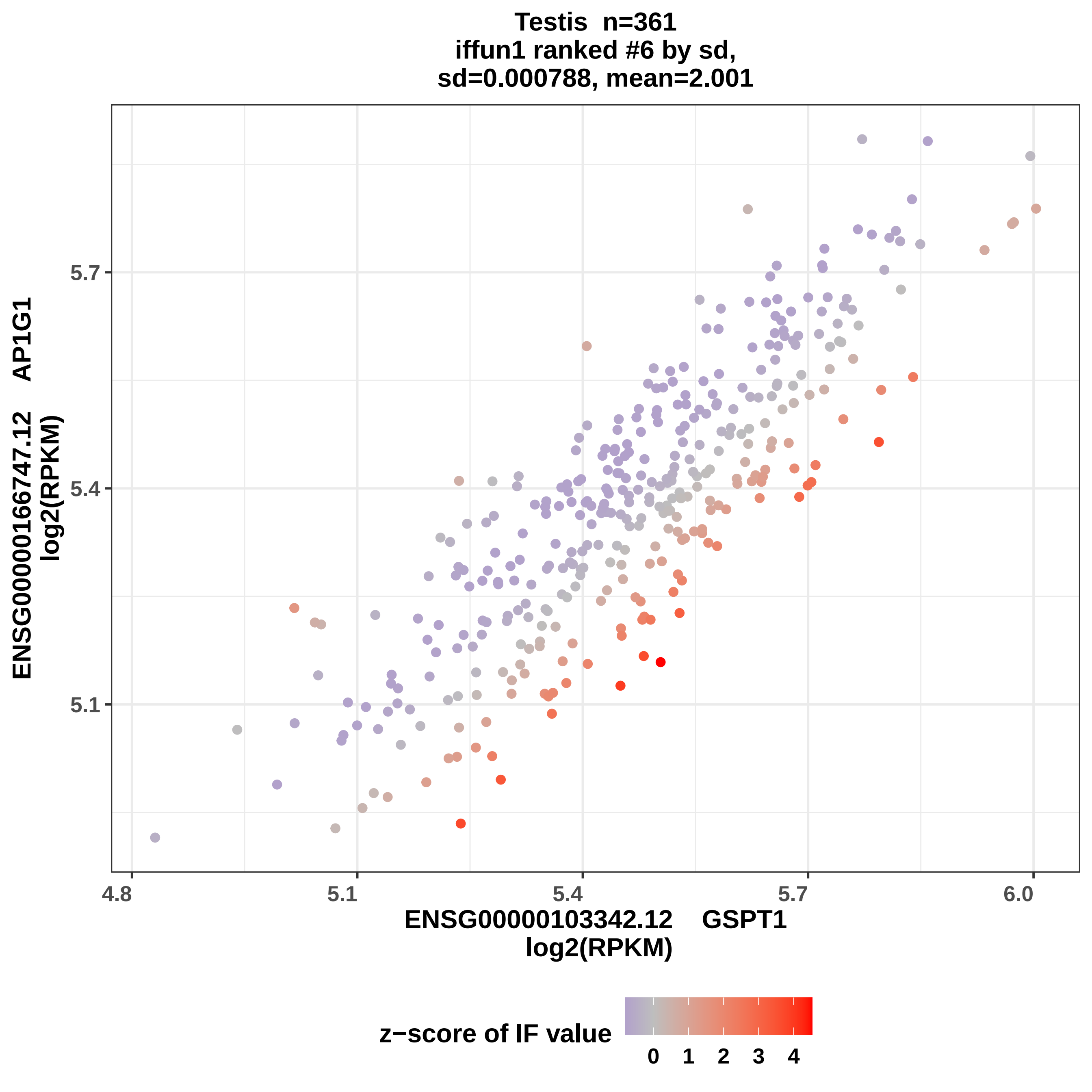
Testis n=361
iffun1 ranked #6 by sd,
sd=0.000788, mean=2.001
ENSG00000166747.12 AP1G1
log2(RPKM)
5.1
5.4
5.7
4.8
5.1
5.4
5.7
6.0
ENSG00000103342.12 GSPT1
log2(RPKM)
z−score of IF value
0
1
2
3
4

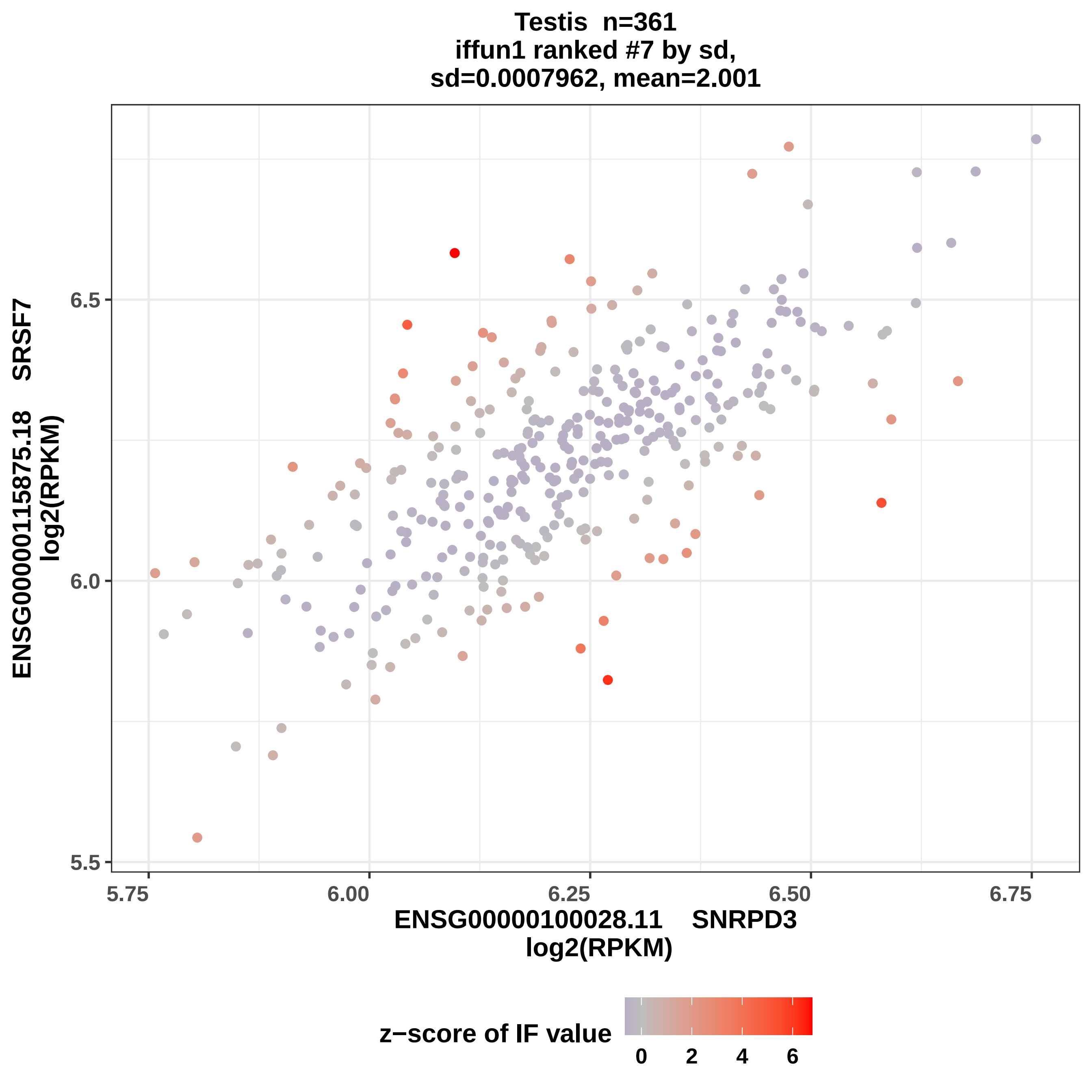
Testis n=361
iffun1 ranked #7 by sd,
sd=0.0007962, mean=2.001
ENSG00000115875.18 SRSF7
log2(RPKM)
6.5
6.0
5.5
5.75
6.00
6.25
6.50
6.75
ENSG00000100028.11 SNRPD3
log2(RPKM)
z−score of IF value
0
2
4
6

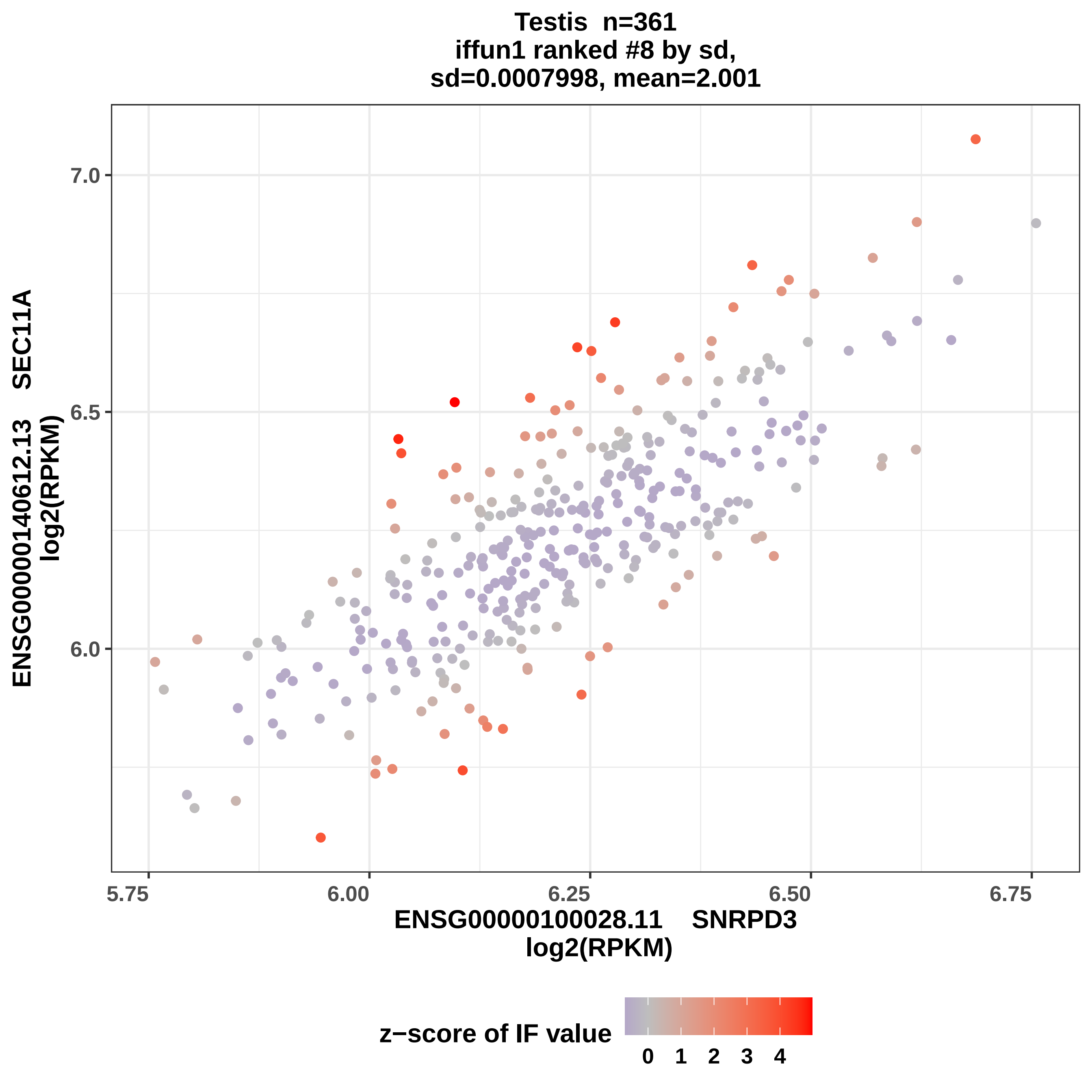

Testis n=361
iffun1 ranked #8 by sd,
sd=0.0007998, mean=2.001
ENSG00000140612.13 SEC11A
log2(RPKM)
7.0
6.5
6.0
5.75
6.00
6.25
6.50
6.75
ENSG00000100028.11 SNRPD3
log2(RPKM)
z−score of IF value
0
1
2
3
4

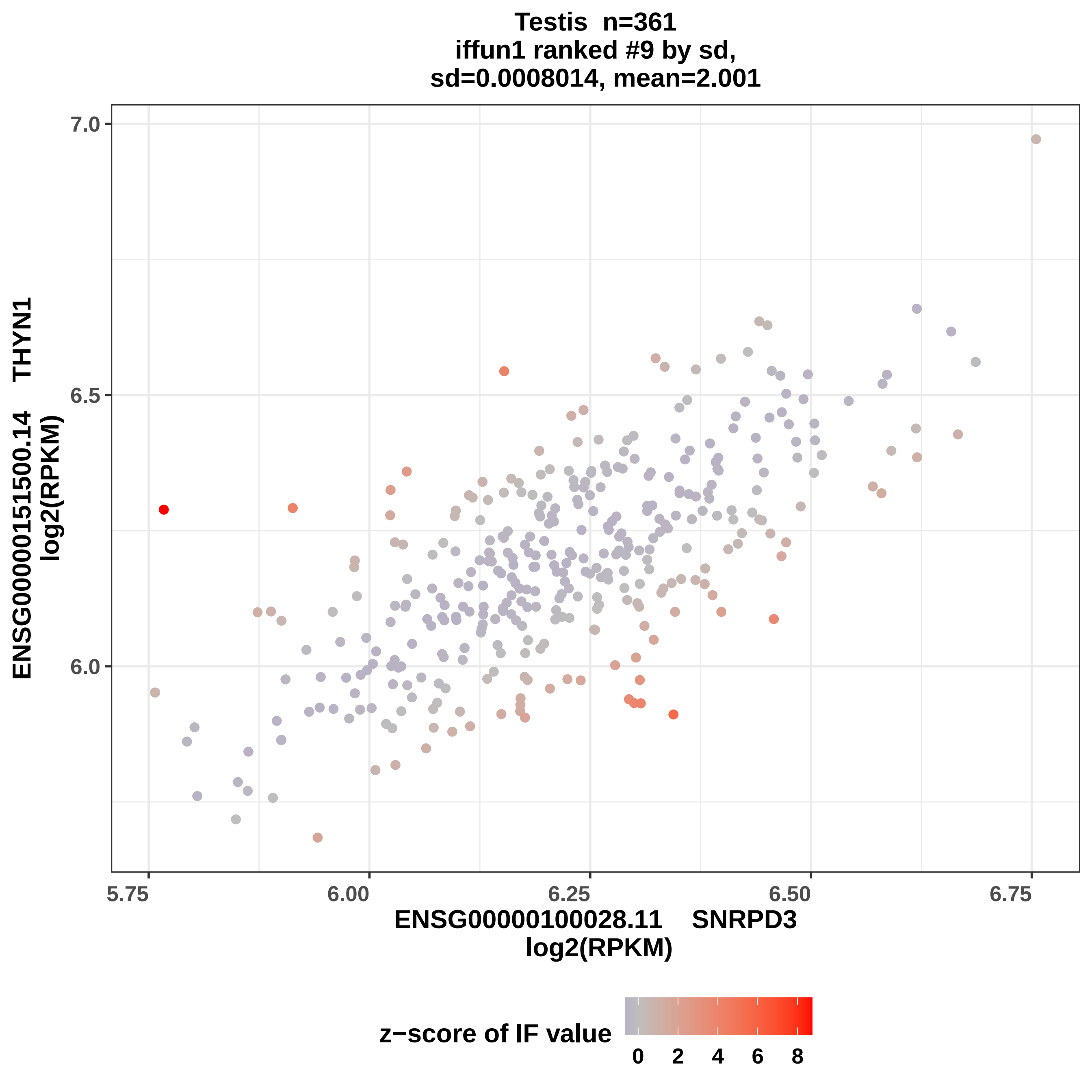

Testis n=361
iffun1 ranked #9 by sd,
sd=0.0008014, mean=2.001
7.0
6.5
6.0
5.75
6.00
6.25
6.50
6.75
ENSG00000151500.14 THYN1
log2(RPKM)
ENSG00000100028.11 SNRPD3
log2(RPKM)
z−score of IF value
0
2
4
6
8

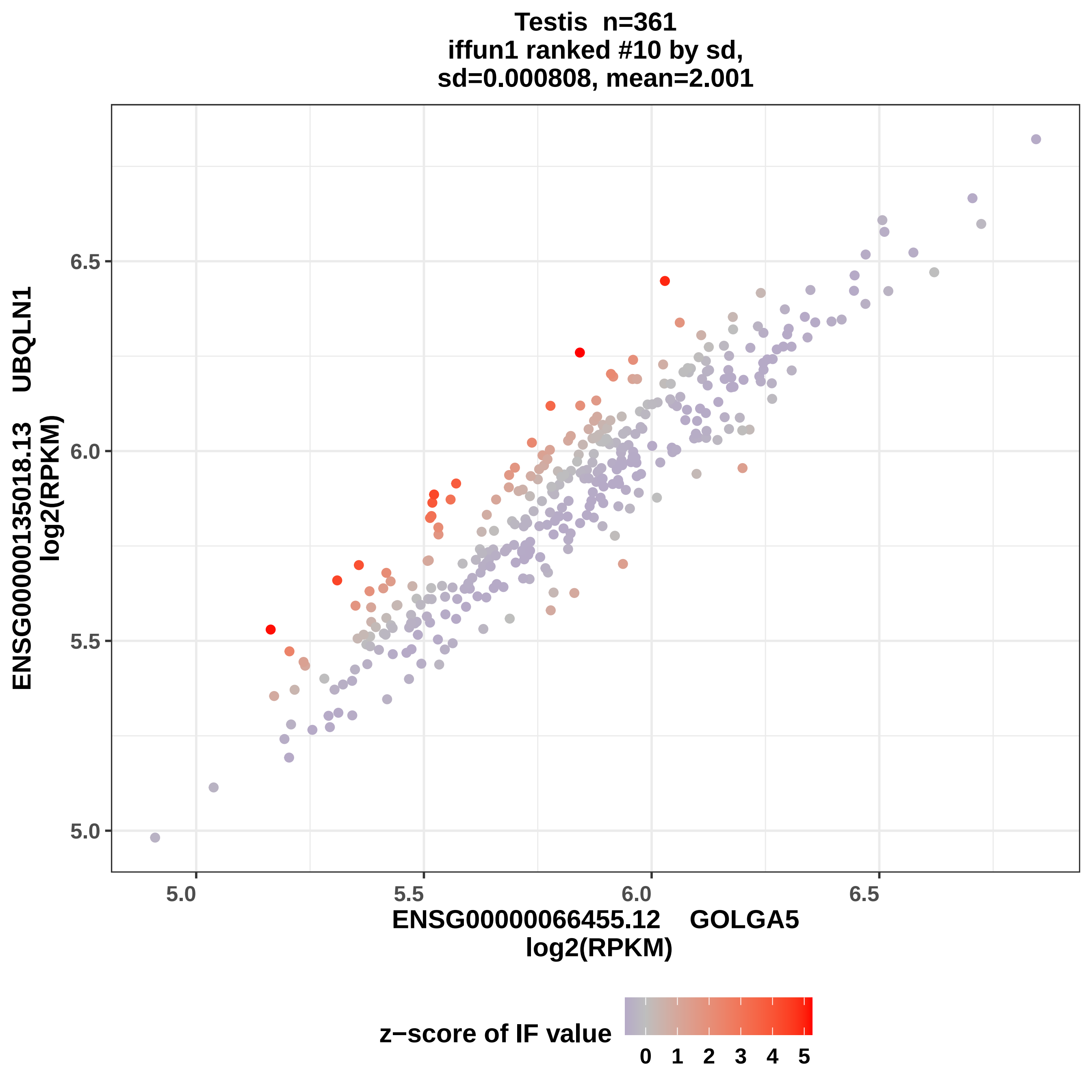
Testis n=361
iffun1 ranked #10 by sd,
sd=0.000808, mean=2.001
ENSG00000135018.13 UBQLN1
log2(RPKM)
6.5
6.0
5.5
5.0
5.0
5.5
6.0
6.5
ENSG00000066455.12 GOLGA5
log2(RPKM)
z−score of IF value
0 1 2 3 4 5

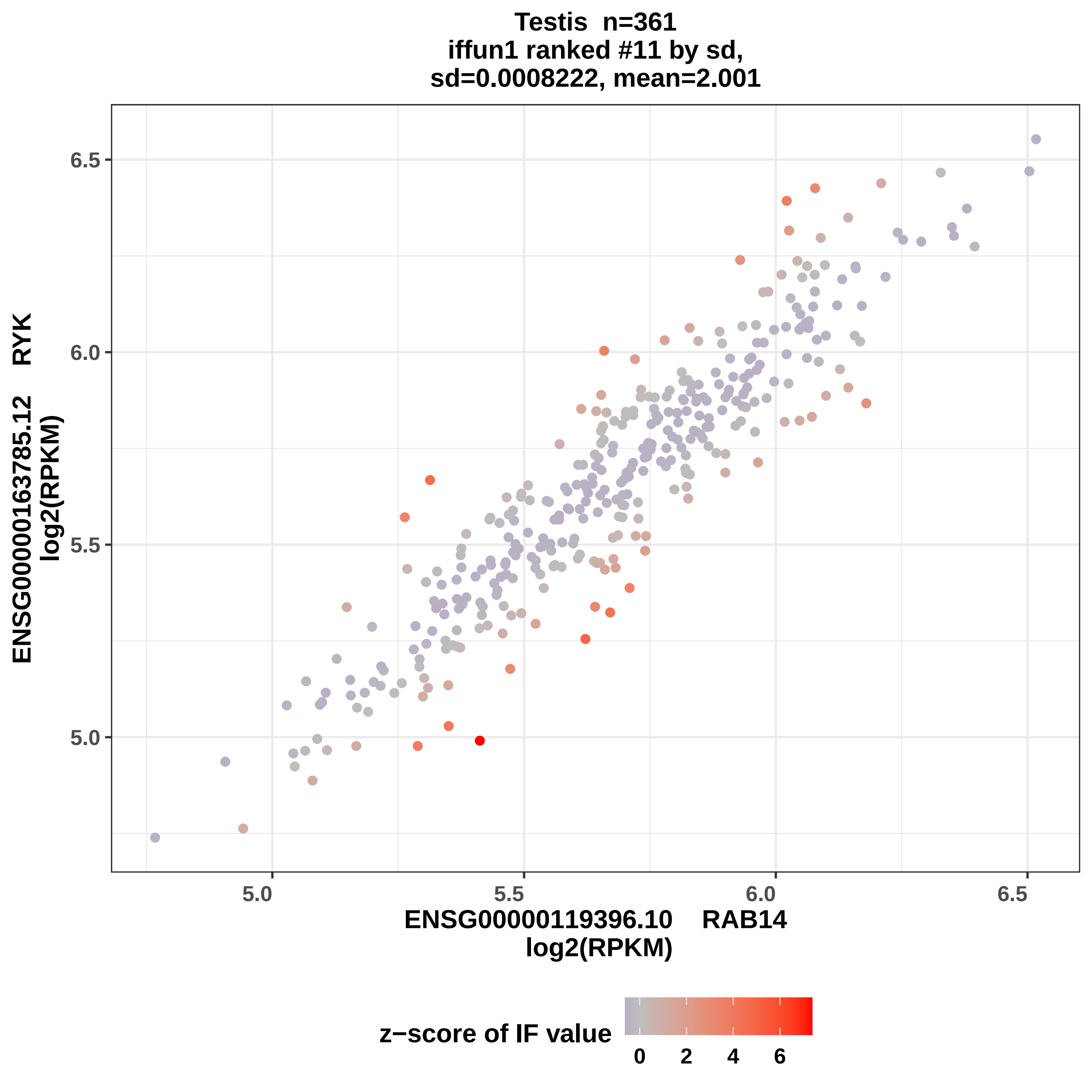

Testis n=361
iffun1 ranked #11 by sd,
sd=0.0008222, mean=2.001
ENSG00000163785.12 RYK
log2(RPKM)
6.5
6.0
5.5
5.0
5.0
5.5
6.0
6.5
ENSG00000119396.10 RAB14
log2(RPKM)
z−score of IF value
0
2
4
6

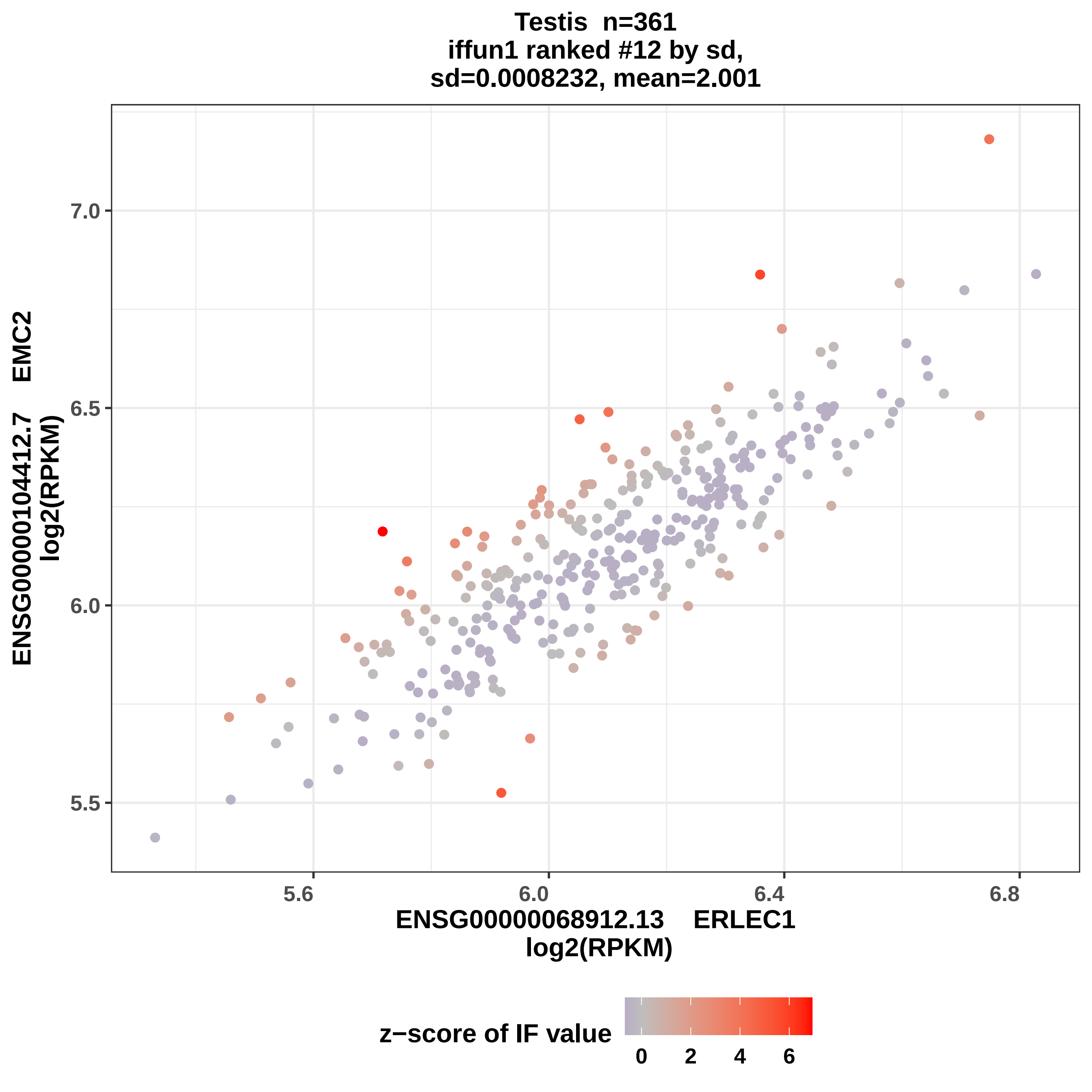
Testis n=361
iffun1 ranked #12 by sd,
sd=0.0008232, mean=2.001
ENSG00000104412.7 EMC2
log2(RPKM)
7.0
6.5
6.0
5.5
5.6
6.0
6.4
6.8
ENSG00000068912.13 ERLEC1
log2(RPKM)
z−score of IF value
0
2
4
6

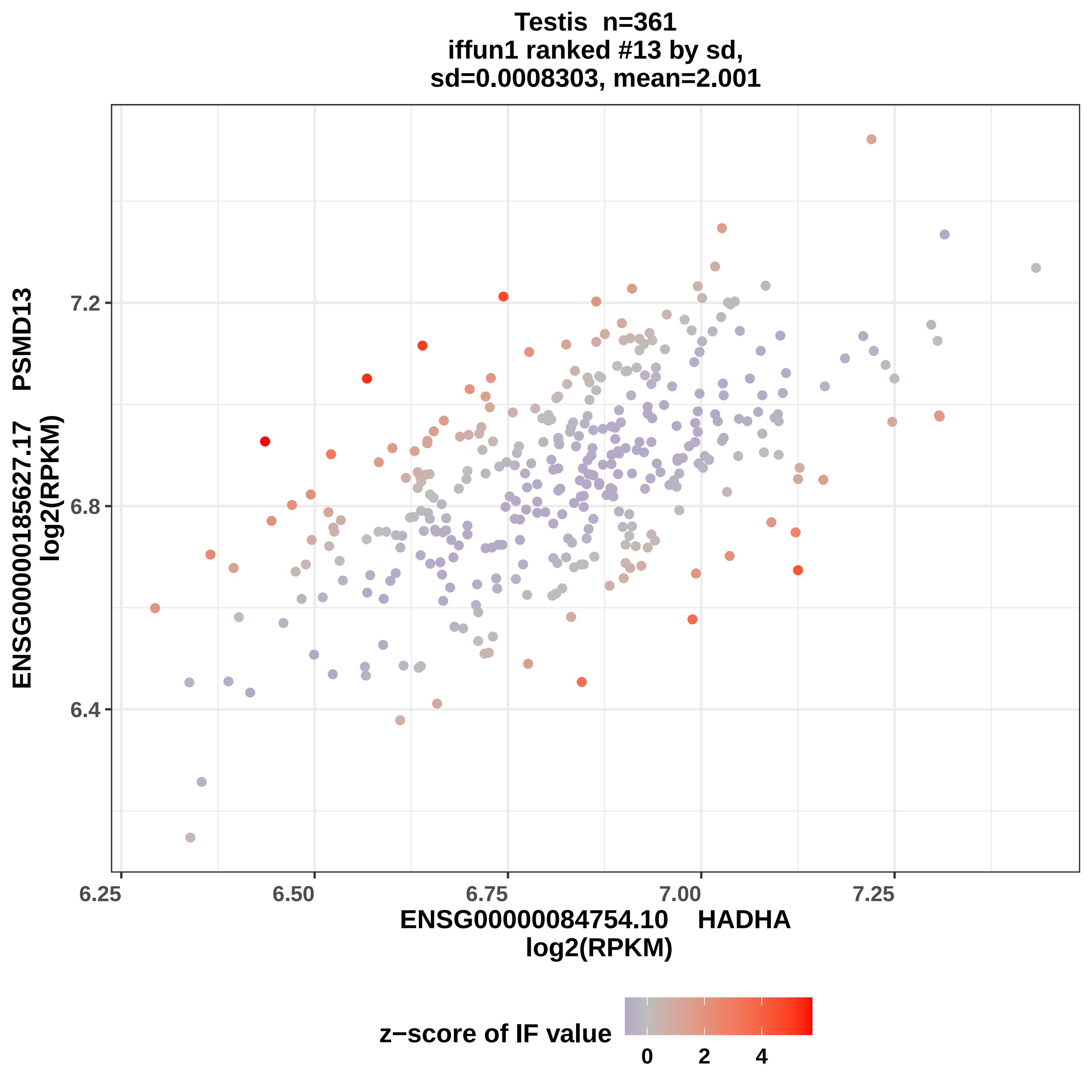

Testis n=361
iffun1 ranked #13 by sd,
sd=0.0008303, mean=2.001
ENSG00000185627.17 PSMD13
log2(RPKM)
7.2
6.8
6.4
6.25
6.50
6.75
7.00
7.25
ENSG00000084754.10 HADHA
log2(RPKM)
z−score of IF value
0
2
4

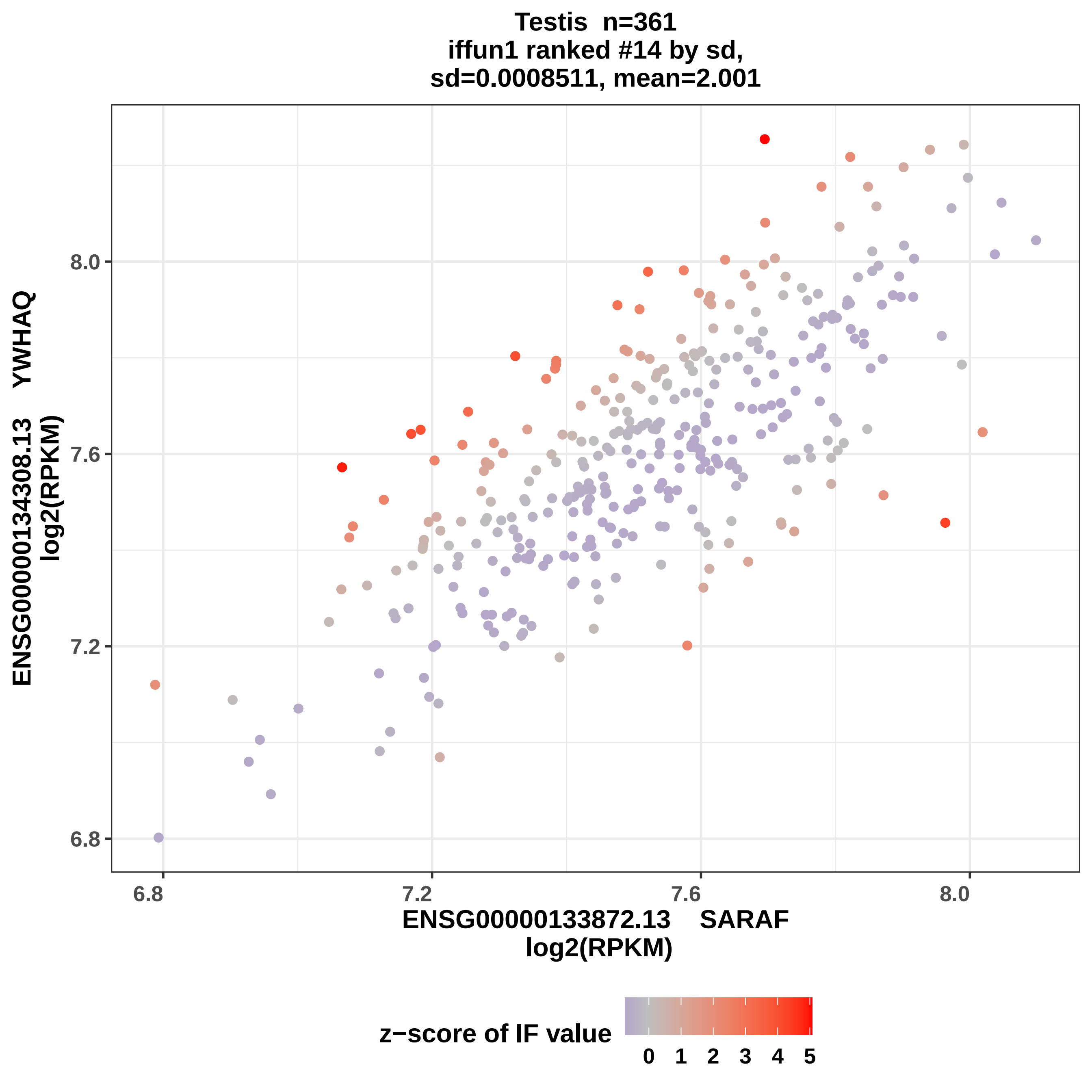
Testis n=361
iffun1 ranked #14 by sd,
sd=0.0008511, mean=2.001
ENSG00000134308.13 YWHAQ
log2(RPKM)
8.0
7.6
7.2
6.8
6.8
7.2
7.6
8.0
ENSG00000133872.13 SARAF
log2(RPKM)
z−score of IF value
0 1 2 3 4 5

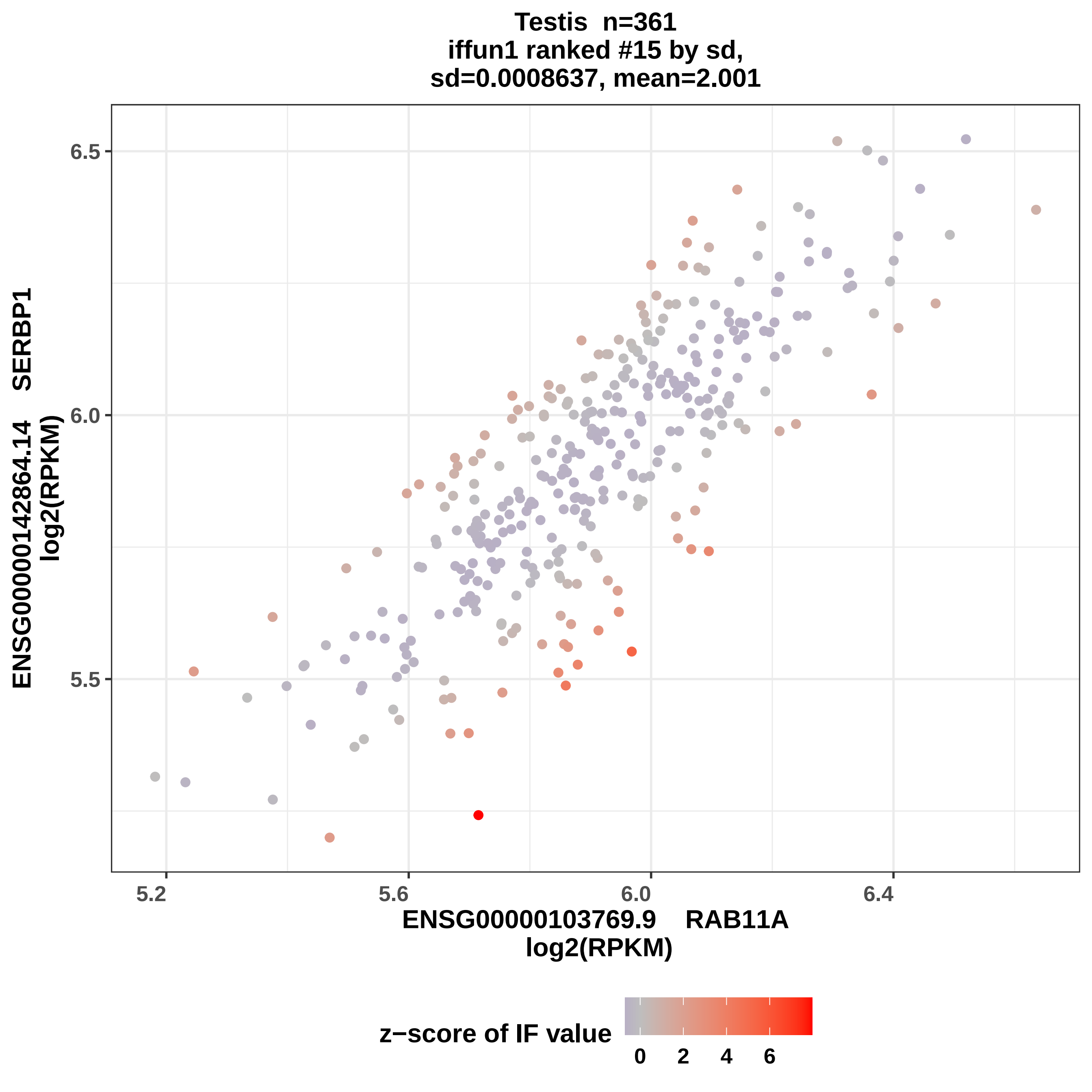

Testis n=361
iffun1 ranked #15 by sd,
sd=0.0008637, mean=2.001
ENSG00000142864.14 SERBP1
log2(RPKM)
6.5
6.0
5.5
5.2
5.6
6.0
6.4
ENSG00000103769.9 RAB11A
log2(RPKM)
z−score of IF value
0
2
4
6

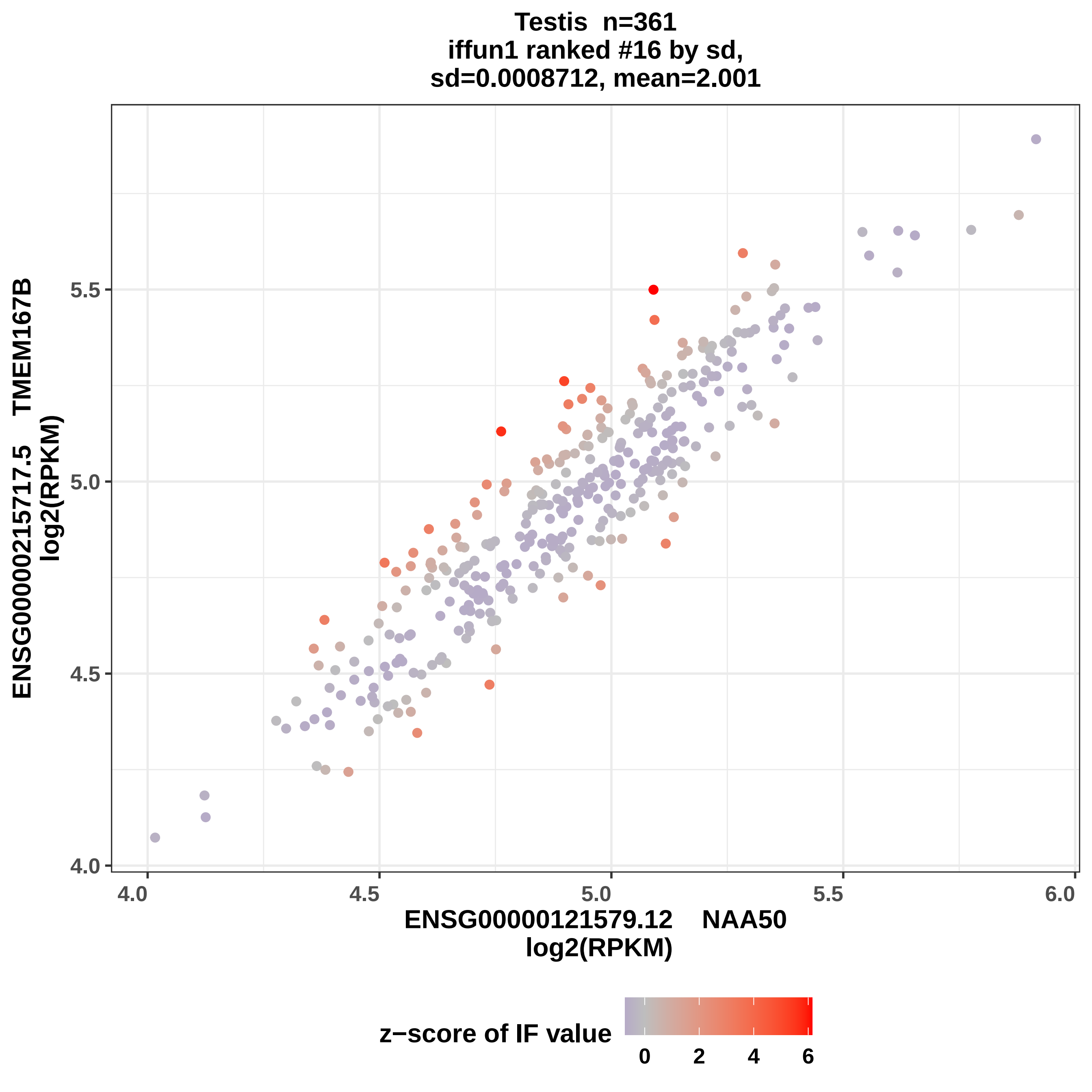
Testis n=361
iffun1 ranked #16 by sd,
sd=0.0008712, mean=2.001
ENSG00000215717.5 TMEM167B
log2(RPKM)
5.5
5.0
4.5
4.0
4.0
4.5
5.0
5.5
6.0
ENSG00000121579.12 NAA50
log2(RPKM)
z−score of IF value
0
2
4
6

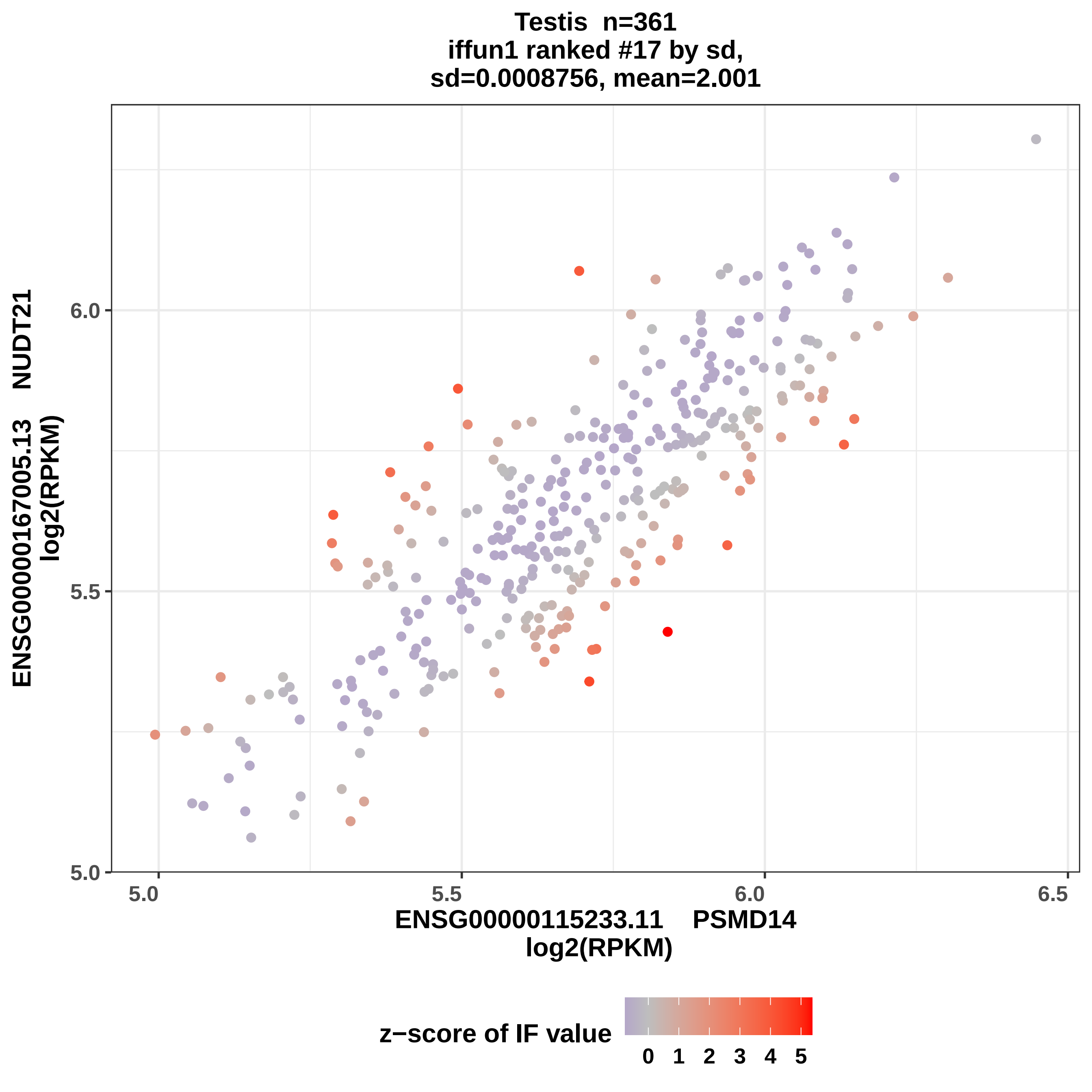

Testis n=361
iffun1 ranked #17 by sd,
sd=0.0008756, mean=2.001
ENSG00000167005.13 NUDT21
log2(RPKM)
6.0
5.5
5.0
5.0
5.5
6.0
6.5
ENSG00000115233.11 PSMD14
log2(RPKM)
z−score of IF value
0 1 2 3 4 5

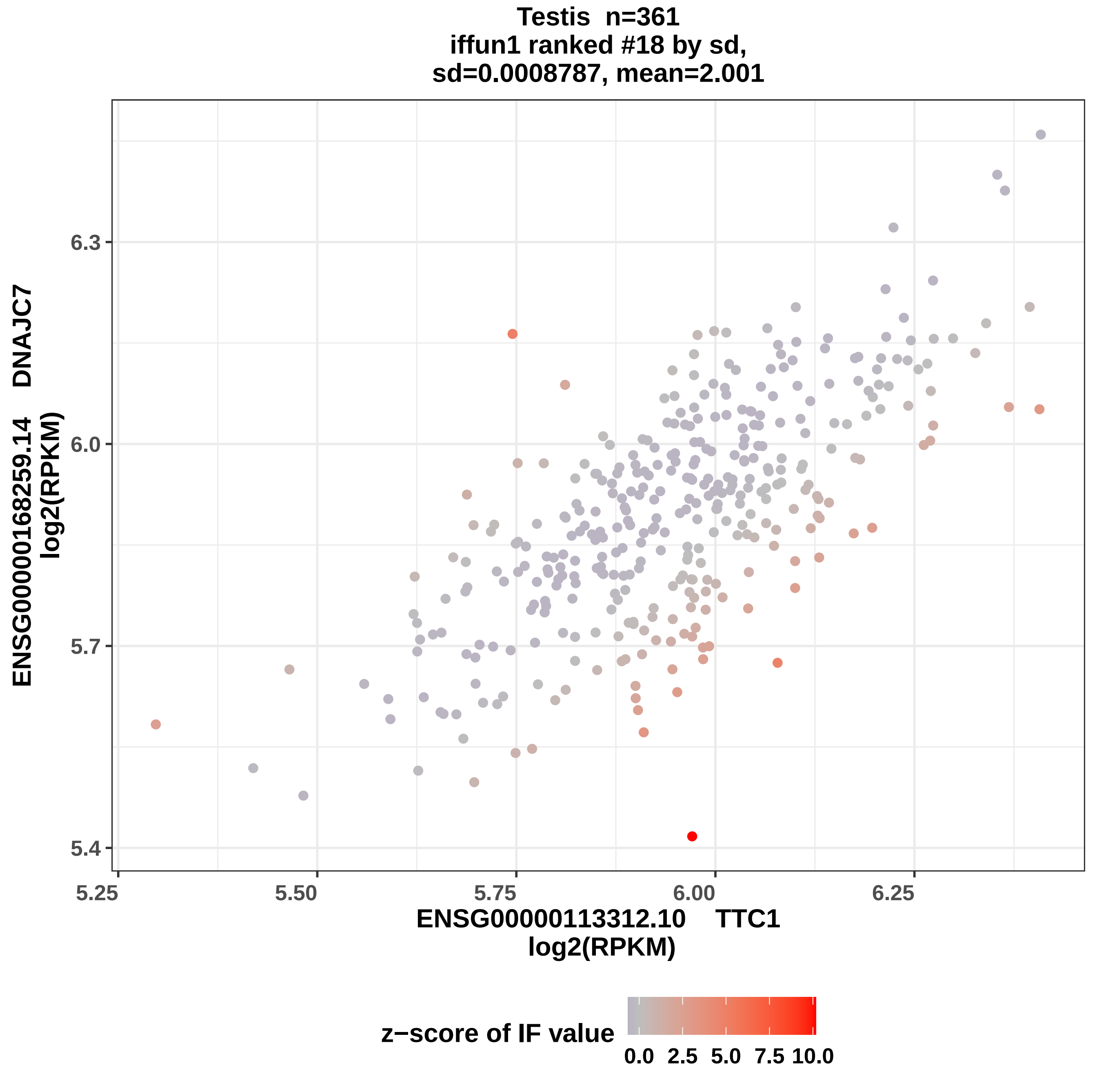
Testis n=361
iffun1 ranked #18 by sd,
sd=0.0008787, mean=2.001
ENSG00000168259.14 DNAJC7
log2(RPKM)
6.3
6.0
5.7
5.4
5.25
5.50
5.75
6.00
6.25
ENSG00000113312.10 TTC1
log2(RPKM)
z−score of IF value
0.0
2.5
5.0
7.5
10.0

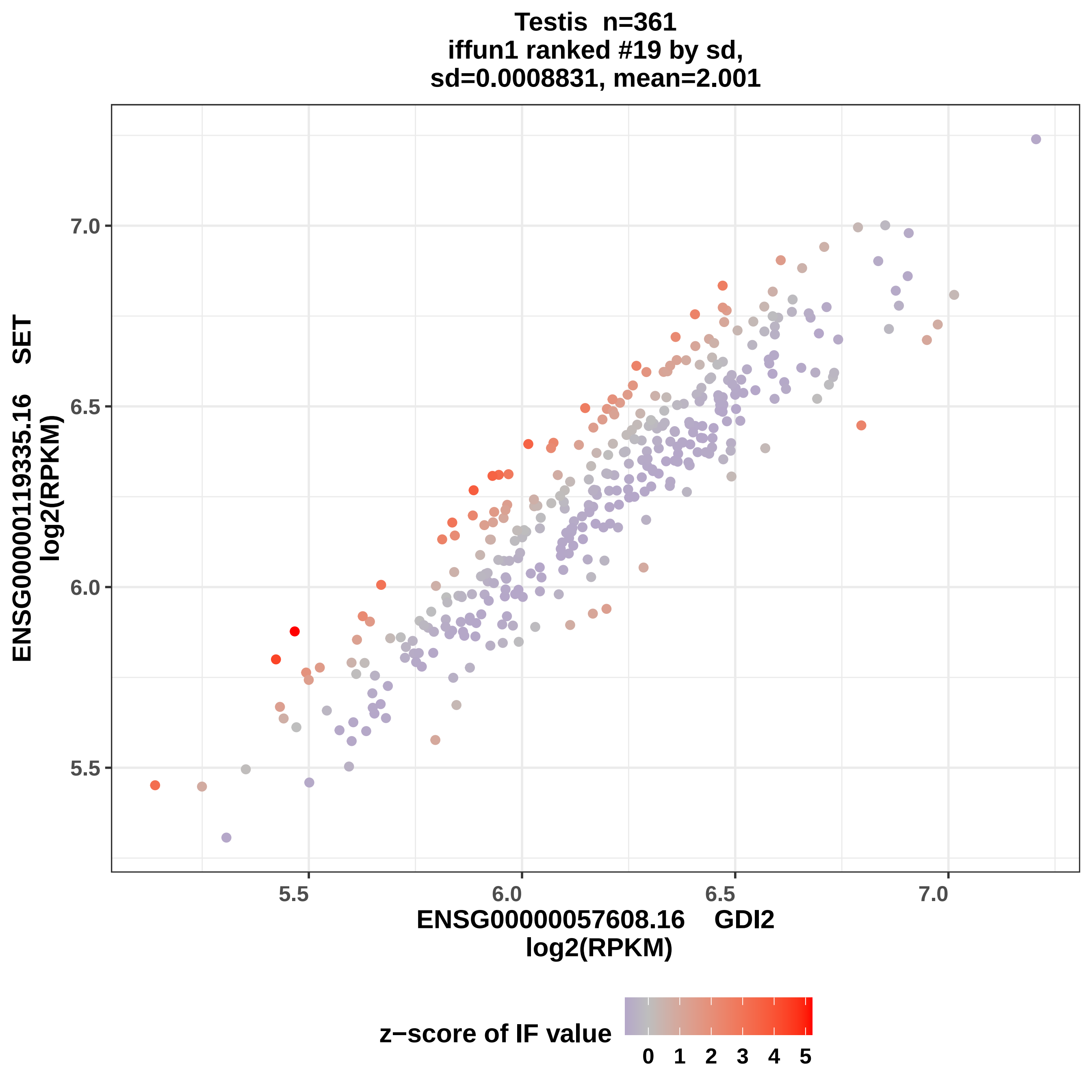

Testis n=361
iffun1 ranked #19 by sd,
sd=0.0008831, mean=2.001
7.0
6.5
6.0
5.5
5.5
6.0
6.5
7.0
ENSG00000119335.16 SET
log2(RPKM)
ENSG00000057608.16 GDI2
log2(RPKM)
z−score of IF value
0 1 2 3 4 5

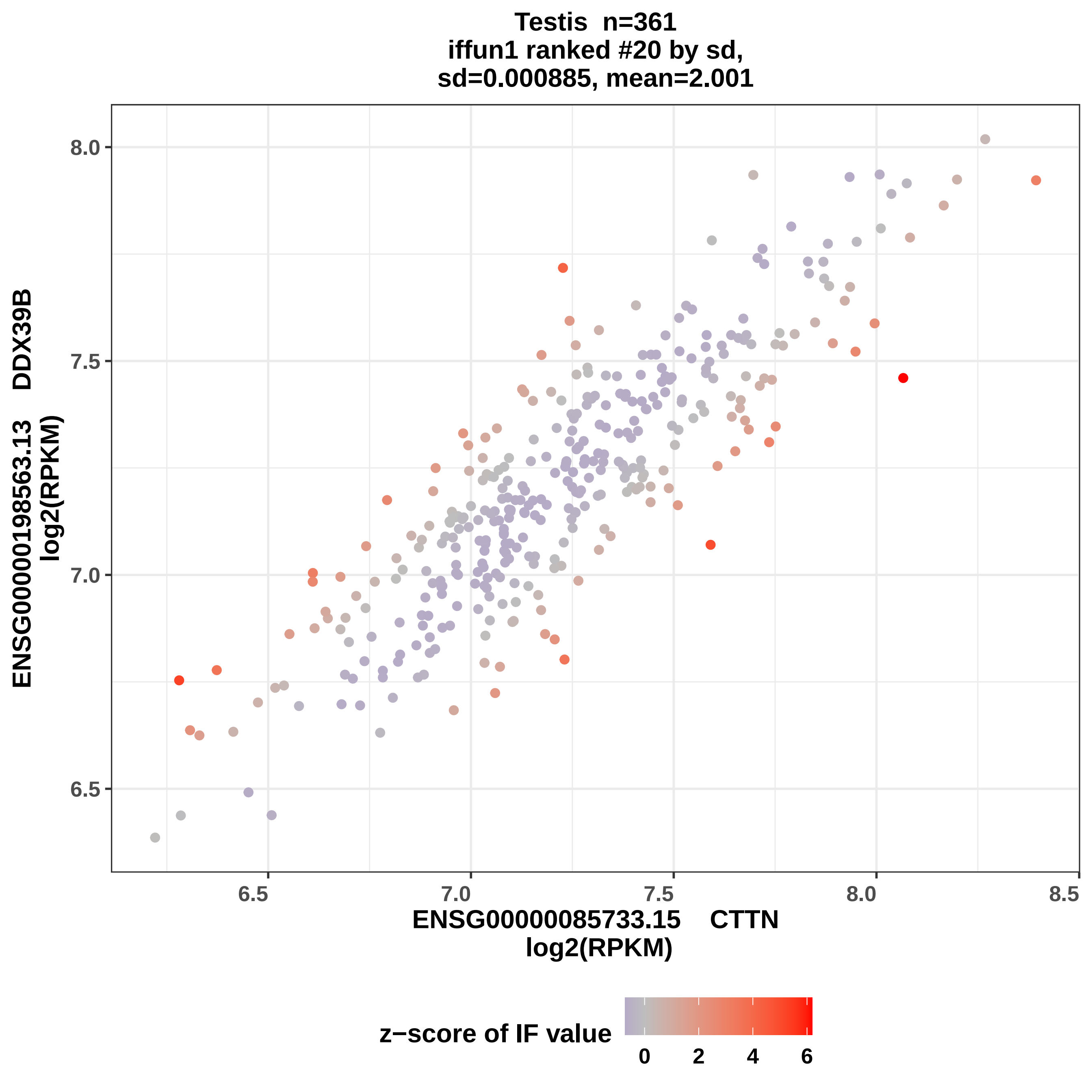

Testis n=361
iffun1 ranked #20 by sd,
sd=0.000885, mean=2.001
8.0
7.5
7.0
6.5
6.5
7.0
7.5
8.0
8.5
ENSG00000198563.13 DDX39B
log2(RPKM)
ENSG00000085733.15 CTTN
log2(RPKM)
z−score of IF value
0
2
4
6

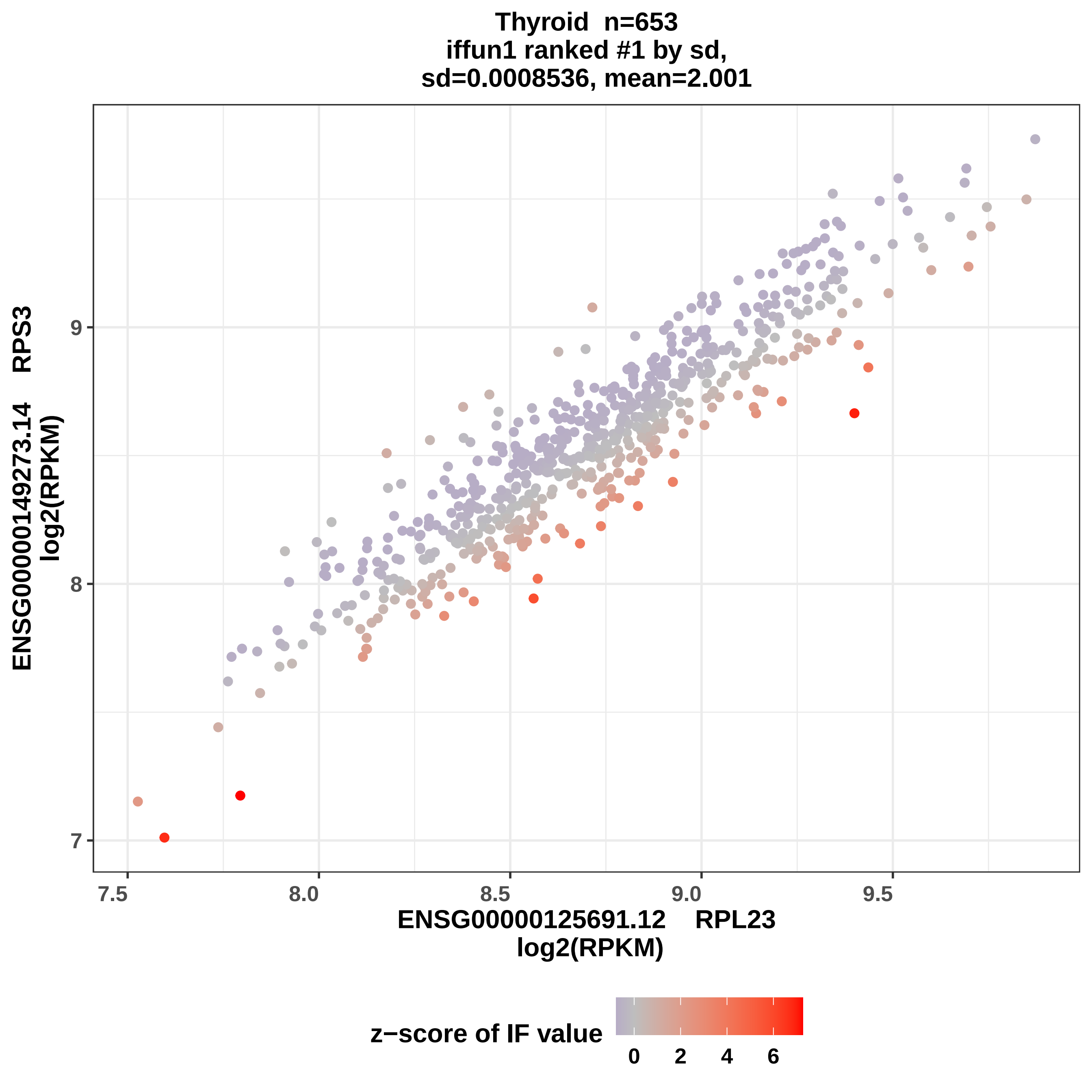

Thyroid n=653
iffun1 ranked #1 by sd,
sd=0.0008536, mean=2.001
ENSG00000149273.14 RPS3
log2(RPKM)
9
8
7
7.5
8.0
8.5
9.0
9.5
ENSG00000125691.12 RPL23
log2(RPKM)
z−score of IF value
0
2
4
6

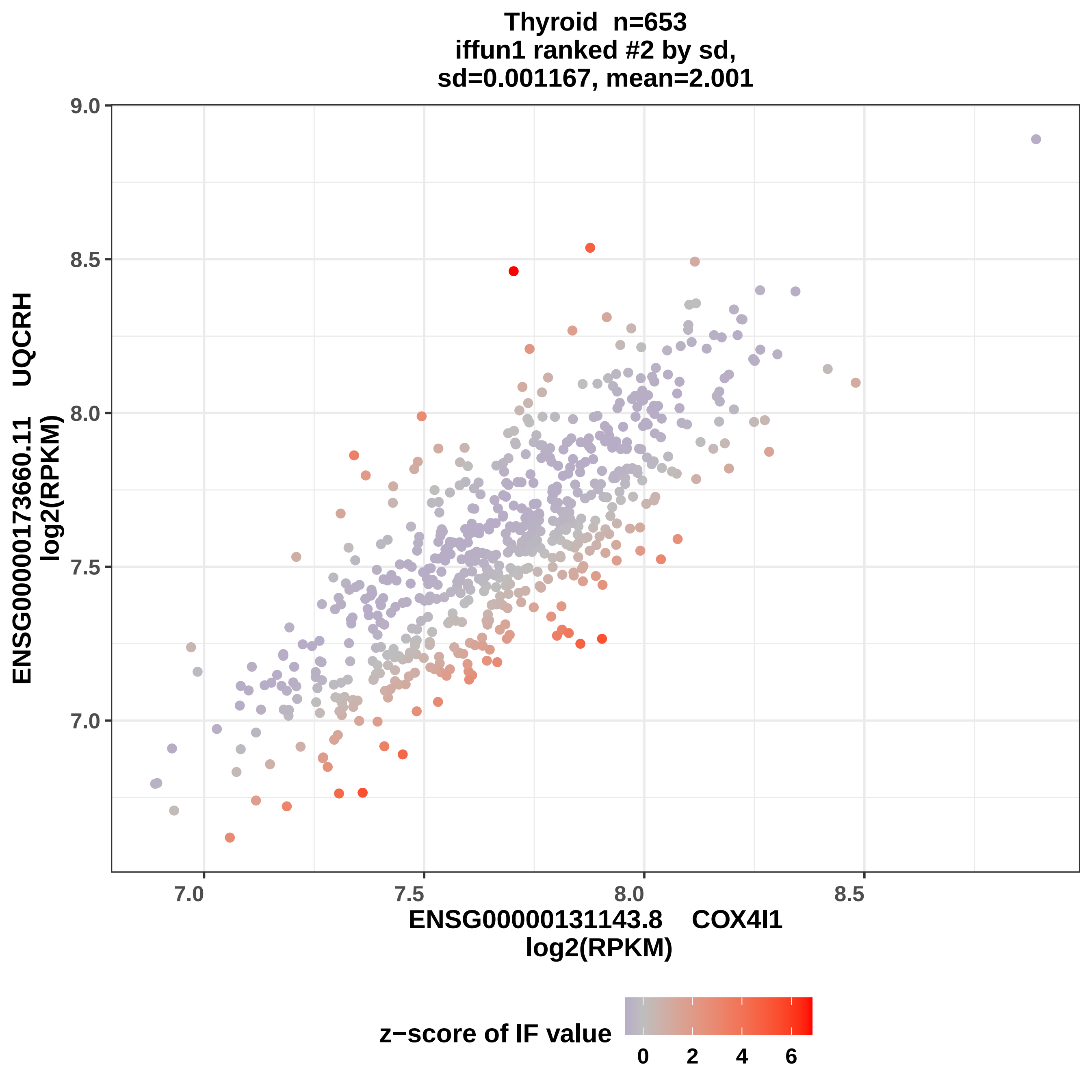
Thyroid n=653
iffun1 ranked #2 by sd,
sd=0.001167, mean=2.001
9.0
8.5
8.0
7.5
7.0
7.0
7.5
8.0
8.5
ENSG00000173660.11 UQCRH
log2(RPKM)
ENSG00000131143.8 COX4I1
log2(RPKM)
z−score of IF value
0
2
4
6

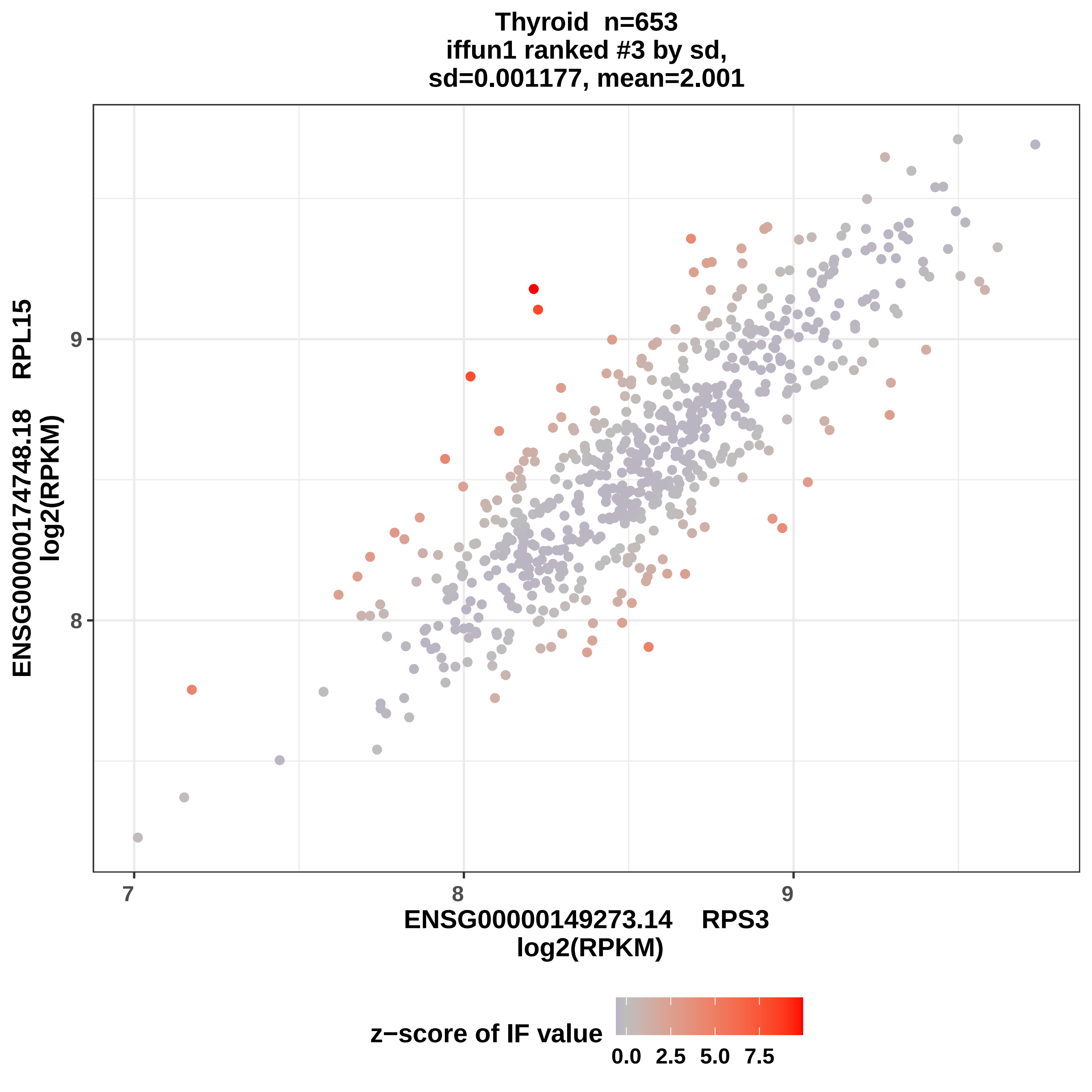

Thyroid n=653
iffun1 ranked #3 by sd,
sd=0.001177, mean=2.001
ENSG00000174748.18 RPL15
log2(RPKM)
9
8
7
8
9
ENSG00000149273.14 RPS3
log2(RPKM)
z−score of IF value
0.0
2.5
5.0
7.5

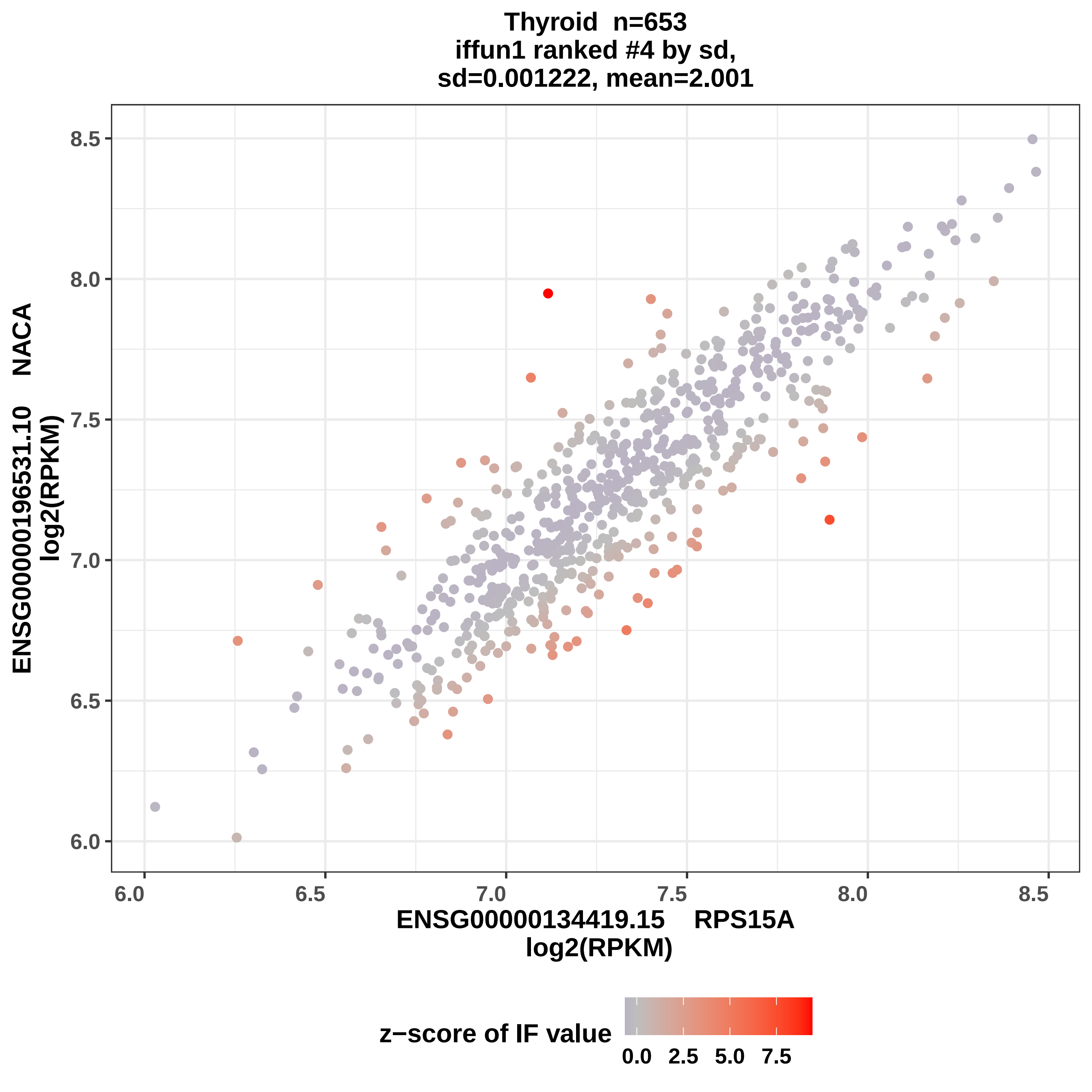

Thyroid n=653
iffun1 ranked #4 by sd,
sd=0.001222, mean=2.001
ENSG00000196531.10 NACA
log2(RPKM)
ENSG00000134419.15 RPS15A
log2(RPKM)
6.0
6.5
7.0
7.5
8.0
8.5
z−score of IF value
0.0
2.5
5.0
7.5

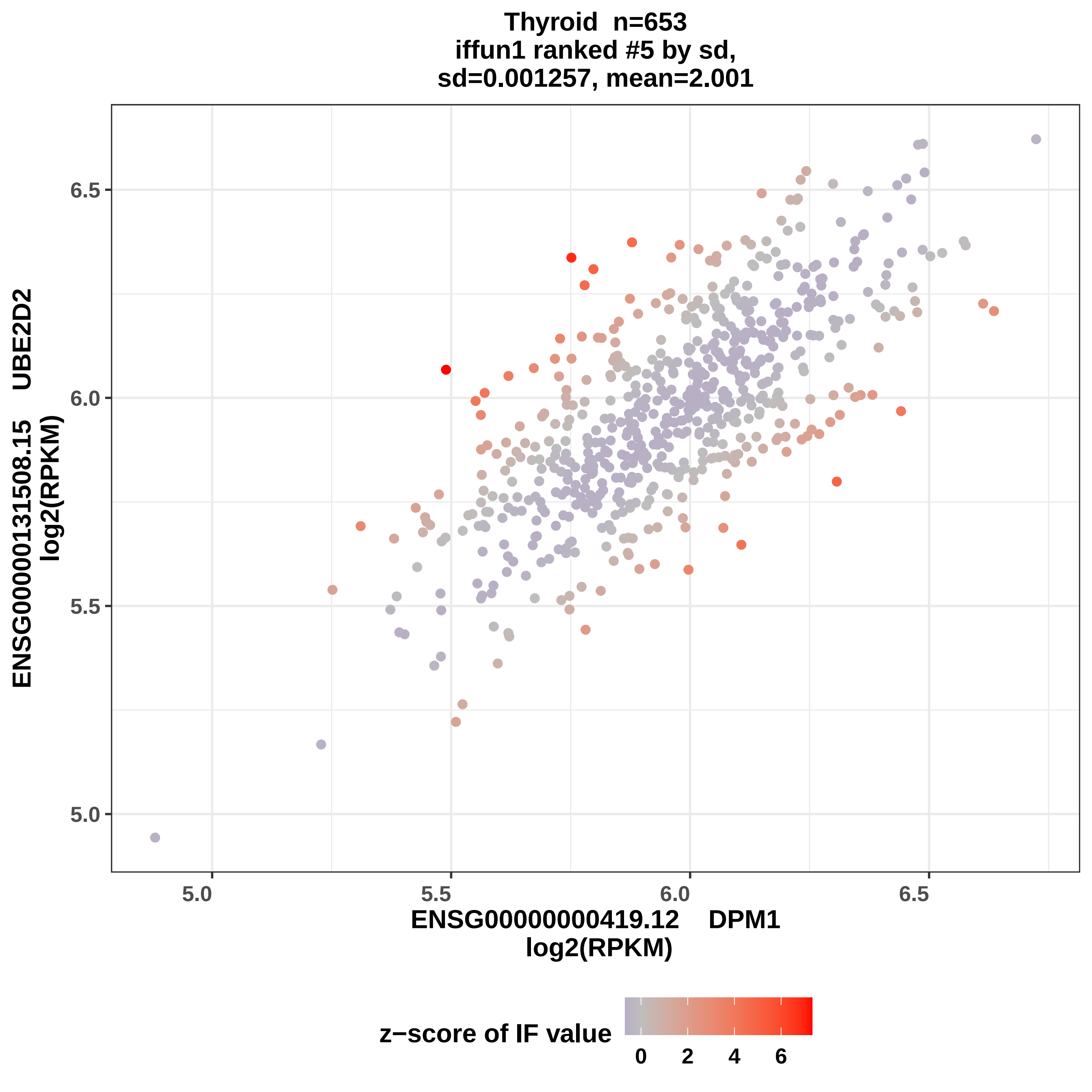
Thyroid n=653
iffun1 ranked #5 by sd,
sd=0.001257, mean=2.001
6.5
6.0
5.5
5.0
ENSG00000131508.15 UBE2D2
log2(RPKM)
5.0
5.5
6.0
6.5
ENSG00000000419.12 DPM1
log2(RPKM)
z−score of IF value
0
2
4
6

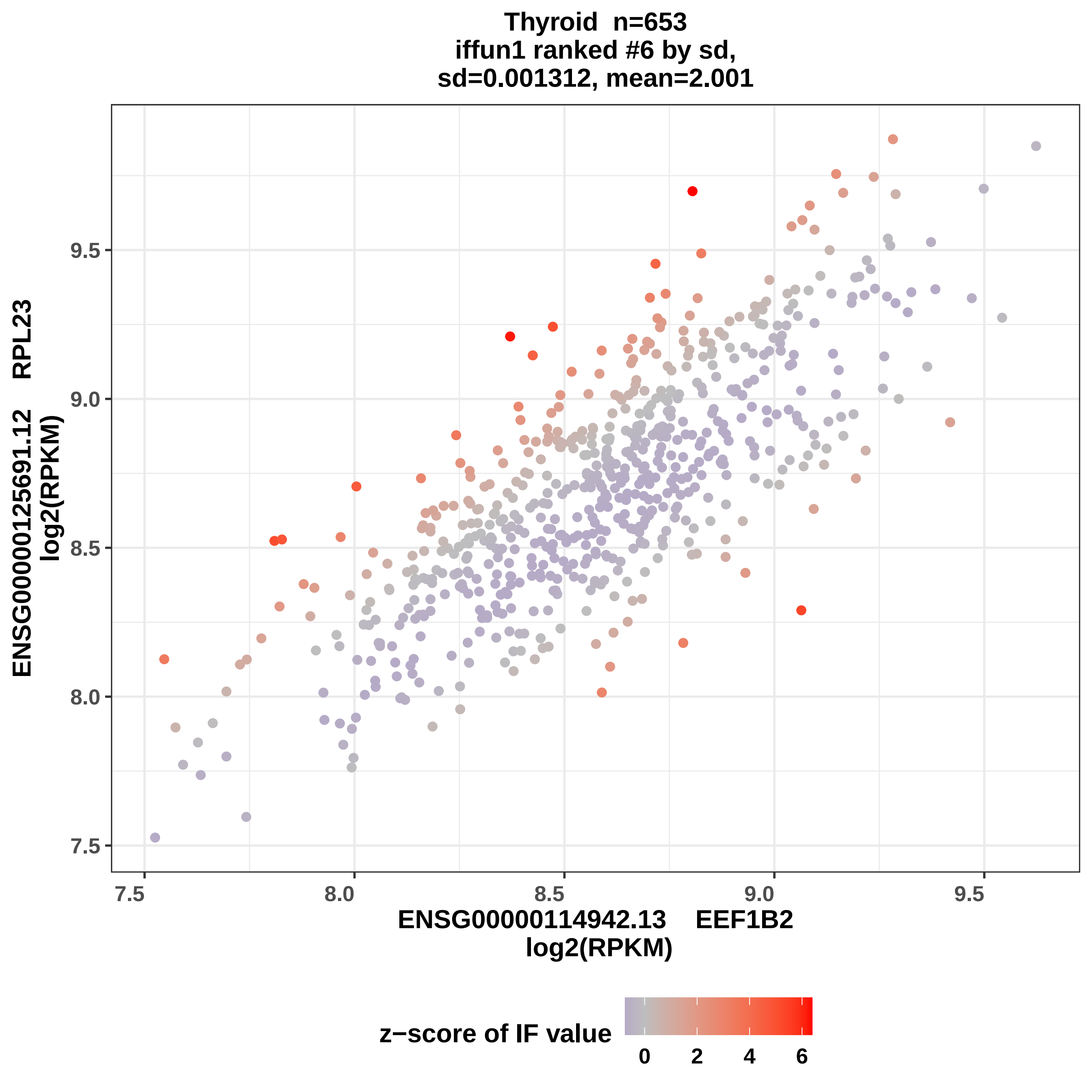
Thyroid n=653
iffun1 ranked #6 by sd,
sd=0.001312, mean=2.001
ENSG00000125691.12 RPL23
log2(RPKM)
9.5
9.0
8.5
8.0
7.5
7.5
8.0
8.5
9.0
9.5
ENSG00000114942.13 EEF1B2
log2(RPKM)
z−score of IF value
0
2
4
6

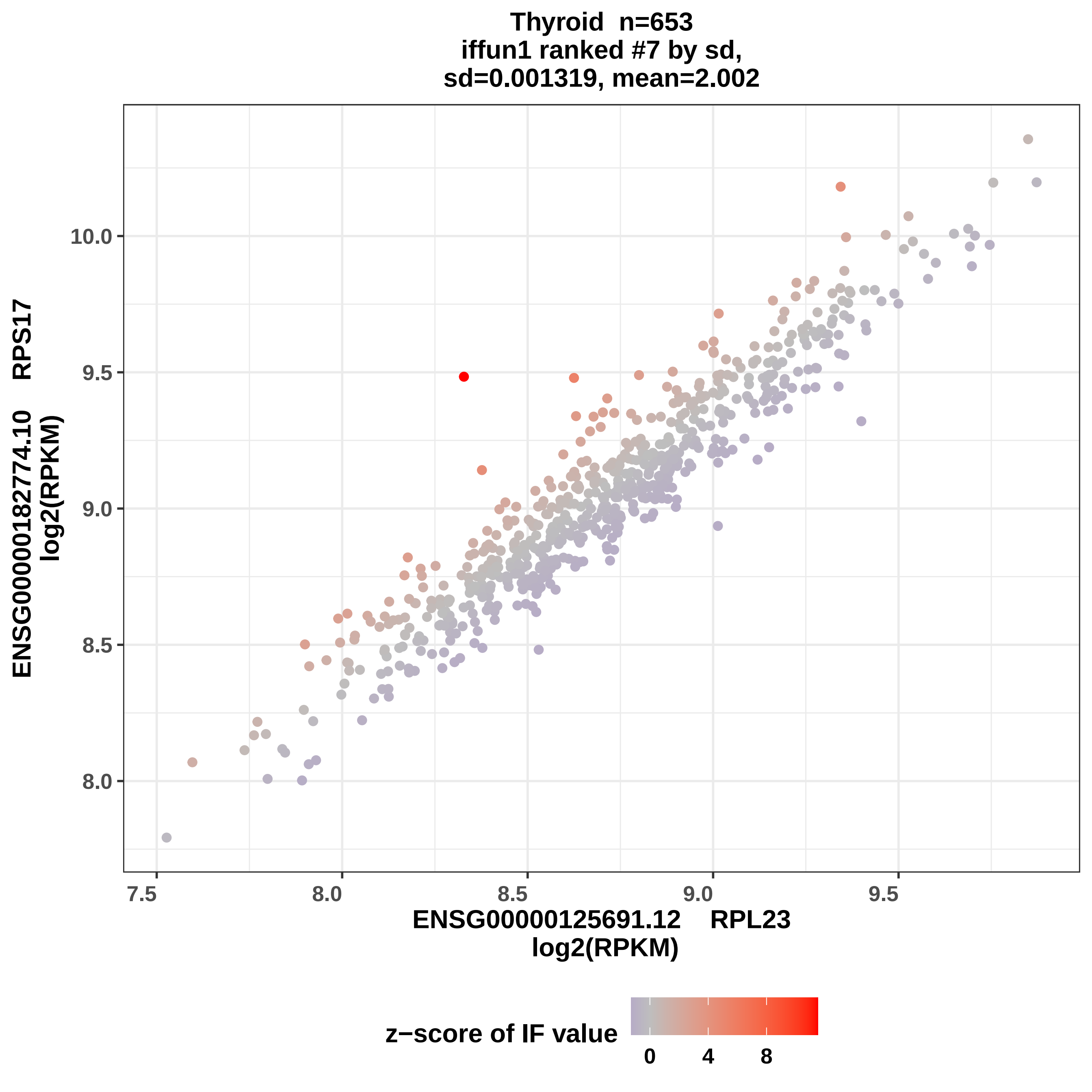
Thyroid n=653
iffun1 ranked #7 by sd,
sd=0.001319, mean=2.002
ENSG00000182774.10 RPS17
log2(RPKM)
10.0
9.5
9.0
8.5
8.0
7.5
8.0
8.5
9.0
9.5
ENSG00000125691.12 RPL23
log2(RPKM)
z−score of IF value
0
4
8

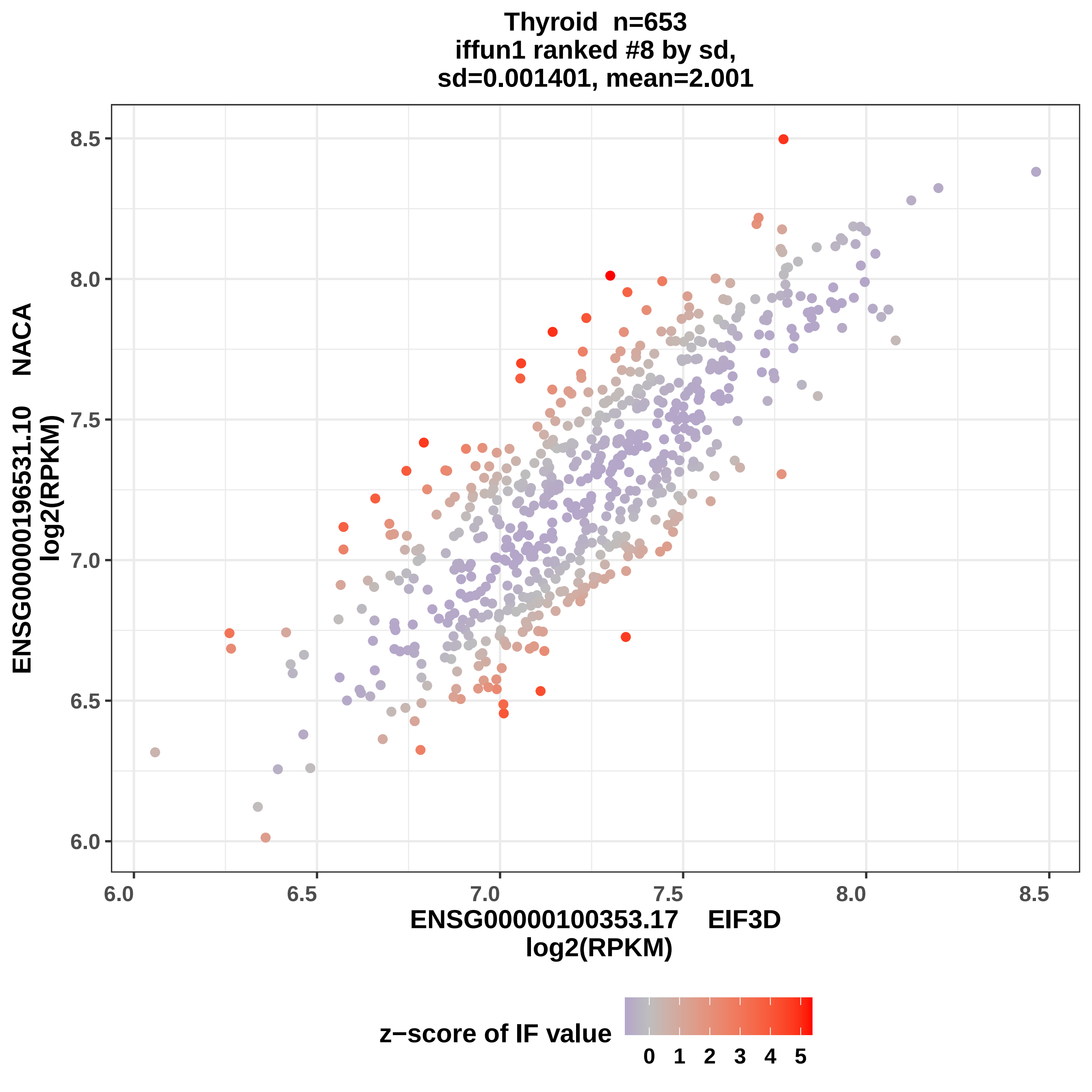

Thyroid n=653
iffun1 ranked #8 by sd,
sd=0.001401, mean=2.001
ENSG00000196531.10 NACA
log2(RPKM)
8.5
8.0
7.5
7.0
6.5
6.0
6.0
6.5
7.0
7.5
8.0
8.5
ENSG00000100353.17 EIF3D
log2(RPKM)
z−score of IF value
0
1
2
3
4
5

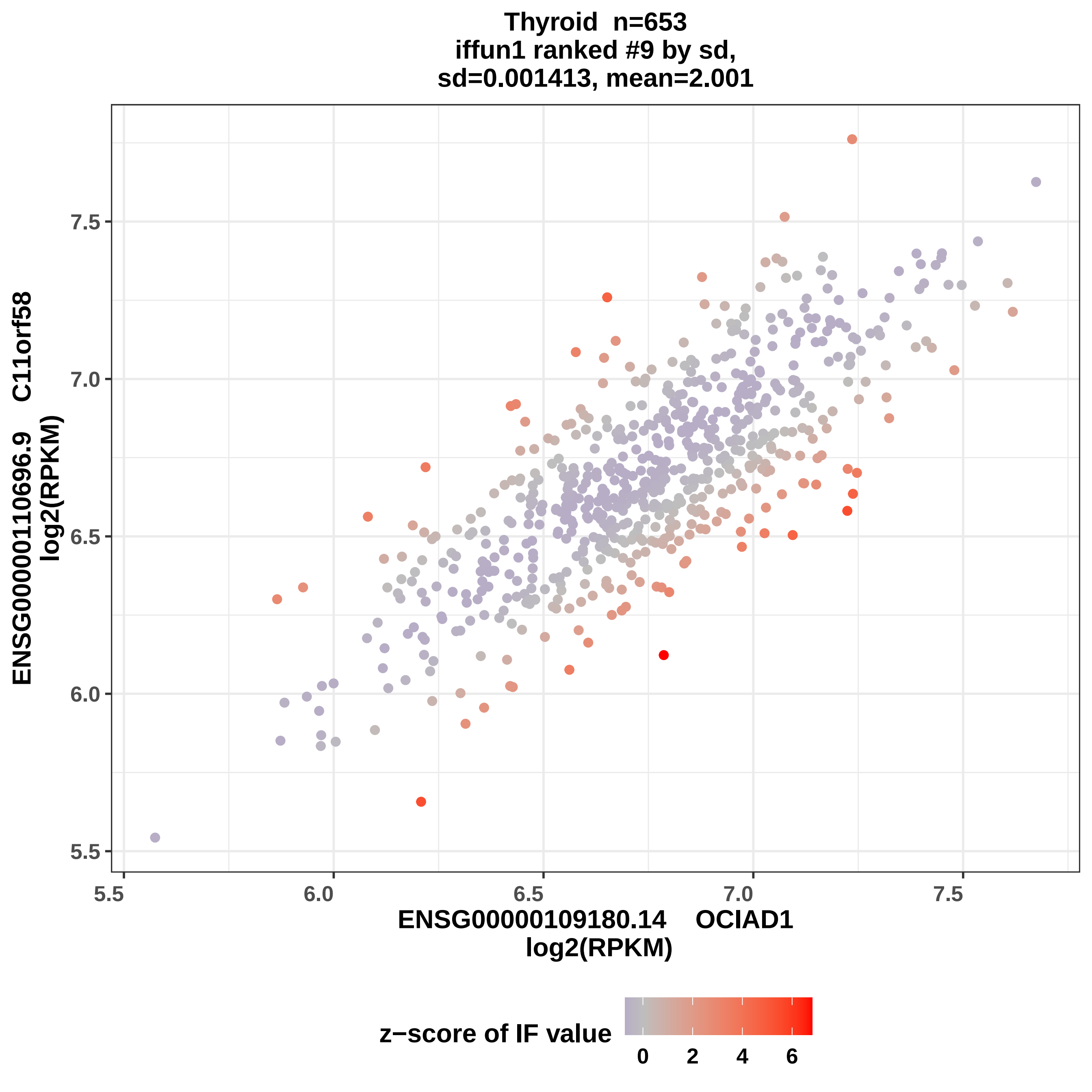
Thyroid n=653
iffun1 ranked #9 by sd,
sd=0.001413, mean=2.001
ENSG00000110696.9 C11orf58
log2(RPKM)
7.5
7.0
6.5
6.0
5.5
5.5
6.0
6.5
7.0
7.5
ENSG00000109180.14 OCIAD1
log2(RPKM)
z−score of IF value
0
2
4
6

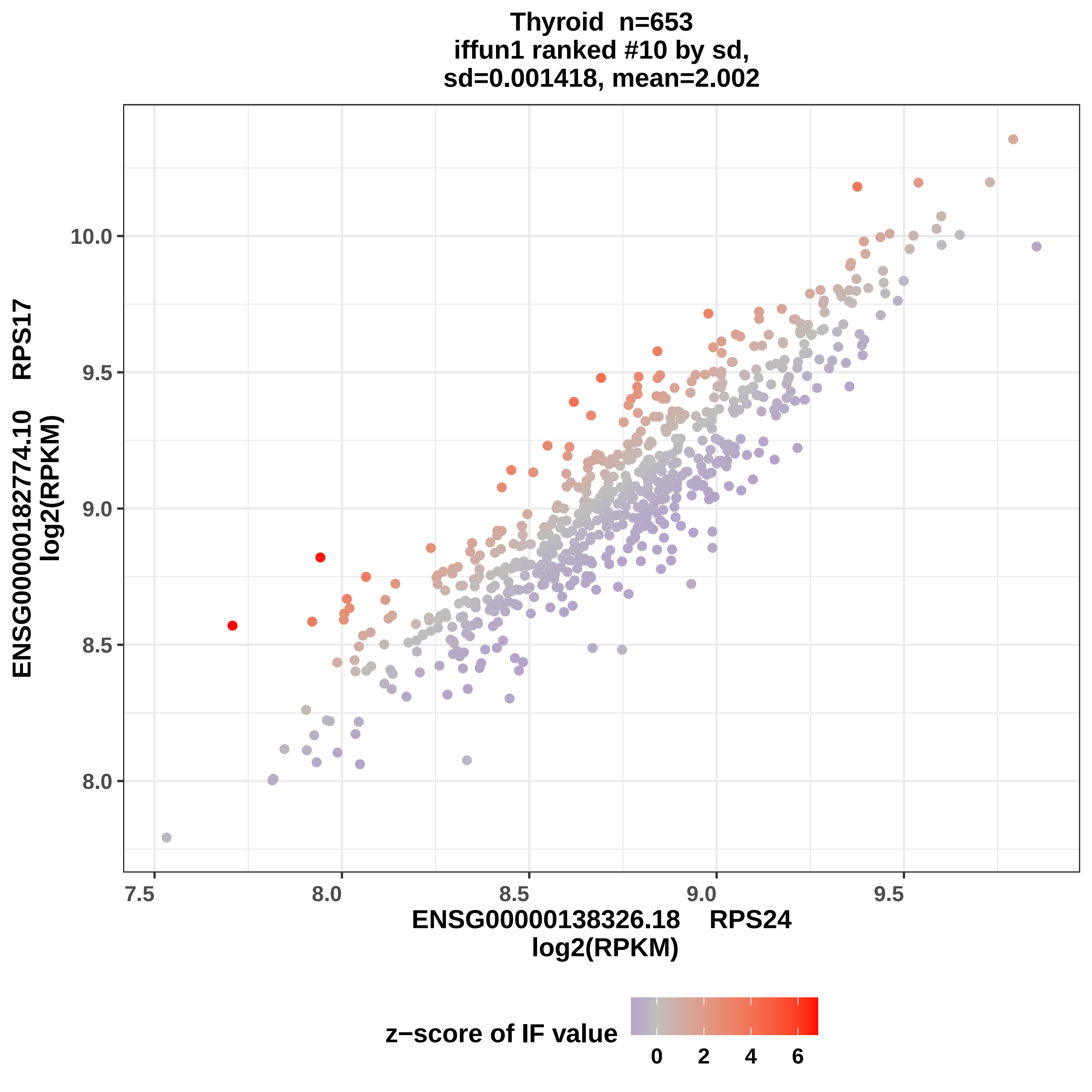
Thyroid n=653
iffun1 ranked #10 by sd,
sd=0.001418, mean=2.002
ENSG00000182774.10 RPS17
log2(RPKM)
10.0
9.5
9.0
8.5
8.0
7.5
8.0
8.5
9.0
9.5
ENSG00000138326.18 RPS24
log2(RPKM)
z−score of IF value
0
2
4
6

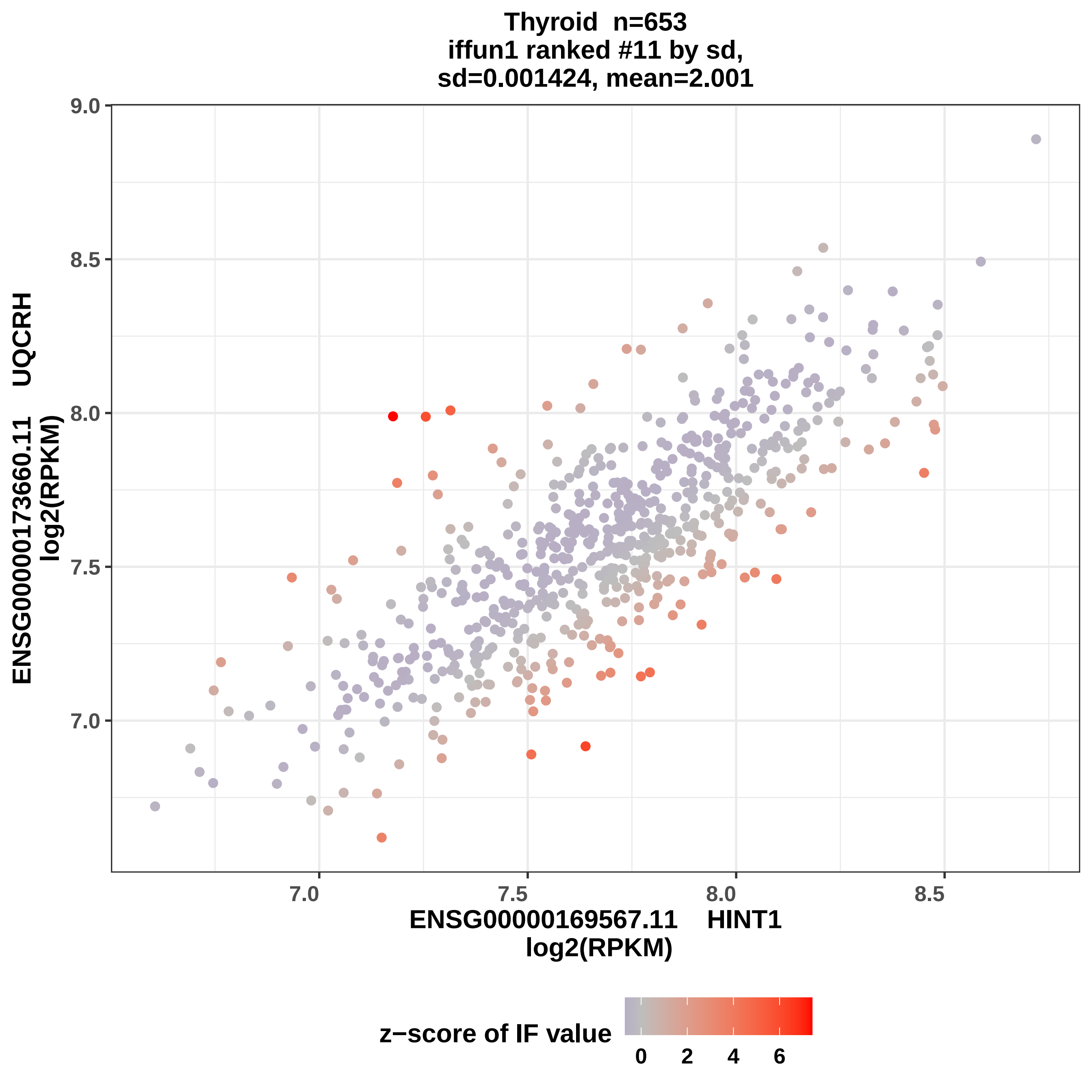

Thyroid n=653
iffun1 ranked #11 by sd,
sd=0.001424, mean=2.001
9.0
8.5
8.0
7.5
7.0
ENSG00000173660.11 UQCRH
log2(RPKM)
7.0
7.5
8.0
8.5
ENSG00000169567.11 HINT1
log2(RPKM)
z−score of IF value
0
2
4
6

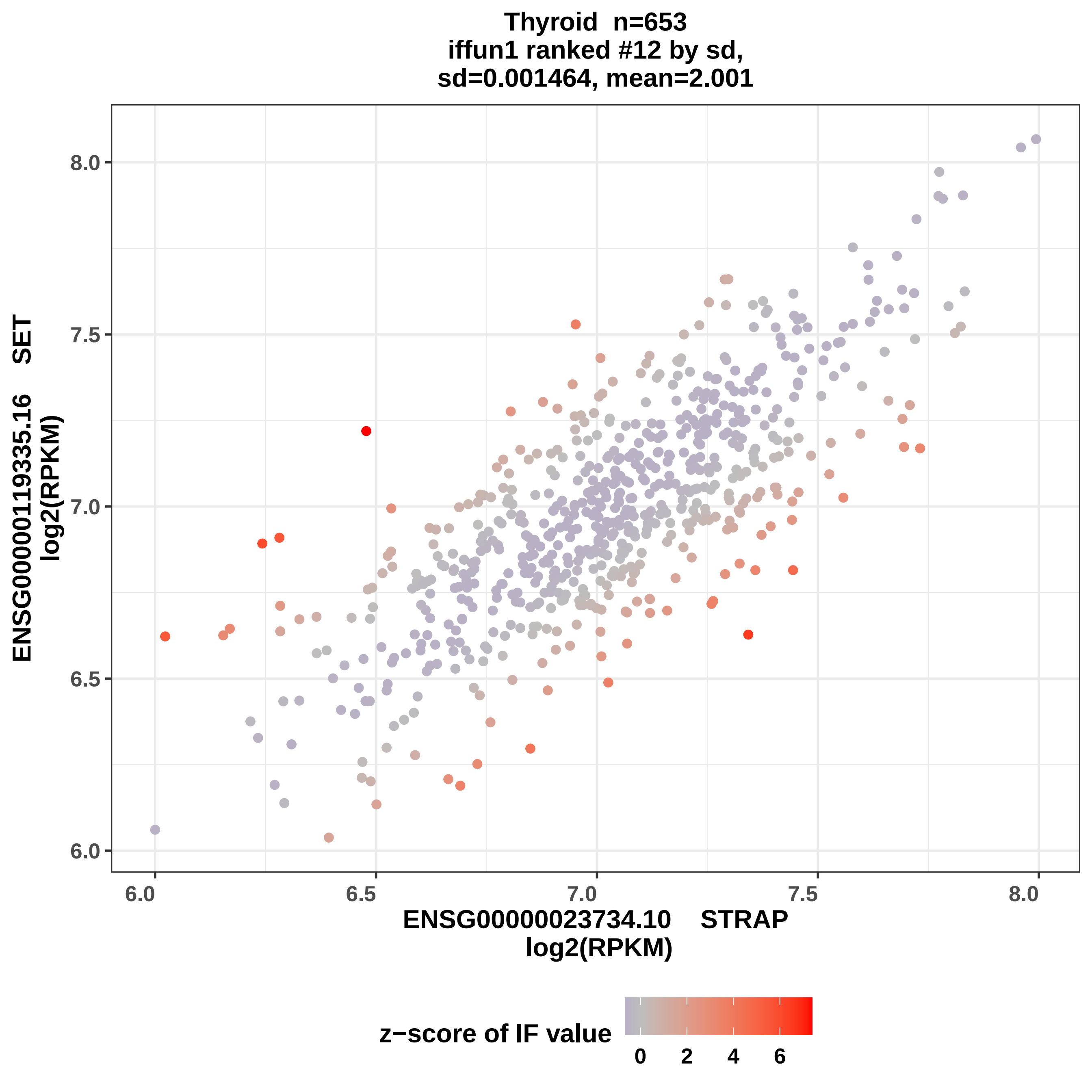
Thyroid n=653
iffun1 ranked #12 by sd,
sd=0.001464, mean=2.001
ENSG00000119335.16 SET
log2(RPKM)
8.0
7.5
7.0
6.5
6.0
6.0
6.5
7.0
7.5
8.0
ENSG00000023734.10 STRAP
log2(RPKM)
z−score of IF value
0
2
4
6

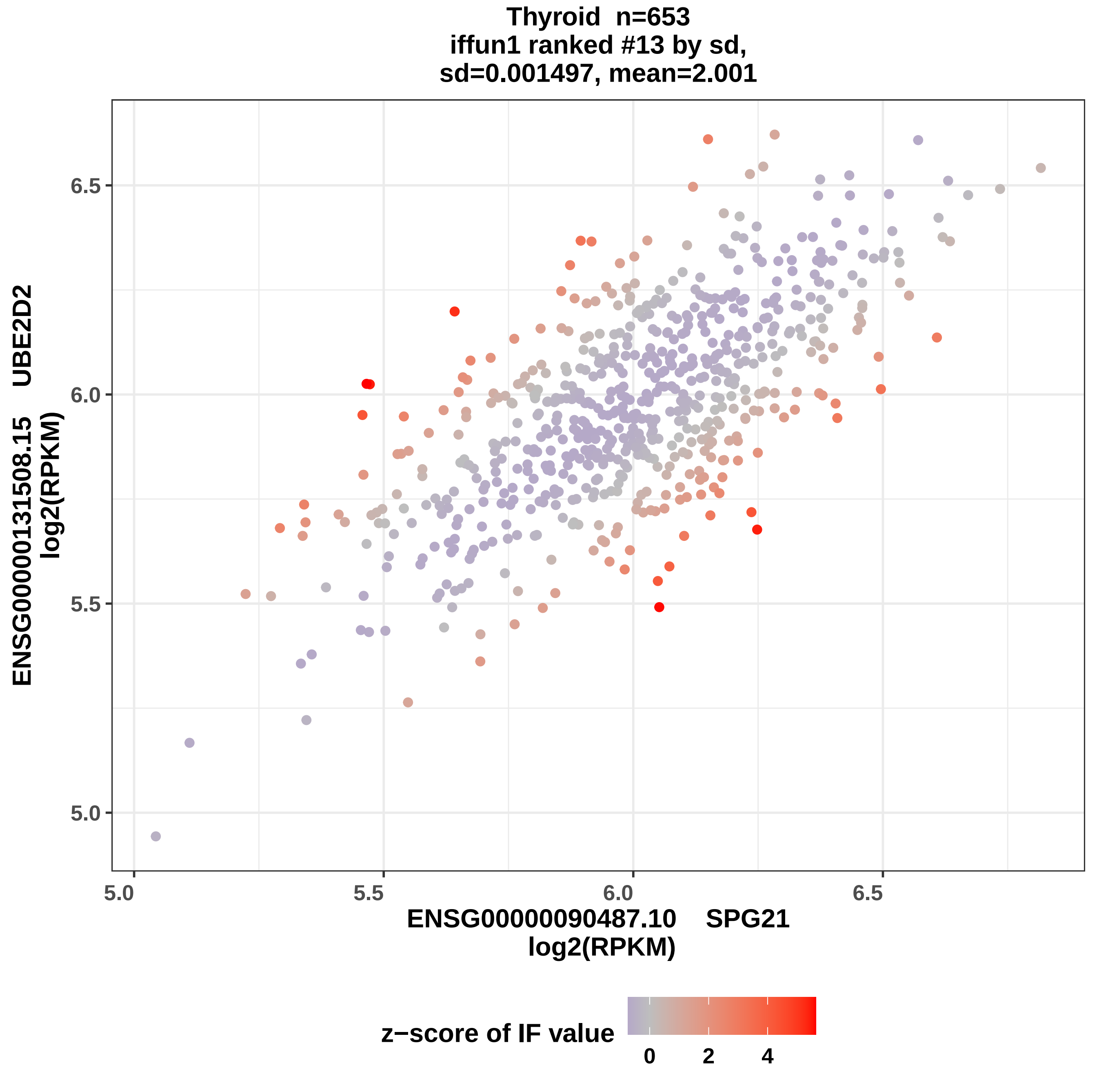
Thyroid n=653
iffun1 ranked #13 by sd,
sd=0.001497, mean=2.001
ENSG00000131508.15 UBE2D2
log2(RPKM)
6.5
6.0
5.5
5.0
5.0
5.5
6.0
6.5
ENSG00000090487.10 SPG21
log2(RPKM)
z−score of IF value
0
2
4

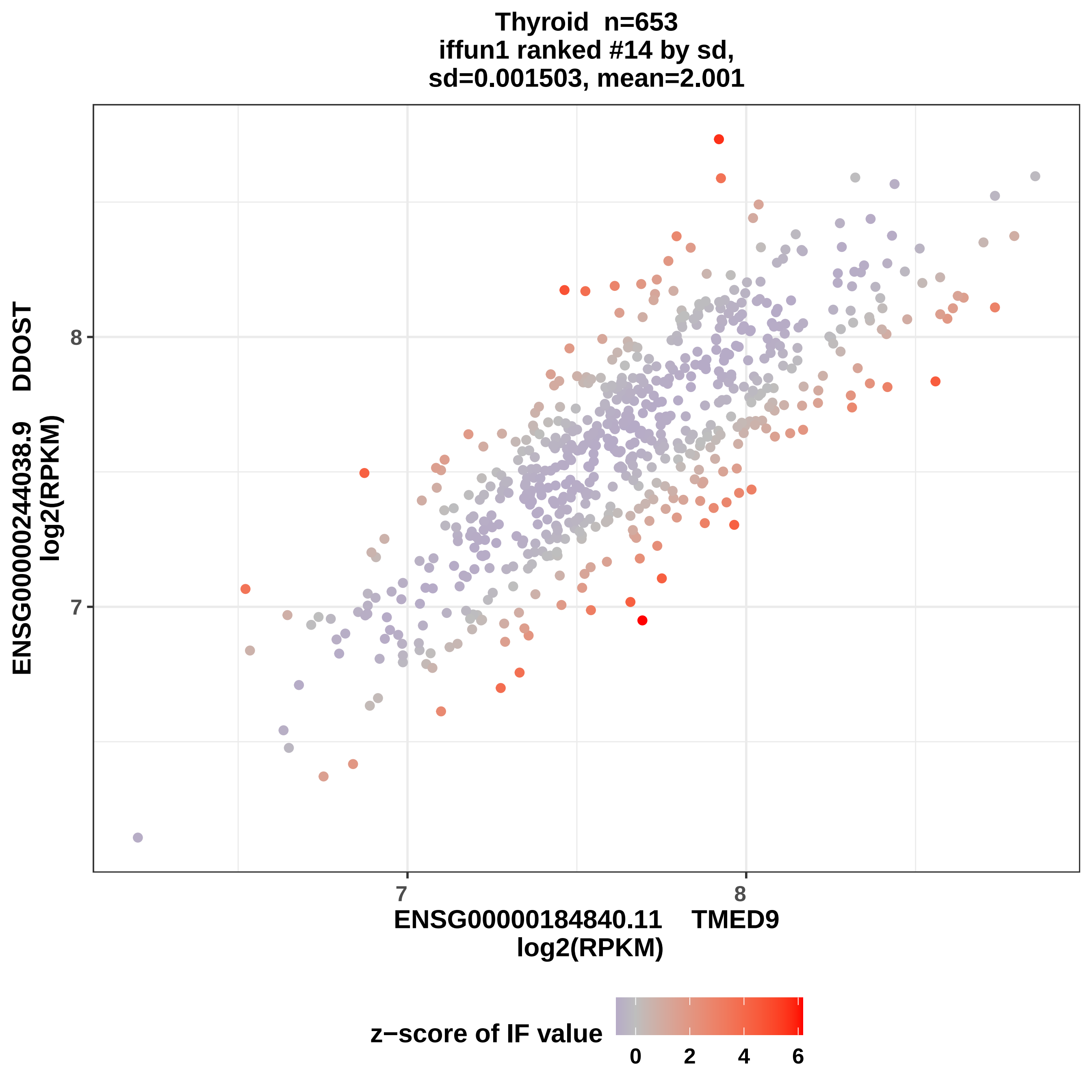
Thyroid n=653
iffun1 ranked #14 by sd,
sd=0.001503, mean=2.001
ENSG00000244038.9 DDOST
log2(RPKM)
8
7
7
8
ENSG00000184840.11 TMED9
log2(RPKM)
z−score of IF value
0
2
4
6

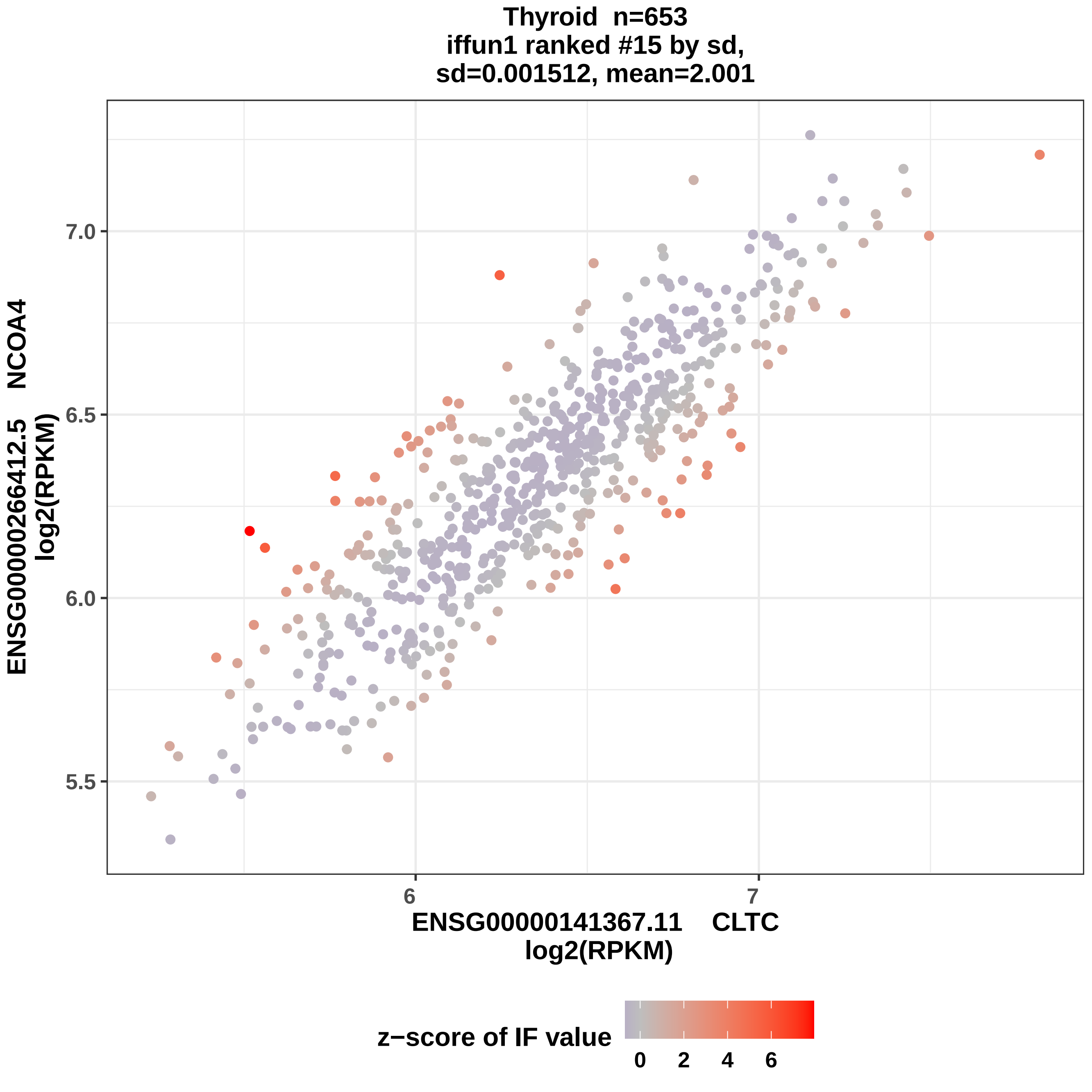
Thyroid n=653
iffun1 ranked #15 by sd,
sd=0.001512, mean=2.001
ENSG00000266412.5 NCOA4
log2(RPKM)
7.0
6.5
6.0
5.5
6
7
ENSG00000141367.11 CLTC
log2(RPKM)
z−score of IF value
0
2
4
6

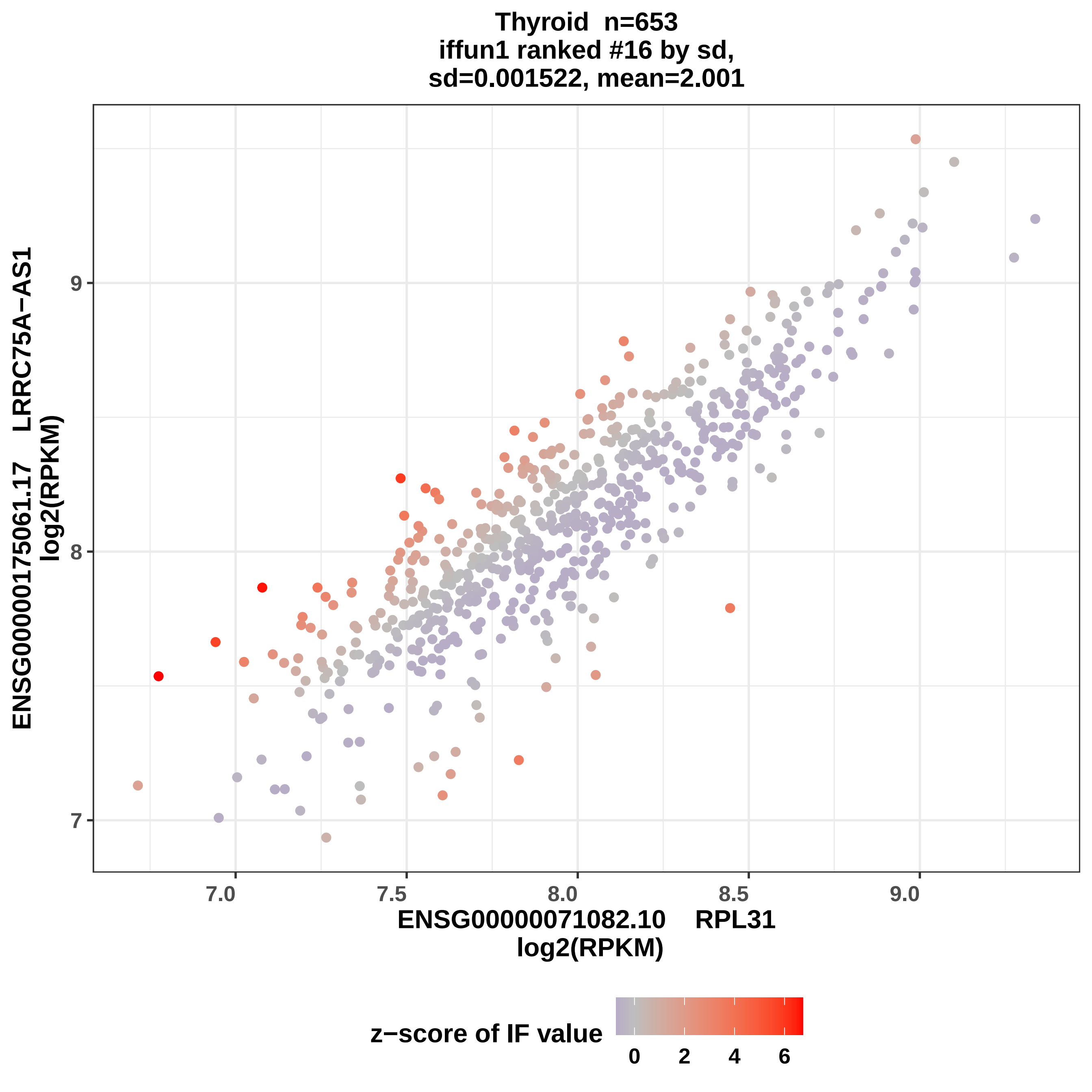
Thyroid n=653
iffun1 ranked #16 by sd,
sd=0.001522, mean=2.001
ENSG00000175061.17 LRRC75A−AS1
log2(RPKM)
9
8
7
7.0
7.5
8.0
8.5
9.0
ENSG00000071082.10 RPL31
log2(RPKM)
z−score of IF value
0
2
4
6

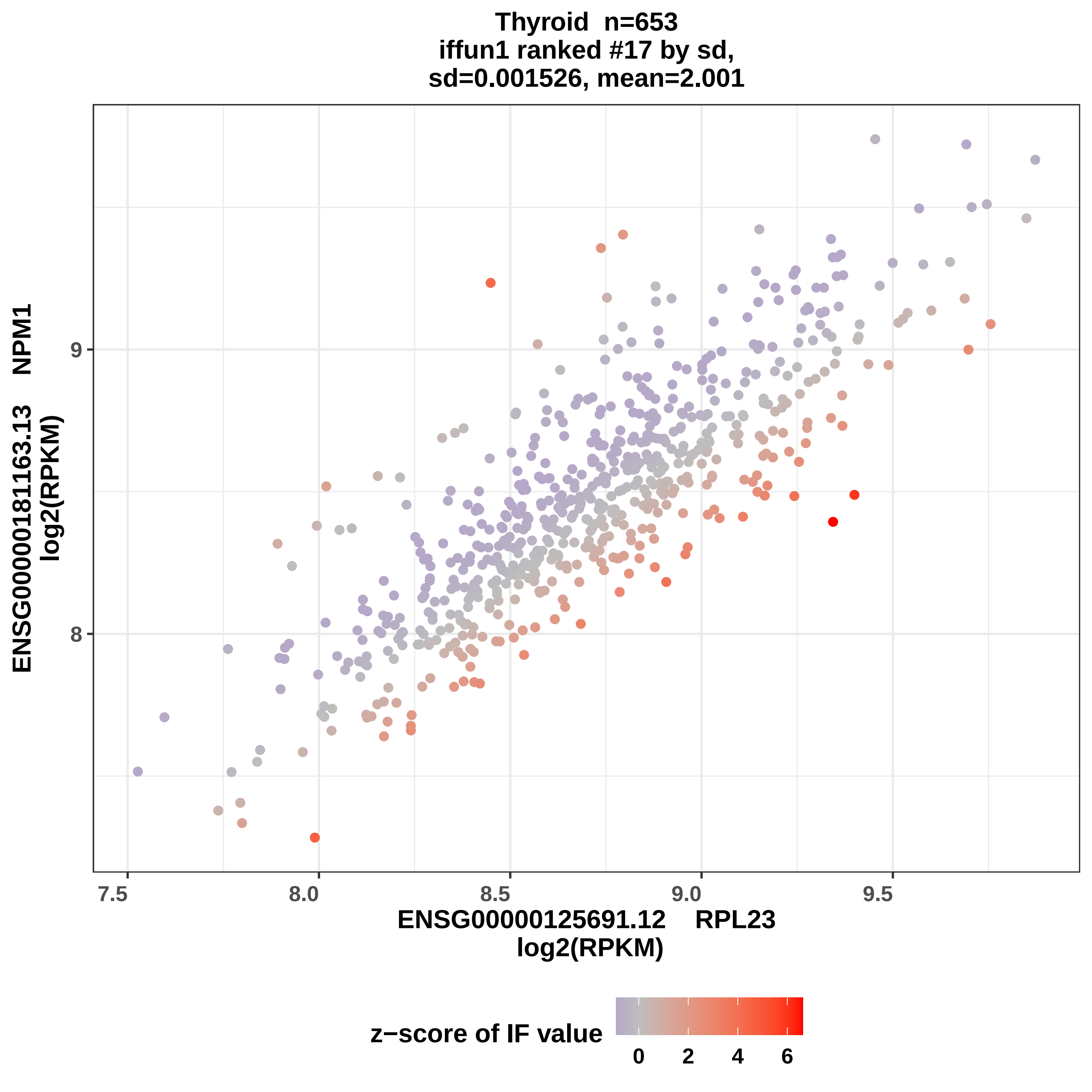

Thyroid n=653
iffun1 ranked #17 by sd,
sd=0.001526, mean=2.001
ENSG00000181163.13 NPM1
log2(RPKM)
9
8
7.5
8.0
8.5
9.0
9.5
ENSG00000125691.12 RPL23
log2(RPKM)
z−score of IF value
0
2
4
6

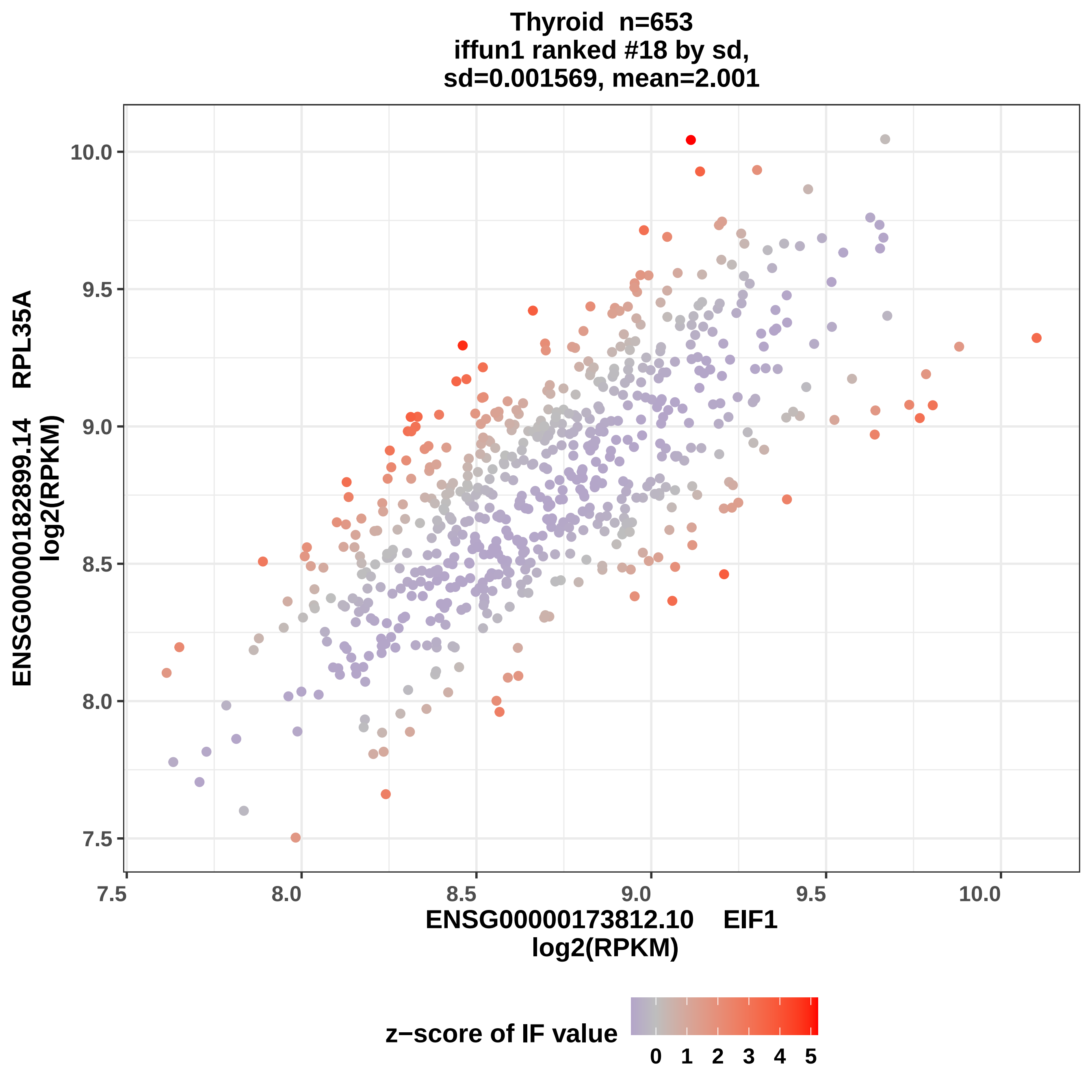

Thyroid n=653
iffun1 ranked #18 by sd,
sd=0.001569, mean=2.001
ENSG00000182899.14 RPL35A
log2(RPKM)
ENSG00000173812.10 EIF1
log2(RPKM)
10.0
9.5
9.0
8.5
8.0
7.5
7.5
8.0
8.5
9.0
9.5
10.0
z−score of IF value
0 1 2 3 4 5

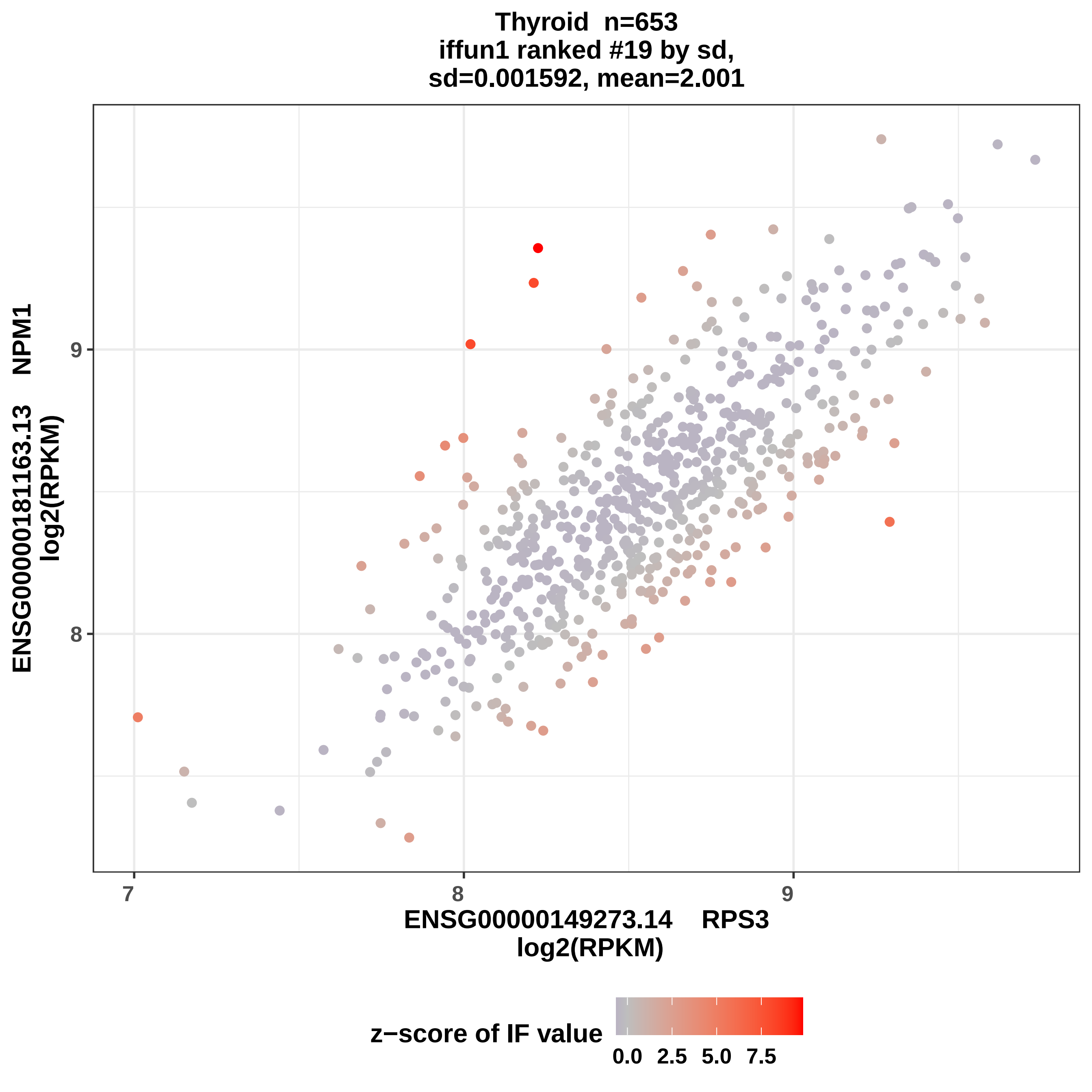
Thyroid n=653
iffun1 ranked #19 by sd,
sd=0.001592, mean=2.001
ENSG00000181163.13 NPM1
log2(RPKM)
9
8
7
8
9
ENSG00000149273.14 RPS3
log2(RPKM)
z−score of IF value
0.0
2.5
5.0
7.5

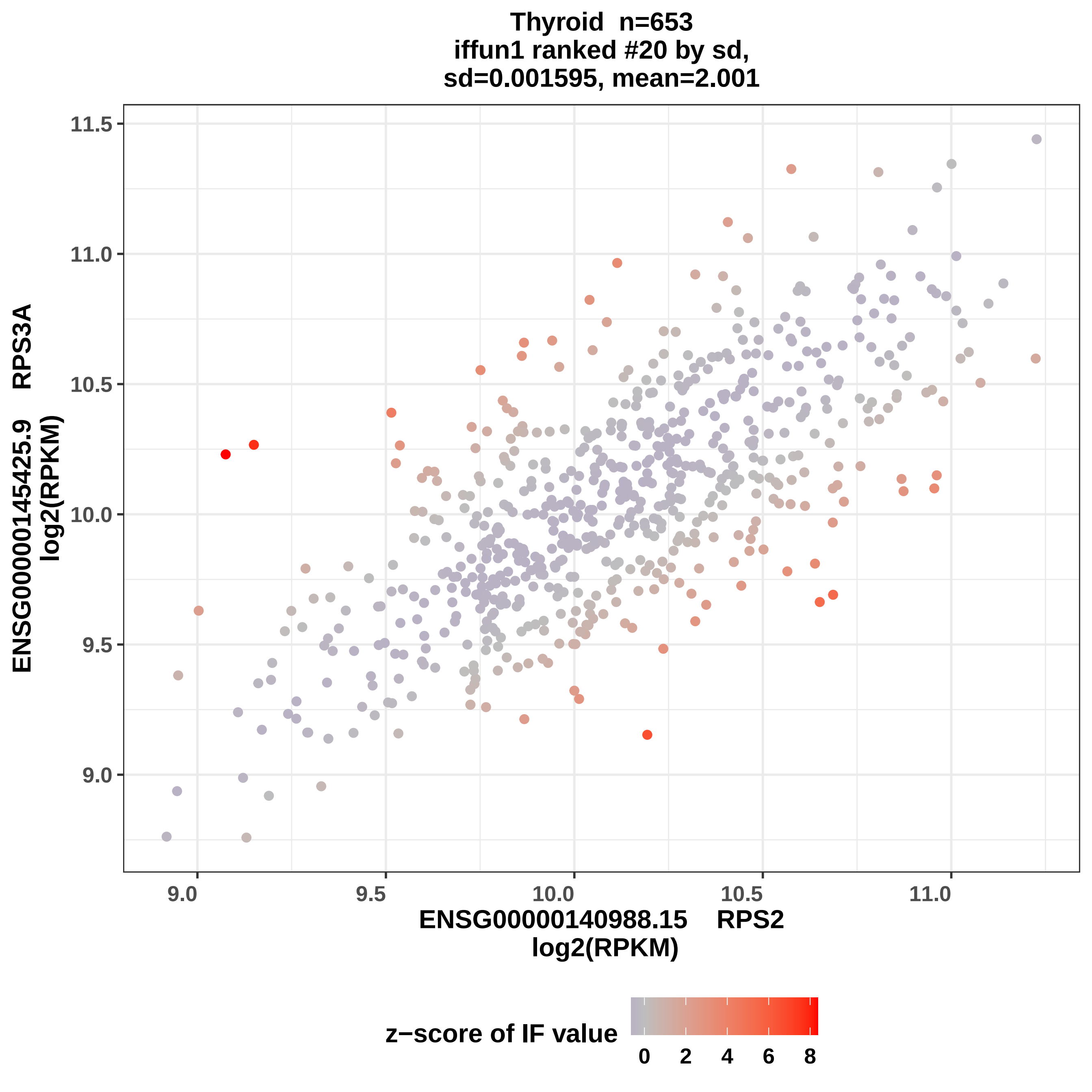

Thyroid n=653
iffun1 ranked #20 by sd,
sd=0.001595, mean=2.001
ENSG00000145425.9 RPS3A
log2(RPKM)
11.5
11.0
10.5
10.0
9.5
9.0
9.0
9.5
10.0
10.5
11.0
ENSG00000140988.15 RPS2
log2(RPKM)
z−score of IF value
0
2
4
6
8

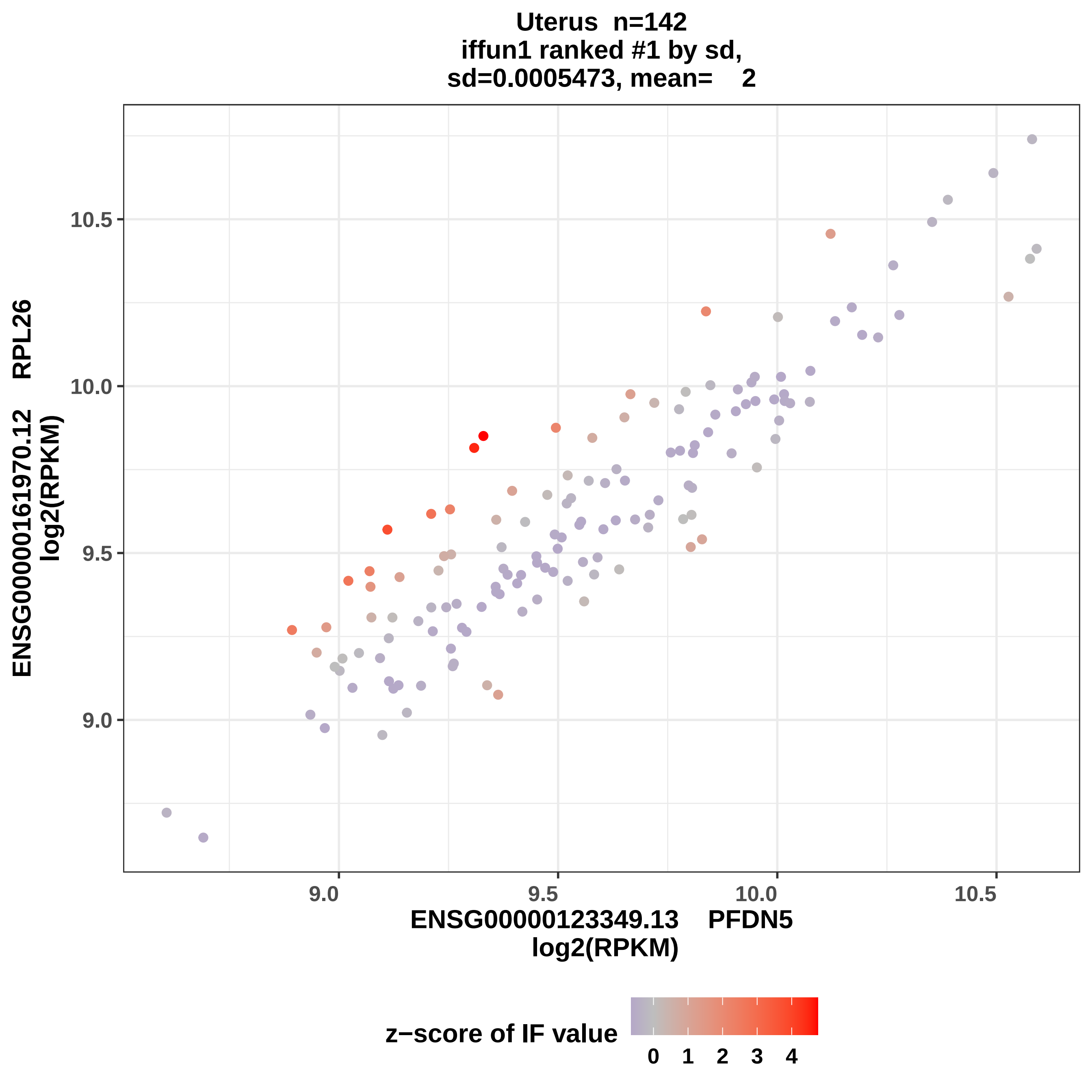
Uterus n=142
iffun1 ranked #1 by sd,
sd=0.0005473, mean= 2
ENSG00000161970.12 RPL26
log2(RPKM)
10.5
10.0
9.5
9.0
9.0
9.5
10.0
10.5
ENSG00000123349.13 PFDN5
log2(RPKM)
z−score of IF value
0
1
2
3
4

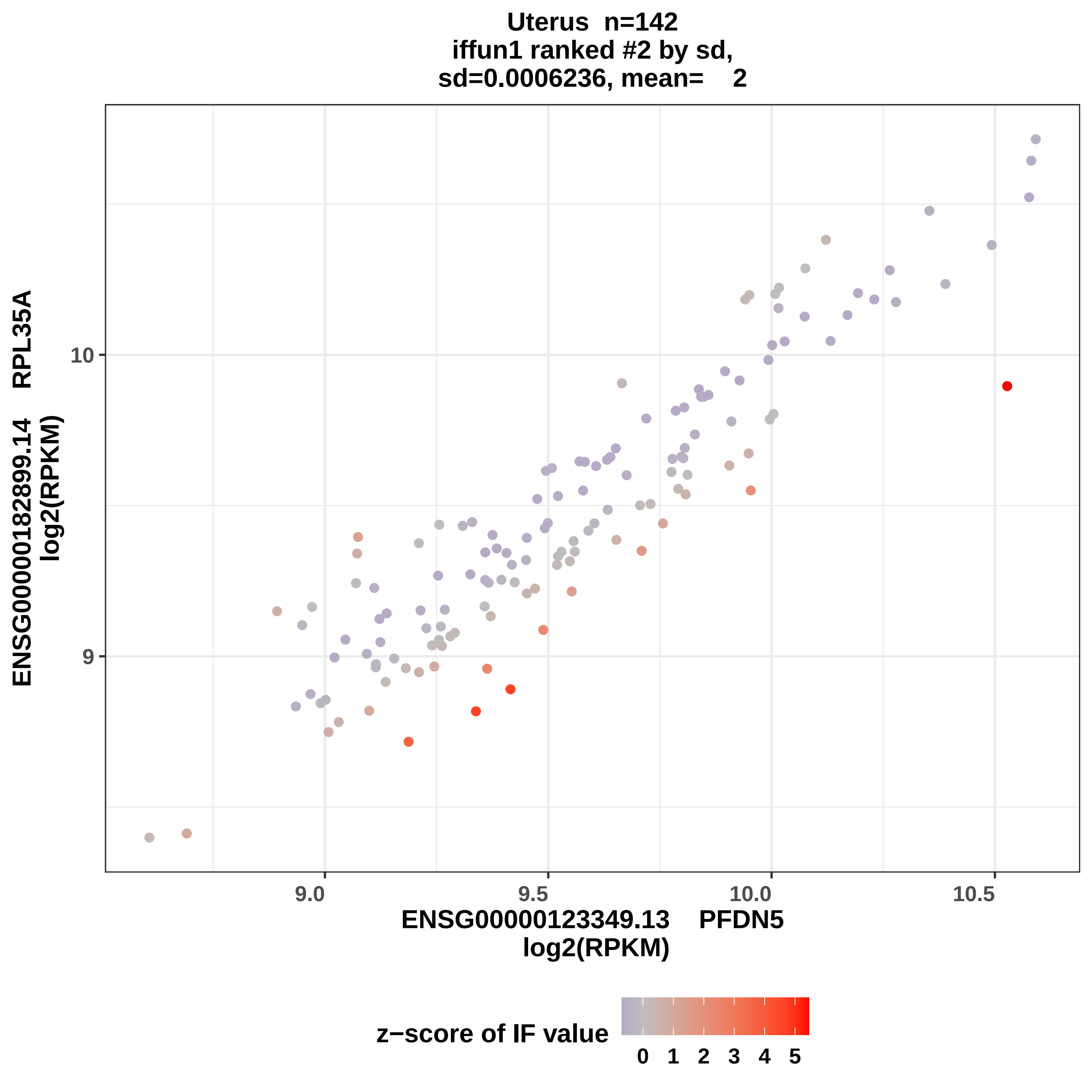
Uterus n=142
iffun1 ranked #2 by sd,
sd=0.0006236, mean= 2
ENSG00000182899.14 RPL35A
log2(RPKM)
10
9
9.0
9.5
10.0
10.5
ENSG00000123349.13 PFDN5
log2(RPKM)
z−score of IF value
0
1
2
3
4
5

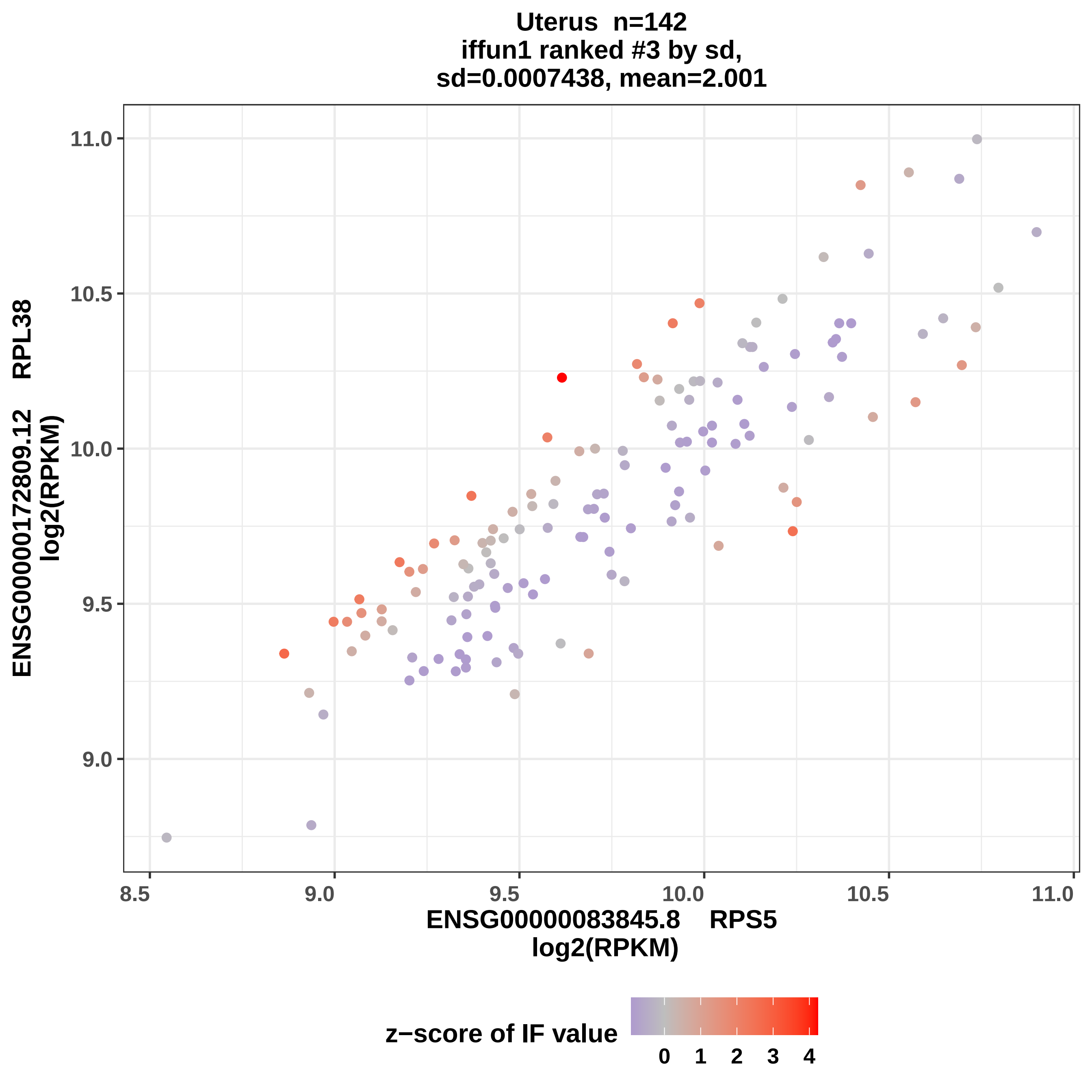
Uterus n=142
iffun1 ranked #3 by sd,
sd=0.0007438, mean=2.001
ENSG00000172809.12 RPL38
log2(RPKM)
11.0
10.5
10.0
9.5
9.0
8.5
9.0
9.5
10.0
10.5
11.0
ENSG00000083845.8 RPS5
log2(RPKM)
z−score of IF value
0
1
2
3
4

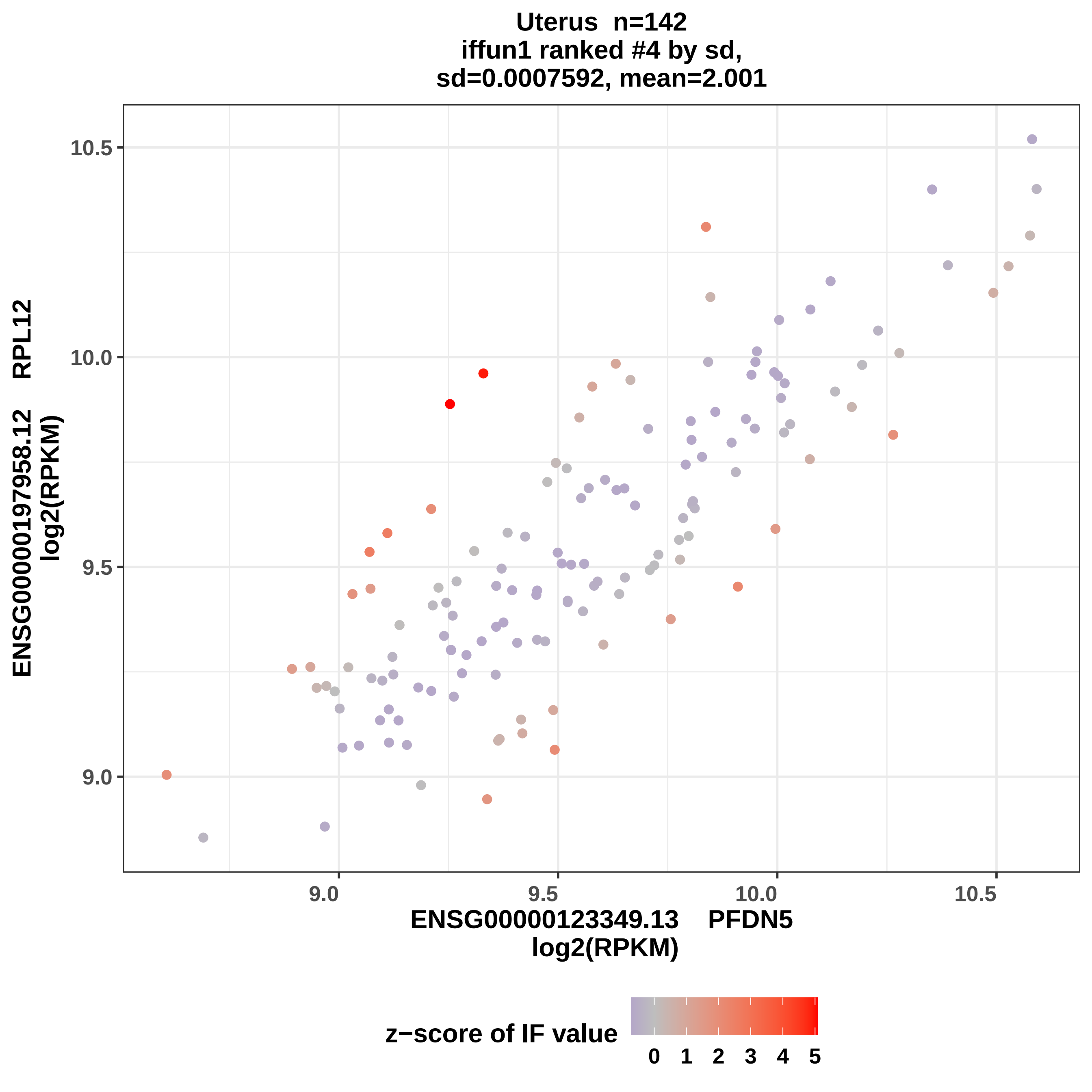

Uterus n=142
iffun1 ranked #4 by sd,
sd=0.0007592, mean=2.001
ENSG00000197958.12 RPL12
log2(RPKM)
10.5
10.0
9.5
9.0
ENSG00000123349.13 PFDN5
log2(RPKM)
z−score of IF value
0 1 2 3 4 5

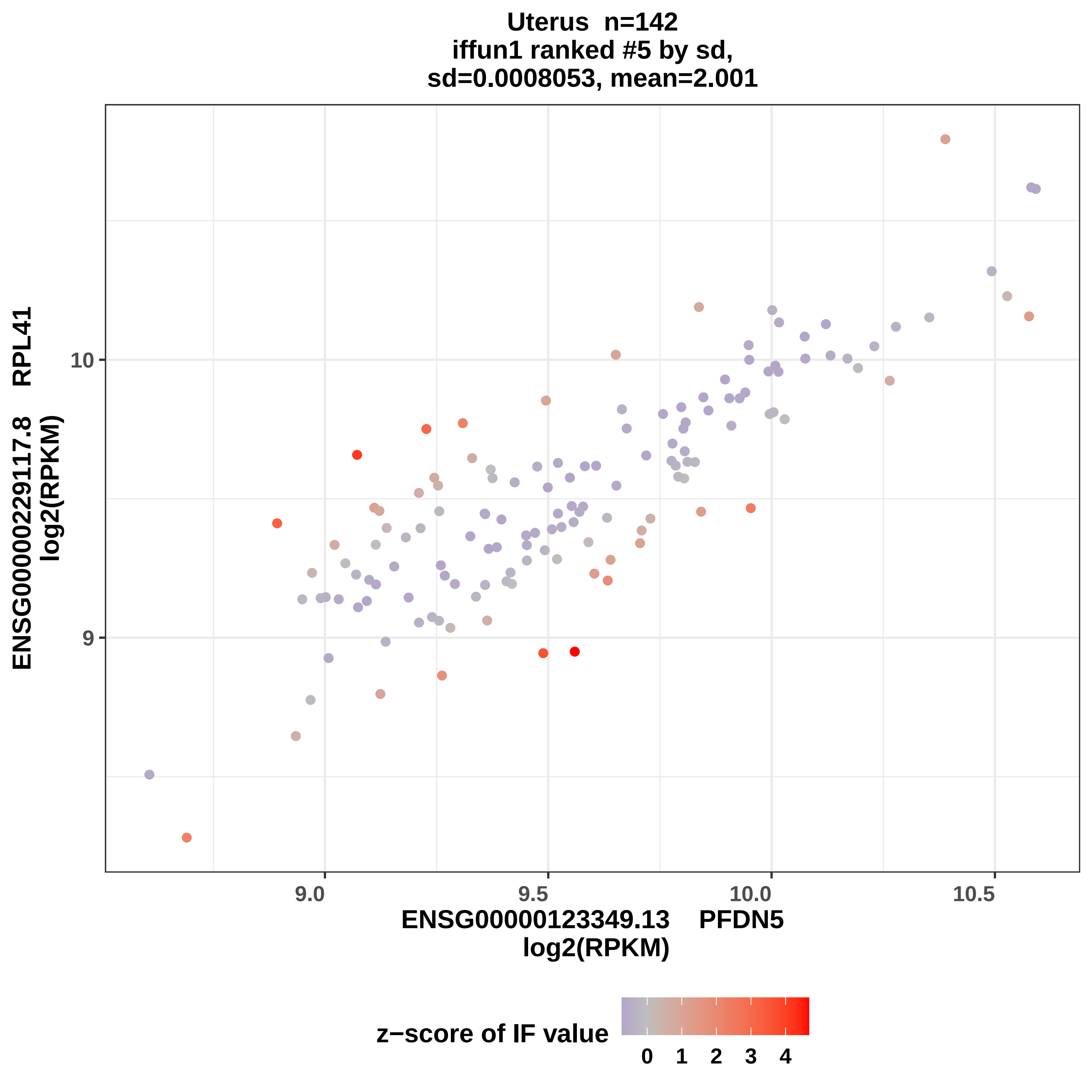
Uterus n=142
iffun1 ranked #5 by sd,
sd=0.0008053, mean=2.001
ENSG00000229117.8 RPL41
log2(RPKM)
10
9
9.0
9.5
10.0
10.5
ENSG00000123349.13 PFDN5
log2(RPKM)
z−score of IF value
0
1
2
3
4

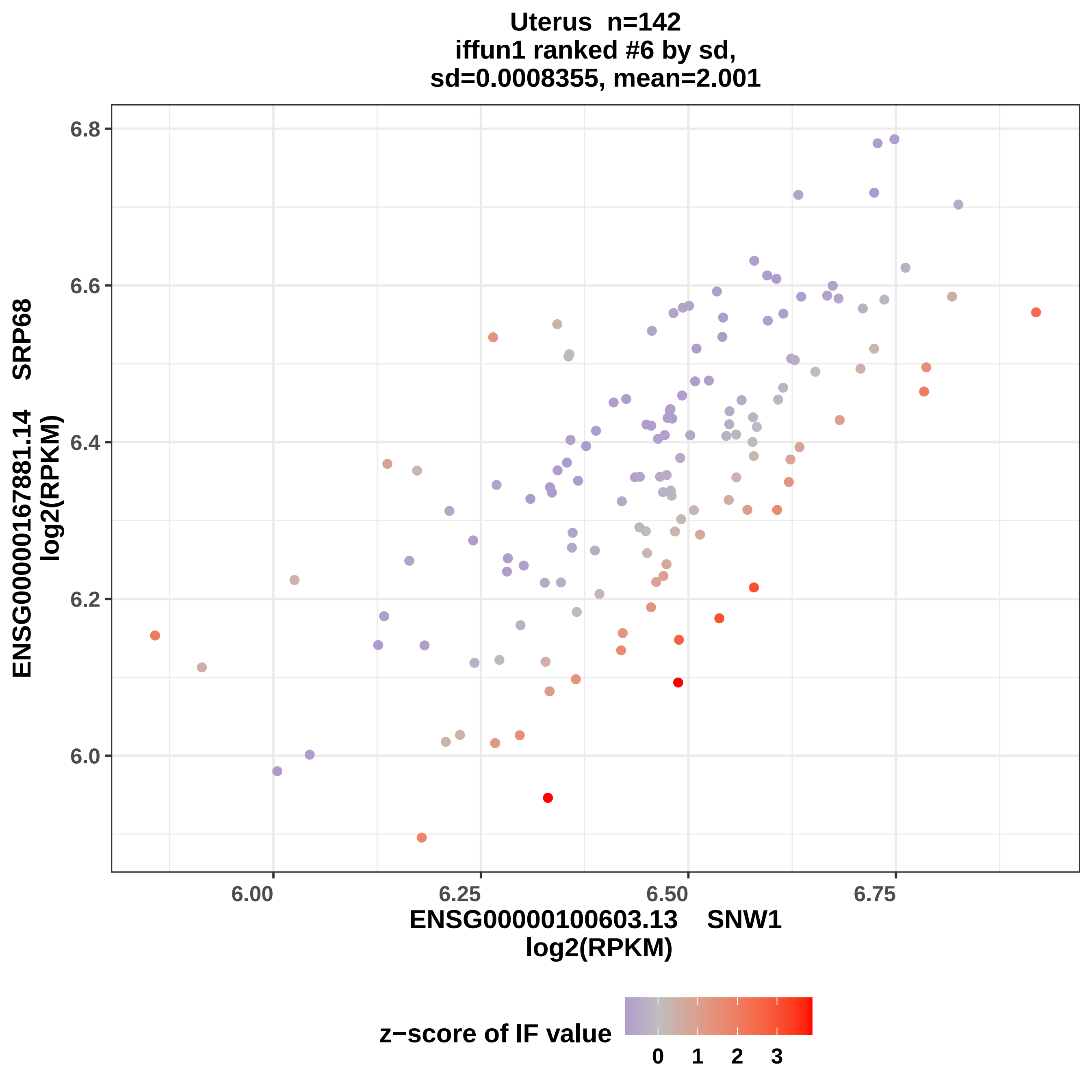
Uterus n=142
iffun1 ranked #6 by sd,
sd=0.0008355, mean=2.001
ENSG00000167881.14 SRP68
log2(RPKM)
6.8
6.6
6.4
6.2
6.0
6.00
6.25
6.50
6.75
ENSG00000100603.13 SNW1
log2(RPKM)
z−score of IF value
0
1
2
3

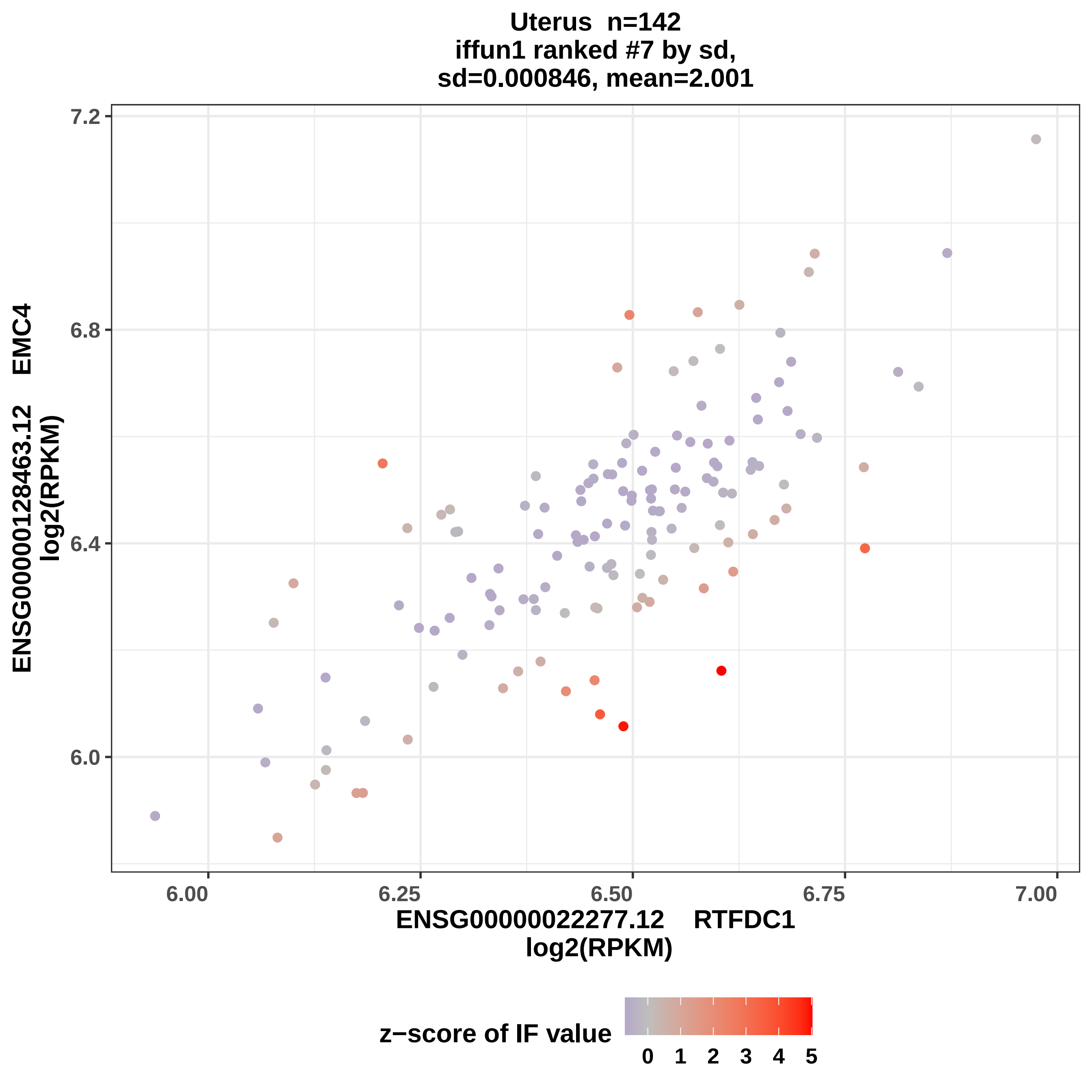

Uterus n=142
iffun1 ranked #7 by sd,
sd=0.000846, mean=2.001
7.2
6.8
6.4
6.0
6.00
6.25
6.50
6.75
7.00
ENSG00000128463.12 EMC4
log2(RPKM)
ENSG00000022277.12 RTFDC1
log2(RPKM)
z−score of IF value
0 1 2 3 4 5

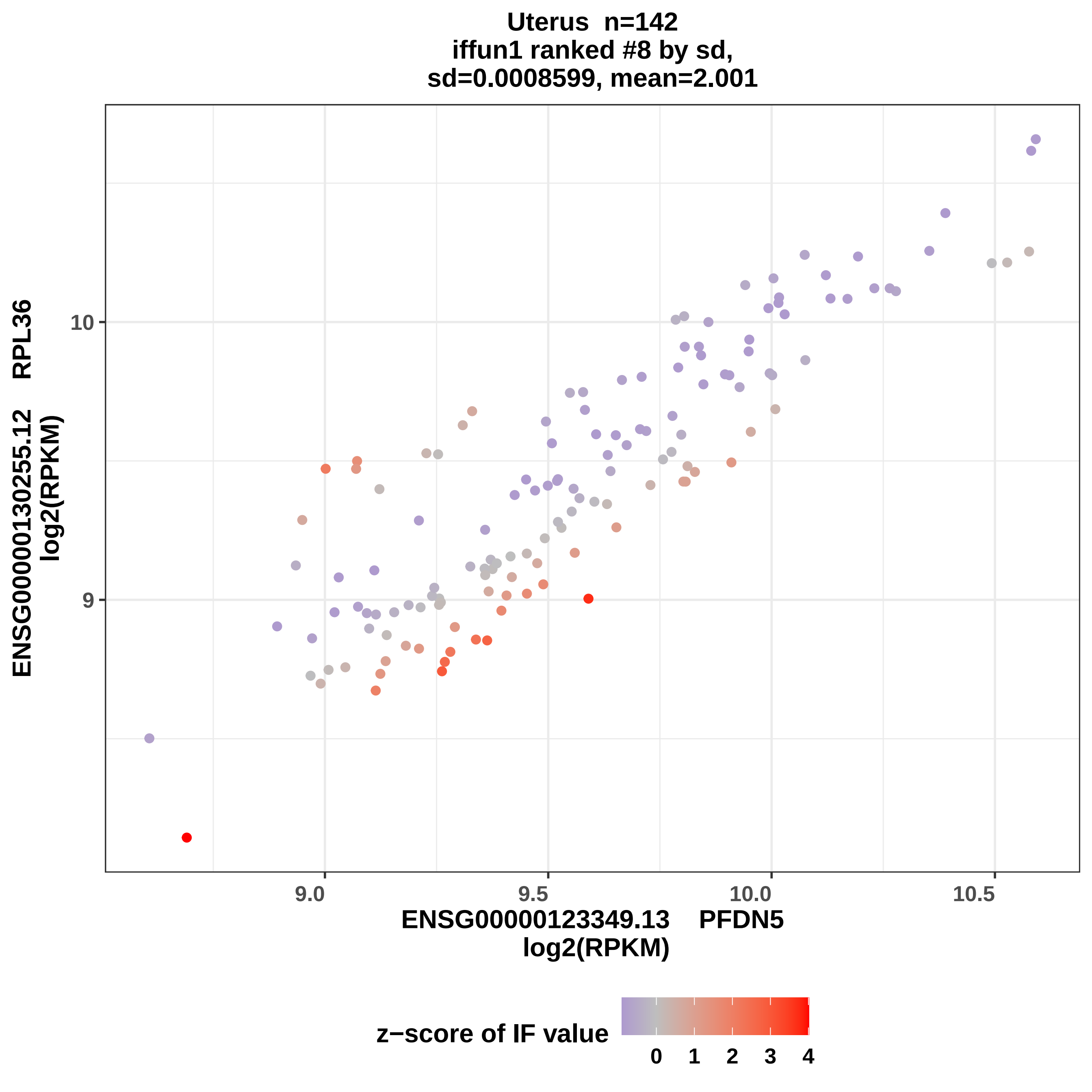

Uterus n=142
iffun1 ranked #8 by sd,
sd=0.0008599, mean=2.001
ENSG00000130255.12 RPL36
log2(RPKM)
10
9
9.0
9.5
10.0
10.5
ENSG00000123349.13 PFDN5
log2(RPKM)
z−score of IF value
0
1
2
3
4

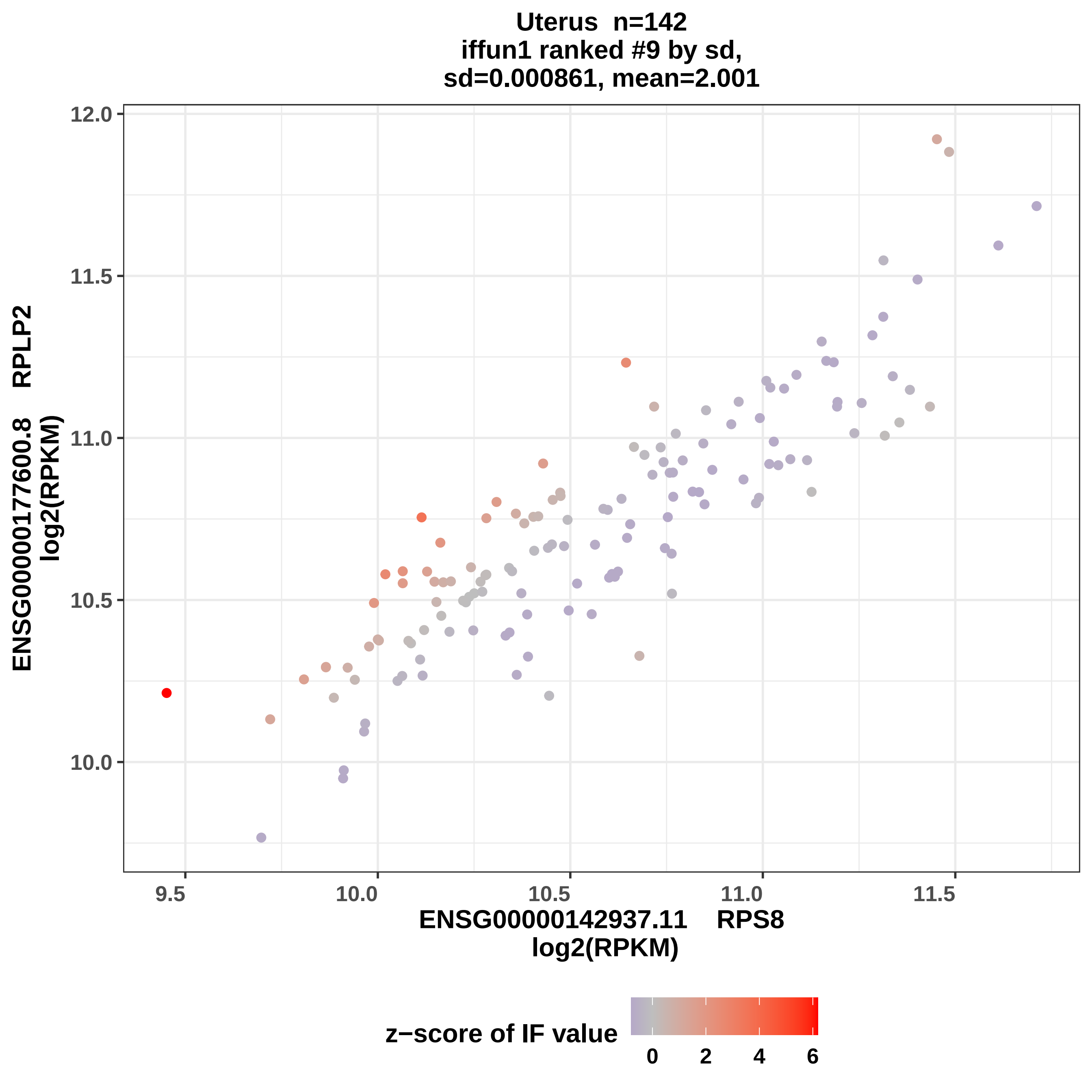
Uterus n=142
iffun1 ranked #9 by sd,
sd=0.000861, mean=2.001
ENSG00000177600.8 RPLP2
log2(RPKM)
ENSG00000142937.11 RPS8
log2(RPKM)
12.0
11.5
11.0
10.5
10.0
9.5
10.0
10.5
11.0
11.5
z−score of IF value
0
2
4
6

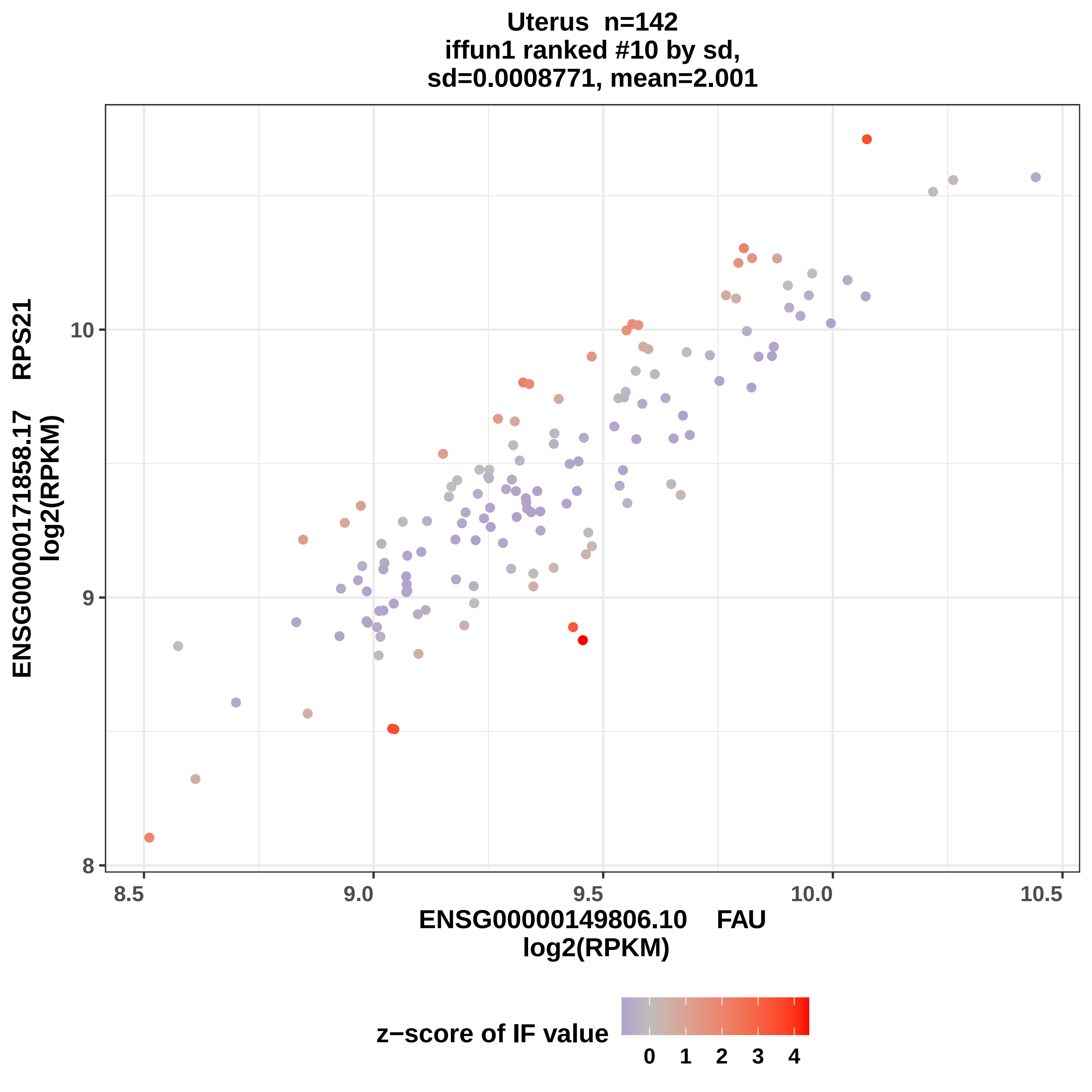
Uterus n=142
iffun1 ranked #10 by sd,
sd=0.0008771, mean=2.001
ENSG00000171858.17 RPS21
log2(RPKM)
10
9
8
8.5
9.0
9.5
10.0
10.5
ENSG00000149806.10 FAU
log2(RPKM)
z−score of IF value
0
1
2
3
4

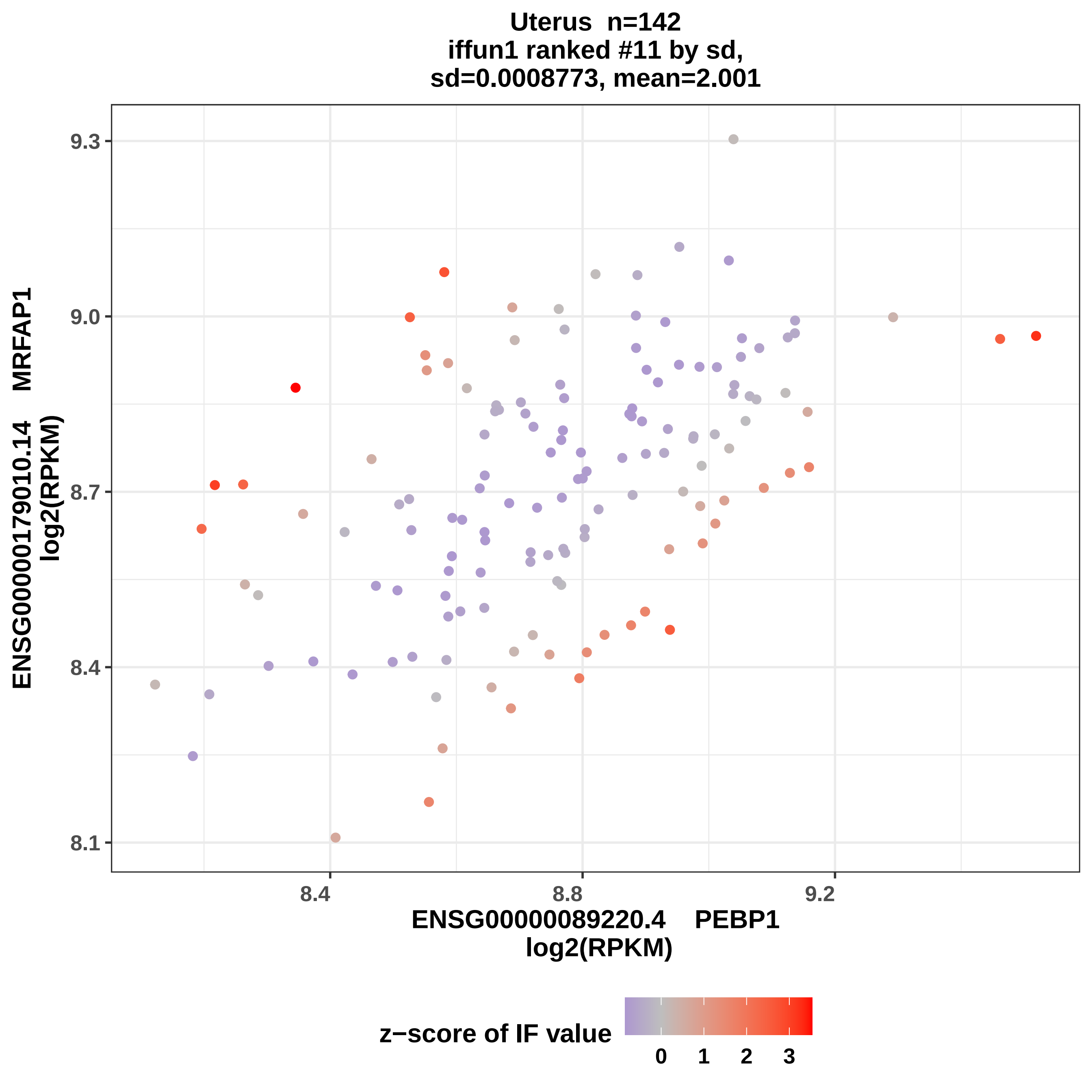

Uterus n=142
iffun1 ranked #11 by sd,
sd=0.0008773, mean=2.001
9.3
9.0
8.7
8.4
8.1
ENSG00000179010.14 MRFAP1
log2(RPKM)
8.4
8.8
9.2
ENSG00000089220.4 PEBP1
log2(RPKM)
z−score of IF value
0
1
2
3

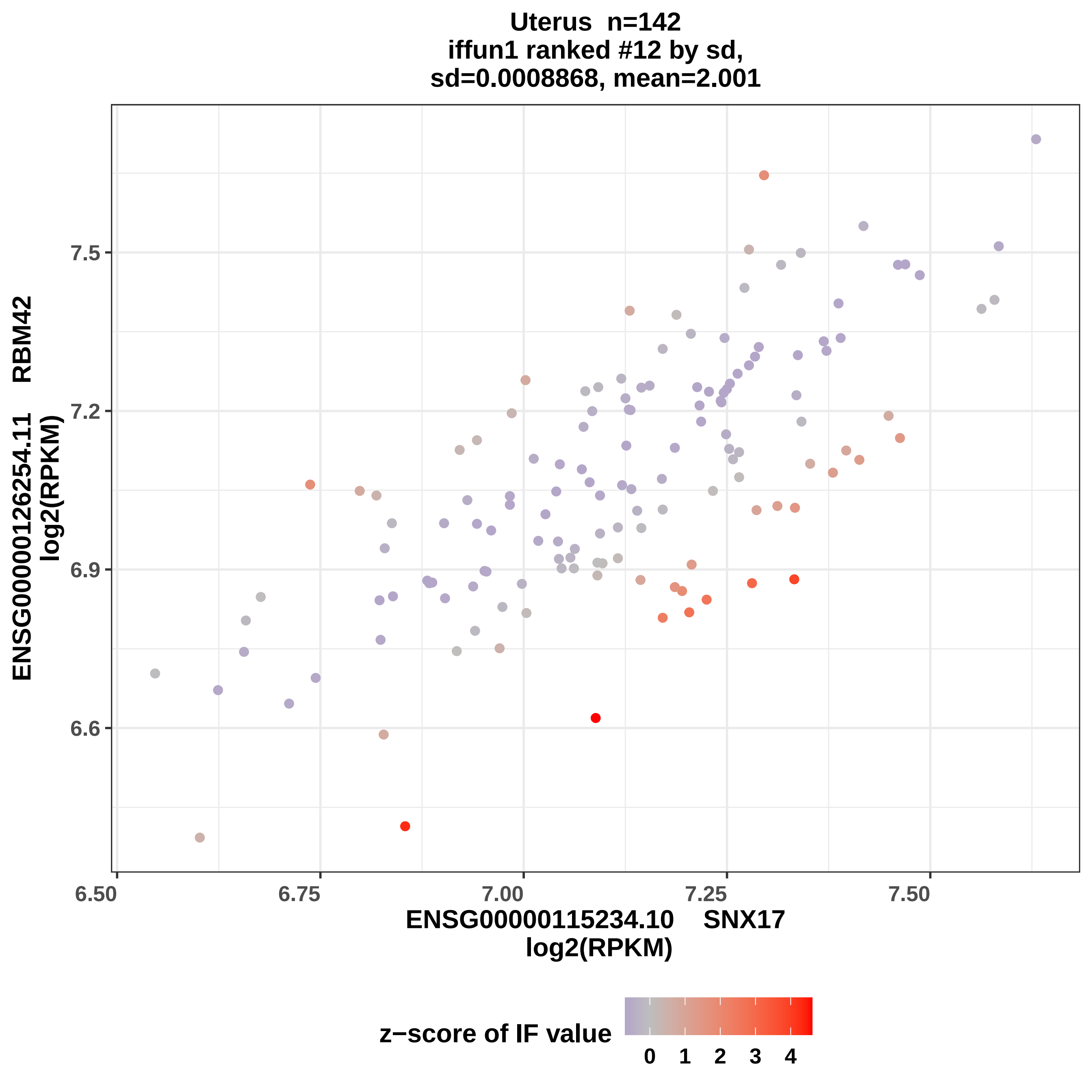
Uterus n=142
iffun1 ranked #12 by sd,
sd=0.0008868, mean=2.001
ENSG00000126254.11 RBM42
log2(RPKM)
7.5
7.2
6.9
6.6
6.50
6.75
7.00
7.25
7.50
ENSG00000115234.10 SNX17
log2(RPKM)
z−score of IF value
0
1
2
3
4

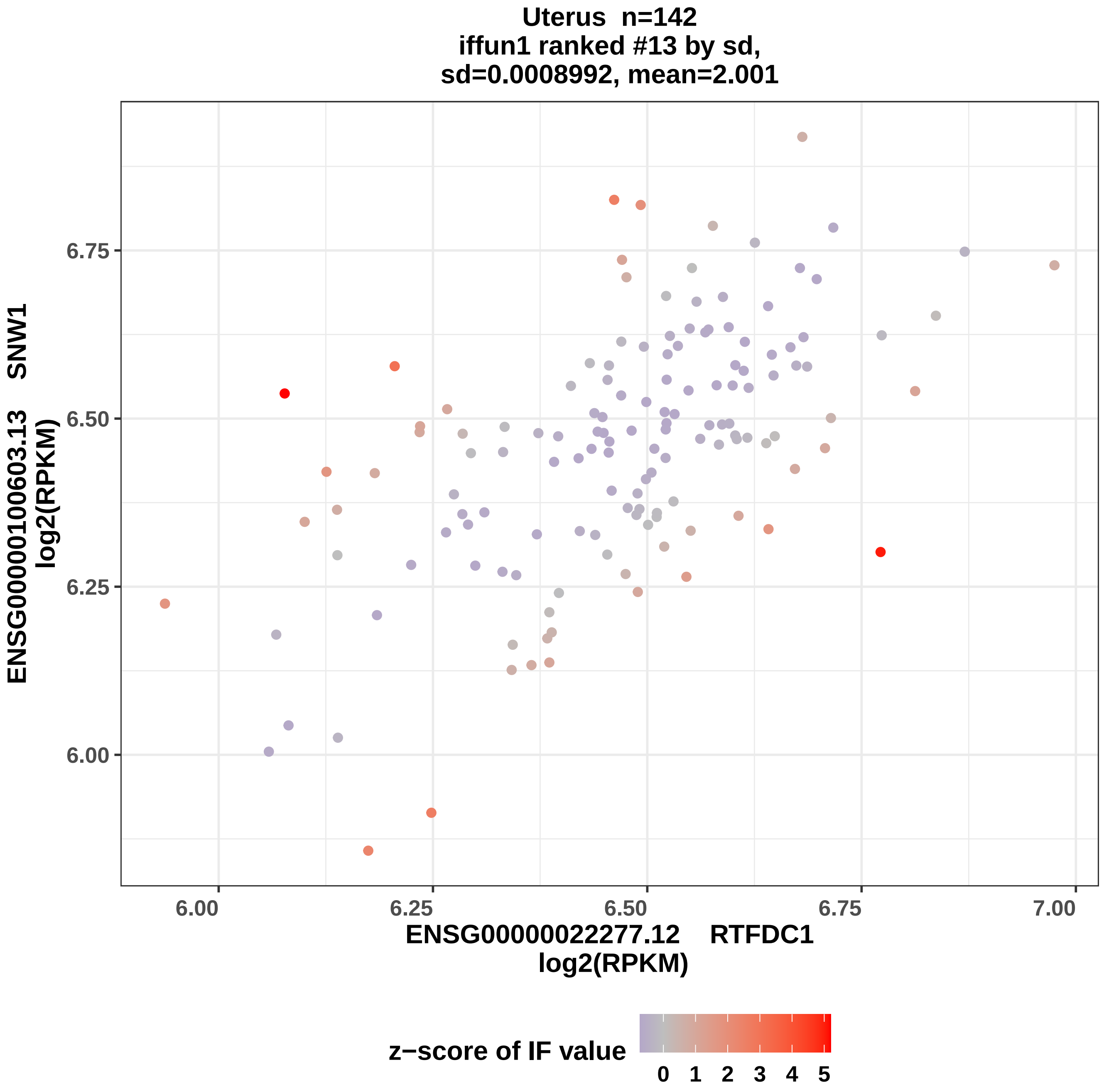
Uterus n=142
iffun1 ranked #13 by sd,
sd=0.0008992, mean=2.001
ENSG00000100603.13 SNW1
log2(RPKM)
6.75
6.50
6.25
6.00
6.00
6.25
6.50
6.75
7.00
ENSG00000022277.12 RTFDC1
log2(RPKM)
z−score of IF value
0 1 2 3 4 5

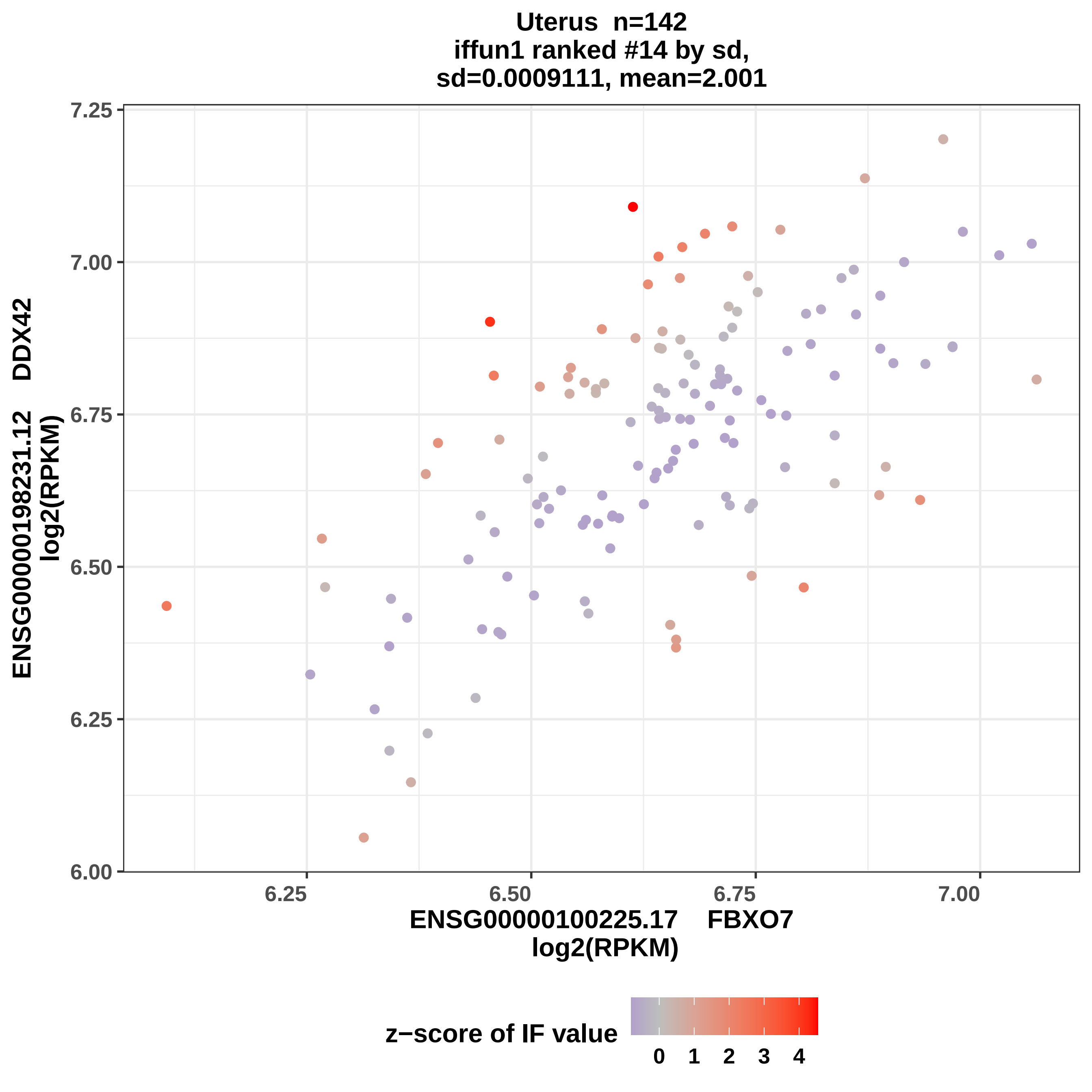
Uterus n=142
iffun1 ranked #14 by sd,
sd=0.0009111, mean=2.001
ENSG00000198231.12 DDX42
log2(RPKM)
ENSG00000100225.17 FBXO7
log2(RPKM)
7.25
7.00
6.75
6.50
6.25
6.00
6.25
6.50
6.75
7.00
z−score of IF value
0
1
2
3
4

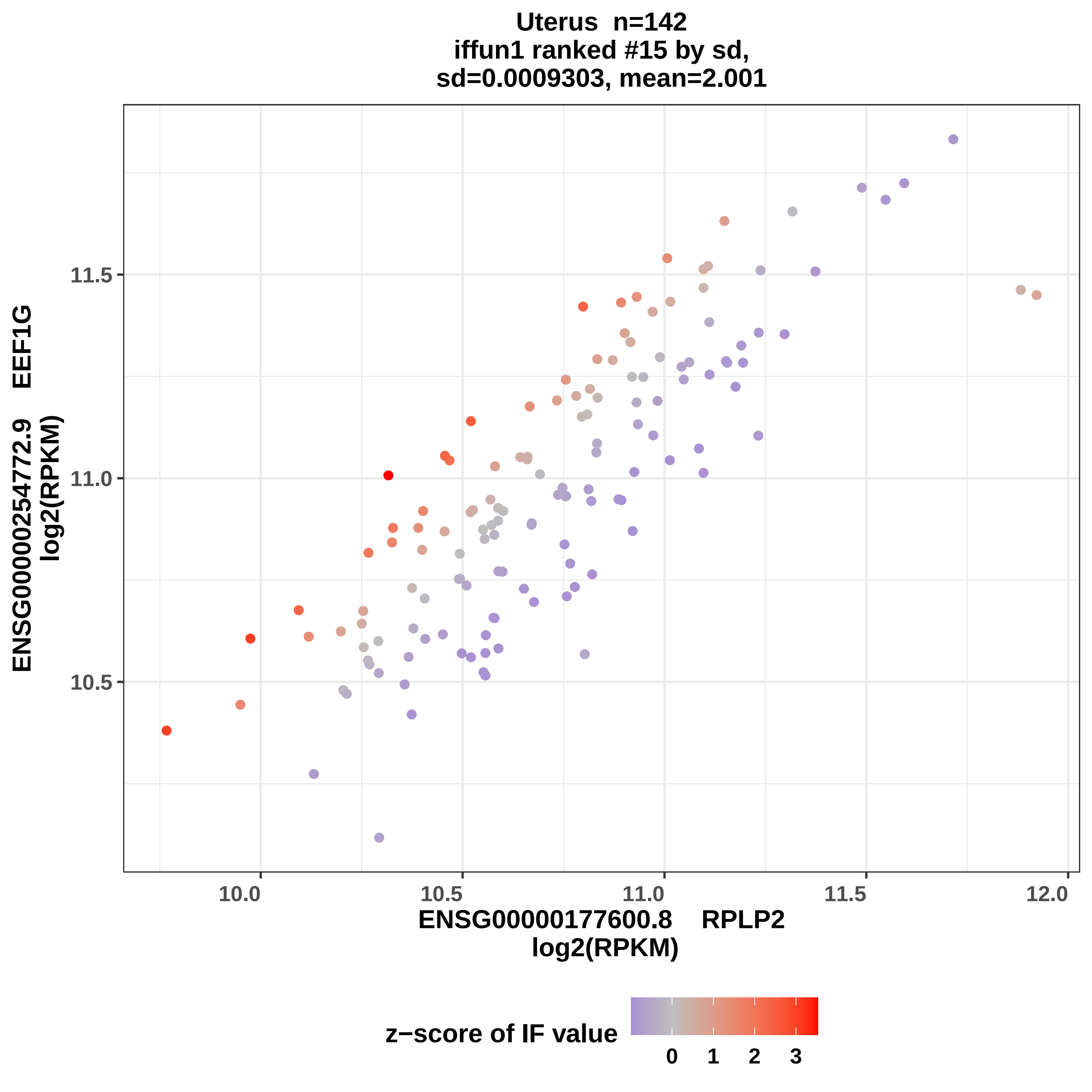
Uterus n=142
iffun1 ranked #15 by sd,
sd=0.0009303, mean=2.001
ENSG00000254772.9 EEF1G
log2(RPKM)
11.5
11.0
10.5
10.0
10.5
11.0
11.5
12.0
ENSG00000177600.8 RPLP2
log2(RPKM)
z−score of IF value
0
1
2
3

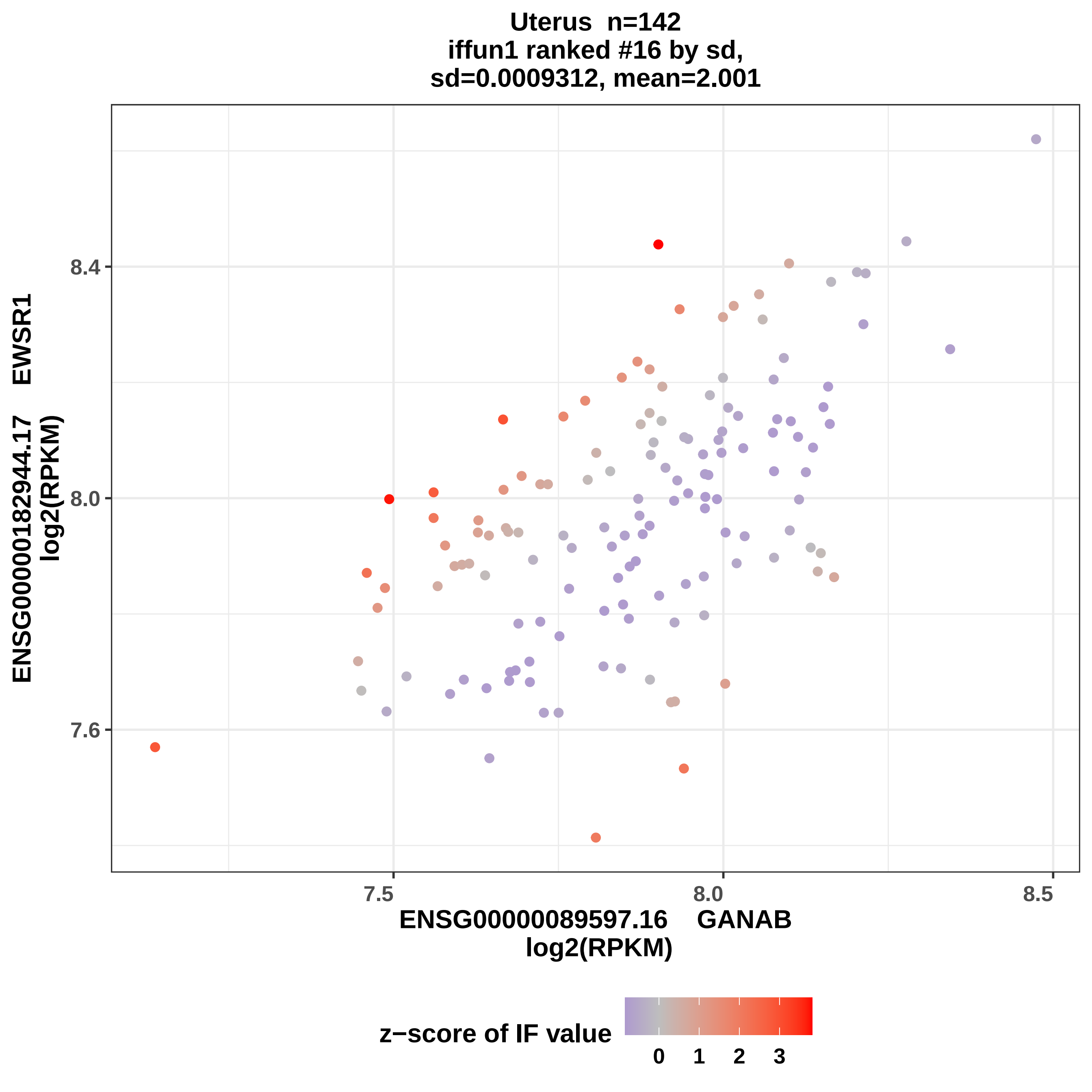

Uterus n=142
iffun1 ranked #16 by sd,
sd=0.0009312, mean=2.001
ENSG00000182944.17 EWSR1
log2(RPKM)
8.4
8.0
7.6
7.5
8.0
8.5
ENSG00000089597.16 GANAB
log2(RPKM)
z−score of IF value
0
1
2
3

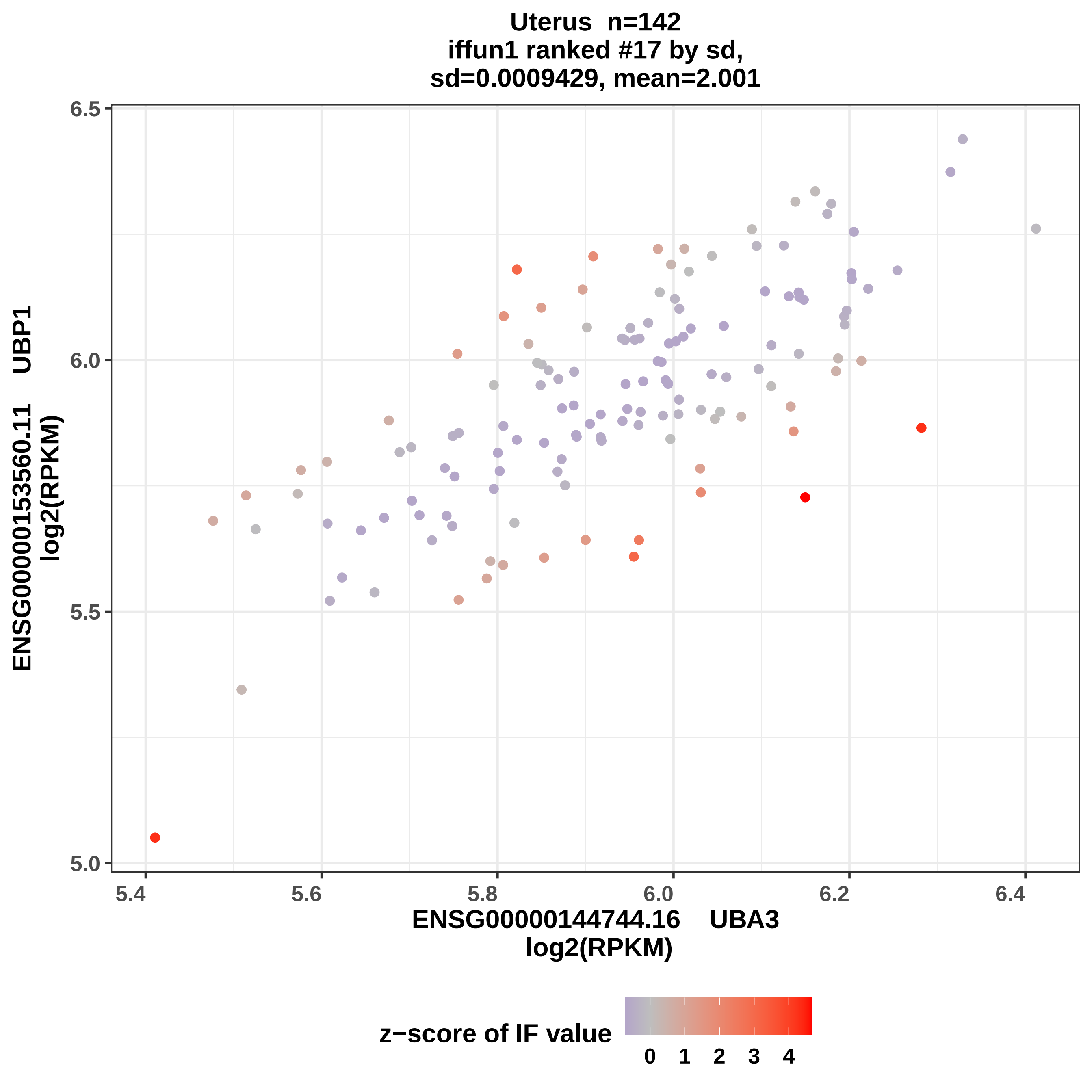
Uterus n=142
iffun1 ranked #17 by sd,
sd=0.0009429, mean=2.001
6.5
6.0
5.5
5.0
5.4
5.6
5.8
6.0
6.2
6.4
ENSG00000153560.11 UBP1
log2(RPKM)
ENSG00000144744.16 UBA3
log2(RPKM)
z−score of IF value
0
1
2
3
4

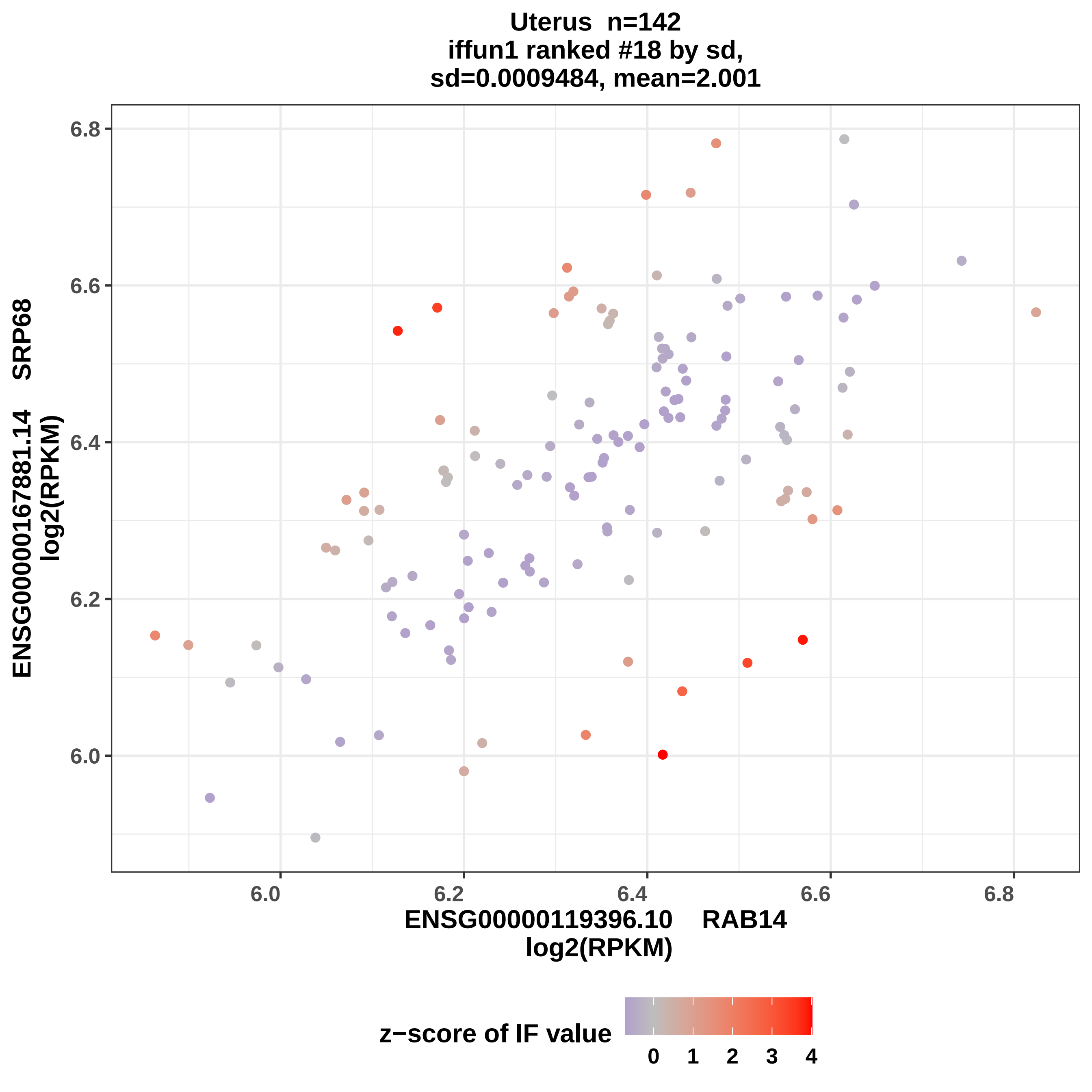
Uterus n=142
iffun1 ranked #18 by sd,
sd=0.0009484, mean=2.001
ENSG00000167881.14 SRP68
log2(RPKM)
6.0
6.2
6.4
6.6
6.8
ENSG00000119396.10 RAB14
log2(RPKM)
z−score of IF value
0
1
2
3
4

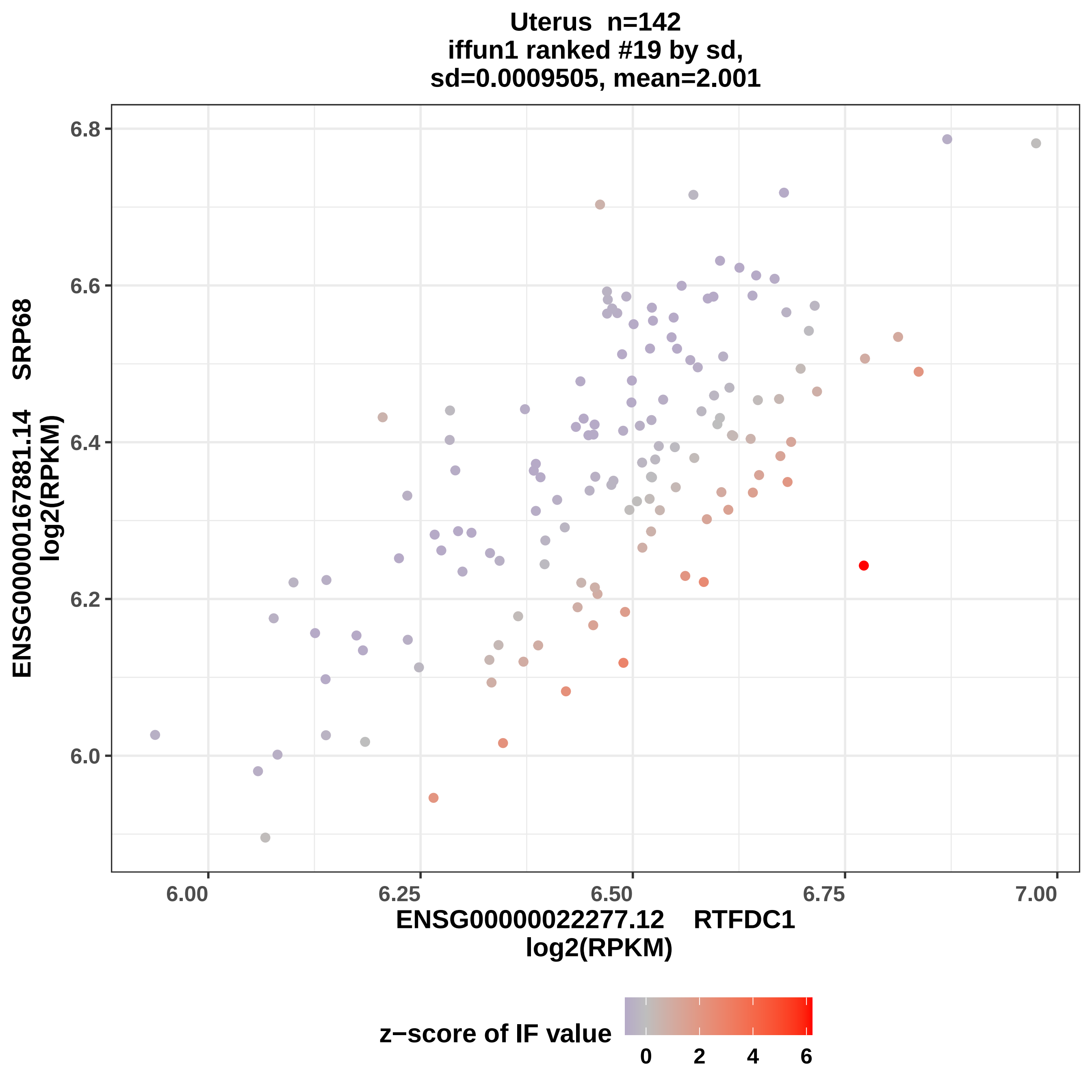
Uterus n=142
iffun1 ranked #19 by sd,
sd=0.0009505, mean=2.001
ENSG00000167881.14 SRP68
log2(RPKM)
6.8
6.6
6.4
6.2
6.0
6.00
6.25
6.50
6.75
7.00
ENSG00000022277.12 RTFDC1
log2(RPKM)
z−score of IF value
0
2
4
6

Uterus n=142
iffun1 ranked #20 by sd,
sd=0.0009546, mean=2.001

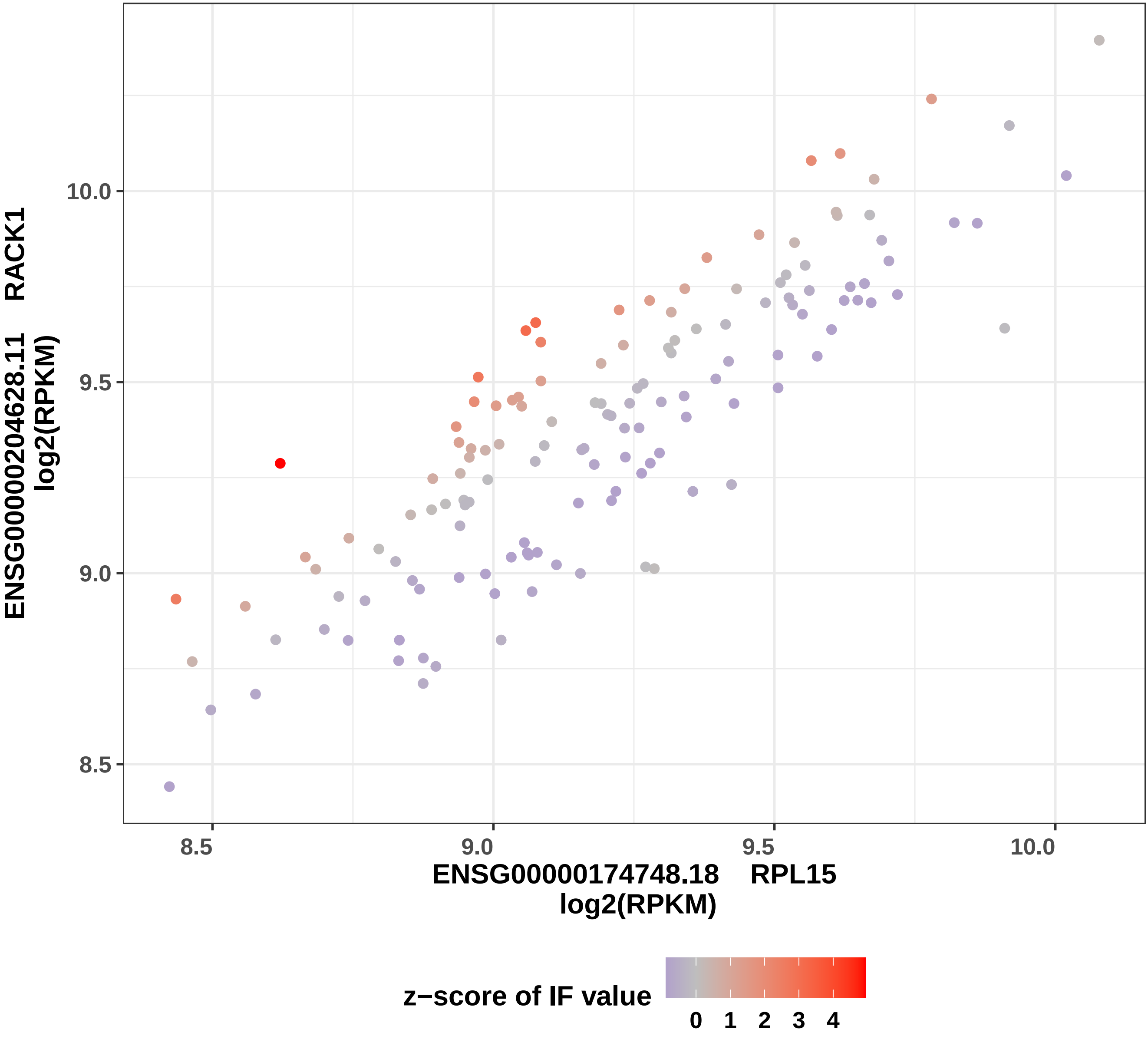

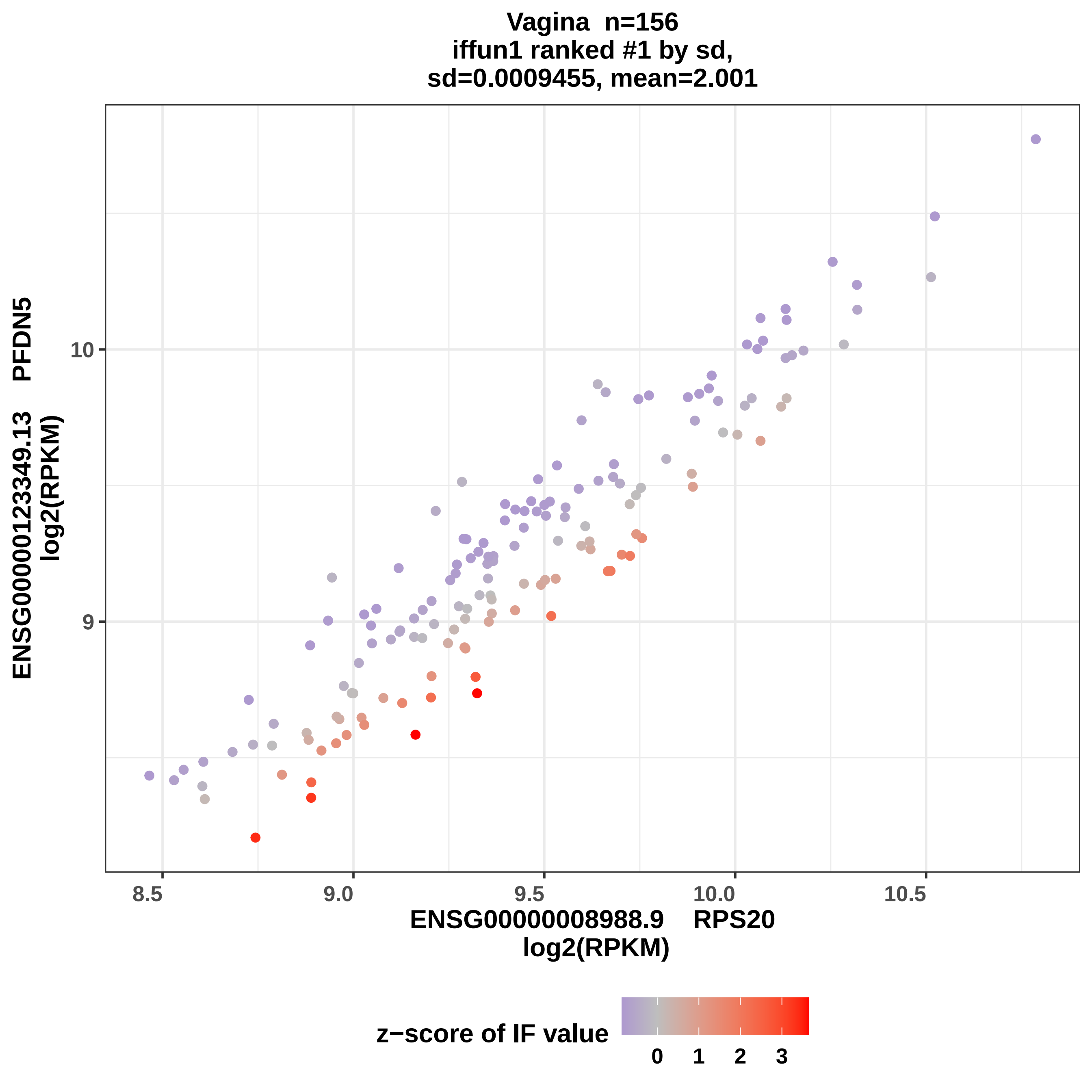
Vagina n=156
iffun1 ranked #1 by sd,
sd=0.0009455, mean=2.001
ENSG00000123349.13 PFDN5
log2(RPKM)
10
9
8.5
9.0
9.5
10.0
10.5
ENSG00000008988.9 RPS20
log2(RPKM)
z−score of IF value
0
1
2
3

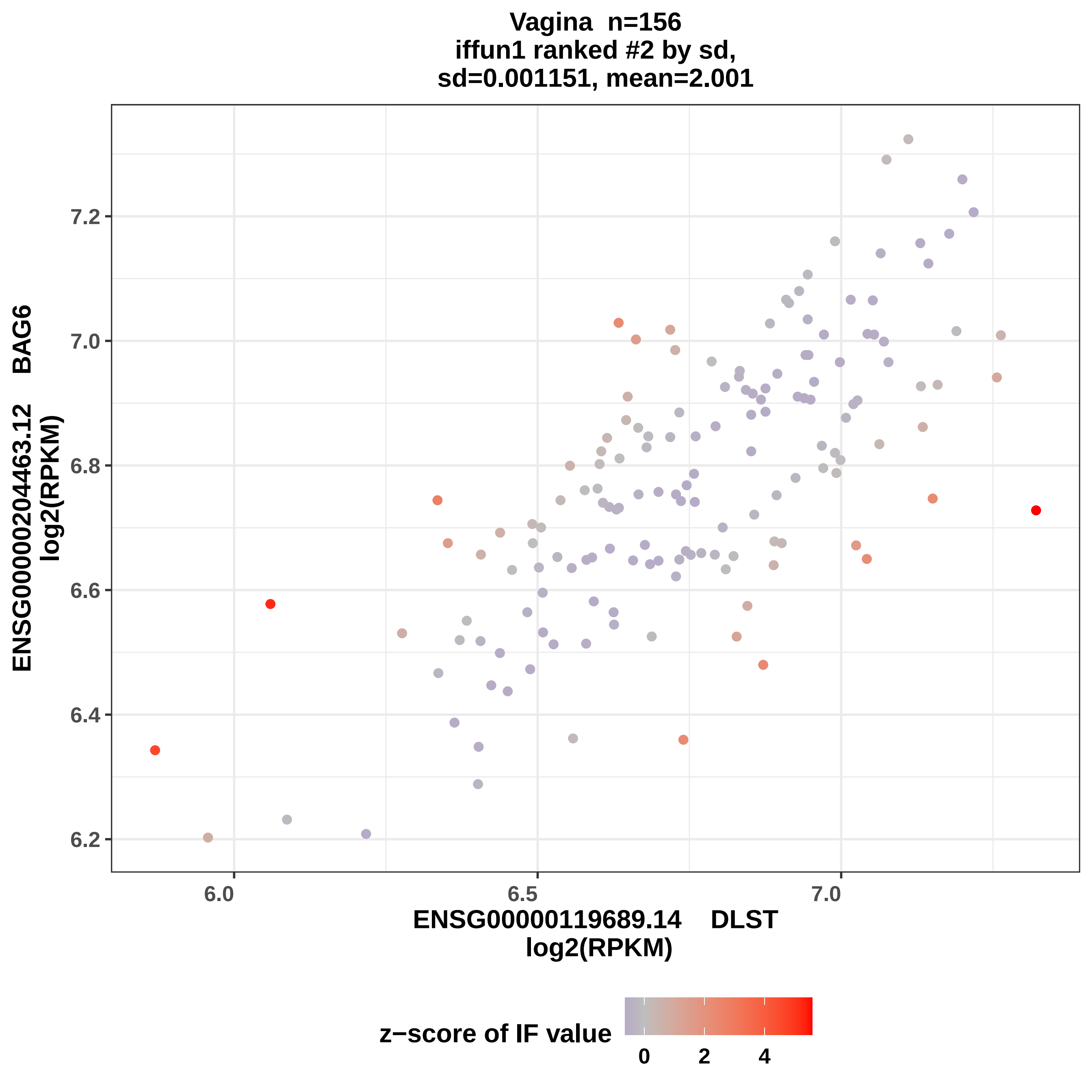
Vagina n=156
iffun1 ranked #2 by sd,
sd=0.001151, mean=2.001
ENSG00000204463.12 BAG6
log2(RPKM)
7.2
7.0
6.8
6.6
6.4
6.2
6.0
6.5
7.0
ENSG00000119689.14 DLST
log2(RPKM)
z−score of IF value
0
2
4

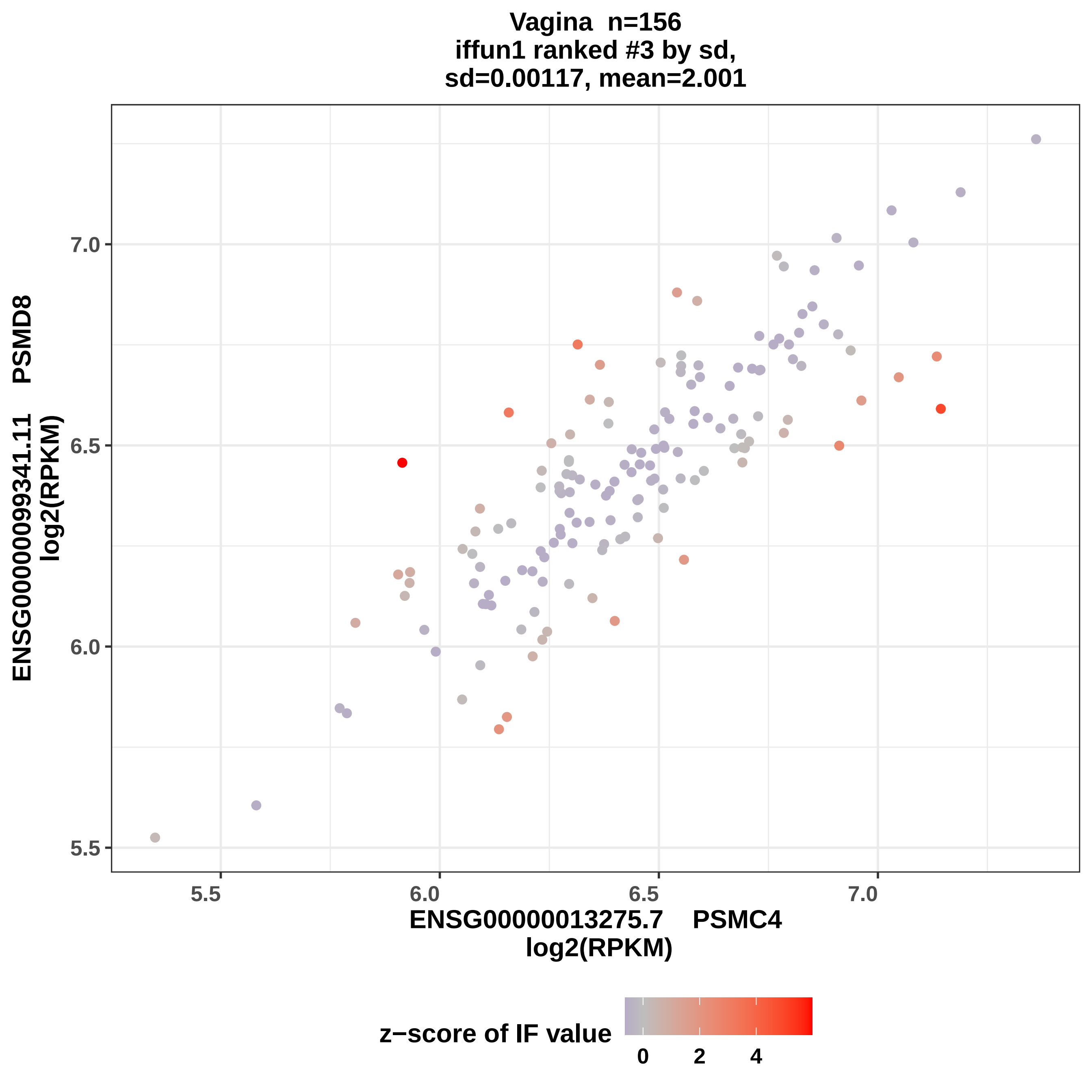
Vagina n=156
iffun1 ranked #3 by sd,
sd=0.00117, mean=2.001
ENSG00000099341.11 PSMD8
log2(RPKM)
7.0
6.5
6.0
5.5
5.5
6.0
6.5
7.0
ENSG00000013275.7 PSMC4
log2(RPKM)
z−score of IF value
0
2
4

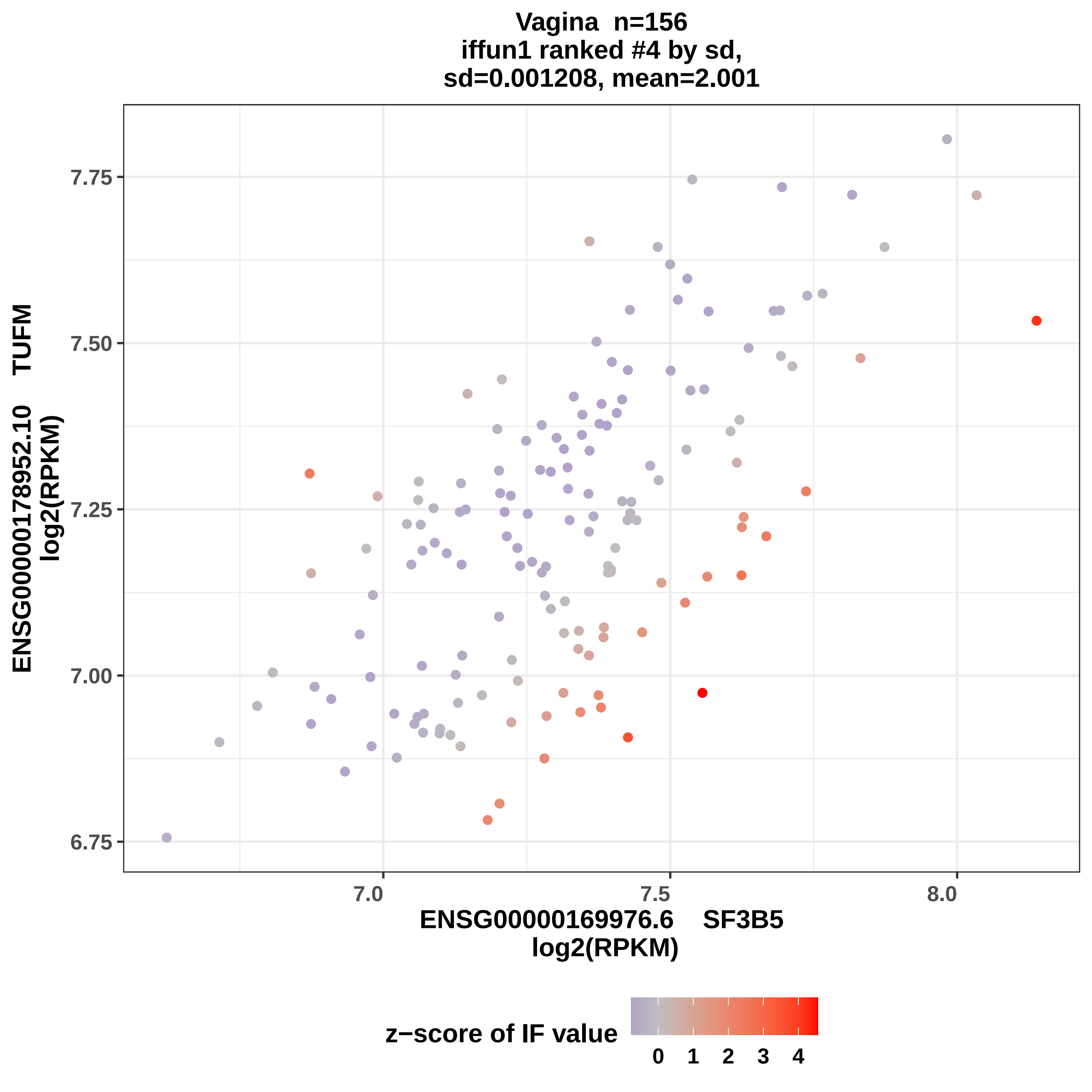

Vagina n=156
iffun1 ranked #4 by sd,
sd=0.001208, mean=2.001
ENSG00000178952.10 TUFM
log2(RPKM)
7.75
7.50
7.25
7.00
6.75
7.0
7.5
8.0
ENSG00000169976.6 SF3B5
log2(RPKM)
z−score of IF value
0
1
2
3
4

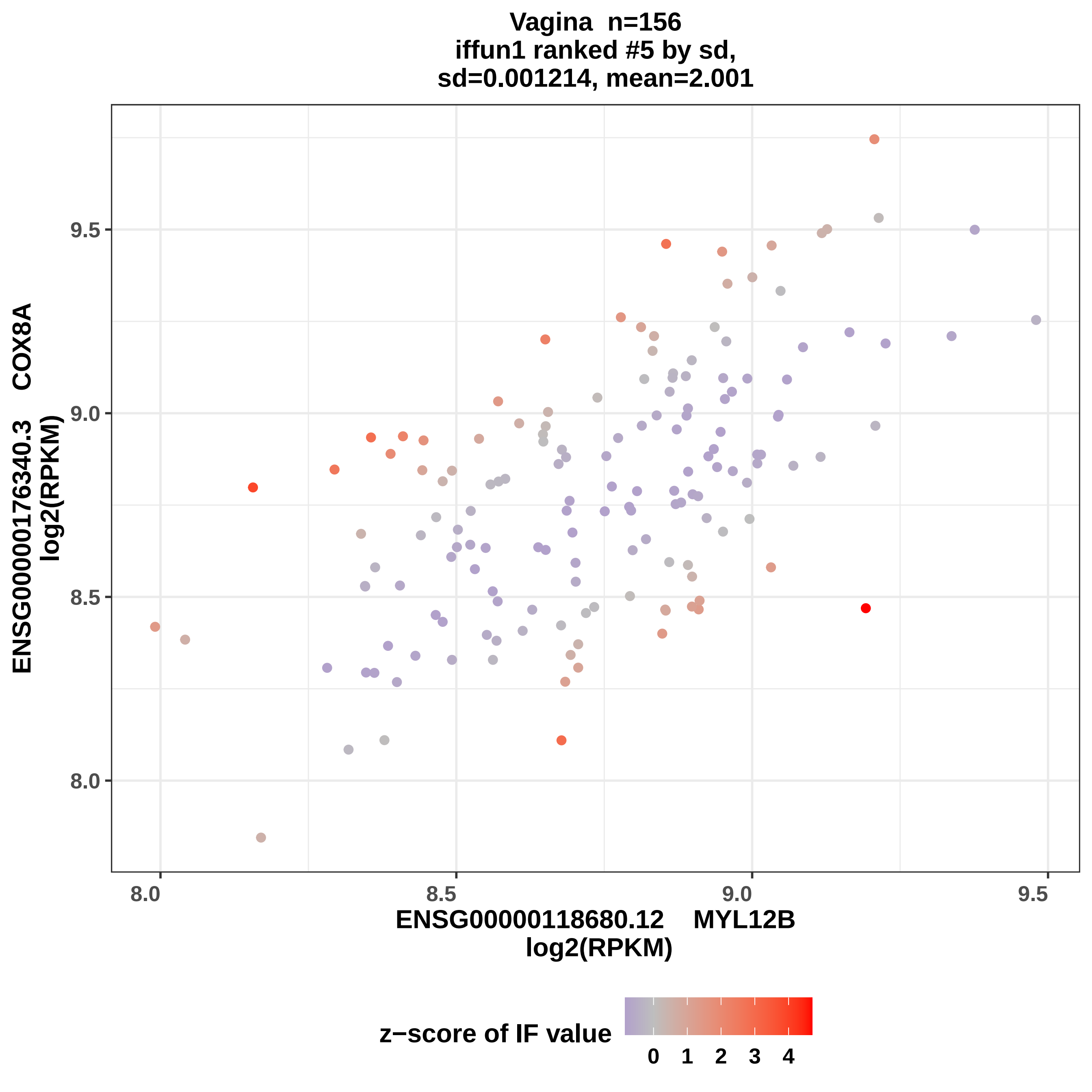

Vagina n=156
iffun1 ranked #5 by sd,
sd=0.001214, mean=2.001
ENSG00000176340.3 COX8A
log2(RPKM)
9.5
9.0
8.5
8.0
8.0
8.5
9.0
9.5
ENSG00000118680.12 MYL12B
log2(RPKM)
z−score of IF value
0
1
2
3
4

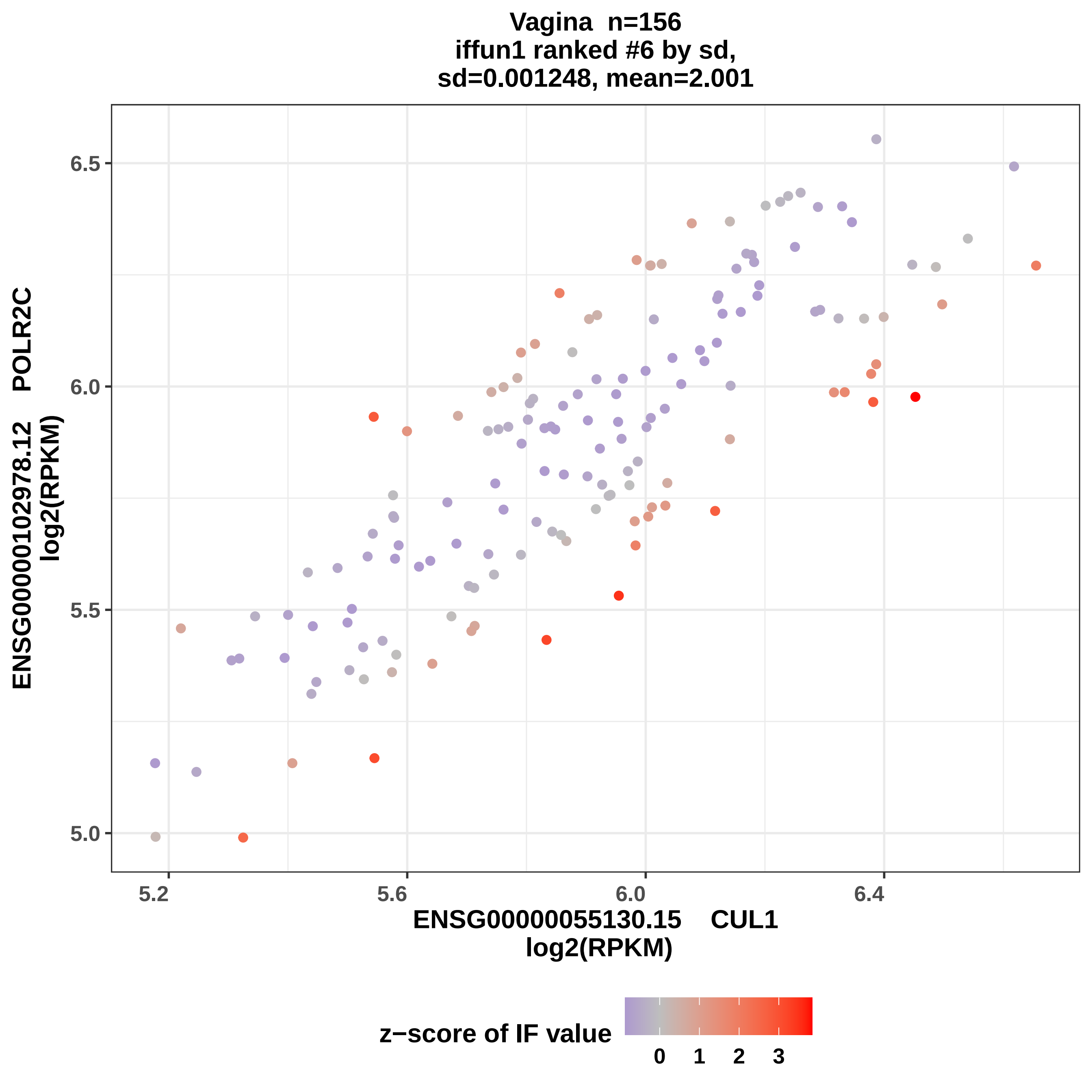

Vagina n=156
iffun1 ranked #6 by sd,
sd=0.001248, mean=2.001
ENSG00000102978.12 POLR2C
log2(RPKM)
6.5
6.0
5.5
5.0
5.2
5.6
6.0
6.4
ENSG00000055130.15 CUL1
log2(RPKM)
z−score of IF value
0
1
2
3

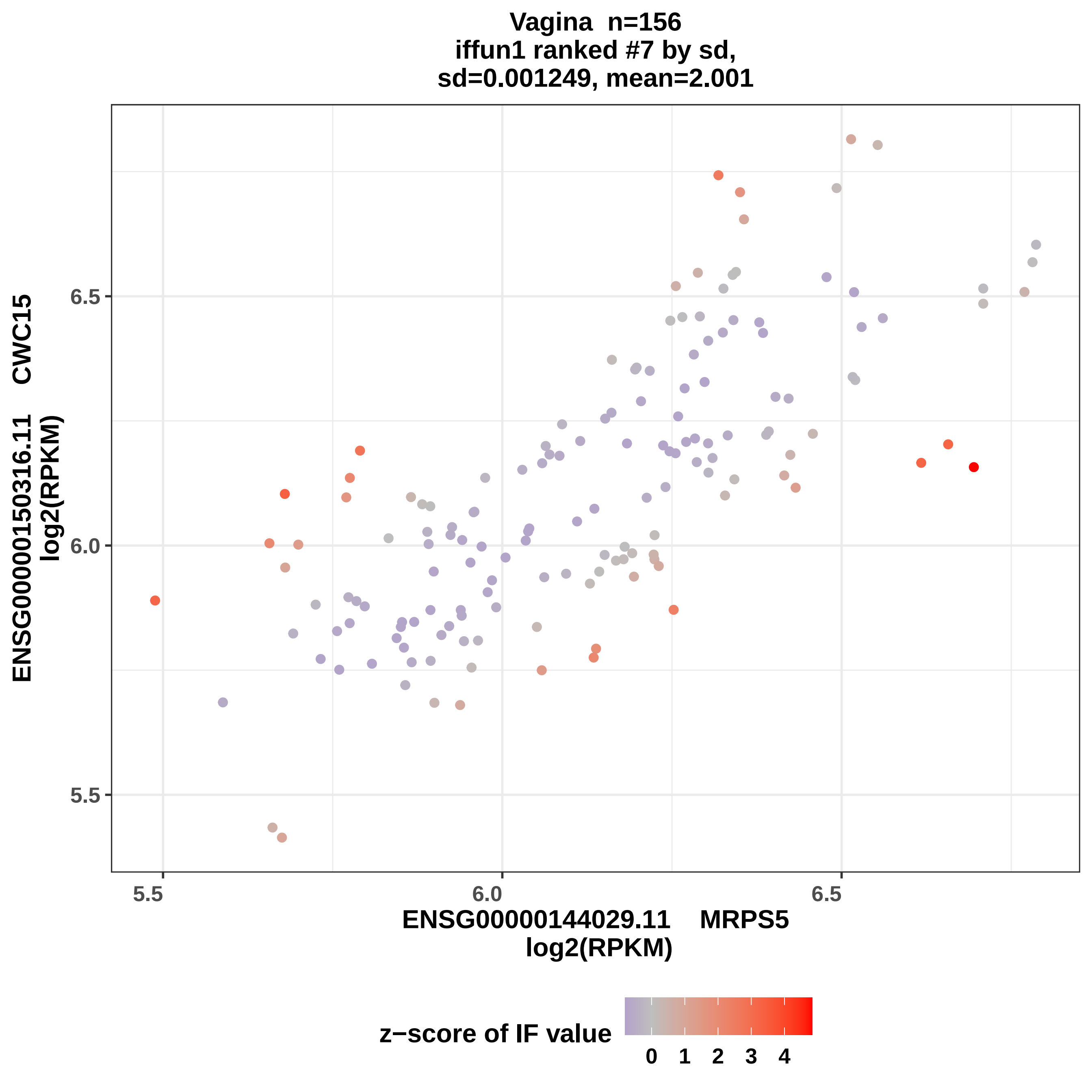

Vagina n=156
iffun1 ranked #7 by sd,
sd=0.001249, mean=2.001
ENSG00000150316.11 CWC15
log2(RPKM)
6.5
6.0
5.5
5.5
6.0
6.5
ENSG00000144029.11 MRPS5
log2(RPKM)
z−score of IF value
0
1
2
3
4

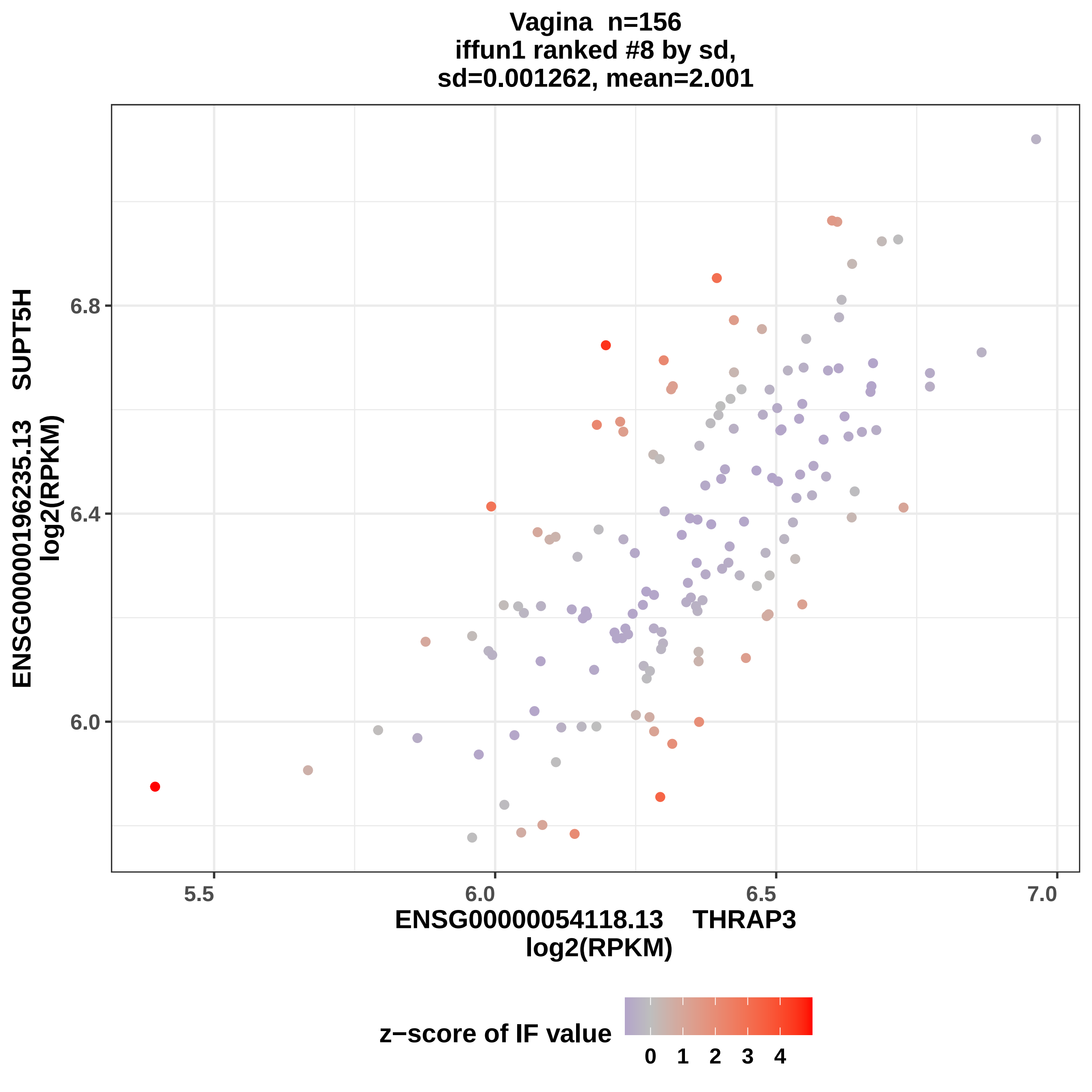

Vagina n=156
iffun1 ranked #8 by sd,
sd=0.001262, mean=2.001
ENSG00000196235.13 SUPT5H
log2(RPKM)
6.8
6.4
6.0
5.5
6.0
6.5
7.0
ENSG00000054118.13 THRAP3
log2(RPKM)
z−score of IF value
0
1
2
3
4

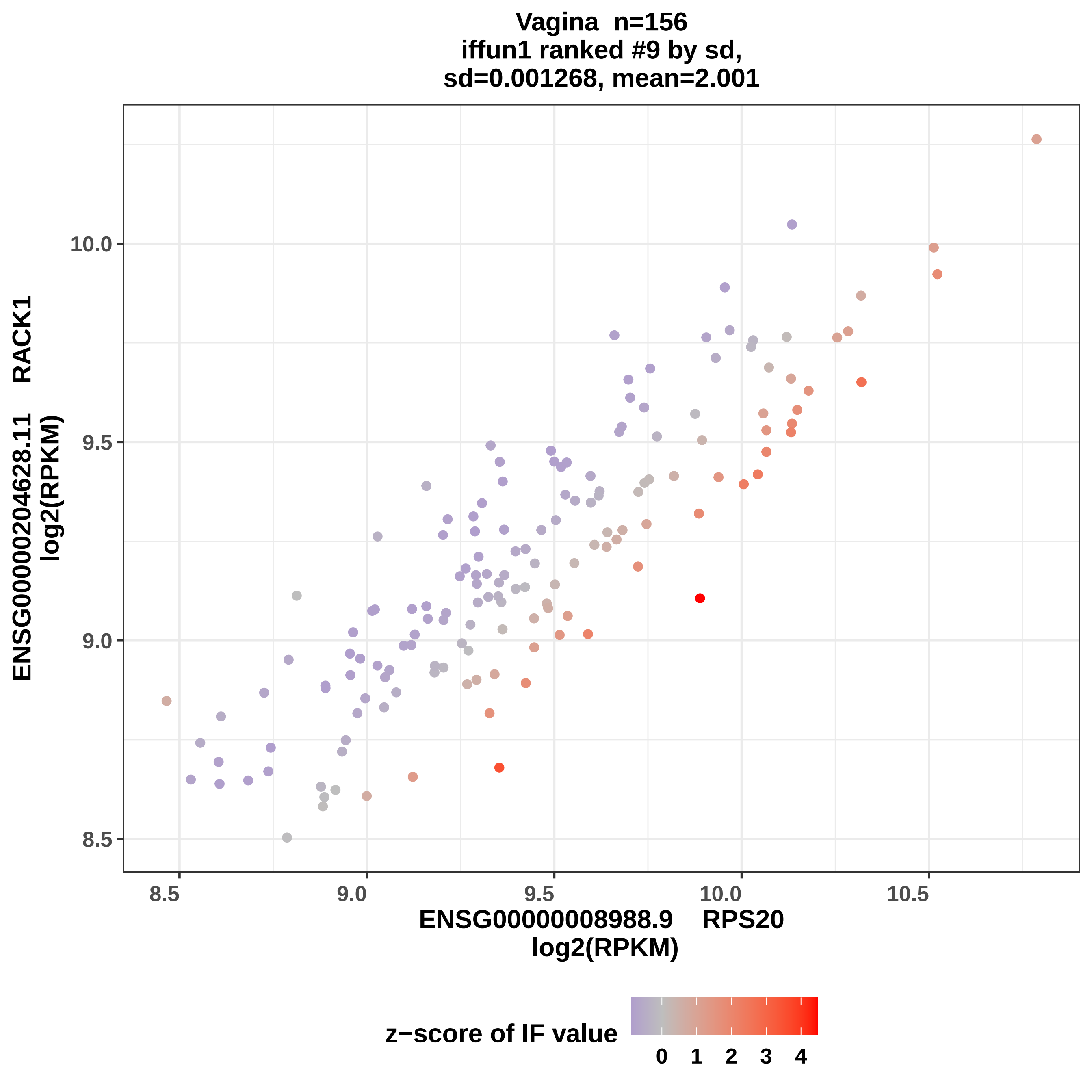
Vagina n=156
iffun1 ranked #9 by sd,
sd=0.001268, mean=2.001
ENSG00000204628.11 RACK1
log2(RPKM)
10.0
9.5
9.0
8.5
8.5
9.0
9.5
10.0
10.5
ENSG00000008988.9 RPS20
log2(RPKM)
z−score of IF value
0
1
2
3
4

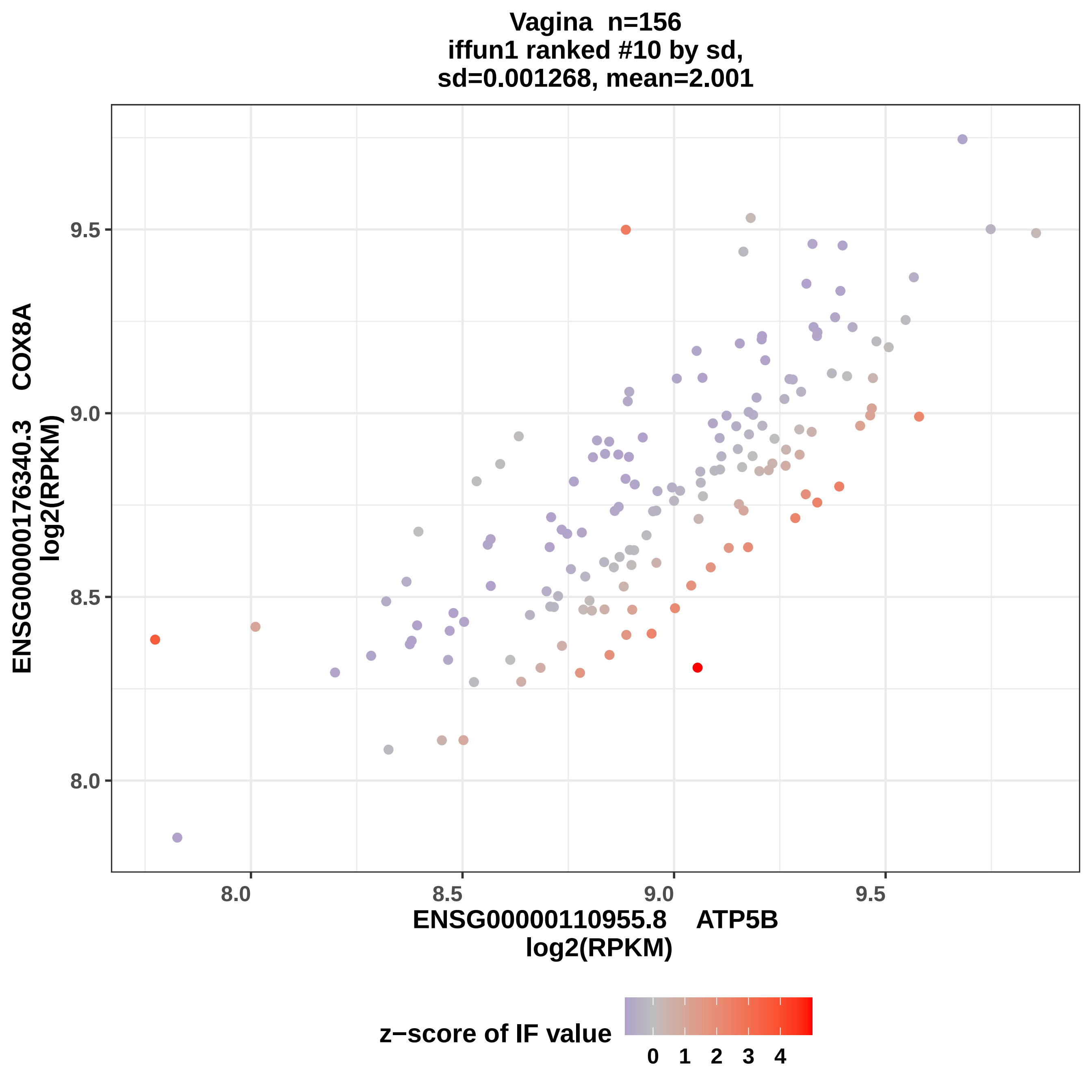

Vagina n=156
iffun1 ranked #10 by sd,
sd=0.001268, mean=2.001
ENSG00000176340.3 COX8A
log2(RPKM)
9.5
9.0
8.5
8.0
ENSG00000110955.8 ATP5B
log2(RPKM)
8.0
8.5
9.0
9.5
z−score of IF value
0
1
2
3
4

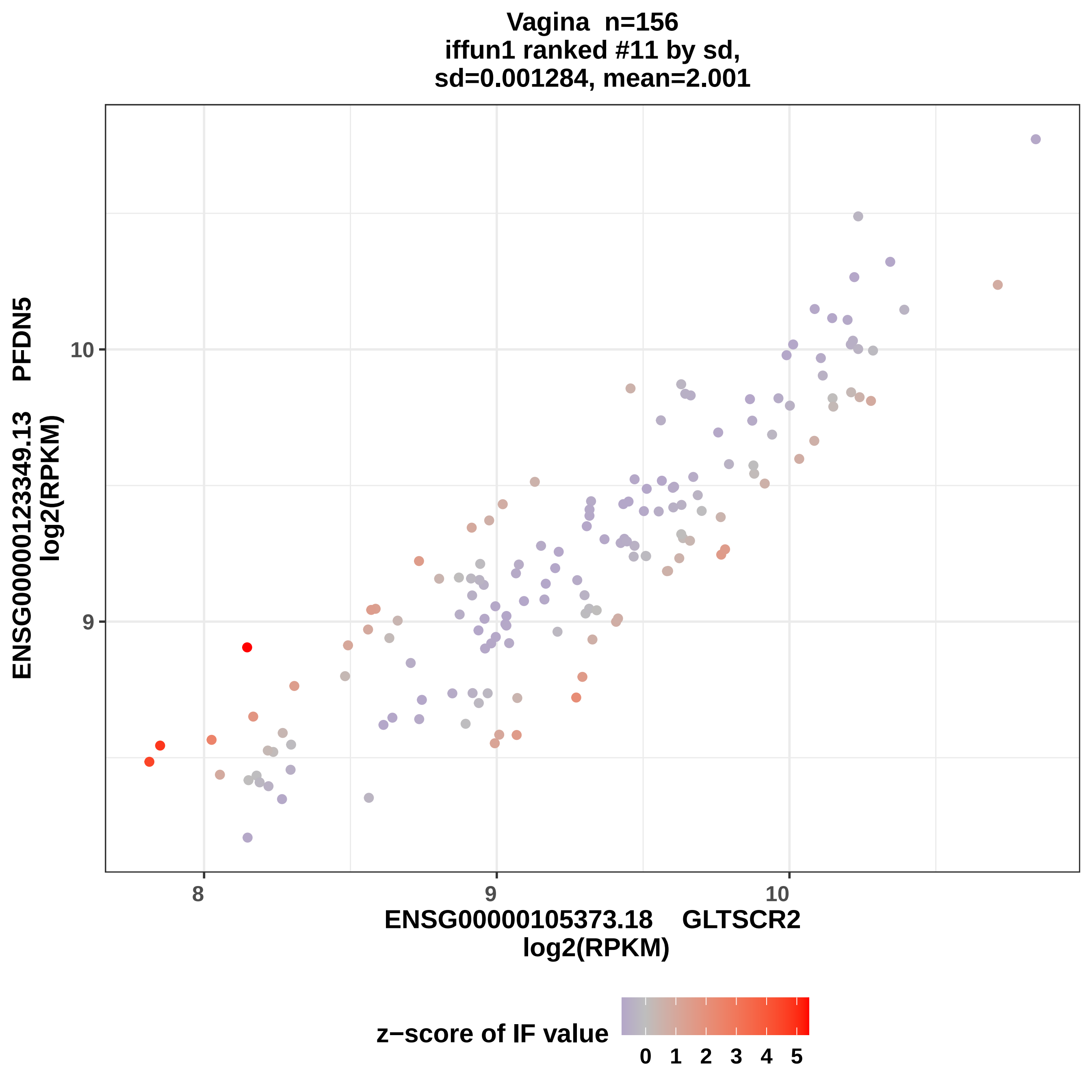

Vagina n=156
iffun1 ranked #11 by sd,
sd=0.001284, mean=2.001
ENSG00000123349.13 PFDN5
log2(RPKM)
10
9
8
9
10
ENSG00000105373.18 GLTSCR2
log2(RPKM)
z−score of IF value
0 1 2 3 4 5

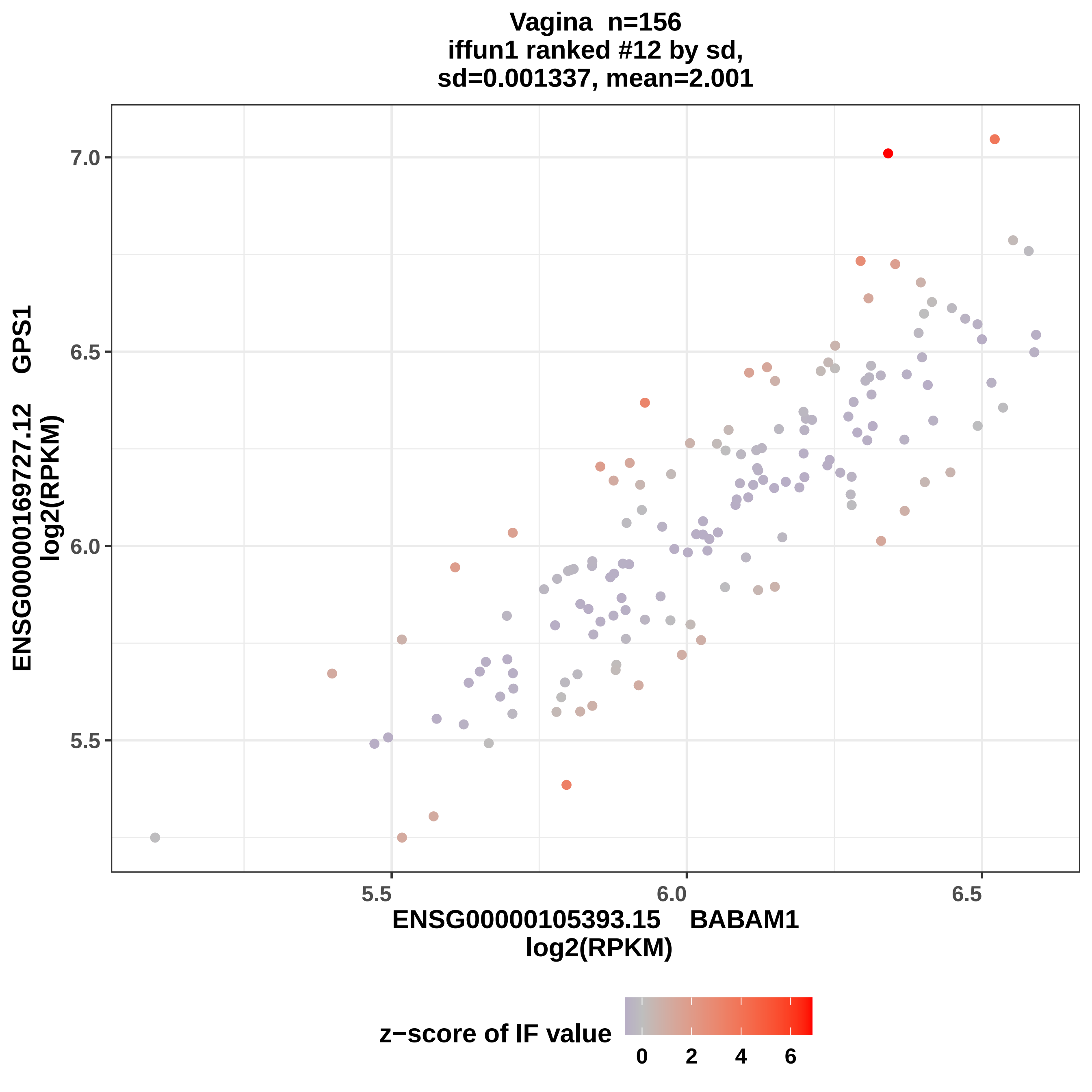

Vagina n=156
iffun1 ranked #12 by sd,
sd=0.001337, mean=2.001
ENSG00000169727.12 GPS1
log2(RPKM)
7.0
6.5
6.0
5.5
5.5
6.0
6.5
ENSG00000105393.15 BABAM1
log2(RPKM)
z−score of IF value
0
2
4
6

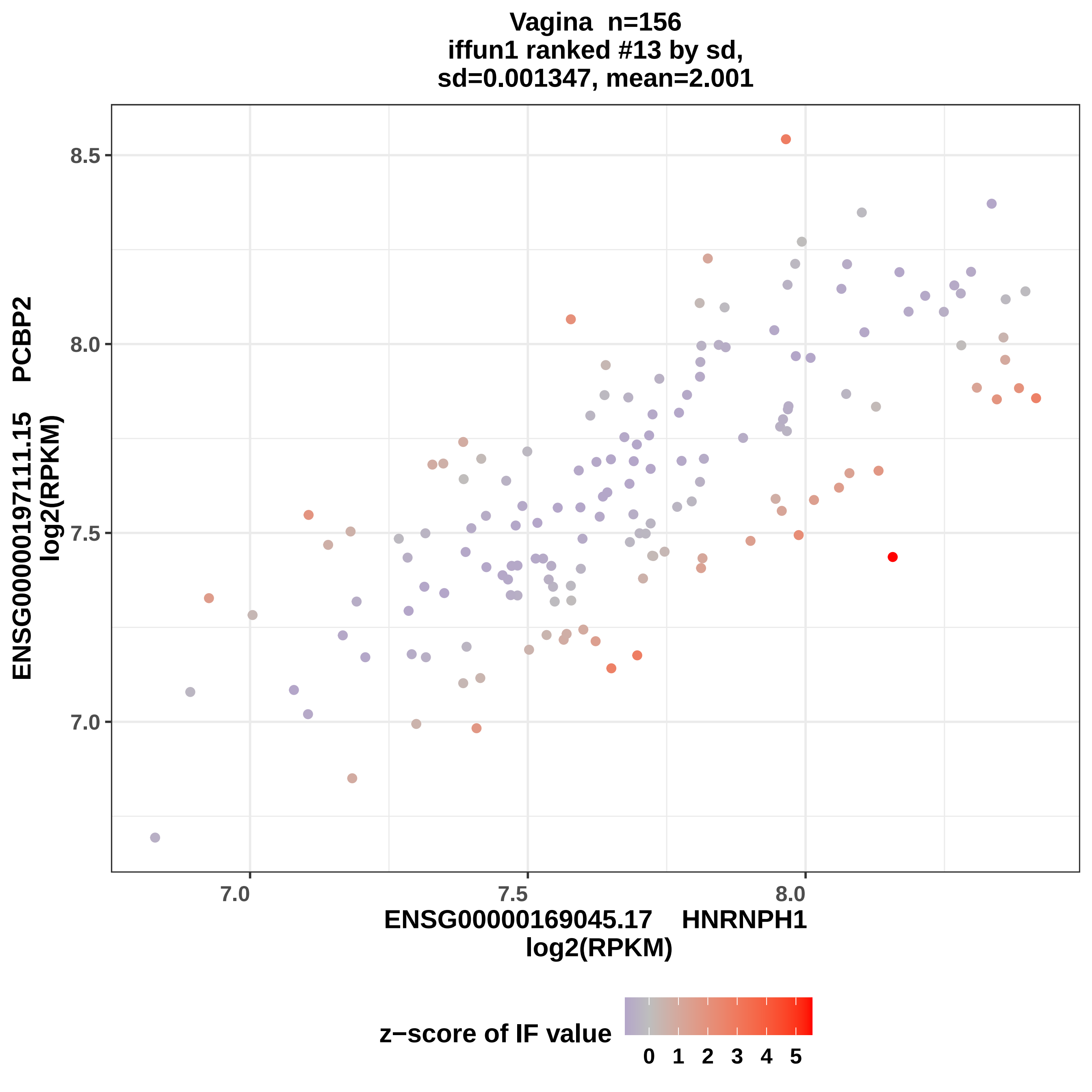
Vagina n=156
iffun1 ranked #13 by sd,
sd=0.001347, mean=2.001
ENSG00000197111.15 PCBP2
log2(RPKM)
8.5
8.0
7.5
7.0
7.0
7.5
8.0
ENSG00000169045.17 HNRNPH1
log2(RPKM)
z−score of IF value
0 1 2 3 4 5

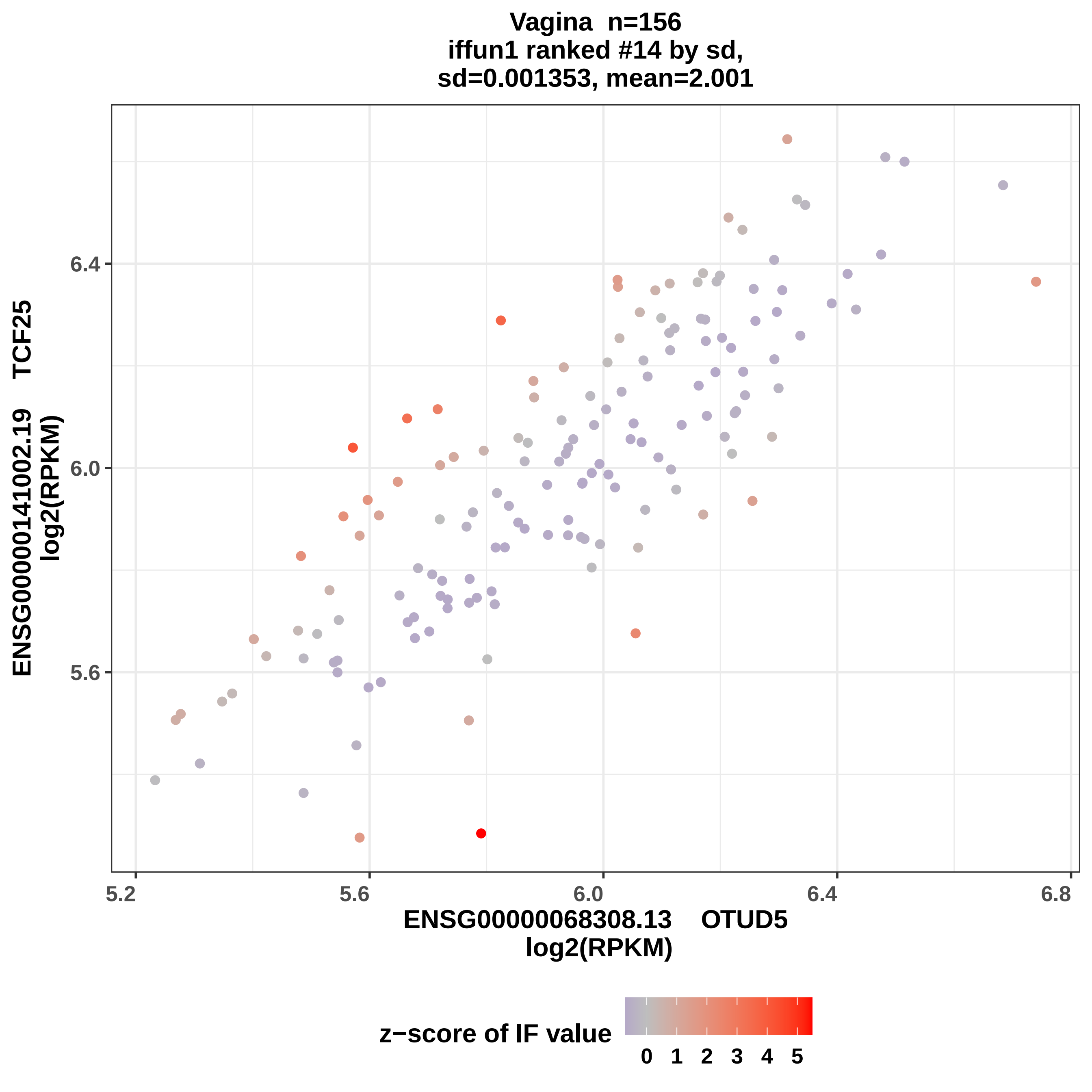
Vagina n=156
iffun1 ranked #14 by sd,
sd=0.001353, mean=2.001
ENSG00000141002.19 TCF25
log2(RPKM)
6.4
6.0
5.6
5.2
5.6
6.0
6.4
6.8
ENSG00000068308.13 OTUD5
log2(RPKM)
z−score of IF value
0 1 2 3 4 5

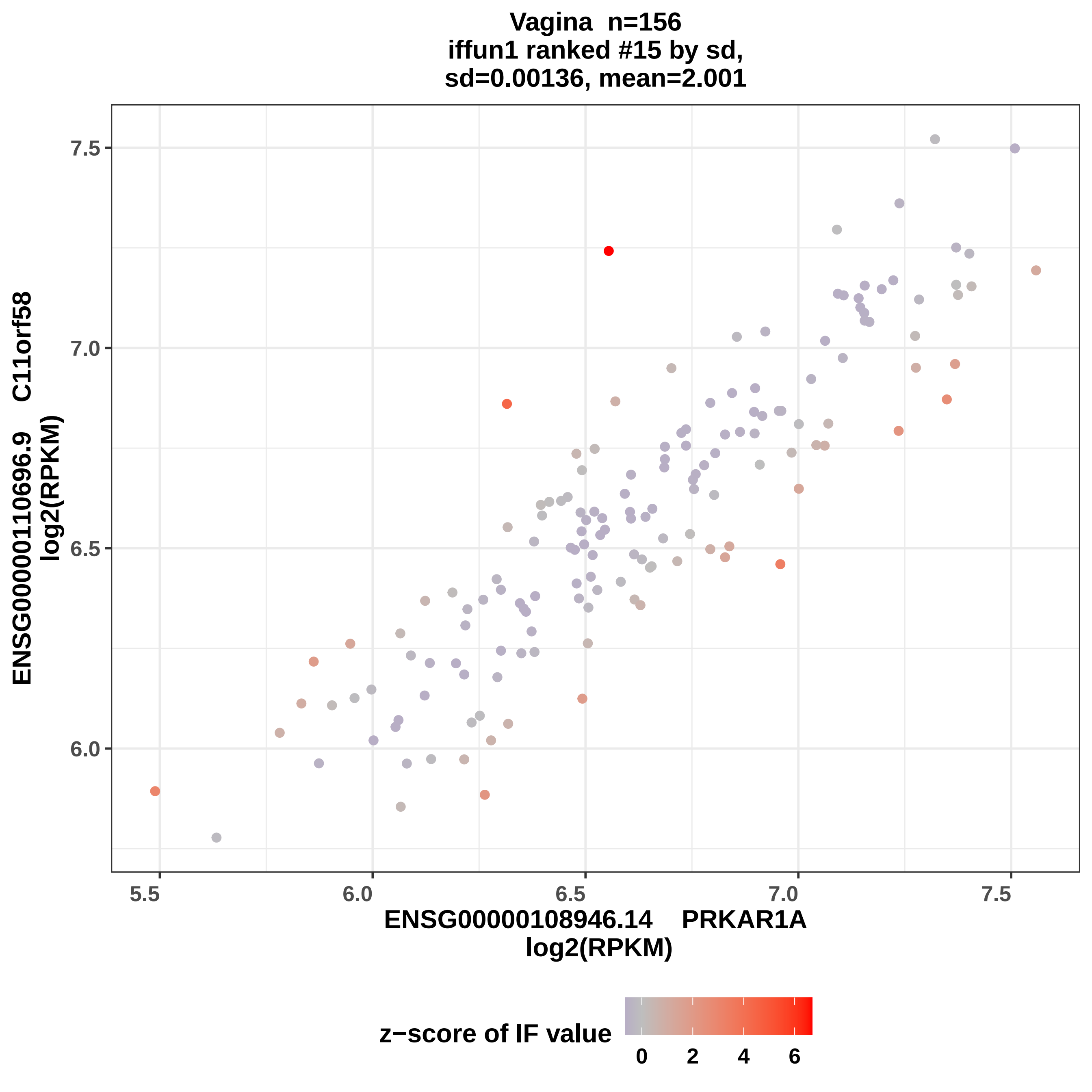

Vagina n=156
iffun1 ranked #15 by sd,
sd=0.00136, mean=2.001
ENSG00000110696.9 C11orf58
log2(RPKM)
7.5
7.0
6.5
6.0
5.5
6.0
6.5
7.0
7.5
ENSG00000108946.14 PRKAR1A
log2(RPKM)
z−score of IF value
0
2
4
6

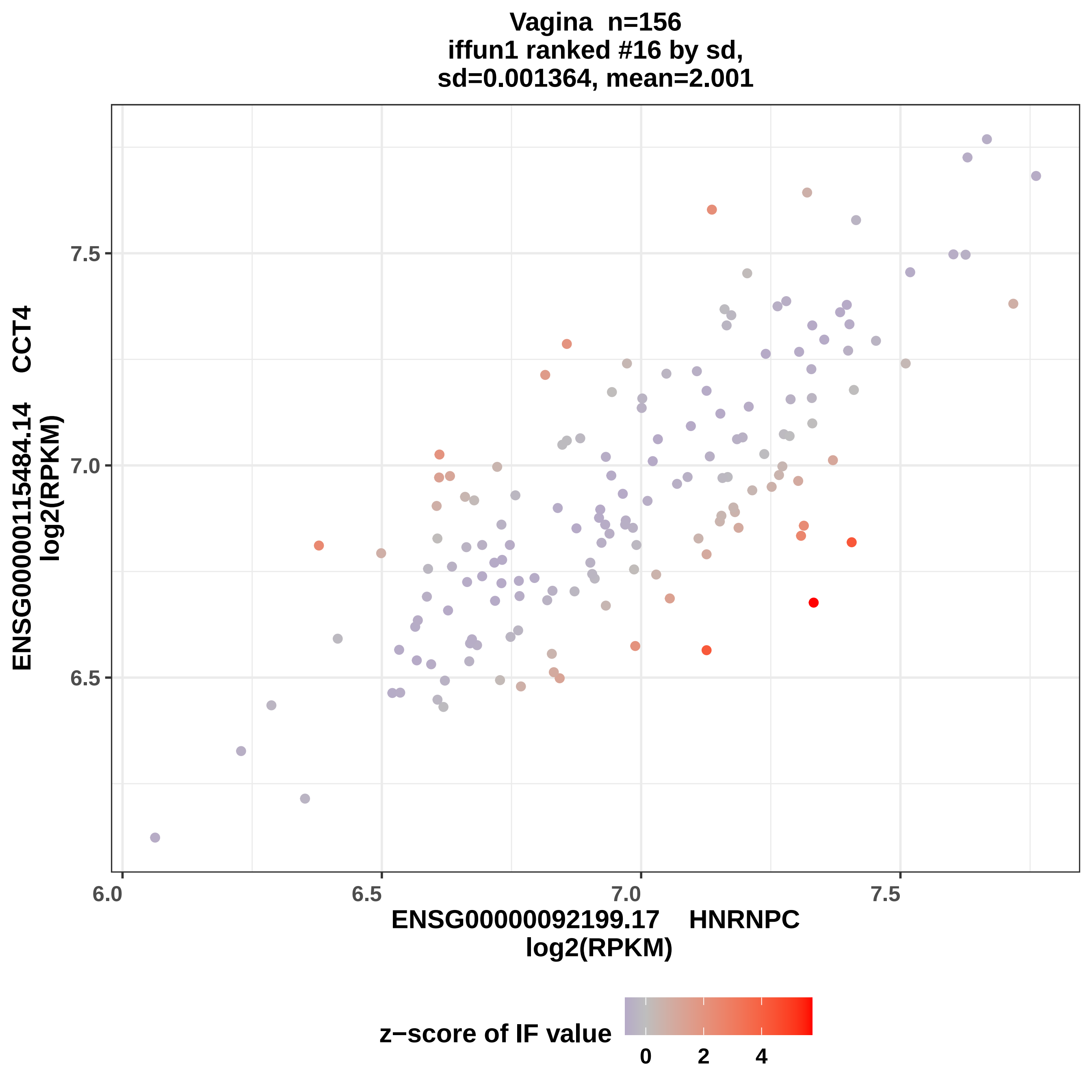
Vagina n=156
iffun1 ranked #16 by sd,
sd=0.001364, mean=2.001
ENSG00000115484.14 CCT4
log2(RPKM)
7.5
7.0
6.5
6.0
6.5
7.0
7.5
ENSG00000092199.17 HNRNPC
log2(RPKM)
z−score of IF value
0
2
4

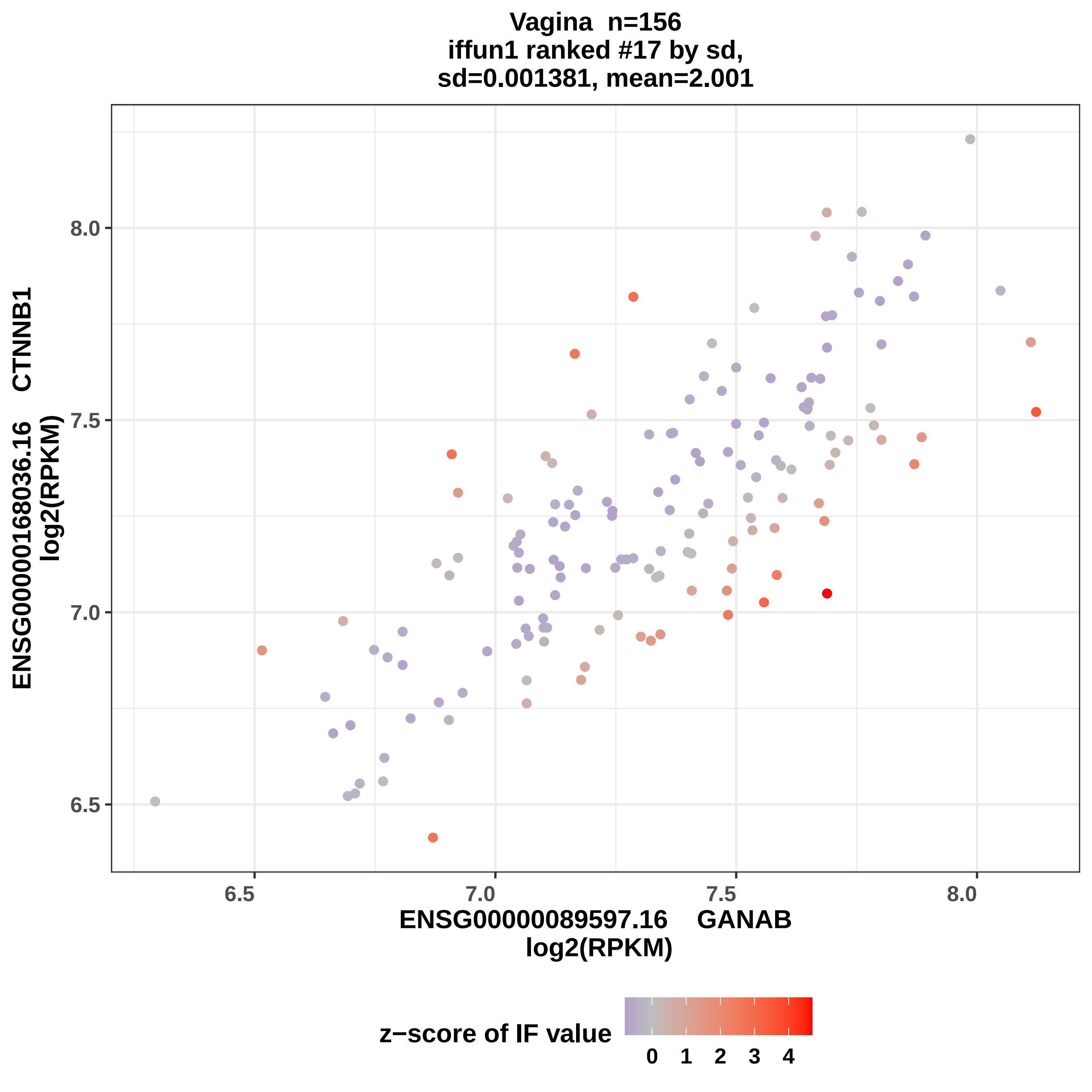

Vagina n=156
iffun1 ranked #17 by sd,
sd=0.001381, mean=2.001
ENSG00000168036.16 CTNNB1
log2(RPKM)
8.0
7.5
7.0
6.5
6.5
7.0
7.5
8.0
ENSG00000089597.16 GANAB
log2(RPKM)
z−score of IF value
0 1 2 3 4

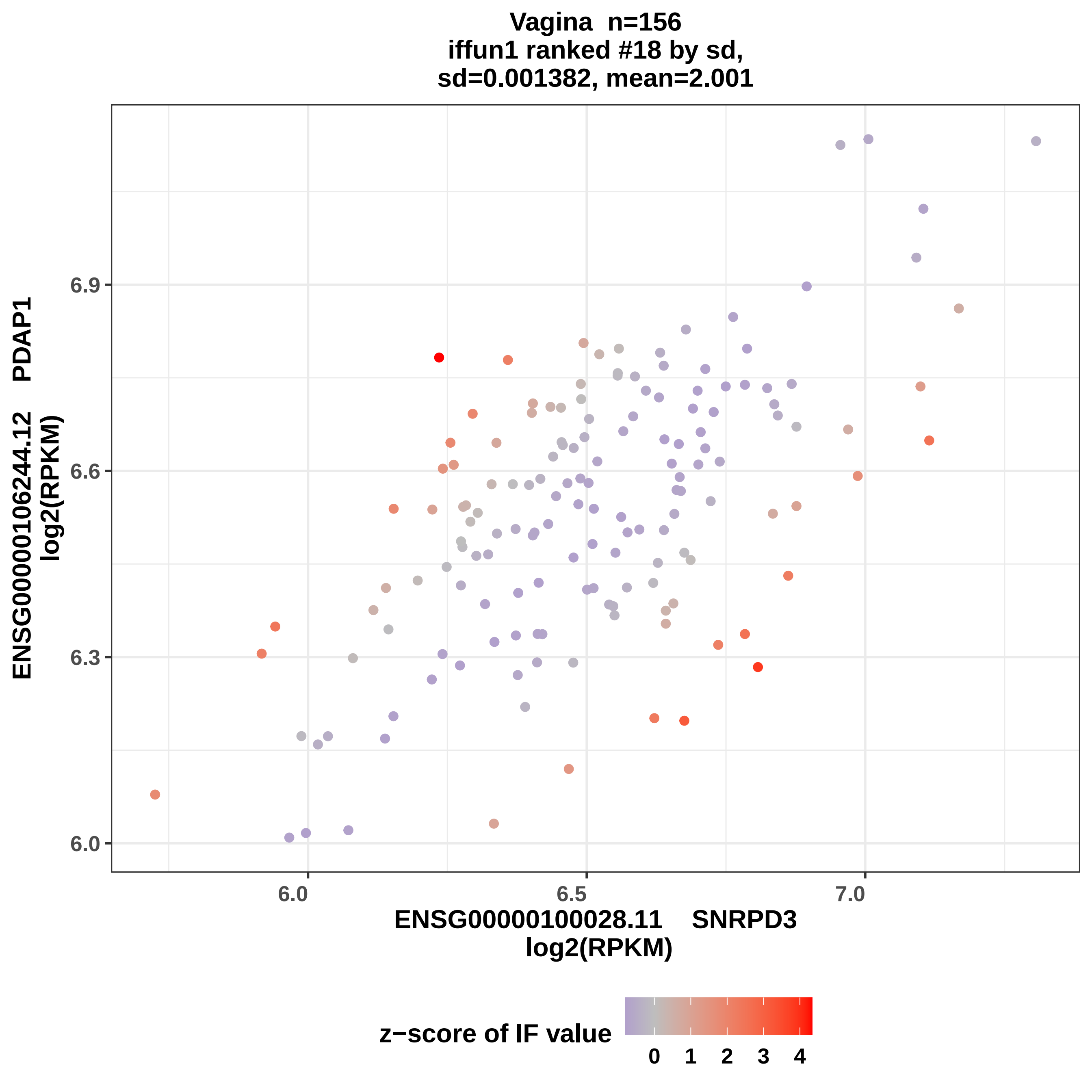

Vagina n=156
iffun1 ranked #18 by sd,
sd=0.001382, mean=2.001
ENSG00000106244.12 PDAP1
log2(RPKM)
6.0
6.3
6.6
6.9
6.0
6.5
7.0
ENSG00000100028.11 SNRPD3
log2(RPKM)
z−score of IF value
0
1
2
3
4

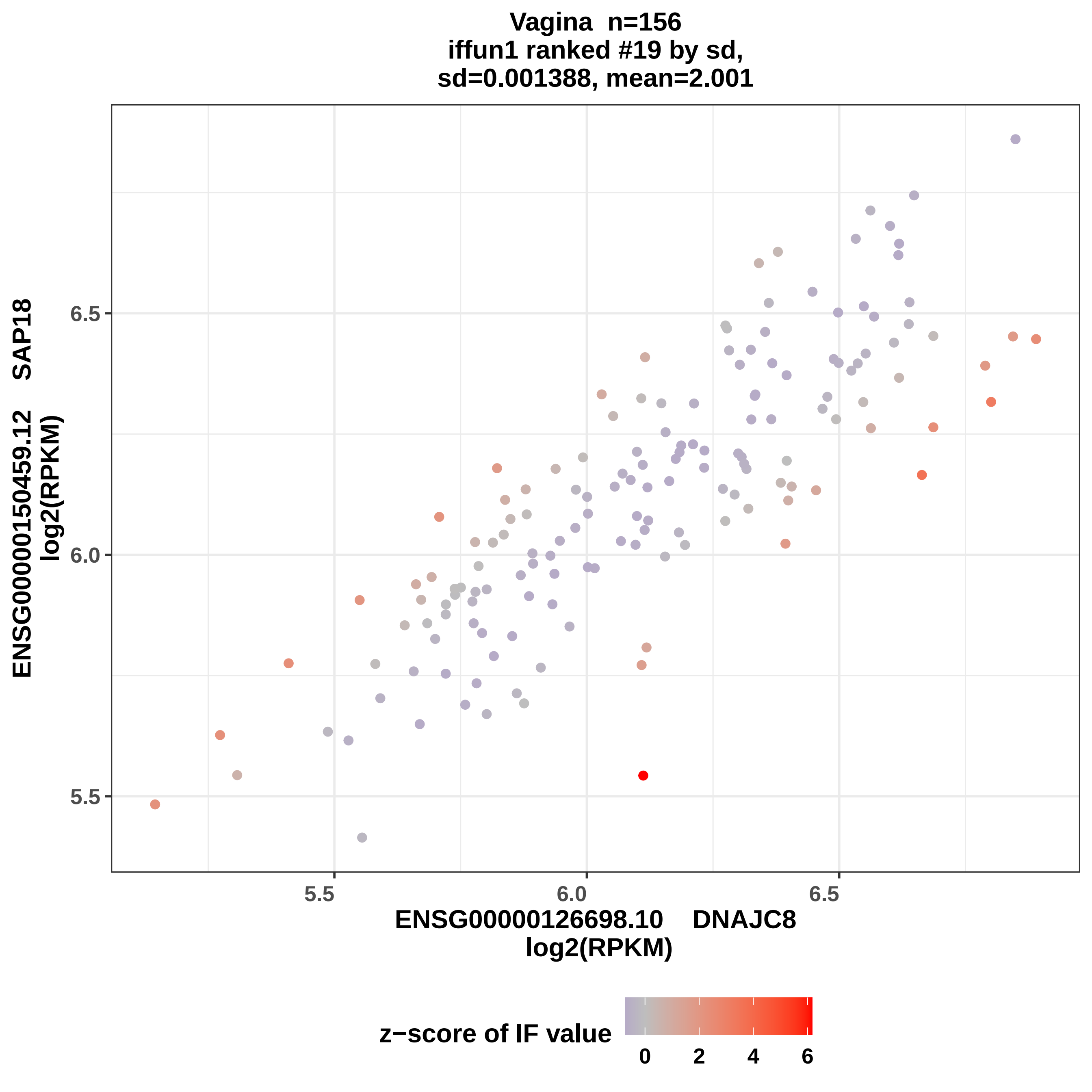

Vagina n=156
iffun1 ranked #19 by sd,
sd=0.001388, mean=2.001
ENSG00000150459.12 SAP18
log2(RPKM)
6.5
6.0
5.5
5.5
6.0
6.5
ENSG00000126698.10 DNAJC8
log2(RPKM)
z−score of IF value
0
2
4
6

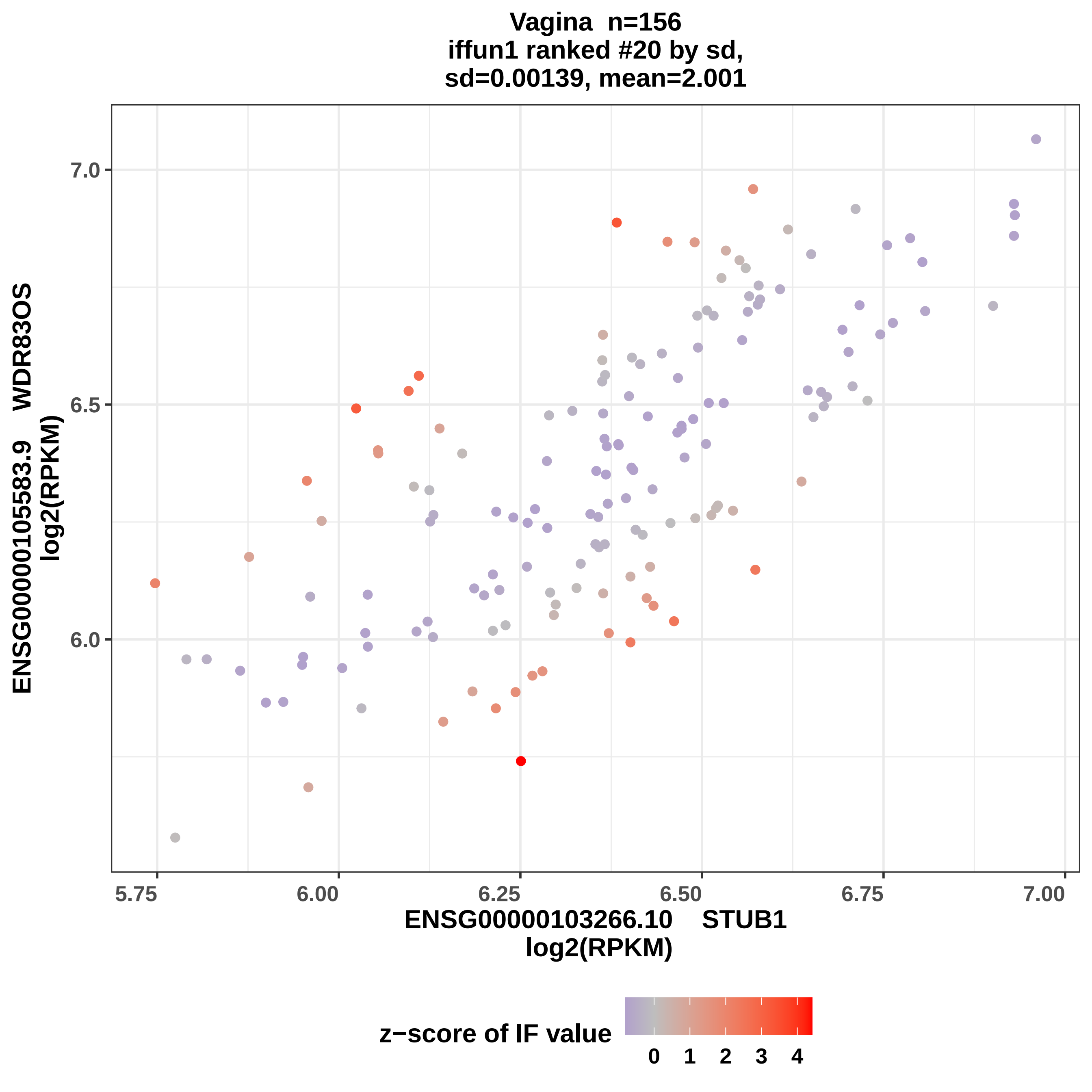

Vagina n=156
iffun1 ranked #20 by sd,
sd=0.00139, mean=2.001
ENSG00000105583.9 WDR83OS
log2(RPKM)
7.0
6.5
6.0
5.75
6.00
6.25
6.50
6.75
7.00
ENSG00000103266.10 STUB1
log2(RPKM)
z−score of IF value
0
1
2
3
4
