## Supplemental Data 7 for "Symmetry as a Fundamental Principle in Defining Gene Expression and Phenotypic Traits"

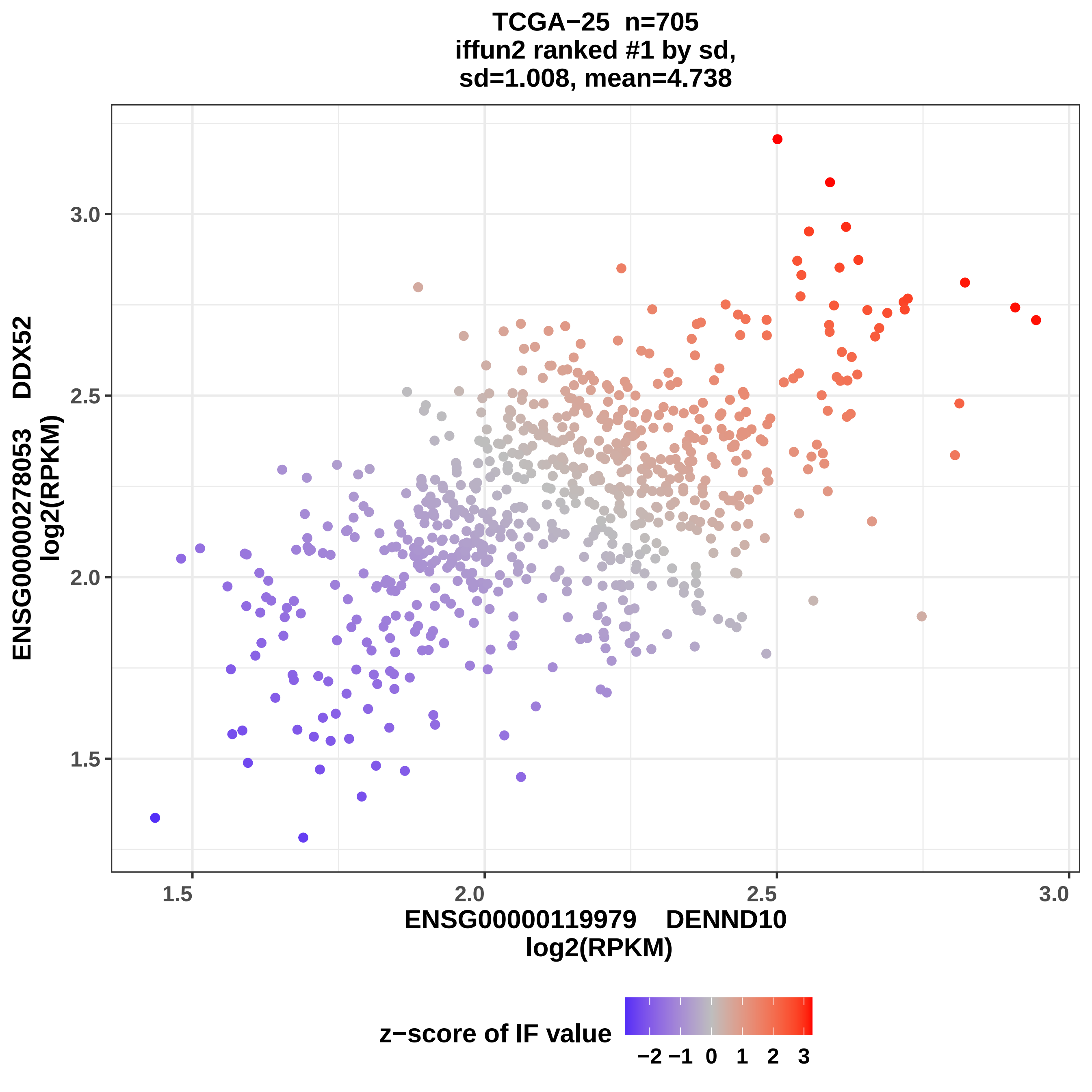

TCGA−25 n=705
iffun2 ranked #1 by sd,
sd=1.008, mean=4.738
ENSG00000278053 DDX52
log2(RPKM)
3.0
2.5
2.0
1.5
1.5
2.0
2.5
3.0
ENSG00000119979 DENND10
log2(RPKM)
z−score of IF value
−2 −1 0 1 2 3

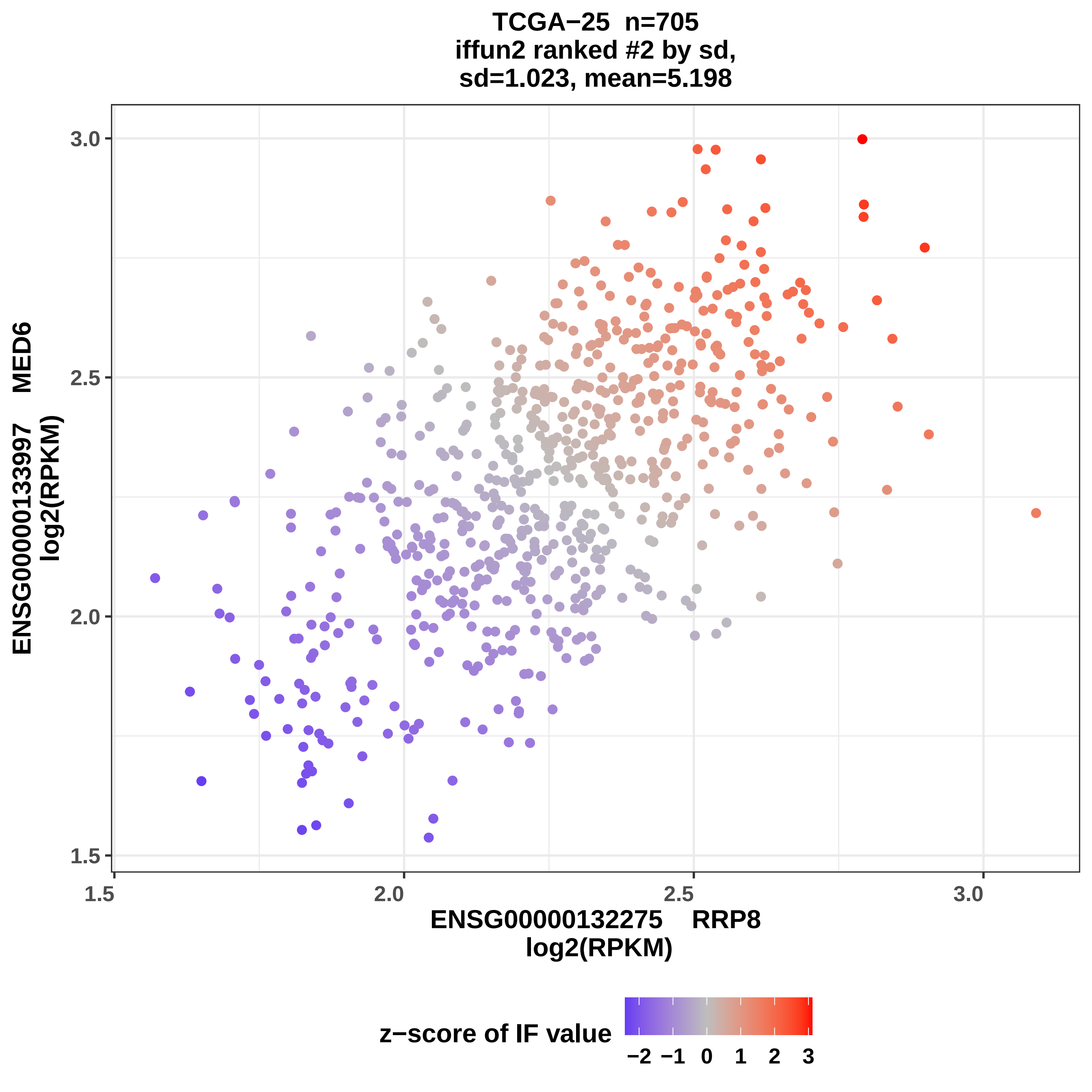
TCGA−25 n=705
iffun2 ranked #2 by sd,
sd=1.023, mean=5.198
ENSG00000133997 MED6
log2(RPKM)
ENSG00000132275 RRP8
log2(RPKM)
3.0
2.5
2.0
1.5
1.5
2.0
2.5
3.0
z−score of IF value
−2 −1 0 1 2 3

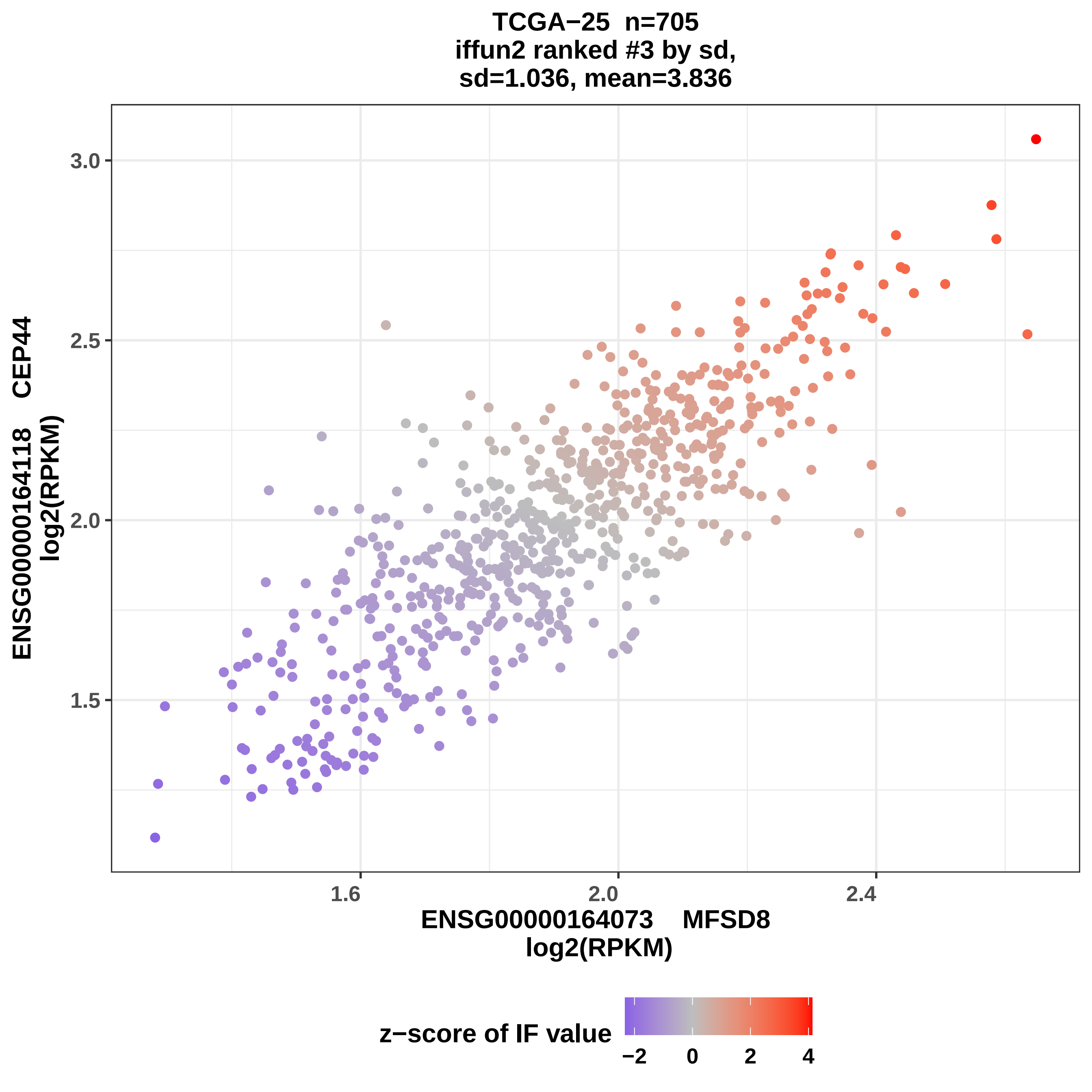
TCGA−25 n=705
iffun2 ranked #3 by sd,
sd=1.036, mean=3.836
ENSG00000164118 CEP44
log2(RPKM)
3.0
2.5
2.0
1.5
1.6
2.0
2.4
ENSG00000164073 MFSD8
log2(RPKM)
z−score of IF value
−2
0
2
4

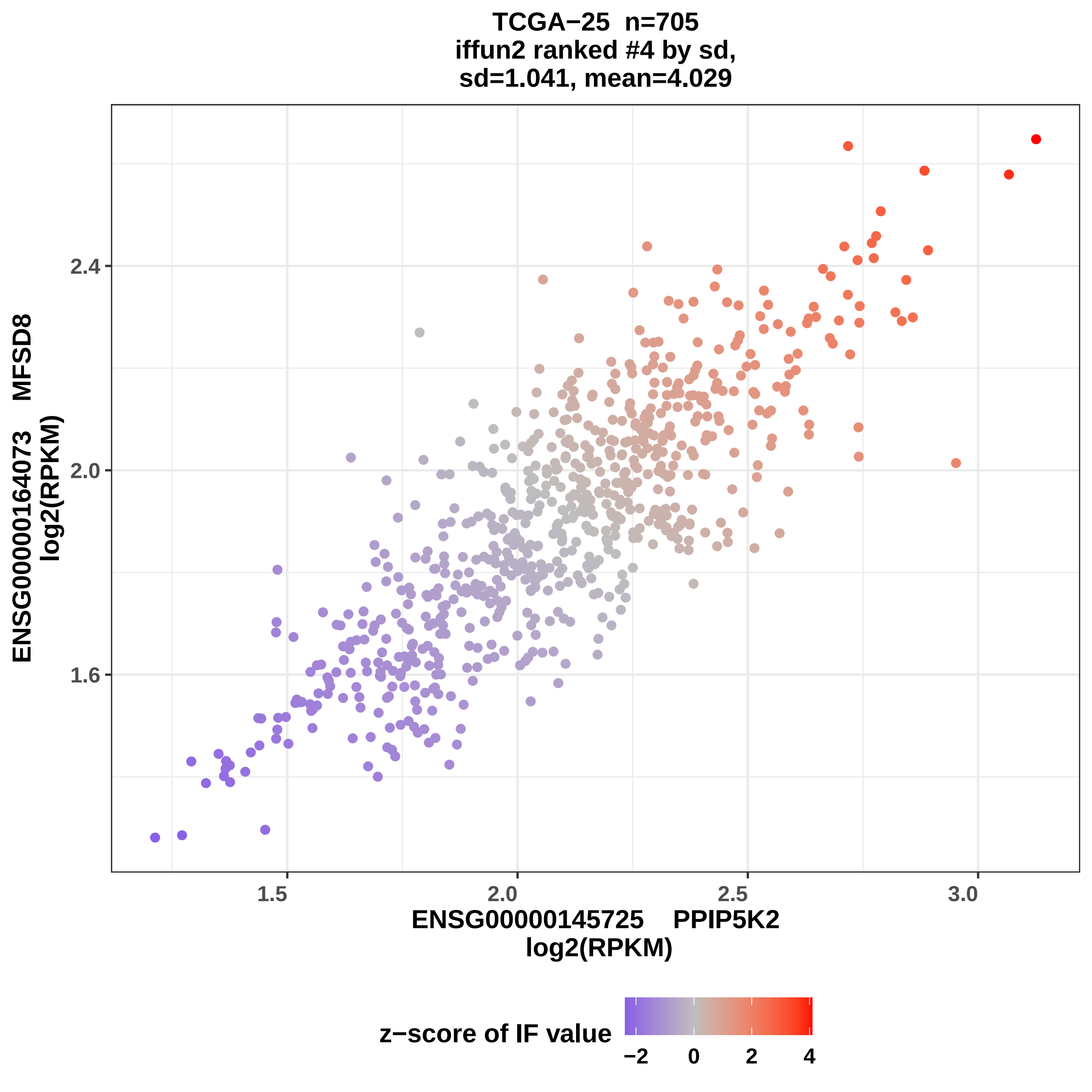

TCGA−25 n=705
iffun2 ranked #4 by sd,
sd=1.041, mean=4.029
ENSG00000164073 MFSD8
log2(RPKM)
2.4
2.0
1.6
1.5
2.0
2.5
3.0
ENSG00000145725 PPIP5K2
log2(RPKM)
z−score of IF value
−2
0
2
4

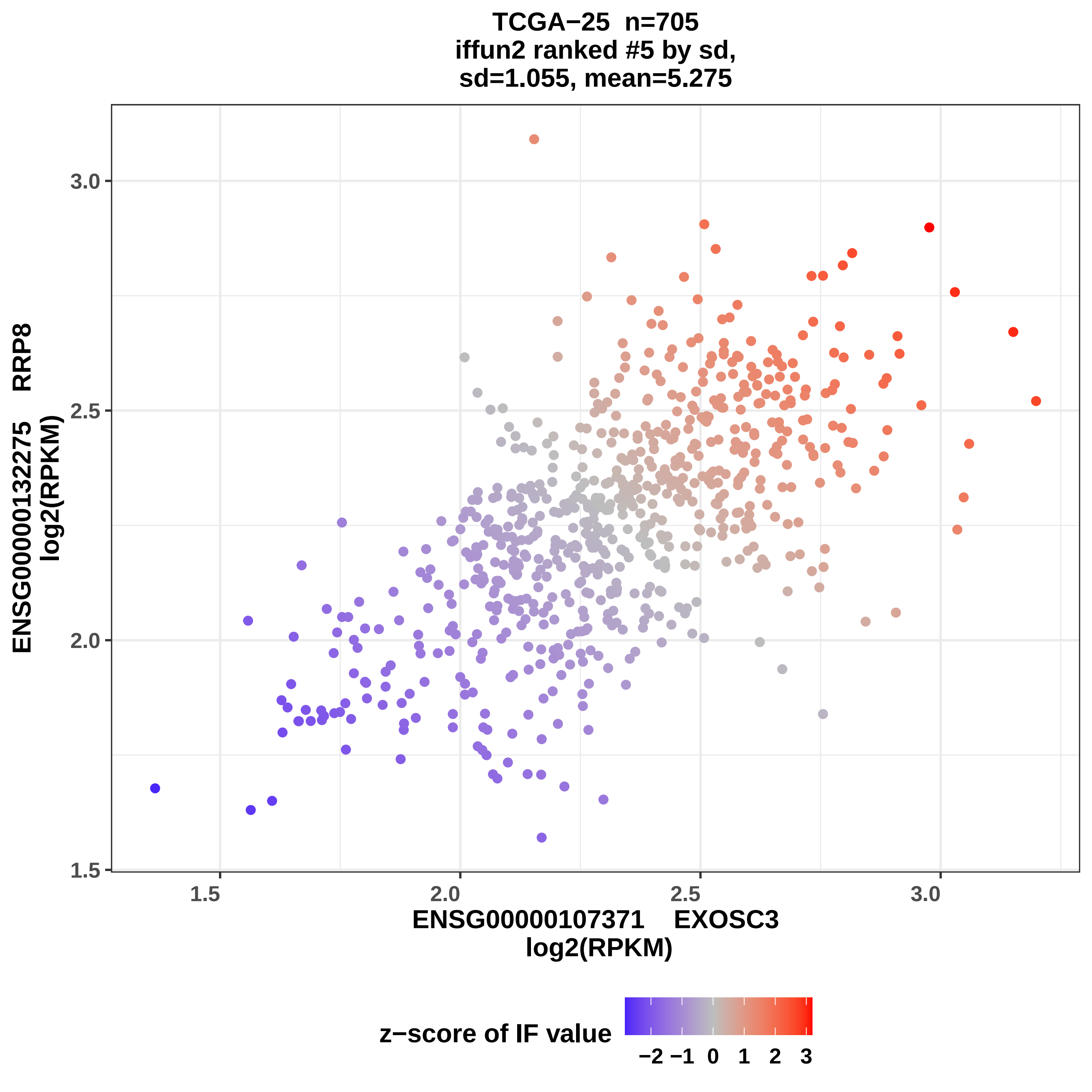
TCGA−25 n=705
iffun2 ranked #5 by sd,
sd=1.055, mean=5.275
ENSG00000132275 RRP8
log2(RPKM)
3.0
2.5
2.0
1.5
1.5
2.0
2.5
3.0
ENSG00000107371 EXOSC3
log2(RPKM)
z−score of IF value
−2 −1 0 1 2 3

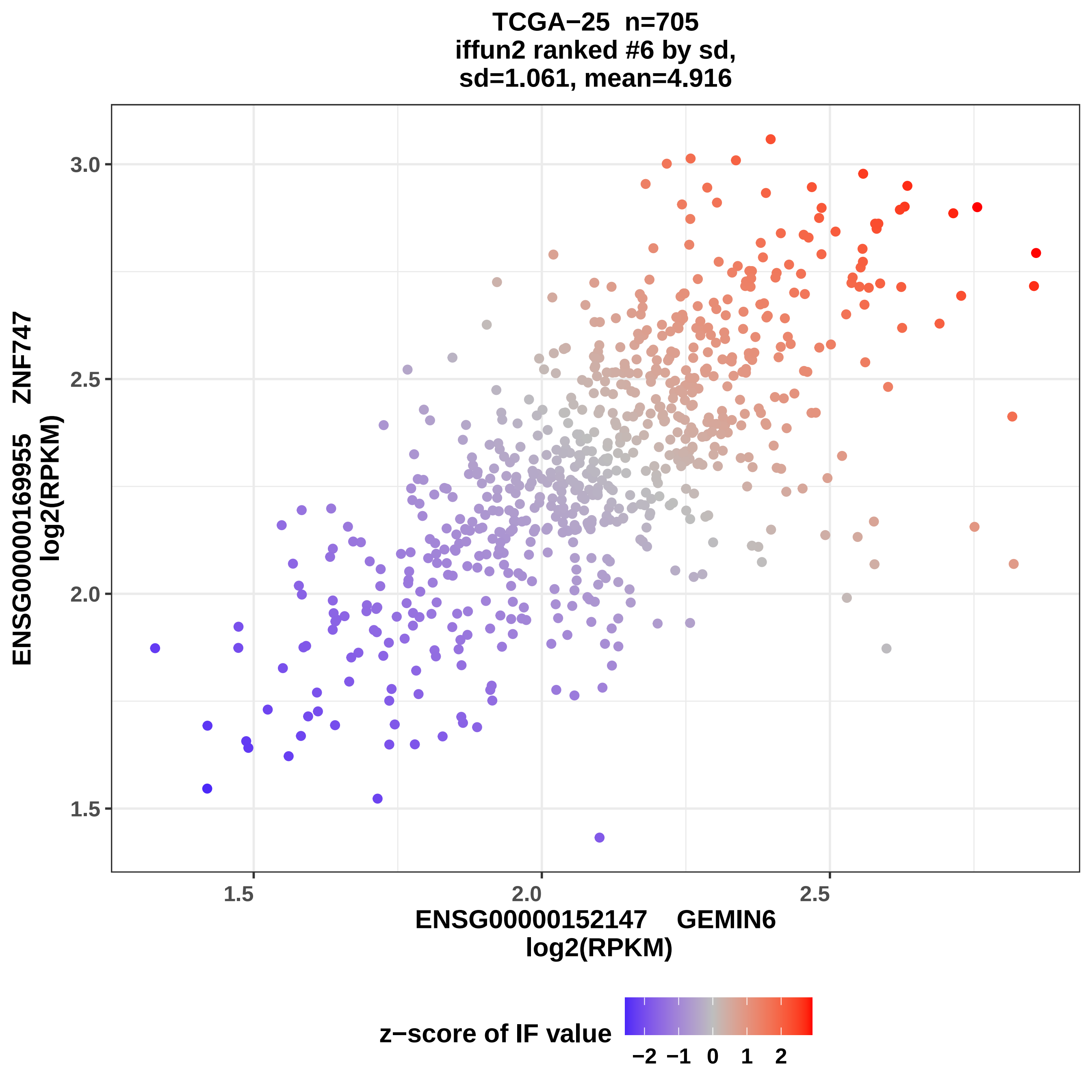
TCGA−25 n=705
iffun2 ranked #6 by sd,
sd=1.061, mean=4.916
ENSG00000169955 ZNF747
log2(RPKM)
3.0
2.5
2.0
1.5
1.5
2.0
2.5
ENSG00000152147 GEMIN6
log2(RPKM)
z−score of IF value
−2 −1 0 1 2

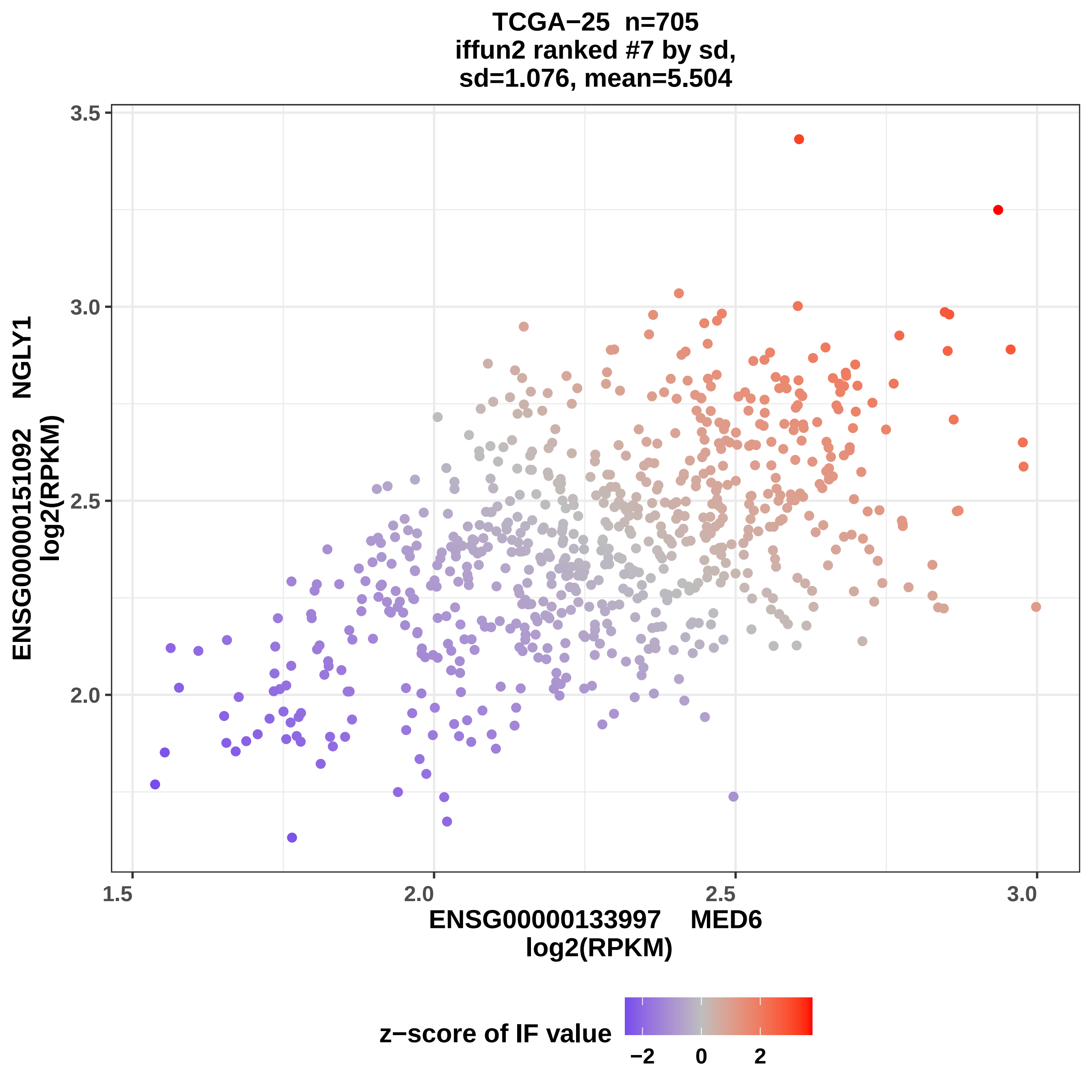

TCGA−25 n=705
iffun2 ranked #7 by sd,
sd=1.076, mean=5.504
ENSG00000151092 NGLY1
log2(RPKM)
ENSG00000133997 MED6
log2(RPKM)
3.5
3.0
2.5
2.0
1.5
2.0
2.5
3.0
z−score of IF value
−2
0
2

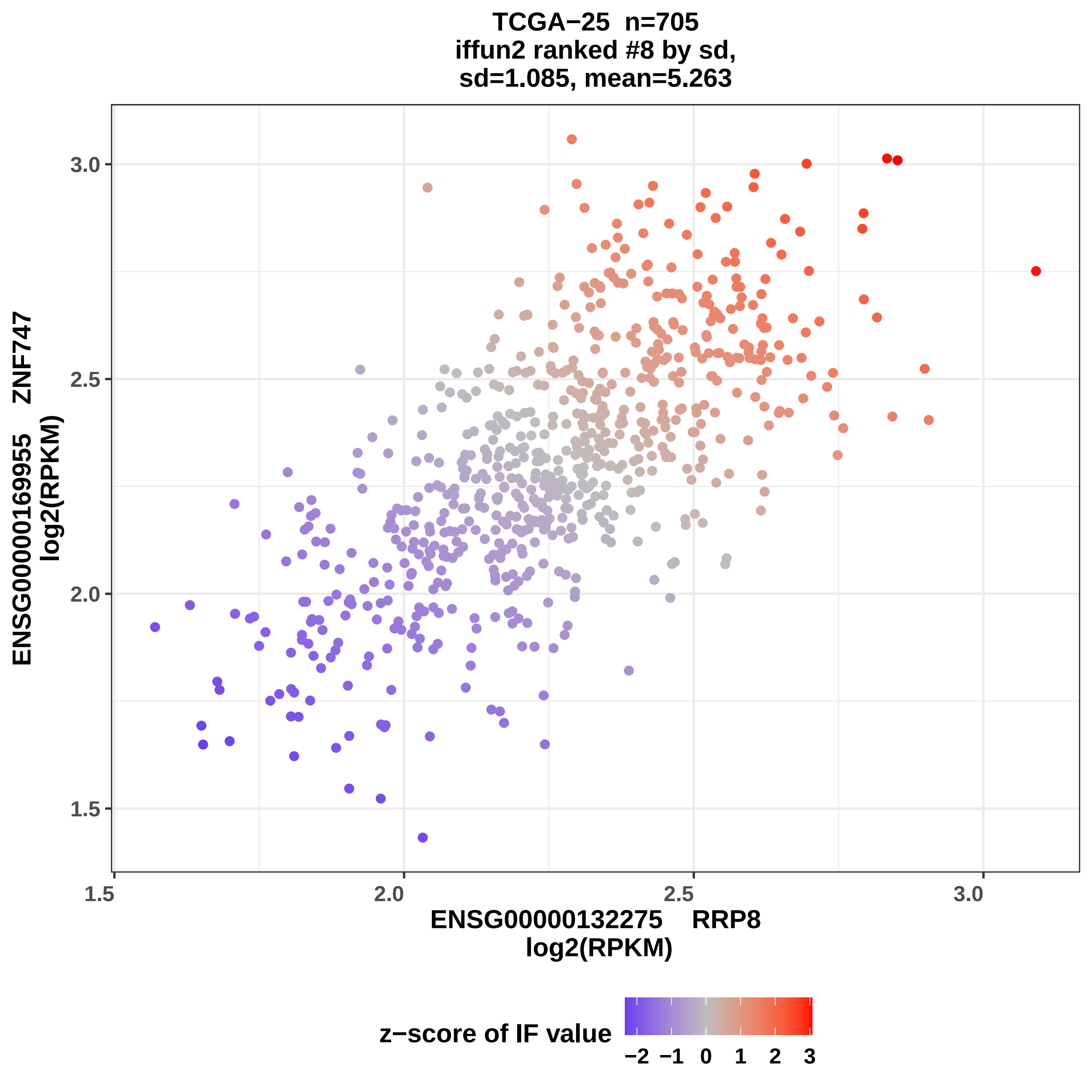

TCGA−25 n=705
iffun2 ranked #8 by sd,
sd=1.085, mean=5.263
3.0
2.5
2.0
1.5
1.5
2.0
2.5
3.0
ENSG00000169955 ZNF747
log2(RPKM)
ENSG00000132275 RRP8
log2(RPKM)
z−score of IF value
−2 −1 0 1 2 3

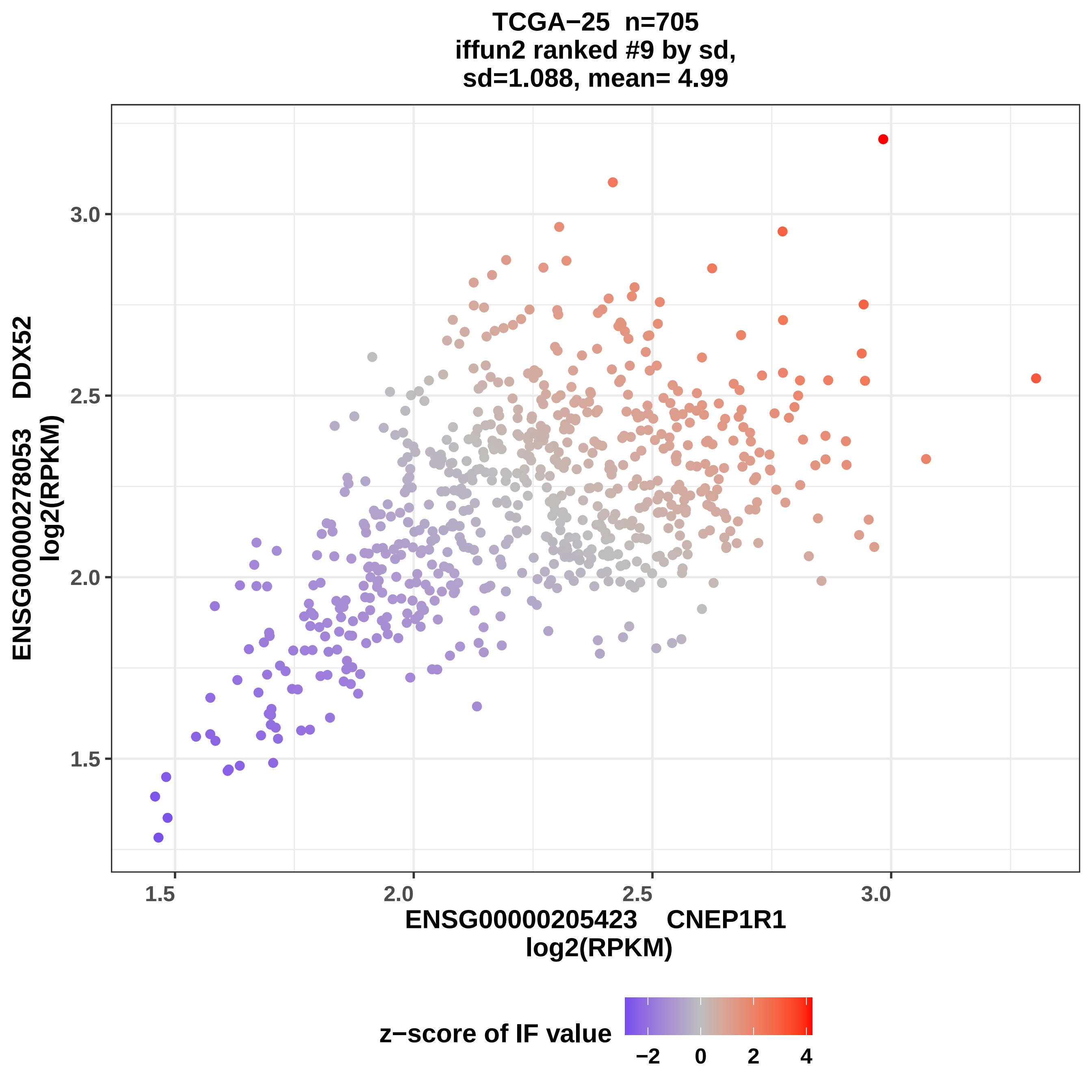
TCGA−25 n=705
iffun2 ranked #9 by sd,
sd=1.088, mean= 4.99
ENSG00000278053 DDX52
log2(RPKM)
ENSG00000205423 CNEP1R1
log2(RPKM)
1.5
2.0
2.5
3.0
z−score of IF value
−2
0
2
4

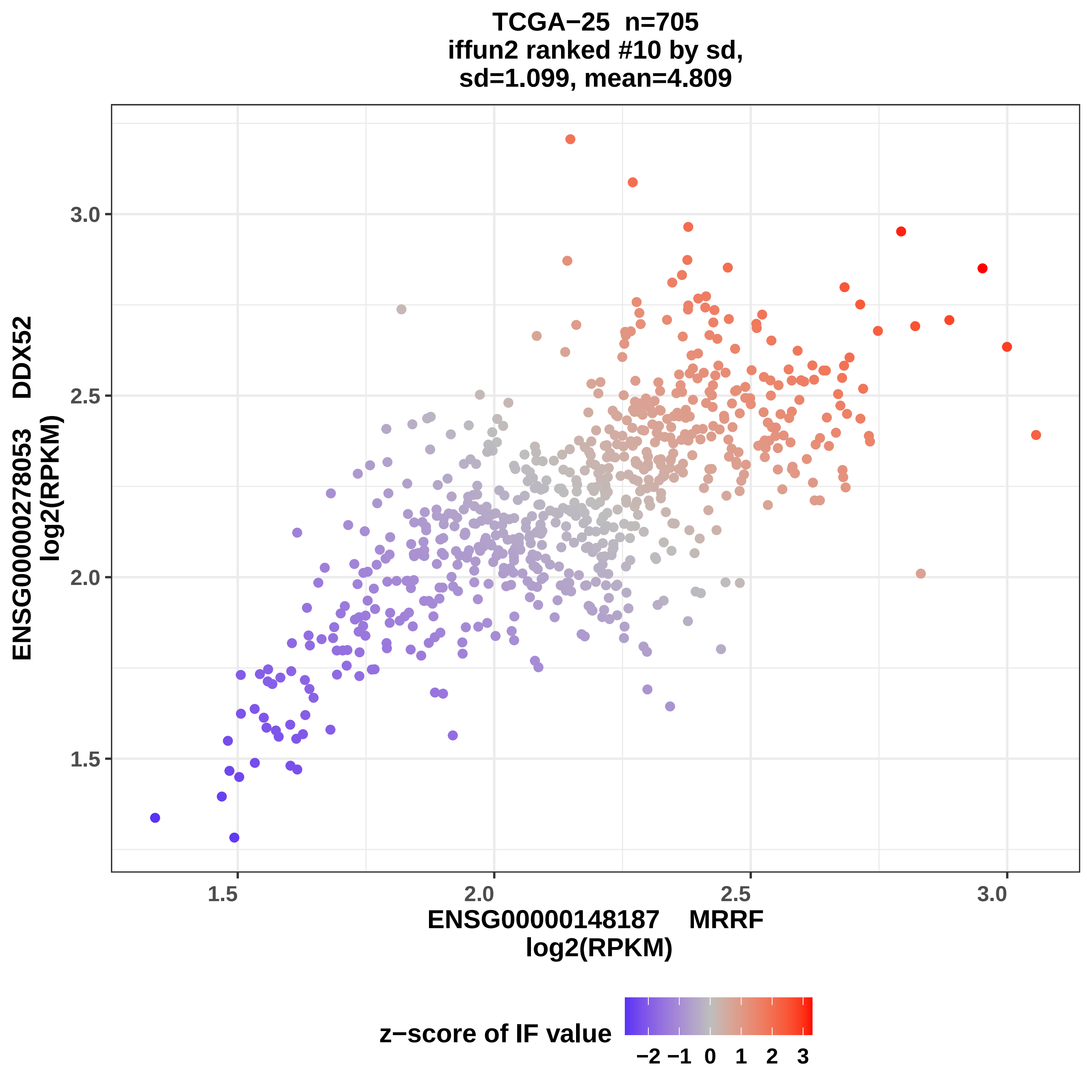

TCGA−25 n=705
iffun2 ranked #10 by sd,
sd=1.099, mean=4.809
ENSG00000278053 DDX52
log2(RPKM)
3.0
2.5
2.0
1.5
1.5
2.0
2.5
3.0
ENSG00000148187 MRRF
log2(RPKM)
z−score of IF value
−2 −1 0 1 2 3

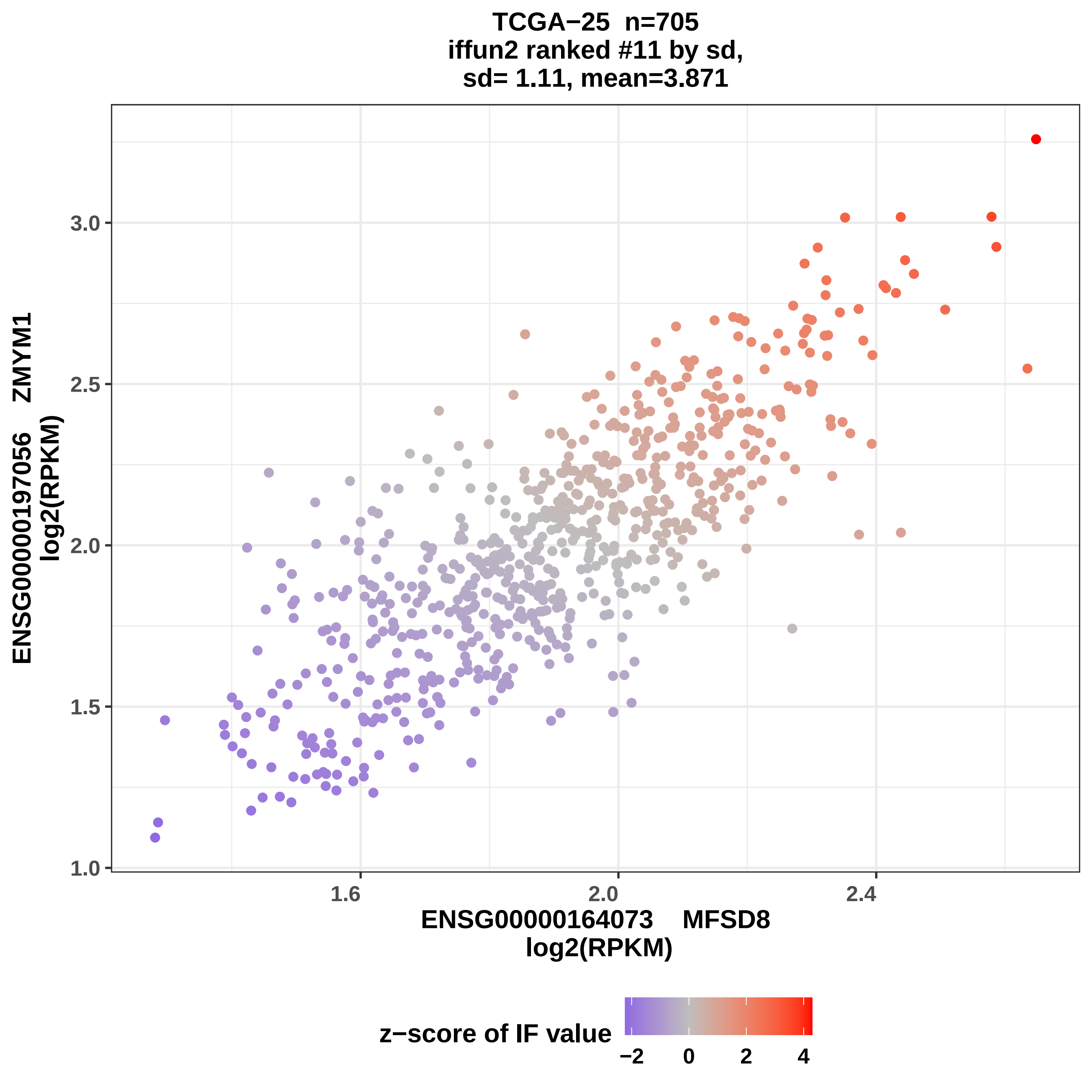

TCGA−25 n=705
iffun2 ranked #11 by sd,
sd= 1.11, mean=3.871
ENSG00000197056 ZMYM1
log2(RPKM)
3.0
2.5
2.0
1.5
1.0
1.6
2.0
2.4
ENSG00000164073 MFSD8
log2(RPKM)
z−score of IF value
−2
0
2
4

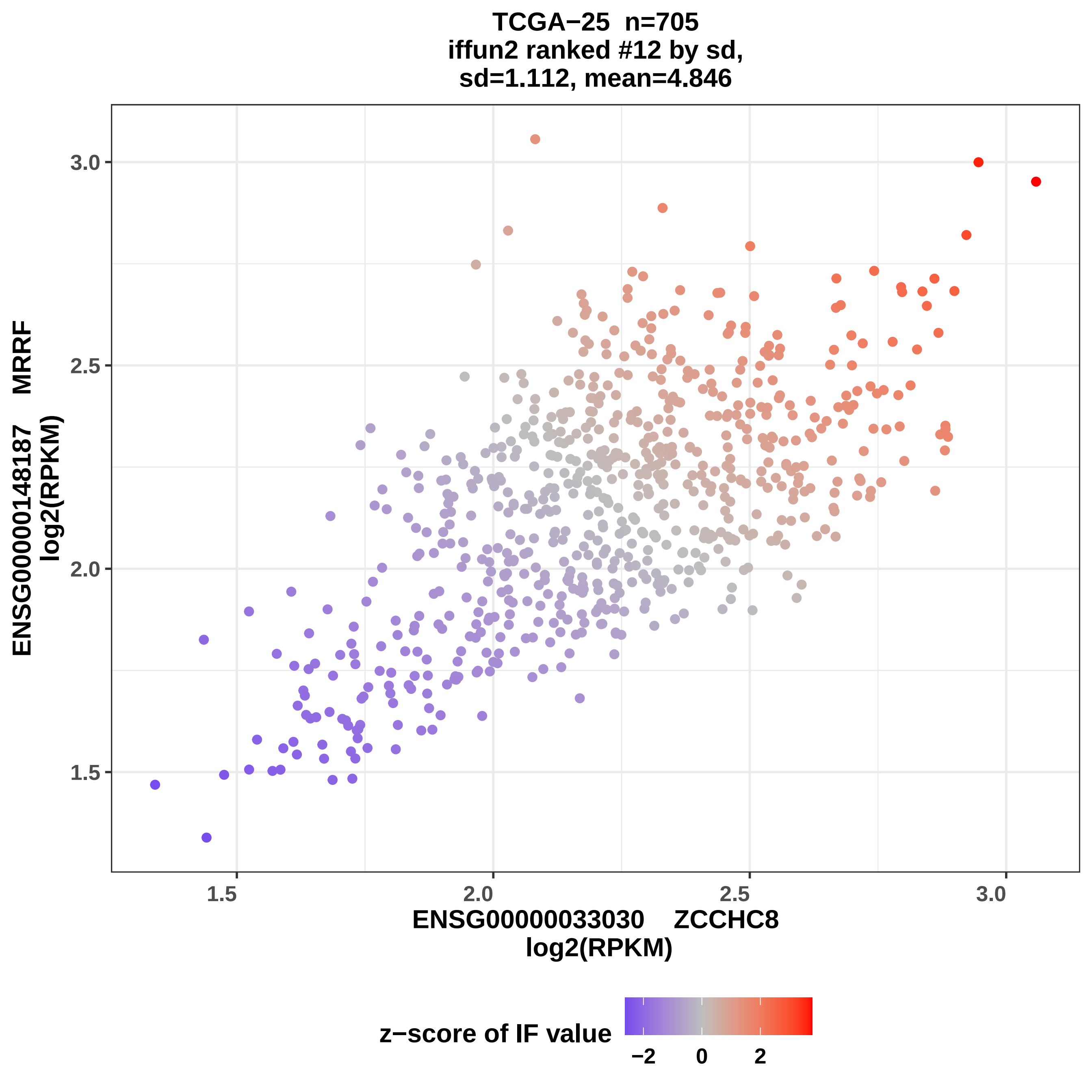
TCGA−25 n=705
iffun2 ranked #12 by sd,
sd=1.112, mean=4.846
ENSG00000148187 MRRF
log2(RPKM)
3.0
2.5
2.0
1.5
1.5
2.0
2.5
3.0
ENSG0000033030 ZCCHC8
log2(RPKM)
z−score of IF value
−2
0
2

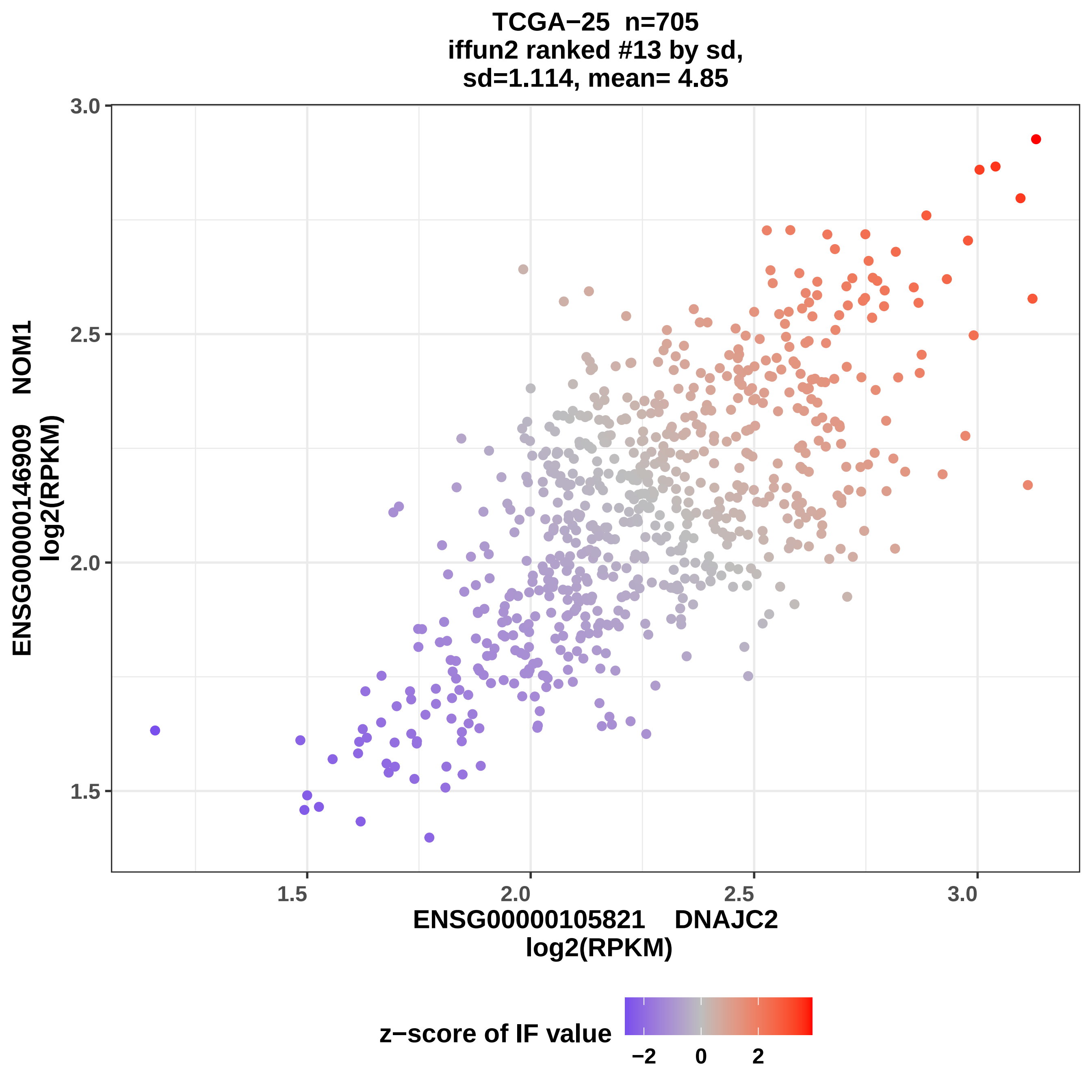
TCGA−25 n=705
iffun2 ranked #13 by sd,
sd=1.114, mean= 4.85
3.0
2.5
2.0
1.5
ENSG00000146909 NOM1
log2(RPKM)
1.5
2.0
2.5
3.0
ENSG00000105821 DNAJC2
log2(RPKM)
z−score of IF value
−2
0
2

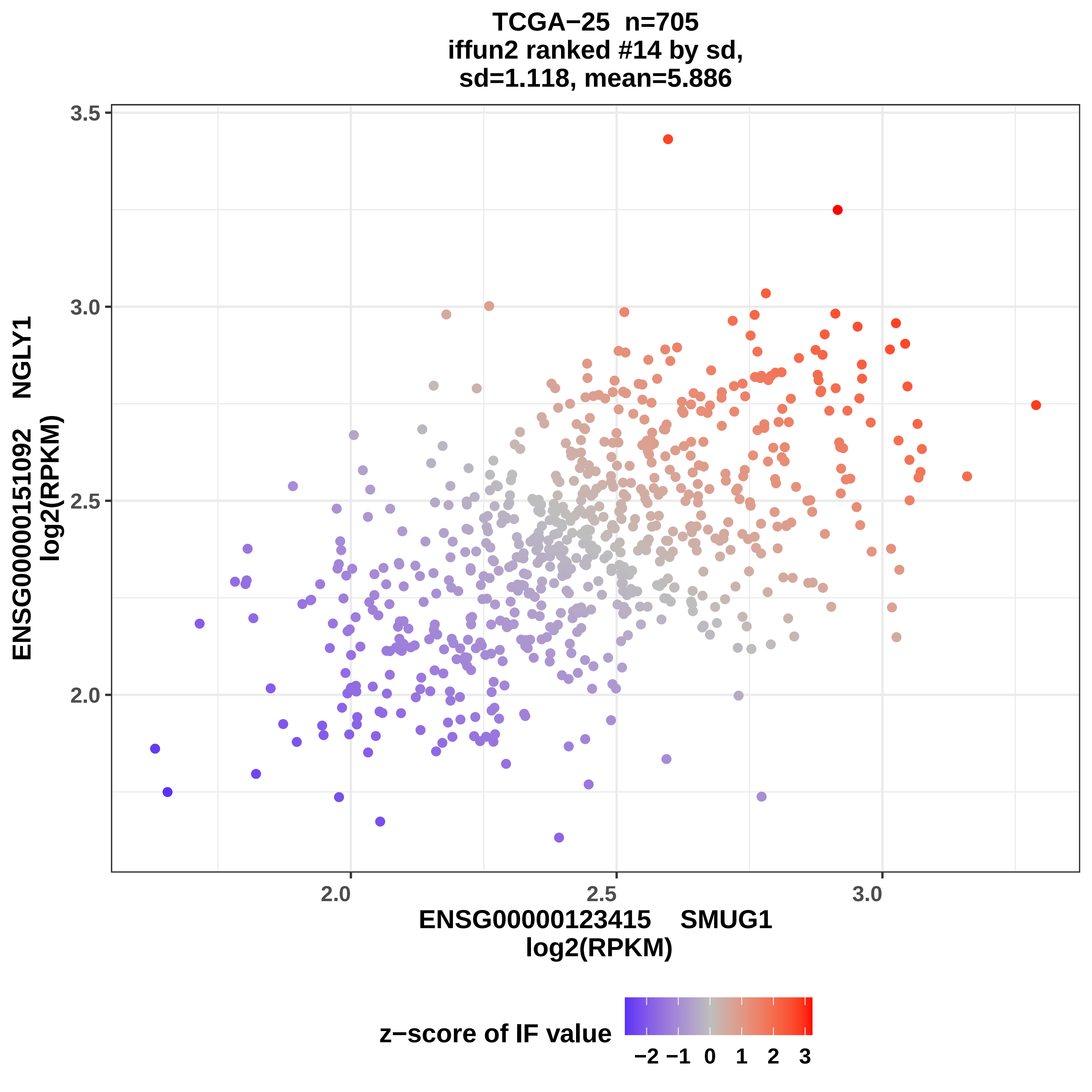

TCGA−25 n=705
iffun2 ranked #14 by sd,
sd=1.118, mean=5.886
3.5
3.0
2.5
2.0
ENSG00000151092 NGLY1
log2(RPKM)
2.0
2.5
3.0
ENSG00000123415 SMUG1
log2(RPKM)
z−score of IF value
−2 −1 0 1 2 3

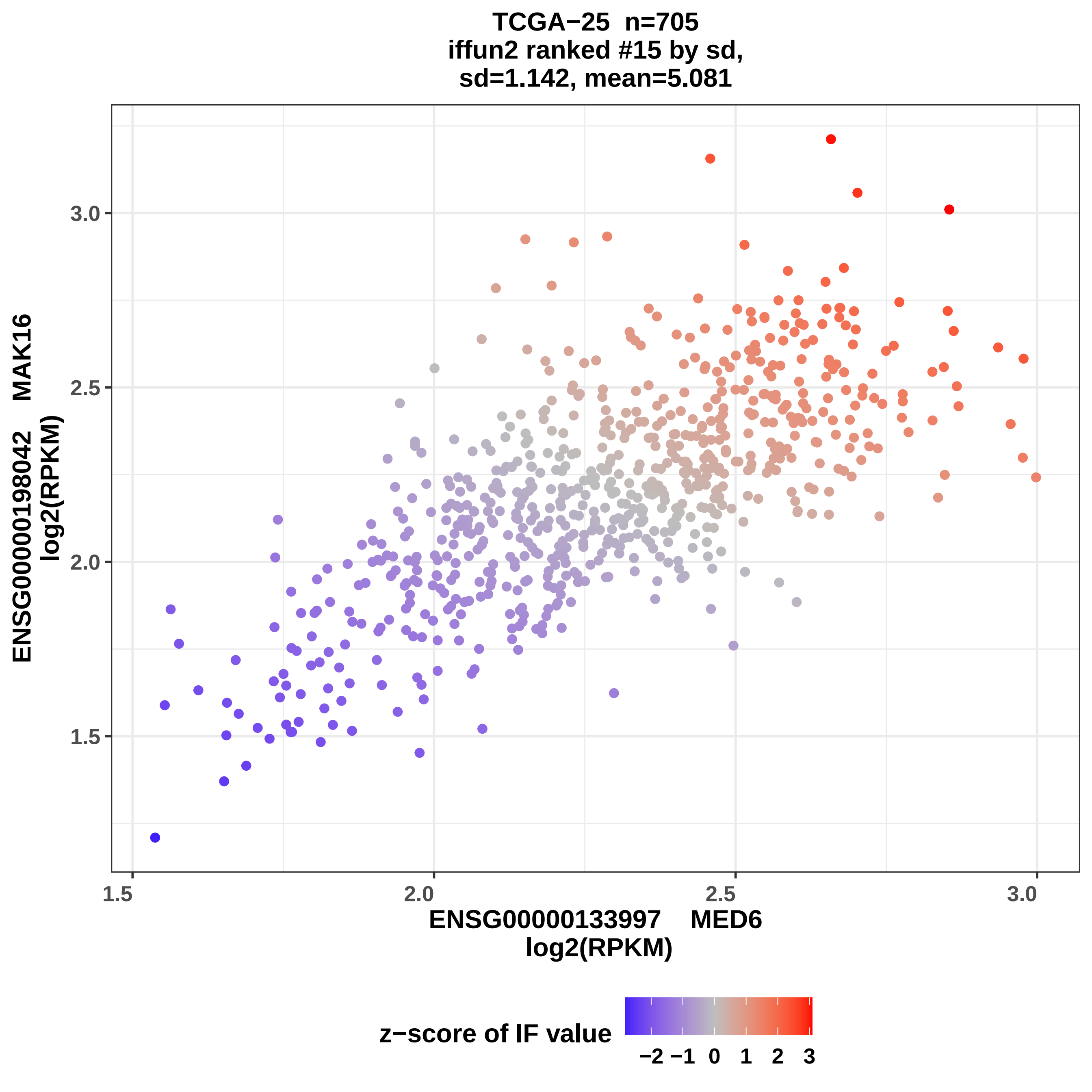
TCGA−25 n=705
iffun2 ranked #15 by sd,
sd=1.142, mean=5.081
ENSG00000198042 MAK16
log2(RPKM)
3.0
2.5
2.0
1.5
1.5
2.0
2.5
3.0
ENSG00000133997 MED6
log2(RPKM)
z−score of IF value
−2 −1 0 1 2 3

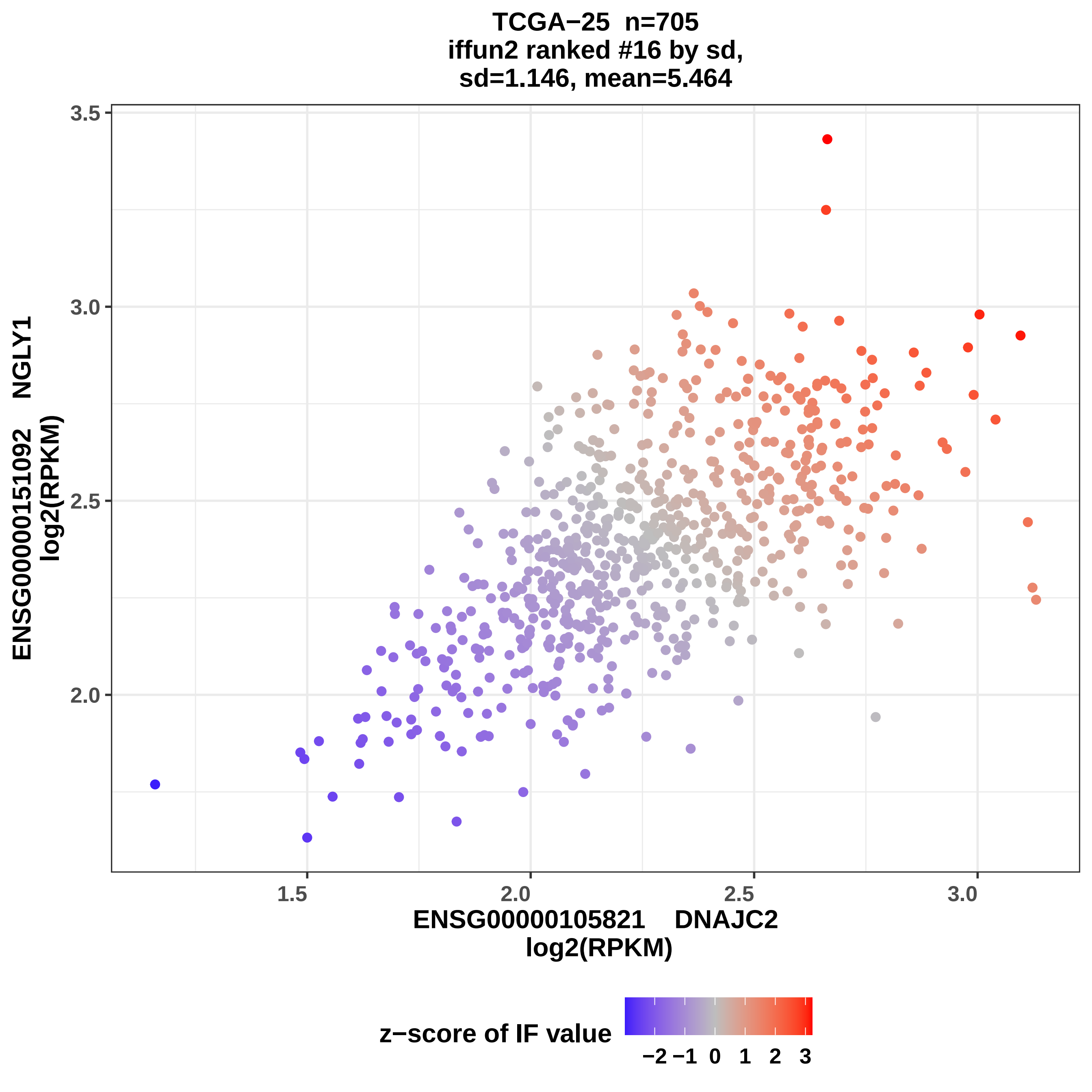
TCGA−25 n=705
iffun2 ranked #16 by sd,
sd=1.146, mean=5.464
ENSG00000151092 NGLY1
log2(RPKM)
3.5
3.0
2.5
2.0
1.5
2.0
2.5
3.0
ENSG00000105821 DNAJC2
log2(RPKM)
z−score of IF value
−2 −1 0 1 2 3

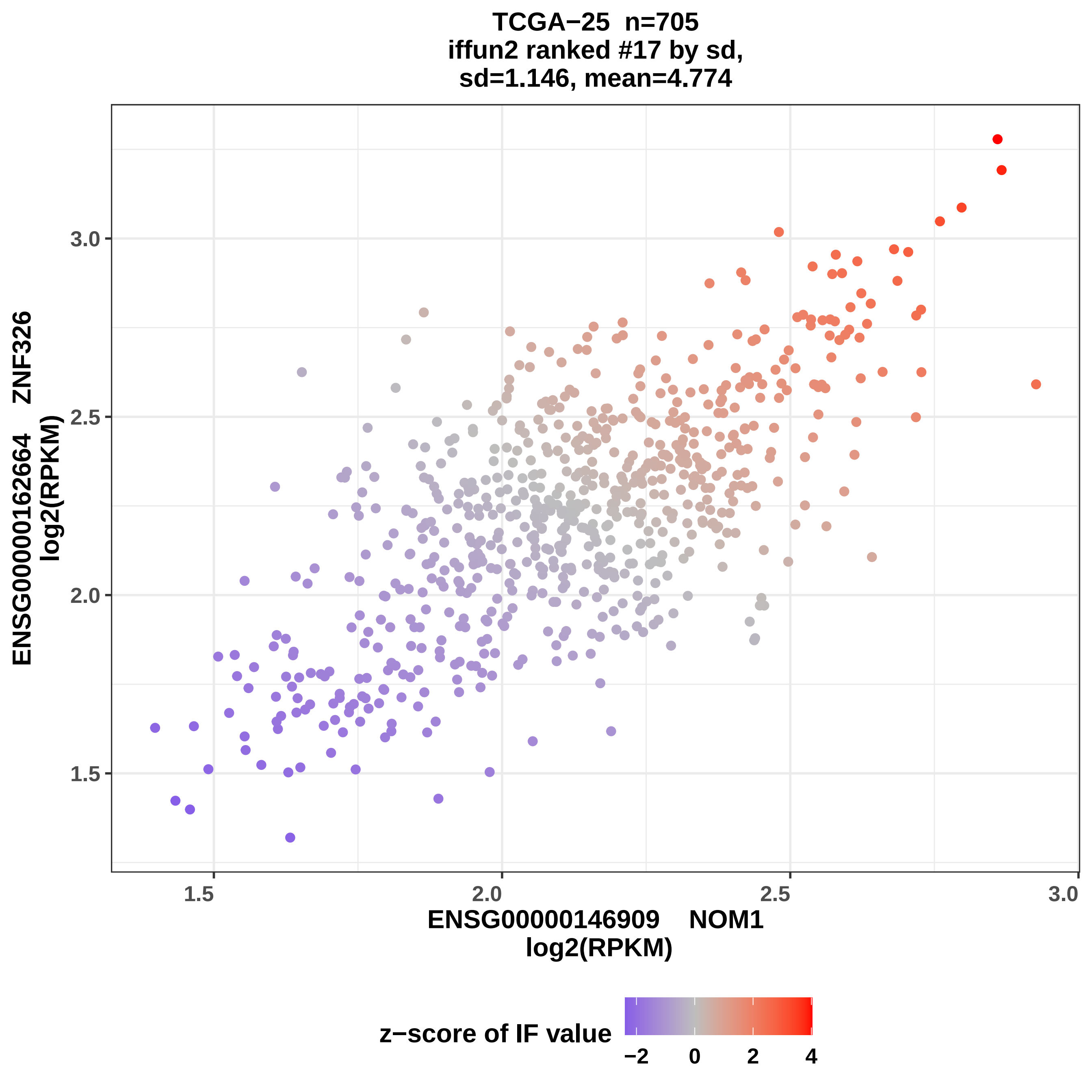
TCGA−25 n=705
iffun2 ranked #17 by sd,
sd=1.146, mean=4.774
ENSG00000162664 ZNF326
log2(RPKM)
3.0
2.5
2.0
1.5
ENSG00000146909 NOM1
log2(RPKM)
1.5
2.0
2.5
3.0
z−score of IF value
−2
0
2
4

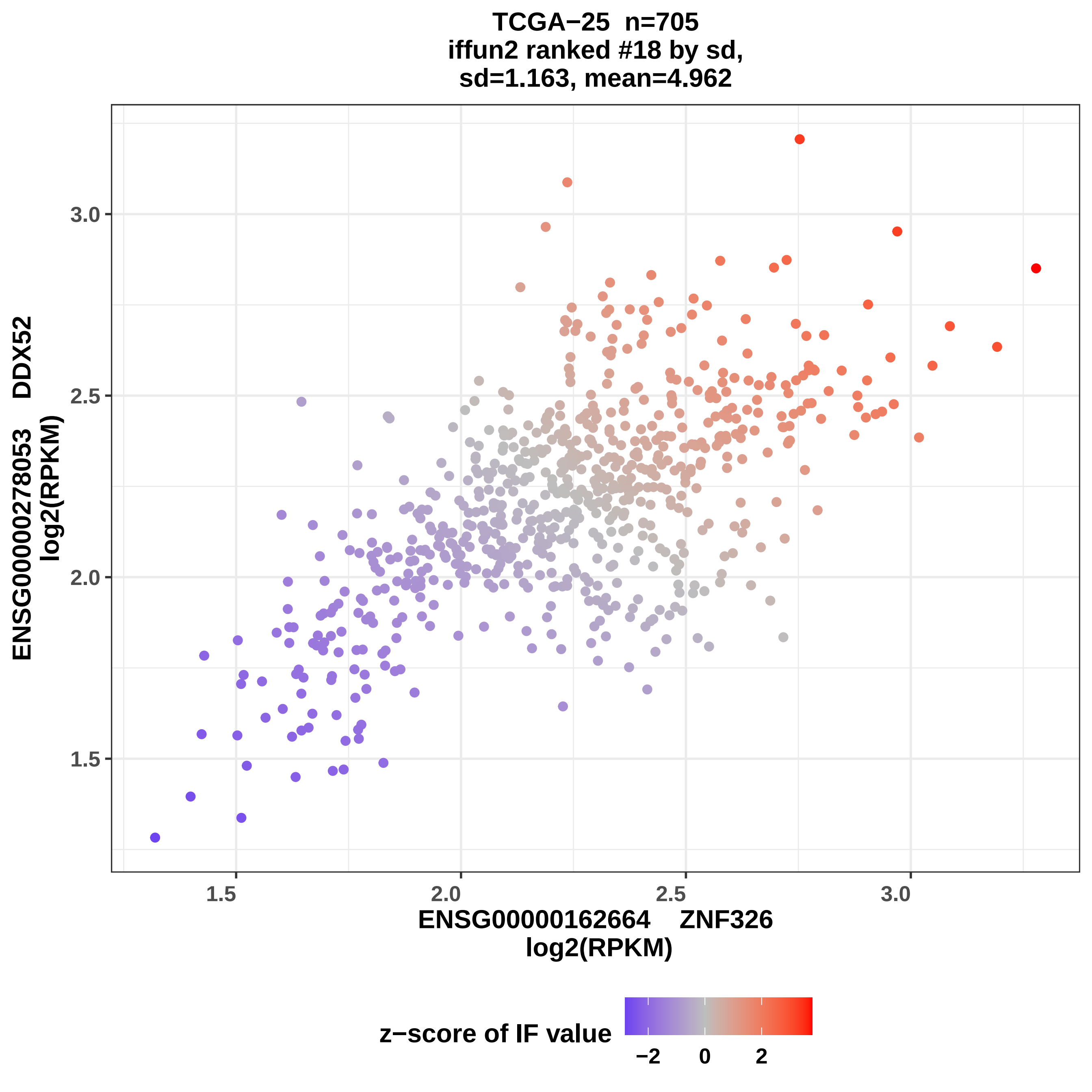

TCGA−25 n=705
iffun2 ranked #18 by sd,
sd=1.163, mean=4.962
3.0
2.5
2.0
1.5
ENSG00000278053 DDX52
log2(RPKM)
1.5
2.0
2.5
3.0
ENSG00000162664 ZNF326
log2(RPKM)
z−score of IF value
−2
0
2

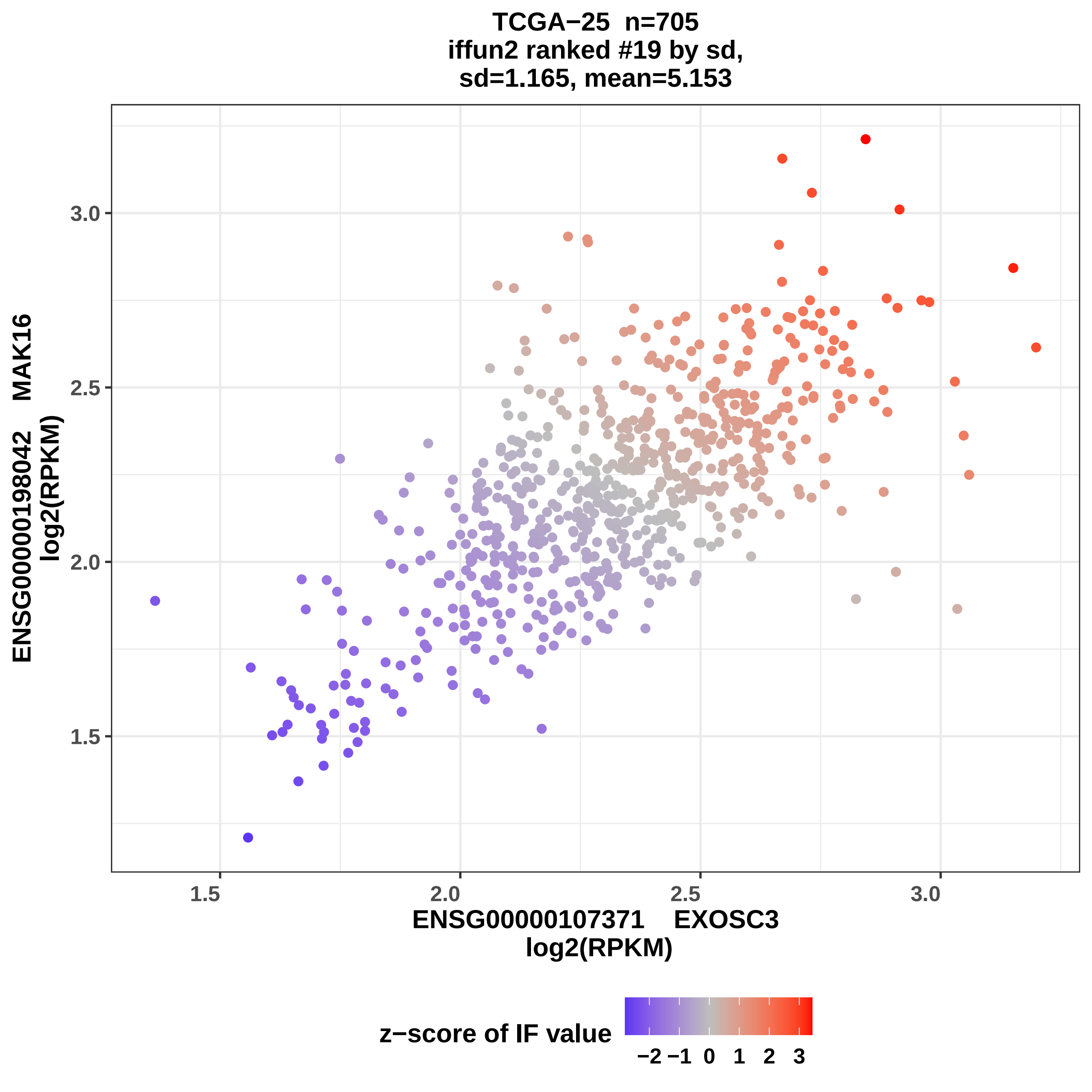

TCGA−25 n=705
iffun2 ranked #19 by sd,
sd=1.165, mean=5.153
ENSG00000198042 MAK16
log2(RPKM)
3.0
2.5
2.0
1.5
1.5
2.0
2.5
3.0
ENSG00000107371 EXOSC3
log2(RPKM)
z−score of IF value
−2 −1 0 1 2 3

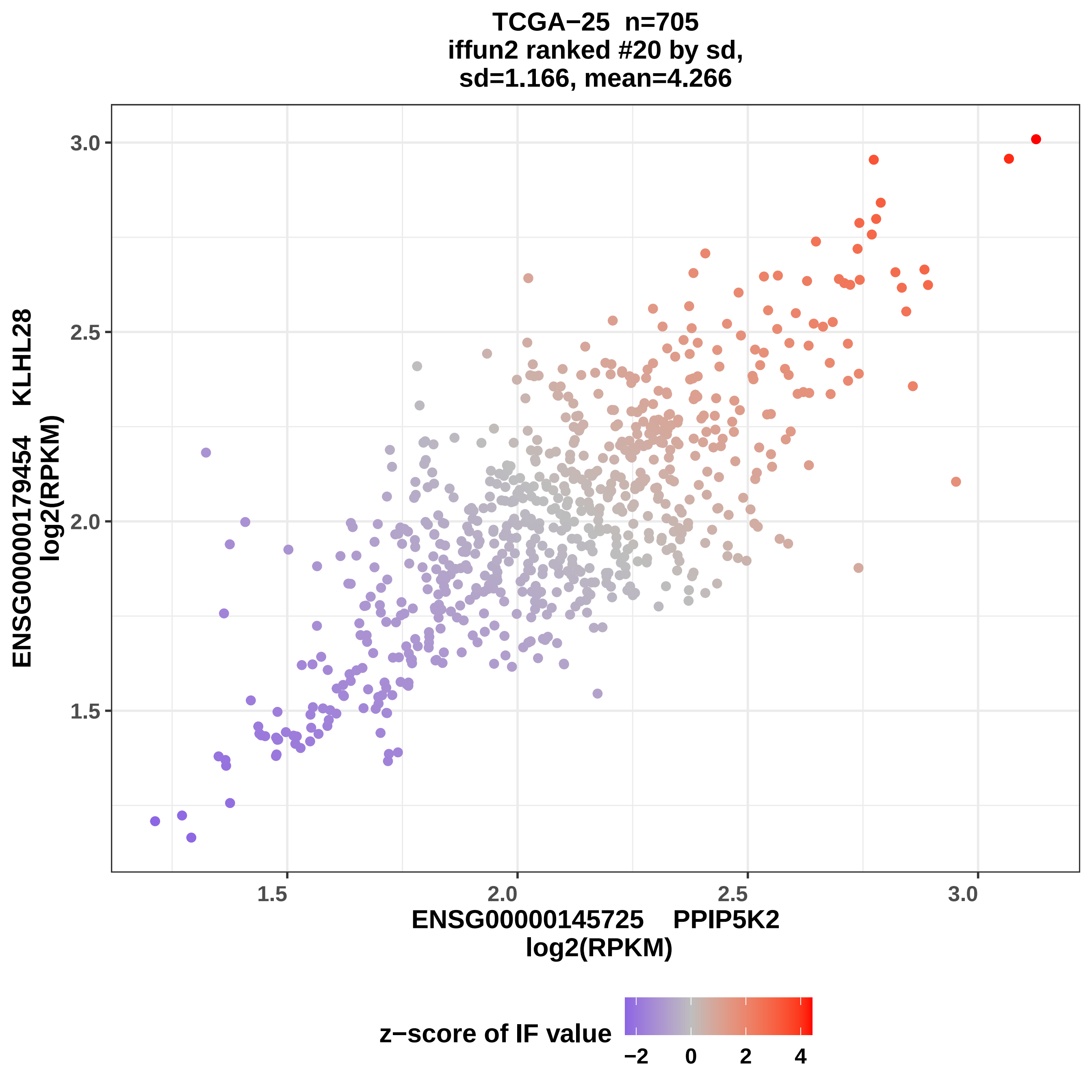
TCGA−25 n=705
iffun2 ranked #20 by sd,
sd=1.166, mean=4.266
ENSG00000179454 KLHL28
log2(RPKM)
3.0
2.5
2.0
1.5
1.5
2.0
2.5
3.0
ENSG00000145725 PPIP5K2
log2(RPKM)
z−score of IF value
−2
0
2
4

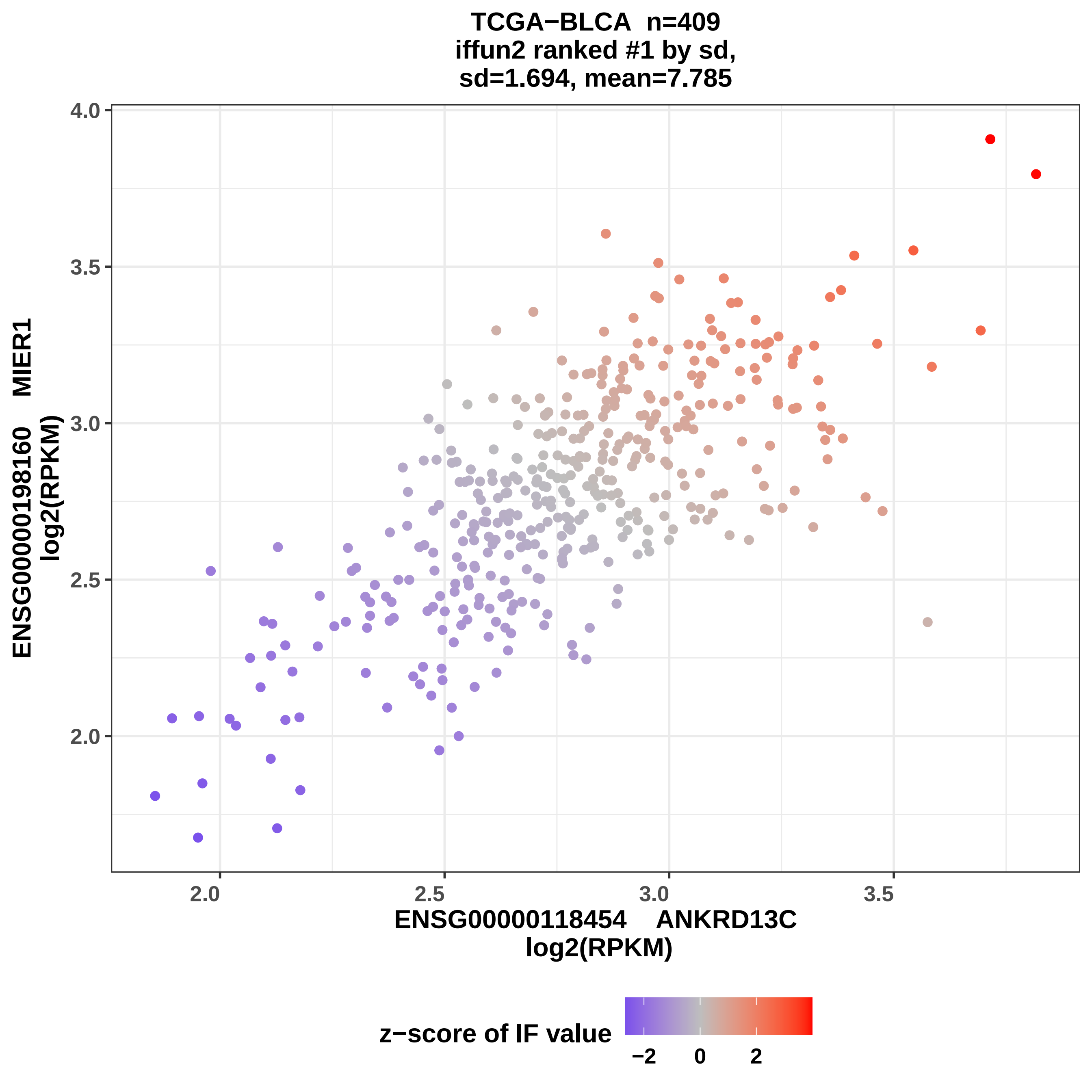
TCGA−BLCA n=409
iffun2 ranked #1 by sd,
sd=1.694, mean=7.785
ENSG00000198160 MIER1
log2(RPKM)
ENSG00000118454 ANKRD13C
log2(RPKM)
4.0
3.5
3.0
2.5
2.0
2.0
2.5
3.0
3.5
z−score of IF value
−2
0
2

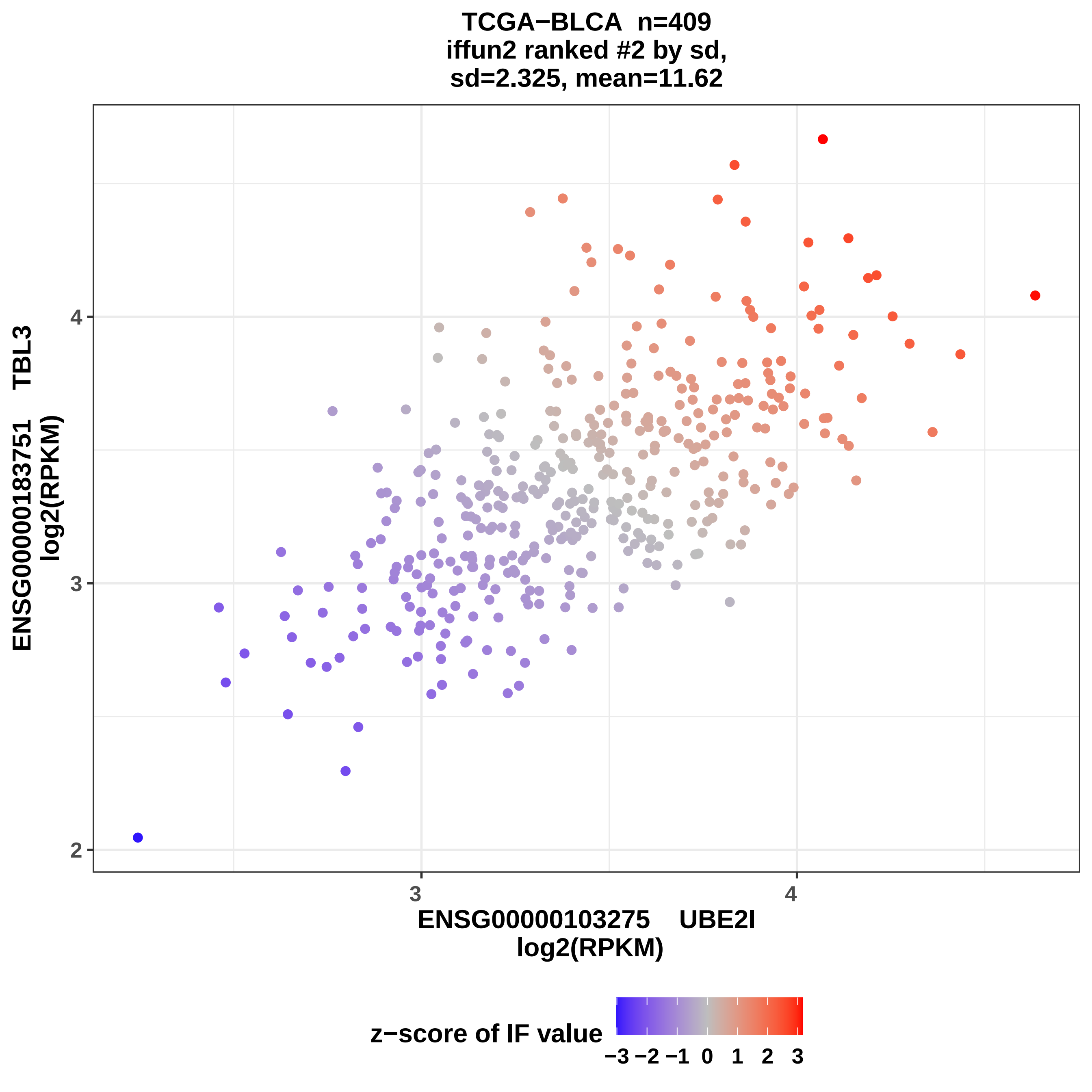
TCGA−BLCA n=409
iffun2 ranked #2 by sd,
sd=2.325, mean=11.62
ENSG00000183751 TBL3
log2(RPKM)
4
3
2
3
4
ENSG00000103275 UBE2I
log2(RPKM)
z−score of IF value
−3 −2 −1 0 1 2 3

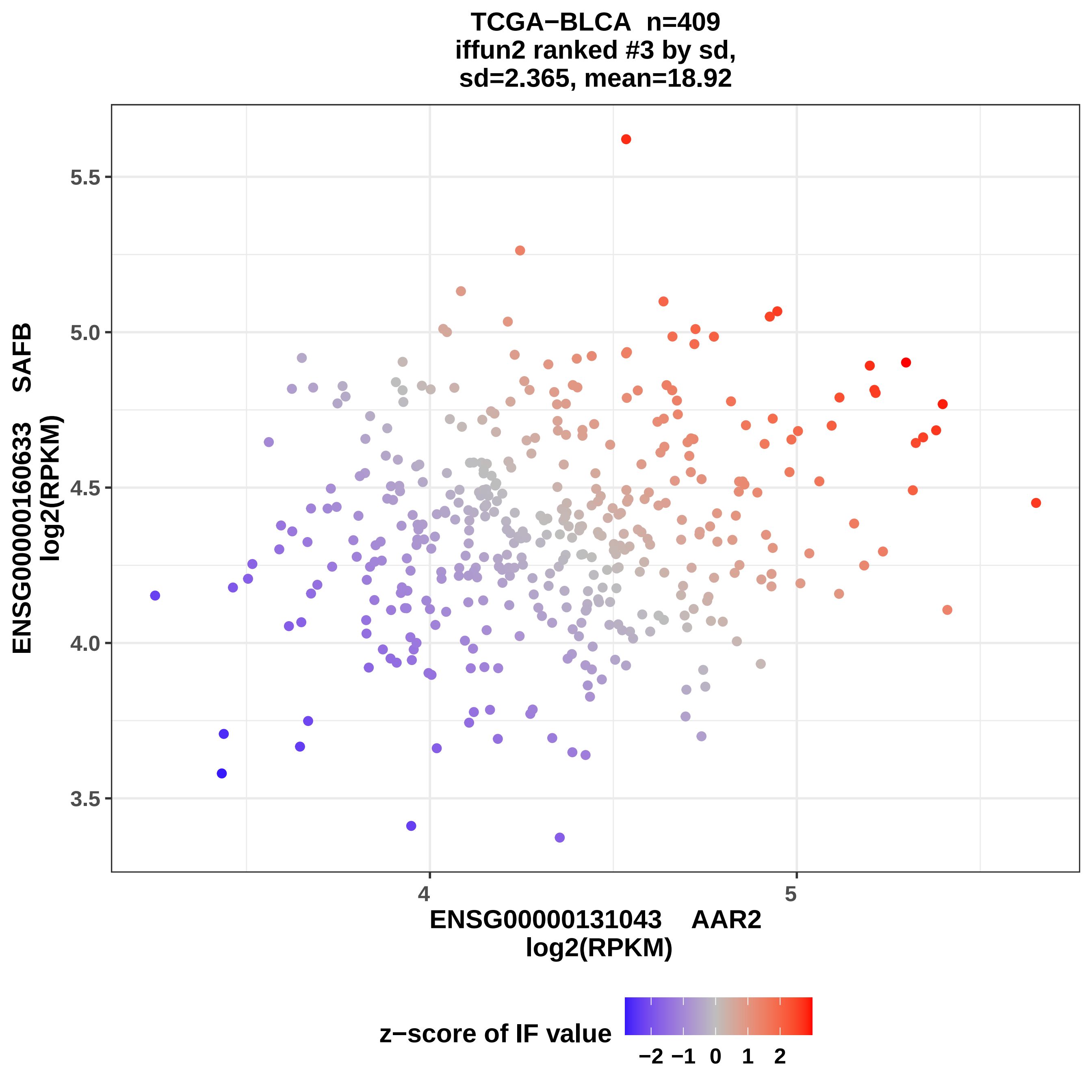
TCGA−BLCA n=409
iffun2 ranked #3 by sd,
sd=2.365, mean=18.92
ENSG00000160633 SAFB
log2(RPKM)
5.5
5.0
4.5
4.0
3.5
4
5
ENSG00000131043 AAR2
log2(RPKM)
z−score of IF value
−2 −1 0 1 2

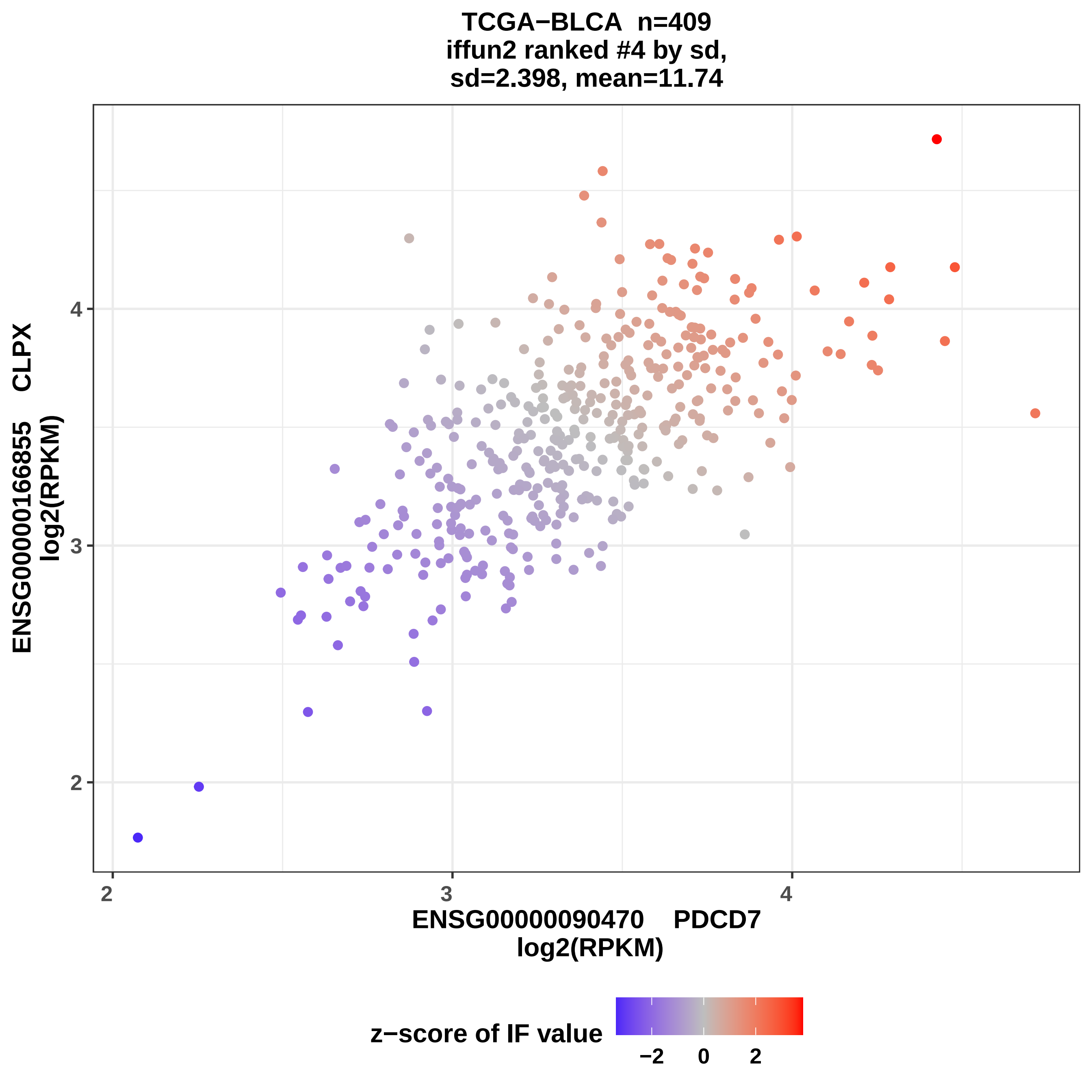
TCGA−BLCA n=409
iffun2 ranked #4 by sd,
sd=2.398, mean=11.74
ENSG00000166855 CLPX
log2(RPKM)
ENSG00000090470 PDCD7
log2(RPKM)
z−score of IF value
−2
0
2
2
3
4

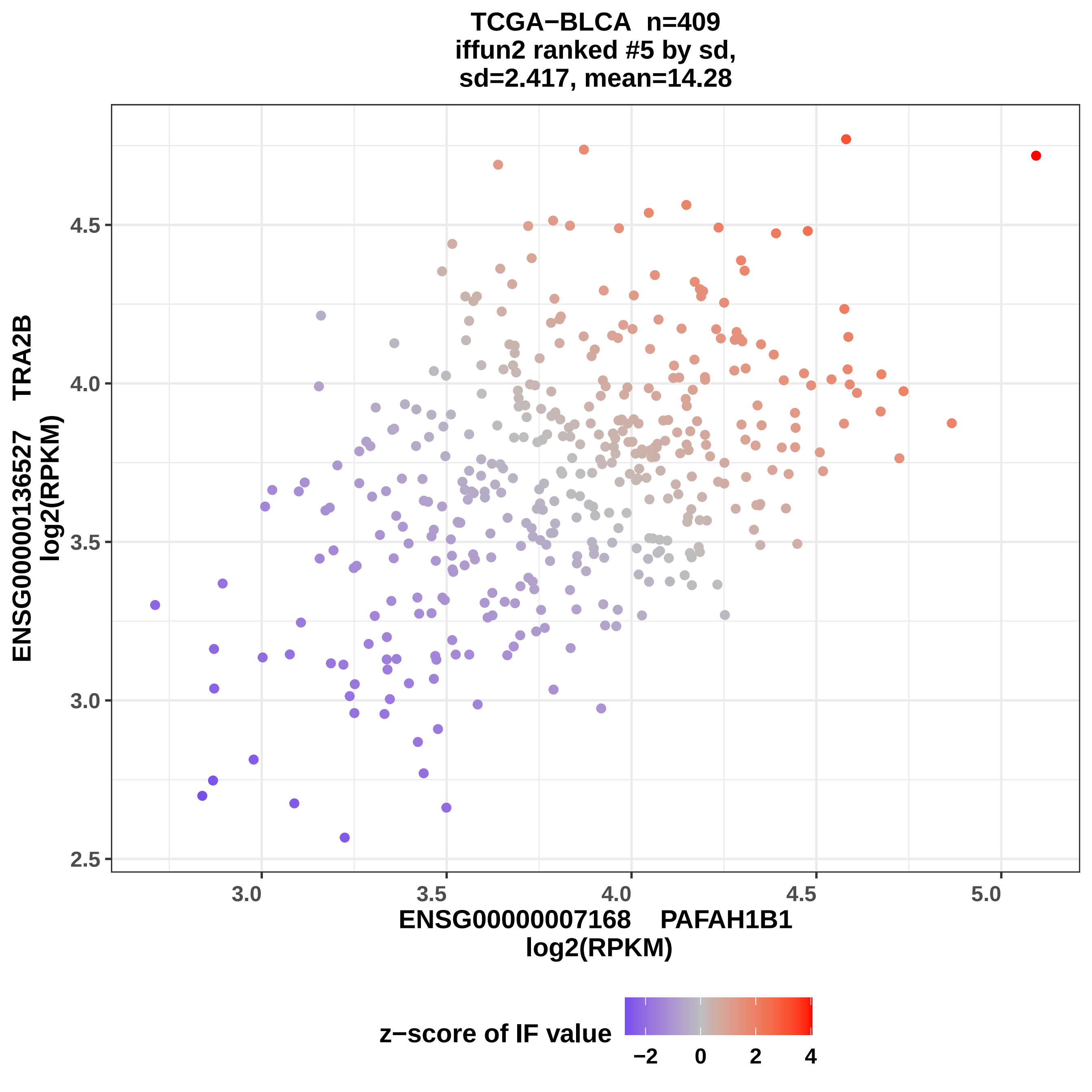
TCGA−BLCA n=409
iffun2 ranked #5 by sd,
sd=2.417, mean=14.28
ENSG00000136527 TRA2B
log2(RPKM)
4.5
4.0
3.5
3.0
2.5
3.0
3.5
4.0
4.5
5.0
ENSG00000007168 PAFAH1B1
log2(RPKM)
z−score of IF value
−2
0
2
4

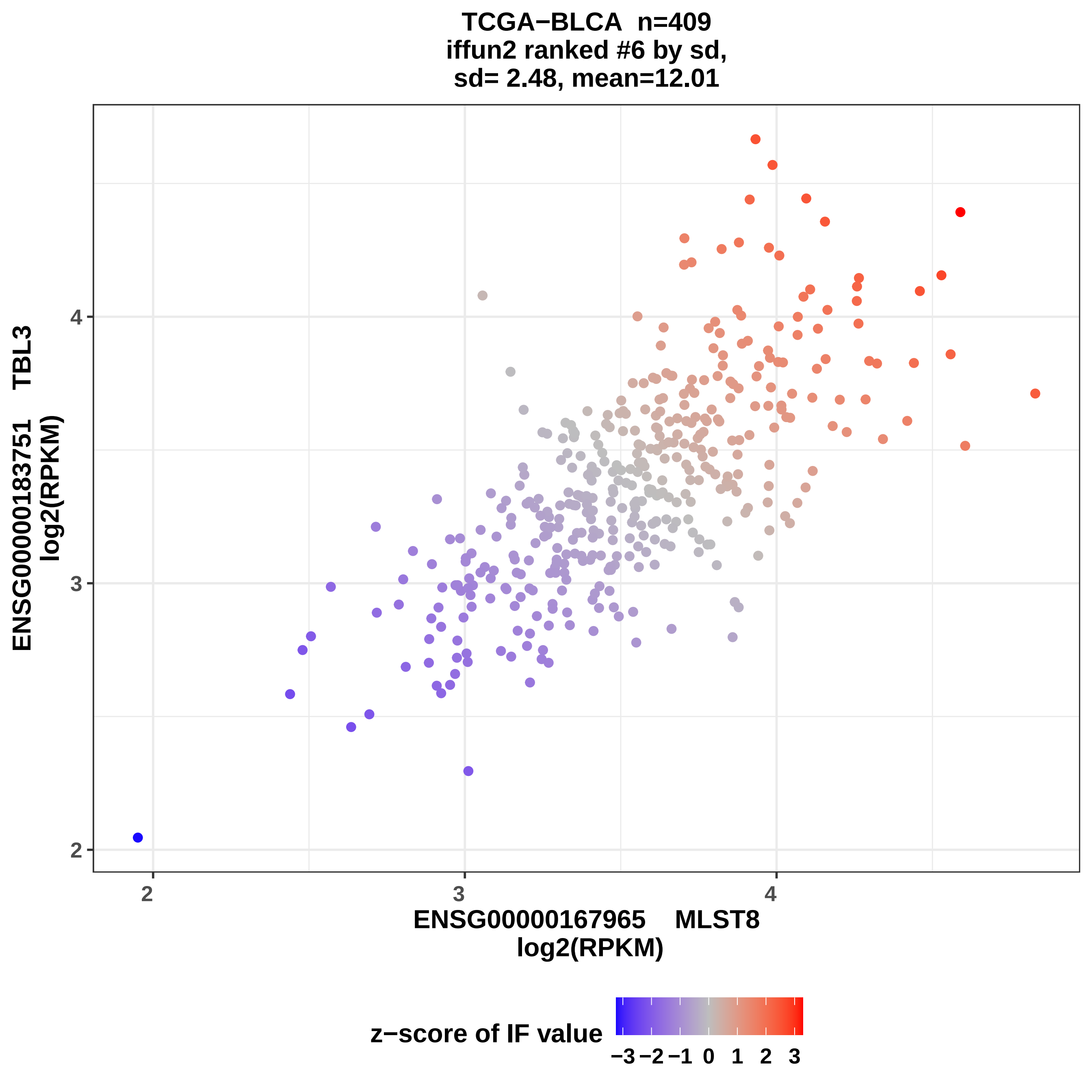

TCGA−BLCA n=409
iffun2 ranked #6 by sd,
sd= 2.48, mean=12.01
ENSG00000183751 TBL3
log2(RPKM)
4
3
2
2
3
4
ENSG00000167965 MLST8
log2(RPKM)
z−score of IF value
−3 −2 −1 0 1 2 3

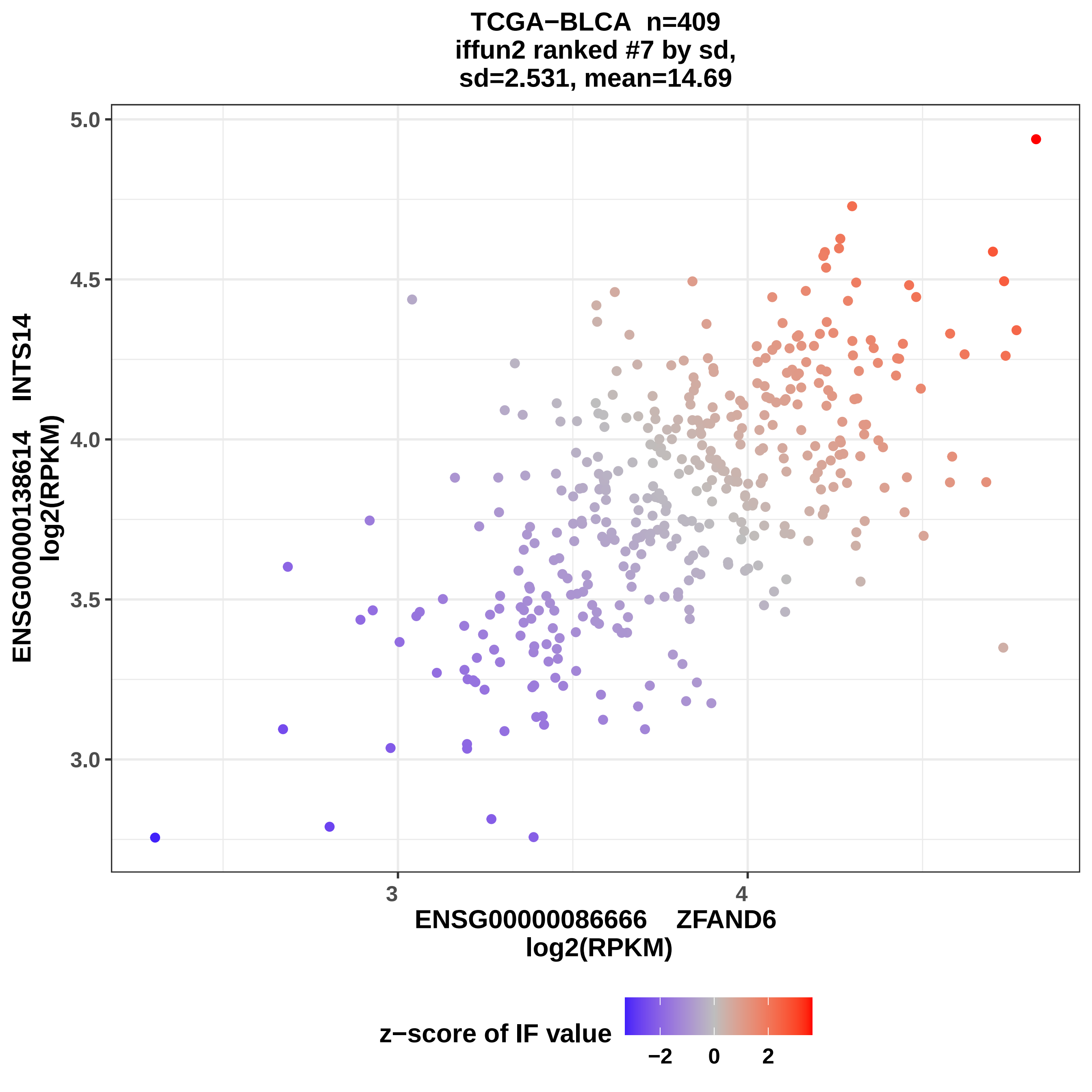
TCGA−BLCA n=409
iffun2 ranked #7 by sd,
sd=2.531, mean=14.69
ENSG00000138614 INTS14
log2(RPKM)
5.0
4.5
4.0
3.5
3.0
3
4
ENSG00000086666 ZFAND6
log2(RPKM)
z−score of IF value
−2
0
2

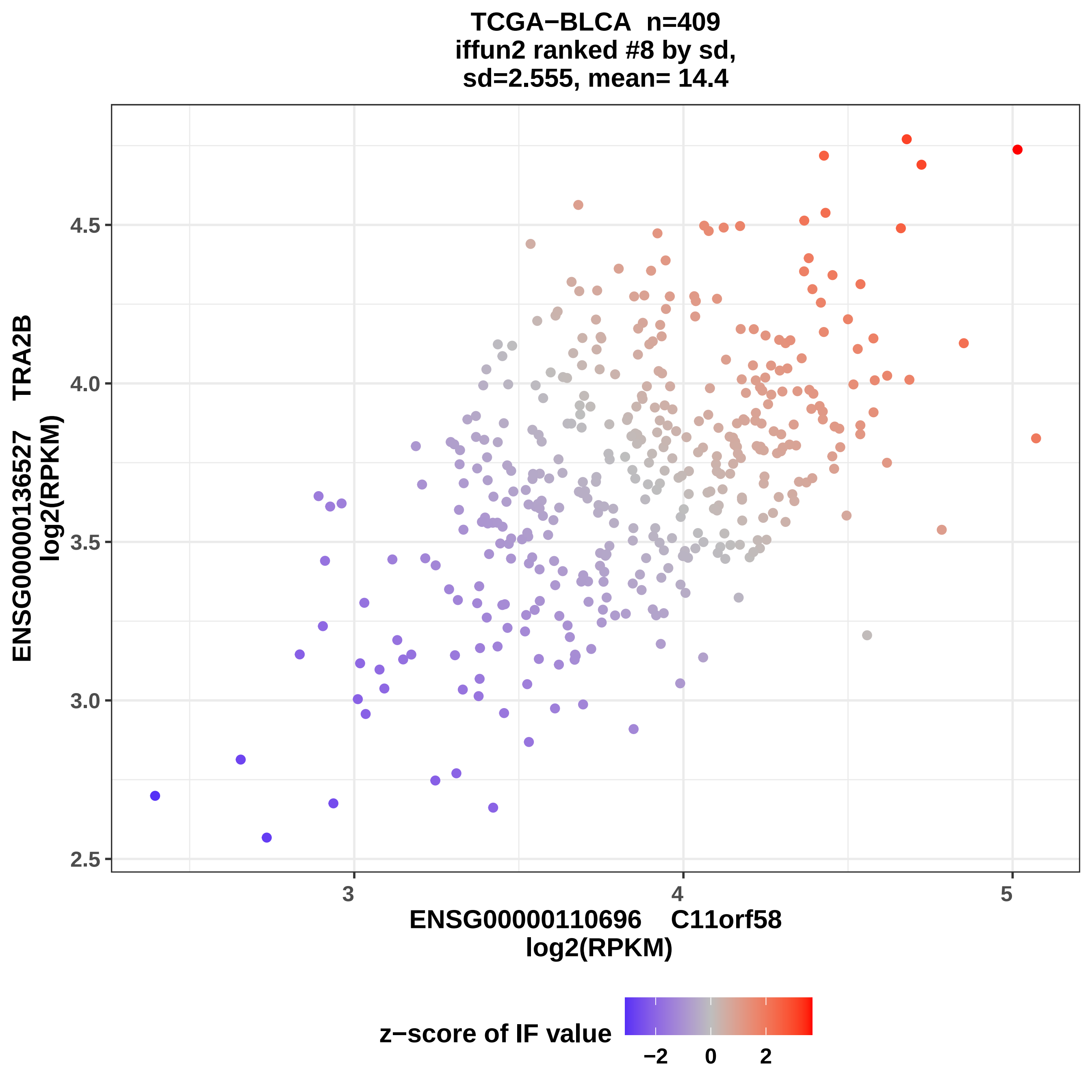
TCGA−BLCA n=409
iffun2 ranked #8 by sd,
sd=2.555, mean= 14.4
ENSG00000136527 TRA2B
log2(RPKM)
4.5
4.0
3.5
3.0
2.5
3
4
5
ENSG00000110696 C11orf58
log2(RPKM)
z−score of IF value
−2
0
2

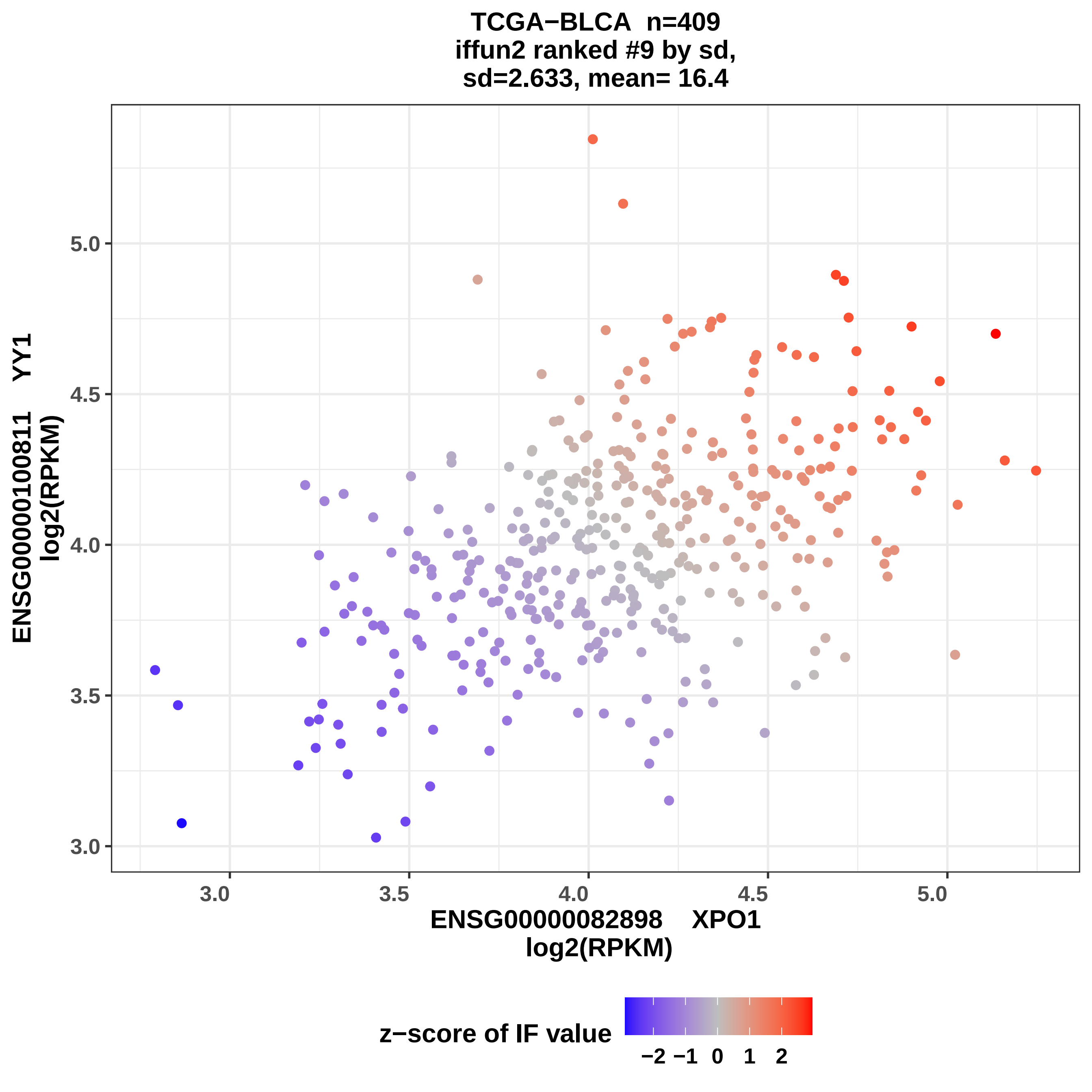

TCGA−BLCA n=409
iffun2 ranked #9 by sd,
sd=2.633, mean= 16.4
ENSG00000100811 YY1
log2(RPKM)
ENSG00000082898 XPO1
log2(RPKM)
3.0
3.5
4.0
4.5
5.0
z−score of IF value
−2 −1 0 1 2

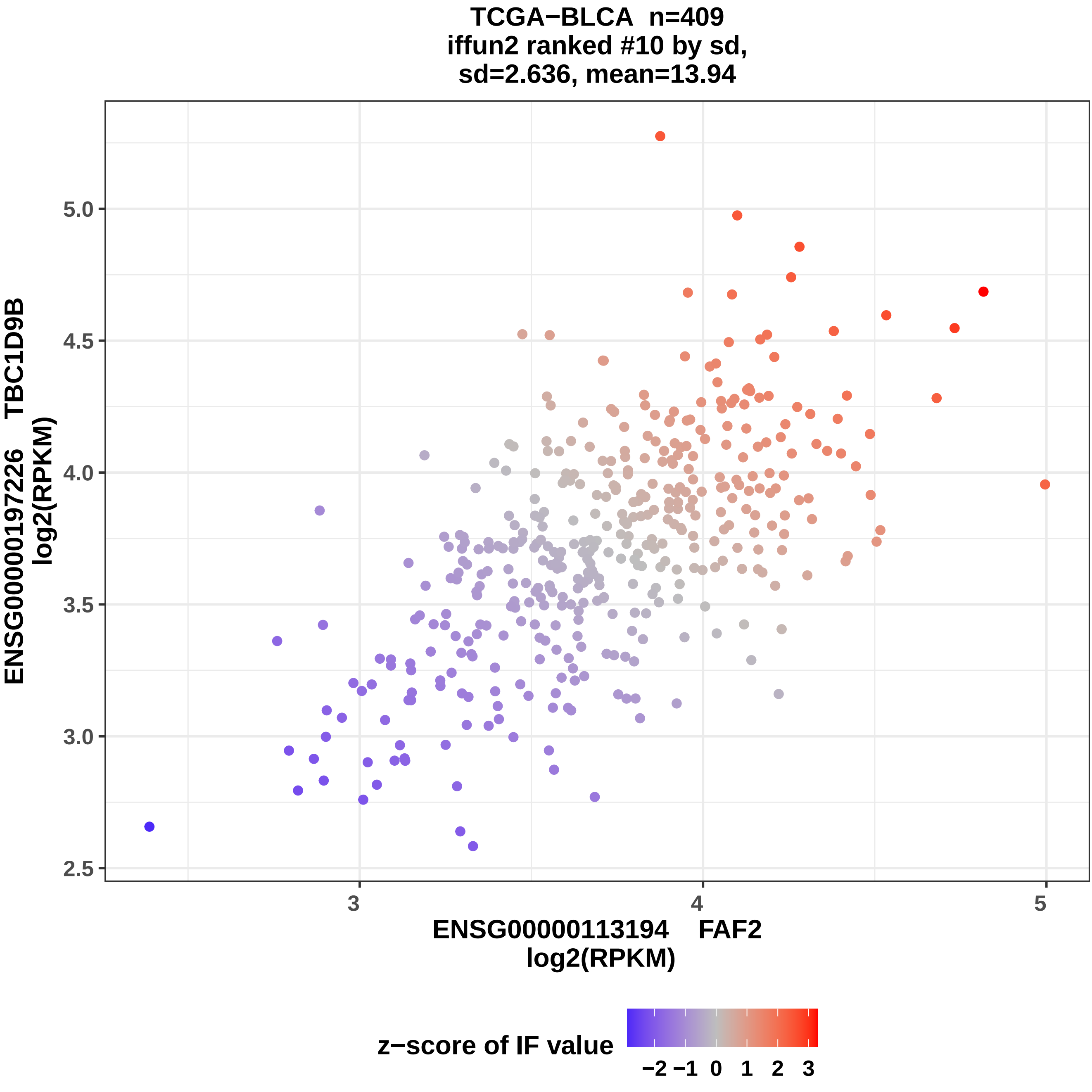

TCGA−BLCA n=409
iffun2 ranked #10 by sd,
sd=2.636, mean=13.94
ENSG00000197226 TBC1D9B
log2(RPKM)
5.0
4.5
4.0
3.5
3.0
2.5
3
4
5
ENSG00000113194 FAF2
log2(RPKM)
z−score of IF value
−2
−1
0
1
2
3

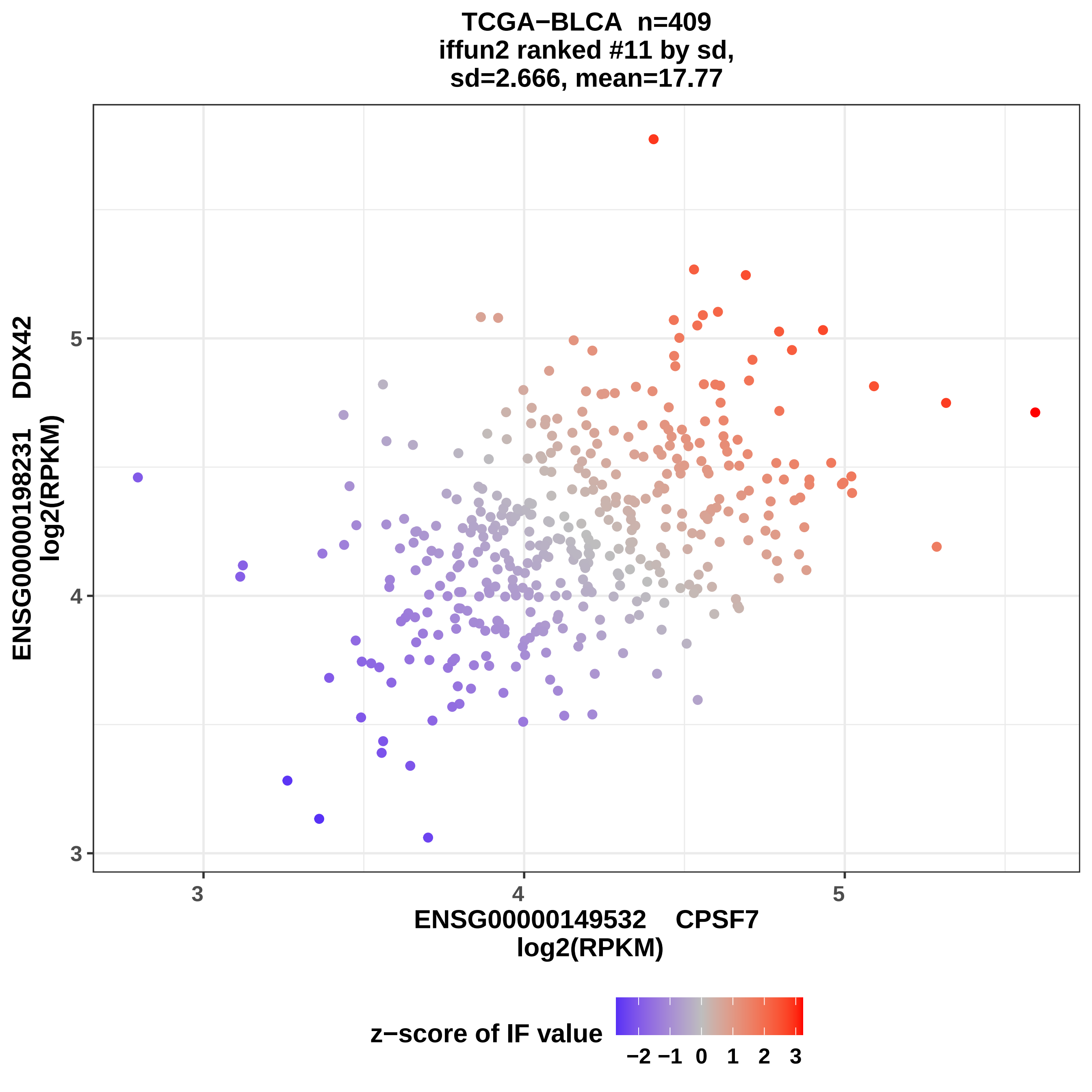
TCGA−BLCA n=409
iffun2 ranked #11 by sd,
sd=2.666, mean=17.77
ENSG00000198231 DDX42
log2(RPKM)
5
4
3
3
4
5
ENSG00000149532 CPSF7
log2(RPKM)
z−score of IF value
−2 −1 0 1 2 3

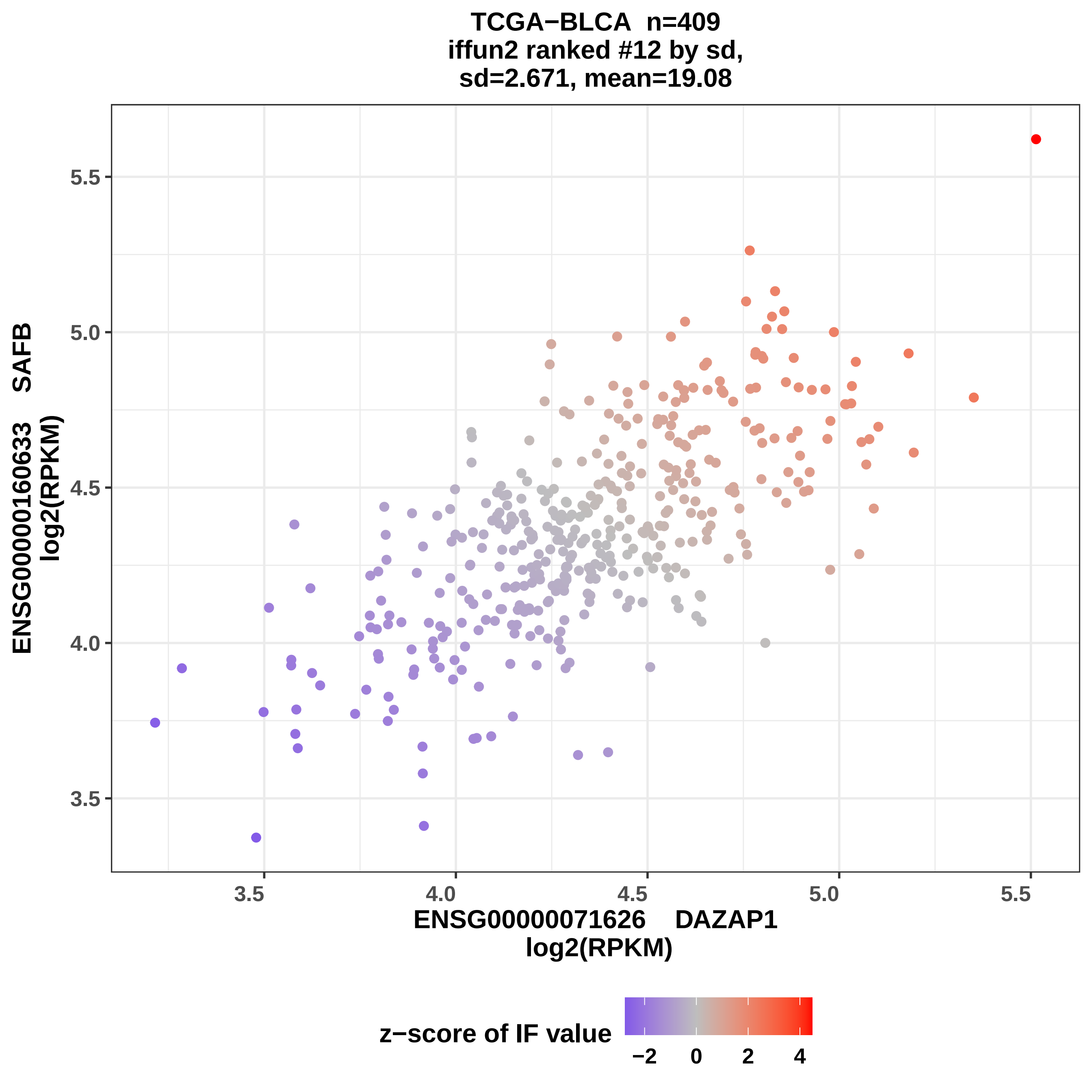
TCGA−BLCA n=409
iffun2 ranked #12 by sd,
sd=2.671, mean=19.08
ENSG00000160633 SAFB
log2(RPKM)
5.5
5.0
4.5
4.0
3.5
3.5
4.0
4.5
5.0
5.5
ENSG00000071626 DAZAP1
log2(RPKM)
z−score of IF value
−2
0
2
4

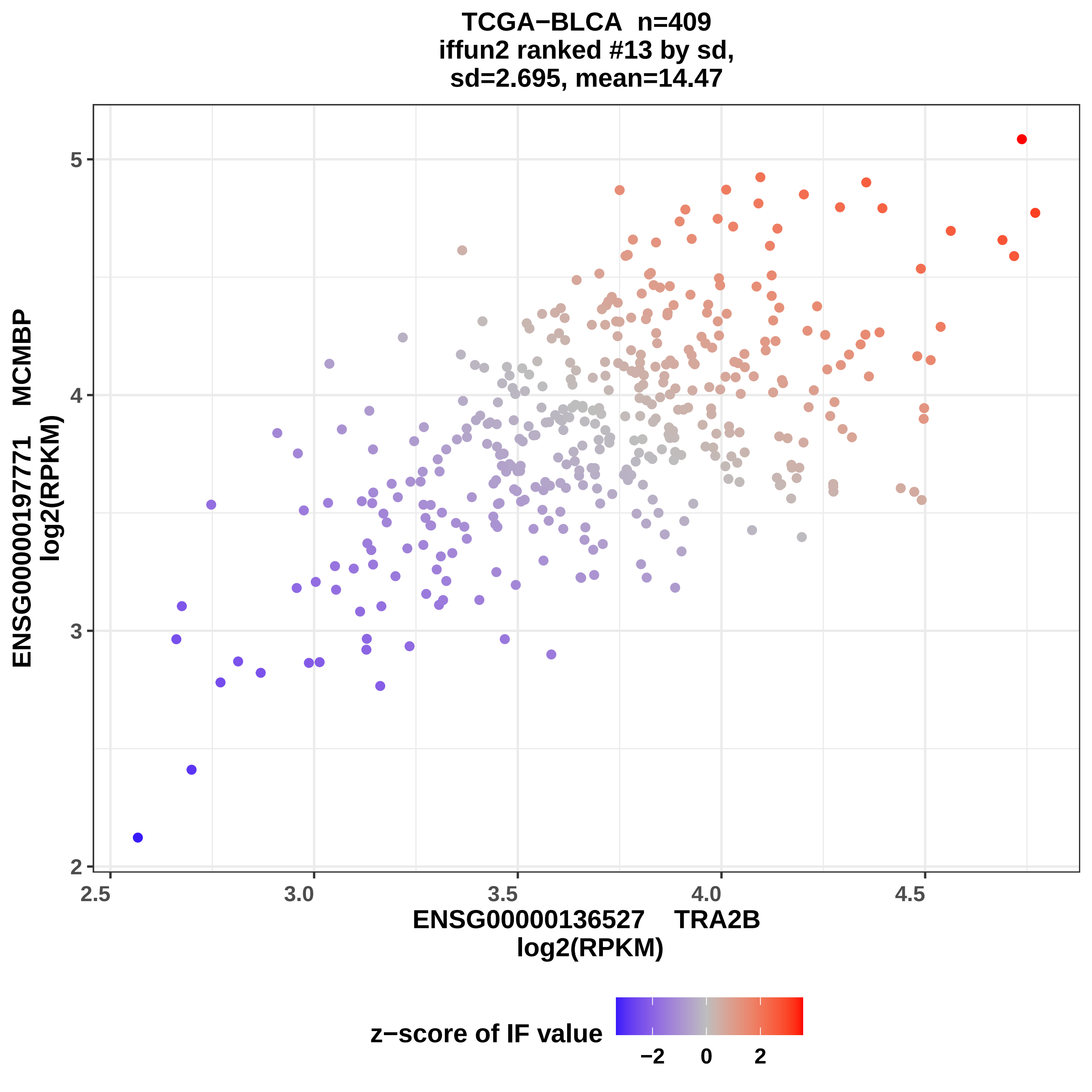
TCGA−BLCA n=409
iffun2 ranked #13 by sd,
sd=2.695, mean=14.47
ENSG00000197771 MCMBP
log2(RPKM)
5
4
3
2
2.5
3.0
3.5
4.0
4.5
ENSG00000136527 TRA2B
log2(RPKM)
z−score of IF value
−2
0
2

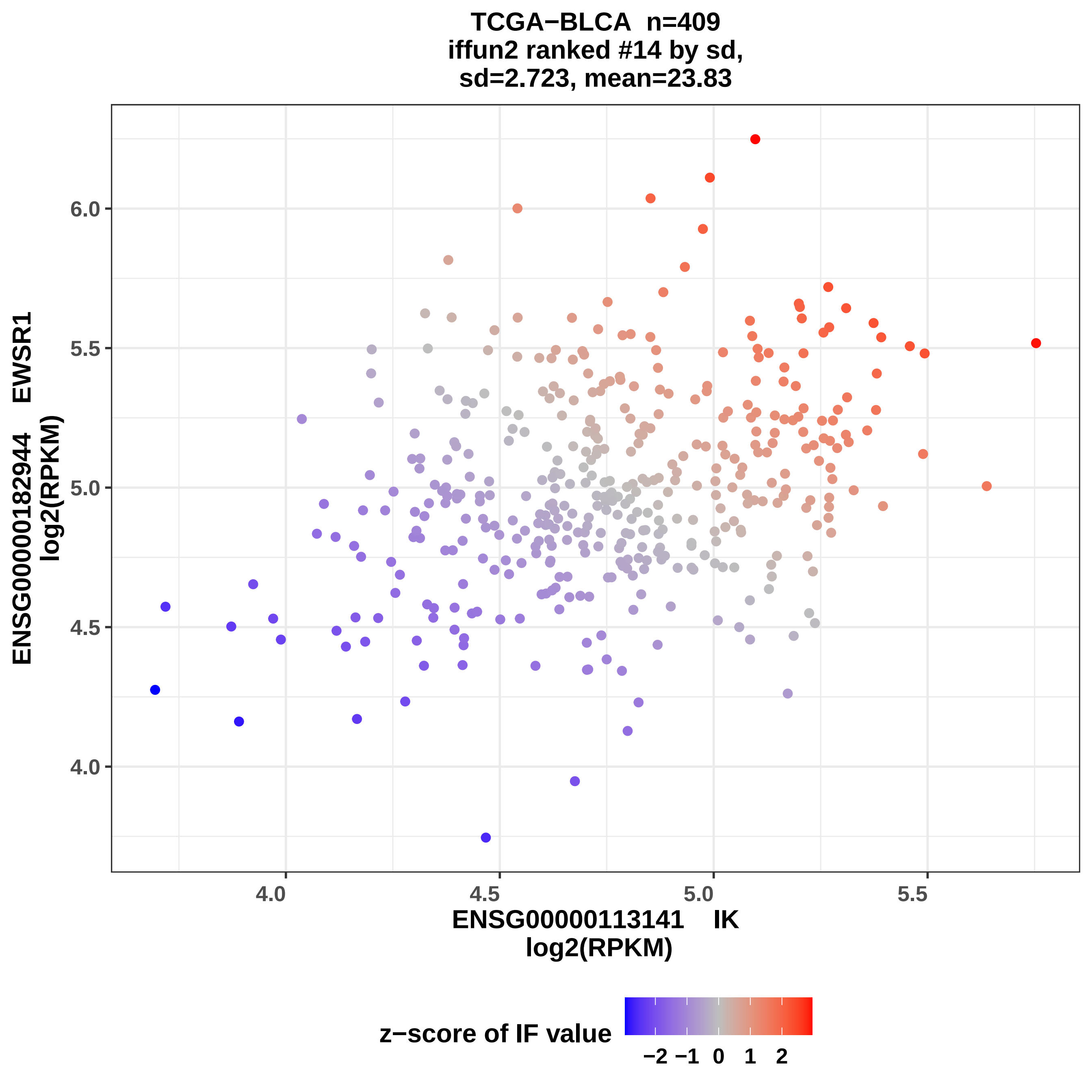

TCGA−BLCA n=409
iffun2 ranked #14 by sd,
sd=2.723, mean=23.83
ENSG00000182944 EWSR1
log2(RPKM)
6.0
5.5
5.0
4.5
4.0
4.0
4.5
5.0
5.5
ENSG00000113141 IK
log2(RPKM)
z−score of IF value
−2 −1 0 1 2

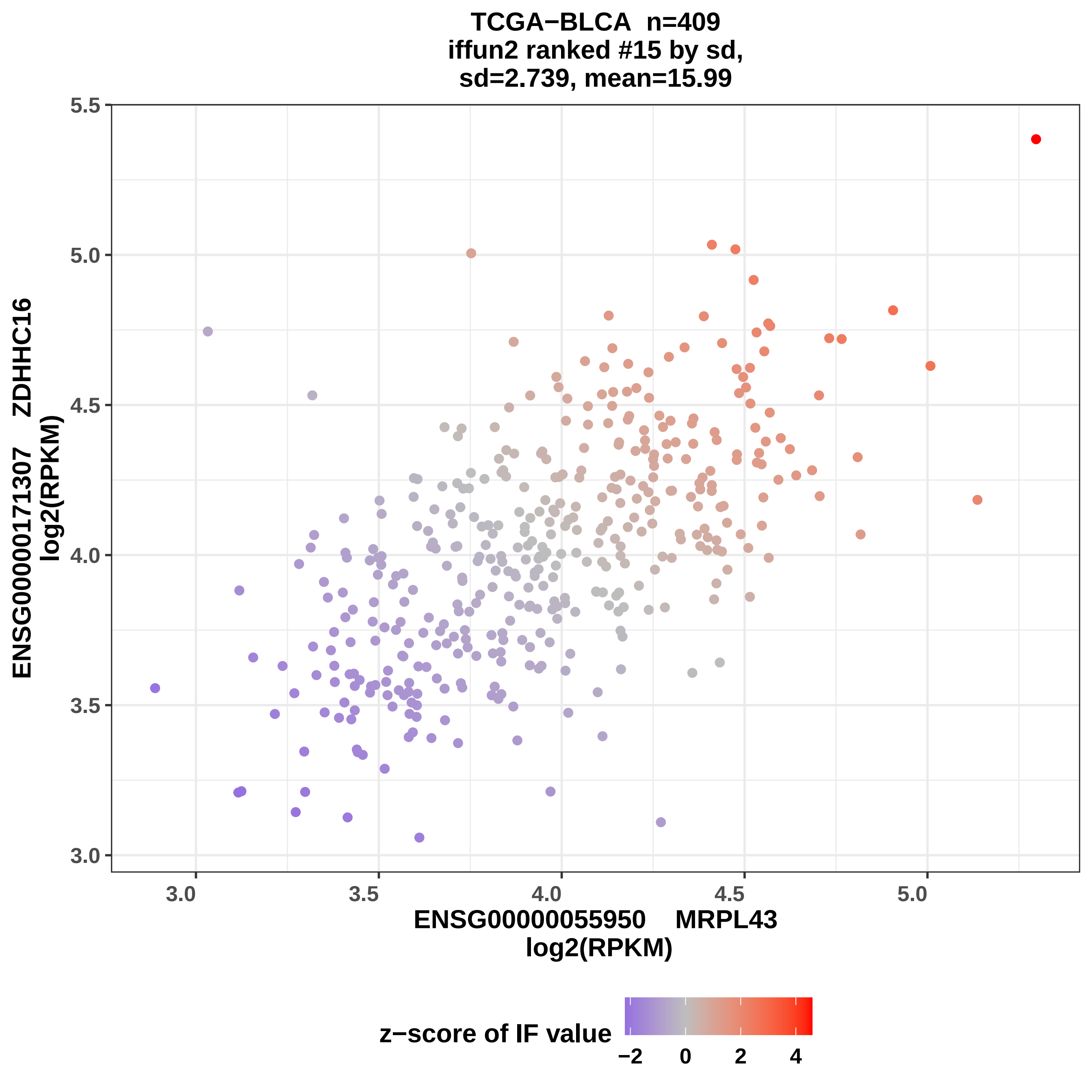
TCGA−BLCA n=409
iffun2 ranked #15 by sd,
sd=2.739, mean=15.99
ENSG00000171307 ZDHHC16
log2(RPKM)
5.5
5.0
4.5
4.0
3.5
3.0
3.0
3.5
4.0
4.5
5.0
ENSG00000055950 MRPL43
log2(RPKM)
z−score of IF value
−2
0
2
4

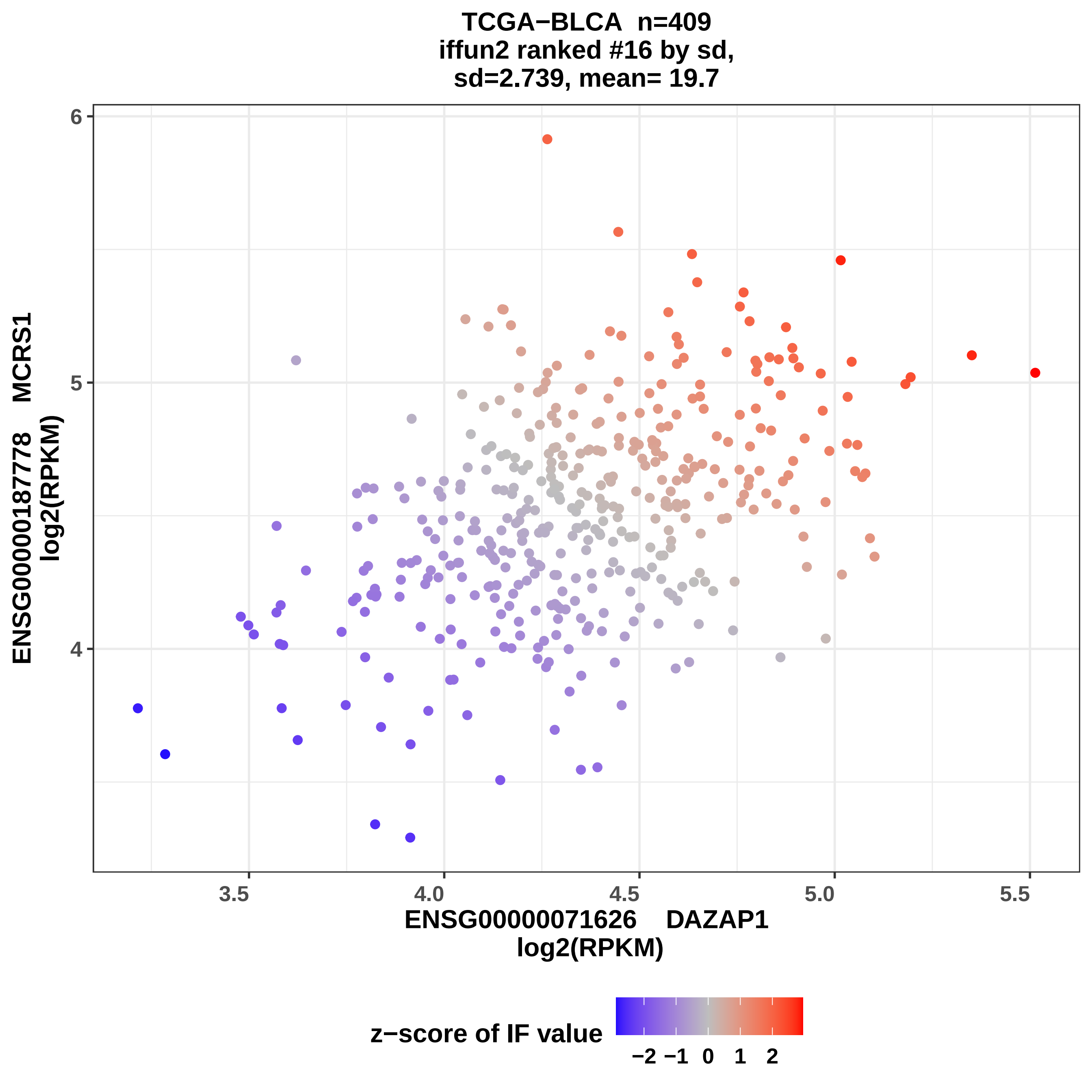
TCGA−BLCA n=409
iffun2 ranked #16 by sd,
sd=2.739, mean= 19.7
ENSG00000187778 MCRS1
log2(RPKM)
6
5
4
3.5
4.0
4.5
5.0
5.5
ENSG00000071626 DAZAP1
log2(RPKM)
z−score of IF value
−2 −1 0 1 2

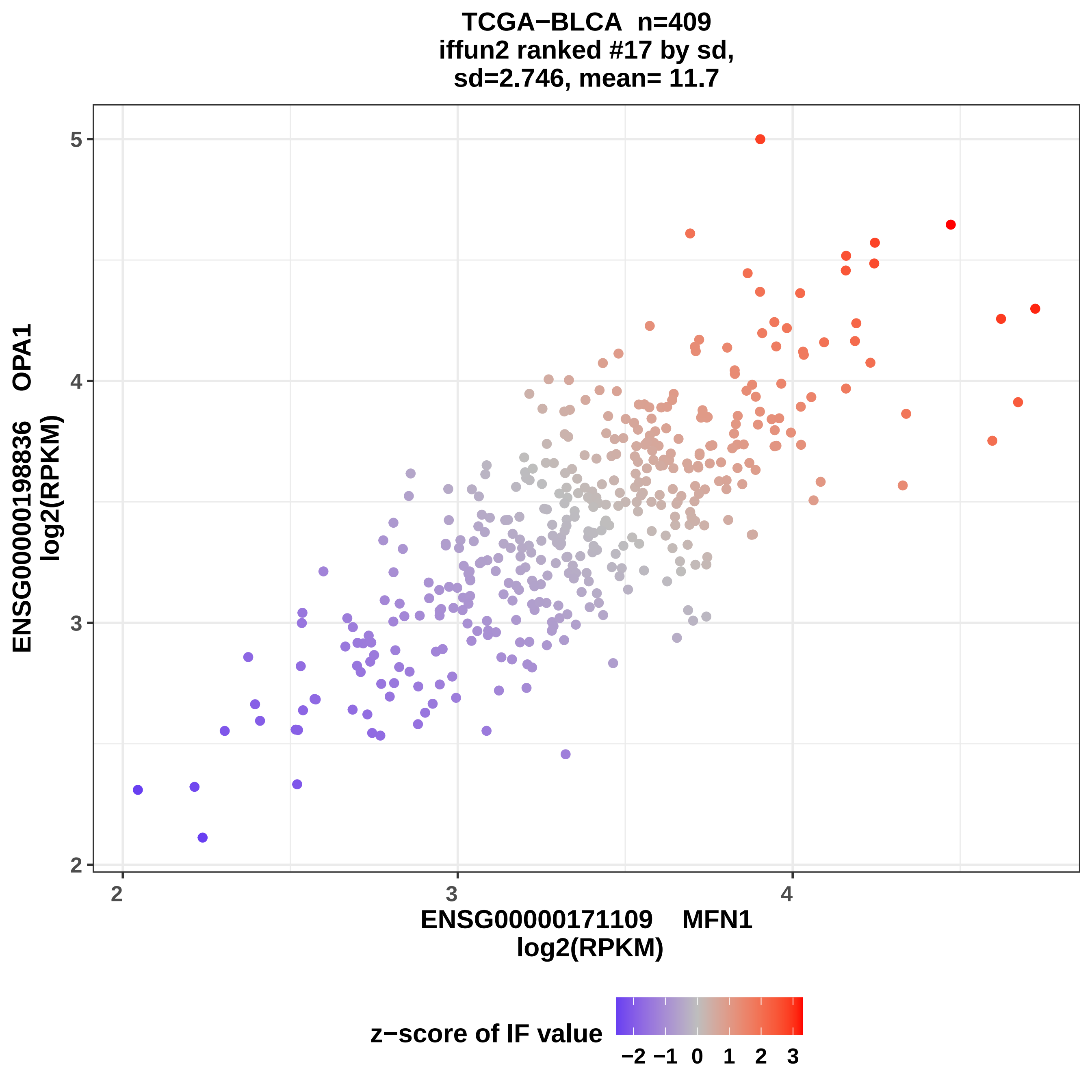
TCGA−BLCA n=409
iffun2 ranked #17 by sd,
sd=2.746, mean= 11.7
ENSG00000198836 OPA1
log2(RPKM)
5
4
3
2
2
3
4
ENSG00000171109 MFN1
log2(RPKM)
z−score of IF value
−2 −1 0 1 2 3

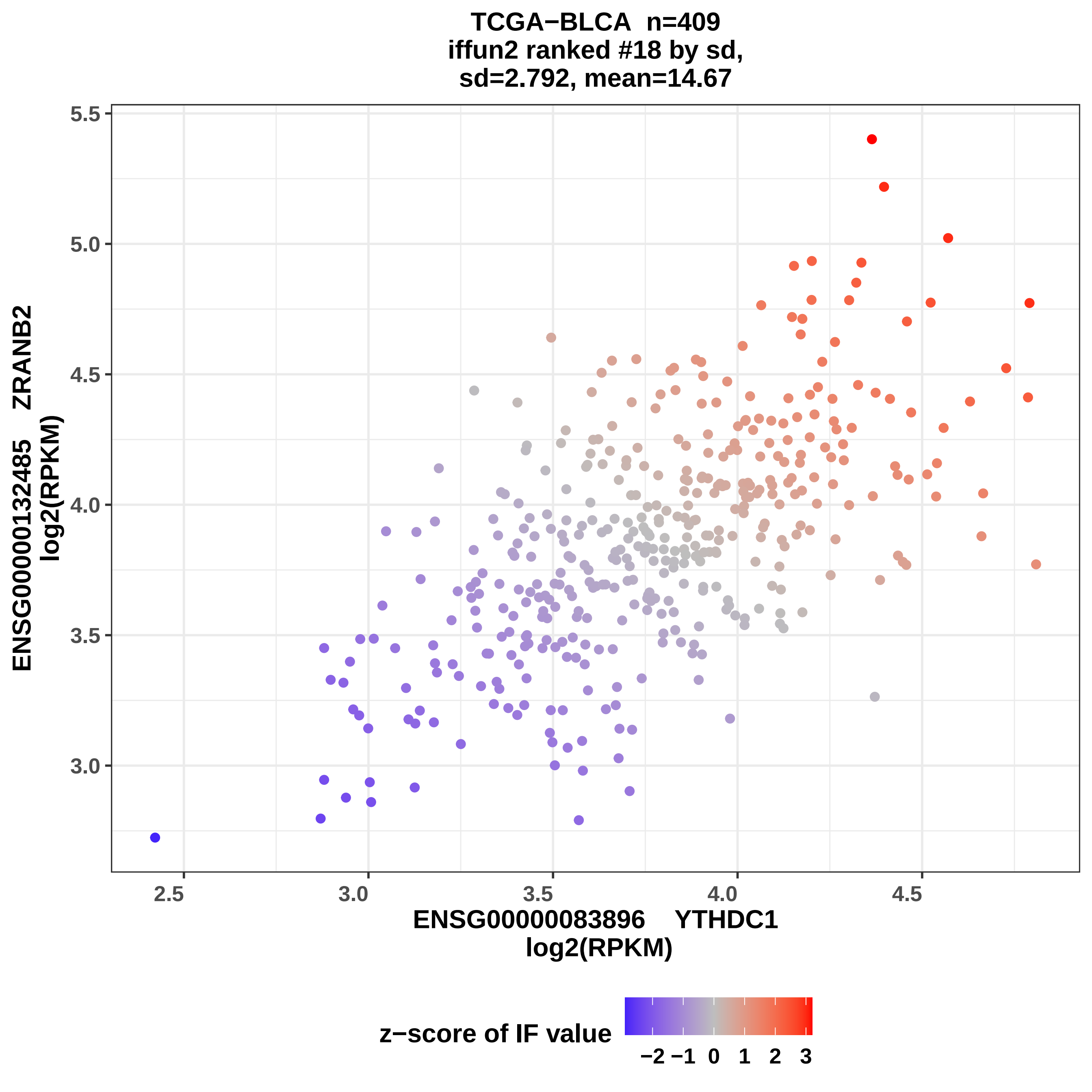

TCGA−BLCA n=409
iffun2 ranked #18 by sd,
sd=2.792, mean=14.67
ENSG00000132485 ZRANB2
log2(RPKM)
5.5
5.0
4.5
4.0
3.5
3.0
2.5
3.0
3.5
4.0
4.5
ENSG00000083896 YTHDC1
log2(RPKM)
z−score of IF value
−2 −1 0 1 2 3

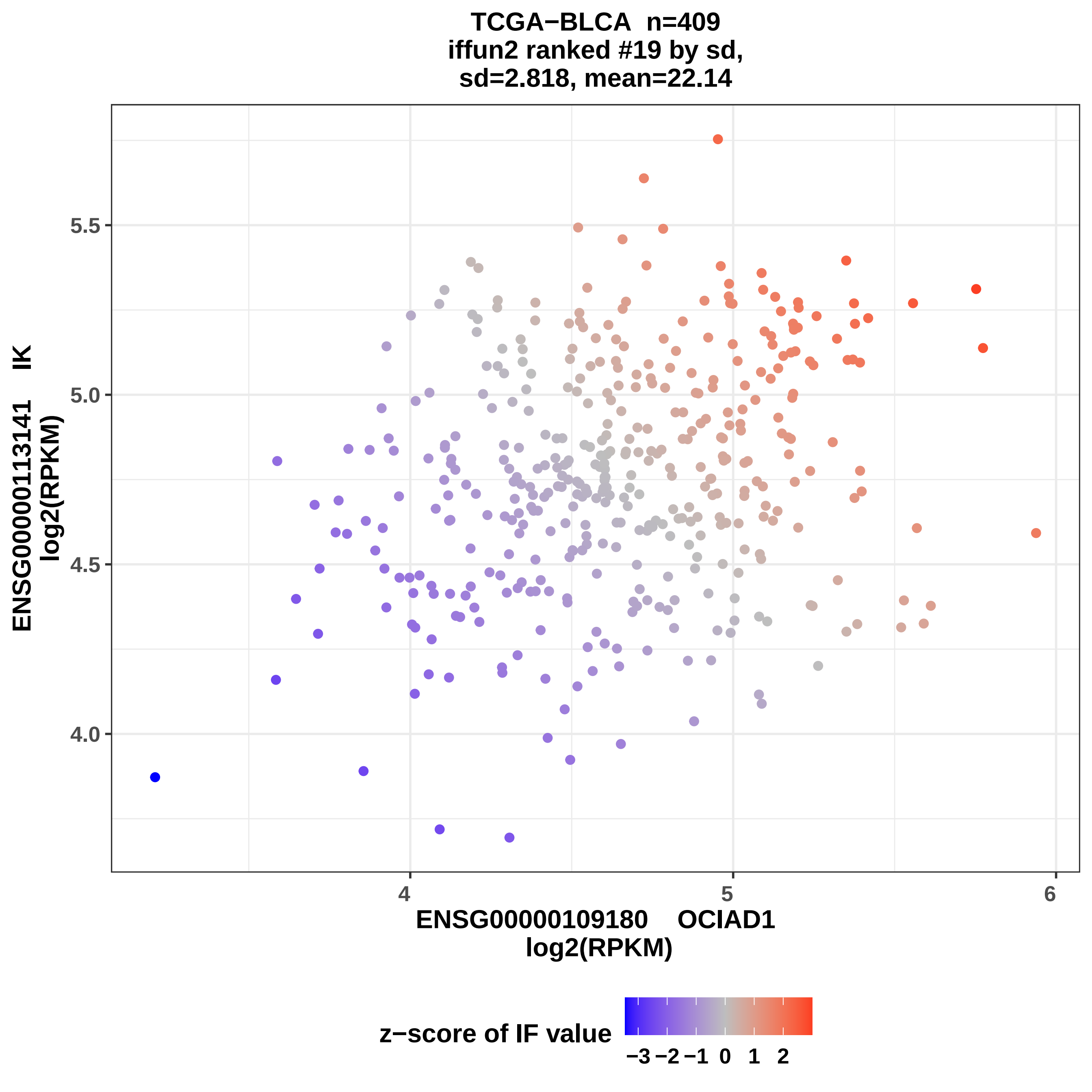
TCGA−BLCA n=409
iffun2 ranked #19 by sd,
sd=2.818, mean=22.14
ENSG00000113141 IK
log2(RPKM)
5.5
5.0
4.5
4.0
4
5
6
ENSG00000109180 OCIAD1
log2(RPKM)
z−score of IF value
−3 −2 −1 0 1 2

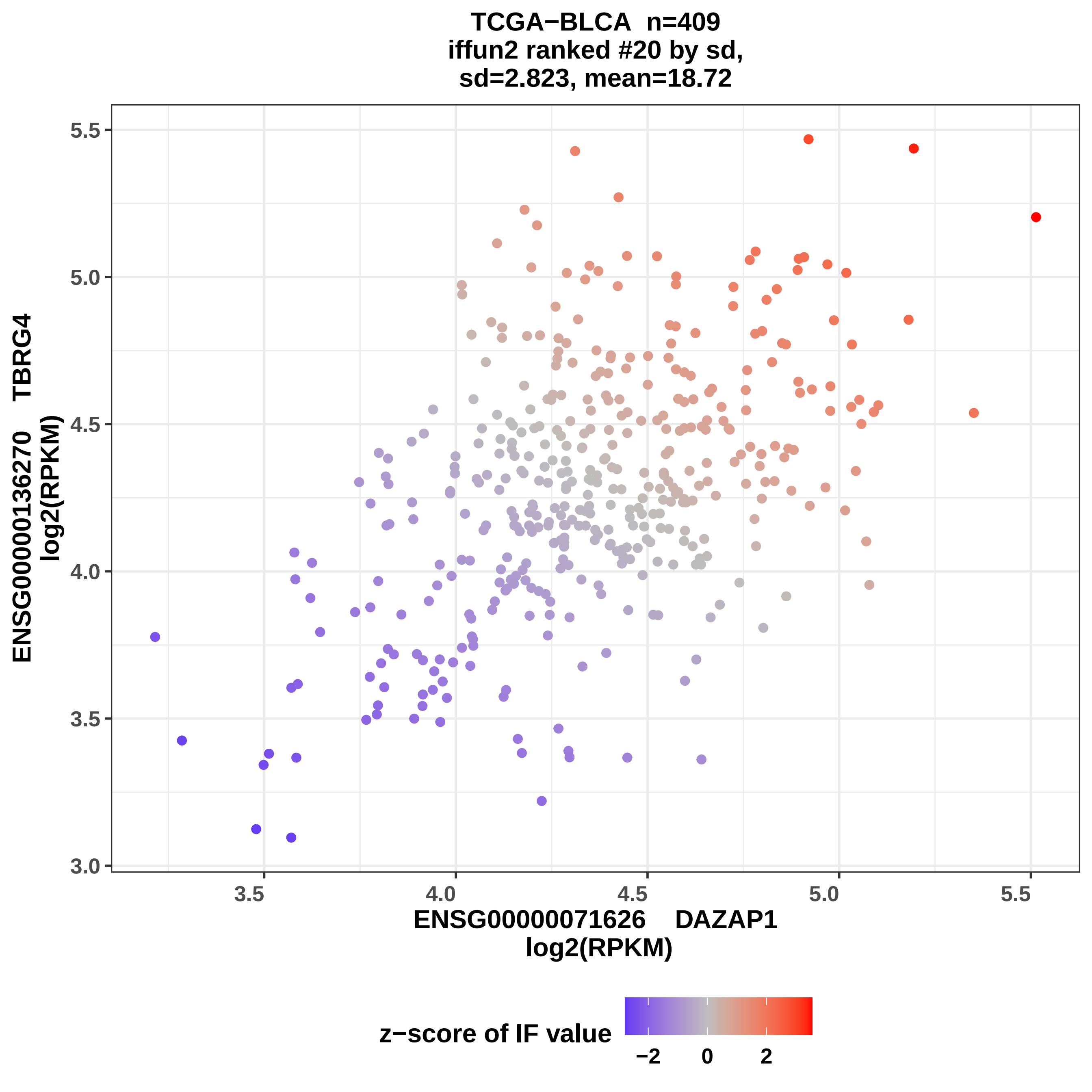

TCGA−BLCA n=409
iffun2 ranked #20 by sd,
sd=2.823, mean=18.72
ENSG00000136270 TBRG4
log2(RPKM)
ENSG00000071626 DAZAP1
log2(RPKM)
3.0
3.5
4.0
4.5
5.0
5.5
z−score of IF value
−2
0
2

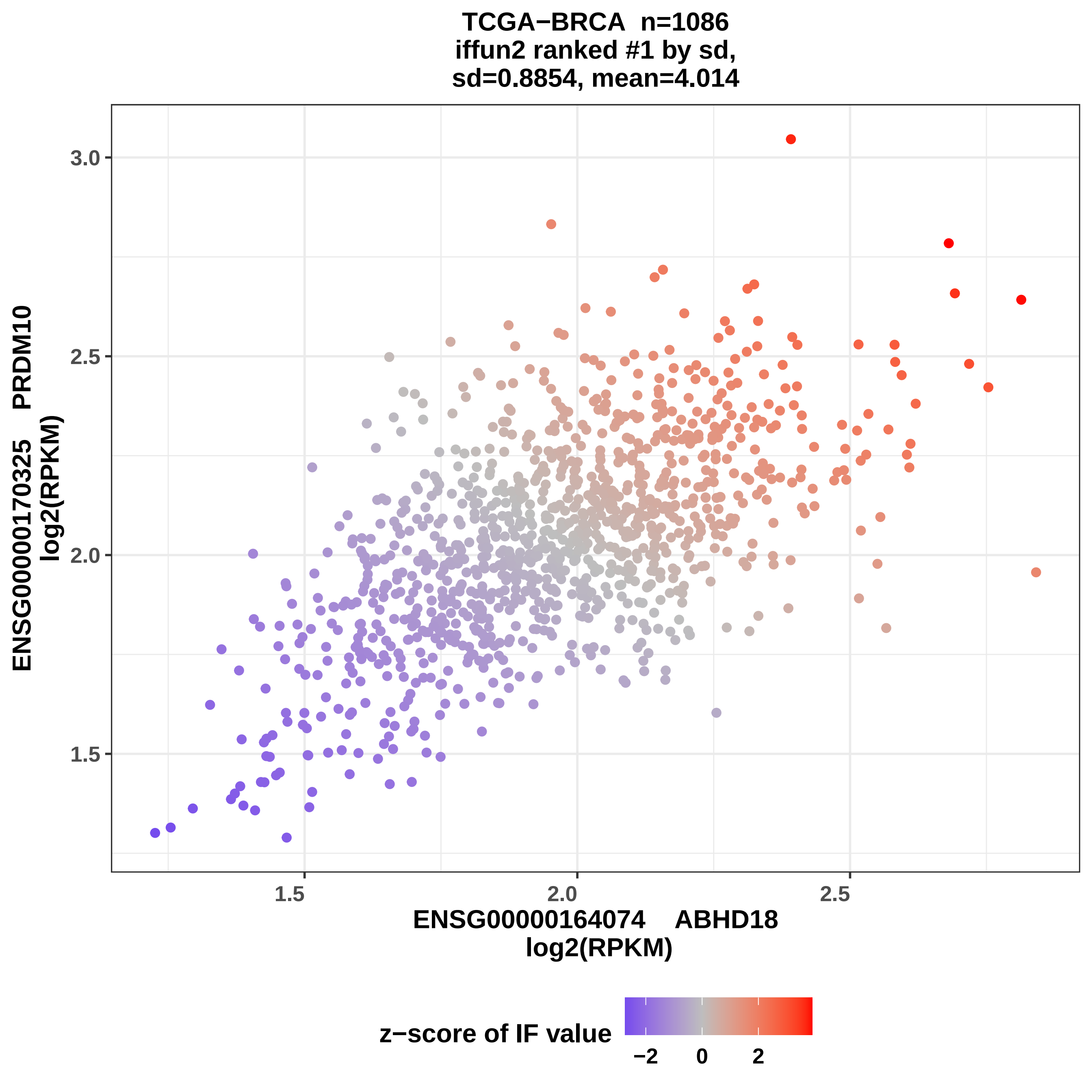
TCGA−BRCA n=1086
iffun2 ranked #1 by sd,
sd=0.8854, mean=4.014
ENSG00000170325 PRDM10
log2(RPKM)
3.0
2.5
2.0
1.5
1.5
2.0
2.5
ENSG00000164074 ABHD18
log2(RPKM)
z−score of IF value
−2
0
2

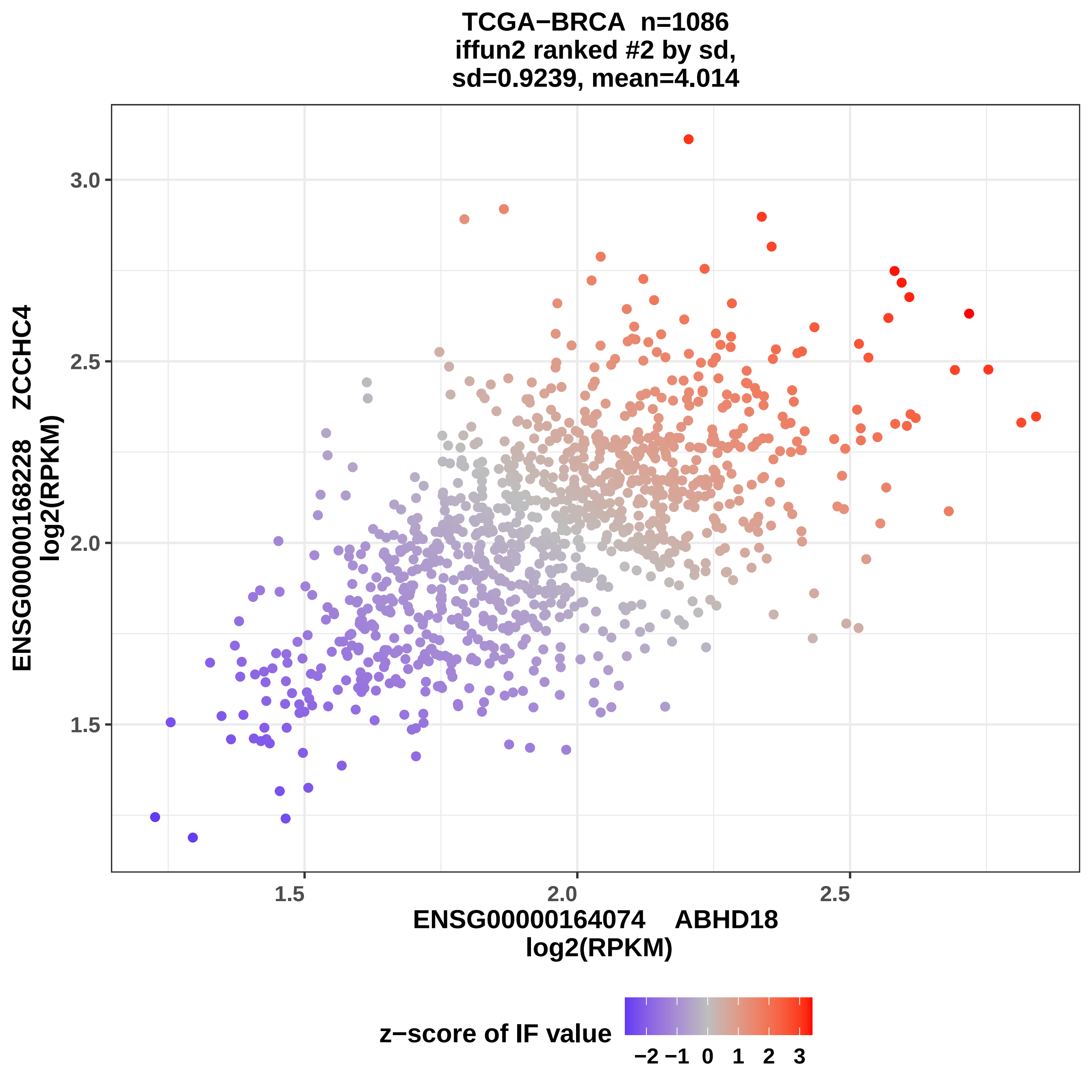
TCGA−BRCA n=1086
iffun2 ranked #2 by sd,
sd=0.9239, mean=4.014
ENSG00000168228 ZCCHC4
log2(RPKM)
3.0
2.5
2.0
1.5
1.5
2.0
2.5
ENSG00000164074 ABHD18
log2(RPKM)
z−score of IF value
−2 −1 0 1 2 3

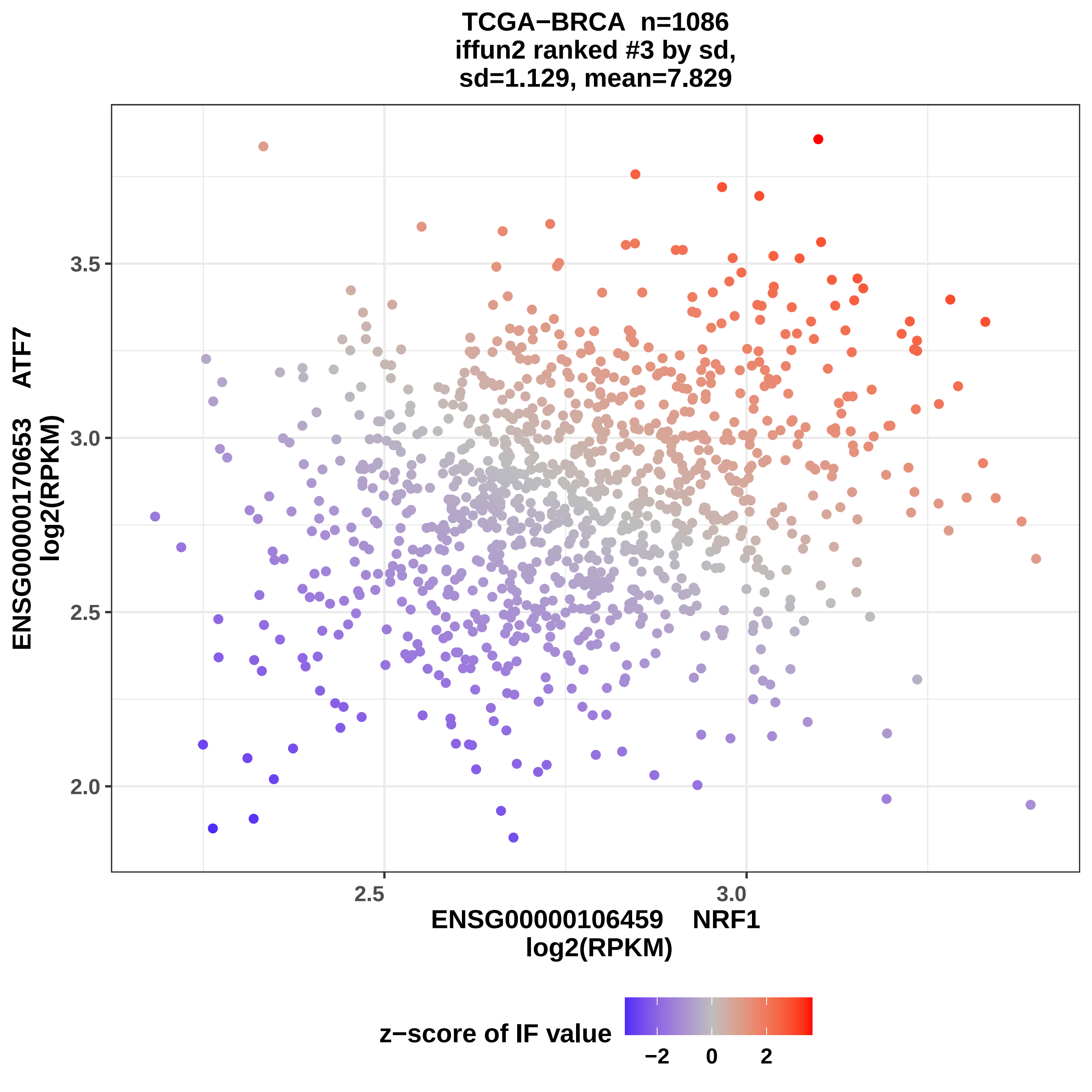
TCGA−BRCA n=1086
iffun2 ranked #3 by sd,
sd=1.129, mean=7.829
ENSG00000170653 ATF7
log2(RPKM)
3.5
3.0
2.5
2.0
2.5
3.0
ENSG00000106459 NRF1
log2(RPKM)
z−score of IF value
−2
0
2

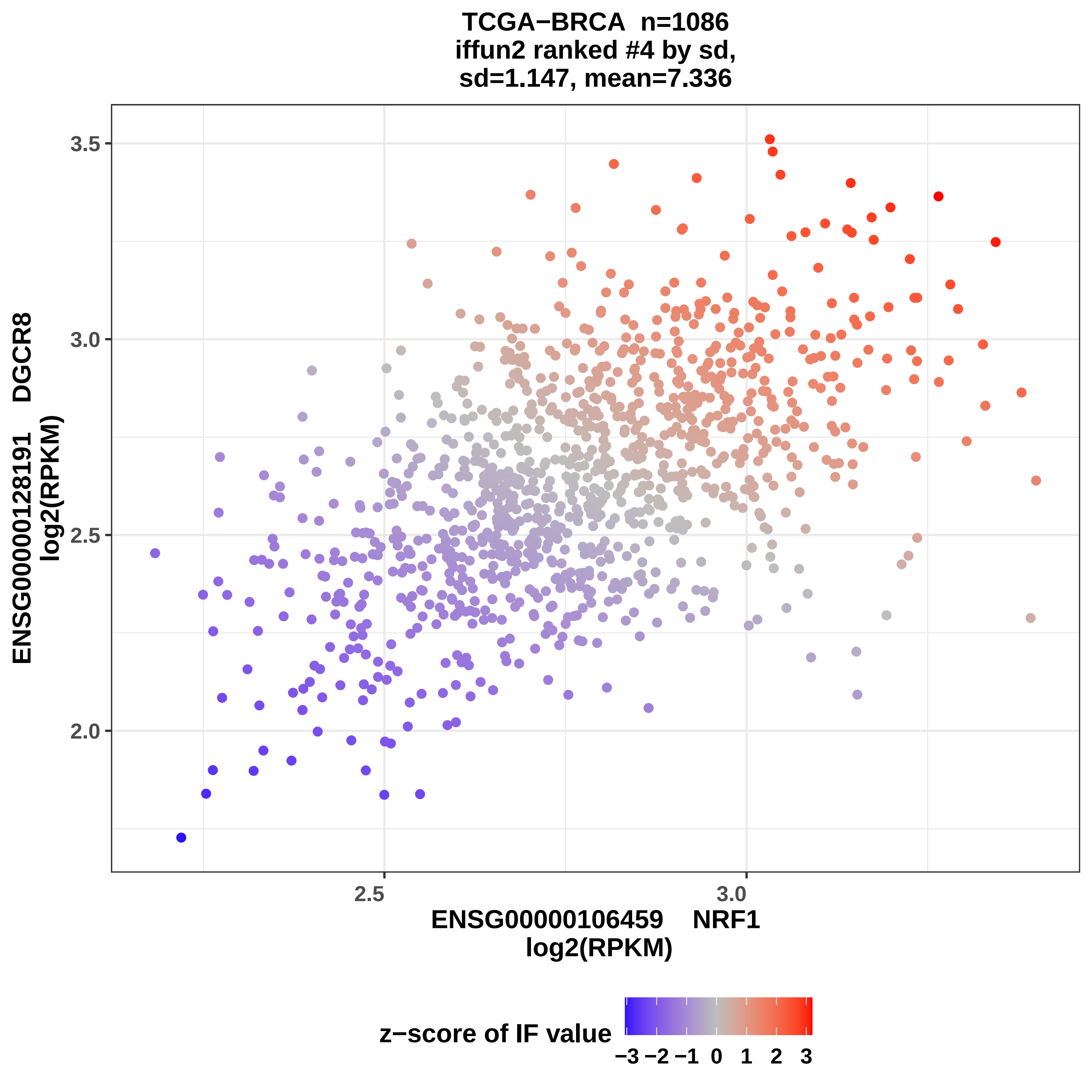
TCGA−BRCA n=1086
iffun2 ranked #4 by sd,
sd=1.147, mean=7.336
ENSG00000128191 DGCR8
log2(RPKM)
3.5
3.0
2.5
2.0
2.5
3.0
ENSG00000106459 NRF1
log2(RPKM)
z−score of IF value
−3 −2 −1 0 1 2 3

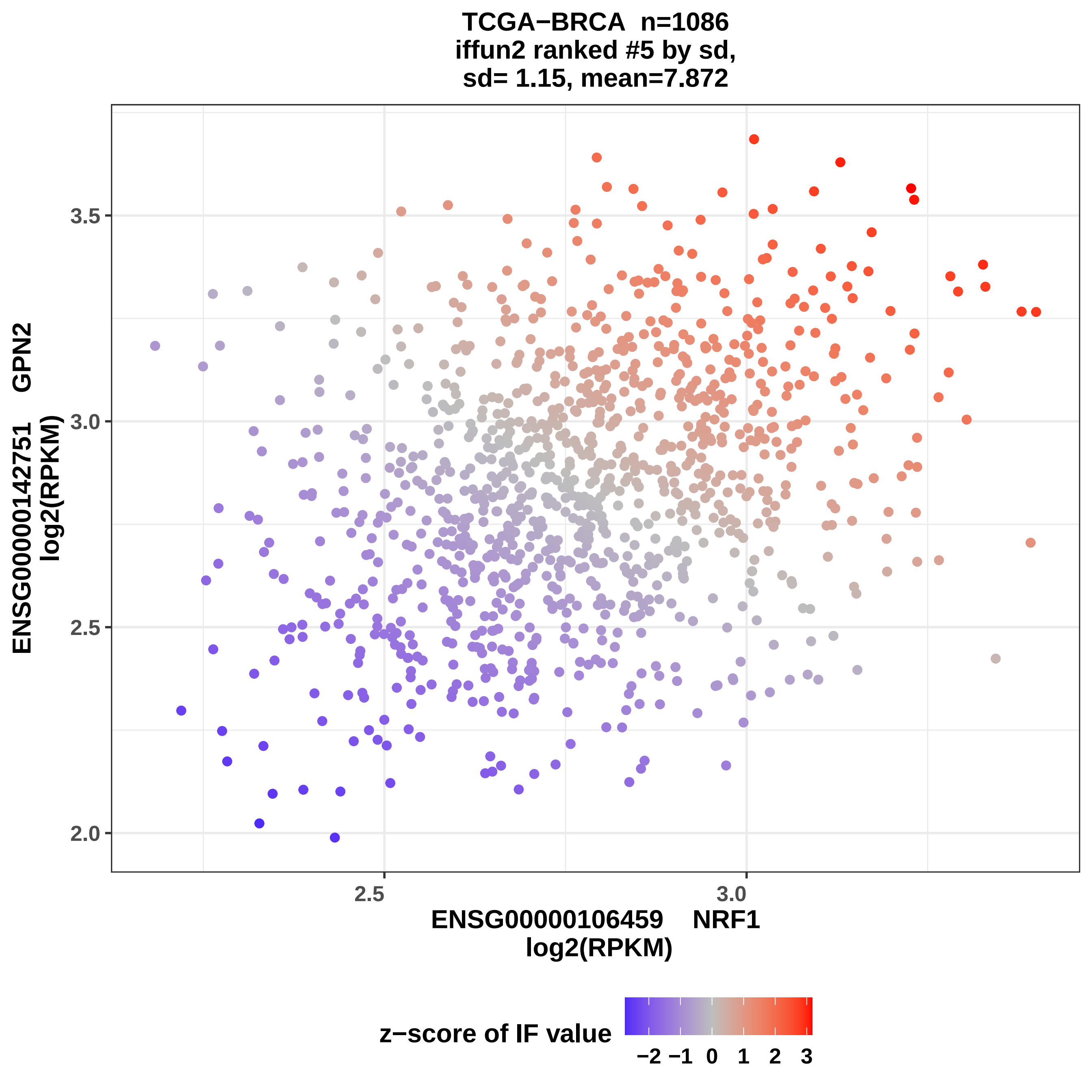

TCGA−BRCA n=1086
iffun2 ranked #5 by sd,
sd= 1.15, mean=7.872
ENSG00000142751 GPN2
log2(RPKM)
3.5
3.0
2.5
2.0
2.5
3.0
ENSG00000106459 NRF1
log2(RPKM)
z−score of IF value
−2 −1 0 1 2 3

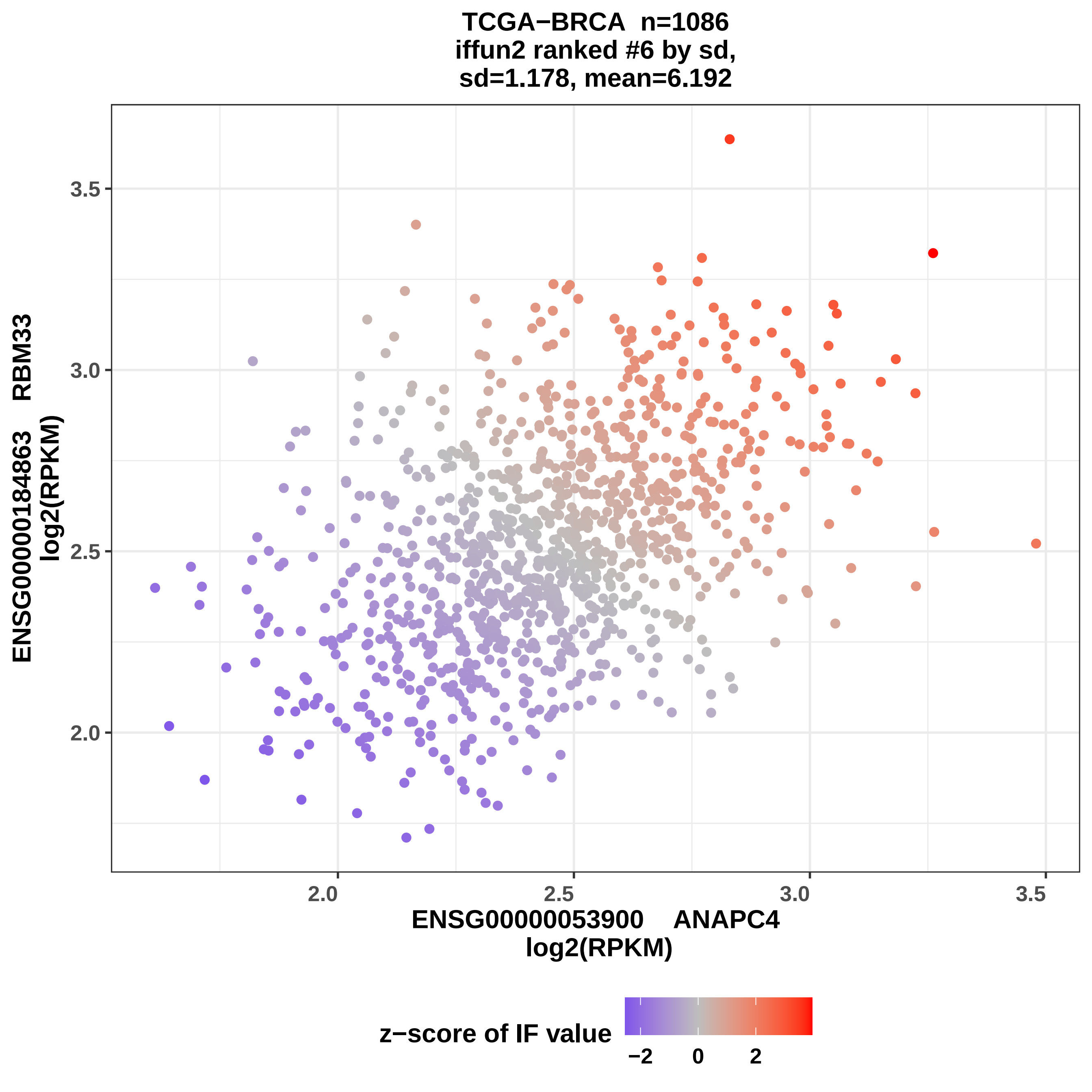

TCGA−BRCA n=1086
iffun2 ranked #6 by sd,
sd=1.178, mean=6.192
3.5
3.0
2.5
2.0
ENSG00000184863 RBM33
log2(RPKM)
2.0
2.5
3.0
3.5
ENSG00000053900 ANAPC4
log2(RPKM)
z−score of IF value
−2
0
2

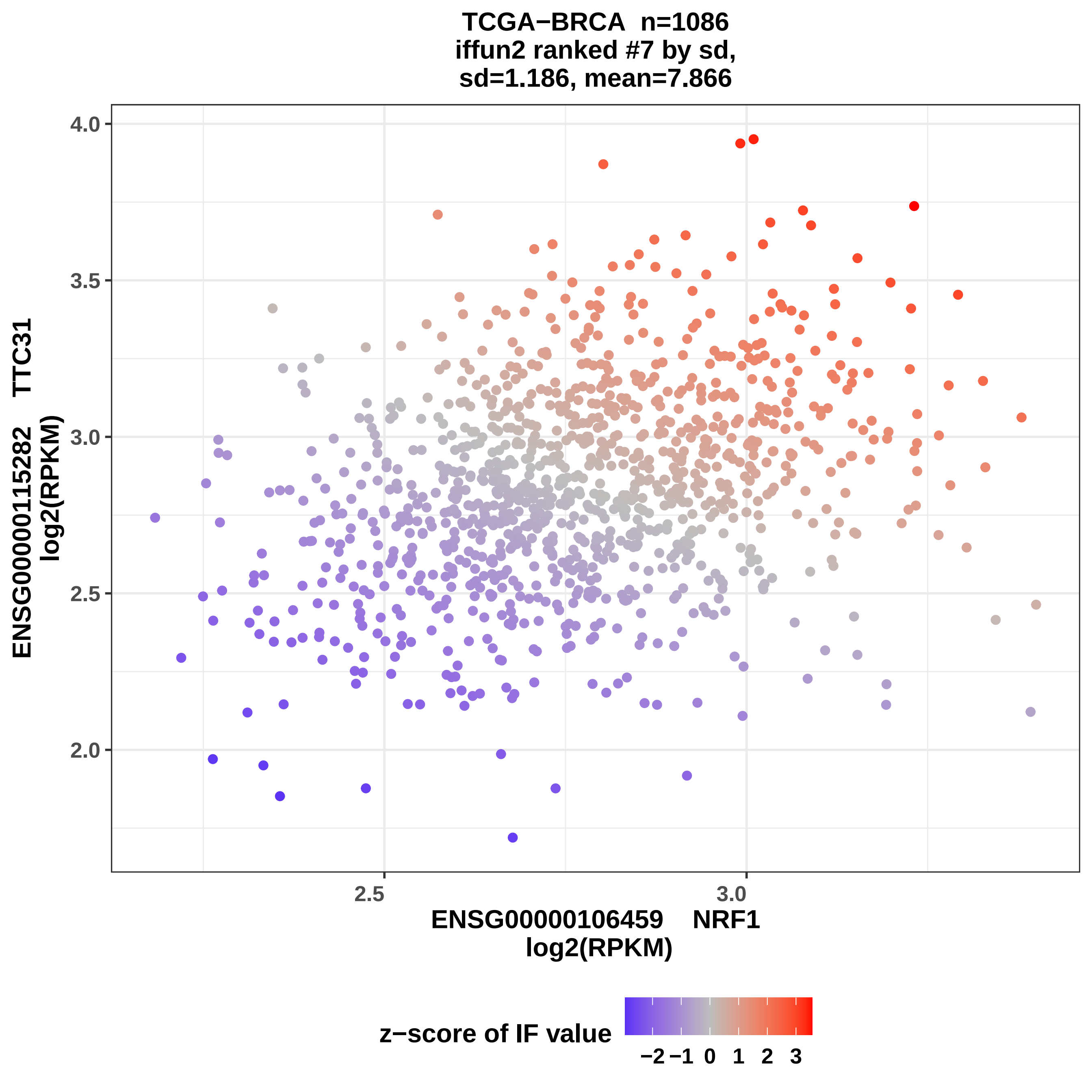
TCGA−BRCA n=1086
iffun2 ranked #7 by sd,
sd=1.186, mean=7.866
4.0
3.5
3.0
2.5
2.0
2.5
3.0
ENSG00000115282 TTC31
log2(RPKM)
ENSG00000106459 NRF1
log2(RPKM)
z−score of IF value
−2 −1 0 1 2 3

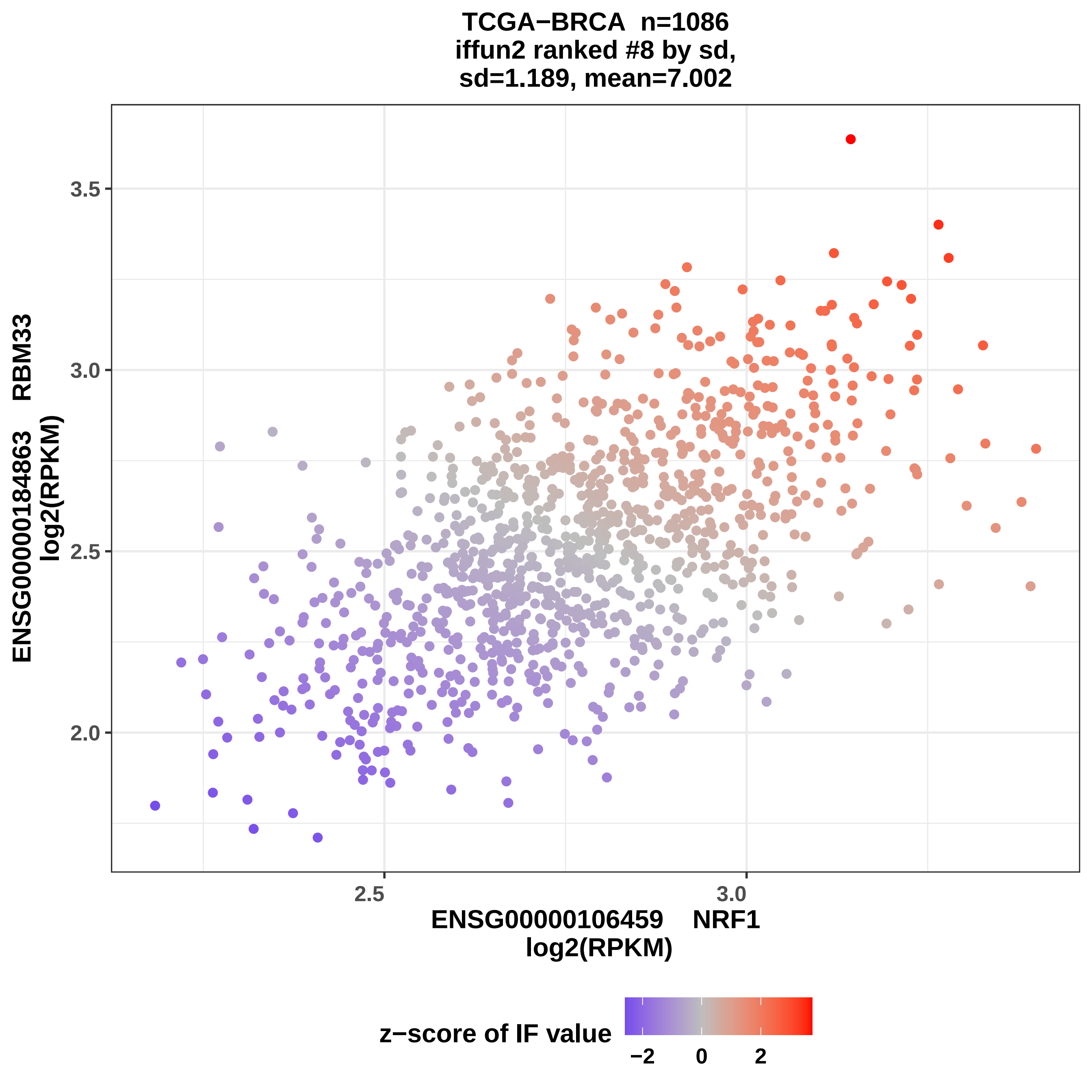
TCGA−BRCA n=1086
iffun2 ranked #8 by sd,
sd=1.189, mean=7.002
3.5
3.0
2.5
2.0
2.5
3.0
ENSG00000106459 NRF1
log2(RPKM)
ENSG00000184863 RBM33
log2(RPKM)
z−score of IF value
−2
0
2

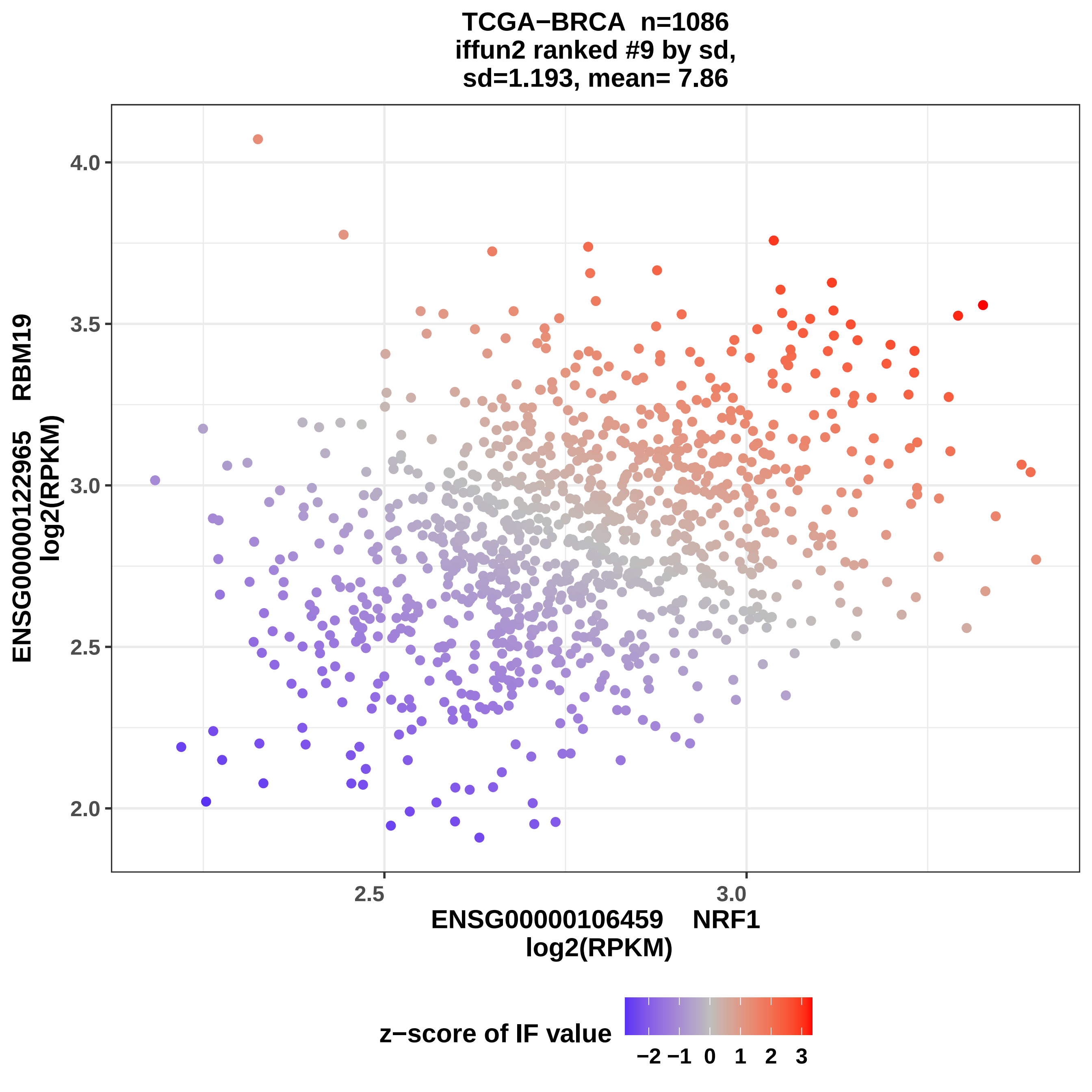
TCGA−BRCA n=1086
iffun2 ranked #9 by sd,
sd=1.193, mean= 7.86
ENSG00000122965 RBM19
log2(RPKM)
4.0
3.5
3.0
2.5
2.0
2.5
3.0
ENSG00000106459 NRF1
log2(RPKM)
z−score of IF value
−2 −1 0 1 2 3

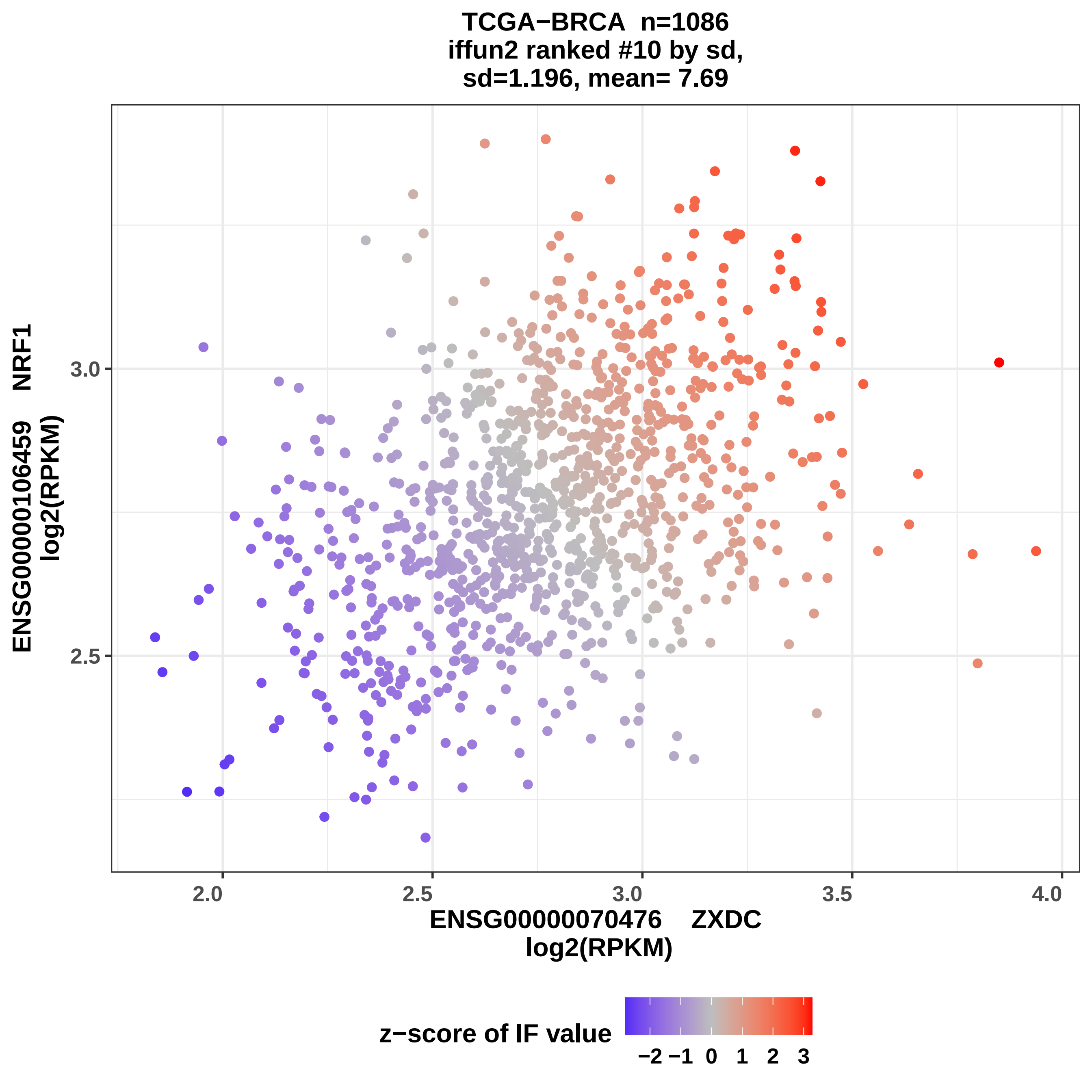
TCGA−BRCA n=1086
iffun2 ranked #10 by sd,
sd=1.196, mean= 7.69
ENSG00000106459 NRF1
log2(RPKM)
3.0
2.5
2.0
2.5
3.0
3.5
4.0
ENSG00000070476 ZXDC
log2(RPKM)
z−score of IF value
−2
−1
0
1
2
3

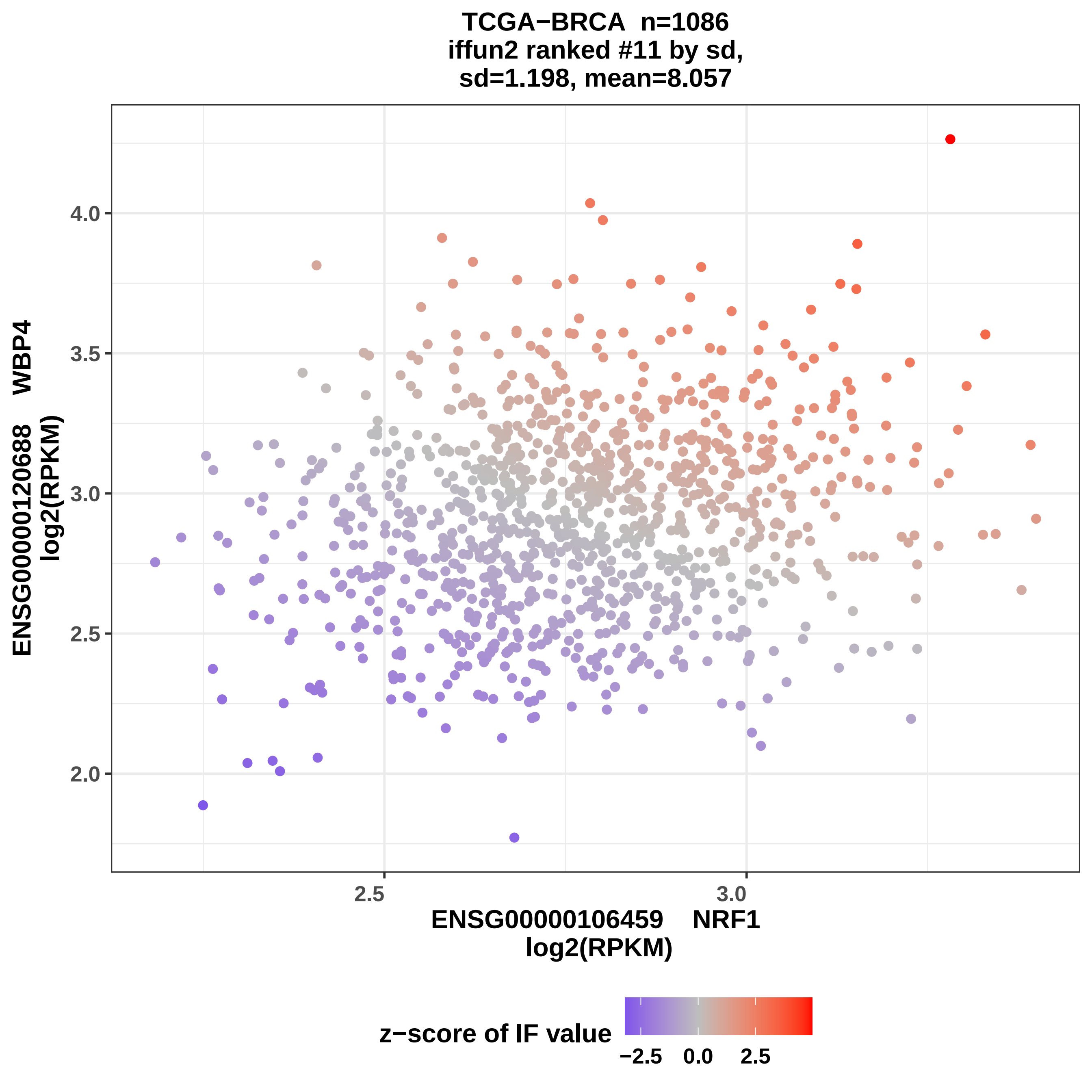
TCGA−BRCA n=1086
iffun2 ranked #11 by sd,
sd=1.198, mean=8.057
ENSG00000120688 WBP4
log2(RPKM)
4.0
3.5
3.0
2.5
2.0
2.5
3.0
ENSG00000106459 NRF1
log2(RPKM)
z−score of IF value
−2.5
0.0
2.5

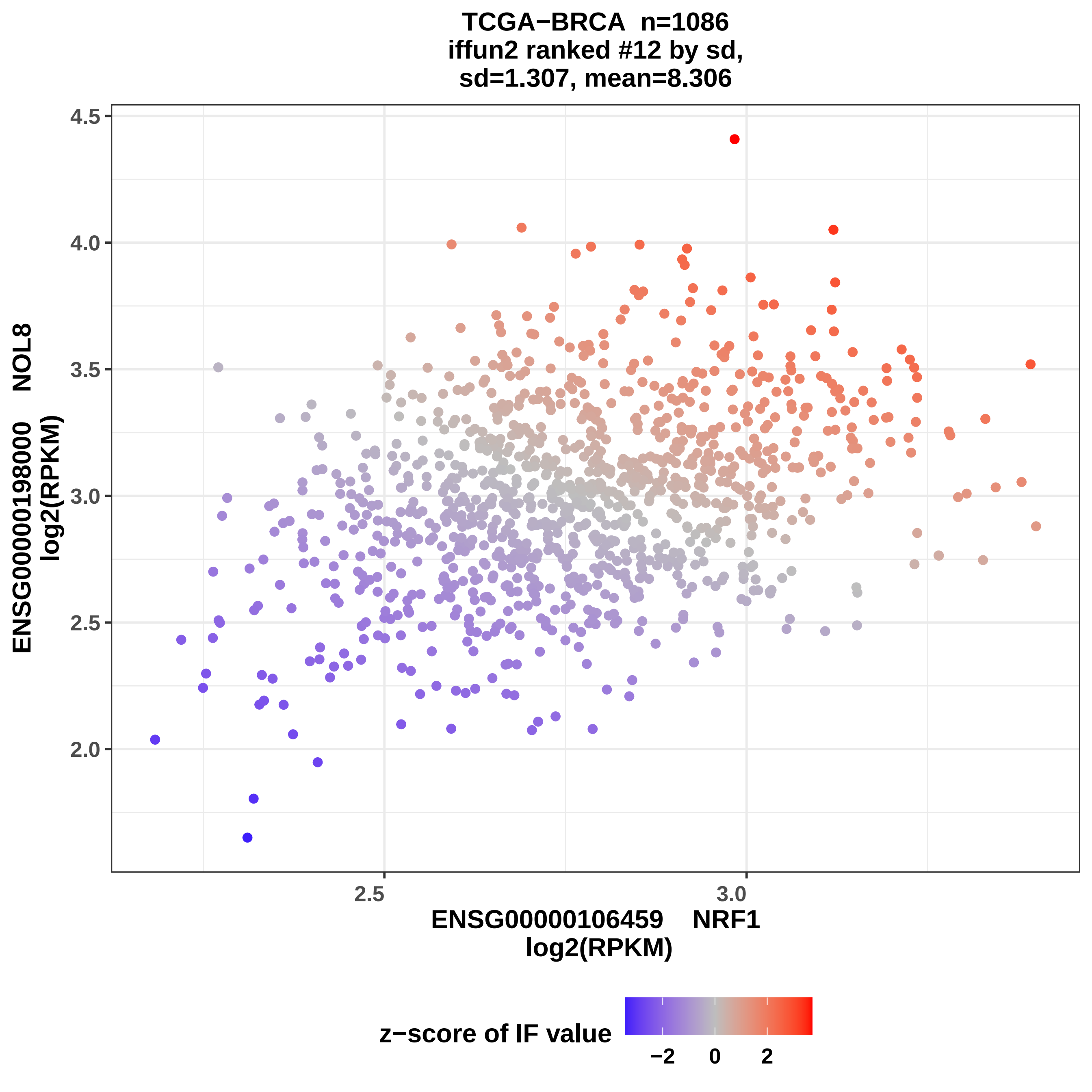

TCGA−BRCA n=1086
iffun2 ranked #12 by sd,
sd=1.307, mean=8.306
ENSG00000198000 NOL8
log2(RPKM)
4.5
4.0
3.5
3.0
2.5
2.0
2.5
3.0
ENSG00000106459 NRF1
log2(RPKM)
z−score of IF value
−2
0
2

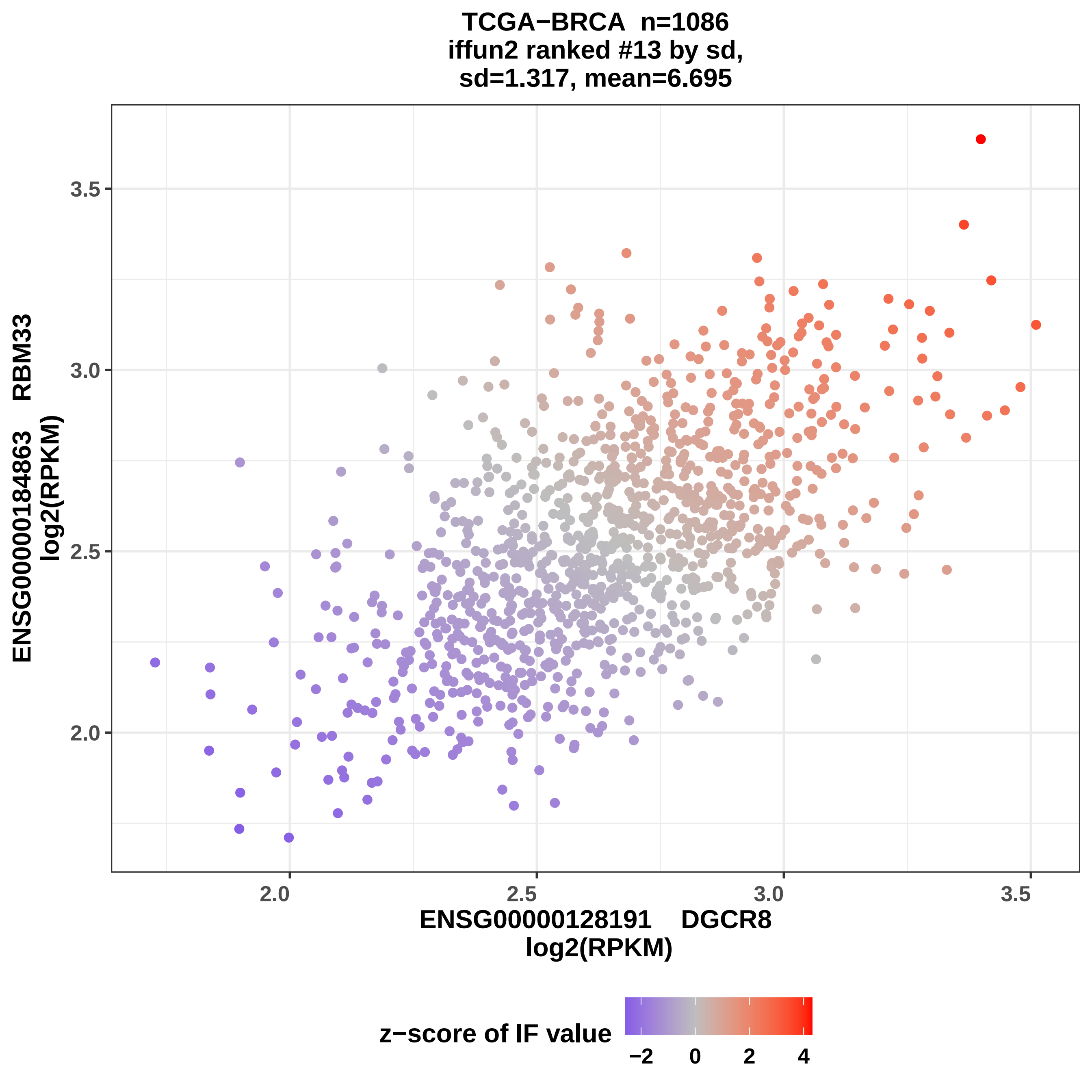

TCGA−BRCA n=1086
iffun2 ranked #13 by sd,
sd=1.317, mean=6.695
ENSG00000184863 RBM33
log2(RPKM)
3.5
3.0
2.5
2.0
ENSG00000128191 DGCR8
log2(RPKM)
z−score of IF value
−2
0
2
4

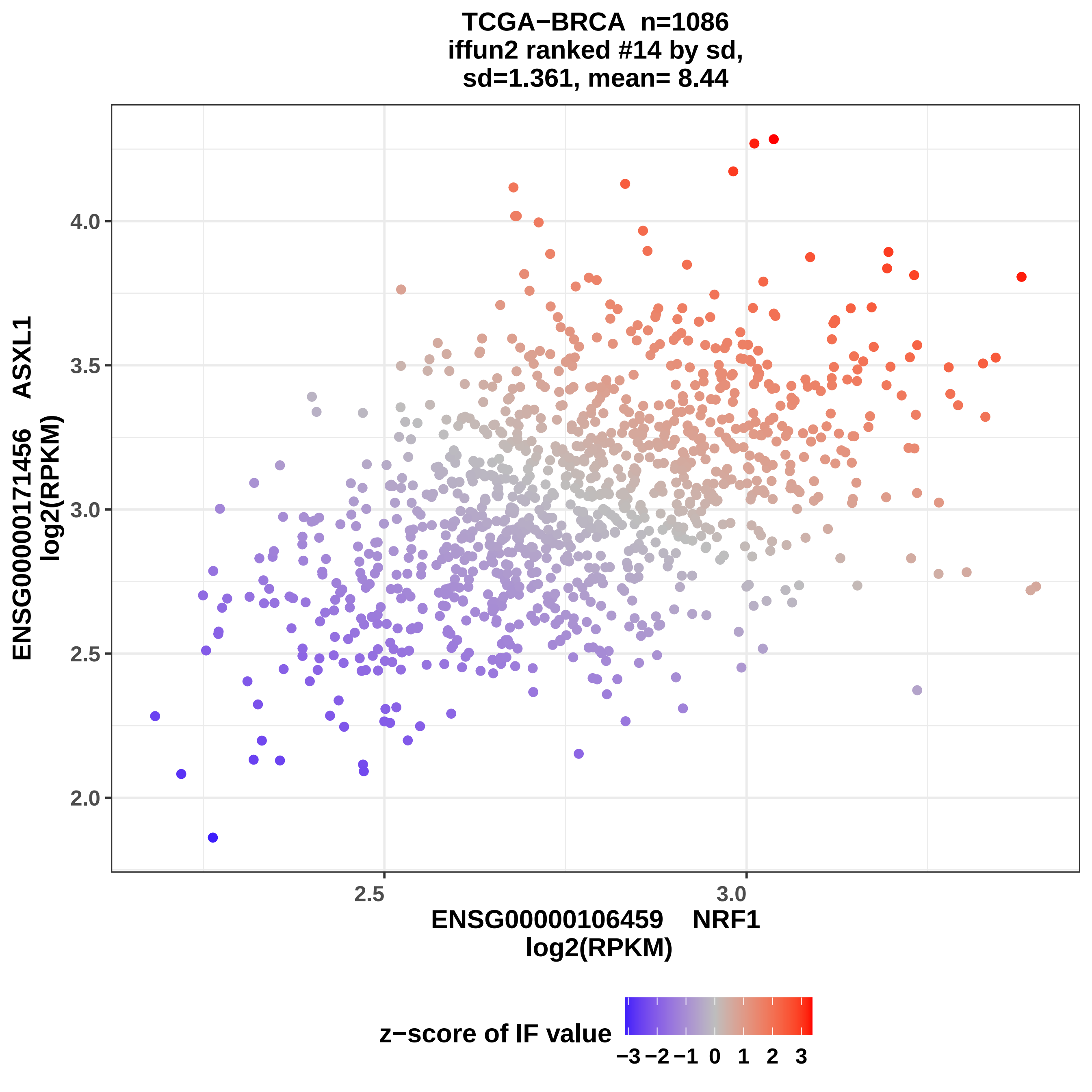
TCGA−BRCA n=1086
iffun2 ranked #14 by sd,
sd=1.361, mean= 8.44
ENSG00000171456 ASXL1
log2(RPKM)
4.0
3.5
3.0
2.5
2.0
2.5
3.0
ENSG00000106459 NRF1
log2(RPKM)
z−score of IF value
−3 −2 −1 0 1 2 3

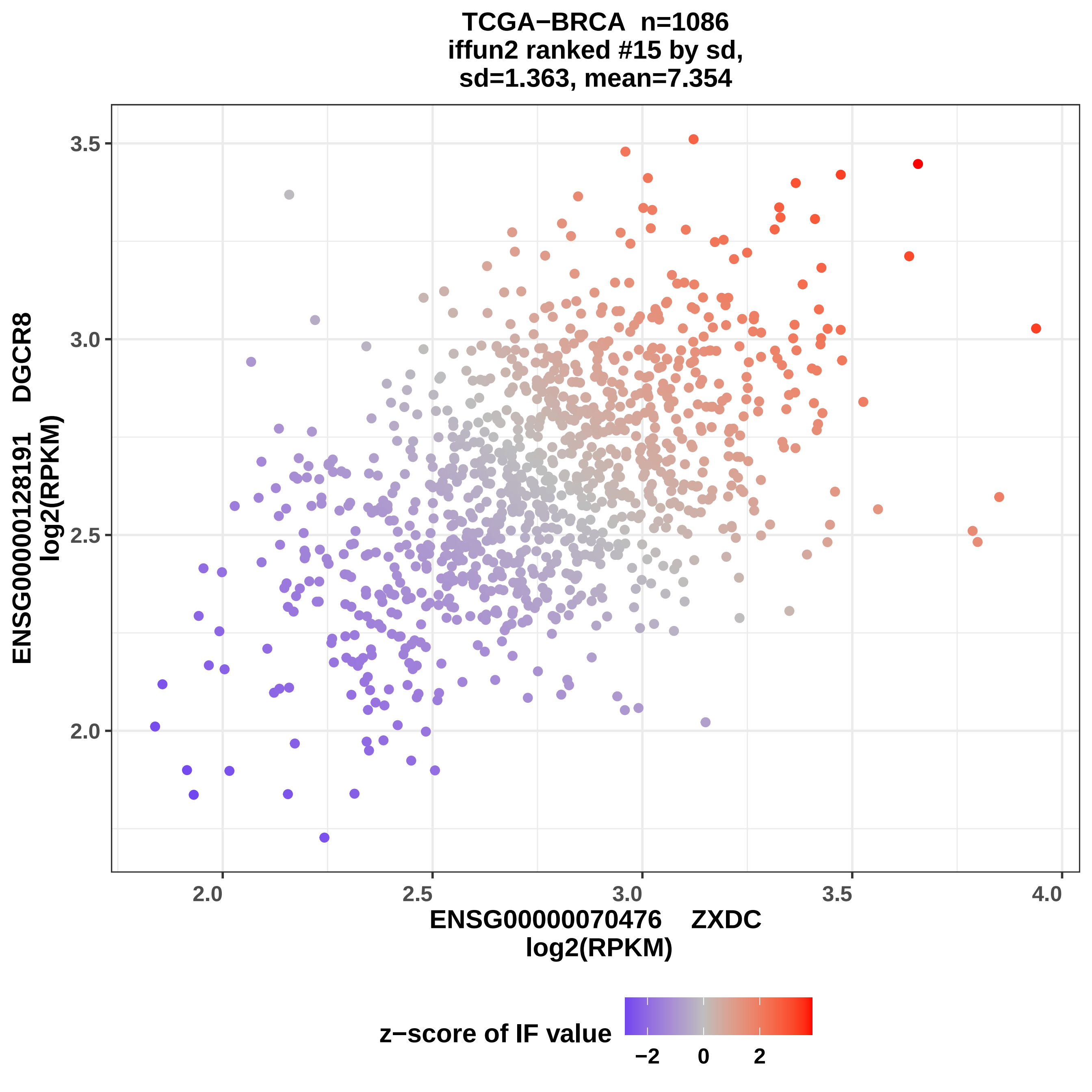
TCGA−BRCA n=1086
iffun2 ranked #15 by sd,
sd=1.363, mean=7.354
ENSG00000128191 DGCR8
log2(RPKM)
3.5
3.0
2.5
2.0
2.0
2.5
3.0
3.5
4.0
ENSG00000070476 ZXDC
log2(RPKM)
z−score of IF value
−2
0
2

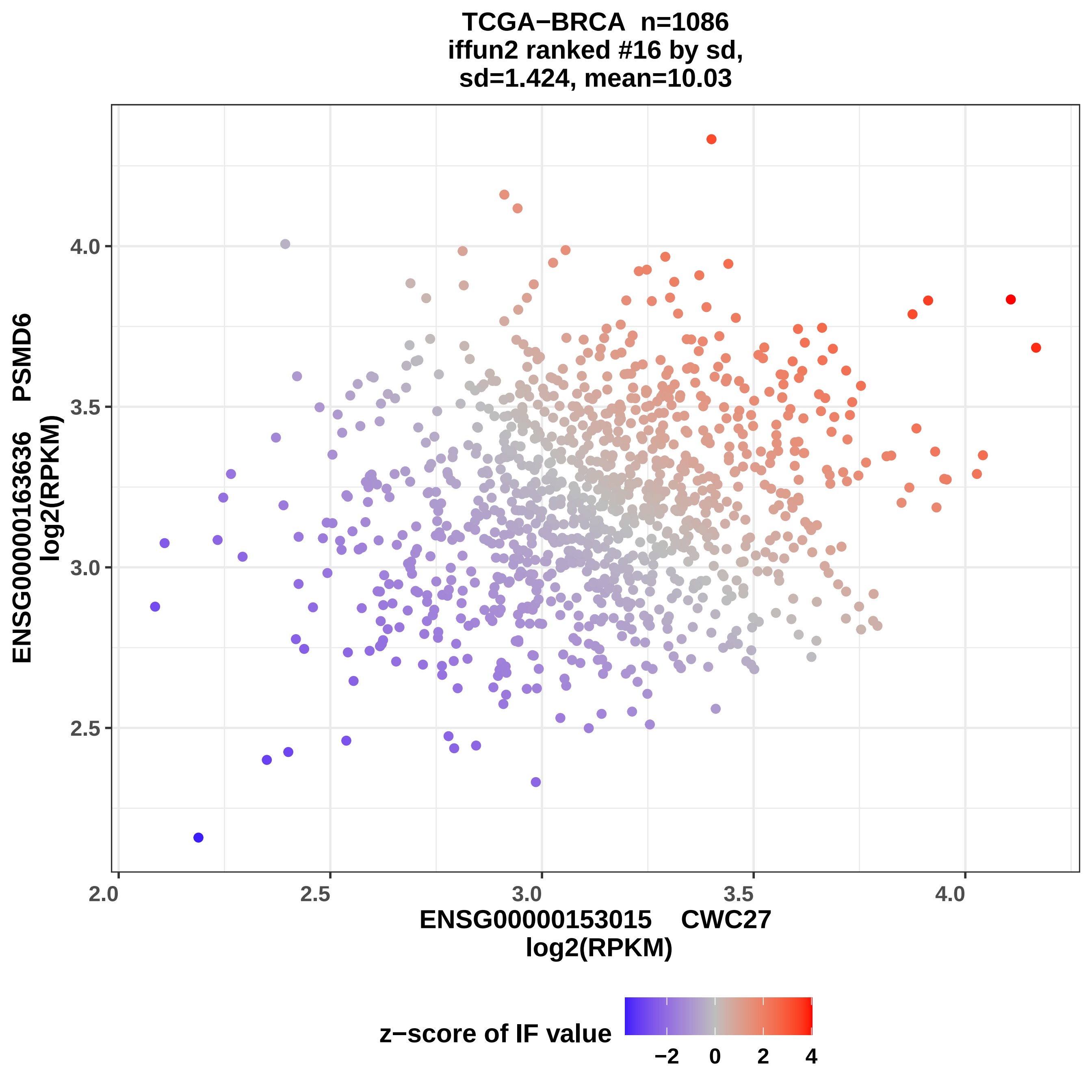

TCGA−BRCA n=1086
iffun2 ranked #16 by sd,
sd=1.424, mean=10.03
ENSG00000163636 PSMD6
log2(RPKM)
4.0
3.5
3.0
2.5
2.0
2.5
3.0
3.5
4.0
ENSG00000153015 CWC27
log2(RPKM)
z−score of IF value
−2
0
2
4

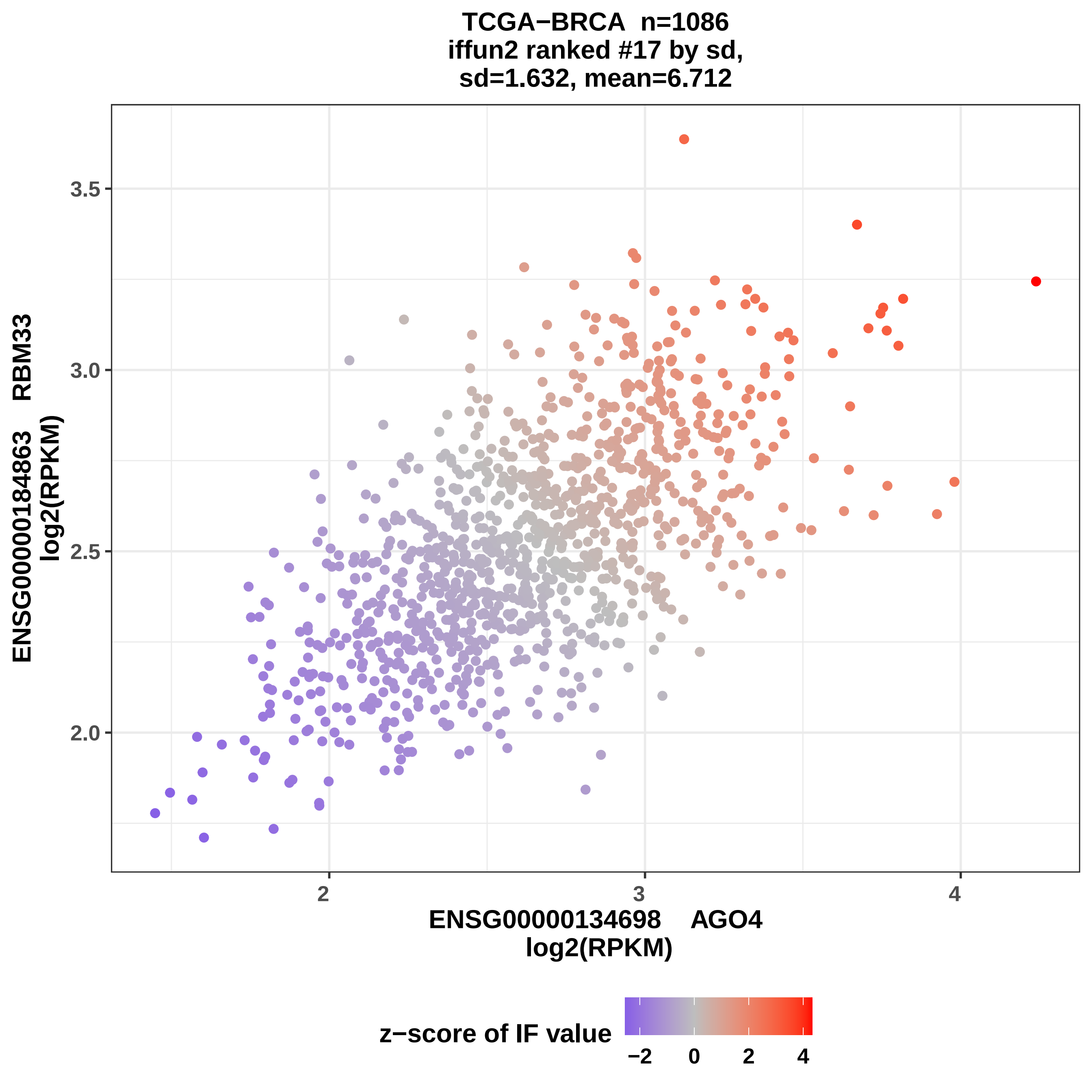

TCGA−BRCA n=1086
iffun2 ranked #17 by sd,
sd=1.632, mean=6.712
ENSG00000184863 RBM33
log2(RPKM)
3.5
3.0
2.5
2.0
2
3
4
ENSG00000134698 AGO4
log2(RPKM)
z−score of IF value
−2
0
2
4

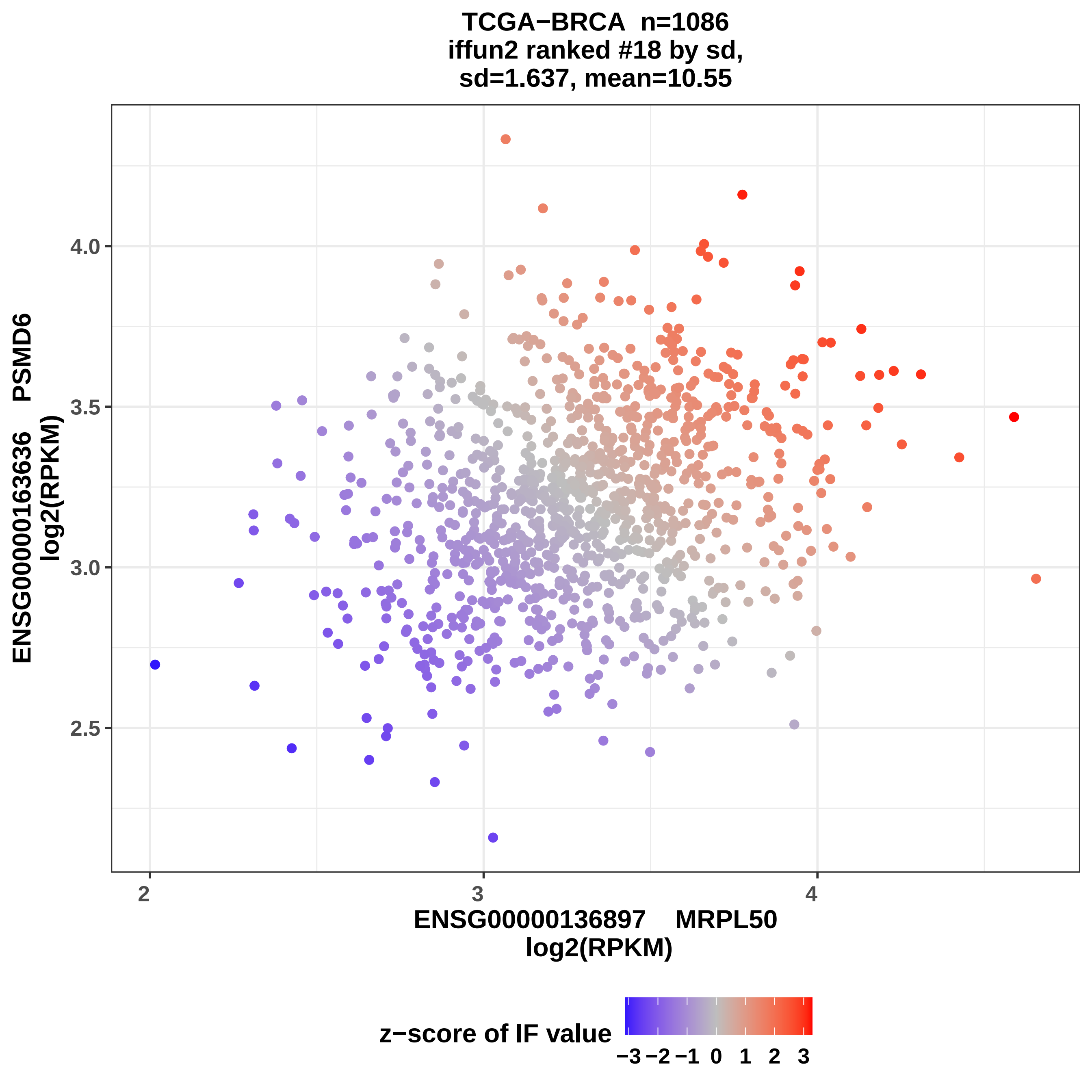
TCGA−BRCA n=1086
iffun2 ranked #18 by sd,
sd=1.637, mean=10.55
ENSG00000163636 PSMD6
log2(RPKM)
4.0
3.5
3.0
2.5
2
3
4
ENSG00000136897 MRPL50
log2(RPKM)
z−score of IF value
−3 −2 −1 0 1 2 3

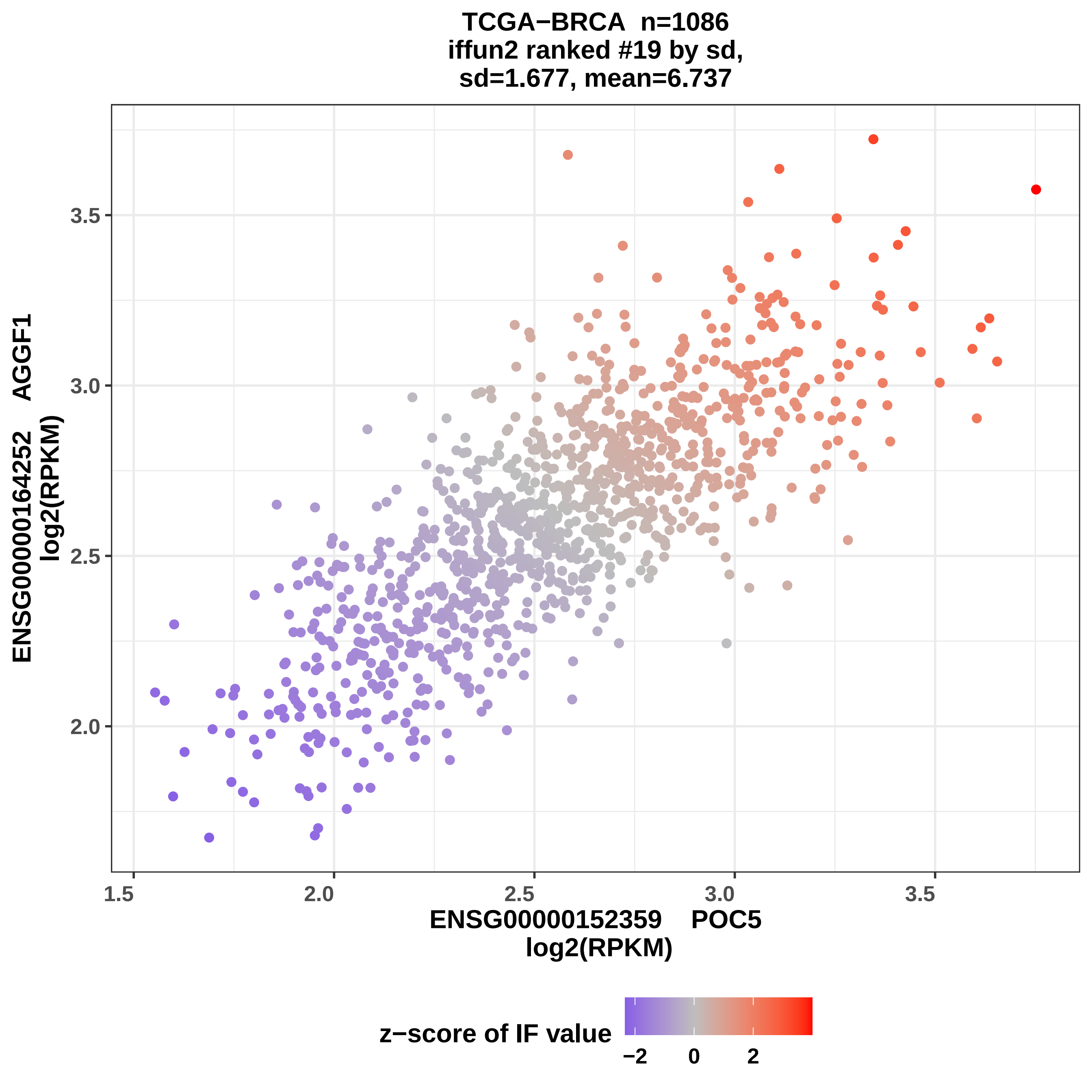

TCGA−BRCA n=1086
iffun2 ranked #19 by sd,
sd=1.677, mean=6.737
ENSG00000164252 AGGF1
log2(RPKM)
3.5
3.0
2.5
2.0
1.5
2.0
2.5
3.0
3.5
ENSG00000152359 POC5
log2(RPKM)
z−score of IF value
−2
0
2

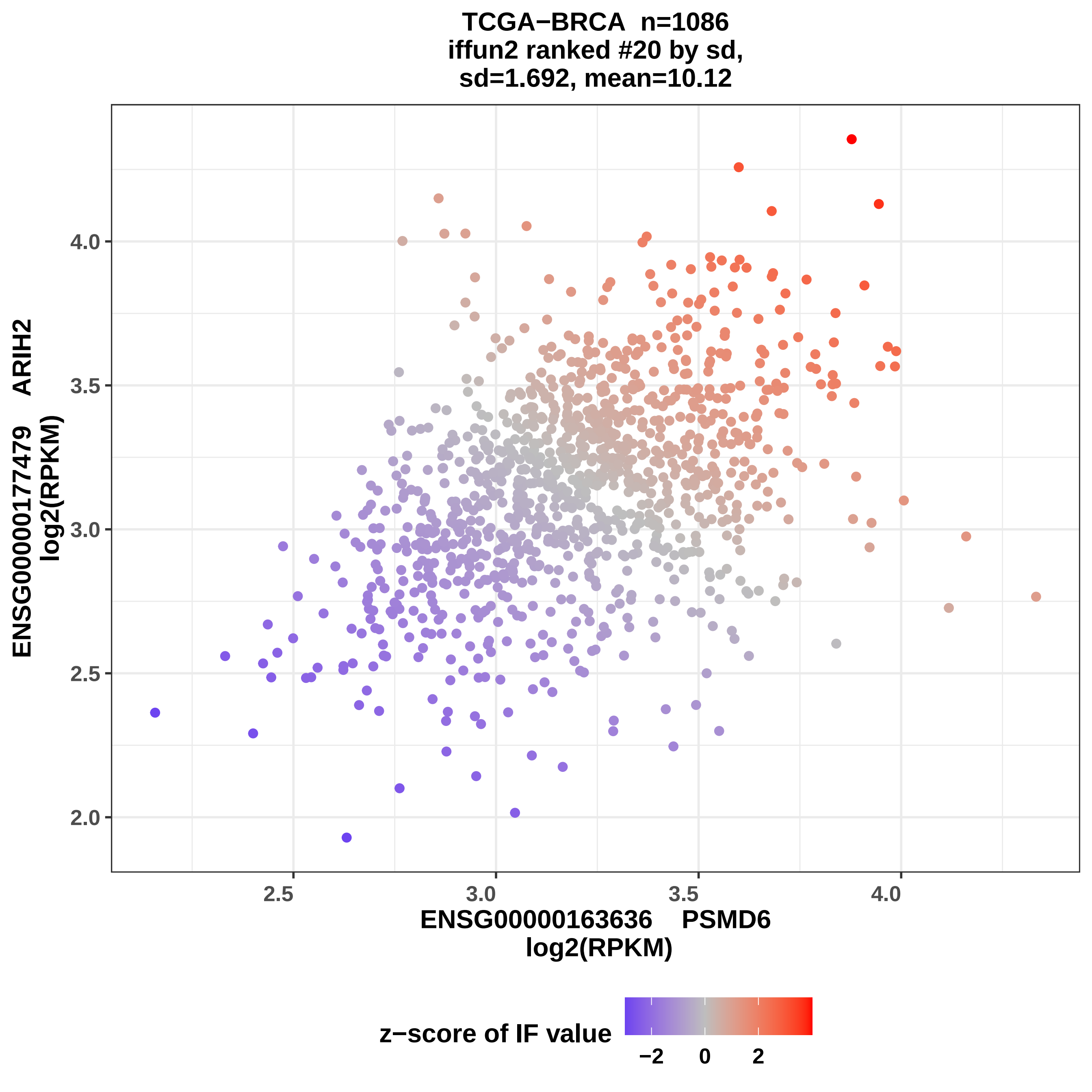
TCGA−BRCA n=1086
iffun2 ranked #20 by sd,
sd=1.692, mean=10.12
ENSG00000177479 ARIH2
log2(RPKM)
4.0
3.5
3.0
2.5
2.0
2.5
3.0
3.5
4.0
ENSG00000163636 PSMD6
log2(RPKM)
z−score of IF value
−2
0
2

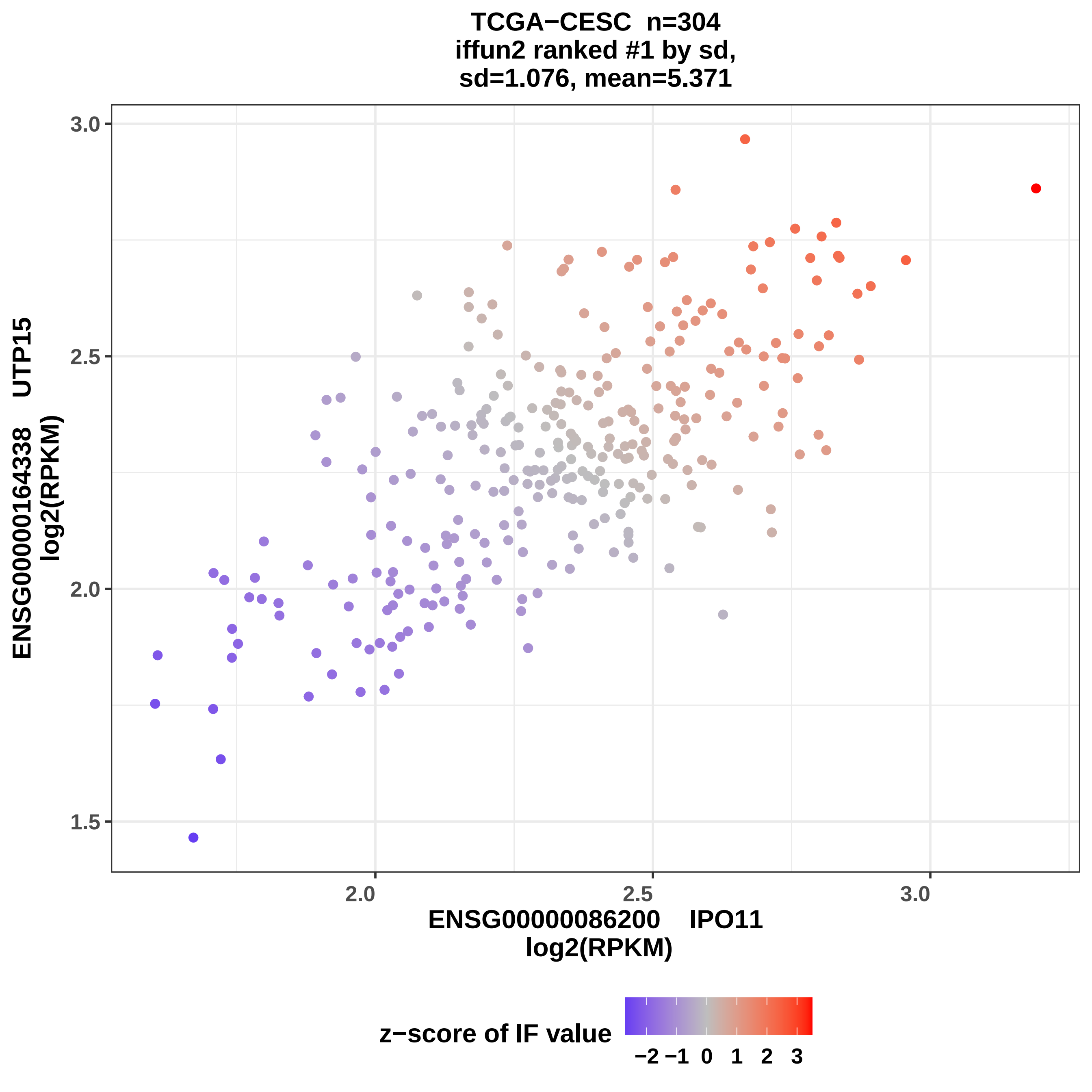
TCGA−CESC n=304
iffun2 ranked #1 by sd,
sd=1.076, mean=5.371
ENSG00000164338 UTP15
log2(RPKM)
ENSG00000086200 IPO11
log2(RPKM)
3.0
2.5
2.0
1.5
2.0
2.5
3.0
z−score of IF value
−2 −1 0 1 2 3

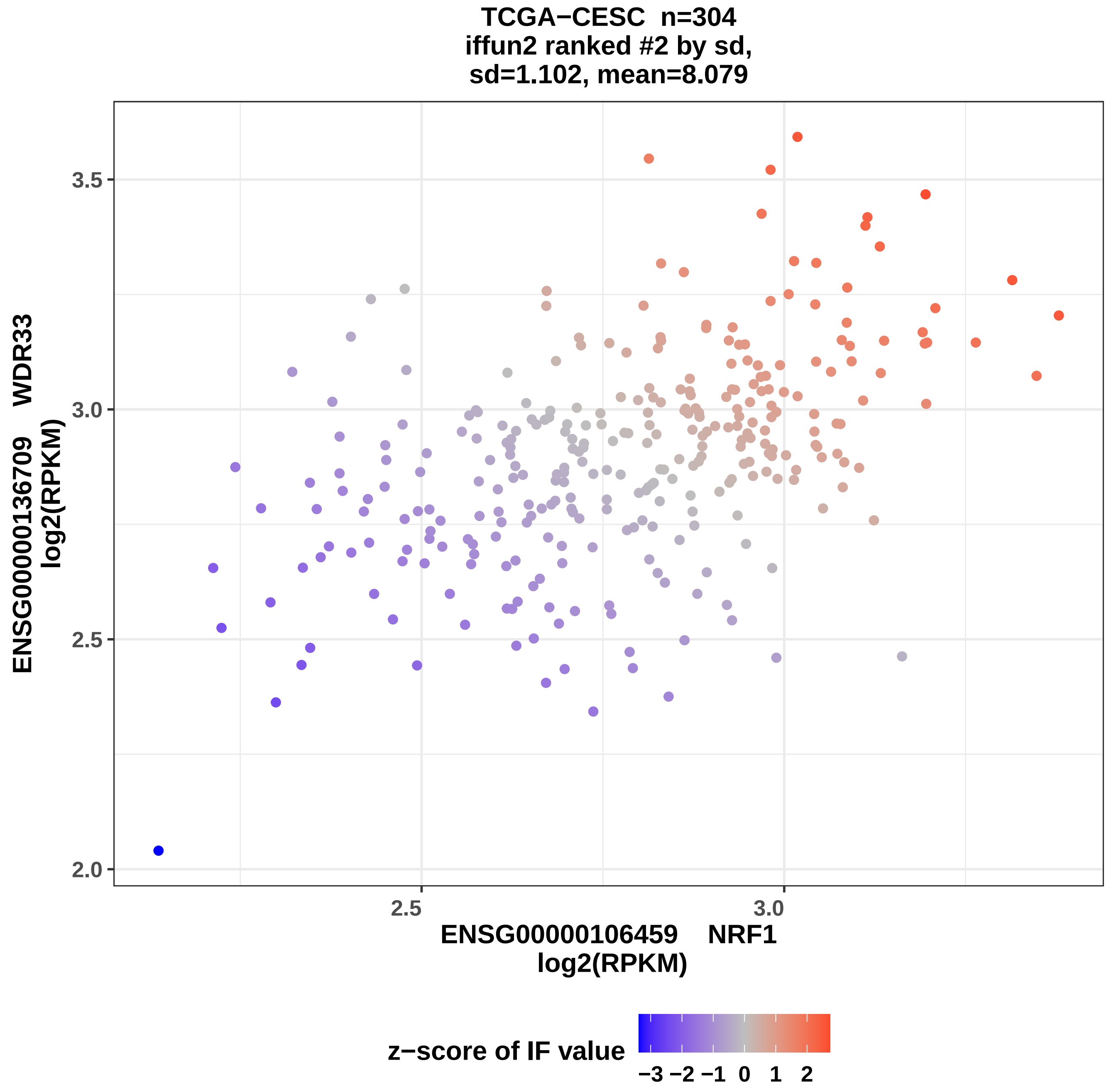
TCGA−CESC n=304
iffun2 ranked #2 by sd,
sd=1.102, mean=8.079
ENSG00000136709 WDR33
log2(RPKM)
3.5
3.0
2.5
2.0
2.5
3.0
ENSG00000106459 NRF1
log2(RPKM)
z−score of IF value
−3 −2 −1 0 1 2

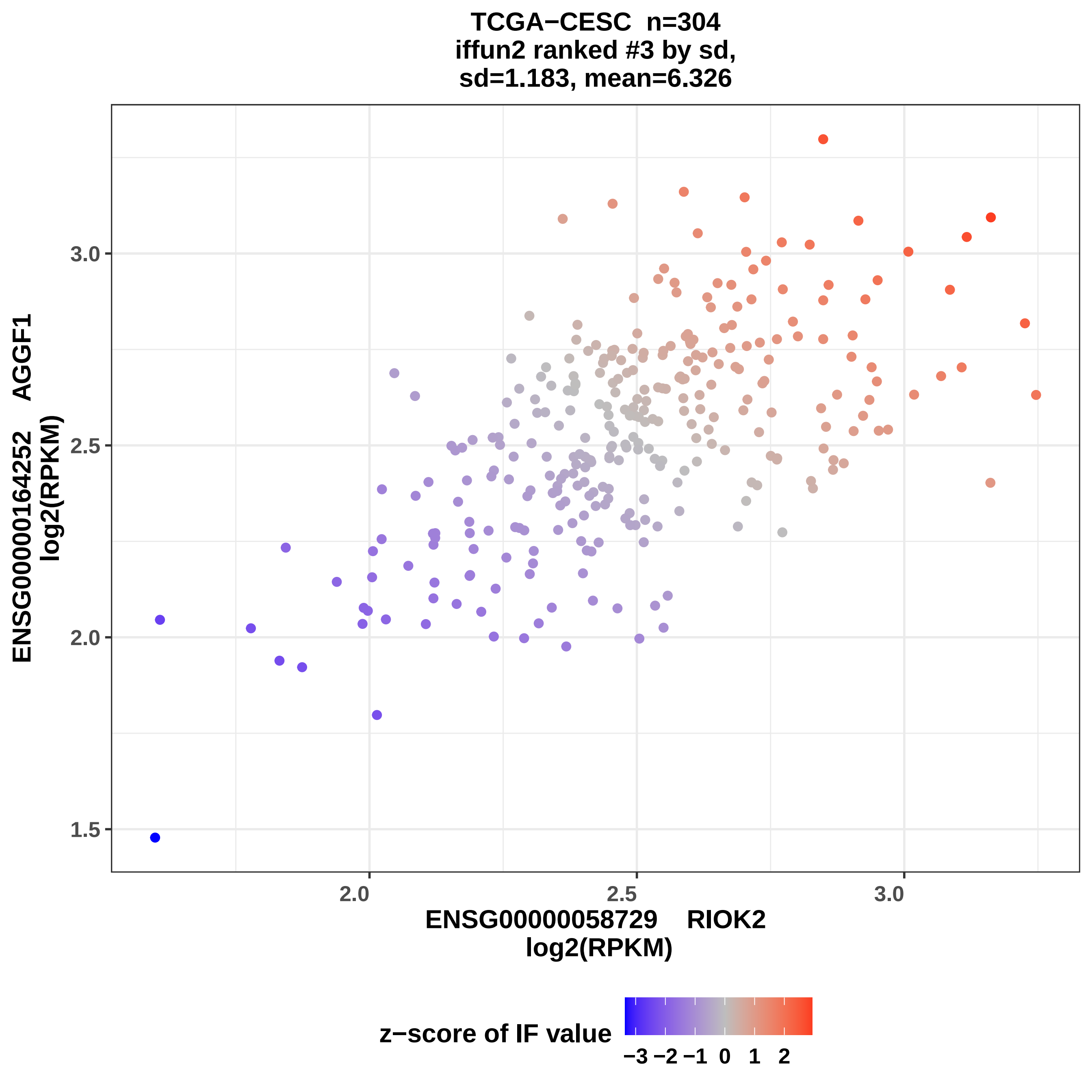
TCGA−CESC n=304
iffun2 ranked #3 by sd,
sd=1.183, mean=6.326
ENSG00000164252 AGGF1
log2(RPKM)
1.5
2.0
2.5
3.0
2.0
2.5
3.0
ENSG00000058729 RIOK2
log2(RPKM)
z−score of IF value
−3 −2 −1 0 1 2

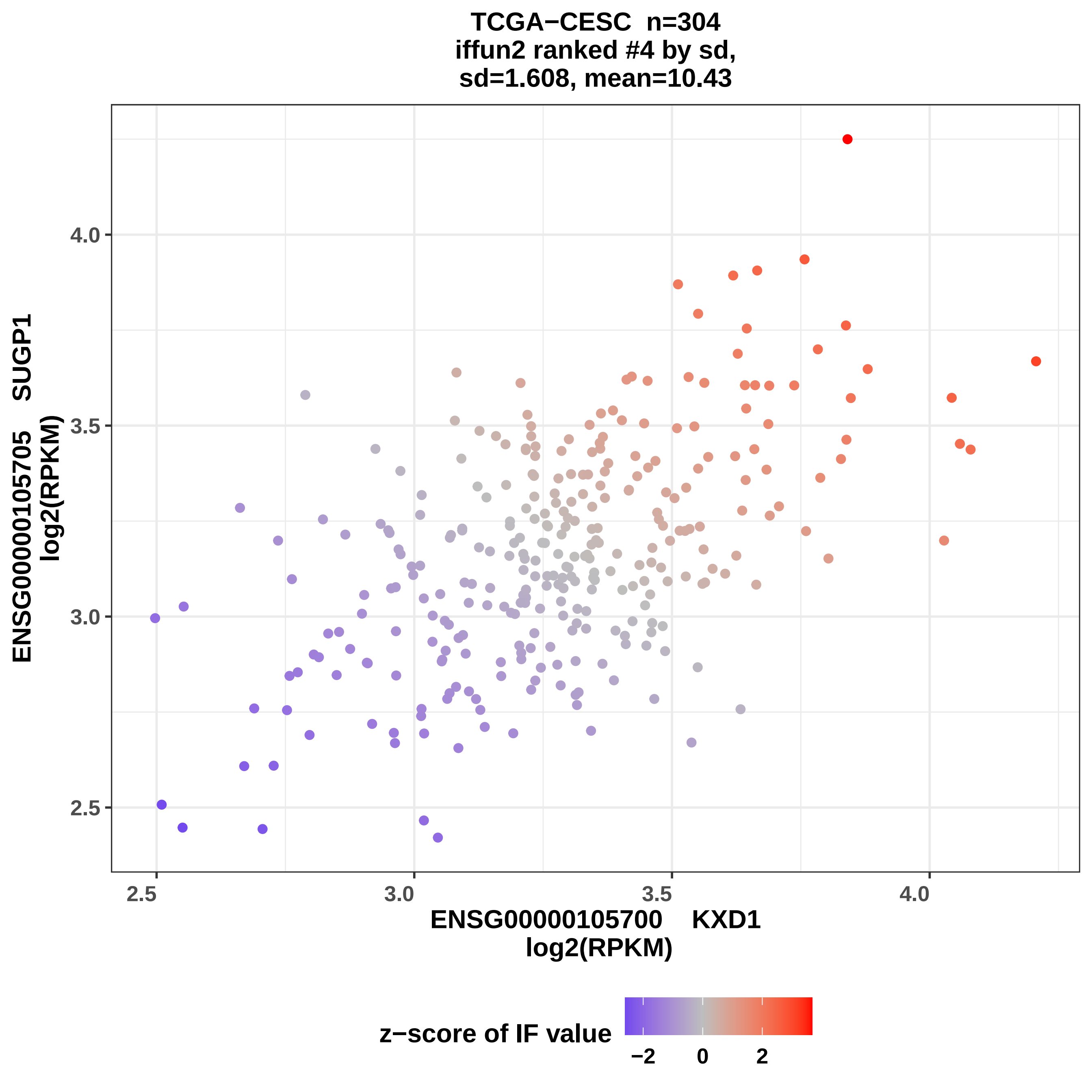

TCGA−CESC n=304
iffun2 ranked #4 by sd,
sd=1.608, mean=10.43
ENSG00000105705 SUGP1
log2(RPKM)
ENSG00000105700 KXD1
log2(RPKM)
2.5
3.0
3.5
4.0
z−score of IF value
−2
0
2

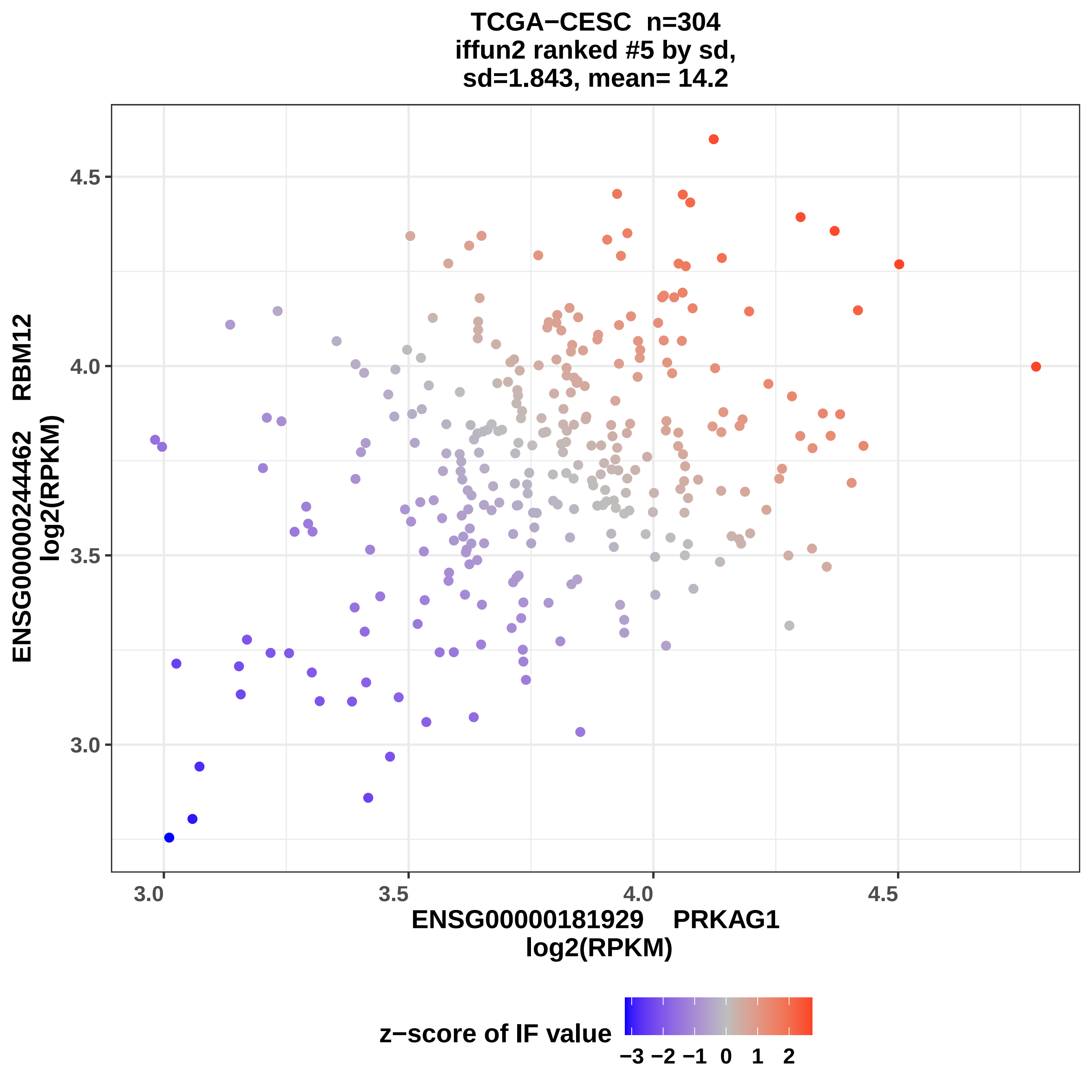
TCGA−CESC n=304
iffun2 ranked #5 by sd,
sd=1.843, mean= 14.2
ENSG00000244462 RBM12
log2(RPKM)
ENSG00000181929 PRKAG1
log2(RPKM)
z−score of IF value
3.0
3.5
4.0
4.5
−3 −2 −1 0 1 2

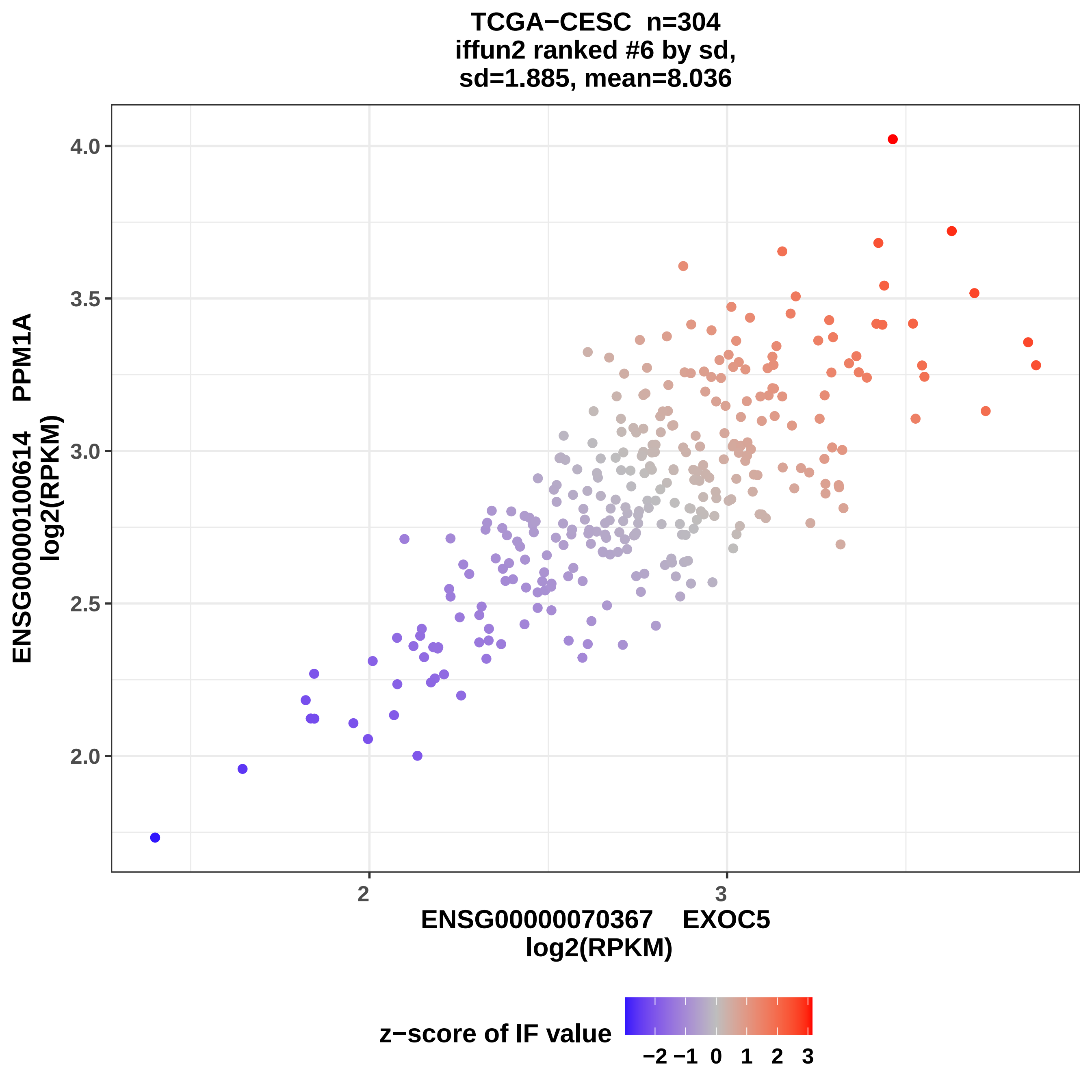

TCGA−CESC n=304
iffun2 ranked #6 by sd,
sd=1.885, mean=8.036
ENSG00000100614 PPM1A
log2(RPKM)
4.0
3.5
3.0
2.5
2.0
2
3
ENSG00000070367 EXOC5
log2(RPKM)
z−score of IF value
−2 −1 0 1 2 3

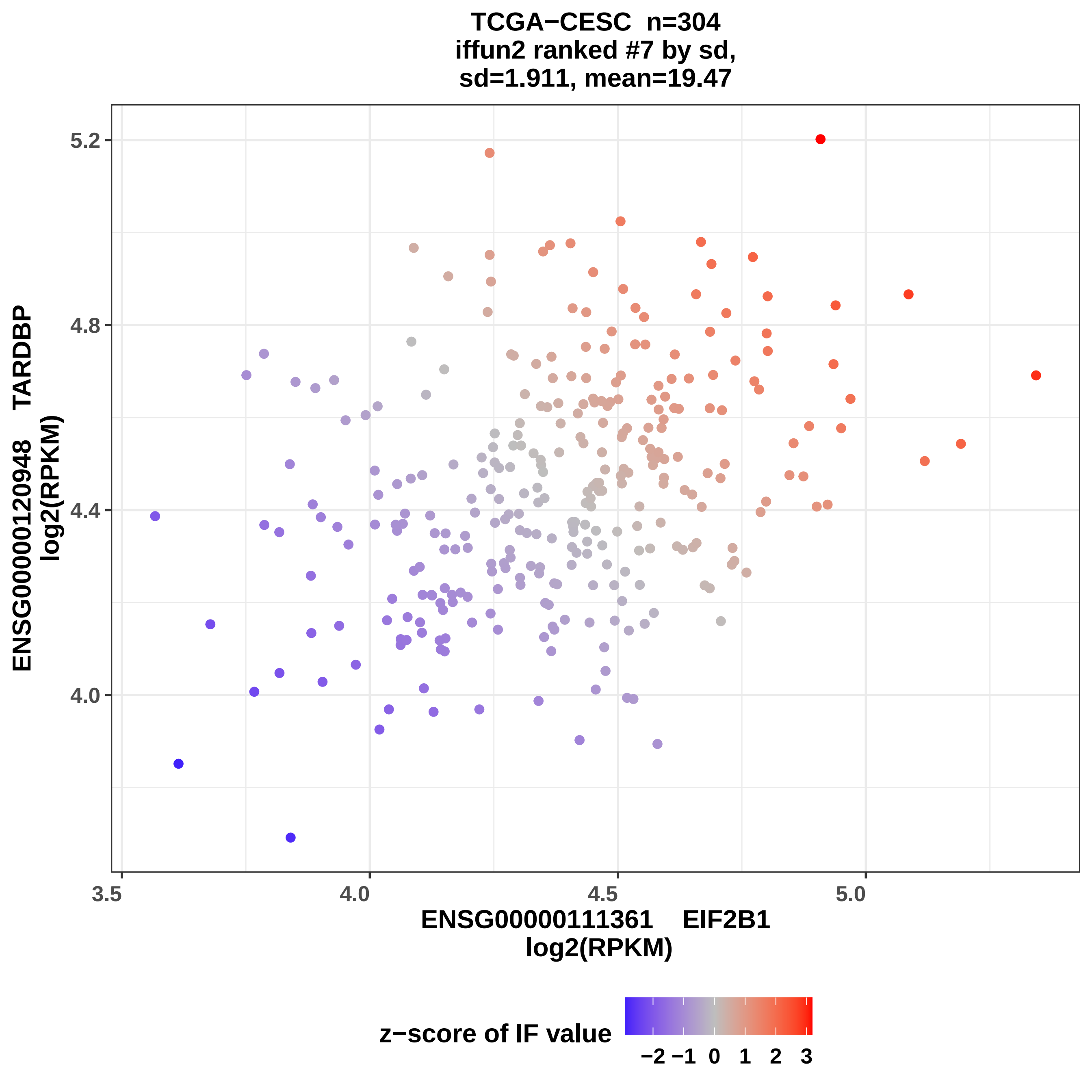
TCGA−CESC n=304
iffun2 ranked #7 by sd,
sd=1.911, mean=19.47
ENSG00000120948 TARDBP
log2(RPKM)
5.2
4.8
4.4
4.0
3.5
4.0
4.5
5.0
ENSG00000111361 EIF2B1
log2(RPKM)
z−score of IF value
−2 −1 0 1 2 3

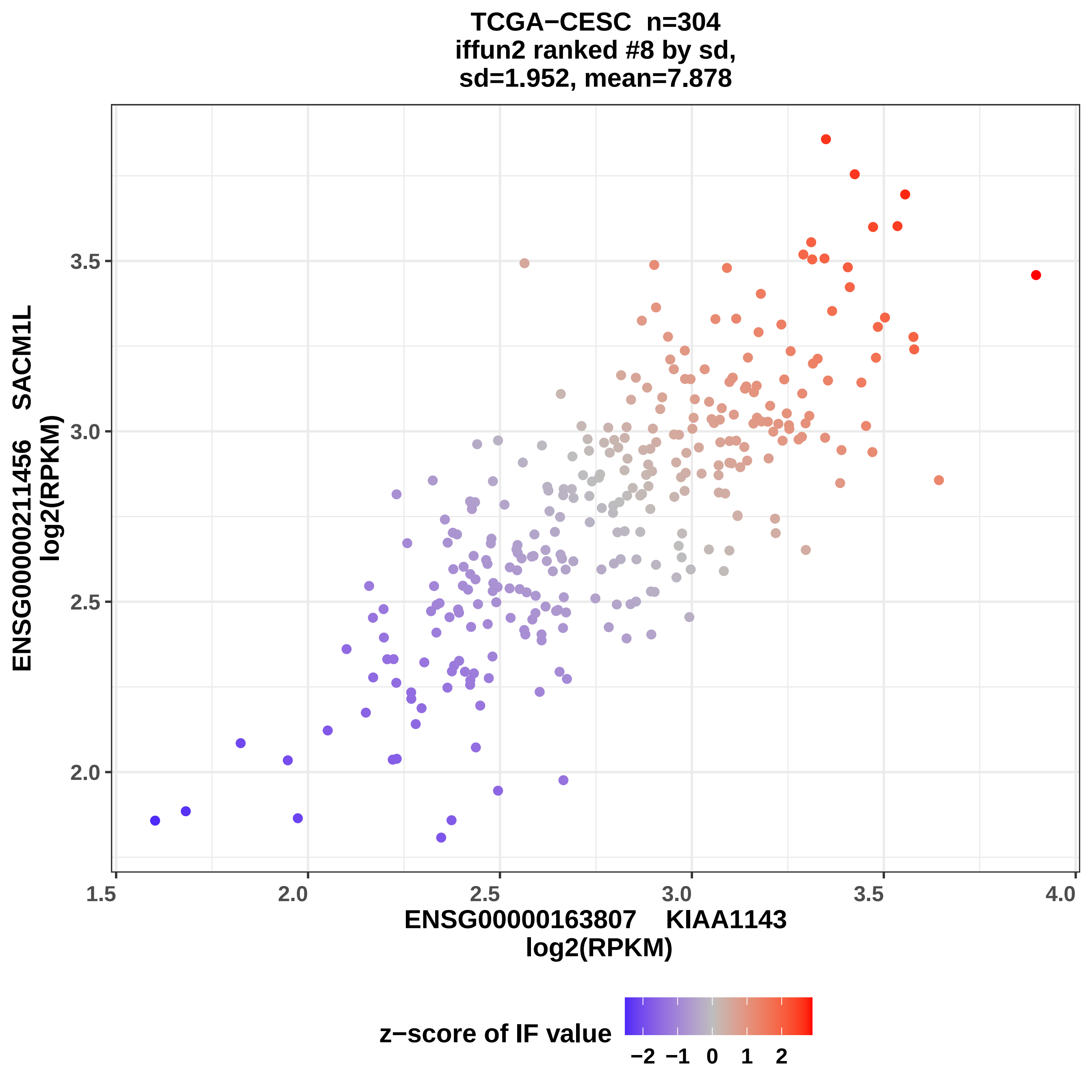
TCGA−CESC n=304
iffun2 ranked #8 by sd,
sd=1.952, mean=7.878
ENSG00000211456 SACM1L
log2(RPKM)
3.5
3.0
2.5
2.0
1.5
2.0
2.5
3.0
3.5
4.0
ENSG00000163807 KIAA1143
log2(RPKM)
z−score of IF value
−2
−1
0
1
2

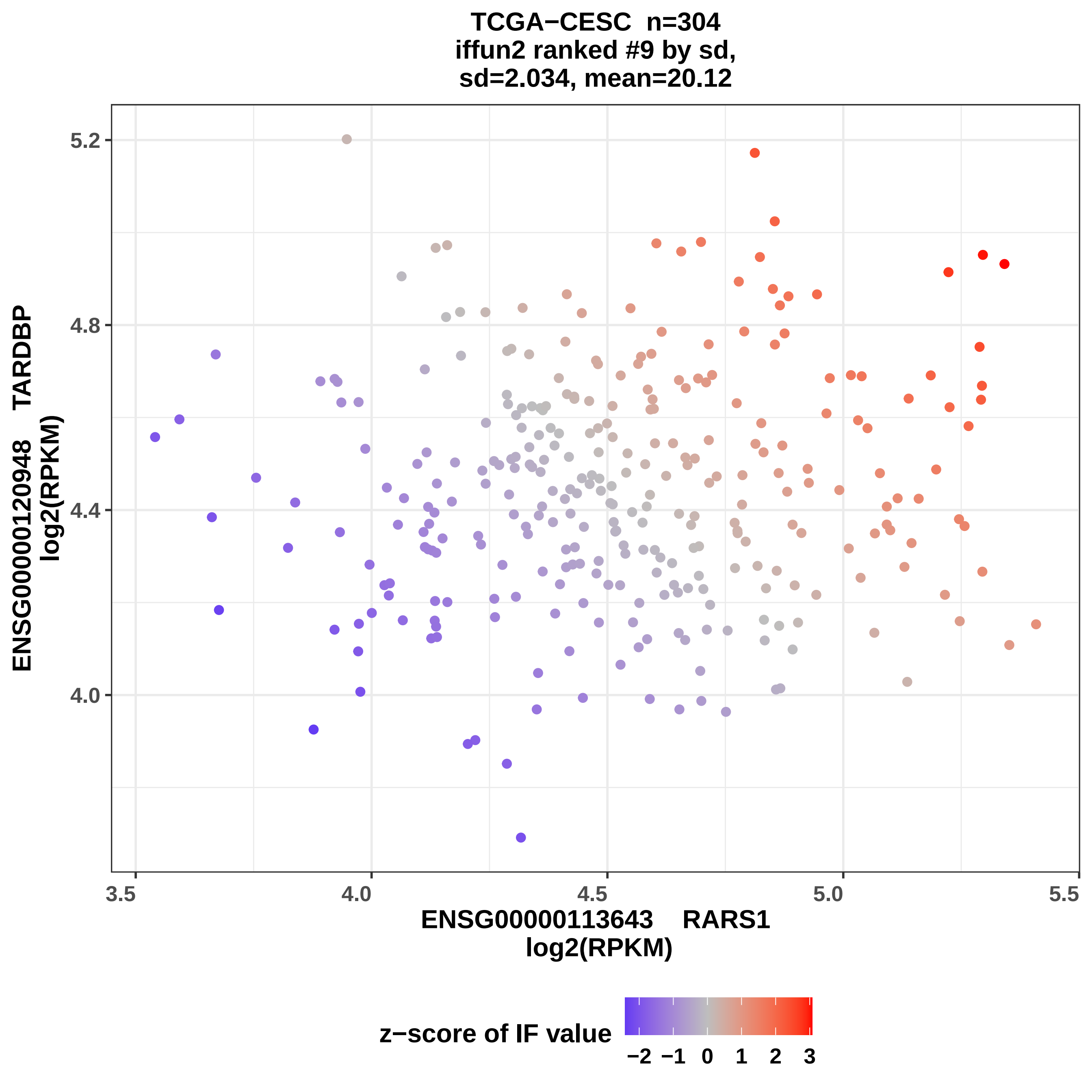

TCGA−CESC n=304
iffun2 ranked #9 by sd,
sd=2.034, mean=20.12
5.2
4.8
4.4
4.0
ENSG00000120948 TARDBP
log2(RPKM)
3.5
4.0
4.5
5.0
5.5
ENSG00000113643 RARS1
log2(RPKM)
z−score of IF value
−2 −1 0 1 2 3

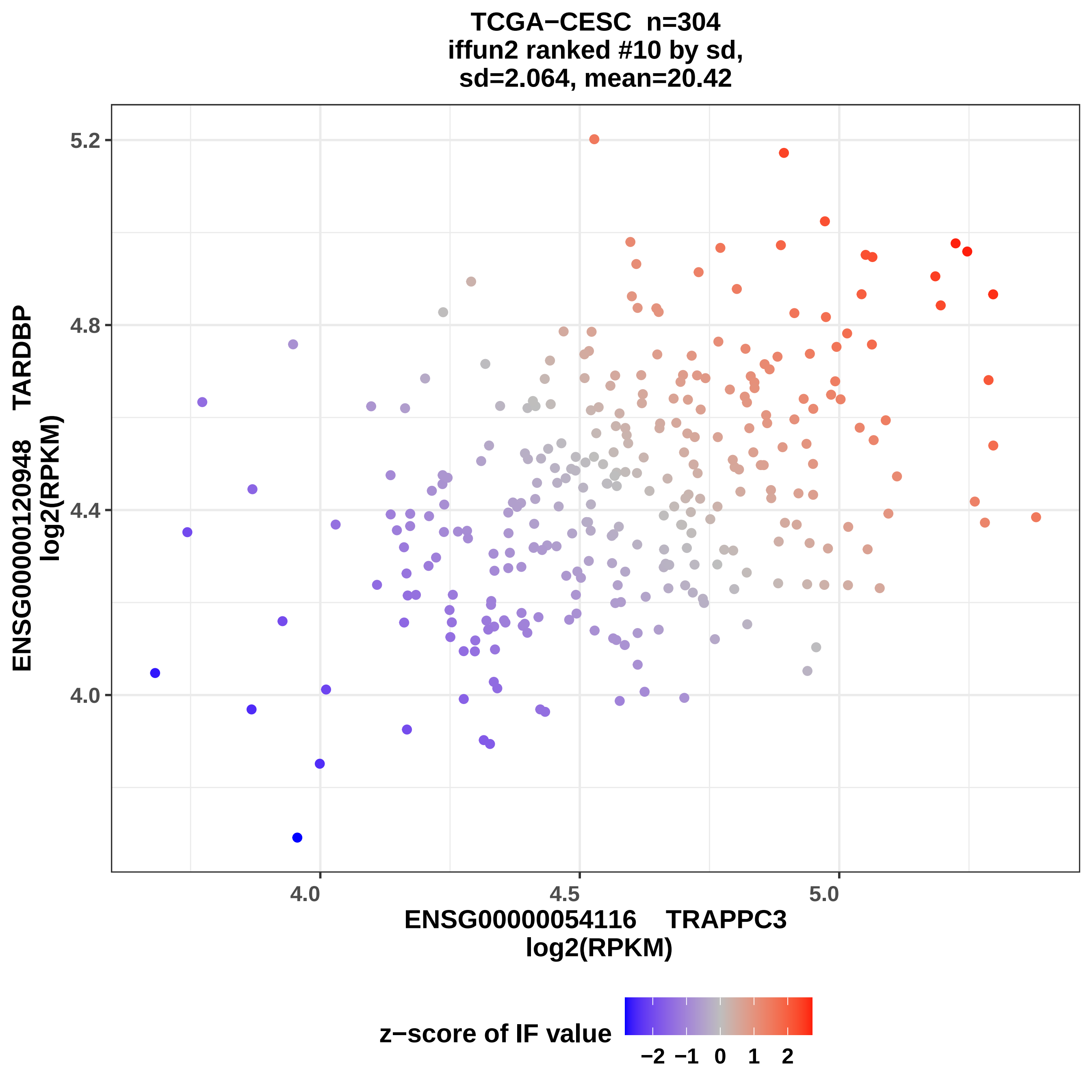
TCGA−CESC n=304
iffun2 ranked #10 by sd,
sd=2.064, mean=20.42
5.2
4.8
4.4
4.0
4.0
4.5
5.0
ENSG00000120948 TARDBP
log2(RPKM)
ENSG00000054116 TRAPPC3
log2(RPKM)
z−score of IF value
−2
−1
0
1
2

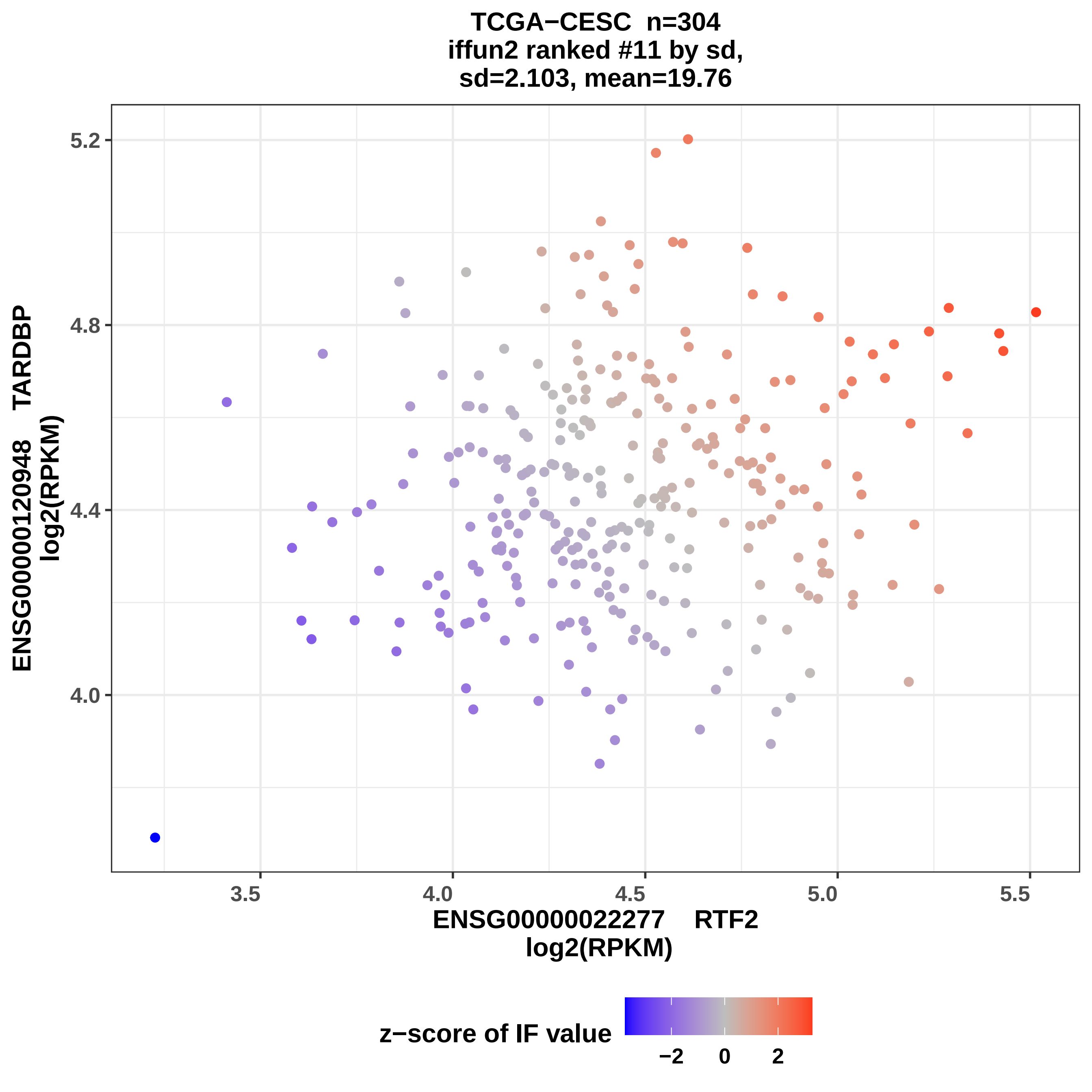

TCGA−CESC n=304
iffun2 ranked #11 by sd,
sd=2.103, mean=19.76
ENSG00000120948 TARDBP
log2(RPKM)
5.2
4.8
4.4
4.0
3.5
4.0
4.5
5.0
5.5
ENSG00000022277 RTF2
log2(RPKM)
z−score of IF value
−2
0
2

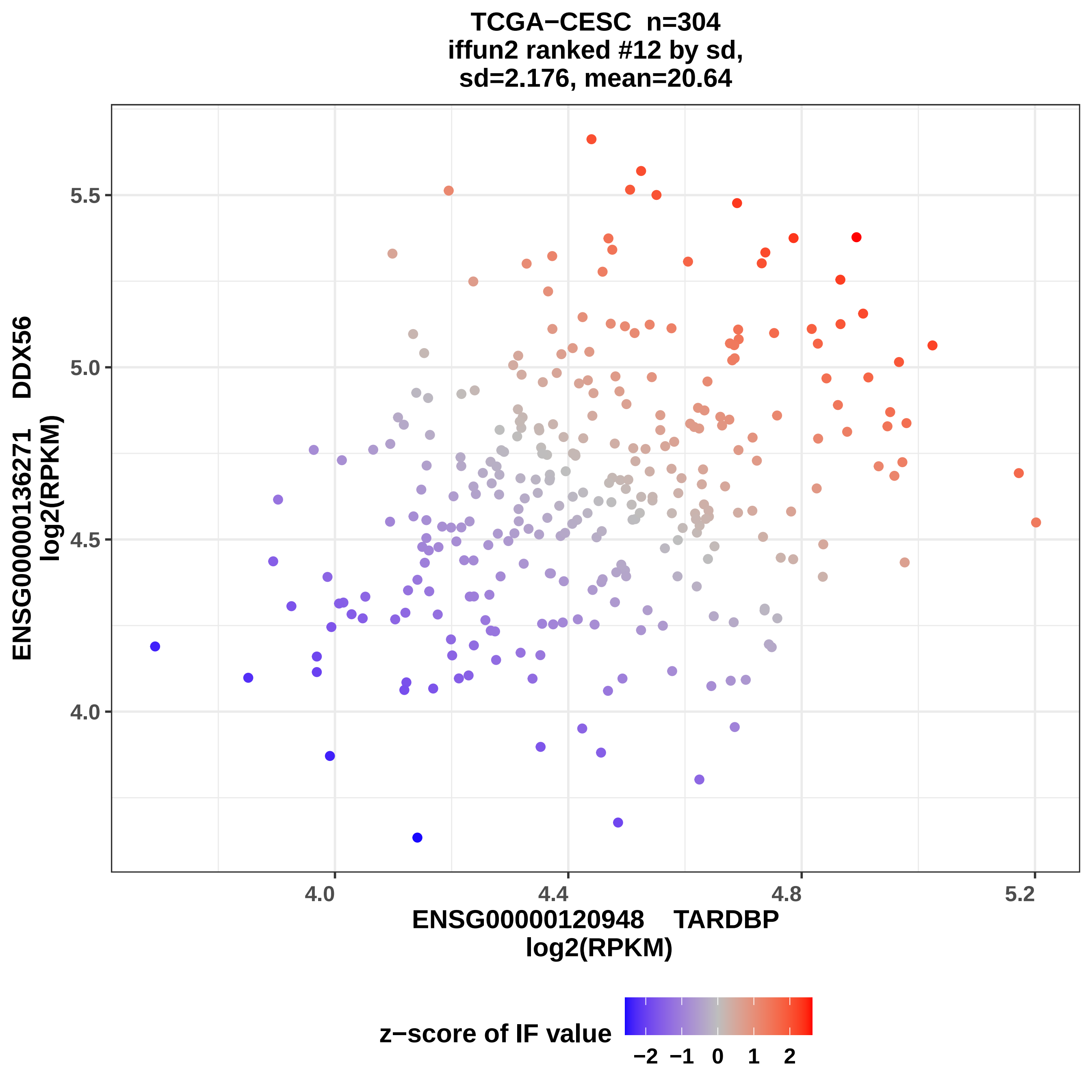

TCGA−CESC n=304
iffun2 ranked #12 by sd,
sd=2.176, mean=20.64
ENSG00000136271 DDX56
log2(RPKM)
5.5
5.0
4.5
4.0
4.0
4.4
4.8
5.2
ENSG00000120948 TARDBP
log2(RPKM)
z−score of IF value
−2 −1 0 1 2

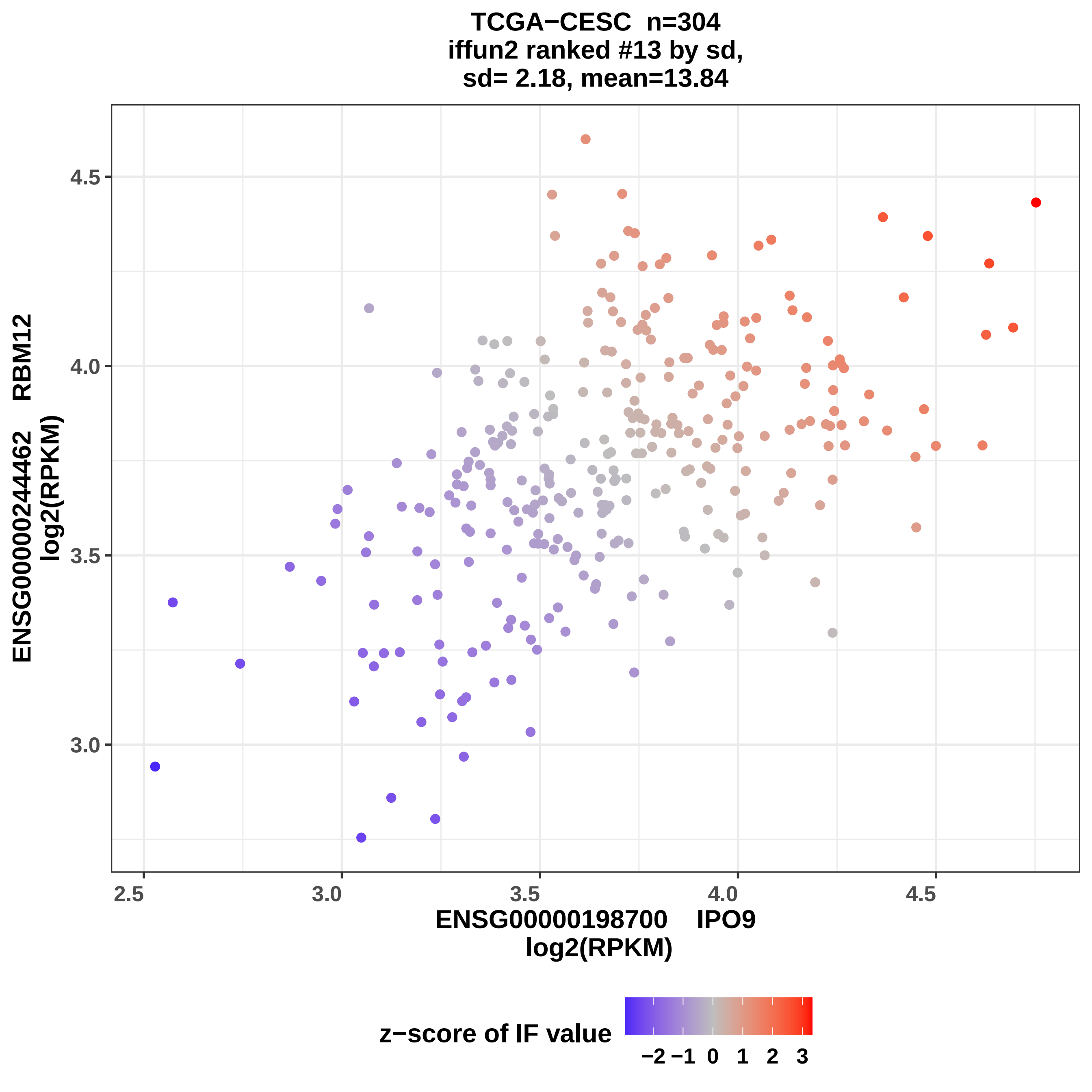
TCGA−CESC n=304
iffun2 ranked #13 by sd,
sd= 2.18, mean=13.84
ENSG00000244462 RBM12
log2(RPKM)
4.5
4.0
3.5
3.0
2.5
3.0
3.5
4.0
4.5
ENSG00000198700 IPO9
log2(RPKM)
z−score of IF value
−2 −1 0 1 2 3

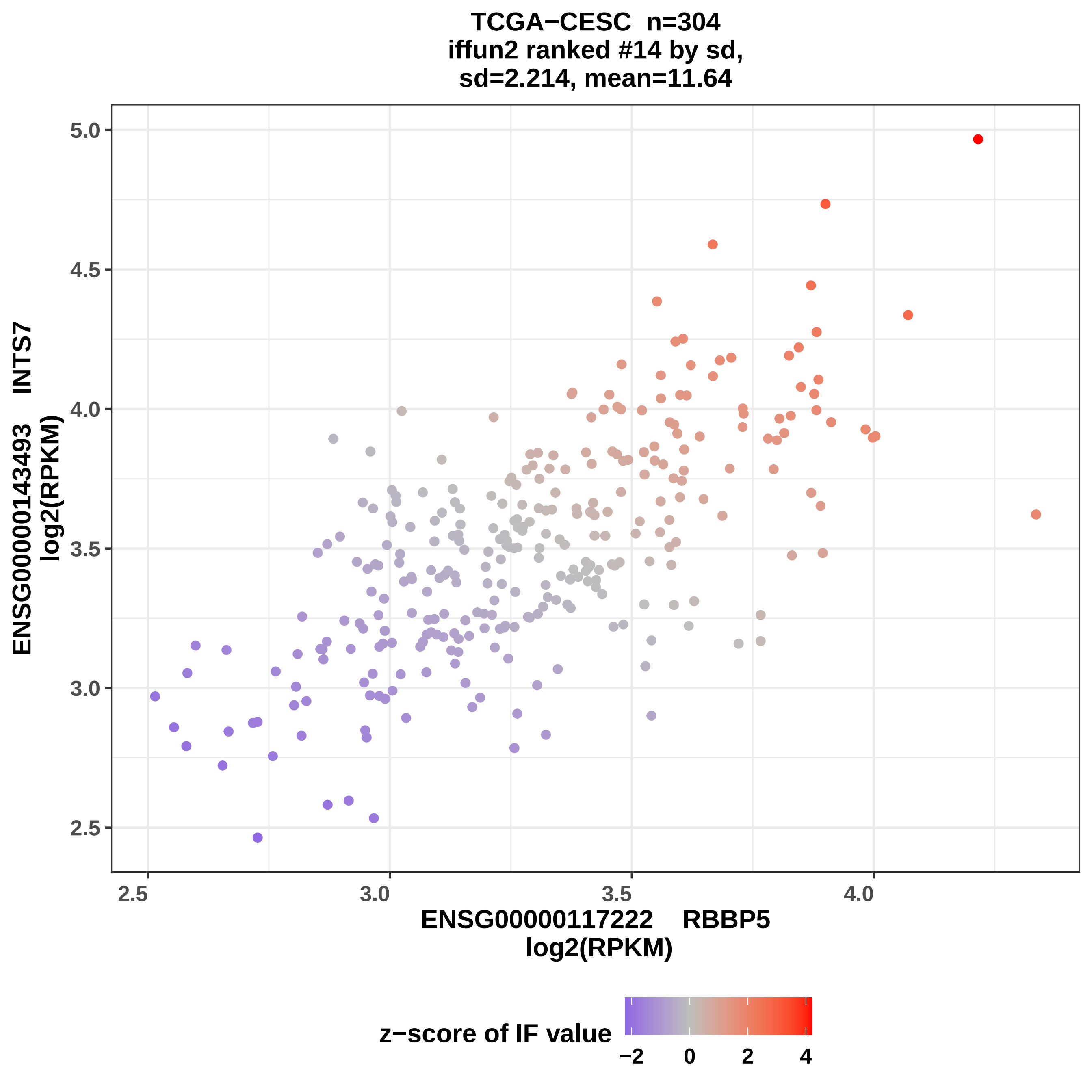
TCGA−CESC n=304
iffun2 ranked #14 by sd,
sd=2.214, mean=11.64
ENSG00000143493 INTS7
log2(RPKM)
5.0
4.5
4.0
3.5
3.0
2.5
2.5
3.0
3.5
4.0
ENSG00000117222 RBBP5
log2(RPKM)
z−score of IF value
−2
0
2
4

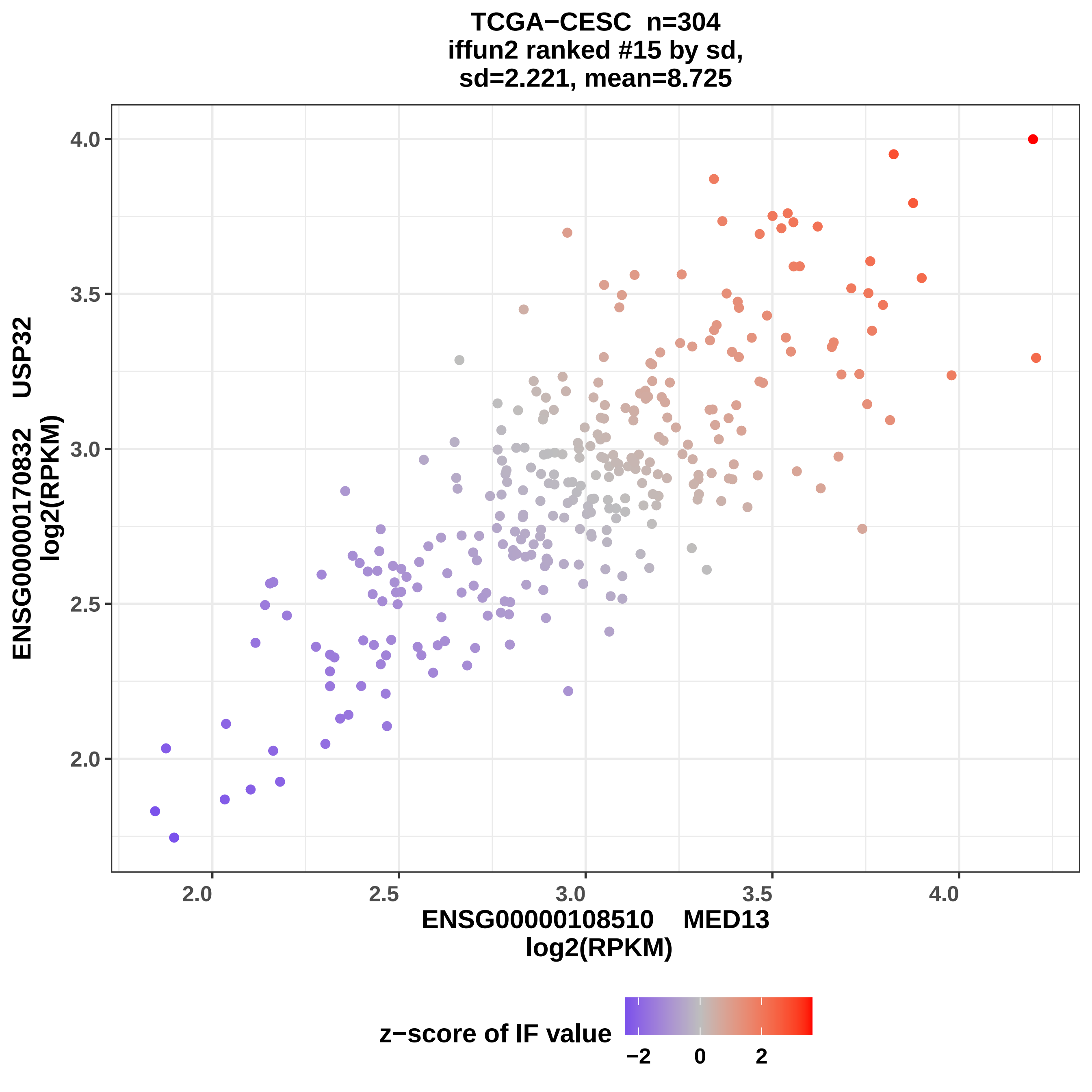
TCGA−CESC n=304
iffun2 ranked #15 by sd,
sd=2.221, mean=8.725
ENSG00000170832 USP32
log2(RPKM)
4.0
3.5
3.0
2.5
2.0
2.0
2.5
3.0
3.5
4.0
ENSG00000108510 MED13
log2(RPKM)
z−score of IF value
−2
0
2

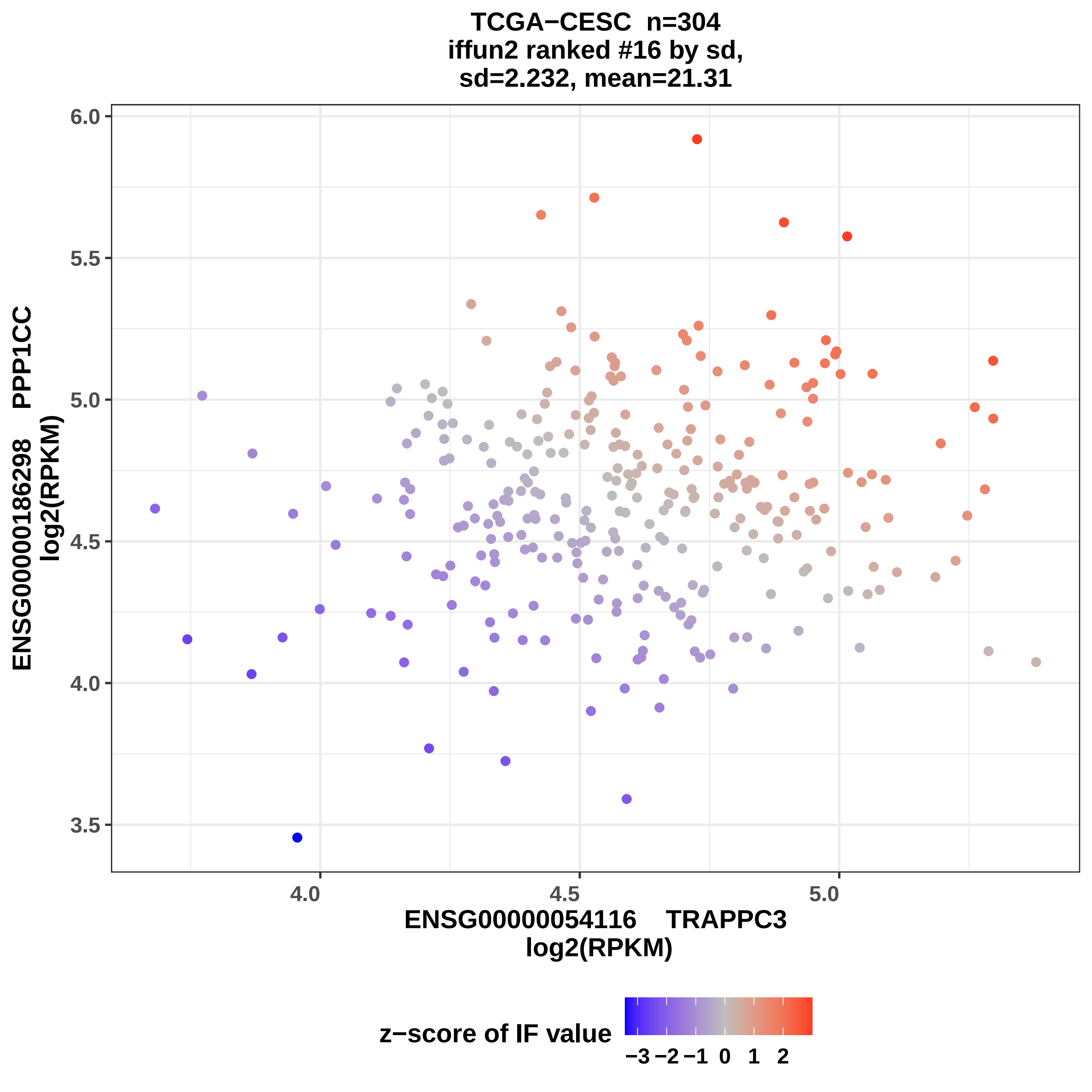
TCGA−CESC n=304
iffun2 ranked #16 by sd,
sd=2.232, mean=21.31
ENSG00000186298 PPP1CC
log2(RPKM)
6.0
5.5
5.0
4.5
4.0
3.5
4.0
4.5
5.0
ENSG00000054116 TRAPPC3
log2(RPKM)
z−score of IF value
−3 −2 −1 0 1 2

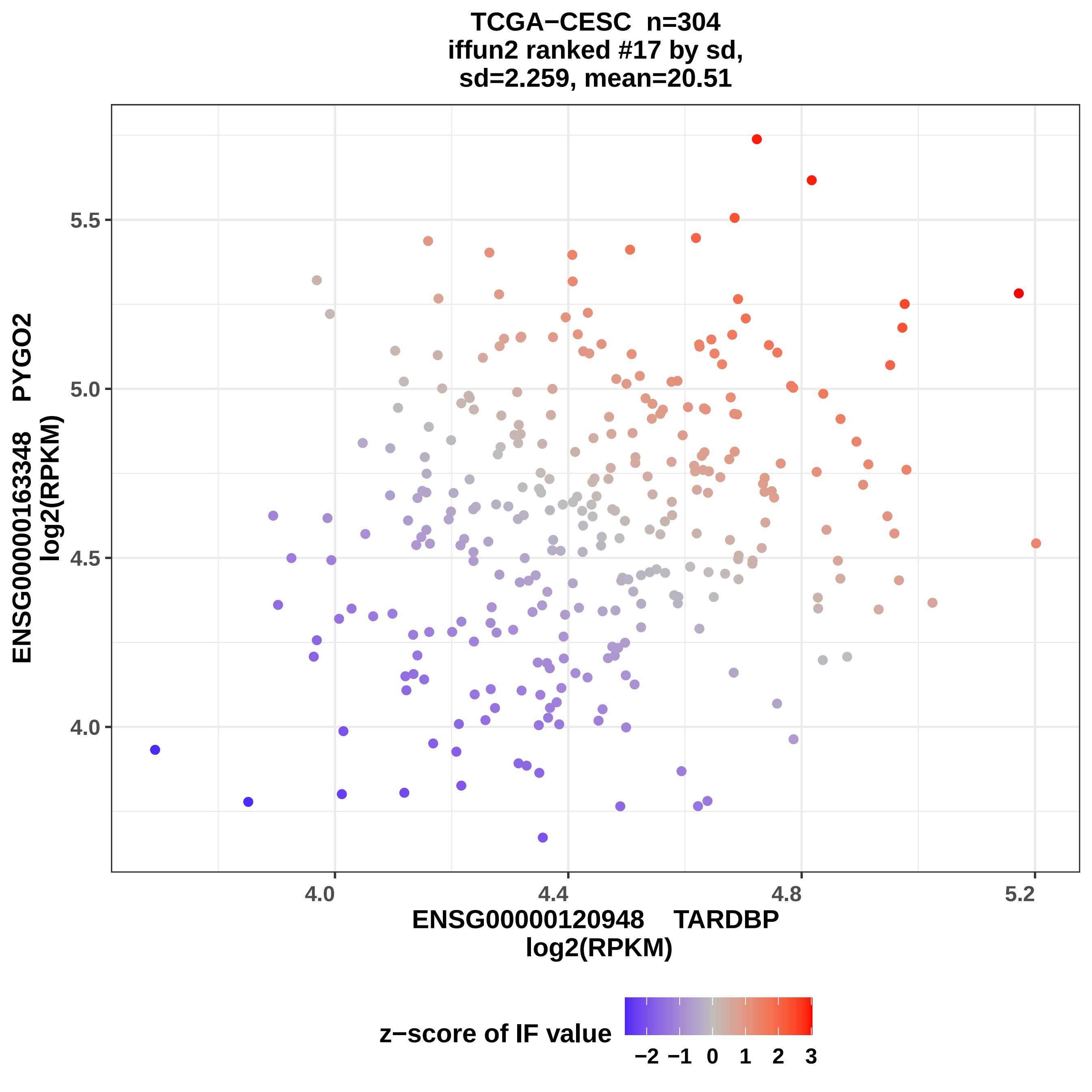
TCGA−CESC n=304
iffun2 ranked #17 by sd,
sd=2.259, mean=20.51
ENSG00000163348 PYGO2
log2(RPKM)
5.5
5.0
4.5
4.0
4.0
4.4
4.8
5.2
ENSG00000120948 TARDBP
log2(RPKM)
z−score of IF value
−2 −1 0 1 2 3

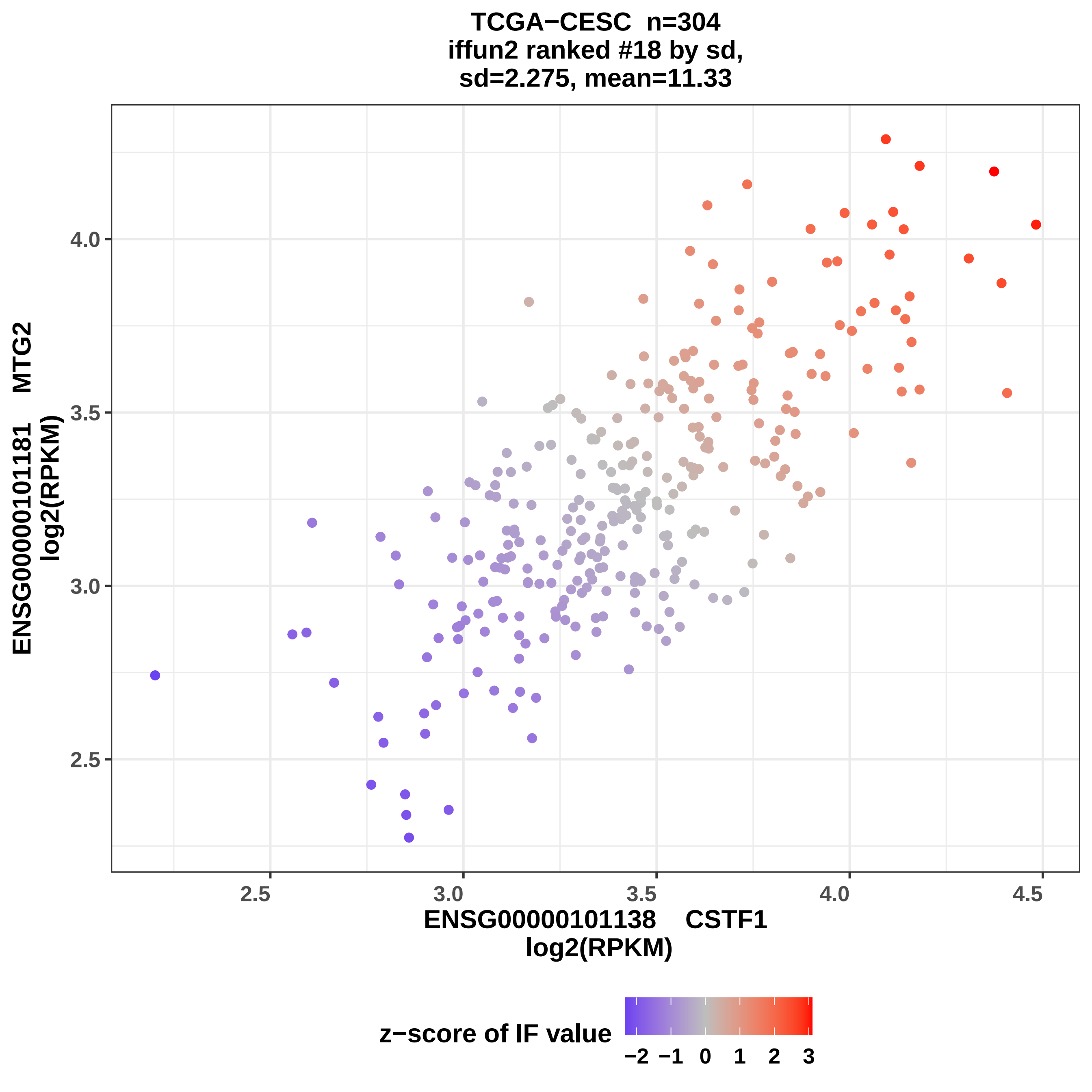
TCGA−CESC n=304
iffun2 ranked #18 by sd,
sd=2.275, mean=11.33
ENSG00000101181 MTG2
log2(RPKM)
4.0
3.5
3.0
2.5
2.5
3.0
3.5
4.0
4.5
ENSG00000101138 CSTF1
log2(RPKM)
z−score of IF value
−2 −1 0 1 2 3

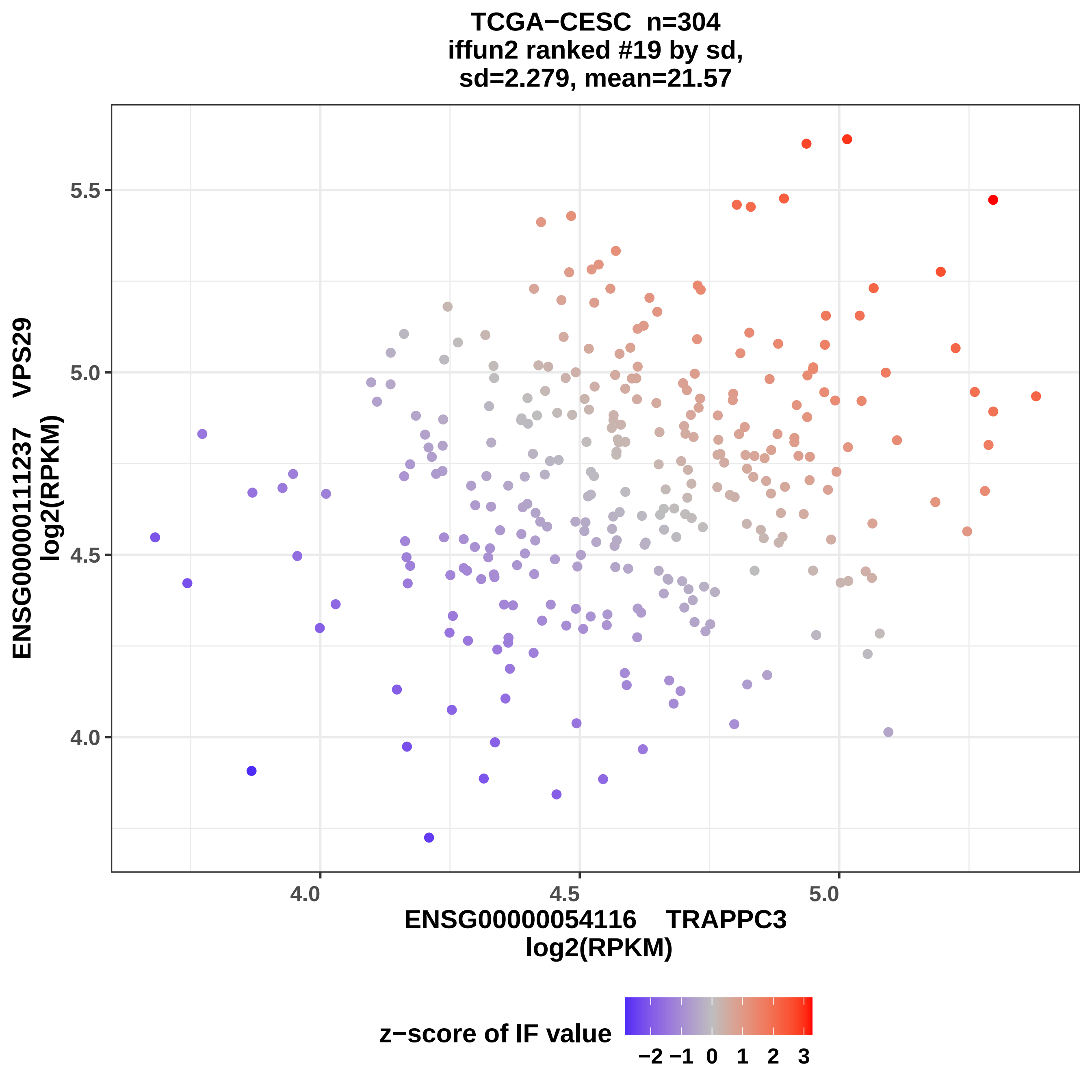

TCGA−CESC n=304
iffun2 ranked #19 by sd,
sd=2.279, mean=21.57
ENSG00000111237 VPS29
log2(RPKM)
5.5
5.0
4.5
4.0
4.0
4.5
5.0
ENSG00000054116 TRAPPC3
log2(RPKM)
z−score of IF value
−2 −1 0 1 2 3

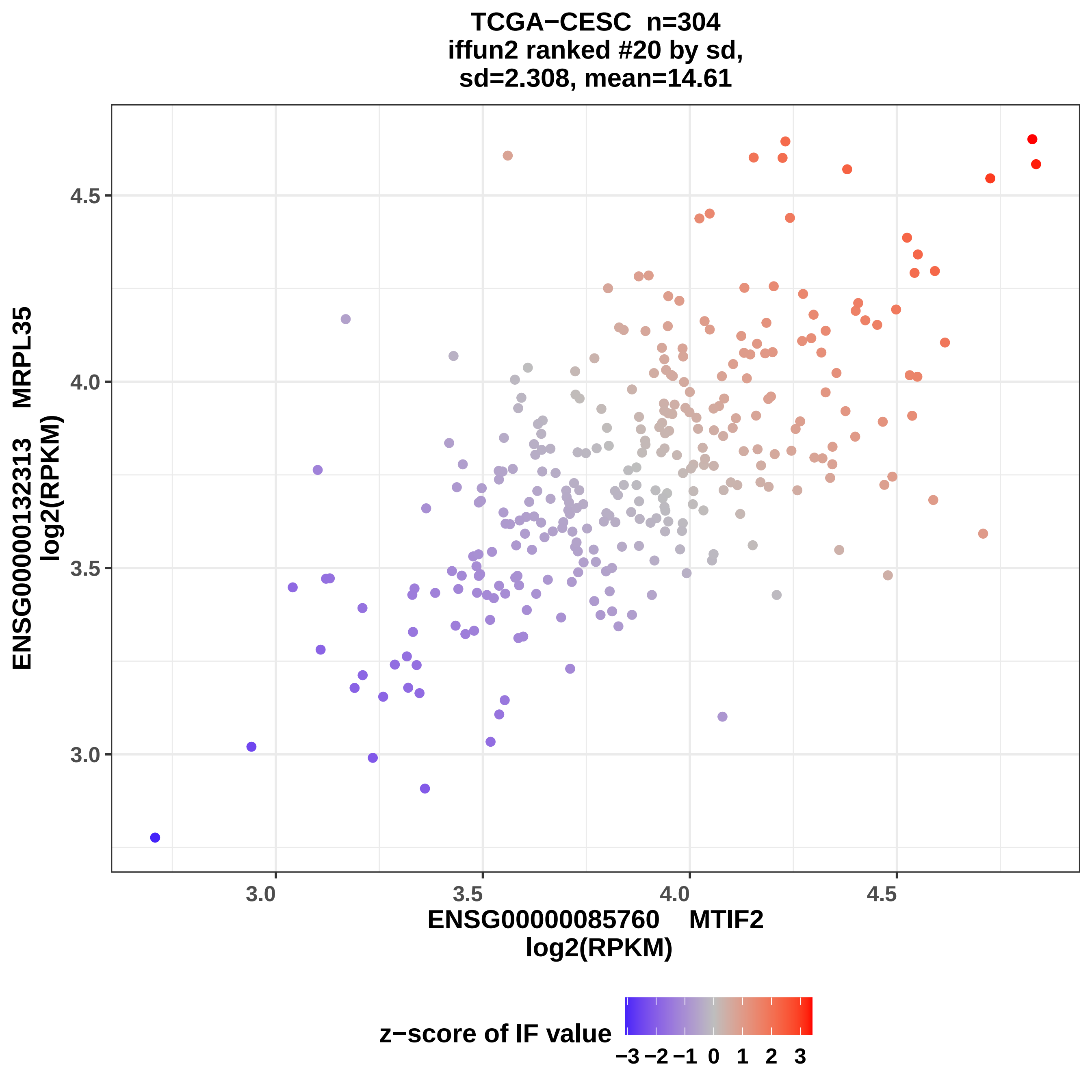
TCGA−CESC n=304
iffun2 ranked #20 by sd,
sd=2.308, mean=14.61
ENSG00000132313 MRPL35
log2(RPKM)
4.5
4.0
3.5
3.0
3.0
3.5
4.0
4.5
ENSG00000085760 MTIF2
log2(RPKM)
z−score of IF value
−3 −2 −1 0 1 2 3

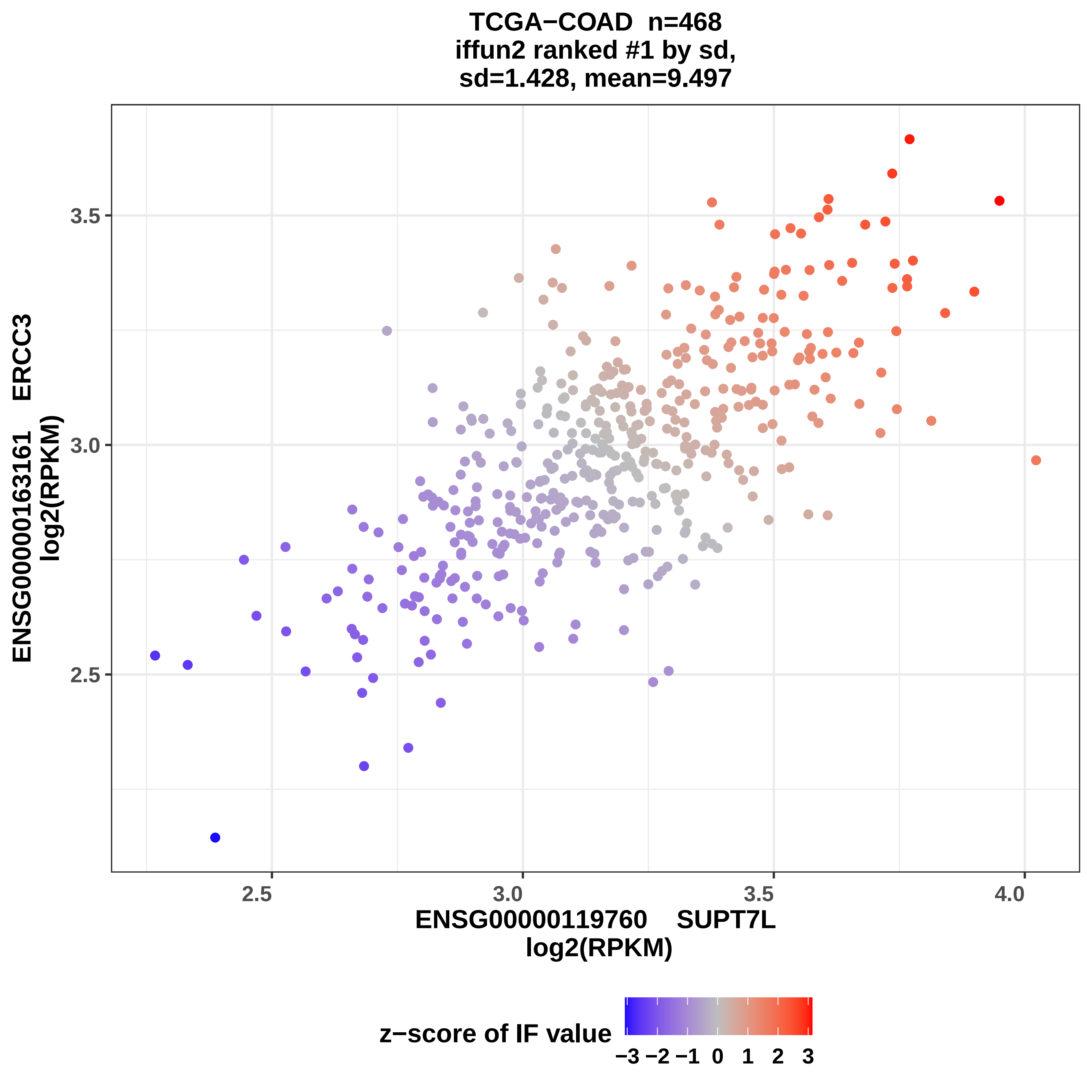

TCGA−COAD n=468
iffun2 ranked #1 by sd,
sd=1.428, mean=9.497
ENSG00000163161 ERCC3
log2(RPKM)
3.5
3.0
2.5
2.5
3.0
3.5
4.0
ENSG00000119760 SUPT7L
log2(RPKM)
z−score of IF value
−3 −2 −1 0 1 2 3

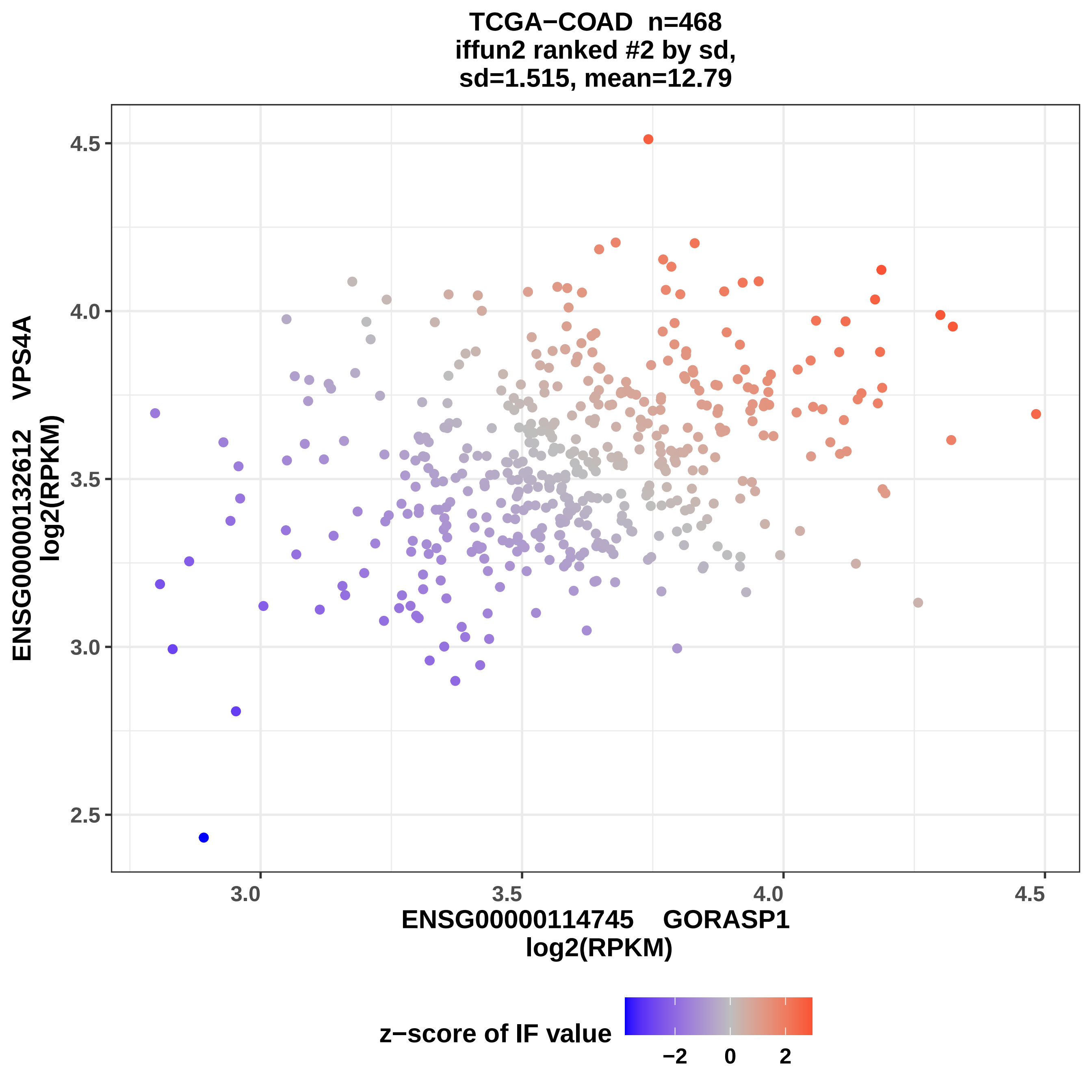
TCGA−COAD n=468
iffun2 ranked #2 by sd,
sd=1.515, mean=12.79
ENSG00000132612 VPS4A
log2(RPKM)
4.5
4.0
3.5
3.0
2.5
3.0
3.5
4.0
4.5
ENSG00000114745 GORASP1
log2(RPKM)
z−score of IF value
−2
0
2

TCGA−COAD  n=468
iffun2 ranked #3 by sd,
sd=1.531, mean=16.75

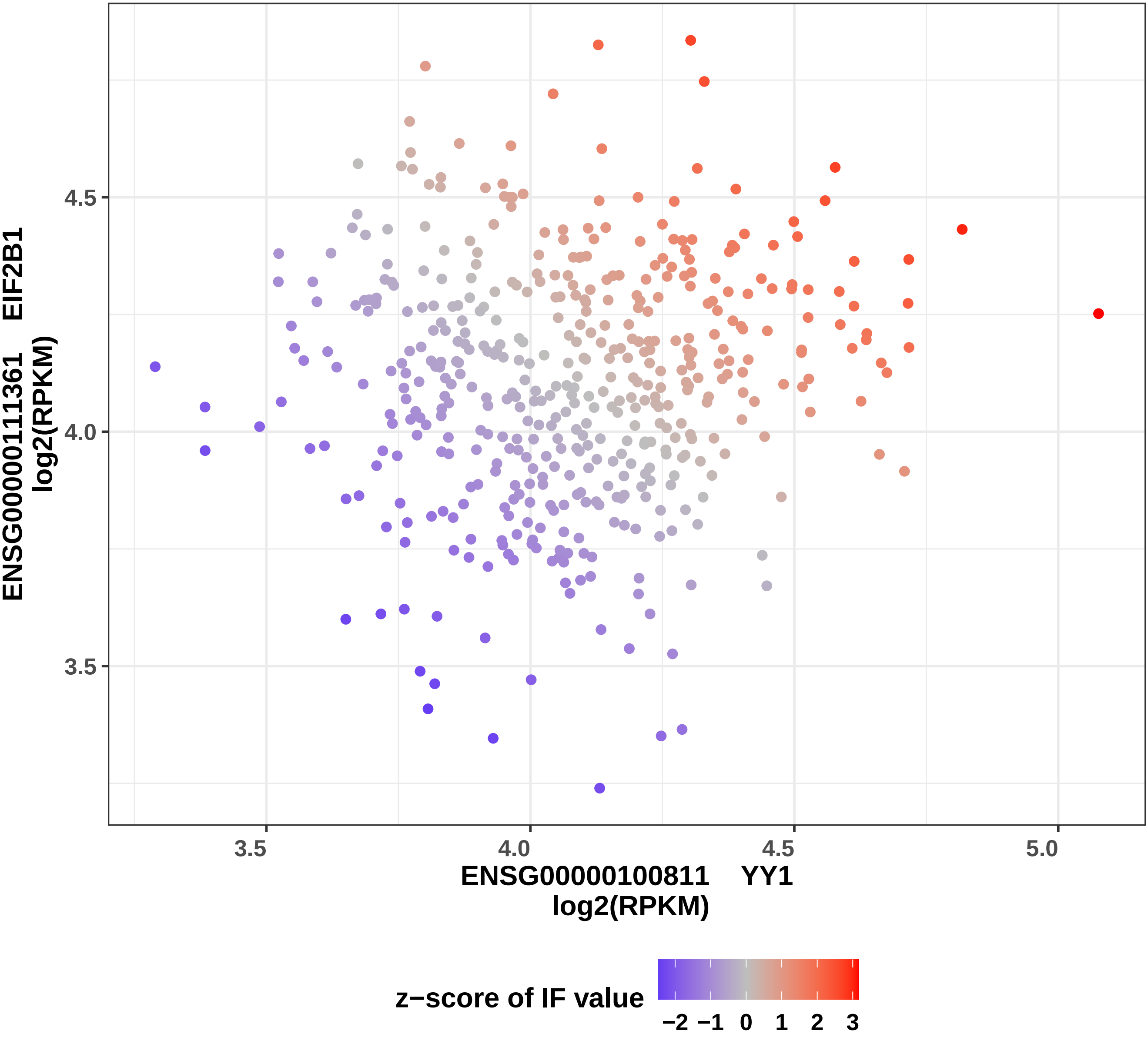

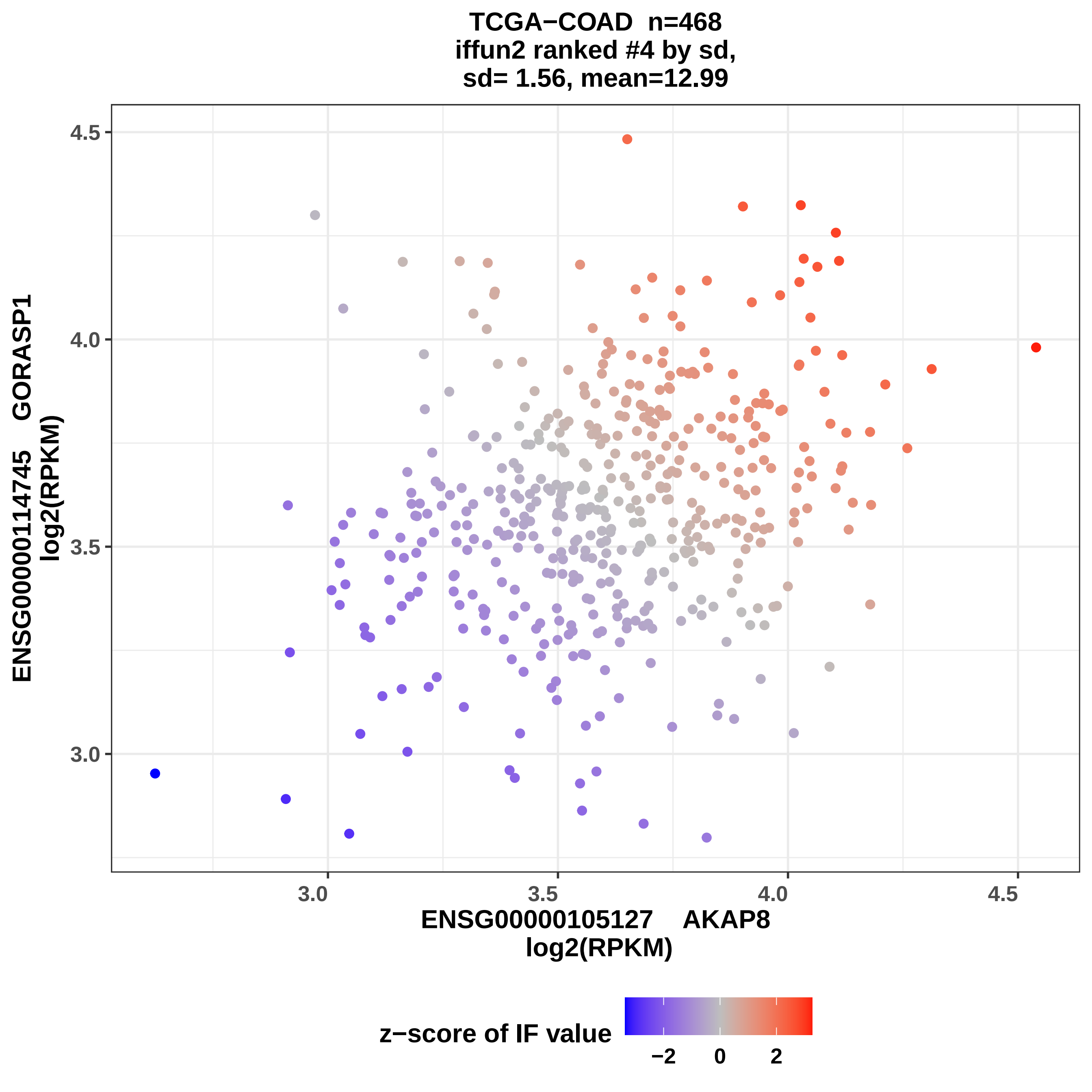
TCGA−COAD n=468
iffun2 ranked #4 by sd,
sd= 1.56, mean=12.99
ENSG00000114745 GORASP1
log2(RPKM)
4.5
4.0
3.5
3.0
3.0
3.5
4.0
4.5
ENSG00000105127 AKAP8
log2(RPKM)
z−score of IF value
−2
0
2

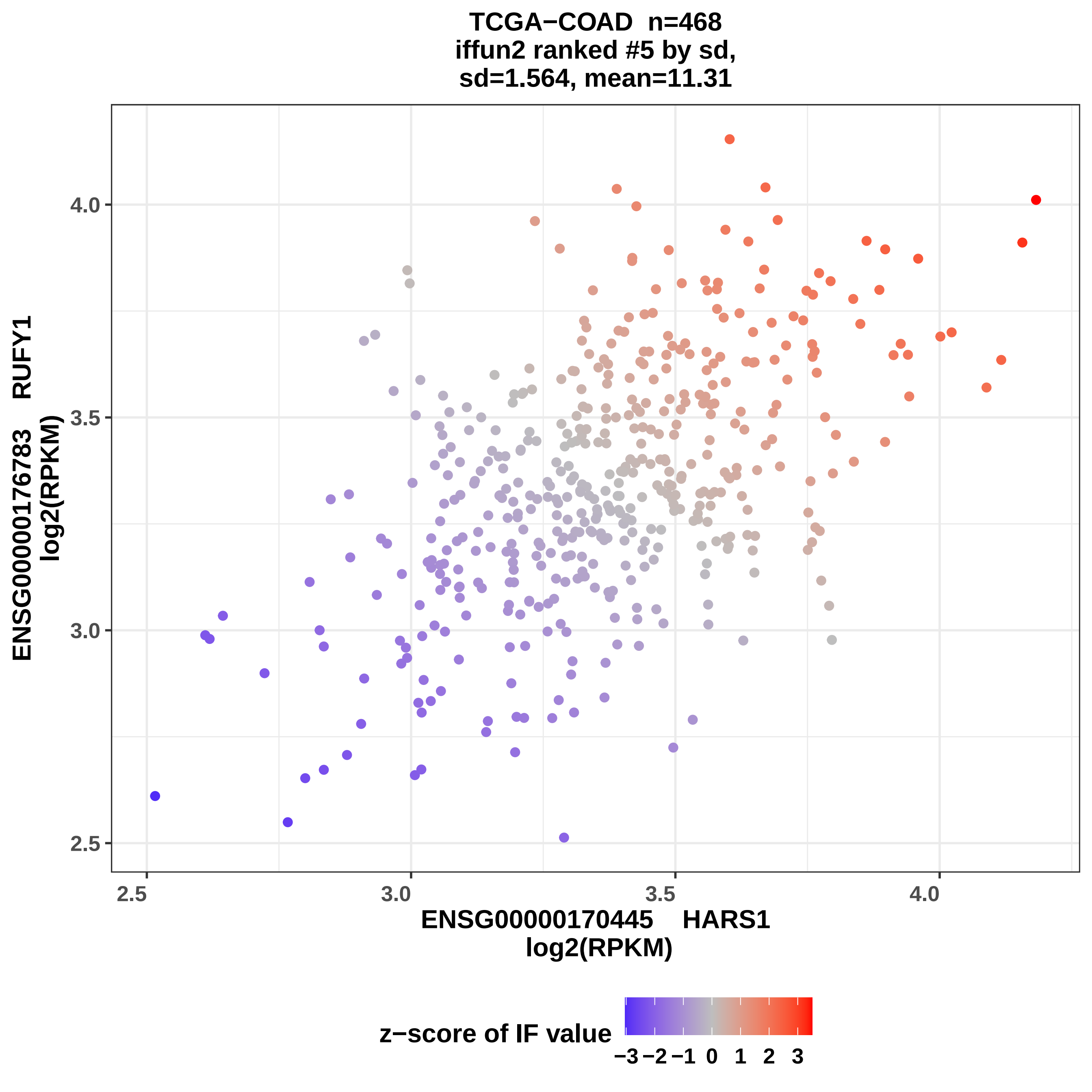
TCGA−COAD n=468
iffun2 ranked #5 by sd,
sd=1.564, mean=11.31
ENSG00000176783 RUFY1
log2(RPKM)
ENSG00000170445 HARS1
log2(RPKM)
4.0
3.5
3.0
2.5
2.5
3.0
3.5
4.0
z−score of IF value
−3 −2 −1 0 1 2 3

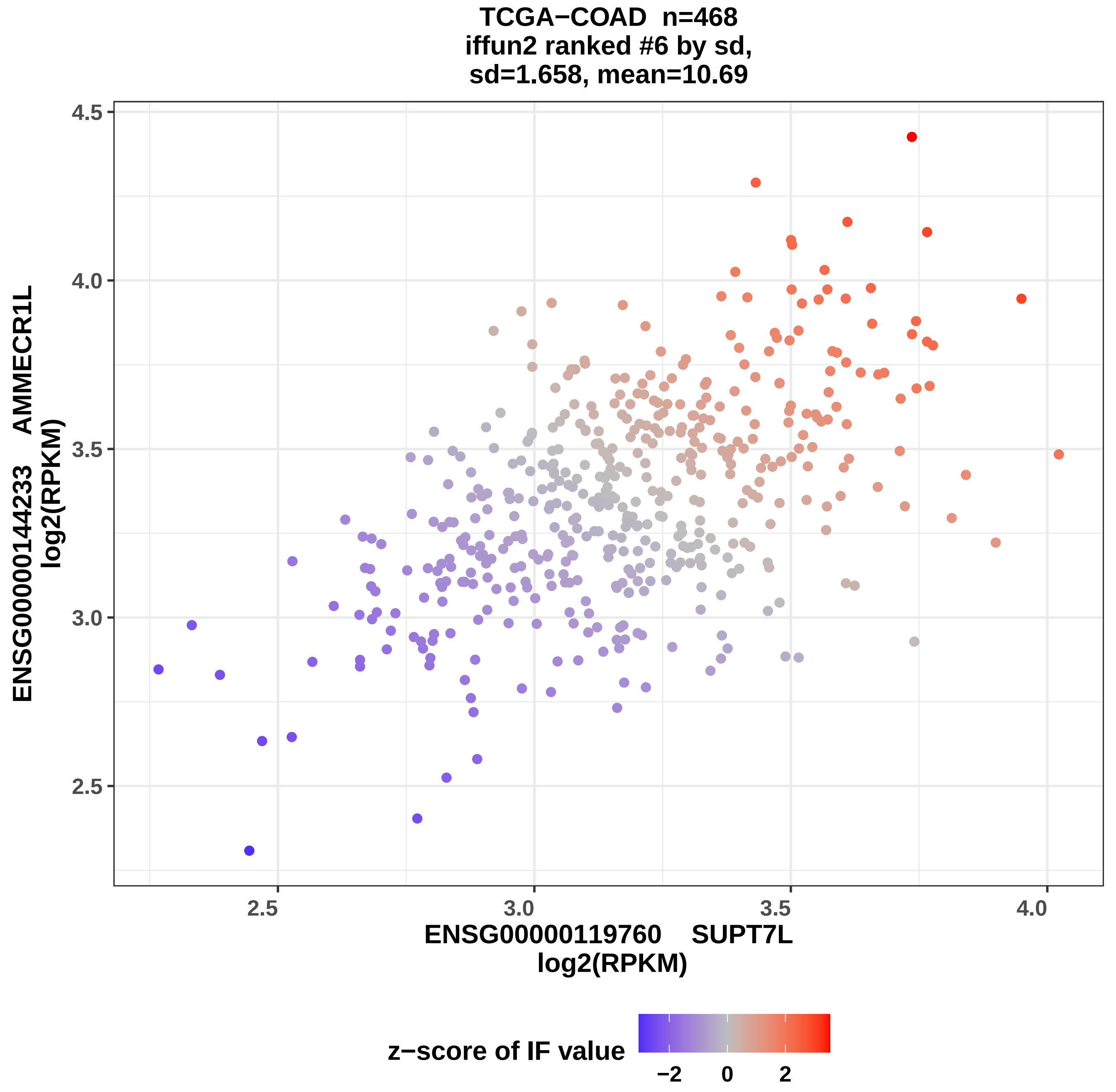
TCGA−COAD n=468
iffun2 ranked #6 by sd,
sd=1.658, mean=10.69
ENSG00000144233 AMMECR1L
log2(RPKM)
4.5
4.0
3.5
3.0
2.5
2.5
3.0
3.5
4.0
ENSG00000119760 SUPT7L
log2(RPKM)
z−score of IF value
−2
0
2

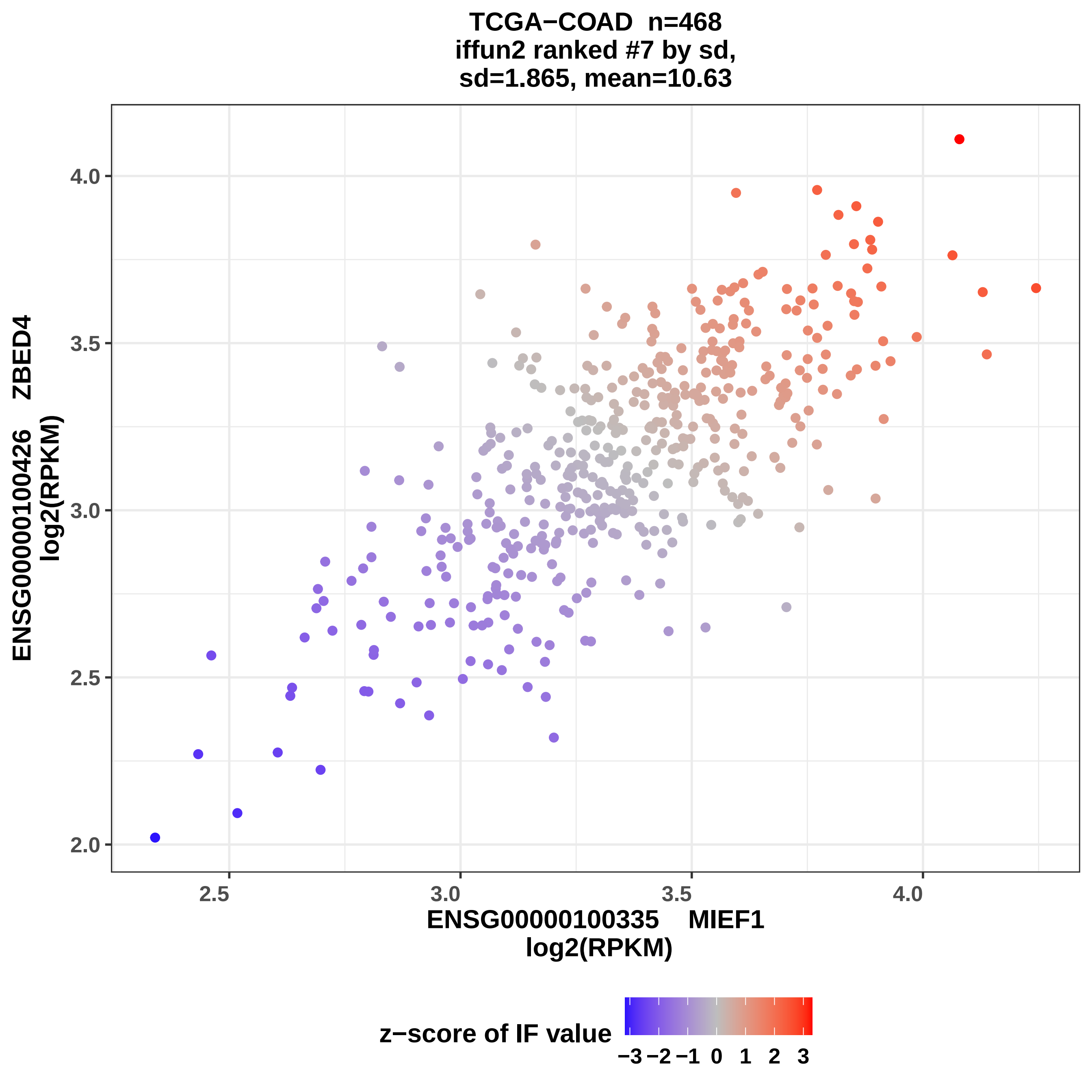
TCGA−COAD n=468
iffun2 ranked #7 by sd,
sd=1.865, mean=10.63
ENSG00000100426 ZBED4
log2(RPKM)
4.0
3.5
3.0
2.5
2.0
2.5
3.0
3.5
4.0
ENSG00000100335 MIEF1
log2(RPKM)
z−score of IF value
−3 −2 −1 0 1 2 3

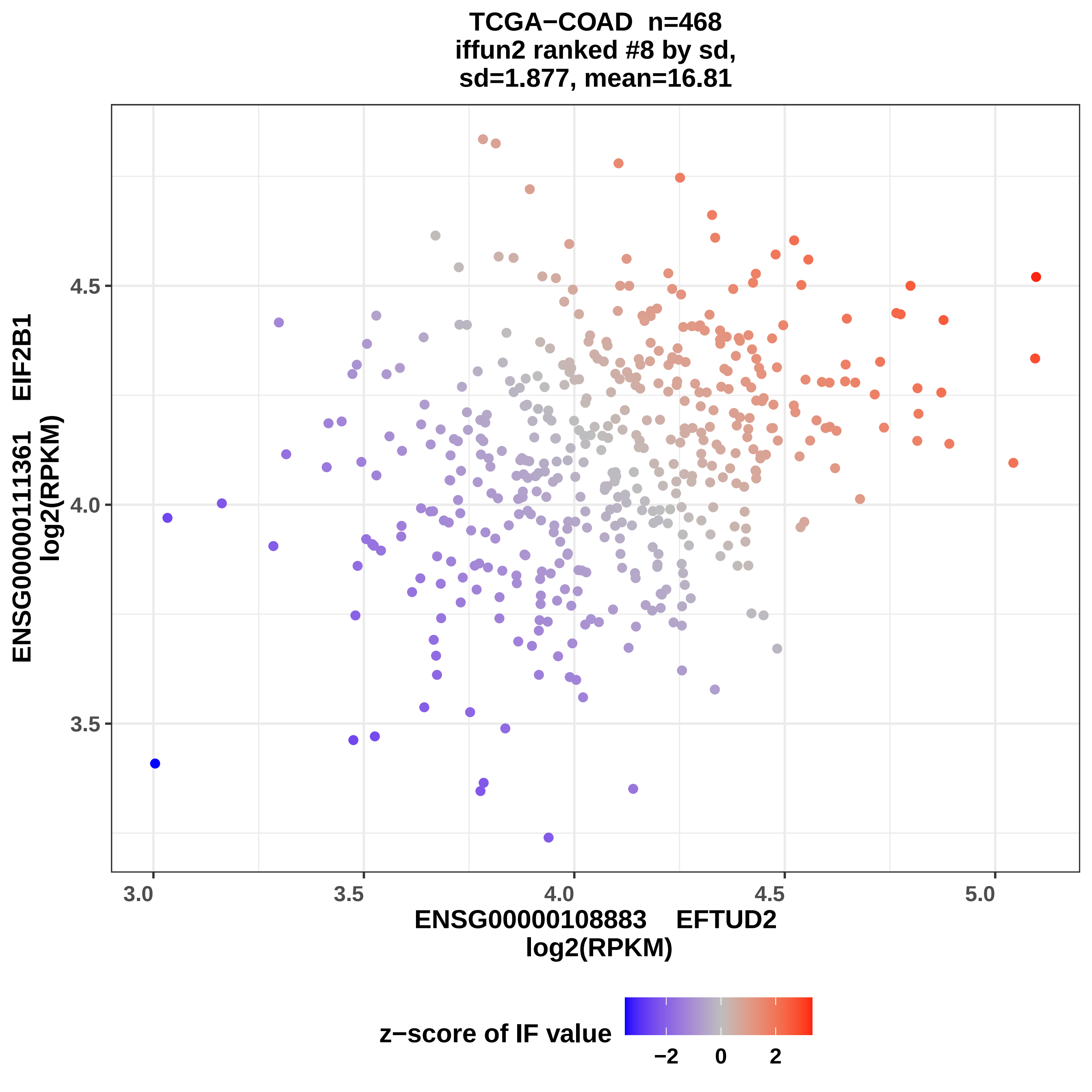
TCGA−COAD n=468
iffun2 ranked #8 by sd,
sd=1.877, mean=16.81
ENSG00000111361 EIF2B1
log2(RPKM)
4.5
4.0
3.5
3.0
3.5
4.0
4.5
5.0
ENSG00000108883 EFTUD2
log2(RPKM)
z−score of IF value
−2
0
2

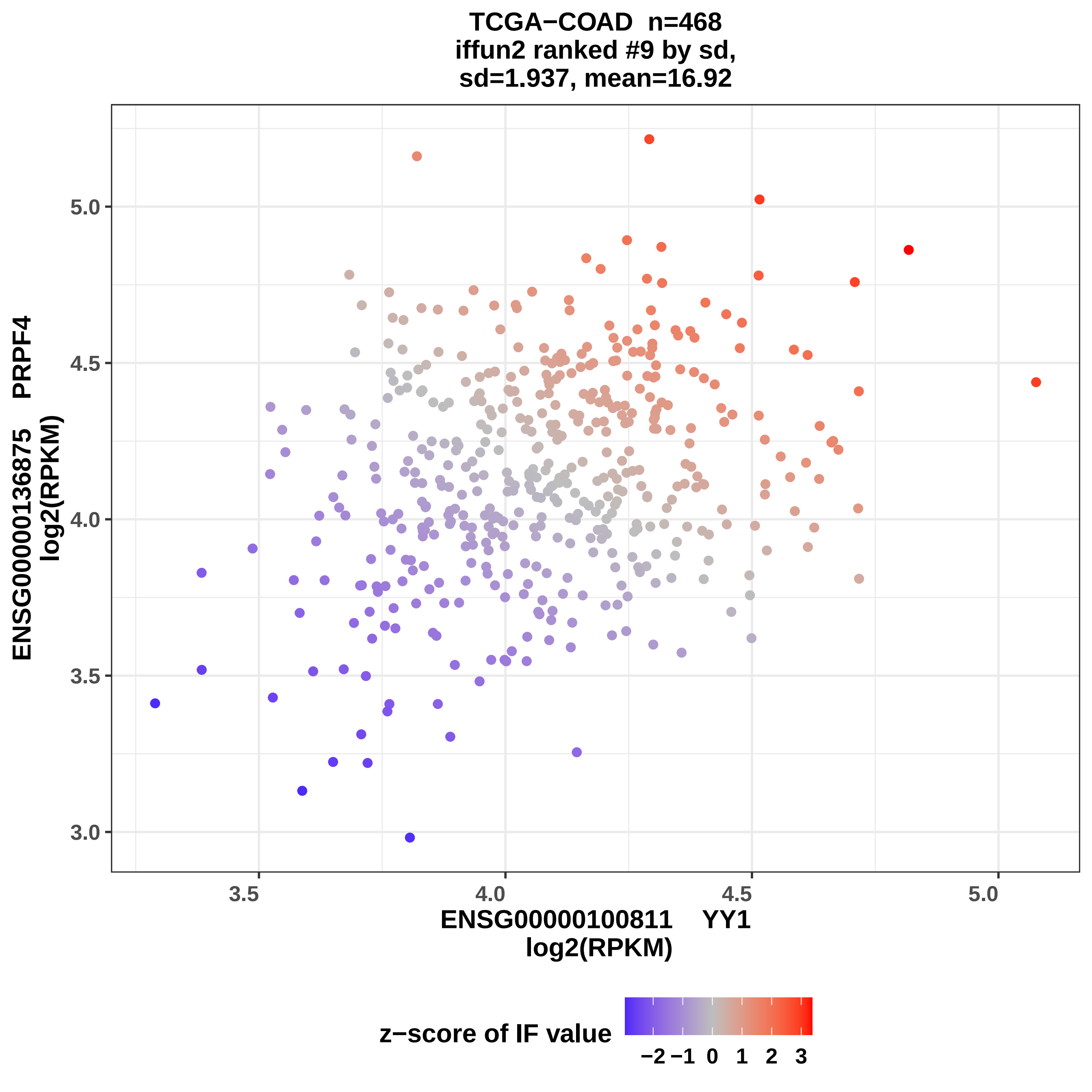
TCGA−COAD n=468
iffun2 ranked #9 by sd,
sd=1.937, mean=16.92
ENSG00000136875 PRPF4
log2(RPKM)
5.0
4.5
4.0
3.5
3.0
3.5
4.0
4.5
5.0
ENSG00000100811 YY1
log2(RPKM)
z−score of IF value
−2
−1
0
1
2
3

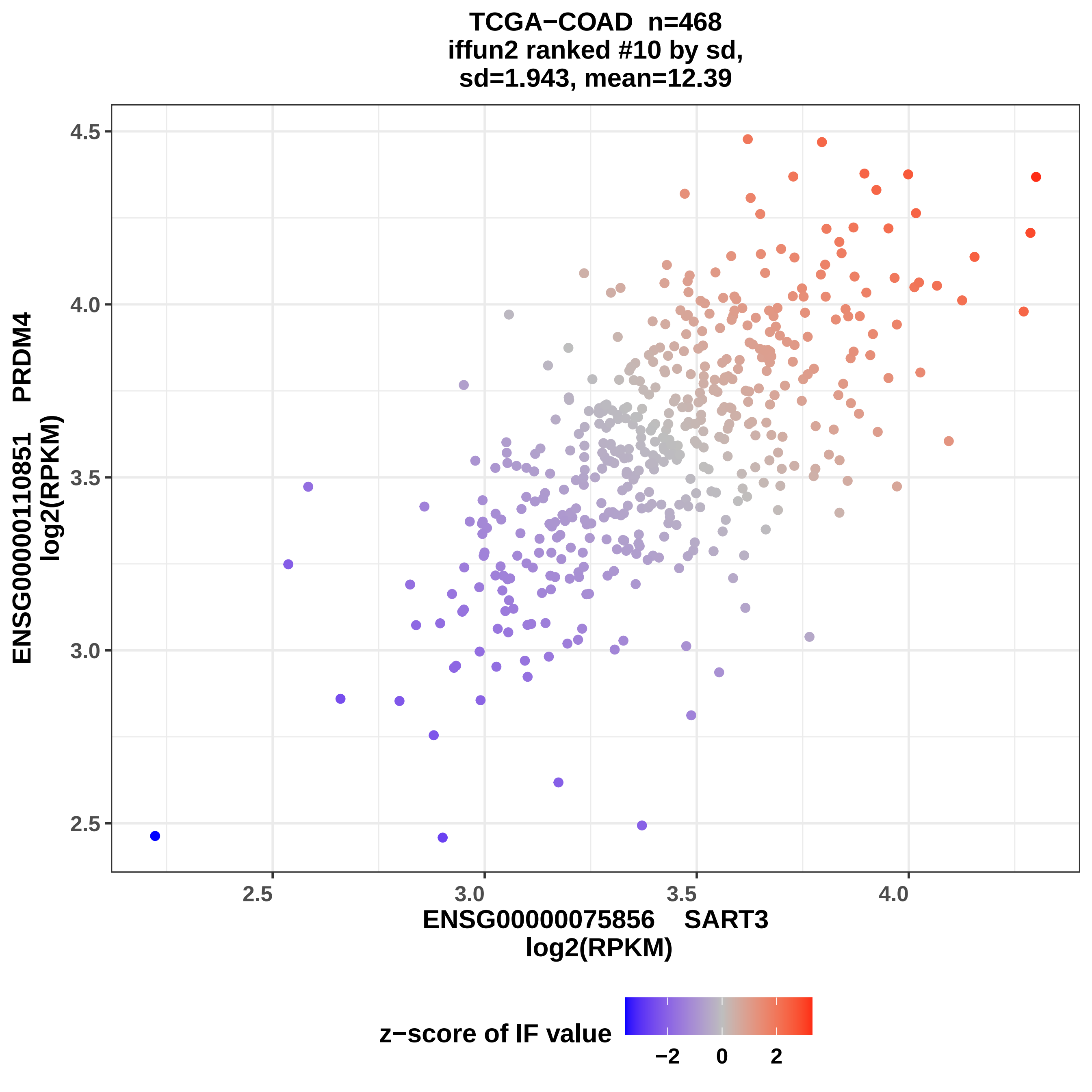
TCGA−COAD n=468
iffun2 ranked #10 by sd,
sd=1.943, mean=12.39
ENSG00000110851 PRDM4
log2(RPKM)
4.5
4.0
3.5
3.0
2.5
2.5
3.0
3.5
4.0
ENSG00000075856 SART3
log2(RPKM)
z−score of IF value
−2
0
2

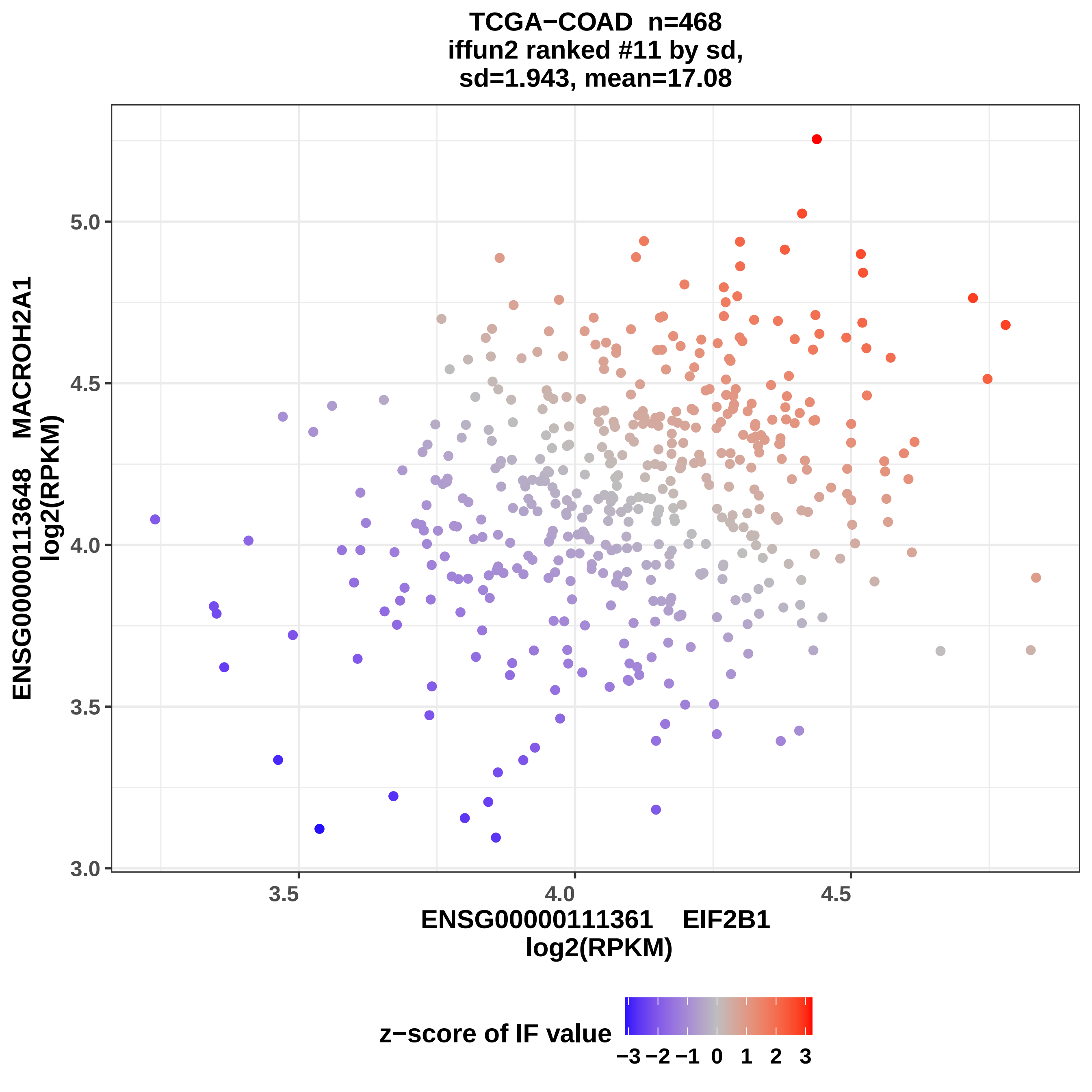
TCGA−COAD n=468
iffun2 ranked #11 by sd,
sd=1.943, mean=17.08
ENSG00000113648 MACROH2A1
log2(RPKM)
3.0
3.5
4.0
4.5
5.0
3.5
4.0
4.5
ENSG00000111361 EIF2B1
log2(RPKM)
z−score of IF value
−3 −2 −1 0 1 2 3

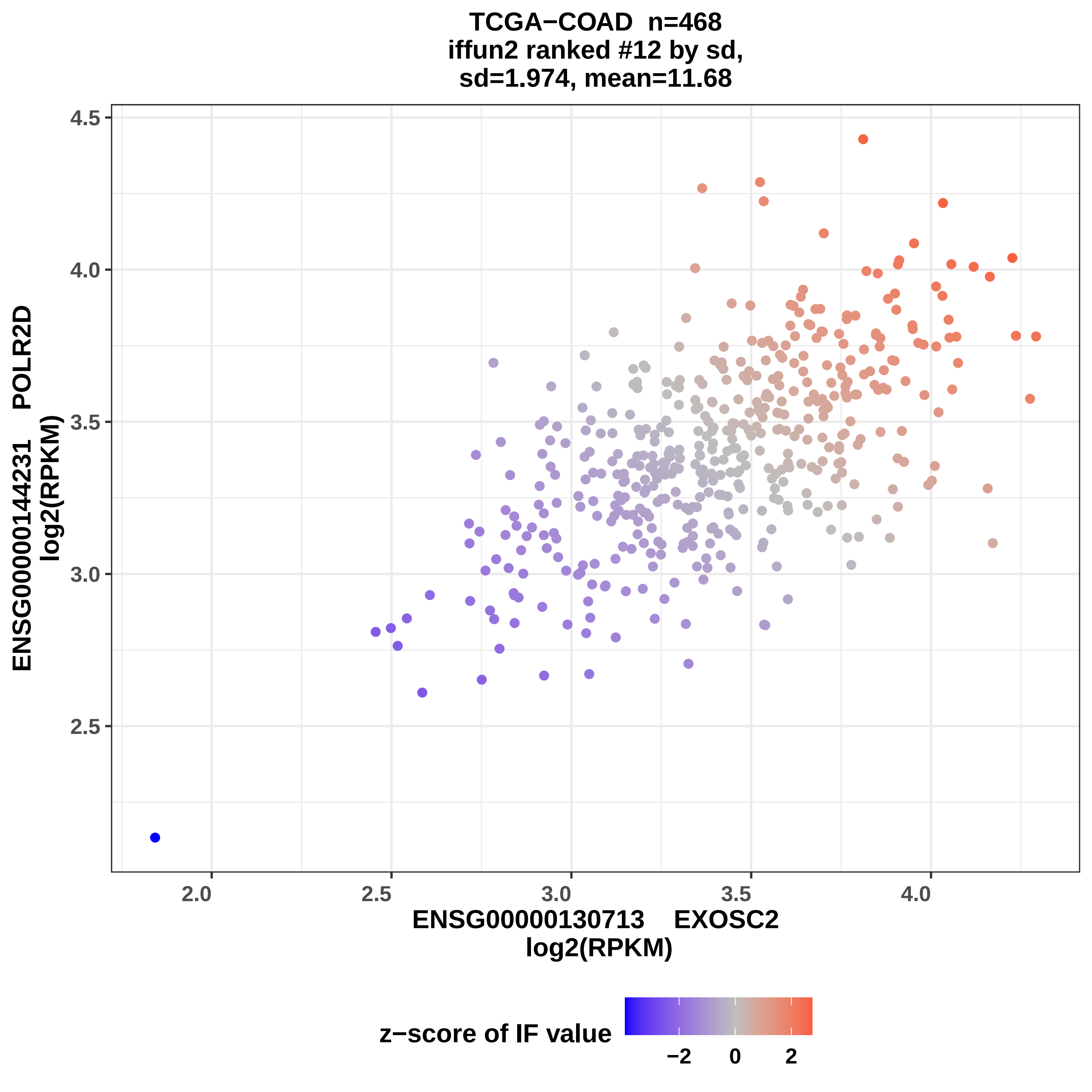

TCGA−COAD n=468
iffun2 ranked #12 by sd,
sd=1.974, mean=11.68
ENSG00000144231 POLR2D
log2(RPKM)
4.5
4.0
3.5
3.0
2.5
2.0
2.5
3.0
3.5
4.0
ENSG00000130713 EXOSC2
log2(RPKM)
z−score of IF value
−2
0
2

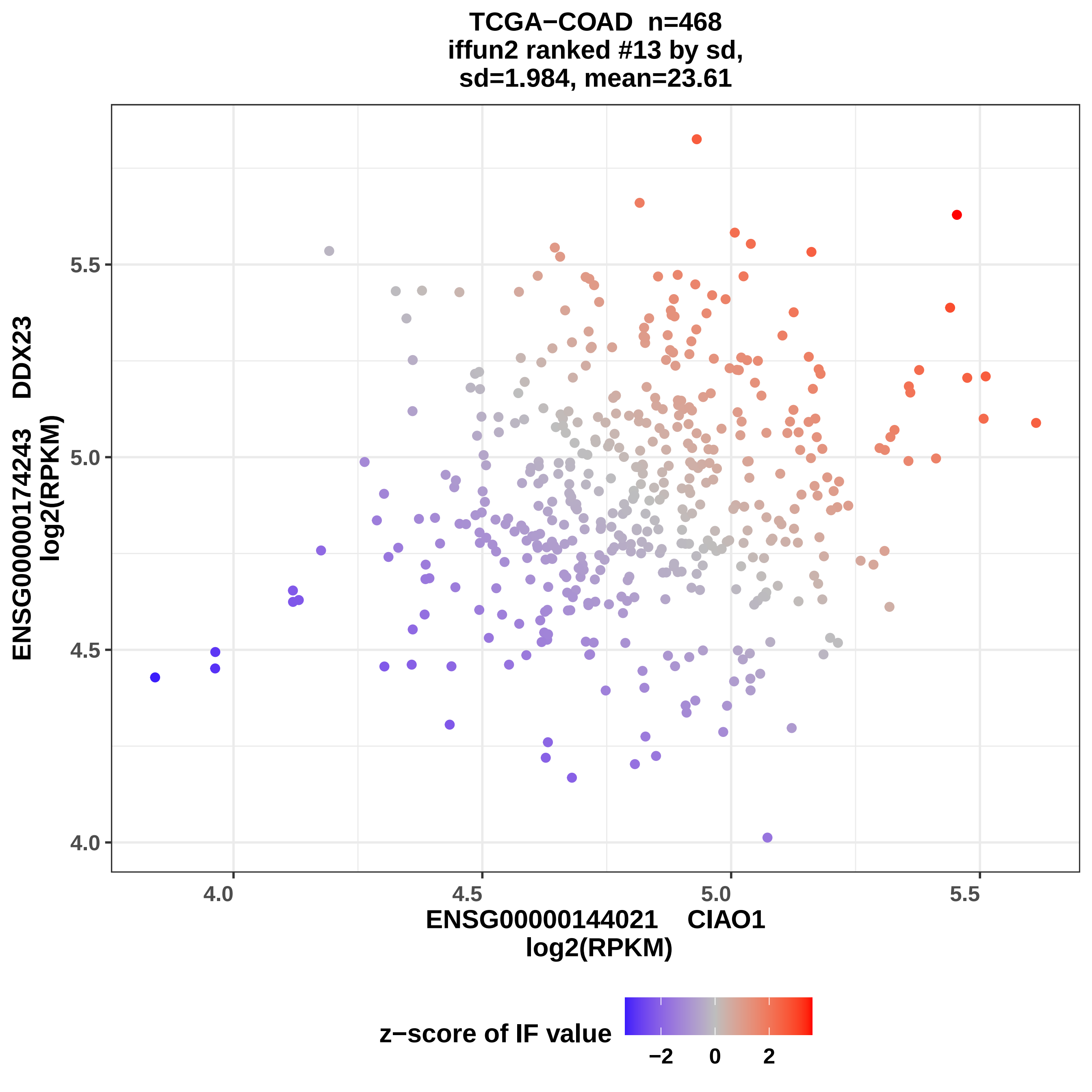

TCGA−COAD n=468
iffun2 ranked #13 by sd,
sd=1.984, mean=23.61
ENSG00000174243 DDX23
log2(RPKM)
4.0
4.5
5.0
5.5
ENSG00000144021 CIAO1
log2(RPKM)
z−score of IF value
−2
0
2

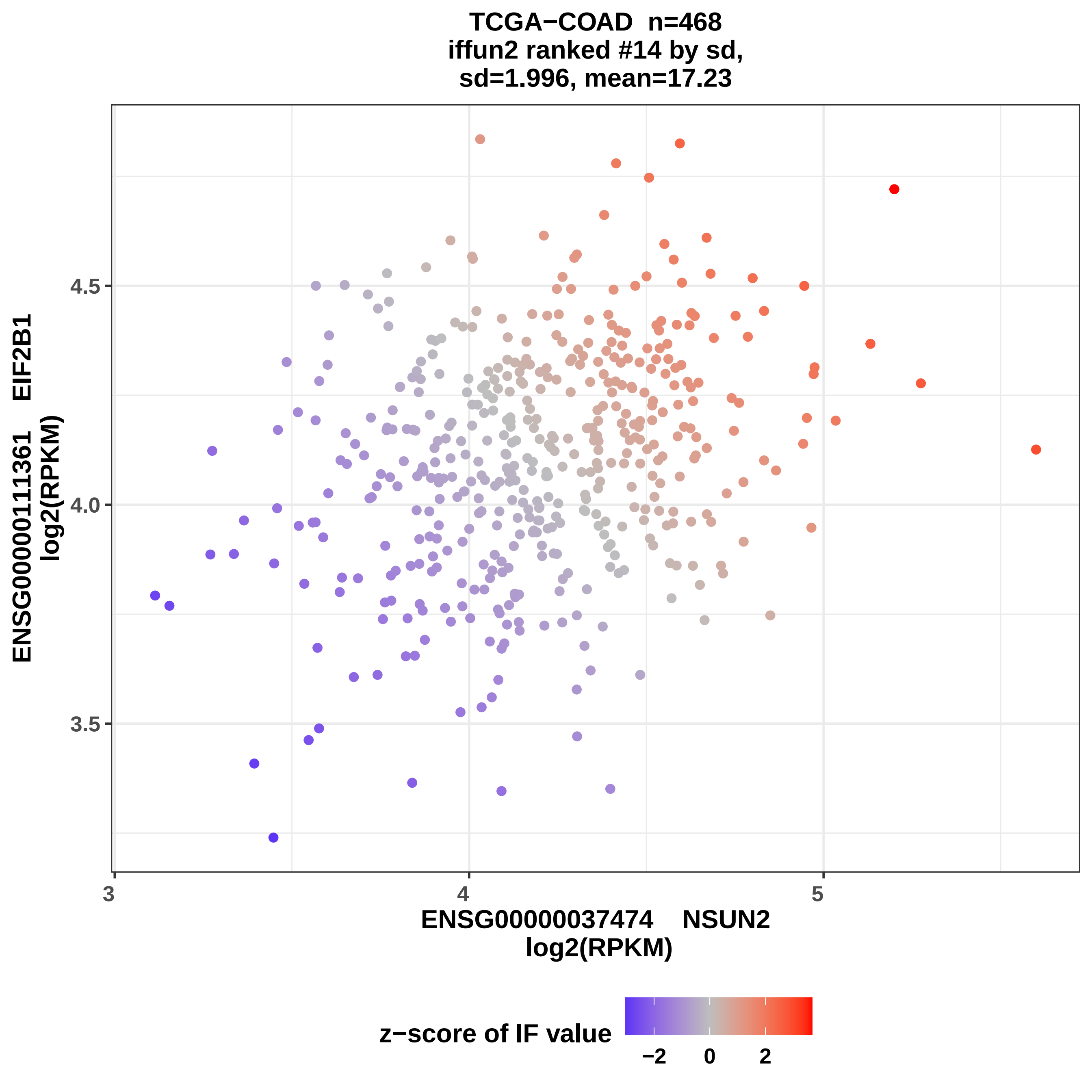
TCGA−COAD n=468
iffun2 ranked #14 by sd,
sd=1.996, mean=17.23
ENSG00000111361 EIF2B1
log2(RPKM)
4.5
4.0
3.5
3
4
5
ENSG00000037474 NSUN2
log2(RPKM)
z−score of IF value
−2
0
2

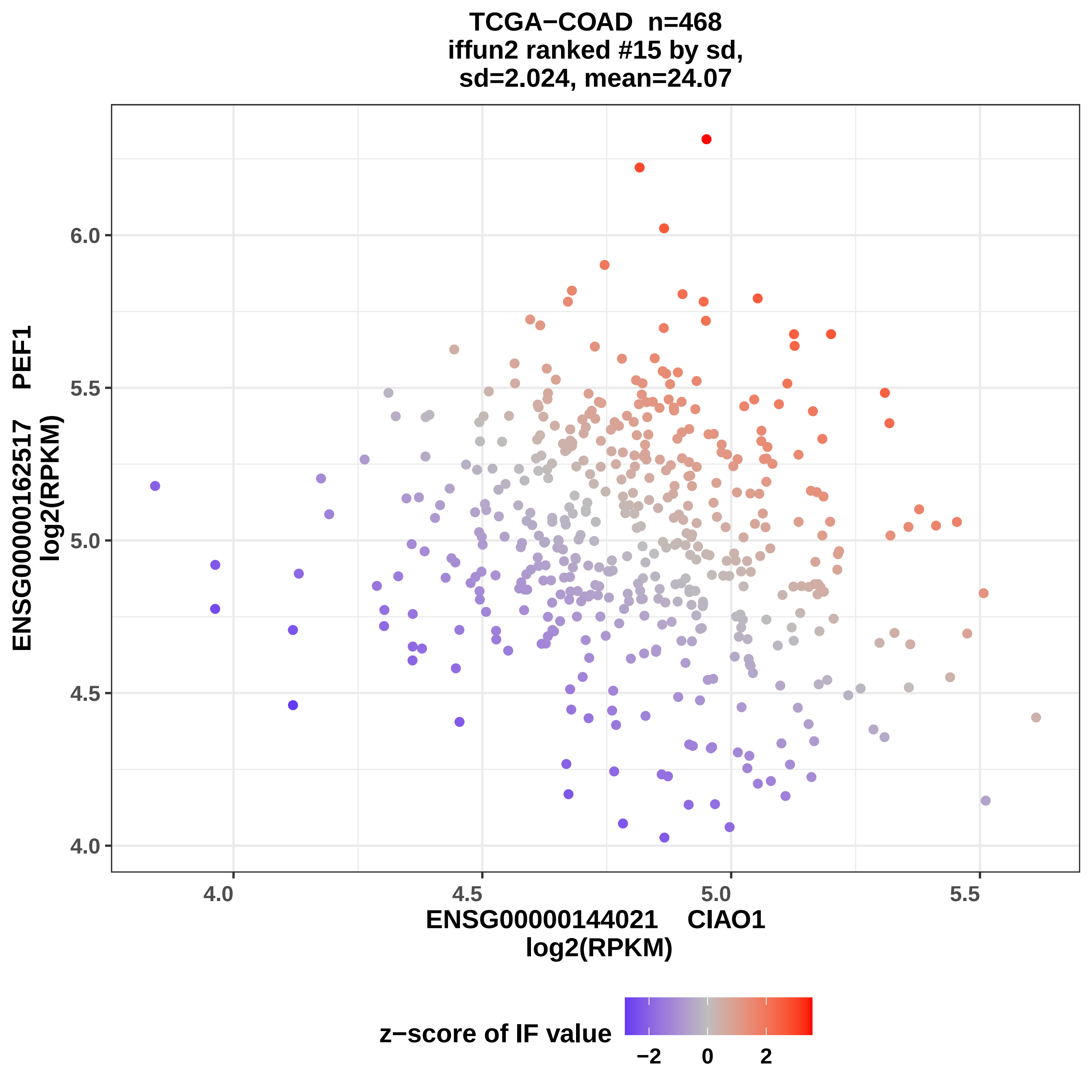
TCGA−COAD n=468
iffun2 ranked #15 by sd,
sd=2.024, mean=24.07
ENSG00000162517 PEF1
log2(RPKM)
6.0
5.5
5.0
4.5
4.0
4.0
4.5
5.0
5.5
ENSG00000144021 CIAO1
log2(RPKM)
z−score of IF value
−2
0
2

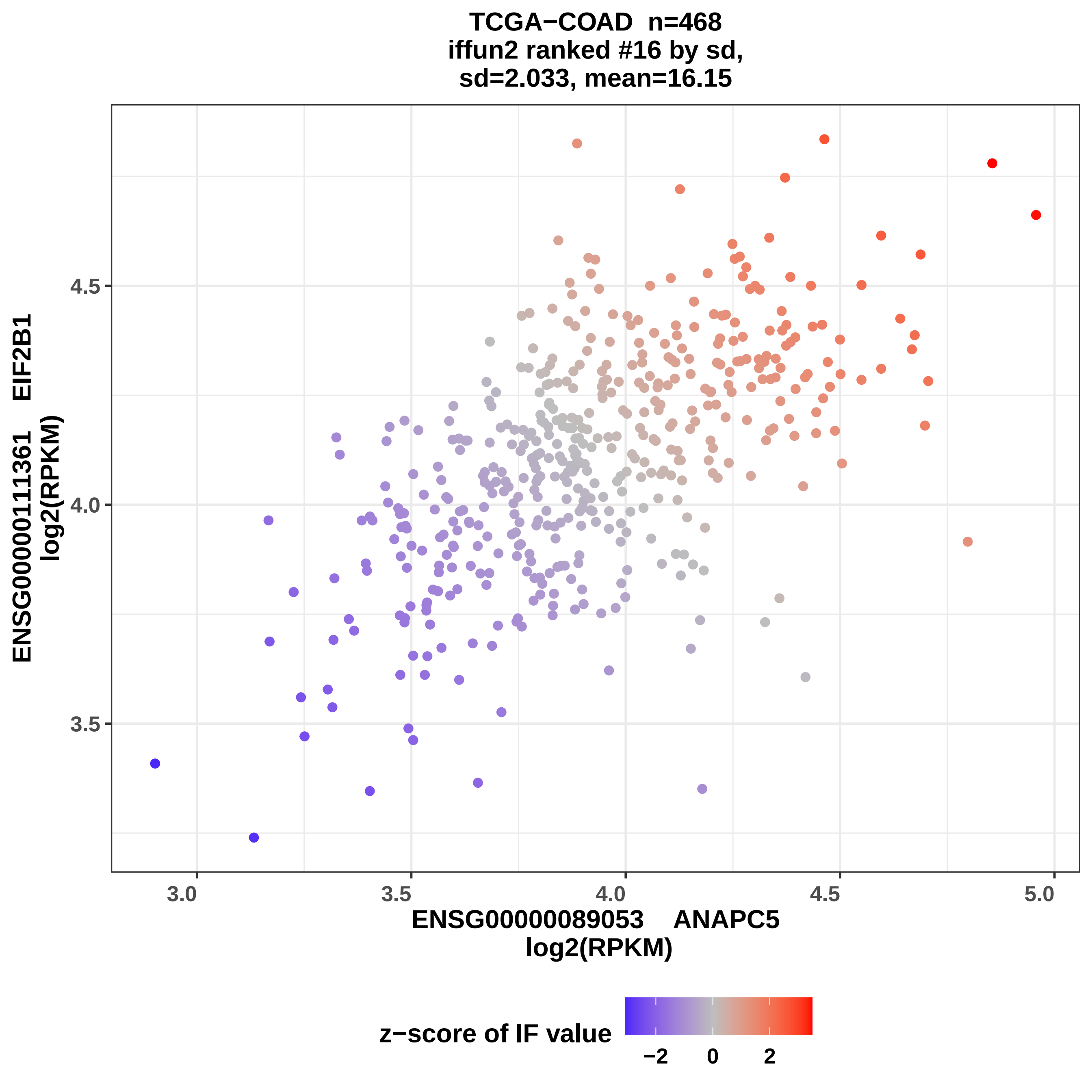
TCGA−COAD n=468
iffun2 ranked #16 by sd,
sd=2.033, mean=16.15
ENSG00000111361 EIF2B1
log2(RPKM)
4.5
4.0
3.5
3.0
3.5
4.0
4.5
5.0
ENSG00000089053 ANAPC5
log2(RPKM)
z−score of IF value
−2
0
2

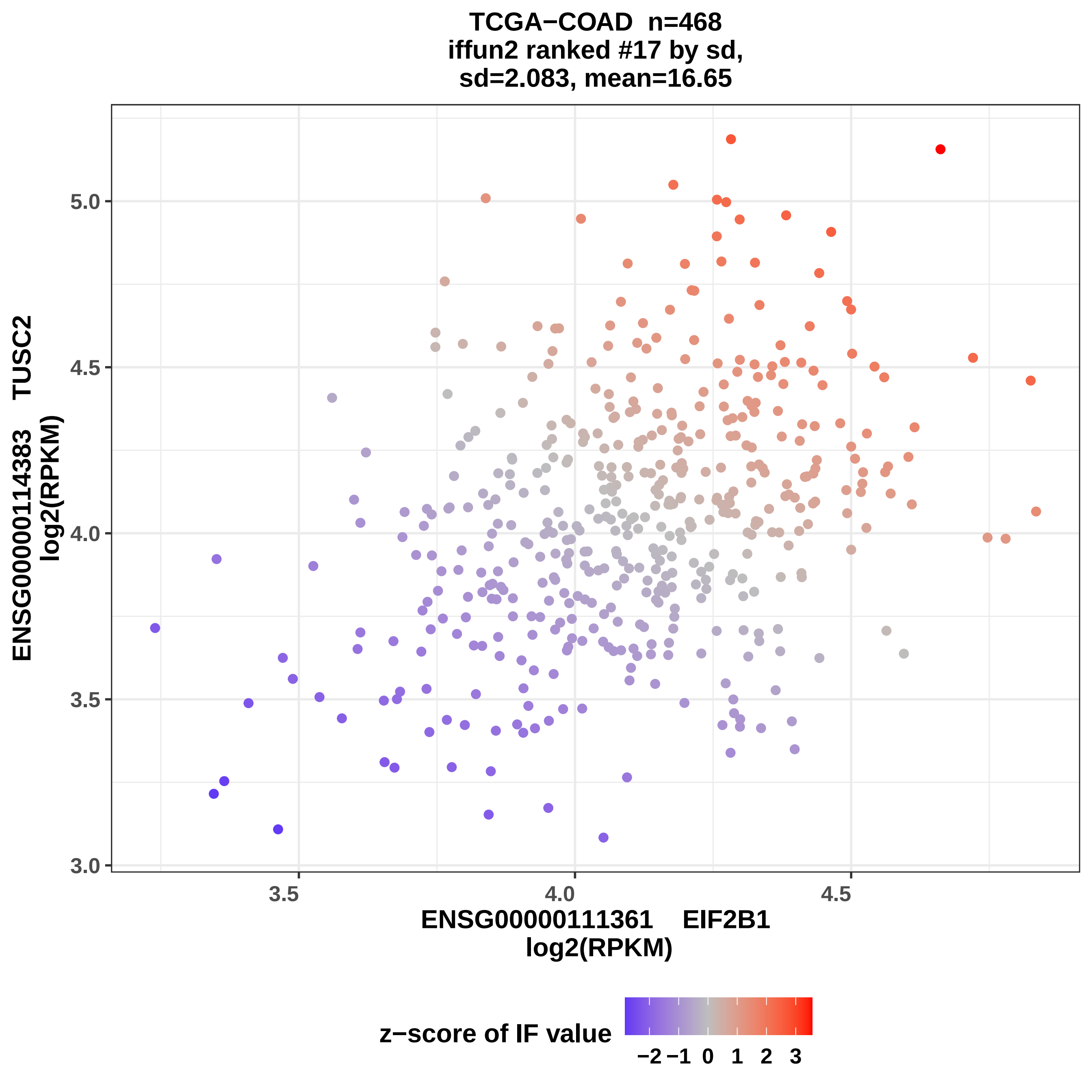

TCGA−COAD n=468
iffun2 ranked #17 by sd,
sd=2.083, mean=16.65
ENSG00000114383 TUSC2
log2(RPKM)
5.0
4.5
4.0
3.5
3.0
3.5
4.0
4.5
ENSG00000111361 EIF2B1
log2(RPKM)
z−score of IF value
−2
−1
0
1
2
3

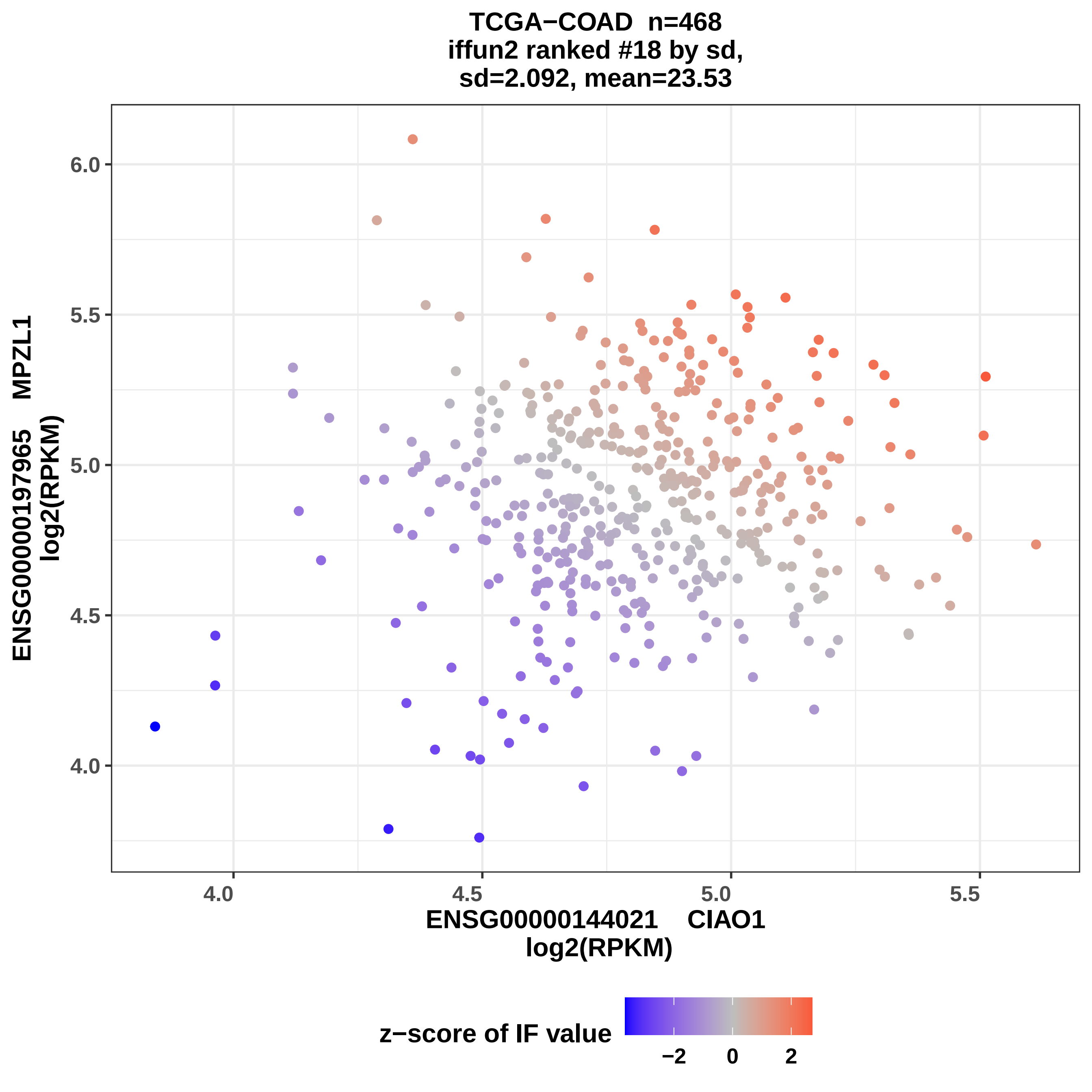

TCGA−COAD n=468
iffun2 ranked #18 by sd,
sd=2.092, mean=23.53
ENSG00000197965 MPZL1
log2(RPKM)
6.0
5.5
5.0
4.5
4.0
4.0
4.5
5.0
5.5
ENSG00000144021 CIAO1
log2(RPKM)
z−score of IF value
−2
0
2

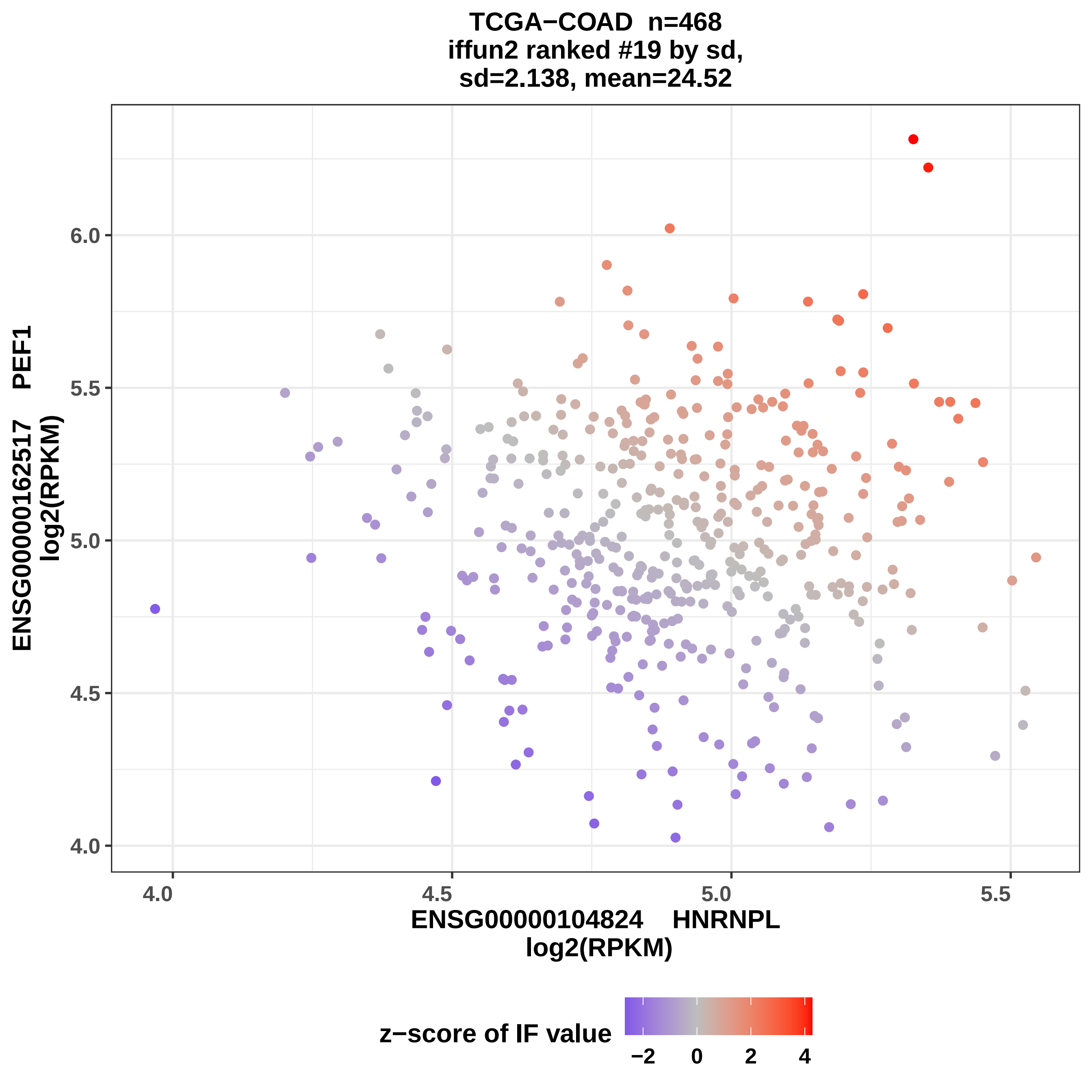
TCGA−COAD n=468
iffun2 ranked #19 by sd,
sd=2.138, mean=24.52
ENSG00000162517 PEF1
log2(RPKM)
ENSG00000104824 HNRNPL
log2(RPKM)
4.0
4.5
5.0
5.5
6.0
z−score of IF value
−2
0
2
4

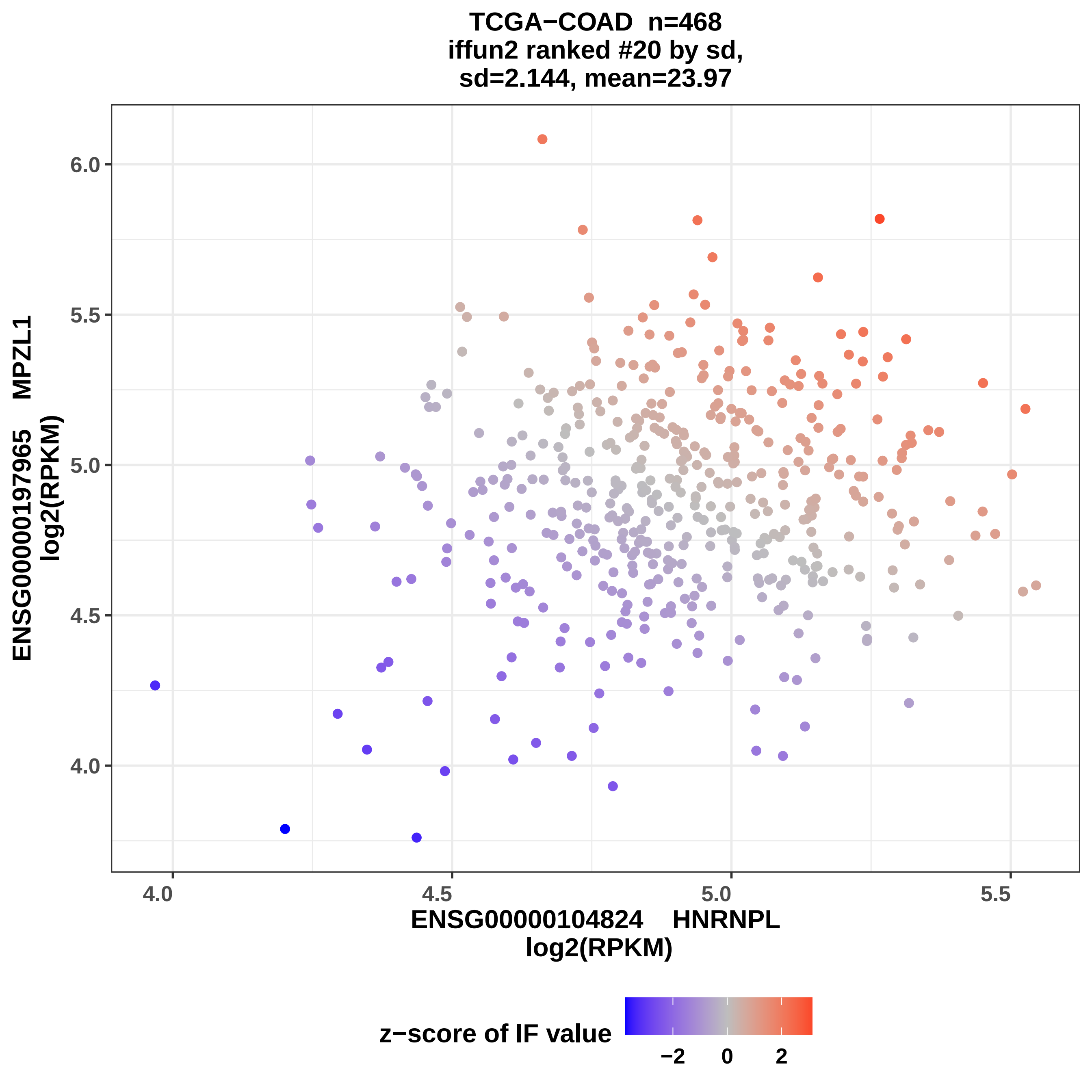

TCGA−COAD n=468
iffun2 ranked #20 by sd,
sd=2.144, mean=23.97
ENSG00000197965 MPZL1
log2(RPKM)
6.0
5.5
5.0
4.5
4.0
4.0
4.5
5.0
5.5
ENSG00000104824 HNRNPL
log2(RPKM)
z−score of IF value
−2
0
2

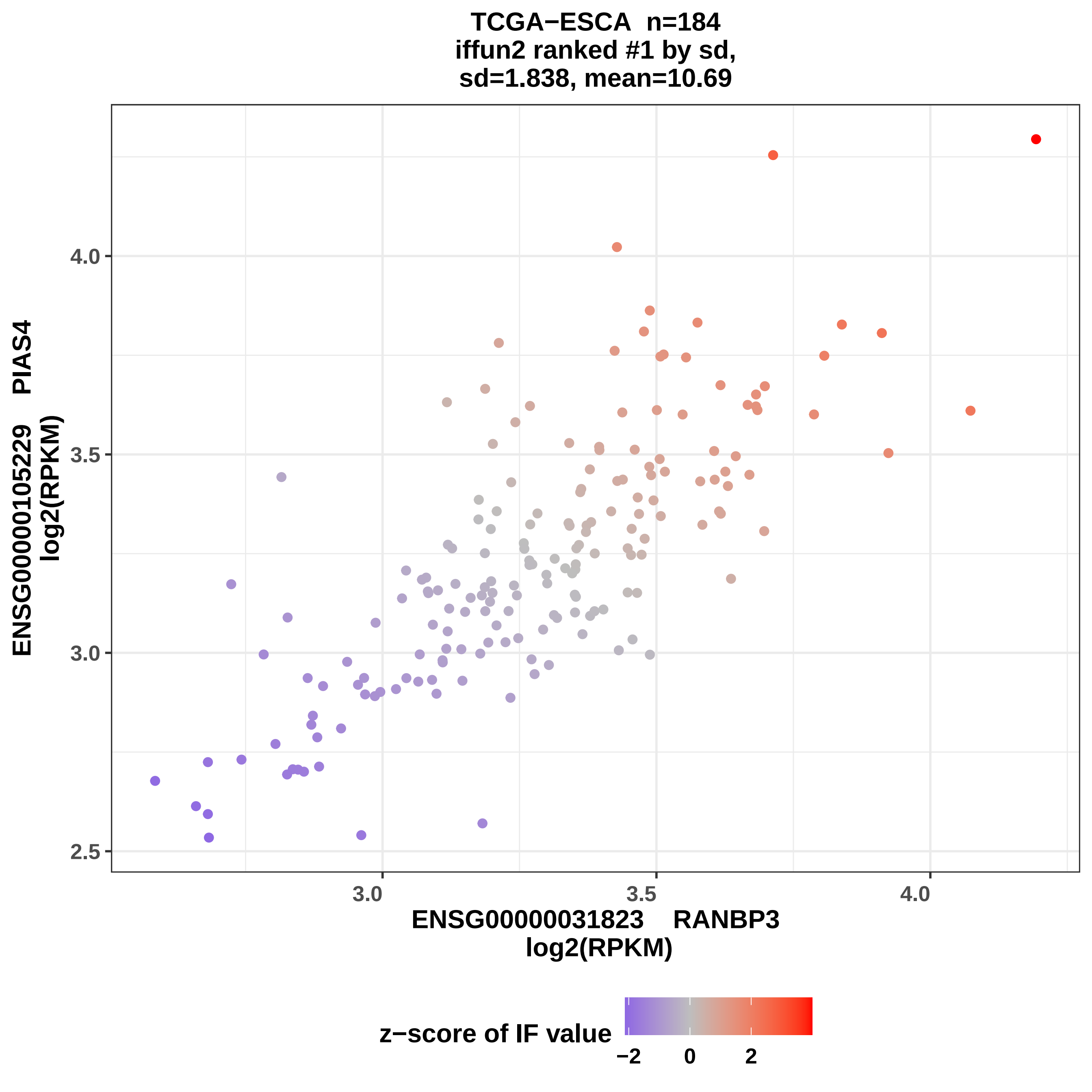
TCGA−ESCA n=184
iffun2 ranked #1 by sd,
sd=1.838, mean=10.69
ENSG00000105229 PIAS4
log2(RPKM)
4.0
3.5
3.0
2.5
3.0
3.5
4.0
ENSG00000031823 RANBP3
log2(RPKM)
z−score of IF value
−2
0
2

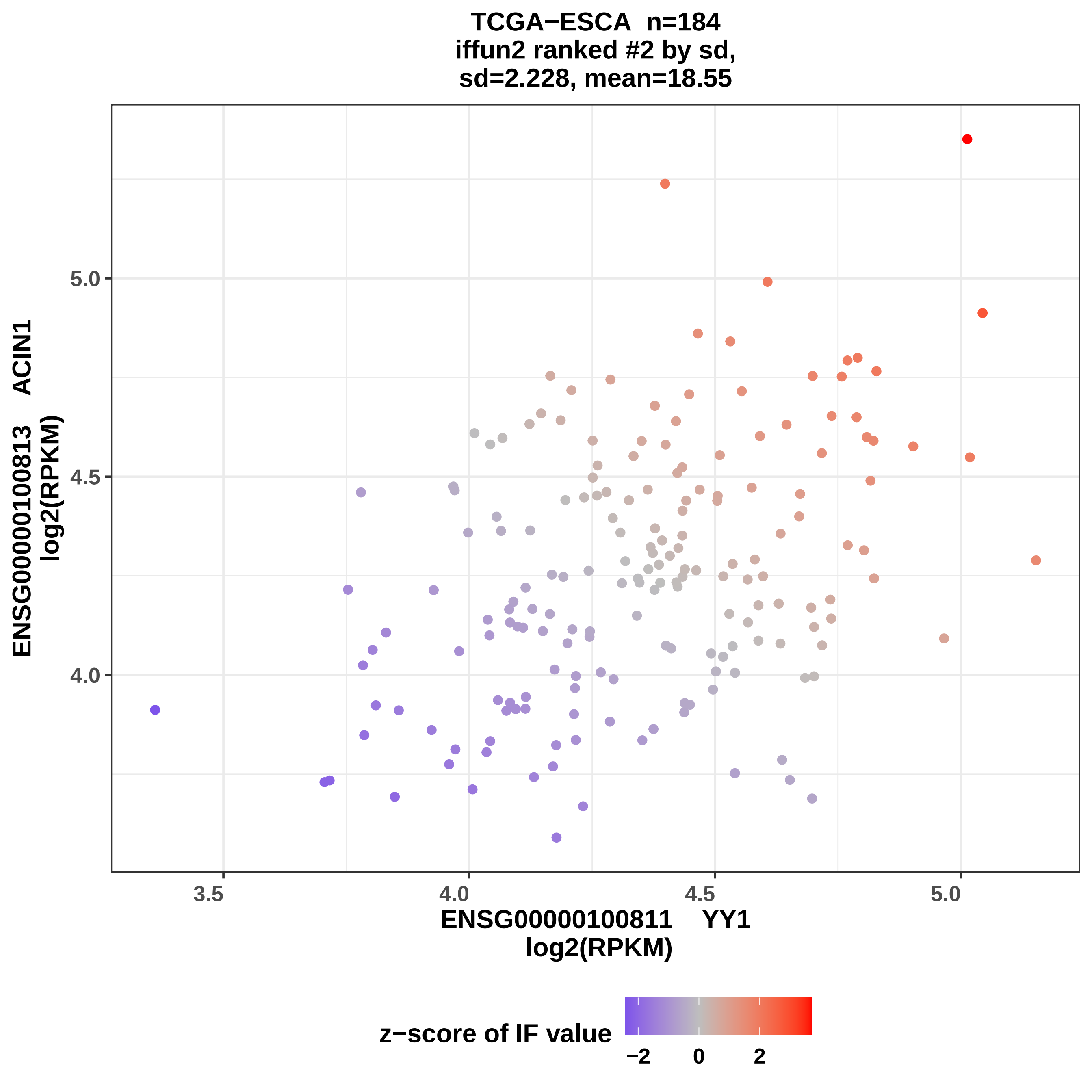

TCGA−ESCA n=184
iffun2 ranked #2 by sd,
sd=2.228, mean=18.55
ENSG00000100813 ACIN1
log2(RPKM)
5.0
4.5
4.0
3.5
4.0
4.5
5.0
ENSG00000100811 YY1
log2(RPKM)
z−score of IF value
−2
0
2

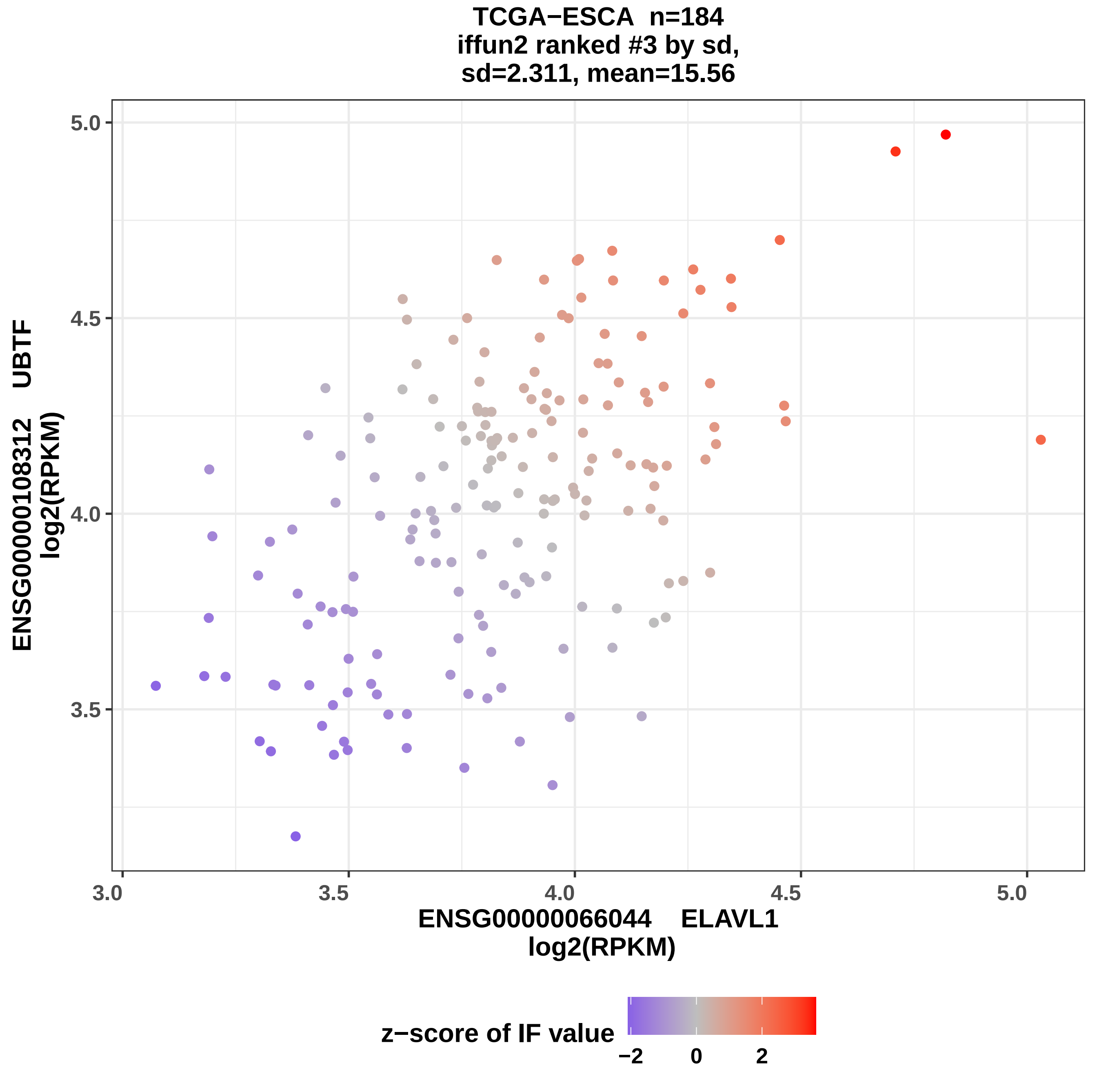

TCGA−ESCA n=184
iffun2 ranked #3 by sd,
sd=2.311, mean=15.56
ENSG00000108312 UBTF
log2(RPKM)
5.0
4.5
4.0
3.5
3.0
3.5
4.0
4.5
5.0
ENSG00000066044 ELAVL1
log2(RPKM)
z−score of IF value
−2
0
2

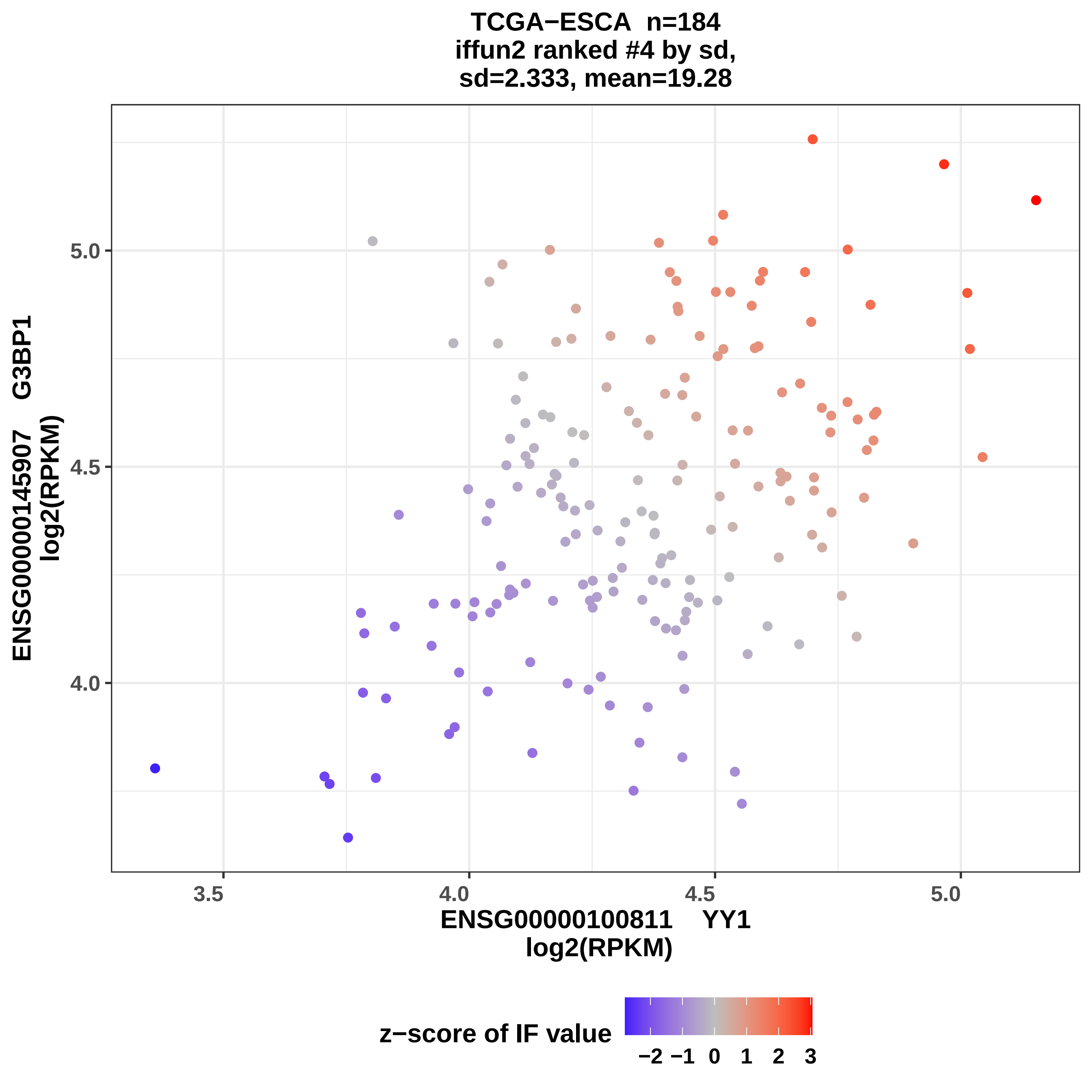
TCGA−ESCA n=184
iffun2 ranked #4 by sd,
sd=2.333, mean=19.28
ENSG00000145907 G3BP1
log2(RPKM)
ENSG00000100811 YY1
log2(RPKM)
3.5
4.0
4.5
5.0
4.0
4.5
5.0
z−score of IF value
−2 −1 0 1 2 3

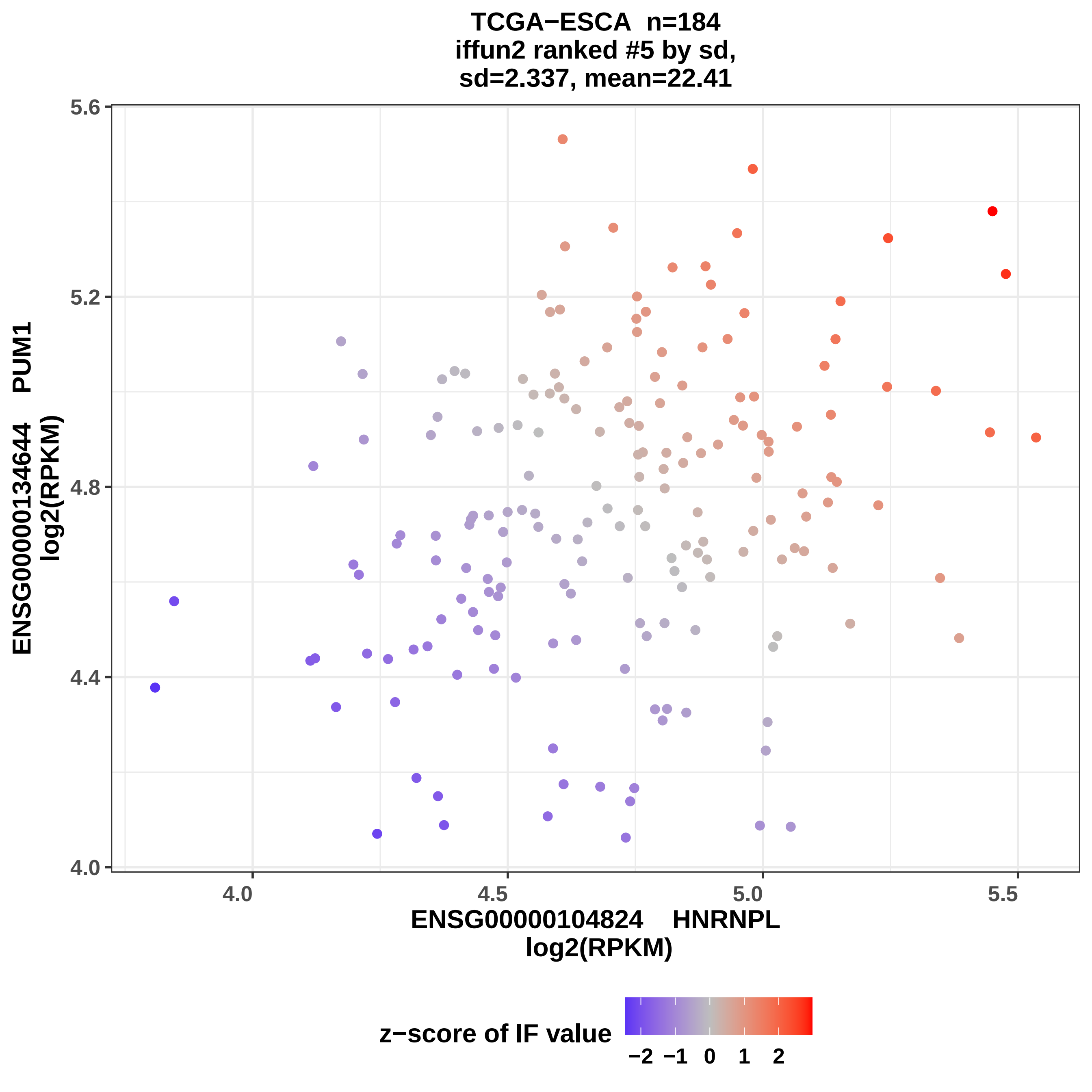
TCGA−ESCA n=184
iffun2 ranked #5 by sd,
sd=2.337, mean=22.41
5.6
5.2
4.8
4.4
4.0
4.0
4.5
5.0
5.5
ENSG00000134644 PUM1
log2(RPKM)
ENSG00000104824 HNRNPL
log2(RPKM)
z−score of IF value
−2
−1
0
1
2

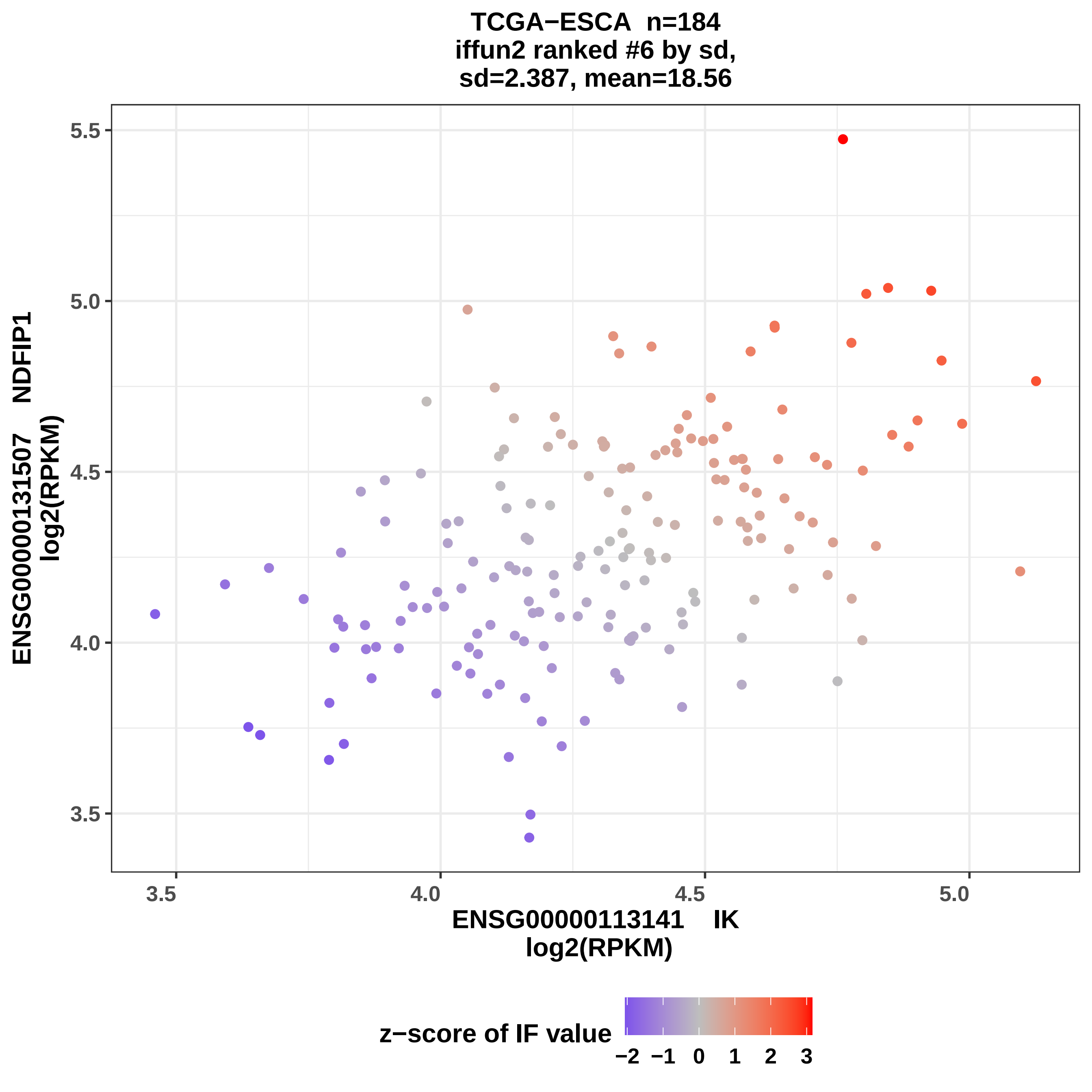

TCGA−ESCA n=184
iffun2 ranked #6 by sd,
sd=2.387, mean=18.56
ENSG00000131507 NDFIP1
log2(RPKM)
5.5
5.0
4.5
4.0
3.5
3.5
4.0
4.5
5.0
ENSG00000113141 IK
log2(RPKM)
z−score of IF value
−2
−1
0
1
2
3

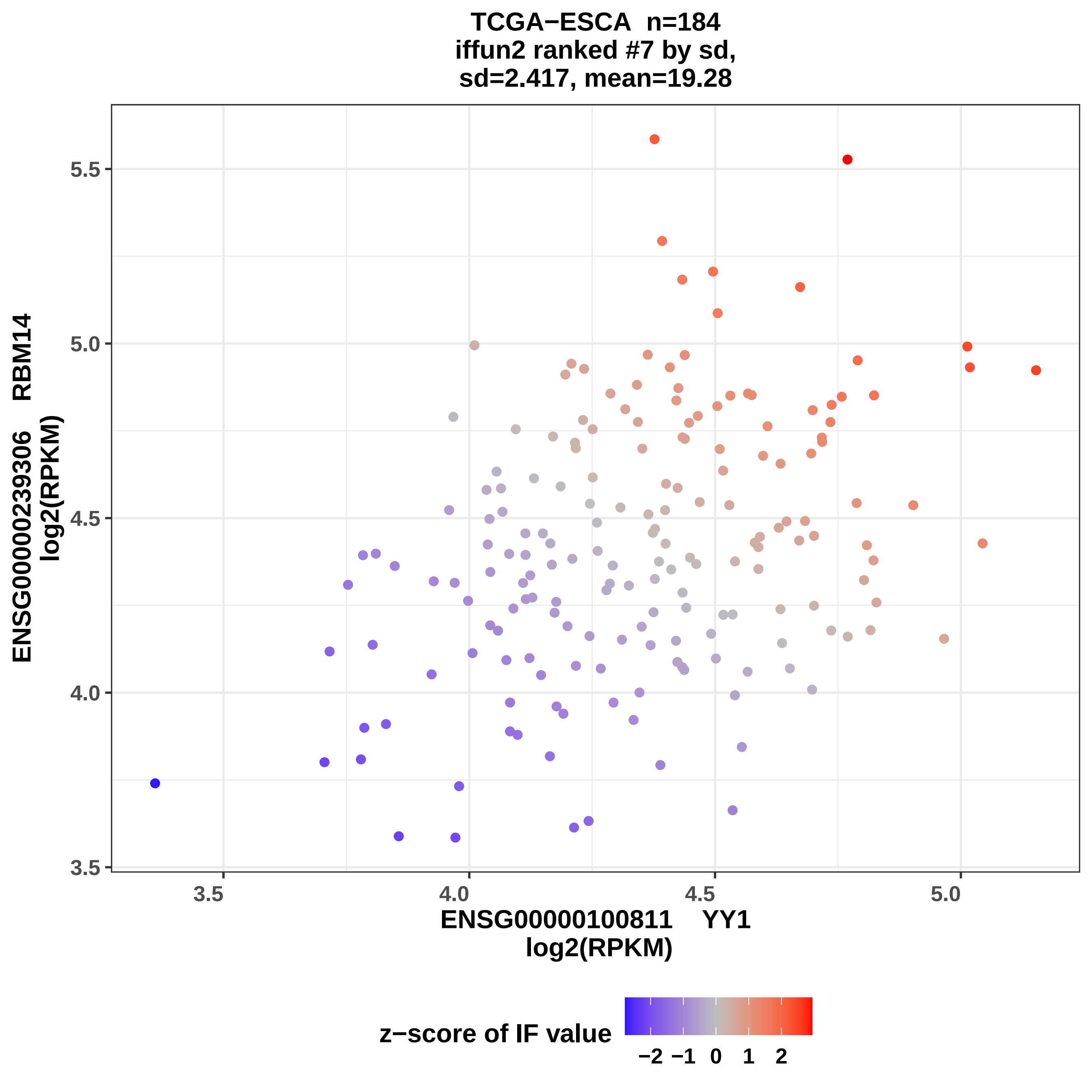
TCGA−ESCA n=184
iffun2 ranked #7 by sd,
sd=2.417, mean=19.28
ENSG00000239306 RBM14
log2(RPKM)
5.5
5.0
4.5
4.0
3.5
3.5
4.0
4.5
5.0
ENSG00000100811 YY1
log2(RPKM)
z−score of IF value
−2 −1 0 1 2

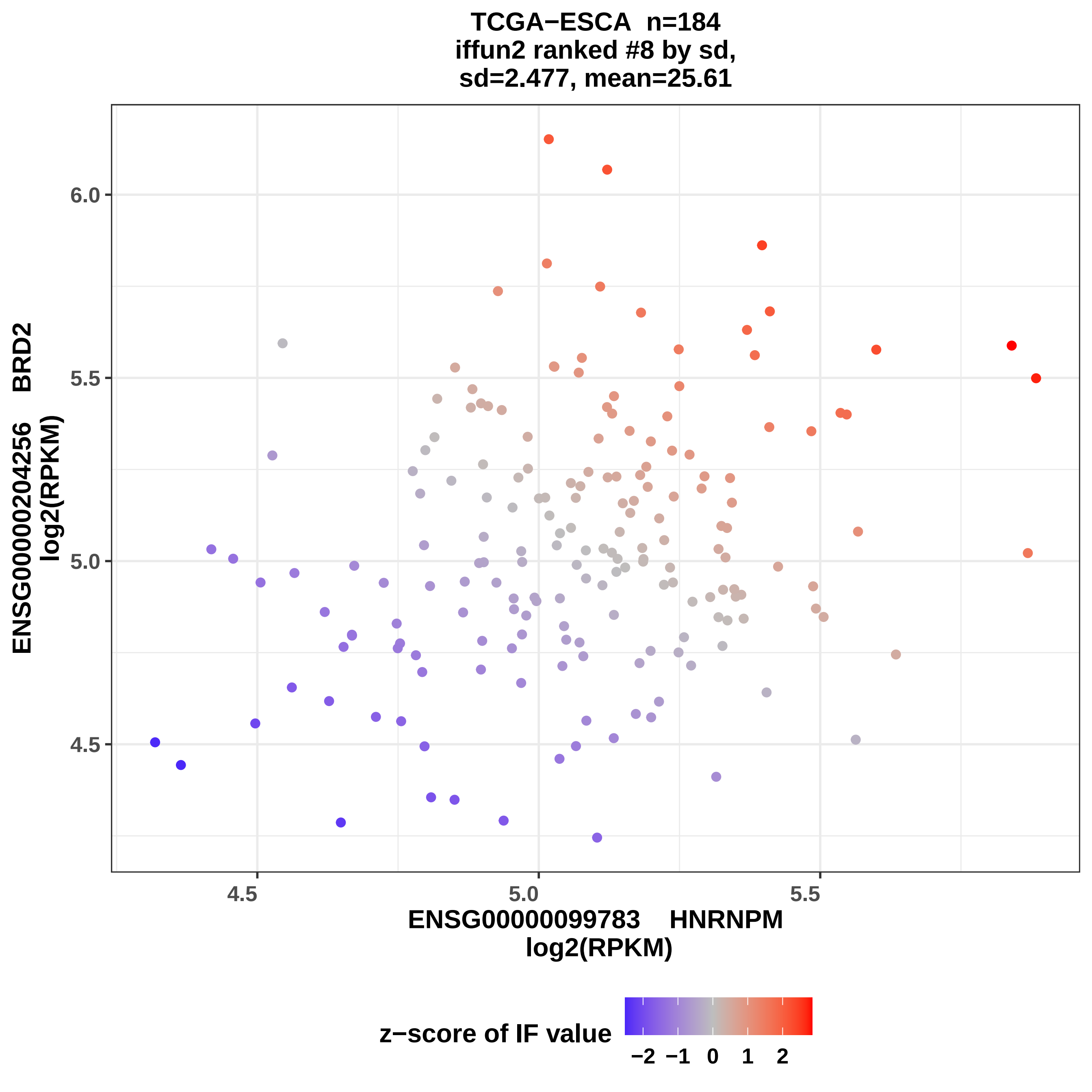

TCGA−ESCA n=184
iffun2 ranked #8 by sd,
sd=2.477, mean=25.61
6.0
5.5
5.0
4.5
4.5
5.0
5.5
ENSG00000204256 BRD2
log2(RPKM)
ENSG00000099783 HNRNPM
log2(RPKM)
z−score of IF value
−2 −1 0 1 2

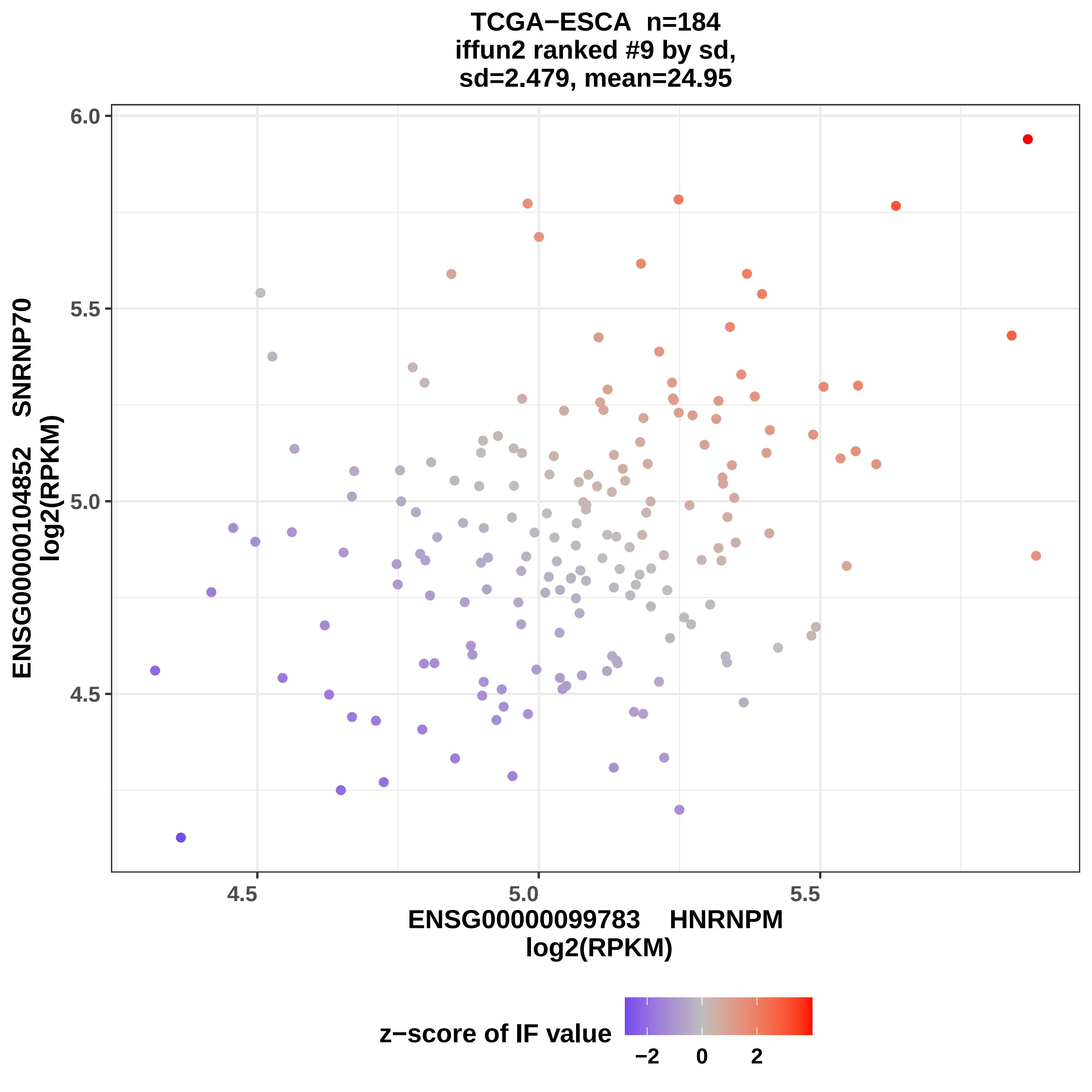
TCGA−ESCA n=184
iffun2 ranked #9 by sd,
sd=2.479, mean=24.95
ENSG00000104852 SNRNP70
log2(RPKM)
ENSG00000099783 HNRNPM
log2(RPKM)
6.0
5.5
5.0
4.5
4.5
5.0
5.5
z−score of IF value
−2
0
2

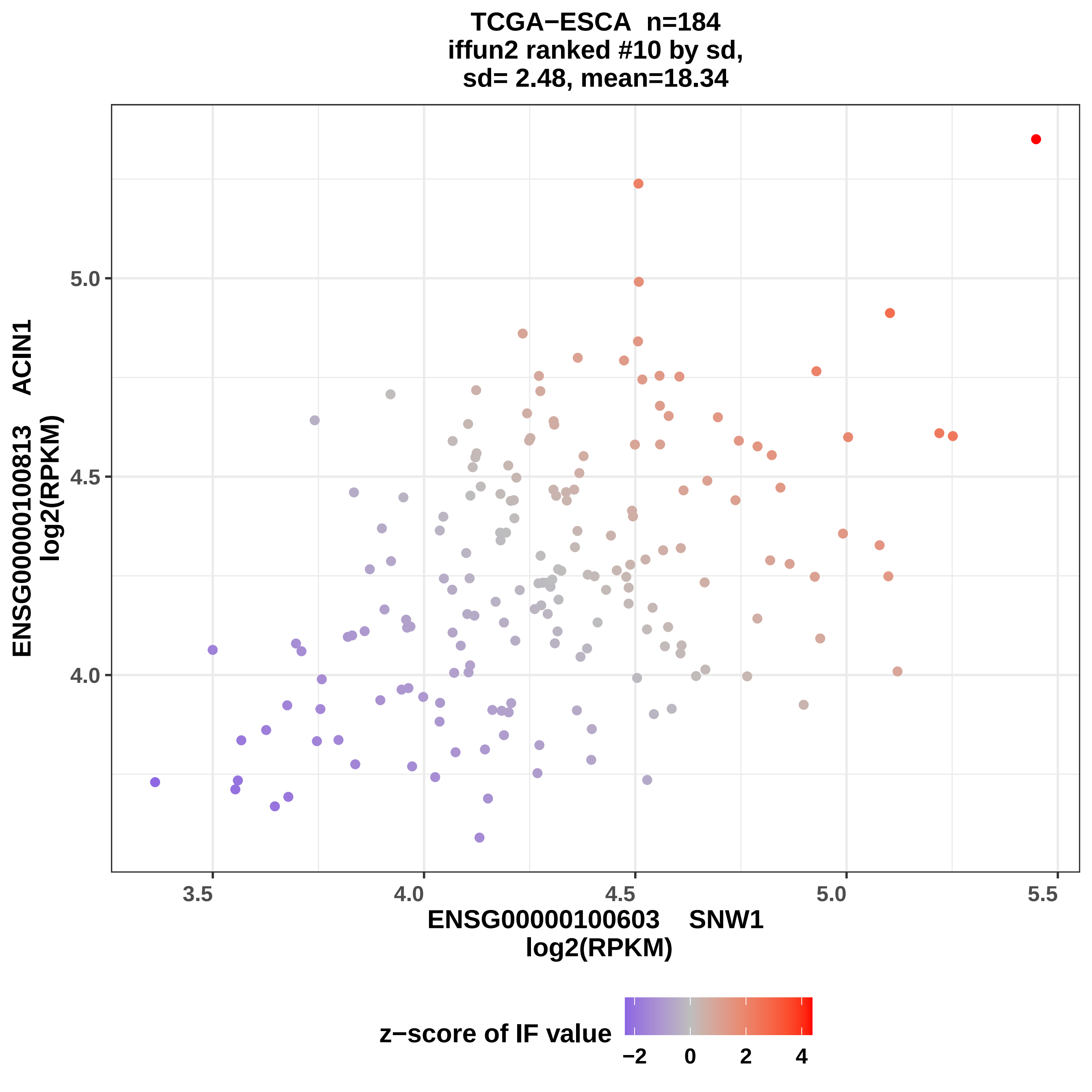
TCGA−ESCA n=184
iffun2 ranked #10 by sd,
sd= 2.48, mean=18.34
ENSG00000100813 ACIN1
log2(RPKM)
5.0
4.5
4.0
3.5
4.0
4.5
5.0
5.5
ENSG00000100603 SNW1
log2(RPKM)
z−score of IF value
−2
0
2
4

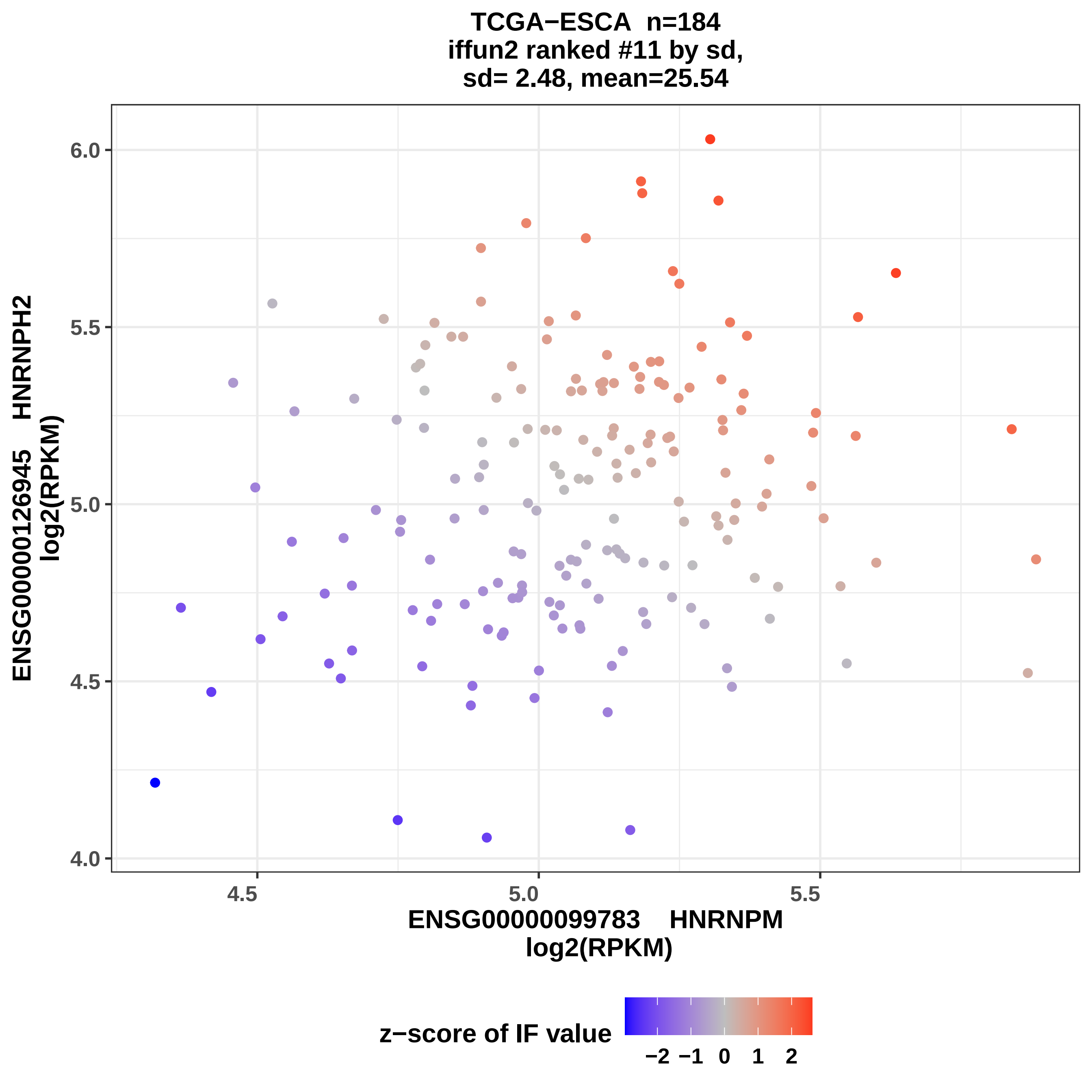
TCGA−ESCA n=184
iffun2 ranked #11 by sd,
sd= 2.48, mean=25.54
6.0
5.5
5.0
4.5
4.0
ENSG00000126945 HNRNPH2
log2(RPKM)
4.5
5.0
5.5
ENSG00000099783 HNRNPM
log2(RPKM)
z−score of IF value
−2 −1 0 1 2

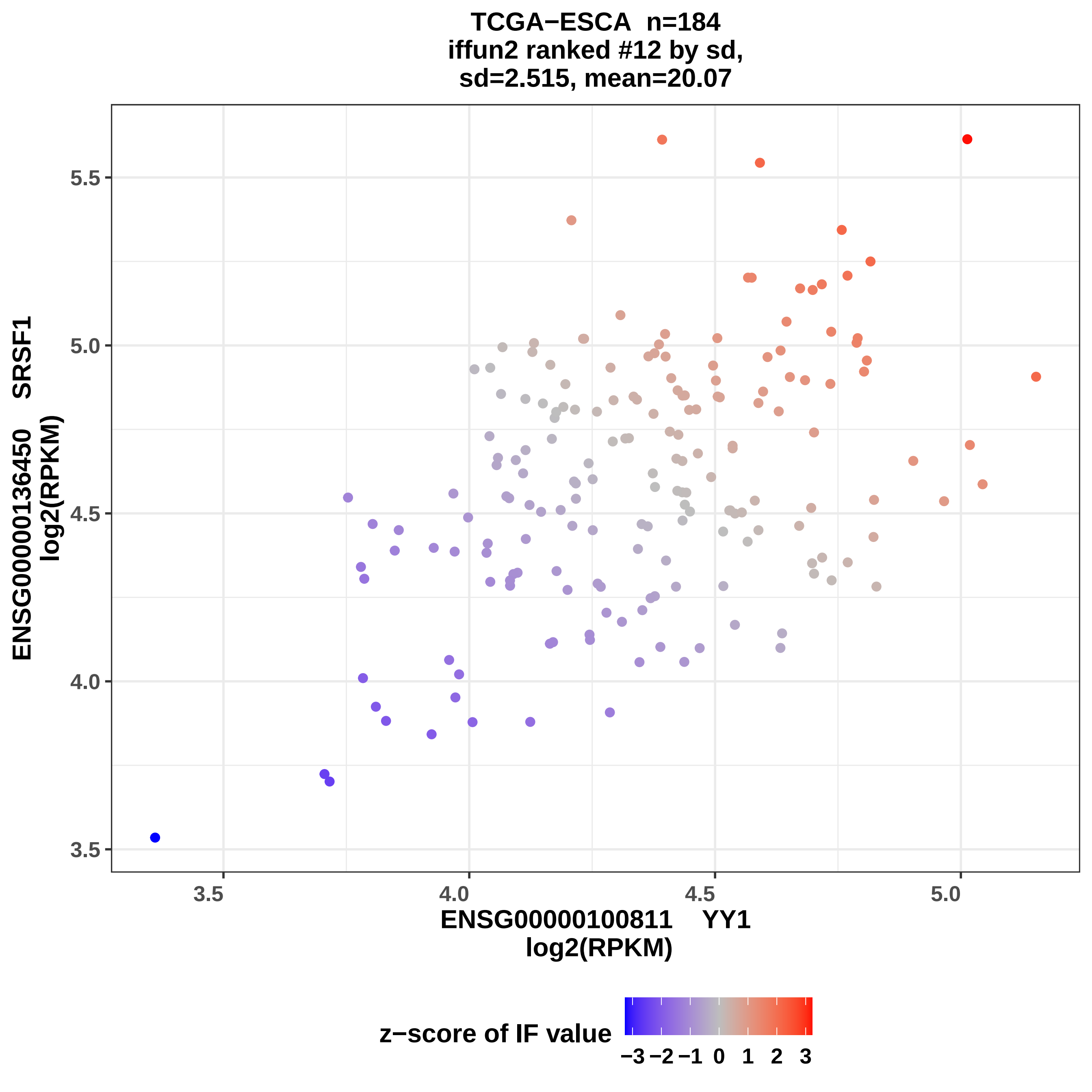
TCGA−ESCA n=184
iffun2 ranked #12 by sd,
sd=2.515, mean=20.07
ENSG00000136450 SRSF1
log2(RPKM)
5.5
5.0
4.5
4.0
3.5
3.5
4.0
4.5
5.0
ENSG00000100811 YY1
log2(RPKM)
z−score of IF value
−3 −2 −1 0 1 2 3

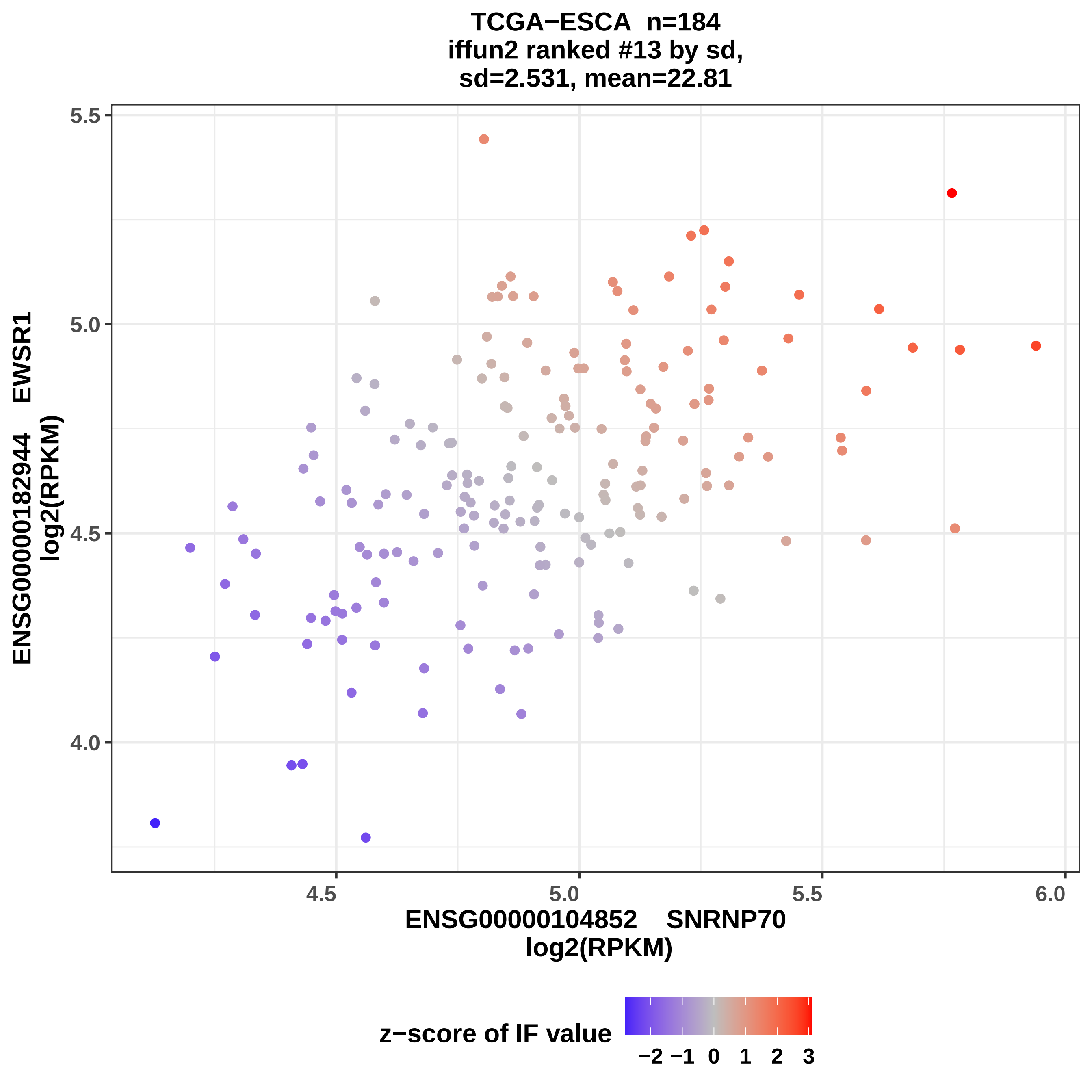

TCGA−ESCA n=184
iffun2 ranked #13 by sd,
sd=2.531, mean=22.81
ENSG00000182944 EWSR1
log2(RPKM)
ENSG00000104852 SNRNP70
log2(RPKM)
5.5
5.0
4.5
4.0
6.0
z−score of IF value
−2 −1 0 1 2 3

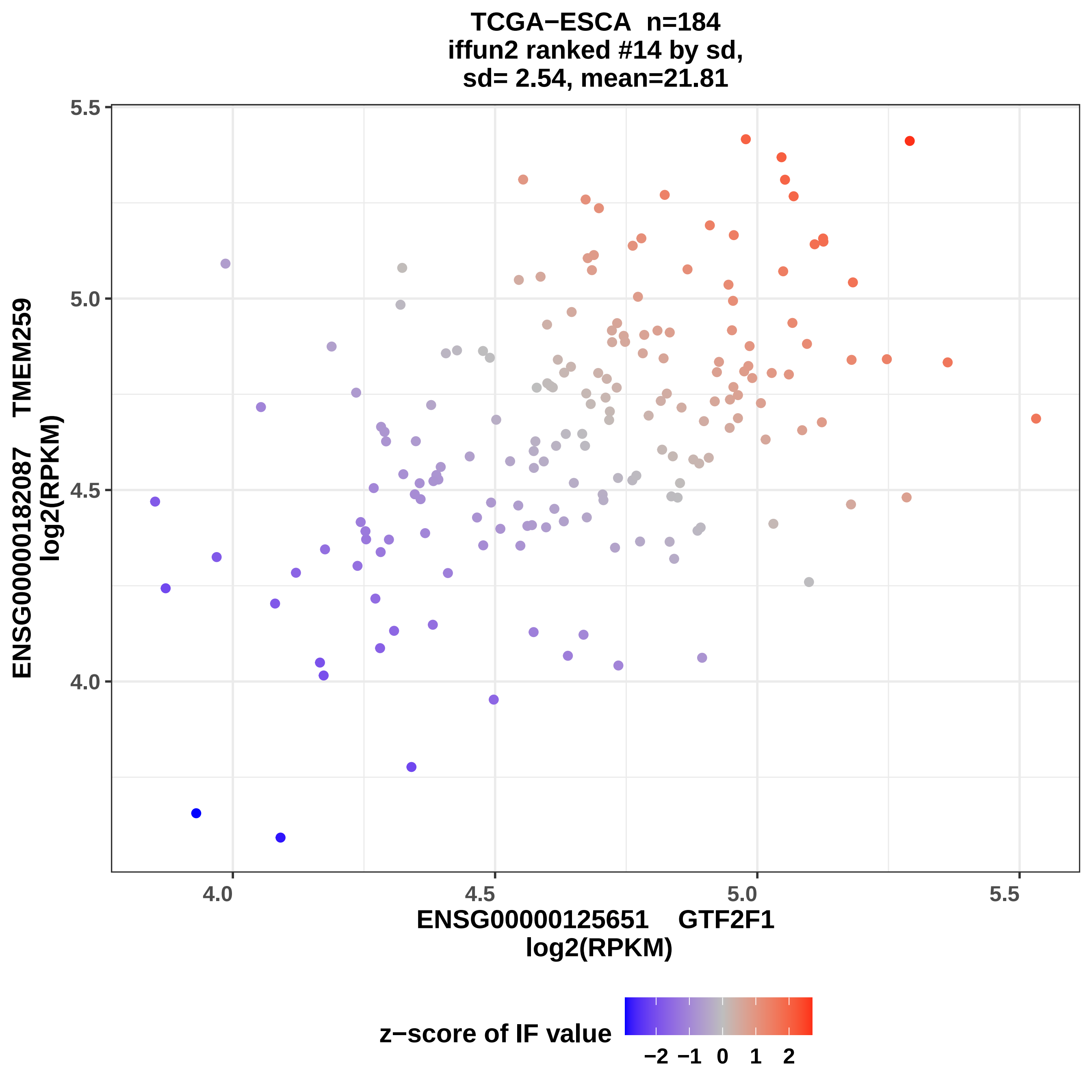

TCGA−ESCA n=184
iffun2 ranked #14 by sd,
sd= 2.54, mean=21.81
ENSG0000182087 TMEM259
log2(RPKM)
5.5
5.0
4.5
4.0
4.0
4.5
5.0
5.5
ENSG0000125651 GTF2F1
log2(RPKM)
z−score of IF value
−2 −1 0 1 2

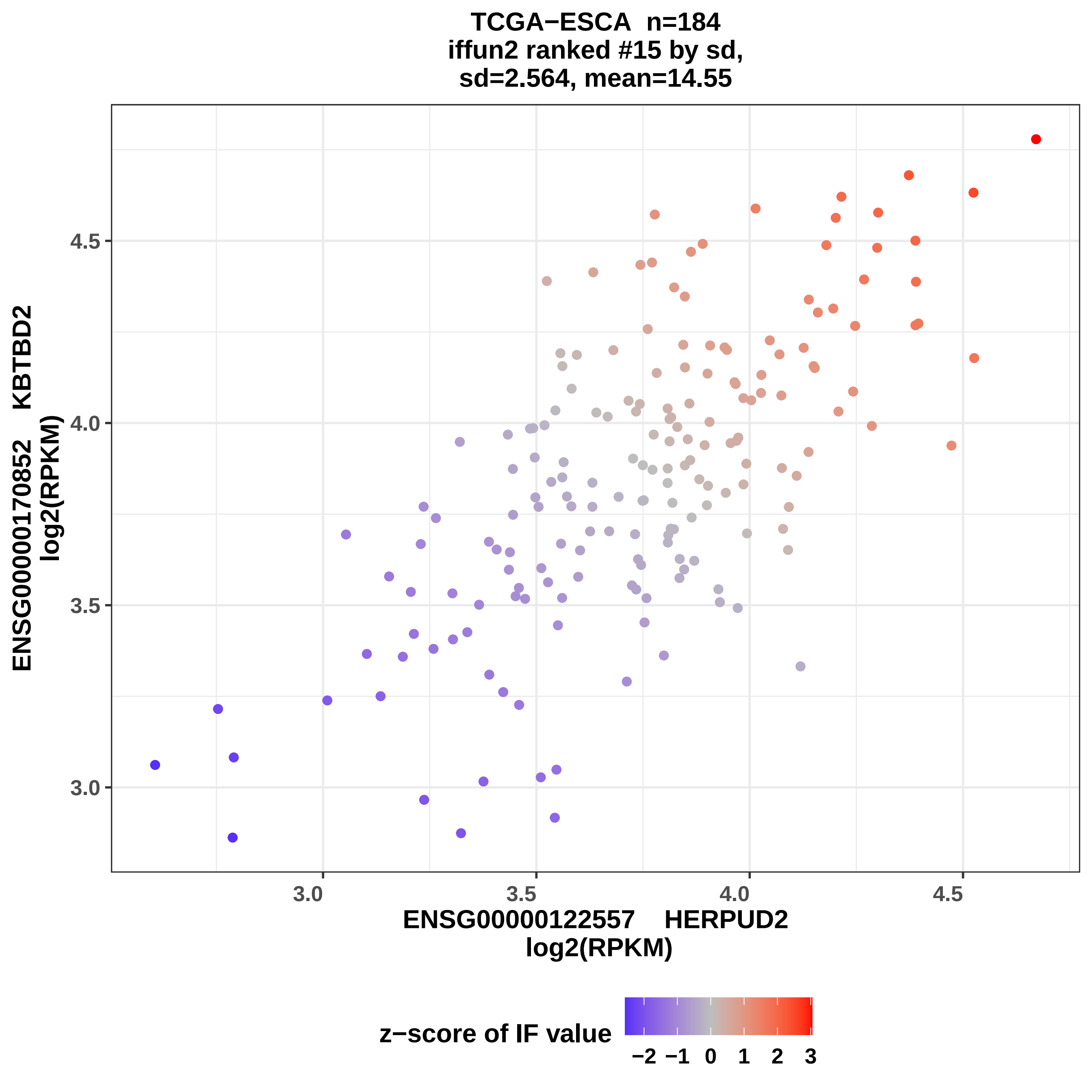
TCGA−ESCA n=184
iffun2 ranked #15 by sd,
sd=2.564, mean=14.55
ENSG00000170852 KBTBD2
log2(RPKM)
4.5
4.0
3.5
3.0
3.0
3.5
4.0
4.5
ENSG00000122557 HERPUD2
log2(RPKM)
z−score of IF value
−2
−1
0
1
2
3

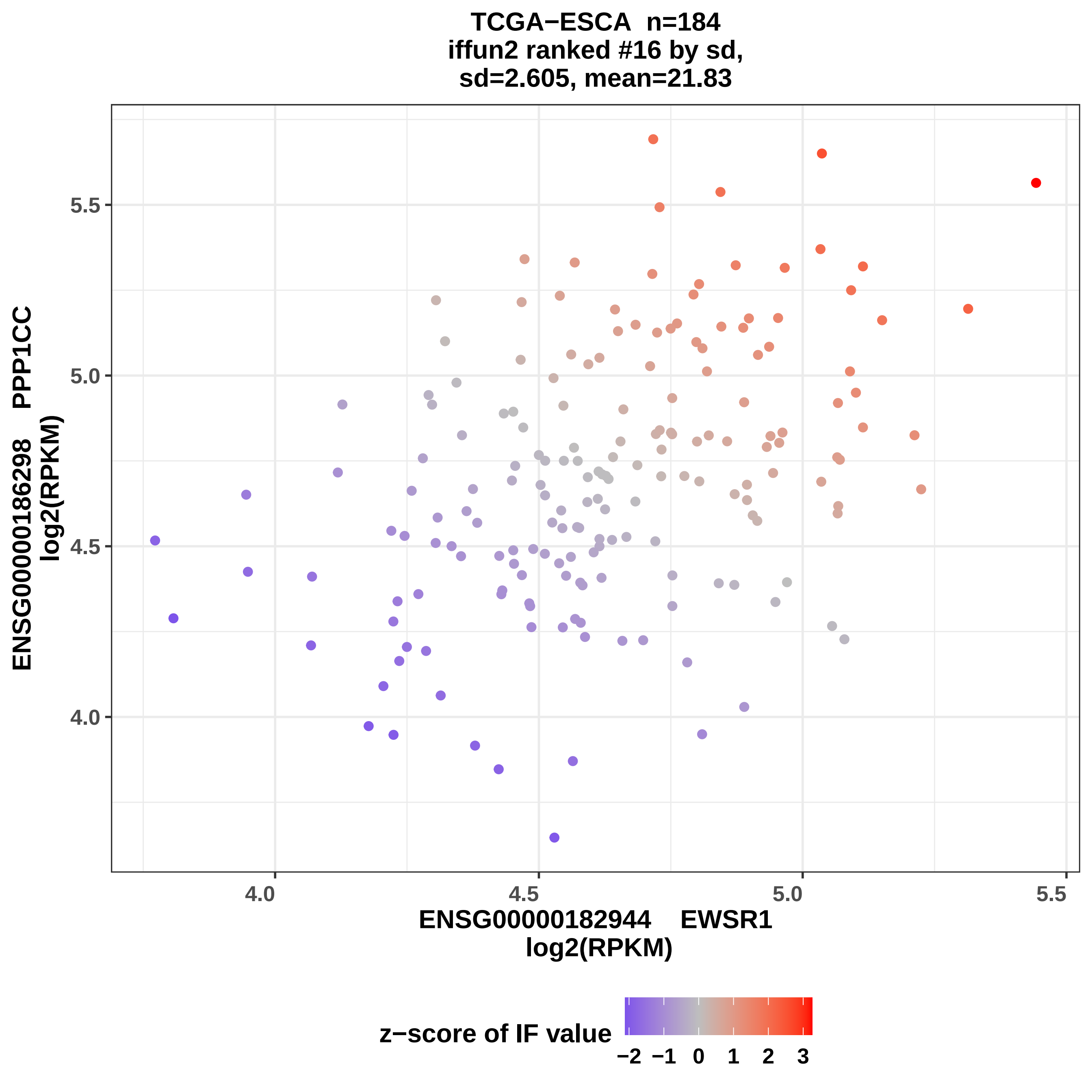

TCGA−ESCA n=184
iffun2 ranked #16 by sd,
sd=2.605, mean=21.83
ENSG00000186298 PPP1CC
log2(RPKM)
5.5
5.0
4.5
4.0
4.0
4.5
5.0
5.5
ENSG00000182944 EWSR1
log2(RPKM)
z−score of IF value
−2
−1
0
1
2
3

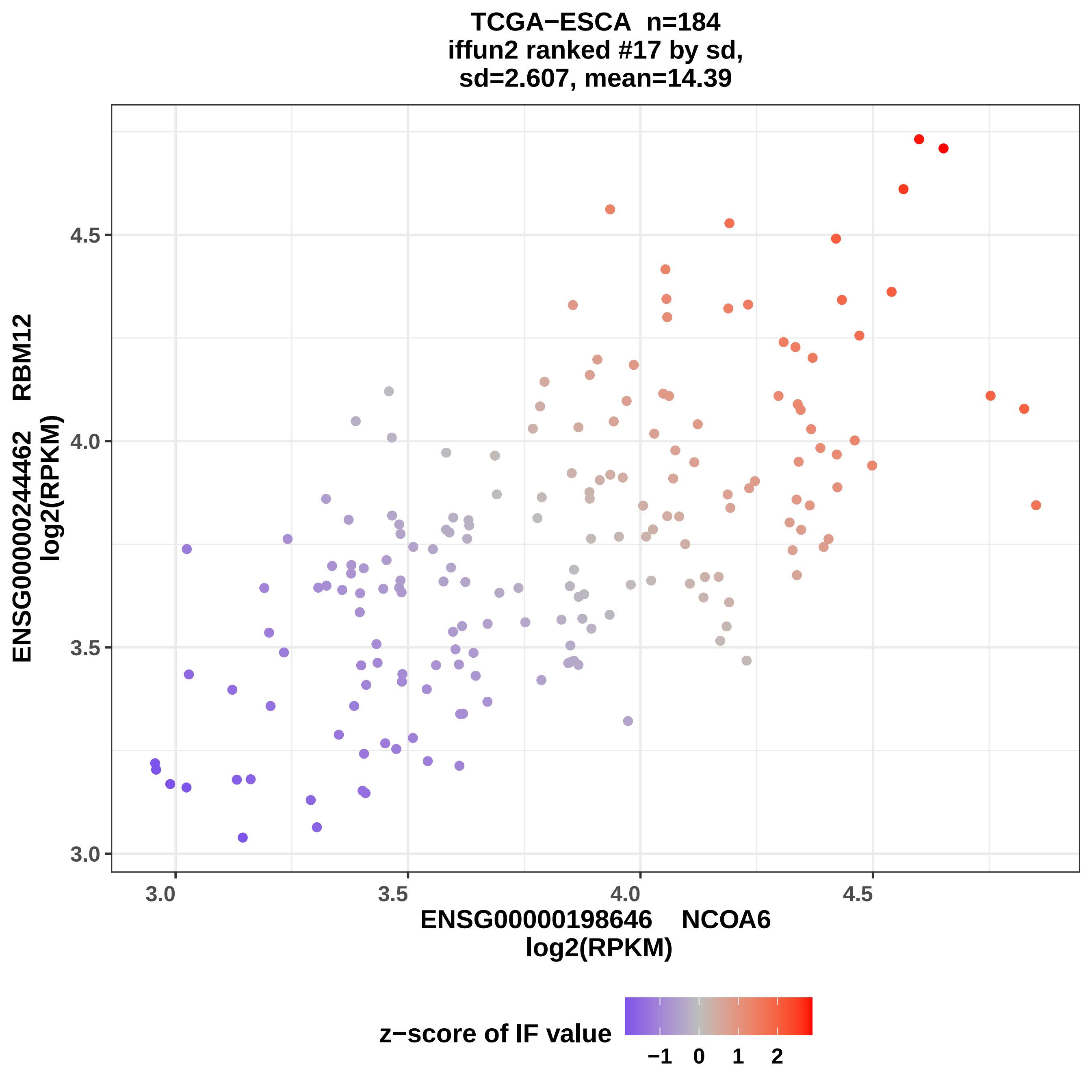
TCGA−ESCA n=184
iffun2 ranked #17 by sd,
sd=2.607, mean=14.39
ENSG00000244462 RBM12
log2(RPKM)
4.5
4.0
3.5
3.0
3.0
3.5
4.0
4.5
ENSG00000198646 NCOA6
log2(RPKM)
z−score of IF value
−1 0 1 2

### TCGA−ESCA n=184
### iffun2 ranked #18 by sd,
### sd=2.628, mean=22.31

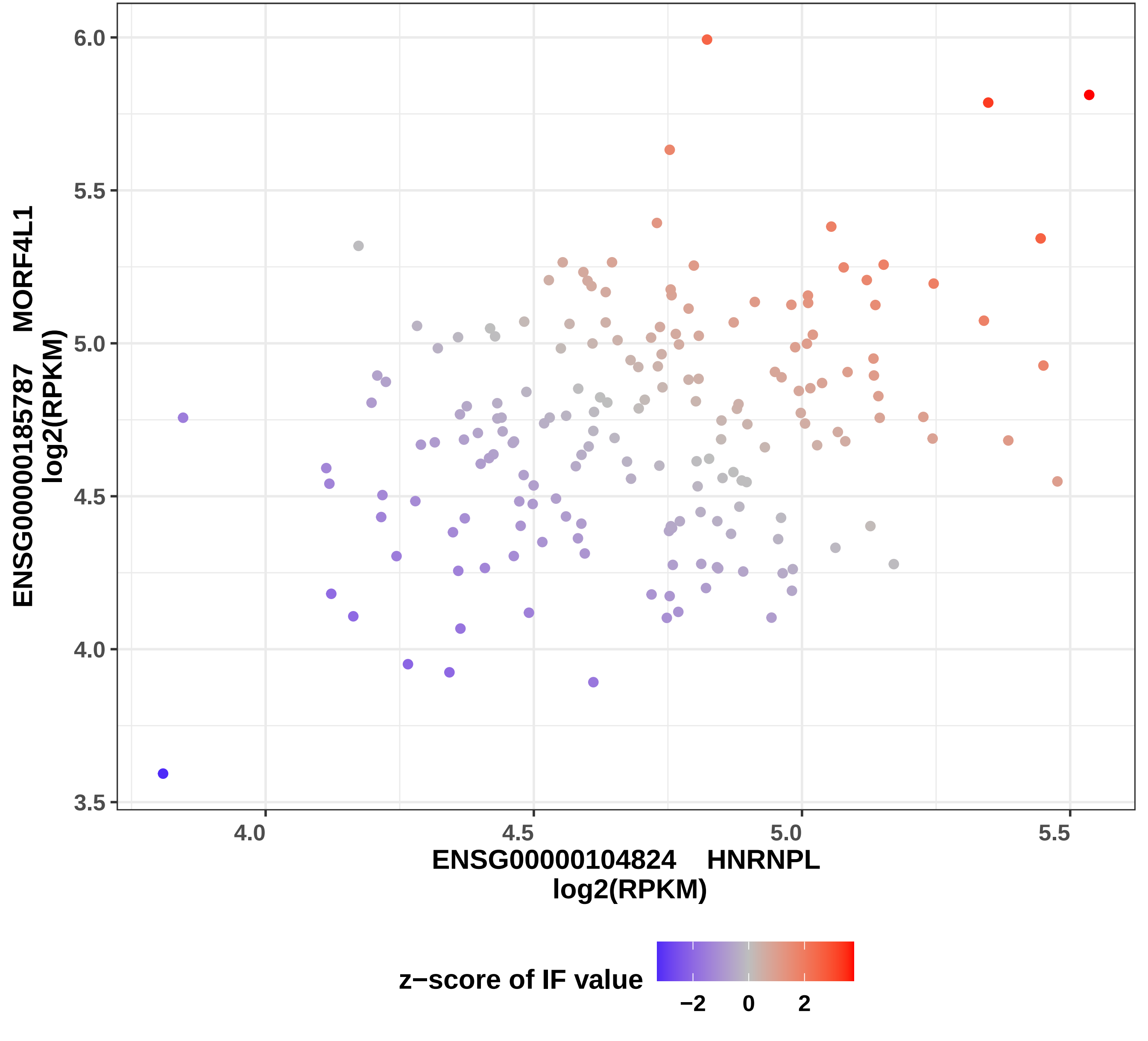

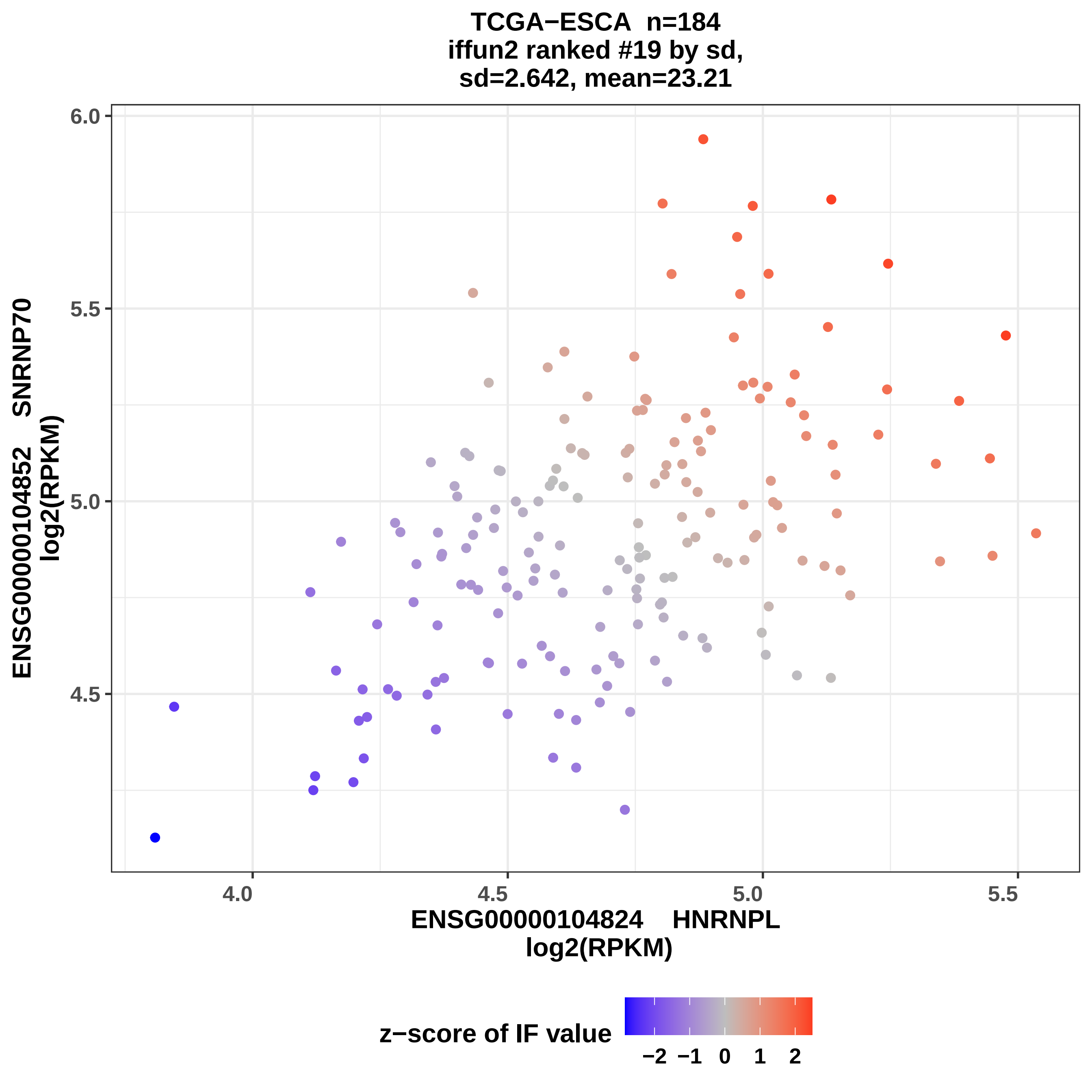
TCGA−ESCA n=184
iffun2 ranked #19 by sd,
sd=2.642, mean=23.21
ENSG00000104852 SNRNP70
log2(RPKM)
6.0
5.5
5.0
4.5
4.0
4.5
5.0
5.5
ENSG00000104824 HNRNPL
log2(RPKM)
z−score of IF value
−2 −1 0 1 2

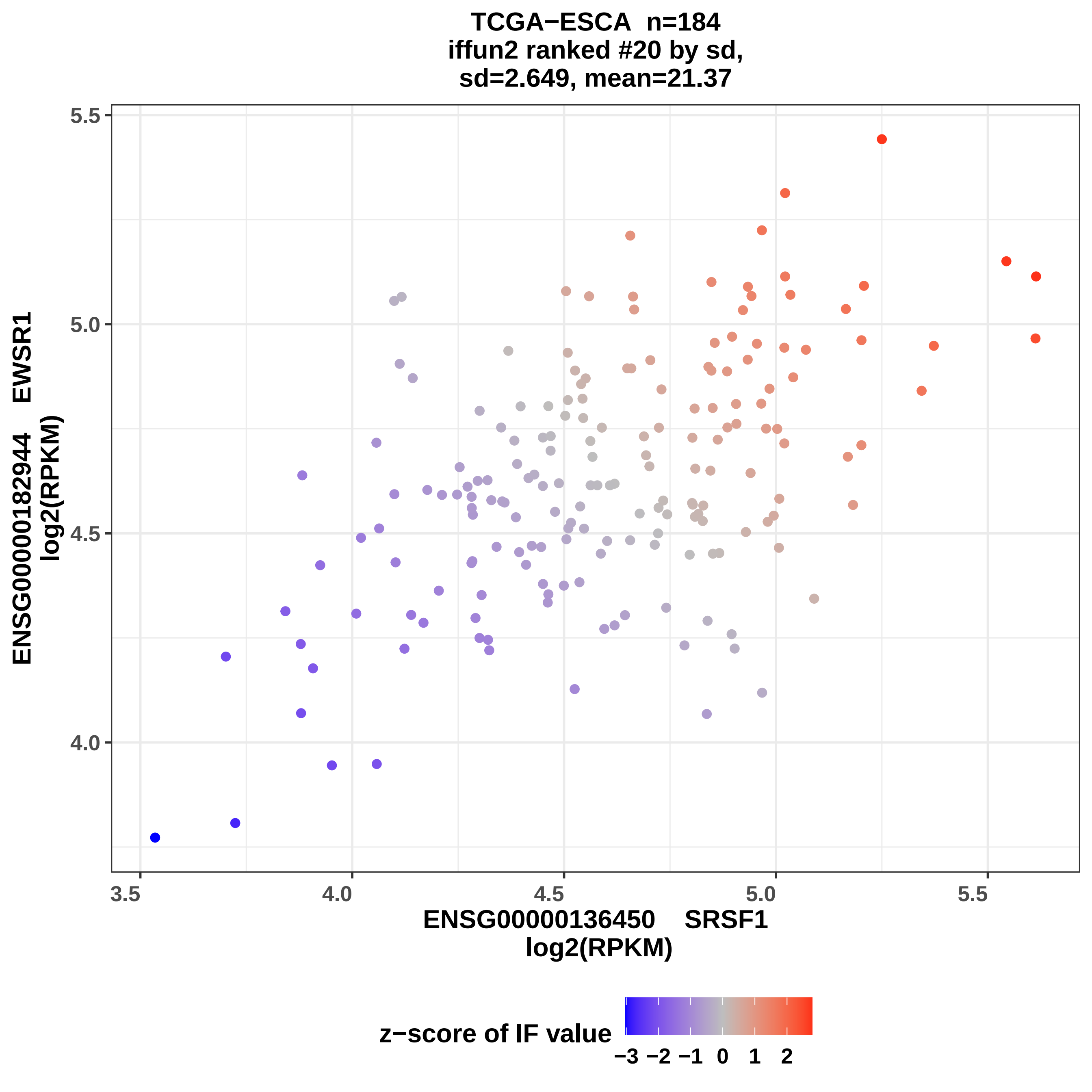

TCGA−ESCA n=184
iffun2 ranked #20 by sd,
sd=2.649, mean=21.37
ENSG00000182944 EWSR1
log2(RPKM)
5.5
5.0
4.5
4.0
3.5
4.0
4.5
5.0
5.5
ENSG00000136450 SRSF1
log2(RPKM)
z−score of IF value
−3 −2 −1 0 1 2

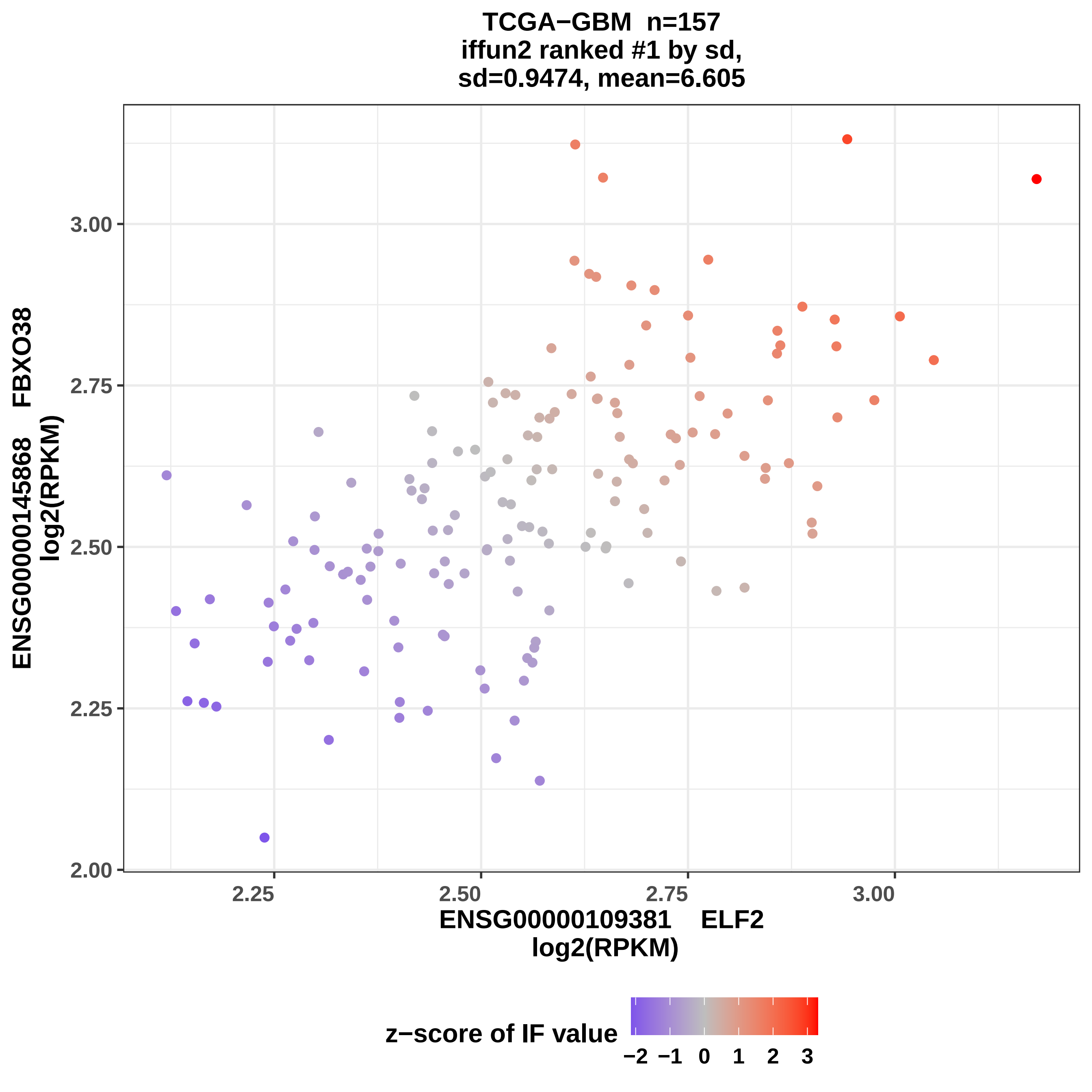
TCGA−GBM n=157
iffun2 ranked #1 by sd,
sd=0.9474, mean=6.605
ENSG00000145868 FBXO38
log2(RPKM)
3.00
2.75
2.50
2.25
2.00
2.25
2.50
2.75
3.00
ENSG00000109381 ELF2
log2(RPKM)
z−score of IF value
−2 −1 0 1 2 3

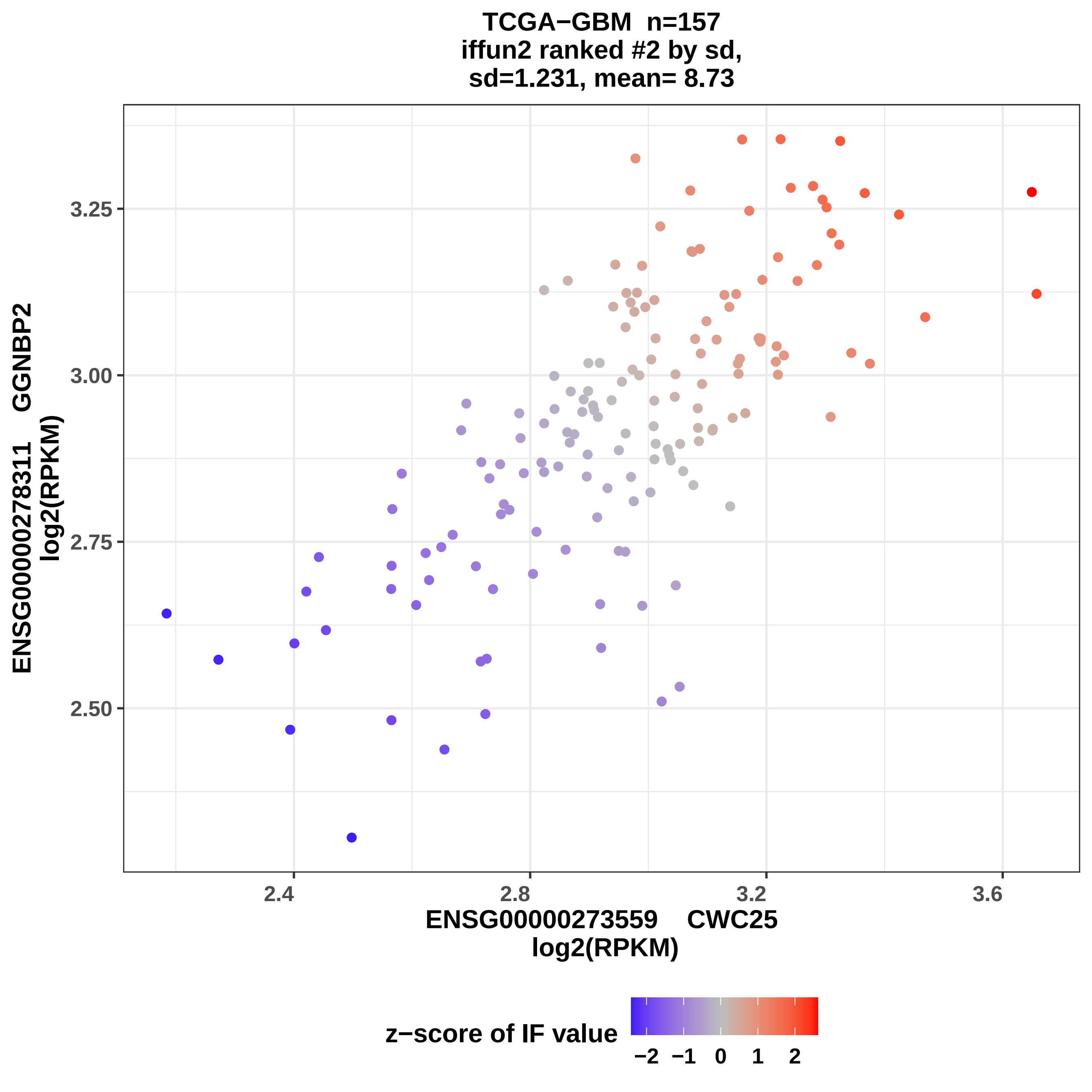
TCGA−GBM n=157
iffun2 ranked #2 by sd,
sd=1.231, mean= 8.73
ENSG00000278311 GGNBP2
log2(RPKM)
3.25
3.00
2.75
2.50
2.4
2.8
3.2
3.6
ENSG00000273559 CWC25
log2(RPKM)
z−score of IF value
−2
−1
0
1
2

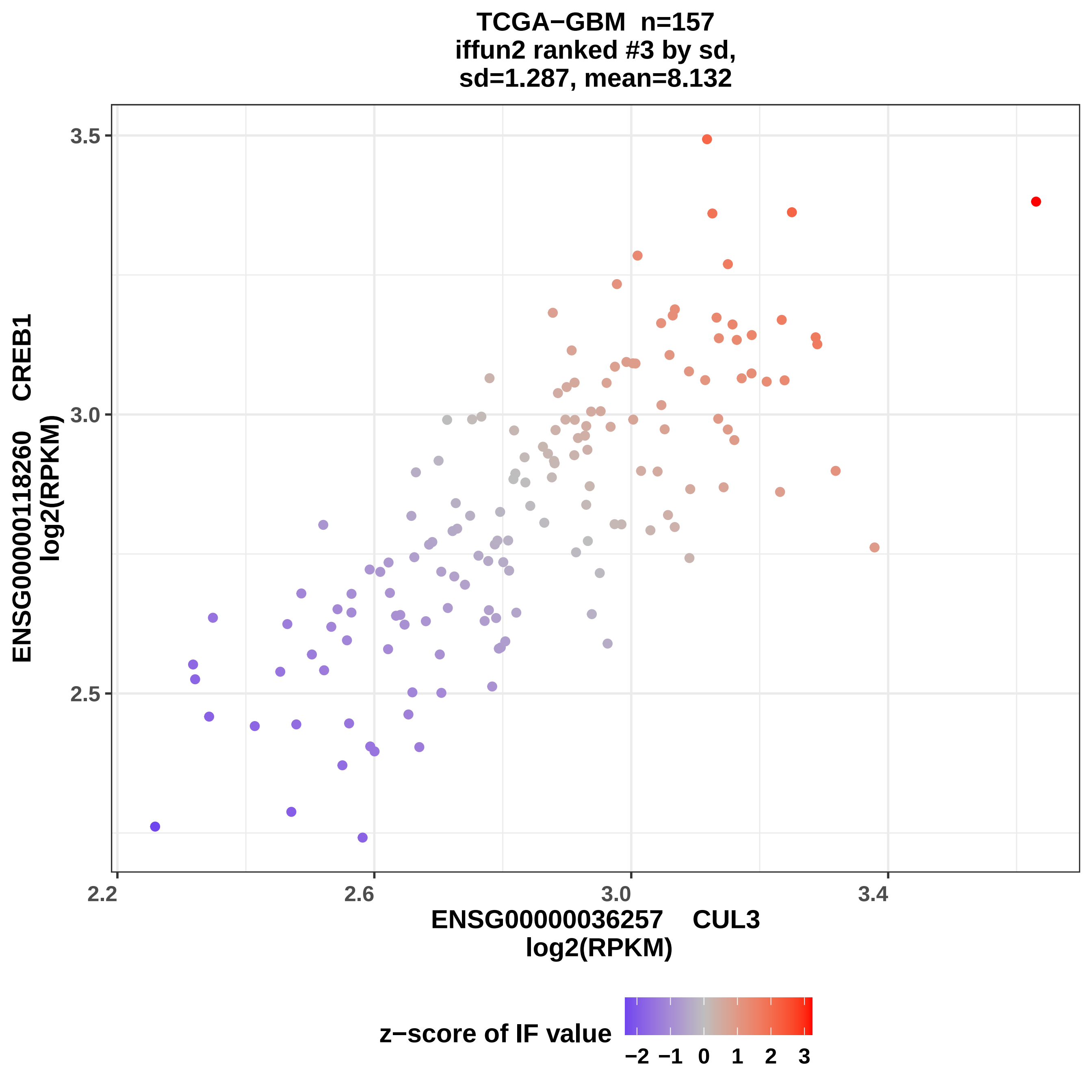
TCGA−GBM n=157
iffun2 ranked #3 by sd,
sd=1.287, mean=8.132
ENSG00000118260 CREB1
log2(RPKM)
3.5
3.0
2.5
2.2
2.6
3.0
3.4
ENSG00000036257 CUL3
log2(RPKM)
z−score of IF value
−2 −1 0 1 2 3

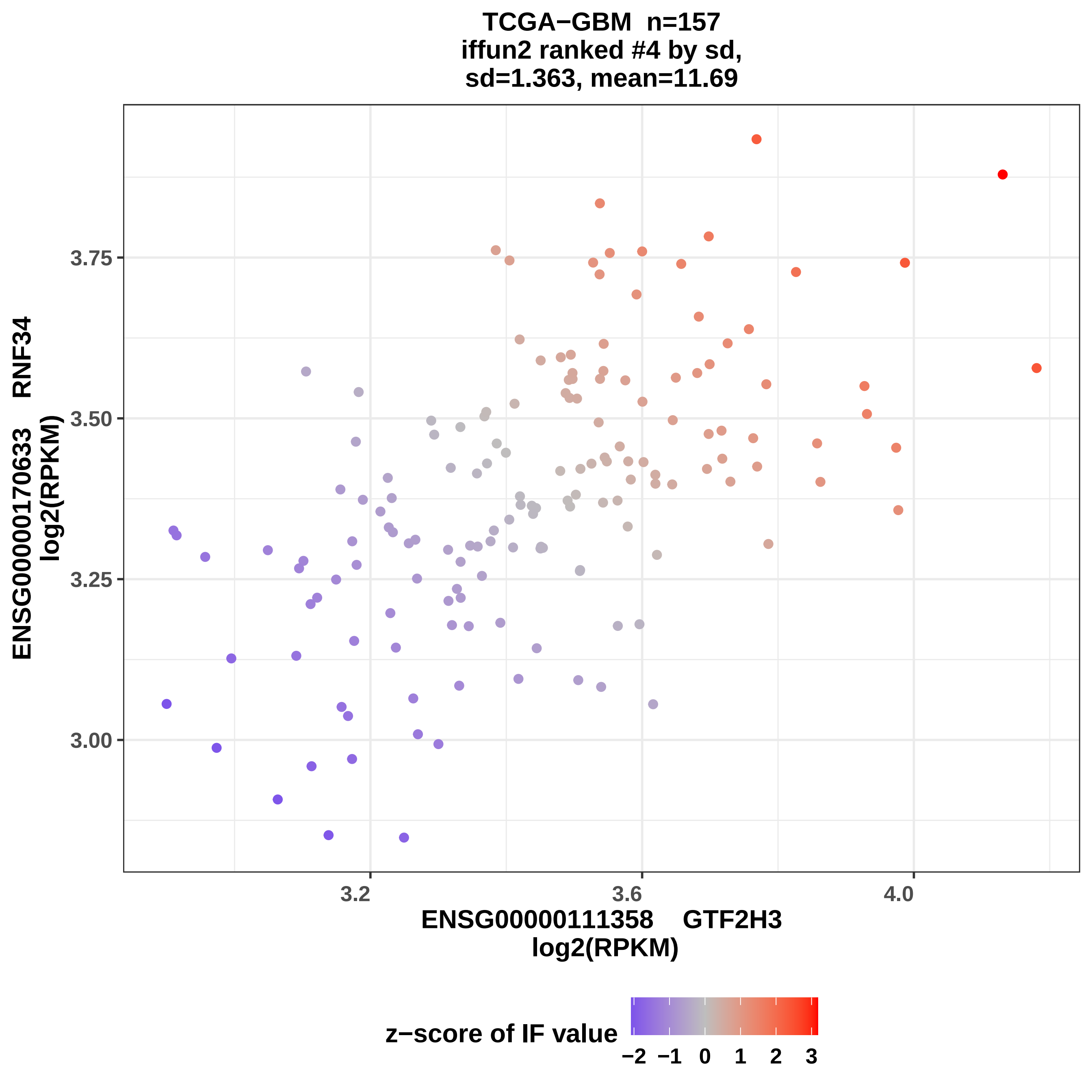
TCGA−GBM n=157
iffun2 ranked #4 by sd,
sd=1.363, mean=11.69
ENSG00000170633 RNF34
log2(RPKM)
3.75
3.50
3.25
3.00
3.2
3.6
4.0
ENSG00000111358 GTF2H3
log2(RPKM)
z−score of IF value
−2 −1 0 1 2 3

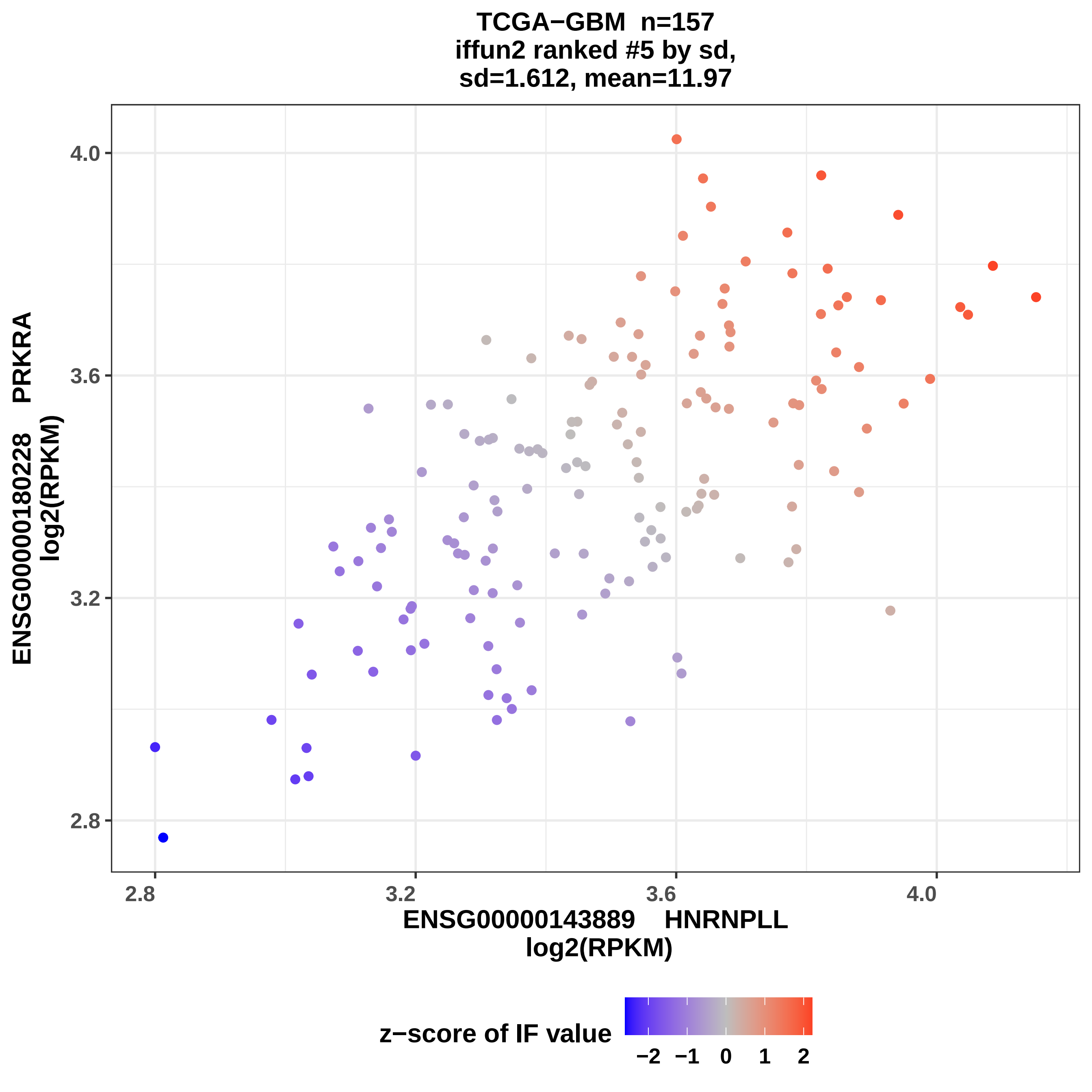

TCGA−GBM n=157
iffun2 ranked #5 by sd,
sd=1.612, mean=11.97
ENSG00000180228 PRKRA
log2(RPKM)
4.0
3.6
3.2
2.8
2.8
3.2
3.6
4.0
ENSG00000143889 HNRNPLL
log2(RPKM)
z−score of IF value
−2
−1
0
1
2

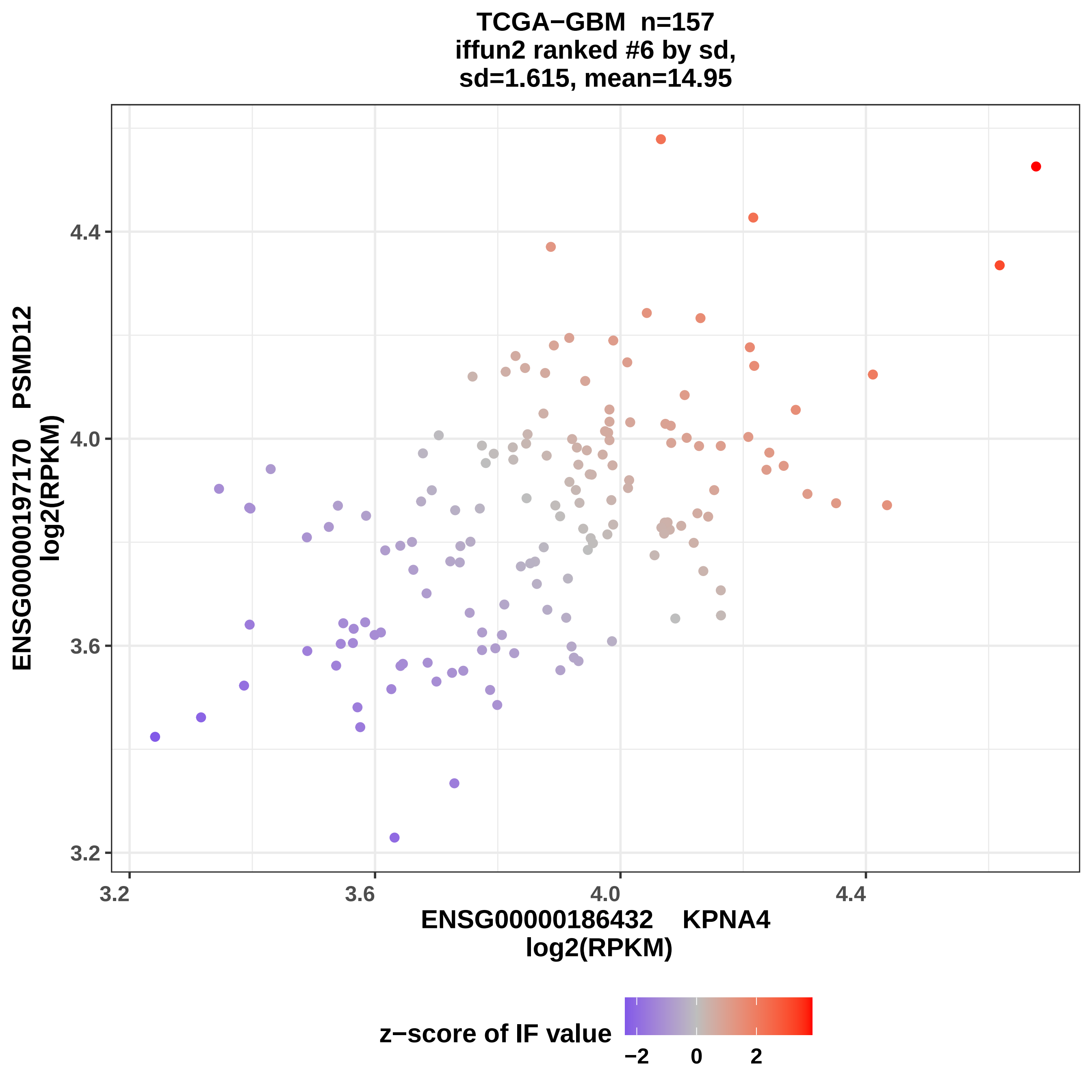
TCGA−GBM n=157
iffun2 ranked #6 by sd,
sd=1.615, mean=14.95
ENSG00000197170 PSMD12
log2(RPKM)
4.4
4.0
3.6
3.2
3.2
3.6
4.0
4.4
ENSG00000186432 KPNA4
log2(RPKM)
z−score of IF value
−2
0
2

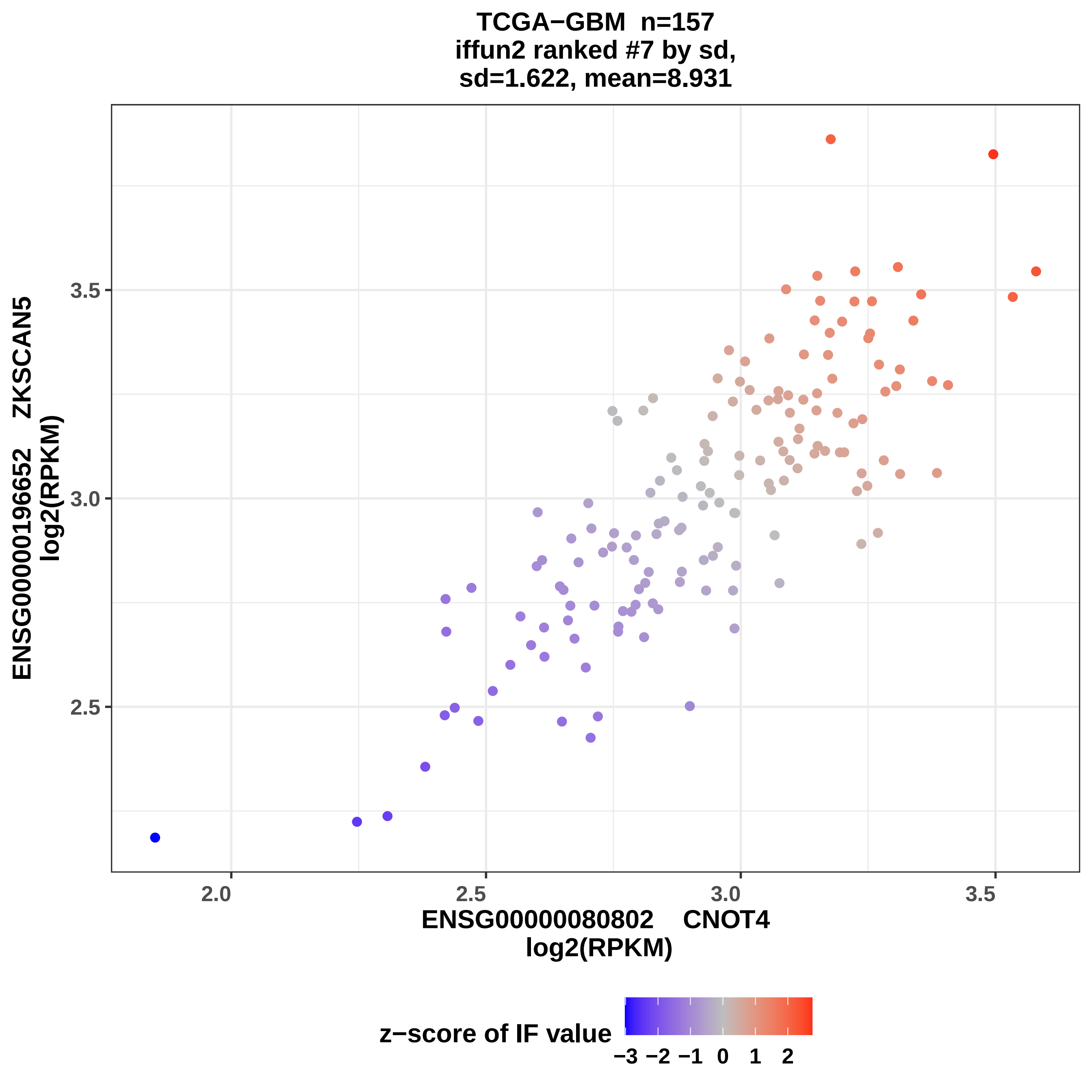
TCGA−GBM n=157
iffun2 ranked #7 by sd,
sd=1.622, mean=8.931
ENSG00000196652 ZKSCAN5
log2(RPKM)
3.5
3.0
2.5
2.0
2.5
3.0
3.5
ENSG00000080802 CNOT4
log2(RPKM)
z−score of IF value
−3 −2 −1 0 1 2

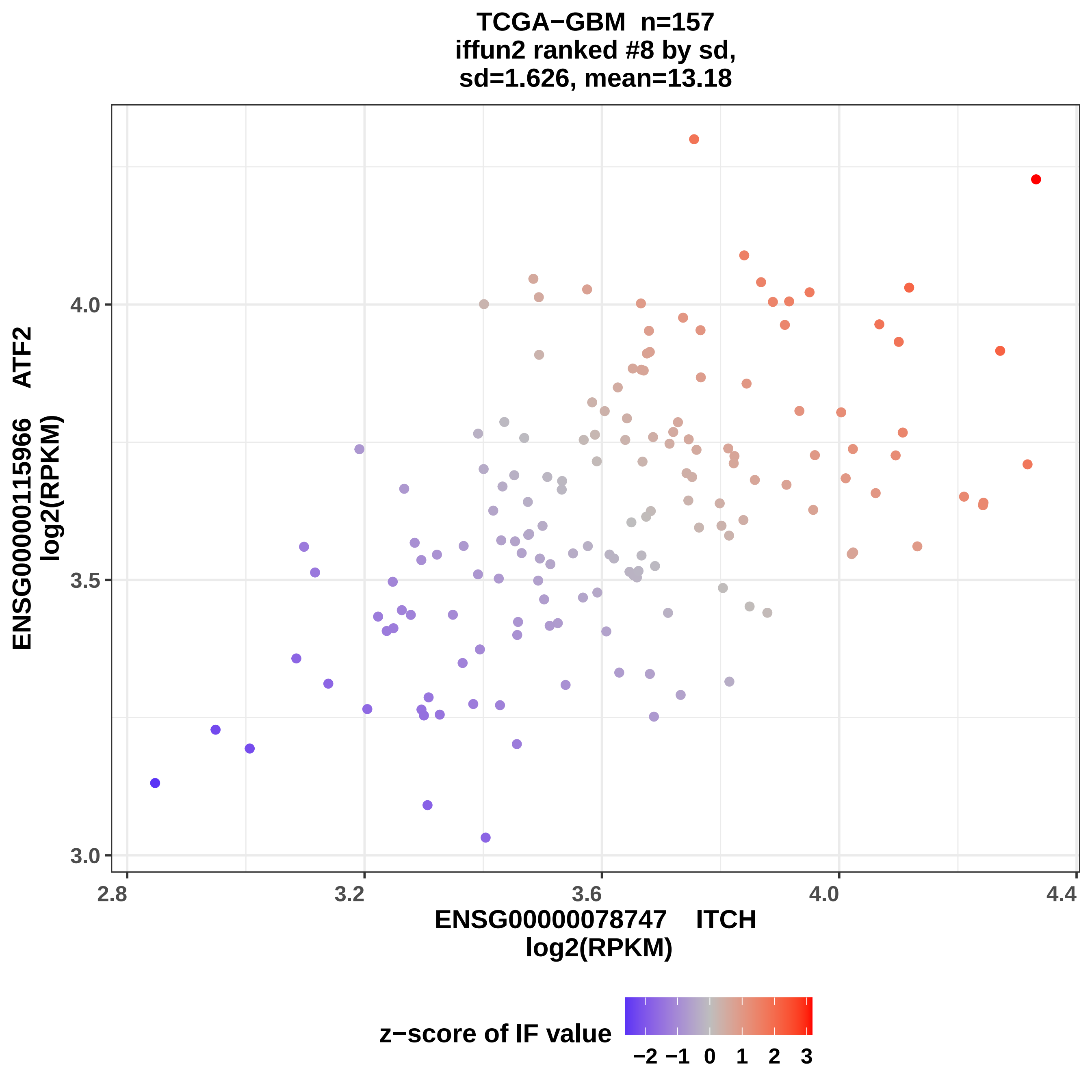
TCGA−GBM n=157
iffun2 ranked #8 by sd,
sd=1.626, mean=13.18
ENSG00000115966 ATF2
log2(RPKM)
4.0
3.5
3.0
2.8
3.2
3.6
4.0
4.4
ENSG00000078747 ITCH
log2(RPKM)
z−score of IF value
−2 −1 0 1 2 3

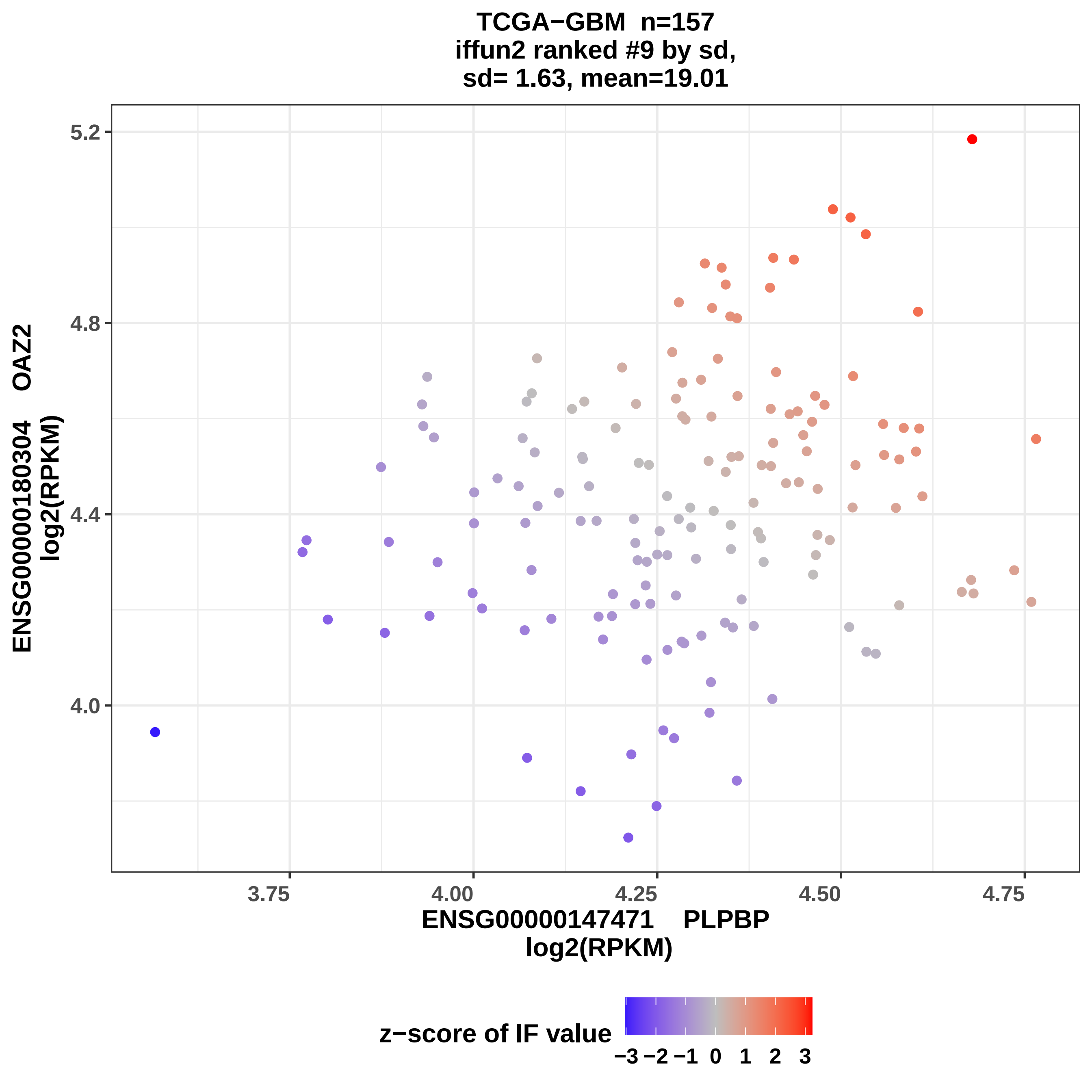

TCGA−GBM n=157
iffun2 ranked #9 by sd,
sd= 1.63, mean=19.01
ENSG00000180304 OAZ2
log2(RPKM)
5.2
4.8
4.4
4.0
3.75
4.00
4.25
4.50
4.75
ENSG00000147471 PLPBP
log2(RPKM)
z−score of IF value
−3 −2 −1 0 1 2 3

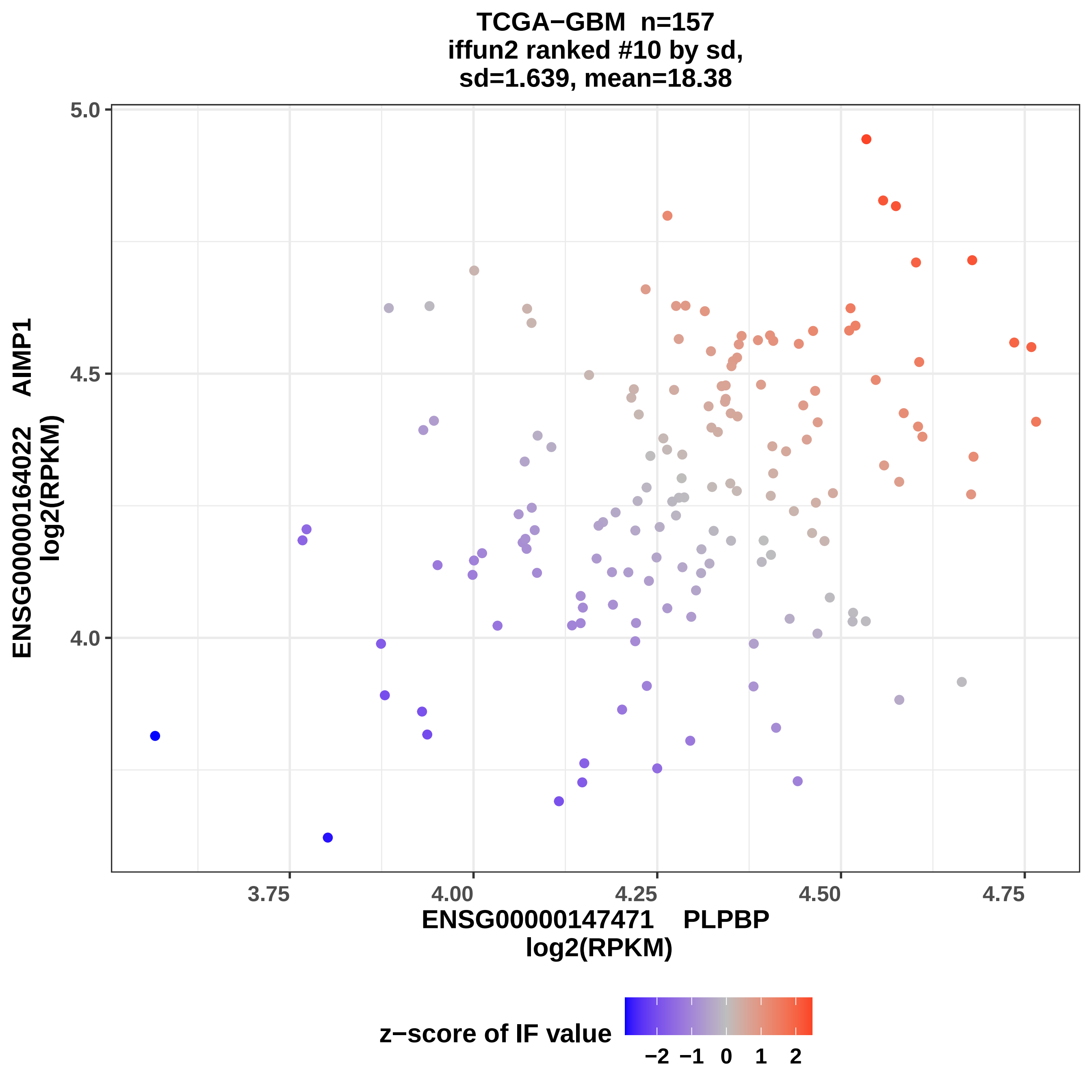
TCGA−GBM n=157
iffun2 ranked #10 by sd,
sd=1.639, mean=18.38
5.0
4.5
4.0
3.75
4.00
4.25
4.50
4.75
ENSG00000147471 PLPBP
log2(RPKM)
ENSG00000164022 AIMP1
log2(RPKM)
z−score of IF value
−2
−1
0
1
2

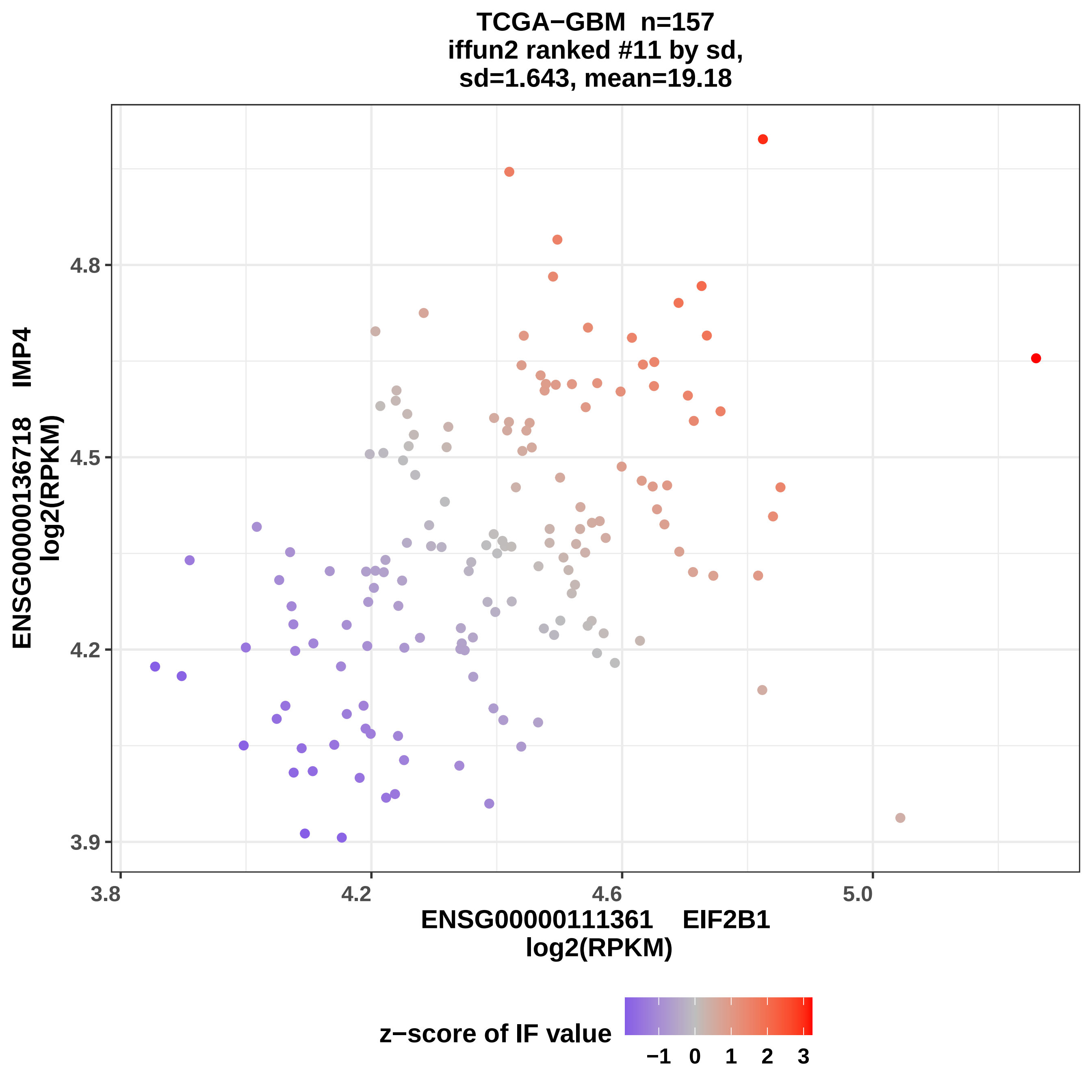

TCGA−GBM n=157
iffun2 ranked #11 by sd,
sd=1.643, mean=19.18
ENSG00000136718 IMP4
log2(RPKM)
4.8
4.5
4.2
3.9
3.8
4.2
4.6
5.0
ENSG00000111361 EIF2B1
log2(RPKM)
z−score of IF value
−1
0
1
2
3

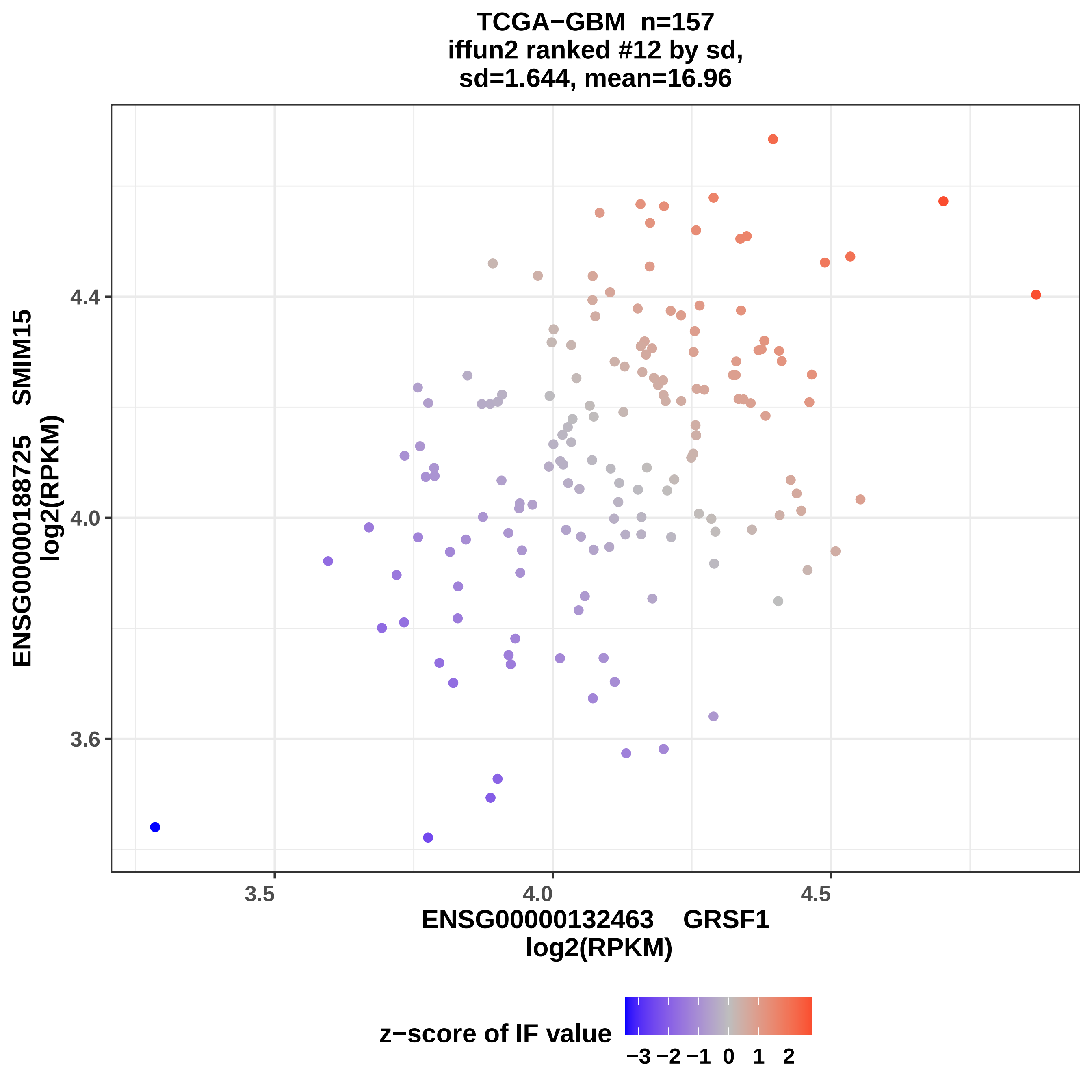

TCGA−GBM n=157
iffun2 ranked #12 by sd,
sd=1.644, mean=16.96
ENSG00000188725 SMIM15
log2(RPKM)
4.4
4.0
3.6
3.5
4.0
4.5
ENSG00000132463 GRSF1
log2(RPKM)
z−score of IF value
−3 −2 −1 0 1 2

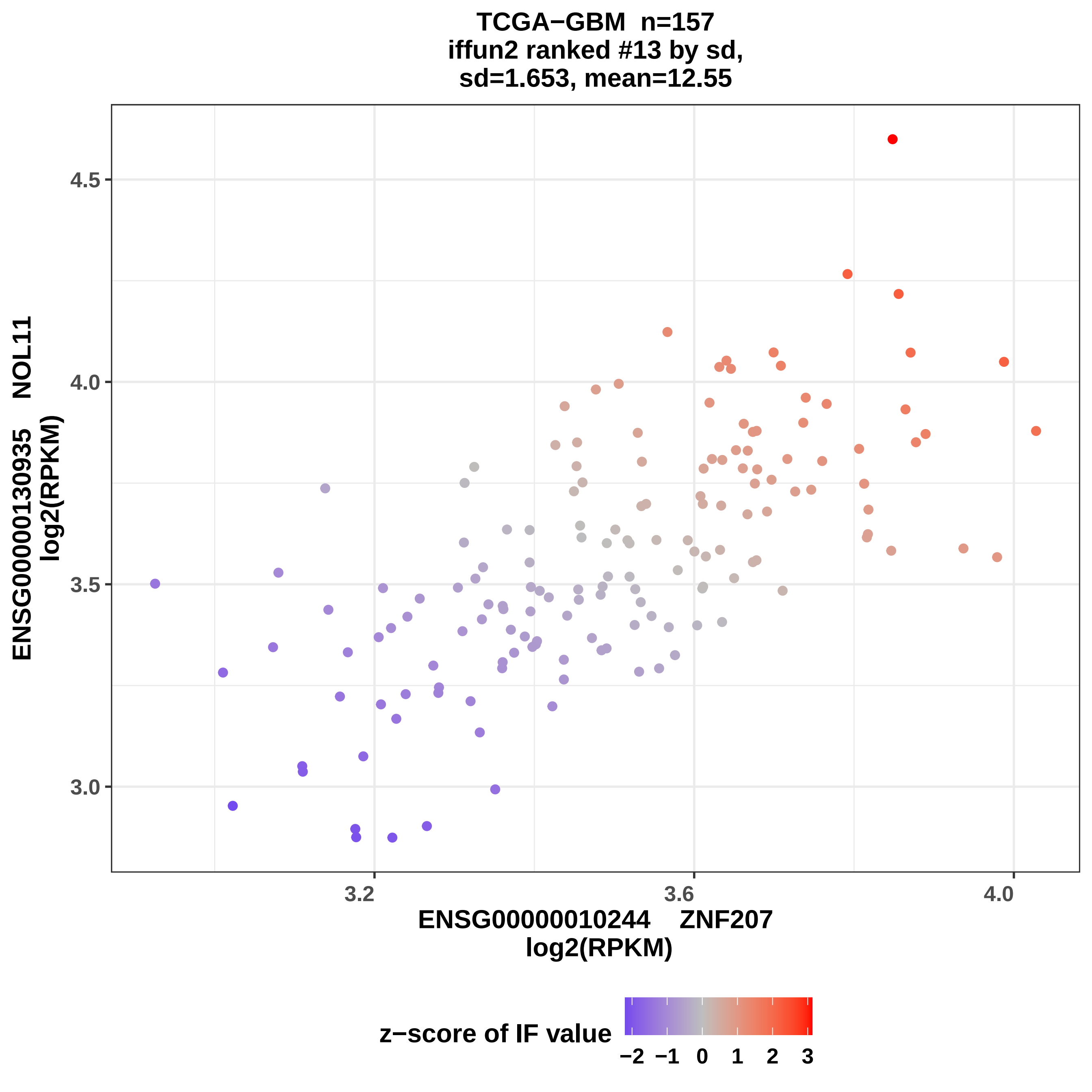
TCGA−GBM n=157
iffun2 ranked #13 by sd,
sd=1.653, mean=12.55
ENSG00000130935 NOL11
log2(RPKM)
4.5
4.0
3.5
3.0
3.2
3.6
4.0
ENSG00000010244 ZNF207
log2(RPKM)
z−score of IF value
−2 −1 0 1 2 3

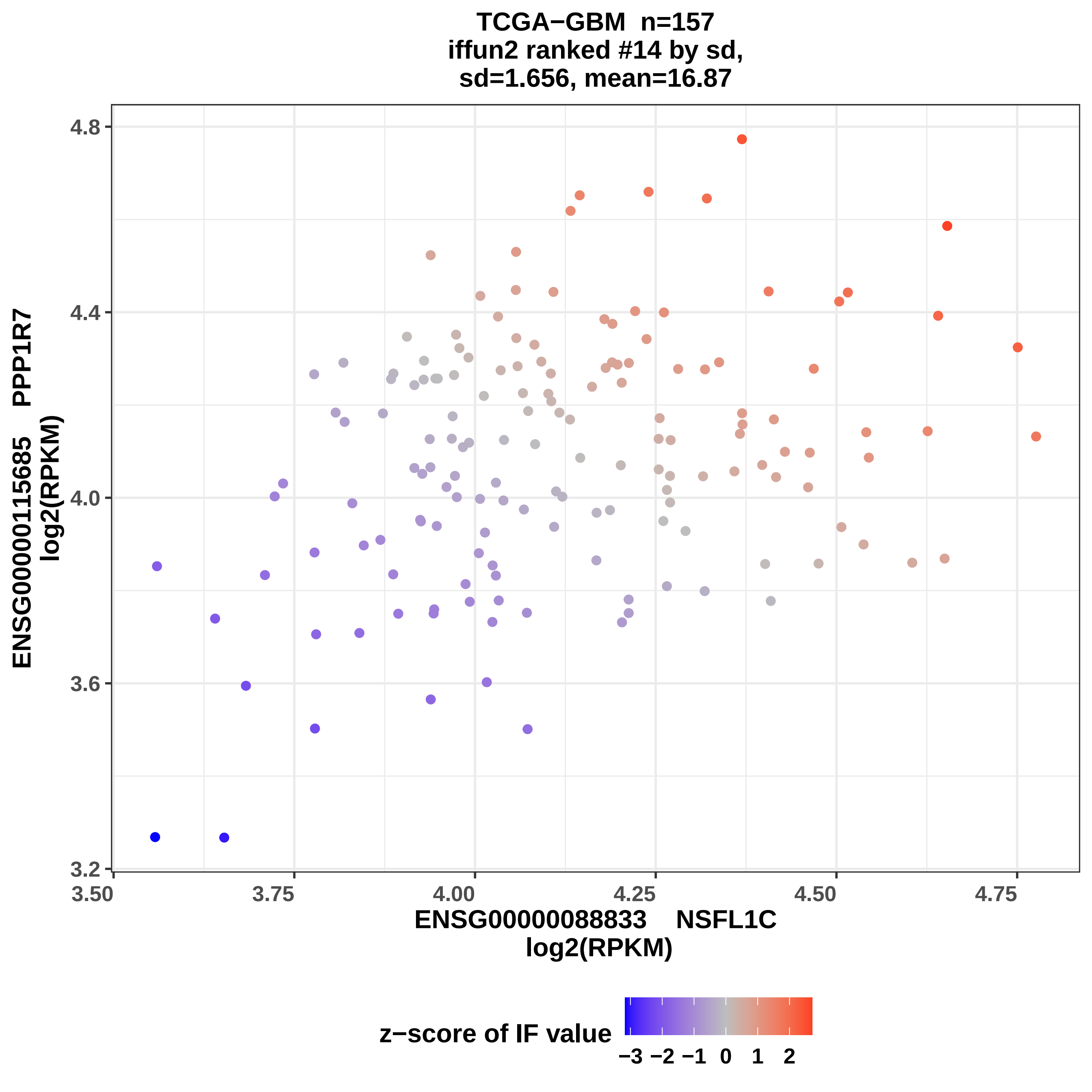
TCGA−GBM n=157
iffun2 ranked #14 by sd,
sd=1.656, mean=16.87
ENSG00000115685 PPP1R7
log2(RPKM)
4.8
4.4
4.0
3.6
3.2
3.50
3.75
4.00
4.25
4.50
4.75
ENSG00000088833 NSFL1C
log2(RPKM)
z−score of IF value
−3 −2 −1 0 1 2

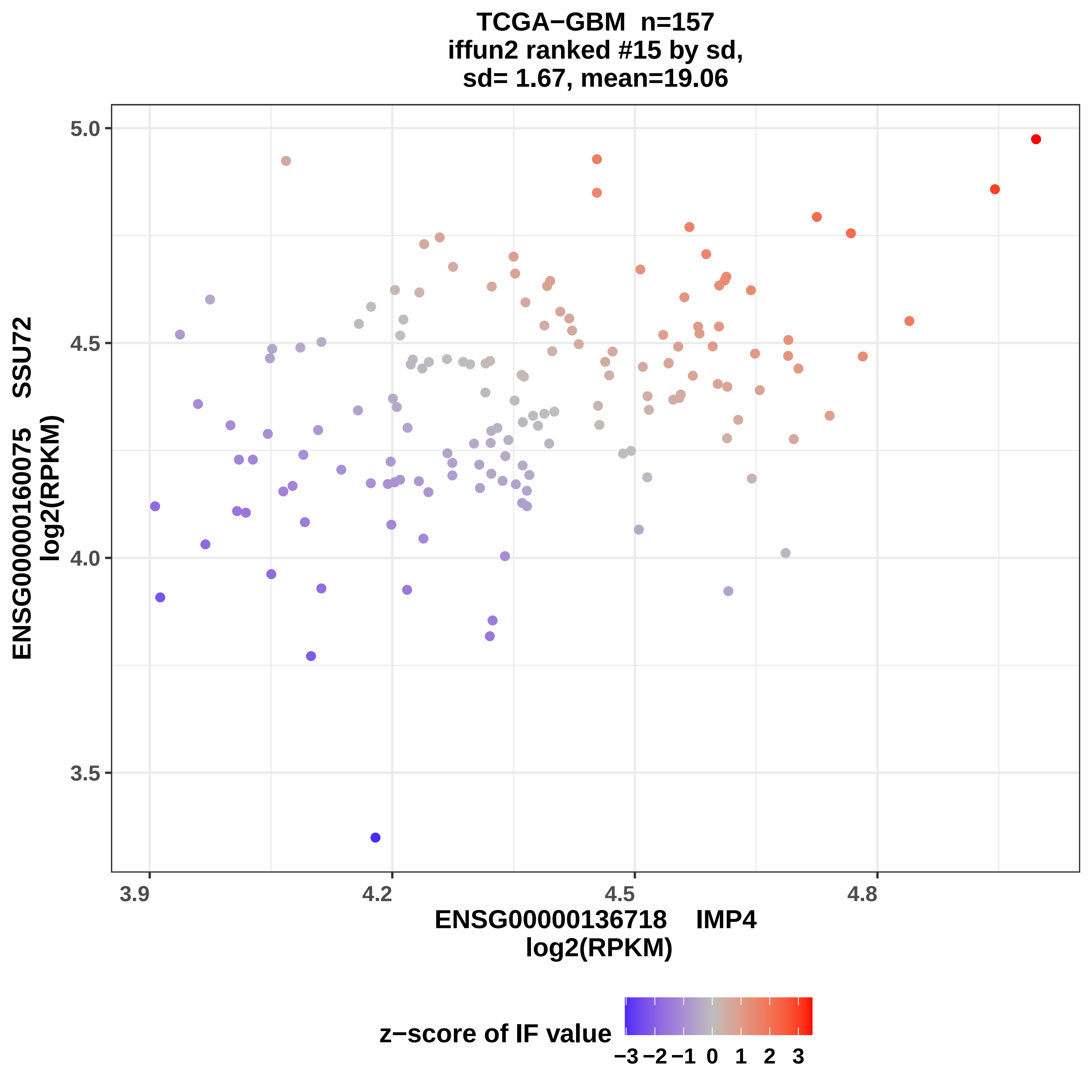

TCGA−GBM n=157
iffun2 ranked #15 by sd,
sd= 1.67, mean=19.06
ENSG00000160075 SSU72
log2(RPKM)
5.0
4.5
4.0
3.5
3.9
4.2
4.5
4.8
ENSG00000136718 IMP4
log2(RPKM)
z−score of IF value
−3 −2 −1 0 1 2 3

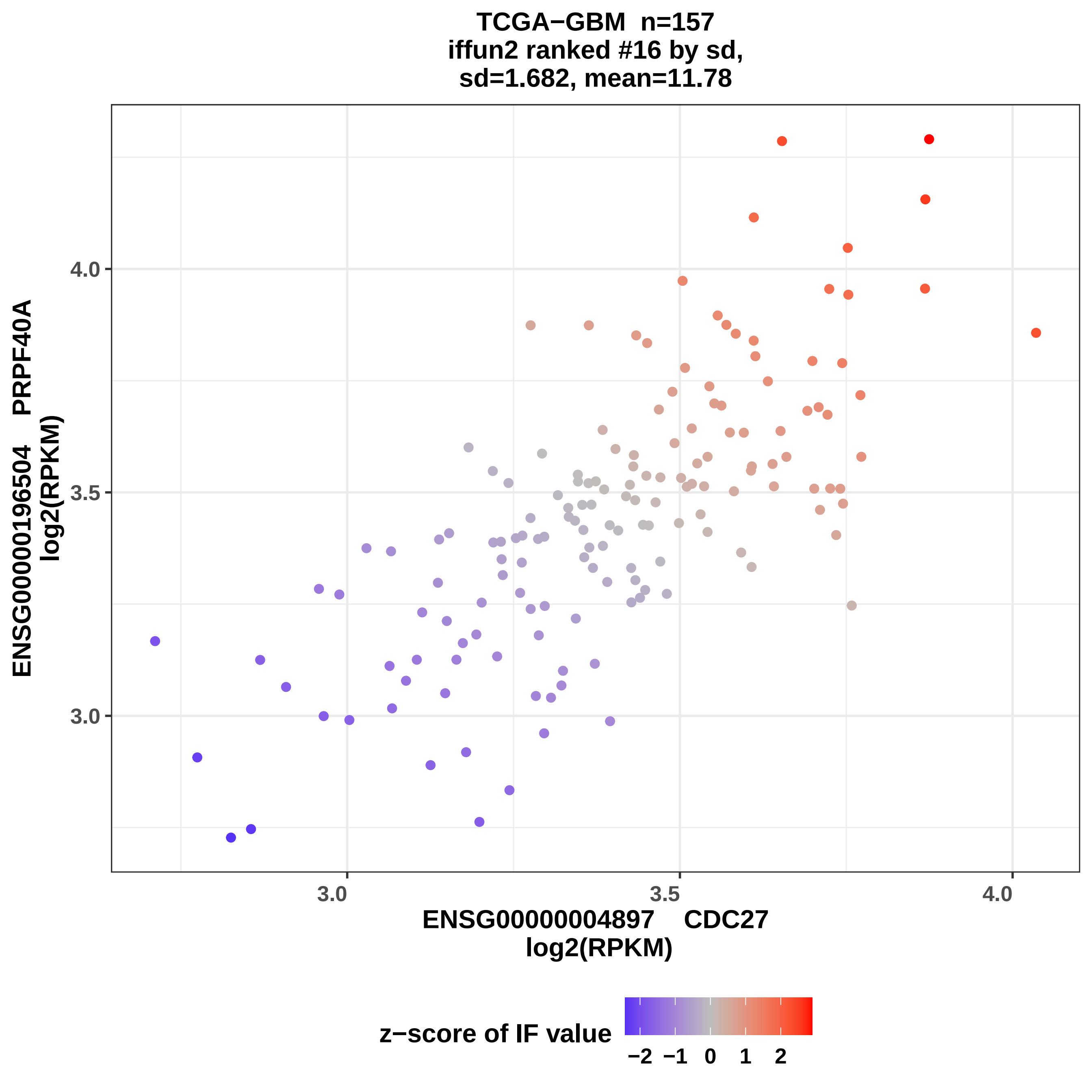
TCGA−GBM n=157
iffun2 ranked #16 by sd,
sd=1.682, mean=11.78
ENSG00000196504 PRPF40A
log2(RPKM)
4.0
3.5
3.0
3.0
3.5
4.0
ENSG00000004897 CDC27
log2(RPKM)
z−score of IF value
−2 −1 0 1 2

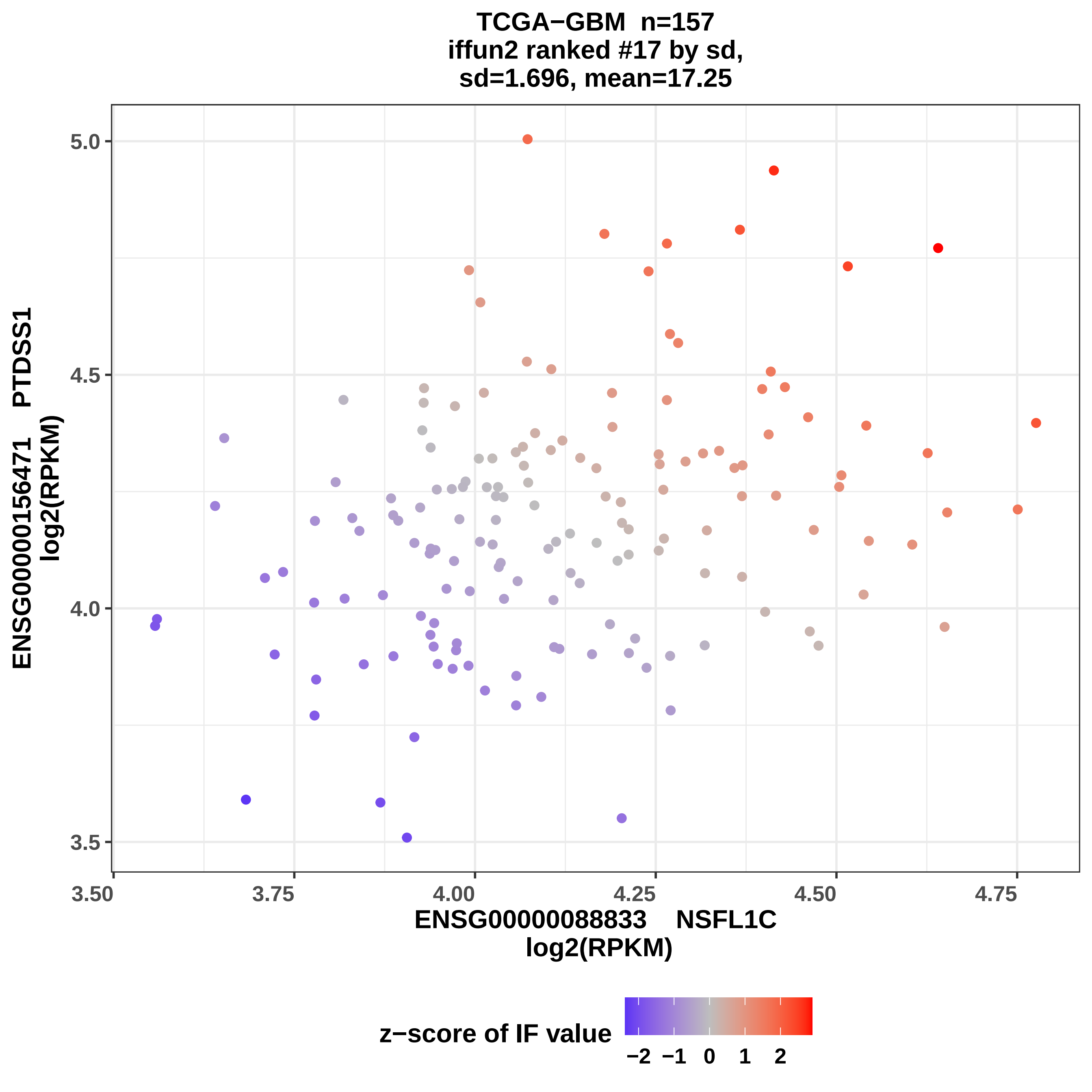

TCGA−GBM n=157
iffun2 ranked #17 by sd,
sd=1.696, mean=17.25
5.0
4.5
4.0
3.5
ENSG00000156471 PTDSS1
log2(RPKM)
3.50
3.75
4.00
4.25
4.50
4.75
ENSG00000088833 NSFL1C
log2(RPKM)
z−score of IF value
−2 −1 0 1 2

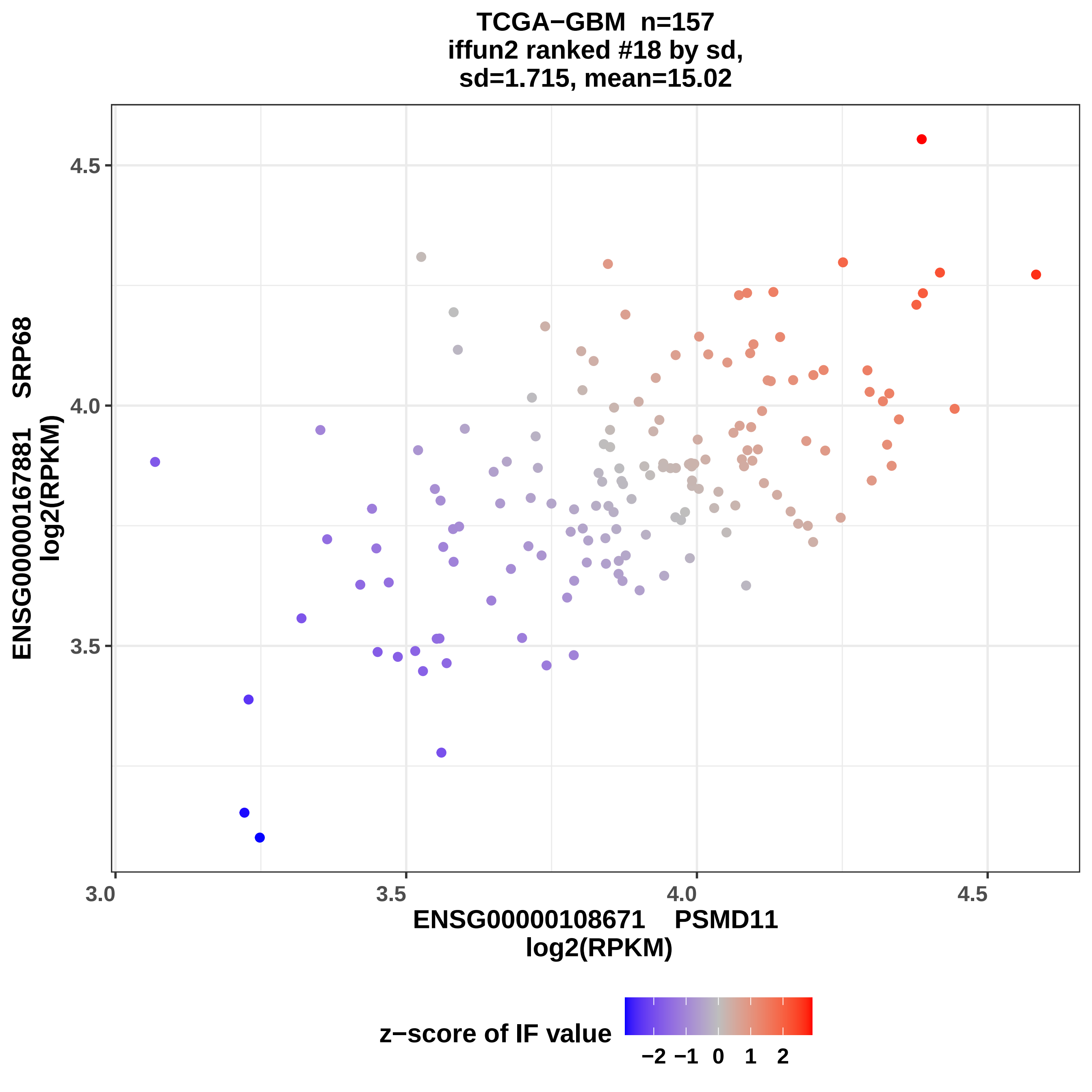
TCGA−GBM n=157
iffun2 ranked #18 by sd,
sd=1.715, mean=15.02
ENSG00000167881 SRP68
log2(RPKM)
4.5
4.0
3.5
3.0
3.5
4.0
4.5
ENSG00000108671 PSMD11
log2(RPKM)
z−score of IF value
−2 −1 0 1 2

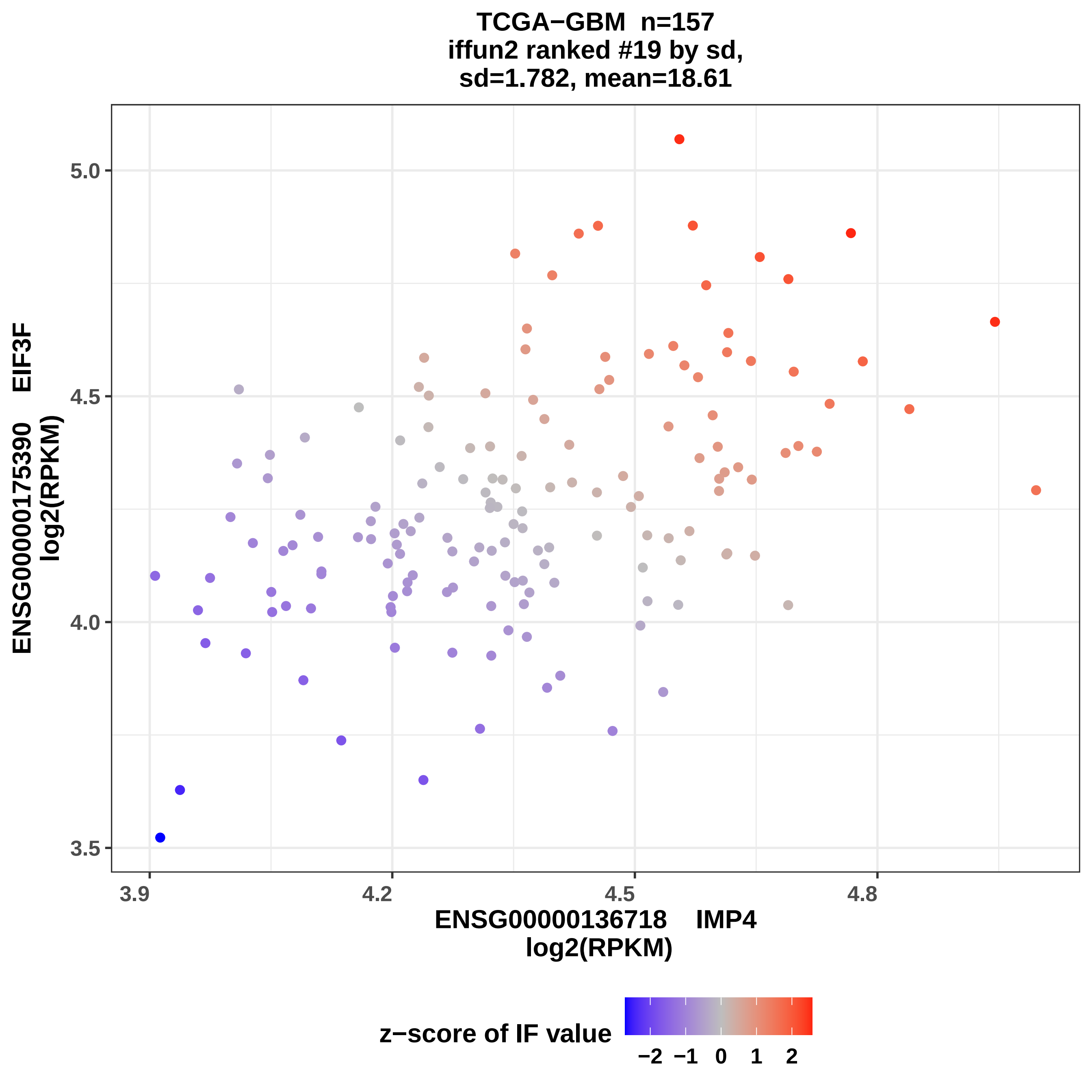
TCGA−GBM n=157
iffun2 ranked #19 by sd,
sd=1.782, mean=18.61
ENSG00000175390 EIF3F
log2(RPKM)
5.0
4.5
4.0
3.5
3.9
4.2
4.5
4.8
ENSG00000136718 IMP4
log2(RPKM)
z−score of IF value
−2 −1 0 1 2

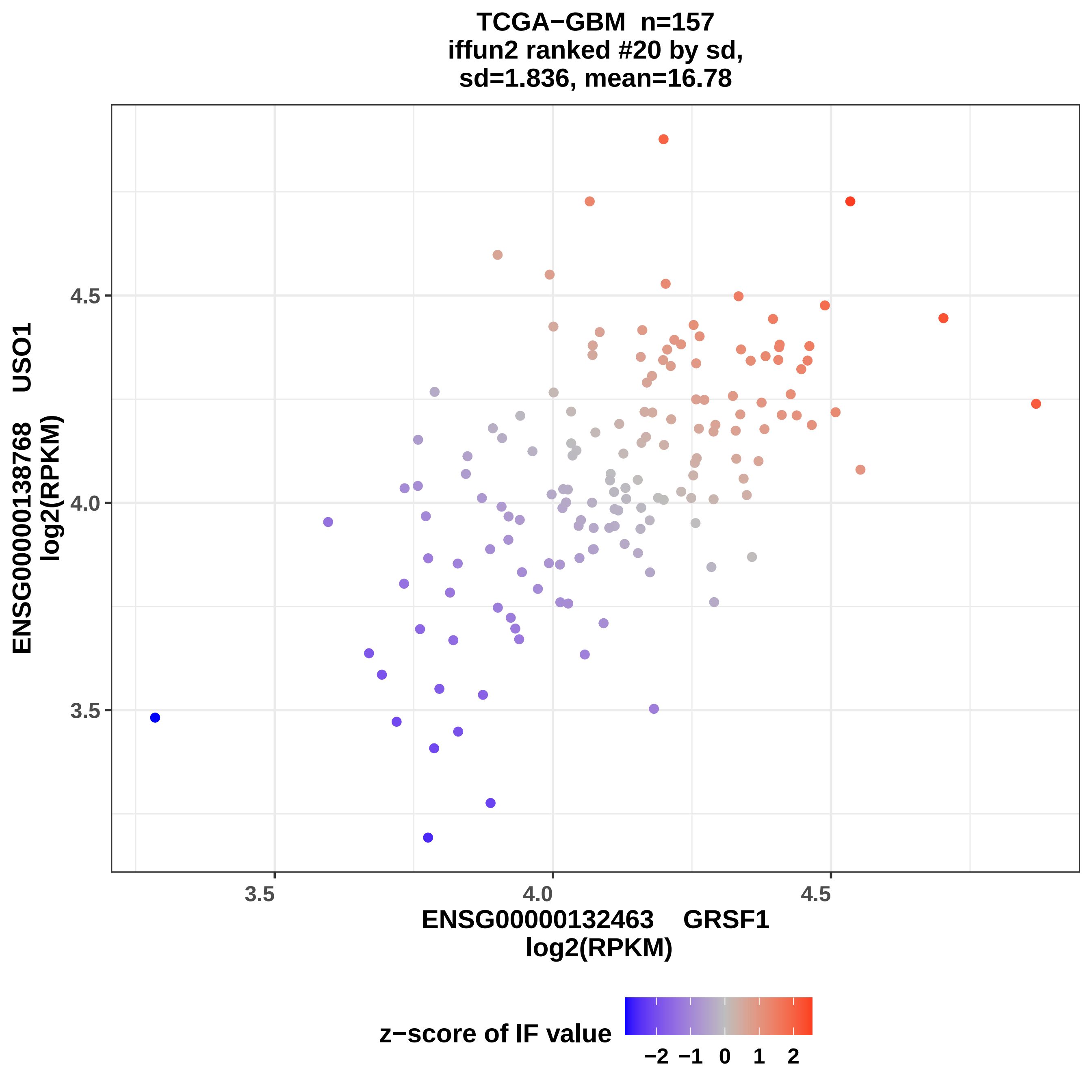
TCGA−GBM n=157
iffun2 ranked #20 by sd,
sd=1.836, mean=16.78
ENSG00000138768 USO1
log2(RPKM)
3.5
4.0
4.5
ENSG00000132463 GRSF1
log2(RPKM)
3.5
4.0
4.5
z−score of IF value
−2
−1
0
1
2

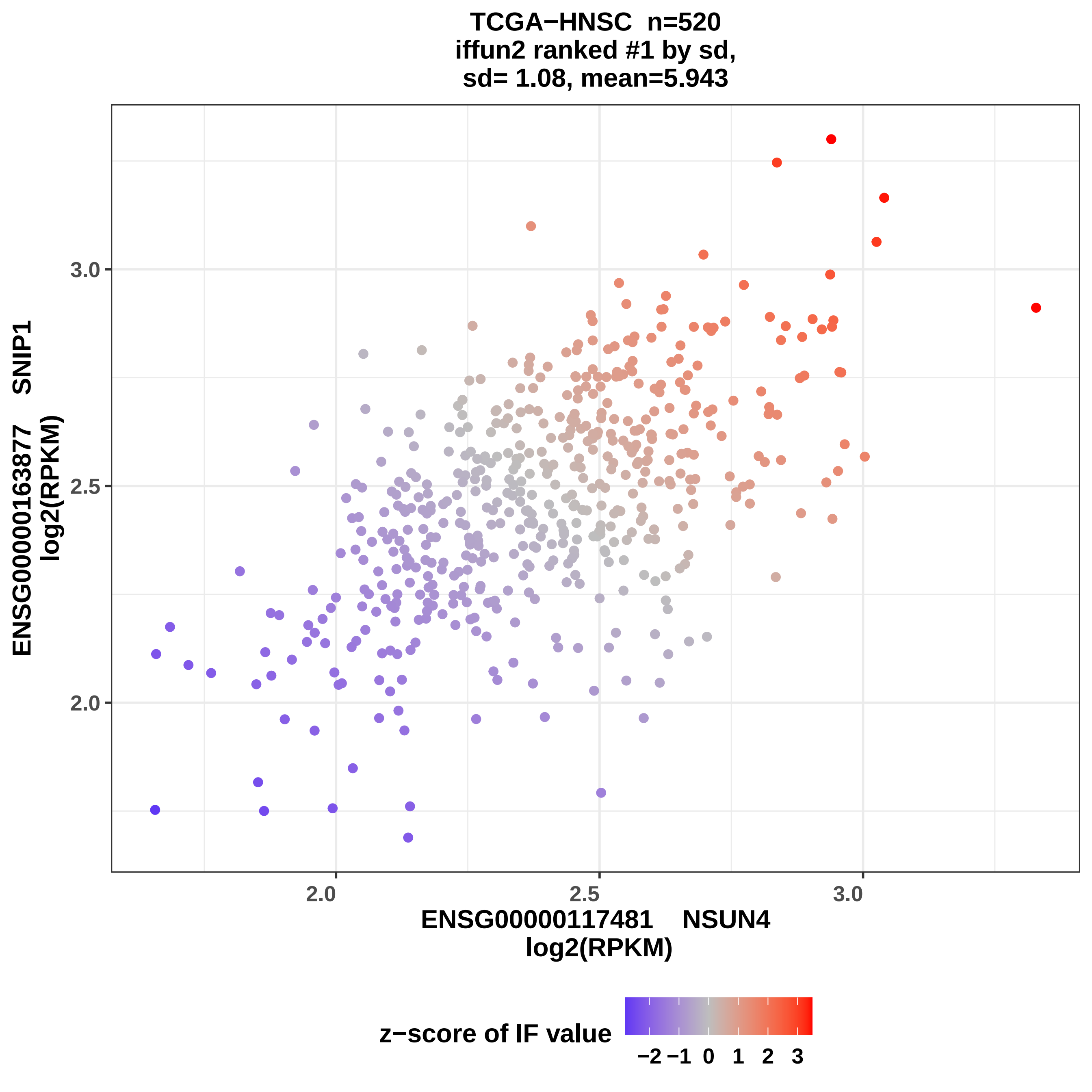

TCGA−HNSC n=520
iffun2 ranked #1 by sd,
sd= 1.08, mean=5.943
ENSG00000163877 SNIP1
log2(RPKM)
3.0
2.5
2.0
2.0
2.5
3.0
ENSG00000117481 NSUN4
log2(RPKM)
z−score of IF value
−2 −1 0 1 2 3

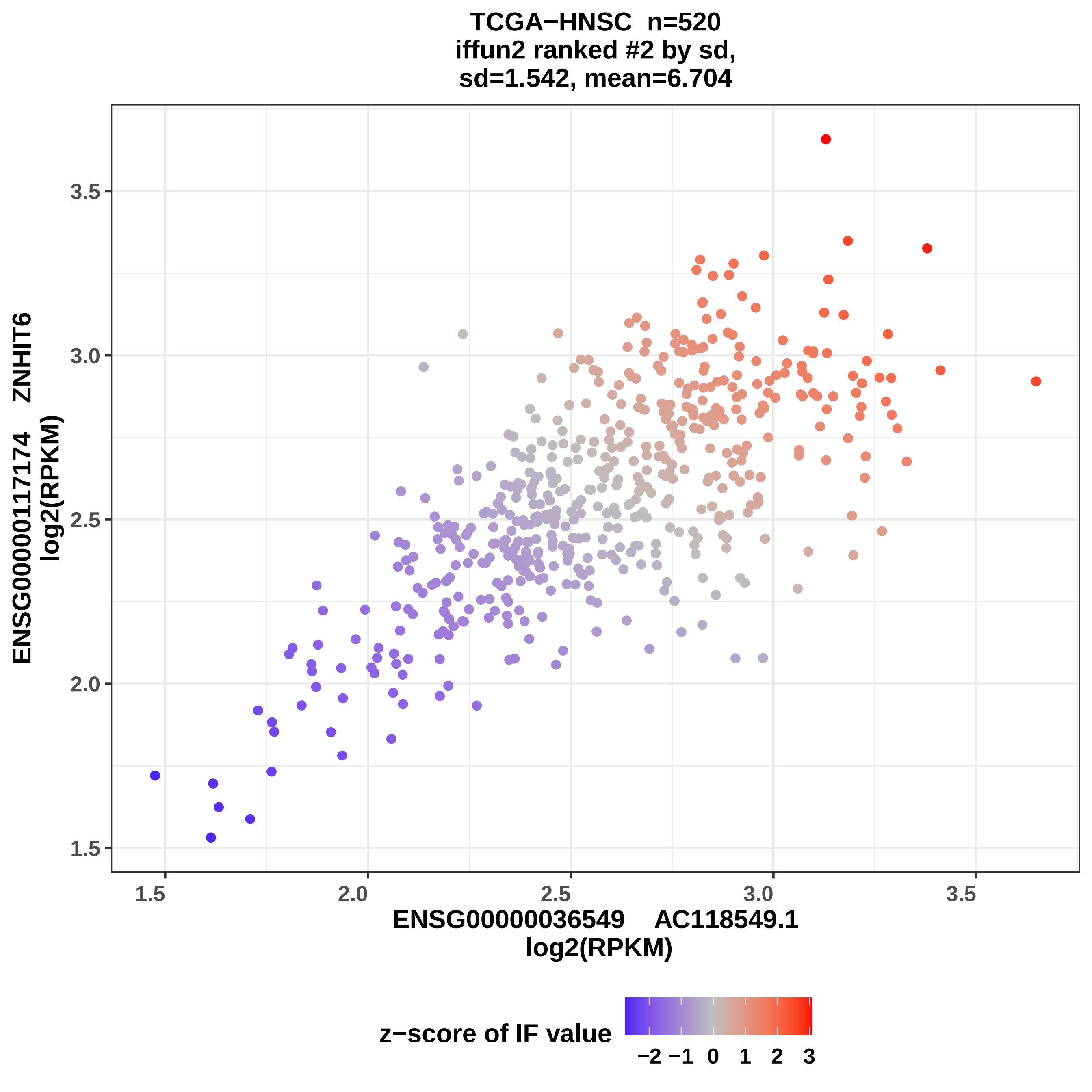
TCGA−HNSC n=520
iffun2 ranked #2 by sd,
sd=1.542, mean=6.704
ENSG00000117174 ZNHIT6
log2(RPKM)
3.5
3.0
2.5
2.0
1.5
1.5
2.0
2.5
3.0
3.5
ENSG00000036549 AC118549.1
log2(RPKM)
z−score of IF value
−2 −1 0 1 2 3

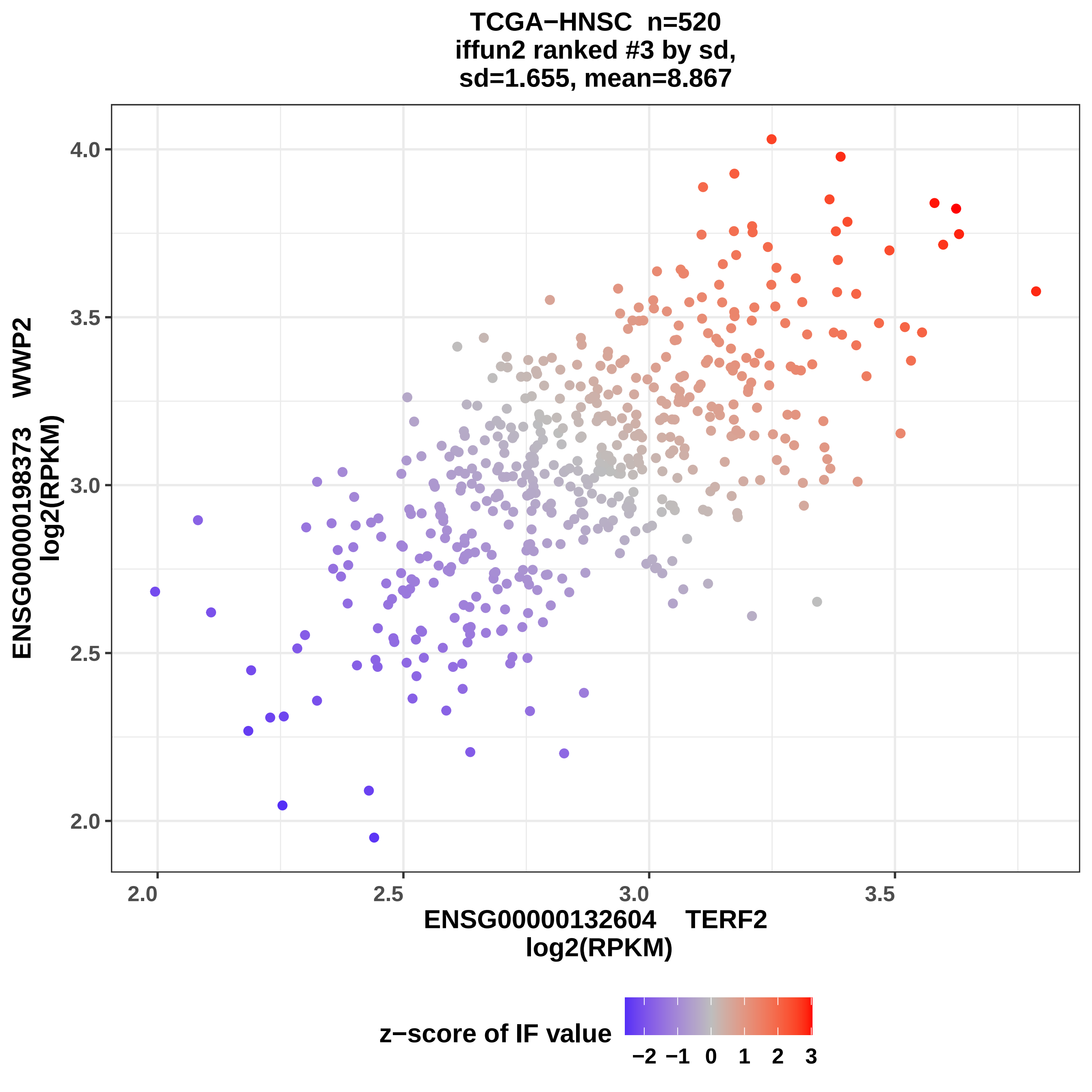
TCGA−HNSC n=520
iffun2 ranked #3 by sd,
sd=1.655, mean=8.867
ENSG00000198373 WWP2
log2(RPKM)
4.0
3.5
3.0
2.5
2.0
2.0
2.5
3.0
3.5
ENSG00000132604 TERF2
log2(RPKM)
z−score of IF value
−2 −1 0 1 2 3

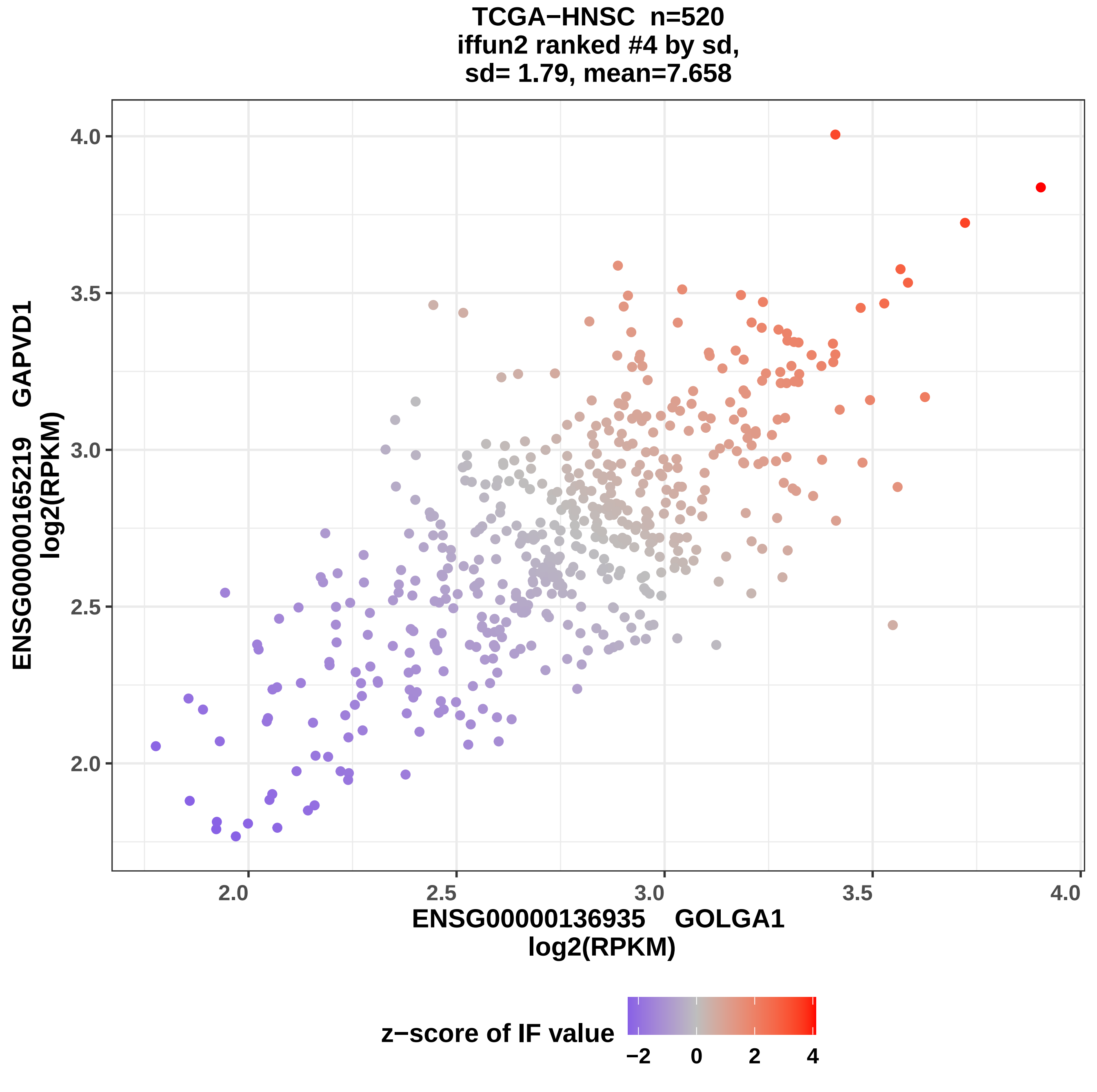
TCGA−HNSC n=520
iffun2 ranked #4 by sd,
sd= 1.79, mean=7.658
ENSG00000165219 GAPVD1
log2(RPKM)
4.0
3.5
3.0
2.5
2.0
2.0
2.5
3.0
3.5
4.0
ENSG00000136935 GOLGA1
log2(RPKM)
z−score of IF value
−2
0
2
4

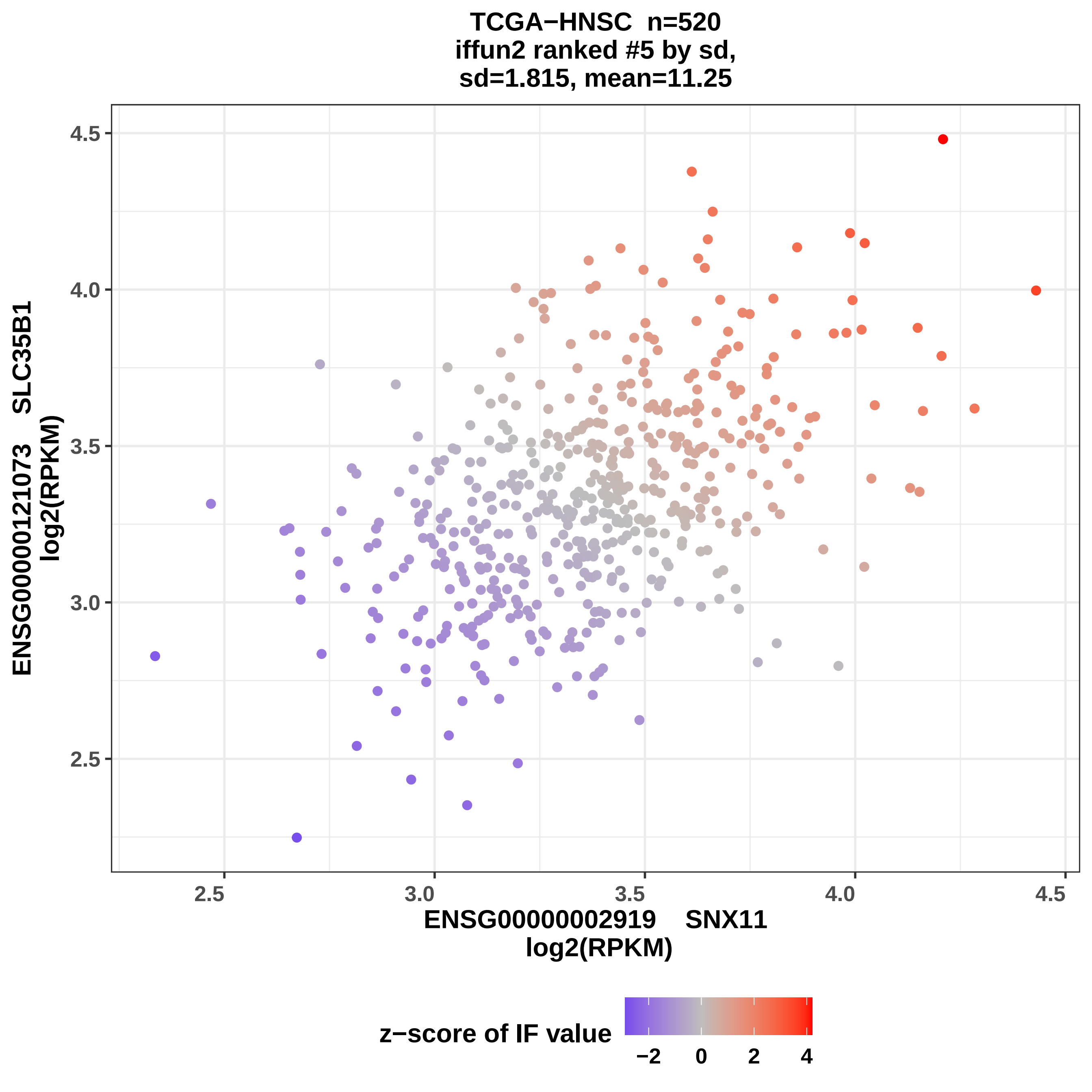
TCGA−HNSC n=520
iffun2 ranked #5 by sd,
sd=1.815, mean=11.25
ENSG00000121073 SLC35B1
log2(RPKM)
4.5
4.0
3.5
3.0
2.5
2.5
3.0
3.5
4.0
4.5
ENSG00000002919 SNX11
log2(RPKM)
z−score of IF value
−2
0
2
4

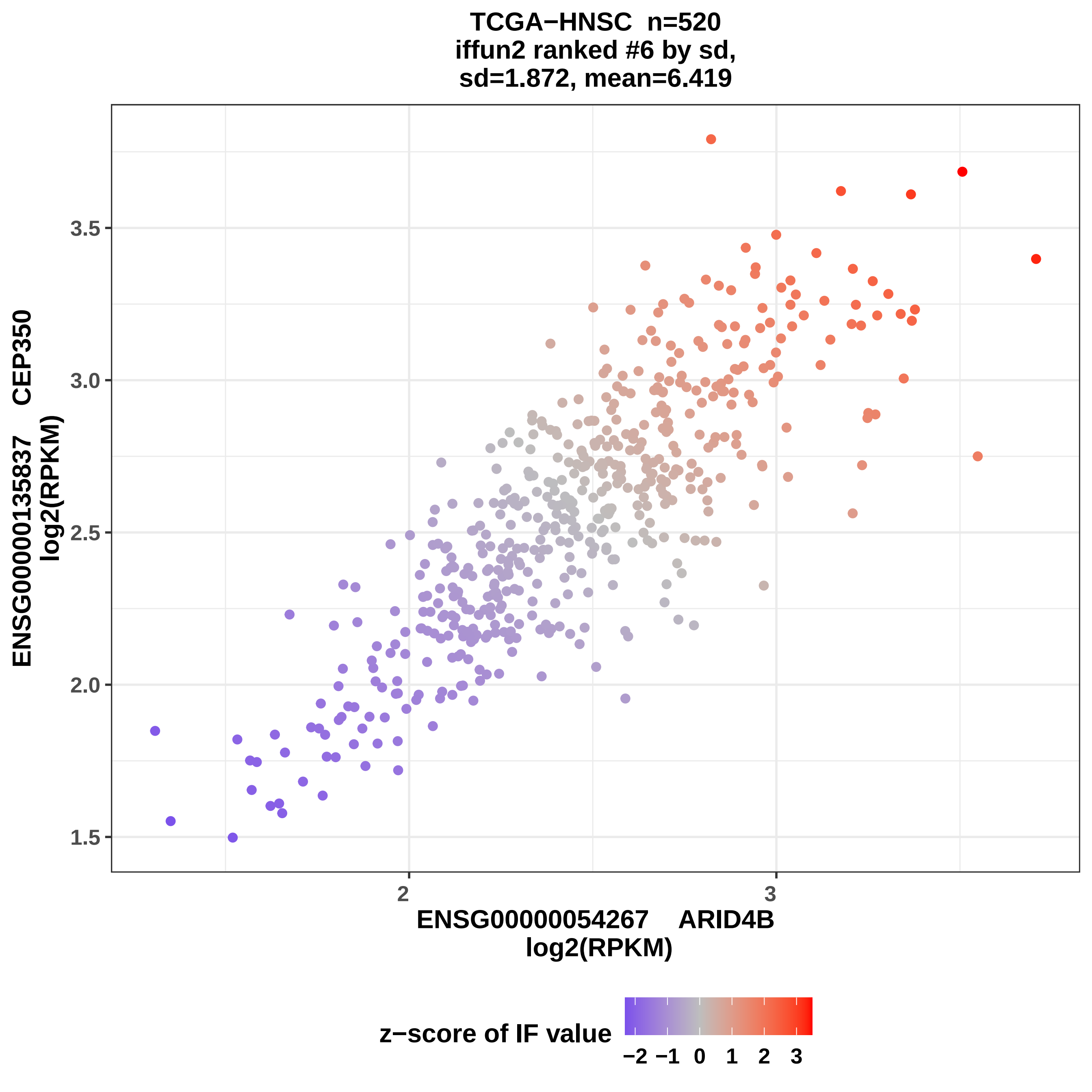
TCGA−HNSC n=520
iffun2 ranked #6 by sd,
sd=1.872, mean=6.419
ENSG00000135837 CEP350
log2(RPKM)
3.5
3.0
2.5
2.0
1.5
2
3
ENSG00000054267 ARID4B
log2(RPKM)
z−score of IF value
−2 −1 0 1 2 3

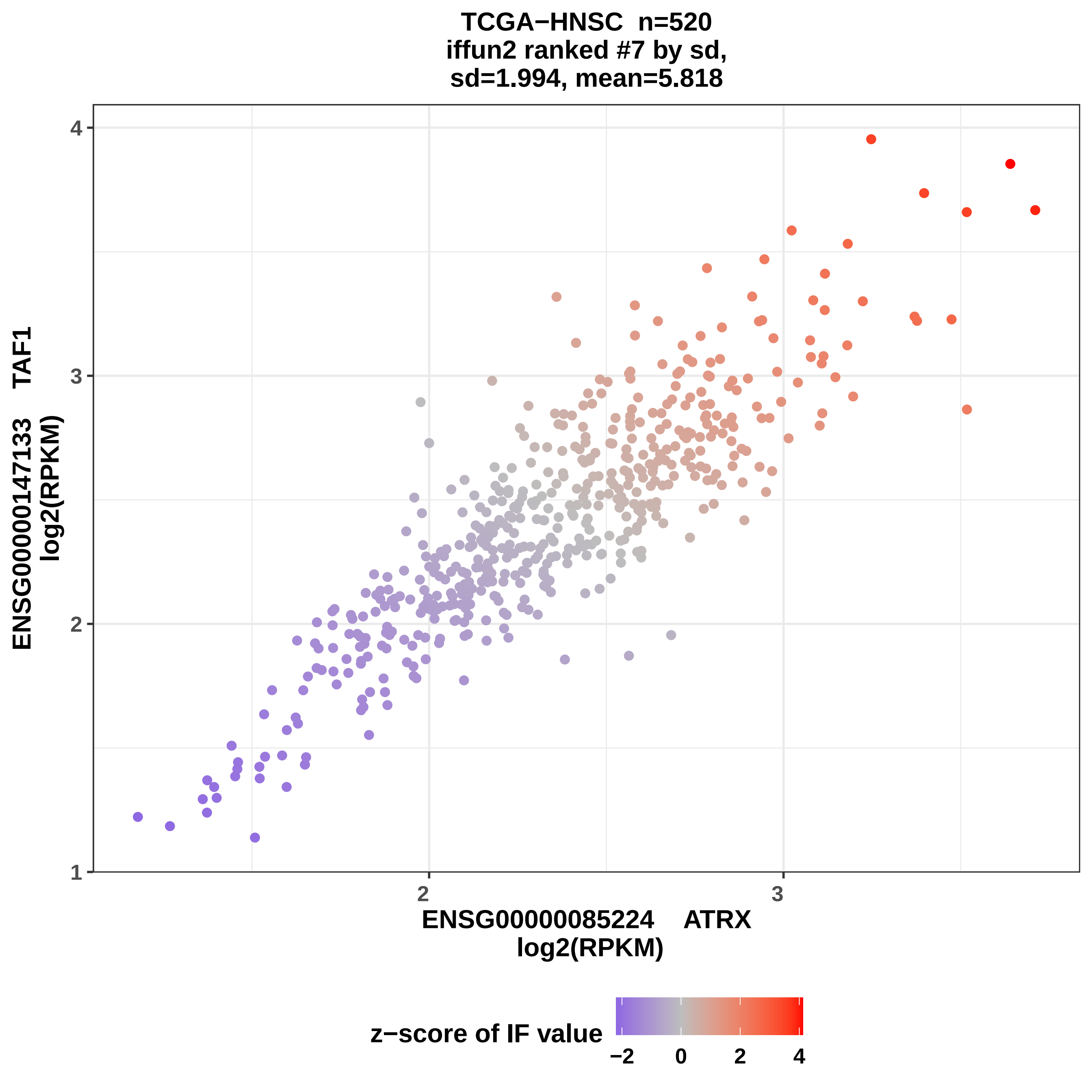

TCGA−HNSC n=520
iffun2 ranked #7 by sd,
sd=1.994, mean=5.818
ENSG00000147133 TAF1
log2(RPKM)
ENSG00000085224 ATRX
log2(RPKM)
z−score of IF value
−2
0
2
4
1
2
3
4

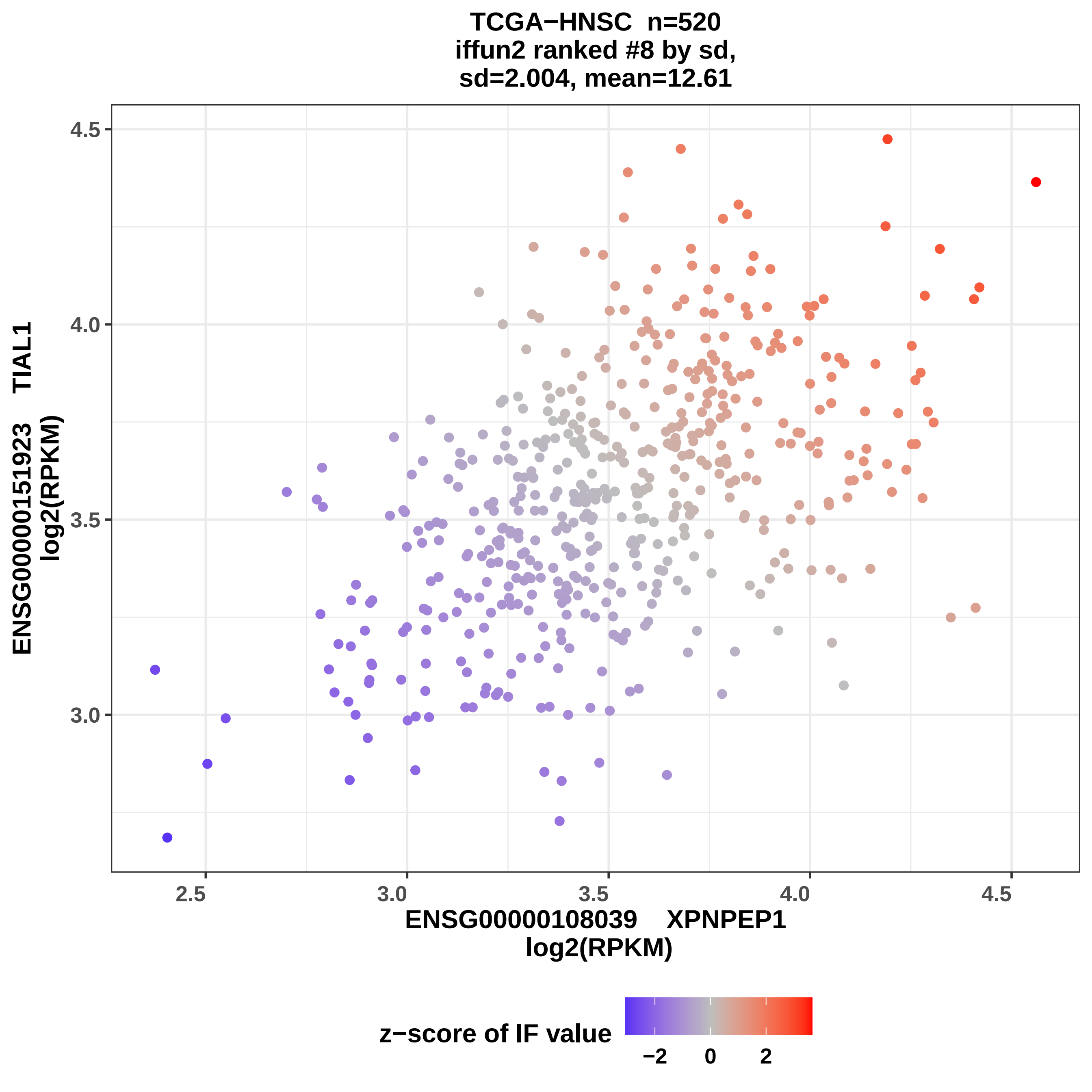

TCGA−HNSC n=520
iffun2 ranked #8 by sd,
sd=2.004, mean=12.61
ENSG00000151923 TIAL1
log2(RPKM)
4.5
4.0
3.5
3.0
2.5
3.0
3.5
4.0
4.5
ENSG00000108039 XPNPEP1
log2(RPKM)
z−score of IF value
−2
0
2

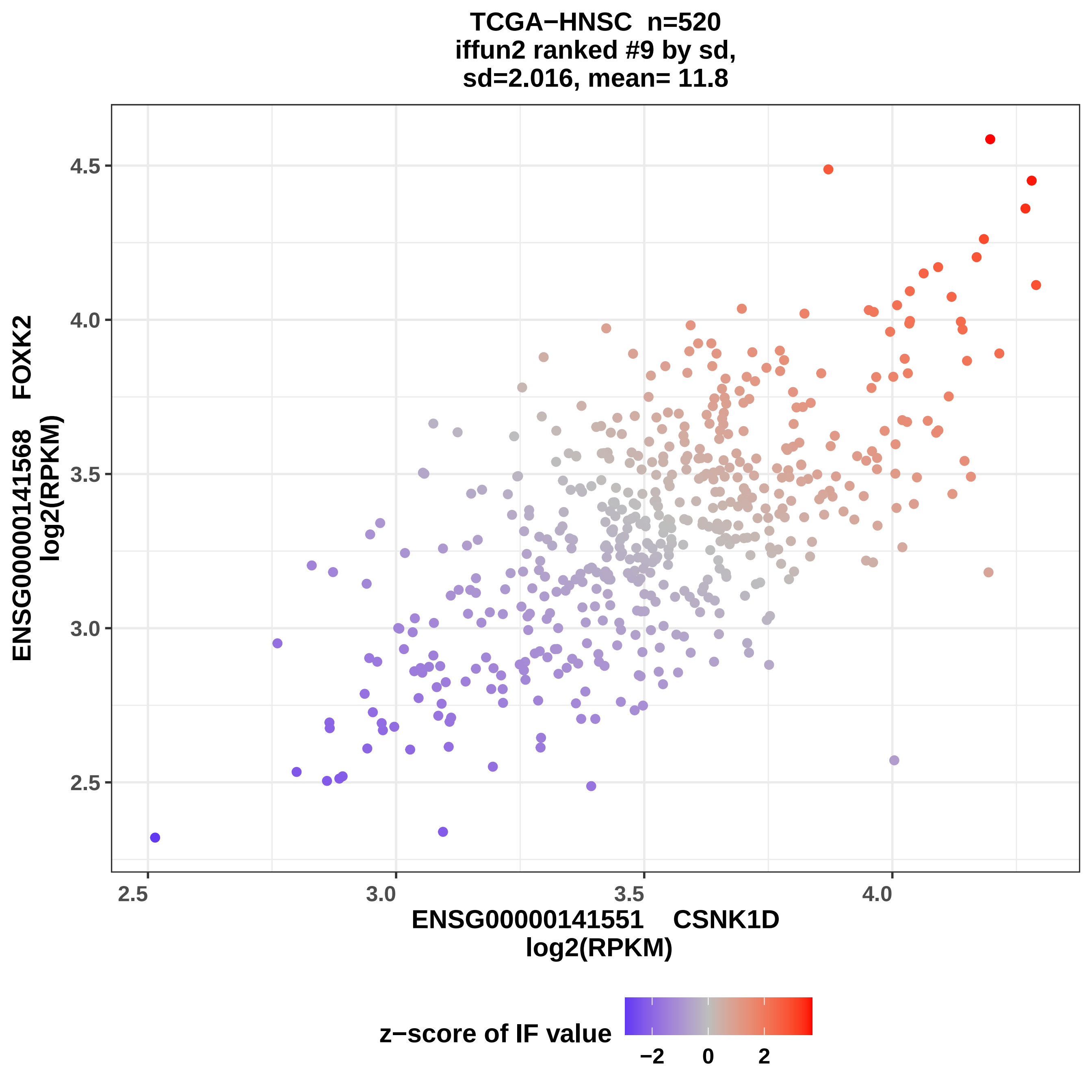

TCGA−HNSC n=520
iffun2 ranked #9 by sd,
sd=2.016, mean= 11.8
ENSG00000141568 FOXK2
log2(RPKM)
4.5
4.0
3.5
3.0
2.5
2.5
3.0
3.5
4.0
ENSG00000141551 CSNK1D
log2(RPKM)
z−score of IF value
−2
0
2

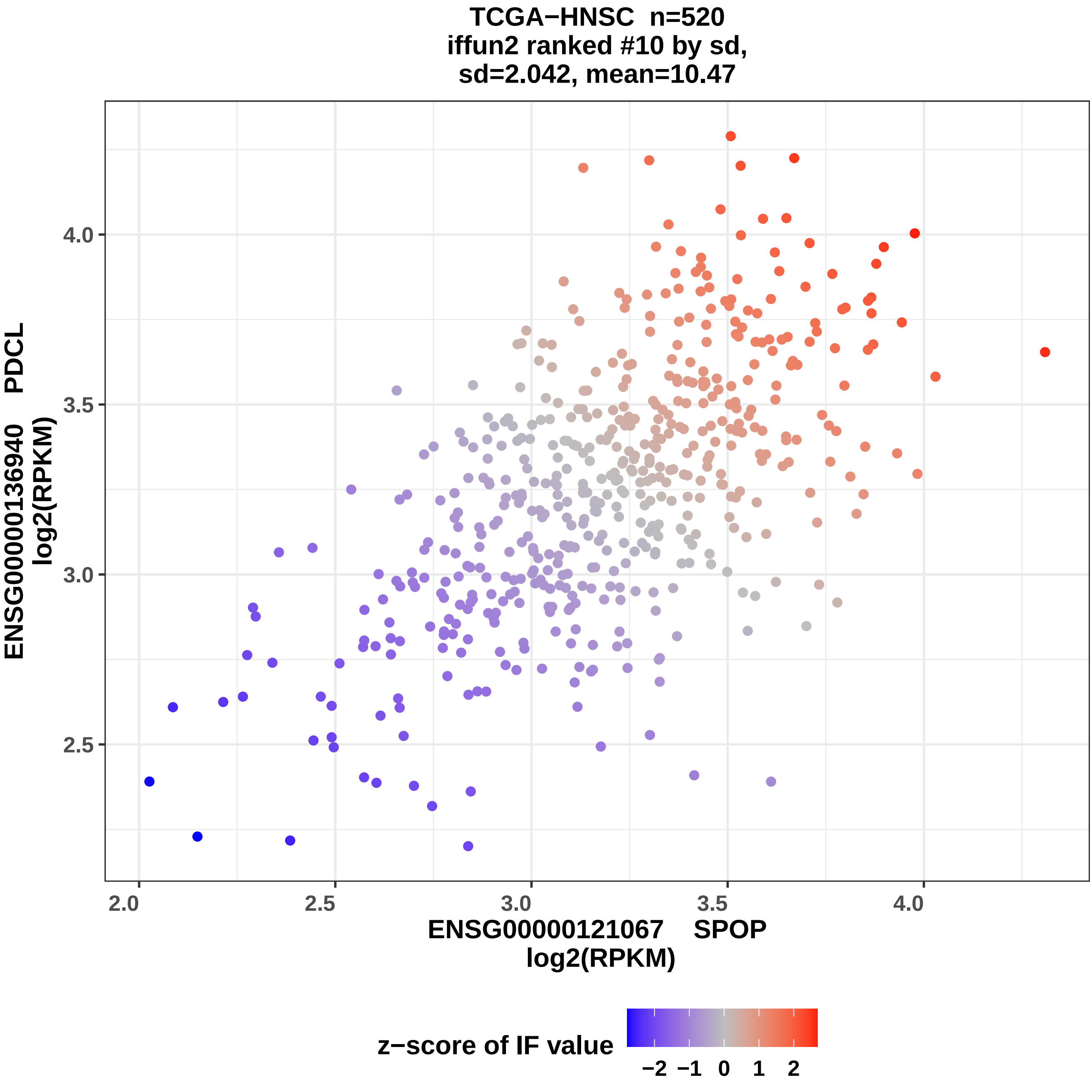
TCGA−HNSC n=520
iffun2 ranked #10 by sd,
sd=2.042, mean=10.47
ENSG00000136940 PDCL
log2(RPKM)
4.0
3.5
3.0
2.5
2.0
2.5
3.0
3.5
4.0
ENSG00000121067 SPOP
log2(RPKM)
z−score of IF value
−2 −1 0 1 2

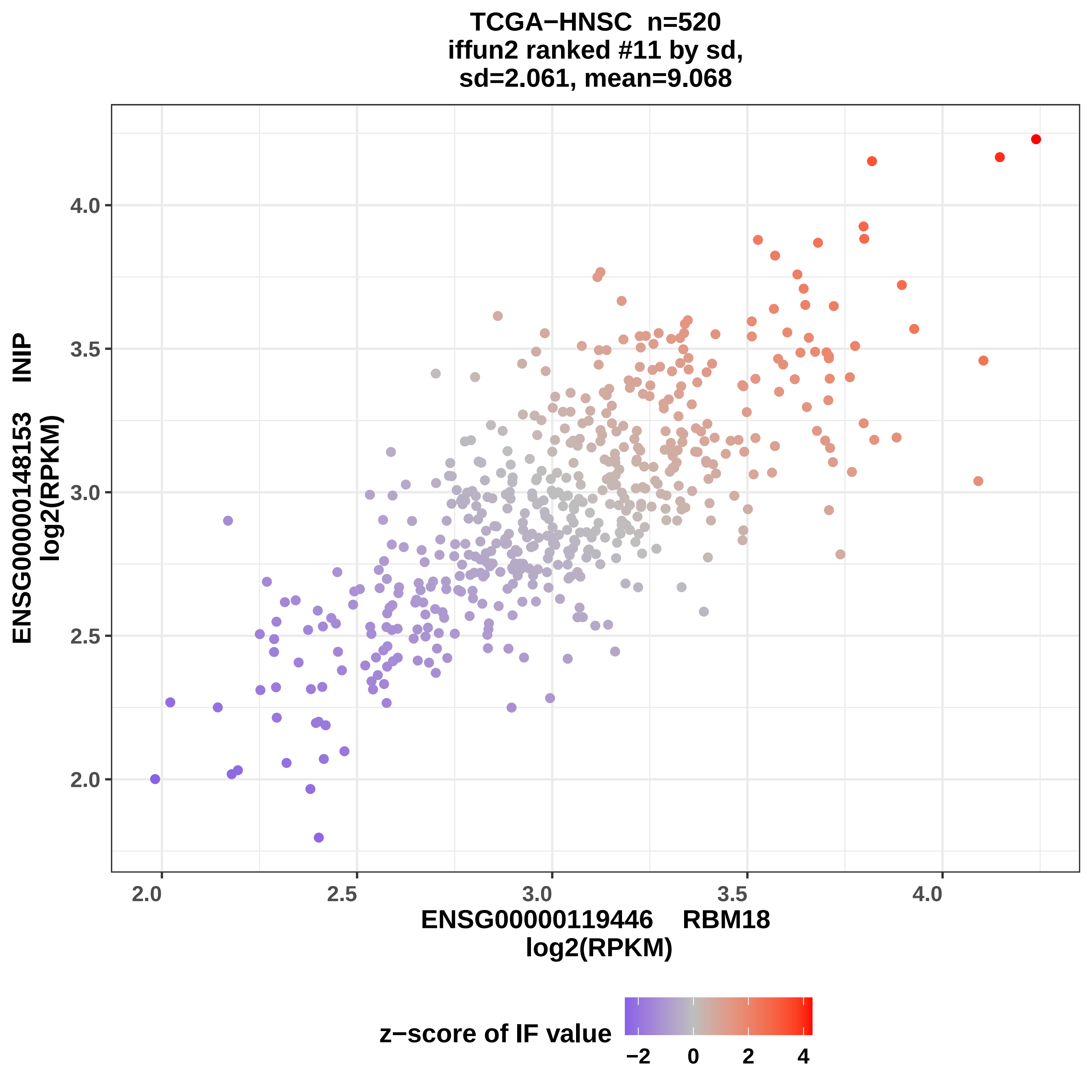
TCGA−HNSC n=520
iffun2 ranked #11 by sd,
sd=2.061, mean=9.068
ENSG00000148153 INIP
log2(RPKM)
ENSG00000119446 RBM18
log2(RPKM)
2.0
2.5
3.0
3.5
4.0
z−score of IF value
−2
0
2
4

TCGA−HNSC n=520
iffun2 ranked #12 by sd,
sd=2.068, mean=14.31

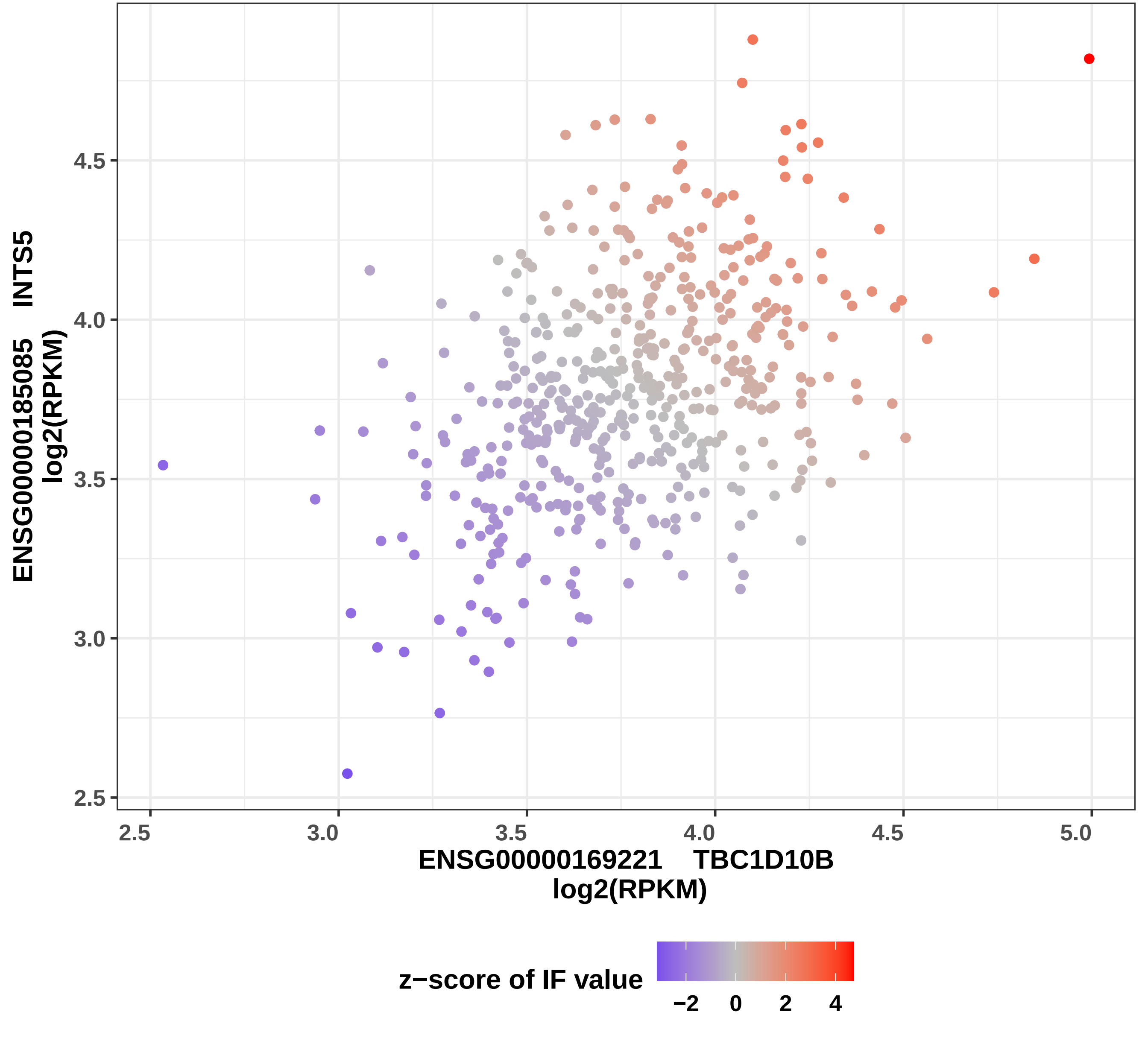

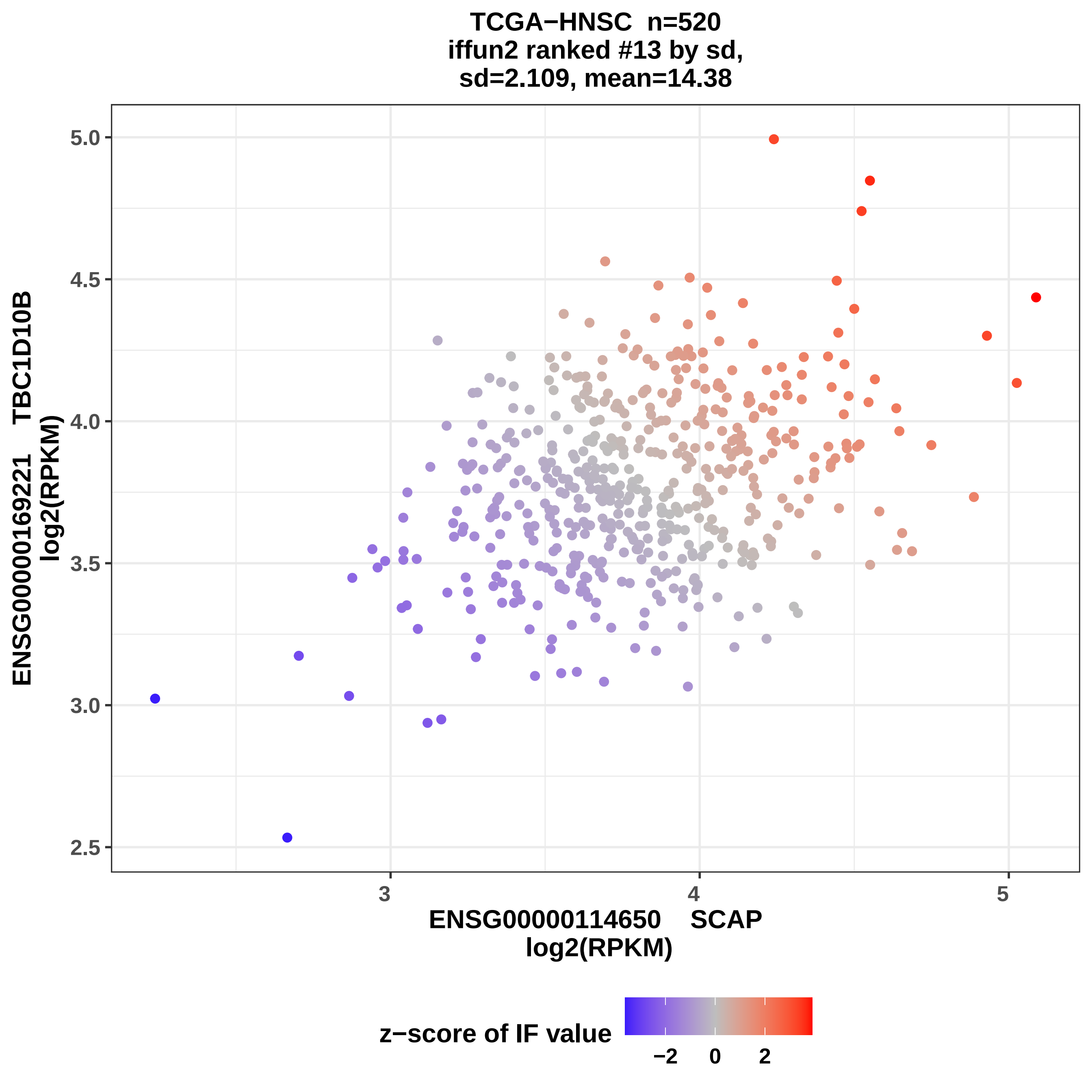

TCGA−HNSC n=520
iffun2 ranked #13 by sd,
sd=2.109, mean=14.38
ENSG00000169221 TBC1D10B
log2(RPKM)
5.0
4.5
4.0
3.5
3.0
2.5
3
4
5
ENSG00000114650 SCAP
log2(RPKM)
z−score of IF value
−2
0
2

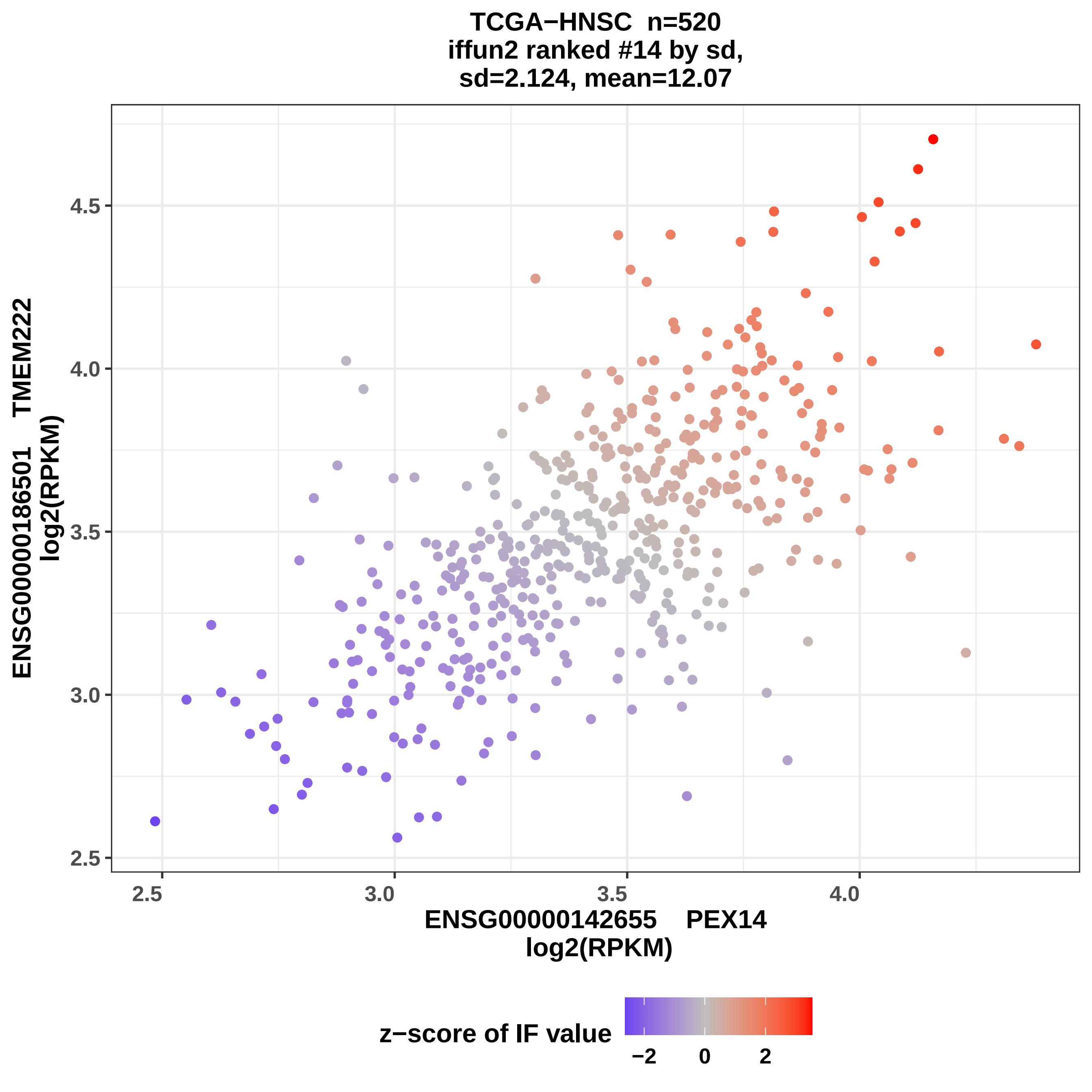
TCGA−HNSC n=520
iffun2 ranked #14 by sd,
sd=2.124, mean=12.07
ENSG00000186501 TMEM222
log2(RPKM)
4.5
4.0
3.5
3.0
2.5
2.5
3.0
3.5
4.0
ENSG00000142655 PEX14
log2(RPKM)
z−score of IF value
−2
0
2

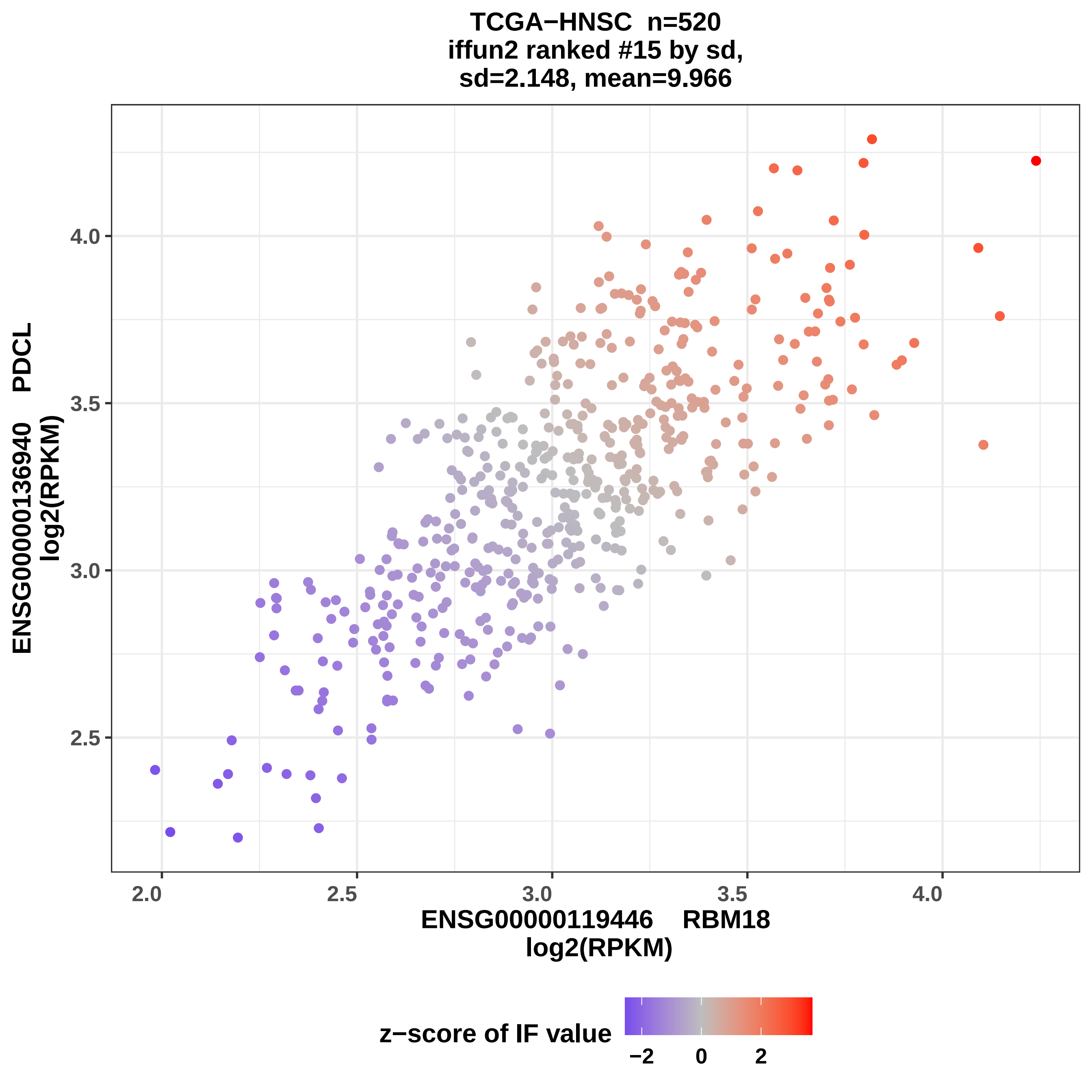

TCGA−HNSC n=520
iffun2 ranked #15 by sd,
sd=2.148, mean=9.966
ENSG00000136940 PDCL
log2(RPKM)
4.0
3.5
3.0
2.5
2.0
2.5
3.0
3.5
4.0
ENSG00000119446 RBM18
log2(RPKM)
z−score of IF value
−2
0
2

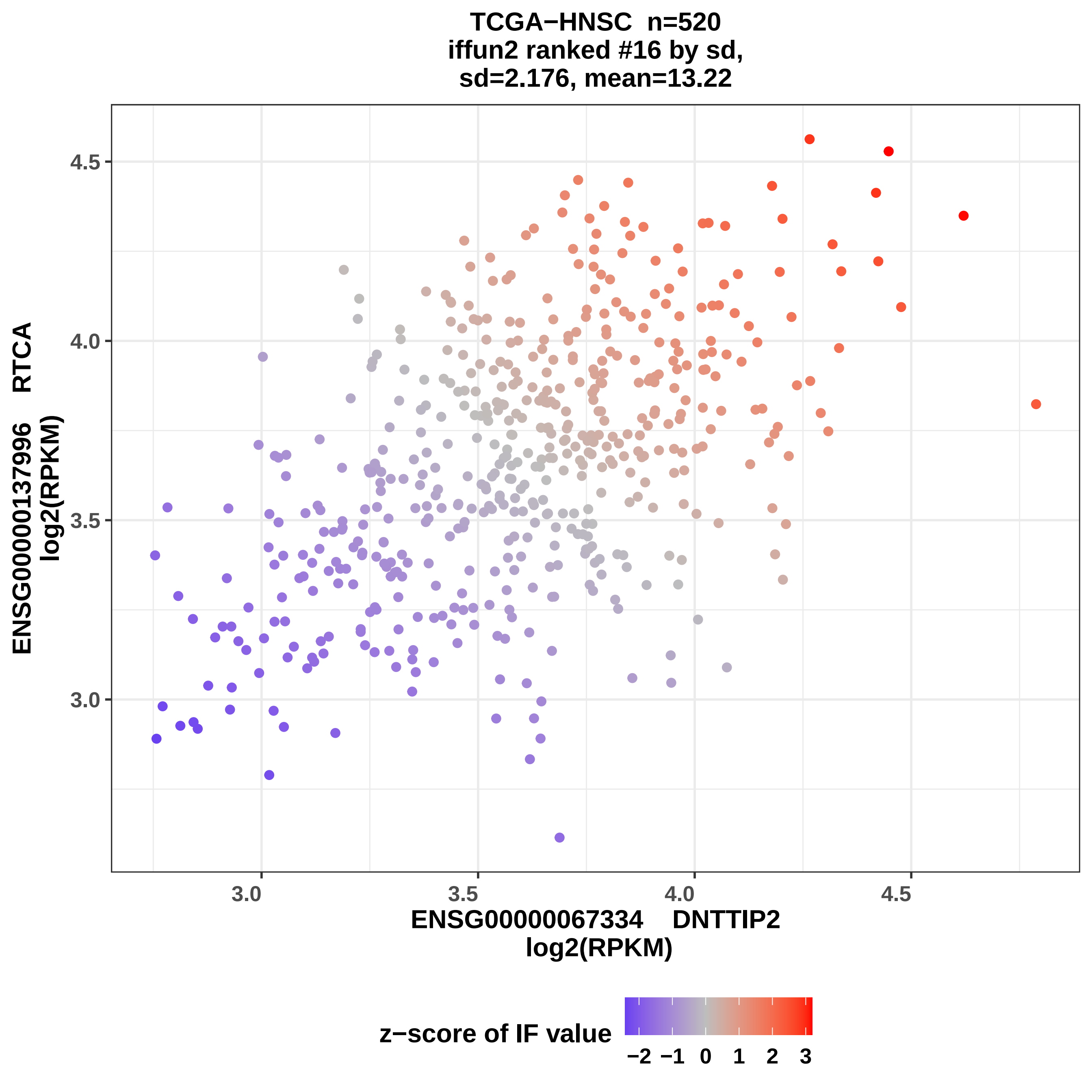
TCGA−HNSC n=520
iffun2 ranked #16 by sd,
sd=2.176, mean=13.22
4.5
4.0
3.5
3.0
3.0
3.5
4.0
4.5
ENSG00000137996 RTCA
log2(RPKM)
ENSG00000067334 DNTTIP2
log2(RPKM)
z−score of IF value
−2 −1 0 1 2 3

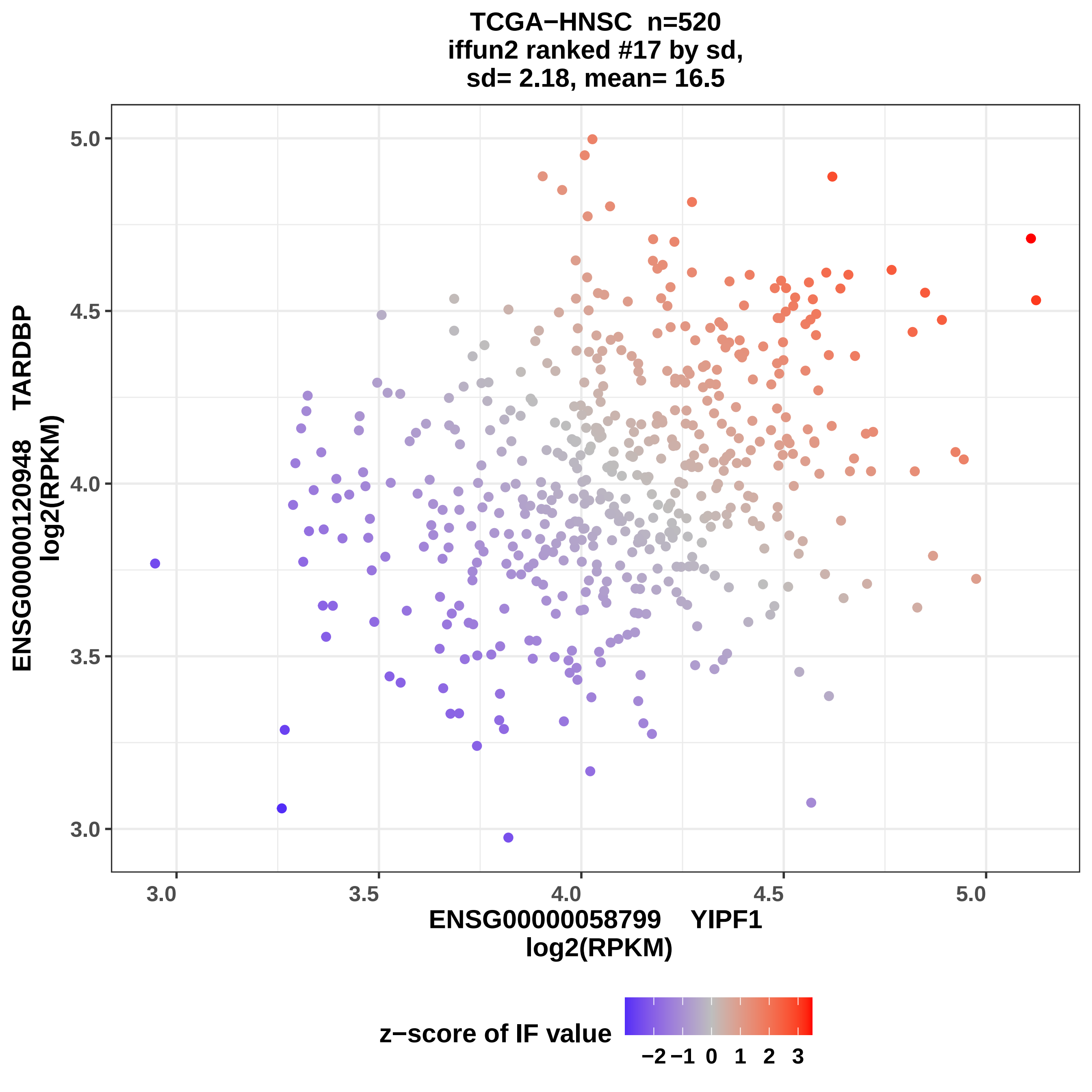

TCGA−HNSC n=520
iffun2 ranked #17 by sd,
sd= 2.18, mean= 16.5
ENSG00000120948 TARDBP
log2(RPKM)
5.0
4.5
4.0
3.5
3.0
3.0
3.5
4.0
4.5
5.0
ENSG00000058799 YIPF1
log2(RPKM)
z−score of IF value
−2
−1
0
1
2
3

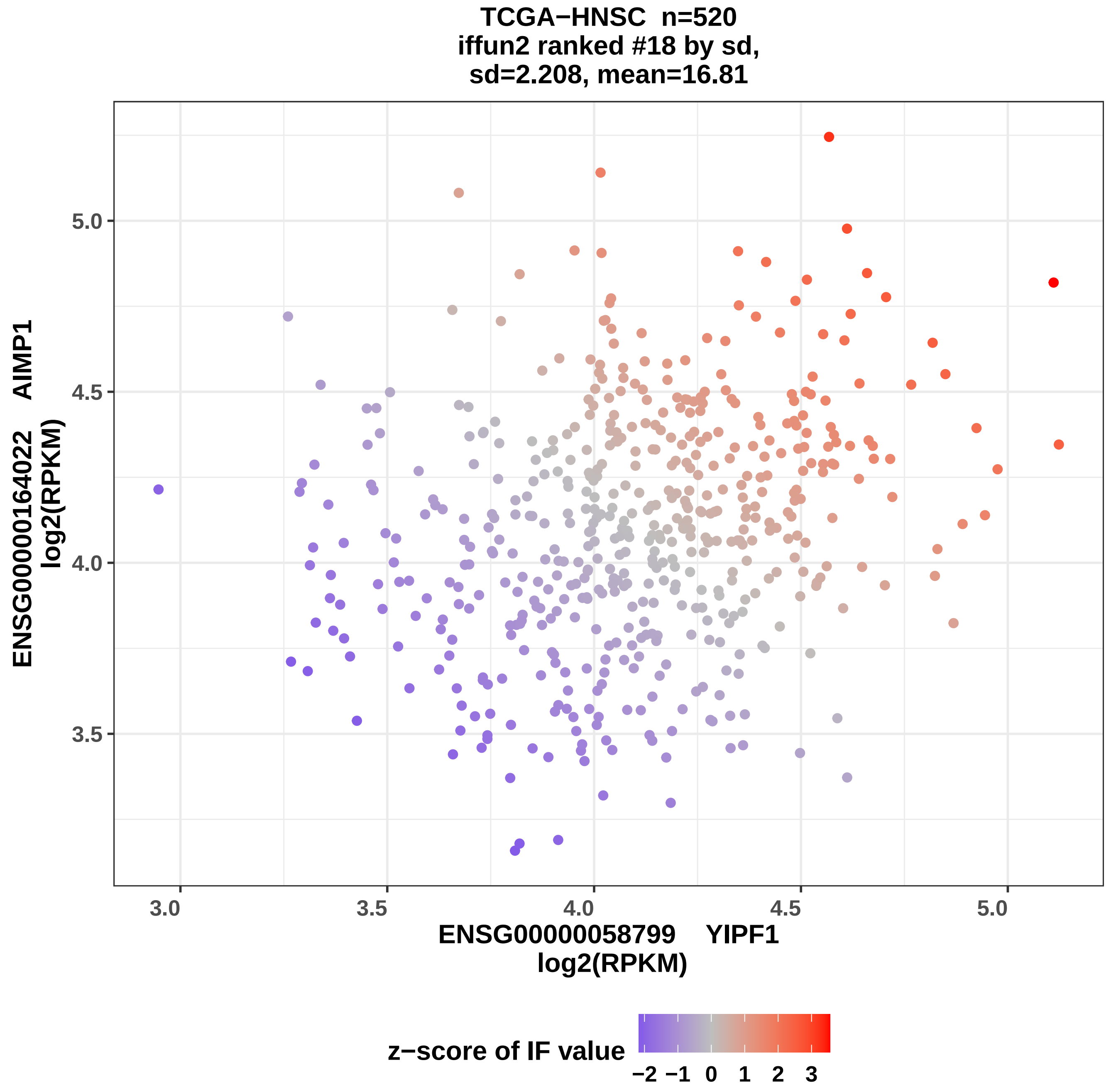

TCGA−HNSC n=520
iffun2 ranked #18 by sd,
sd=2.208, mean=16.81
ENSG00000164022 AIMP1
log2(RPKM)
5.0
4.5
4.0
3.5
3.0
3.5
4.0
4.5
5.0
ENSG00000058799 YIPF1
log2(RPKM)
z−score of IF value
−2 −1 0 1 2 3

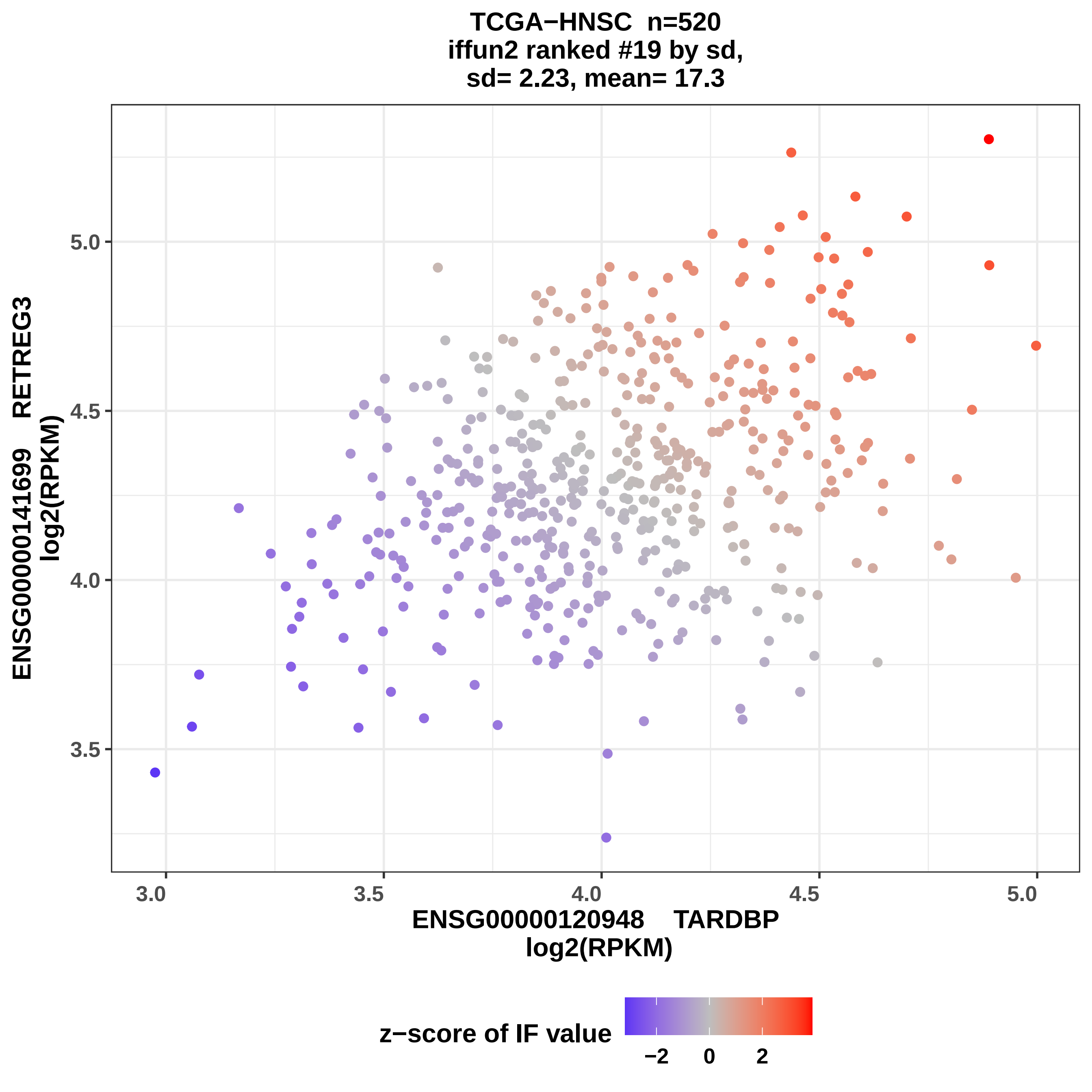
TCGA−HNSC n=520
iffun2 ranked #19 by sd,
sd= 2.23, mean= 17.3
ENSG00000141699 RETREG3
log2(RPKM)
5.0
4.5
4.0
3.5
3.0
3.5
4.0
4.5
5.0
ENSG00000120948 TARDBP
log2(RPKM)
z−score of IF value
−2
0
2

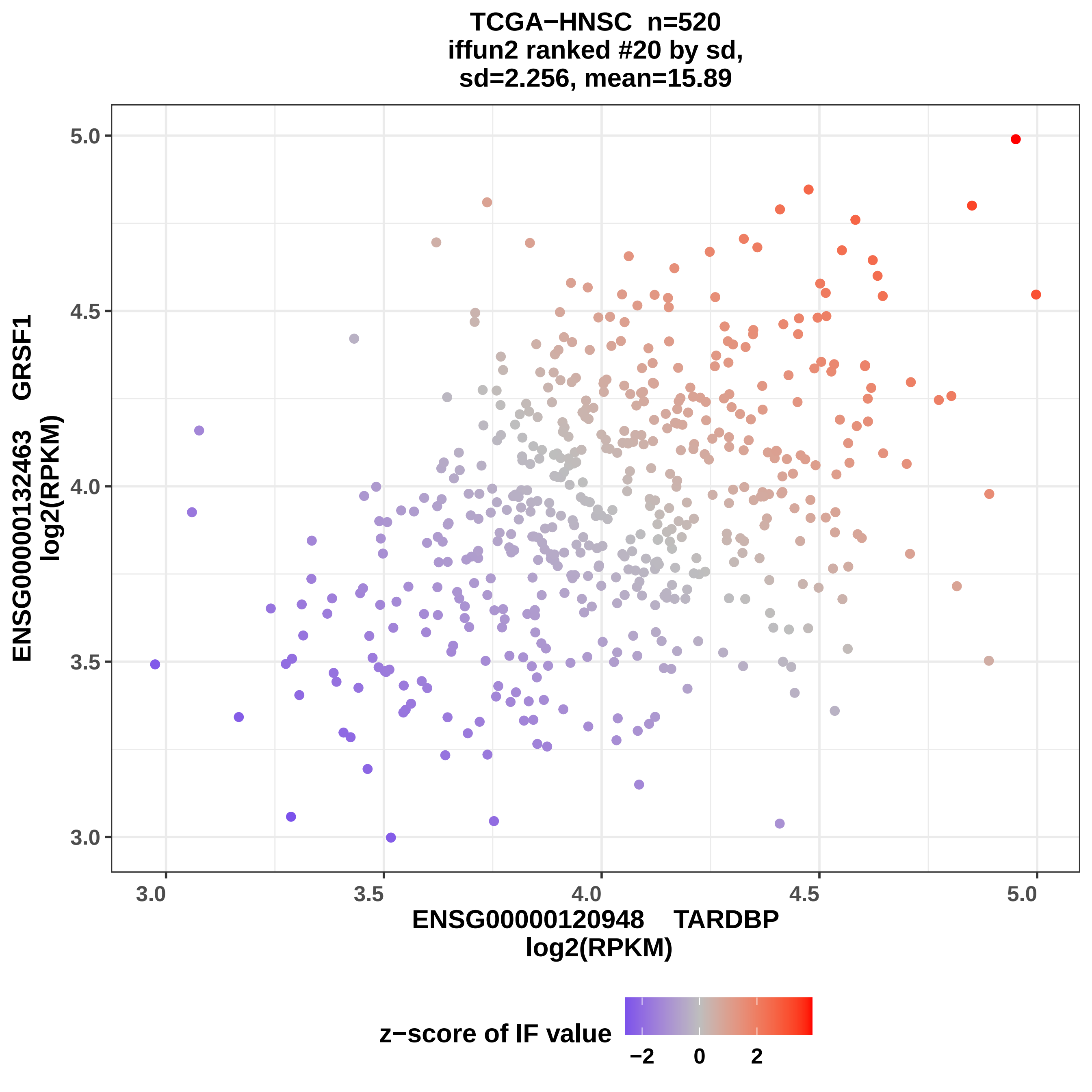
TCGA−HNSC n=520
iffun2 ranked #20 by sd,
sd=2.256, mean=15.89
ENSG00000132463 GRSF1
log2(RPKM)
5.0
4.5
4.0
3.5
3.0
3.0
3.5
4.0
4.5
5.0
ENSG00000120948 TARDBP
log2(RPKM)
z−score of IF value
−2
0
2

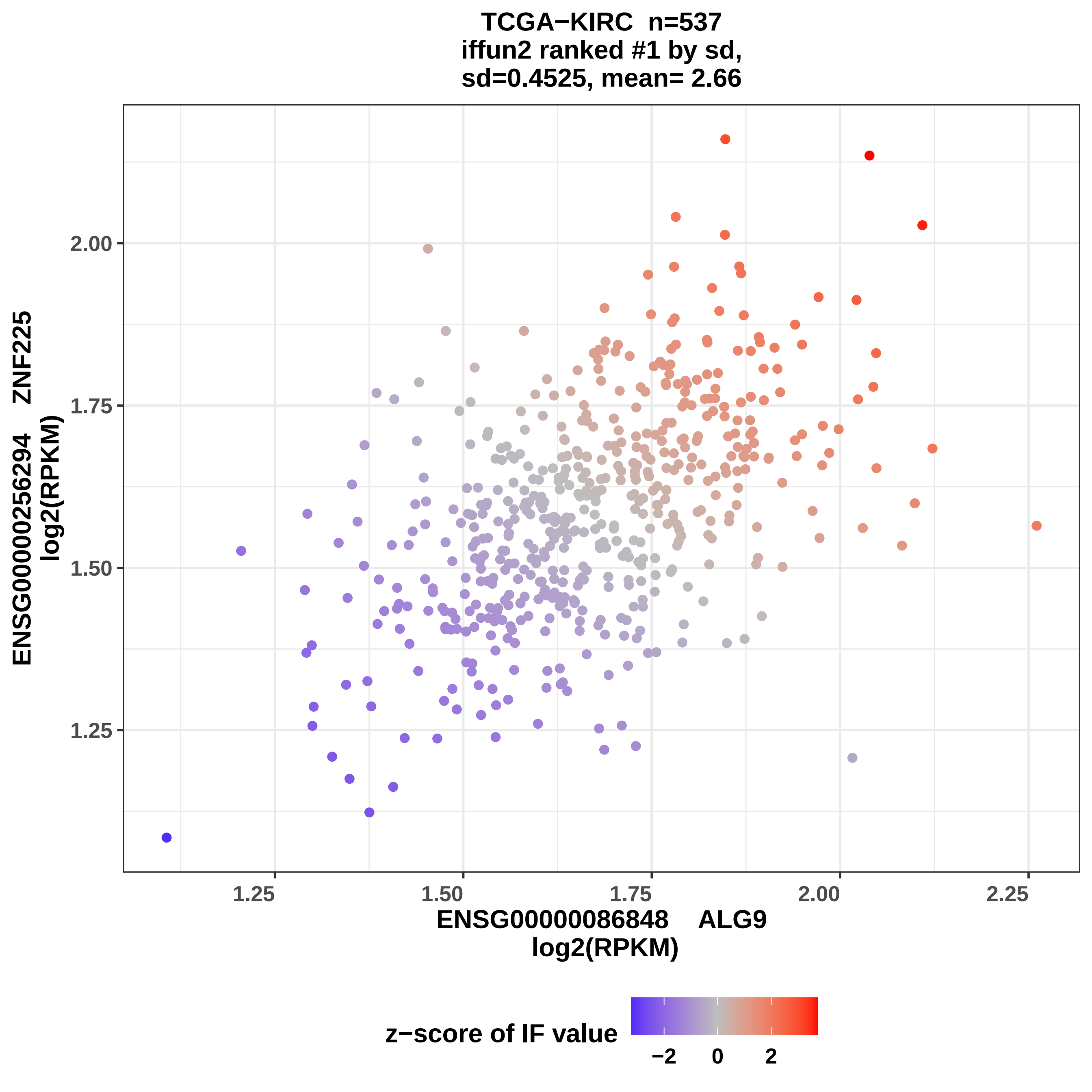
TCGA−KIRC n=537
iffun2 ranked #1 by sd,
sd=0.4525, mean= 2.66
ENSG00000256294 ZNF225
log2(RPKM)
2.00
1.75
1.50
1.25
1.25
1.50
1.75
2.00
2.25
ENSG00000086848 ALG9
log2(RPKM)
z−score of IF value
−2
0
2

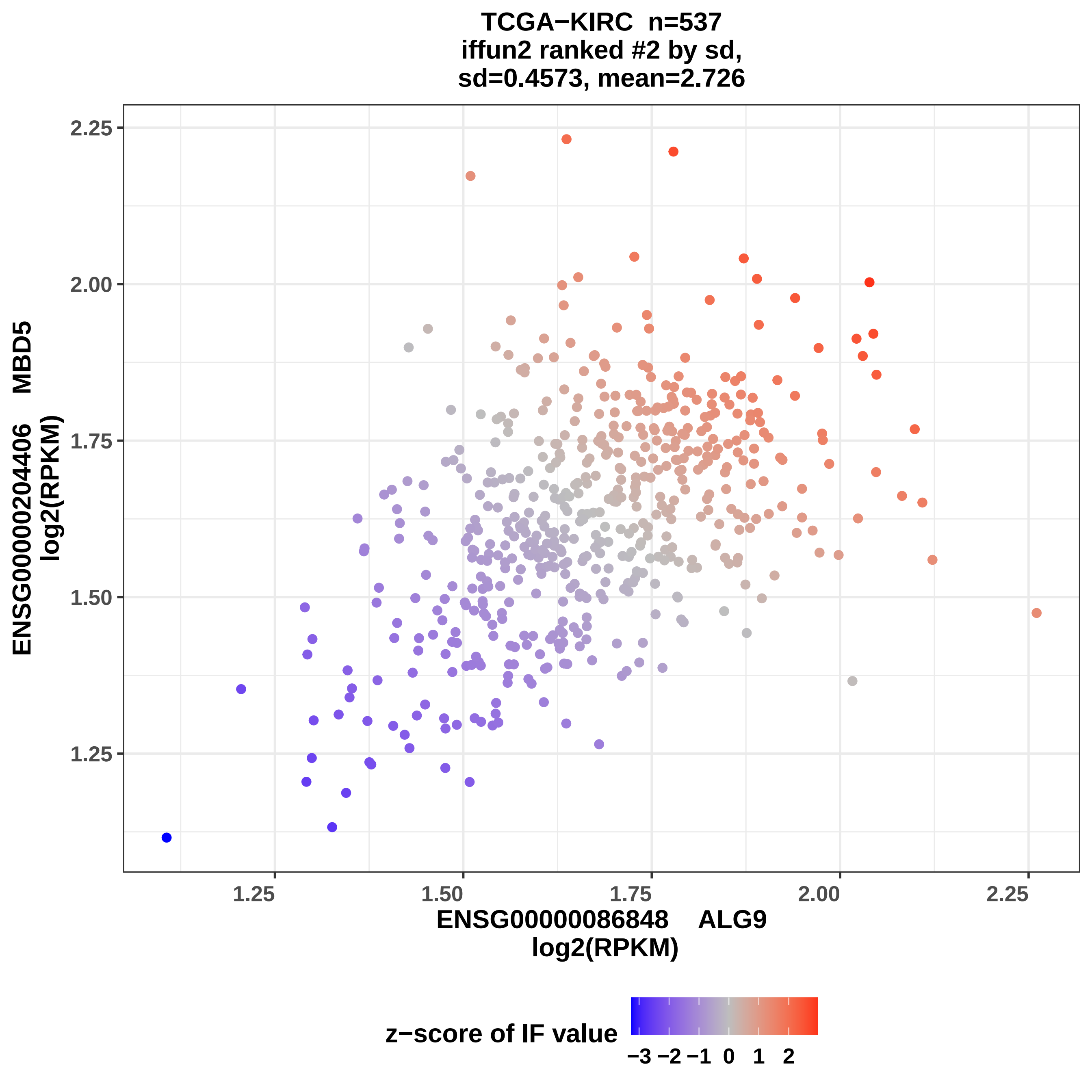
TCGA−KIRC n=537
iffun2 ranked #2 by sd,
sd=0.4573, mean=2.726
2.25
2.00
1.75
1.50
1.25
ENSG00000204406 MBD5
log2(RPKM)
1.25
1.50
1.75
2.00
2.25
ENSG00000086848 ALG9
log2(RPKM)
z−score of IF value
−3 −2 −1 0 1 2

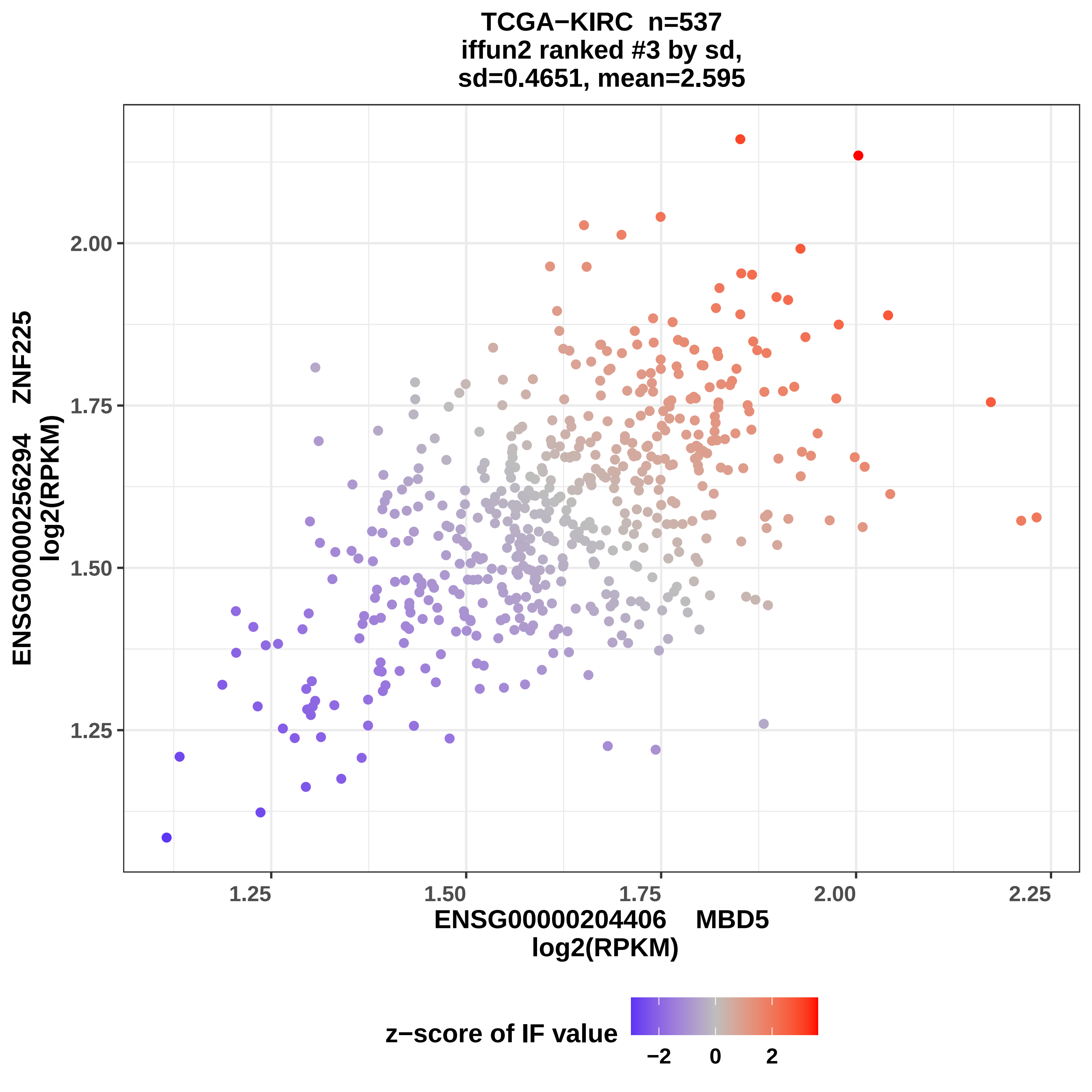
TCGA−KIRC n=537
iffun2 ranked #3 by sd,
sd=0.4651, mean=2.595
ENSG00000256294 ZNF225
log2(RPKM)
2.00
1.75
1.50
1.25
1.25
1.50
1.75
2.00
2.25
ENSG00000204406 MBD5
log2(RPKM)
z−score of IF value
−2
0
2

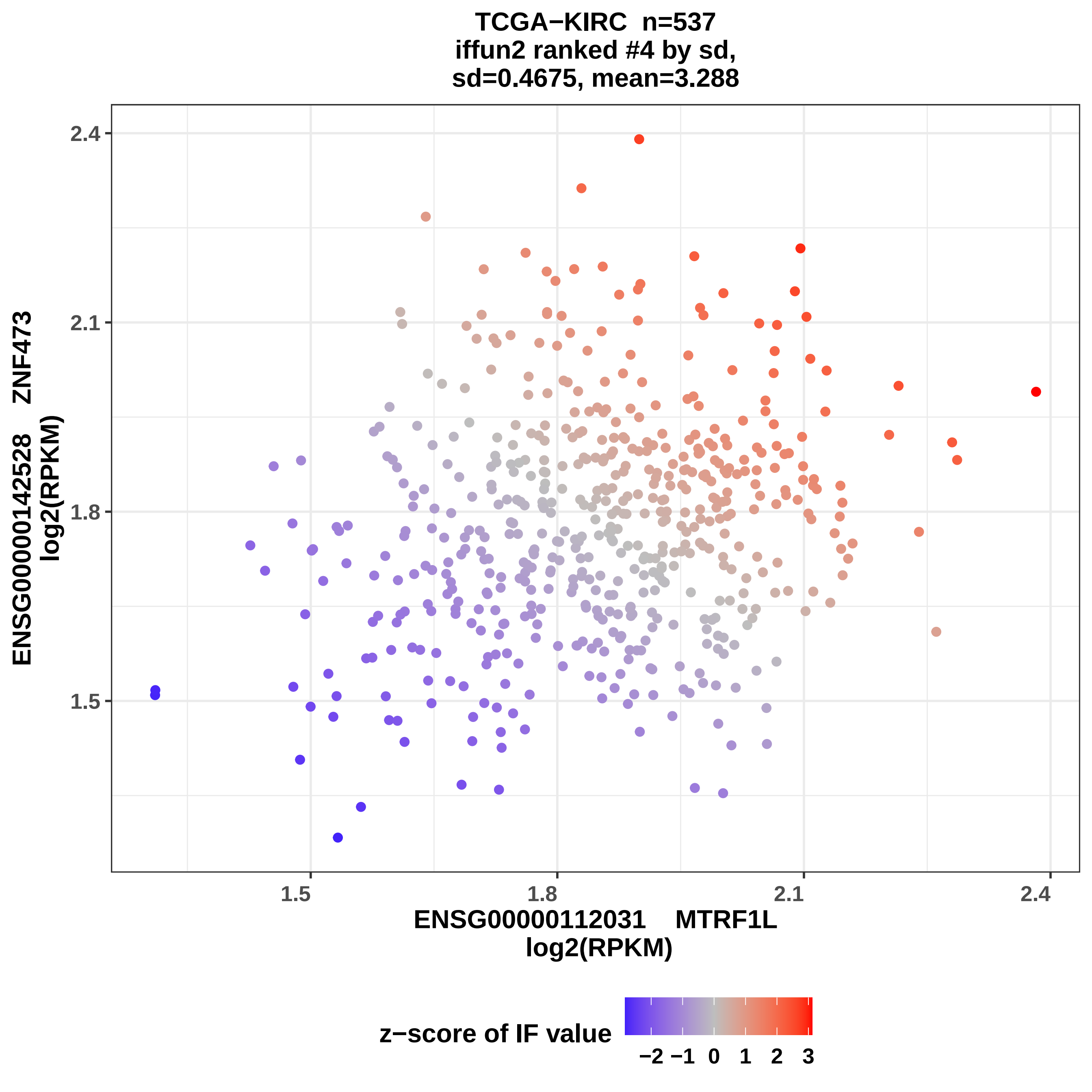
TCGA−KIRC n=537
iffun2 ranked #4 by sd,
sd=0.4675, mean=3.288
ENSG00000142528 ZNF473
log2(RPKM)
2.4
2.1
1.8
1.5
ENSG00000112031 MTRF1L
log2(RPKM)
1.5
1.8
2.1
2.4
z−score of IF value
−2 −1 0 1 2 3

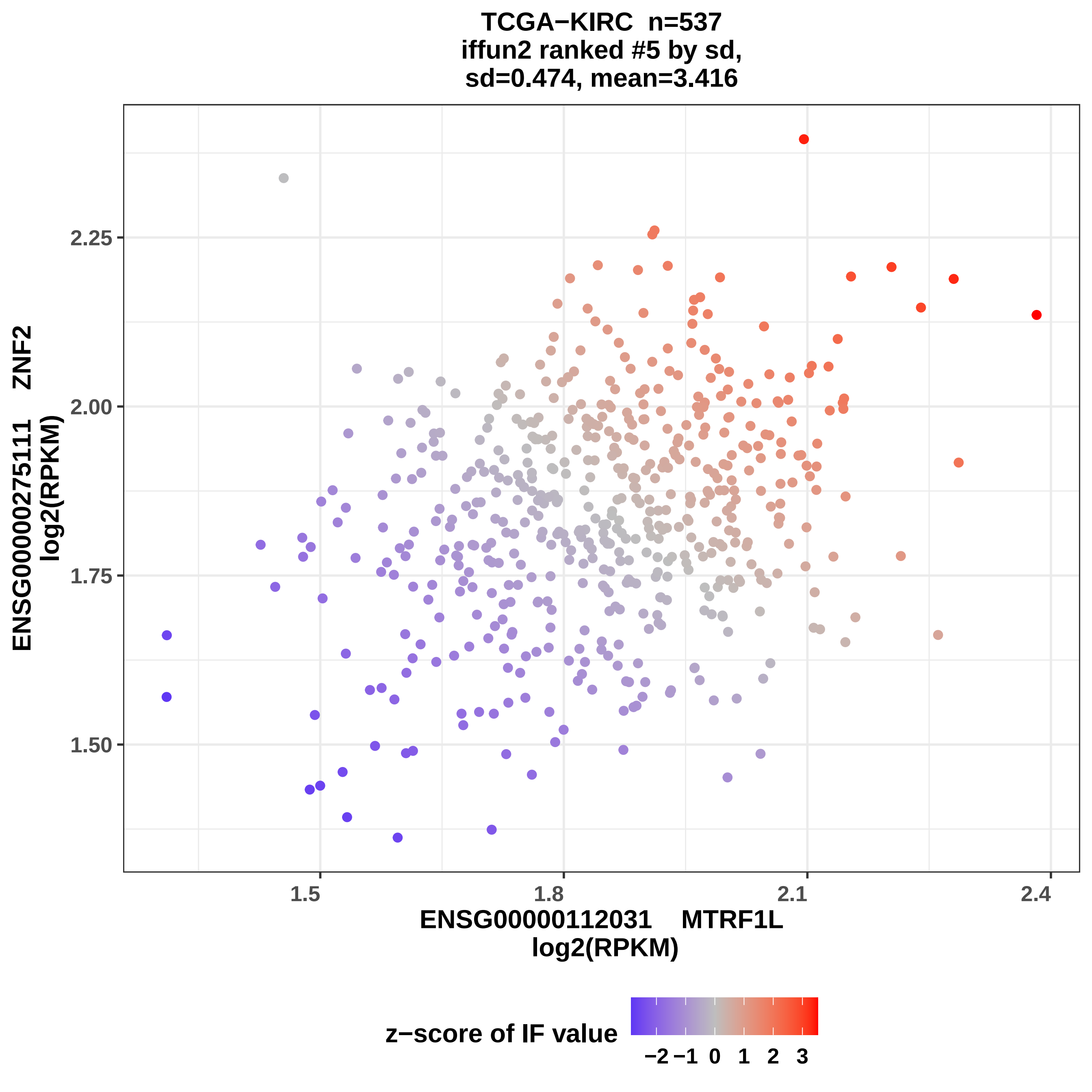

TCGA−KIRC n=537
iffun2 ranked #5 by sd,
sd=0.474, mean=3.416
ENSG00000275111 ZNF2
log2(RPKM)
2.25
2.00
1.75
1.50
1.5
1.8
2.1
2.4
ENSG00000112031 MTRF1L
log2(RPKM)
z−score of IF value
−2
−1
0
1
2
3

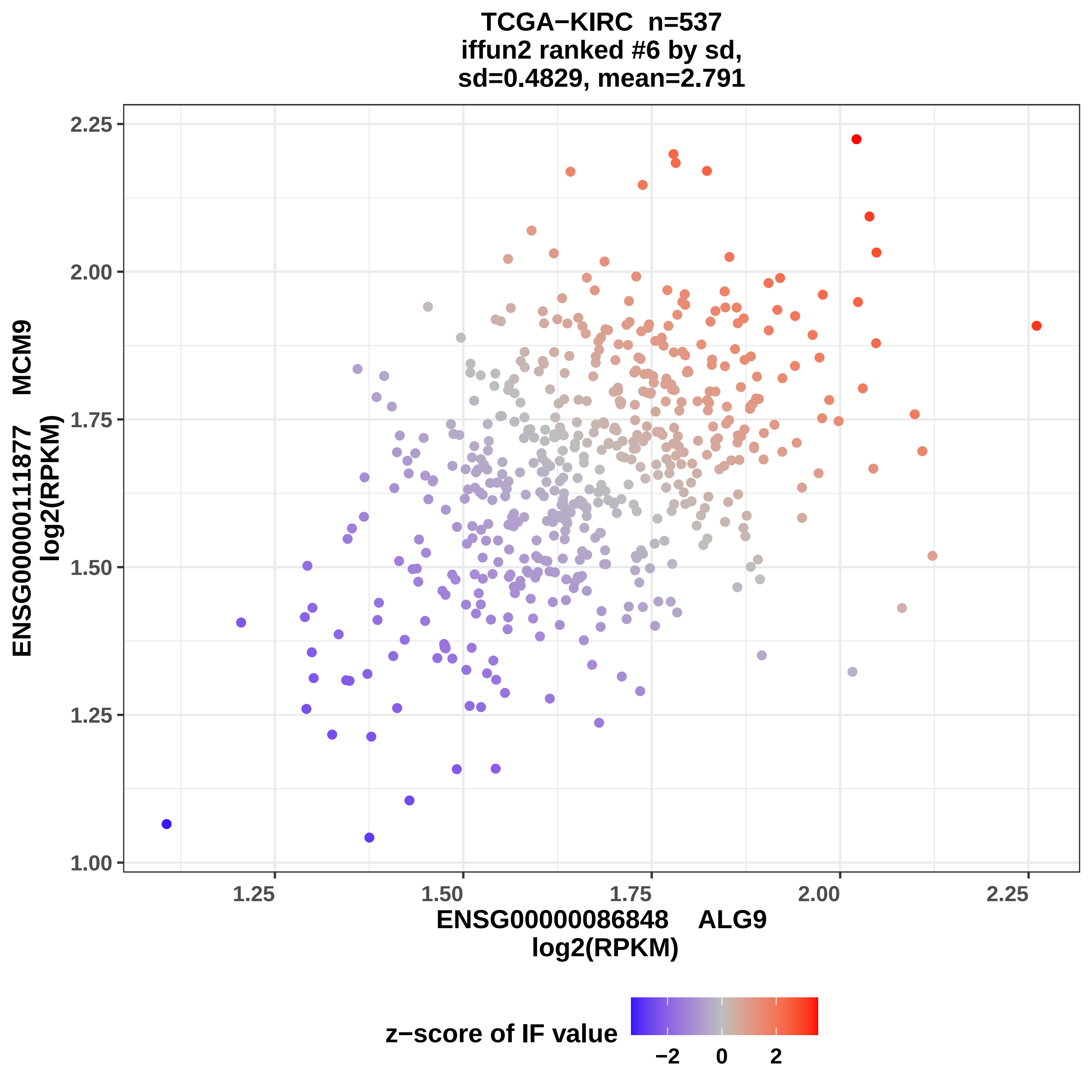
TCGA−KIRC n=537
iffun2 ranked #6 by sd,
sd=0.4829, mean=2.791
ENSG00000111877 MCM9
log2(RPKM)
ENSG00000086848 ALG9
log2(RPKM)
1.00
1.25
1.50
1.75
2.00
2.25
z−score of IF value
−2
0
2

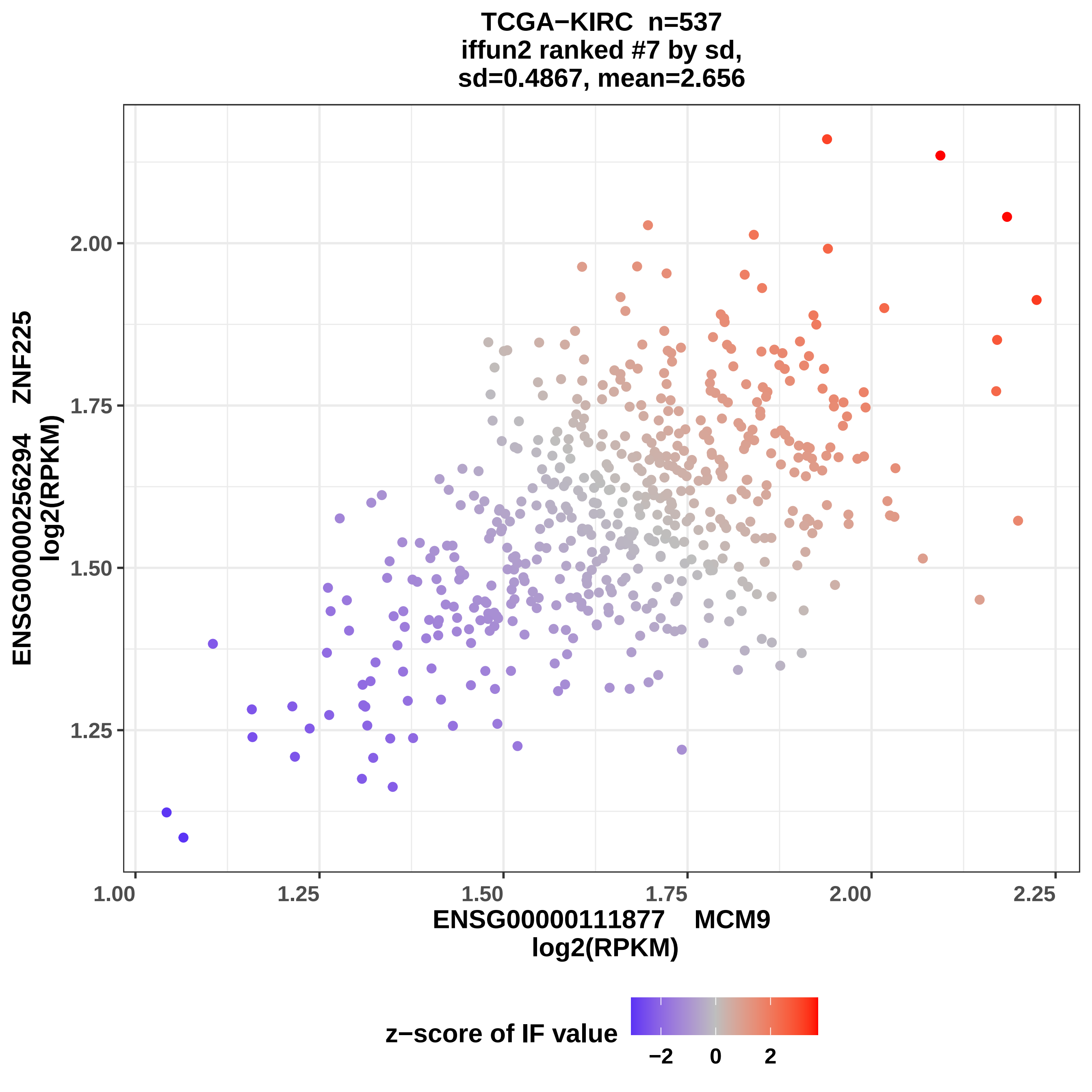
TCGA−KIRC n=537
iffun2 ranked #7 by sd,
sd=0.4867, mean=2.656
ENSG00000256294 ZNF225
log2(RPKM)
2.00
1.75
1.50
1.25
1.00
1.25
1.50
1.75
2.00
2.25
ENSG00000111877 MCM9
log2(RPKM)
z−score of IF value
−2
0
2

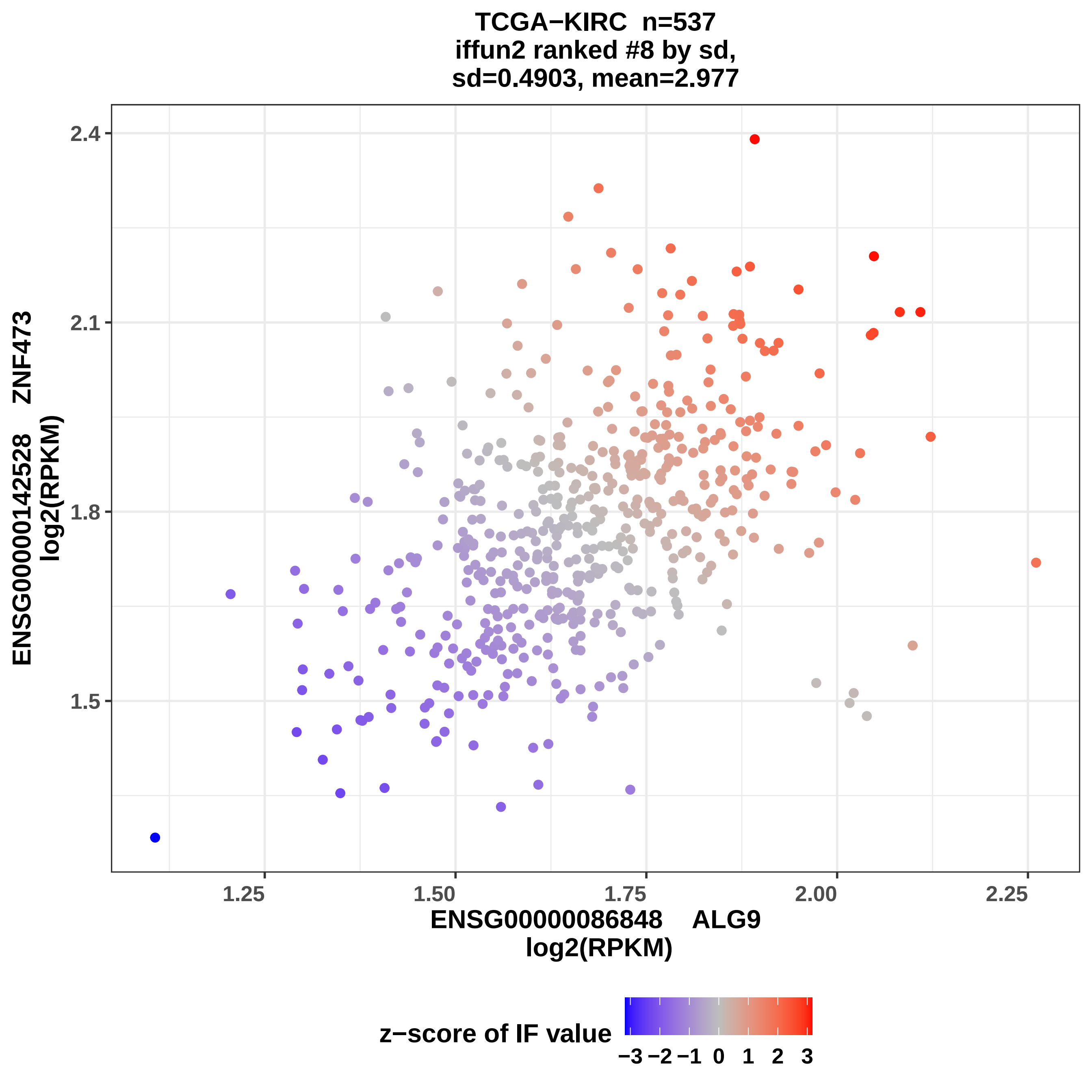

TCGA−KIRC n=537
iffun2 ranked #8 by sd,
sd=0.4903, mean=2.977
ENSG00000142528 ZNF473
log2(RPKM)
2.4
2.1
1.8
1.5
1.25
1.50
1.75
2.00
2.25
ENSG00000086848 ALG9
log2(RPKM)
z−score of IF value
−3 −2 −1 0 1 2 3

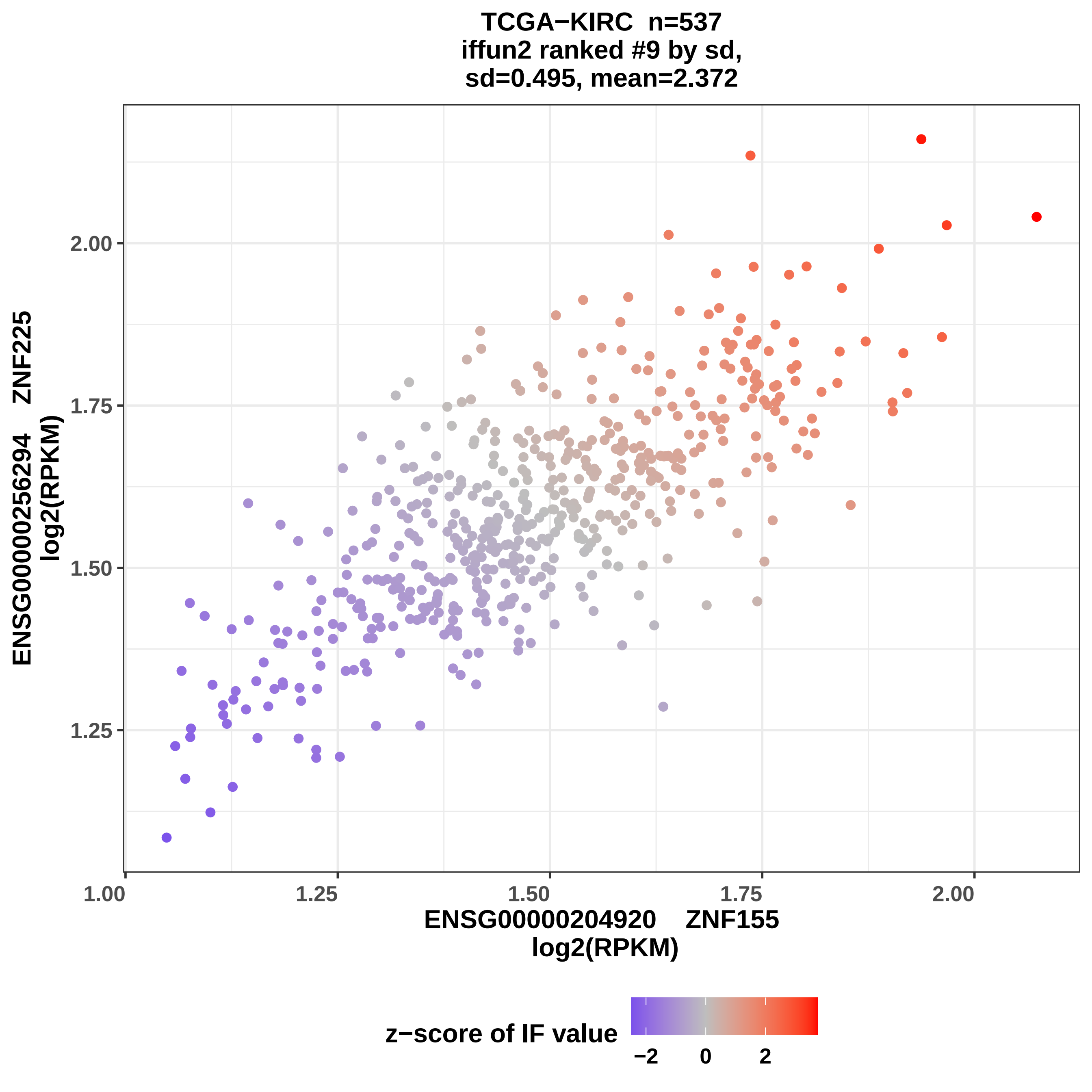

TCGA−KIRC n=537
iffun2 ranked #9 by sd,
sd=0.495, mean=2.372
ENSG00000256294 ZNF225
log2(RPKM)
2.00
1.75
1.50
1.25
1.00
1.25
1.50
1.75
2.00
ENSG00000204920 ZNF155
log2(RPKM)
z−score of IF value
−2
0
2

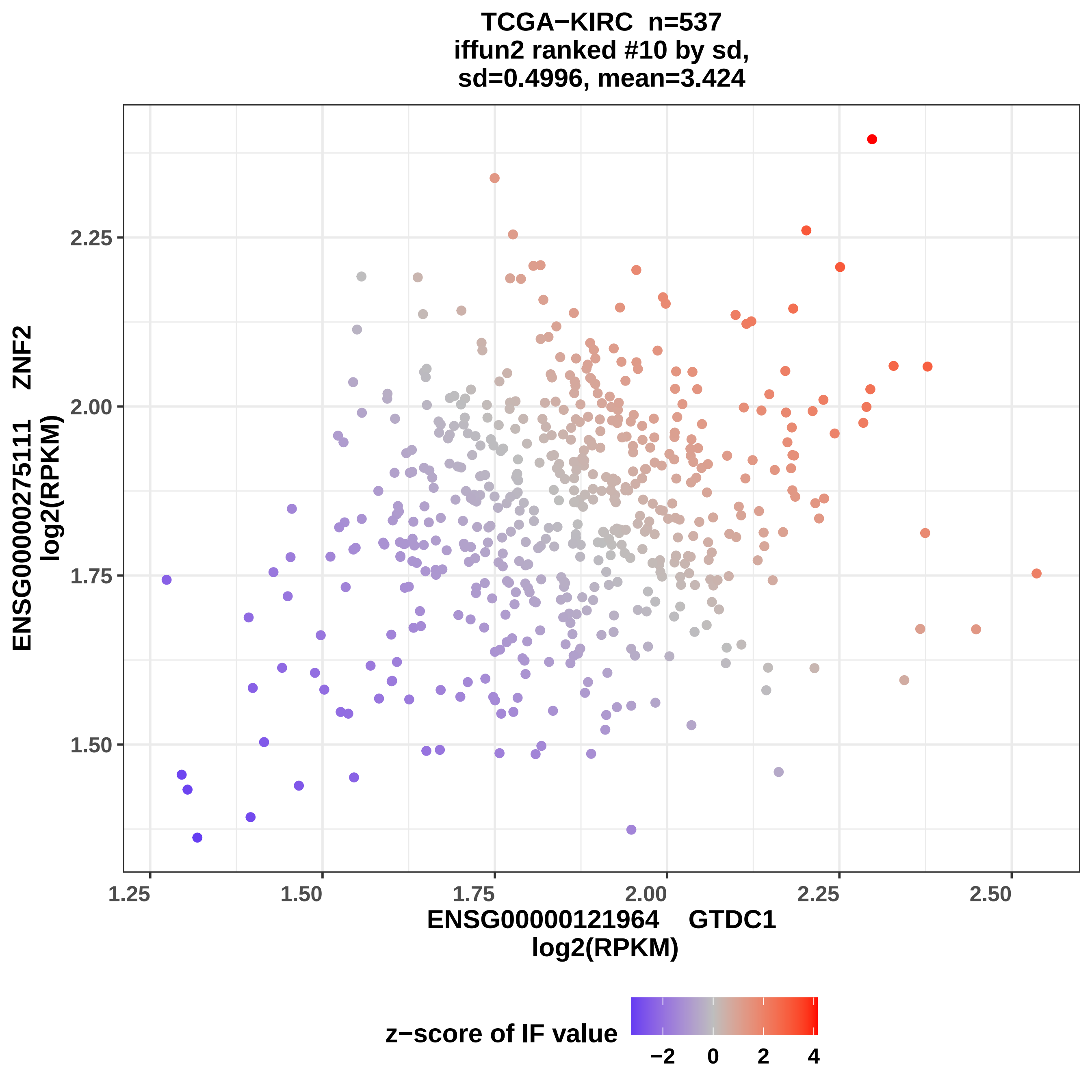

TCGA−KIRC n=537
iffun2 ranked #10 by sd,
sd=0.4996, mean=3.424
ENSG00000275111 ZNF2
log2(RPKM)
ENSG00000121964 GTDC1
log2(RPKM)
1.25
1.50
1.75
2.00
2.25
2.50
z−score of IF value
−2
0
2
4

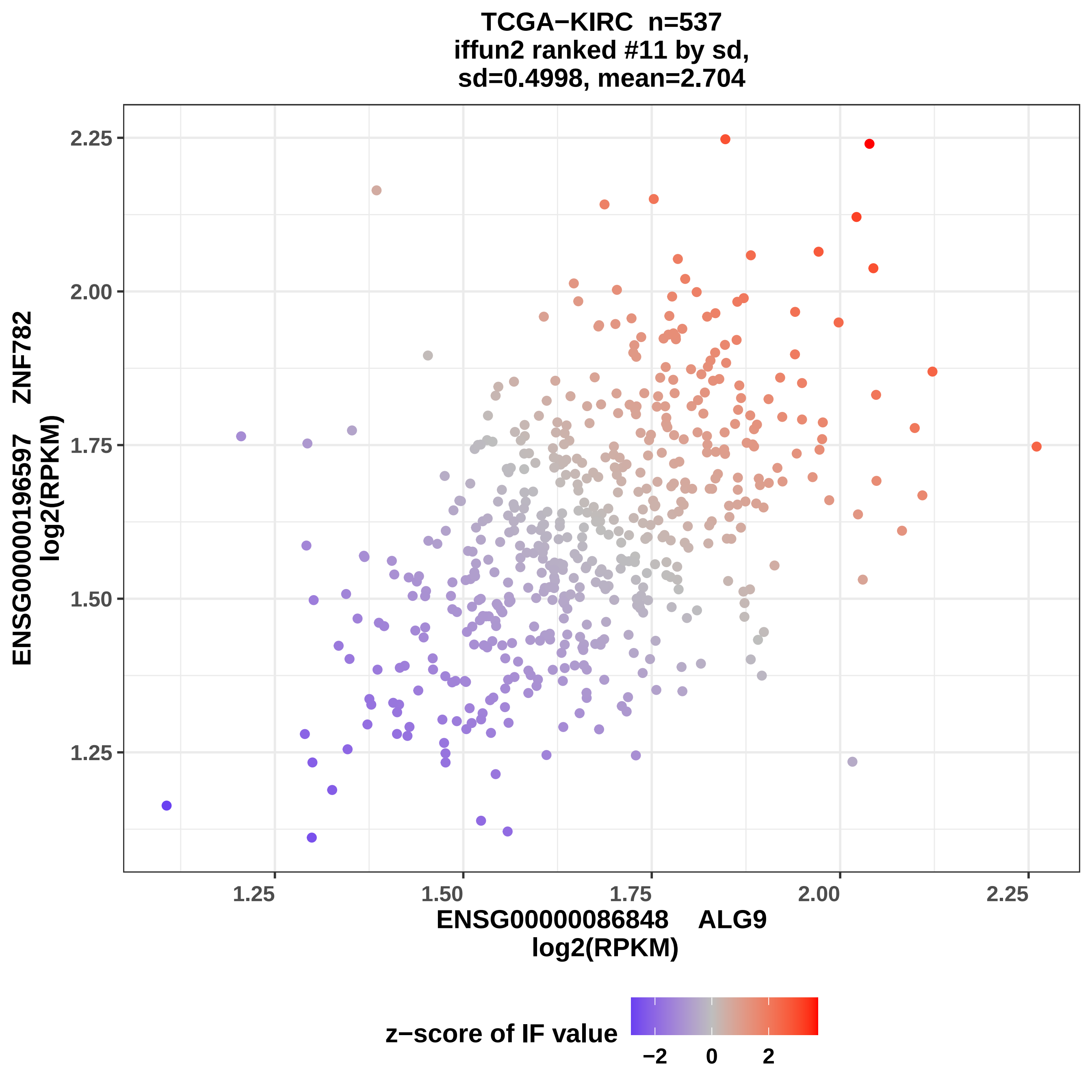
TCGA−KIRC n=537
iffun2 ranked #11 by sd,
sd=0.4998, mean=2.704
ENSG00000196597 ZNF782
log2(RPKM)
ENSG00000086848 ALG9
log2(RPKM)
2.25
2.00
1.75
1.50
1.25
1.25
1.50
1.75
2.00
2.25
z−score of IF value
−2
0
2

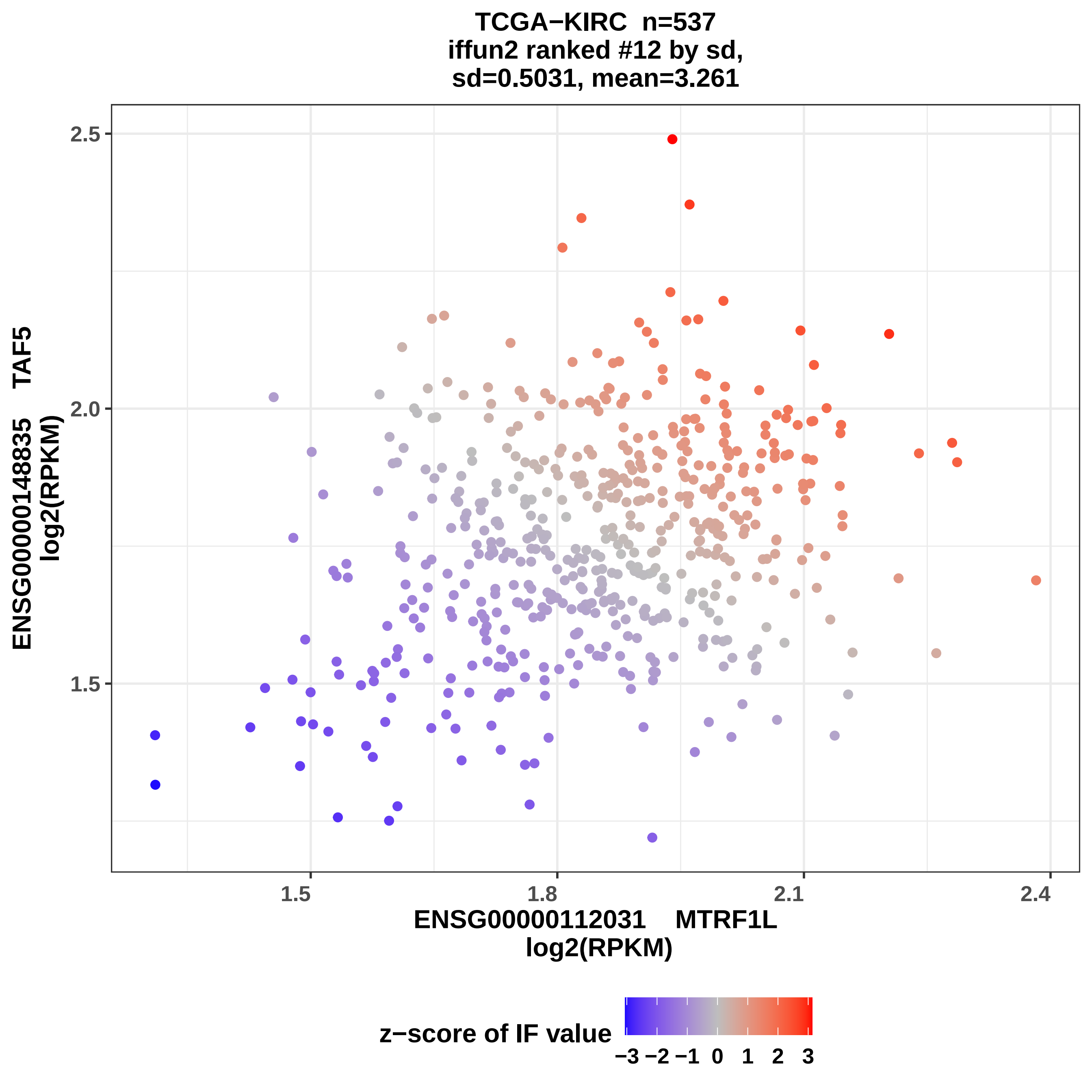
TCGA−KIRC n=537
iffun2 ranked #12 by sd,
sd=0.5031, mean=3.261
ENSG00000148835 TAF5
log2(RPKM)
2.5
2.0
1.5
1.5
1.8
2.1
2.4
ENSG00000112031 MTRF1L
log2(RPKM)
z−score of IF value
−3 −2 −1 0 1 2 3

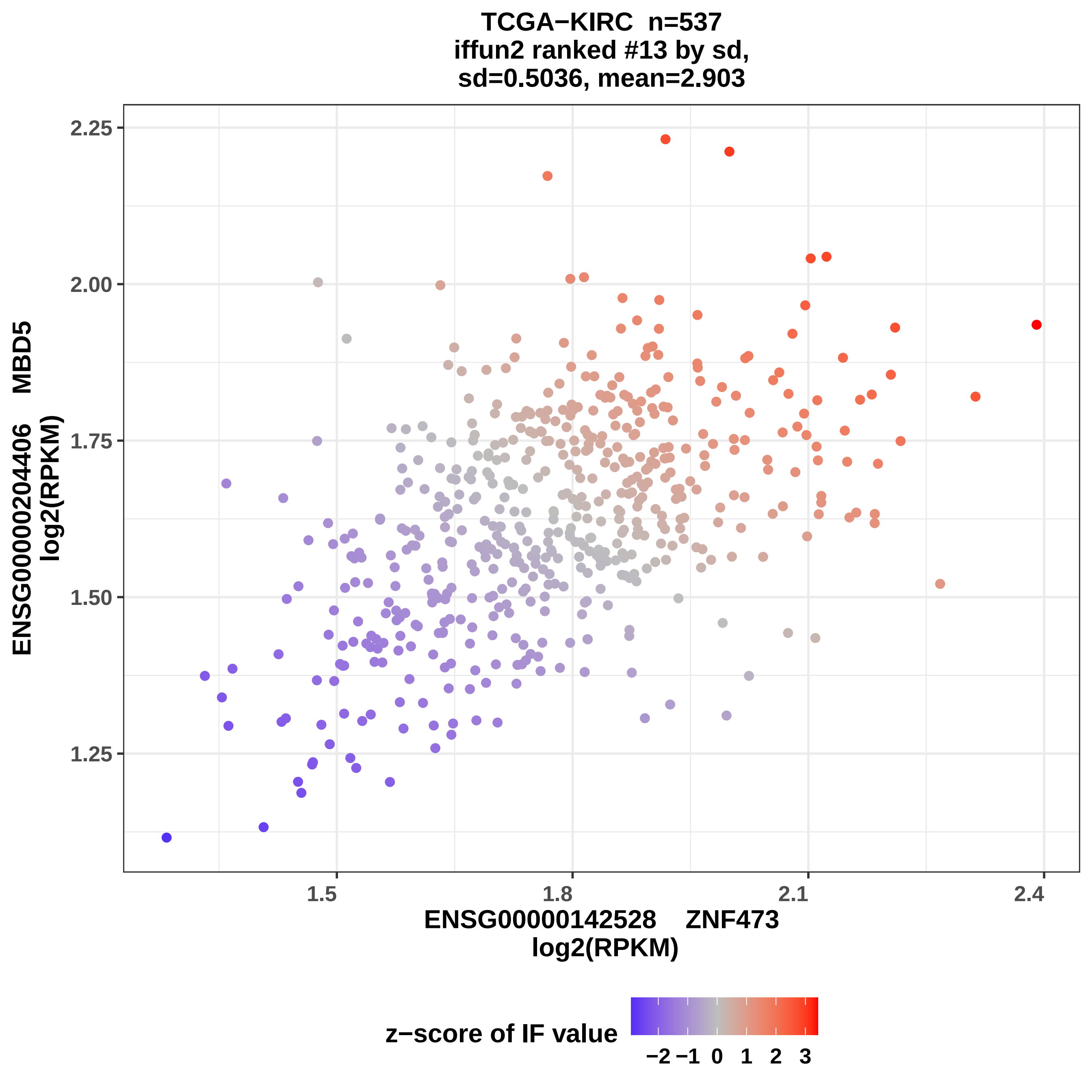

TCGA−KIRC n=537
iffun2 ranked #13 by sd,
sd=0.5036, mean=2.903
ENSG00000204406 MBD5
log2(RPKM)
2.25
2.00
1.75
1.50
1.25
1.5
1.8
2.1
2.4
ENSG00000142528 ZNF473
log2(RPKM)
z−score of IF value
−2 −1 0 1 2 3

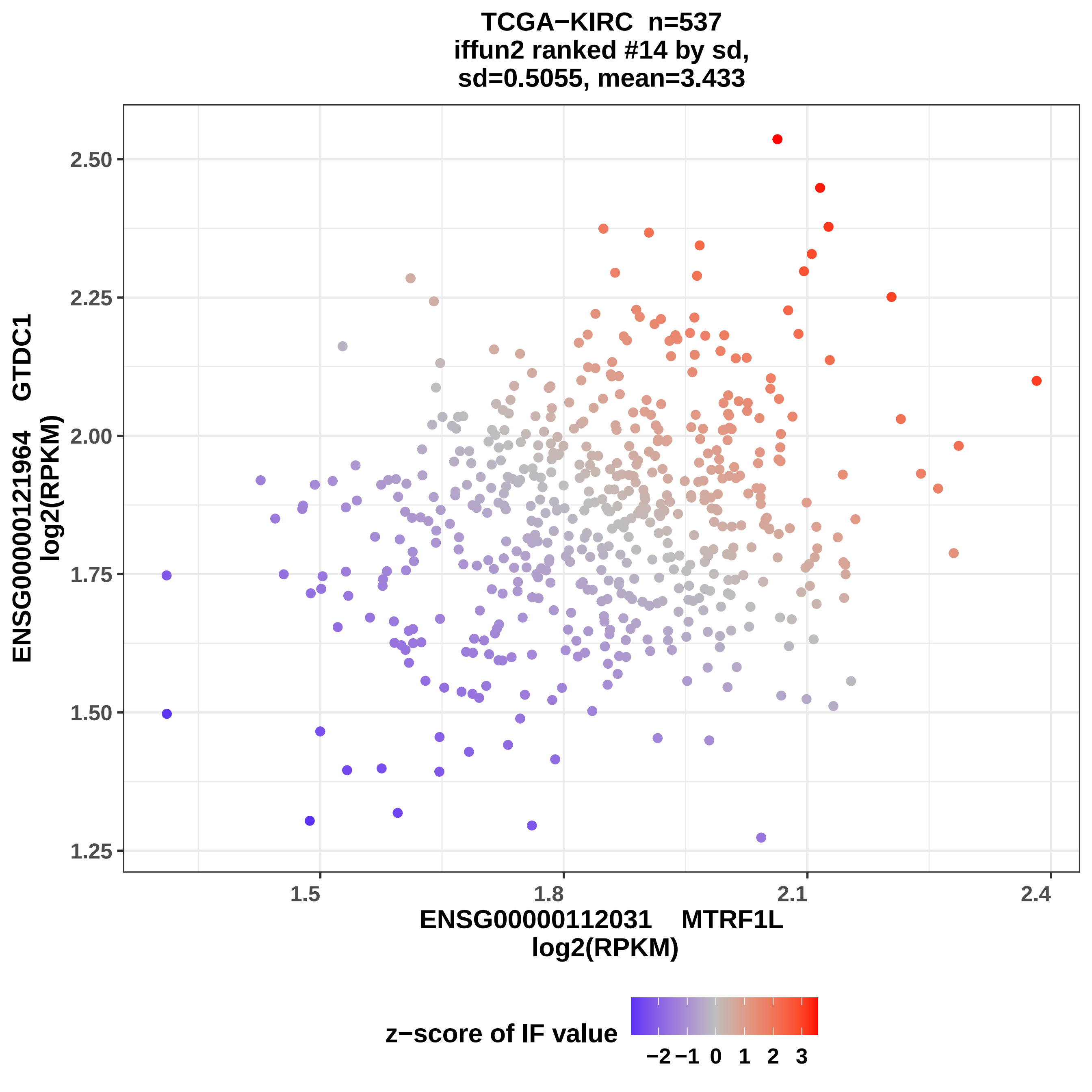

TCGA−KIRC n=537
iffun2 ranked #14 by sd,
sd=0.5055, mean=3.433
2.50
2.25
2.00
1.75
1.50
1.25
1.5
1.8
2.1
2.4
ENSG00000121964 GTDC1
log2(RPKM)
ENSG00000112031 MTRF1L
log2(RPKM)
z−score of IF value
−2 −1 0 1 2 3

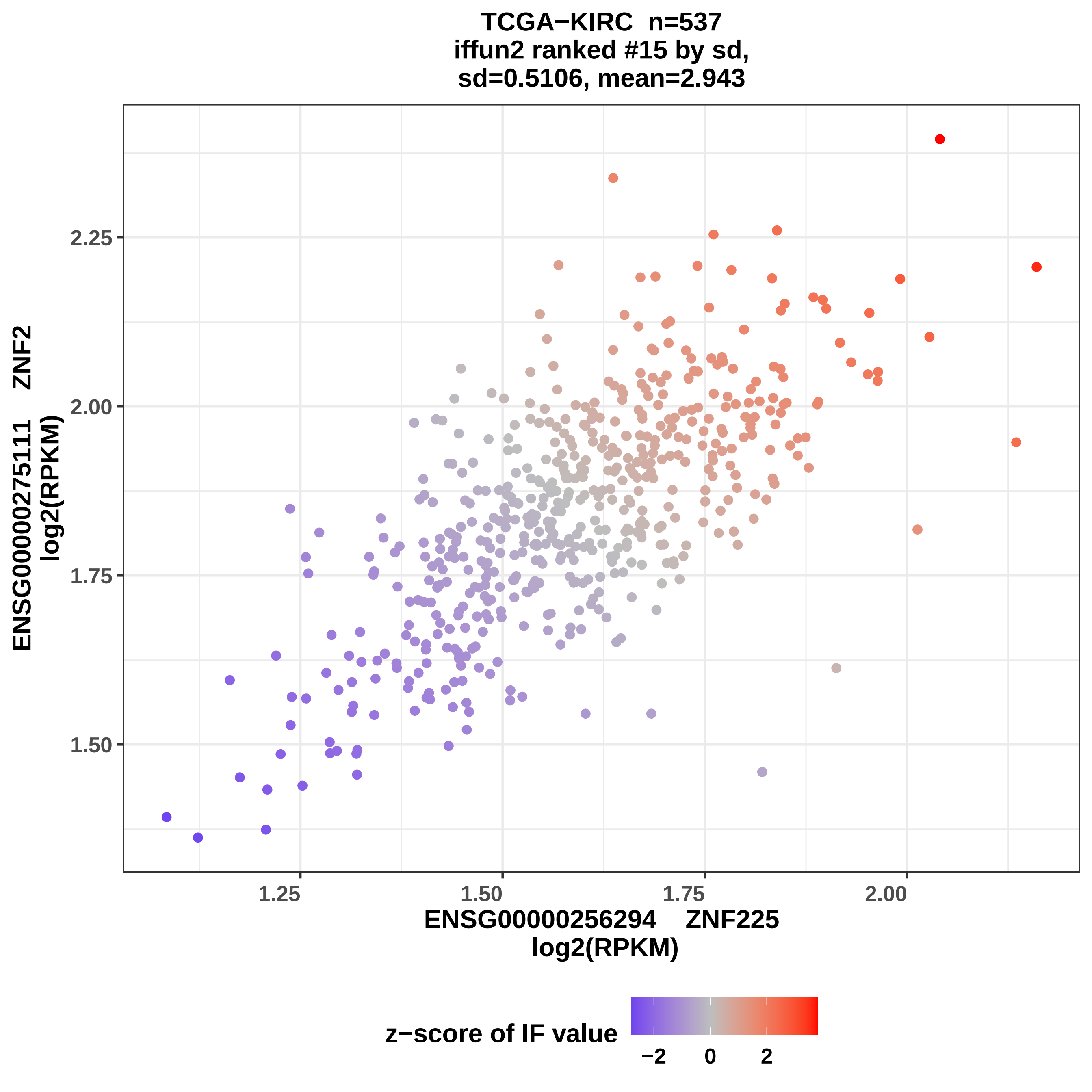

TCGA−KIRC n=537
iffun2 ranked #15 by sd,
sd=0.5106, mean=2.943
ENSG00000275111 ZNF2
log2(RPKM)
2.25
2.00
1.75
1.50
1.25
1.50
1.75
2.00
ENSG00000256294 ZNF225
log2(RPKM)
z−score of IF value
−2
0
2

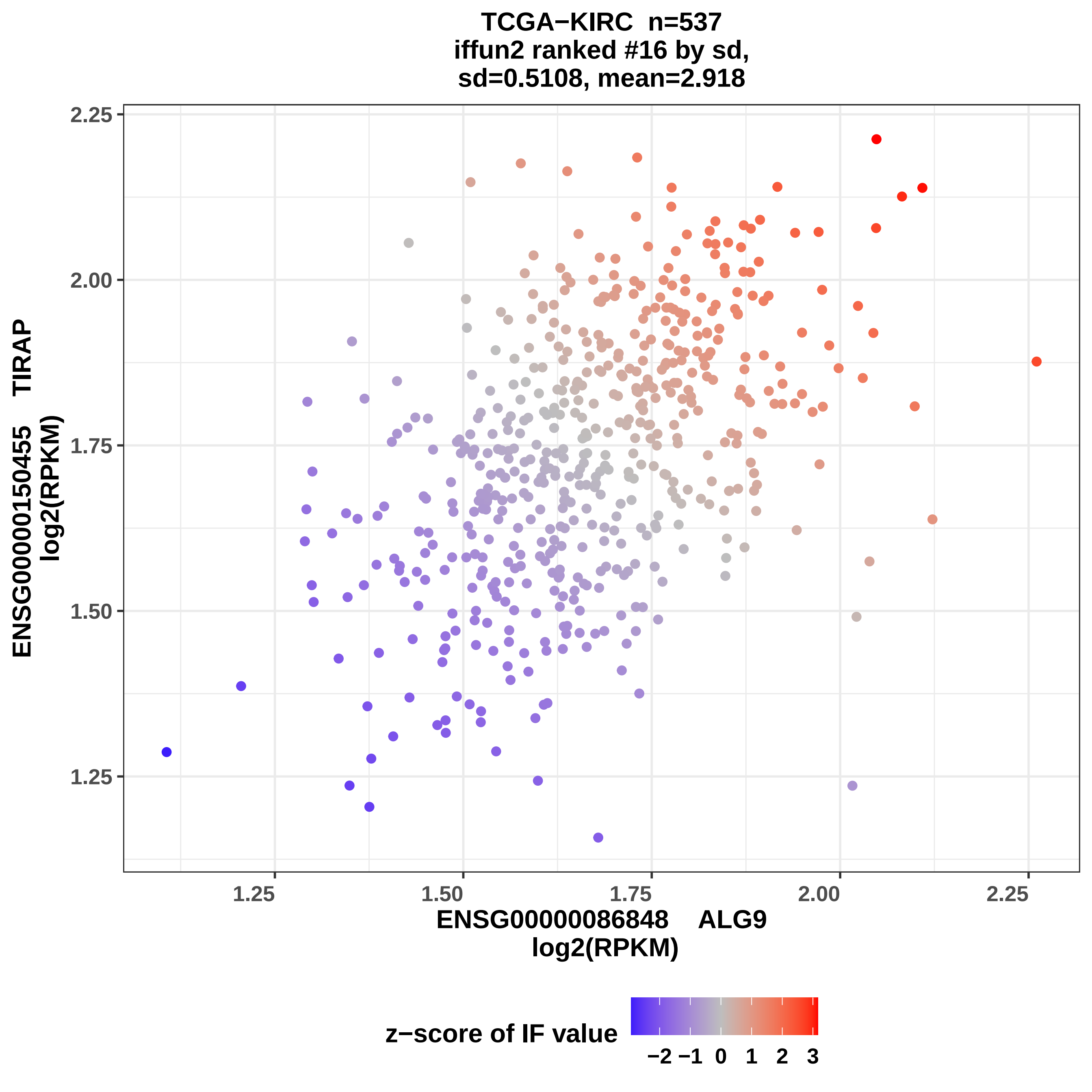

TCGA−KIRC n=537
iffun2 ranked #16 by sd,
sd=0.5108, mean=2.918
ENSG00000150455 TIRAP
log2(RPKM)
ENSG00000086848 ALG9
log2(RPKM)
2.25
2.00
1.75
1.50
1.25
1.25
1.50
1.75
2.00
2.25
z−score of IF value
−2 −1 0 1 2 3

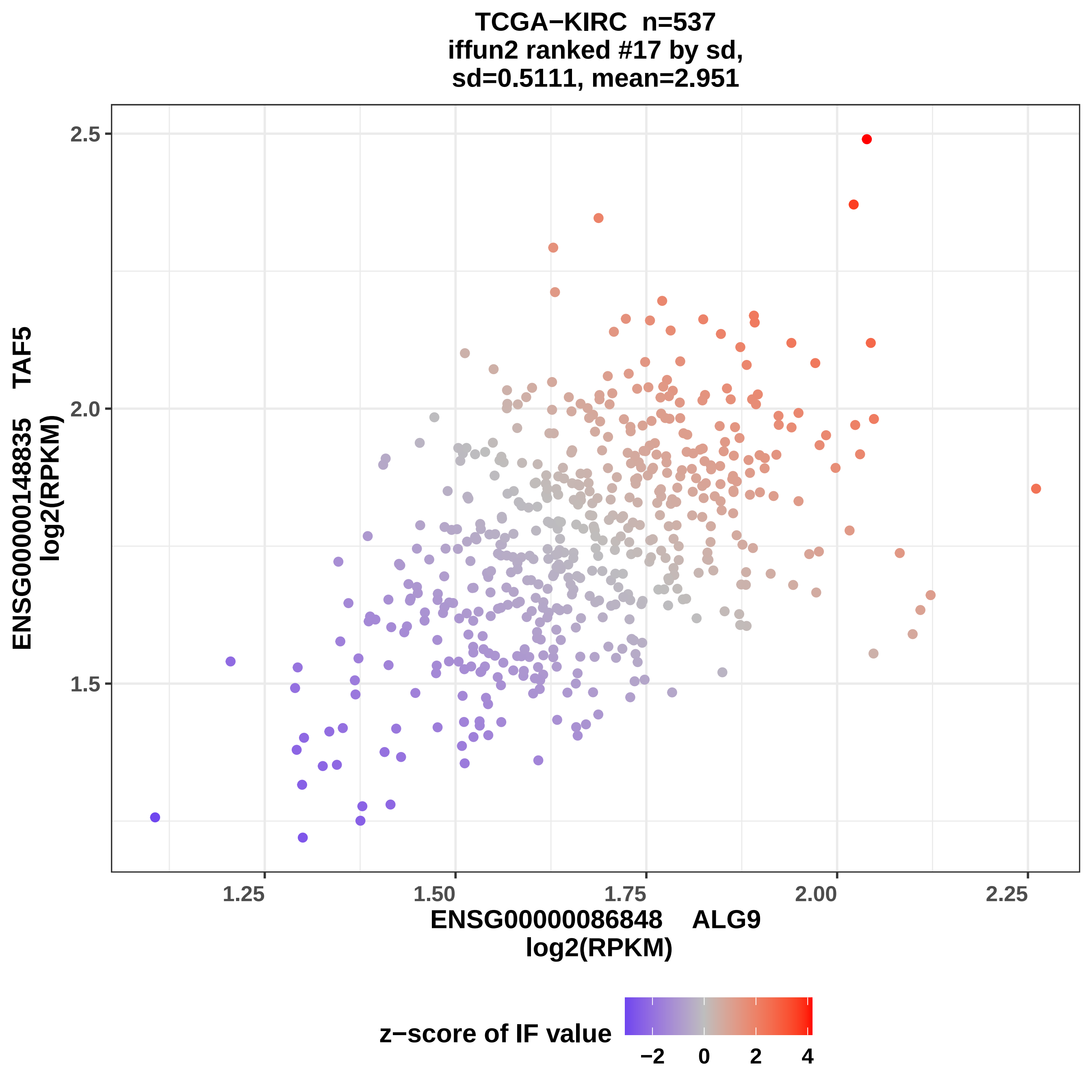
TCGA−KIRC n=537
iffun2 ranked #17 by sd,
sd=0.5111, mean=2.951
ENSG00000148835 TAF5
log2(RPKM)
2.5
2.0
1.5
1.25
1.50
1.75
2.00
2.25
ENSG00000086848 ALG9
log2(RPKM)
z−score of IF value
−2
0
2
4

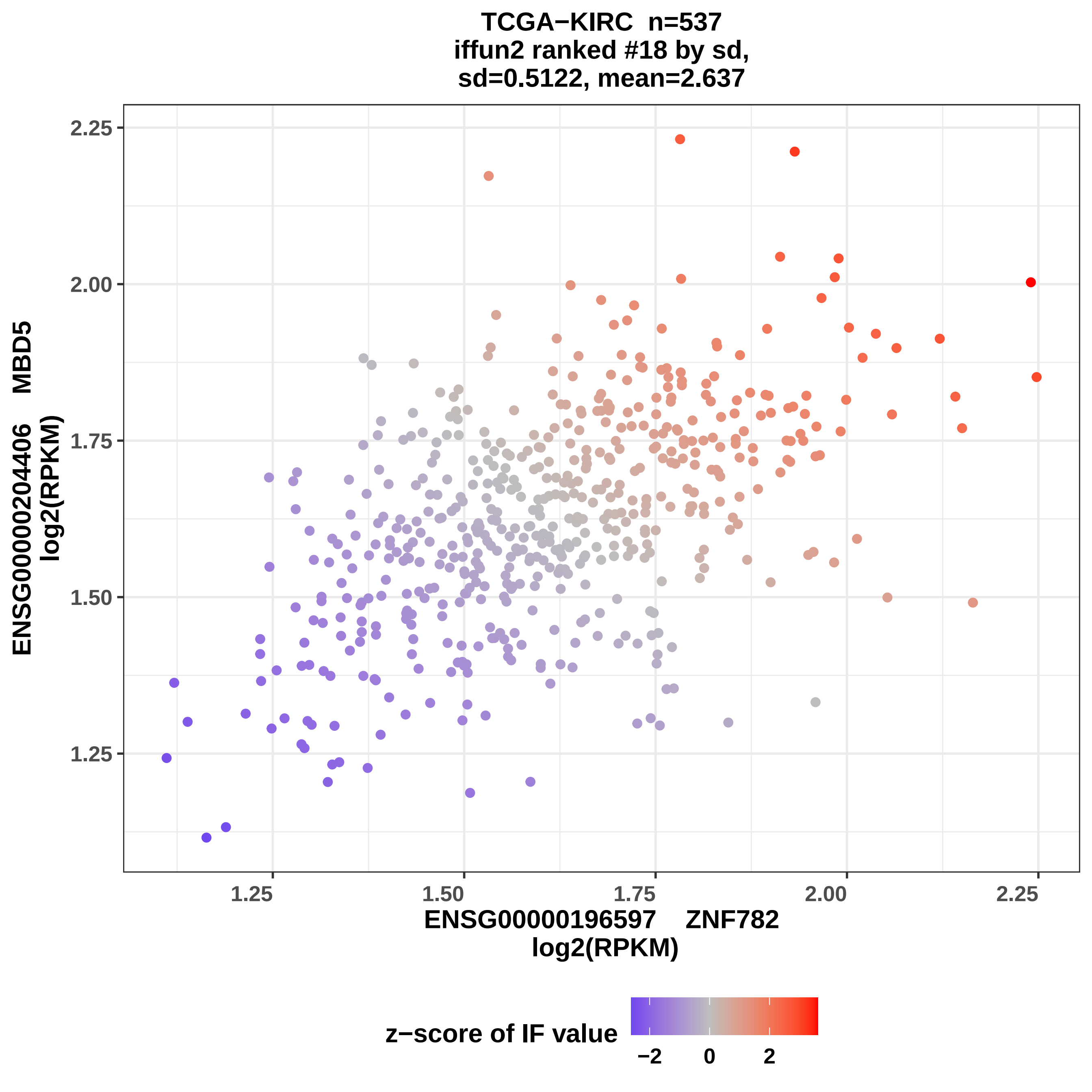

TCGA−KIRC n=537
iffun2 ranked #18 by sd,
sd=0.5122, mean=2.637
ENSG00000204406 MBD5
log2(RPKM)
2.25
2.00
1.75
1.50
1.25
1.25
1.50
1.75
2.00
2.25
ENSG00000196597 ZNF782
log2(RPKM)
z−score of IF value
−2
0
2

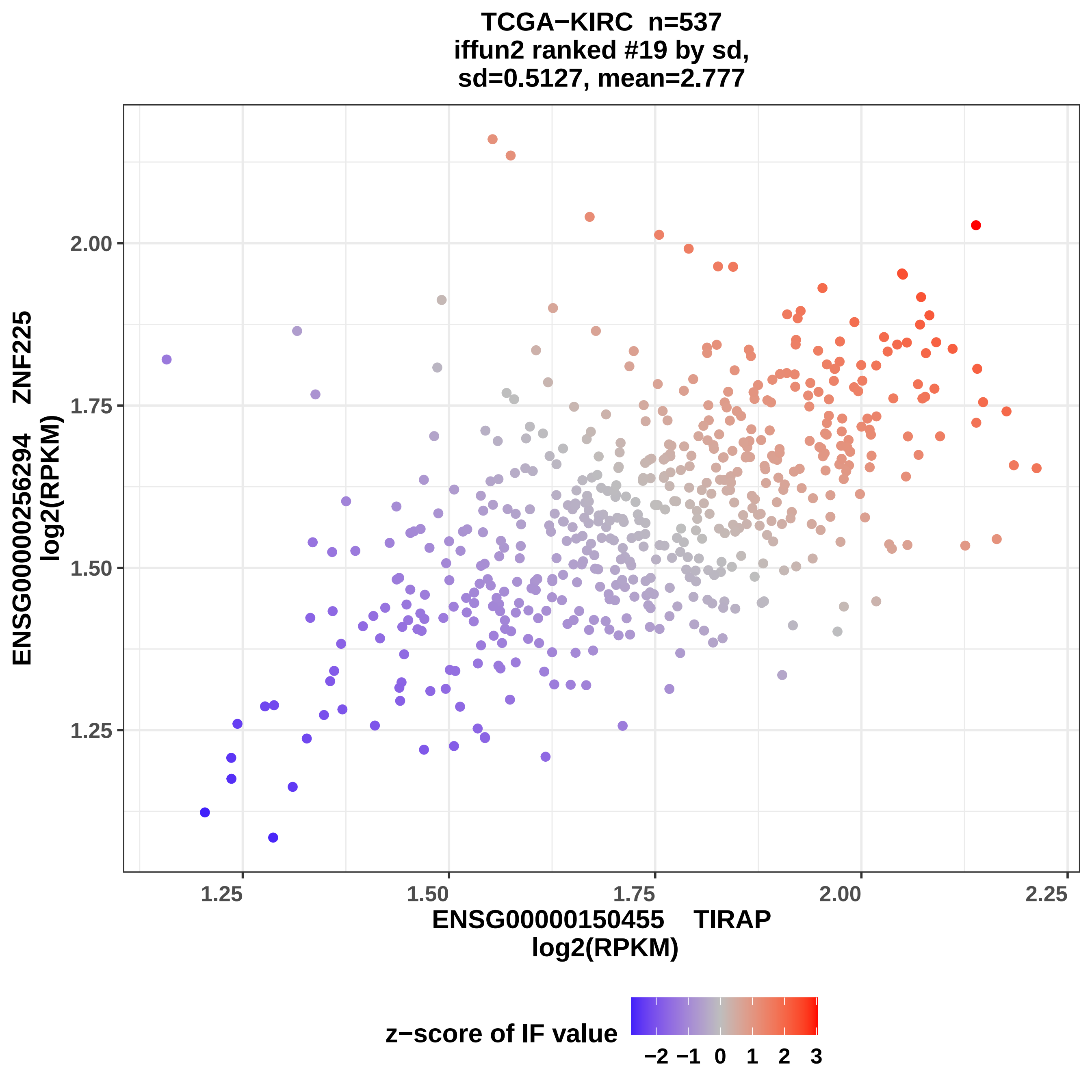

TCGA−KIRC n=537
iffun2 ranked #19 by sd,
sd=0.5127, mean=2.777
ENSG00000256294 ZNF225
log2(RPKM)
2.00
1.75
1.50
1.25
1.25
1.50
1.75
2.00
2.25
ENSG00000150455 TIRAP
log2(RPKM)
z−score of IF value
−2 −1 0 1 2 3

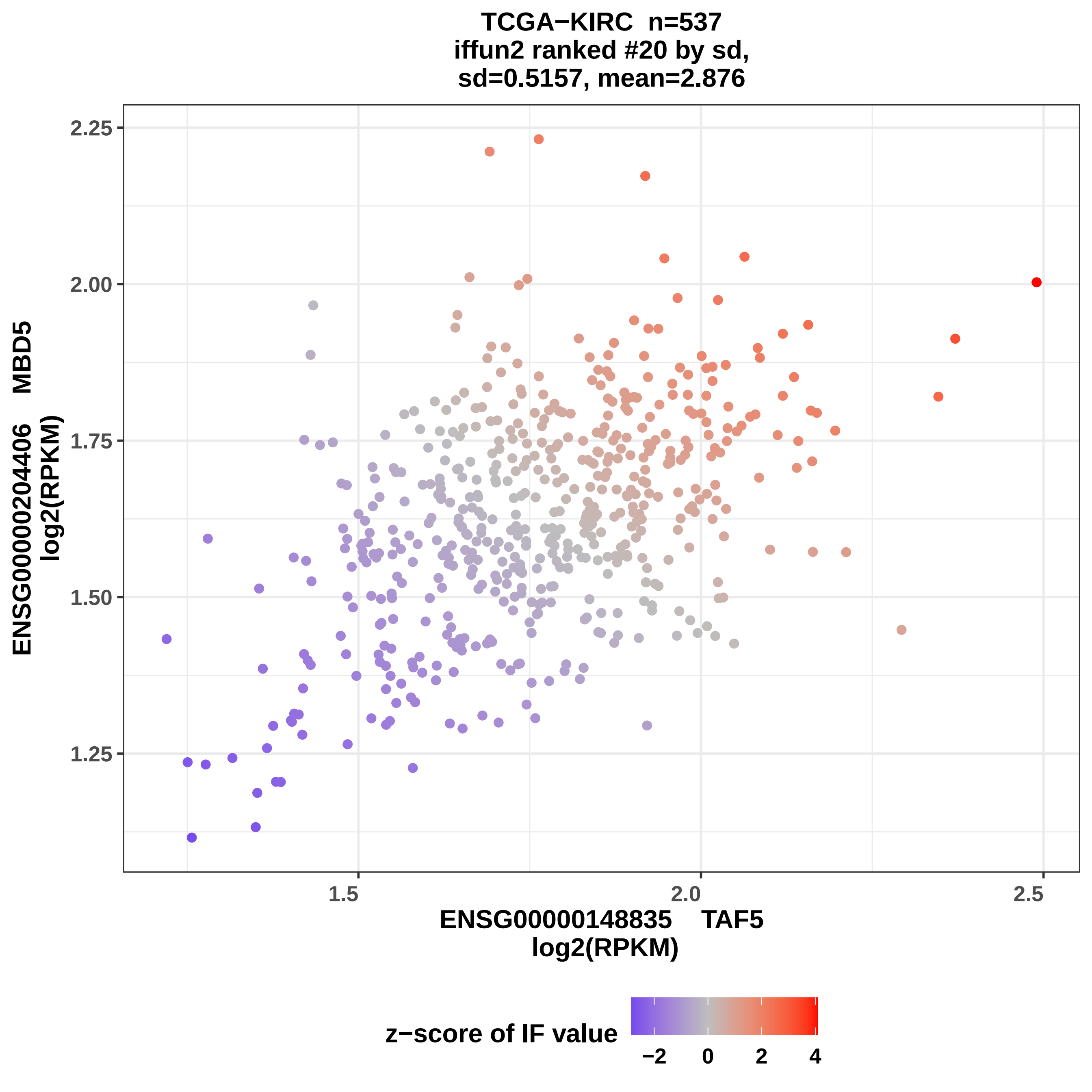
TCGA−KIRC n=537
iffun2 ranked #20 by sd,
sd=0.5157, mean=2.876
ENSG00000204406 MBD5
log2(RPKM)
2.25
2.00
1.75
1.50
1.25
1.5
2.0
2.5
ENSG00000148835 TAF5
log2(RPKM)
z−score of IF value
−2
0
2
4

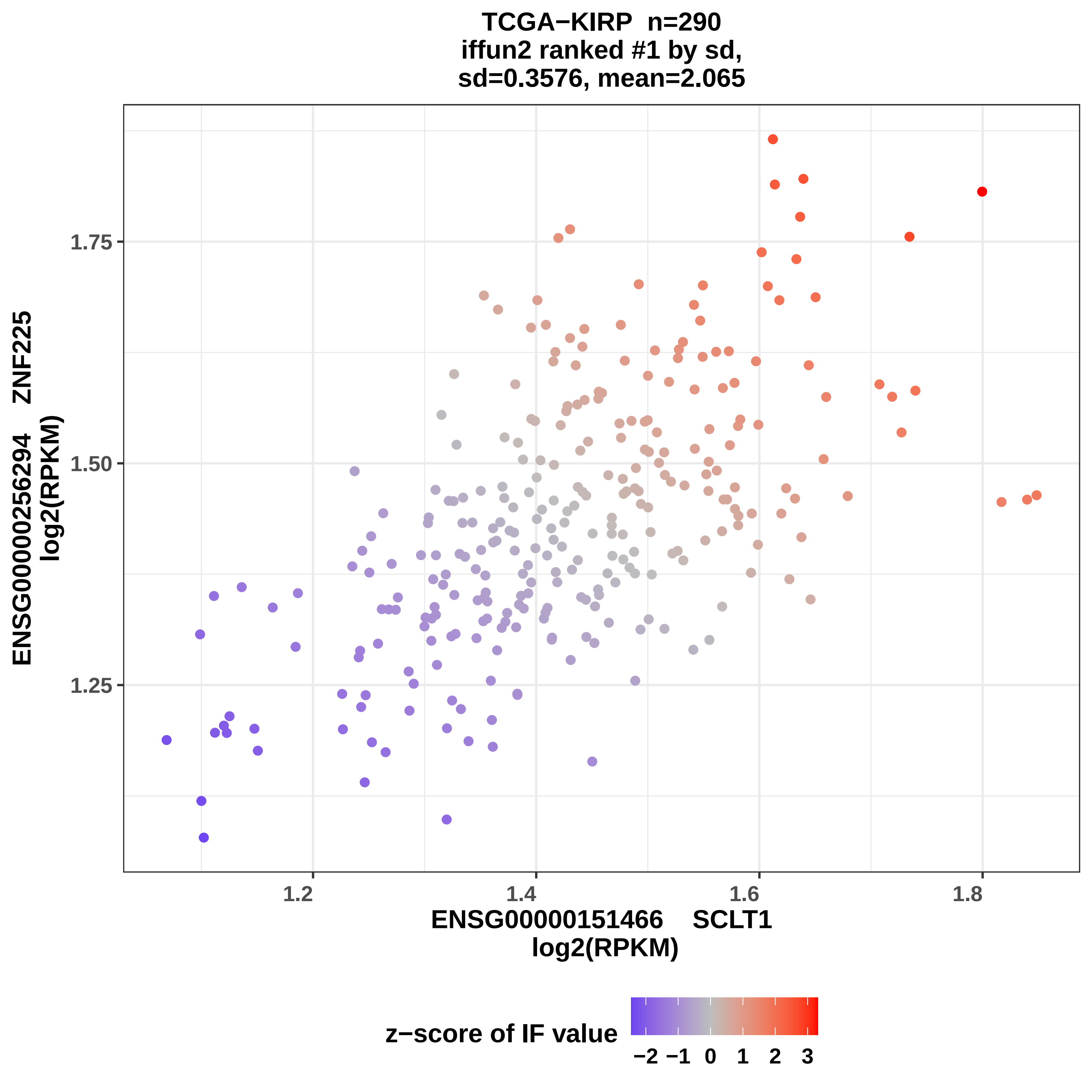
TCGA−KIRP n=290
iffun2 ranked #1 by sd,
sd=0.3576, mean=2.065
ENSG00000256294 ZNF225
log2(RPKM)
1.75
1.50
1.25
1.2
1.4
1.6
1.8
ENSG00000151466 SCLT1
log2(RPKM)
z−score of IF value
−2 −1 0 1 2 3

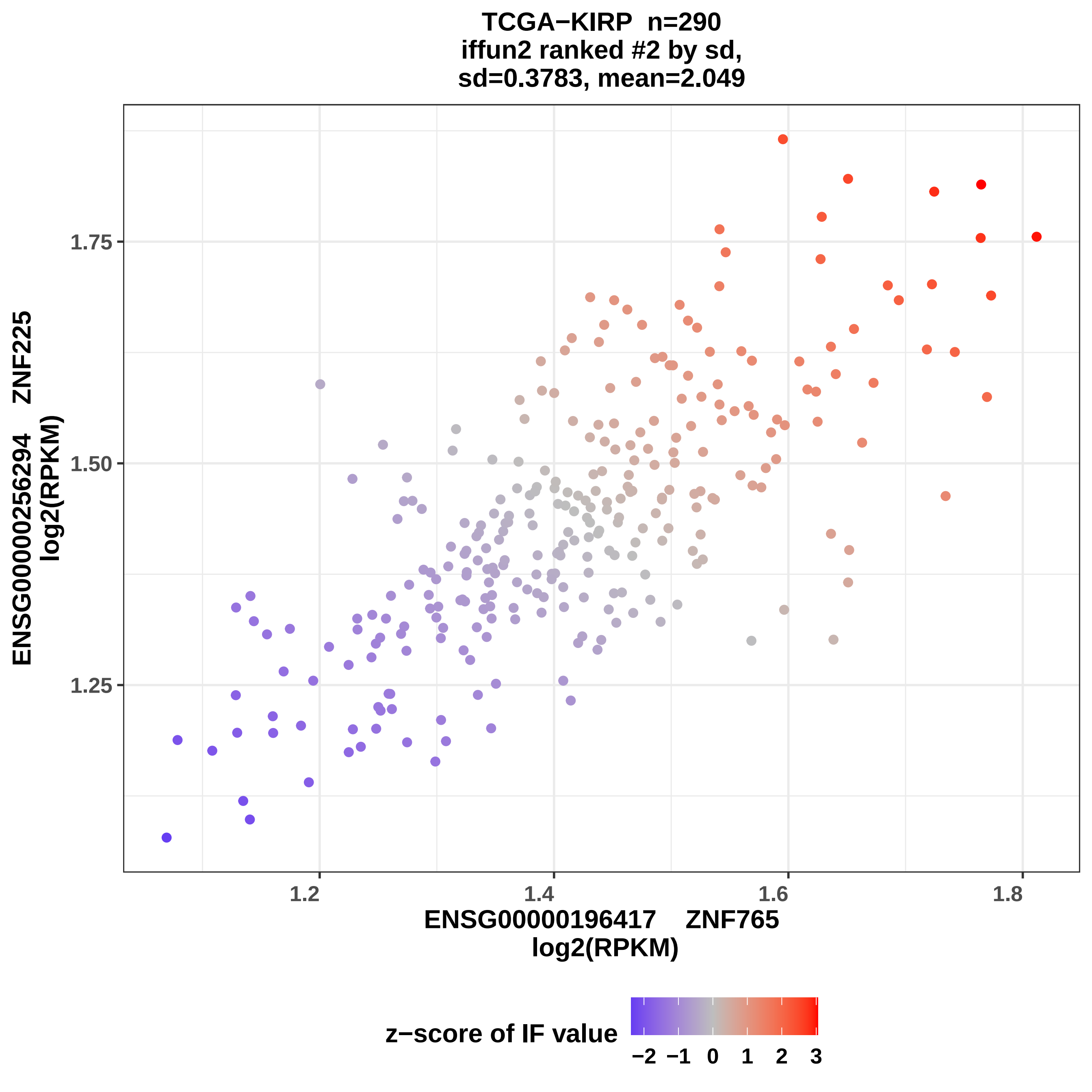

TCGA−KIRP n=290
iffun2 ranked #2 by sd,
sd=0.3783, mean=2.049
1.75
1.50
1.25
1.2
1.4
1.6
1.8
ENSG00000256294 ZNF225
log2(RPKM)
ENSG00000196417 ZNF765
log2(RPKM)
z−score of IF value
−2 −1 0 1 2 3

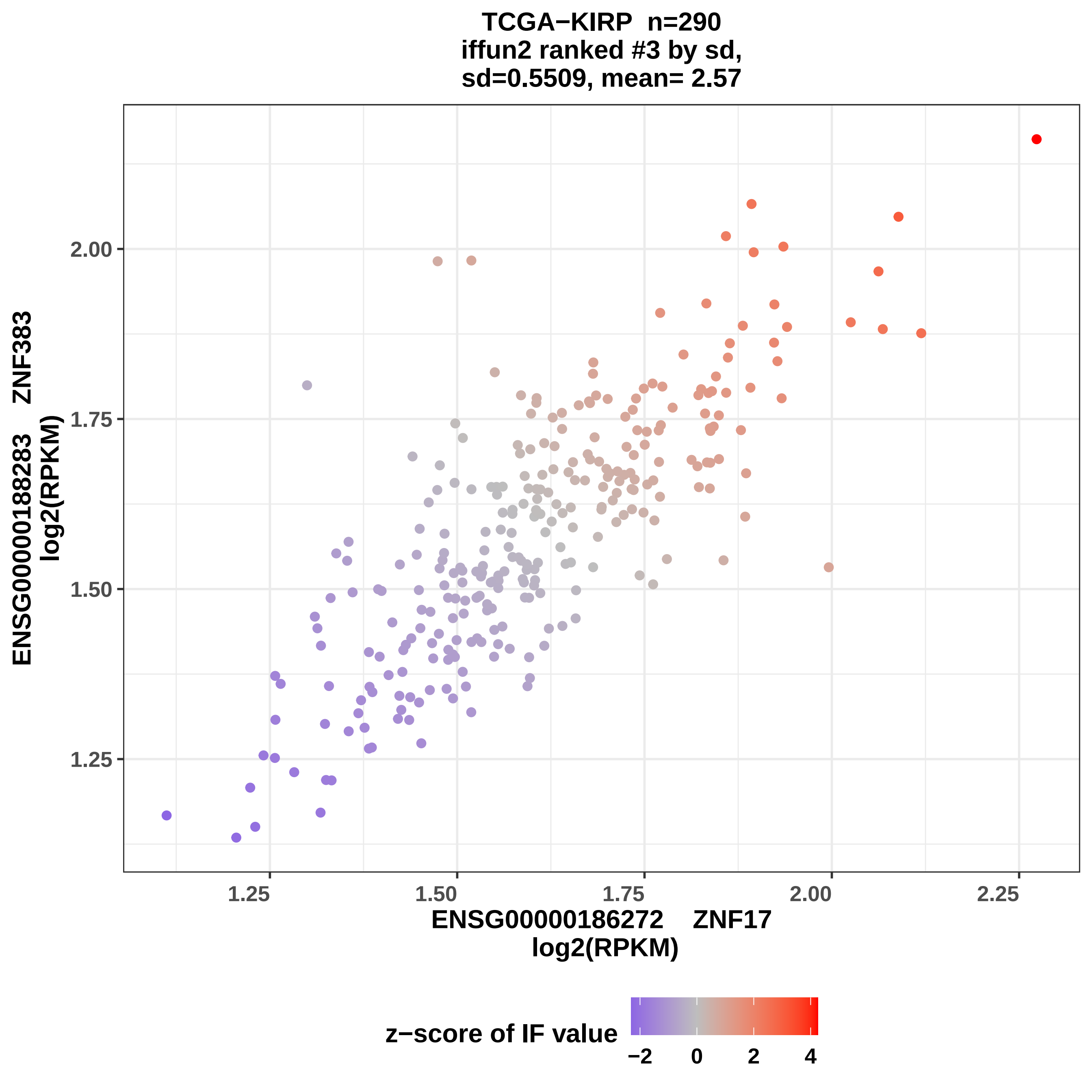
TCGA−KIRP n=290
iffun2 ranked #3 by sd,
sd=0.5509, mean= 2.57
ENSG00000188283 ZNF383
log2(RPKM)
1.25
1.50
1.75
2.00
1.25
1.50
1.75
2.00
2.25
ENSG00000186272 ZNF17
log2(RPKM)
z−score of IF value
−2
0
2
4

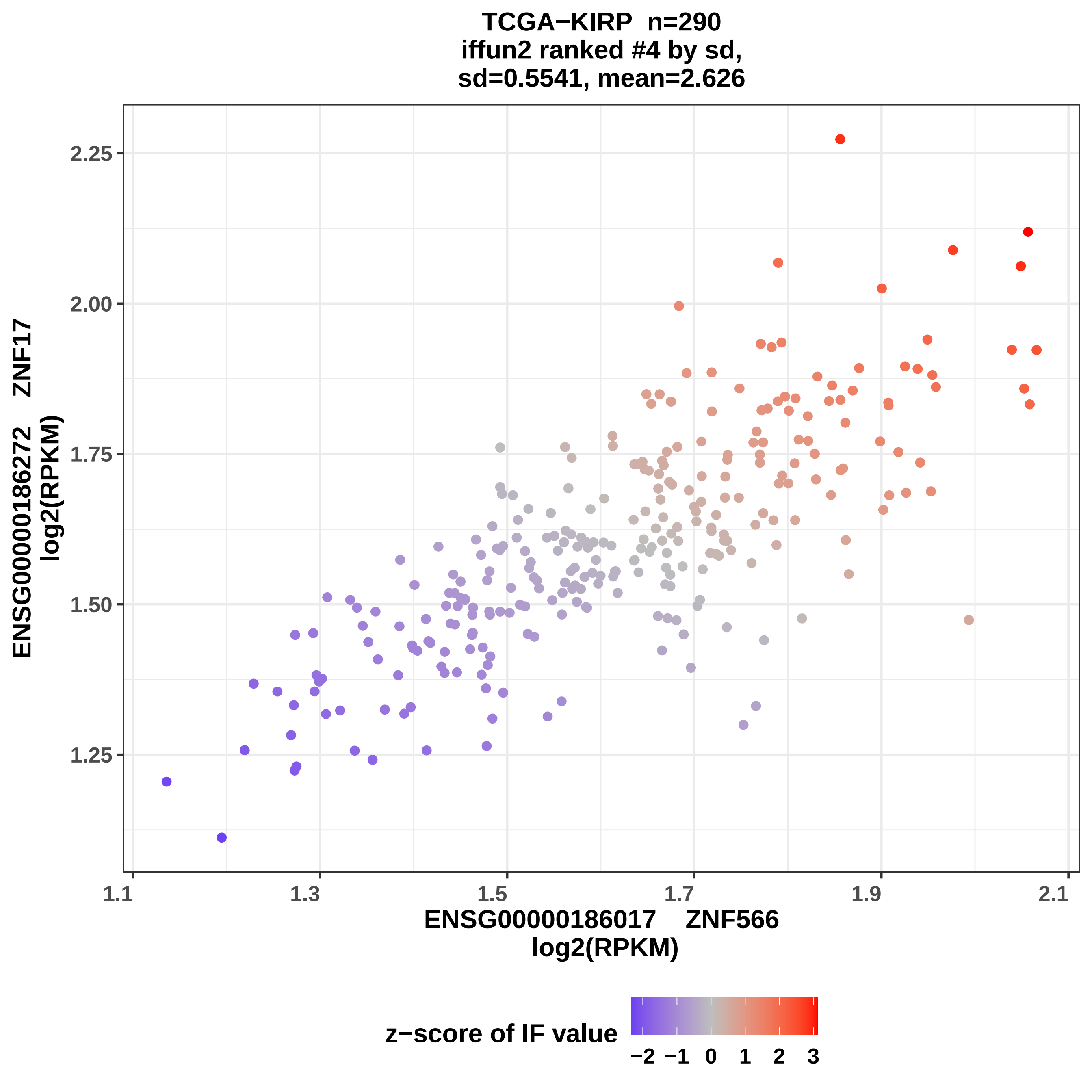
TCGA−KIRP n=290
iffun2 ranked #4 by sd,
sd=0.5541, mean=2.626
ENSG00000186272 ZNF17
log2(RPKM)
2.25
2.00
1.75
1.50
1.25
1.1
1.3
1.5
1.7
1.9
2.1
ENSG00000186017 ZNF566
log2(RPKM)
z−score of IF value
−2 −1 0 1 2 3

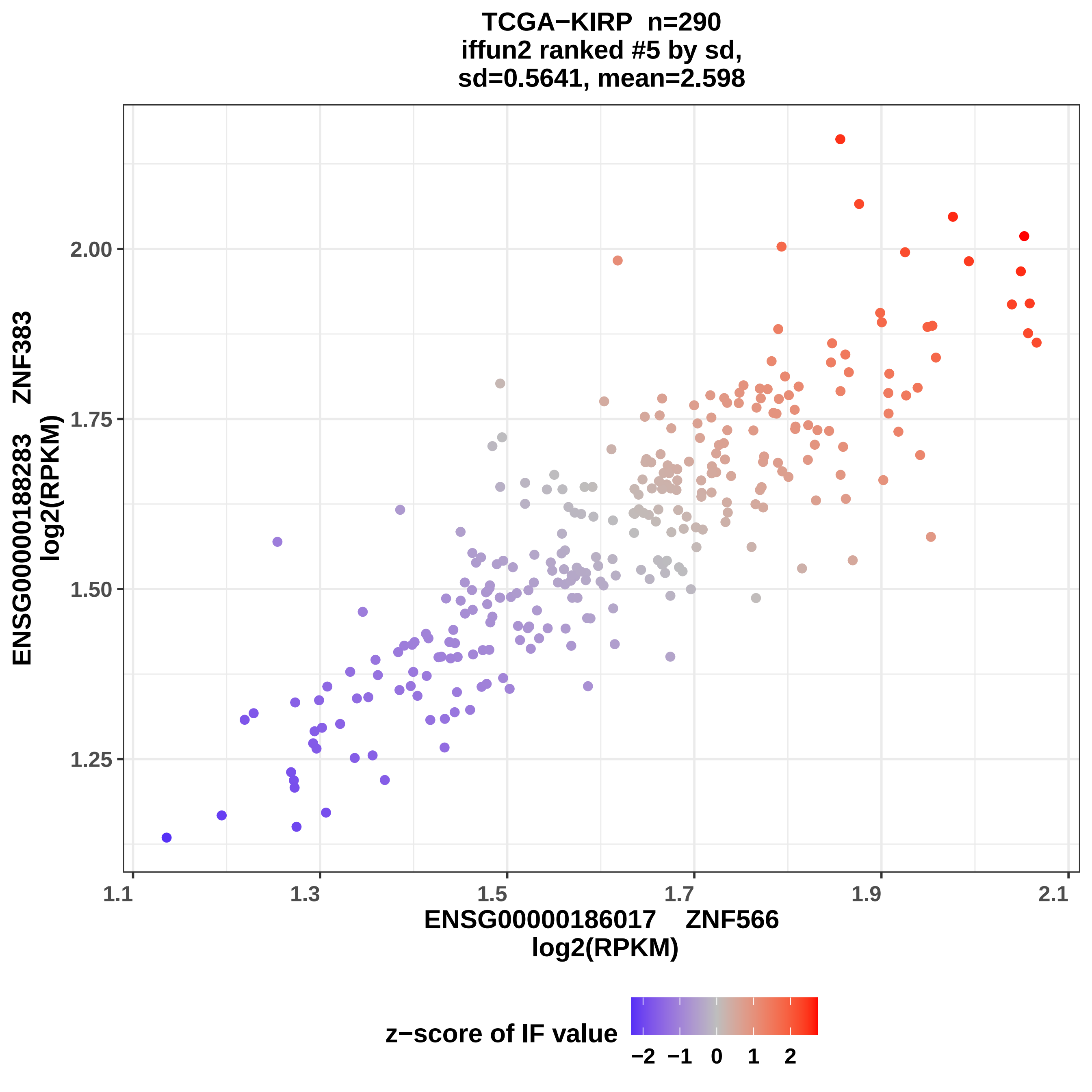

TCGA−KIRP n=290
iffun2 ranked #5 by sd,
sd=0.5641, mean=2.598
ENSG00000188283 ZNF383
log2(RPKM)
2.00
1.75
1.50
1.25
1.1
1.3
1.5
1.7
1.9
2.1
ENSG00000186017 ZNF566
log2(RPKM)
z−score of IF value
−2
−1
0
1
2

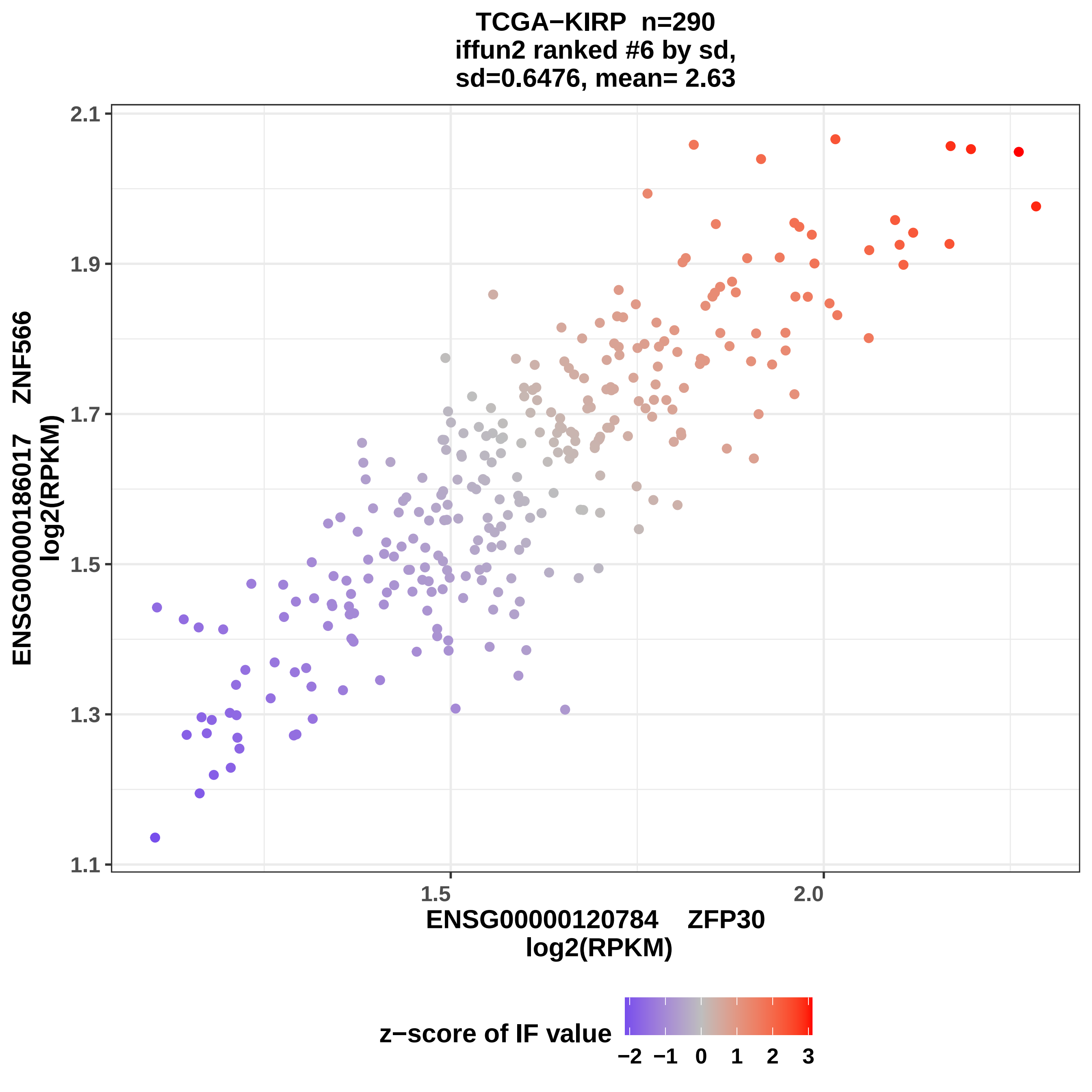

TCGA−KIRP n=290
iffun2 ranked #6 by sd,
sd=0.6476, mean= 2.63
ENSG00000186017 ZNF566
log2(RPKM)
2.1
1.9
1.7
1.5
1.3
1.1
1.5
2.0
ENSG00000120784 ZFP30
log2(RPKM)
z−score of IF value
−2 −1 0 1 2 3

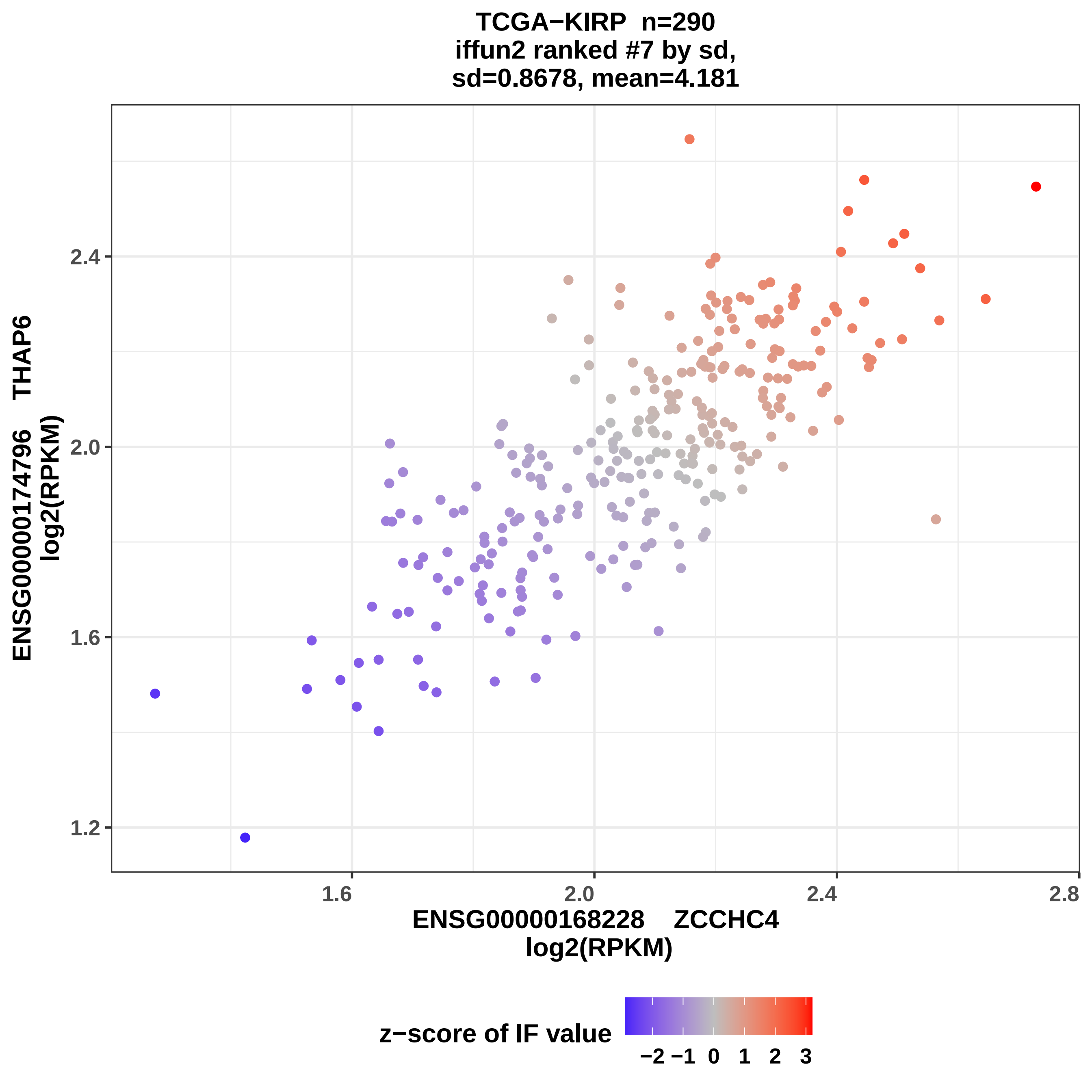

TCGA−KIRP n=290
iffun2 ranked #7 by sd,
sd=0.8678, mean=4.181
ENSG00000174796 THAP6
log2(RPKM)
2.4
2.0
1.6
1.2
1.6
2.0
2.4
2.8
ENSG00000168228 ZCCHC4
log2(RPKM)
z−score of IF value
−2 −1 0 1 2 3

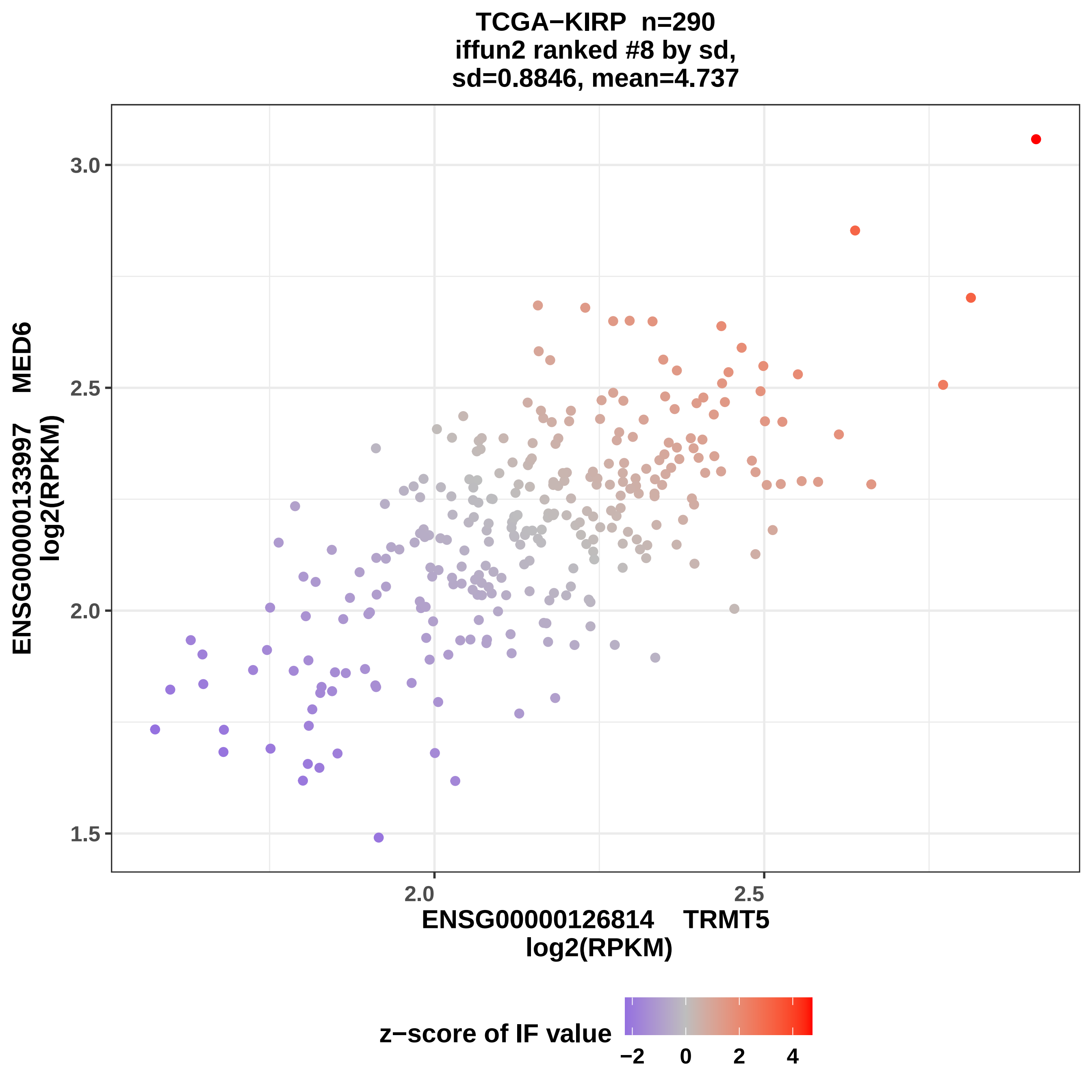

TCGA−KIRP n=290
iffun2 ranked #8 by sd,
sd=0.8846, mean=4.737
ENSG00000133997 MED6
log2(RPKM)
3.0
2.5
2.0
1.5
2.0
2.5
ENSG00000126814 TRMT5
log2(RPKM)
z−score of IF value
−2
0
2
4

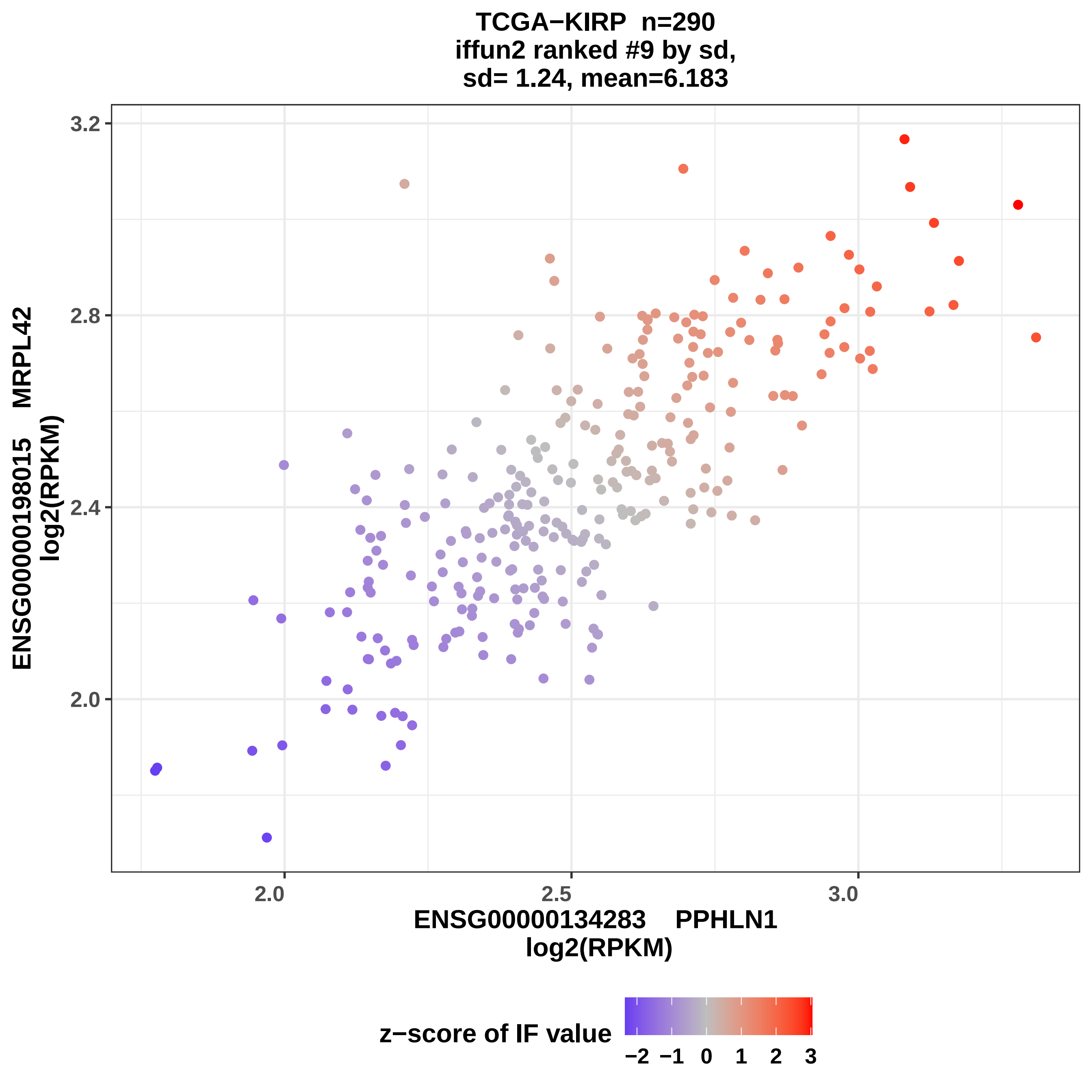
TCGA−KIRP n=290
iffun2 ranked #9 by sd,
sd= 1.24, mean=6.183
ENSG00000198015 MRPL42
log2(RPKM)
3.2
2.8
2.4
2.0
2.0
2.5
3.0
ENSG00000134283 PPHLN1
log2(RPKM)
z−score of IF value
−2
−1
0
1
2
3

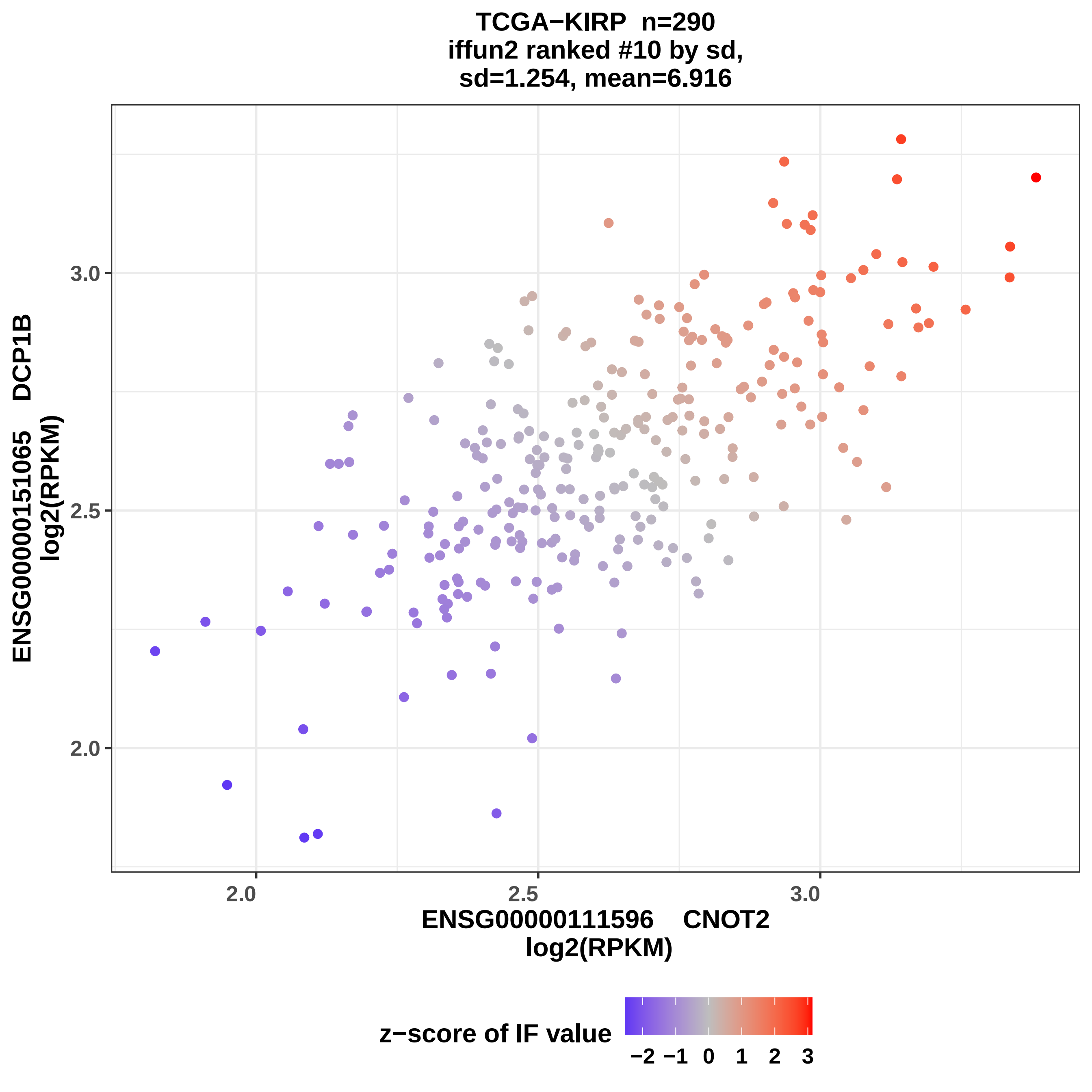
TCGA−KIRP n=290
iffun2 ranked #10 by sd,
sd=1.254, mean=6.916
ENSG00000151065 DCP1B
log2(RPKM)
3.0
2.5
2.0
2.0
2.5
3.0
ENSG00000111596 CNOT2
log2(RPKM)
z−score of IF value
−2 −1 0 1 2 3

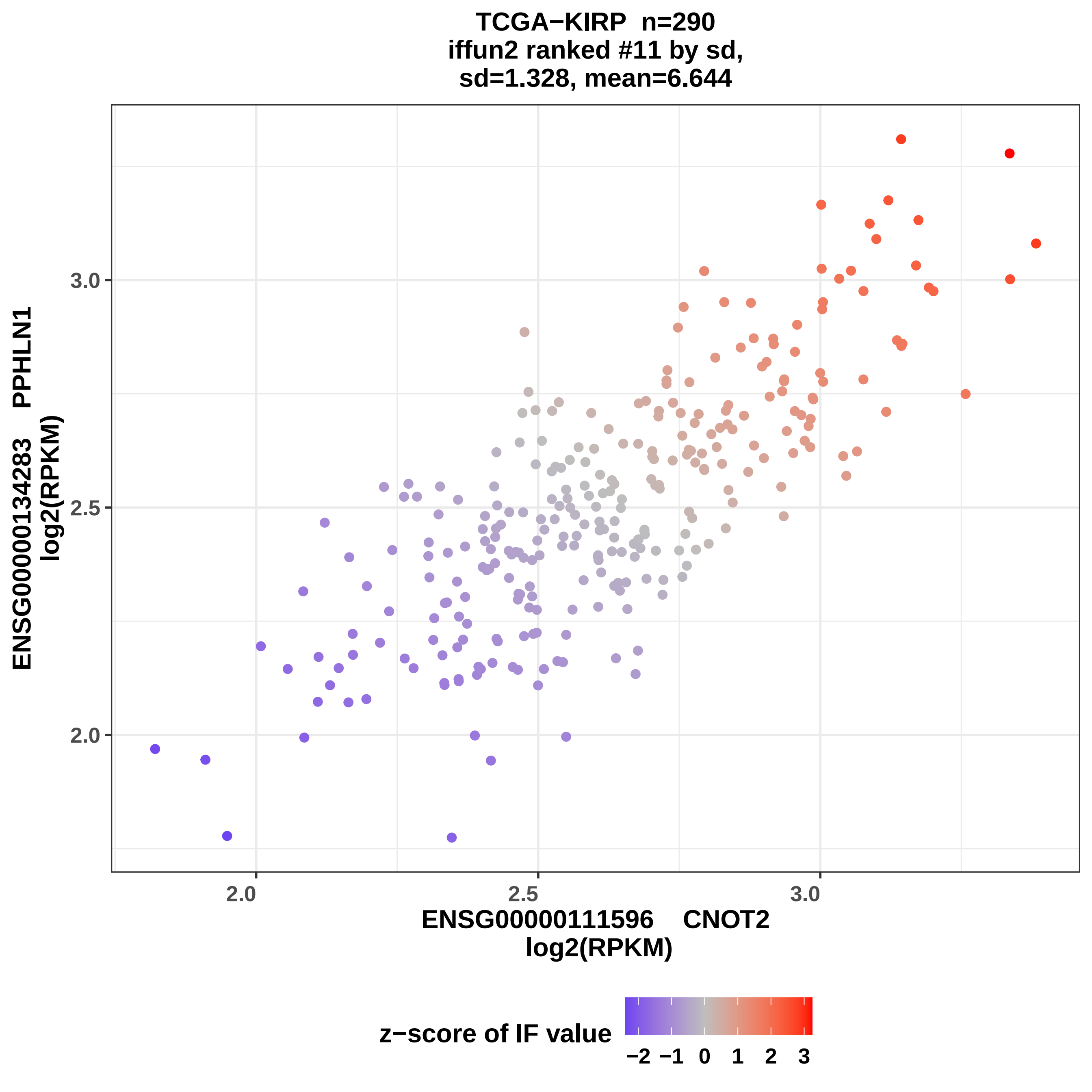

TCGA−KIRP n=290
iffun2 ranked #11 by sd,
sd=1.328, mean=6.644
ENSG00000134283 PPHLN1
log2(RPKM)
3.0
2.5
2.0
2.0
2.5
3.0
ENSG00000111596 CNOT2
log2(RPKM)
z−score of IF value
−2 −1 0 1 2 3

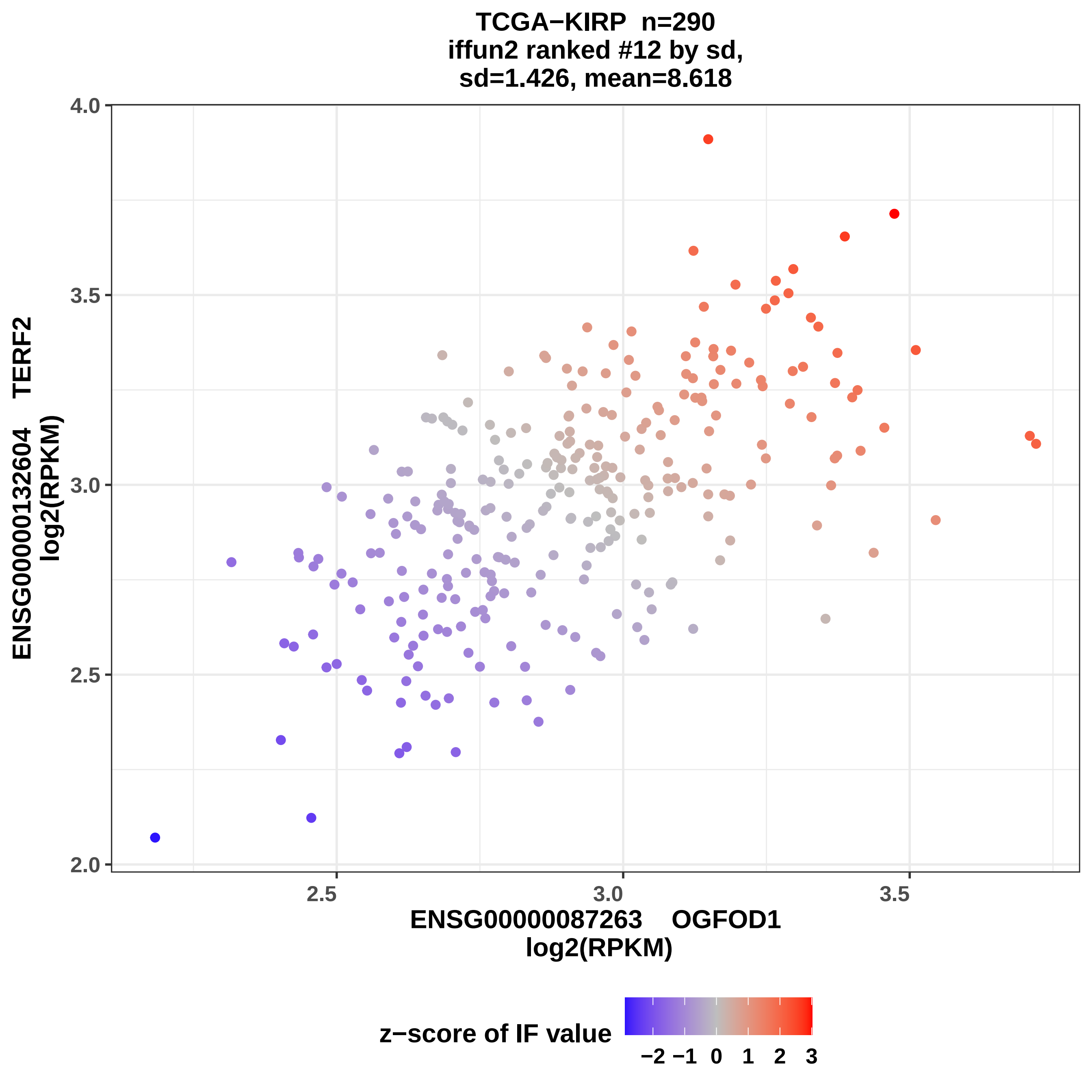
TCGA−KIRP n=290
iffun2 ranked #12 by sd,
sd=1.426, mean=8.618
ENSG00000132604 TERF2
log2(RPKM)
4.0
3.5
3.0
2.5
2.0
2.5
3.0
3.5
ENSG00000087263 OGFOD1
log2(RPKM)
z−score of IF value
−2 −1 0 1 2 3

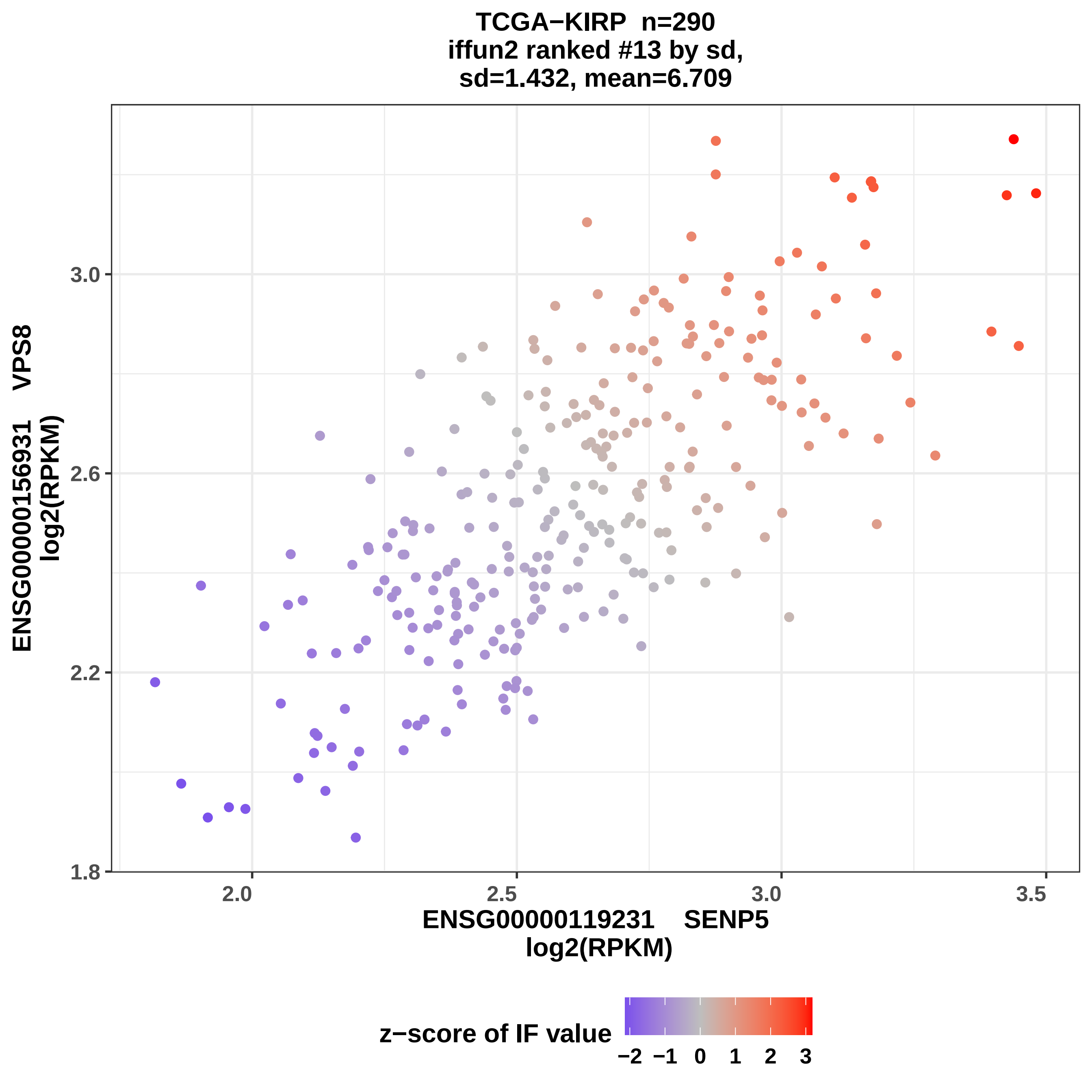

TCGA−KIRP n=290
iffun2 ranked #13 by sd,
sd=1.432, mean=6.709
ENSG00000156931 VPS8
log2(RPKM)
3.0
2.6
2.2
1.8
2.0
2.5
3.0
3.5
ENSG00000119231 SENP5
log2(RPKM)
z−score of IF value
−2 −1 0 1 2 3

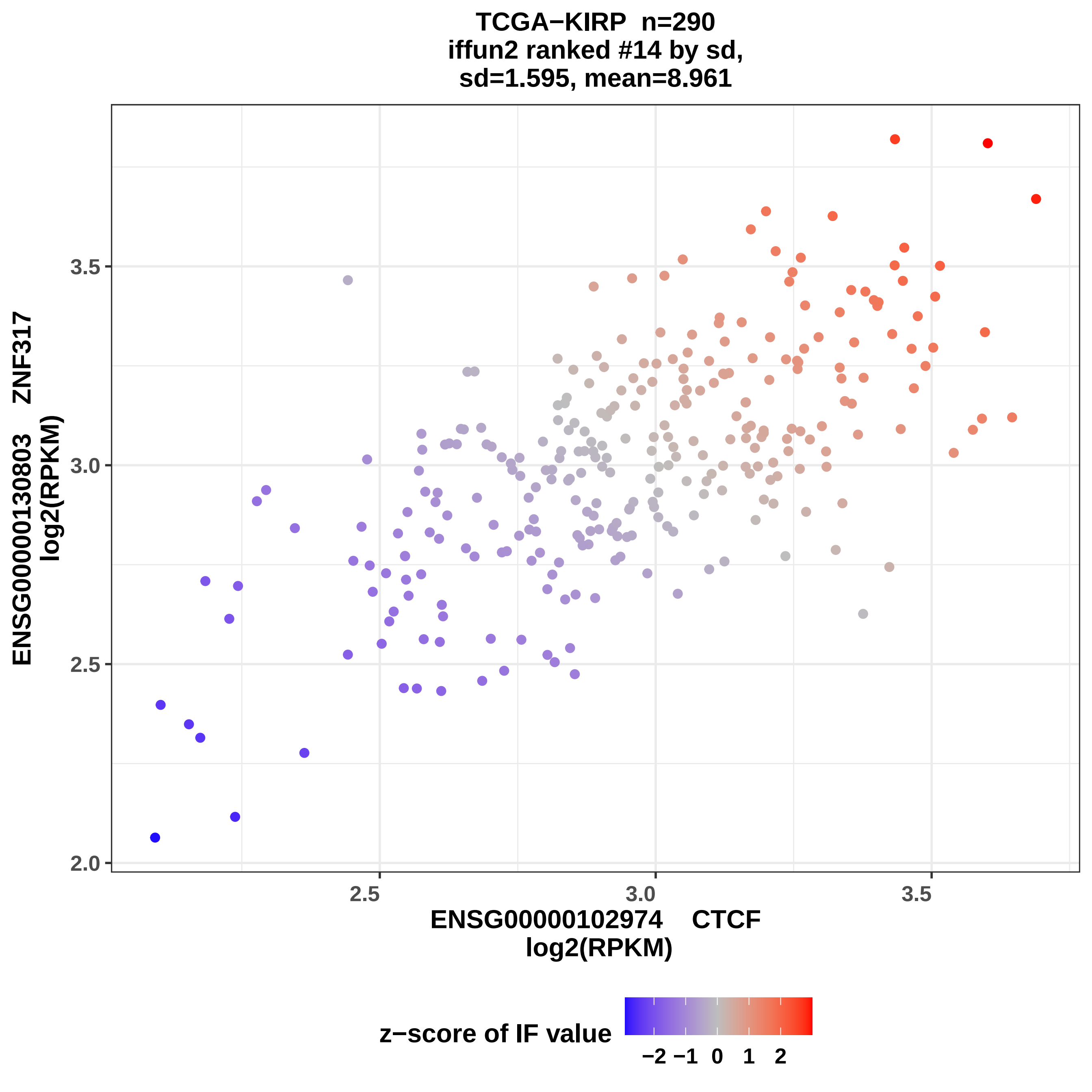
TCGA−KIRP n=290
iffun2 ranked #14 by sd,
sd=1.595, mean=8.961
ENSG00000130803 ZNF317
log2(RPKM)
2.0
2.5
3.0
3.5
ENSG00000102974 CTCF
log2(RPKM)
2.5
3.0
3.5
z−score of IF value
−2 −1 0 1 2

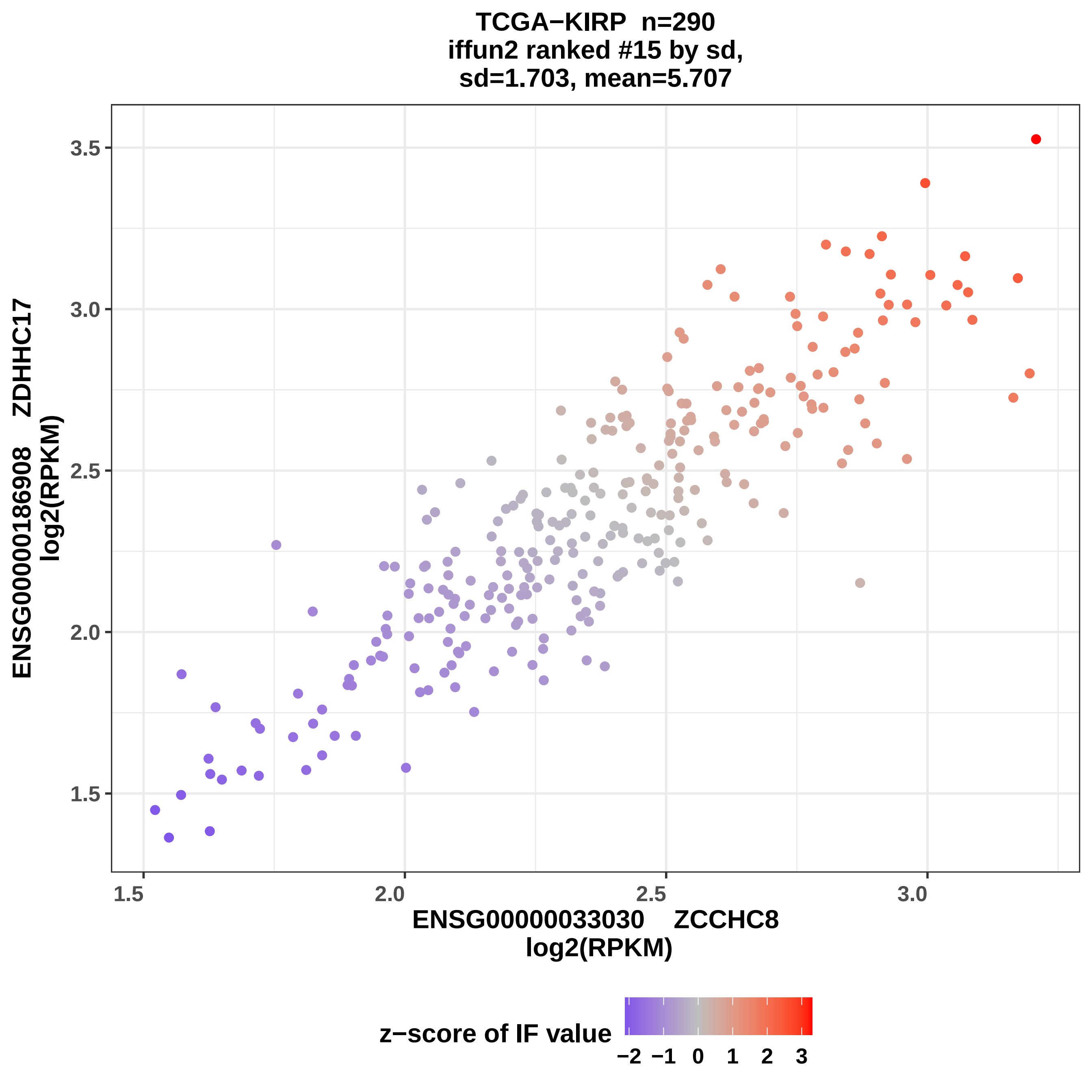
TCGA−KIRP n=290
iffun2 ranked #15 by sd,
sd=1.703, mean=5.707
ENSG00000186908 ZDHHC17
log2(RPKM)
ENSG00000033030 ZCCHC8
log2(RPKM)
1.5
2.0
2.5
3.0
3.5
z−score of IF value
−2 −1 0 1 2 3

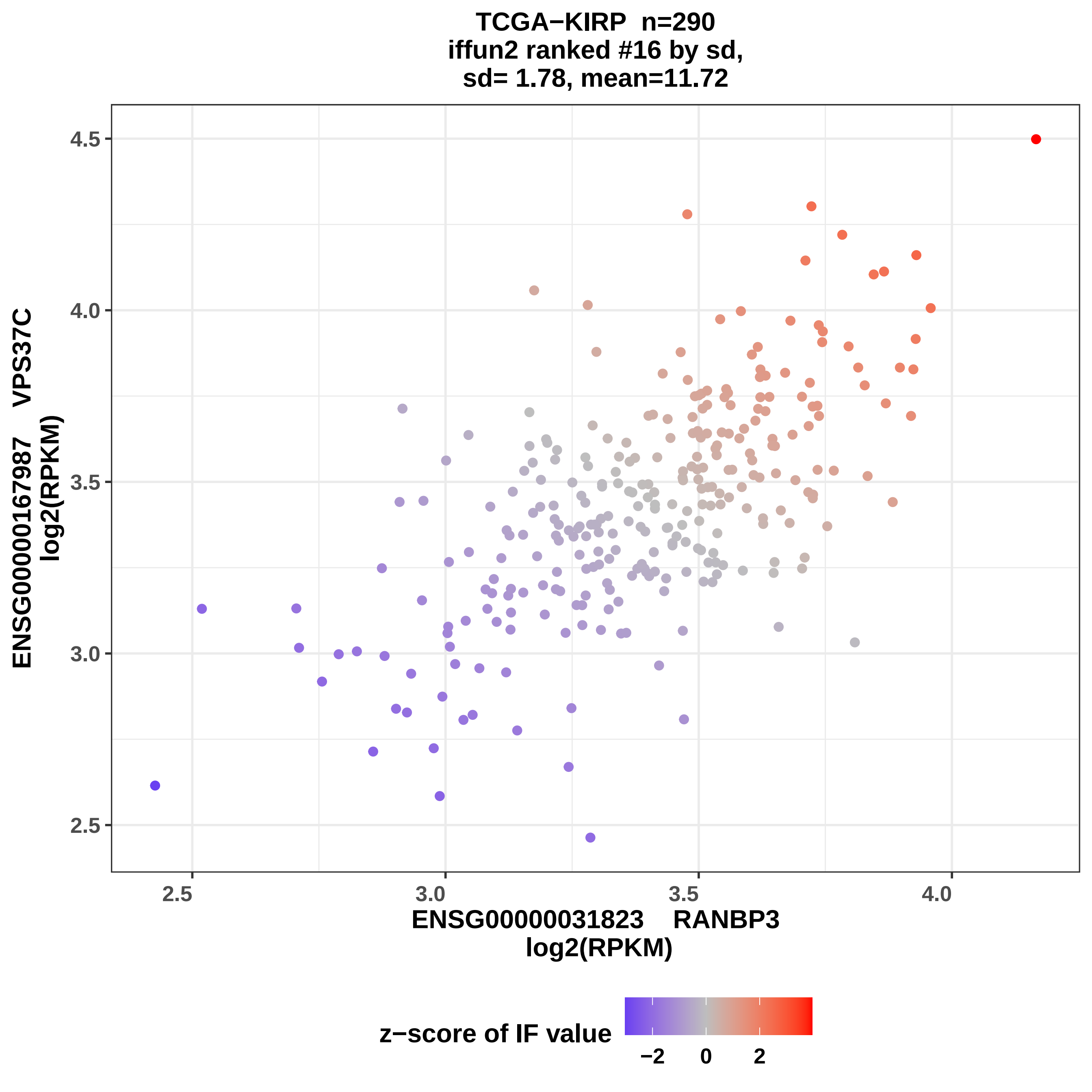

TCGA−KIRP n=290
iffun2 ranked #16 by sd,
sd= 1.78, mean=11.72
ENSG00000167987 VPS37C
log2(RPKM)
4.5
4.0
3.5
3.0
2.5
2.5
3.0
3.5
4.0
ENSG00000031823 RANBP3
log2(RPKM)
z−score of IF value
−2
0
2

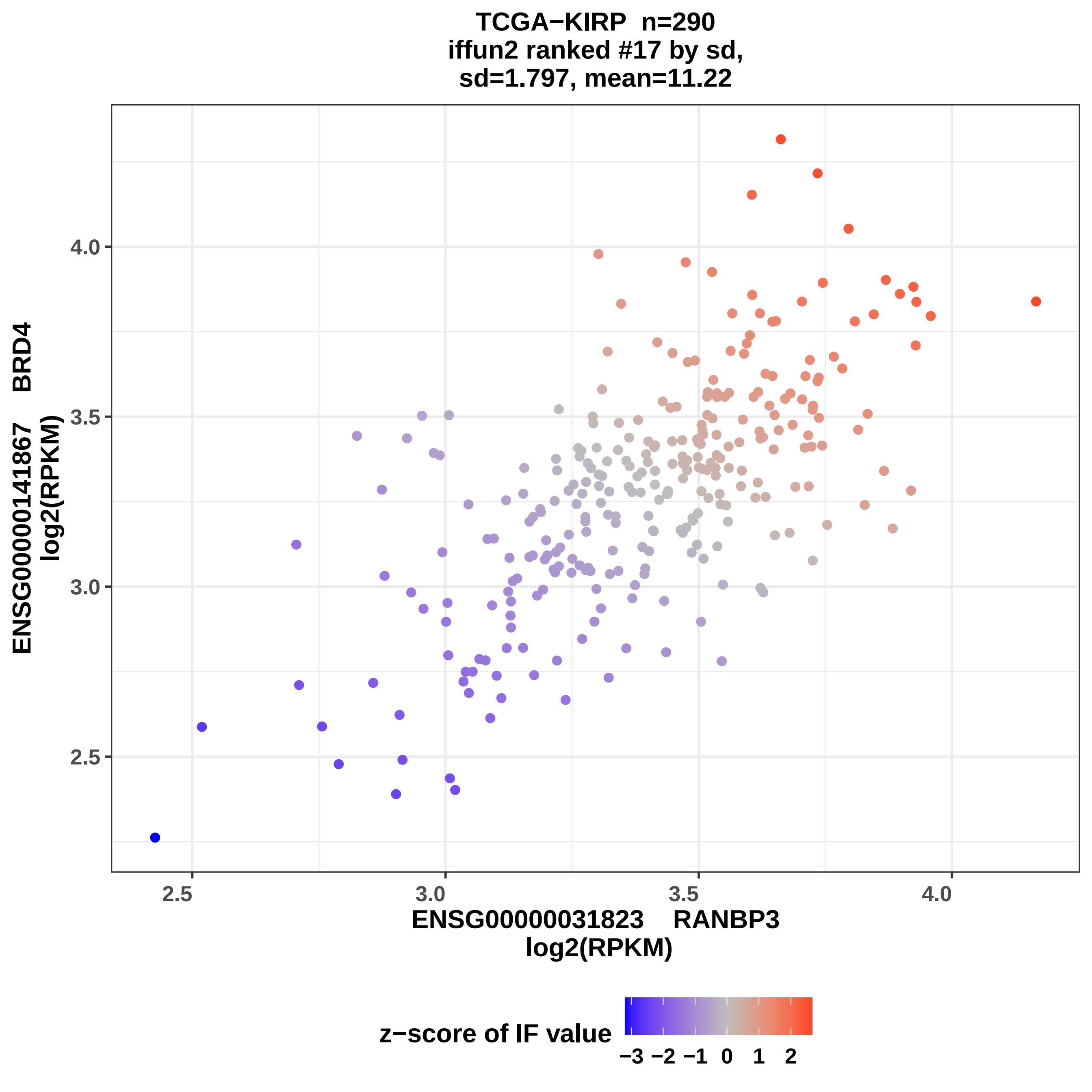

TCGA−KIRP n=290
iffun2 ranked #17 by sd,
sd=1.797, mean=11.22
ENSG00000141867 BRD4
log2(RPKM)
ENSG00000031823 RANBP3
log2(RPKM)
2.5
3.0
3.5
4.0
2.5
3.0
3.5
4.0
z−score of IF value
−3 −2 −1 0 1 2

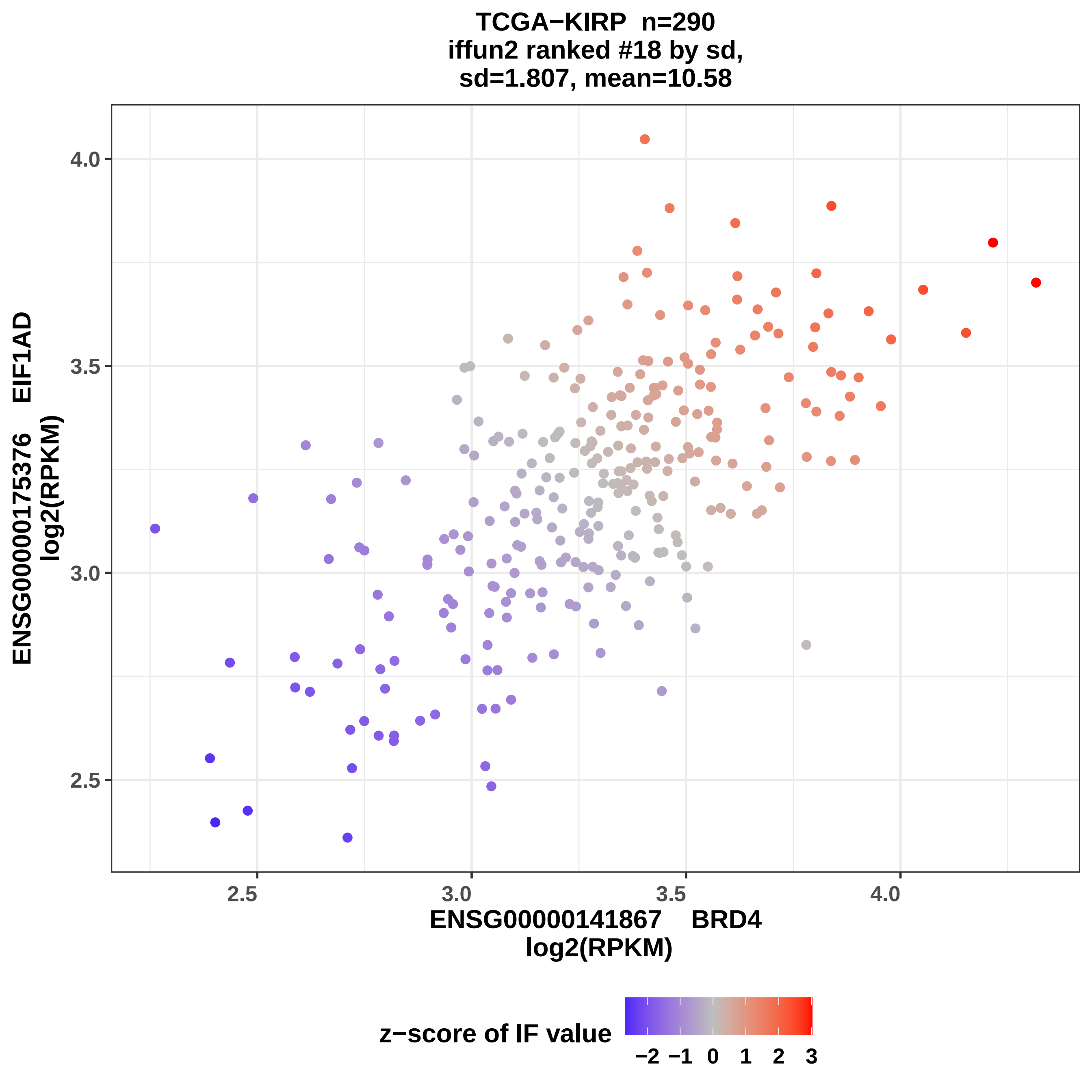
TCGA−KIRP n=290
iffun2 ranked #18 by sd,
sd=1.807, mean=10.58
ENSG00000175376 EIF1AD
log2(RPKM)
4.0
3.5
3.0
2.5
2.5
3.0
3.5
4.0
ENSG00000141867 BRD4
log2(RPKM)
z−score of IF value
−2 −1 0 1 2 3

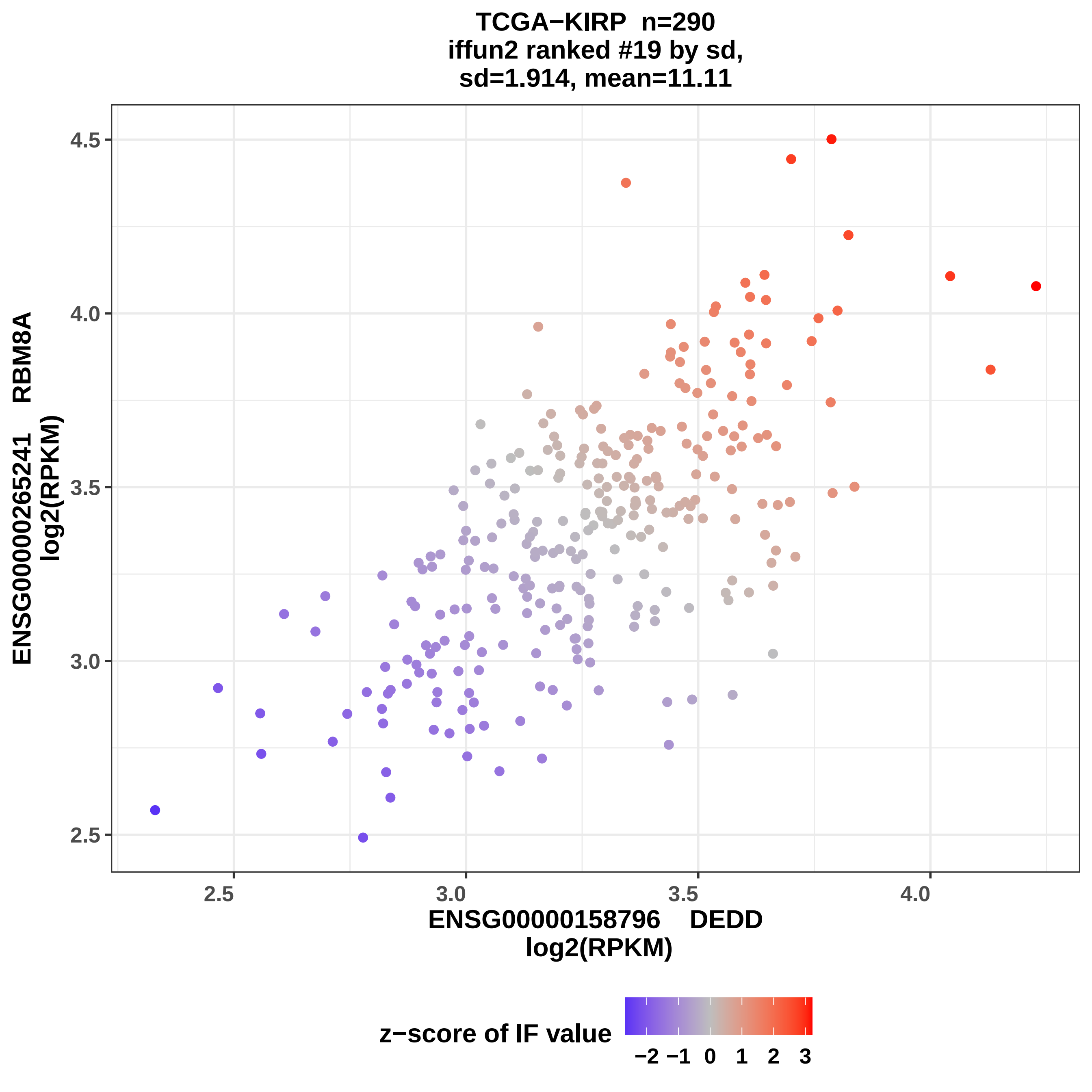

TCGA−KIRP n=290
iffun2 ranked #19 by sd,
sd=1.914, mean=11.11
ENSG00000265241 RBM8A
log2(RPKM)
4.5
4.0
3.5
3.0
2.5
2.5
3.0
3.5
4.0
ENSG00000158796 DEDD
log2(RPKM)
z−score of IF value
−2 −1 0 1 2 3

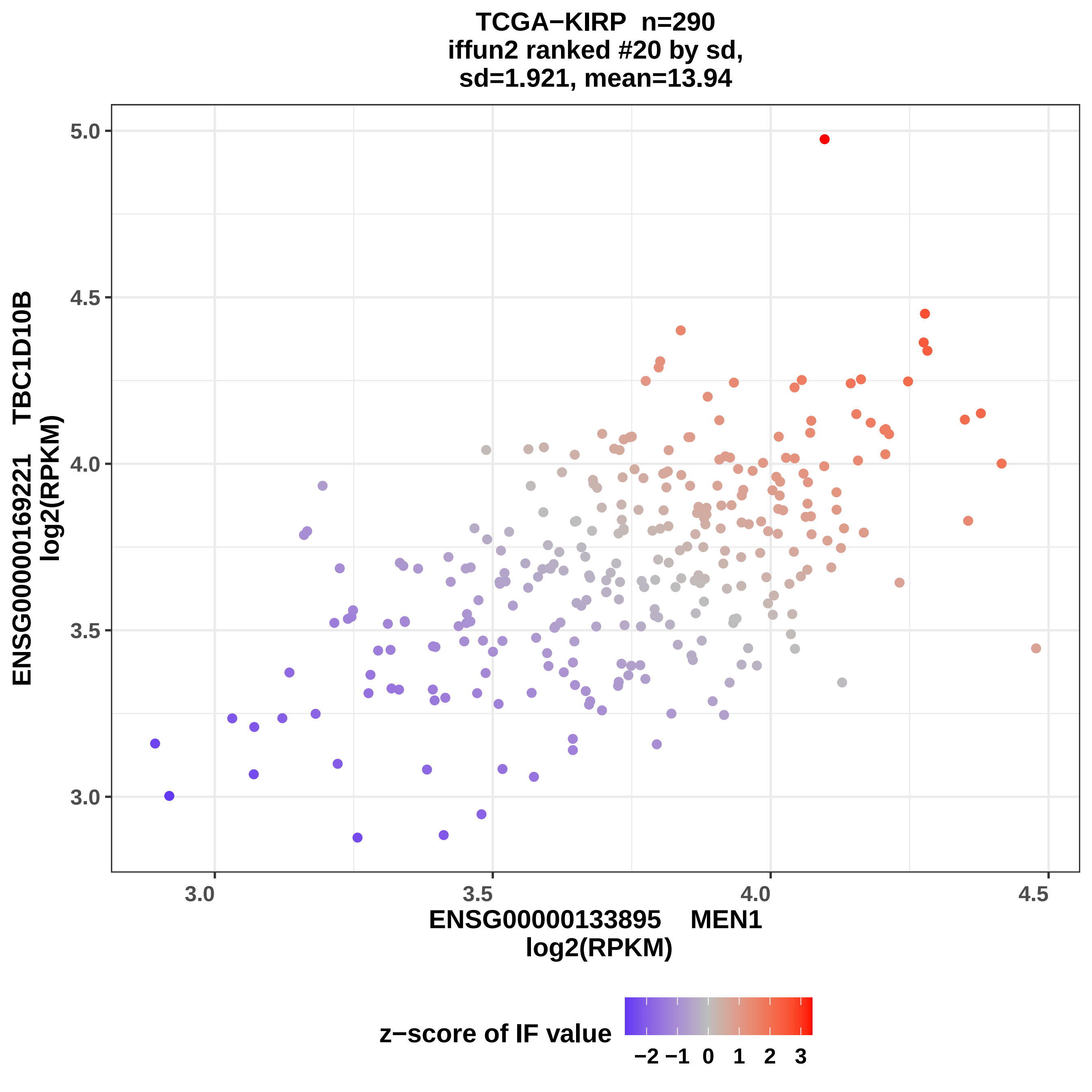

TCGA−KIRP n=290
iffun2 ranked #20 by sd,
sd=1.921, mean=13.94
ENSG00000169221 TBC1D10B
log2(RPKM)
5.0
4.5
4.0
3.5
3.0
3.0
3.5
4.0
4.5
ENSG00000133895 MEN1
log2(RPKM)
z−score of IF value
−2 −1 0 1 2 3

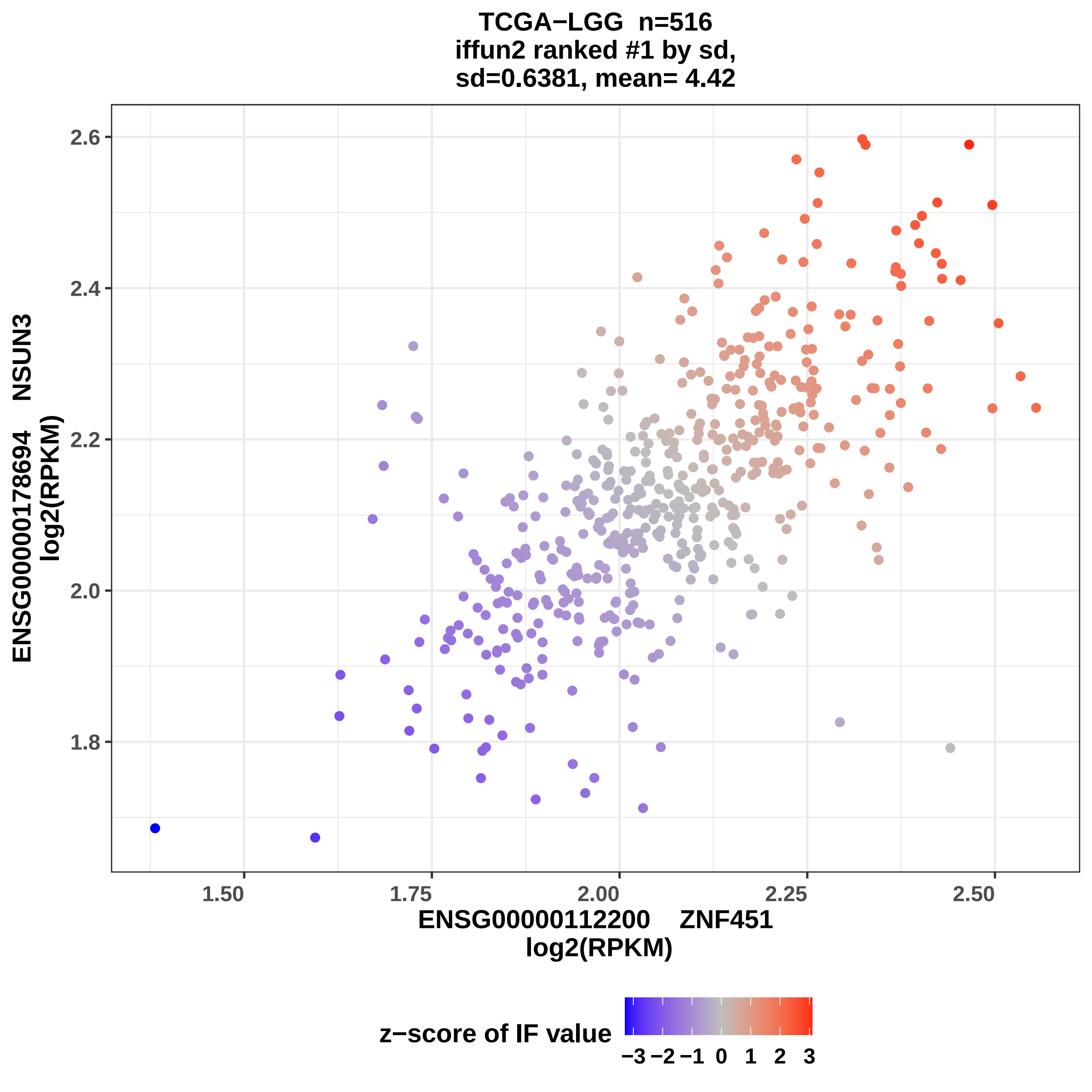
TCGA−LGG n=516
iffun2 ranked #1 by sd,
sd=0.6381, mean= 4.42
2.6
2.4
2.2
2.0
1.8
1.50
1.75
2.00
2.25
2.50
ENSG00000178694 NSUN3
log2(RPKM)
ENSG00000112200 ZNF451
log2(RPKM)
z−score of IF value
−3 −2 −1 0 1 2 3

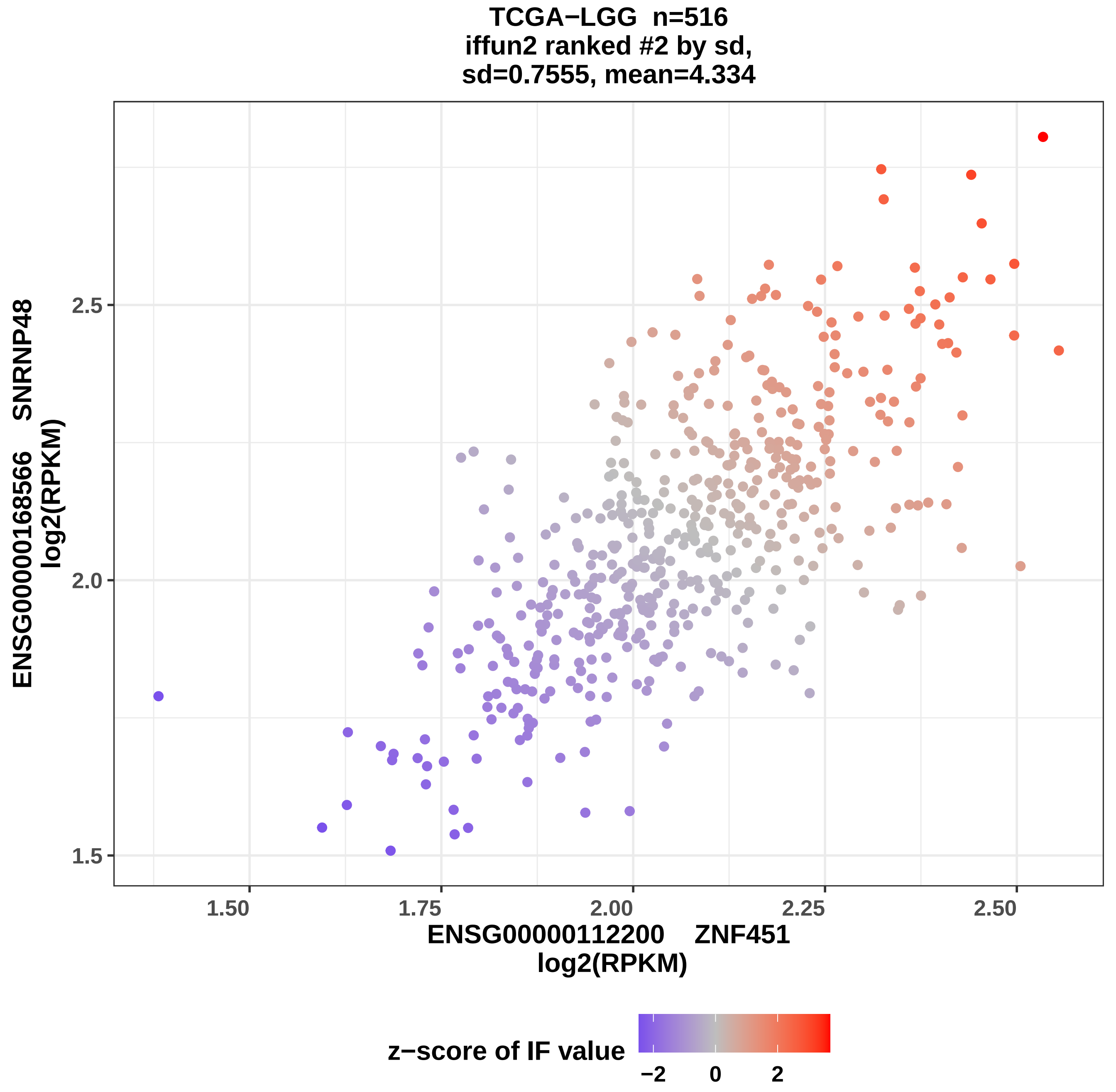

TCGA−LGG n=516
iffun2 ranked #2 by sd,
sd=0.7555, mean=4.334
ENSG00000168566 SNRNP48
log2(RPKM)
2.5
2.0
1.5
1.50
1.75
2.00
2.25
2.50
ENSG00000112200 ZNF451
log2(RPKM)
z−score of IF value
−2
0
2

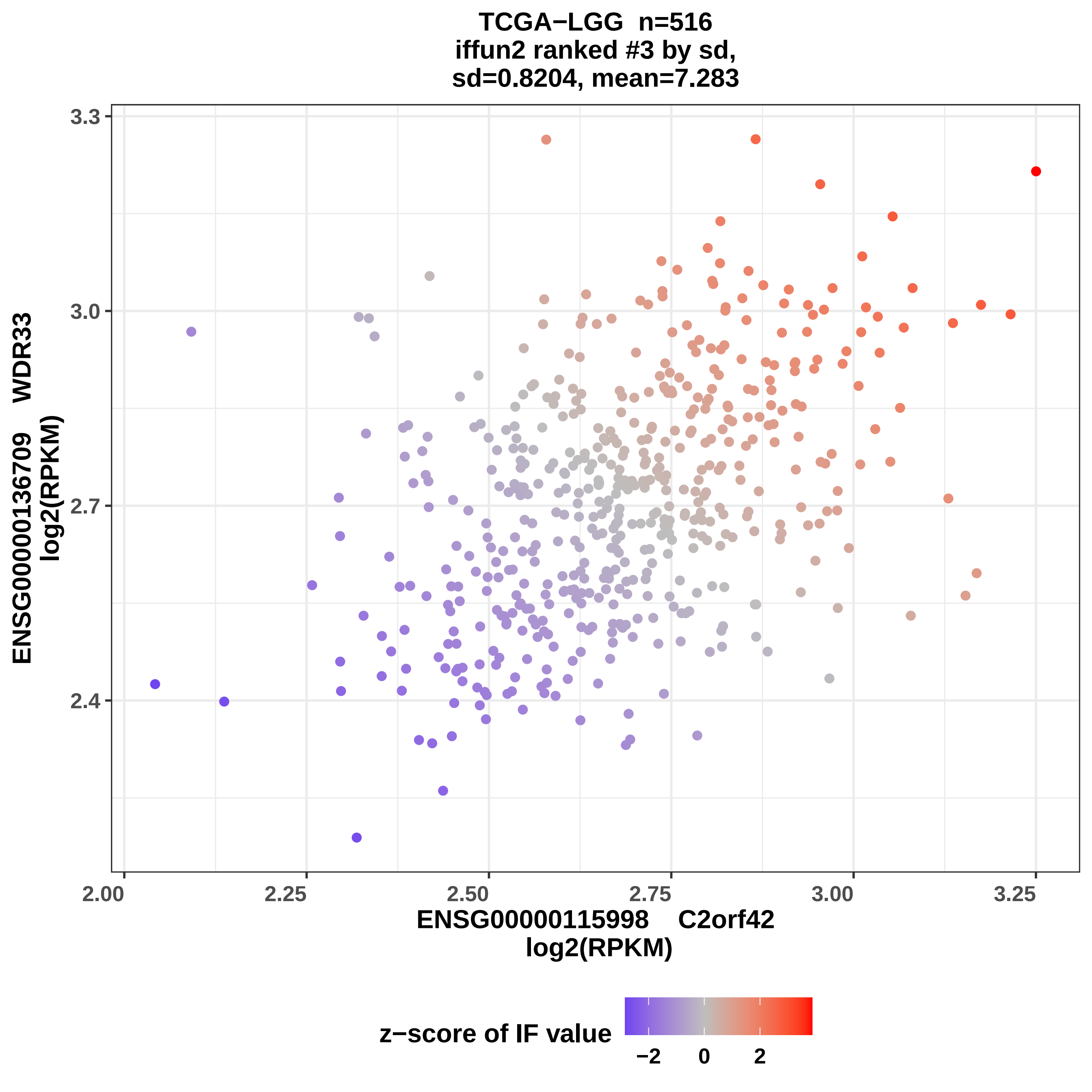
TCGA−LGG n=516
iffun2 ranked #3 by sd,
sd=0.8204, mean=7.283
ENSG00000136709 WDR33
log2(RPKM)
ENSG00000115998 C2orf42
log2(RPKM)
3.3
3.0
2.7
2.4
2.00
2.25
2.50
2.75
3.00
3.25
z−score of IF value
−2
0
2

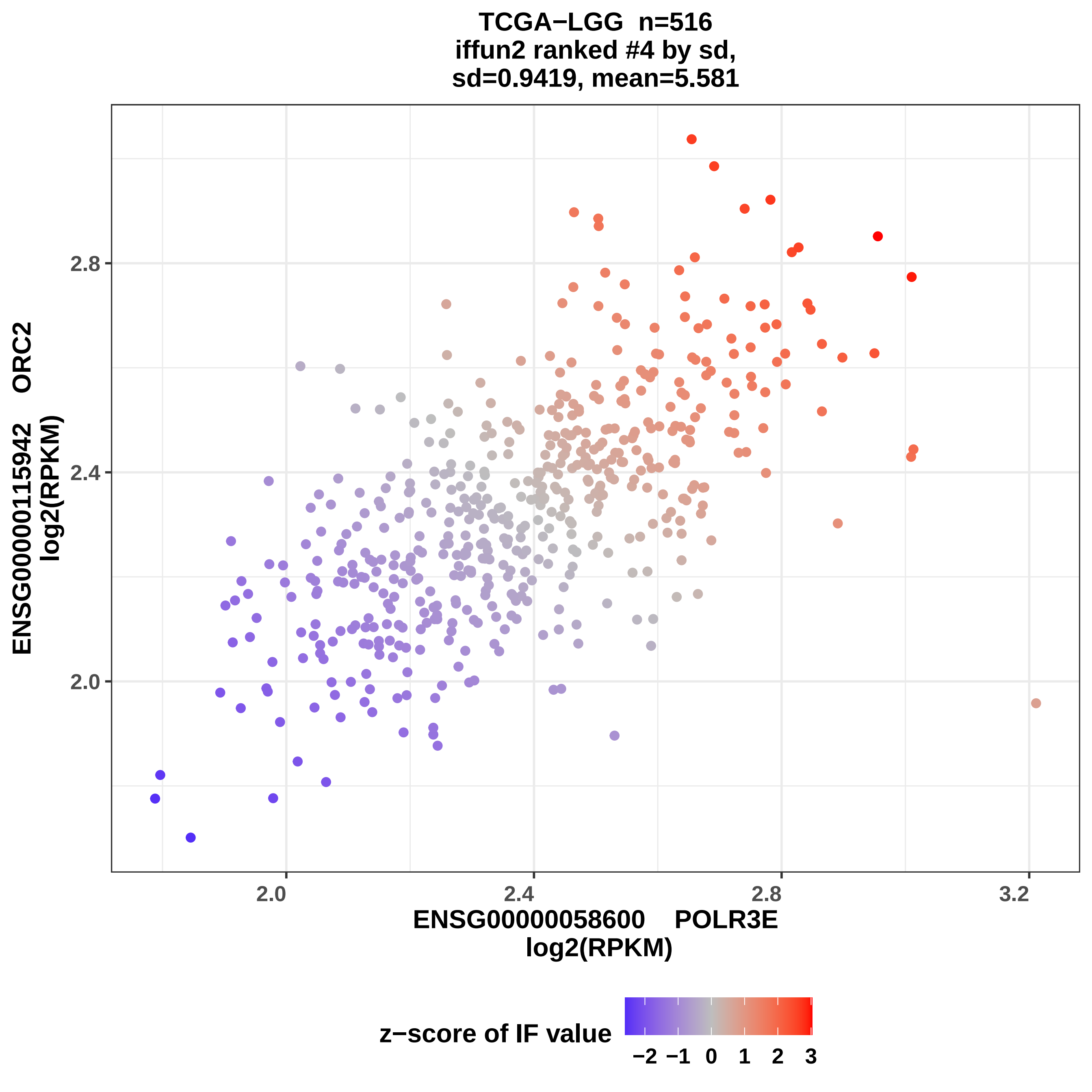
TCGA−LGG n=516
iffun2 ranked #4 by sd,
sd=0.9419, mean=5.581
ENSG00000115942 ORC2
log2(RPKM)
ENSG00000058600 POLR3E
log2(RPKM)
2.0
2.4
2.8
3.2
z−score of IF value
−2 −1 0 1 2 3

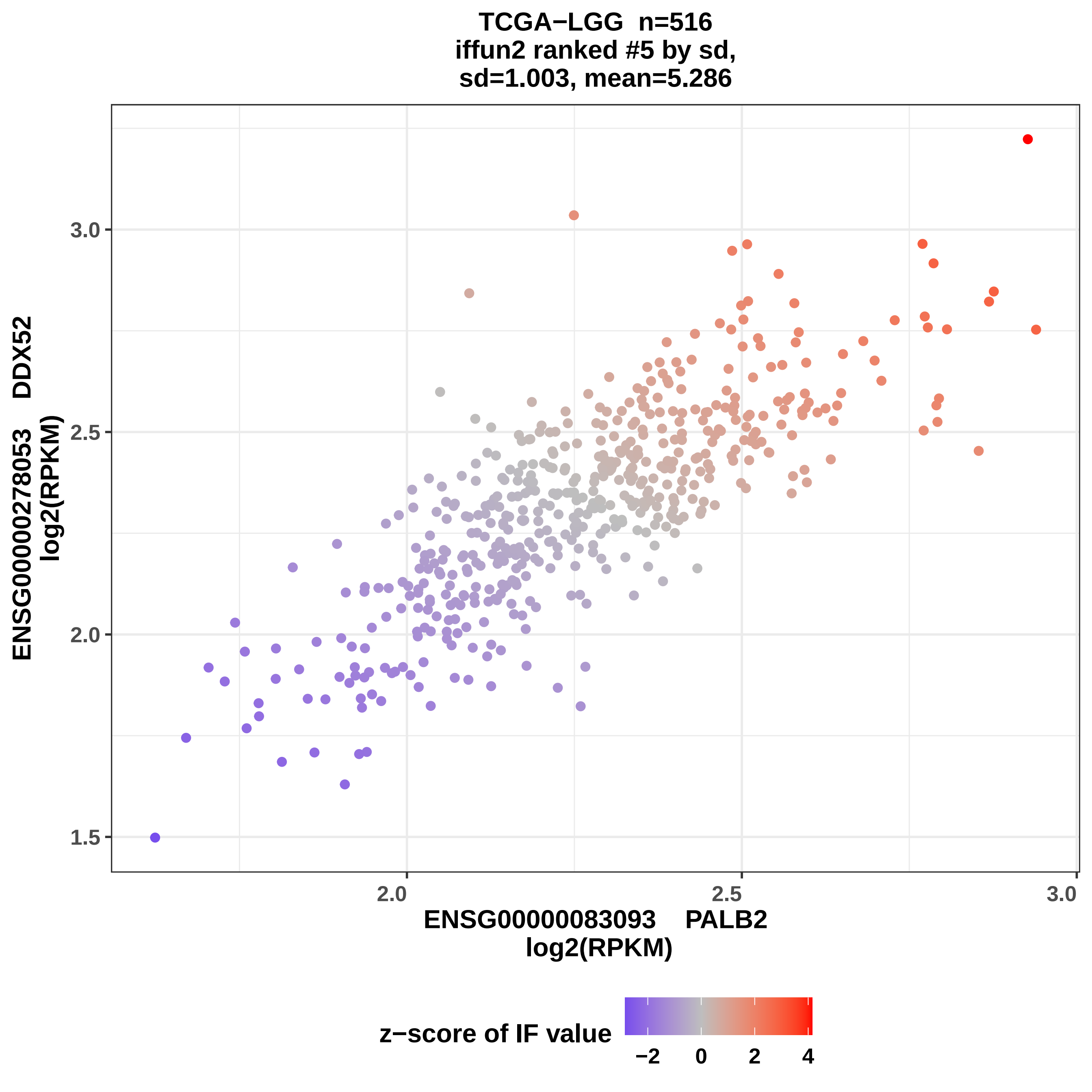

TCGA−LGG n=516
iffun2 ranked #5 by sd,
sd=1.003, mean=5.286
ENSG00000278053 DDX52
log2(RPKM)
1.5
2.0
2.5
3.0
ENSG00000083093 PALB2
log2(RPKM)
z−score of IF value
−2
0
2
4

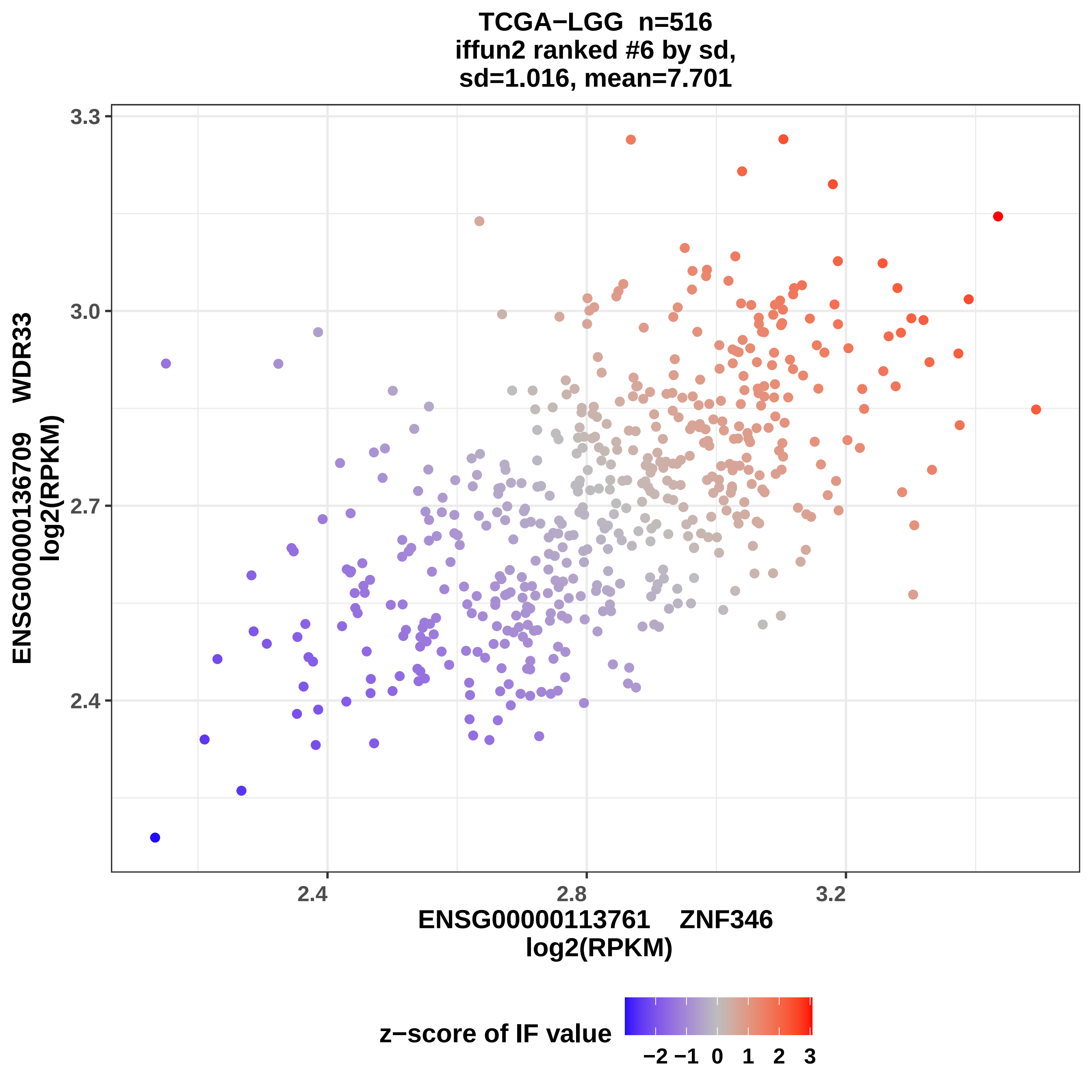
TCGA−LGG n=516
iffun2 ranked #6 by sd,
sd=1.016, mean=7.701
3.3
3.0
2.7
2.4
2.4
2.8
3.2
ENSG00000136709 WDR33
log2(RPKM)
ENSG00000113761 ZNF346
log2(RPKM)
z−score of IF value
−2 −1 0 1 2 3

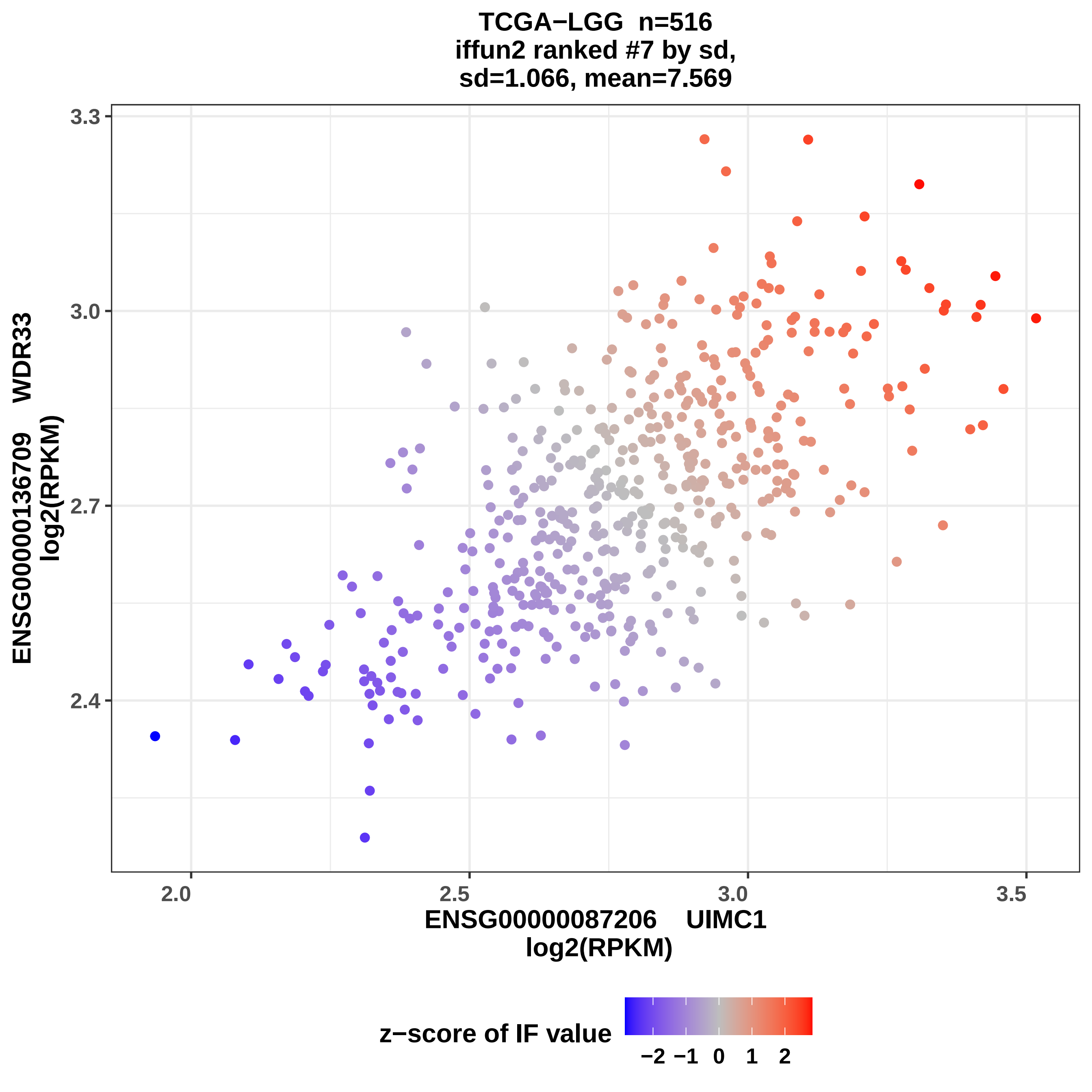

TCGA−LGG n=516
iffun2 ranked #7 by sd,
sd=1.066, mean=7.569
ENSG00000136709 WDR33
log2(RPKM)
3.3
3.0
2.7
2.4
2.0
2.5
3.0
3.5
ENSG00000087206 UIMC1
log2(RPKM)
z−score of IF value
−2 −1 0 1 2

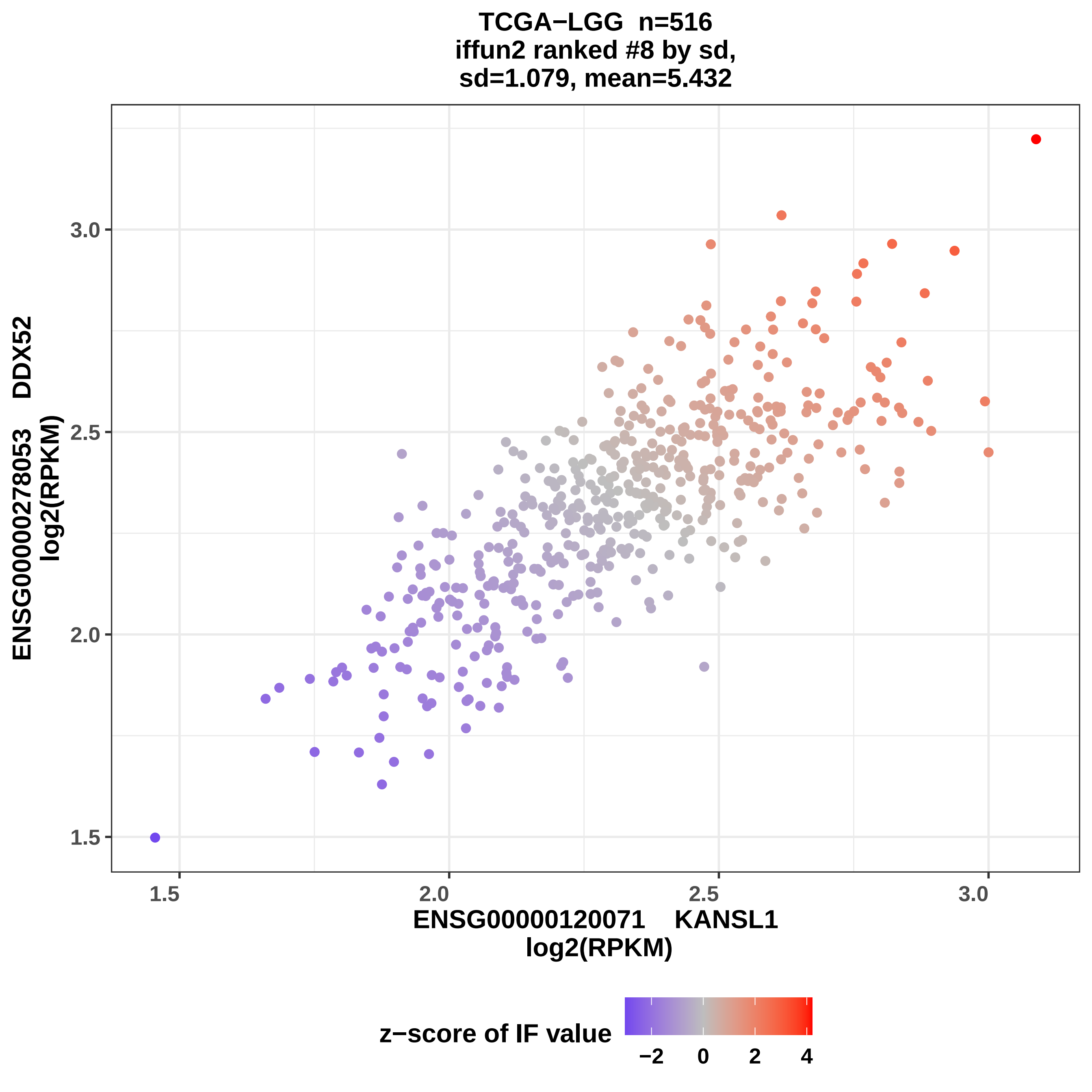
TCGA−LGG n=516
iffun2 ranked #8 by sd,
sd=1.079, mean=5.432
ENSG00000278053 DDX52
log2(RPKM)
ENSG00000120071 KANSL1
log2(RPKM)
1.5
2.0
2.5
3.0
z−score of IF value
−2
0
2
4

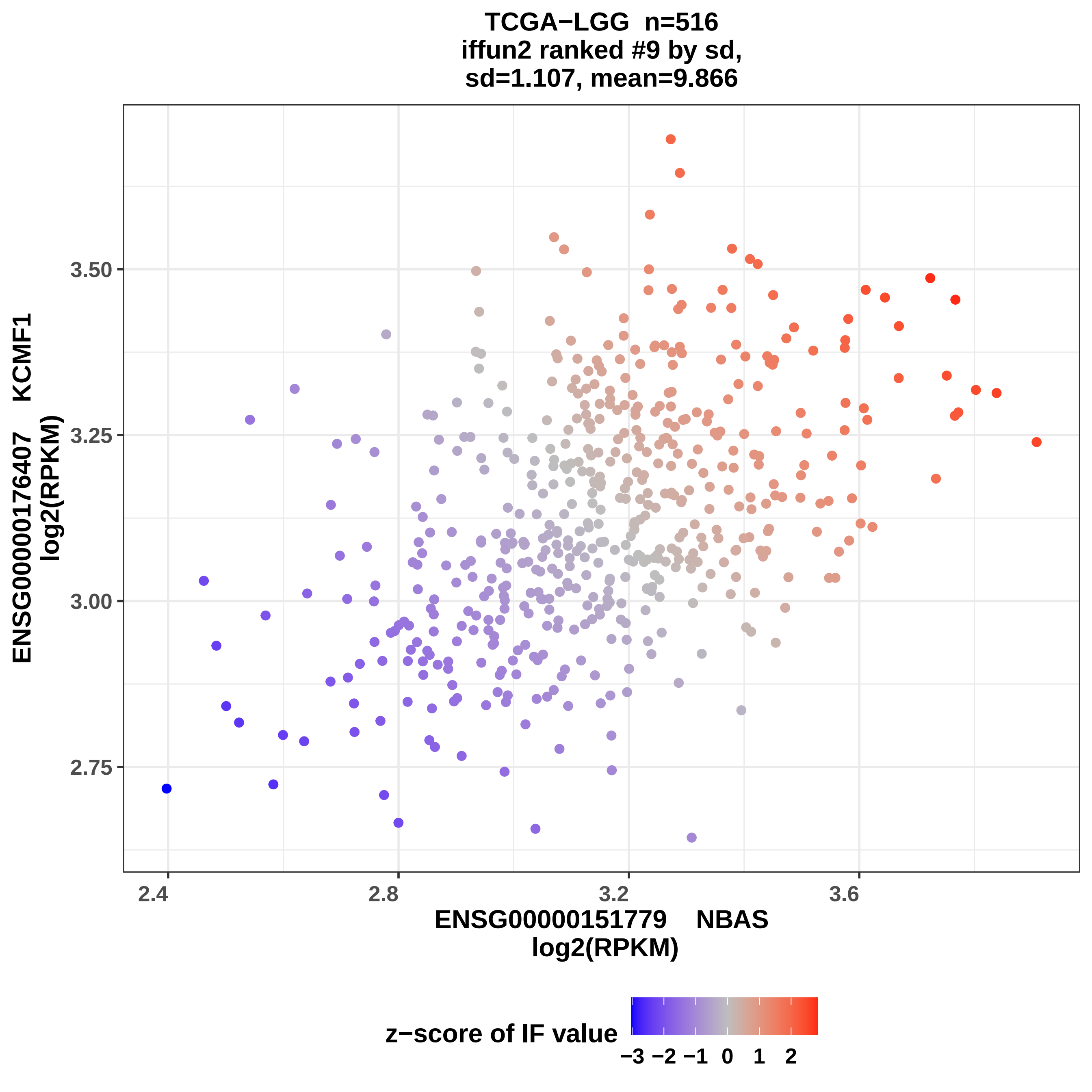
TCGA−LGG n=516
iffun2 ranked #9 by sd,
sd=1.107, mean=9.866
ENSG00000176407 KCMF1
log2(RPKM)
3.50
3.25
3.00
2.75
2.4
2.8
3.2
3.6
ENSG00000151779 NBAS
log2(RPKM)
z−score of IF value
−3 −2 −1 0 1 2

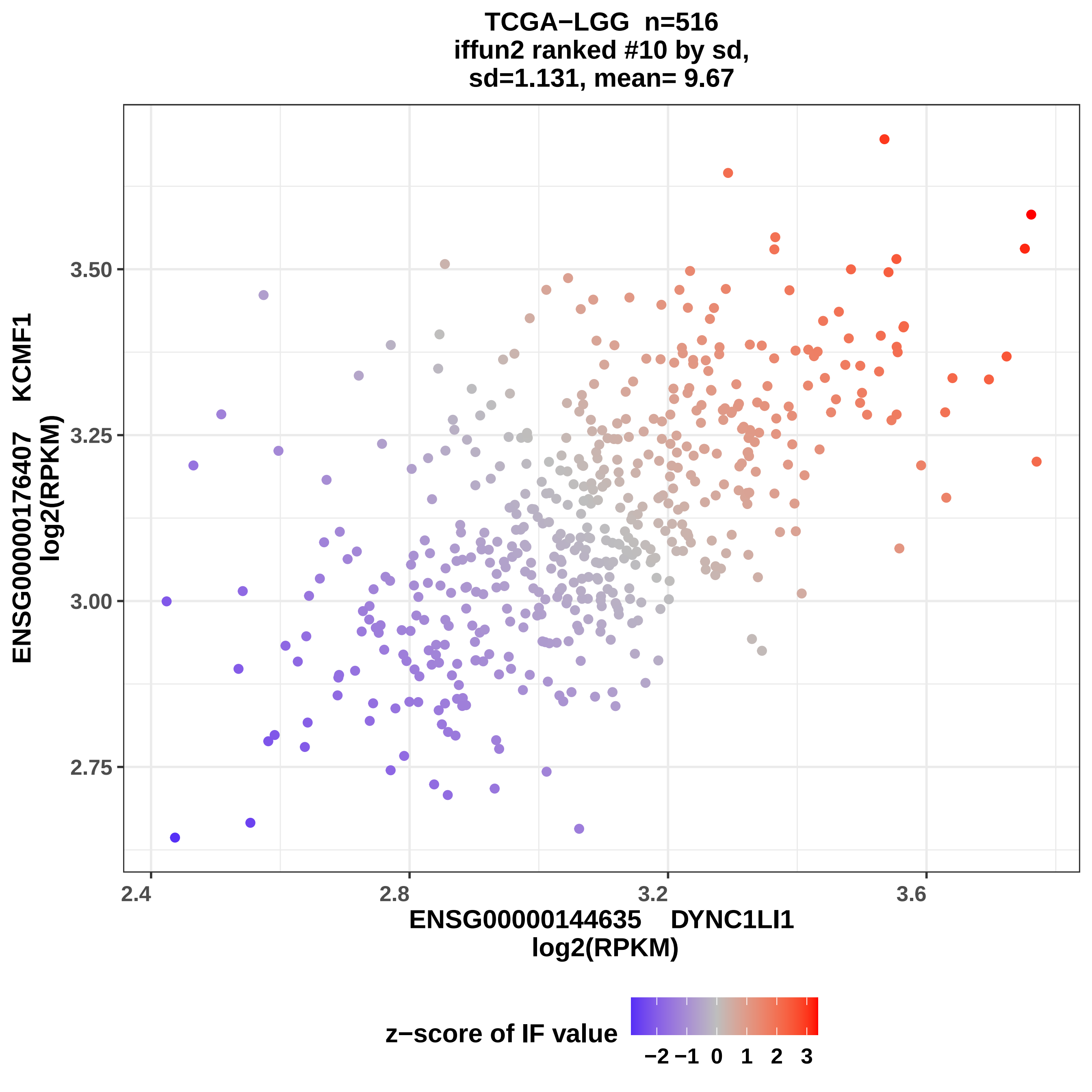

TCGA−LGG n=516
iffun2 ranked #10 by sd,
sd=1.131, mean= 9.67
ENSG00000176407 KCMF1
log2(RPKM)
3.50
3.25
3.00
2.75
2.4
2.8
3.2
3.6
ENSG00000144635 DYNC1LI1
log2(RPKM)
z−score of IF value
−2 −1 0 1 2 3

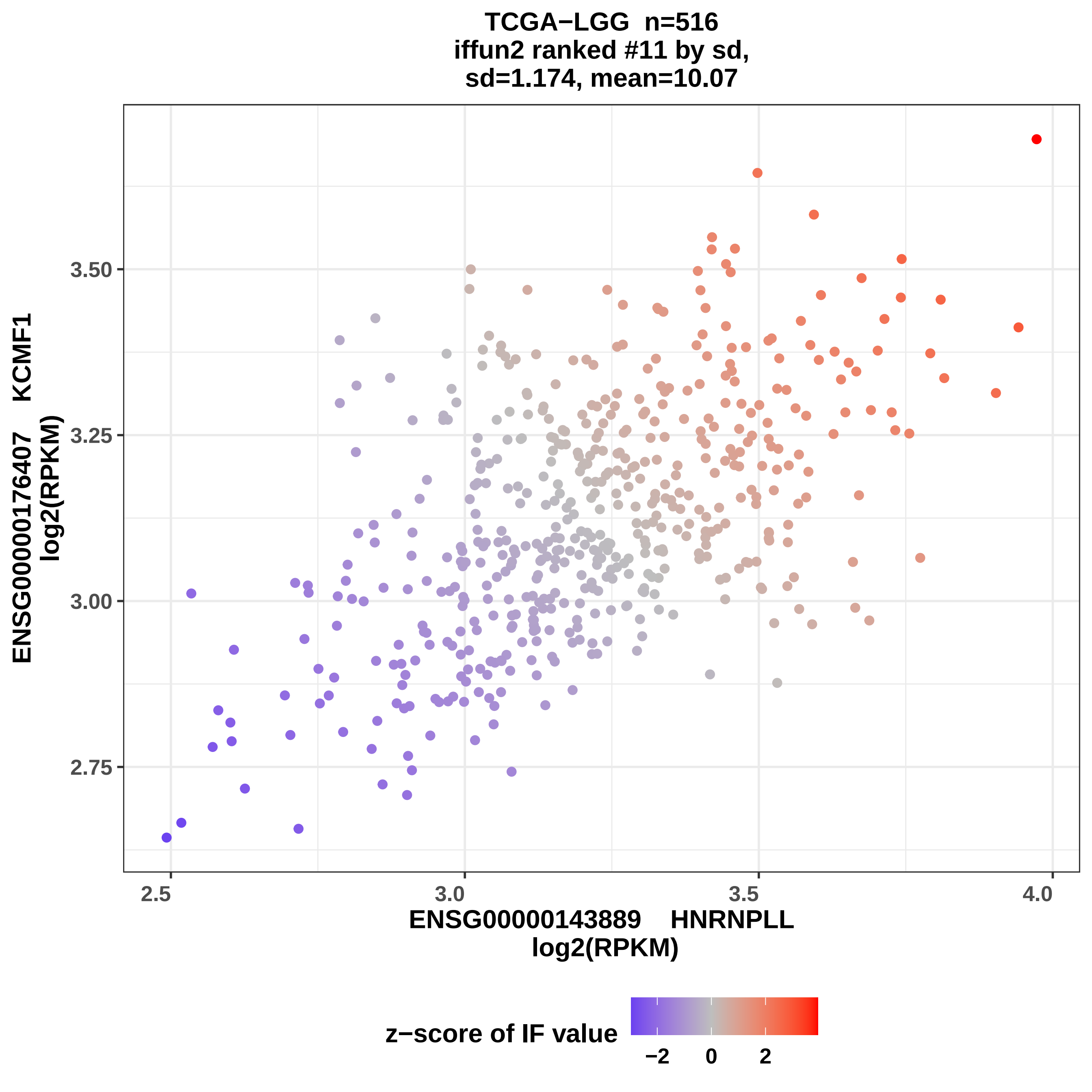

TCGA−LGG n=516
iffun2 ranked #11 by sd,
sd=1.174, mean=10.07
ENSG00000176407 KCMF1
log2(RPKM)
3.50
3.25
3.00
2.75
2.5
3.0
3.5
4.0
ENSG00000143889 HNRNPLL
log2(RPKM)
z−score of IF value
−2
0
2

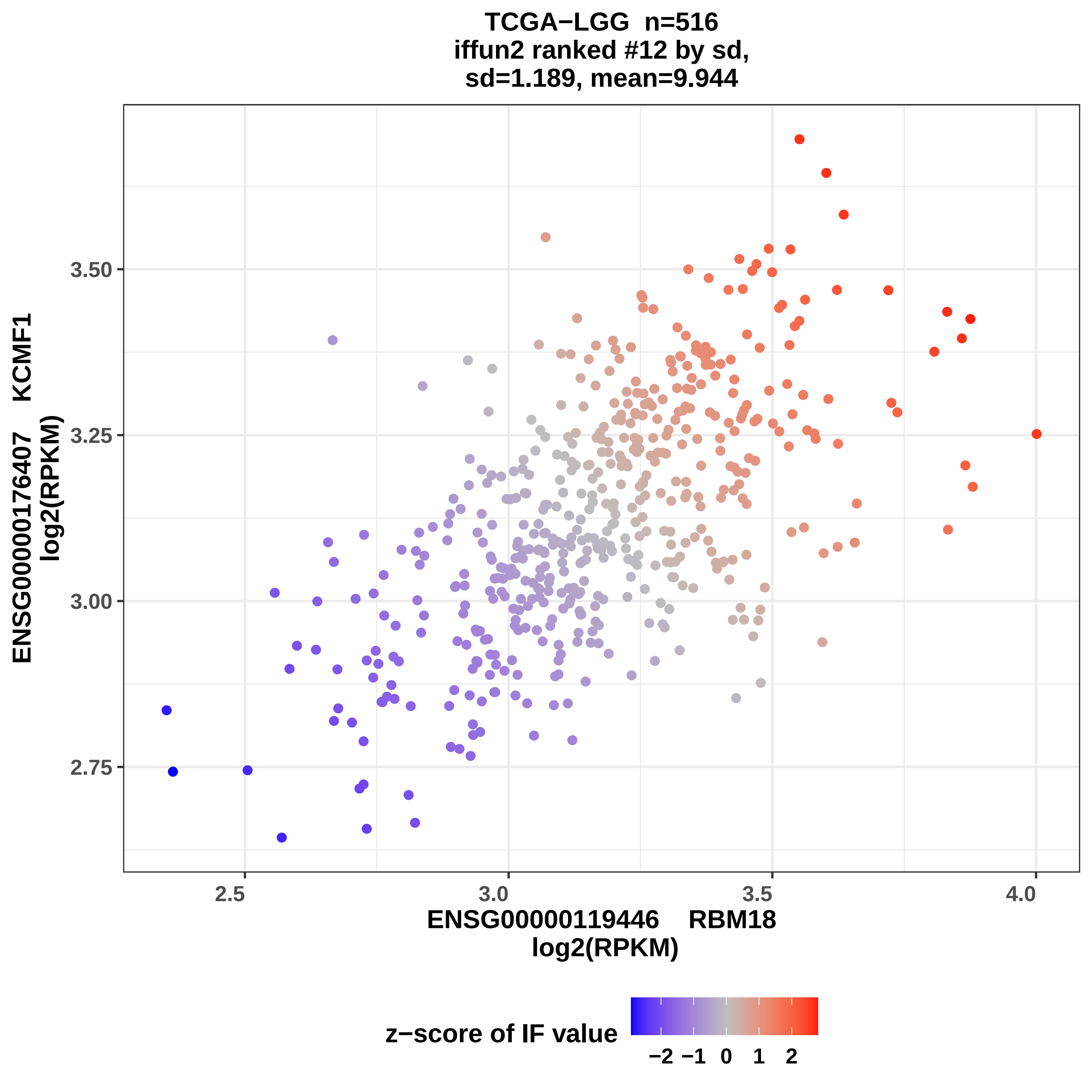

TCGA−LGG n=516
iffun2 ranked #12 by sd,
sd=1.189, mean=9.944
ENSG00000176407 KCMF1
log2(RPKM)
3.50
3.25
3.00
2.75
2.5
3.0
3.5
4.0
ENSG00000119446 RBM18
log2(RPKM)
z−score of IF value
−2
−1
0
1
2

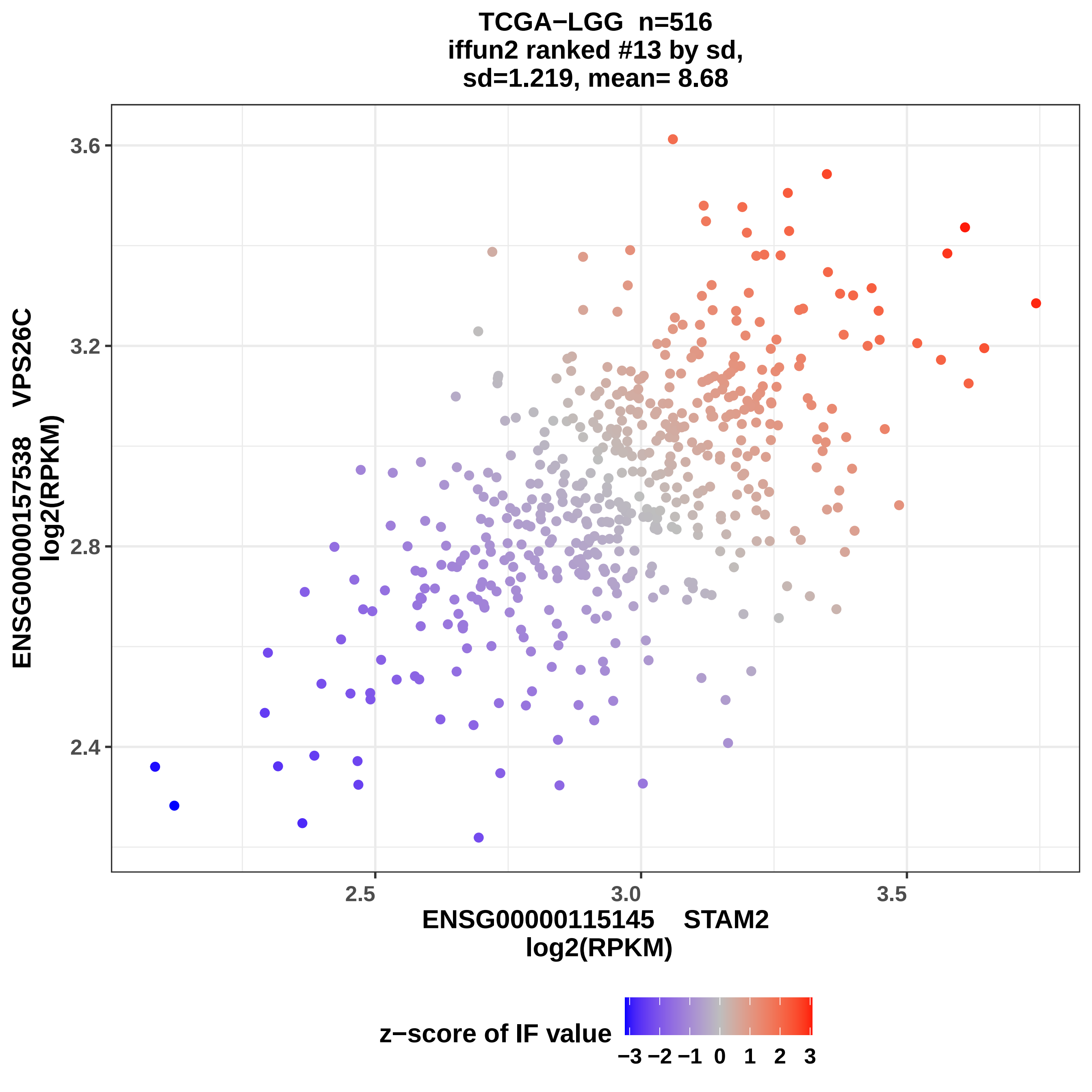

TCGA−LGG n=516
iffun2 ranked #13 by sd,
sd=1.219, mean= 8.68
3.6
3.2
2.8
2.4
2.5
3.0
3.5
ENSG00000115145 STAM2
log2(RPKM)
ENSG00000157538 VPS26C
log2(RPKM)
z−score of IF value
−3 −2 −1 0 1 2 3

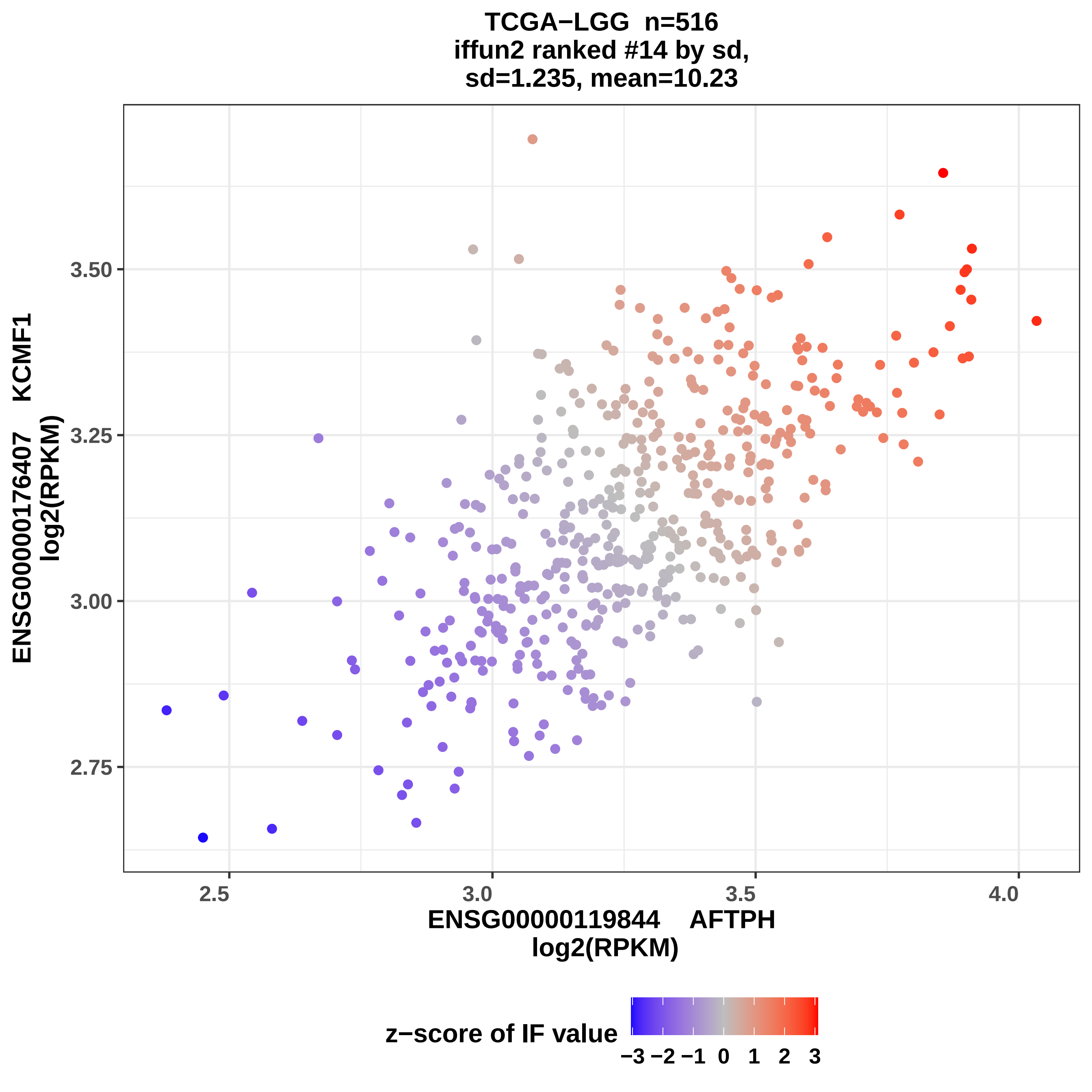
TCGA−LGG n=516
iffun2 ranked #14 by sd,
sd=1.235, mean=10.23
ENSG00000176407 KCMF1
log2(RPKM)
3.50
3.25
3.00
2.75
2.5
3.0
3.5
4.0
ENSG00000119844 AFTPH
log2(RPKM)
z−score of IF value
−3 −2 −1 0 1 2 3

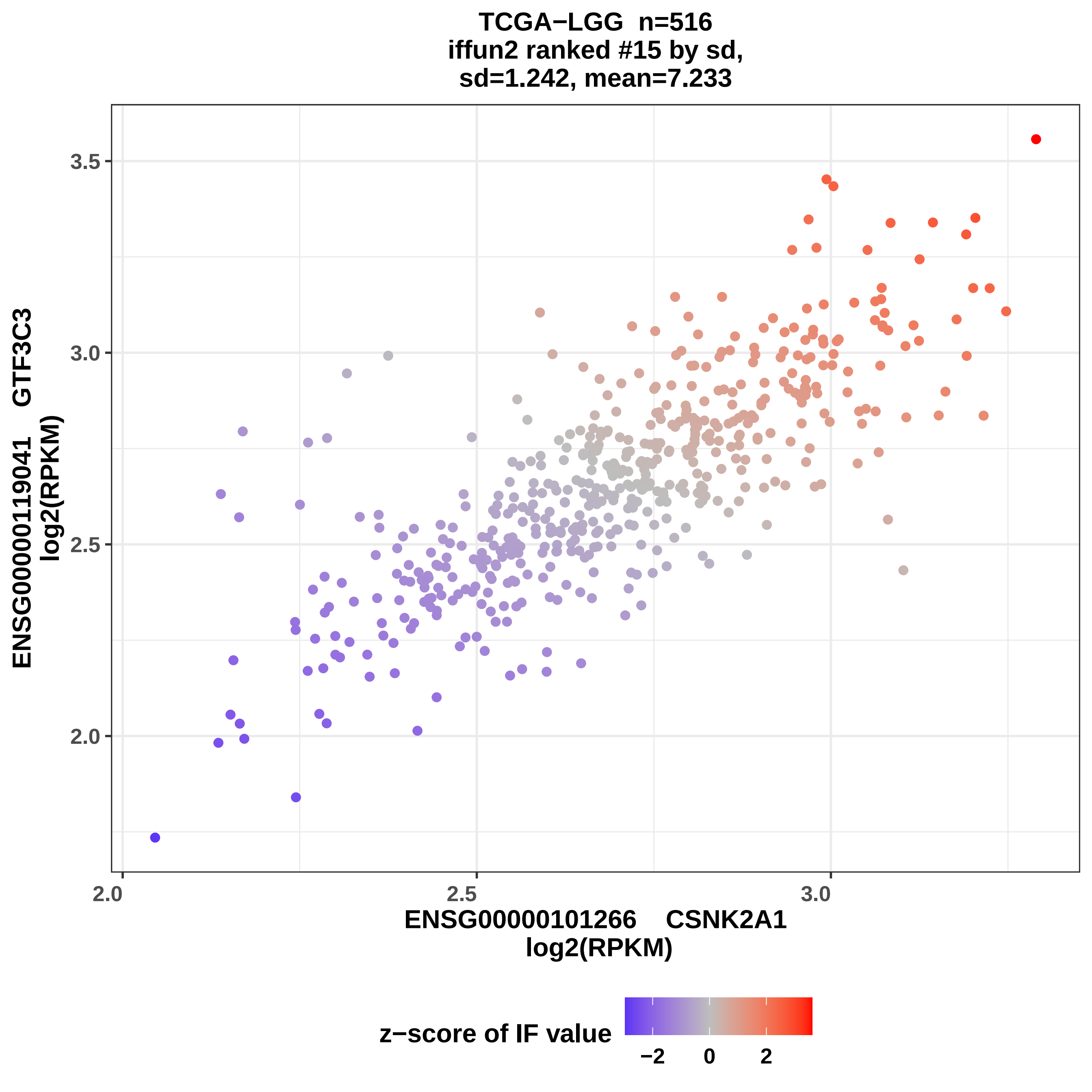
TCGA−LGG n=516
iffun2 ranked #15 by sd,
sd=1.242, mean=7.233
ENSG00000119041 GTF3C3
log2(RPKM)
3.5
3.0
2.5
2.0
2.0
2.5
3.0
ENSG00000101266 CSNK2A1
log2(RPKM)
z−score of IF value
−2
0
2

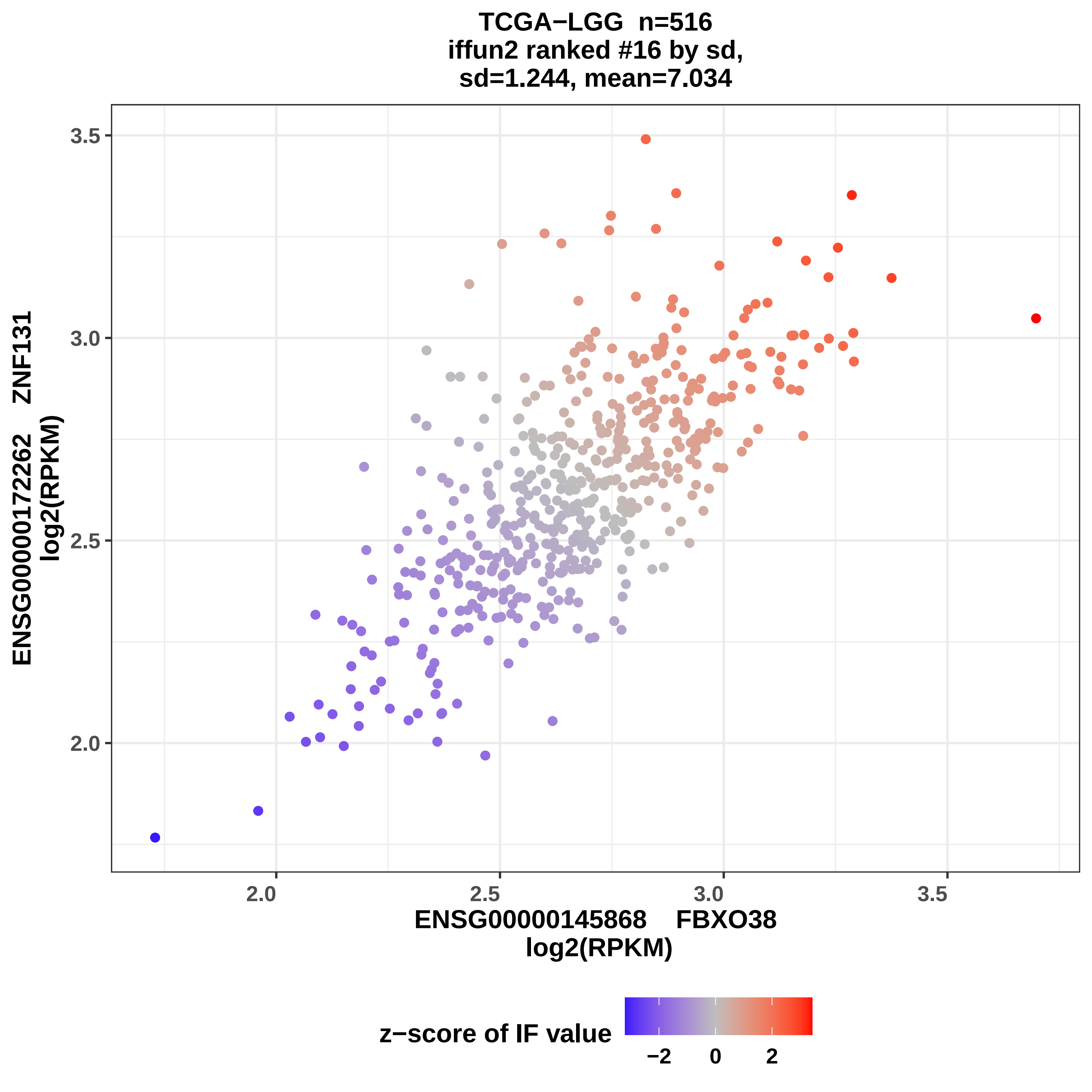
TCGA−LGG n=516
iffun2 ranked #16 by sd,
sd=1.244, mean=7.034
ENSG00000172262 ZNF131
log2(RPKM)
3.5
3.0
2.5
2.0
2.0
2.5
3.0
3.5
ENSG00000145868 FBXO38
log2(RPKM)
z−score of IF value
−2
0
2

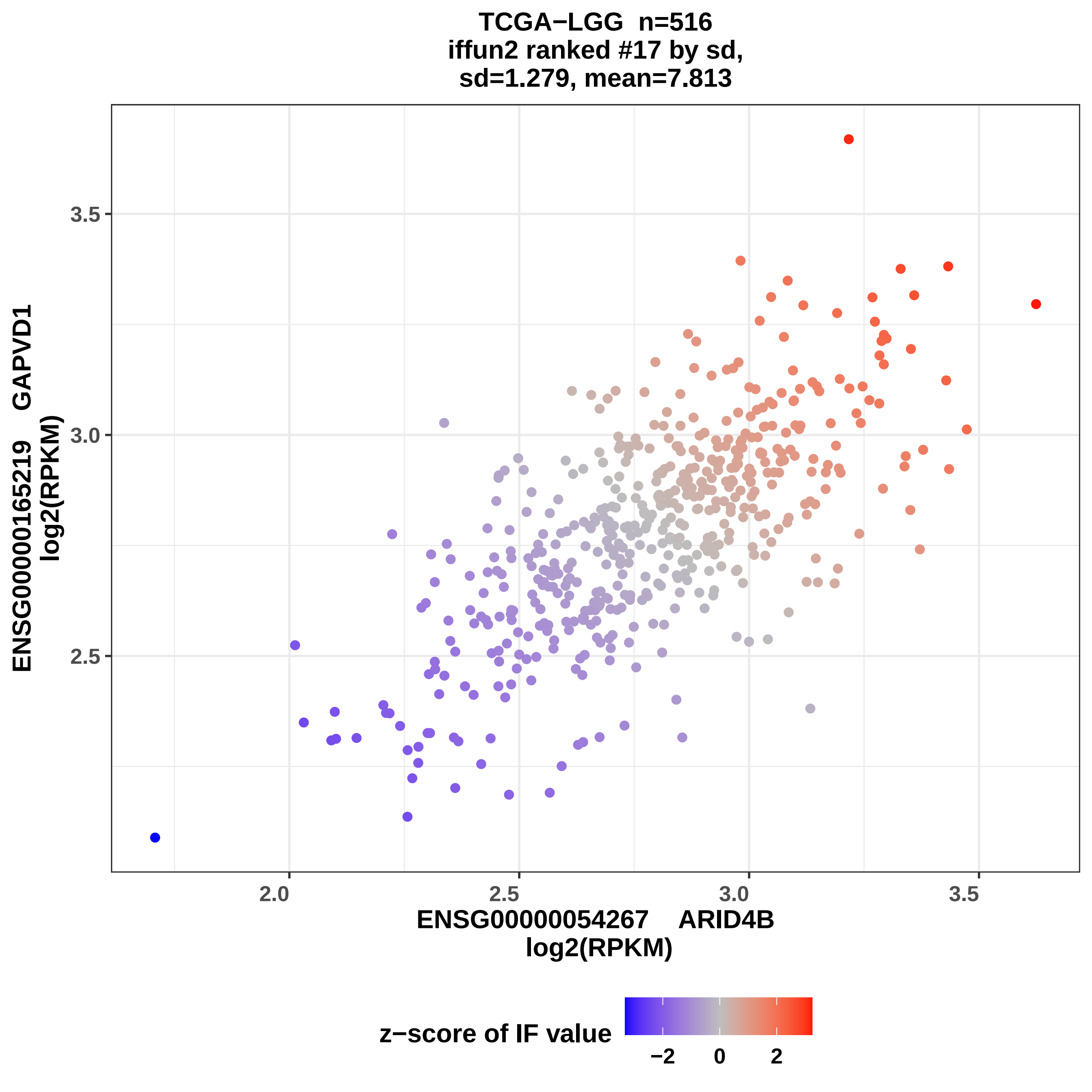
TCGA−LGG n=516
iffun2 ranked #17 by sd,
sd=1.279, mean=7.813
ENSG00000165219 GAPVD1
log2(RPKM)
3.5
3.0
2.5
2.0
2.5
3.0
3.5
ENSG00000054267 ARID4B
log2(RPKM)
z−score of IF value
−2
0
2

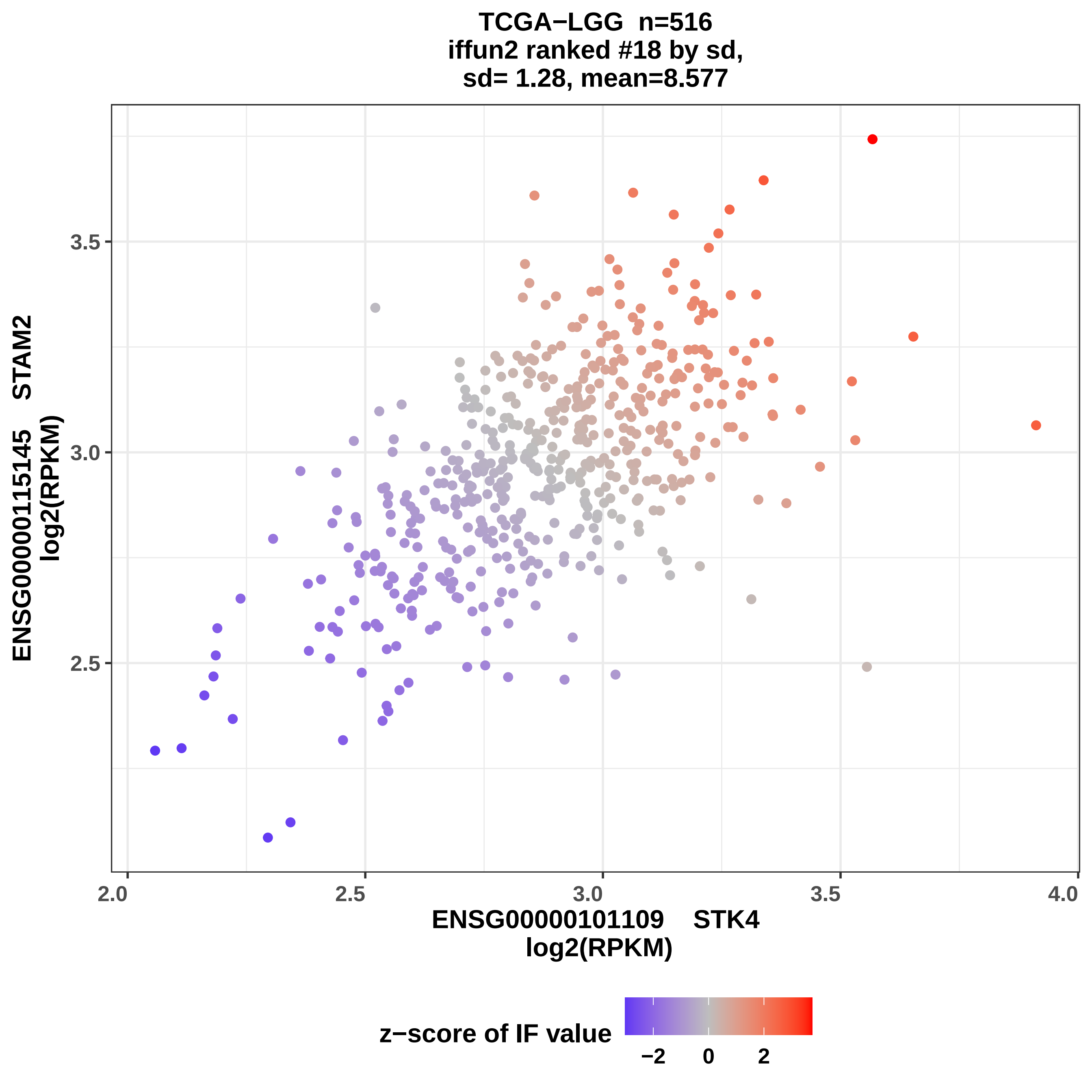

TCGA−LGG n=516
iffun2 ranked #18 by sd,
sd= 1.28, mean=8.577
ENSG00000115145 STAM2
log2(RPKM)
3.5
3.0
2.5
2.0
2.5
3.0
3.5
4.0
ENSG00000101109 STK4
log2(RPKM)
z−score of IF value
−2
0
2

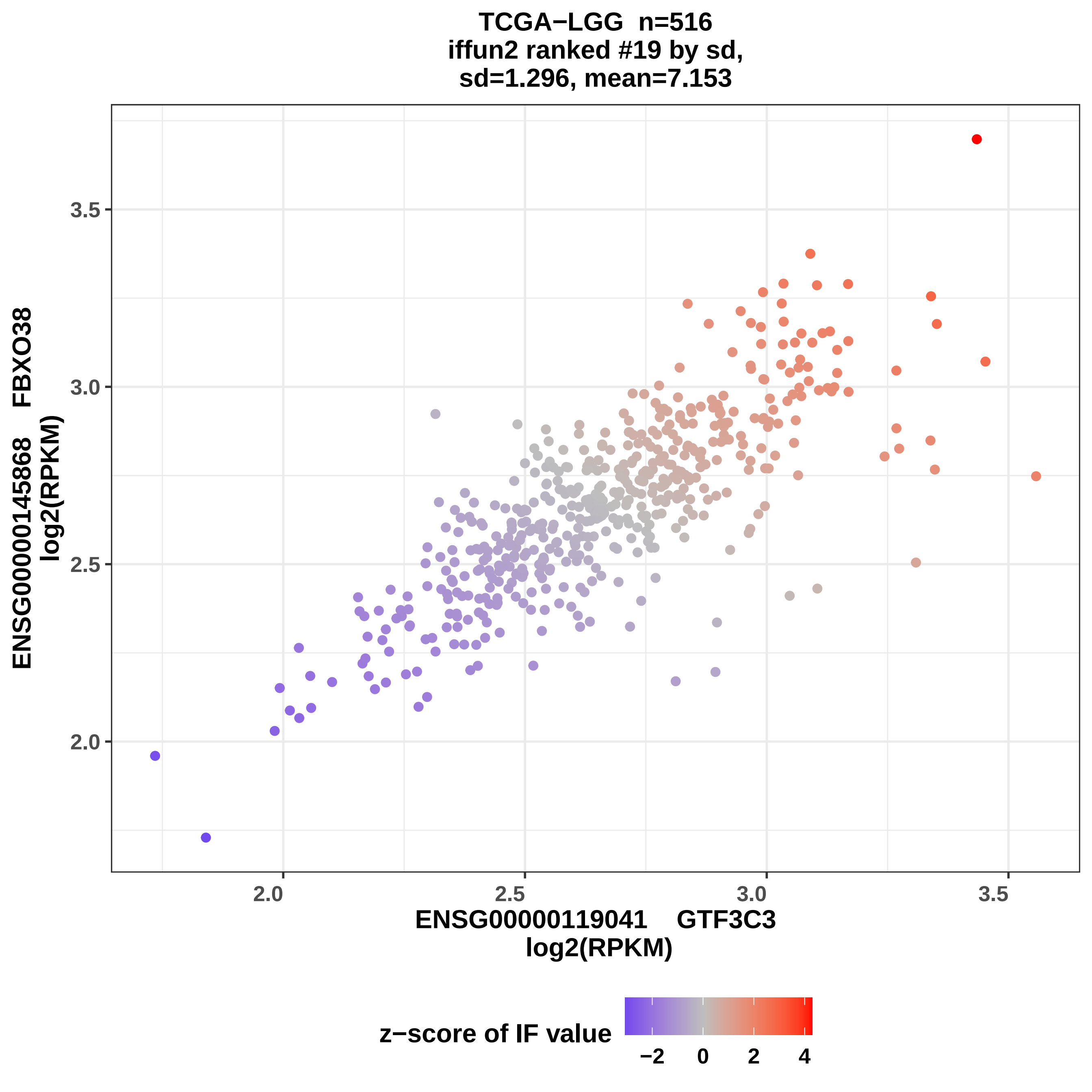

TCGA−LGG n=516
iffun2 ranked #19 by sd,
sd=1.296, mean=7.153
ENSG00000145868 FBXO38
log2(RPKM)
3.5
3.0
2.5
2.0
2.0
2.5
3.0
3.5
ENSG00000119041 GTF3C3
log2(RPKM)
z−score of IF value
−2
0
2
4

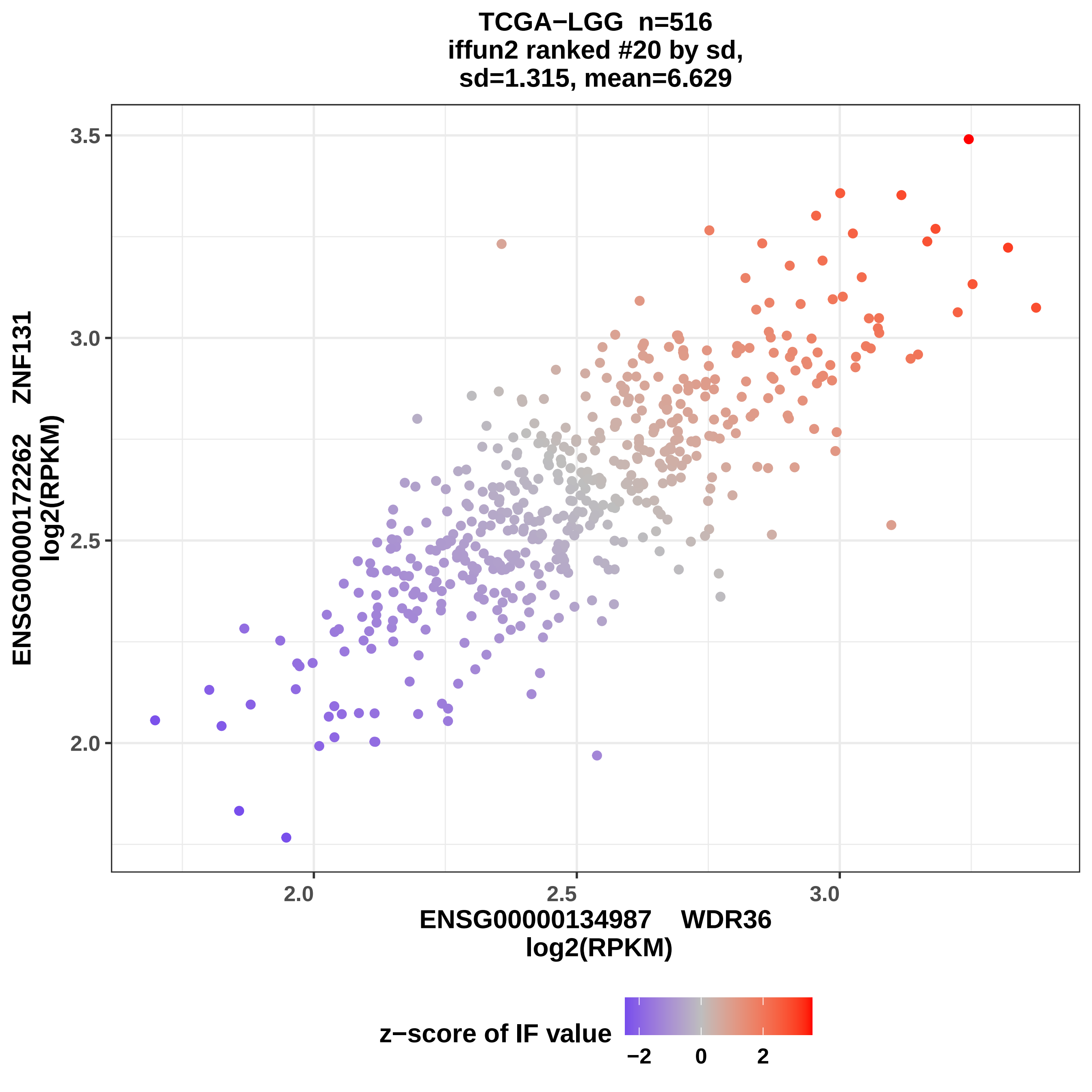

TCGA−LGG n=516
iffun2 ranked #20 by sd,
sd=1.315, mean=6.629
3.5
3.0
2.5
2.0
ENSG00000172262 ZNF131
log2(RPKM)
2.0
2.5
3.0
ENSG00000134987 WDR36
log2(RPKM)
z−score of IF value
−2
0
2

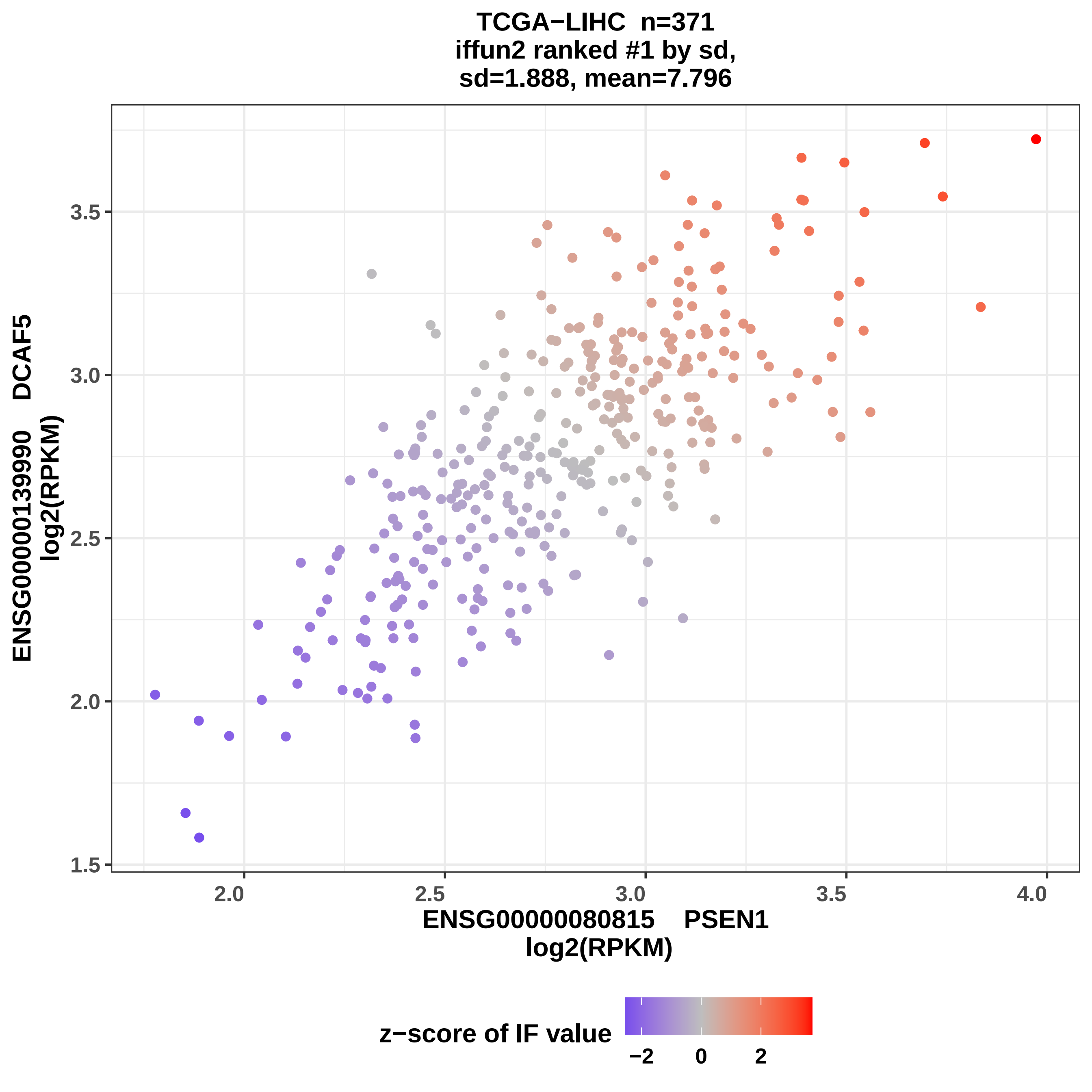

TCGA−LIHC n=371
iffun2 ranked #1 by sd,
sd=1.888, mean=7.796
ENSG00000139990 DCAF5
log2(RPKM)
3.5
3.0
2.5
2.0
1.5
2.0
2.5
3.0
3.5
4.0
ENSG00000080815 PSEN1
log2(RPKM)
z−score of IF value
−2
0
2

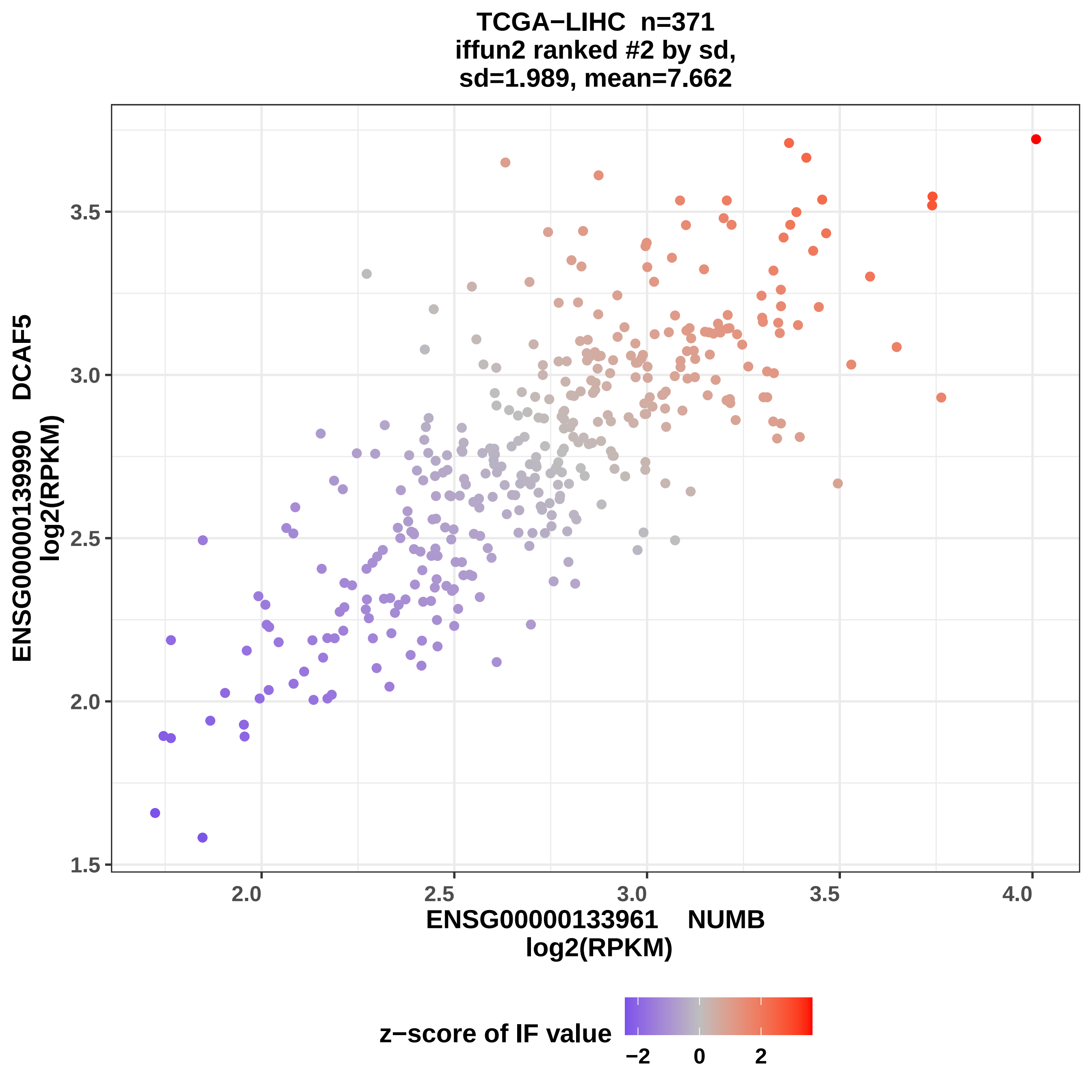

TCGA−LIHC n=371
iffun2 ranked #2 by sd,
sd=1.989, mean=7.662
ENSG00000139990 DCAF5
log2(RPKM)
3.5
3.0
2.5
2.0
1.5
2.0
2.5
3.0
3.5
4.0
ENSG00000133961 NUMB
log2(RPKM)
z−score of IF value
−2
0
2

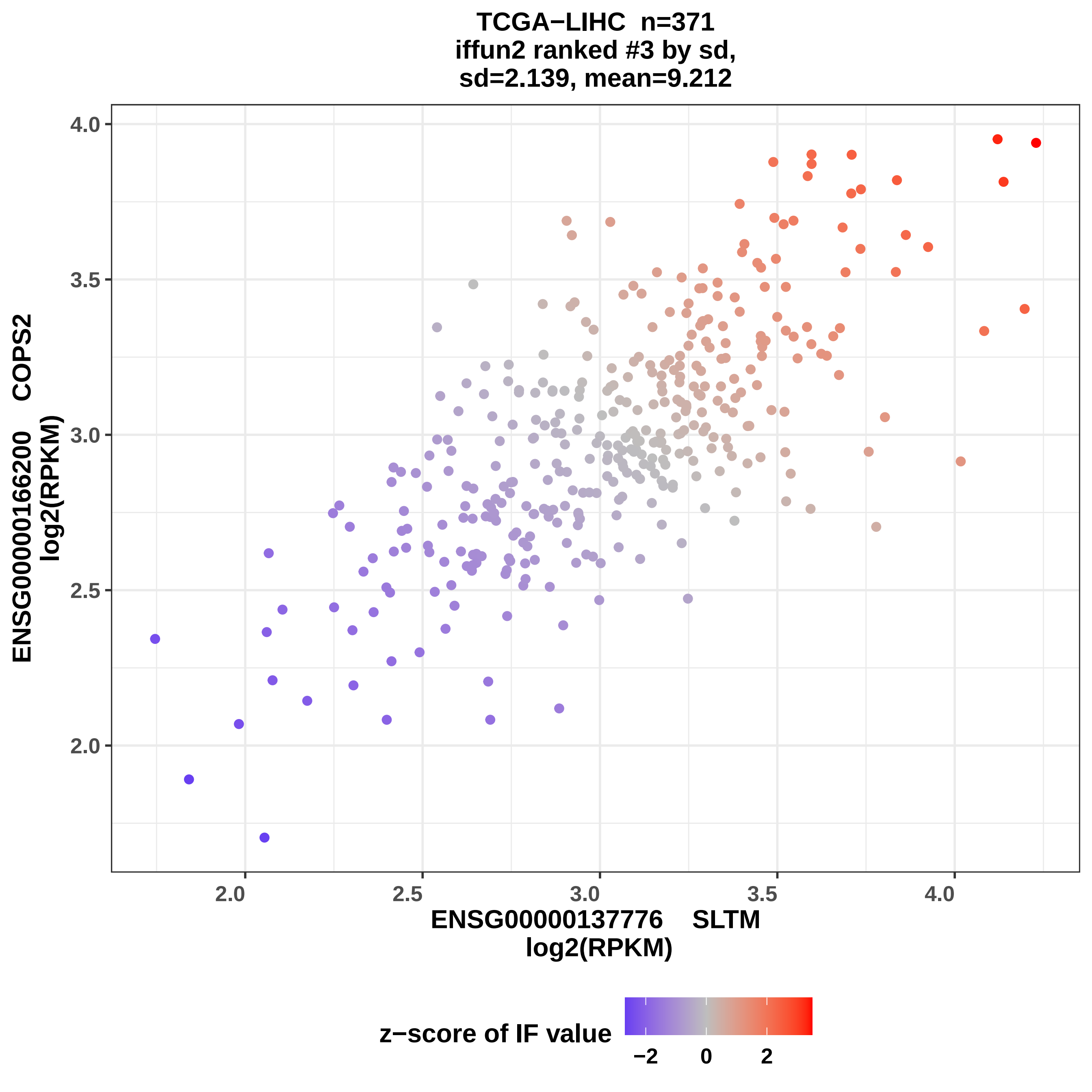
TCGA−LIHC n=371
iffun2 ranked #3 by sd,
sd=2.139, mean=9.212
ENSG00000166200 COPS2
log2(RPKM)
4.0
3.5
3.0
2.5
2.0
2.0
2.5
3.0
3.5
4.0
ENSG00000137776 SLTM
log2(RPKM)
z−score of IF value
−2
0
2

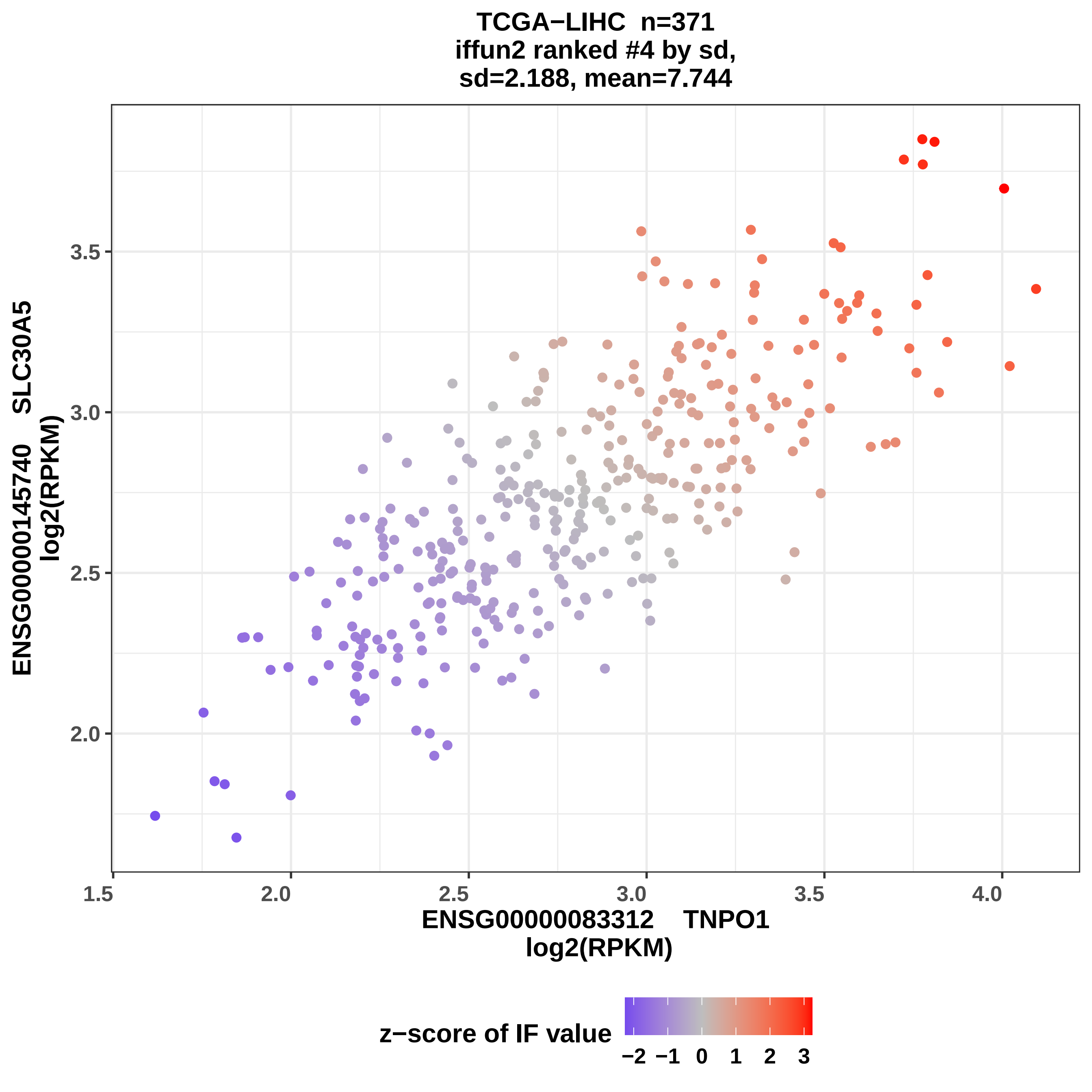

TCGA−LIHC n=371
iffun2 ranked #4 by sd,
sd=2.188, mean=7.744
ENSG00000145740 SLC30A5
log2(RPKM)
3.5
3.0
2.5
2.0
1.5
2.0
2.5
3.0
3.5
4.0
ENSG00000083312 TNPO1
log2(RPKM)
z−score of IF value
−2
−1
0
1
2
3

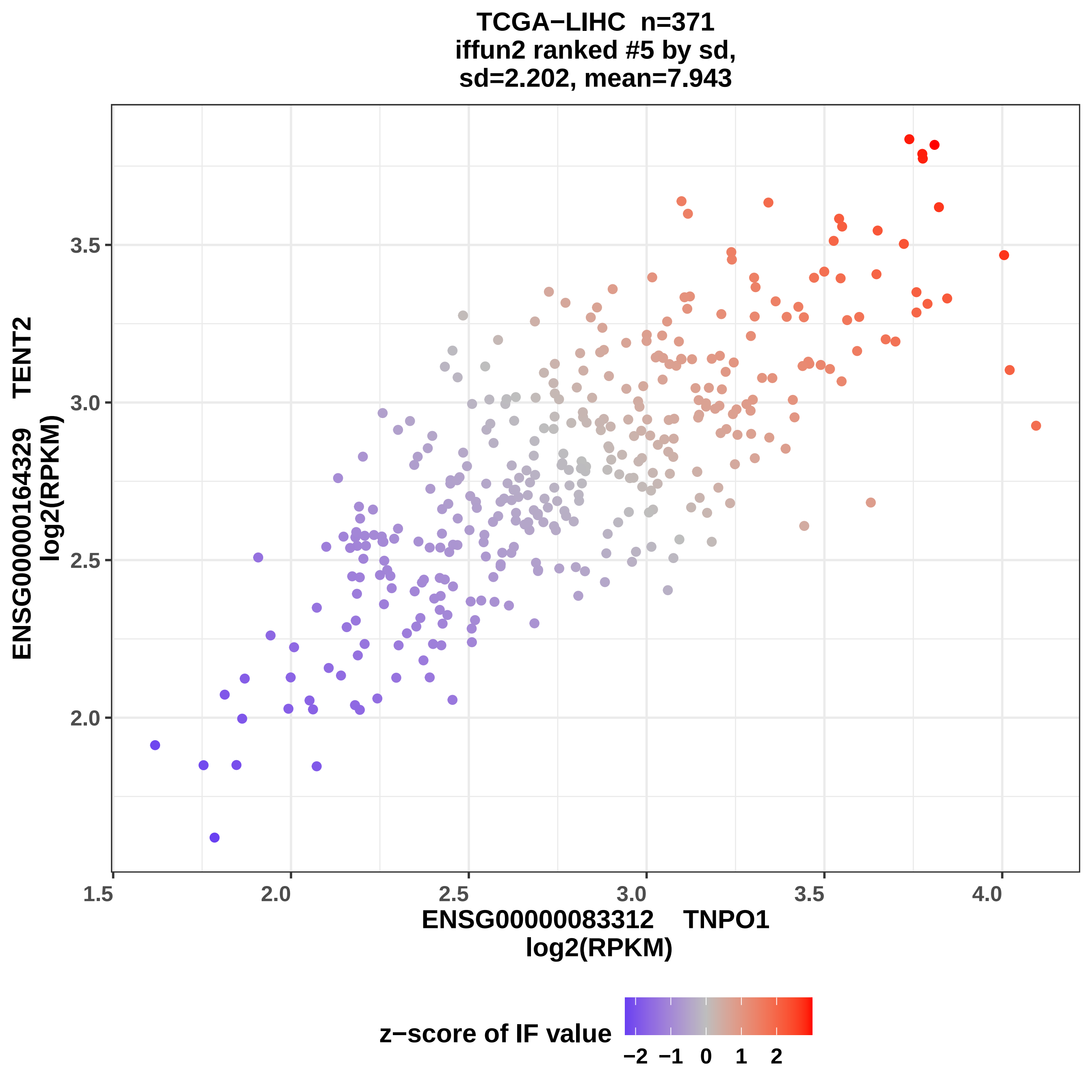

TCGA−LIHC n=371
iffun2 ranked #5 by sd,
sd=2.202, mean=7.943
ENSG00000164329 TENT2
log2(RPKM)
ENSG00000083312 TNPO1
log2(RPKM)
2.0
2.5
3.0
3.5
1.5
2.0
2.5
3.0
3.5
4.0
z−score of IF value
−2 −1 0 1 2

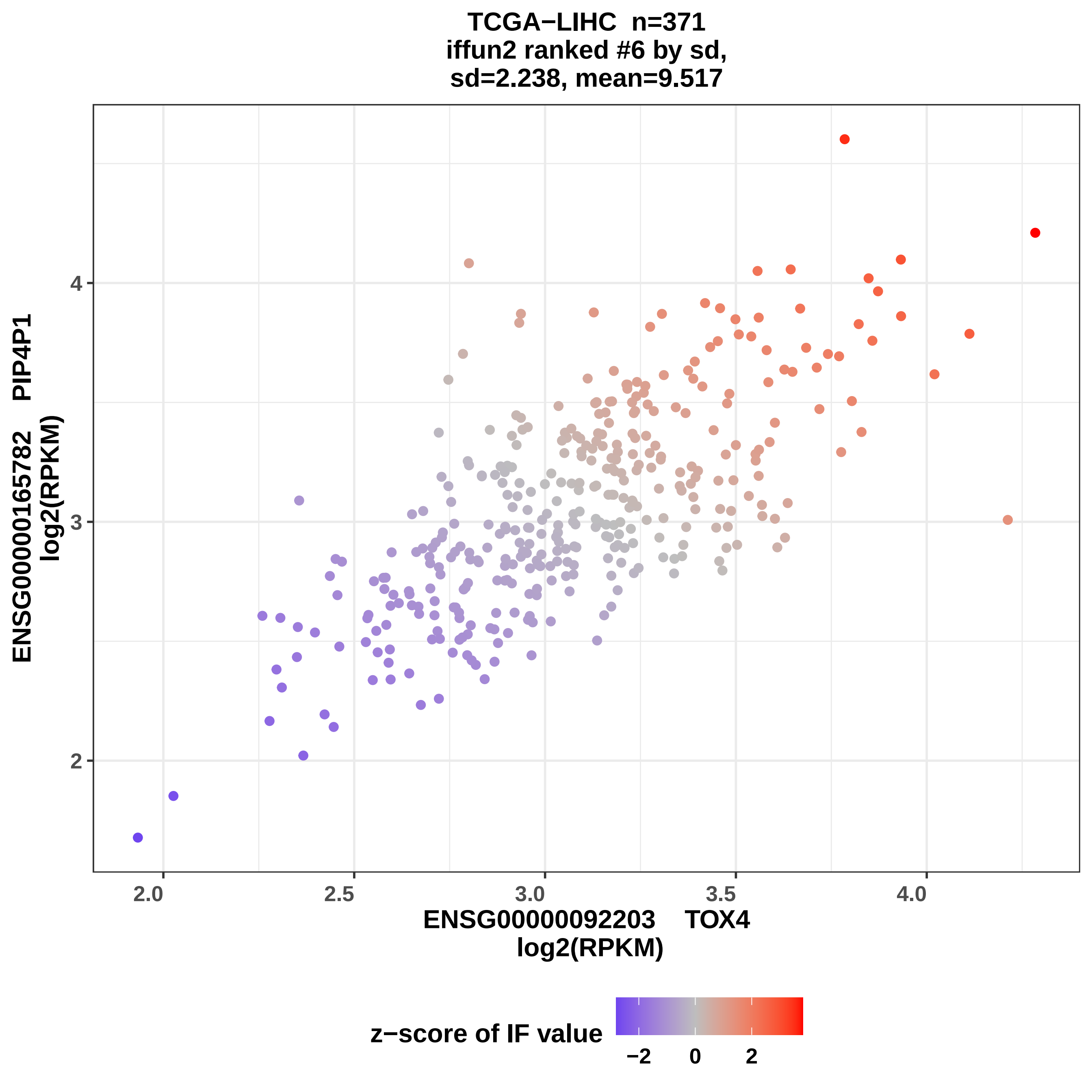
TCGA−LIHC n=371
iffun2 ranked #6 by sd,
sd=2.238, mean=9.517
ENSG00000165782 PIP4P1
log2(RPKM)
ENSG00000092203 TOX4
log2(RPKM)
z−score of IF value
2.0
2.5
3.0
3.5
4.0
2
3
4
−2
0
2

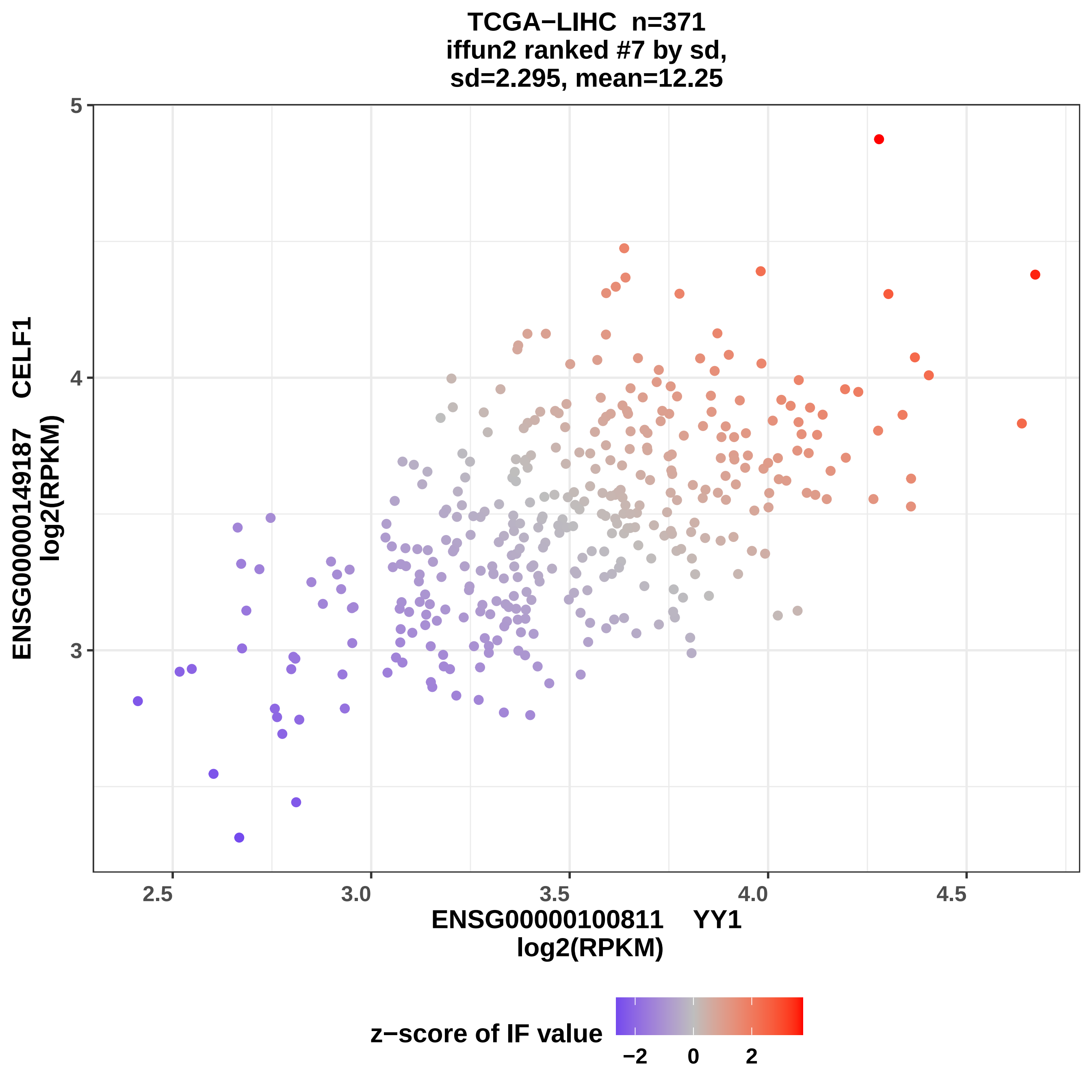
TCGA−LIHC n=371
iffun2 ranked #7 by sd,
sd=2.295, mean=12.25
ENSG00000149187 CELF1
log2(RPKM)
5
4
3
2.5
3.0
3.5
4.0
4.5
ENSG00000100811 YY1
log2(RPKM)
z−score of IF value
−2
0
2

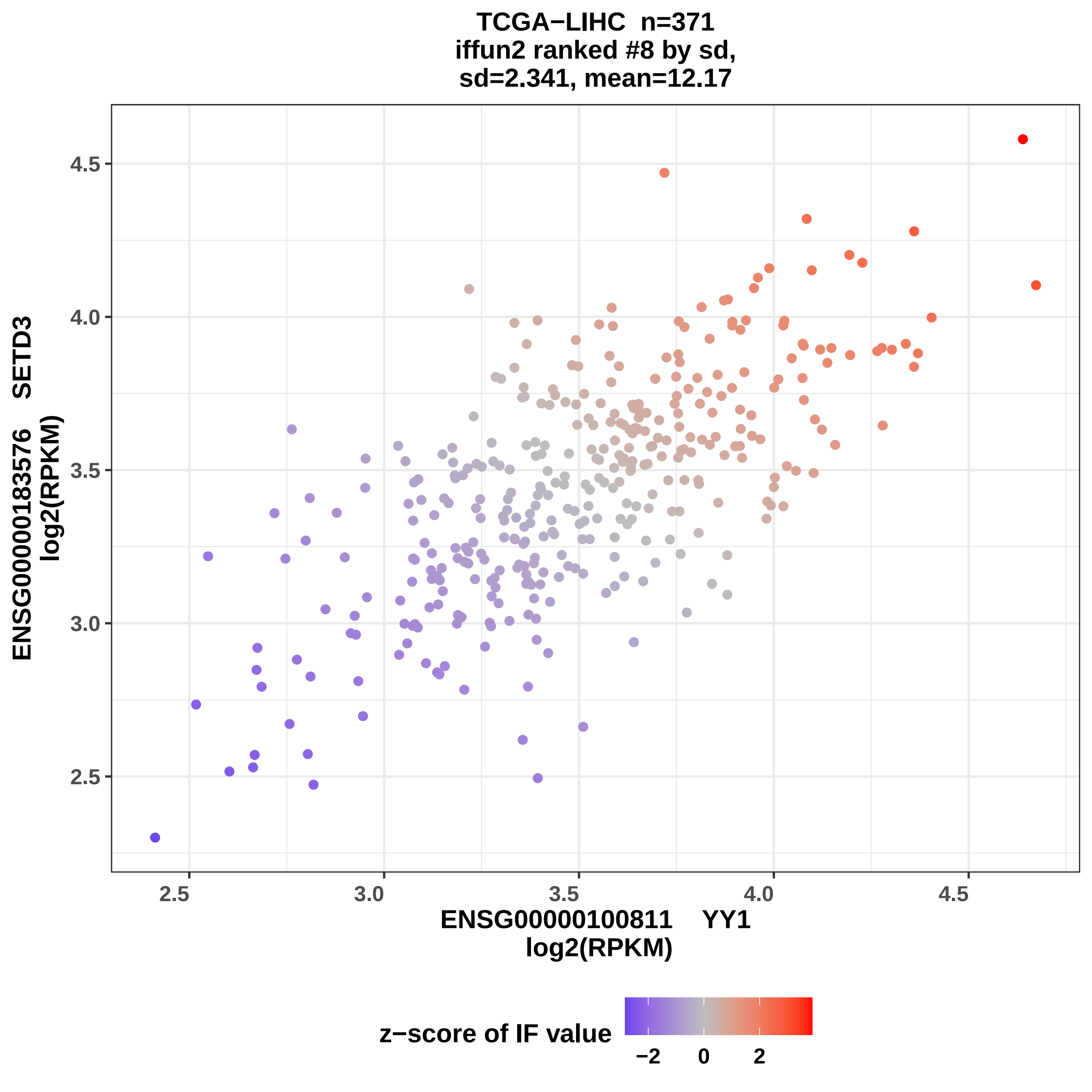

TCGA−LIHC n=371
iffun2 ranked #8 by sd,
sd=2.341, mean=12.17
ENSG00000183576 SETD3
log2(RPKM)
4.5
4.0
3.5
3.0
2.5
2.5
3.0
3.5
4.0
4.5
ENSG00000100811 YY1
log2(RPKM)
z−score of IF value
−2
0
2

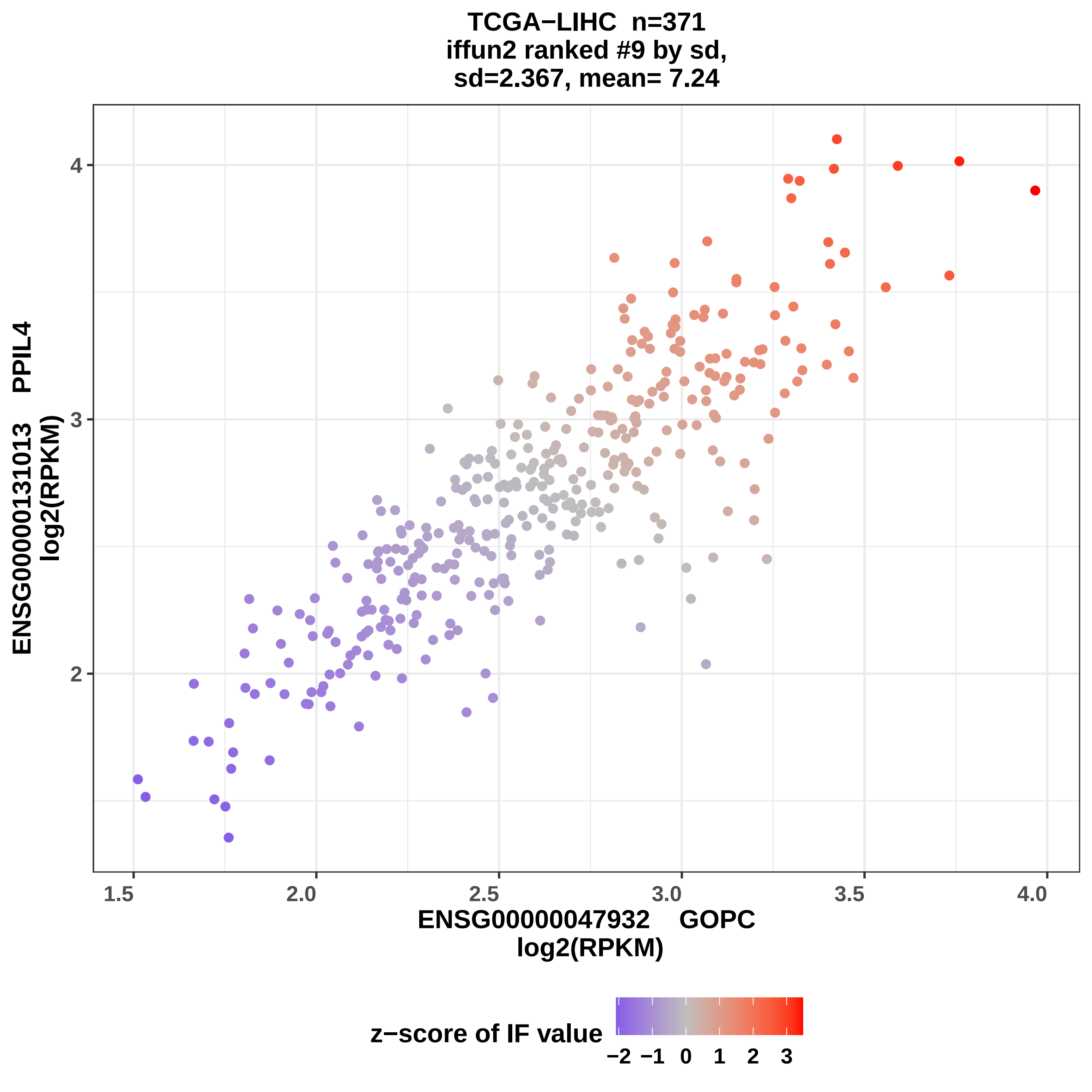

TCGA−LIHC n=371
iffun2 ranked #9 by sd,
sd=2.367, mean= 7.24
ENSG00000131013 PPIL4
log2(RPKM)
ENSG00000047932 GOPC
log2(RPKM)
z−score of IF value
−2 −1 0 1 2 3
1.5
2.0
2.5
3.0
3.5
4.0
2
3
4

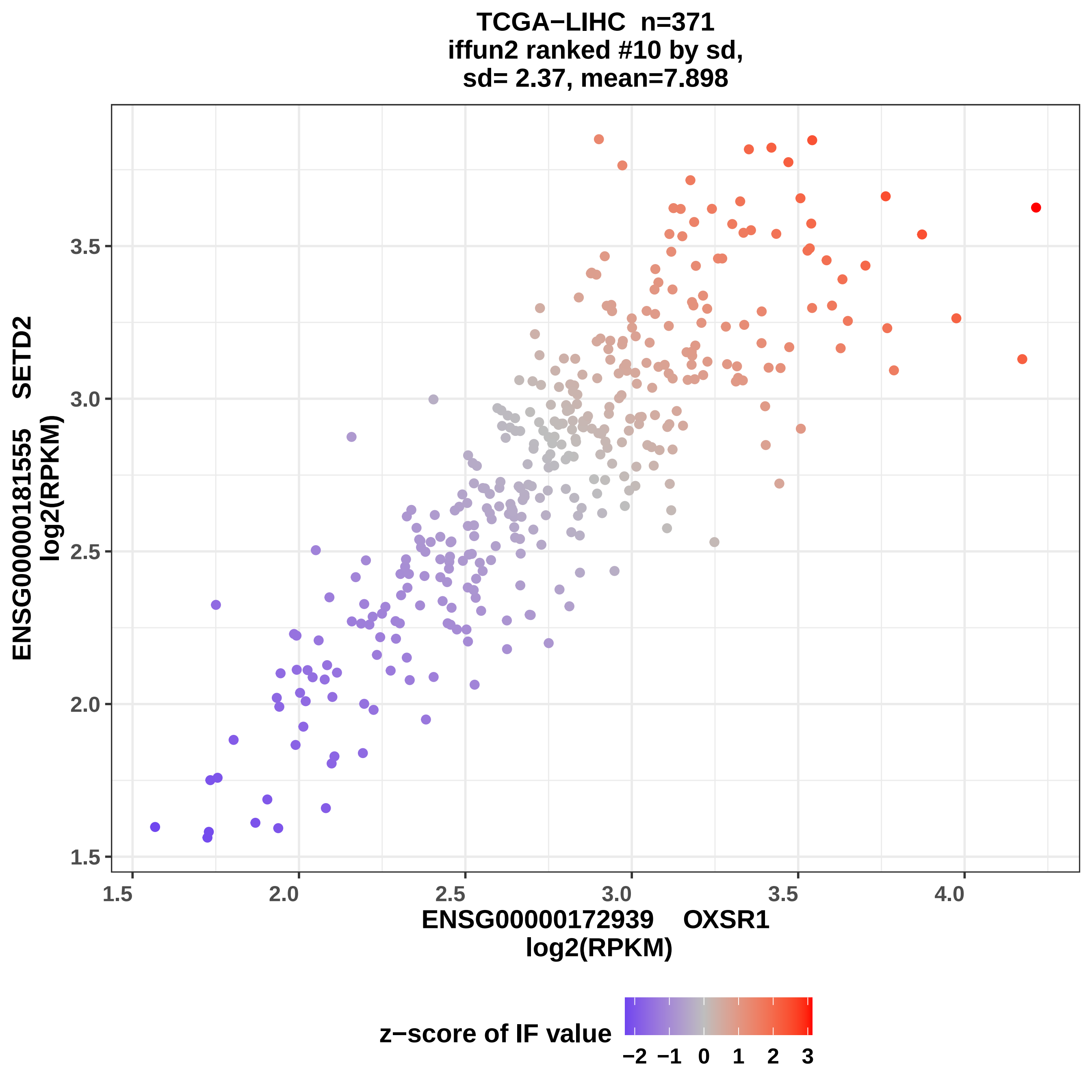
TCGA−LIHC n=371
iffun2 ranked #10 by sd,
sd= 2.37, mean=7.898
ENSG00000181555 SETD2
log2(RPKM)
3.5
3.0
2.5
2.0
1.5
1.5
2.0
2.5
3.0
3.5
4.0
ENSG00000172939 OXSR1
log2(RPKM)
z−score of IF value
−2 −1 0 1 2 3

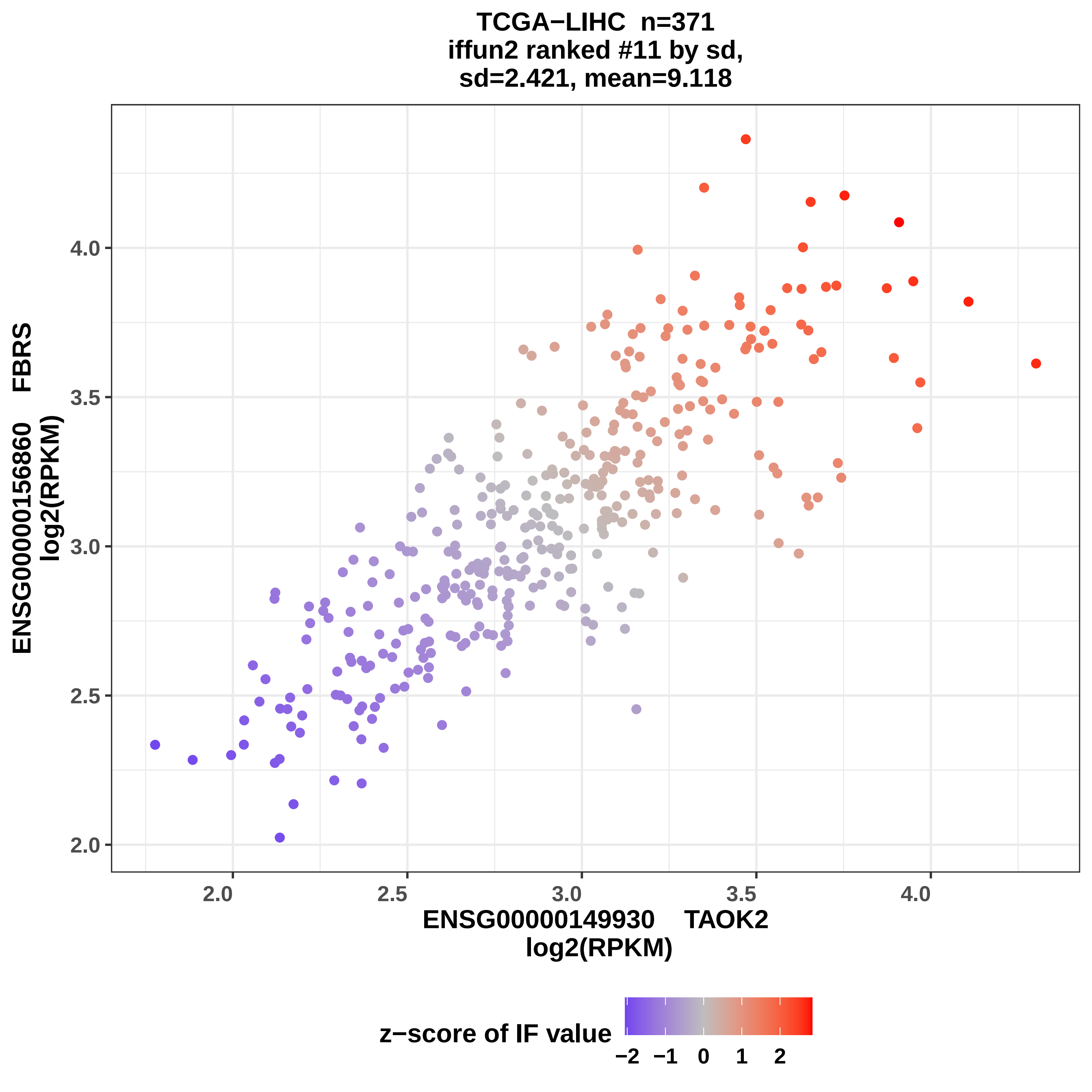
TCGA−LIHC n=371
iffun2 ranked #11 by sd,
sd=2.421, mean=9.118
ENSG00000156860 FBRS
log2(RPKM)
2.0
2.5
3.0
3.5
4.0
ENSG00000149930 TAOK2
log2(RPKM)
z−score of IF value
−2 −1 0 1 2

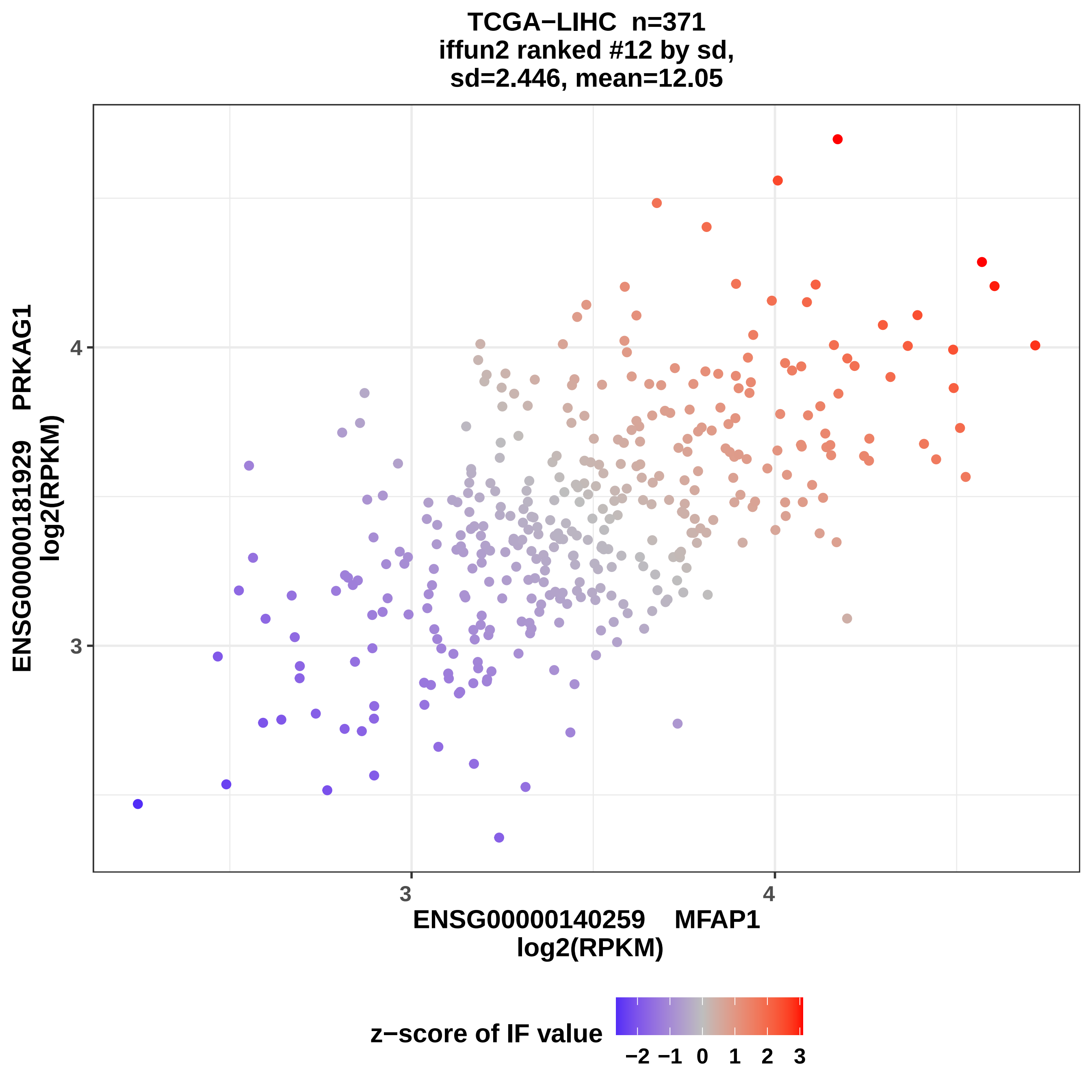
TCGA−LIHC n=371
iffun2 ranked #12 by sd,
sd=2.446, mean=12.05
ENSG00000181929 PRKAG1
log2(RPKM)
4
3
ENSG00000140259 MFAP1
log2(RPKM)
z−score of IF value
−2 −1 0 1 2 3

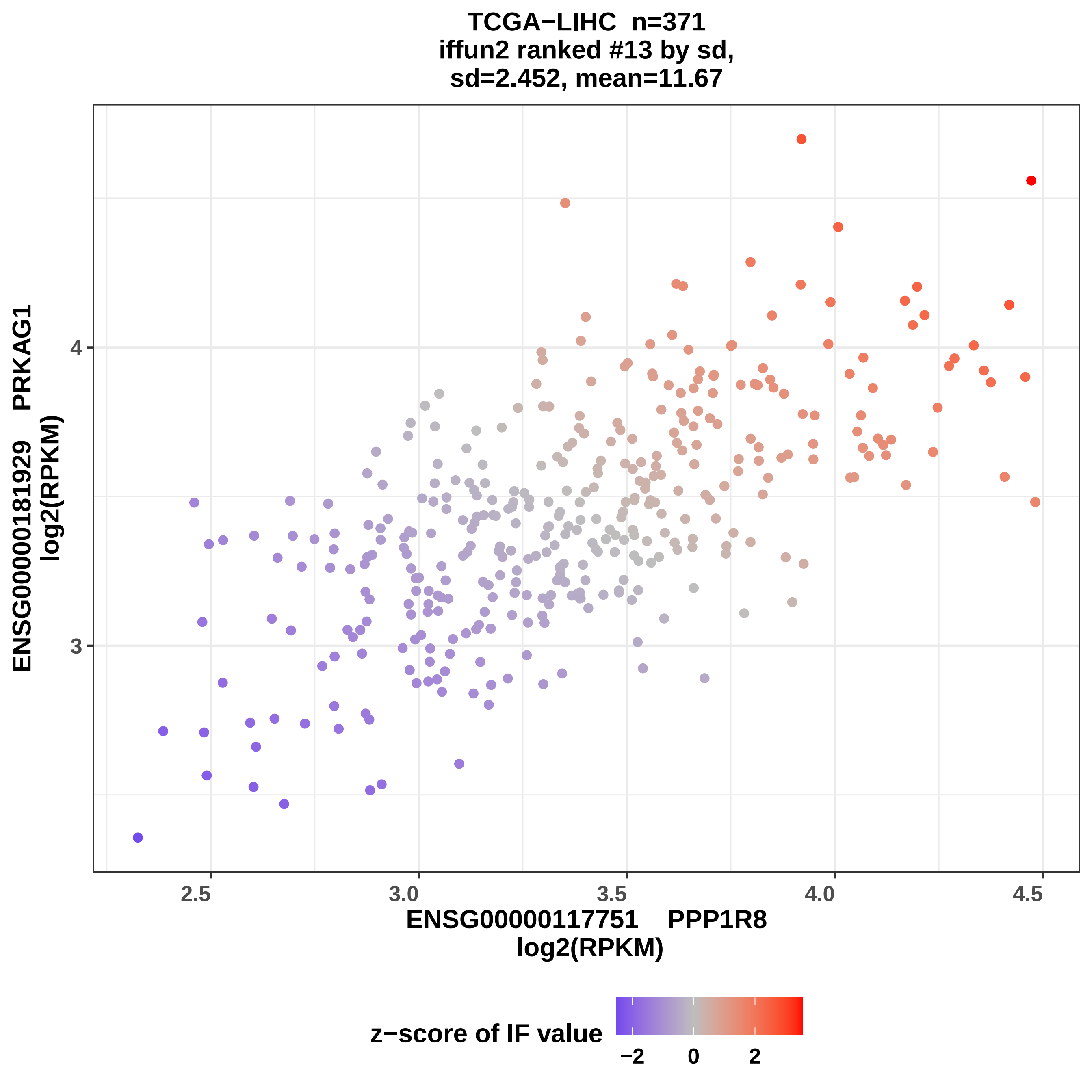
TCGA−LIHC n=371
iffun2 ranked #13 by sd,
sd=2.452, mean=11.67
ENSG00000181929 PRKAG1
log2(RPKM)
4
3
2.5
3.0
3.5
4.0
4.5
ENSG00000117751 PPP1R8
log2(RPKM)
z−score of IF value
−2
0
2

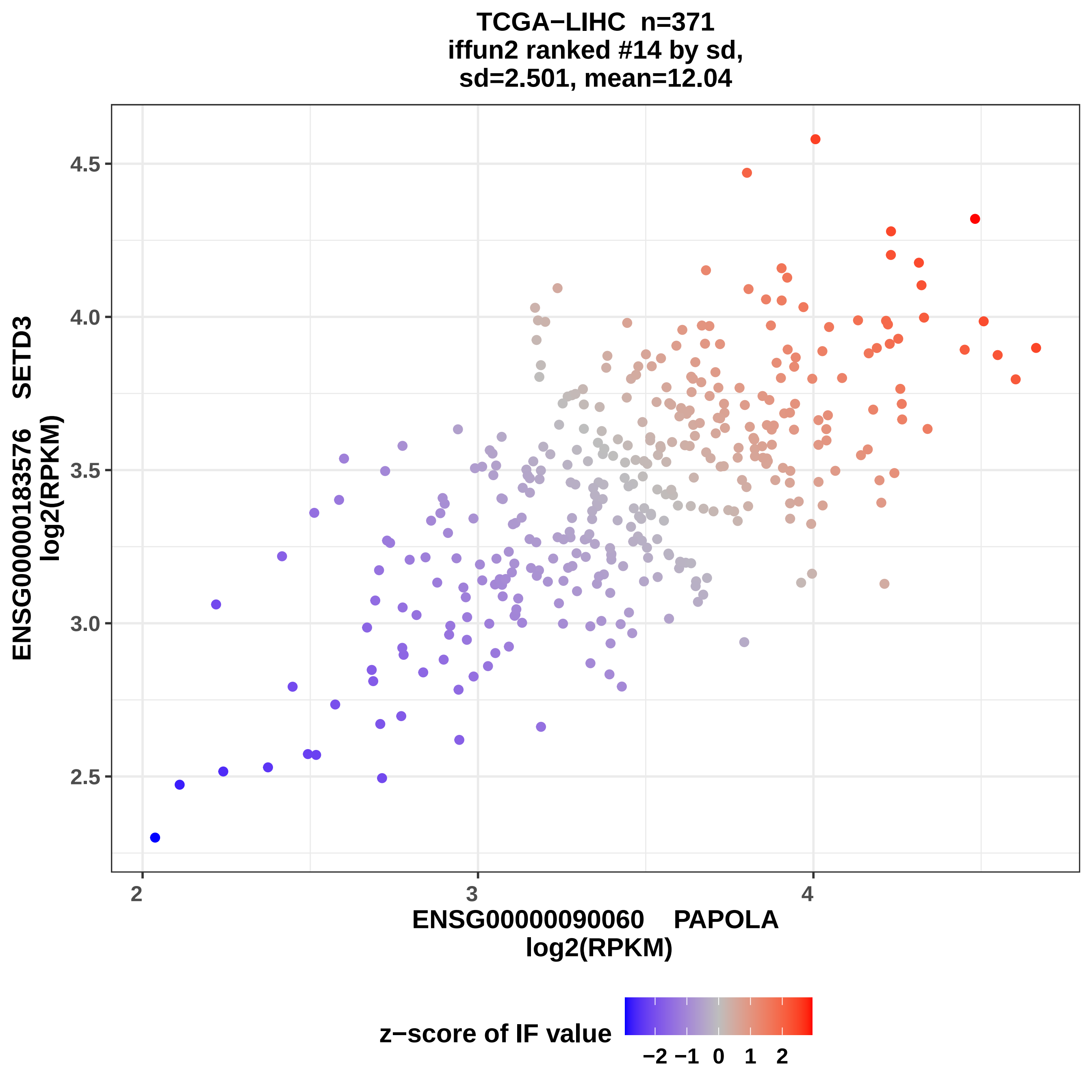
TCGA−LIHC n=371
iffun2 ranked #14 by sd,
sd=2.501, mean=12.04
ENSG00000183576 SETD3
log2(RPKM)
4.5
4.0
3.5
3.0
2.5
2
3
4
ENSG00000090060 PAPOLA
log2(RPKM)
z−score of IF value
−2 −1 0 1 2

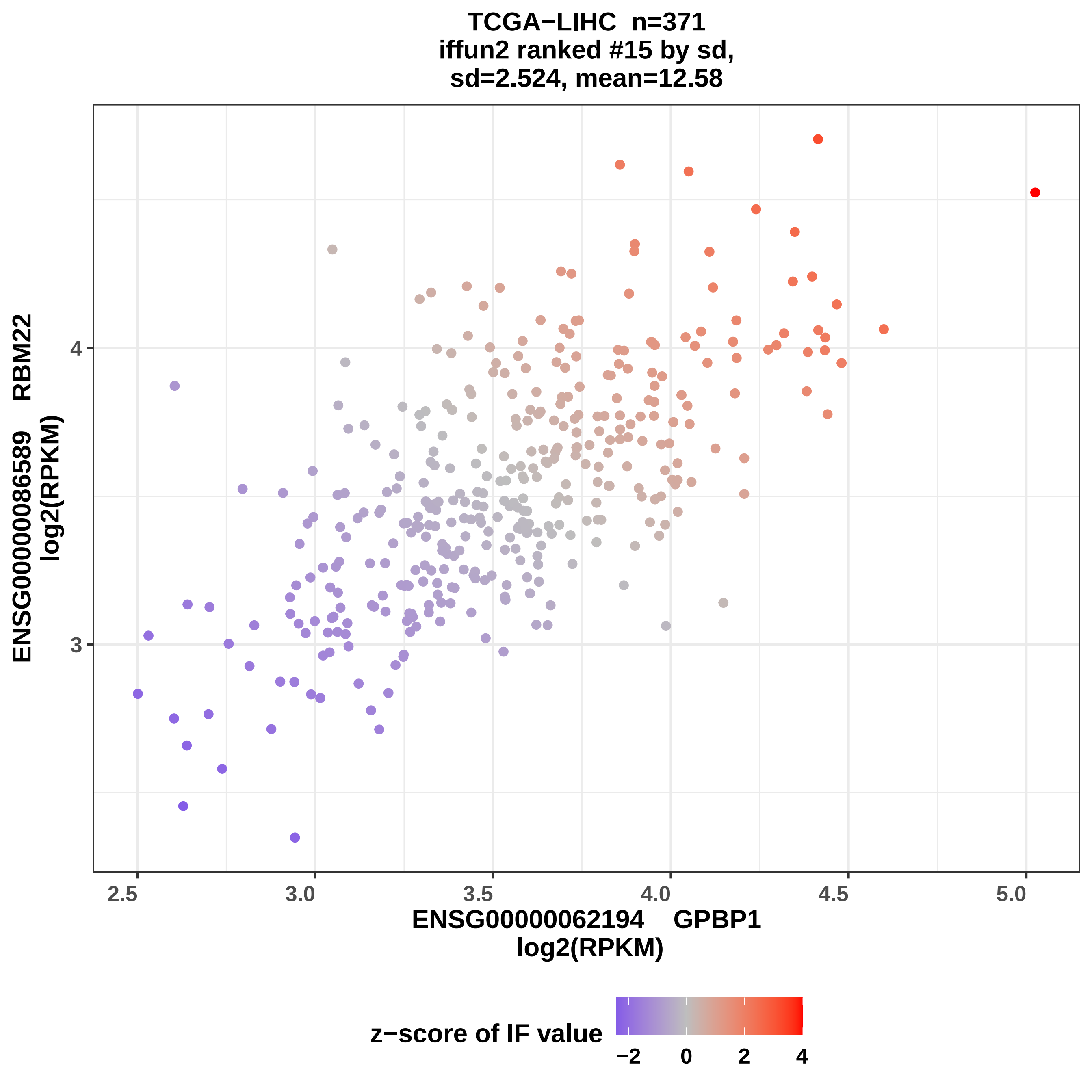

TCGA−LIHC n=371
iffun2 ranked #15 by sd,
sd=2.524, mean=12.58
ENSG00000086589 RBM22
log2(RPKM)
4
3
2.5
3.0
3.5
4.0
4.5
5.0
ENSG00000062194 GPBP1
log2(RPKM)
z−score of IF value
−2
0
2
4

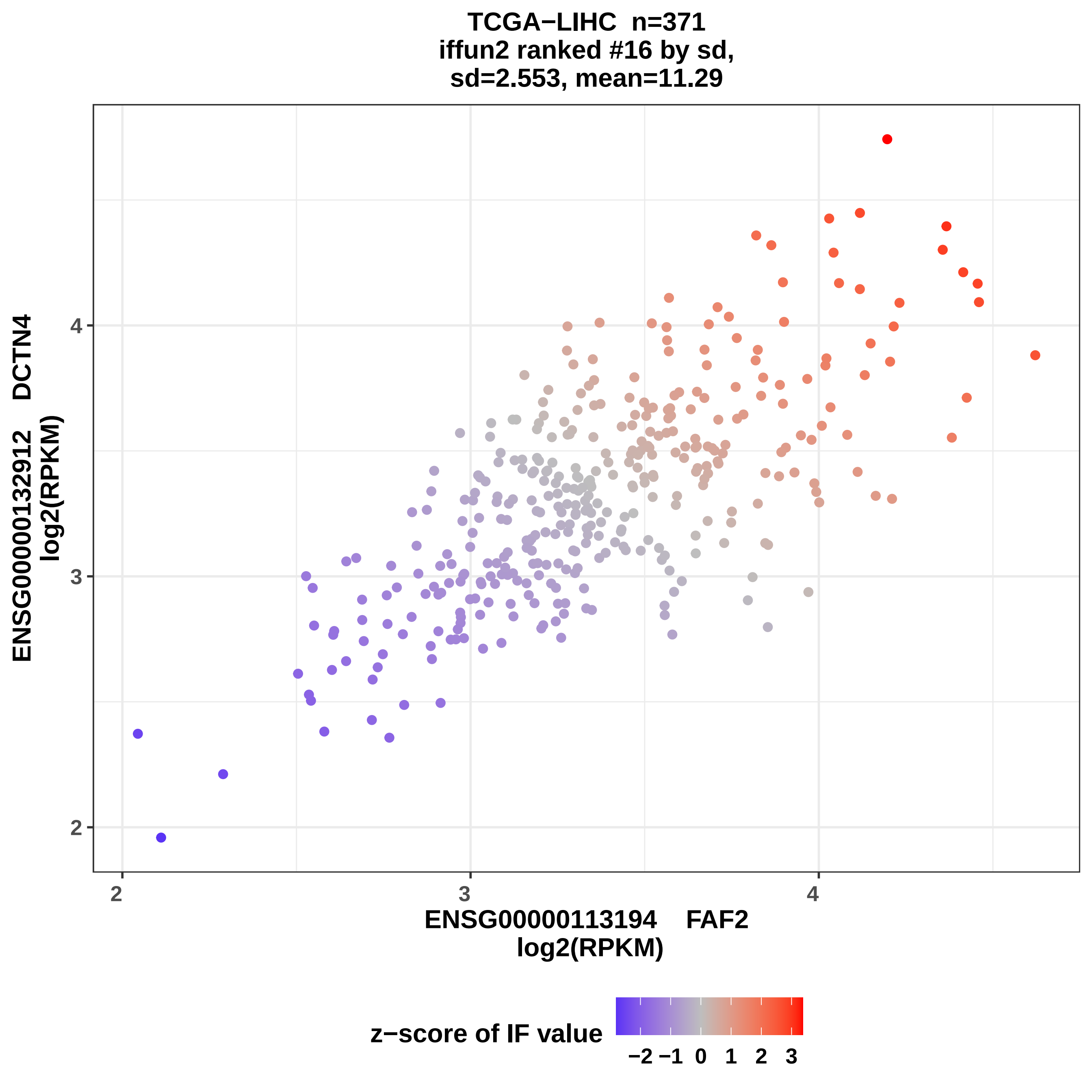
TCGA−LIHC n=371
iffun2 ranked #16 by sd,
sd=2.553, mean=11.29
ENSG00000132912 DCTN4
log2(RPKM)
ENSG00000113194 FAF2
log2(RPKM)
z−score of IF value
−2 −1 0 1 2 3
2
3
4

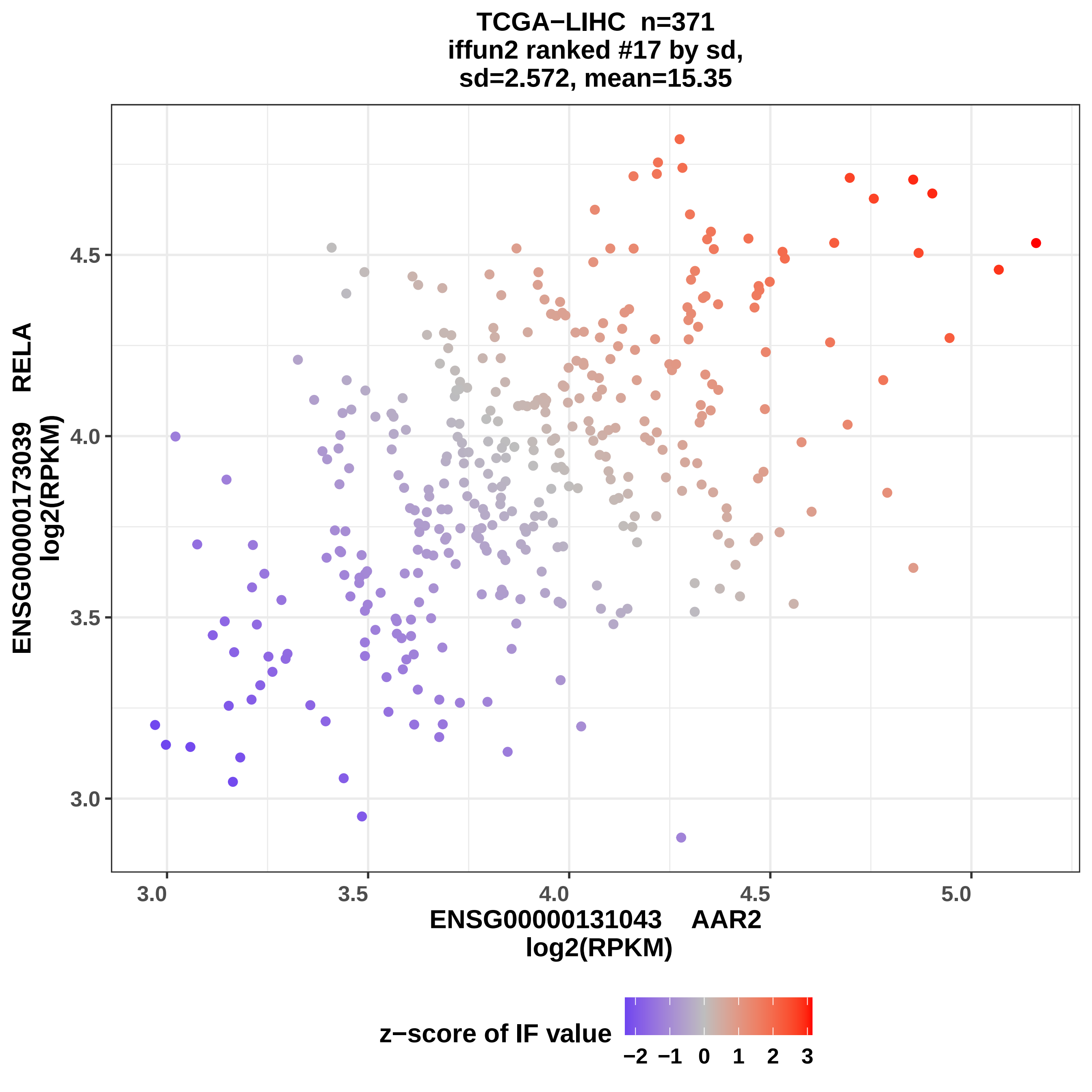
TCGA−LIHC n=371
iffun2 ranked #17 by sd,
sd=2.572, mean=15.35
ENSG00000173039 RELA
log2(RPKM)
4.5
4.0
3.5
3.0
3.0
3.5
4.0
4.5
5.0
ENSG00000131043 AAR2
log2(RPKM)
z−score of IF value
−2 −1 0 1 2 3

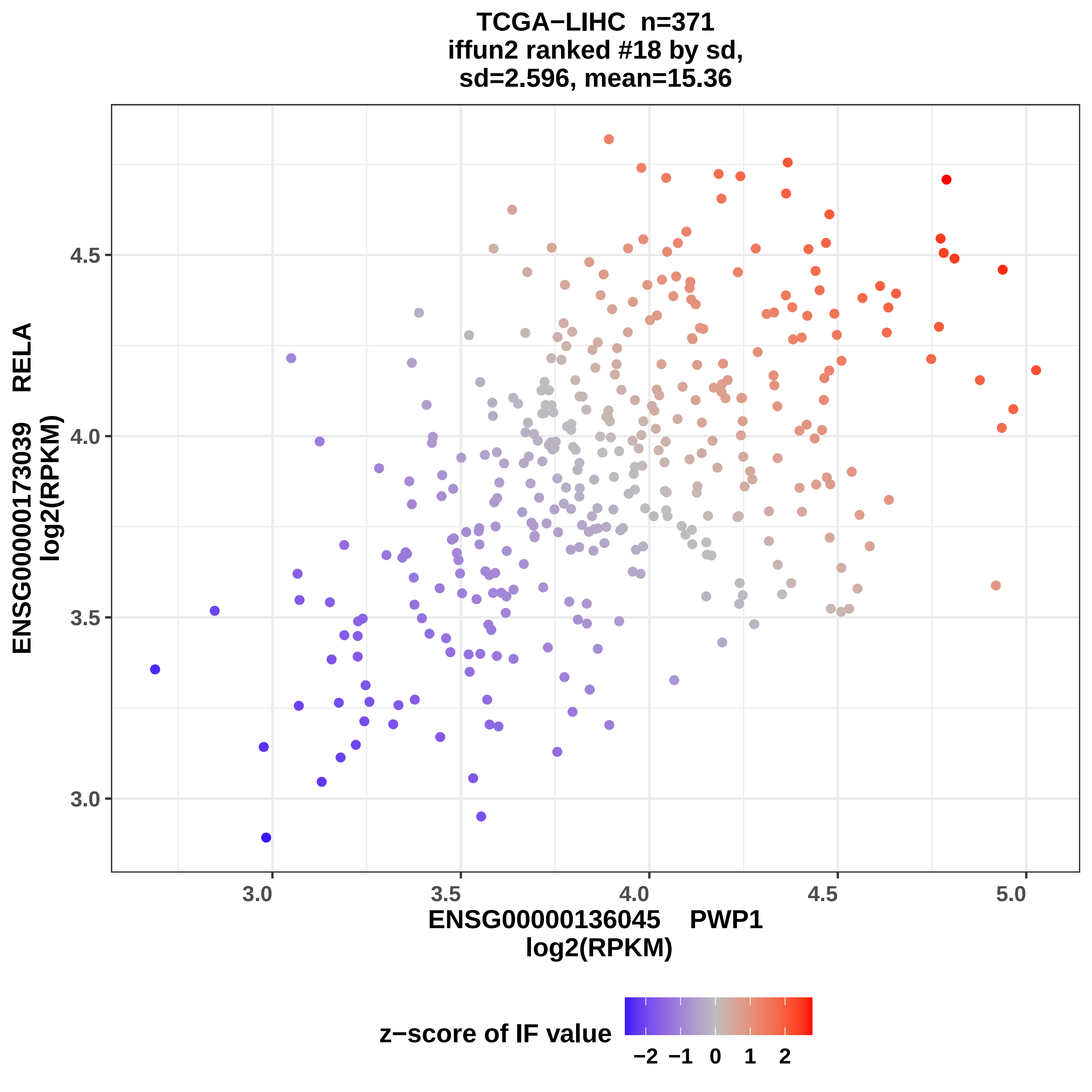

TCGA−LIHC n=371
iffun2 ranked #18 by sd,
sd=2.596, mean=15.36
ENSG00000173039 RELA
log2(RPKM)
4.5
4.0
3.5
3.0
3.0
3.5
4.0
4.5
5.0
ENSG00000136045 PWP1
log2(RPKM)
z−score of IF value
−2 −1 0 1 2

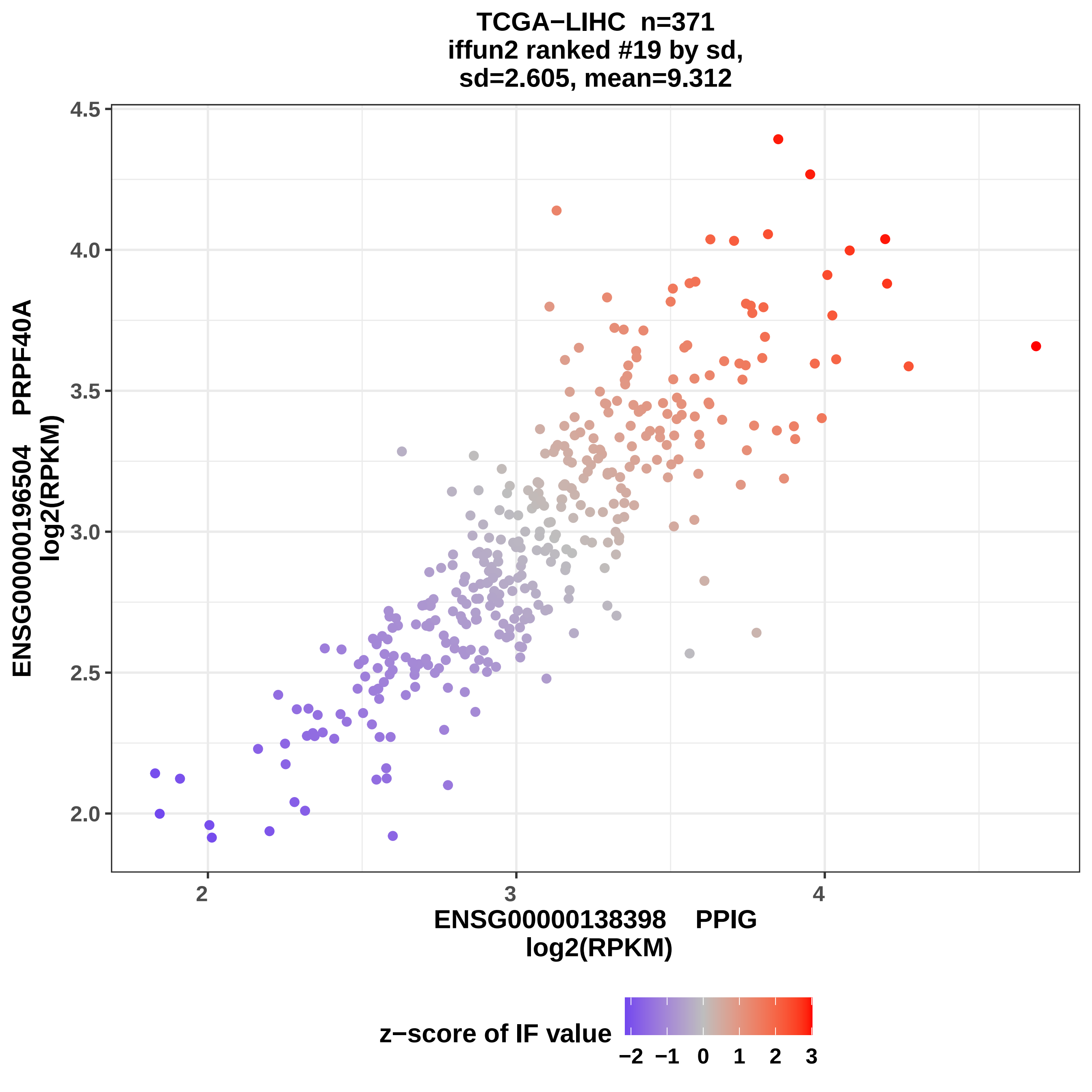
TCGA−LIHC n=371
iffun2 ranked #19 by sd,
sd=2.605, mean=9.312
ENSG00000196504 PRPF40A
log2(RPKM)
ENSG00000138398 PPIG
log2(RPKM)
2.0
2.5
3.0
3.5
4.0
4.5
2
3
4
z−score of IF value
−2 −1 0 1 2 3

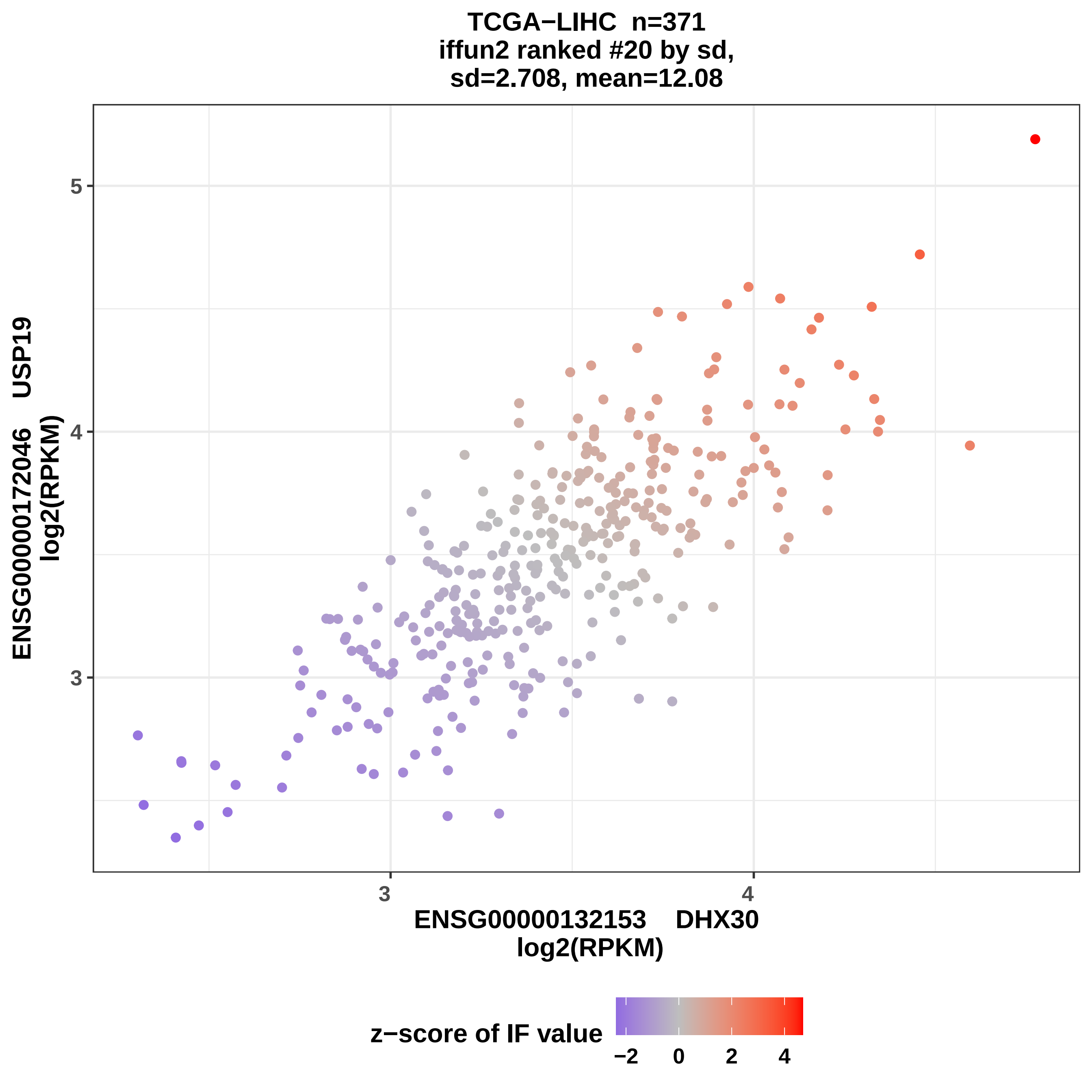
TCGA−LIHC n=371
iffun2 ranked #20 by sd,
sd=2.708, mean=12.08
ENSG00000172046 USP19
log2(RPKM)
5
4
3
3
4
ENSG00000132153 DHX30
log2(RPKM)
z−score of IF value
−2
0
2
4

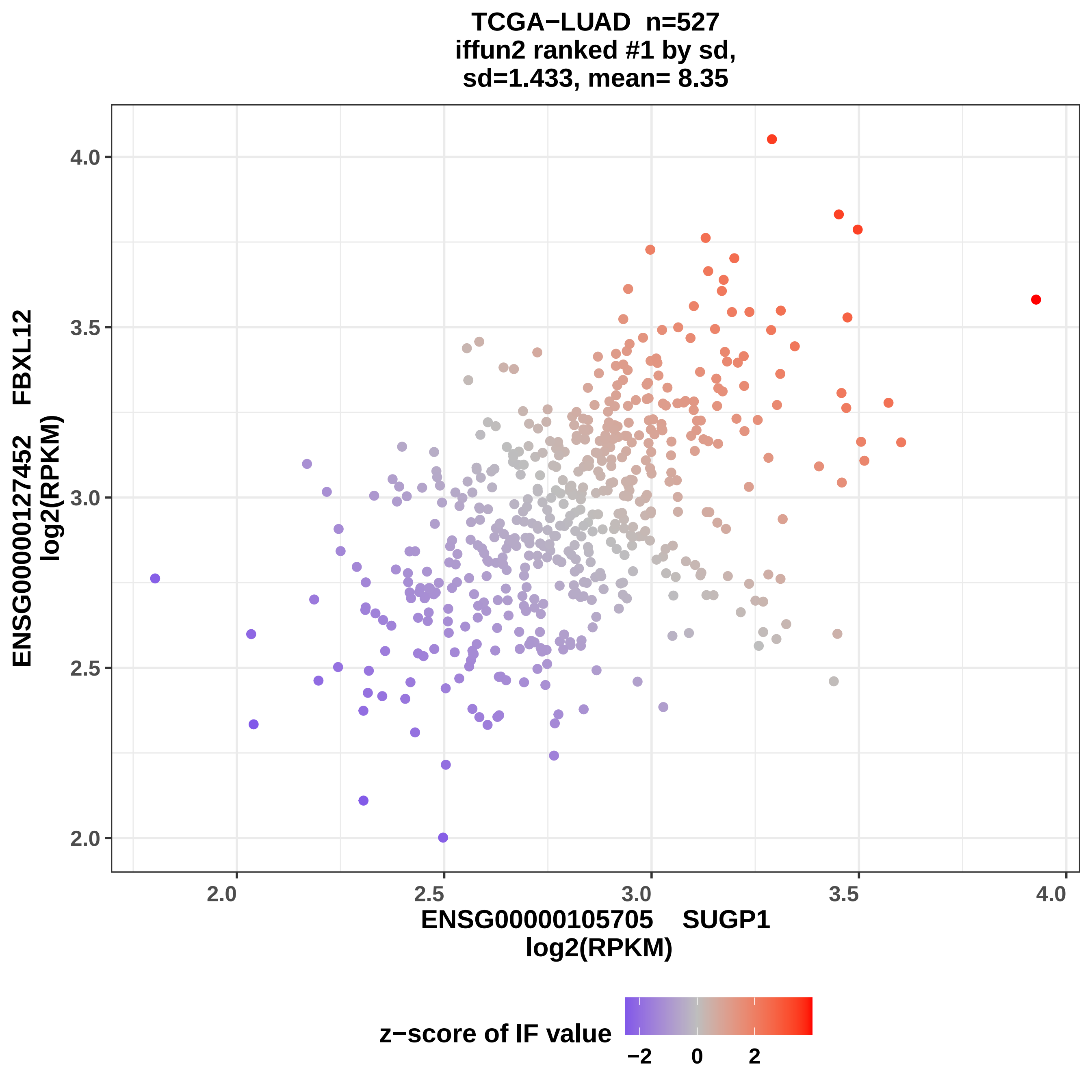

TCGA−LUAD n=527
iffun2 ranked #1 by sd,
sd=1.433, mean= 8.35
ENSG00000127452 FBXL12
log2(RPKM)
4.0
3.5
3.0
2.5
2.0
2.0
2.5
3.0
3.5
4.0
ENSG00000105705 SUGP1
log2(RPKM)
z−score of IF value
−2
0
2

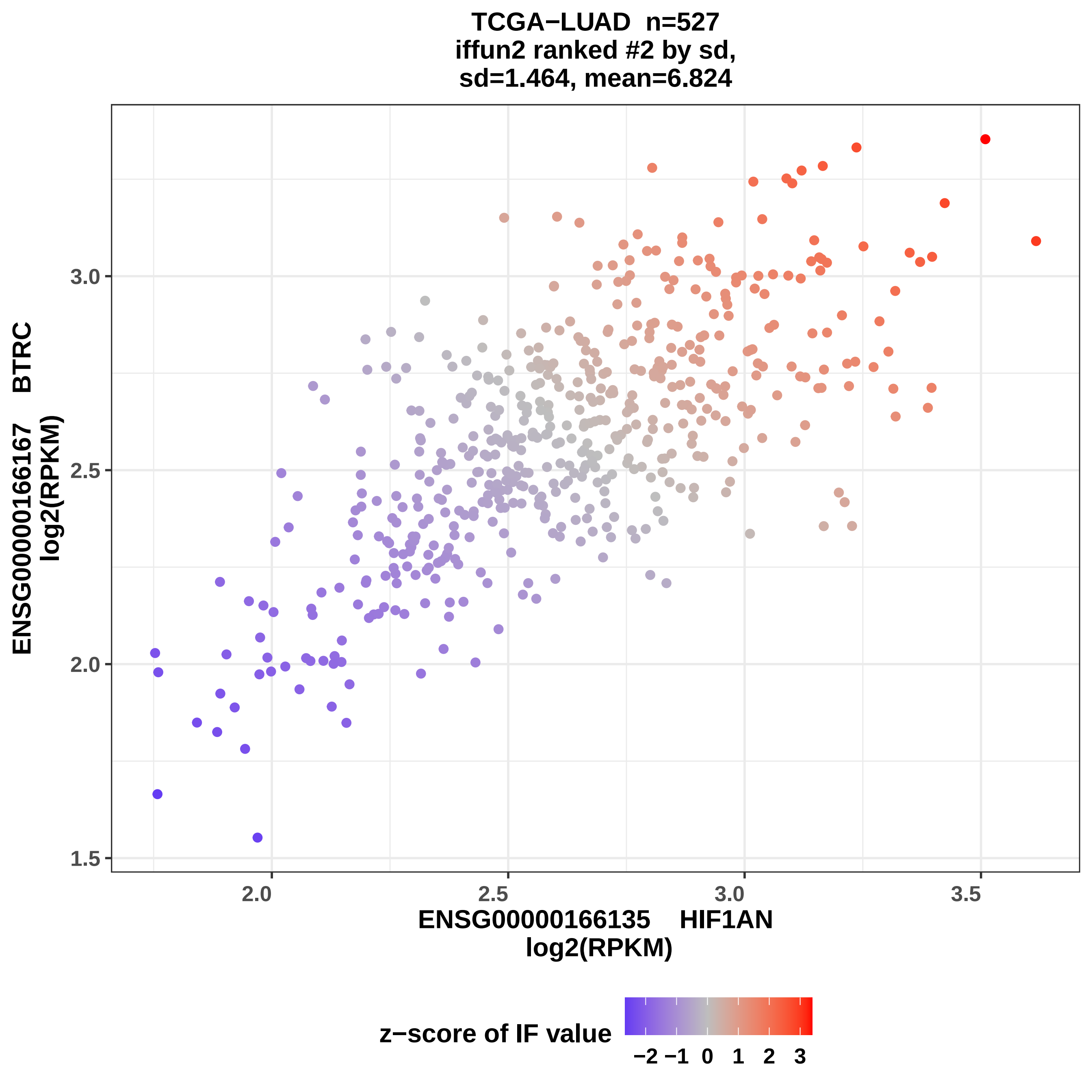

TCGA−LUAD n=527
iffun2 ranked #2 by sd,
sd=1.464, mean=6.824
ENSG00000166167 BTRC
log2(RPKM)
ENSG00000166135 HIF1AN
log2(RPKM)
1.5
2.0
2.5
3.0
3.5
z−score of IF value
−2 −1 0 1 2 3

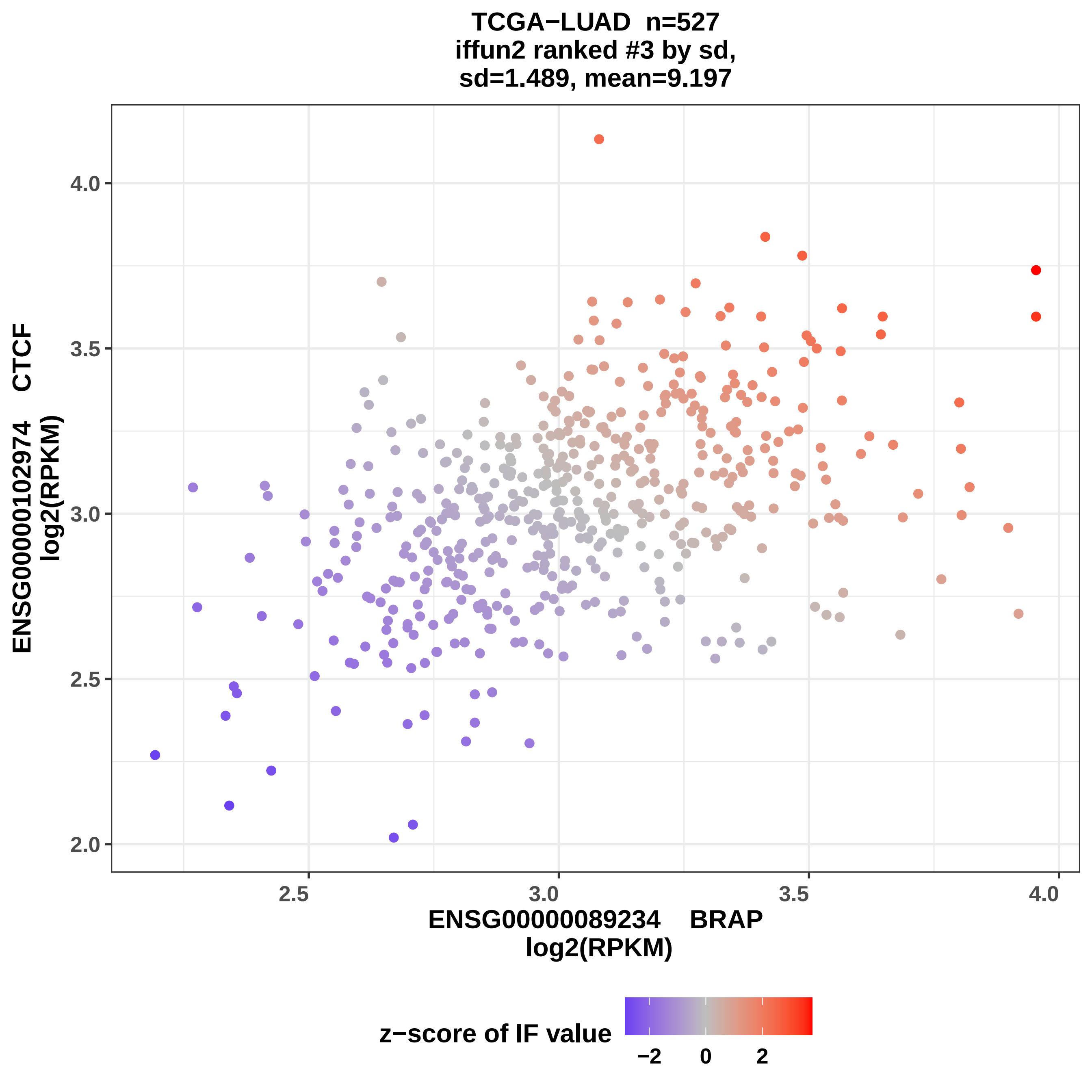
TCGA−LUAD n=527
iffun2 ranked #3 by sd,
sd=1.489, mean=9.197
ENSG00000102974 CTCF
log2(RPKM)
ENSG00000089234 BRAP
log2(RPKM)
2.0
2.5
3.0
3.5
4.0
2.5
3.0
3.5
4.0
z−score of IF value
−2
0
2

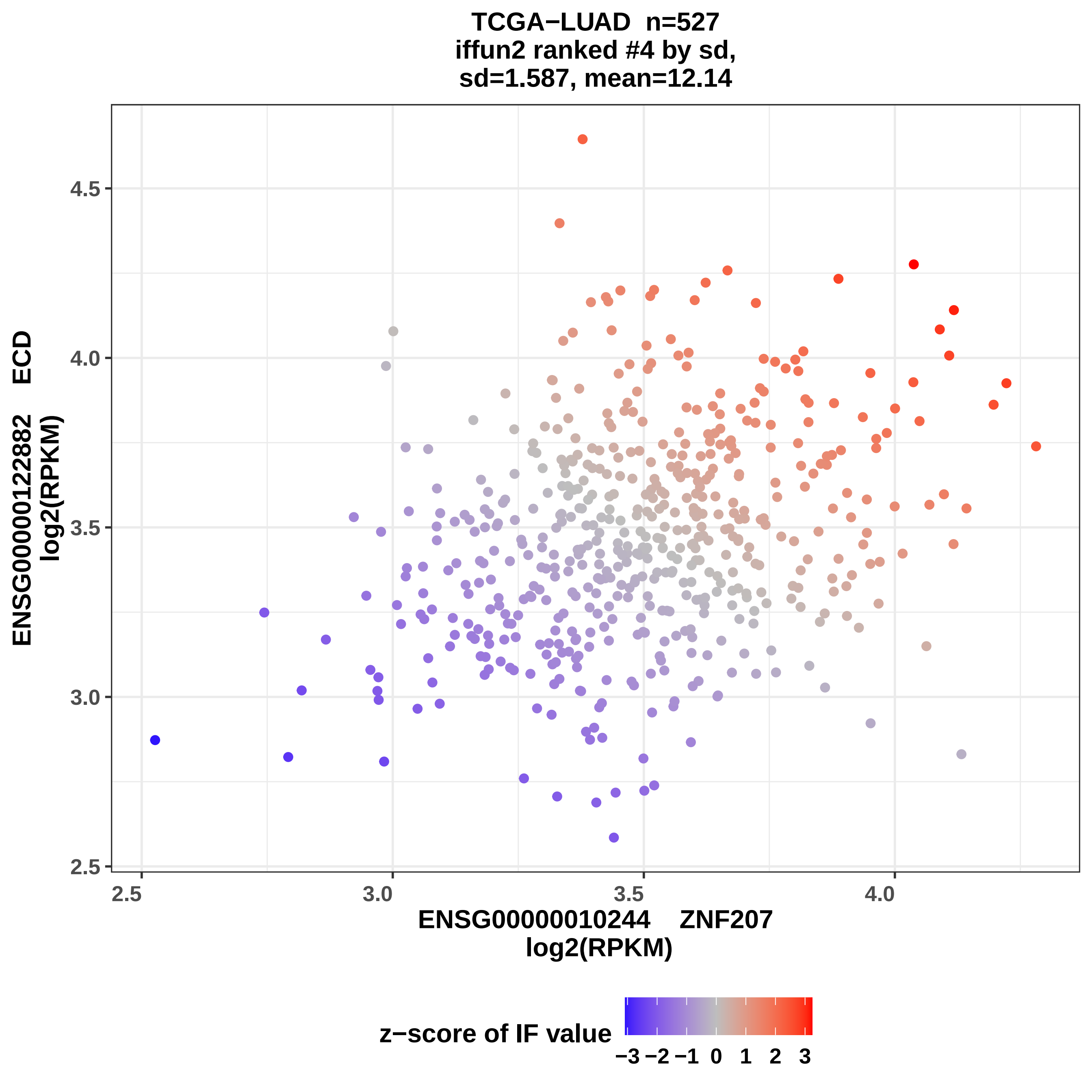

TCGA−LUAD n=527
iffun2 ranked #4 by sd,
sd=1.587, mean=12.14
ENSG00000122882 ECD
log2(RPKM)
4.5
4.0
3.5
3.0
2.5
2.5
3.0
3.5
4.0
ENSG00000010244 ZNF207
log2(RPKM)
z−score of IF value
−3 −2 −1 0 1 2 3

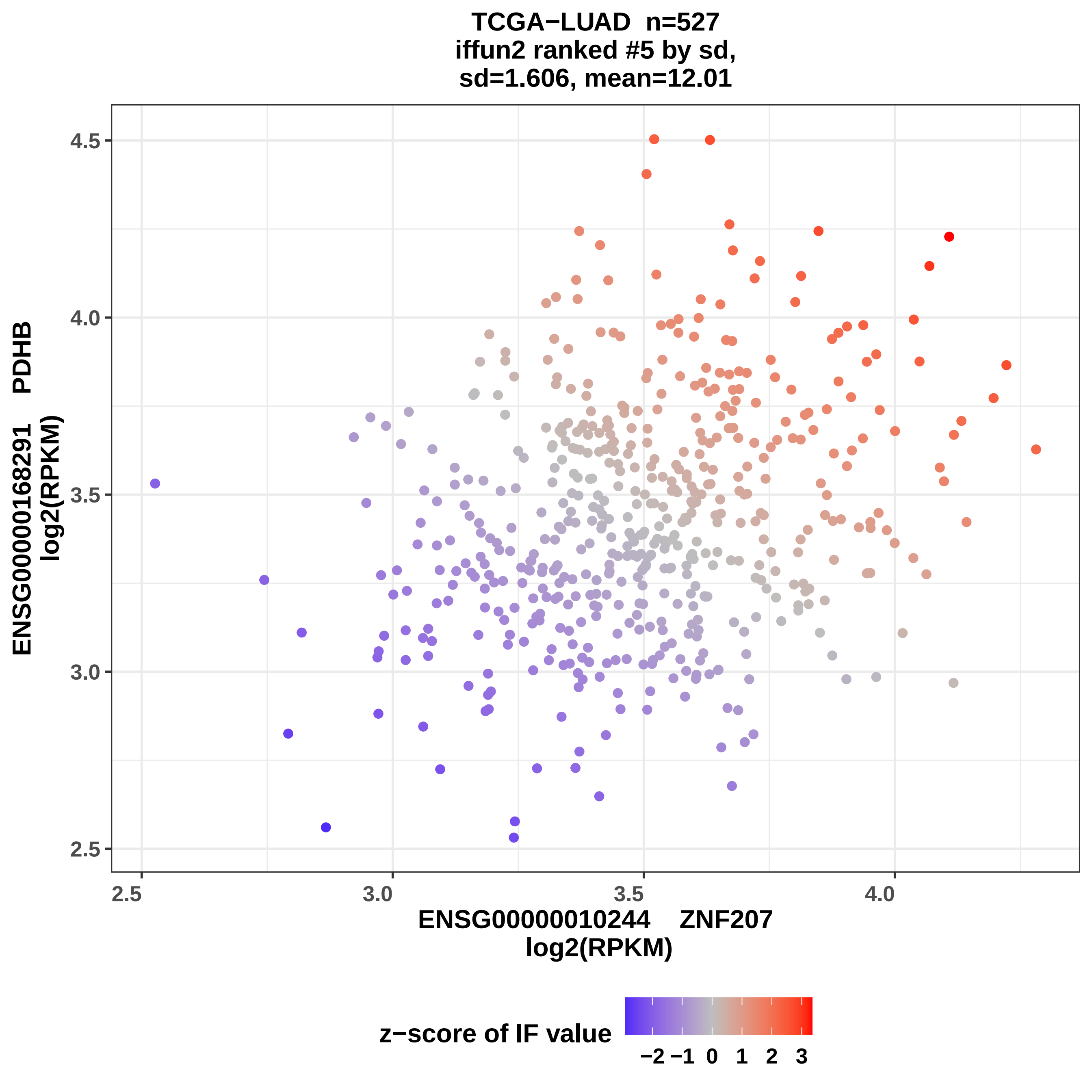
TCGA−LUAD n=527
iffun2 ranked #5 by sd,
sd=1.606, mean=12.01
ENSG00000168291 PDHB
log2(RPKM)
ENSG00000010244 ZNF207
log2(RPKM)
4.5
4.0
3.5
3.0
2.5
2.5
3.0
3.5
4.0
z−score of IF value
−2 −1 0 1 2 3

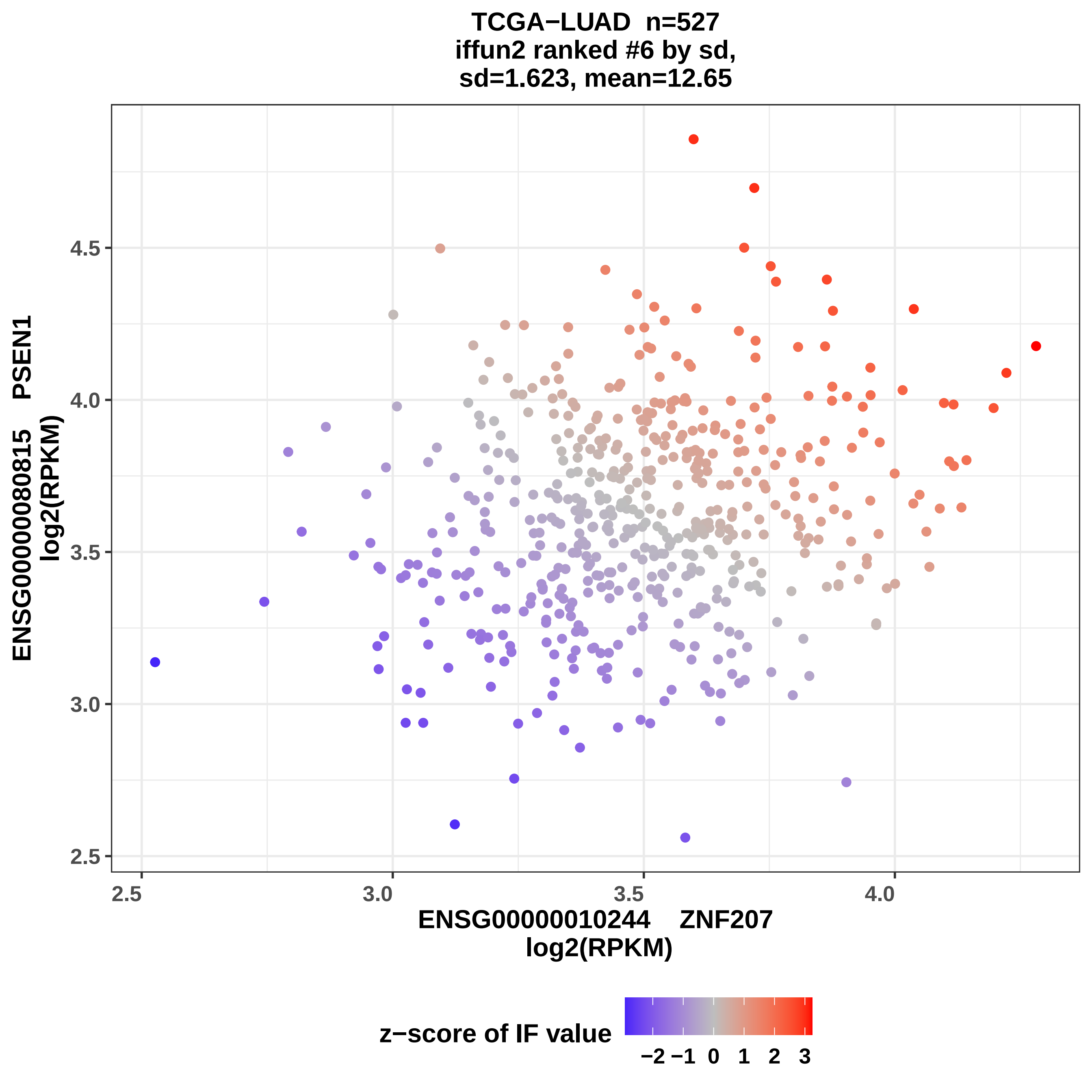
TCGA−LUAD n=527
iffun2 ranked #6 by sd,
sd=1.623, mean=12.65
ENSG00000080815 PSEN1
log2(RPKM)
4.5
4.0
3.5
3.0
2.5
2.5
3.0
3.5
4.0
ENSG00000010244 ZNF207
log2(RPKM)
z−score of IF value
−2 −1 0 1 2 3

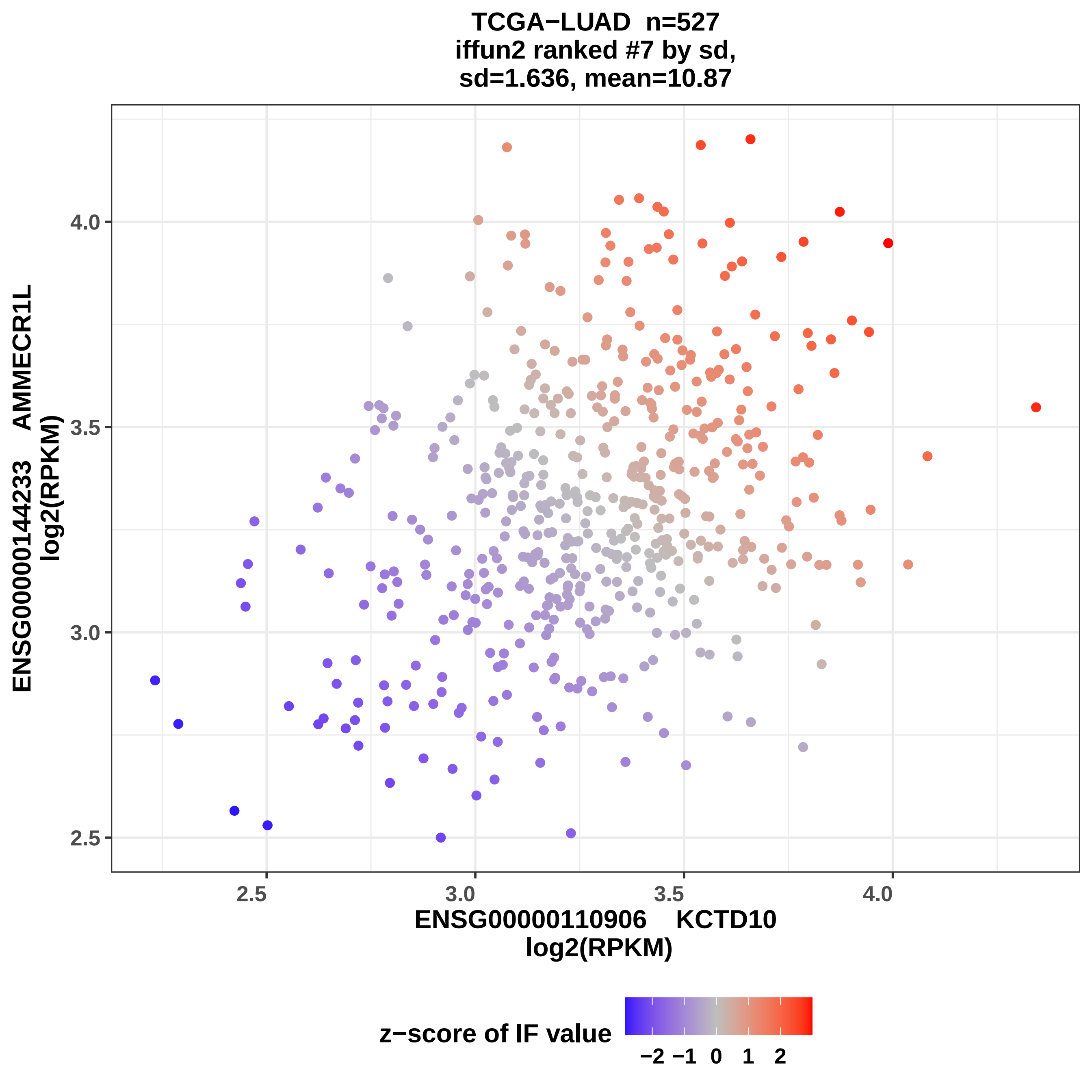

TCGA−LUAD n=527
iffun2 ranked #7 by sd,
sd=1.636, mean=10.87
ENSG00000144233 AMMECR1L
log2(RPKM)
4.0
3.5
3.0
2.5
2.5
3.0
3.5
4.0
ENSG00000110906 KCTD10
log2(RPKM)
z−score of IF value
−2 −1 0 1 2

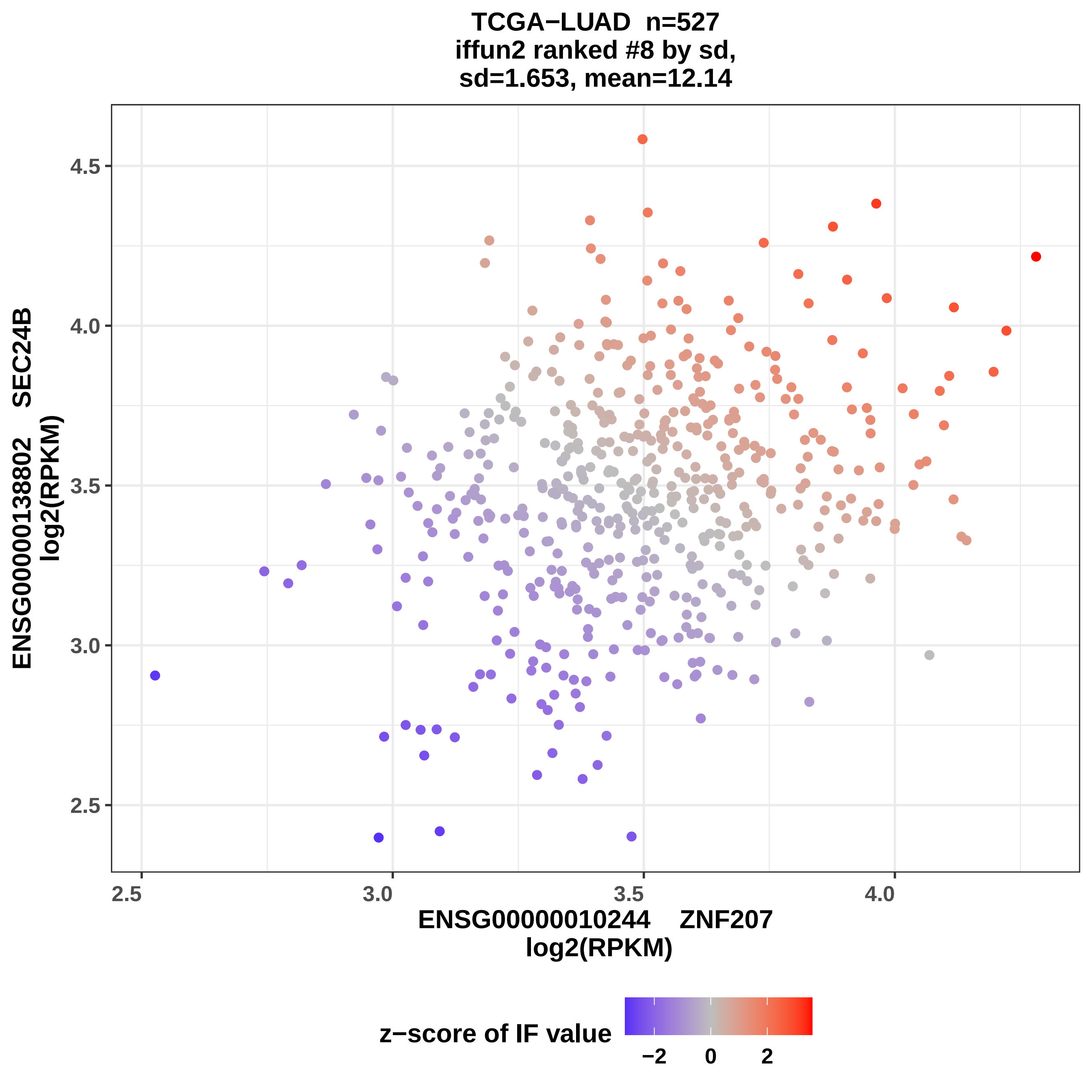
TCGA−LUAD n=527
iffun2 ranked #8 by sd,
sd=1.653, mean=12.14
ENSG00000138802 SEC24B
log2(RPKM)
4.5
4.0
3.5
3.0
2.5
2.5
3.0
3.5
4.0
ENSG00000010244 ZNF207
log2(RPKM)
z−score of IF value
−2
0
2

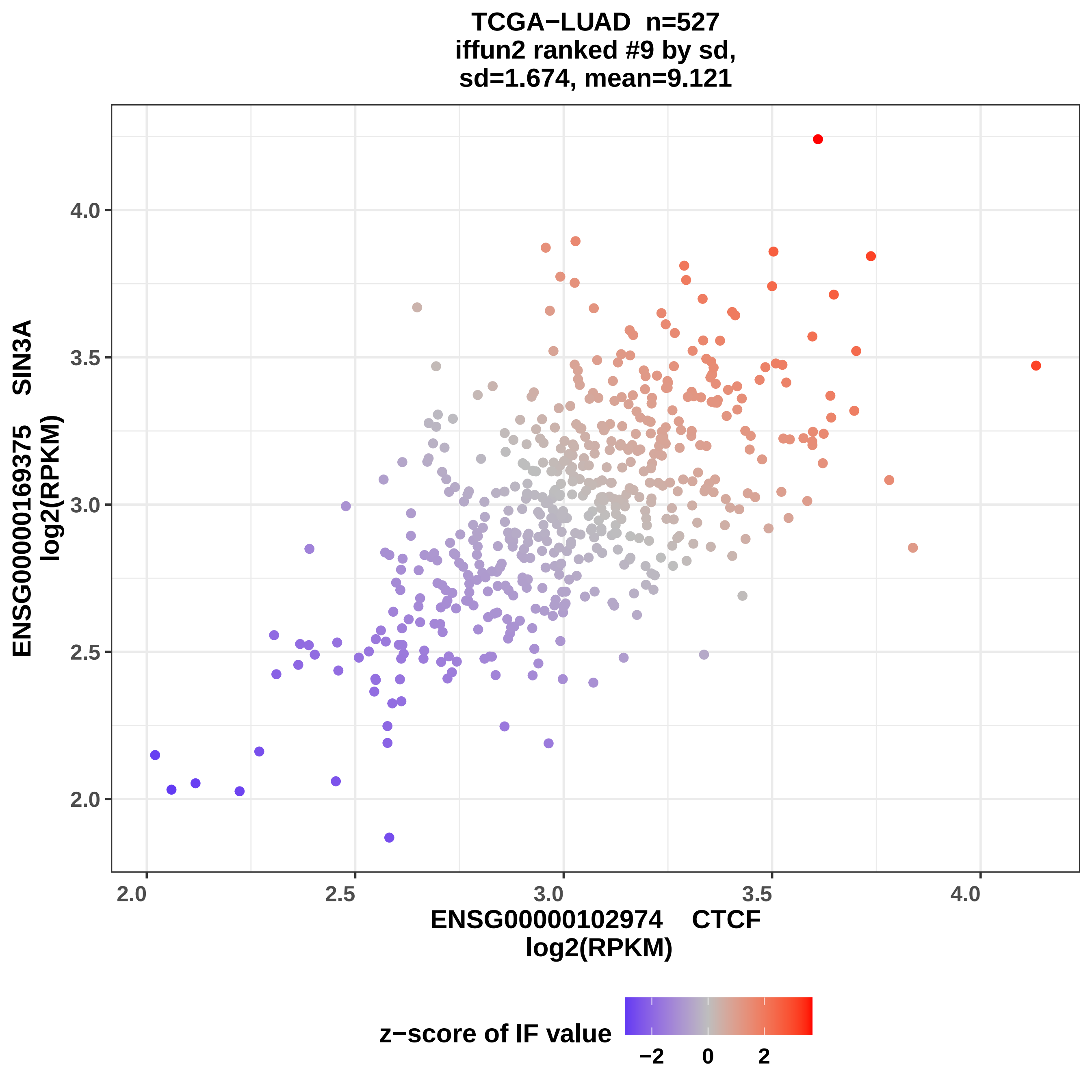
TCGA−LUAD n=527
iffun2 ranked #9 by sd,
sd=1.674, mean=9.121
ENSG00000169375 SIN3A
log2(RPKM)
4.0
3.5
3.0
2.5
2.0
2.0
2.5
3.0
3.5
4.0
ENSG00000102974 CTCF
log2(RPKM)
z−score of IF value
−2
0
2

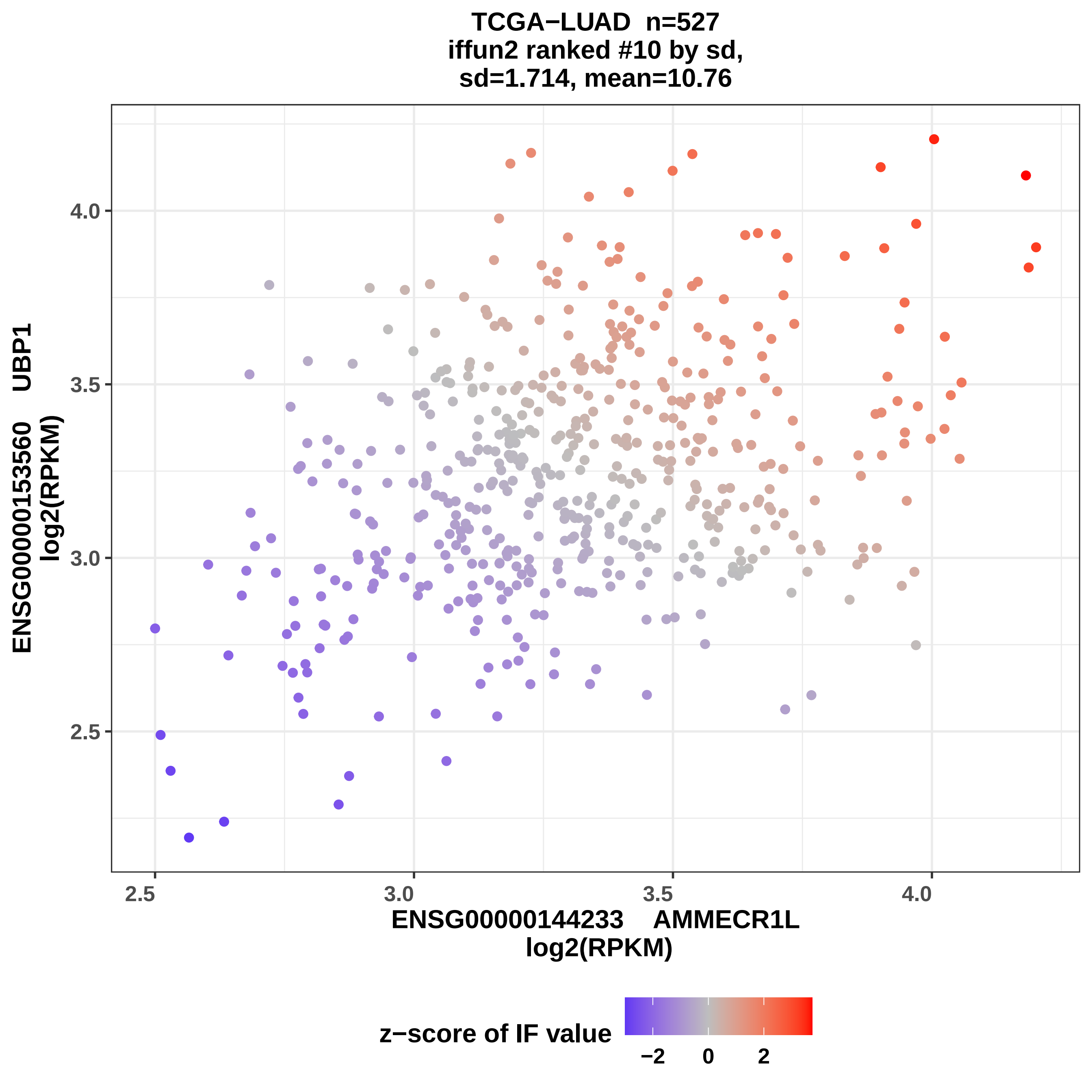
TCGA−LUAD n=527
iffun2 ranked #10 by sd,
sd=1.714, mean=10.76
ENSG00000153560 UBP1
log2(RPKM)
4.0
3.5
3.0
2.5
2.5
3.0
3.5
4.0
ENSG00000144233 AMMECR1L
log2(RPKM)
z−score of IF value
−2
0
2

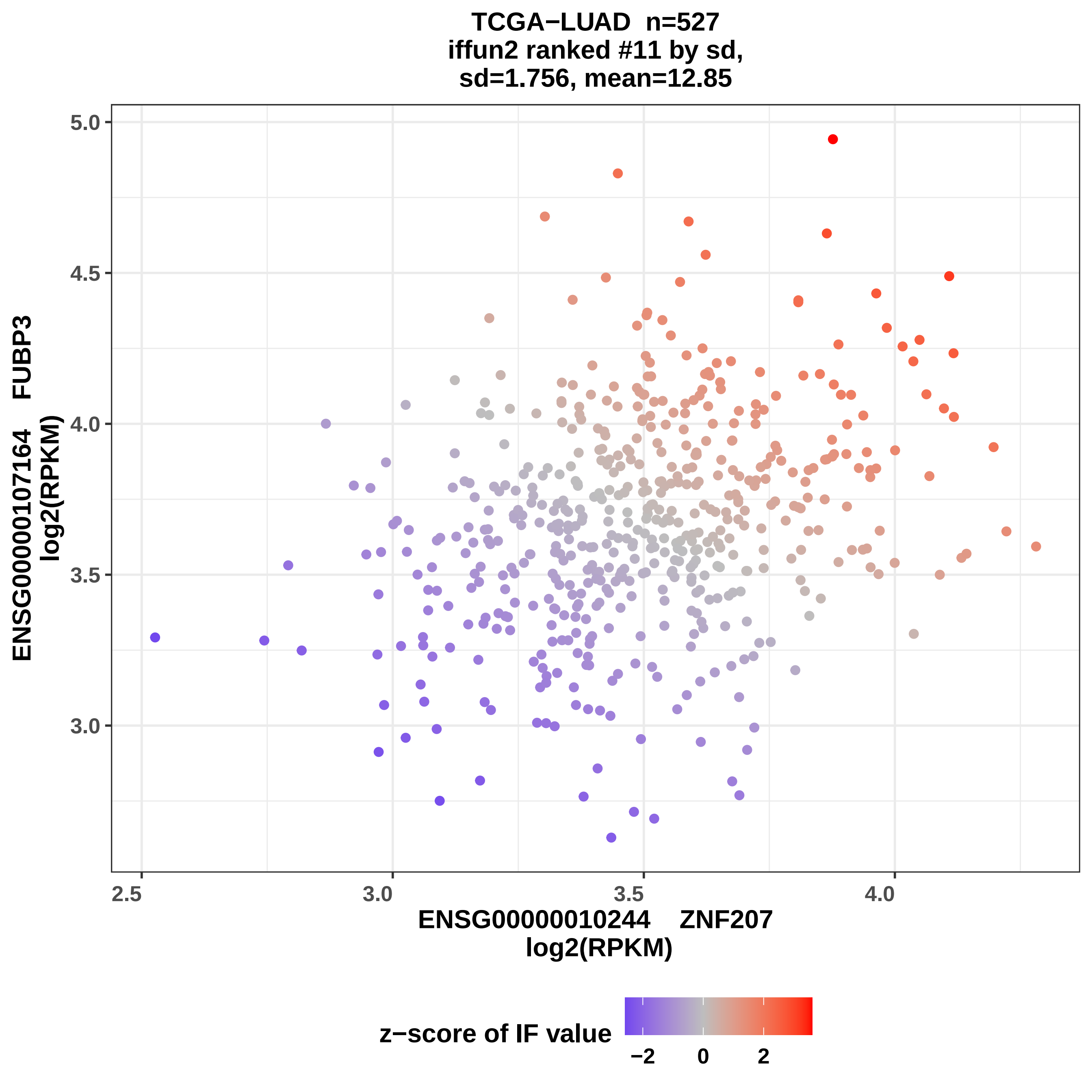
TCGA−LUAD n=527
iffun2 ranked #11 by sd,
sd=1.756, mean=12.85
ENSG00000107164 FUBP3
log2(RPKM)
5.0
4.5
4.0
3.5
3.0
2.5
3.0
3.5
4.0
ENSG00000010244 ZNF207
log2(RPKM)
z−score of IF value
−2
0
2

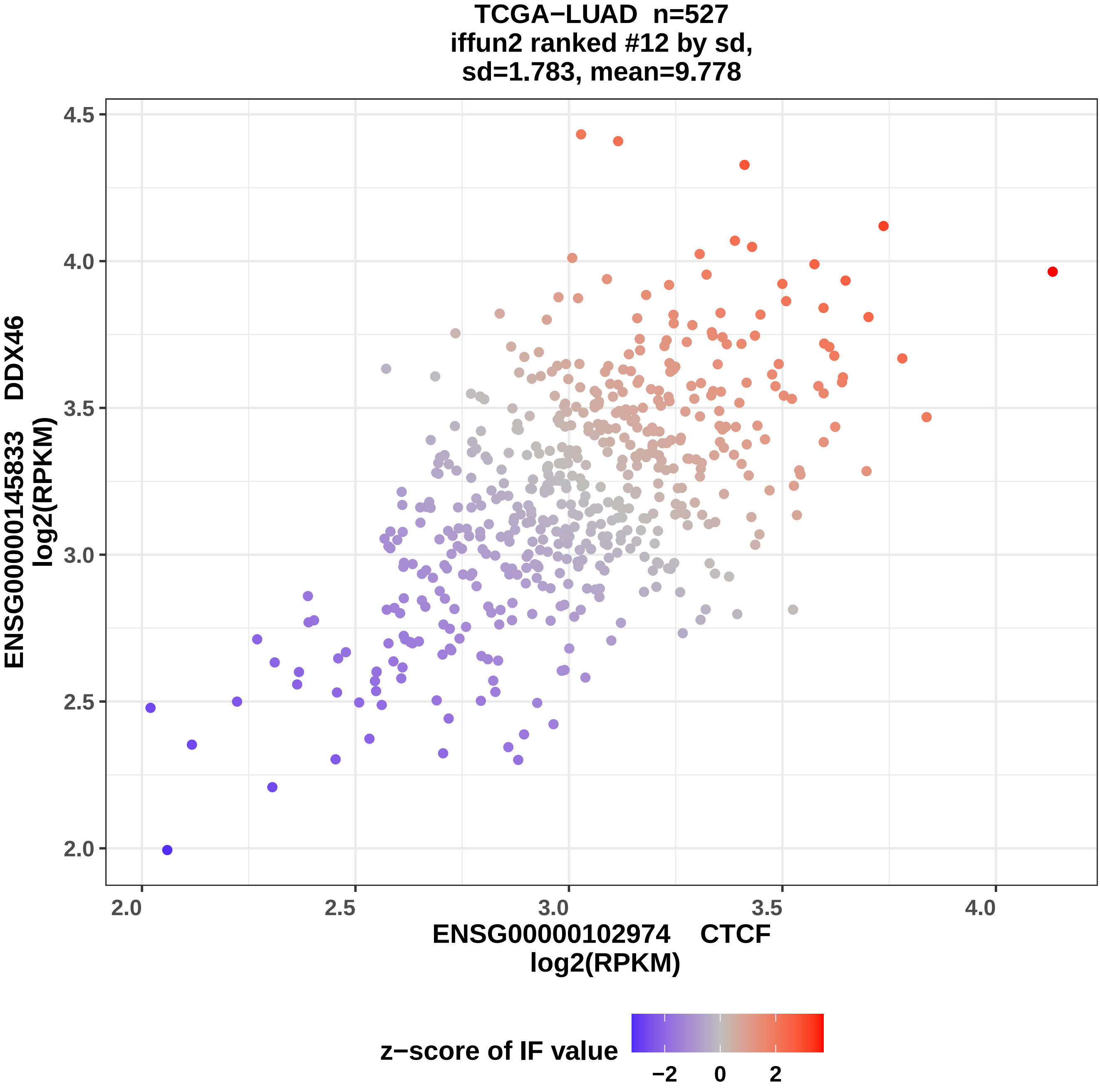
TCGA−LUAD n=527
iffun2 ranked #12 by sd,
sd=1.783, mean=9.778
ENSG00000145833 DDX46
log2(RPKM)
4.5
4.0
3.5
3.0
2.5
2.0
2.0
2.5
3.0
3.5
4.0
ENSG00000102974 CTCF
log2(RPKM)
z−score of IF value
−2
0
2

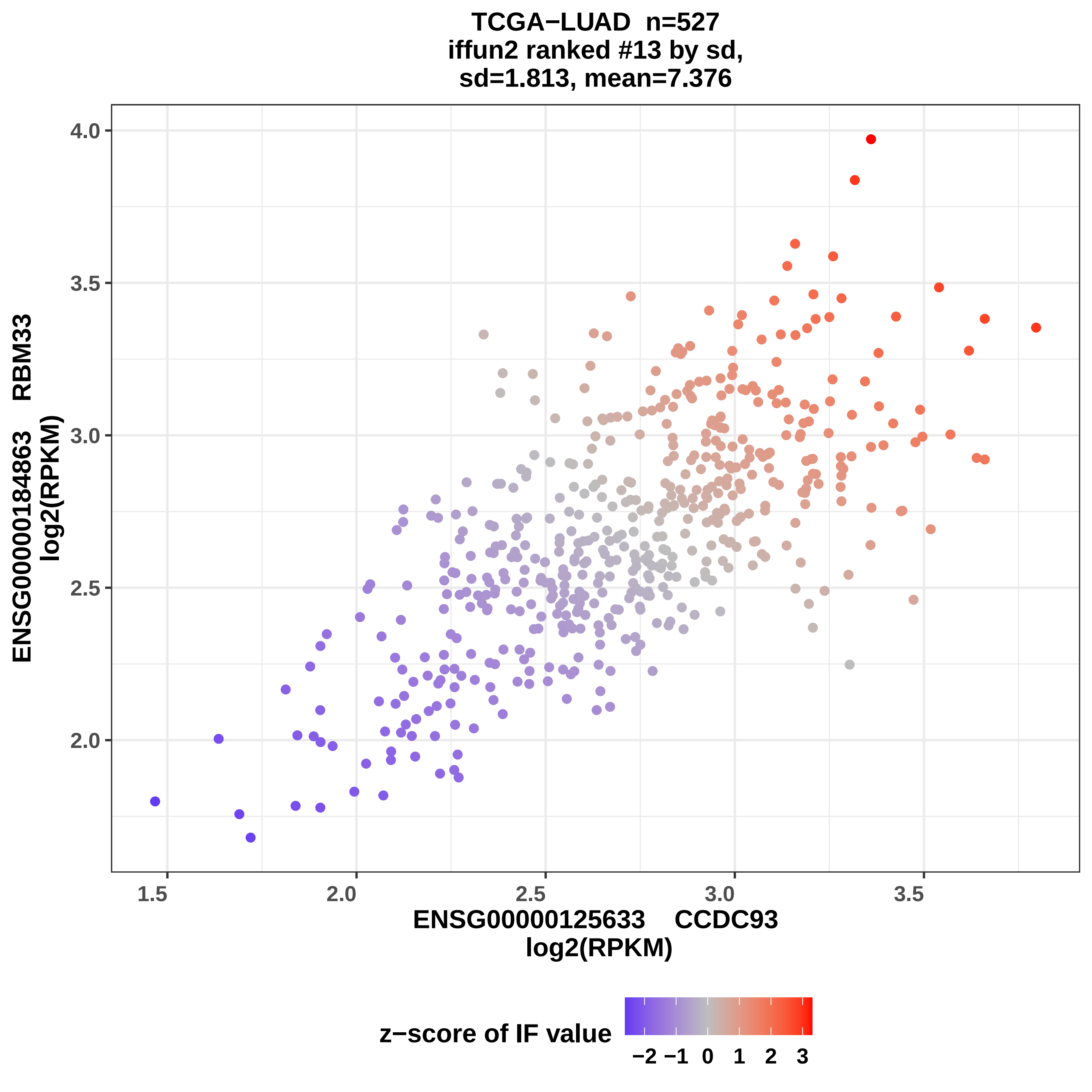
TCGA−LUAD n=527
iffun2 ranked #13 by sd,
sd=1.813, mean=7.376
ENSG00000184863 RBM33
log2(RPKM)
4.0
3.5
3.0
2.5
2.0
1.5
2.0
2.5
3.0
3.5
ENSG00000125633 CCDC93
log2(RPKM)
z−score of IF value
−2 −1 0 1 2 3

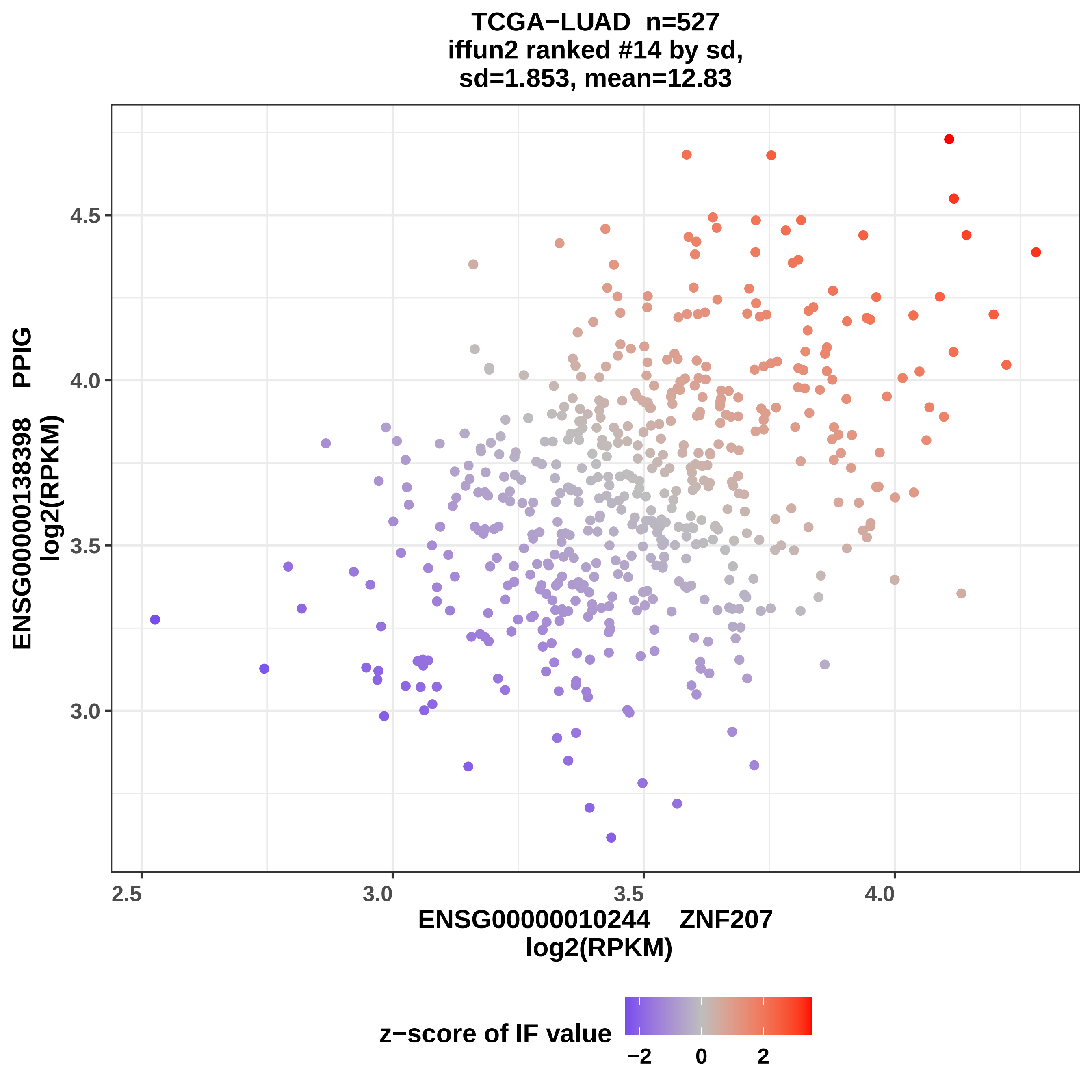
TCGA−LUAD n=527
iffun2 ranked #14 by sd,
sd=1.853, mean=12.83
ENSG00000138398 PPIG
log2(RPKM)
4.5
4.0
3.5
3.0
2.5
3.0
3.5
4.0
ENSG00000010244 ZNF207
log2(RPKM)
z−score of IF value
−2
0
2

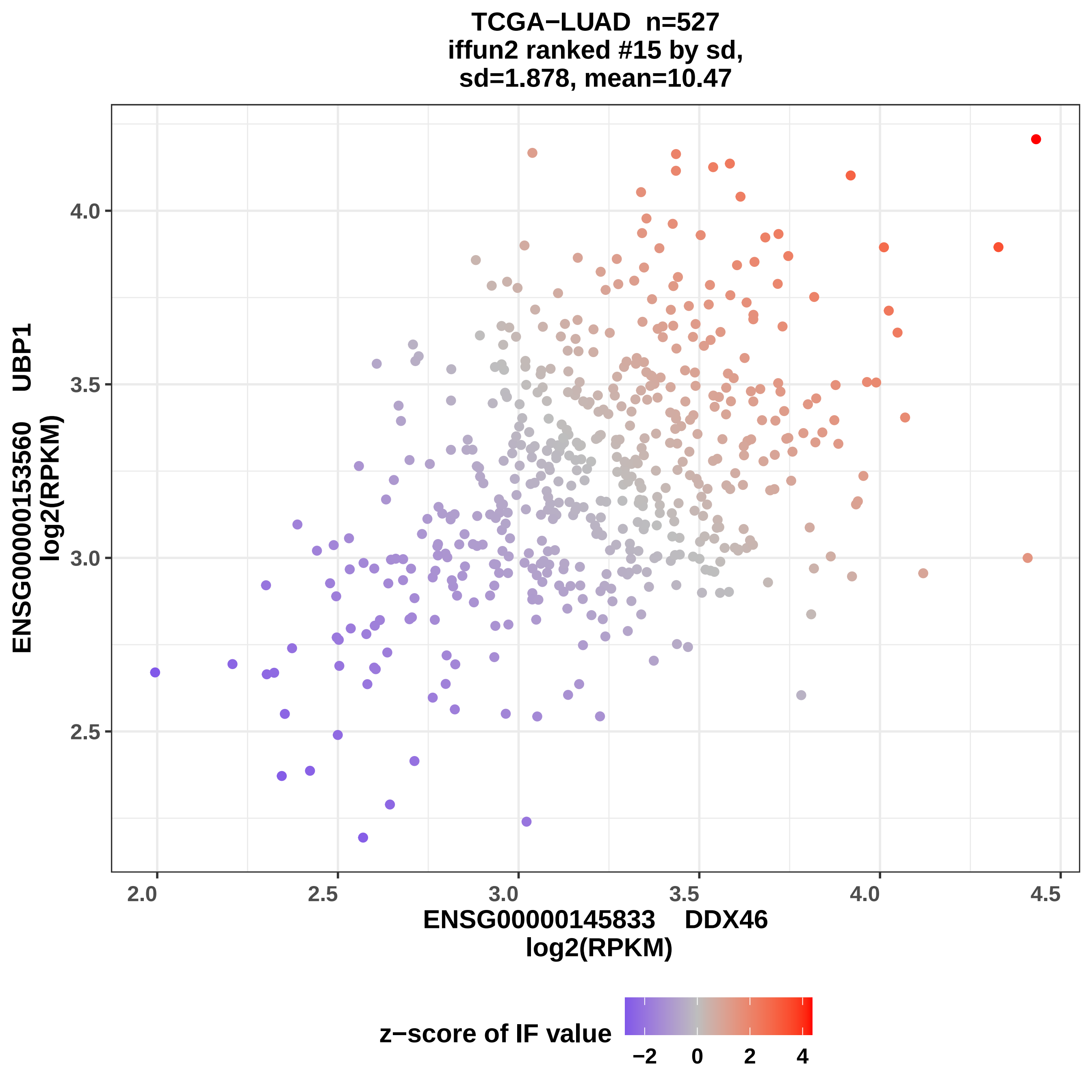

TCGA−LUAD n=527
iffun2 ranked #15 by sd,
sd=1.878, mean=10.47
ENSG00000153560 UBP1
log2(RPKM)
4.0
3.5
3.0
2.5
2.0
2.5
3.0
3.5
4.0
4.5
ENSG00000145833 DDX46
log2(RPKM)
z−score of IF value
−2
0
2
4

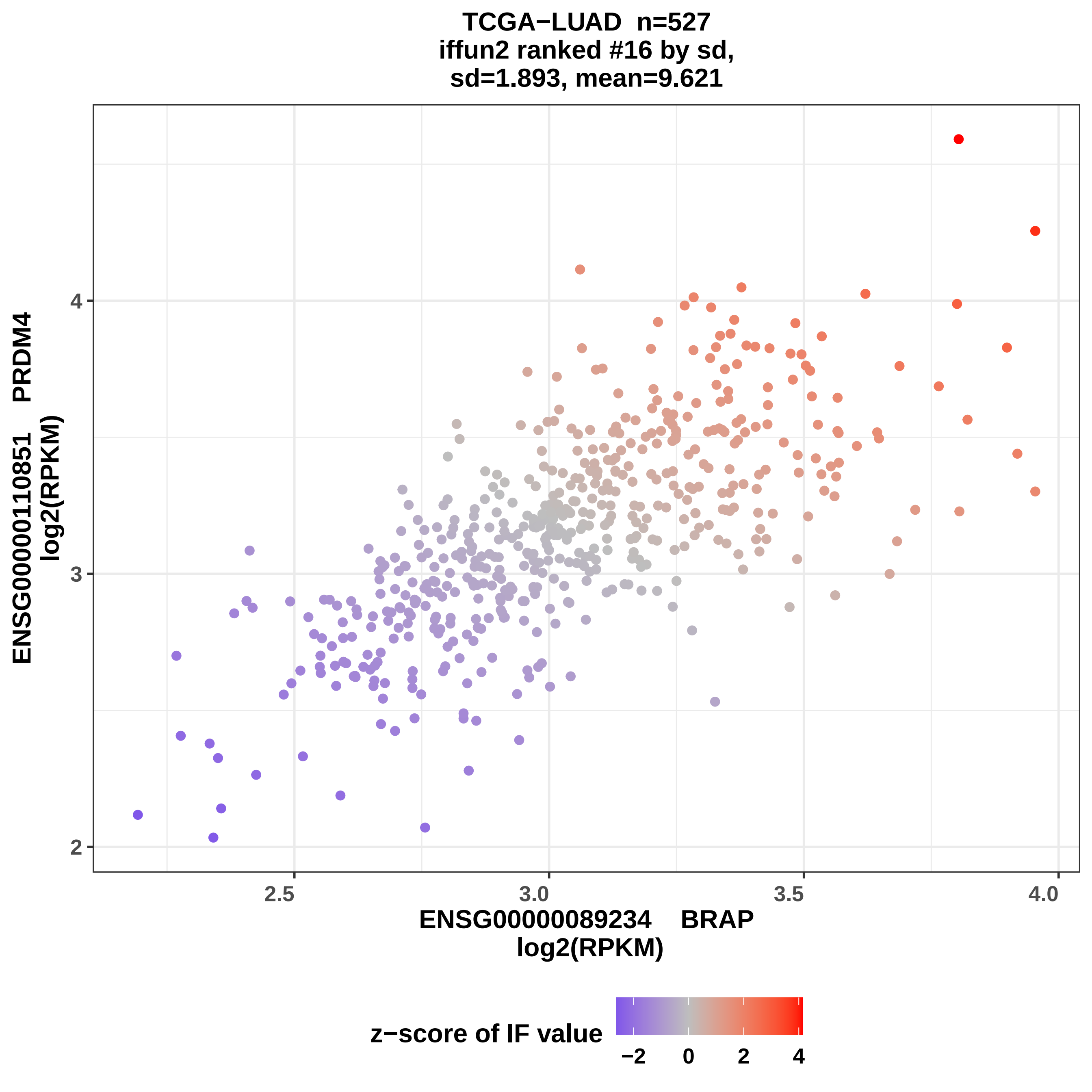

TCGA−LUAD n=527
iffun2 ranked #16 by sd,
sd=1.893, mean=9.621
ENSG00000110851 PRDM4
log2(RPKM)
4
3
2
2.5
3.0
3.5
4.0
ENSG00000089234 BRAP
log2(RPKM)
z−score of IF value
−2
0
2
4

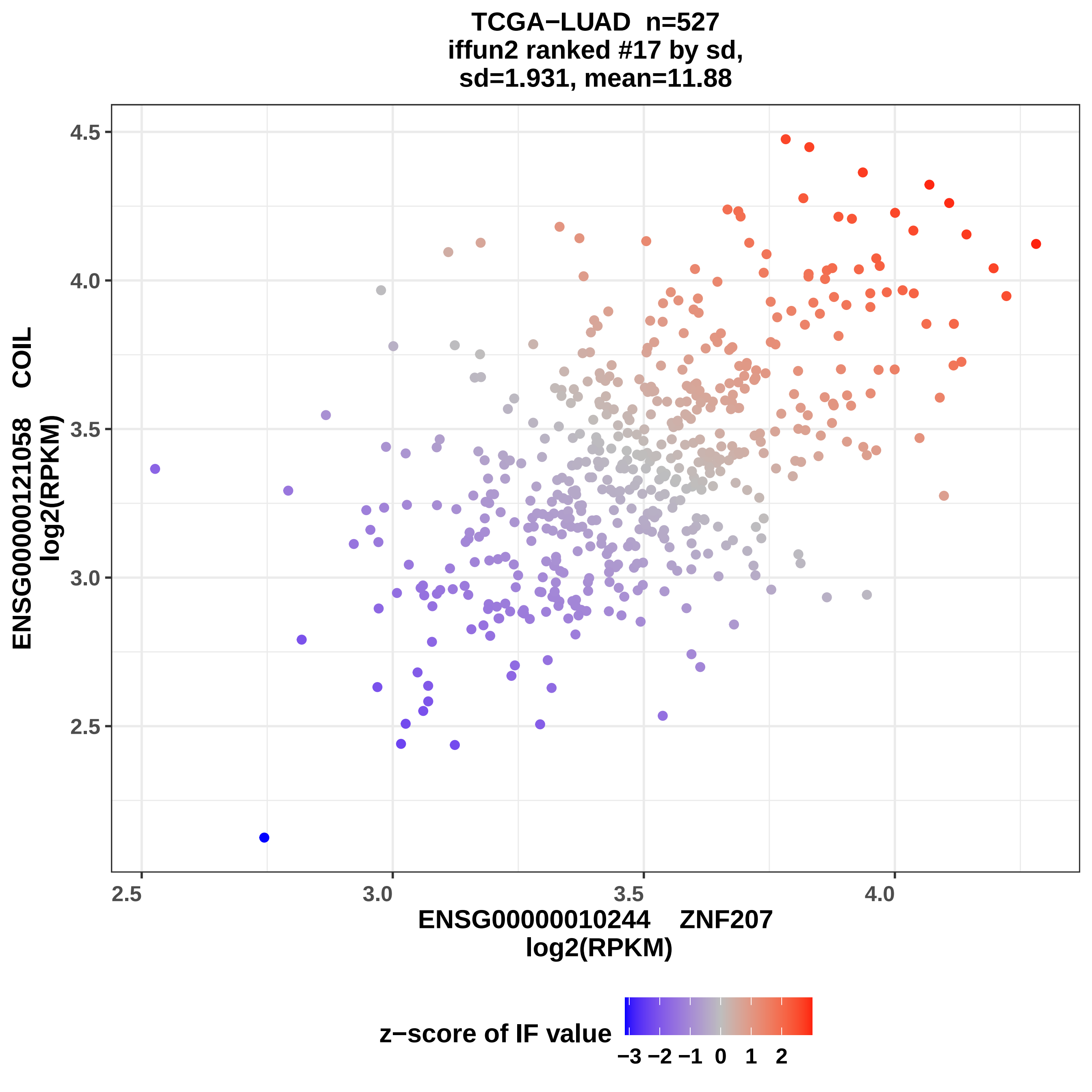
TCGA−LUAD n=527
iffun2 ranked #17 by sd,
sd=1.931, mean=11.88
ENSG00000121058 COIL
log2(RPKM)
ENSG00000010244 ZNF207
log2(RPKM)
z−score of IF value
−3 −2 −1 0 1 2
2.5
3.0
3.5
4.0
4.5

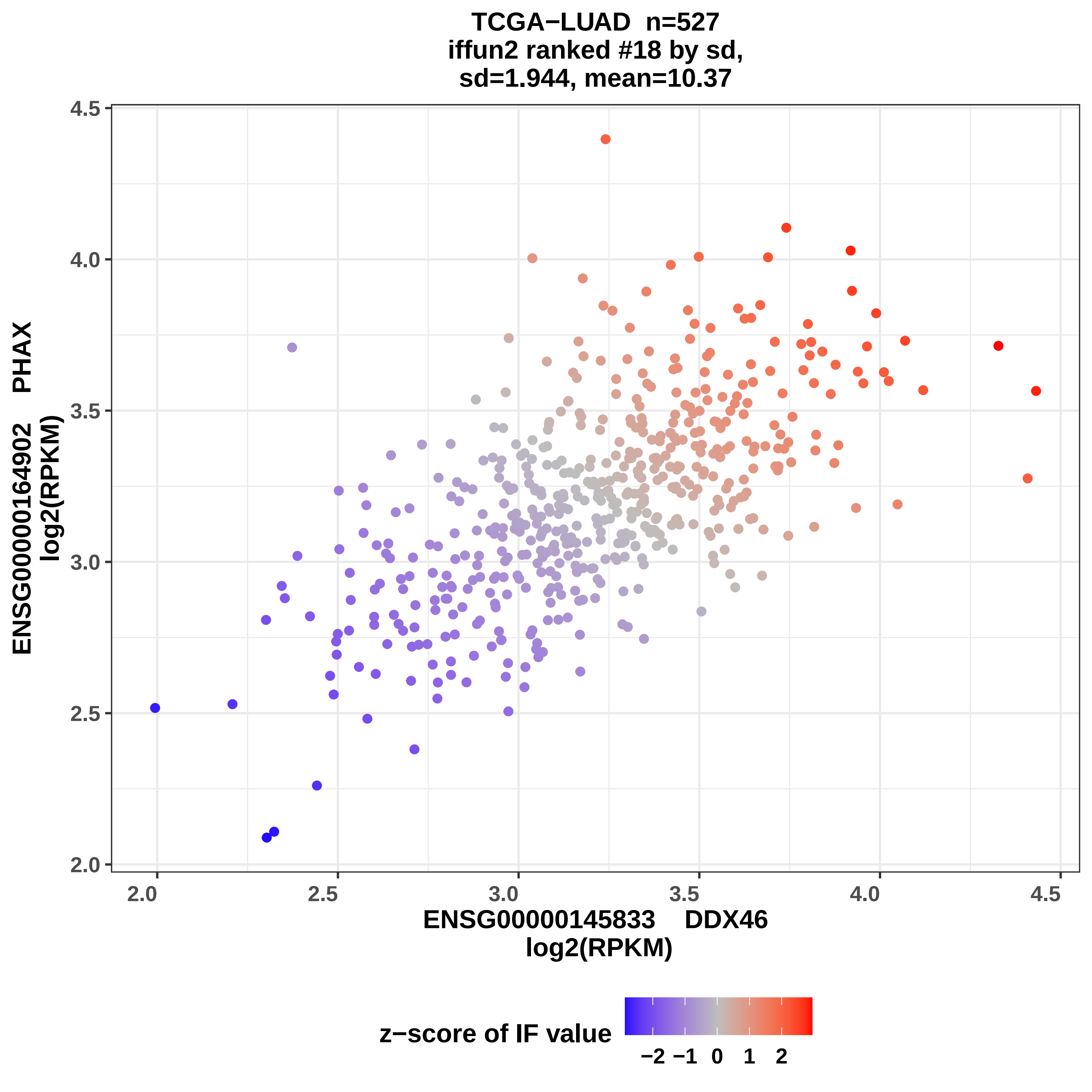

TCGA−LUAD n=527
iffun2 ranked #18 by sd,
sd=1.944, mean=10.37
ENSG00000164902 PHAX
log2(RPKM)
ENSG00000145833 DDX46
log2(RPKM)
4.5
4.0
3.5
3.0
2.5
2.0
2.0
2.5
3.0
3.5
4.0
4.5
z−score of IF value
−2 −1 0 1 2

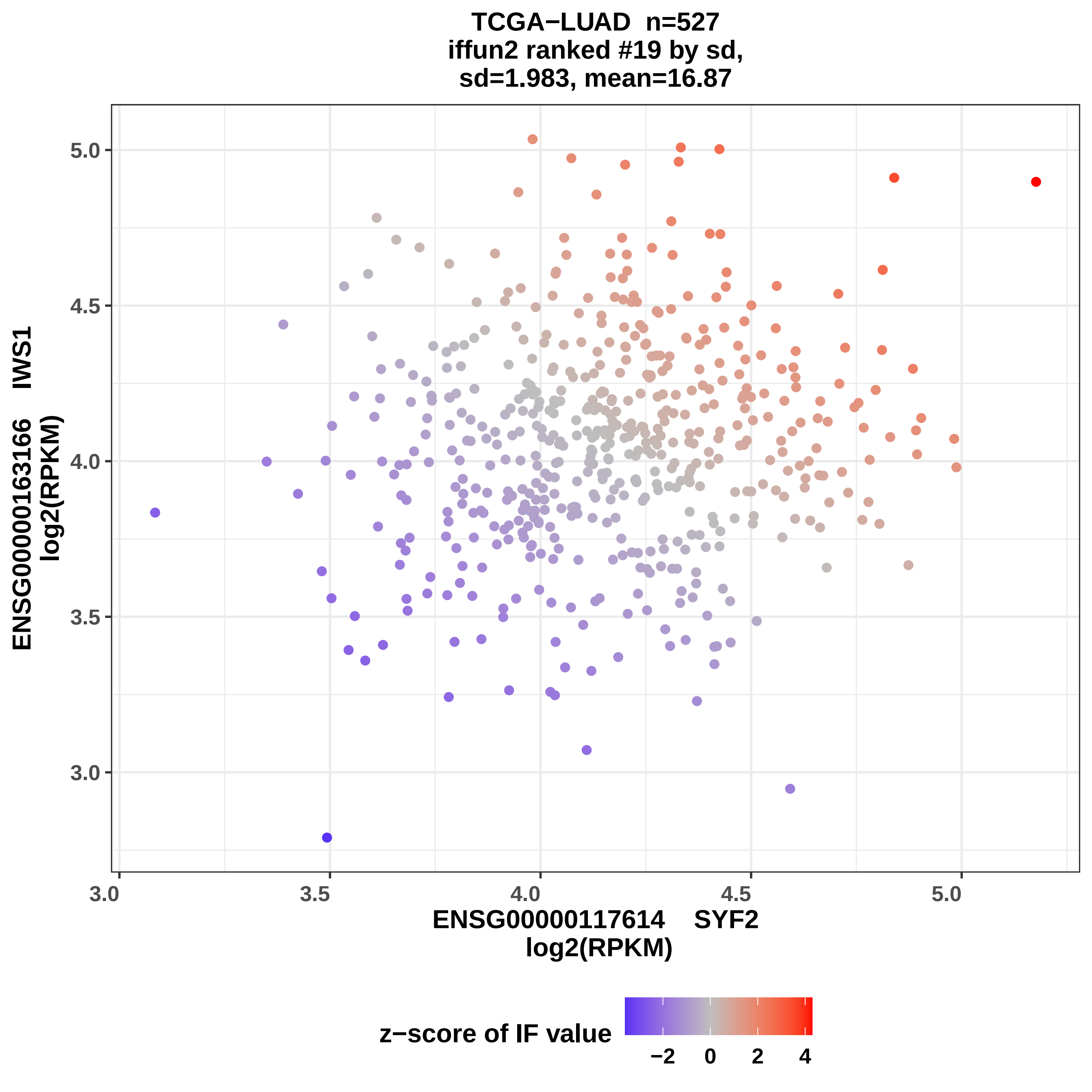
TCGA−LUAD n=527
iffun2 ranked #19 by sd,
sd=1.983, mean=16.87
ENSG00000163166 IWS1
log2(RPKM)
5.0
4.5
4.0
3.5
3.0
3.0
3.5
4.0
4.5
5.0
ENSG00000117614 SYF2
log2(RPKM)
z−score of IF value
−2
0
2
4

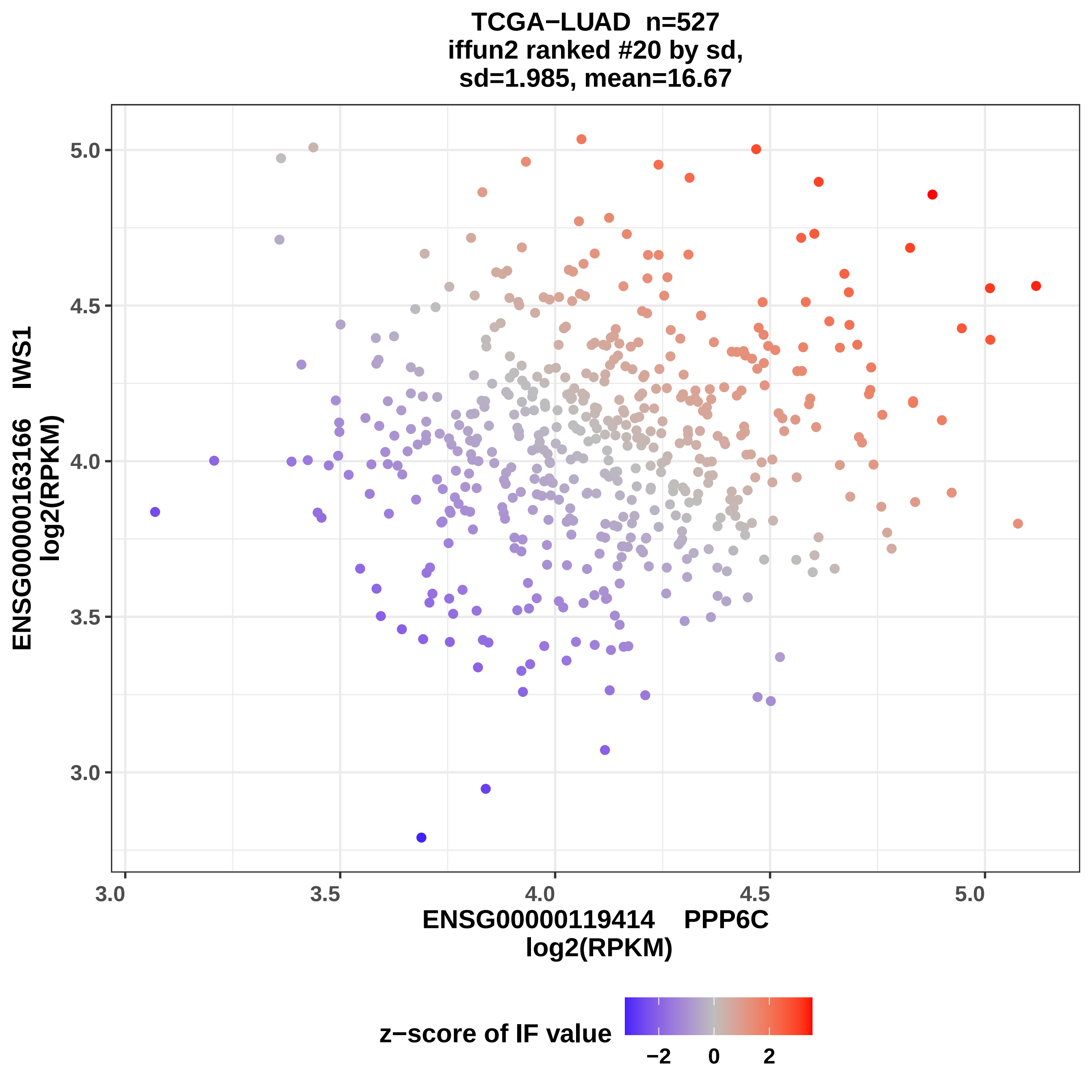
TCGA−LUAD n=527
iffun2 ranked #20 by sd,
sd=1.985, mean=16.67
ENSG00000163166 IWS1
log2(RPKM)
5.0
4.5
4.0
3.5
3.0
3.0
3.5
4.0
4.5
5.0
ENSG00000119414 PPP6C
log2(RPKM)
z−score of IF value
−2
0
2

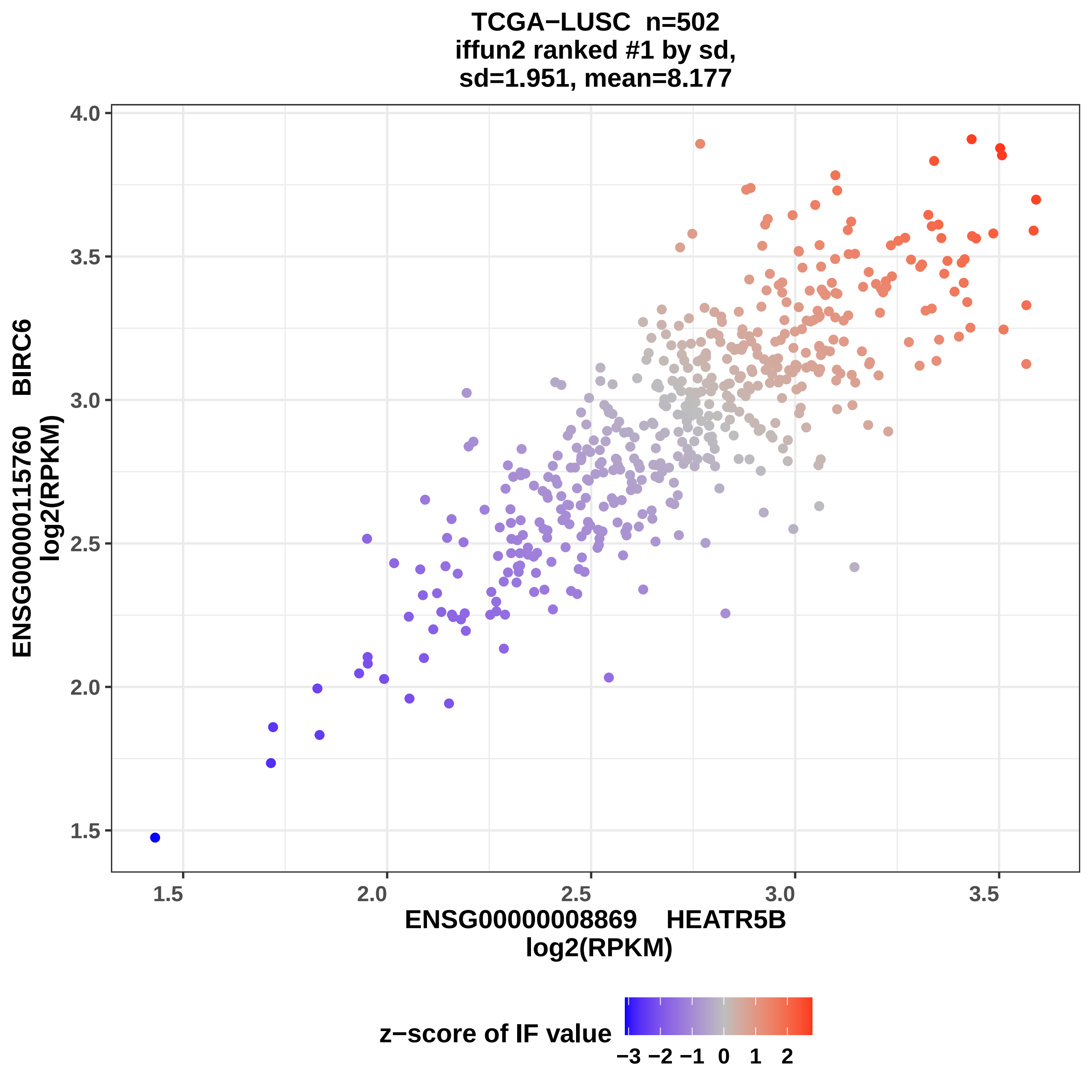
TCGA−LUSC n=502
iffun2 ranked #1 by sd,
sd=1.951, mean=8.177
ENSG00000115760 BIRC6
log2(RPKM)
4.0
3.5
3.0
2.5
2.0
1.5
1.5
2.0
2.5
3.0
3.5
ENSG00000008869 HEATR5B
log2(RPKM)
z−score of IF value
−3 −2 −1 0 1 2

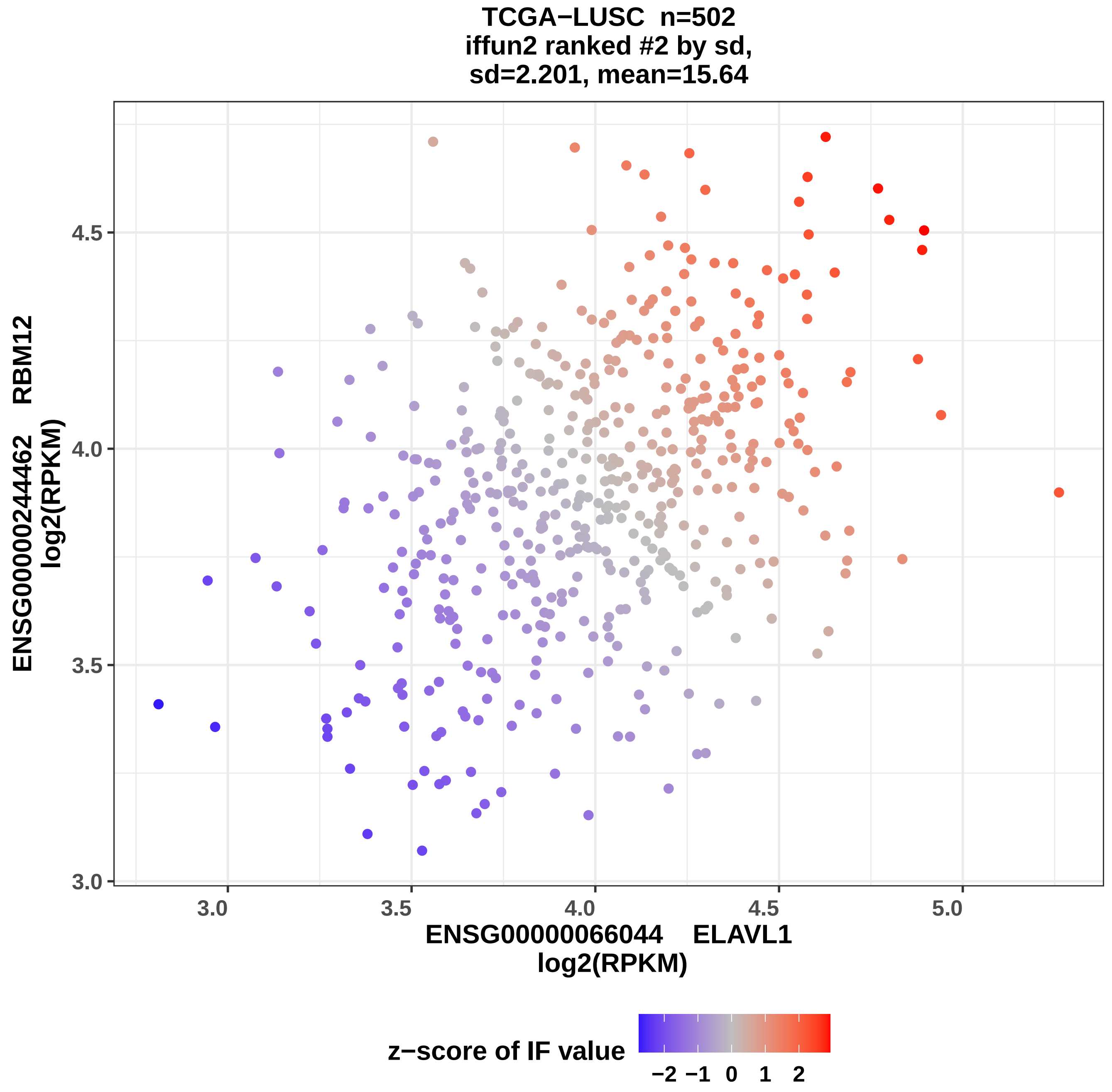
TCGA−LUSC n=502
iffun2 ranked #2 by sd,
sd=2.201, mean=15.64
ENSG00000244462 RBM12
log2(RPKM)
ENSG00000066044 ELAVL1
log2(RPKM)
3.0
3.5
4.0
4.5
5.0
z−score of IF value
−2
−1
0
1
2

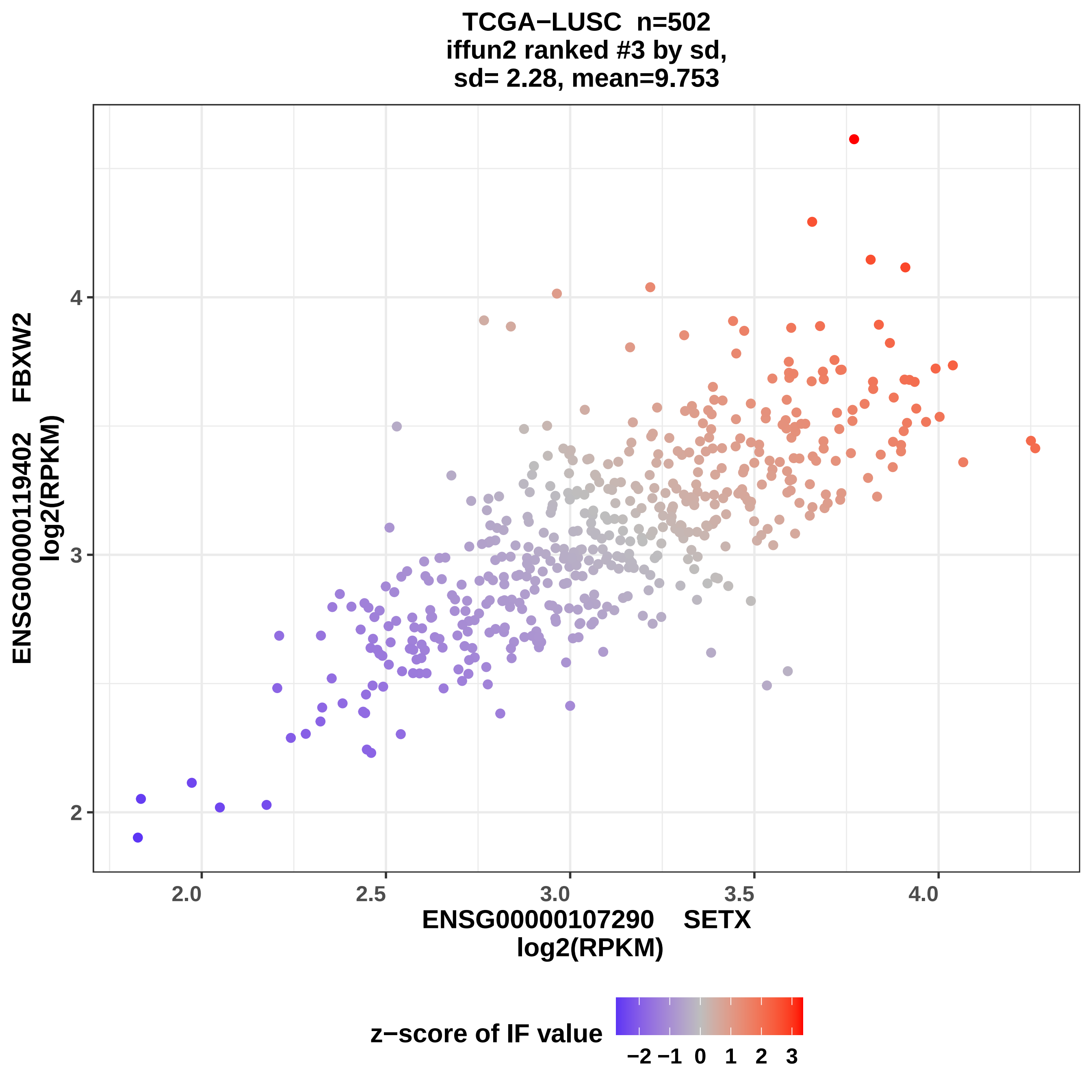
TCGA−LUSC n=502
iffun2 ranked #3 by sd,
sd= 2.28, mean=9.753
ENSG00000119402 FBXW2
log2(RPKM)
2
3
4
2.0
2.5
3.0
3.5
4.0
ENSG00000107290 SETX
log2(RPKM)
z−score of IF value
−2 −1 0 1 2 3

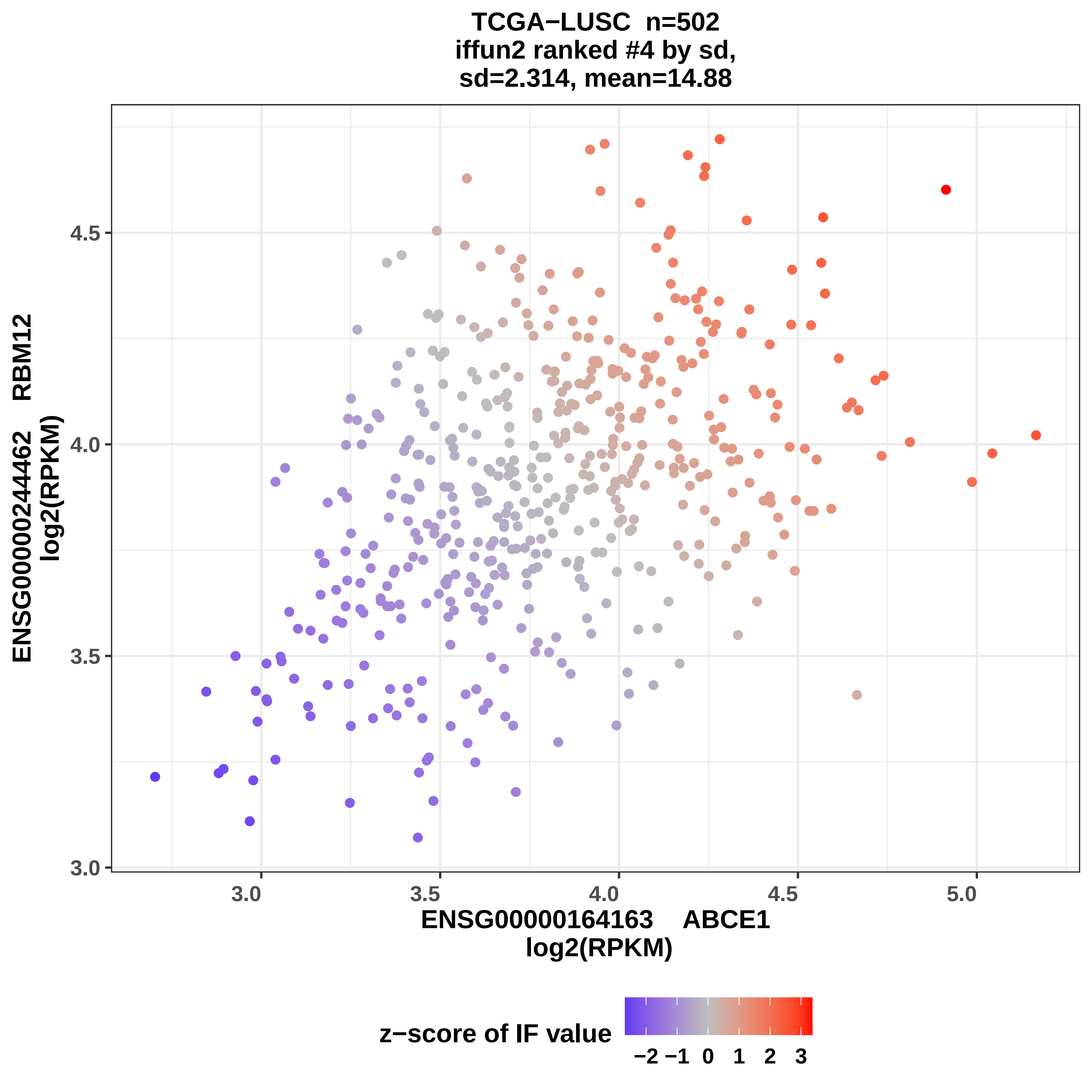

TCGA−LUSC n=502
iffun2 ranked #4 by sd,
sd=2.314, mean=14.88
ENSG00000244462 RBM12
log2(RPKM)
4.5
4.0
3.5
3.0
3.0
3.5
4.0
4.5
5.0
ENSG00000164163 ABCE1
log2(RPKM)
z−score of IF value
−2 −1 0 1 2 3

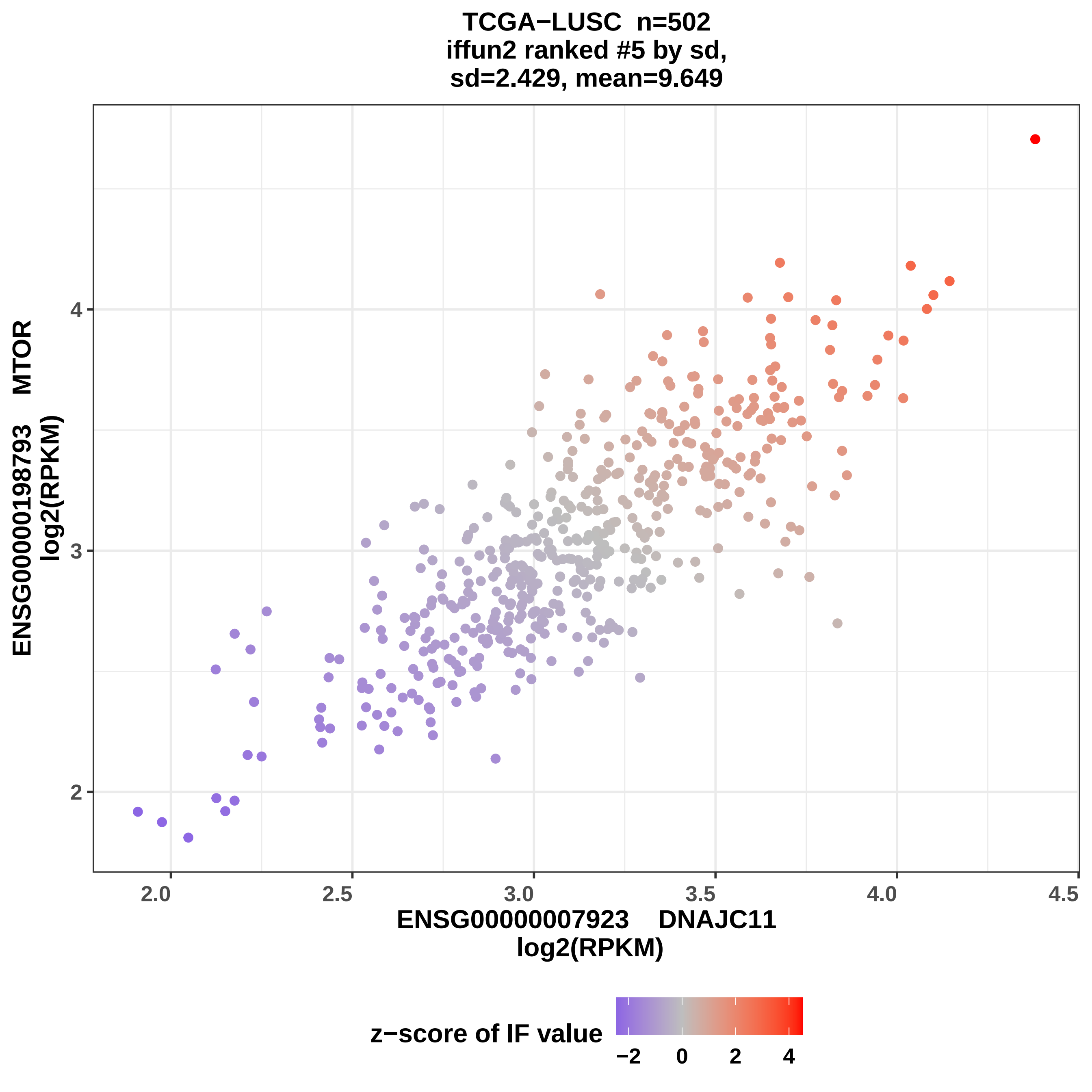

TCGA−LUSC n=502
iffun2 ranked #5 by sd,
sd=2.429, mean=9.649
ENSG00000198793 MTOR
log2(RPKM)
ENSG00000007923 DNAJC11
log2(RPKM)
z−score of IF value
2
3
4
2.0
2.5
3.0
3.5
4.0
4.5
−2
0
2
4

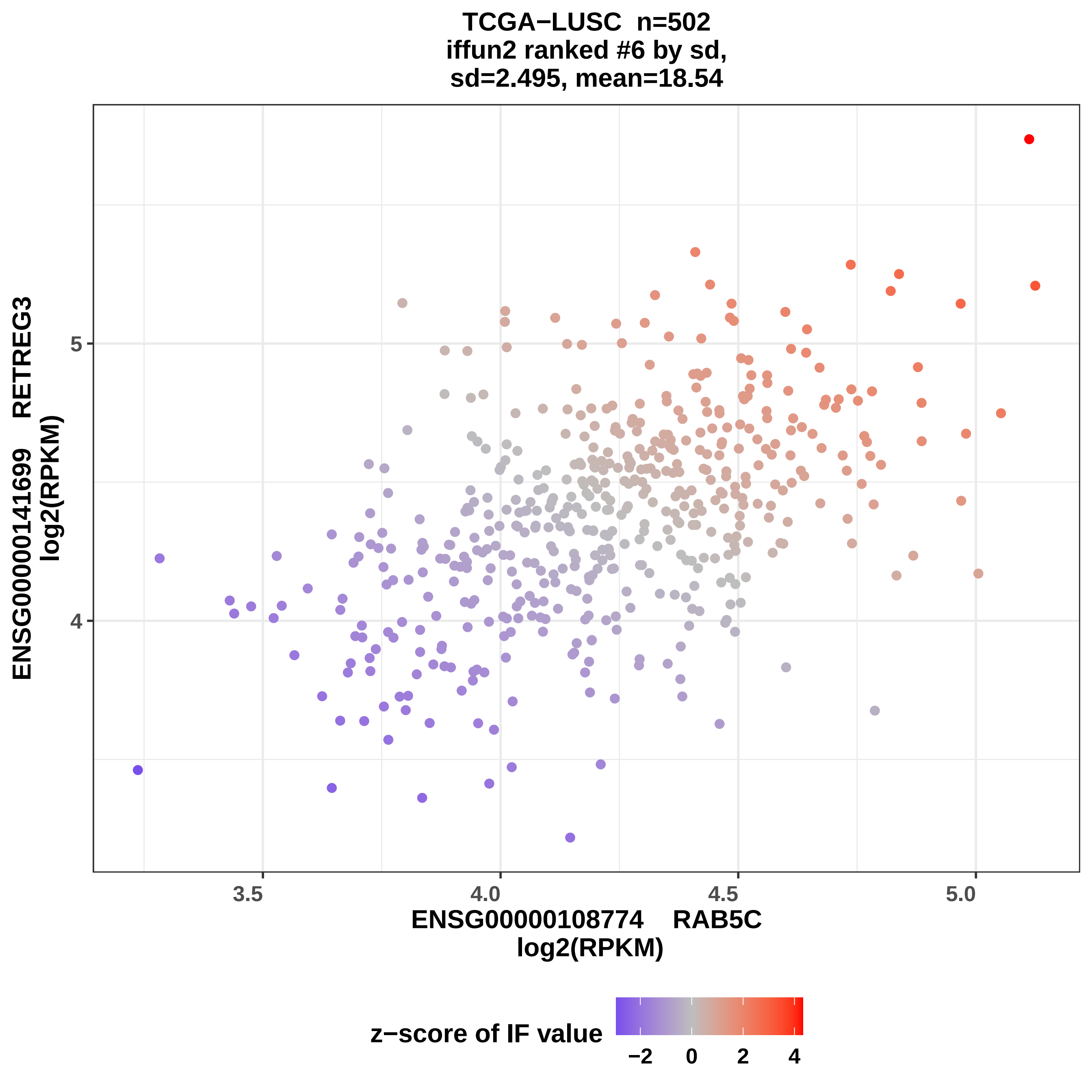
TCGA−LUSC n=502
iffun2 ranked #6 by sd,
sd=2.495, mean=18.54
ENSG00000141699 RETREG3
log2(RPKM)
5
4
3.5
4.0
4.5
5.0
ENSG00000108774 RAB5C
log2(RPKM)
z−score of IF value
−2
0
2
4

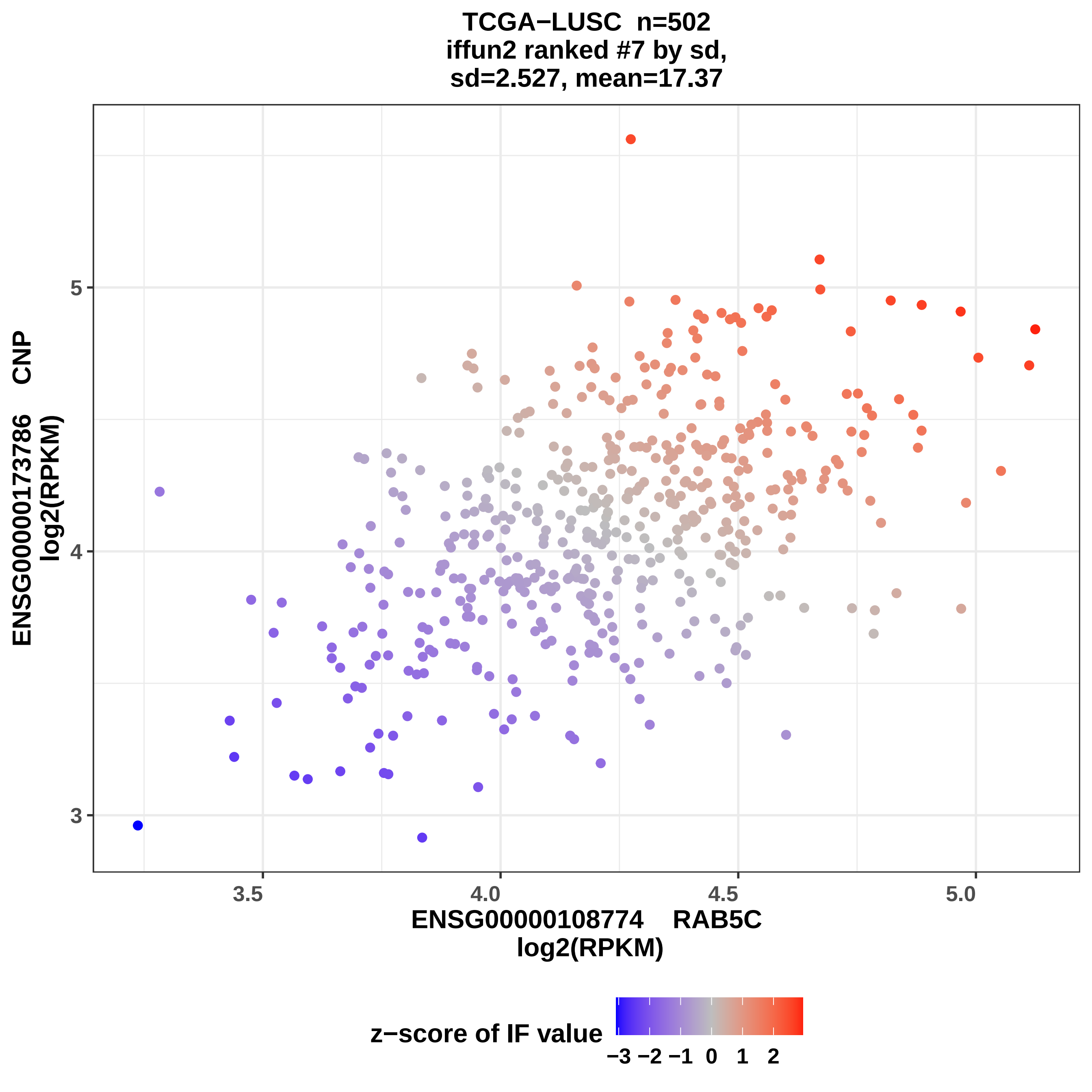
TCGA−LUSC n=502
iffun2 ranked #7 by sd,
sd=2.527, mean=17.37
ENSG00000173786 CNP
log2(RPKM)
5
4
3
3.5
4.0
4.5
5.0
ENSG00000108774 RAB5C
log2(RPKM)
z−score of IF value
−3 −2 −1 0 1 2

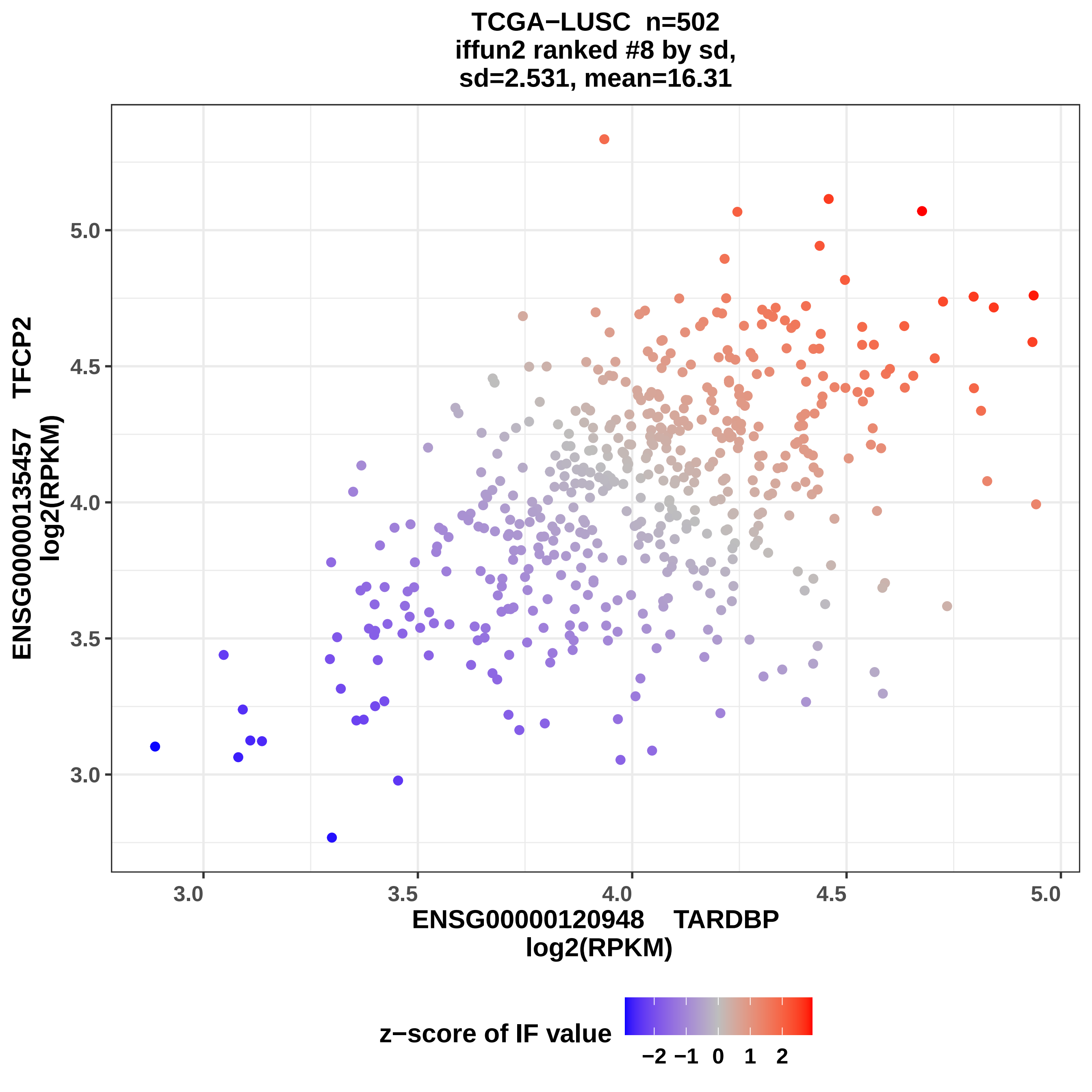
TCGA−LUSC n=502
iffun2 ranked #8 by sd,
sd=2.531, mean=16.31
ENSG00000135457 TFCP2
log2(RPKM)
5.0
4.5
4.0
3.5
3.0
3.0
3.5
4.0
4.5
5.0
ENSG00000120948 TARDBP
log2(RPKM)
z−score of IF value
−2 −1 0 1 2

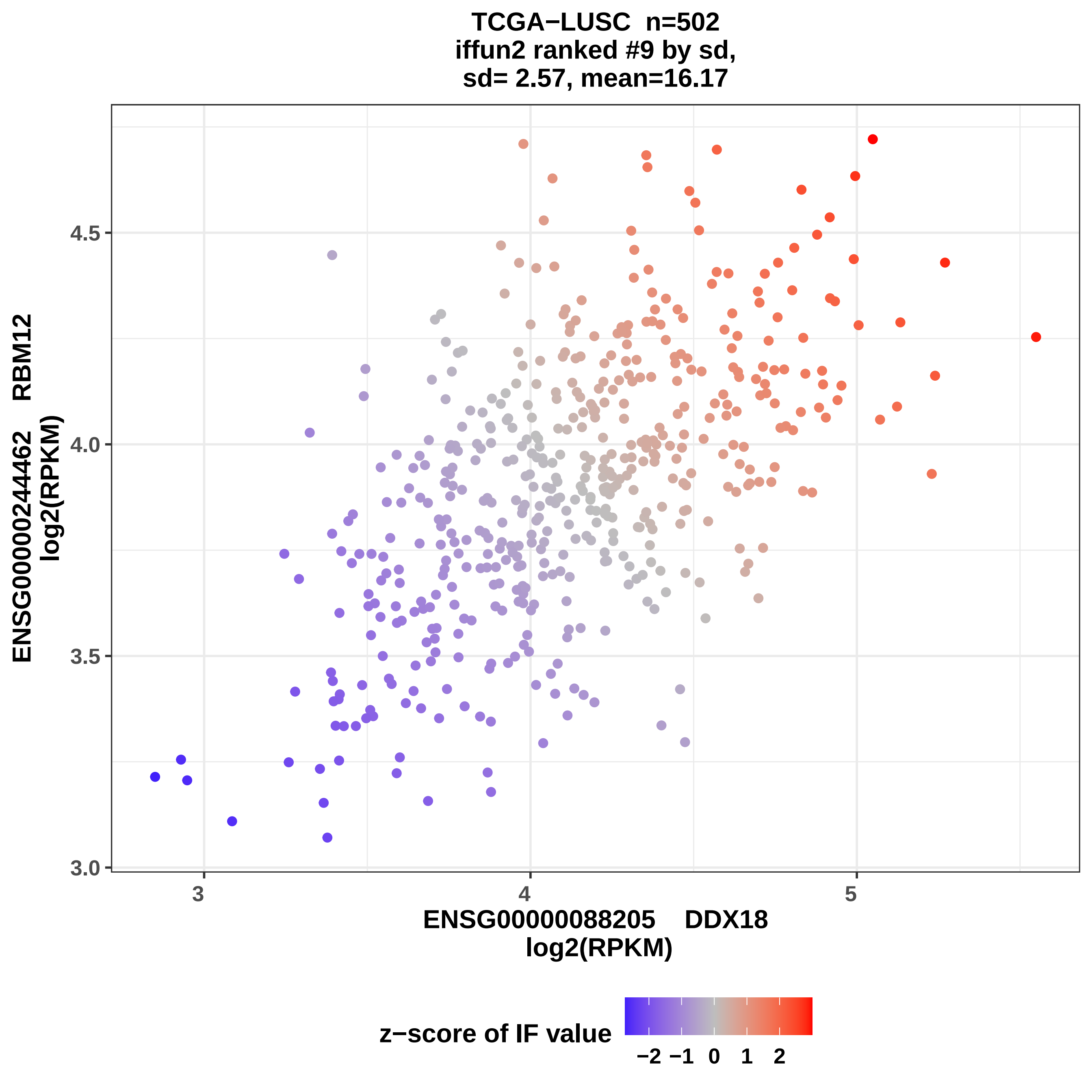

TCGA−LUSC n=502
iffun2 ranked #9 by sd,
sd= 2.57, mean=16.17
ENSG00000244462 RBM12
log2(RPKM)
3.0
3.5
4.0
4.5
3
4
5
ENSG00000088205 DDX18
log2(RPKM)
z−score of IF value
−2 −1 0 1 2

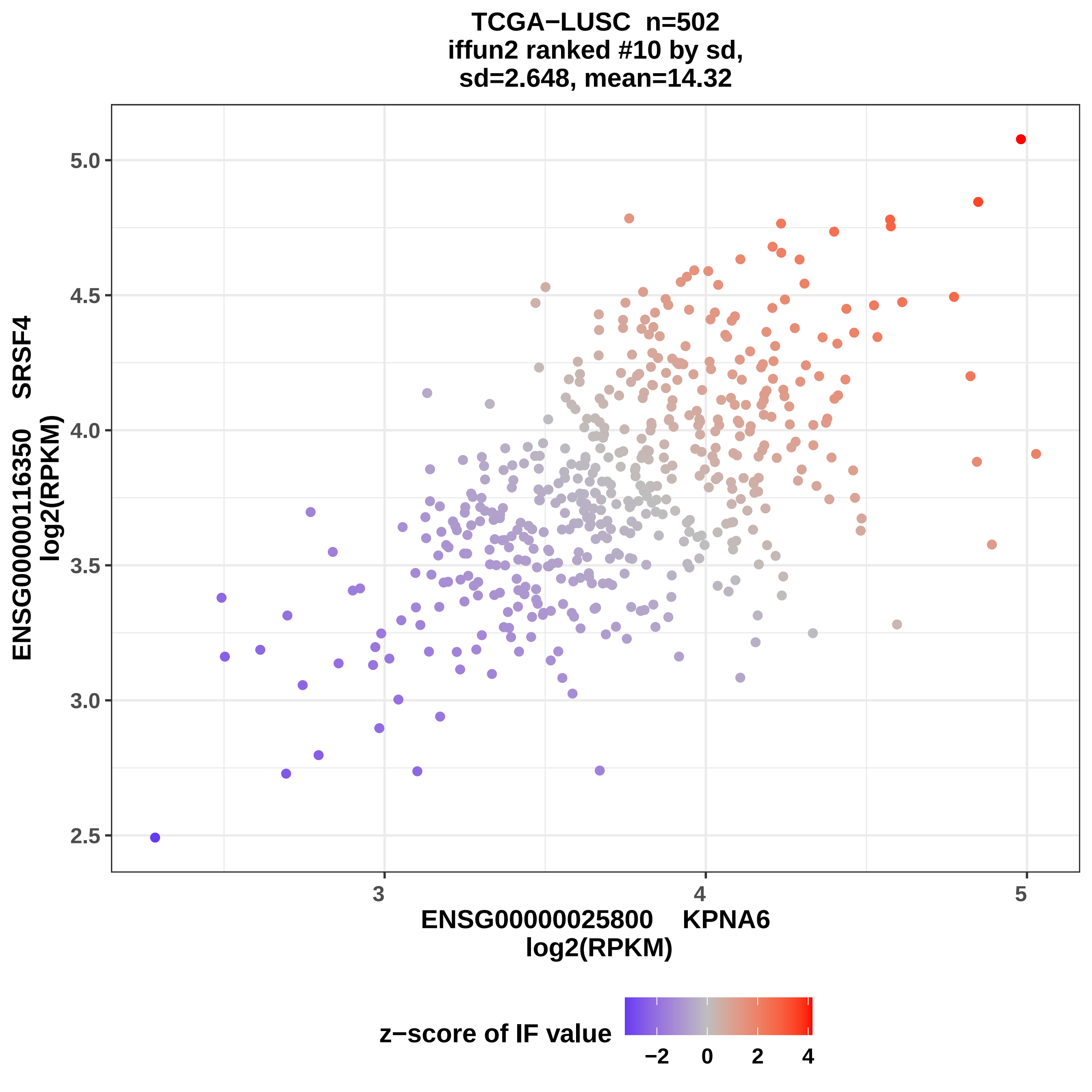

TCGA−LUSC n=502
iffun2 ranked #10 by sd,
sd=2.648, mean=14.32
ENSG00000116350 SRSF4
log2(RPKM)
5.0
4.5
4.0
3.5
3.0
2.5
3
4
5
ENSG00000025800 KPNA6
log2(RPKM)
z−score of IF value
−2
0
2
4

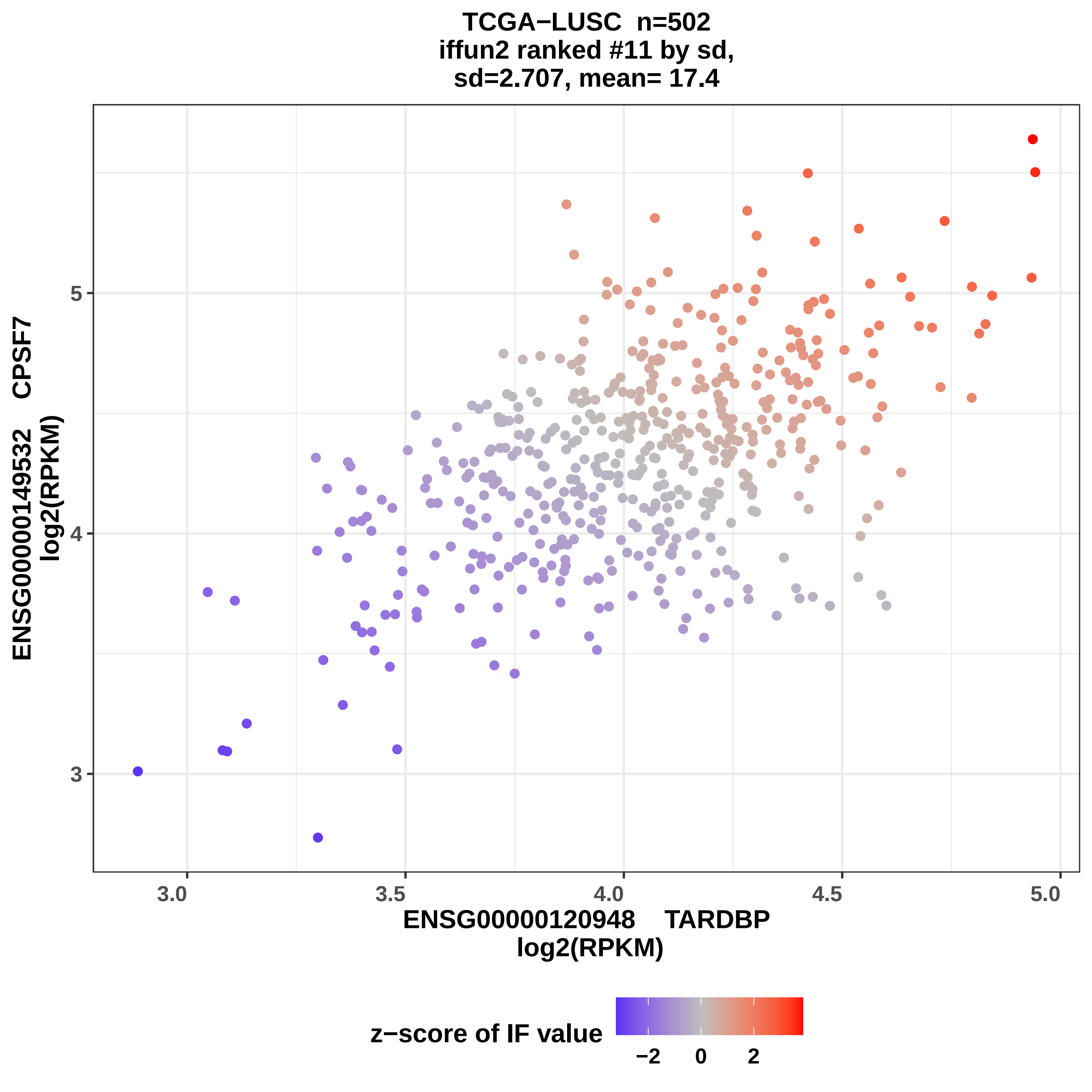
TCGA−LUSC n=502
iffun2 ranked #11 by sd,
sd=2.707, mean= 17.4
ENSG00000149532 CPSF7
log2(RPKM)
3
4
5
3.0
3.5
4.0
4.5
5.0
ENSG00000120948 TARDBP
log2(RPKM)
z−score of IF value
−2
0
2

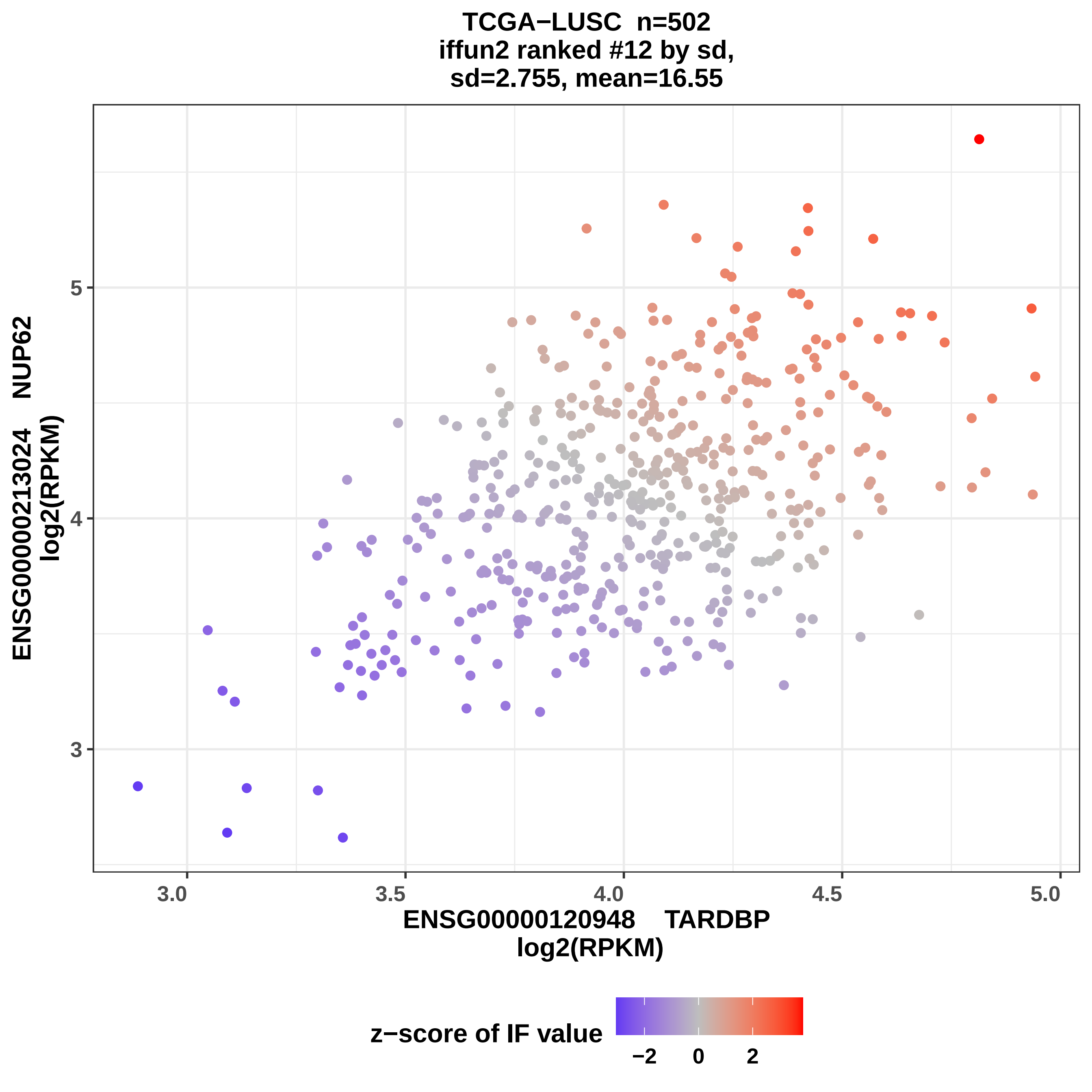
TCGA−LUSC n=502
iffun2 ranked #12 by sd,
sd=2.755, mean=16.55
ENSG00000213024 NUP62
log2(RPKM)
5
4
3
3.0
3.5
4.0
4.5
5.0
ENSG00000120948 TARDBP
log2(RPKM)
z−score of IF value
−2
0
2

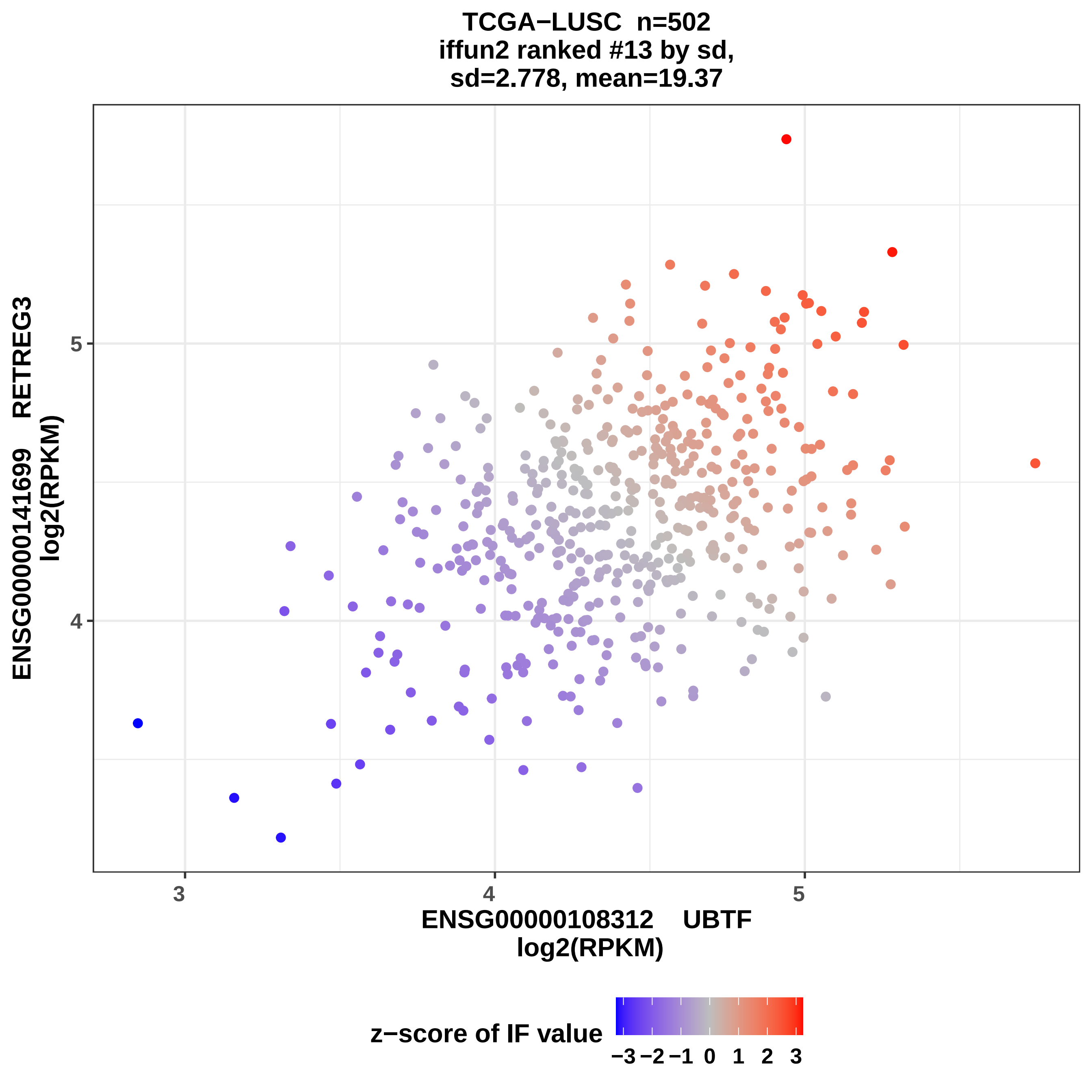
TCGA−LUSC n=502
iffun2 ranked #13 by sd,
sd=2.778, mean=19.37
ENSG00000141699 RETREG3
log2(RPKM)
ENSG00000108312 UBTF
log2(RPKM)
z−score of IF value
−3 −2 −1 0 1 2 3
3
4
5
4
5

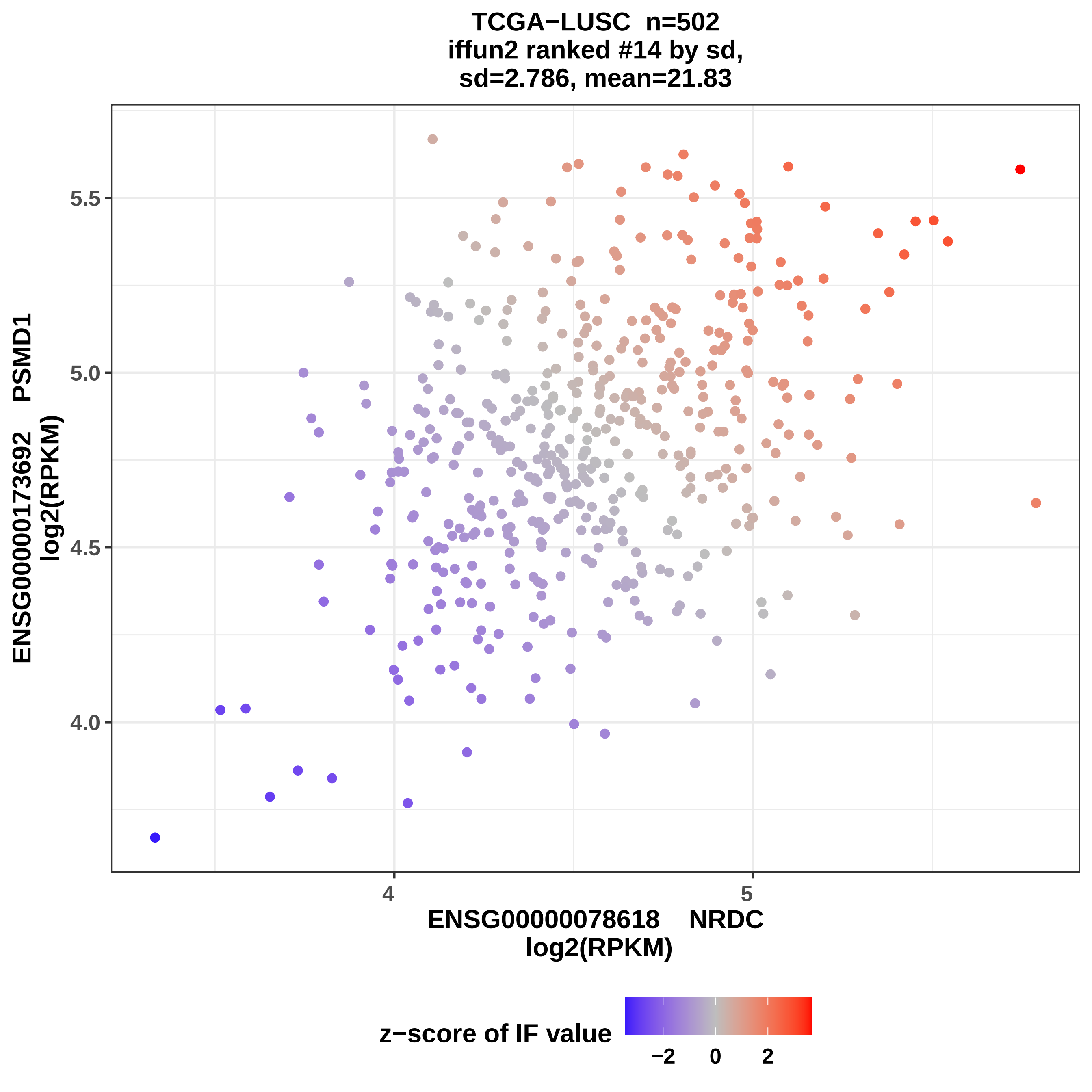
TCGA−LUSC n=502
iffun2 ranked #14 by sd,
sd=2.786, mean=21.83
ENSG00000173692 PSMD1
log2(RPKM)
5.5
5.0
4.5
4.0
4
5
ENSG00000078618 NRDC
log2(RPKM)
z−score of IF value
−2
0
2

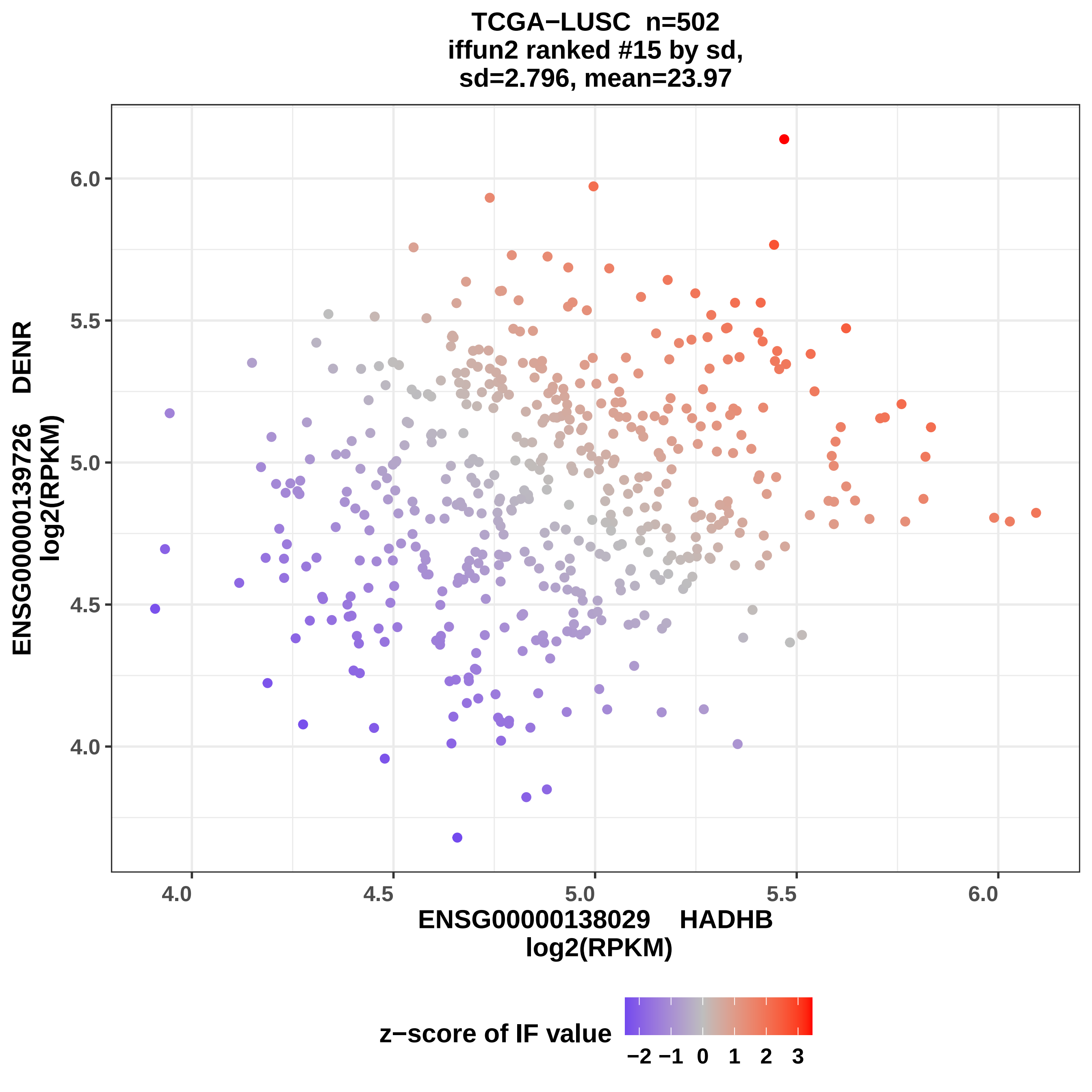

TCGA−LUSC n=502
iffun2 ranked #15 by sd,
sd=2.796, mean=23.97
ENSG00000139726 DENR
log2(RPKM)
6.0
5.5
5.0
4.5
4.0
4.0
4.5
5.0
5.5
6.0
ENSG00000138029 HADHB
log2(RPKM)
z−score of IF value
−2 −1 0 1 2 3

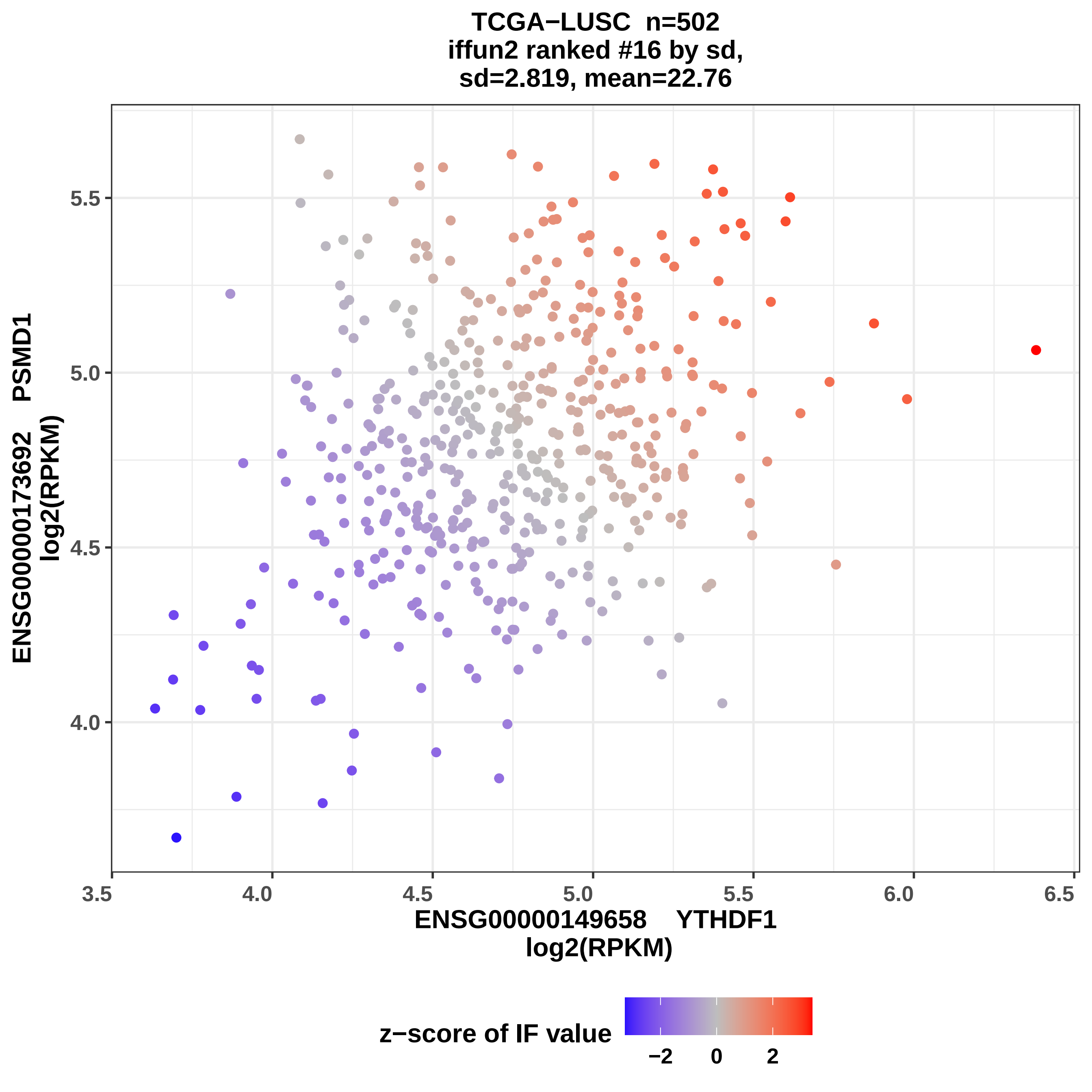

TCGA−LUSC n=502
iffun2 ranked #16 by sd,
sd=2.819, mean=22.76
ENSG00000173692 PSMD1
log2(RPKM)
5.5
5.0
4.5
4.0
3.5
4.0
4.5
5.0
5.5
6.0
6.5
ENSG00000149658 YTHDF1
log2(RPKM)
z−score of IF value
−2
0
2

TCGA−LUSC n=502
iffun2 ranked #17 by sd,
sd= 2.83, mean=20.83

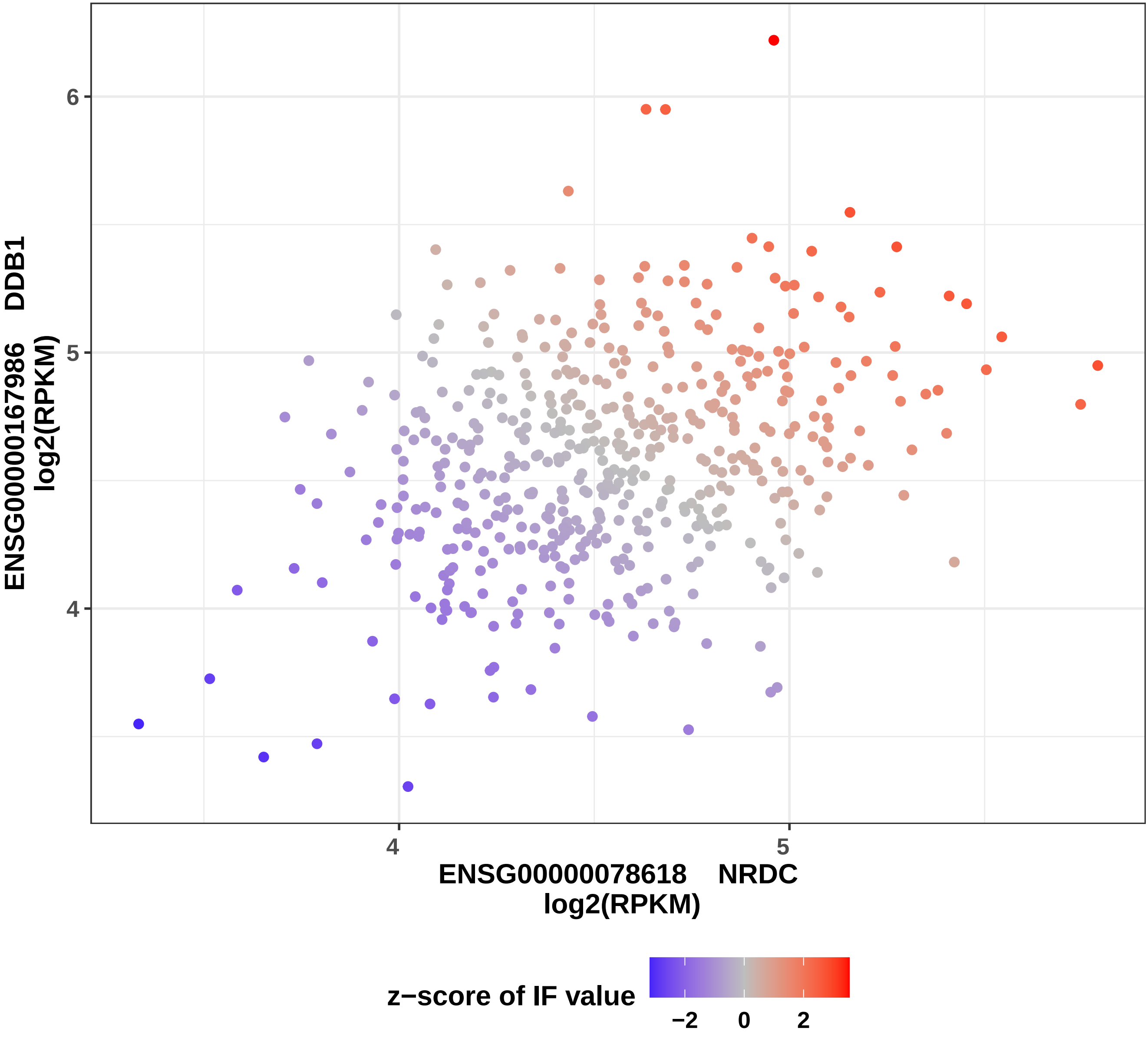

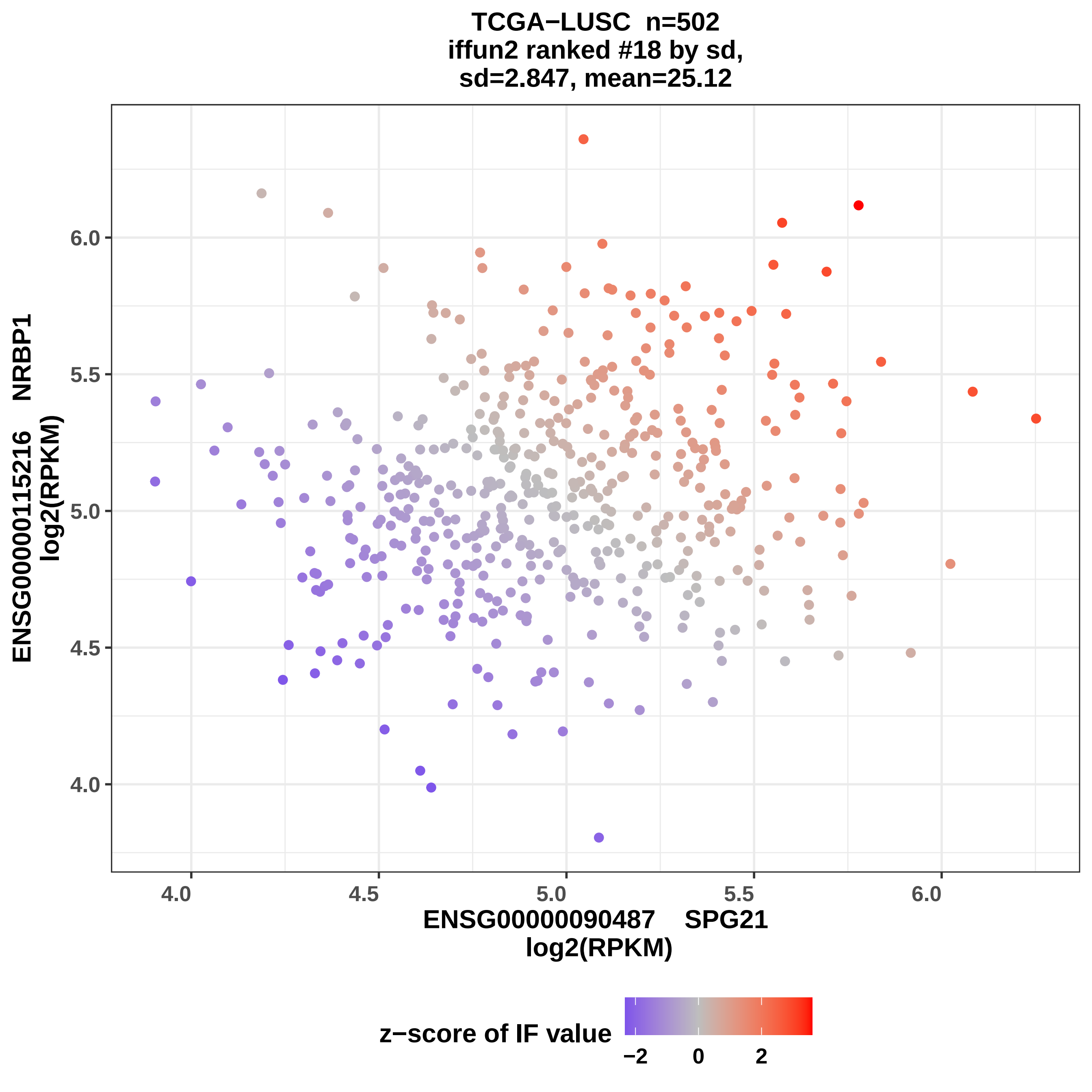
TCGA−LUSC n=502
iffun2 ranked #18 by sd,
sd=2.847, mean=25.12
ENSG00000115216 NRBP1
log2(RPKM)
4.0
4.5
5.0
5.5
6.0
ENSG00000090487 SPG21
log2(RPKM)
z−score of IF value
−2
0
2

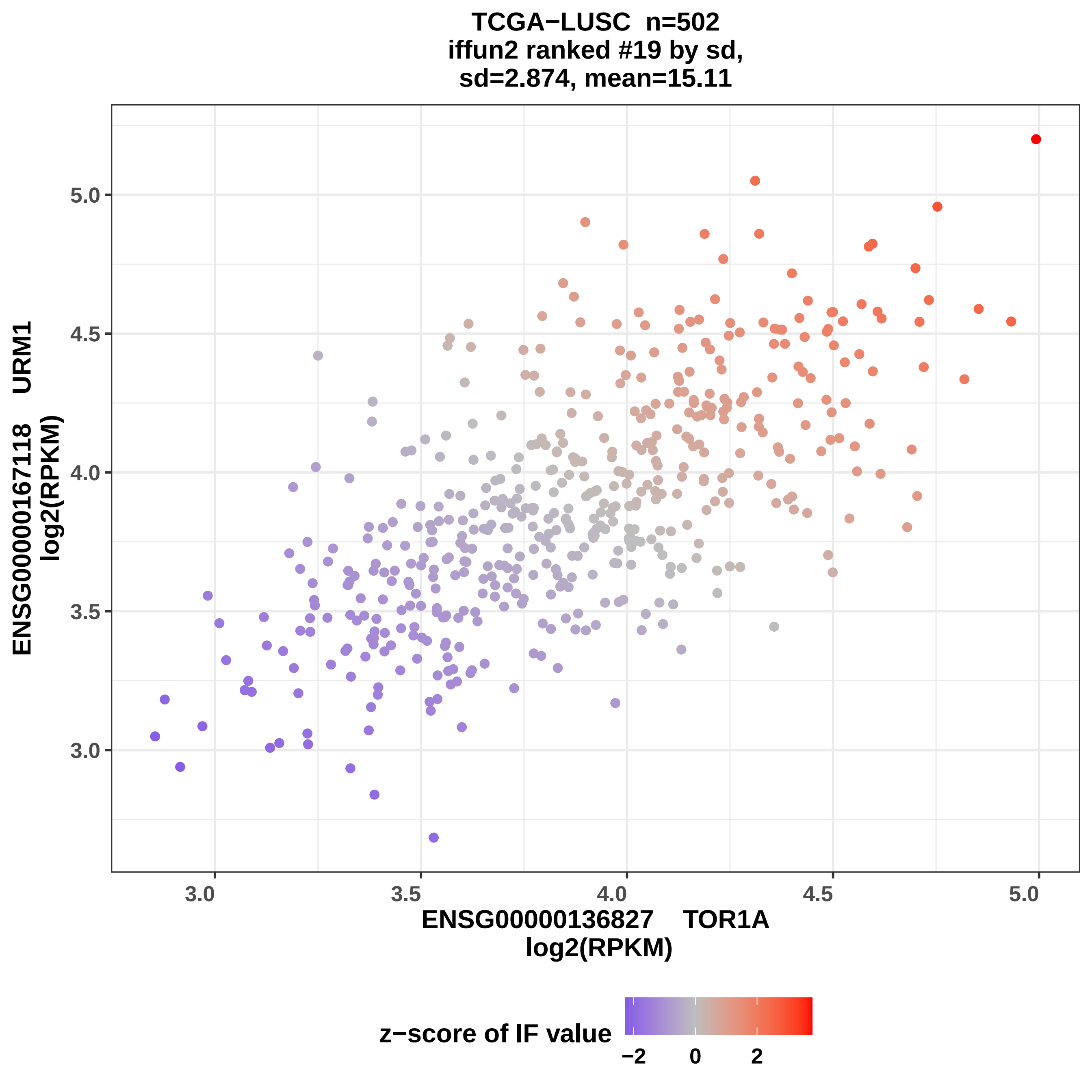
TCGA−LUSC n=502
iffun2 ranked #19 by sd,
sd=2.874, mean=15.11
ENSG00000167118 URM1
log2(RPKM)
ENSG00000136827 TOR1A
log2(RPKM)
3.0
3.5
4.0
4.5
5.0
z−score of IF value
−2
0
2

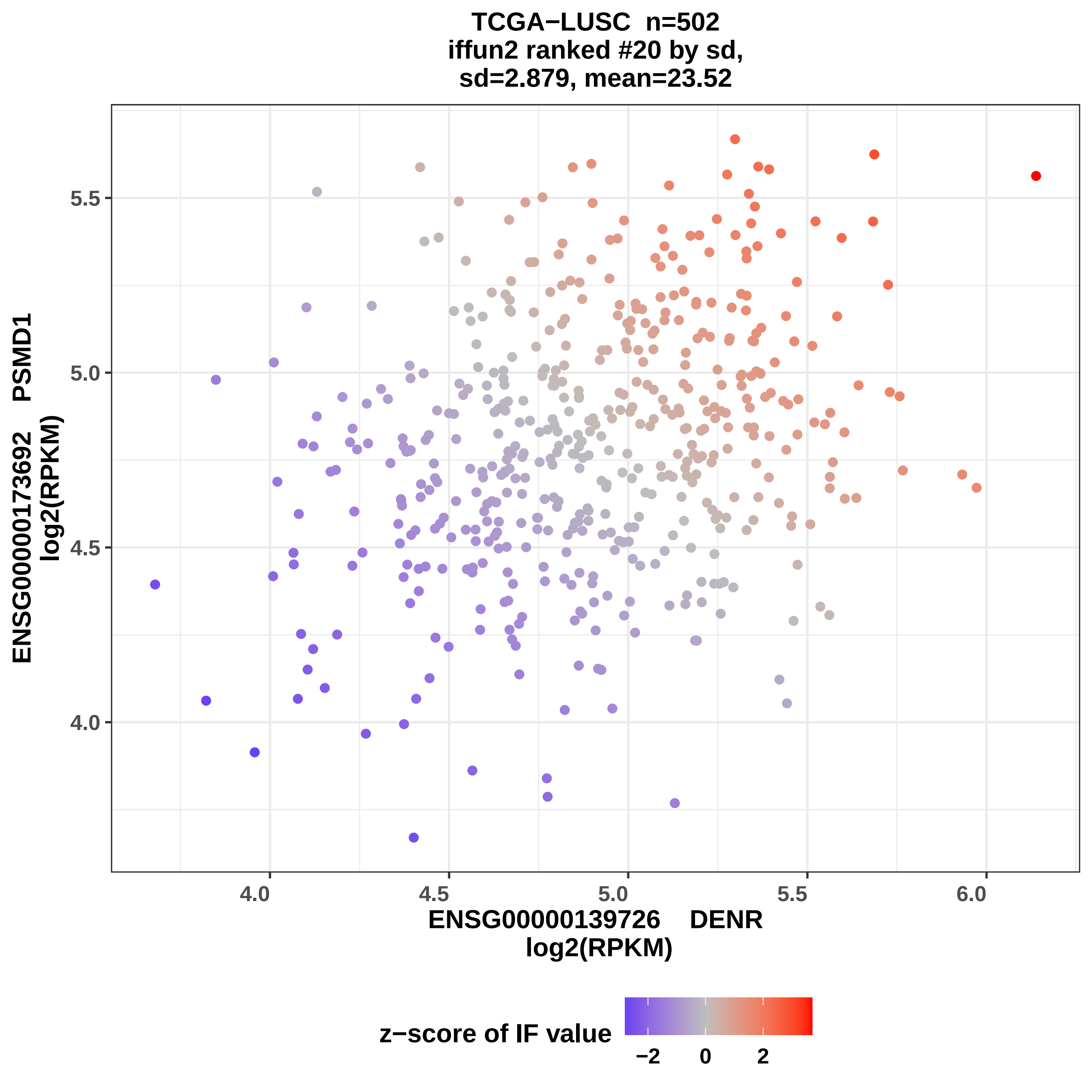
TCGA−LUSC n=502
iffun2 ranked #20 by sd,
sd=2.879, mean=23.52
ENSG00000173692 PSMD1
log2(RPKM)
5.5
5.0
4.5
4.0
4.0
4.5
5.0
5.5
6.0
ENSG00000139726 DENR
log2(RPKM)
z−score of IF value
−2
0
2

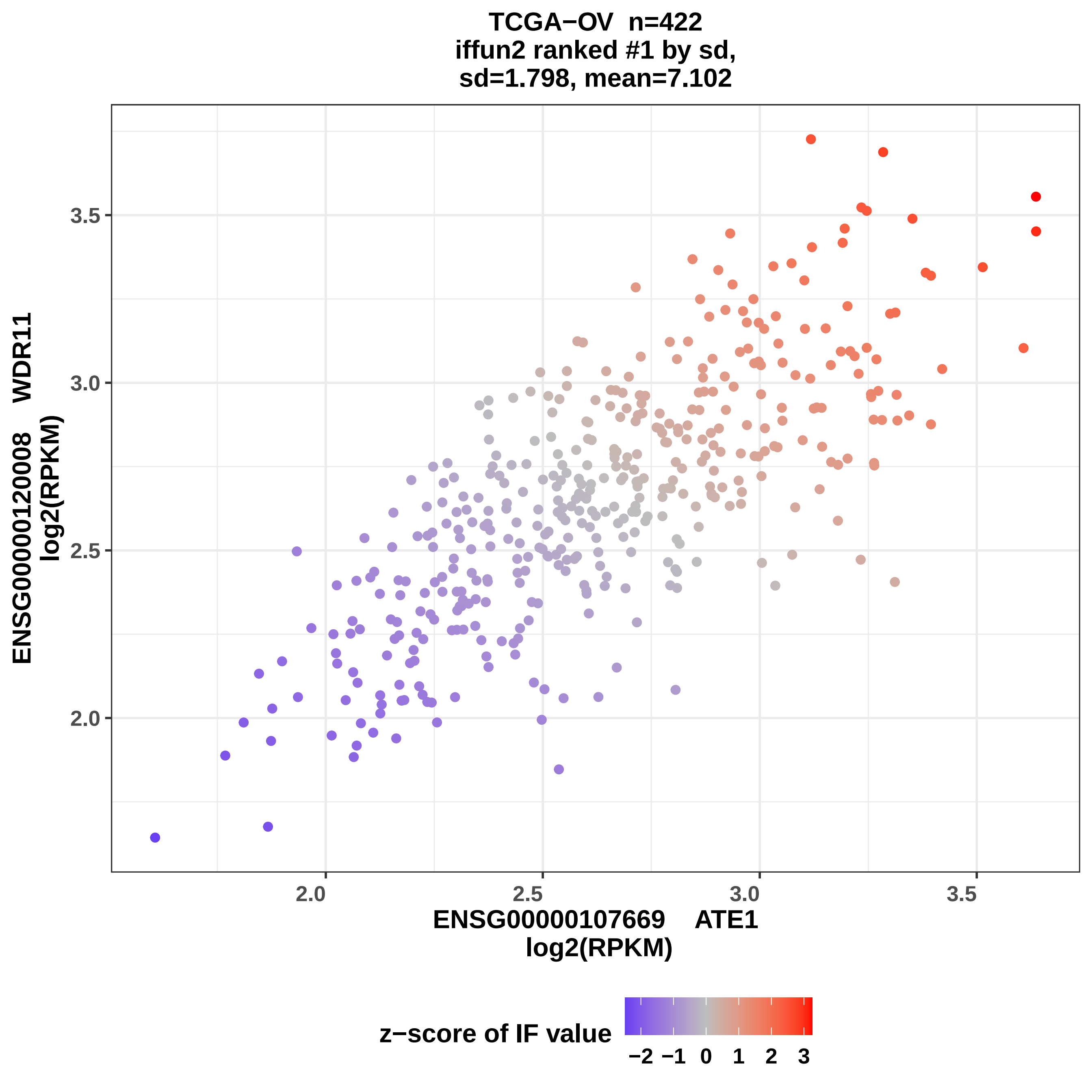
TCGA−OV n=422
iffun2 ranked #1 by sd,
sd=1.798, mean=7.102
ENSG00000120008 WDR11
log2(RPKM)
3.5
3.0
2.5
2.0
2.0
2.5
3.0
3.5
ENSG00000107669 ATE1
log2(RPKM)
z−score of IF value
−2 −1 0 1 2 3

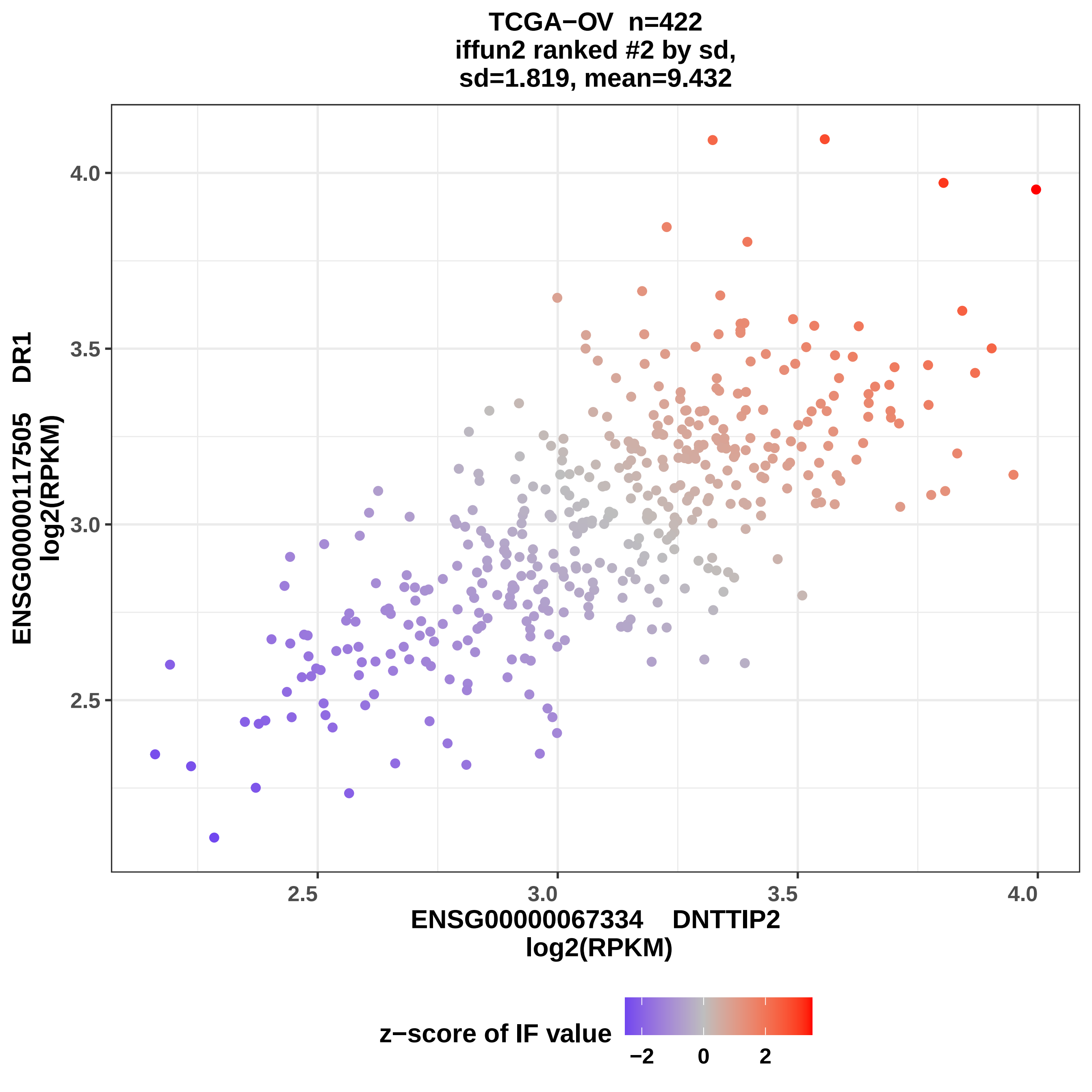

TCGA−OV n=422
iffun2 ranked #2 by sd,
sd=1.819, mean=9.432
ENSG00000117505 DR1
log2(RPKM)
ENSG00000067334 DNTTIP2
log2(RPKM)
2.5
3.0
3.5
4.0
z−score of IF value
−2
0
2

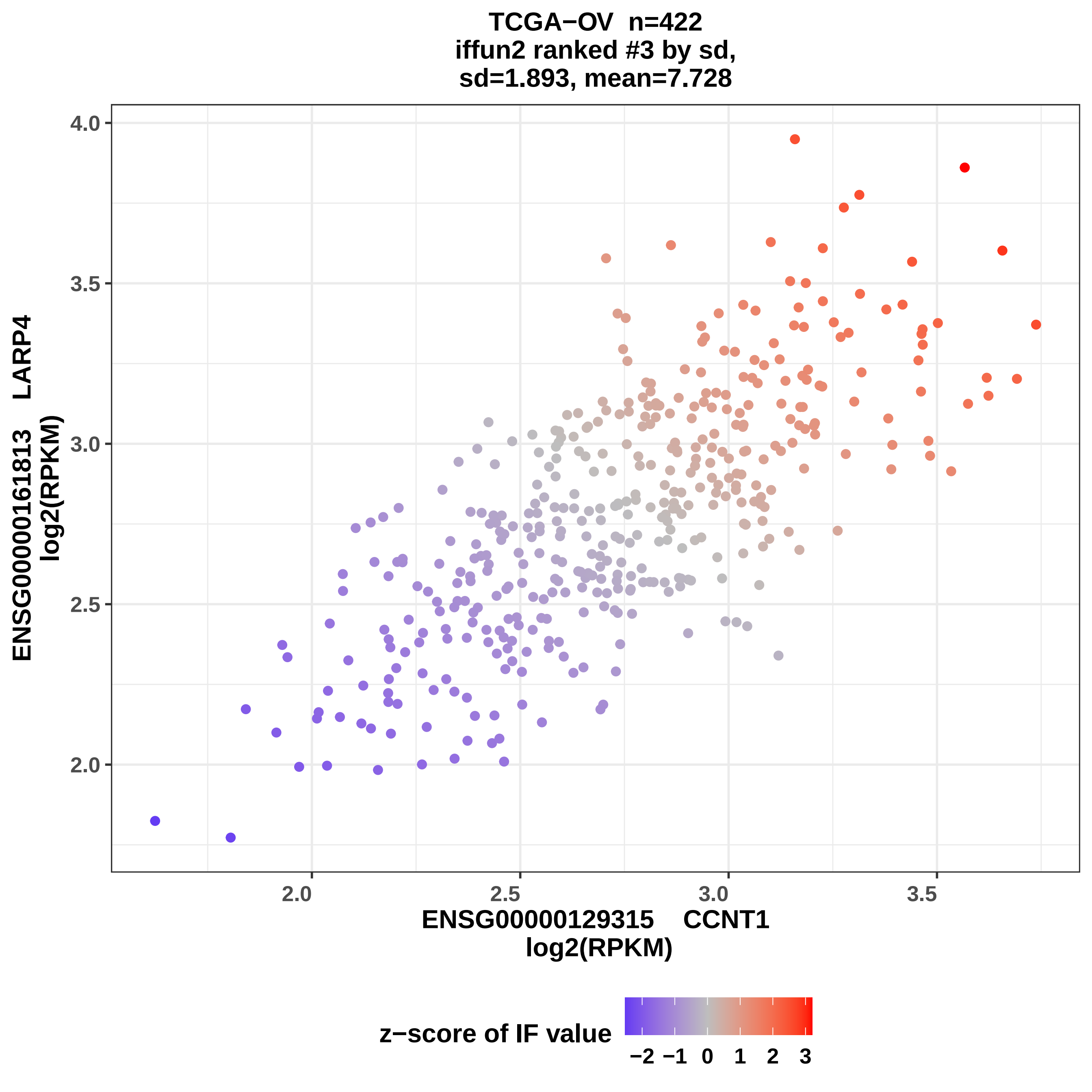

TCGA−OV n=422
iffun2 ranked #3 by sd,
sd=1.893, mean=7.728
ENSG00000161813 LARP4
log2(RPKM)
4.0
3.5
3.0
2.5
2.0
2.0
2.5
3.0
3.5
ENSG00000129315 CCNT1
log2(RPKM)
z−score of IF value
−2 −1 0 1 2 3

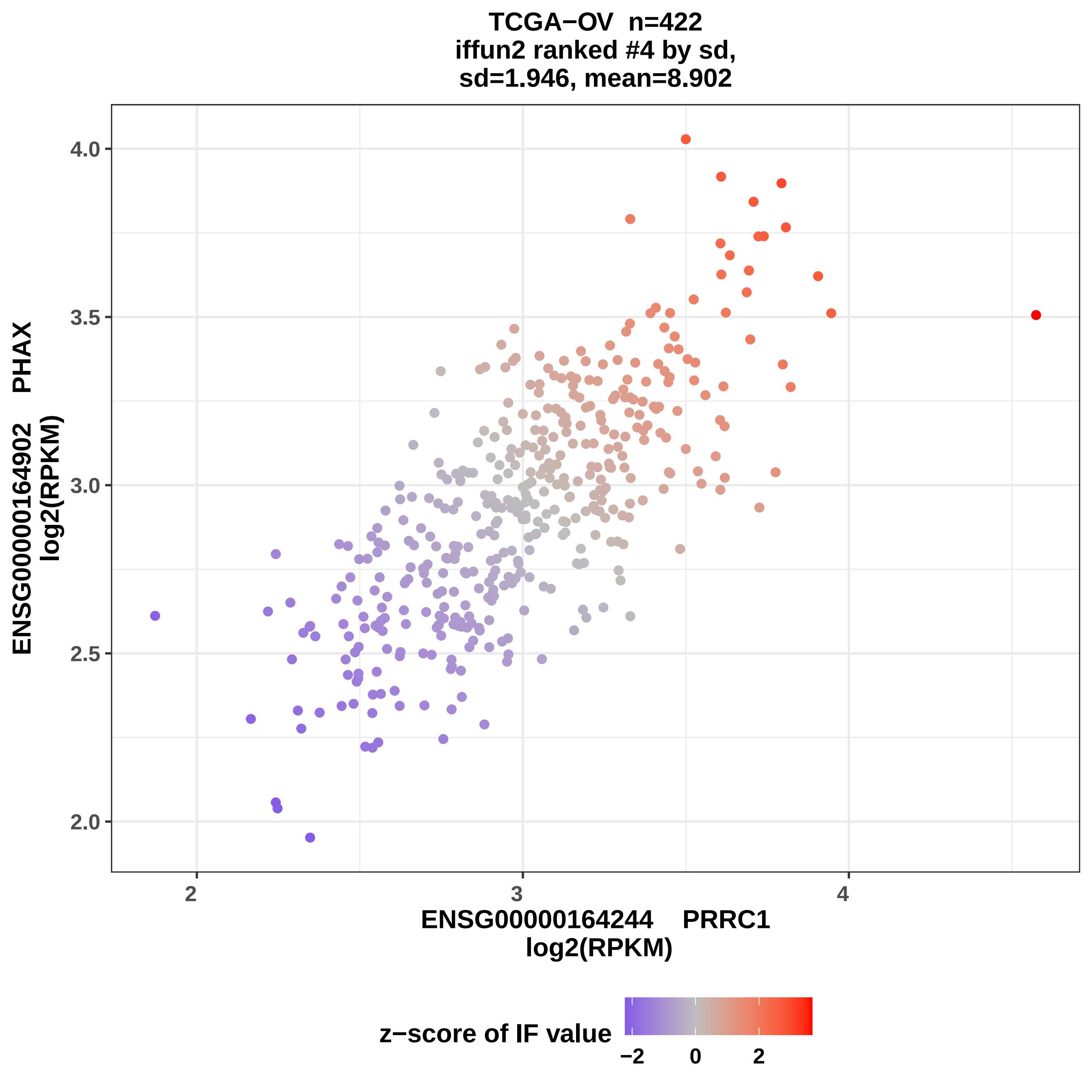
TCGA−OV n=422
iffun2 ranked #4 by sd,
sd=1.946, mean=8.902
ENSG00000164902 PHAX
log2(RPKM)
4.0
3.5
3.0
2.5
2.0
2
3
4
ENSG00000164244 PRRC1
log2(RPKM)
z−score of IF value
−2
0
2

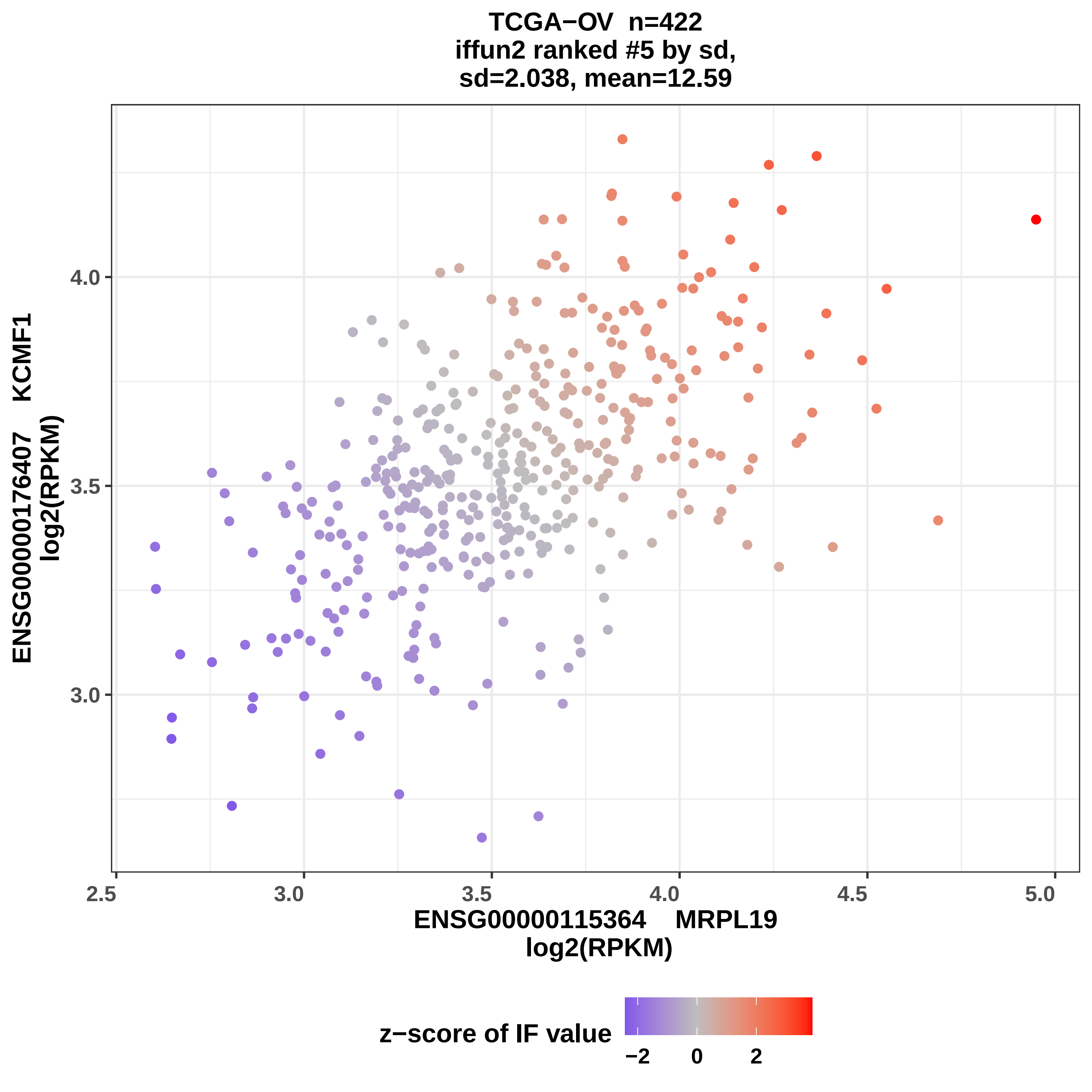
TCGA−OV n=422
iffun2 ranked #5 by sd,
sd=2.038, mean=12.59
ENSG00000176407 KCMF1
log2(RPKM)
ENSG00000115364 MRPL19
log2(RPKM)
z−score of IF value
−2
0
2
4.0
3.5
3.0
2.5
3.0
3.5
4.0
4.5
5.0

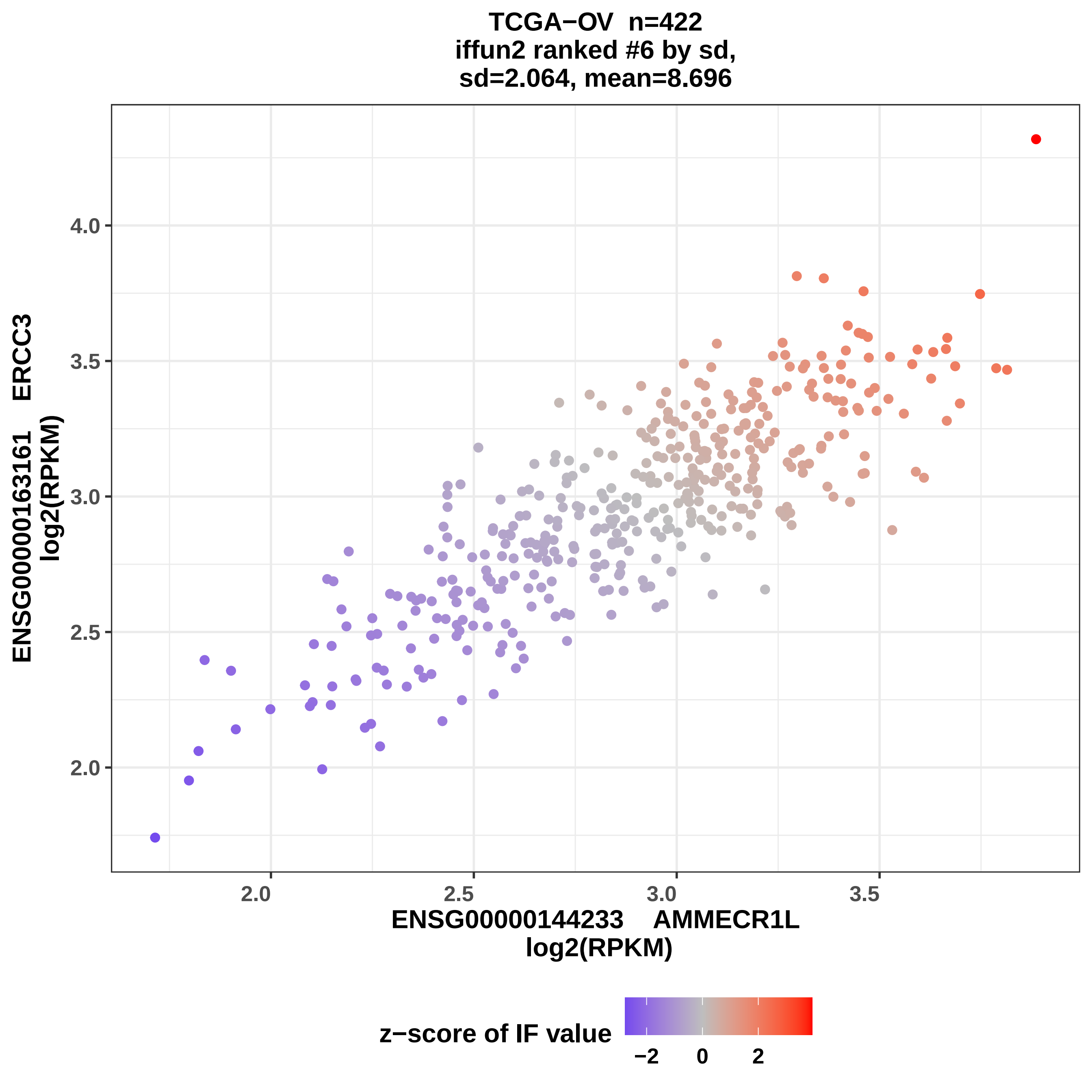

TCGA−OV n=422
iffun2 ranked #6 by sd,
sd=2.064, mean=8.696
ENSG00000163161 ERCC3
log2(RPKM)
4.0
3.5
3.0
2.5
2.0
2.0
2.5
3.0
3.5
ENSG00000144233 AMMECR1L
log2(RPKM)
z−score of IF value
−2
0
2

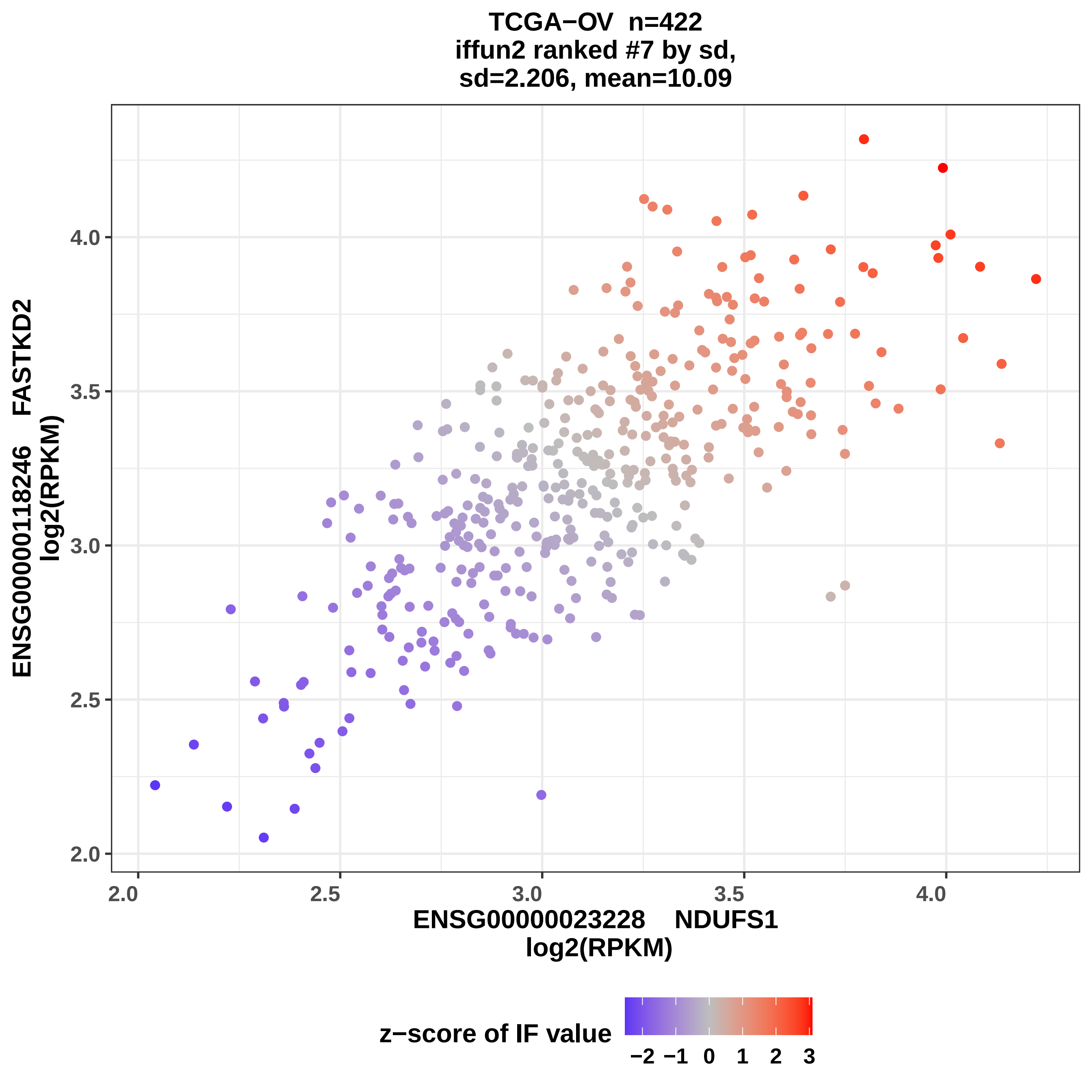
TCGA−OV n=422
iffun2 ranked #7 by sd,
sd=2.206, mean=10.09
ENSG00000118246 FASTKD2
log2(RPKM)
ENSG00000023228 NDUFS1
log2(RPKM)
2.0
2.5
3.0
3.5
4.0
z−score of IF value
−2 −1 0 1 2 3

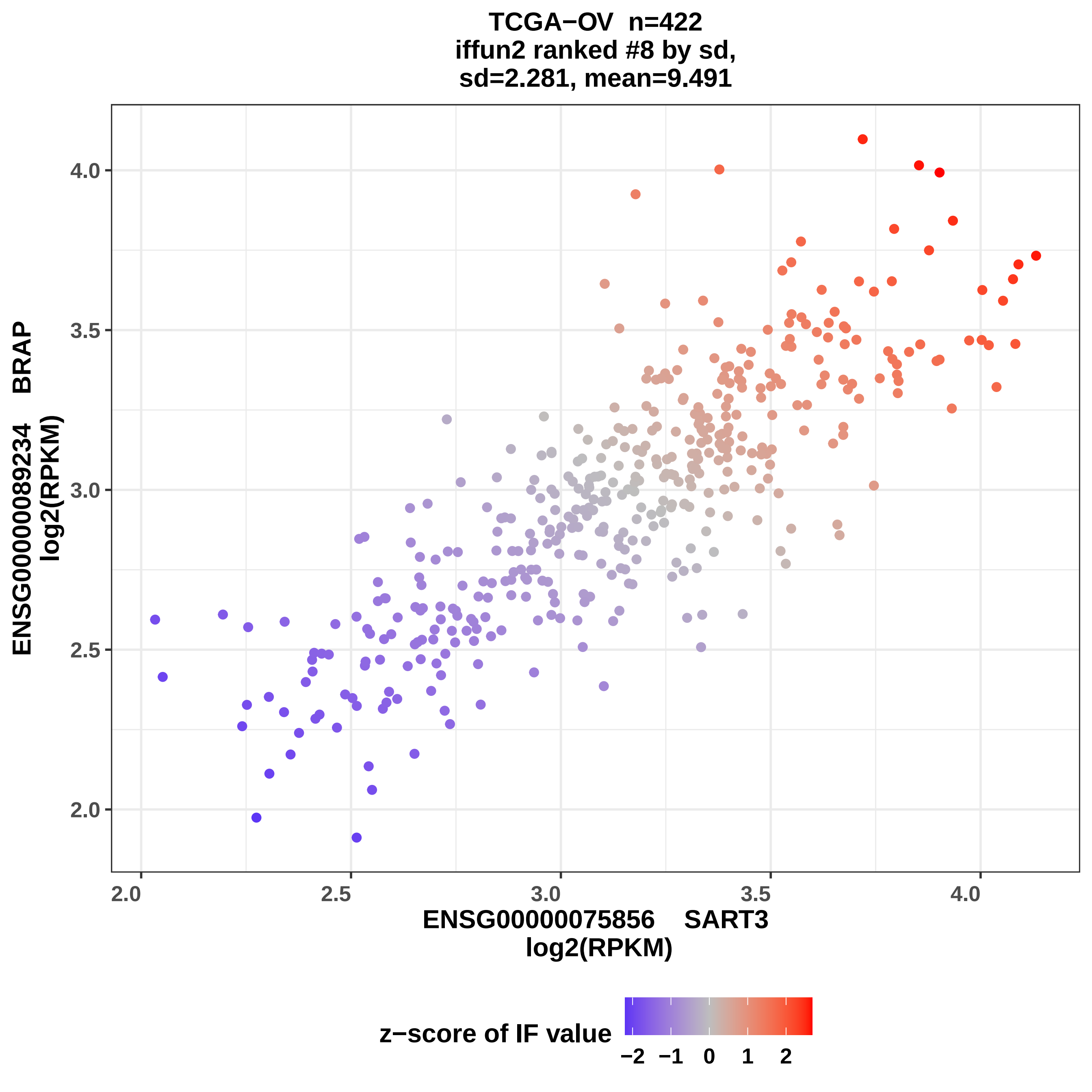
TCGA−OV n=422
iffun2 ranked #8 by sd,
sd=2.281, mean=9.491
ENSG00000089234 BRAP
log2(RPKM)
4.0
3.5
3.0
2.5
2.0
2.0
2.5
3.0
3.5
4.0
ENSG00000075856 SART3
log2(RPKM)
z−score of IF value
−2
−1
0
1
2

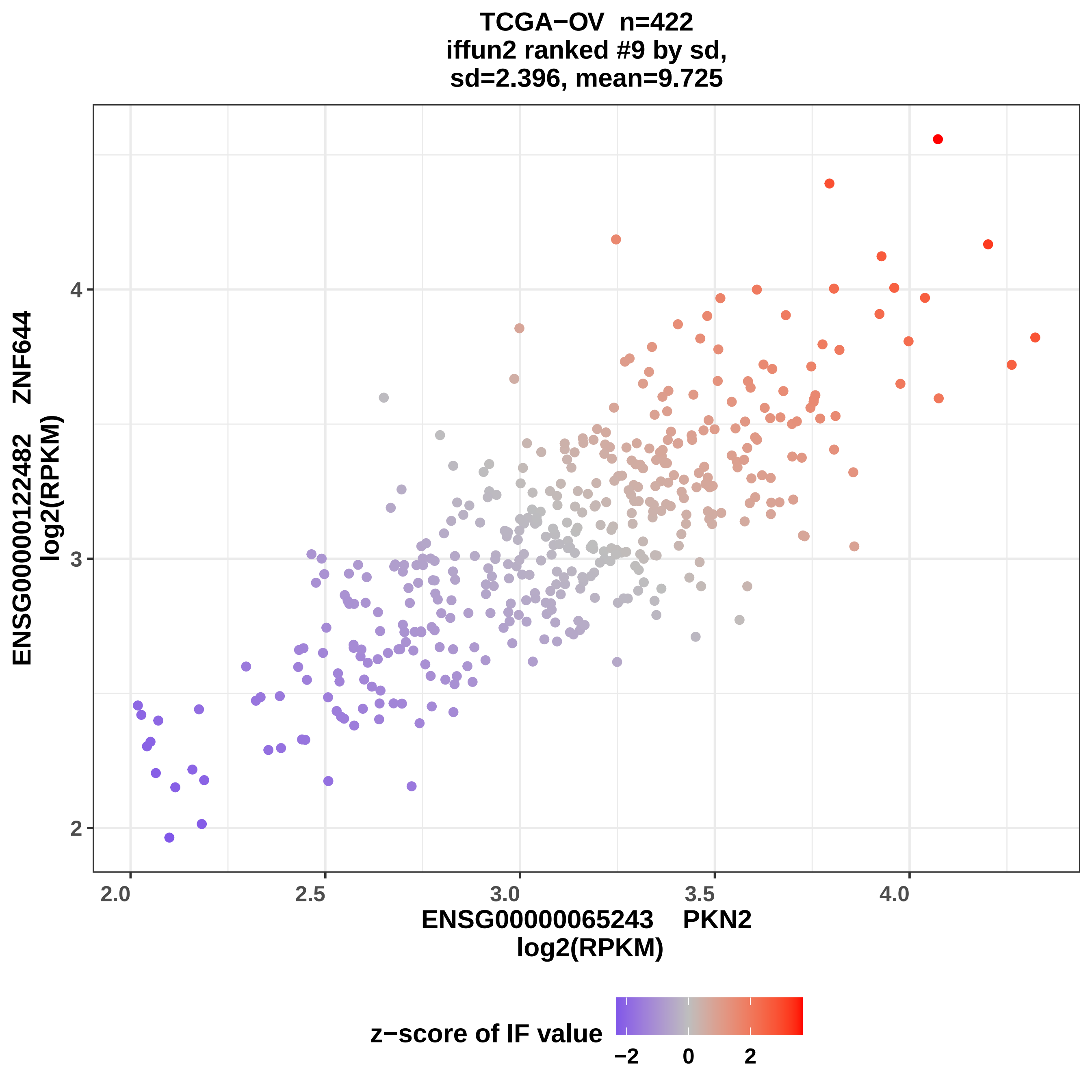
TCGA−OV n=422
iffun2 ranked #9 by sd,
sd=2.396, mean=9.725
ENSG00000122482 ZNF644
log2(RPKM)
2
3
4
2.0
2.5
3.0
3.5
4.0
ENSG00000065243 PKN2
log2(RPKM)
z−score of IF value
−2
0
2

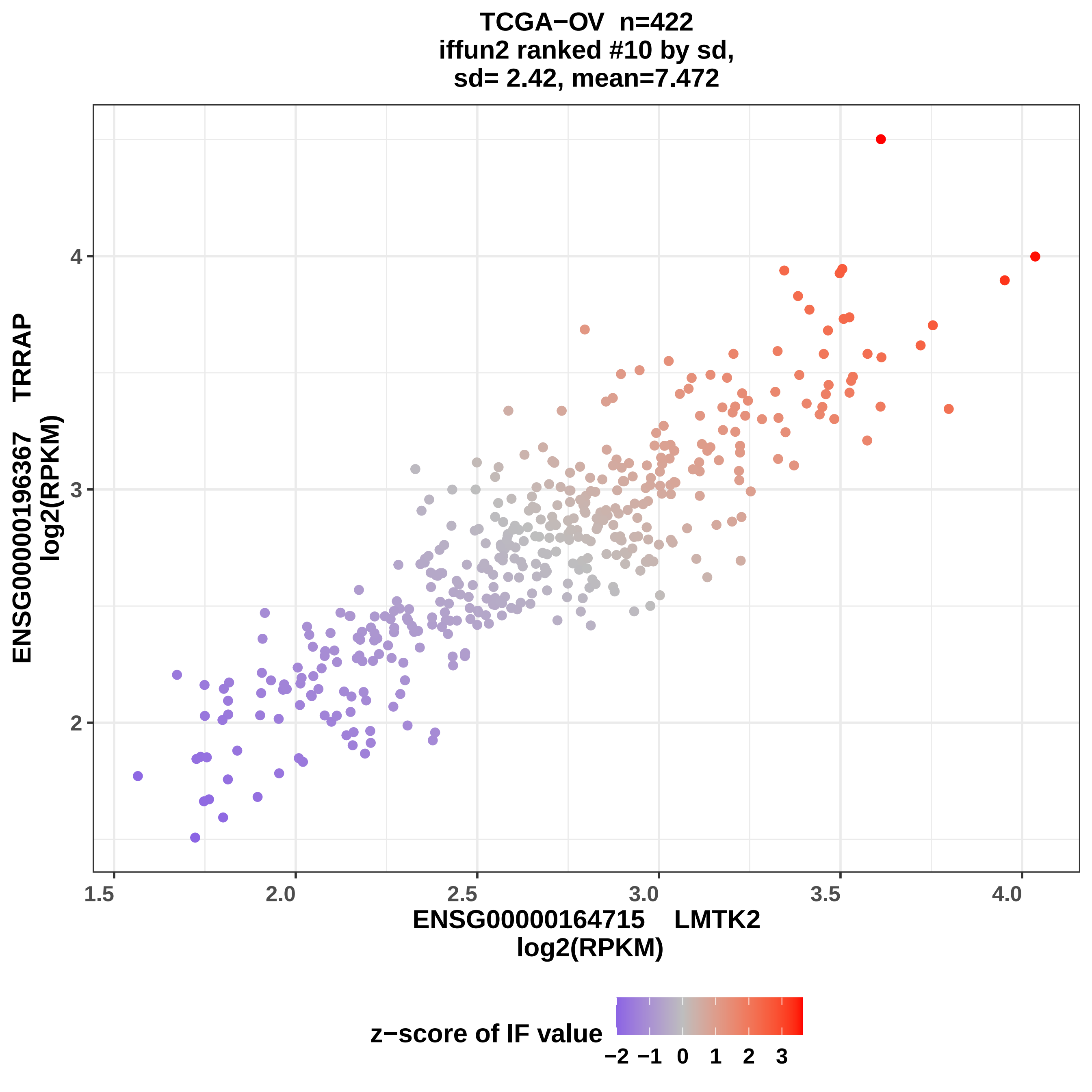
TCGA−OV n=422
iffun2 ranked #10 by sd,
sd= 2.42, mean=7.472
ENSG00000196367 TRRAP
log2(RPKM)
2
3
4
1.5
2.0
2.5
3.0
3.5
4.0
ENSG00000164715 LMTK2
log2(RPKM)
z−score of IF value
−2
−1
0
1
2
3

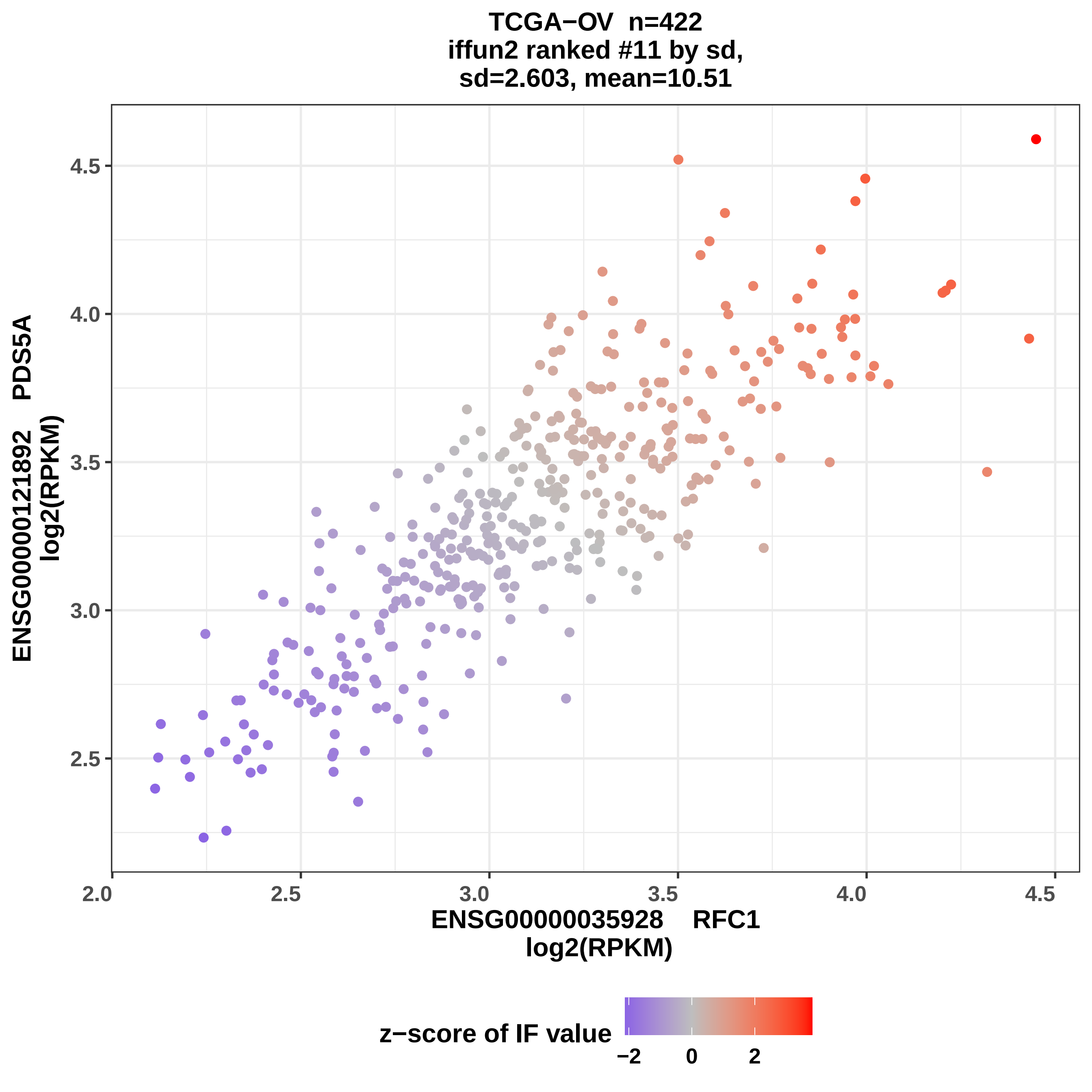
TCGA−OV n=422
iffun2 ranked #11 by sd,
sd=2.603, mean=10.51
ENSG00000121892 PDS5A
log2(RPKM)
2.5
3.0
3.5
4.0
4.5
2.0
2.5
3.0
3.5
4.0
4.5
ENSG00000035928 RFC1
log2(RPKM)
z−score of IF value
−2
0
2

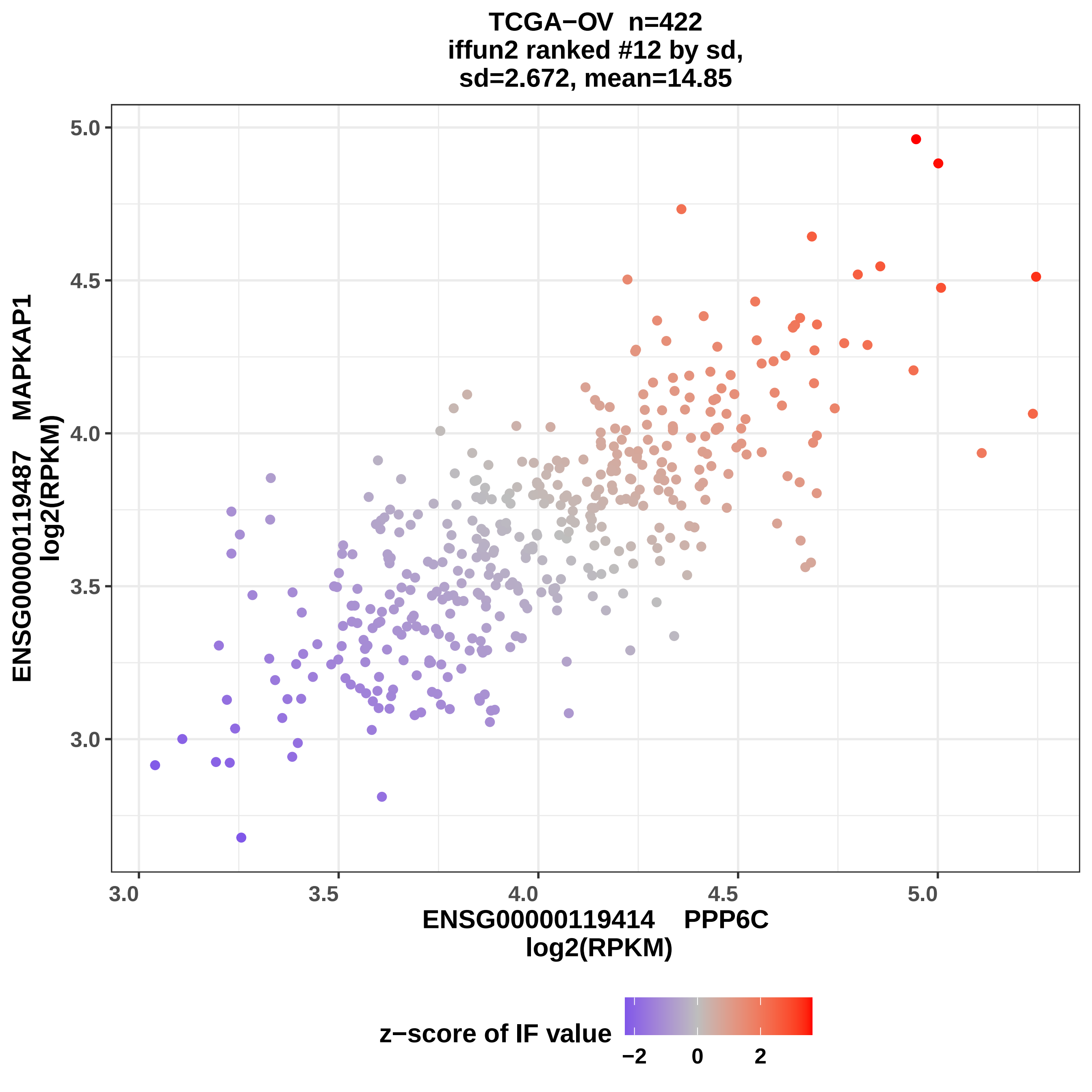

TCGA−OV n=422
iffun2 ranked #12 by sd,
sd=2.672, mean=14.85
ENSG00000119487 MAPKAP1
log2(RPKM)
5.0
4.5
4.0
3.5
3.0
3.0
3.5
4.0
4.5
5.0
ENSG00000119414 PPP6C
log2(RPKM)
z−score of IF value
−2
0
2

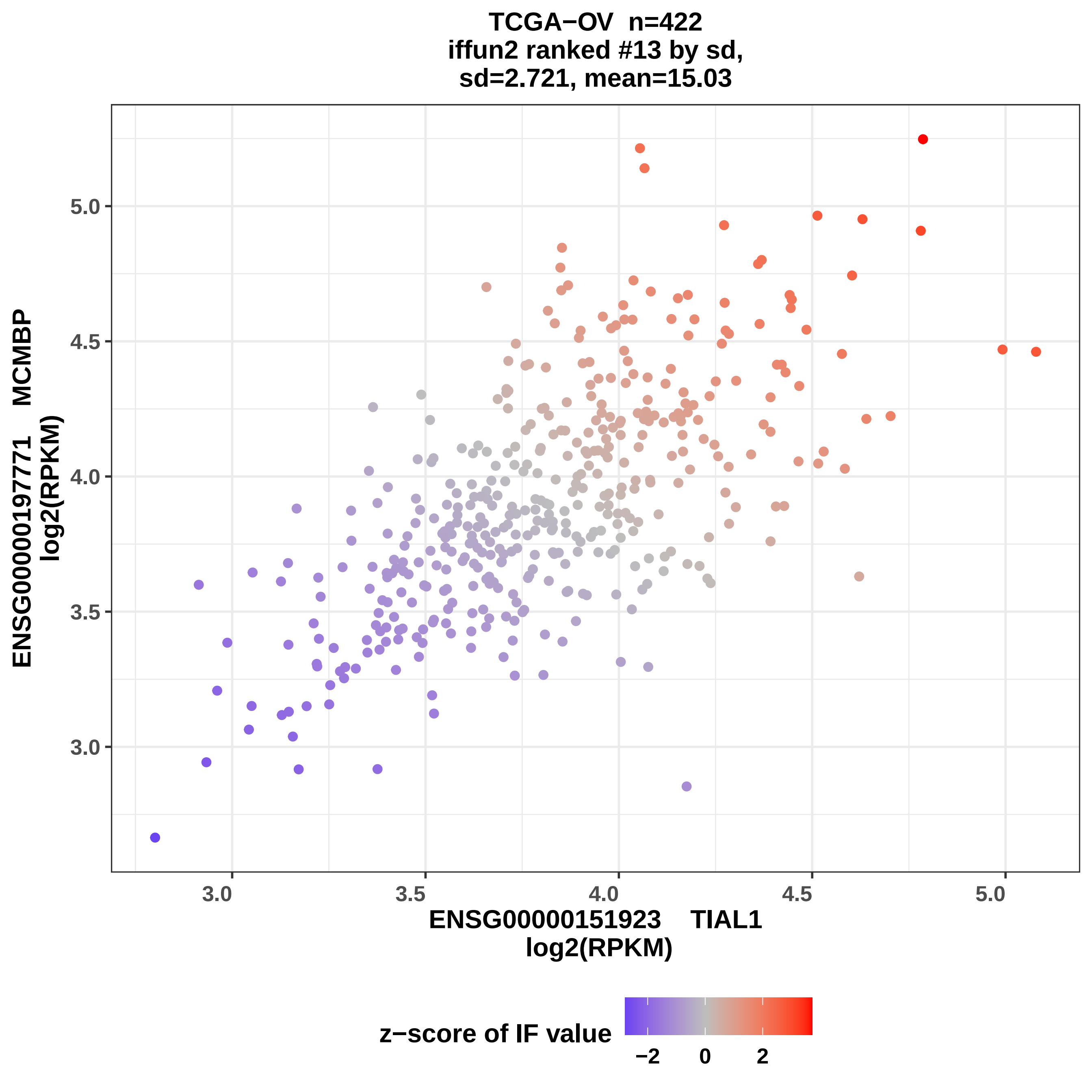
TCGA−OV n=422
iffun2 ranked #13 by sd,
sd=2.721, mean=15.03
ENSG00000197771 MCMBP
log2(RPKM)
5.0
4.5
4.0
3.5
3.0
3.0
3.5
4.0
4.5
5.0
ENSG00000151923 TIAL1
log2(RPKM)
z−score of IF value
−2
0
2

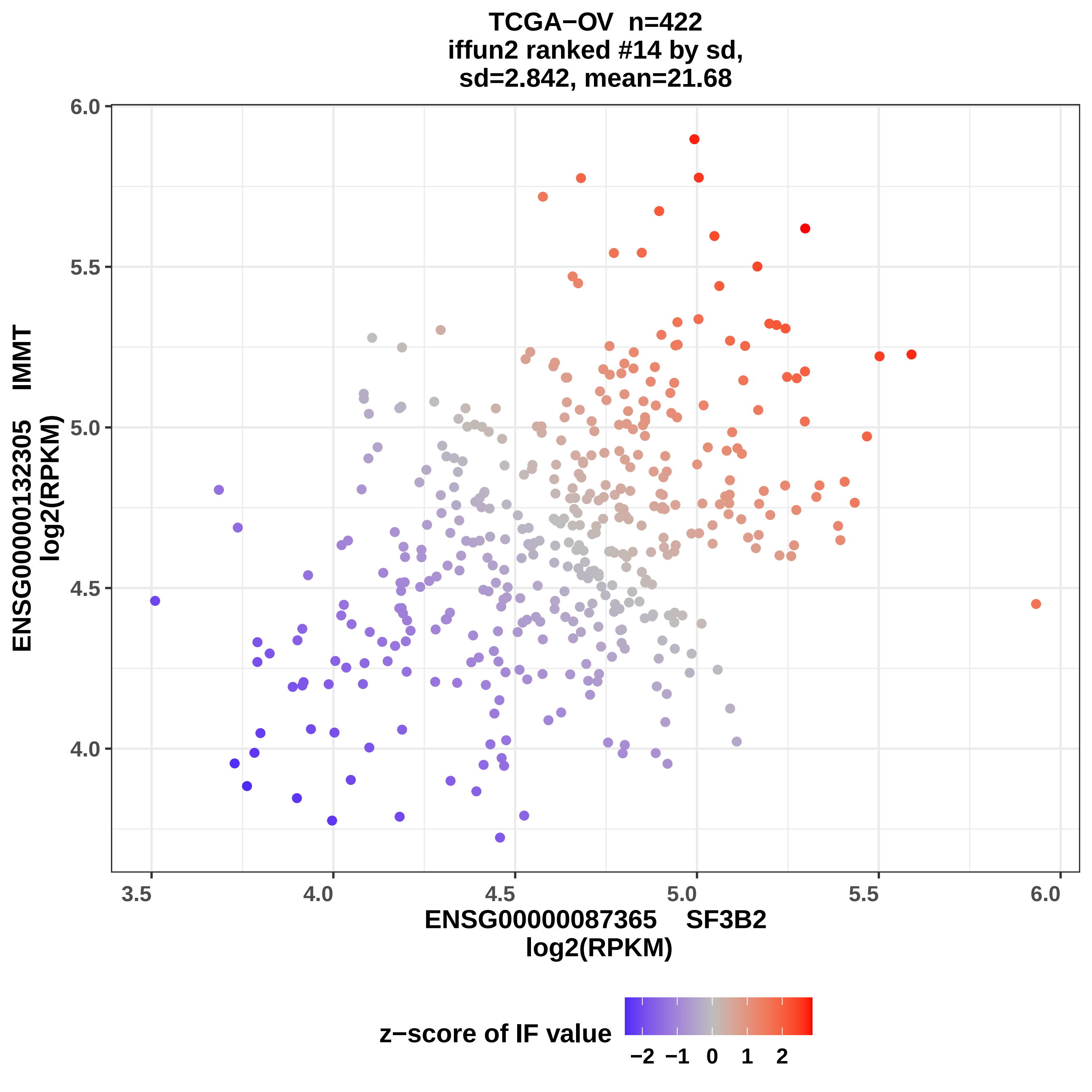
TCGA−OV n=422
iffun2 ranked #14 by sd,
sd=2.842, mean=21.68
ENSG00000132305 IMMT
log2(RPKM)
ENSG00000087365 SF3B2
log2(RPKM)
z−score of IF value
−2 −1 0 1 2
3.5
4.0
4.5
5.0
5.5
6.0

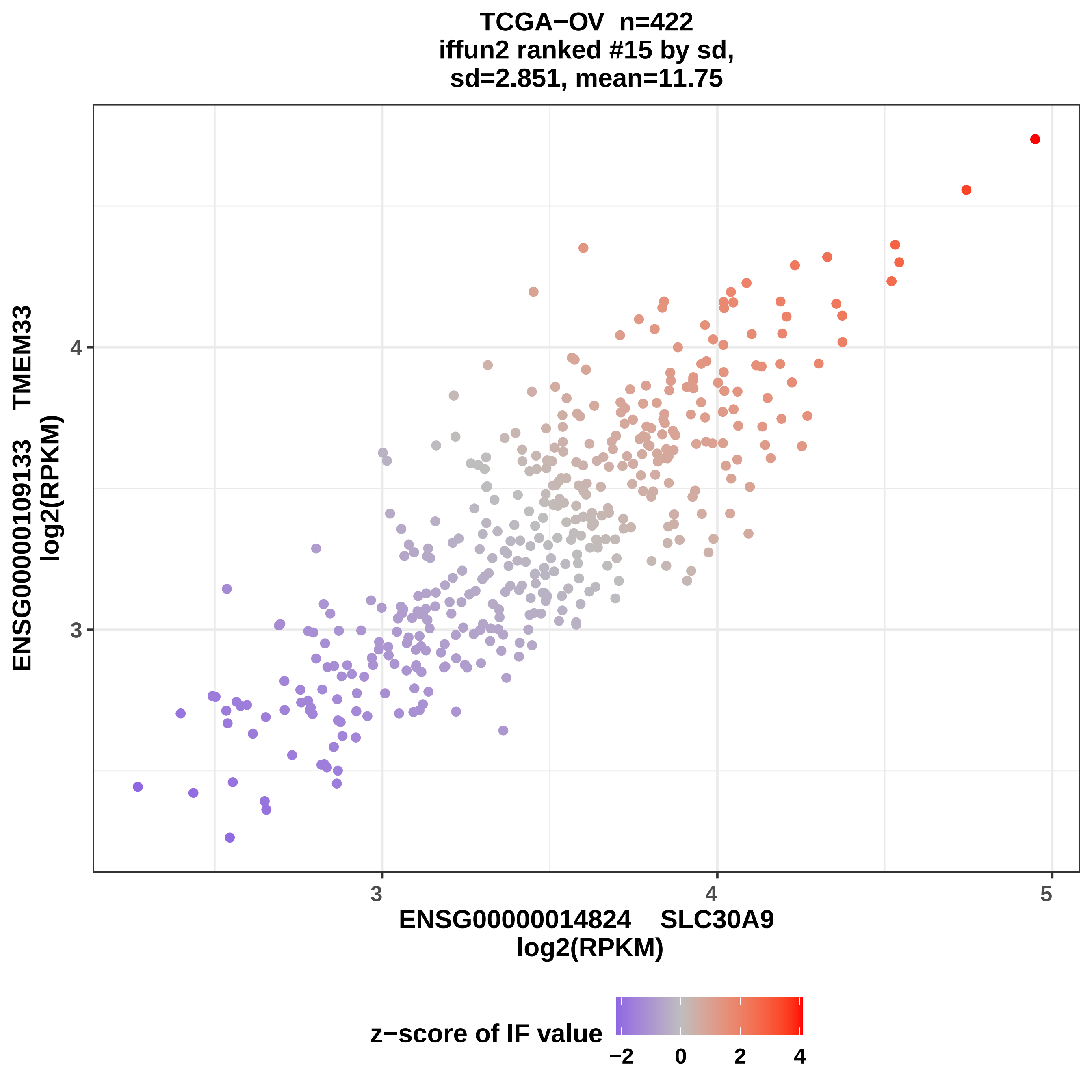

TCGA−OV n=422
iffun2 ranked #15 by sd,
sd=2.851, mean=11.75
ENSG00000109133 TMEM33
log2(RPKM)
4
3
3
4
5
ENSG00000014824 SLC30A9
log2(RPKM)
z−score of IF value
−2
0
2
4

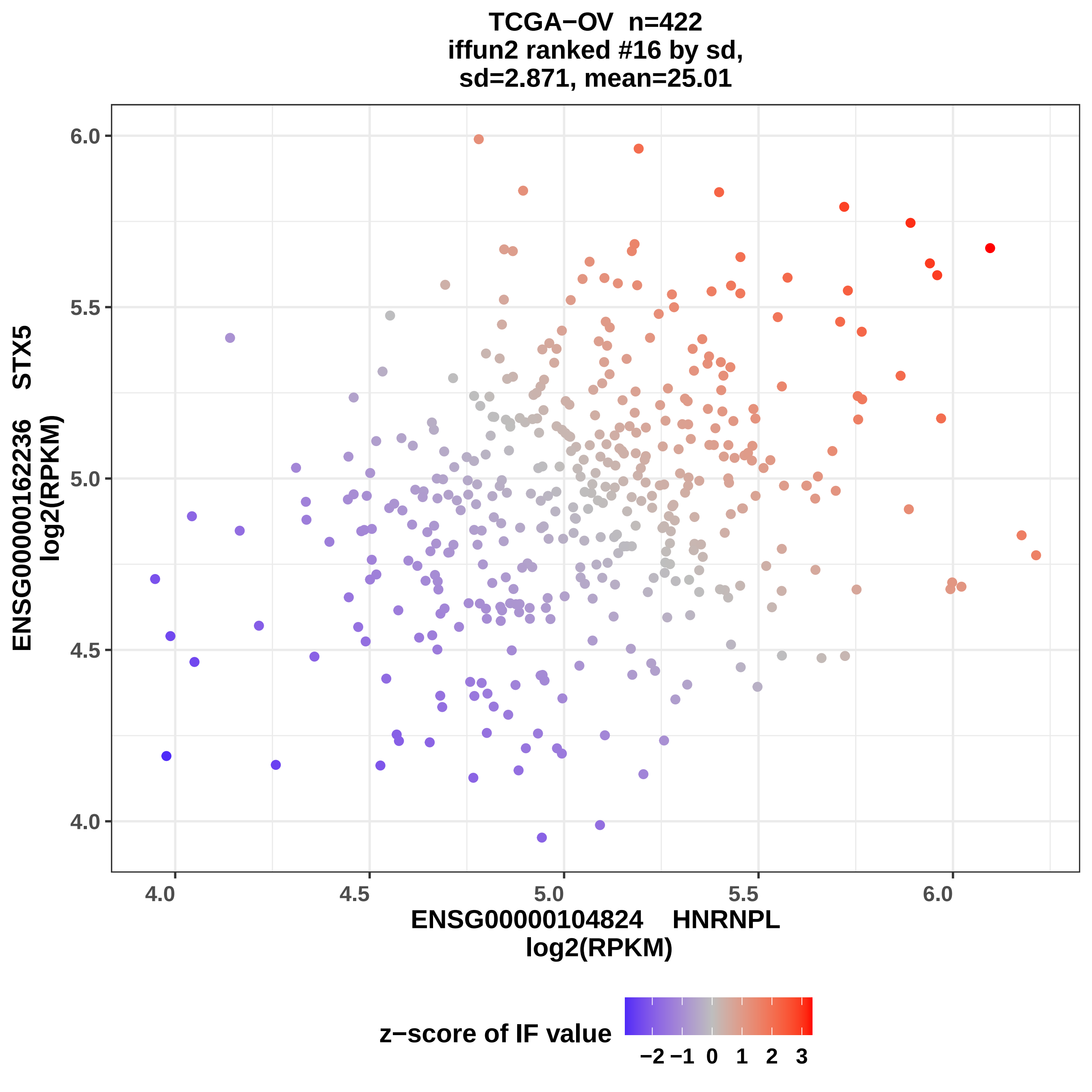
TCGA−OV n=422
iffun2 ranked #16 by sd,
sd=2.871, mean=25.01
ENSG00000162236 STX5
log2(RPKM)
6.0
5.5
5.0
4.5
4.0
4.0
4.5
5.0
5.5
6.0
ENSG00000104824 HNRNPL
log2(RPKM)
z−score of IF value
−2 −1 0 1 2 3

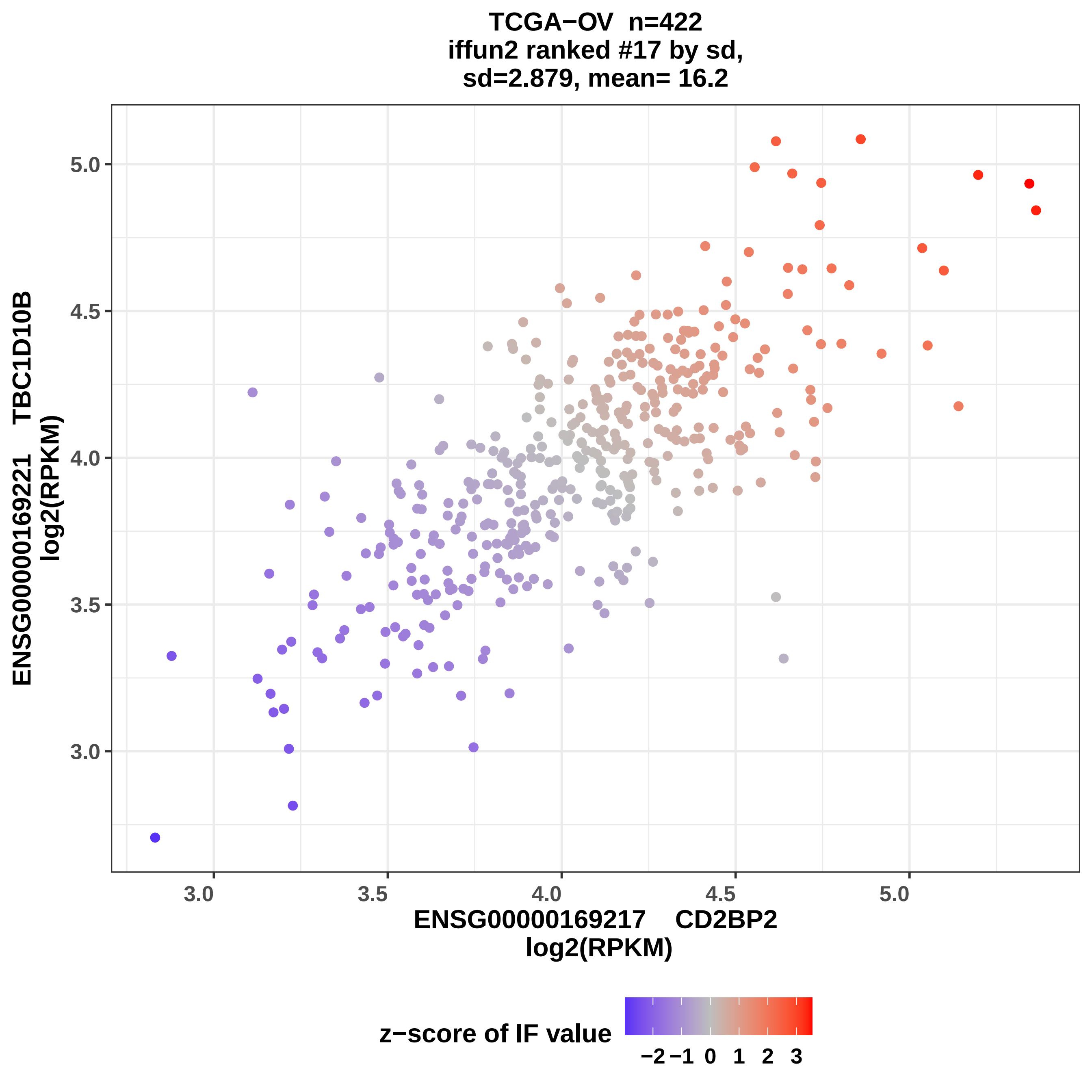

TCGA−OV n=422
iffun2 ranked #17 by sd,
sd=2.879, mean= 16.2
ENSG00000169221 TBC1D10B
log2(RPKM)
5.0
4.5
4.0
3.5
3.0
3.0
3.5
4.0
4.5
5.0
ENSG00000169217 CD2BP2
log2(RPKM)
z−score of IF value
−2
−1
0
1
2
3

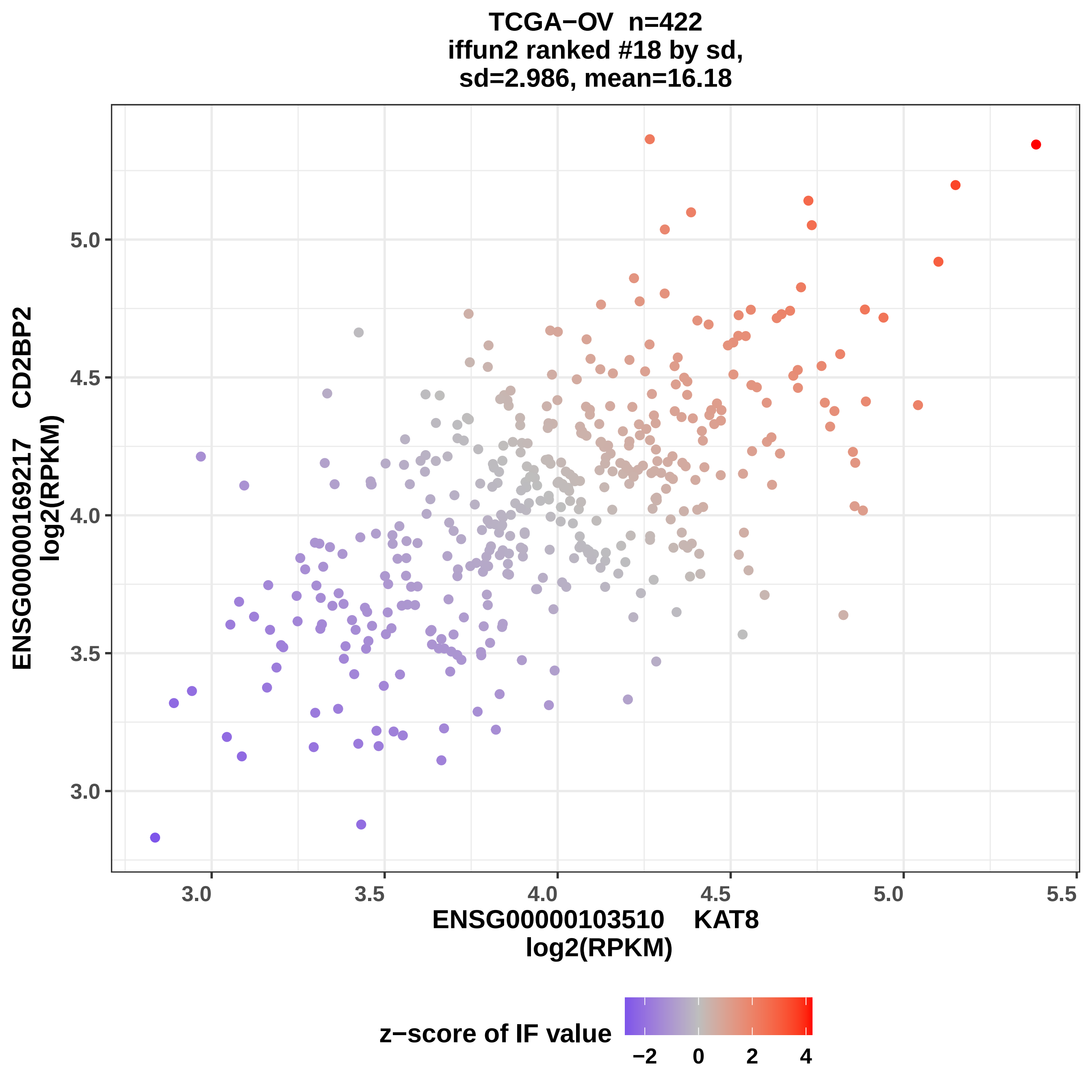

TCGA−OV n=422
iffun2 ranked #18 by sd,
sd=2.986, mean=16.18
ENSG00000169217 CD2BP2
log2(RPKM)
5.0
4.5
4.0
3.5
3.0
3.0
3.5
4.0
4.5
5.0
5.5
ENSG00000103510 KAT8
log2(RPKM)
z−score of IF value
−2
0
2
4

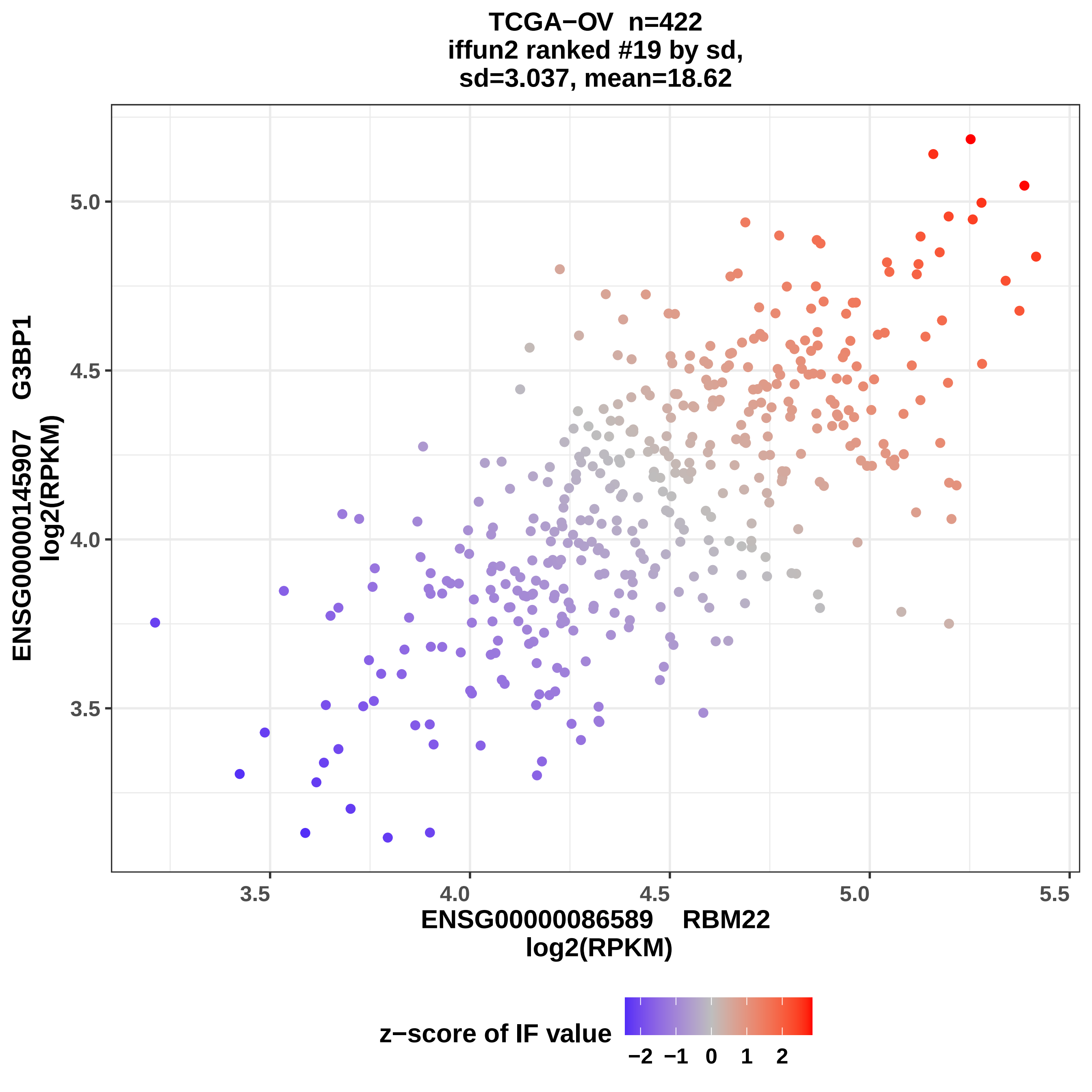
TCGA−OV n=422
iffun2 ranked #19 by sd,
sd=3.037, mean=18.62
ENSG00000145907 G3BP1
log2(RPKM)
5.0
4.5
4.0
3.5
3.5
4.0
4.5
5.0
5.5
ENSG00000086589 RBM22
log2(RPKM)
z−score of IF value
−2 −1 0 1 2

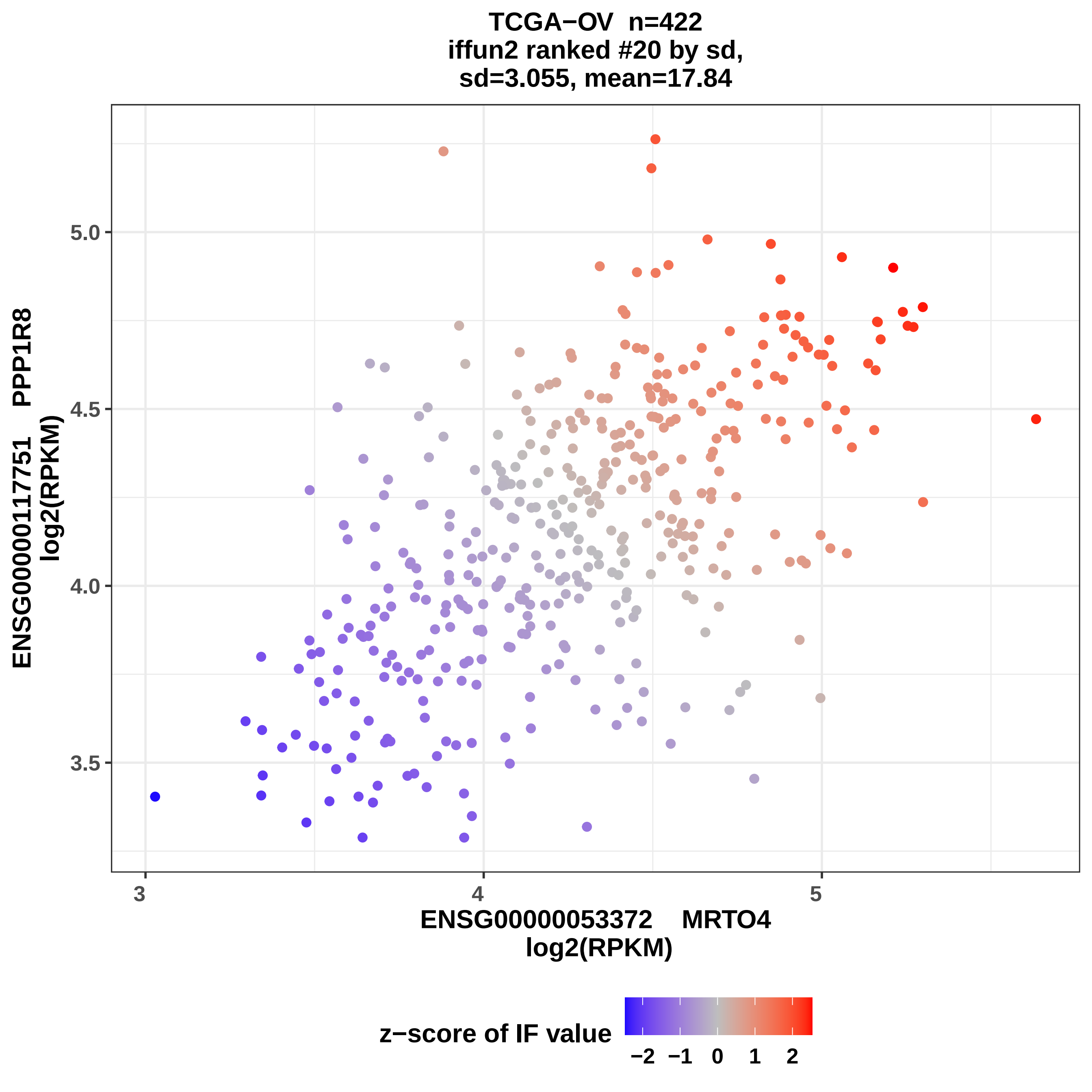
TCGA−OV n=422
iffun2 ranked #20 by sd,
sd=3.055, mean=17.84
ENSG00000117751 PPP1R8
log2(RPKM)
5.0
4.5
4.0
3.5
3
4
5
ENSG00000053372 MRTO4
log2(RPKM)
z−score of IF value
−2
−1
0
1
2

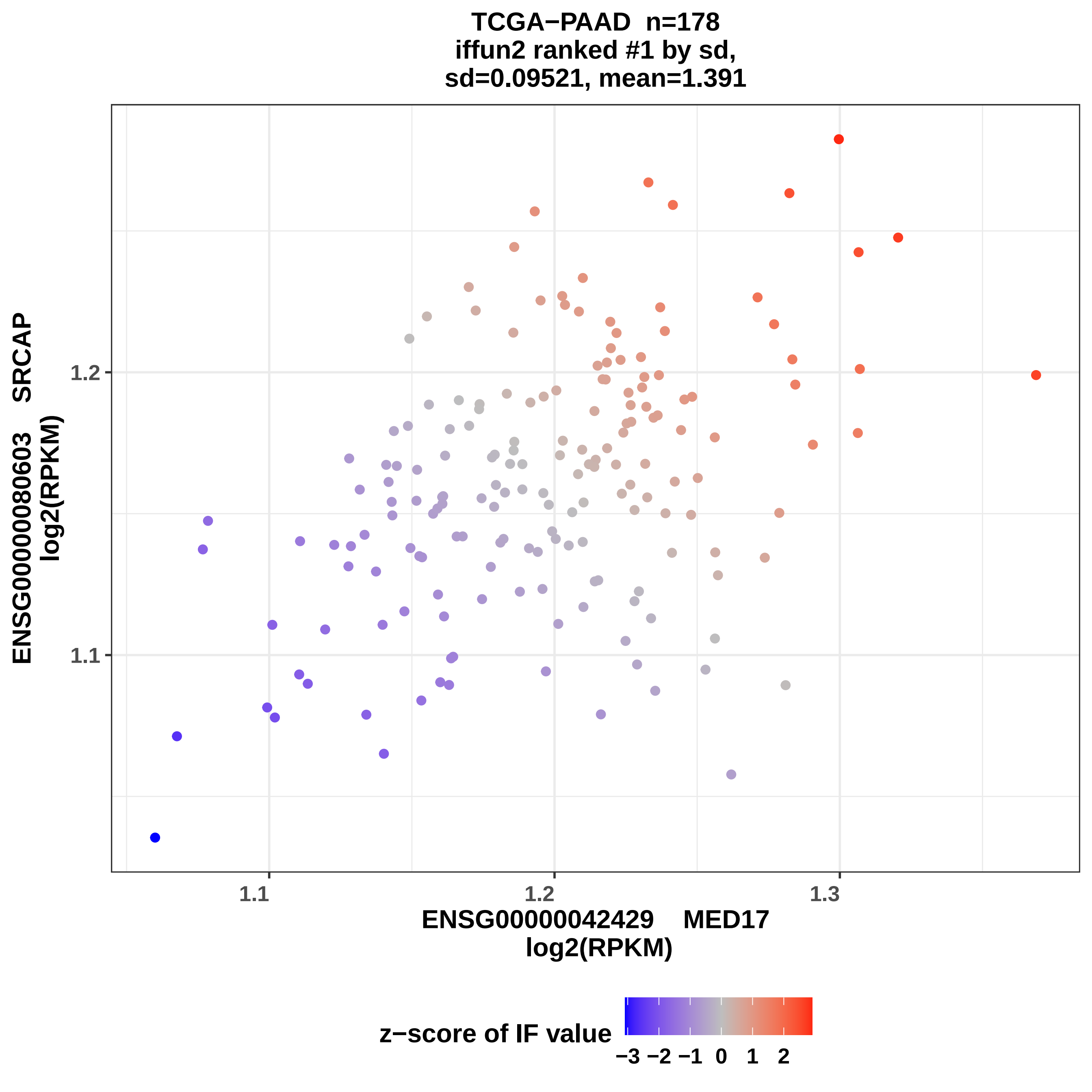
TCGA−PAAD n=178
iffun2 ranked #1 by sd,
sd=0.09521, mean=1.391
ENSG00000080603 SRCAP
log2(RPKM)
1.2
1.1
1.1
1.2
1.3
ENSG00000042429 MED17
log2(RPKM)
z−score of IF value
−3 −2 −1 0 1 2

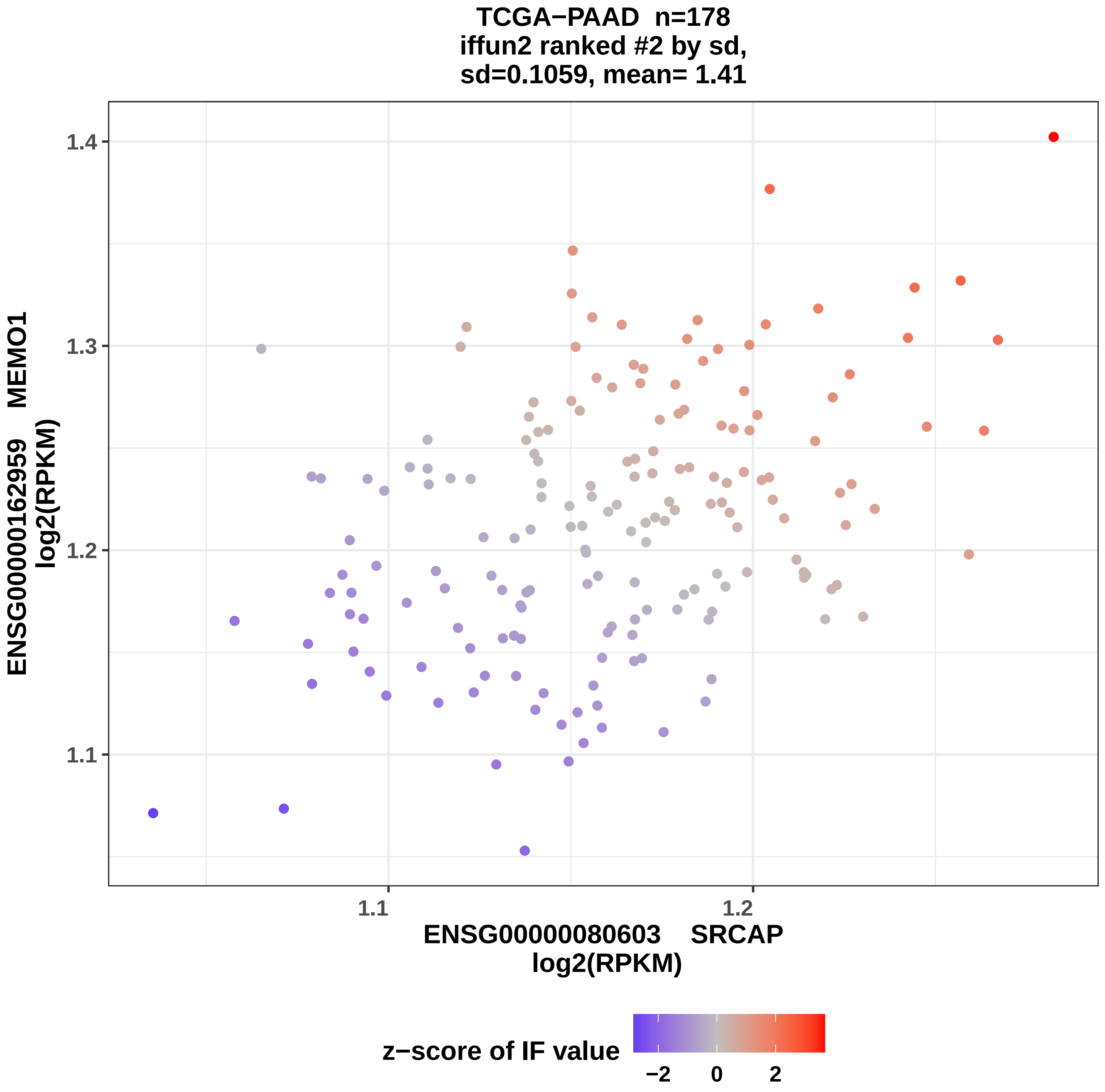
TCGA−PAAD n=178
iffun2 ranked #2 by sd,
sd=0.1059, mean= 1.41
ENSG00000162959 MEMO1
log2(RPKM)
1.4
1.3
1.2
1.1
1.1
1.2
ENSG00000080603 SRCAP
log2(RPKM)
z−score of IF value
−2
0
2

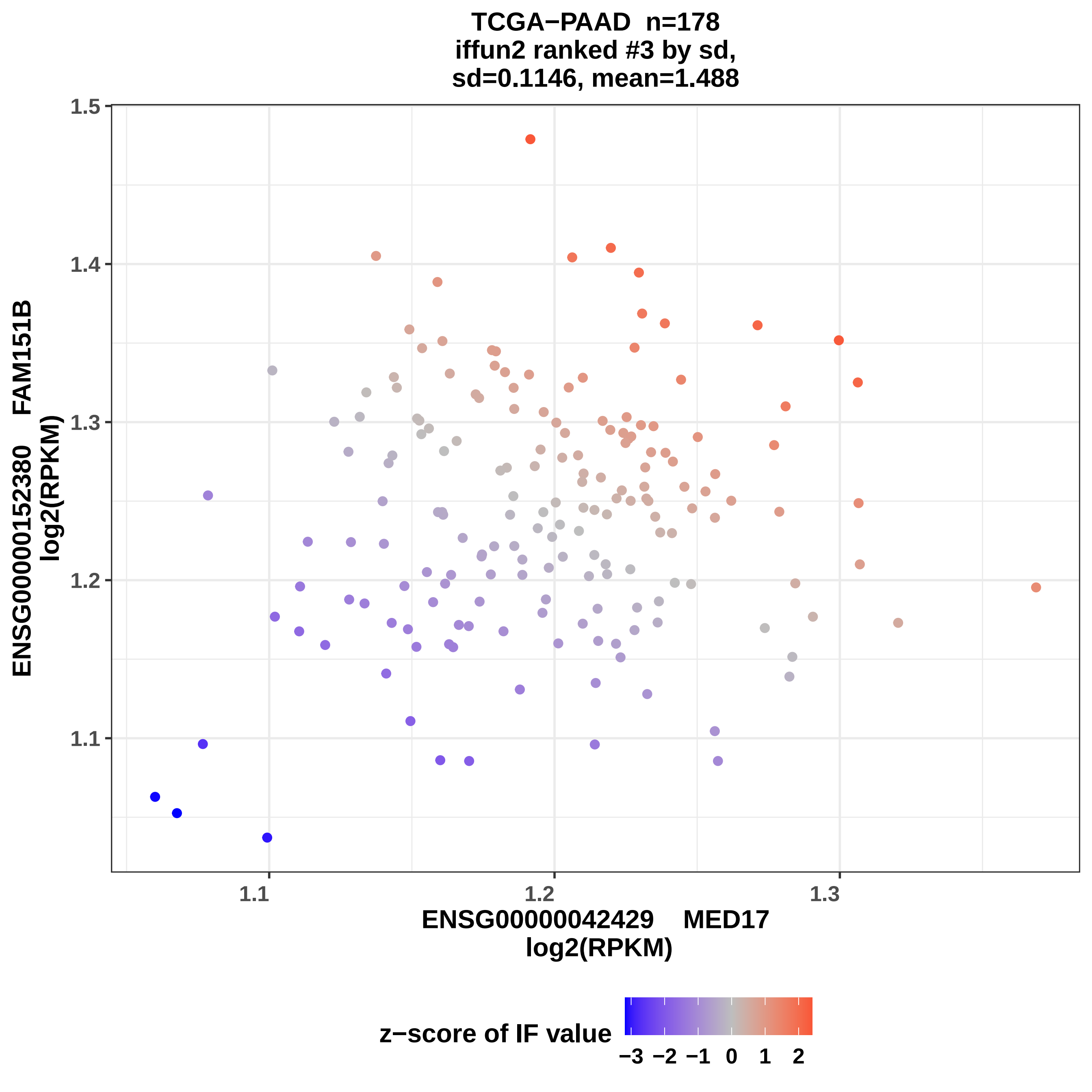
TCGA−PAAD n=178
iffun2 ranked #3 by sd,
sd=0.1146, mean=1.488
ENSG00000152380 FAM151B
log2(RPKM)
1.5
1.4
1.3
1.2
1.1
1.1
1.2
1.3
ENSG00000042429 MED17
log2(RPKM)
z−score of IF value
−3 −2 −1 0 1 2

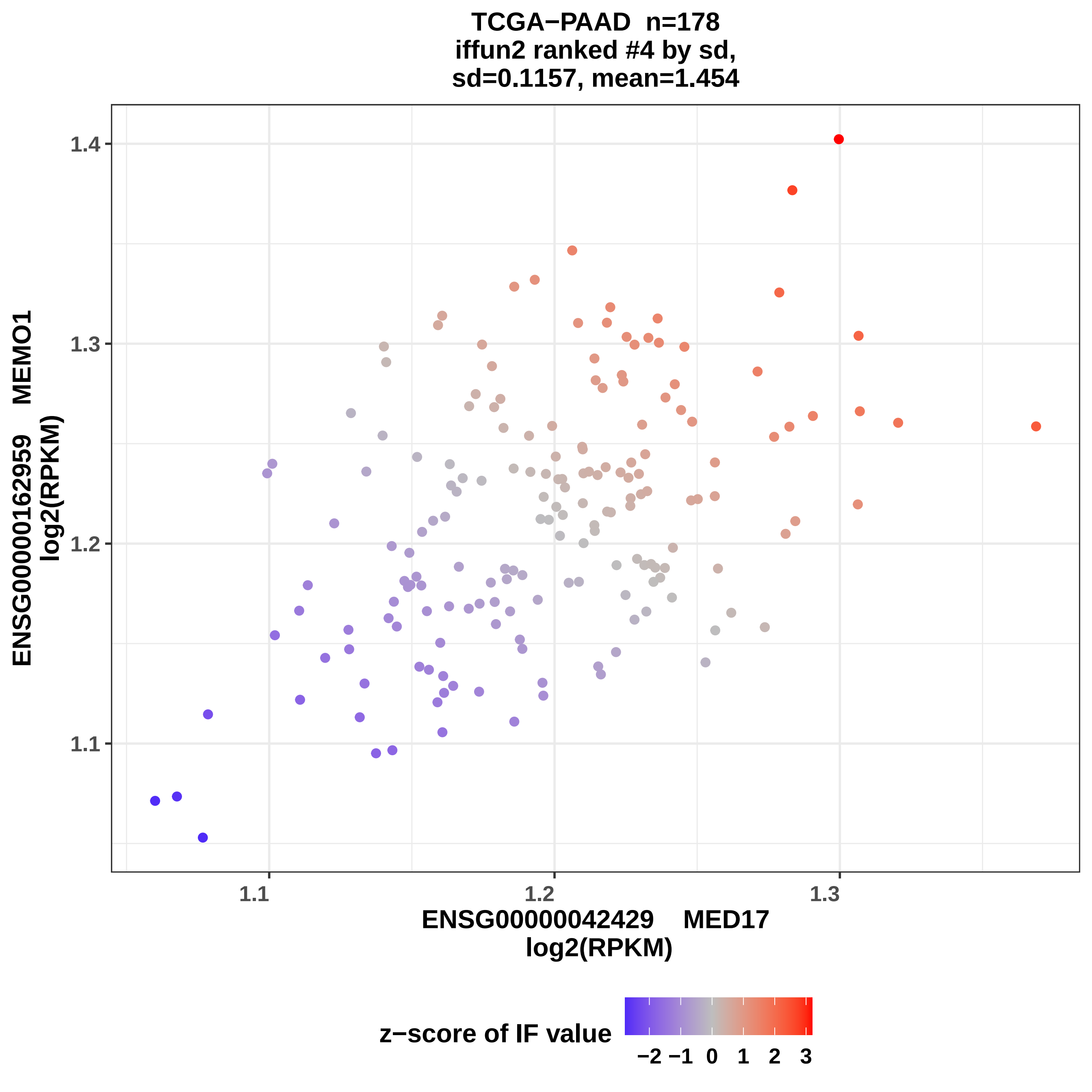
TCGA−PAAD n=178
iffun2 ranked #4 by sd,
sd=0.1157, mean=1.454
1.4
1.3
1.2
1.1
1.1
1.2
1.3
ENSG00000162959 MEMO1
log2(RPKM)
ENSG00000042429 MED17
log2(RPKM)
z−score of IF value
−2 −1 0 1 2 3

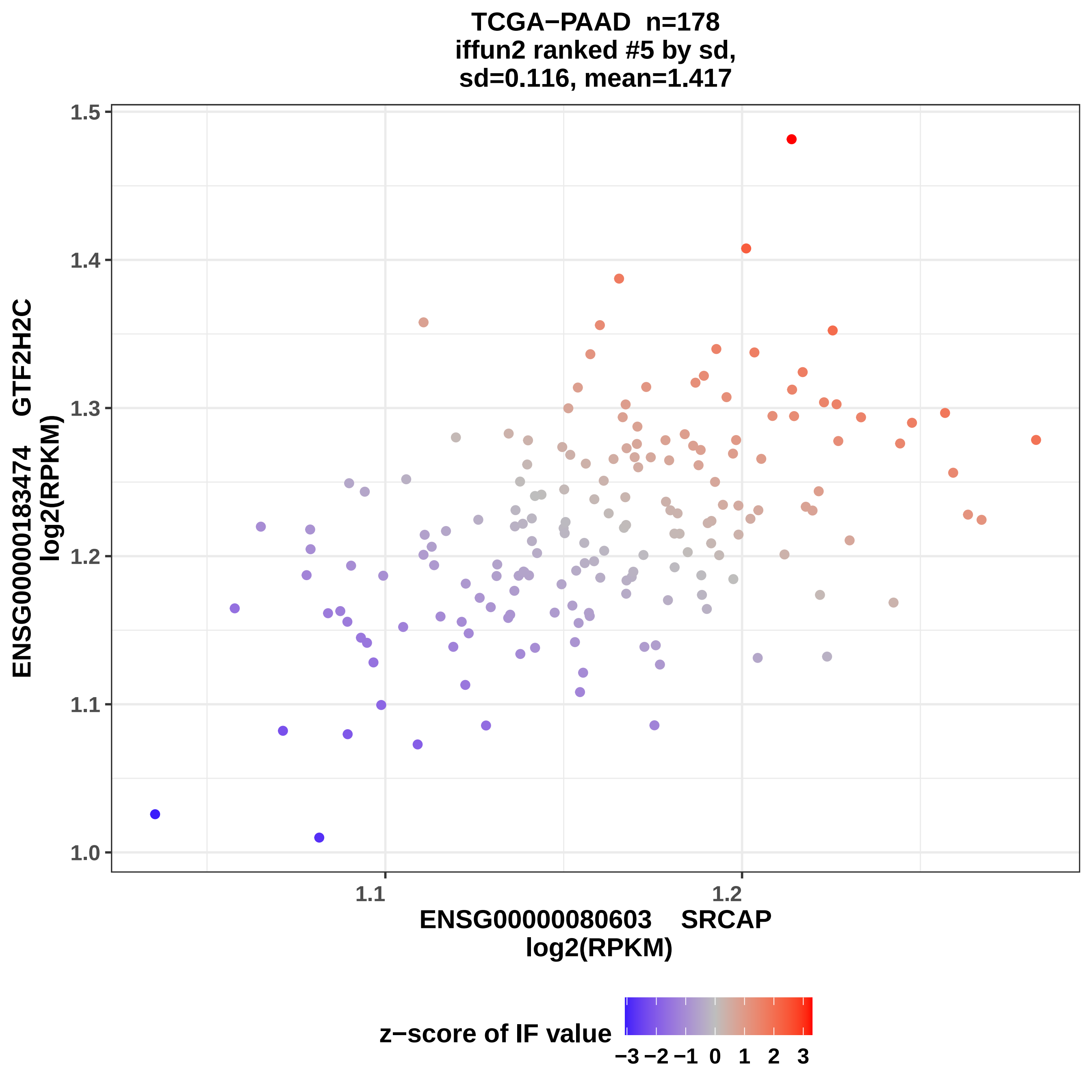
TCGA−PAAD n=178
iffun2 ranked #5 by sd,
sd=0.116, mean=1.417
ENSG00000183474 GTF2H2C
log2(RPKM)
1.0
1.1
1.2
1.3
1.4
1.5
1.1
1.2
ENSG00000080603 SRCAP
log2(RPKM)
z−score of IF value
−3 −2 −1 0 1 2 3

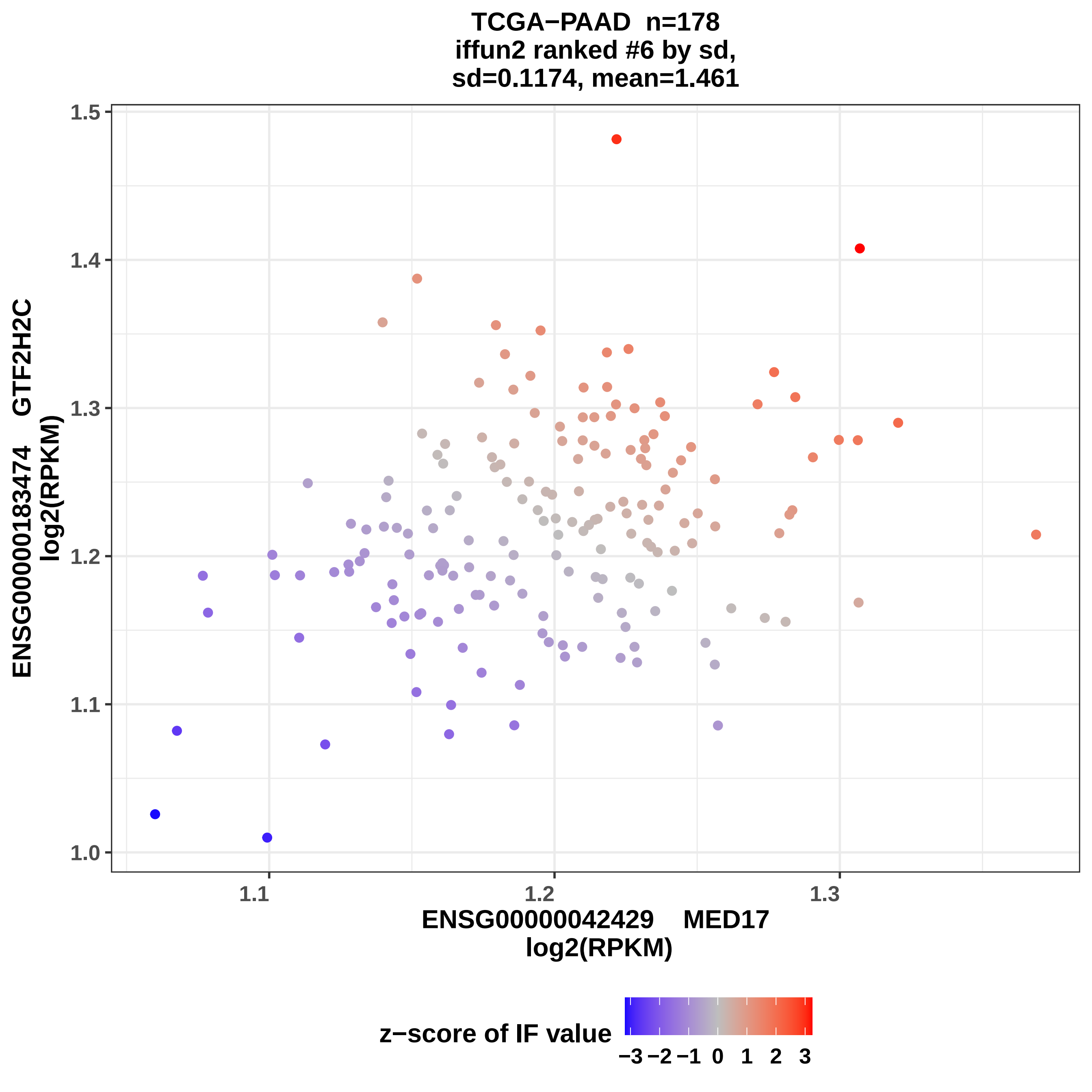
TCGA−PAAD n=178
iffun2 ranked #6 by sd,
sd=0.1174, mean=1.461
ENSG00000183474 GTF2H2C
log2(RPKM)
1.5
1.4
1.3
1.2
1.1
1.0
1.1
1.2
1.3
ENSG00000042429 MED17
log2(RPKM)
z−score of IF value
−3 −2 −1 0 1 2 3

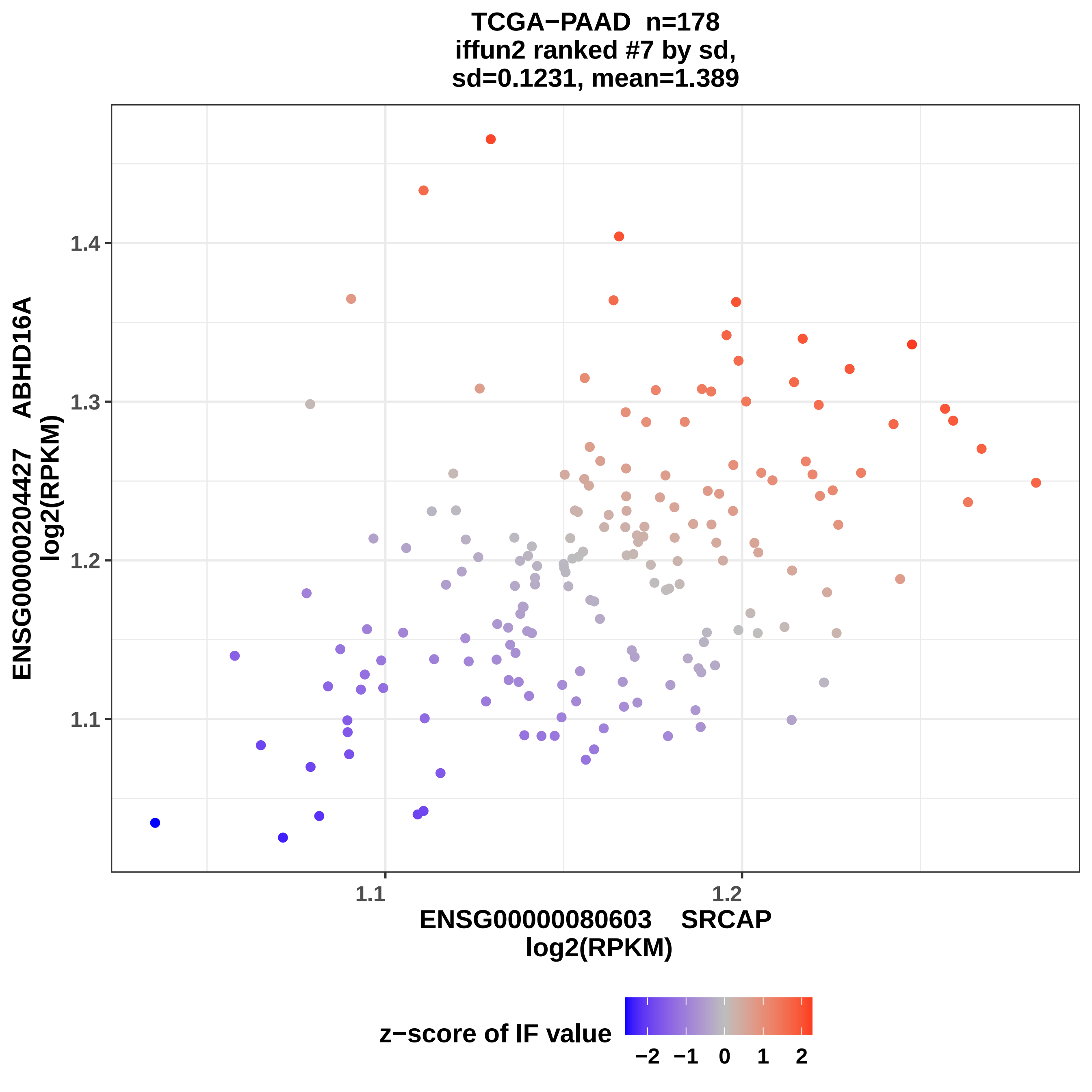

TCGA−PAAD n=178
iffun2 ranked #7 by sd,
sd=0.1231, mean=1.389
ENSG00000204427 ABHD16A
log2(RPKM)
1.4
1.3
1.2
1.1
1.1
1.2
ENSG00000080603 SRCAP
log2(RPKM)
z−score of IF value
−2
−1
0
1
2

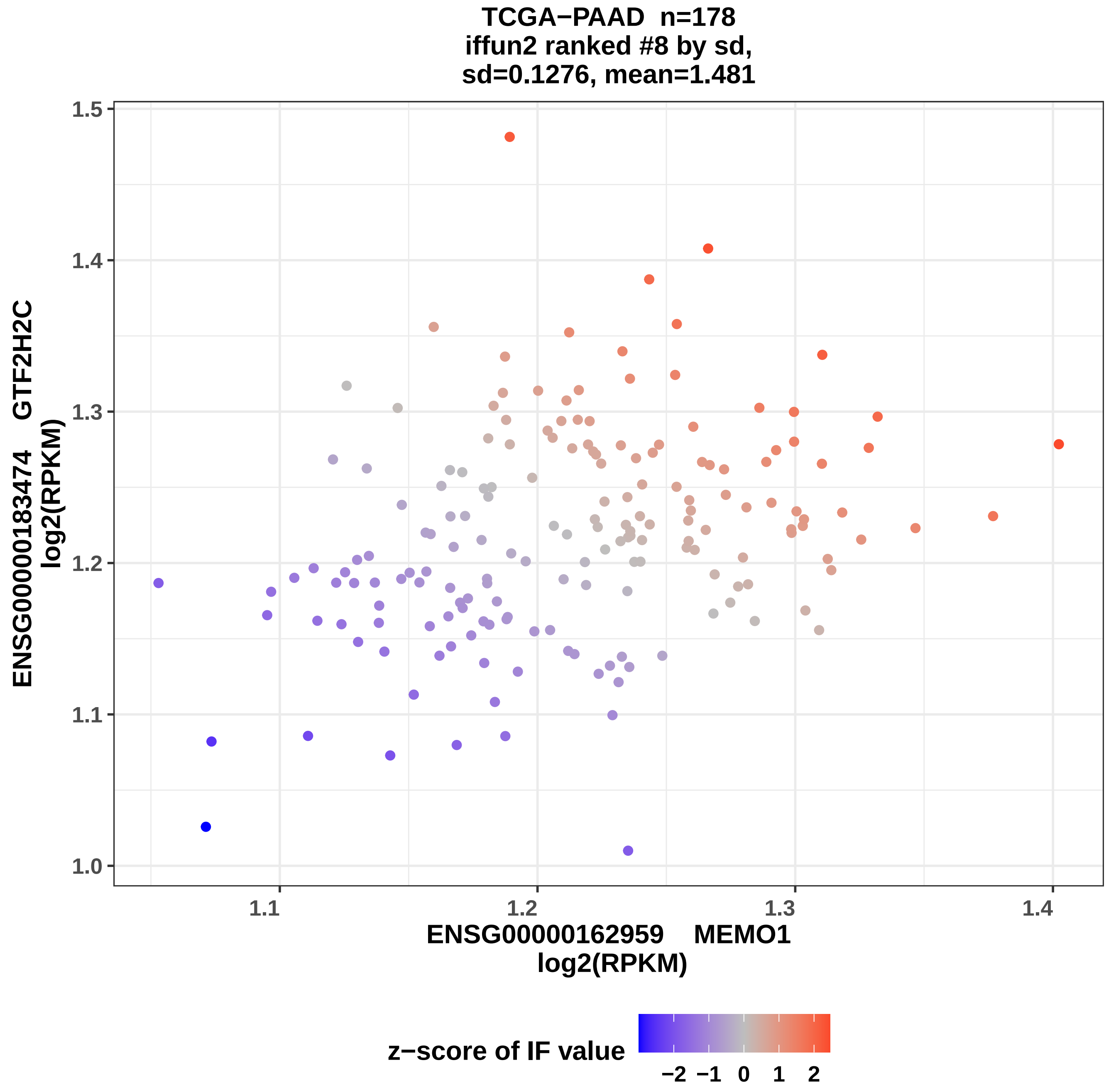
TCGA−PAAD n=178
iffun2 ranked #8 by sd,
sd=0.1276, mean=1.481
ENSG00000183474 GTF2H2C
log2(RPKM)
1.0
1.1
1.2
1.3
1.4
1.5
1.1
1.2
1.3
1.4
ENSG00000162959 MEMO1
log2(RPKM)
z−score of IF value
−2 −1 0 1 2

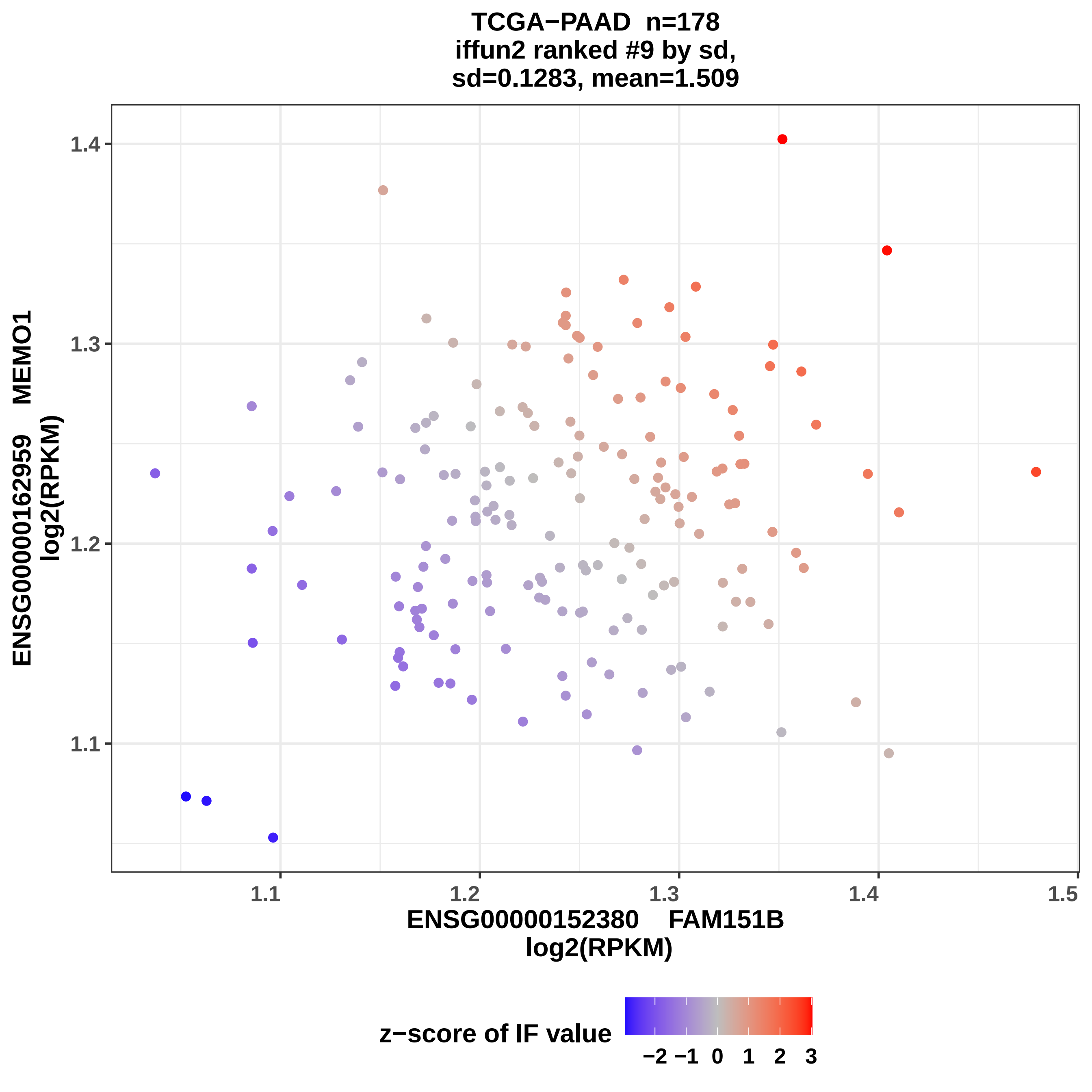
TCGA−PAAD n=178
iffun2 ranked #9 by sd,
sd=0.1283, mean=1.509
ENSG00000162959 MEMO1
log2(RPKM)
1.4
1.3
1.2
1.1
1.1
1.2
1.3
1.4
1.5
ENSG00000152380 FAM151B
log2(RPKM)
z−score of IF value
−2 −1 0 1 2 3

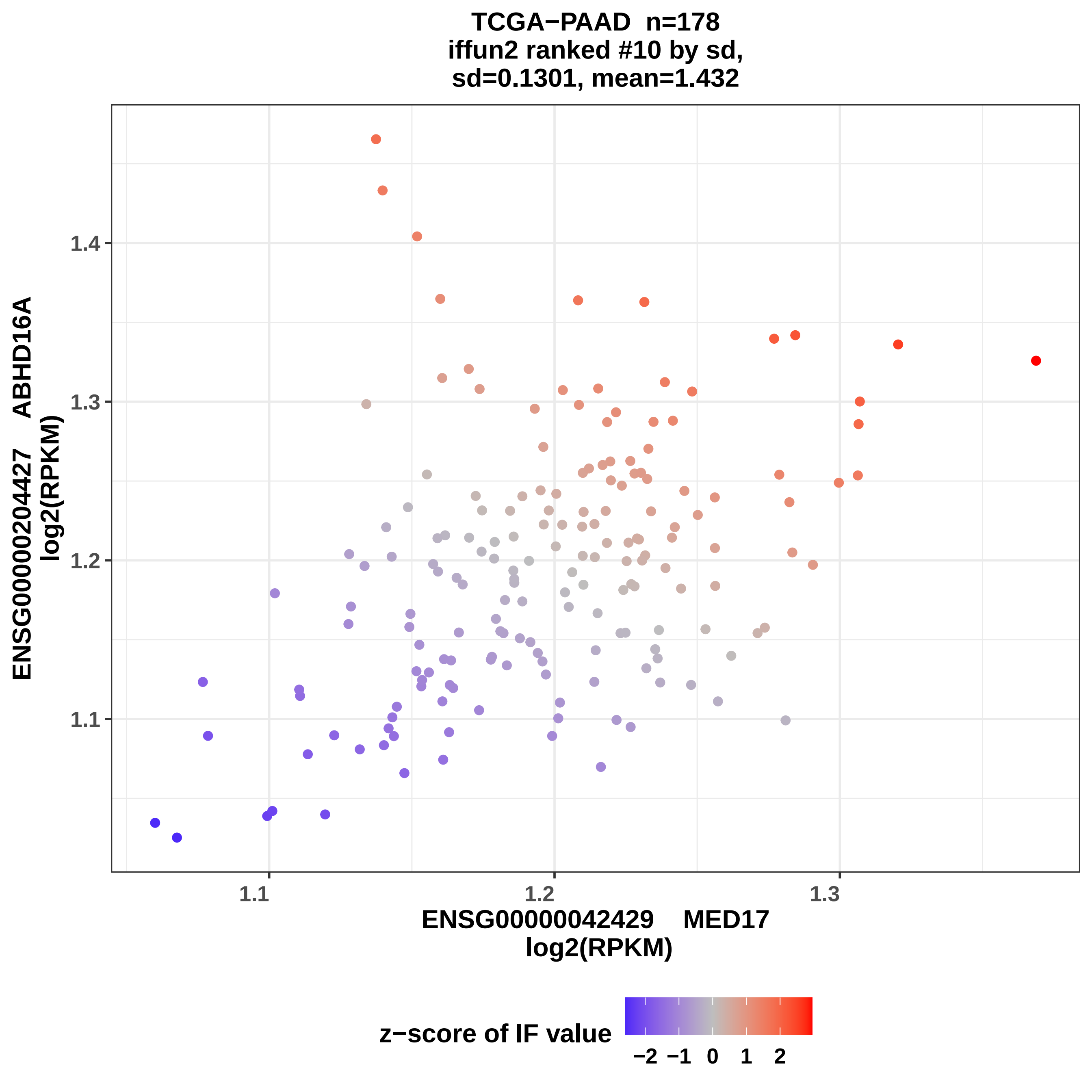
TCGA−PAAD n=178
iffun2 ranked #10 by sd,
sd=0.1301, mean=1.432
ENSG00000204427 ABHD16A
log2(RPKM)
1.4
1.3
1.2
1.1
1.1
1.2
1.3
ENSG00000042429 MED17
log2(RPKM)
z−score of IF value
−2 −1 0 1 2

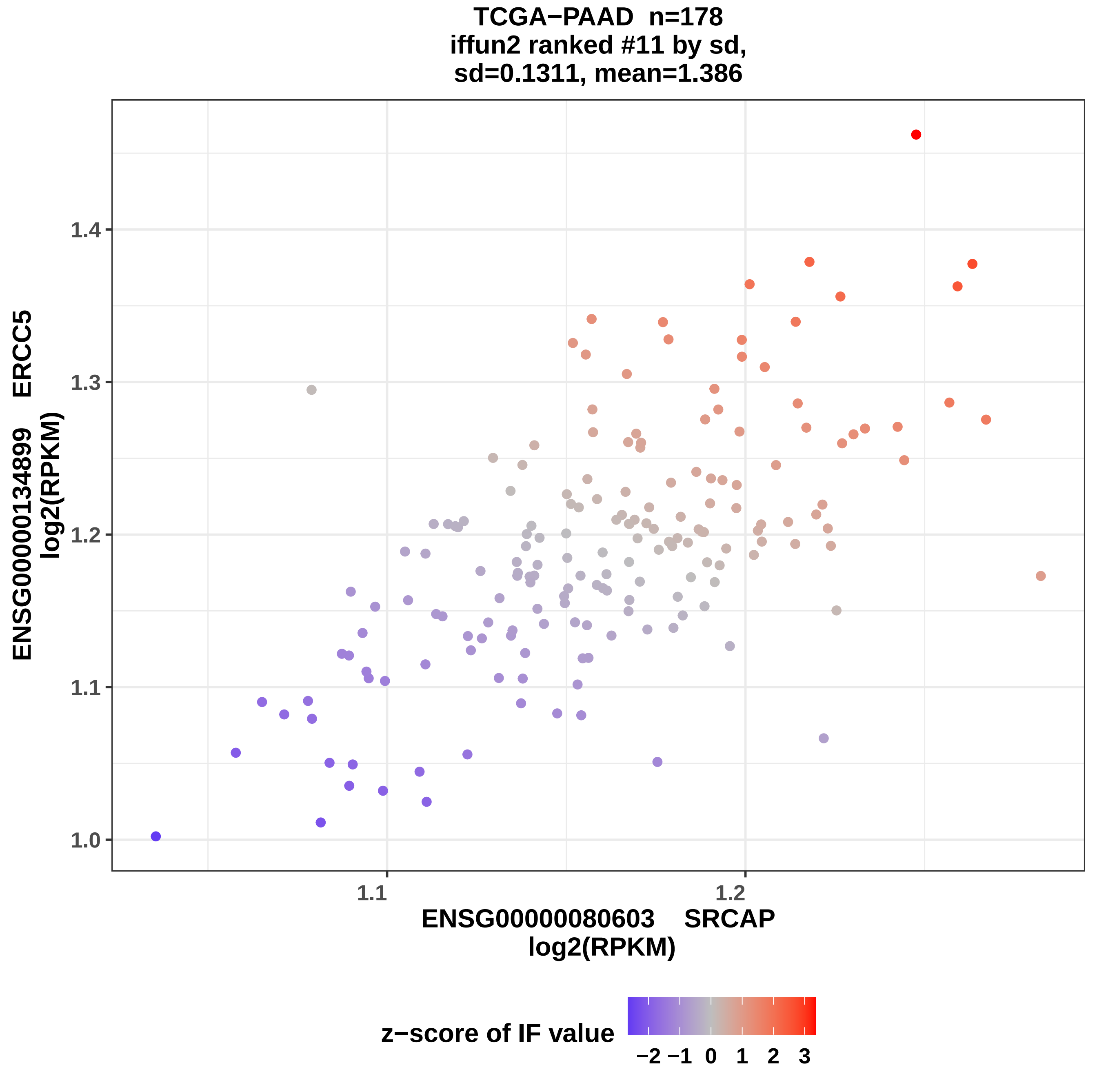
TCGA−PAAD n=178
iffun2 ranked #11 by sd,
sd=0.1311, mean=1.386
ENSG00000134899 ERCC5
log2(RPKM)
1.0
1.1
1.2
1.3
1.4
1.1
1.2
ENSG00000080603 SRCAP
log2(RPKM)
z−score of IF value
−2 −1 0 1 2 3

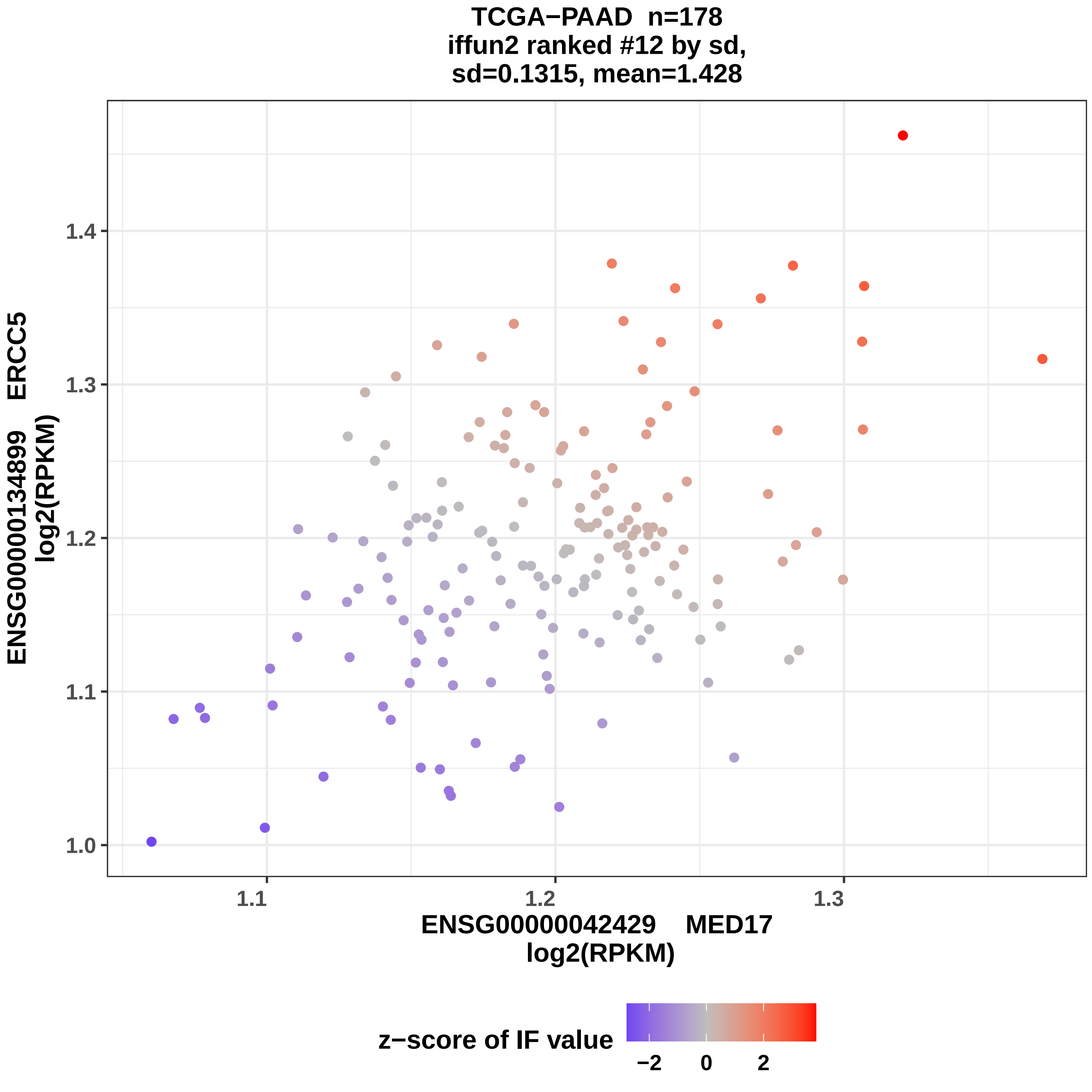
TCGA−PAAD n=178
iffun2 ranked #12 by sd,
sd=0.1315, mean=1.428
ENSG00000134899 ERCC5
log2(RPKM)
1.4
1.3
1.2
1.1
1.0
1.1
1.2
1.3
ENSG00000042429 MED17
log2(RPKM)
z−score of IF value
−2
0
2

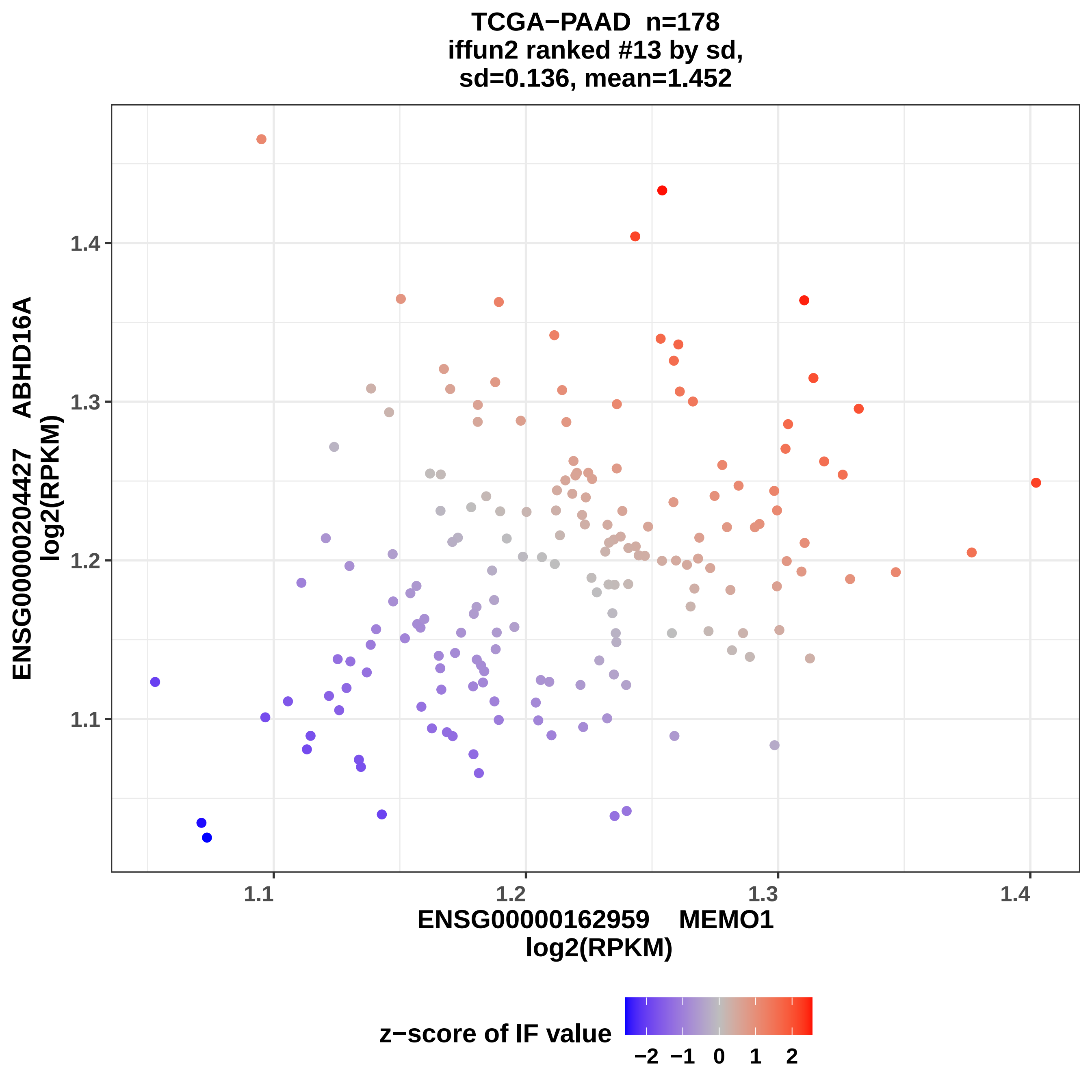

TCGA−PAAD n=178
iffun2 ranked #13 by sd,
sd=0.136, mean=1.452
ENSG00000204427 ABHD16A
log2(RPKM)
1.1
1.2
1.3
1.4
ENSG00000162959 MEMO1
log2(RPKM)
1.1
1.2
1.3
1.4
z−score of IF value
−2
−1
0
1
2

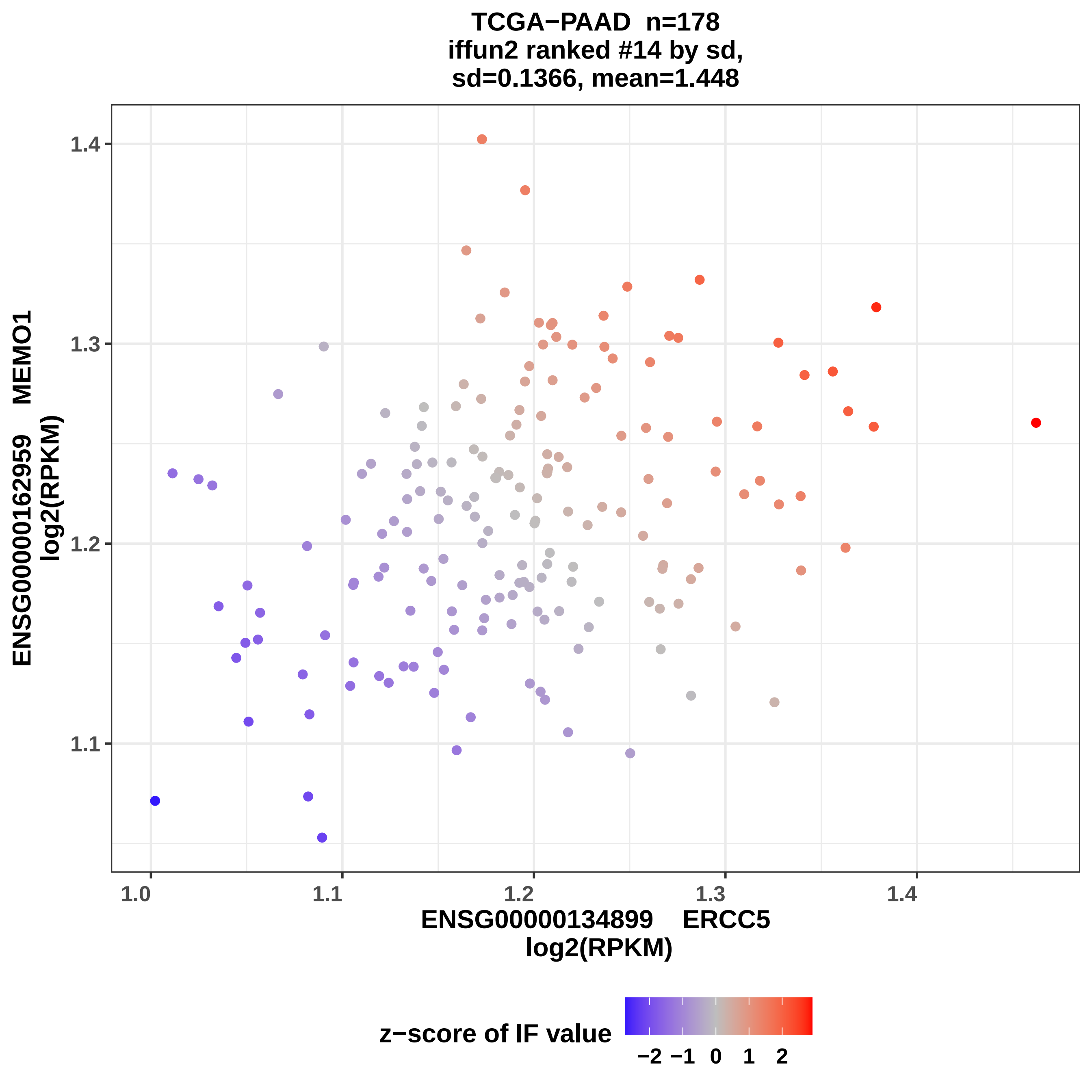
TCGA−PAAD n=178
iffun2 ranked #14 by sd,
sd=0.1366, mean=1.448
ENSG00000162959 MEMO1
log2(RPKM)
1.4
1.3
1.2
1.1
1.0
1.1
1.2
1.3
1.4
ENSG00000134899 ERCC5
log2(RPKM)
z−score of IF value
−2 −1 0 1 2

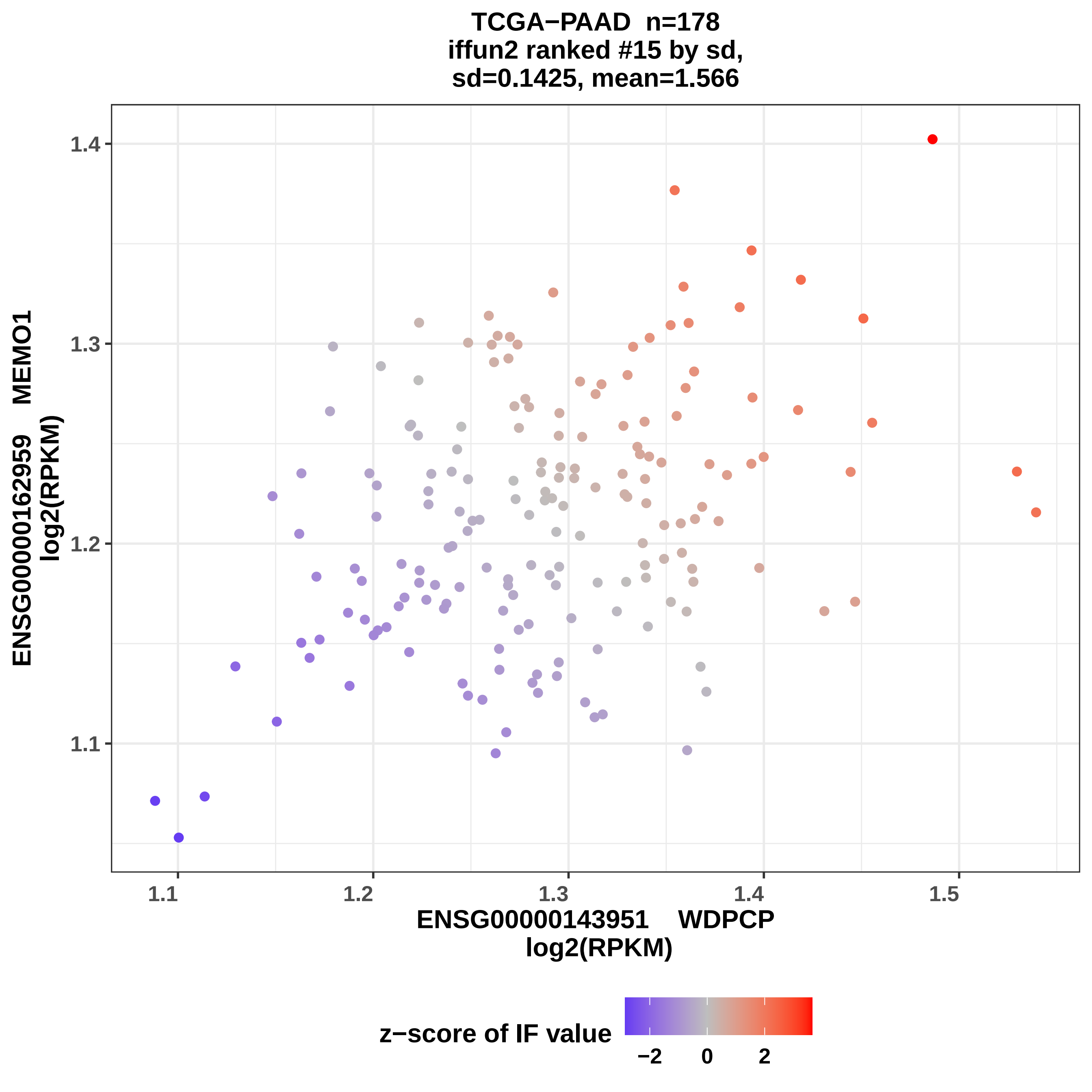

TCGA−PAAD n=178
iffun2 ranked #15 by sd,
sd=0.1425, mean=1.566
ENSG00000162959 MEMO1
log2(RPKM)
1.4
1.3
1.2
1.1
1.1
1.2
1.3
1.4
1.5
ENSG00000143951 WDPCP
log2(RPKM)
z−score of IF value
−2
0
2

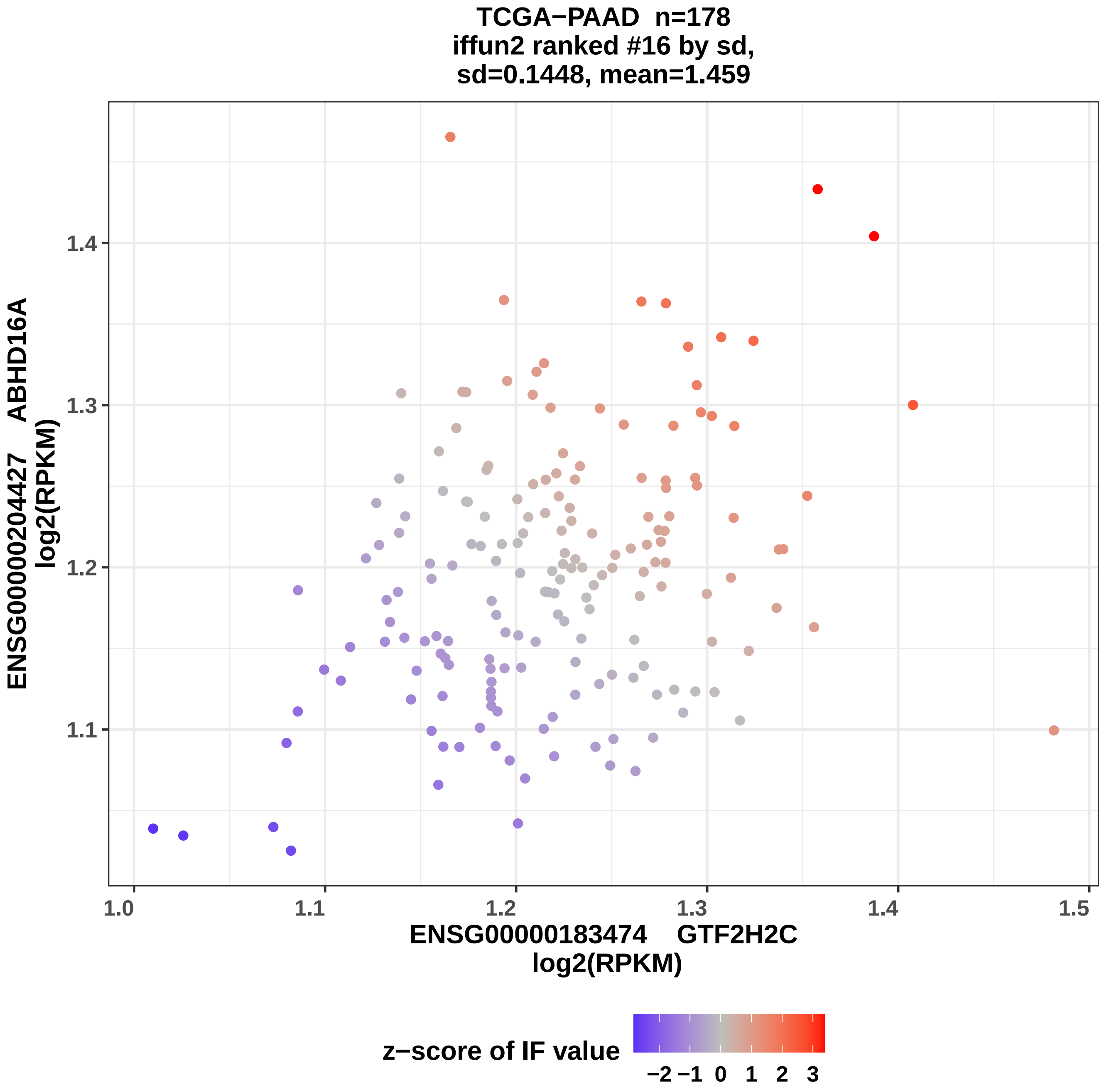
TCGA−PAAD n=178
iffun2 ranked #16 by sd,
sd=0.1448, mean=1.459
ENSG00000204427 ABHD16A
log2(RPKM)
1.1
1.2
1.3
1.4
1.0
1.1
1.2
1.3
1.4
1.5
ENSG00000183474 GTF2H2C
log2(RPKM)
z−score of IF value
−2 −1 0 1 2 3

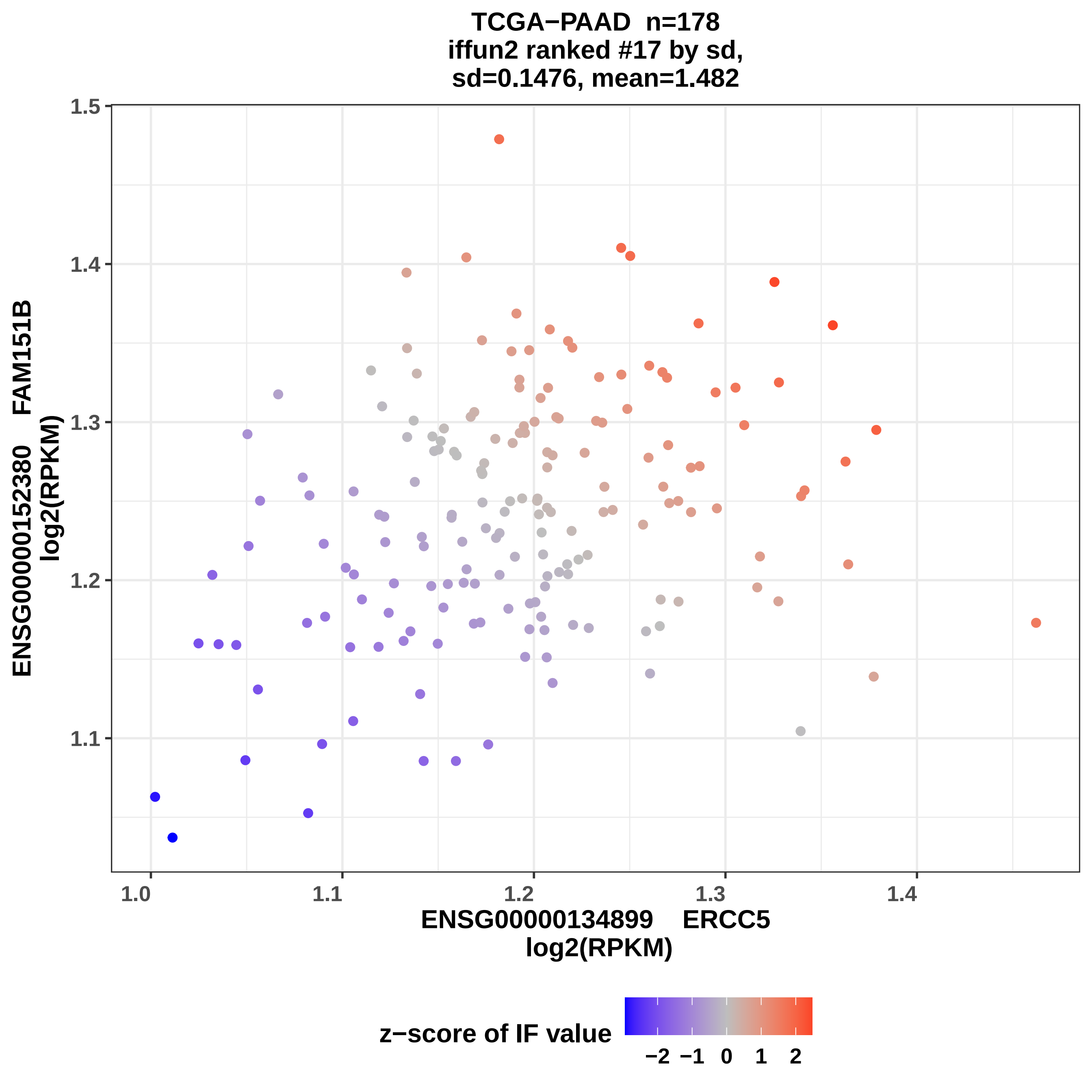

TCGA−PAAD n=178
iffun2 ranked #17 by sd,
sd=0.1476, mean=1.482
ENSG00000152380 FAM151B
log2(RPKM)
1.5
1.4
1.3
1.2
1.1
1.0
1.1
1.2
1.3
1.4
ENSG00000134899 ERCC5
log2(RPKM)
z−score of IF value
−2 −1 0 1 2

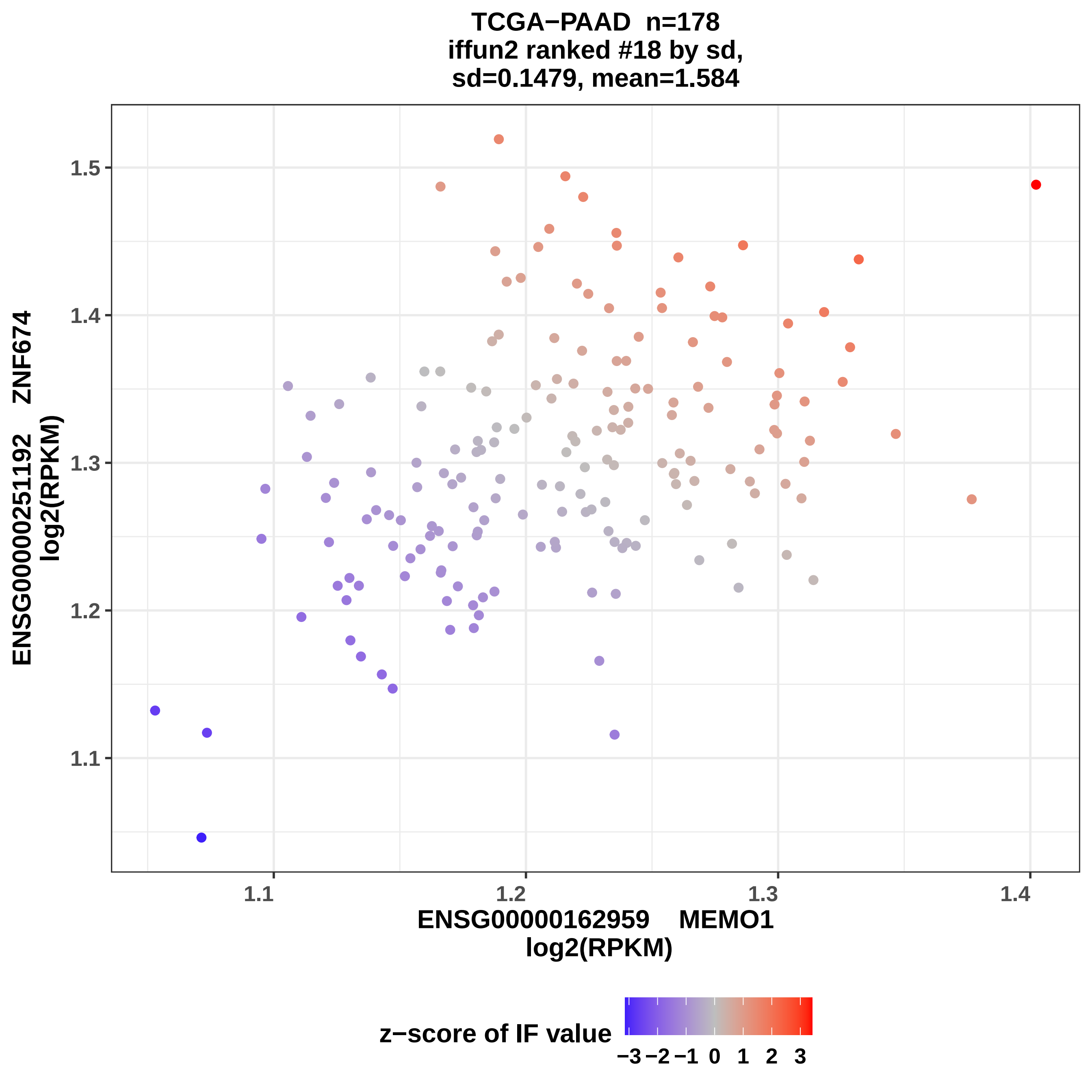
TCGA−PAAD n=178
iffun2 ranked #18 by sd,
sd=0.1479, mean=1.584
1.5
1.4
1.3
1.2
1.1
ENSG00000251192 ZNF674
log2(RPKM)
1.1
1.2
1.3
1.4
ENSG00000162959 MEMO1
log2(RPKM)
z−score of IF value
−3 −2 −1 0 1 2 3

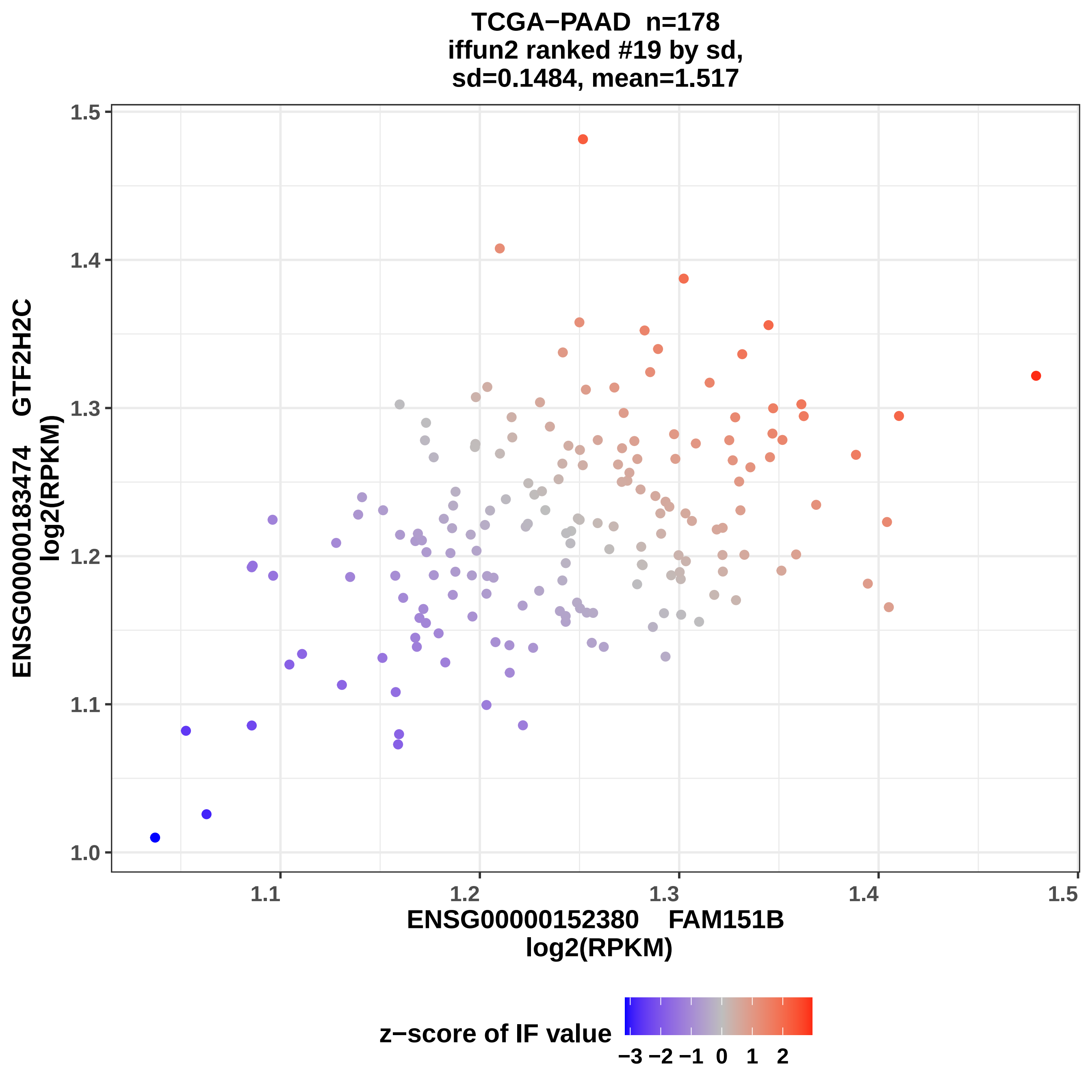
TCGA−PAAD n=178
iffun2 ranked #19 by sd,
sd=0.1484, mean=1.517
ENSG00000183474 GTF2H2C
log2(RPKM)
1.0
1.1
1.2
1.3
1.4
1.5
ENSG00000152380 FAM151B
log2(RPKM)
z−score of IF value
−3 −2 −1 0 1 2

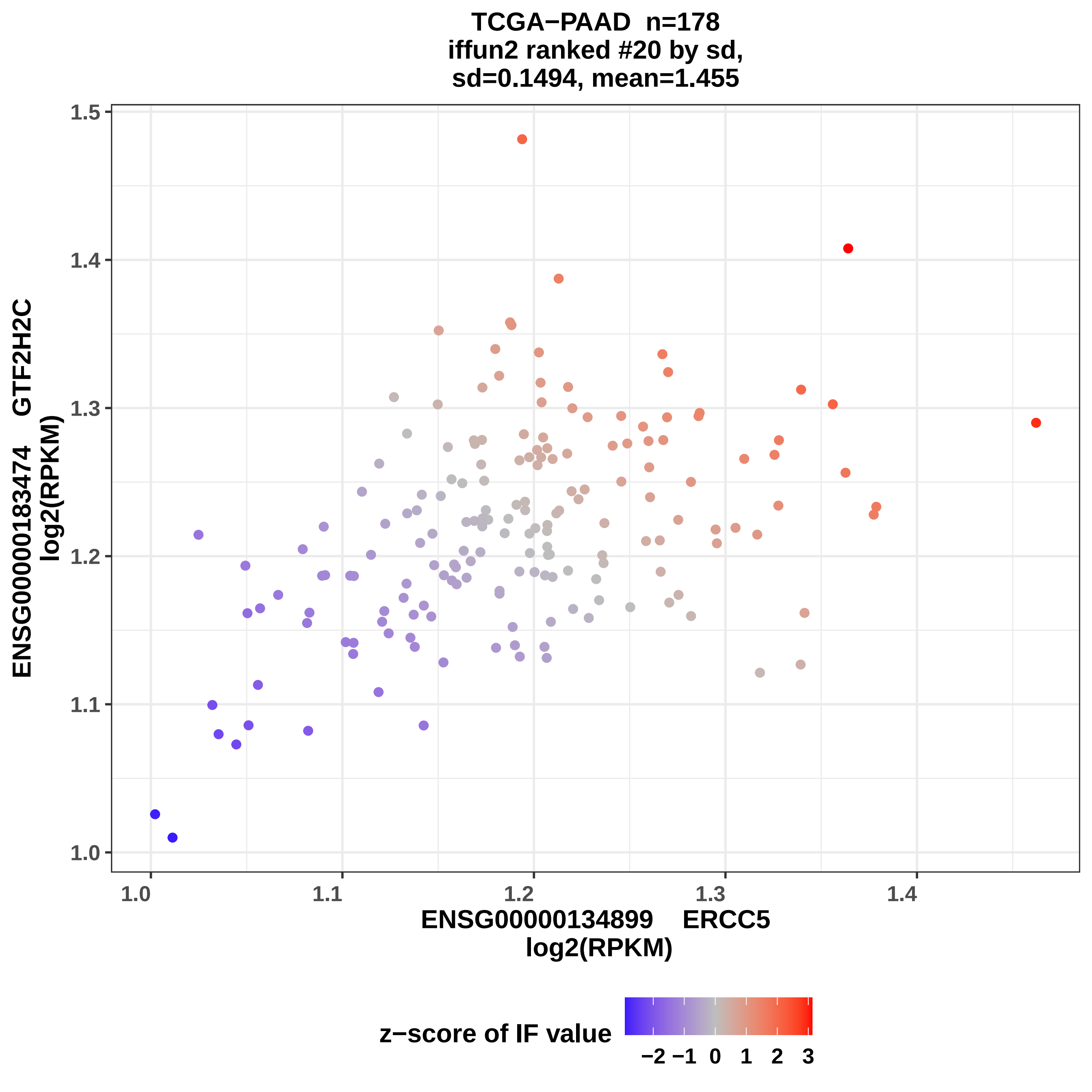
TCGA−PAAD n=178
iffun2 ranked #20 by sd,
sd=0.1494, mean=1.455
ENSG00000183474 GTF2H2C
log2(RPKM)
ENSG00000134899 ERCC5
log2(RPKM)
1.0
1.1
1.2
1.3
1.4
1.5
z−score of IF value
−2 −1 0 1 2 3

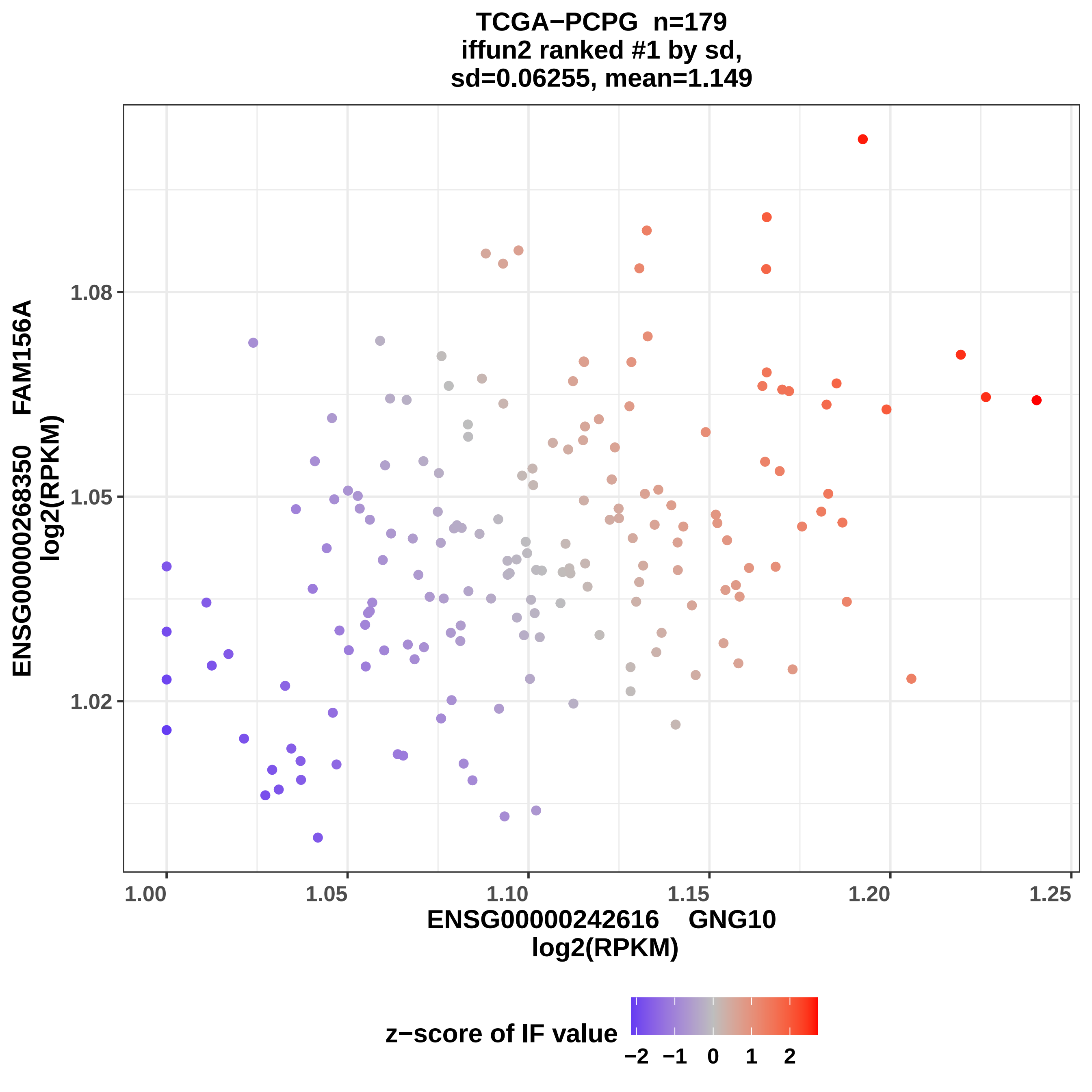

TCGA−PCPG n=179
iffun2 ranked #1 by sd,
sd=0.06255, mean=1.149
ENSG00000268350 FAM156A
log2(RPKM)
1.08
1.05
1.02
1.00
1.05
1.10
1.15
1.20
1.25
ENSG00000242616 GNG10
log2(RPKM)
z−score of IF value
−2
−1
0
1
2

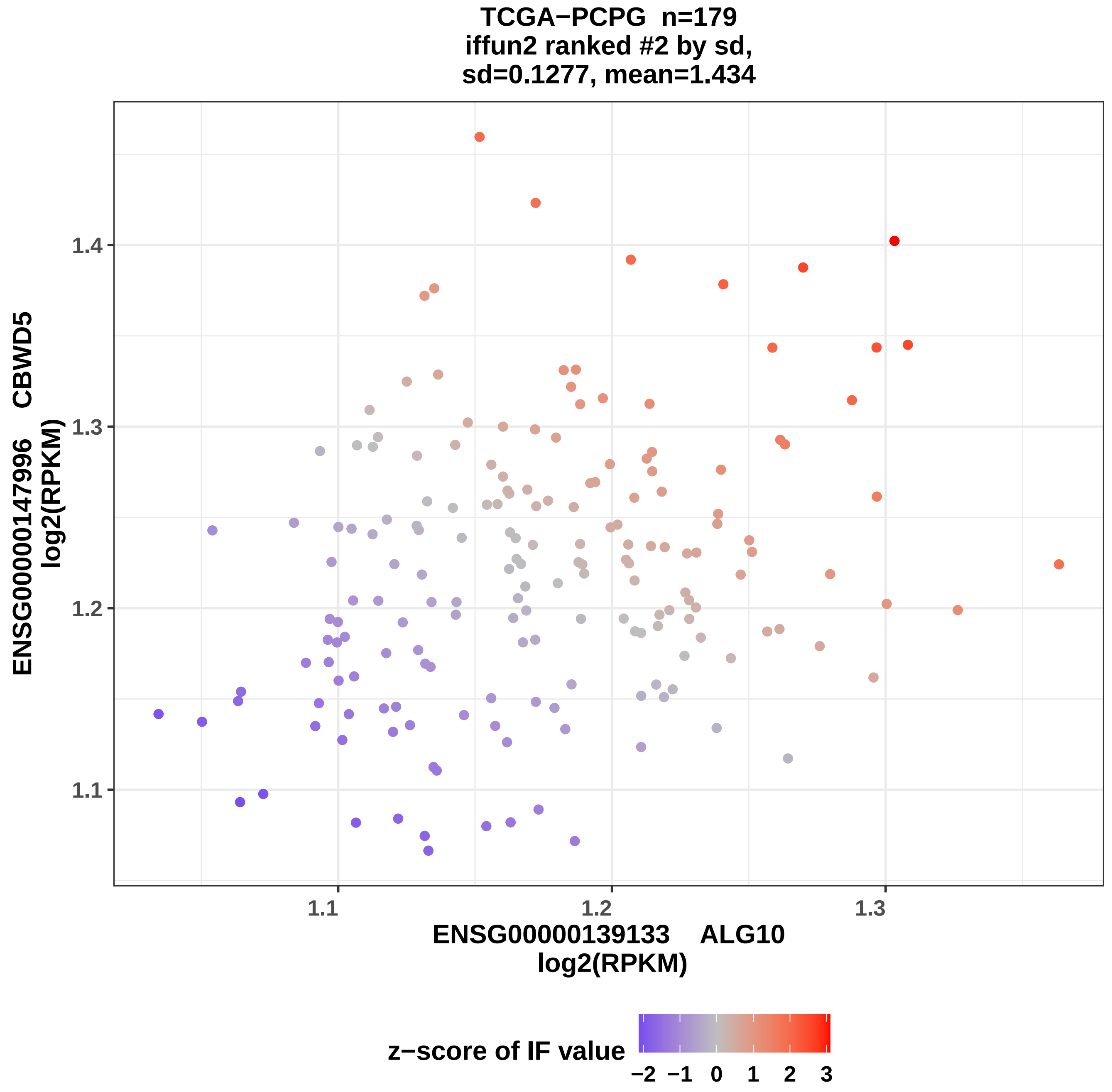
TCGA−PCPG n=179
iffun2 ranked #2 by sd,
sd=0.1277, mean=1.434
ENSG00000147996 CBWD5
log2(RPKM)
1.4
1.3
1.2
1.1
ENSG00000139133 ALG10
log2(RPKM)
1.1
1.2
1.3
z−score of IF value
−2 −1 0 1 2 3

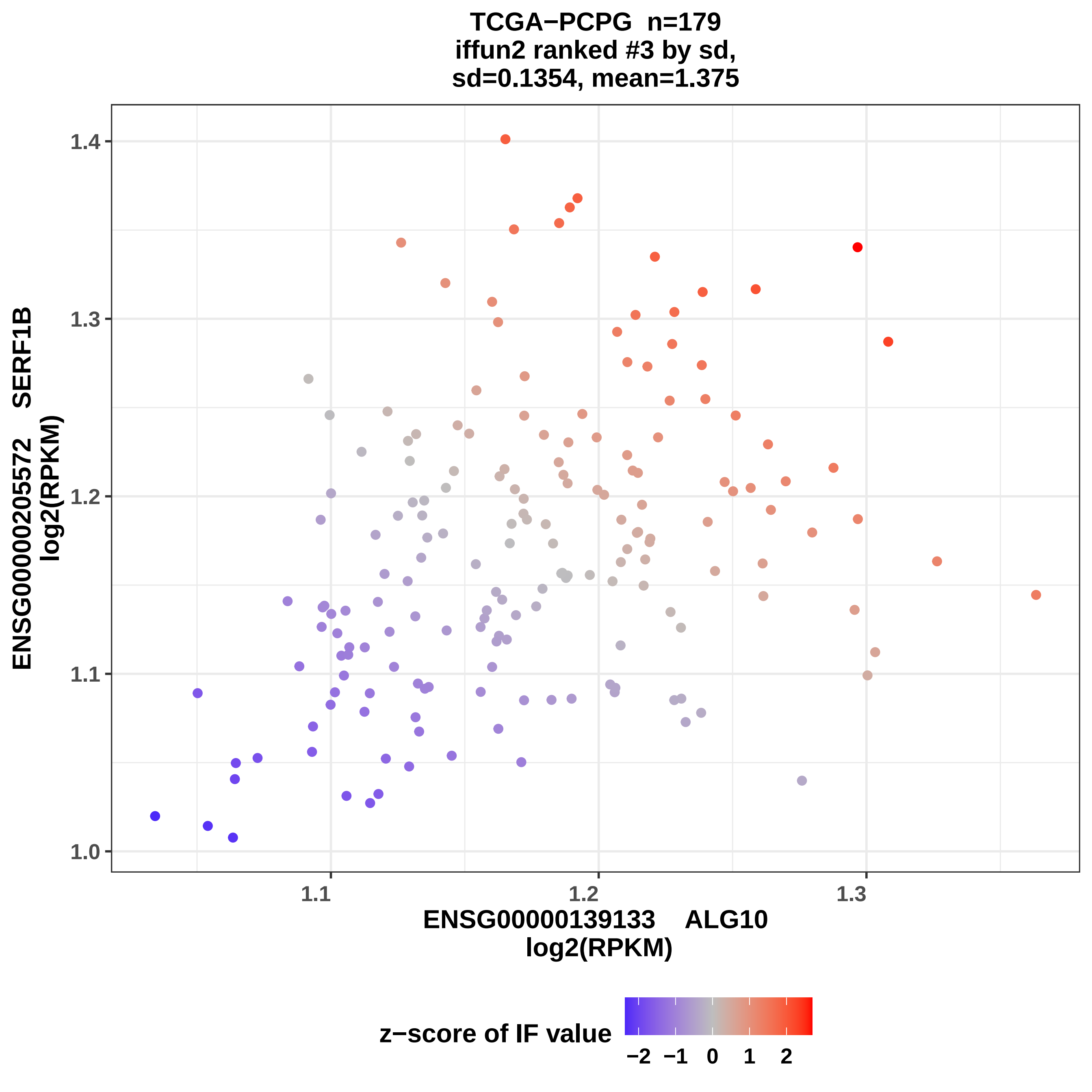
TCGA−PCPG n=179
iffun2 ranked #3 by sd,
sd=0.1354, mean=1.375
ENSG00000205572 SERF1B
log2(RPKM)
1.0
1.1
1.2
1.3
1.4
1.1
1.2
1.3
ENSG00000139133 ALG10
log2(RPKM)
z−score of IF value
−2 −1 0 1 2

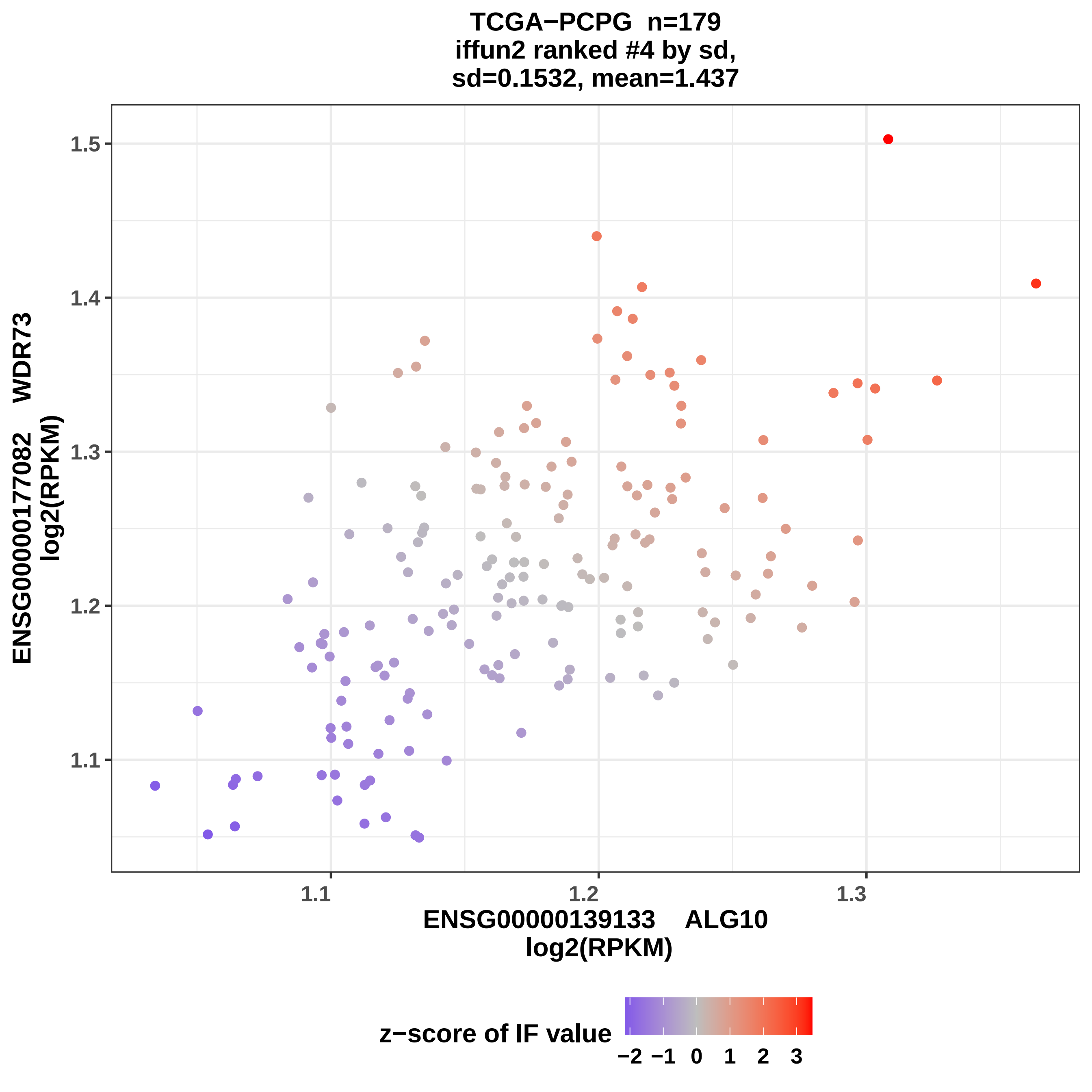

TCGA−PCPG n=179
iffun2 ranked #4 by sd,
sd=0.1532, mean=1.437
1.5
1.4
1.3
1.2
1.1
ENSG00000177082 WDR73
log2(RPKM)
1.1
1.2
1.3
ENSG00000139133 ALG10
log2(RPKM)
z−score of IF value
−2 −1 0 1 2 3

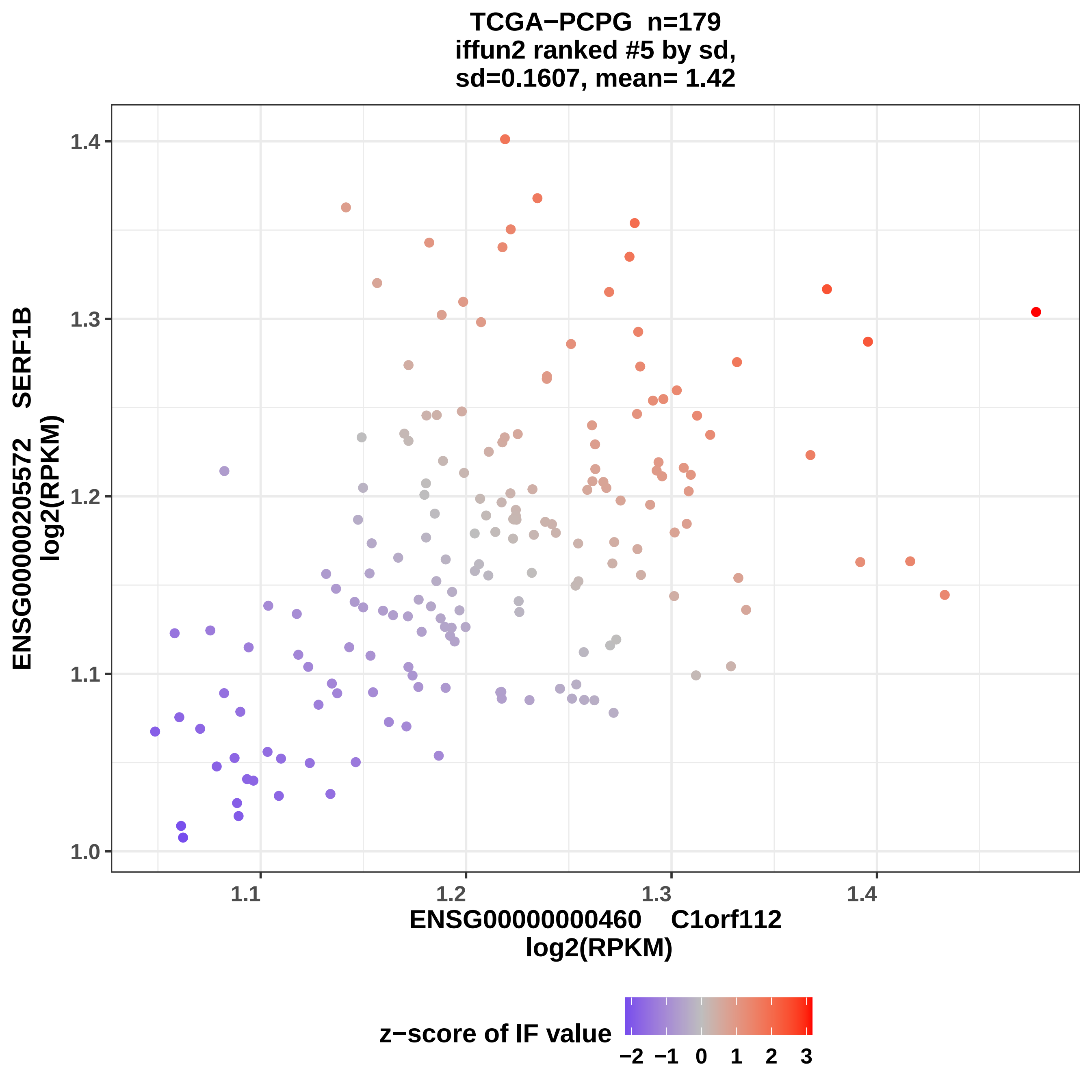
TCGA−PCPG n=179
iffun2 ranked #5 by sd,
sd=0.1607, mean= 1.42
ENSG00000205572 SERF1B
log2(RPKM)
1.0
1.1
1.2
1.3
1.4
ENSG00000000460 C1orf112
log2(RPKM)
z−score of IF value
−2 −1 0 1 2 3

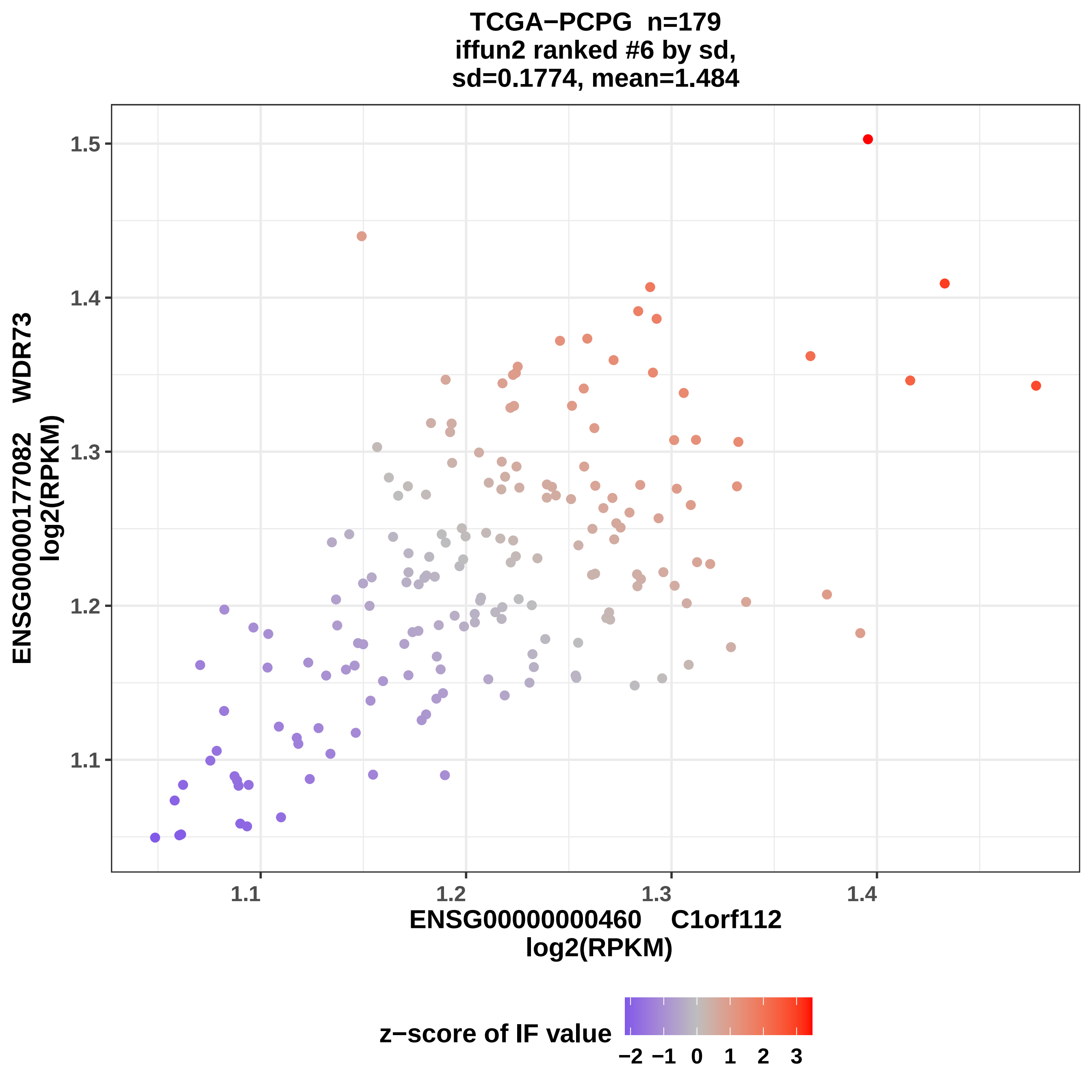

TCGA−PCPG n=179
iffun2 ranked #6 by sd,
sd=0.1774, mean=1.484
ENSG00000177082 WDR73
log2(RPKM)
1.5
1.4
1.3
1.2
1.1
1.1
1.2
1.3
1.4
ENSG00000000460 C1orf112
log2(RPKM)
z−score of IF value
−2
−1
0
1
2
3

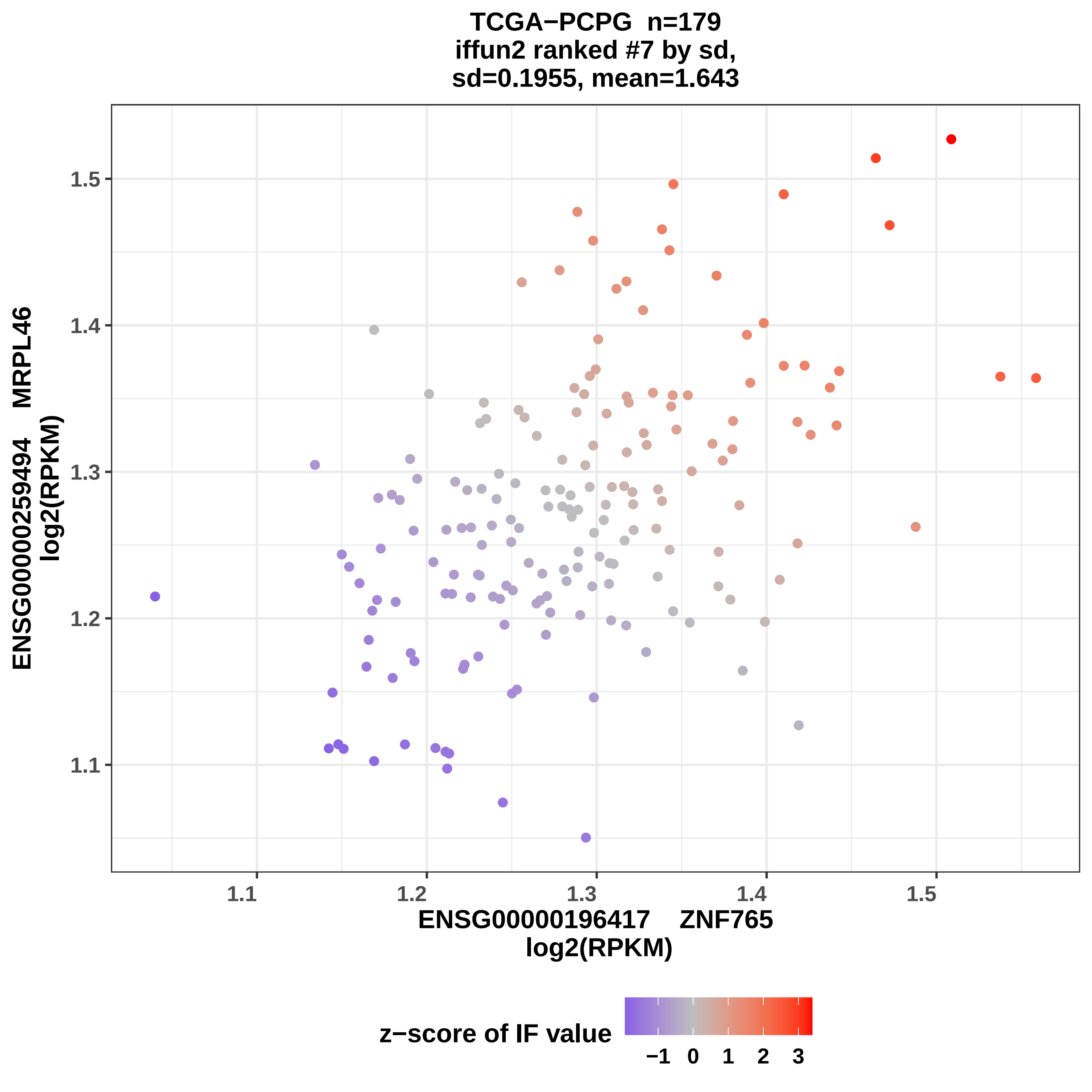
TCGA−PCPG n=179
iffun2 ranked #7 by sd,
sd=0.1955, mean=1.643
ENSG00000259494 MRPL46
log2(RPKM)
1.5
1.4
1.3
1.2
1.1
ENSG00000196417 ZNF765
log2(RPKM)
z−score of IF value
−1
0
1
2
3

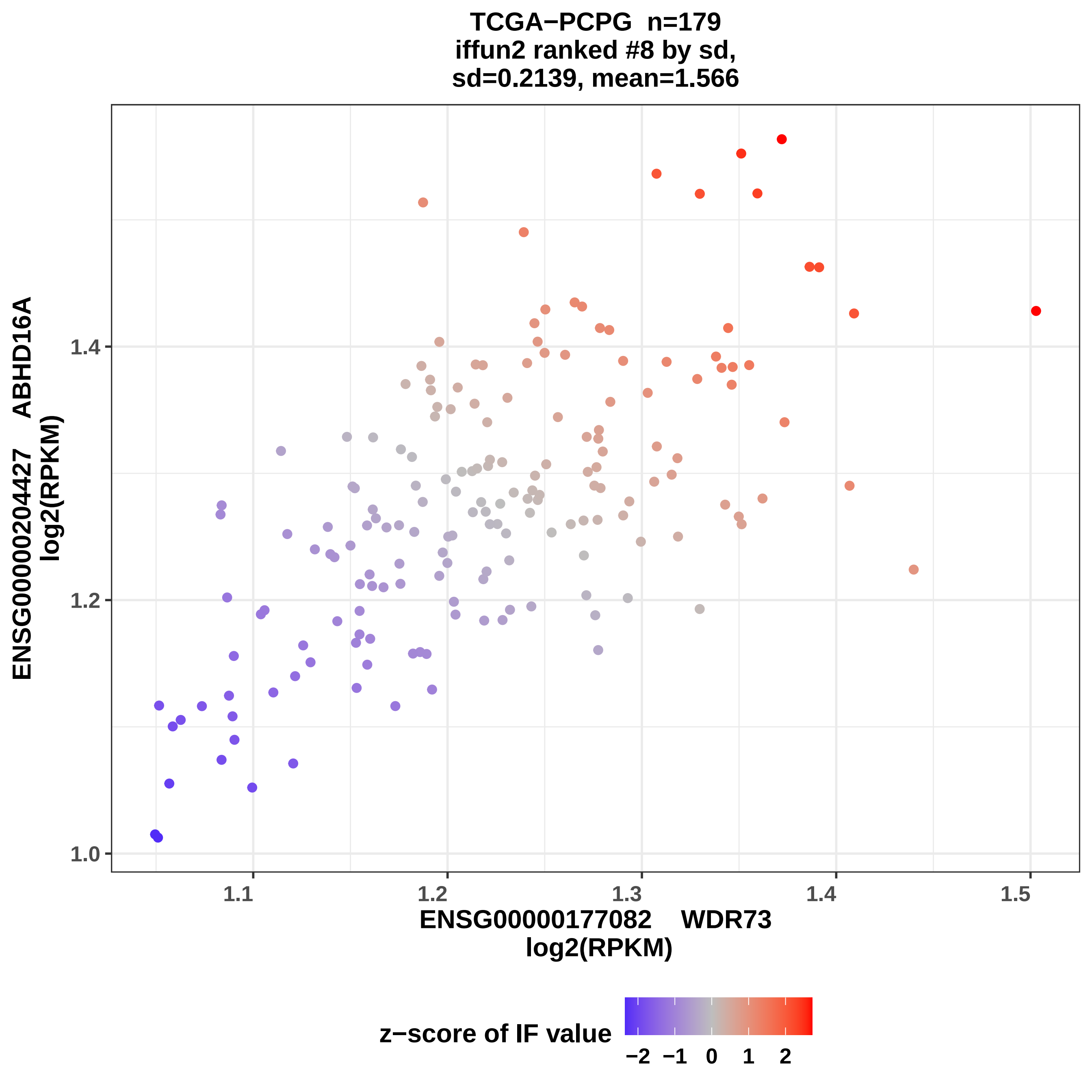
TCGA−PCPG n=179
iffun2 ranked #8 by sd,
sd=0.2139, mean=1.566
ENSG00000204427 ABHD16A
log2(RPKM)
1.0
1.2
1.4
1.1
1.2
1.3
1.4
1.5
ENSG00000177082 WDR73
log2(RPKM)
z−score of IF value
−2
−1
0
1
2

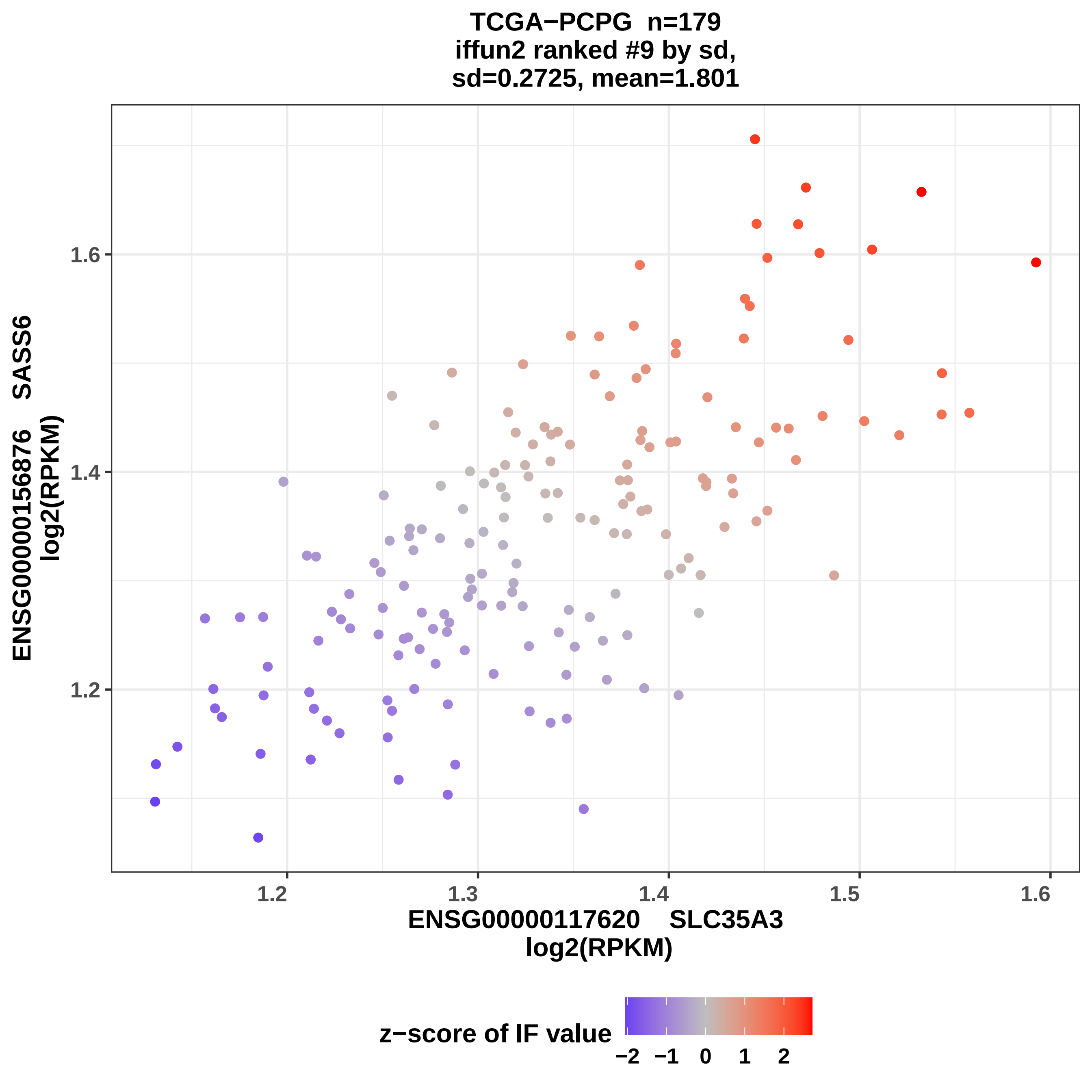

TCGA−PCPG n=179
iffun2 ranked #9 by sd,
sd=0.2725, mean=1.801
ENSG00000156876 SASS6
log2(RPKM)
1.6
1.4
1.2
1.2
1.3
1.4
1.5
1.6
ENSG00000117620 SLC35A3
log2(RPKM)
z−score of IF value
−2
−1
0
1
2

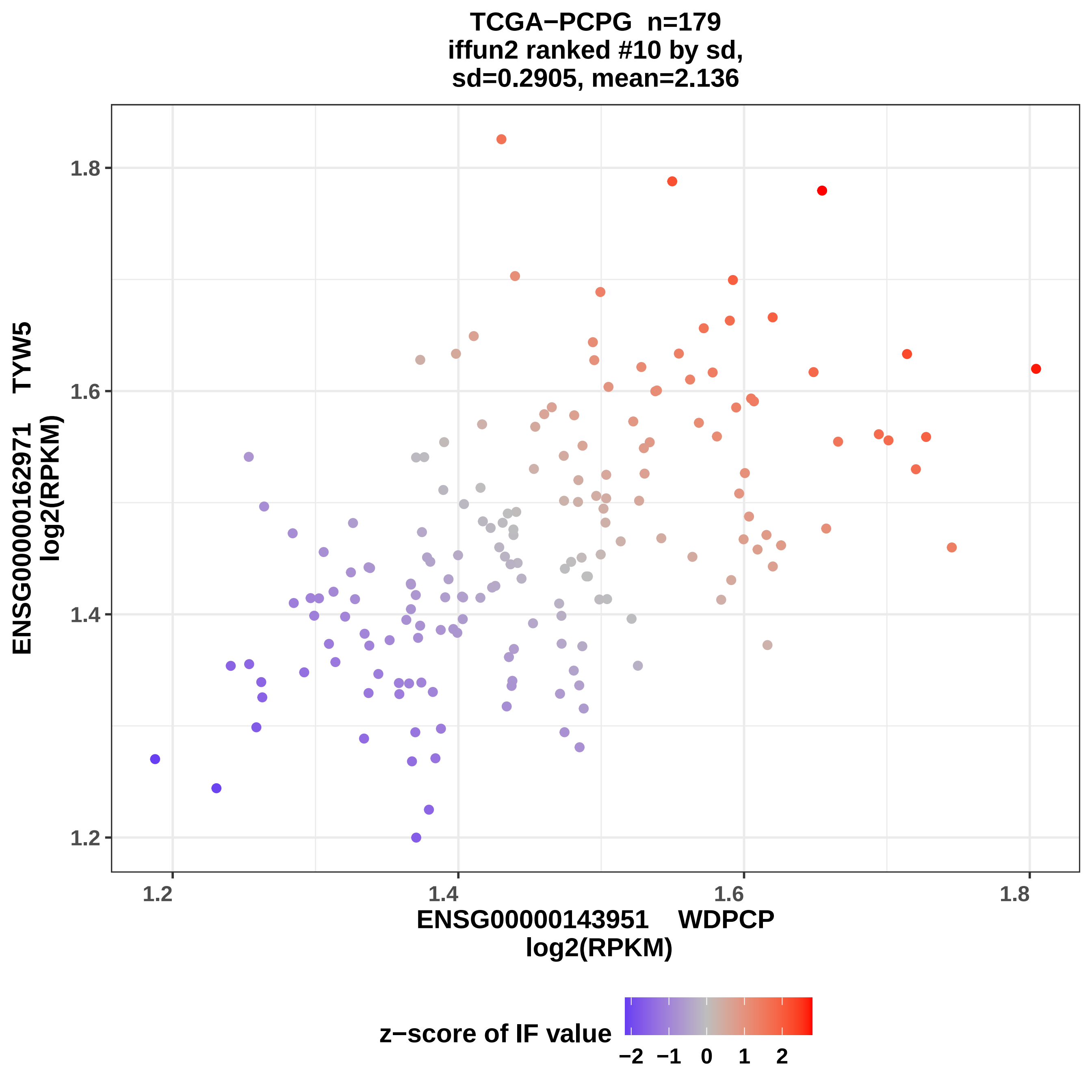
TCGA−PCPG n=179
iffun2 ranked #10 by sd,
sd=0.2905, mean=2.136
ENSG00000162971 TYW5
log2(RPKM)
1.8
1.6
1.4
1.2
1.2
1.4
1.6
1.8
ENSG00000143951 WDPCP
log2(RPKM)
z−score of IF value
−2
−1
0
1
2

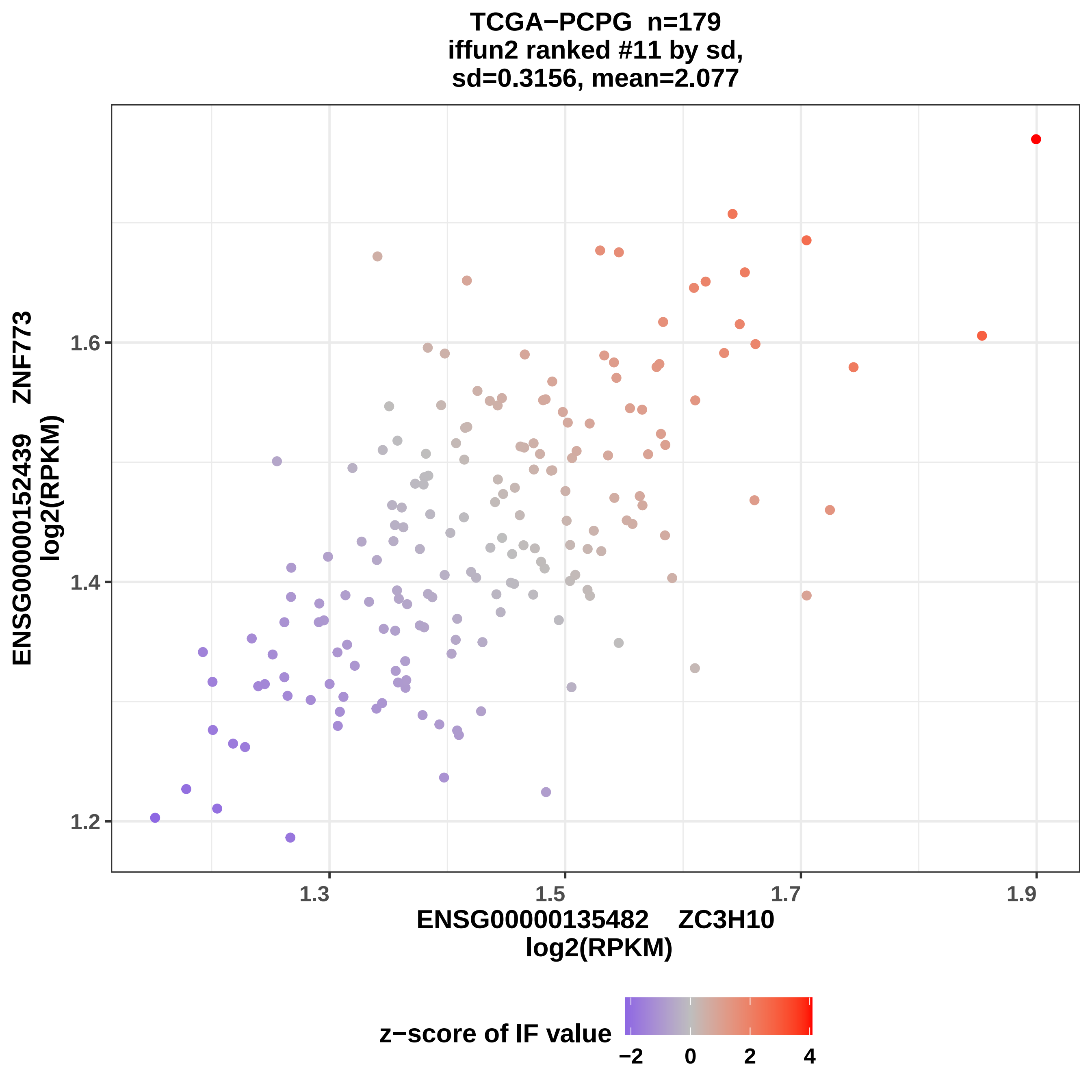
TCGA−PCPG n=179
iffun2 ranked #11 by sd,
sd=0.3156, mean=2.077
ENSG00000152439 ZNF773
log2(RPKM)
1.2
1.4
1.6
1.3
1.5
1.7
1.9
ENSG00000135482 ZC3H10
log2(RPKM)
z−score of IF value
−2
0
2
4

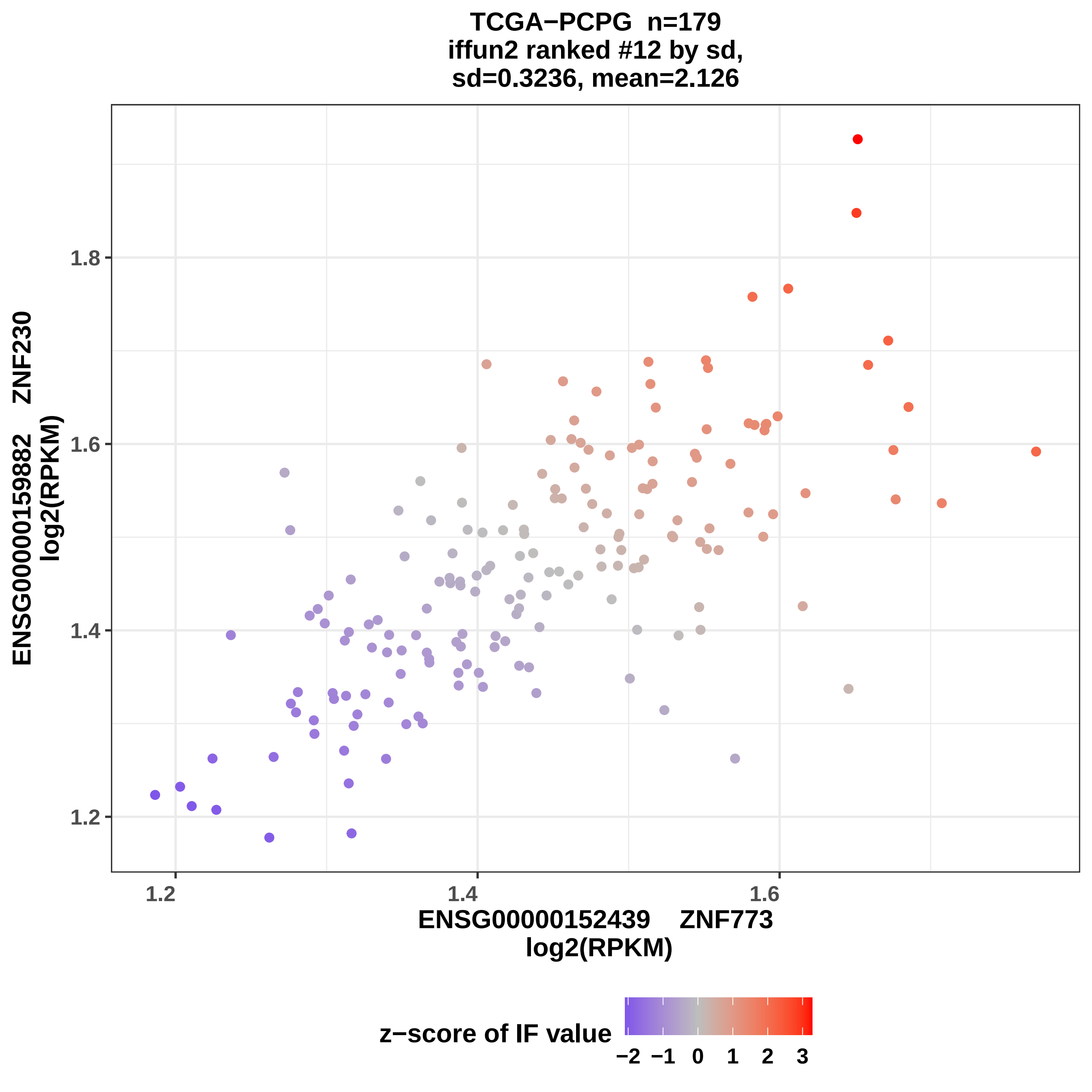
TCGA−PCPG n=179
iffun2 ranked #12 by sd,
sd=0.3236, mean=2.126
ENSG00000159882 ZNF230
log2(RPKM)
1.8
1.6
1.4
1.2
1.2
1.4
1.6
ENSG00000152439 ZNF773
log2(RPKM)
z−score of IF value
−2 −1 0 1 2 3

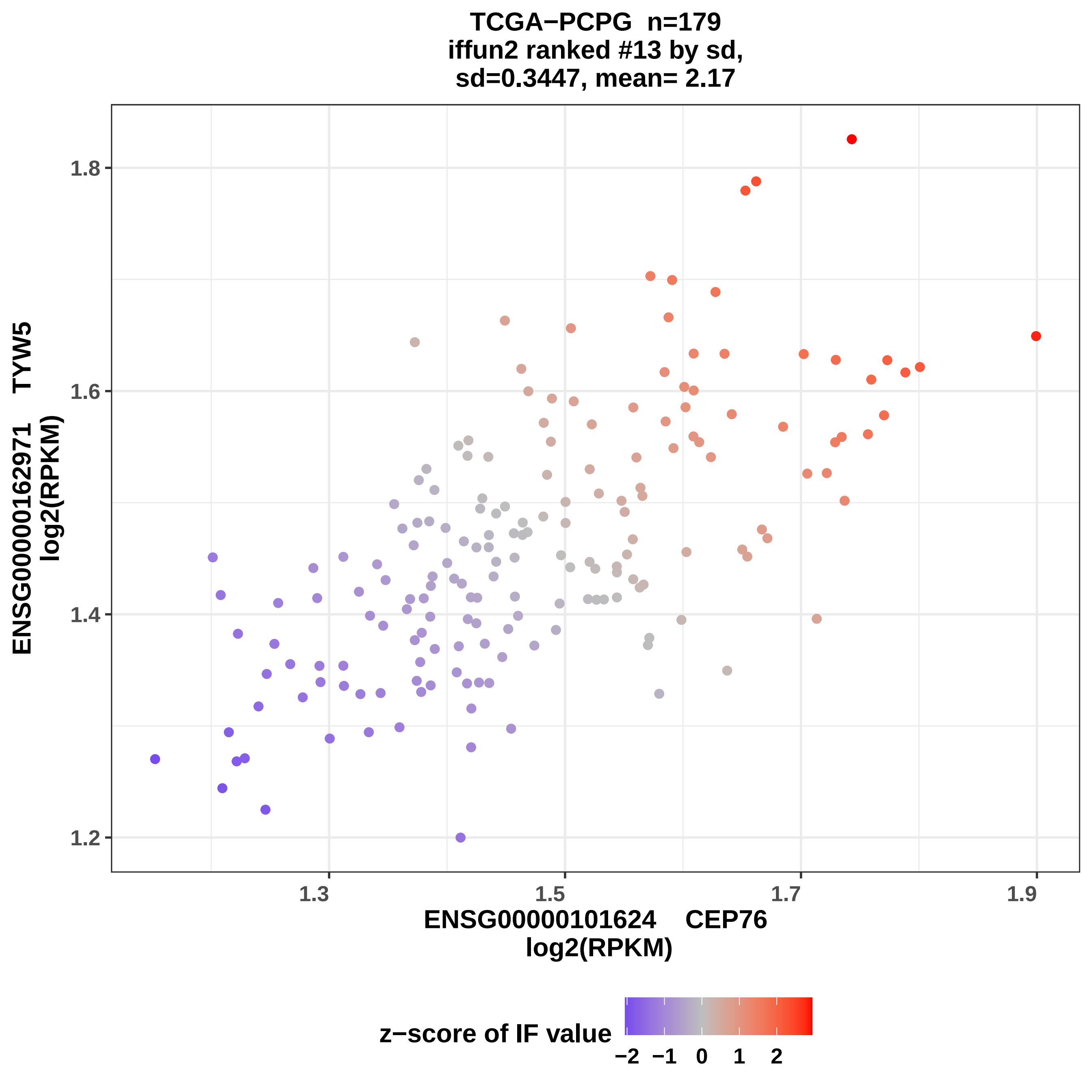
TCGA−PCPG n=179
iffun2 ranked #13 by sd,
sd=0.3447, mean= 2.17
ENSG00000162971 TYW5
log2(RPKM)
1.8
1.6
1.4
1.2
1.3
1.5
1.7
1.9
ENSG00000101624 CEP76
log2(RPKM)
z−score of IF value
−2 −1 0 1 2

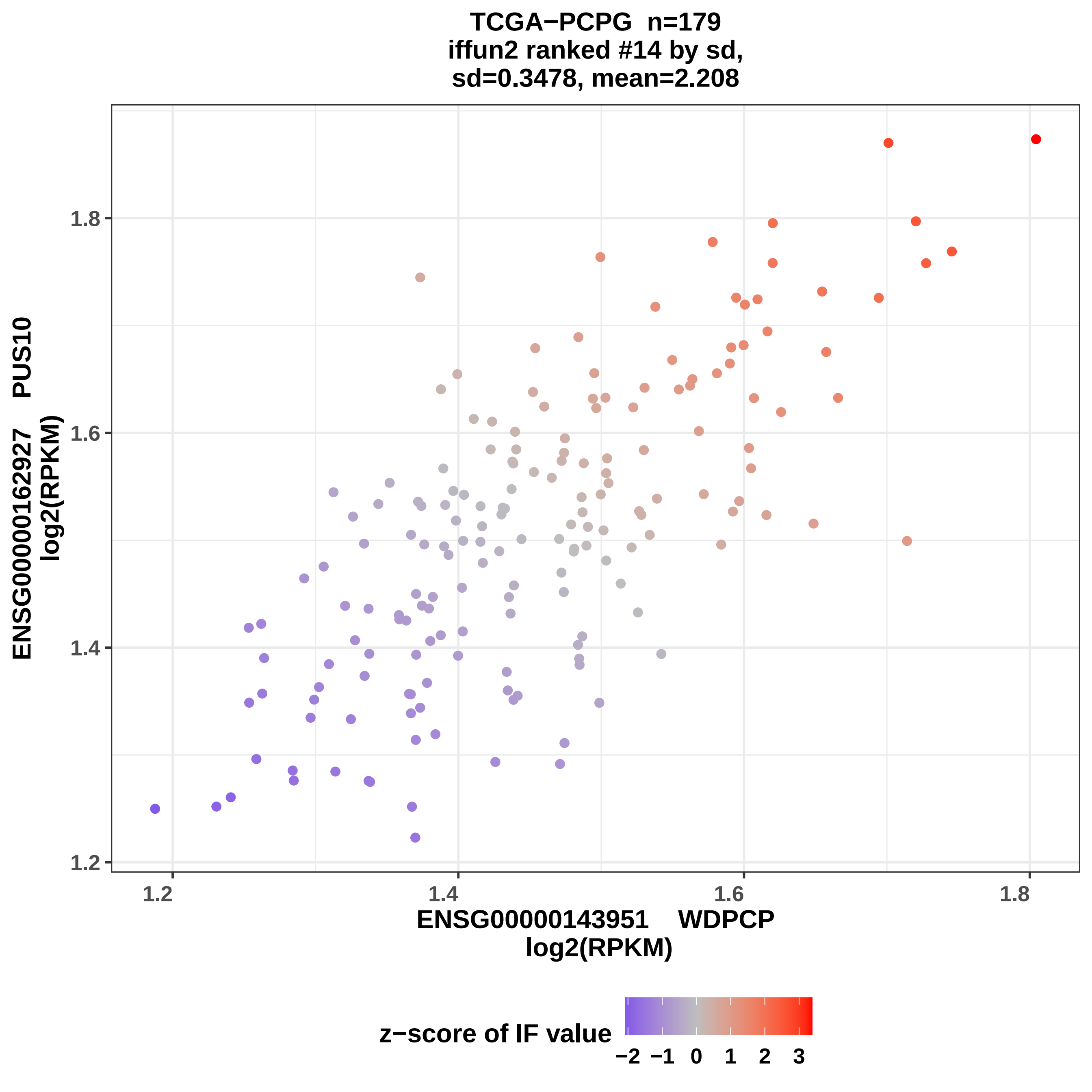
TCGA−PCPG n=179
iffun2 ranked #14 by sd,
sd=0.3478, mean=2.208
ENSG00000162927 PUS10
log2(RPKM)
1.2
1.4
1.6
1.8
ENSG00000143951 WDPCP
log2(RPKM)
z−score of IF value
−2 −1 0 1 2 3

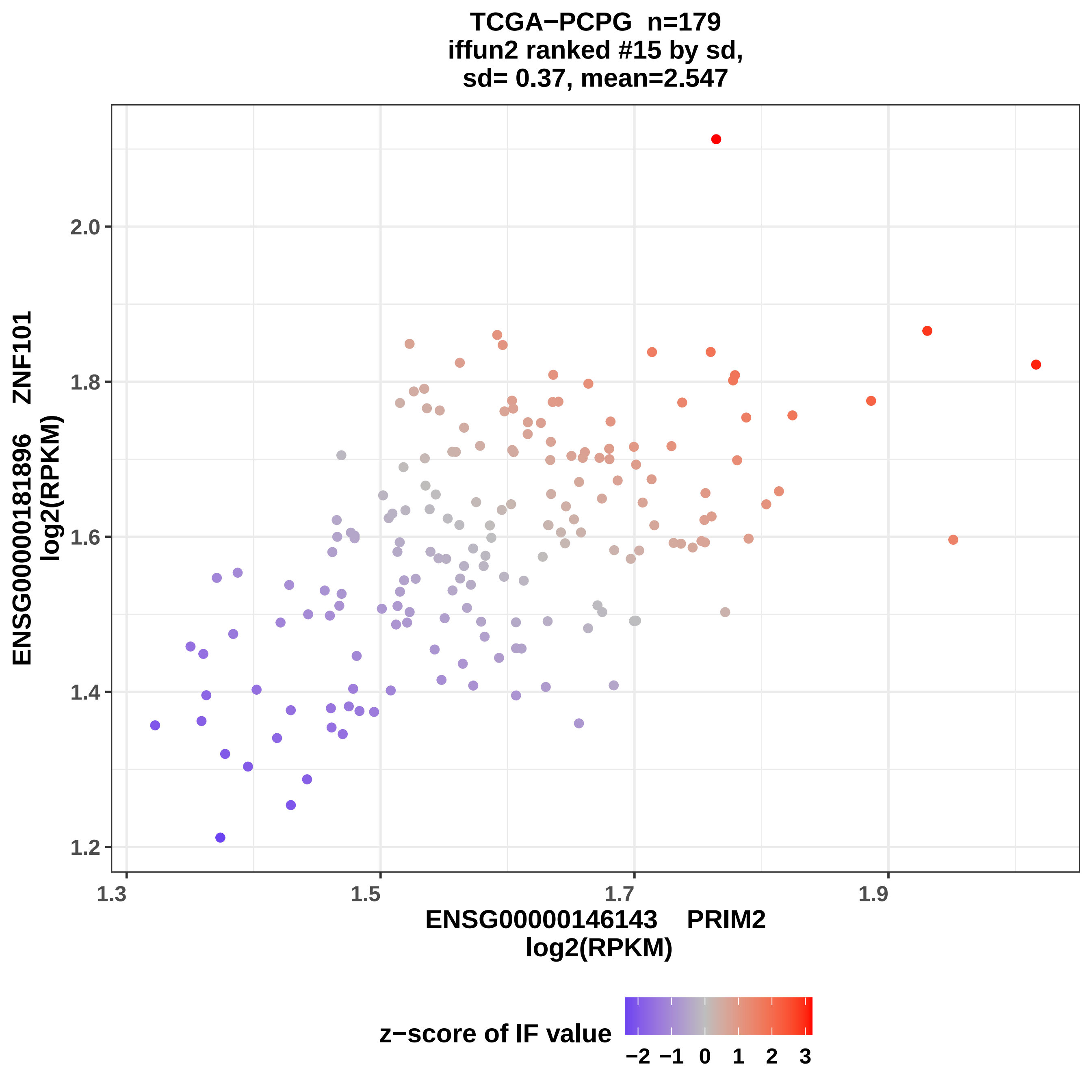
TCGA−PCPG n=179
iffun2 ranked #15 by sd,
sd= 0.37, mean=2.547
ENSG00000181896 ZNF101
log2(RPKM)
2.0
1.8
1.6
1.4
1.2
1.3
1.5
1.7
1.9
ENSG00000146143 PRIM2
log2(RPKM)
z−score of IF value
−2 −1 0 1 2 3

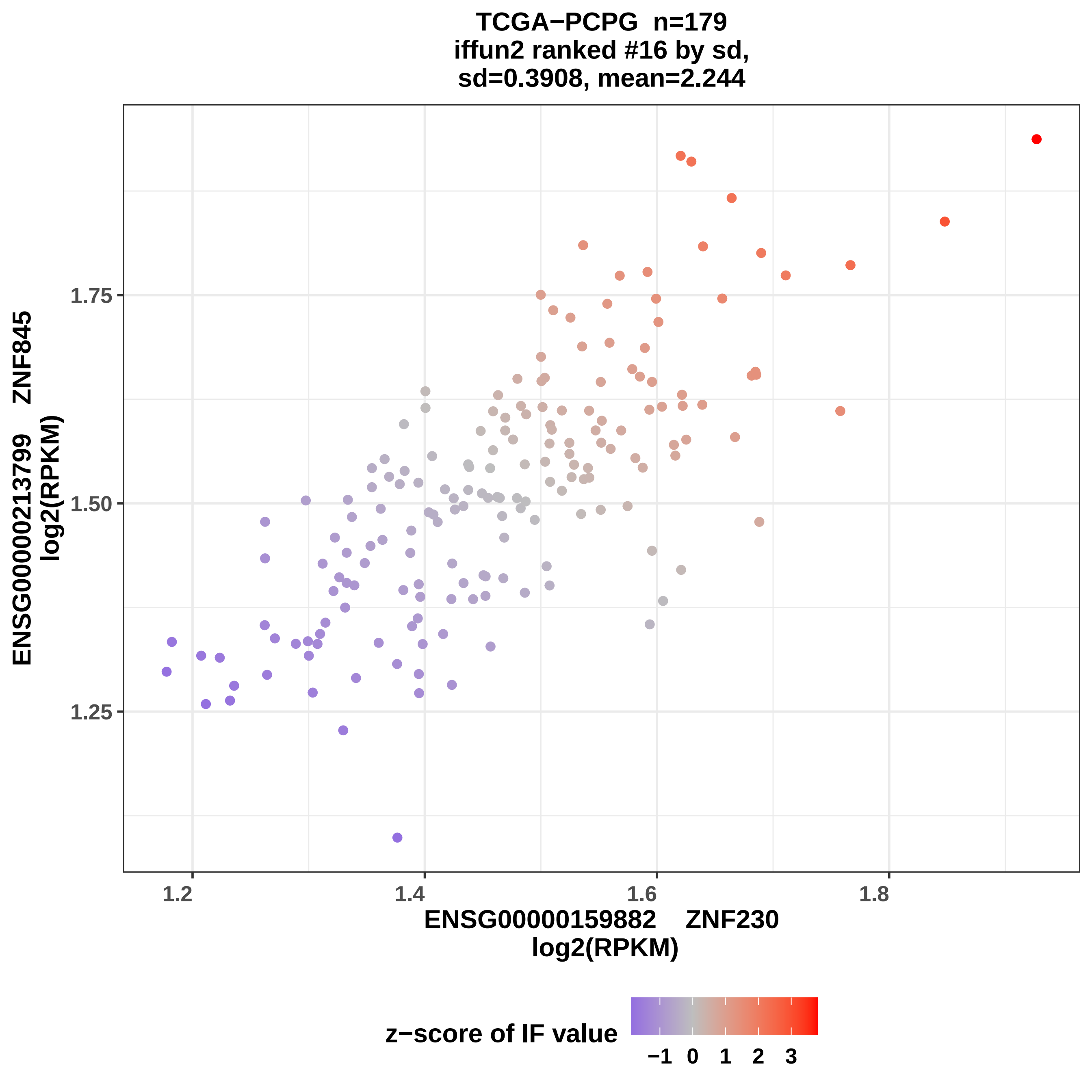
TCGA−PCPG n=179
iffun2 ranked #16 by sd,
sd=0.3908, mean=2.244
ENSG00000213799 ZNF845
log2(RPKM)
1.75
1.50
1.25
1.2
1.4
1.6
1.8
ENSG00000159882 ZNF230
log2(RPKM)
z−score of IF value
−1
0
1
2
3

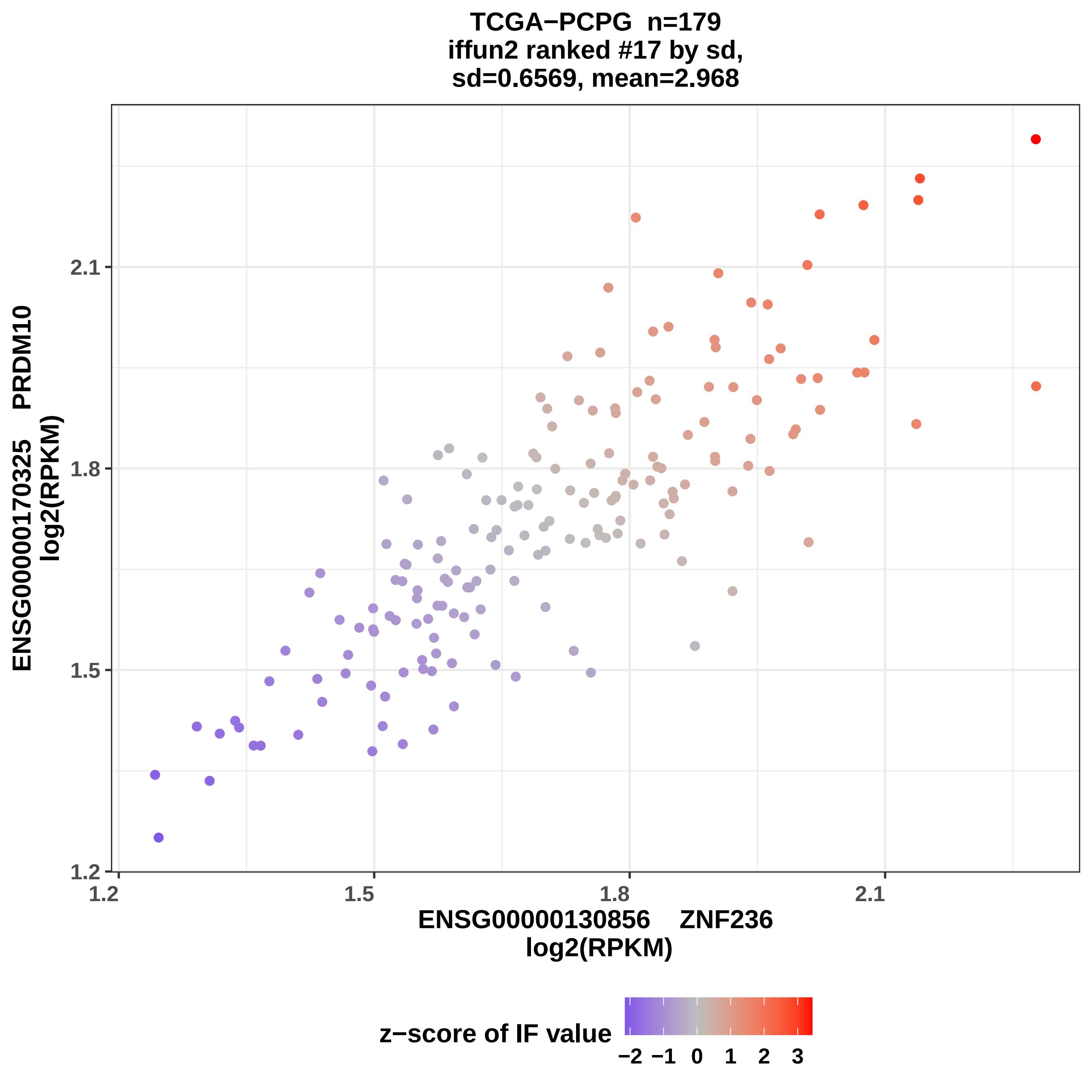

TCGA−PCPG n=179
iffun2 ranked #17 by sd,
sd=0.6569, mean=2.968
ENSG00000170325 PRDM10
log2(RPKM)
ENSG00000130856 ZNF236
log2(RPKM)
1.2
1.5
1.8
2.1
z−score of IF value
−2 −1 0 1 2 3

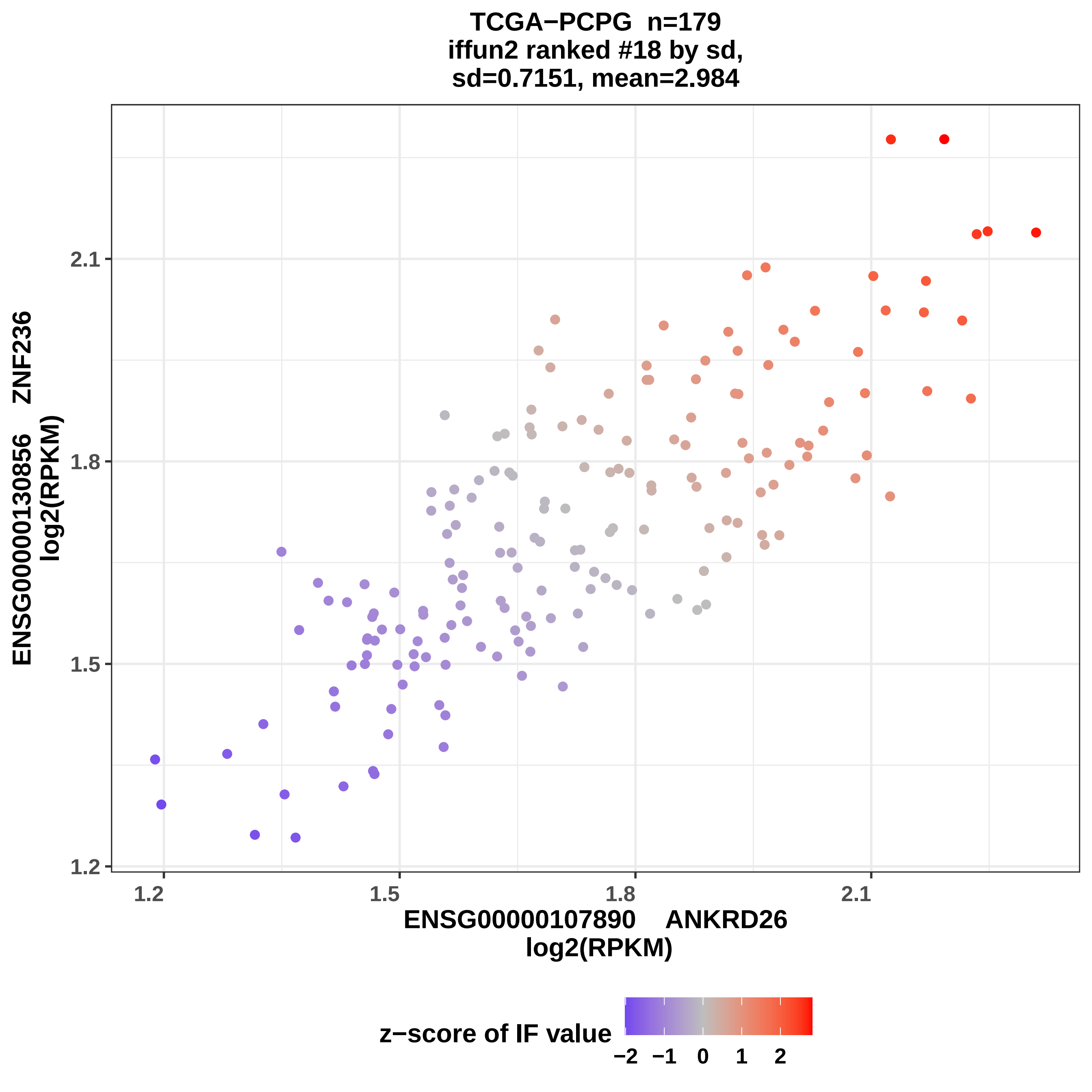

TCGA−PCPG n=179
iffun2 ranked #18 by sd,
sd=0.7151, mean=2.984
ENSG00000130856 ZNF236
log2(RPKM)
2.1
1.8
1.5
1.2
1.2
1.5
1.8
2.1
ENSG00000107890 ANKRD26
log2(RPKM)
z−score of IF value
−2 −1 0 1 2

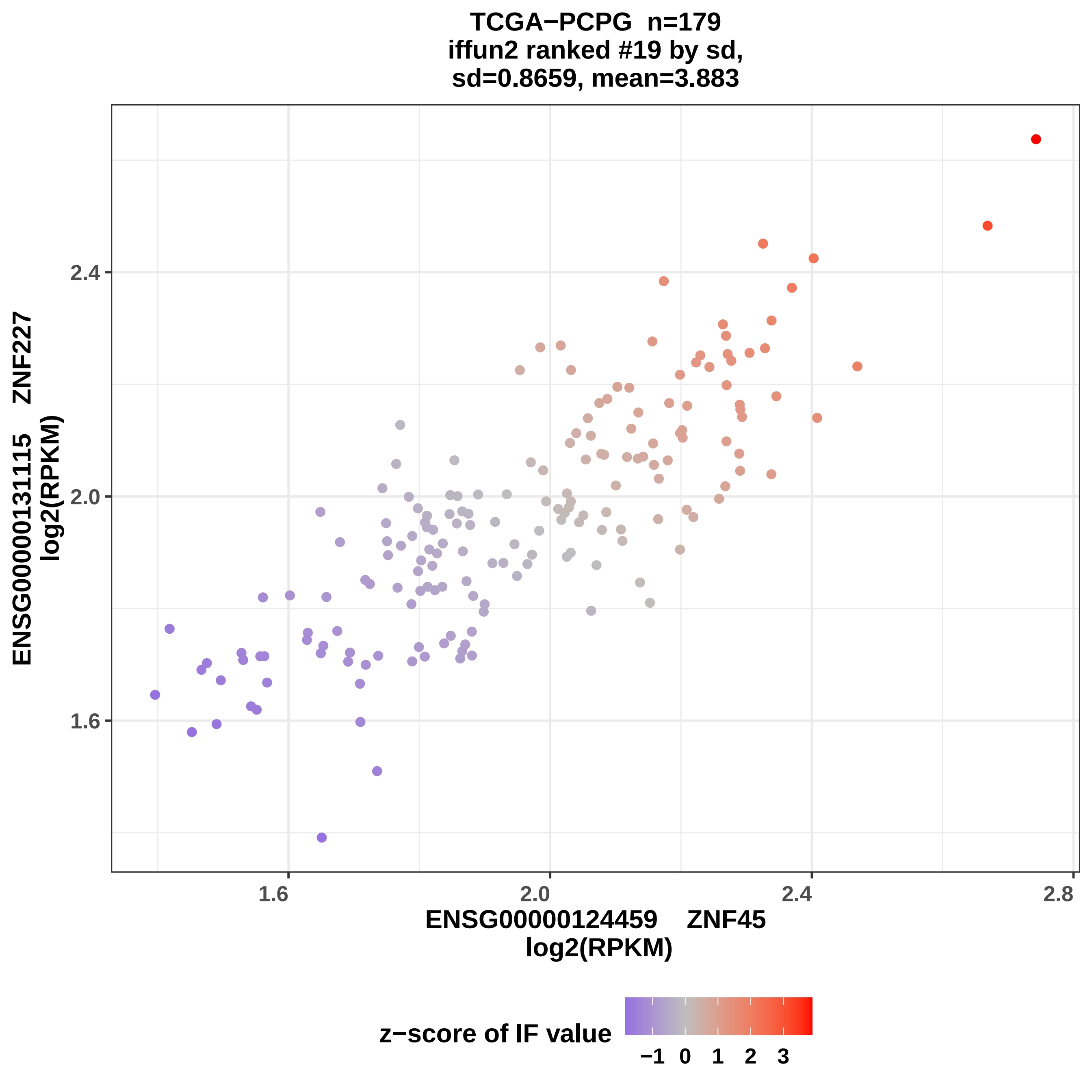

TCGA−PCPG n=179
iffun2 ranked #19 by sd,
sd=0.8659, mean=3.883
ENSG00000131115 ZNF227
log2(RPKM)
1.6
2.0
2.4
ENSG00000124459 ZNF45
log2(RPKM)
1.6
2.0
2.4
2.8
z−score of IF value
−1 0 1 2 3

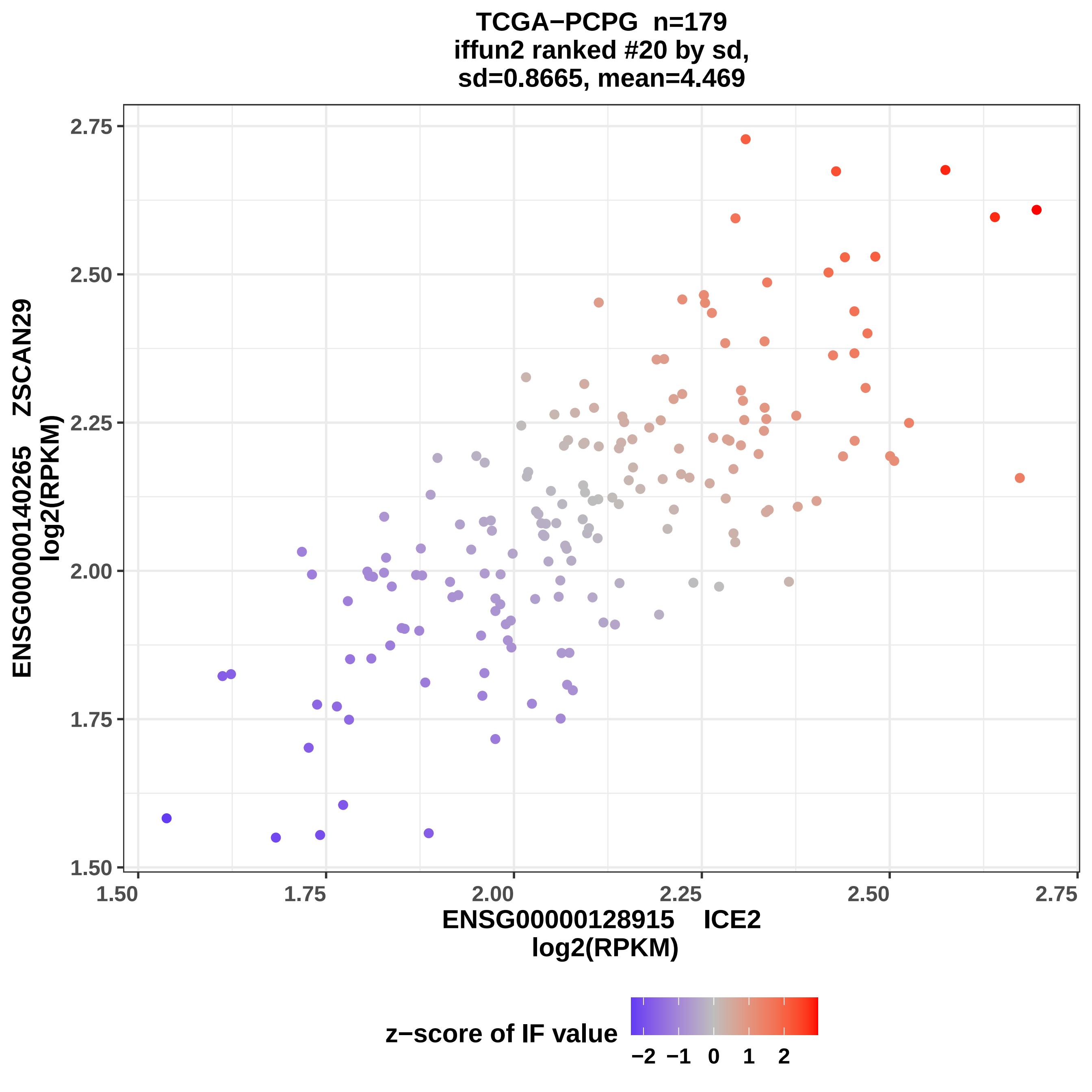
TCGA−PCPG n=179
iffun2 ranked #20 by sd,
sd=0.8665, mean=4.469
ENSG00000140265 ZSCAN29
log2(RPKM)
1.50
1.75
2.00
2.25
2.50
2.75
ENSG00000128915 ICE2
log2(RPKM)
z−score of IF value
−2 −1 0 1 2

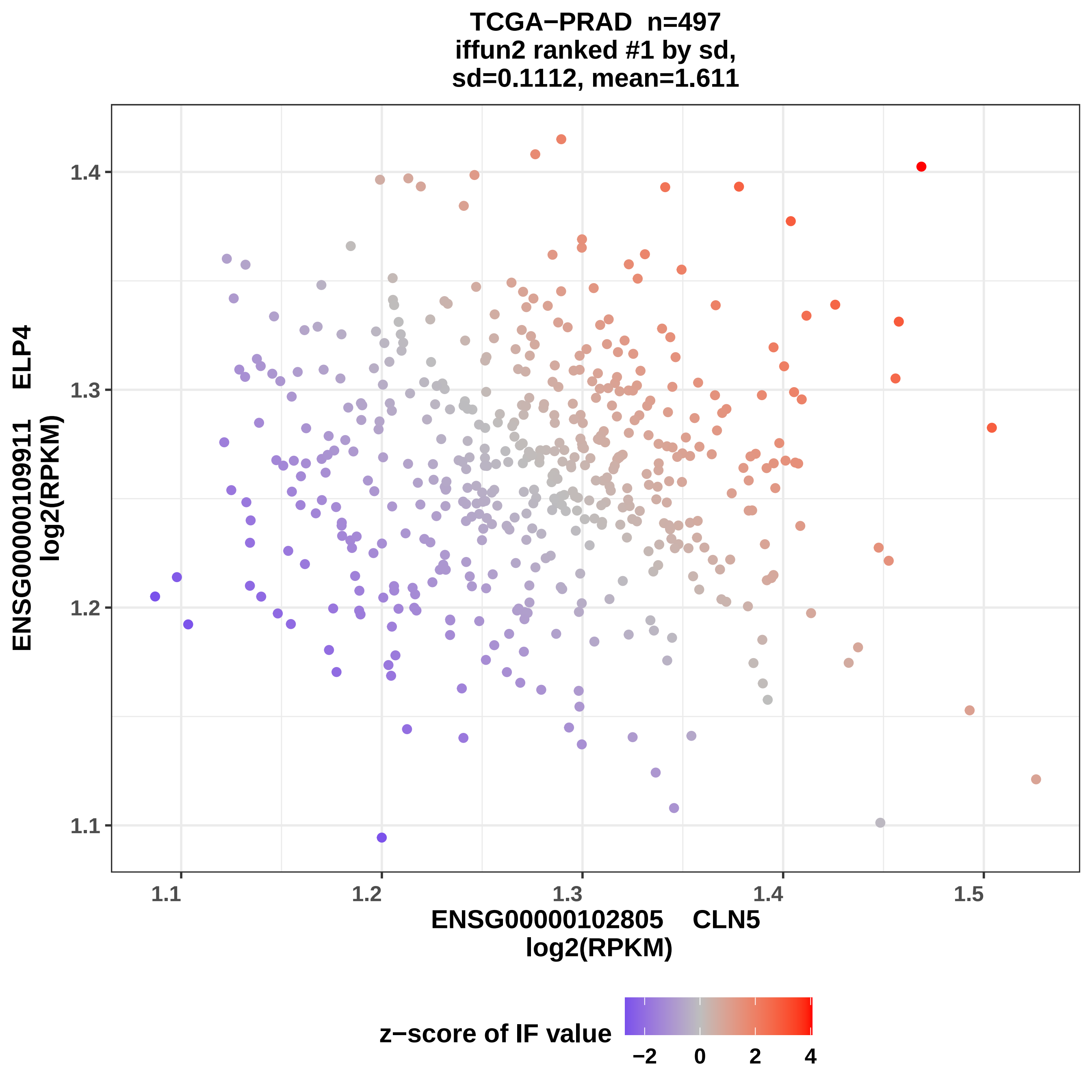

TCGA−PRAD n=497
iffun2 ranked #1 by sd,
sd=0.1112, mean=1.611
ENSG00000109911 ELP4
log2(RPKM)
1.4
1.3
1.2
1.1
1.1
1.2
1.3
1.4
1.5
ENSG00000102805 CLN5
log2(RPKM)
z−score of IF value
−2
0
2
4

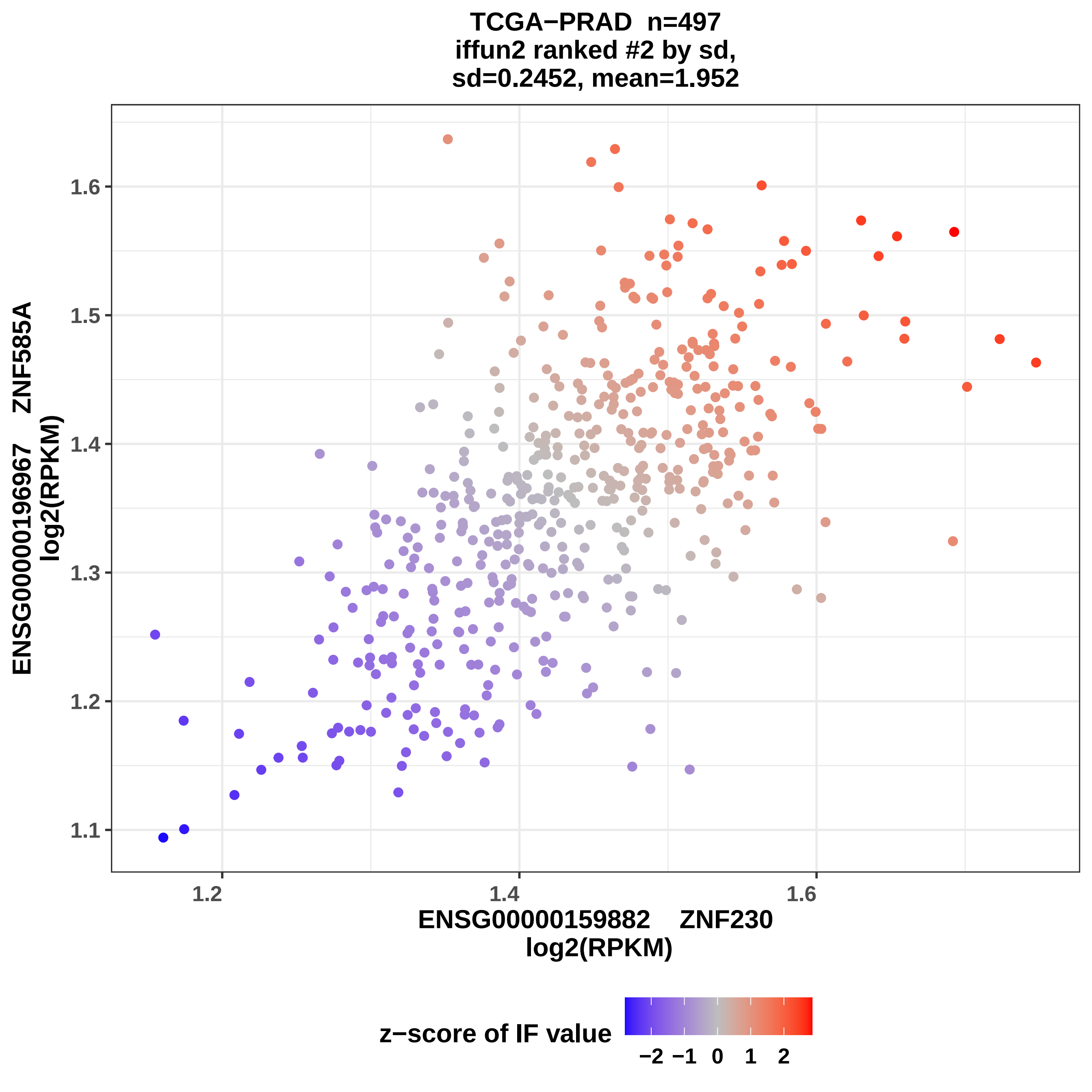

TCGA−PRAD n=497
iffun2 ranked #2 by sd,
sd=0.2452, mean=1.952
ENSG00000196967 ZNF585A
log2(RPKM)
1.1
1.2
1.3
1.4
1.5
1.6
1.2
1.4
1.6
ENSG00000159882 ZNF230
log2(RPKM)
z−score of IF value
−2
−1
0
1
2

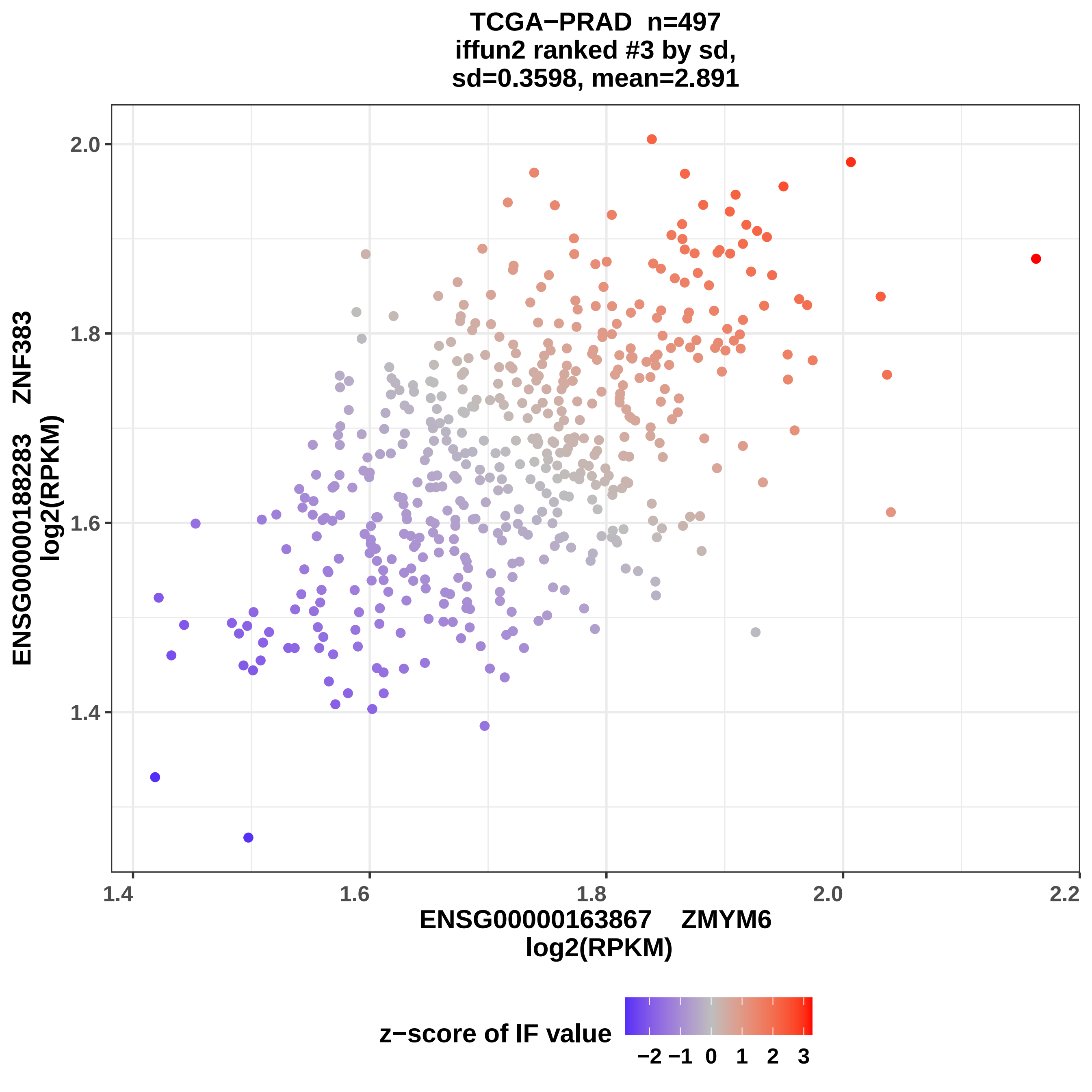
TCGA−PRAD n=497
iffun2 ranked #3 by sd,
sd=0.3598, mean=2.891
ENSG00000188283 ZNF383
log2(RPKM)
2.0
1.8
1.6
1.4
1.4
1.6
1.8
2.0
2.2
ENSG00000163867 ZMYM6
log2(RPKM)
z−score of IF value
−2 −1 0 1 2 3

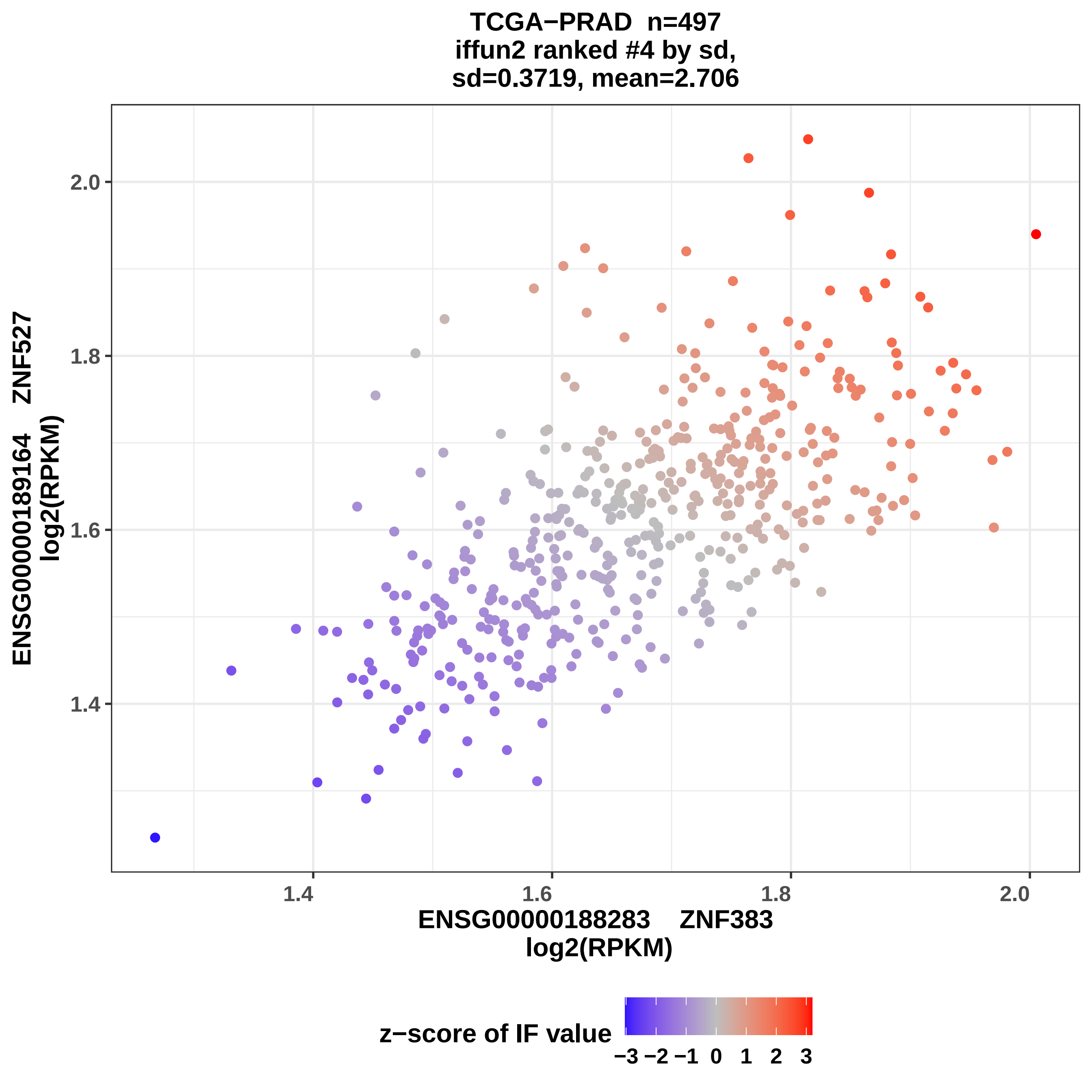

TCGA−PRAD n=497
iffun2 ranked #4 by sd,
sd=0.3719, mean=2.706
ENSG00000189164 ZNF527
log2(RPKM)
2.0
1.8
1.6
1.4
1.4
1.6
1.8
2.0
ENSG00000188283 ZNF383
log2(RPKM)
z−score of IF value
−3 −2 −1 0 1 2 3

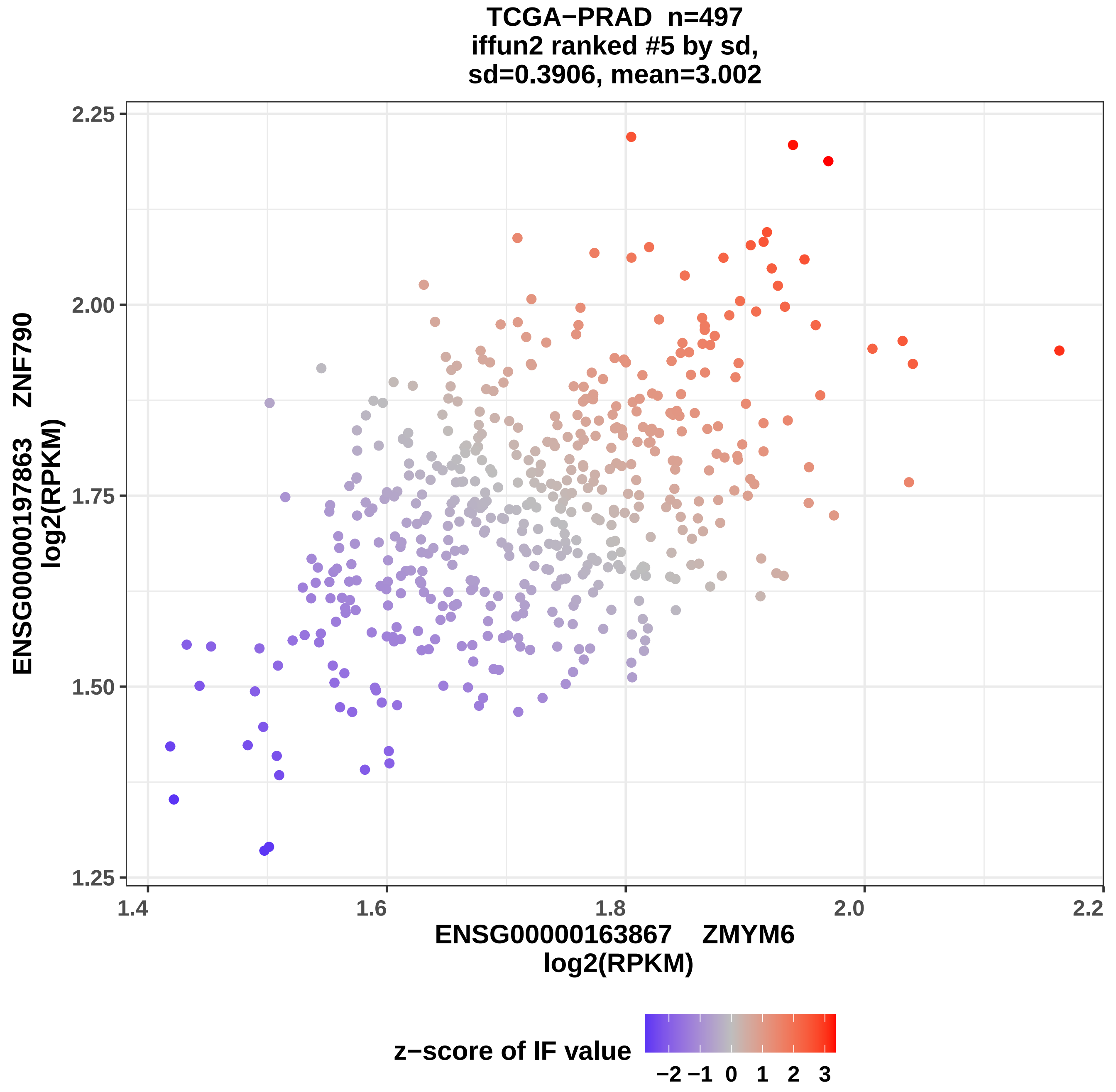
TCGA−PRAD n=497
iffun2 ranked #5 by sd,
sd=0.3906, mean=3.002
ENSG00000197863 ZNF790
log2(RPKM)
2.25
2.00
1.75
1.50
1.25
1.4
1.6
1.8
2.0
2.2
ENSG00000163867 ZMYM6
log2(RPKM)
z−score of IF value
−2 −1 0 1 2 3

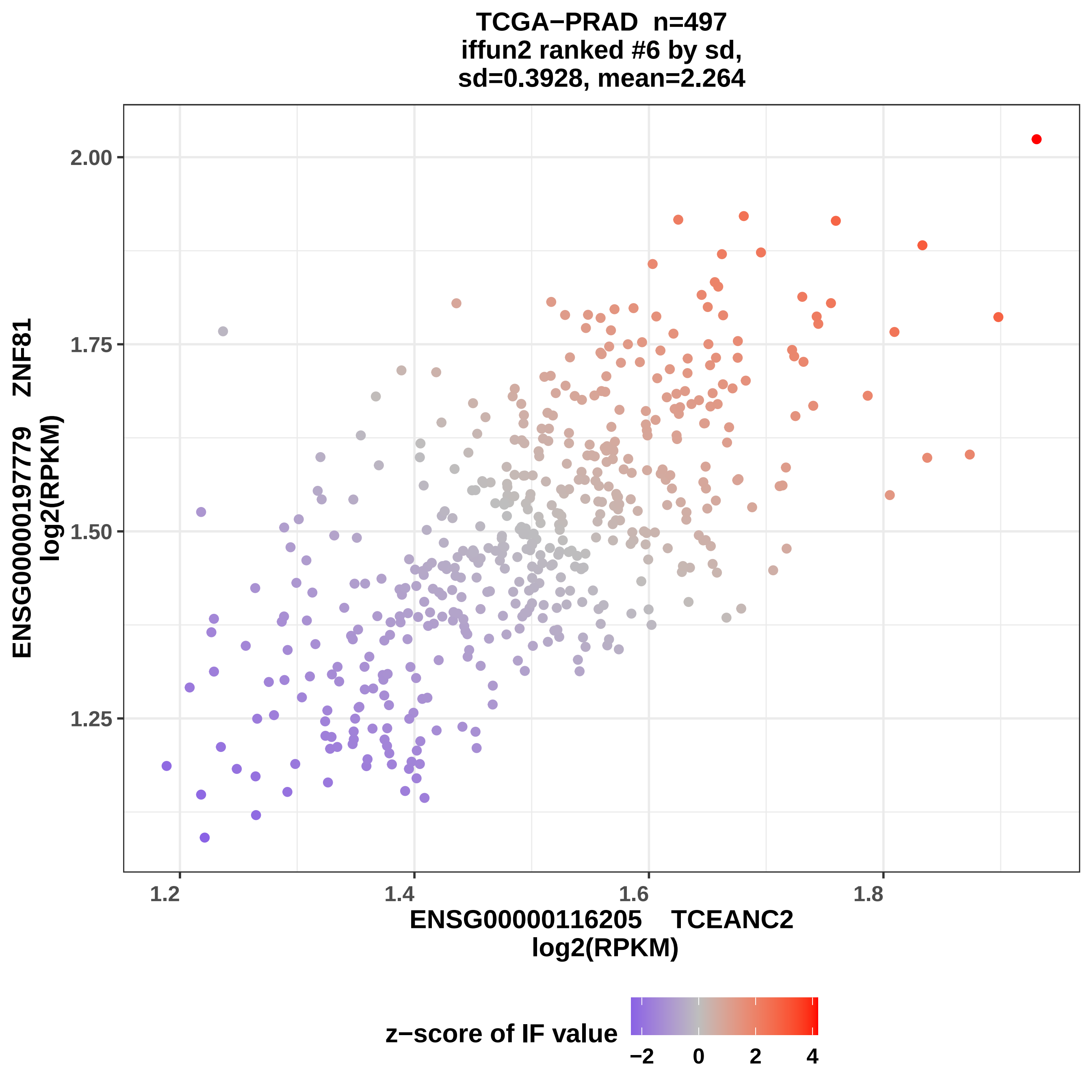

TCGA−PRAD n=497
iffun2 ranked #6 by sd,
sd=0.3928, mean=2.264
ENSG00000197779 ZNF81
log2(RPKM)
2.00
1.75
1.50
1.25
1.2
1.4
1.6
1.8
ENSG00000116205 TCEANC2
log2(RPKM)
z−score of IF value
−2
0
2
4

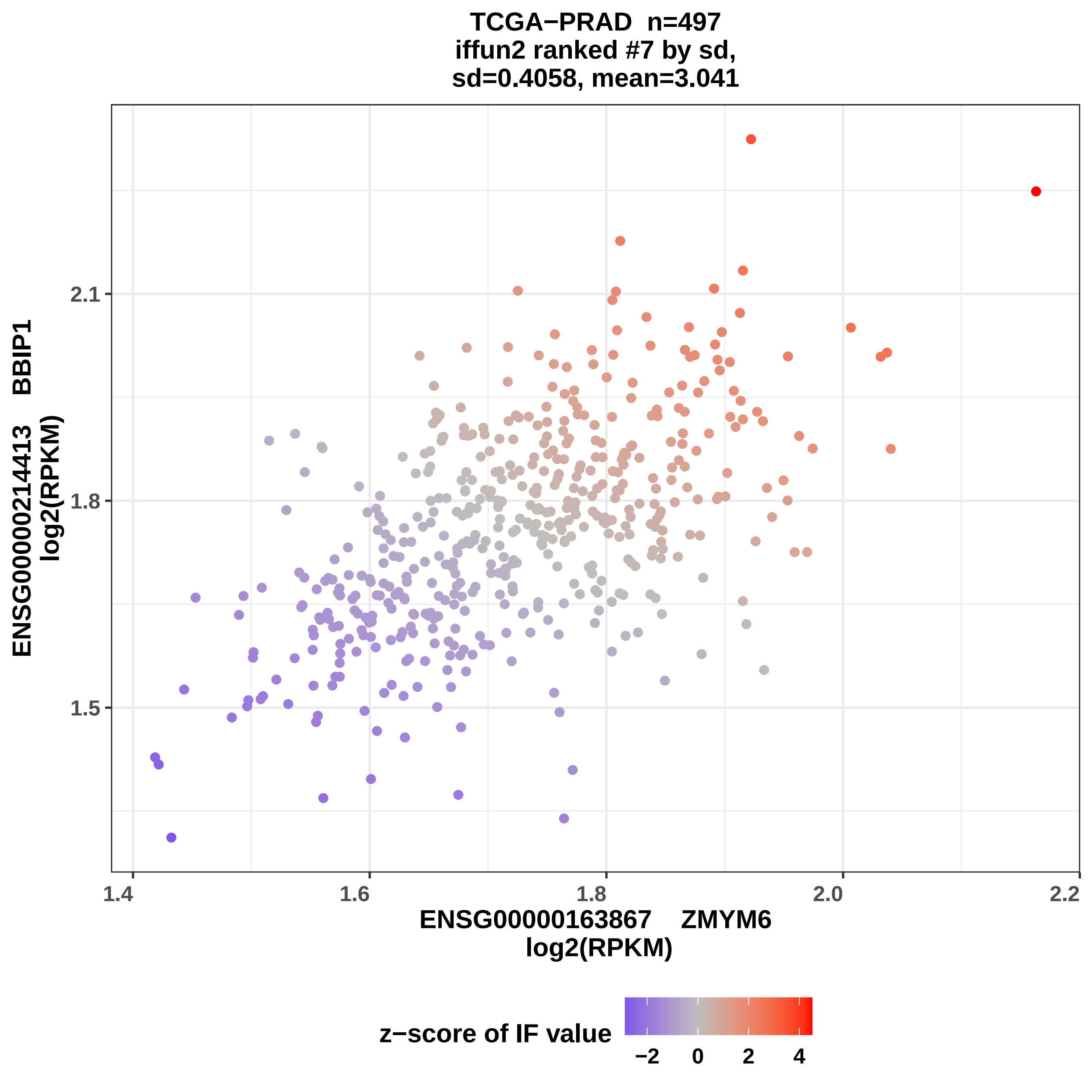
TCGA−PRAD n=497
iffun2 ranked #7 by sd,
sd=0.4058, mean=3.041
ENSG00000214413 BBIP1
log2(RPKM)
2.1
1.8
1.5
1.4
1.6
1.8
2.0
2.2
ENSG00000163867 ZMYM6
log2(RPKM)
z−score of IF value
−2
0
2
4

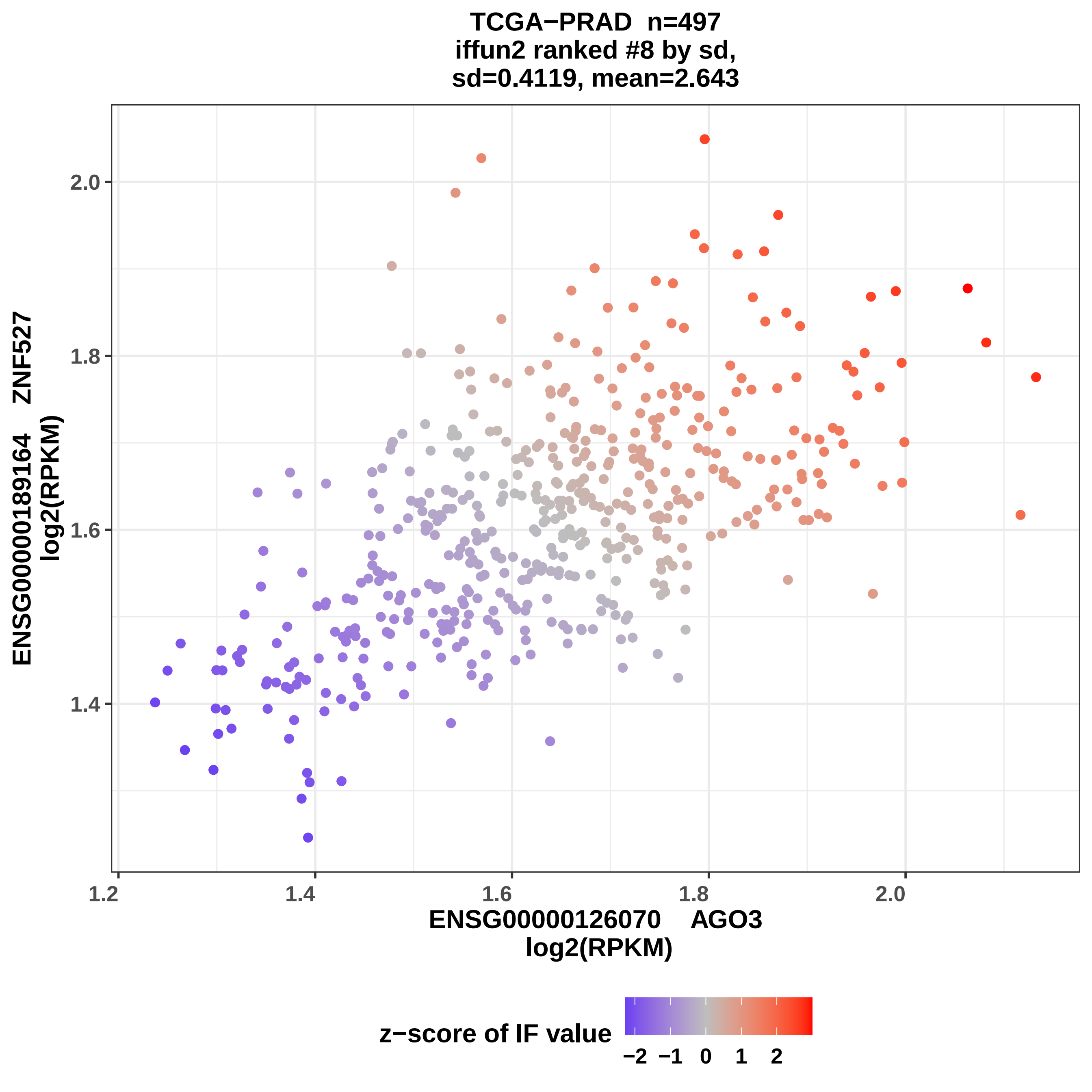
TCGA−PRAD n=497
iffun2 ranked #8 by sd,
sd=0.4119, mean=2.643
ENSG00000189164 ZNF527
log2(RPKM)
2.0
1.8
1.6
1.4
1.2
1.4
1.6
1.8
2.0
ENSG00000126070 AGO3
log2(RPKM)
z−score of IF value
−2
−1
0
1
2

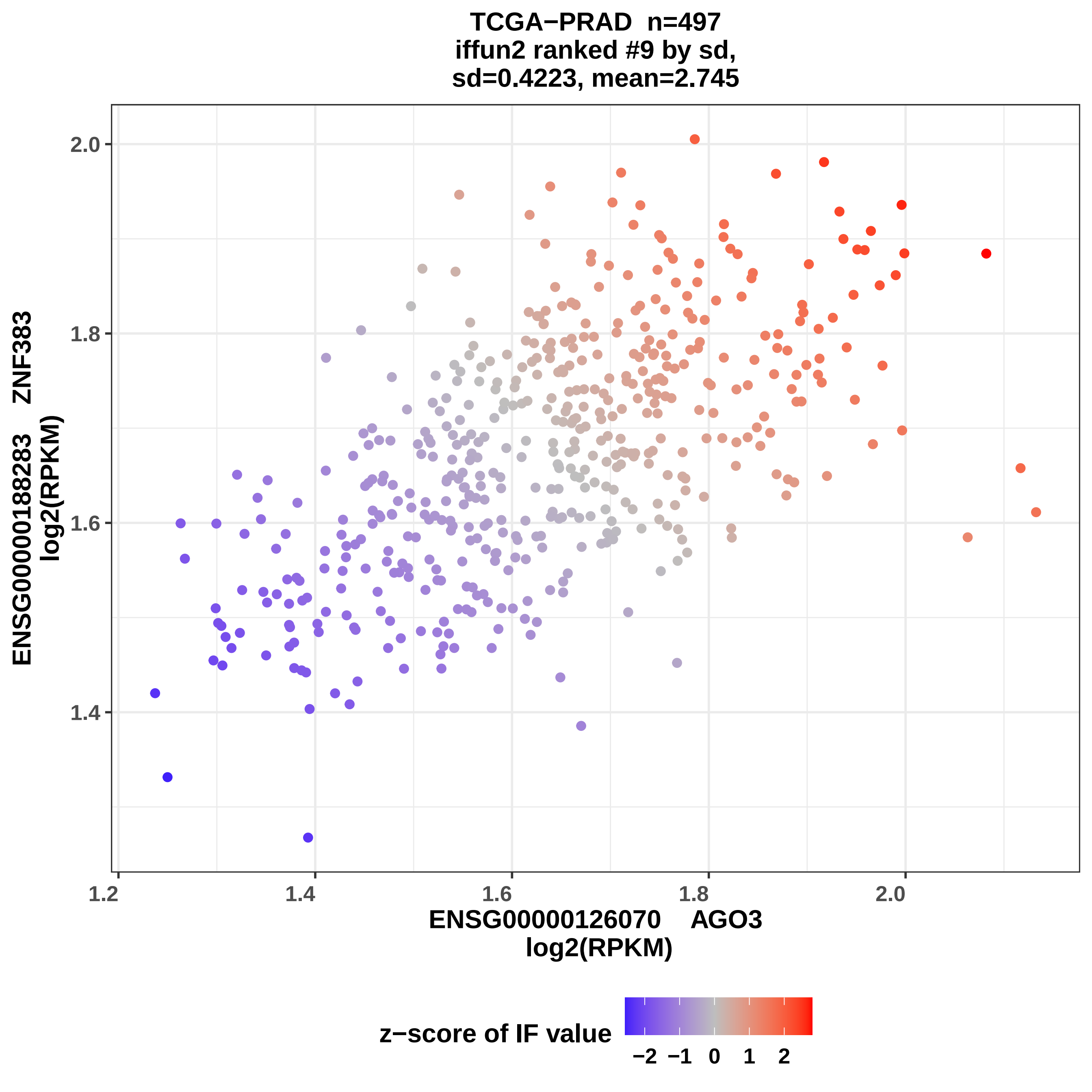
TCGA−PRAD n=497
iffun2 ranked #9 by sd,
sd=0.4223, mean=2.745
ENSG00000188283 ZNF383
log2(RPKM)
2.0
1.8
1.6
1.4
1.2
1.4
1.6
1.8
2.0
ENSG00000126070 AGO3
log2(RPKM)
z−score of IF value
−2
−1
0
1
2

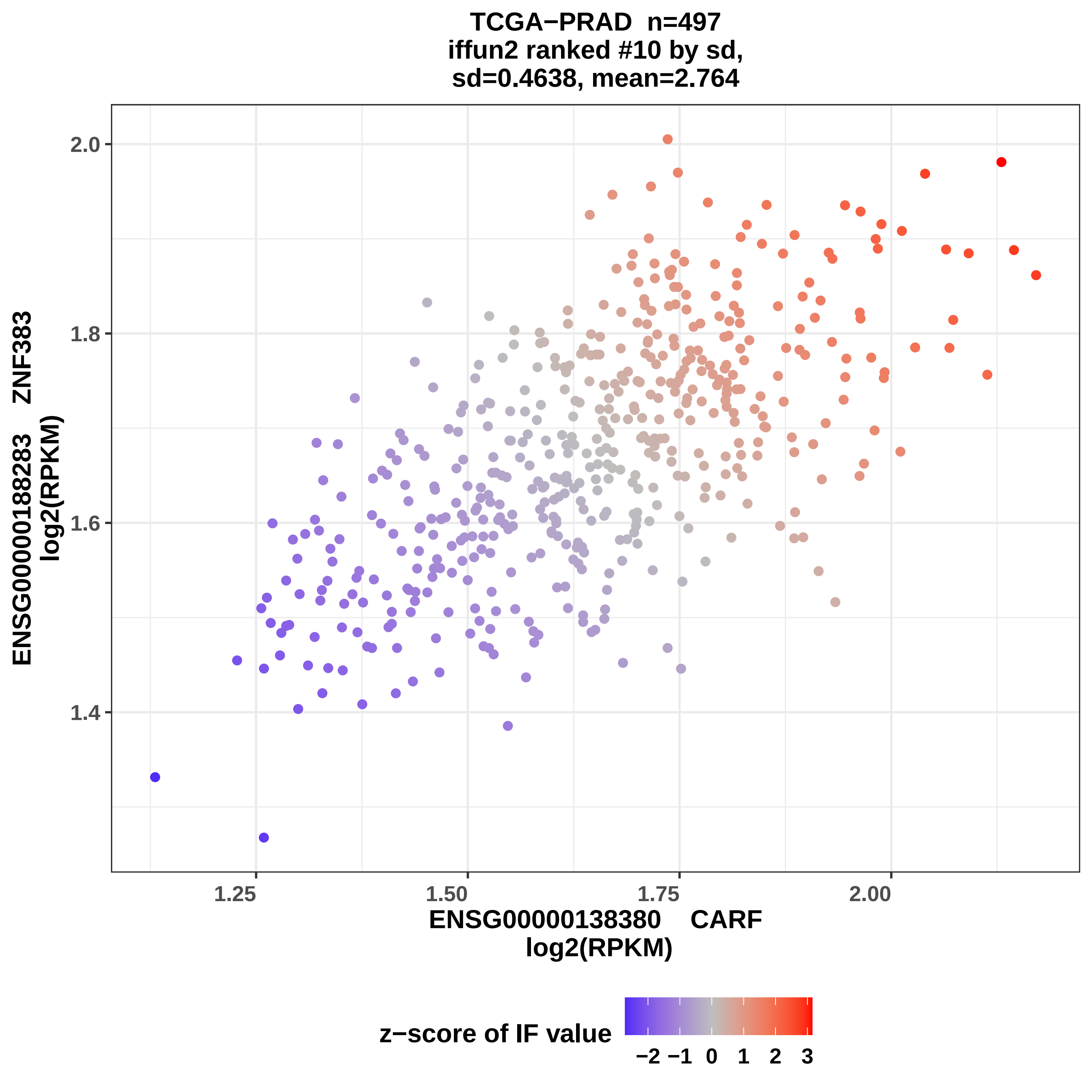

TCGA−PRAD n=497
iffun2 ranked #10 by sd,
sd=0.4638, mean=2.764
ENSG00000188283 ZNF383
log2(RPKM)
2.0
1.8
1.6
1.4
1.25
1.50
1.75
2.00
ENSG00000138380 CARF
log2(RPKM)
z−score of IF value
−2 −1 0 1 2 3

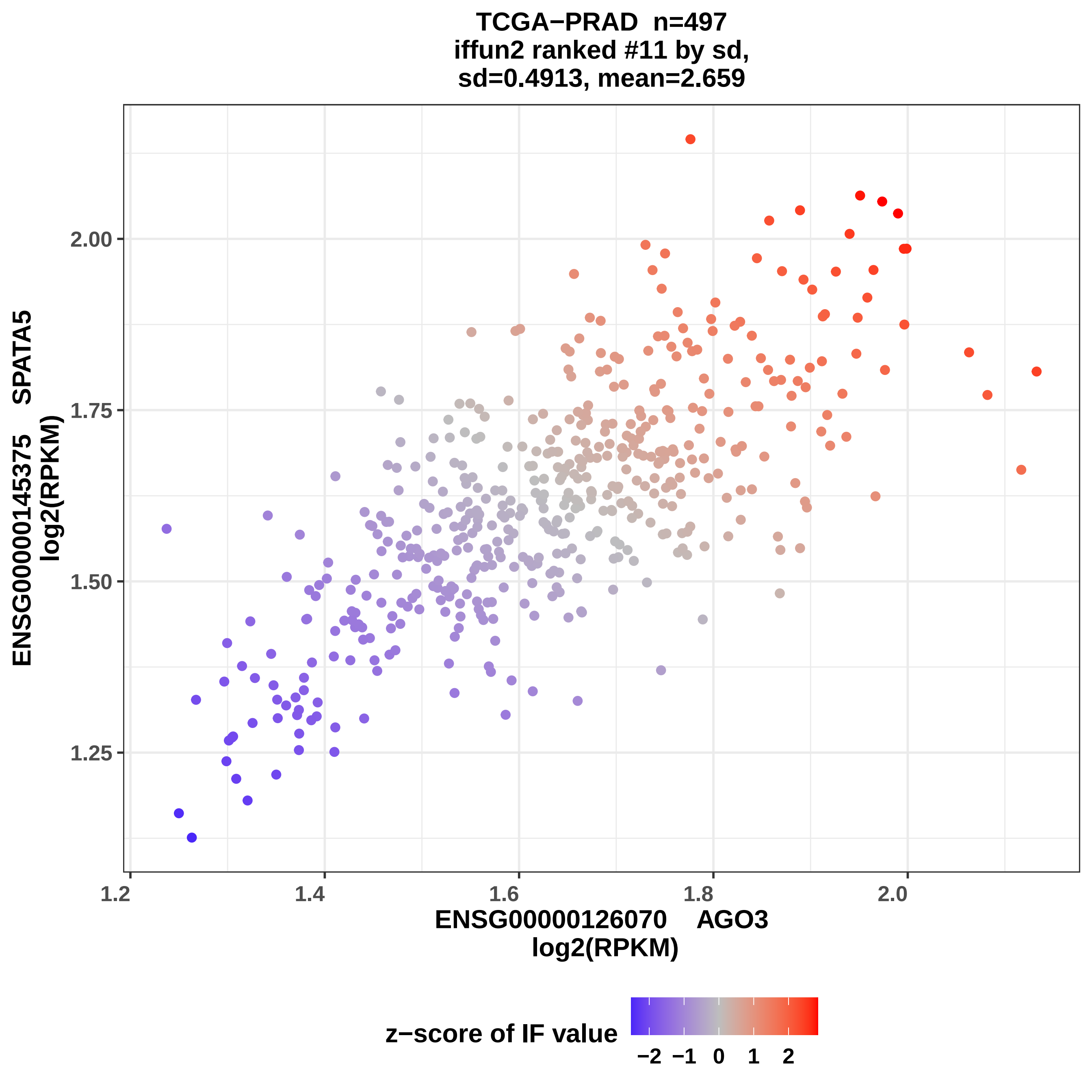
TCGA−PRAD n=497
iffun2 ranked #11 by sd,
sd=0.4913, mean=2.659
ENSG00000145375 SPATA5
log2(RPKM)
1.25
1.50
1.75
2.00
1.2
1.4
1.6
1.8
2.0
ENSG00000126070 AGO3
log2(RPKM)
z−score of IF value
−2
−1
0
1
2

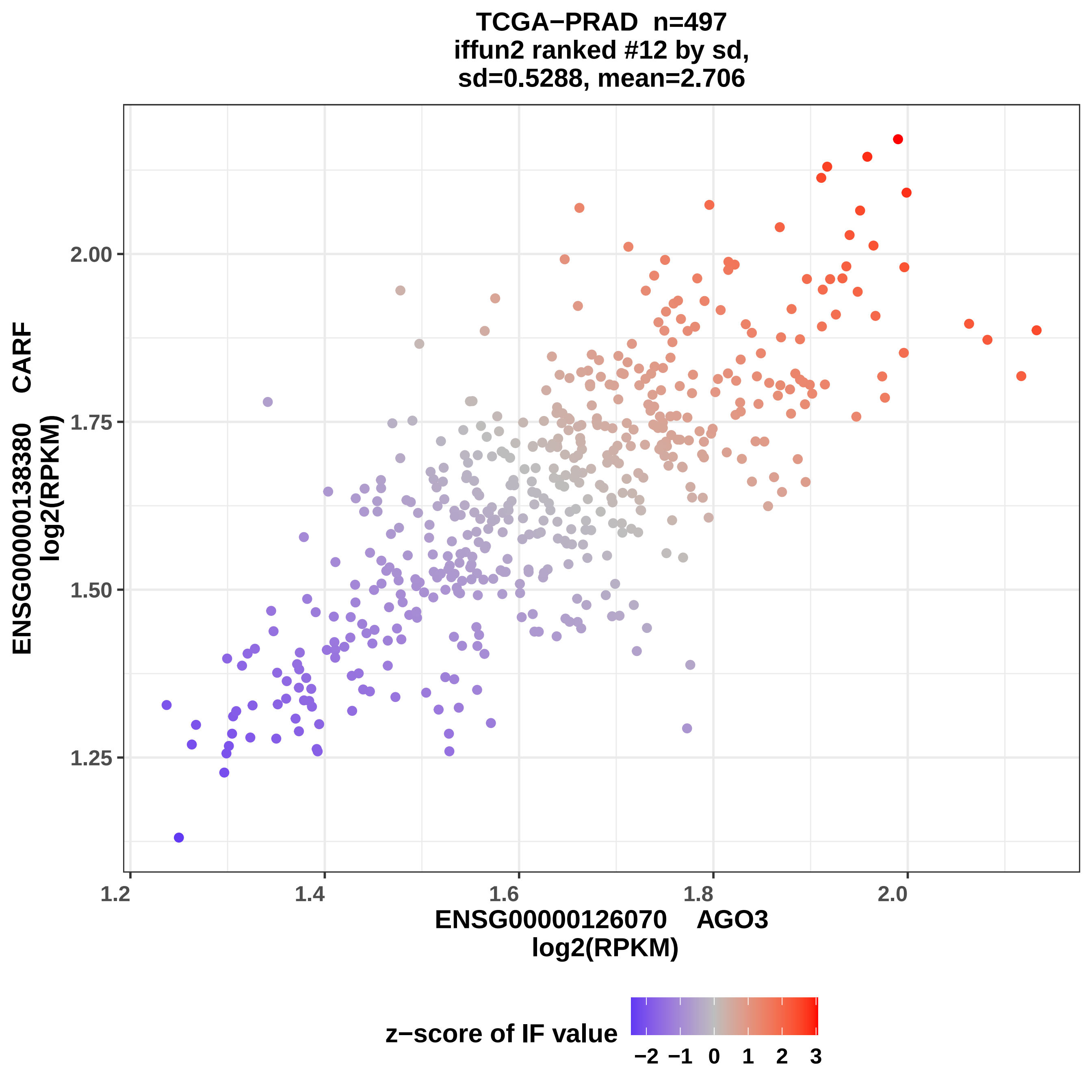
TCGA−PRAD n=497
iffun2 ranked #12 by sd,
sd=0.5288, mean=2.706
ENSG00000138380 CARF
log2(RPKM)
2.00
1.75
1.50
1.25
1.2
1.4
1.6
1.8
2.0
ENSG00000126070 AGO3
log2(RPKM)
z−score of IF value
−2 −1 0 1 2 3

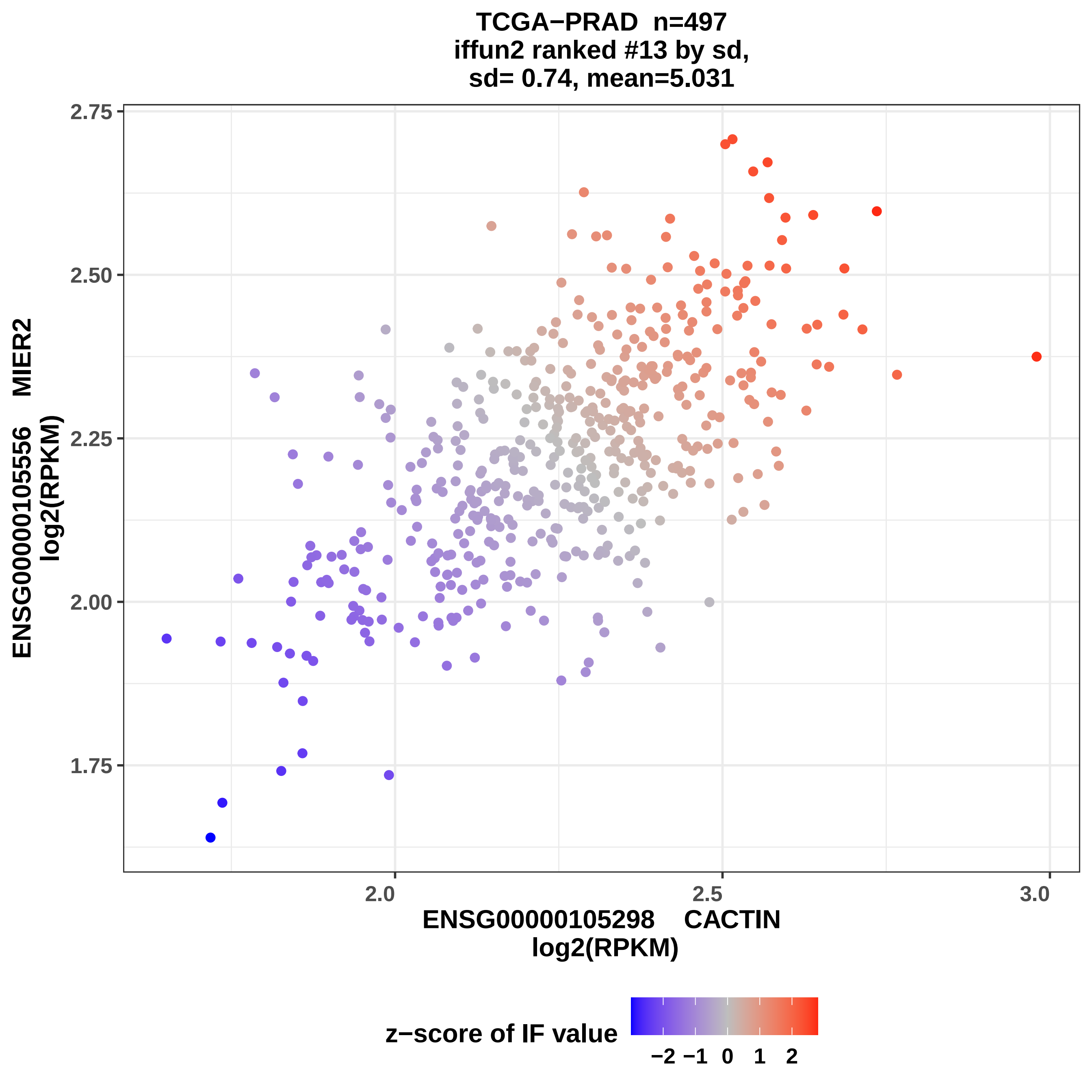
TCGA−PRAD n=497
iffun2 ranked #13 by sd,
sd= 0.74, mean=5.031
ENSG00000105556 MIER2
log2(RPKM)
2.75
2.50
2.25
2.00
1.75
2.0
2.5
3.0
ENSG00000105298 CACTIN
log2(RPKM)
z−score of IF value
−2 −1 0 1 2

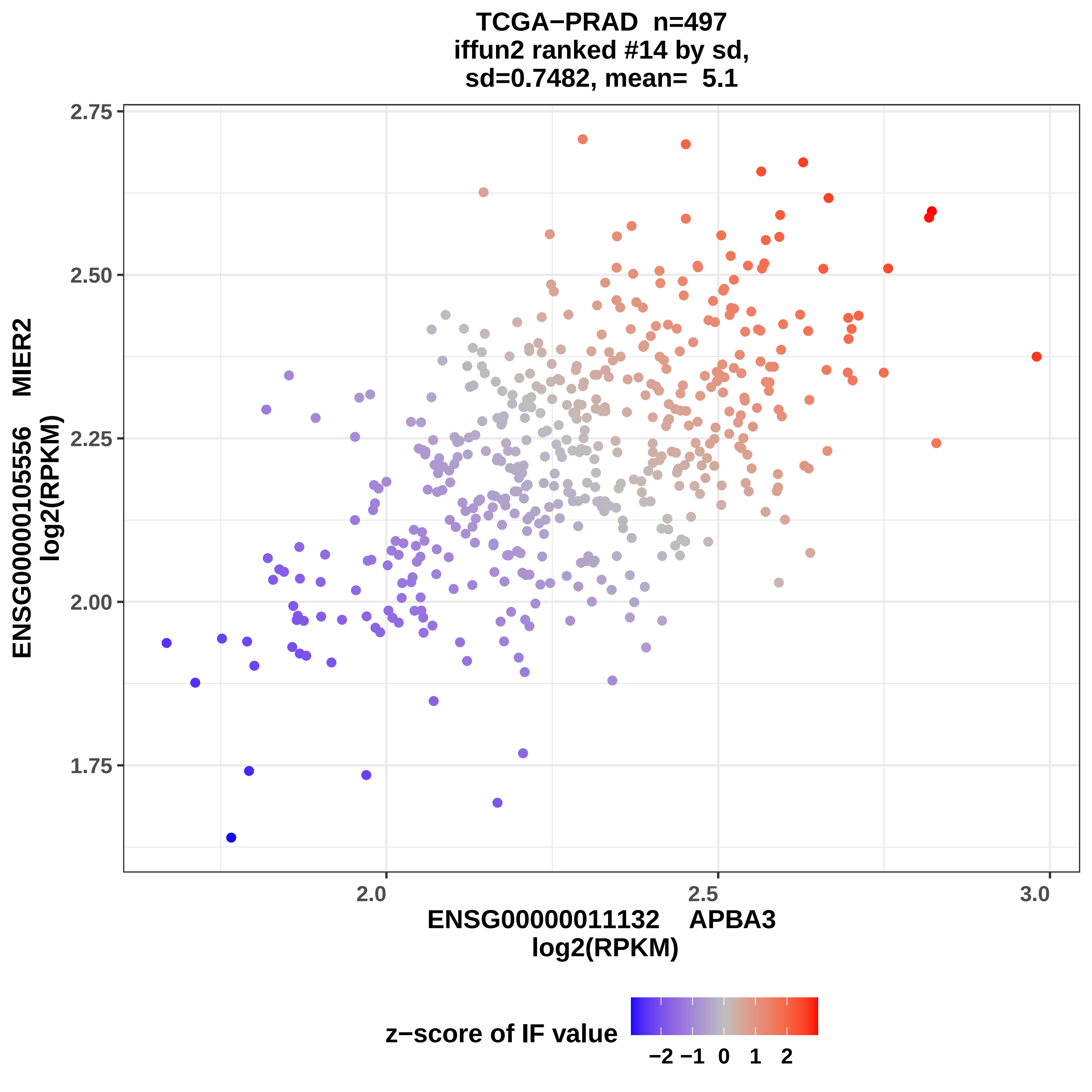

TCGA−PRAD n=497
iffun2 ranked #14 by sd,
sd=0.7482, mean= 5.1
ENSG00000105556 MIER2
log2(RPKM)
2.75
2.50
2.25
2.00
1.75
2.0
2.5
3.0
ENSG00000011132 APBA3
log2(RPKM)
z−score of IF value
−2 −1 0 1 2

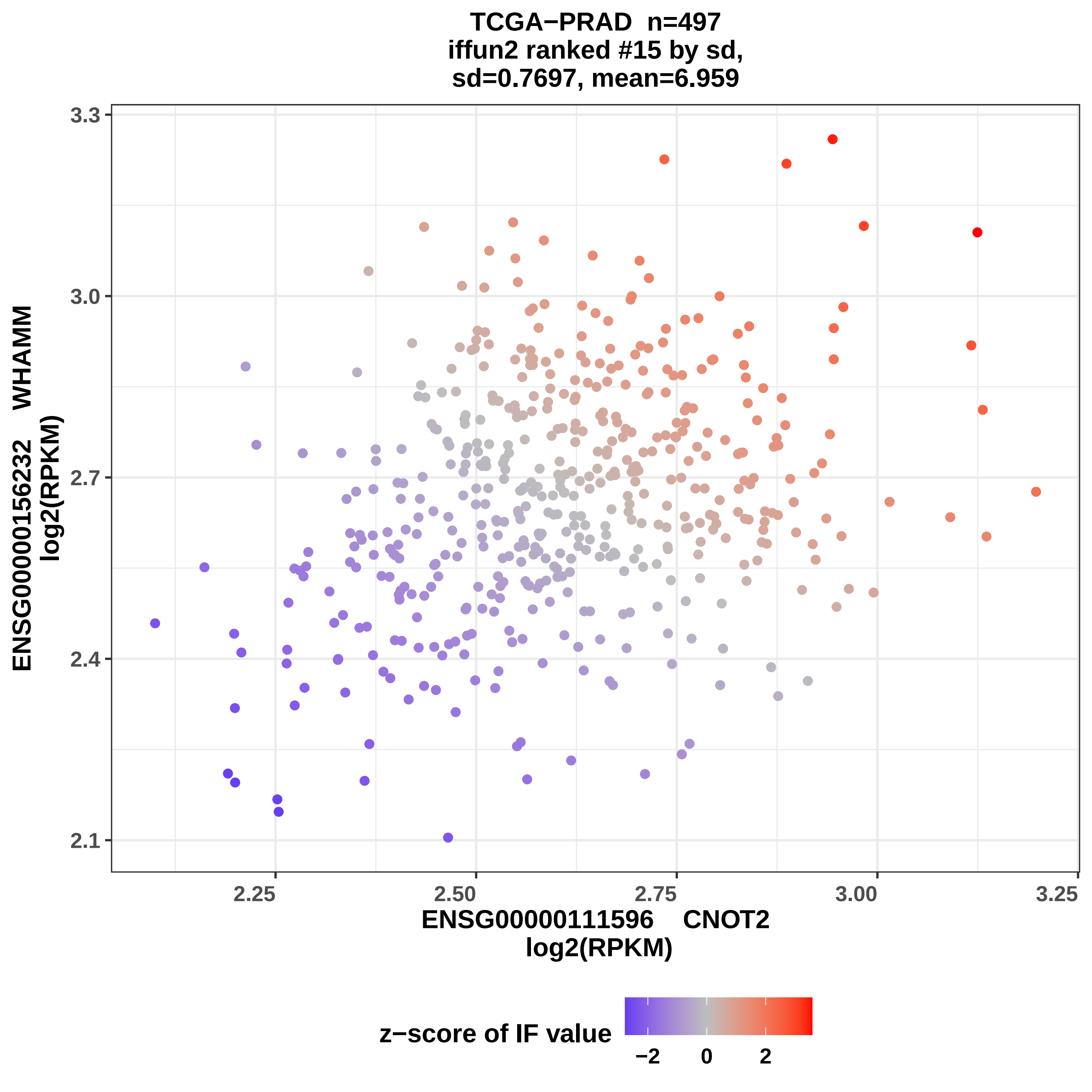

TCGA−PRAD n=497
iffun2 ranked #15 by sd,
sd=0.7697, mean=6.959
ENSG00000156232 WHAMM
log2(RPKM)
3.3
3.0
2.7
2.4
2.1
2.25
2.50
2.75
3.00
3.25
ENSG00000111596 CNOT2
log2(RPKM)
z−score of IF value
−2
0
2

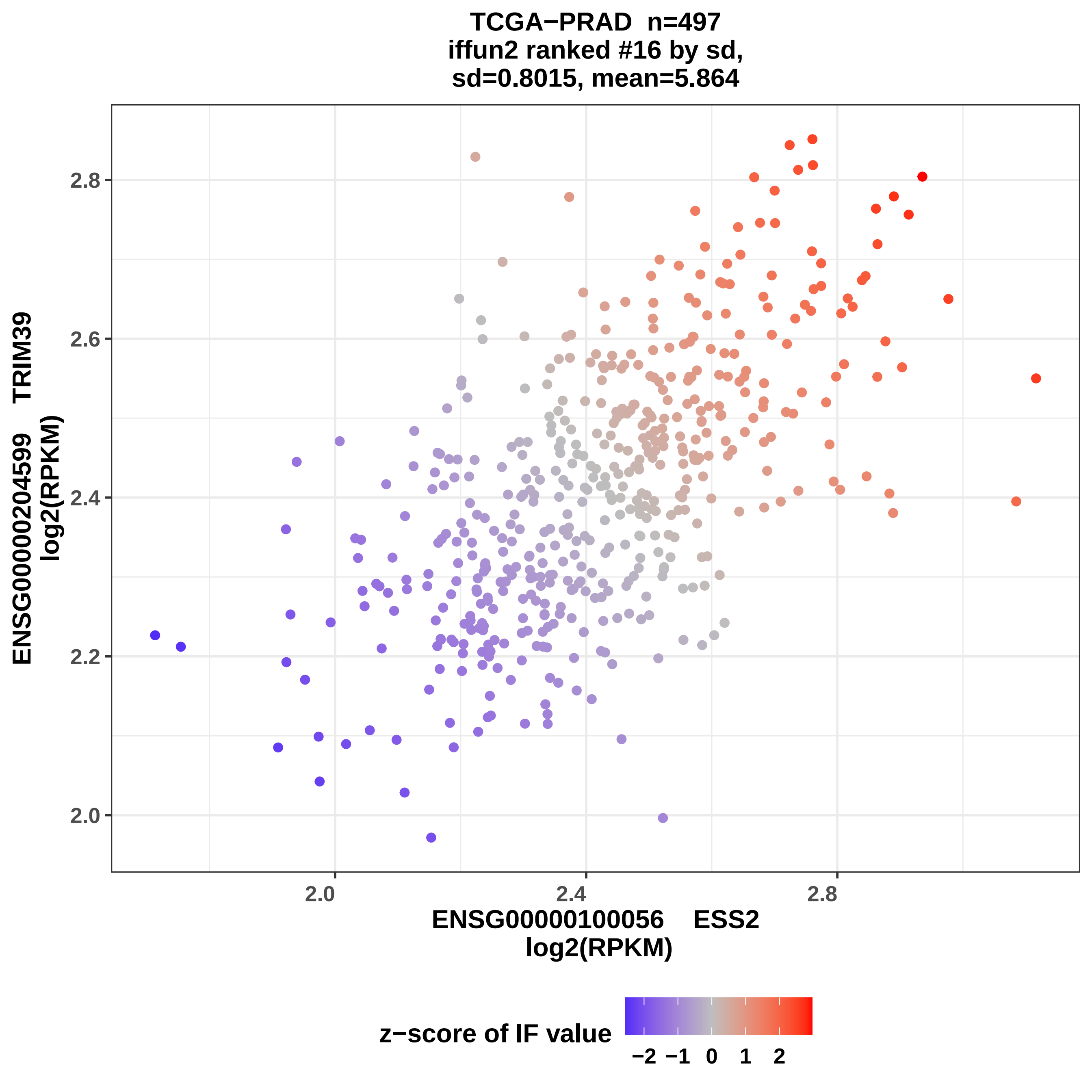

TCGA−PRAD n=497
iffun2 ranked #16 by sd,
sd=0.8015, mean=5.864
ENSG00000204599 TRIM39
log2(RPKM)
2.0
2.2
2.4
2.6
2.8
2.0
2.4
2.8
ENSG00000100056 ESS2
log2(RPKM)
z−score of IF value
−2
−1
0
1
2

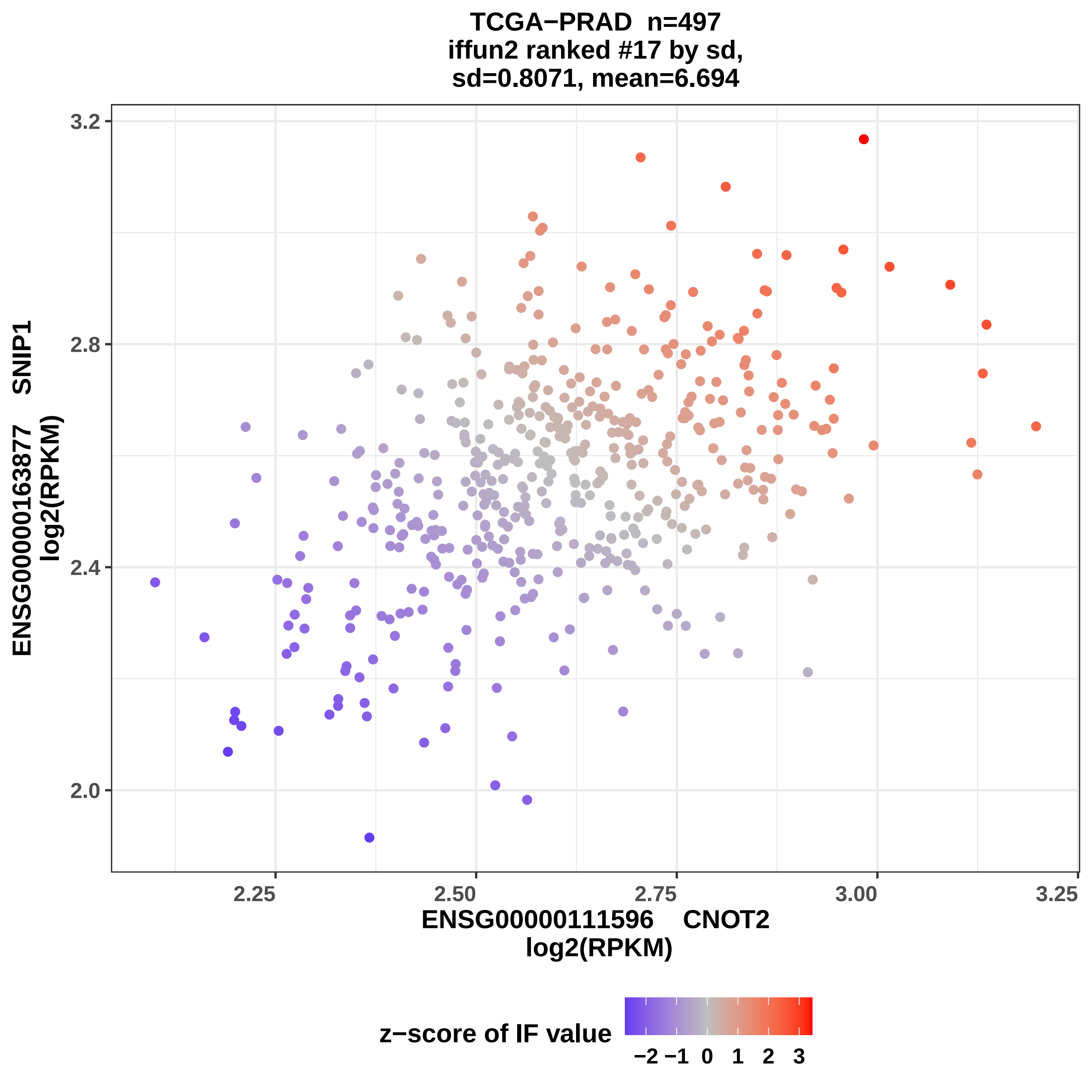

TCGA−PRAD n=497
iffun2 ranked #17 by sd,
sd=0.8071, mean=6.694
ENSG00000163877 SNIP1
log2(RPKM)
3.2
2.8
2.4
2.0
2.25
2.50
2.75
3.00
3.25
ENSG00000111596 CNOT2
log2(RPKM)
z−score of IF value
−2 −1 0 1 2 3

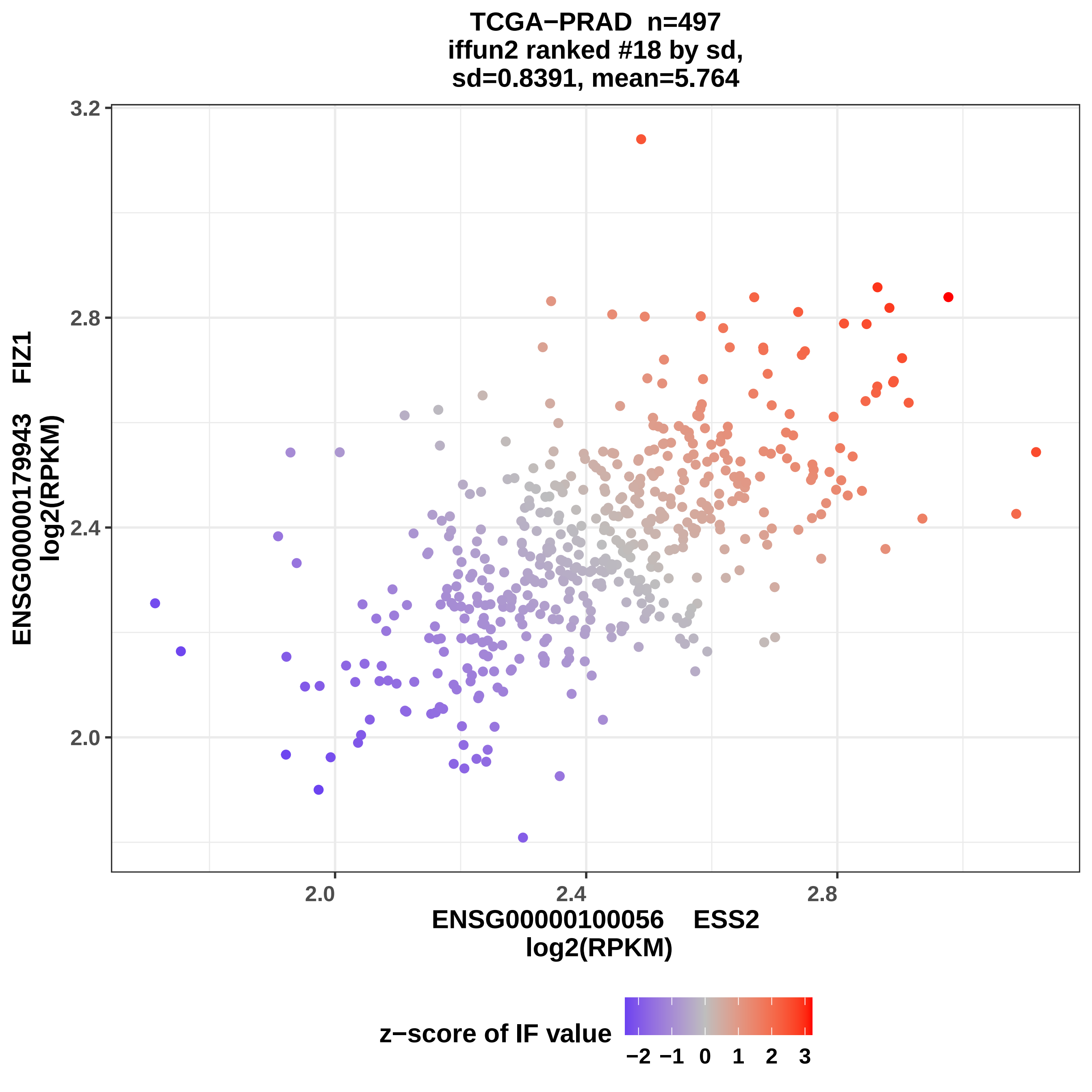
TCGA−PRAD n=497
iffun2 ranked #18 by sd,
sd=0.8391, mean=5.764
3.2
2.8
2.4
2.0
2.0
2.4
2.8
ENSG00000179943 FIZ1
log2(RPKM)
ENSG00000100056 ESS2
log2(RPKM)
z−score of IF value
−2 −1 0 1 2 3

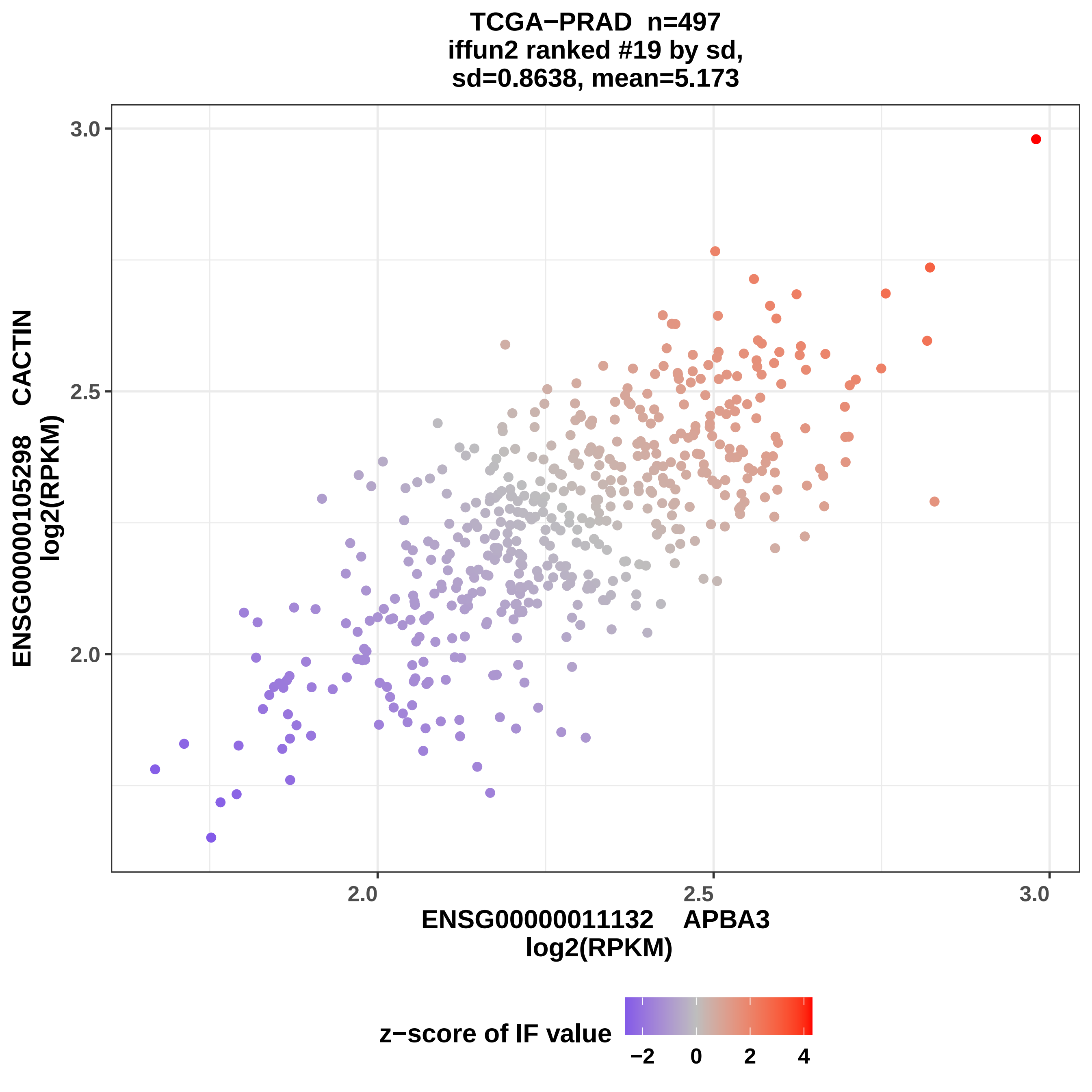
TCGA−PRAD n=497
iffun2 ranked #19 by sd,
sd=0.8638, mean=5.173
ENSG00000105298 CACTIN
log2(RPKM)
3.0
2.5
2.0
2.0
2.5
3.0
ENSG00000011132 APBA3
log2(RPKM)
z−score of IF value
−2
0
2
4

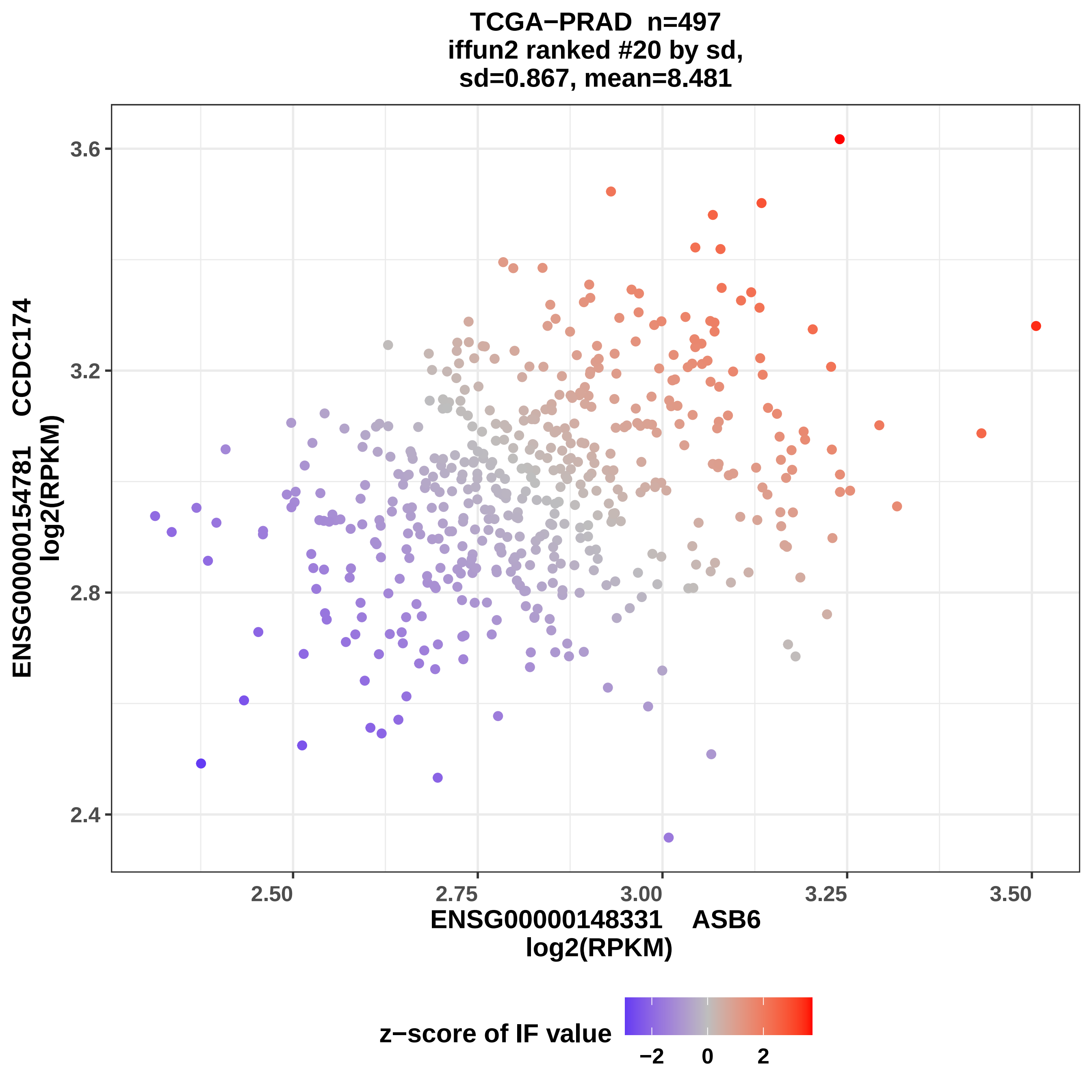
TCGA−PRAD n=497
iffun2 ranked #20 by sd,
sd=0.867, mean=8.481
ENSG00000154781 CCDC174
log2(RPKM)
3.6
3.2
2.8
2.4
2.50
2.75
3.00
3.25
3.50
ENSG00000148331 ASB6
log2(RPKM)
z−score of IF value
−2
0
2

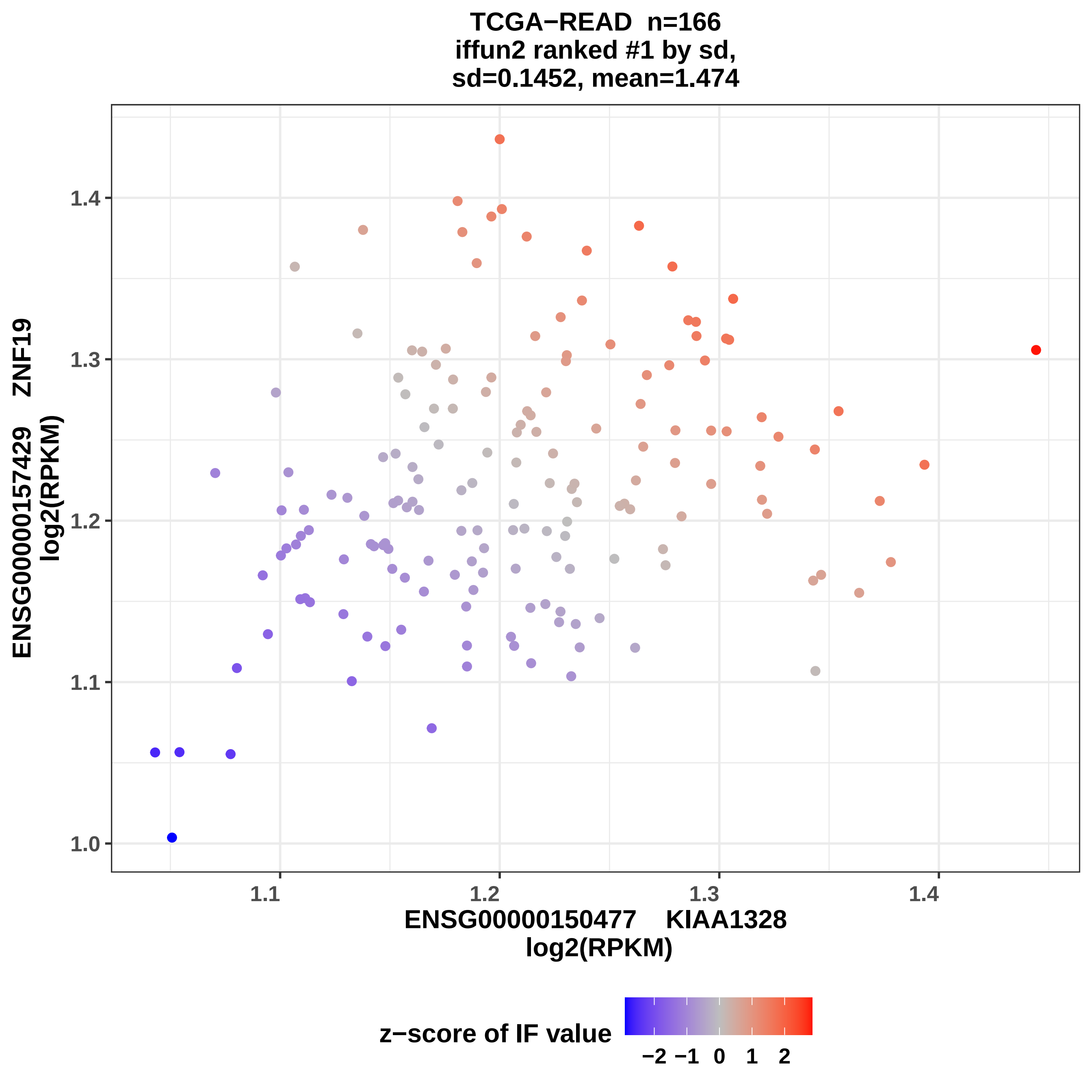
TCGA−READ n=166
iffun2 ranked #1 by sd,
sd=0.1452, mean=1.474
ENSG00000157429 ZNF19
log2(RPKM)
1.0
1.1
1.2
1.3
1.4
ENSG00000150477 KIAA1328
log2(RPKM)
z−score of IF value
−2 −1 0 1 2

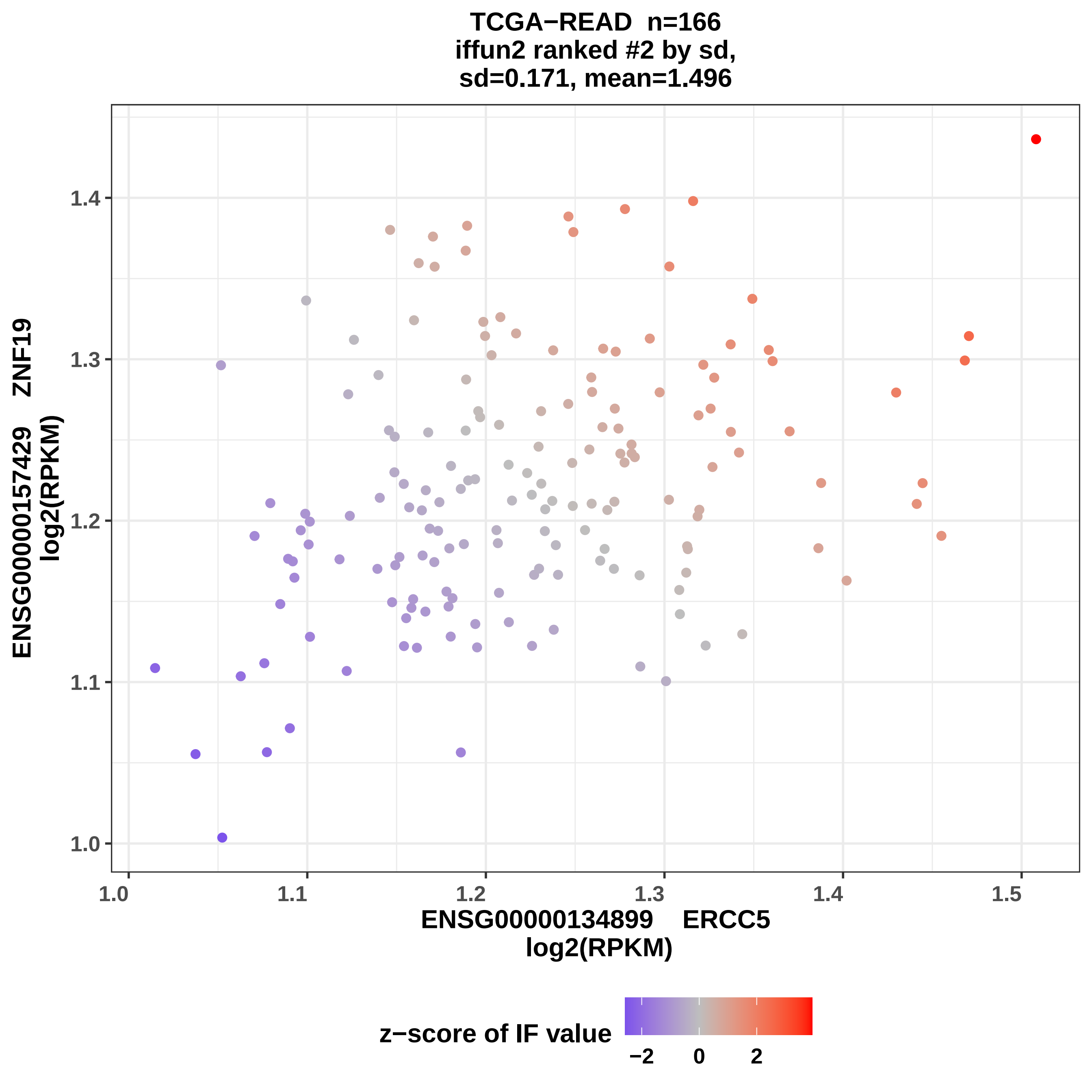

TCGA−READ n=166
iffun2 ranked #2 by sd,
sd=0.171, mean=1.496
ENSG00000157429 ZNF19
log2(RPKM)
ENSG00000134899 ERCC5
log2(RPKM)
1.0
1.1
1.2
1.3
1.4
1.5
z−score of IF value
−2
0
2

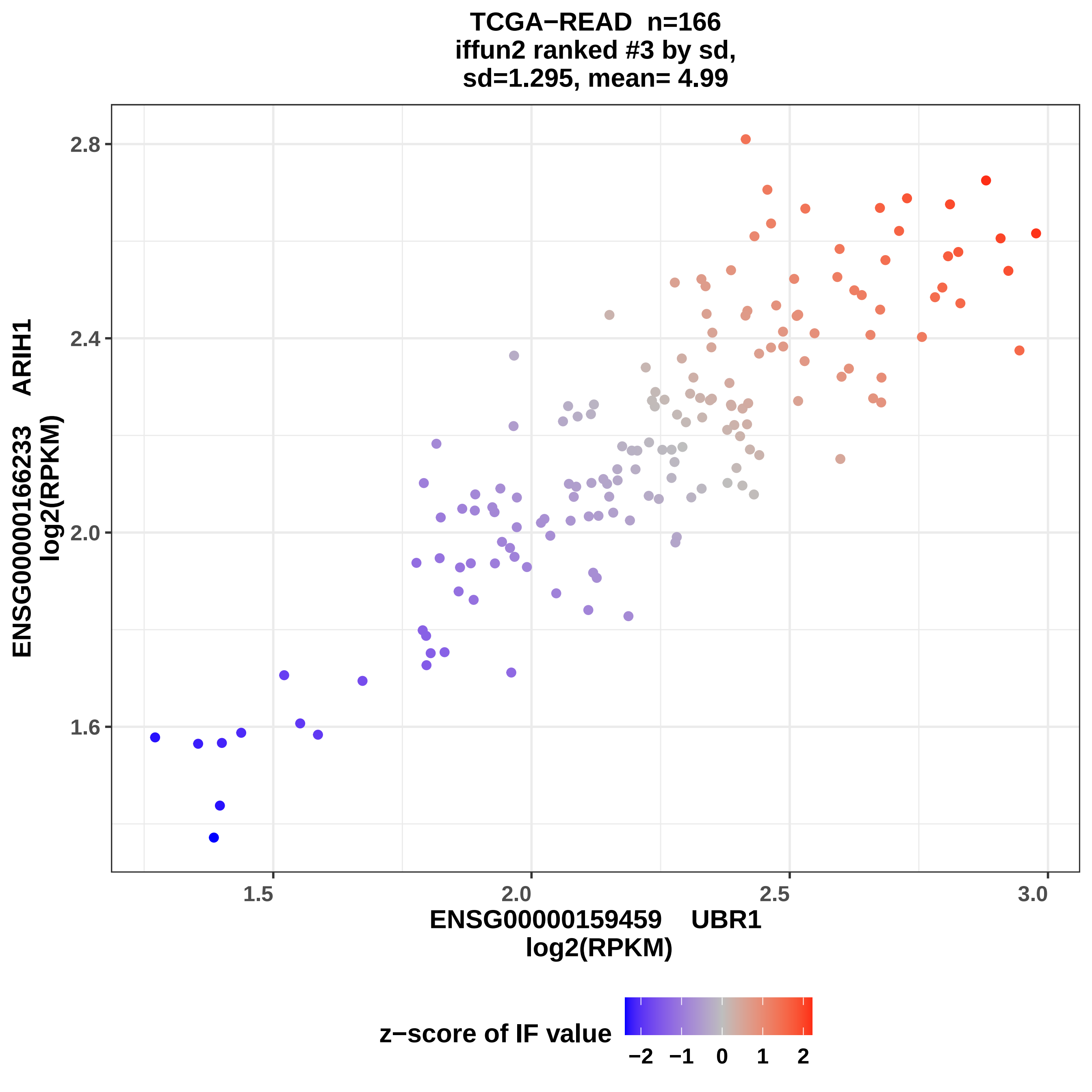

TCGA−READ n=166
iffun2 ranked #3 by sd,
sd=1.295, mean= 4.99
ENSG00000166233 ARIH1
log2(RPKM)
2.8
2.4
2.0
1.6
1.5
2.0
2.5
3.0
ENSG00000159459 UBR1
log2(RPKM)
z−score of IF value
−2
−1
0
1
2

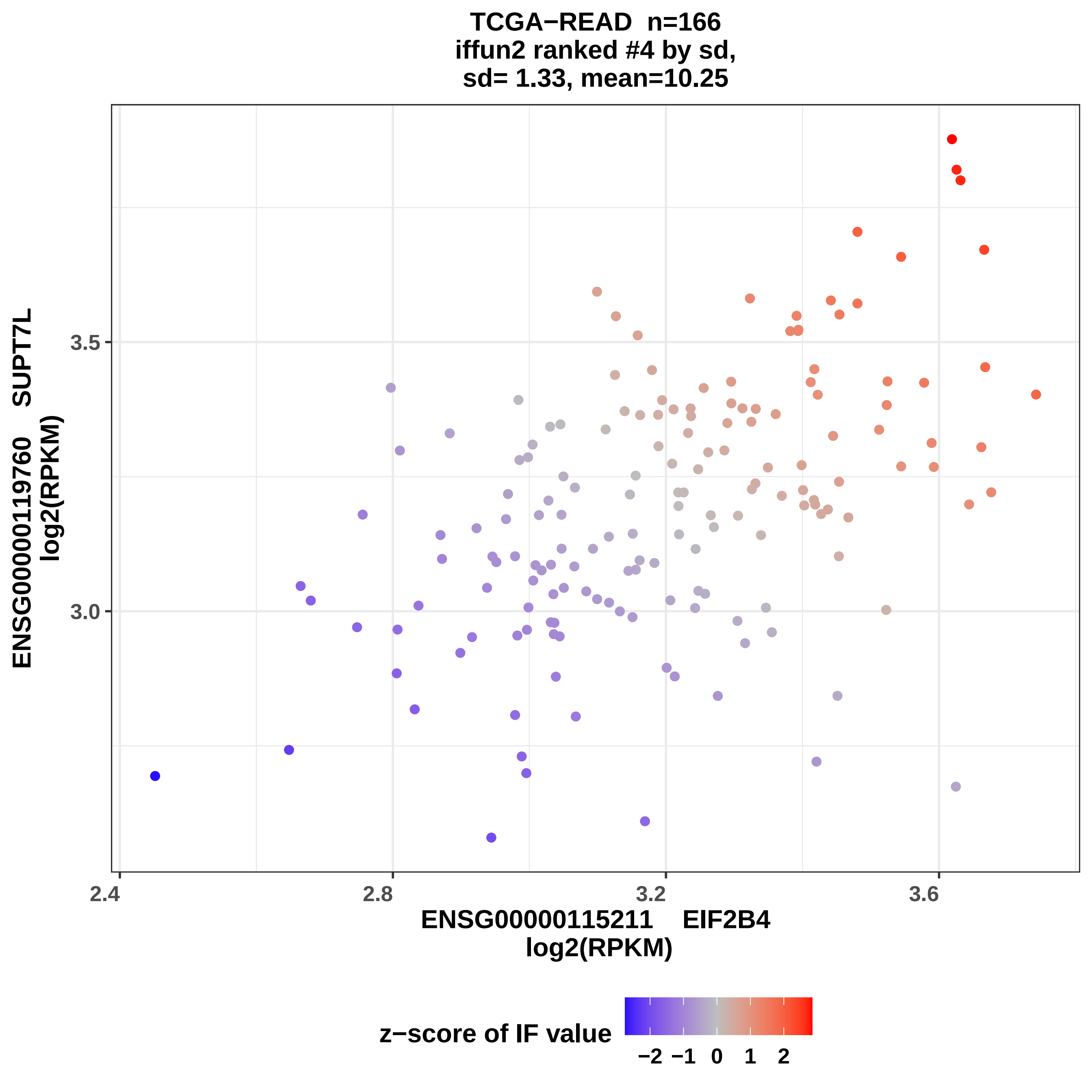
TCGA−READ n=166
iffun2 ranked #4 by sd,
sd= 1.33, mean=10.25
ENSG00000119760 SUPT7L
log2(RPKM)
3.0
3.5
2.4
2.8
3.2
3.6
ENSG00000115211 EIF2B4
log2(RPKM)
z−score of IF value
−2 −1 0 1 2

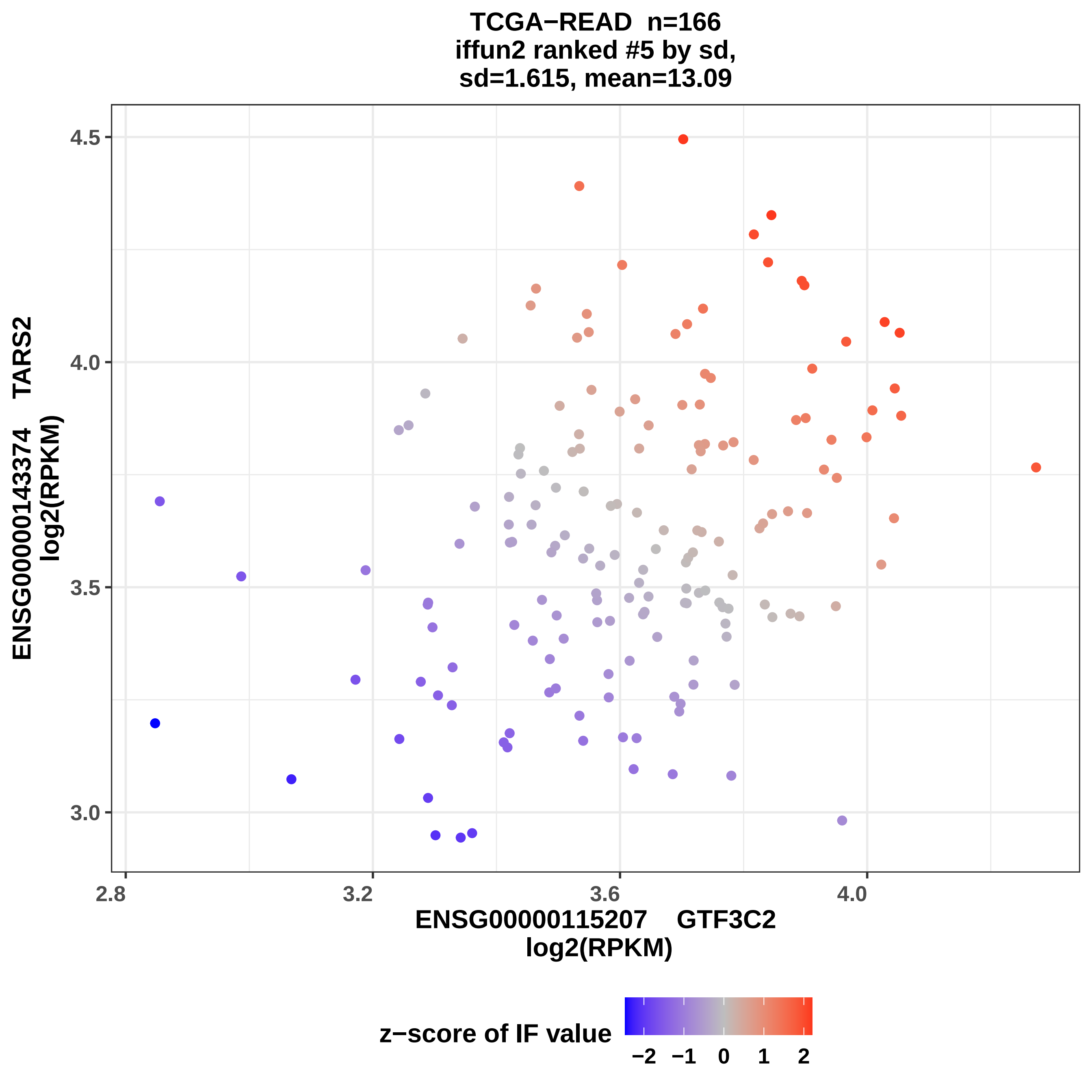
TCGA−READ n=166
iffun2 ranked #5 by sd,
sd=1.615, mean=13.09
ENSG00000143374 TARS2
log2(RPKM)
4.5
4.0
3.5
3.0
2.8
3.2
3.6
4.0
ENSG00000115207 GTF3C2
log2(RPKM)
z−score of IF value
−2
−1
0
1
2

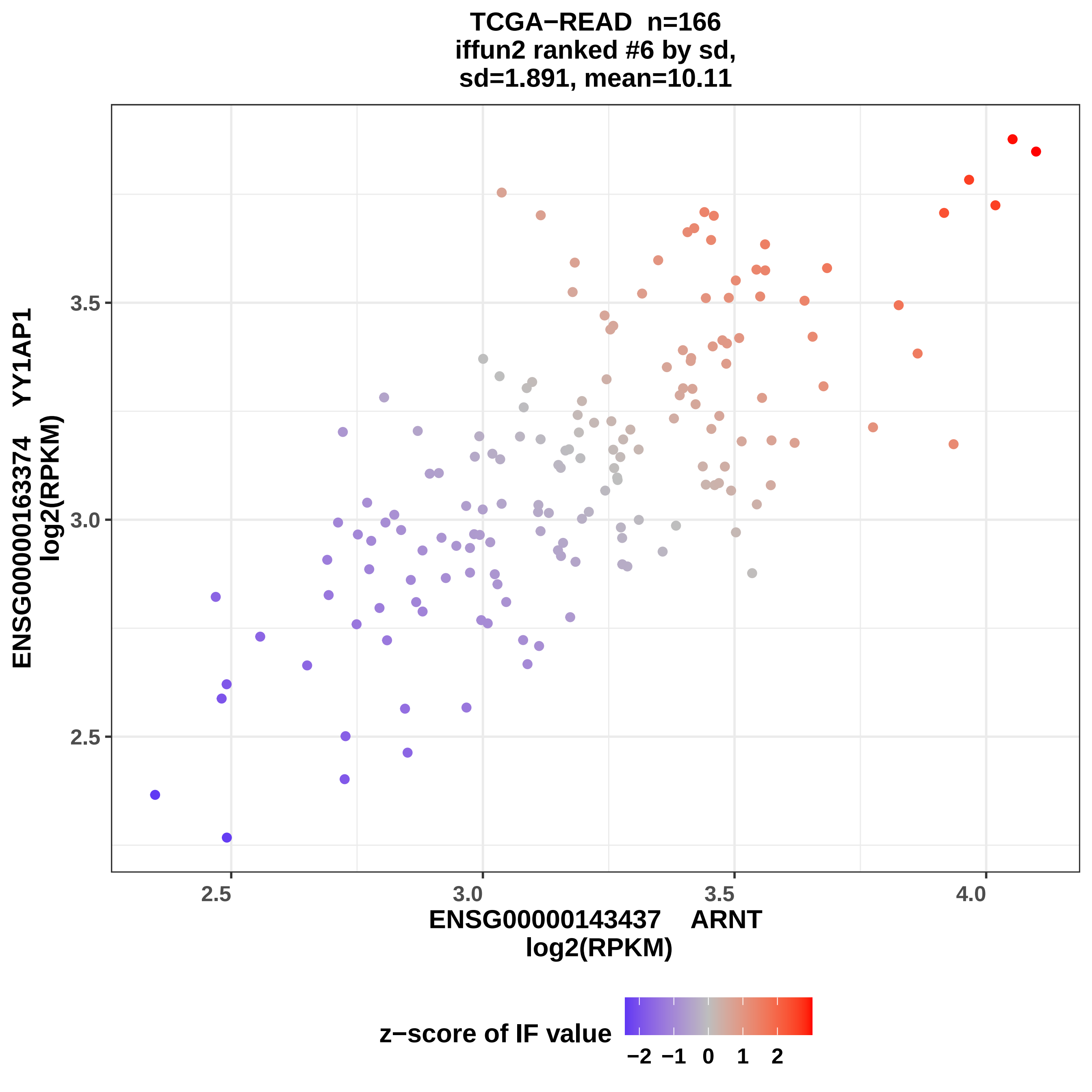

TCGA−READ n=166
iffun2 ranked #6 by sd,
sd=1.891, mean=10.11
ENSG00000163374 YY1AP1
log2(RPKM)
3.5
3.0
2.5
2.5
3.0
3.5
4.0
ENSG00000143437 ARNT
log2(RPKM)
z−score of IF value
−2 −1 0 1 2

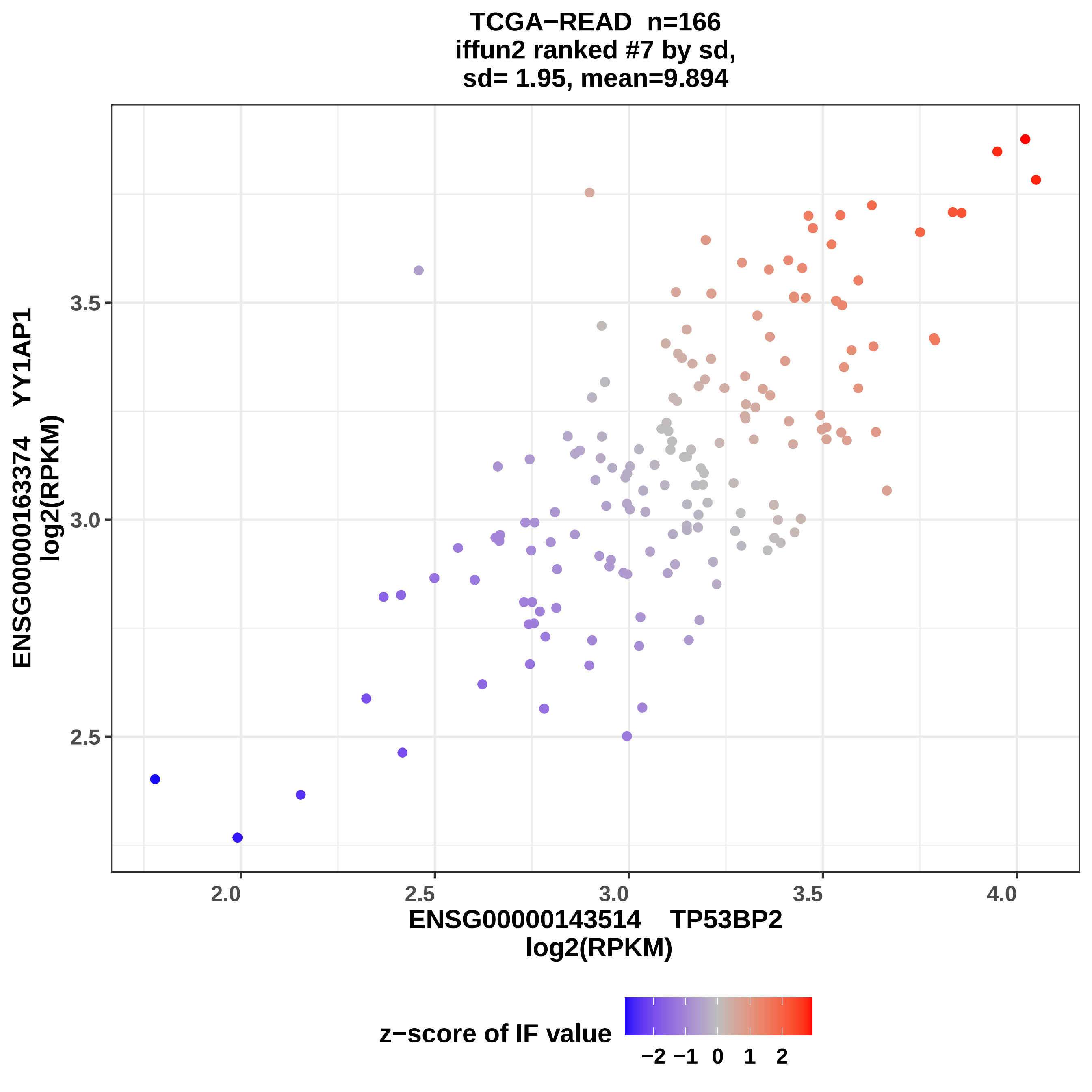
TCGA−READ n=166
iffun2 ranked #7 by sd,
sd= 1.95, mean=9.894
ENSG00000163374 YY1AP1
log2(RPKM)
3.5
3.0
2.5
2.0
2.5
3.0
3.5
4.0
ENSG00000143514 TP53BP2
log2(RPKM)
z−score of IF value
−2 −1 0 1 2

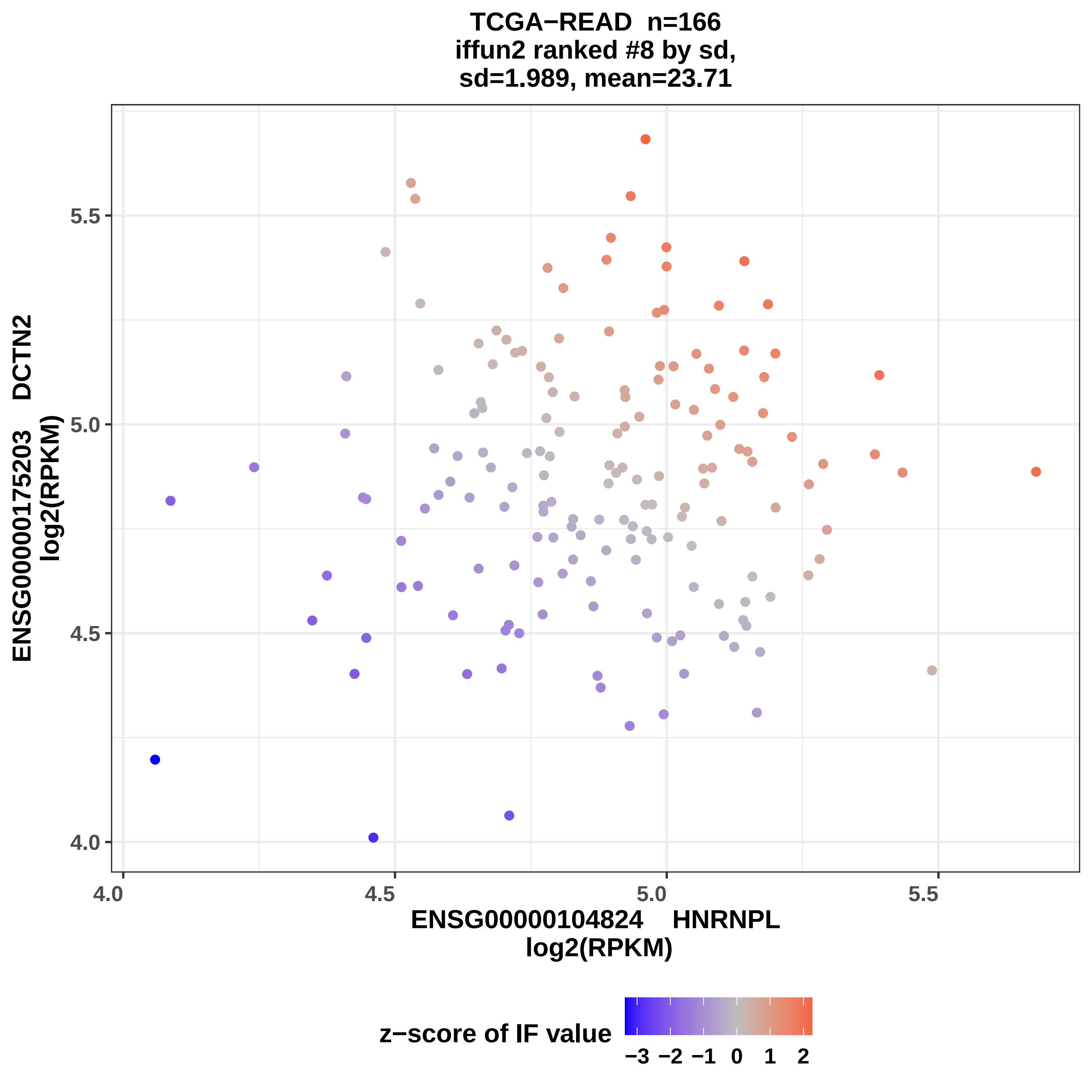
TCGA−READ n=166
iffun2 ranked #8 by sd,
sd=1.989, mean=23.71
ENSG00000175203 DCTN2
log2(RPKM)
5.5
5.0
4.5
4.0
4.0
4.5
5.0
5.5
ENSG00000104824 HNRNPL
log2(RPKM)
z−score of IF value
−3 −2 −1 0 1 2

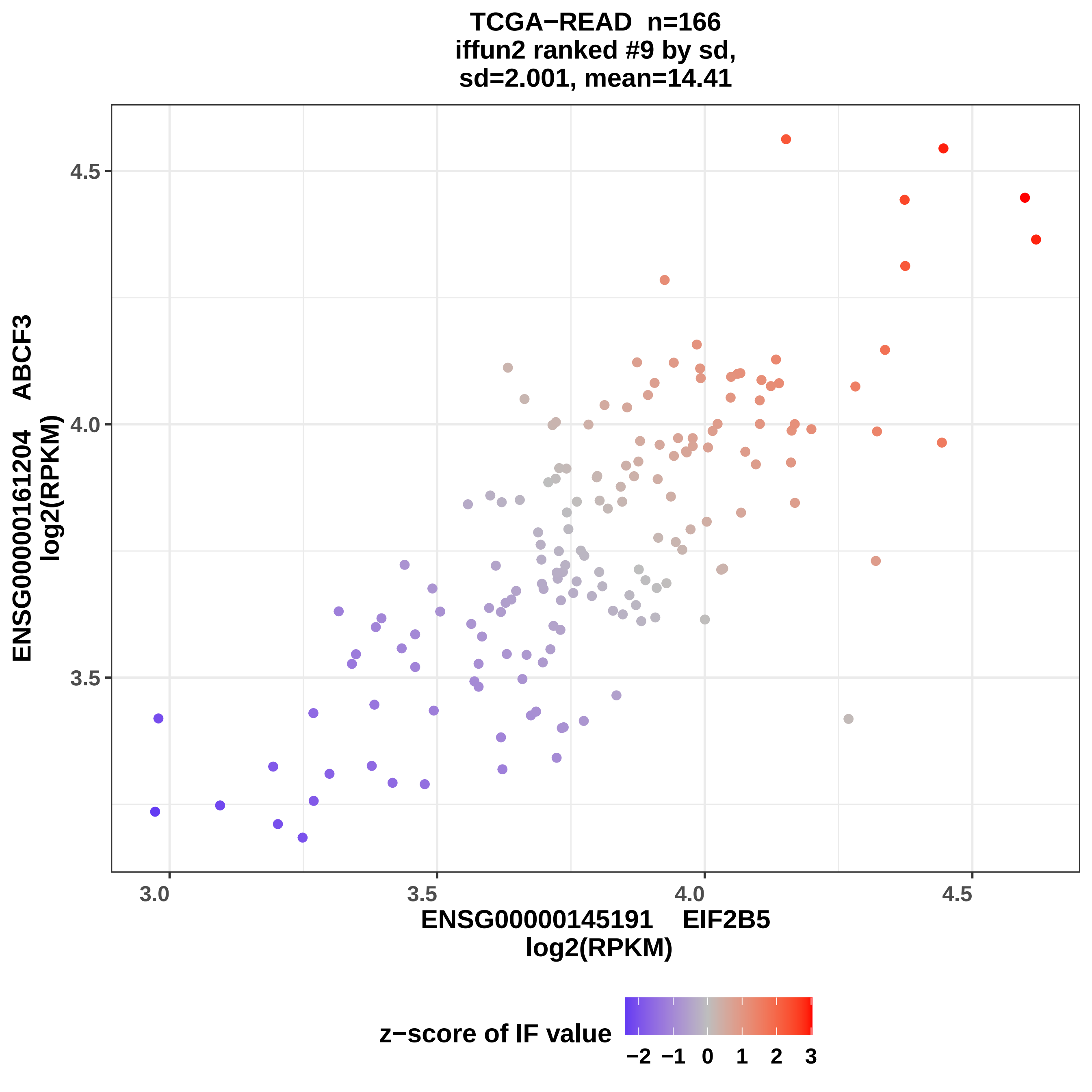

TCGA−READ n=166
iffun2 ranked #9 by sd,
sd=2.001, mean=14.41
4.5
4.0
3.5
3.0
3.5
4.0
4.5
ENSG00000161204 ABCF3
log2(RPKM)
ENSG00000145191 EIF2B5
log2(RPKM)
z−score of IF value
−2 −1 0 1 2 3

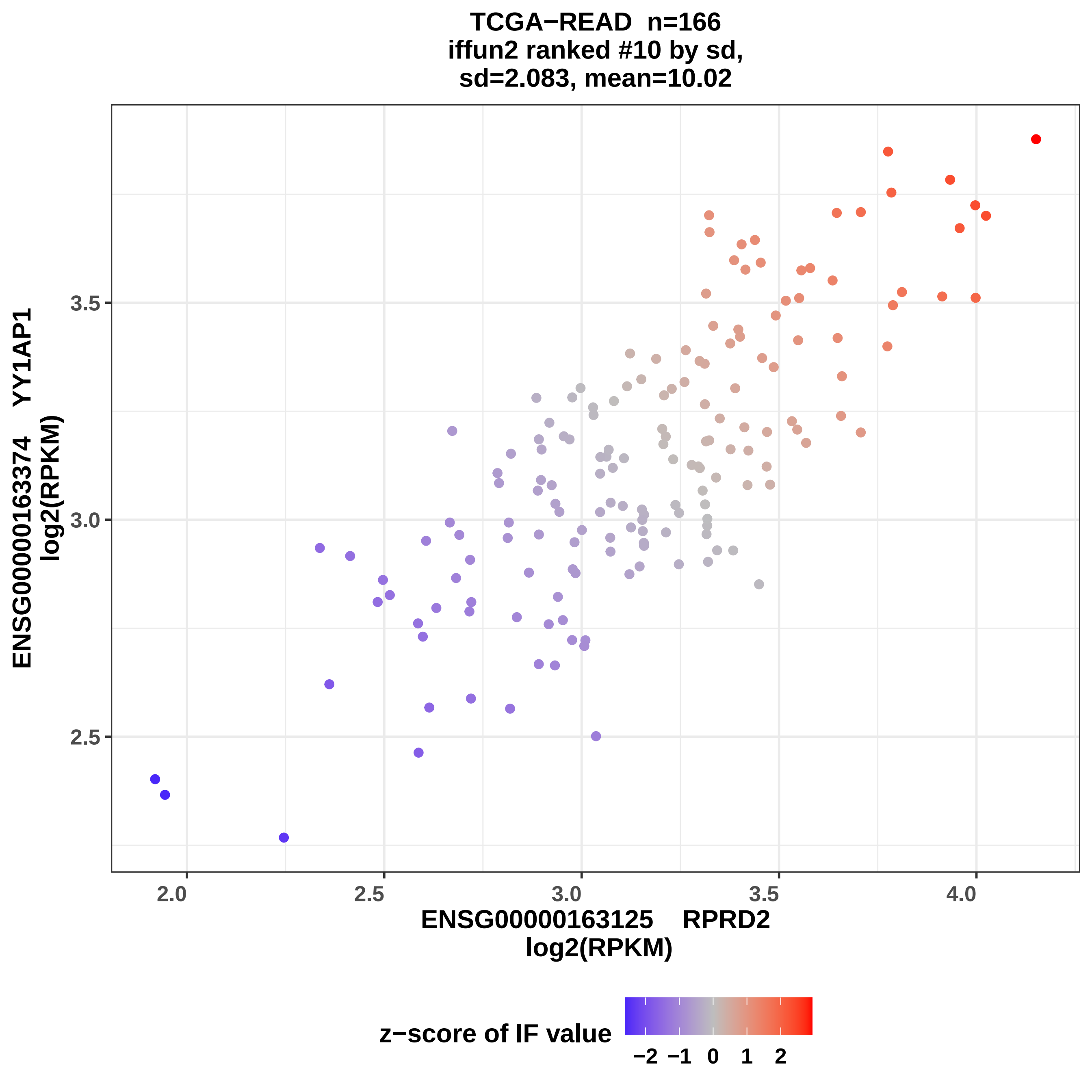
TCGA−READ n=166
iffun2 ranked #10 by sd,
sd=2.083, mean=10.02
ENSG00000163374 YY1AP1
log2(RPKM)
ENSG00000163125 RPRD2
log2(RPKM)
2.5
3.0
3.5
2.0
2.5
3.0
3.5
4.0
z−score of IF value
−2 −1 0 1 2

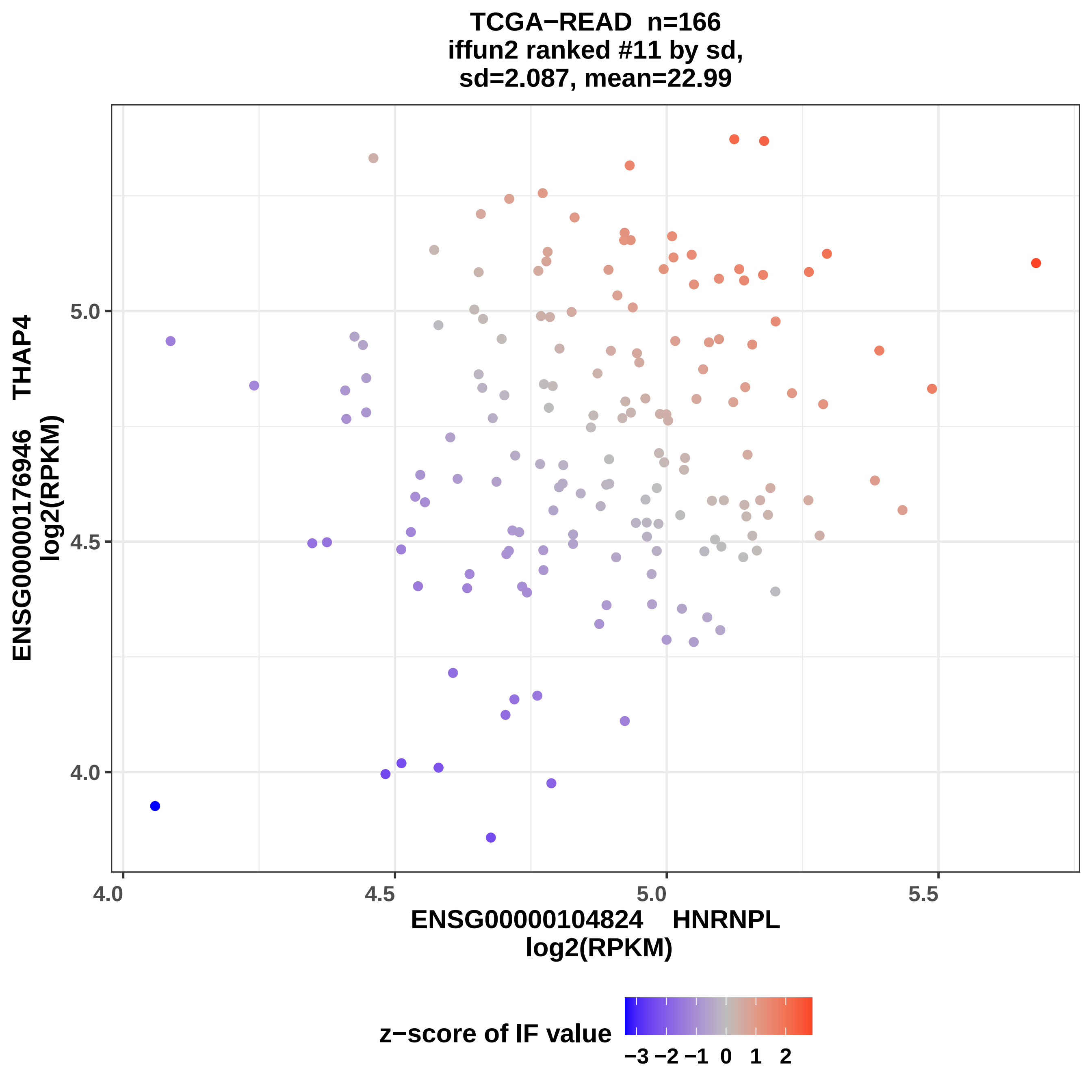
TCGA−READ n=166
iffun2 ranked #11 by sd,
sd=2.087, mean=22.99
ENSG00000176946 THAP4
log2(RPKM)
5.0
4.5
4.0
4.0
4.5
5.0
5.5
ENSG00000104824 HNRNPL
log2(RPKM)
z−score of IF value
−3 −2 −1 0 1 2

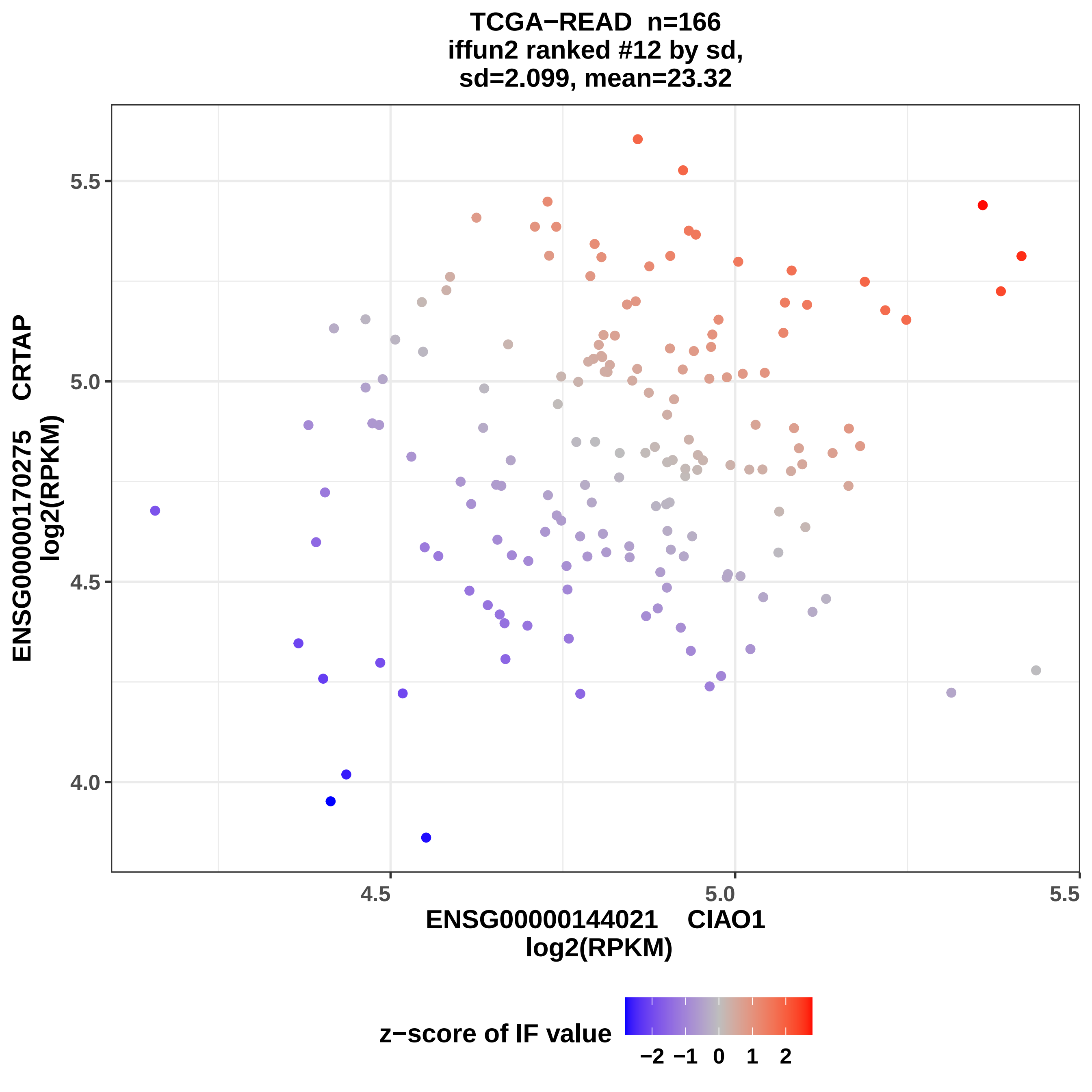
TCGA−READ n=166
iffun2 ranked #12 by sd,
sd=2.099, mean=23.32
ENSG00000170275 CRTAP
log2(RPKM)
5.5
5.0
4.5
4.0
4.5
5.0
5.5
ENSG00000144021 CIAO1
log2(RPKM)
z−score of IF value
−2
−1
0
1
2

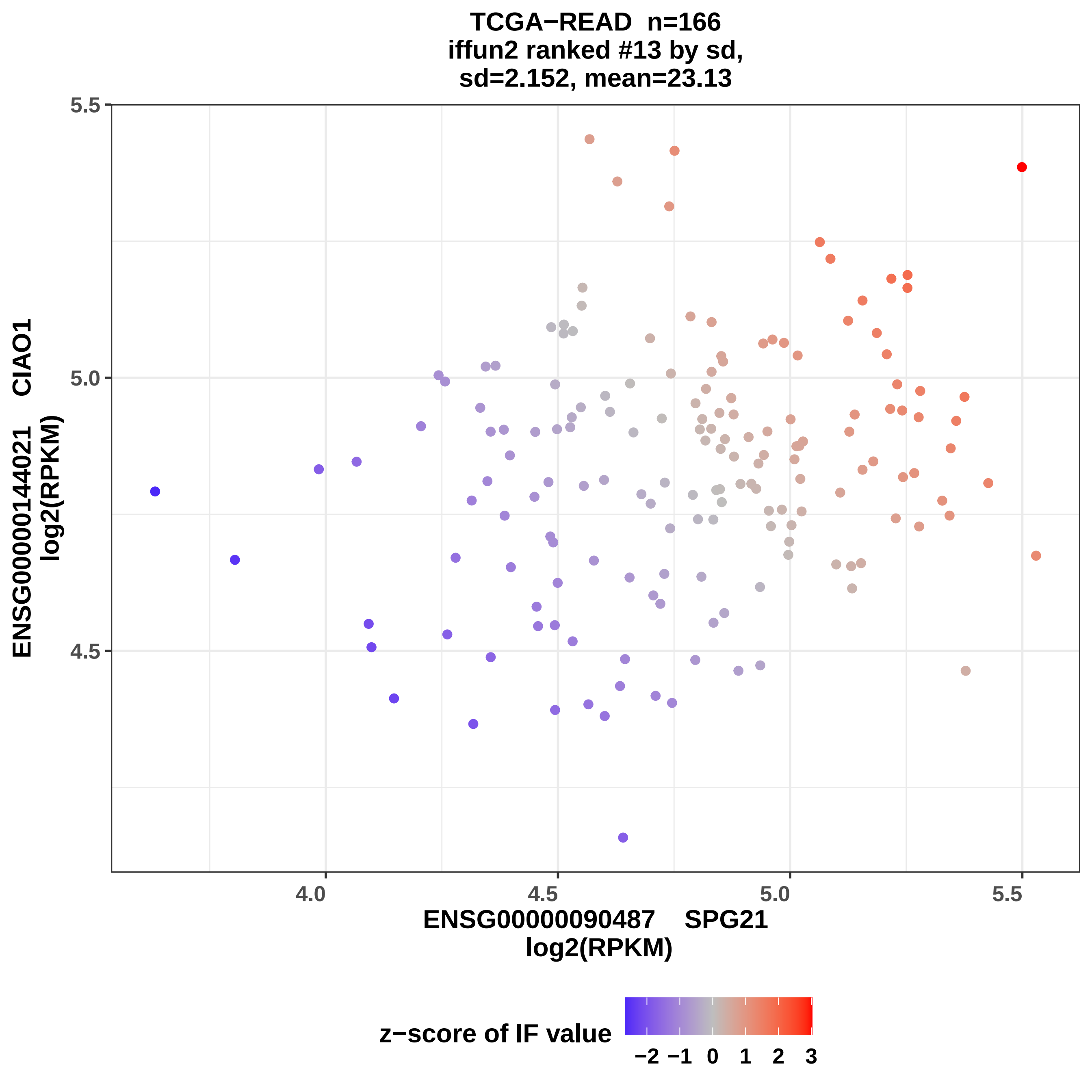

TCGA−READ n=166
iffun2 ranked #13 by sd,
sd=2.152, mean=23.13
5.5
5.0
4.5
4.0
4.5
5.0
5.5
ENSG00000144021 CIAO1
log2(RPKM)
ENSG00000090487 SPG21
log2(RPKM)
z−score of IF value
−2 −1 0 1 2 3

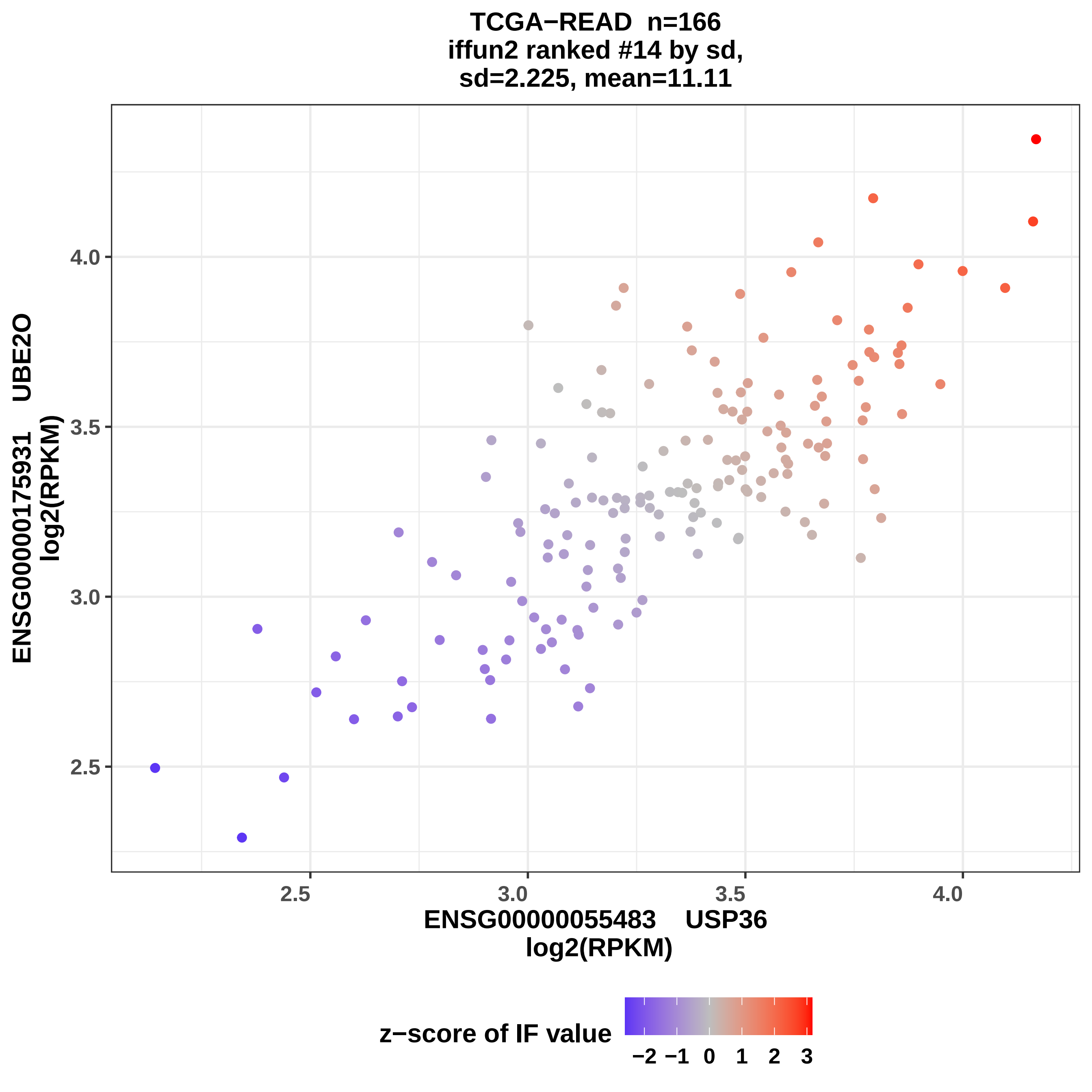

TCGA−READ n=166
iffun2 ranked #14 by sd,
sd=2.225, mean=11.11
ENSG00000175931 UBE2O
log2(RPKM)
4.0
3.5
3.0
2.5
2.5
3.0
3.5
4.0
ENSG00000055483 USP36
log2(RPKM)
z−score of IF value
−2 −1 0 1 2 3

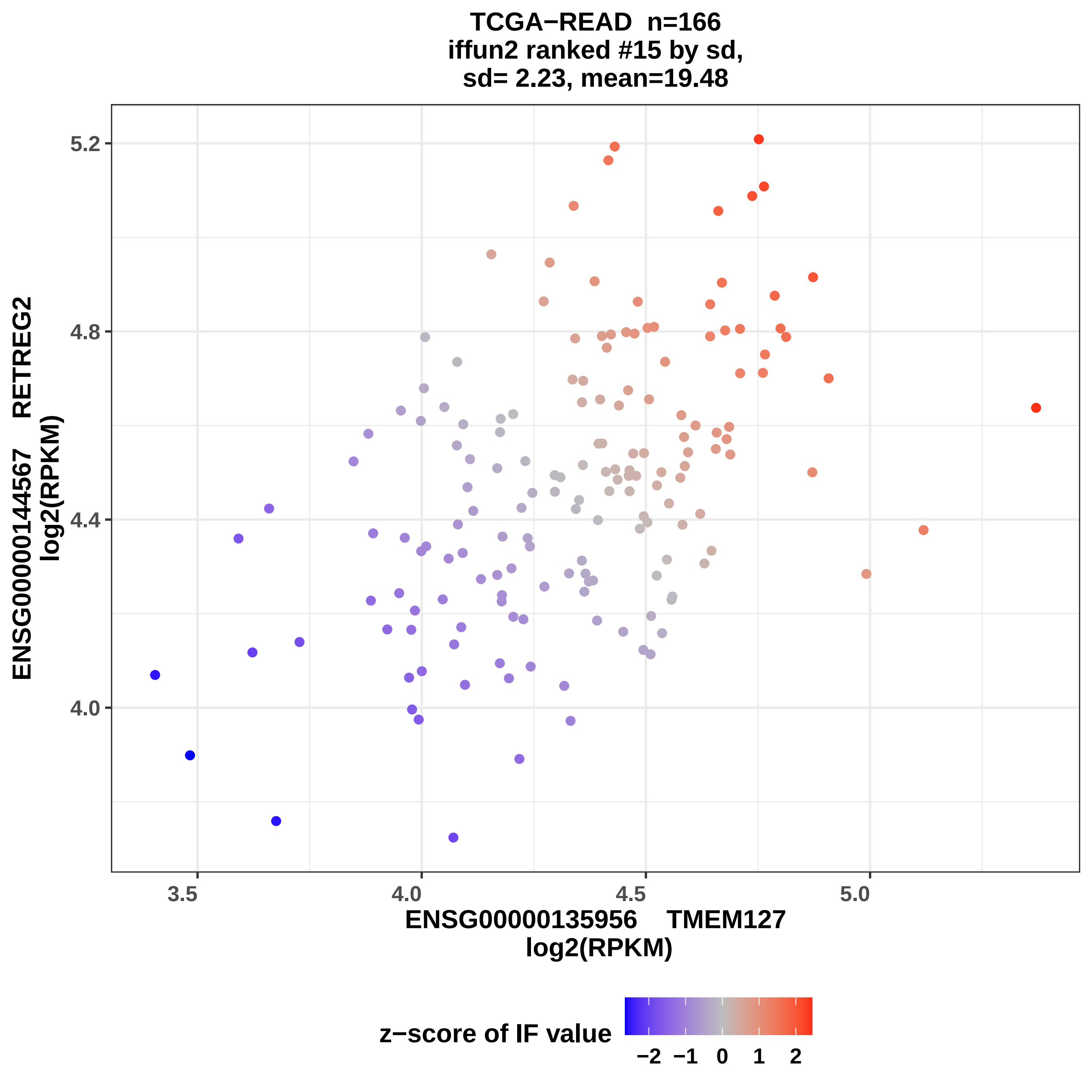
TCGA−READ n=166
iffun2 ranked #15 by sd,
sd= 2.23, mean=19.48
ENSG00000144567 RETREG2
log2(RPKM)
5.2
4.8
4.4
4.0
3.5
4.0
4.5
5.0
ENSG00000135956 TMEM127
log2(RPKM)
z−score of IF value
−2 −1 0 1 2

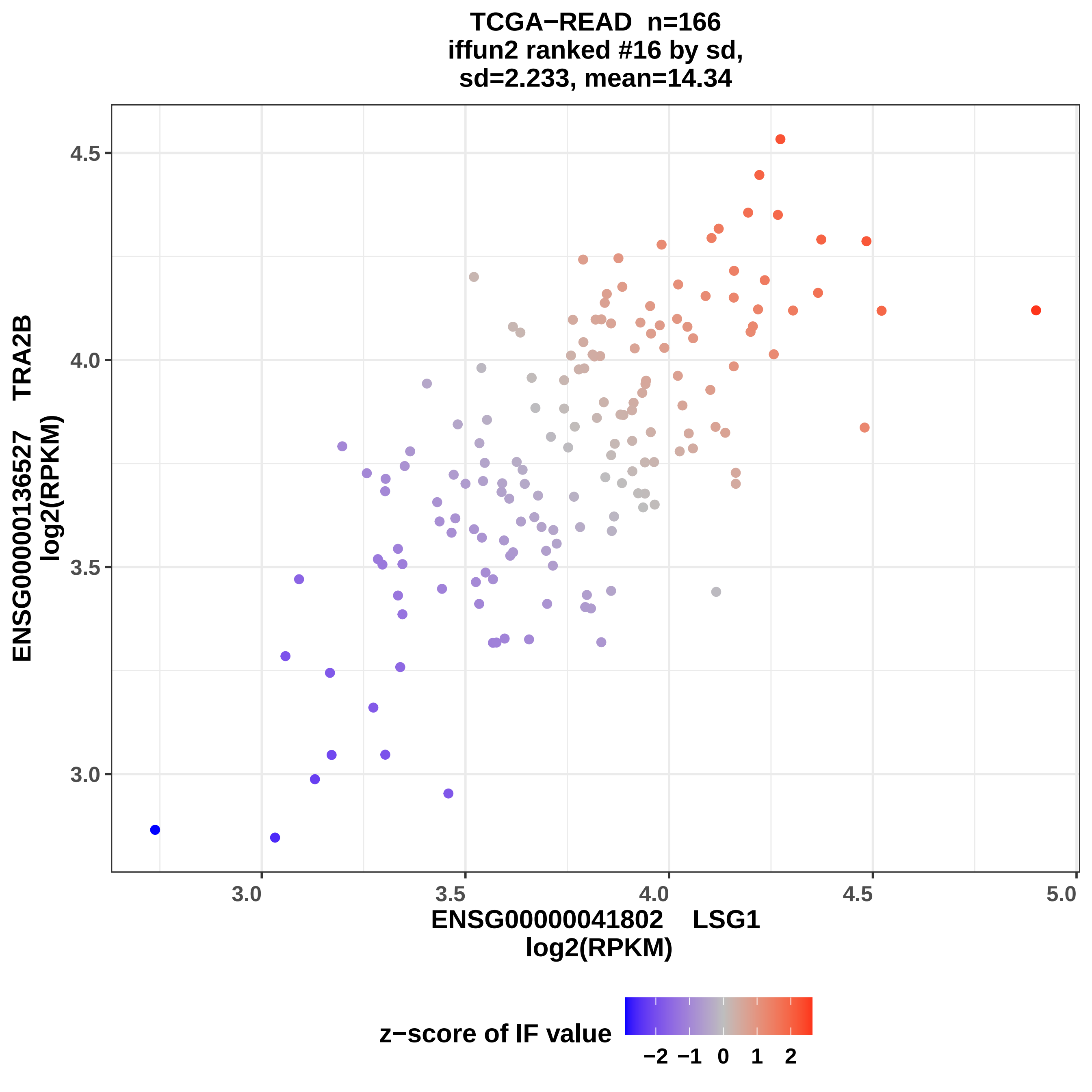

TCGA−READ n=166
iffun2 ranked #16 by sd,
sd=2.233, mean=14.34
ENSG00000136527 TRA2B
log2(RPKM)
4.5
4.0
3.5
3.0
3.0
3.5
4.0
4.5
5.0
ENSG00000041802 LSG1
log2(RPKM)
z−score of IF value
−2
−1
0
1
2

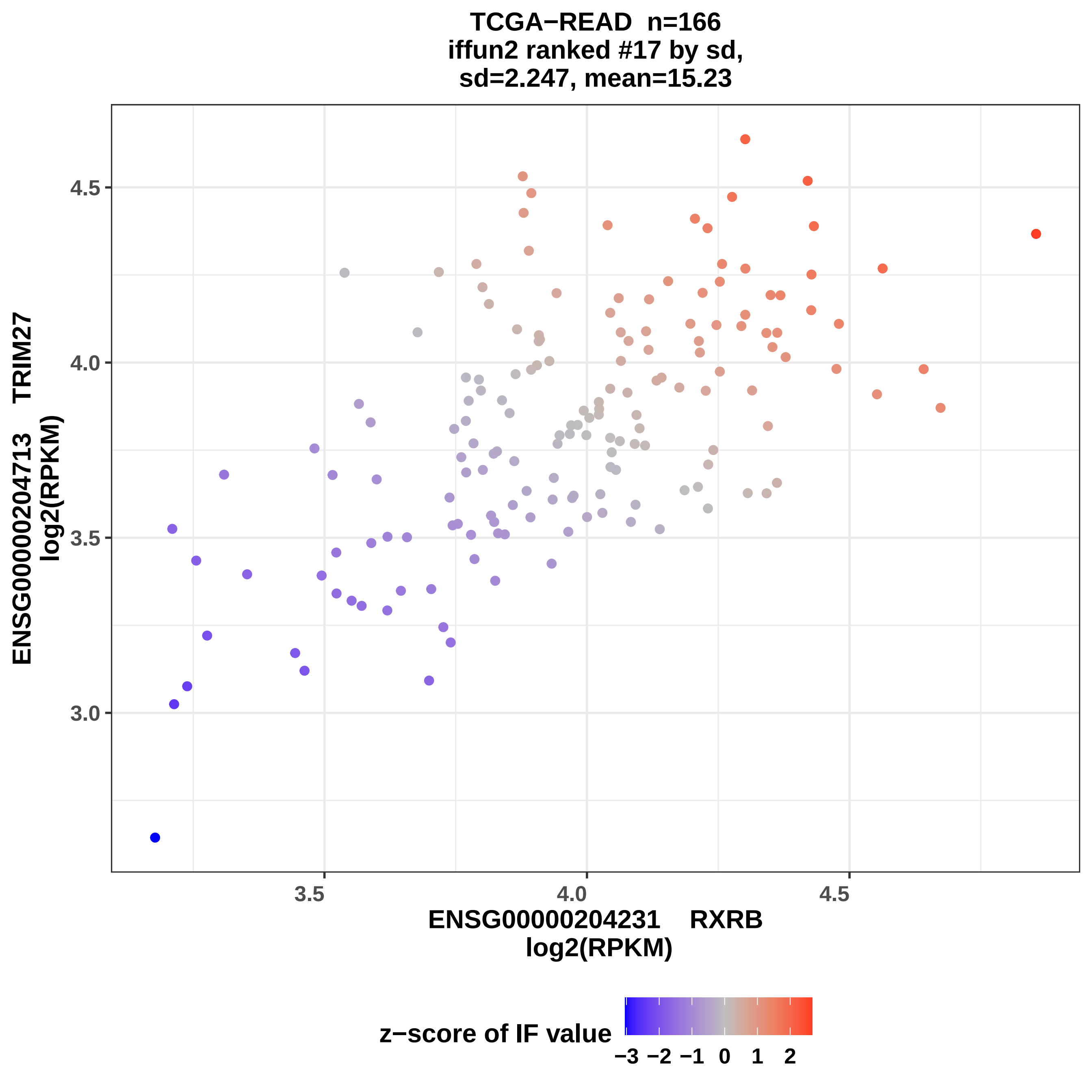

TCGA−READ n=166
iffun2 ranked #17 by sd,
sd=2.247, mean=15.23
4.5
4.0
3.5
3.0
ENSG00000204713 TRIM27
log2(RPKM)
3.5
4.0
4.5
ENSG00000204231 RXRB
log2(RPKM)
z−score of IF value
−3 −2 −1 0 1 2

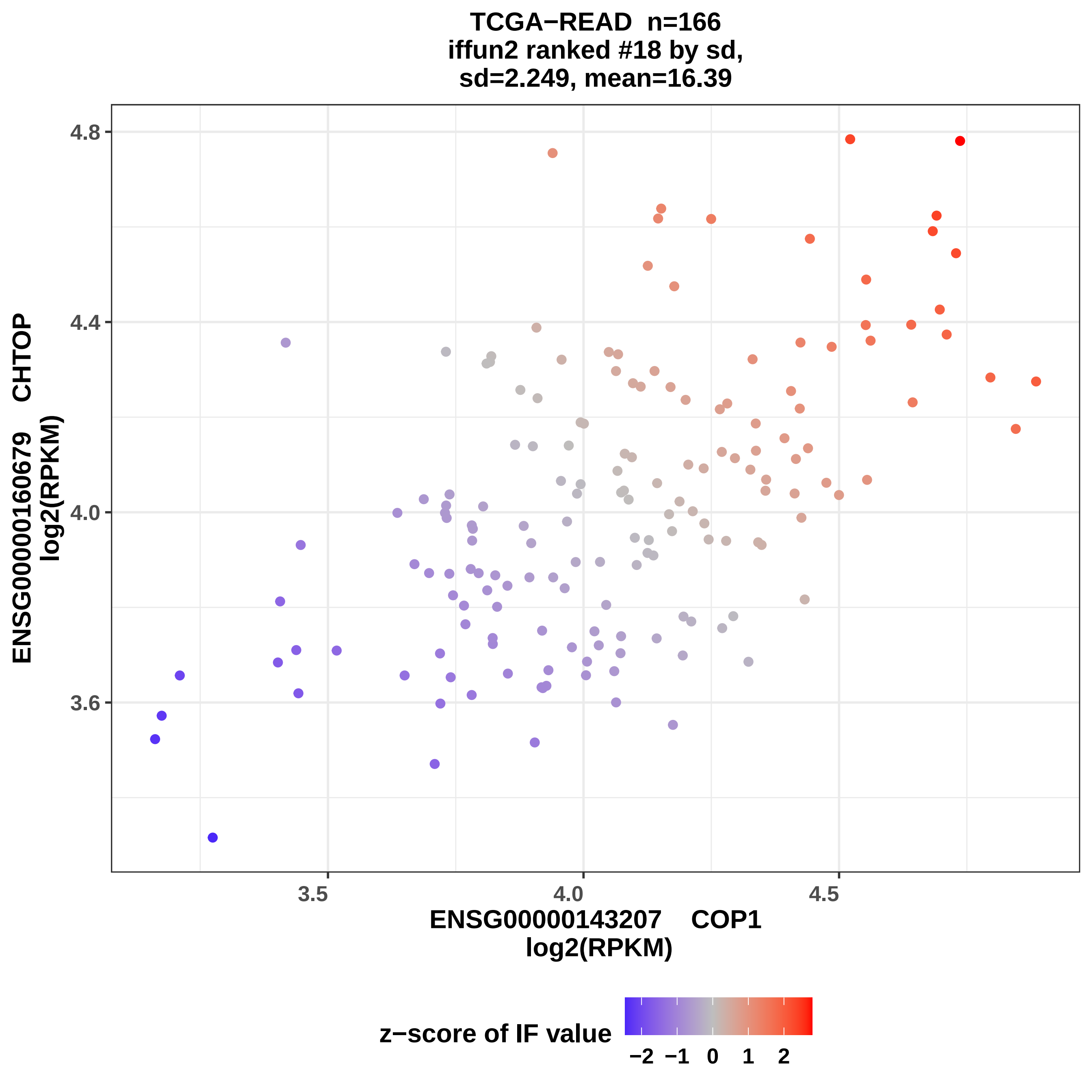

TCGA−READ n=166
iffun2 ranked #18 by sd,
sd=2.249, mean=16.39
ENSG00000160679 CHTOP
log2(RPKM)
4.8
4.4
4.0
3.6
3.5
4.0
4.5
ENSG00000143207 COP1
log2(RPKM)
z−score of IF value
−2 −1 0 1 2

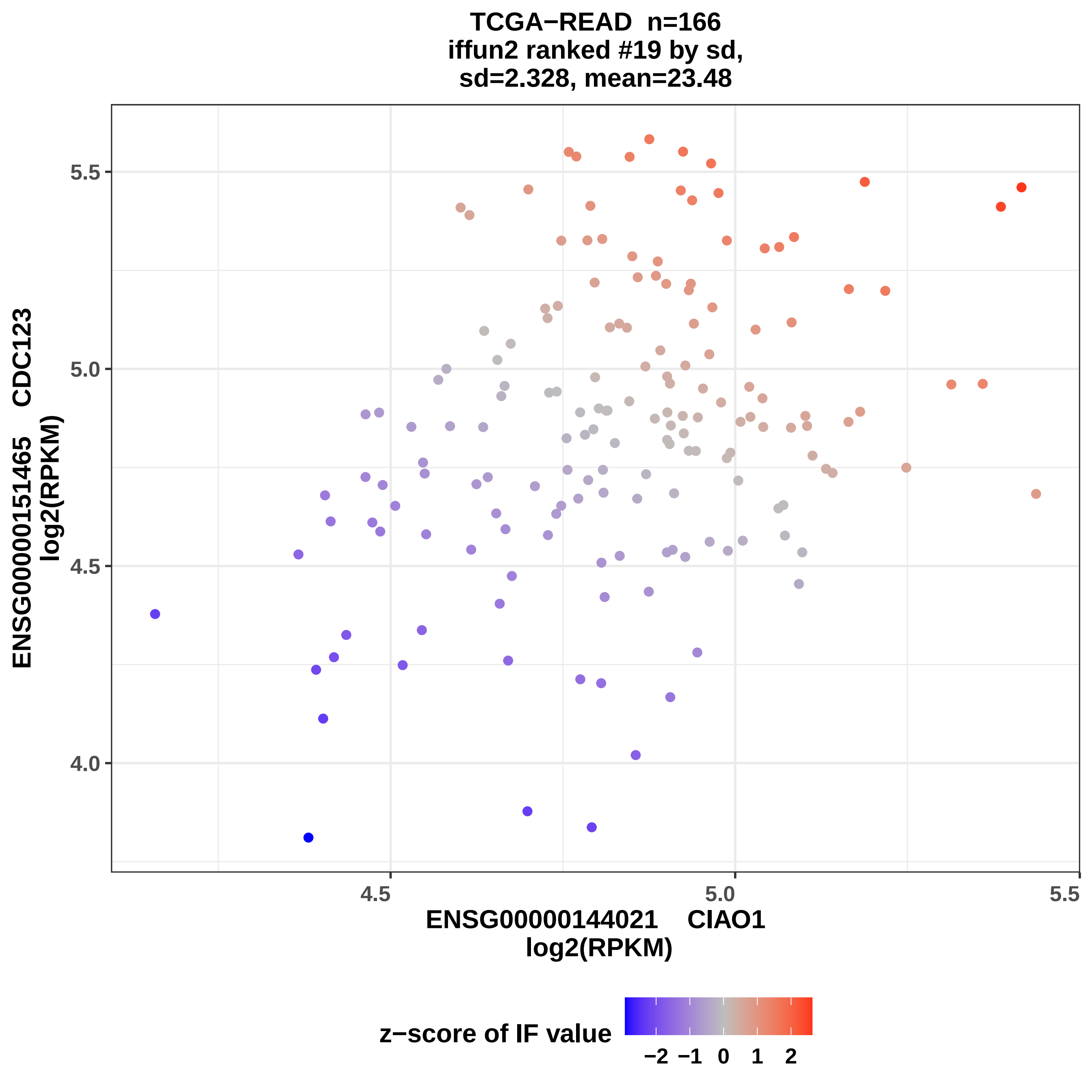

TCGA−READ n=166
iffun2 ranked #19 by sd,
sd=2.328, mean=23.48
ENSG00000151465 CDC123
log2(RPKM)
ENSG00000144021 CIAO1
log2(RPKM)
5.5
5.0
4.5
4.0
4.5
5.0
5.5
z−score of IF value
−2
−1
0
1
2

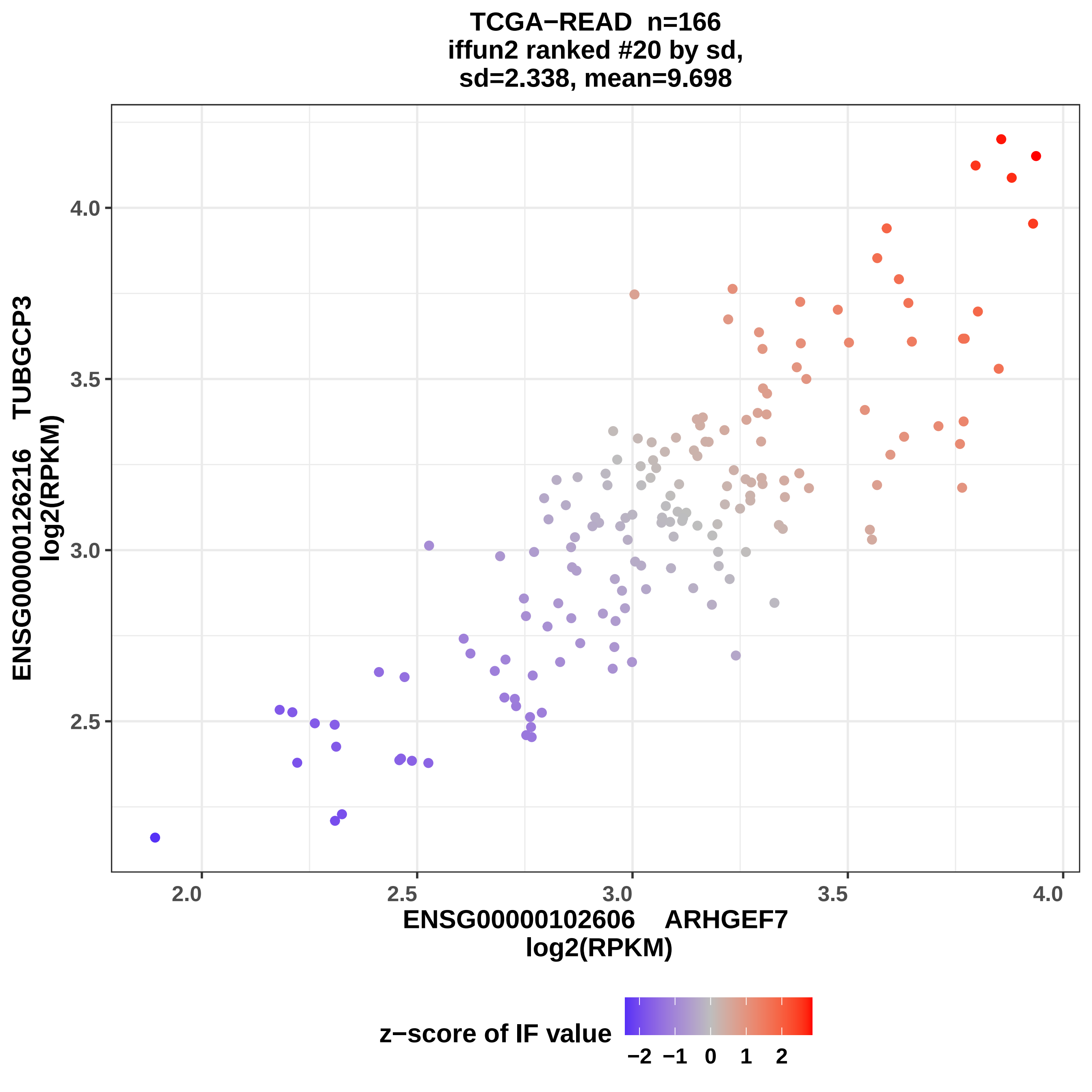
TCGA−READ n=166
iffun2 ranked #20 by sd,
sd=2.338, mean=9.698
ENSG00000126216 TUBGCP3
log2(RPKM)
4.0
3.5
3.0
2.5
2.0
2.5
3.0
3.5
4.0
ENSG00000102606 ARHGEF7
log2(RPKM)
z−score of IF value
−2 −1 0 1 2

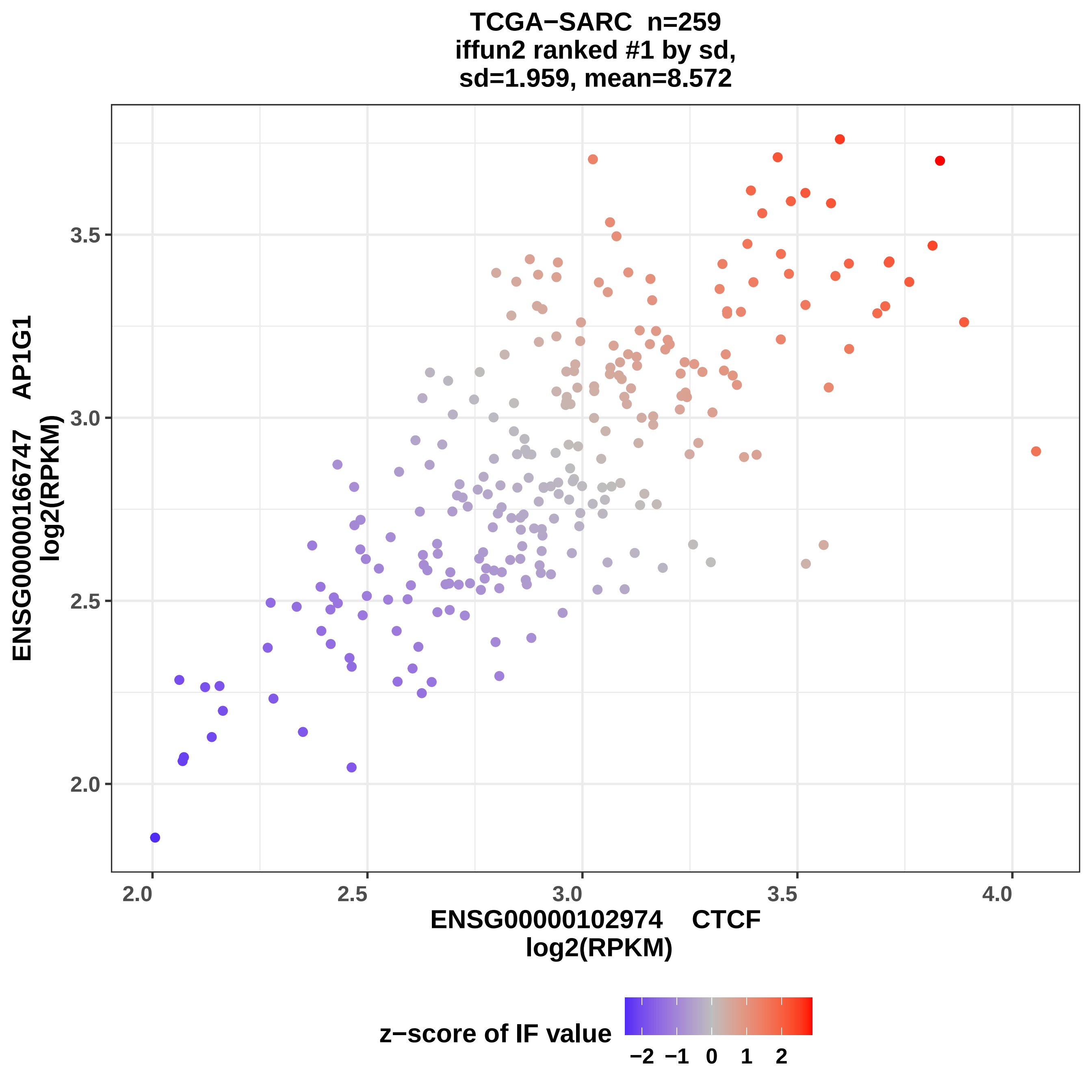
TCGA−SARC n=259
iffun2 ranked #1 by sd,
sd=1.959, mean=8.572
ENSG00000166747 AP1G1
log2(RPKM)
3.5
3.0
2.5
2.0
2.0
2.5
3.0
3.5
4.0
ENSG00000102974 CTCF
log2(RPKM)
z−score of IF value
−2 −1 0 1 2

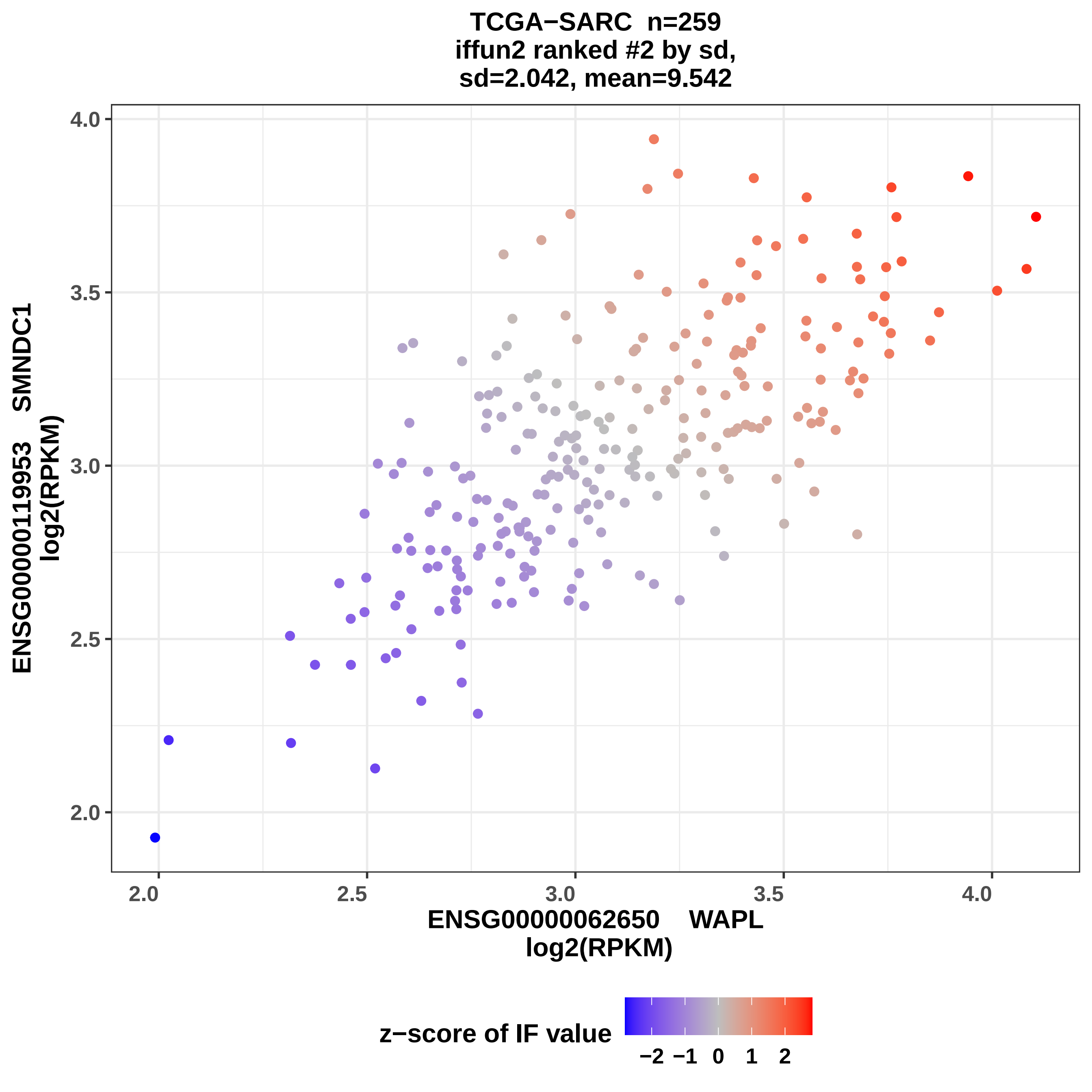
TCGA−SARC n=259
iffun2 ranked #2 by sd,
sd=2.042, mean=9.542
ENSG00000119953 SMNDC1
log2(RPKM)
4.0
3.5
3.0
2.5
2.0
2.0
2.5
3.0
3.5
4.0
ENSG00000062650 WAPL
log2(RPKM)
z−score of IF value
−2
−1
0
1
2

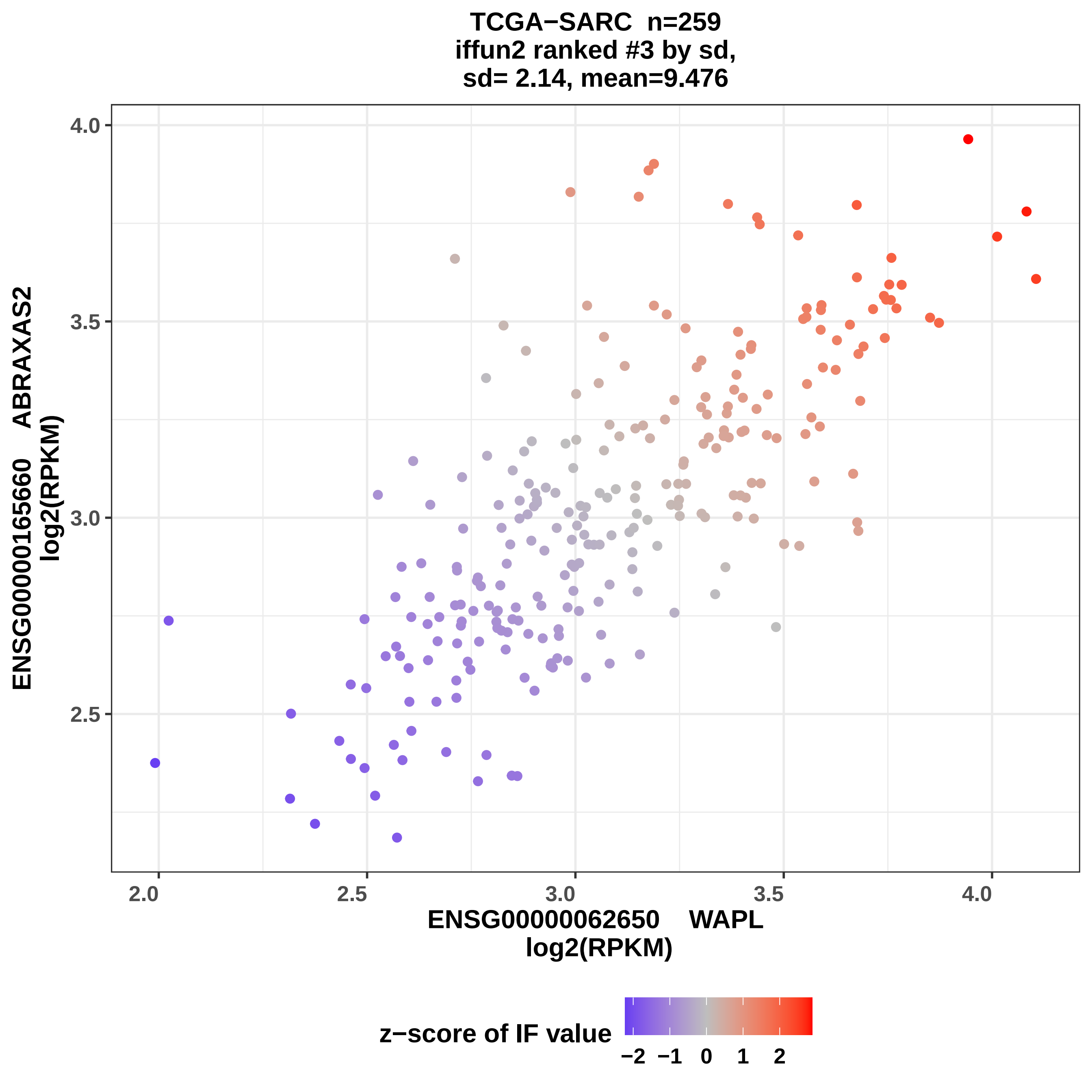
TCGA−SARC n=259
iffun2 ranked #3 by sd,
sd= 2.14, mean=9.476
ENSG00000165660 ABRAXAS2
log2(RPKM)
4.0
3.5
3.0
2.5
2.0
2.5
3.0
3.5
4.0
ENSG00000062650 WAPL
log2(RPKM)
z−score of IF value
−2
−1
0
1
2

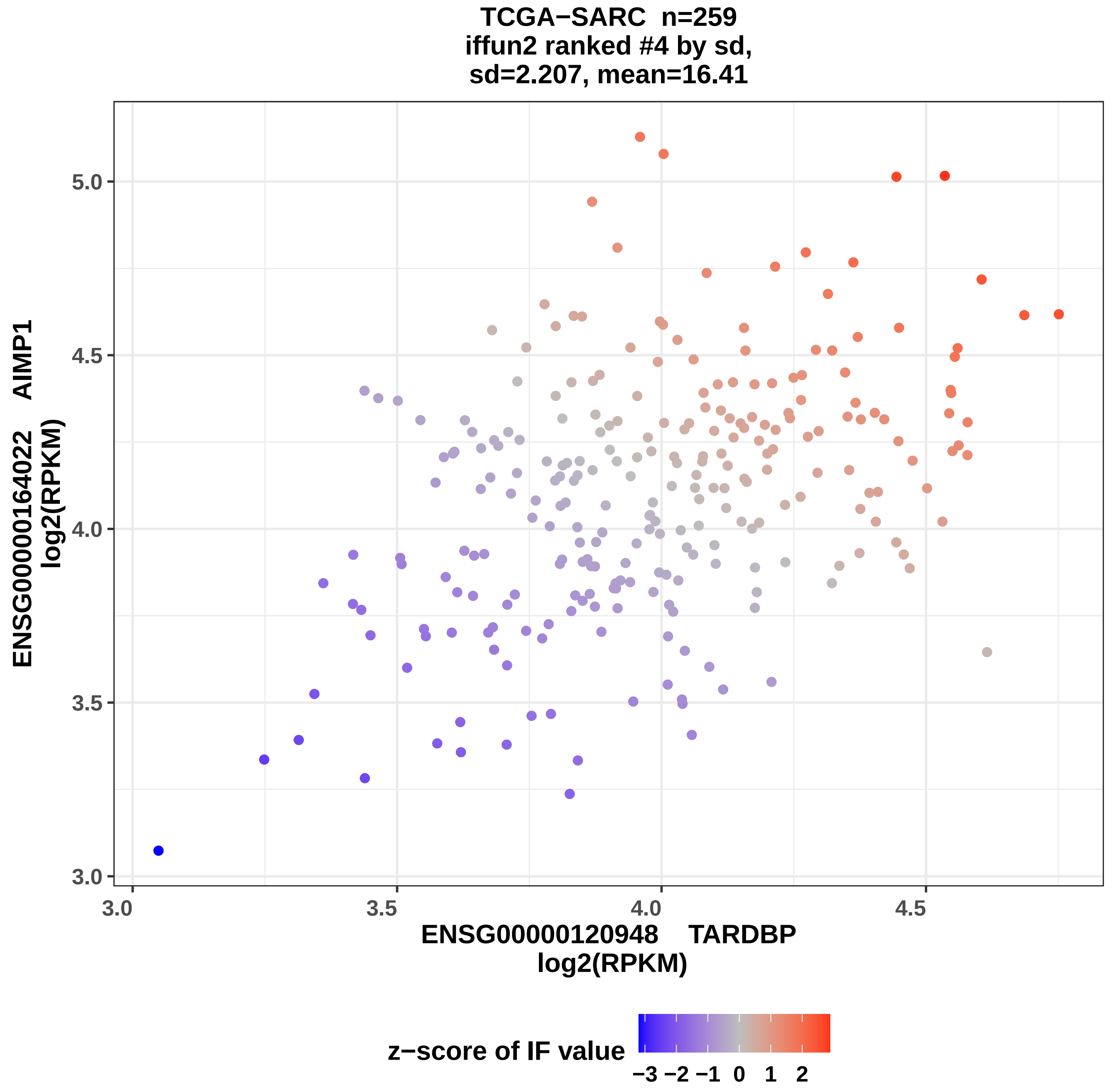
TCGA−SARC n=259
iffun2 ranked #4 by sd,
sd=2.207, mean=16.41
ENSG00000164022 AIMP1
log2(RPKM)
5.0
4.5
4.0
3.5
3.0
3.0
3.5
4.0
4.5
ENSG00000120948 TARDBP
log2(RPKM)
z−score of IF value
−3 −2 −1 0 1 2

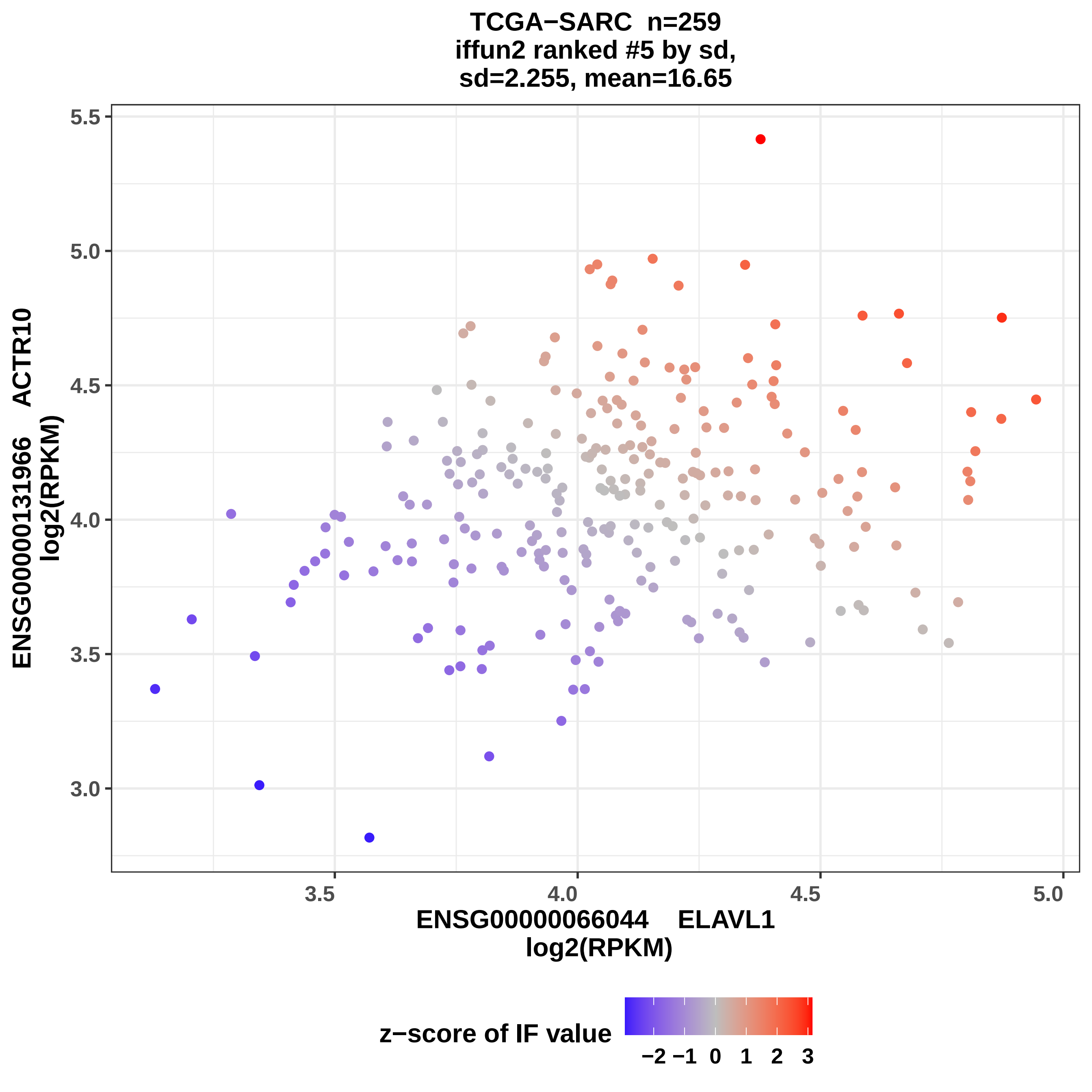
TCGA−SARC n=259
iffun2 ranked #5 by sd,
sd=2.255, mean=16.65
ENSG00000131966 ACTR10
log2(RPKM)
ENSG00000066044 ELAVL1
log2(RPKM)
z−score of IF value
−2 −1 0 1 2 3

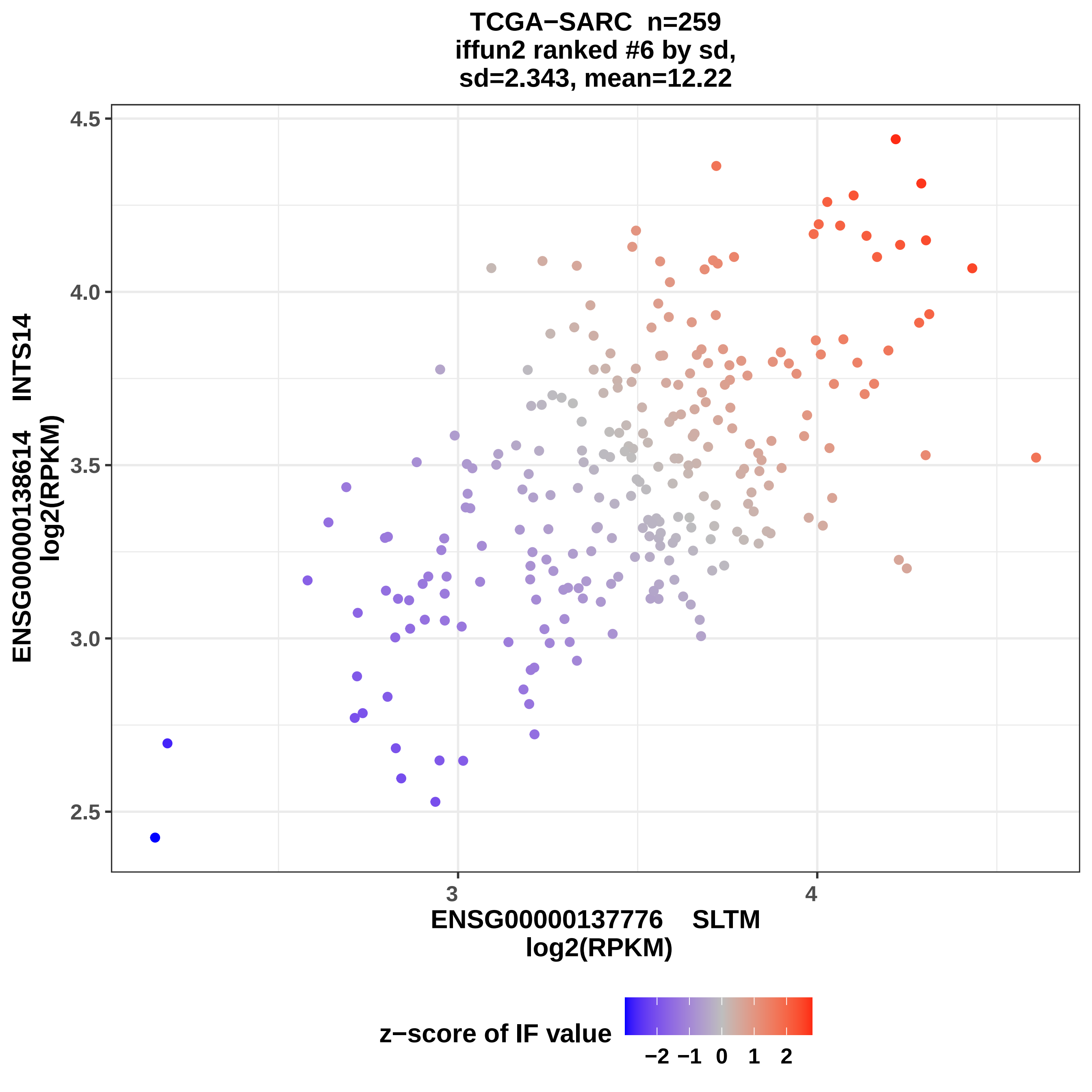
TCGA−SARC n=259
iffun2 ranked #6 by sd,
sd=2.343, mean=12.22
4.5
4.0
3.5
3.0
2.5
3
4
ENSG00000137776 SLTM
log2(RPKM)
ENSG00000138614 INTS14
log2(RPKM)
z−score of IF value
−2 −1 0 1 2

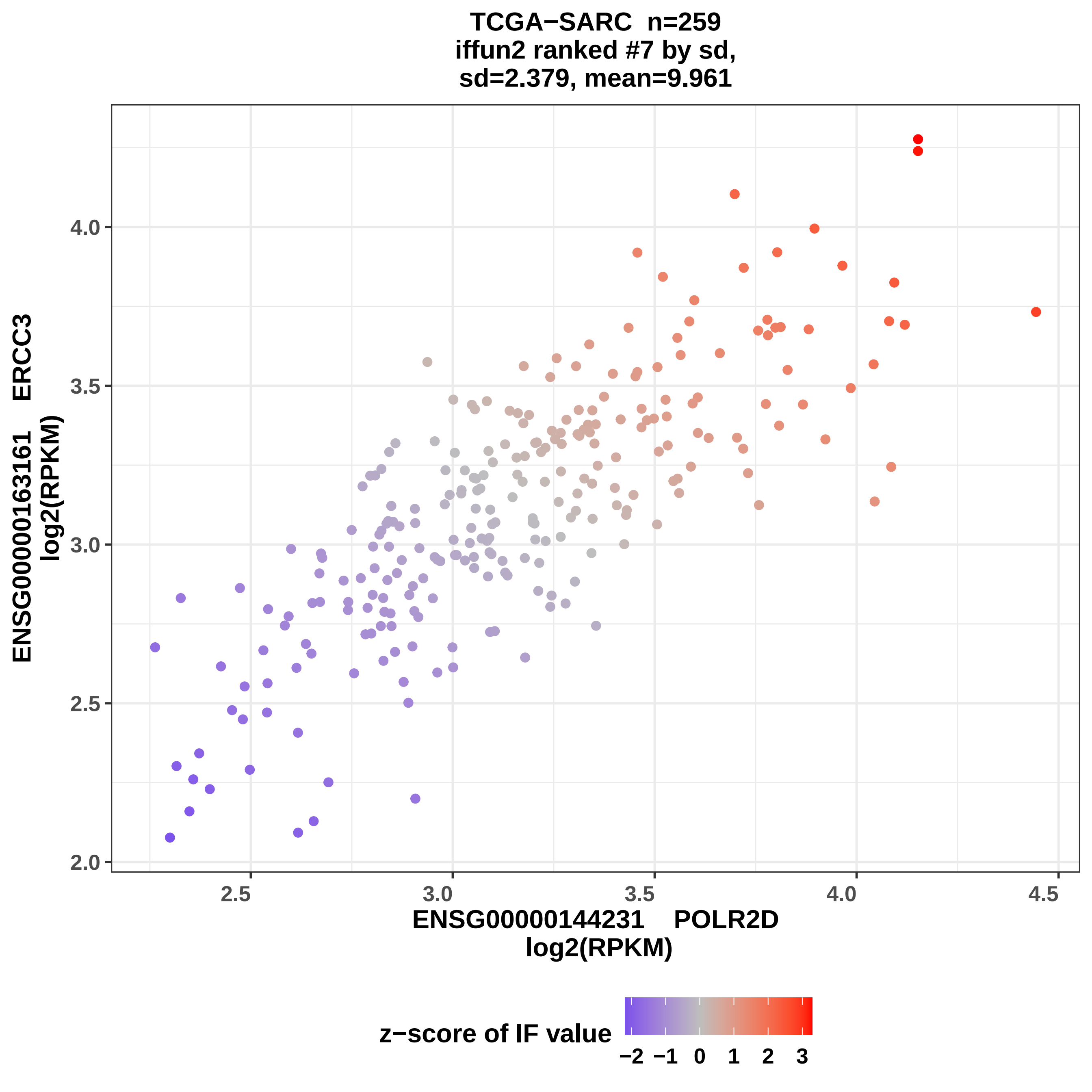
TCGA−SARC n=259
iffun2 ranked #7 by sd,
sd=2.379, mean=9.961
ENSG00000163161 ERCC3
log2(RPKM)
4.0
3.5
3.0
2.5
2.0
2.5
3.0
3.5
4.0
4.5
ENSG00000144231 POLR2D
log2(RPKM)
z−score of IF value
−2 −1 0 1 2 3

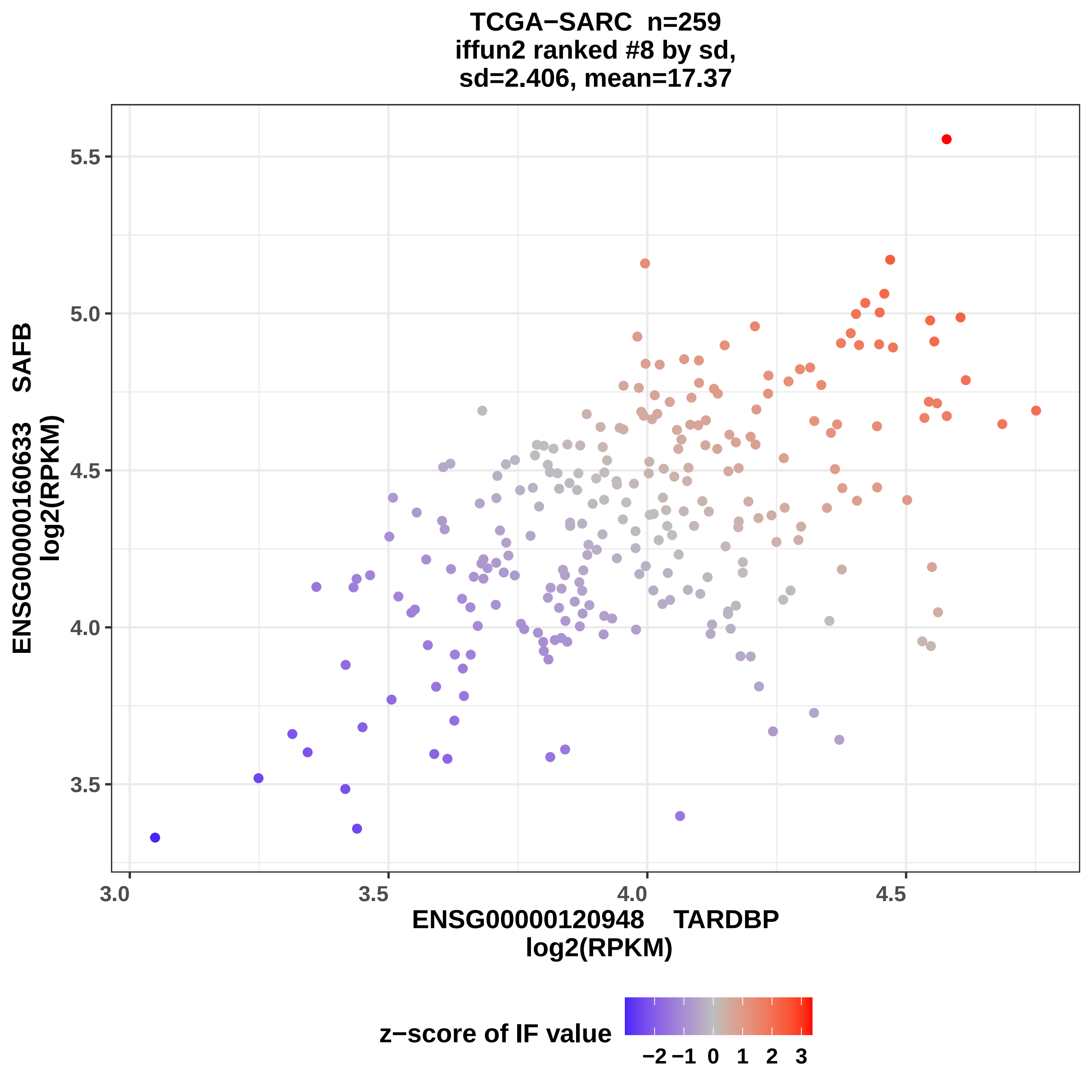
TCGA−SARC n=259
iffun2 ranked #8 by sd,
sd=2.406, mean=17.37
ENSG00000160633 SAFB
log2(RPKM)
ENSG00000120948 TARDBP
log2(RPKM)
5.5
5.0
4.5
4.0
3.5
3.0
3.5
4.0
4.5
z−score of IF value
−2 −1 0 1 2 3

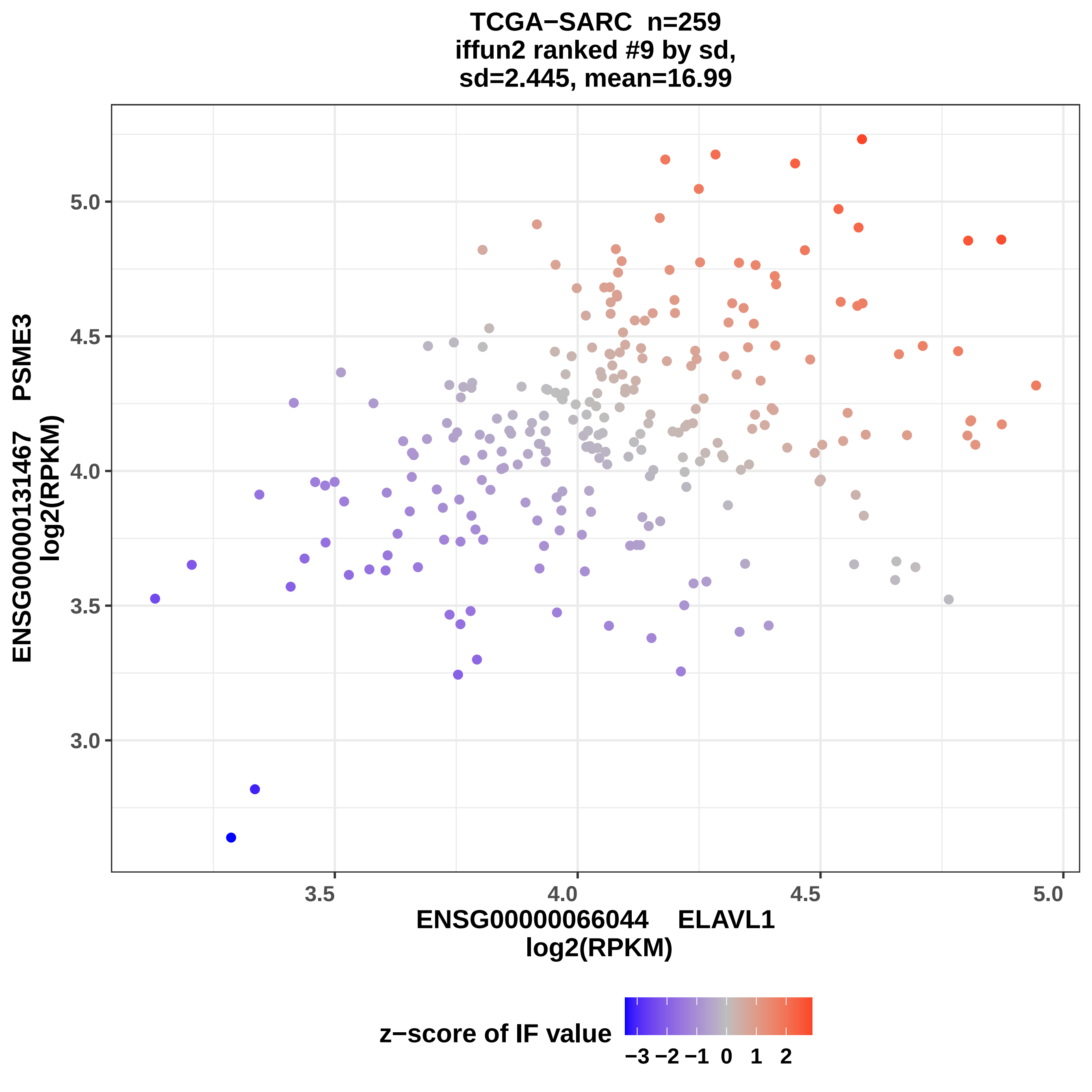
TCGA−SARC n=259
iffun2 ranked #9 by sd,
sd=2.445, mean=16.99
ENSG00000131467 PSME3
log2(RPKM)
5.0
4.5
4.0
3.5
3.0
3.5
4.0
4.5
5.0
ENSG00000066044 ELAVL1
log2(RPKM)
z−score of IF value
−3 −2 −1 0 1 2

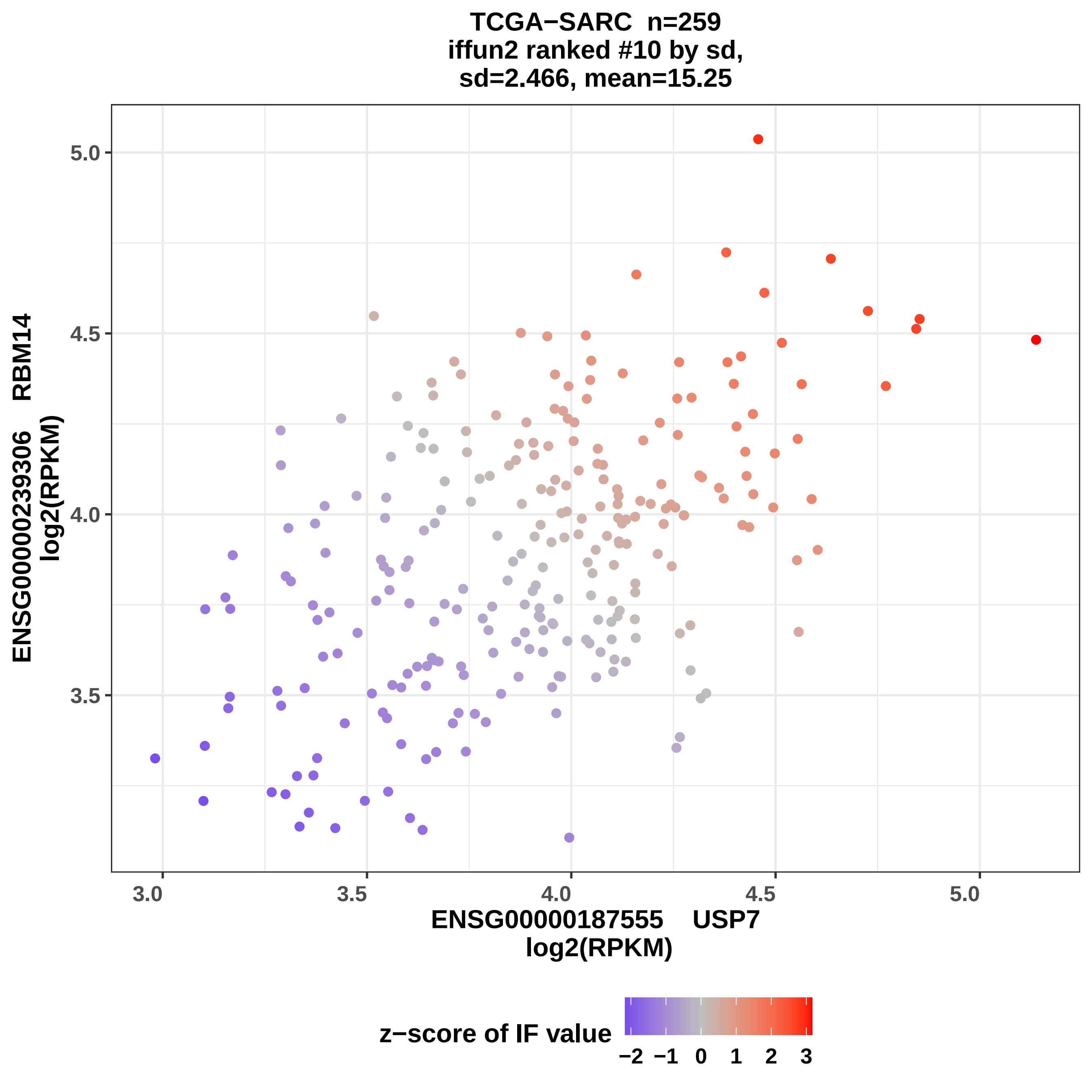
TCGA−SARC n=259
iffun2 ranked #10 by sd,
sd=2.466, mean=15.25
ENSG00000239306 RBM14
log2(RPKM)
5.0
4.5
4.0
3.5
3.0
3.5
4.0
4.5
5.0
ENSG00000187555 USP7
log2(RPKM)
z−score of IF value
−2 −1 0 1 2 3

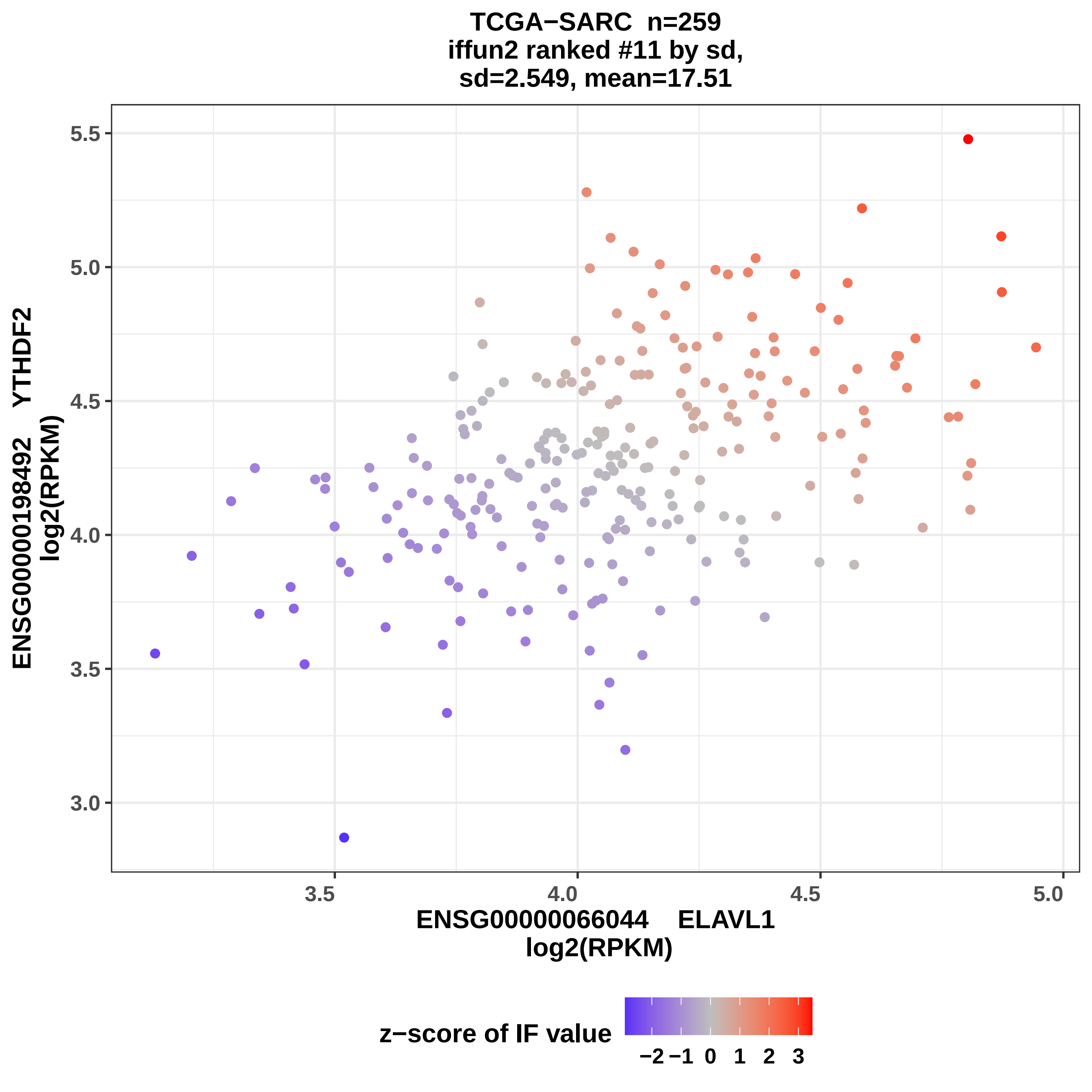

TCGA−SARC n=259
iffun2 ranked #11 by sd,
sd=2.549, mean=17.51
ENSG00000198492 YTHDF2
log2(RPKM)
5.5
5.0
4.5
4.0
3.5
3.0
3.5
4.0
4.5
5.0
ENSG00000066044 ELAVL1
log2(RPKM)
z−score of IF value
−2
−1
0
1
2
3

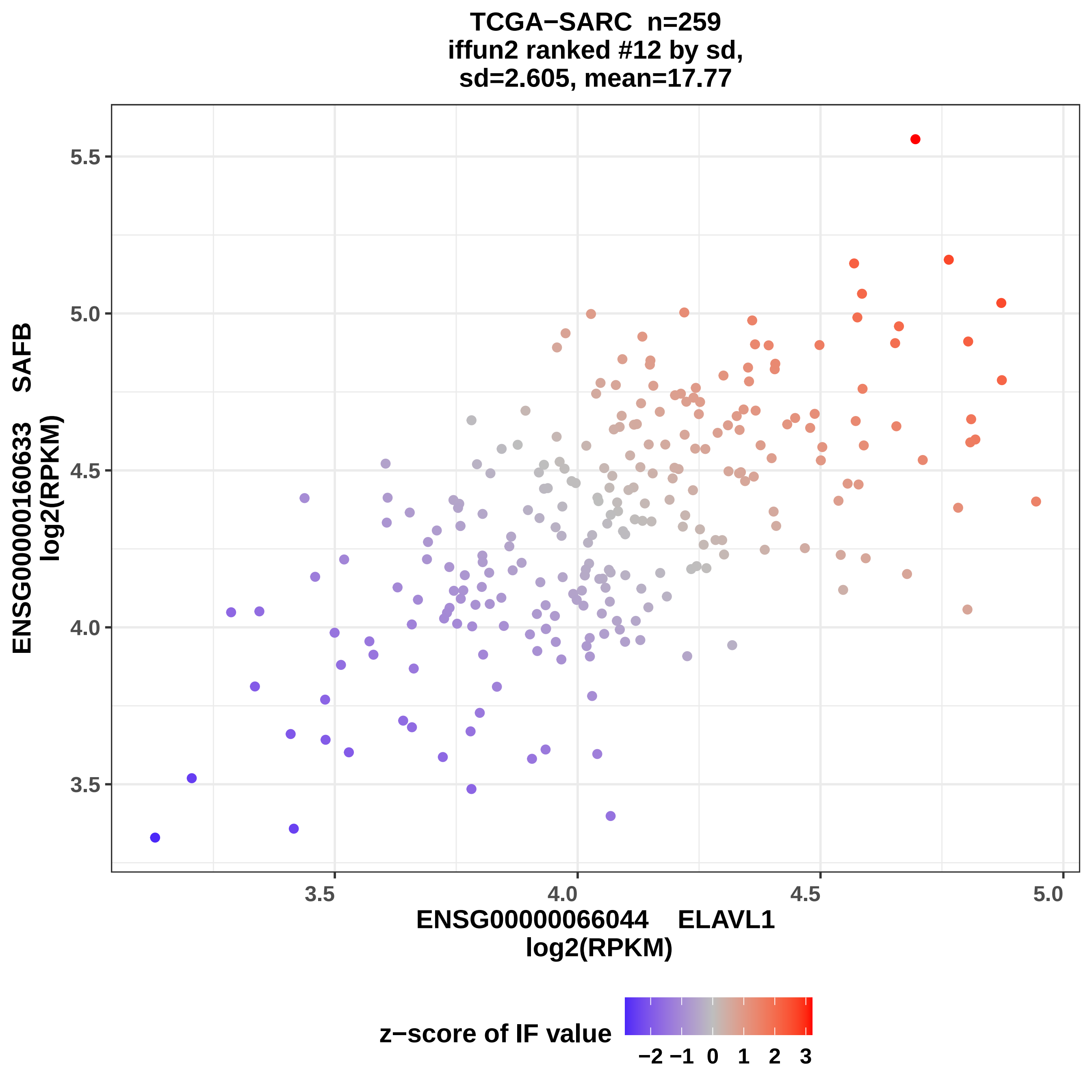

TCGA−SARC n=259
iffun2 ranked #12 by sd,
sd=2.605, mean=17.77
ENSG00000160633 SAFB
log2(RPKM)
5.5
5.0
4.5
4.0
3.5
3.5
4.0
4.5
5.0
ENSG00000066044 ELAVL1
log2(RPKM)
z−score of IF value
−2 −1 0 1 2 3

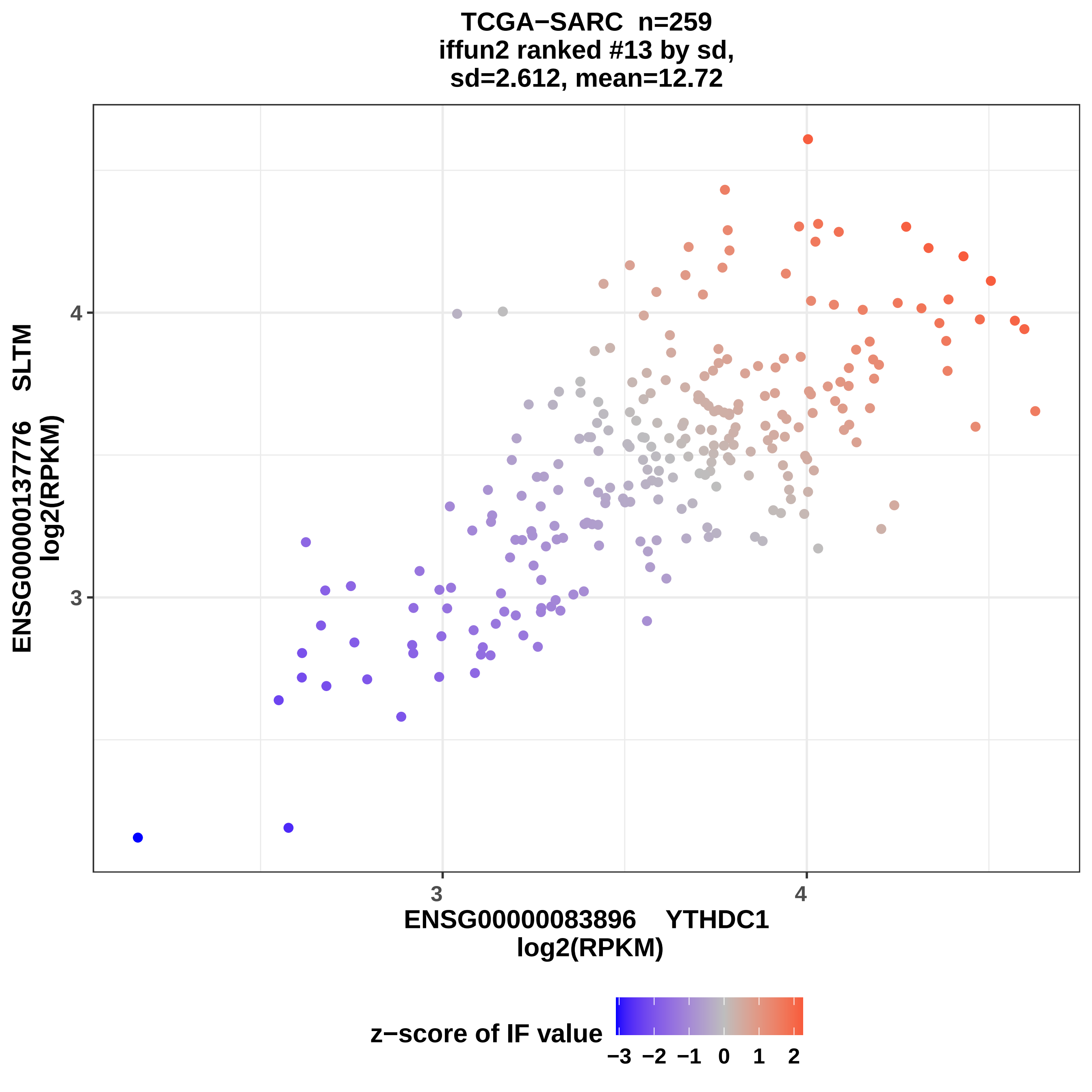
TCGA−SARC n=259
iffun2 ranked #13 by sd,
sd=2.612, mean=12.72
ENSG00000137776 SLTM
log2(RPKM)
4
3
3
4
ENSG00000083896 YTHDC1
log2(RPKM)
z−score of IF value
−3 −2 −1 0 1 2

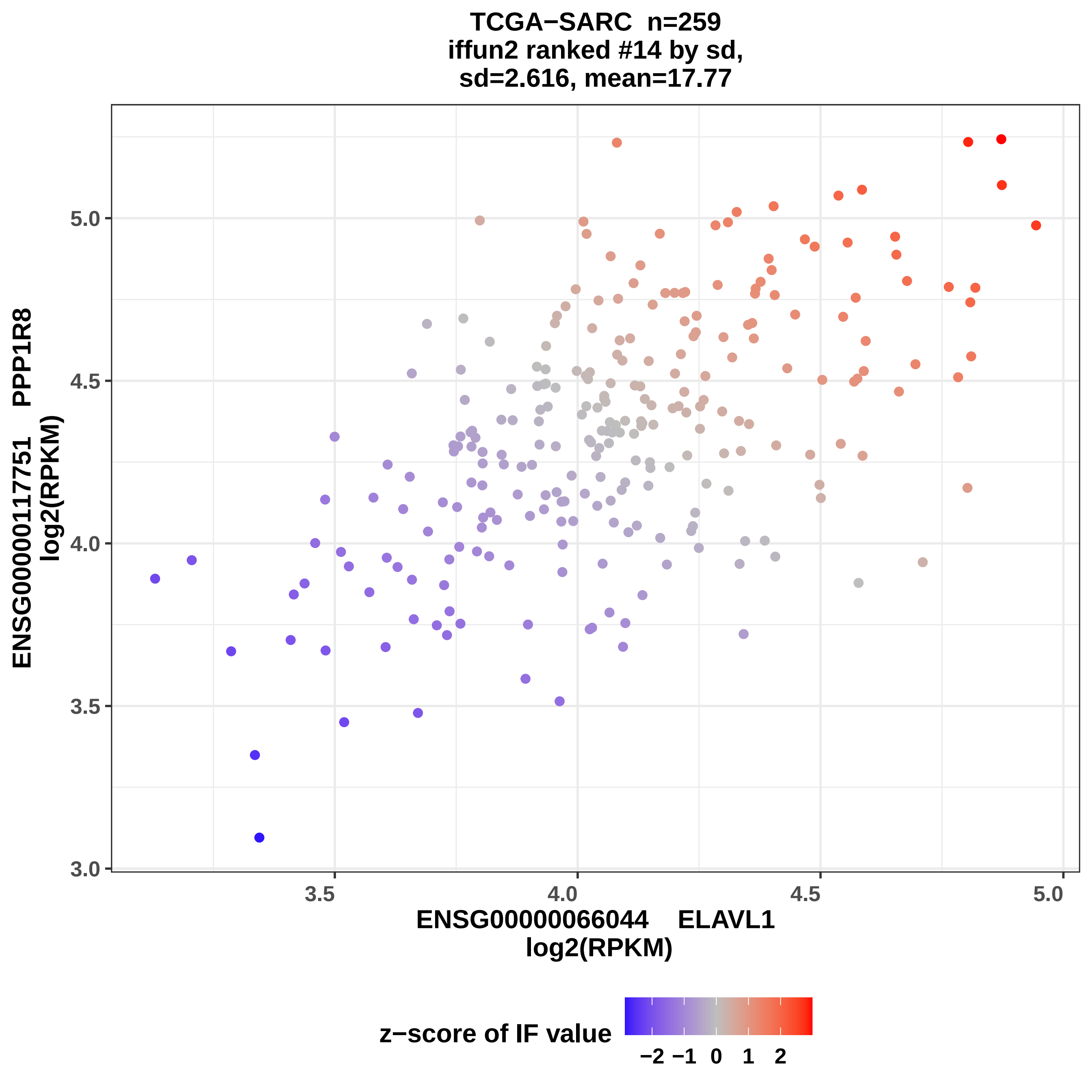
TCGA−SARC n=259
iffun2 ranked #14 by sd,
sd=2.616, mean=17.77
ENSG00000117751 PPP1R8
log2(RPKM)
5.0
4.5
4.0
3.5
3.0
3.5
4.0
4.5
5.0
ENSG00000066044 ELAVL1
log2(RPKM)
z−score of IF value
−2 −1 0 1 2

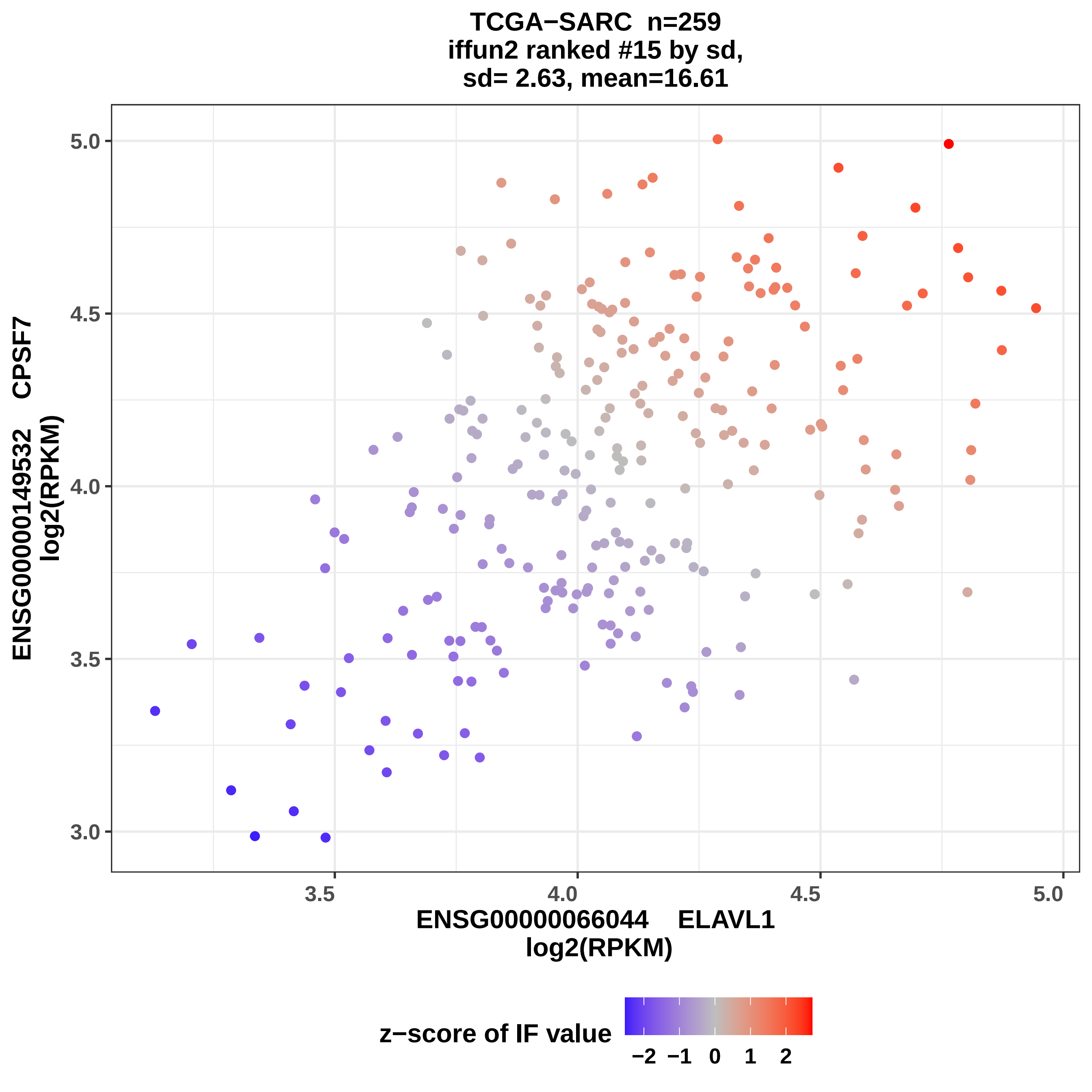
TCGA−SARC n=259
iffun2 ranked #15 by sd,
sd= 2.63, mean=16.61
ENSG00000149532 CPSF7
log2(RPKM)
5.0
4.5
4.0
3.5
3.0
3.5
4.0
4.5
5.0
ENSG00000066044 ELAVL1
log2(RPKM)
z−score of IF value
−2 −1 0 1 2

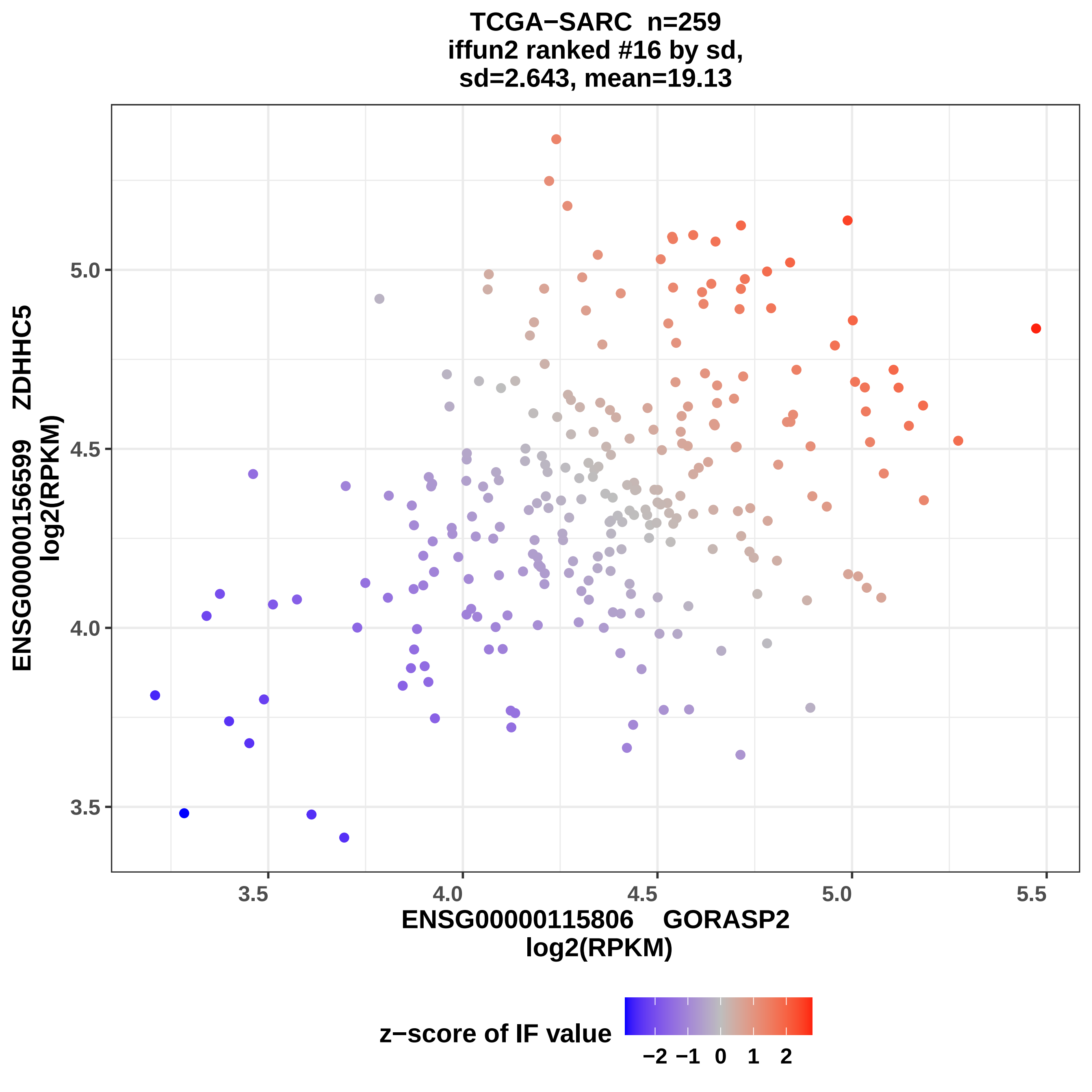
TCGA−SARC n=259
iffun2 ranked #16 by sd,
sd=2.643, mean=19.13
ENSG00000156599 ZDHHC5
log2(RPKM)
5.0
4.5
4.0
3.5
3.5
4.0
4.5
5.0
5.5
ENSG00000115806 GORASP2
log2(RPKM)
z−score of IF value
−2 −1 0 1 2

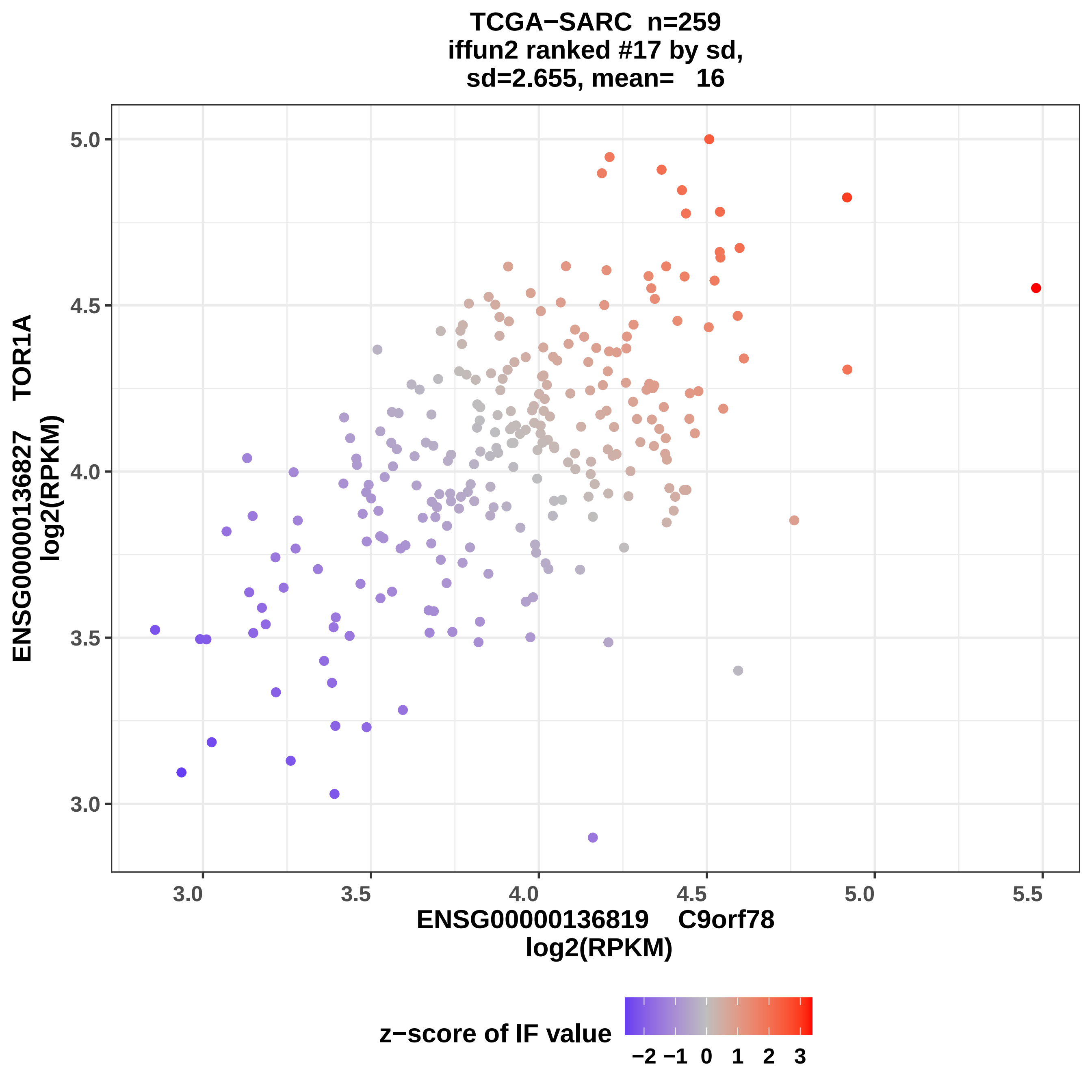

TCGA−SARC n=259
iffun2 ranked #17 by sd,
sd=2.655, mean= 16
ENSG00000136827 TOR1A
log2(RPKM)
5.0
4.5
4.0
3.5
3.0
3.0
3.5
4.0
4.5
5.0
5.5
ENSG00000136819 C9orf78
log2(RPKM)
z−score of IF value
−2 −1 0 1 2 3

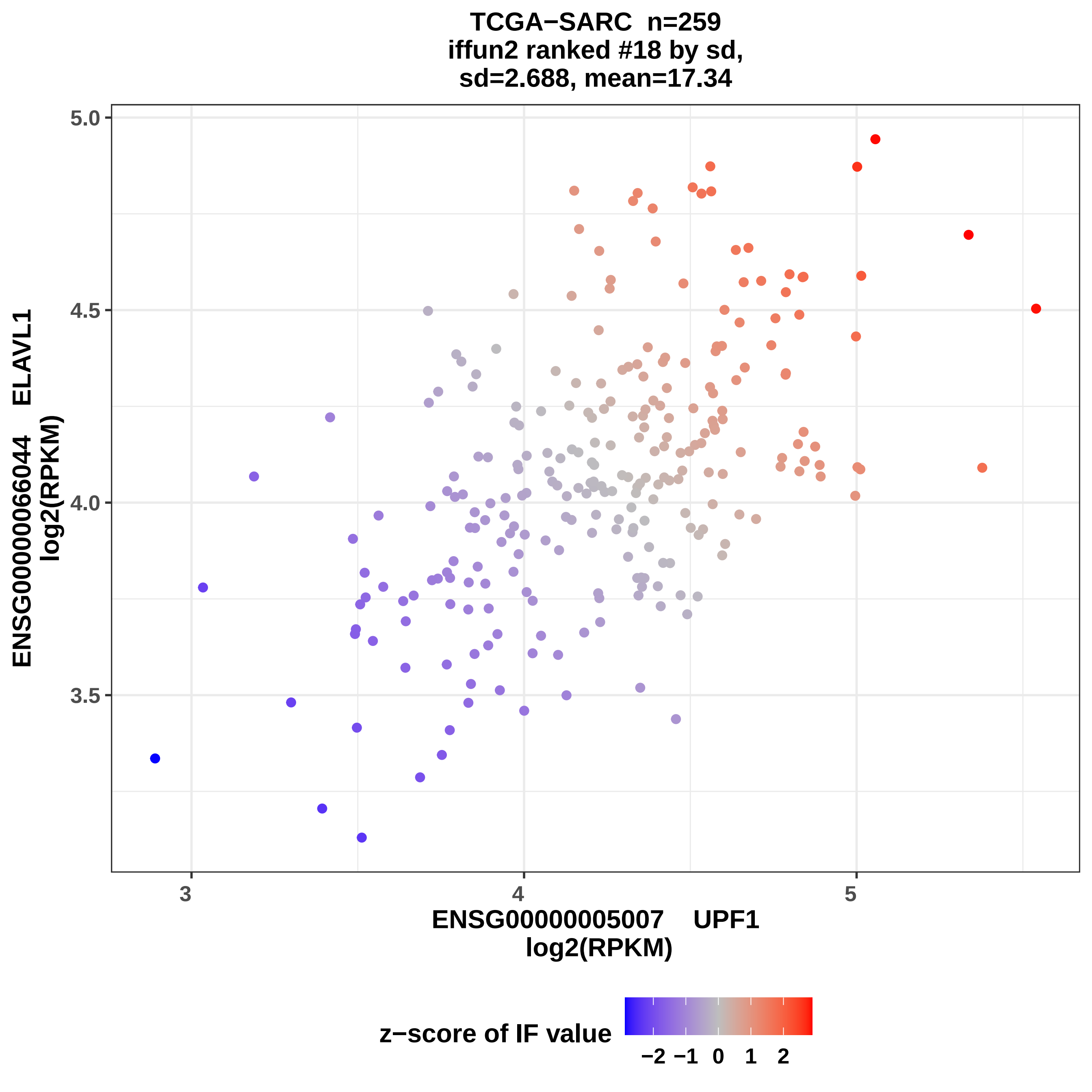

TCGA−SARC n=259
iffun2 ranked #18 by sd,
sd=2.688, mean=17.34
ENSG00000066044 ELAVL1
log2(RPKM)
5.0
4.5
4.0
3.5
3
4
5
ENSG00000005007 UPF1
log2(RPKM)
z−score of IF value
−2
−1
0
1
2

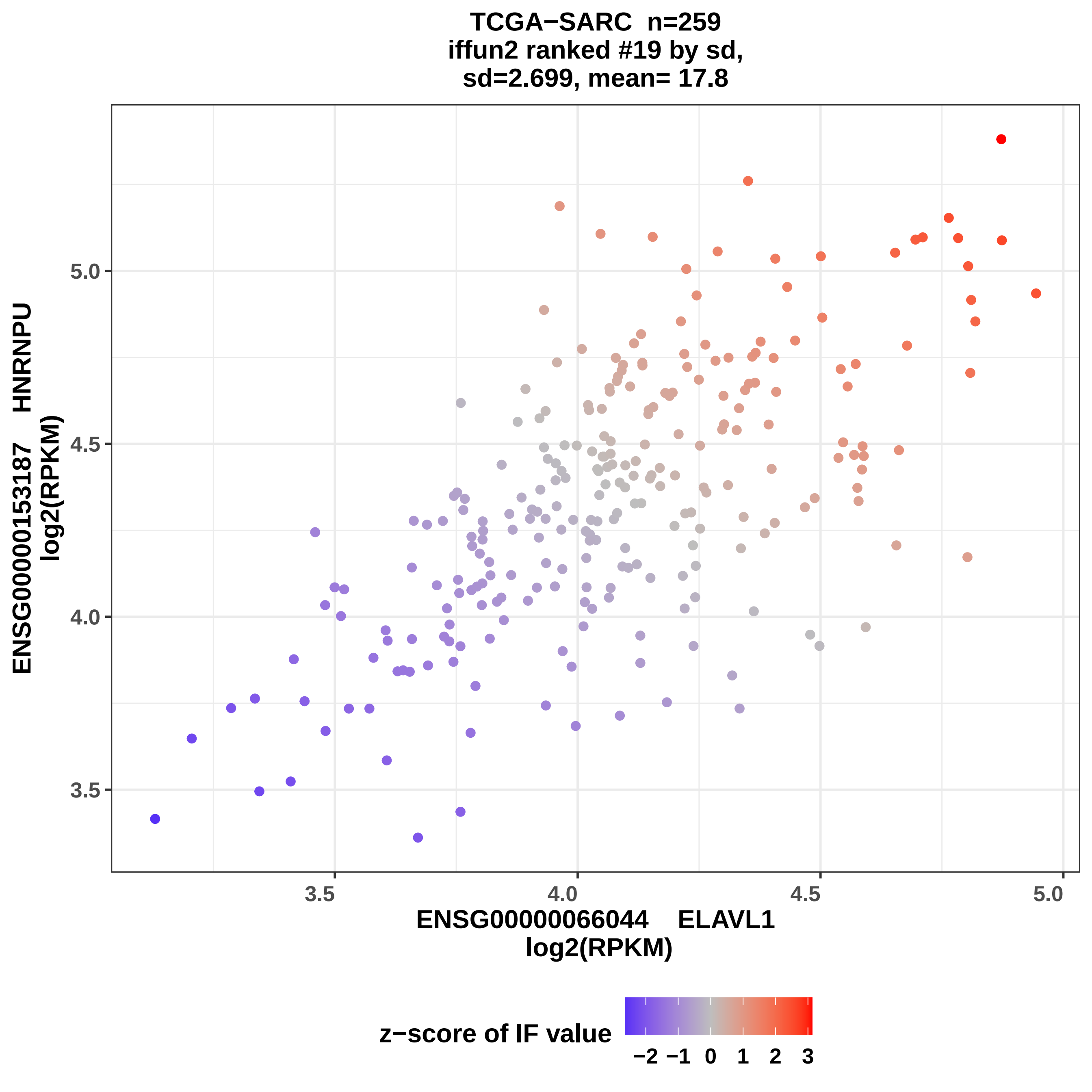
TCGA−SARC n=259
iffun2 ranked #19 by sd,
sd=2.699, mean= 17.8
ENSG00000153187 HNRNPU
log2(RPKM)
5.0
4.5
4.0
3.5
3.5
4.0
4.5
5.0
ENSG00000066044 ELAVL1
log2(RPKM)
z−score of IF value
−2 −1 0 1 2 3

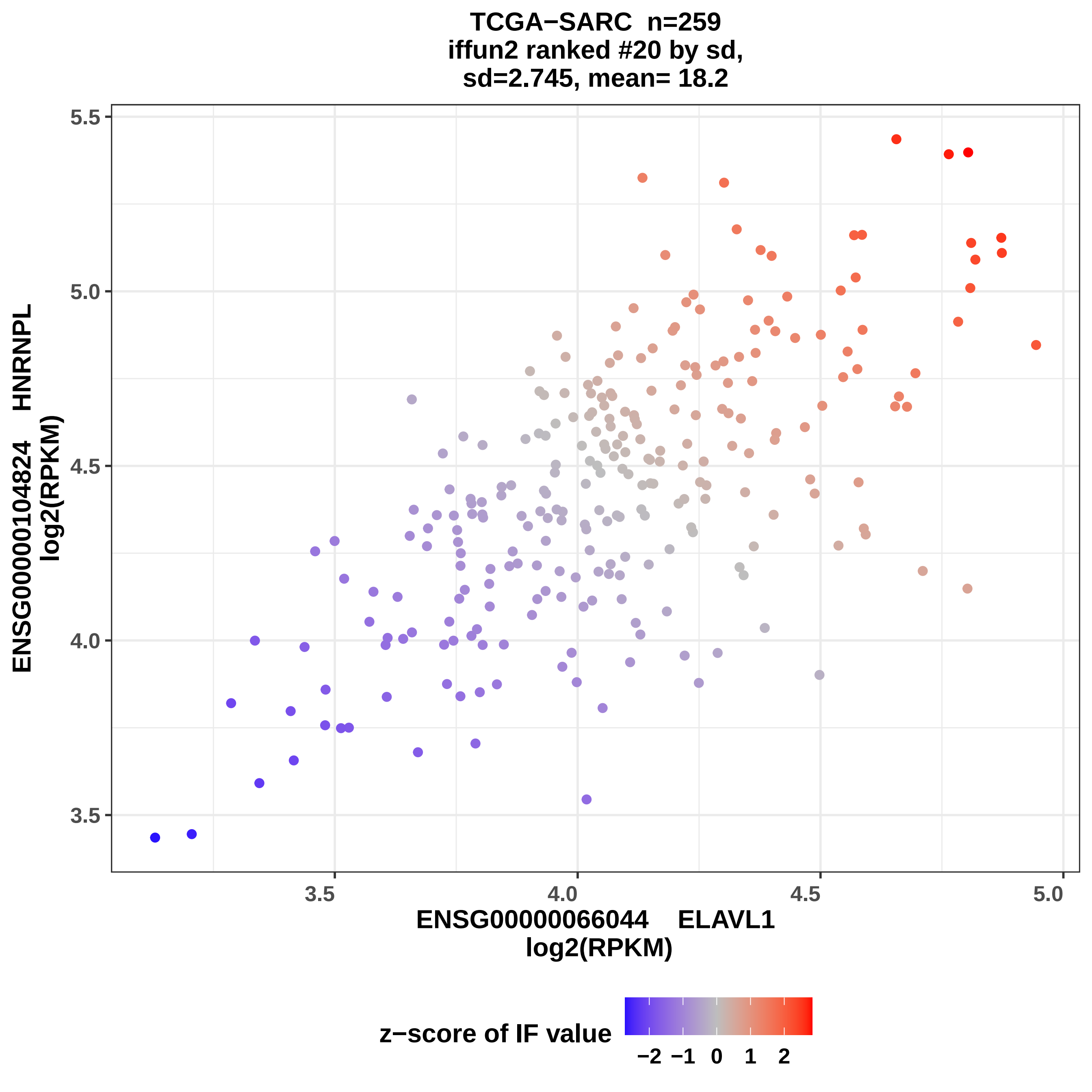

TCGA−SARC n=259
iffun2 ranked #20 by sd,
sd=2.745, mean= 18.2
5.5
5.0
4.5
4.0
3.5
ENSG00000104824 HNRNPL
log2(RPKM)
3.5
4.0
4.5
5.0
ENSG00000066044 ELAVL1
log2(RPKM)
z−score of IF value
−2 −1 0 1 2

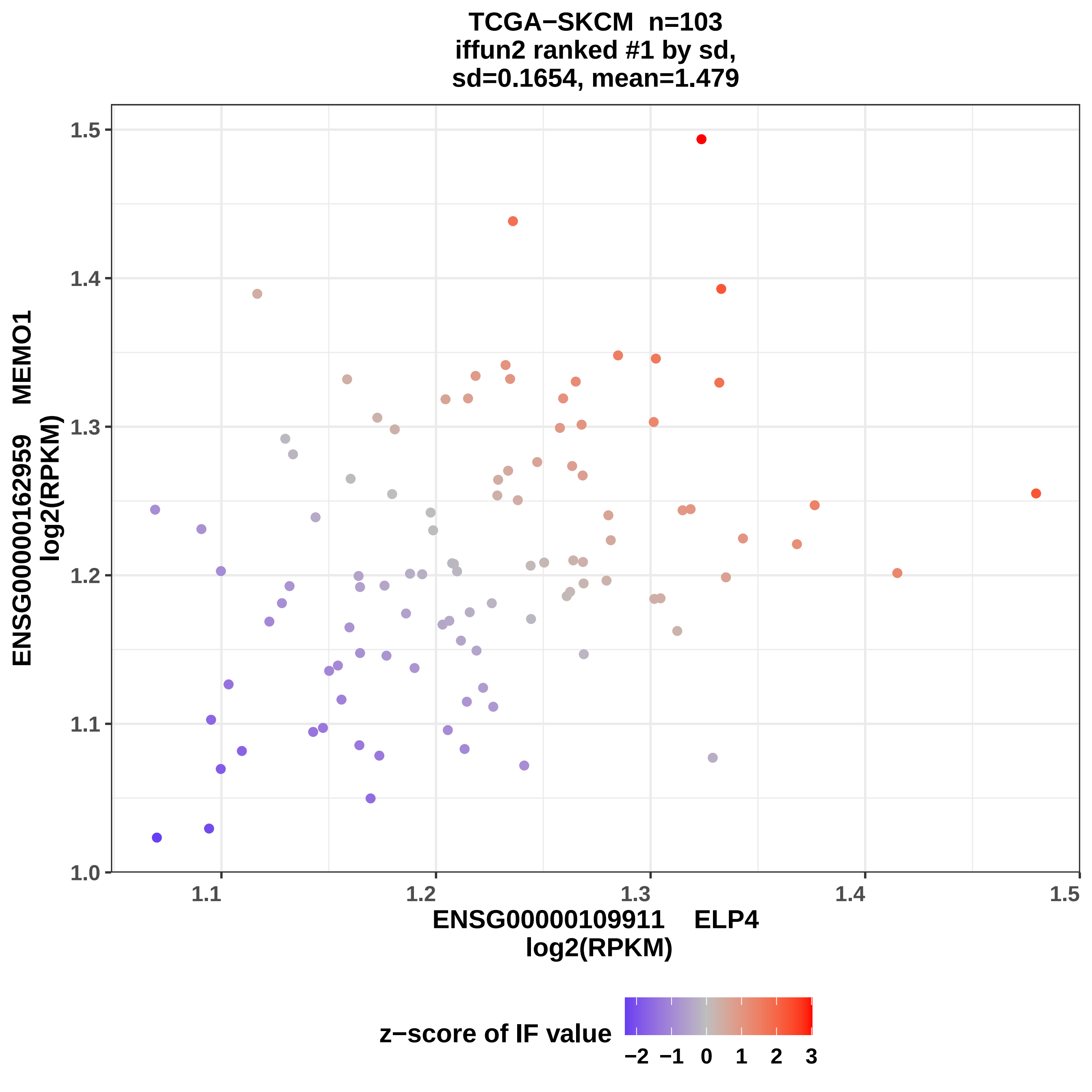

TCGA−SKCM n=103
iffun2 ranked #1 by sd,
sd=0.1654, mean=1.479
ENSG00000162959 MEMO1
log2(RPKM)
1.5
1.4
1.3
1.2
1.1
1.0
1.1
1.2
1.3
1.4
1.5
ENSG00000109911 ELP4
log2(RPKM)
z−score of IF value
−2 −1 0 1 2 3

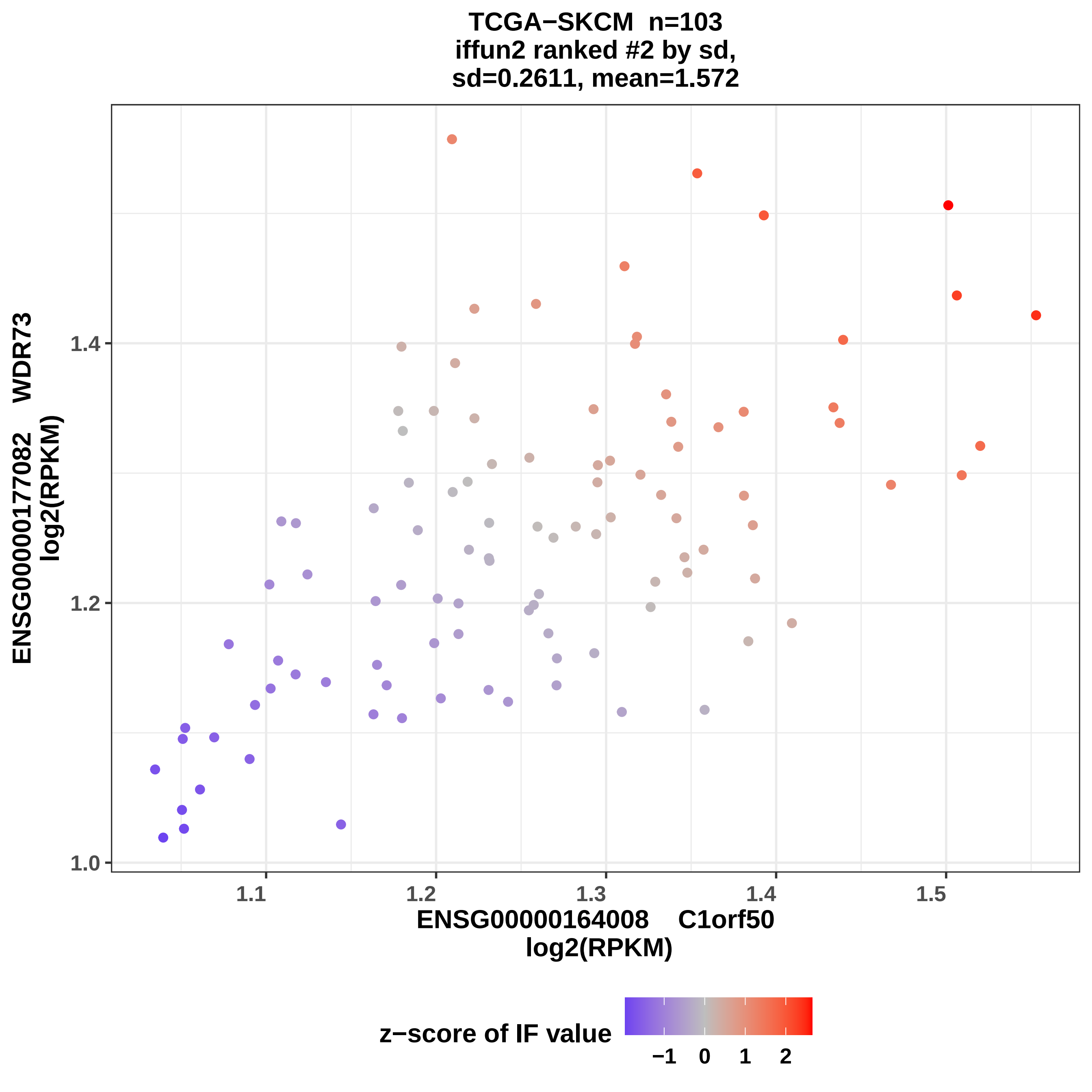
TCGA−SKCM n=103
iffun2 ranked #2 by sd,
sd=0.2611, mean=1.572
ENSG00000177082 WDR73
log2(RPKM)
1.0
1.2
1.4
1.1
1.2
1.3
1.4
1.5
ENSG00000164008 C1orf50
log2(RPKM)
z−score of IF value
−1
0
1
2

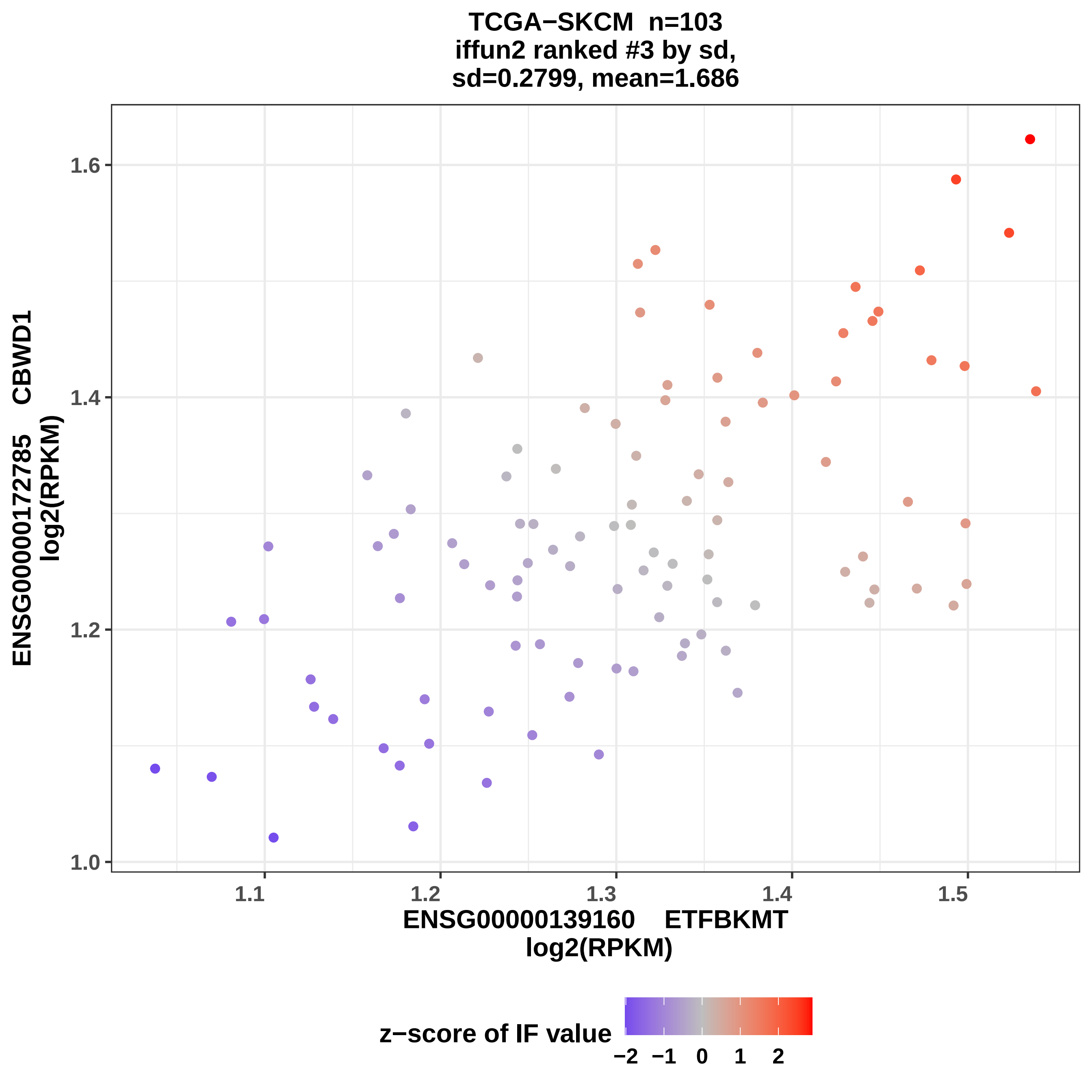

TCGA−SKCM n=103
iffun2 ranked #3 by sd,
sd=0.2799, mean=1.686
ENSG00000172785 CBWD1
log2(RPKM)
1.0
1.2
1.4
1.6
1.1
1.2
1.3
1.4
1.5
ENSG00000139160 ETFBKMT
log2(RPKM)
z−score of IF value
−2 −1 0 1 2

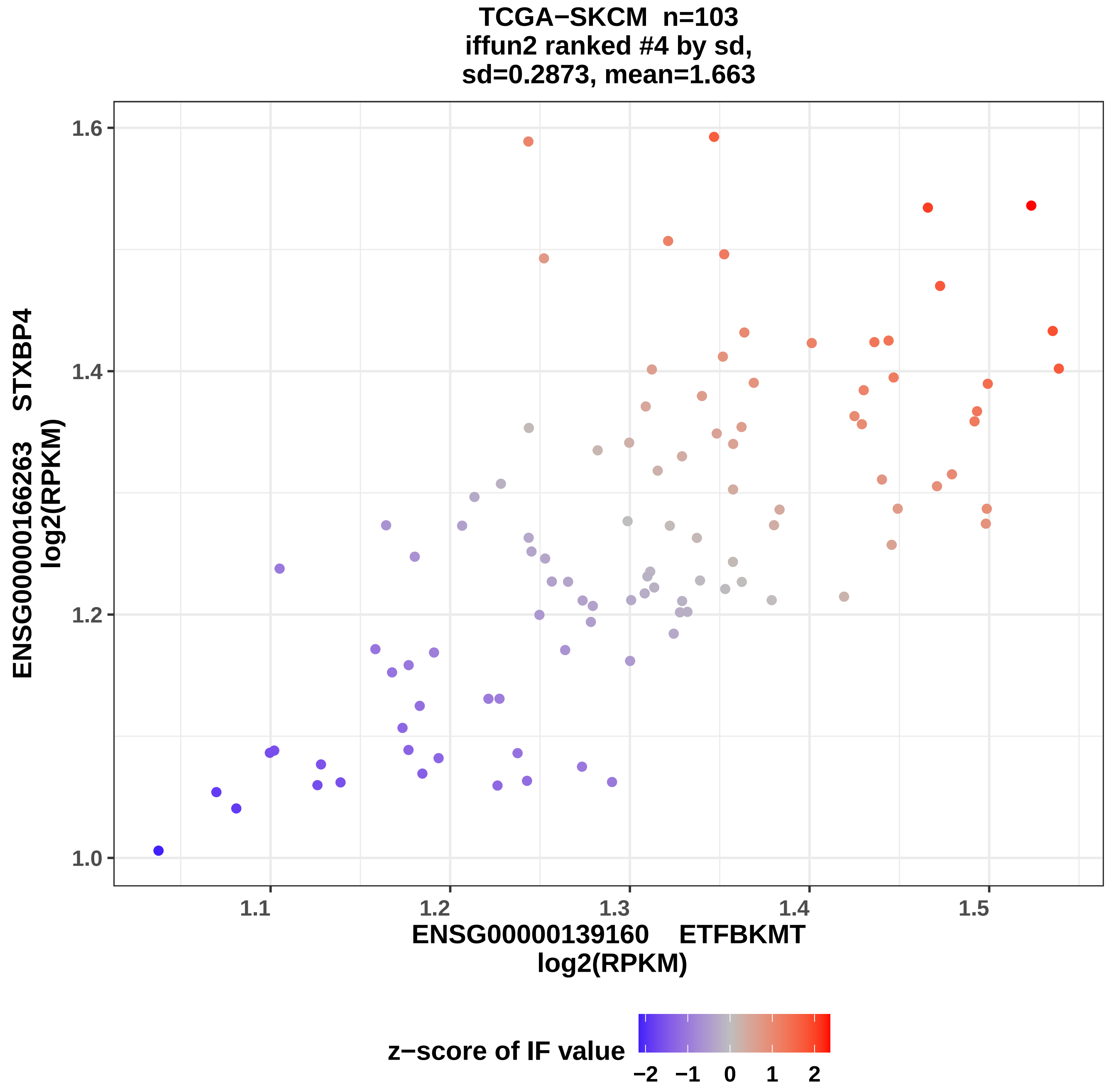

TCGA−SKCM n=103
iffun2 ranked #4 by sd,
sd=0.2873, mean=1.663
ENSG00000166263 STXBP4
log2(RPKM)
1.0
1.2
1.4
1.6
1.1
1.2
1.3
1.4
1.5
ENSG00000139160 ETFBKMT
log2(RPKM)
z−score of IF value
−2
−1
0
1
2

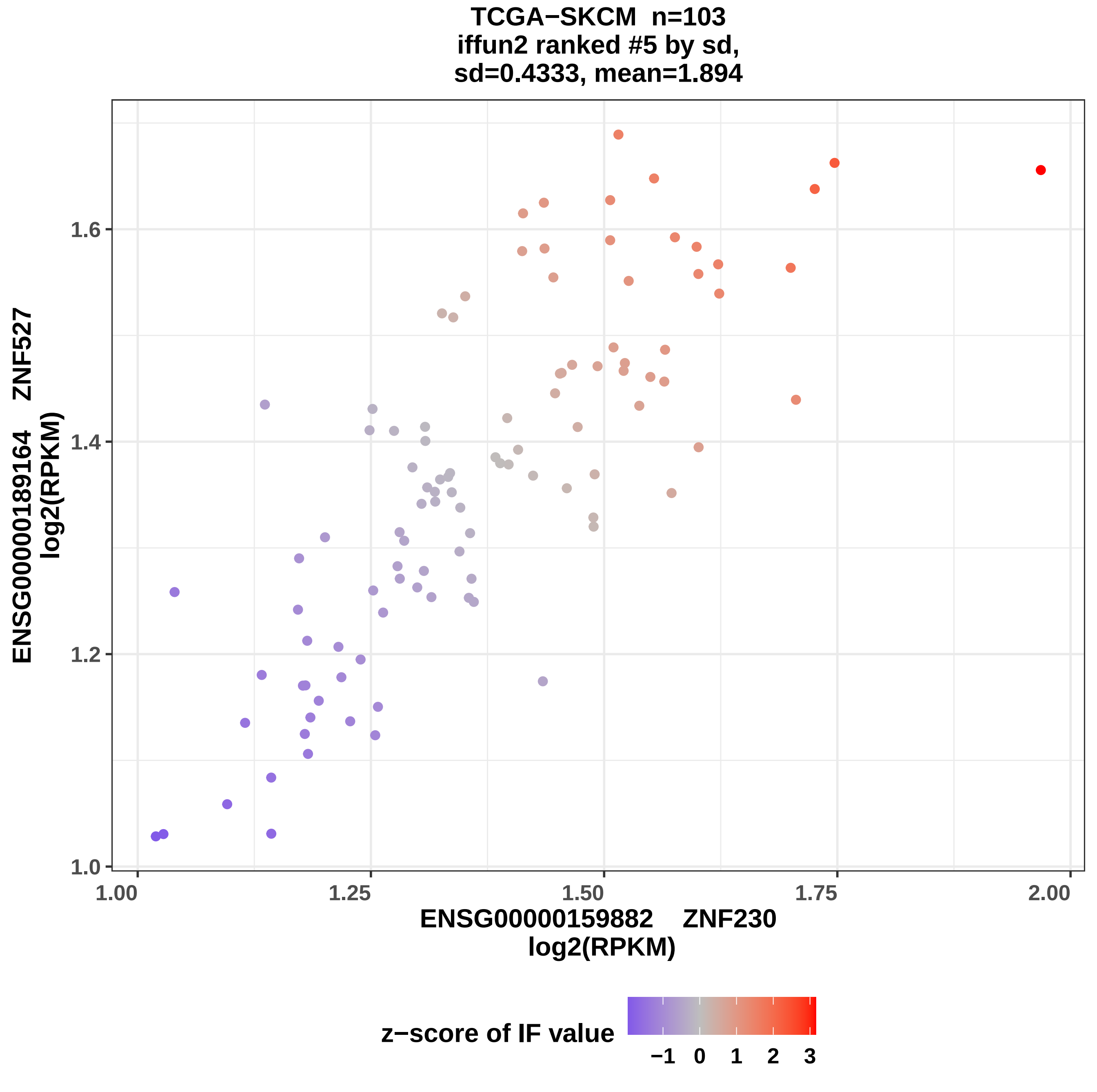
TCGA−SKCM n=103
iffun2 ranked #5 by sd,
sd=0.4333, mean=1.894
ENSG00000189164 ZNF527
log2(RPKM)
1.0
1.2
1.4
1.6
1.00
1.25
1.50
1.75
2.00
ENSG00000159882 ZNF230
log2(RPKM)
z−score of IF value
−1
0
1
2
3

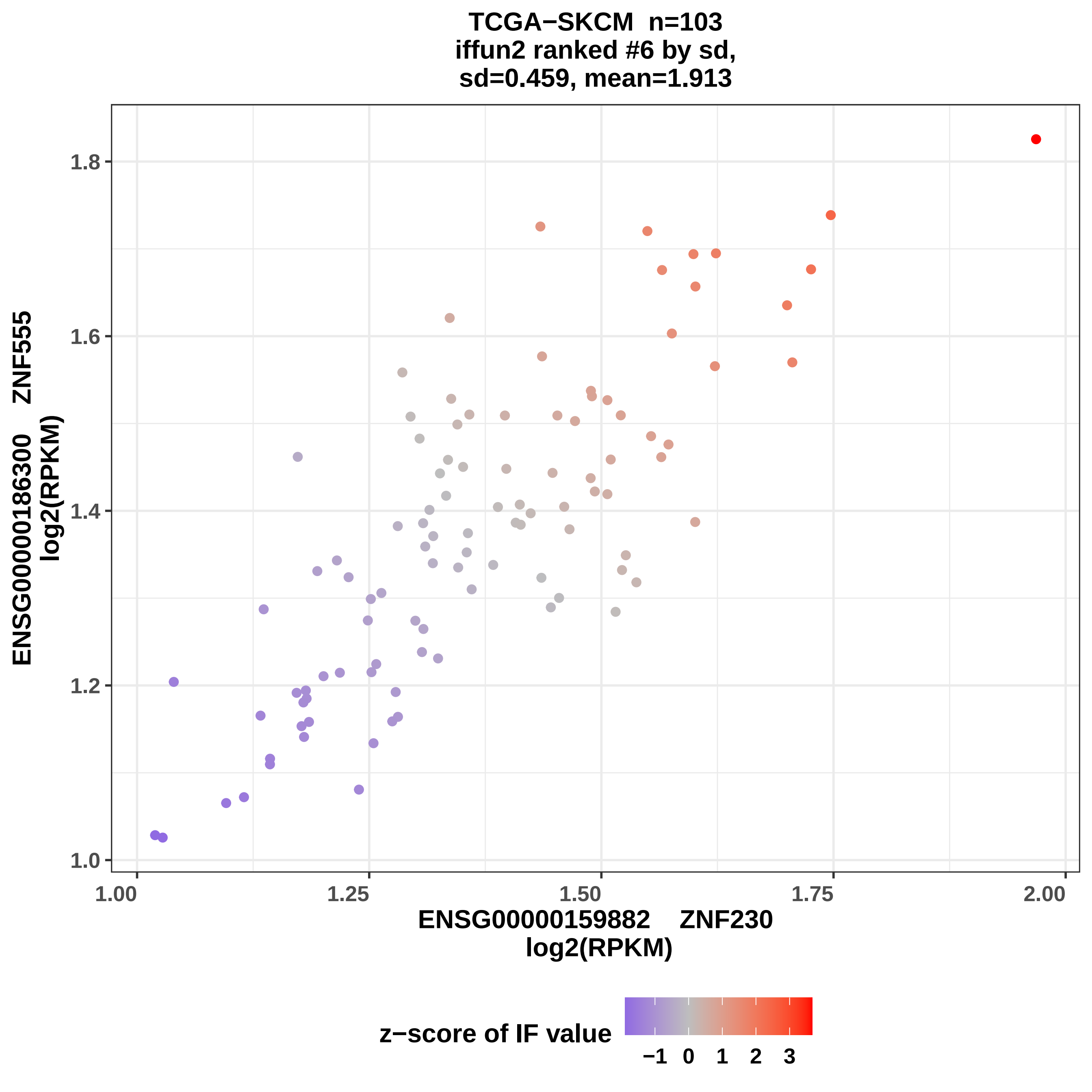
TCGA−SKCM n=103
iffun2 ranked #6 by sd,
sd=0.459, mean=1.913
ENSG00000186300 ZNF555
log2(RPKM)
1.0
1.2
1.4
1.6
1.8
1.00
1.25
1.50
1.75
2.00
ENSG00000159882 ZNF230
log2(RPKM)
z−score of IF value
−1
0
1
2
3

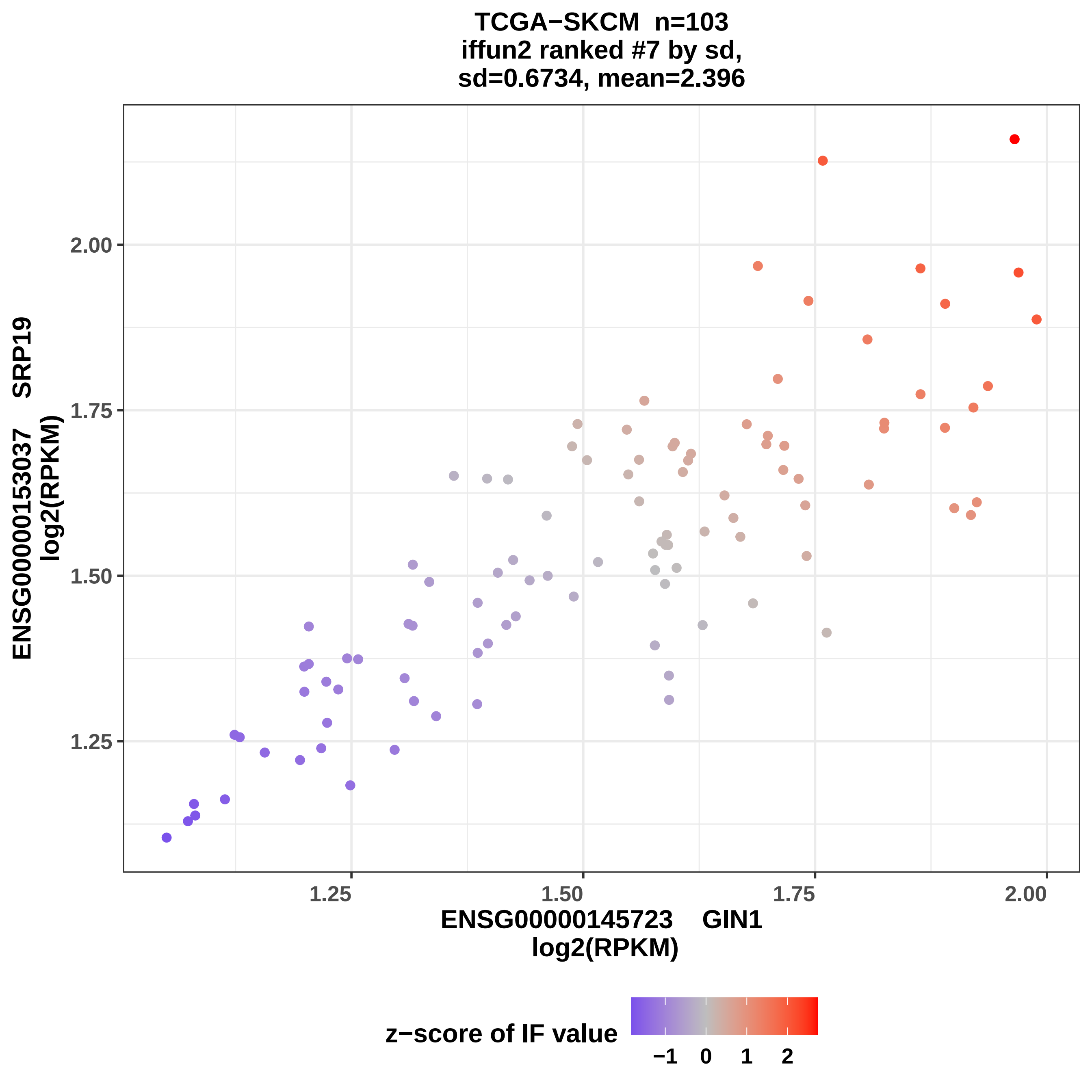
TCGA−SKCM n=103
iffun2 ranked #7 by sd,
sd=0.6734, mean=2.396
ENSG00000153037 SRP19
log2(RPKM)
2.00
1.75
1.50
1.25
1.25
1.50
1.75
2.00
ENSG00000145723 GIN1
log2(RPKM)
z−score of IF value
−1
0
1
2

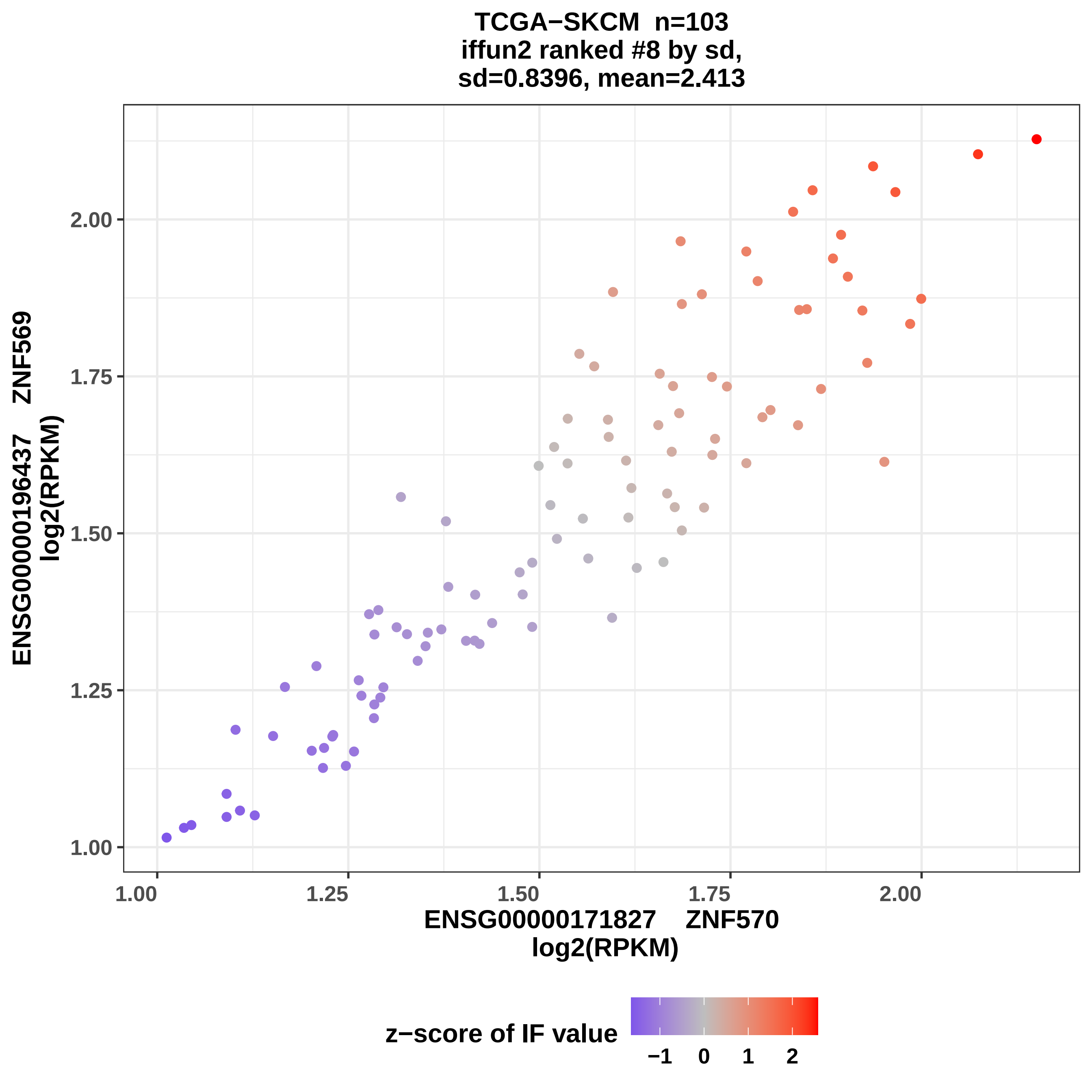
TCGA−SKCM n=103
iffun2 ranked #8 by sd,
sd=0.8396, mean=2.413
ENSG00000196437 ZNF569
log2(RPKM)
1.00
1.25
1.50
1.75
2.00
ENSG00000171827 ZNF570
log2(RPKM)
z−score of IF value
−1
0
1
2

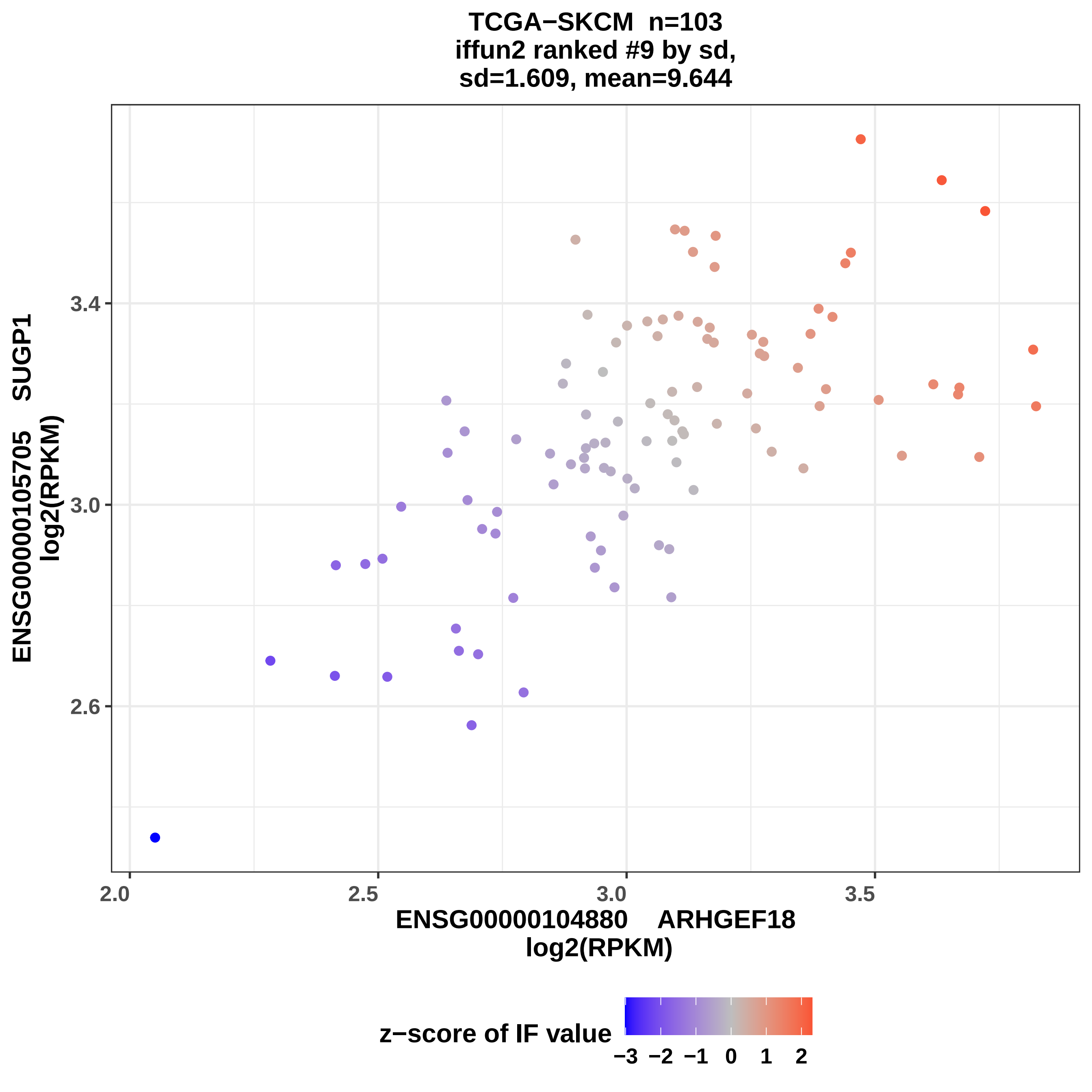

TCGA−SKCM n=103
iffun2 ranked #9 by sd,
sd=1.609, mean=9.644
ENSG00000105705 SUGP1
log2(RPKM)
3.4
3.0
2.6
2.0
2.5
3.0
3.5
ENSG00000104880 ARHGEF18
log2(RPKM)
z−score of IF value
−3 −2 −1 0 1 2

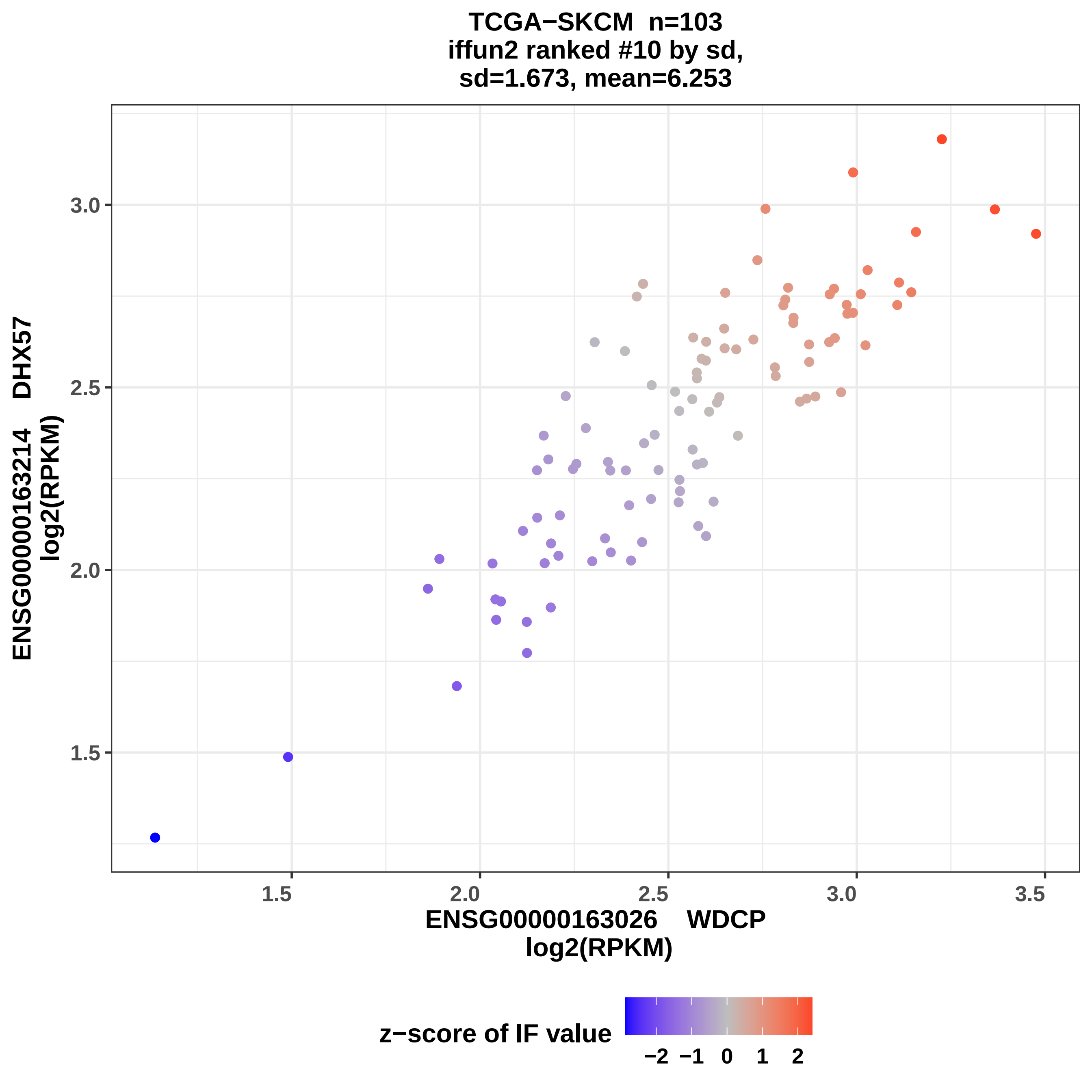

TCGA−SKCM n=103
iffun2 ranked #10 by sd,
sd=1.673, mean=6.253
ENSG00000163214 DHX57
log2(RPKM)
3.0
2.5
2.0
1.5
1.5
2.0
2.5
3.0
3.5
ENSG00000163026 WDCP
log2(RPKM)
z−score of IF value
−2 −1 0 1 2

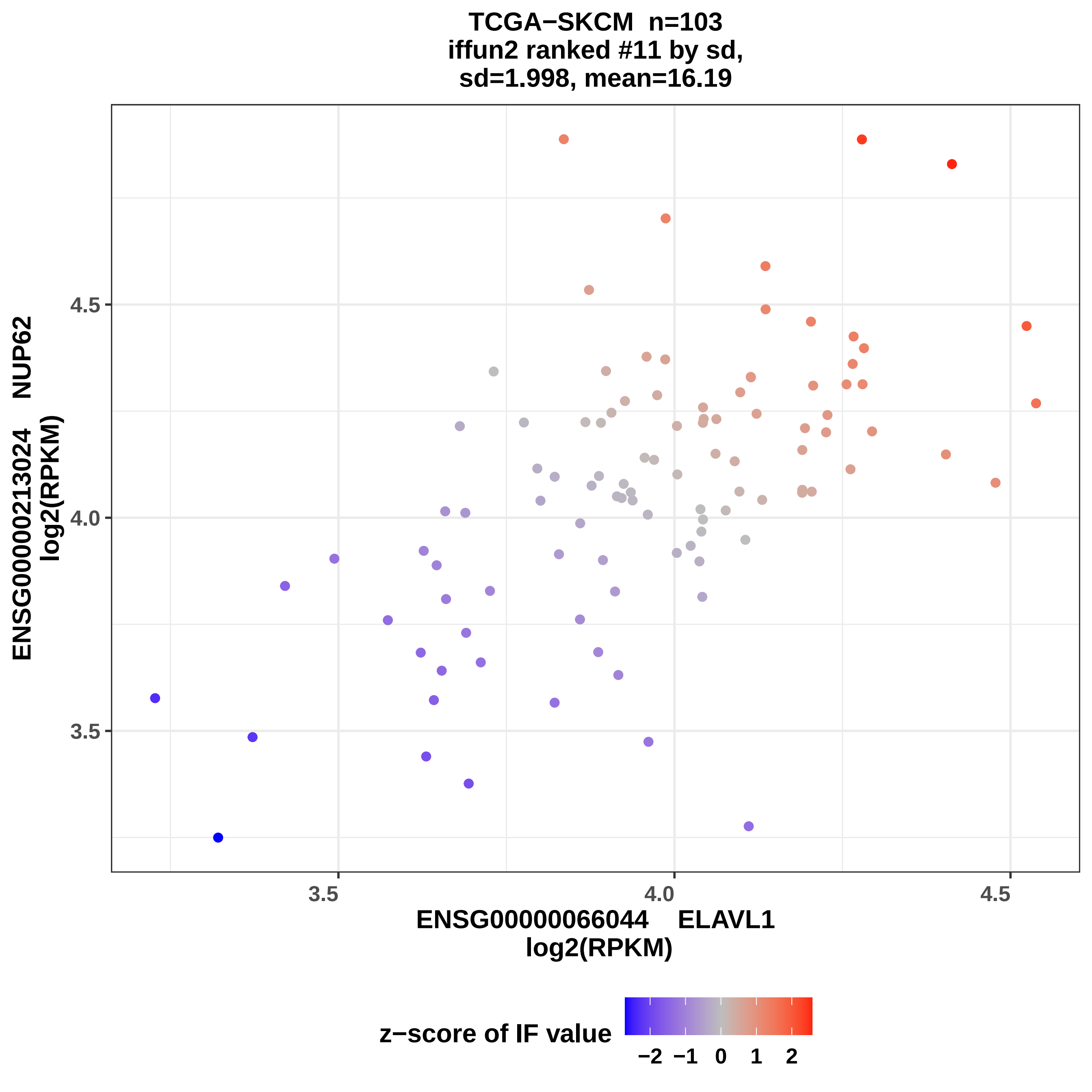

TCGA−SKCM n=103
iffun2 ranked #11 by sd,
sd=1.998, mean=16.19
ENSG00000213024 NUP62
log2(RPKM)
4.5
4.0
3.5
3.5
4.0
4.5
ENSG00000066044 ELAVL1
log2(RPKM)
z−score of IF value
−2
−1
0
1
2

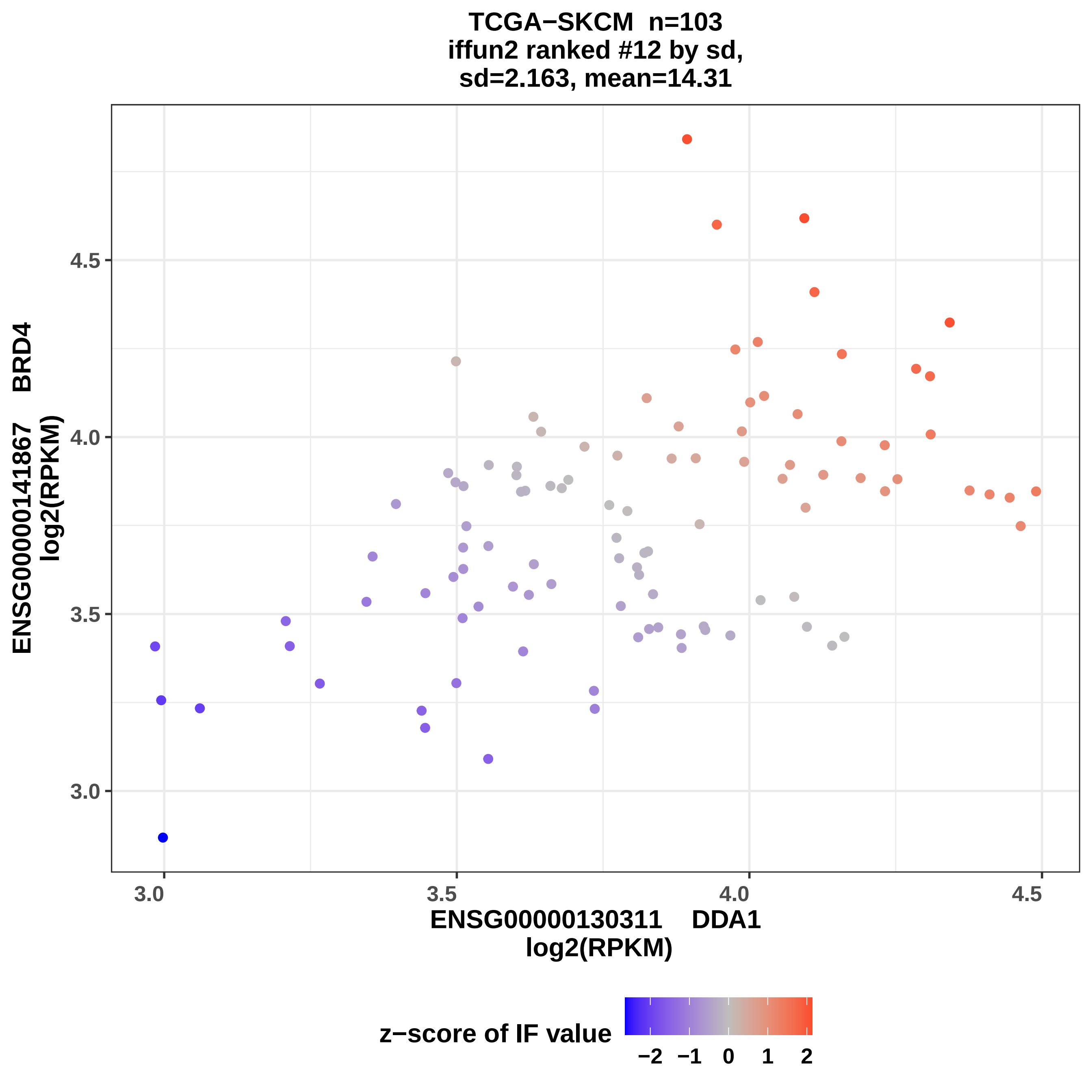
TCGA−SKCM n=103
iffun2 ranked #12 by sd,
sd=2.163, mean=14.31
ENSG00000141867 BRD4
log2(RPKM)
4.5
4.0
3.5
3.0
3.0
3.5
4.0
4.5
ENSG00000130311 DDA1
log2(RPKM)
z−score of IF value
−2 −1 0 1 2

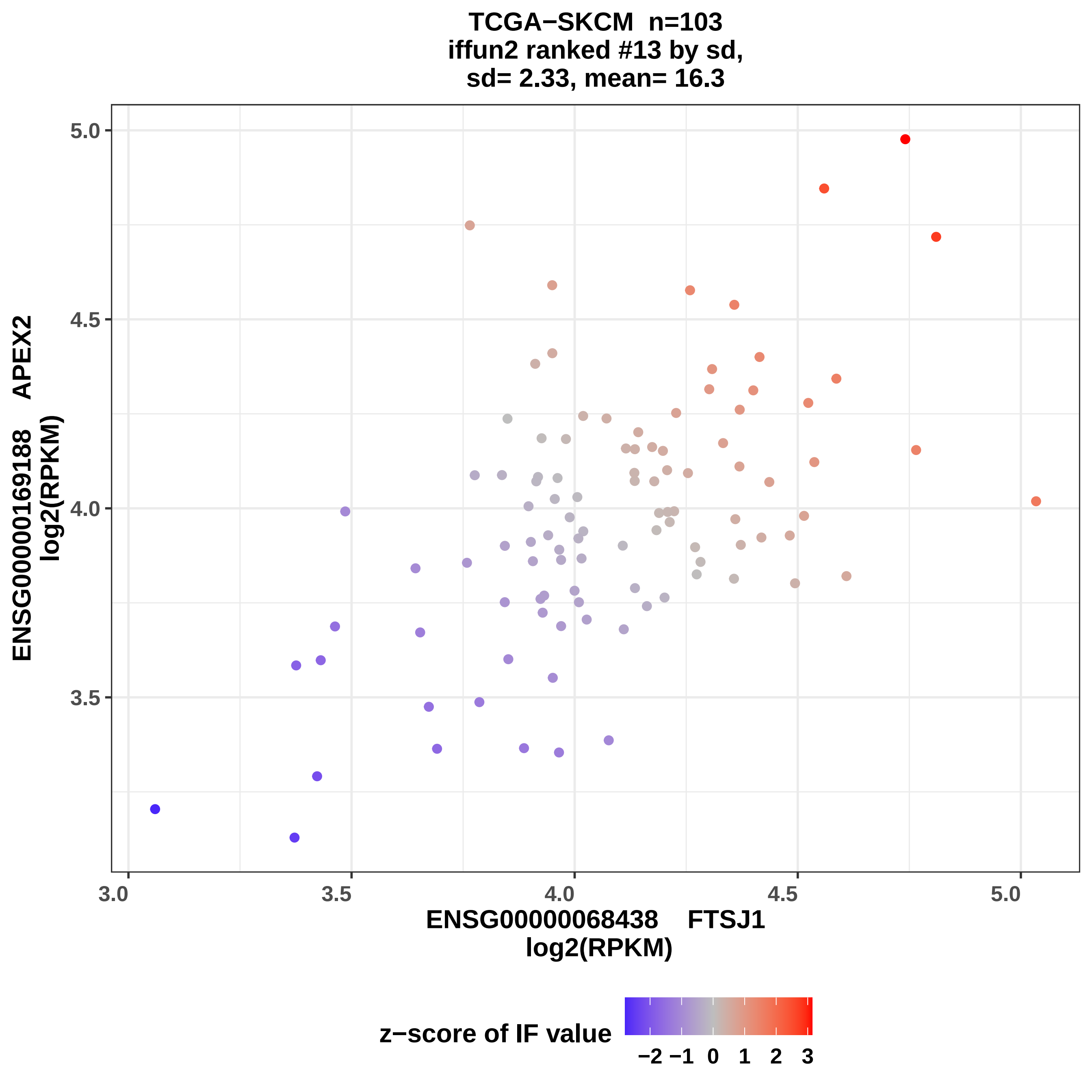

TCGA−SKCM n=103
iffun2 ranked #13 by sd,
sd= 2.33, mean= 16.3
ENSG00000169188 APEX2
log2(RPKM)
5.0
4.5
4.0
3.5
3.0
3.5
4.0
4.5
5.0
ENSG00000068438 FTSJ1
log2(RPKM)
z−score of IF value
−2 −1 0 1 2 3

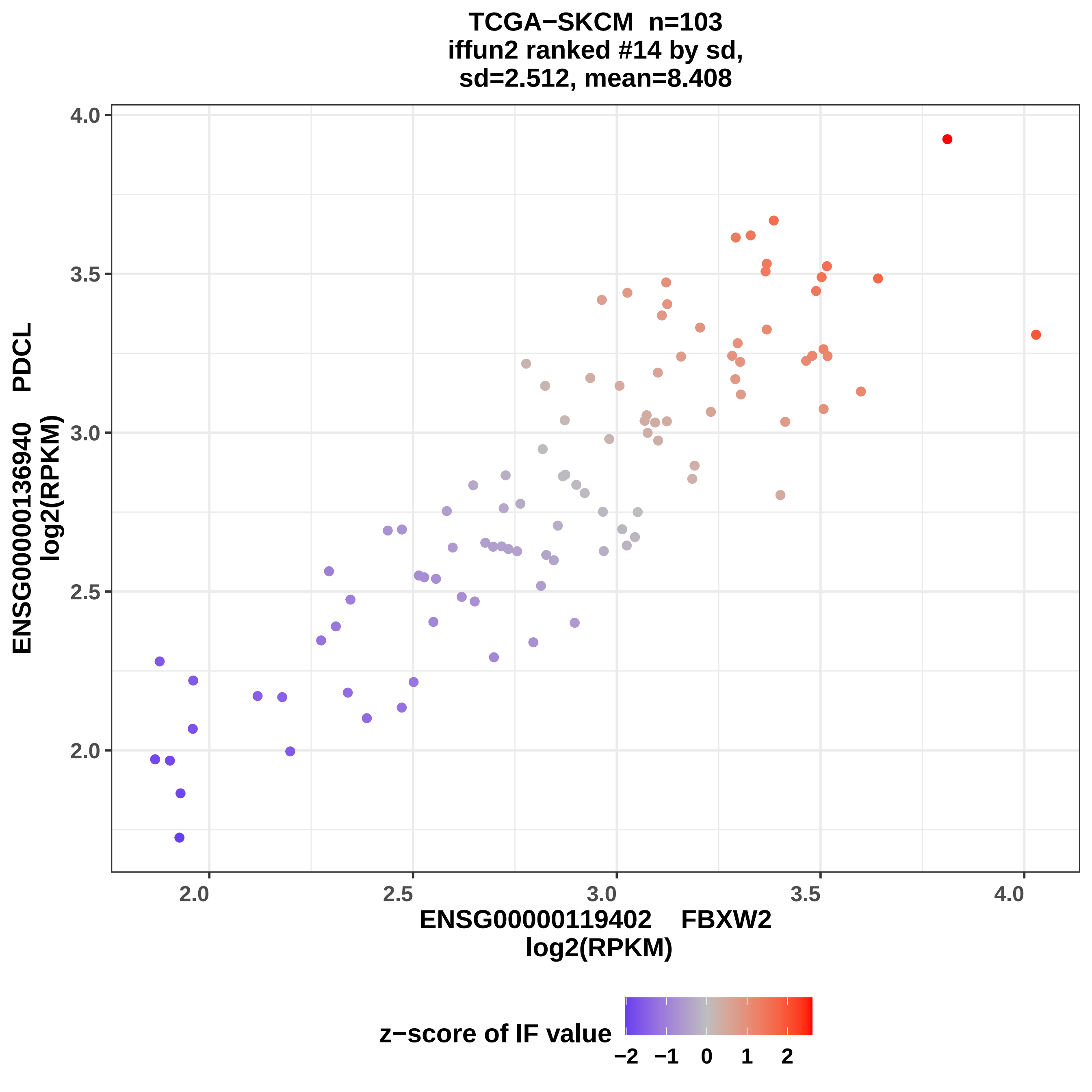

TCGA−SKCM n=103
iffun2 ranked #14 by sd,
sd=2.512, mean=8.408
ENSG00000136940 PDCL
log2(RPKM)
4.0
3.5
3.0
2.5
2.0
2.0
2.5
3.0
3.5
4.0
ENSG00000119402 FBXW2
log2(RPKM)
z−score of IF value
−2
−1
0
1
2

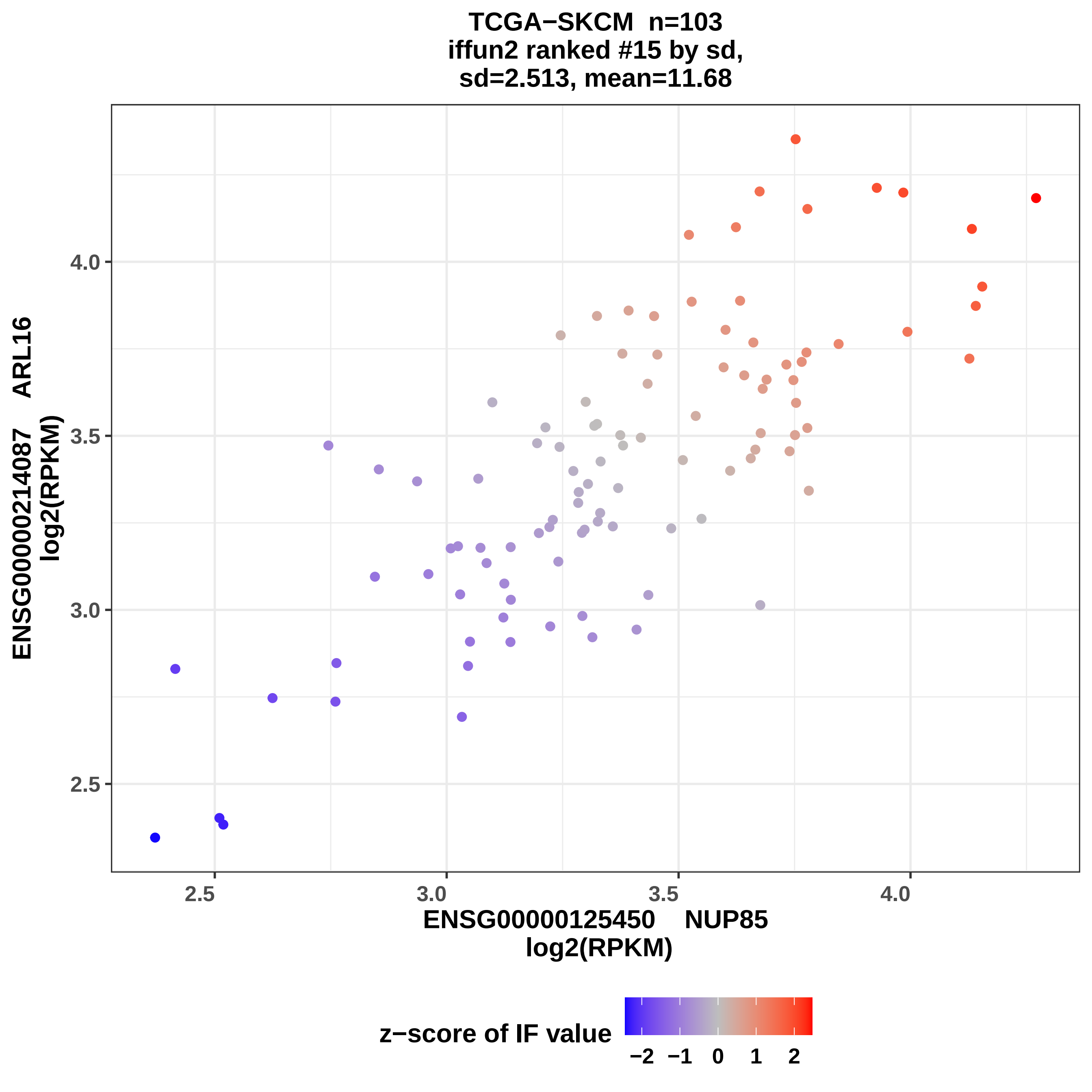
TCGA−SKCM n=103
iffun2 ranked #15 by sd,
sd=2.513, mean=11.68
ENSG00000214087 ARL16
log2(RPKM)
4.0
3.5
3.0
2.5
2.5
3.0
3.5
4.0
ENSG00000125450 NUP85
log2(RPKM)
z−score of IF value
−2
−1
0
1
2

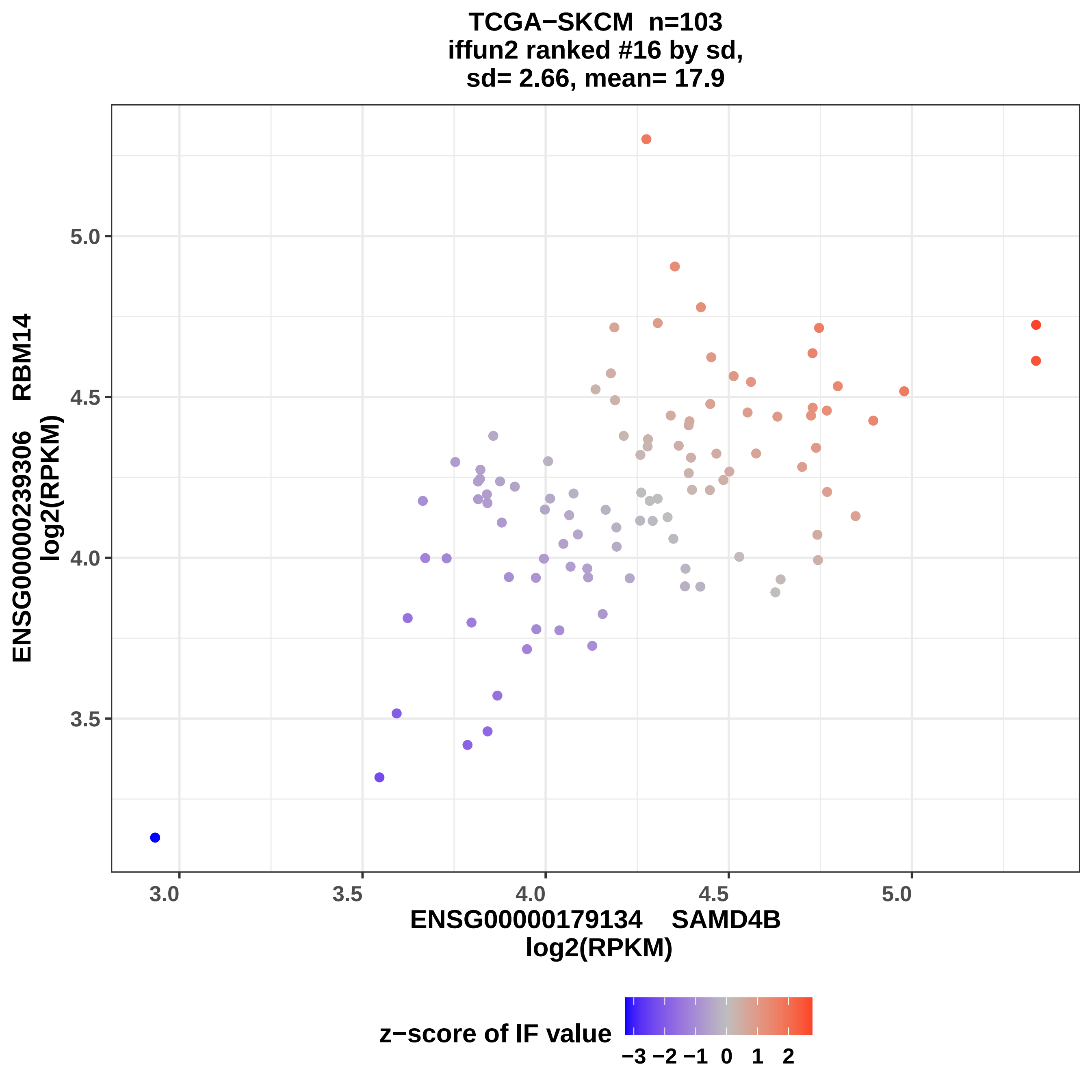
TCGA−SKCM n=103
iffun2 ranked #16 by sd,
sd= 2.66, mean= 17.9
ENSG00000239306 RBM14
log2(RPKM)
5.0
4.5
4.0
3.5
3.0
3.5
4.0
4.5
5.0
ENSG00000179134 SAMD4B
log2(RPKM)
z−score of IF value
−3 −2 −1 0 1 2

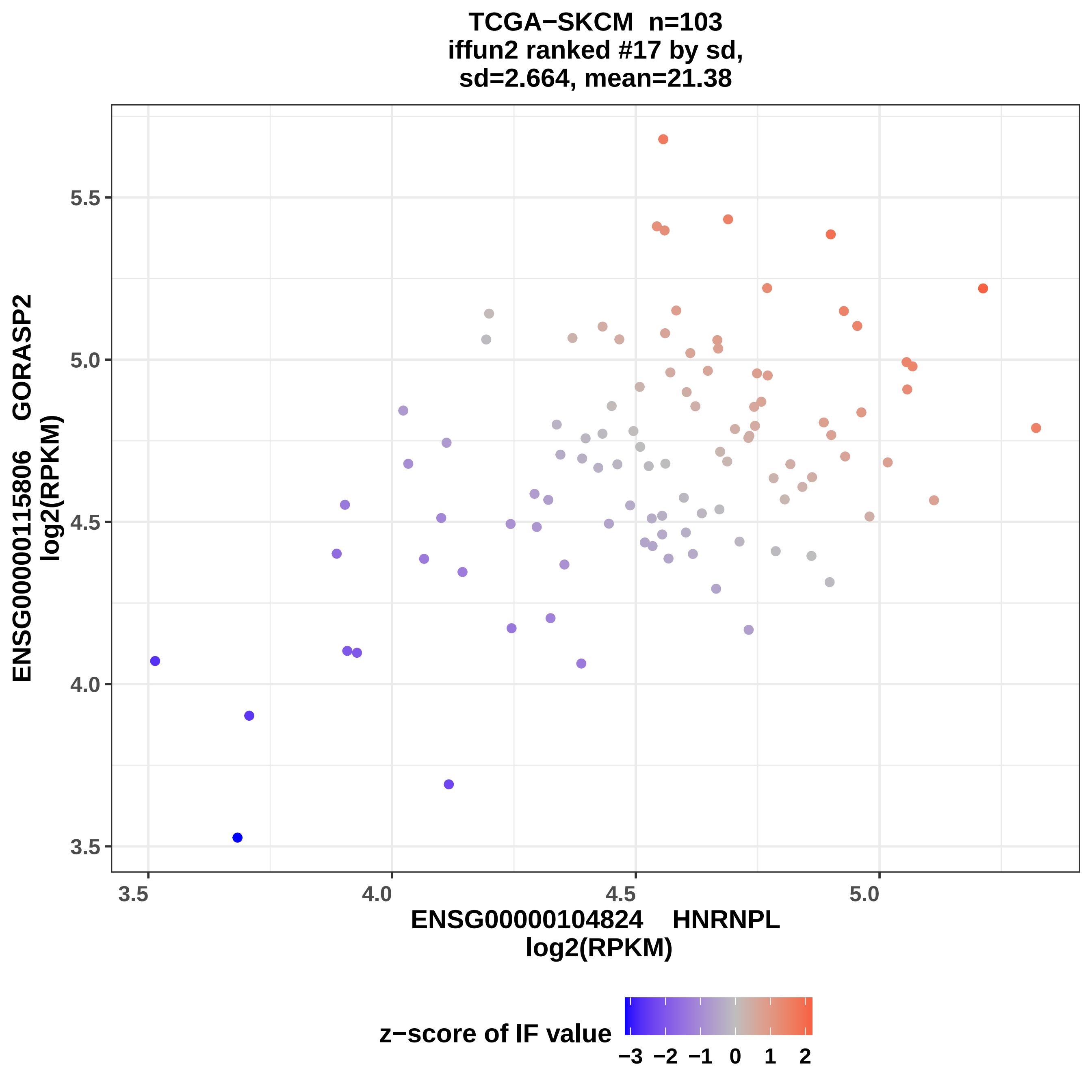
TCGA−SKCM n=103
iffun2 ranked #17 by sd,
sd=2.664, mean=21.38
ENSG00000115806 GORASP2
log2(RPKM)
5.5
5.0
4.5
4.0
3.5
3.5
4.0
4.5
5.0
ENSG00000104824 HNRNPL
log2(RPKM)
z−score of IF value
−3 −2 −1 0 1 2

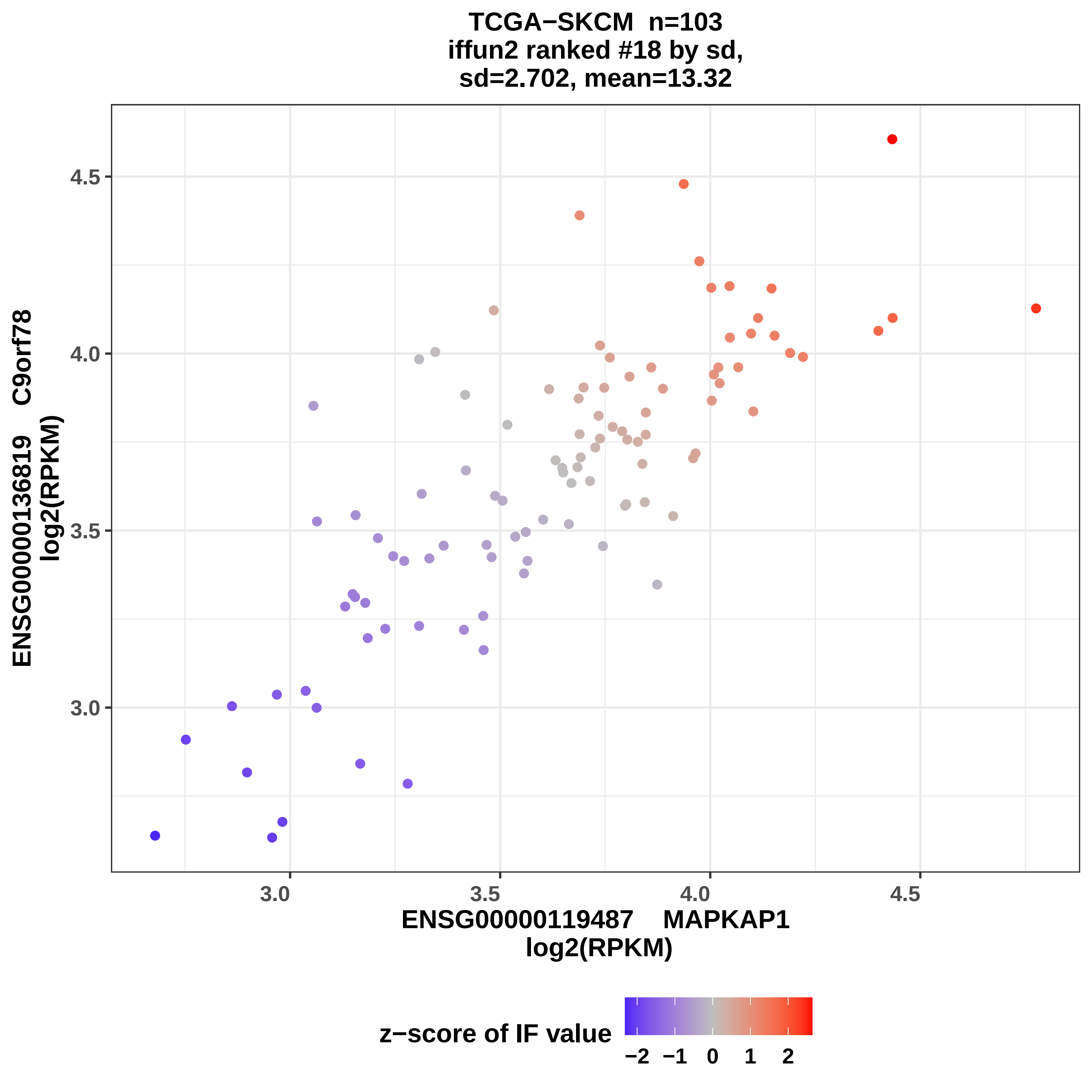
TCGA−SKCM n=103
iffun2 ranked #18 by sd,
sd=2.702, mean=13.32
ENSG00000136819 C9orf78
log2(RPKM)
4.5
4.0
3.5
3.0
3.0
3.5
4.0
4.5
ENSG00000119487 MAPKAP1
log2(RPKM)
z−score of IF value
−2
−1
0
1
2

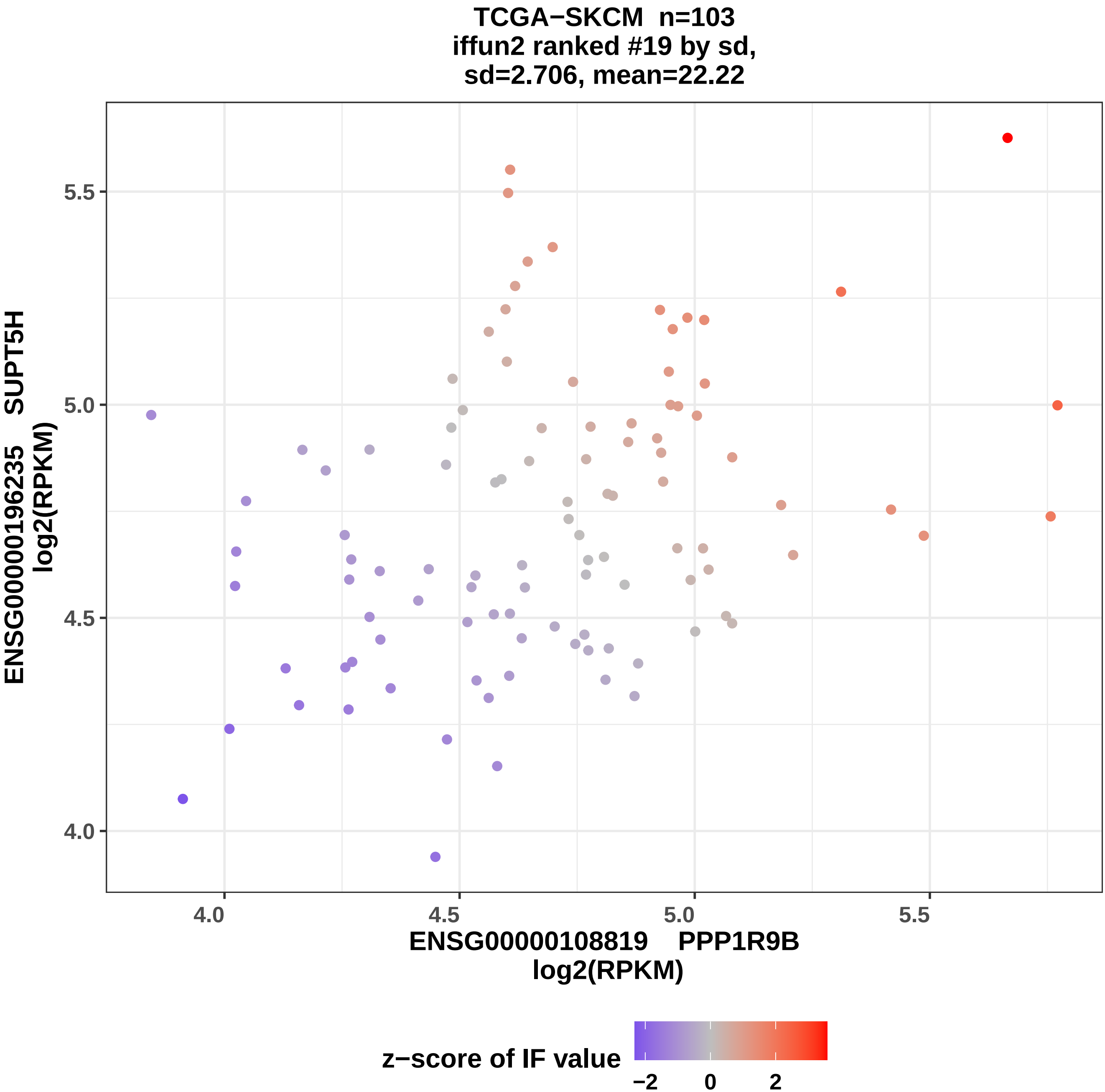

TCGA−SKCM n=103
iffun2 ranked #19 by sd,
sd=2.706, mean=22.22
ENSG00000196235 SUPT5H
log2(RPKM)
5.5
5.0
4.5
4.0
4.0
4.5
5.0
5.5
ENSG00000108819 PPP1R9B
log2(RPKM)
z−score of IF value
−2
0
2

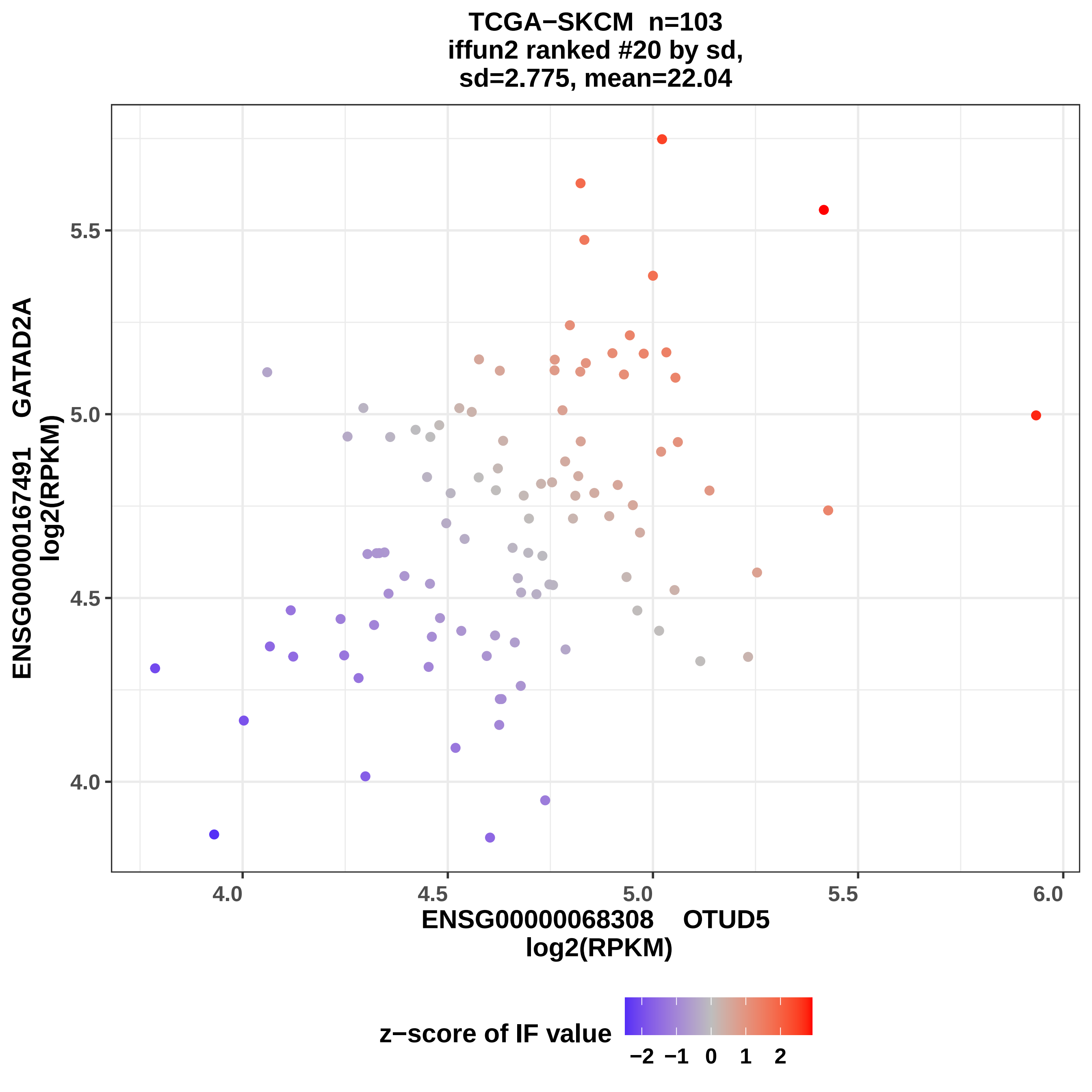

TCGA−SKCM n=103
iffun2 ranked #20 by sd,
sd=2.775, mean=22.04
ENSG00000167491 GATAD2A
log2(RPKM)
5.5
5.0
4.5
4.0
4.0
4.5
5.0
5.5
6.0
ENSG00000068308 OTUD5
log2(RPKM)
z−score of IF value
−2
−1
0
1
2

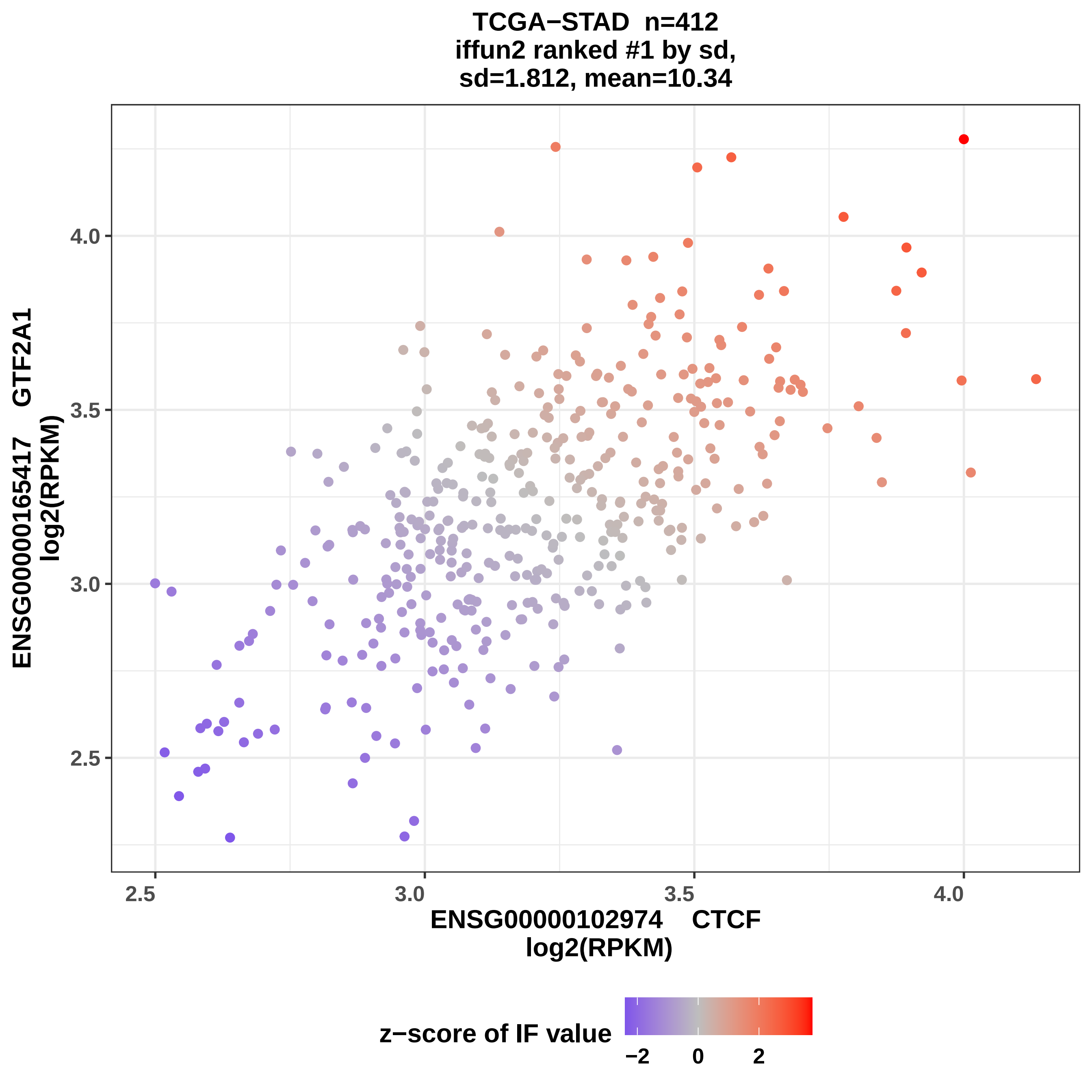

TCGA−STAD n=412
iffun2 ranked #1 by sd,
sd=1.812, mean=10.34
ENSG00000165417 GTF2A1
log2(RPKM)
4.0
3.5
3.0
2.5
2.5
3.0
3.5
4.0
ENSG00000102974 CTCF
log2(RPKM)
z−score of IF value
−2
0
2

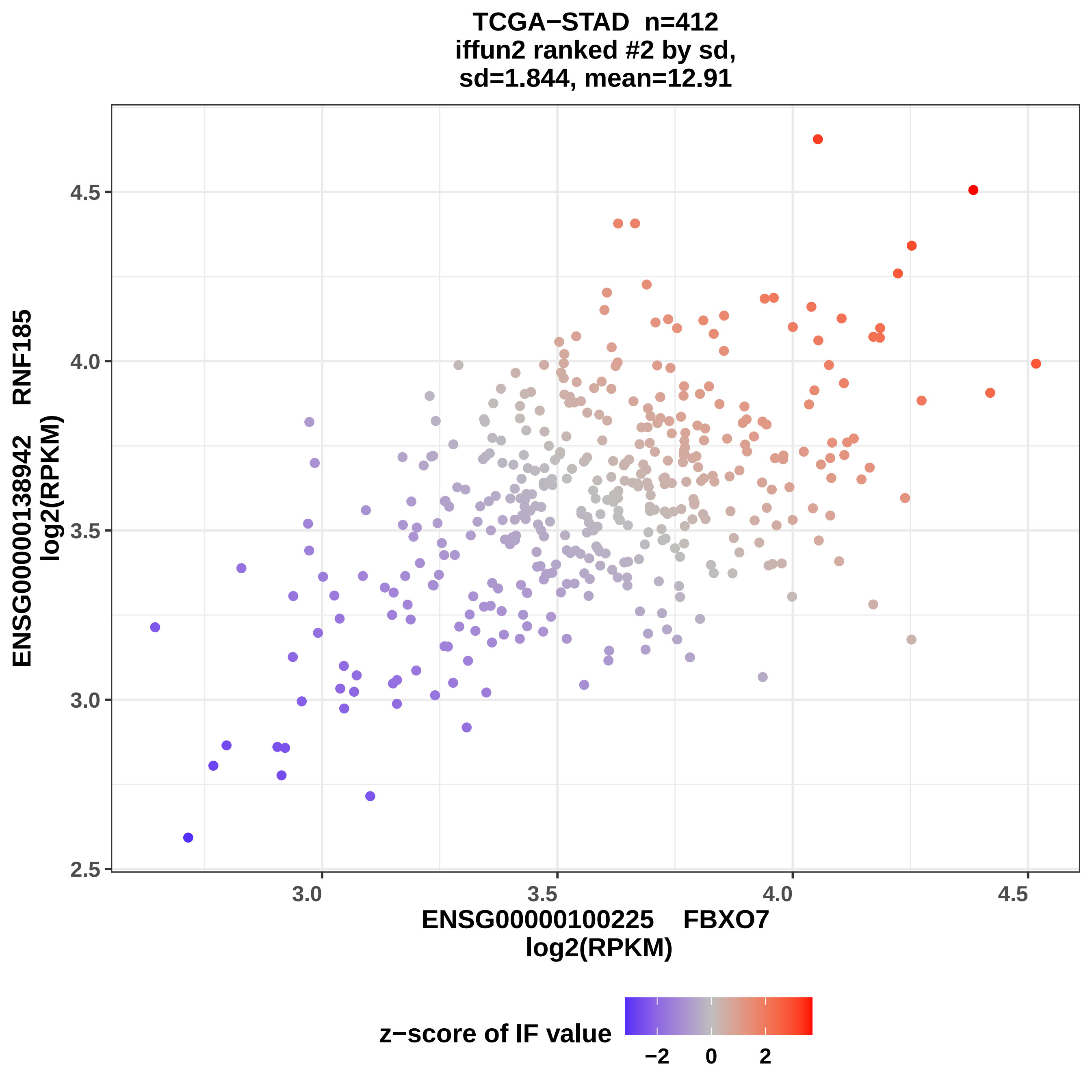
TCGA−STAD n=412
iffun2 ranked #2 by sd,
sd=1.844, mean=12.91
ENSG00000138942 RNF185
log2(RPKM)
4.5
4.0
3.5
3.0
2.5
3.0
3.5
4.0
4.5
ENSG00000100225 FBXO7
log2(RPKM)
z−score of IF value
−2
0
2

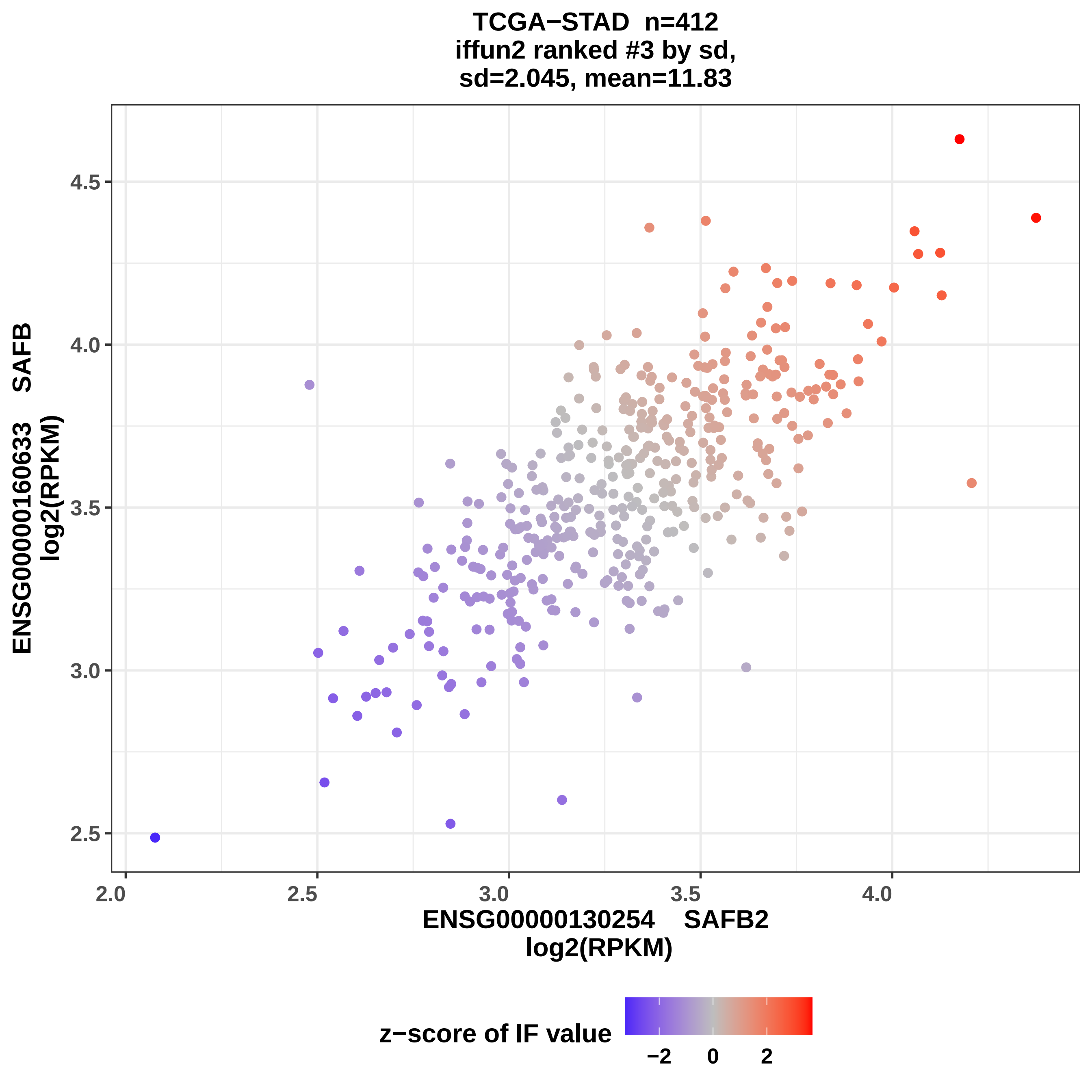
TCGA−STAD n=412
iffun2 ranked #3 by sd,
sd=2.045, mean=11.83
ENSG00000160633 SAFB
log2(RPKM)
4.5
4.0
3.5
3.0
2.5
2.0
2.5
3.0
3.5
4.0
ENSG00000130254 SAFB2
log2(RPKM)
z−score of IF value
−2
0
2

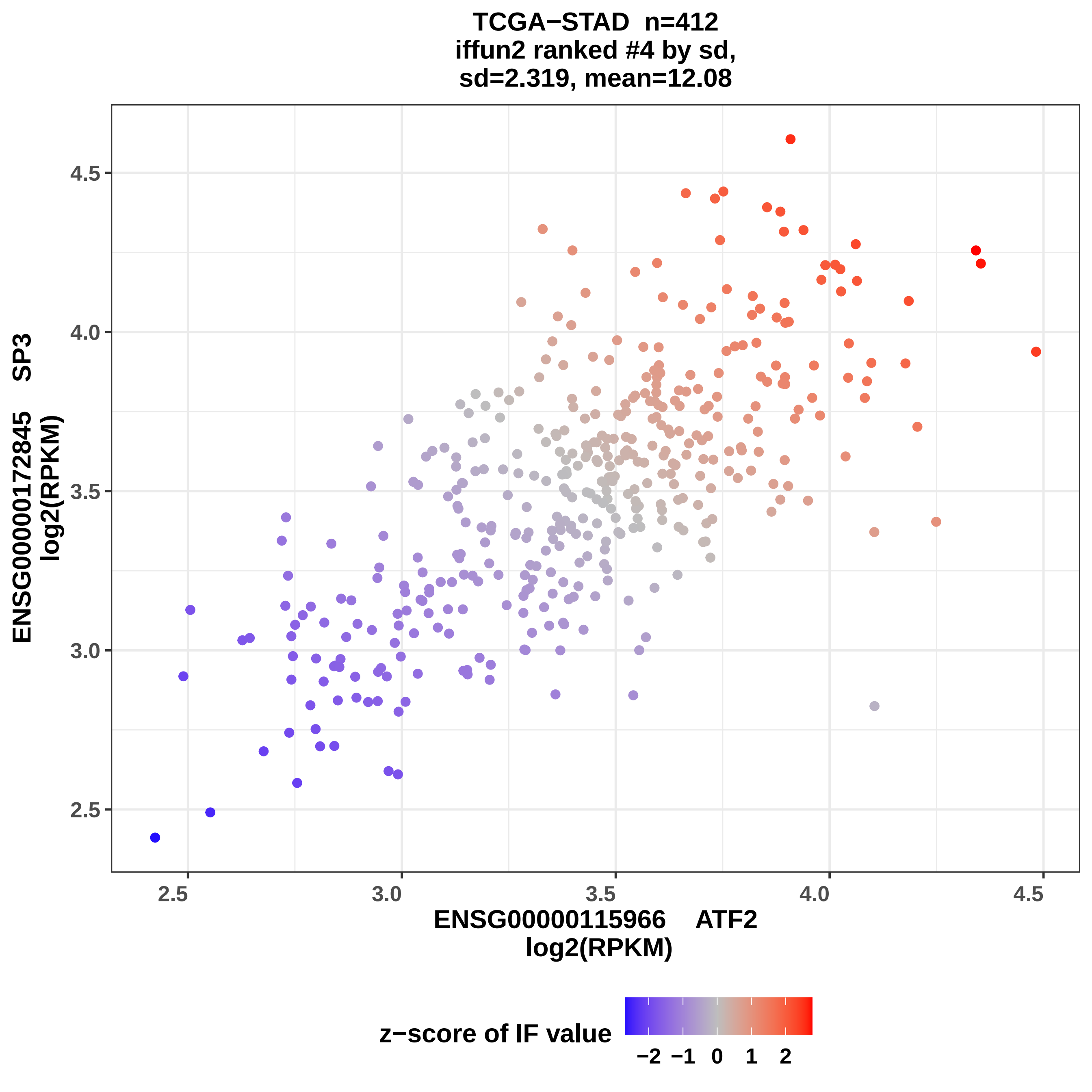
TCGA−STAD n=412
iffun2 ranked #4 by sd,
sd=2.319, mean=12.08
ENSG00000172845 SP3
log2(RPKM)
ENSG00000115966 ATF2
log2(RPKM)
2.5
3.0
3.5
4.0
4.5
z−score of IF value
−2 −1 0 1 2

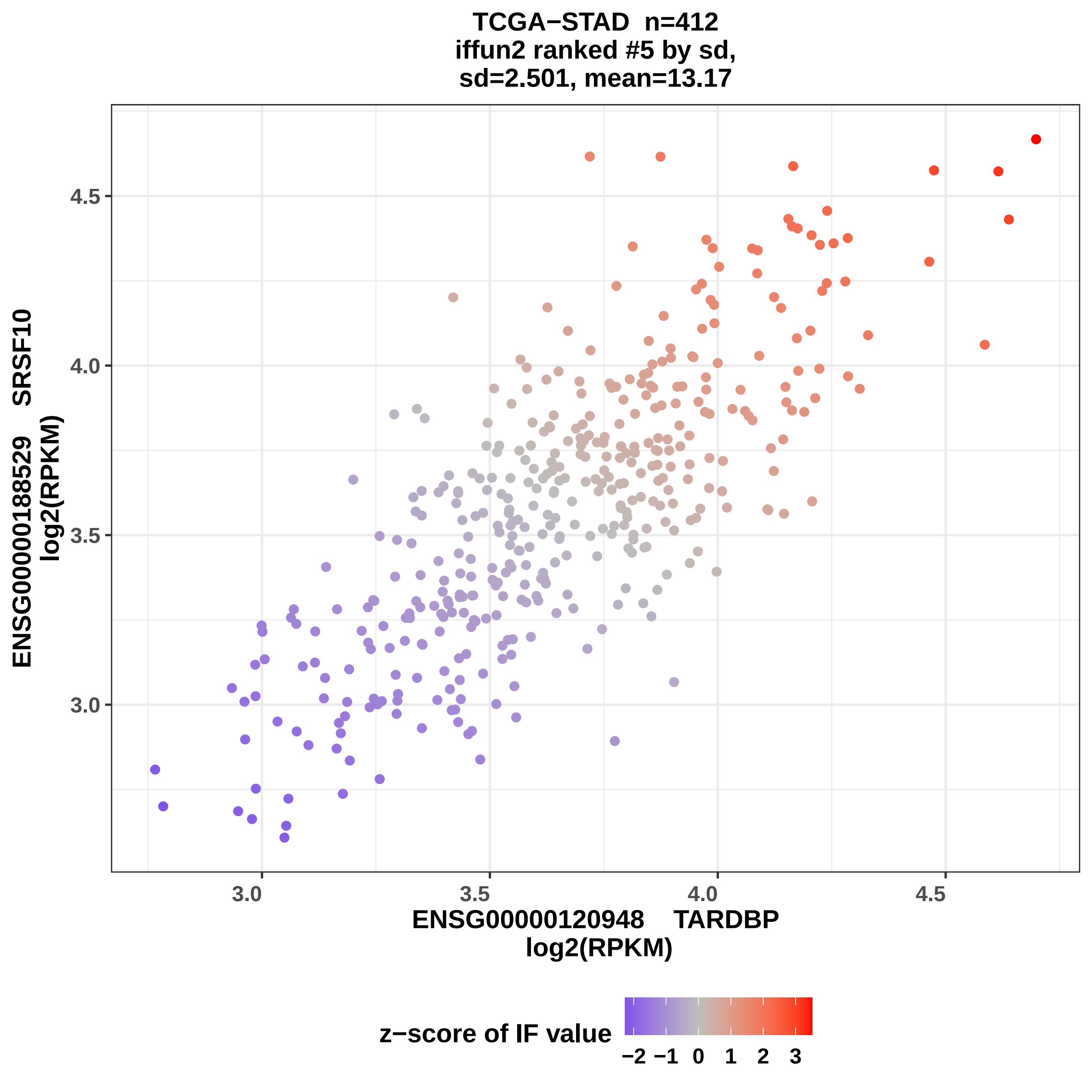
TCGA−STAD n=412
iffun2 ranked #5 by sd,
sd=2.501, mean=13.17
ENSG00000188529 SRSF10
log2(RPKM)
4.5
4.0
3.5
3.0
3.0
3.5
4.0
4.5
ENSG00000120948 TARDBP
log2(RPKM)
z−score of IF value
−2
−1
0
1
2
3

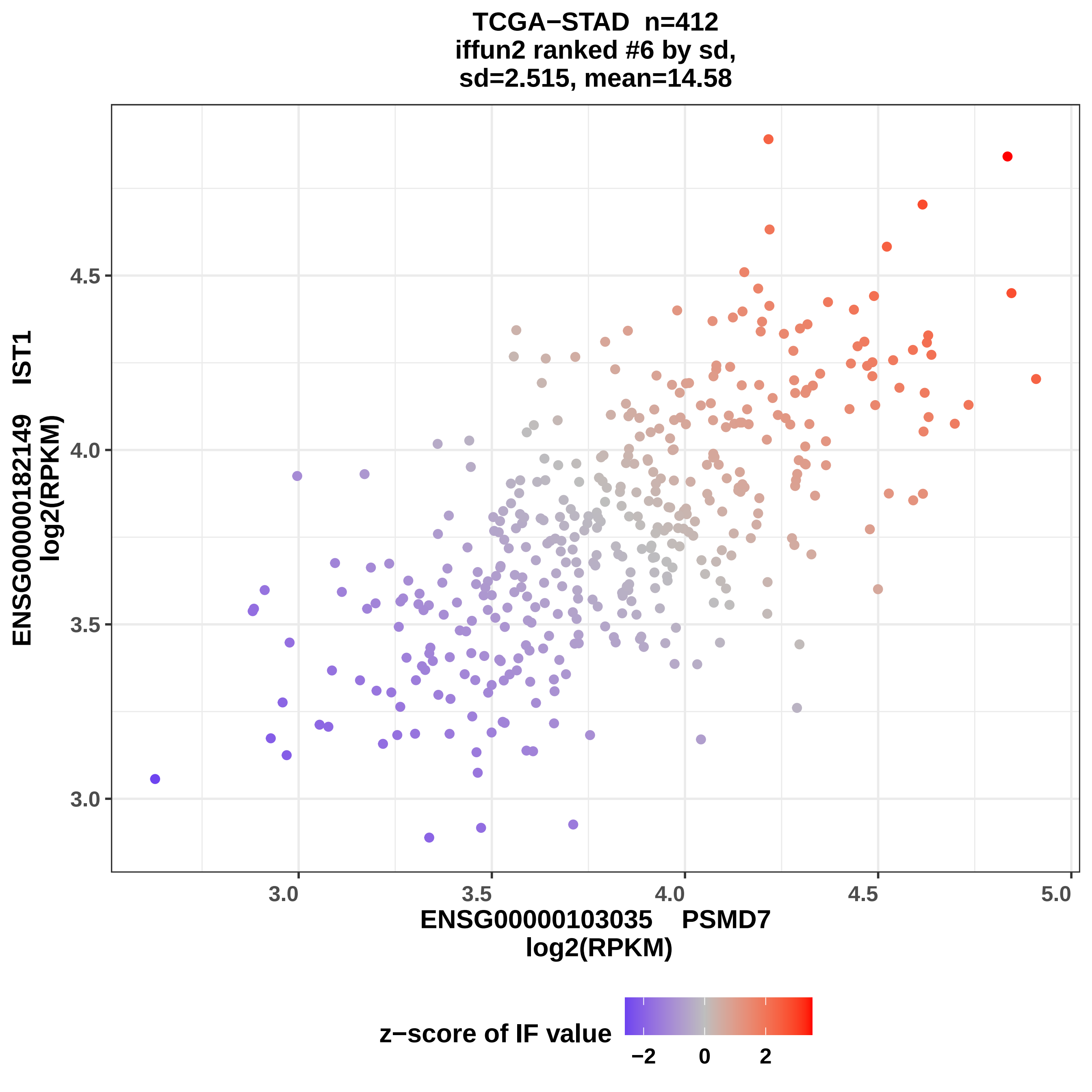
TCGA−STAD n=412
iffun2 ranked #6 by sd,
sd=2.515, mean=14.58
ENSG00000182149 IST1
log2(RPKM)
4.5
4.0
3.5
3.0
5.0
ENSG00000103035 PSMD7
log2(RPKM)
z−score of IF value
−2
0
2

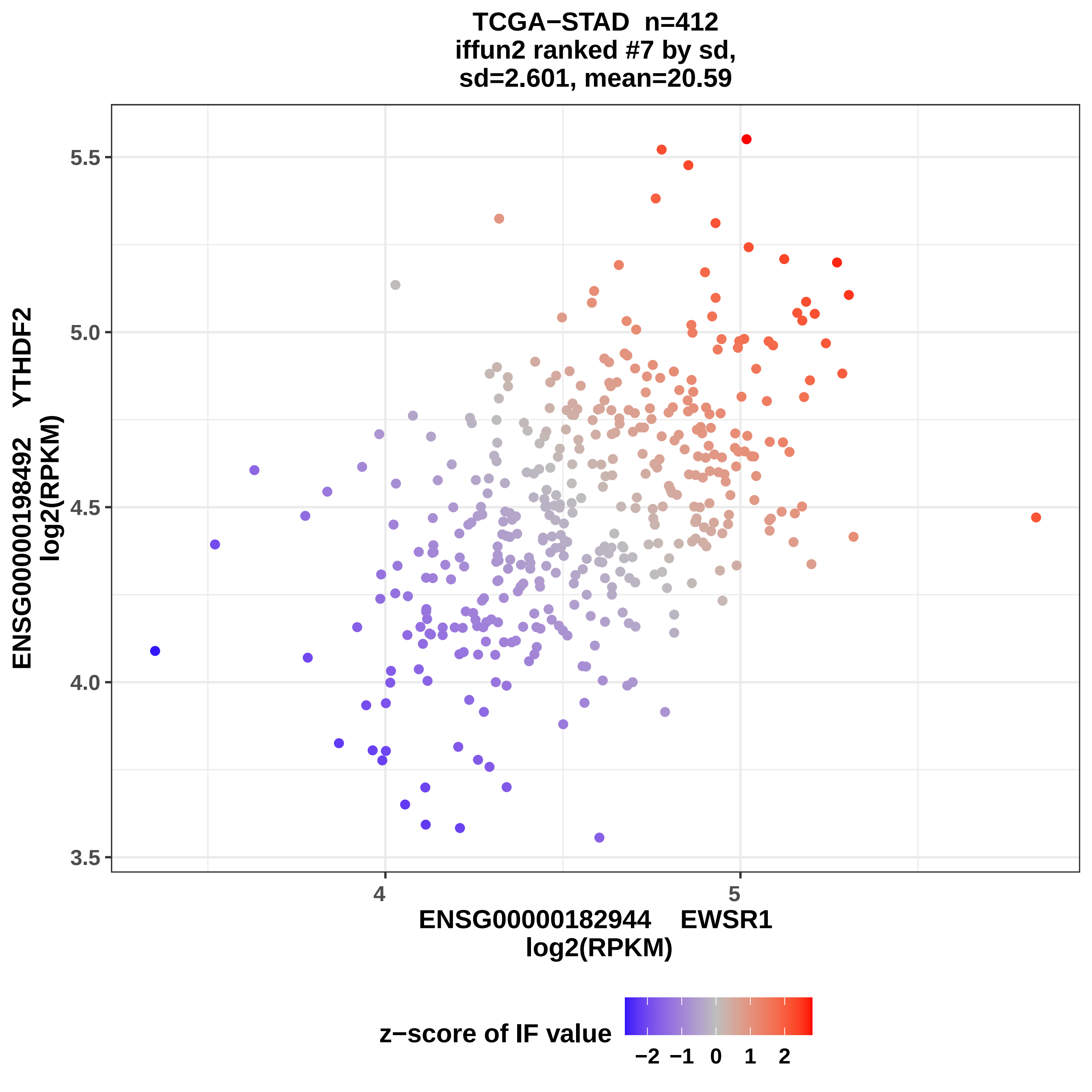
TCGA−STAD n=412
iffun2 ranked #7 by sd,
sd=2.601, mean=20.59
5.5
5.0
4.5
4.0
3.5
ENSG00000198492 YTHDF2
log2(RPKM)
4
5
ENSG00000182944 EWSR1
log2(RPKM)
z−score of IF value
−2 −1 0 1 2

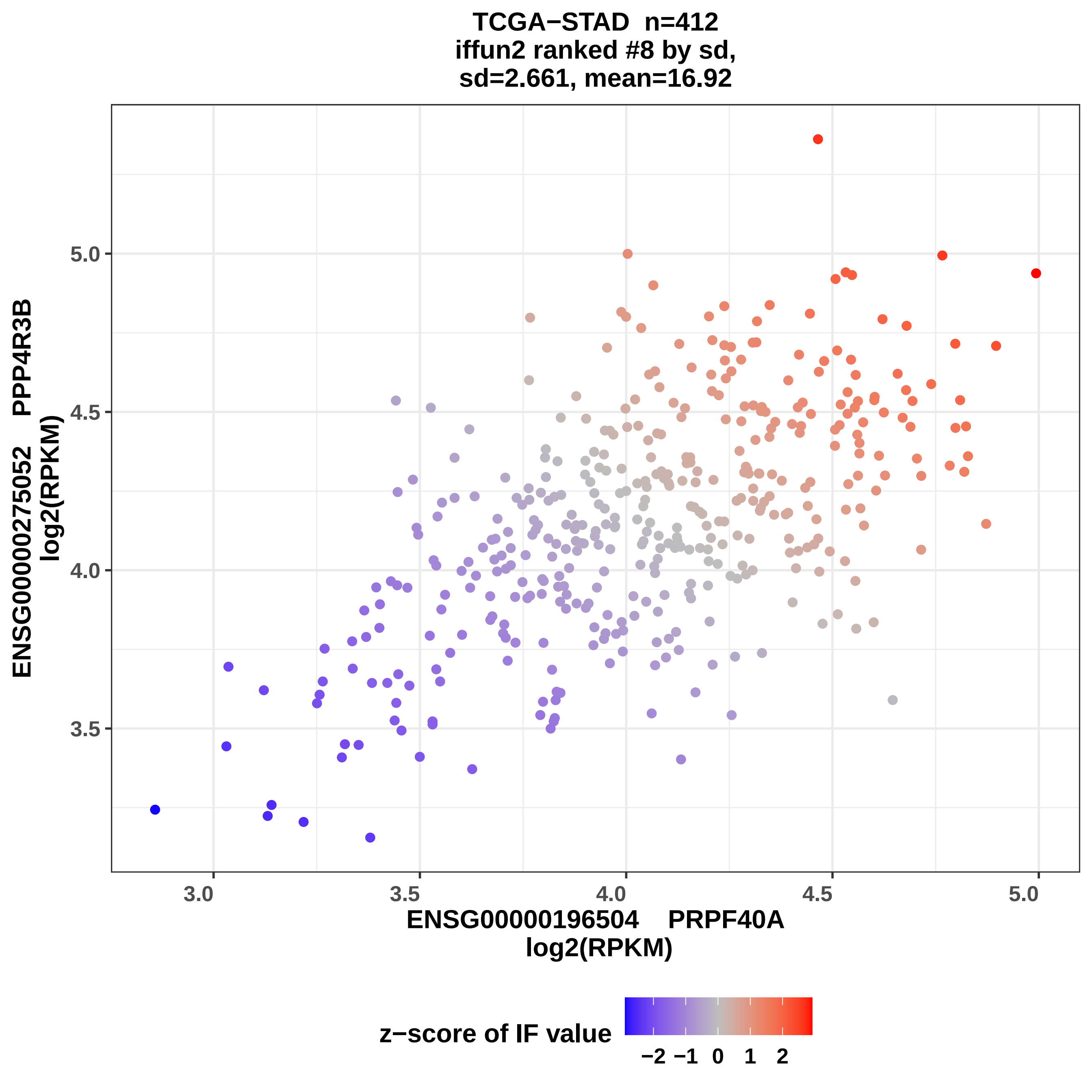
TCGA−STAD n=412
iffun2 ranked #8 by sd,
sd=2.661, mean=16.92
ENSG00000275052 PPP4R3B
log2(RPKM)
5.0
4.5
4.0
3.5
3.0
3.5
4.0
4.5
5.0
ENSG00000196504 PRPF40A
log2(RPKM)
z−score of IF value
−2 −1 0 1 2

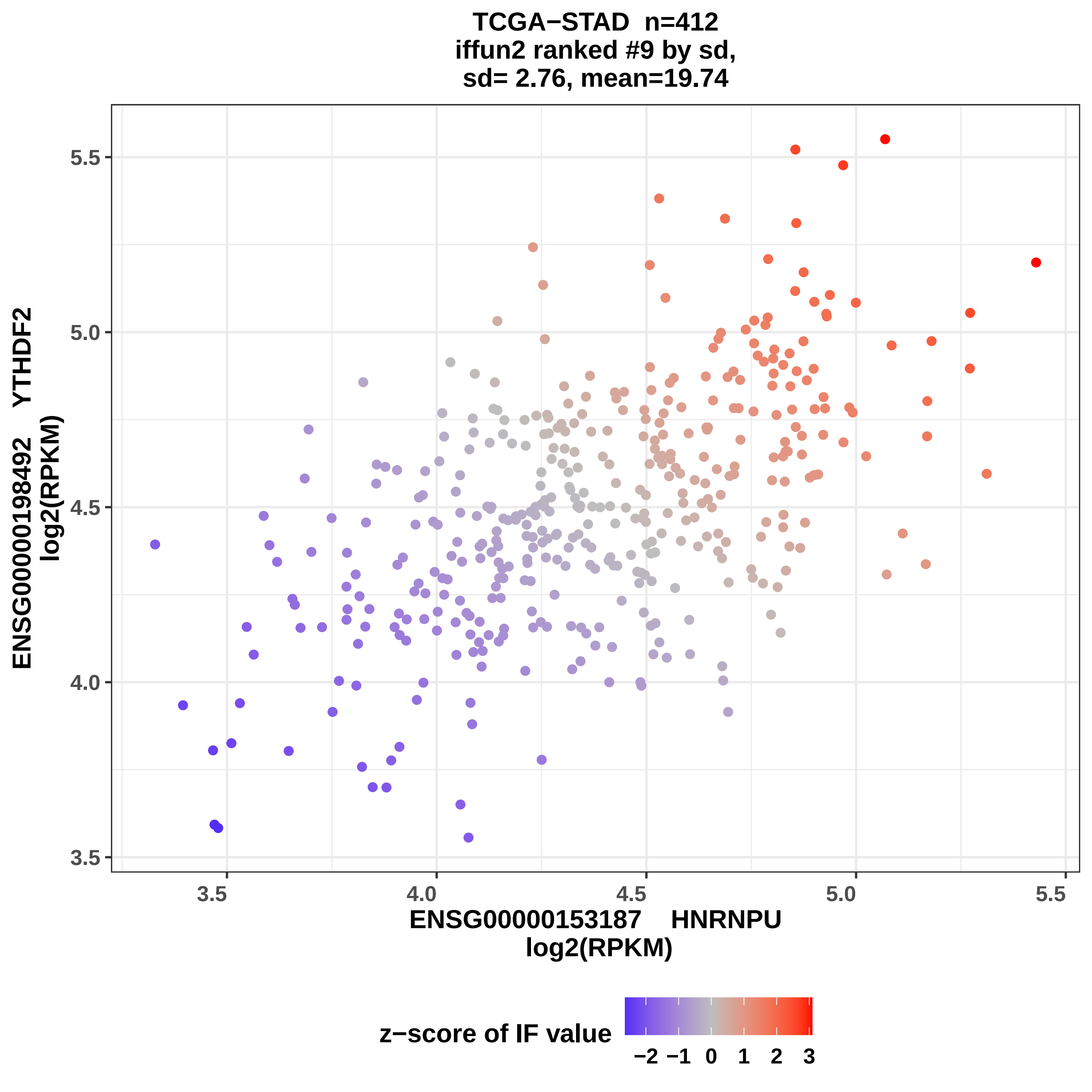
TCGA−STAD n=412
iffun2 ranked #9 by sd,
sd= 2.76, mean=19.74
ENSG00000198492 YTHDF2
log2(RPKM)
5.5
5.0
4.5
4.0
3.5
3.5
4.0
4.5
5.0
5.5
ENSG00000153187 HNRNPU
log2(RPKM)
z−score of IF value
−2 −1 0 1 2 3

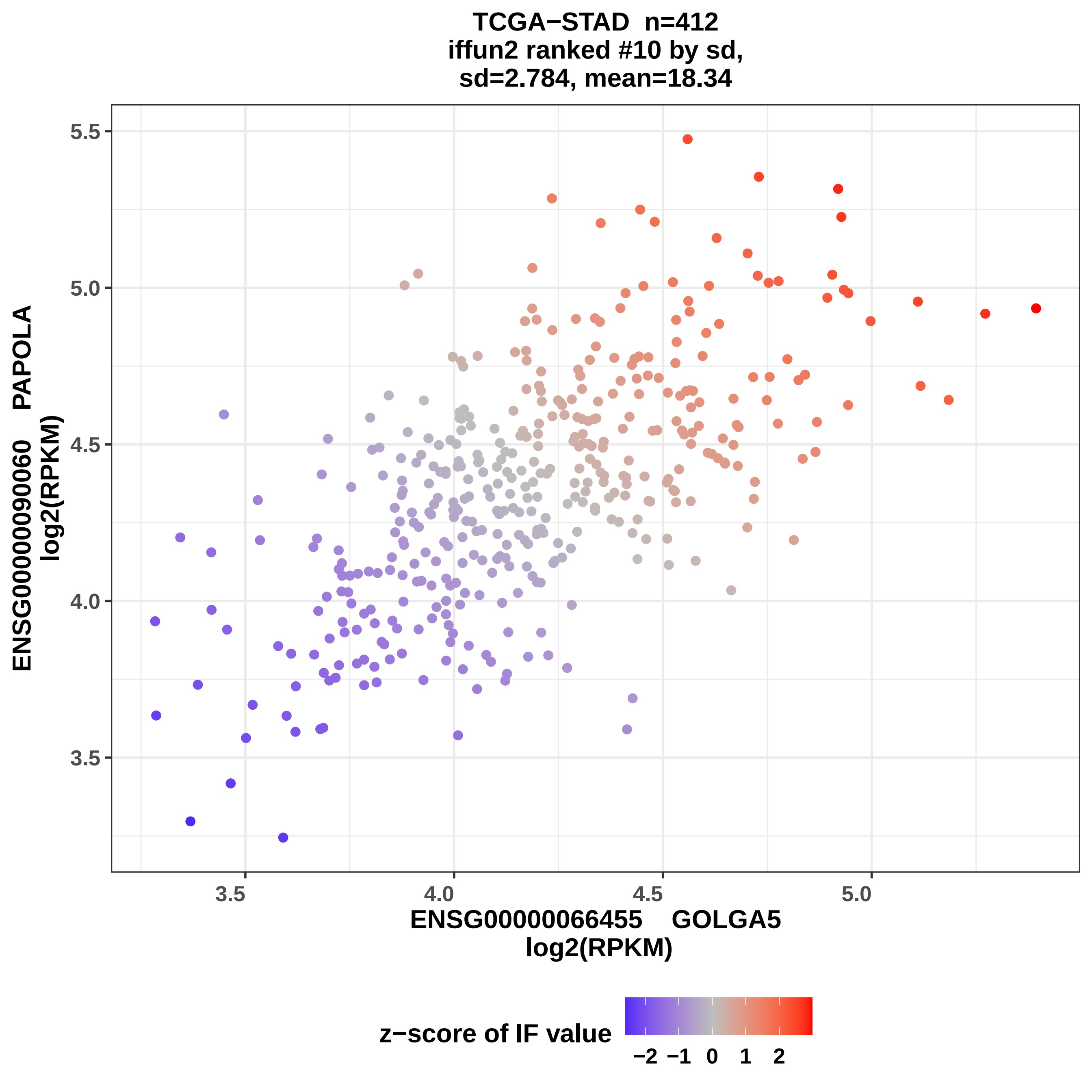
TCGA−STAD n=412
iffun2 ranked #10 by sd,
sd=2.784, mean=18.34
ENSG00000090060 PAPOLA
log2(RPKM)
5.5
5.0
4.5
4.0
3.5
3.5
4.0
4.5
5.0
ENSG00000066455 GOLGA5
log2(RPKM)
z−score of IF value
−2 −1 0 1 2

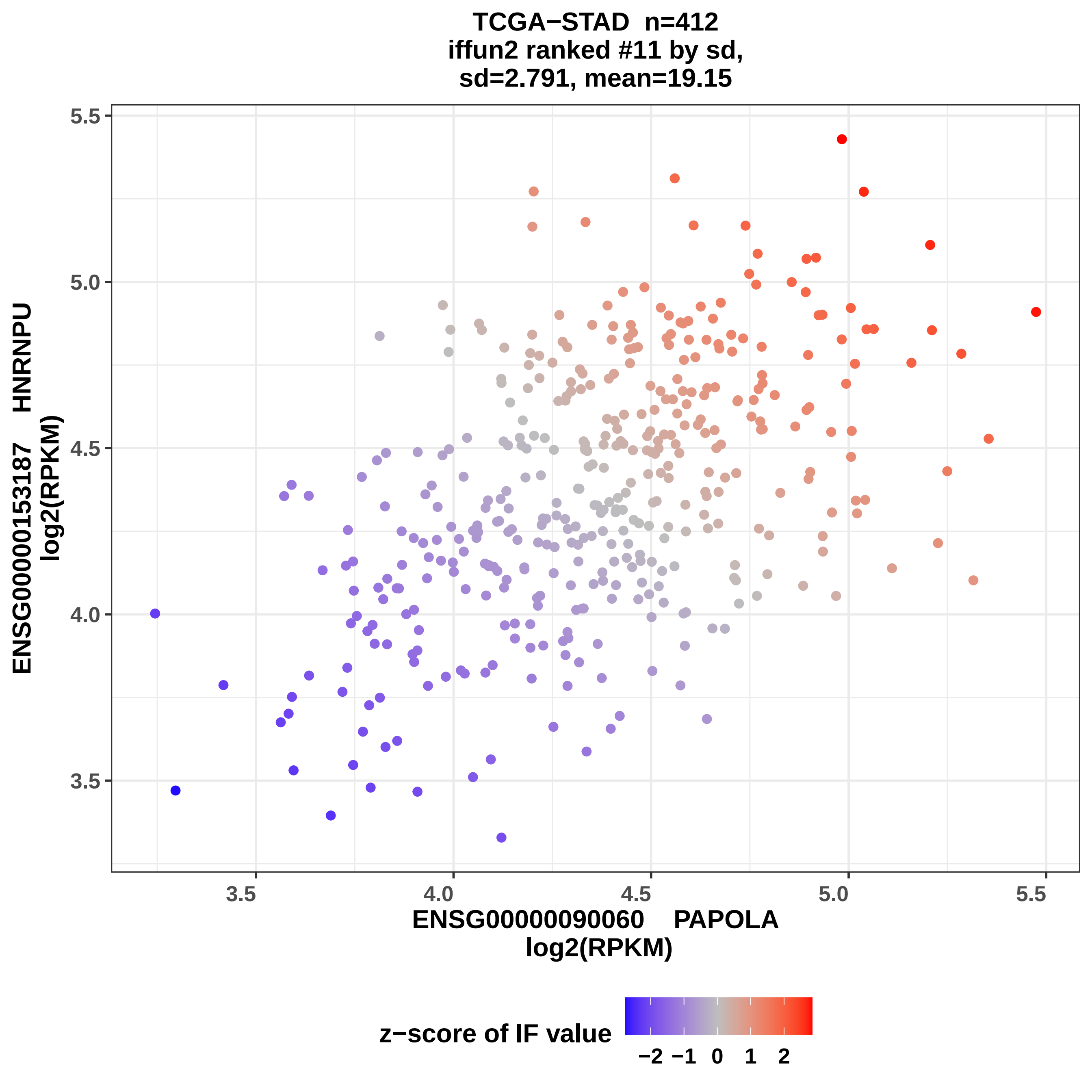

TCGA−STAD n=412
iffun2 ranked #11 by sd,
sd=2.791, mean=19.15
ENSG00000153187 HNRNPU
log2(RPKM)
5.5
5.0
4.5
4.0
3.5
3.5
4.0
4.5
5.0
5.5
ENSG00000090060 PAPOLA
log2(RPKM)
z−score of IF value
−2
−1
0
1
2

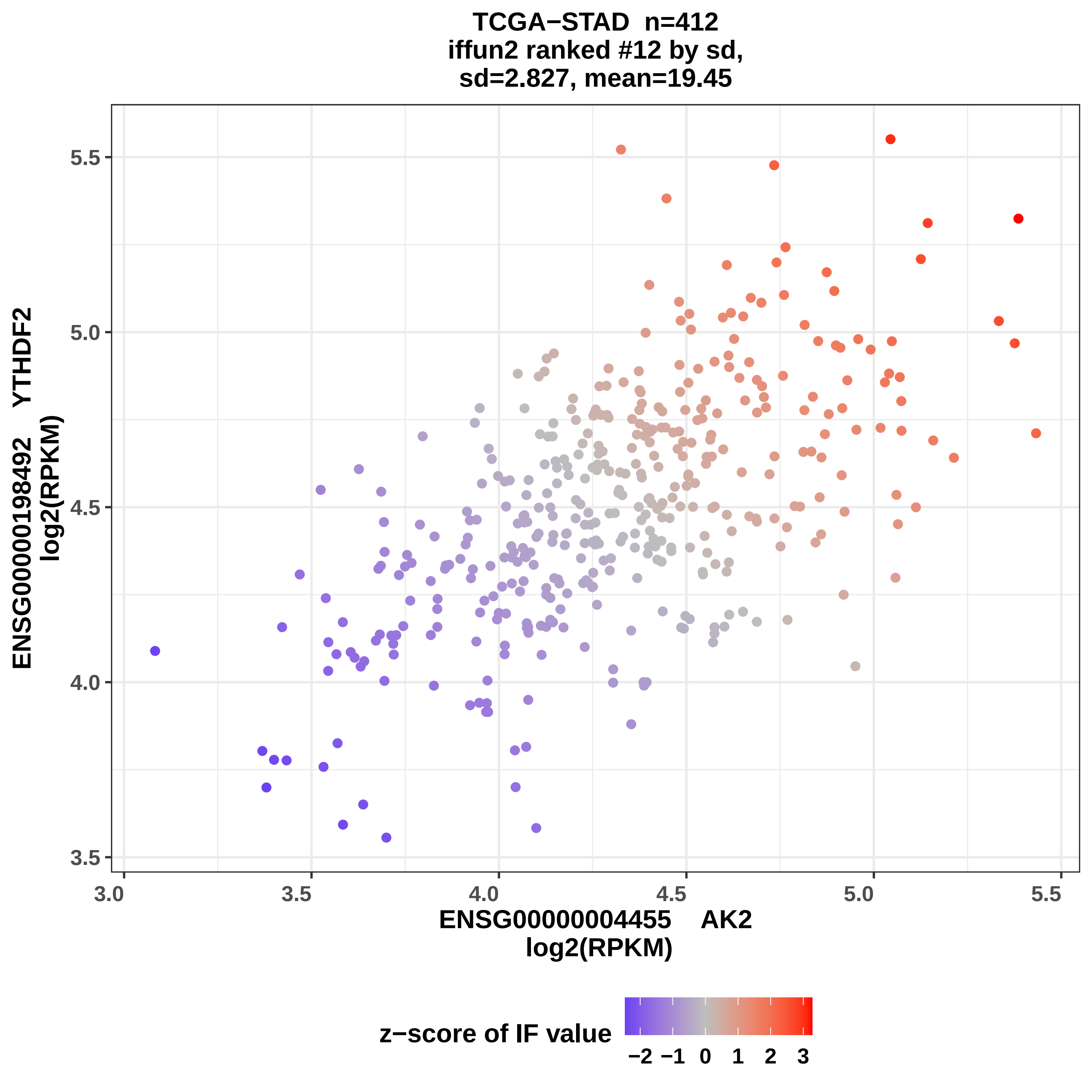
TCGA−STAD n=412
iffun2 ranked #12 by sd,
sd=2.827, mean=19.45
ENSG00000198492 YTHDF2
log2(RPKM)
5.5
5.0
4.5
4.0
3.5
3.0
3.5
4.0
4.5
5.0
5.5
ENSG00000004455 AK2
log2(RPKM)
z−score of IF value
−2 −1 0 1 2 3

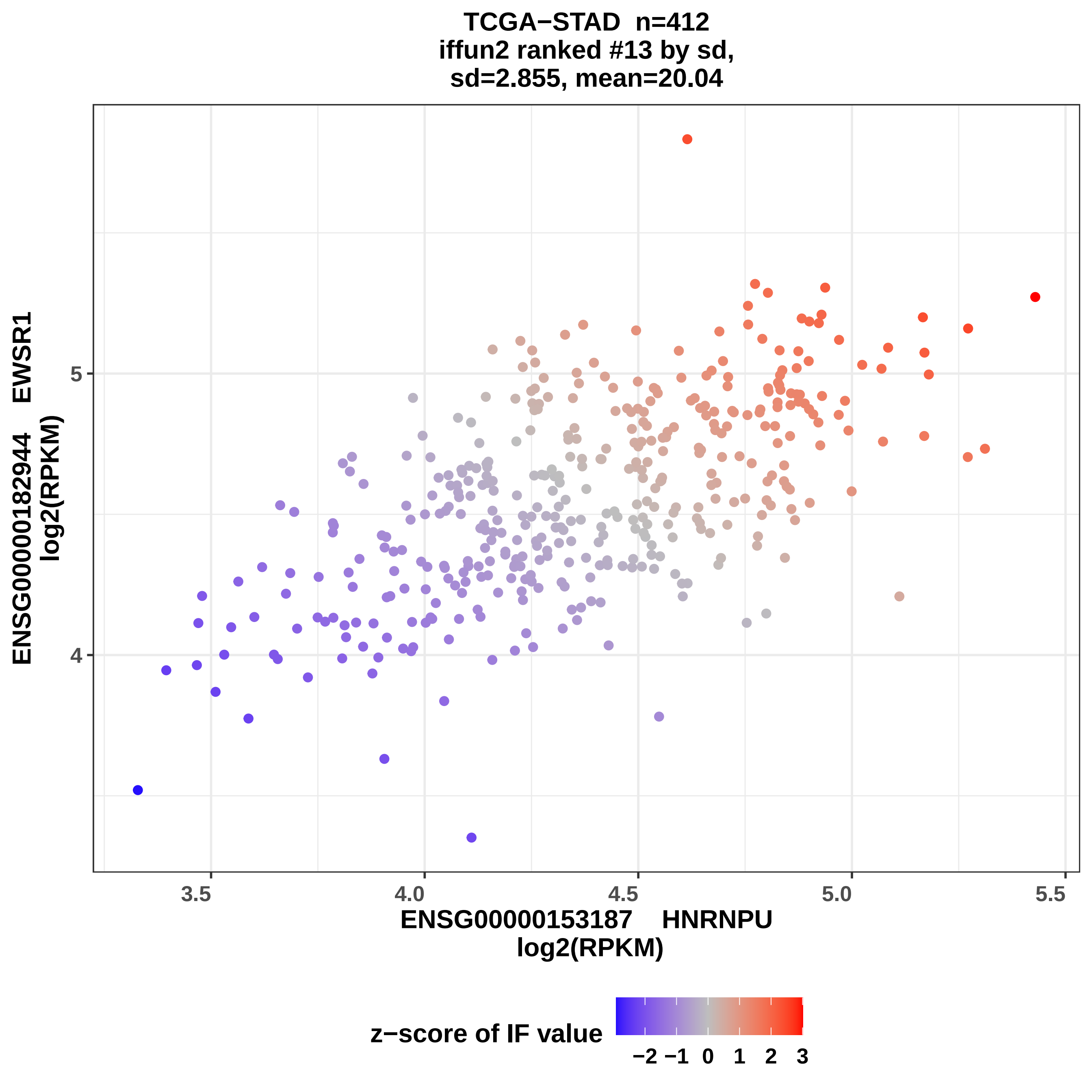
TCGA−STAD n=412
iffun2 ranked #13 by sd,
sd=2.855, mean=20.04
ENSG00000182944 EWSR1
log2(RPKM)
5
4
3.5
4.0
4.5
5.0
5.5
ENSG00000153187 HNRNPU
log2(RPKM)
z−score of IF value
−2
−1
0
1
2
3

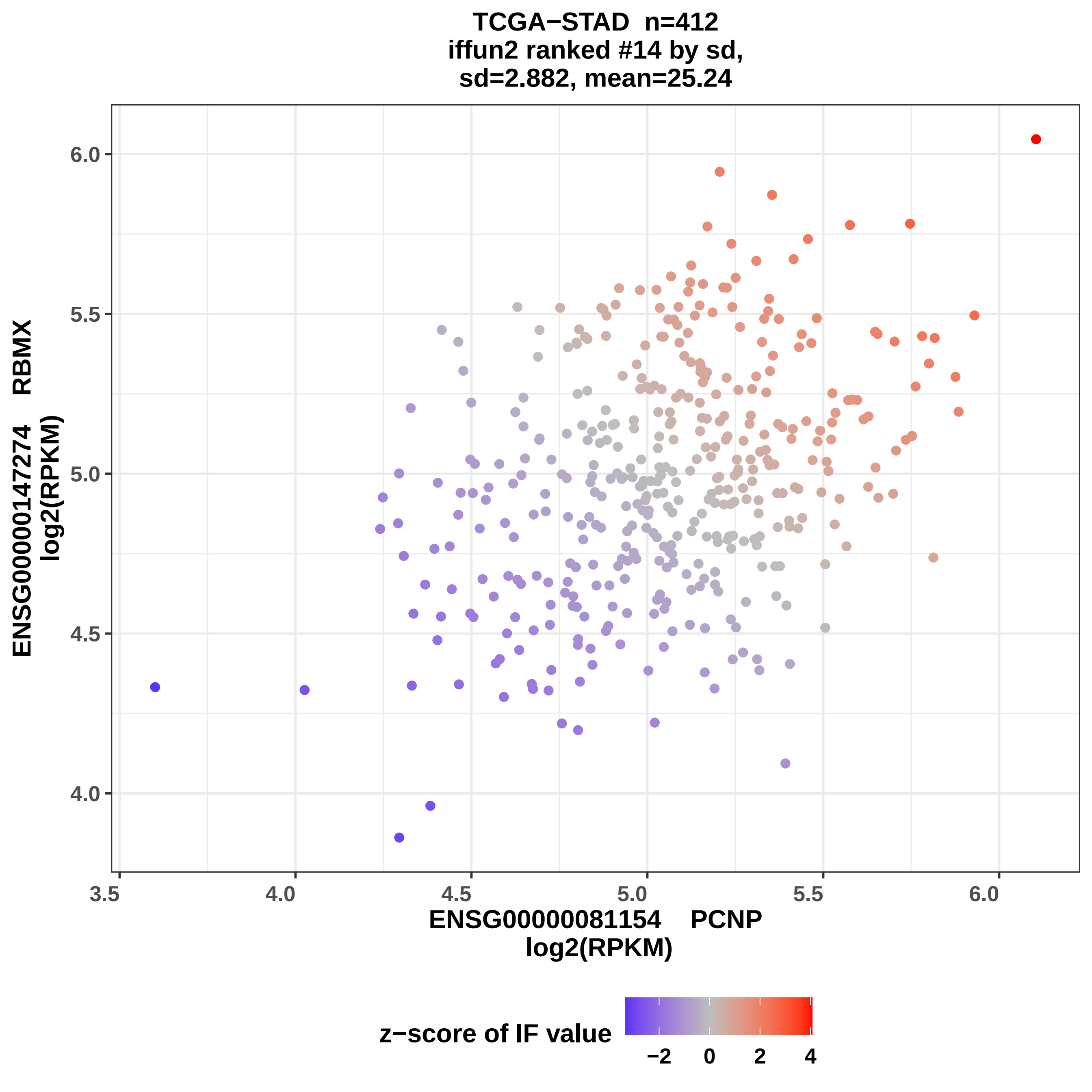
TCGA−STAD n=412
iffun2 ranked #14 by sd,
sd=2.882, mean=25.24
ENSG00000147274 RBMX
log2(RPKM)
6.0
5.5
5.0
4.5
4.0
3.5
4.0
4.5
5.0
5.5
6.0
ENSG00000081154 PCNP
log2(RPKM)
z−score of IF value
−2
0
2
4

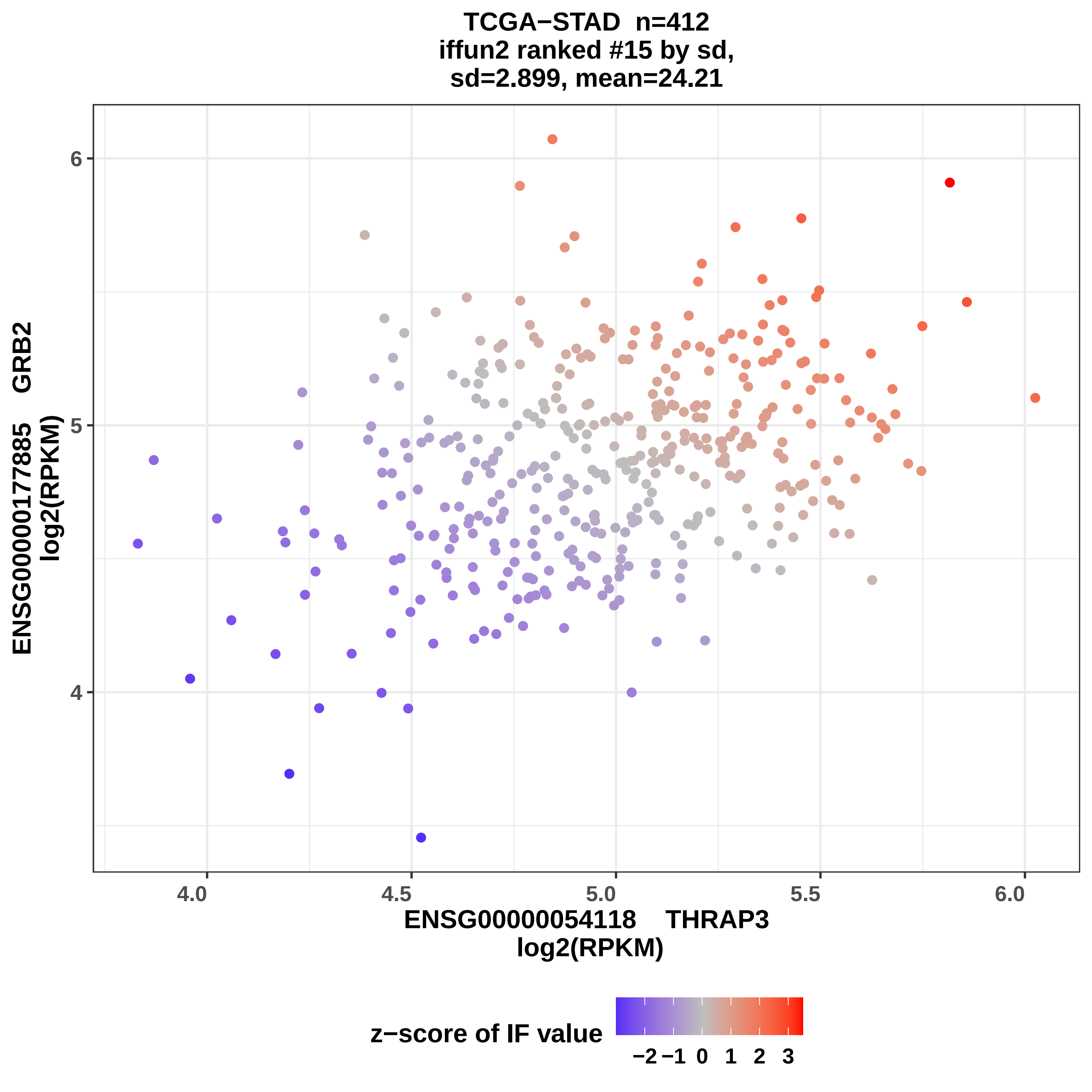
TCGA−STAD n=412
iffun2 ranked #15 by sd,
sd=2.899, mean=24.21
ENSG00000177885 GRB2
log2(RPKM)
ENSG00000054118 THRAP3
log2(RPKM)
4.0
4.5
5.0
5.5
6.0
4
5
6
z−score of IF value
−2
−1
0
1
2
3

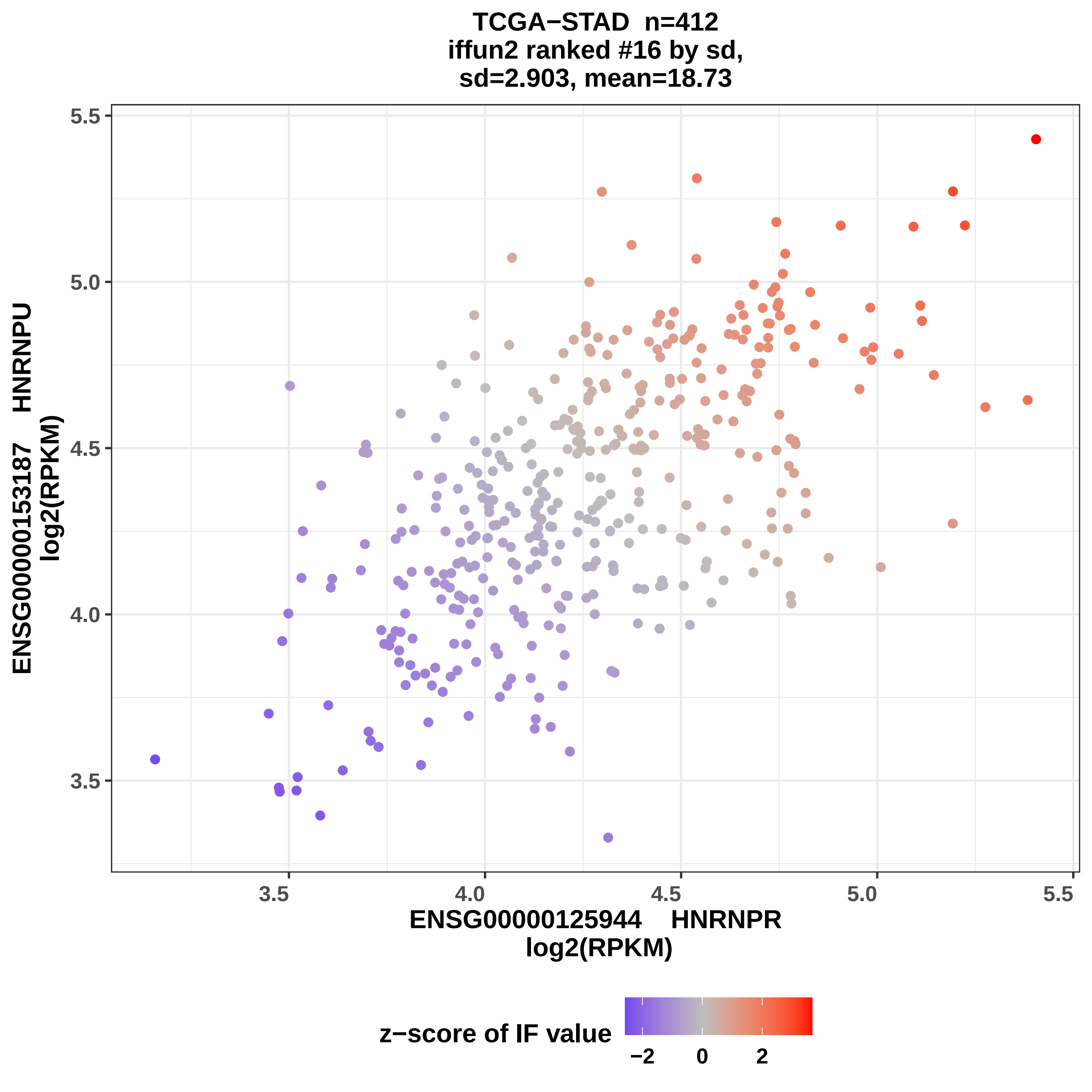
TCGA−STAD n=412
iffun2 ranked #16 by sd,
sd=2.903, mean=18.73
ENSG00000153187 HNRNPU
log2(RPKM)
ENSG00000125944 HNRNPR
log2(RPKM)
5.5
5.0
4.5
4.0
3.5
3.5
4.0
4.5
5.0
5.5
z−score of IF value
−2
0
2

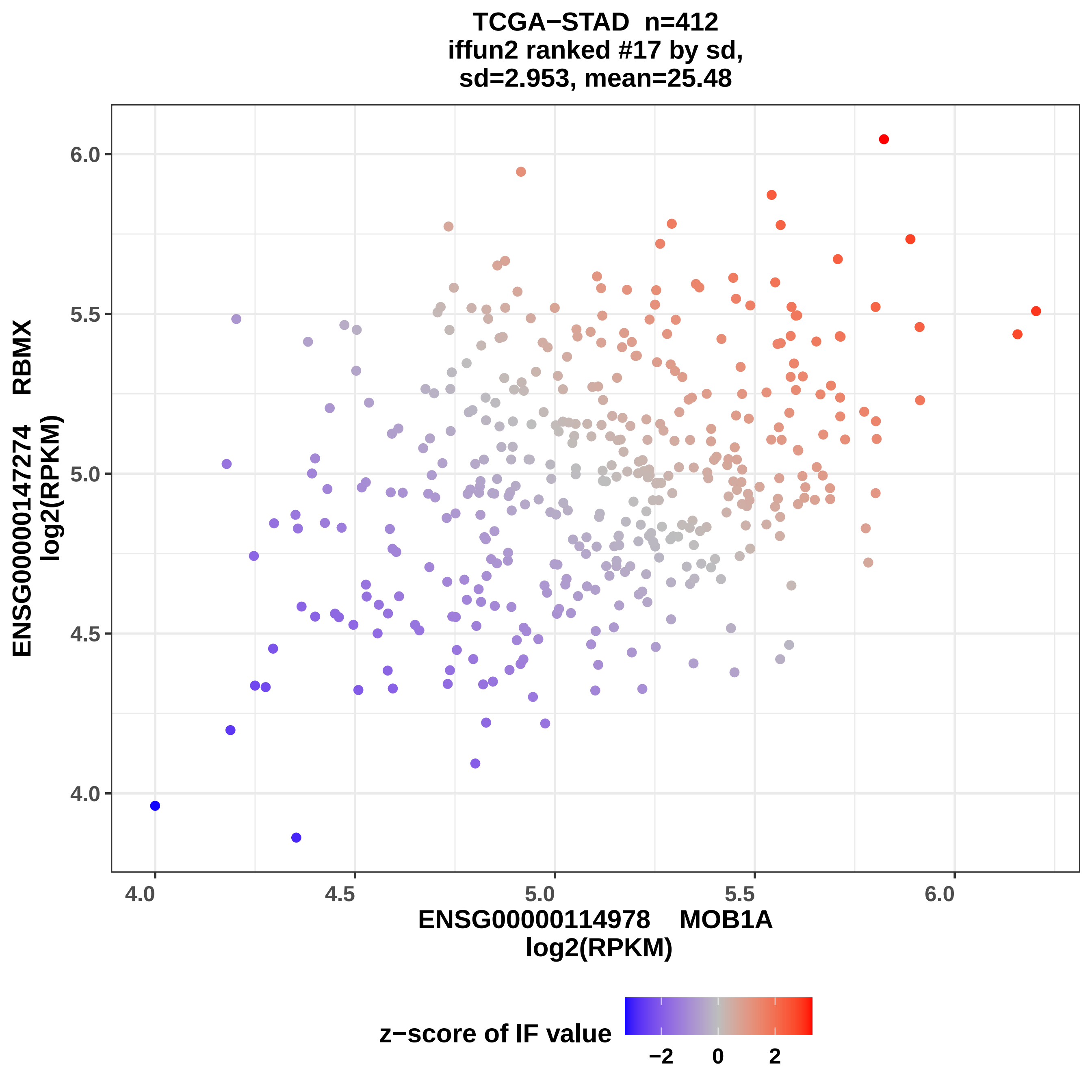

TCGA−STAD n=412
iffun2 ranked #17 by sd,
sd=2.953, mean=25.48
ENSG00000147274 RBMX
log2(RPKM)
6.0
5.5
5.0
4.5
4.0
4.0
4.5
5.0
5.5
6.0
ENSG00000114978 MOB1A
log2(RPKM)
z−score of IF value
−2
0
2

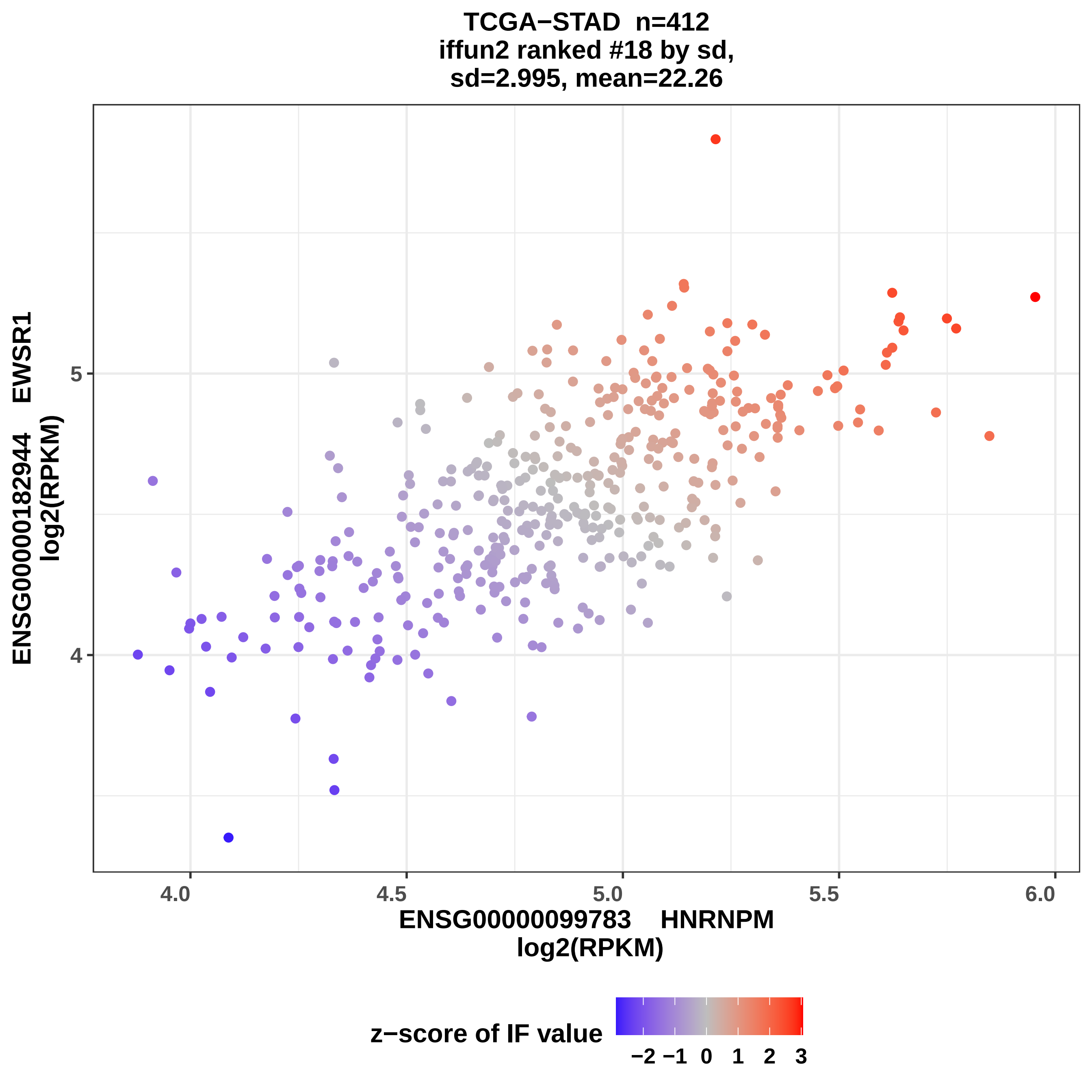
TCGA−STAD n=412
iffun2 ranked #18 by sd,
sd=2.995, mean=22.26
ENSG00000182944 EWSR1
log2(RPKM)
5
4
4.0
4.5
5.0
5.5
6.0
ENSG00000099783 HNRNPM
log2(RPKM)
z−score of IF value
−2 −1 0 1 2 3

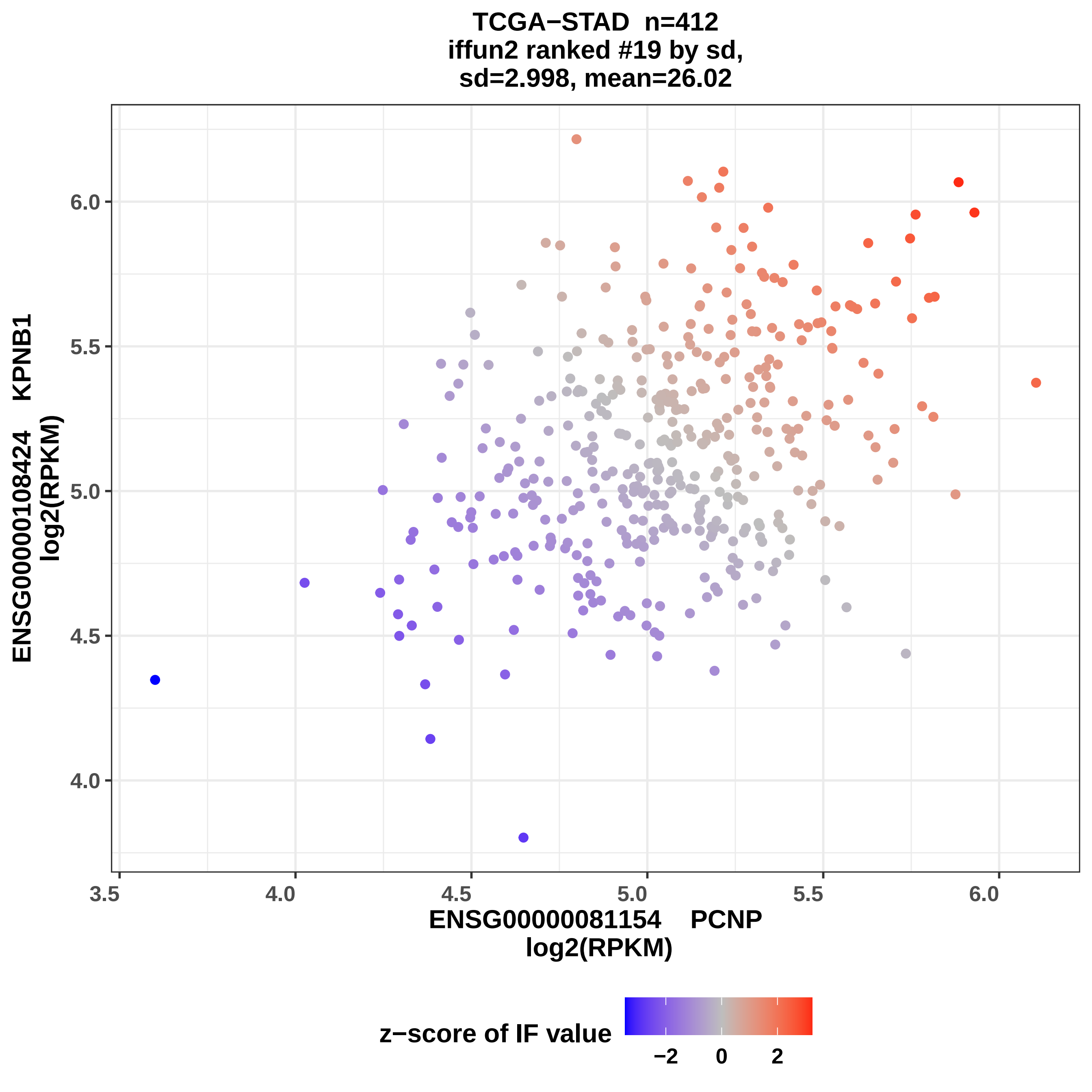

TCGA−STAD n=412
iffun2 ranked #19 by sd,
sd=2.998, mean=26.02
ENSG00000108424 KPNB1
log2(RPKM)
6.0
5.5
5.0
4.5
4.0
3.5
4.0
4.5
5.0
5.5
6.0
ENSG00000081154 PCNP
log2(RPKM)
z−score of IF value
−2
0
2

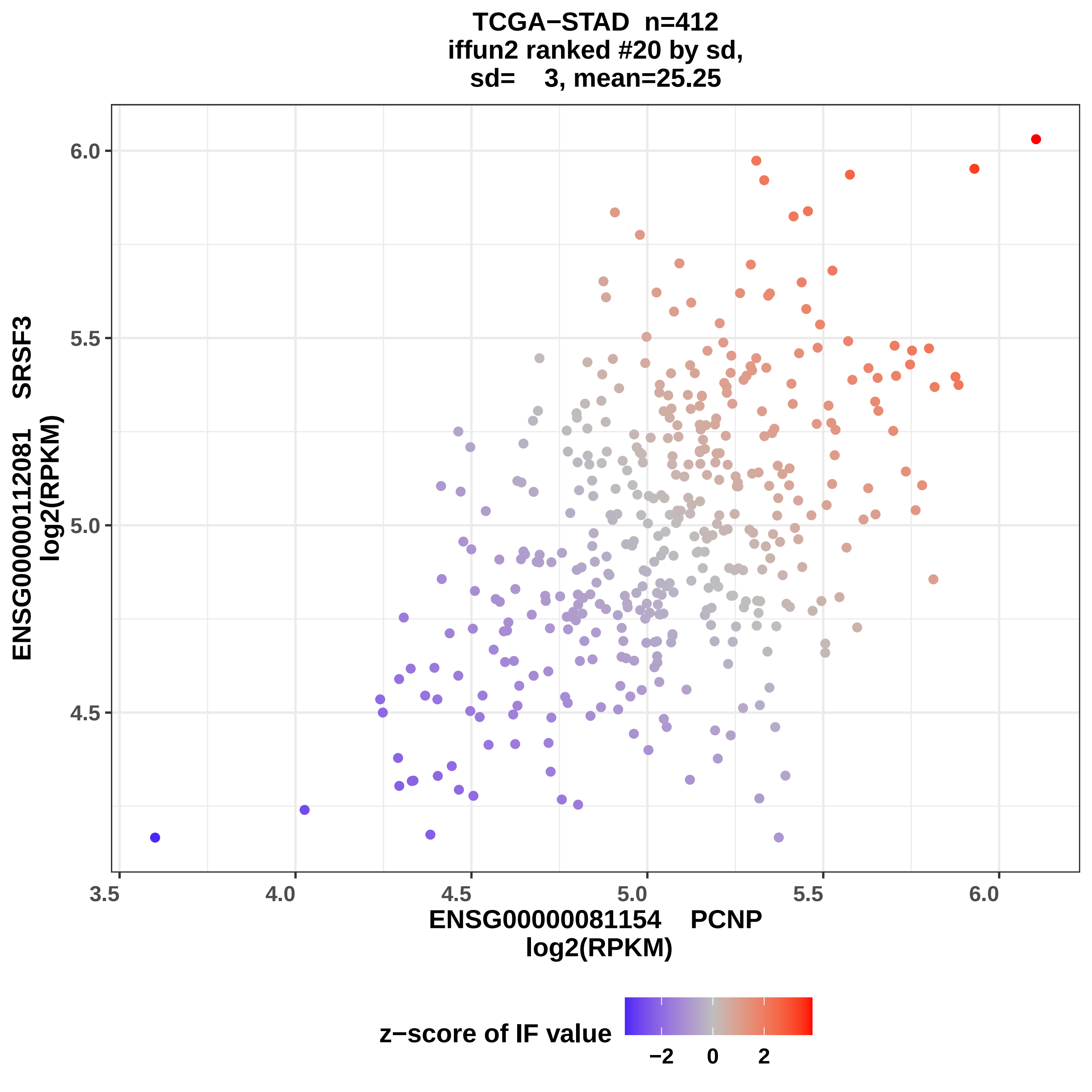

TCGA−STAD n=412
iffun2 ranked #20 by sd,
sd= 3, mean=25.25
ENSG00000112081 SRSF3
log2(RPKM)
6.0
5.5
5.0
4.5
3.5
4.0
4.5
5.0
5.5
6.0
ENSG00000081154 PCNP
log2(RPKM)
z−score of IF value
−2
0
2

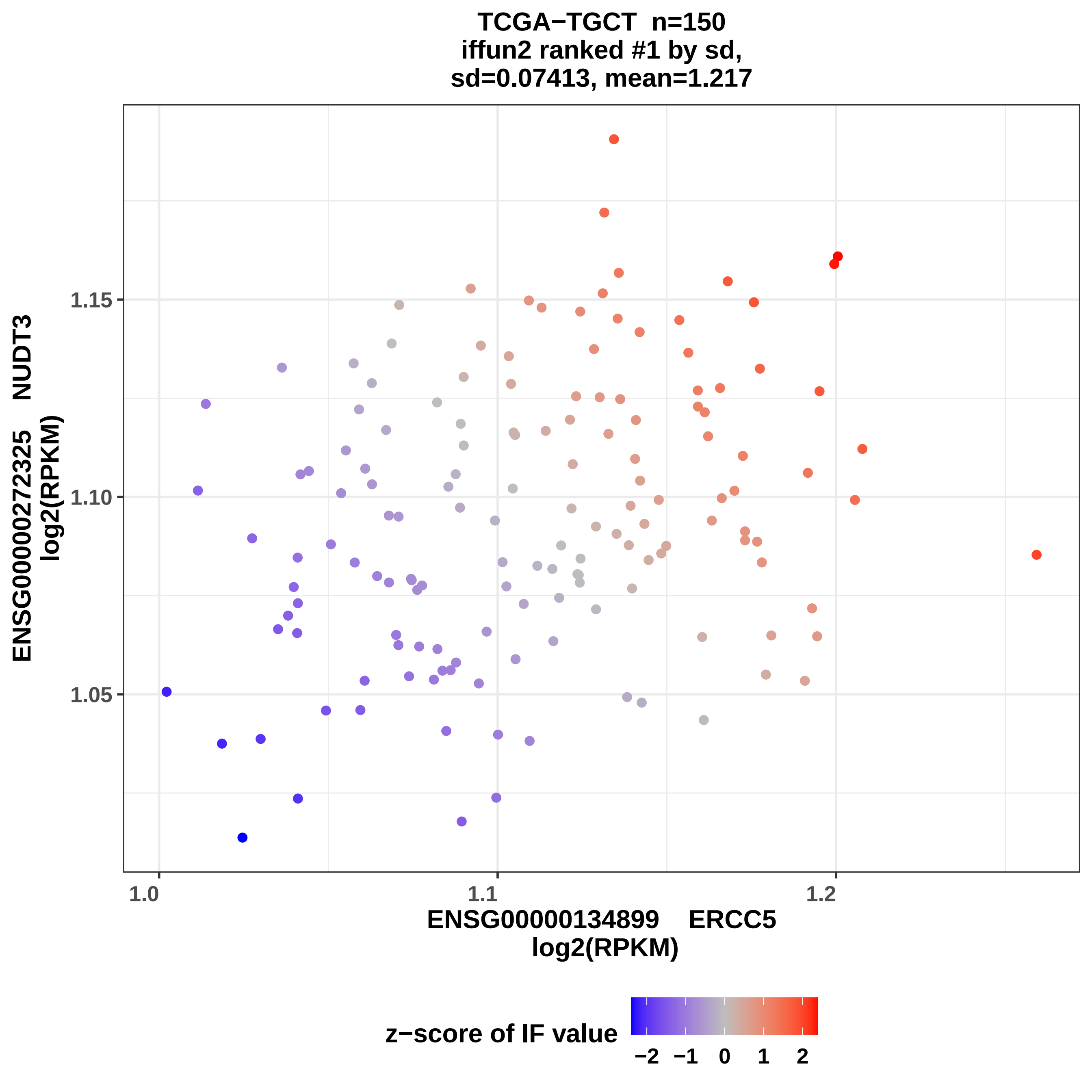

TCGA−TGCT n=150
iffun2 ranked #1 by sd,
sd=0.07413, mean=1.217
ENSG00000272325 NUDT3
log2(RPKM)
1.15
1.10
1.05
1.0
1.1
1.2
ENSG00000134899 ERCC5
log2(RPKM)
z−score of IF value
−2
−1
0
1
2

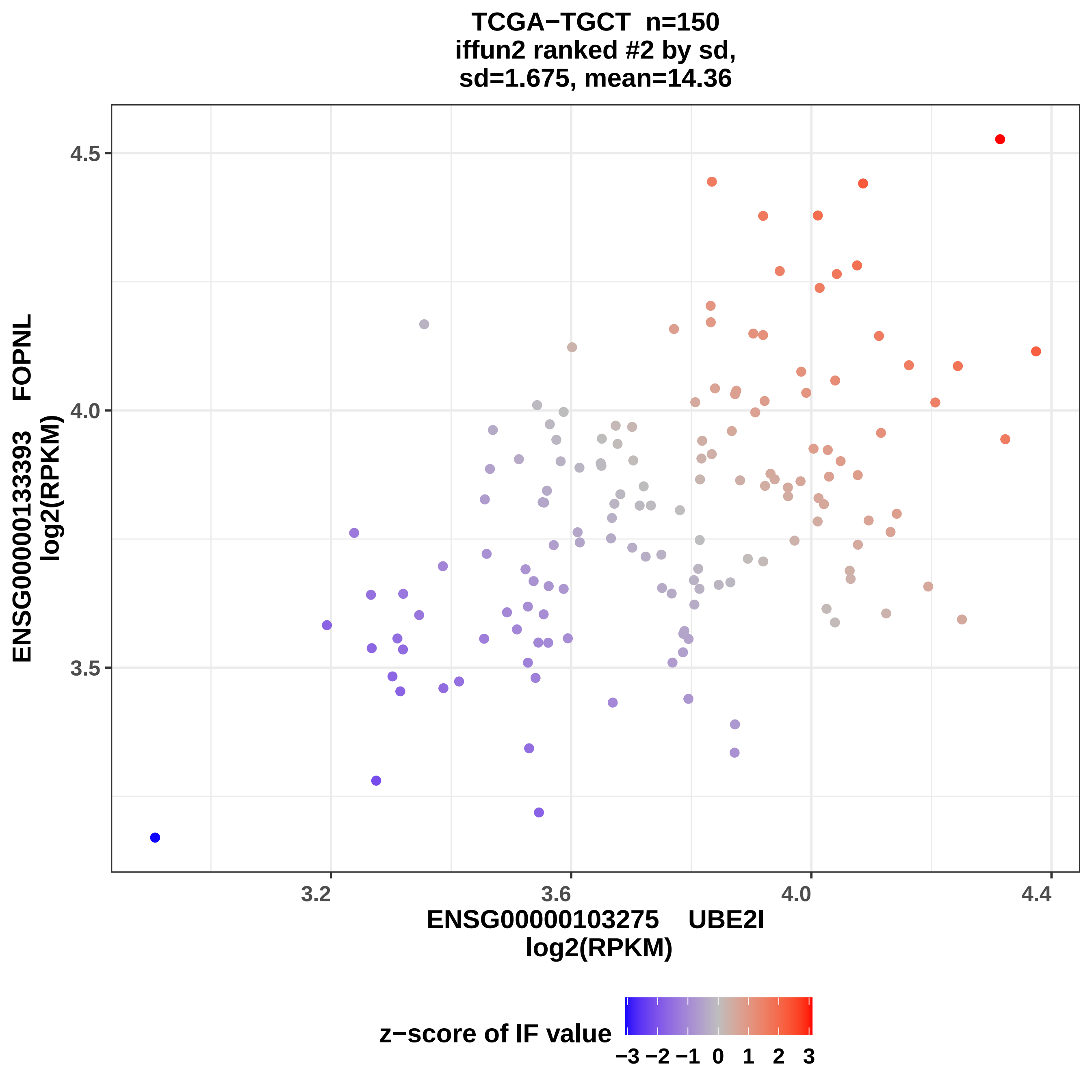

TCGA−TGCT n=150
iffun2 ranked #2 by sd,
sd=1.675, mean=14.36
ENSG00000133393 FOPNL
log2(RPKM)
4.5
4.0
3.5
3.2
3.6
4.0
4.4
ENSG00000103275 UBE2I
log2(RPKM)
z−score of IF value
−3 −2 −1 0 1 2 3

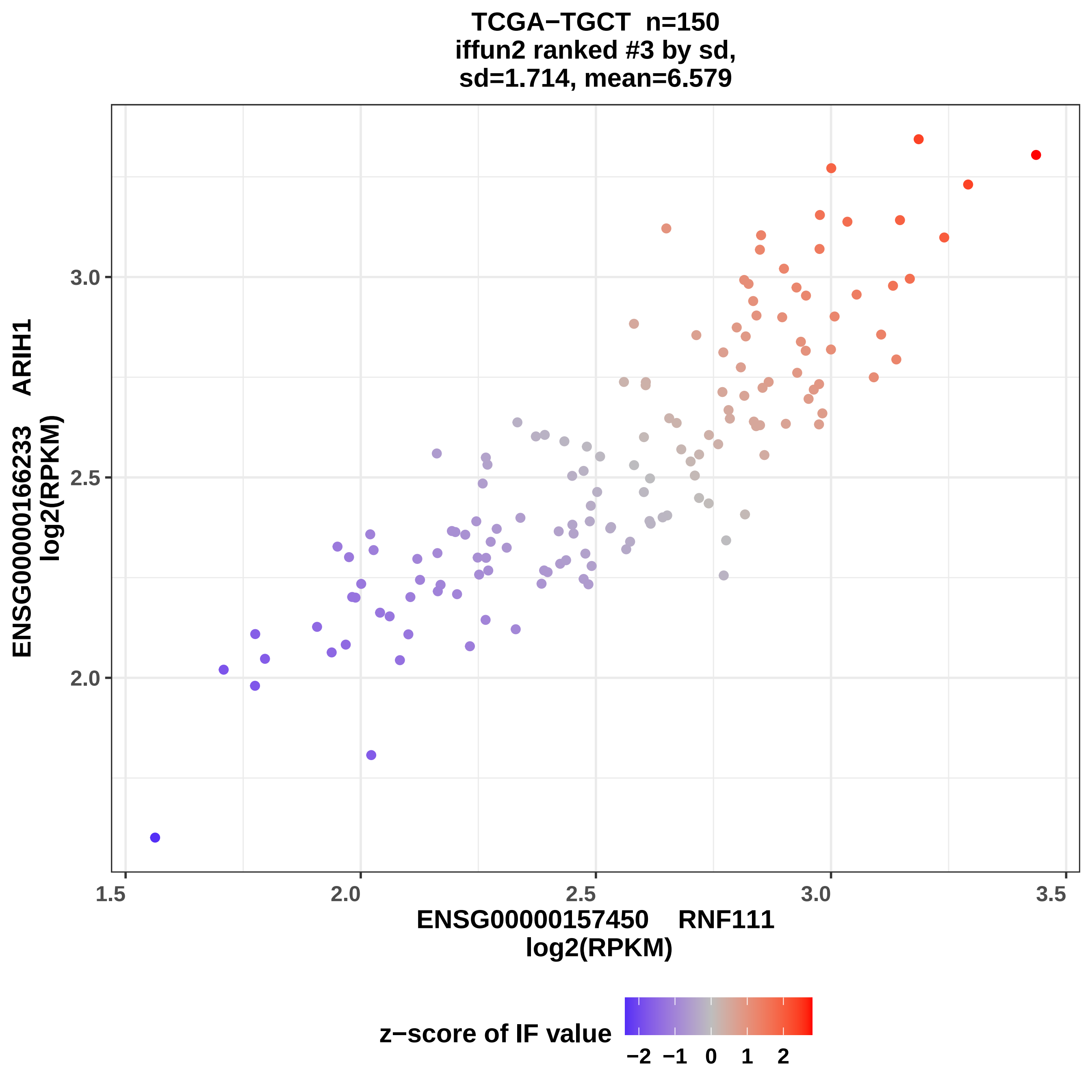
TCGA−TGCT n=150
iffun2 ranked #3 by sd,
sd=1.714, mean=6.579
ENSG00000166233 ARIH1
log2(RPKM)
3.0
2.5
2.0
1.5
2.0
2.5
3.0
3.5
ENSG00000157450 RNF111
log2(RPKM)
z−score of IF value
−2
−1
0
1
2

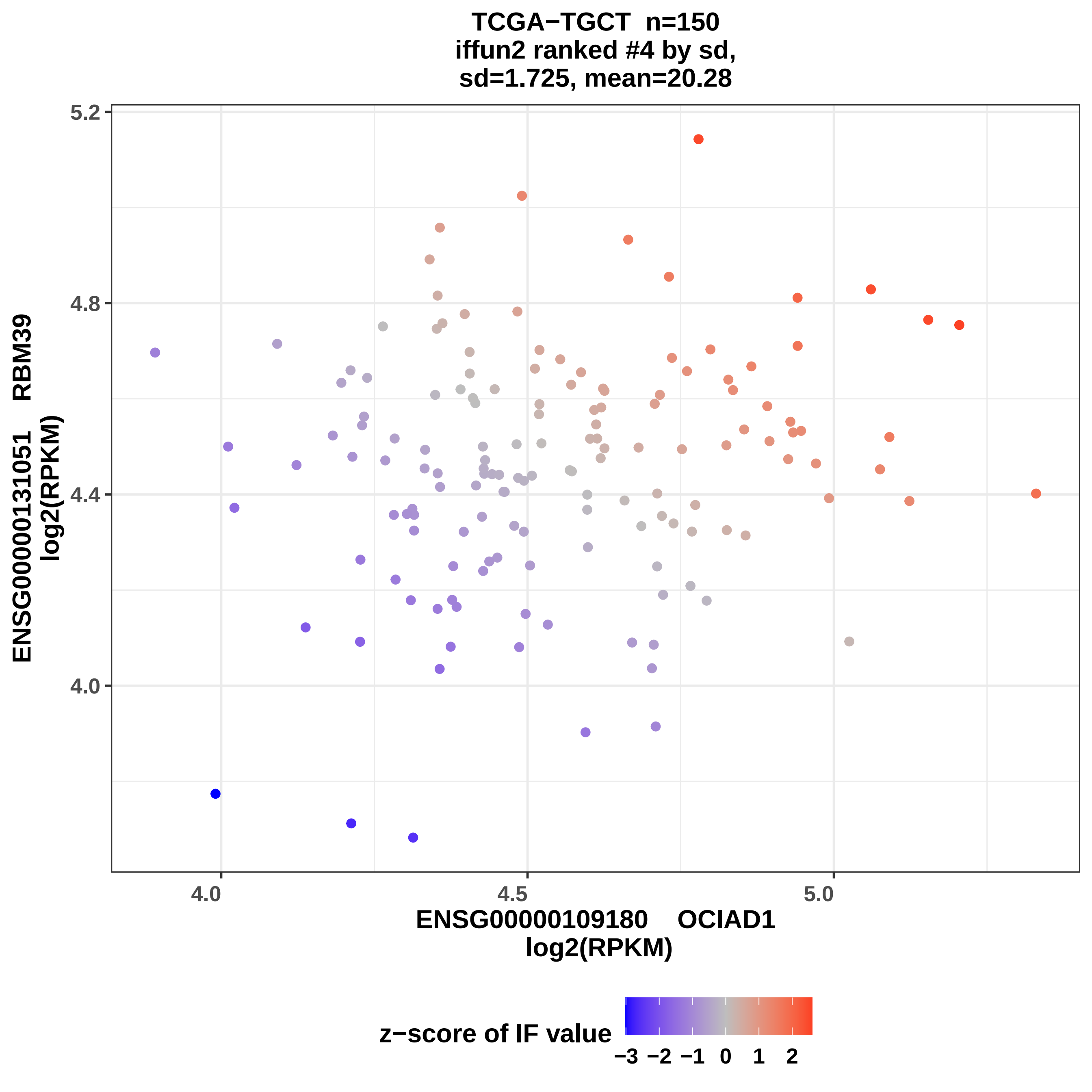
TCGA−TGCT n=150
iffun2 ranked #4 by sd,
sd=1.725, mean=20.28
ENSG00000131051 RBM39
log2(RPKM)
ENSG00000109180 OCIAD1
log2(RPKM)
5.2
4.8
4.4
4.0
4.0
4.5
5.0
z−score of IF value
−3 −2 −1 0 1 2

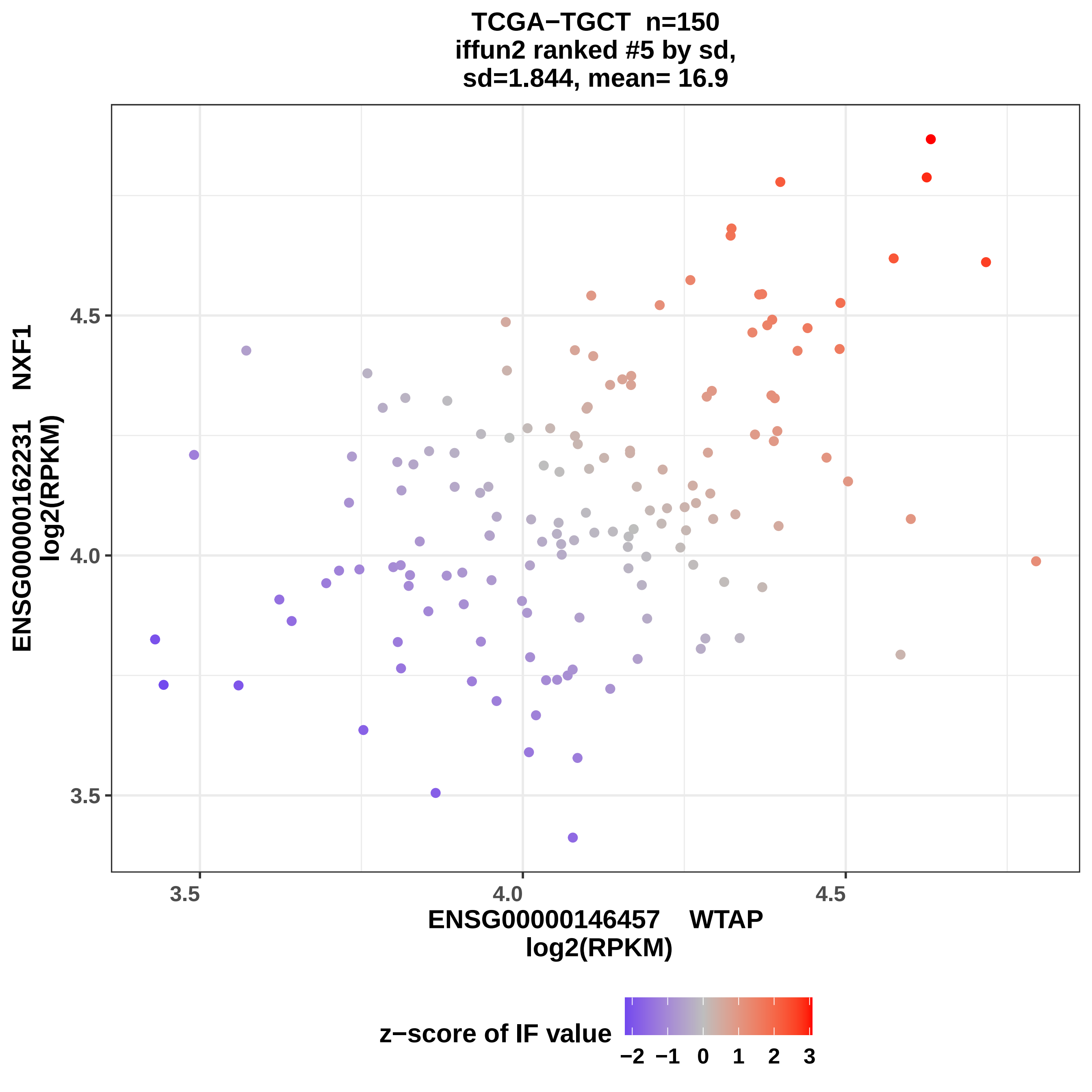

TCGA−TGCT n=150
iffun2 ranked #5 by sd,
sd=1.844, mean= 16.9
ENSG00000162231 NXF1
log2(RPKM)
4.5
4.0
3.5
3.5
4.0
4.5
ENSG00000146457 WTAP
log2(RPKM)
z−score of IF value
−2 −1 0 1 2 3

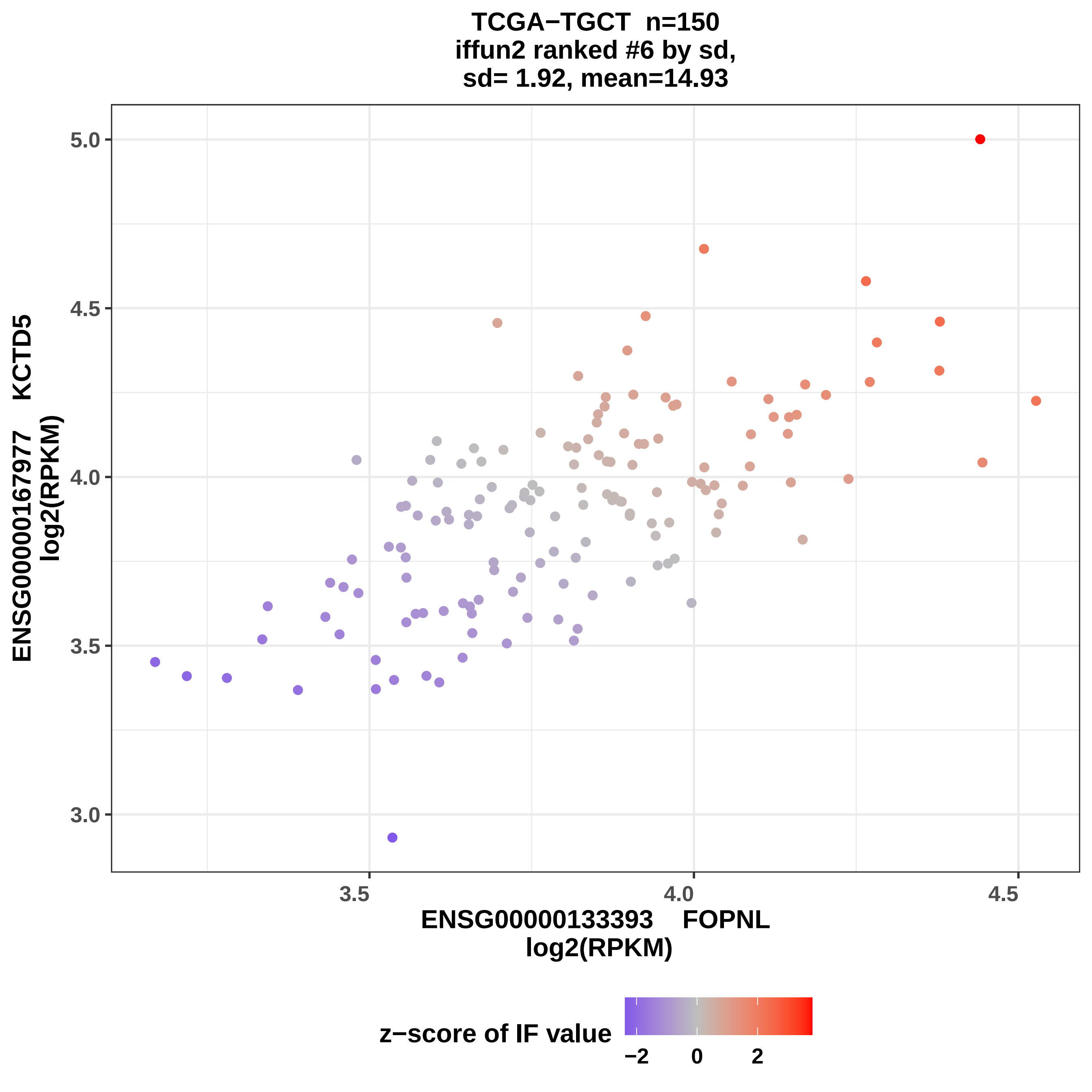

TCGA−TGCT n=150
iffun2 ranked #6 by sd,
sd= 1.92, mean=14.93
5.0
4.5
4.0
3.5
3.0
3.5
4.0
4.5
ENSG00000167977 KCTD5
log2(RPKM)
ENSG00000133393 FOPNL
log2(RPKM)
z−score of IF value
−2
0
2

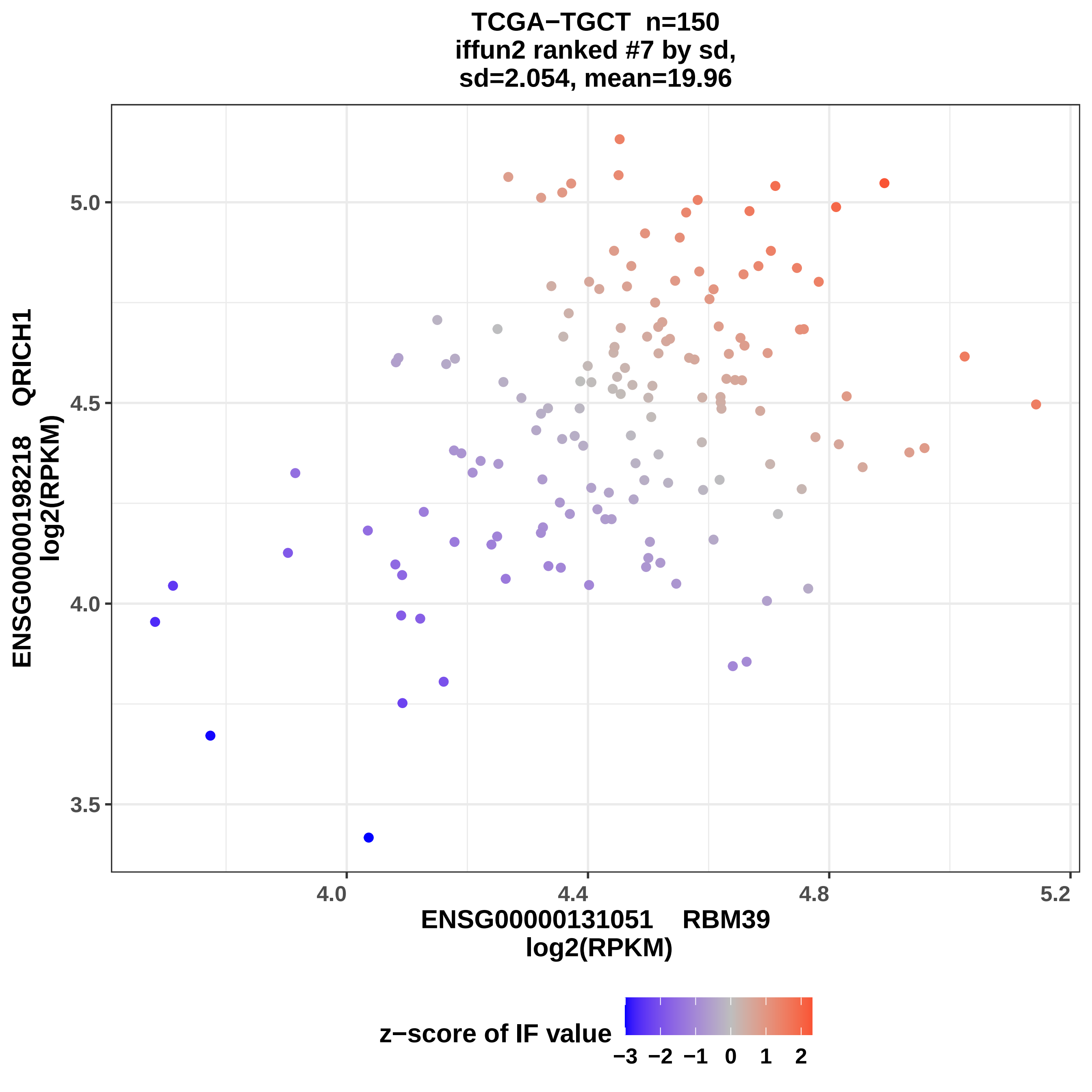

TCGA−TGCT n=150
iffun2 ranked #7 by sd,
sd=2.054, mean=19.96
ENSG00000198218 QRICH1
log2(RPKM)
5.0
4.5
4.0
3.5
4.0
4.4
4.8
5.2
ENSG00000131051 RBM39
log2(RPKM)
z−score of IF value
−3
−2
−1
0
1
2

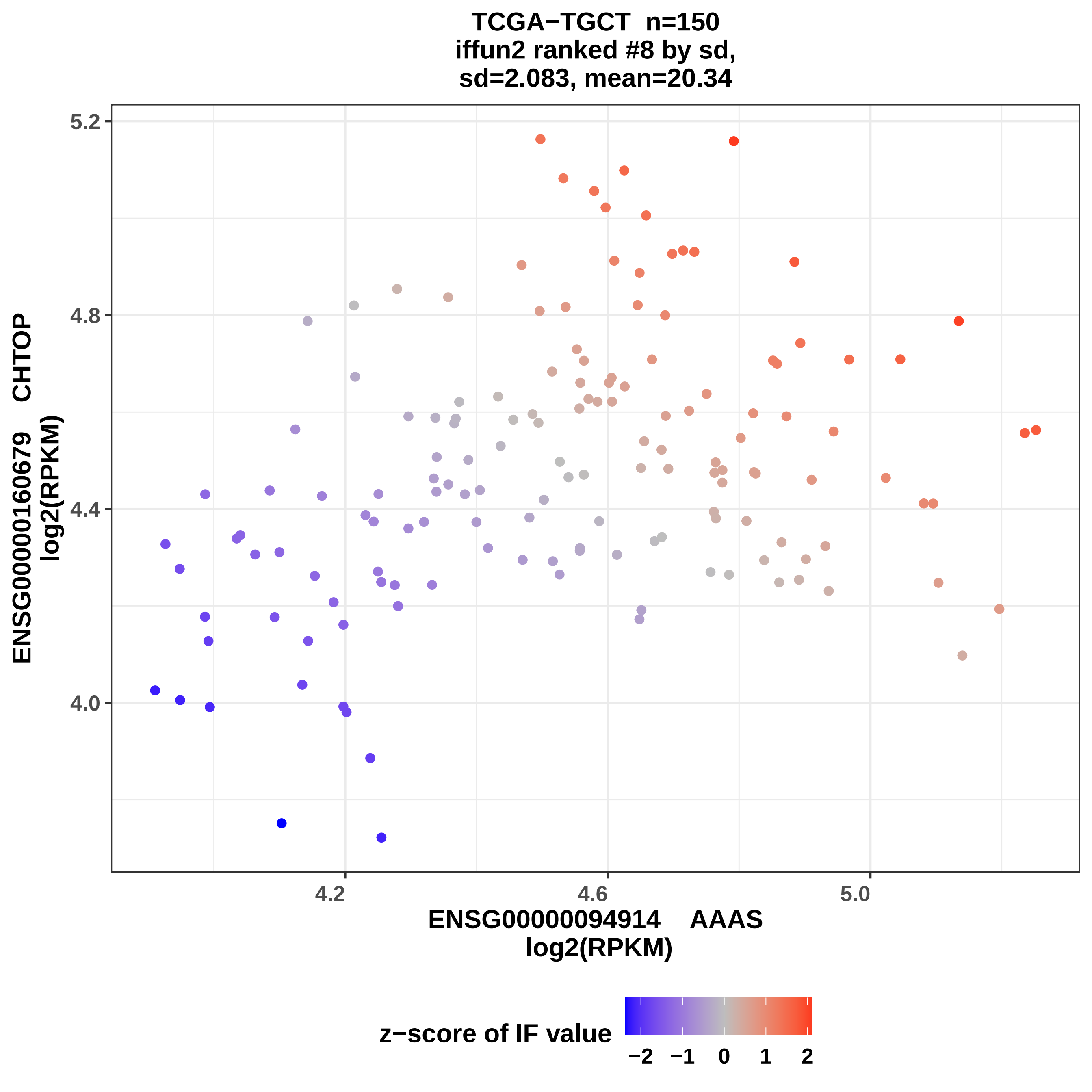

TCGA−TGCT n=150
iffun2 ranked #8 by sd,
sd=2.083, mean=20.34
ENSG00000160679 CHTOP
log2(RPKM)
5.2
4.8
4.4
4.0
4.2
4.6
5.0
ENSG00000094914 AAAS
log2(RPKM)
z−score of IF value
−2
−1
0
1
2

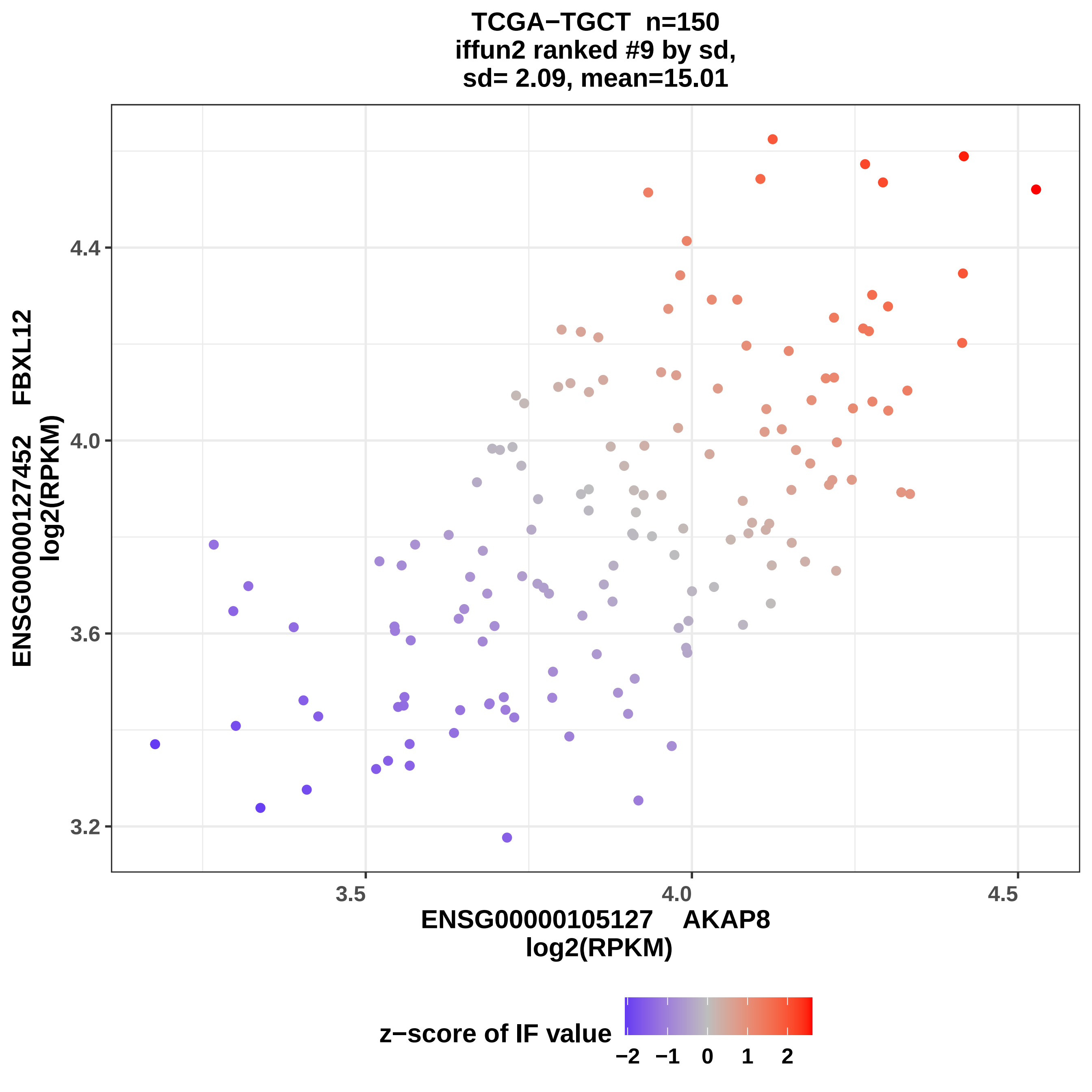

TCGA−TGCT n=150
iffun2 ranked #9 by sd,
sd= 2.09, mean=15.01
ENSG00000127452 FBXL12
log2(RPKM)
4.4
4.0
3.6
3.2
3.5
4.0
4.5
ENSG00000105127 AKAP8
log2(RPKM)
z−score of IF value
−2
−1
0
1
2

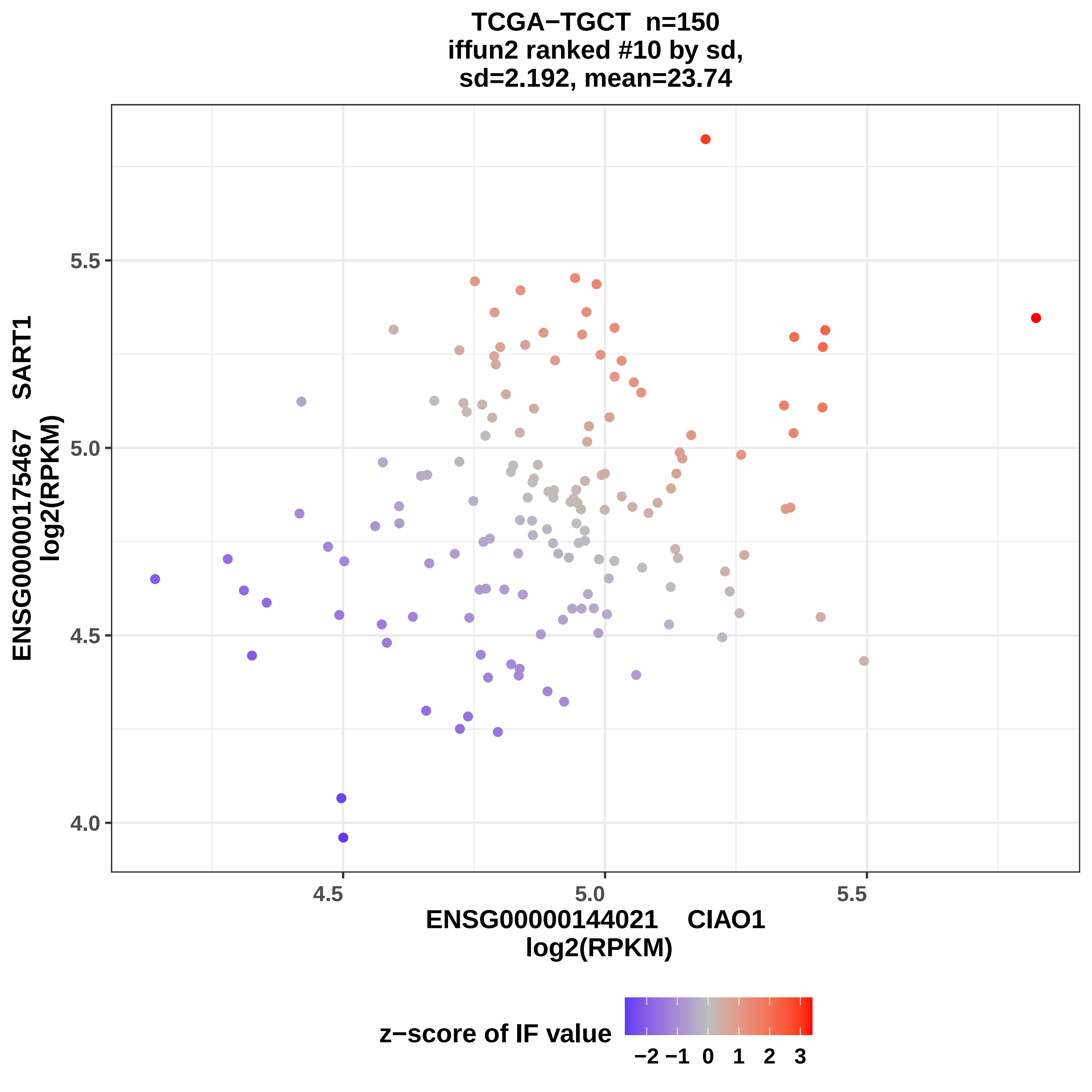
TCGA−TGCT n=150
iffun2 ranked #10 by sd,
sd=2.192, mean=23.74
ENSG00000175467 SART1
log2(RPKM)
5.5
5.0
4.5
4.0
4.5
5.0
5.5
ENSG00000144021 CIAO1
log2(RPKM)
z−score of IF value
−2 −1 0 1 2 3

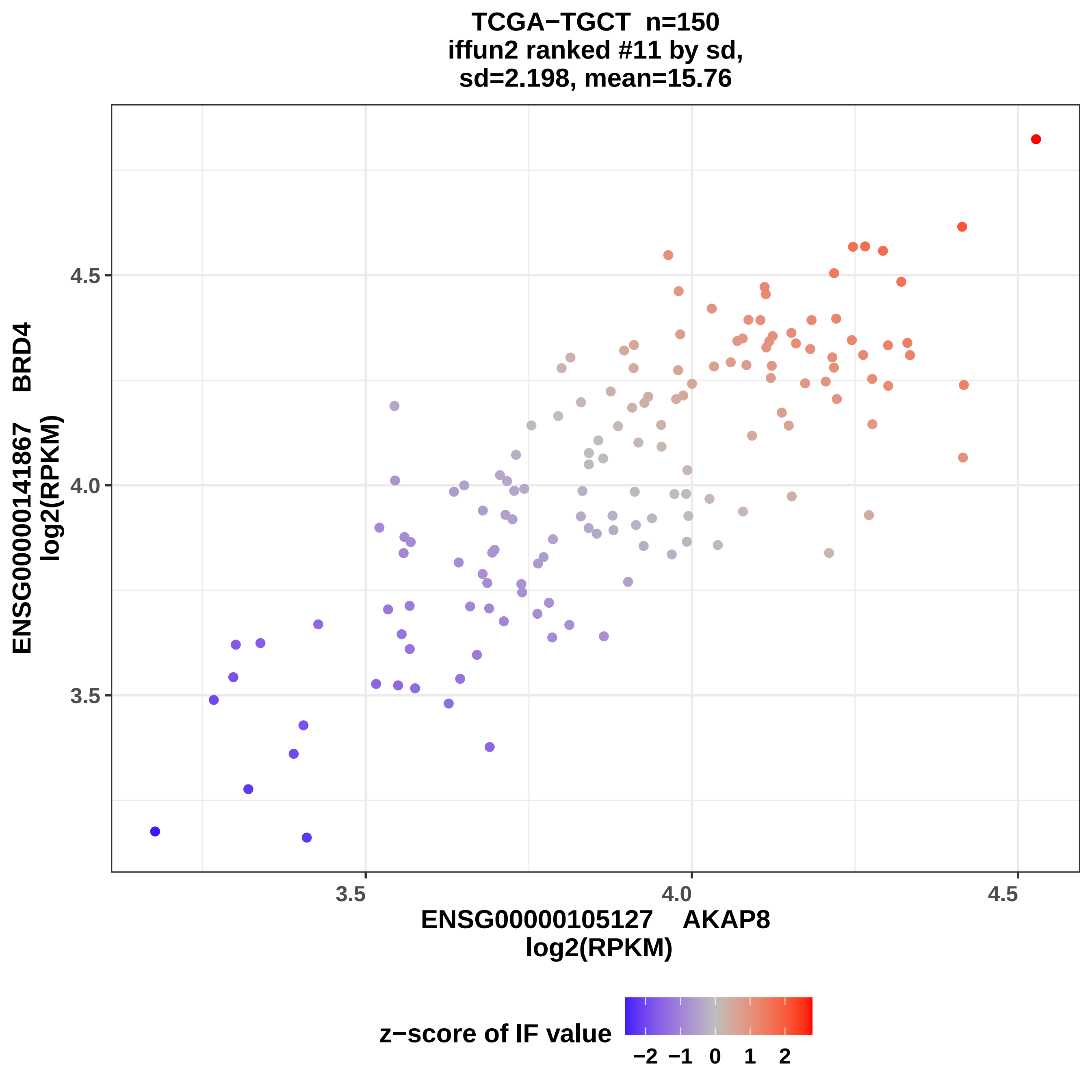

TCGA−TGCT n=150
iffun2 ranked #11 by sd,
sd=2.198, mean=15.76
ENSG00000141867 BRD4
log2(RPKM)
4.5
4.0
3.5
3.5
4.0
4.5
ENSG00000105127 AKAP8
log2(RPKM)
z−score of IF value
−2 −1 0 1 2

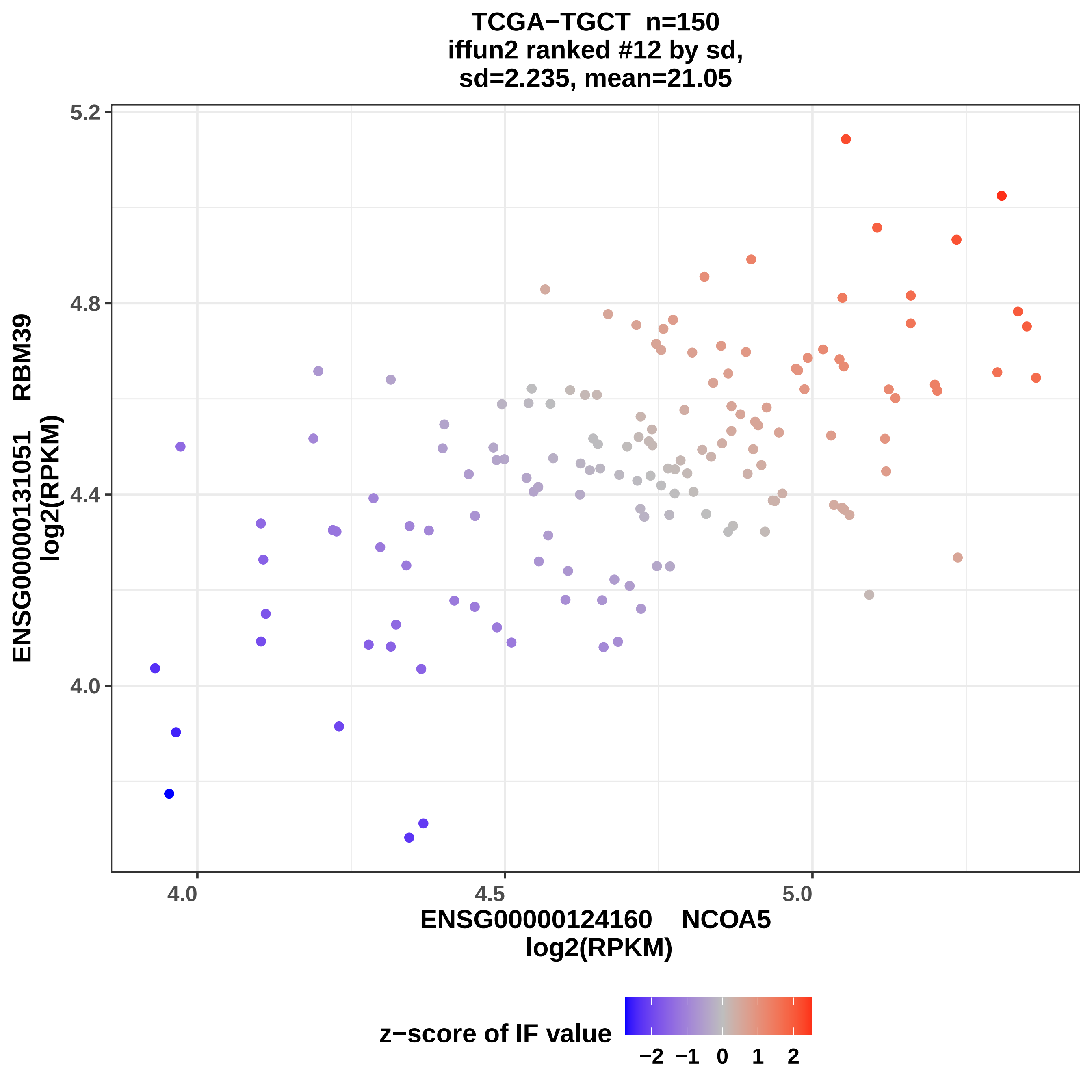
TCGA−TGCT n=150
iffun2 ranked #12 by sd,
sd=2.235, mean=21.05
ENSG00000131051 RBM39
log2(RPKM)
ENSG00000124160 NCOA5
log2(RPKM)
5.2
4.8
4.4
4.0
4.0
4.5
5.0
z−score of IF value
−2
−1
0
1
2

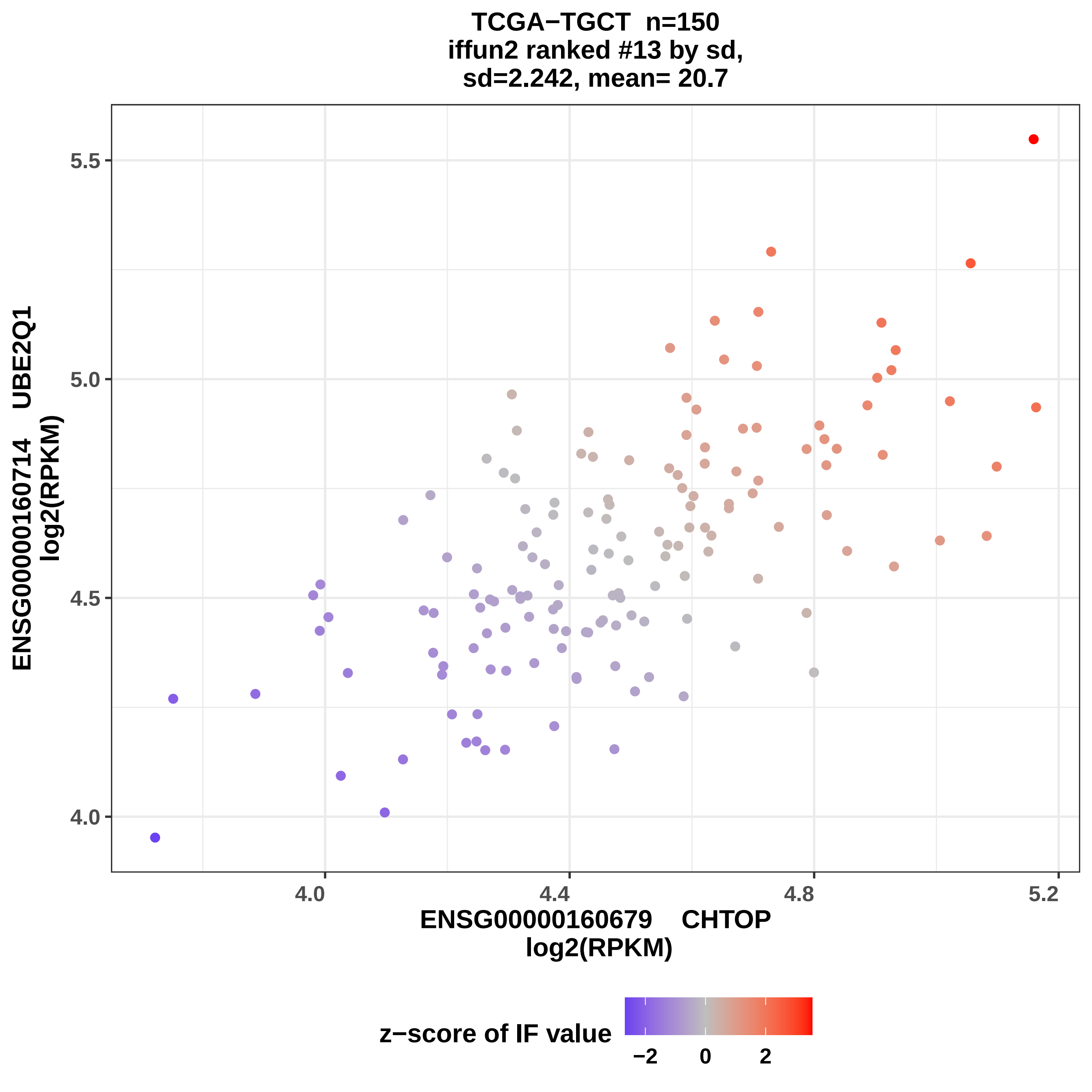

TCGA−TGCT n=150
iffun2 ranked #13 by sd,
sd=2.242, mean= 20.7
ENSG00000160714 UBE2Q1
log2(RPKM)
5.5
5.0
4.5
4.0
4.0
4.4
4.8
5.2
ENSG00000160679 CHTOP
log2(RPKM)
z−score of IF value
−2
0
2

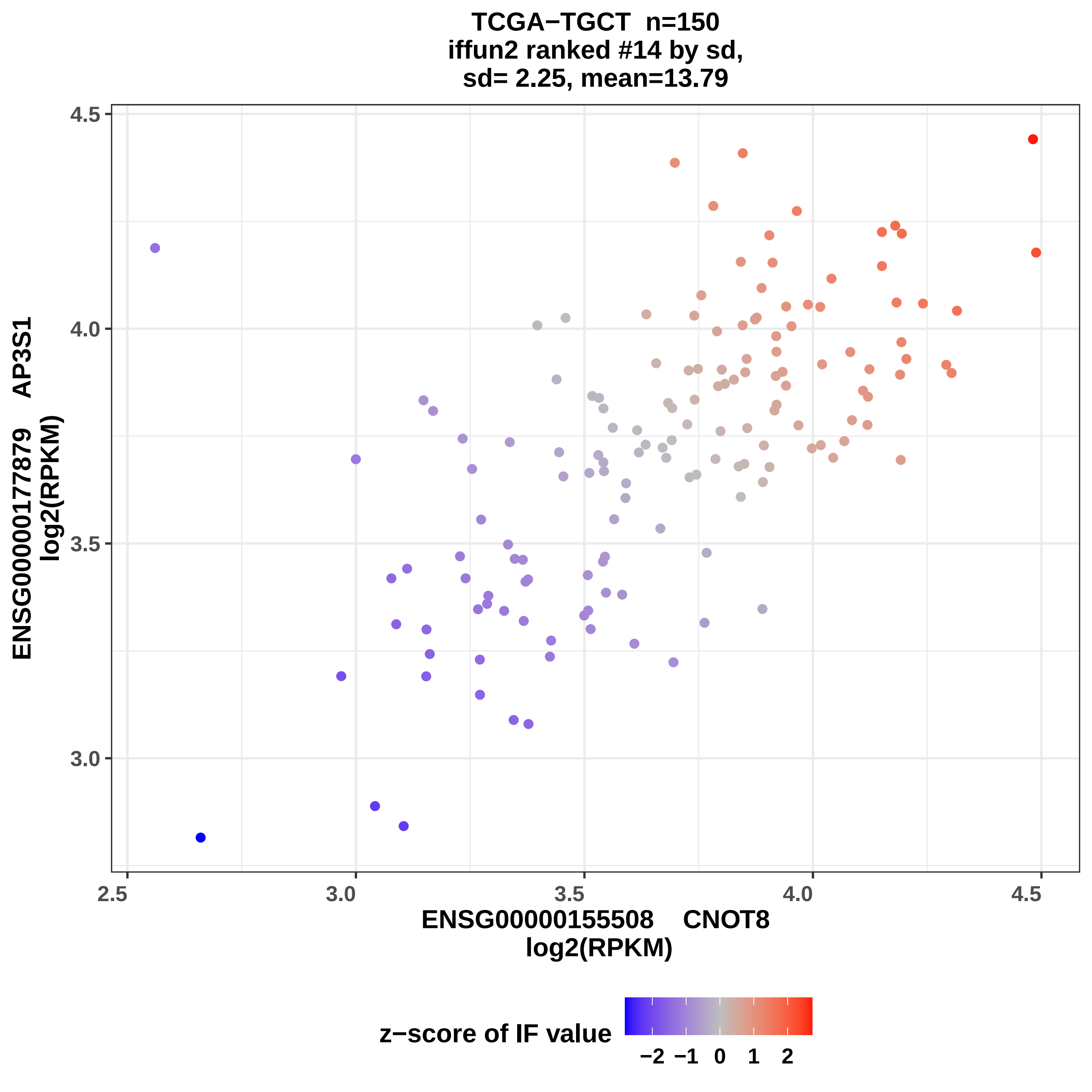
TCGA−TGCT n=150
iffun2 ranked #14 by sd,
sd= 2.25, mean=13.79
ENSG00000177879 AP3S1
log2(RPKM)
4.5
4.0
3.5
3.0
2.5
3.0
3.5
4.0
4.5
ENSG00000155508 CNOT8
log2(RPKM)
z−score of IF value
−2 −1 0 1 2

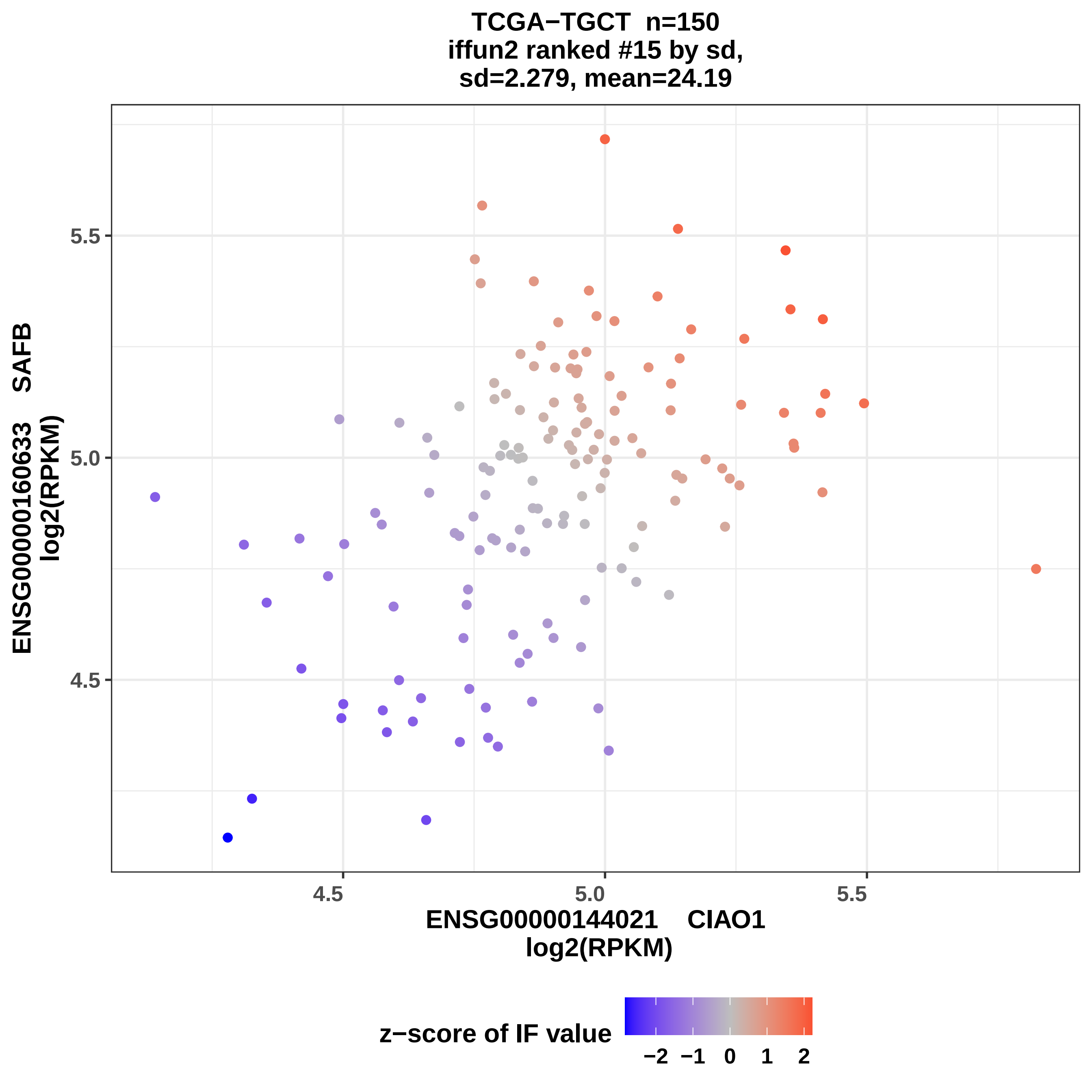
TCGA−TGCT n=150
iffun2 ranked #15 by sd,
sd=2.279, mean=24.19
ENSG00000160633 SAFB
log2(RPKM)
5.5
5.0
4.5
4.5
5.0
5.5
ENSG00000144021 CIAO1
log2(RPKM)
z−score of IF value
−2 −1 0 1 2

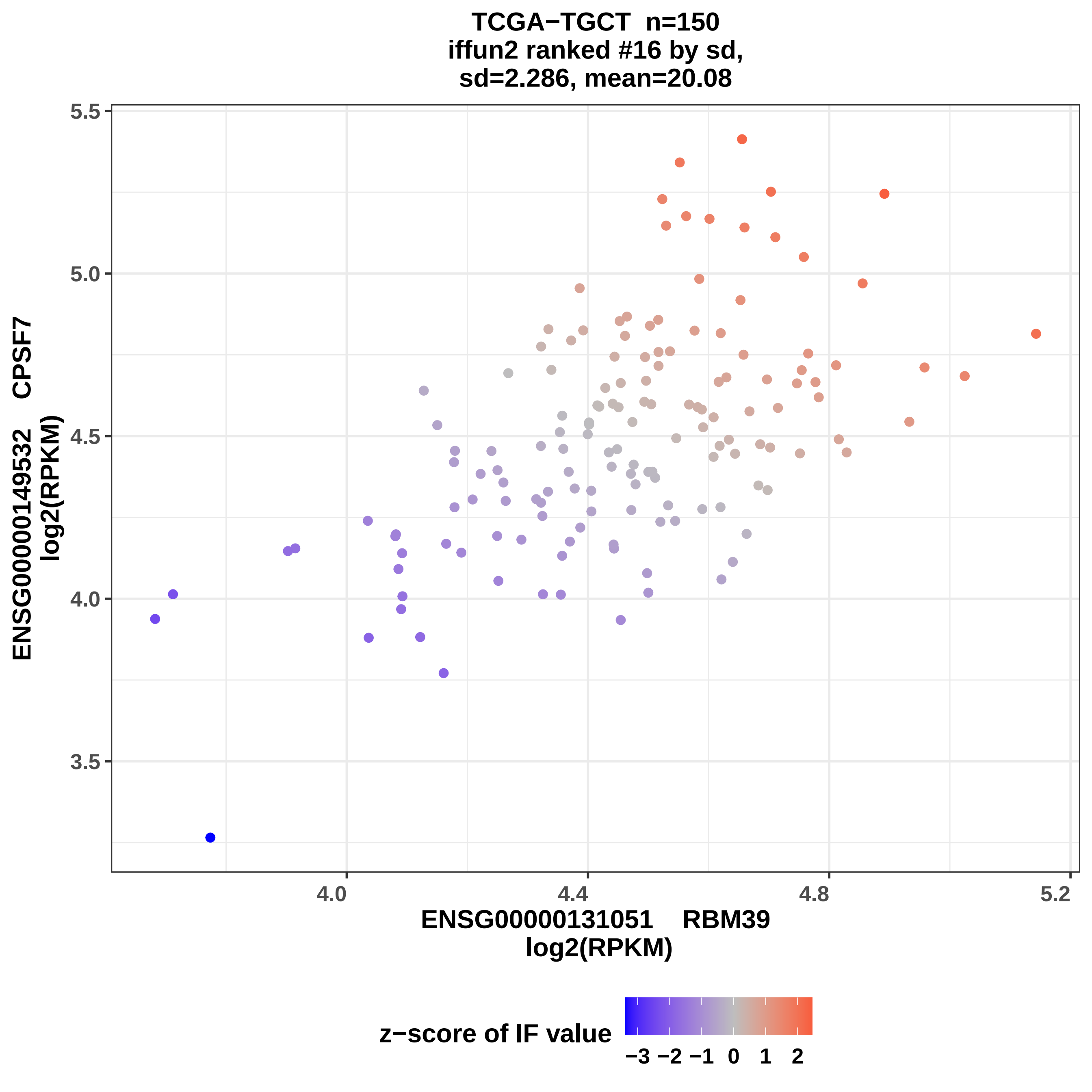

TCGA−TGCT n=150
iffun2 ranked #16 by sd,
sd=2.286, mean=20.08
ENSG00000149532 CPSF7
log2(RPKM)
5.5
5.0
4.5
4.0
3.5
4.0
4.4
4.8
5.2
ENSG00000131051 RBM39
log2(RPKM)
z−score of IF value
−3 −2 −1 0 1 2

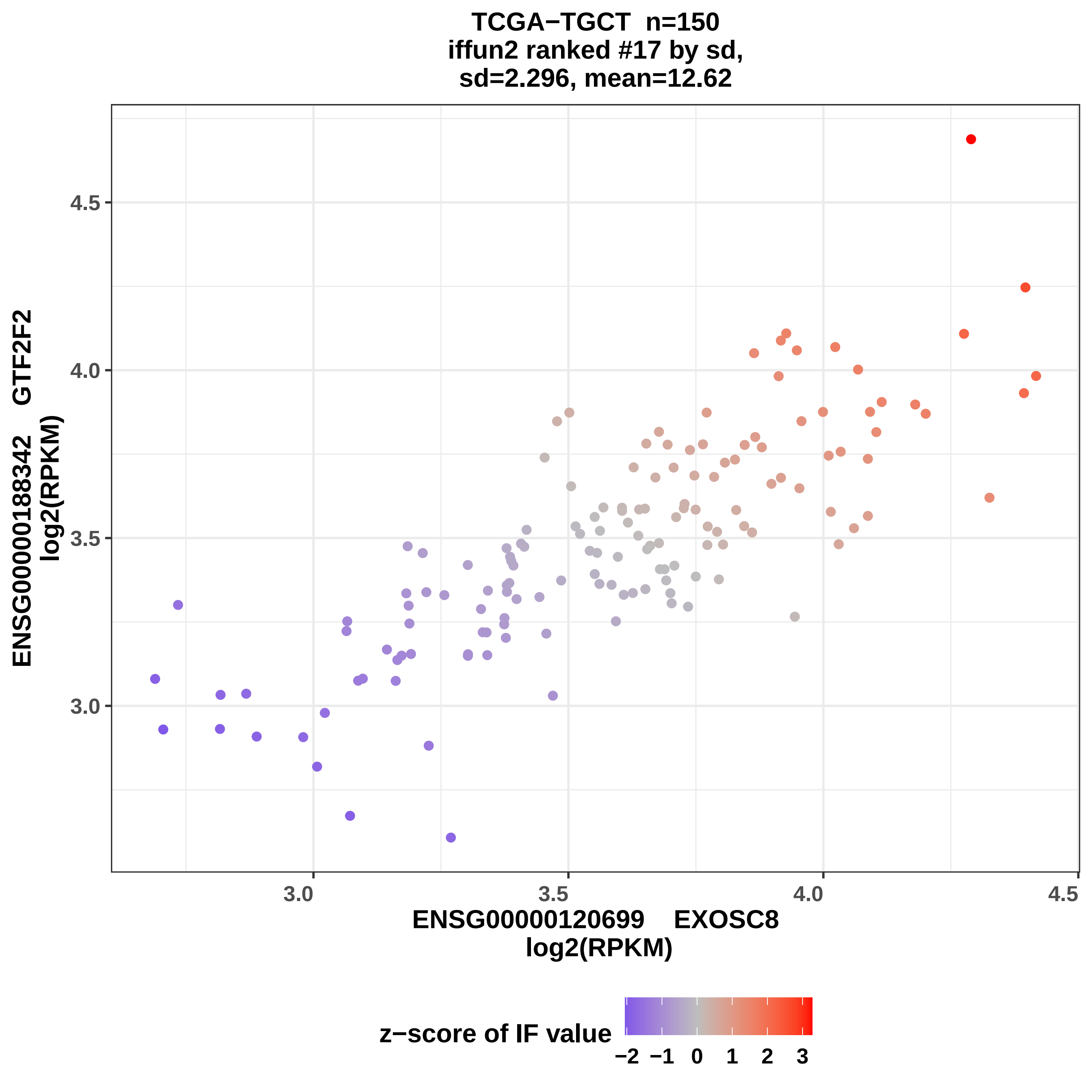
TCGA−TGCT n=150
iffun2 ranked #17 by sd,
sd=2.296, mean=12.62
ENSG00000188342 GTF2F2
log2(RPKM)
4.5
4.0
3.5
3.0
3.0
3.5
4.0
4.5
ENSG00000120699 EXOSC8
log2(RPKM)
z−score of IF value
−2 −1 0 1 2 3

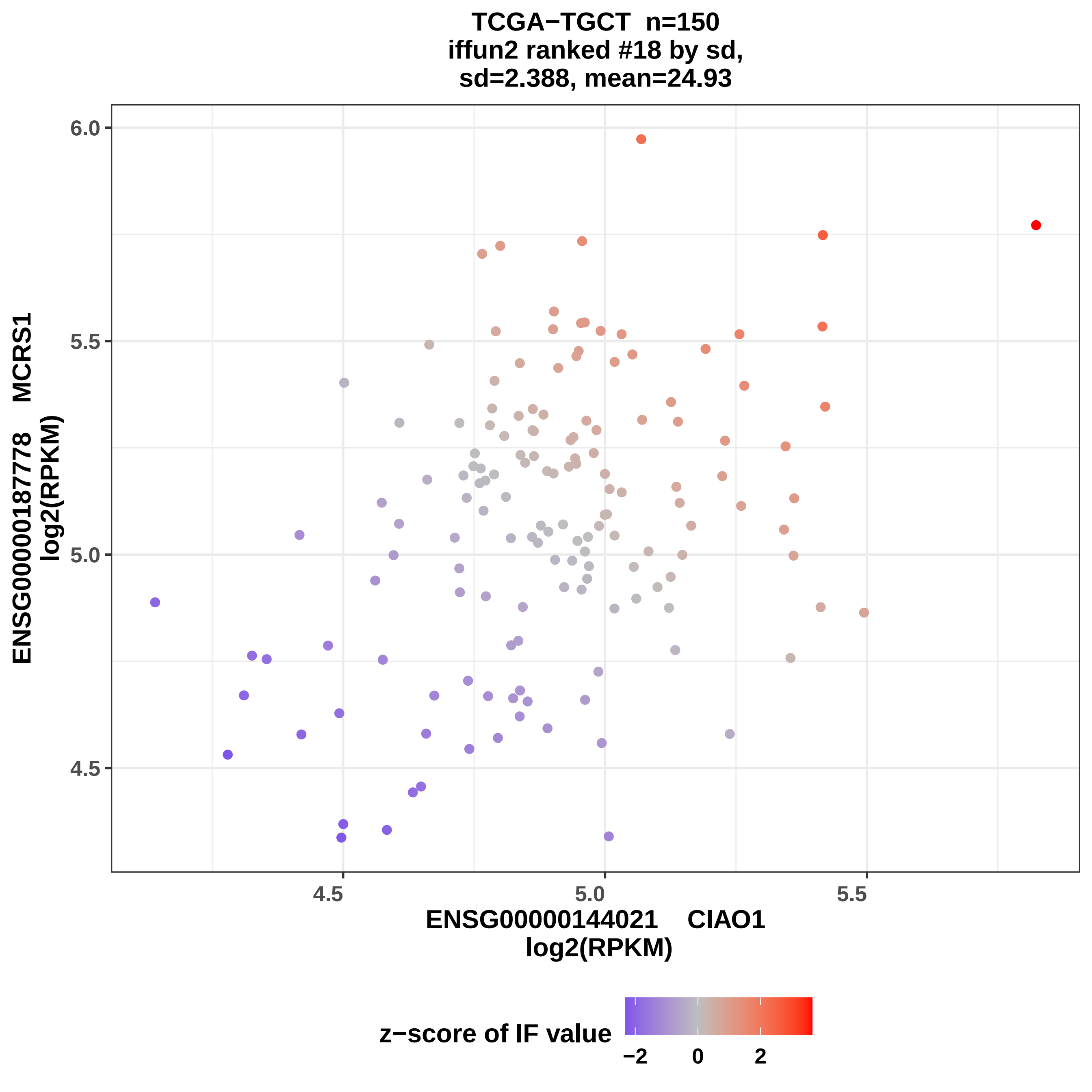
TCGA−TGCT n=150
iffun2 ranked #18 by sd,
sd=2.388, mean=24.93
ENSG00000187778 MCRS1
log2(RPKM)
6.0
5.5
5.0
4.5
ENSG00000144021 CIAO1
log2(RPKM)
z−score of IF value
−2
0
2

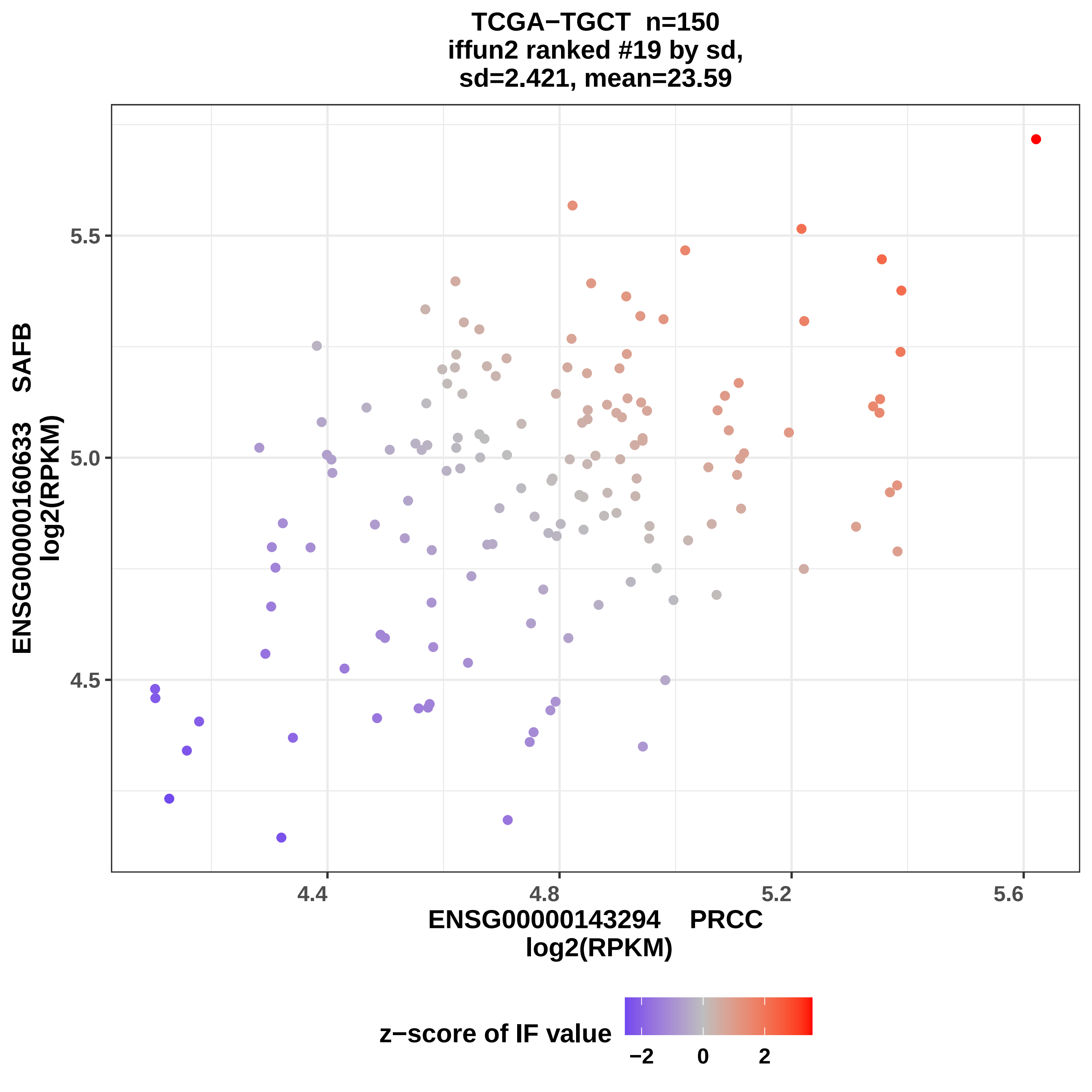
TCGA−TGCT n=150
iffun2 ranked #19 by sd,
sd=2.421, mean=23.59
ENSG00000160633 SAFB
log2(RPKM)
5.5
5.0
4.5
4.4
4.8
5.2
5.6
ENSG00000143294 PRCC
log2(RPKM)
z−score of IF value
−2
0
2

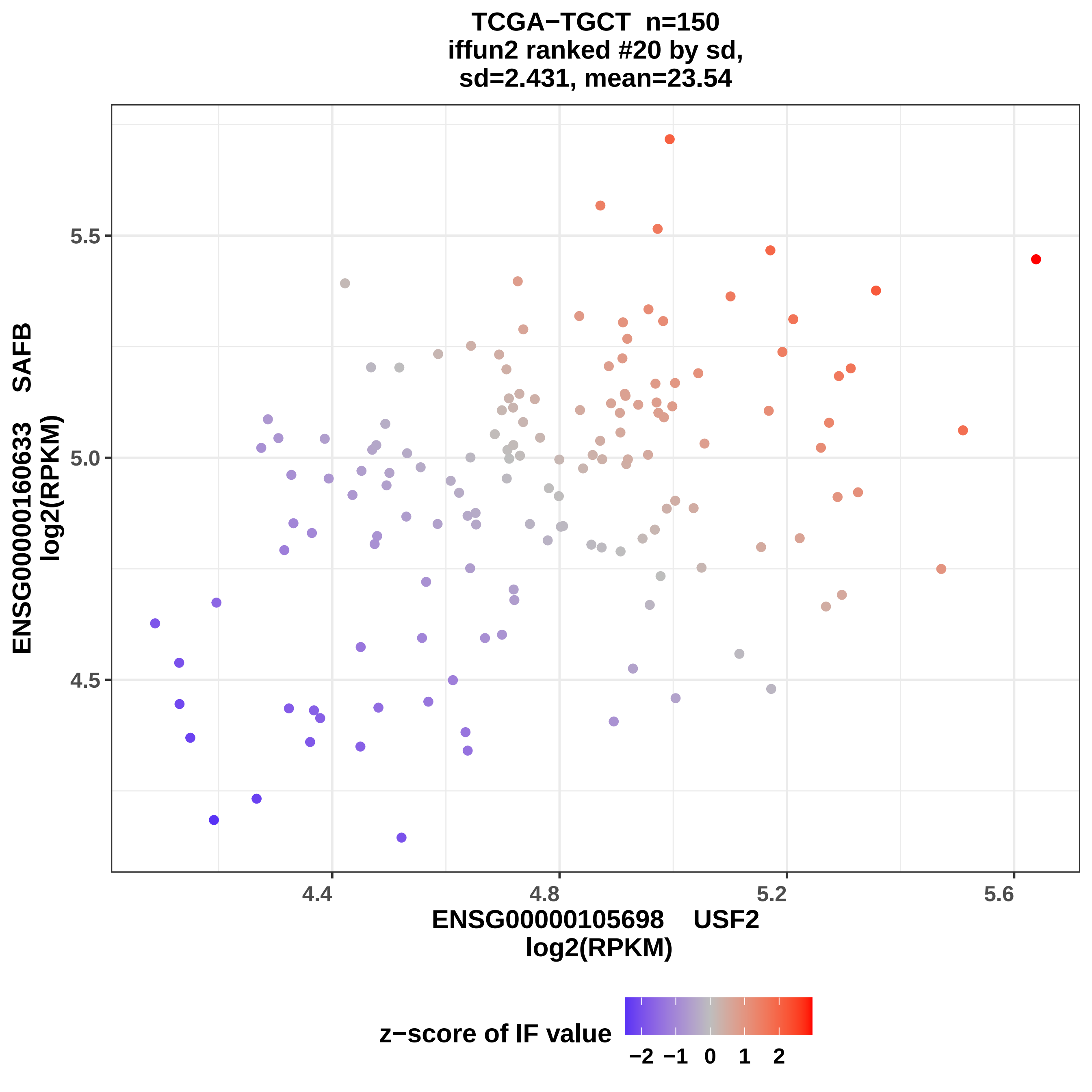
TCGA−TGCT n=150
iffun2 ranked #20 by sd,
sd=2.431, mean=23.54
ENSG00000160633 SAFB
log2(RPKM)
5.5
5.0
4.5
4.4
4.8
5.2
5.6
ENSG00000105698 USF2
log2(RPKM)
z−score of IF value
−2 −1 0 1 2

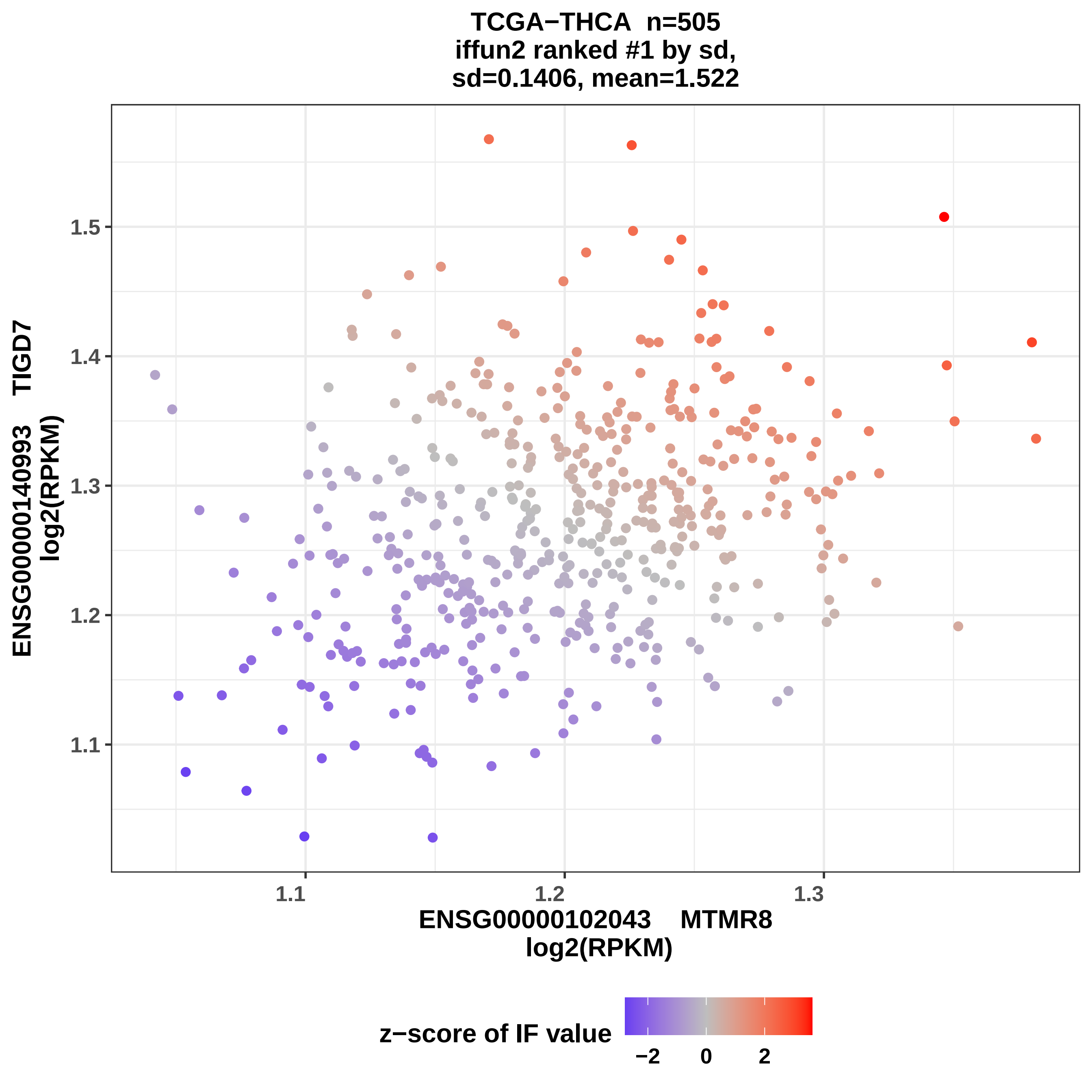

TCGA−THCA n=505
iffun2 ranked #1 by sd,
sd=0.1406, mean=1.522
ENSG00000140993 TIGD7
log2(RPKM)
1.5
1.4
1.3
1.2
1.1
1.1
1.2
1.3
ENSG00000102043 MTMR8
log2(RPKM)
z−score of IF value
−2
0
2

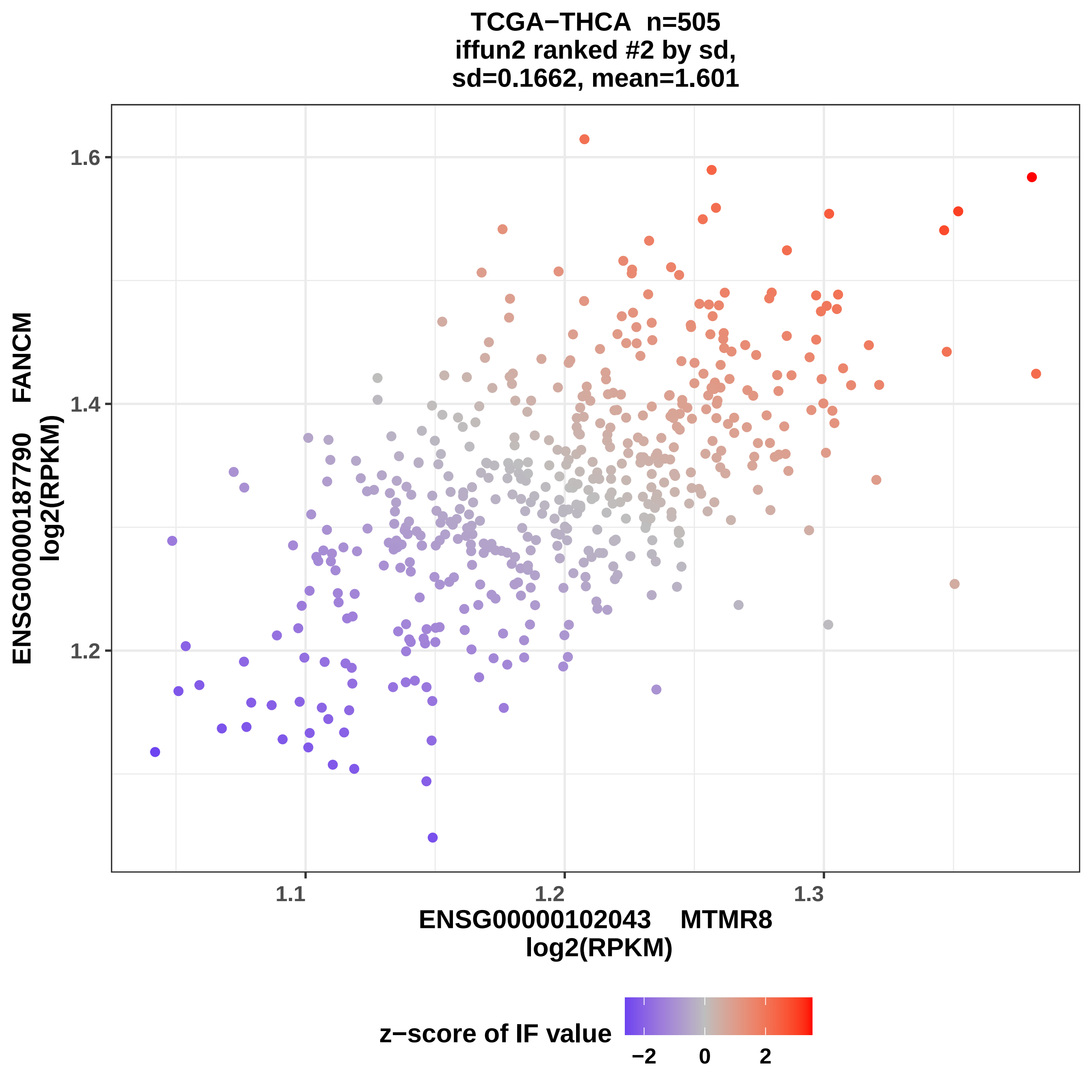
TCGA−THCA n=505
iffun2 ranked #2 by sd,
sd=0.1662, mean=1.601
1.6
1.4
1.2
1.1
1.2
1.3
ENSG00000187790 FANCM
log2(RPKM)
ENSG00000102043 MTMR8
log2(RPKM)
z−score of IF value
−2
0
2

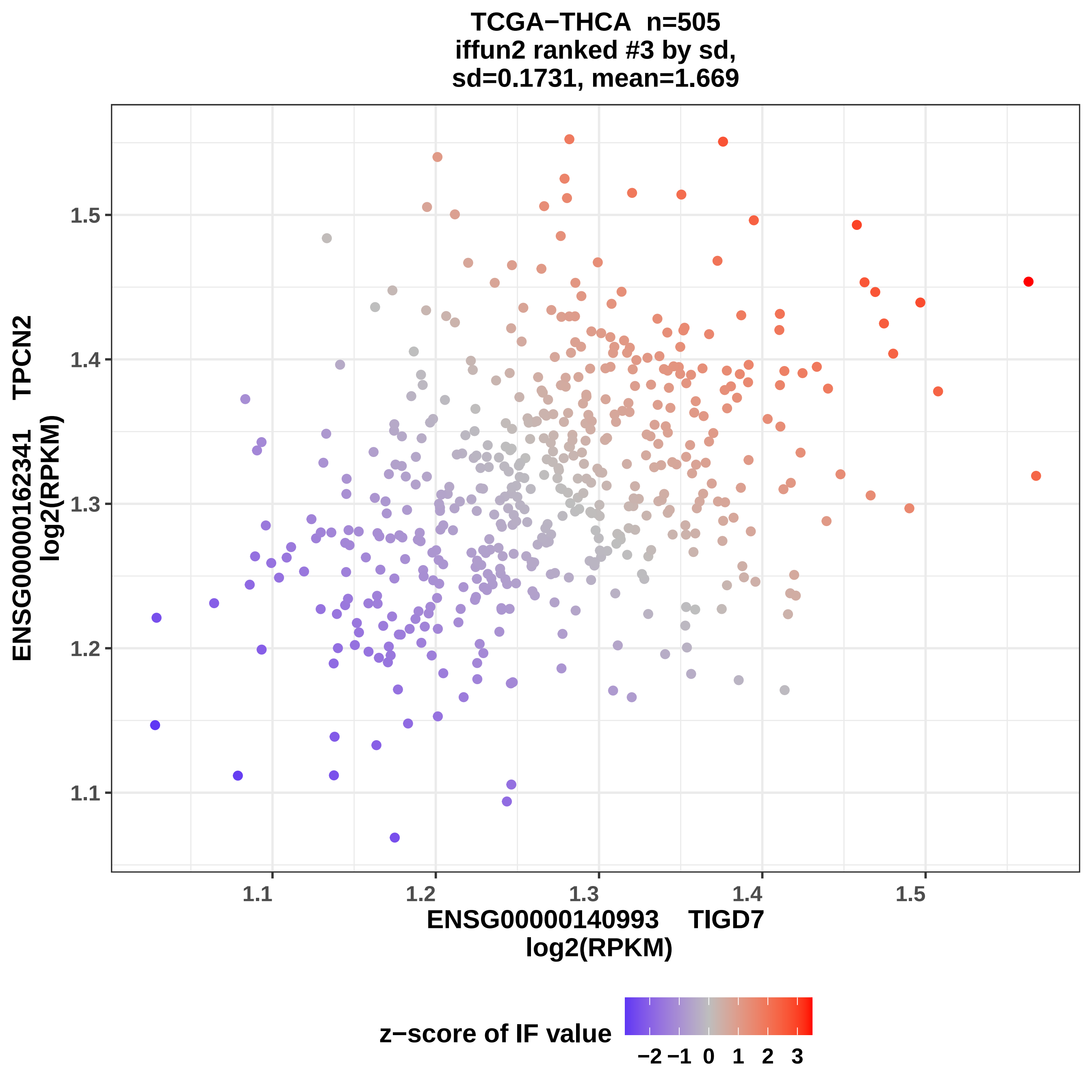
TCGA−THCA n=505
iffun2 ranked #3 by sd,
sd=0.1731, mean=1.669
ENSG00000162341 TPCN2
log2(RPKM)
1.1
1.2
1.3
1.4
1.5
ENSG00000140993 TIGD7
log2(RPKM)
z−score of IF value
−2 −1 0 1 2 3

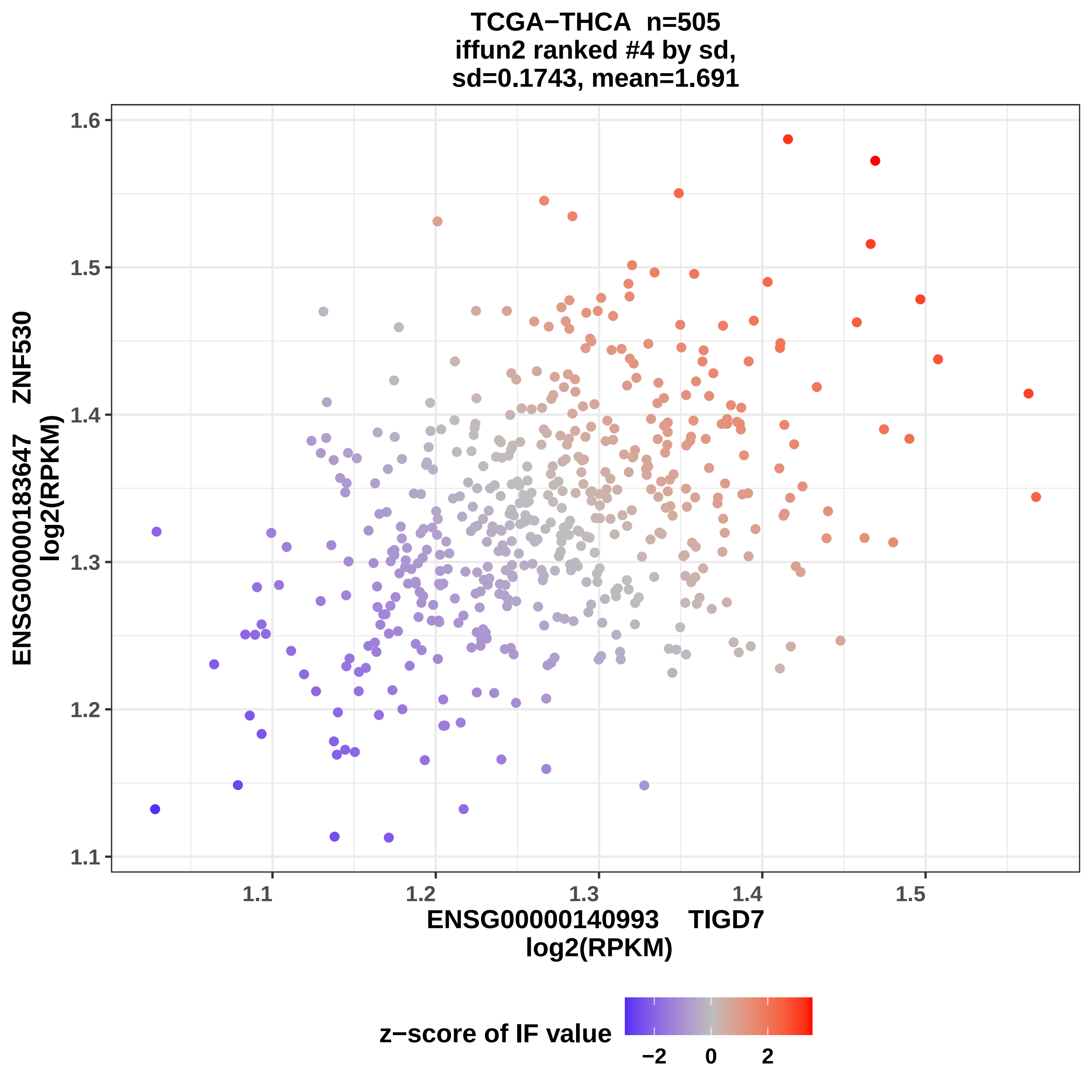
TCGA−THCA n=505
iffun2 ranked #4 by sd,
sd=0.1743, mean=1.691
ENSG00000183647 ZNF530
log2(RPKM)
ENSG00000140993 TIGD7
log2(RPKM)
1.1
1.2
1.3
1.4
1.5
1.6
z−score of IF value
−2
0
2

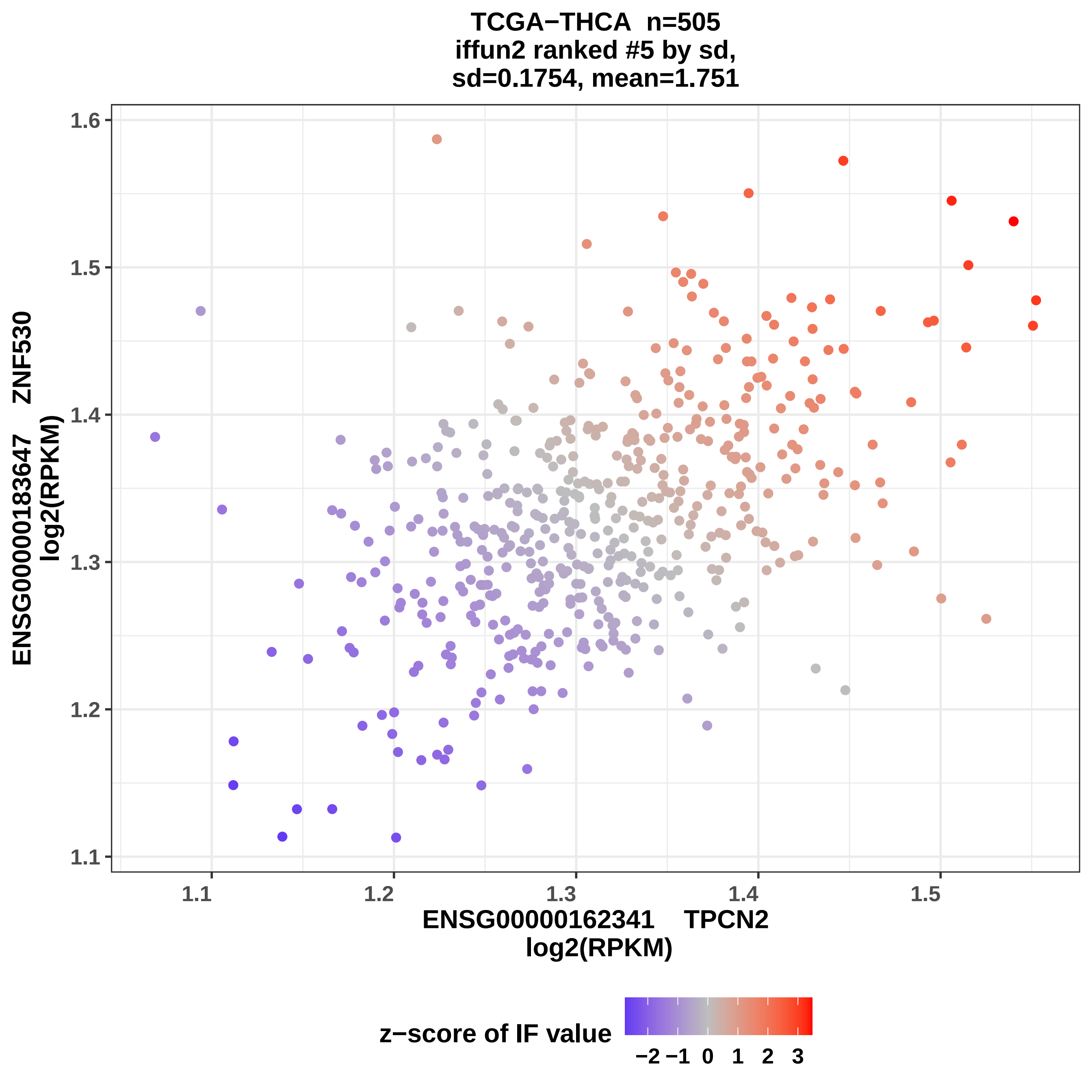
TCGA−THCA n=505
iffun2 ranked #5 by sd,
sd=0.1754, mean=1.751
ENSG00000183647 ZNF530
log2(RPKM)
1.6
1.5
1.4
1.3
1.2
1.1
1.1
1.2
1.3
1.4
1.5
ENSG00000162341 TPCN2
log2(RPKM)
z−score of IF value
−2
−1
0
1
2
3

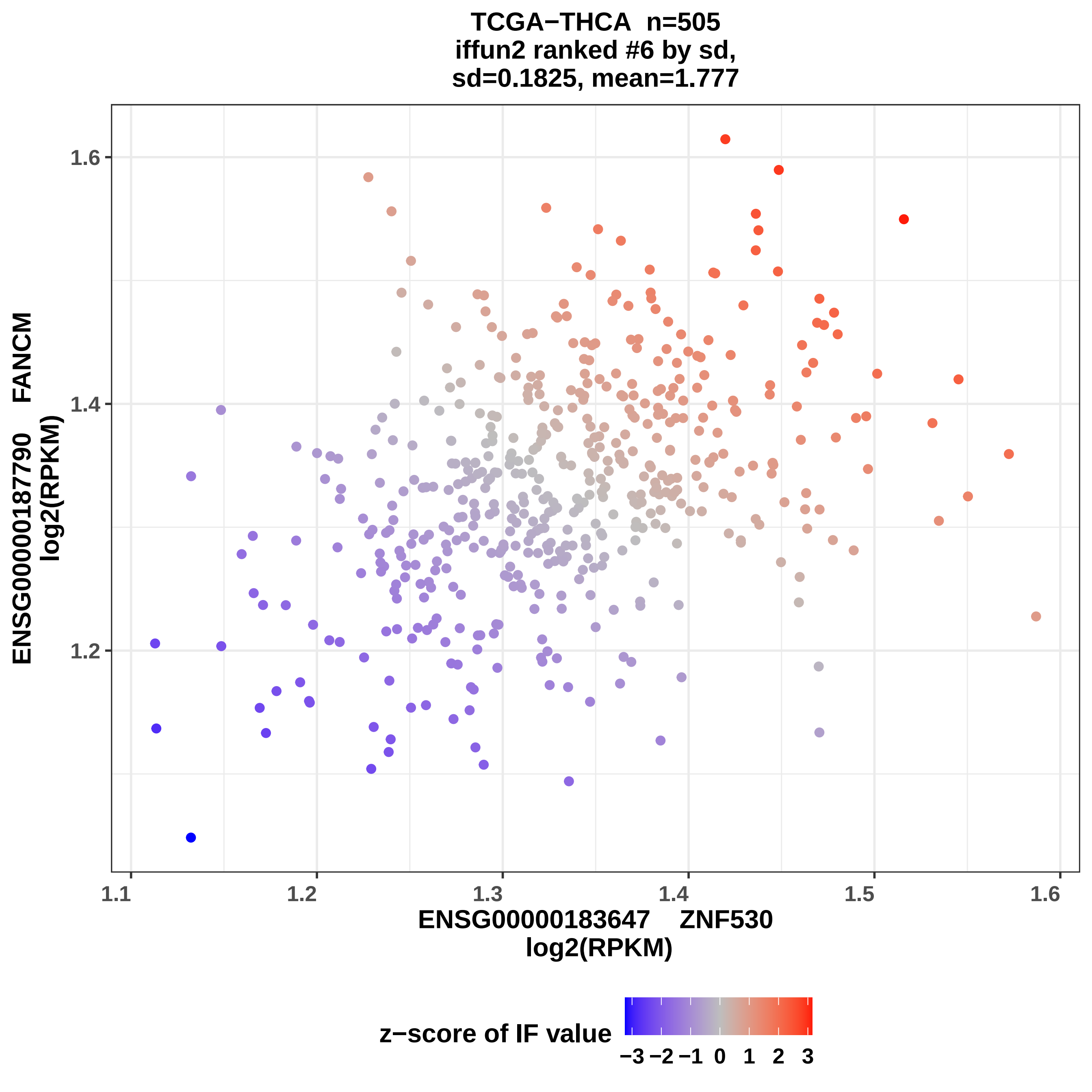

TCGA−THCA n=505
iffun2 ranked #6 by sd,
sd=0.1825, mean=1.777
ENSG00000187790 FANCM
log2(RPKM)
1.6
1.4
1.2
1.1
1.2
1.3
1.4
1.5
1.6
ENSG00000183647 ZNF530
log2(RPKM)
z−score of IF value
−3 −2 −1 0 1 2 3

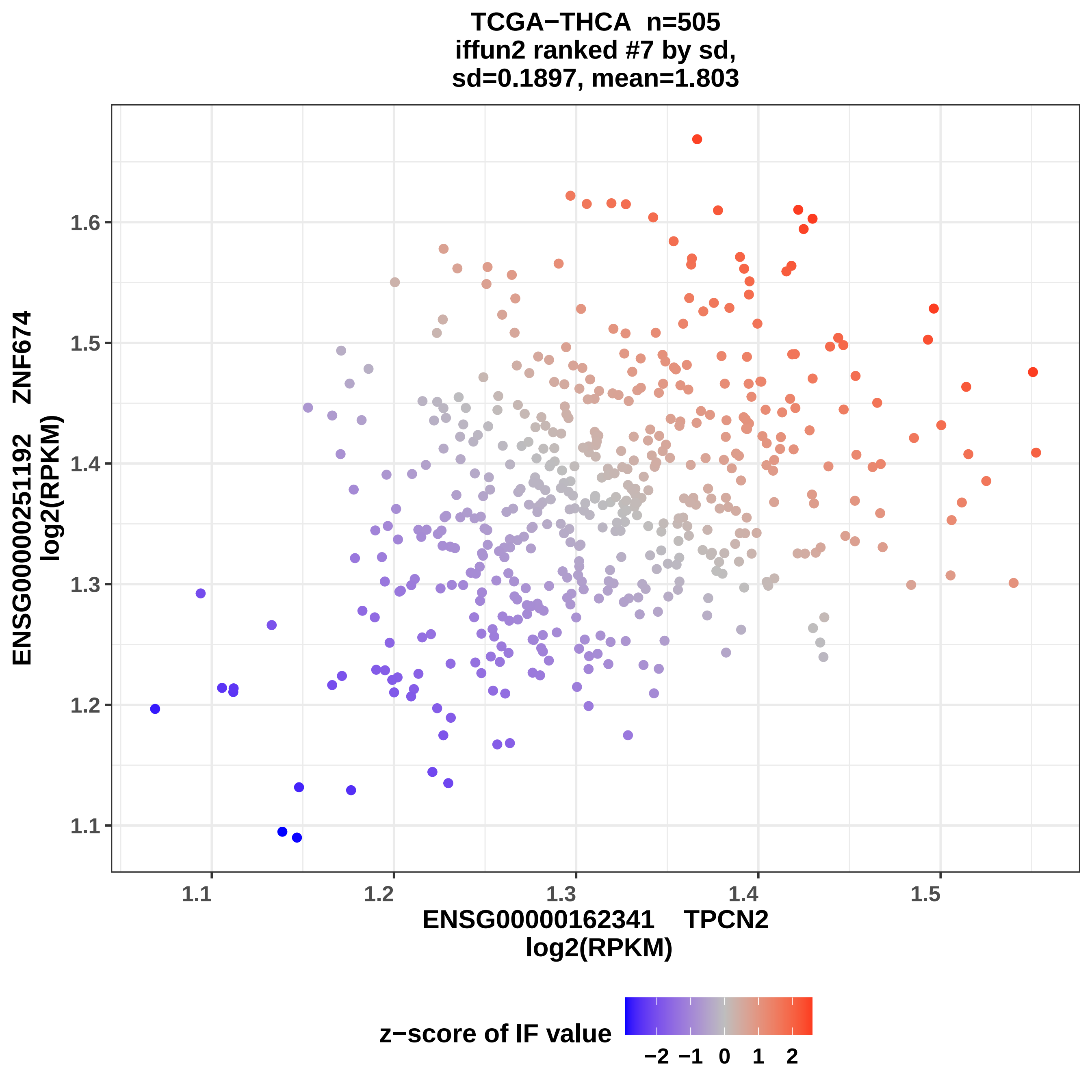
TCGA−THCA n=505
iffun2 ranked #7 by sd,
sd=0.1897, mean=1.803
ENSG00000251192 ZNF674
log2(RPKM)
1.1
1.2
1.3
1.4
1.5
1.6
ENSG00000162341 TPCN2
log2(RPKM)
z−score of IF value
−2
−1
0
1
2

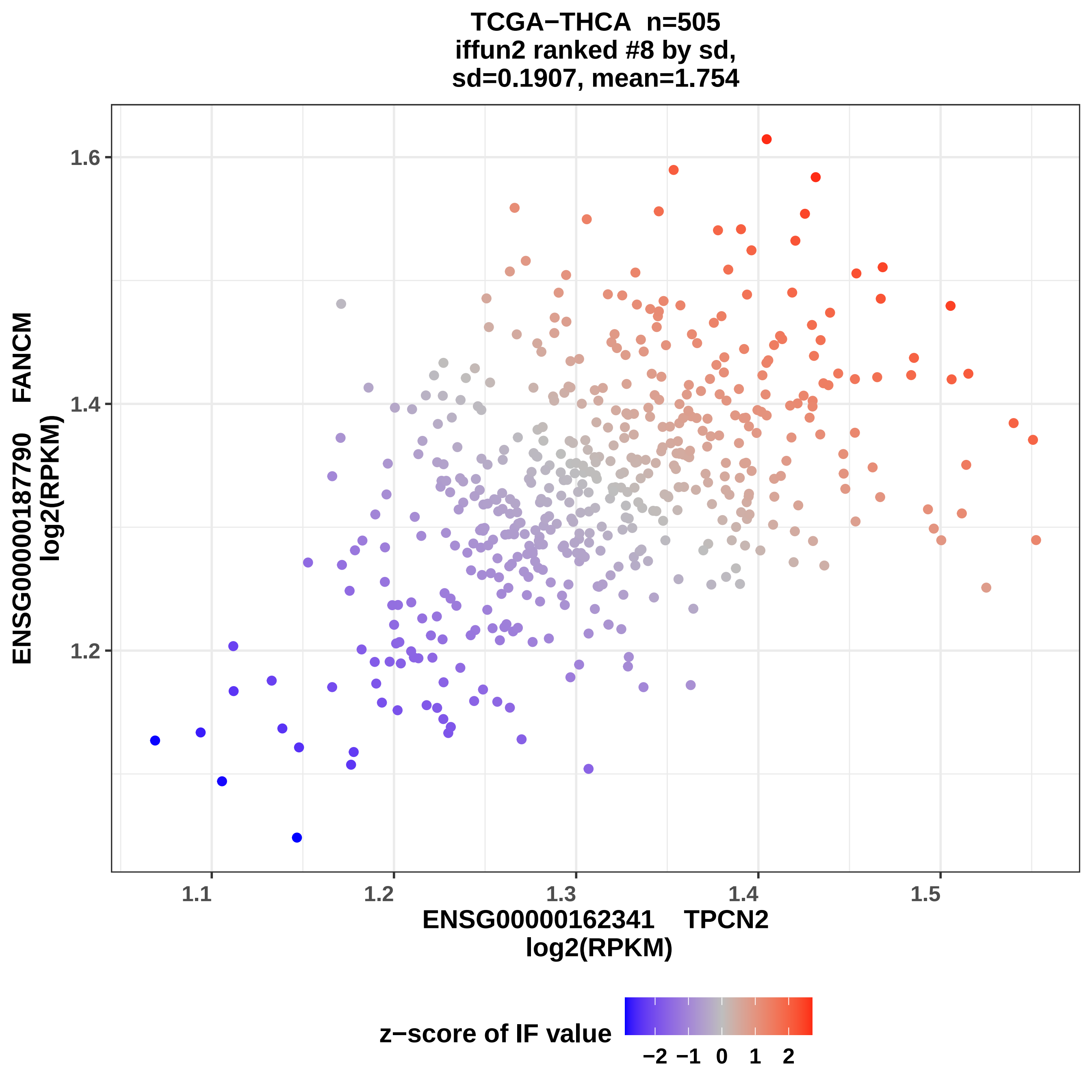

TCGA−THCA n=505
iffun2 ranked #8 by sd,
sd=0.1907, mean=1.754
ENSG00000187790 FANCM
log2(RPKM)
1.6
1.4
1.2
1.1
1.2
1.3
1.4
1.5
ENSG00000162341 TPCN2
log2(RPKM)
z−score of IF value
−2
−1
0
1
2

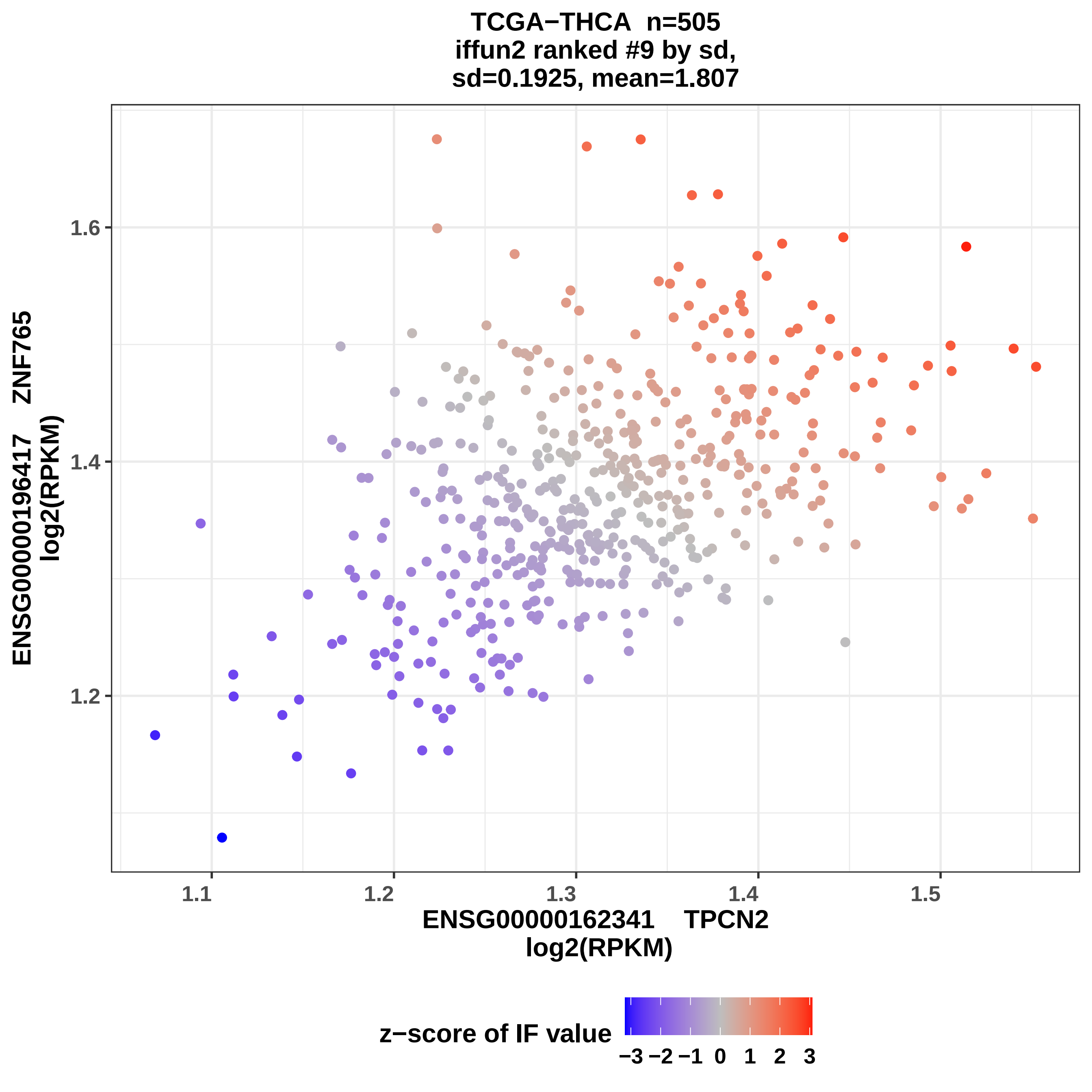
TCGA−THCA n=505
iffun2 ranked #9 by sd,
sd=0.1925, mean=1.807
ENSG00000196417 ZNF765
log2(RPKM)
1.2
1.4
1.6
1.1
1.2
1.3
1.4
1.5
ENSG00000162341 TPCN2
log2(RPKM)
z−score of IF value
−3 −2 −1 0 1 2 3

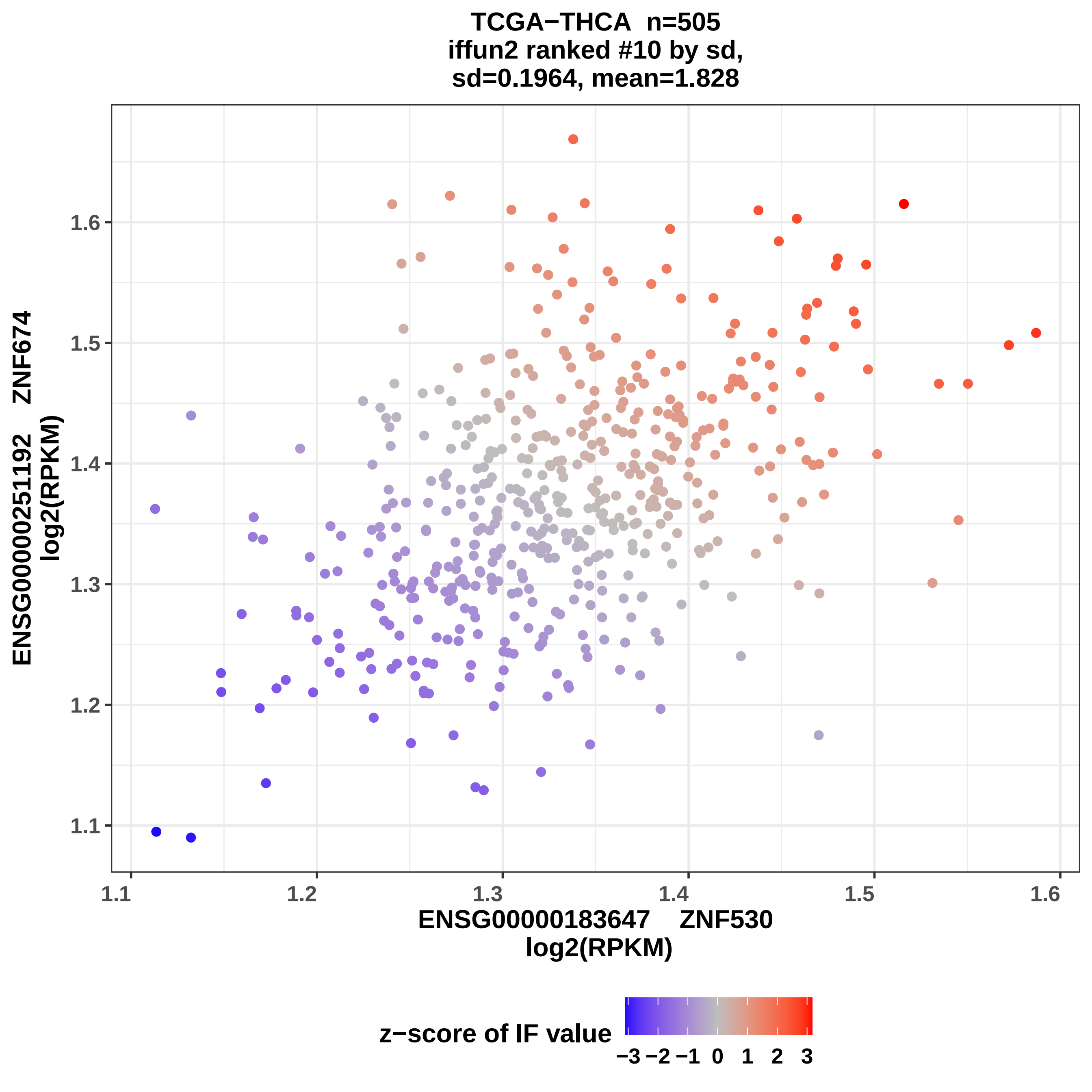
TCGA−THCA n=505
iffun2 ranked #10 by sd,
sd=0.1964, mean=1.828
ENSG00000251192 ZNF674
log2(RPKM)
1.1
1.2
1.3
1.4
1.5
1.6
ENSG00000183647 ZNF530
log2(RPKM)
z−score of IF value
−3 −2 −1 0 1 2 3

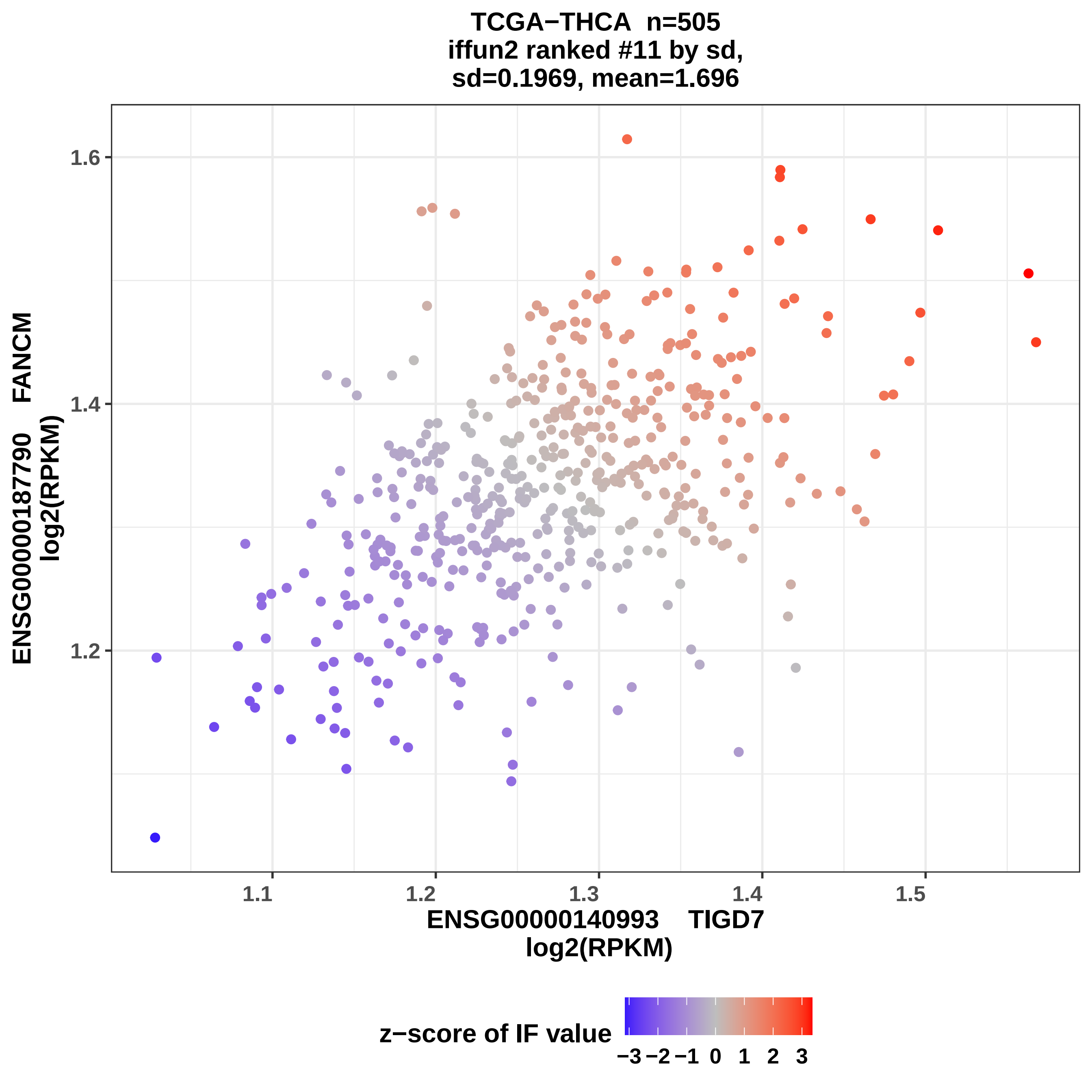

TCGA−THCA n=505
iffun2 ranked #11 by sd,
sd=0.1969, mean=1.696
ENSG00000187790 FANCM
log2(RPKM)
1.6
1.4
1.2
1.1
1.2
1.3
1.4
1.5
ENSG00000140993 TIGD7
log2(RPKM)
z−score of IF value
−3 −2 −1 0 1 2 3

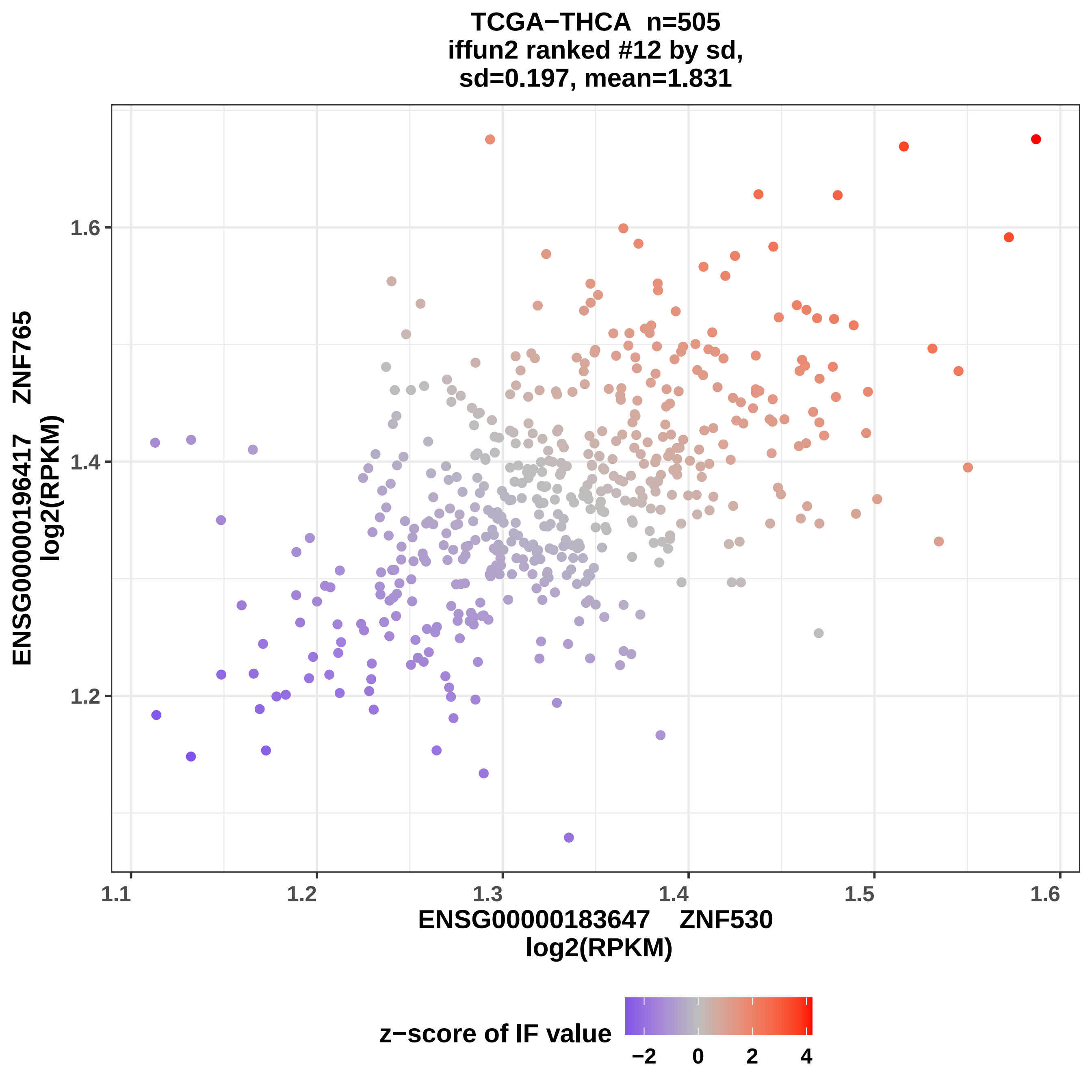

TCGA−THCA n=505
iffun2 ranked #12 by sd,
sd=0.197, mean=1.831
ENSG00000196417 ZNF765
log2(RPKM)
1.6
1.4
1.2
1.1
1.2
1.3
1.4
1.5
1.6
ENSG00000183647 ZNF530
log2(RPKM)
z−score of IF value
−2
0
2
4

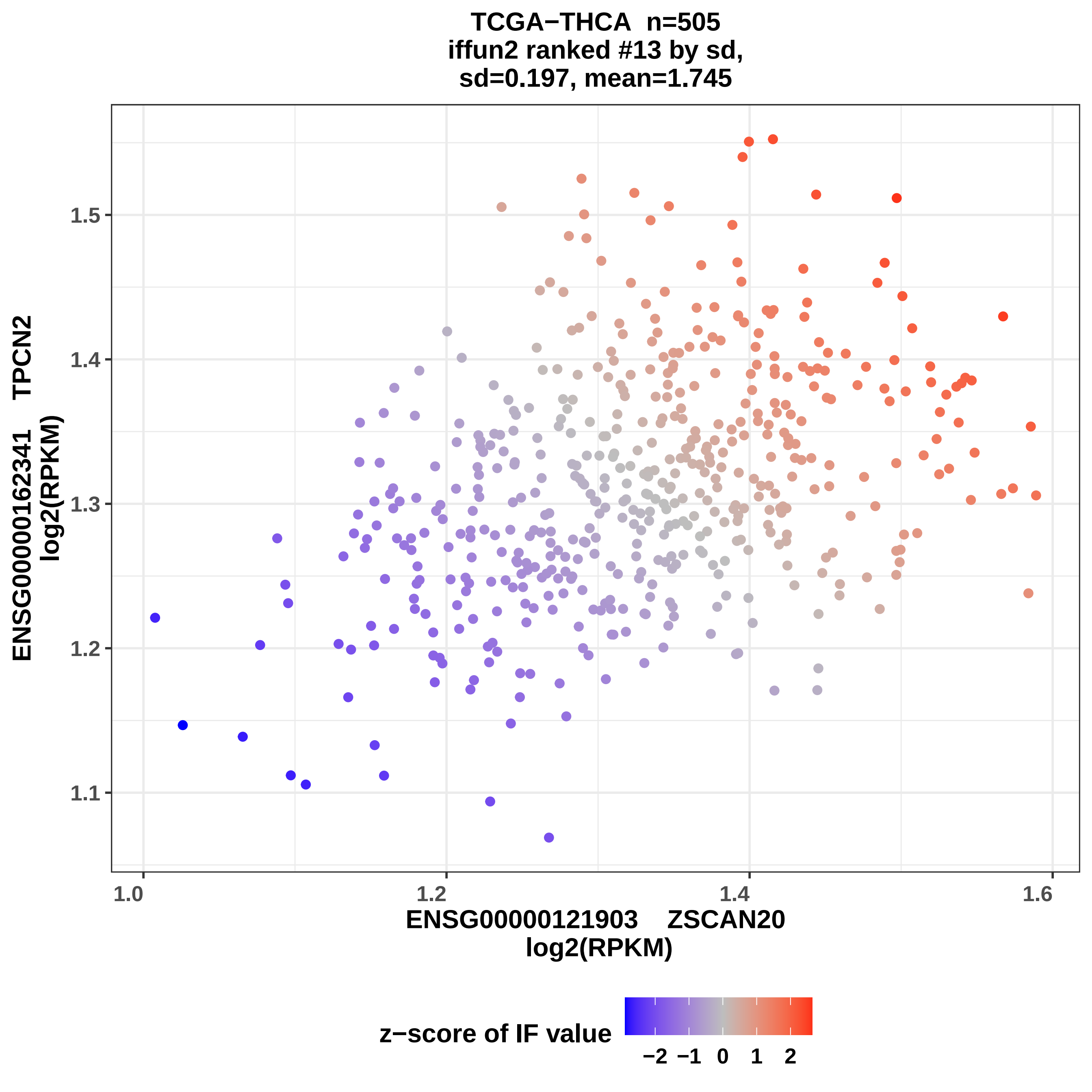
TCGA−THCA n=505
iffun2 ranked #13 by sd,
sd=0.197, mean=1.745
ENSG00000162341 TPCN2
log2(RPKM)
1.5
1.4
1.3
1.2
1.1
1.0
1.2
1.4
1.6
ENSG00000121903 ZSCAN20
log2(RPKM)
z−score of IF value
−2 −1 0 1 2

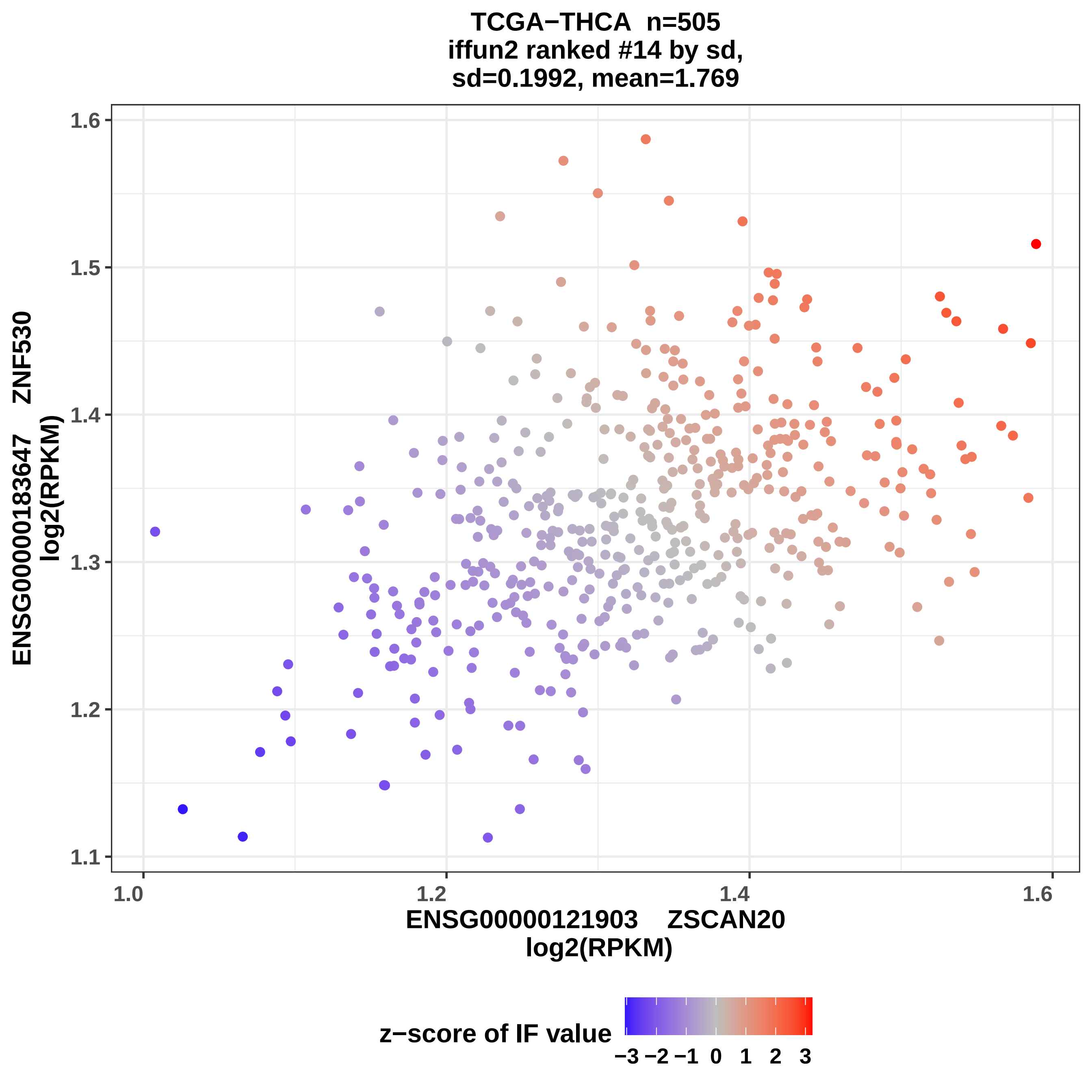
TCGA−THCA n=505
iffun2 ranked #14 by sd,
sd=0.1992, mean=1.769
ENSG00000183647 ZNF530
log2(RPKM)
1.1
1.2
1.3
1.4
1.5
1.6
1.0
1.2
1.4
1.6
ENSG00000121903 ZSCAN20
log2(RPKM)
z−score of IF value
−3 −2 −1 0 1 2 3

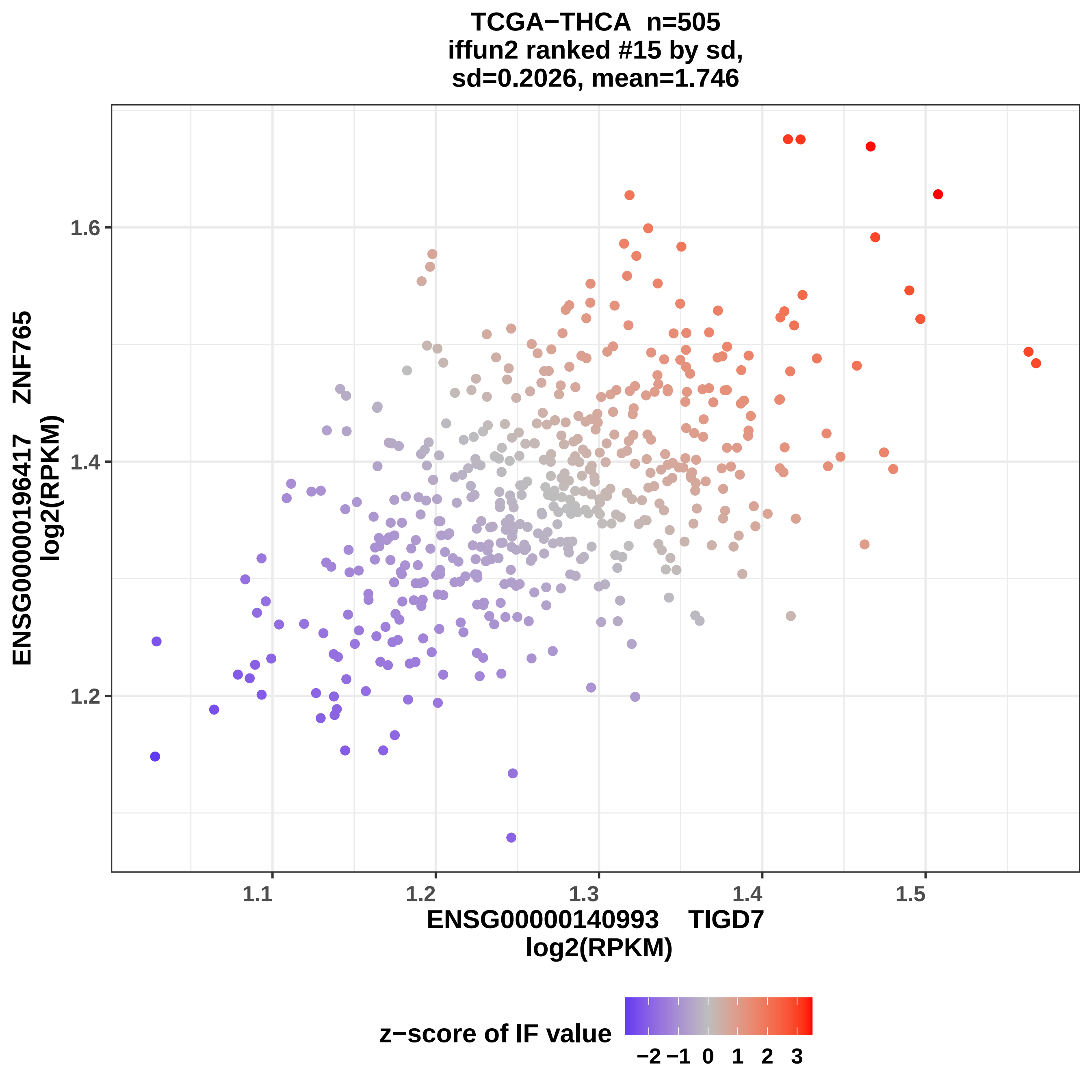

TCGA−THCA n=505
iffun2 ranked #15 by sd,
sd=0.2026, mean=1.746
ENSG00000196417 ZNF765
log2(RPKM)
1.2
1.4
1.6
1.1
1.2
1.3
1.4
1.5
ENSG00000140993 TIGD7
log2(RPKM)
z−score of IF value
−2
−1
0
1
2
3

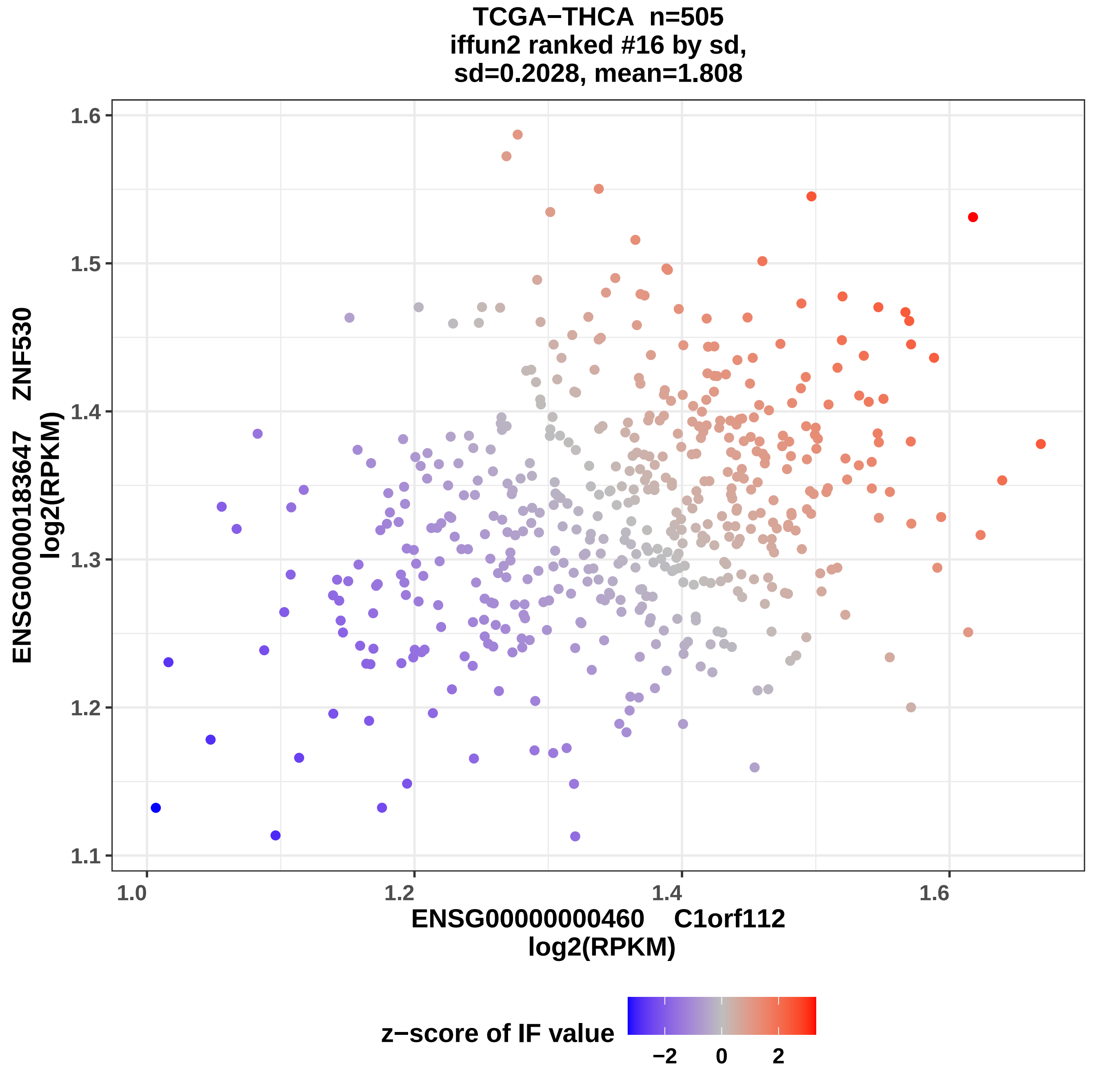

TCGA−THCA n=505
iffun2 ranked #16 by sd,
sd=0.2028, mean=1.808
ENSG00000183647 ZNF530
log2(RPKM)
1.1
1.2
1.3
1.4
1.5
1.6
1.0
1.2
1.4
1.6
ENSG00000000460 C1orf112
log2(RPKM)
z−score of IF value
−2
0
2

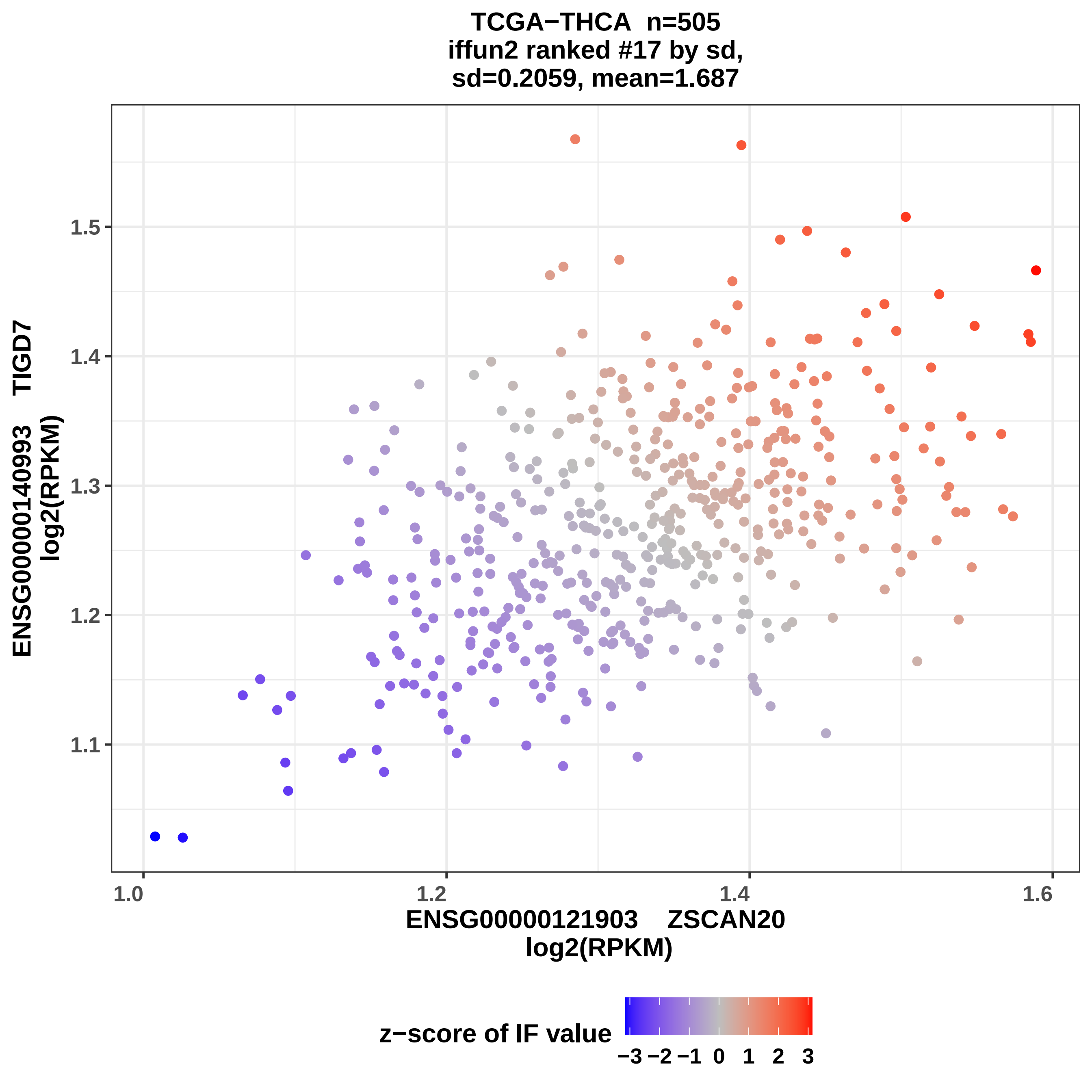
TCGA−THCA n=505
iffun2 ranked #17 by sd,
sd=0.2059, mean=1.687
ENSG00000140993 TIGD7
log2(RPKM)
ENSG00000121903 ZSCAN20
log2(RPKM)
1.0
1.2
1.4
1.6
1.1
1.2
1.3
1.4
1.5
z−score of IF value
−3 −2 −1 0 1 2 3

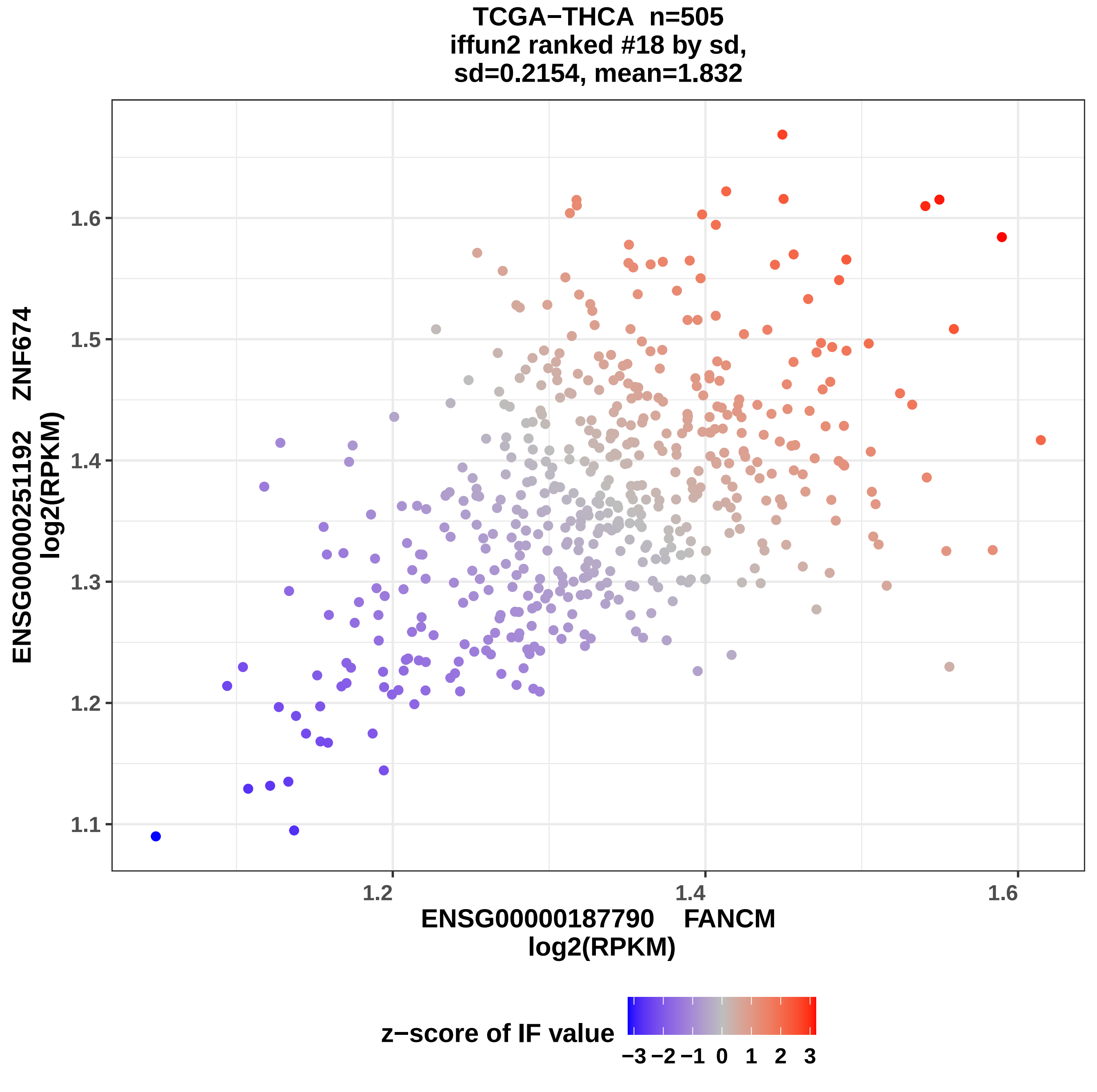
TCGA−THCA n=505
iffun2 ranked #18 by sd,
sd=0.2154, mean=1.832
ENSG00000251192 ZNF674
log2(RPKM)
1.1
1.2
1.3
1.4
1.5
1.6
1.2
1.4
1.6
ENSG00000187790 FANCM
log2(RPKM)
z−score of IF value
−3 −2 −1 0 1 2 3

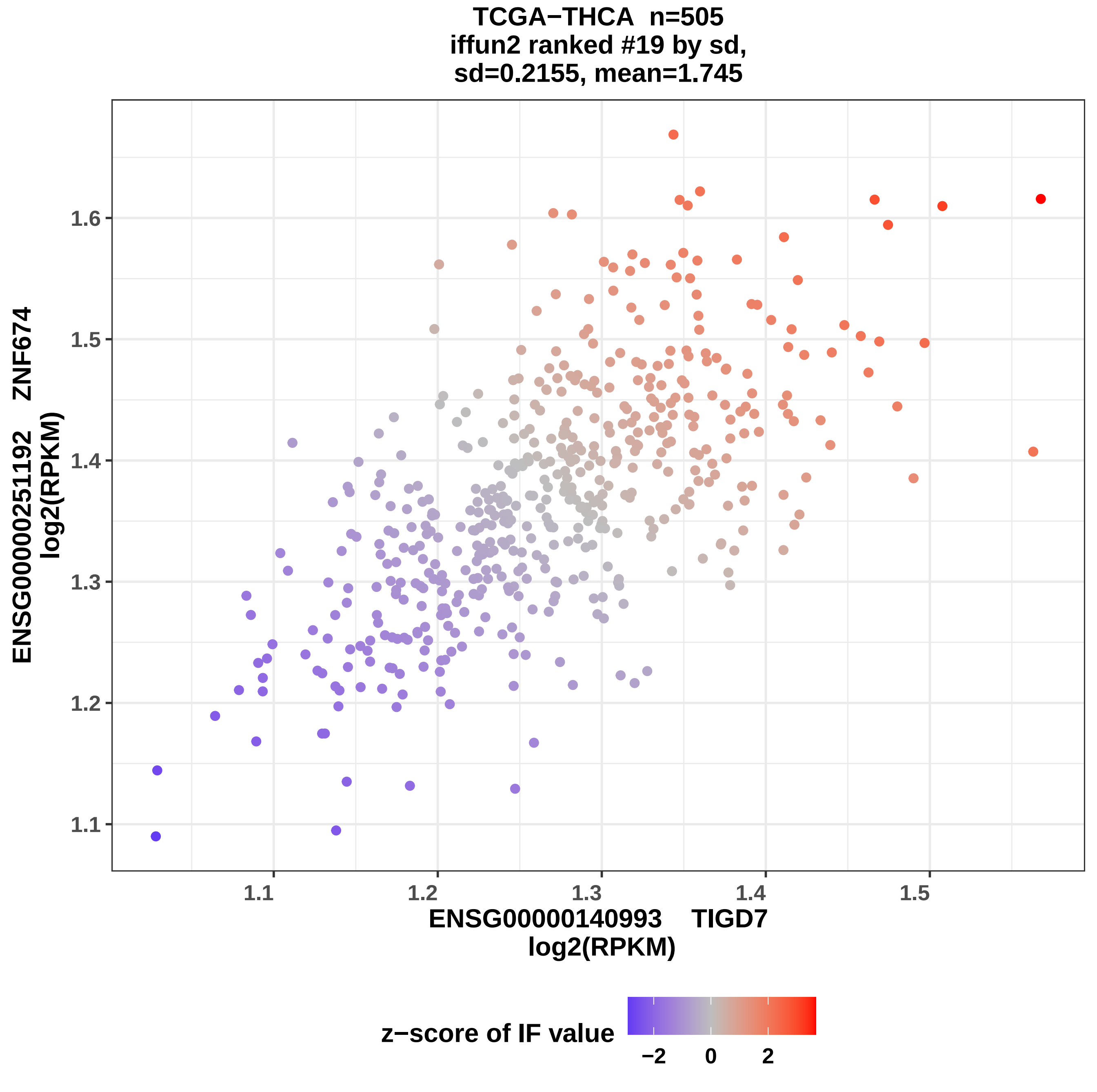
TCGA−THCA n=505
iffun2 ranked #19 by sd,
sd=0.2155, mean=1.745
ENSG00000251192 ZNF674
log2(RPKM)
1.1
1.2
1.3
1.4
1.5
1.6
ENSG00000140993 TIGD7
log2(RPKM)
z−score of IF value
−2
0
2

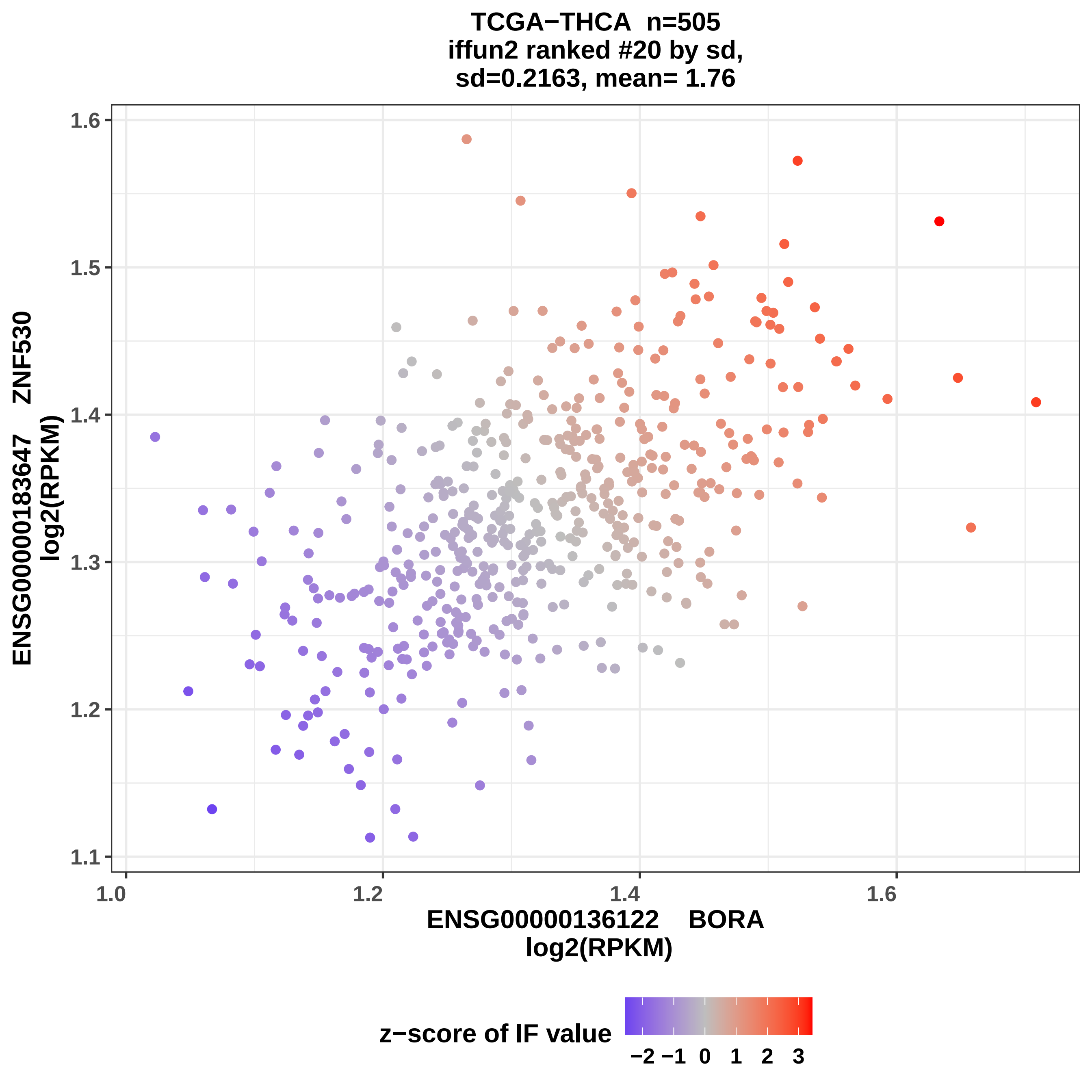
TCGA−THCA n=505
iffun2 ranked #20 by sd,
sd=0.2163, mean= 1.76
ENSG00000183647 ZNF530
log2(RPKM)
ENSG00000136122 BORA
log2(RPKM)
1.1
1.2
1.3
1.4
1.5
1.6
1.0
1.2
1.4
1.6
z−score of IF value
−2
−1
0
1
2
3

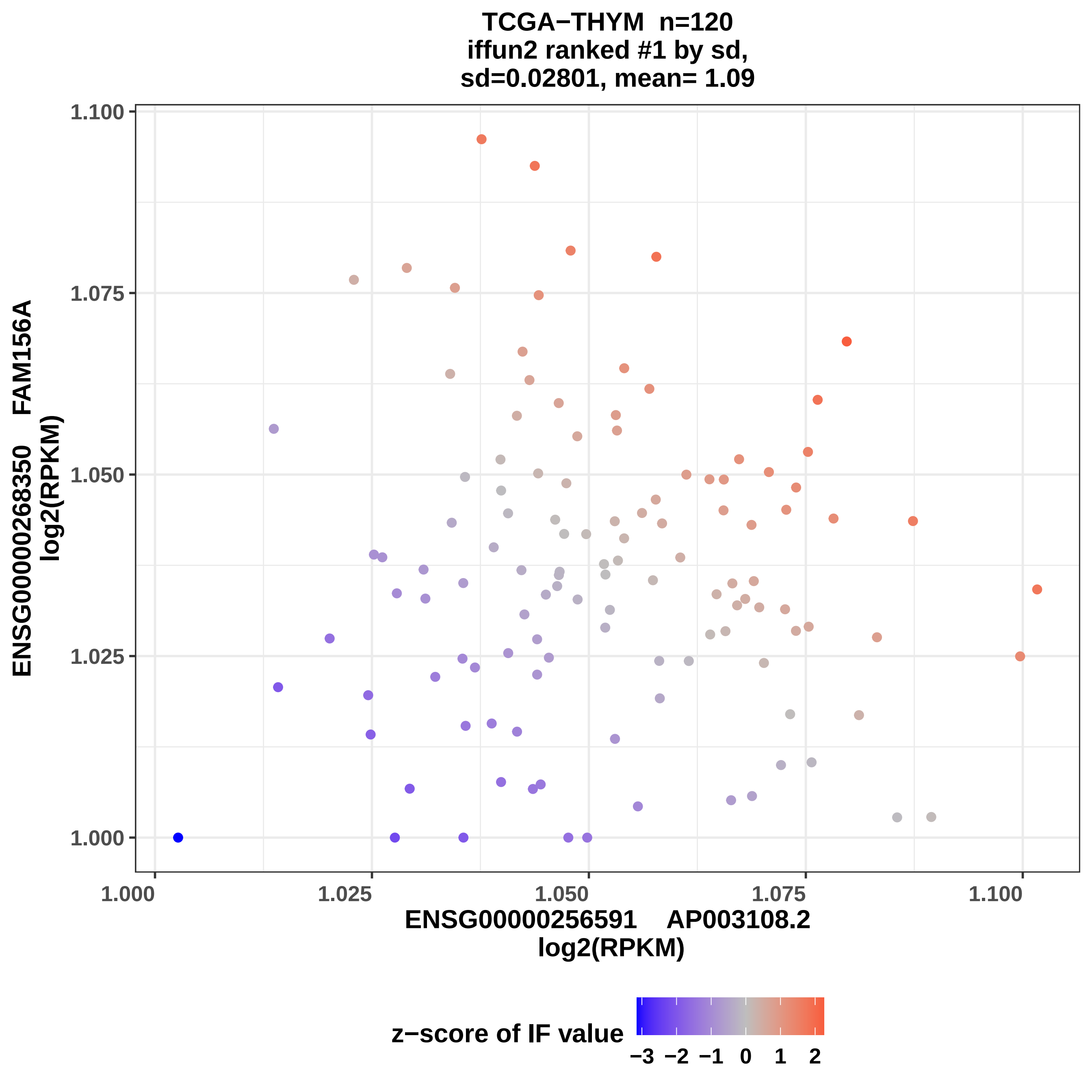
TCGA−THYM n=120
iffun2 ranked #1 by sd,
sd=0.02801, mean= 1.09
ENSG00000268350 FAM156A
log2(RPKM)
1.100
1.075
1.050
1.025
1.000
1.000
1.025
1.050
1.075
1.100
ENSG00000256591 AP003108.2
log2(RPKM)
z−score of IF value
−3 −2 −1 0 1 2

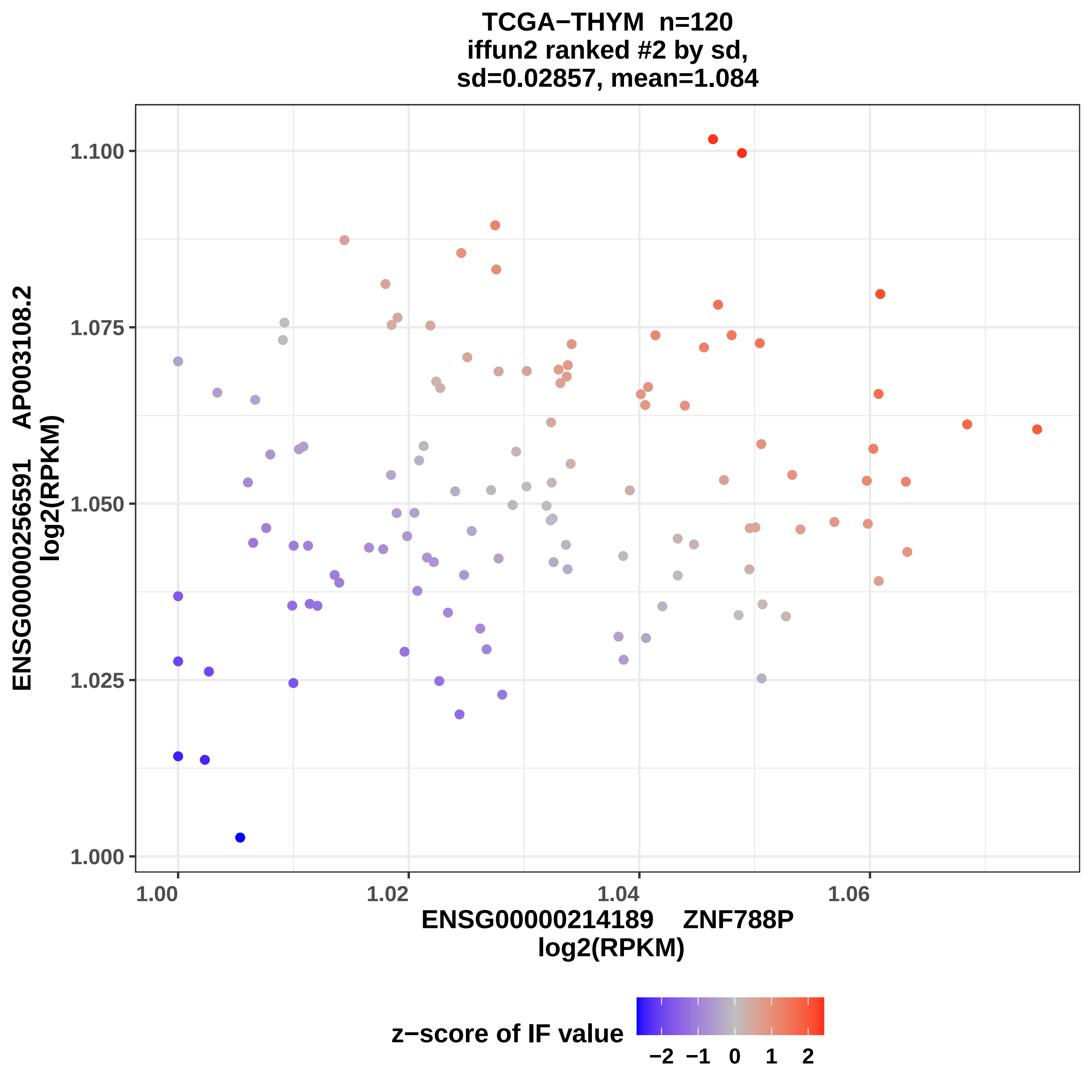
TCGA−THYM n=120
iffun2 ranked #2 by sd,
sd=0.02857, mean=1.084
ENSG00000256591 AP003108.2
log2(RPKM)
1.100
1.075
1.050
1.025
1.000
1.00
1.02
1.04
1.06
ENSG00000214189 ZNF788P
log2(RPKM)
z−score of IF value
−2
−1
0
1
2

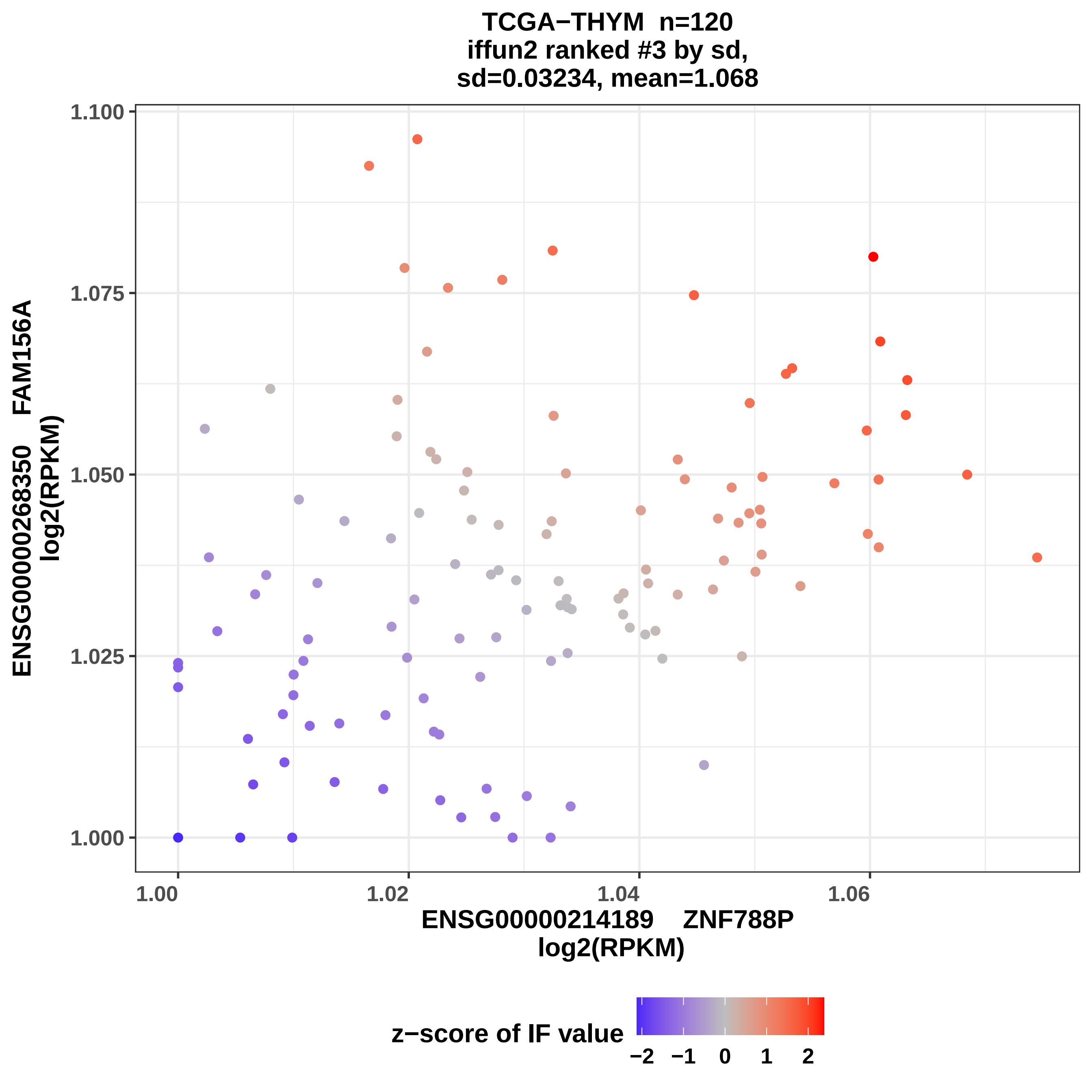
TCGA−THYM n=120
iffun2 ranked #3 by sd,
sd=0.03234, mean=1.068
ENSG00000268350 FAM156A
log2(RPKM)
1.100
1.075
1.050
1.025
1.000
1.00
1.02
1.04
1.06
ENSG00000214189 ZNF788P
log2(RPKM)
z−score of IF value
−2
−1
0
1
2

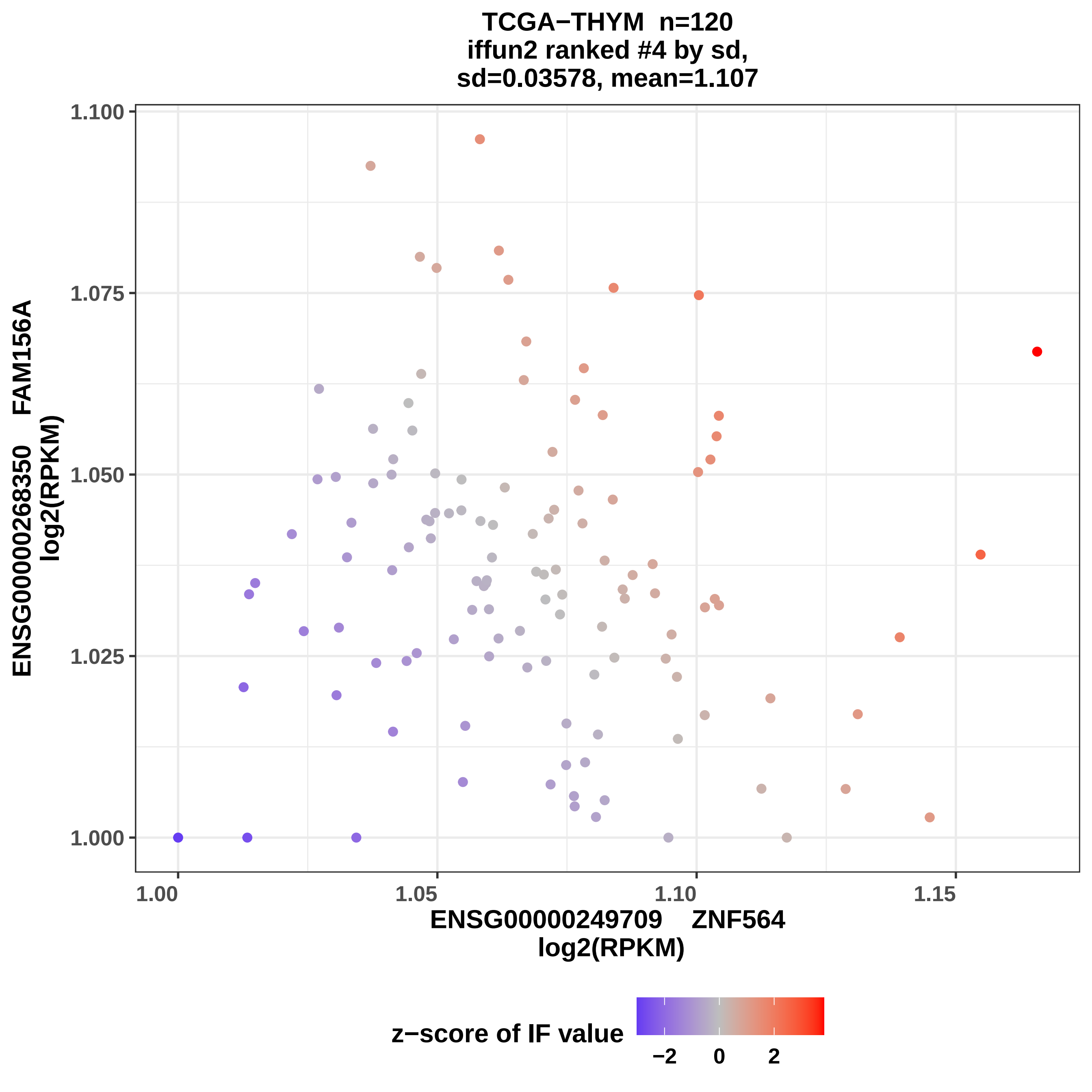
TCGA−THYM n=120
iffun2 ranked #4 by sd,
sd=0.03578, mean=1.107
ENSG00000268350 FAM156A
log2(RPKM)
1.100
1.075
1.050
1.025
1.000
1.00
1.05
1.10
1.15
ENSG00000249709 ZNF564
log2(RPKM)
z−score of IF value
−2
0
2

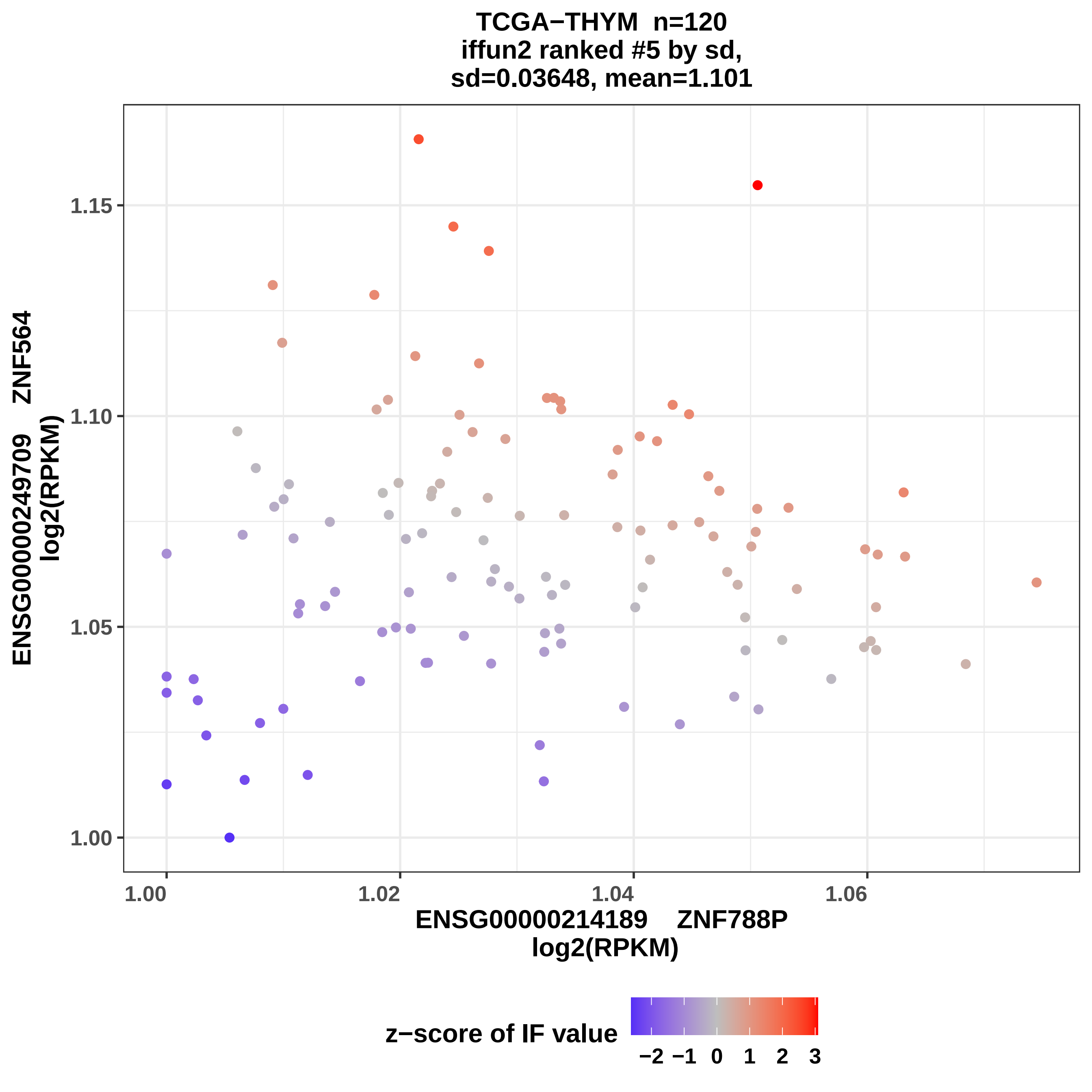
TCGA−THYM n=120
iffun2 ranked #5 by sd,
sd=0.03648, mean=1.101
ENSG00000249709 ZNF564
log2(RPKM)
1.15
1.10
1.05
1.00
1.00
1.02
1.04
1.06
ENSG00000214189 ZNF788P
log2(RPKM)
z−score of IF value
−2 −1 0 1 2 3

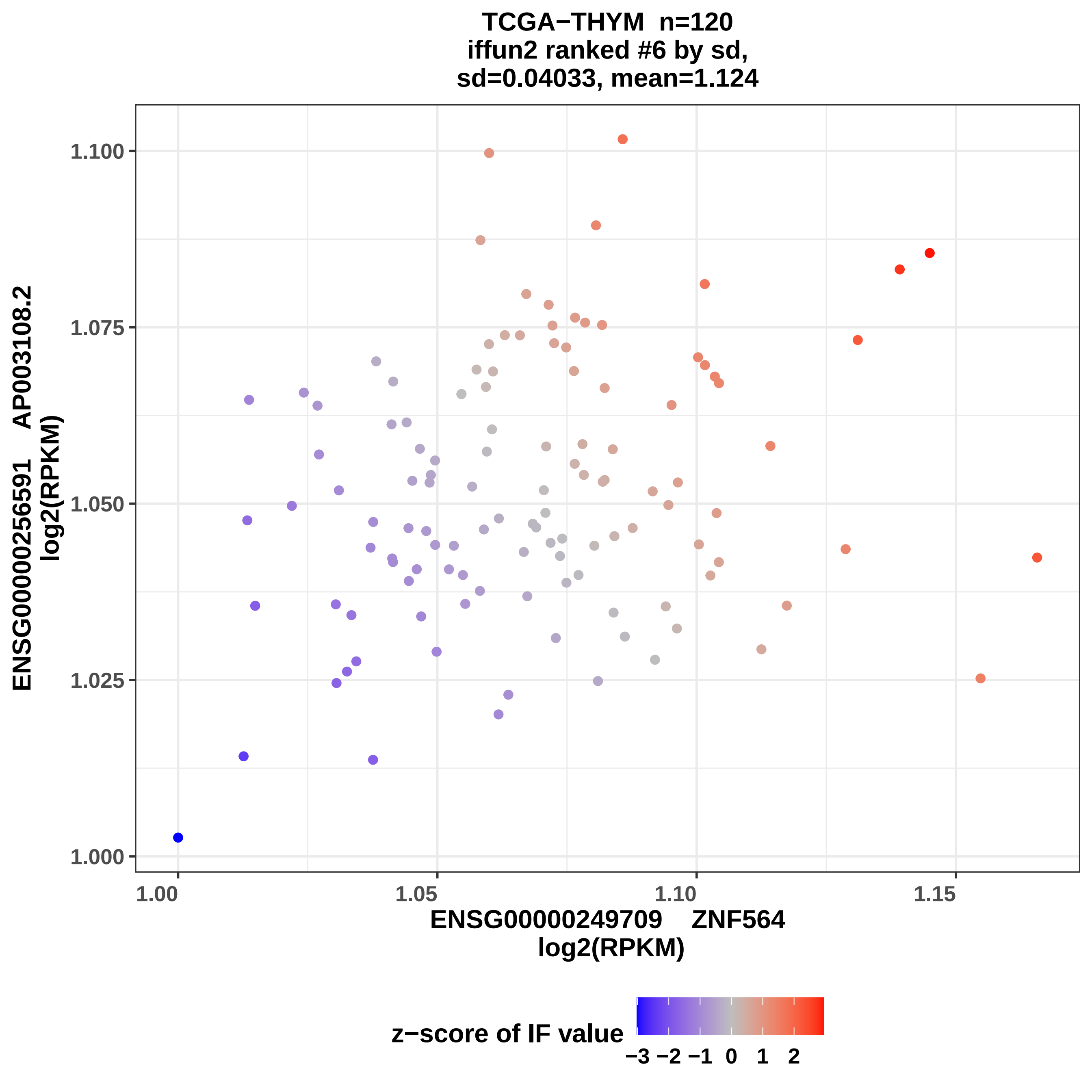

TCGA−THYM n=120
iffun2 ranked #6 by sd,
sd=0.04033, mean=1.124
ENSG00000256591 AP003108.2
log2(RPKM)
1.100
1.075
1.050
1.025
1.000
1.00
1.05
1.10
1.15
ENSG00000249709 ZNF564
log2(RPKM)
z−score of IF value
−3 −2 −1 0 1 2

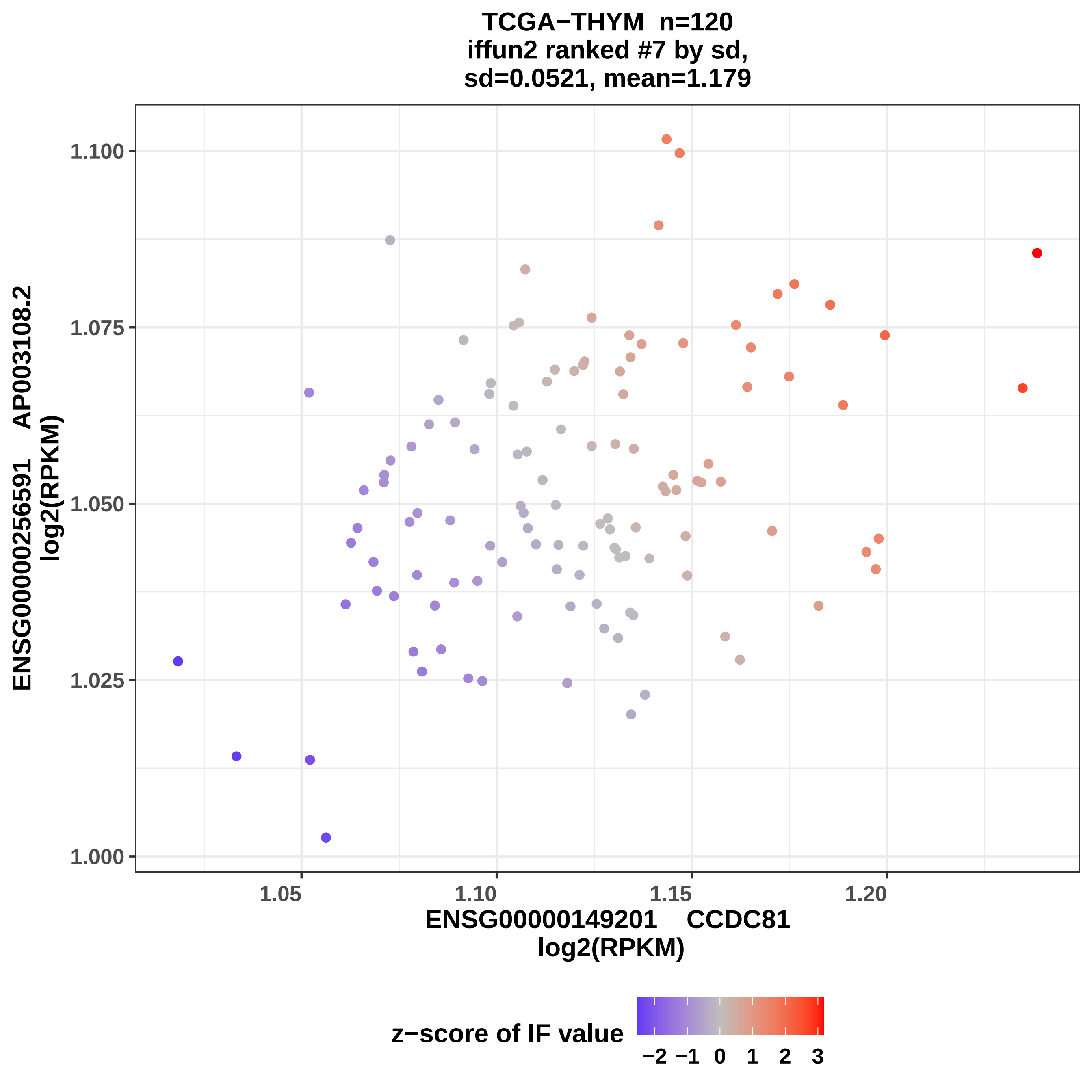
TCGA−THYM n=120
iffun2 ranked #7 by sd,
sd=0.0521, mean=1.179
ENSG00000256591 AP003108.2
log2(RPKM)
1.100
1.075
1.050
1.025
1.000
1.05
1.10
1.15
1.20
ENSG00000149201 CCDC81
log2(RPKM)
z−score of IF value
−2 −1 0 1 2 3

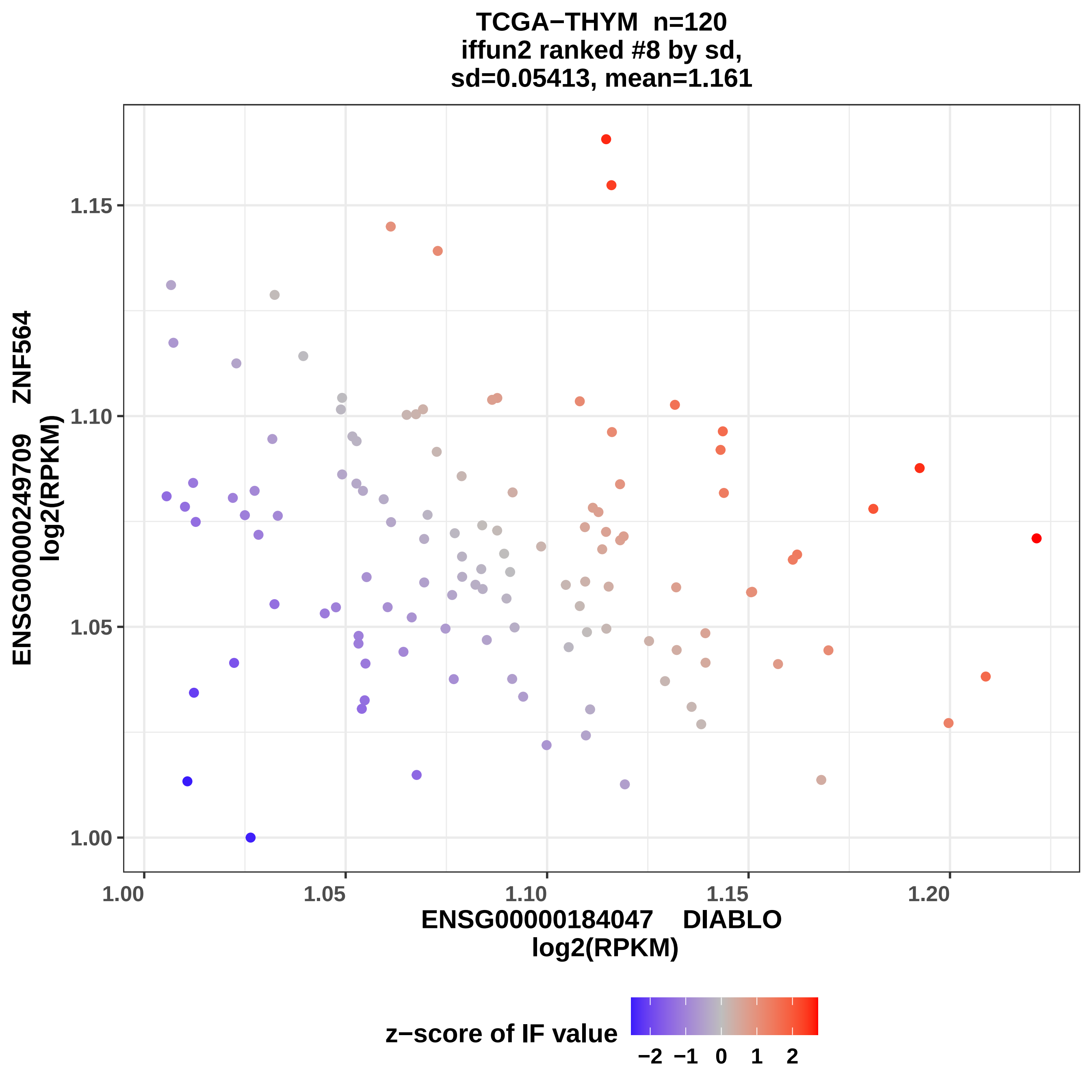
TCGA−THYM n=120
iffun2 ranked #8 by sd,
sd=0.05413, mean=1.161
ENSG00000249709 ZNF564
log2(RPKM)
1.15
1.10
1.05
1.00
1.00
1.05
1.10
1.15
1.20
ENSG00000184047 DIABLO
log2(RPKM)
z−score of IF value
−2
−1
0
1
2

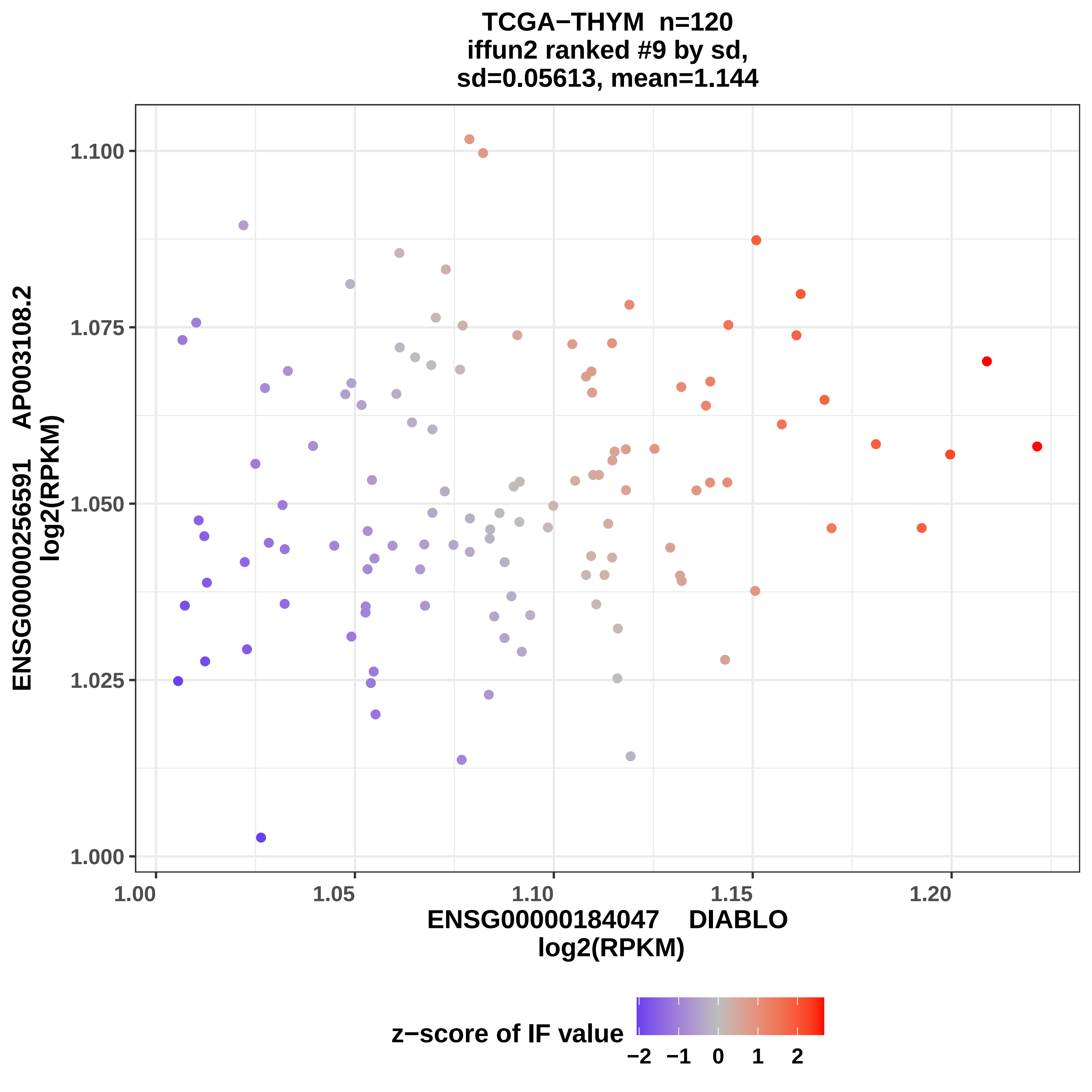

TCGA−THYM n=120
iffun2 ranked #9 by sd,
sd=0.05613, mean=1.144
ENSG00000256591 AP003108.2
log2(RPKM)
1.100
1.075
1.050
1.025
1.000
1.00
1.05
1.10
1.15
1.20
ENSG00000184047 DIABLO
log2(RPKM)
z−score of IF value
−2
−1
0
1
2

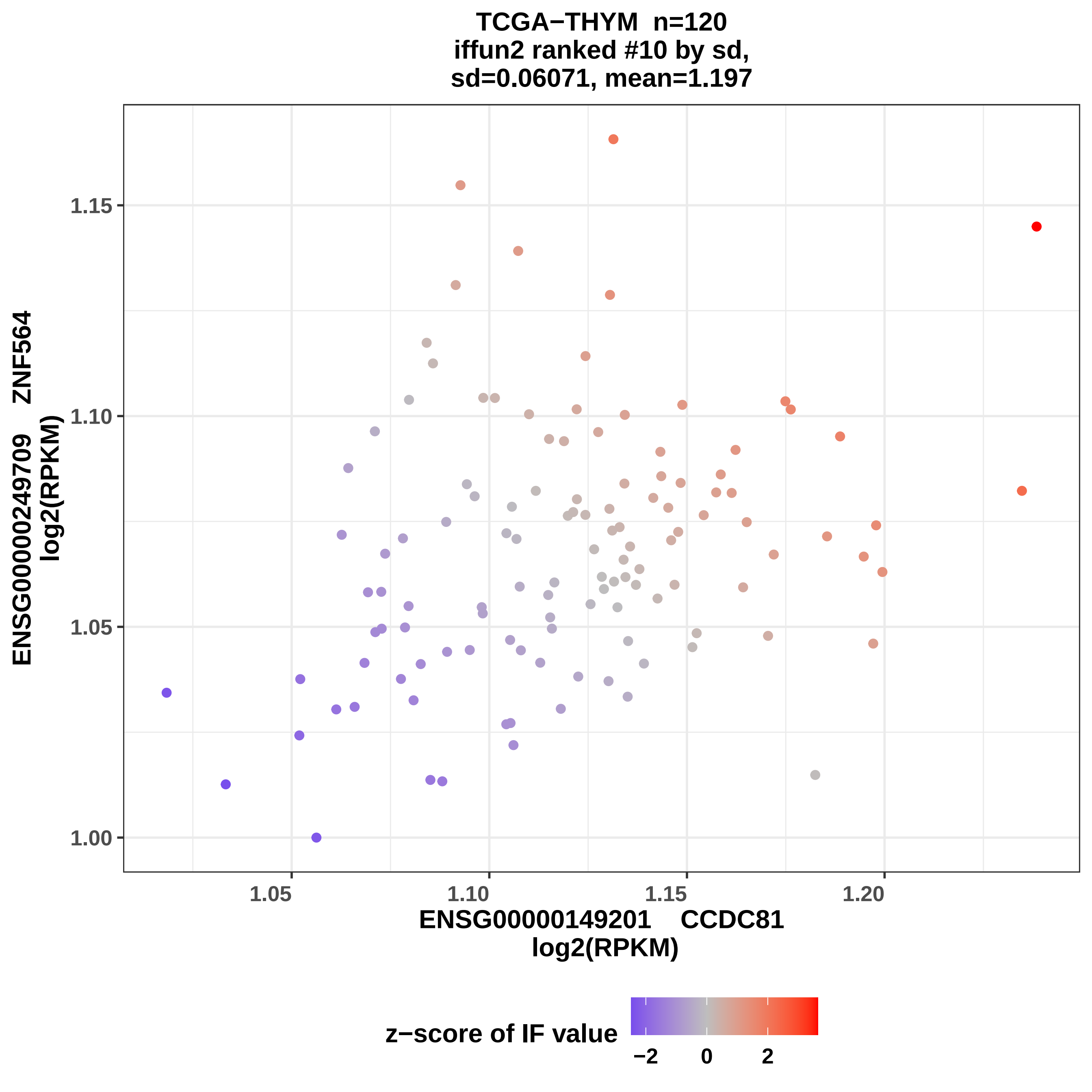
TCGA−THYM n=120
iffun2 ranked #10 by sd,
sd=0.06071, mean=1.197
ENSG00000249709 ZNF564
log2(RPKM)
1.15
1.10
1.05
1.00
1.05
1.10
1.15
1.20
ENSG00000149201 CCDC81
log2(RPKM)
z−score of IF value
−2
0
2

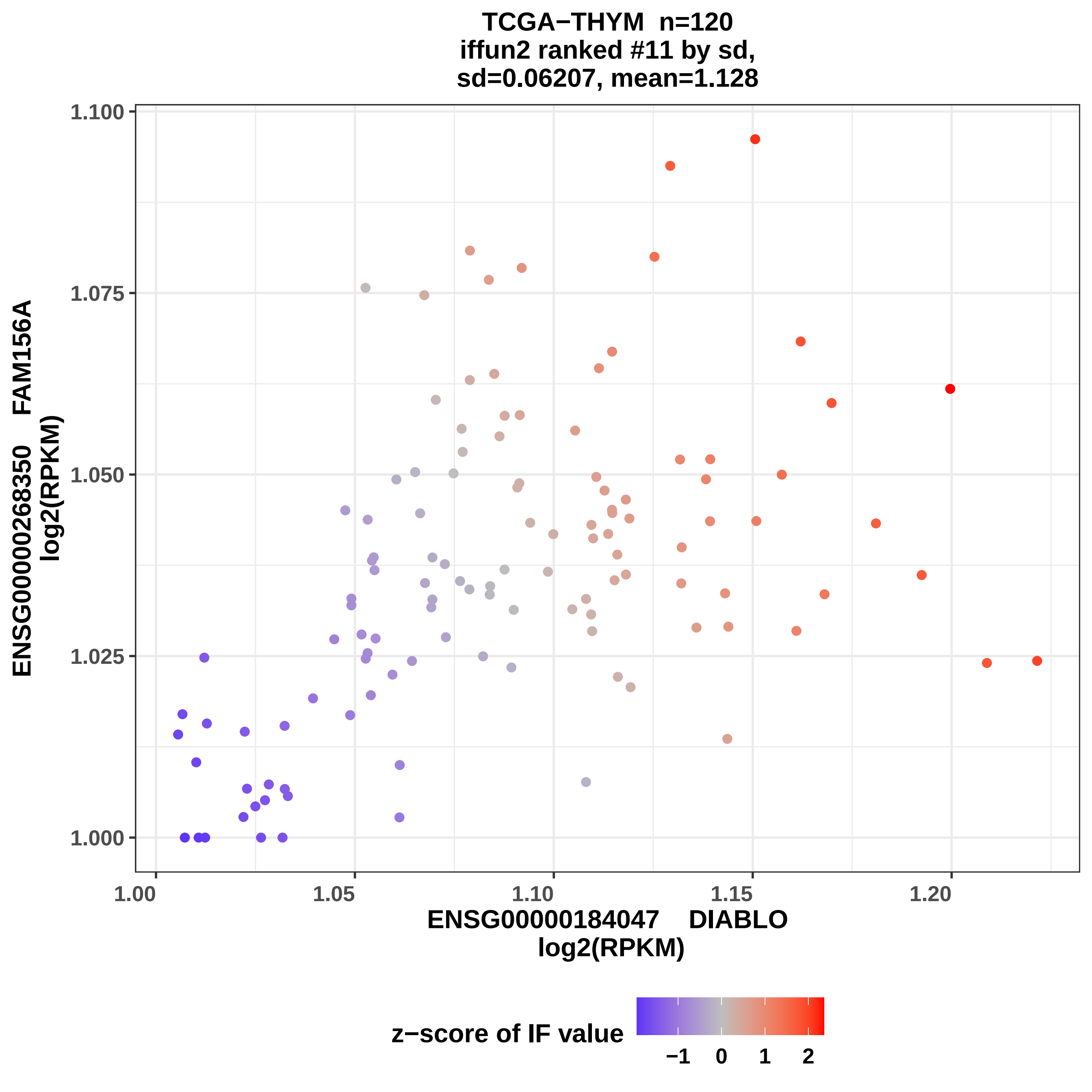

TCGA−THYM n=120
iffun2 ranked #11 by sd,
sd=0.06207, mean=1.128
ENSG00000268350 FAM156A
log2(RPKM)
1.100
1.075
1.050
1.025
1.000
1.00
1.05
1.10
1.15
1.20
ENSG00000184047 DIABLO
log2(RPKM)
z−score of IF value
−1
0
1
2

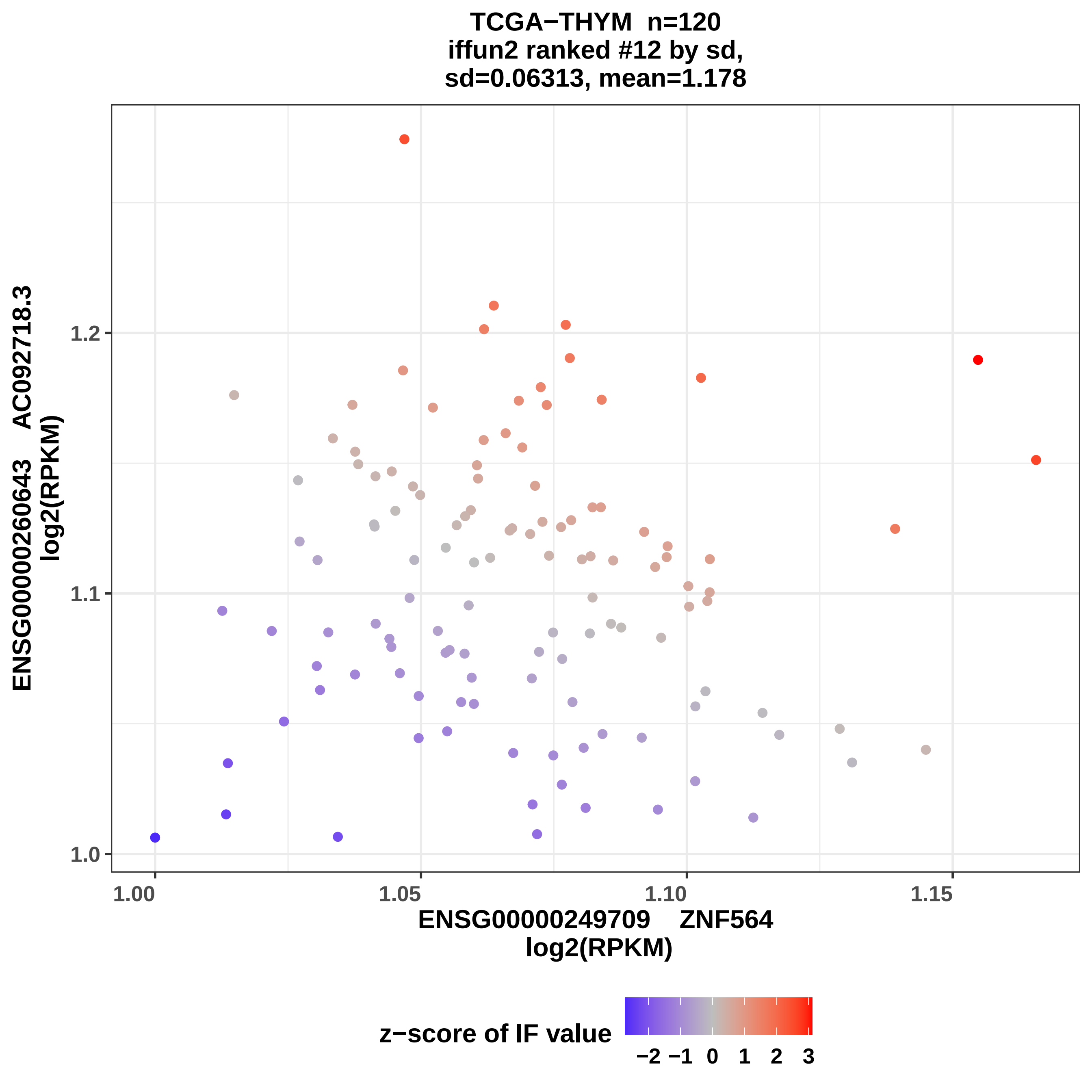
TCGA−THYM n=120
iffun2 ranked #12 by sd,
sd=0.06313, mean=1.178
ENSG00000260643 AC092718.3
log2(RPKM)
1.0
1.1
1.2
1.00
1.05
1.10
1.15
ENSG00000249709 ZNF564
log2(RPKM)
z−score of IF value
−2 −1 0 1 2 3

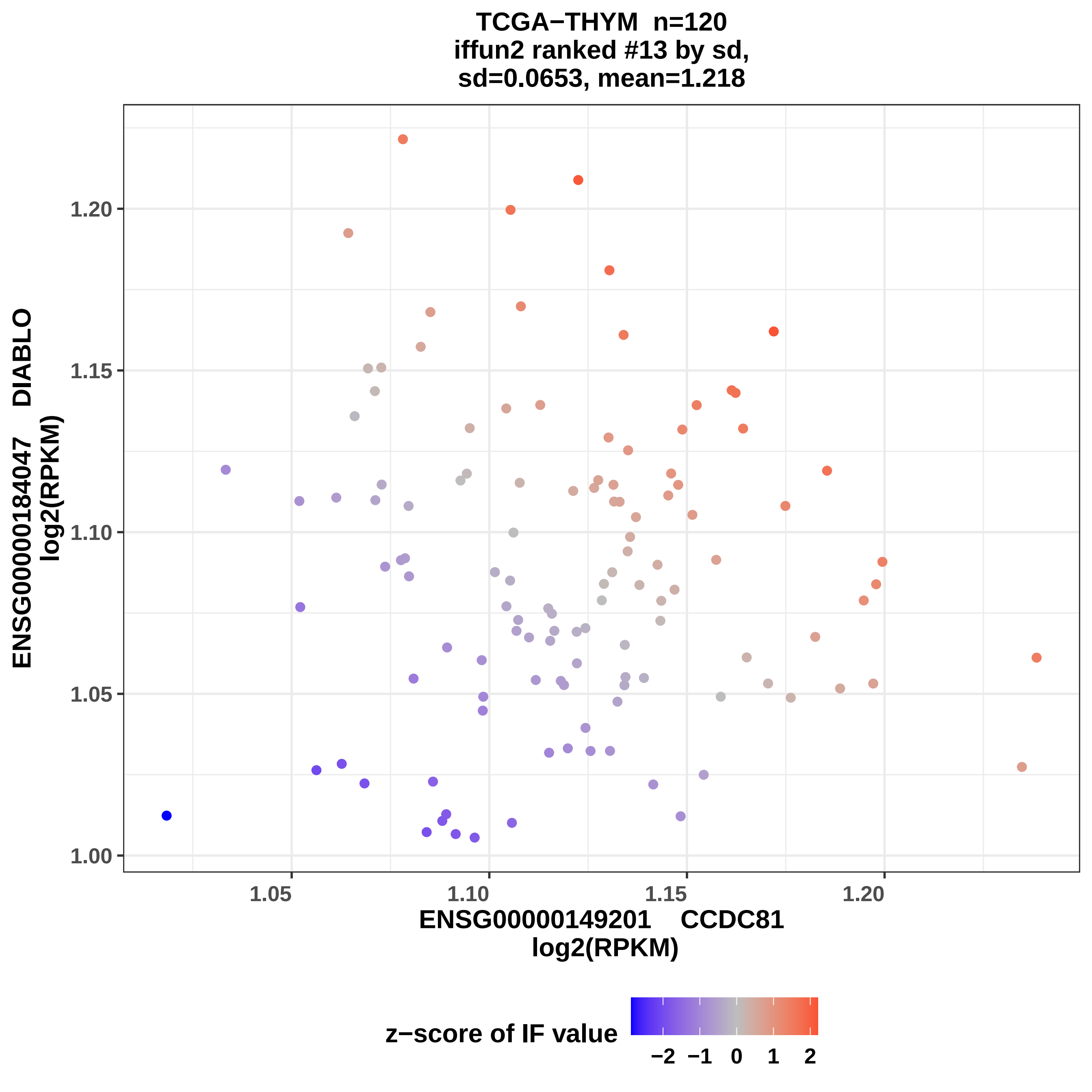
TCGA−THYM n=120
iffun2 ranked #13 by sd,
sd=0.0653, mean=1.218
ENSG00000184047 DIABLO
log2(RPKM)
1.20
1.15
1.10
1.05
1.00
1.05
1.10
1.15
1.20
ENSG00000149201 CCDC81
log2(RPKM)
z−score of IF value
−2
−1
0
1
2

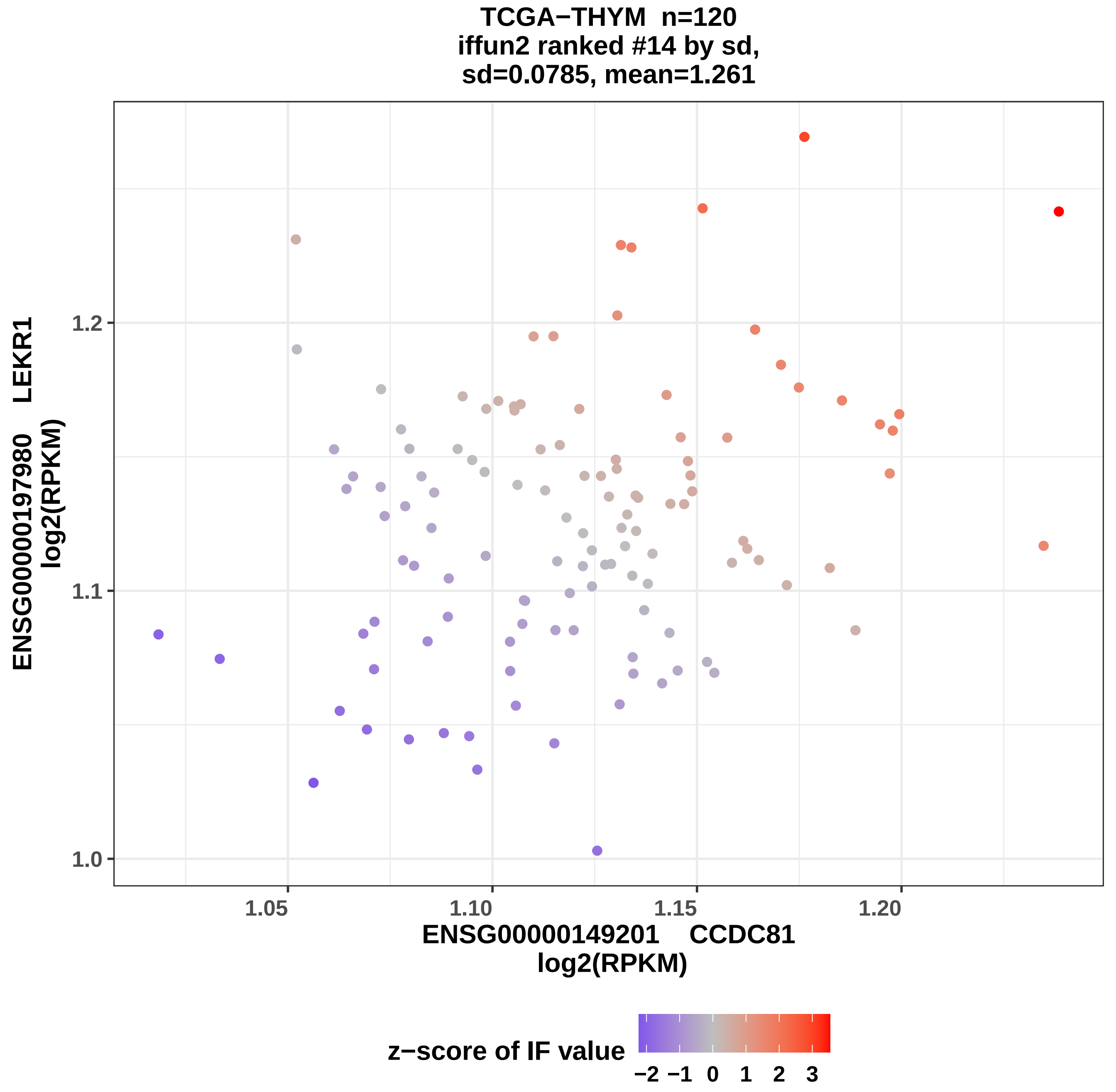
TCGA−THYM n=120
iffun2 ranked #14 by sd,
sd=0.0785, mean=1.261
ENSG00000197980 LEKR1
log2(RPKM)
1.2
1.1
1.0
1.05
1.10
1.15
1.20
ENSG00000149201 CCDC81
log2(RPKM)
z−score of IF value
−2 −1 0 1 2 3

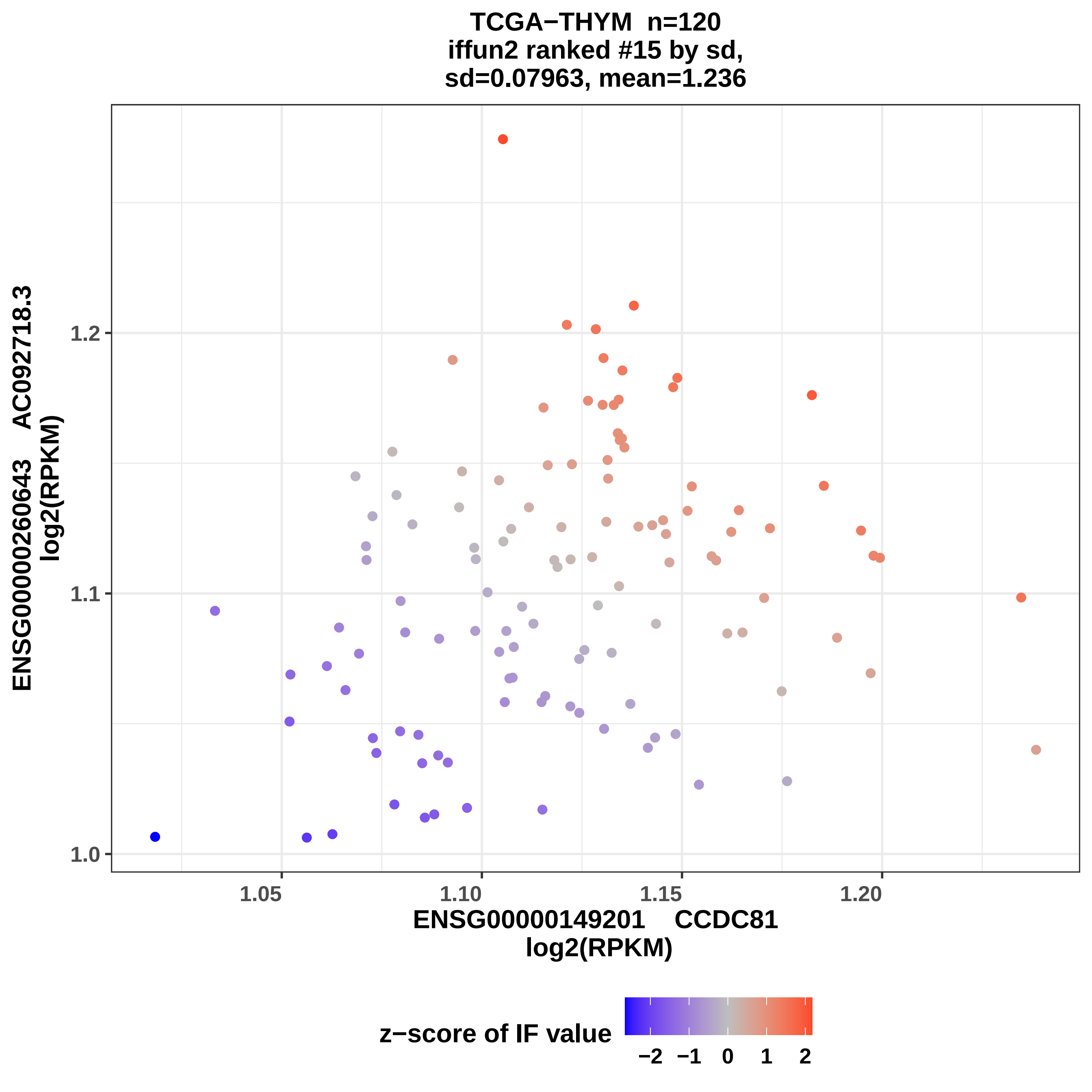
TCGA−THYM n=120
iffun2 ranked #15 by sd,
sd=0.07963, mean=1.236
ENSG00000260643 AC092718.3
log2(RPKM)
1.0
1.1
1.2
1.05
1.10
1.15
1.20
ENSG00000149201 CCDC81
log2(RPKM)
z−score of IF value
−2
−1
0
1
2

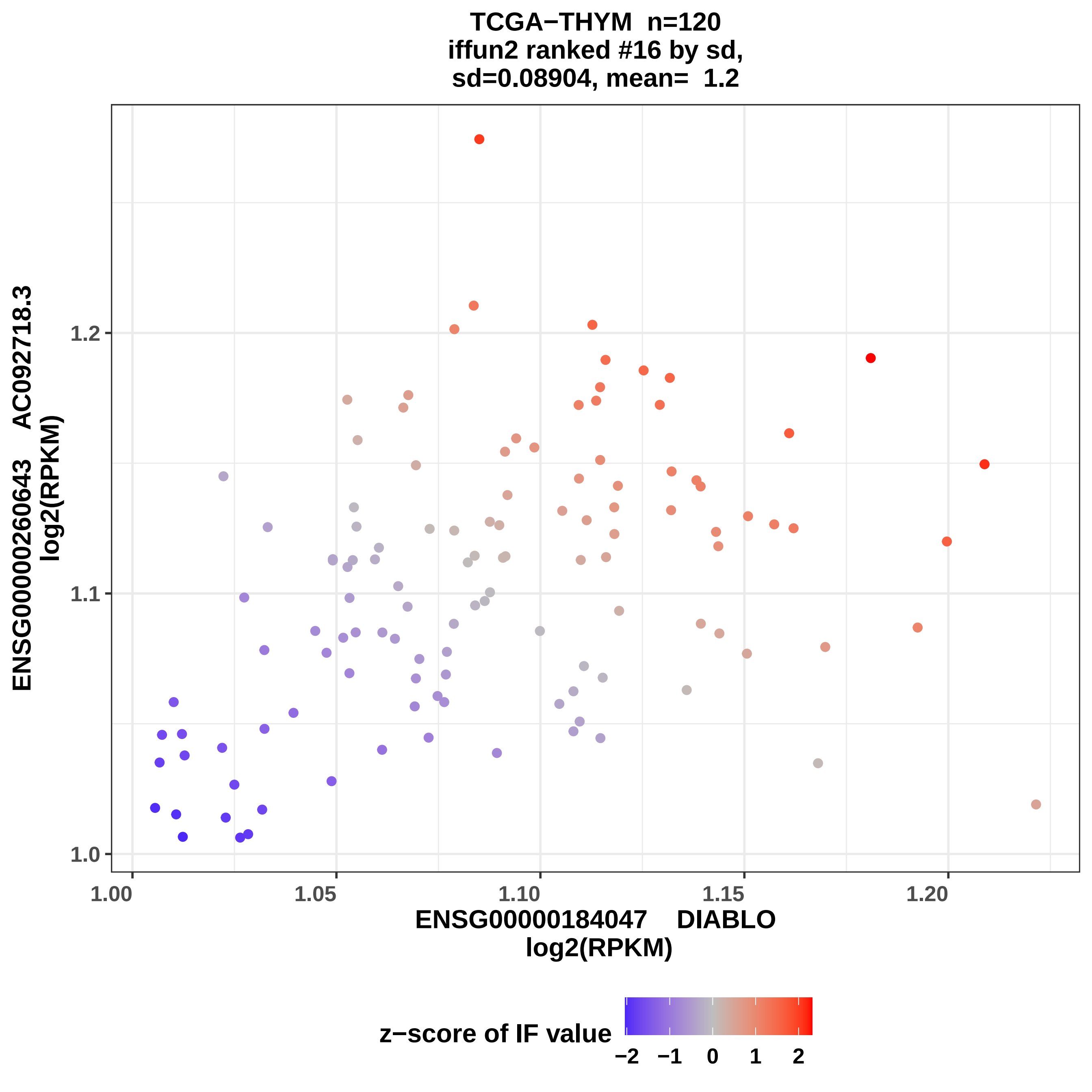
TCGA−THYM n=120
iffun2 ranked #16 by sd,
sd=0.08904, mean= 1.2
ENSG00000260643 AC092718.3
log2(RPKM)
1.0
1.1
1.2
1.00
1.05
1.10
1.15
1.20
ENSG00000184047 DIABLO
log2(RPKM)
z−score of IF value
−2
−1
0
1
2

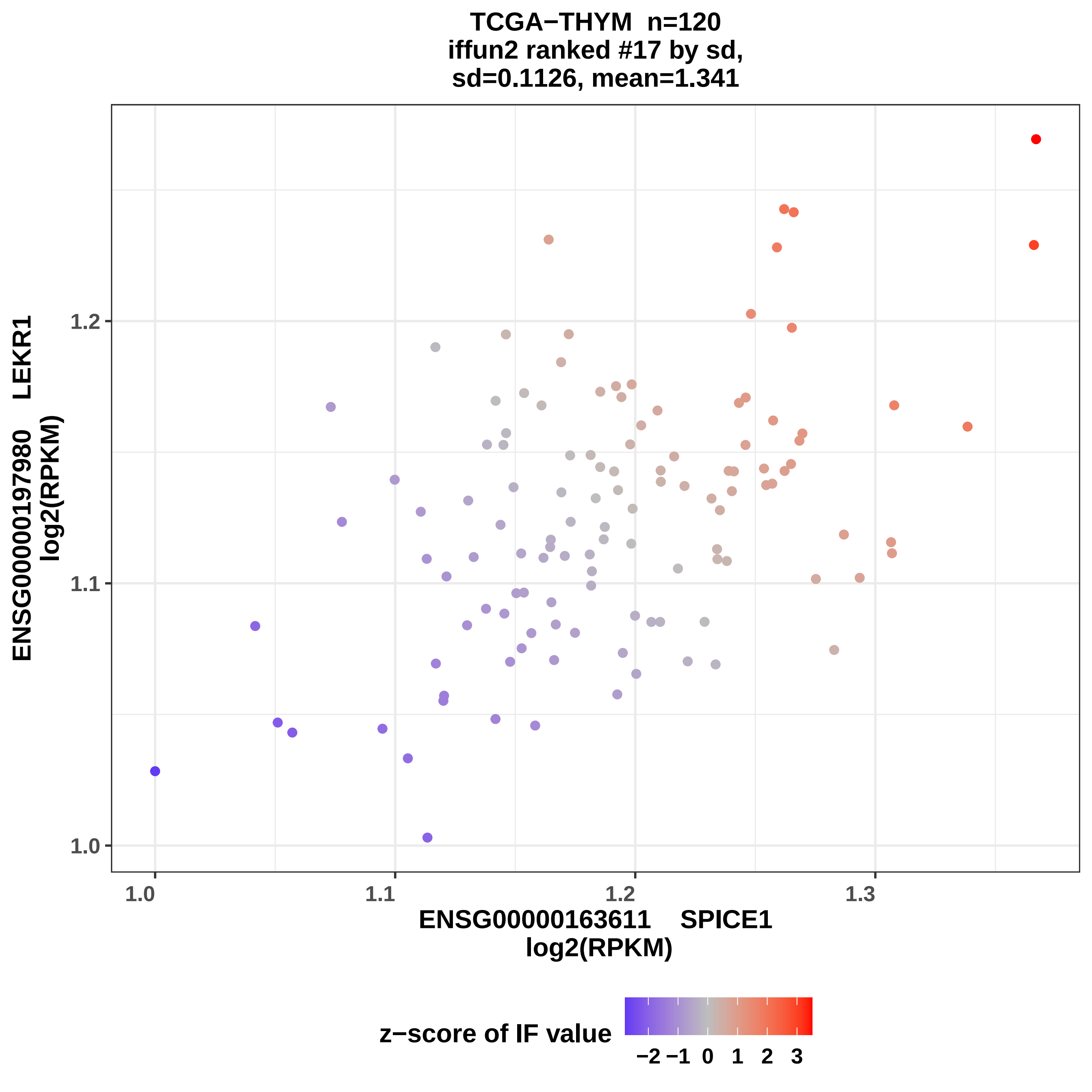
TCGA−THYM n=120
iffun2 ranked #17 by sd,
sd=0.1126, mean=1.341
ENSG00000197980 LEKR1
log2(RPKM)
1.0
1.1
1.2
1.3
ENSG00000163611 SPICE1
log2(RPKM)
z−score of IF value
−2
−1
0
1
2
3

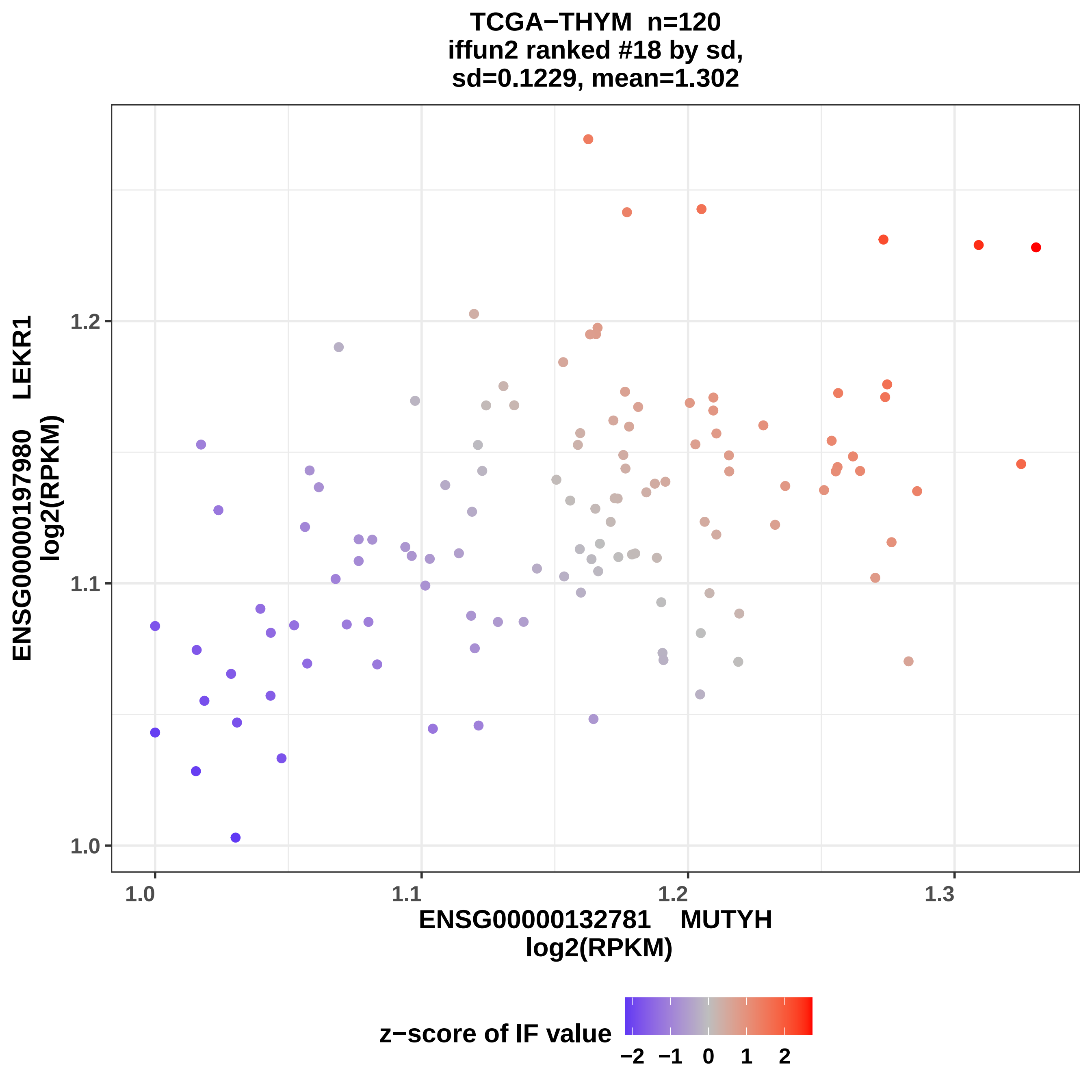

TCGA−THYM n=120
iffun2 ranked #18 by sd,
sd=0.1229, mean=1.302
ENSG00000197980 LEKR1
log2(RPKM)
1.2
1.1
1.0
1.0
1.1
1.2
1.3
ENSG00000132781 MUTYH
log2(RPKM)
z−score of IF value
−2
−1
0
1
2

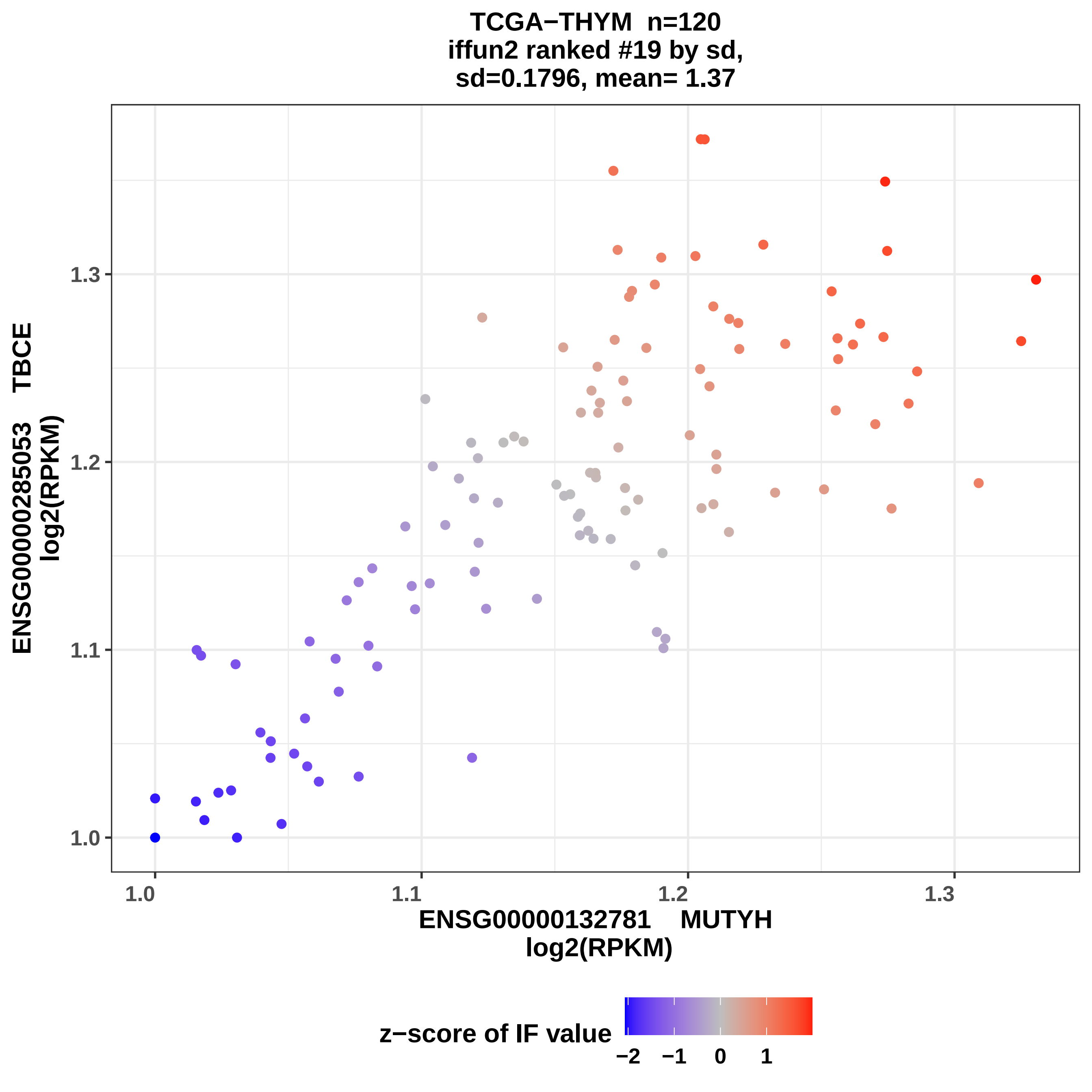

TCGA−THYM n=120
iffun2 ranked #19 by sd,
sd=0.1796, mean= 1.37
ENSG00000285053 TBCE
log2(RPKM)
1.0
1.1
1.2
1.3
ENSG00000132781 MUTYH
log2(RPKM)
z−score of IF value
−2
−1
0
1

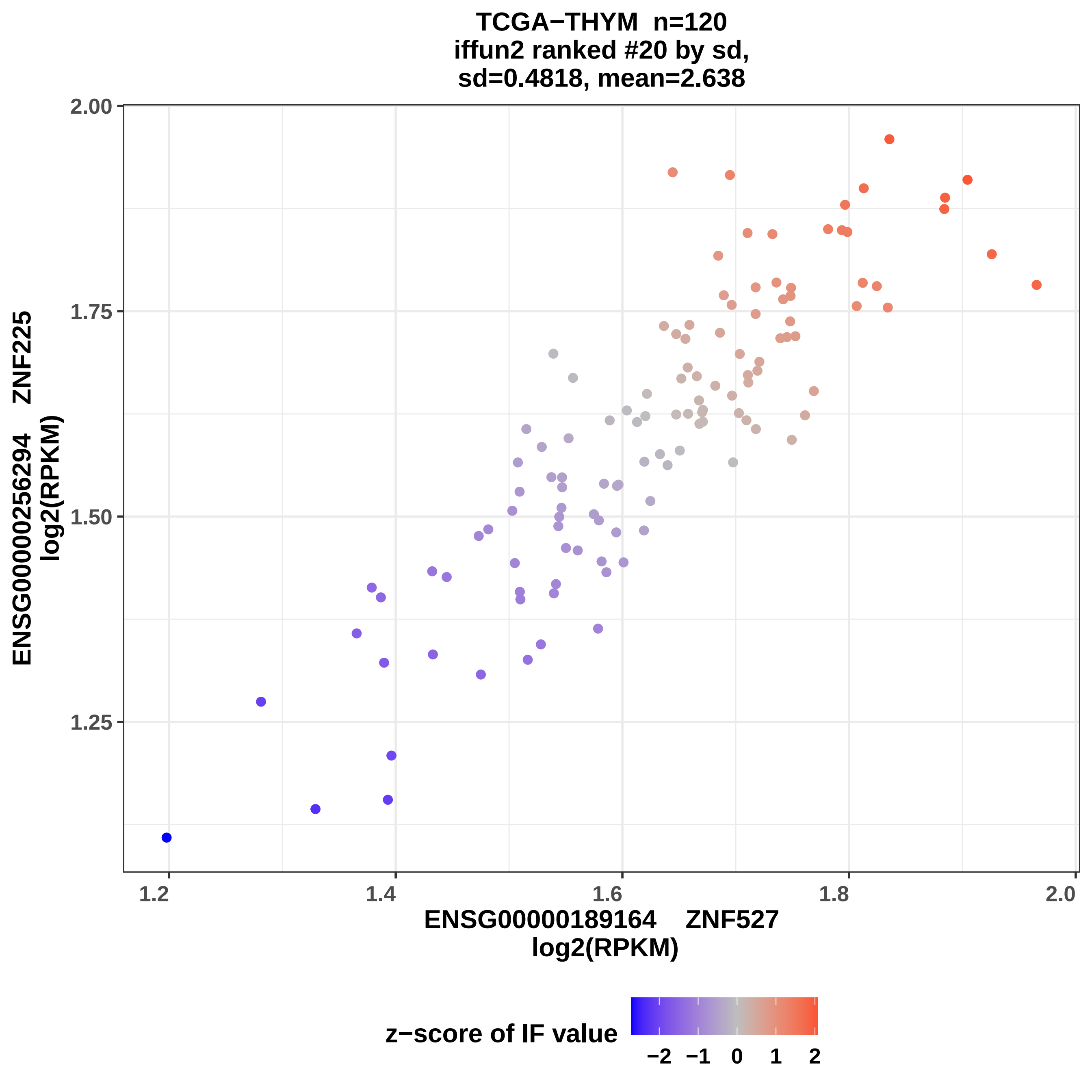

TCGA−THYM n=120
iffun2 ranked #20 by sd,
sd=0.4818, mean=2.638
2.00
1.75
1.50
1.25
1.2
1.4
1.6
1.8
2.0
ENSG00000256294 ZNF225
log2(RPKM)
ENSG00000189164 ZNF527
log2(RPKM)
z−score of IF value
−2
−1
0
1
2

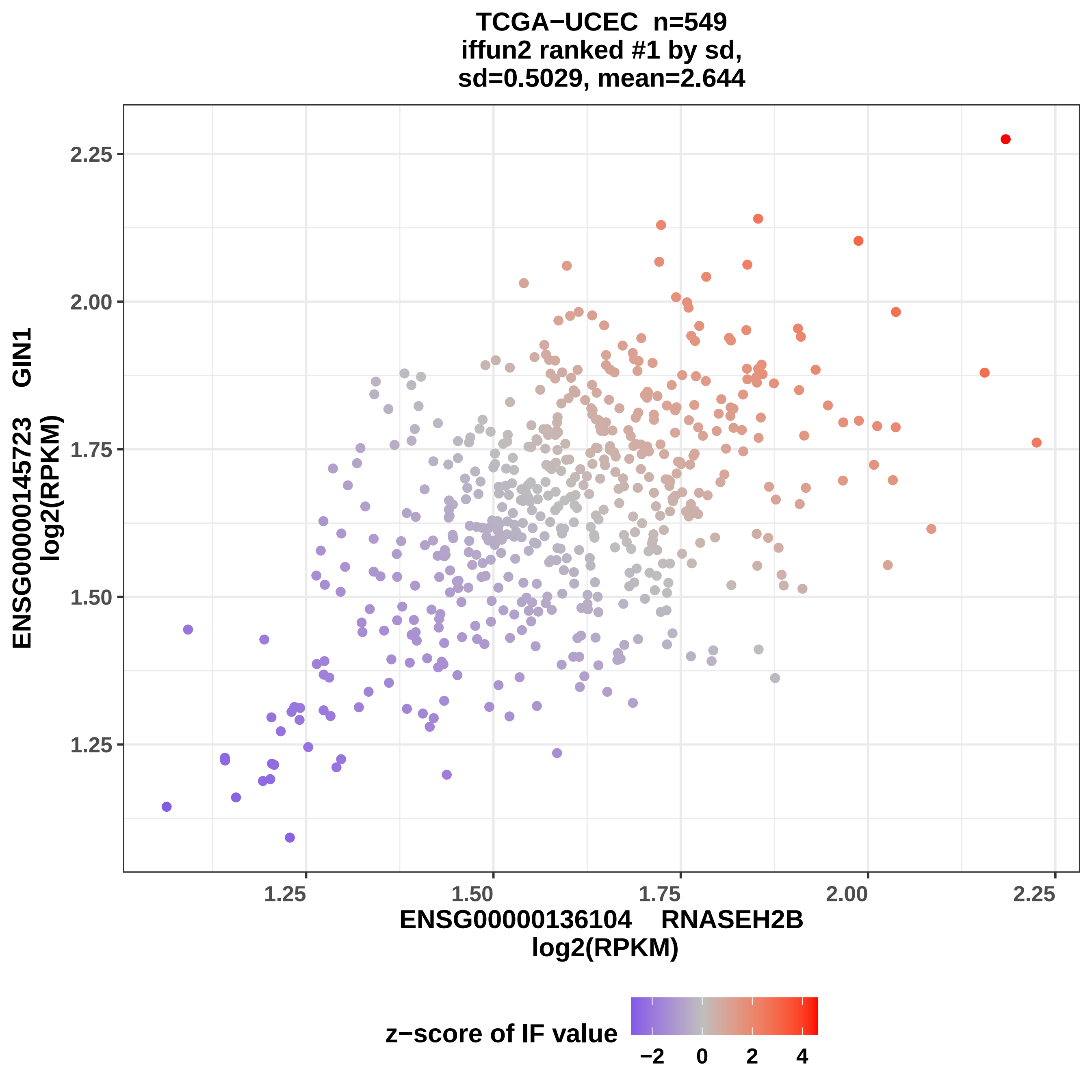
TCGA−UCEC n=549
iffun2 ranked #1 by sd,
sd=0.5029, mean=2.644
ENSG00000145723 GIN1
log2(RPKM)
2.25
2.00
1.75
1.50
1.25
ENSG00000136104 RNASEH2B
log2(RPKM)
1.25
1.50
1.75
2.00
2.25
z−score of IF value
−2
0
2
4

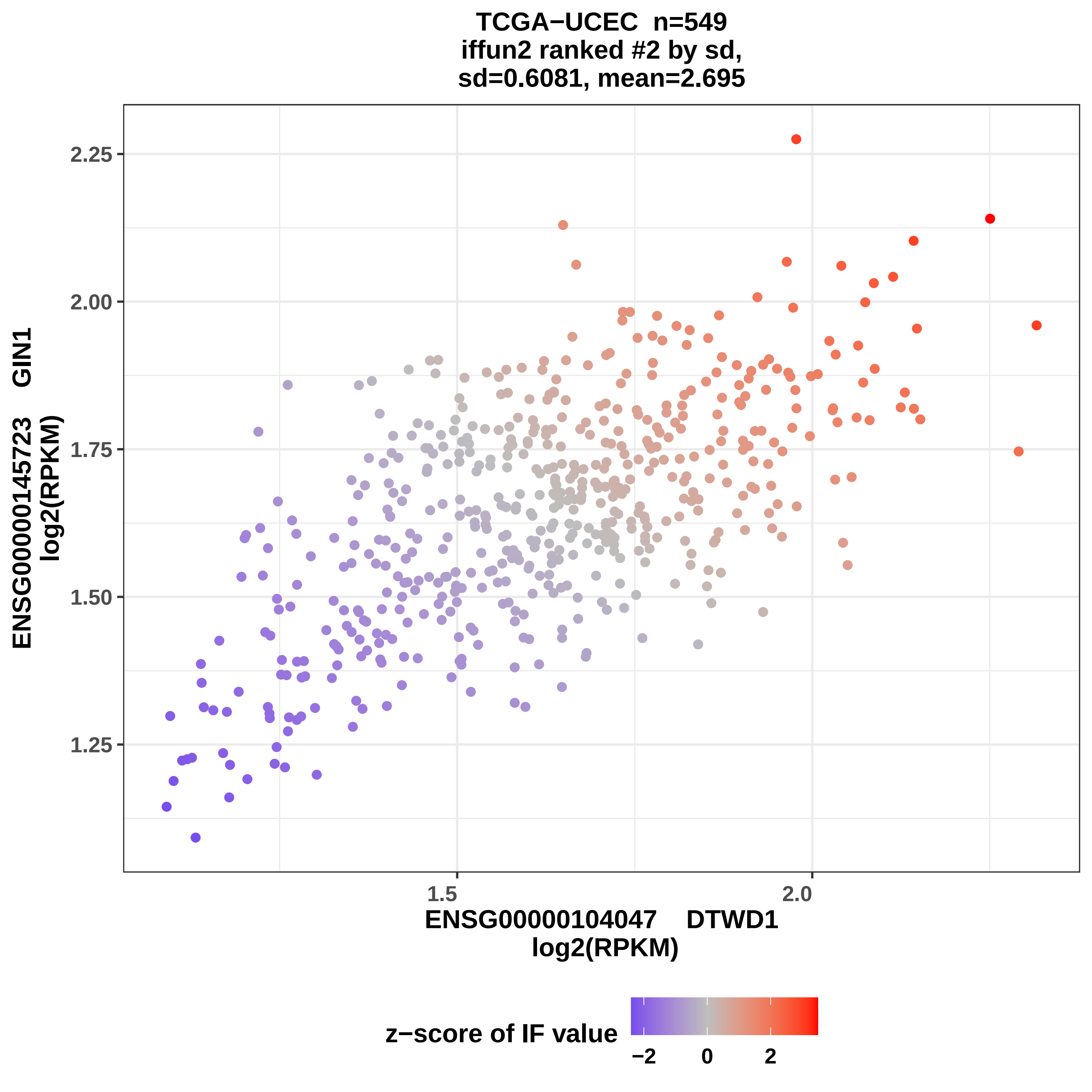

TCGA−UCEC n=549
iffun2 ranked #2 by sd,
sd=0.6081, mean=2.695
2.25
2.00
1.75
1.50
1.25
1.5
2.0
ENSG00000145723 GIN1
log2(RPKM)
ENSG00000104047 DTWD1
log2(RPKM)
z−score of IF value
−2
0
2

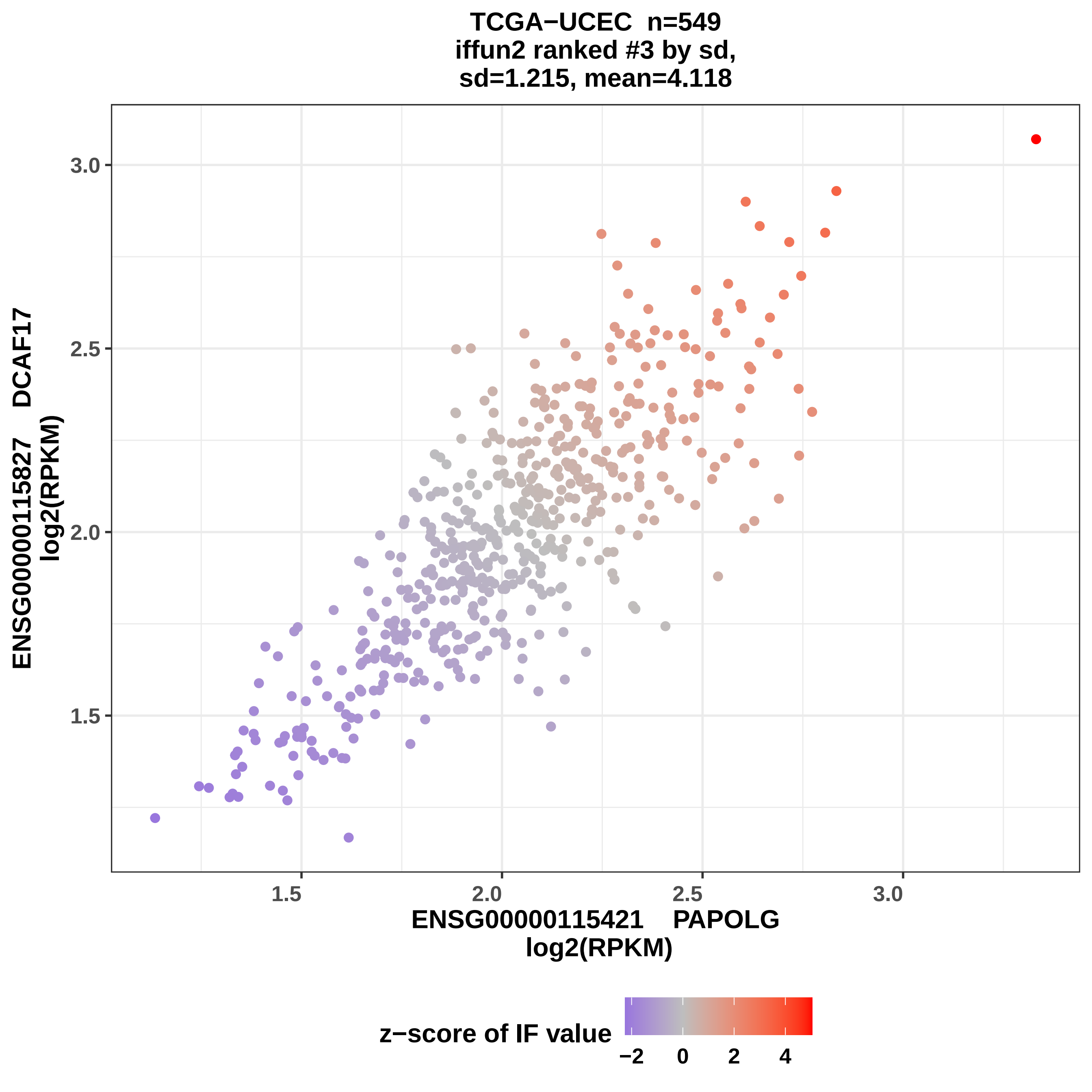
TCGA−UCEC n=549
iffun2 ranked #3 by sd,
sd=1.215, mean=4.118
3.0
2.5
2.0
1.5
1.5
2.0
2.5
3.0
ENSG00000115827 DCAF17
log2(RPKM)
ENSG00000115421 PAPOLG
log2(RPKM)
z−score of IF value
−2
0
2
4

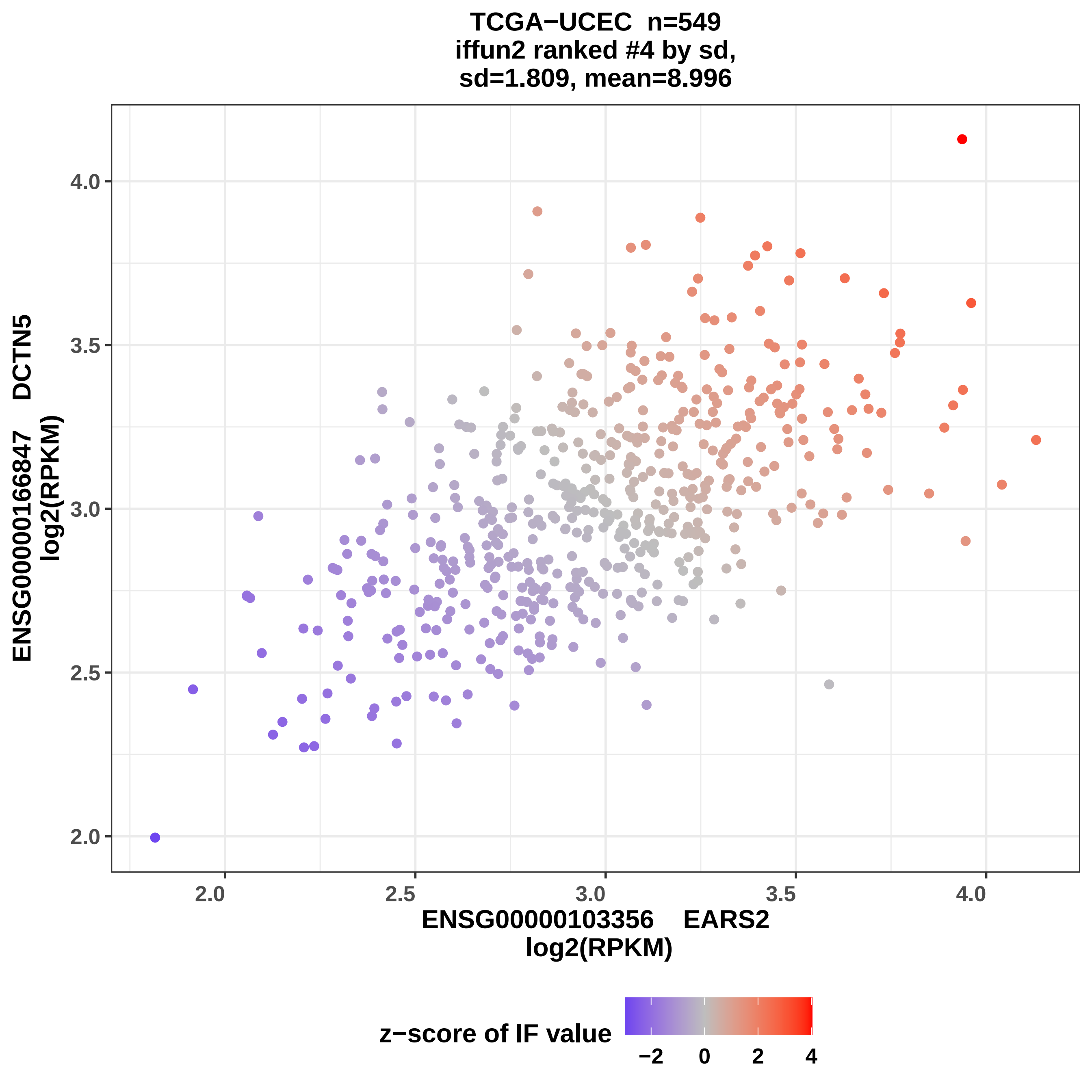

TCGA−UCEC n=549
iffun2 ranked #4 by sd,
sd=1.809, mean=8.996
ENSG00000166847 DCTN5
log2(RPKM)
4.0
3.5
3.0
2.5
2.0
2.0
2.5
3.0
3.5
4.0
ENSG00000103356 EARS2
log2(RPKM)
z−score of IF value
−2
0
2
4

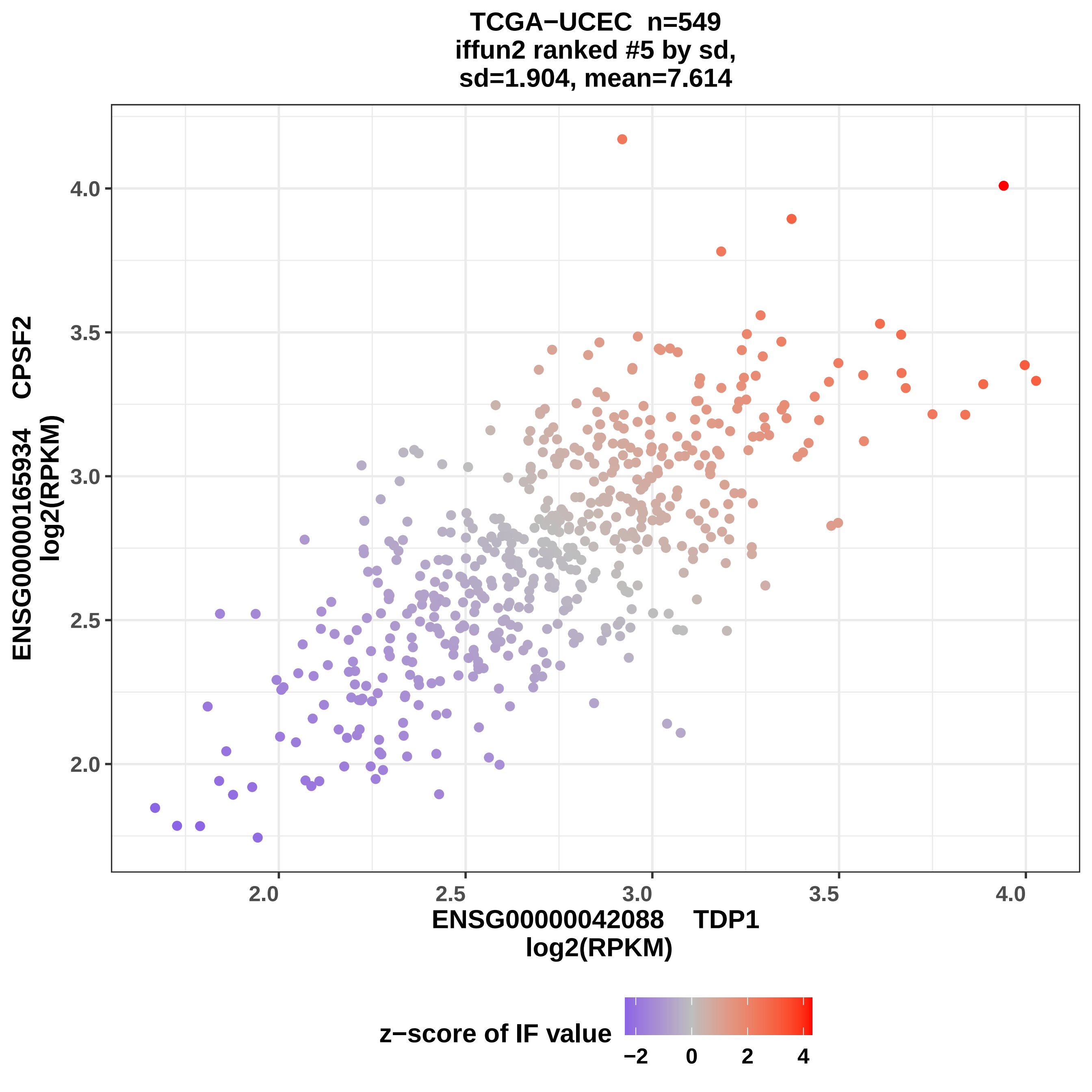
TCGA−UCEC n=549
iffun2 ranked #5 by sd,
sd=1.904, mean=7.614
ENSG00000165934 CPSF2
log2(RPKM)
4.0
3.5
3.0
2.5
2.0
2.0
2.5
3.0
3.5
4.0
ENSG00000042088 TDP1
log2(RPKM)
z−score of IF value
−2
0
2
4

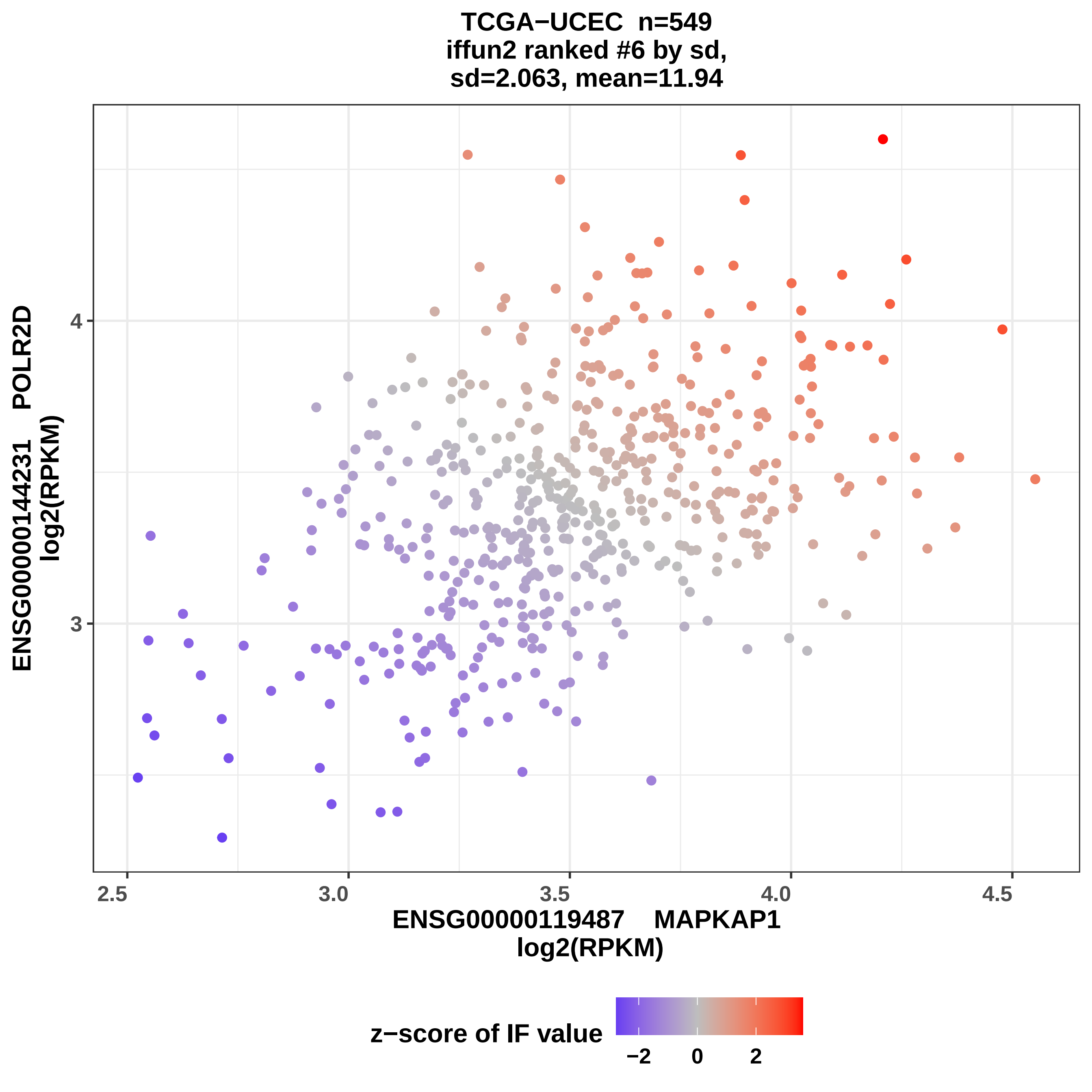
TCGA−UCEC n=549
iffun2 ranked #6 by sd,
sd=2.063, mean=11.94
ENSG00000144231 POLR2D
log2(RPKM)
4
3
2.5
3.0
3.5
4.0
4.5
ENSG00000119487 MAPKAP1
log2(RPKM)
z−score of IF value
−2
0
2

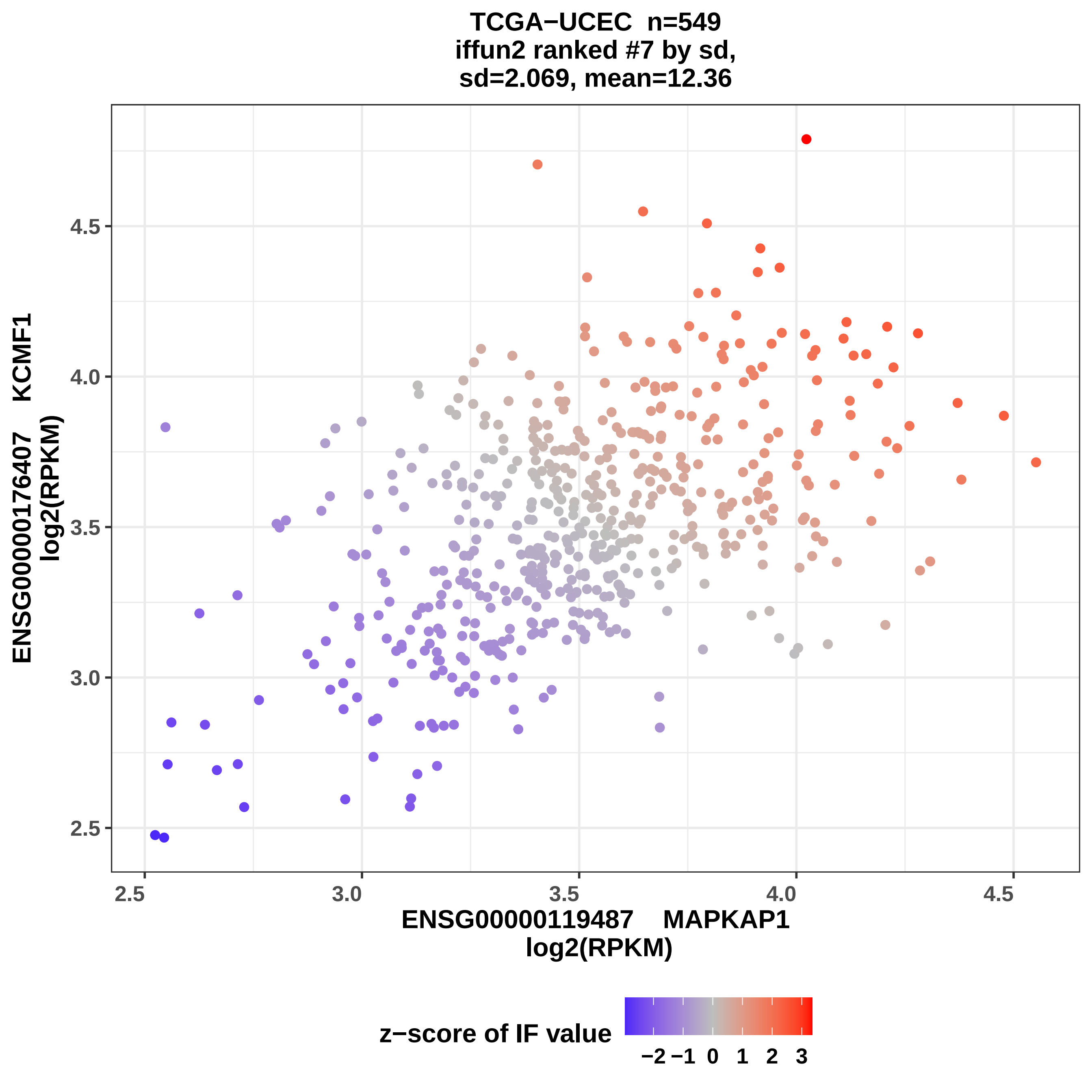

TCGA−UCEC n=549
iffun2 ranked #7 by sd,
sd=2.069, mean=12.36
ENSG00000176407 KCMF1
log2(RPKM)
ENSG00000119487 MAPKAP1
log2(RPKM)
2.5
3.0
3.5
4.0
4.5
z−score of IF value
−2 −1 0 1 2 3

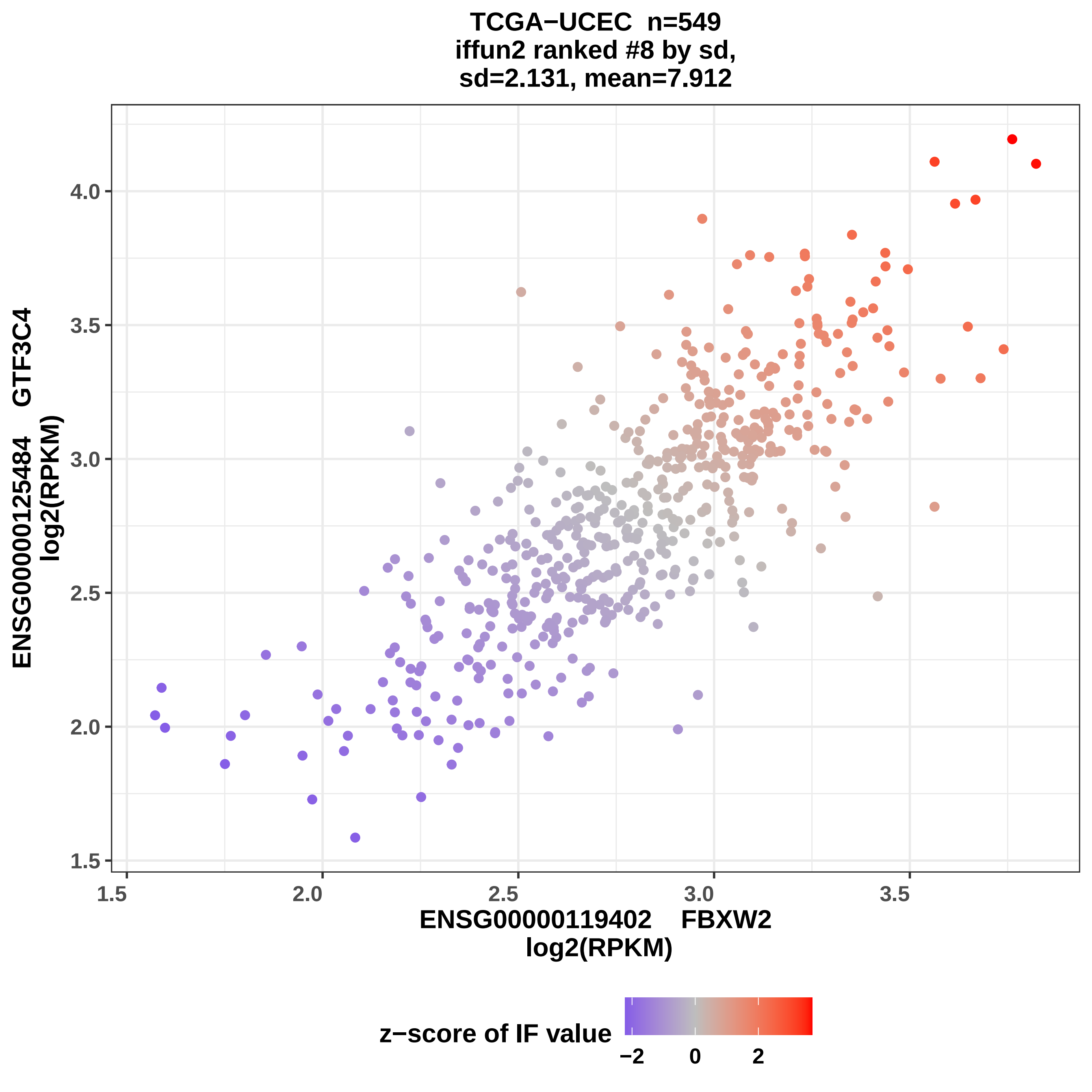
TCGA−UCEC n=549
iffun2 ranked #8 by sd,
sd=2.131, mean=7.912
ENSG00000125484 GTF3C4
log2(RPKM)
4.0
3.5
3.0
2.5
2.0
1.5
1.5
2.0
2.5
3.0
3.5
ENSG00000119402 FBXW2
log2(RPKM)
z−score of IF value
−2
0
2

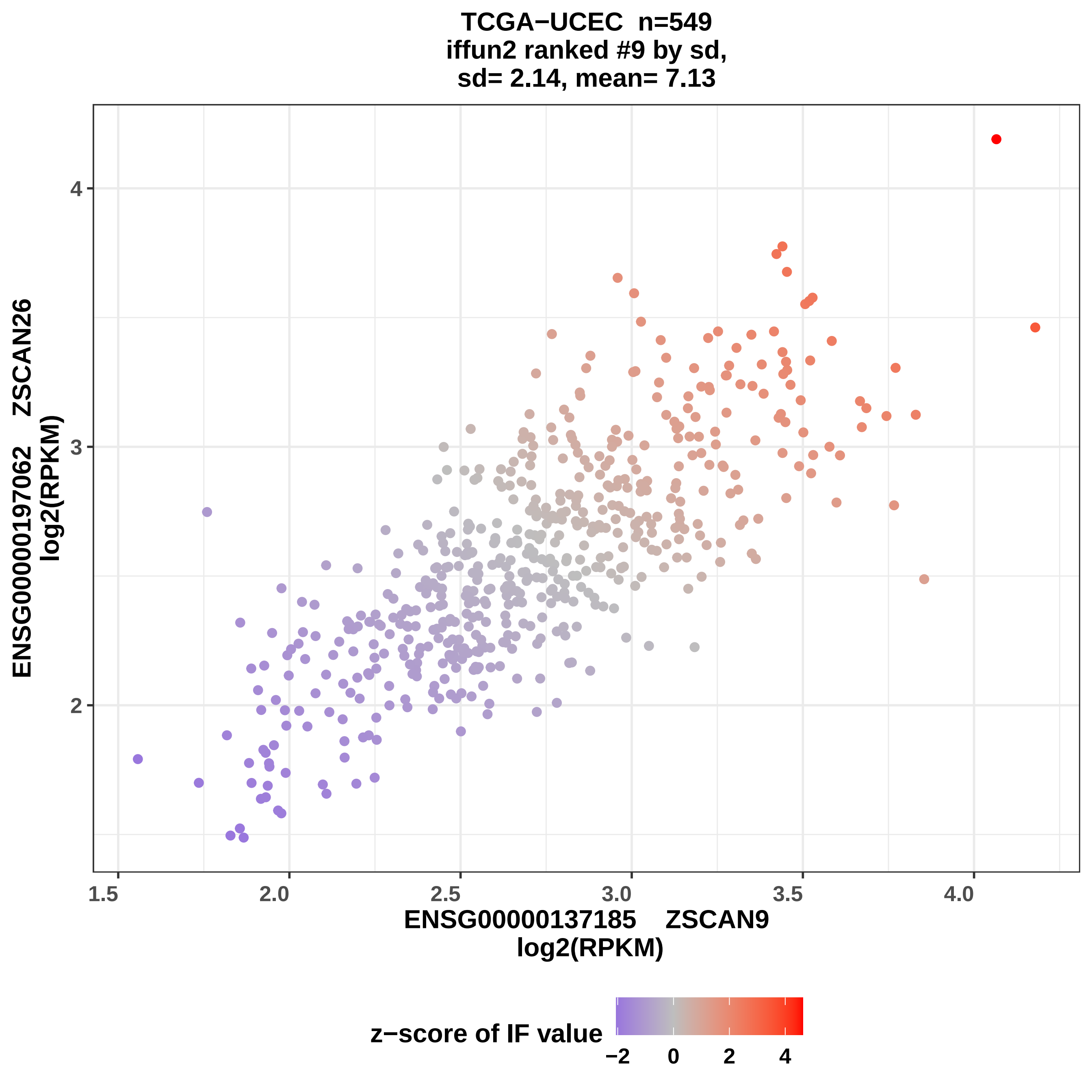

TCGA−UCEC n=549
iffun2 ranked #9 by sd,
sd= 2.14, mean= 7.13
ENSG00000197062 ZSCAN26
log2(RPKM)
4
3
2
1.5
2.0
2.5
3.0
3.5
4.0
ENSG00000137185 ZSCAN9
log2(RPKM)
z−score of IF value
−2
0
2
4

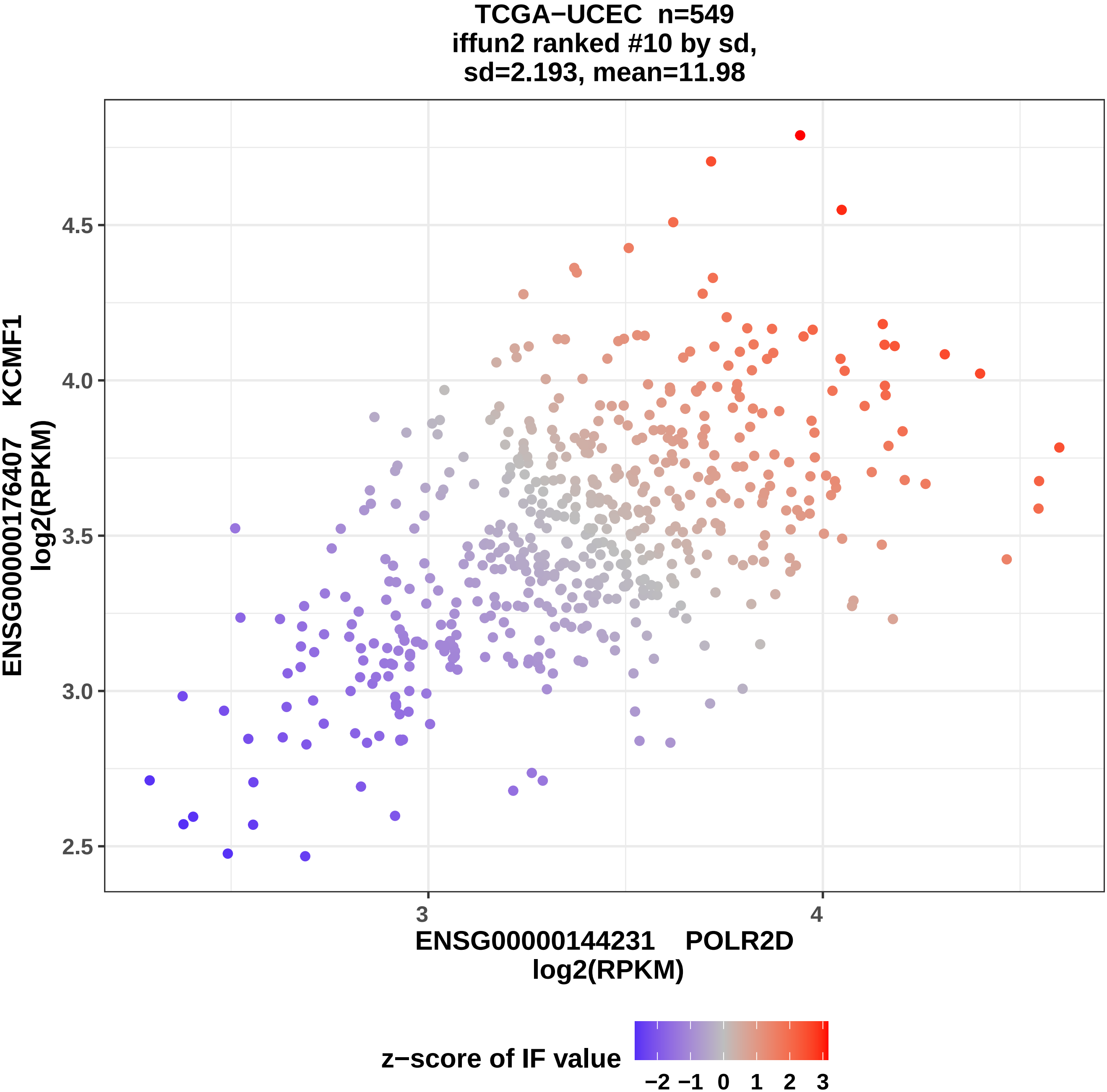

TCGA−UCEC n=549
iffun2 ranked #10 by sd,
sd=2.193, mean=11.98
ENSG00000176407 KCMF1
log2(RPKM)
4.5
4.0
3.5
3.0
2.5
3
4
ENSG00000144231 POLR2D
log2(RPKM)
z−score of IF value
−2
−1
0
1
2
3

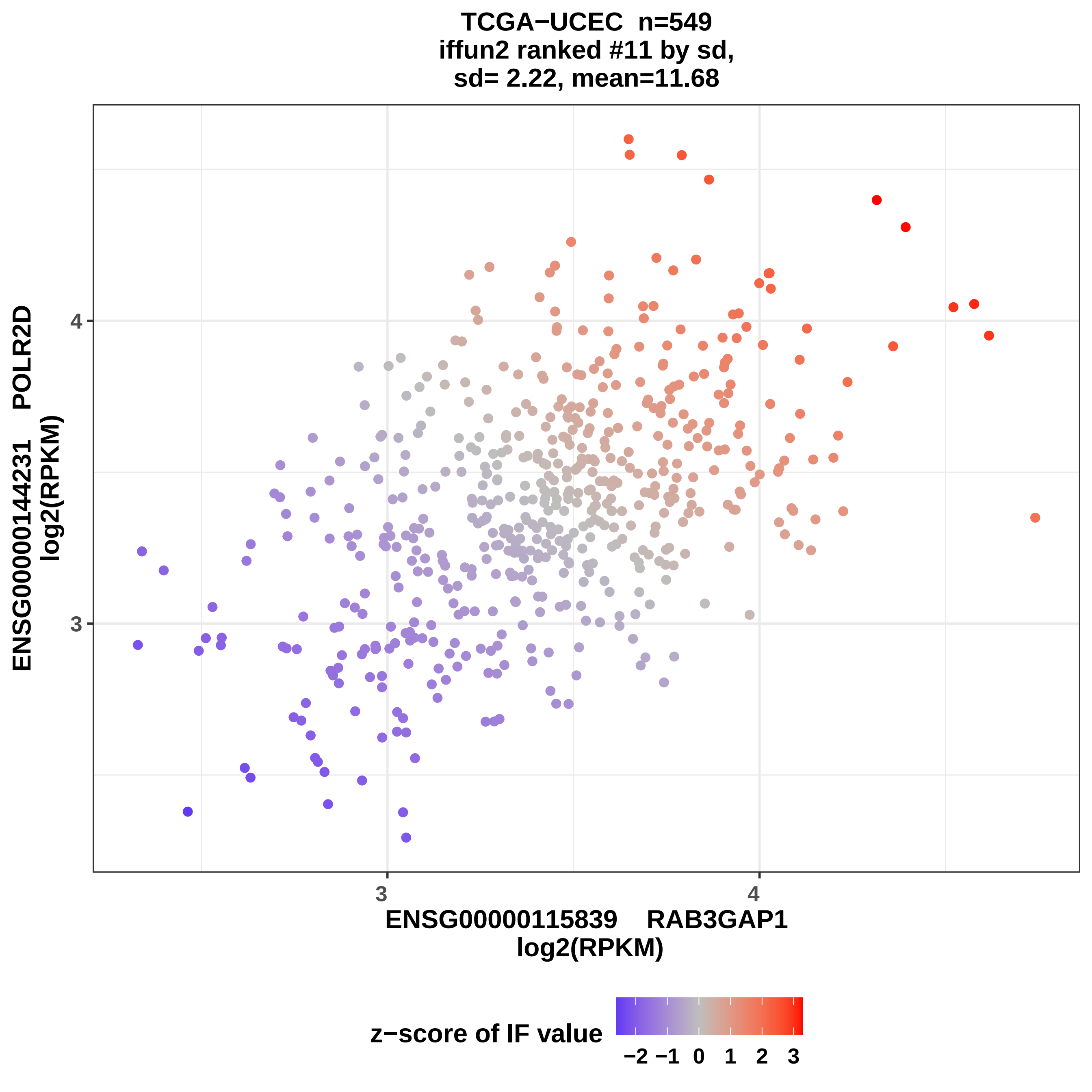

TCGA−UCEC n=549
iffun2 ranked #11 by sd,
sd= 2.22, mean=11.68
ENSG00000144231 POLR2D
log2(RPKM)
4
3
ENSG00000115839 RAB3GAP1
log2(RPKM)
z−score of IF value
−2 −1 0 1 2 3

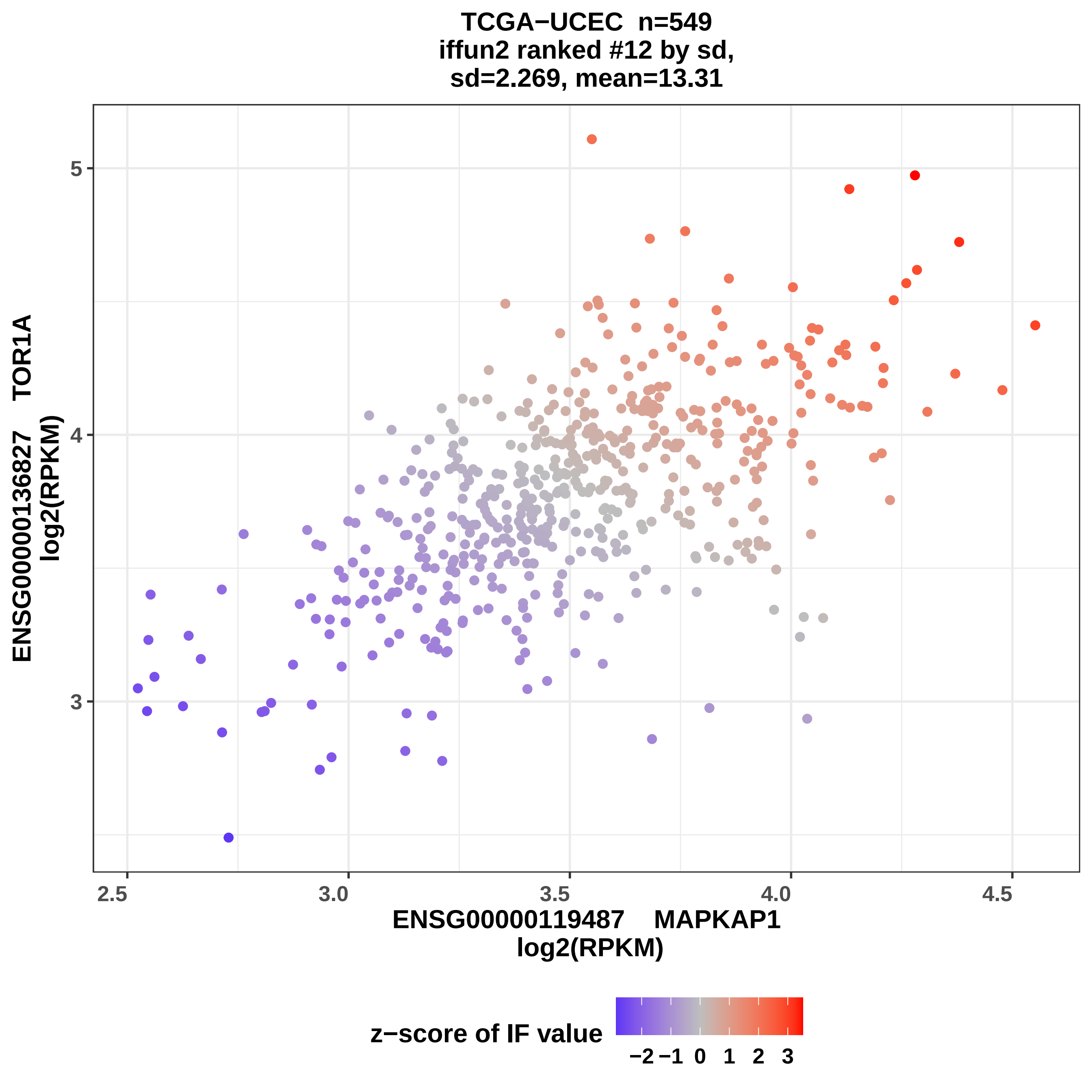

TCGA−UCEC n=549
iffun2 ranked #12 by sd,
sd=2.269, mean=13.31
5
4
3
2.5
3.0
3.5
4.0
4.5
ENSG00000136827 TOR1A
log2(RPKM)
ENSG00000119487 MAPKAP1
log2(RPKM)
z−score of IF value
−2
−1
0
1
2
3

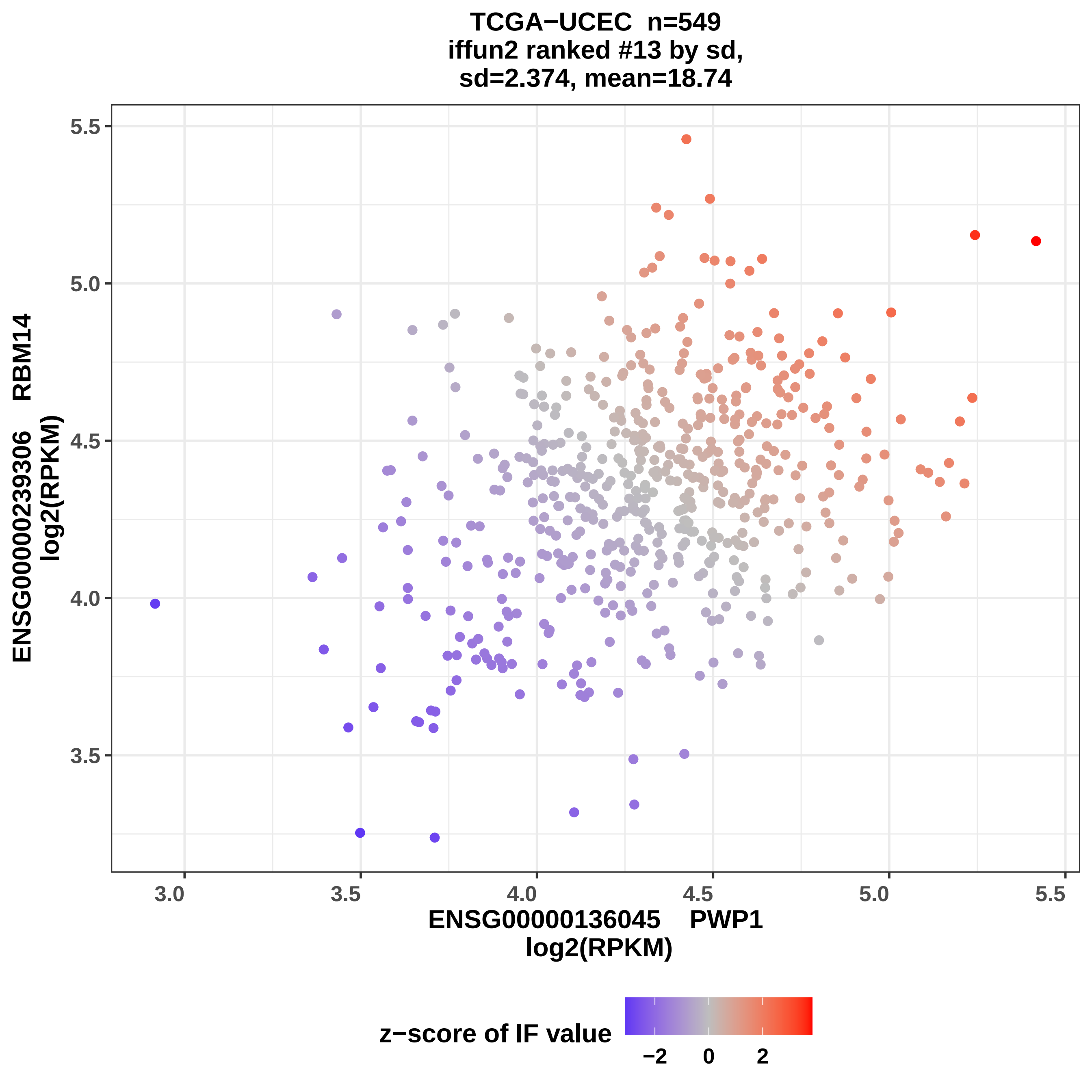

TCGA−UCEC n=549
iffun2 ranked #13 by sd,
sd=2.374, mean=18.74
ENSG00000239306 RBM14
log2(RPKM)
5.5
5.0
4.5
4.0
3.5
3.0
3.5
4.0
4.5
5.0
5.5
ENSG00000136045 PWP1
log2(RPKM)
z−score of IF value
−2
0
2

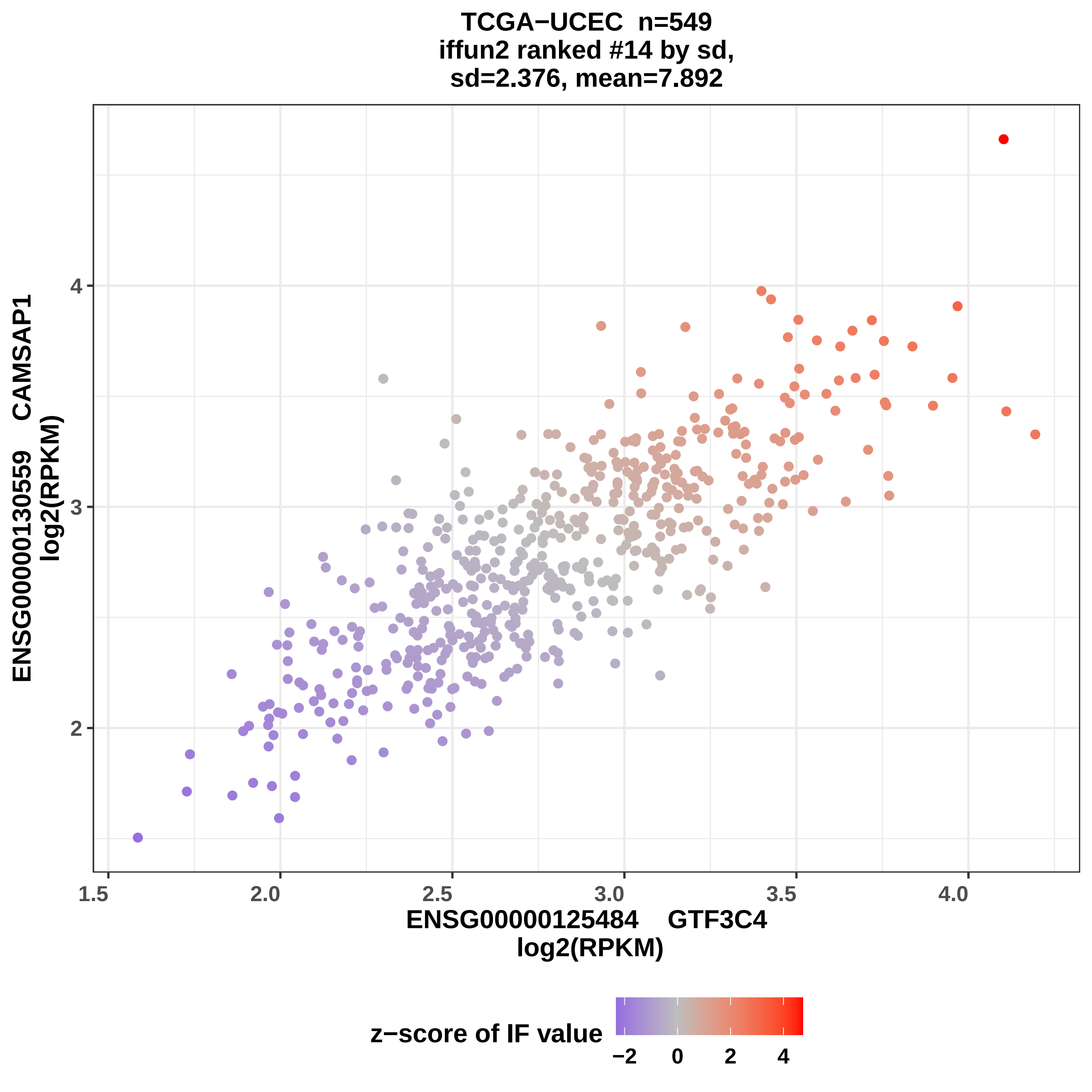
TCGA−UCEC n=549
iffun2 ranked #14 by sd,
sd=2.376, mean=7.892
ENSG00000130559 CAMSAP1
log2(RPKM)
ENSG00000125484 GTF3C4
log2(RPKM)
z−score of IF value
−2
0
2
4
1.5
2.0
2.5
3.0
3.5
4.0
2
3
4

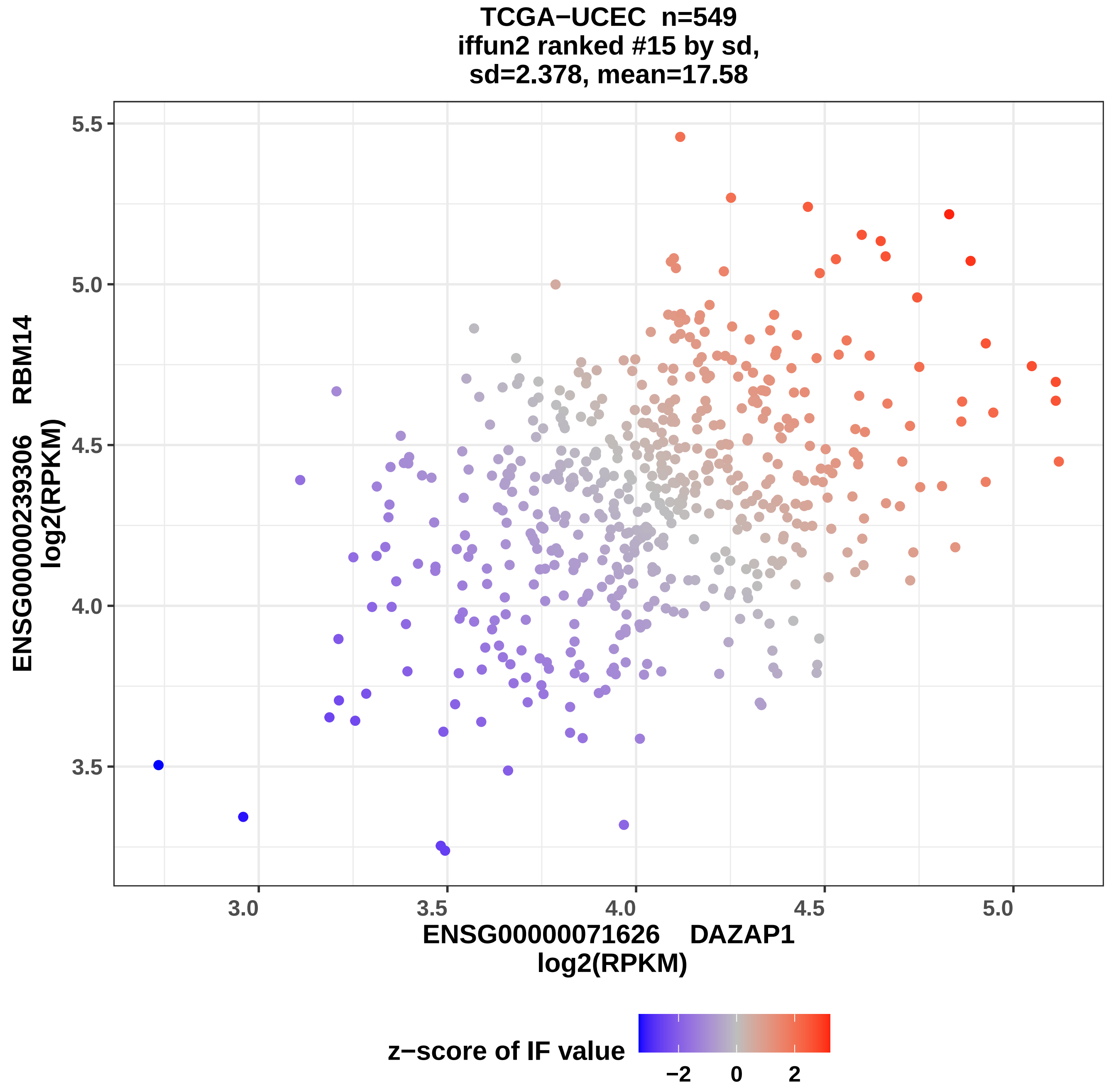

TCGA−UCEC n=549
iffun2 ranked #15 by sd,
sd=2.378, mean=17.58
5.5
5.0
4.5
4.0
3.5
3.0
3.5
4.0
4.5
5.0
ENSG00000239306 RBM14
log2(RPKM)
ENSG00000071626 DAZAP1
log2(RPKM)
z−score of IF value
−2
0
2

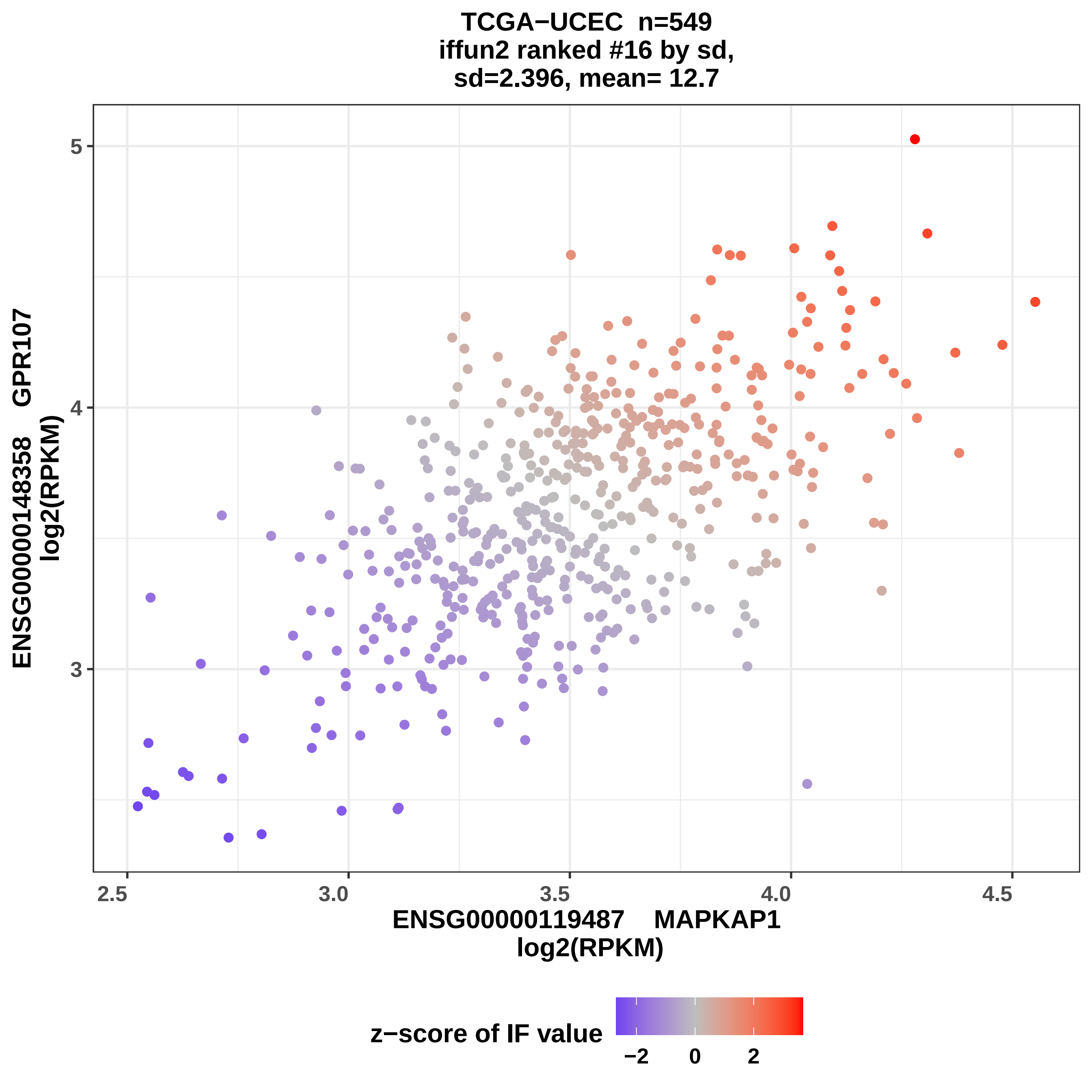

TCGA−UCEC n=549
iffun2 ranked #16 by sd,
sd=2.396, mean= 12.7
ENSG00000148358 GPR107
log2(RPKM)
5
4
3
2.5
3.0
3.5
4.0
4.5
ENSG00000119487 MAPKAP1
log2(RPKM)
z−score of IF value
−2
0
2

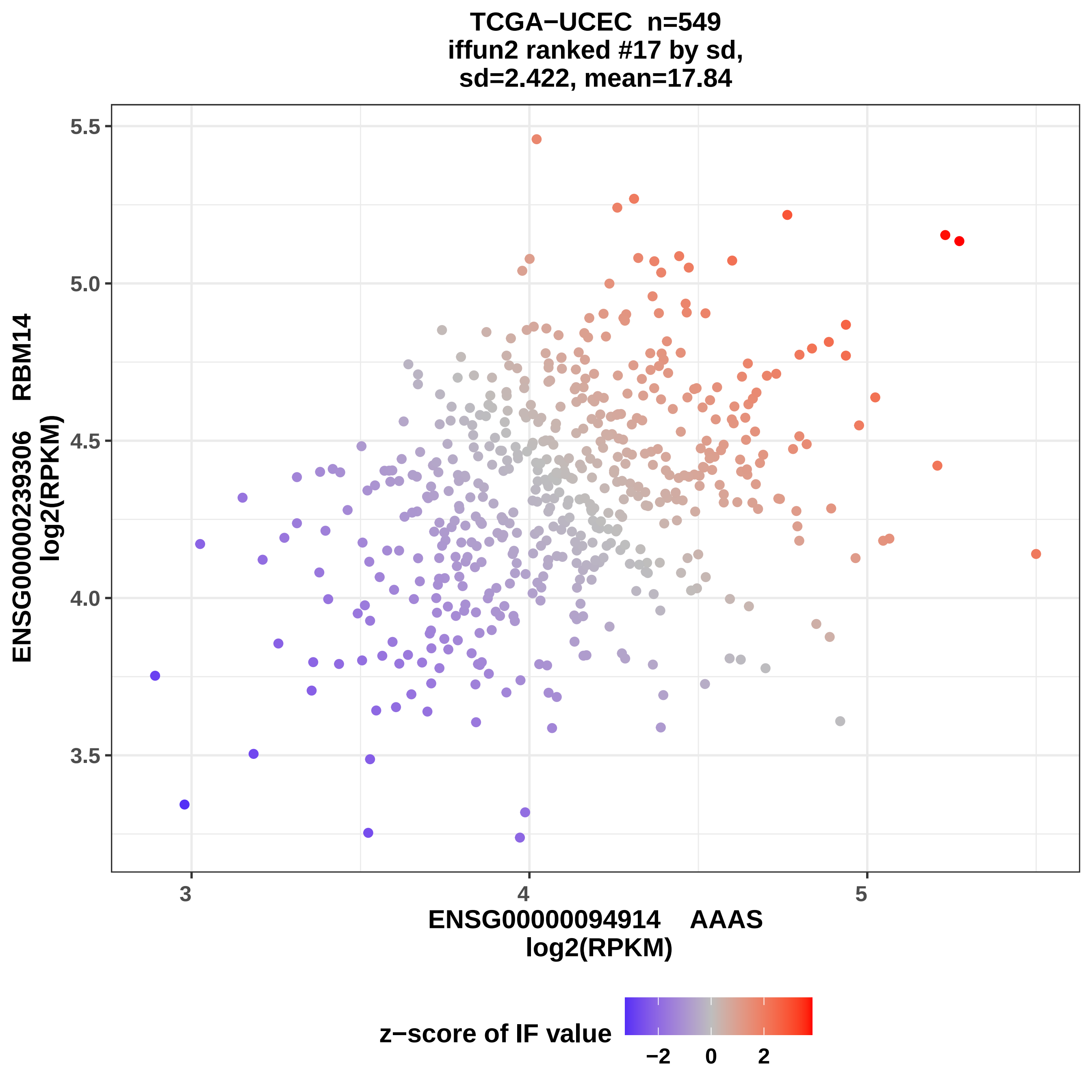
TCGA−UCEC n=549
iffun2 ranked #17 by sd,
sd=2.422, mean=17.84
ENSG00000239306 RBM14
log2(RPKM)
5.5
5.0
4.5
4.0
3.5
3
4
5
ENSG00000094914 AAAS
log2(RPKM)
z−score of IF value
−2
0
2

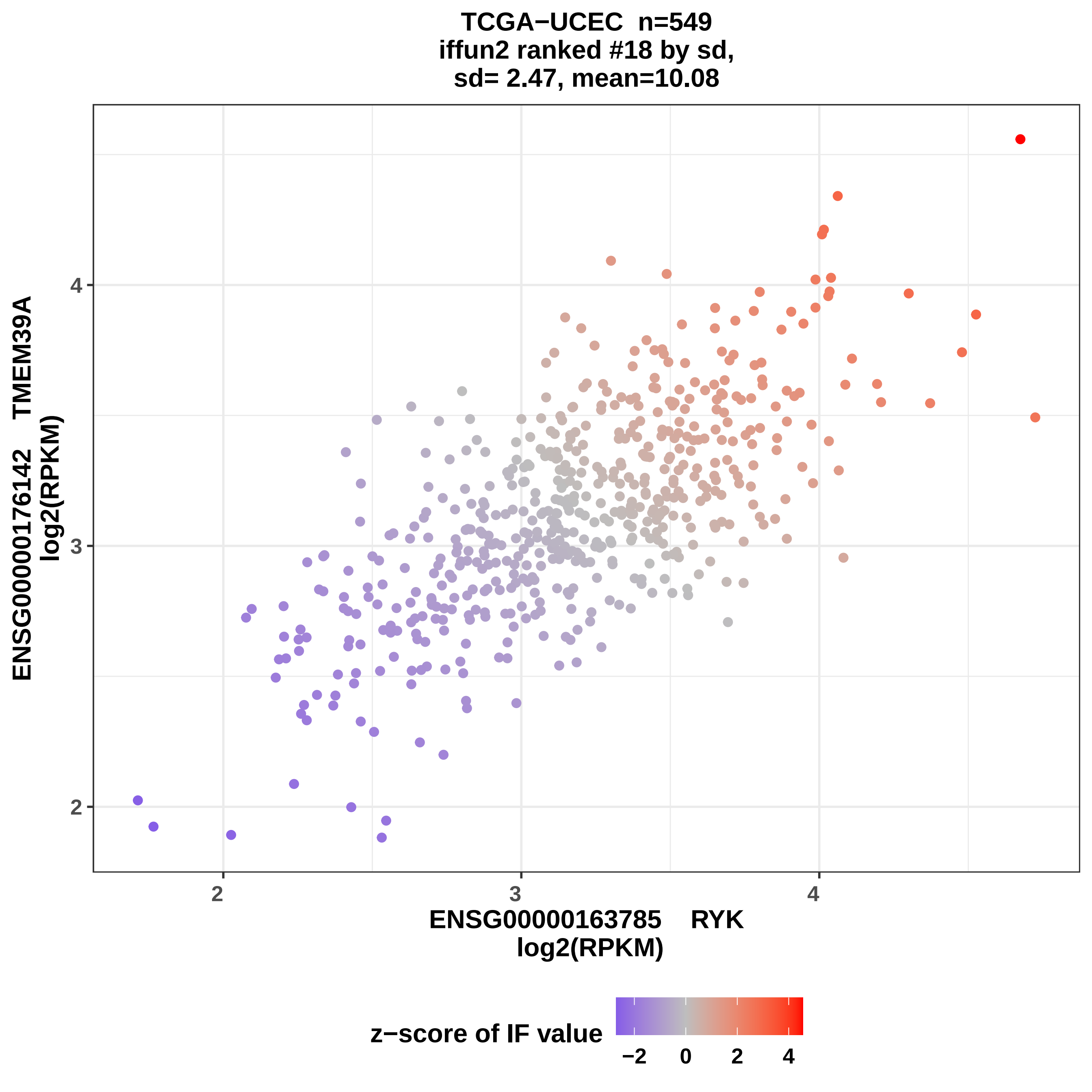
TCGA−UCEC n=549
iffun2 ranked #18 by sd,
sd= 2.47, mean=10.08
ENSG00000176142 TMEM39A
log2(RPKM)
4
3
2
2
3
4
ENSG00000163785 RYK
log2(RPKM)
z−score of IF value
−2
0
2
4

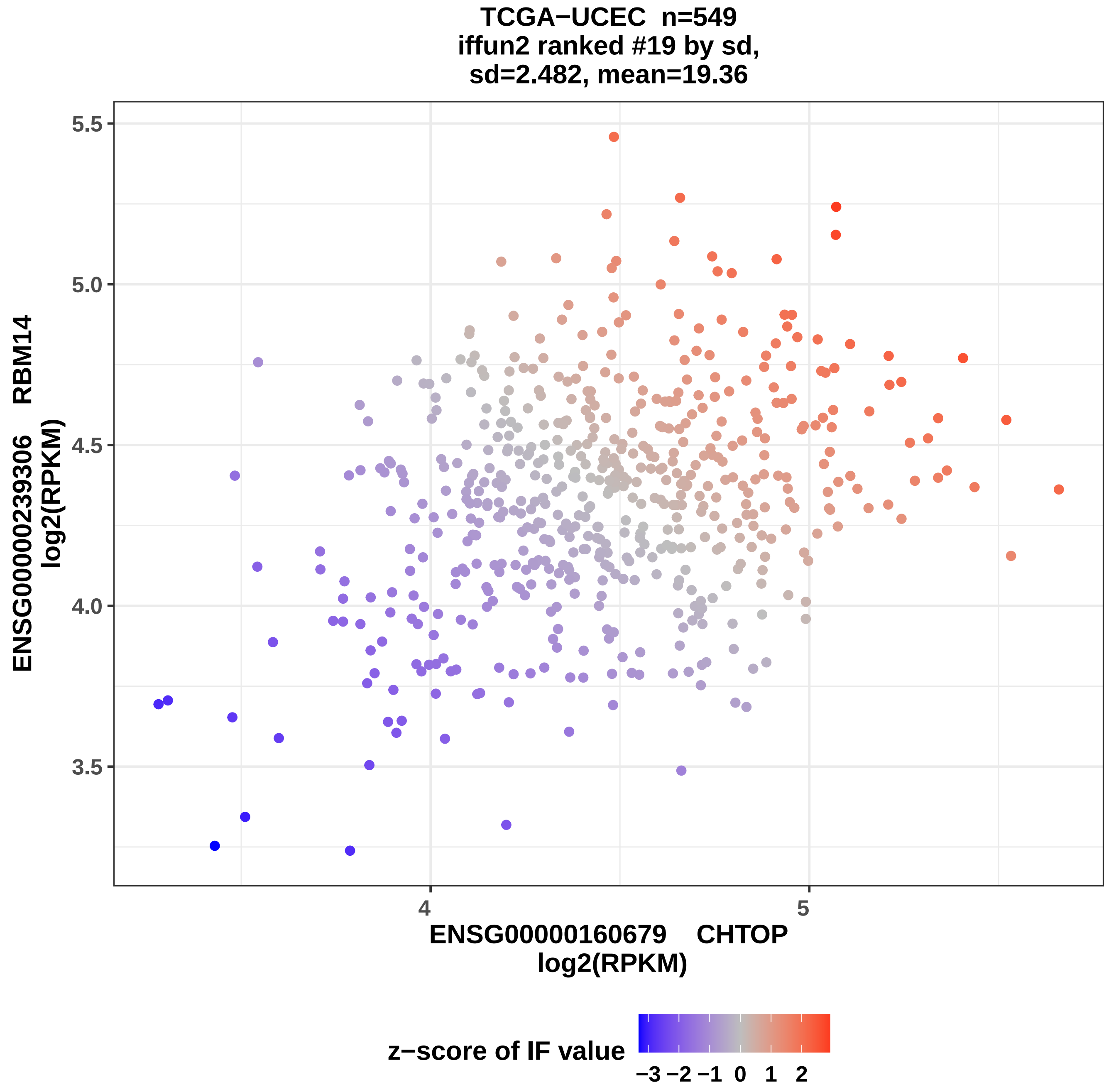
TCGA−UCEC n=549
iffun2 ranked #19 by sd,
sd=2.482, mean=19.36
ENSG00000239306 RBM14
log2(RPKM)
5.5
5.0
4.5
4.0
3.5
4
5
ENSG00000160679 CHTOP
log2(RPKM)
z−score of IF value
−3 −2 −1 0 1 2

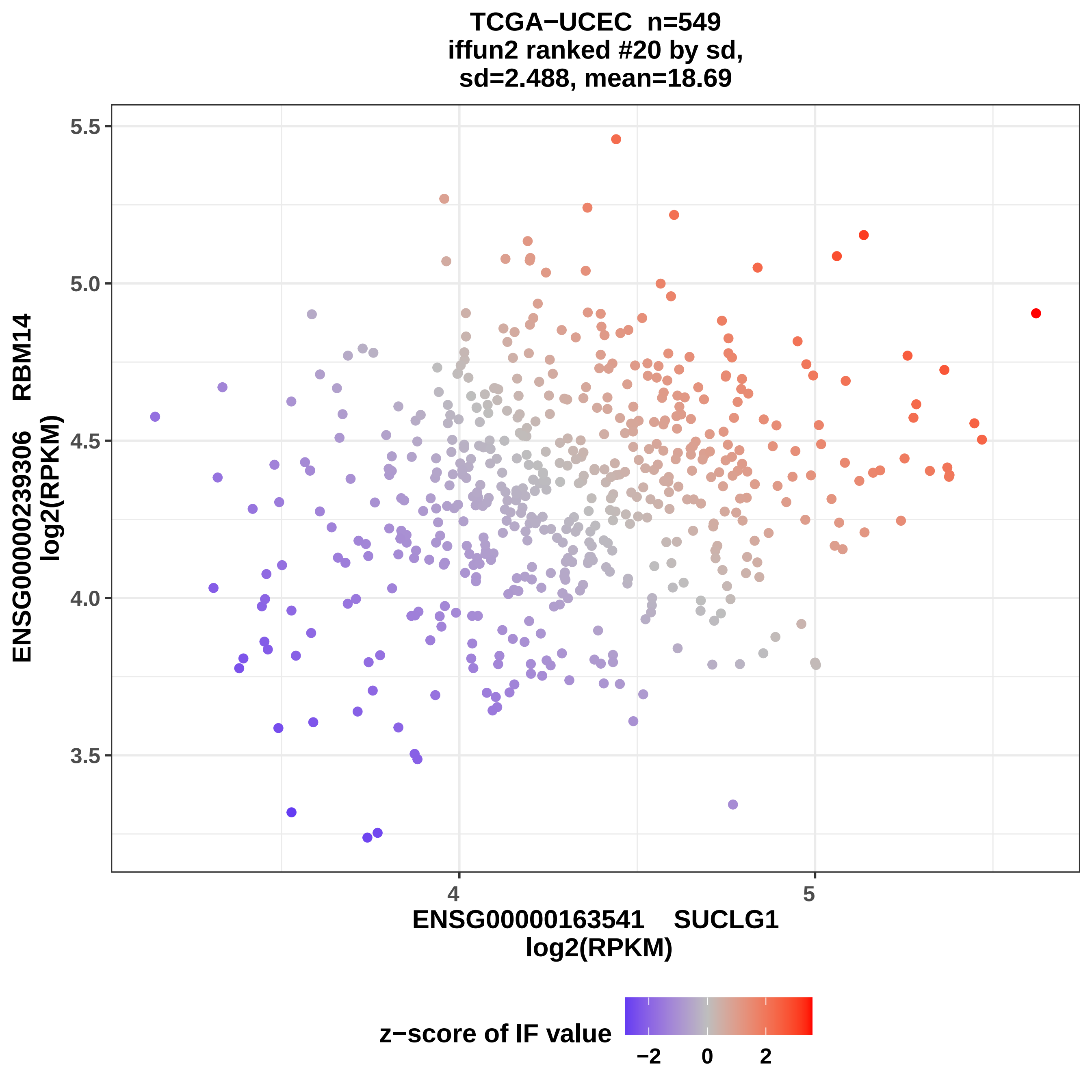

TCGA−UCEC n=549
iffun2 ranked #20 by sd,
sd=2.488, mean=18.69
ENSG00000239306 RBM14
log2(RPKM)
5.5
5.0
4.5
4.0
3.5
4
5
ENSG00000163541 SUCLG1
log2(RPKM)
z−score of IF value
−2
0
2
