## Supplemental Data 6 for "Symmetry as a Fundamental Principle in Defining Gene Expression and Phenotypic Traits"

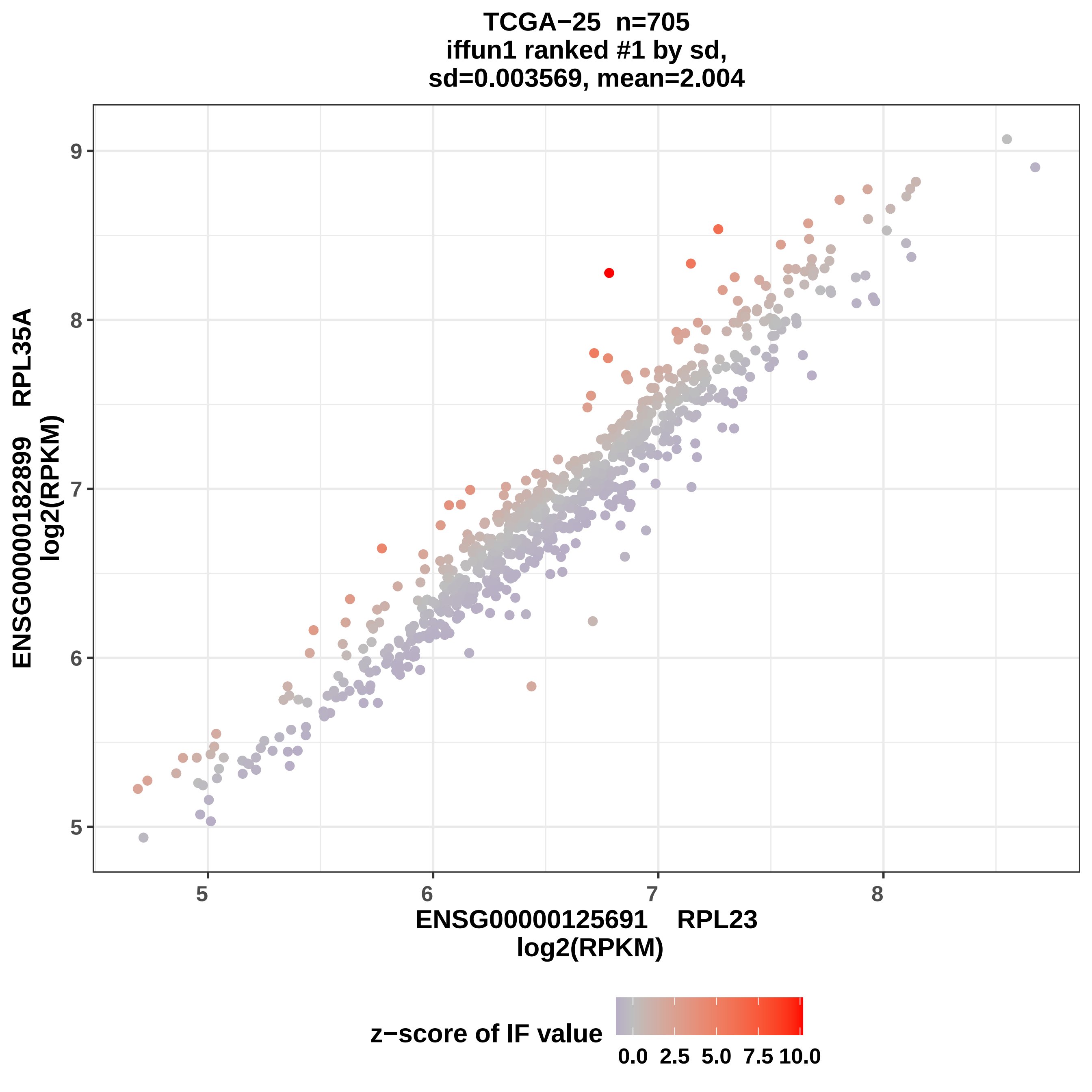
TCGA−25 n=705
iffun1 ranked #1 by sd,
sd=0.003569, mean=2.004
ENSG00000182899 RPL35A
log2(RPKM)
9
8
7
6
5
5
6
7
8
ENSG00000125691 RPL23
log2(RPKM)
z−score of IF value
0.0
2.5
5.0
7.5
10.0

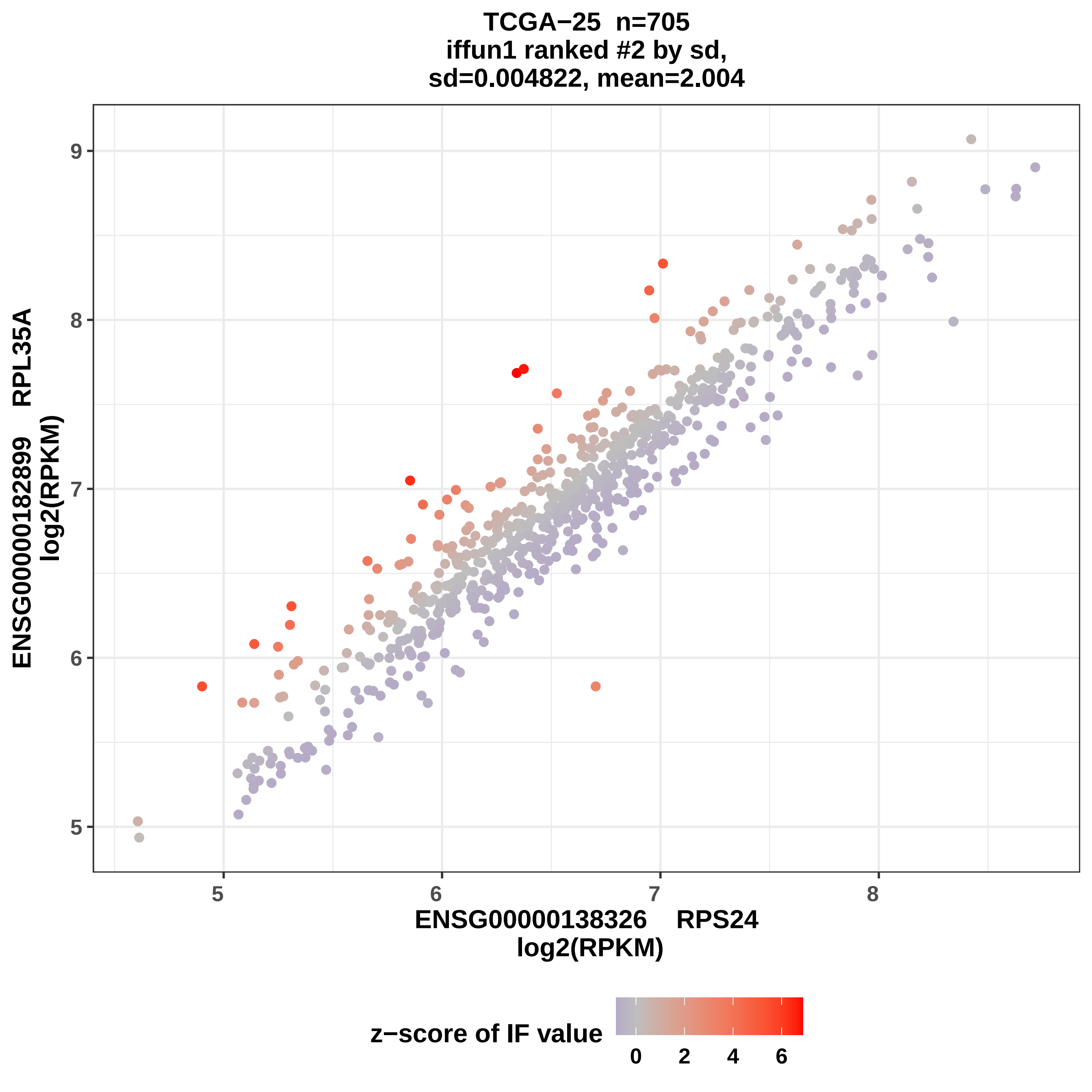
TCGA−25 n=705
iffun1 ranked #2 by sd,
sd=0.004822, mean=2.004
ENSG00000182899 RPL35A
log2(RPKM)
ENSG00000138326 RPS24
log2(RPKM)
z−score of IF value
5
6
7
8
9
0
2
4
6

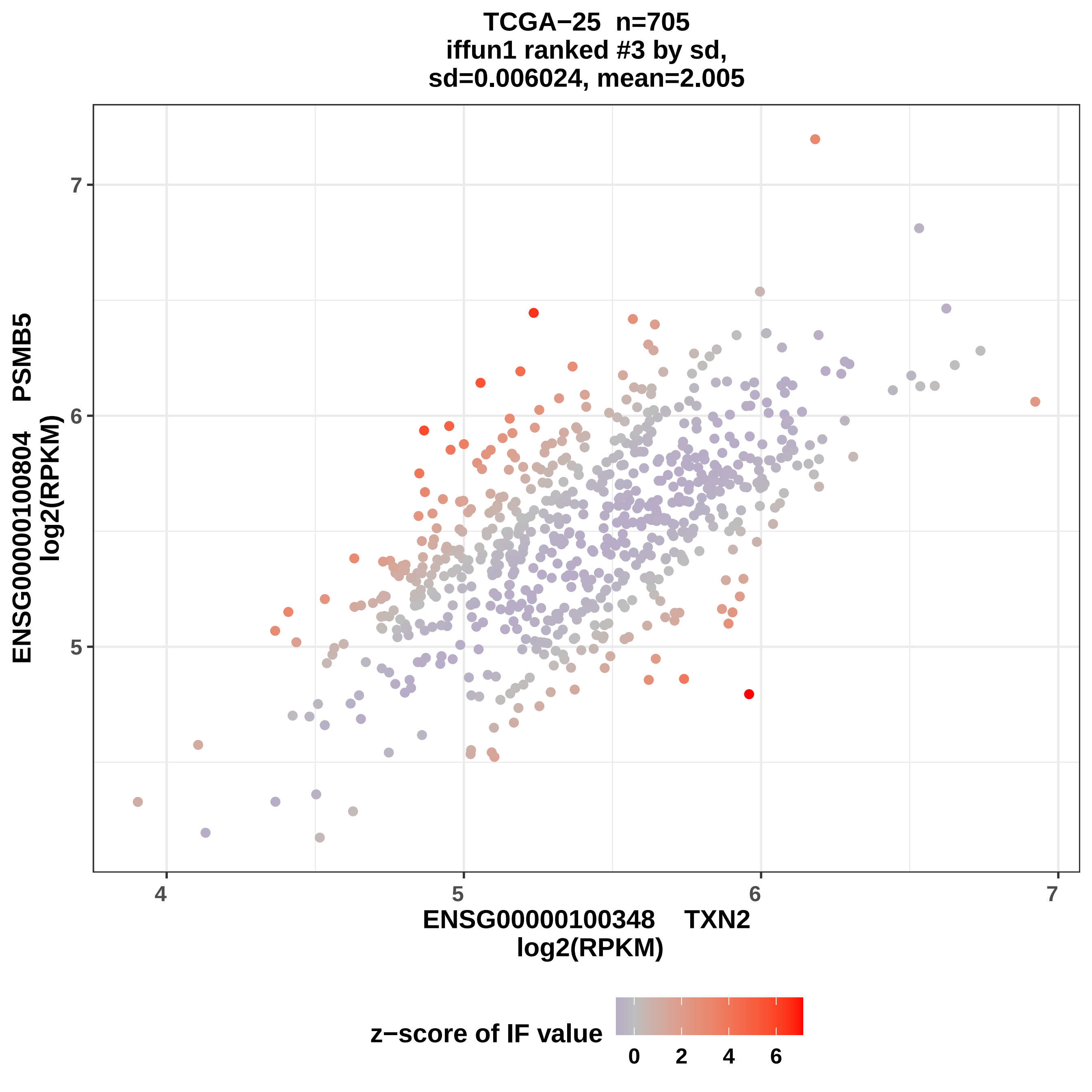
TCGA−25 n=705
iffun1 ranked #3 by sd,
sd=0.006024, mean=2.005
ENSG00000100804 PSMB5
log2(RPKM)
7
6
5
4
5
6
7
ENSG00000100348 TXN2
log2(RPKM)
z−score of IF value
0
2
4
6

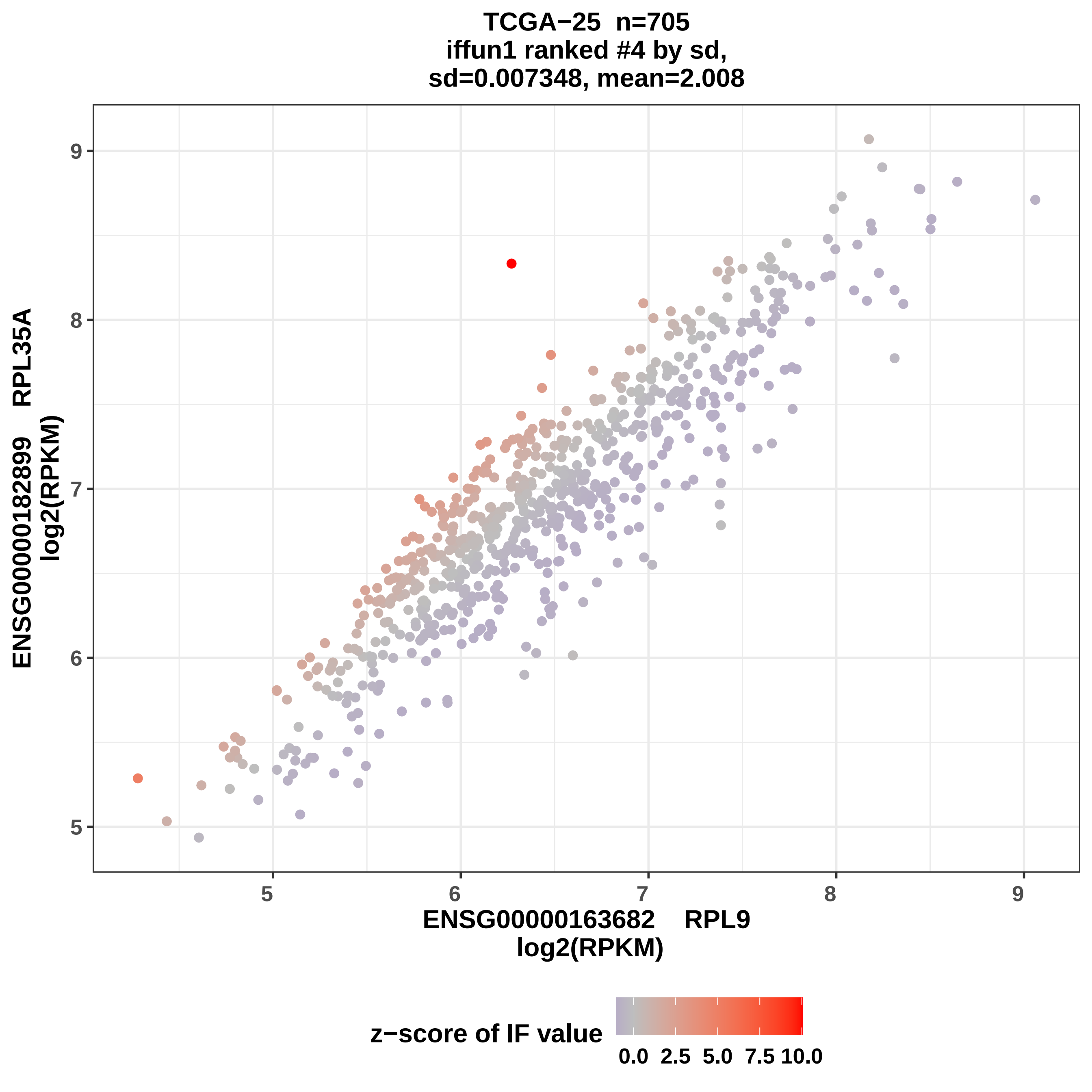

TCGA−25 n=705
iffun1 ranked #4 by sd,
sd=0.007348, mean=2.008
ENSG00000182899 RPL35A
log2(RPKM)
9
8
7
6
5
ENSG00000163682 RPL9
log2(RPKM)
z−score of IF value
0.0
2.5
5.0
7.5
10.0

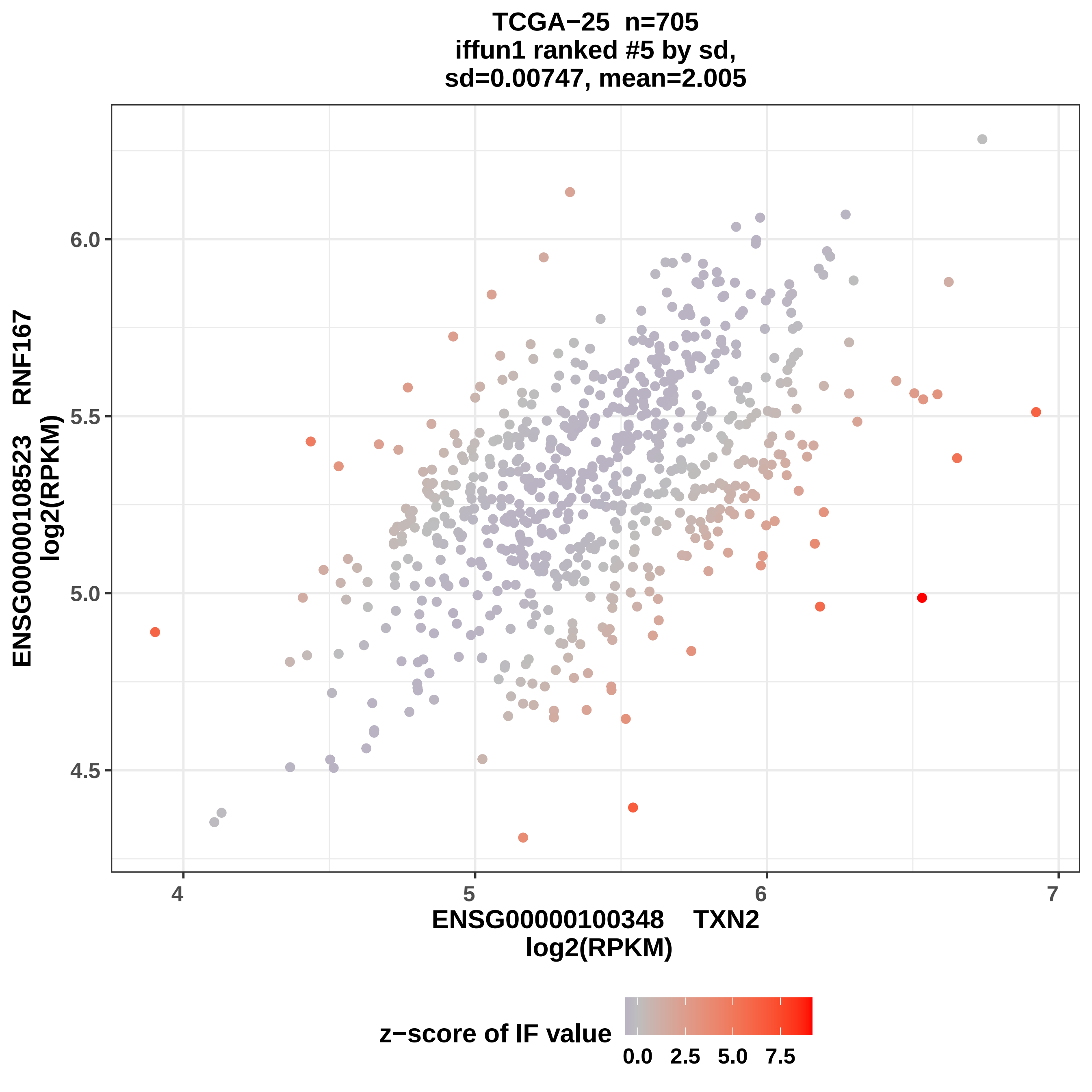

TCGA−25 n=705
iffun1 ranked #5 by sd,
sd=0.00747, mean=2.005
ENSG00000108523 RNF167
log2(RPKM)
6.0
5.5
5.0
4.5
4
5
6
7
ENSG00000100348 TXN2
log2(RPKM)
z−score of IF value
0.0
2.5
5.0
7.5

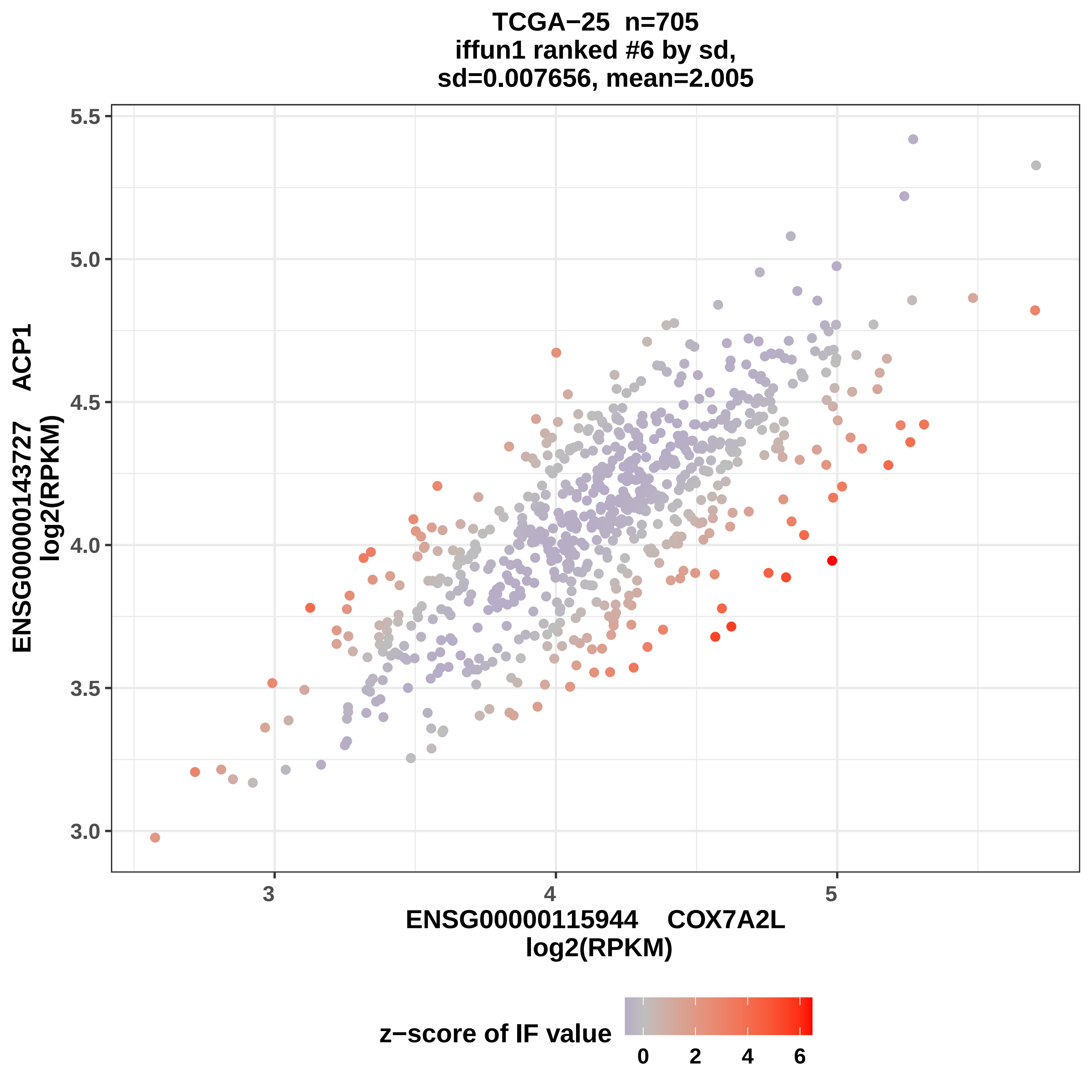

TCGA−25 n=705
iffun1 ranked #6 by sd,
sd=0.007656, mean=2.005
ENSG00000143727 ACP1
log2(RPKM)
5.5
5.0
4.5
4.0
3.5
3.0
3
4
5
ENSG00000115944 COX7A2L
log2(RPKM)
z−score of IF value
0
2
4
6

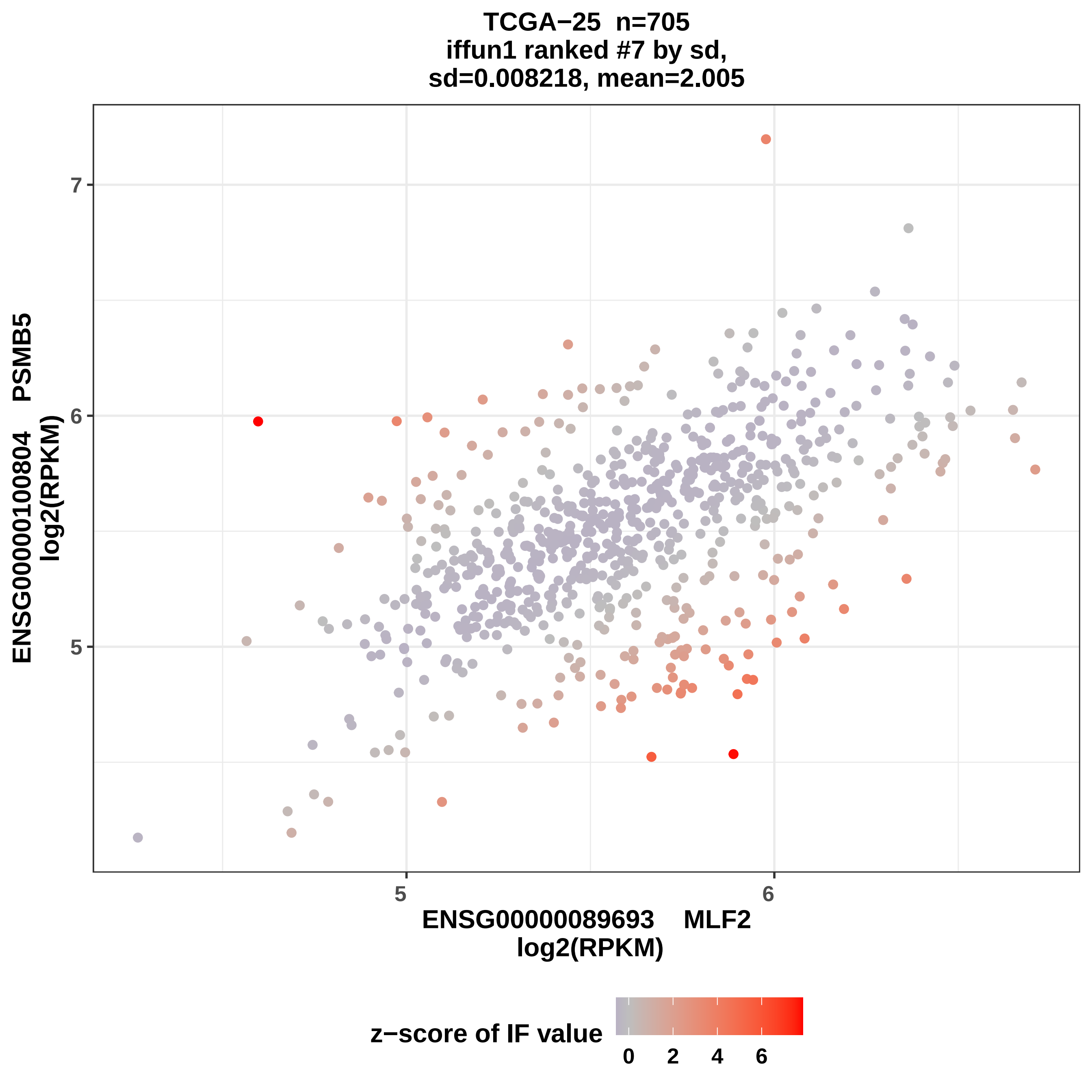

TCGA−25 n=705
iffun1 ranked #7 by sd,
sd=0.008218, mean=2.005
ENSG00000100804 PSMB5
log2(RPKM)
7
6
5
5
6
ENSG00000089693 MLF2
log2(RPKM)
z−score of IF value
0
2
4
6

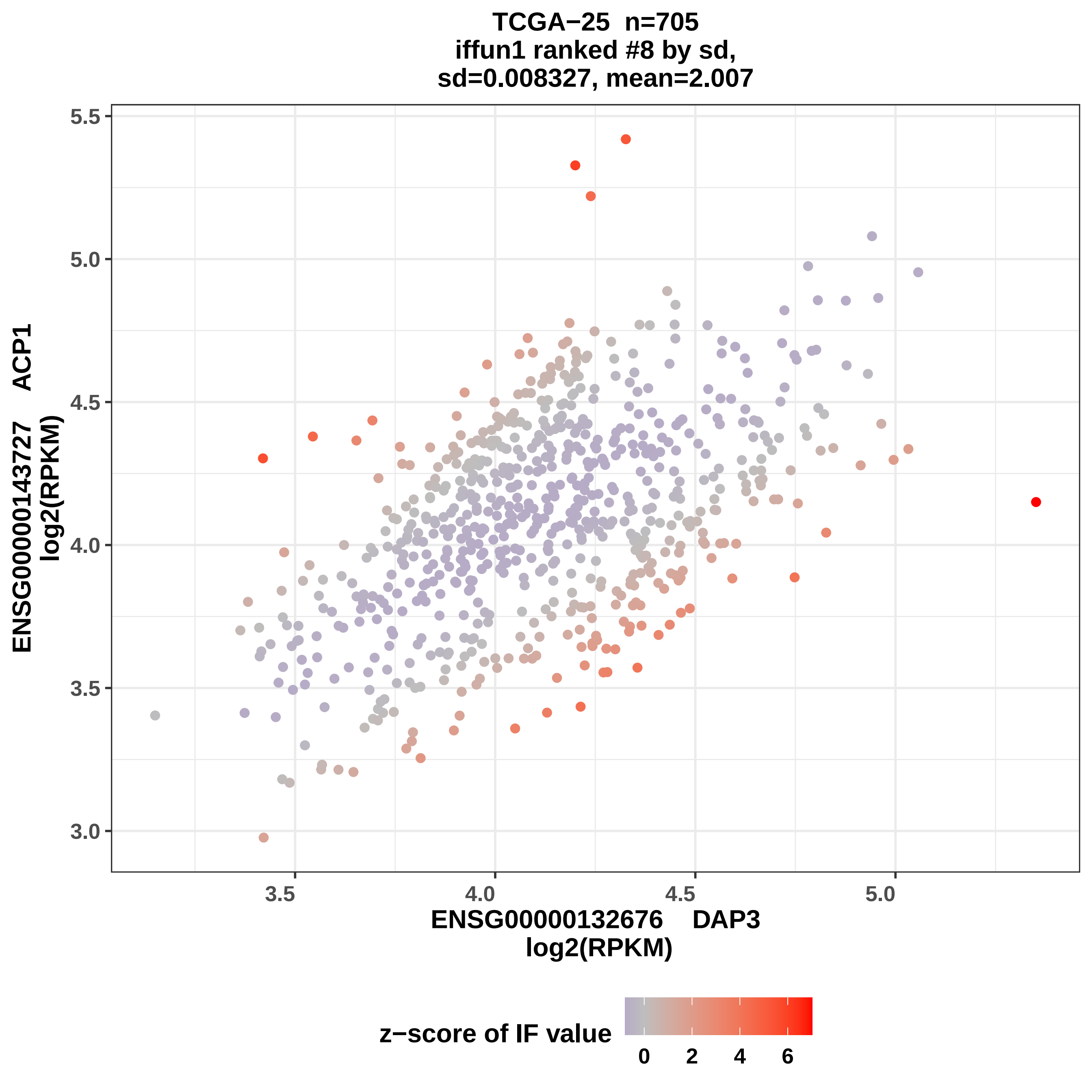

TCGA−25 n=705
iffun1 ranked #8 by sd,
sd=0.008327, mean=2.007
ENSG00000143727 ACP1
log2(RPKM)
5.5
5.0
4.5
4.0
3.5
3.0
3.5
4.0
4.5
5.0
ENSG00000132676 DAP3
log2(RPKM)
z−score of IF value
0
2
4
6

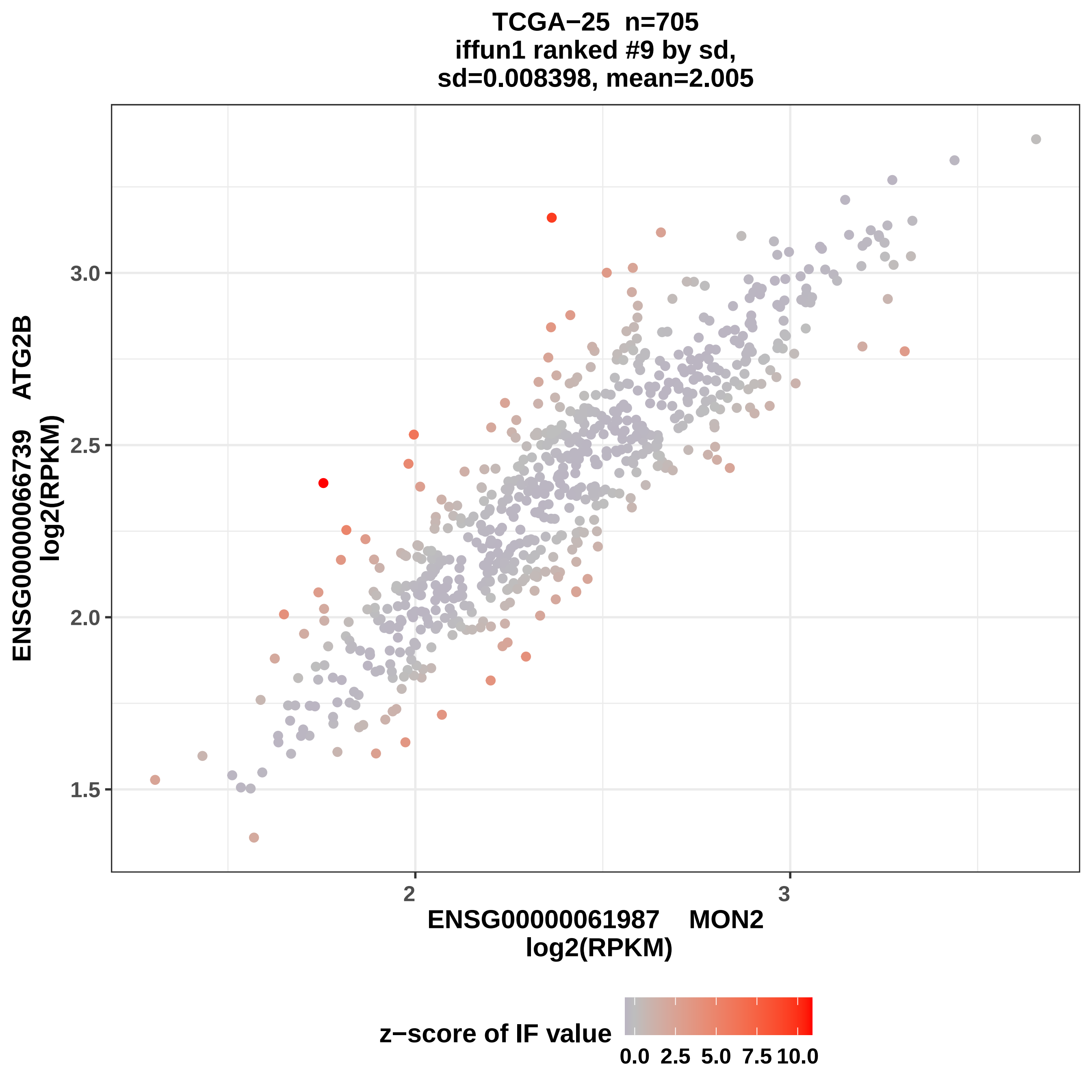
TCGA−25 n=705
iffun1 ranked #9 by sd,
sd=0.008398, mean=2.005
ENSG00000066739 ATG2B
log2(RPKM)
3.0
2.5
2.0
1.5
2
3
ENSG00000061987 MON2
log2(RPKM)
z−score of IF value
0.0
2.5
5.0
7.5
10.0

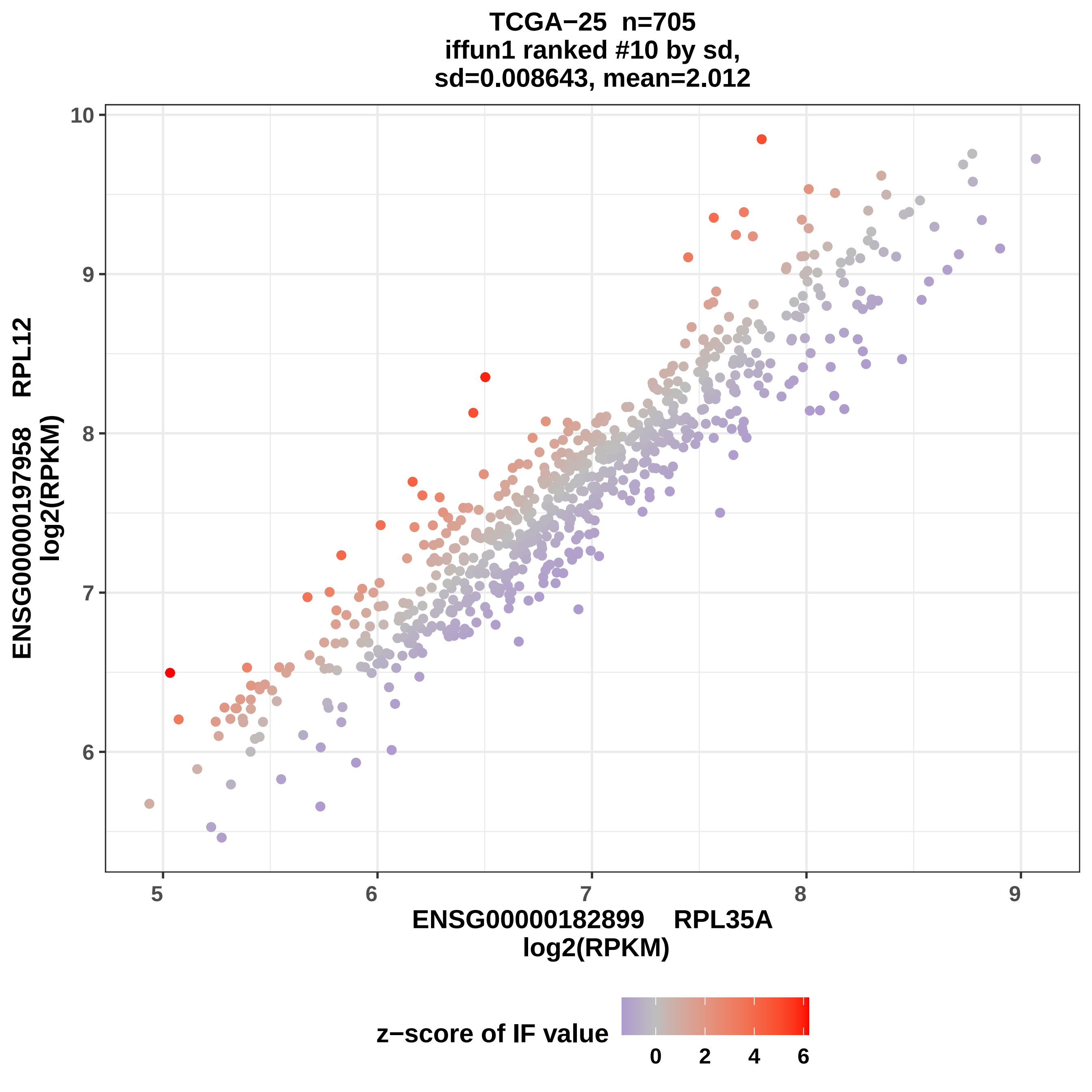

TCGA−25 n=705
iffun1 ranked #10 by sd,
sd=0.008643, mean=2.012
ENSG00000197958 RPL12
log2(RPKM)
10
9
8
7
6
5
6
7
8
9
ENSG00000182899 RPL35A
log2(RPKM)
z−score of IF value
0
2
4
6

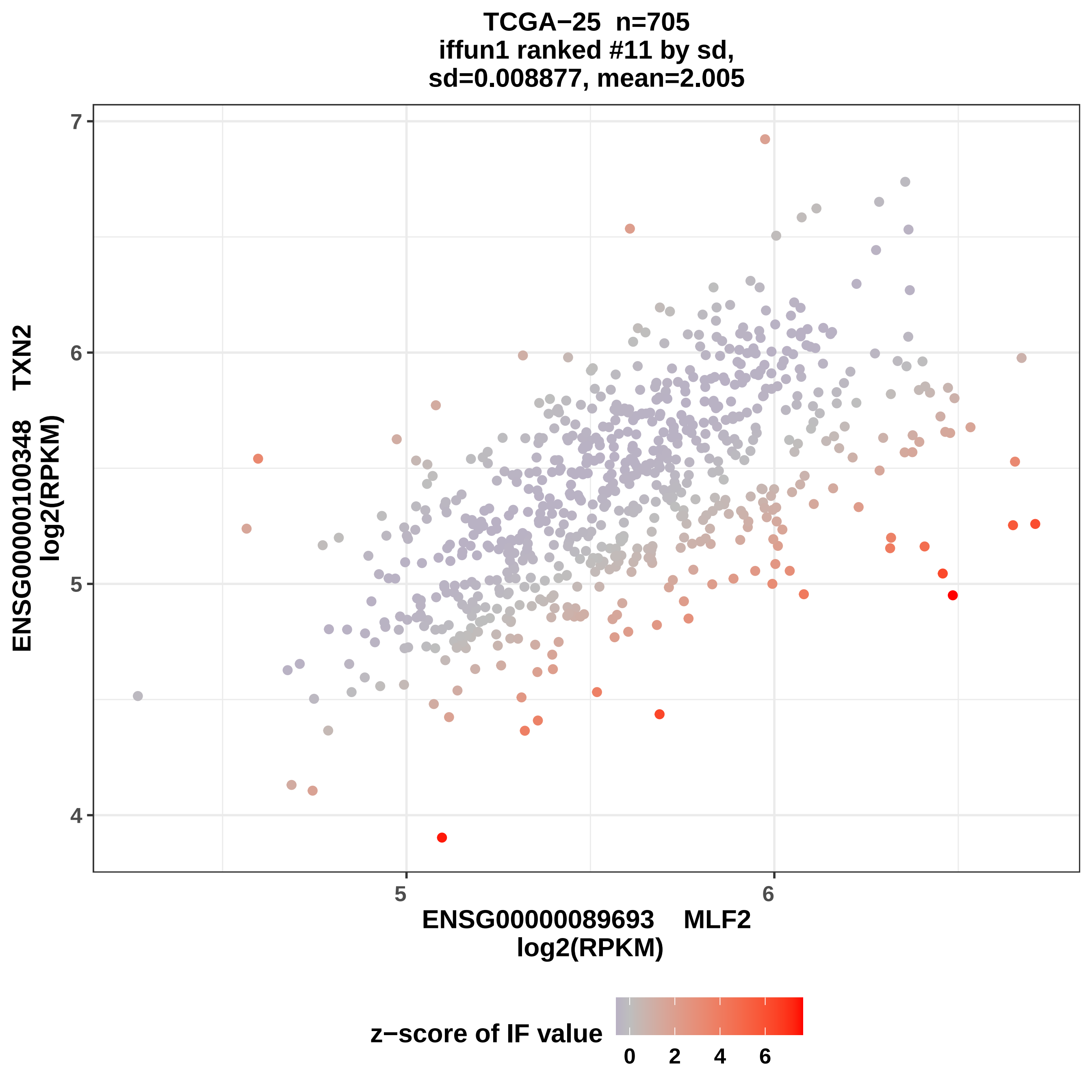
TCGA−25 n=705
iffun1 ranked #11 by sd,
sd=0.008877, mean=2.005
ENSG00000100348 TXN2
log2(RPKM)
ENSG00000089693 MLF2
log2(RPKM)
7
6
5
4
5
6
z−score of IF value
0
2
4
6

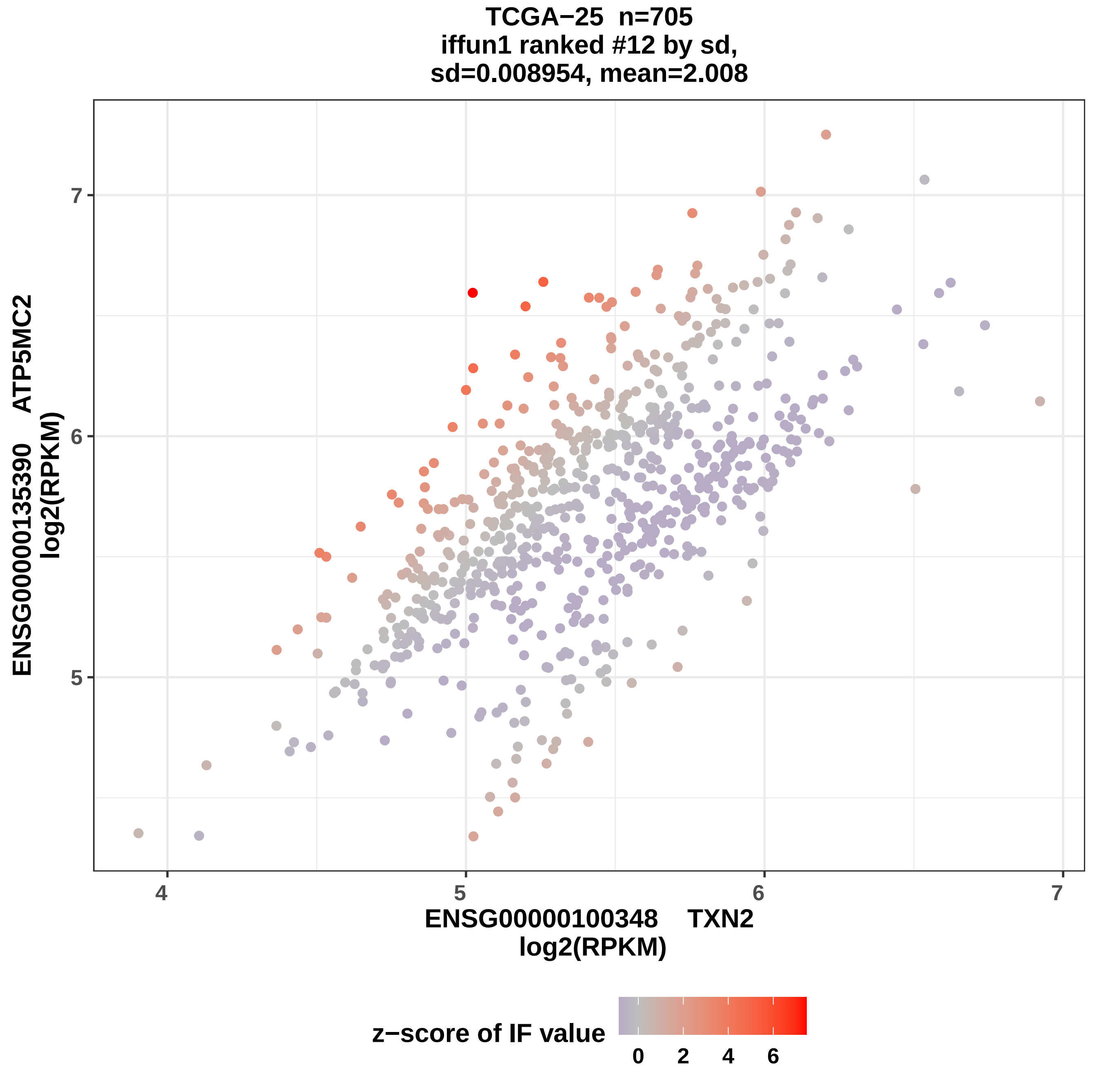
TCGA−25 n=705
iffun1 ranked #12 by sd,
sd=0.008954, mean=2.008
ENSG00000135390 ATP5MC2
log2(RPKM)
7
6
5
4
5
6
7
ENSG00000100348 TXN2
log2(RPKM)
z−score of IF value
0
2
4
6

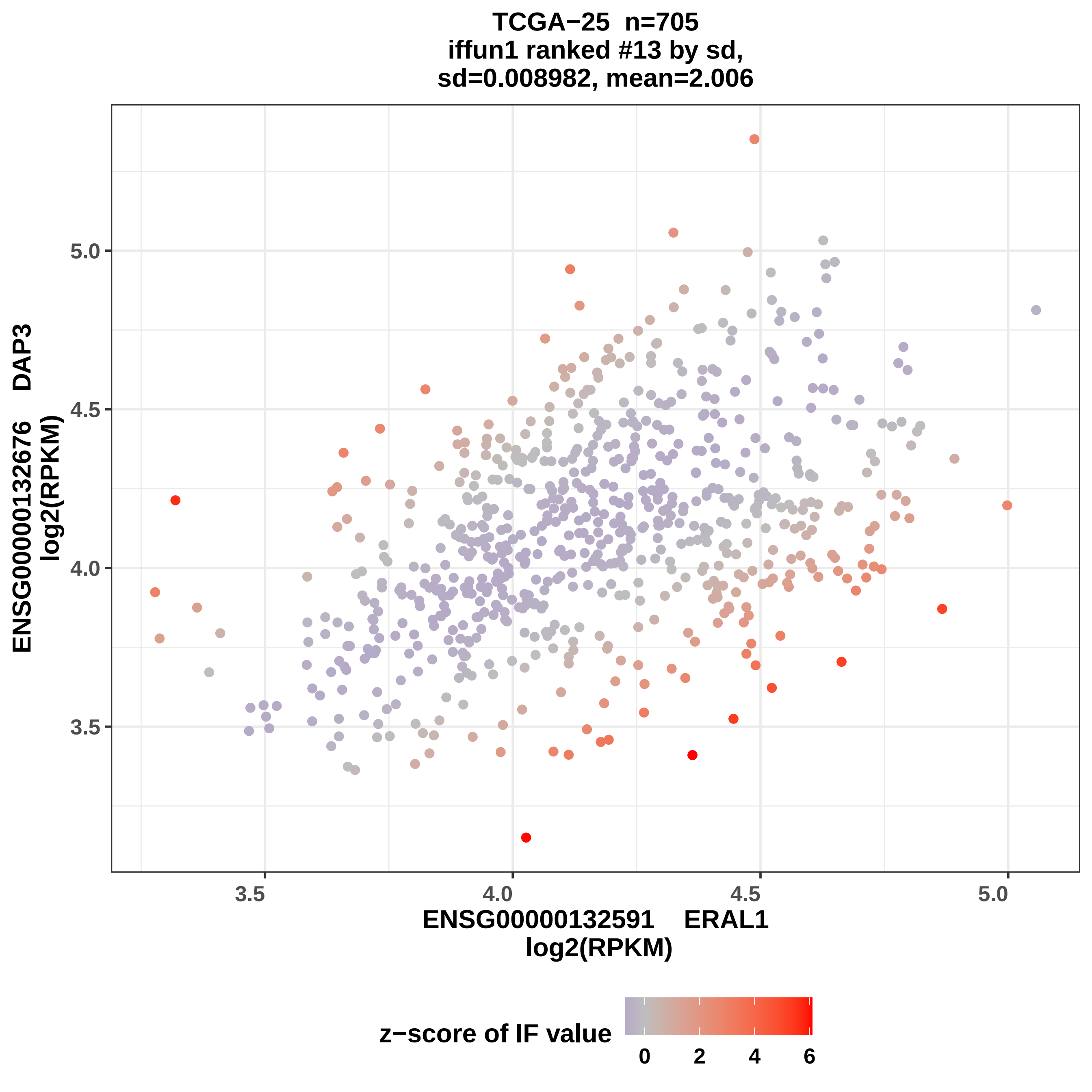
TCGA−25 n=705
iffun1 ranked #13 by sd,
sd=0.008982, mean=2.006
ENSG00000132676 DAP3
log2(RPKM)
5.0
4.5
4.0
3.5
3.5
4.0
4.5
5.0
ENSG00000132591 ERAL1
log2(RPKM)
z−score of IF value
0
2
4
6

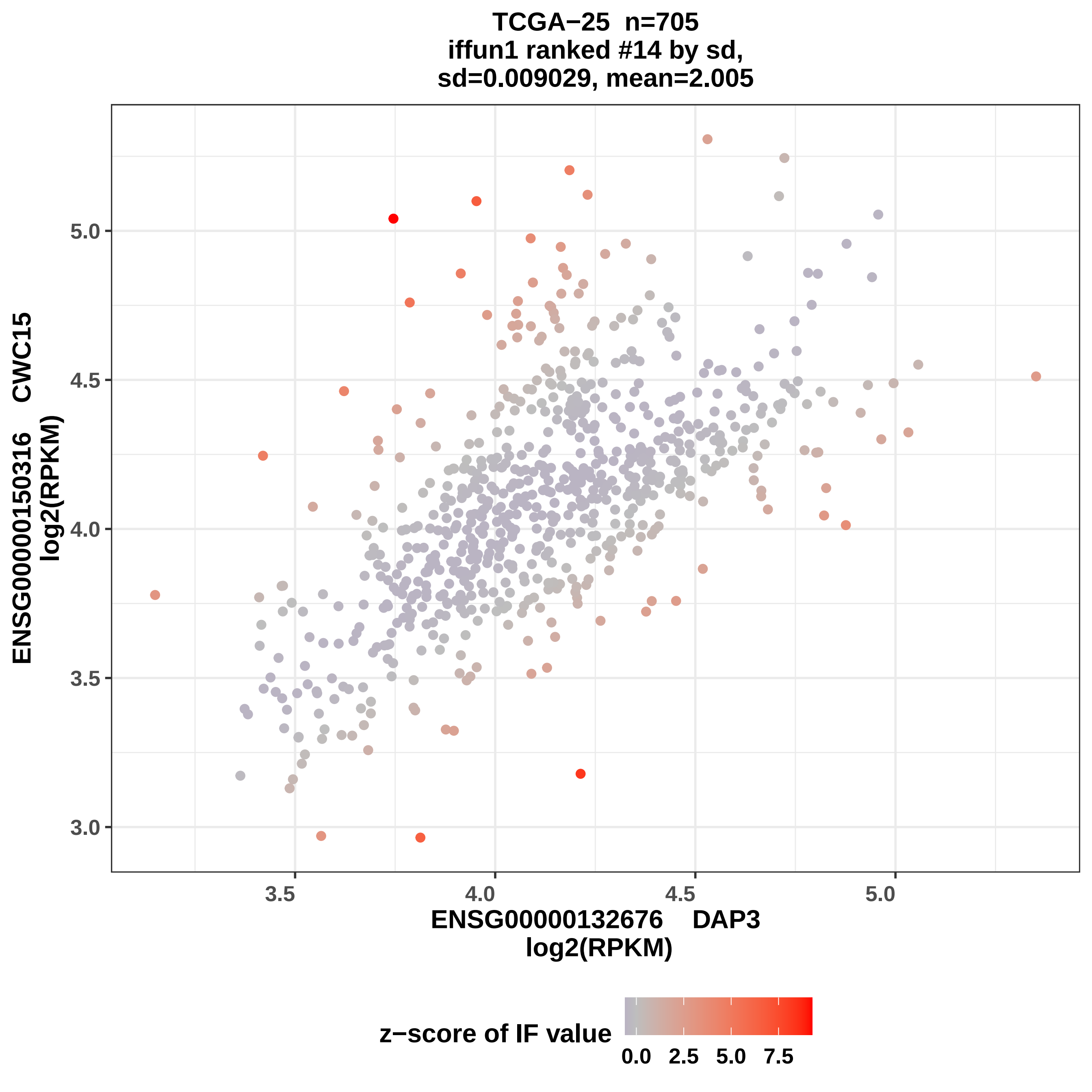
TCGA−25 n=705
iffun1 ranked #14 by sd,
sd=0.009029, mean=2.005
ENSG00000150316 CWC15
log2(RPKM)
5.0
4.5
4.0
3.5
3.0
3.5
4.0
4.5
5.0
ENSG00000132676 DAP3
log2(RPKM)
z−score of IF value
0.0
2.5
5.0
7.5

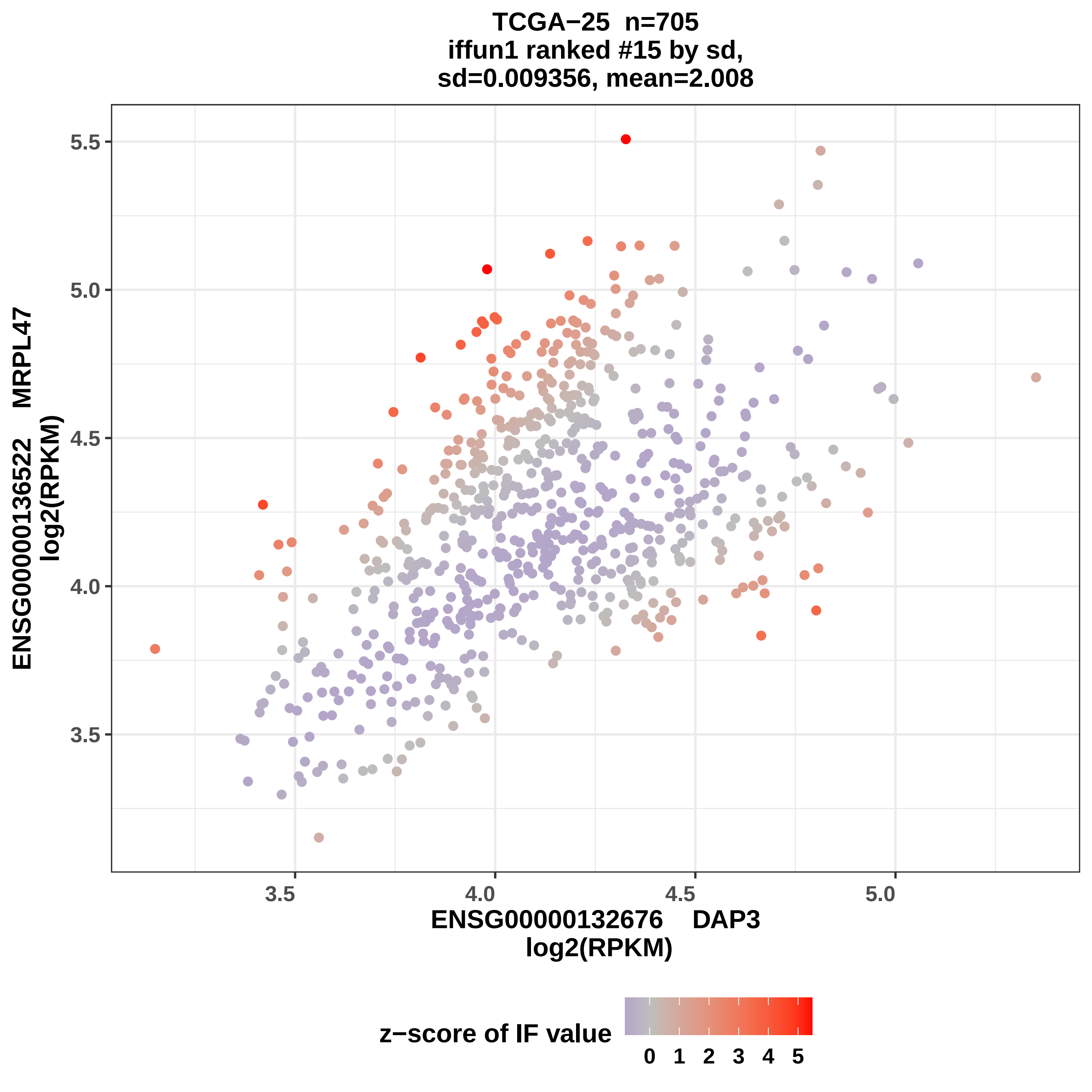

TCGA−25 n=705
iffun1 ranked #15 by sd,
sd=0.009356, mean=2.008
5.5
5.0
4.5
4.0
3.5
ENSG00000136522 MRPL47
log2(RPKM)
3.5
4.0
4.5
5.0
ENSG00000132676 DAP3
log2(RPKM)
z−score of IF value
0 1 2 3 4 5

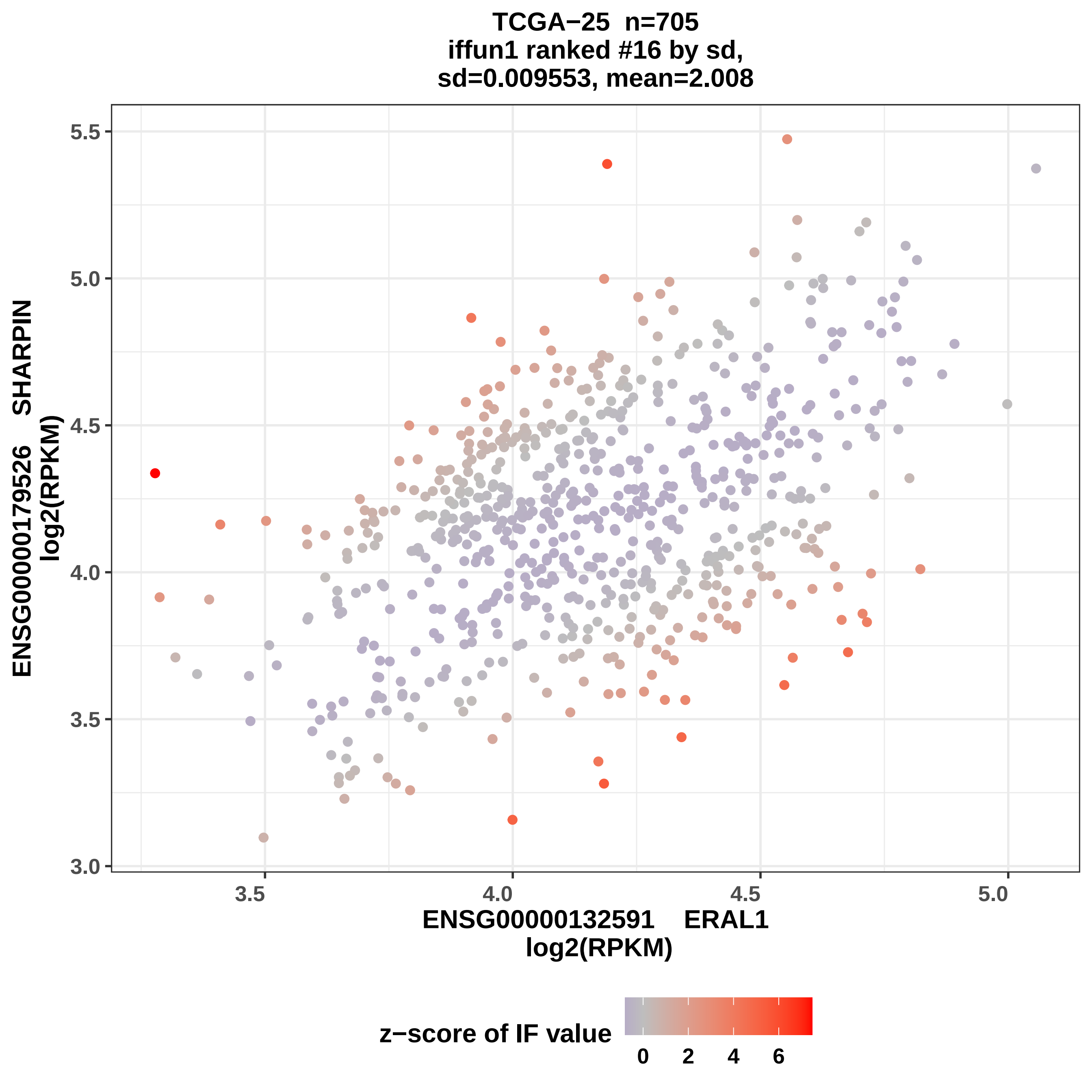

TCGA−25 n=705
iffun1 ranked #16 by sd,
sd=0.009553, mean=2.008
ENSG00000179526 SHARPIN
log2(RPKM)
5.5
5.0
4.5
4.0
3.5
3.0
3.5
4.0
4.5
5.0
ENSG00000132591 ERAL1
log2(RPKM)
z−score of IF value
0
2
4
6

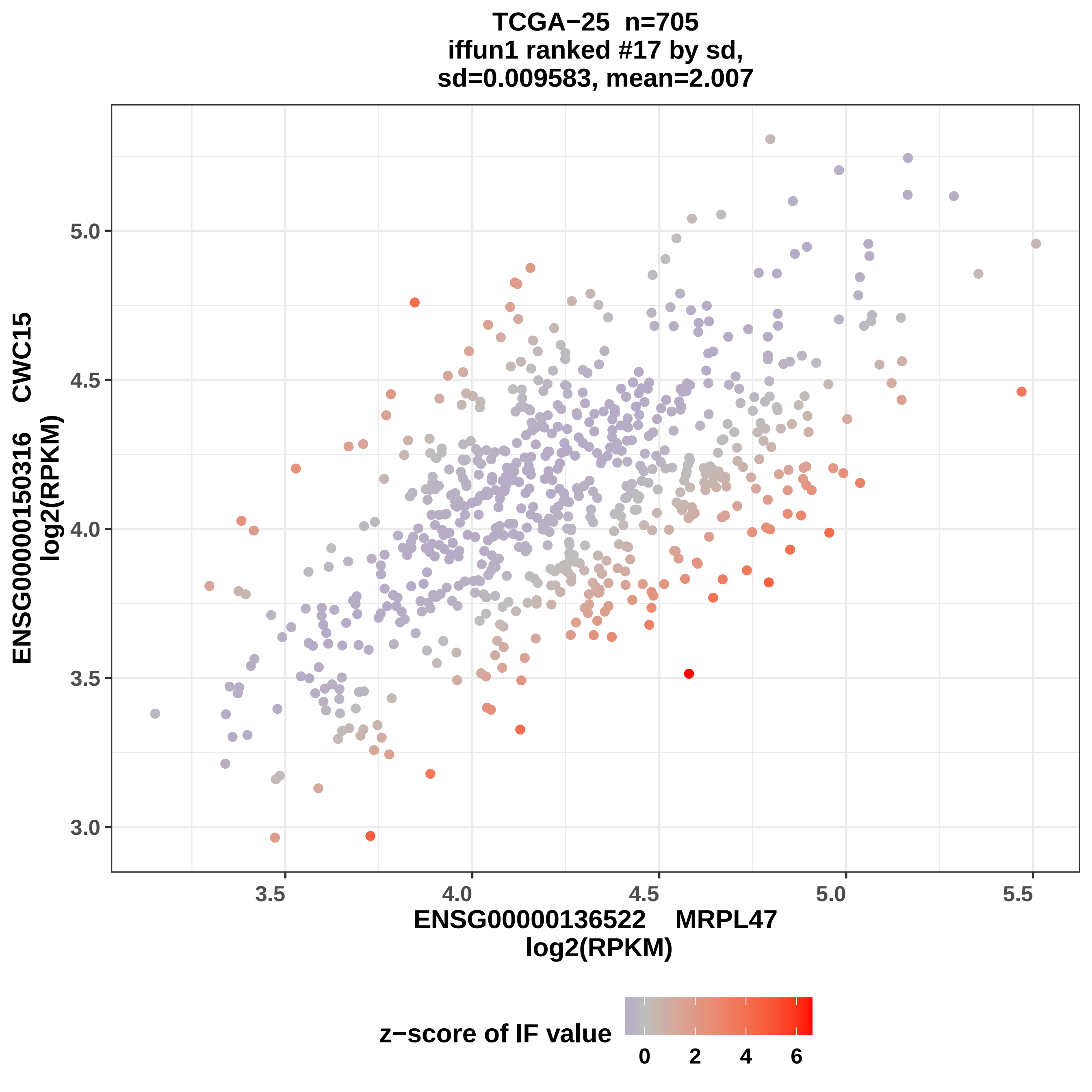

TCGA−25 n=705
iffun1 ranked #17 by sd,
sd=0.009583, mean=2.007
ENSG00000150316 CWC15
log2(RPKM)
ENSG00000136522 MRPL47
log2(RPKM)
5.0
4.5
4.0
3.5
3.0
3.5
4.0
4.5
5.0
5.5
z−score of IF value
0
2
4
6

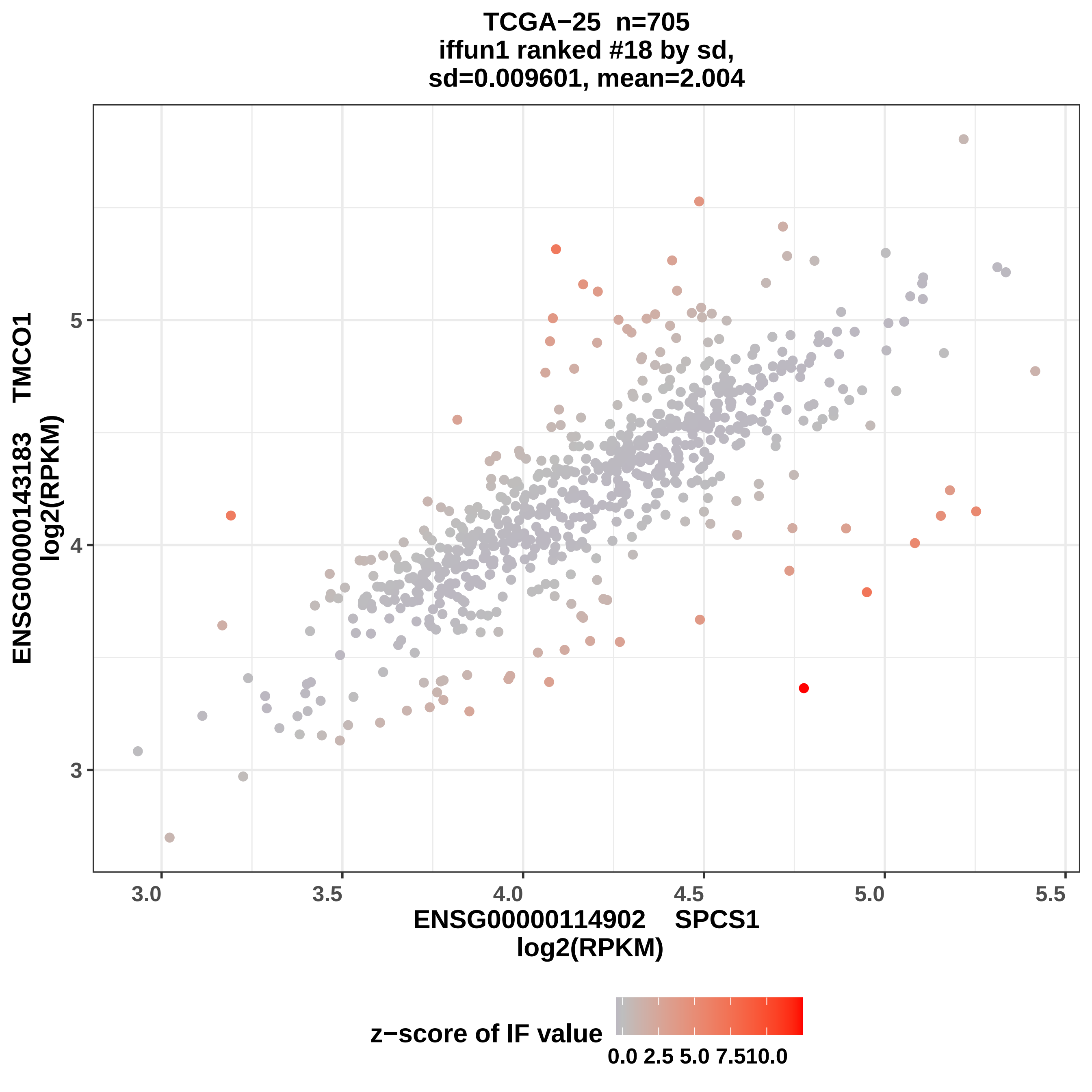

TCGA−25 n=705
iffun1 ranked #18 by sd,
sd=0.009601, mean=2.004
ENSG00000143183 TMCO1
log2(RPKM)
3
4
5
3.0
3.5
4.0
4.5
5.0
5.5
ENSG00000114902 SPCS1
log2(RPKM)
z−score of IF value
0.0
2.5
5.0
7.5
10.0

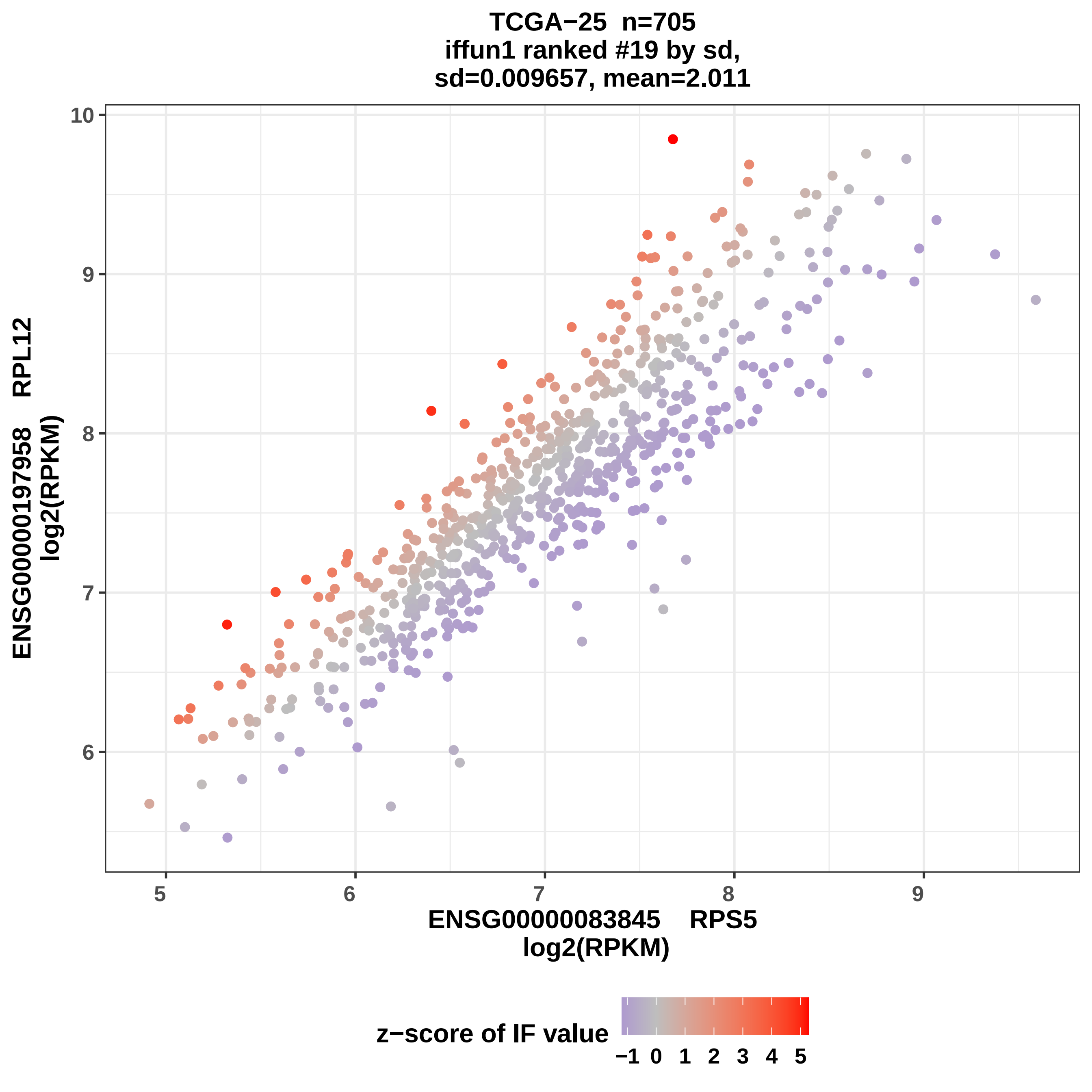
TCGA−25 n=705
iffun1 ranked #19 by sd,
sd=0.009657, mean=2.011
ENSG00000197958 RPL12
log2(RPKM)
10
9
8
7
6
5
6
7
8
9
ENSG00000083845 RPS5
log2(RPKM)
z−score of IF value
−1 0 1 2 3 4 5

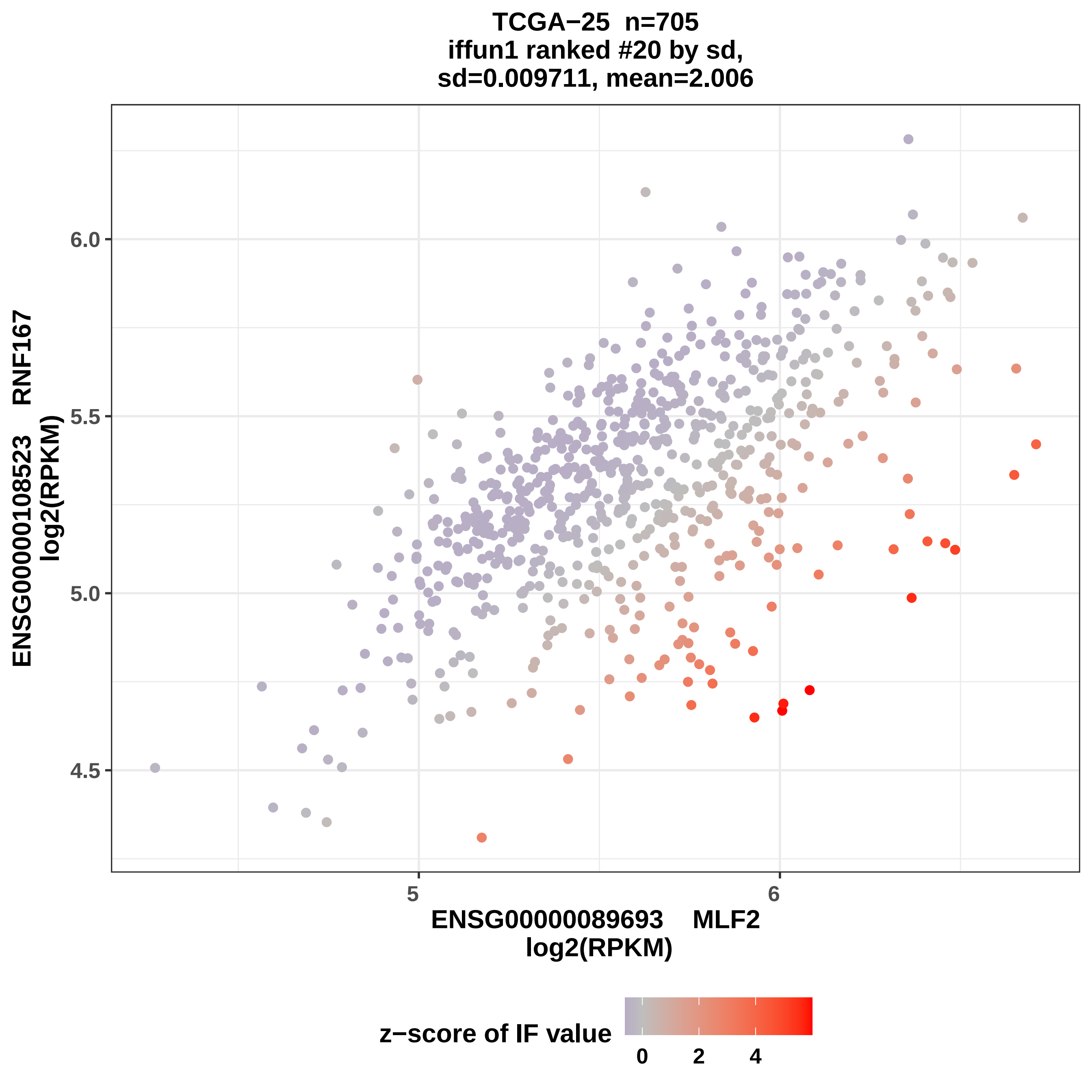

TCGA−25 n=705
iffun1 ranked #20 by sd,
sd=0.009711, mean=2.006
ENSG00000108523 RNF167
log2(RPKM)
6.0
5.5
5.0
4.5
5
6
ENSG00000089693 MLF2
log2(RPKM)
z−score of IF value
0
2
4

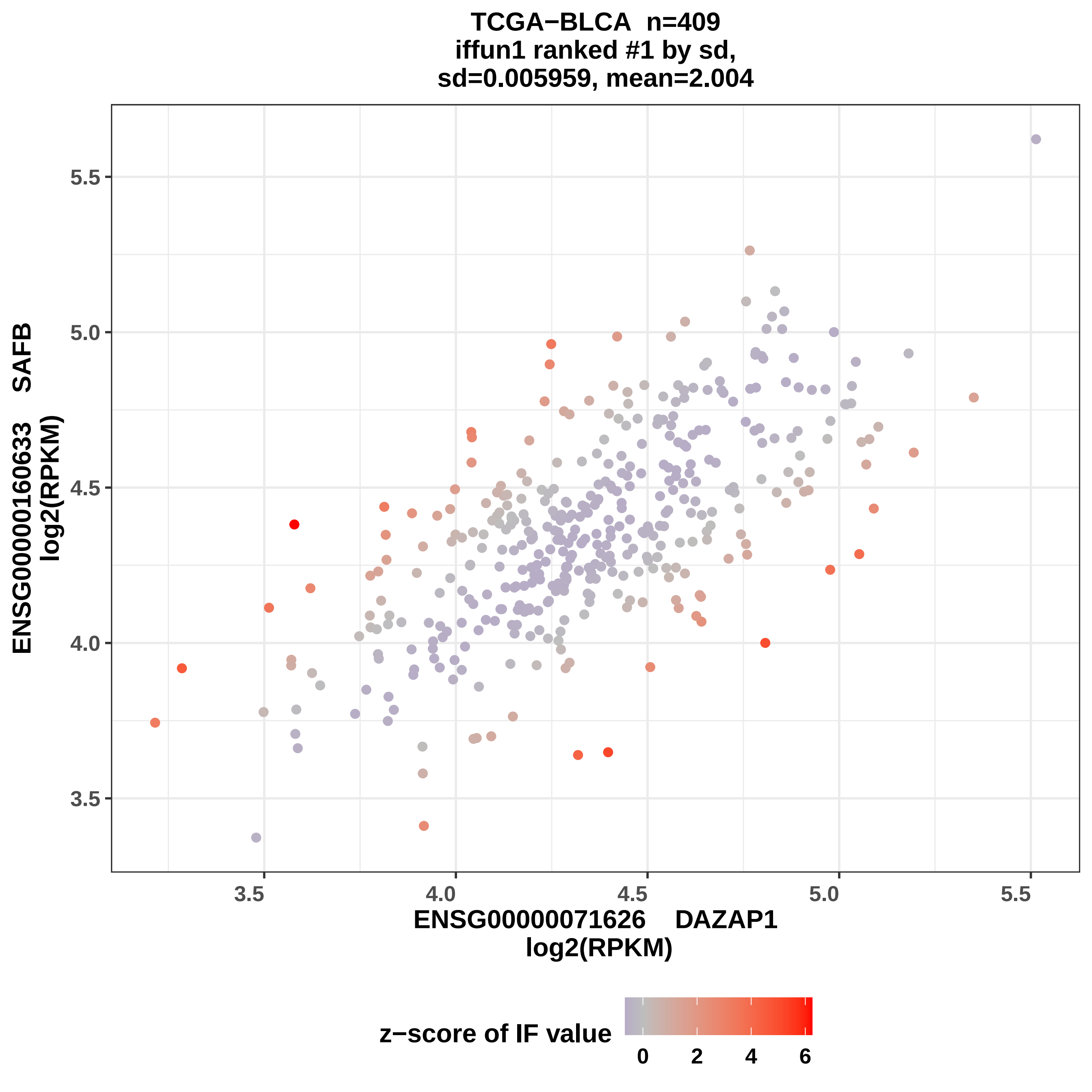

TCGA−BLCA n=409
iffun1 ranked #1 by sd,
sd=0.005959, mean=2.004
ENSG00000160633 SAFB
log2(RPKM)
5.5
5.0
4.5
4.0
3.5
3.5
4.0
4.5
5.0
5.5
ENSG00000071626 DAZAP1
log2(RPKM)
z−score of IF value
0
2
4
6

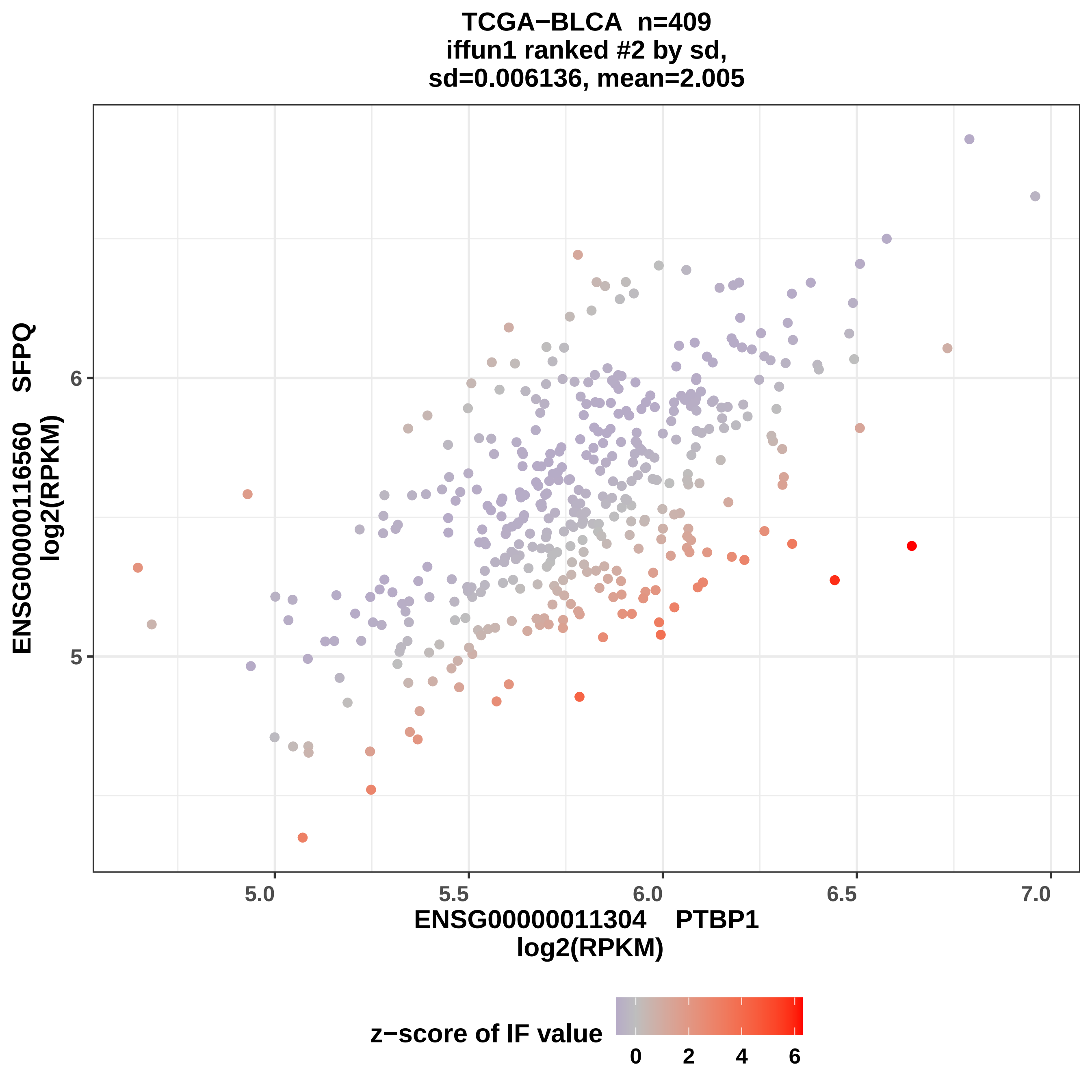
TCGA−BLCA n=409
iffun1 ranked #2 by sd,
sd=0.006136, mean=2.005
ENSG00000116560 SFPQ
log2(RPKM)
6
5
5.0
5.5
6.0
6.5
7.0
ENSG00000011304 PTBP1
log2(RPKM)
z−score of IF value
0
2
4
6

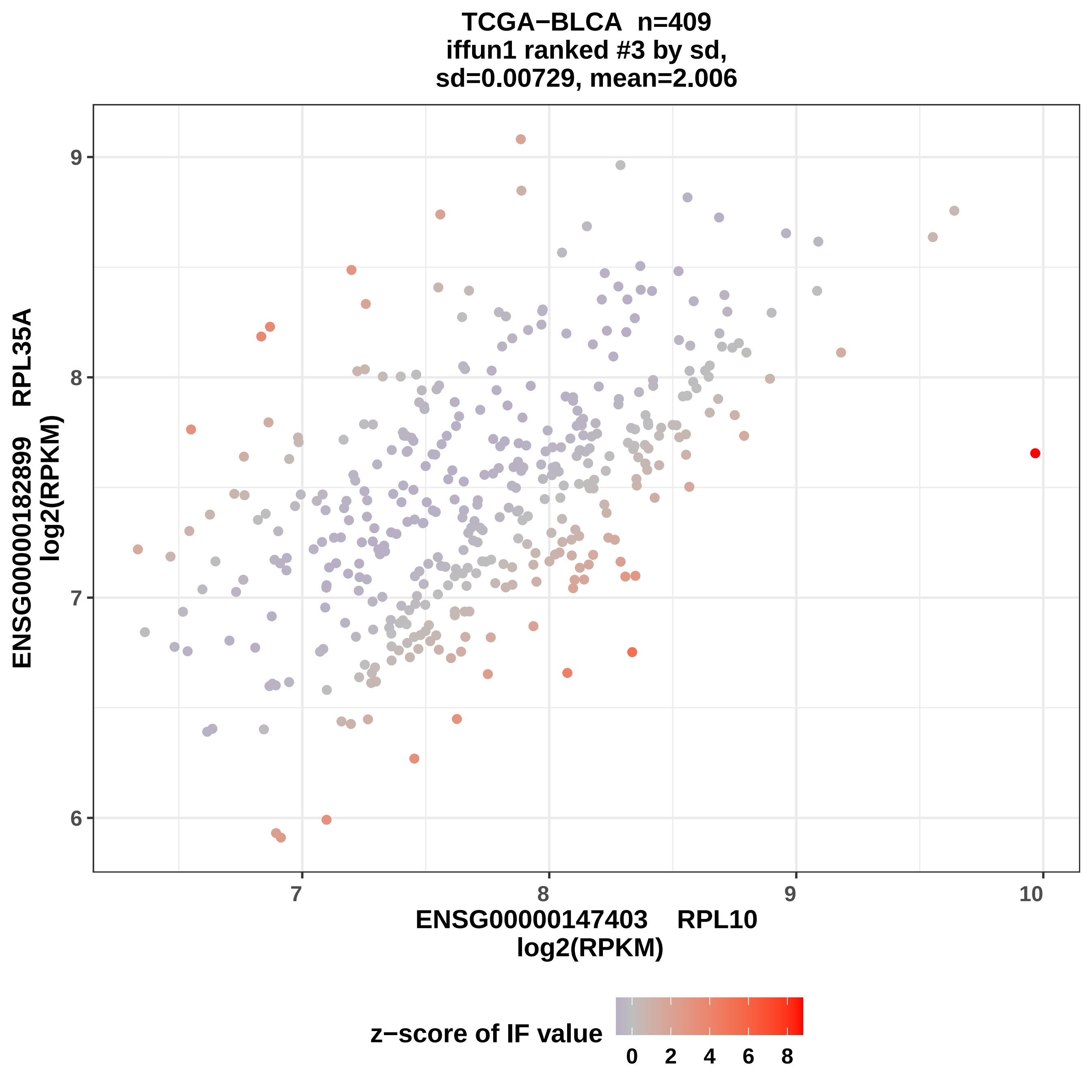

TCGA−BLCA n=409
iffun1 ranked #3 by sd,
sd=0.00729, mean=2.006
ENSG00000182899 RPL35A
log2(RPKM)
ENSG00000147403 RPL10
log2(RPKM)
z−score of IF value
6
7
8
9
7
8
9
10
0
2
4
6
8

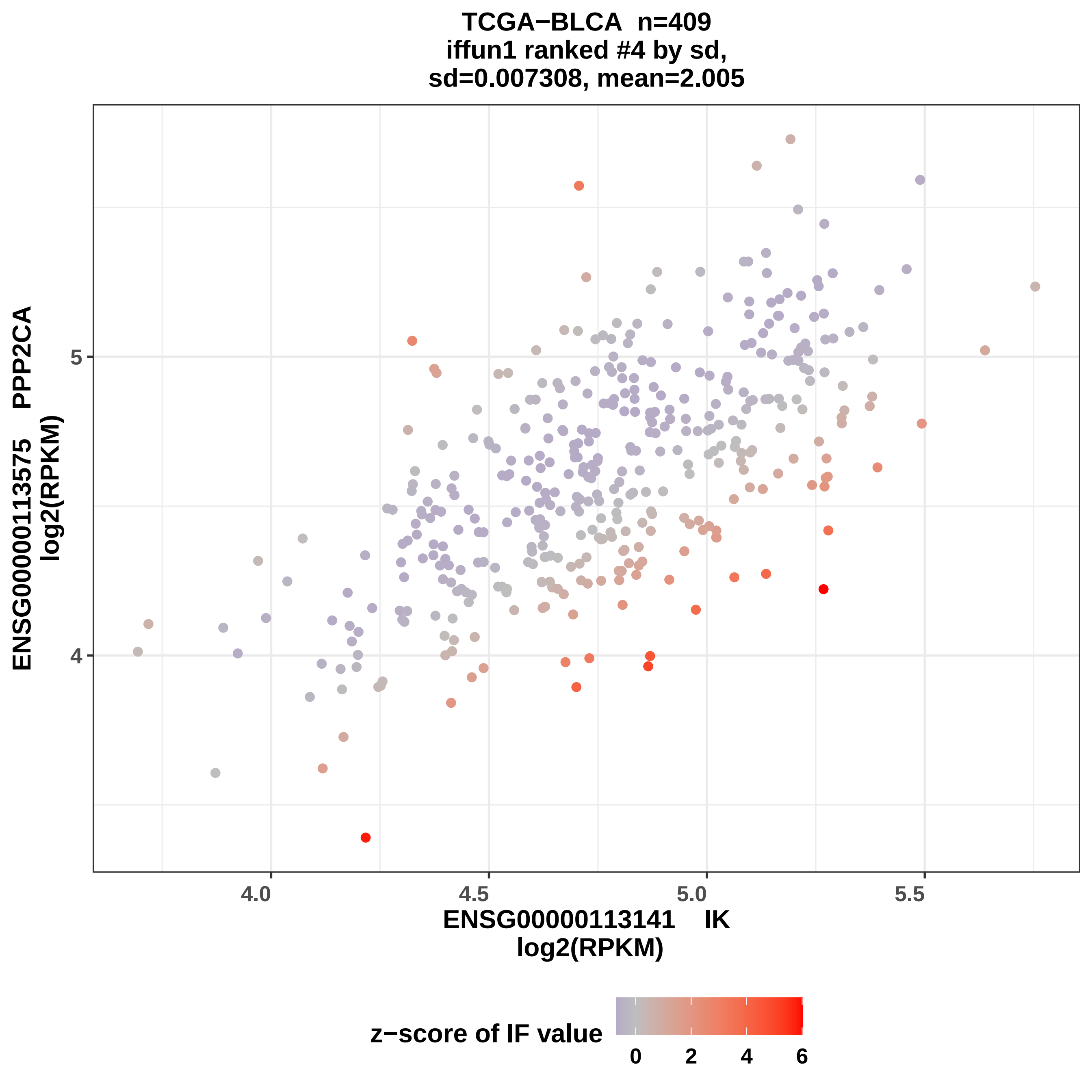
TCGA−BLCA n=409
iffun1 ranked #4 by sd,
sd=0.007308, mean=2.005
ENSG00000113575 PPP2CA
log2(RPKM)
5
4
4.0
4.5
5.0
5.5
ENSG00000113141 IK
log2(RPKM)
z−score of IF value
0
2
4
6

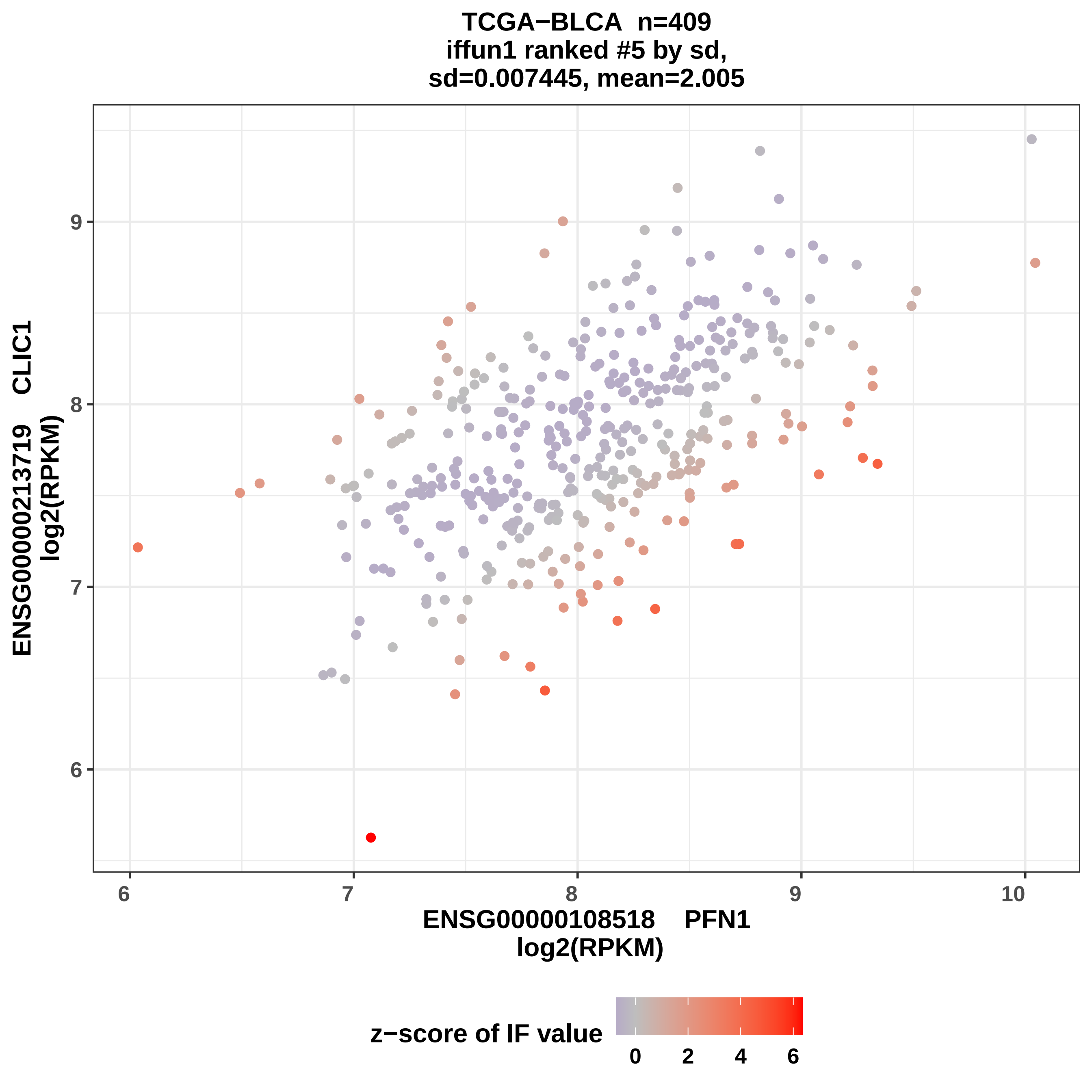

TCGA−BLCA n=409
iffun1 ranked #5 by sd,
sd=0.007445, mean=2.005
ENSG00000213719 CLIC1
log2(RPKM)
ENSG00000108518 PFN1
log2(RPKM)
6
7
8
9
10
z−score of IF value
0
2
4
6

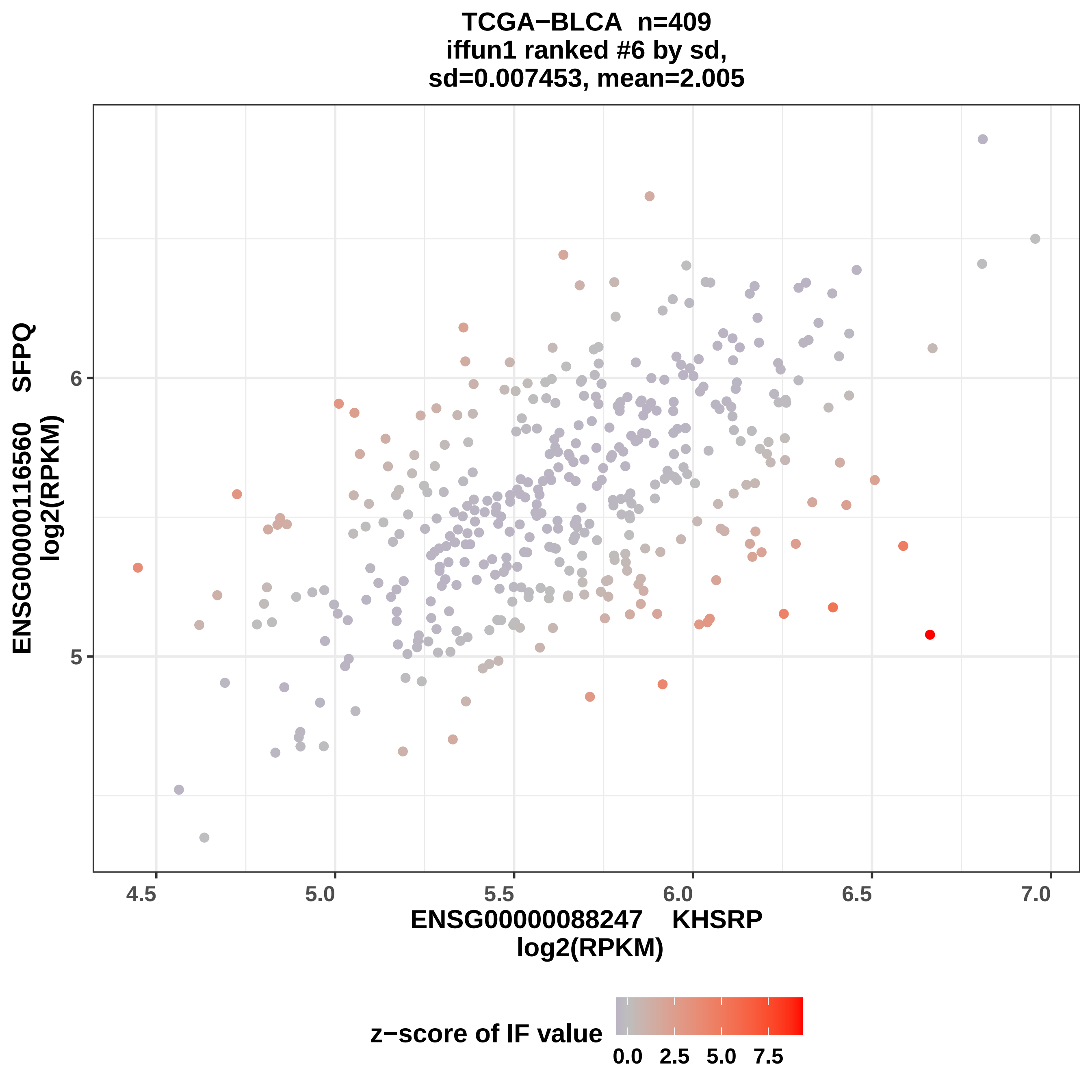

TCGA−BLCA n=409
iffun1 ranked #6 by sd,
sd=0.007453, mean=2.005
ENSG00000116560 SFPQ
log2(RPKM)
6
5
4.5
5.0
5.5
6.0
6.5
7.0
ENSG00000088247 KHSRP
log2(RPKM)
z−score of IF value
0.0
2.5
5.0
7.5

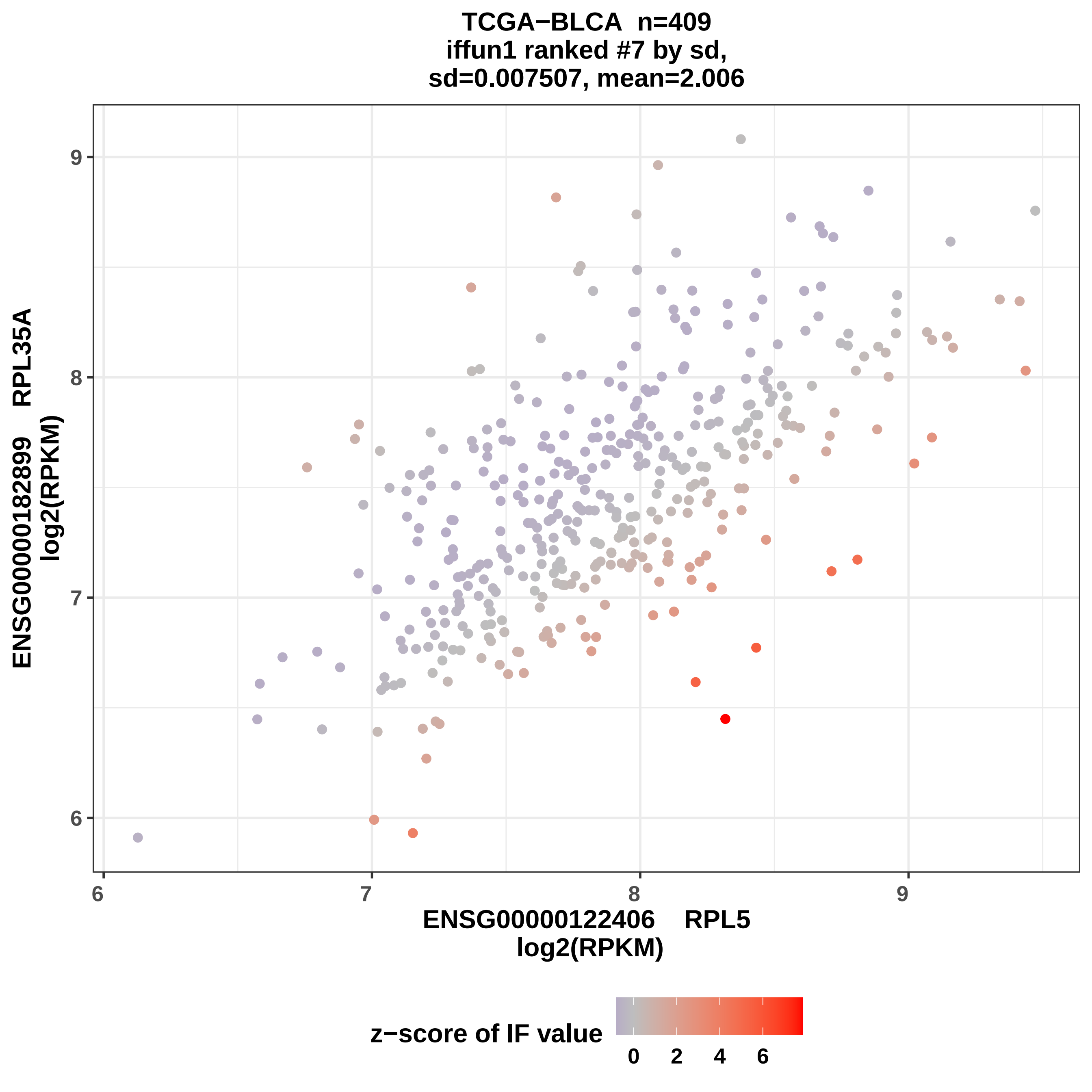
TCGA−BLCA n=409
iffun1 ranked #7 by sd,
sd=0.007507, mean=2.006
ENSG00000182899 RPL35A
log2(RPKM)
ENSG00000122406 RPL5
log2(RPKM)
z−score of IF value
6
7
8
9
0
2
4
6

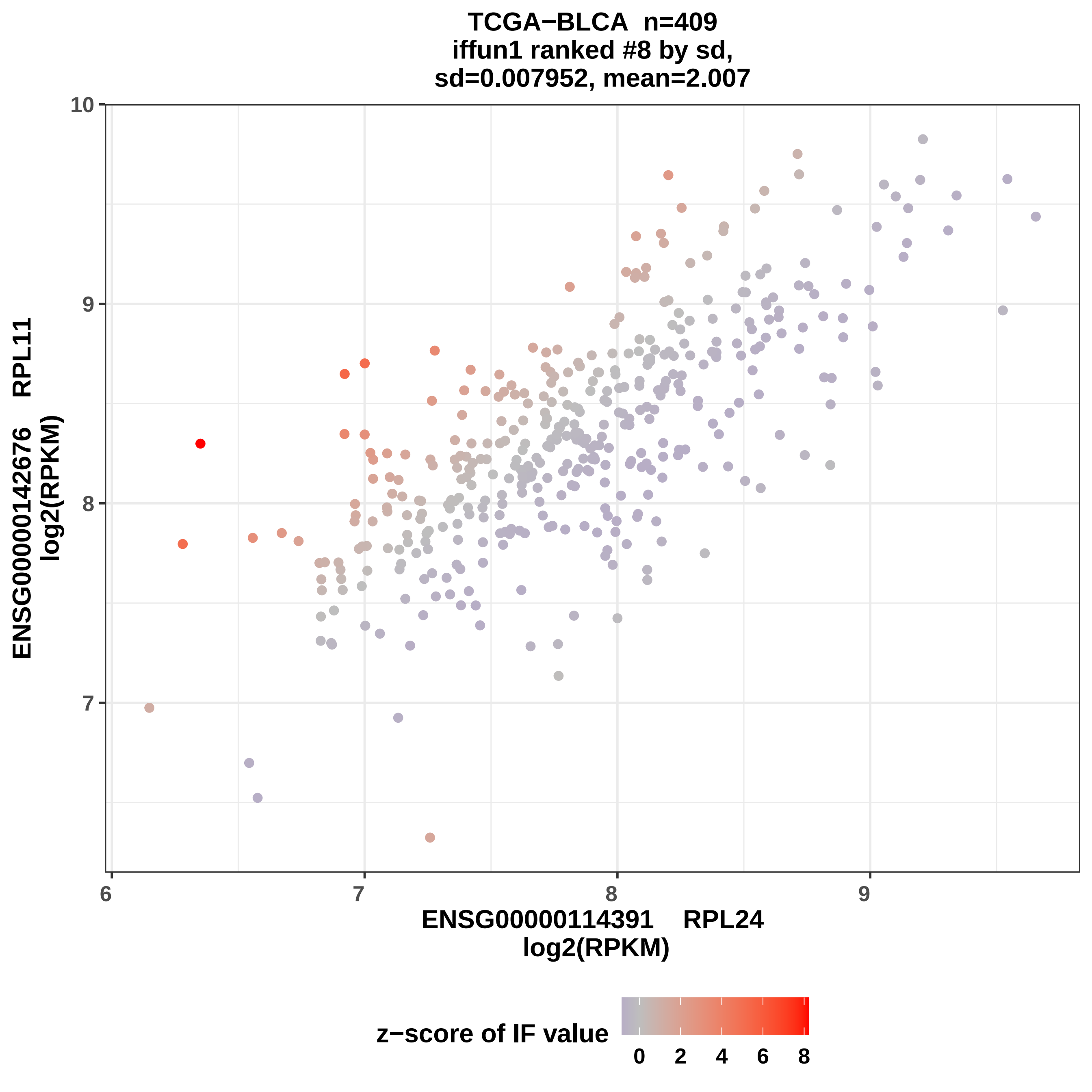

TCGA−BLCA n=409
iffun1 ranked #8 by sd,
sd=0.007952, mean=2.007
ENSG00000142676 RPL11
log2(RPKM)
ENSG00000114391 RPL24
log2(RPKM)
z−score of IF value
10
9
8
7
6
0
2
4
6
8

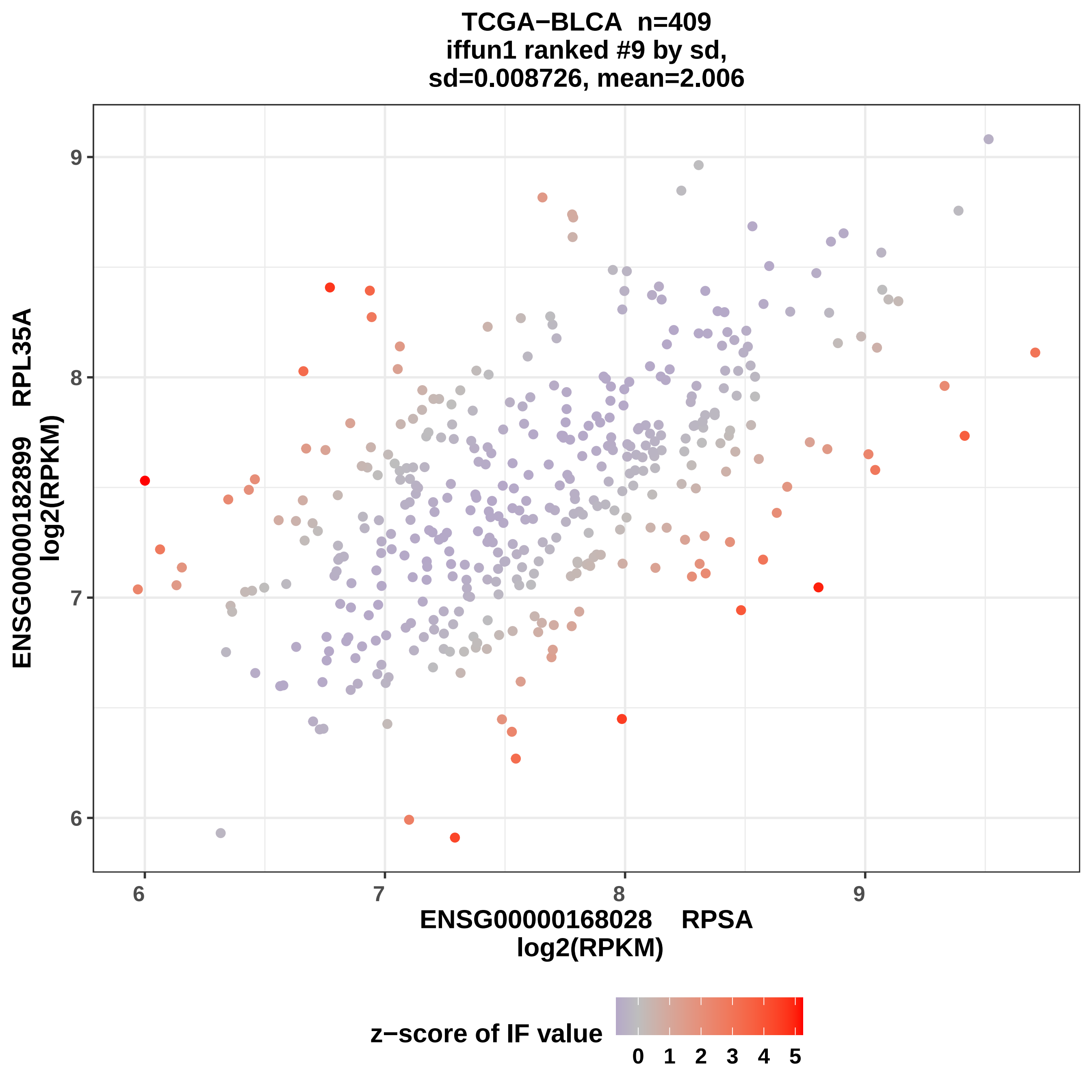
TCGA−BLCA n=409
iffun1 ranked #9 by sd,
sd=0.008726, mean=2.006
ENSG00000182899 RPL35A
log2(RPKM)
9
8
7
6
ENSG00000168028 RPSA
log2(RPKM)
z−score of IF value
0 1 2 3 4 5

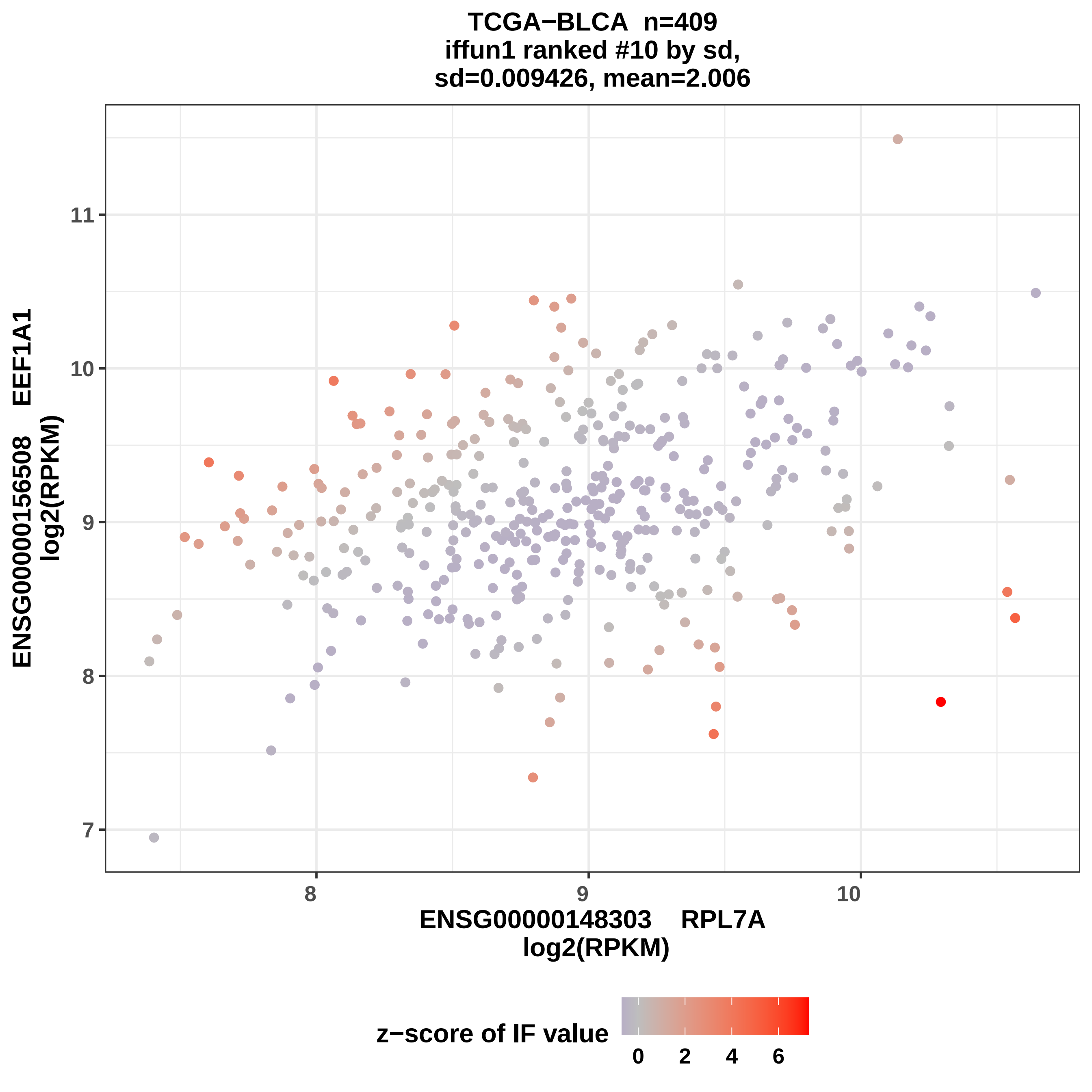
TCGA−BLCA n=409
iffun1 ranked #10 by sd,
sd=0.009426, mean=2.006
ENSG00000156508 EEF1A1
log2(RPKM)
ENSG00000148303 RPL7A
log2(RPKM)
7
8
9
10
11
8
9
10
z−score of IF value
0
2
4
6

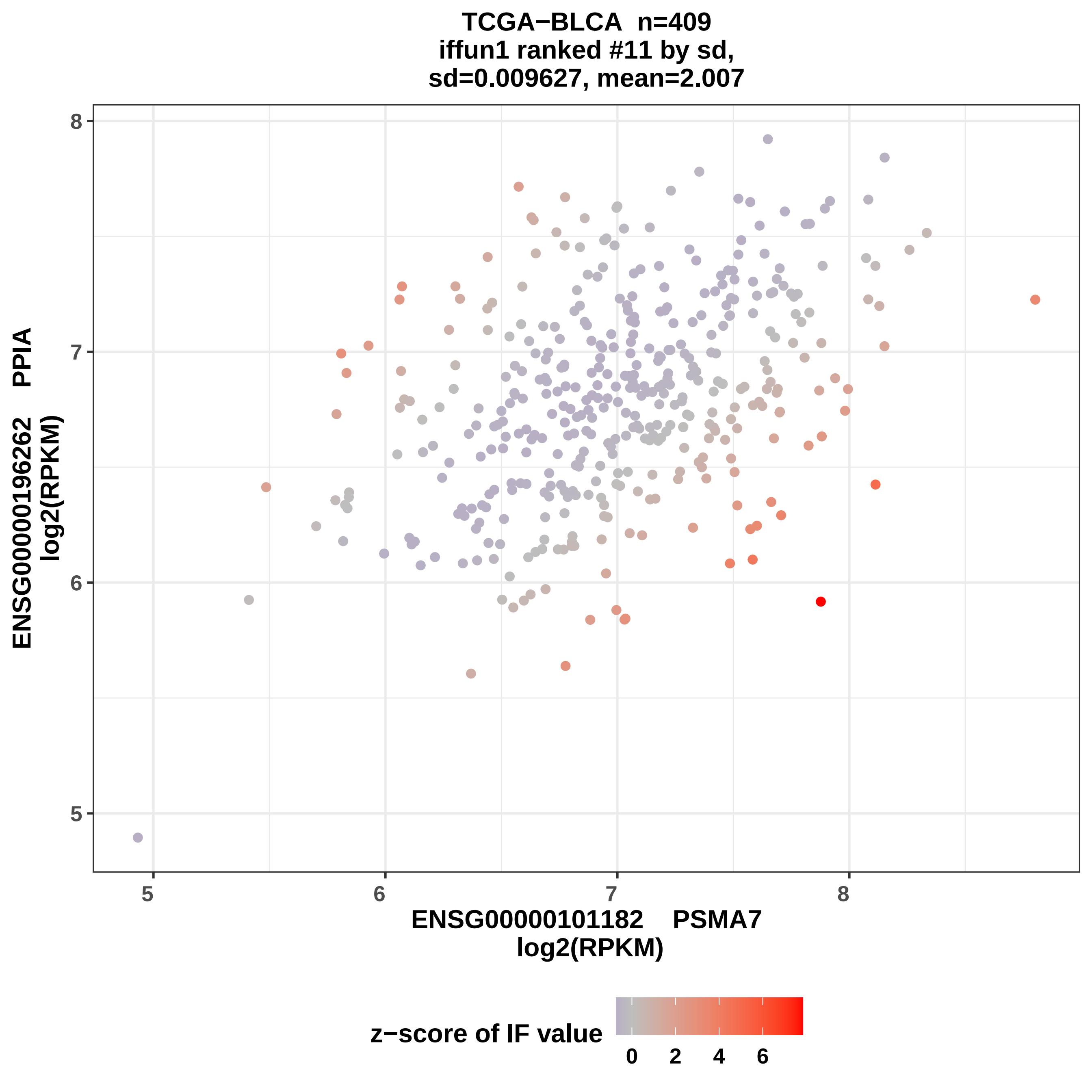

TCGA−BLCA n=409
iffun1 ranked #11 by sd,
sd=0.009627, mean=2.007
ENSG00000196262 PPIA
log2(RPKM)
ENSG00000101182 PSMA7
log2(RPKM)
5
6
7
8
z−score of IF value
0
2
4
6

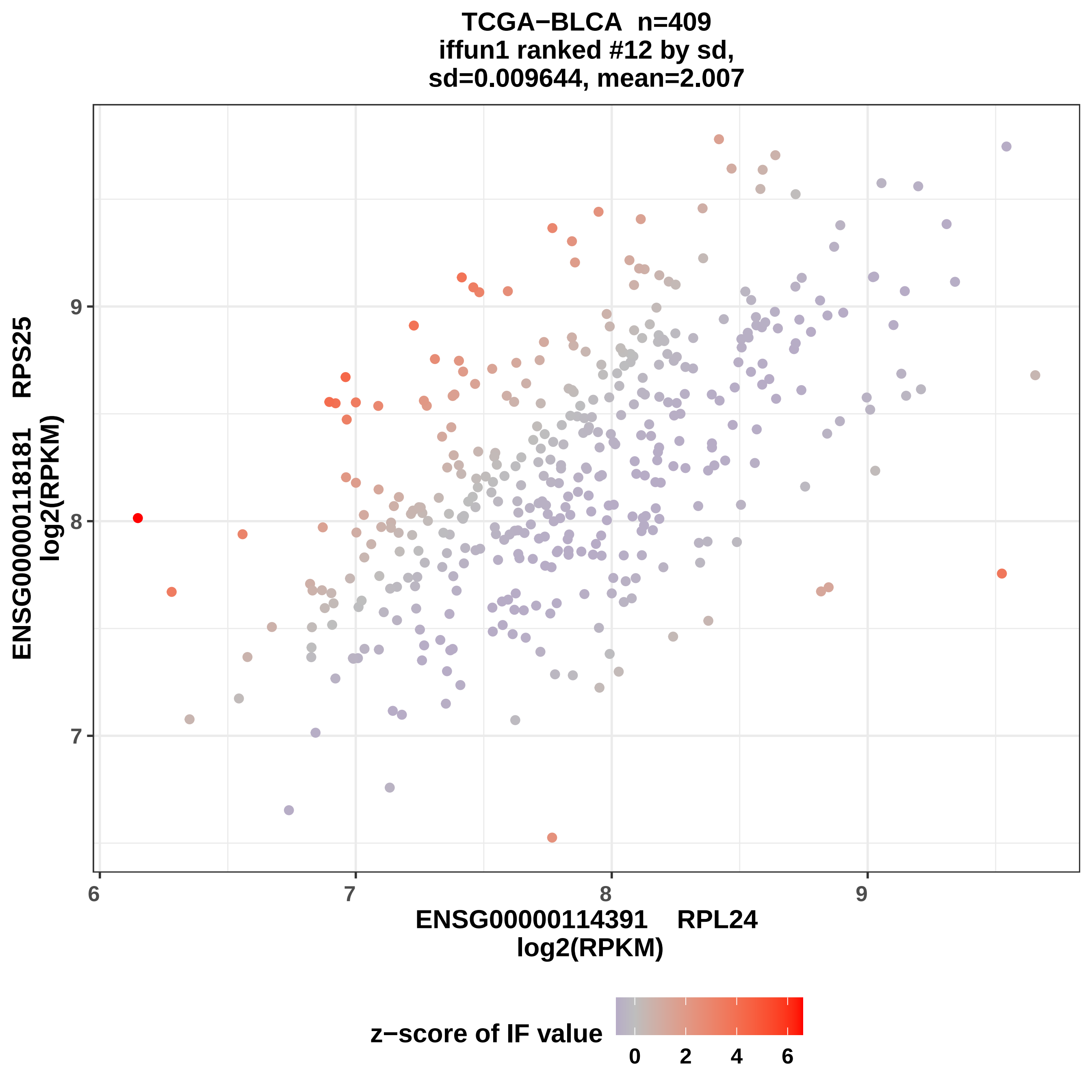
TCGA−BLCA n=409
iffun1 ranked #12 by sd,
sd=0.009644, mean=2.007
ENSG00000118181 RPS25
log2(RPKM)
ENSG00000114391 RPL24
log2(RPKM)
9
8
7
6
7
8
9
z−score of IF value
0
2
4
6

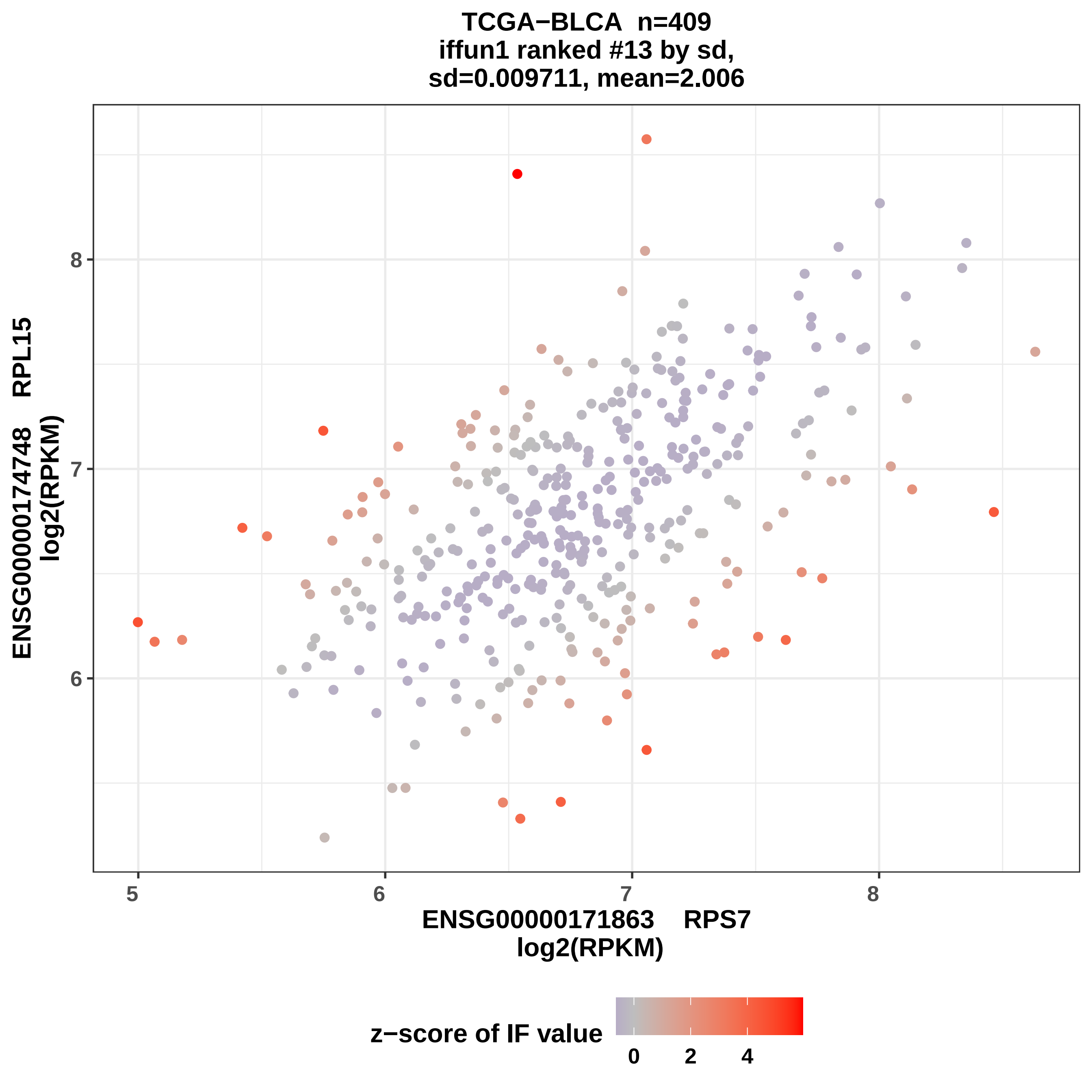

TCGA−BLCA n=409
iffun1 ranked #13 by sd,
sd=0.009711, mean=2.006
ENSG00000174748 RPL15
log2(RPKM)
8
7
6
5
6
7
8
ENSG00000171863 RPS7
log2(RPKM)
z−score of IF value
0
2
4

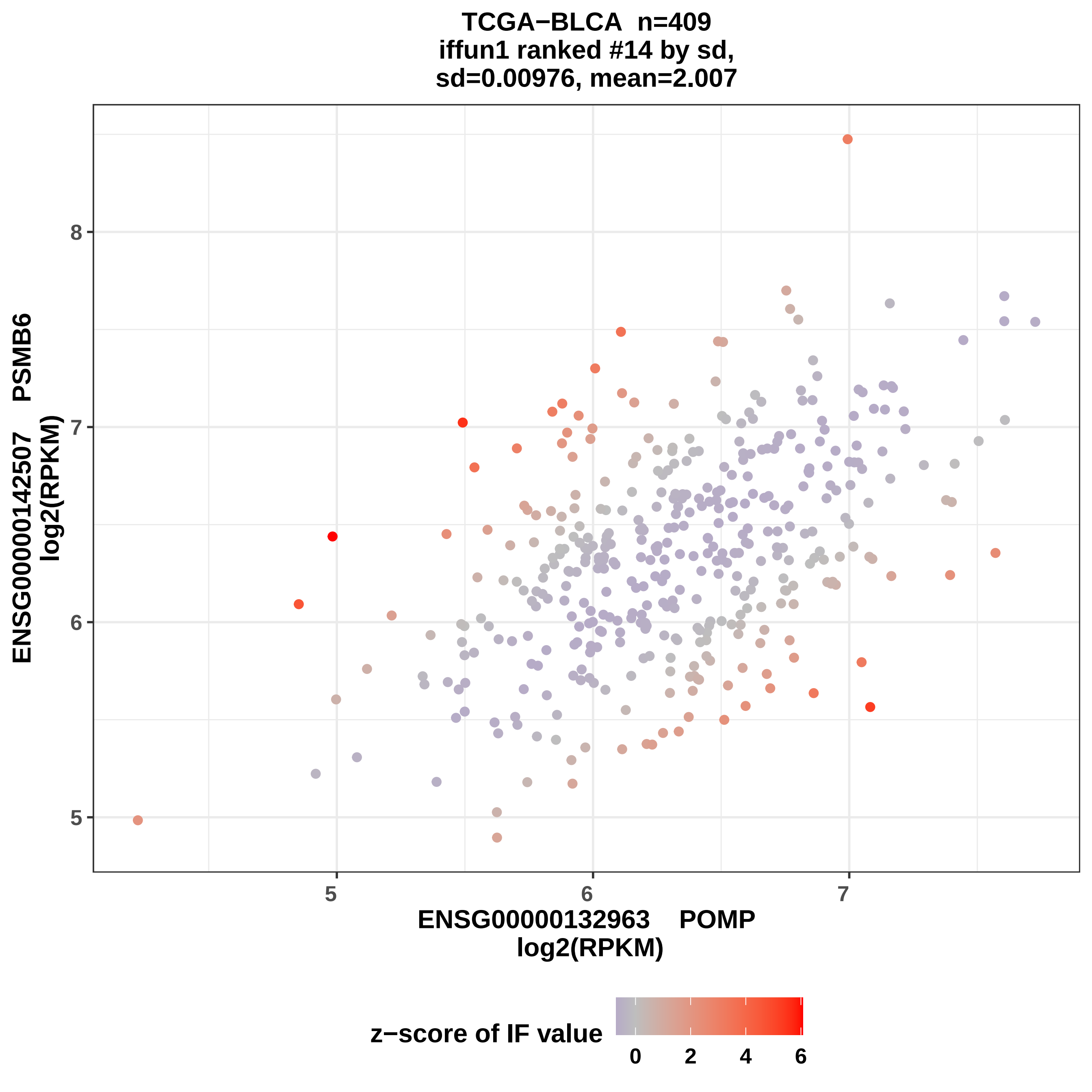
TCGA−BLCA n=409
iffun1 ranked #14 by sd,
sd=0.00976, mean=2.007
ENSG00000142507 PSMB6
log2(RPKM)
ENSG00000132963 POMP
log2(RPKM)
z−score of IF value
5
6
7
8
0
2
4
6

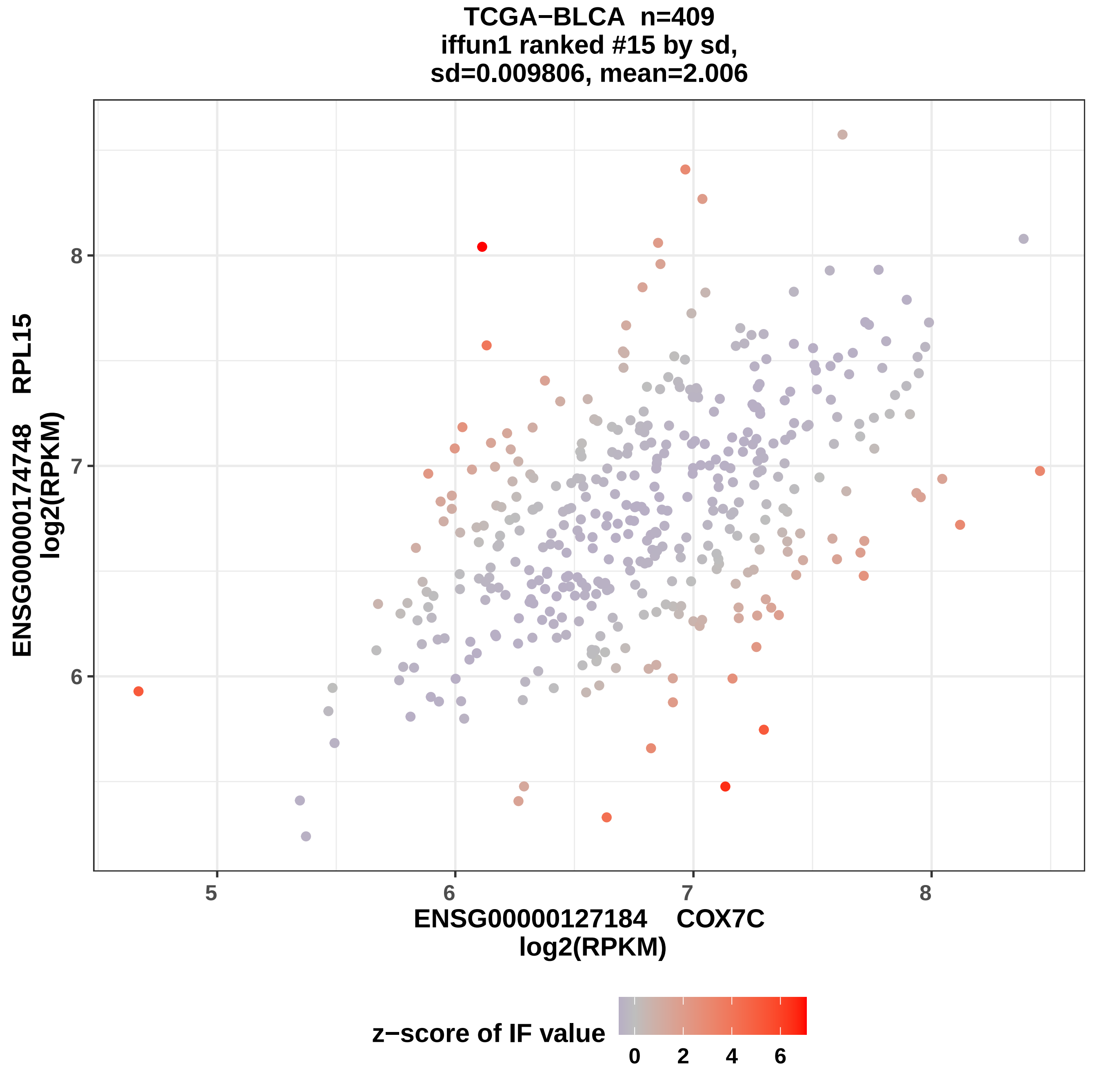

TCGA−BLCA n=409
iffun1 ranked #15 by sd,
sd=0.009806, mean=2.006
ENSG00000174748 RPL15
log2(RPKM)
8
7
6
5
6
7
8
ENSG00000127184 COX7C
log2(RPKM)
z−score of IF value
0
2
4
6

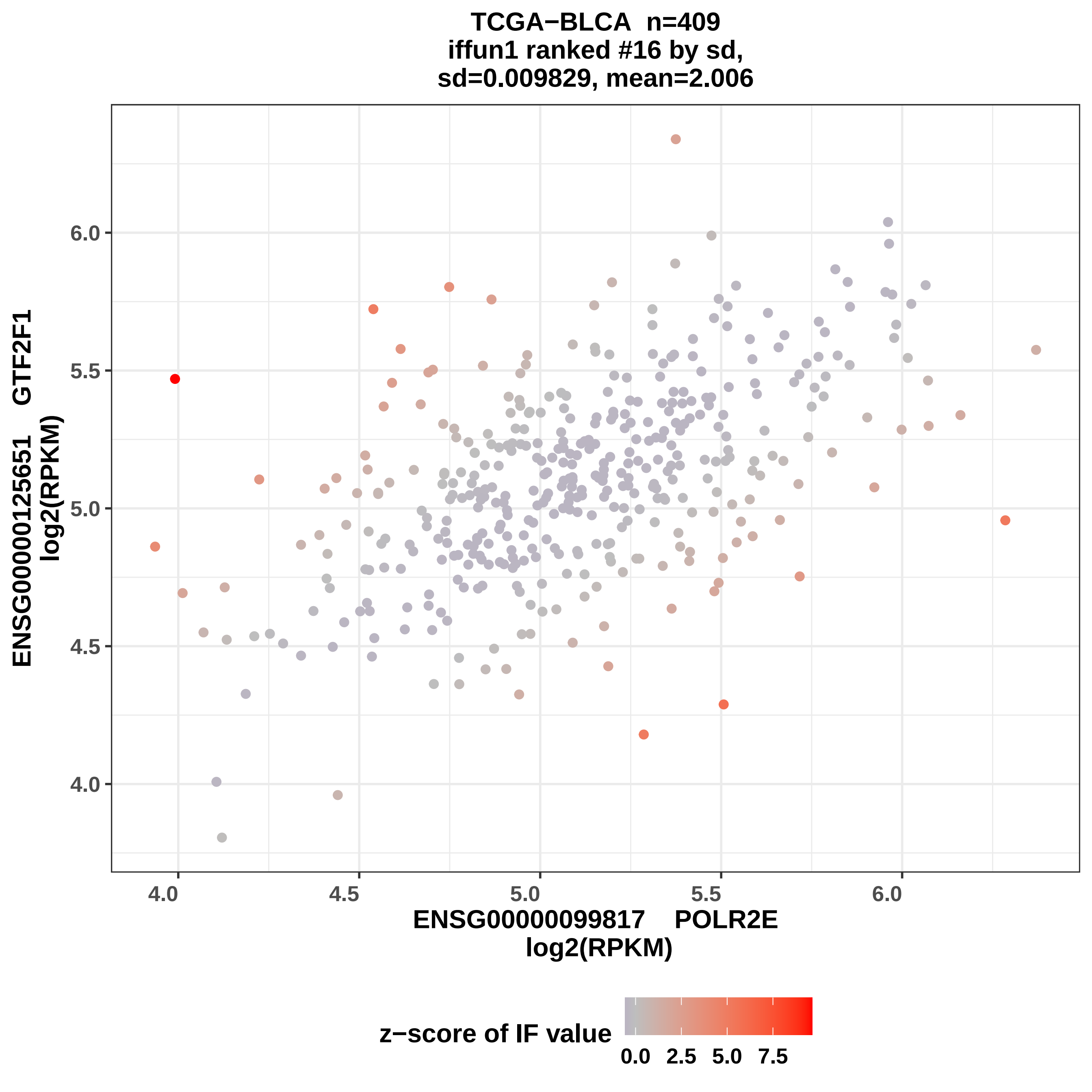
TCGA−BLCA n=409
iffun1 ranked #16 by sd,
sd=0.009829, mean=2.006
ENSG00000125651 GTF2F1
log2(RPKM)
6.0
5.5
5.0
4.5
4.0
4.0
4.5
5.0
5.5
6.0
ENSG00000099817 POLR2E
log2(RPKM)
z−score of IF value
0.0
2.5
5.0
7.5

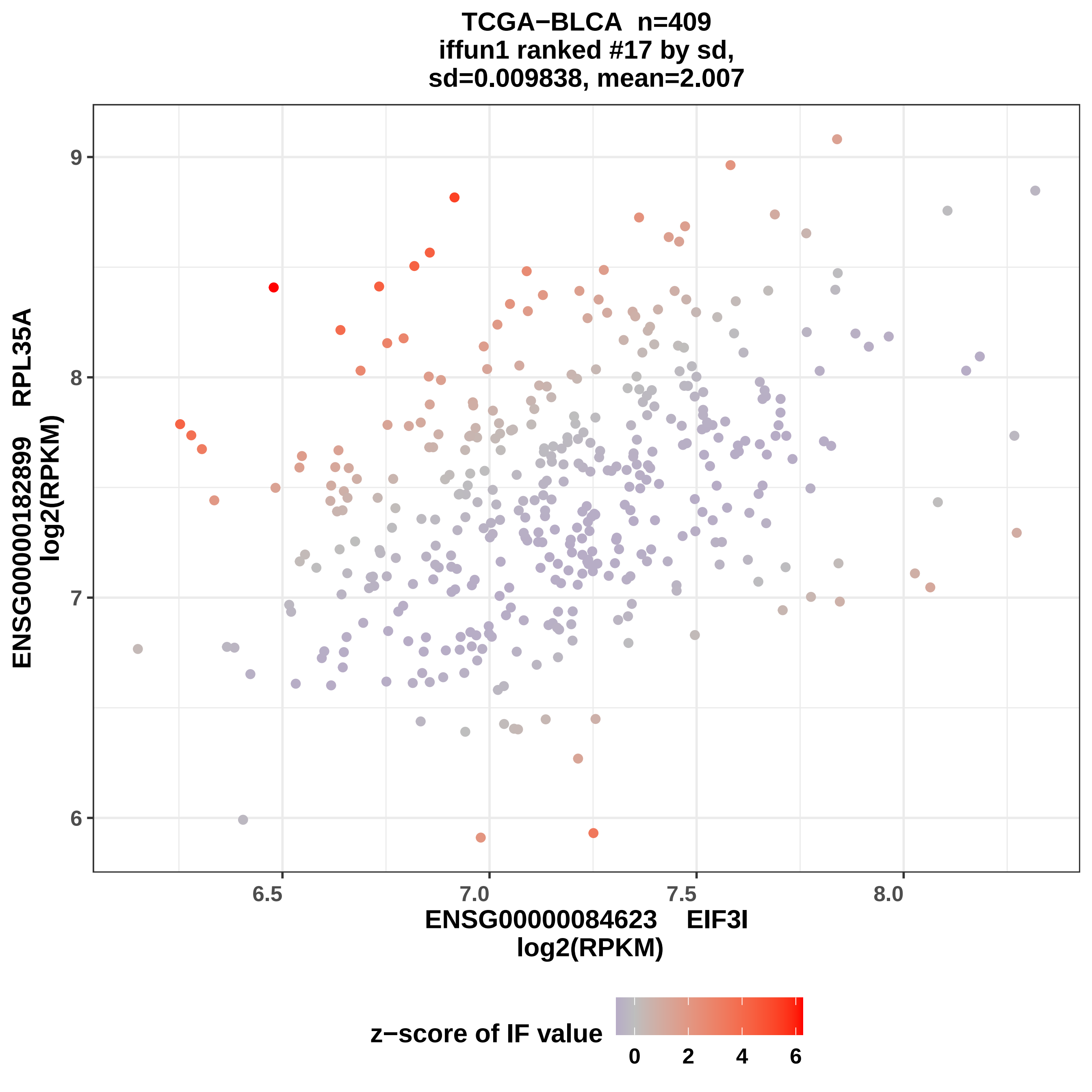
TCGA−BLCA n=409
iffun1 ranked #17 by sd,
sd=0.009838, mean=2.007
ENSG00000182899 RPL35A
log2(RPKM)
ENSG00000084623 EIF3I
log2(RPKM)
z−score of IF value
9
8
7
6
6.5
7.0
7.5
8.0
0
2
4
6

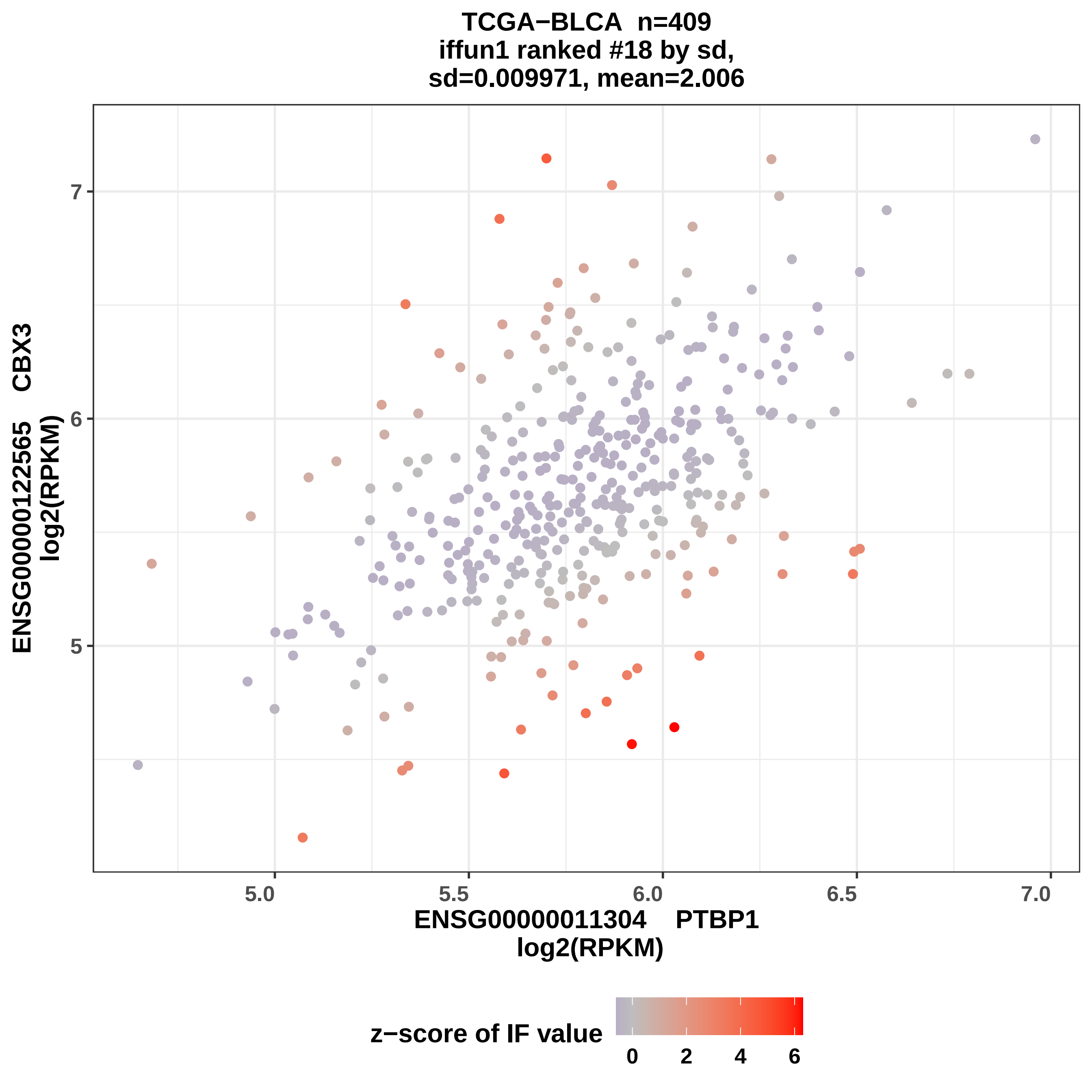
TCGA−BLCA n=409
iffun1 ranked #18 by sd,
sd=0.009971, mean=2.006
ENSG00000122565 CBX3
log2(RPKM)
7
6
5
5.0
5.5
6.0
6.5
7.0
ENSG00000011304 PTBP1
log2(RPKM)
z−score of IF value
0
2
4
6

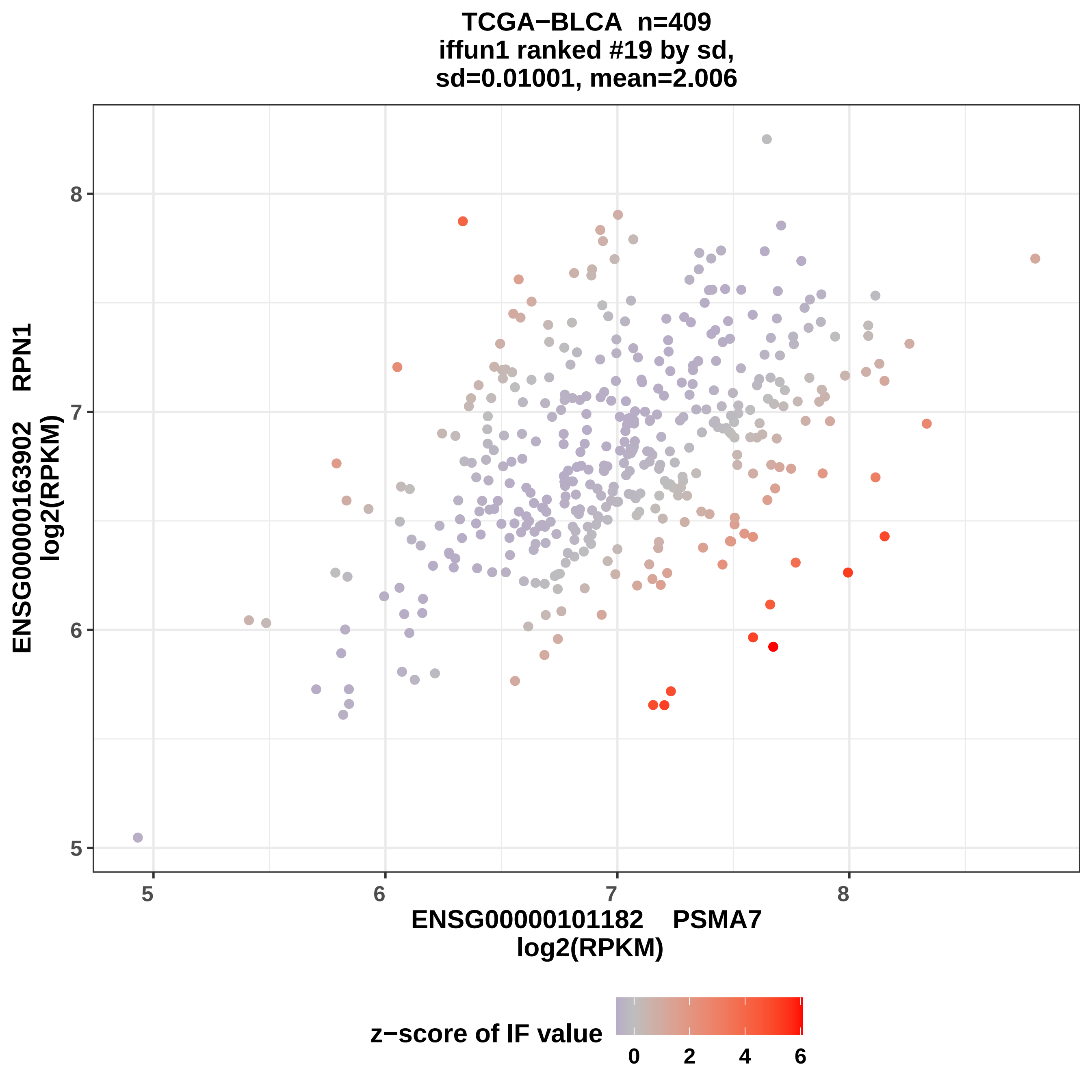

TCGA−BLCA n=409
iffun1 ranked #19 by sd,
sd=0.01001, mean=2.006
ENSG00000163902 RPN1
log2(RPKM)
ENSG00000101182 PSMA7
log2(RPKM)
z−score of IF value
0
2
4
6
5
6
7
8

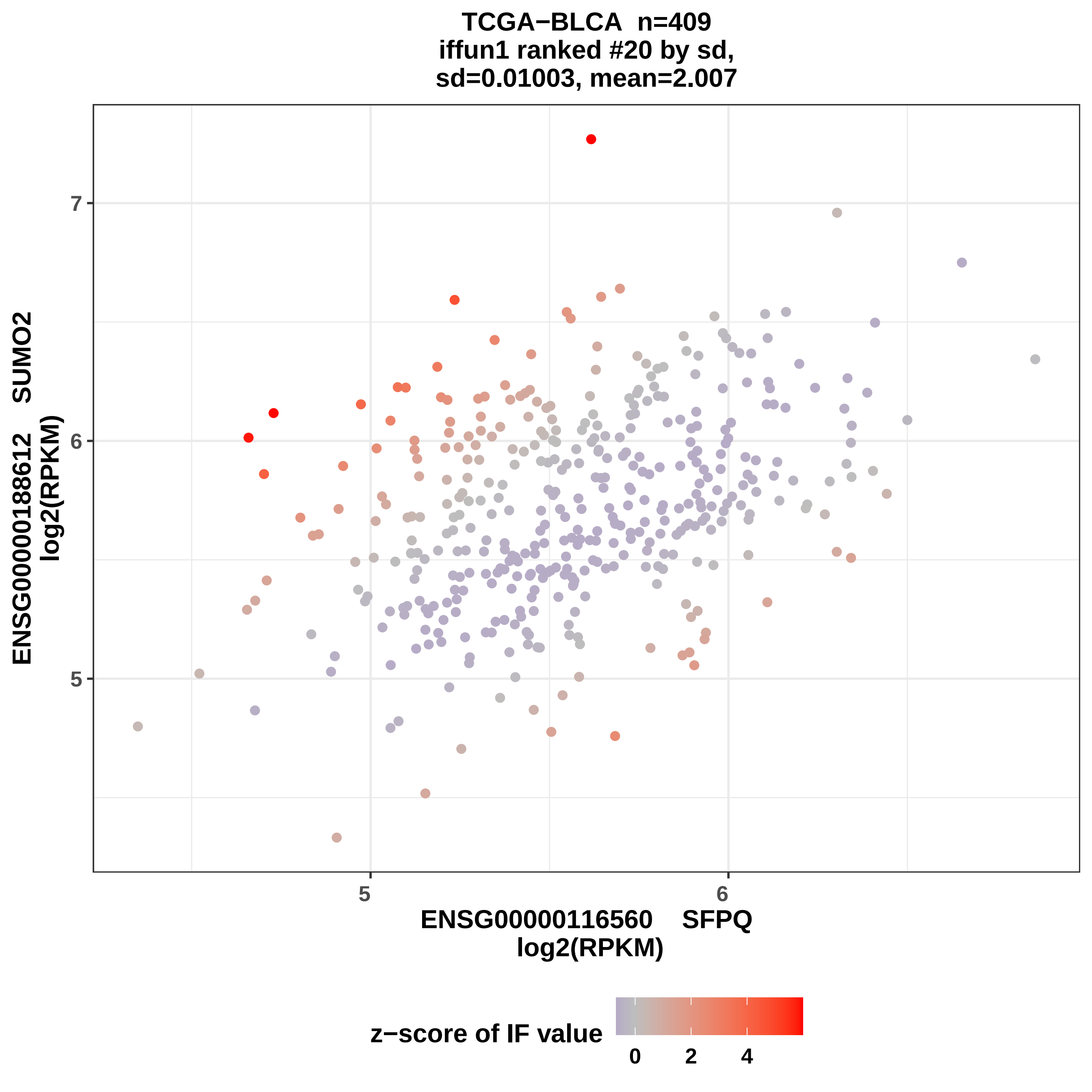

TCGA−BLCA n=409
iffun1 ranked #20 by sd,
sd=0.01003, mean=2.007
ENSG00000188612 SUMO2
log2(RPKM)
7
6
5
5
6
ENSG00000116560 SFPQ
log2(RPKM)
z−score of IF value
0
2
4

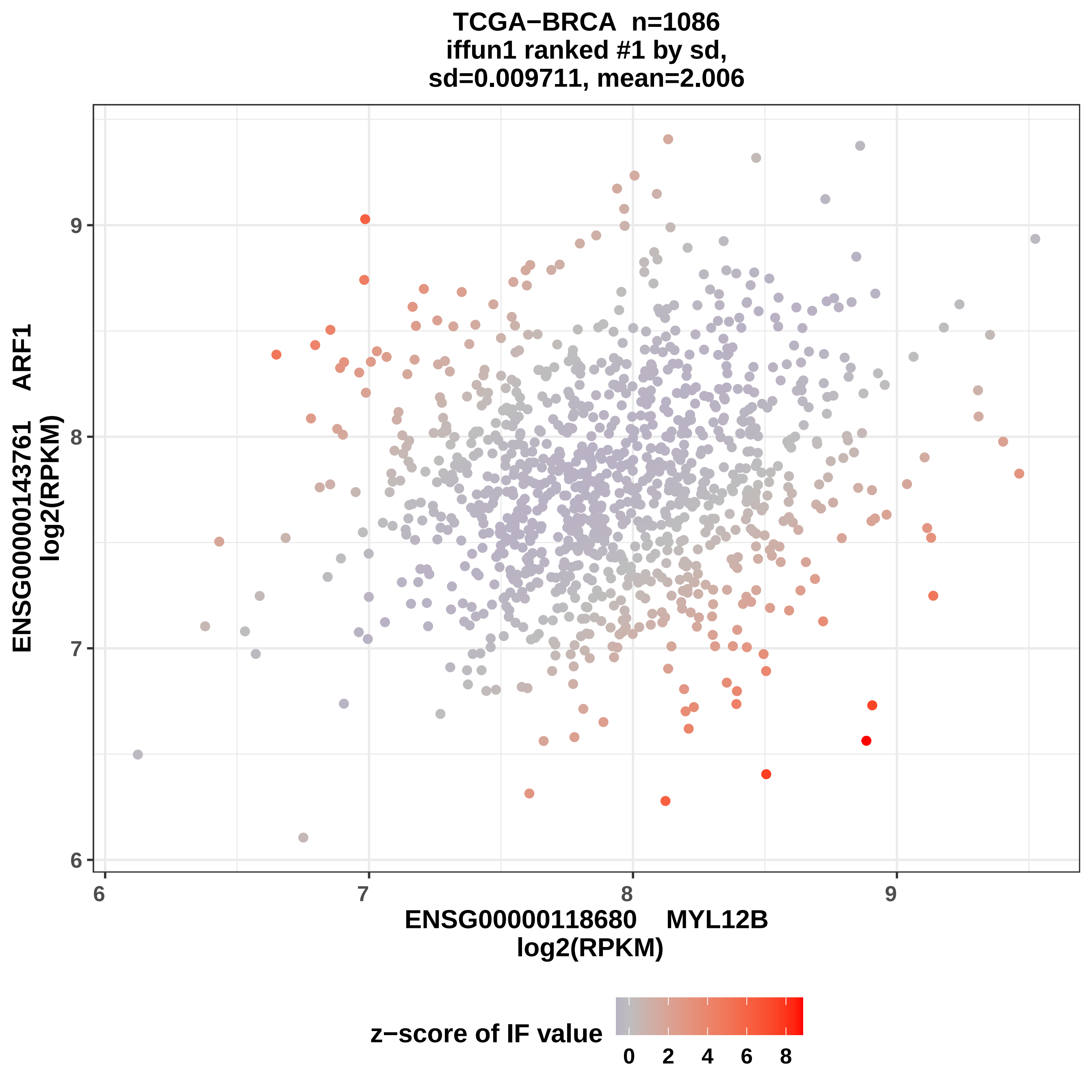
TCGA−BRCA n=1086
iffun1 ranked #1 by sd,
sd=0.009711, mean=2.006
ENSG00000143761 ARF1
log2(RPKM)
9
8
7
6
6
7
8
9
ENSG00000118680 MYL12B
log2(RPKM)
z−score of IF value
0
2
4
6
8

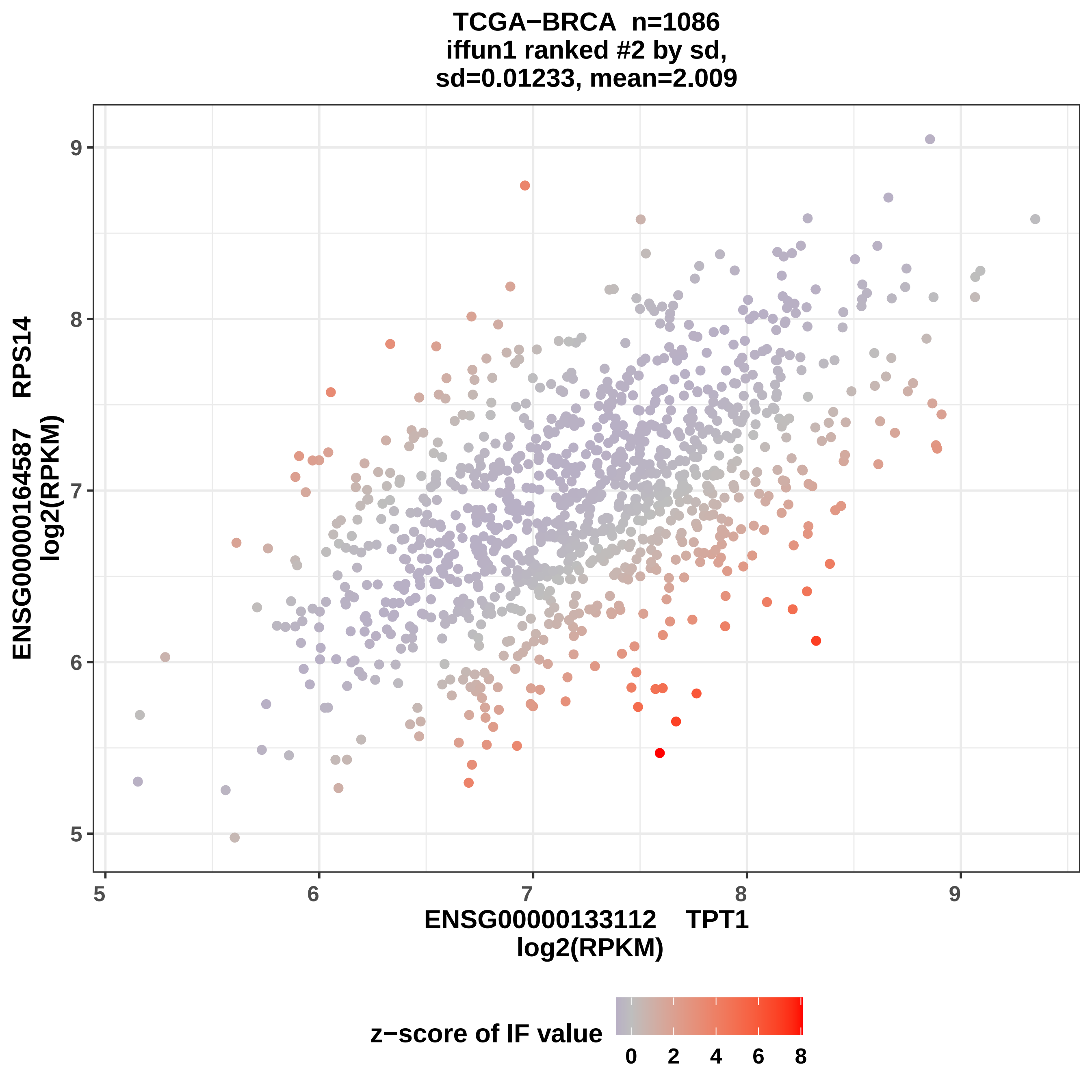
TCGA−BRCA n=1086
iffun1 ranked #2 by sd,
sd=0.01233, mean=2.009
ENSG00000164587 RPS14
log2(RPKM)
9
8
7
6
5
ENSG00000133112 TPT1
log2(RPKM)
5
6
7
8
9
z−score of IF value
0
2
4
6
8

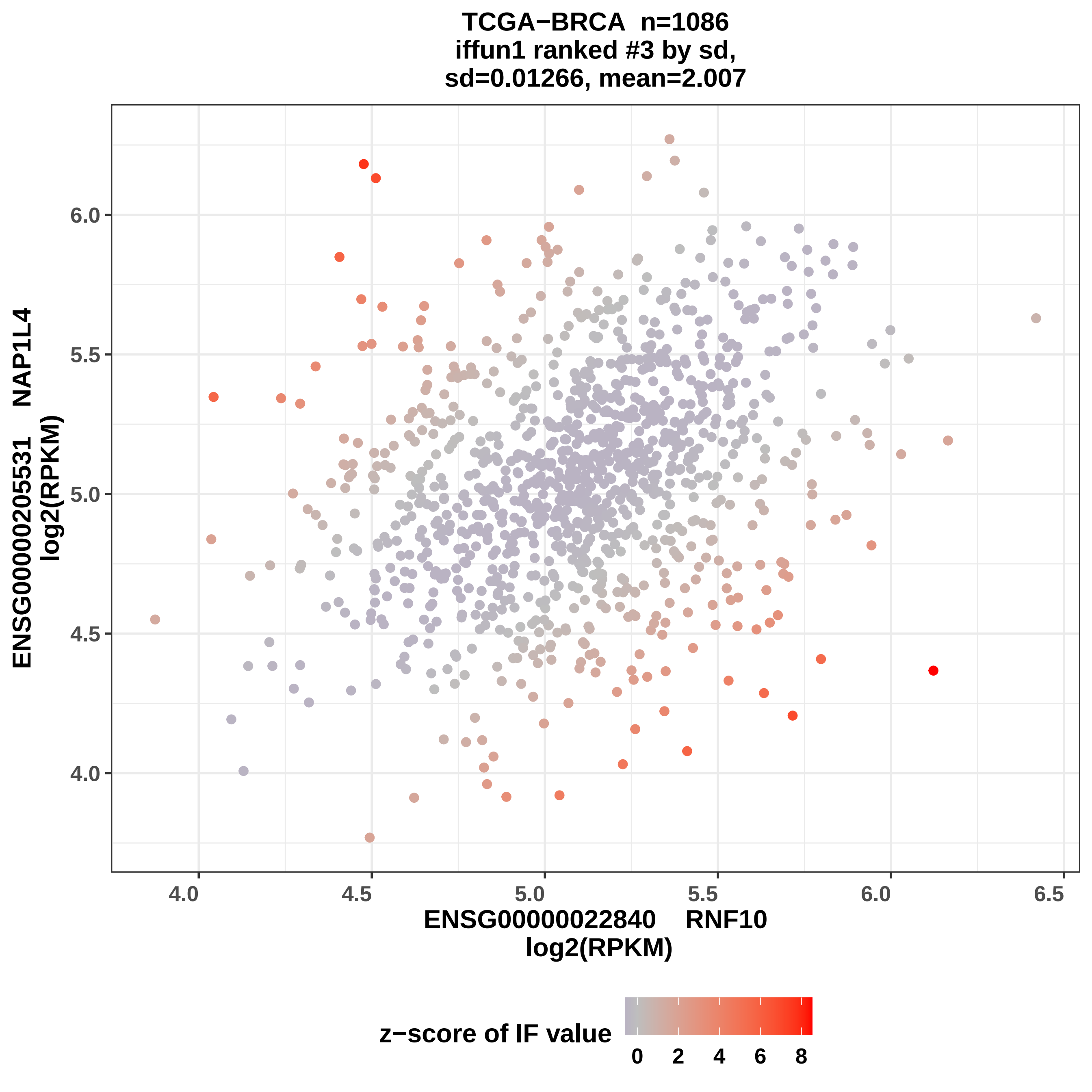

TCGA−BRCA n=1086
iffun1 ranked #3 by sd,
sd=0.01266, mean=2.007
ENSG00000205531 NAP1L4
log2(RPKM)
6.0
5.5
5.0
4.5
4.0
4.0
4.5
5.0
5.5
6.0
6.5
ENSG00000022840 RNF10
log2(RPKM)
z−score of IF value
0
2
4
6
8

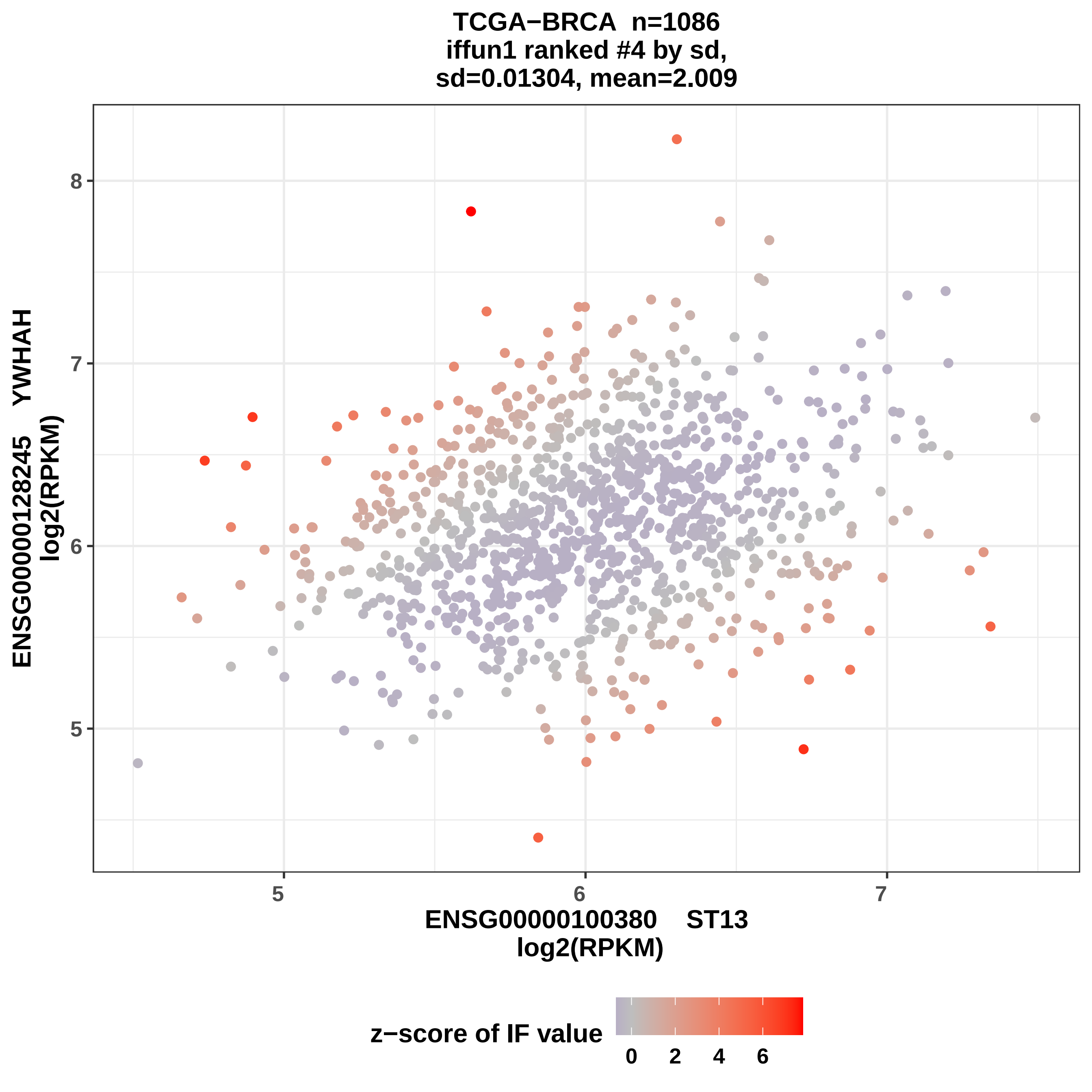
TCGA−BRCA n=1086
iffun1 ranked #4 by sd,
sd=0.01304, mean=2.009
ENSG00000128245 YWHAH
log2(RPKM)
ENSG00000100380 ST13
log2(RPKM)
5
6
7
8
z−score of IF value
0
2
4
6

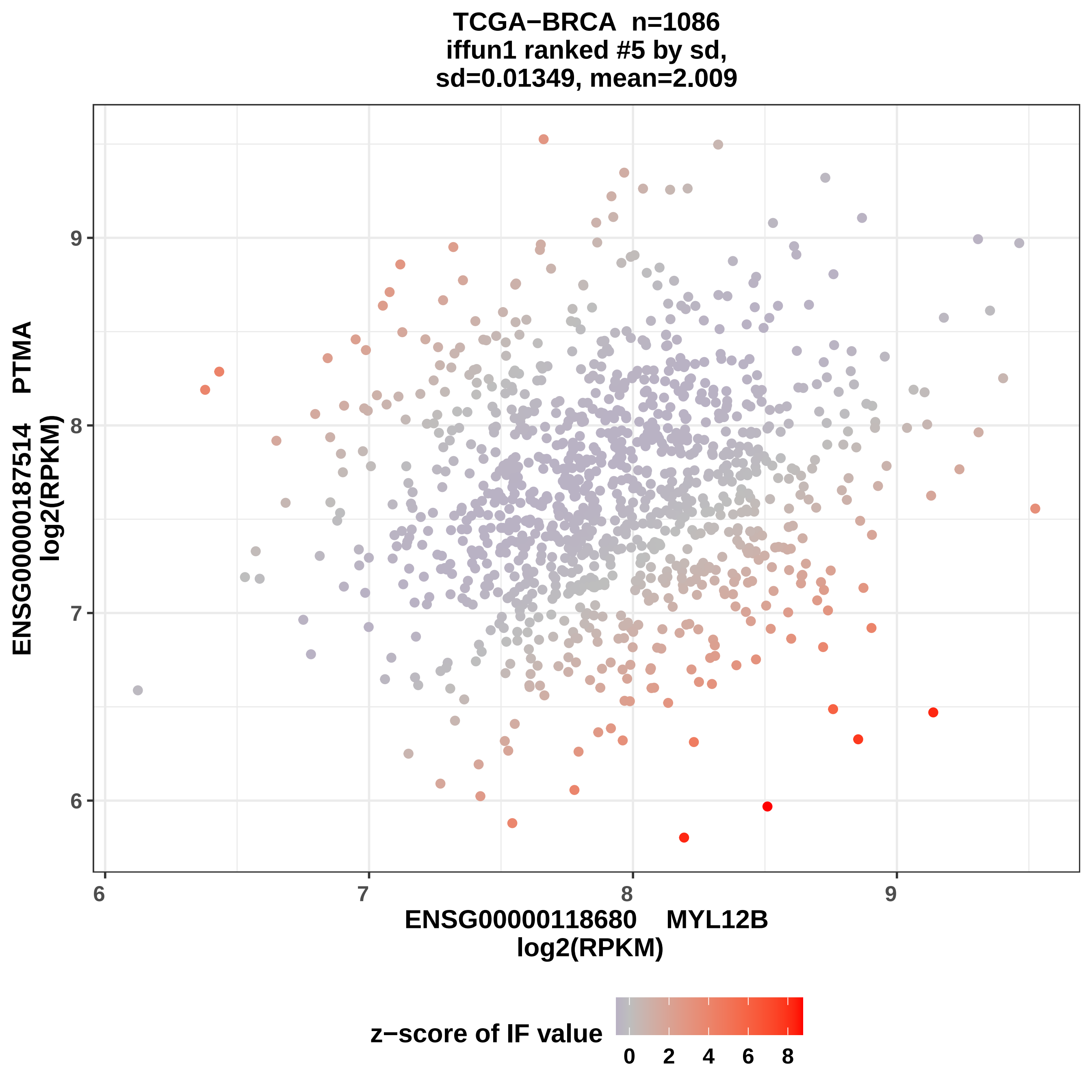

TCGA−BRCA n=1086
iffun1 ranked #5 by sd,
sd=0.01349, mean=2.009
ENSG00000187514 PTMA
log2(RPKM)
ENSG00000118680 MYL12B
log2(RPKM)
z−score of IF value
0
2
4
6
8
6
7
8
9

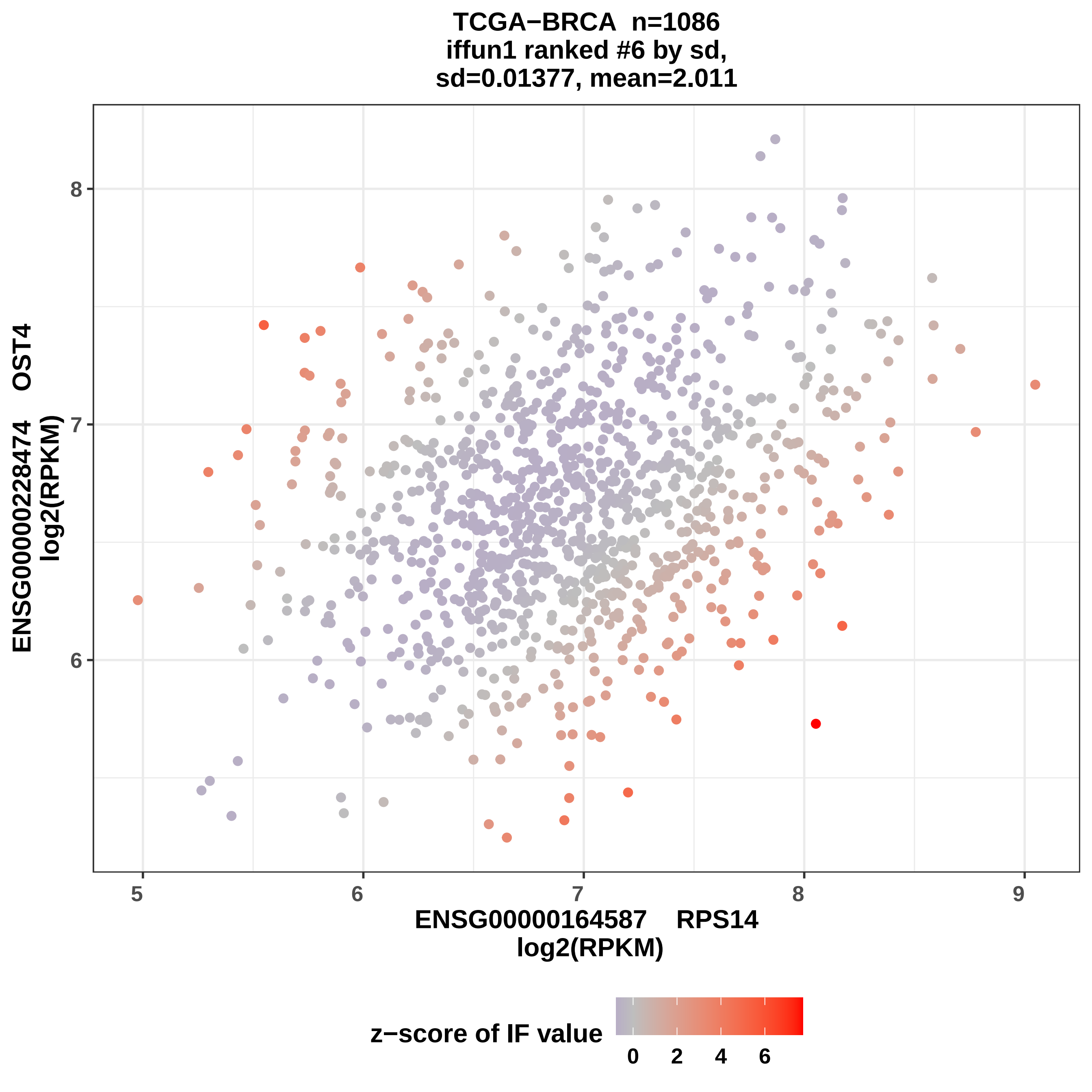
TCGA−BRCA n=1086
iffun1 ranked #6 by sd,
sd=0.01377, mean=2.011
ENSG00000228474 OST4
log2(RPKM)
8
7
6
5
6
7
8
9
ENSG00000164587 RPS14
log2(RPKM)
z−score of IF value
0
2
4
6

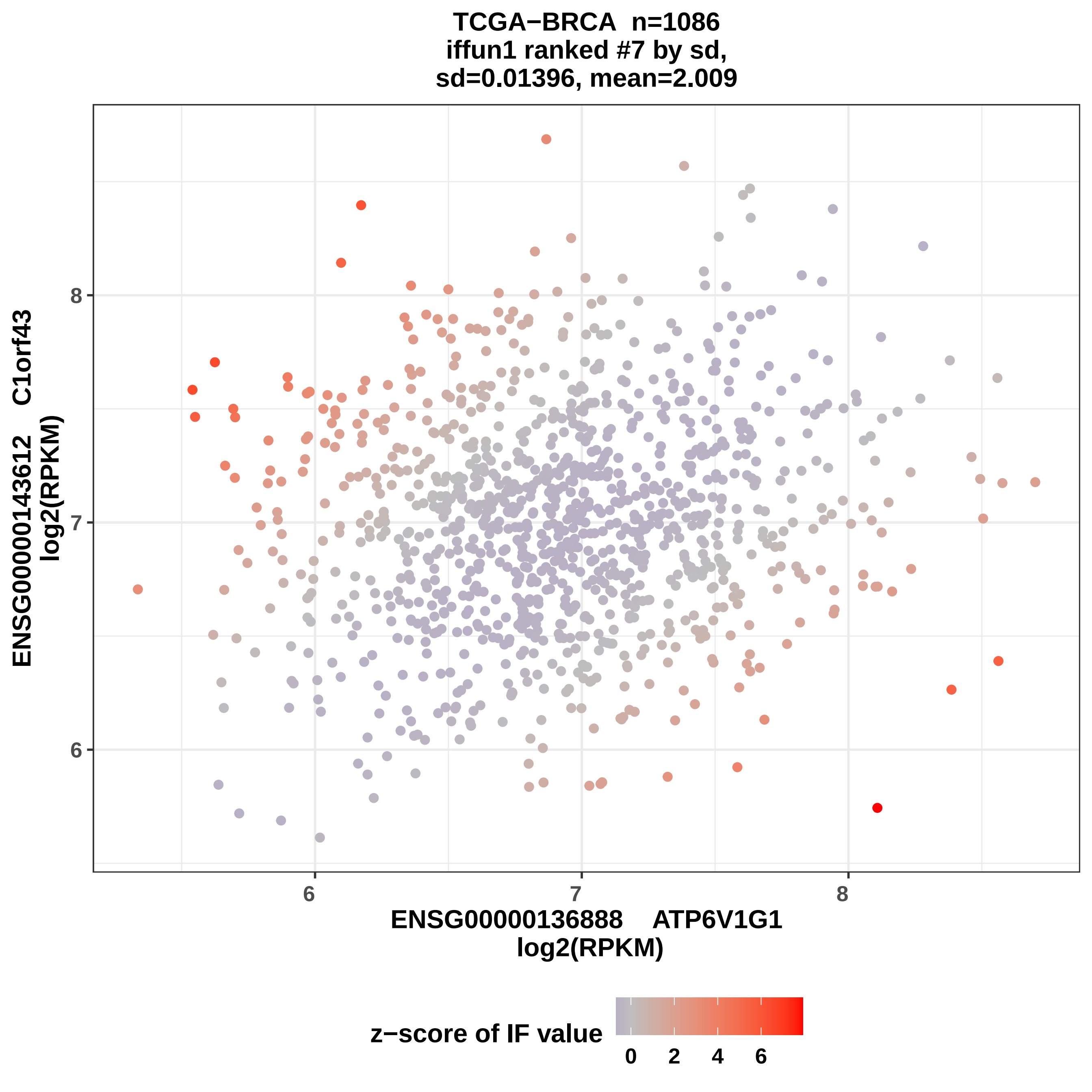

TCGA−BRCA n=1086
iffun1 ranked #7 by sd,
sd=0.01396, mean=2.009
ENSG00000143612 C1orf43
log2(RPKM)
6
7
8
ENSG00000136888 ATP6V1G1
log2(RPKM)
z−score of IF value
0
2
4
6

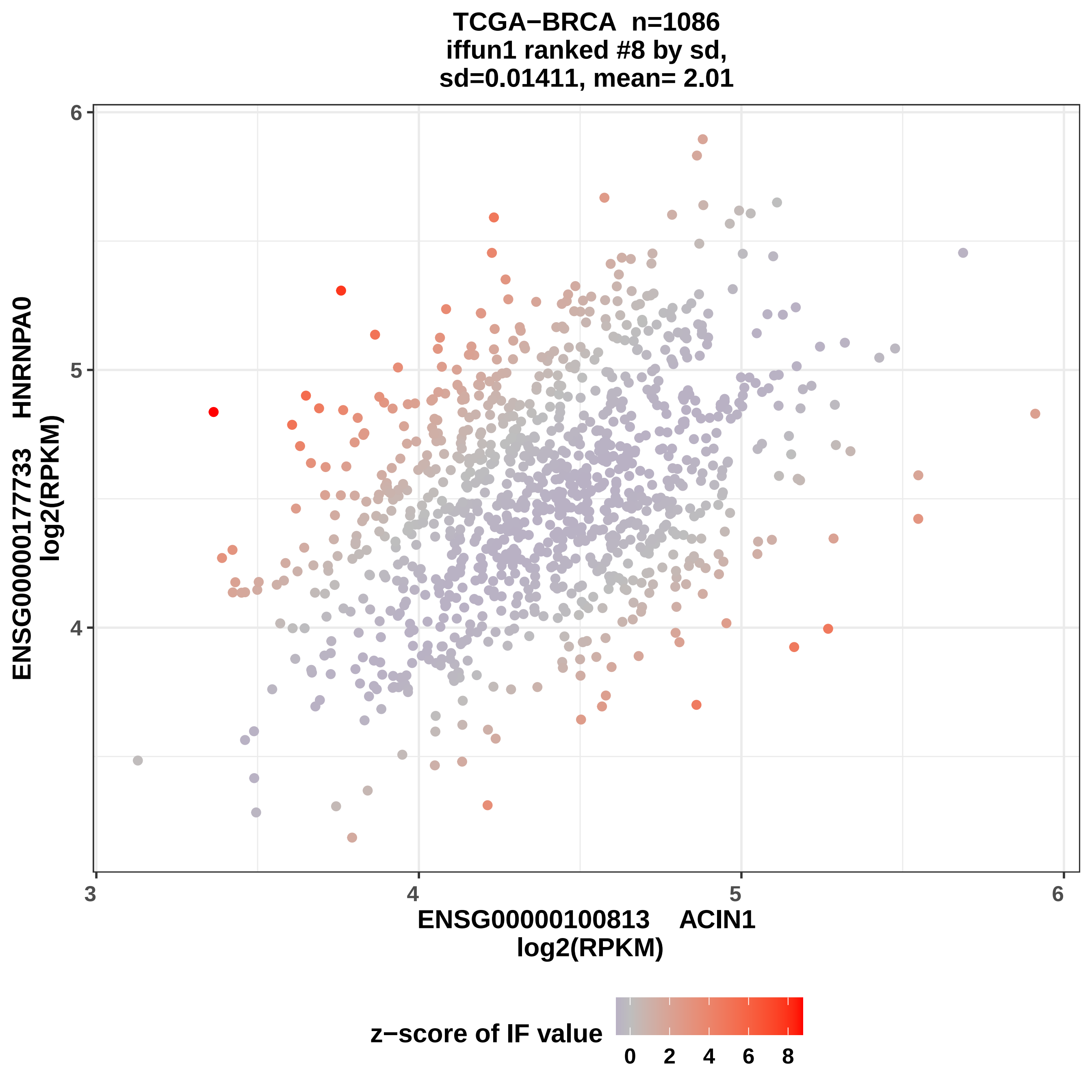
TCGA−BRCA n=1086
iffun1 ranked #8 by sd,
sd=0.01411, mean= 2.01
ENSG00000177733 HNRNPA0
log2(RPKM)
6
5
4
3
4
5
6
ENSG00000100813 ACIN1
log2(RPKM)
z−score of IF value
0
2
4
6
8

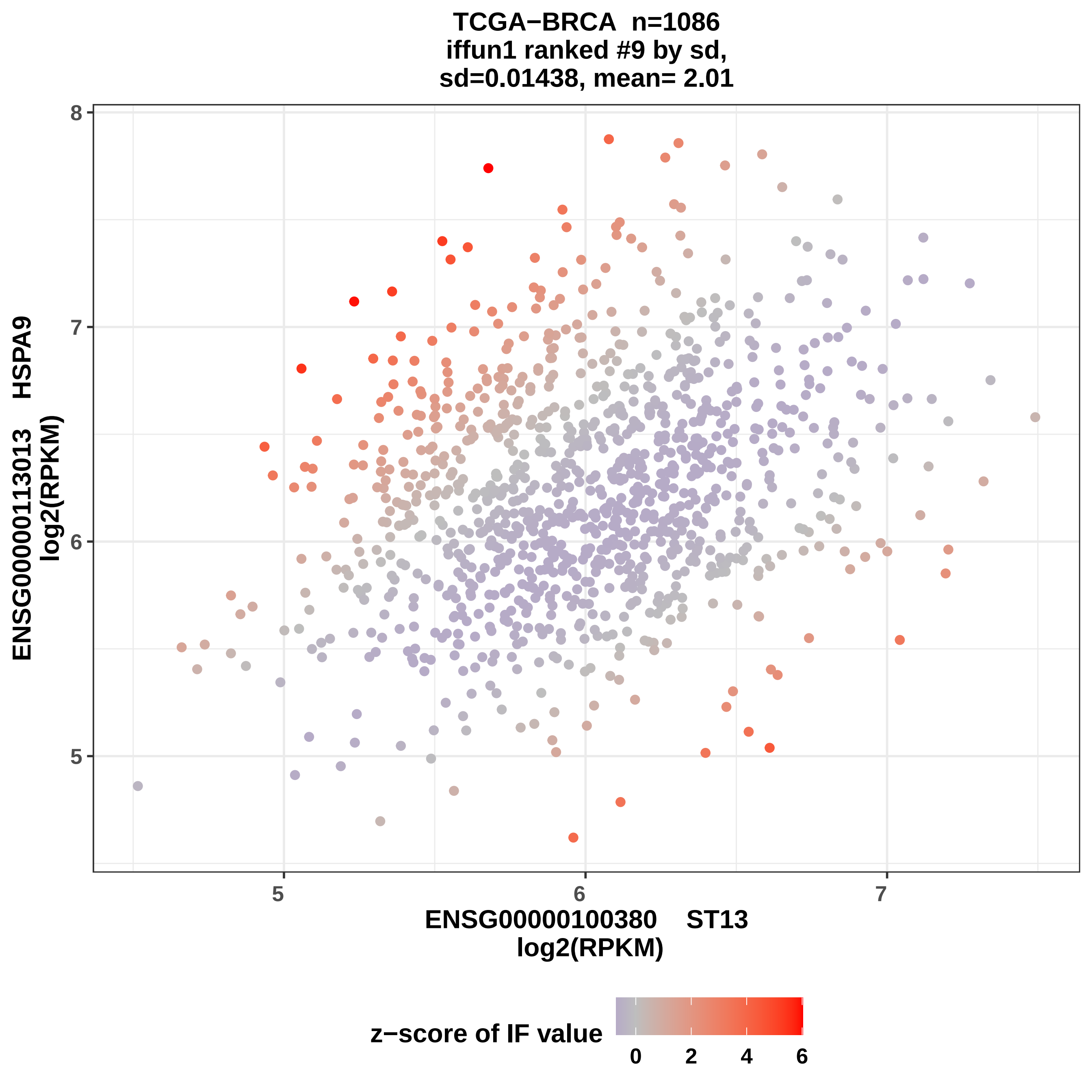

TCGA−BRCA n=1086
iffun1 ranked #9 by sd,
sd=0.01438, mean= 2.01
ENSG00000113013 HSPA9
log2(RPKM)
ENSG00000100380 ST13
log2(RPKM)
8
7
6
5
5
6
7
z−score of IF value
0
2
4
6

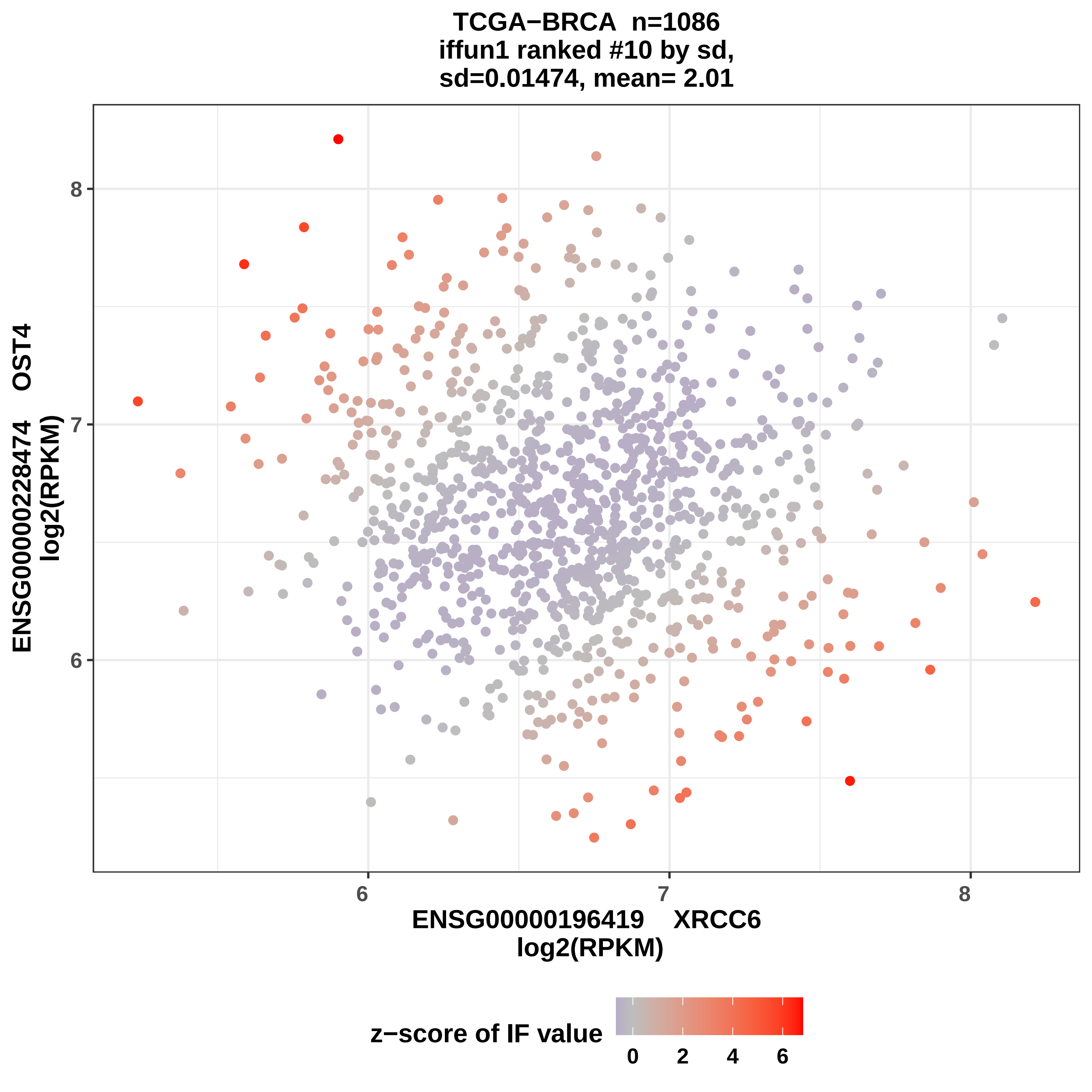
TCGA−BRCA n=1086
iffun1 ranked #10 by sd,
sd=0.01474, mean= 2.01
ENSG00000228474 OST4
log2(RPKM)
8
7
6
ENSG00000196419 XRCC6
log2(RPKM)
6
7
8
z−score of IF value
0
2
4
6

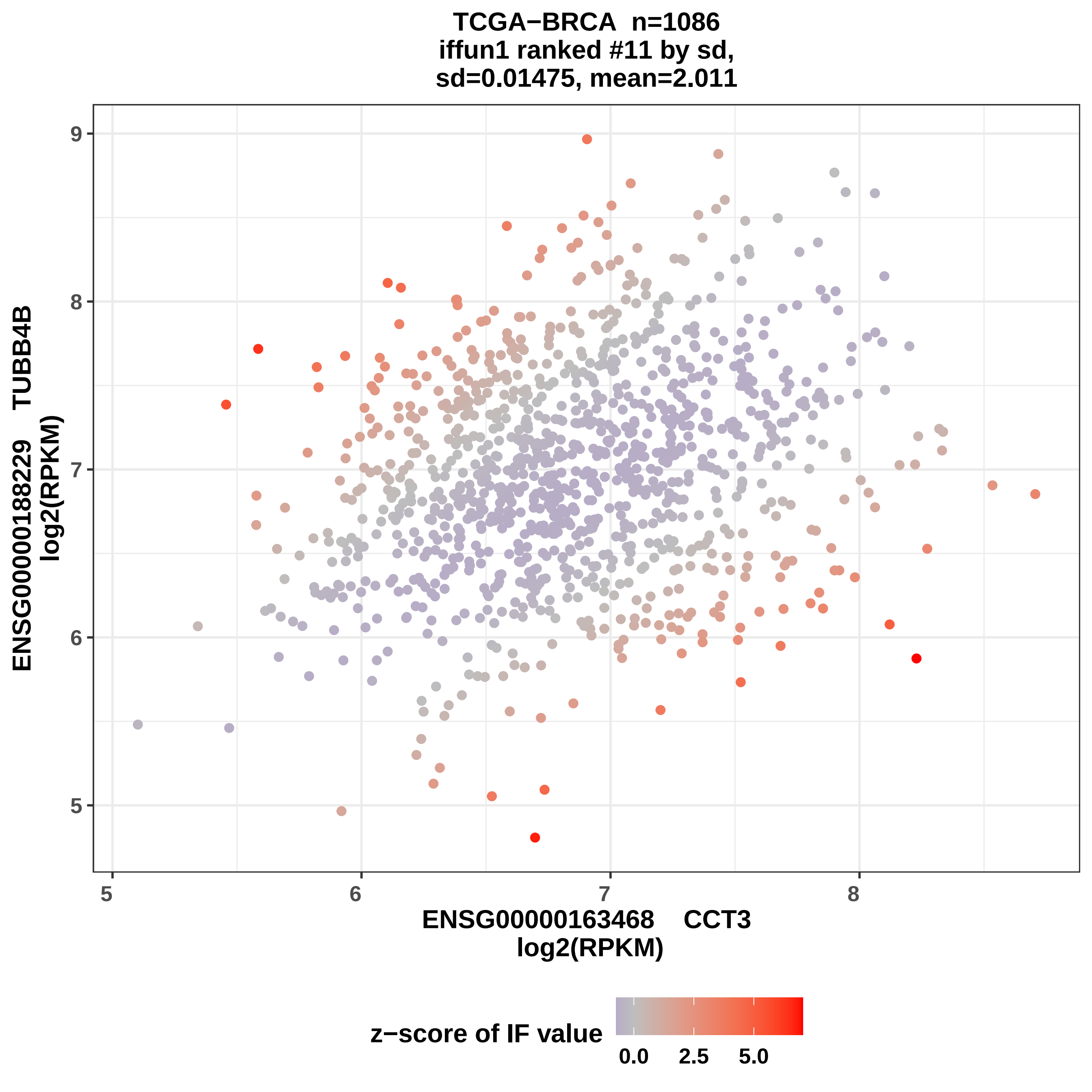
TCGA−BRCA n=1086
iffun1 ranked #11 by sd,
sd=0.01475, mean=2.011
ENSG00000188229 TUBB4B
log2(RPKM)
ENSG00000163468 CCT3
log2(RPKM)
9
8
7
6
5
5
6
7
8
z−score of IF value
0.0
2.5
5.0

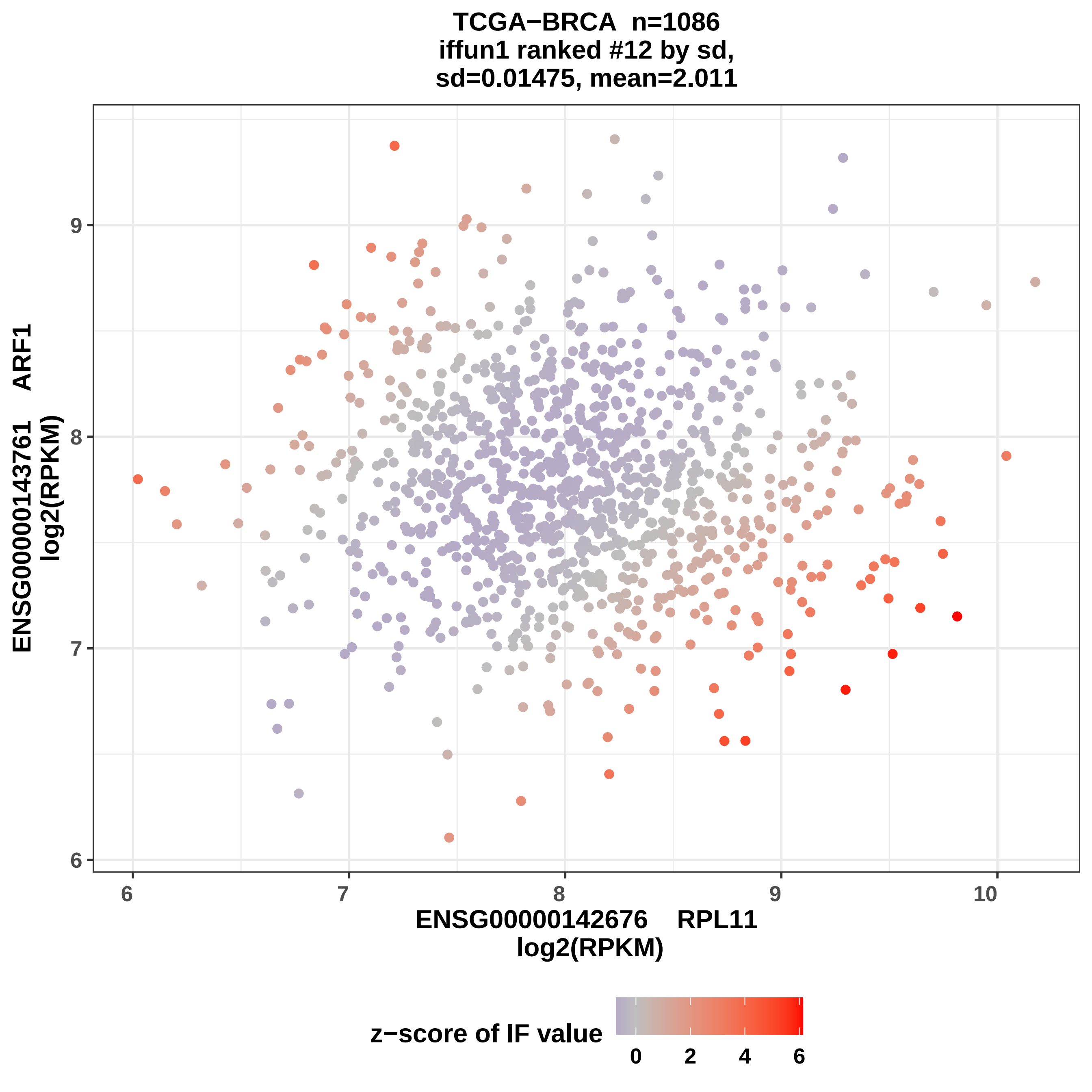

TCGA−BRCA n=1086
iffun1 ranked #12 by sd,
sd=0.01475, mean=2.011
ENSG00000143761 ARF1
log2(RPKM)
ENSG00000142676 RPL11
log2(RPKM)
z−score of IF value
6
7
8
9
10
0
2
4
6

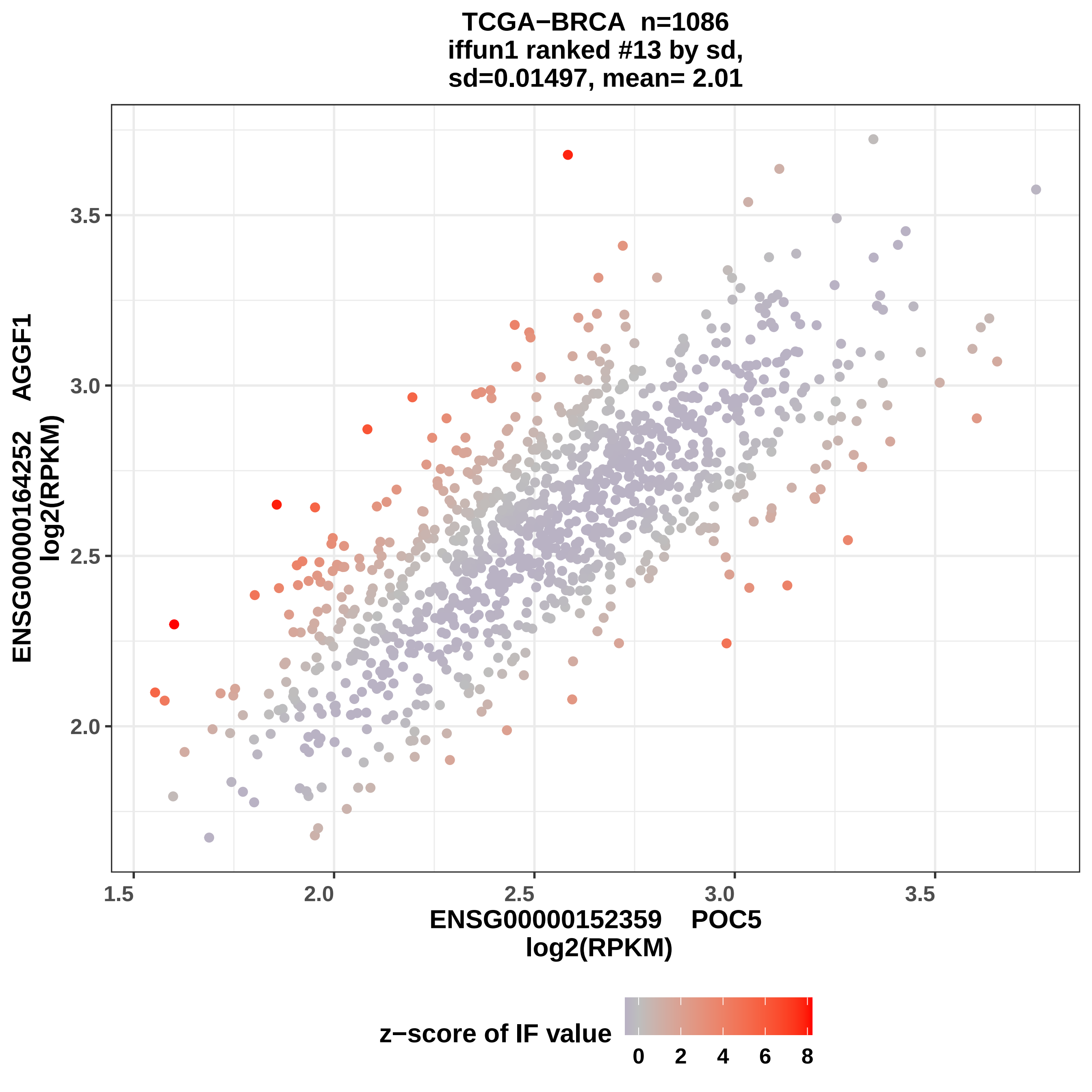
TCGA−BRCA n=1086
iffun1 ranked #13 by sd,
sd=0.01497, mean= 2.01
ENSG00000164252 AGGF1
log2(RPKM)
3.5
3.0
2.5
2.0
1.5
2.0
2.5
3.0
3.5
ENSG00000152359 POC5
log2(RPKM)
z−score of IF value
0
2
4
6
8

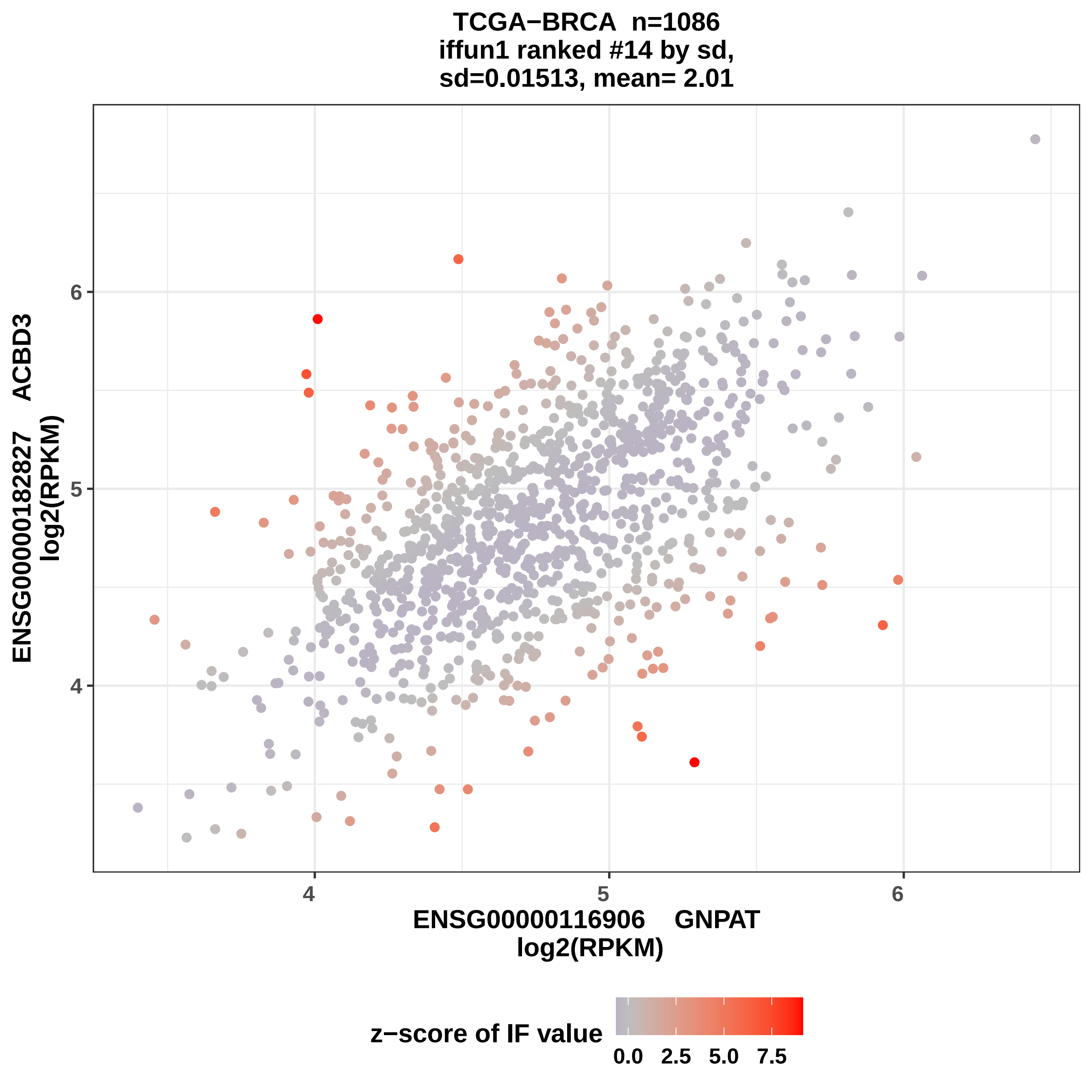
TCGA−BRCA n=1086
iffun1 ranked #14 by sd,
sd=0.01513, mean= 2.01
ENSG00000182827 ACBD3
log2(RPKM)
6
5
4
4
5
6
ENSG00000116906 GNPAT
log2(RPKM)
z−score of IF value
0.0
2.5
5.0
7.5

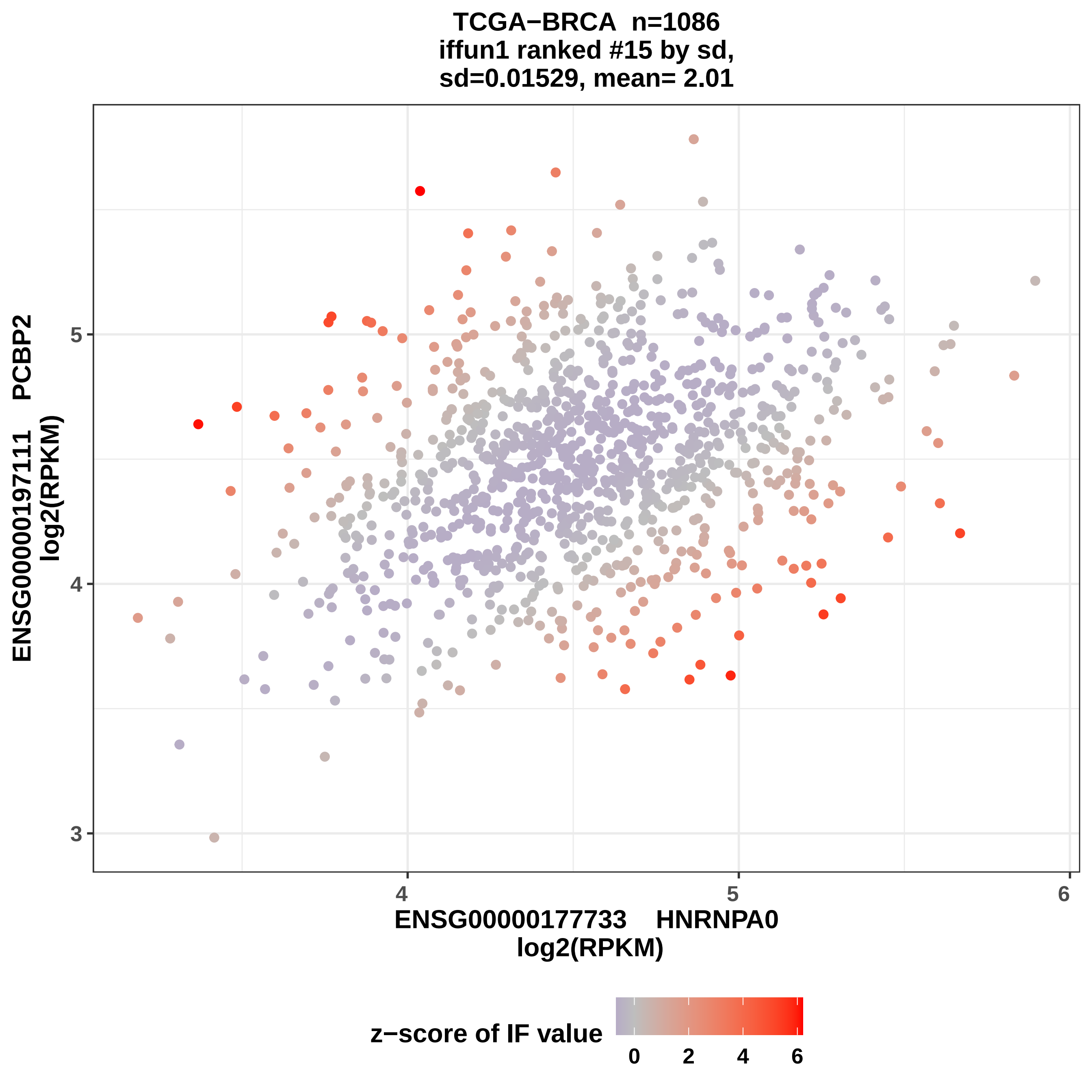
TCGA−BRCA n=1086
iffun1 ranked #15 by sd,
sd=0.01529, mean= 2.01
ENSG00000197111 PCBP2
log2(RPKM)
5
4
3
4
5
6
ENSG00000177733 HNRNPA0
log2(RPKM)
z−score of IF value
0
2
4
6

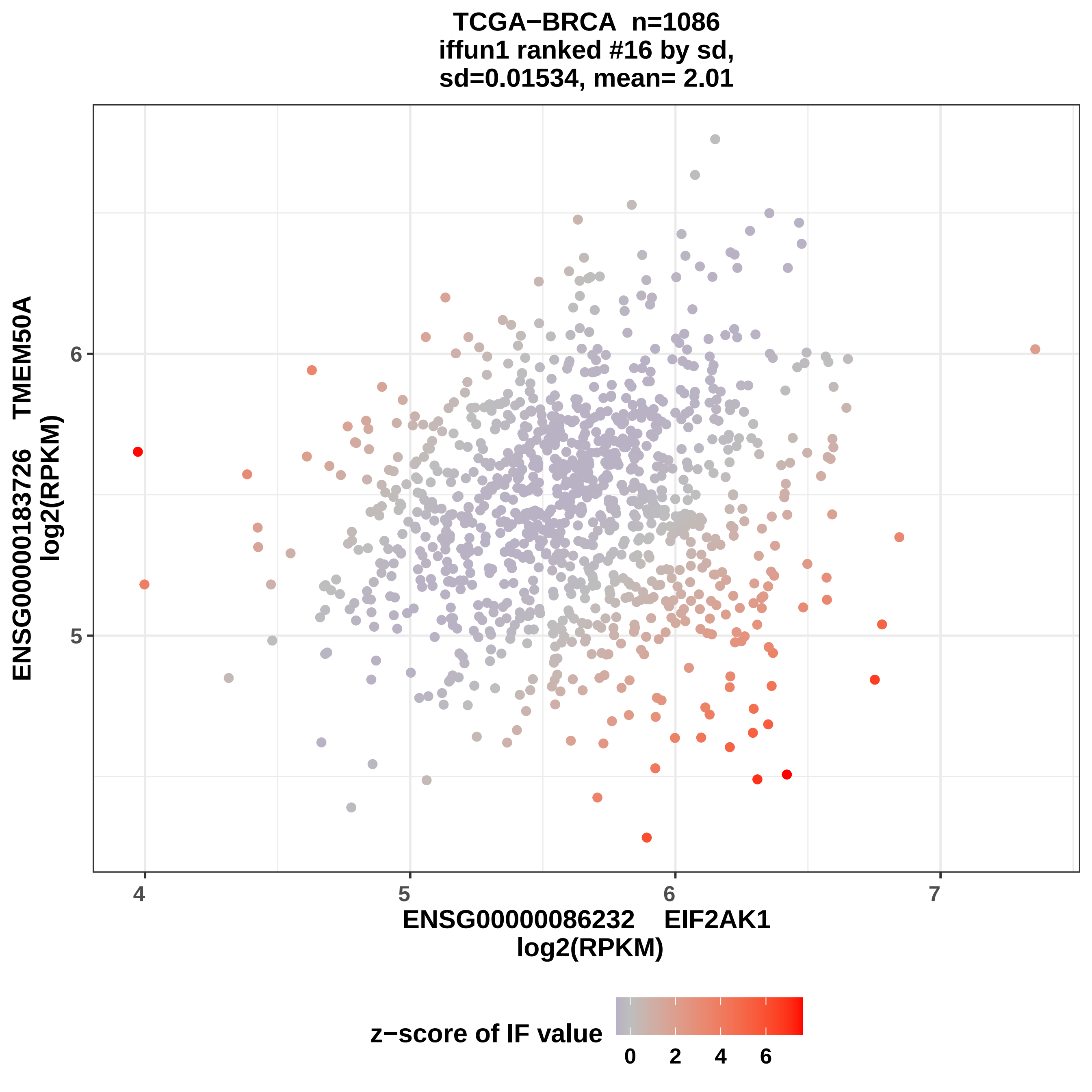

TCGA−BRCA n=1086
iffun1 ranked #16 by sd,
sd=0.01534, mean= 2.01
ENSG00000183726 TMEM50A
log2(RPKM)
6
5
4
5
6
7
ENSG00000086232 EIF2AK1
log2(RPKM)
z−score of IF value
0
2
4
6

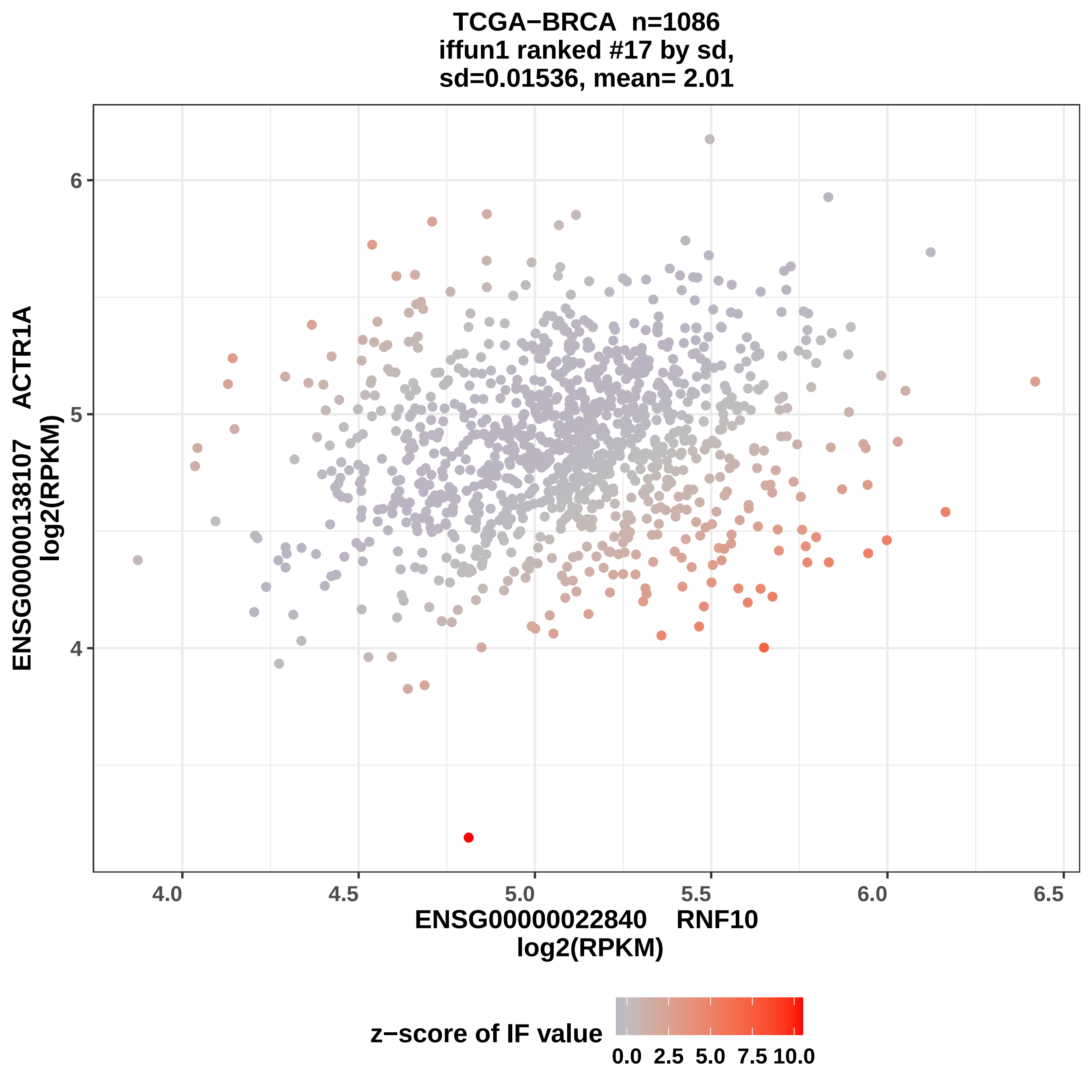
TCGA−BRCA n=1086
iffun1 ranked #17 by sd,
sd=0.01536, mean= 2.01
ENSG00000138107 ACTR1A
log2(RPKM)
4
5
6
4.0
4.5
5.0
5.5
6.0
6.5
ENSG00000022840 RNF10
log2(RPKM)
z−score of IF value
0.0
2.5
5.0
7.5
10.0

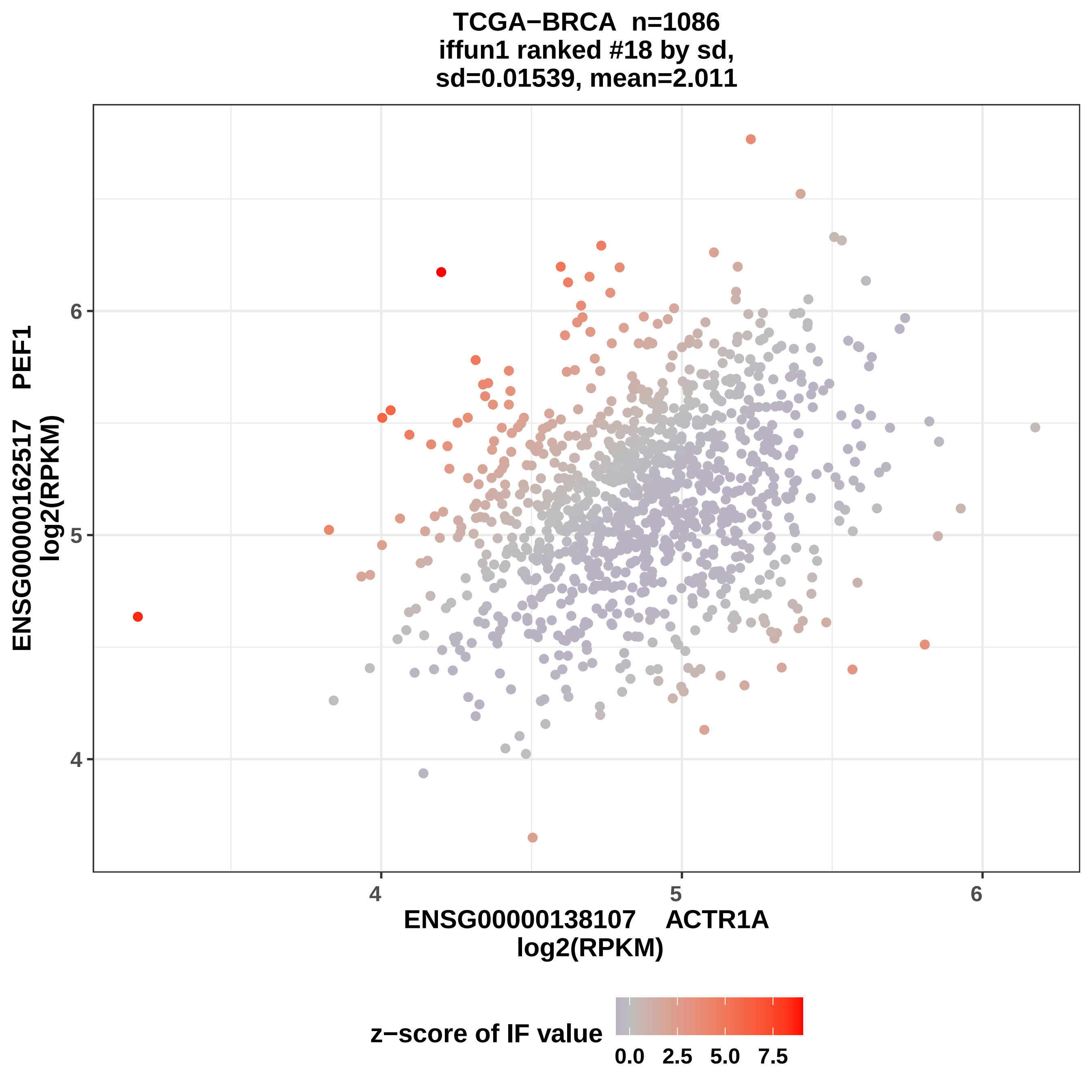
TCGA−BRCA n=1086
iffun1 ranked #18 by sd,
sd=0.01539, mean=2.011
ENSG00000162517 PEF1
log2(RPKM)
4
5
6
ENSG00000138107 ACTR1A
log2(RPKM)
z−score of IF value
0.0
2.5
5.0
7.5

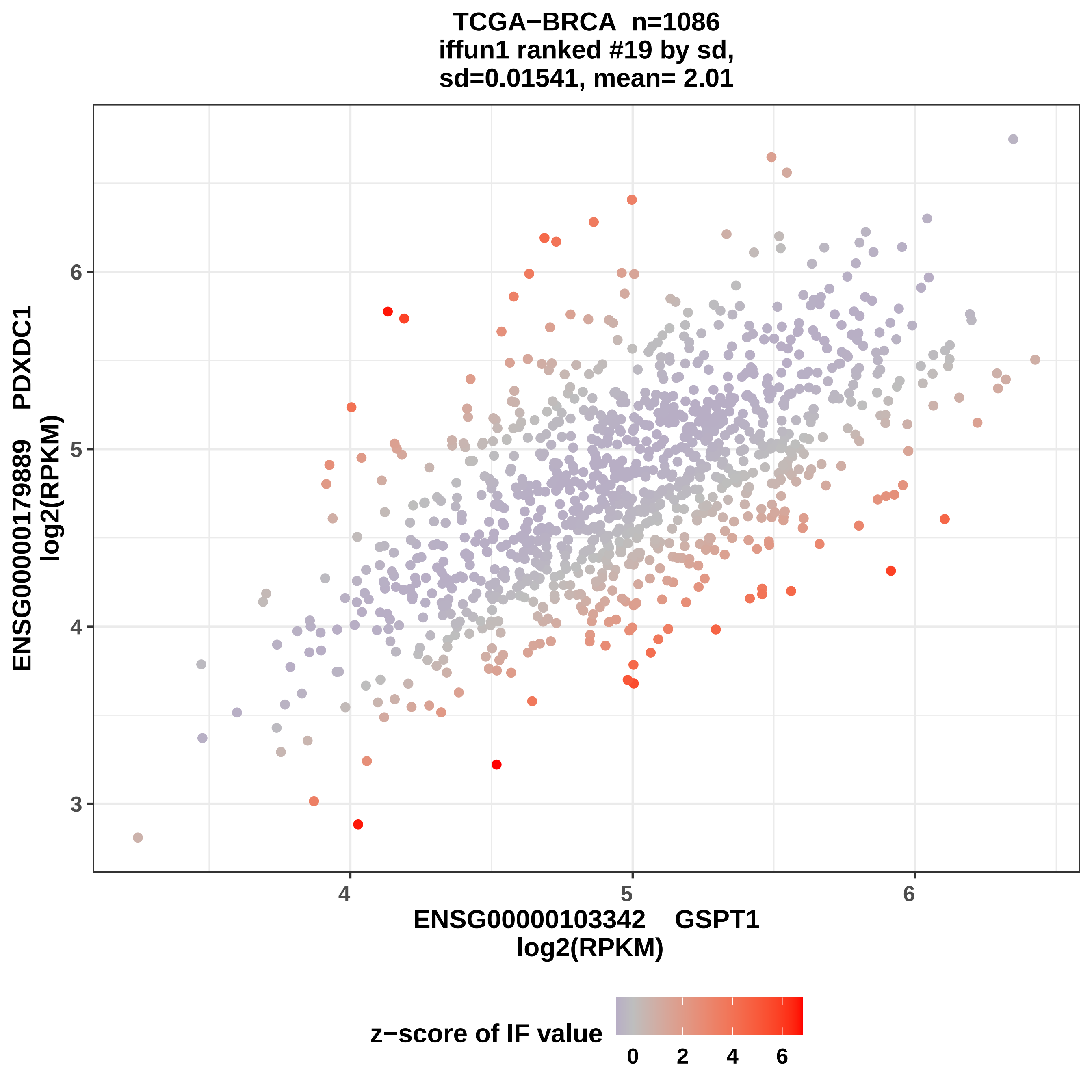

TCGA−BRCA n=1086
iffun1 ranked #19 by sd,
sd=0.01541, mean= 2.01
ENSG00000179889 PDXDC1
log2(RPKM)
6
5
4
3
4
5
6
ENSG00000103342 GSPT1
log2(RPKM)
z−score of IF value
0
2
4
6

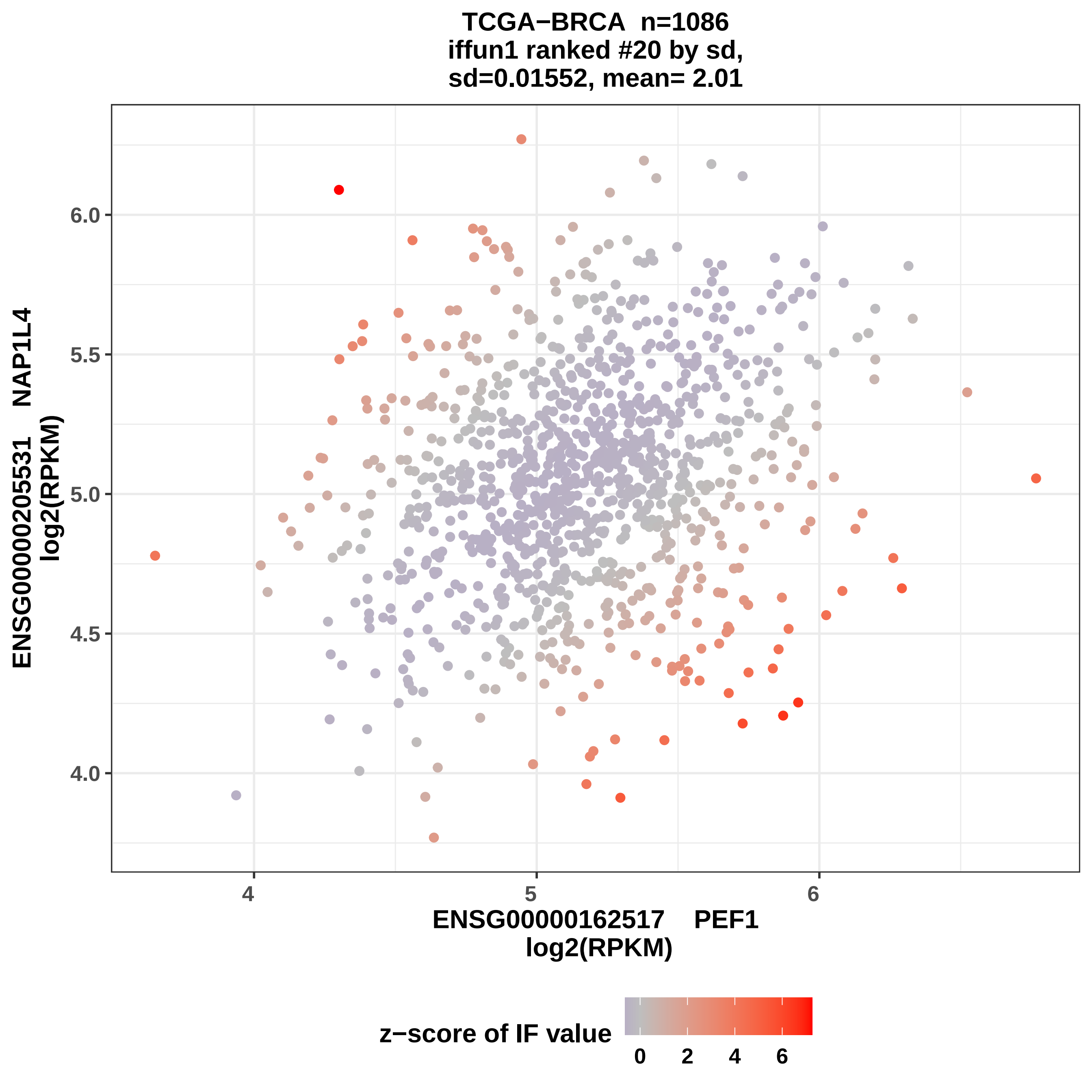

TCGA−BRCA n=1086
iffun1 ranked #20 by sd,
sd=0.01552, mean= 2.01
ENSG00000205531 NAP1L4
log2(RPKM)
6.0
5.5
5.0
4.5
4.0
4
5
6
ENSG00000162517 PEF1
log2(RPKM)
z−score of IF value
0
2
4
6

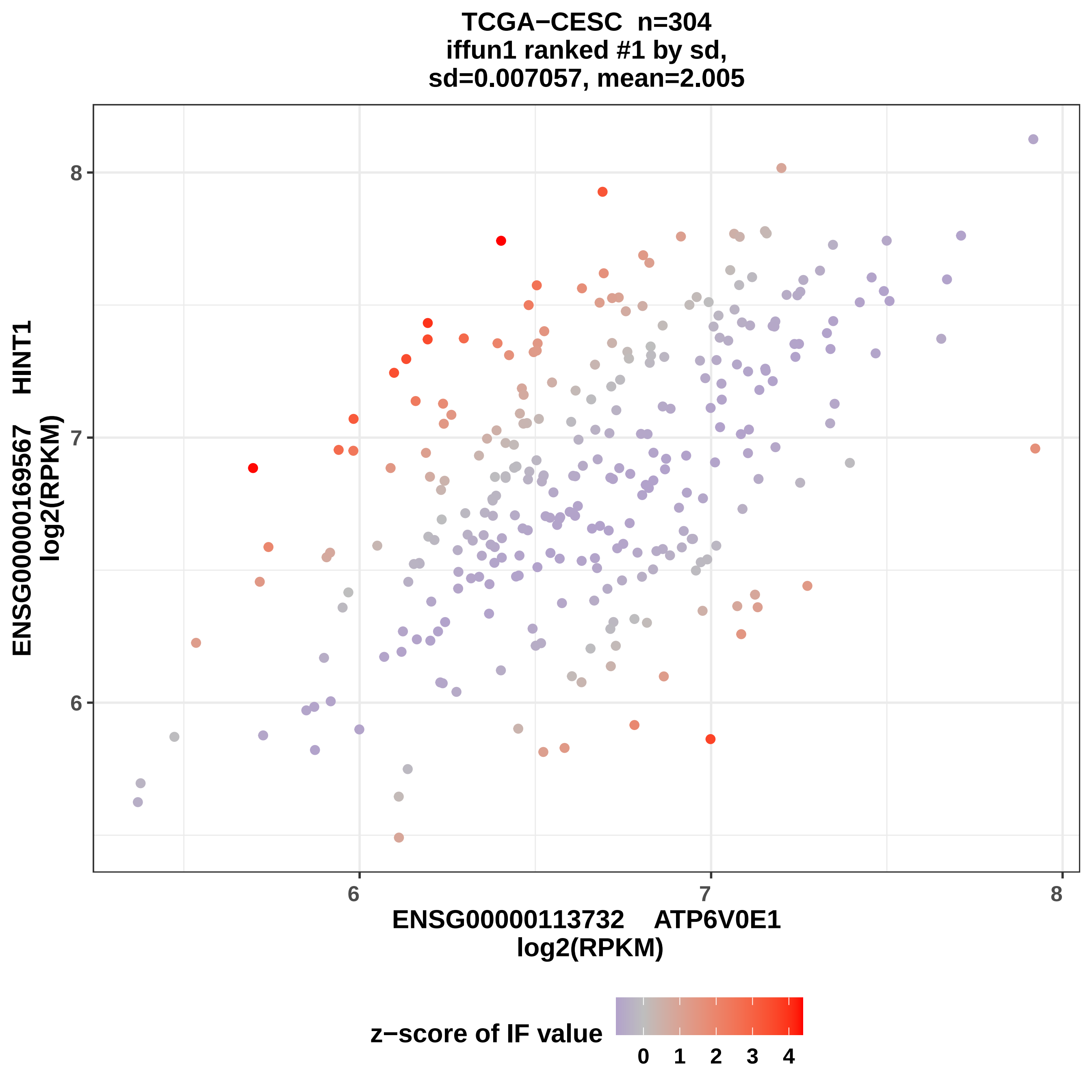
TCGA−CESC n=304
iffun1 ranked #1 by sd,
sd=0.007057, mean=2.005
ENSG00000169567 HINT1
log2(RPKM)
8
7
6
6
7
8
ENSG00000113732 ATP6V0E1
log2(RPKM)
z−score of IF value
0
1
2
3
4

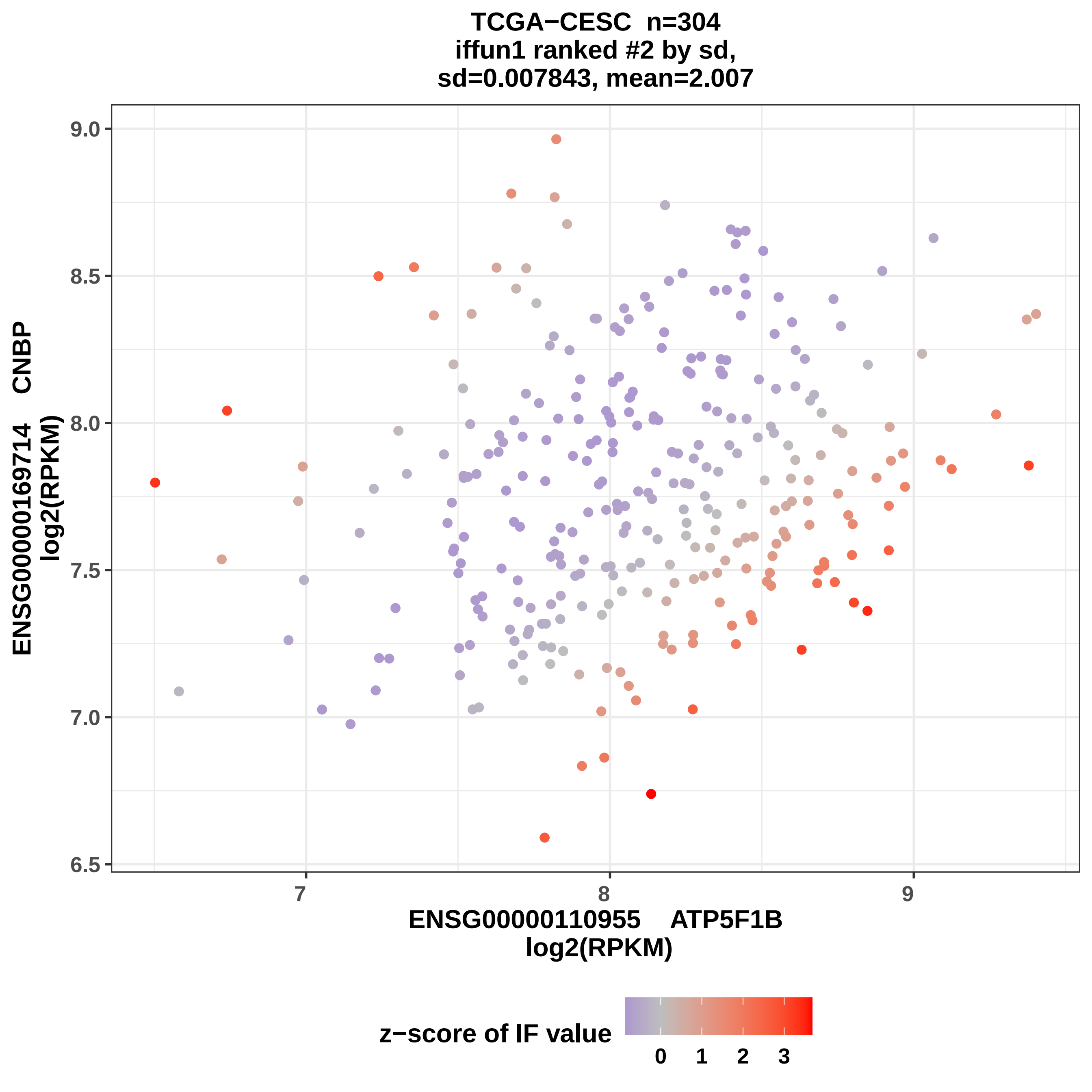

TCGA−CESC n=304
iffun1 ranked #2 by sd,
sd=0.007843, mean=2.007
ENSG00000169714 CNBP
log2(RPKM)
9.0
8.5
8.0
7.5
7.0
6.5
7
8
9
ENSG00000110955 ATP5F1B
log2(RPKM)
z−score of IF value
0
1
2
3

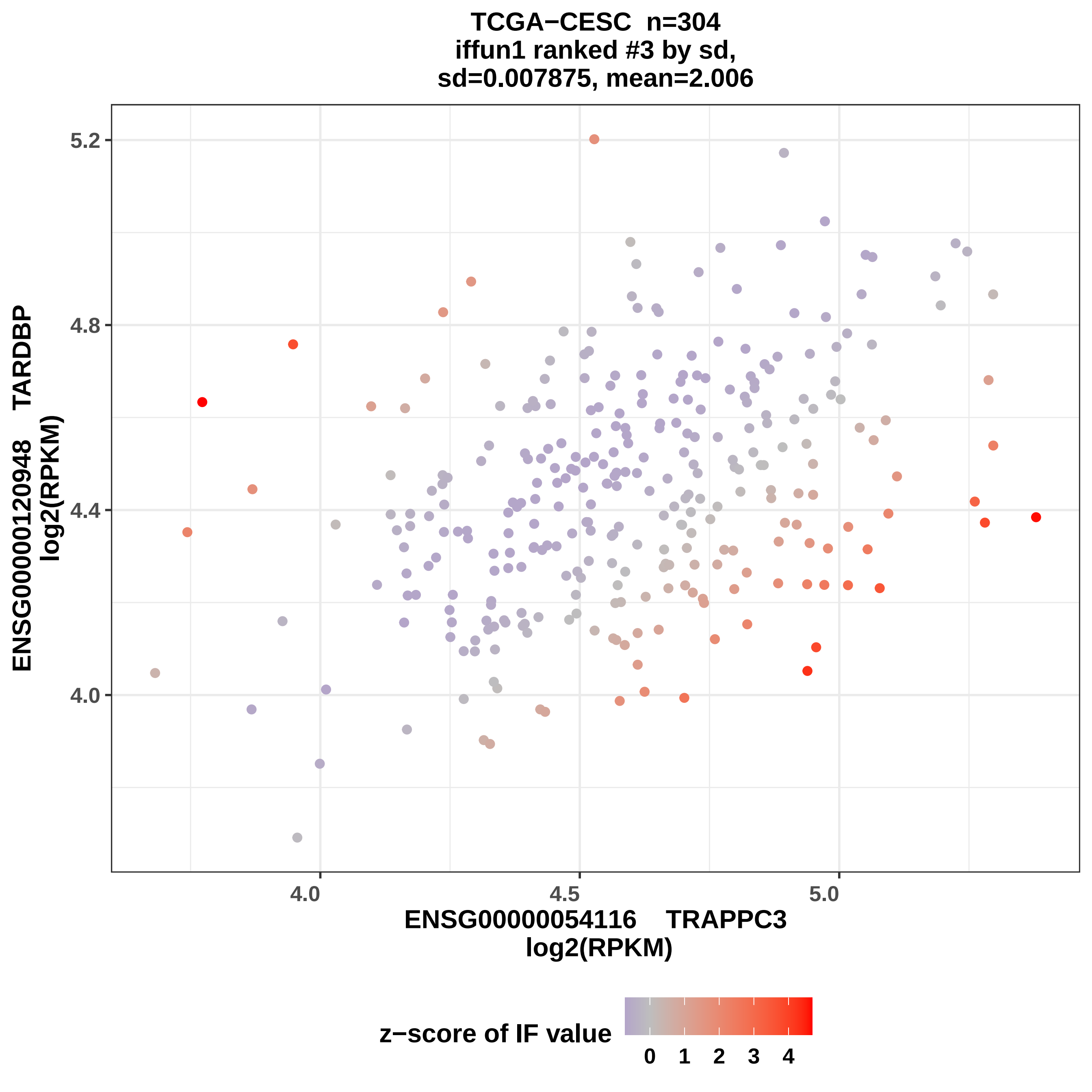

TCGA−CESC n=304
iffun1 ranked #3 by sd,
sd=0.007875, mean=2.006
ENSG00000120948 TARDBP
log2(RPKM)
5.2
4.8
4.4
4.0
4.0
4.5
5.0
ENSG00000054116 TRAPPC3
log2(RPKM)
z−score of IF value
0 1 2 3 4

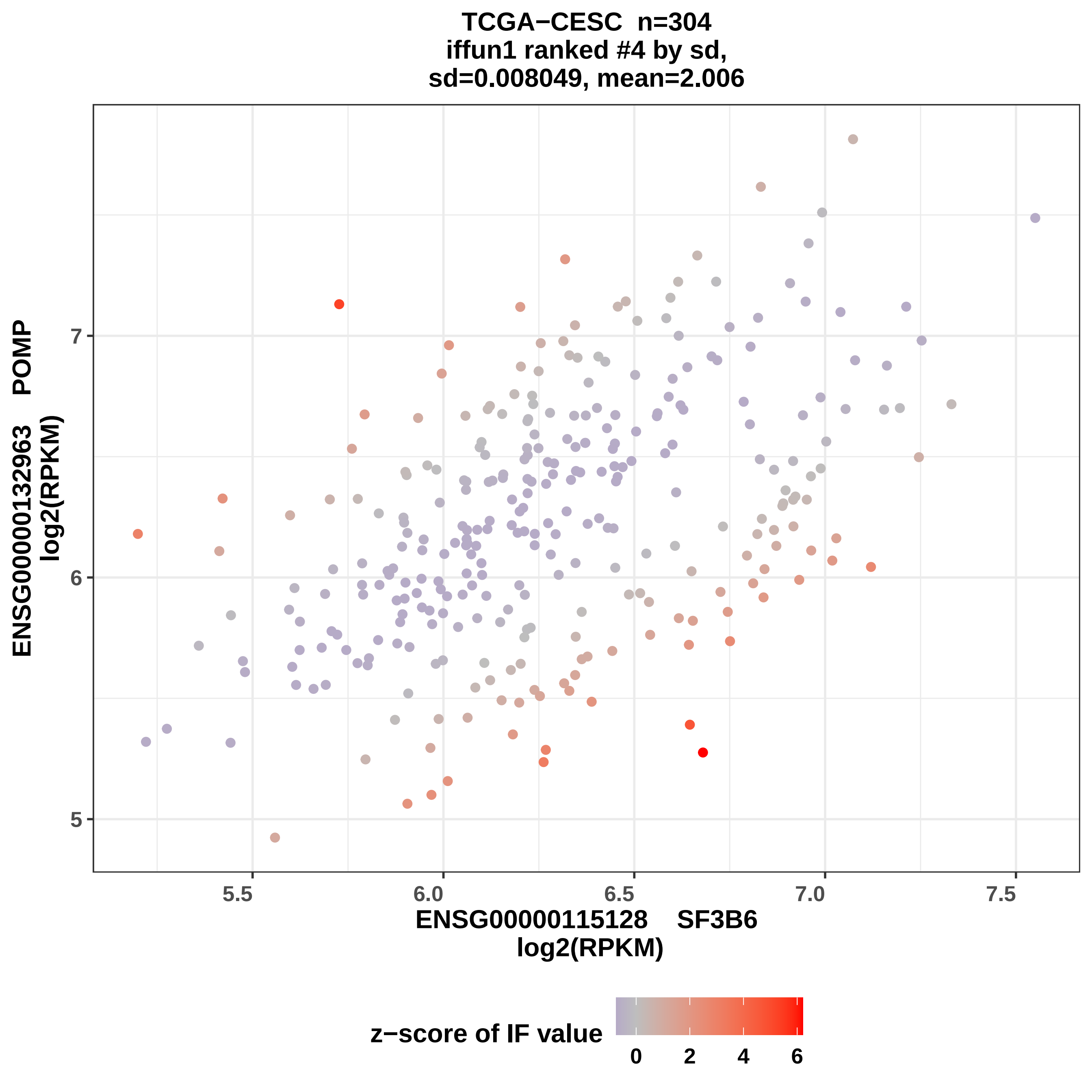

TCGA−CESC n=304
iffun1 ranked #4 by sd,
sd=0.008049, mean=2.006
ENSG00000132963 POMP
log2(RPKM)
7
6
5
5.5
6.0
6.5
7.0
7.5
ENSG00000115128 SF3B6
log2(RPKM)
z−score of IF value
0
2
4
6

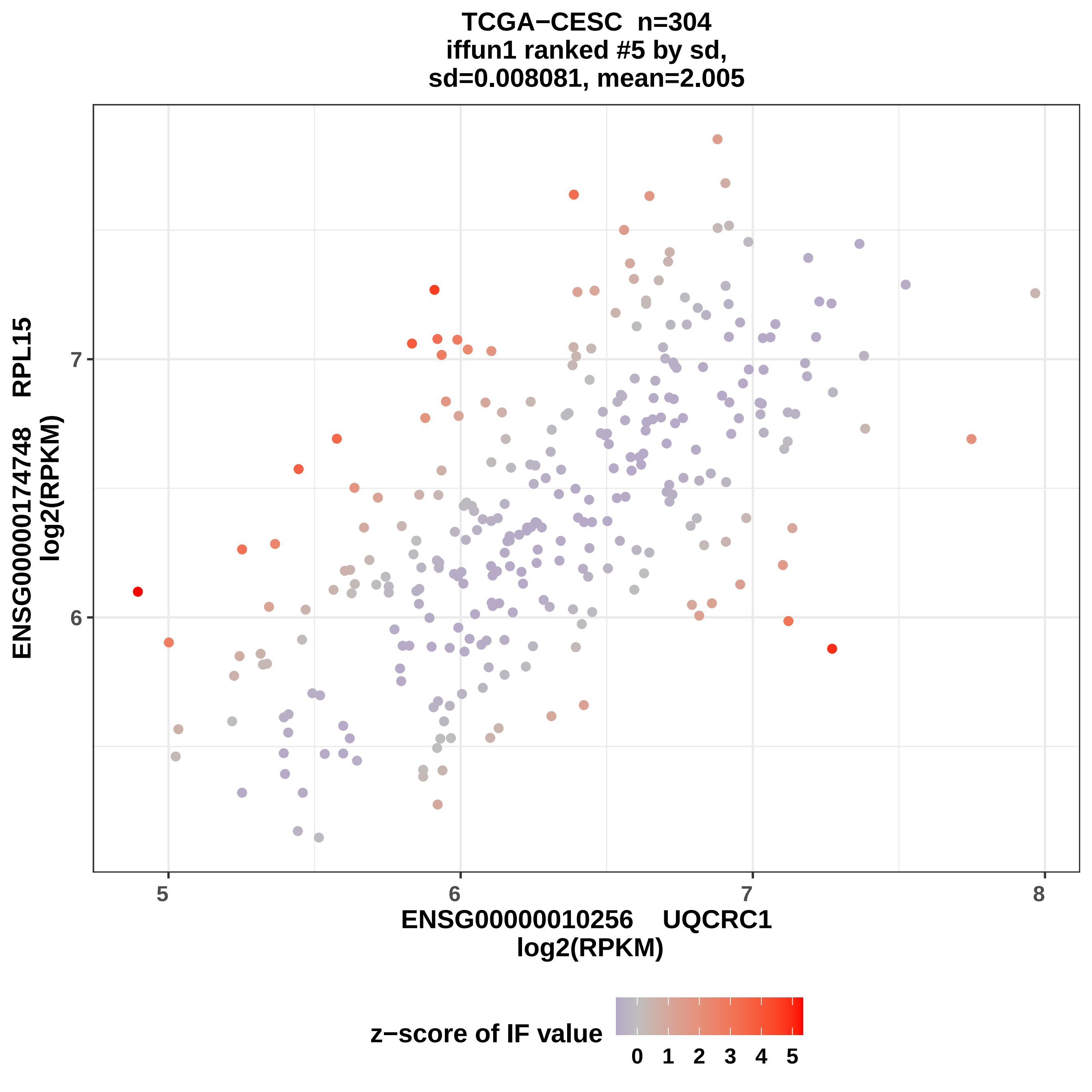
TCGA−CESC n=304
iffun1 ranked #5 by sd,
sd=0.008081, mean=2.005
ENSG00000174748 RPL15
log2(RPKM)
6
7
5
6
7
8
ENSG00000010256 UQCRC1
log2(RPKM)
z−score of IF value
0
1
2
3
4
5

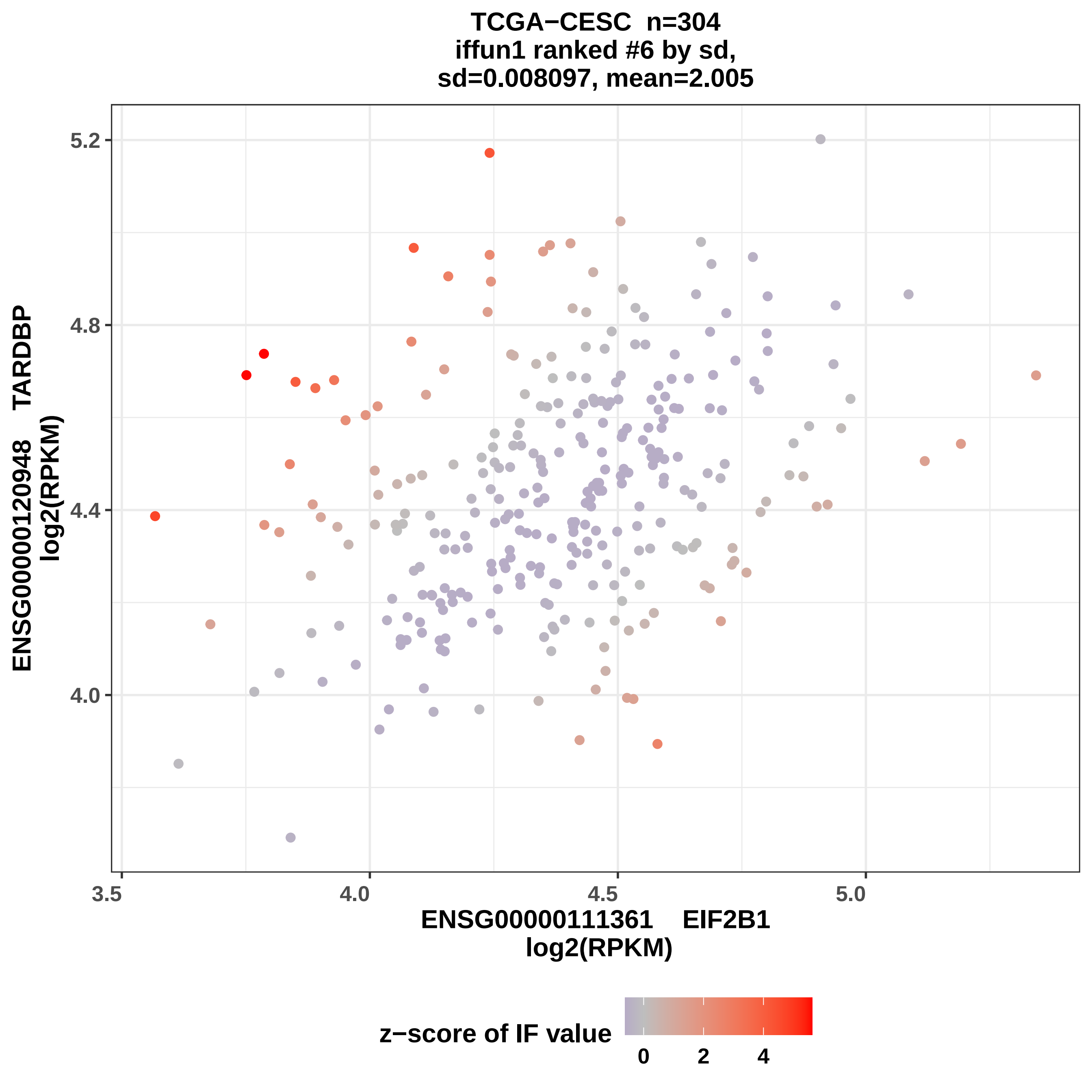
TCGA−CESC n=304
iffun1 ranked #6 by sd,
sd=0.008097, mean=2.005
ENSG00000120948 TARDBP
log2(RPKM)
5.2
4.8
4.4
4.0
3.5
4.0
4.5
5.0
ENSG00000111361 EIF2B1
log2(RPKM)
z−score of IF value
0
2
4

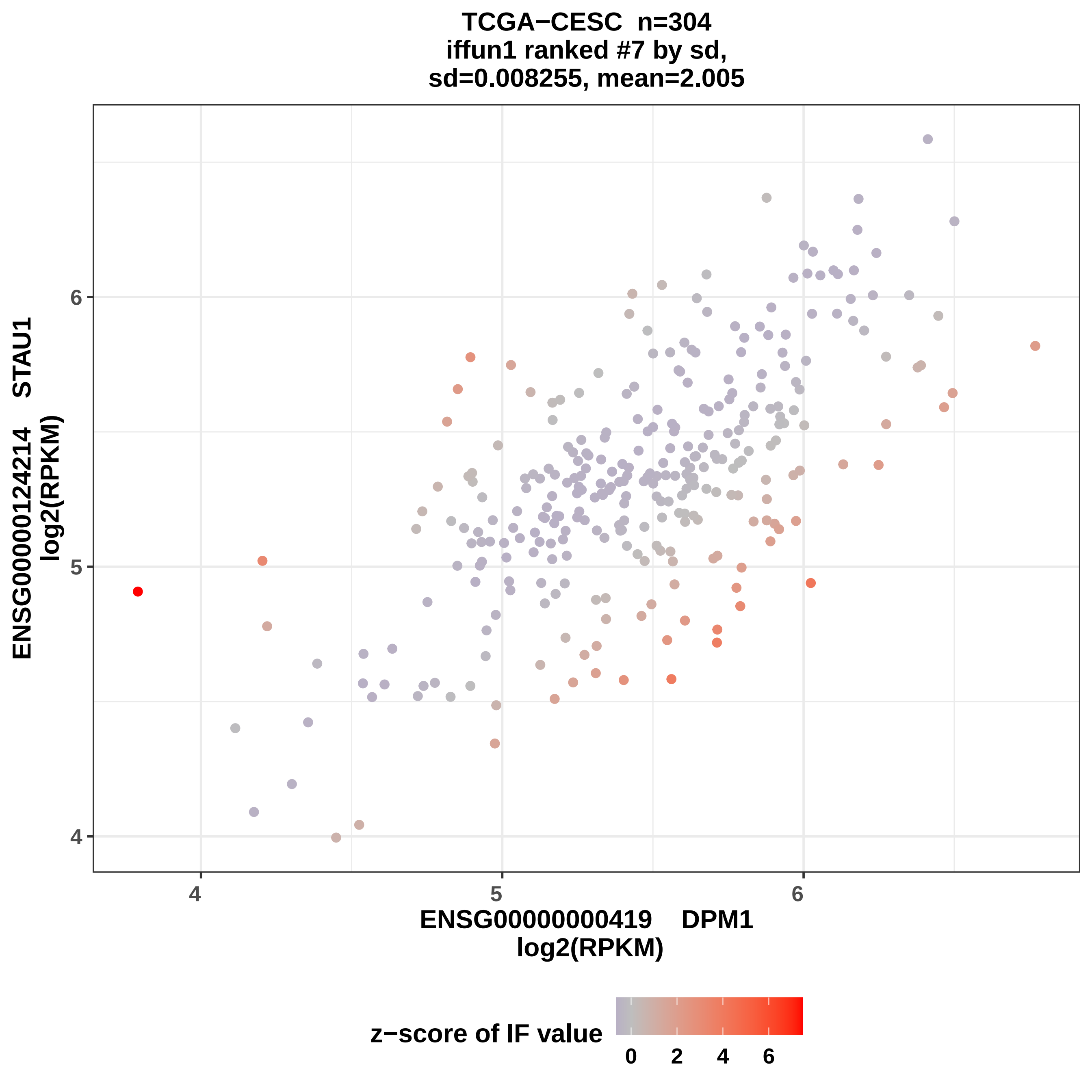
TCGA−CESC n=304
iffun1 ranked #7 by sd,
sd=0.008255, mean=2.005
ENSG00000124214 STAU1
log2(RPKM)
6
5
4
ENSG00000000419 DPM1
log2(RPKM)
z−score of IF value
0
2
4
6

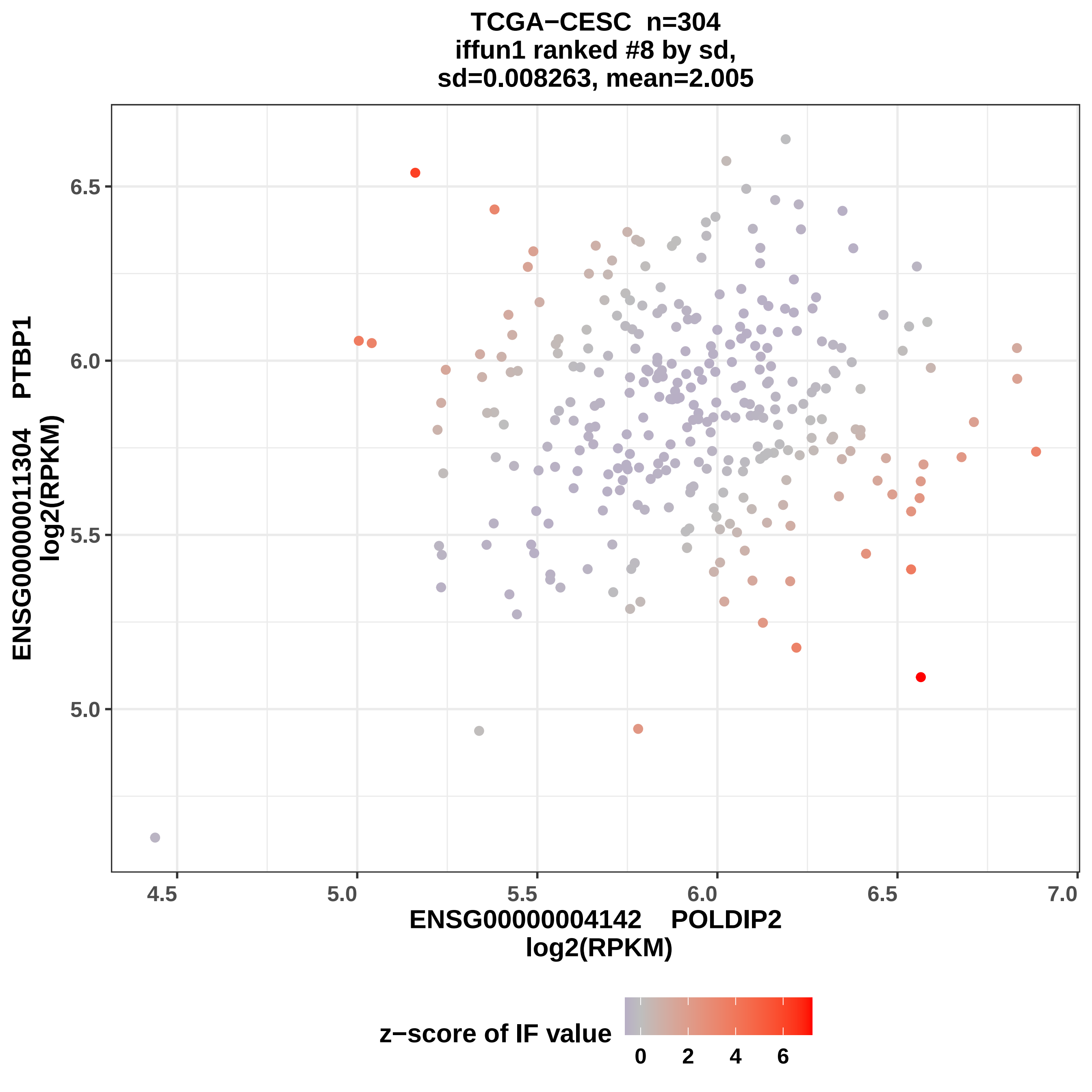
TCGA−CESC n=304
iffun1 ranked #8 by sd,
sd=0.008263, mean=2.005
ENSG00000011304 PTBP1
log2(RPKM)
6.5
6.0
5.5
5.0
4.5
5.0
5.5
6.0
6.5
7.0
ENSG00000004142 POLDIP2
log2(RPKM)
z−score of IF value
0
2
4
6

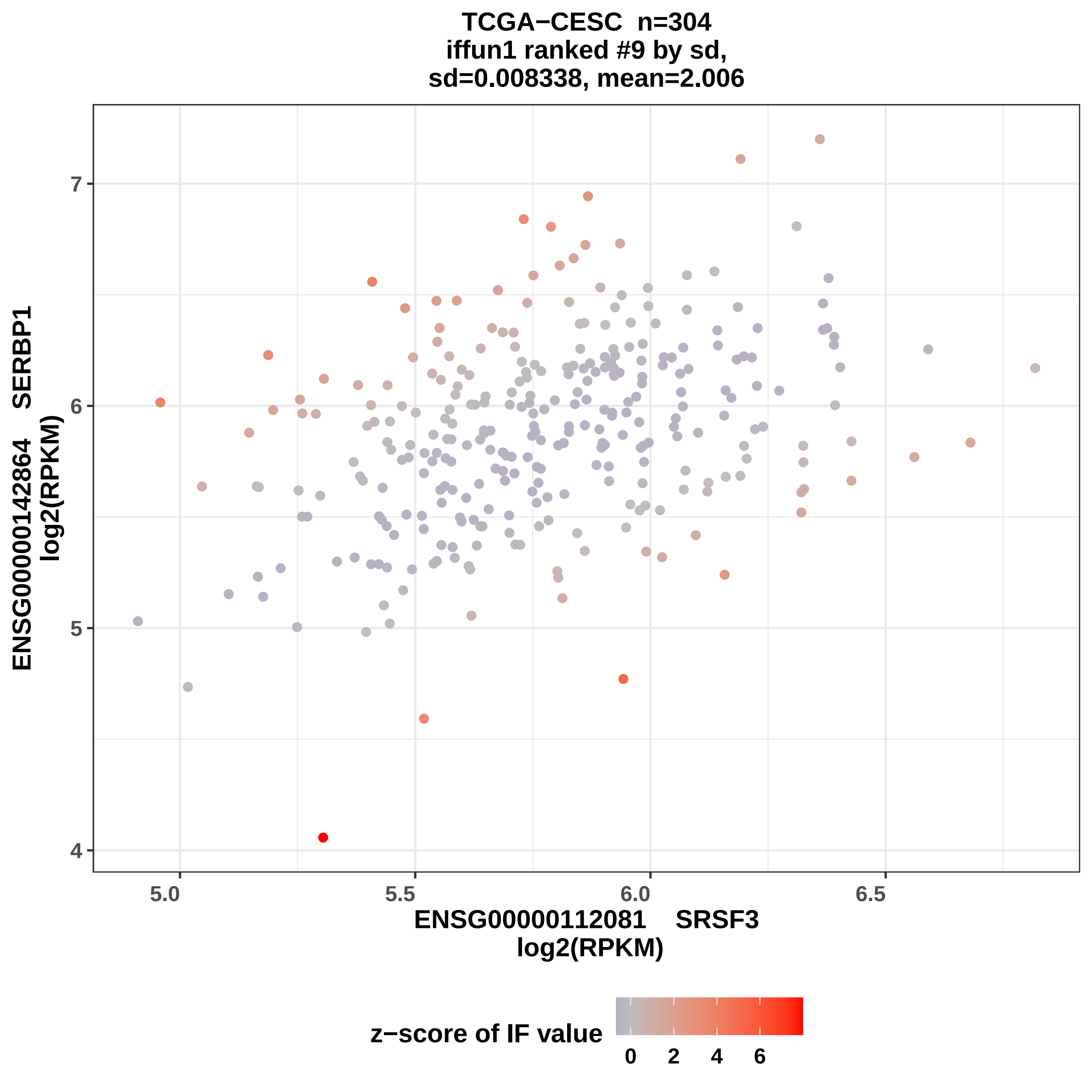
TCGA−CESC n=304
iffun1 ranked #9 by sd,
sd=0.008338, mean=2.006
ENSG00000142864 SERBP1
log2(RPKM)
7
6
5
4
5.0
5.5
6.0
6.5
ENSG00000112081 SRSF3
log2(RPKM)
z−score of IF value
0
2
4
6

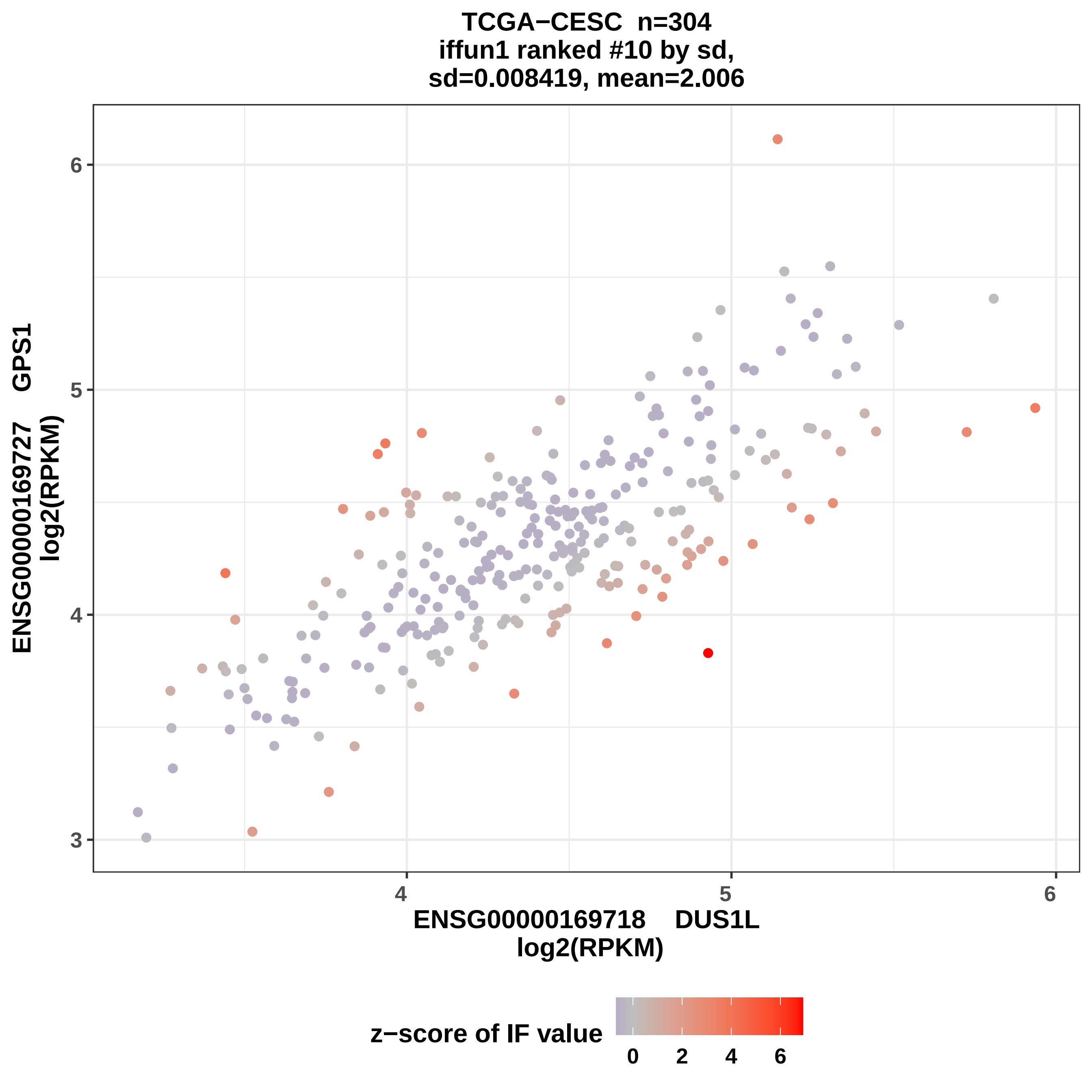
TCGA−CESC n=304
iffun1 ranked #10 by sd,
sd=0.008419, mean=2.006
ENSG00000169727 GPS1
log2(RPKM)
ENSG00000169718 DUS1L
log2(RPKM)
z−score of IF value
0
2
4
6
3
4
5
6

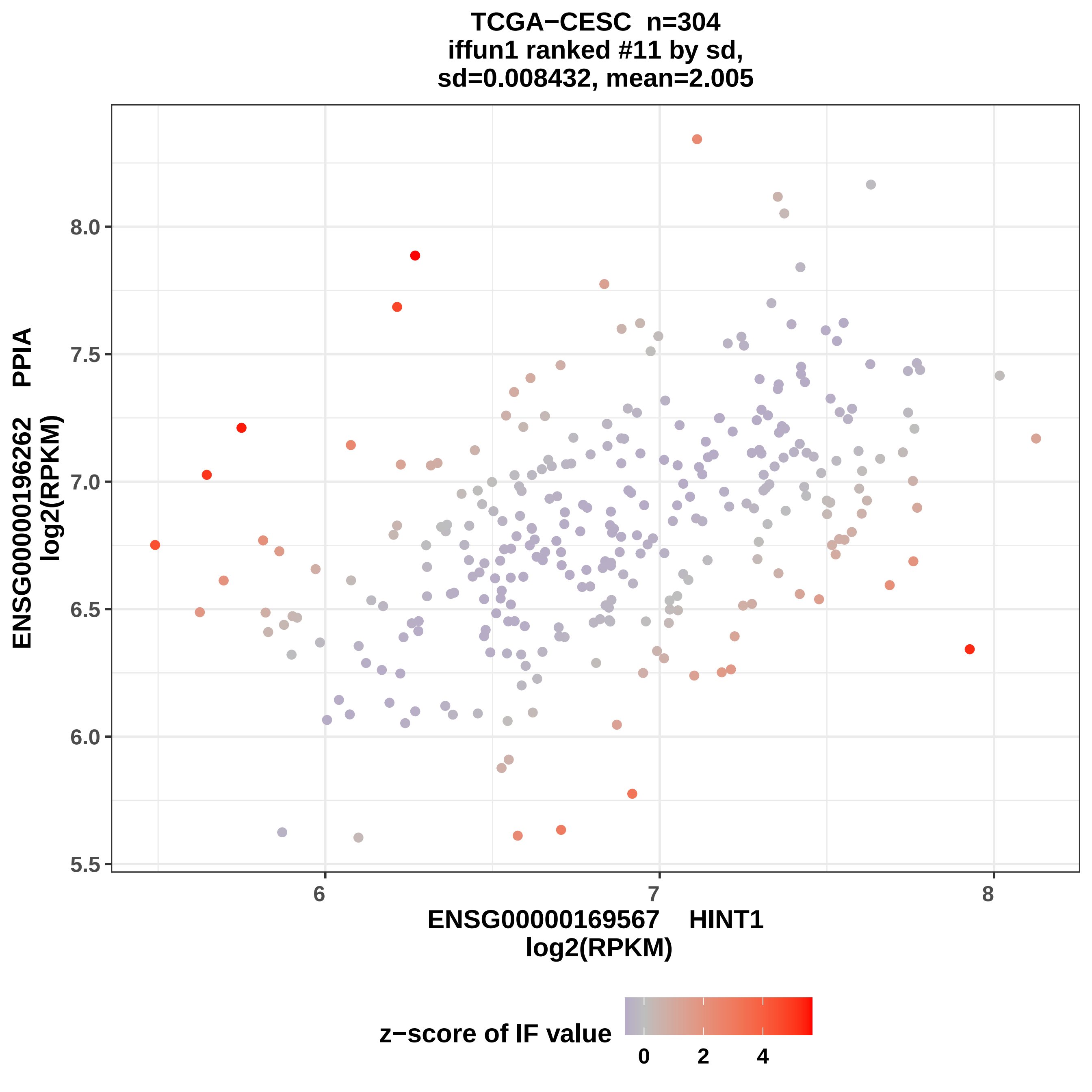

TCGA−CESC n=304
iffun1 ranked #11 by sd,
sd=0.008432, mean=2.005
ENSG00000196262 PPIA
log2(RPKM)
8.0
7.5
7.0
6.5
6.0
5.5
6
7
8
ENSG00000169567 HINT1
log2(RPKM)
z−score of IF value
0
2
4

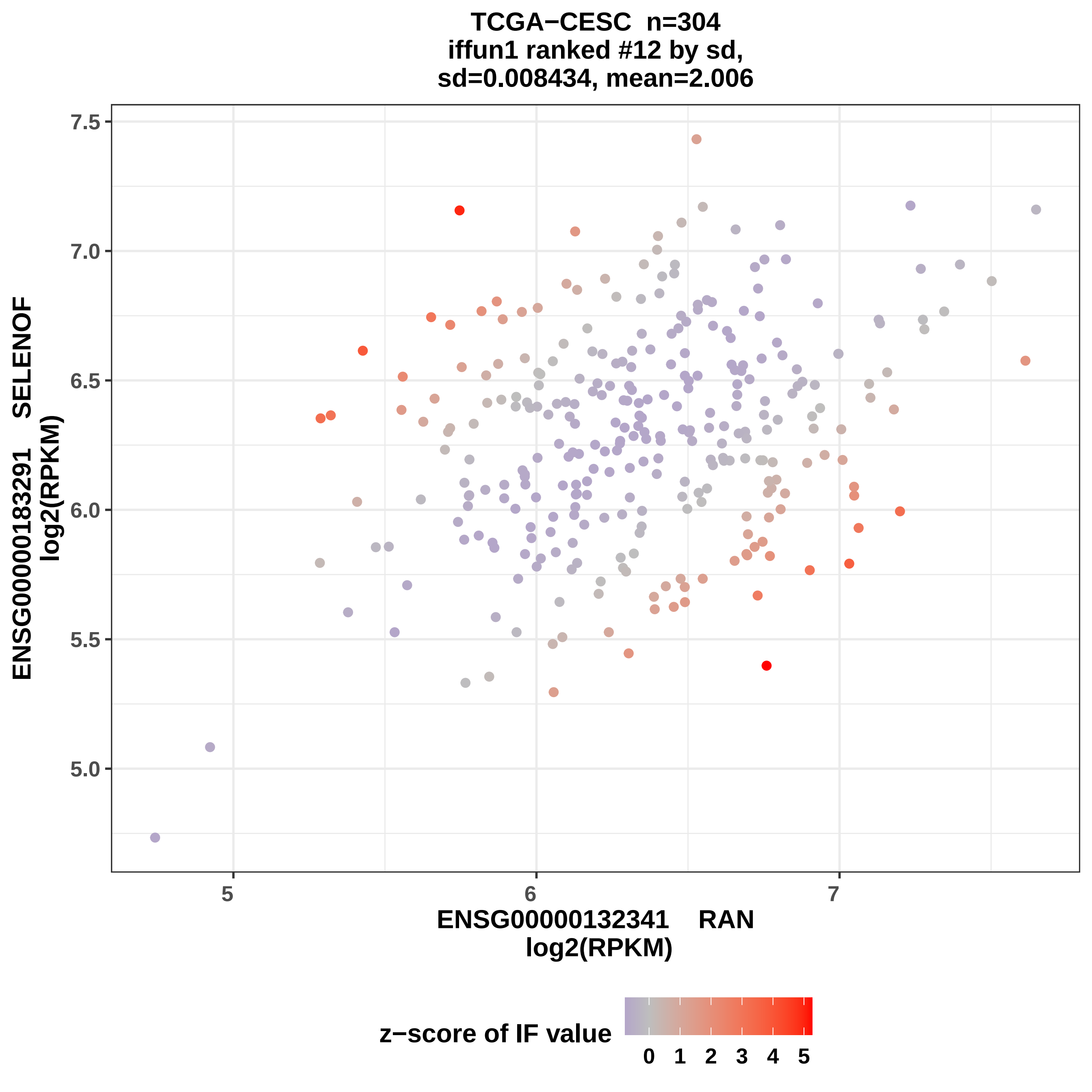
TCGA−CESC n=304
iffun1 ranked #12 by sd,
sd=0.008434, mean=2.006
ENSG00000183291 SELENOF
log2(RPKM)
7.5
7.0
6.5
6.0
5.5
5.0
5
6
7
ENSG00000132341 RAN
log2(RPKM)
z−score of IF value
0 1 2 3 4 5

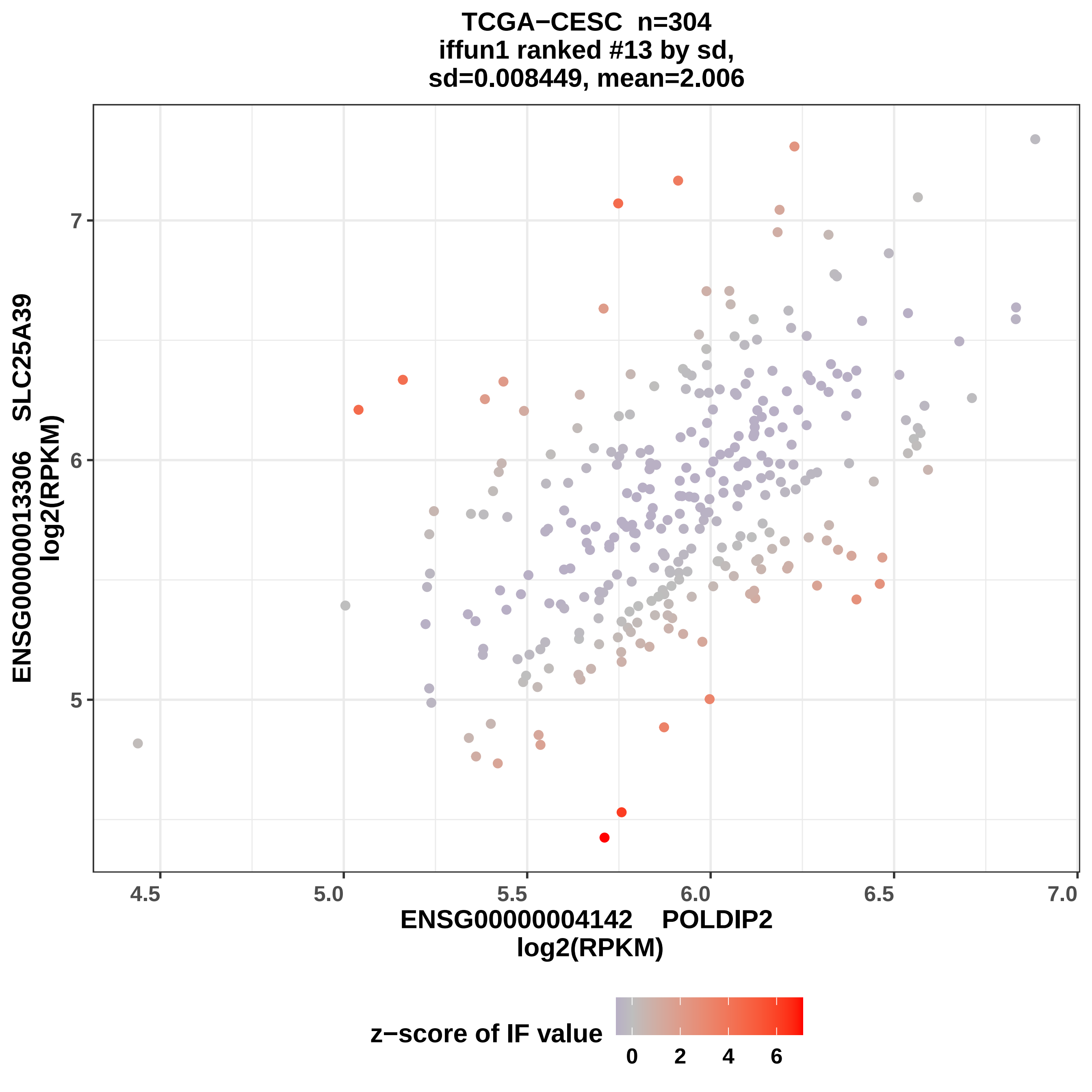

TCGA−CESC n=304
iffun1 ranked #13 by sd,
sd=0.008449, mean=2.006
ENSG00000013306 SLC25A39
log2(RPKM)
7
6
5
4.5
5.0
5.5
6.0
6.5
7.0
ENSG00000004142 POLDIP2
log2(RPKM)
z−score of IF value
0
2
4
6

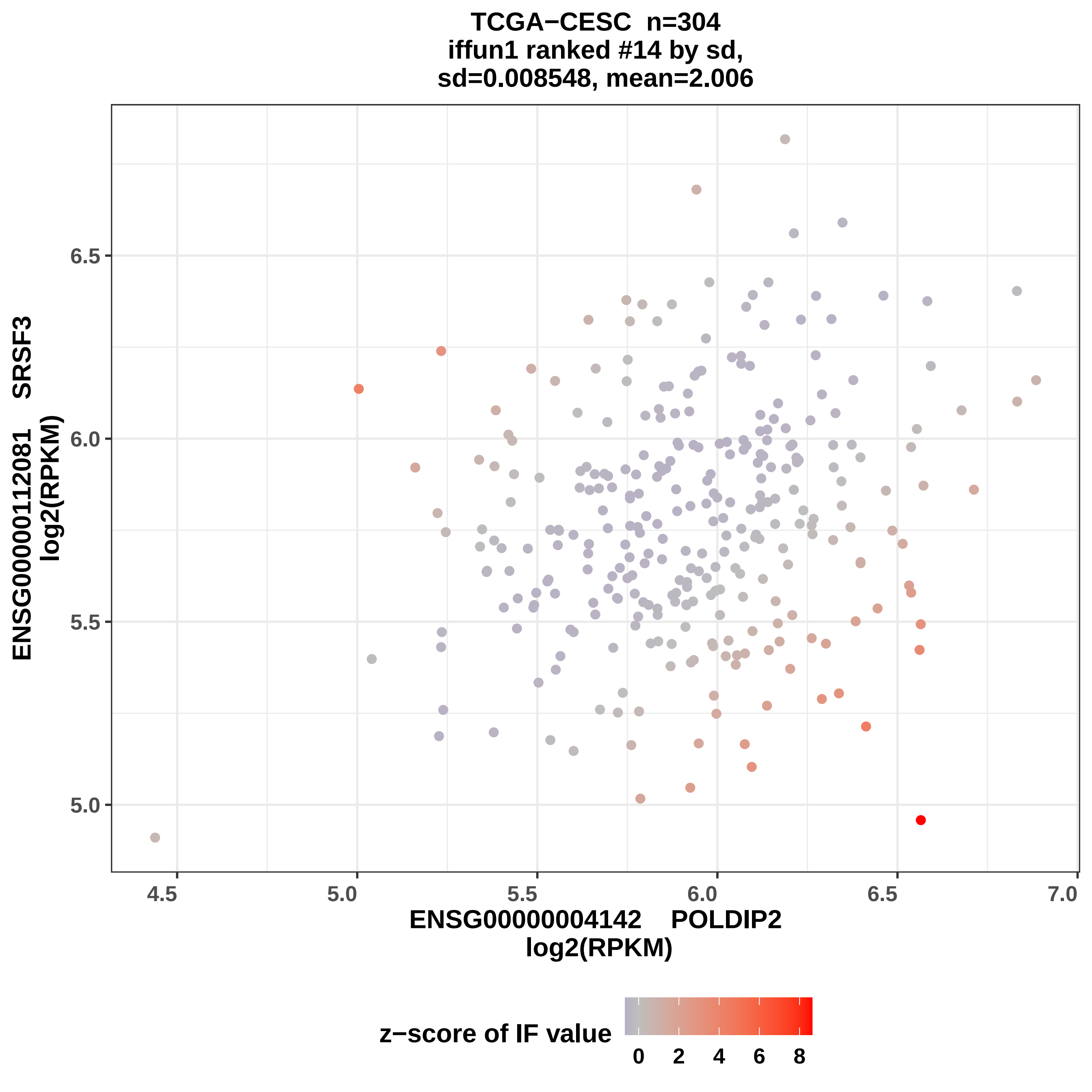
TCGA−CESC n=304
iffun1 ranked #14 by sd,
sd=0.008548, mean=2.006
ENSG00000112081 SRSF3
log2(RPKM)
6.5
6.0
5.5
5.0
4.5
5.0
5.5
6.0
6.5
7.0
ENSG00000004142 POLDIP2
log2(RPKM)
z−score of IF value
0
2
4
6
8

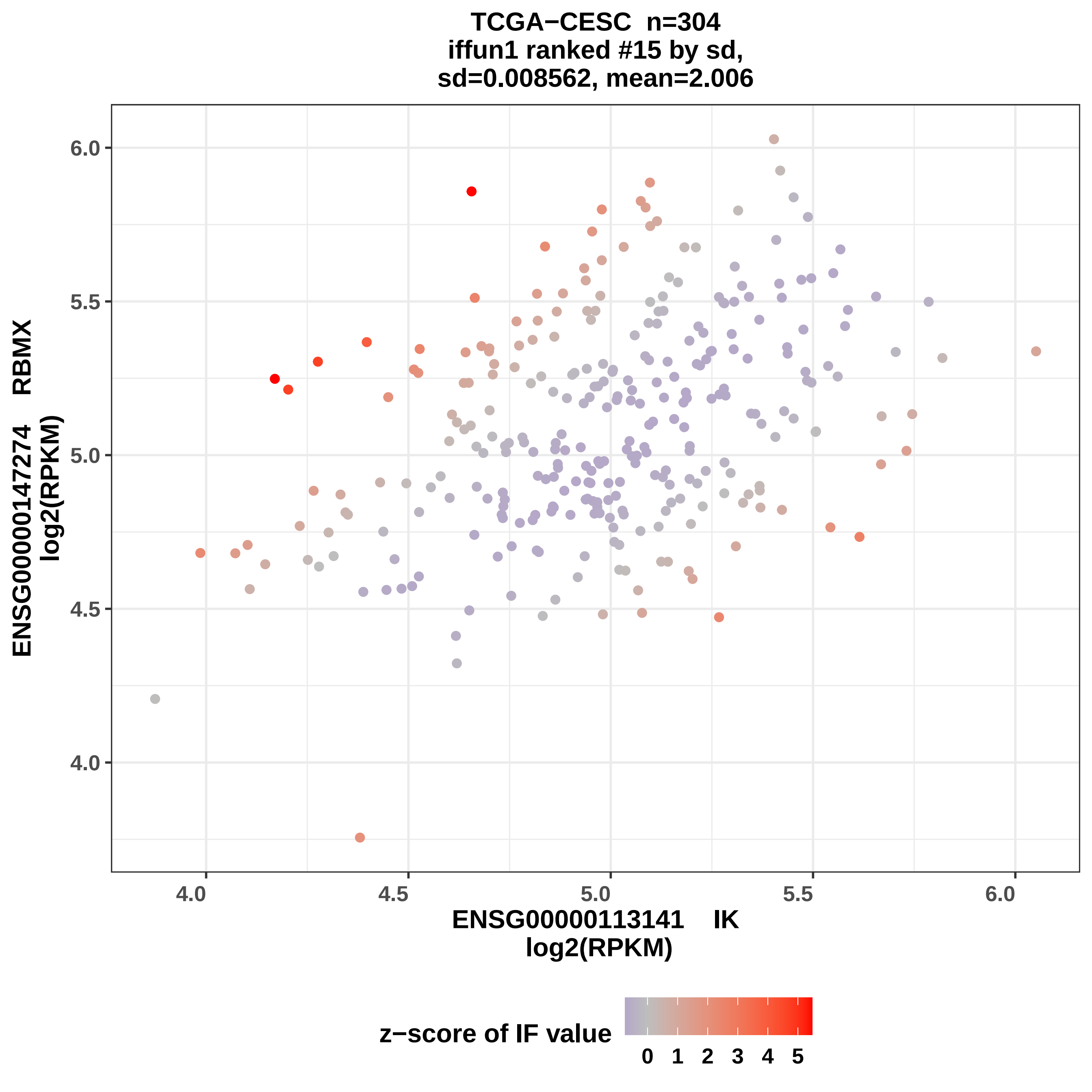
TCGA−CESC n=304
iffun1 ranked #15 by sd,
sd=0.008562, mean=2.006
ENSG00000147274 RBMX
log2(RPKM)
6.0
5.5
5.0
4.5
4.0
4.0
4.5
5.0
5.5
6.0
ENSG00000113141 IK
log2(RPKM)
z−score of IF value
0 1 2 3 4 5

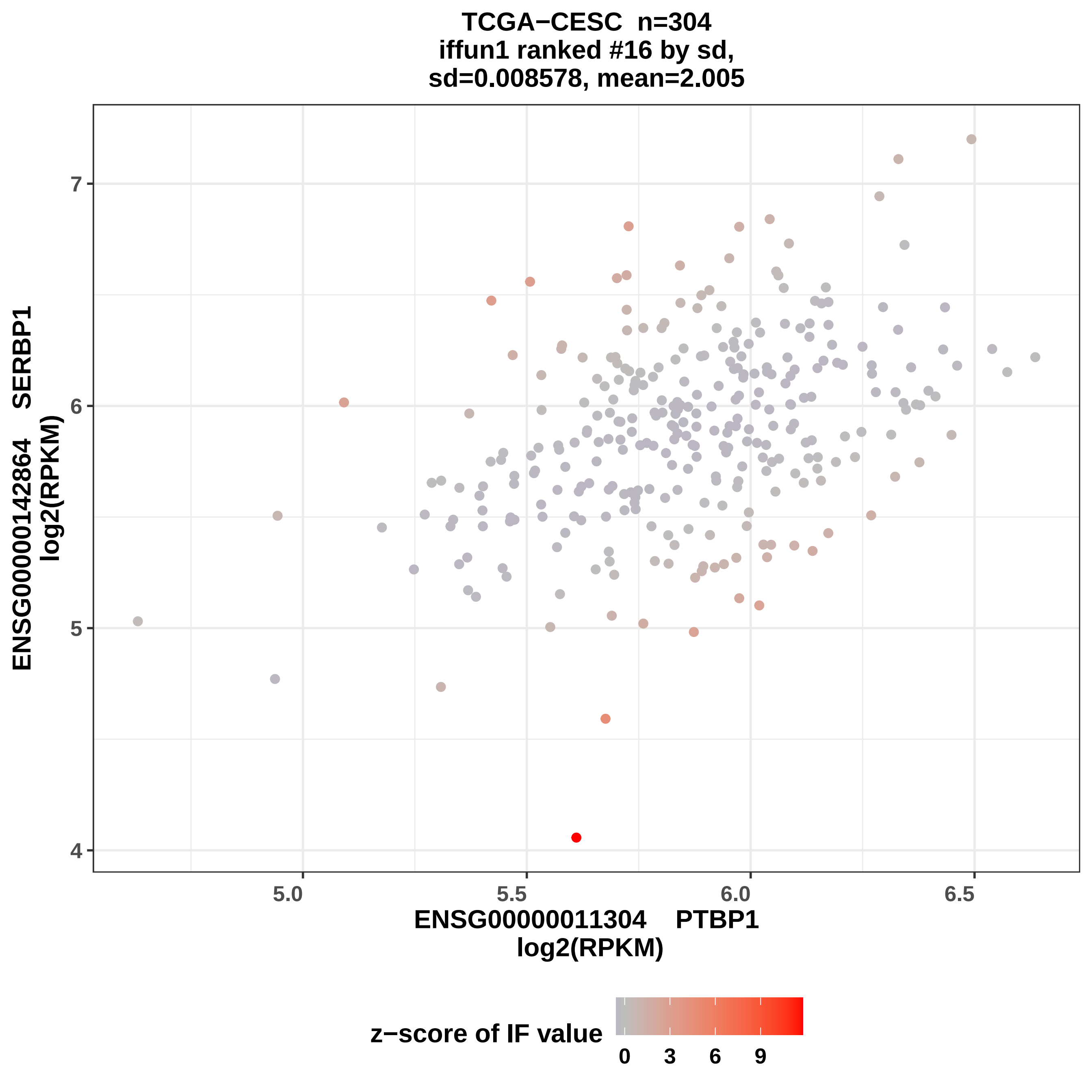
TCGA−CESC n=304
iffun1 ranked #16 by sd,
sd=0.008578, mean=2.005
ENSG00000142864 SERBP1
log2(RPKM)
ENSG00000011304 PTBP1
log2(RPKM)
4
5
6
7
5.0
5.5
6.0
6.5
z−score of IF value
0
3
6
9

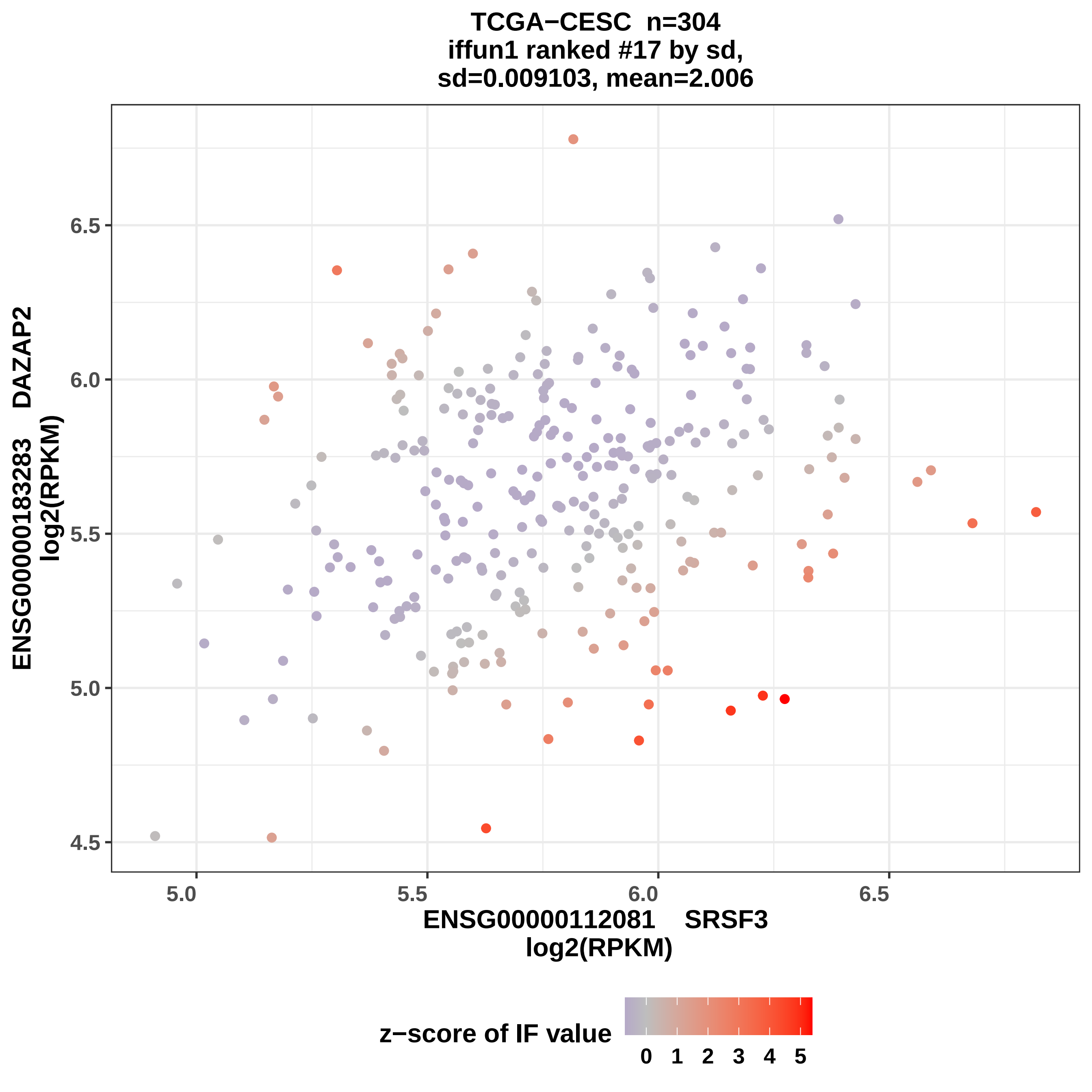

TCGA−CESC n=304
iffun1 ranked #17 by sd,
sd=0.009103, mean=2.006
ENSG00000183283 DAZAP2
log2(RPKM)
6.5
6.0
5.5
5.0
4.5
5.0
5.5
6.0
6.5
ENSG00000112081 SRSF3
log2(RPKM)
z−score of IF value
0 1 2 3 4 5

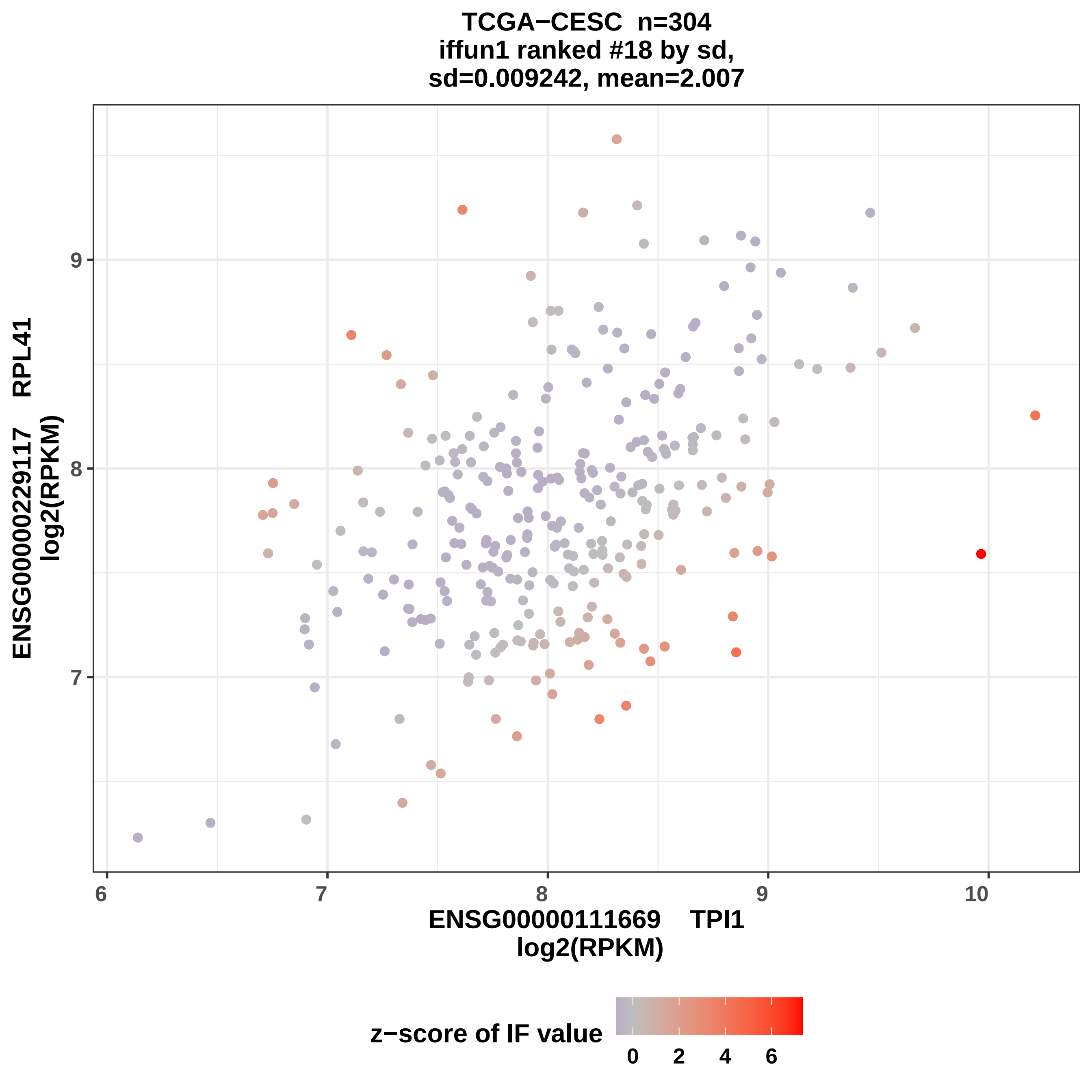
TCGA−CESC n=304
iffun1 ranked #18 by sd,
sd=0.009242, mean=2.007
ENSG00000229117 RPL41
log2(RPKM)
ENSG00000111669 TPI1
log2(RPKM)
z−score of IF value
0
2
4
6
6
7
8
9
10
7
8
9

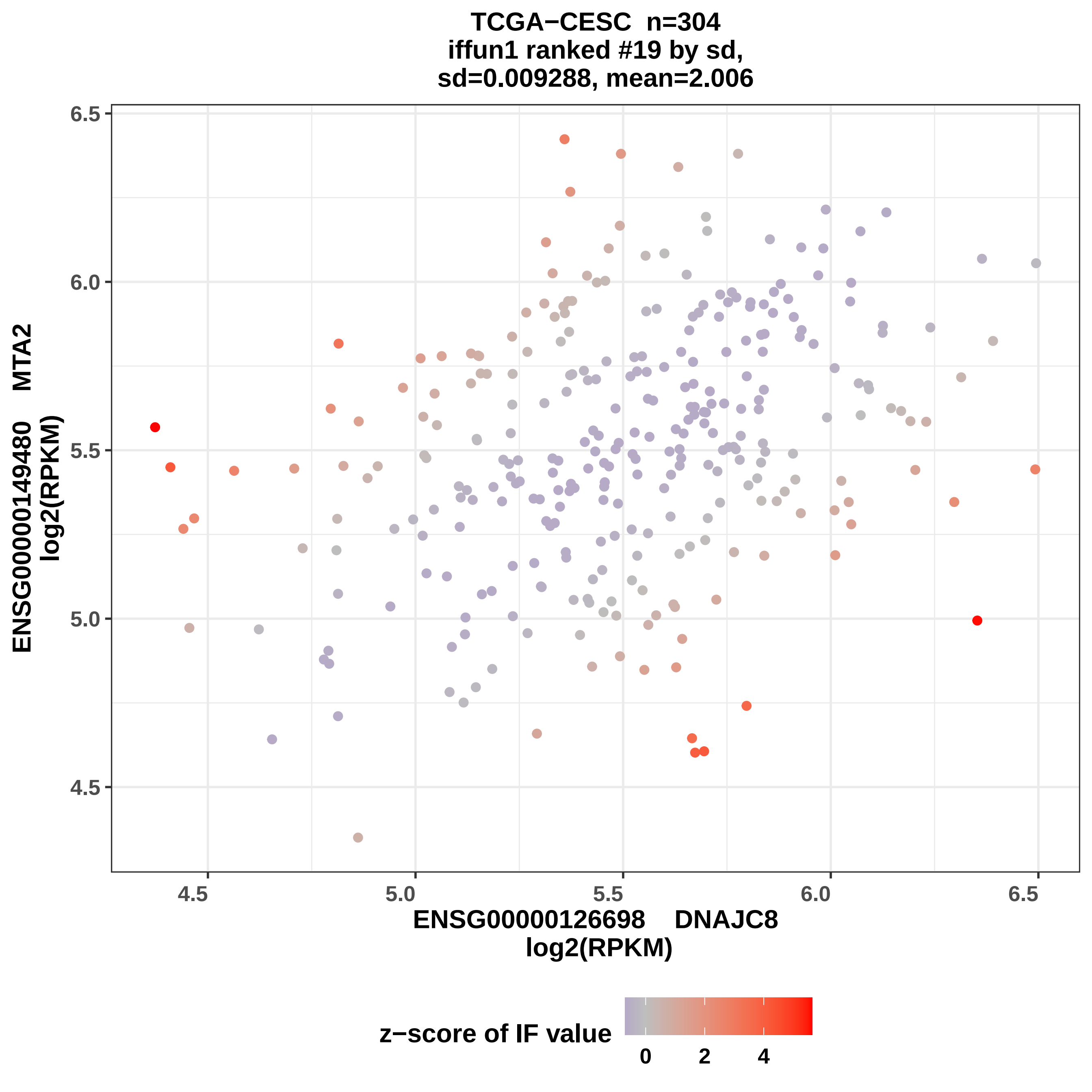

TCGA−CESC n=304
iffun1 ranked #19 by sd,
sd=0.009288, mean=2.006
ENSG00000149480 MTA2
log2(RPKM)
6.5
6.0
5.5
5.0
4.5
4.5
5.0
5.5
6.0
6.5
ENSG00000126698 DNAJC8
log2(RPKM)
z−score of IF value
0
2
4

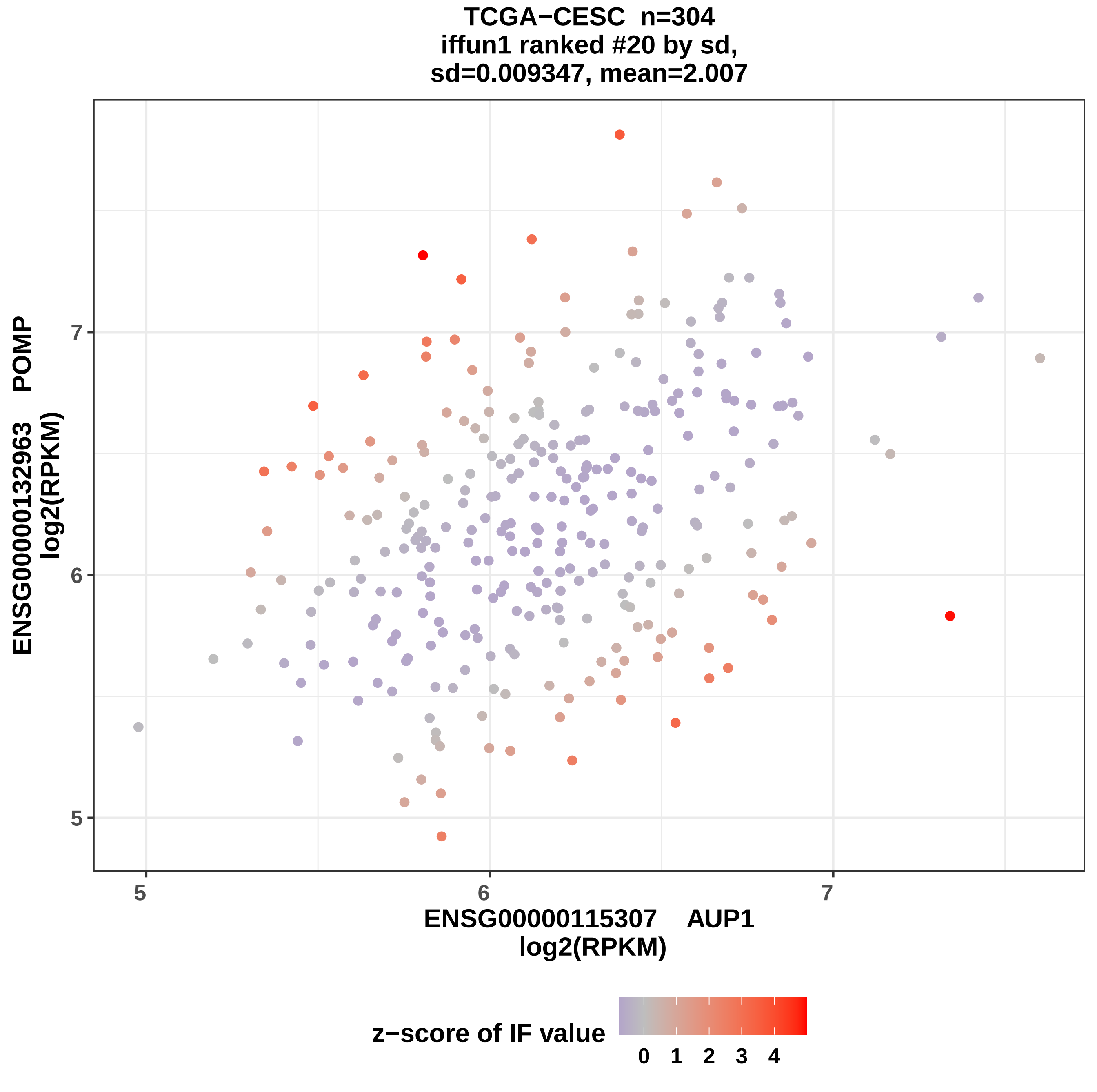
TCGA−CESC n=304
iffun1 ranked #20 by sd,
sd=0.009347, mean=2.007
ENSG00000132963 POMP
log2(RPKM)
5
6
7
ENSG00000115307 AUP1
log2(RPKM)
z−score of IF value
0 1 2 3 4

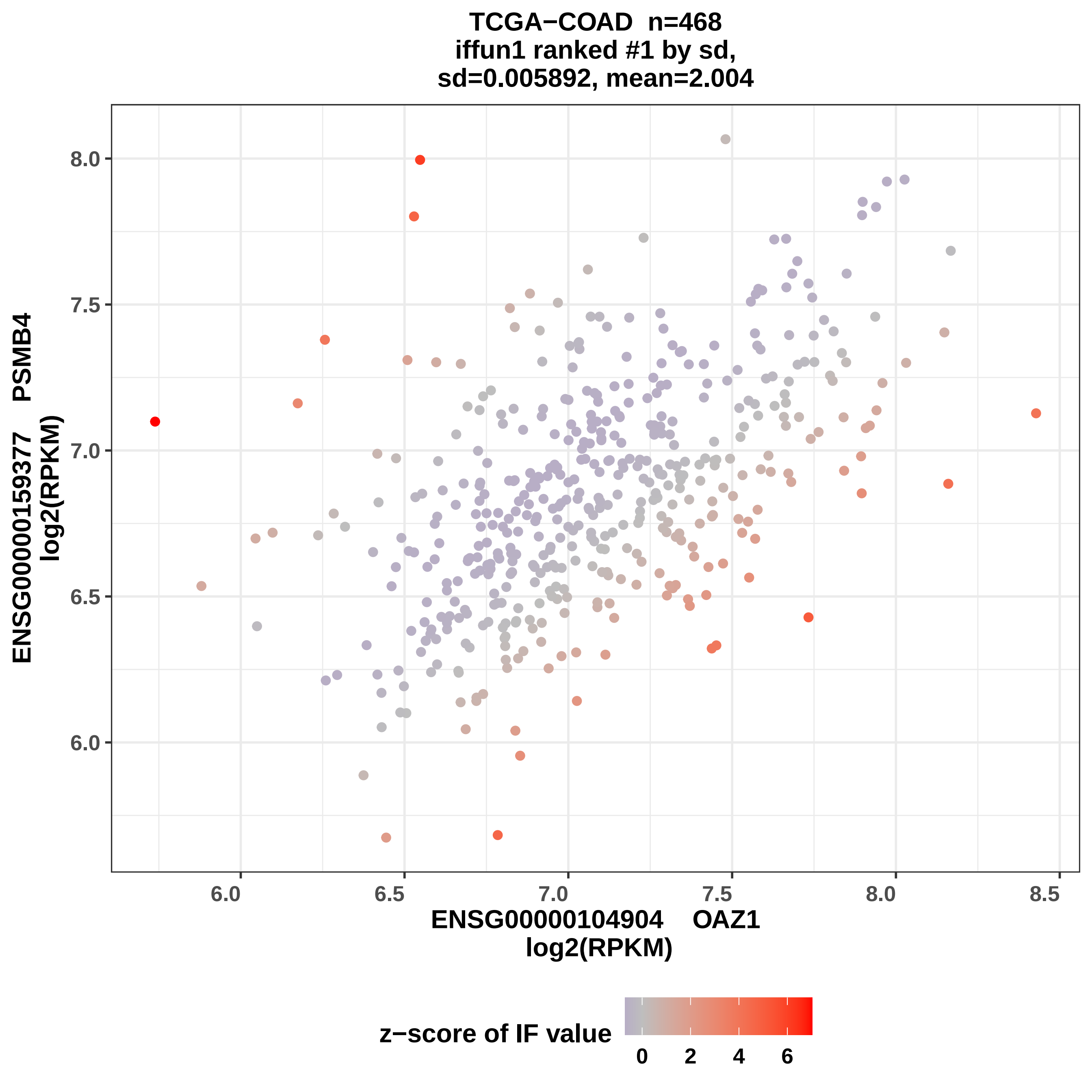
TCGA−COAD n=468
iffun1 ranked #1 by sd,
sd=0.005892, mean=2.004
ENSG00000159377 PSMB4
log2(RPKM)
8.0
7.5
7.0
6.5
6.0
6.0
6.5
7.0
7.5
8.0
8.5
ENSG00000104904 OAZ1
log2(RPKM)
z−score of IF value
0
2
4
6

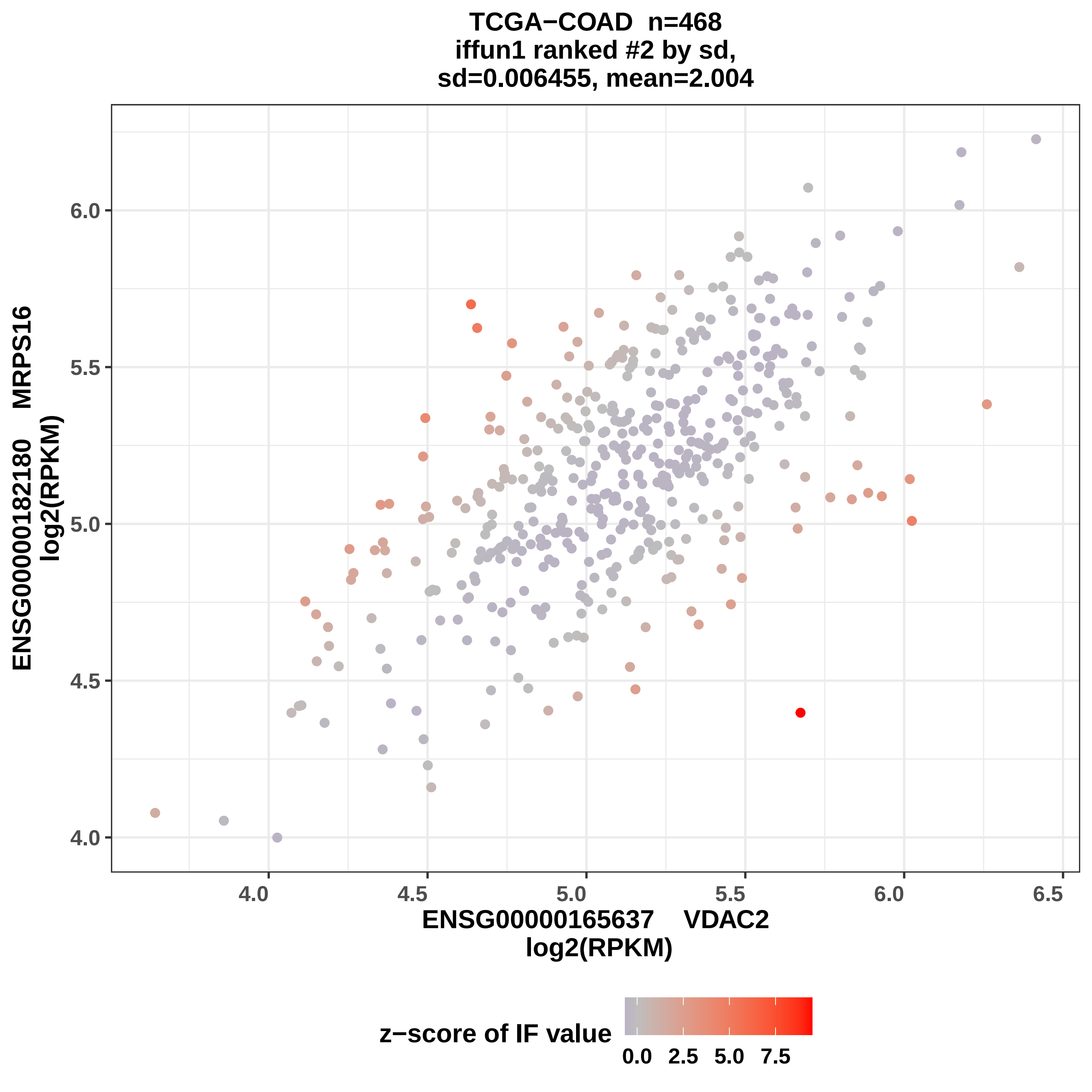

TCGA−COAD n=468
iffun1 ranked #2 by sd,
sd=0.006455, mean=2.004
ENSG00000182180 MRPS16
log2(RPKM)
6.0
5.5
5.0
4.5
4.0
4.0
4.5
5.0
5.5
6.0
6.5
ENSG00000165637 VDAC2
log2(RPKM)
z−score of IF value
0.0
2.5
5.0
7.5

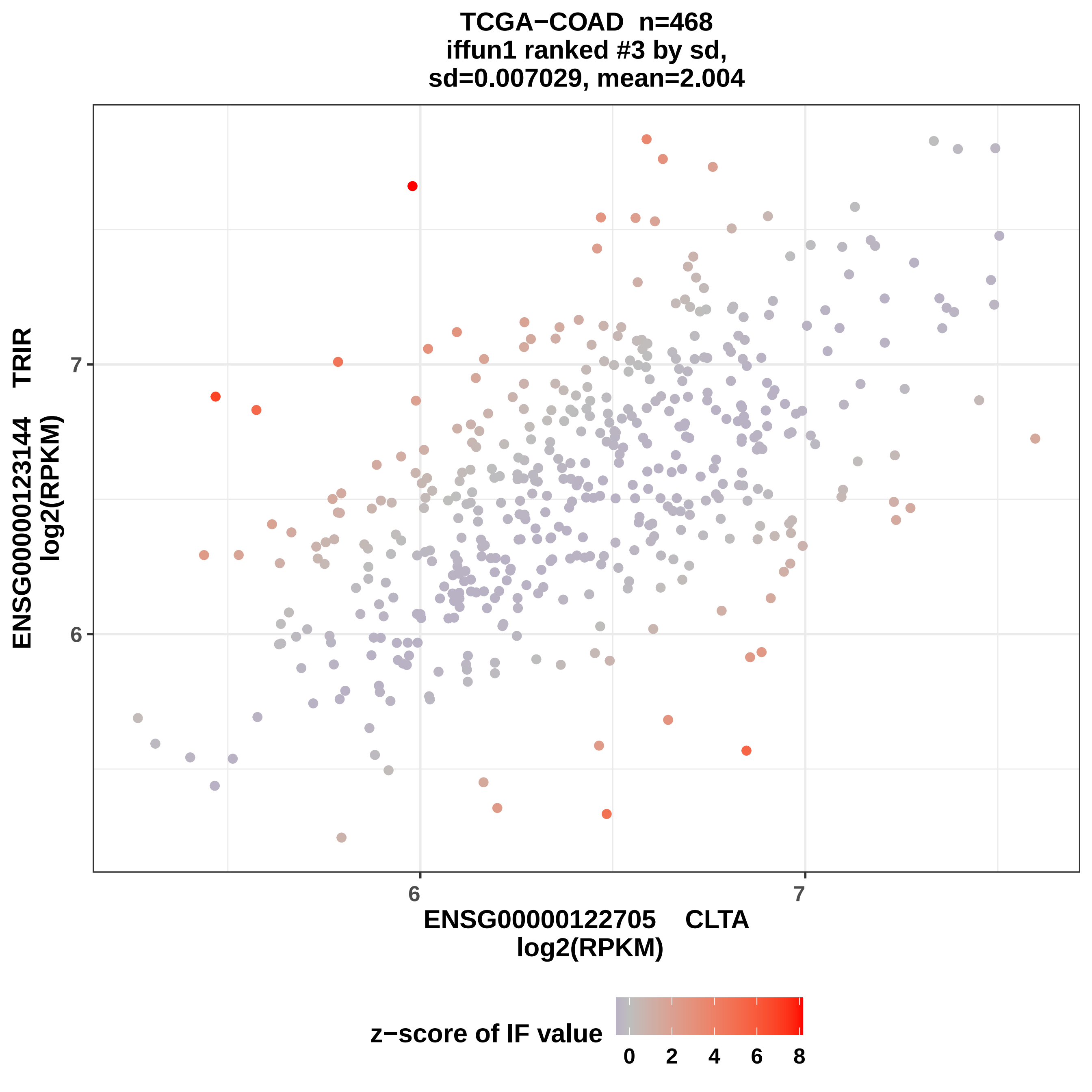
TCGA−COAD n=468
iffun1 ranked #3 by sd,
sd=0.007029, mean=2.004
ENSG00000123144 TRIR
log2(RPKM)
7
6
ENSG00000122705 CLTA
log2(RPKM)
z−score of IF value
0
2
4
6
8

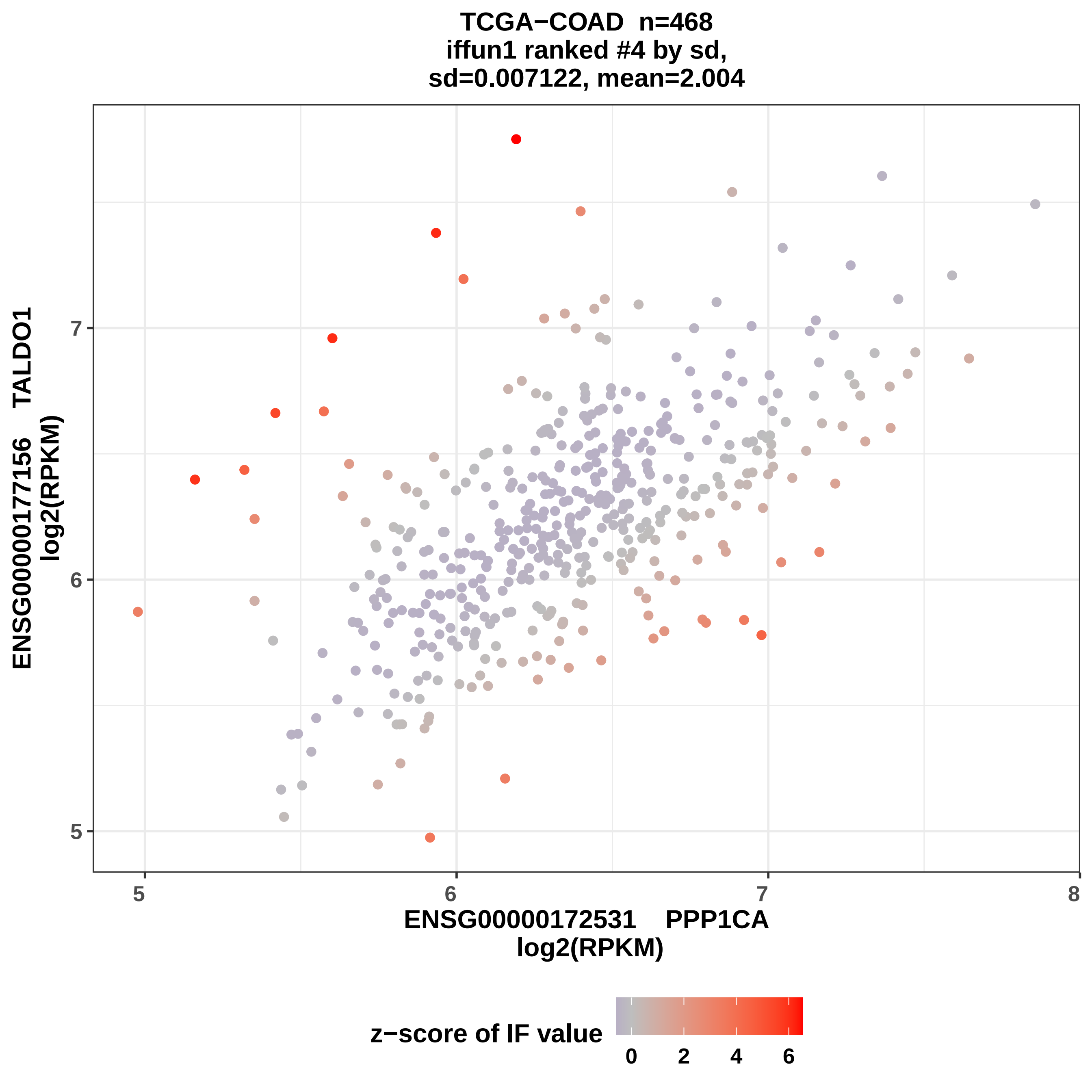

TCGA−COAD n=468
iffun1 ranked #4 by sd,
sd=0.007122, mean=2.004
ENSG00000177156 TALDO1
log2(RPKM)
ENSG00000172531 PPP1CA
log2(RPKM)
5
6
7
8
z−score of IF value
0
2
4
6

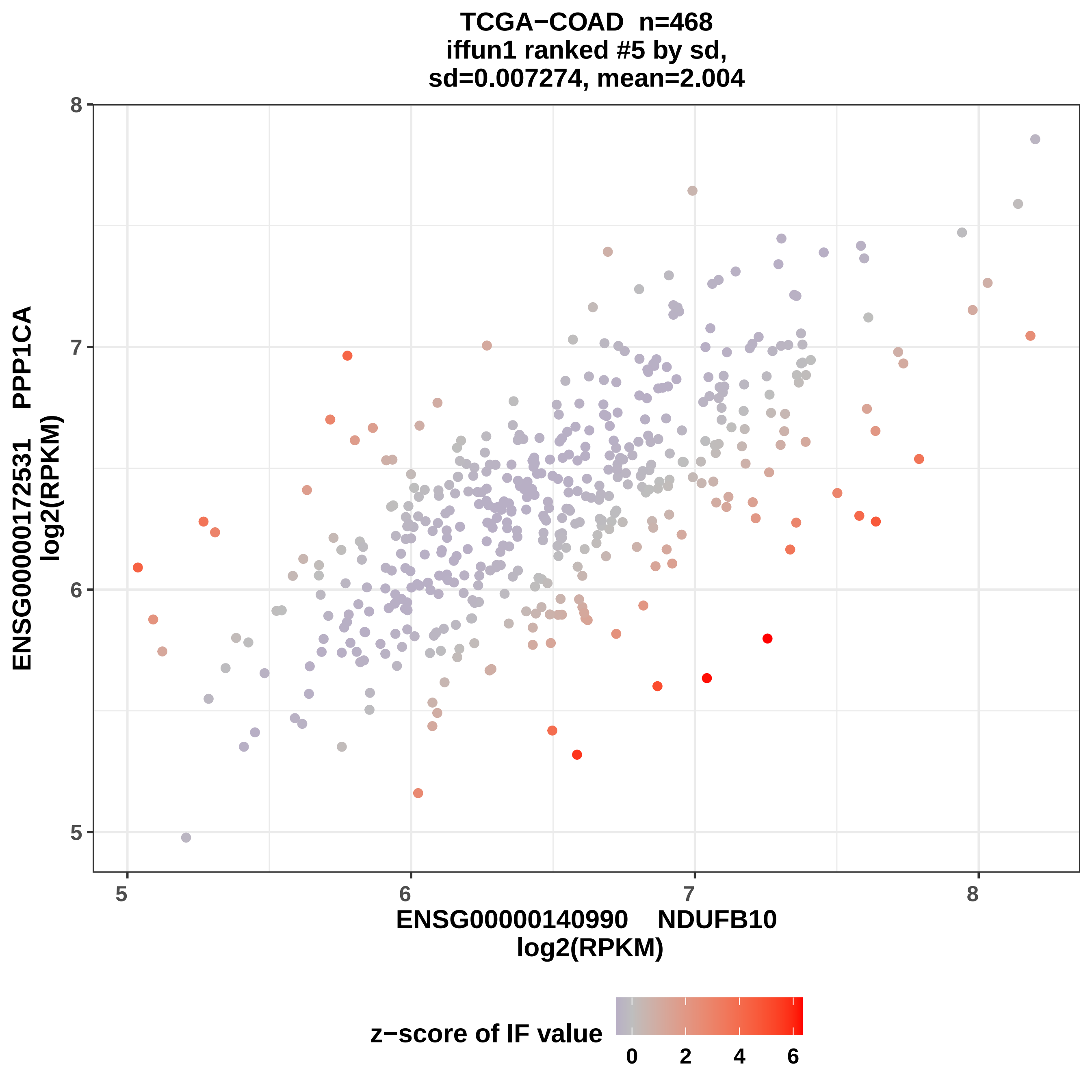
TCGA−COAD n=468
iffun1 ranked #5 by sd,
sd=0.007274, mean=2.004
ENSG00000172531 PPP1CA
log2(RPKM)
ENSG00000140990 NDUFB10
log2(RPKM)
z−score of IF value
5
6
7
8
0
2
4
6

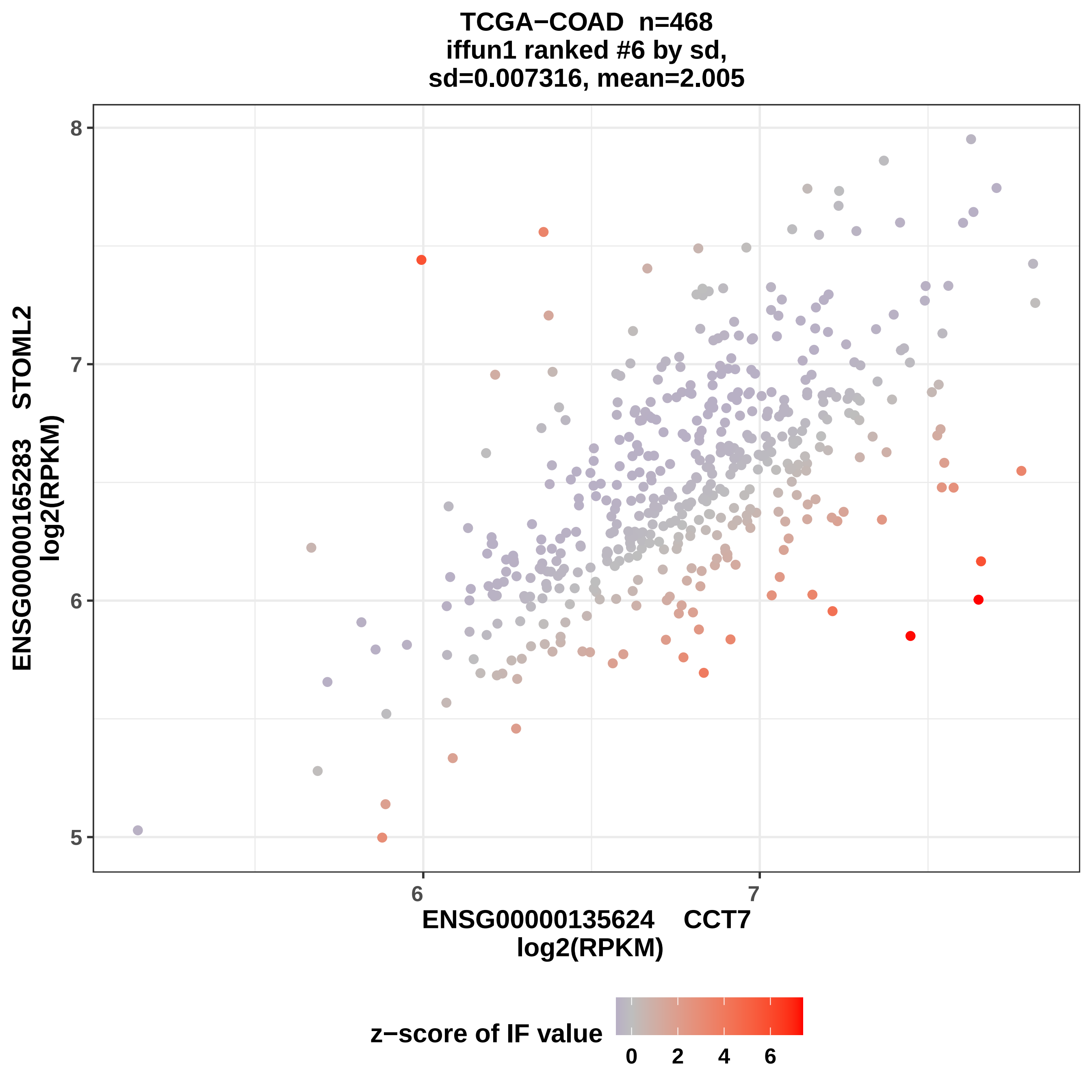
TCGA−COAD n=468
iffun1 ranked #6 by sd,
sd=0.007316, mean=2.005
ENSG00000165283 STOML2
log2(RPKM)
ENSG00000135624 CCT7
log2(RPKM)
8
7
6
5
6
7
z−score of IF value
0
2
4
6

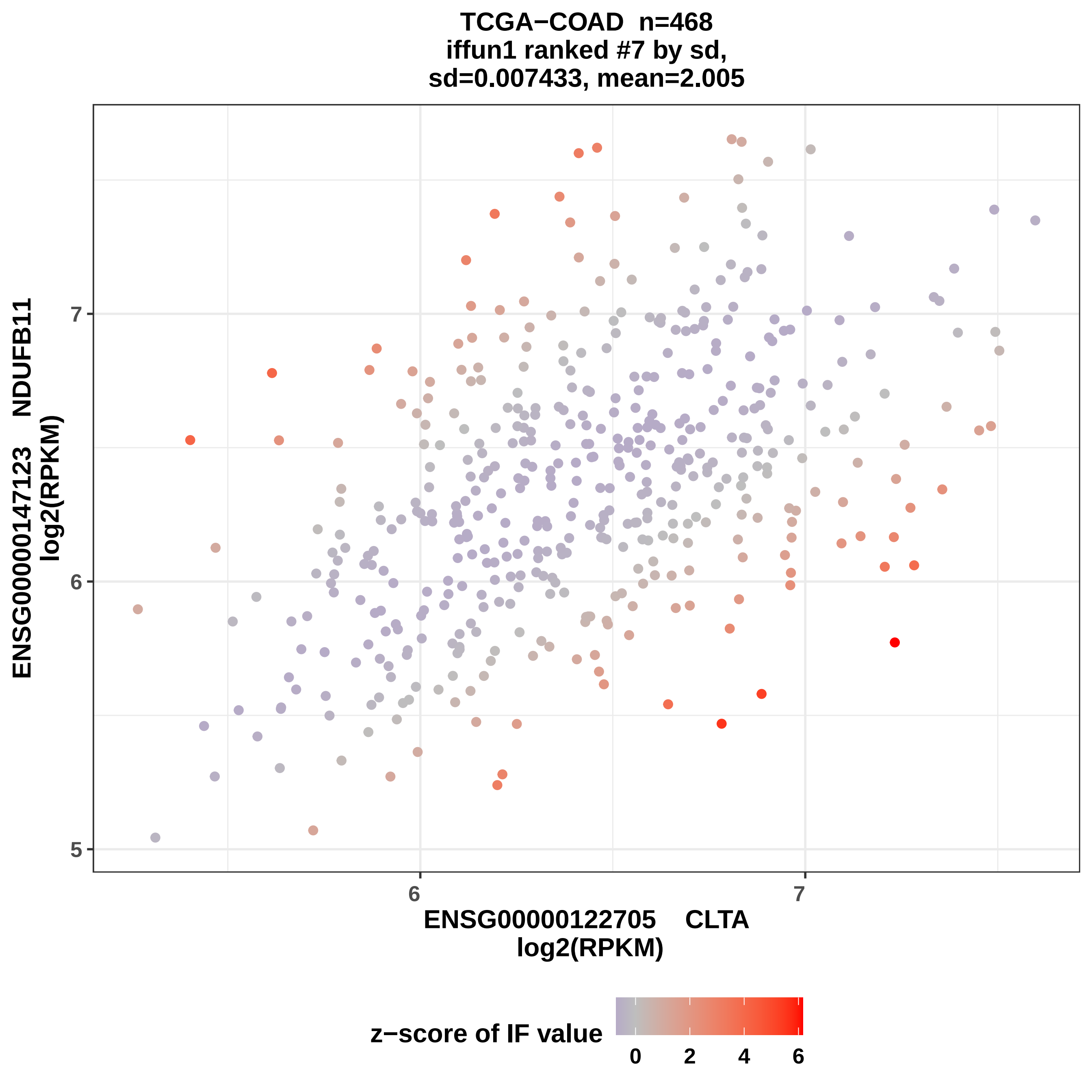
TCGA−COAD n=468
iffun1 ranked #7 by sd,
sd=0.007433, mean=2.005
ENSG00000147123 NDUFB11
log2(RPKM)
7
6
5
6
7
ENSG00000122705 CLTA
log2(RPKM)
z−score of IF value
0
2
4
6

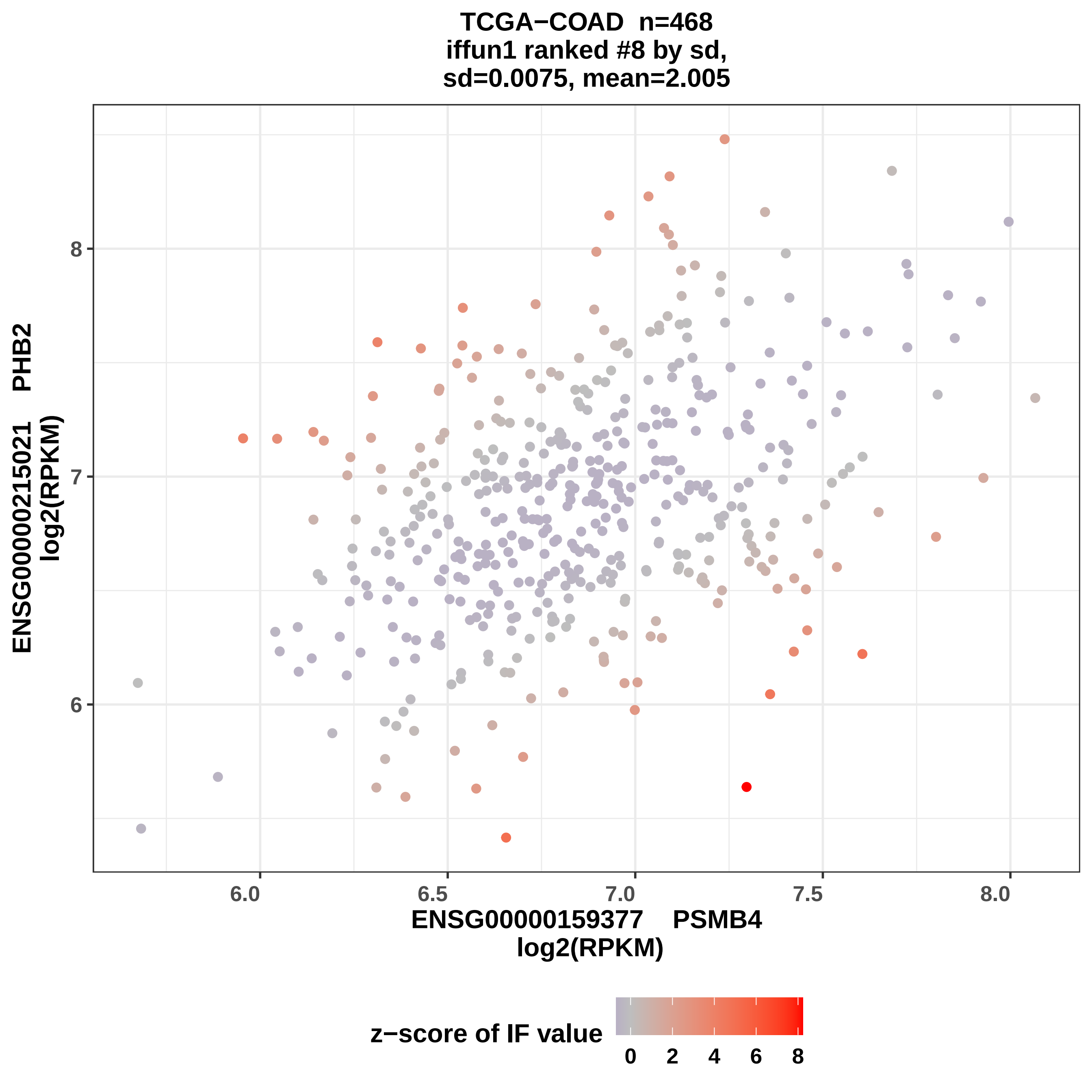
TCGA−COAD n=468
iffun1 ranked #8 by sd,
sd=0.0075, mean=2.005
8
7
6
6.0
6.5
7.0
7.5
8.0
ENSG00000215021 PHB2
log2(RPKM)
ENSG00000159377 PSMB4
log2(RPKM)
z−score of IF value
0
2
4
6
8

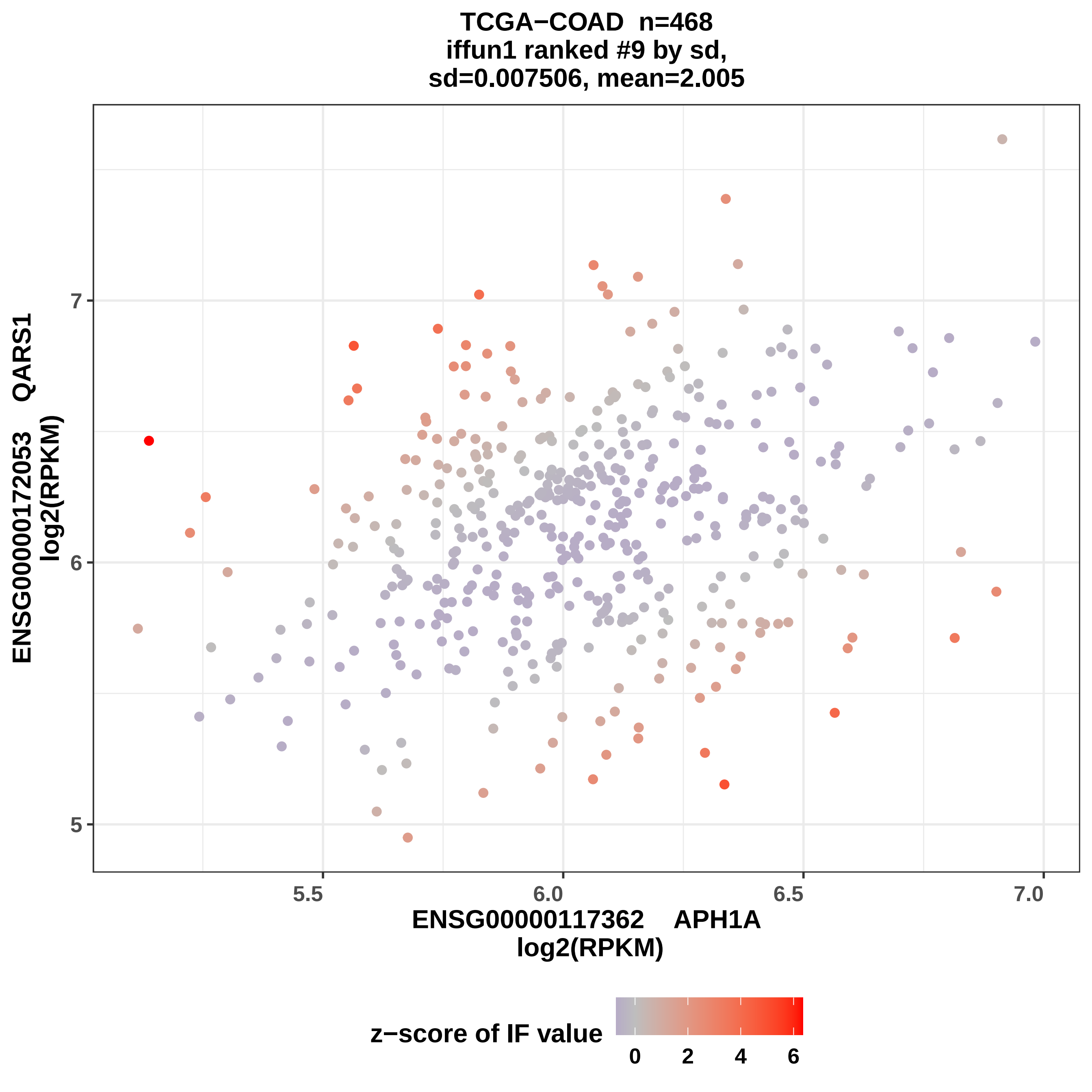
TCGA−COAD n=468
iffun1 ranked #9 by sd,
sd=0.007506, mean=2.005
ENSG00000172053 QARS1
log2(RPKM)
7
6
5
5.5
6.0
6.5
7.0
ENSG00000117362 APH1A
log2(RPKM)
z−score of IF value
0
2
4
6

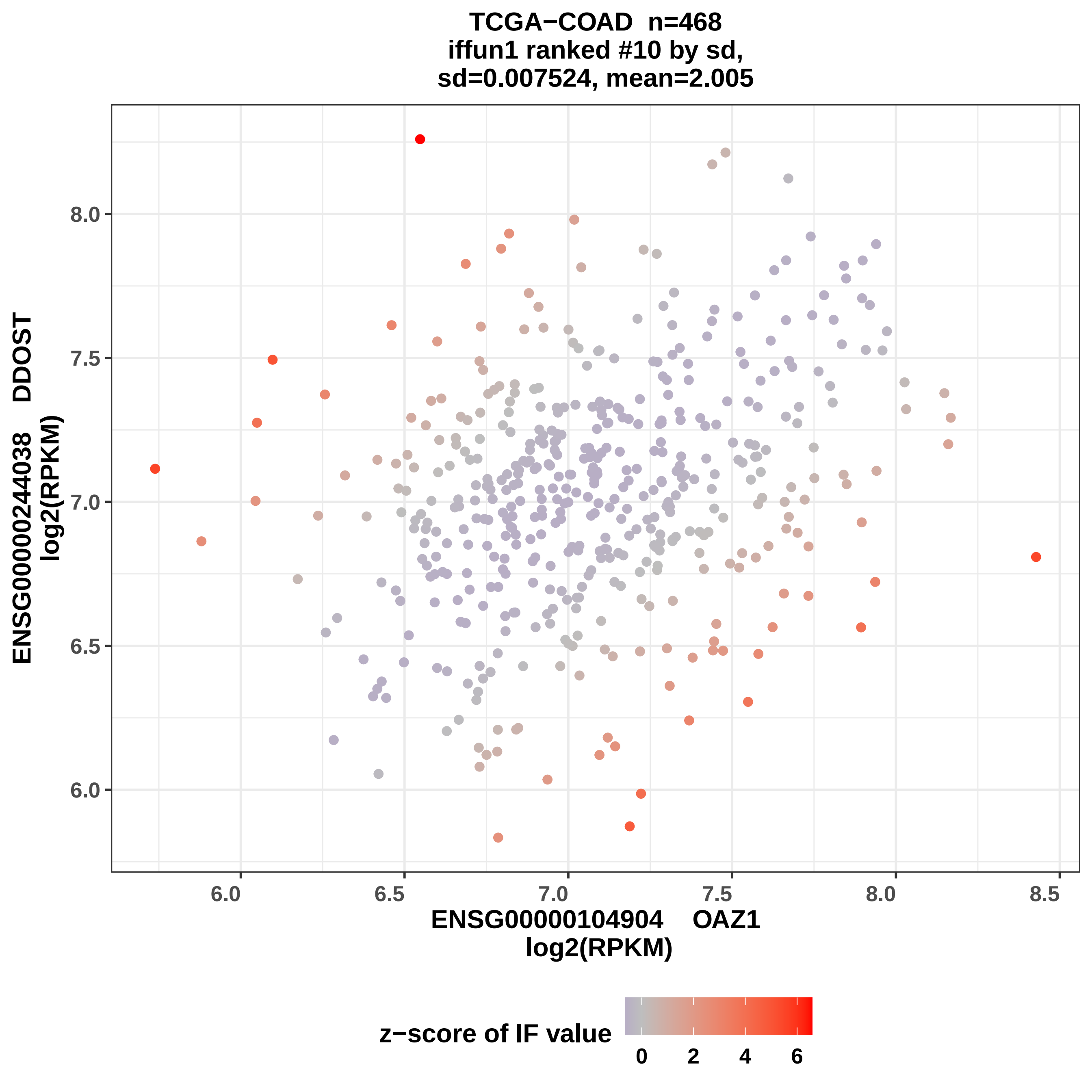

TCGA−COAD n=468
iffun1 ranked #10 by sd,
sd=0.007524, mean=2.005
ENSG00000244038 DDOST
log2(RPKM)
8.0
7.5
7.0
6.5
6.0
6.0
6.5
7.0
7.5
8.0
8.5
ENSG00000104904 OAZ1
log2(RPKM)
z−score of IF value
0
2
4
6

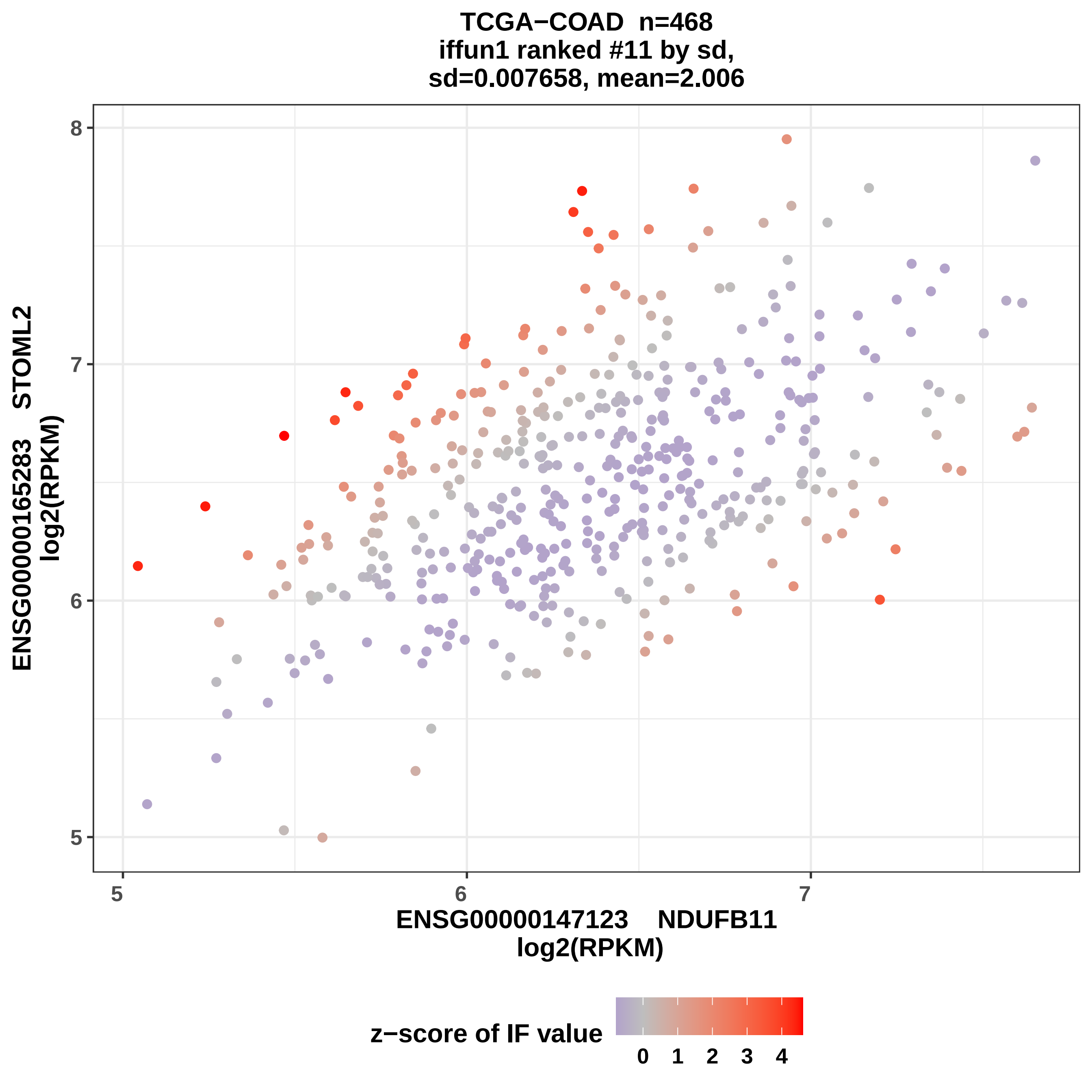

TCGA−COAD n=468
iffun1 ranked #11 by sd,
sd=0.007658, mean=2.006
ENSG00000165283 STOML2
log2(RPKM)
ENSG00000147123 NDUFB11
log2(RPKM)
5
6
7
8
5
6
7
z−score of IF value
0
1
2
3
4

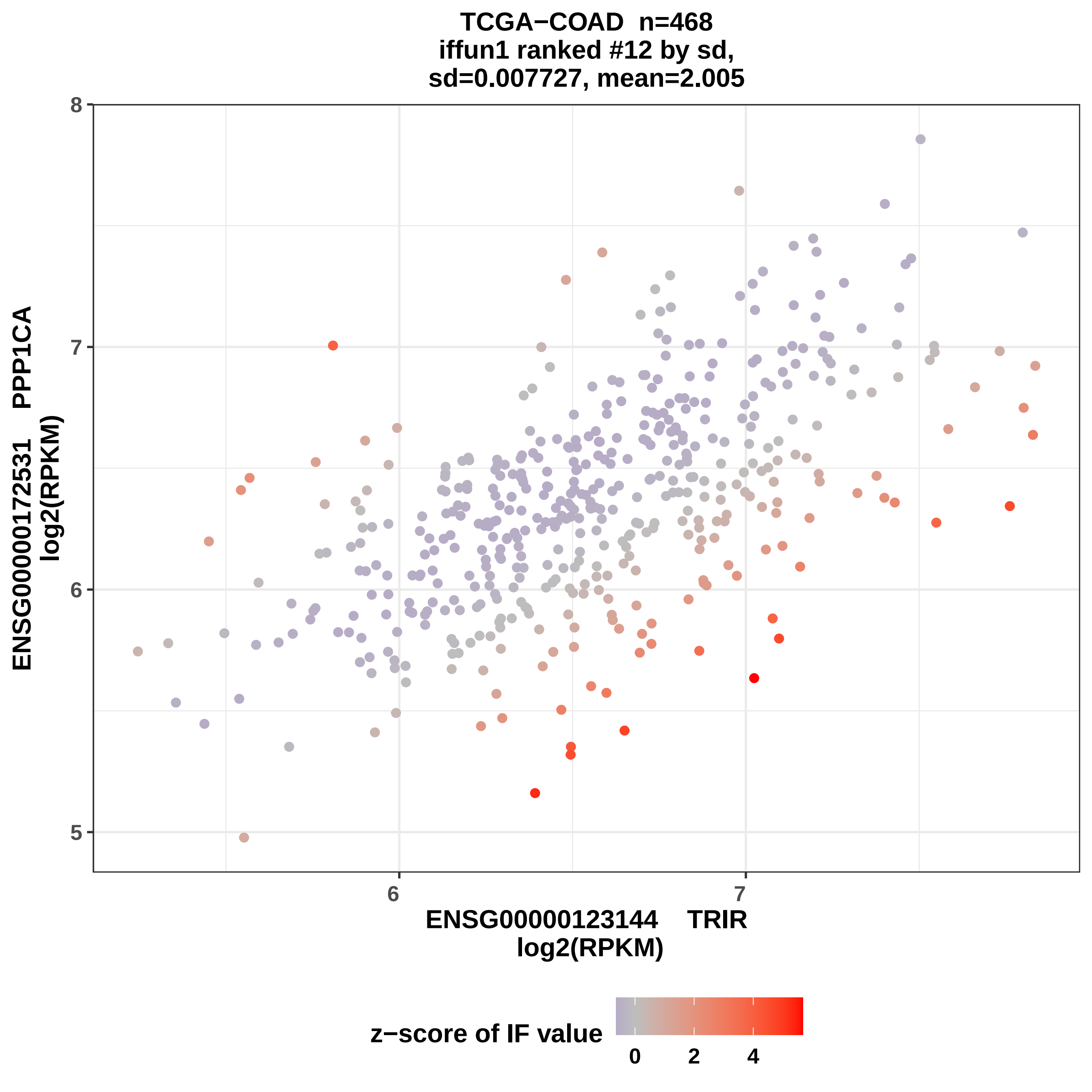
TCGA−COAD n=468
iffun1 ranked #12 by sd,
sd=0.007727, mean=2.005
ENSG00000172531 PPP1CA
log2(RPKM)
8
7
6
5
6
7
ENSG00000123144 TRIR
log2(RPKM)
z−score of IF value
0
2
4

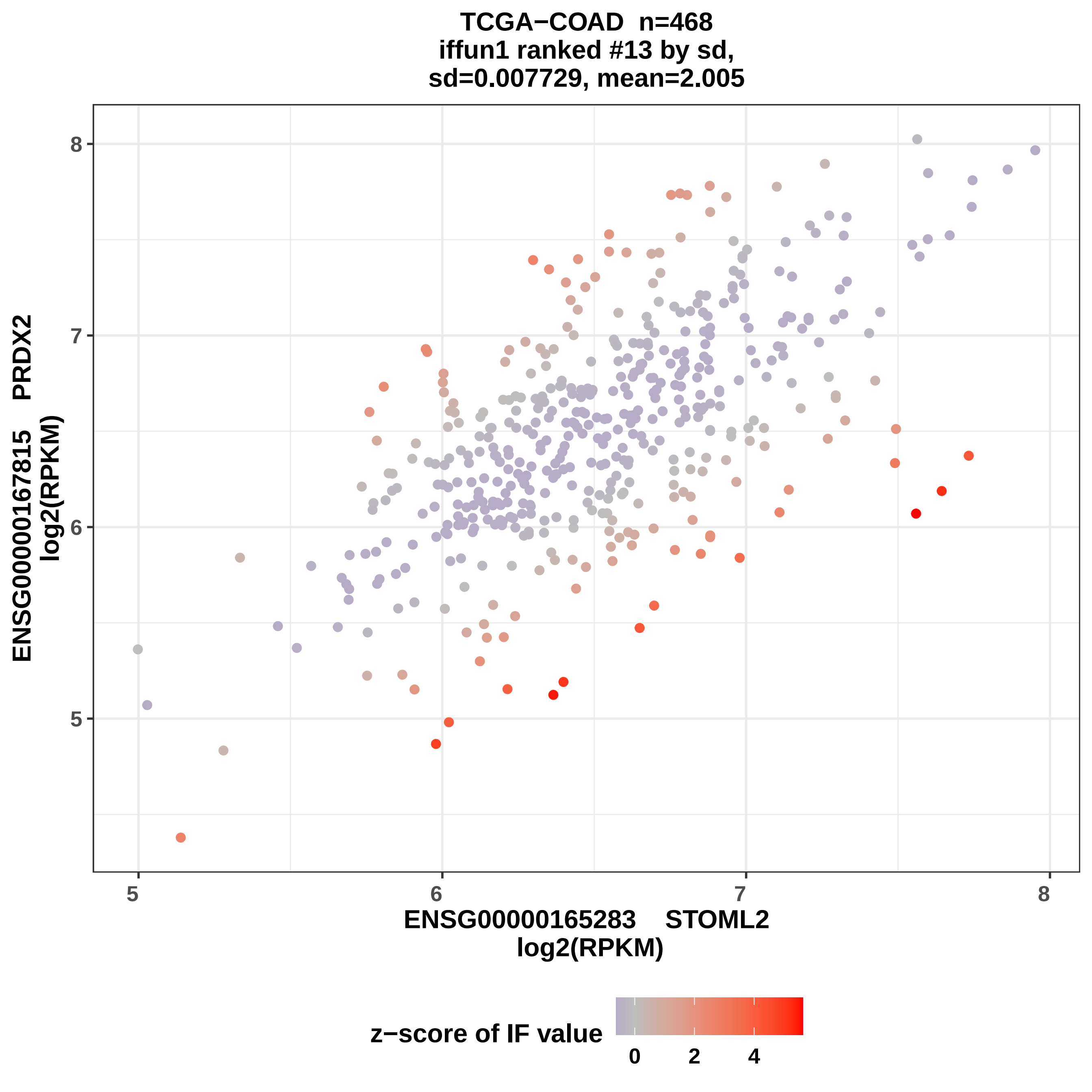
TCGA−COAD n=468
iffun1 ranked #13 by sd,
sd=0.007729, mean=2.005
ENSG00000167815 PRDX2
log2(RPKM)
8
7
6
5
5
6
7
8
ENSG00000165283 STOML2
log2(RPKM)
z−score of IF value
0
2
4

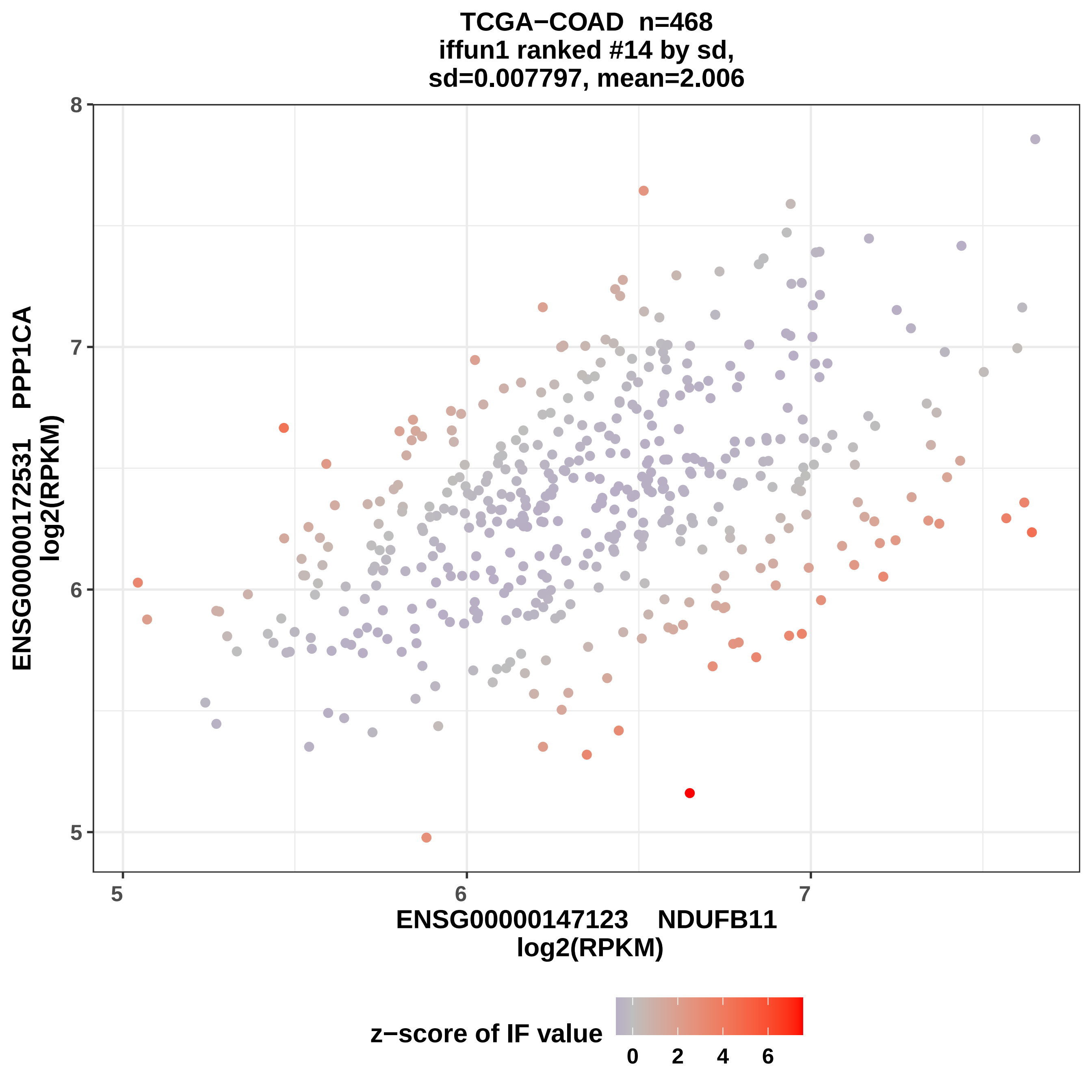

TCGA−COAD n=468
iffun1 ranked #14 by sd,
sd=0.007797, mean=2.006
ENSG00000172531 PPP1CA
log2(RPKM)
8
7
6
5
5
6
7
ENSG00000147123 NDUFB11
log2(RPKM)
z−score of IF value
0
2
4
6

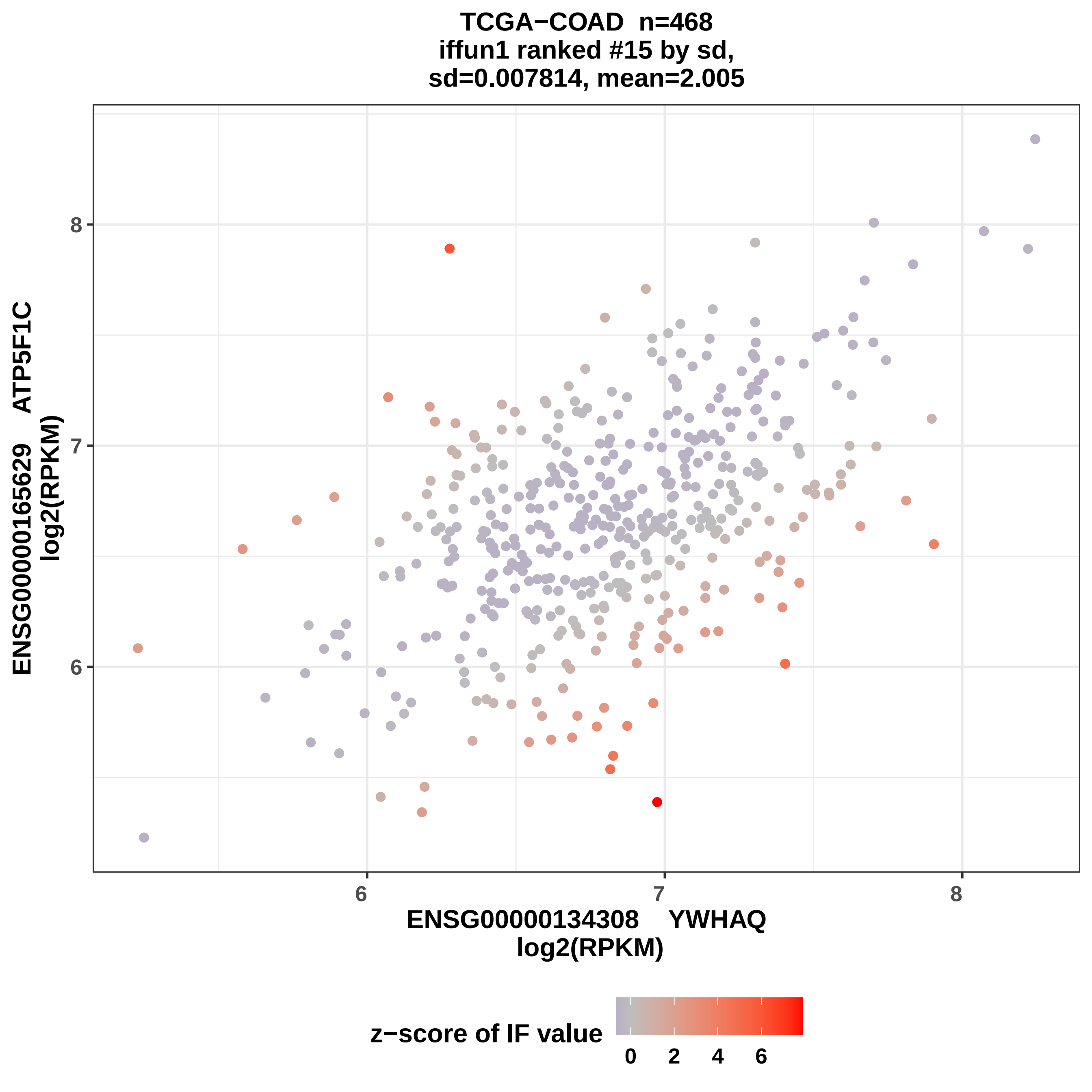

TCGA−COAD n=468
iffun1 ranked #15 by sd,
sd=0.007814, mean=2.005
ENSG00000165629 ATP5F1C
log2(RPKM)
8
7
6
6
7
8
ENSG00000134308 YWHAQ
log2(RPKM)
z−score of IF value
0
2
4
6

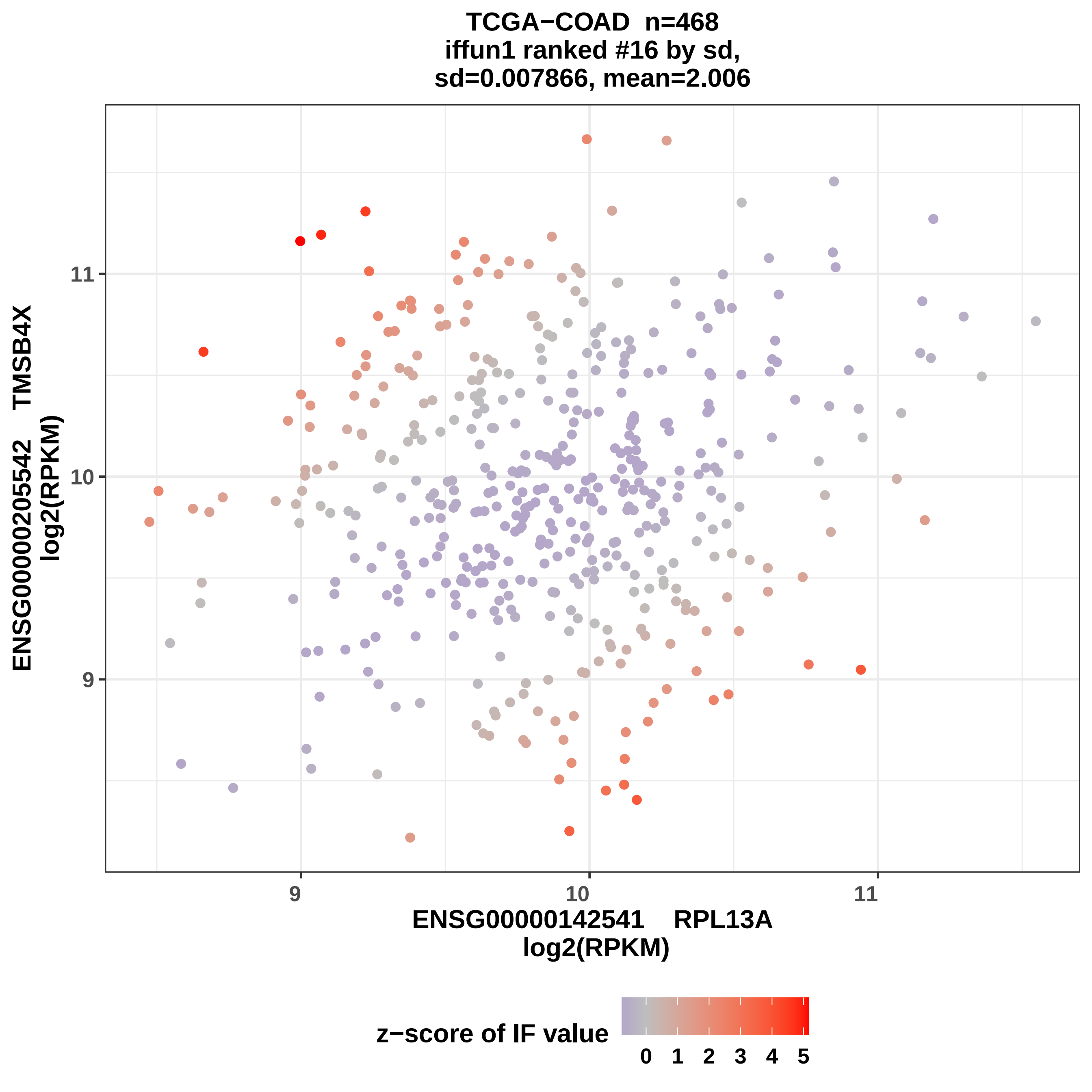

TCGA−COAD n=468
iffun1 ranked #16 by sd,
sd=0.007866, mean=2.006
ENSG00000205542 TMSB4X
log2(RPKM)
11
10
9
ENSG00000142541 RPL13A
log2(RPKM)
z−score of IF value
0 1 2 3 4 5

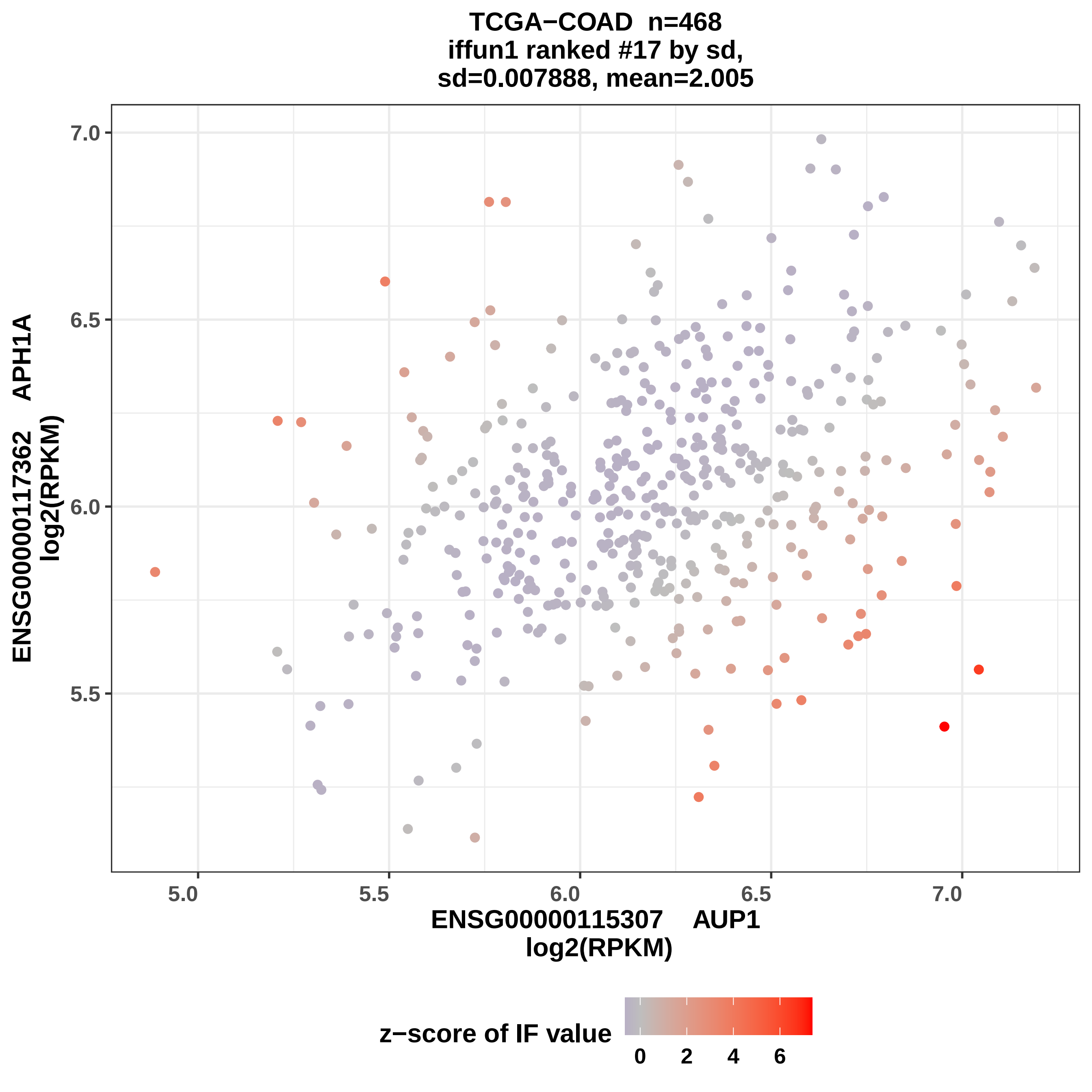

TCGA−COAD n=468
iffun1 ranked #17 by sd,
sd=0.007888, mean=2.005
ENSG00000117362 APH1A
log2(RPKM)
7.0
6.5
6.0
5.5
5.0
5.5
6.0
6.5
7.0
ENSG00000115307 AUP1
log2(RPKM)
z−score of IF value
0
2
4
6

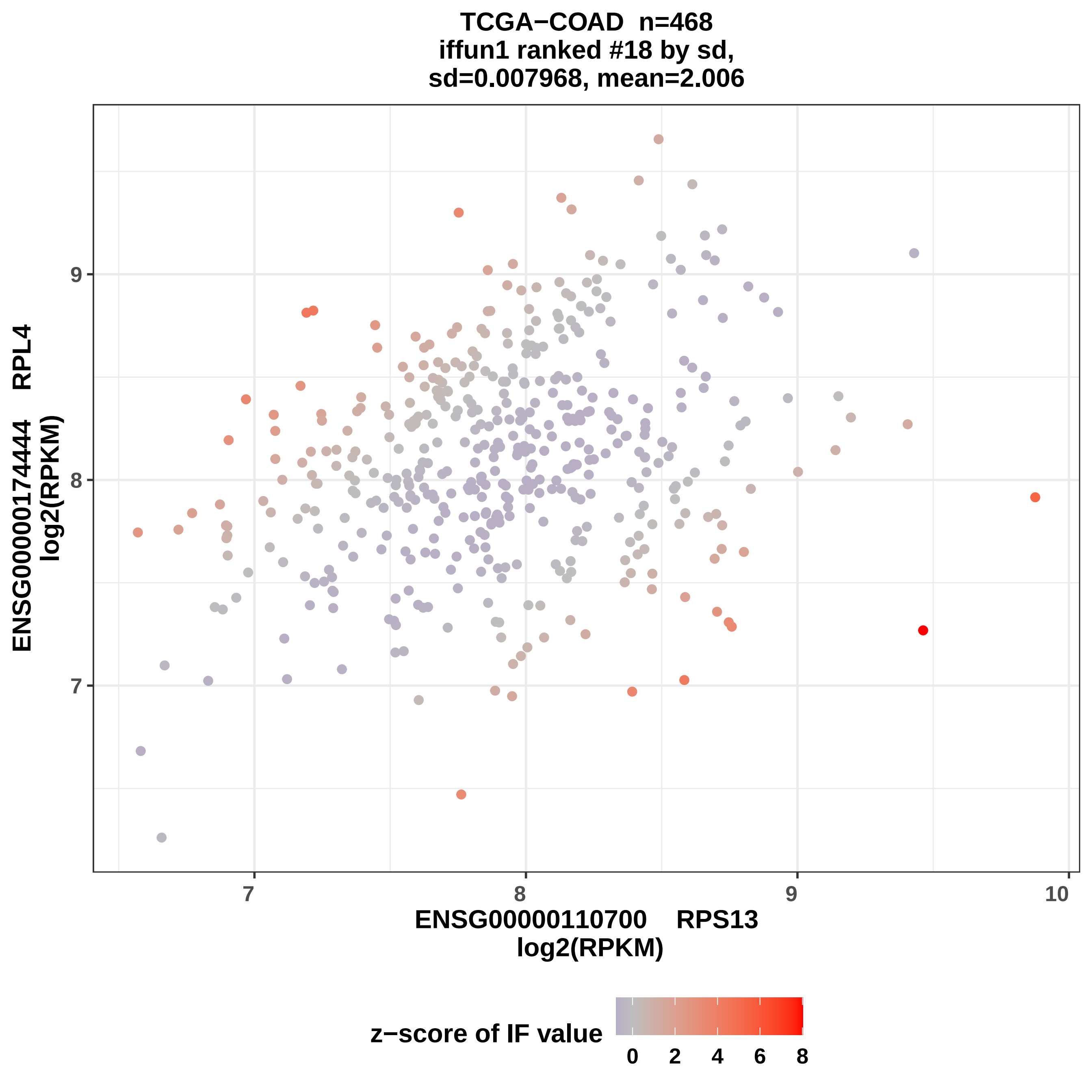
TCGA−COAD n=468
iffun1 ranked #18 by sd,
sd=0.007968, mean=2.006
ENSG00000174444 RPL4
log2(RPKM)
9
8
7
7
8
9
10
ENSG00000110700 RPS13
log2(RPKM)
z−score of IF value
0
2
4
6
8

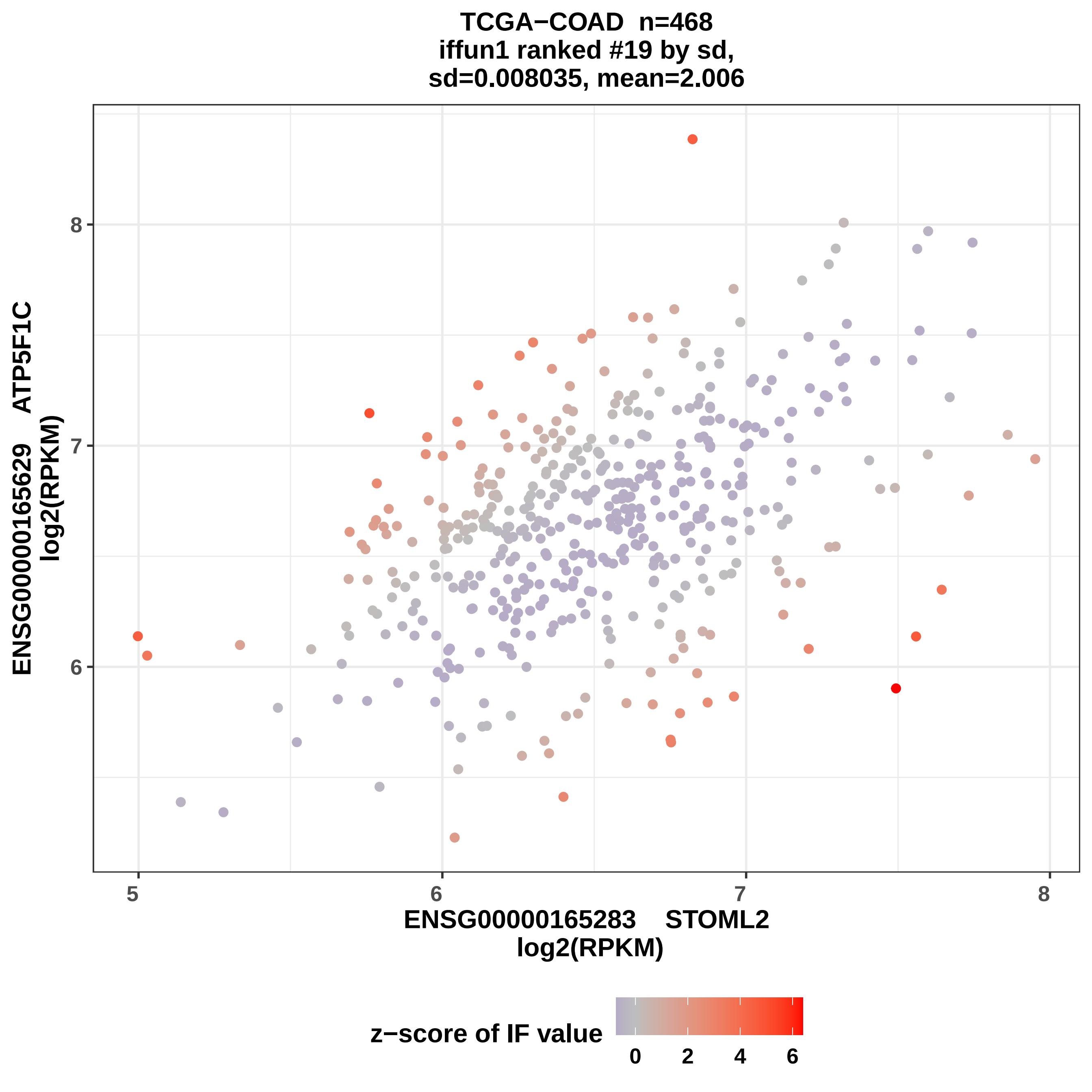
TCGA−COAD n=468
iffun1 ranked #19 by sd,
sd=0.008035, mean=2.006
ENSG00000165629 ATP5F1C
log2(RPKM)
8
7
6
5
6
7
8
ENSG00000165283 STOML2
log2(RPKM)
z−score of IF value
0
2
4
6

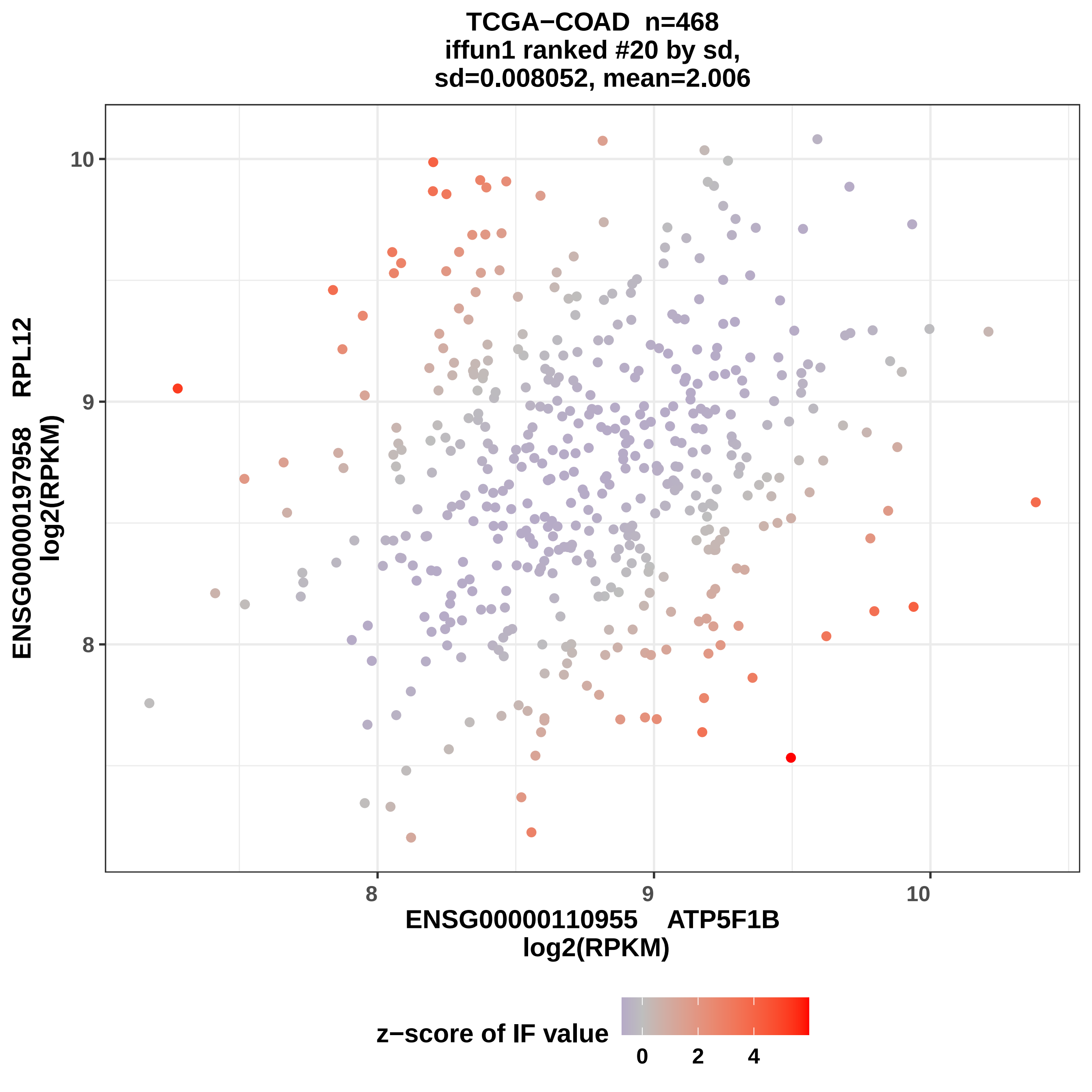
TCGA−COAD n=468
iffun1 ranked #20 by sd,
sd=0.008052, mean=2.006
ENSG00000197958 RPL12
log2(RPKM)
10
9
8
8
9
10
ENSG00000110955 ATP5F1B
log2(RPKM)
z−score of IF value
0
2
4

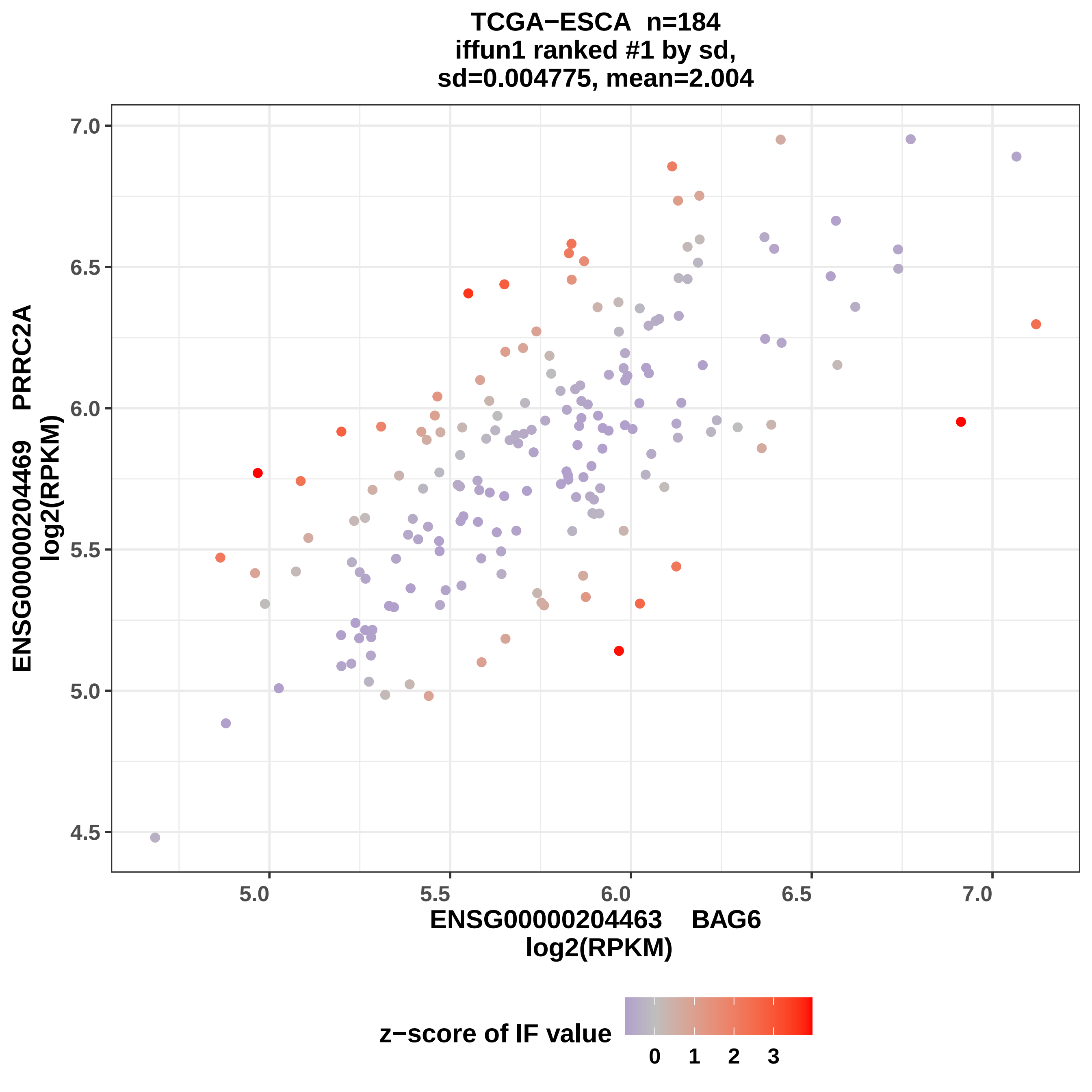

TCGA−ESCA n=184
iffun1 ranked #1 by sd,
sd=0.004775, mean=2.004
ENSG00000204469 PRRC2A
log2(RPKM)
ENSG00000204463 BAG6
log2(RPKM)
7.0
6.5
6.0
5.5
5.0
4.5
5.0
5.5
6.0
6.5
7.0
z−score of IF value
0
1
2
3

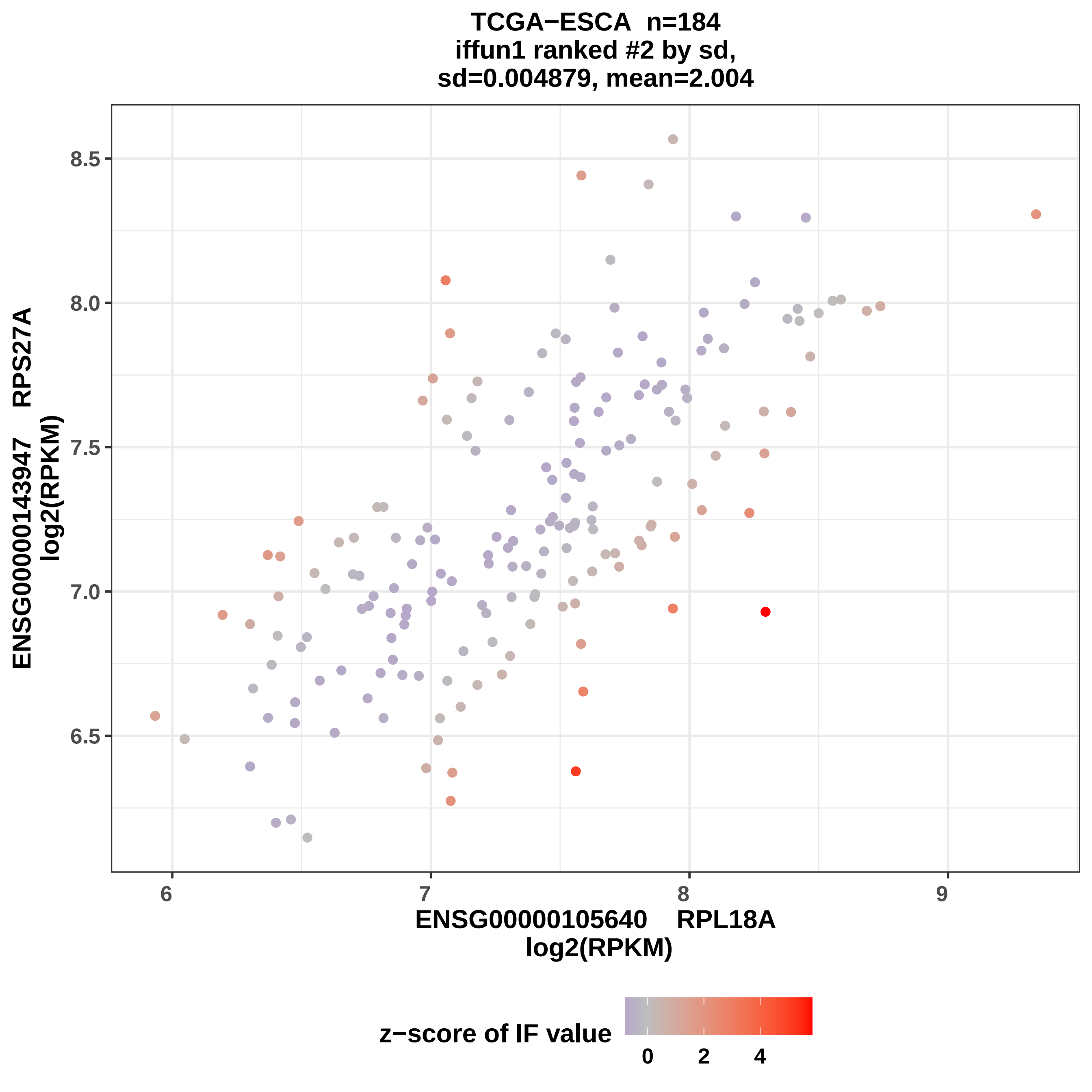
TCGA−ESCA n=184
iffun1 ranked #2 by sd,
sd=0.004879, mean=2.004
ENSG00000143947 RPS27A
log2(RPKM)
8.5
8.0
7.5
7.0
6.5
6
7
8
9
ENSG00000105640 RPL18A
log2(RPKM)
z−score of IF value
0
2
4

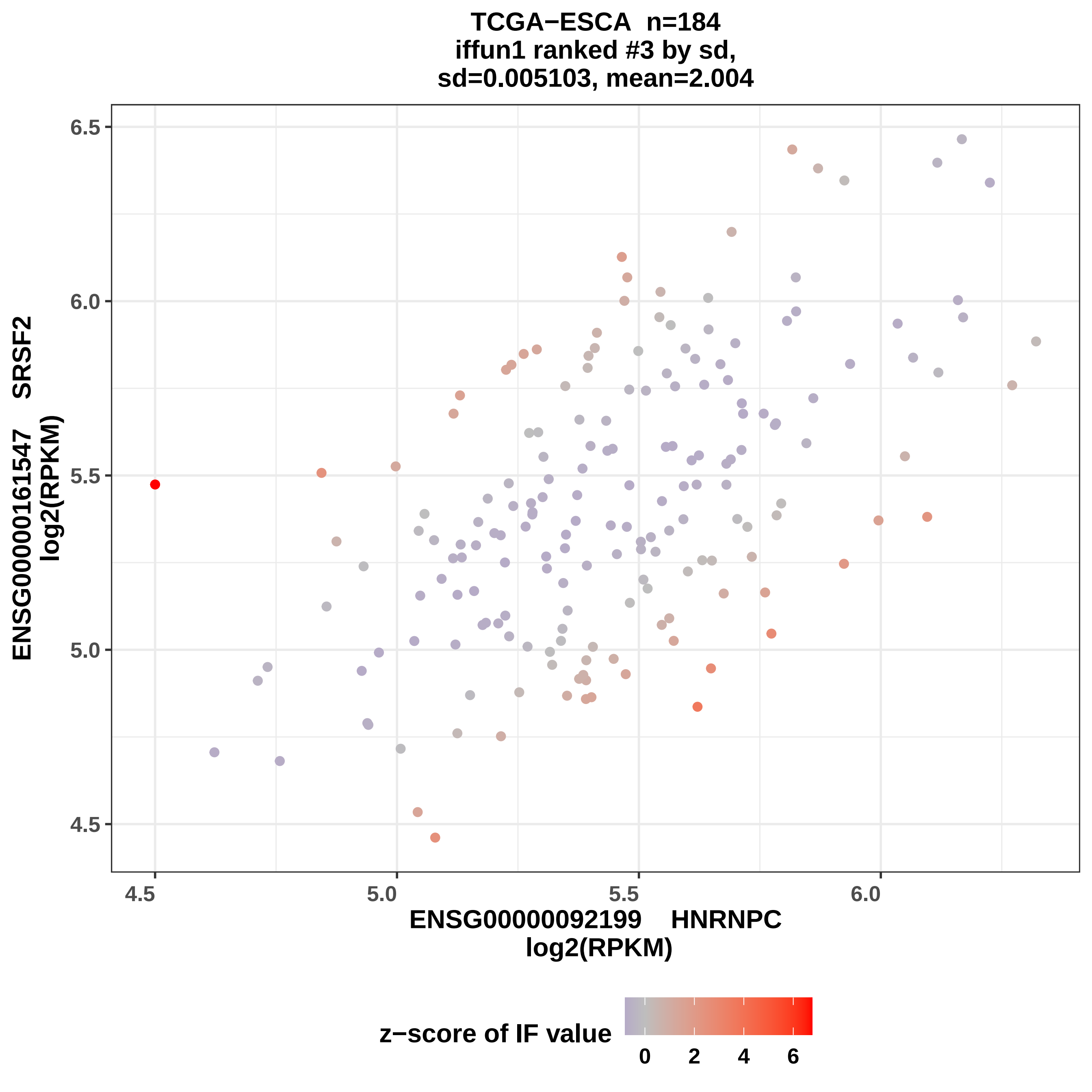
TCGA−ESCA n=184
iffun1 ranked #3 by sd,
sd=0.005103, mean=2.004
ENSG00000161547 SRSF2
log2(RPKM)
6.5
6.0
5.5
5.0
4.5
4.5
5.0
5.5
6.0
ENSG00000092199 HNRNPC
log2(RPKM)
z−score of IF value
0
2
4
6

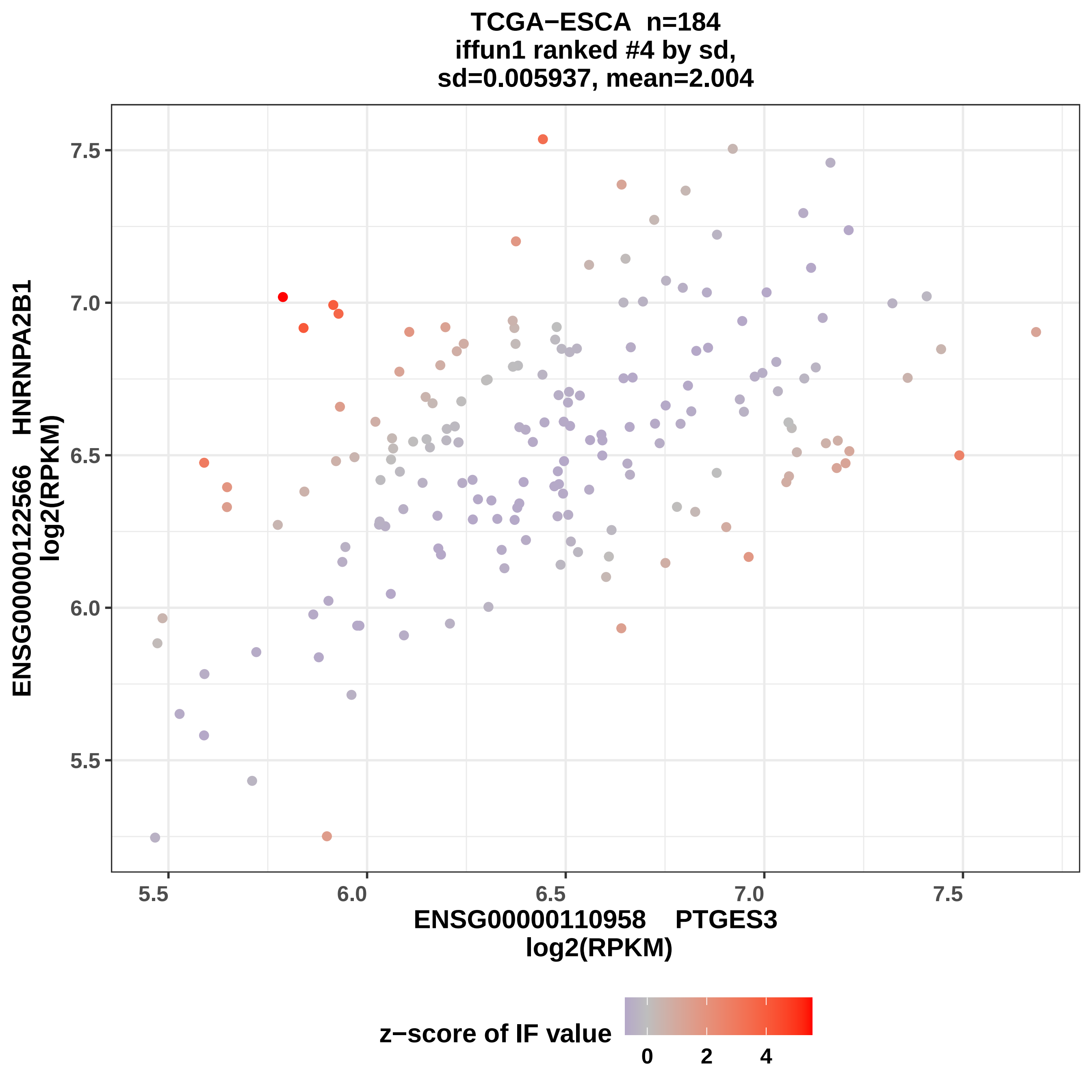
TCGA−ESCA n=184
iffun1 ranked #4 by sd,
sd=0.005937, mean=2.004
ENSG00000122566 HNRNPA2B1
log2(RPKM)
7.5
7.0
6.5
6.0
5.5
5.5
6.0
6.5
7.0
7.5
ENSG00000110958 PTGES3
log2(RPKM)
z−score of IF value
0
2
4

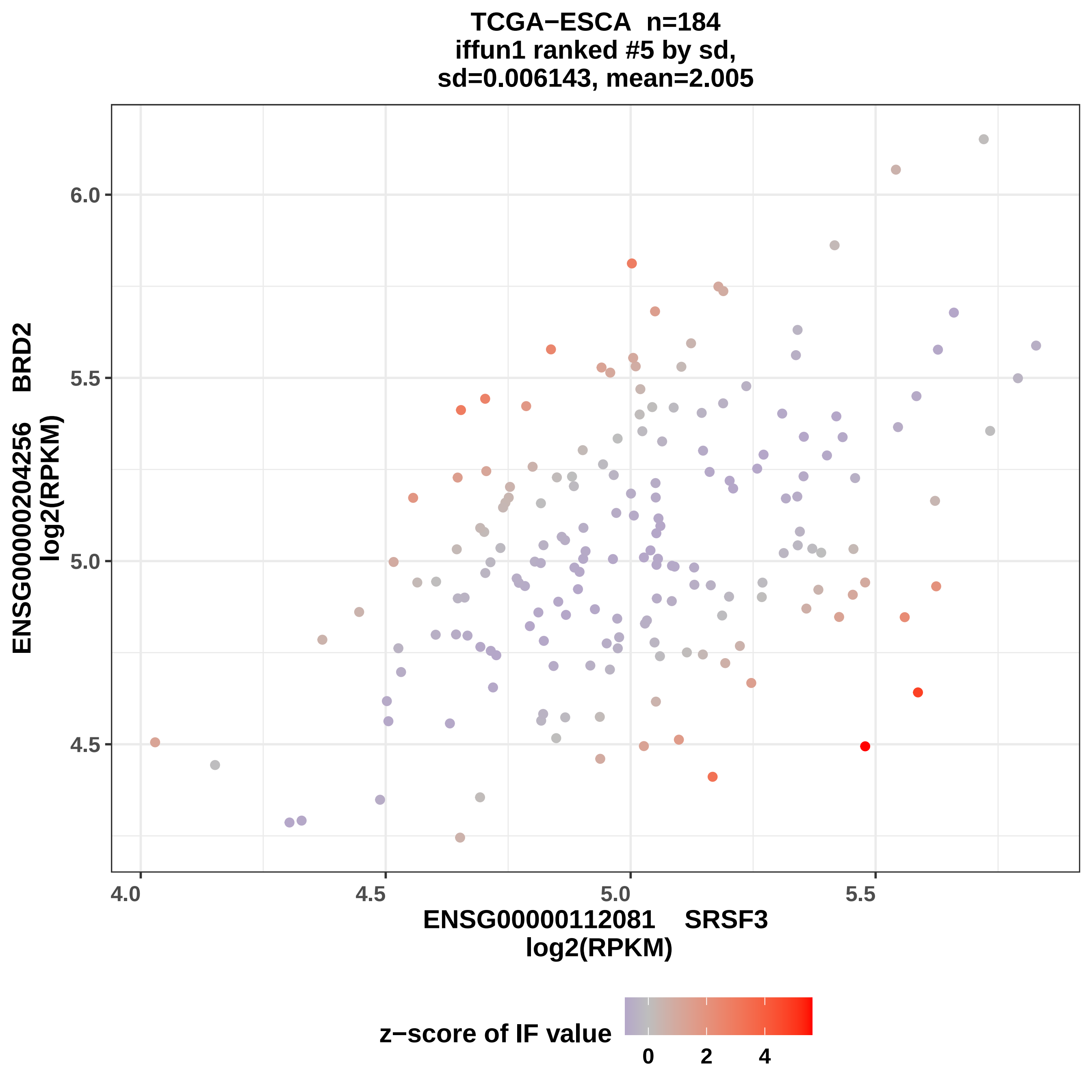

TCGA−ESCA n=184
iffun1 ranked #5 by sd,
sd=0.006143, mean=2.005
ENSG00000204256 BRD2
log2(RPKM)
6.0
5.5
5.0
4.5
4.0
4.5
5.0
5.5
ENSG00000112081 SRSF3
log2(RPKM)
z−score of IF value
0
2
4

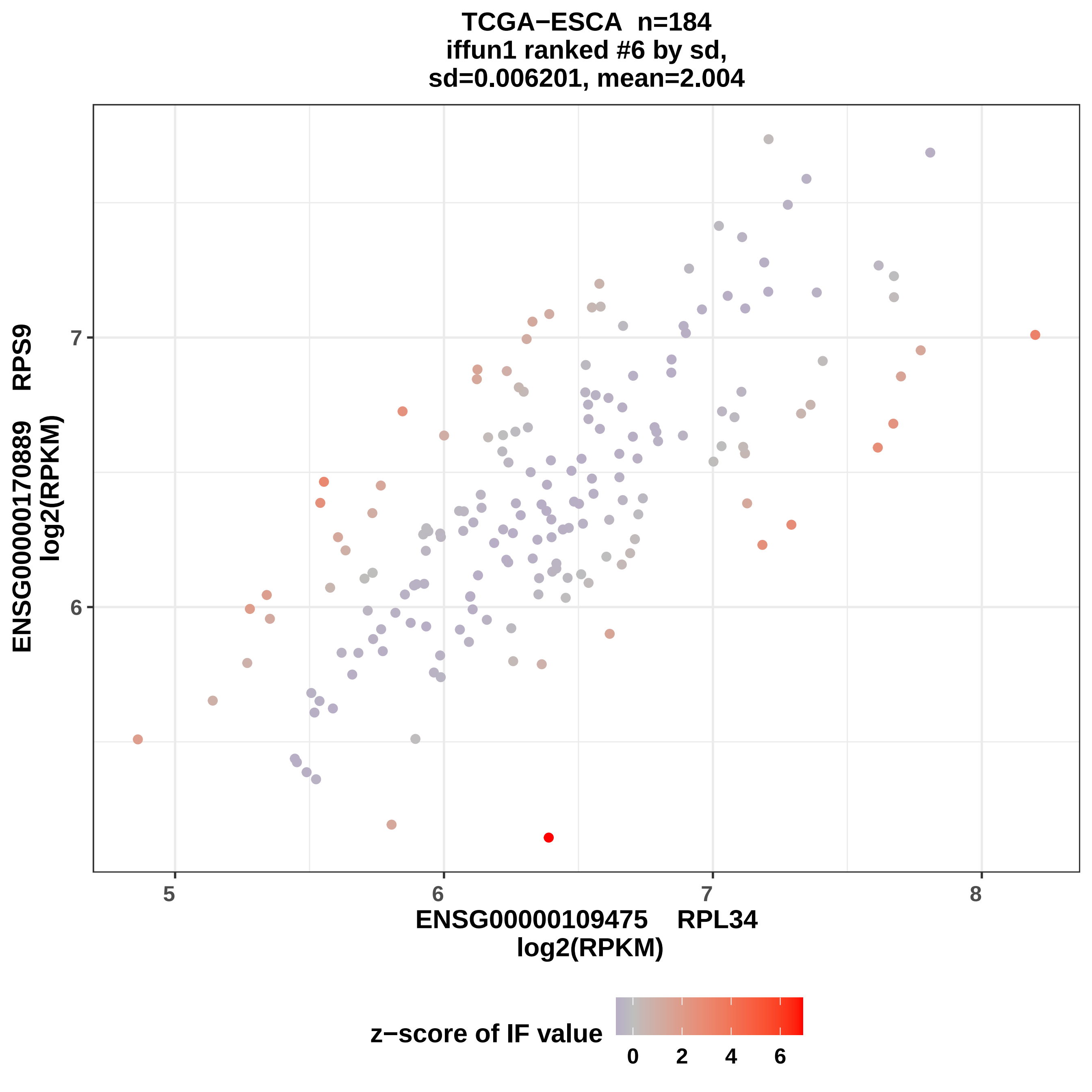

TCGA−ESCA n=184
iffun1 ranked #6 by sd,
sd=0.006201, mean=2.004
ENSG00000170889 RPS9
log2(RPKM)
ENSG00000109475 RPL34
log2(RPKM)
5
6
7
8
z−score of IF value
0
2
4
6

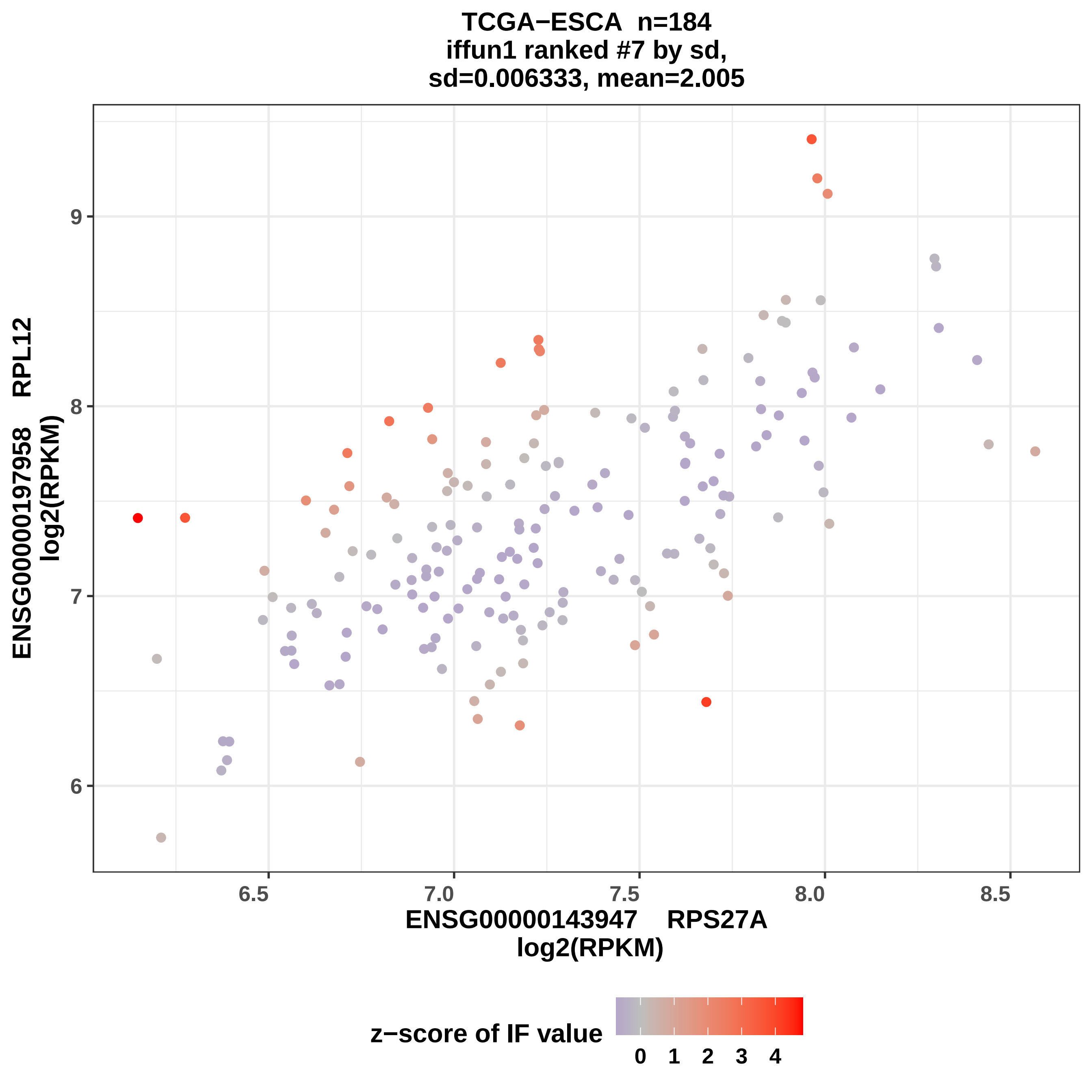

TCGA−ESCA n=184
iffun1 ranked #7 by sd,
sd=0.006333, mean=2.005
ENSG00000197958 RPL12
log2(RPKM)
9
8
7
6
6.5
7.0
7.5
8.0
8.5
ENSG00000143947 RPS27A
log2(RPKM)
z−score of IF value
0
1
2
3
4

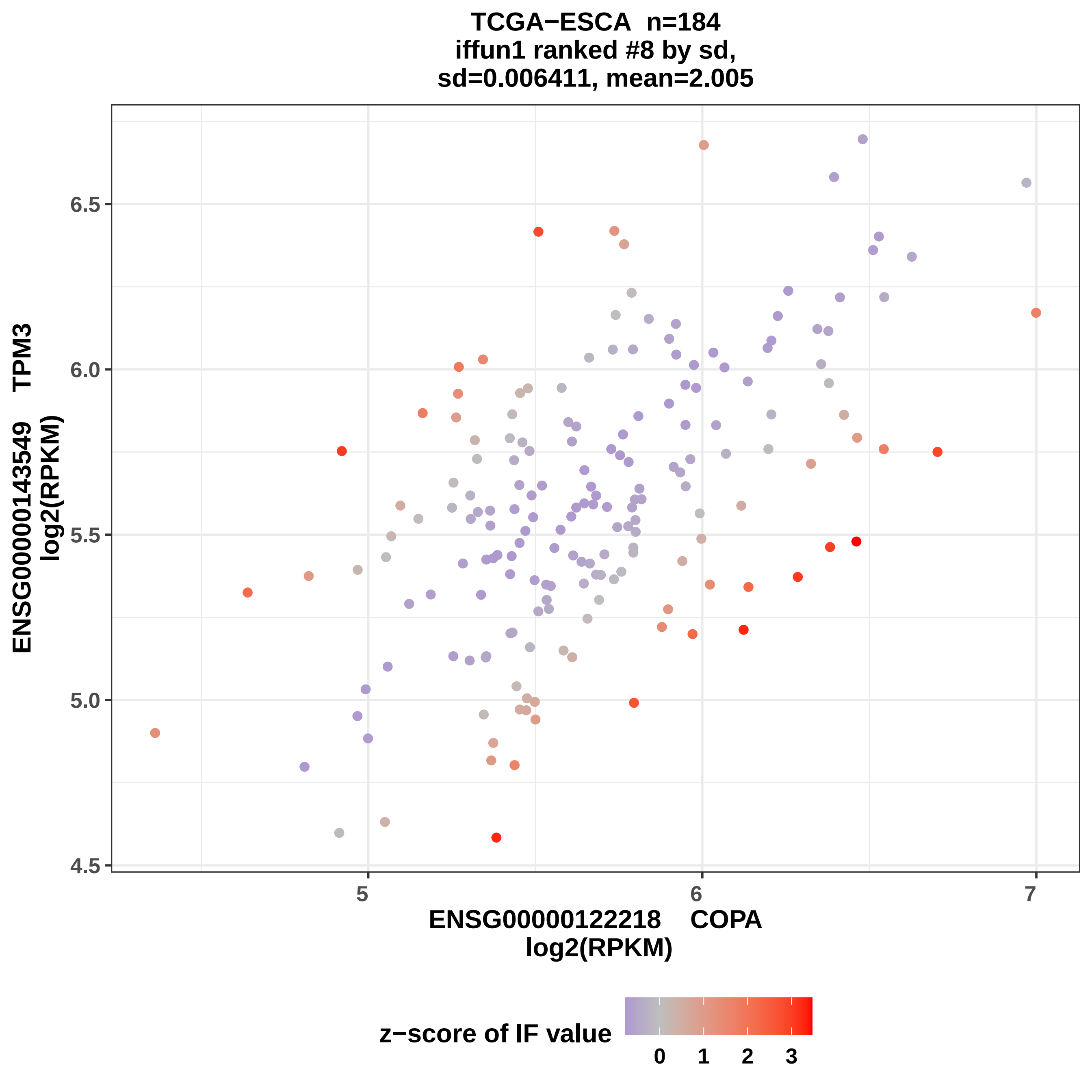

TCGA−ESCA n=184
iffun1 ranked #8 by sd,
sd=0.006411, mean=2.005
ENSG00000143549 TPM3
log2(RPKM)
6.5
6.0
5.5
5.0
4.5
5
6
7
ENSG00000122218 COPA
log2(RPKM)
z−score of IF value
0
1
2
3

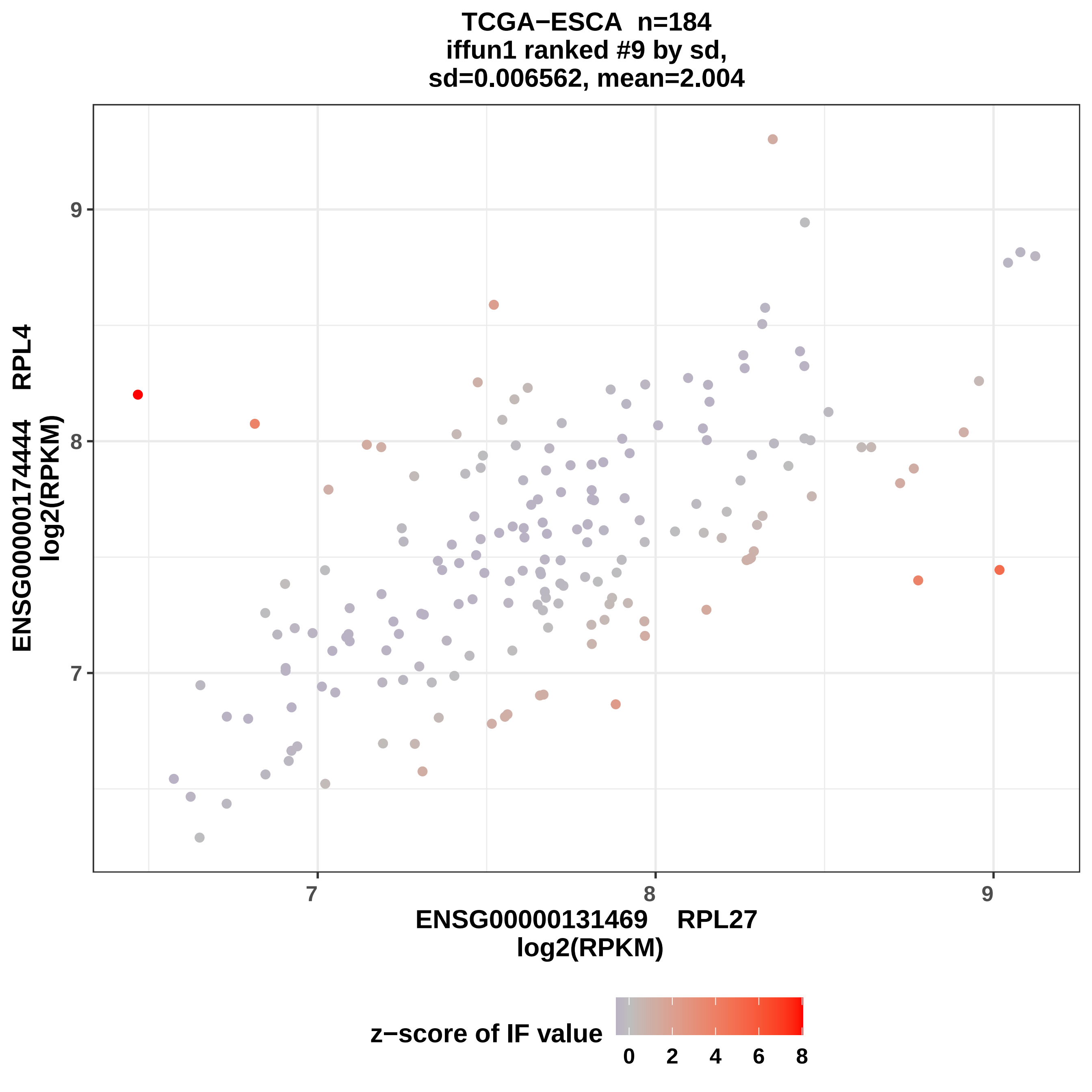
TCGA−ESCA n=184
iffun1 ranked #9 by sd,
sd=0.006562, mean=2.004
ENSG00000174444 RPL4
log2(RPKM)
9
8
7
7
8
9
ENSG00000131469 RPL27
log2(RPKM)
z−score of IF value
0
2
4
6
8

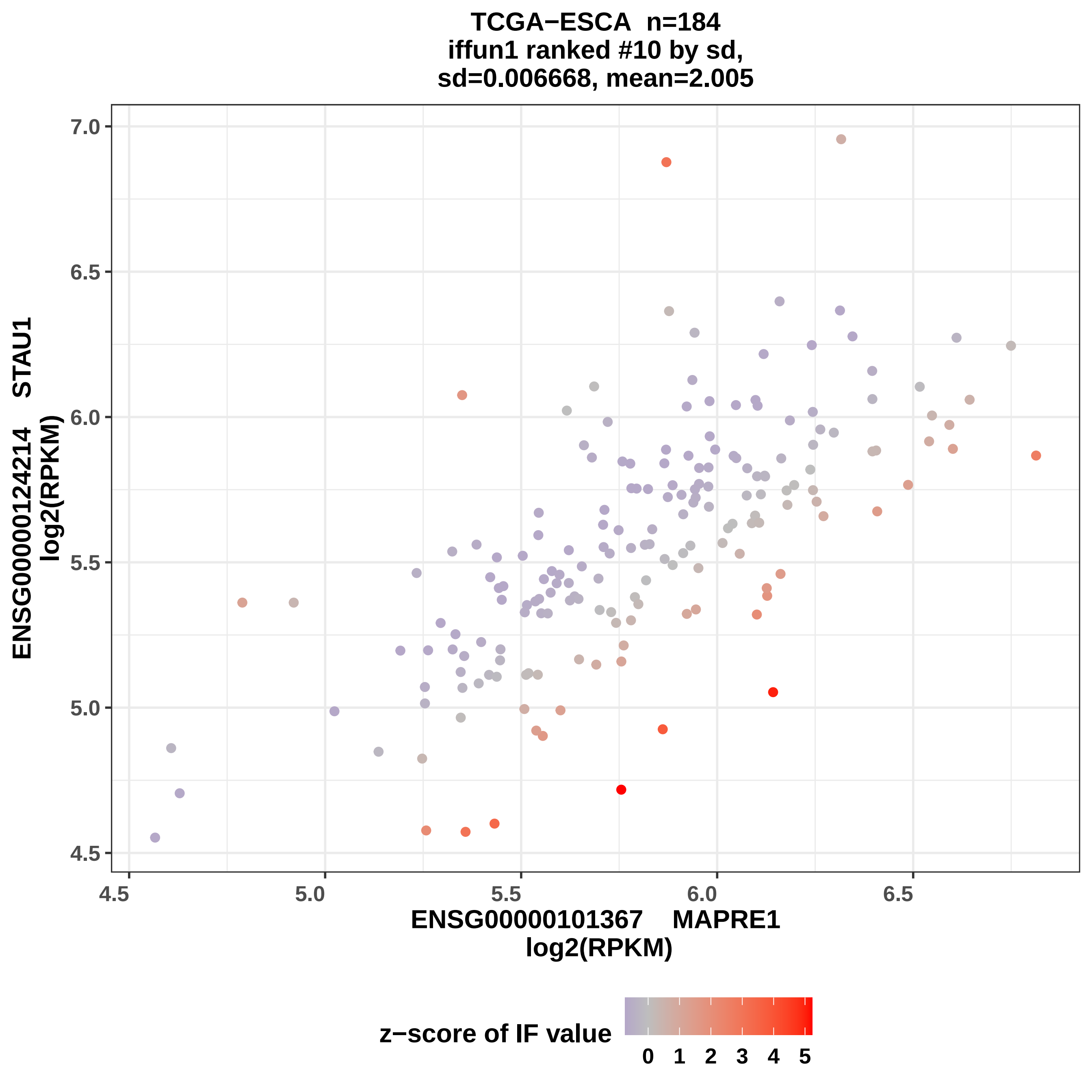

TCGA−ESCA n=184
iffun1 ranked #10 by sd,
sd=0.006668, mean=2.005
ENSG00000124214 STAU1
log2(RPKM)
7.0
6.5
6.0
5.5
5.0
4.5
4.5
5.0
5.5
6.0
6.5
ENSG00000101367 MAPRE1
log2(RPKM)
z−score of IF value
0 1 2 3 4 5

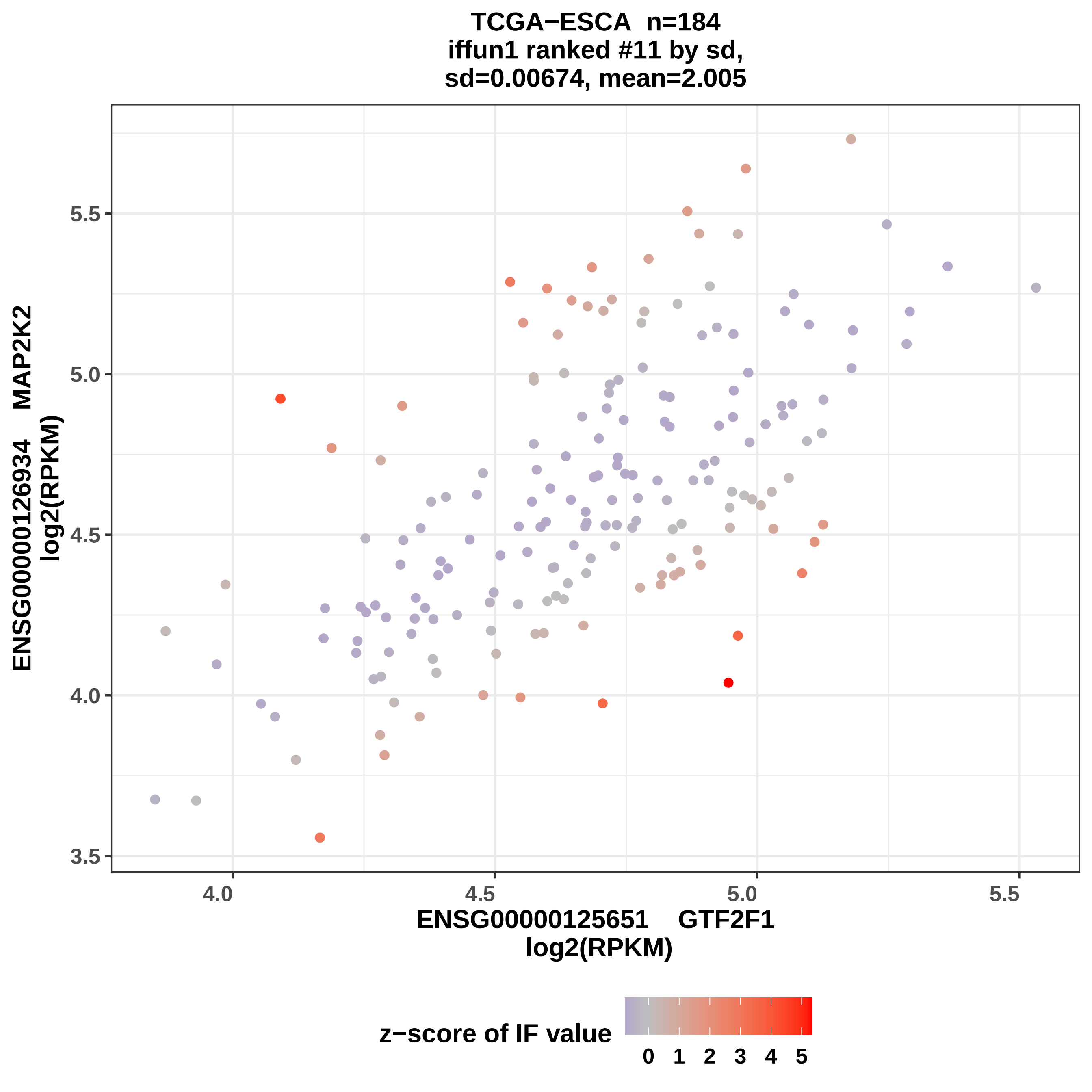
TCGA−ESCA n=184
iffun1 ranked #11 by sd,
sd=0.00674, mean=2.005
ENSG00000126934 MAP2K2
log2(RPKM)
5.5
5.0
4.5
4.0
3.5
4.0
4.5
5.0
5.5
ENSG0000125651 GTF2F1
log2(RPKM)
z−score of IF value
0 1 2 3 4 5

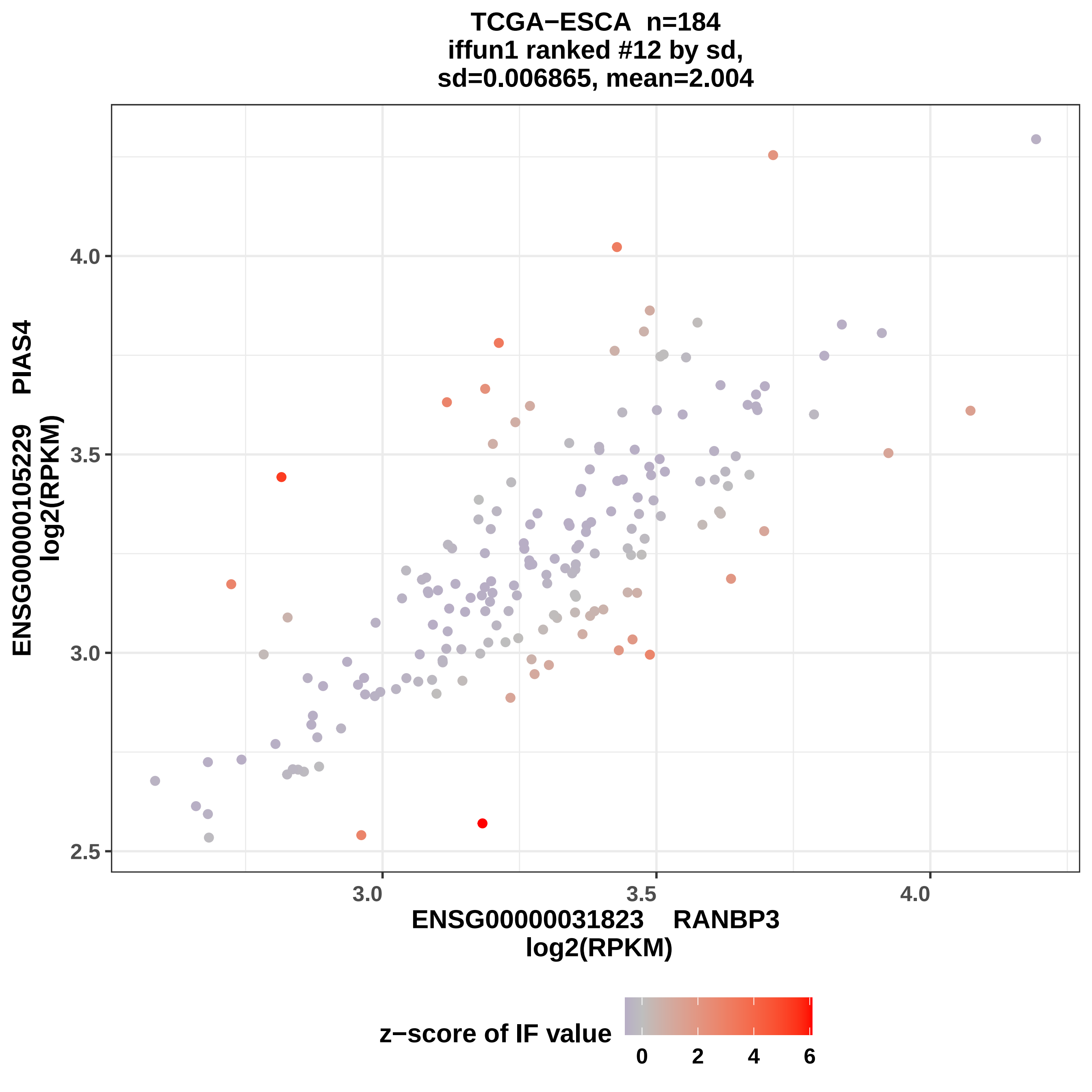
TCGA−ESCA n=184
iffun1 ranked #12 by sd,
sd=0.006865, mean=2.004
ENSG00000105229 PIAS4
log2(RPKM)
4.0
3.5
3.0
2.5
3.0
3.5
4.0
ENSG00000031823 RANBP3
log2(RPKM)
z−score of IF value
0
2
4
6

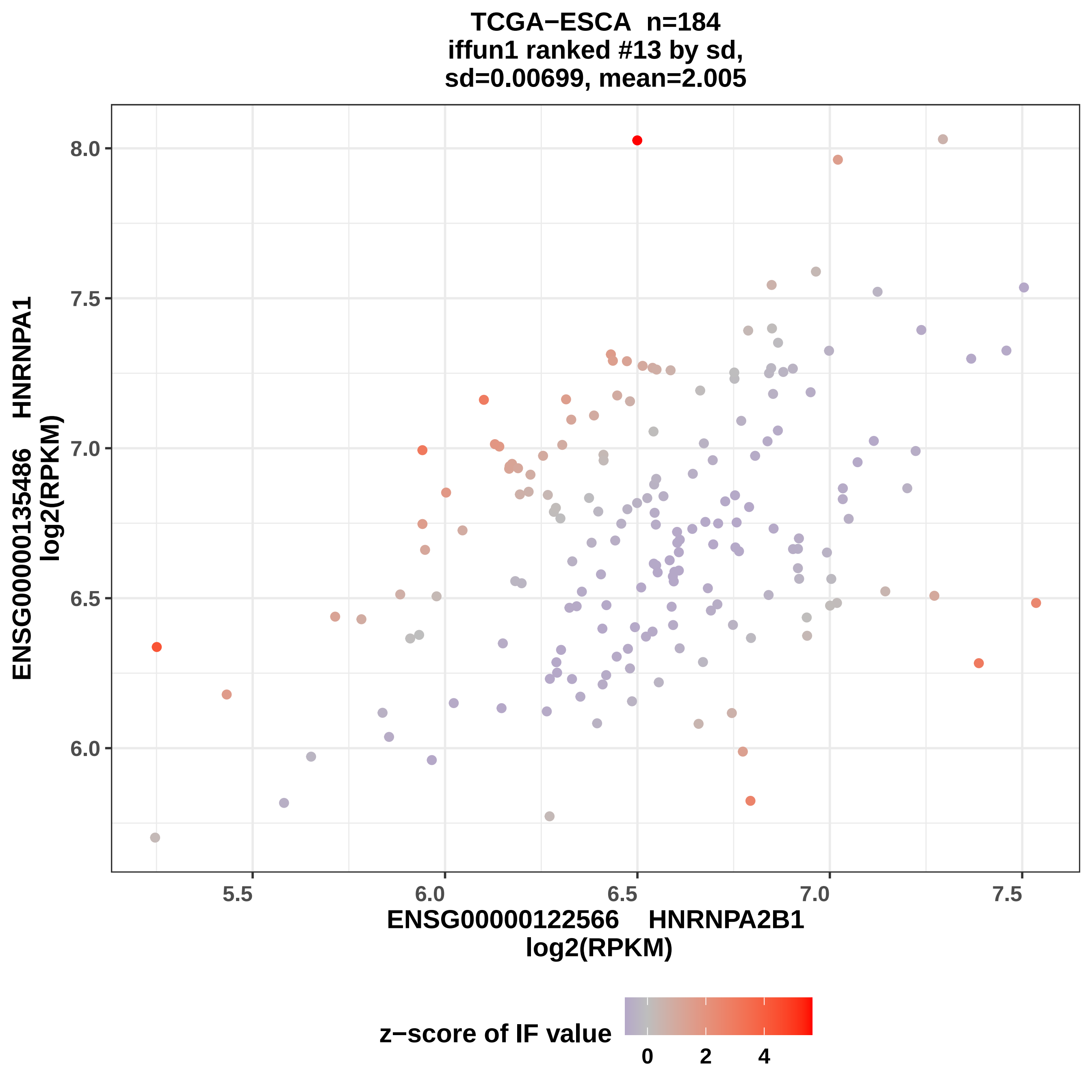
TCGA−ESCA n=184
iffun1 ranked #13 by sd,
sd=0.00699, mean=2.005
ENSG00000135486 HNRNPA1
log2(RPKM)
8.0
7.5
7.0
6.5
6.0
5.5
6.0
6.5
7.0
7.5
ENSG00000122566 HNRNPA2B1
log2(RPKM)
z−score of IF value
0
2
4

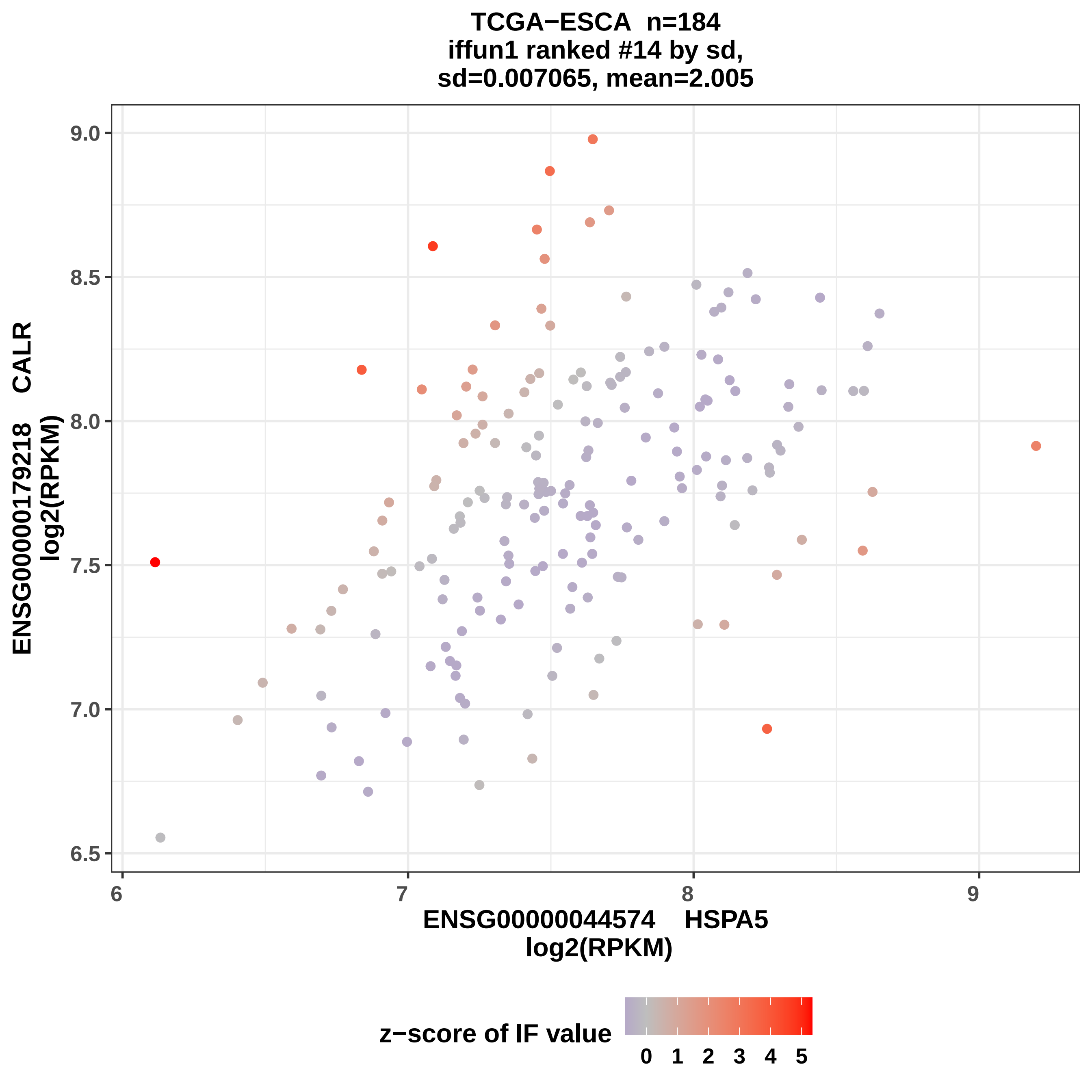
TCGA−ESCA n=184
iffun1 ranked #14 by sd,
sd=0.007065, mean=2.005
ENSG00000179218 CALR
log2(RPKM)
9.0
8.5
8.0
7.5
7.0
6.5
6
7
8
9
ENSG00000044574 HSPA5
log2(RPKM)
z−score of IF value
0 1 2 3 4 5

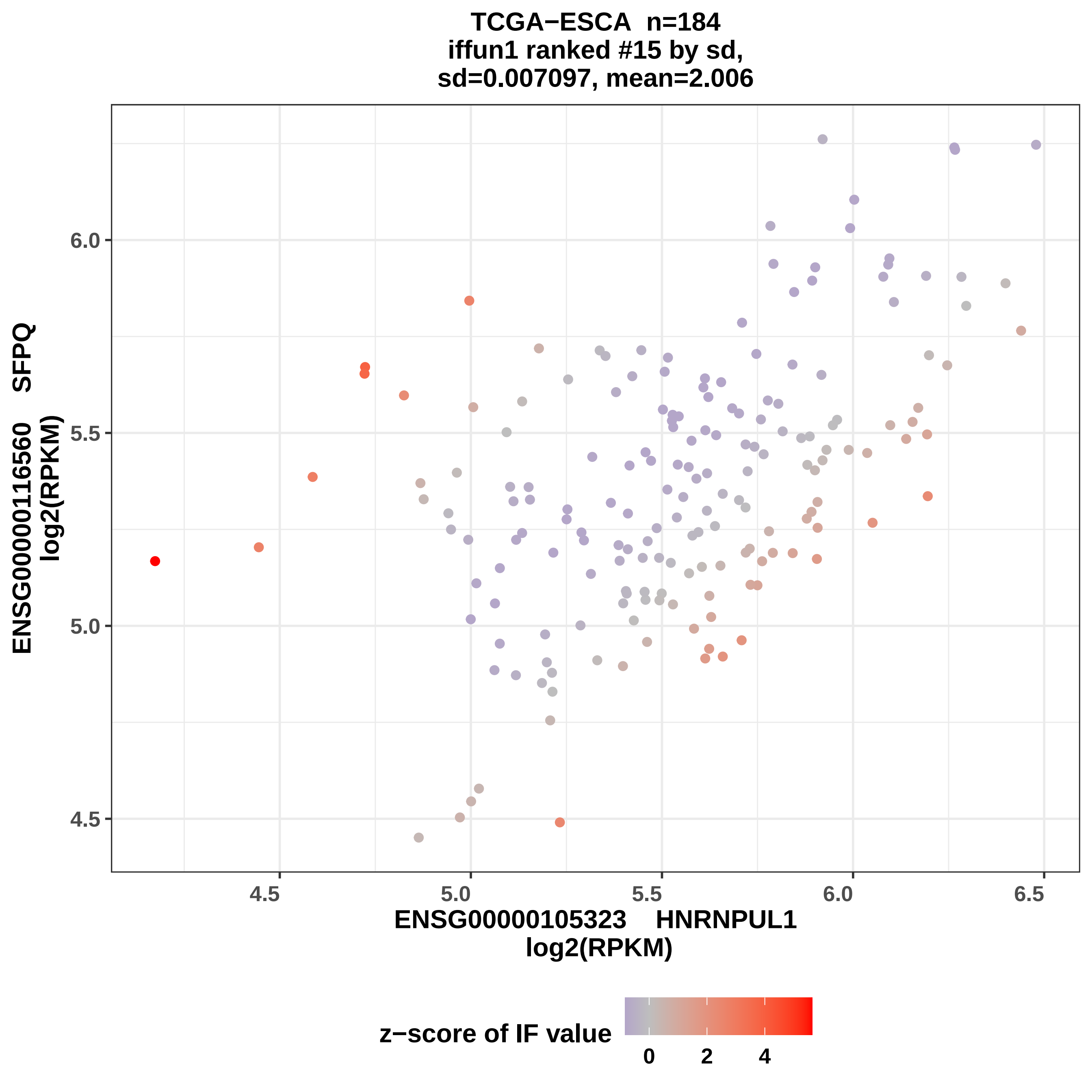
TCGA−ESCA n=184
iffun1 ranked #15 by sd,
sd=0.007097, mean=2.006
ENSG00000116560 SFPQ
log2(RPKM)
6.0
5.5
5.0
4.5
4.5
5.0
5.5
6.0
6.5
ENSG00000105323 HNRNPUL1
log2(RPKM)
z−score of IF value
0
2
4

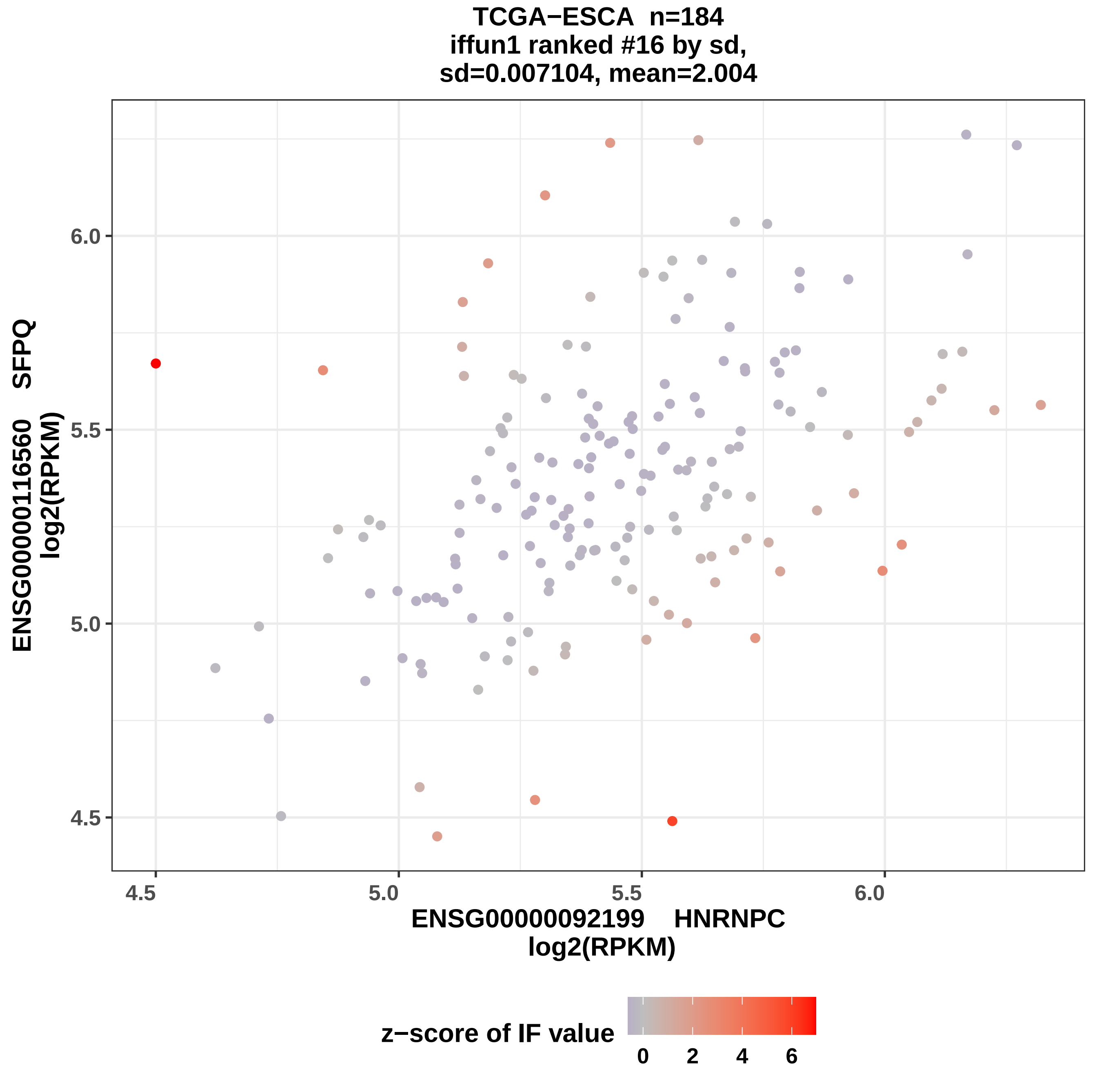

TCGA−ESCA n=184
iffun1 ranked #16 by sd,
sd=0.007104, mean=2.004
ENSG00000116560 SFPQ
log2(RPKM)
6.0
5.5
5.0
4.5
4.5
5.0
5.5
6.0
ENSG00000092199 HNRNPC
log2(RPKM)
z−score of IF value
0
2
4
6

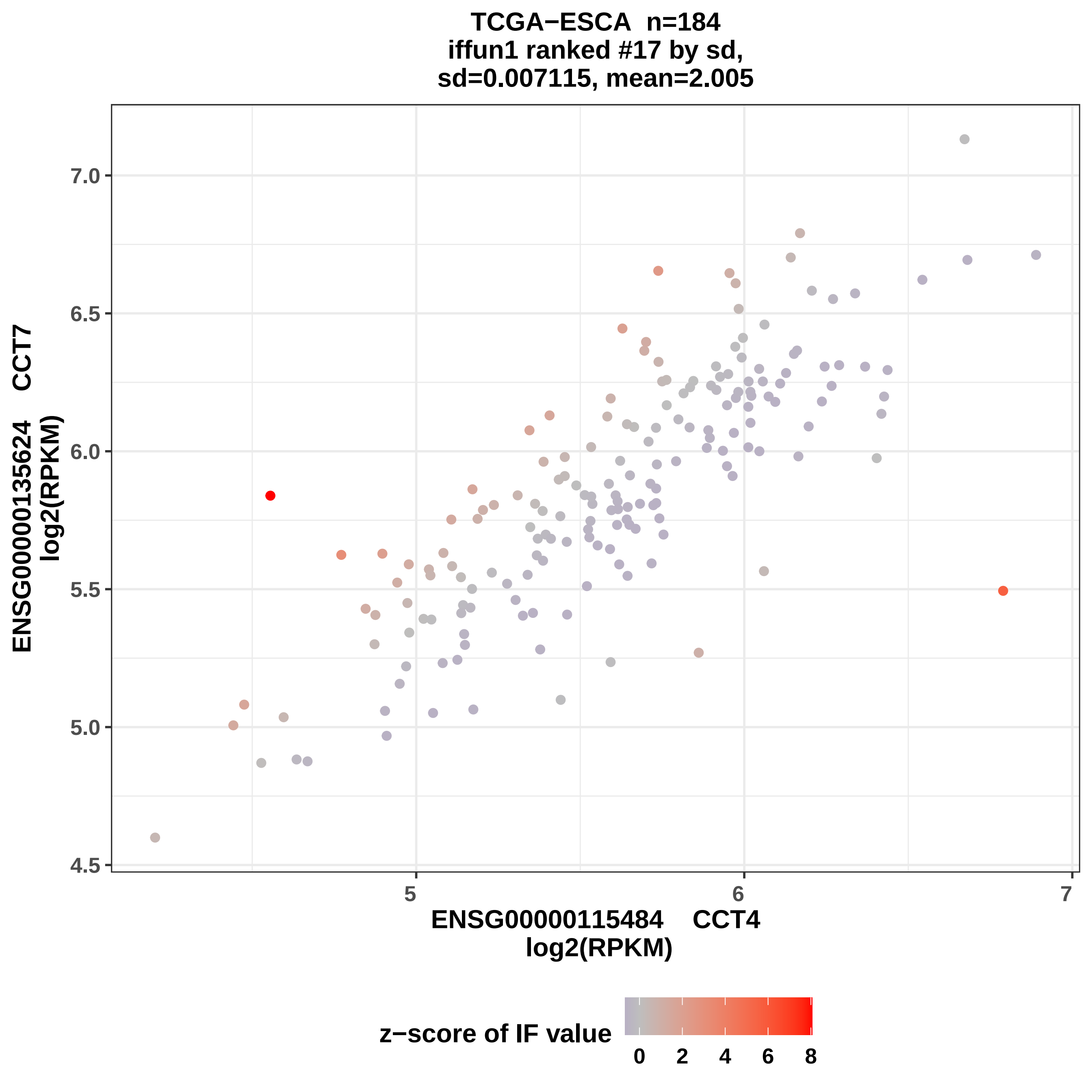

TCGA−ESCA n=184
iffun1 ranked #17 by sd,
sd=0.007115, mean=2.005
ENSG00000135624 CCT7
log2(RPKM)
7.0
6.5
6.0
5.5
5.0
4.5
5
6
7
ENSG00000115484 CCT4
log2(RPKM)
z−score of IF value
0
2
4
6
8

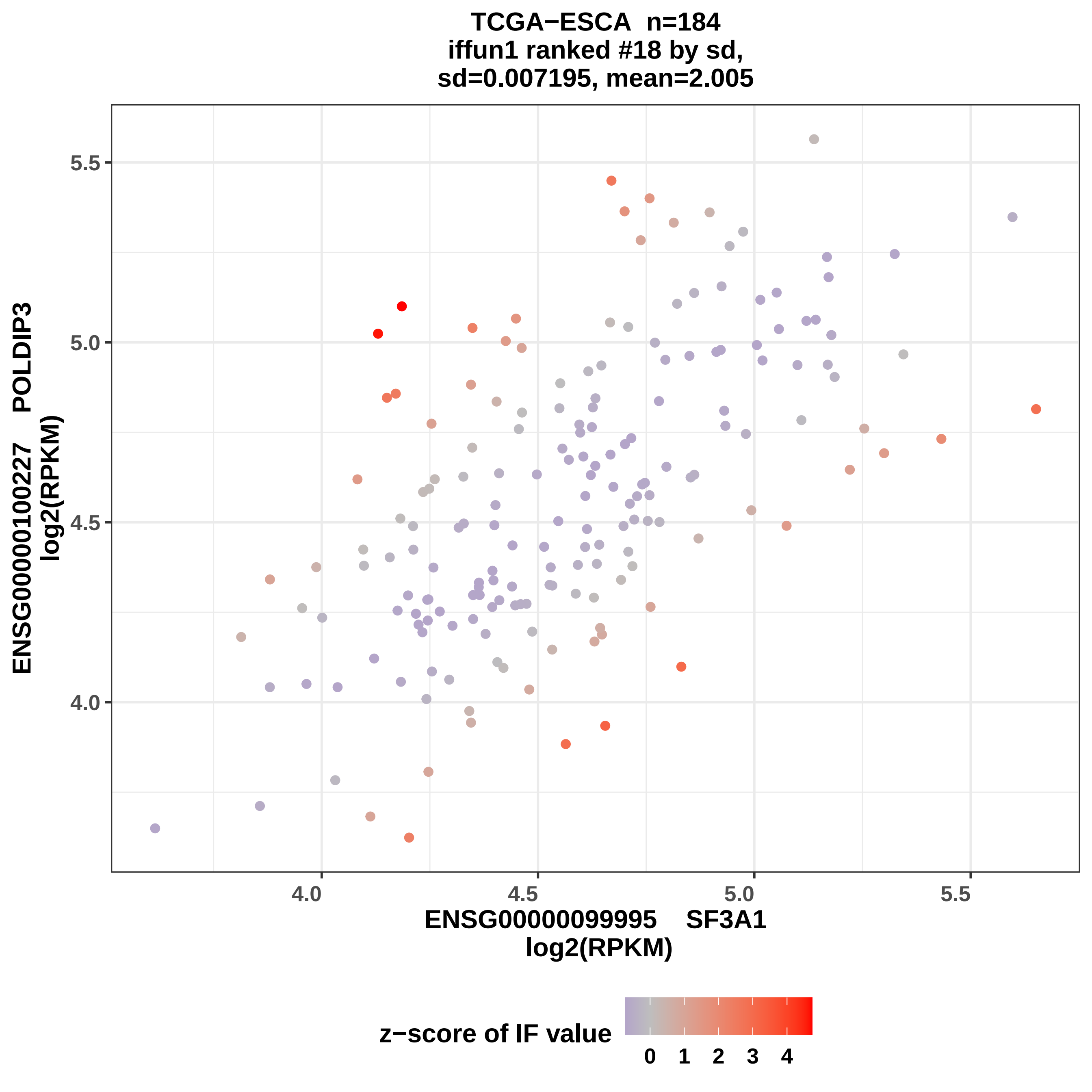
TCGA−ESCA n=184
iffun1 ranked #18 by sd,
sd=0.007195, mean=2.005
ENSG00000100227 POLDIP3
log2(RPKM)
5.5
5.0
4.5
4.0
4.0
4.5
5.0
5.5
ENSG00000099995 SF3A1
log2(RPKM)
z−score of IF value
0
1
2
3
4

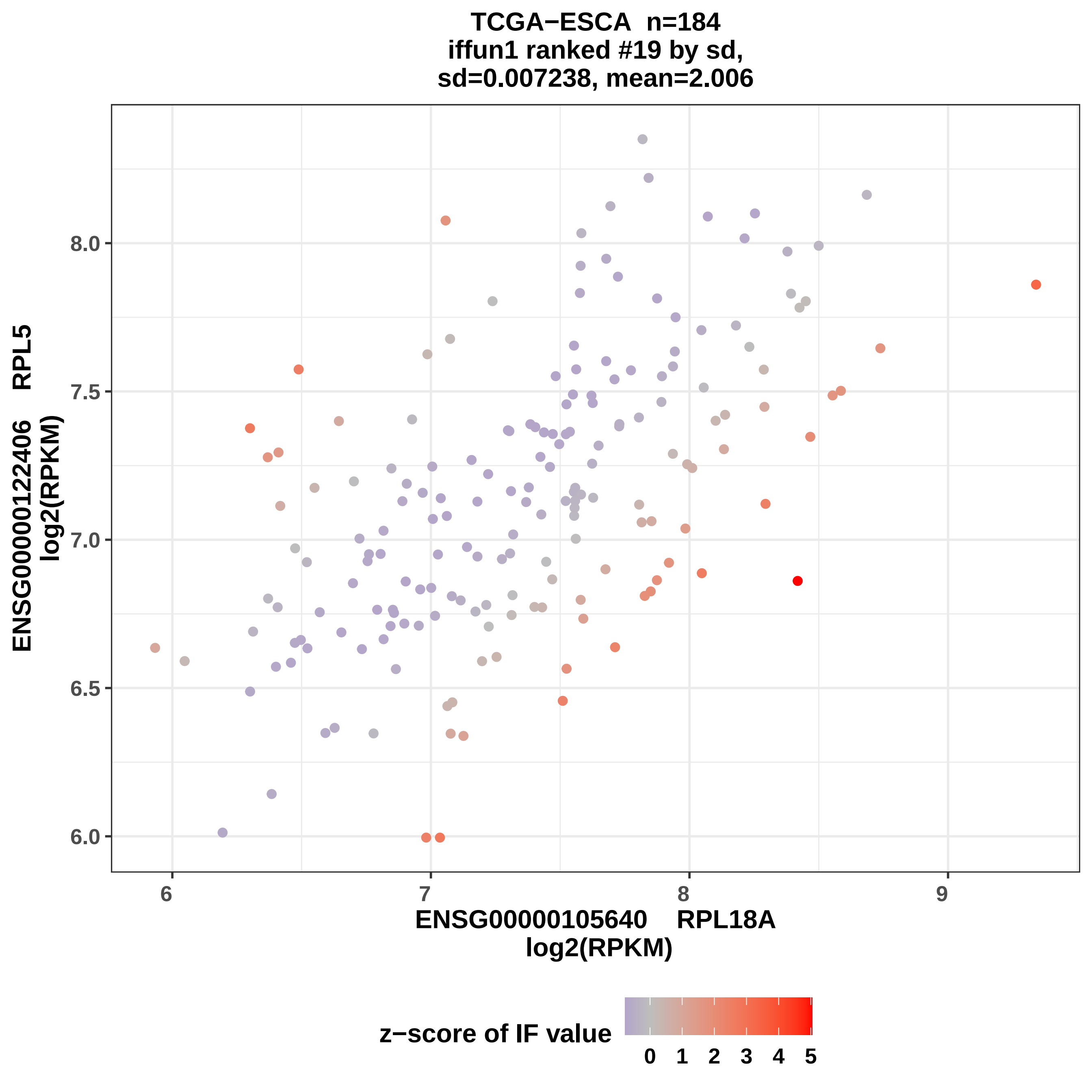

TCGA−ESCA n=184
iffun1 ranked #19 by sd,
sd=0.007238, mean=2.006
ENSG00000122406 RPL5
log2(RPKM)
8.0
7.5
7.0
6.5
6.0
6
7
8
9
ENSG00000105640 RPL18A
log2(RPKM)
z−score of IF value
0
1
2
3
4
5

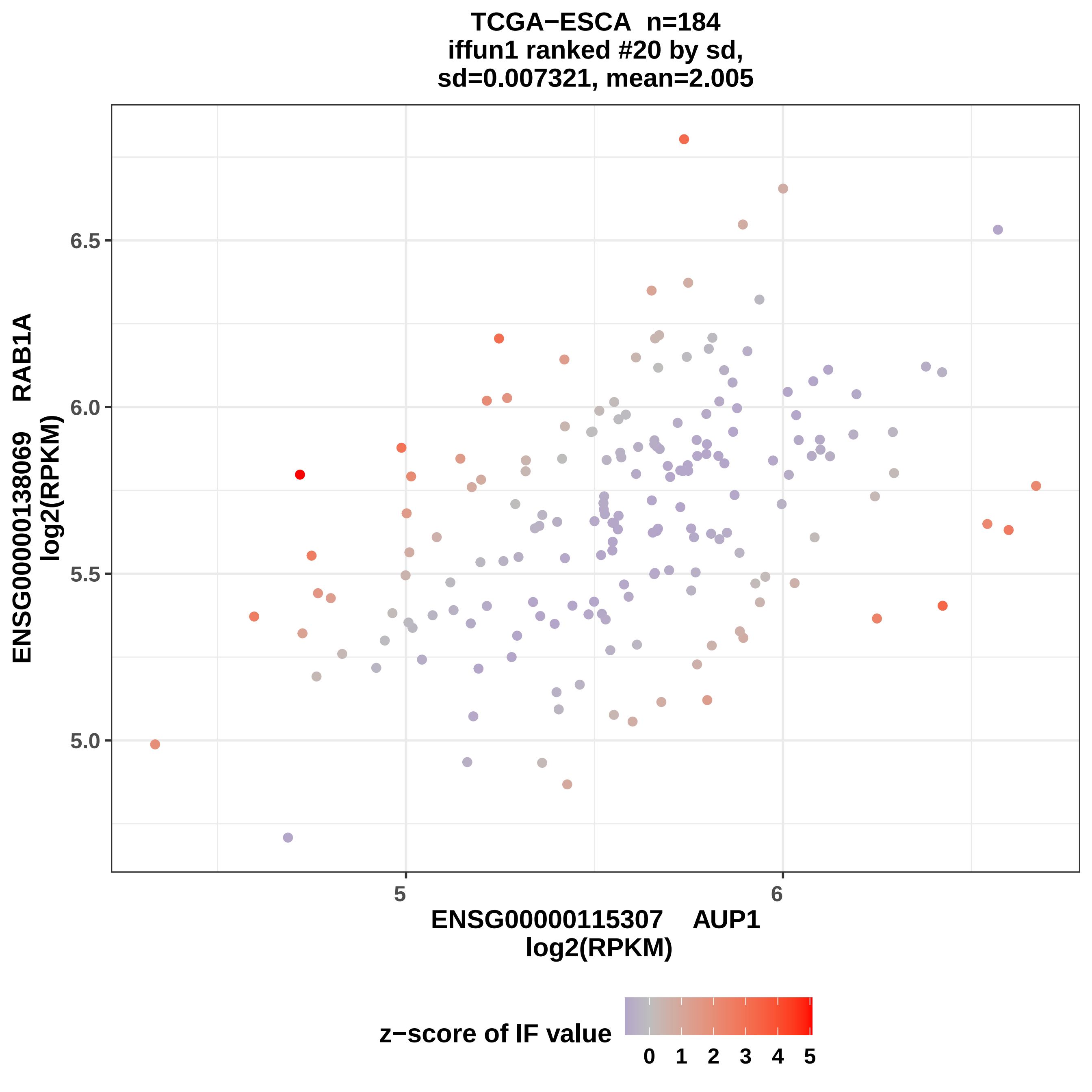
TCGA−ESCA n=184
iffun1 ranked #20 by sd,
sd=0.007321, mean=2.005
ENSG00000138069 RAB1A
log2(RPKM)
6.5
6.0
5.5
5.0
5
6
ENSG00000115307 AUP1
log2(RPKM)
z−score of IF value
0 1 2 3 4 5

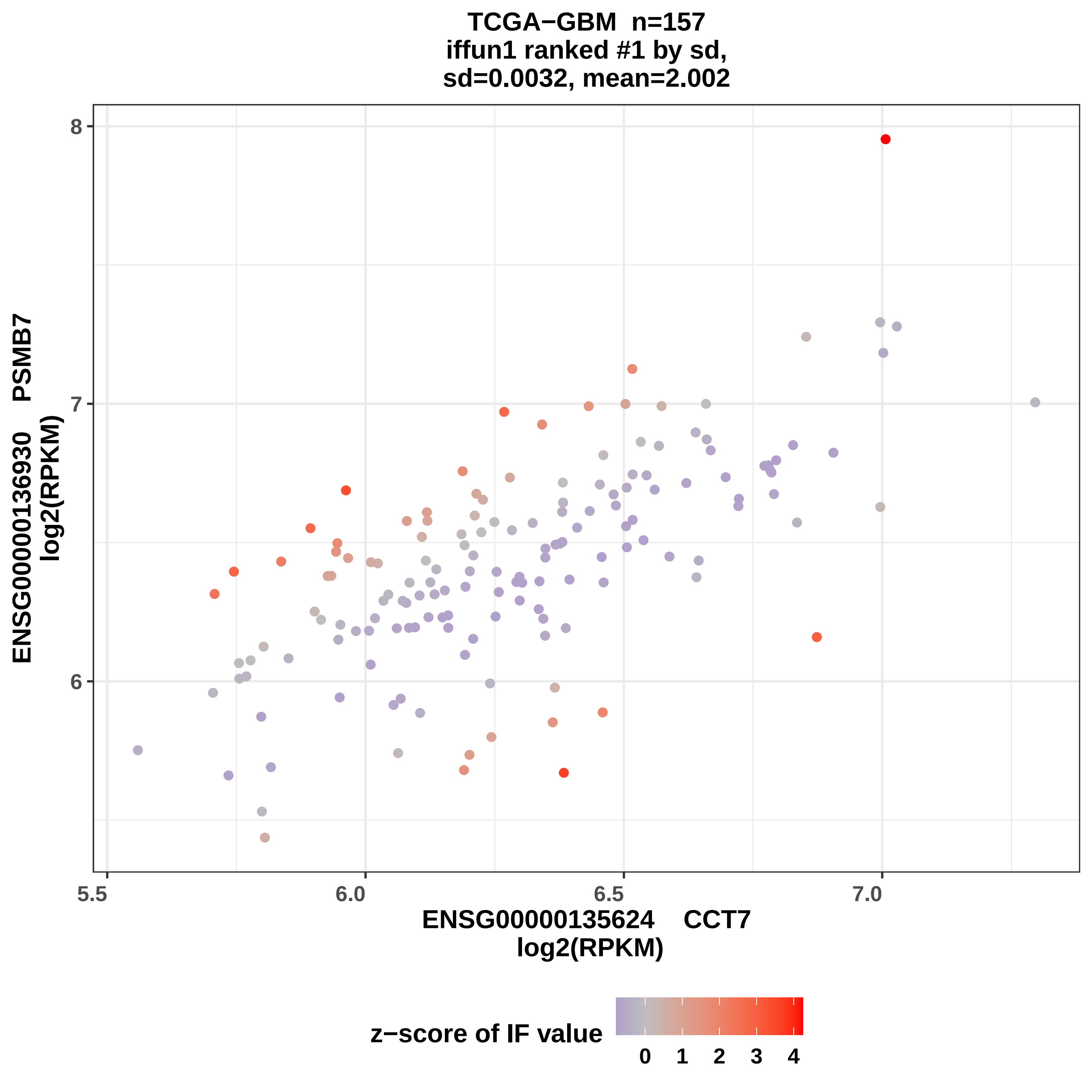

TCGA−GBM n=157
iffun1 ranked #1 by sd,
sd=0.0032, mean=2.002
ENSG00000136930 PSMB7
log2(RPKM)
8
7
6
5.5
6.0
6.5
7.0
ENSG00000135624 CCT7
log2(RPKM)
z−score of IF value
0 1 2 3 4

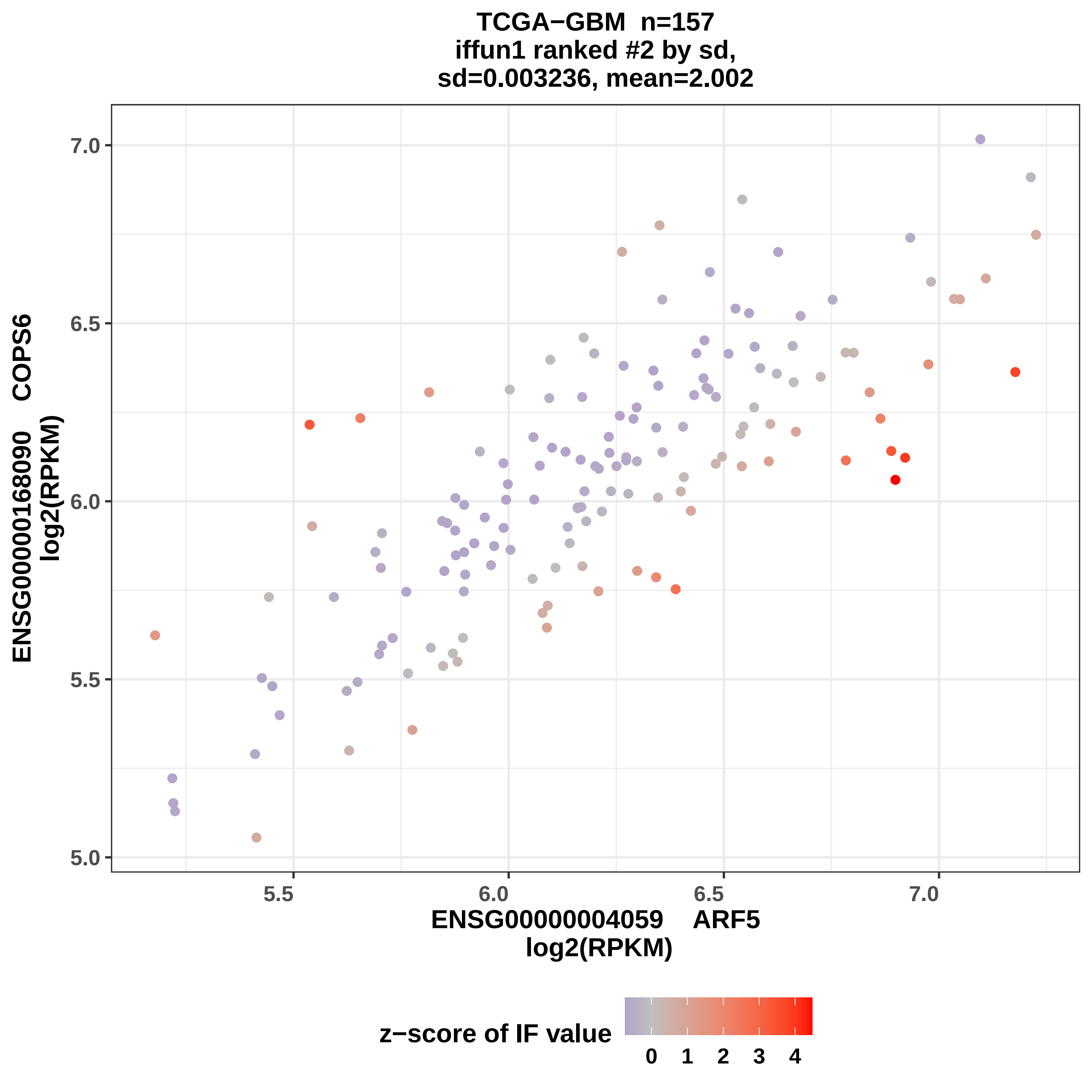
TCGA−GBM n=157
iffun1 ranked #2 by sd,
sd=0.003236, mean=2.002
ENSG00000168090 COPS6
log2(RPKM)
7.0
6.5
6.0
5.5
5.0
5.5
6.0
6.5
7.0
ENSG00000004059 ARF5
log2(RPKM)
z−score of IF value
0 1 2 3 4

TCGA−GBM n=157
iffun1 ranked #3 by sd,
sd=0.003398, mean=2.002

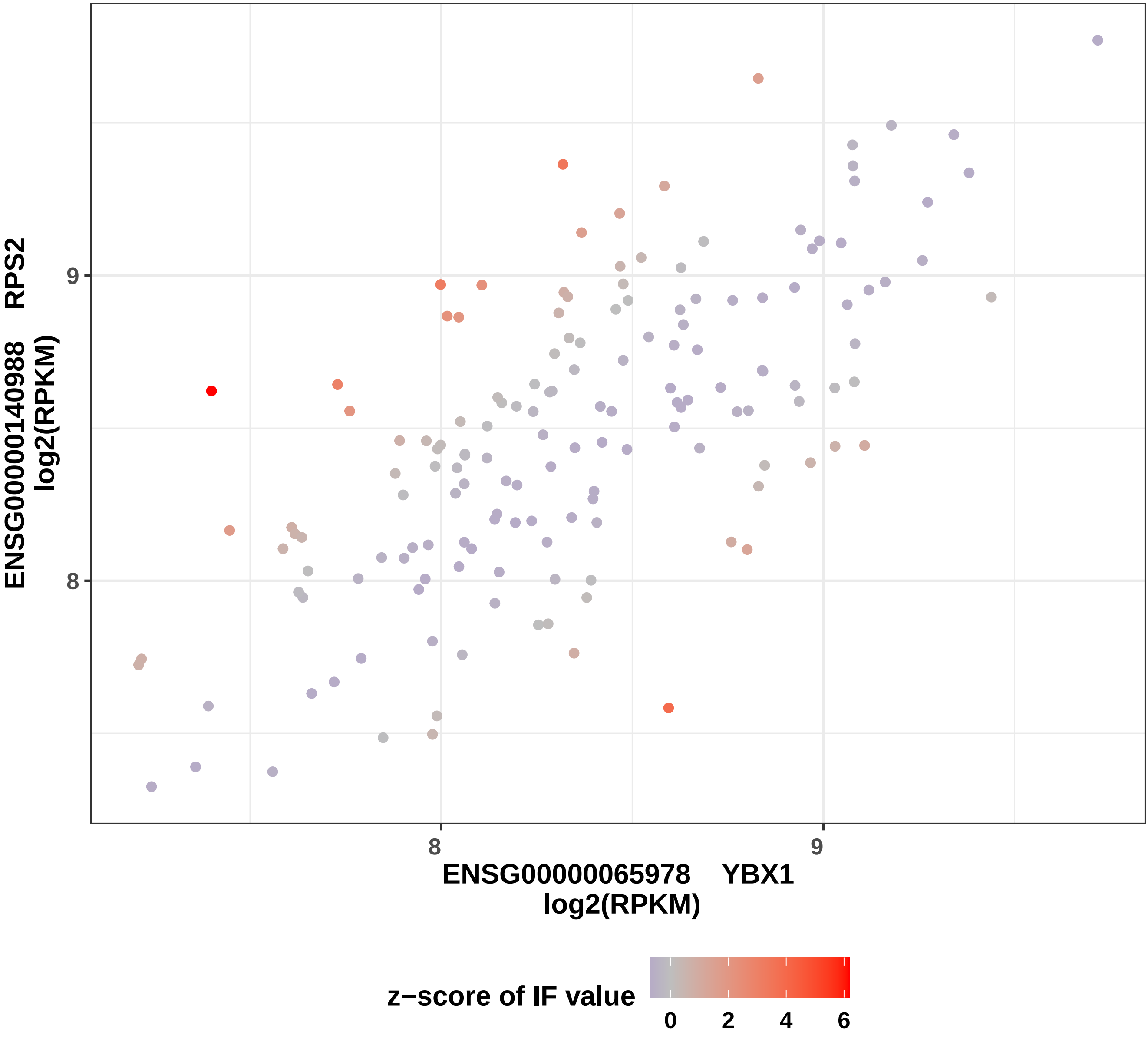

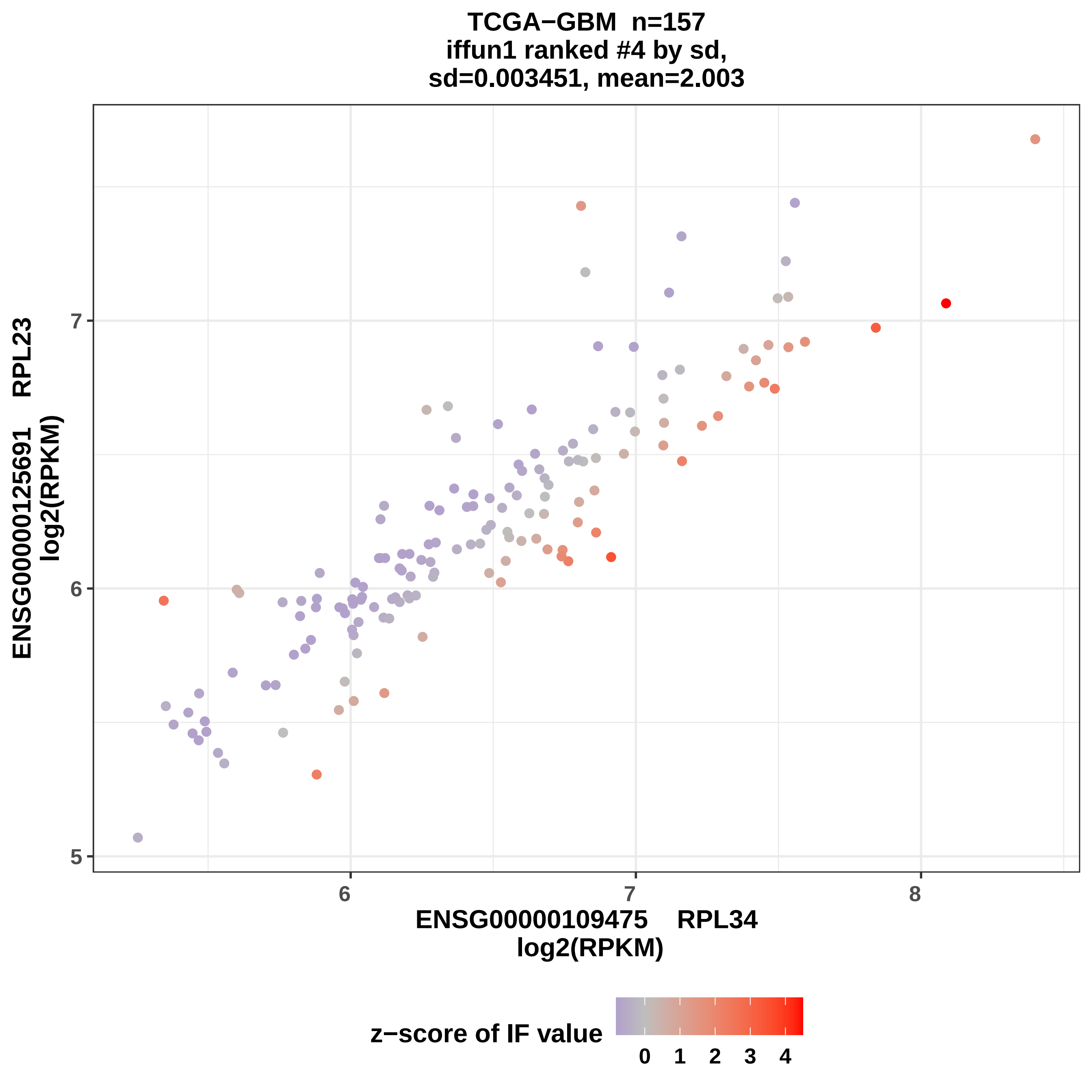
TCGA−GBM n=157
iffun1 ranked #4 by sd,
sd=0.003451, mean=2.003
ENSG00000125691 RPL23
log2(RPKM)
ENSG00000109475 RPL34
log2(RPKM)
5
6
7
8
z−score of IF value
0
1
2
3
4

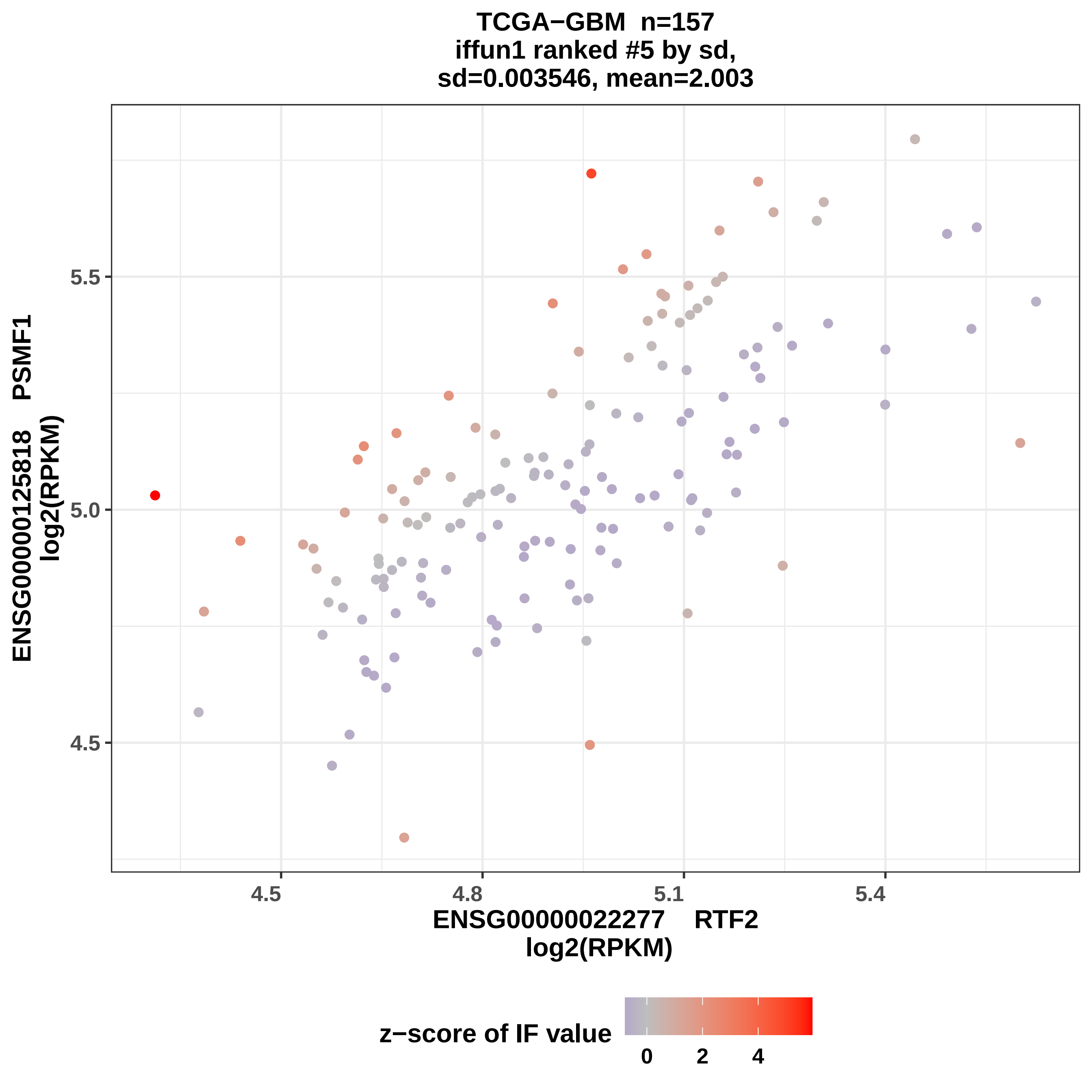
TCGA−GBM n=157
iffun1 ranked #5 by sd,
sd=0.003546, mean=2.003
ENSG00000125818 PSMF1
log2(RPKM)
5.5
5.0
4.5
4.5
4.8
5.1
5.4
ENSG00000022277 RTF2
log2(RPKM)
z−score of IF value
0
2
4

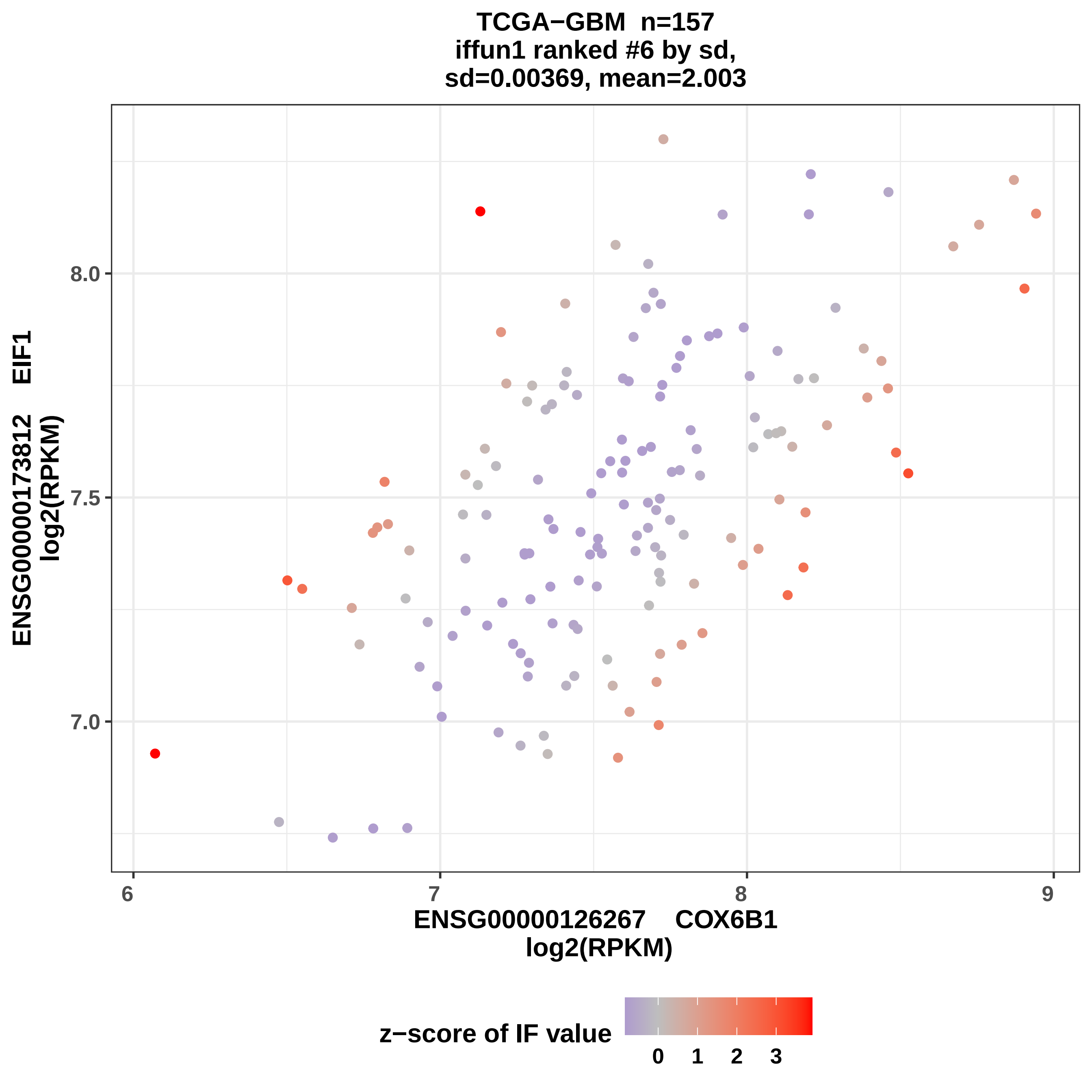
TCGA−GBM n=157
iffun1 ranked #6 by sd,
sd=0.00369, mean=2.003
ENSG00000173812 EIF1
log2(RPKM)
8.0
7.5
7.0
6
7
8
9
ENSG00000126267 COX6B1
log2(RPKM)
z−score of IF value
0
1
2
3

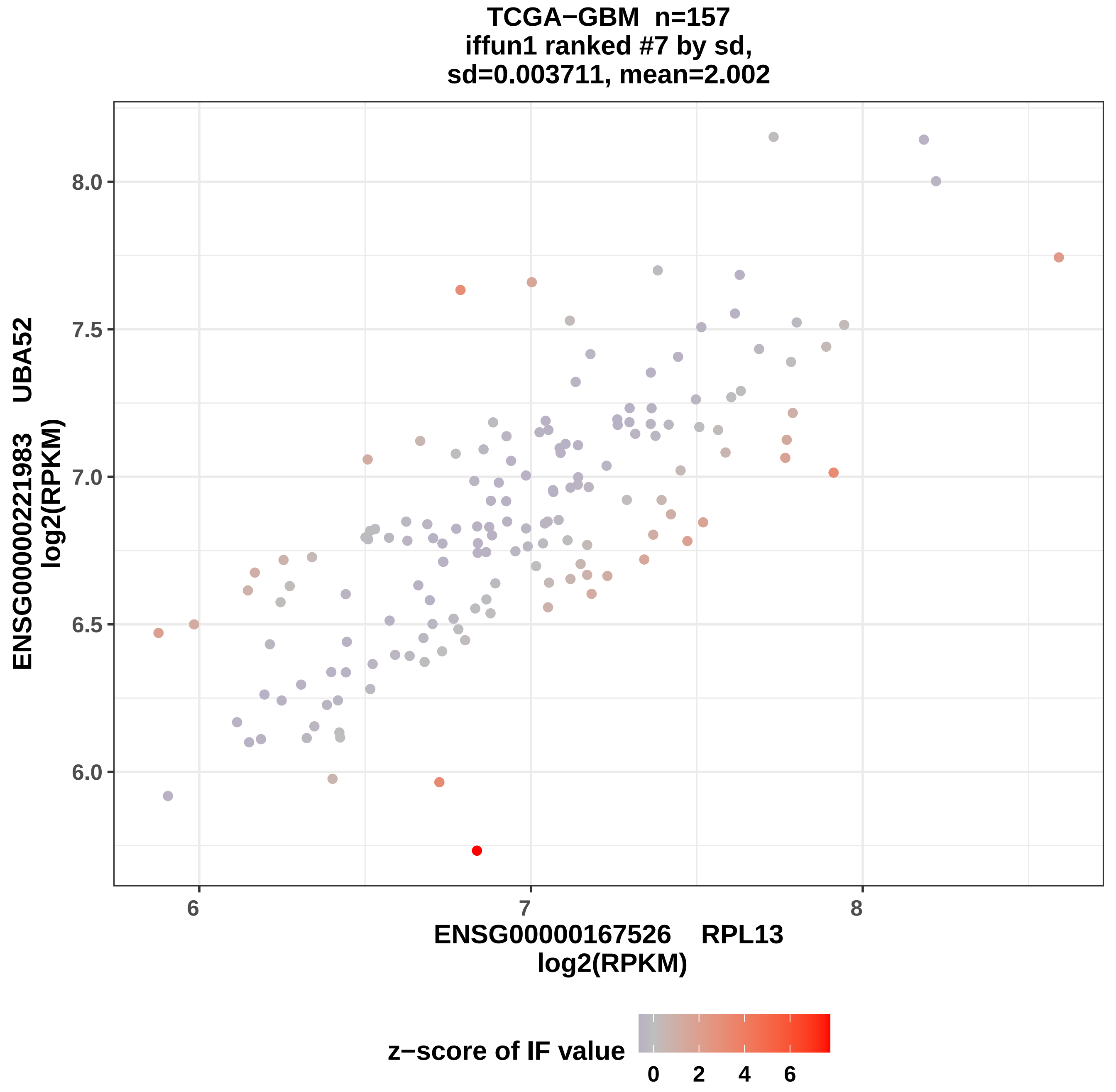

TCGA−GBM n=157
iffun1 ranked #7 by sd,
sd=0.003711, mean=2.002
ENSG00000221983 UBA52
log2(RPKM)
8.0
7.5
7.0
6.5
6.0
6
7
8
ENSG00000167526 RPL13
log2(RPKM)
z−score of IF value
0
2
4
6

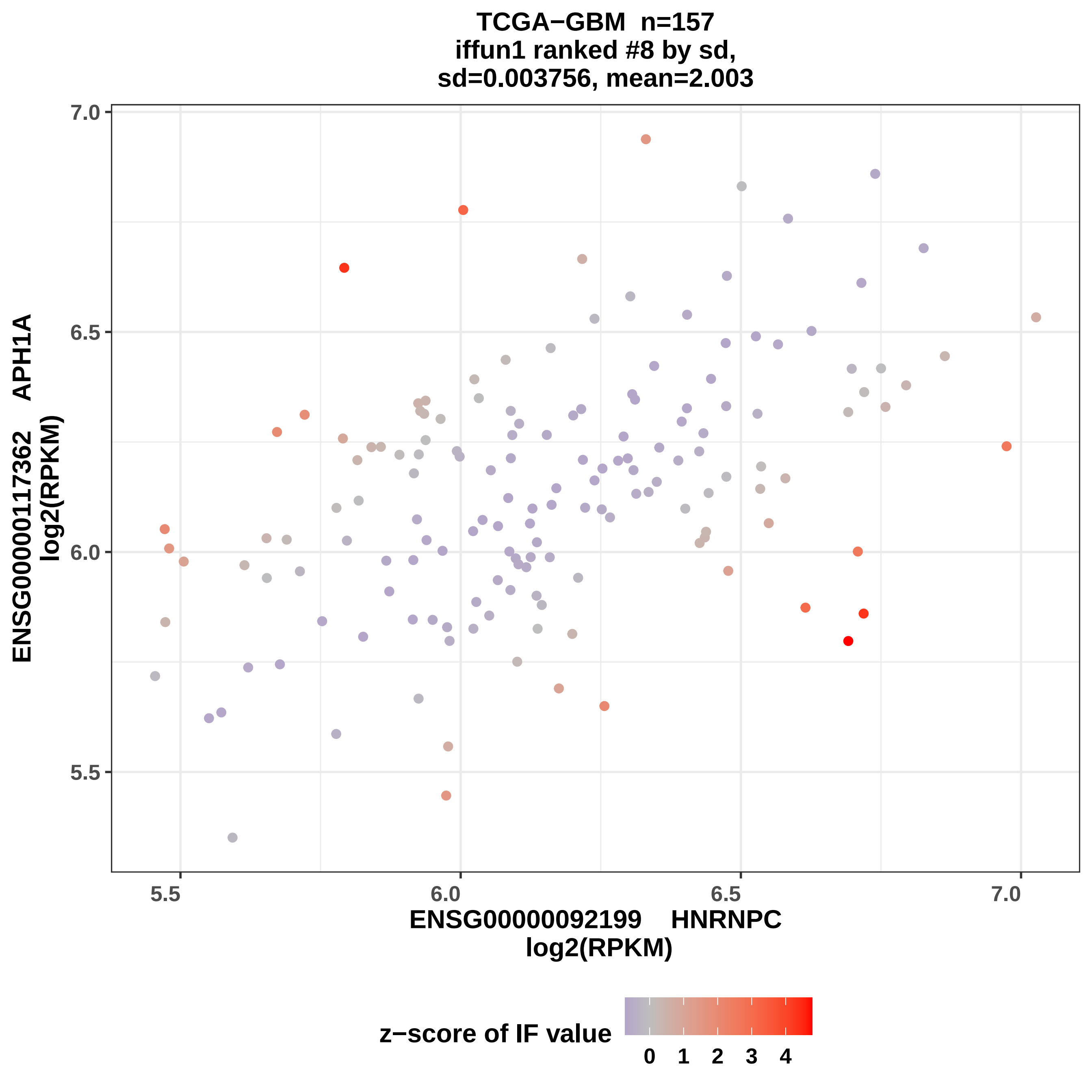
TCGA−GBM n=157
iffun1 ranked #8 by sd,
sd=0.003756, mean=2.003
ENSG00000117362 APH1A
log2(RPKM)
7.0
6.5
6.0
5.5
5.5
6.0
6.5
7.0
ENSG00000092199 HNRNPC
log2(RPKM)
z−score of IF value
0
1
2
3
4

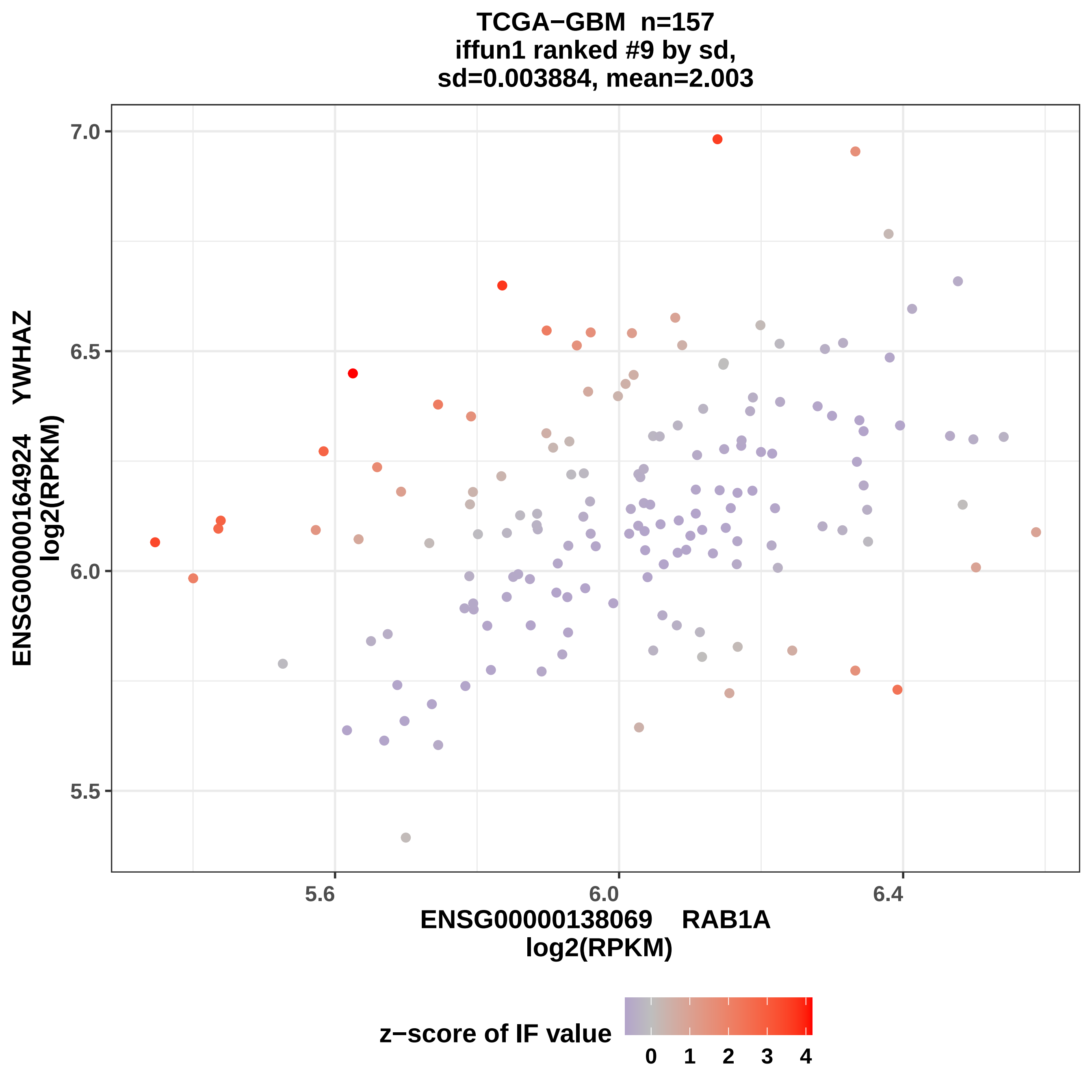
TCGA−GBM n=157
iffun1 ranked #9 by sd,
sd=0.003884, mean=2.003
ENSG00000164924 YWHAZ
log2(RPKM)
7.0
6.5
6.0
5.5
5.6
6.0
6.4
ENSG00000138069 RAB1A
log2(RPKM)
z−score of IF value
0
1
2
3
4

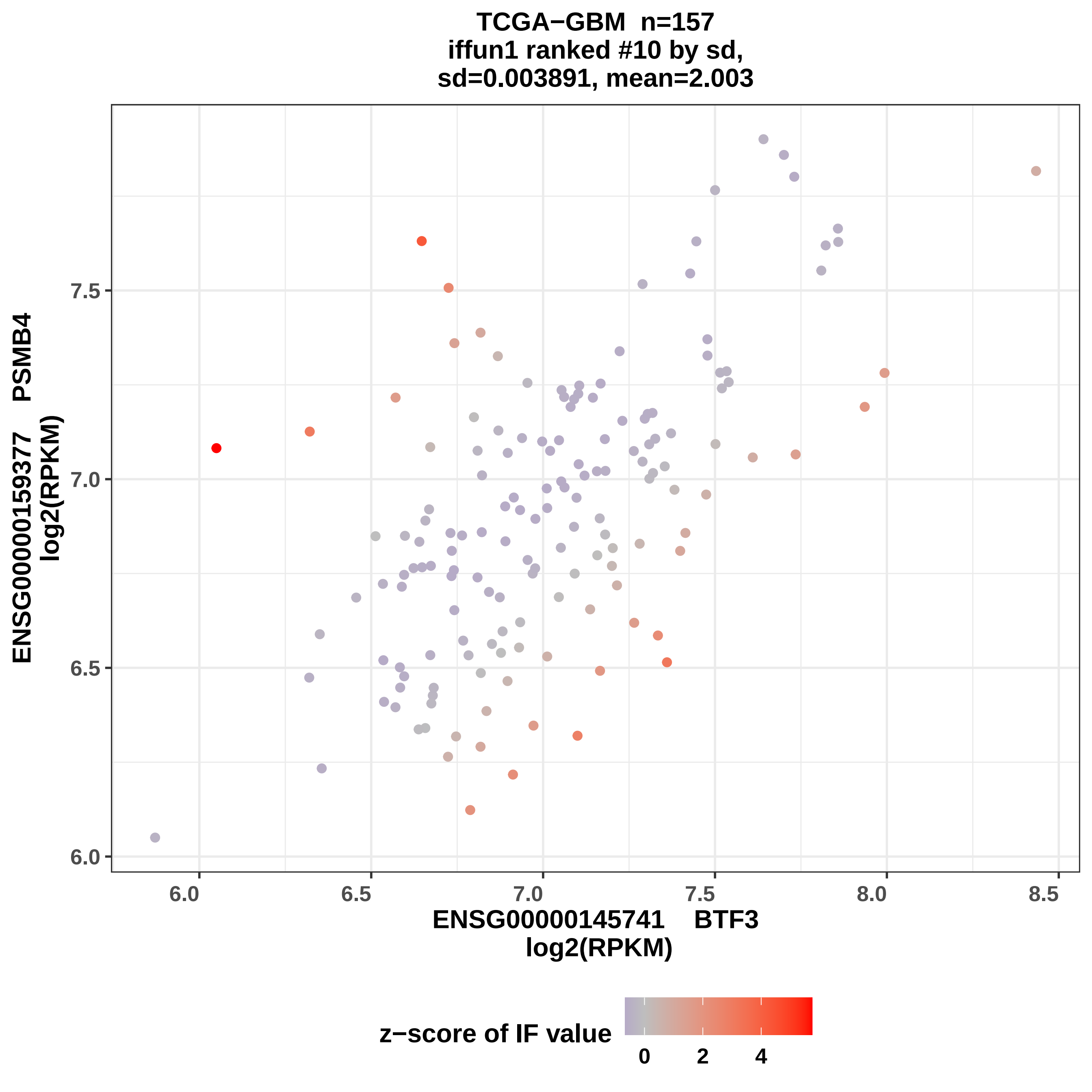
TCGA−GBM n=157
iffun1 ranked #10 by sd,
sd=0.003891, mean=2.003
ENSG00000159377 PSMB4
log2(RPKM)
7.5
7.0
6.5
6.0
6.0
6.5
7.0
7.5
8.0
8.5
ENSG00000145741 BTF3
log2(RPKM)
z−score of IF value
0
2
4

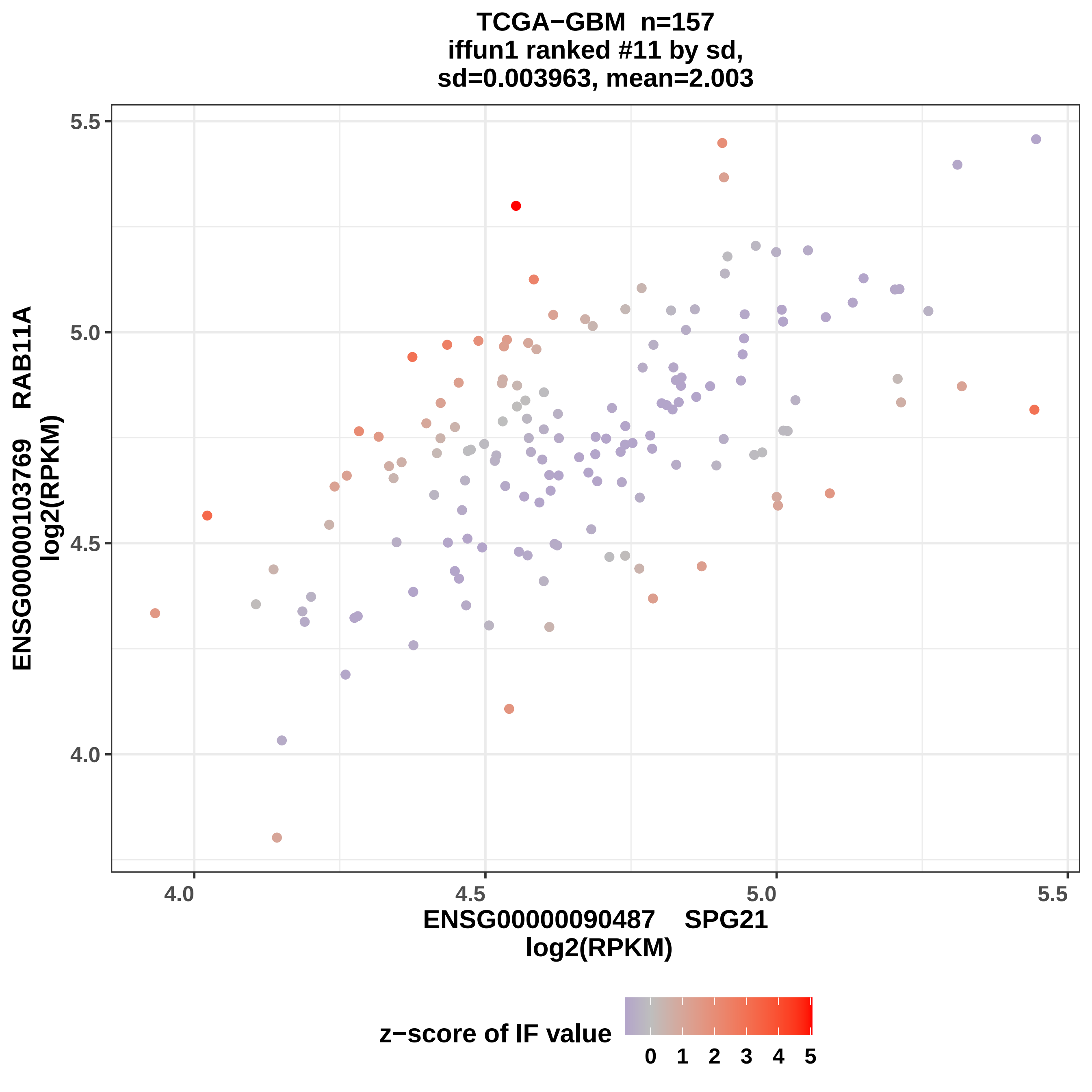
TCGA−GBM n=157
iffun1 ranked #11 by sd,
sd=0.003963, mean=2.003
ENSG00000103769 RAB11A
log2(RPKM)
5.5
5.0
4.5
4.0
4.0
4.5
5.0
5.5
ENSG00000090487 SPG21
log2(RPKM)
z−score of IF value
0 1 2 3 4 5

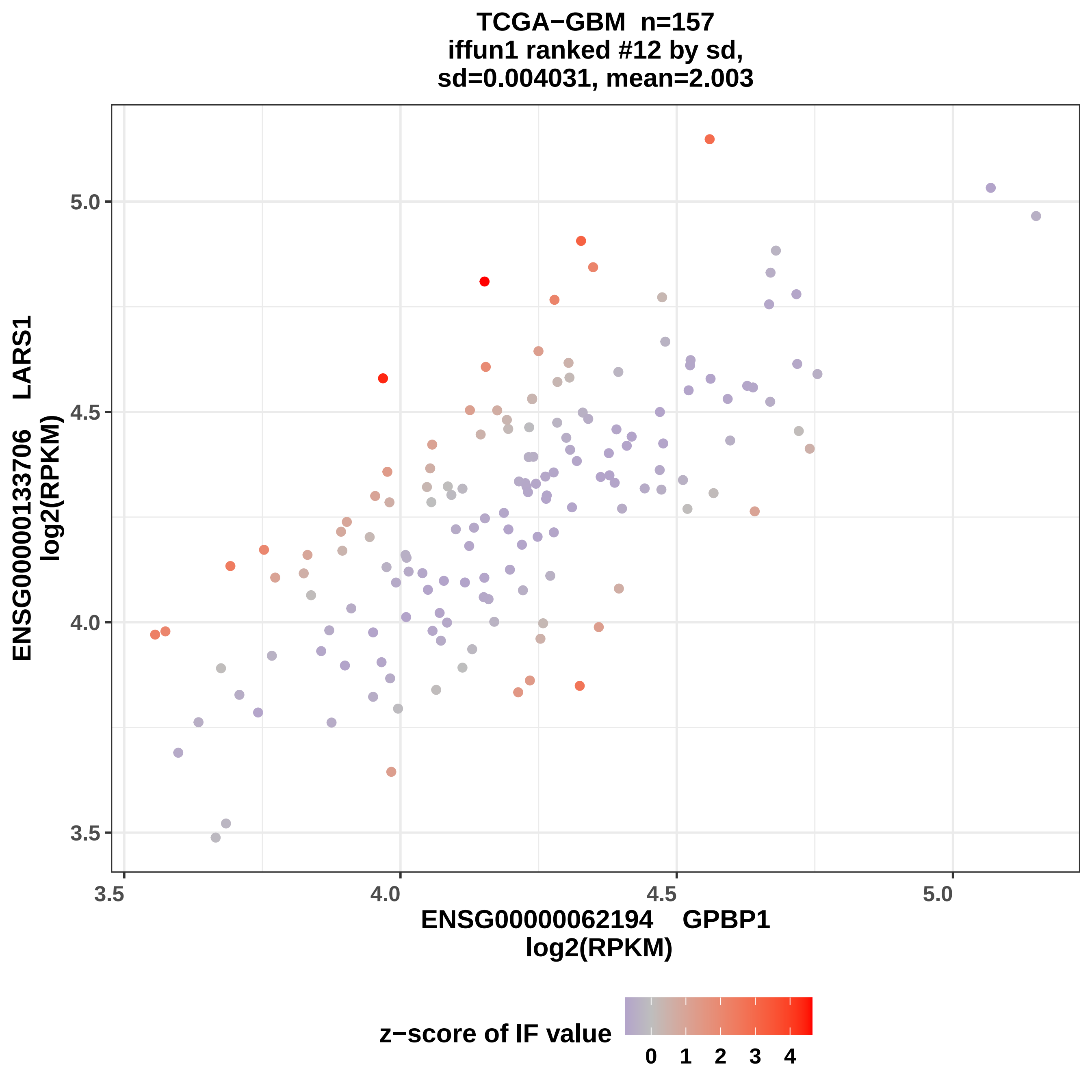
TCGA−GBM n=157
iffun1 ranked #12 by sd,
sd=0.004031, mean=2.003
ENSG00000133706 LARS1
log2(RPKM)
5.0
4.5
4.0
3.5
3.5
4.0
4.5
5.0
ENSG00000062194 GPBP1
log2(RPKM)
z−score of IF value
0
1
2
3
4

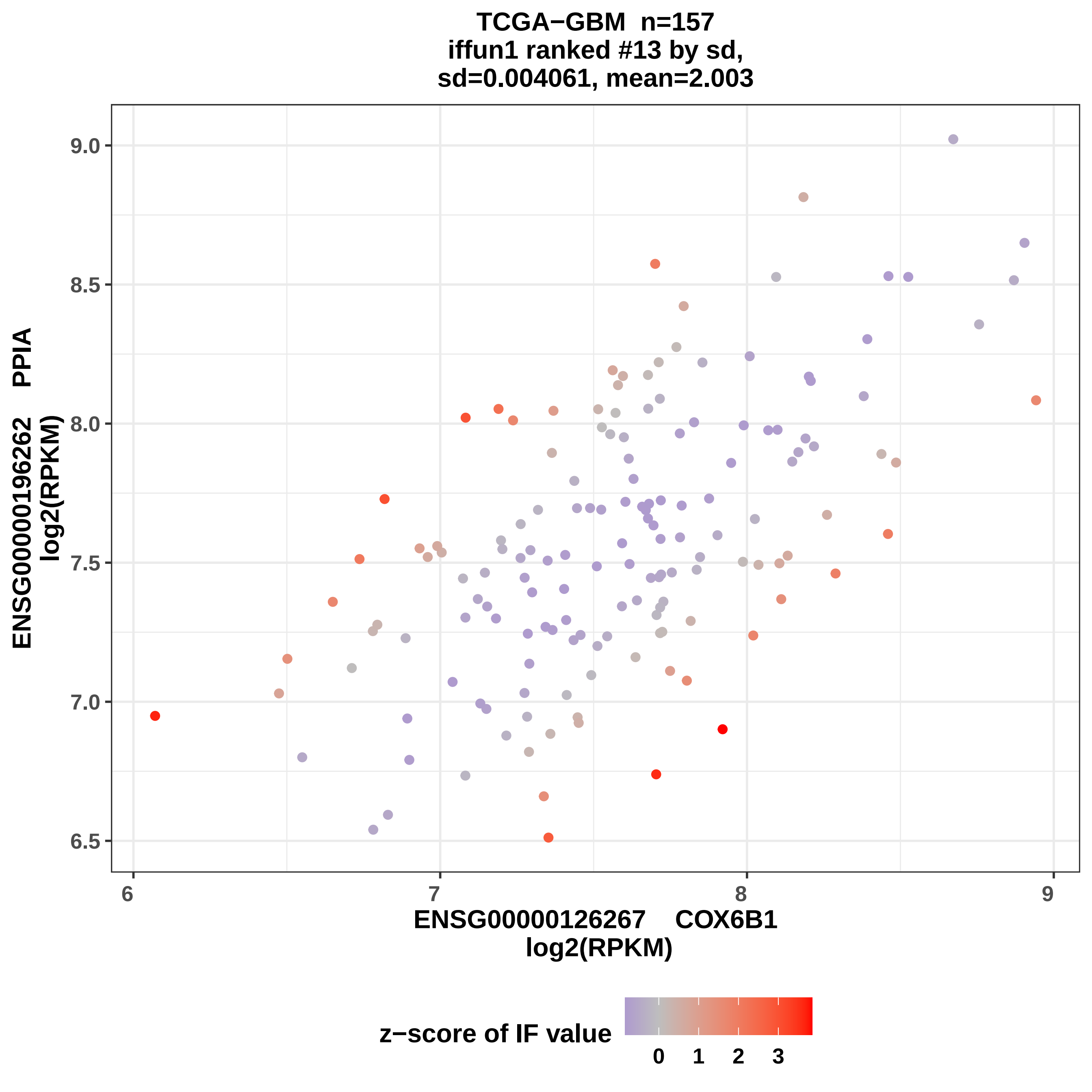

TCGA−GBM n=157
iffun1 ranked #13 by sd,
sd=0.004061, mean=2.003
ENSG00000196262 PPIA
log2(RPKM)
9.0
8.5
8.0
7.5
7.0
6.5
6
7
8
9
ENSG00000126267 COX6B1
log2(RPKM)
z−score of IF value
0
1
2
3

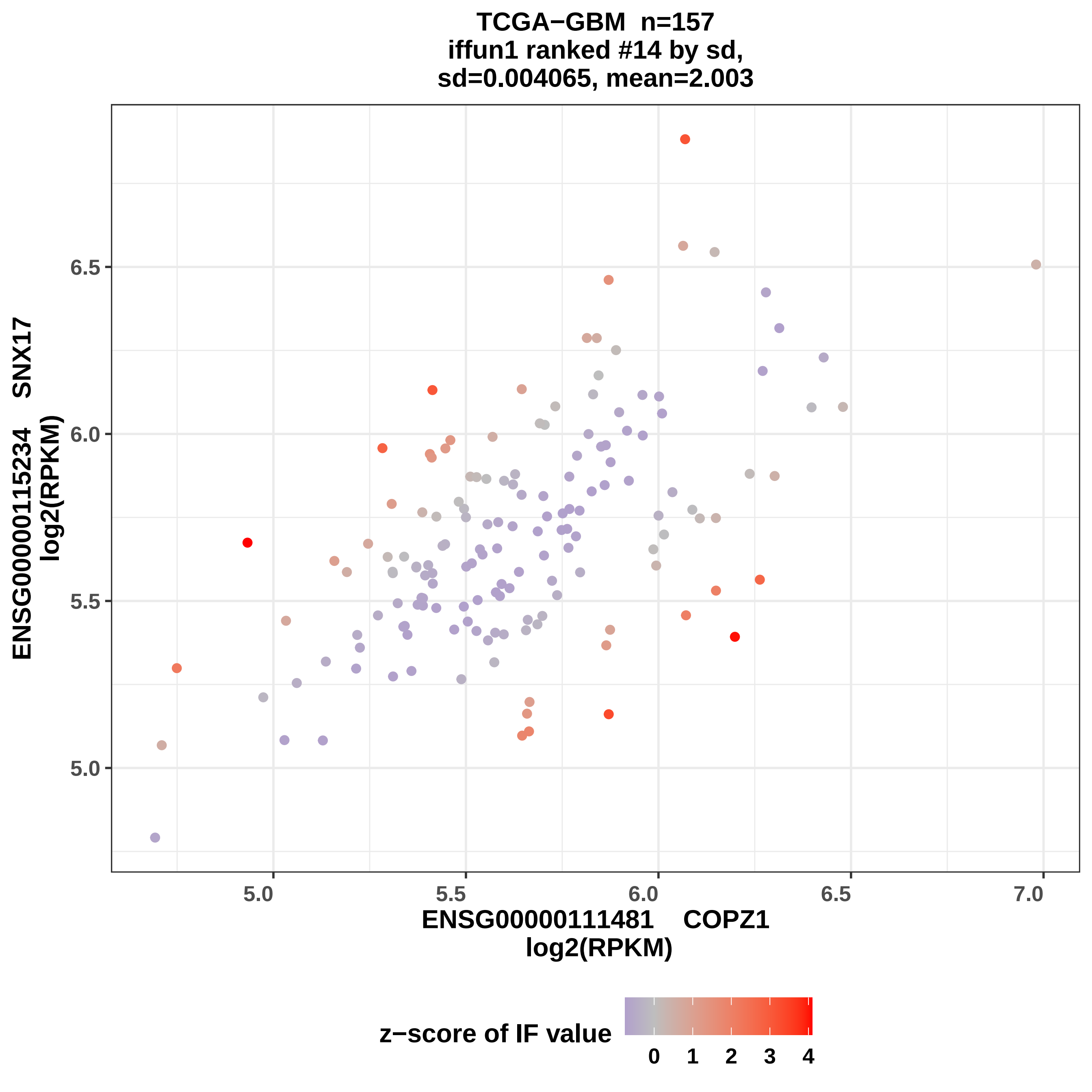
TCGA−GBM n=157
iffun1 ranked #14 by sd,
sd=0.004065, mean=2.003
ENSG00000115234 SNX17
log2(RPKM)
6.5
6.0
5.5
5.0
5.0
5.5
6.0
6.5
7.0
ENSG00000111481 COPZ1
log2(RPKM)
z−score of IF value
0
1
2
3
4

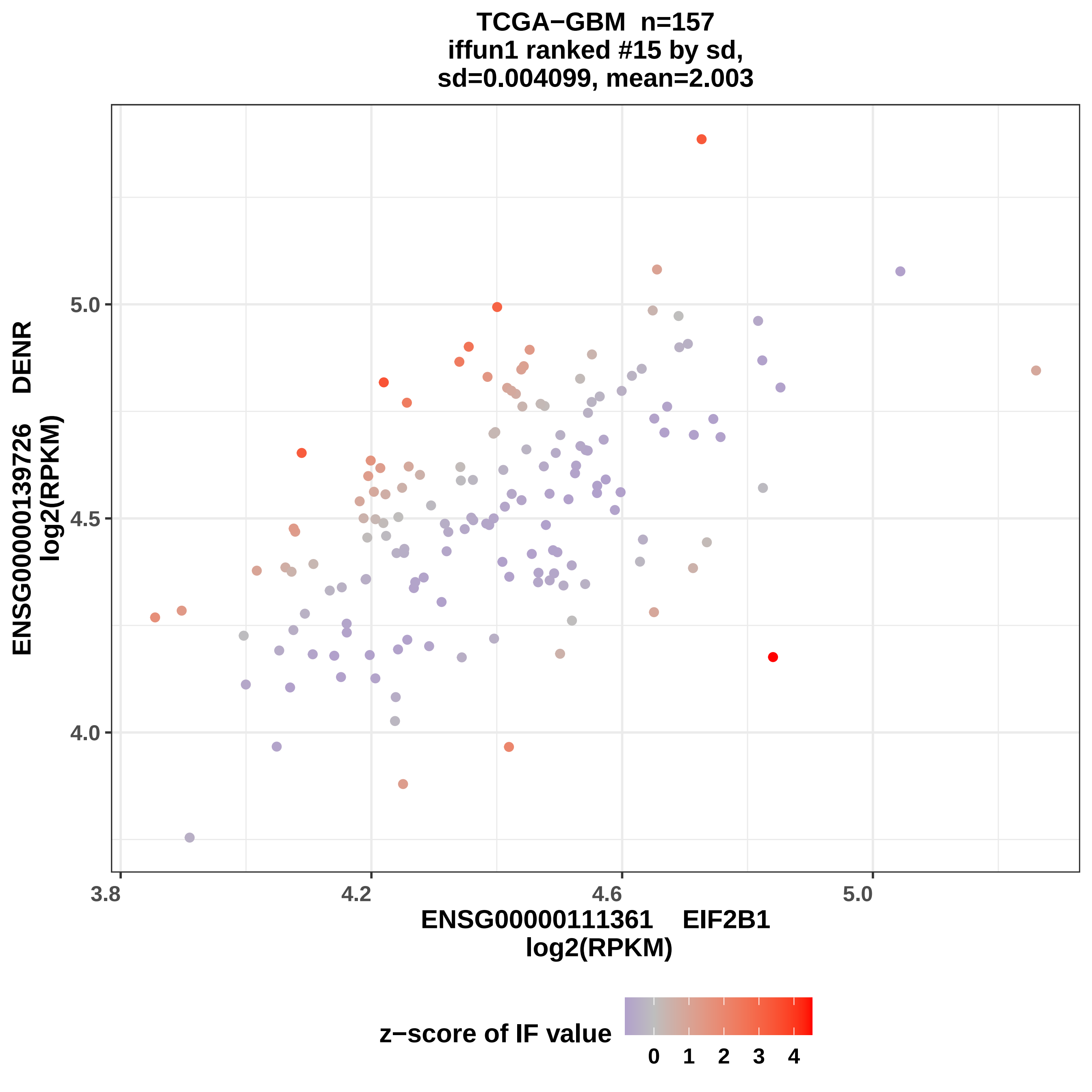
TCGA−GBM n=157
iffun1 ranked #15 by sd,
sd=0.004099, mean=2.003
ENSG00000139726 DENR
log2(RPKM)
5.0
4.5
4.0
3.8
4.2
4.6
5.0
ENSG00000111361 EIF2B1
log2(RPKM)
z−score of IF value
0
1
2
3
4

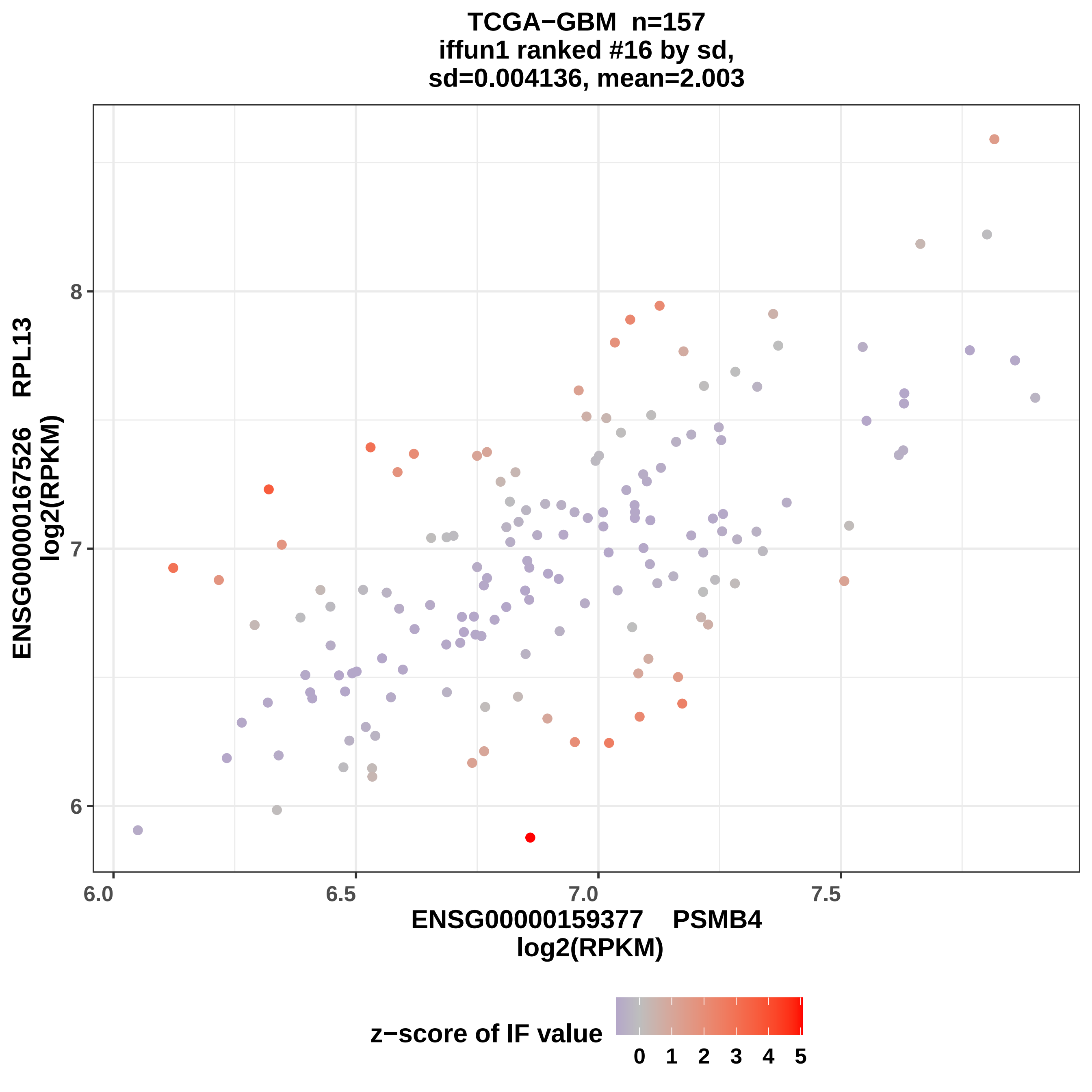

TCGA−GBM n=157
iffun1 ranked #16 by sd,
sd=0.004136, mean=2.003
ENSG00000167526 RPL13
log2(RPKM)
8
7
6
6.0
6.5
7.0
7.5
ENSG00000159377 PSMB4
log2(RPKM)
z−score of IF value
0 1 2 3 4 5

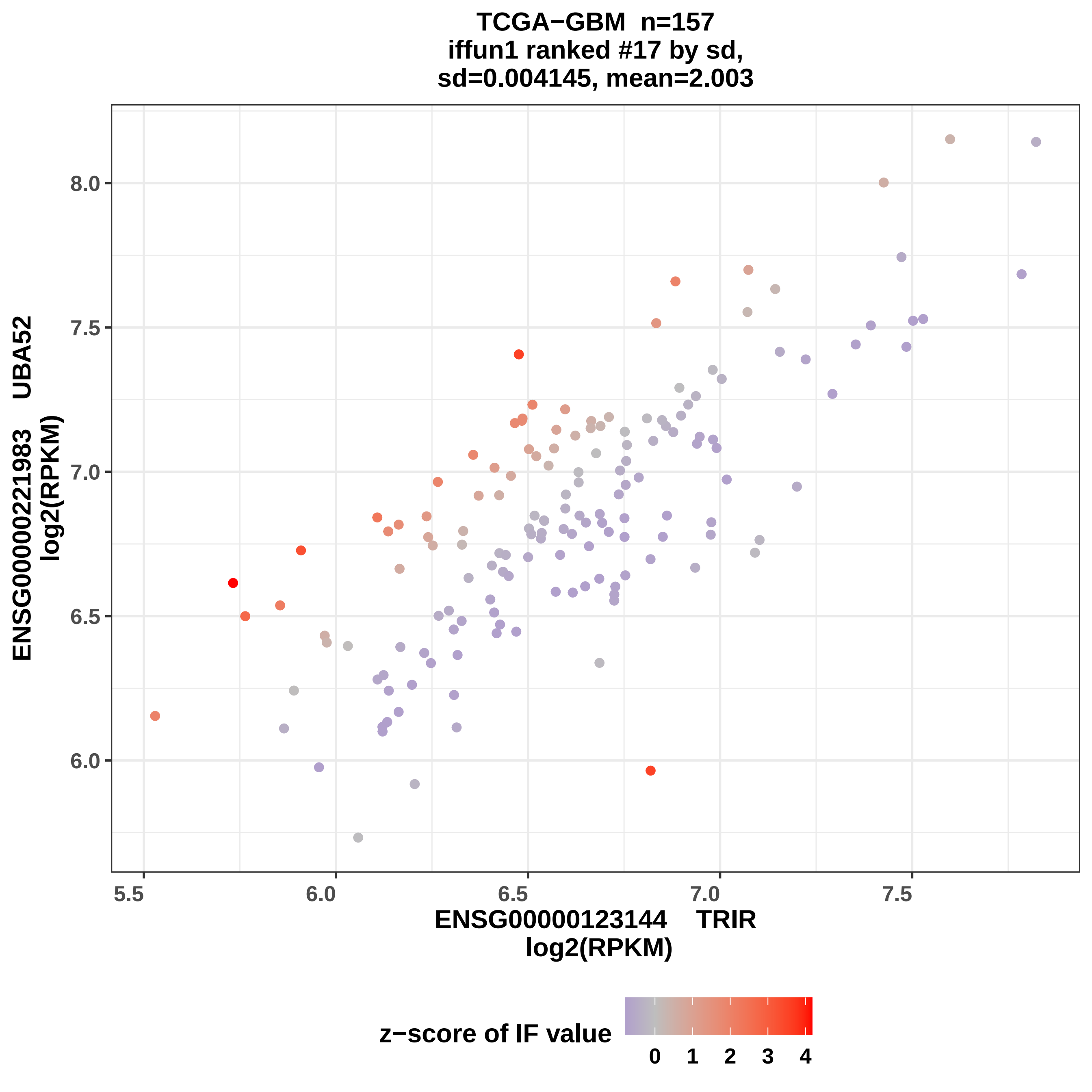

TCGA−GBM n=157
iffun1 ranked #17 by sd,
sd=0.004145, mean=2.003
ENSG00000221983 UBA52
log2(RPKM)
8.0
7.5
7.0
6.5
6.0
5.5
6.0
6.5
7.0
7.5
ENSG00000123144 TRIR
log2(RPKM)
z−score of IF value
0
1
2
3
4

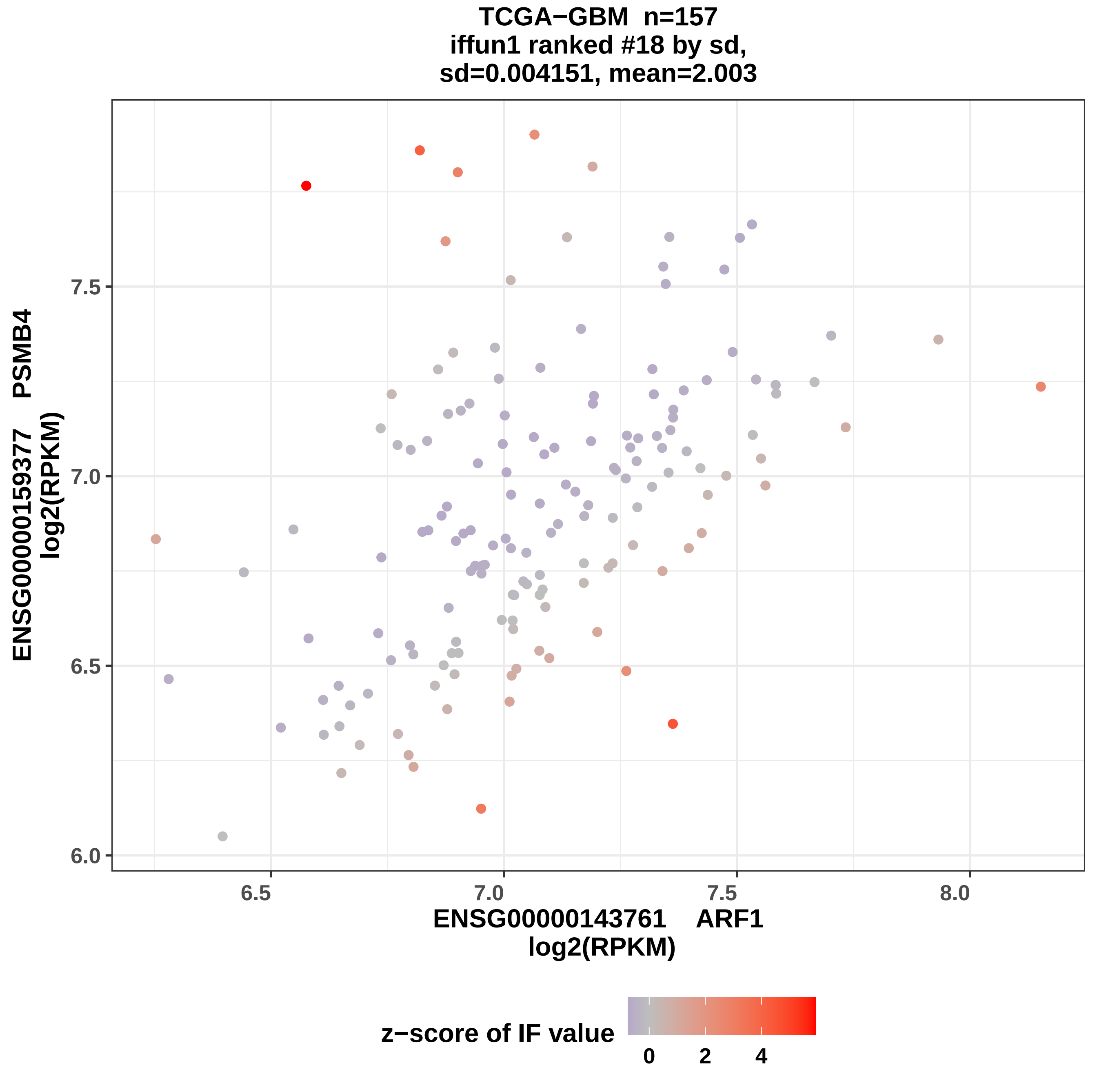
TCGA−GBM n=157
iffun1 ranked #18 by sd,
sd=0.004151, mean=2.003
ENSG00000159377 PSMB4
log2(RPKM)
7.5
7.0
6.5
6.0
6.5
7.0
7.5
8.0
ENSG00000143761 ARF1
log2(RPKM)
z−score of IF value
0
2
4

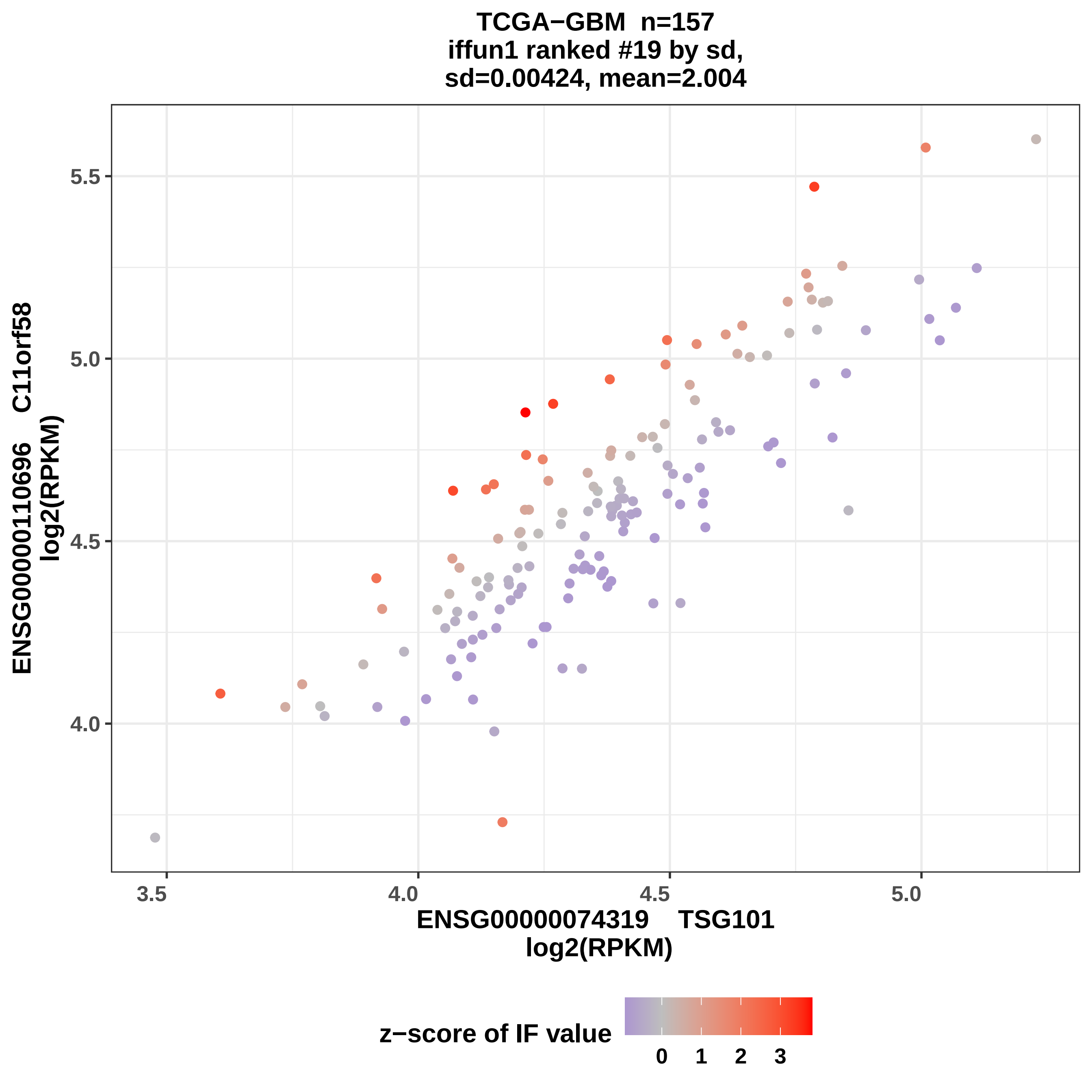

TCGA−GBM n=157
iffun1 ranked #19 by sd,
sd=0.00424, mean=2.004
ENSG00000110696 C11orf58
log2(RPKM)
5.5
5.0
4.5
4.0
3.5
4.0
4.5
5.0
ENSG00000074319 TSG101
log2(RPKM)
z−score of IF value
0
1
2
3

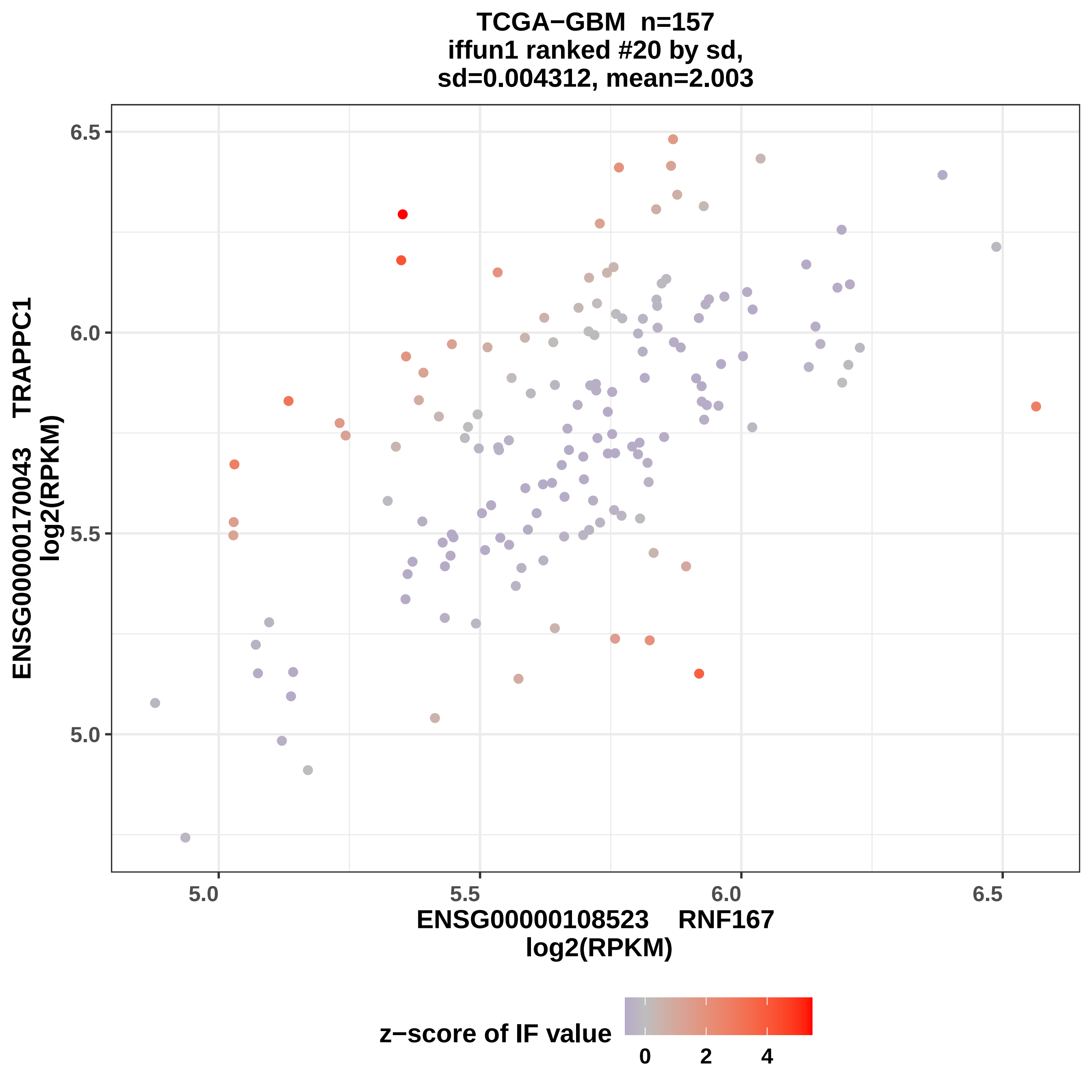

TCGA−GBM n=157
iffun1 ranked #20 by sd,
sd=0.004312, mean=2.003
ENSG00000170043 TRAPPC1
log2(RPKM)
6.5
6.0
5.5
5.0
5.0
5.5
6.0
6.5
ENSG00000108523 RNF167
log2(RPKM)
z−score of IF value
0
2
4

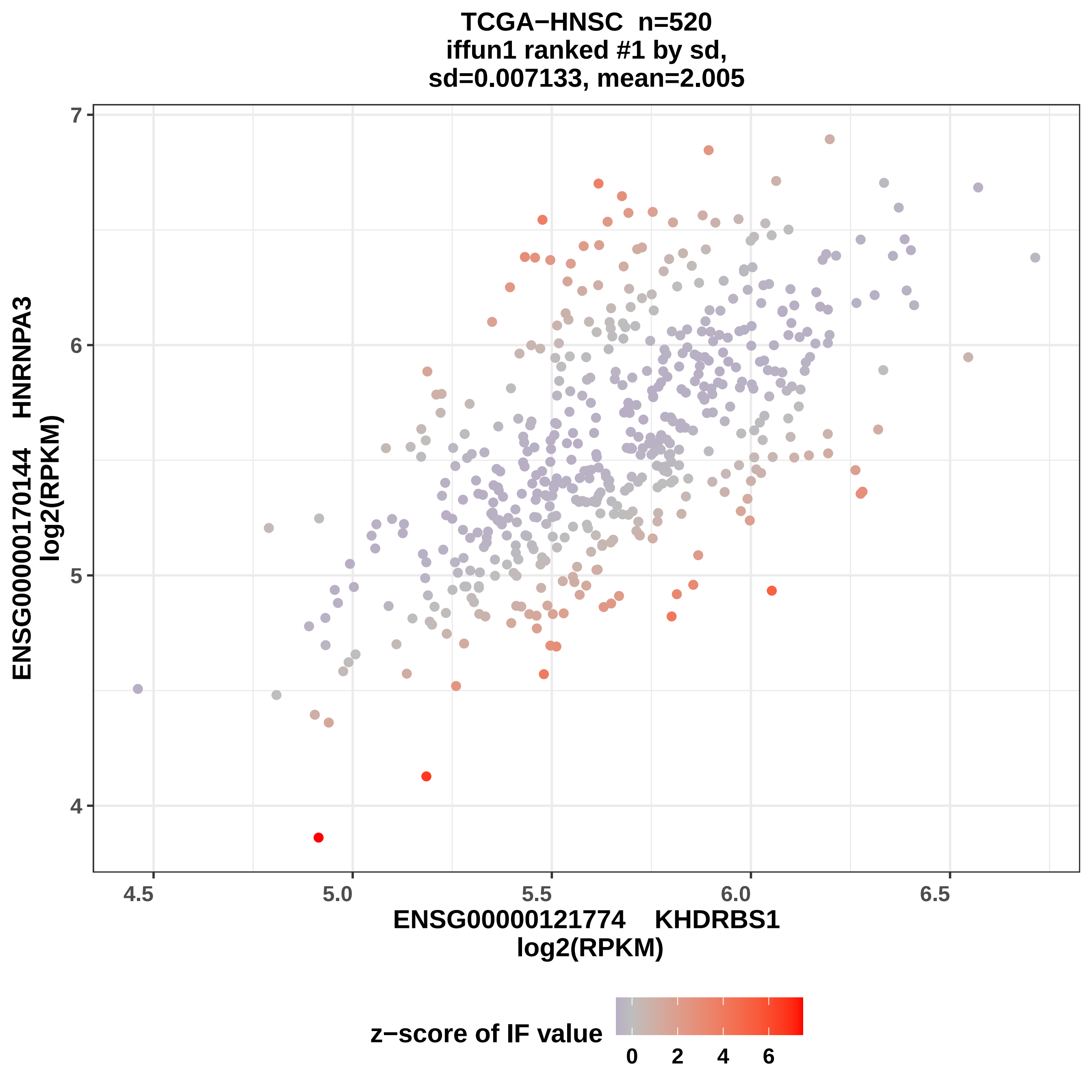
TCGA−HNSC n=520
iffun1 ranked #1 by sd,
sd=0.007133, mean=2.005
ENSG00000170144 HNRNPA3
log2(RPKM)
4
5
6
7
4.5
5.0
5.5
6.0
6.5
ENSG00000121774 KHDRBS1
log2(RPKM)
z−score of IF value
0
2
4
6

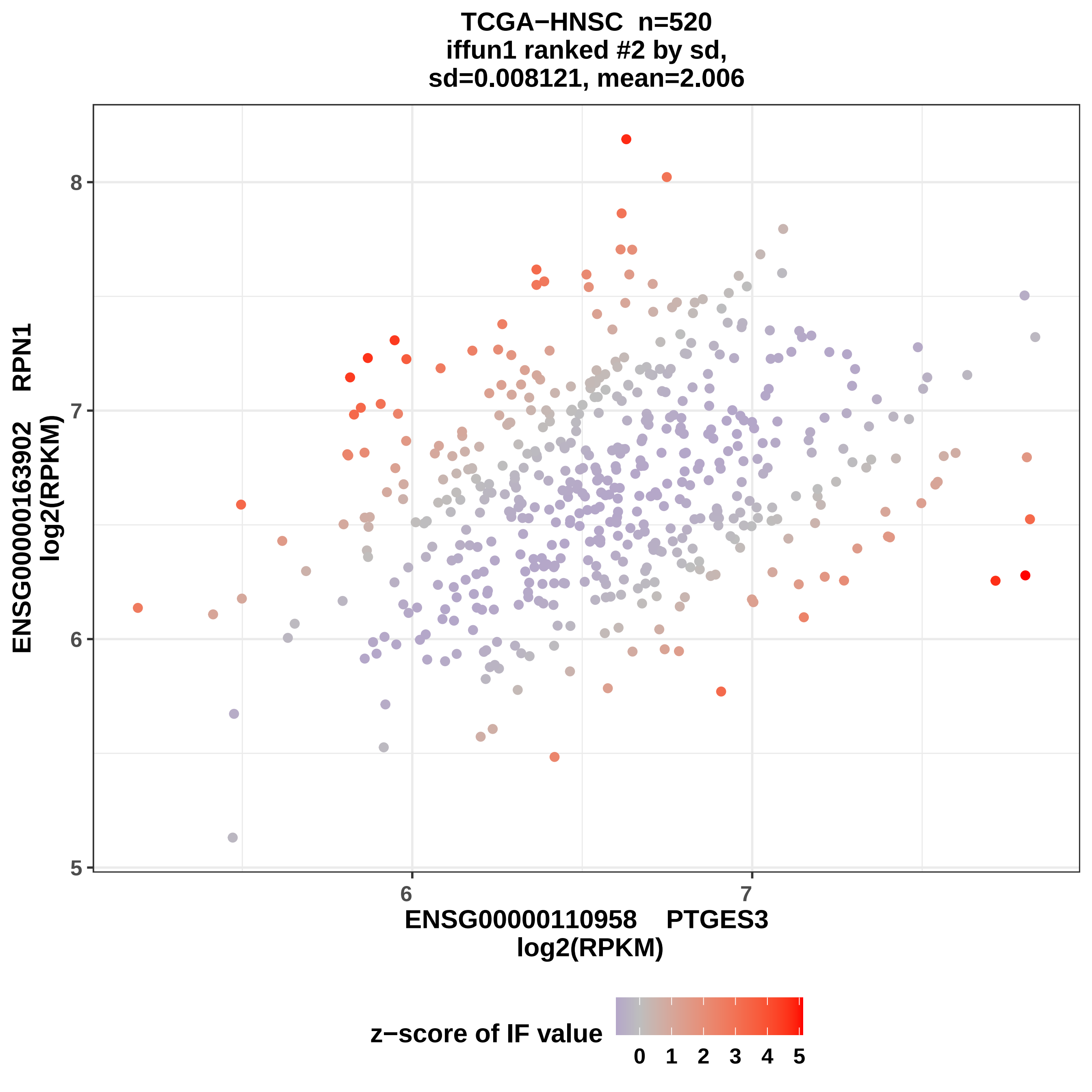

TCGA−HNSC n=520
iffun1 ranked #2 by sd,
sd=0.008121, mean=2.006
ENSG00000163902 RPN1
log2(RPKM)
8
7
6
5
6
7
ENSG0000110958 PTGES3
log2(RPKM)
z−score of IF value
0 1 2 3 4 5

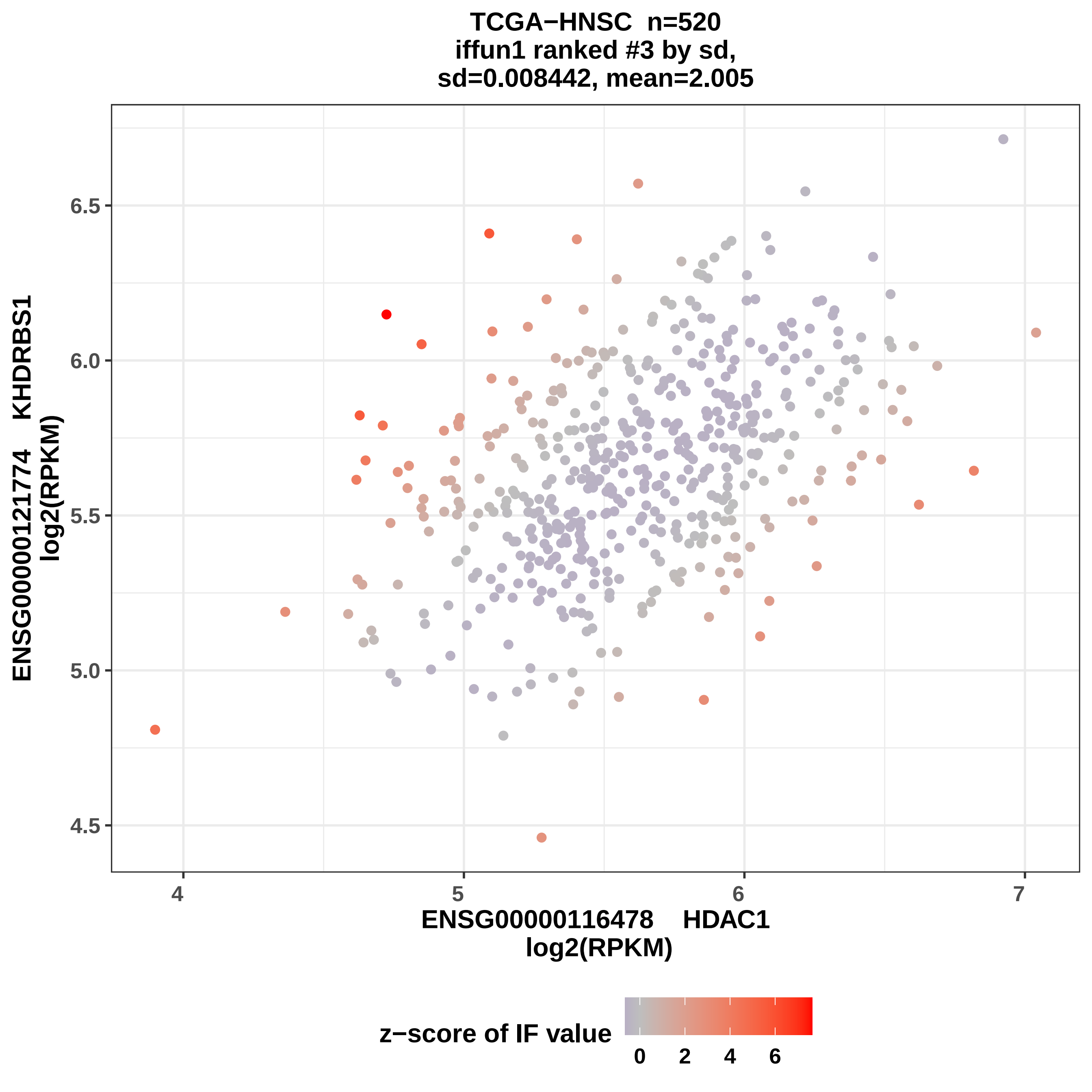

TCGA−HNSC n=520
iffun1 ranked #3 by sd,
sd=0.008442, mean=2.005
ENSG00000121774 KHDRBS1
log2(RPKM)
6.5
6.0
5.5
5.0
4.5
4
5
6
7
ENSG00000116478 HDAC1
log2(RPKM)
z−score of IF value
0
2
4
6

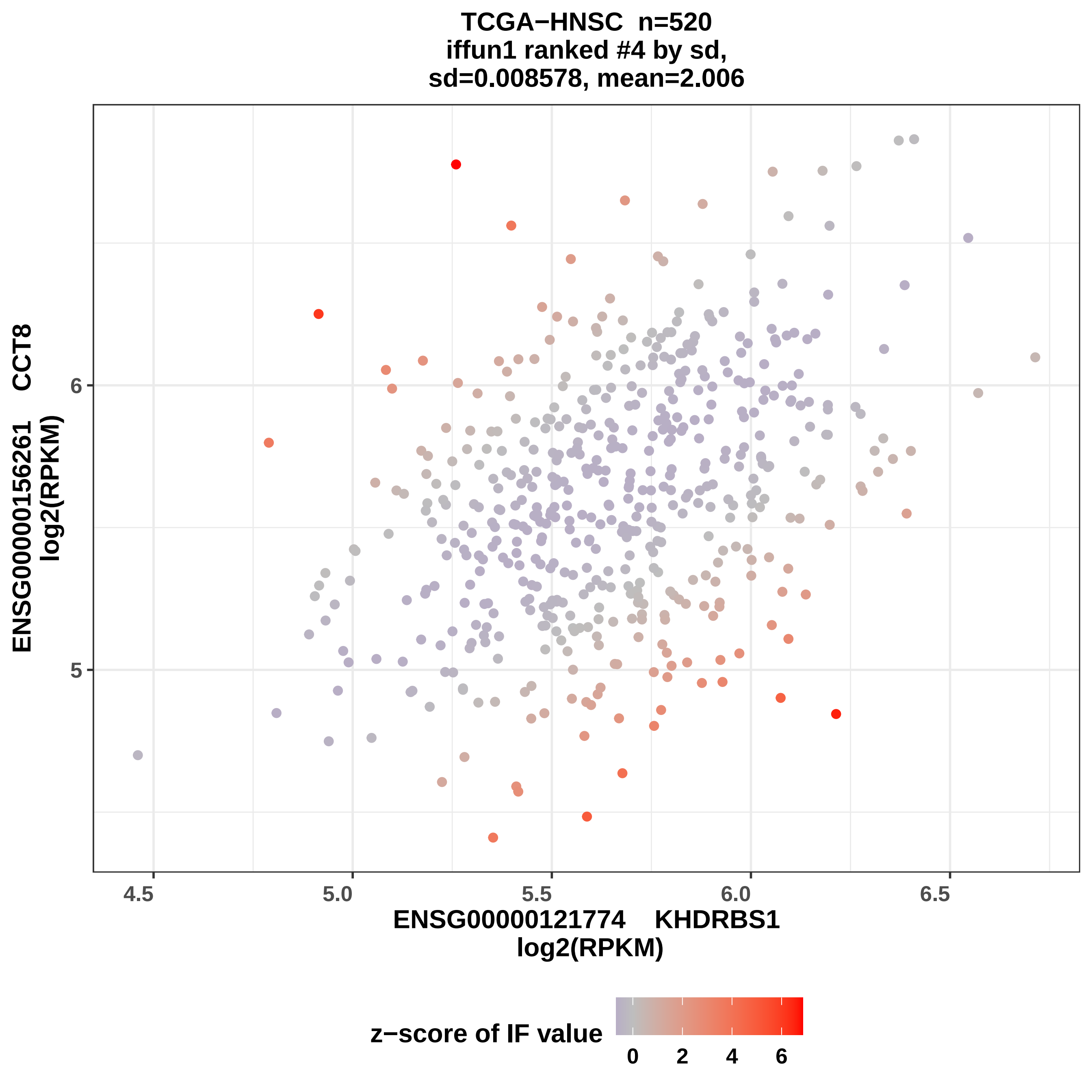
TCGA−HNSC n=520
iffun1 ranked #4 by sd,
sd=0.008578, mean=2.006
ENSG00000156261 CCT8
log2(RPKM)
6
5
4.5
5.0
5.5
6.0
6.5
ENSG00000121774 KHDRBS1
log2(RPKM)
z−score of IF value
0
2
4
6

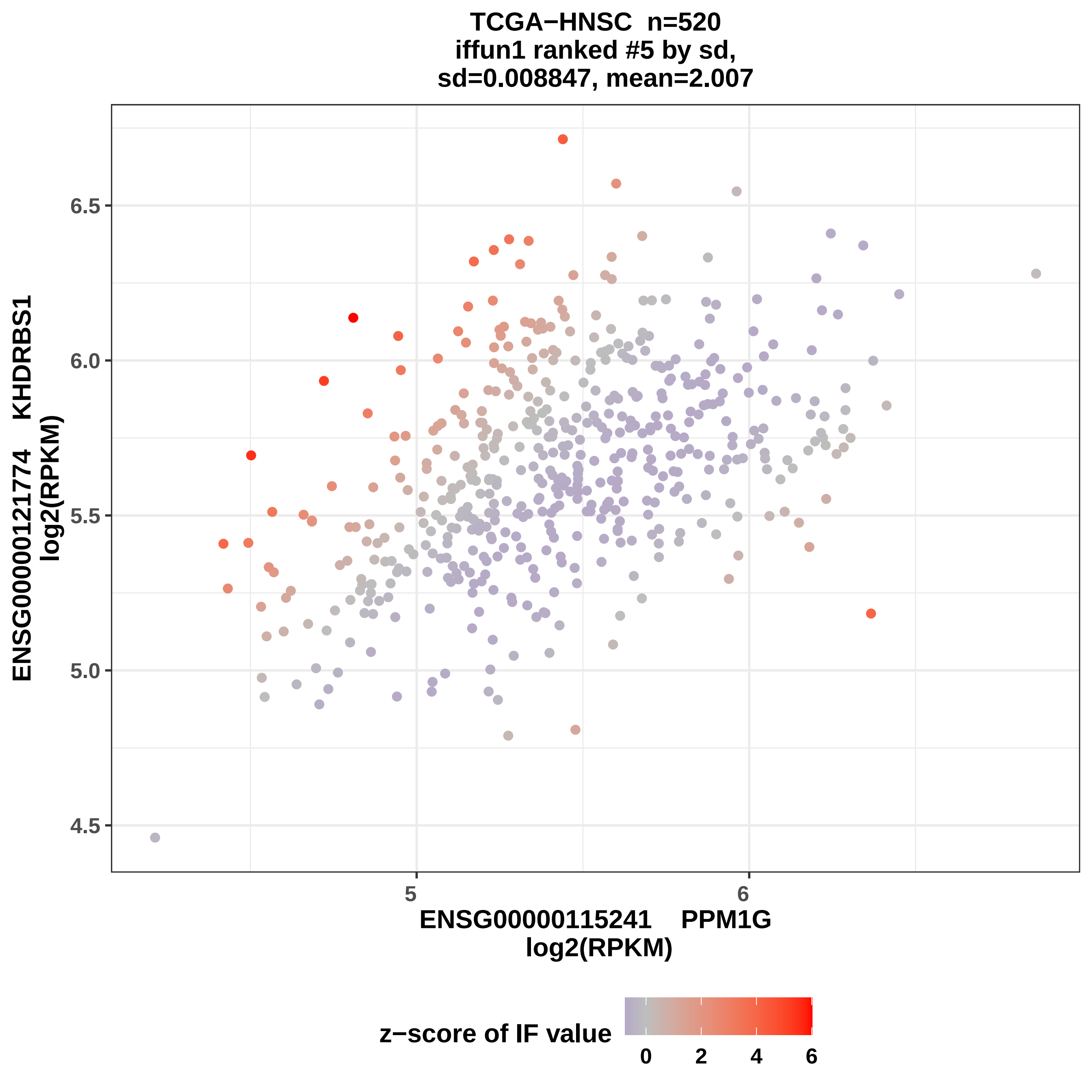
TCGA−HNSC n=520
iffun1 ranked #5 by sd,
sd=0.008847, mean=2.007
ENSG00000121774 KHDRBS1
log2(RPKM)
6.5
6.0
5.5
5.0
4.5
5
6
ENSG00000115241 PPM1G
log2(RPKM)
z−score of IF value
0
2
4
6

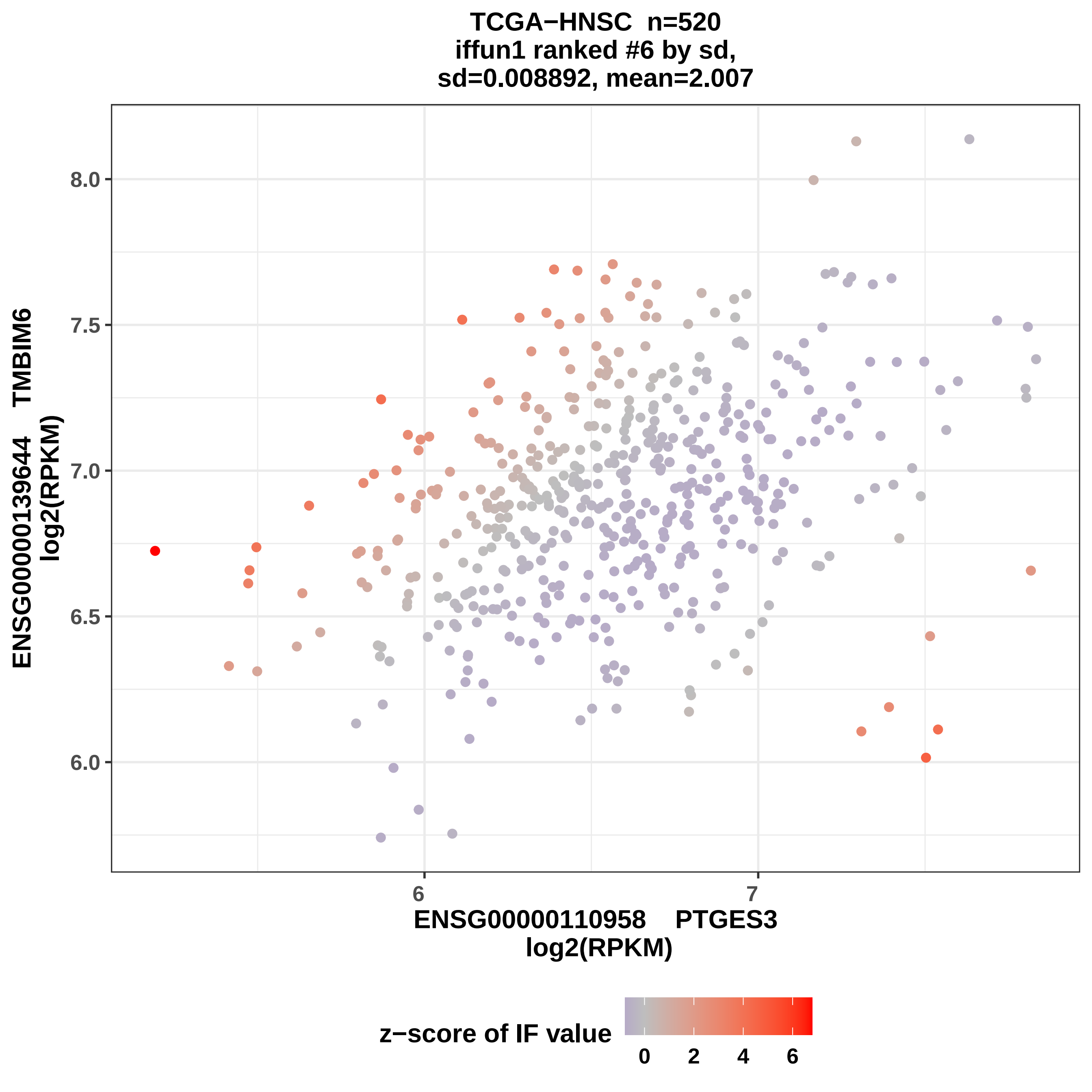
TCGA−HNSC n=520
iffun1 ranked #6 by sd,
sd=0.008892, mean=2.007
ENSG00000139644 TMBIM6
log2(RPKM)
8.0
7.5
7.0
6.5
6.0
6
7
ENSG00000110958 PTGES3
log2(RPKM)
z−score of IF value
0
2
4
6

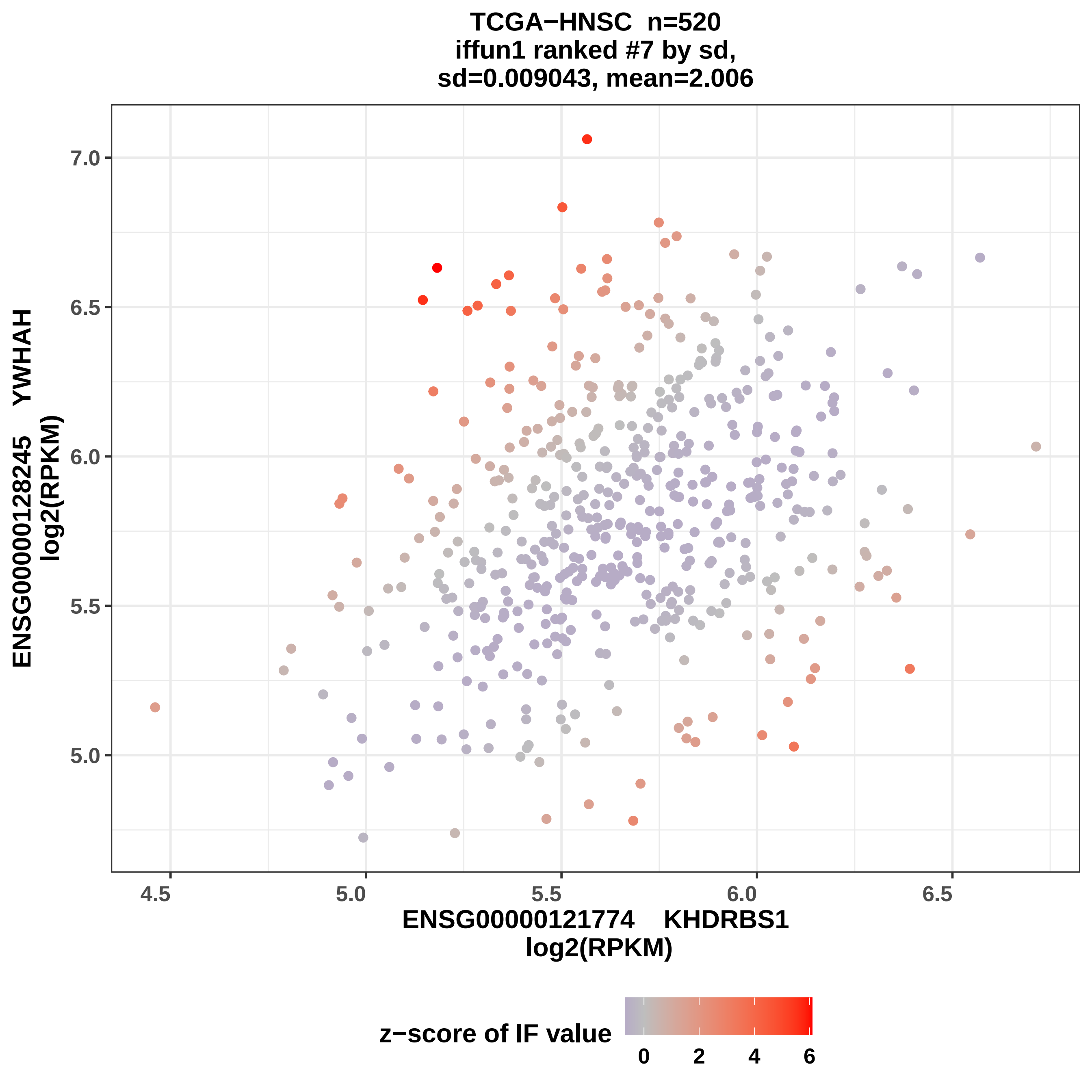
TCGA−HNSC n=520
iffun1 ranked #7 by sd,
sd=0.009043, mean=2.006
ENSG00000128245 YWHAH
log2(RPKM)
7.0
6.5
6.0
5.5
5.0
4.5
5.0
5.5
6.0
6.5
ENSG00000121774 KHDRBS1
log2(RPKM)
z−score of IF value
0
2
4
6

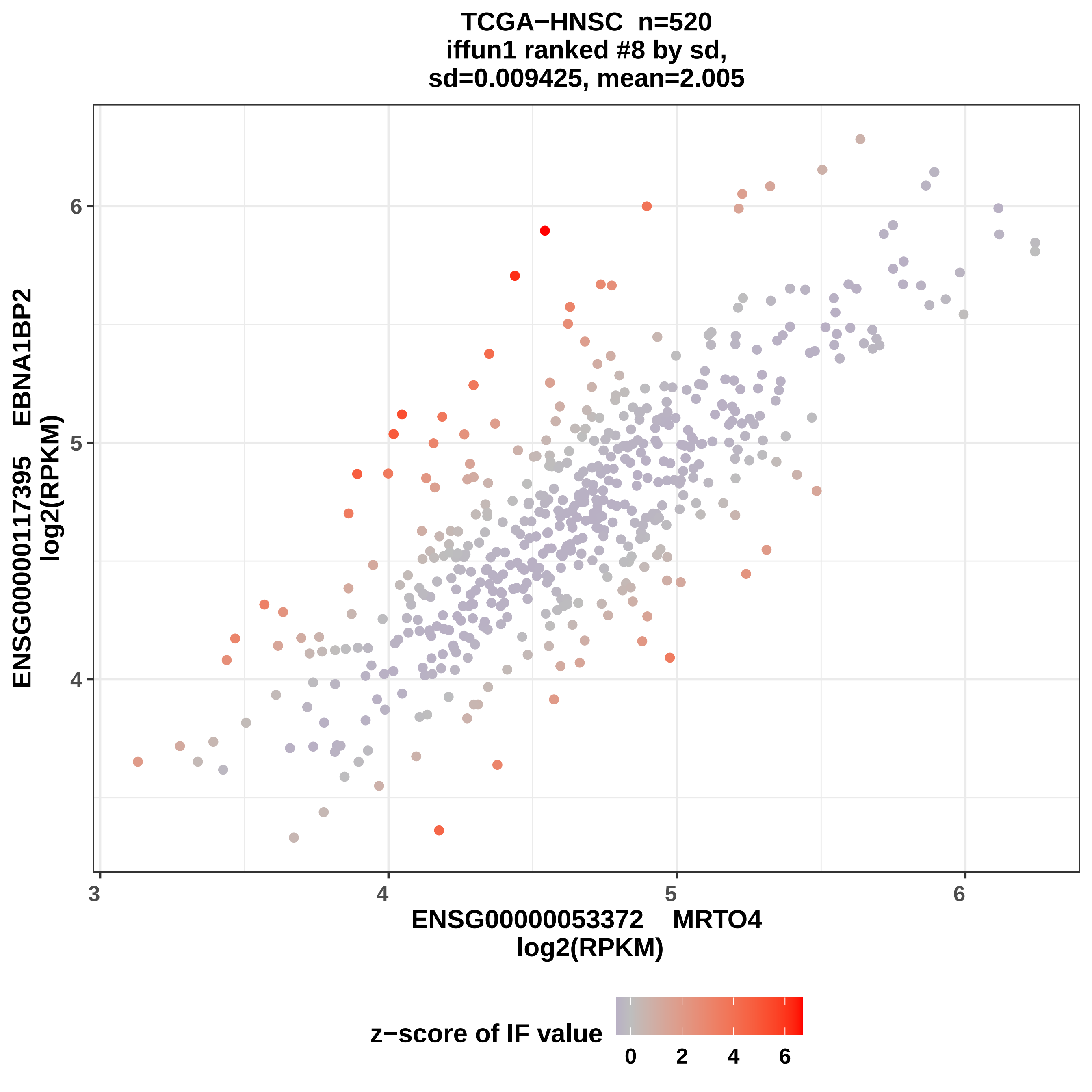

TCGA−HNSC n=520
iffun1 ranked #8 by sd,
sd=0.009425, mean=2.005
ENSG00000117395 EBNA1BP2
log2(RPKM)
6
5
4
3
ENSG00000053372 MRTO4
log2(RPKM)
z−score of IF value
0
2
4
6

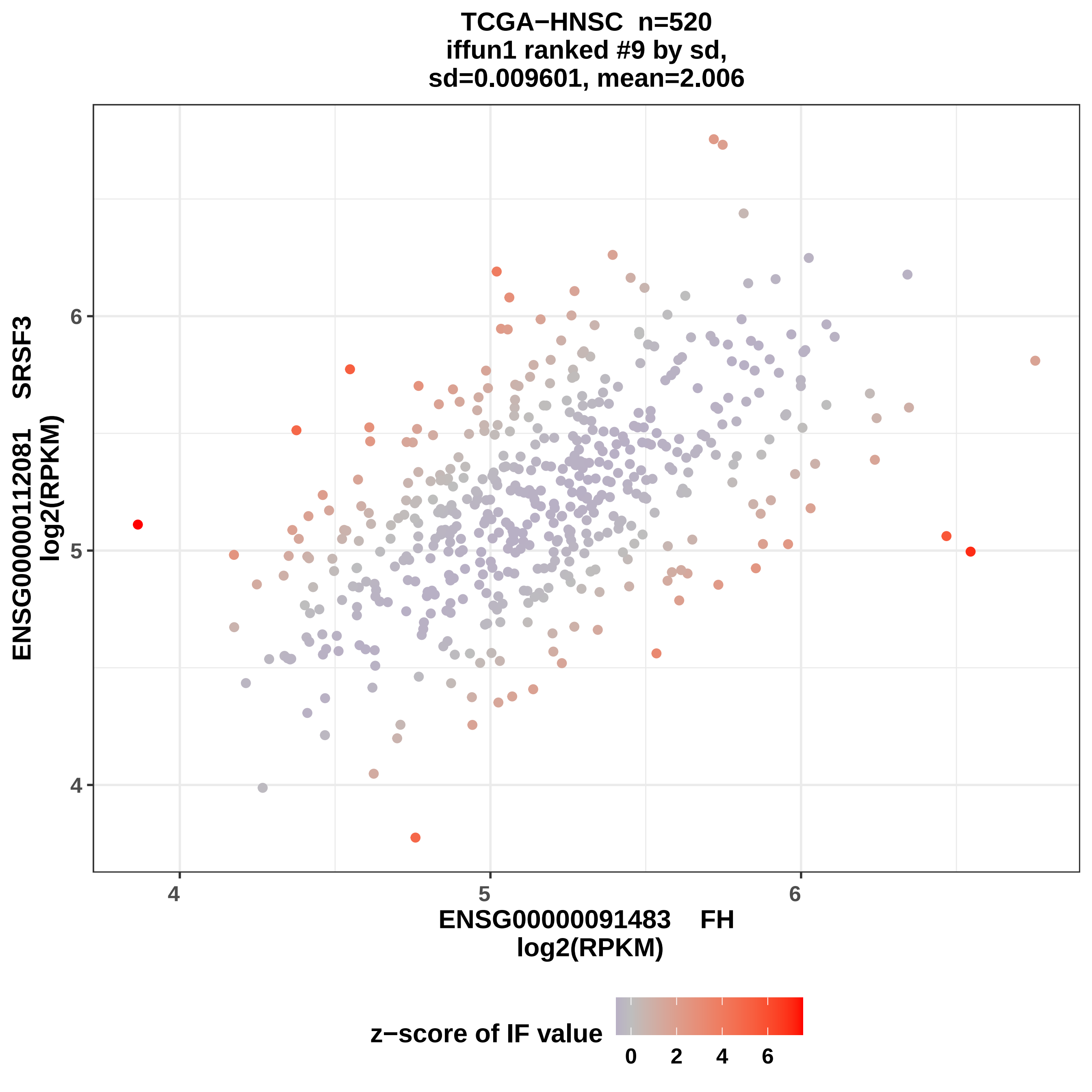

TCGA−HNSC n=520
iffun1 ranked #9 by sd,
sd=0.009601, mean=2.006
ENSG00000112081 SRSF3
log2(RPKM)
6
5
4
4
5
6
ENSG00000091483 FH
log2(RPKM)
z−score of IF value
0
2
4
6

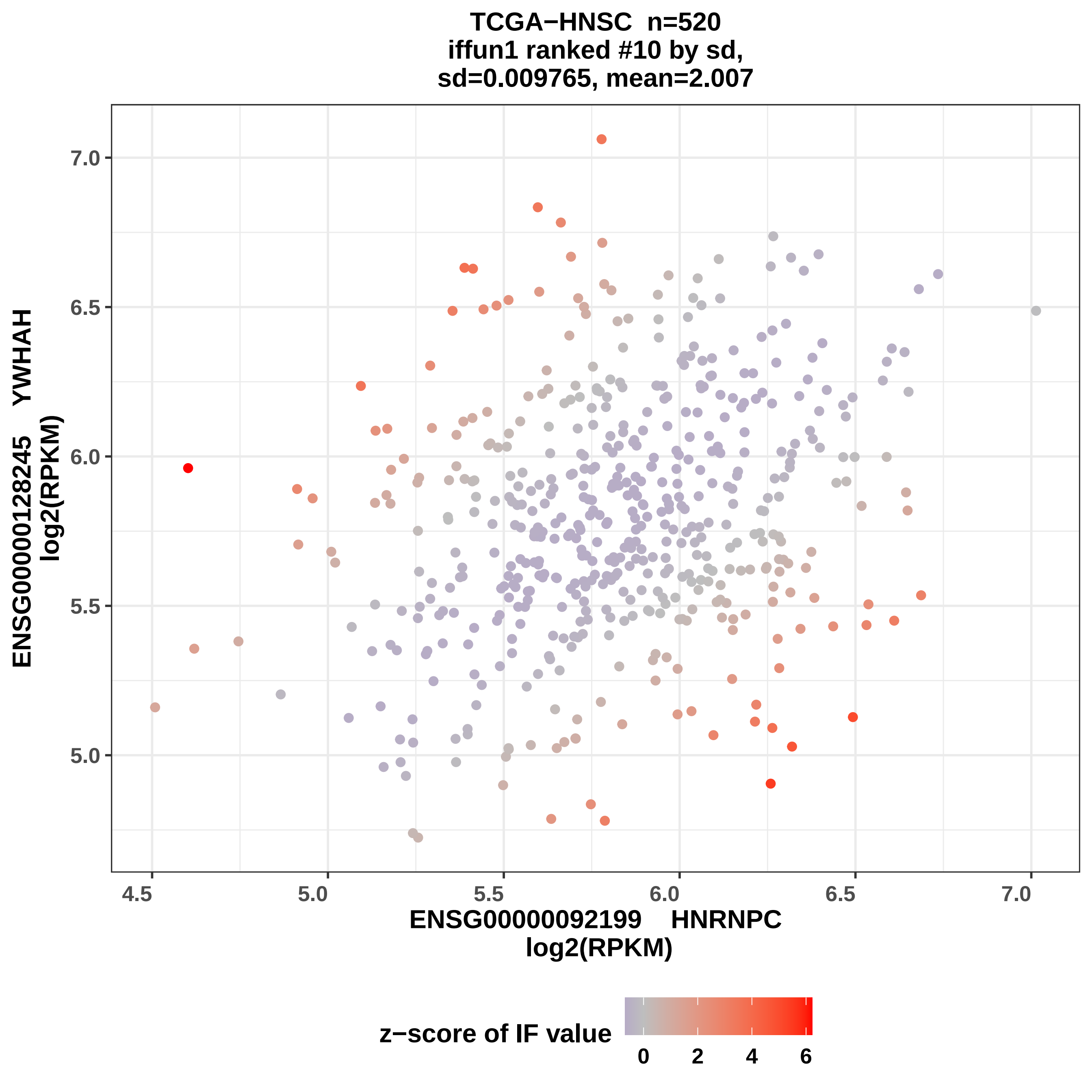
TCGA−HNSC n=520
iffun1 ranked #10 by sd,
sd=0.009765, mean=2.007
ENSG00000128245 YWHAH
log2(RPKM)
7.0
6.5
6.0
5.5
5.0
4.5
5.0
5.5
6.0
6.5
7.0
ENSG00000092199 HNRNPC
log2(RPKM)
z−score of IF value
0
2
4
6

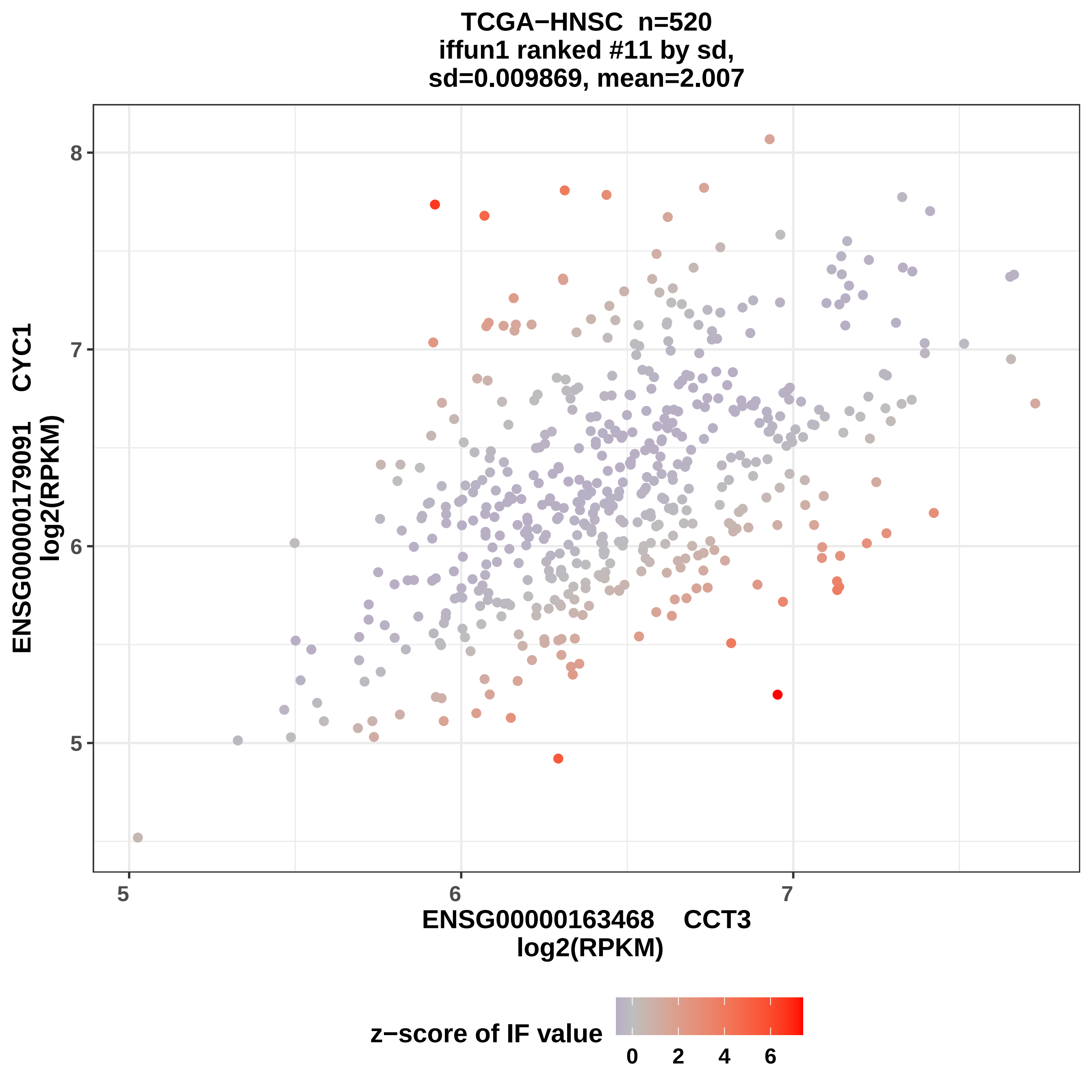
TCGA−HNSC n=520
iffun1 ranked #11 by sd,
sd=0.009869, mean=2.007
ENSG00000179091 CYC1
log2(RPKM)
8
7
6
5
5
6
7
ENSG00000163468 CCT3
log2(RPKM)
z−score of IF value
0
2
4
6

TCGA−HNSC n=520
iffun1 ranked #12 by sd,
sd=0.009961, mean=2.006

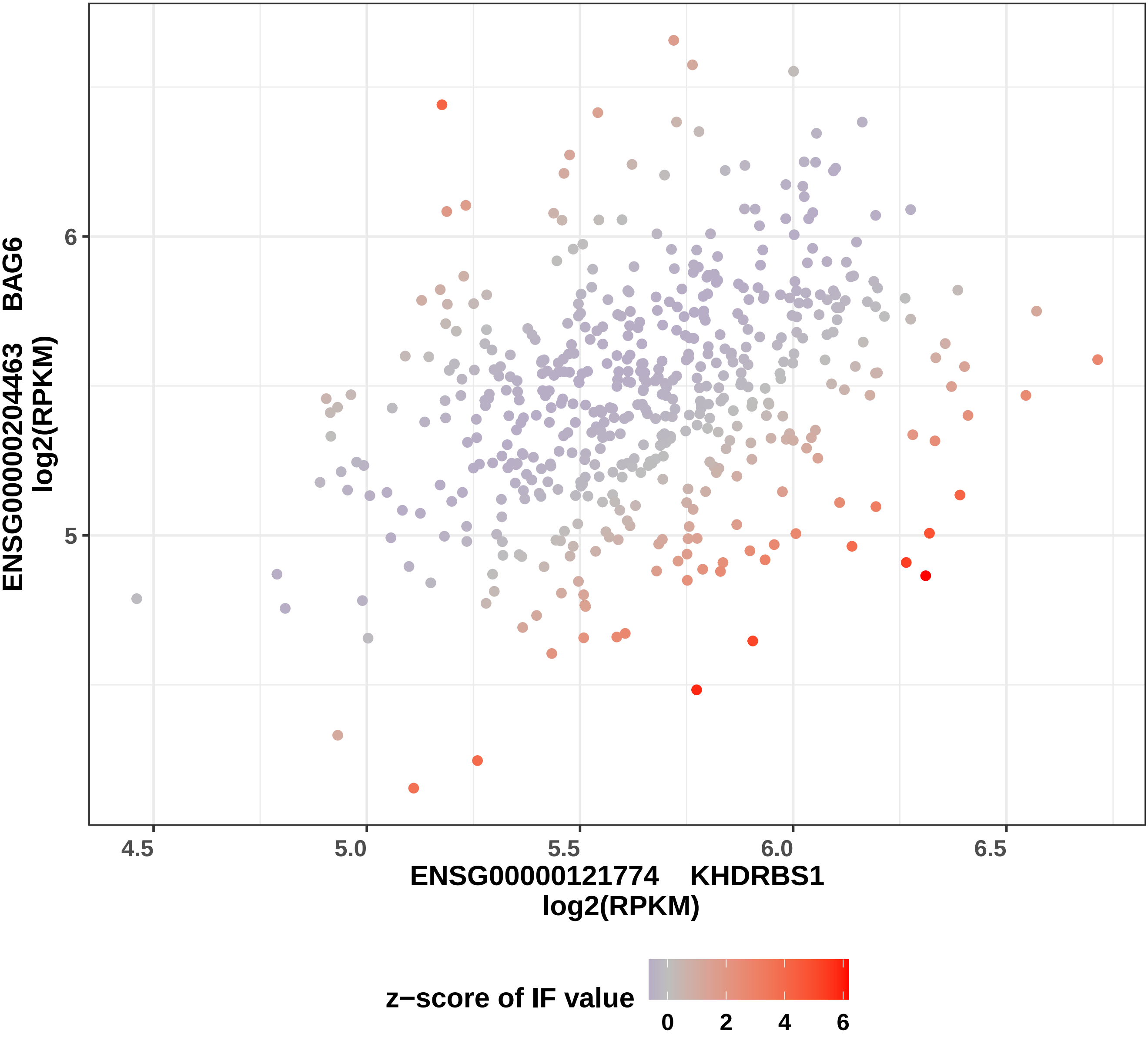

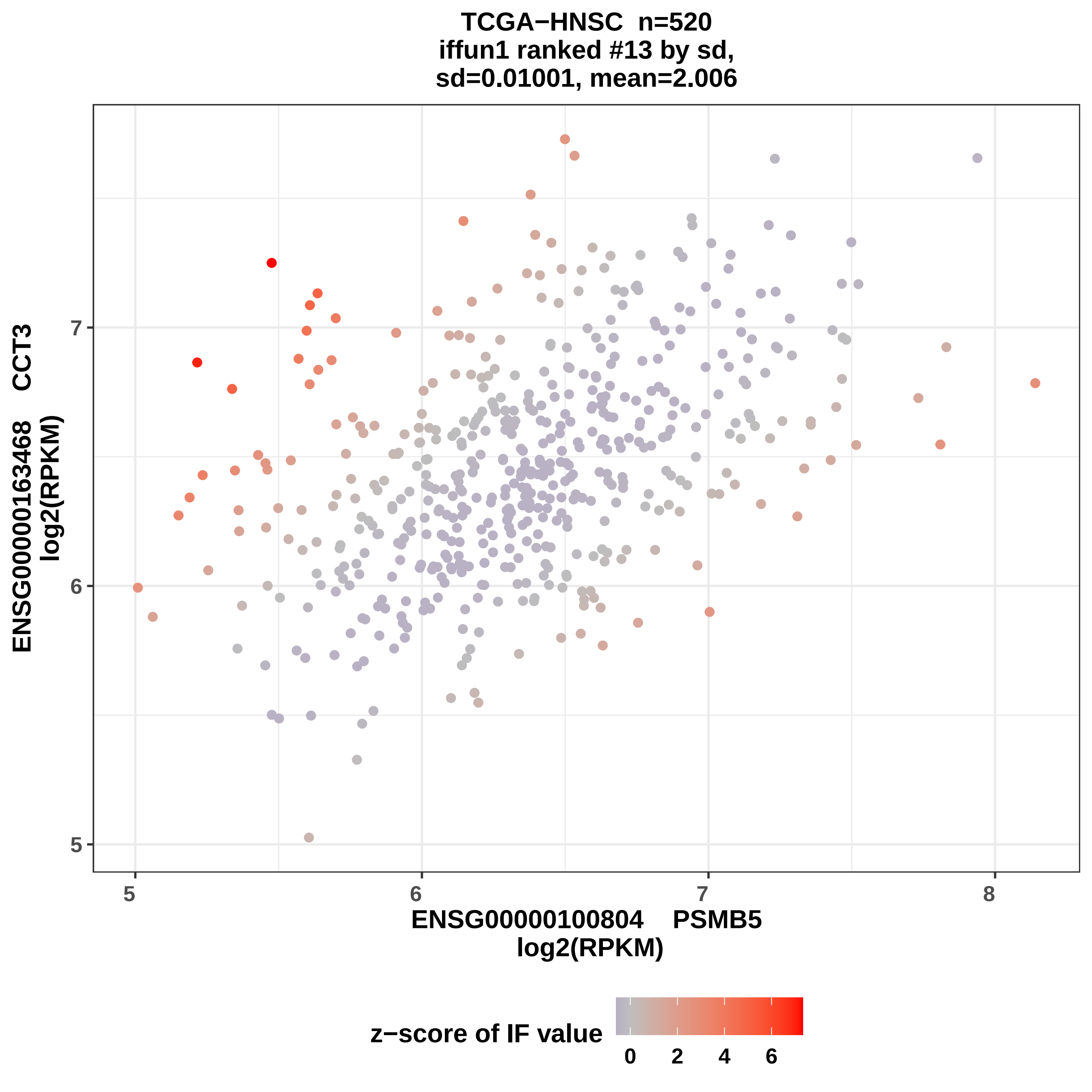

TCGA−HNSC n=520
iffun1 ranked #13 by sd,
sd=0.01001, mean=2.006
ENSG00000163468 CCT3
log2(RPKM)
5
6
7
8
ENSG00000100804 PSMB5
log2(RPKM)
z−score of IF value
0
2
4
6

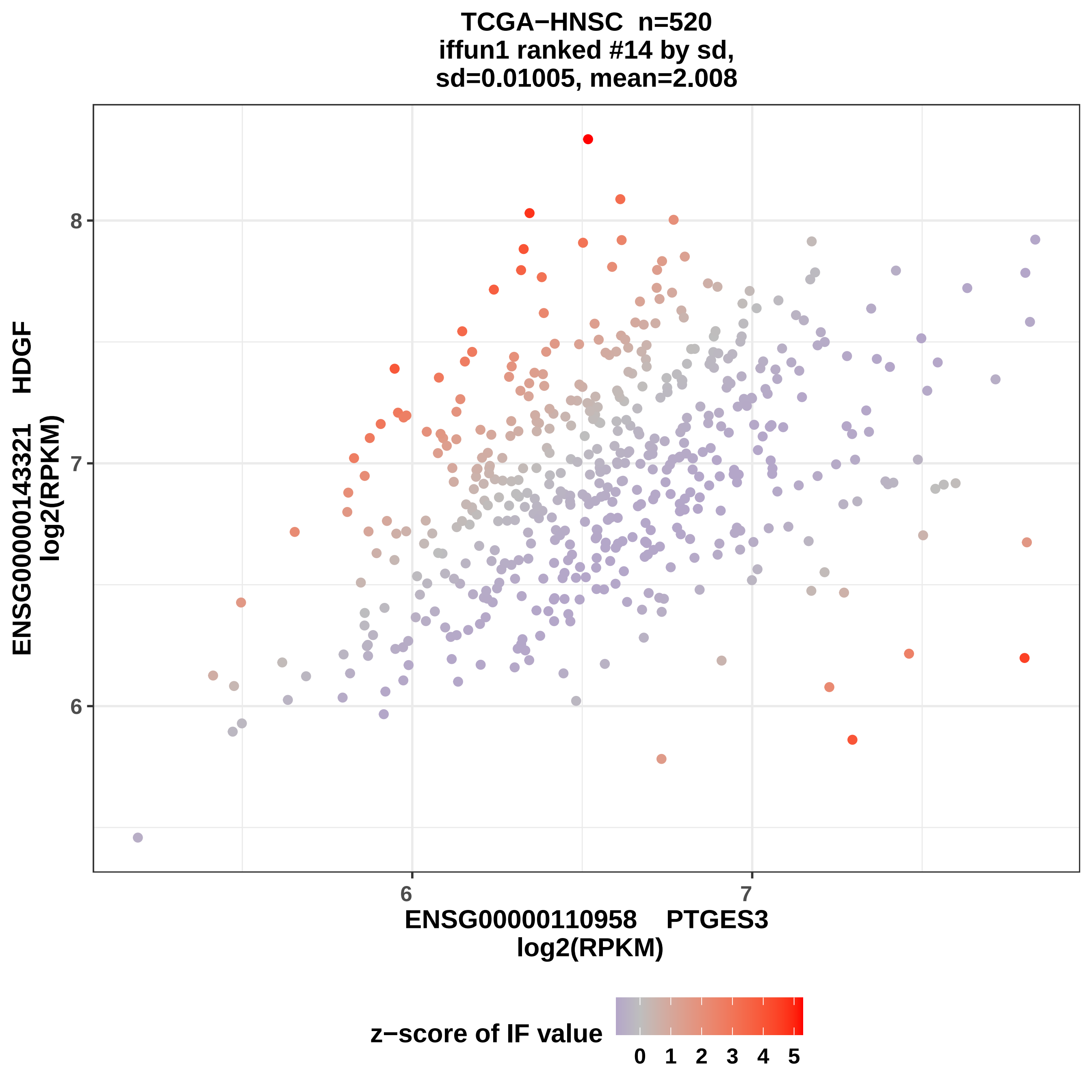
TCGA−HNSC n=520
iffun1 ranked #14 by sd,
sd=0.01005, mean=2.008
8
7
6
ENSG00000143321 HDGF
log2(RPKM)
6
7
ENSG00000110958 PTGES3
log2(RPKM)
z−score of IF value
0 1 2 3 4 5

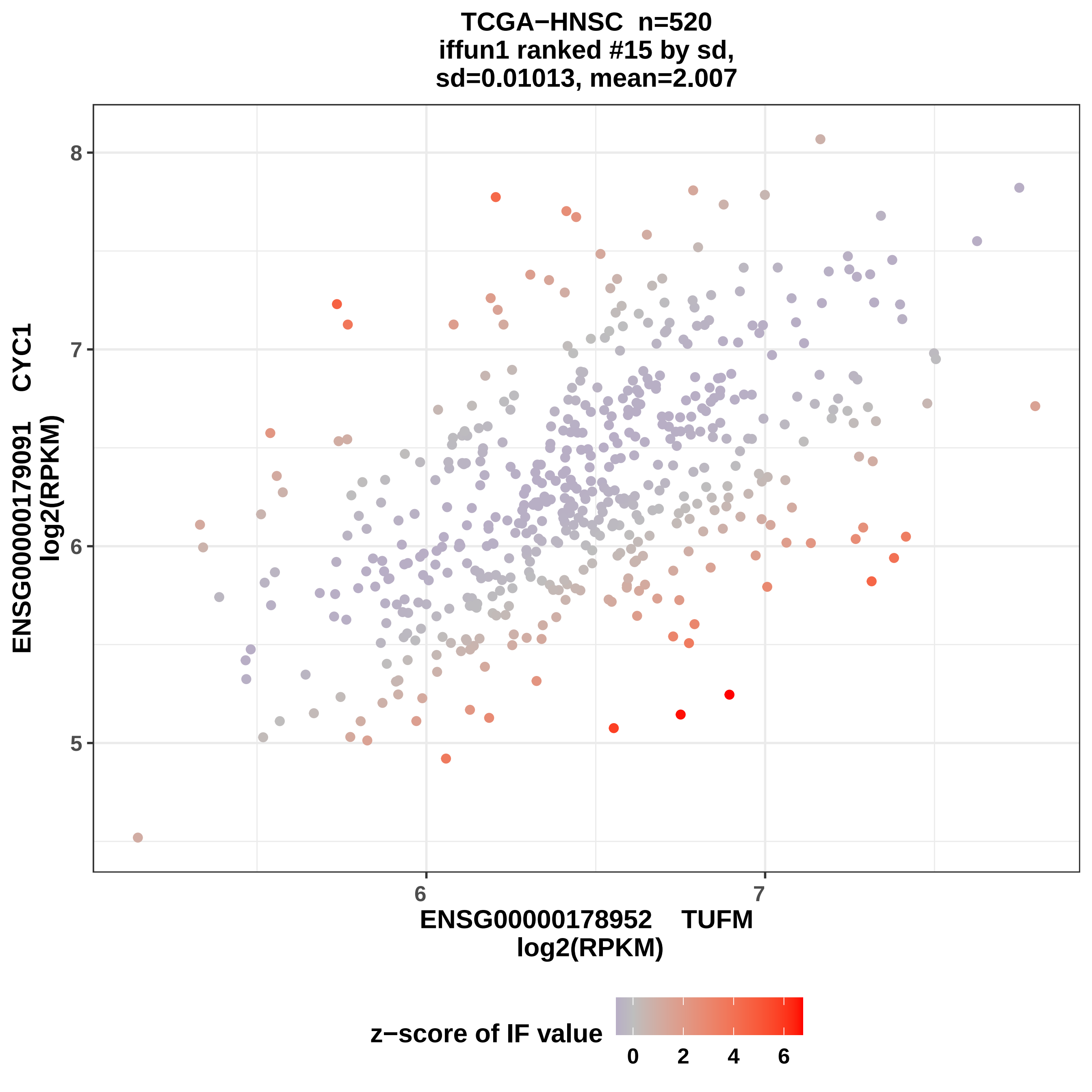
TCGA−HNSC n=520
iffun1 ranked #15 by sd,
sd=0.01013, mean=2.007
ENSG00000179091 CYC1
log2(RPKM)
8
7
6
5
6
7
ENSG0000178952 TUFM
log2(RPKM)
z−score of IF value
0
2
4
6

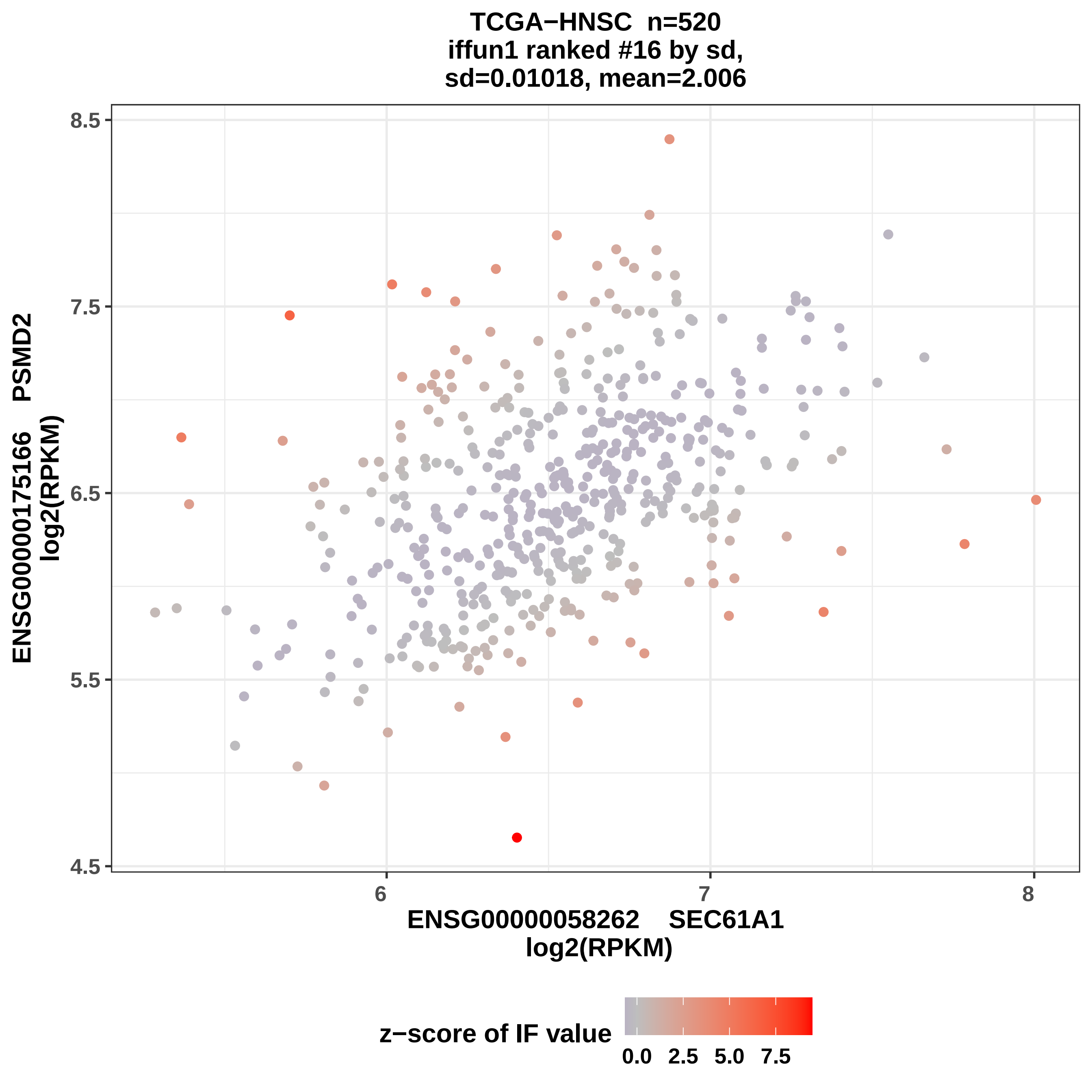

TCGA−HNSC n=520
iffun1 ranked #16 by sd,
sd=0.01018, mean=2.006
ENSG00000175166 PSMD2
log2(RPKM)
8.5
7.5
6.5
5.5
4.5
6
7
8
ENSG00000058262 SEC61A1
log2(RPKM)
z−score of IF value
0.0
2.5
5.0
7.5

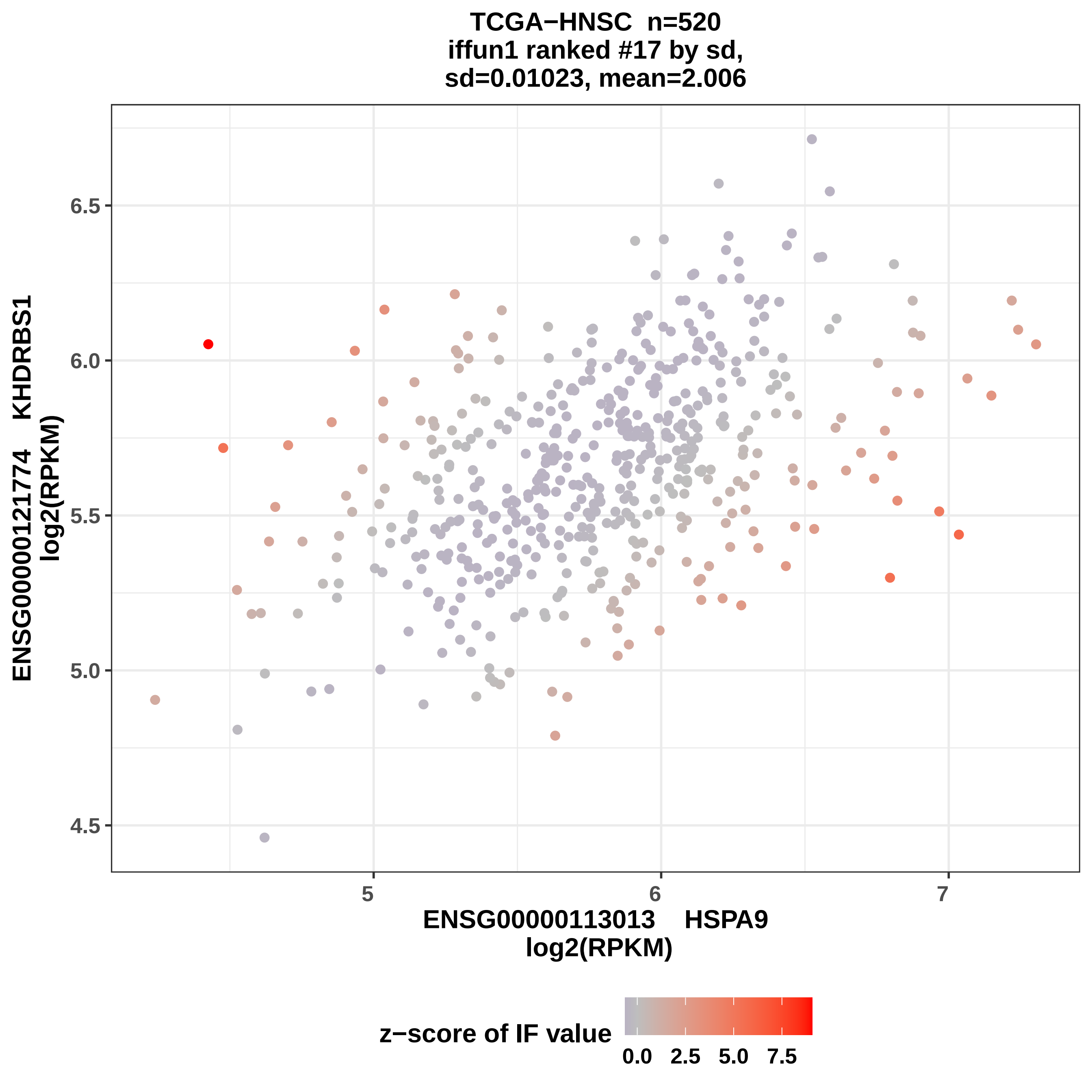
TCGA−HNSC n=520
iffun1 ranked #17 by sd,
sd=0.01023, mean=2.006
ENSG00000121774 KHDRBS1
log2(RPKM)
6.5
6.0
5.5
5.0
4.5
5
6
7
ENSG00000113013 HSPA9
log2(RPKM)
z−score of IF value
0.0
2.5
5.0
7.5

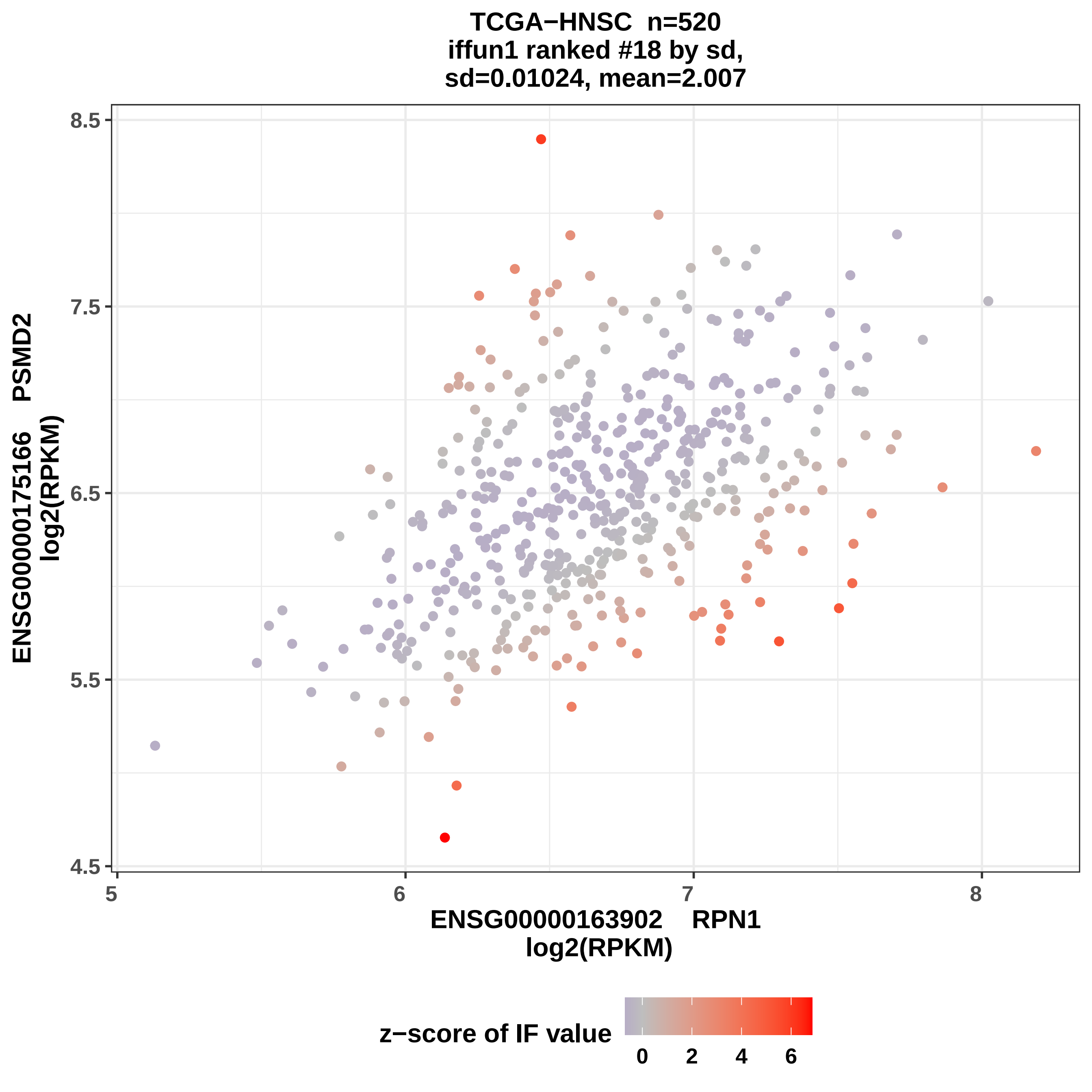

TCGA−HNSC n=520
iffun1 ranked #18 by sd,
sd=0.01024, mean=2.007
ENSG00000175166 PSMD2
log2(RPKM)
8.5
7.5
6.5
5.5
4.5
5
6
7
8
ENSG00000163902 RPN1
log2(RPKM)
z−score of IF value
0
2
4
6

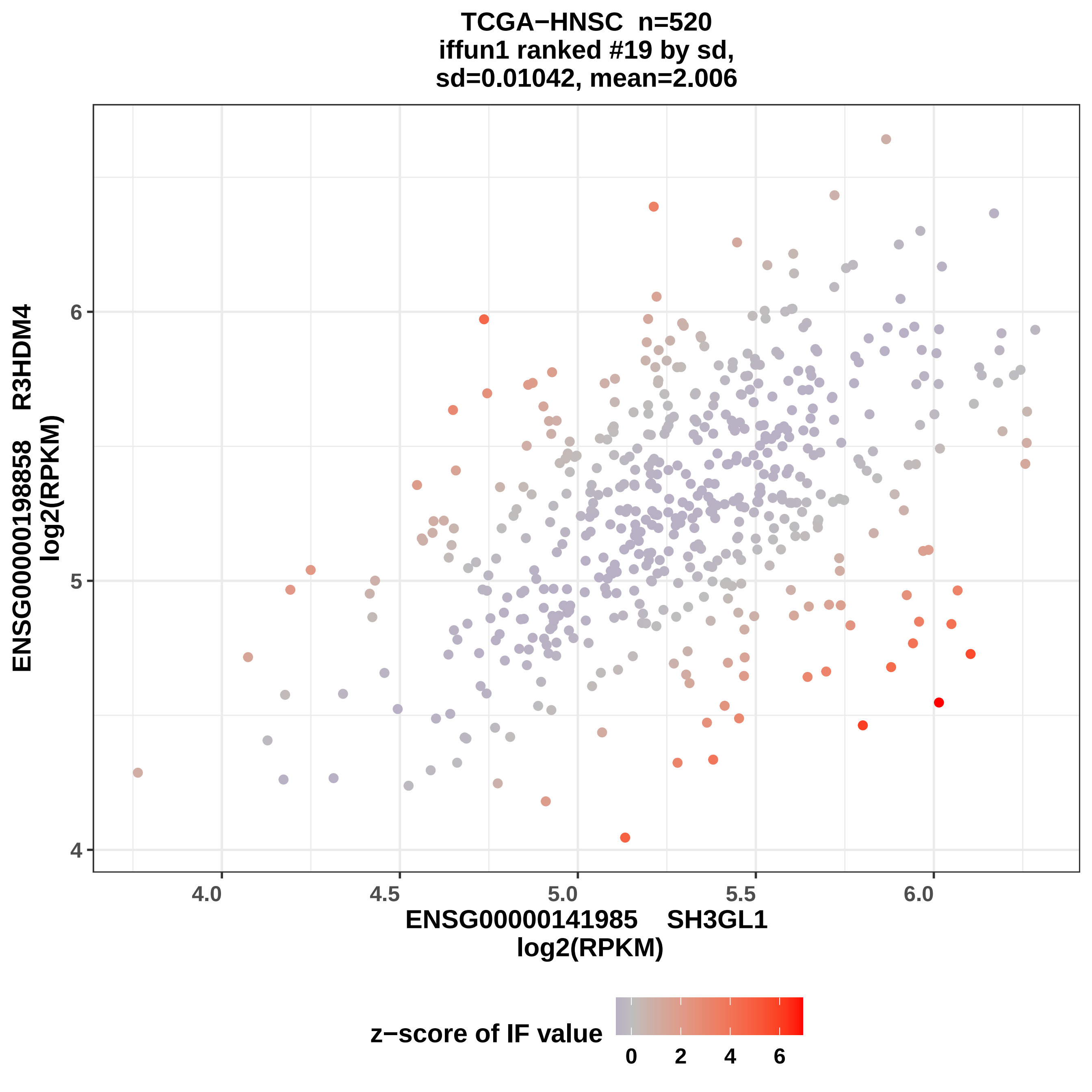

TCGA−HNSC n=520
iffun1 ranked #19 by sd,
sd=0.01042, mean=2.006
ENSG00000198858 R3HDM4
log2(RPKM)
4
5
6
4.0
4.5
5.0
5.5
6.0
ENSG00000141985 SH3GL1
log2(RPKM)
z−score of IF value
0
2
4
6

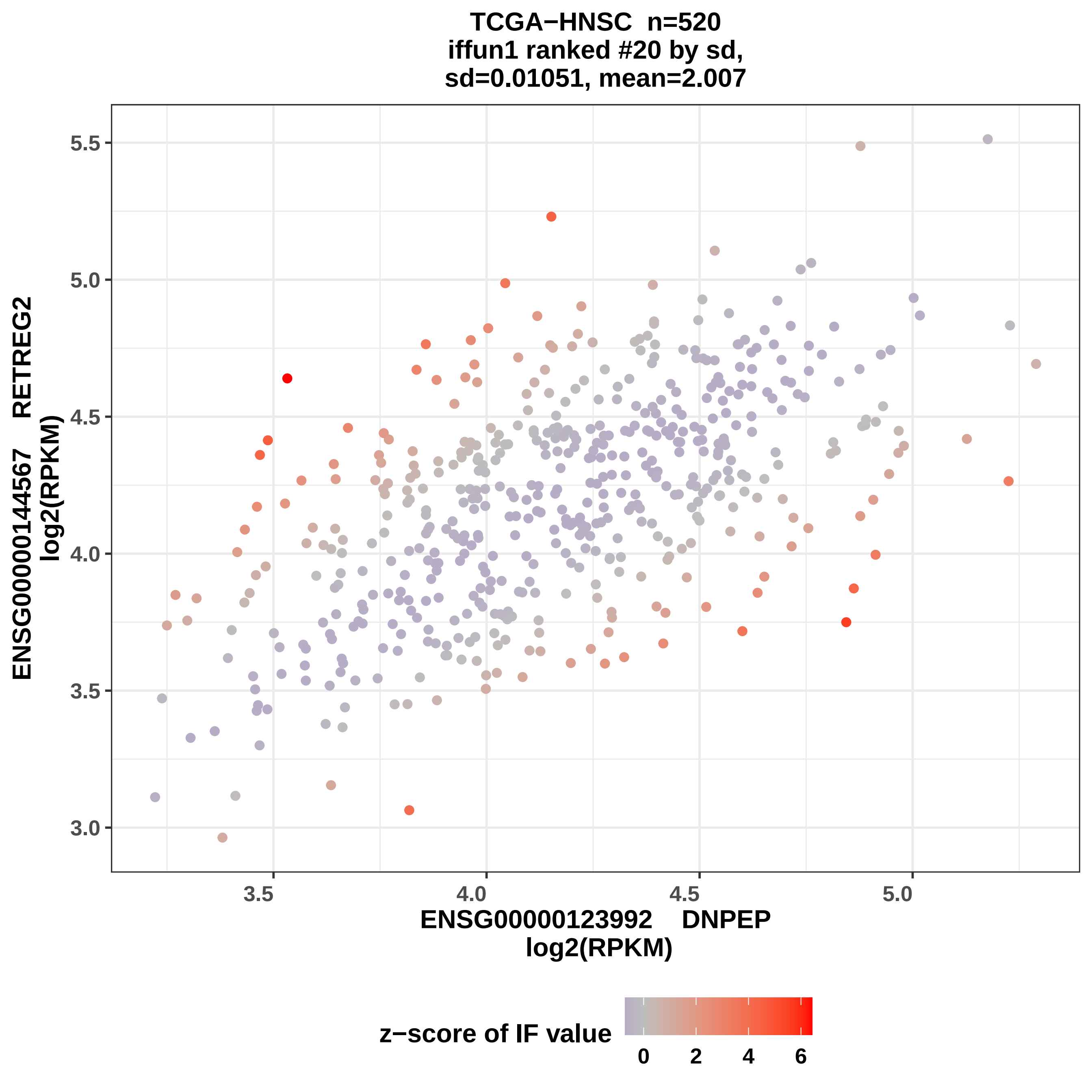
TCGA−HNSC n=520
iffun1 ranked #20 by sd,
sd=0.01051, mean=2.007
ENSG00000144567 RETREG2
log2(RPKM)
ENSG00000123992 DNPEP
log2(RPKM)
3.0
3.5
4.0
4.5
5.0
5.5
z−score of IF value
0
2
4
6

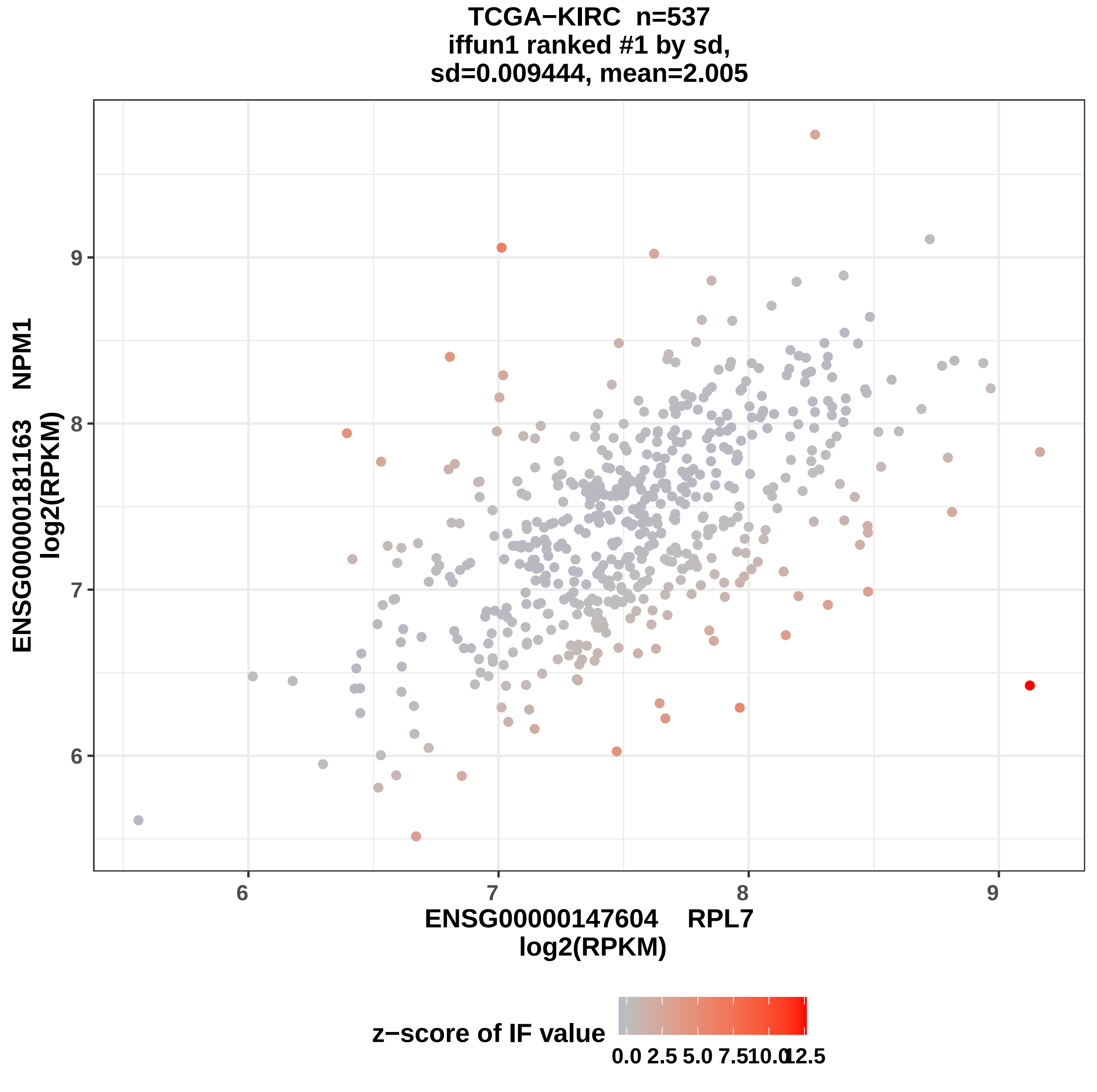
TCGA−KIRC n=537
iffun1 ranked #1 by sd,
sd=0.009444, mean=2.005
ENSG00000181163 NPM1
log2(RPKM)
ENSG00000147604 RPL7
log2(RPKM)
6
7
8
9
z−score of IF value
0.0
2.5
5.0
7.5
10.0
12.5

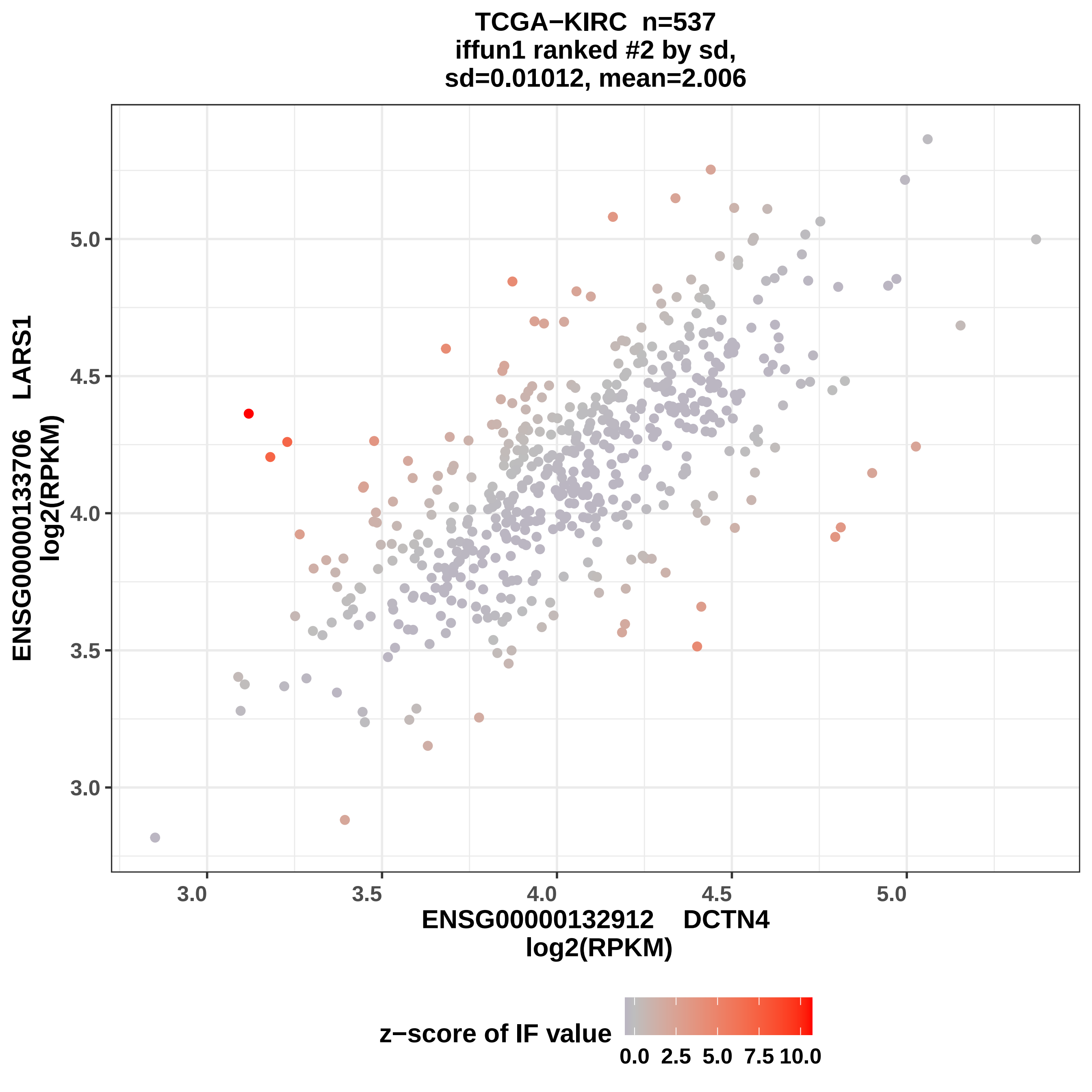

TCGA−KIRC n=537
iffun1 ranked #2 by sd,
sd=0.01012, mean=2.006
ENSG00000133706 LARS1
log2(RPKM)
5.0
4.5
4.0
3.5
3.0
3.0
3.5
4.0
4.5
5.0
ENSG00000132912 DCTN4
log2(RPKM)
z−score of IF value
0.0
2.5
5.0
7.5
10.0

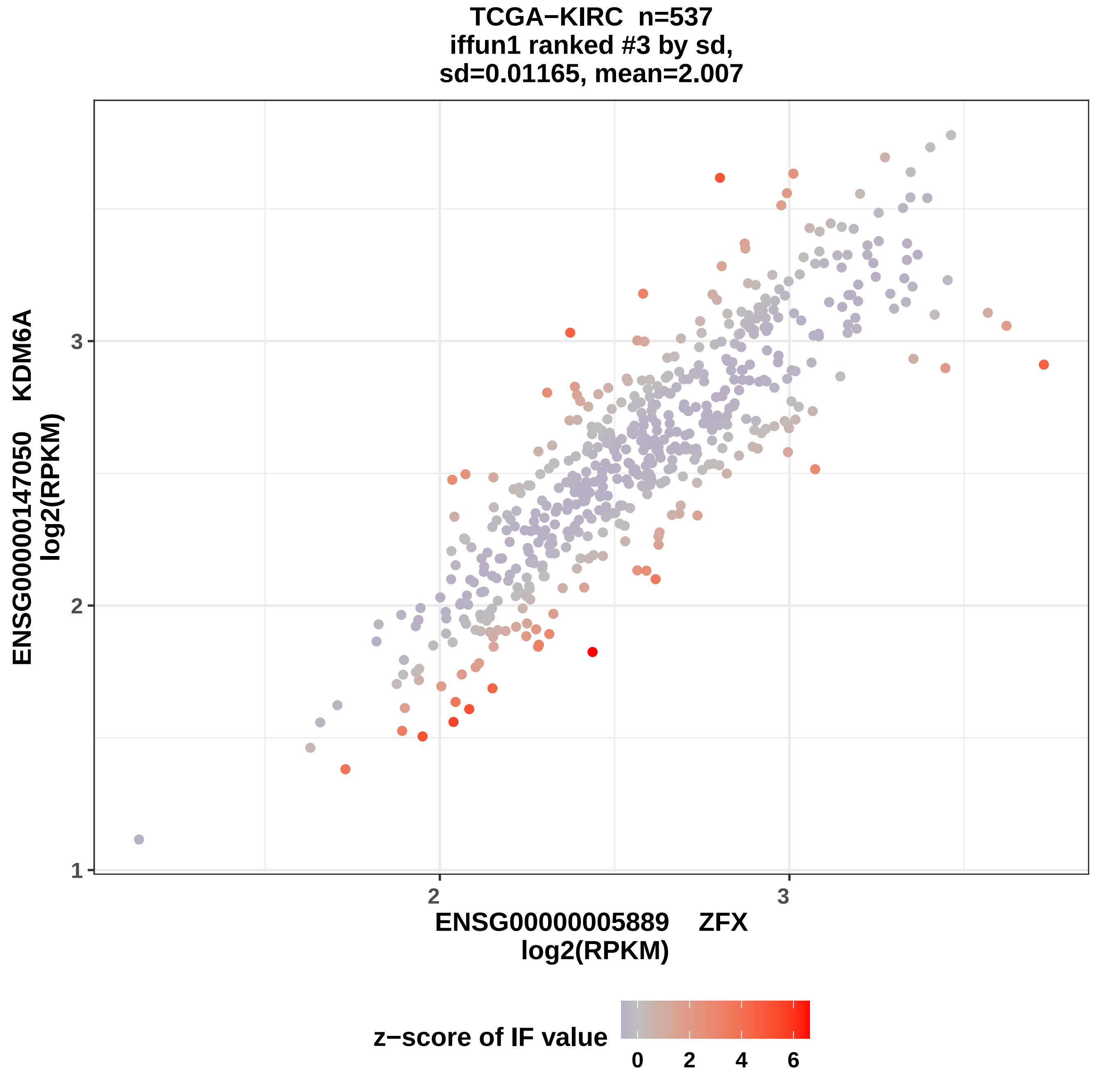

TCGA−KIRC n=537
iffun1 ranked #3 by sd,
sd=0.01165, mean=2.007
ENSG00000147050 KDM6A
log2(RPKM)
1
2
3
2
3
ENSG00000005889 ZFX
log2(RPKM)
z−score of IF value
0
2
4
6

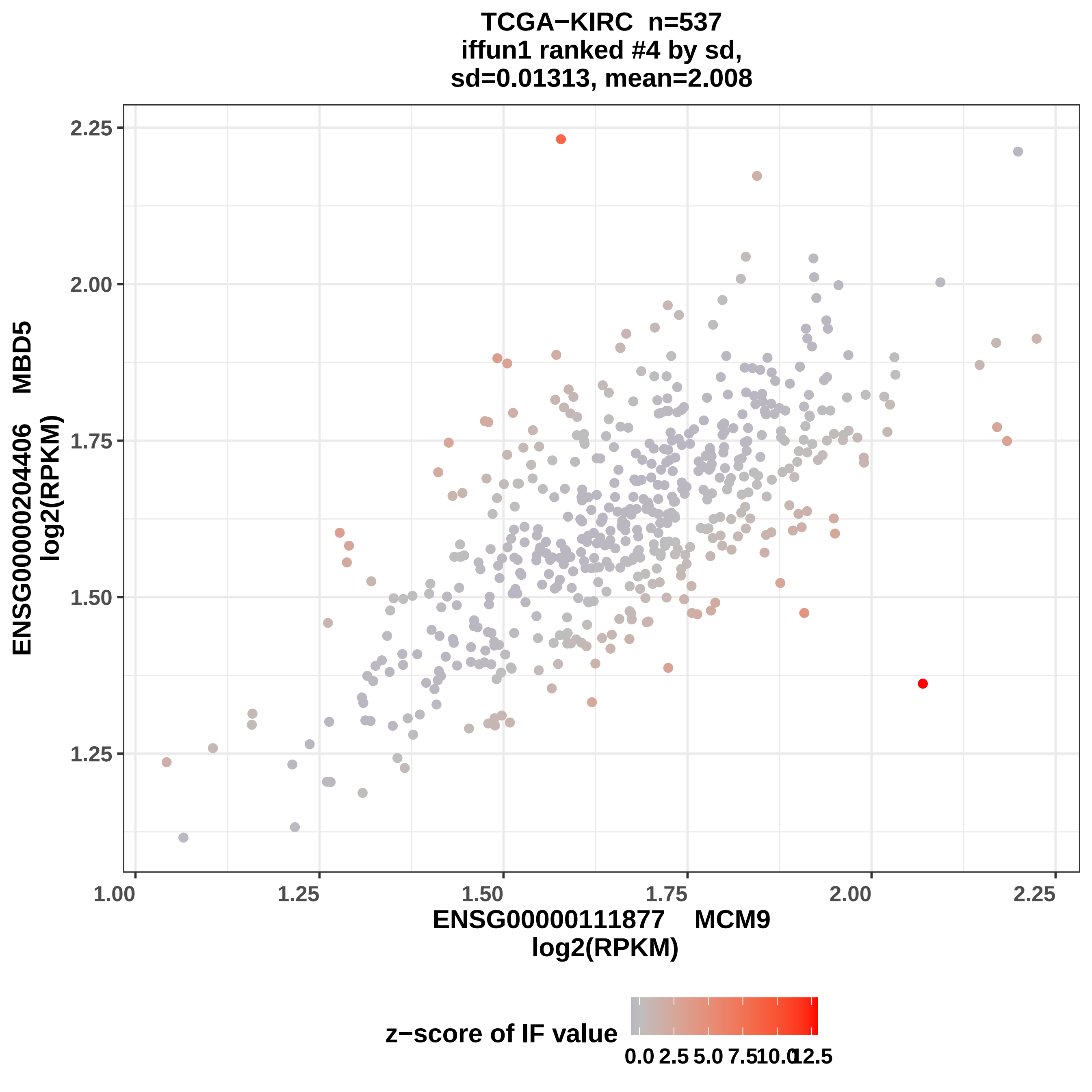
TCGA−KIRC n=537
iffun1 ranked #4 by sd,
sd=0.01313, mean=2.008
ENSG00000204406 MBD5
log2(RPKM)
ENSG00000111877 MCM9
log2(RPKM)
1.00
1.25
1.50
1.75
2.00
2.25
z−score of IF value
0.0
2.5
5.0
7.5
10.0
12.5

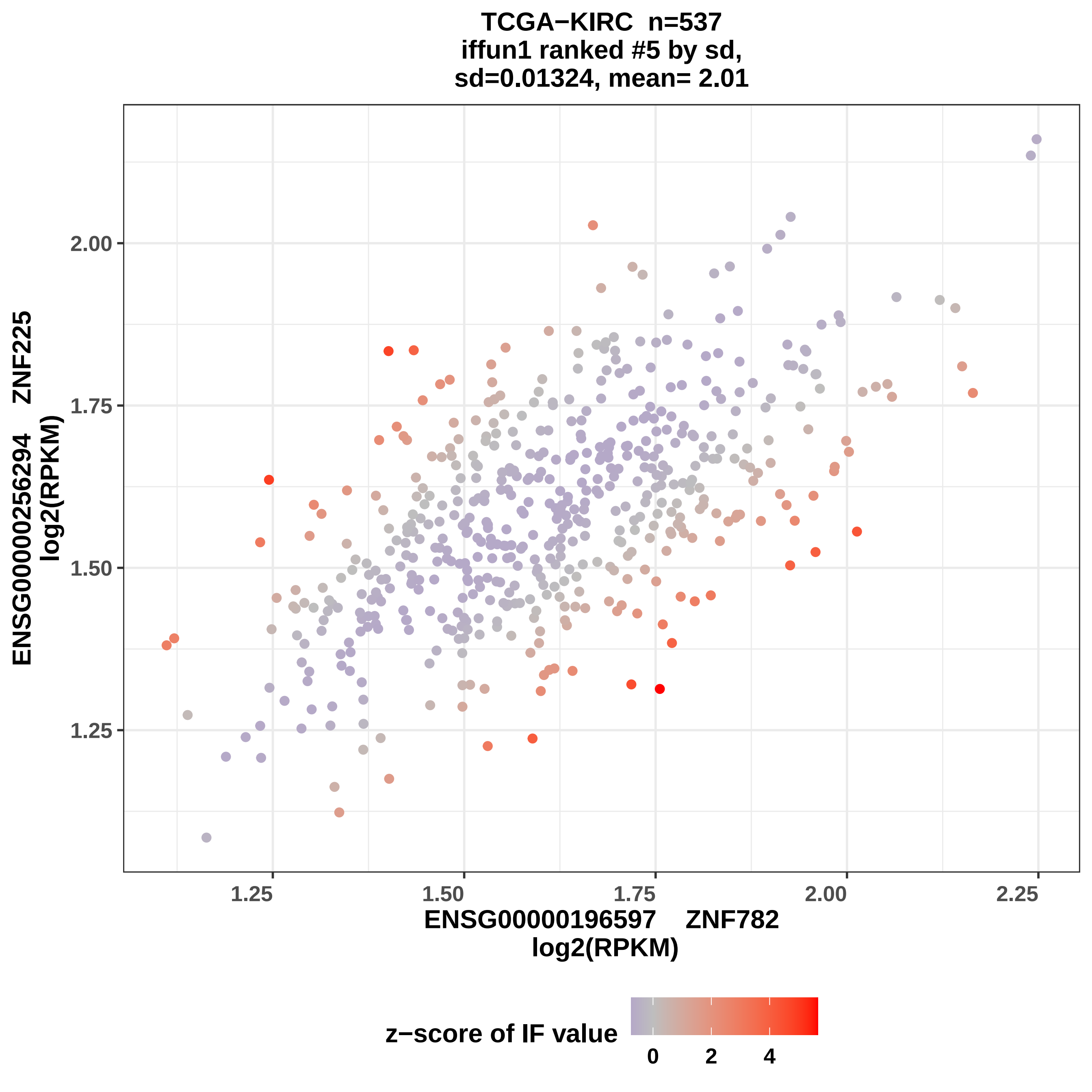
TCGA−KIRC n=537
iffun1 ranked #5 by sd,
sd=0.01324, mean= 2.01
ENSG00000256294 ZNF225
log2(RPKM)
2.00
1.75
1.50
1.25
1.25
1.50
1.75
2.00
2.25
ENSG00000196597 ZNF782
log2(RPKM)
z−score of IF value
0
2
4

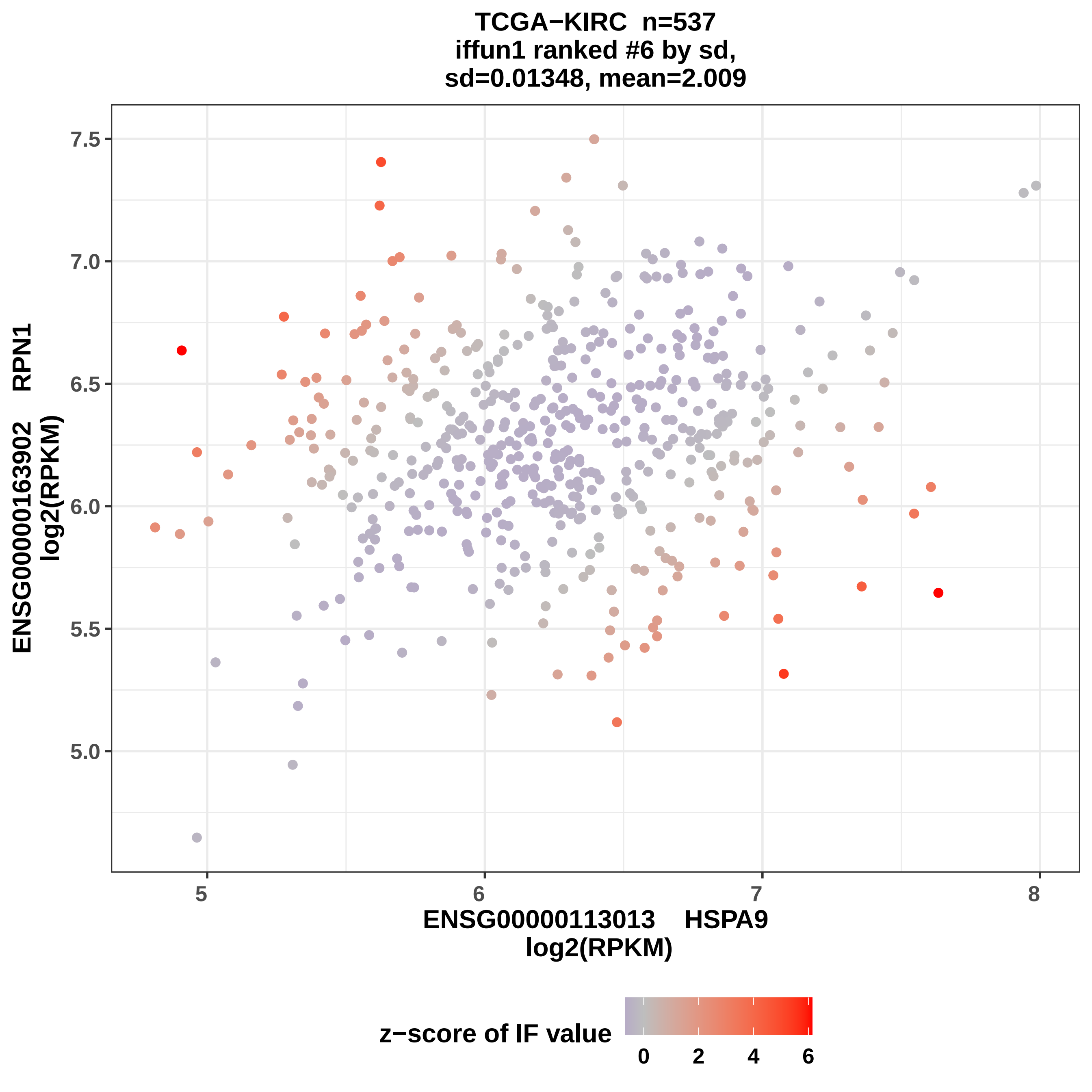
TCGA−KIRC n=537
iffun1 ranked #6 by sd,
sd=0.01348, mean=2.009
ENSG00000163902 RPN1
log2(RPKM)
7.5
7.0
6.5
6.0
5.5
5.0
5
6
7
8
ENSG00000113013 HSPA9
log2(RPKM)
z−score of IF value
0
2
4
6

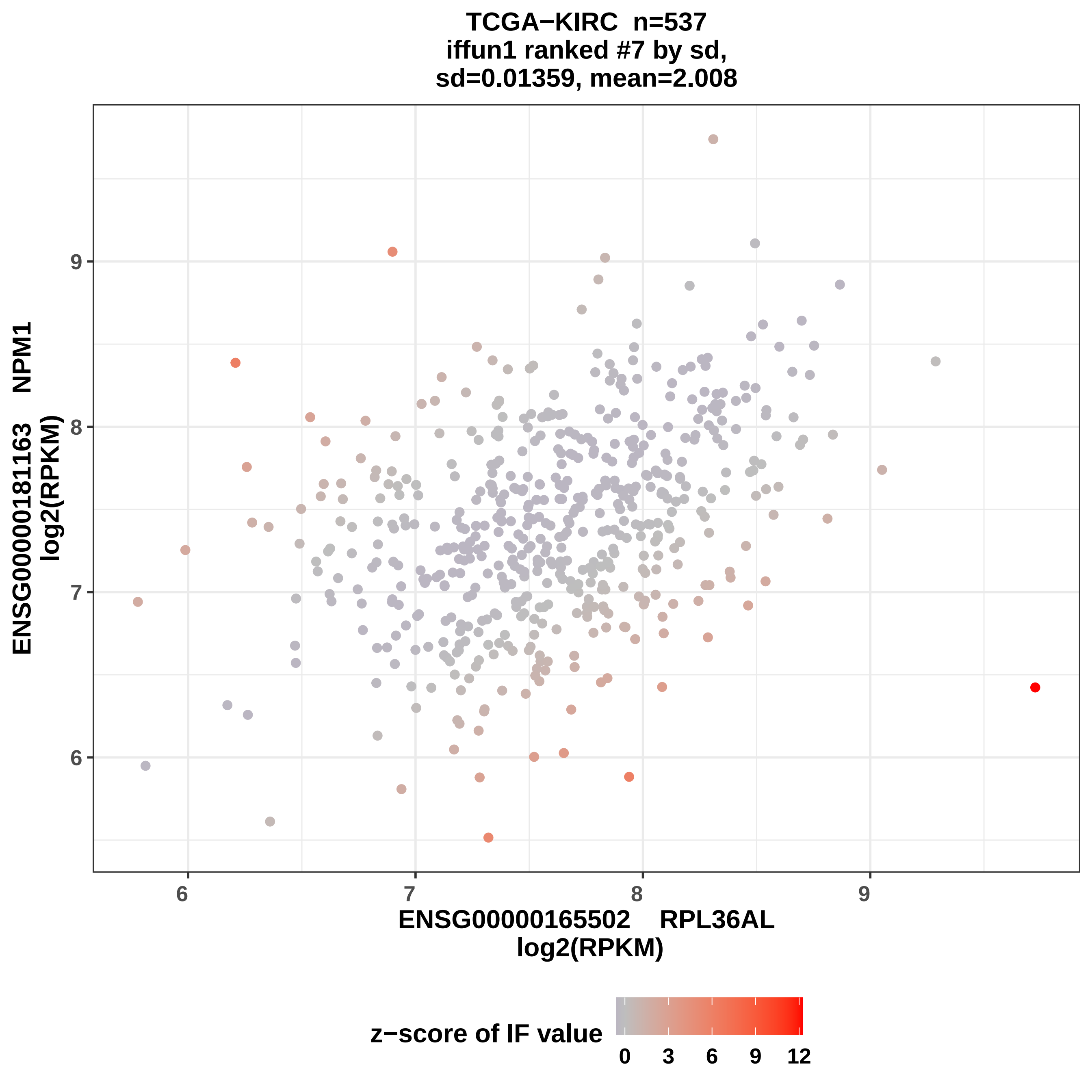
TCGA−KIRC n=537
iffun1 ranked #7 by sd,
sd=0.01359, mean=2.008
ENSG00000181163 NPM1
log2(RPKM)
9
8
7
6
6
7
8
9
ENSG00000165502 RPL36AL
log2(RPKM)
z−score of IF value
0
3
6
9
12

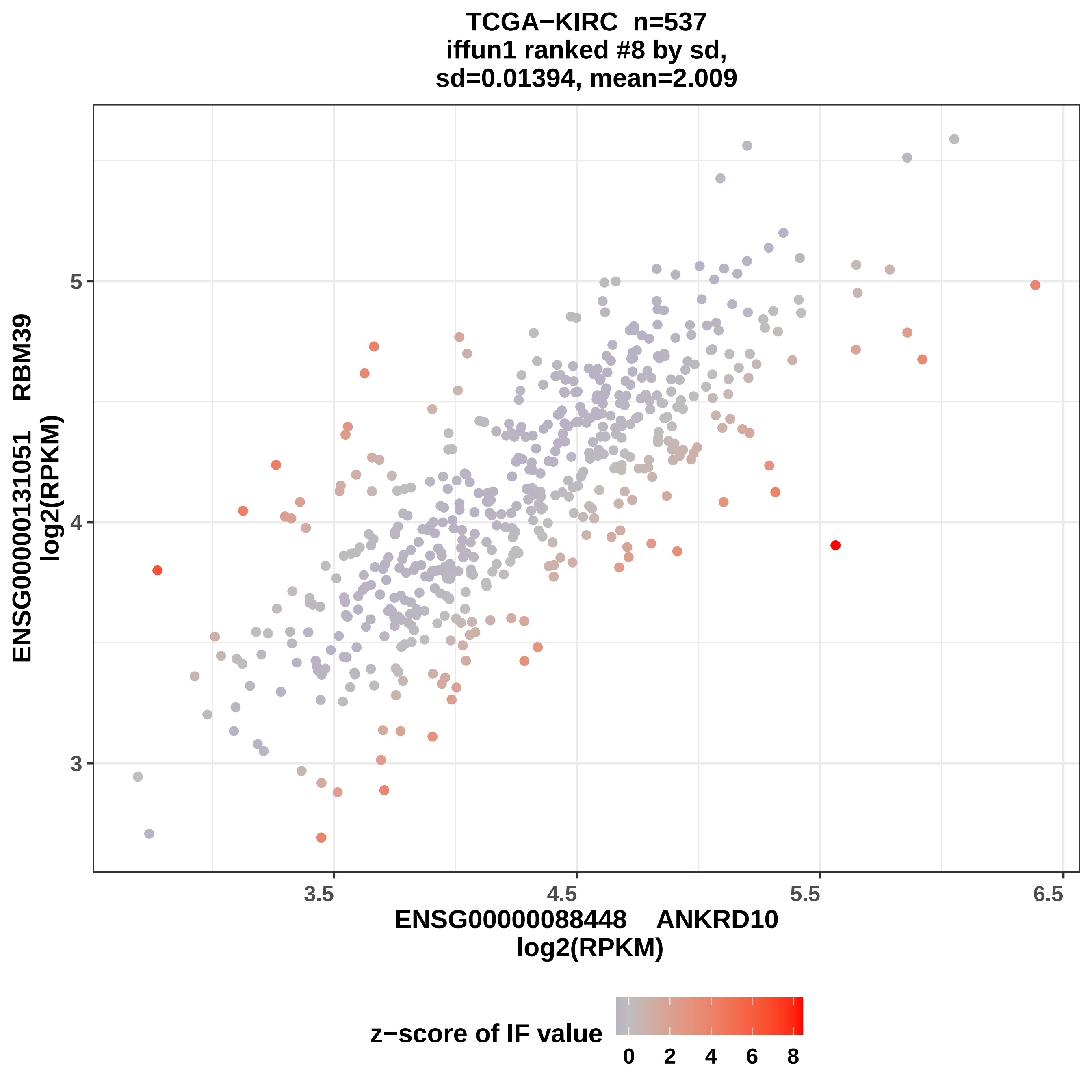
TCGA−KIRC n=537
iffun1 ranked #8 by sd,
sd=0.01394, mean=2.009
ENSG00000131051 RBM39
log2(RPKM)
5
4
3
3.5
4.5
5.5
6.5
ENSG00000088448 ANKRD10
log2(RPKM)
z−score of IF value
0
2
4
6
8

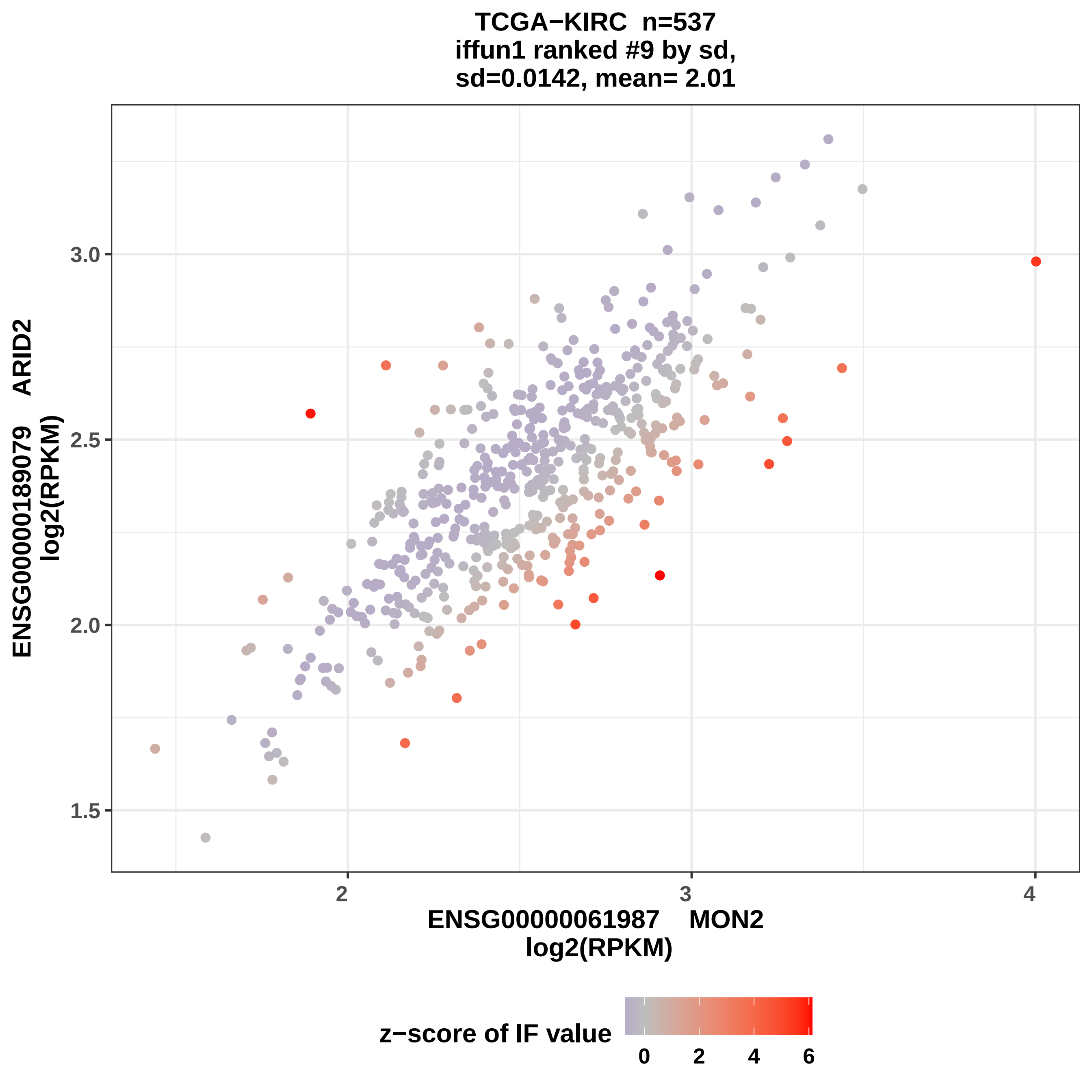

TCGA−KIRC n=537
iffun1 ranked #9 by sd,
sd=0.0142, mean= 2.01
ENSG00000189079 ARID2
log2(RPKM)
1.5
2.0
2.5
3.0
2
3
4
ENSG00000061987 MON2
log2(RPKM)
z−score of IF value
0
2
4
6

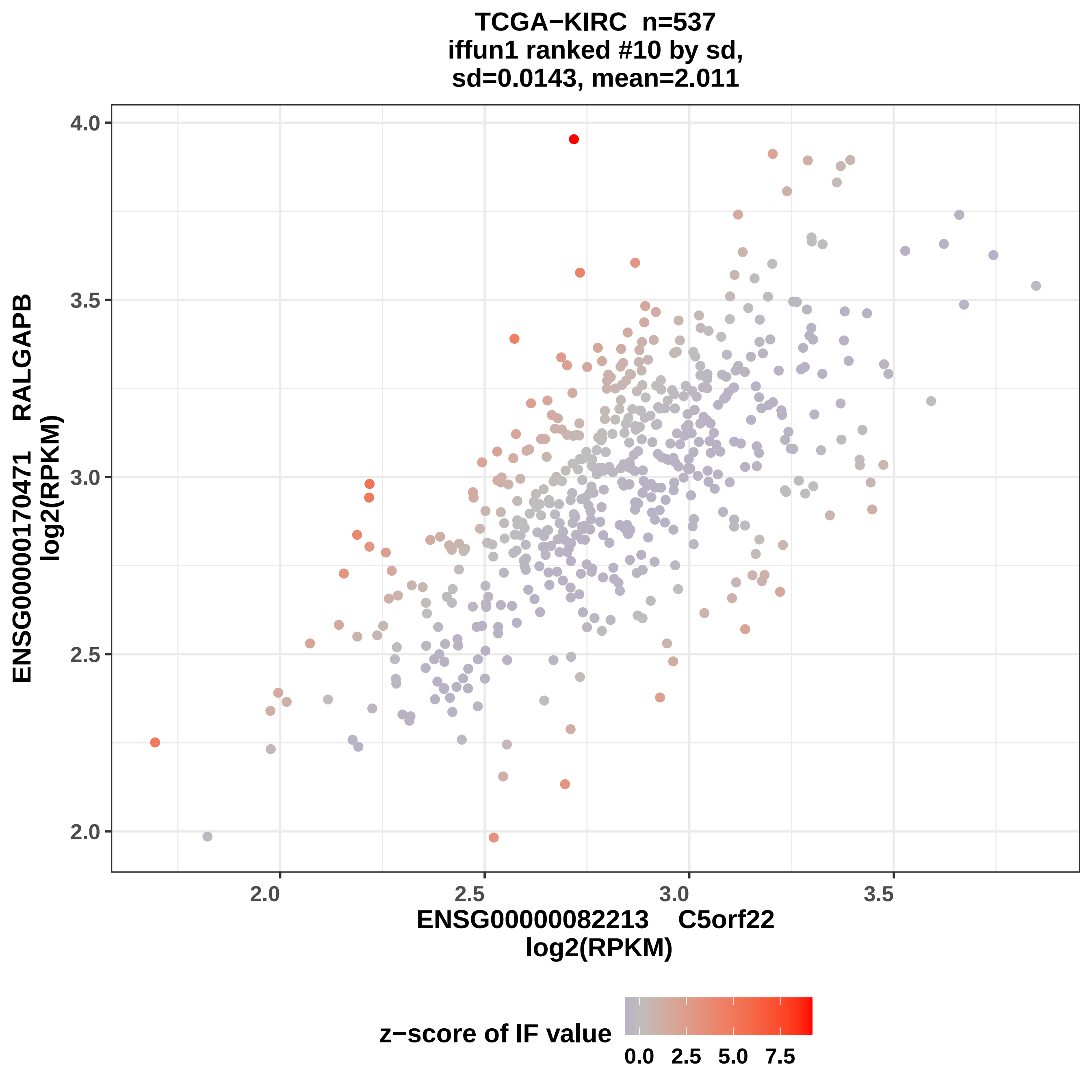

TCGA−KIRC n=537
iffun1 ranked #10 by sd,
sd=0.0143, mean=2.011
ENSG00000170471 RALGAPB
log2(RPKM)
4.0
3.5
3.0
2.5
2.0
2.0
2.5
3.0
3.5
ENSG00000082213 C5orf22
log2(RPKM)
z−score of IF value
0.0
2.5
5.0
7.5

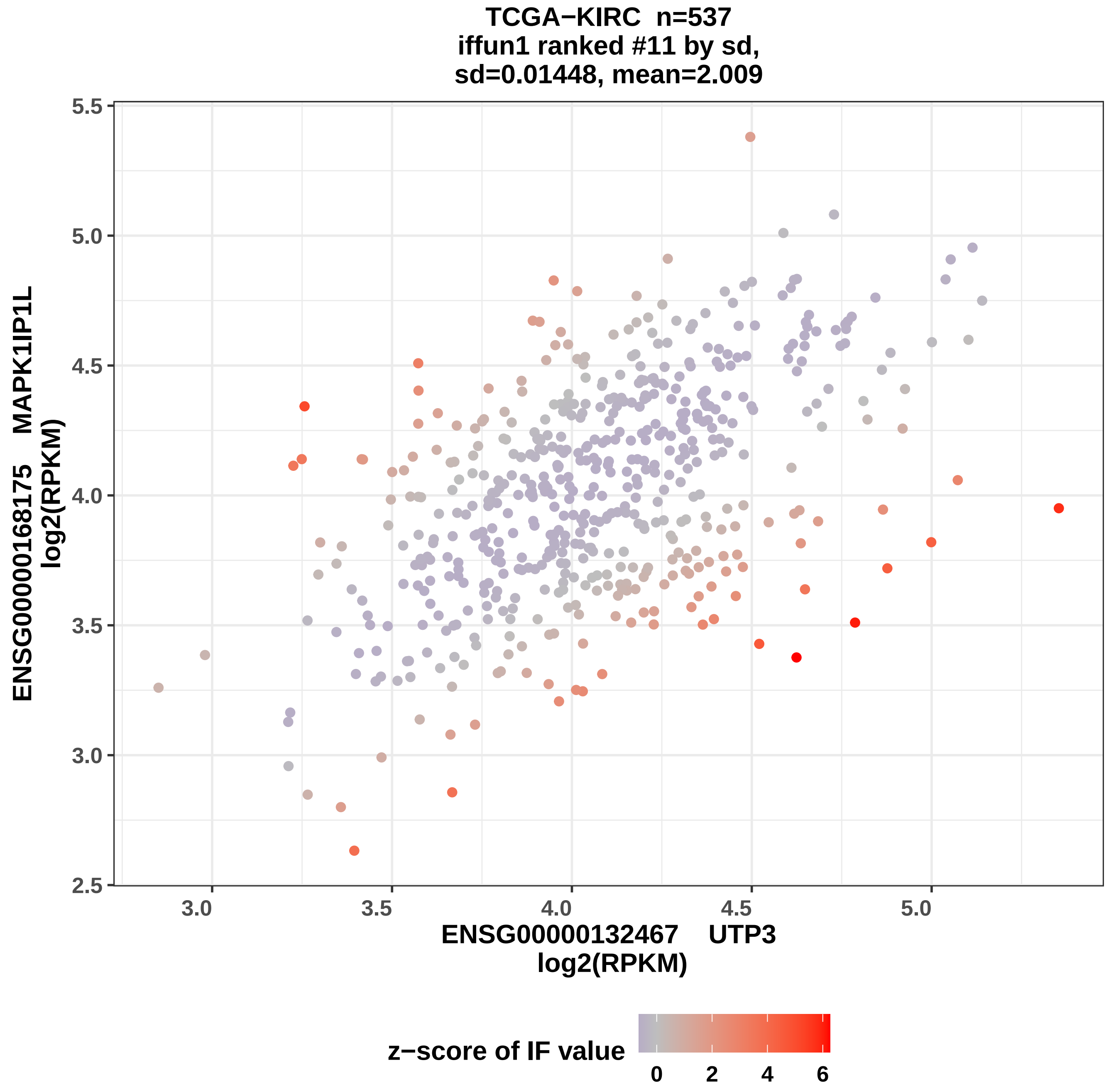

TCGA−KIRC n=537
iffun1 ranked #11 by sd,
sd=0.01448, mean=2.009
ENSG00000168175 MAPK1IP1L
log2(RPKM)
5.5
5.0
4.5
4.0
3.5
3.0
2.5
3.0
3.5
4.0
4.5
5.0
ENSG00000132467 UTP3
log2(RPKM)
z−score of IF value
0
2
4
6

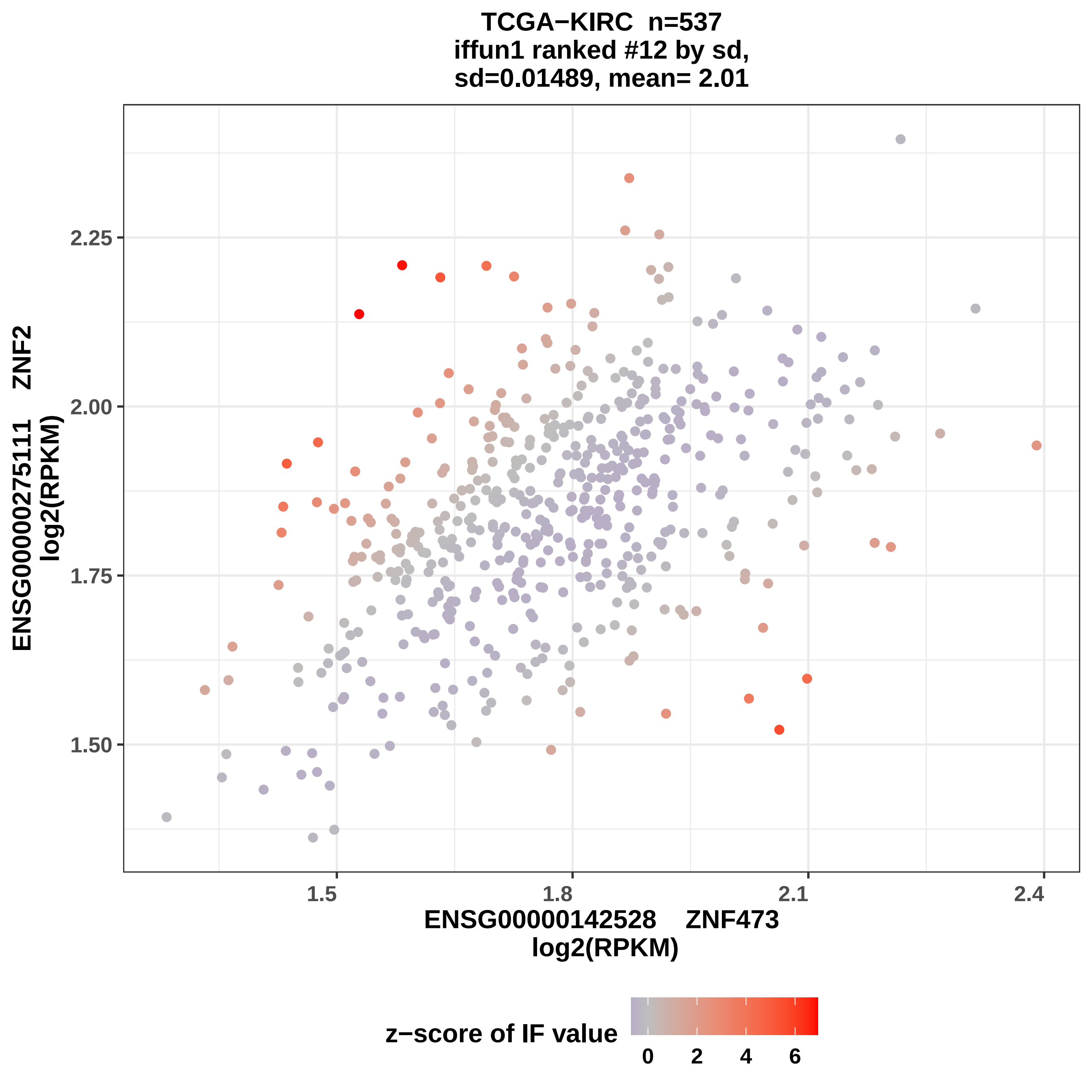
TCGA−KIRC n=537
iffun1 ranked #12 by sd,
sd=0.01489, mean= 2.01
ENSG00000275111 ZNF2
log2(RPKM)
2.25
2.00
1.75
1.50
1.5
1.8
2.1
2.4
ENSG00000142528 ZNF473
log2(RPKM)
z−score of IF value
0
2
4
6

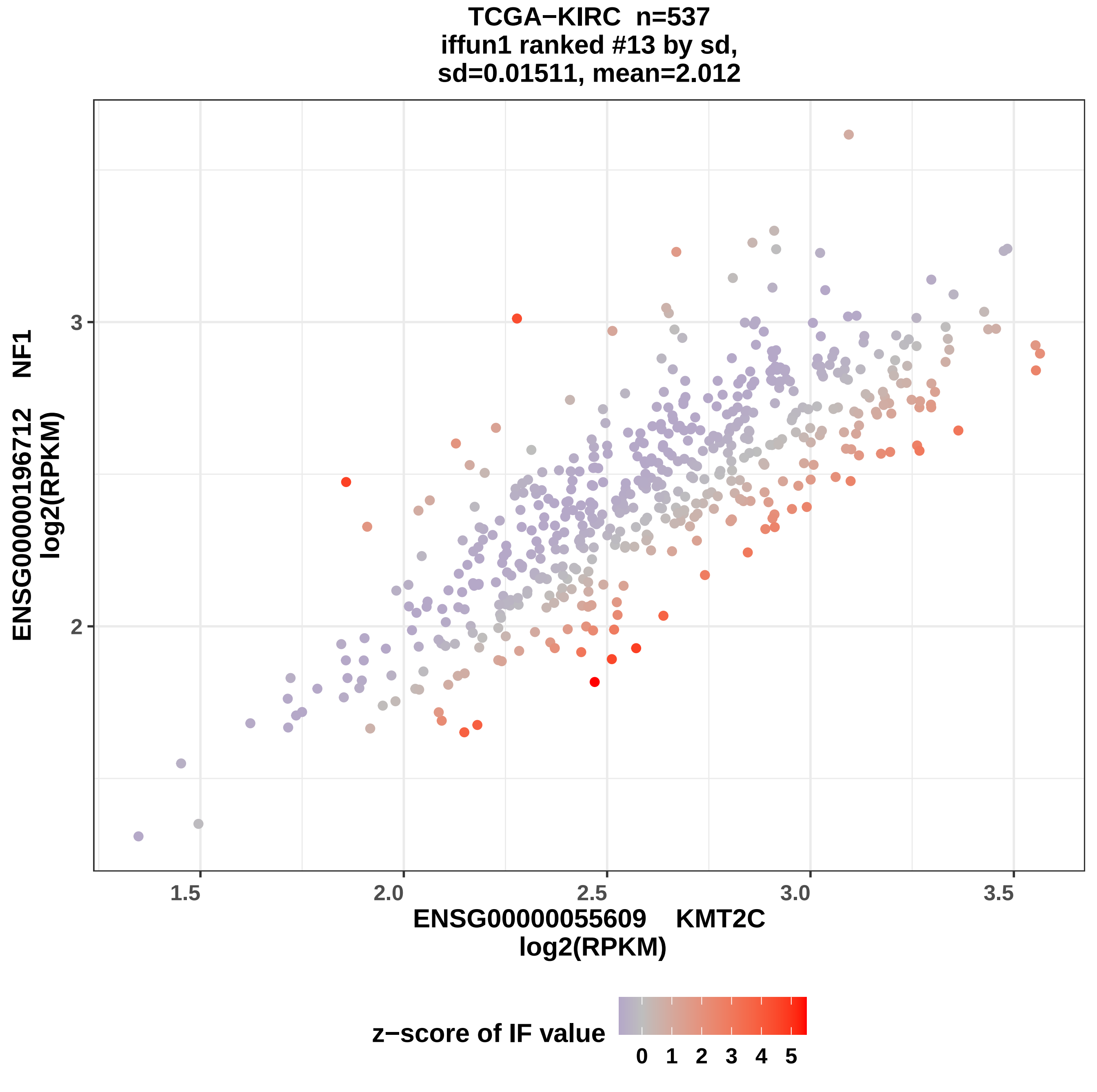
TCGA−KIRC n=537
iffun1 ranked #13 by sd,
sd=0.01511, mean=2.012
ENSG00000196712 NF1
log2(RPKM)
3
2
1.5
2.0
2.5
3.0
3.5
ENSG00000055609 KMT2C
log2(RPKM)
z−score of IF value
0 1 2 3 4 5

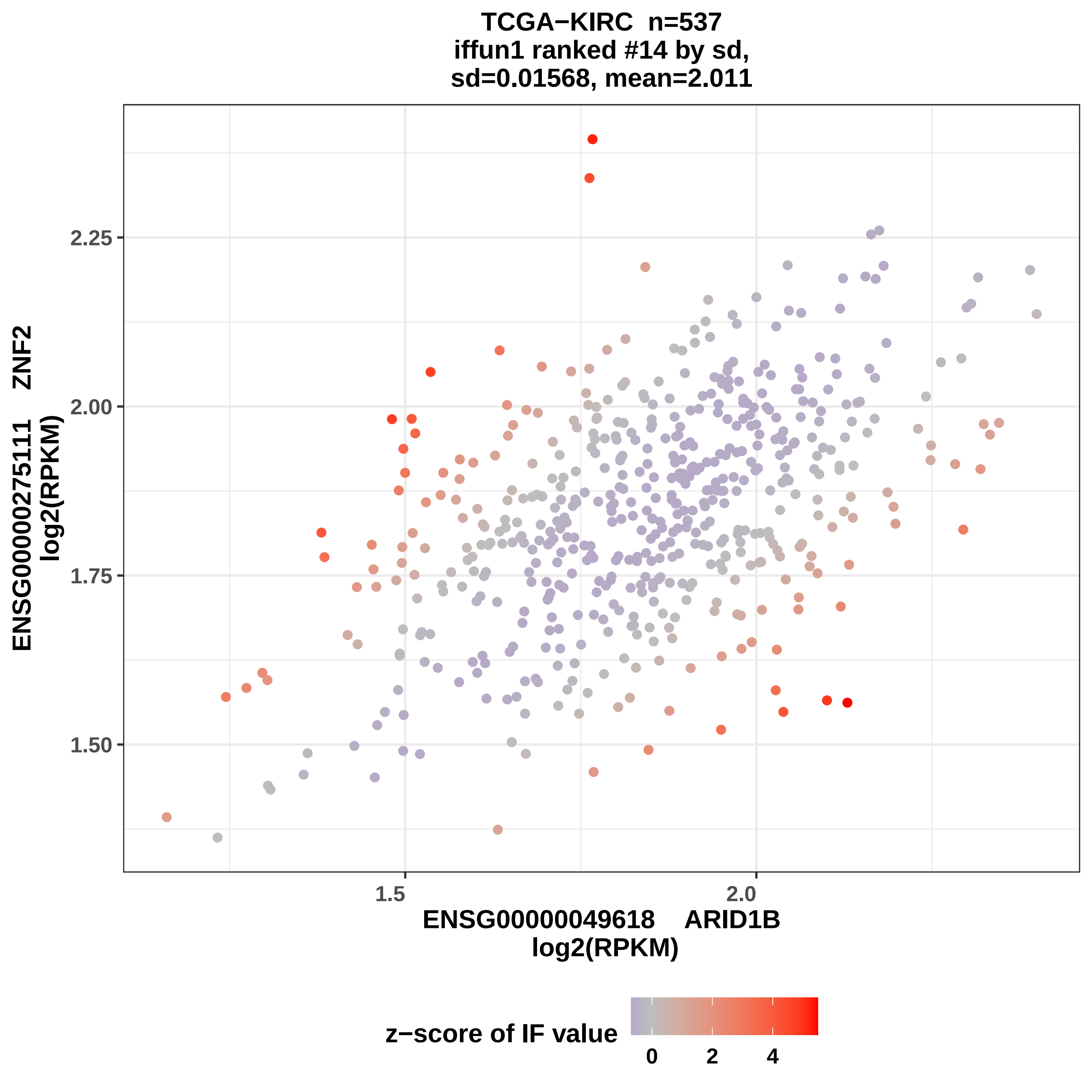

TCGA−KIRC n=537
iffun1 ranked #14 by sd,
sd=0.01568, mean=2.011
ENSG00000275111 ZNF2
log2(RPKM)
2.25
2.00
1.75
1.50
1.5
2.0
ENSG00000049618 ARID1B
log2(RPKM)
z−score of IF value
0
2
4

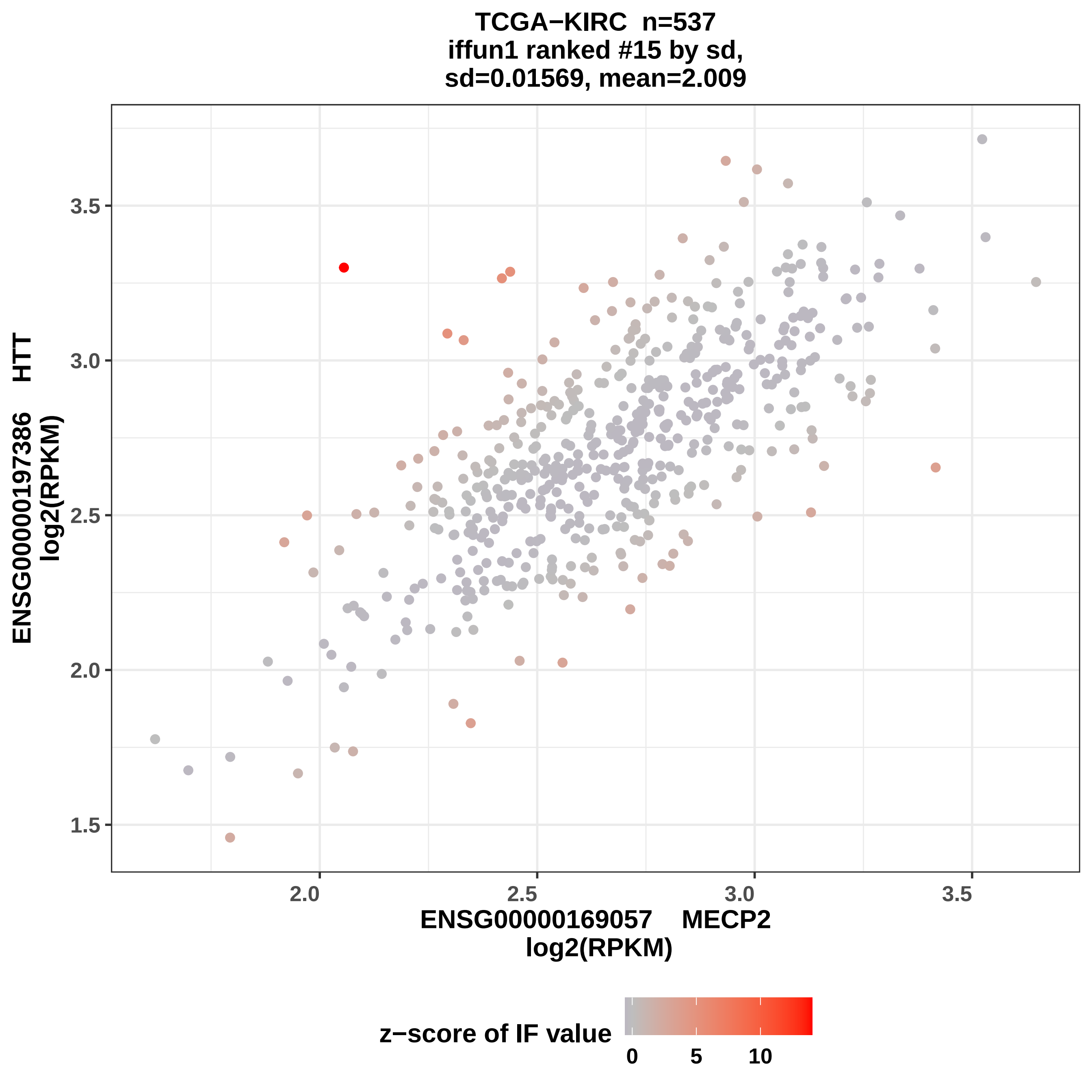
TCGA−KIRC n=537
iffun1 ranked #15 by sd,
sd=0.01569, mean=2.009
ENSG00000197386 HTT
log2(RPKM)
3.5
3.0
2.5
2.0
1.5
2.0
2.5
3.0
3.5
ENSG00000169057 MECP2
log2(RPKM)
z−score of IF value
0
5
10

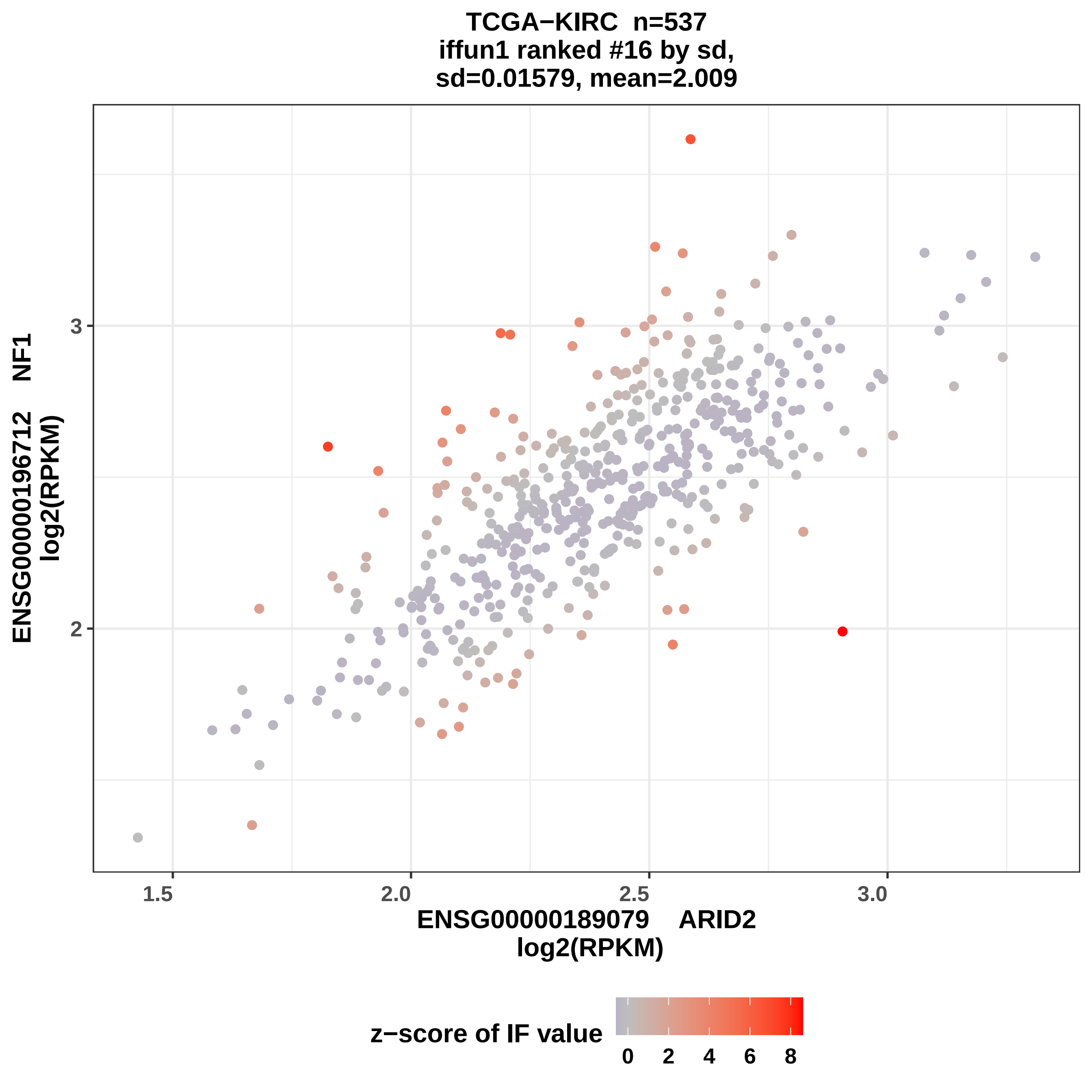
TCGA−KIRC n=537
iffun1 ranked #16 by sd,
sd=0.01579, mean=2.009
ENSG00000196712 NF1
log2(RPKM)
3
2
1.5
2.0
2.5
3.0
ENSG00000189079 ARID2
log2(RPKM)
z−score of IF value
0
2
4
6
8

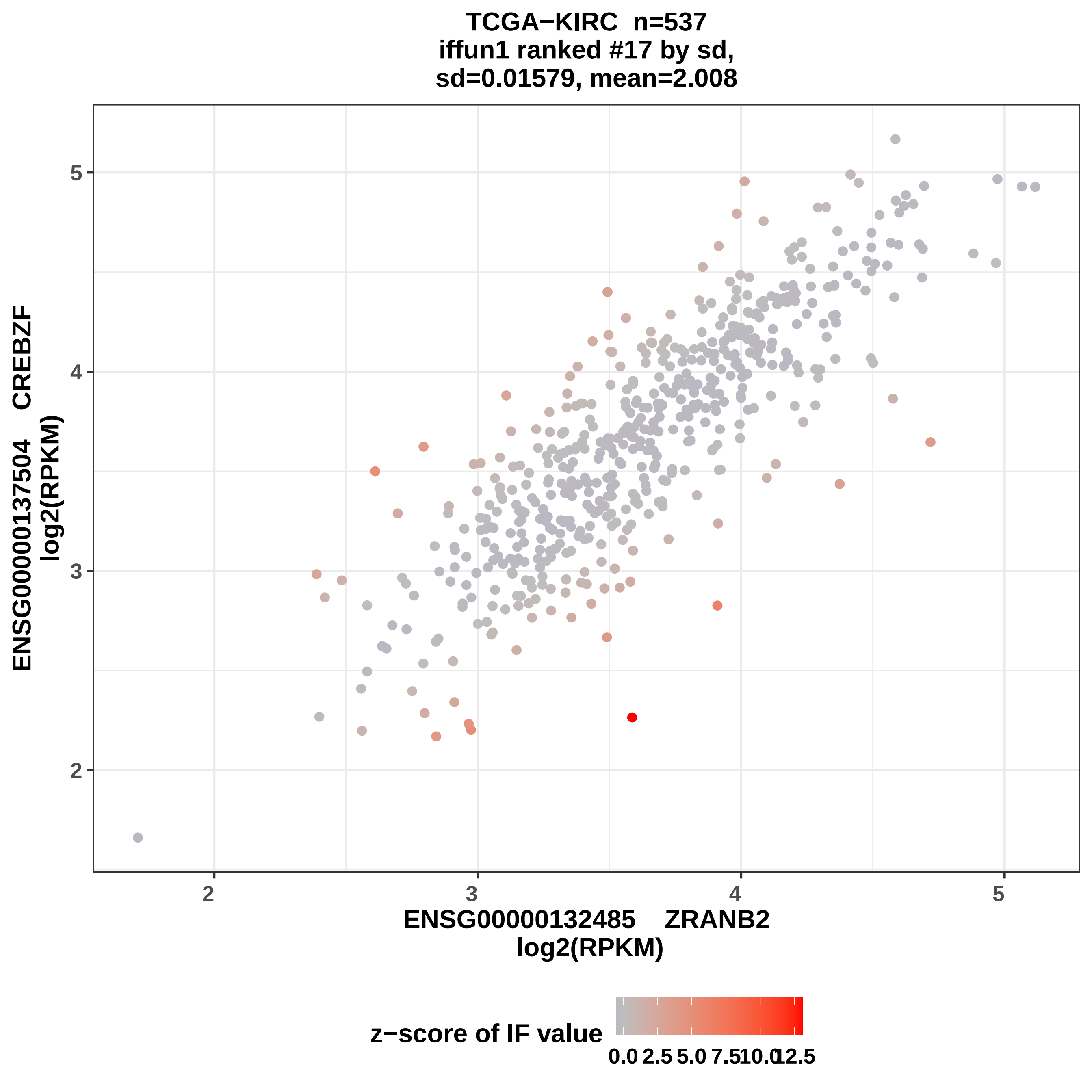

TCGA−KIRC n=537
iffun1 ranked #17 by sd,
sd=0.01579, mean=2.008
ENSG00000137504 CREBZF
log2(RPKM)
ENSG00000132485 ZRANB2
log2(RPKM)
2
3
4
5
2
3
4
5
z−score of IF value
0.0
2.5
5.0
7.5
10.0
12.5

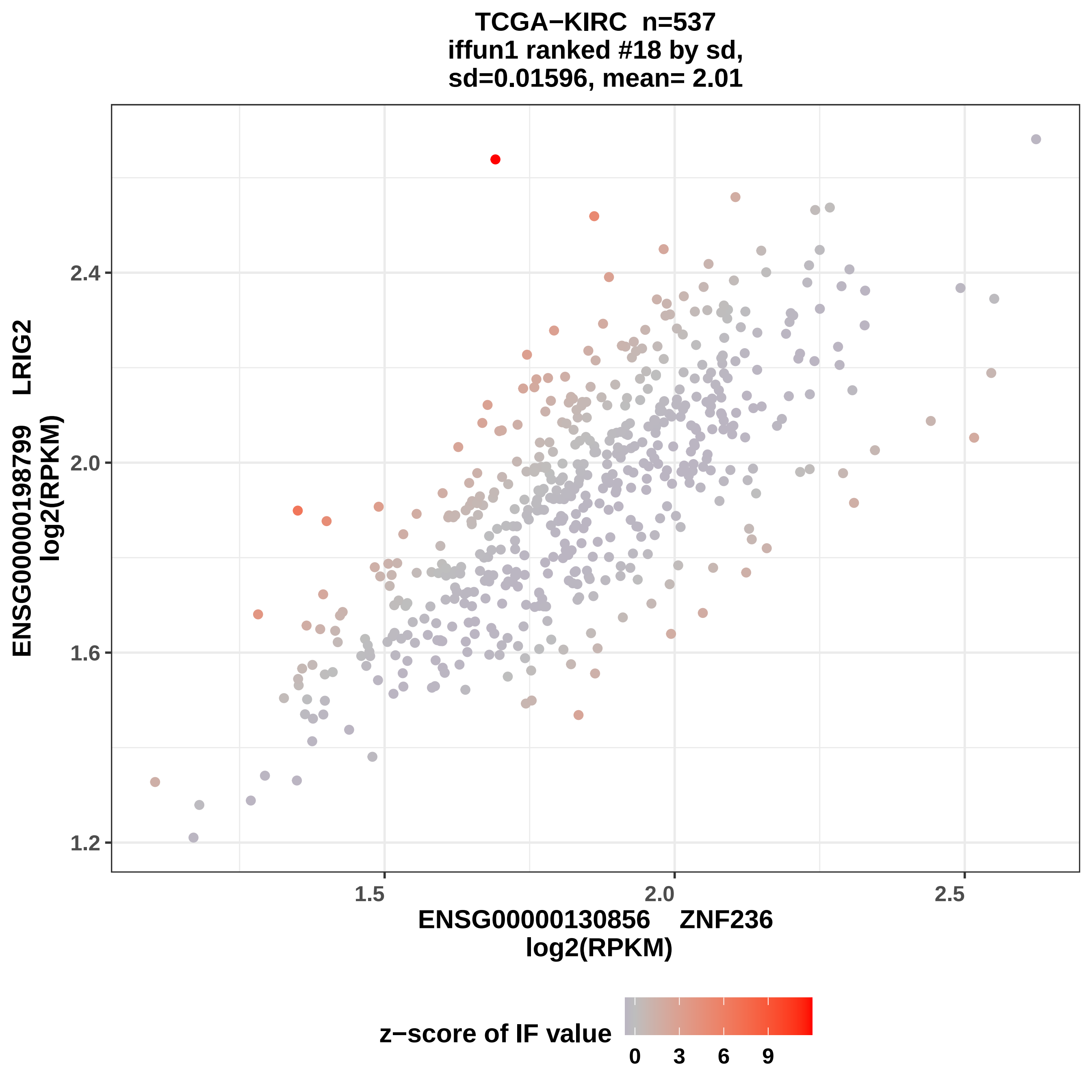

TCGA−KIRC n=537
iffun1 ranked #18 by sd,
sd=0.01596, mean= 2.01
ENSG00000198799 LRIG2
log2(RPKM)
2.4
2.0
1.6
1.2
1.5
2.0
2.5
ENSG00000130856 ZNF236
log2(RPKM)
z−score of IF value
0
3
6
9

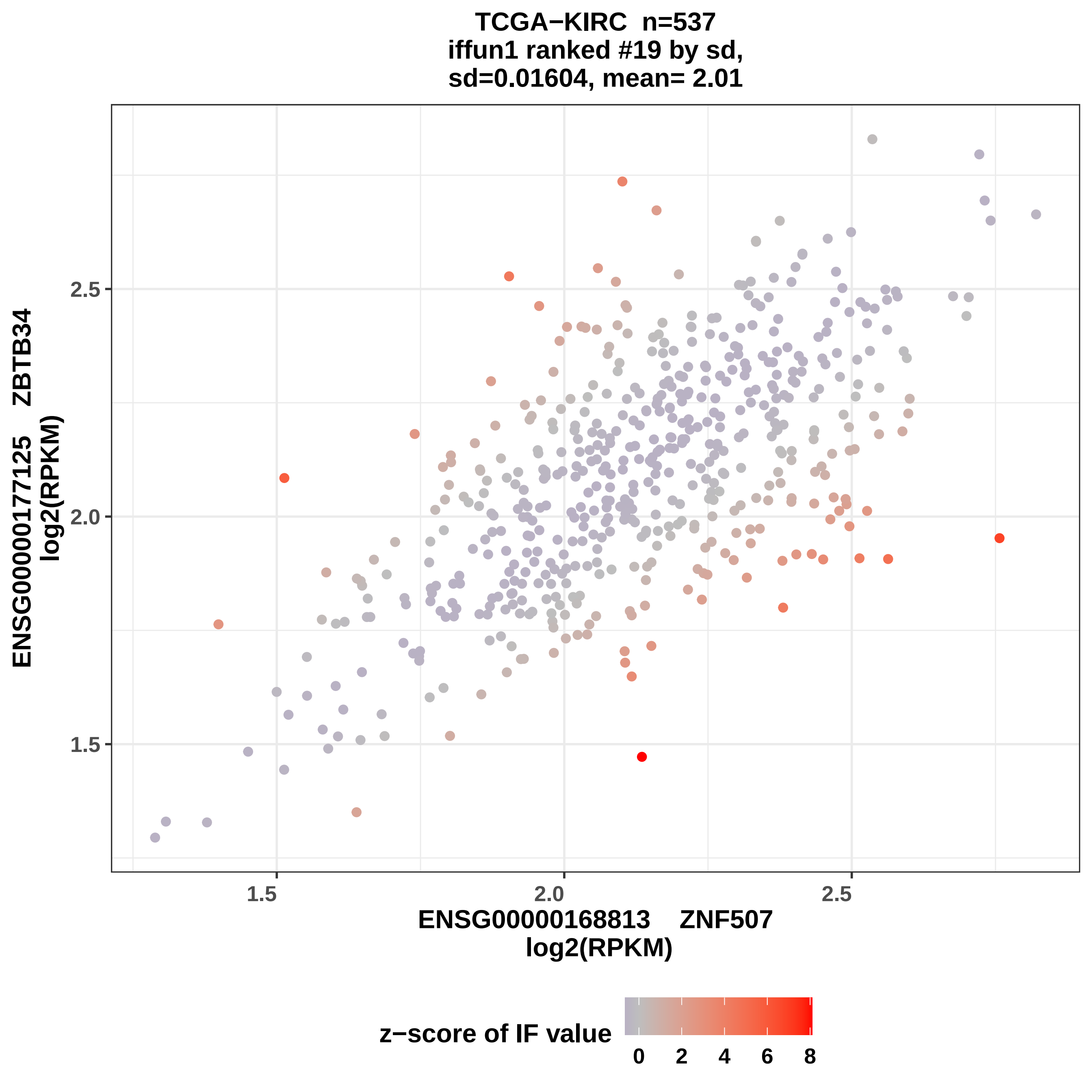

TCGA−KIRC n=537
iffun1 ranked #19 by sd,
sd=0.01604, mean= 2.01
ENSG00000177125 ZBTB34
log2(RPKM)
2.5
2.0
1.5
1.5
2.0
2.5
ENSG00000168813 ZNF507
log2(RPKM)
z−score of IF value
0
2
4
6
8

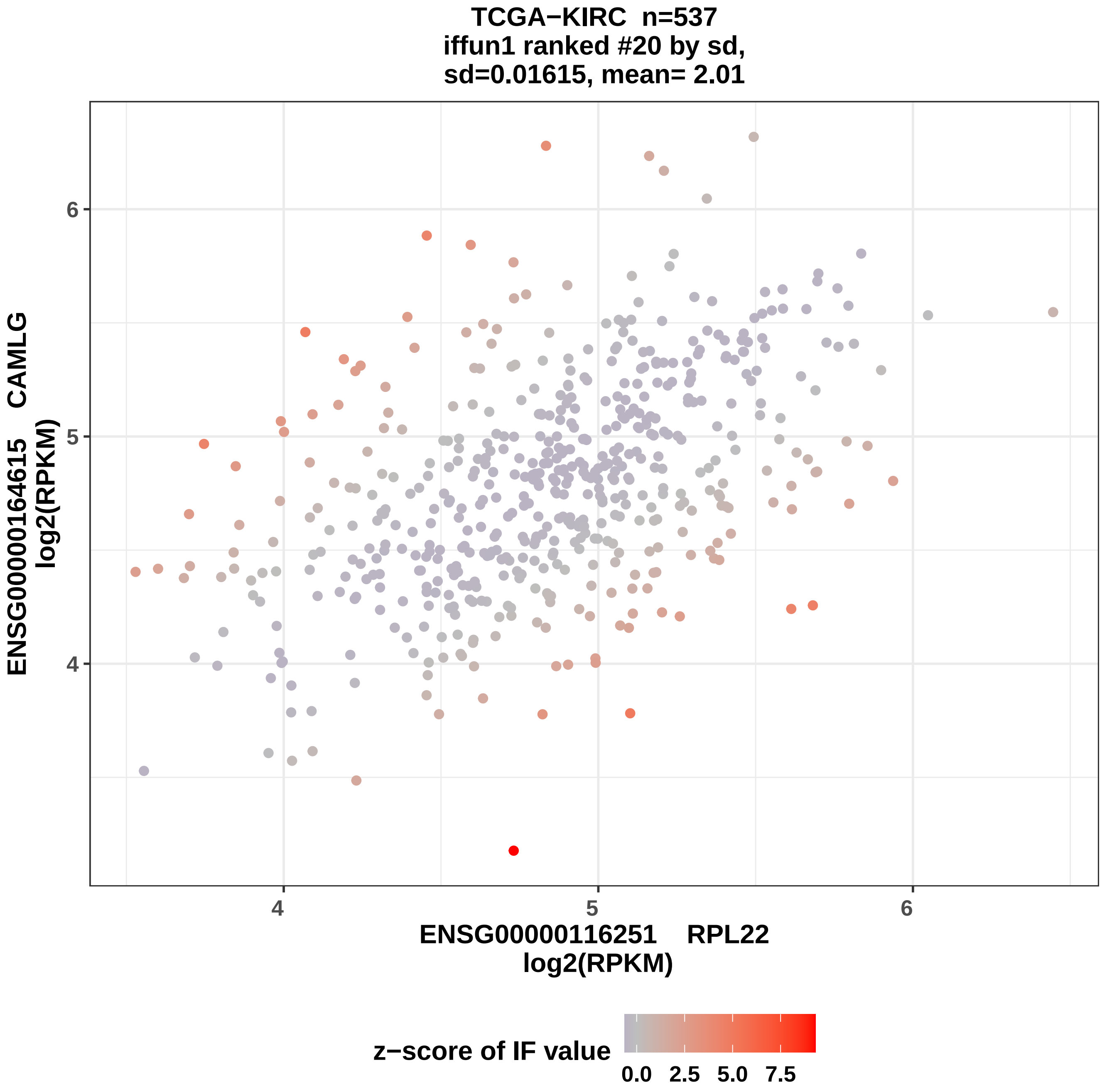
TCGA−KIRC n=537
iffun1 ranked #20 by sd,
sd=0.01615, mean= 2.01
ENSG00000164615 CAMLG
log2(RPKM)
ENSG00000116251 RPL22
log2(RPKM)
4
5
6
4
5
6
z−score of IF value
0.0
2.5
5.0
7.5

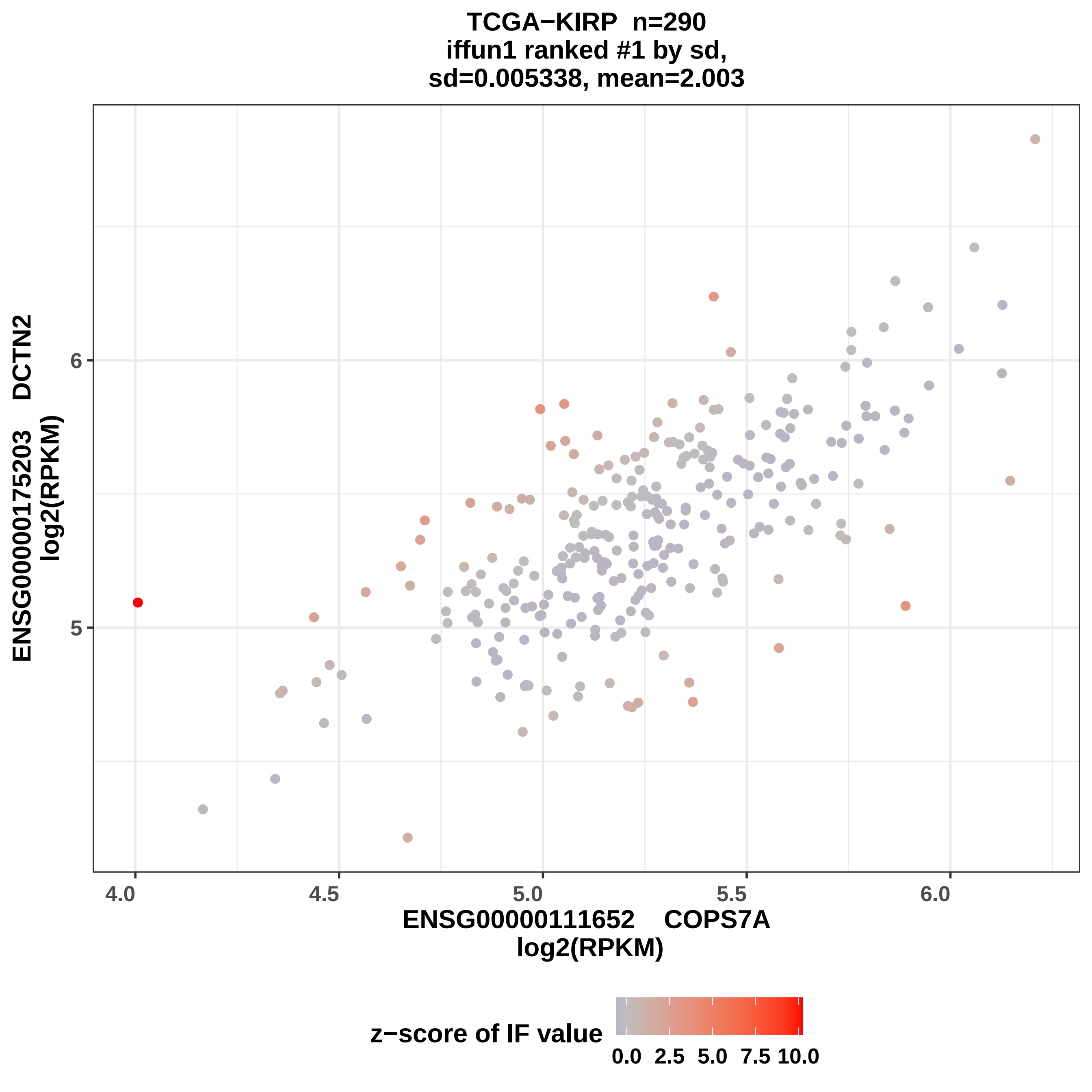
TCGA−KIRP n=290
iffun1 ranked #1 by sd,
sd=0.005338, mean=2.003
ENSG00000175203 DCTN2
log2(RPKM)
6
5
4.0
4.5
5.0
5.5
6.0
ENSG00000111652 COPS7A
log2(RPKM)
z−score of IF value
0.0
2.5
5.0
7.5
10.0

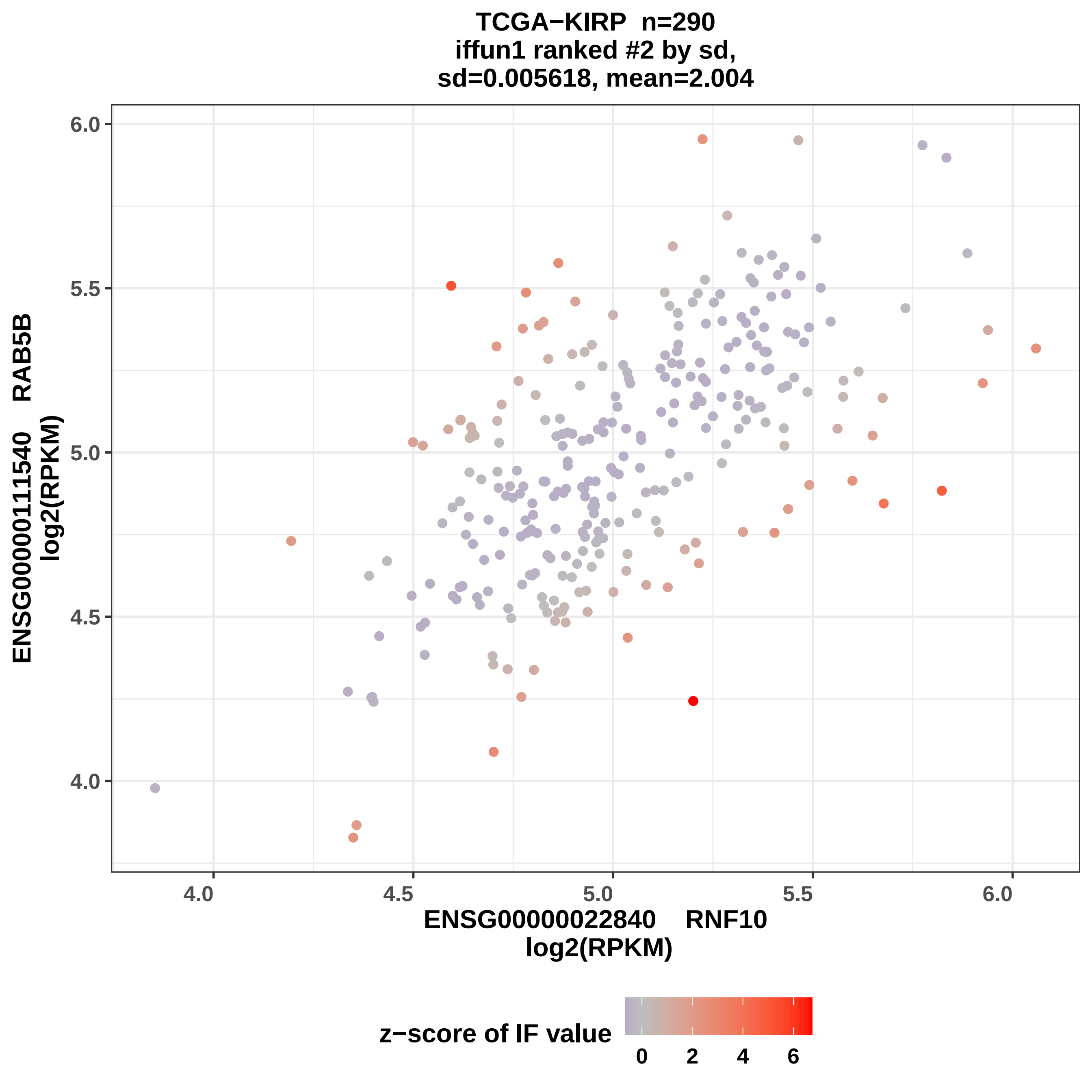
TCGA−KIRP n=290
iffun1 ranked #2 by sd,
sd=0.005618, mean=2.004
ENSG00000111540 RAB5B
log2(RPKM)
6.0
5.5
5.0
4.5
4.0
4.0
4.5
5.0
5.5
6.0
ENSG00000022840 RNF10
log2(RPKM)
z−score of IF value
0
2
4
6

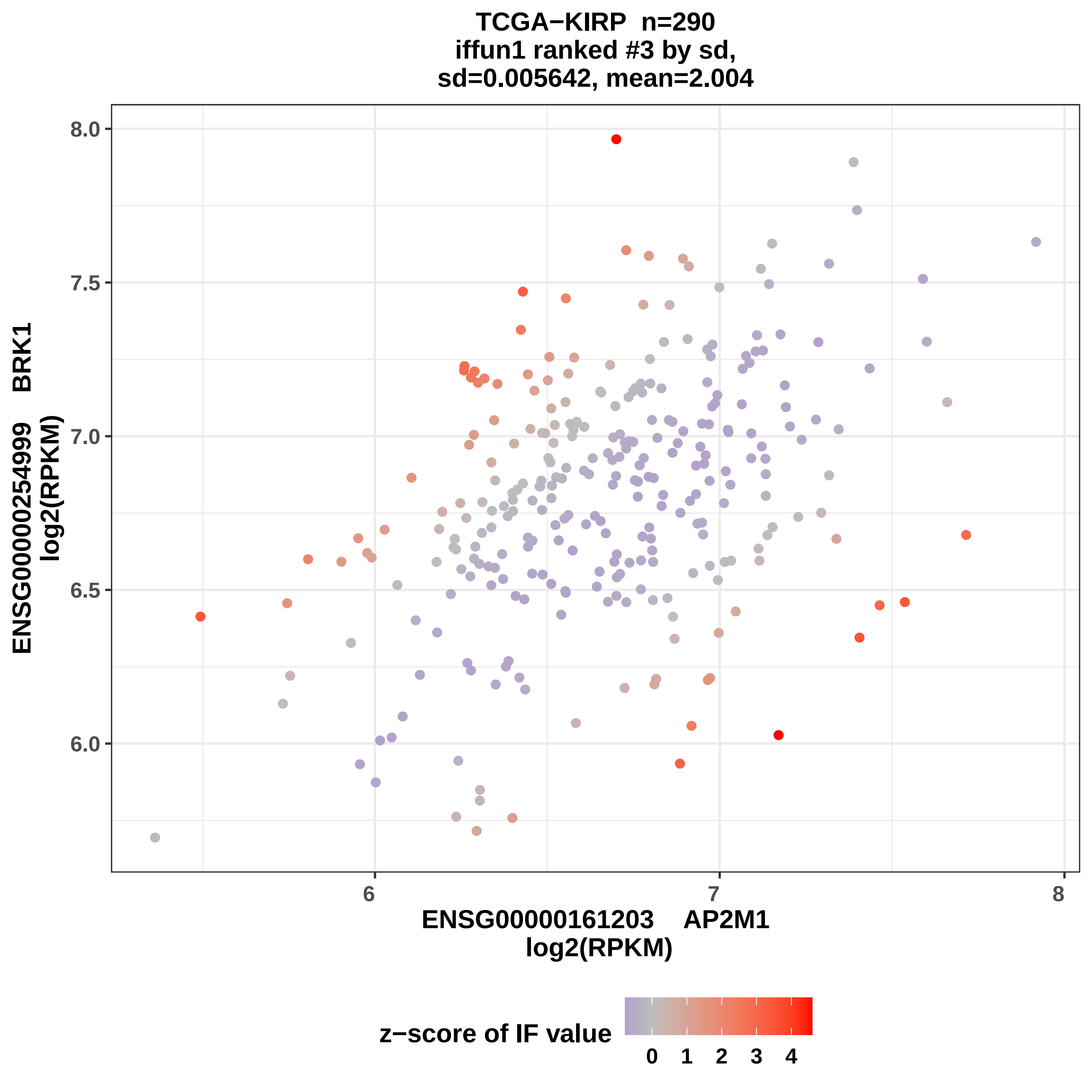
TCGA−KIRP n=290
iffun1 ranked #3 by sd,
sd=0.005642, mean=2.004
ENSG00000254999 BRK1
log2(RPKM)
8.0
7.5
7.0
6.5
6.0
6
7
8
ENSG00000161203 AP2M1
log2(RPKM)
z−score of IF value
0
1
2
3
4

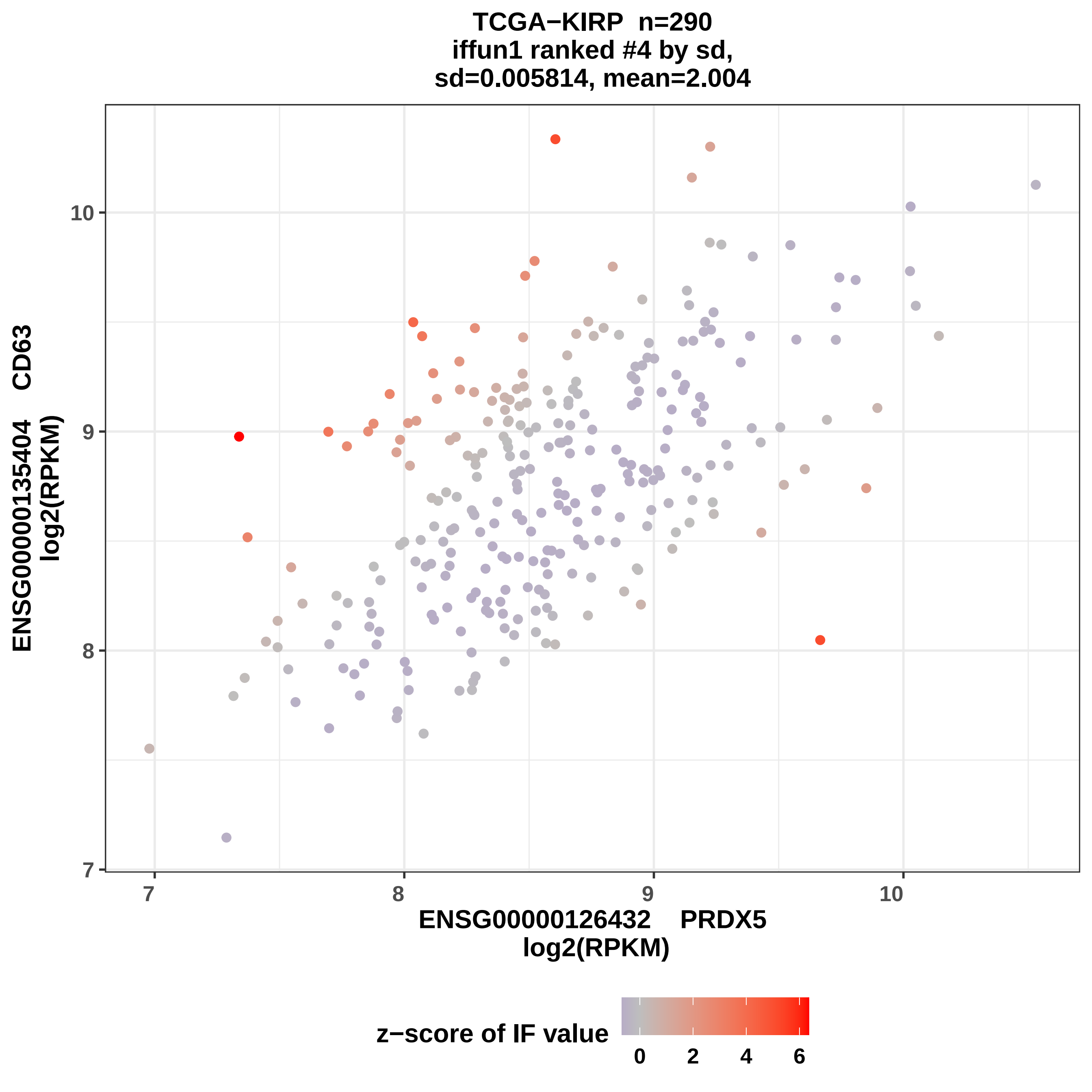
TCGA−KIRP n=290
iffun1 ranked #4 by sd,
sd=0.005814, mean=2.004
ENSG00000135404 CD63
log2(RPKM)
ENSG00000126432 PRDX5
log2(RPKM)
7
8
9
10
z−score of IF value
0
2
4
6

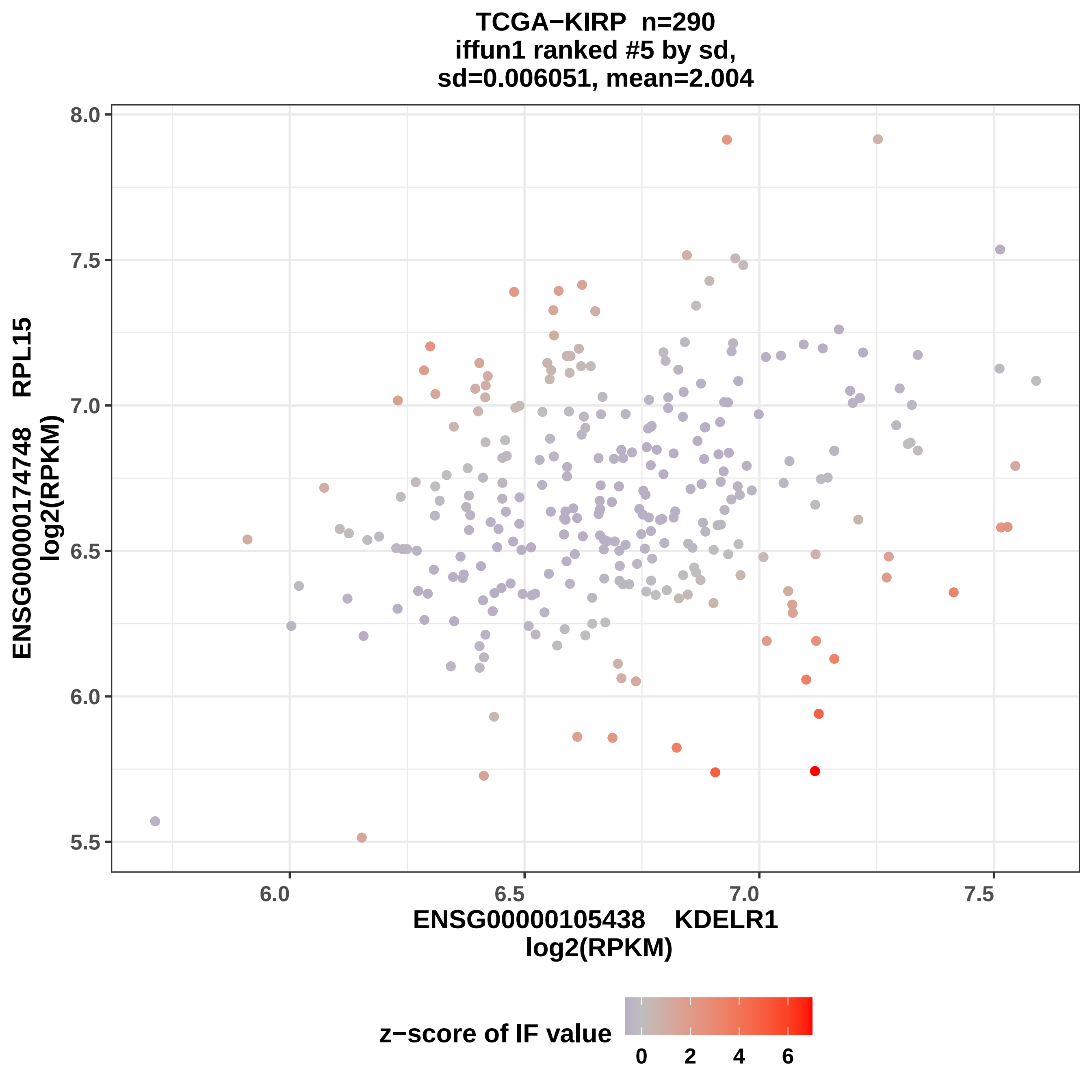
TCGA−KIRP n=290
iffun1 ranked #5 by sd,
sd=0.006051, mean=2.004
ENSG00000174748 RPL15
log2(RPKM)
8.0
7.5
7.0
6.5
6.0
5.5
6.0
6.5
7.0
7.5
ENSG00000105438 KDELR1
log2(RPKM)
z−score of IF value
0
2
4
6

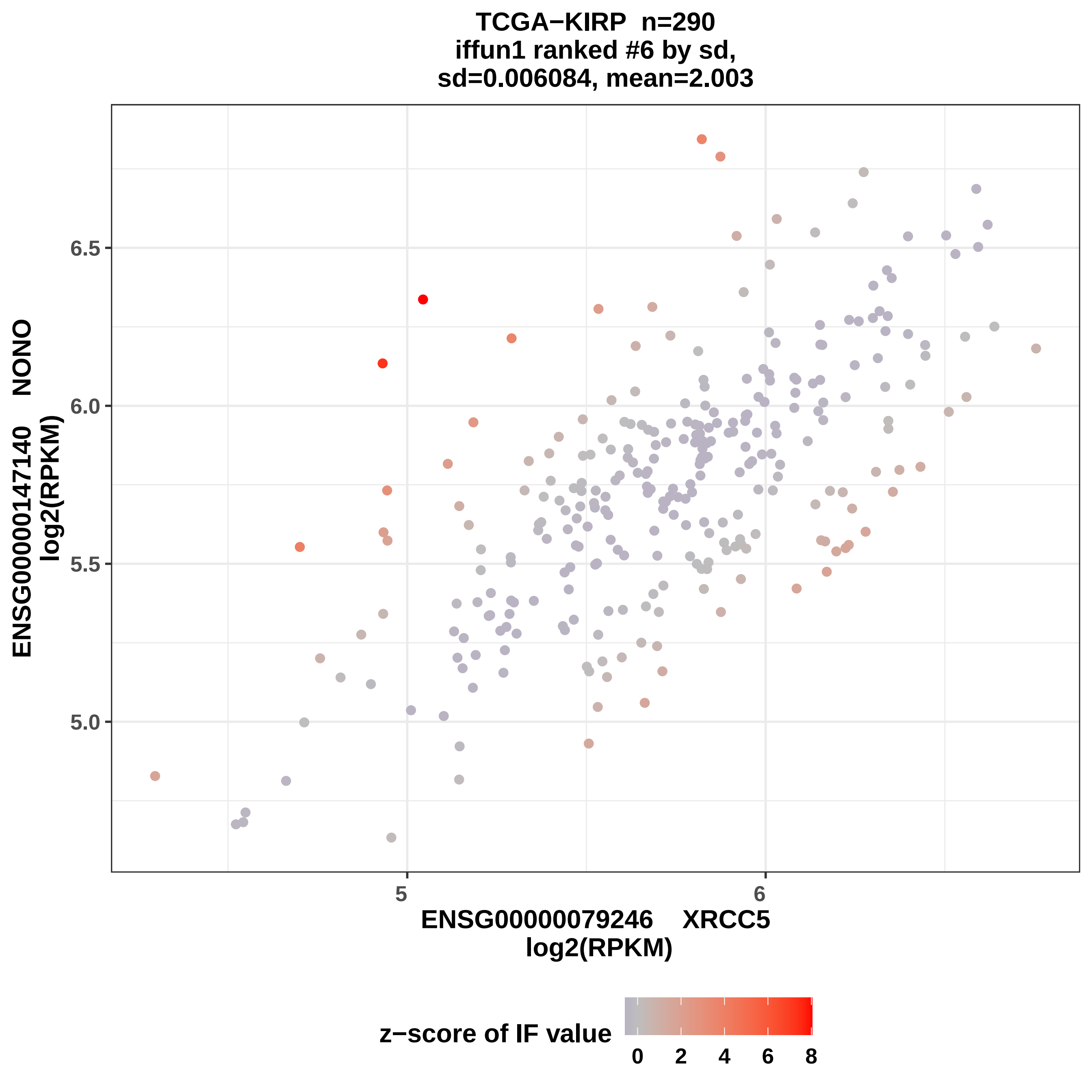

TCGA−KIRP n=290
iffun1 ranked #6 by sd,
sd=0.006084, mean=2.003
6.5
6.0
5.5
5.0
5
6
ENSG00000147140 NONO
log2(RPKM)
ENSG00000079246 XRCC5
log2(RPKM)
z−score of IF value
0
2
4
6
8

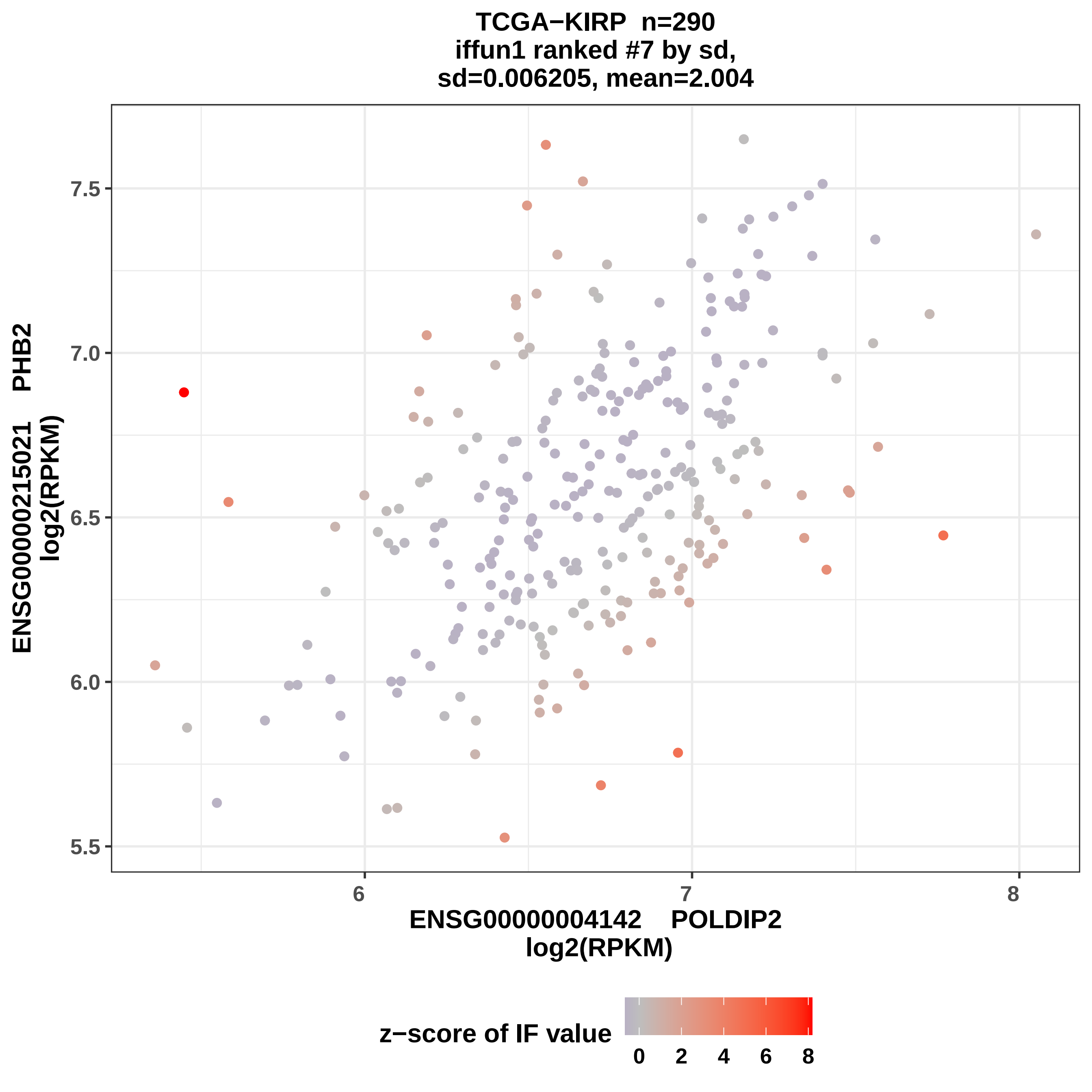

TCGA−KIRP n=290
iffun1 ranked #7 by sd,
sd=0.006205, mean=2.004
ENSG00000215021 PHB2
log2(RPKM)
7.5
7.0
6.5
6.0
5.5
6
7
8
ENSG00000004142 POLDIP2
log2(RPKM)
z−score of IF value
0 2 4 6 8

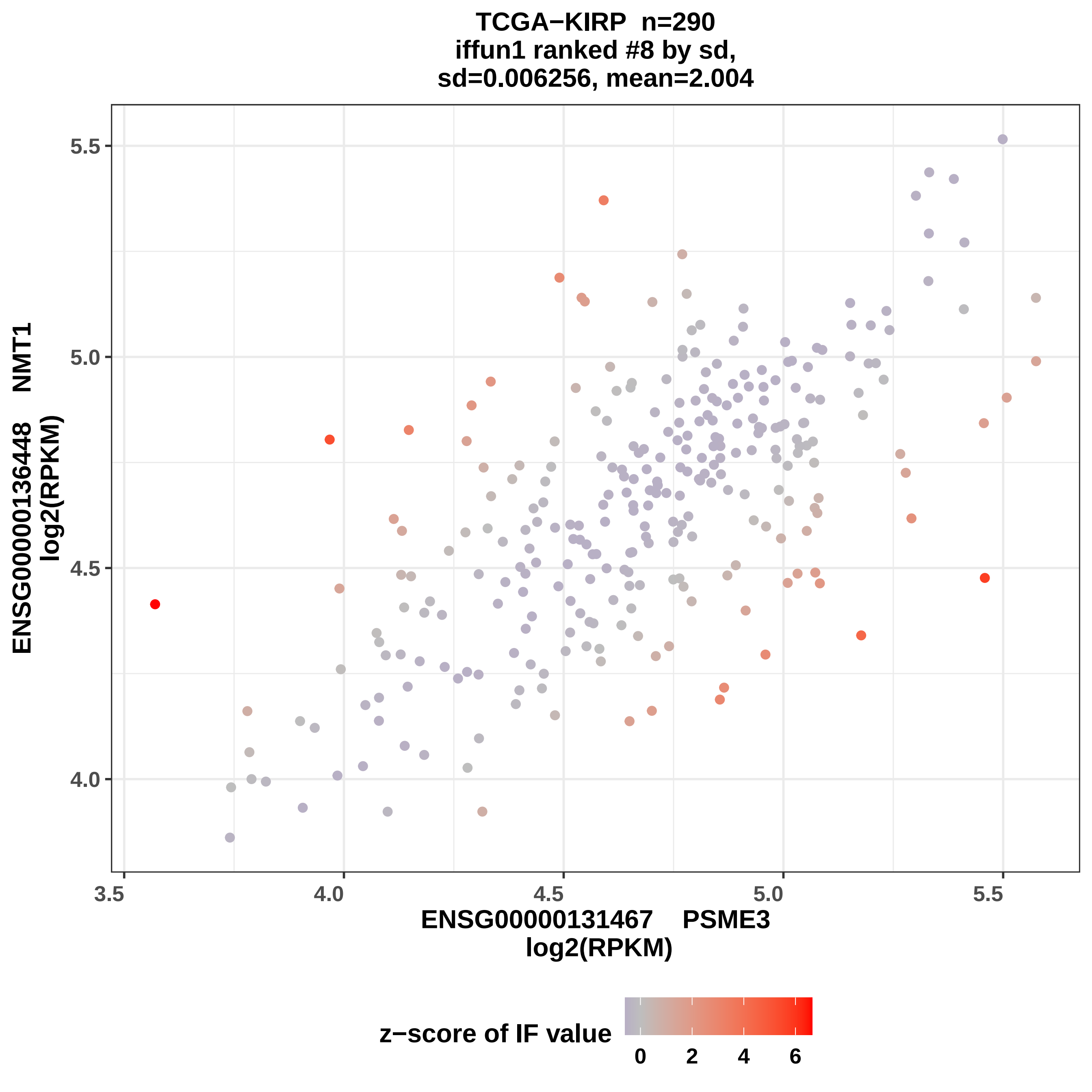

TCGA−KIRP n=290
iffun1 ranked #8 by sd,
sd=0.006256, mean=2.004
ENSG00000136448 NMT1
log2(RPKM)
ENSG00000131467 PSME3
log2(RPKM)
5.5
5.0
4.5
4.0
3.5
4.0
4.5
5.0
5.5
z−score of IF value
0
2
4
6

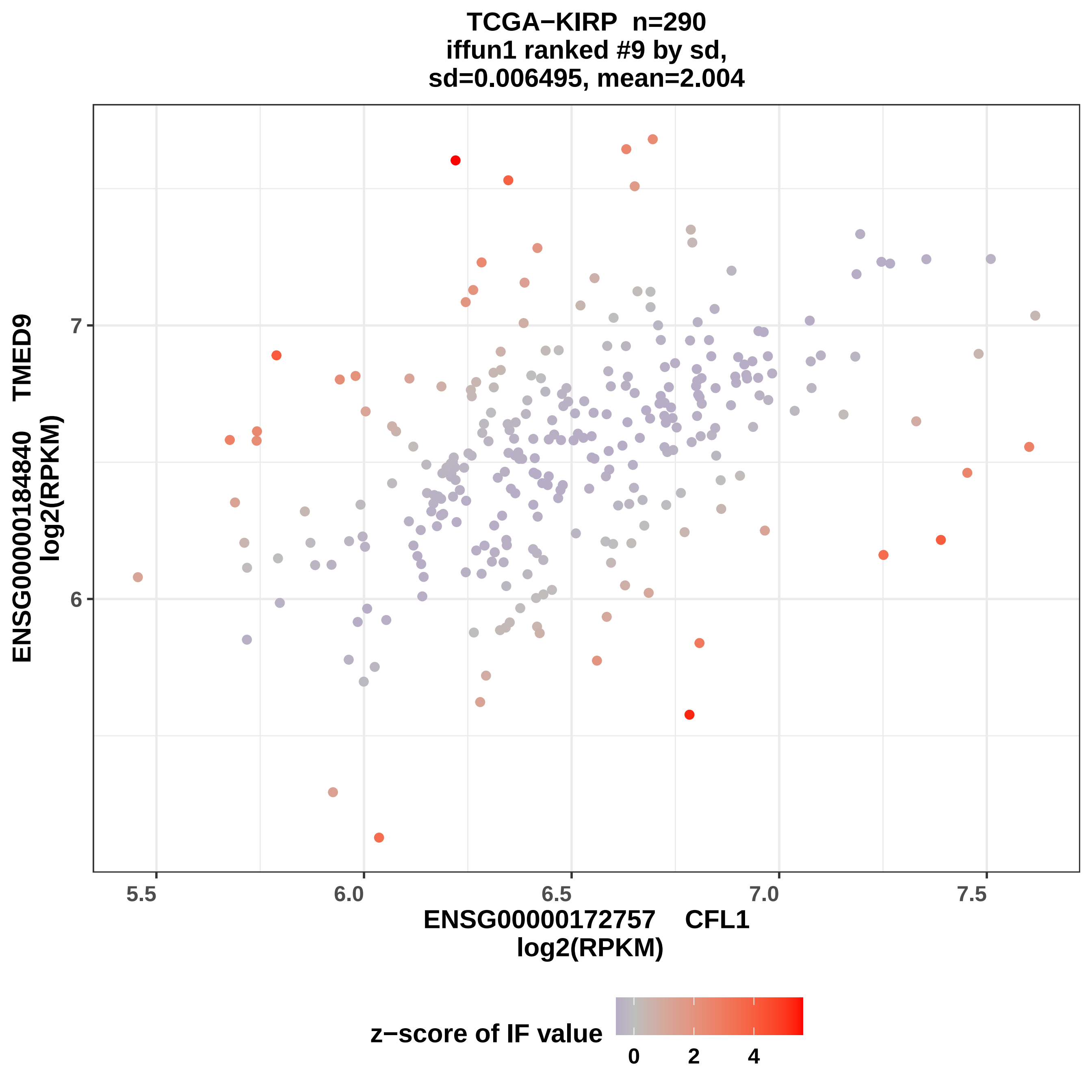

TCGA−KIRP n=290
iffun1 ranked #9 by sd,
sd=0.006495, mean=2.004
ENSG00000184840 TMED9
log2(RPKM)
7
6
5.5
6.0
6.5
7.0
7.5
ENSG00000172757 CFL1
log2(RPKM)
z−score of IF value
0
2
4

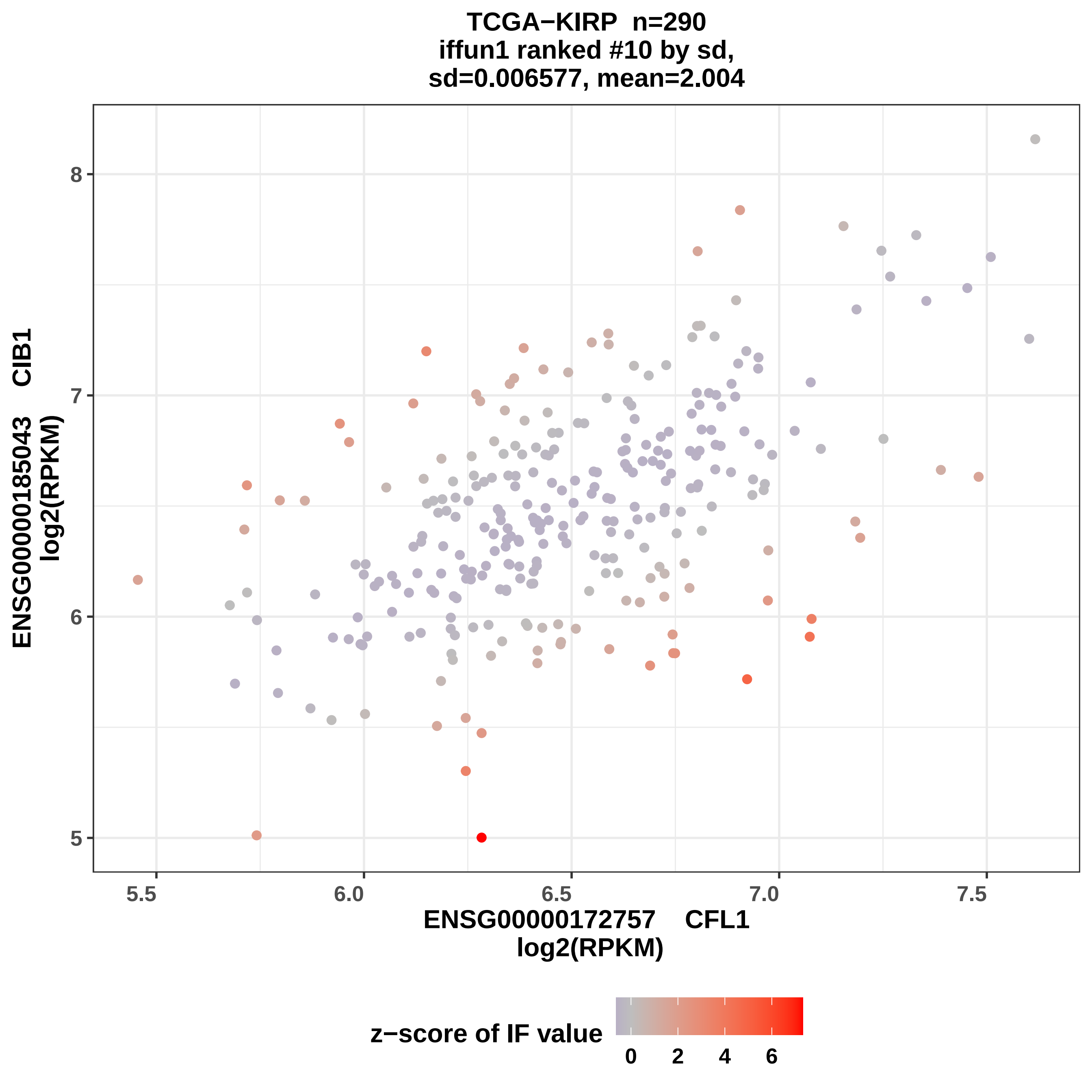
TCGA−KIRP n=290
iffun1 ranked #10 by sd,
sd=0.006577, mean=2.004
ENSG00000185043 CIB1
log2(RPKM)
8
7
6
5
5.5
6.0
6.5
7.0
7.5
ENSG00000172757 CFL1
log2(RPKM)
z−score of IF value
0
2
4
6

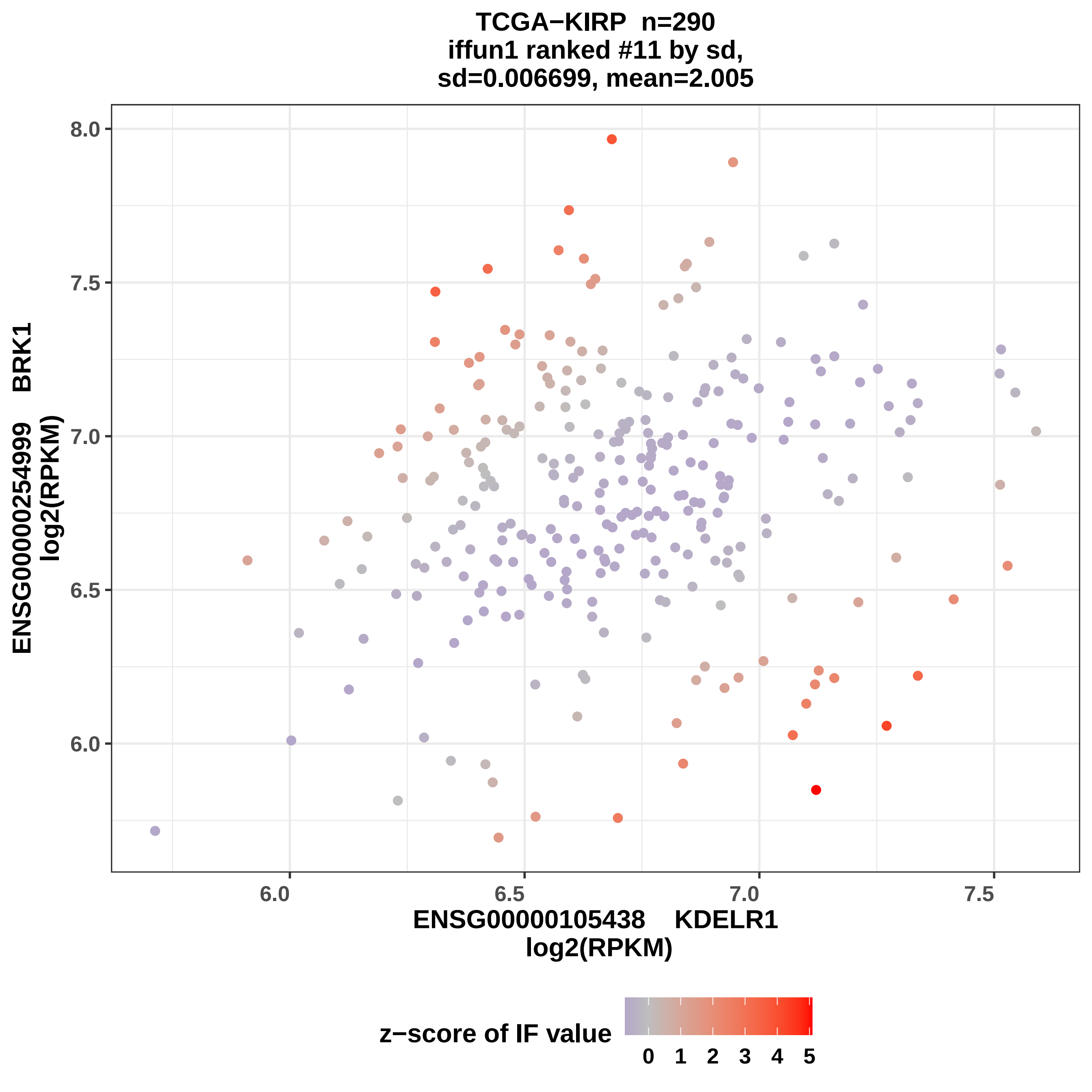
TCGA−KIRP n=290
iffun1 ranked #11 by sd,
sd=0.006699, mean=2.005
ENSG00000254999 BRK1
log2(RPKM)
8.0
7.5
7.0
6.5
6.0
6.0
6.5
7.0
7.5
ENSG00000105438 KDELR1
log2(RPKM)
z−score of IF value
0 1 2 3 4 5

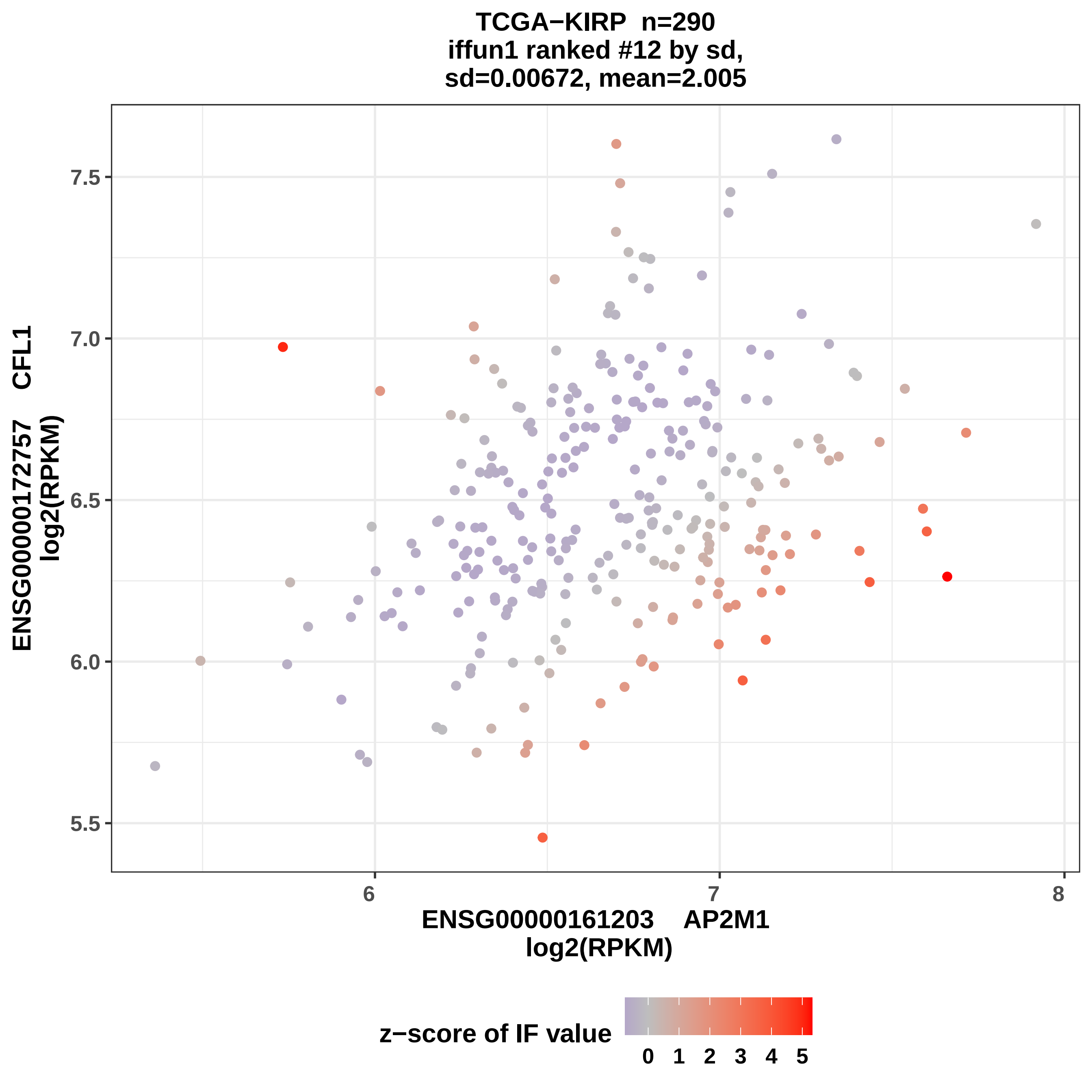
TCGA−KIRP n=290
iffun1 ranked #12 by sd,
sd=0.00672, mean=2.005
ENSG00000172757 CFL1
log2(RPKM)
7.5
7.0
6.5
6.0
5.5
6
7
8
ENSG00000161203 AP2M1
log2(RPKM)
z−score of IF value
0 1 2 3 4 5

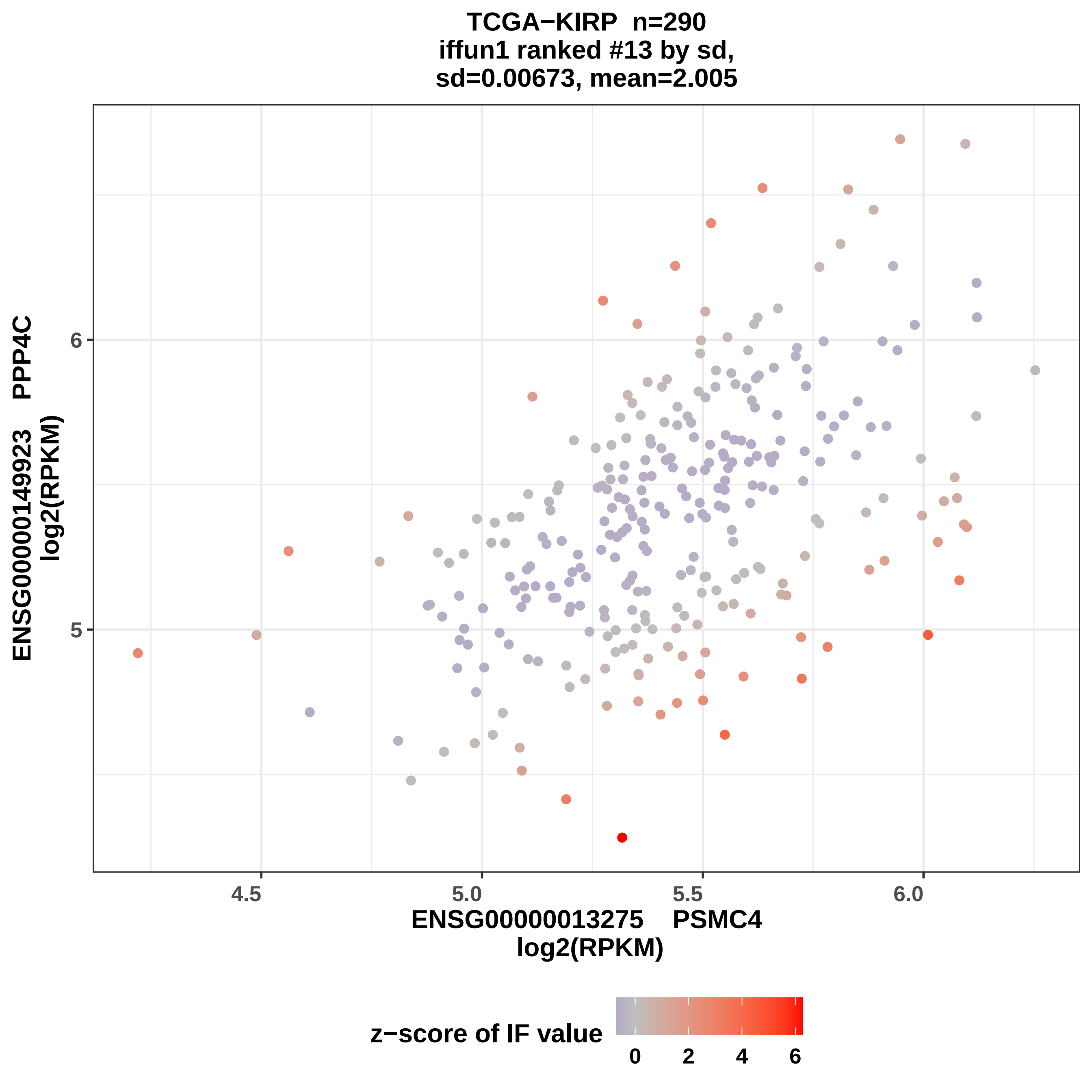

TCGA−KIRP n=290
iffun1 ranked #13 by sd,
sd=0.00673, mean=2.005
ENSG00000149923 PPP4C
log2(RPKM)
6
5
4.5
5.0
5.5
6.0
ENSG00000013275 PSMC4
log2(RPKM)
z−score of IF value
0
2
4
6

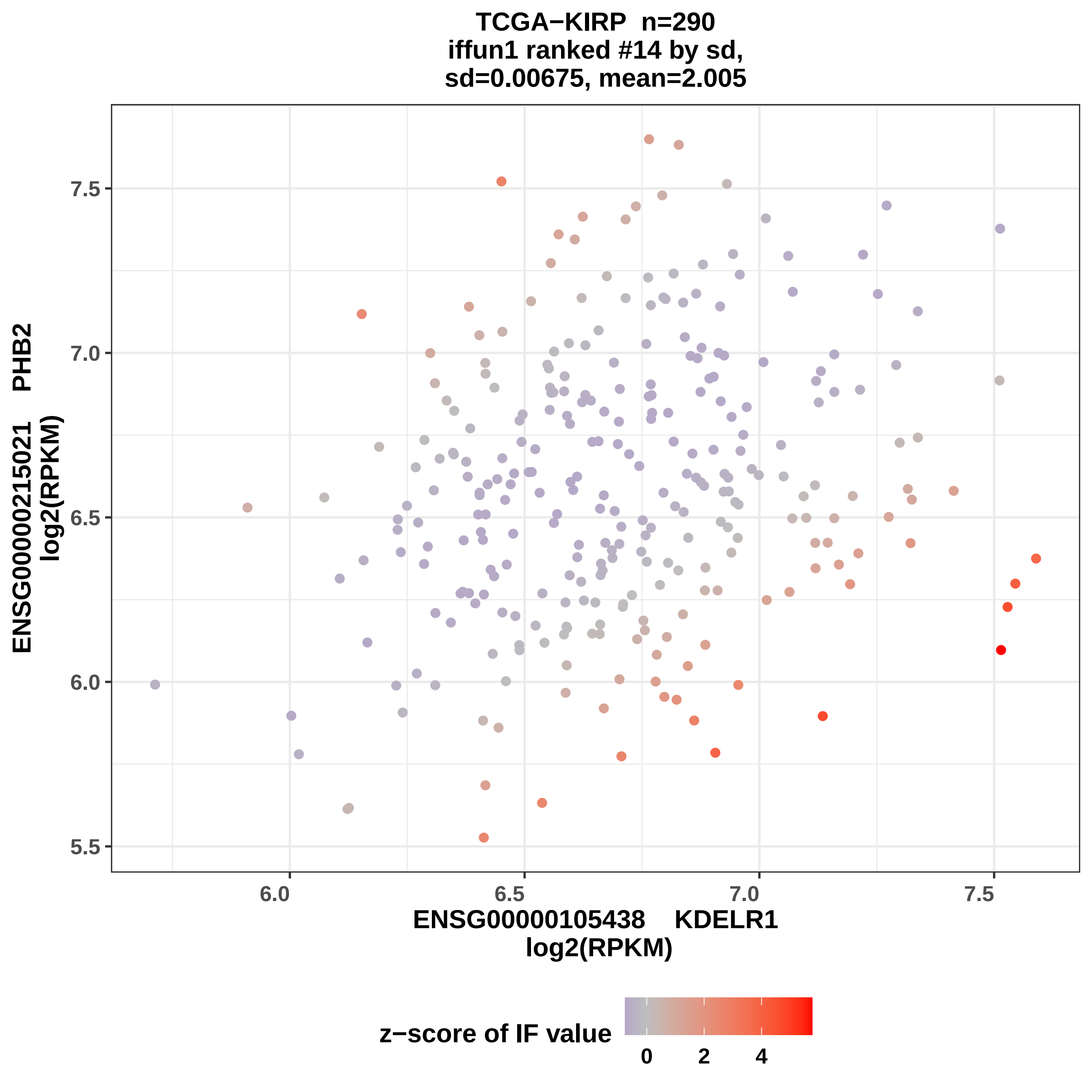

TCGA−KIRP n=290
iffun1 ranked #14 by sd,
sd=0.00675, mean=2.005
ENSG00000215021 PHB2
log2(RPKM)
7.5
7.0
6.5
6.0
5.5
6.0
6.5
7.0
7.5
ENSG00000105438 KDELR1
log2(RPKM)
z−score of IF value
0
2
4

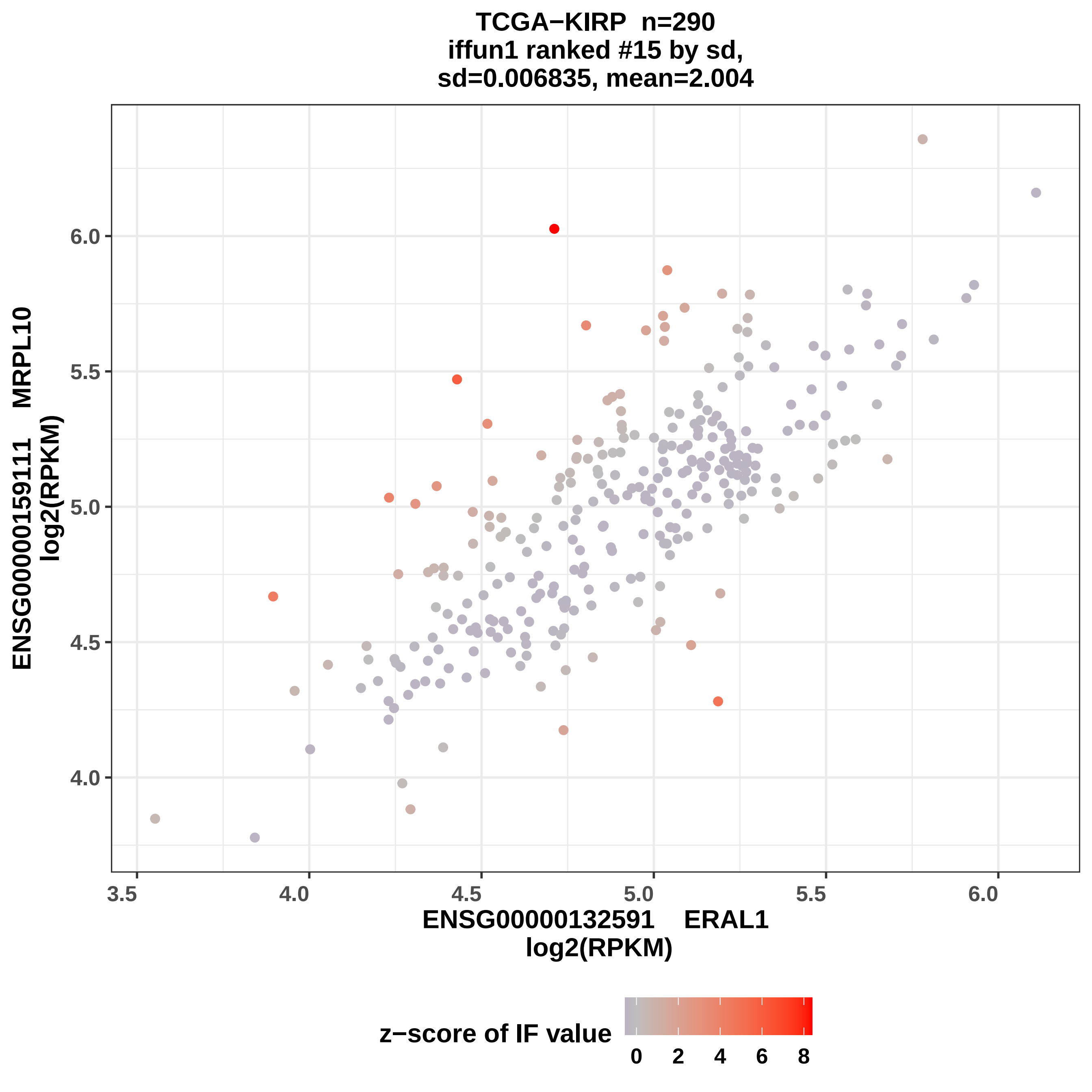

TCGA−KIRP n=290
iffun1 ranked #15 by sd,
sd=0.006835, mean=2.004
ENSG00000159111 MRPL10
log2(RPKM)
6.0
5.5
5.0
4.5
4.0
3.5
4.0
4.5
5.0
5.5
6.0
ENSG00000132591 ERAL1
log2(RPKM)
z−score of IF value
0
2
4
6
8

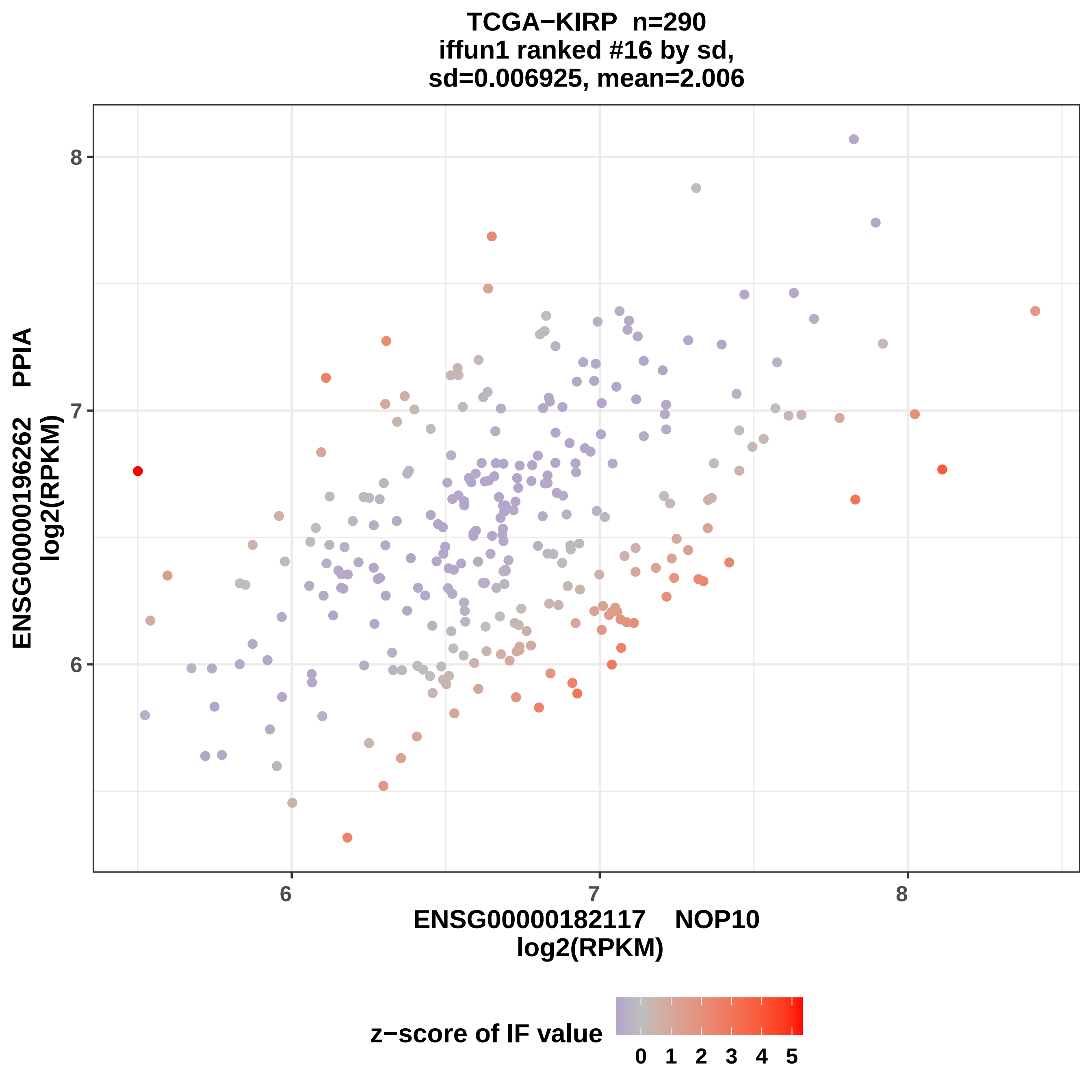

TCGA−KIRP n=290
iffun1 ranked #16 by sd,
sd=0.006925, mean=2.006
ENSG00000196262 PPIA
log2(RPKM)
ENSG00000182117 NOP10
log2(RPKM)
z−score of IF value
0 1 2 3 4 5

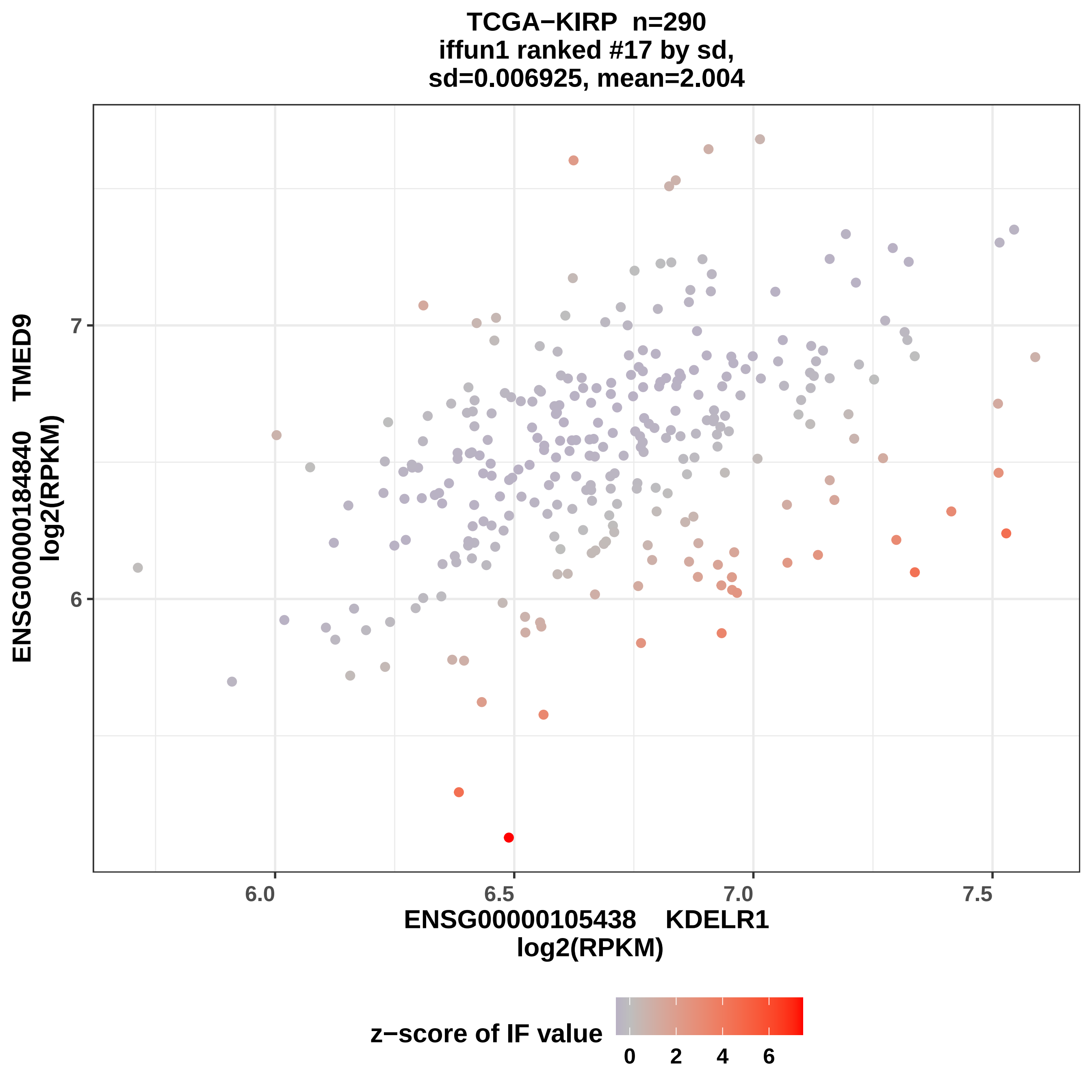

TCGA−KIRP n=290
iffun1 ranked #17 by sd,
sd=0.006925, mean=2.004
ENSG00000184840 TMED9
log2(RPKM)
7
6
6.0
6.5
7.0
7.5
ENSG00000105438 KDELR1
log2(RPKM)
z−score of IF value
0
2
4
6

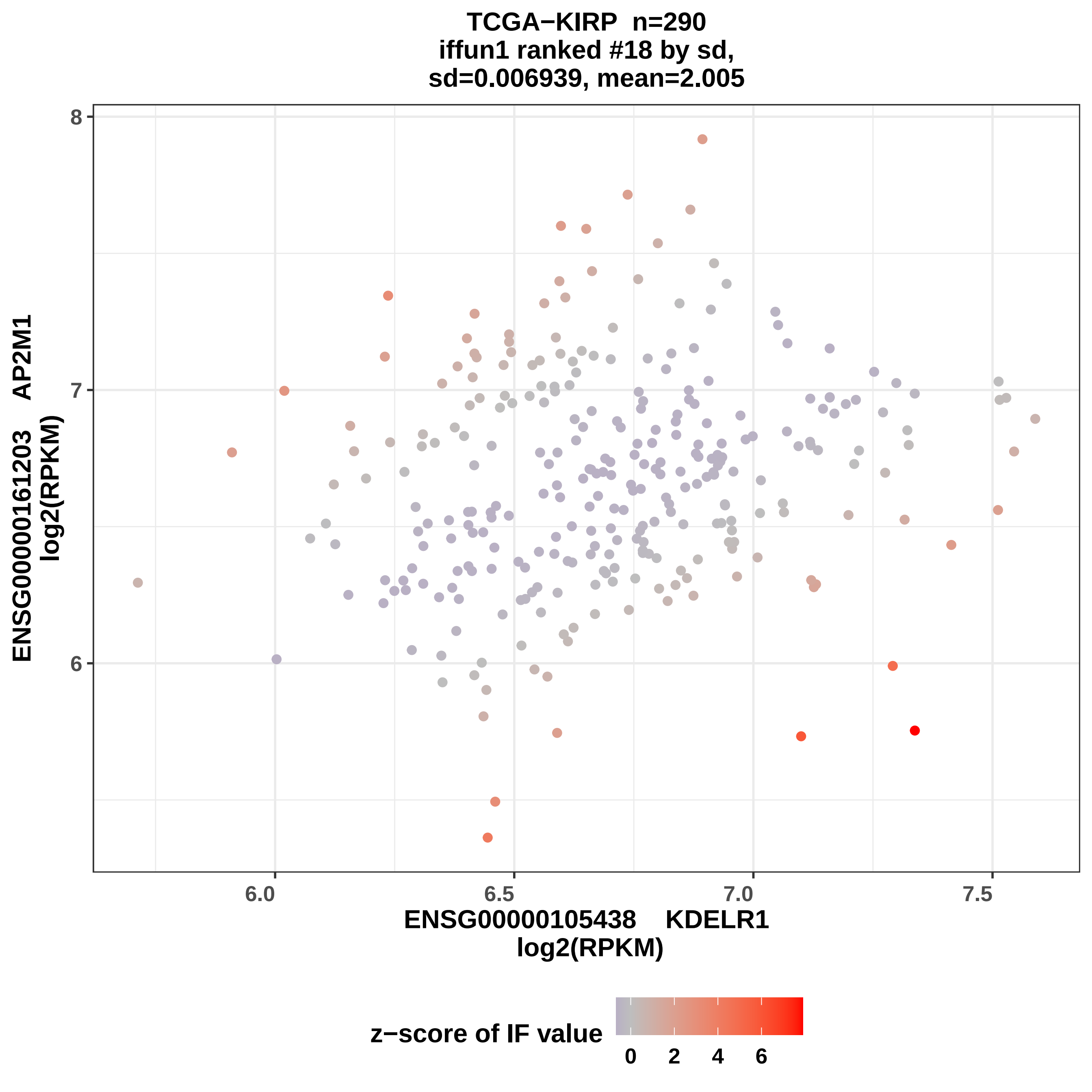

TCGA−KIRP n=290
iffun1 ranked #18 by sd,
sd=0.006939, mean=2.005
8
7
6
ENSG00000161203 AP2M1
log2(RPKM)
6.0
6.5
7.0
7.5
ENSG00000105438 KDELR1
log2(RPKM)
z−score of IF value
0
2
4
6

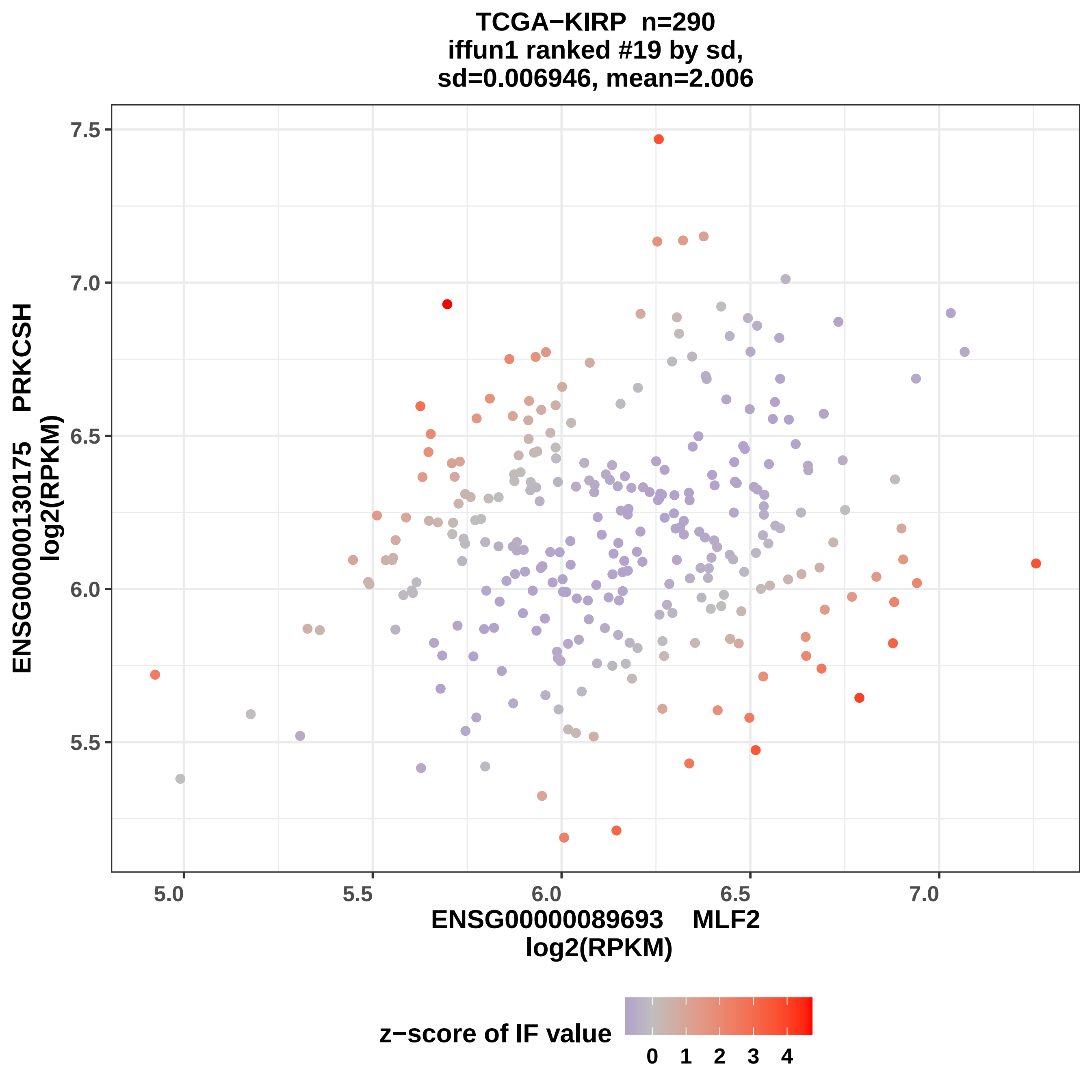

TCGA−KIRP n=290
iffun1 ranked #19 by sd,
sd=0.006946, mean=2.006
7.5
7.0
6.5
6.0
5.5
ENSG00000130175 PRKCSH
log2(RPKM)
5.0
5.5
6.0
6.5
7.0
ENSG00000089693 MLF2
log2(RPKM)
z−score of IF value
0 1 2 3 4

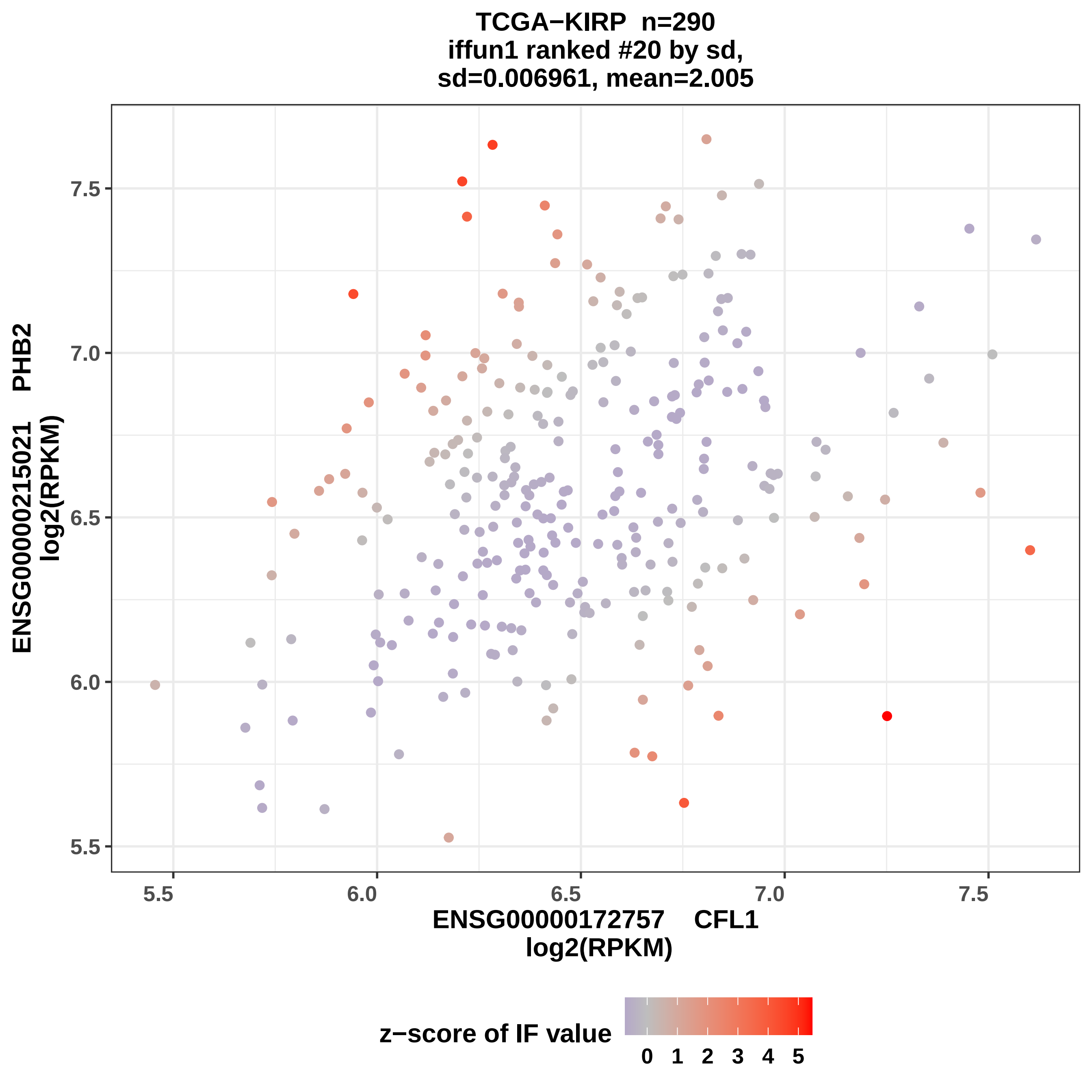
TCGA−KIRP n=290
iffun1 ranked #20 by sd,
sd=0.006961, mean=2.005
ENSG00000215021 PHB2
log2(RPKM)
7.5
7.0
6.5
6.0
5.5
5.5
6.0
6.5
7.0
7.5
ENSG00000172757 CFL1
log2(RPKM)
z−score of IF value
0
1
2
3
4
5

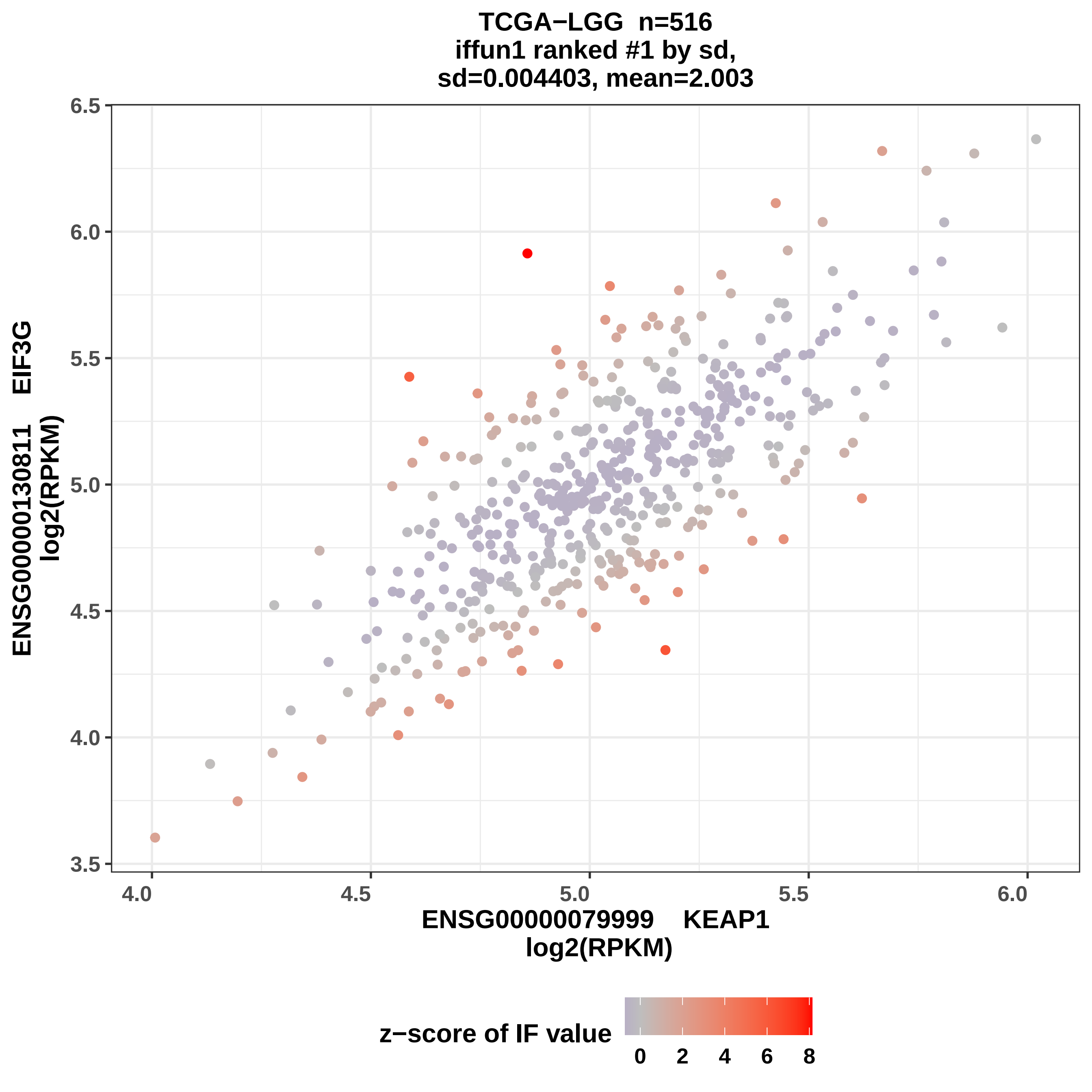
TCGA−LGG n=516
iffun1 ranked #1 by sd,
sd=0.004403, mean=2.003
ENSG00000130811 EIF3G
log2(RPKM)
ENSG00000079999 KEAP1
log2(RPKM)
3.5
4.0
4.5
5.0
5.5
6.0
6.5
4.0
4.5
5.0
5.5
6.0
z−score of IF value
0
2
4
6
8

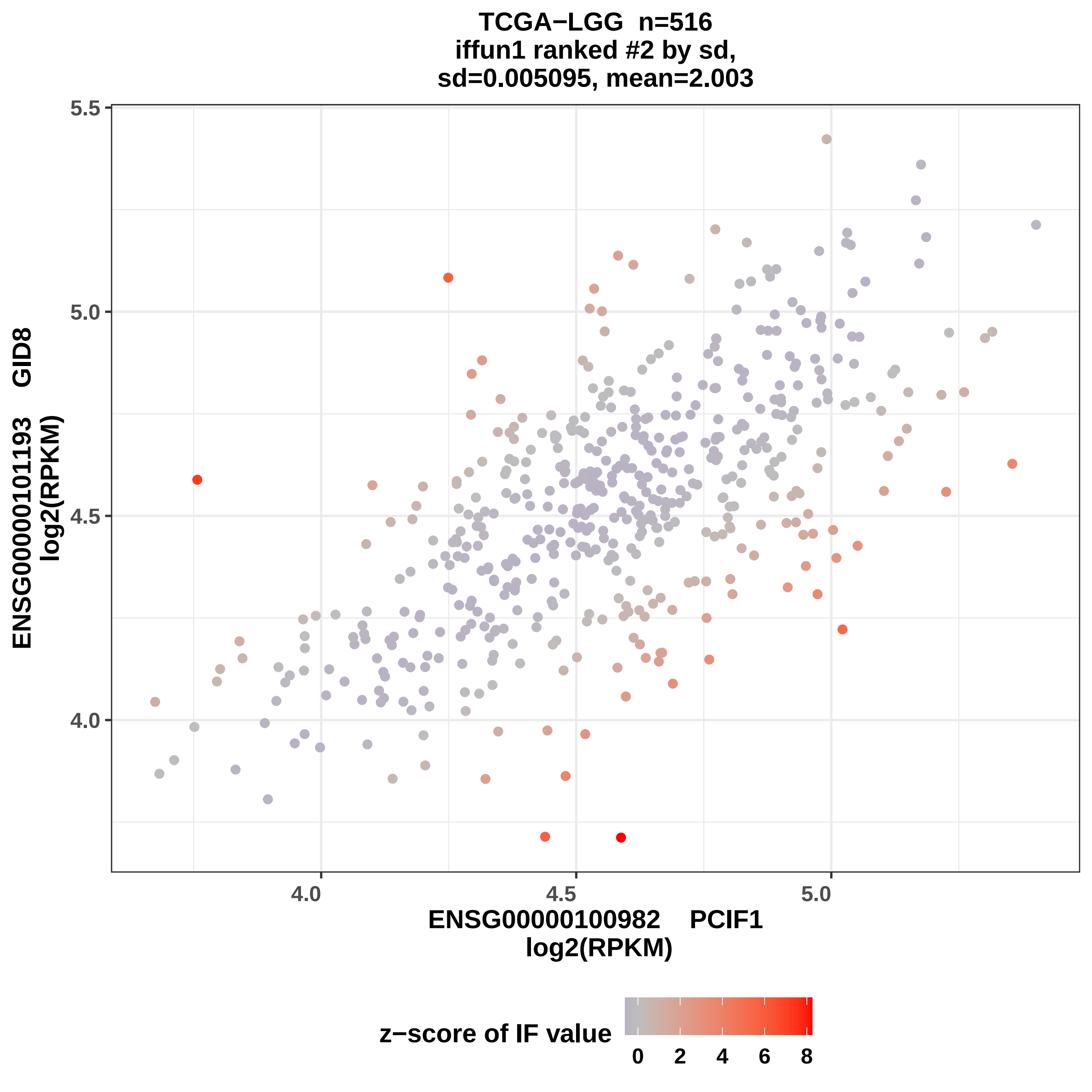

TCGA−LGG n=516
iffun1 ranked #2 by sd,
sd=0.005095, mean=2.003
ENSG00000101193 GID8
log2(RPKM)
5.5
5.0
4.5
4.0
4.0
4.5
5.0
ENSG00000100982 PCIF1
log2(RPKM)
z−score of IF value
0
2
4
6
8

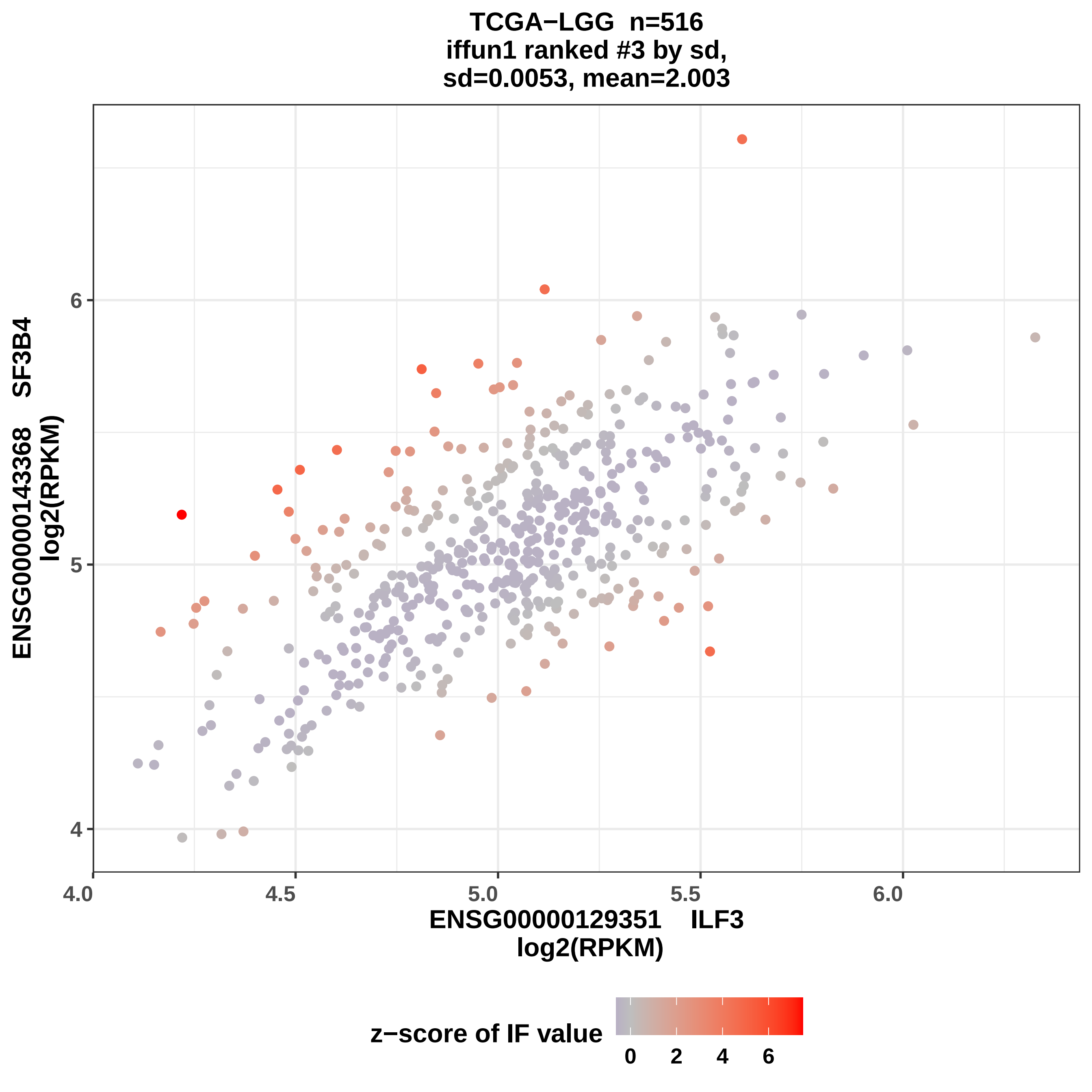

TCGA−LGG n=516
iffun1 ranked #3 by sd,
sd=0.0053, mean=2.003
ENSG00000143368 SF3B4
log2(RPKM)
ENSG00000129351 ILF3
log2(RPKM)
4
5
6
4.0
4.5
5.0
5.5
6.0
z−score of IF value
0
2
4
6

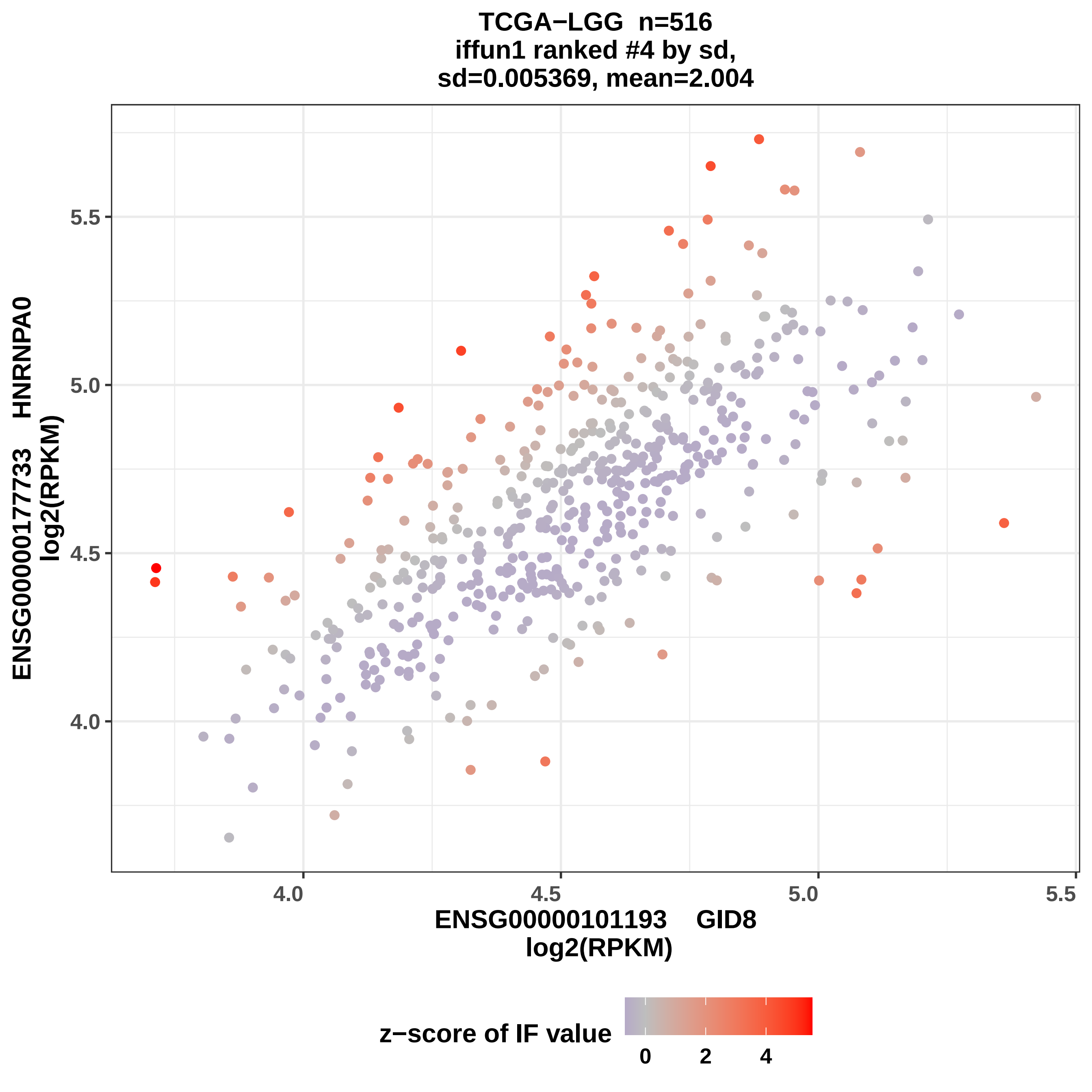
TCGA−LGG n=516
iffun1 ranked #4 by sd,
sd=0.005369, mean=2.004
ENSG00000177733 HNRNPA0
log2(RPKM)
5.5
5.0
4.5
4.0
4.0
4.5
5.0
5.5
ENSG00000101193 GID8
log2(RPKM)
z−score of IF value
0
2
4

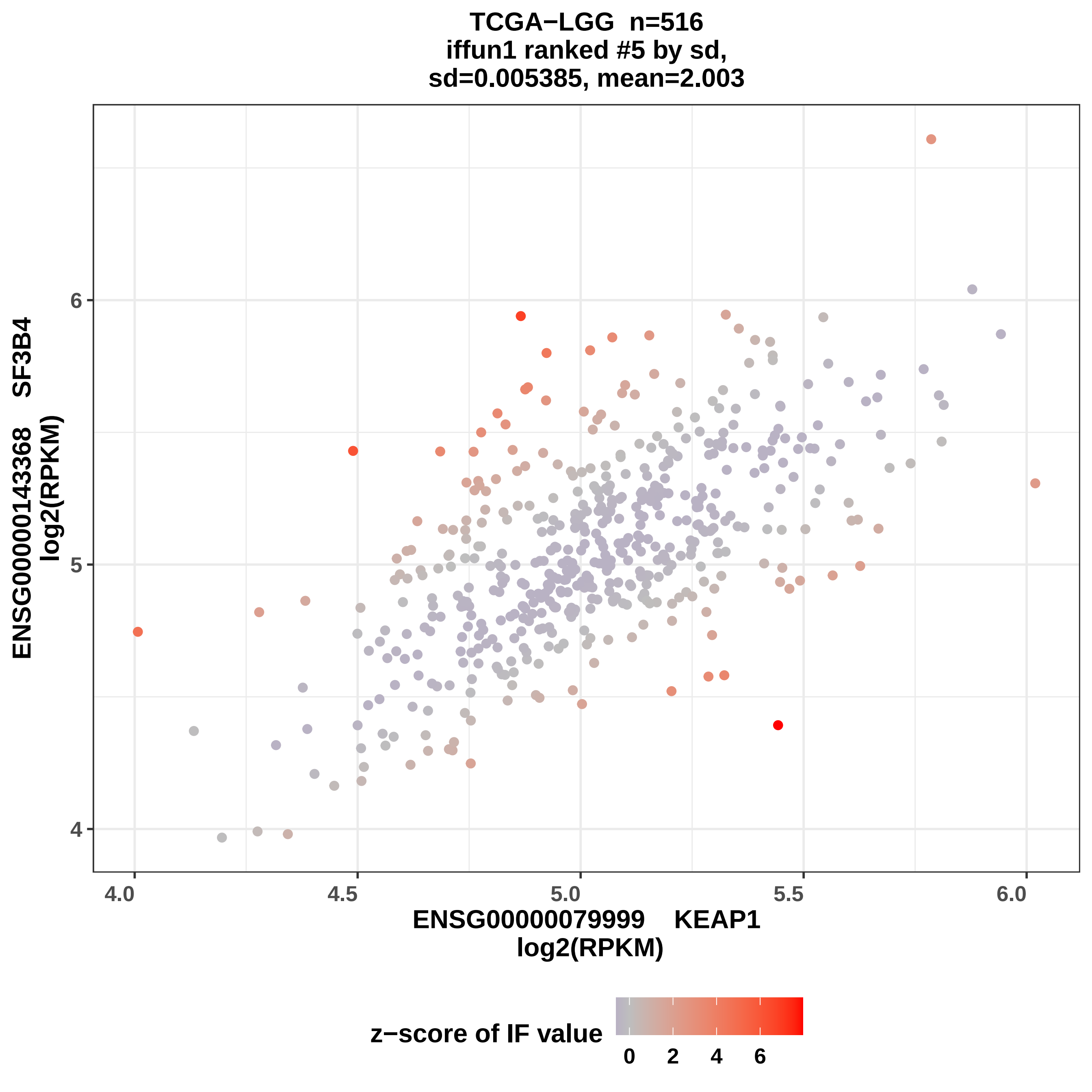
TCGA−LGG n=516
iffun1 ranked #5 by sd,
sd=0.005385, mean=2.003
ENSG00000143368 SF3B4
log2(RPKM)
ENSG00000079999 KEAP1
log2(RPKM)
4
5
6
4.0
4.5
5.0
5.5
6.0
z−score of IF value
0
2
4
6

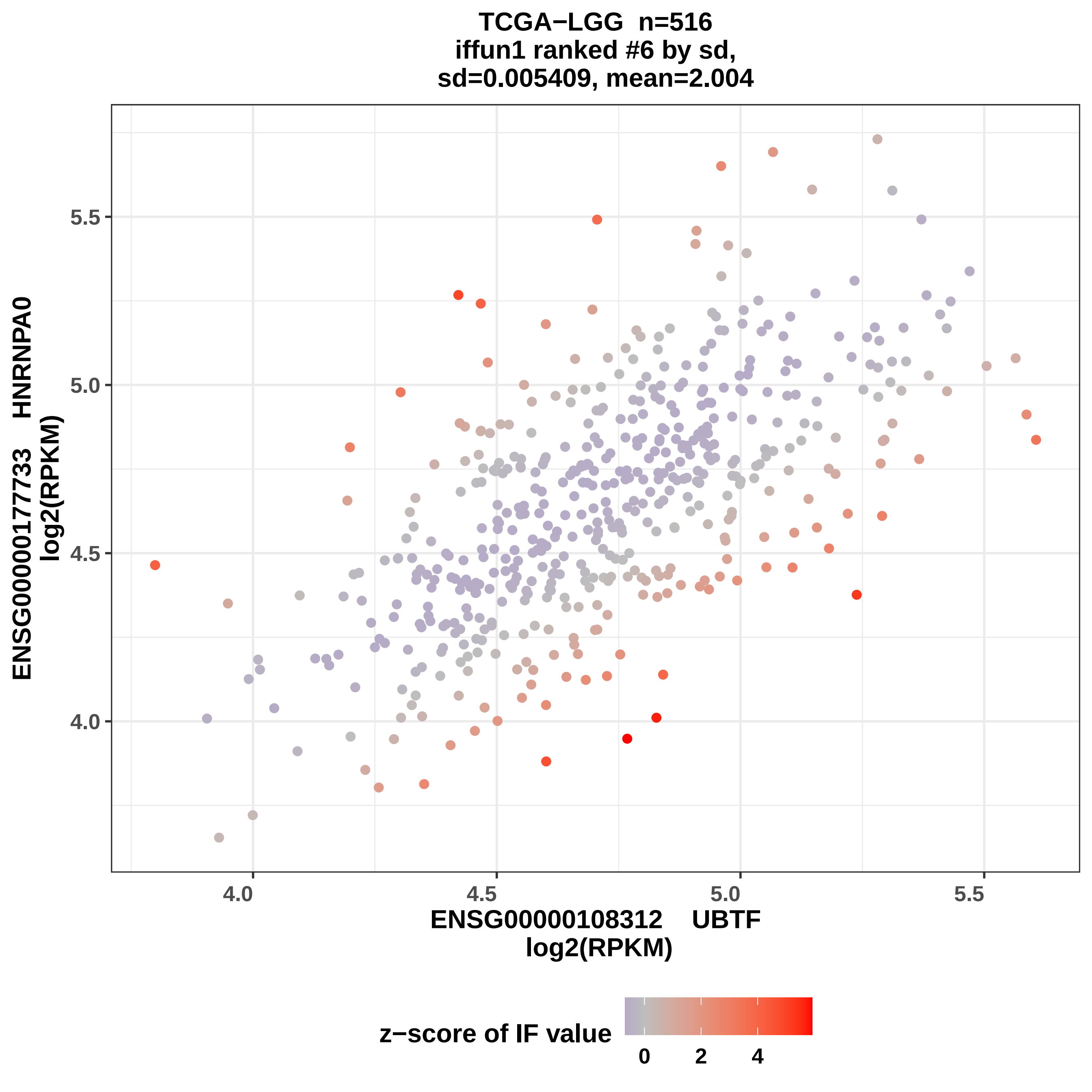

TCGA−LGG n=516
iffun1 ranked #6 by sd,
sd=0.005409, mean=2.004
ENSG00000177733 HNRNPA0
log2(RPKM)
5.5
5.0
4.5
4.0
4.0
4.5
5.0
5.5
ENSG00000108312 UBTF
log2(RPKM)
z−score of IF value
0
2
4

TCGA−LGG  n=516
iffun1 ranked #7 by sd,
sd=0.005547, mean=2.004

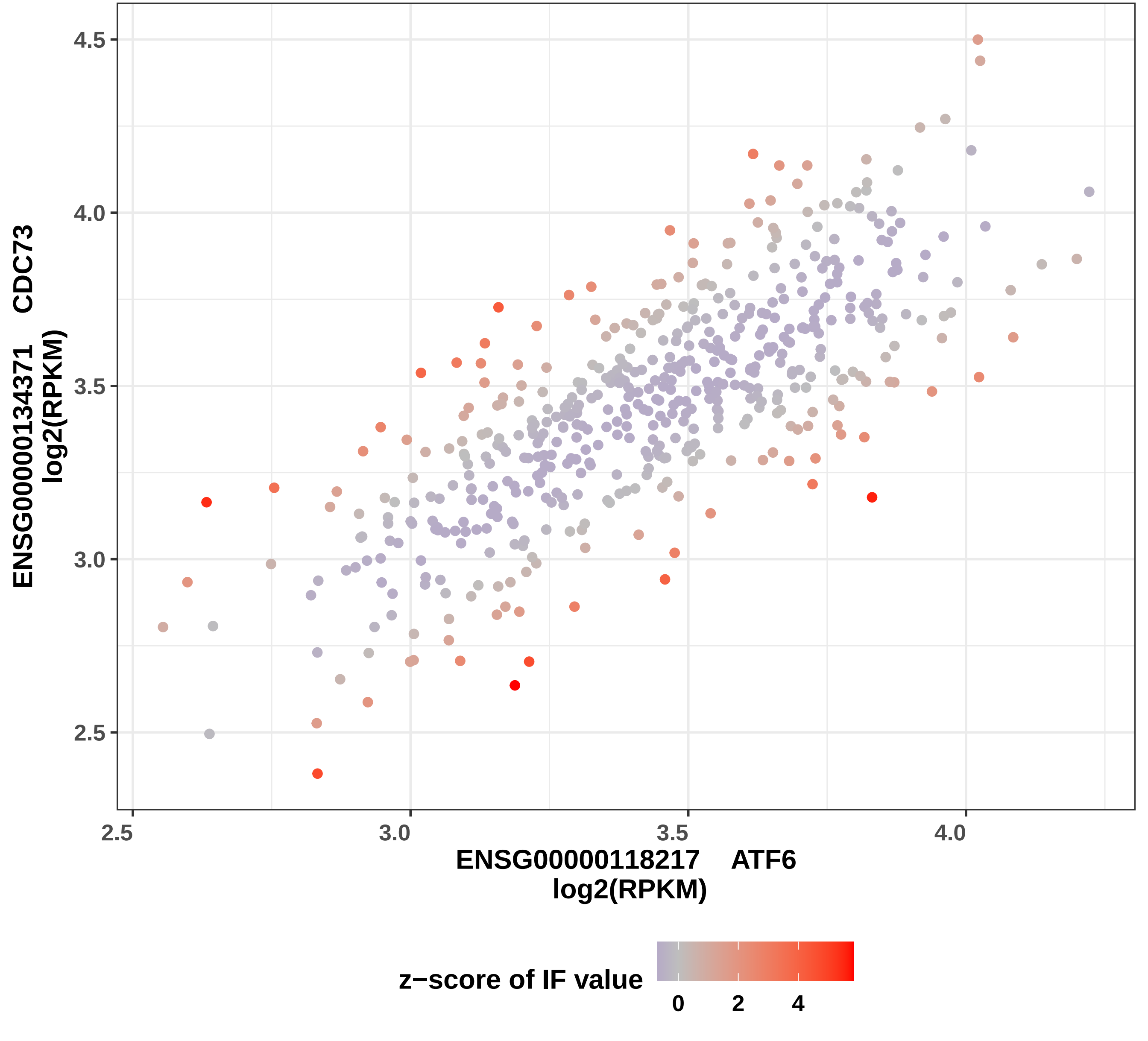

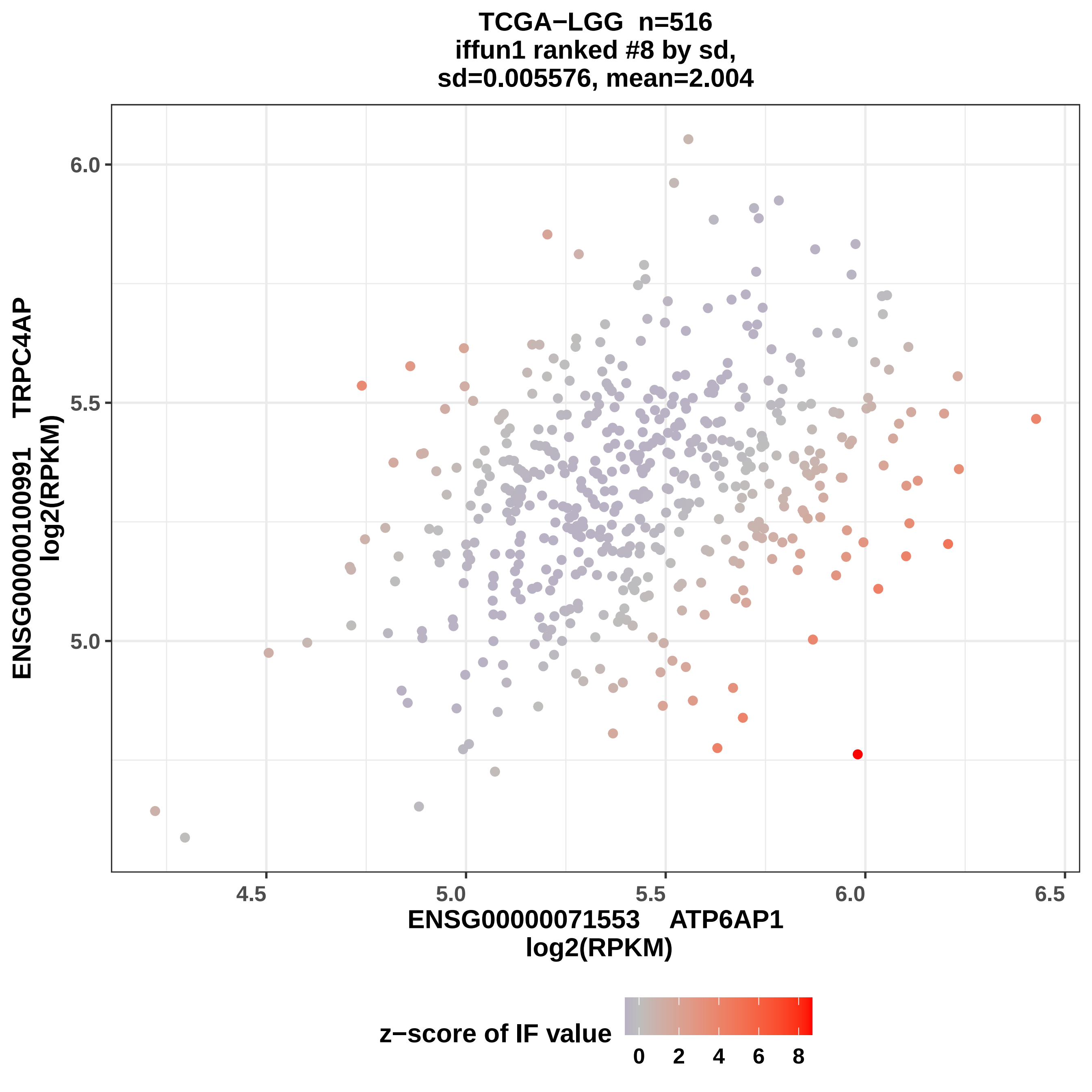
TCGA−LGG n=516
iffun1 ranked #8 by sd,
sd=0.005576, mean=2.004
ENSG00000100991 TRPC4AP
log2(RPKM)
6.0
5.5
5.0
4.5
5.0
5.5
6.0
6.5
ENSG00000071553 ATP6AP1
log2(RPKM)
z−score of IF value
0
2
4
6
8

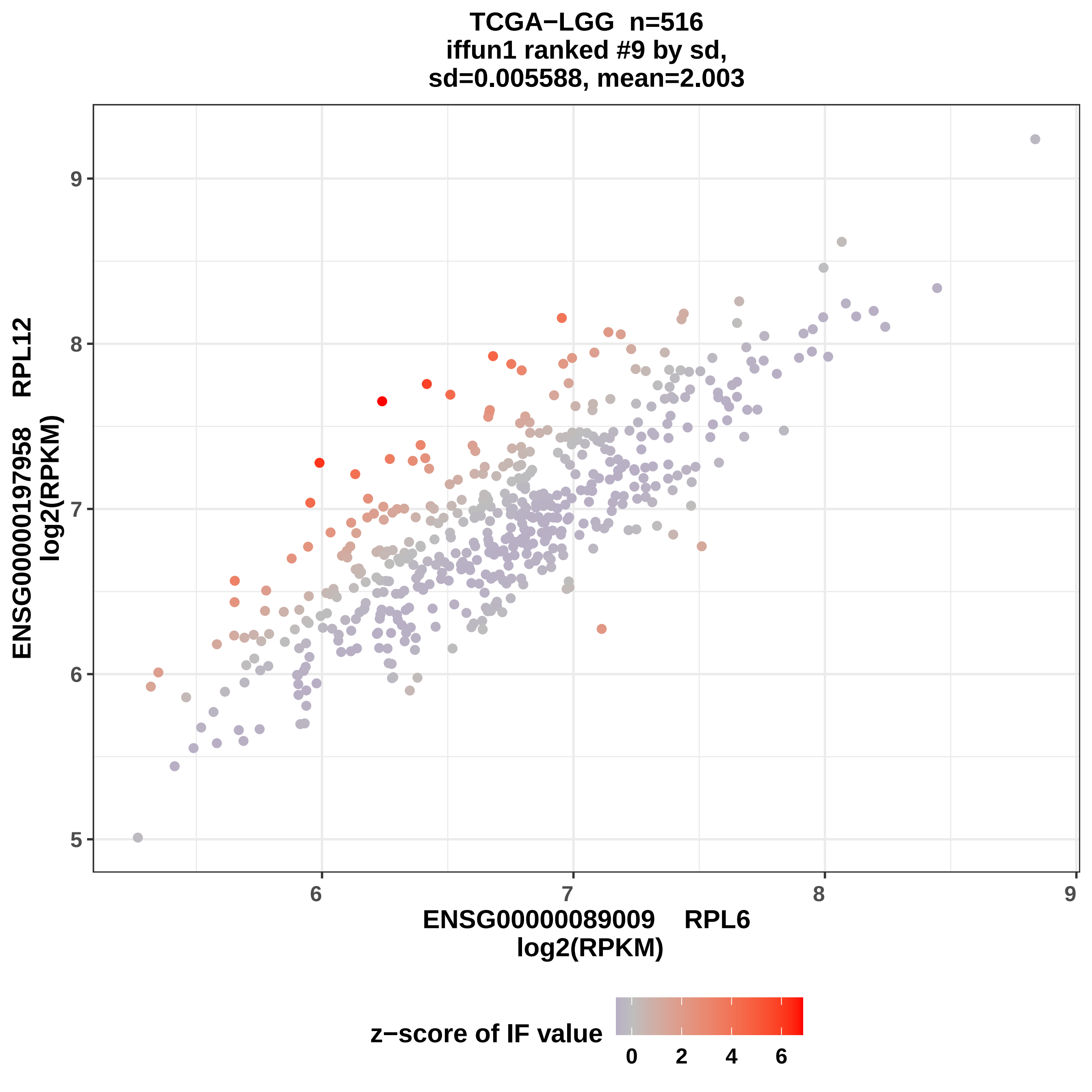

TCGA−LGG n=516
iffun1 ranked #9 by sd,
sd=0.005588, mean=2.003
ENSG00000197958 RPL12
log2(RPKM)
ENSG00000089009 RPL6
log2(RPKM)
z−score of IF value
0
2
4
6
5
6
7
8
9

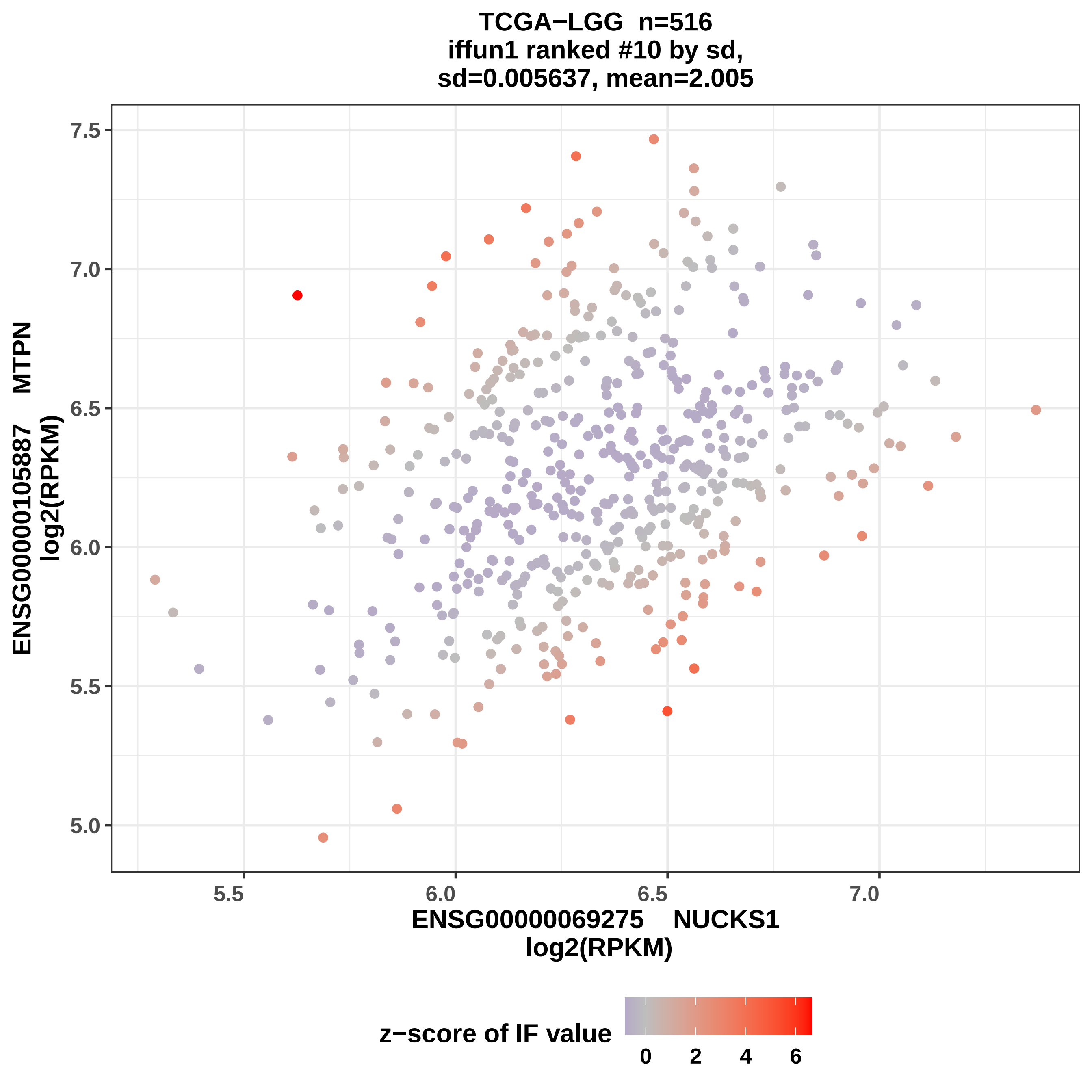
TCGA−LGG n=516
iffun1 ranked #10 by sd,
sd=0.005637, mean=2.005
ENSG00000105887 MTPN
log2(RPKM)
7.5
7.0
6.5
6.0
5.5
5.0
5.5
6.0
6.5
7.0
ENSG00000069275 NUCKS1
log2(RPKM)
z−score of IF value
0
2
4
6

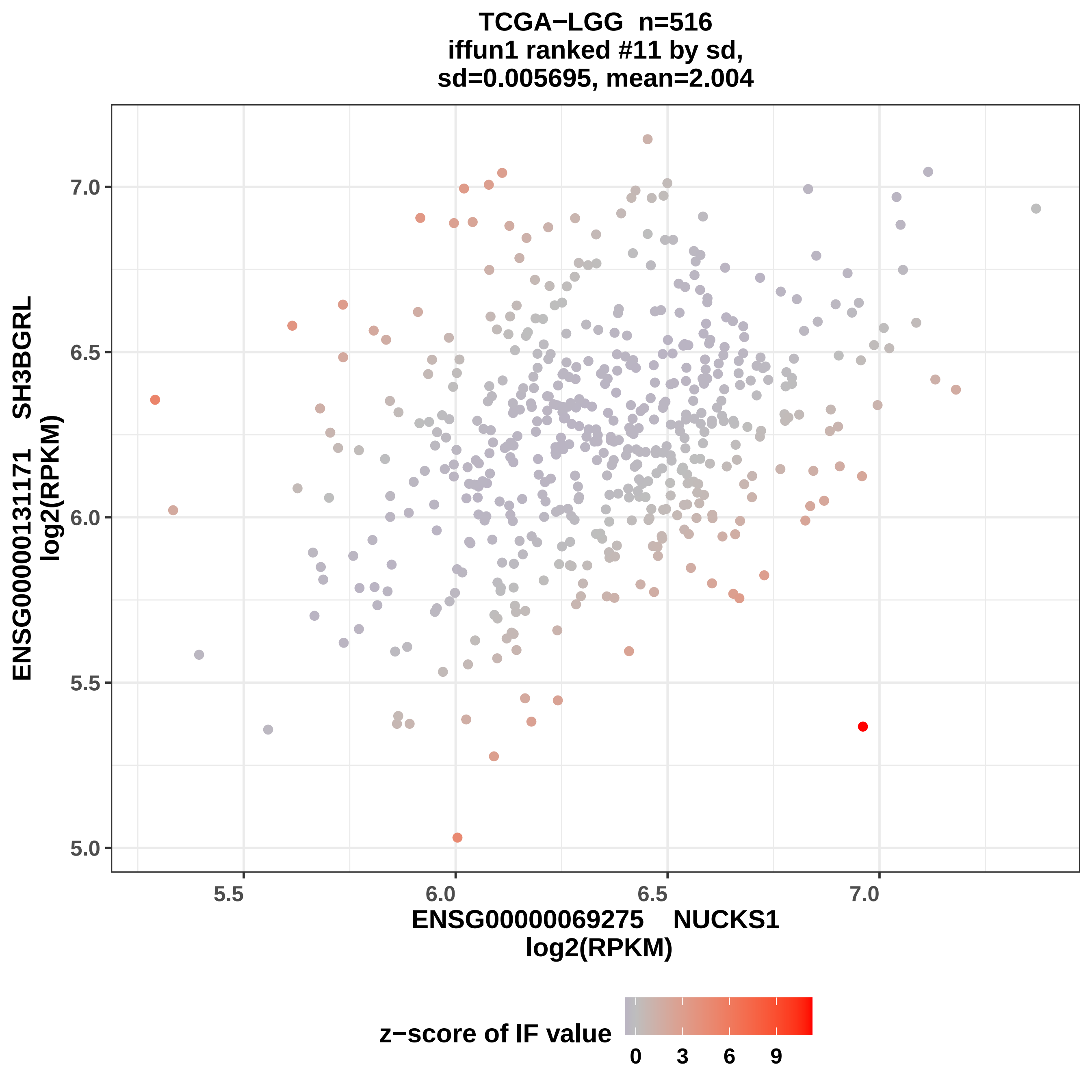
TCGA−LGG n=516
iffun1 ranked #11 by sd,
sd=0.005695, mean=2.004
ENSG00000131171 SH3BGRL
log2(RPKM)
ENSG00000069275 NUCKS1
log2(RPKM)
5.0
5.5
6.0
6.5
7.0
z−score of IF value
0
3
6
9

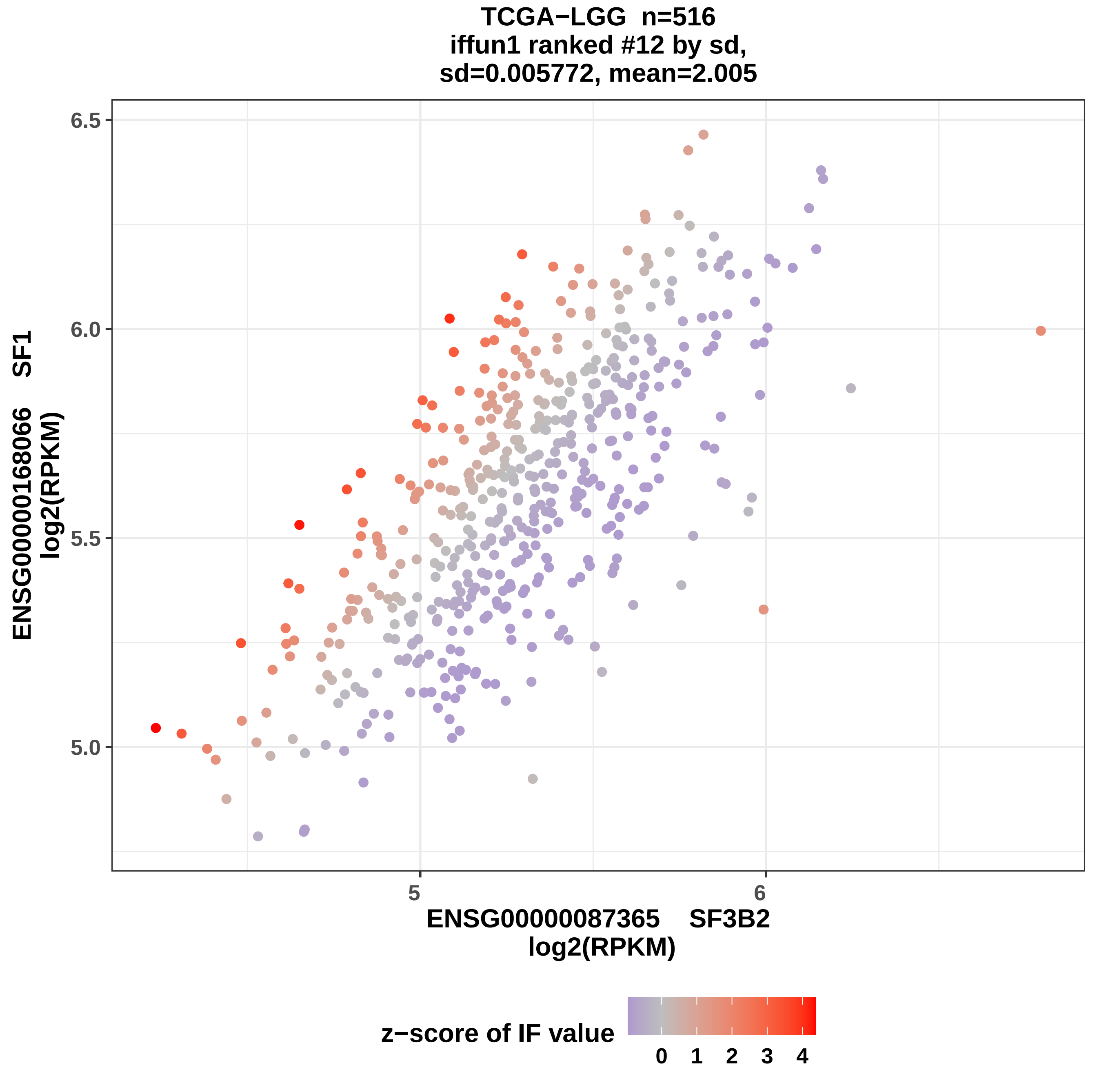
TCGA−LGG n=516
iffun1 ranked #12 by sd,
sd=0.005772, mean=2.005
ENSG00000168066 SF1
log2(RPKM)
6.5
6.0
5.5
5.0
5
6
ENSG00000087365 SF3B2
log2(RPKM)
z−score of IF value
0 1 2 3 4

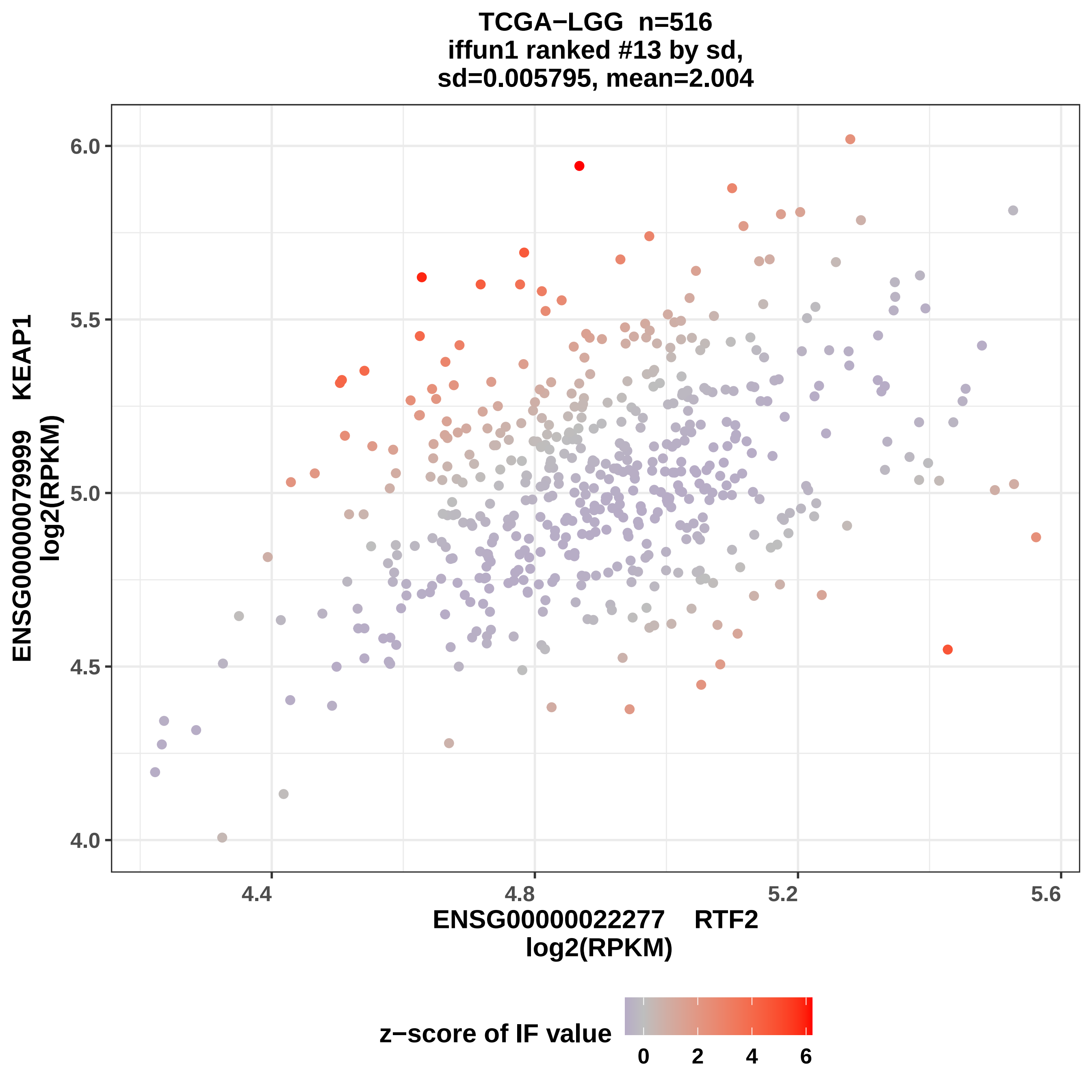
TCGA−LGG n=516
iffun1 ranked #13 by sd,
sd=0.005795, mean=2.004
ENSG00000079999 KEAP1
log2(RPKM)
6.0
5.5
5.0
4.5
4.0
4.4
4.8
5.2
5.6
ENSG00000022277 RTF2
log2(RPKM)
z−score of IF value
0
2
4
6

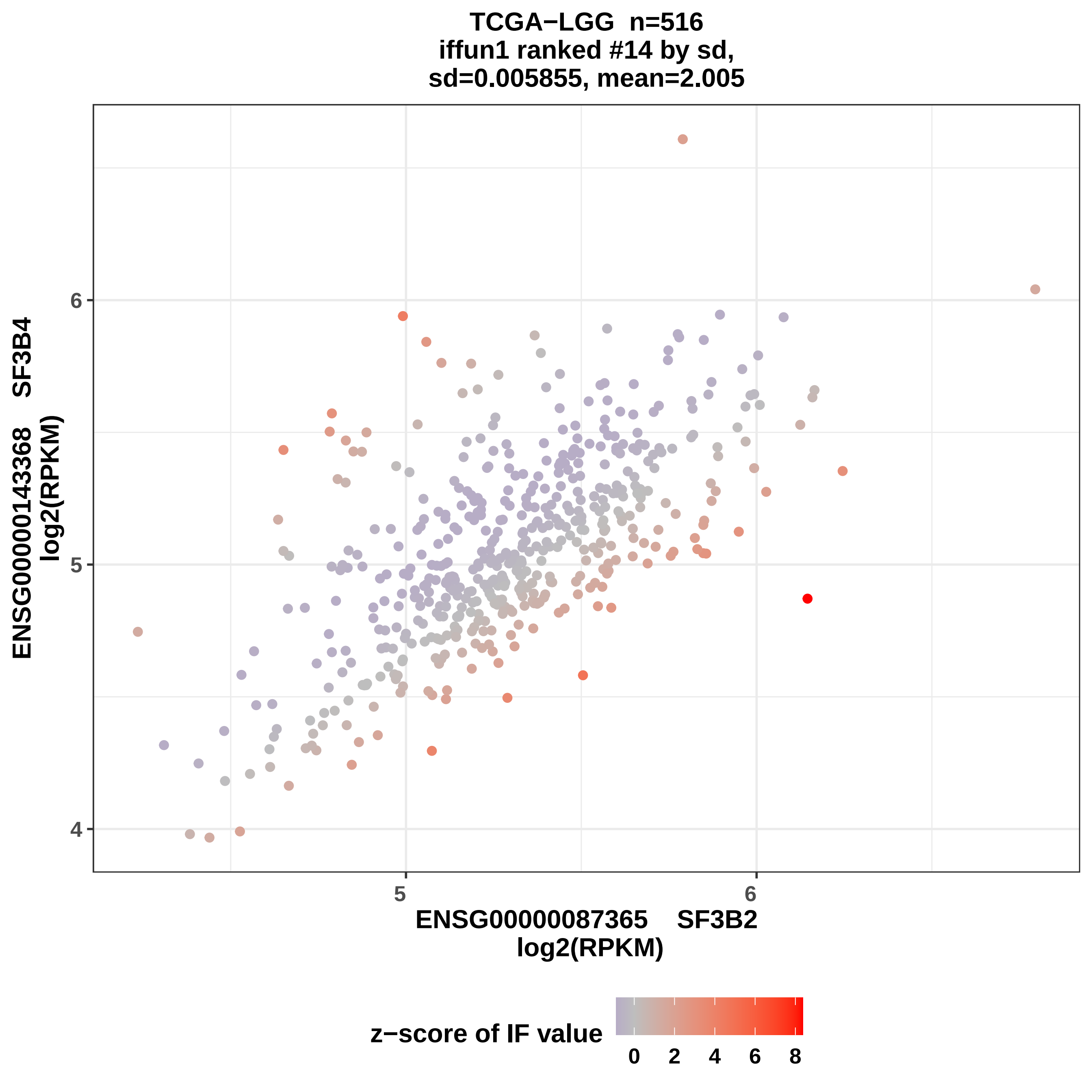
TCGA−LGG n=516
iffun1 ranked #14 by sd,
sd=0.005855, mean=2.005
ENSG00000143368 SF3B4
log2(RPKM)
6
5
4
5
6
ENSG00000087365 SF3B2
log2(RPKM)
z−score of IF value
0
2
4
6
8

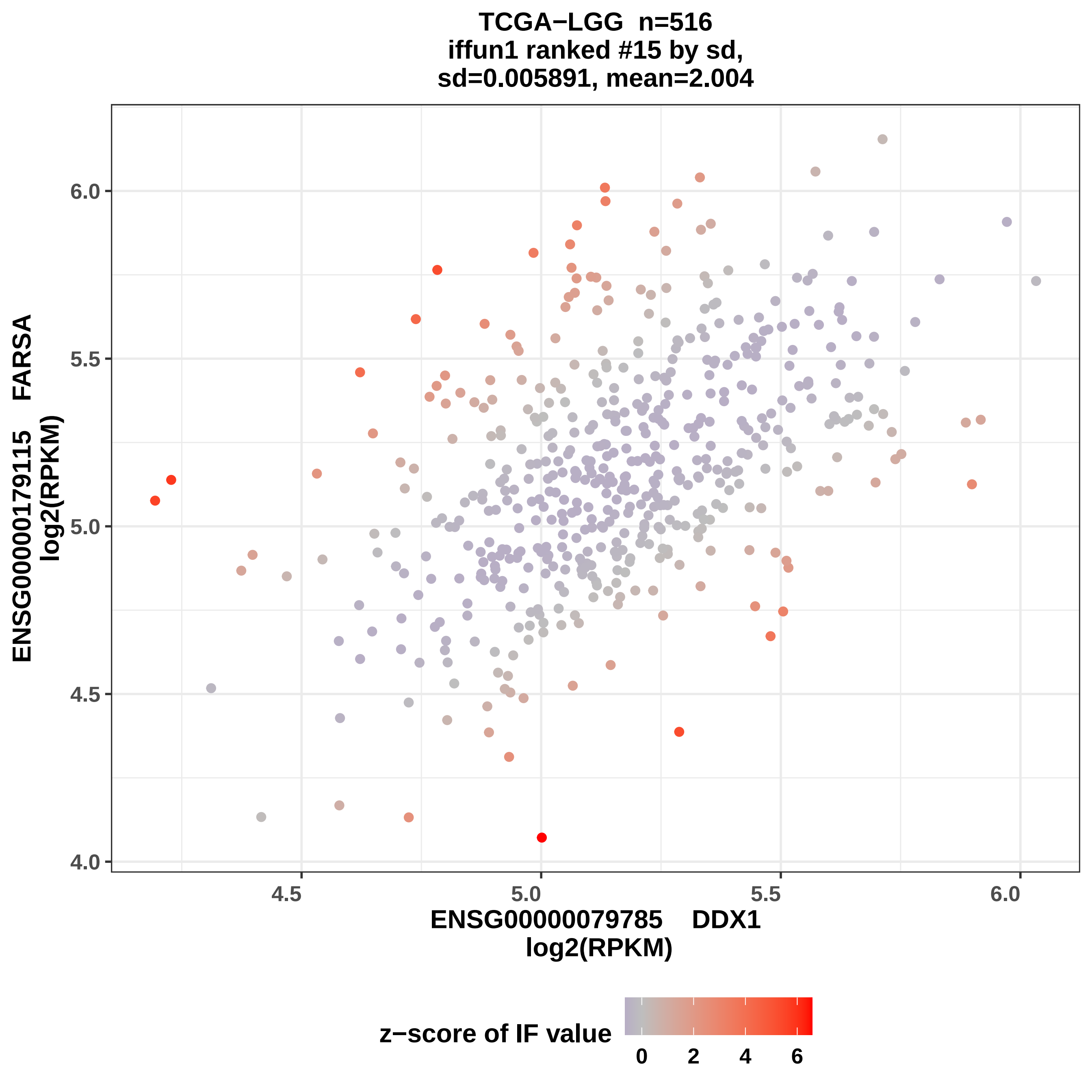
TCGA−LGG n=516
iffun1 ranked #15 by sd,
sd=0.005891, mean=2.004
ENSG00000179115 FARSA
log2(RPKM)
6.0
5.5
5.0
4.5
4.0
4.5
5.0
5.5
6.0
ENSG00000079785 DDX1
log2(RPKM)
z−score of IF value
0
2
4
6

TCGA−LGG n=516
iffun1 ranked #16 by sd,
sd=0.005901, mean=2.004

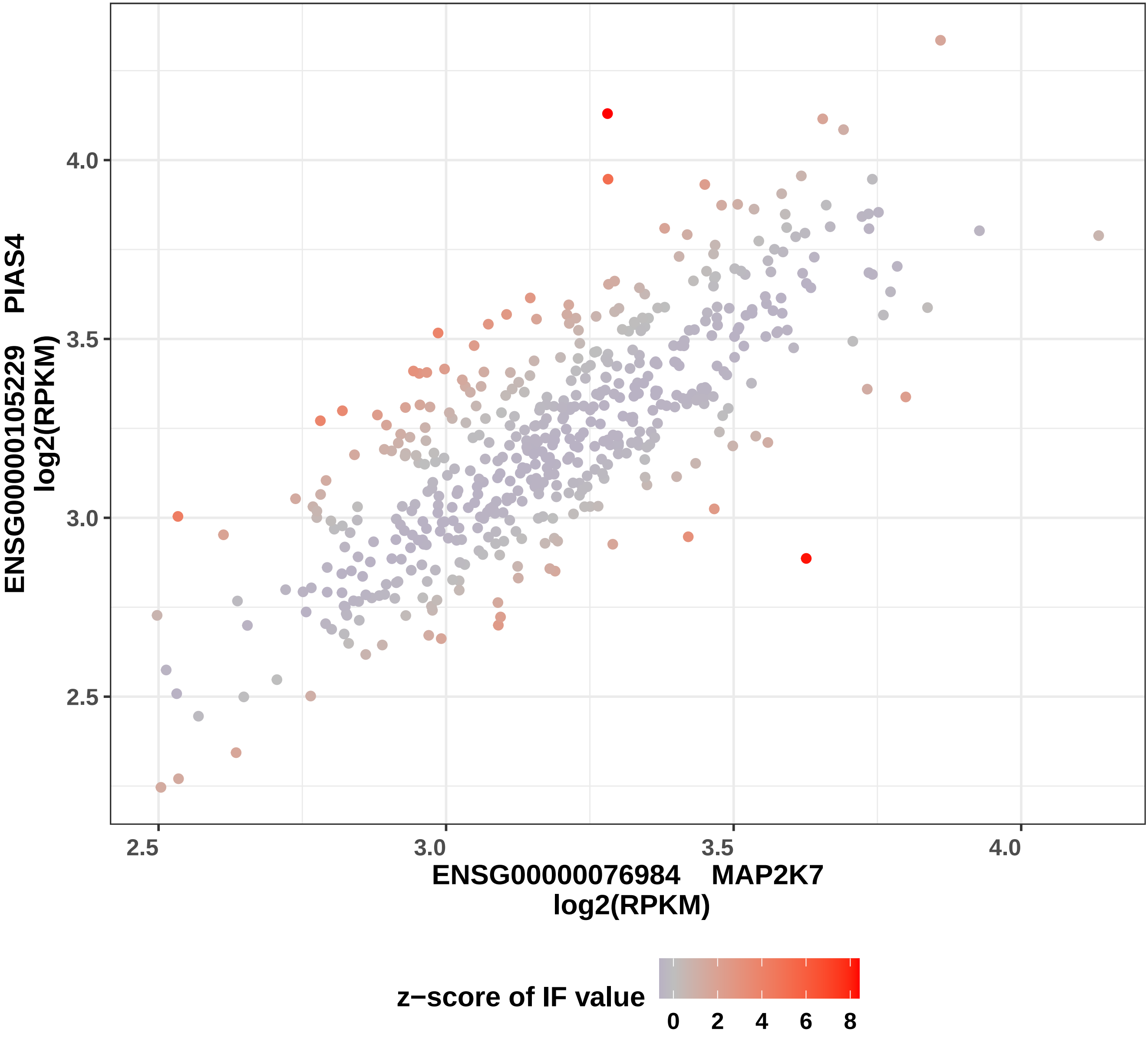

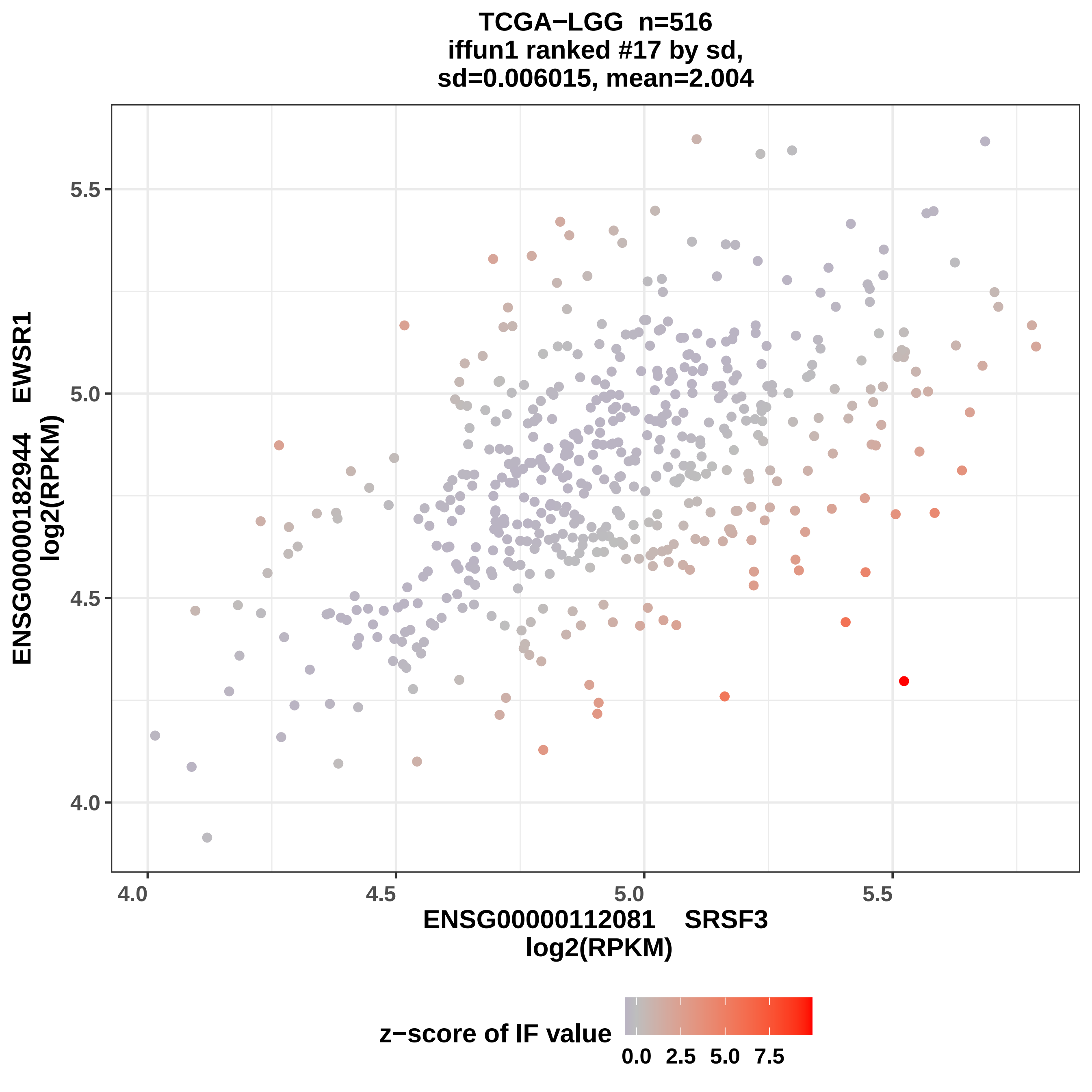
TCGA−LGG n=516
iffun1 ranked #17 by sd,
sd=0.006015, mean=2.004
ENSG00000182944 EWSR1
log2(RPKM)
5.5
5.0
4.5
4.0
4.0
4.5
5.0
5.5
ENSG00000112081 SRSF3
log2(RPKM)
z−score of IF value
0.0 2.5 5.0 7.5

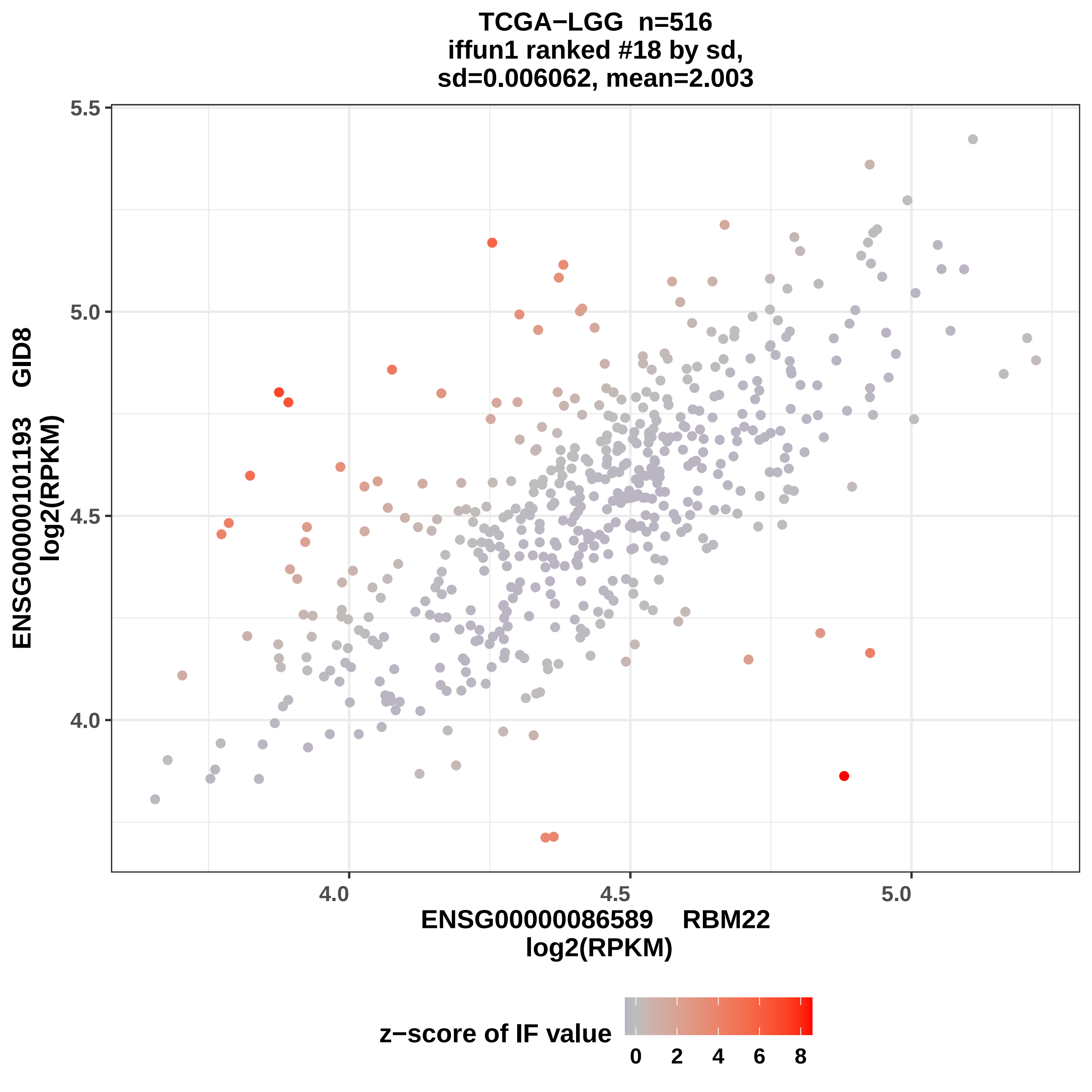
TCGA−LGG n=516
iffun1 ranked #18 by sd,
sd=0.006062, mean=2.003
5.5
5.0
4.5
4.0
ENSG00000101193 GID8
log2(RPKM)
4.0
4.5
5.0
ENSG00000086589 RBM22
log2(RPKM)
z−score of IF value
0
2
4
6
8

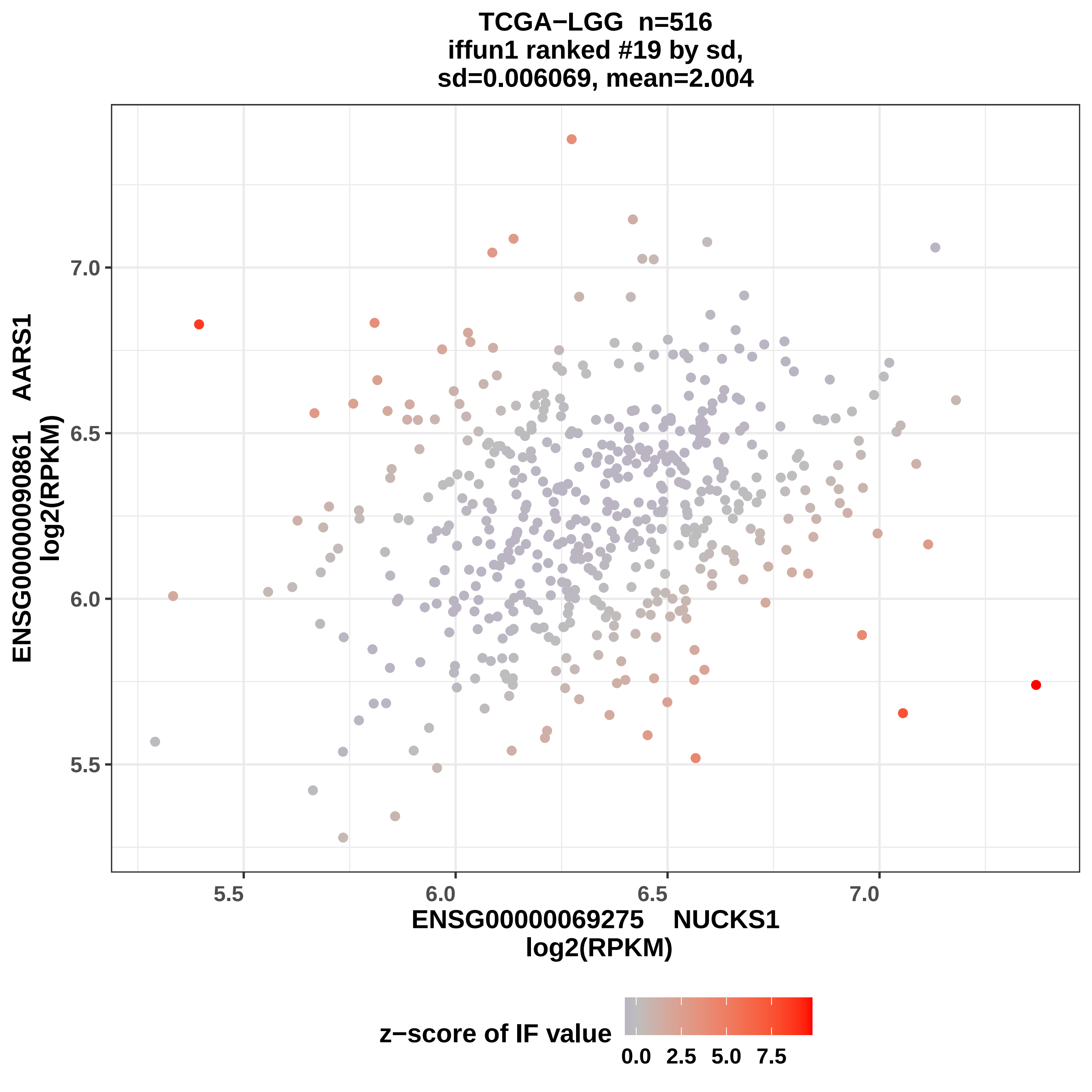

TCGA−LGG n=516
iffun1 ranked #19 by sd,
sd=0.006069, mean=2.004
7.0
6.5
6.0
5.5
5.5
6.0
6.5
7.0
ENSG00000090861 AARS1
log2(RPKM)
ENSG00000069275 NUCKS1
log2(RPKM)
z−score of IF value
0.0
2.5
5.0
7.5

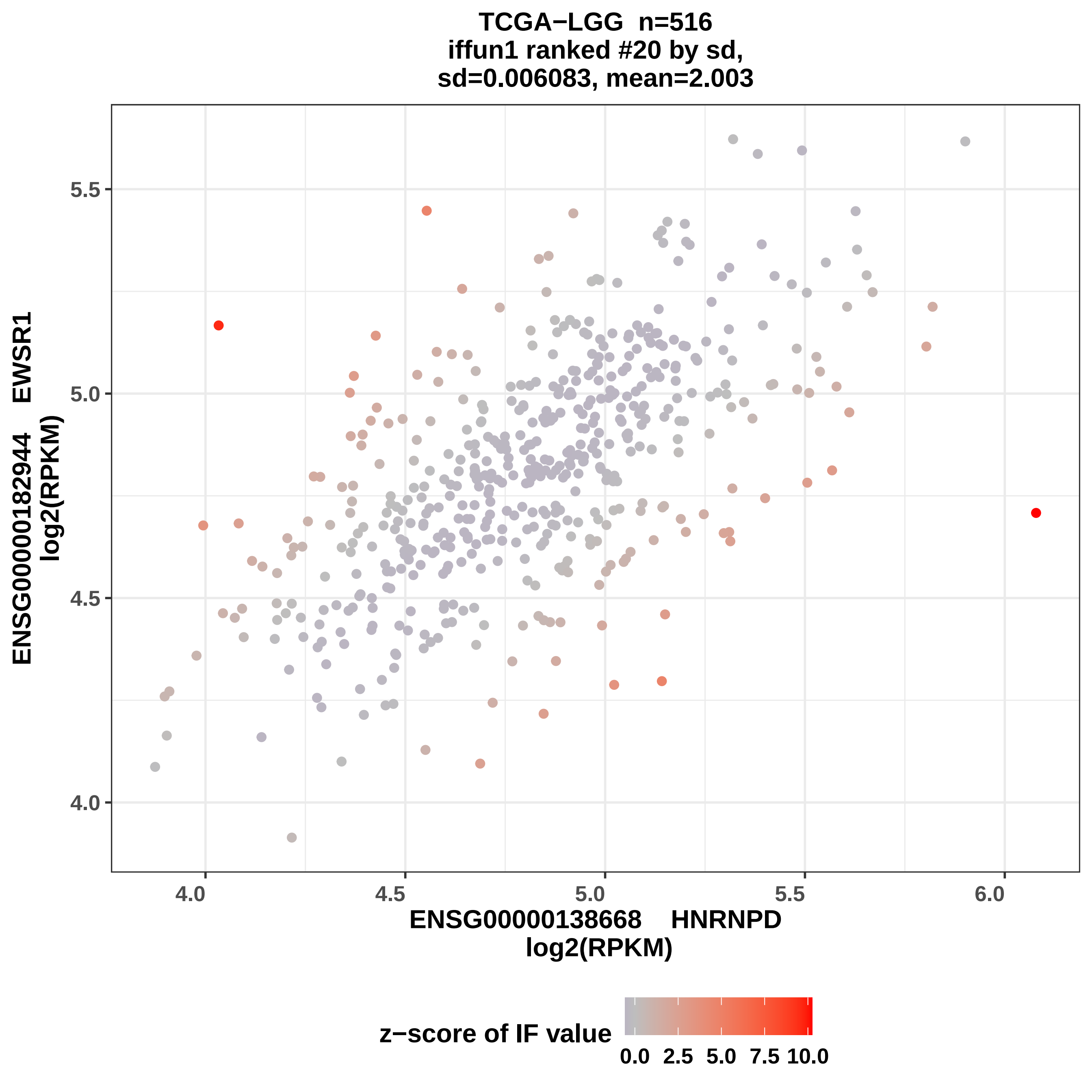
TCGA−LGG n=516
iffun1 ranked #20 by sd,
sd=0.006083, mean=2.003
ENSG00000182944 EWSR1
log2(RPKM)
5.5
5.0
4.5
4.0
4.0
4.5
5.0
5.5
6.0
ENSG00000138668 HNRNPD
log2(RPKM)
z−score of IF value
0.0
2.5
5.0
7.5
10.0

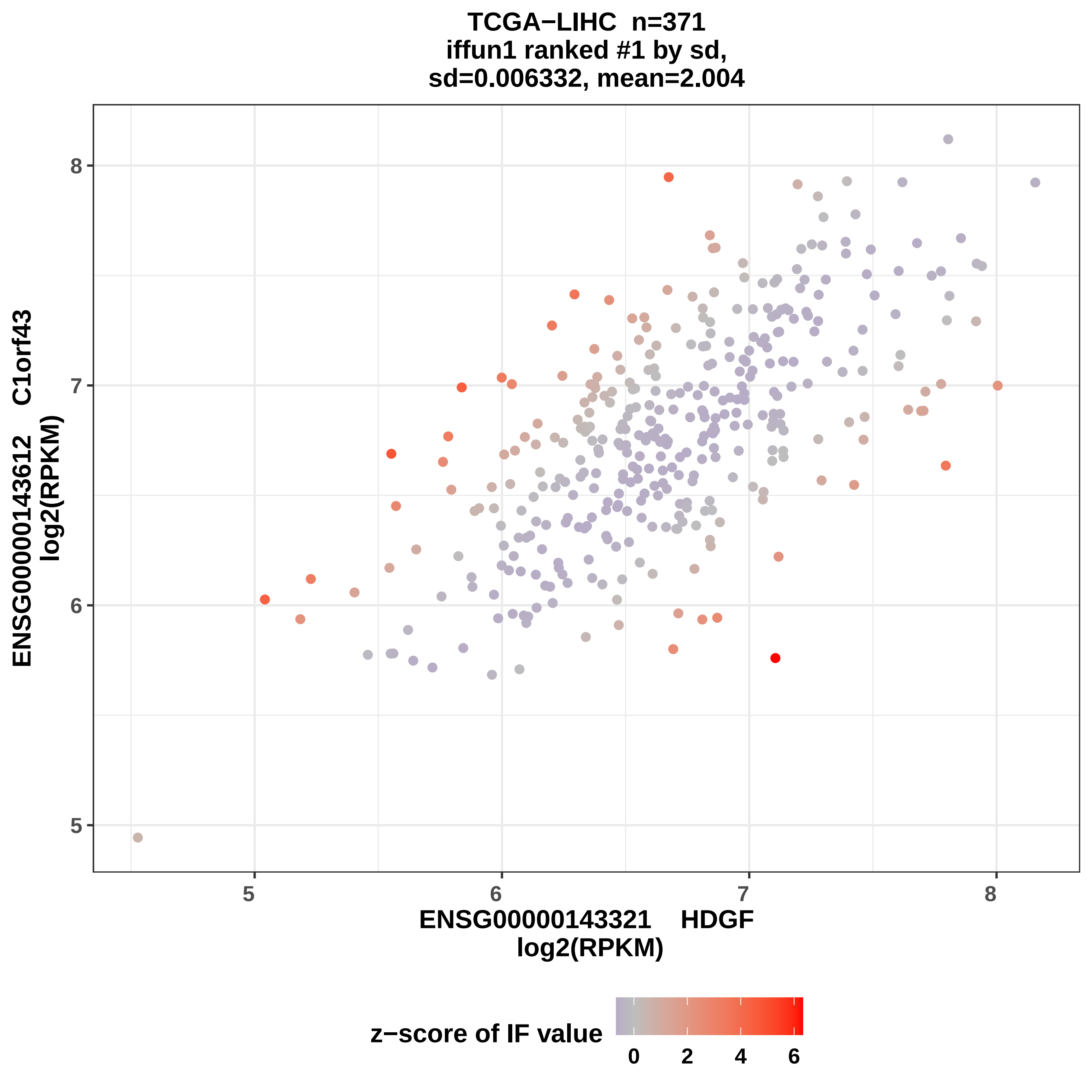
TCGA−LIHC n=371
iffun1 ranked #1 by sd,
sd=0.006332, mean=2.004
ENSG00000143612 C1orf43
log2(RPKM)
8
7
6
5
5
6
7
8
ENSG00000143321 HDGF
log2(RPKM)
z−score of IF value
0
2
4
6

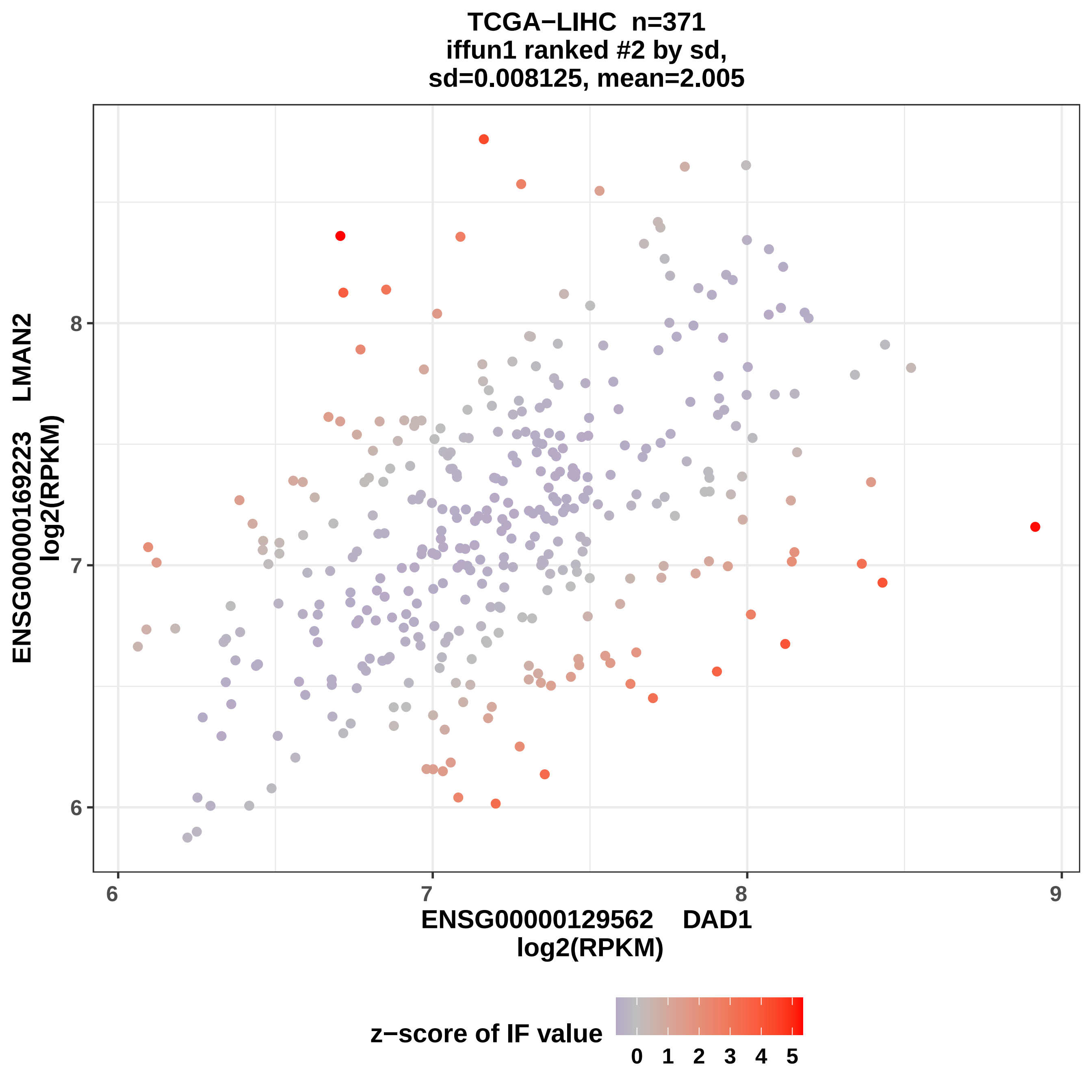

TCGA−LIHC n=371
iffun1 ranked #2 by sd,
sd=0.008125, mean=2.005
ENSG00000169223 LMAN2
log2(RPKM)
8
7
6
6
7
8
9
ENSG00000129562 DAD1
log2(RPKM)
z−score of IF value
0 1 2 3 4 5

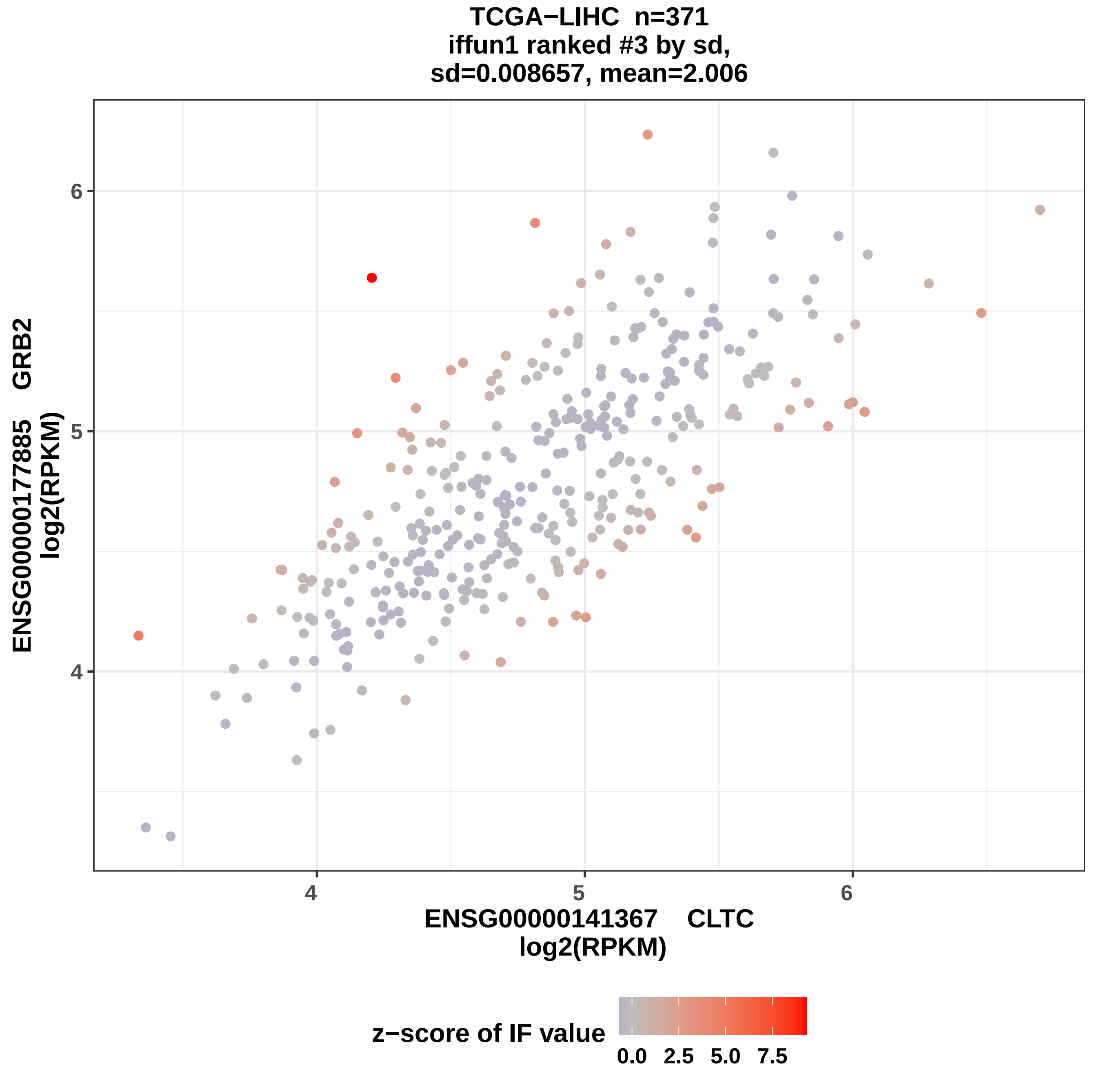
TCGA−LIHC n=371
iffun1 ranked #3 by sd,
sd=0.008657, mean=2.006
ENSG00000177885 GRB2
log2(RPKM)
6
5
4
4
5
6
ENSG00000141367 CLTC
log2(RPKM)
z−score of IF value
0.0
2.5
5.0
7.5

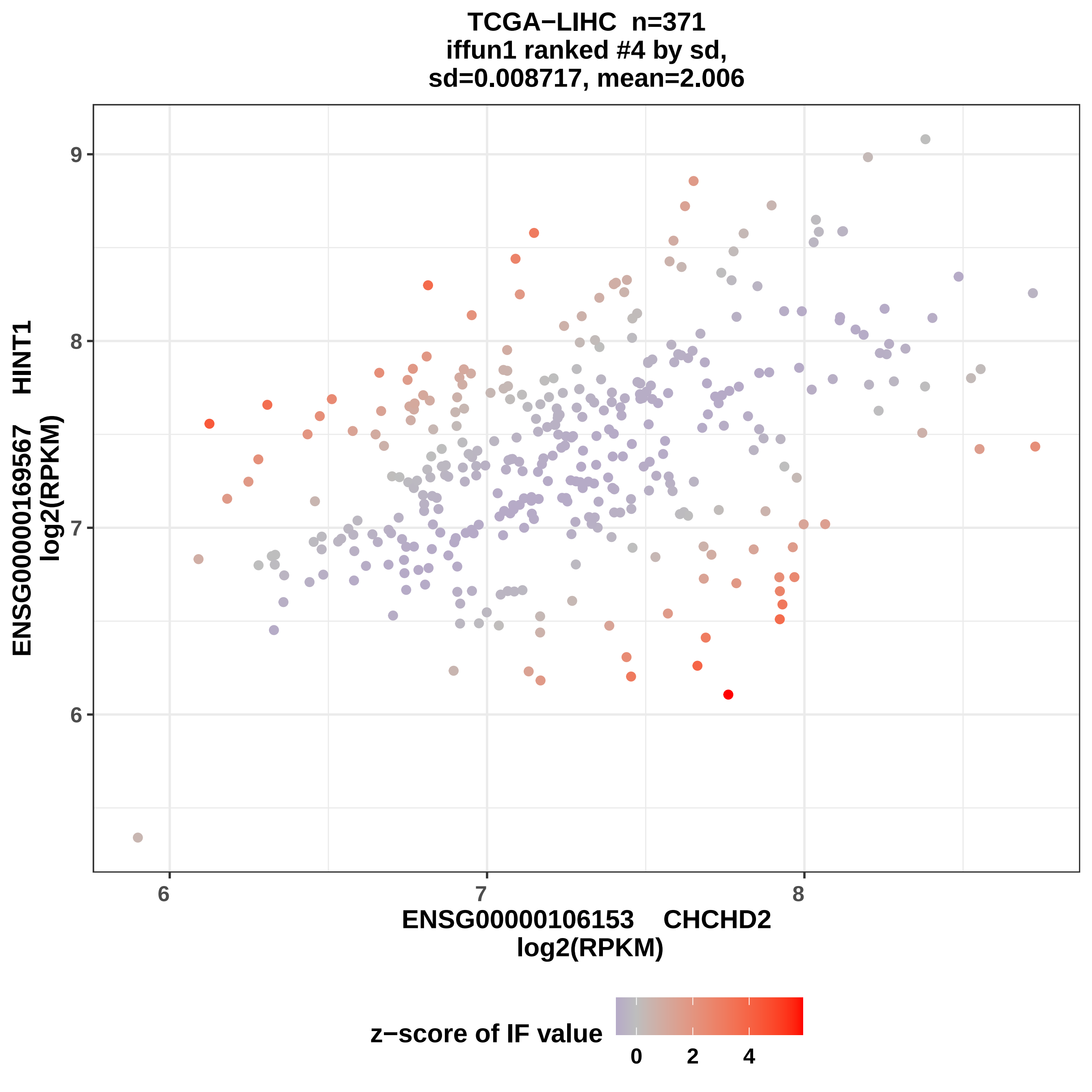

TCGA−LIHC n=371
iffun1 ranked #4 by sd,
sd=0.008717, mean=2.006
ENSG00000169567 HINT1
log2(RPKM)
9
8
7
6
6
7
8
ENSG00000106153 CHCHD2
log2(RPKM)
z−score of IF value
0
2
4

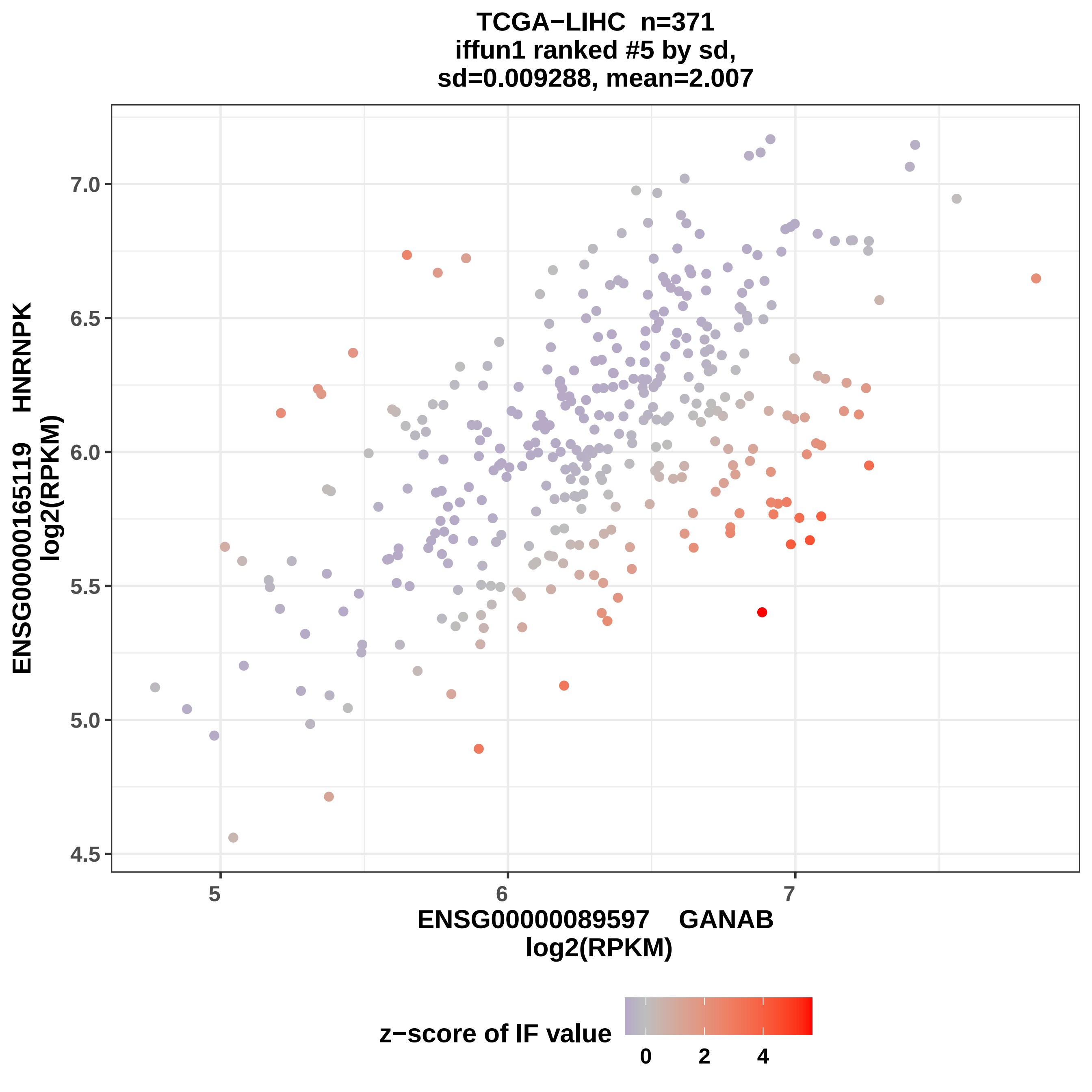
TCGA−LIHC n=371
iffun1 ranked #5 by sd,
sd=0.009288, mean=2.007
ENSG00000165119 HNRNPK
log2(RPKM)
7.0
6.5
6.0
5.5
5.0
4.5
5
6
7
ENSG00000089597 GANAB
log2(RPKM)
z−score of IF value
0
2
4

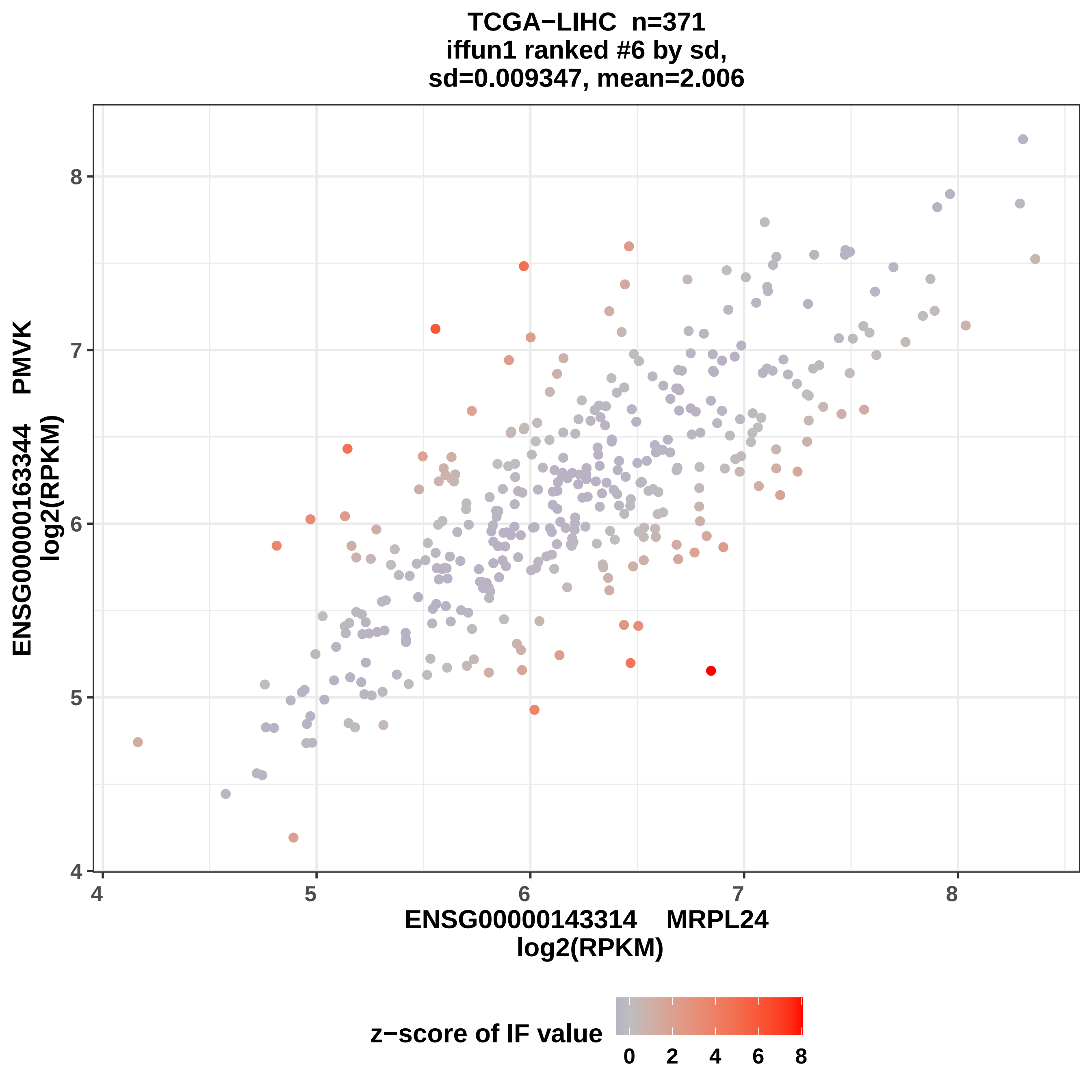

TCGA−LIHC n=371
iffun1 ranked #6 by sd,
sd=0.009347, mean=2.006
ENSG00000163344 PMVK
log2(RPKM)
ENSG00000143314 MRPL24
log2(RPKM)
z−score of IF value
4
5
6
7
8
0
2
4
6
8

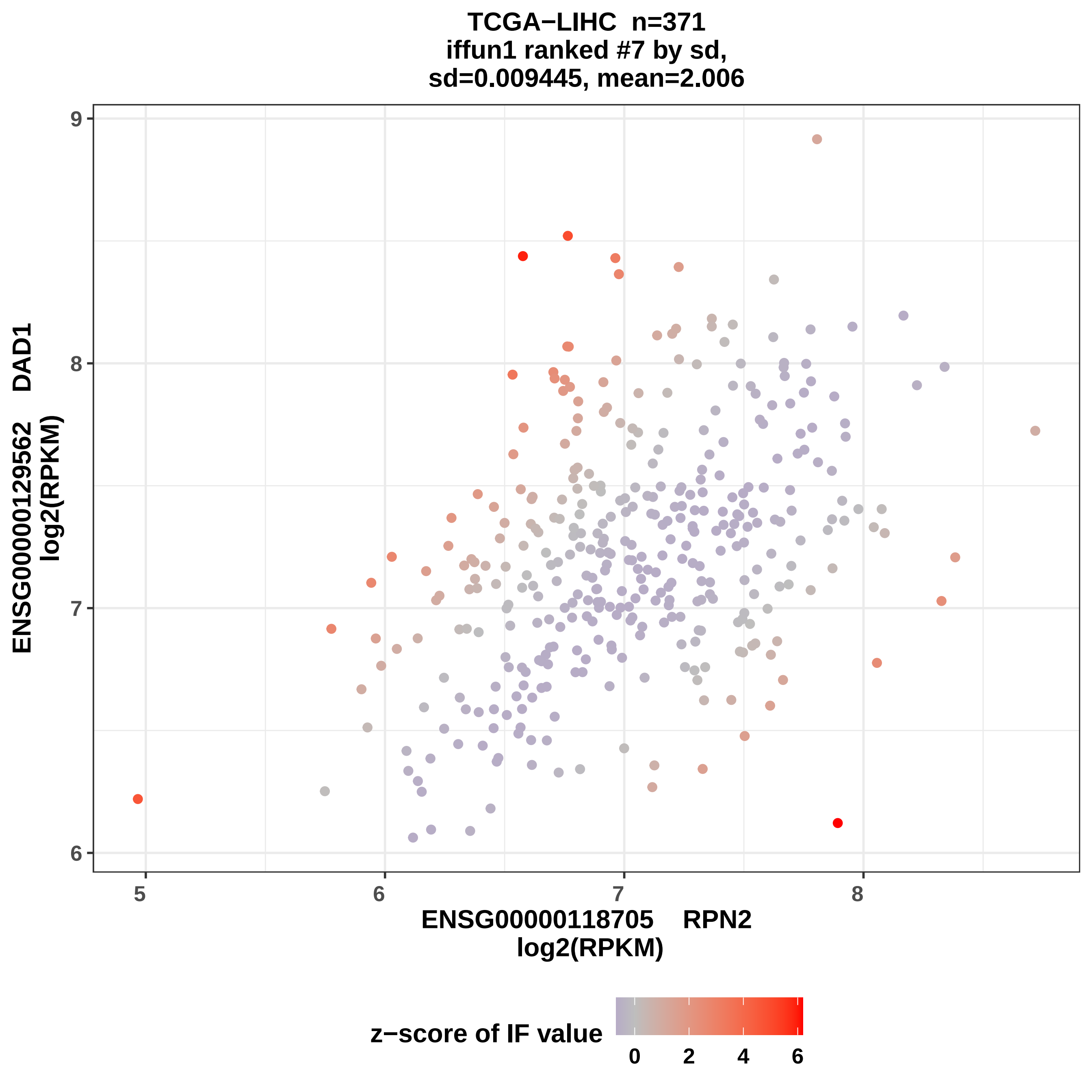
TCGA−LIHC n=371
iffun1 ranked #7 by sd,
sd=0.009445, mean=2.006
ENSG00000129562 DAD1
log2(RPKM)
9
8
7
6
5
6
7
8
ENSG00000118705 RPN2
log2(RPKM)
z−score of IF value
0
2
4
6

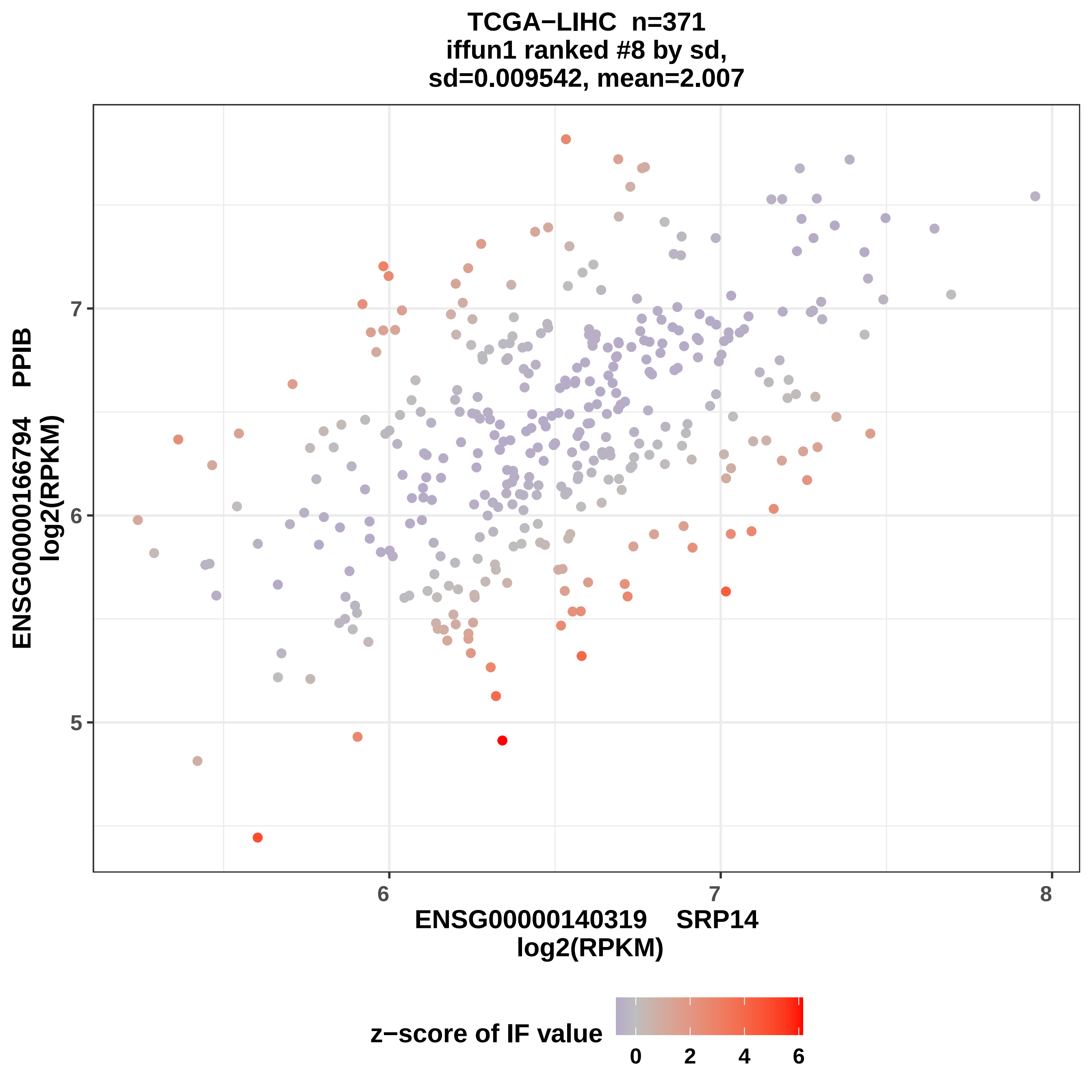
TCGA−LIHC n=371
iffun1 ranked #8 by sd,
sd=0.009542, mean=2.007
ENSG00000166794 PPIB
log2(RPKM)
ENSG00000140319 SRP14
log2(RPKM)
z−score of IF value
0
2
4
6
5
6
7
8

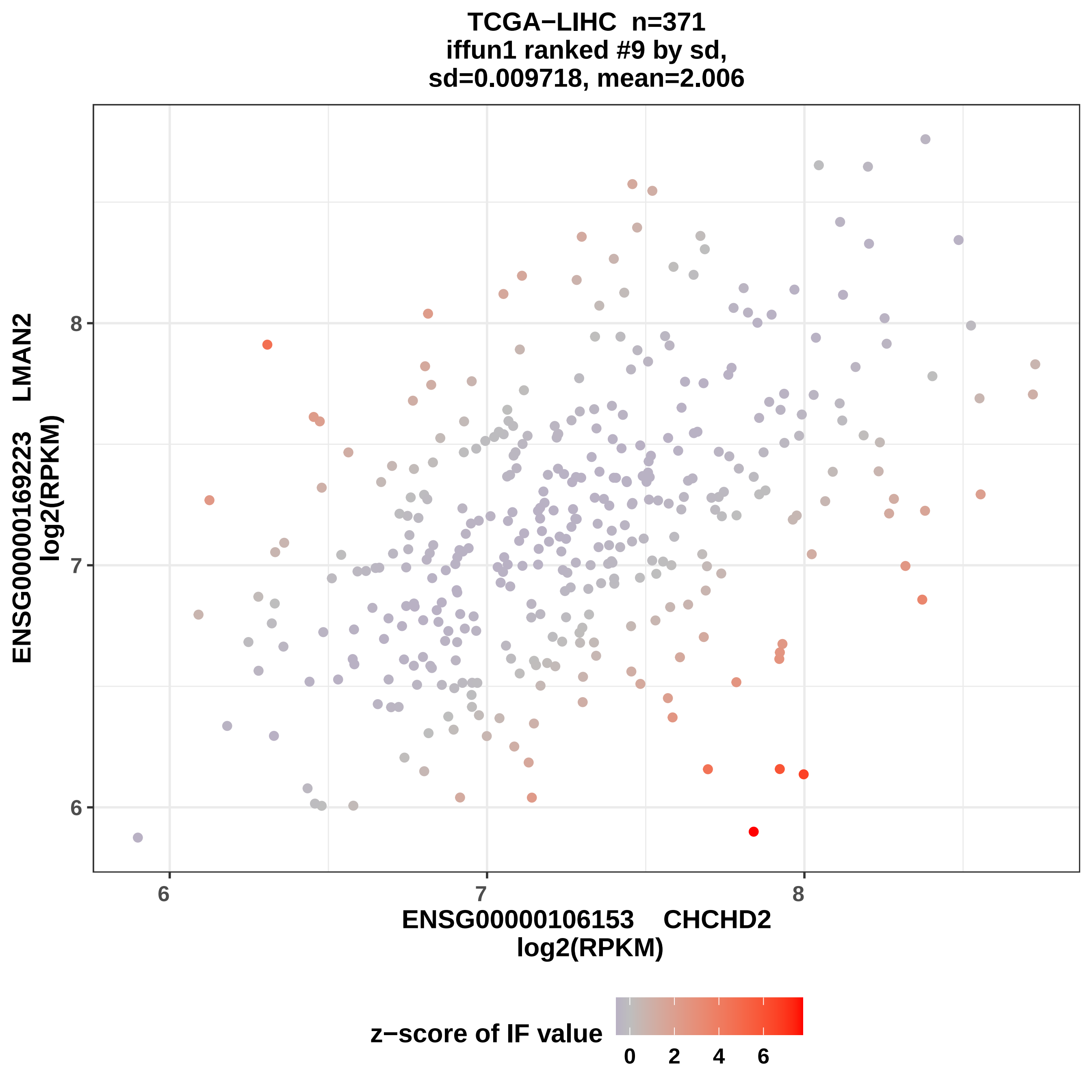

TCGA−LIHC n=371
iffun1 ranked #9 by sd,
sd=0.009718, mean=2.006
ENSG00000169223 LMAN2
log2(RPKM)
8
7
6
ENSG00000106153 CHCHD2
log2(RPKM)
z−score of IF value
0
2
4
6

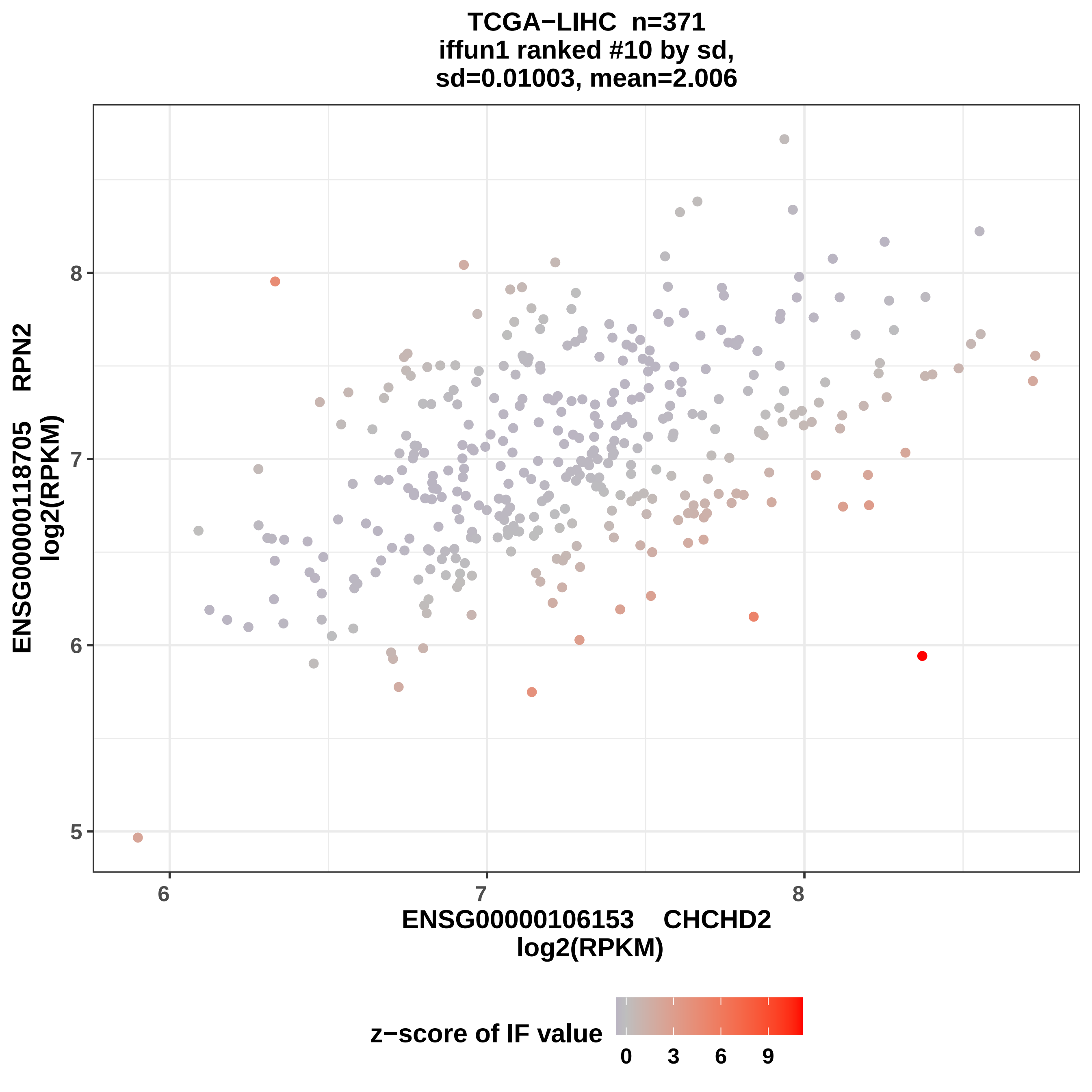

TCGA−LIHC n=371
iffun1 ranked #10 by sd,
sd=0.01003, mean=2.006
ENSG00000118705 RPN2
log2(RPKM)
8
7
6
5
6
7
8
ENSG00000106153 CHCHD2
log2(RPKM)
z−score of IF value
0
3
6
9

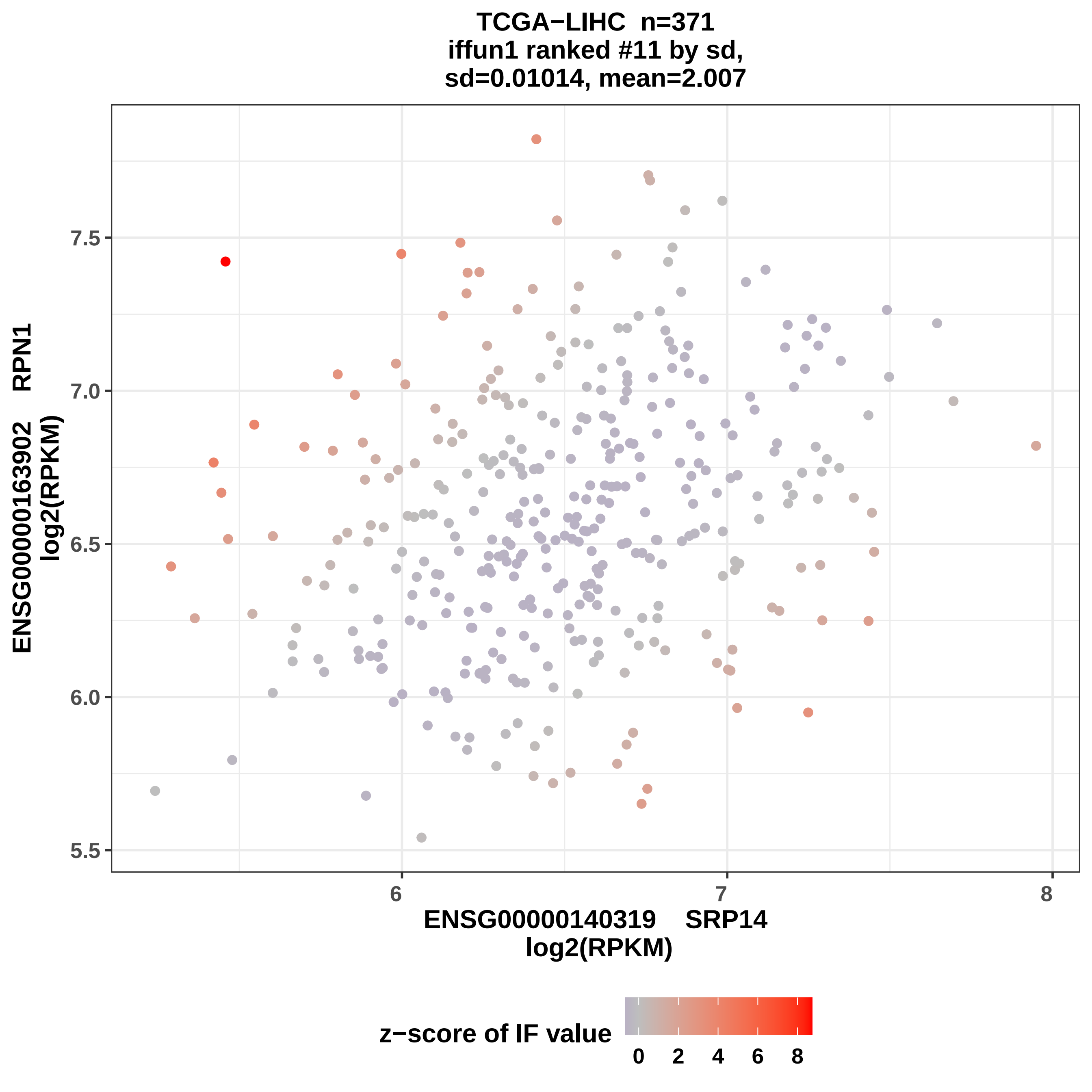

TCGA−LIHC n=371
iffun1 ranked #11 by sd,
sd=0.01014, mean=2.007
ENSG00000163902 RPN1
log2(RPKM)
7.5
7.0
6.5
6.0
5.5
6
7
8
ENSG00000140319 SRP14
log2(RPKM)
z−score of IF value
0
2
4
6
8

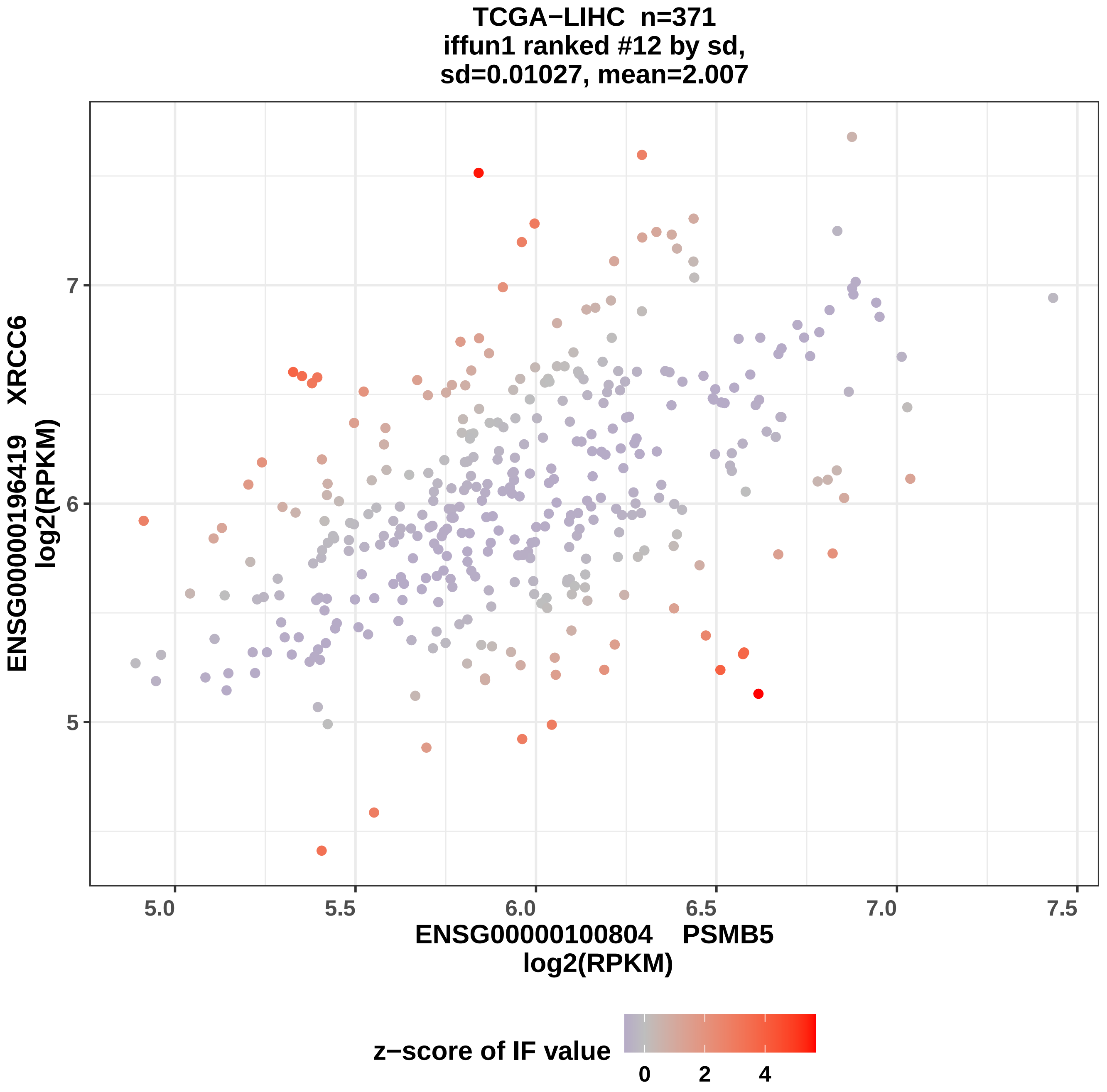

TCGA−LIHC n=371
iffun1 ranked #12 by sd,
sd=0.01027, mean=2.007
ENSG00000196419 XRCC6
log2(RPKM)
7
6
5
5.0
5.5
6.0
6.5
7.0
7.5
ENSG00000100804 PSMB5
log2(RPKM)
z−score of IF value
0
2
4

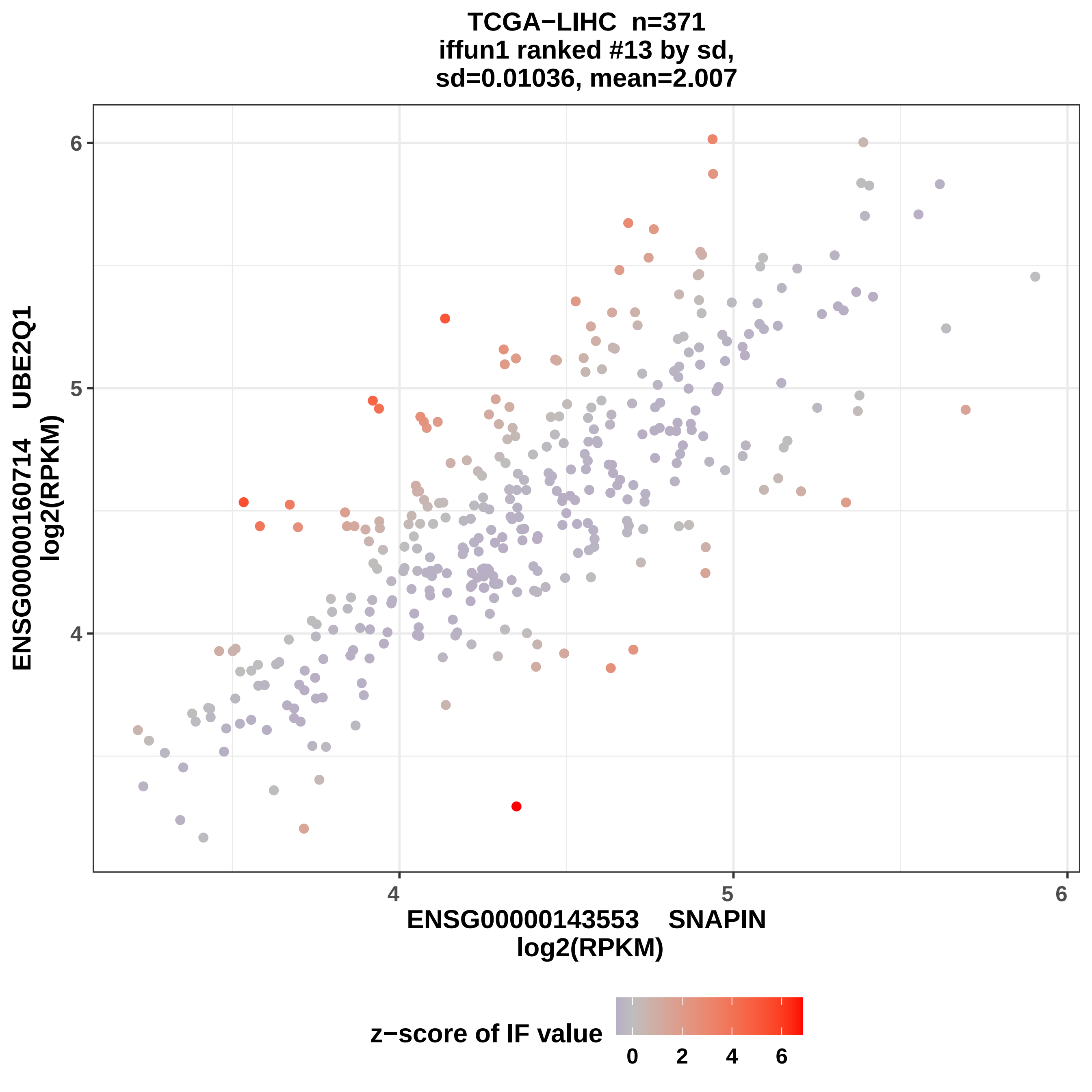
TCGA−LIHC n=371
iffun1 ranked #13 by sd,
sd=0.01036, mean=2.007
ENSG00000160714 UBE2Q1
log2(RPKM)
6
5
4
ENSG00000143553 SNAPIN
log2(RPKM)
4
5
6
z−score of IF value
0
2
4
6

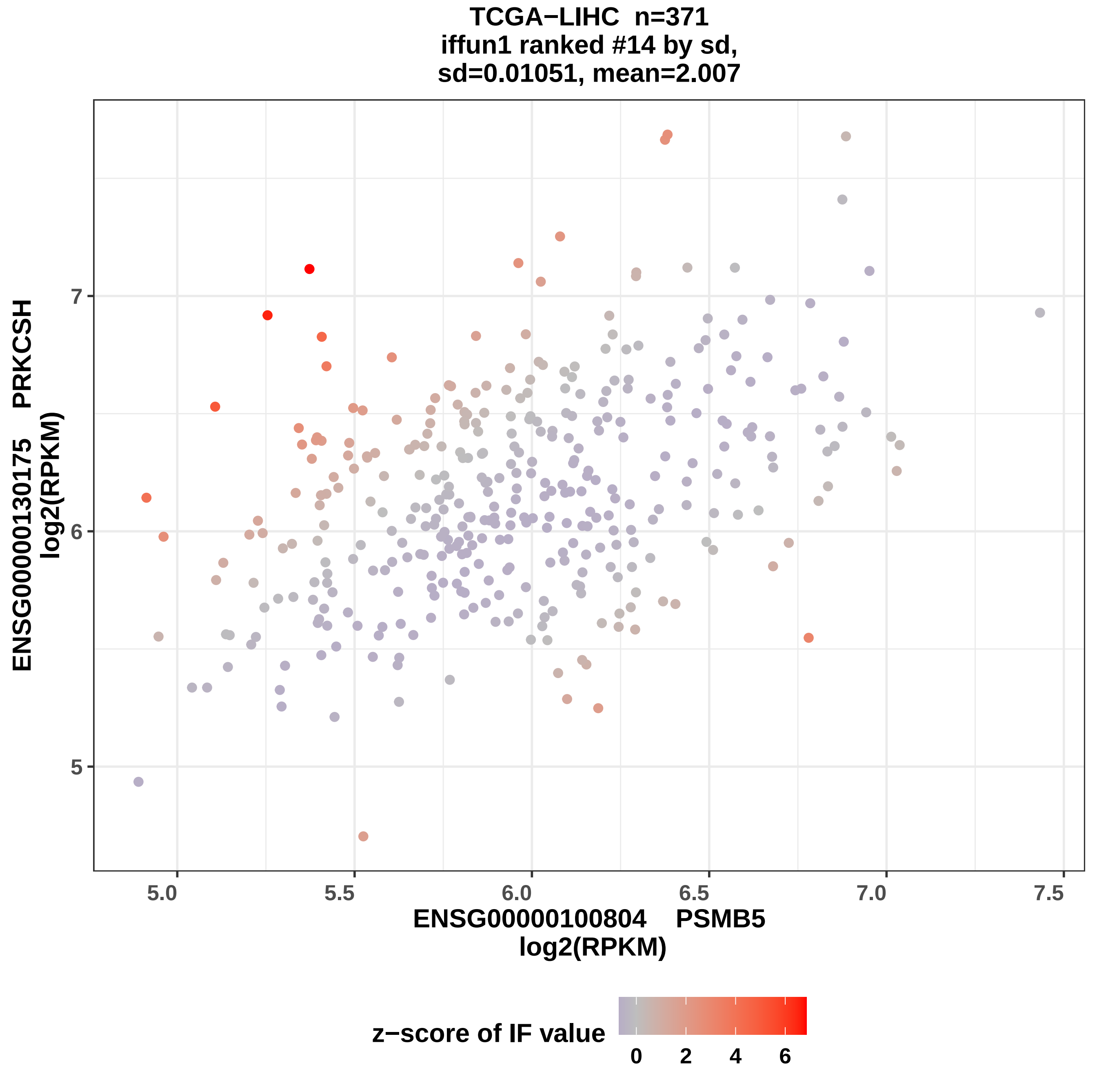
TCGA−LIHC n=371
iffun1 ranked #14 by sd,
sd=0.01051, mean=2.007
ENSG00000130175 PRKCSH
log2(RPKM)
5
6
7
5.0
5.5
6.0
6.5
7.0
7.5
ENSG00000100804 PSMB5
log2(RPKM)
z−score of IF value
0
2
4
6

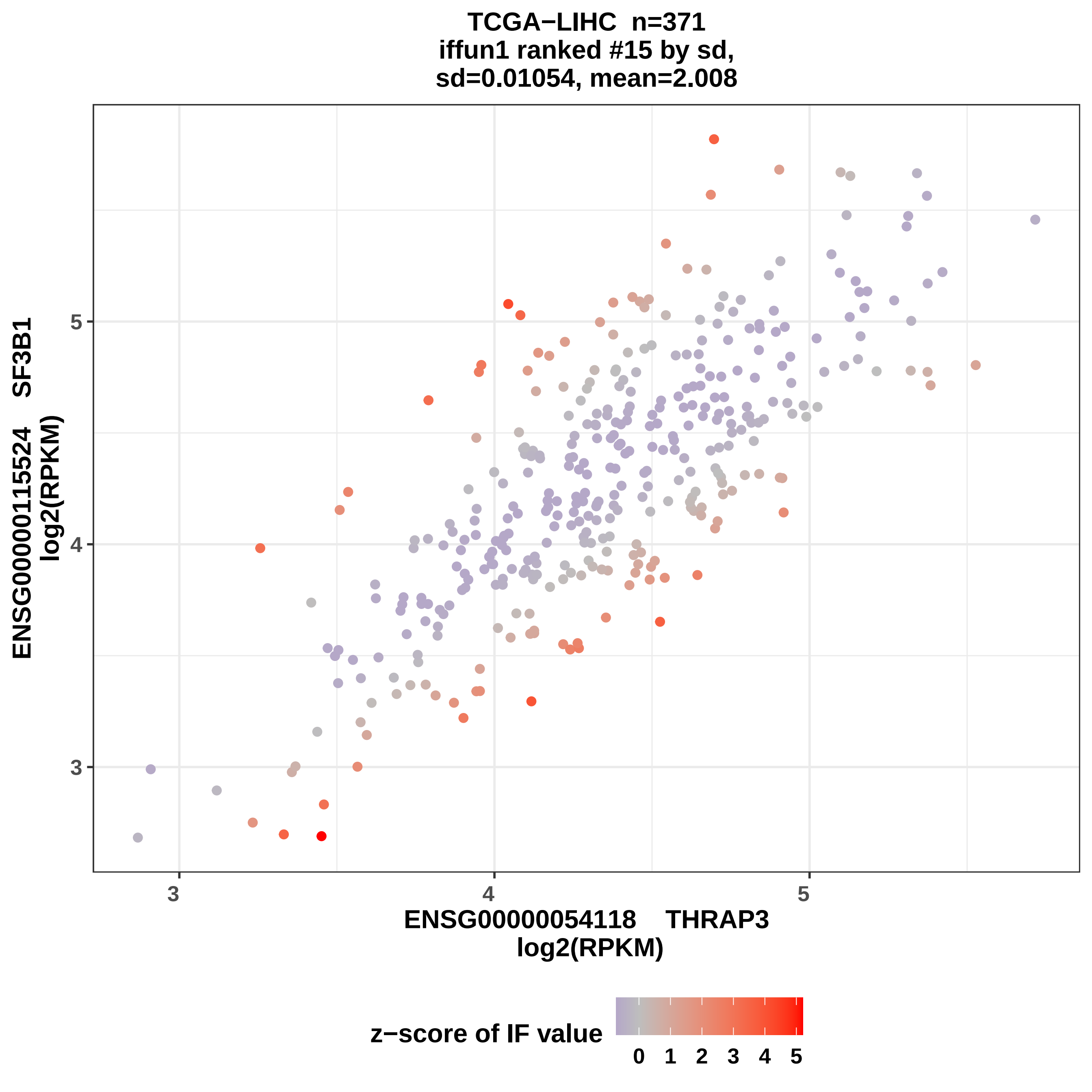

TCGA−LIHC n=371
iffun1 ranked #15 by sd,
sd=0.01054, mean=2.008
ENSG00000115524 SF3B1
log2(RPKM)
ENSG00000054118 THRAP3
log2(RPKM)
3
4
5
3
4
5
z−score of IF value
0
1
2
3
4
5

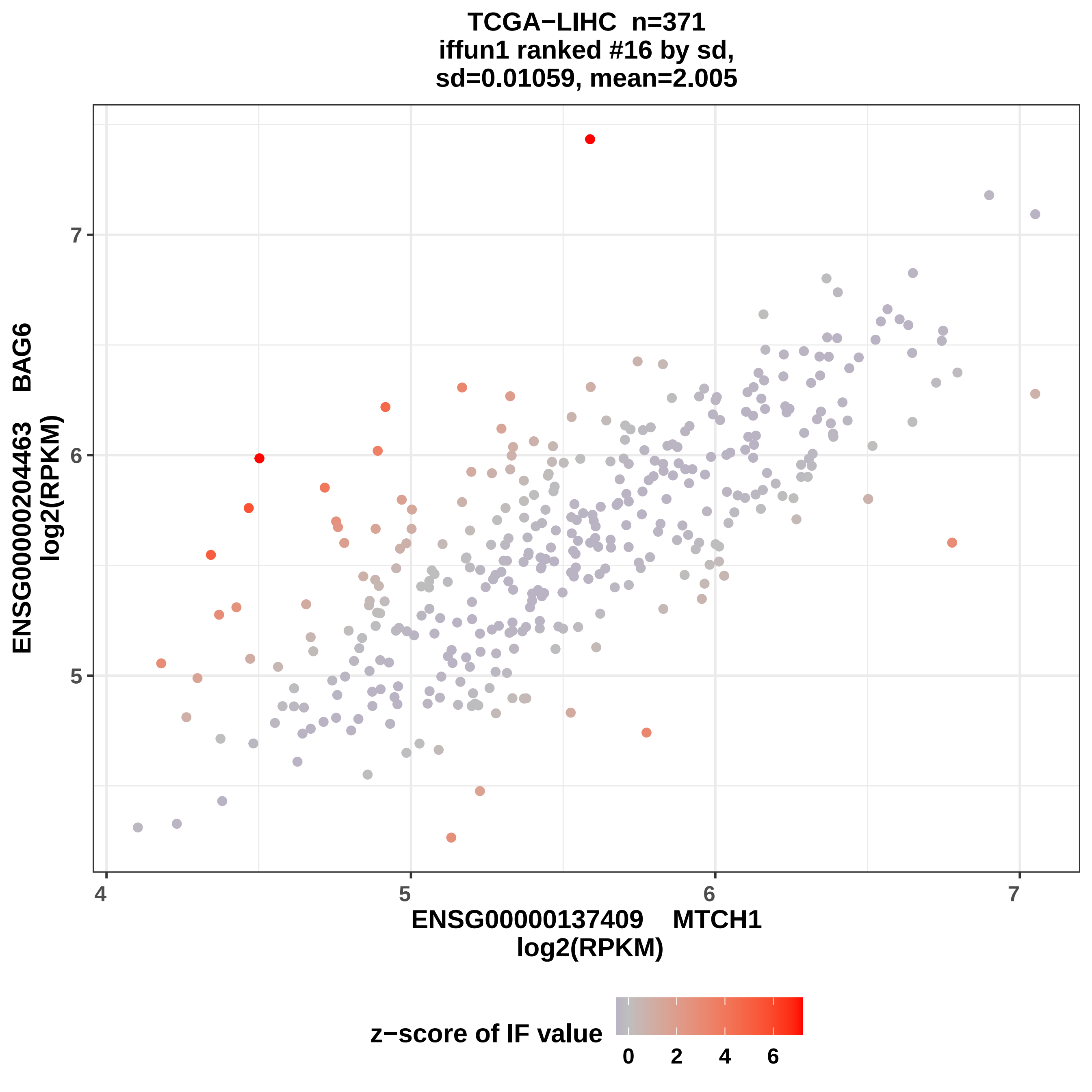
TCGA−LIHC n=371
iffun1 ranked #16 by sd,
sd=0.01059, mean=2.005
ENSG00000204463 BAG6
log2(RPKM)
ENSG00000137409 MTCH1
log2(RPKM)
4
5
6
7
z−score of IF value
0
2
4
6

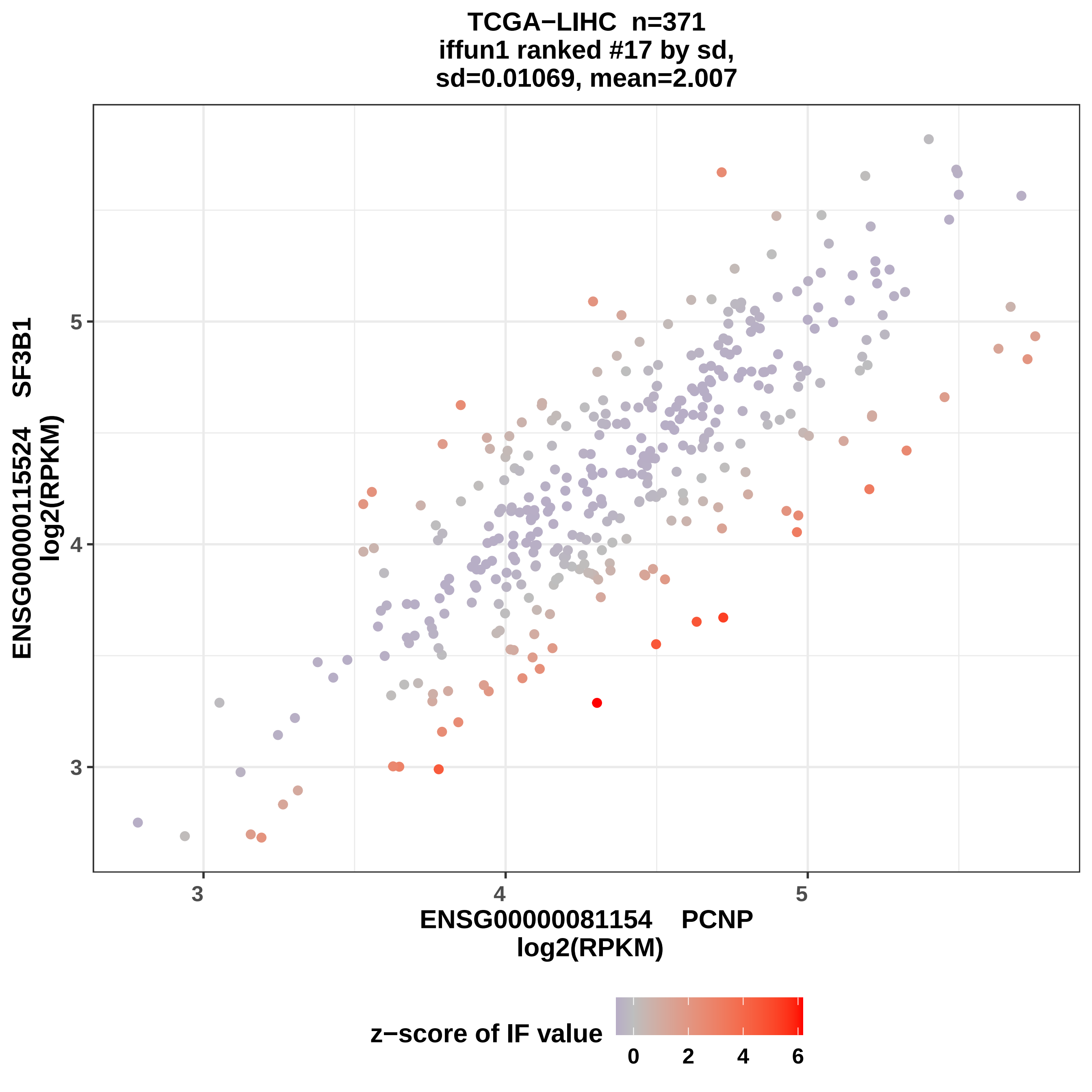

TCGA−LIHC n=371
iffun1 ranked #17 by sd,
sd=0.01069, mean=2.007
ENSG00000115524 SF3B1
log2(RPKM)
ENSG00000081154 PCNP
log2(RPKM)
3
4
5
3
4
5
z−score of IF value
0
2
4
6

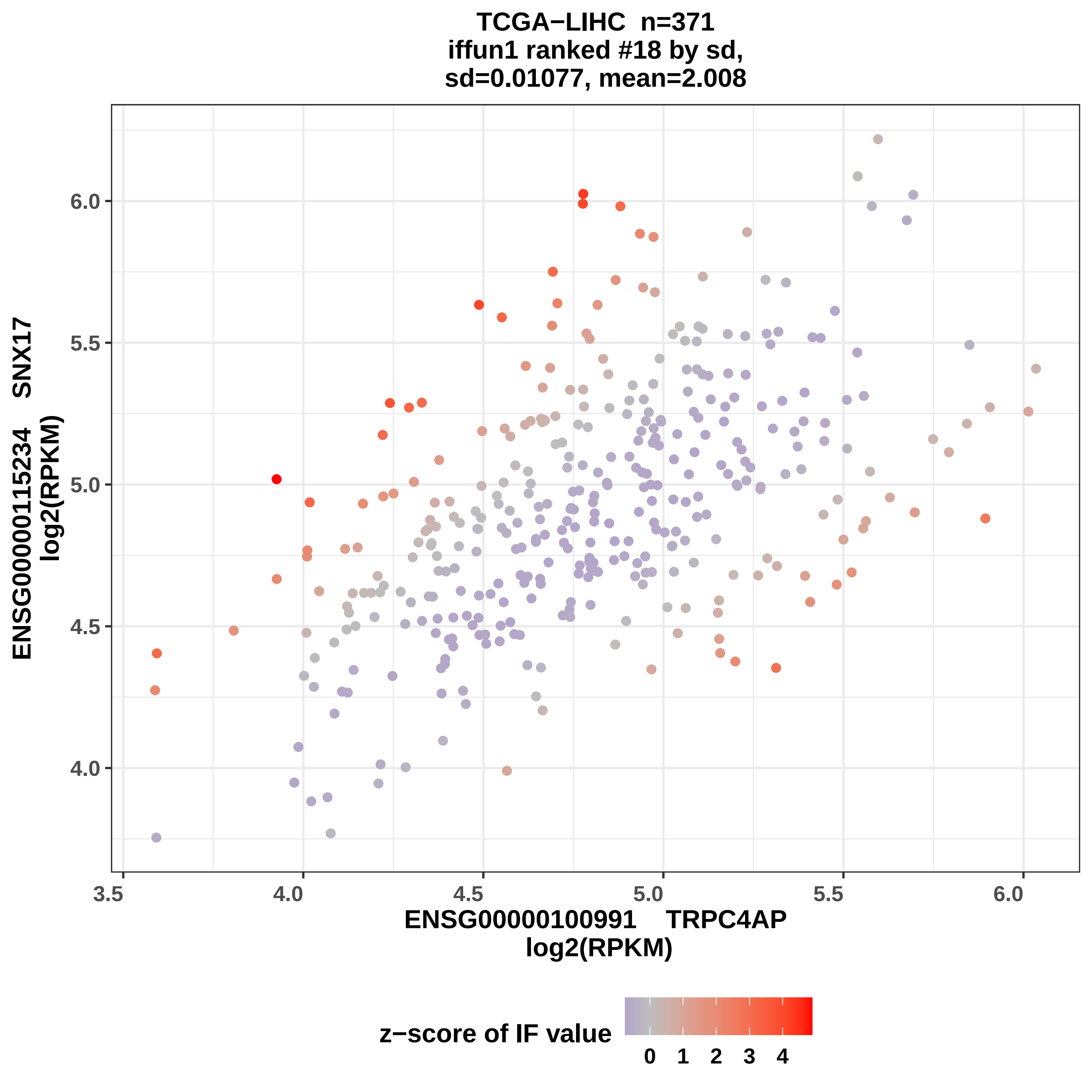
TCGA−LIHC n=371
iffun1 ranked #18 by sd,
sd=0.01077, mean=2.008
ENSG00000115234 SNX17
log2(RPKM)
6.0
5.5
5.0
4.5
4.0
3.5
4.0
4.5
5.0
5.5
6.0
ENSG00000100991 TRPC4AP
log2(RPKM)
z−score of IF value
0
1
2
3
4

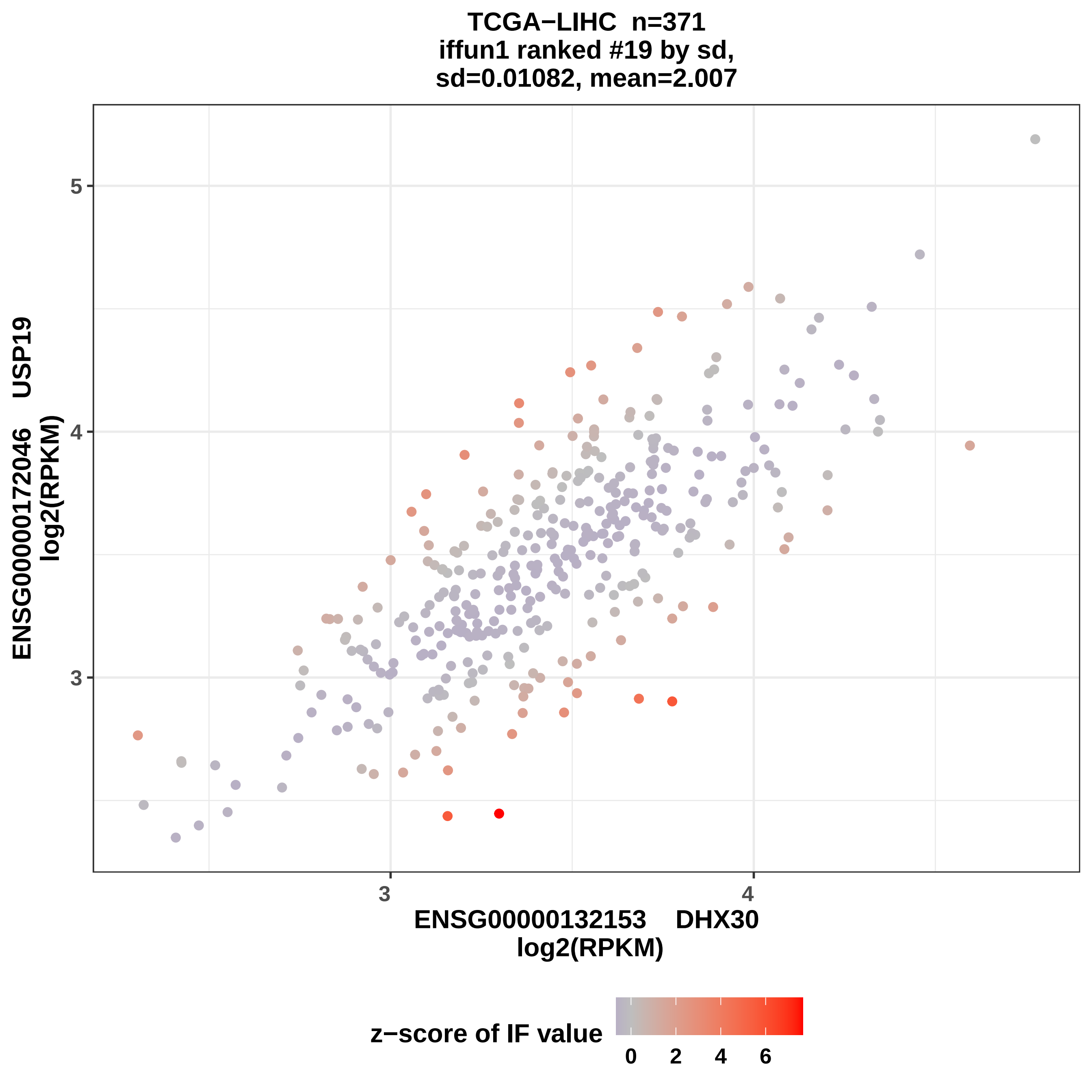

TCGA−LIHC n=371
iffun1 ranked #19 by sd,
sd=0.01082, mean=2.007
ENSG00000172046 USP19
log2(RPKM)
5
4
3
3
4
ENSG00000132153 DHX30
log2(RPKM)
z−score of IF value
0
2
4
6

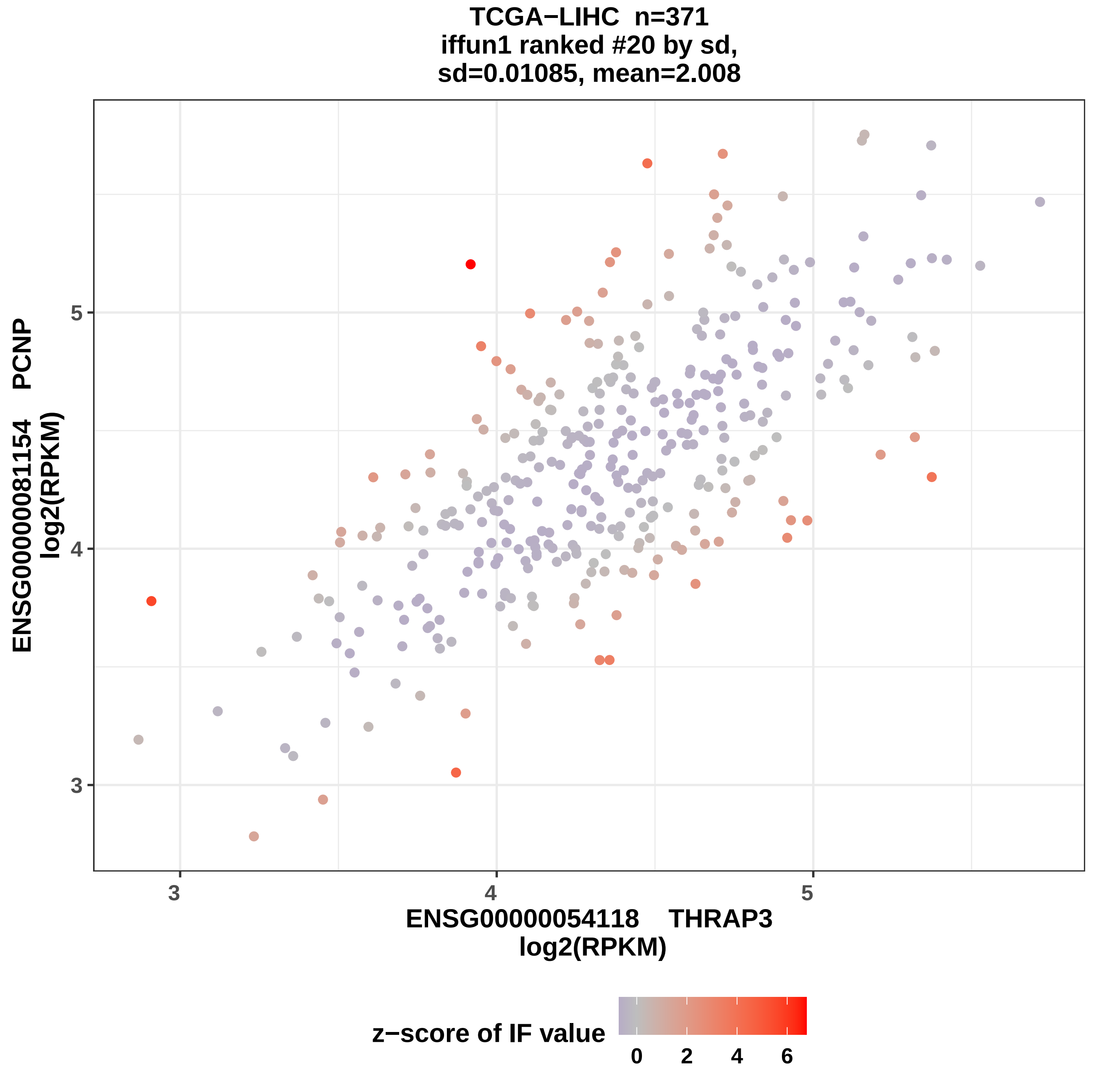
TCGA−LIHC n=371
iffun1 ranked #20 by sd,
sd=0.01085, mean=2.008
ENSG00000081154 PCNP
log2(RPKM)
ENSG00000054118 THRAP3
log2(RPKM)
z−score of IF value
3
4
5
0
2
4
6

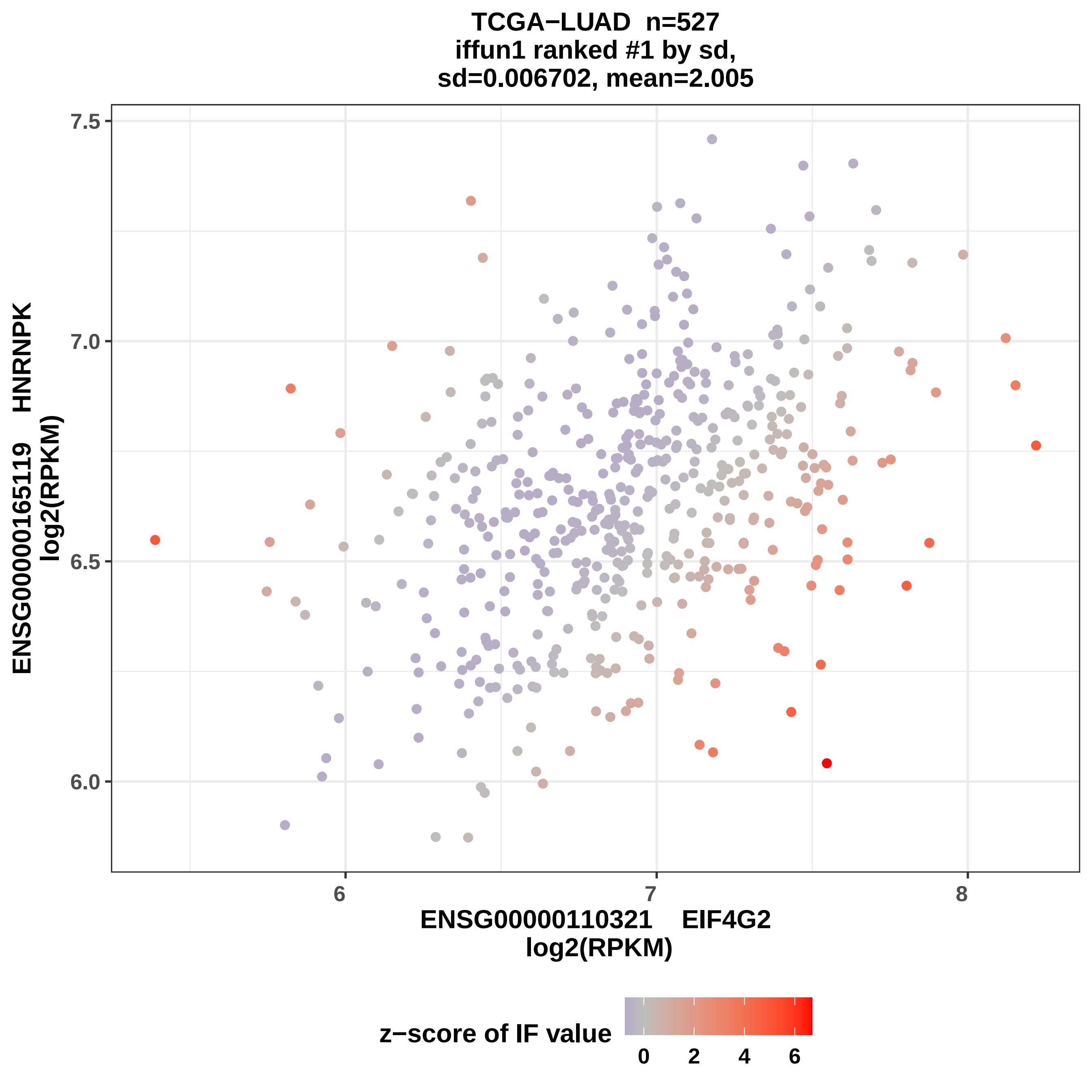

TCGA−LUAD n=527
iffun1 ranked #1 by sd,
sd=0.006702, mean=2.005
ENSG00000165119 HNRNPK
log2(RPKM)
7.5
7.0
6.5
6.0
6
7
8
ENSG00000110321 EIF4G2
log2(RPKM)
z−score of IF value
0
2
4
6

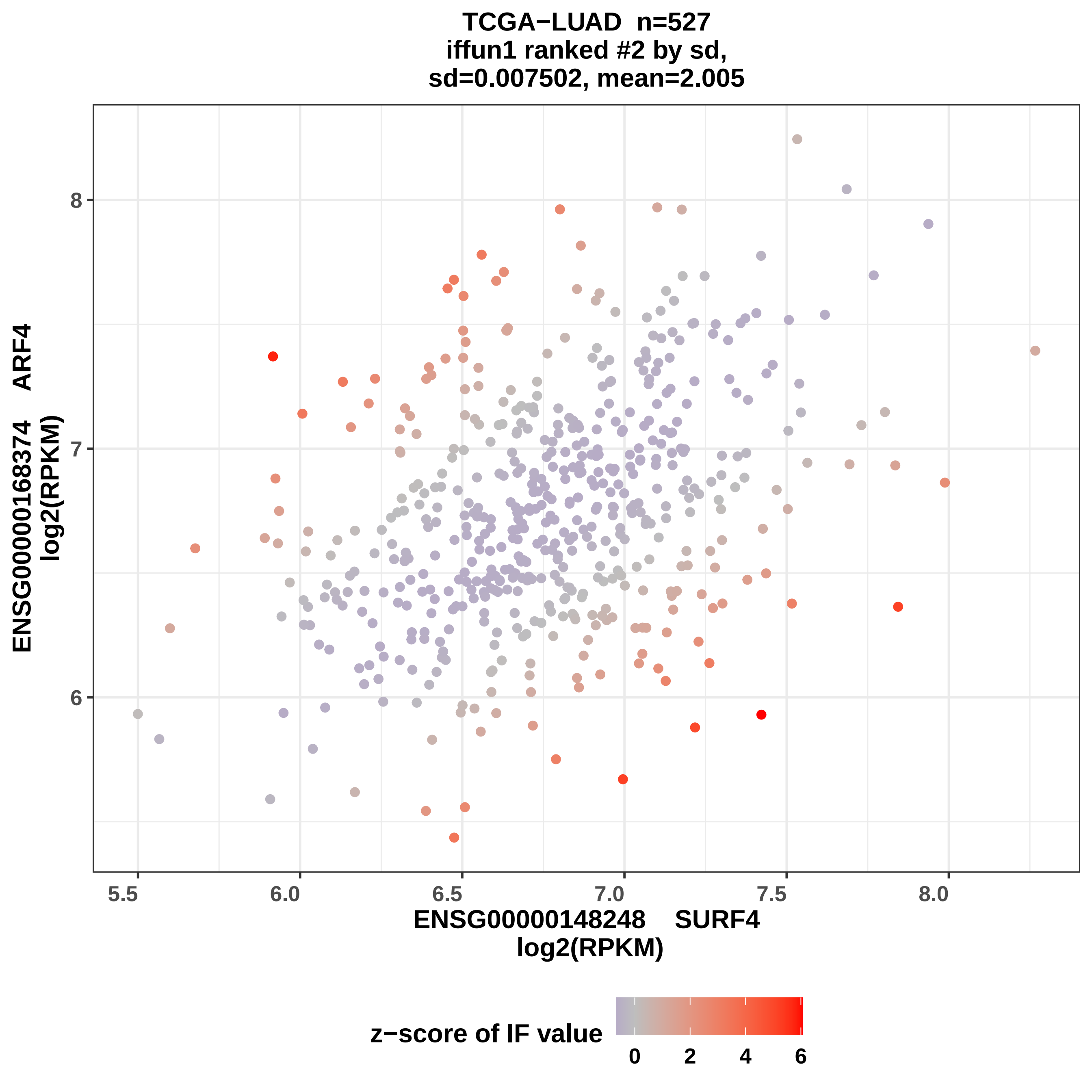

TCGA−LUAD n=527
iffun1 ranked #2 by sd,
sd=0.007502, mean=2.005
ENSG00000168374 ARF4
log2(RPKM)
8
7
6
5.5
6.0
6.5
7.0
7.5
8.0
ENSG00000148248 SURF4
log2(RPKM)
z−score of IF value
0
2
4
6

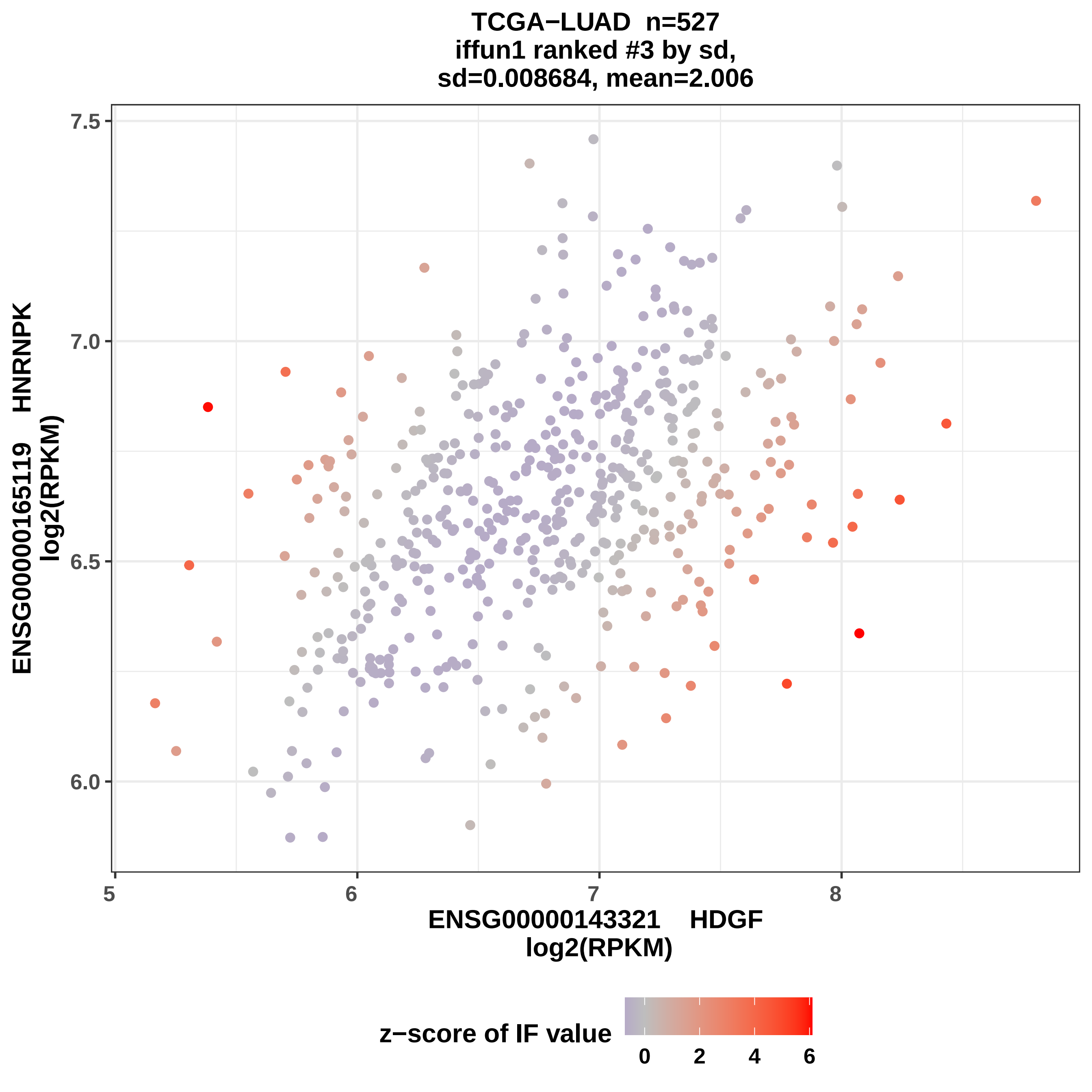

TCGA−LUAD n=527
iffun1 ranked #3 by sd,
sd=0.008684, mean=2.006
ENSG00000165119 HNRNPK
log2(RPKM)
7.5
7.0
6.5
6.0
5
6
7
8
ENSG00000143321 HDGF
log2(RPKM)
z−score of IF value
0
2
4
6

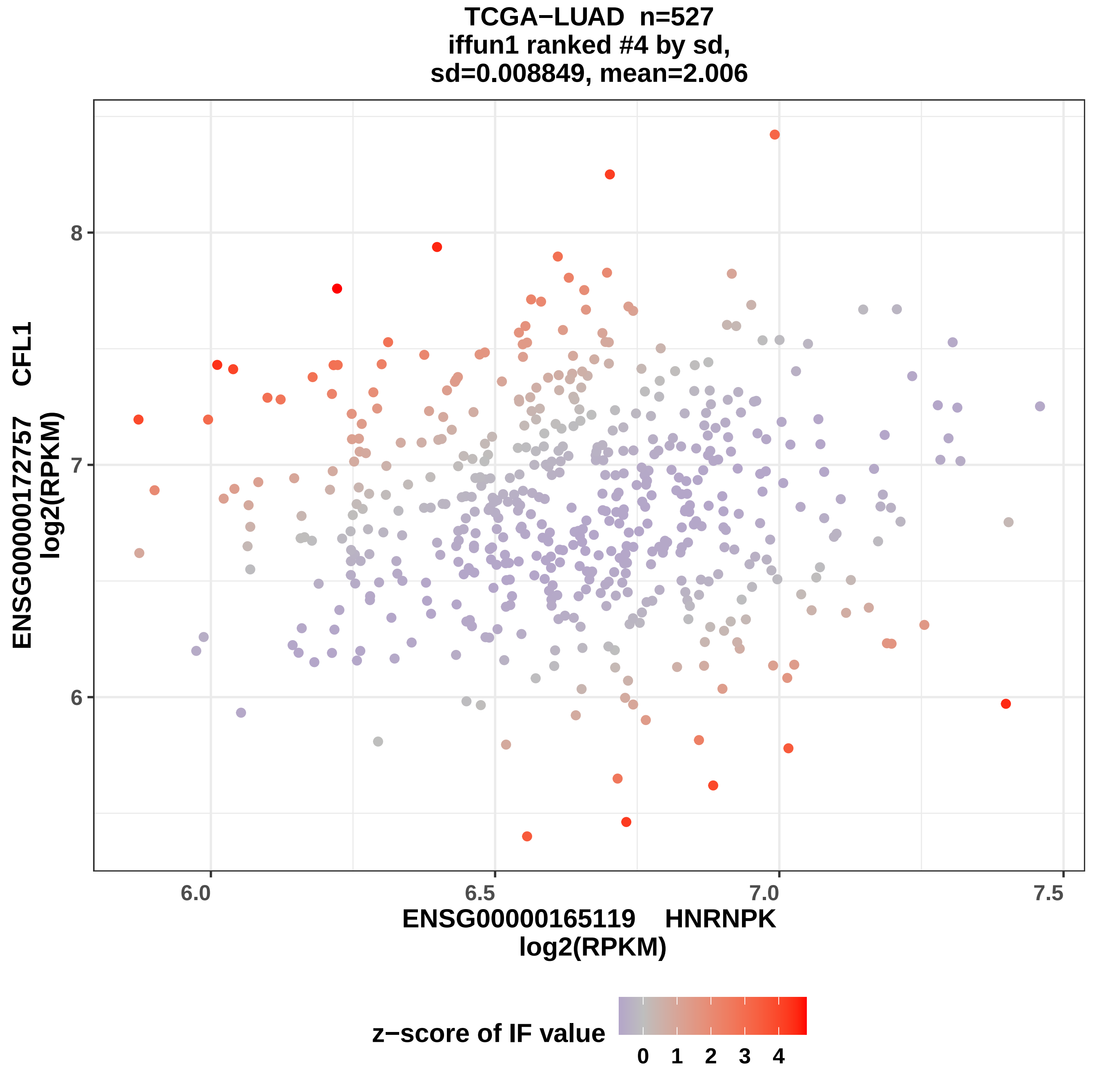
TCGA−LUAD n=527
iffun1 ranked #4 by sd,
sd=0.008849, mean=2.006
ENSG00000172757 CFL1
log2(RPKM)
8
7
6
6.0
6.5
7.0
7.5
ENSG00000165119 HNRNPK
log2(RPKM)
z−score of IF value
0
1
2
3
4

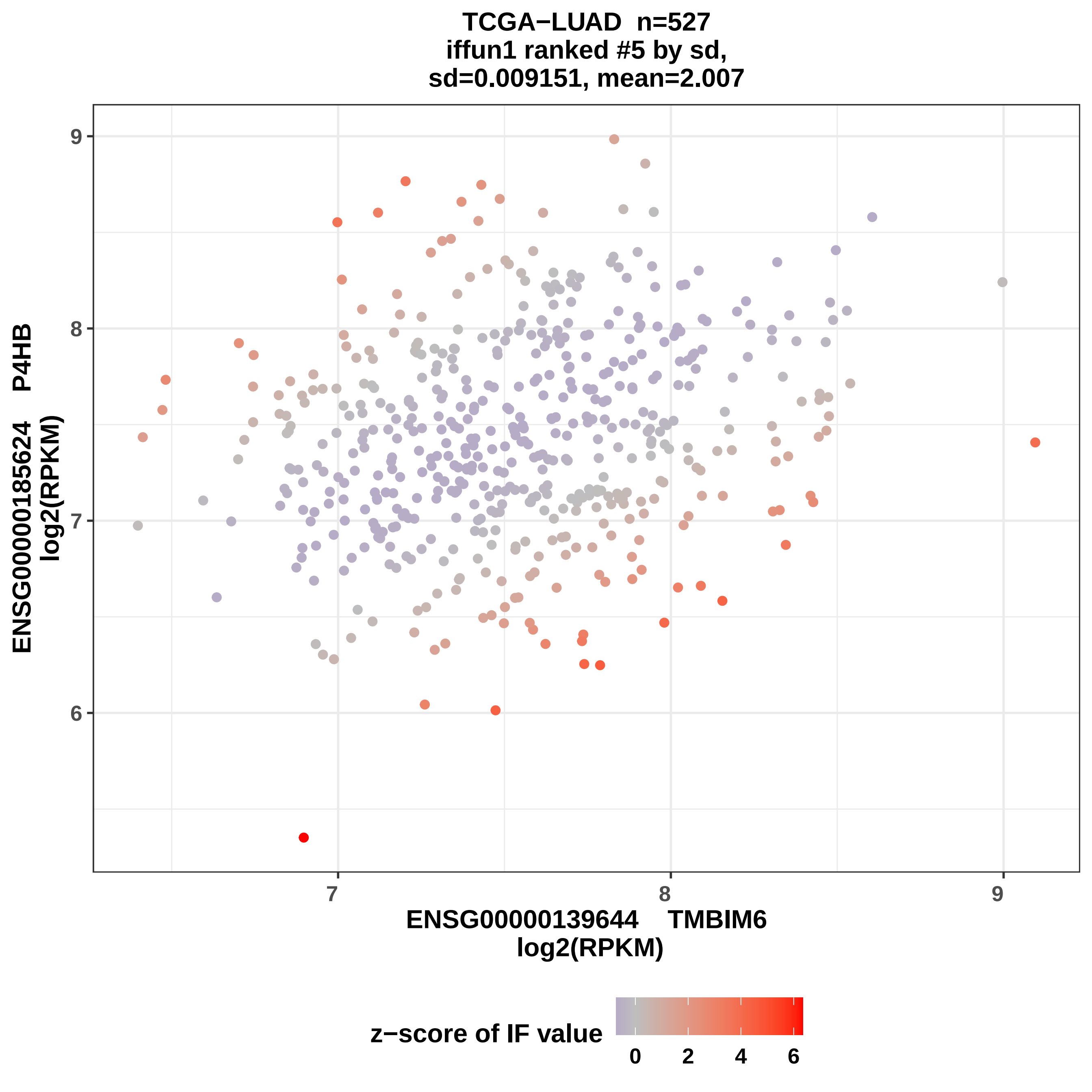
TCGA−LUAD n=527
iffun1 ranked #5 by sd,
sd=0.009151, mean=2.007
ENSG00000185624 P4HB
log2(RPKM)
ENSG00000139644 TMBIM6
log2(RPKM)
z−score of IF value
9
8
7
6
7
8
9
0
2
4
6

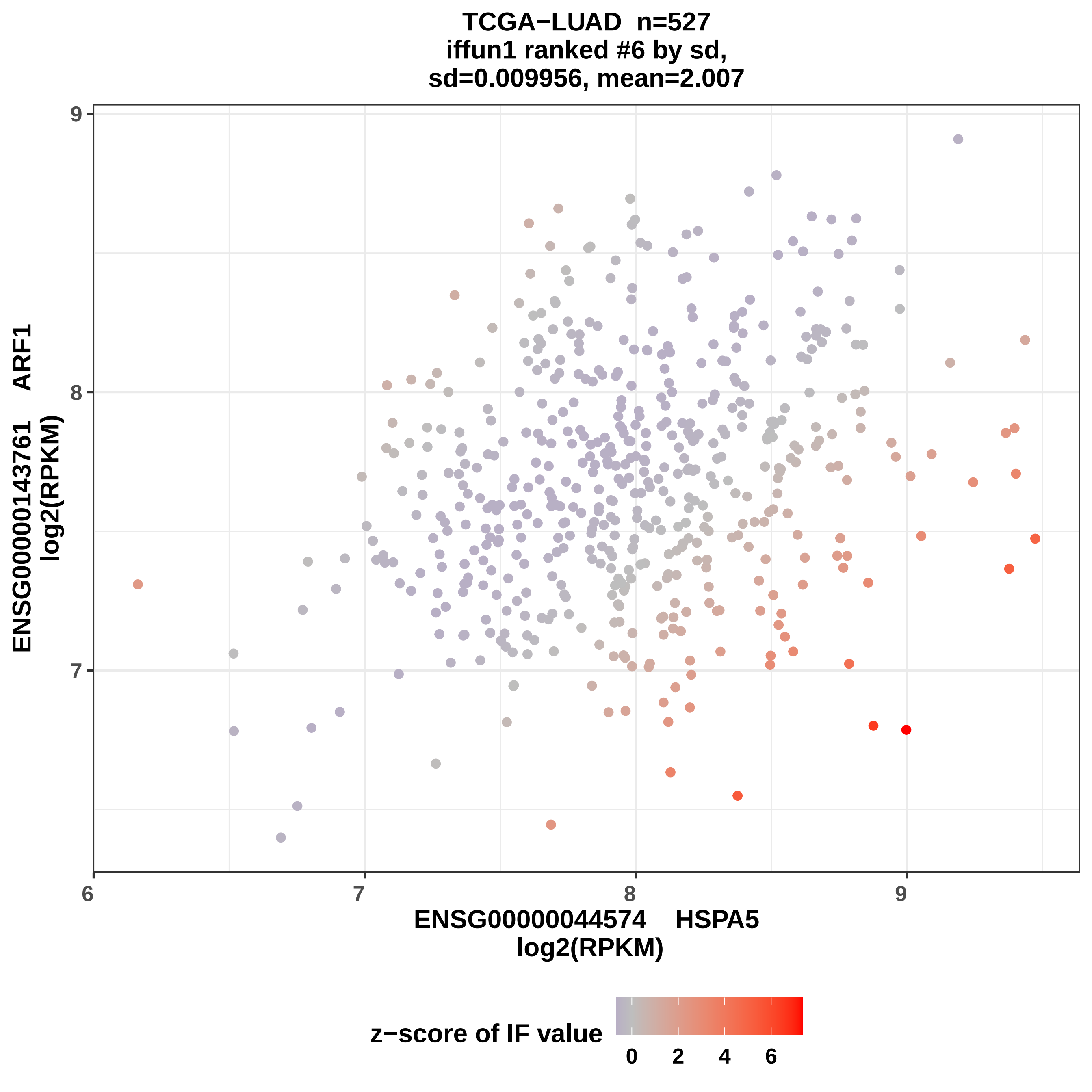

TCGA−LUAD n=527
iffun1 ranked #6 by sd,
sd=0.009956, mean=2.007
ENSG00000143761 ARF1
log2(RPKM)
ENSG00000044574 HSPA5
log2(RPKM)
9
8
7
6
7
8
9
z−score of IF value
0
2
4
6

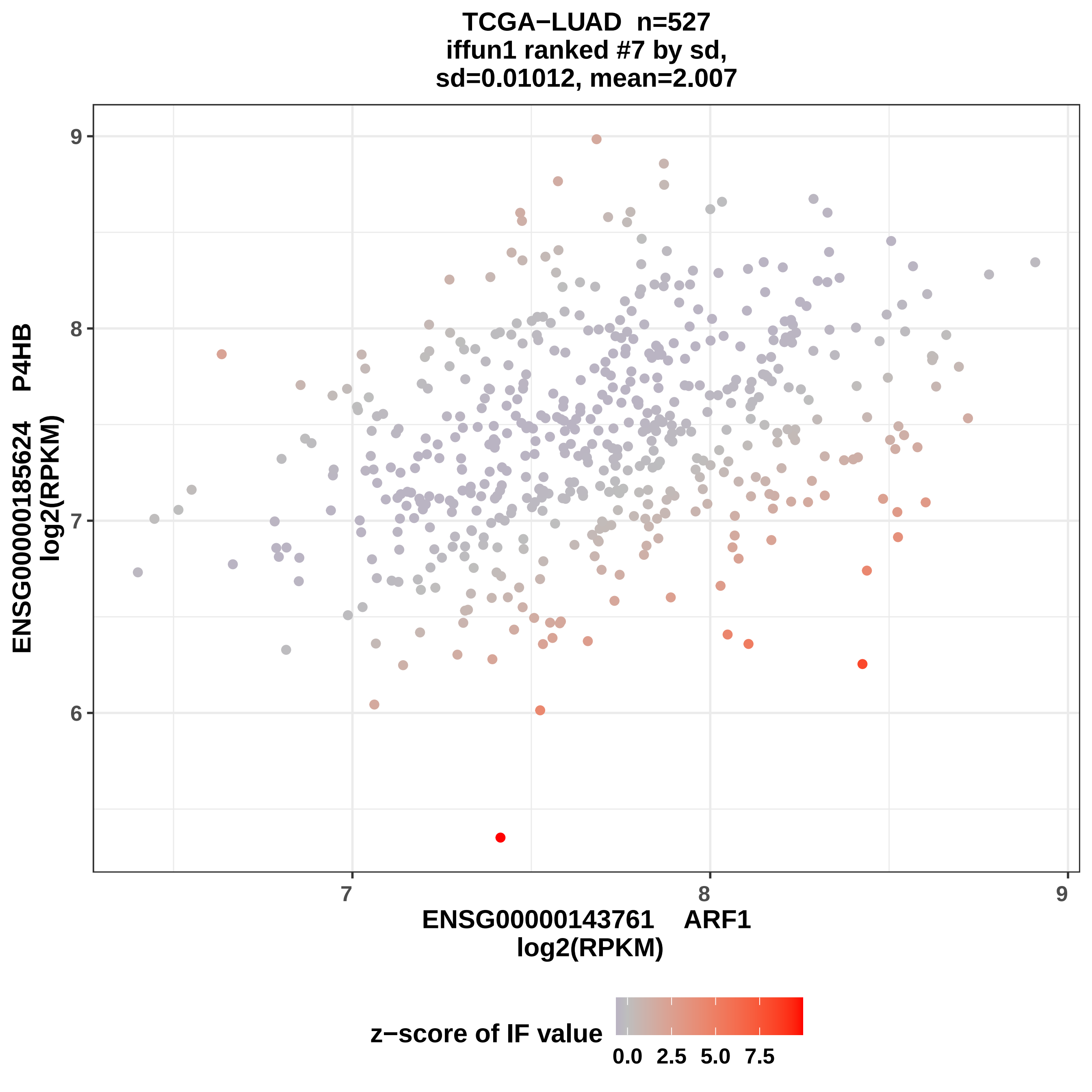
TCGA−LUAD n=527
iffun1 ranked #7 by sd,
sd=0.01012, mean=2.007
ENSG00000185624 P4HB
log2(RPKM)
9
8
7
6
7
8
9
ENSG00000143761 ARF1
log2(RPKM)
z−score of IF value
0.0
2.5
5.0
7.5

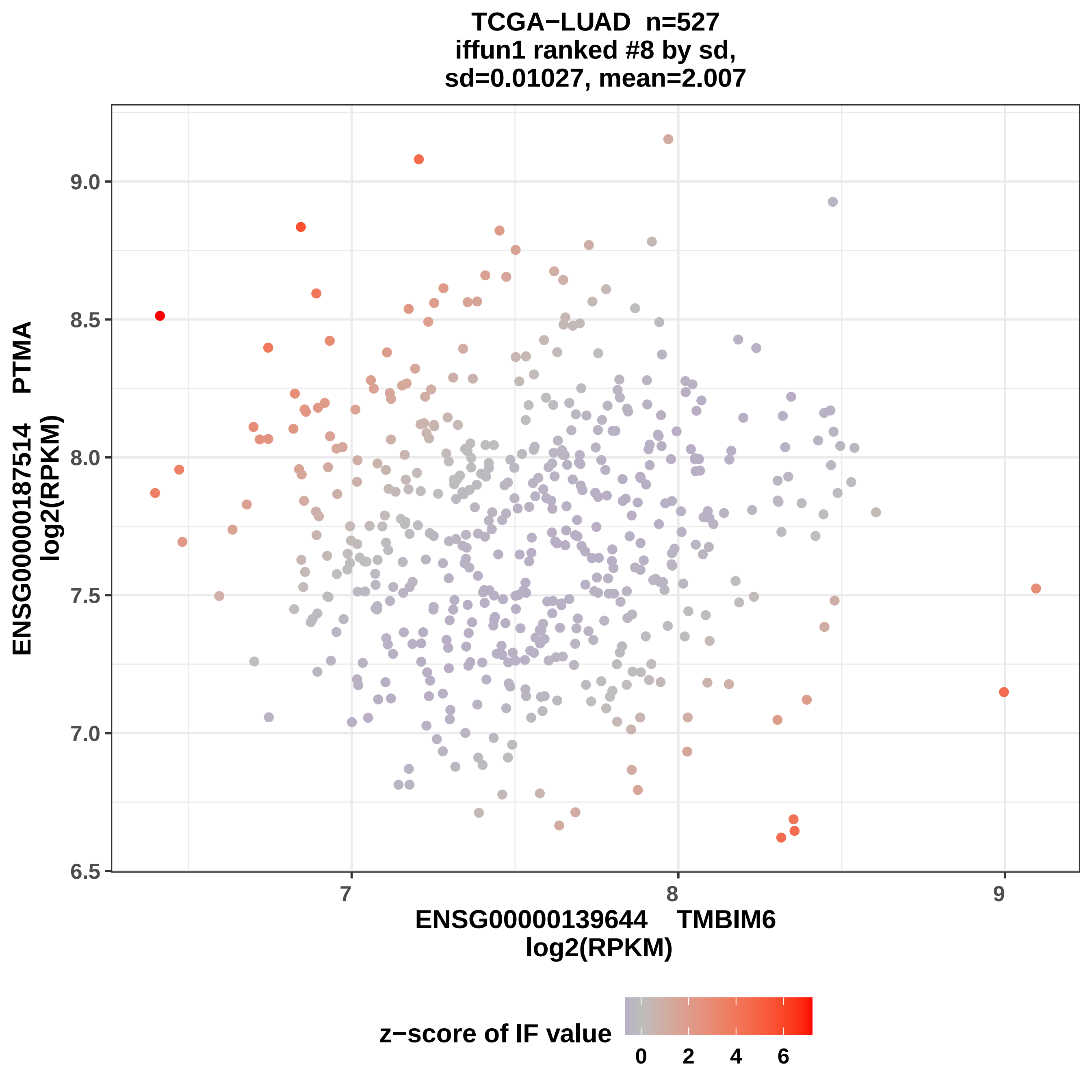

TCGA−LUAD n=527
iffun1 ranked #8 by sd,
sd=0.01027, mean=2.007
ENSG00000187514 PTMA
log2(RPKM)
ENSG00000139644 TMBIM6
log2(RPKM)
9.0
8.5
8.0
7.5
7.0
6.5
7
8
9
z−score of IF value
0
2
4
6

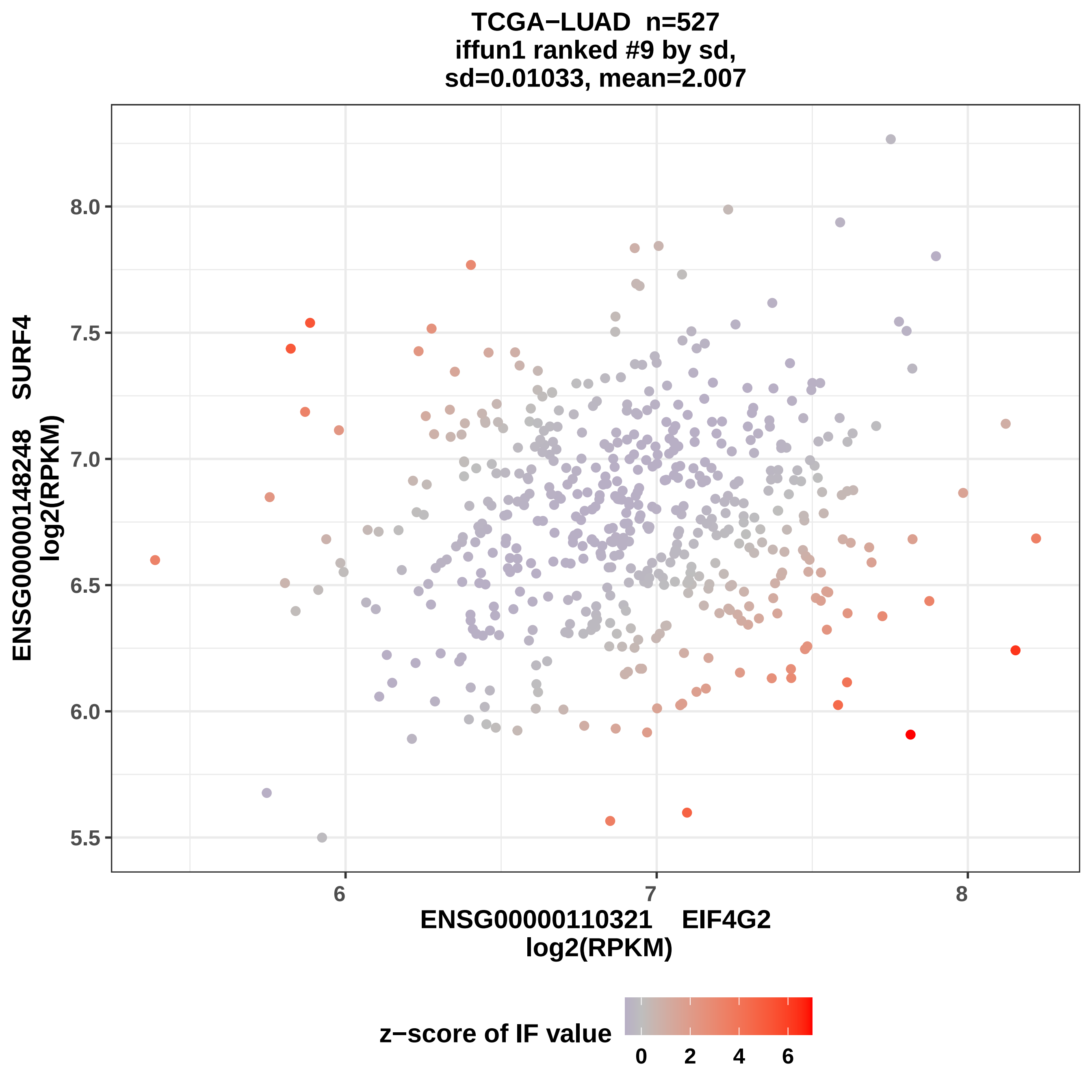
TCGA−LUAD n=527
iffun1 ranked #9 by sd,
sd=0.01033, mean=2.007
ENSG00000148248 SURF4
log2(RPKM)
8.0
7.5
7.0
6.5
6.0
5.5
6
7
8
ENSG00000110321 EIF4G2
log2(RPKM)
z−score of IF value
0
2
4
6

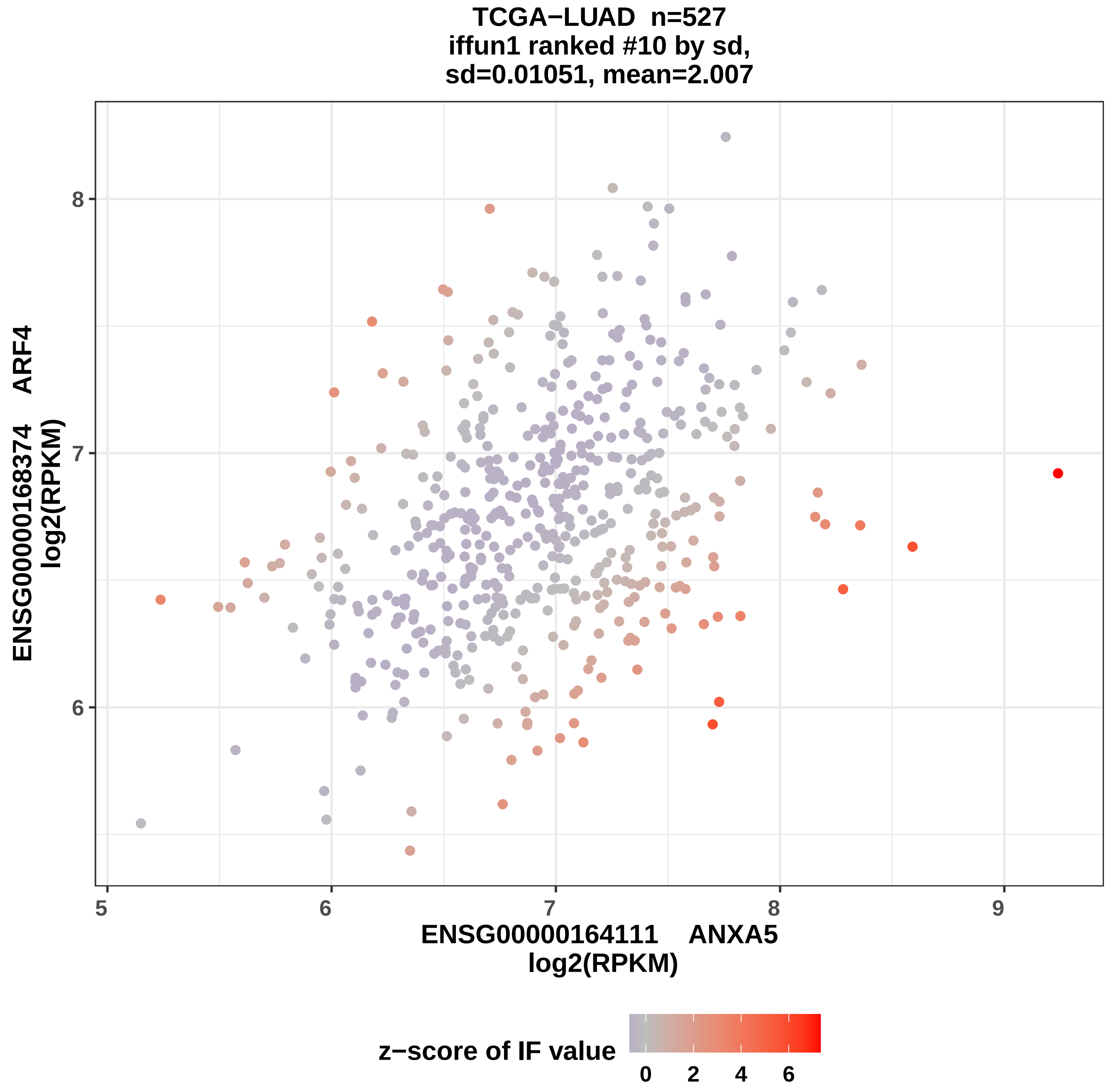

TCGA−LUAD n=527
iffun1 ranked #10 by sd,
sd=0.01051, mean=2.007
ENSG00000168374 ARF4
log2(RPKM)
8
7
6
5
6
7
8
9
ENSG00000164111 ANXA5
log2(RPKM)
z−score of IF value
0
2
4
6

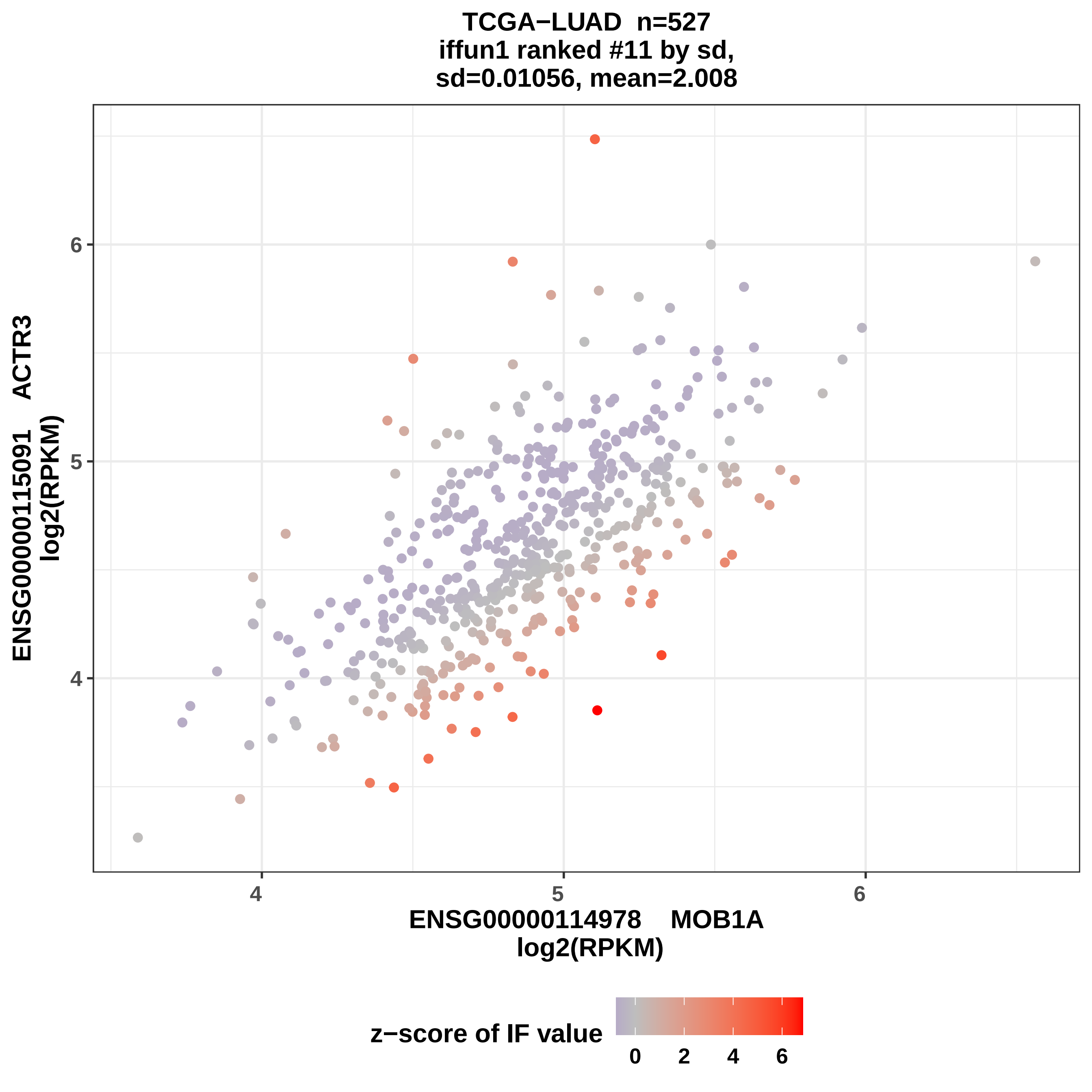

TCGA−LUAD n=527
iffun1 ranked #11 by sd,
sd=0.01056, mean=2.008
ENSG00000115091 ACTR3
log2(RPKM)
ENSG00000114978 MOB1A
log2(RPKM)
z−score of IF value
0
2
4
6

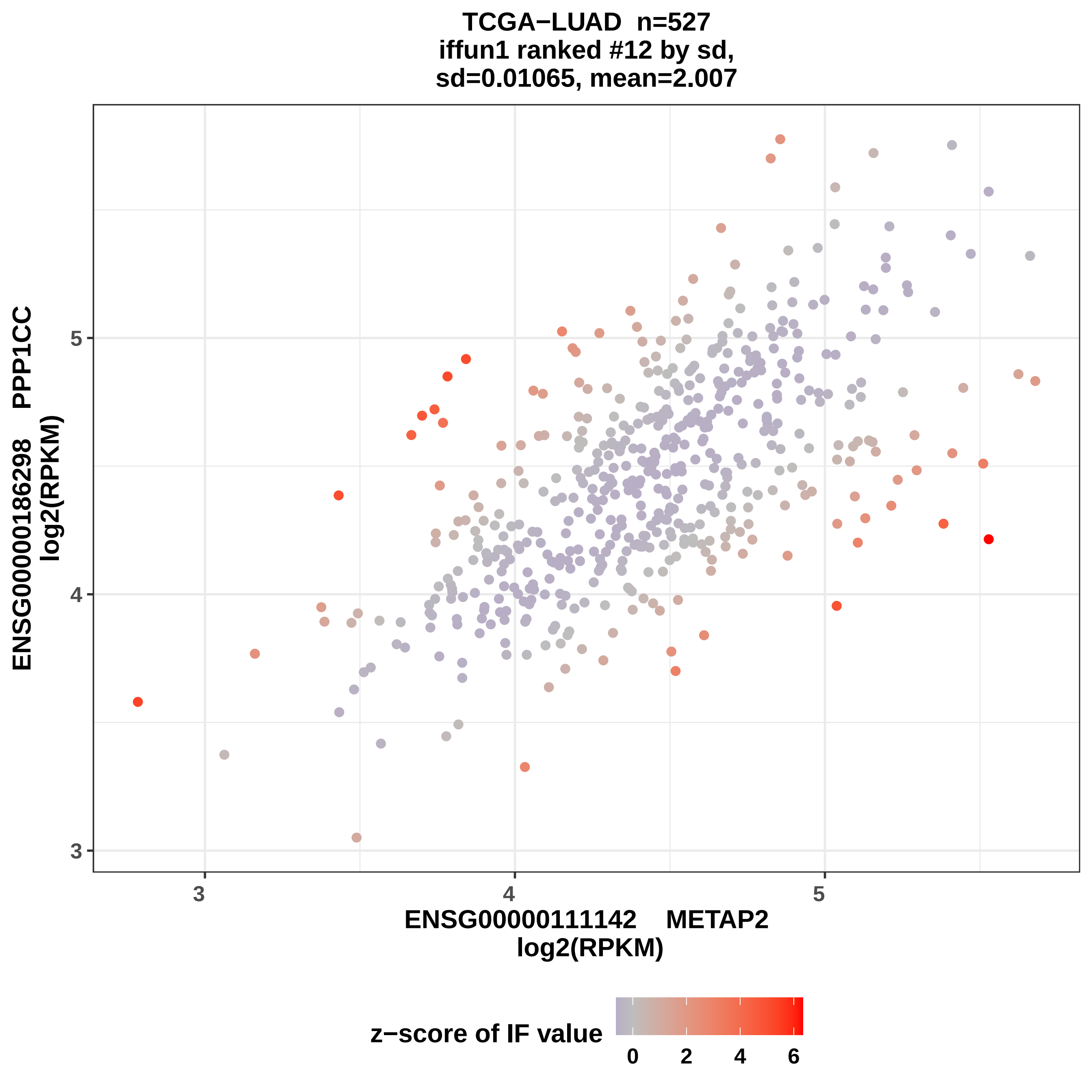
TCGA−LUAD n=527
iffun1 ranked #12 by sd,
sd=0.01065, mean=2.007
ENSG00000186298 PPP1CC
log2(RPKM)
5
4
3
ENSG00000111142 METAP2
log2(RPKM)
3
4
5
z−score of IF value
0
2
4
6

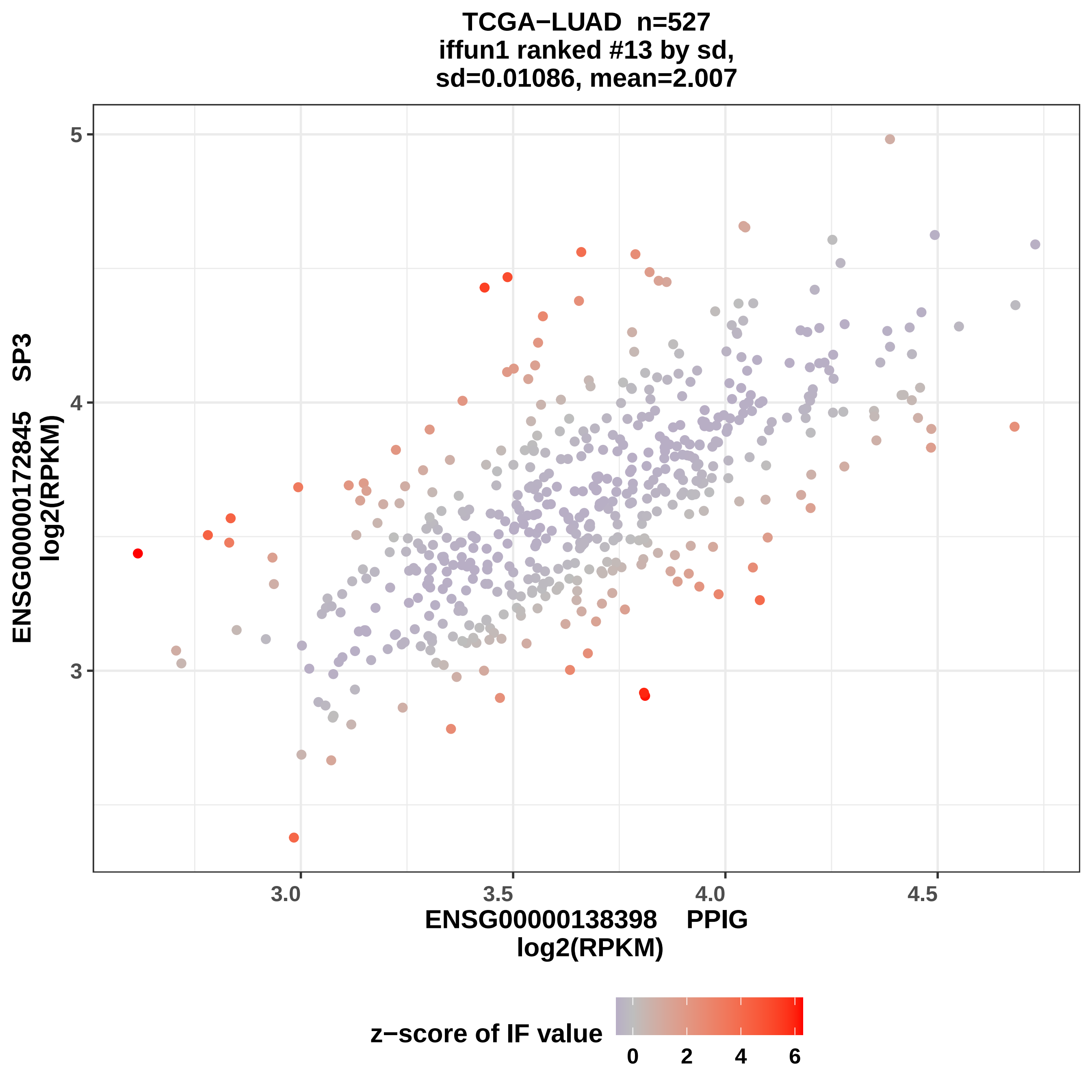

TCGA−LUAD n=527
iffun1 ranked #13 by sd,
sd=0.01086, mean=2.007
ENSG00000172845 SP3
log2(RPKM)
ENSG00000138398 PPIG
log2(RPKM)
z−score of IF value
3.0
3.5
4.0
4.5
3
4
5
0
2
4
6

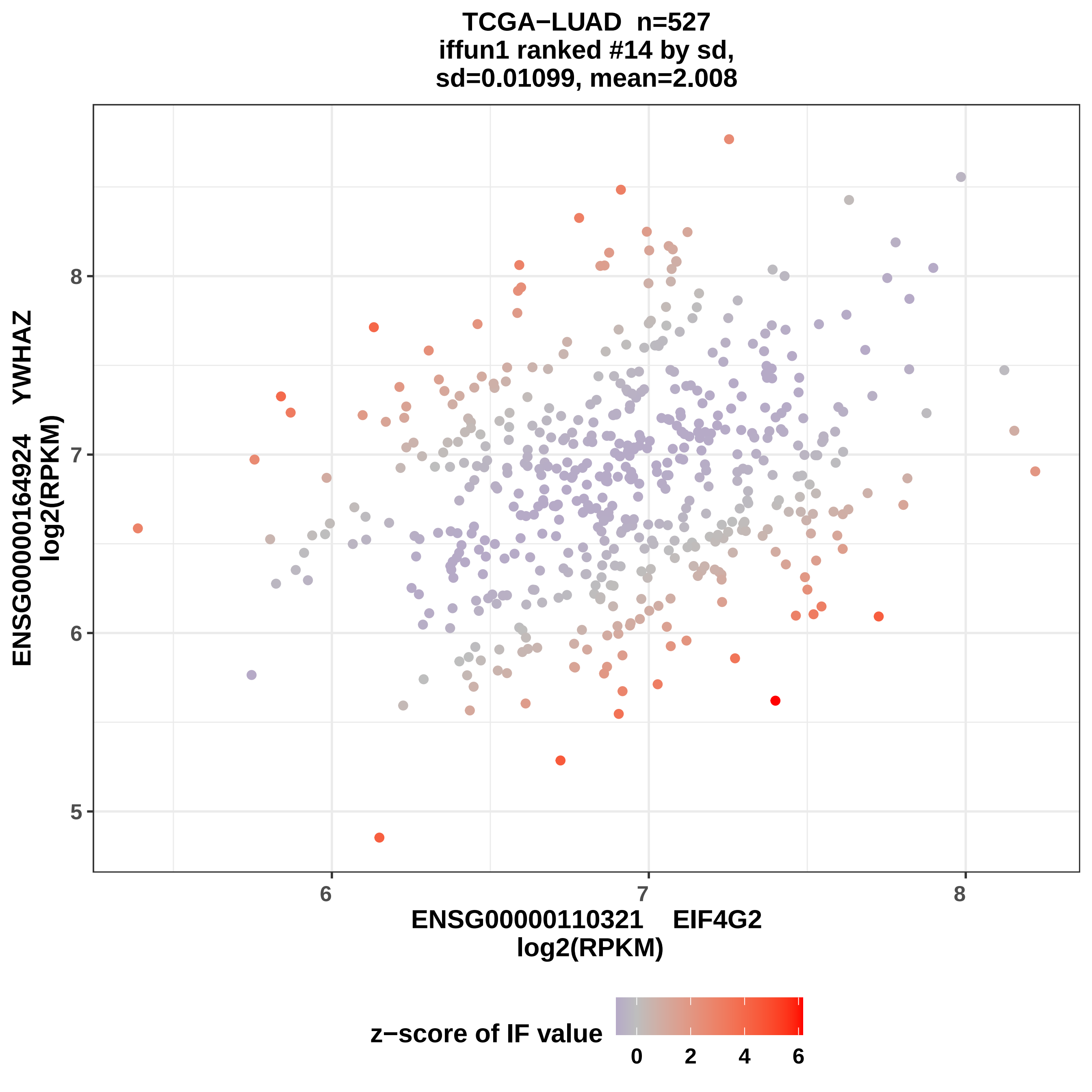
TCGA−LUAD n=527
iffun1 ranked #14 by sd,
sd=0.01099, mean=2.008
ENSG00000164924 YWHAZ
log2(RPKM)
5
6
7
8
ENSG00000110321 EIF4G2
log2(RPKM)
6
7
8
z−score of IF value
0
2
4
6

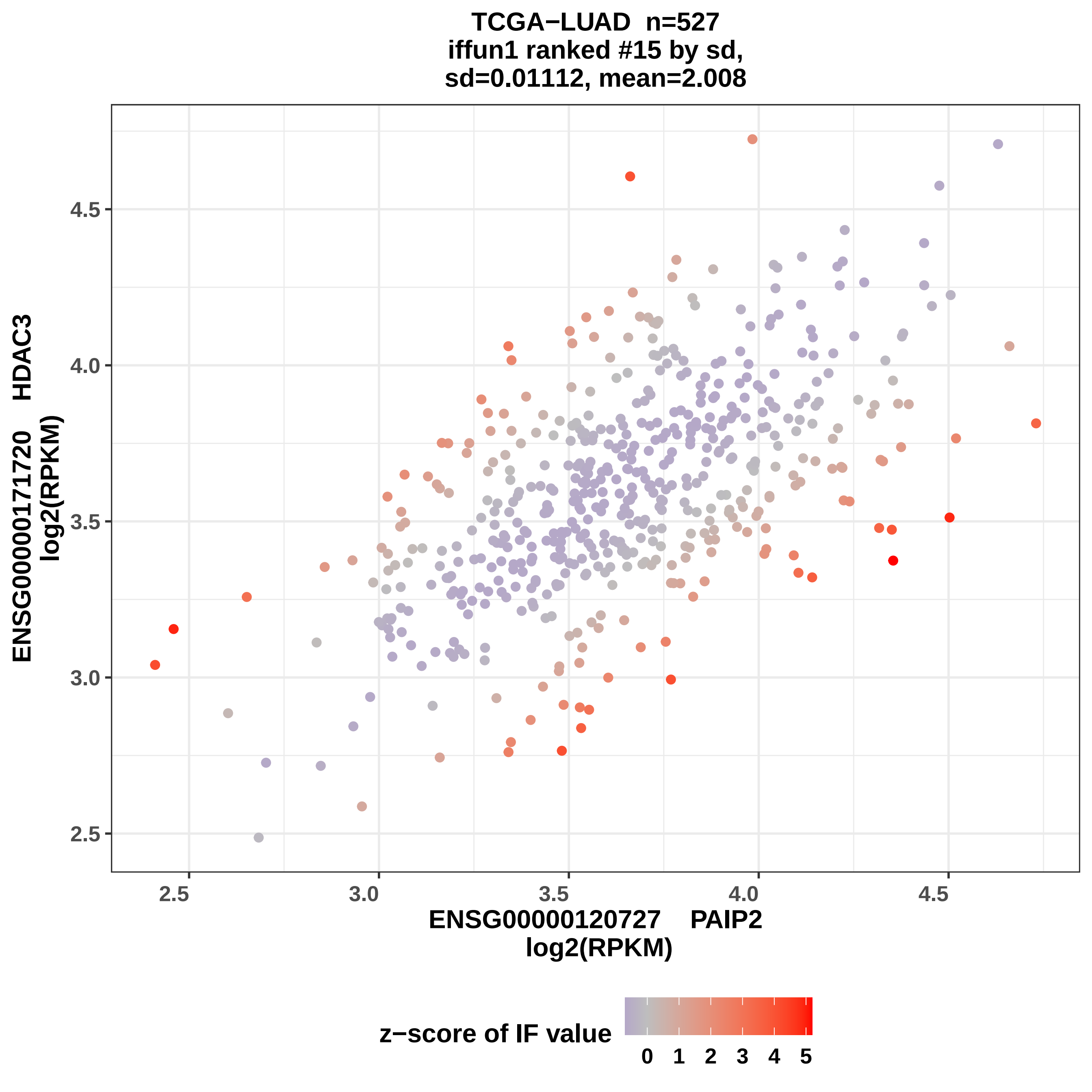

TCGA−LUAD n=527
iffun1 ranked #15 by sd,
sd=0.01112, mean=2.008
ENSG00000171720 HDAC3
log2(RPKM)
4.5
4.0
3.5
3.0
2.5
2.5
3.0
3.5
4.0
4.5
ENSG00000120727 PAIP2
log2(RPKM)
z−score of IF value
0
1
2
3
4
5

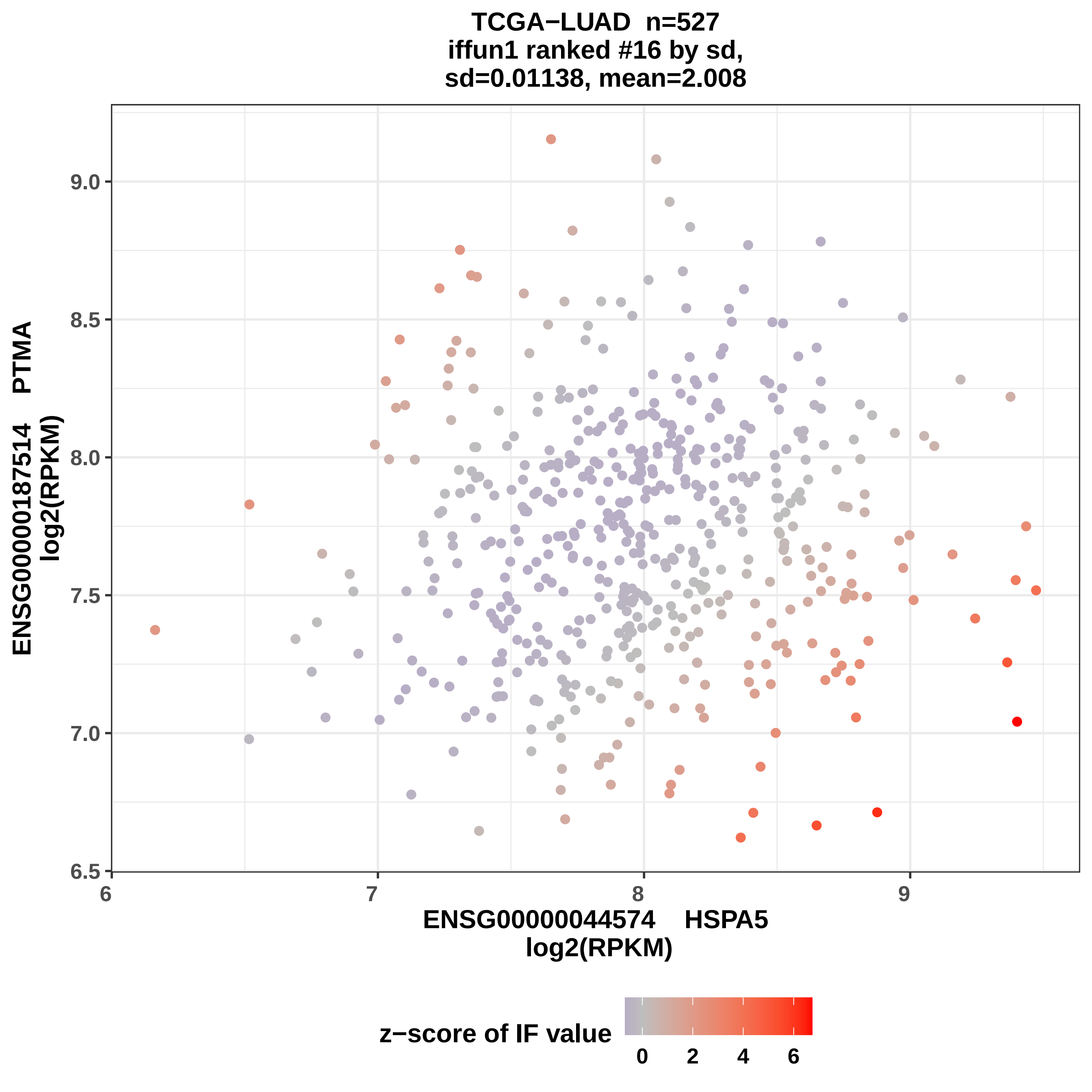
TCGA−LUAD n=527
iffun1 ranked #16 by sd,
sd=0.01138, mean=2.008
ENSG00000187514 PTMA
log2(RPKM)
ENSG00000044574 HSPA5
log2(RPKM)
6.5
7.0
7.5
8.0
8.5
9.0
6
7
8
9
z−score of IF value
0
2
4
6

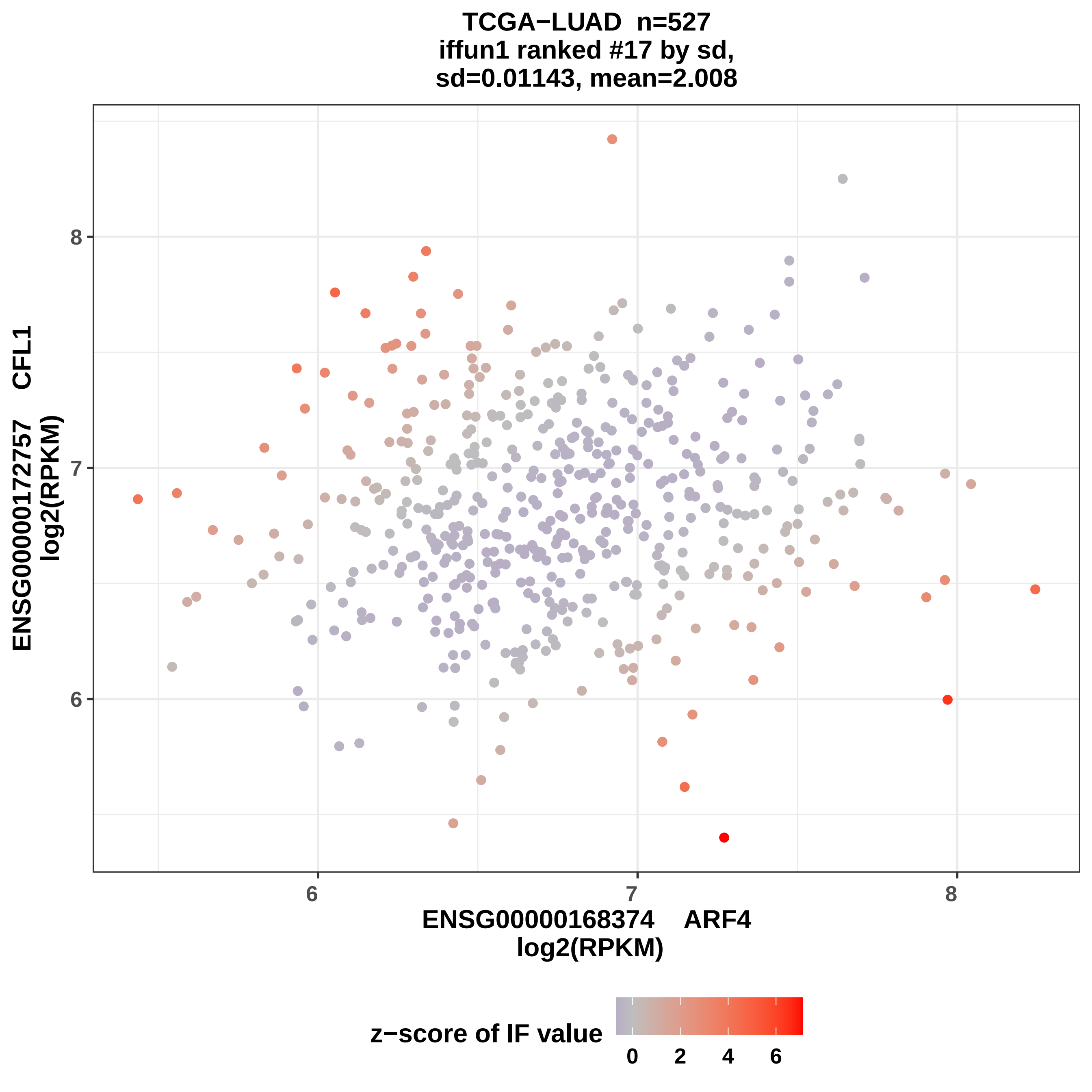

TCGA−LUAD n=527
iffun1 ranked #17 by sd,
sd=0.01143, mean=2.008
ENSG00000172757 CFL1
log2(RPKM)
8
7
6
6
7
8
ENSG00000168374 ARF4
log2(RPKM)
z−score of IF value
0
2
4
6

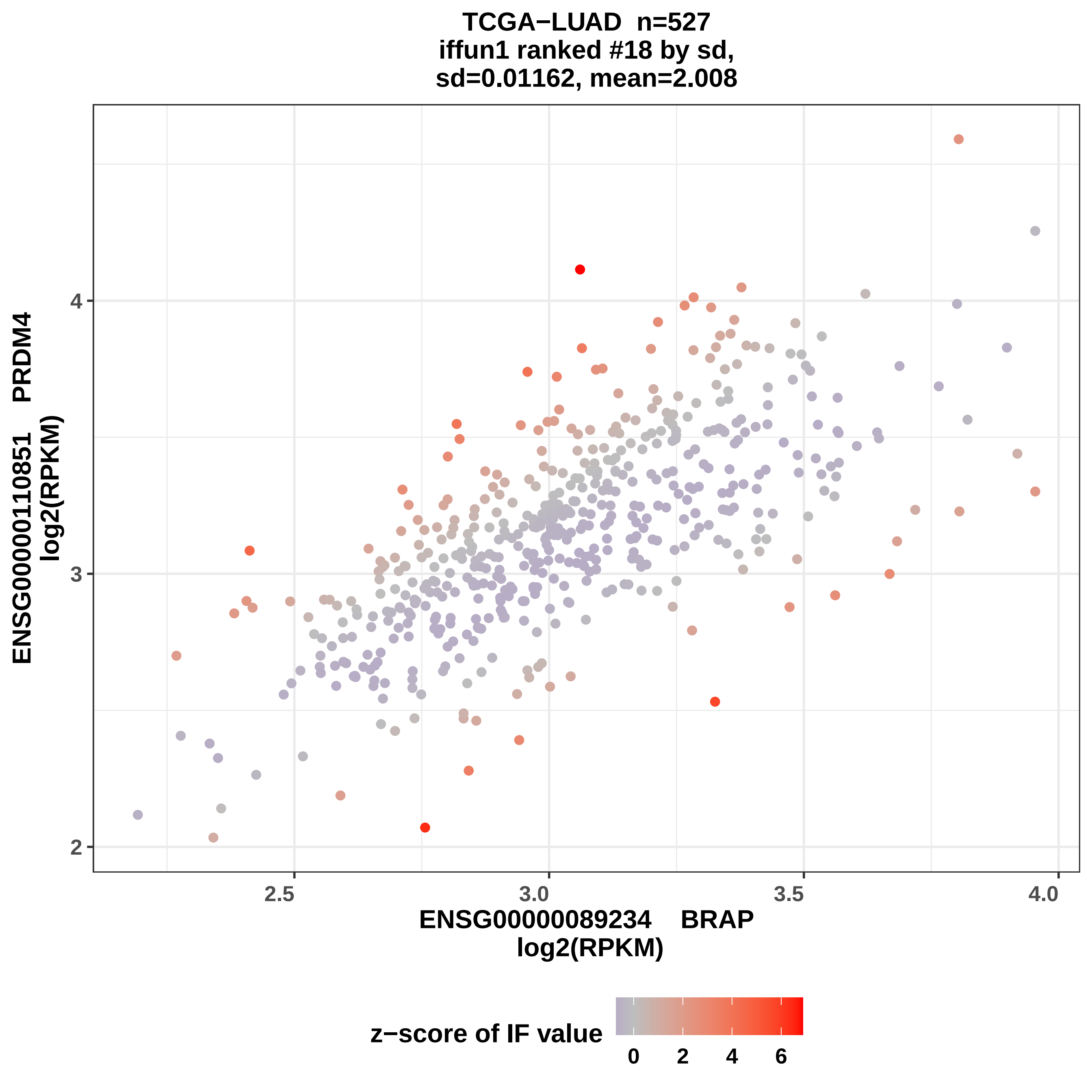

TCGA−LUAD n=527
iffun1 ranked #18 by sd,
sd=0.01162, mean=2.008
ENSG00000110851 PRDM4
log2(RPKM)
2
3
4
2.5
3.0
3.5
4.0
ENSG00000089234 BRAP
log2(RPKM)
z−score of IF value
0
2
4
6

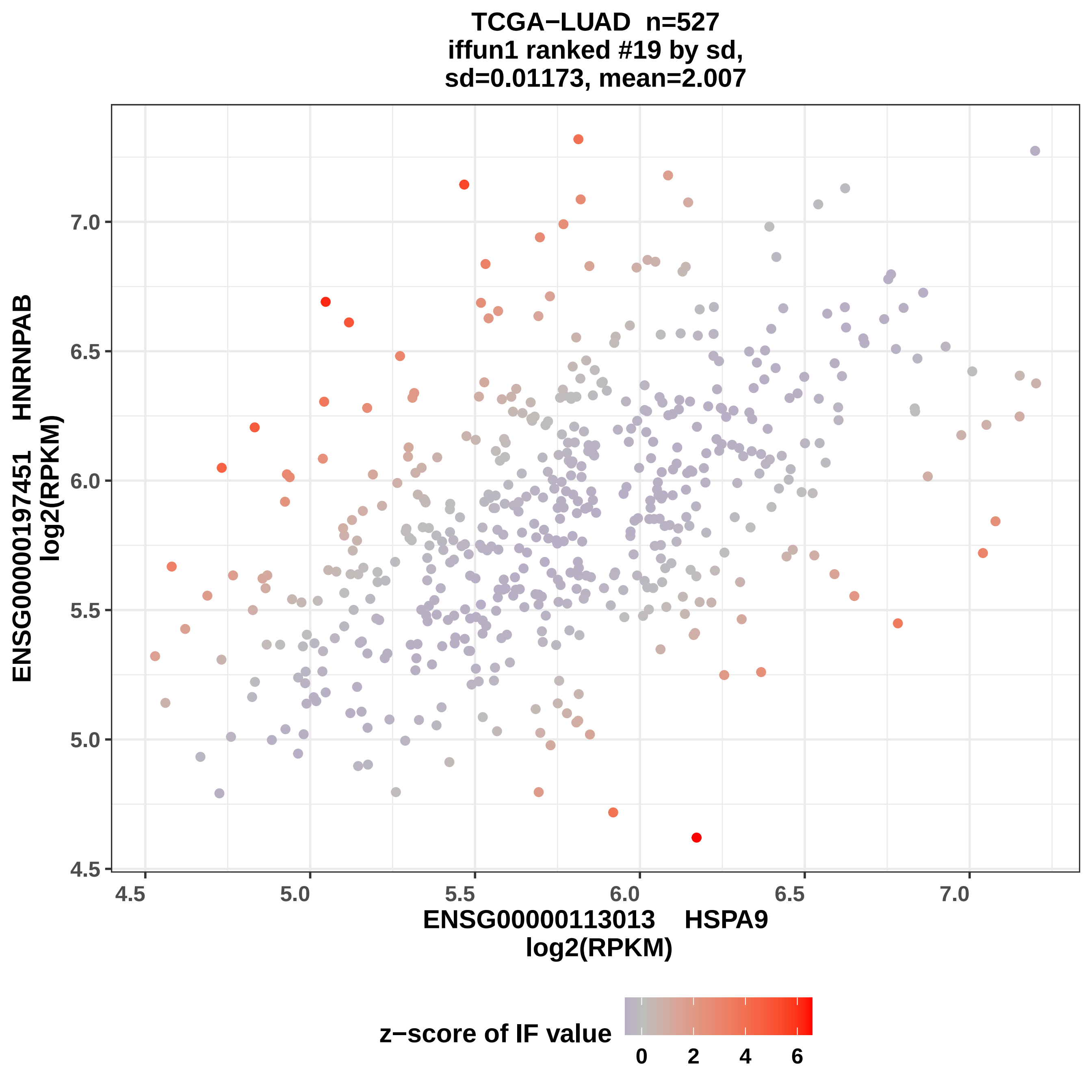
TCGA−LUAD n=527
iffun1 ranked #19 by sd,
sd=0.01173, mean=2.007
ENSG00000197451 HNRNPAB
log2(RPKM)
ENSG00000113013 HSPA9
log2(RPKM)
4.5
5.0
5.5
6.0
6.5
7.0
z−score of IF value
0
2
4
6

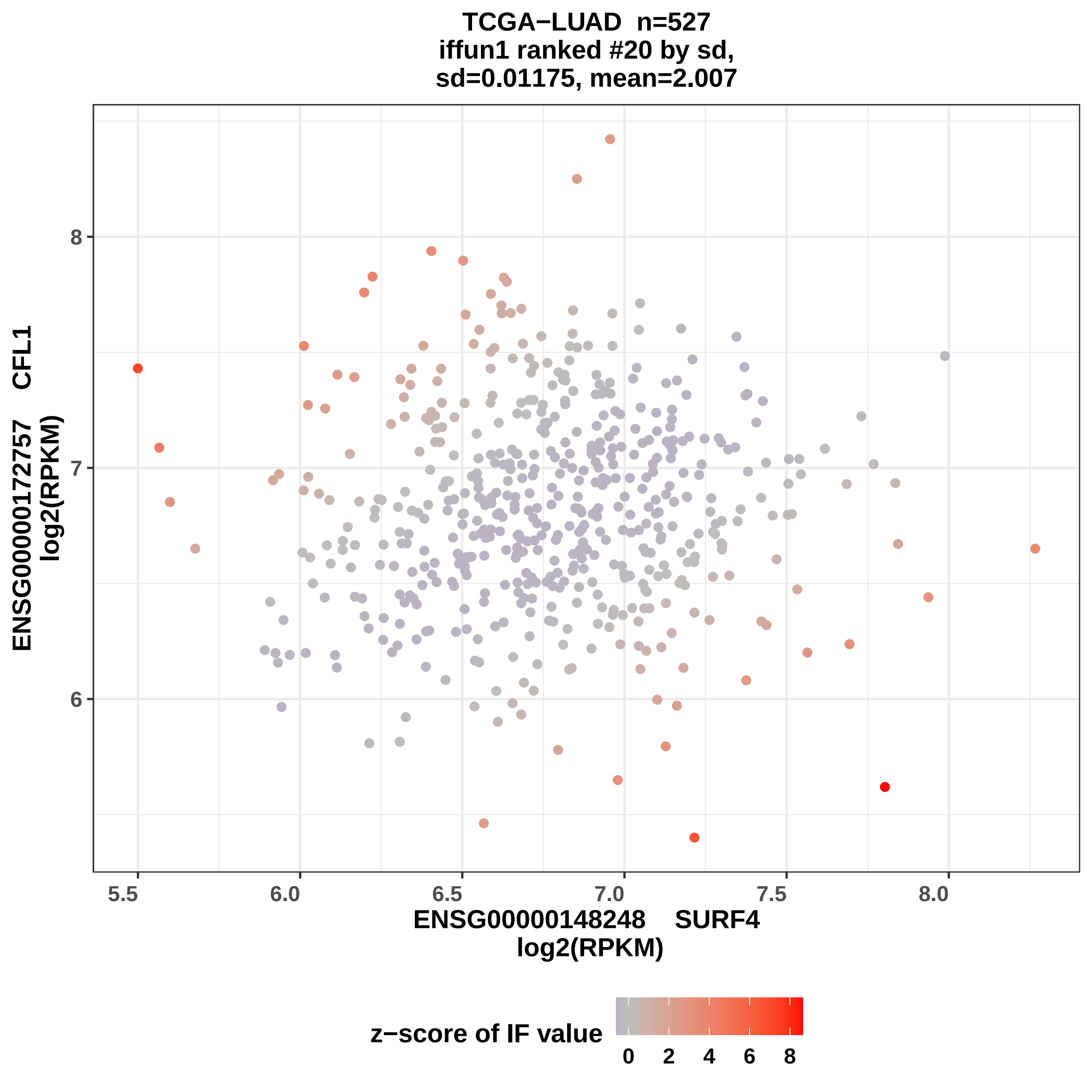

TCGA−LUAD n=527
iffun1 ranked #20 by sd,
sd=0.01175, mean=2.007
ENSG00000172757 CFL1
log2(RPKM)
8
7
6
5.5
6.0
6.5
7.0
7.5
8.0
ENSG00000148248 SURF4
log2(RPKM)
z−score of IF value
0
2
4
6
8

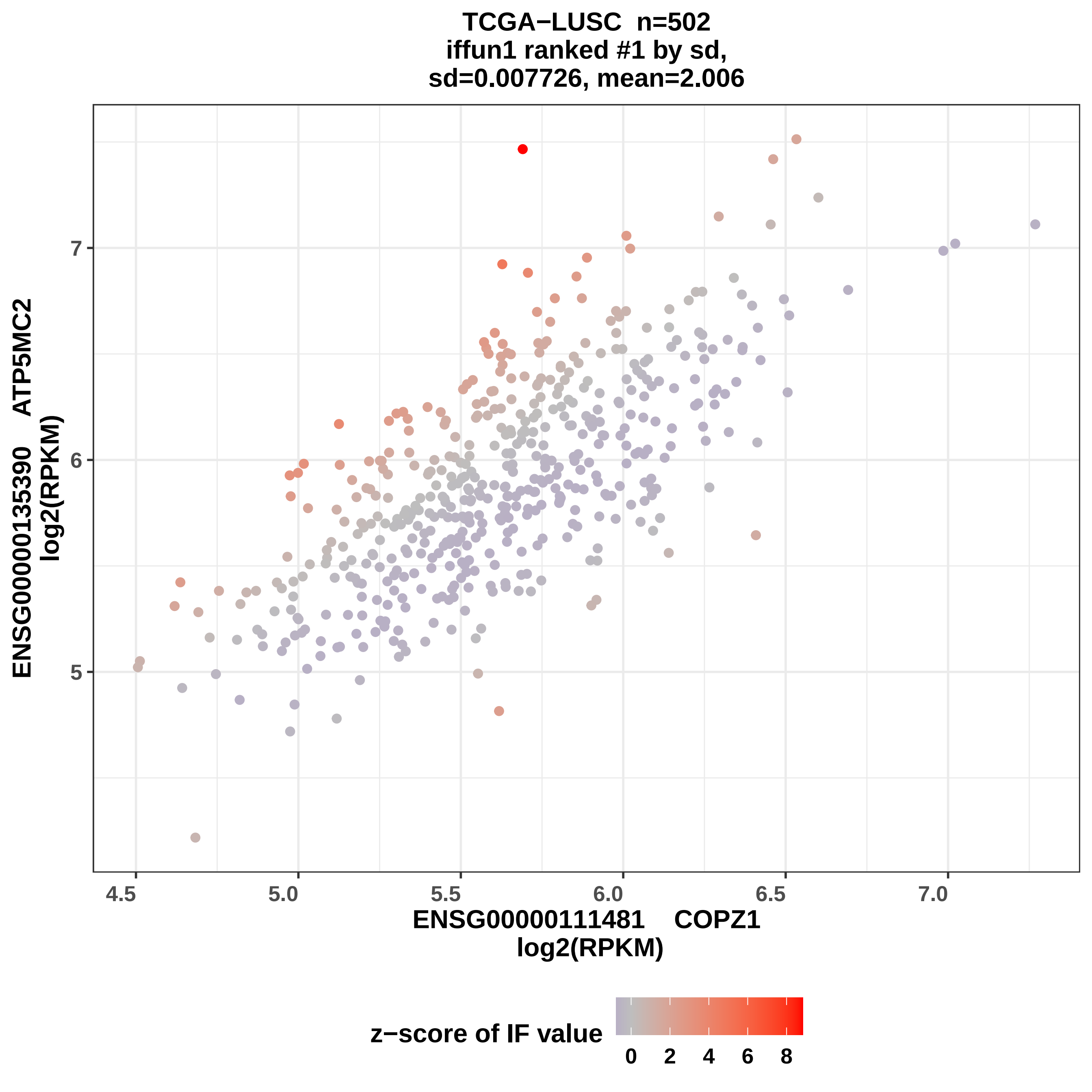

TCGA−LUSC n=502
iffun1 ranked #1 by sd,
sd=0.007726, mean=2.006
ENSG00000135390 ATP5MC2
log2(RPKM)
5
6
7
4.5
5.0
5.5
6.0
6.5
7.0
ENSG00000111481 COPZ1
log2(RPKM)
z−score of IF value
0
2
4
6
8

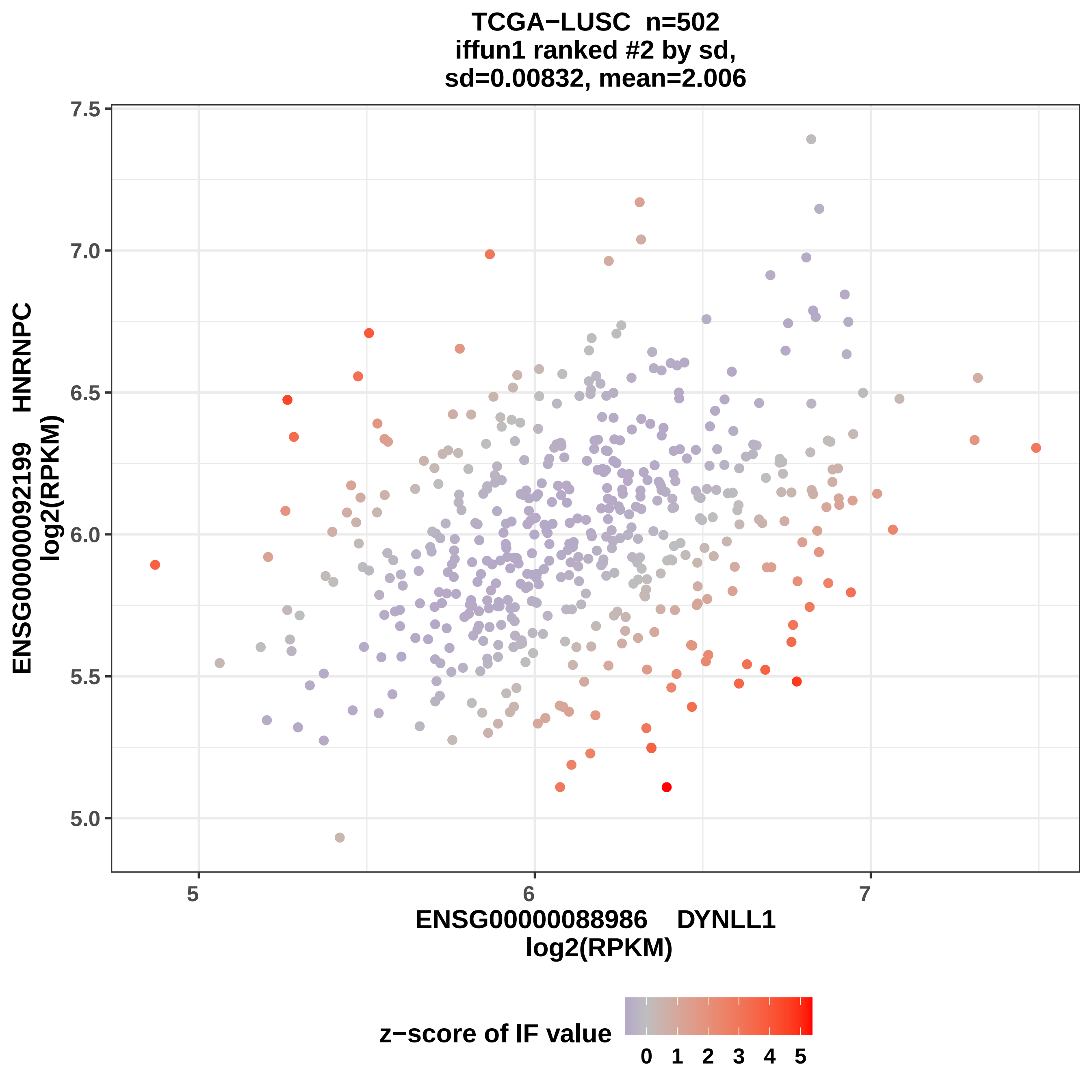

TCGA−LUSC n=502
iffun1 ranked #2 by sd,
sd=0.00832, mean=2.006
ENSG00000092199 HNRNPC
log2(RPKM)
7.5
7.0
6.5
6.0
5.5
5.0
5
6
7
ENSG00000088986 DYNLL1
log2(RPKM)
z−score of IF value
0 1 2 3 4 5

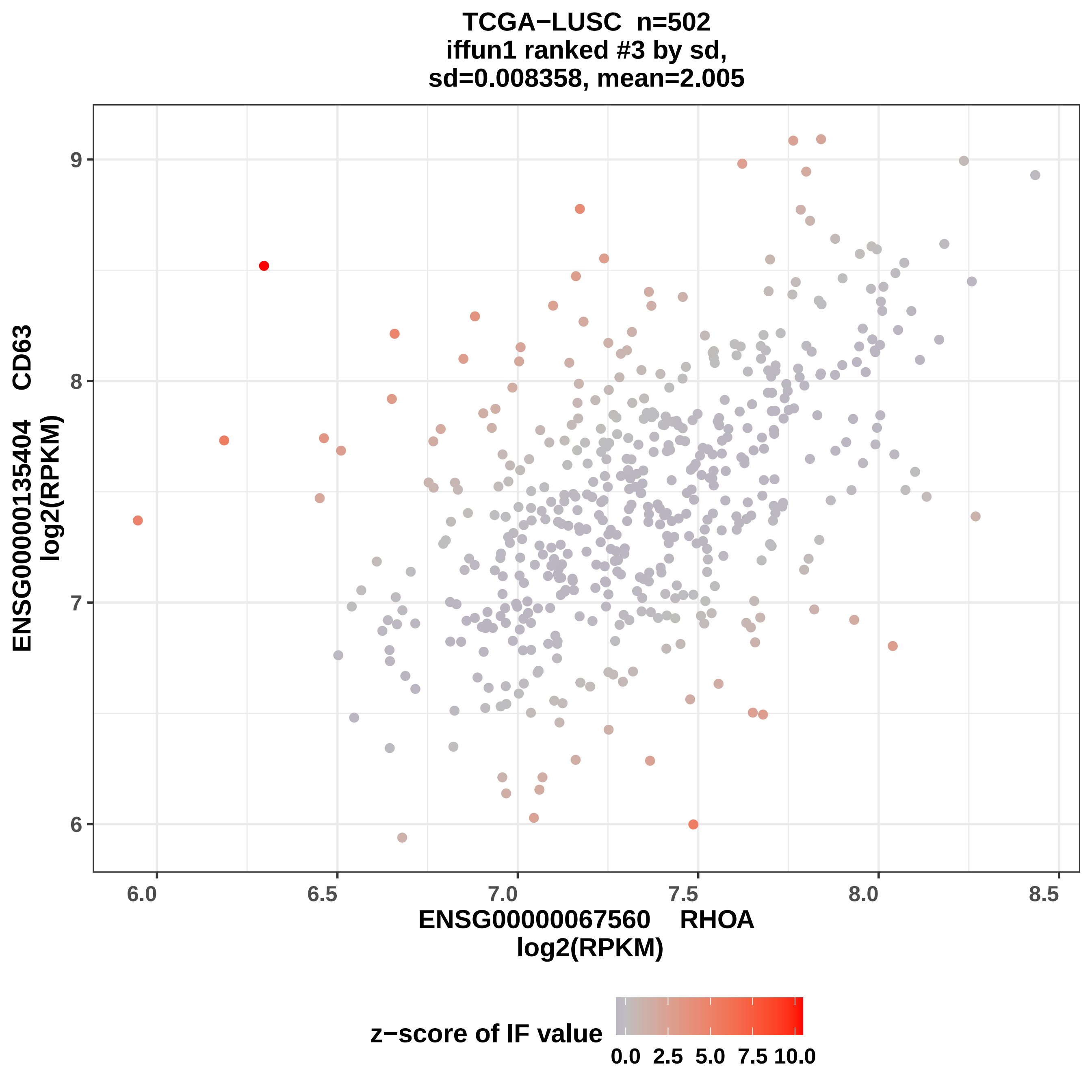
TCGA−LUSC n=502
iffun1 ranked #3 by sd,
sd=0.008358, mean=2.005
ENSG00000135404 CD63
log2(RPKM)
9
8
7
6
6.0
6.5
7.0
7.5
8.0
8.5
ENSG00000067560 RHOA
log2(RPKM)
z−score of IF value
0.0
2.5
5.0
7.5
10.0

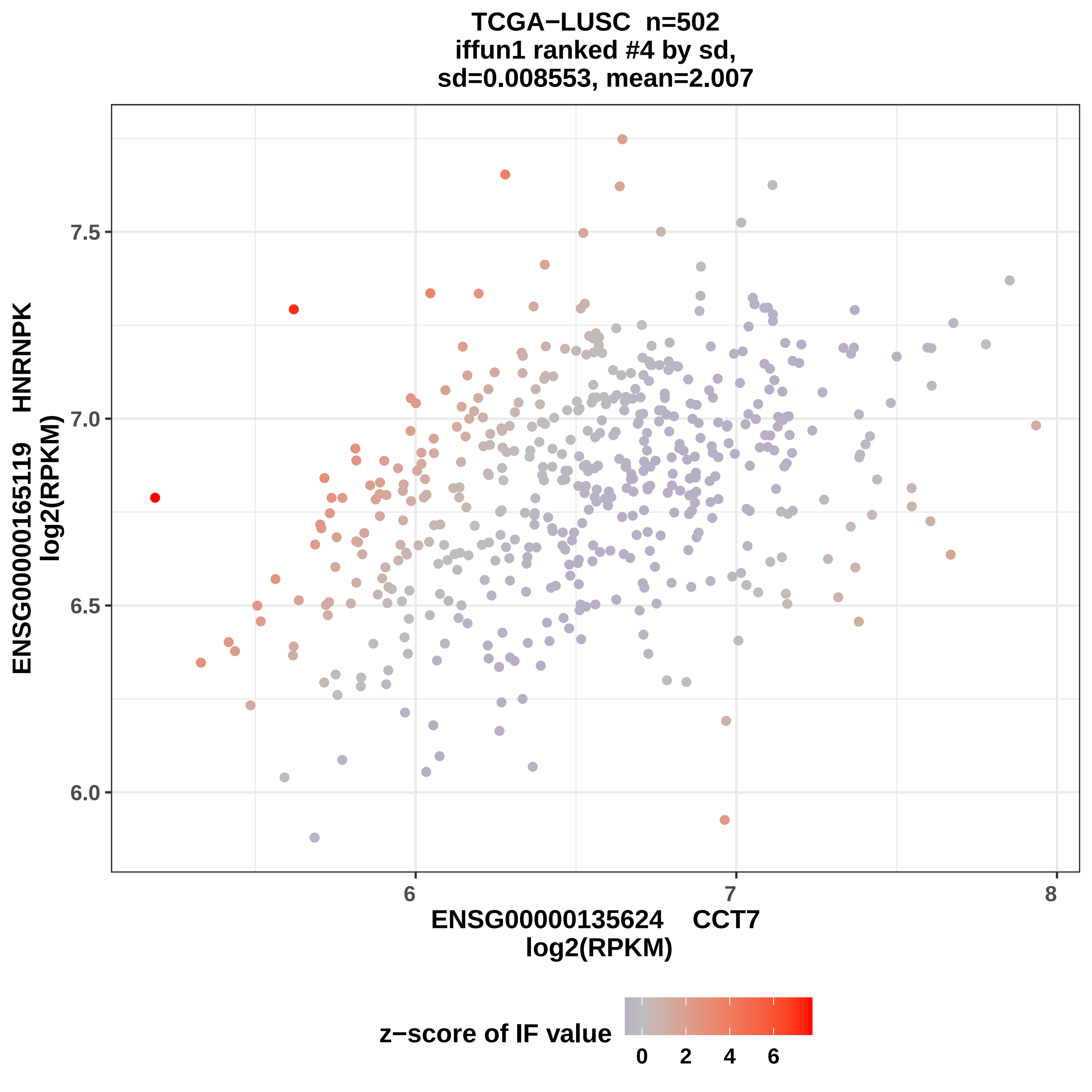

TCGA−LUSC n=502
iffun1 ranked #4 by sd,
sd=0.008553, mean=2.007
ENSG00000165119 HNRNPK
log2(RPKM)
7.5
7.0
6.5
6.0
6
7
8
ENSG00000135624 CCT7
log2(RPKM)
z−score of IF value
0
2
4
6

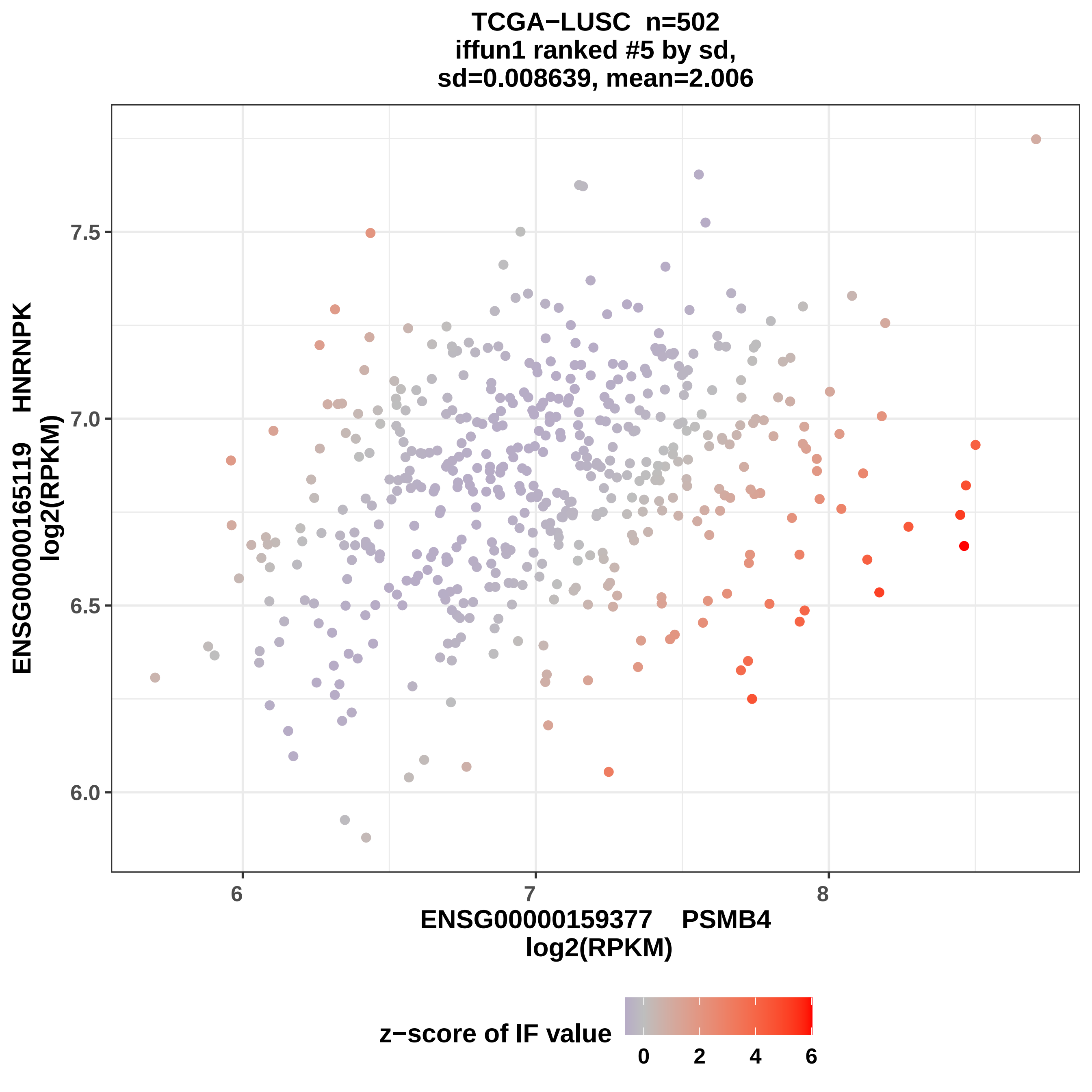

TCGA−LUSC n=502
iffun1 ranked #5 by sd,
sd=0.008639, mean=2.006
ENSG00000165119 HNRNPK
log2(RPKM)
7.5
7.0
6.5
6.0
6
7
8
ENSG00000159377 PSMB4
log2(RPKM)
z−score of IF value
0
2
4
6

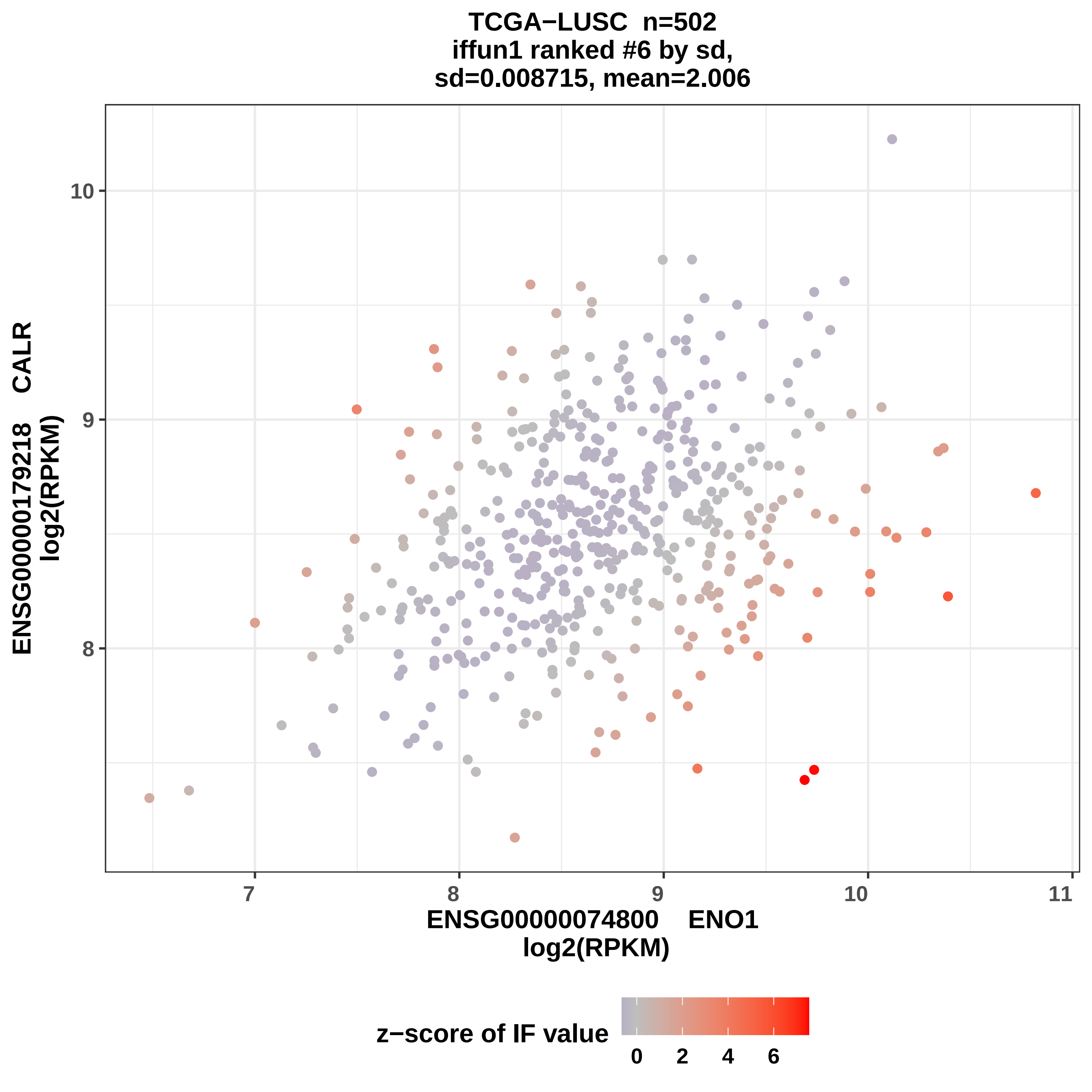

TCGA−LUSC n=502
iffun1 ranked #6 by sd,
sd=0.008715, mean=2.006
ENSG00000179218 CALR
log2(RPKM)
10
9
8
7
8
9
10
11
ENSG00000074800 ENO1
log2(RPKM)
z−score of IF value
0
2
4
6

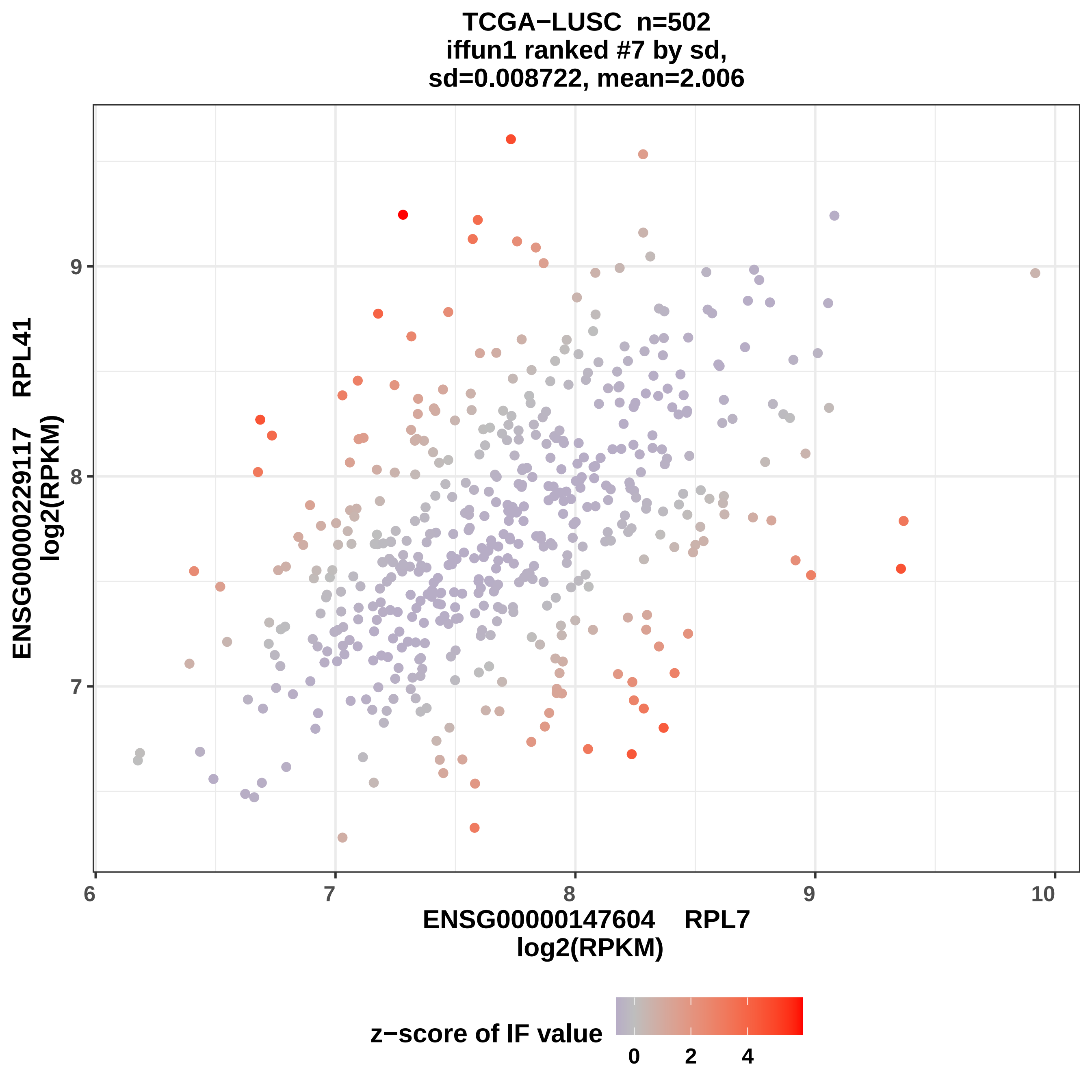

TCGA−LUSC n=502
iffun1 ranked #7 by sd,
sd=0.008722, mean=2.006
ENSG00000229117 RPL41
log2(RPKM)
ENSG00000147604 RPL7
log2(RPKM)
6
7
8
9
10
z−score of IF value
0
2
4

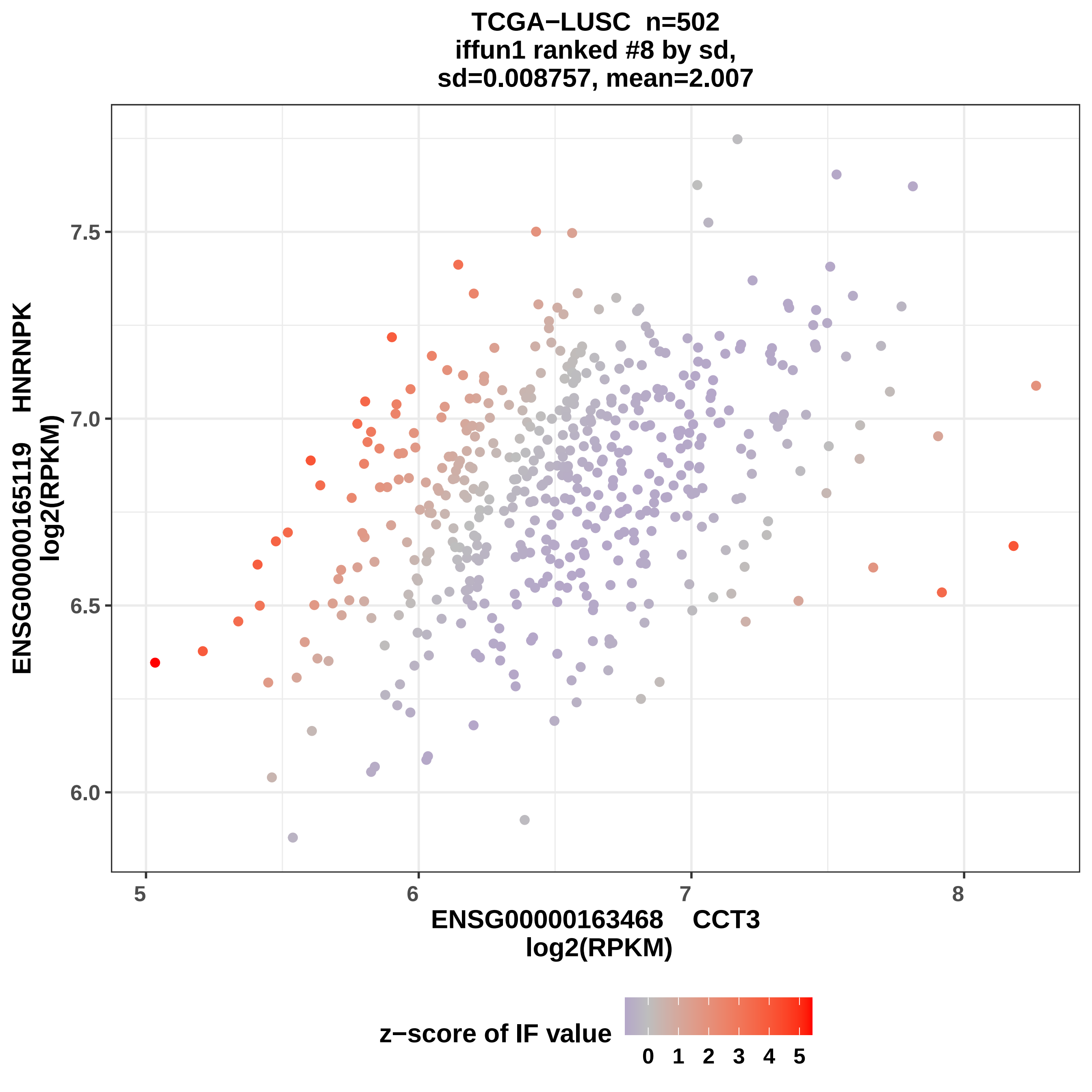

TCGA−LUSC n=502
iffun1 ranked #8 by sd,
sd=0.008757, mean=2.007
ENSG00000165119 HNRNPK
log2(RPKM)
7.5
7.0
6.5
6.0
5
6
7
8
ENSG00000163468 CCT3
log2(RPKM)
z−score of IF value
0 1 2 3 4 5

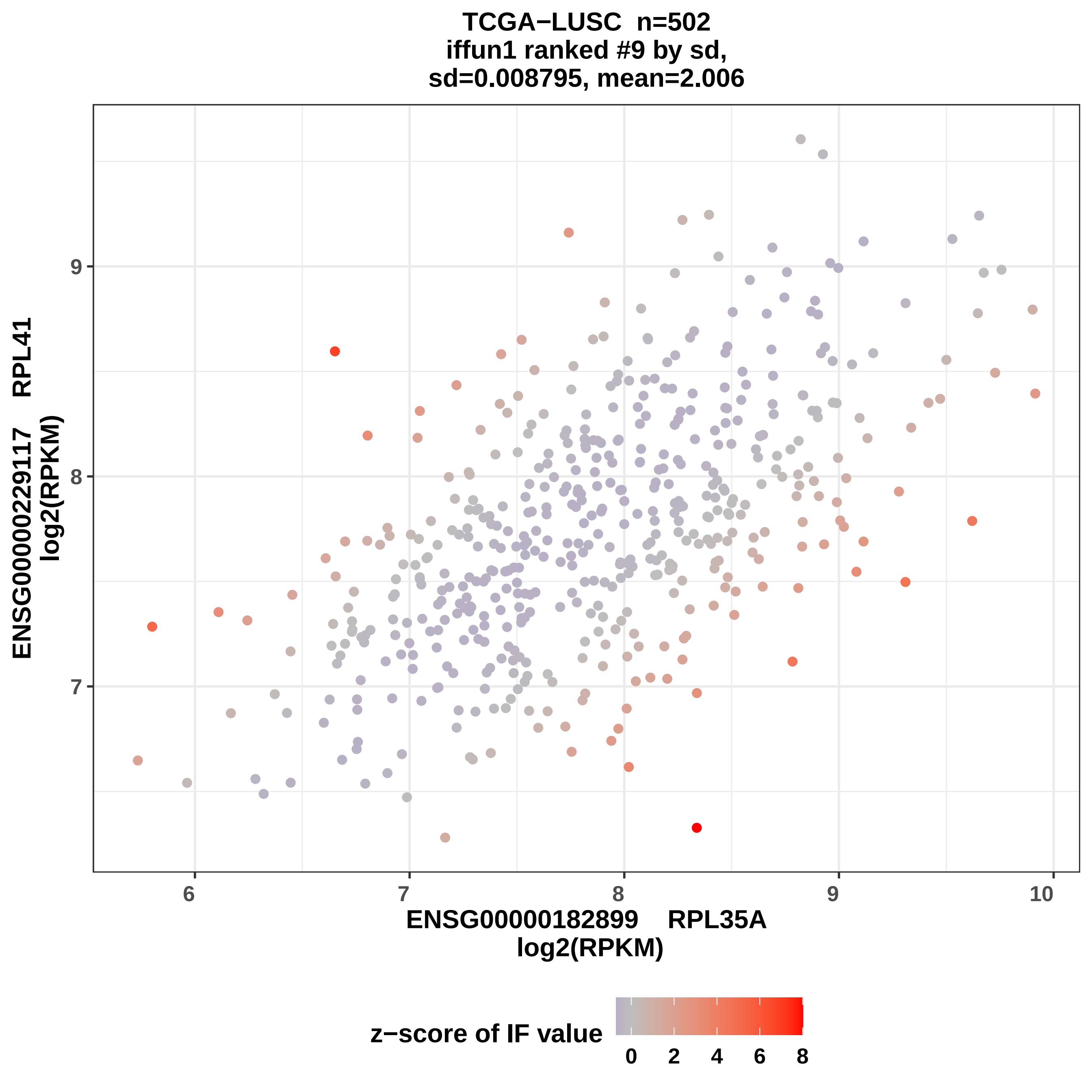
TCGA−LUSC n=502
iffun1 ranked #9 by sd,
sd=0.008795, mean=2.006
ENSG00000229117 RPL41
log2(RPKM)
9
8
7
6
7
8
9
10
ENSG00000182899 RPL35A
log2(RPKM)
z−score of IF value
0
2
4
6
8

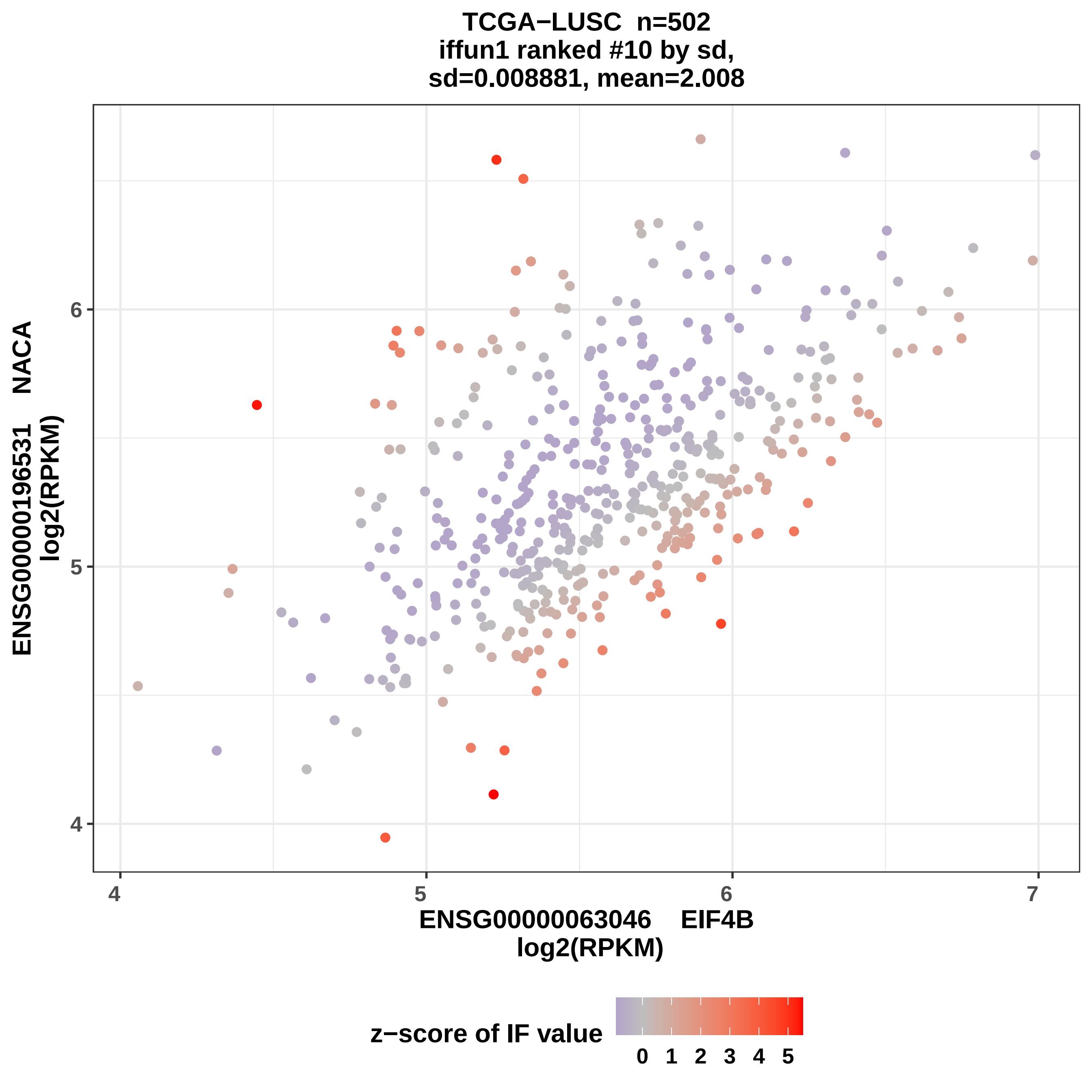

TCGA−LUSC n=502
iffun1 ranked #10 by sd,
sd=0.008881, mean=2.008
ENSG00000196531 NACA
log2(RPKM)
ENSG00000063046 EIF4B
log2(RPKM)
4
5
6
7
z−score of IF value
0 1 2 3 4 5

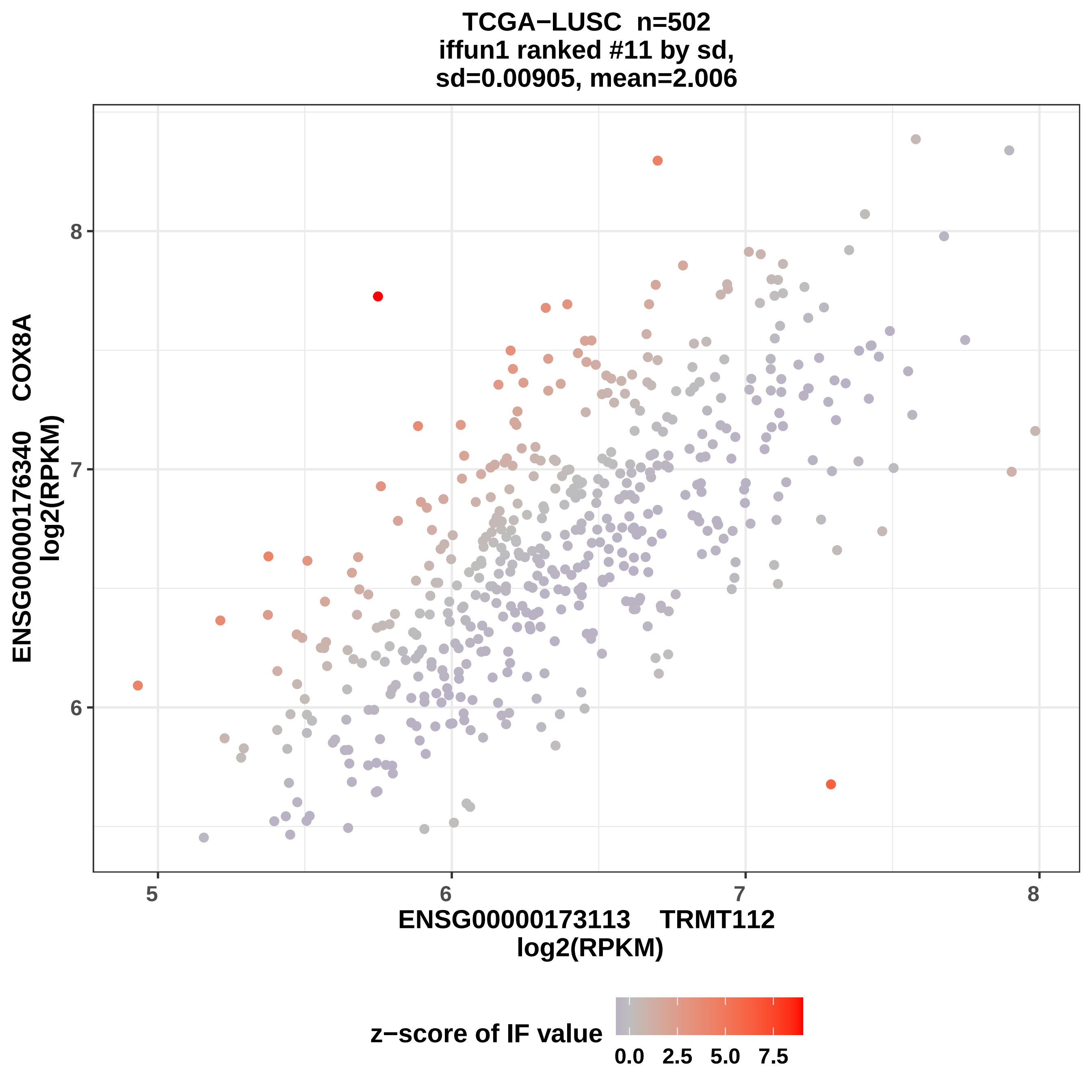
TCGA−LUSC n=502
iffun1 ranked #11 by sd,
sd=0.00905, mean=2.006
ENSG00000176340 COX8A
log2(RPKM)
8
7
6
5
6
7
8
ENSG00000173113 TRMT112
log2(RPKM)
z−score of IF value
0.0
2.5
5.0
7.5

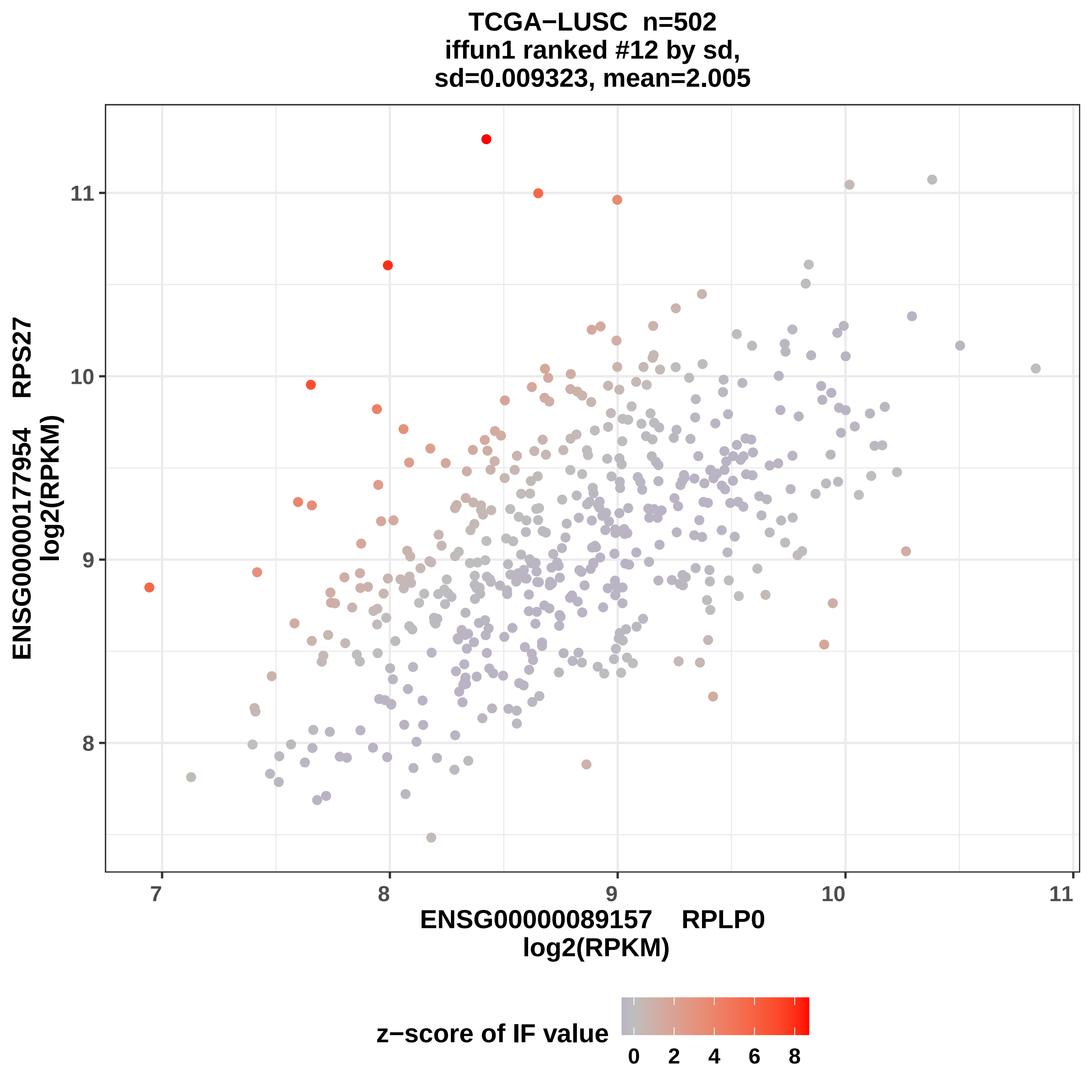

TCGA−LUSC n=502
iffun1 ranked #12 by sd,
sd=0.009323, mean=2.005
ENSG00000177954 RPS27
log2(RPKM)
ENSG00000089157 RPLP0
log2(RPKM)
8
9
10
11
7
8
9
10
11
z−score of IF value
0
2
4
6
8

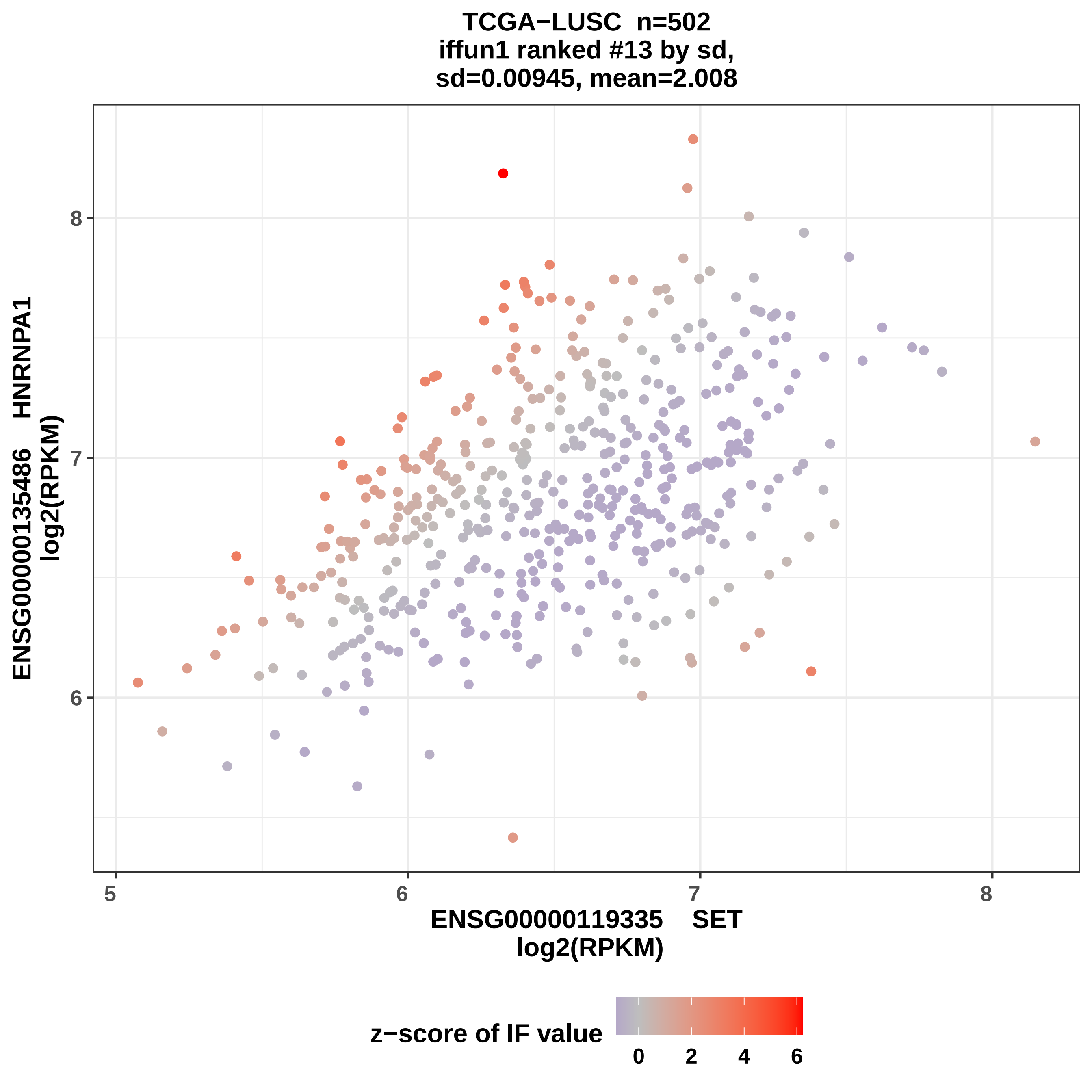
TCGA−LUSC n=502
iffun1 ranked #13 by sd,
sd=0.00945, mean=2.008
ENSG00000135486 HNRNPA1
log2(RPKM)
8
7
6
5
6
7
8
ENSG00000119335 SET
log2(RPKM)
z−score of IF value
0
2
4
6

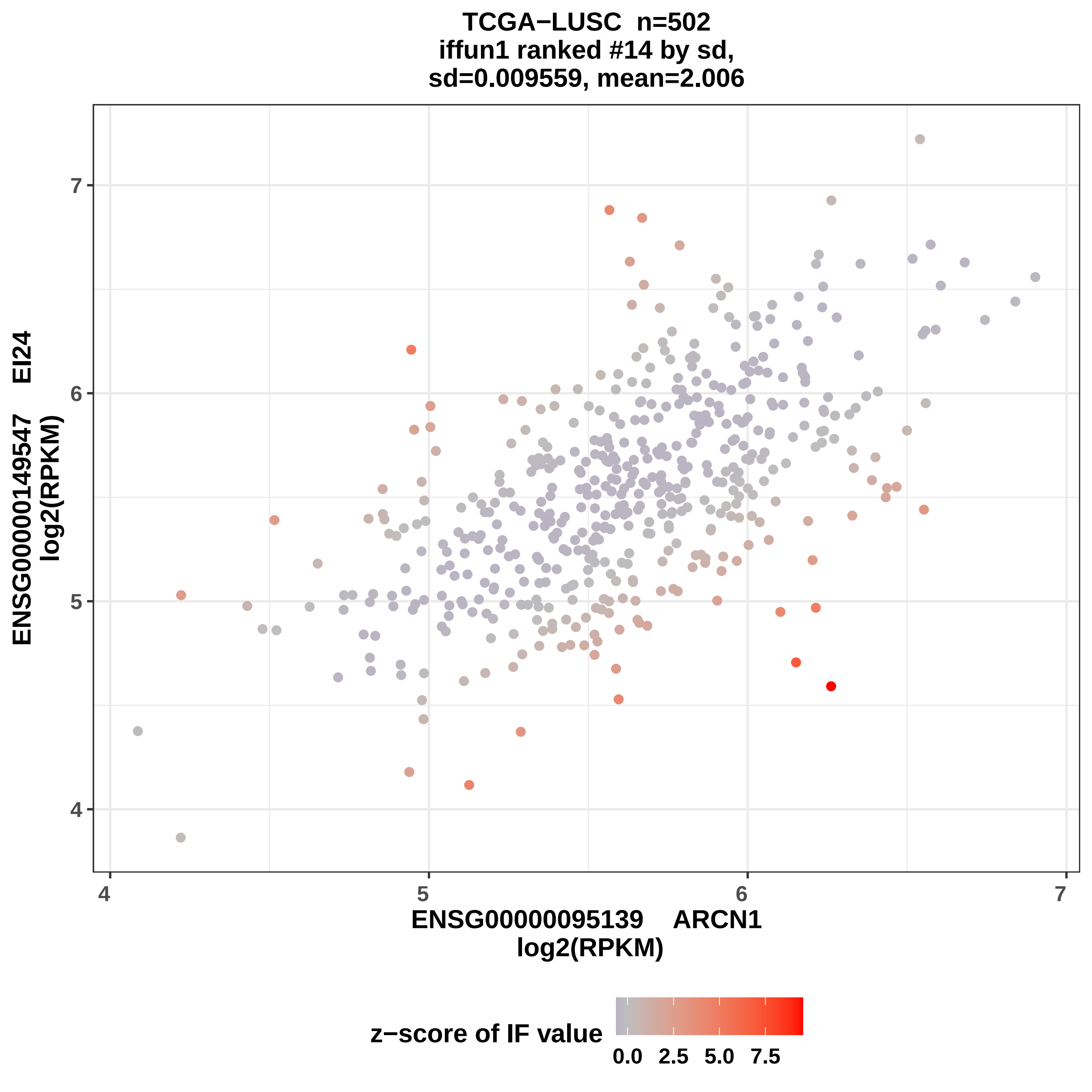
TCGA−LUSC n=502
iffun1 ranked #14 by sd,
sd=0.009559, mean=2.006
ENSG00000149547 EI24
log2(RPKM)
7
6
5
4
4
5
6
7
ENSG00000095139 ARCN1
log2(RPKM)
z−score of IF value
0.0
2.5
5.0
7.5

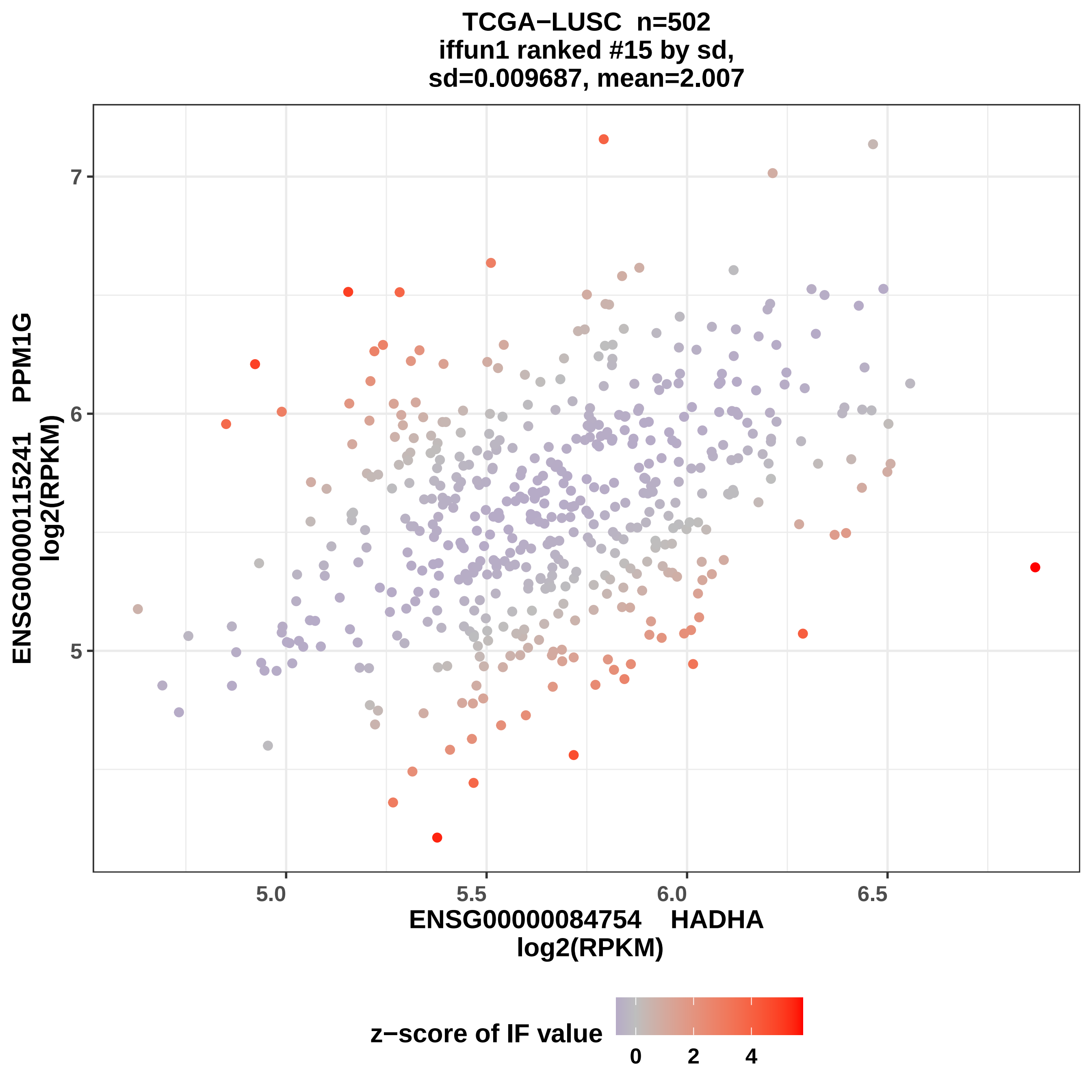

TCGA−LUSC n=502
iffun1 ranked #15 by sd,
sd=0.009687, mean=2.007
ENSG00000115241 PPM1G
log2(RPKM)
7
6
5
5.0
5.5
6.0
6.5
ENSG00000084754 HADHA
log2(RPKM)
z−score of IF value
0
2
4

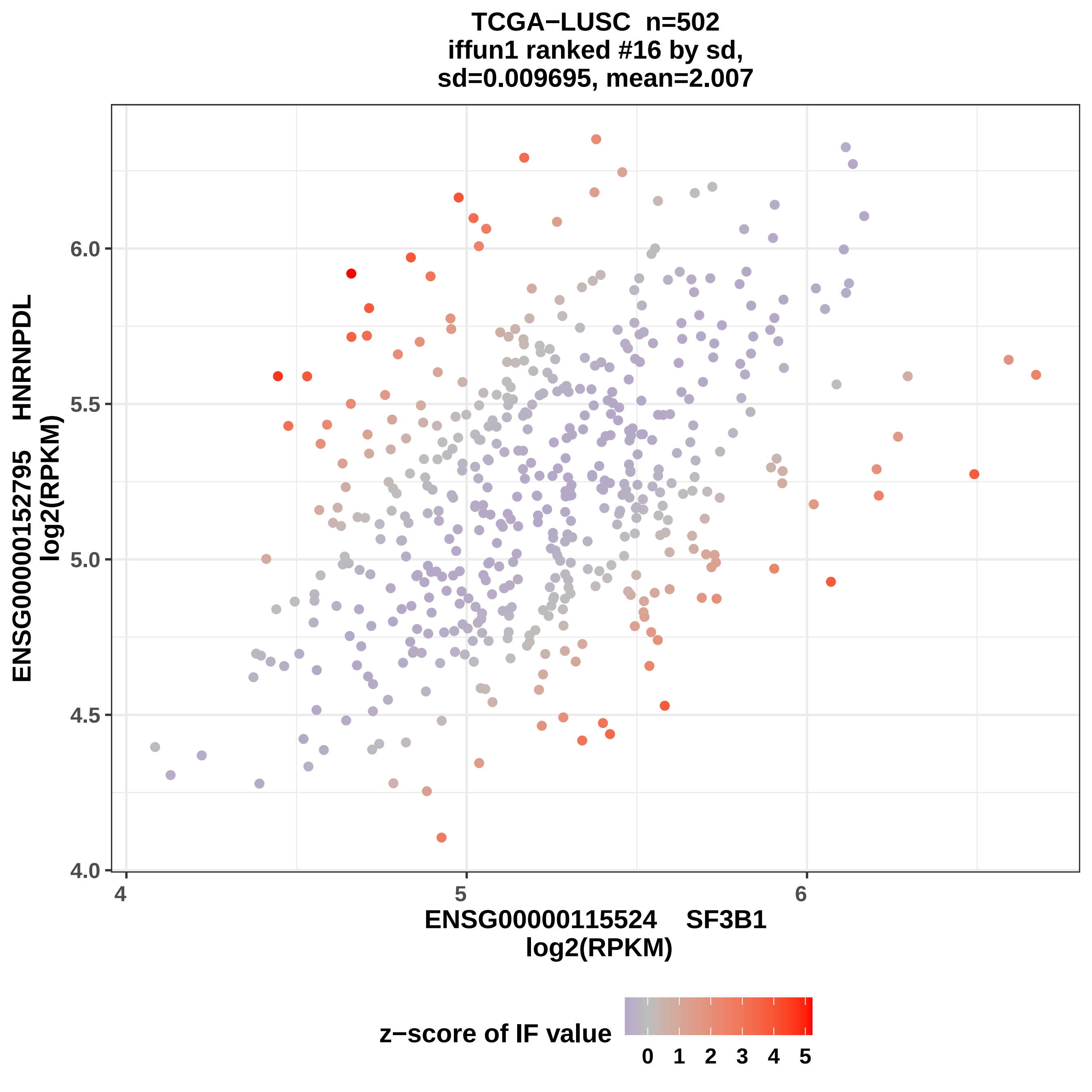

TCGA−LUSC n=502
iffun1 ranked #16 by sd,
sd=0.009695, mean=2.007
ENSG00000152795 HNRNPDL
log2(RPKM)
6.0
5.5
5.0
4.5
4.0
4
5
6
ENSG00000115524 SF3B1
log2(RPKM)
z−score of IF value
0 1 2 3 4 5

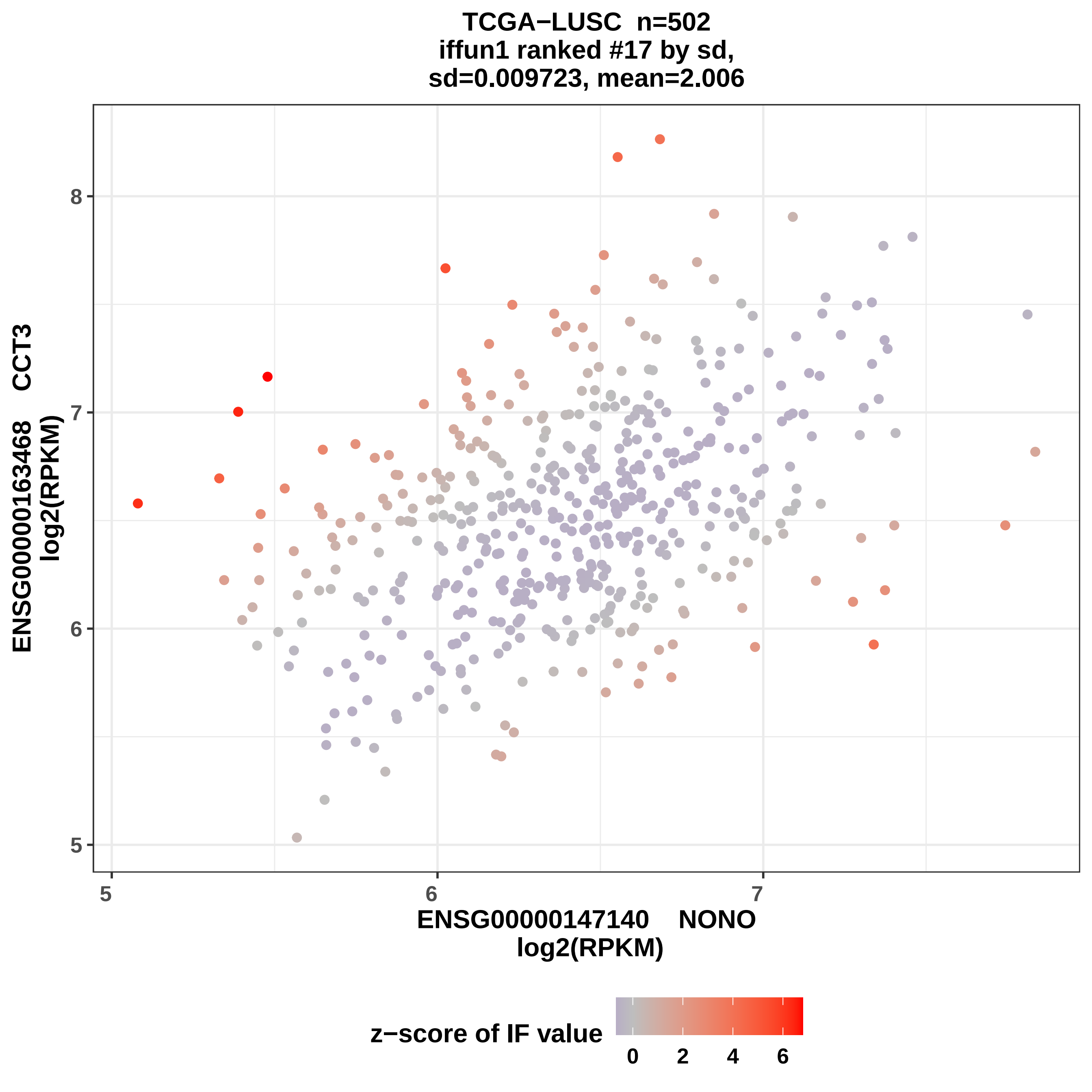

TCGA−LUSC n=502
iffun1 ranked #17 by sd,
sd=0.009723, mean=2.006
ENSG00000163468 CCT3
log2(RPKM)
8
7
6
5
ENSG00000147140 NONO
log2(RPKM)
z−score of IF value
0
2
4
6

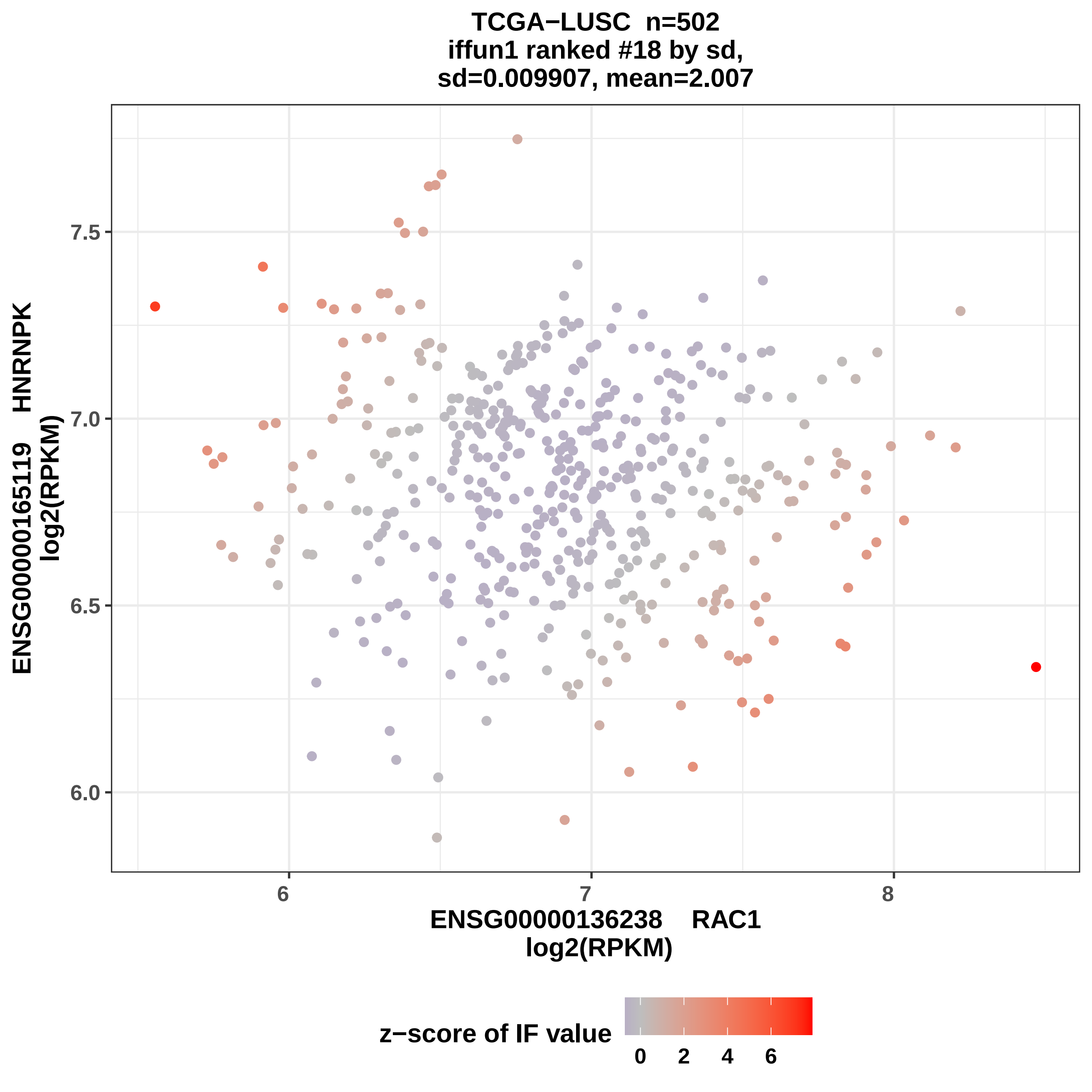
TCGA−LUSC n=502
iffun1 ranked #18 by sd,
sd=0.009907, mean=2.007
ENSG00000165119 HNRNPK
log2(RPKM)
7.5
7.0
6.5
6.0
6
7
8
ENSG00000136238 RAC1
log2(RPKM)
z−score of IF value
0
2
4
6

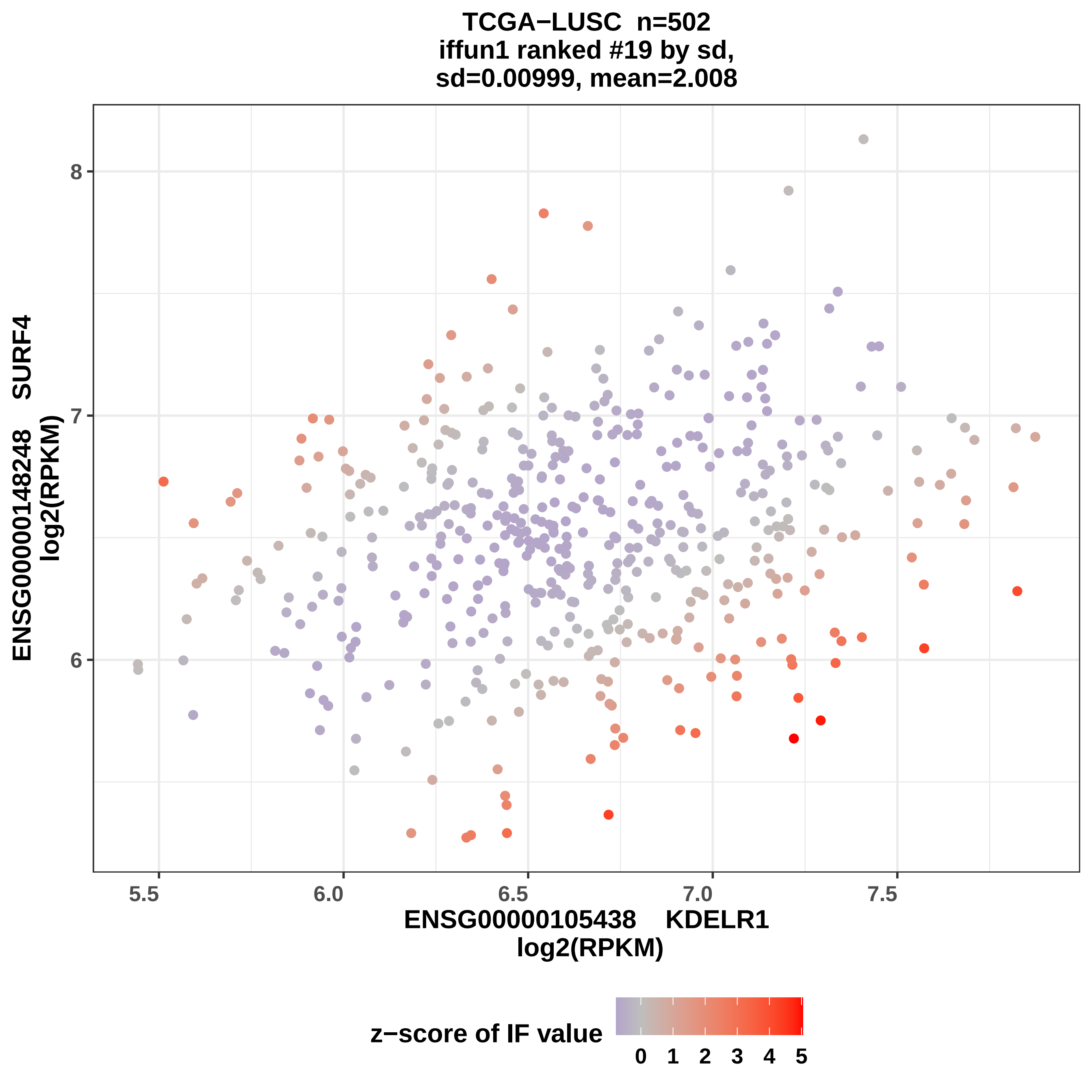

TCGA−LUSC n=502
iffun1 ranked #19 by sd,
sd=0.00999, mean=2.008
ENSG00000148248 SURF4
log2(RPKM)
8
7
6
5.5
6.0
6.5
7.0
7.5
ENSG00000105438 KDELR1
log2(RPKM)
z−score of IF value
0 1 2 3 4 5

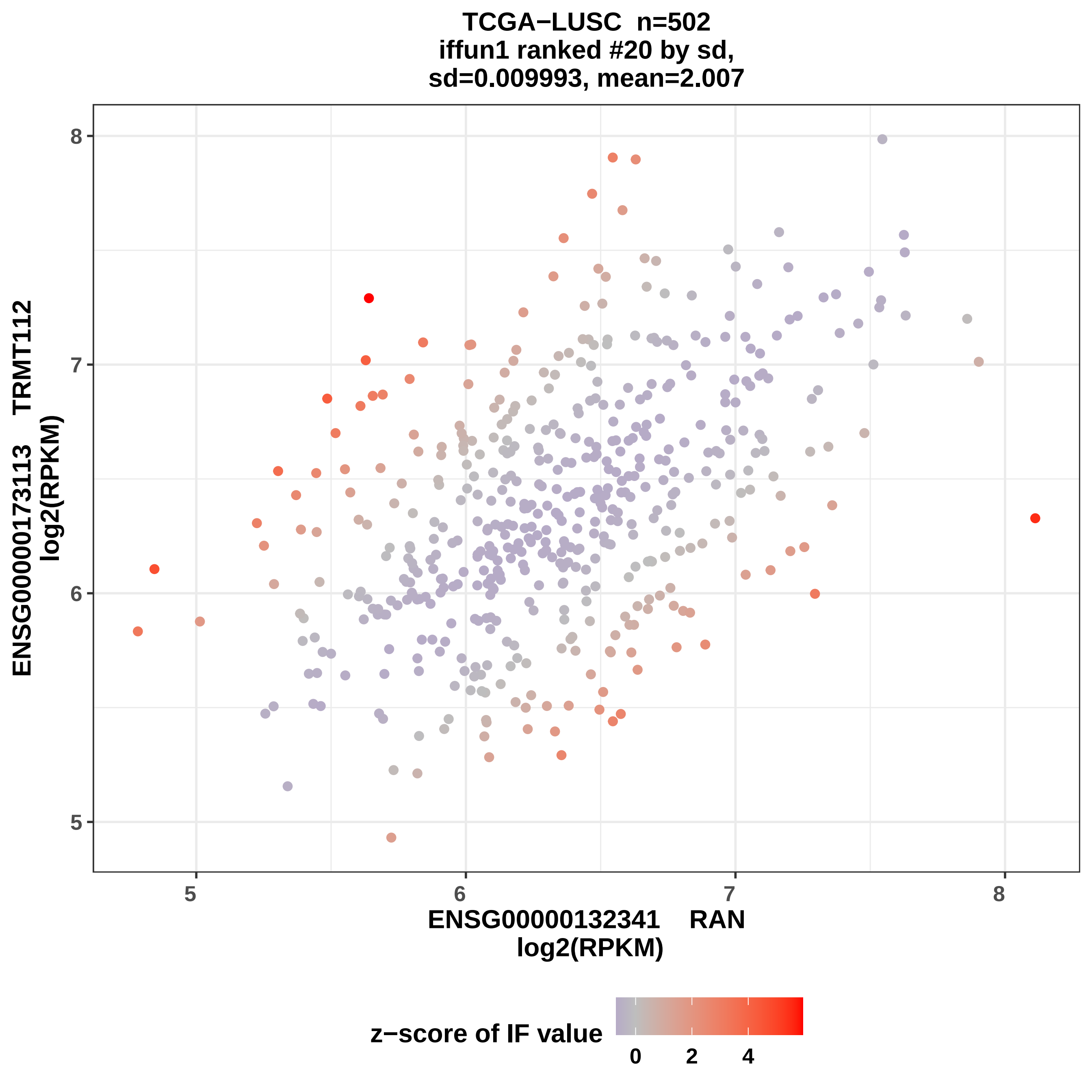

TCGA−LUSC n=502
iffun1 ranked #20 by sd,
sd=0.009993, mean=2.007
ENSG00000173113 TRMT112
log2(RPKM)
8
7
6
5
5
6
7
8
ENSG00000132341 RAN
log2(RPKM)
z−score of IF value
0
2
4

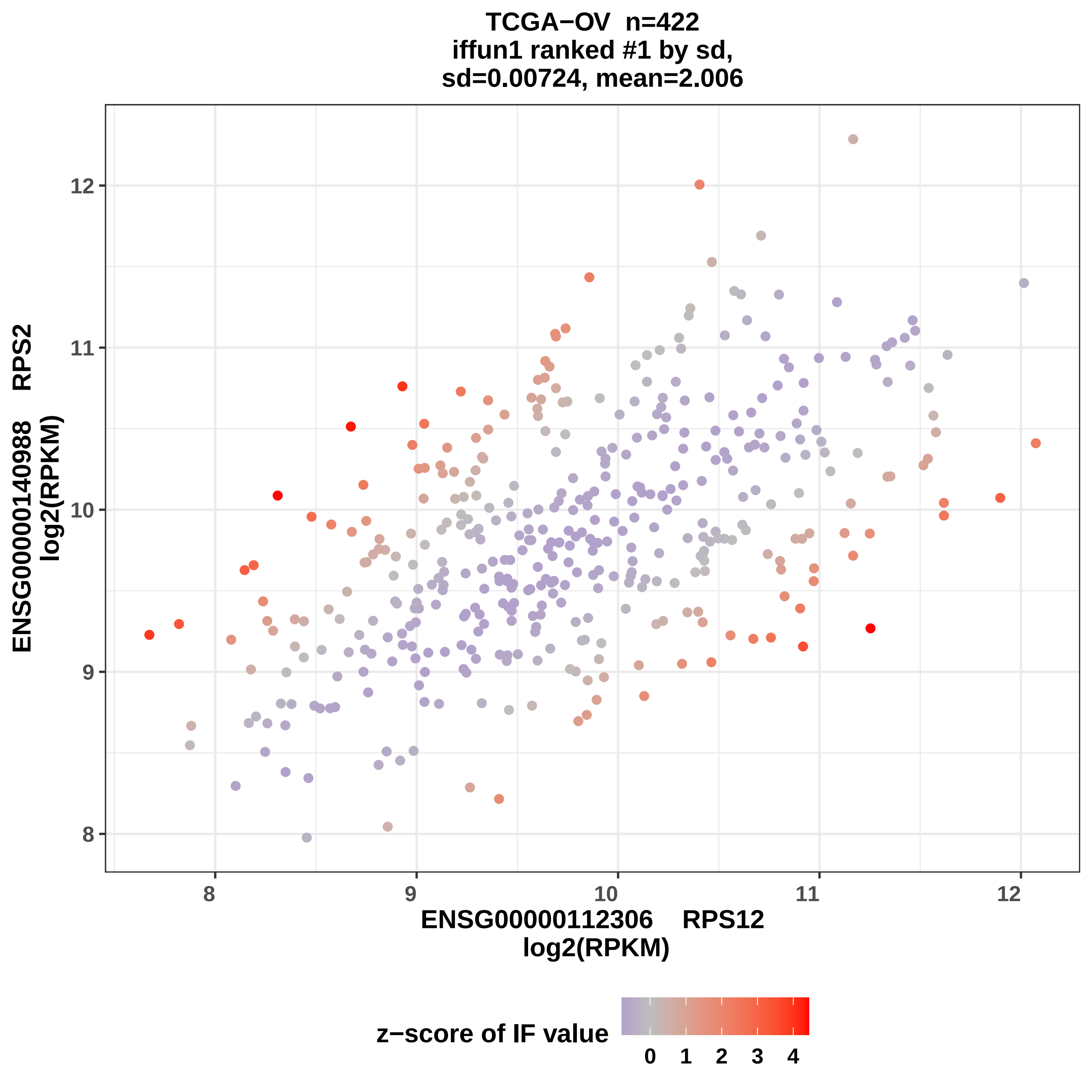
TCGA−OV n=422
iffun1 ranked #1 by sd,
sd=0.00724, mean=2.006
ENSG00000140988 RPS2
log2(RPKM)
ENSG00000112306 RPS12
log2(RPKM)
8
9
10
11
12
z−score of IF value
0
1
2
3
4

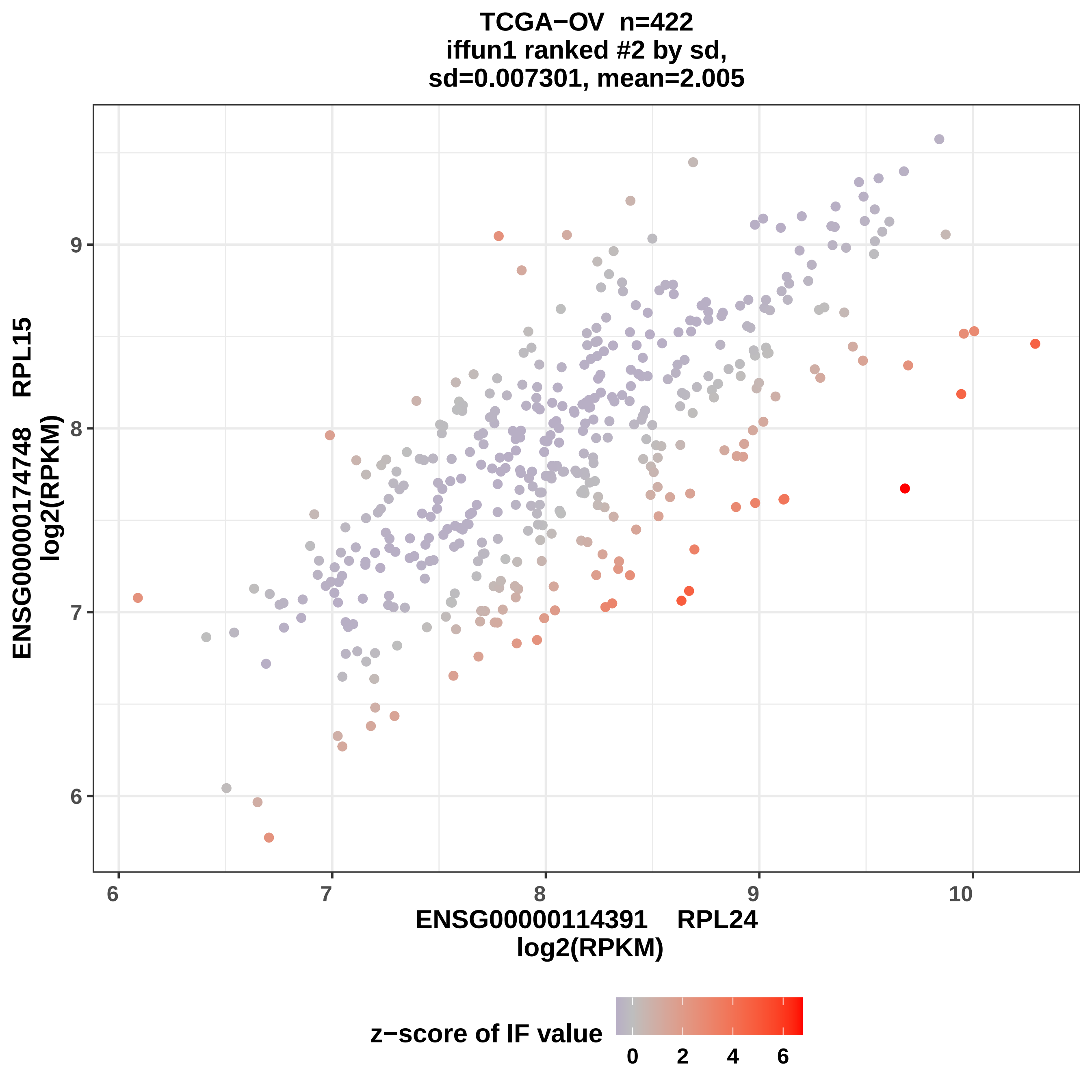

TCGA−OV n=422
iffun1 ranked #2 by sd,
sd=0.007301, mean=2.005
ENSG00000174748 RPL15
log2(RPKM)
9
8
7
6
6
7
8
9
10
ENSG00000114391 RPL24
log2(RPKM)
z−score of IF value
0
2
4
6

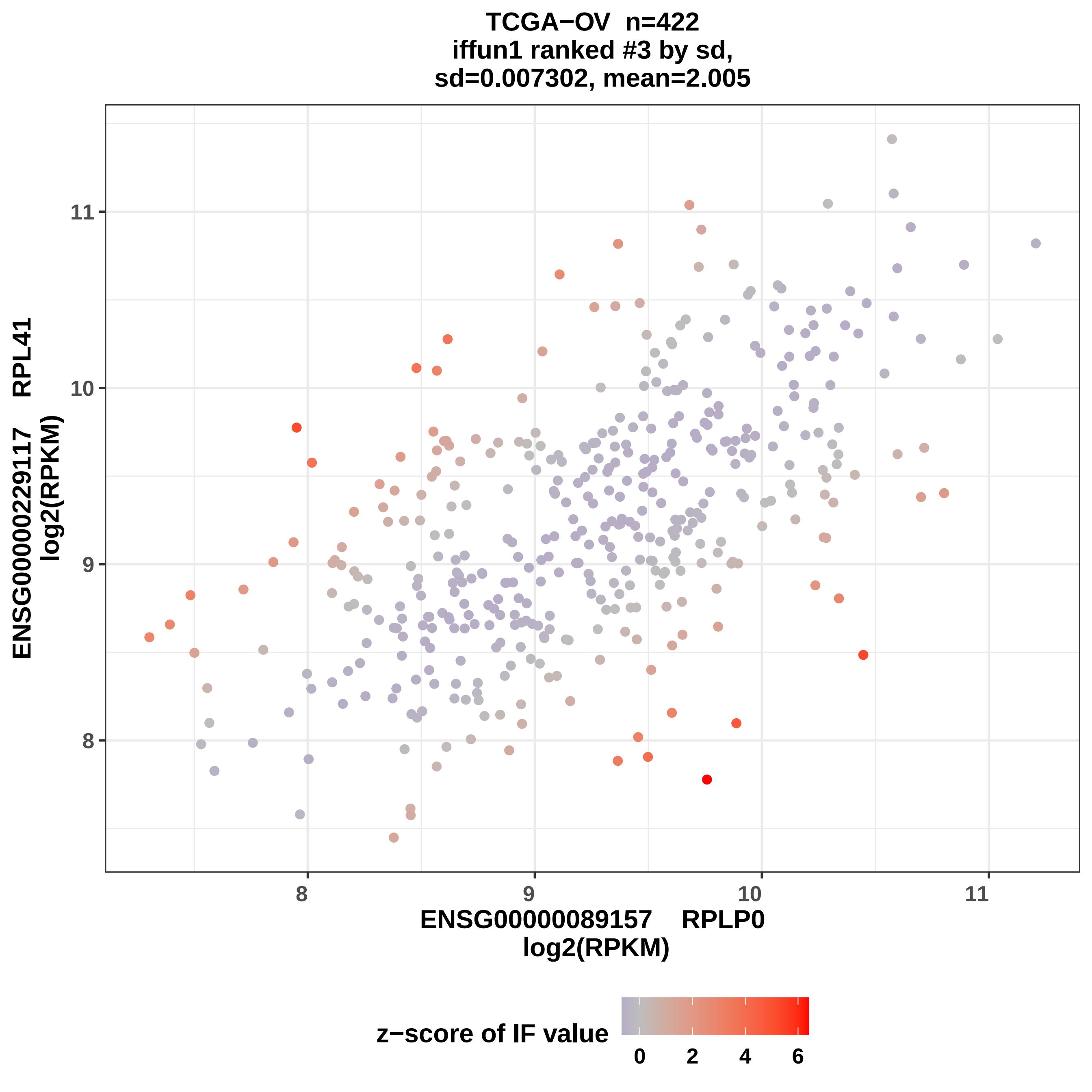
TCGA−OV n=422
iffun1 ranked #3 by sd,
sd=0.007302, mean=2.005
ENSG00000229117 RPL41
log2(RPKM)
11
10
9
8
ENSG00000089157 RPLP0
log2(RPKM)
z−score of IF value
0
2
4
6

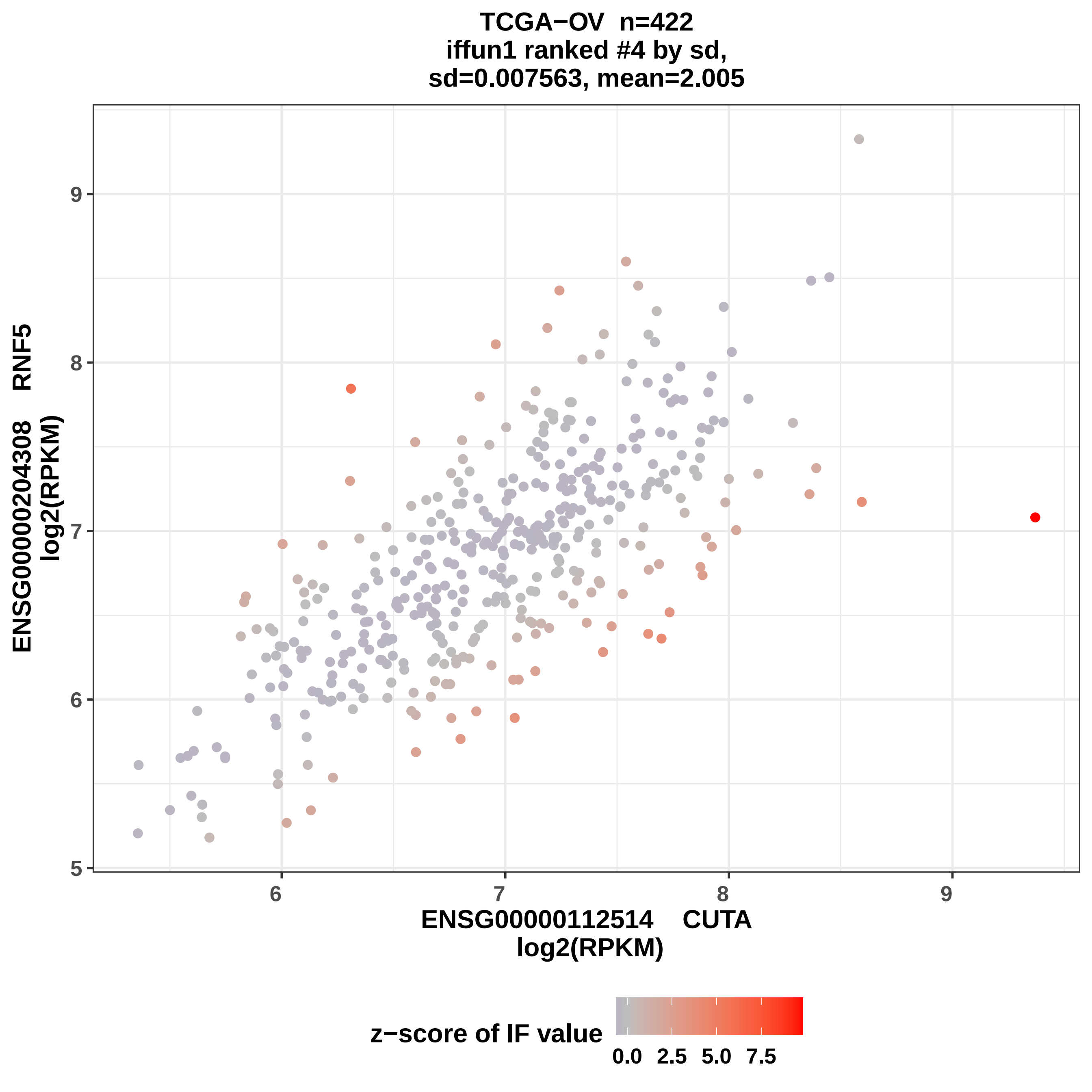

TCGA−OV n=422
iffun1 ranked #4 by sd,
sd=0.007563, mean=2.005
ENSG00000204308 RNF5
log2(RPKM)
ENSG00000112514 CUTA
log2(RPKM)
z−score of IF value
0.0 2.5 5.0 7.5

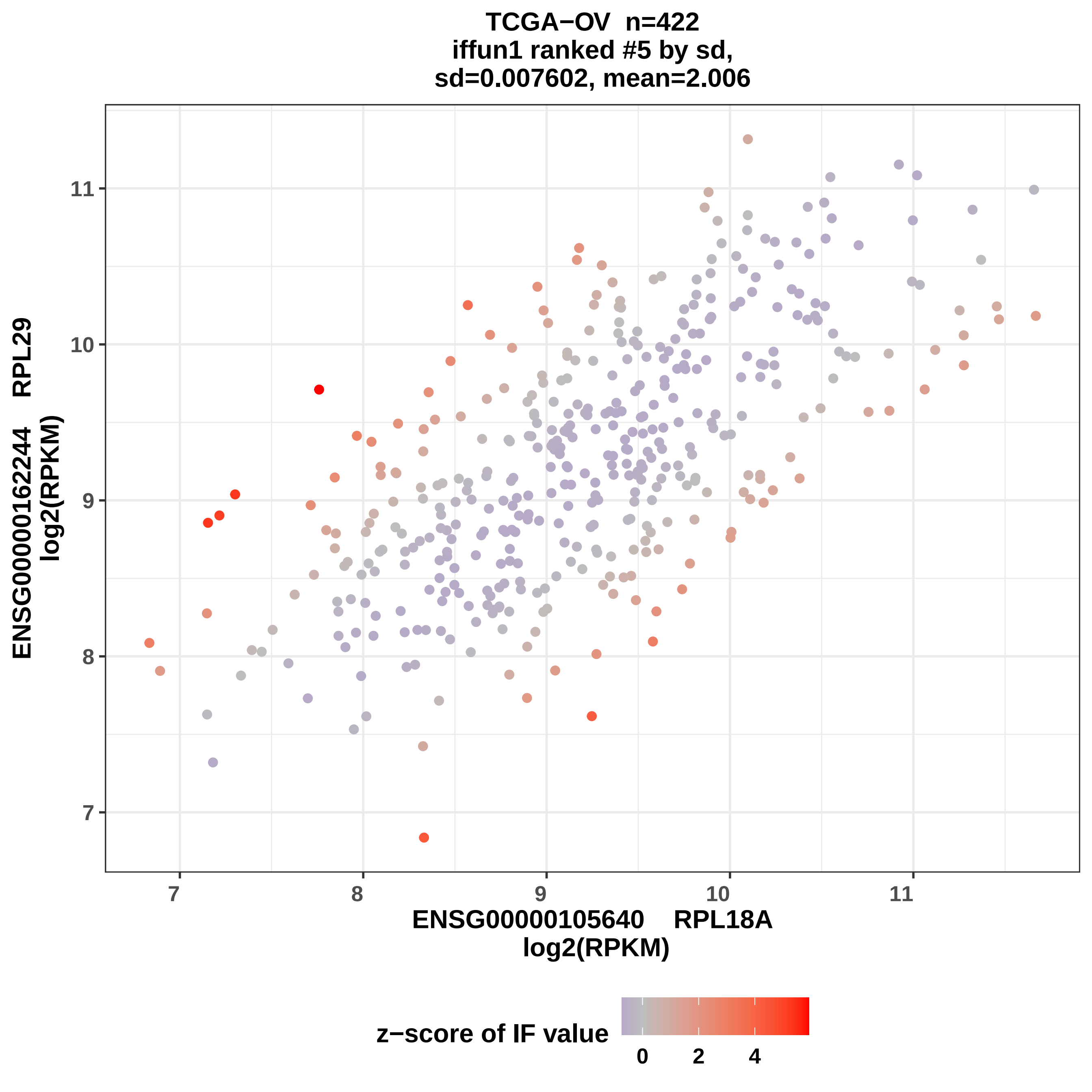
TCGA−OV n=422
iffun1 ranked #5 by sd,
sd=0.007602, mean=2.006
ENSG00000162244 RPL29
log2(RPKM)
11
10
9
8
7
ENSG00000105640 RPL18A
log2(RPKM)
z−score of IF value
0
2
4

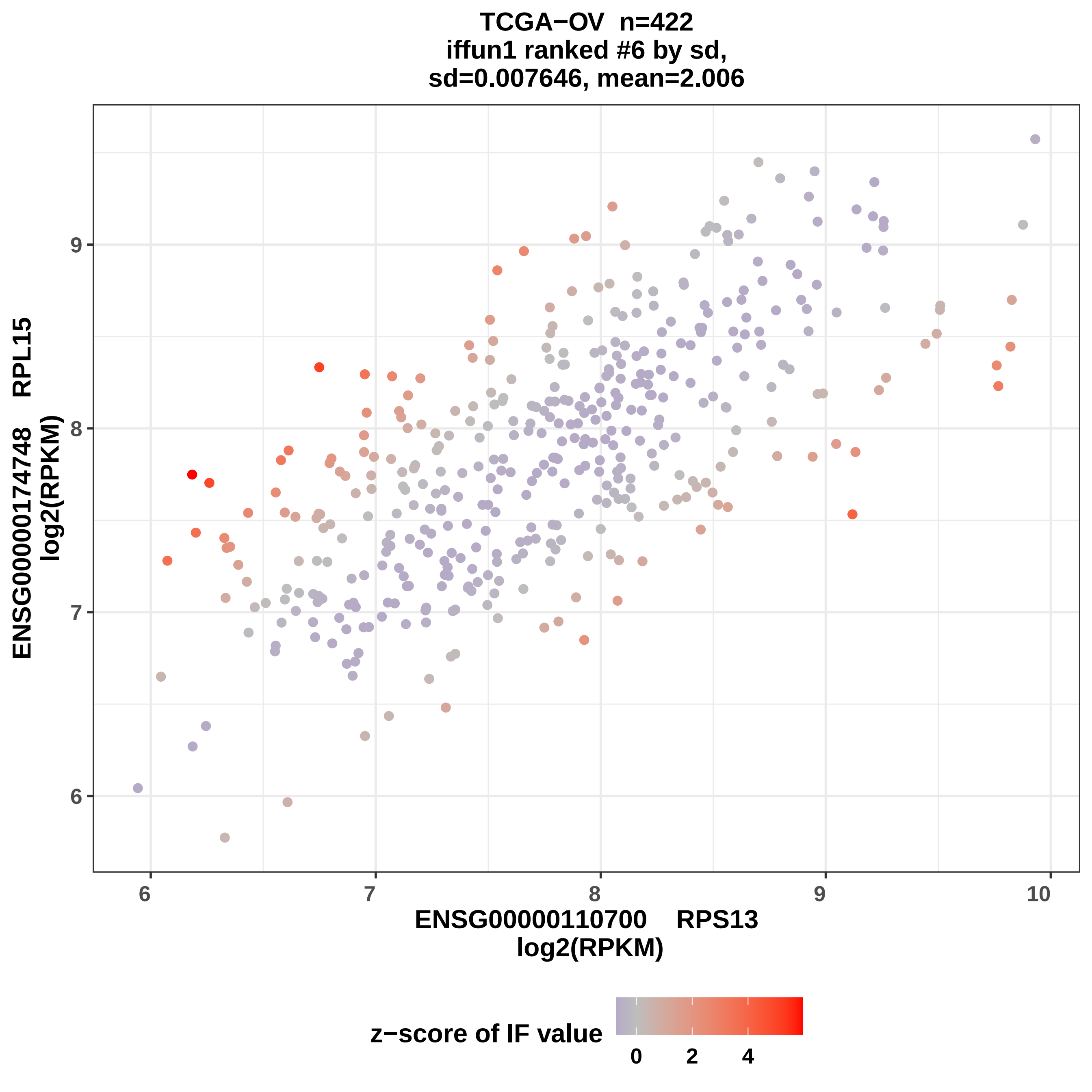
TCGA−OV n=422
iffun1 ranked #6 by sd,
sd=0.007646, mean=2.006
ENSG00000174748 RPL15
log2(RPKM)
ENSG00000110700 RPS13
log2(RPKM)
6
7
8
9
10
z−score of IF value
0
2
4

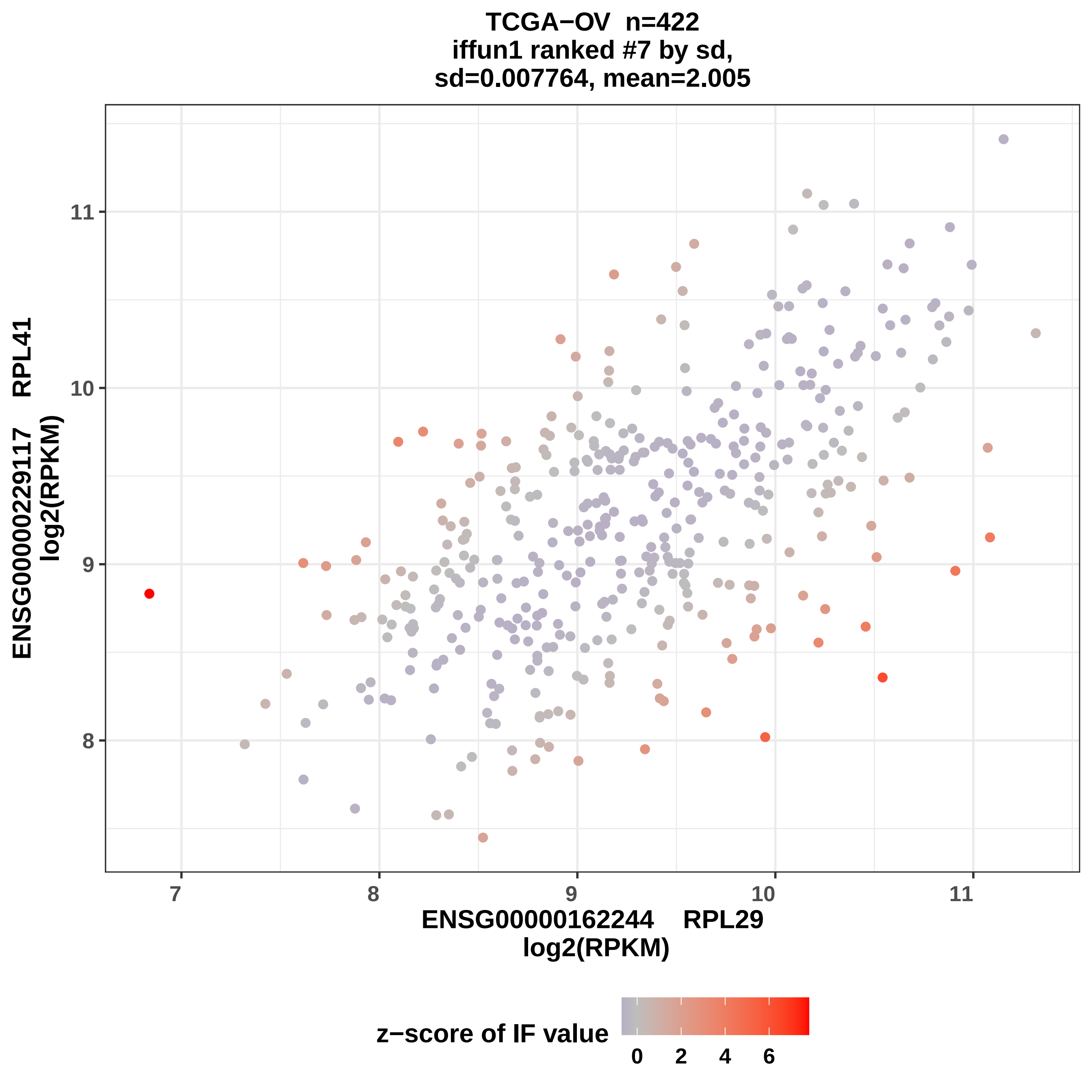

TCGA−OV n=422
iffun1 ranked #7 by sd,
sd=0.007764, mean=2.005
ENSG00000229117 RPL41
log2(RPKM)
ENSG00000162244 RPL29
log2(RPKM)
11
10
9
8
7
8
9
10
11
z−score of IF value
0
2
4
6

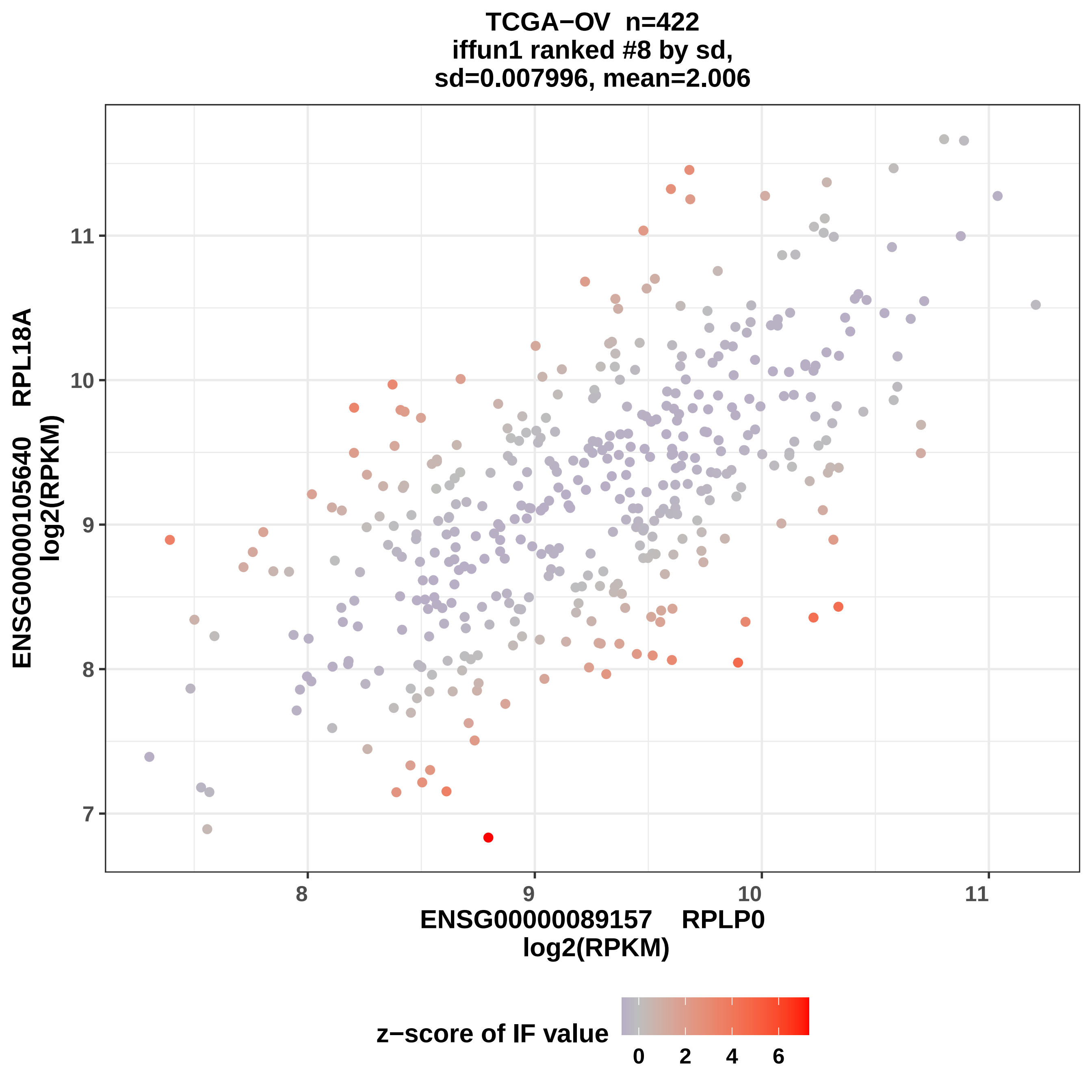
TCGA−OV n=422
iffun1 ranked #8 by sd,
sd=0.007996, mean=2.006
ENSG00000105640 RPL18A
log2(RPKM)
ENSG00000089157 RPLP0
log2(RPKM)
7
8
9
10
11
8
9
10
11
z−score of IF value
0
2
4
6

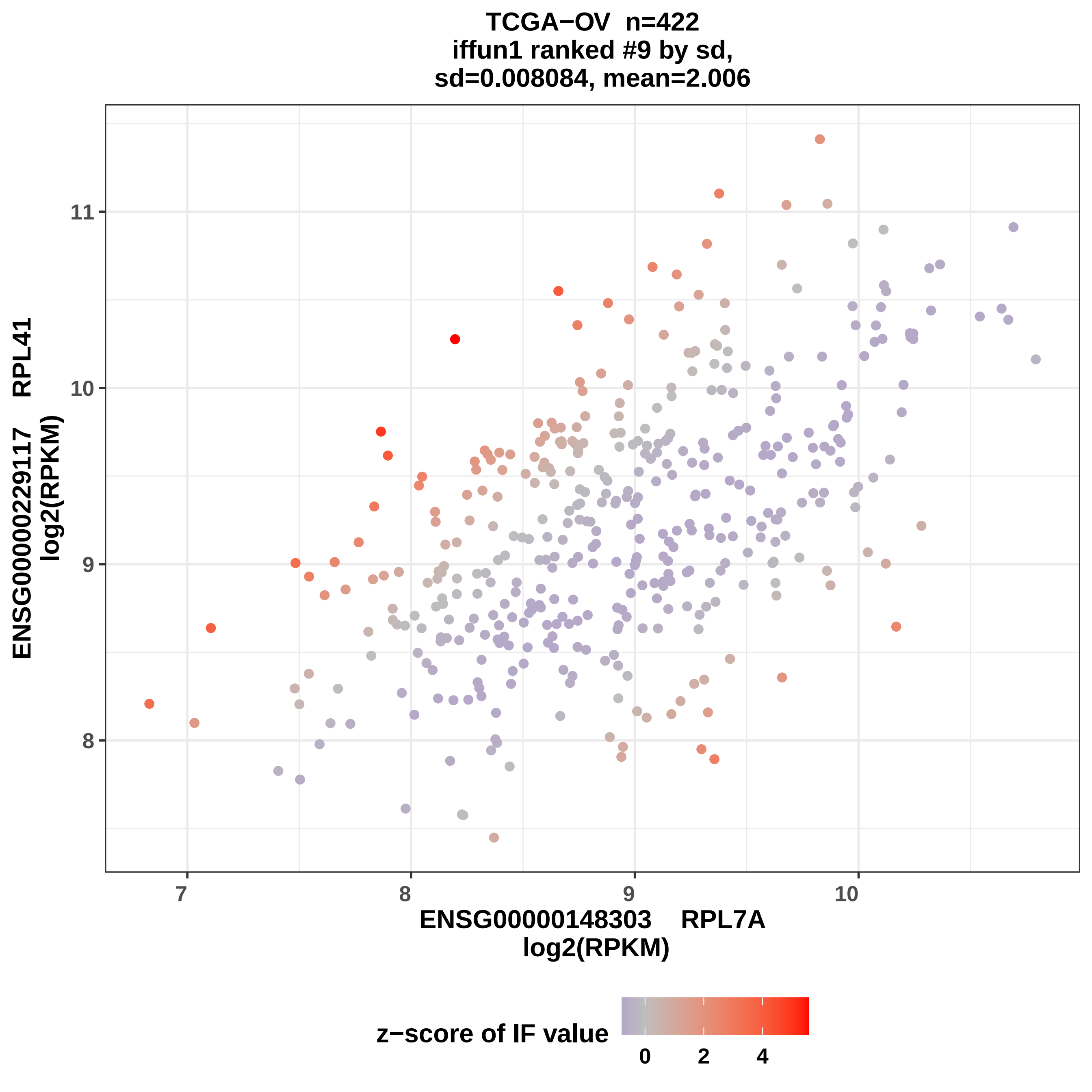
TCGA−OV n=422
iffun1 ranked #9 by sd,
sd=0.008084, mean=2.006
ENSG00000229117 RPL41
log2(RPKM)
11
10
9
8
7
8
9
10
ENSG00000148303 RPL7A
log2(RPKM)
z−score of IF value
0
2
4

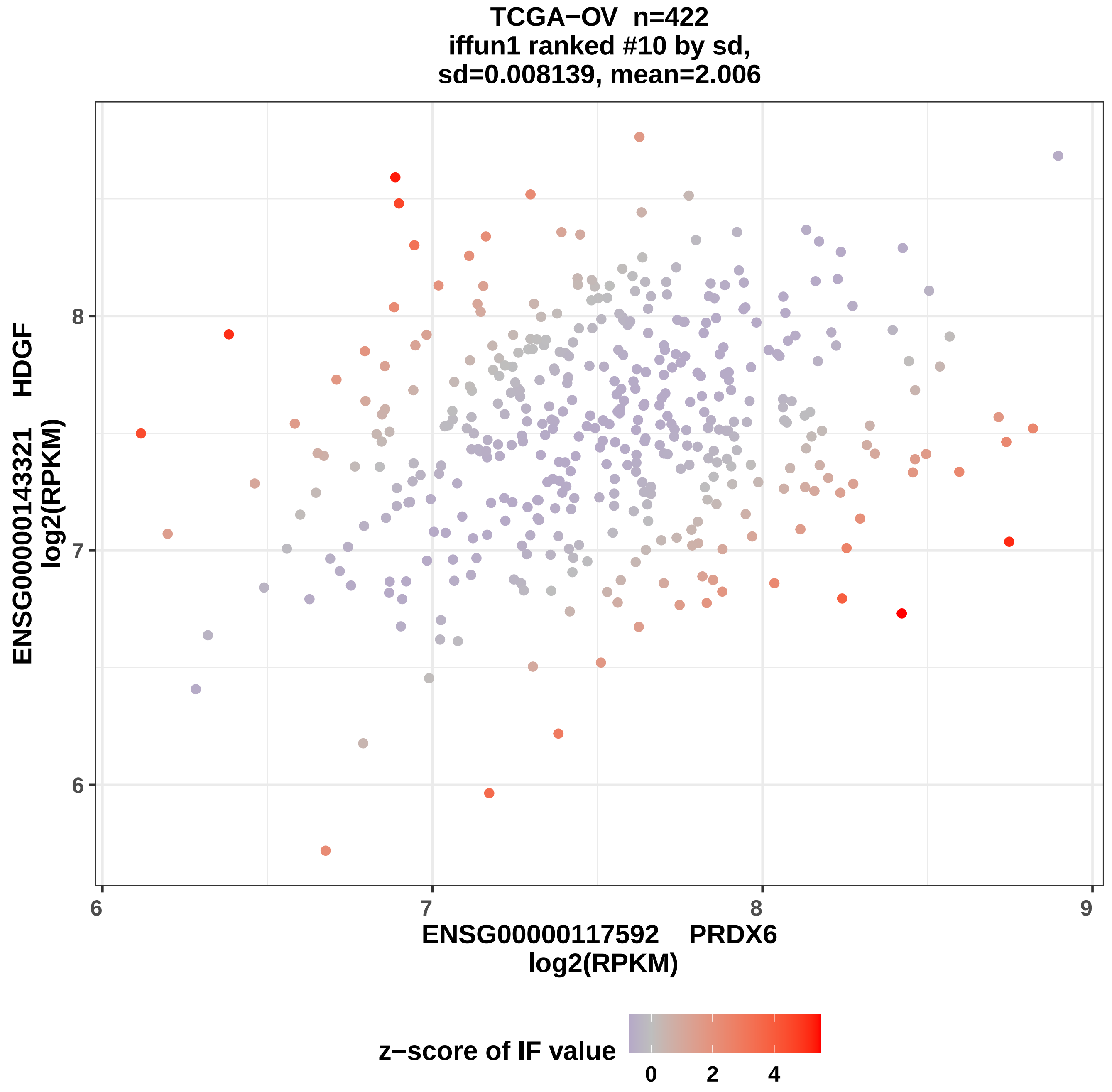

TCGA−OV n=422
iffun1 ranked #10 by sd,
sd=0.008139, mean=2.006
ENSG00000143321 HDGF
log2(RPKM)
ENSG00000117592 PRDX6
log2(RPKM)
6
7
8
9
z−score of IF value
0
2
4

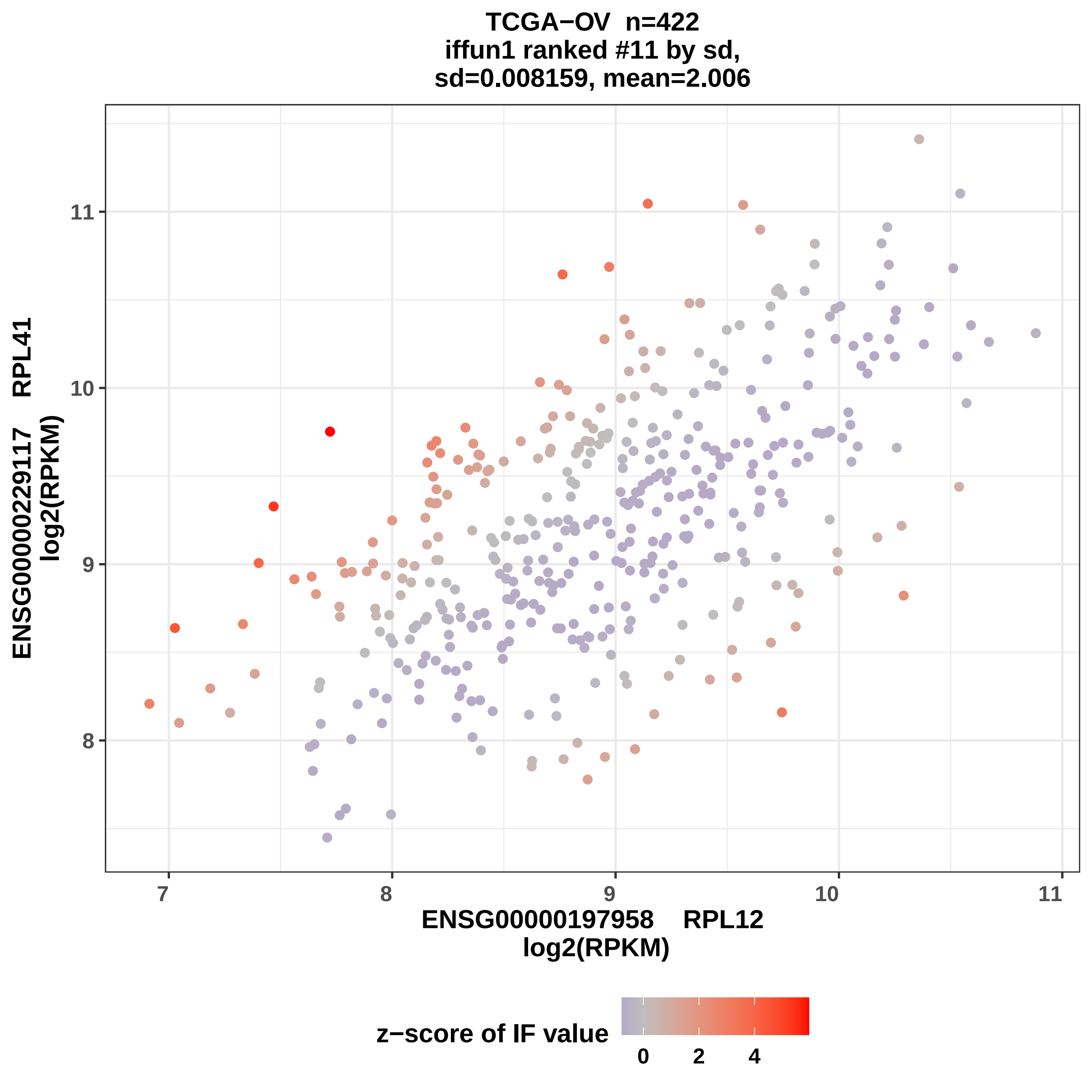

TCGA−OV n=422
iffun1 ranked #11 by sd,
sd=0.008159, mean=2.006
ENSG00000229117 RPL41
log2(RPKM)
ENSG00000197958 RPL12
log2(RPKM)
7
8
9
10
11
8
9
10
11
z−score of IF value
0
2
4

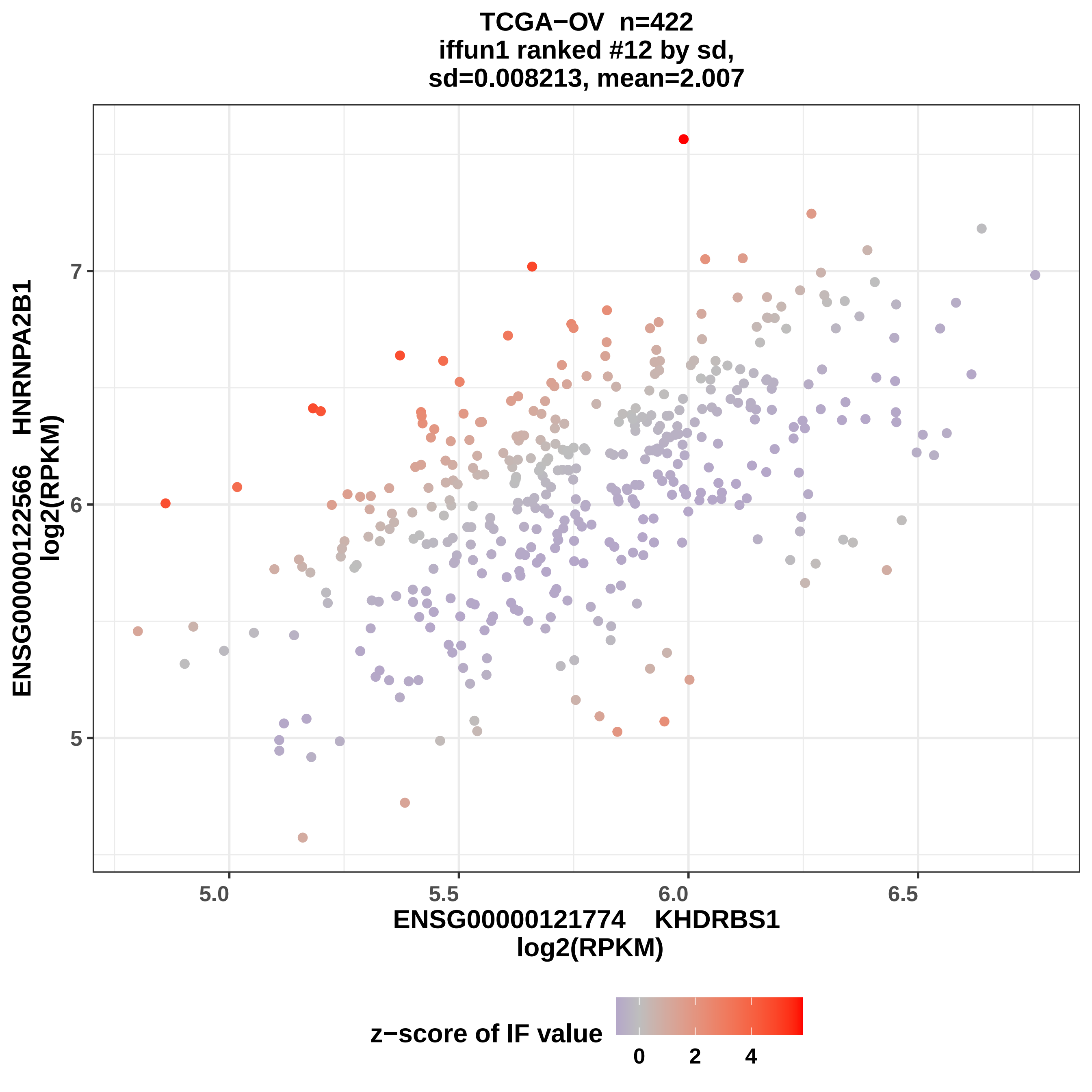
TCGA−OV n=422
iffun1 ranked #12 by sd,
sd=0.008213, mean=2.007
ENSG00000122566 HNRNPA2B1
log2(RPKM)
ENSG00000121774 KHDRBS1
log2(RPKM)
5
6
7
5.0
5.5
6.0
6.5
z−score of IF value
0
2
4

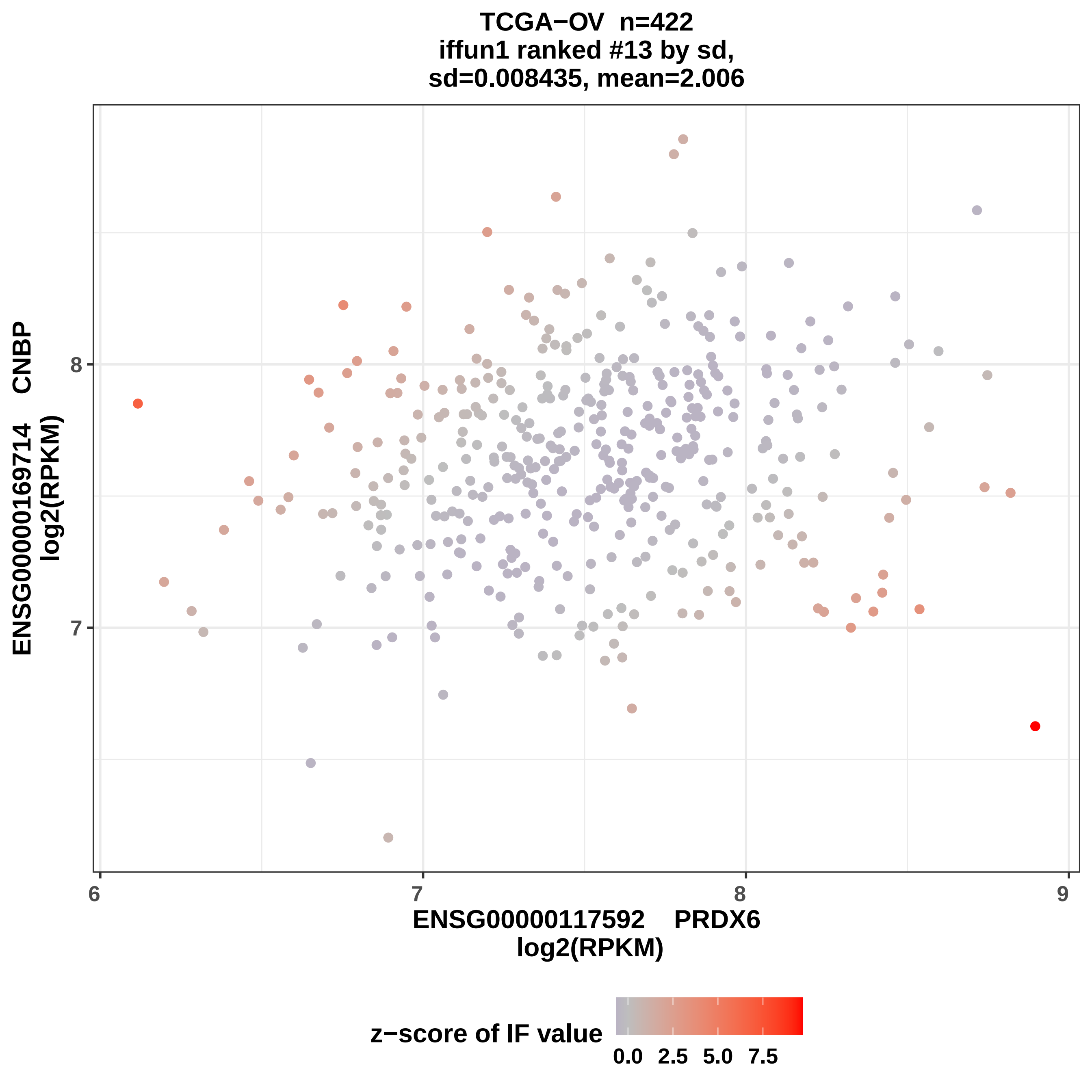
TCGA−OV n=422
iffun1 ranked #13 by sd,
sd=0.008435, mean=2.006
ENSG00000169714 CNBP
log2(RPKM)
ENSG00000117592 PRDX6
log2(RPKM)
6
7
8
9
7
8
z−score of IF value
0.0
2.5
5.0
7.5

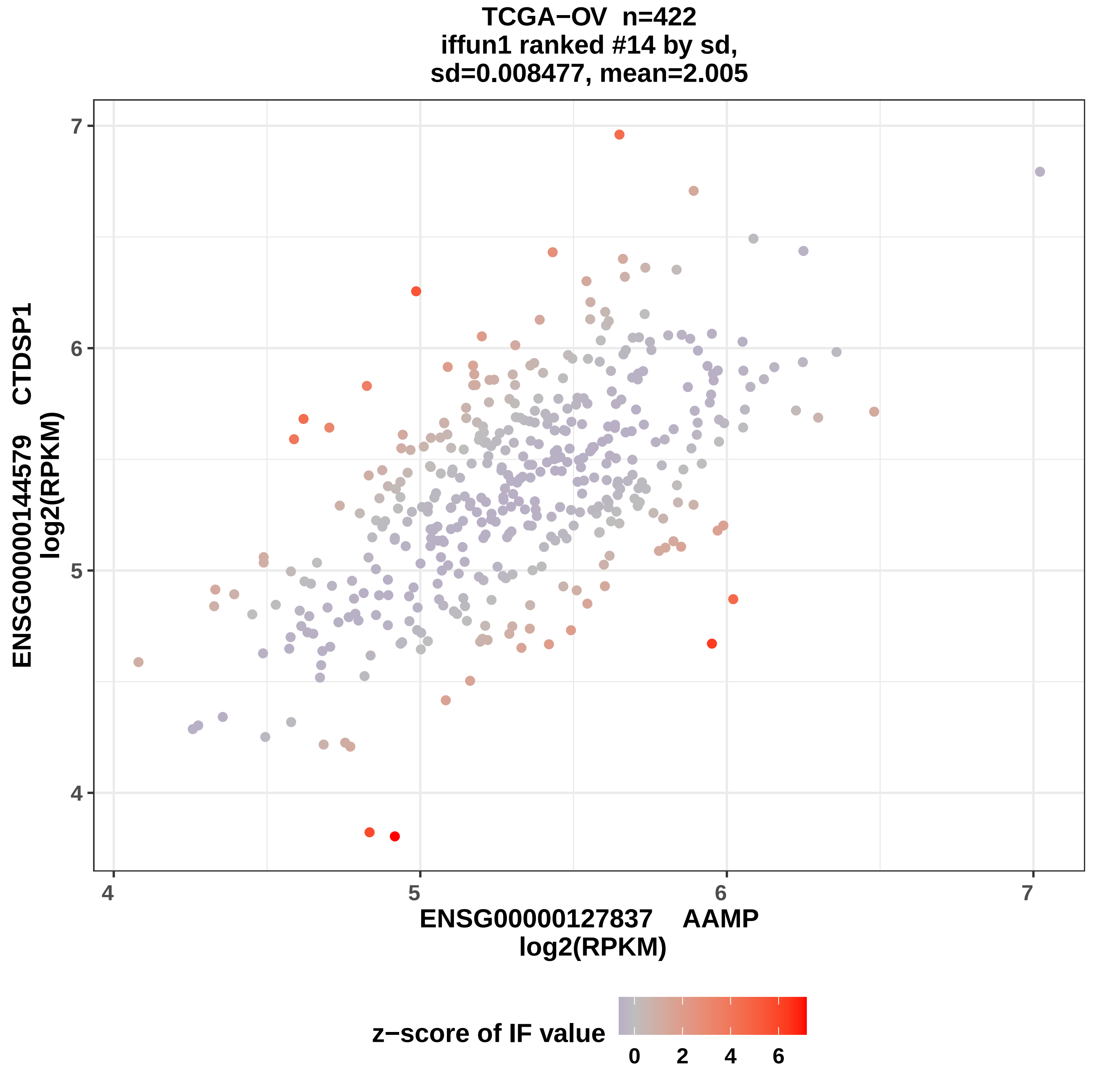
TCGA−OV n=422
iffun1 ranked #14 by sd,
sd=0.008477, mean=2.005
ENSG00000144579 CTDSP1
log2(RPKM)
ENSG00000127837 AAMP
log2(RPKM)
4
5
6
7
4
5
6
7
z−score of IF value
0
2
4
6

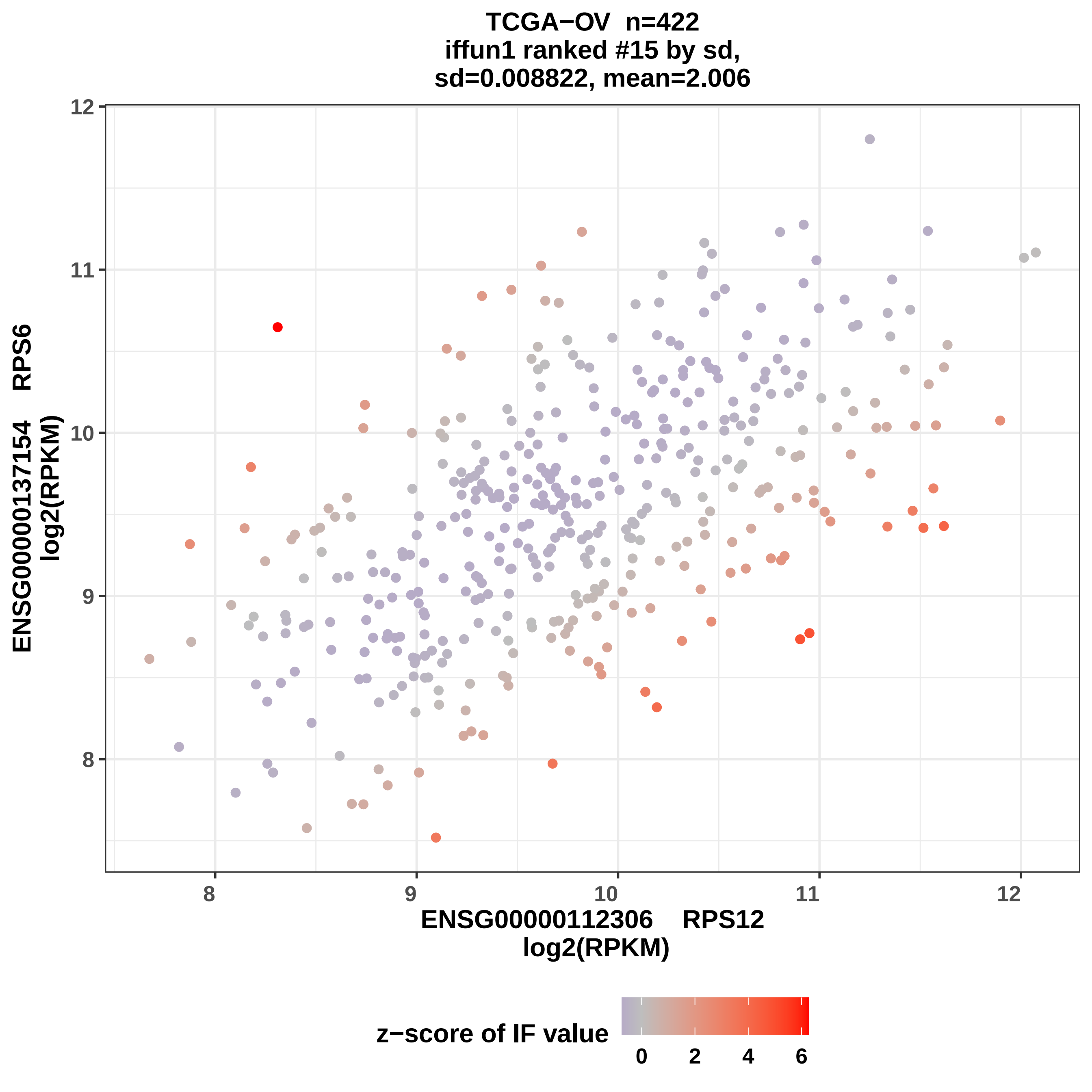

TCGA−OV n=422
iffun1 ranked #15 by sd,
sd=0.008822, mean=2.006
ENSG00000137154 RPS6
log2(RPKM)
ENSG00000112306 RPS12
log2(RPKM)
8
9
10
11
12
z−score of IF value
0
2
4
6

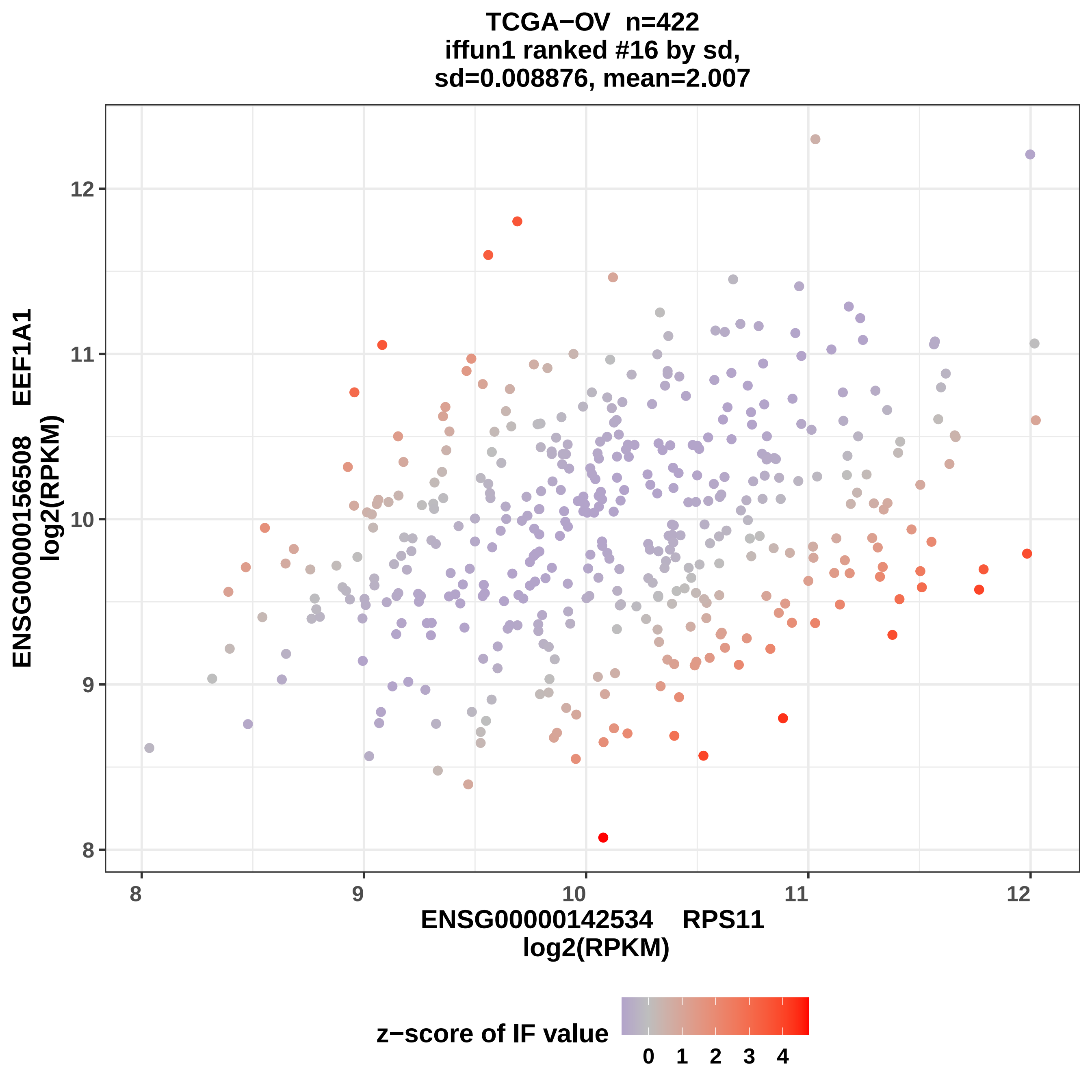

TCGA−OV n=422
iffun1 ranked #16 by sd,
sd=0.008876, mean=2.007
ENSG00000156508 EEF1A1
log2(RPKM)
12
11
10
9
8
ENSG00000142534 RPS11
log2(RPKM)
z−score of IF value
0 1 2 3 4

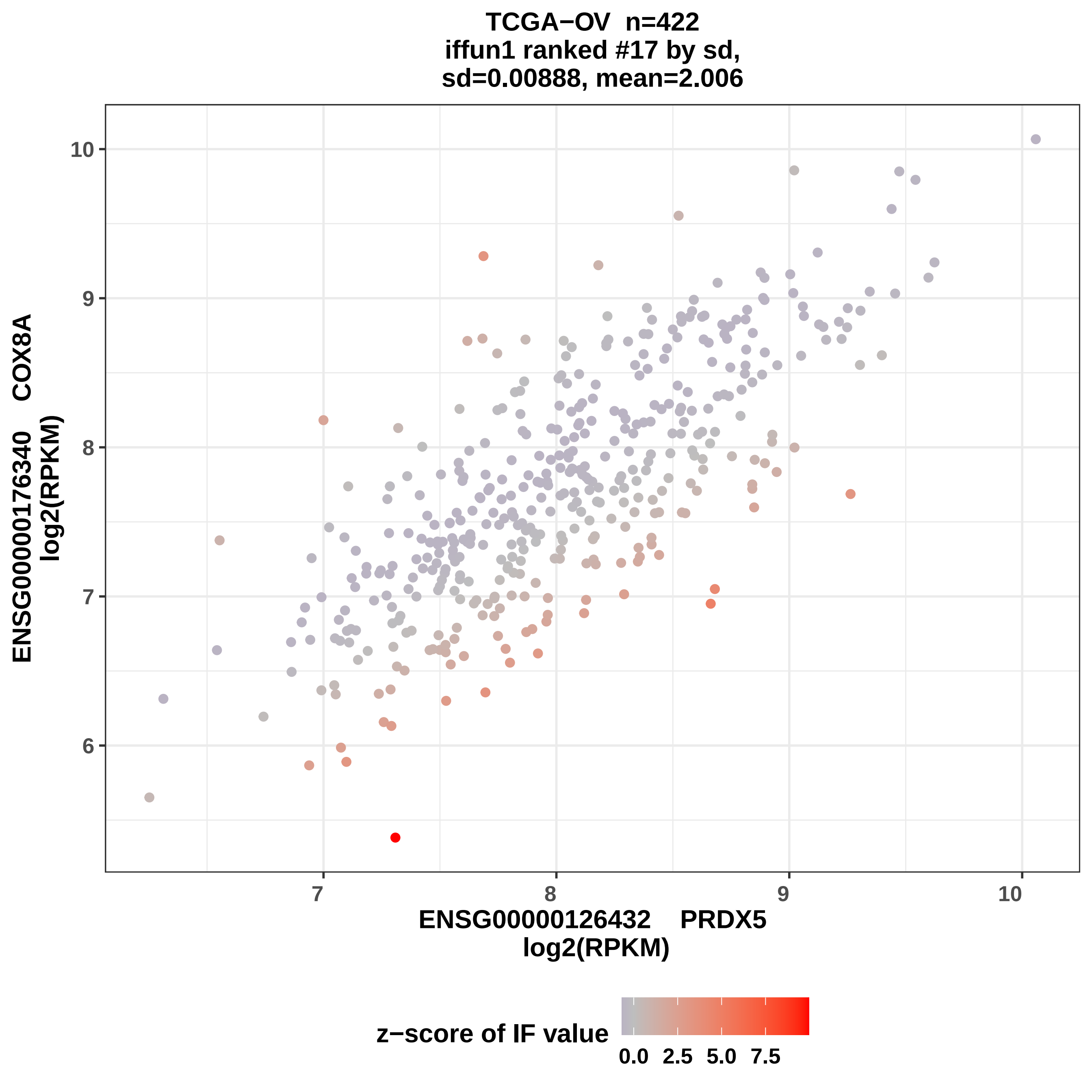

TCGA−OV n=422
iffun1 ranked #17 by sd,
sd=0.00888, mean=2.006
ENSG00000176340 COX8A
log2(RPKM)
10
9
8
7
6
7
8
9
10
ENSG00000126432 PRDX5
log2(RPKM)
z−score of IF value
0.0
2.5
5.0
7.5

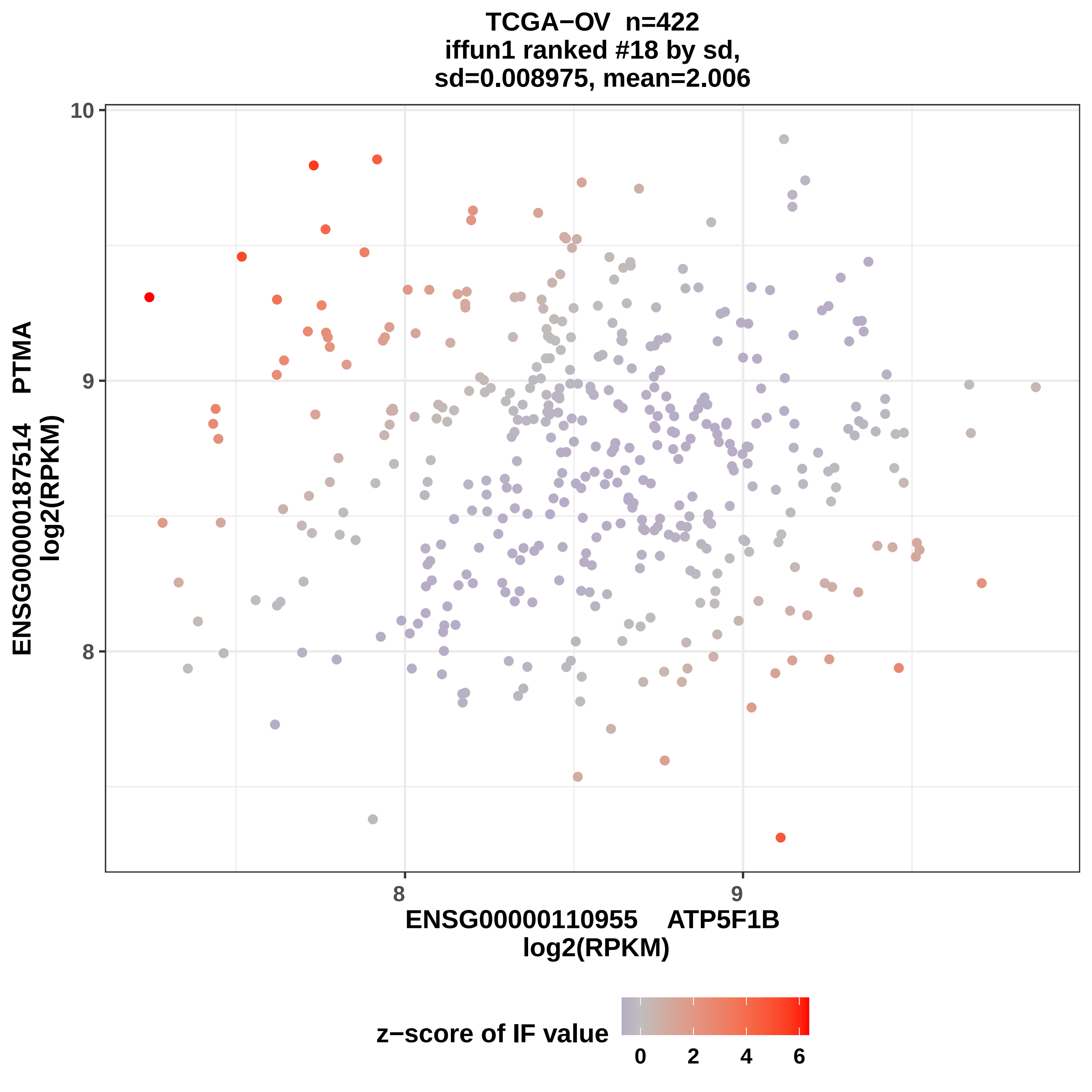
TCGA−OV n=422
iffun1 ranked #18 by sd,
sd=0.008975, mean=2.006
10
9
8
8
9
ENSG00000187514 PTMA
log2(RPKM)
ENSG00000110955 ATP5F1B
log2(RPKM)
z−score of IF value
0
2
4
6

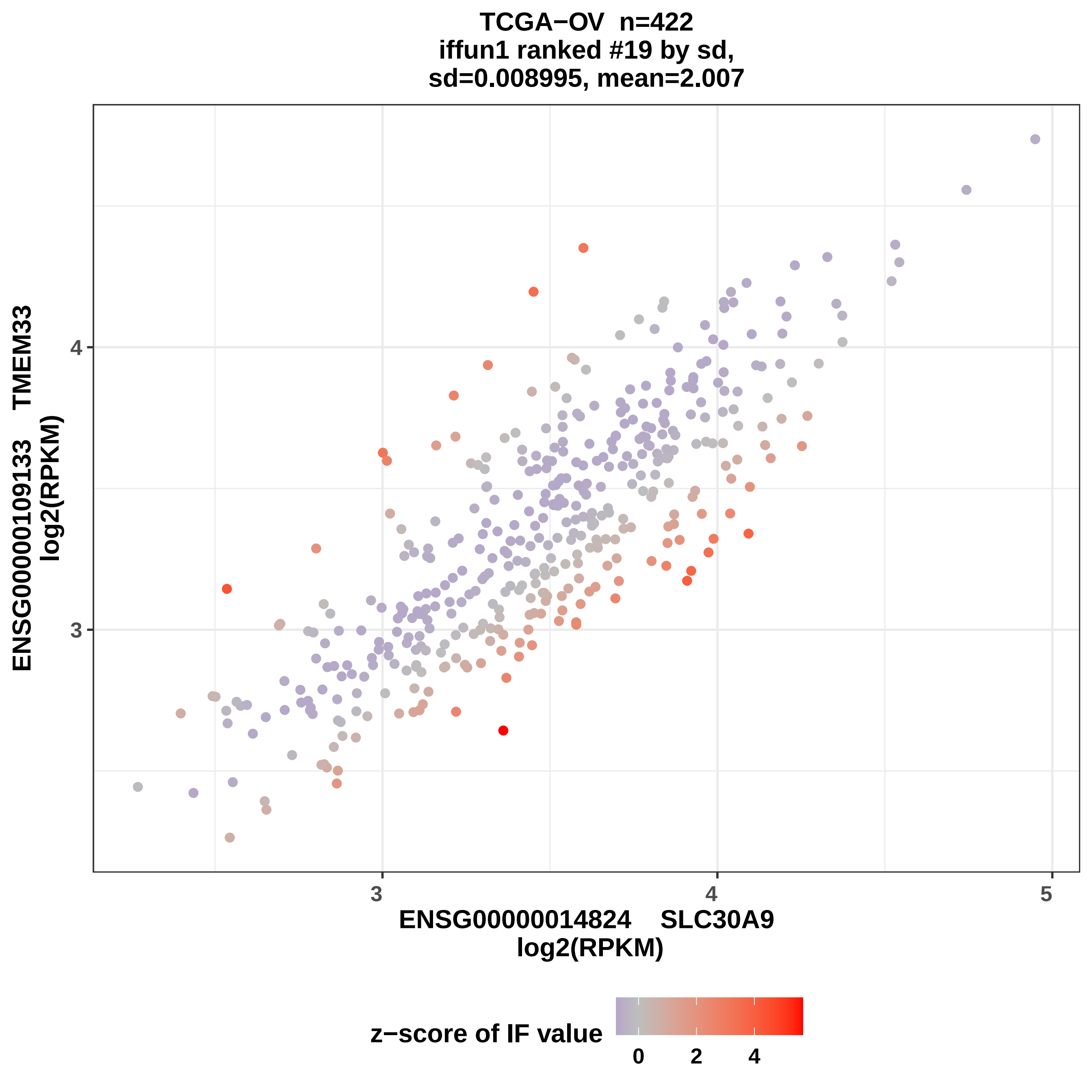

TCGA−OV n=422
iffun1 ranked #19 by sd,
sd=0.008995, mean=2.007
ENSG00000109133 TMEM33
log2(RPKM)
4
3
3
4
5
ENSG00000014824 SLC30A9
log2(RPKM)
z−score of IF value
0
2
4

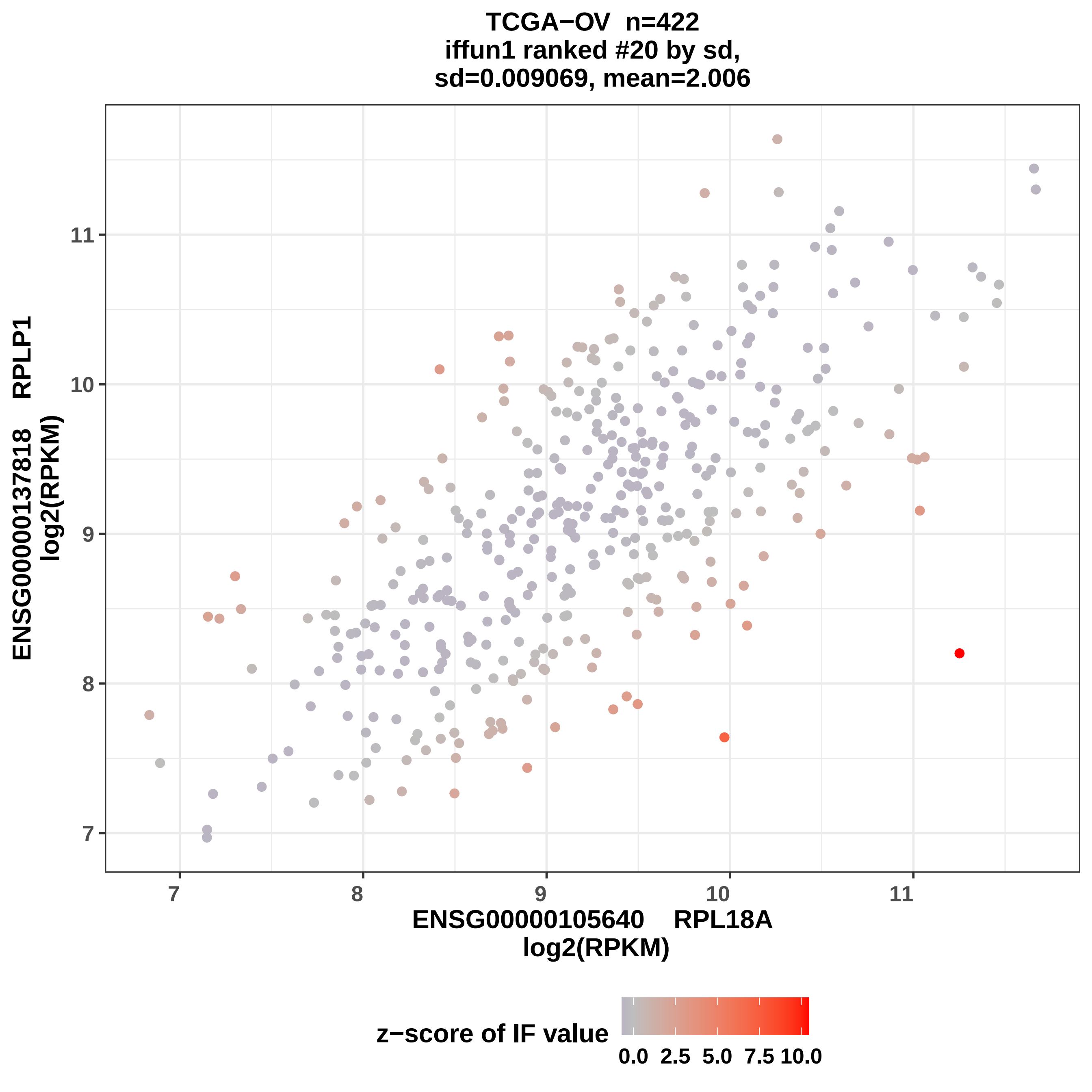
TCGA−OV n=422
iffun1 ranked #20 by sd,
sd=0.009069, mean=2.006
ENSG00000137818 RPLP1
log2(RPKM)
11
10
9
8
7
7
8
9
10
11
ENSG00000105640 RPL18A
log2(RPKM)
z−score of IF value
0.0
2.5
5.0
7.5
10.0

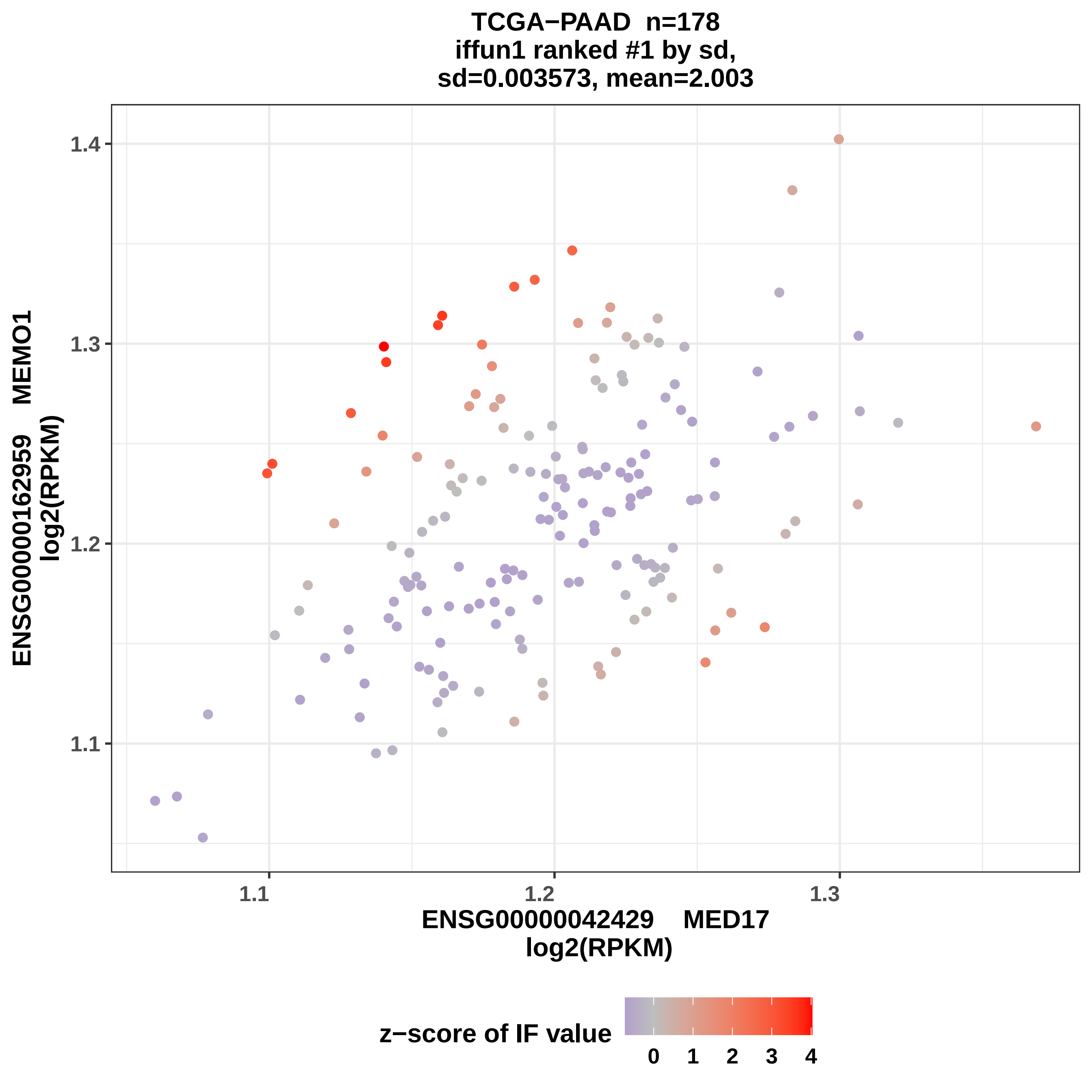

TCGA−PAAD n=178
iffun1 ranked #1 by sd,
sd=0.003573, mean=2.003
ENSG00000162959 MEMO1
log2(RPKM)
1.4
1.3
1.2
1.1
1.1
1.2
1.3
ENSG00000042429 MED17
log2(RPKM)
z−score of IF value
0
1
2
3
4

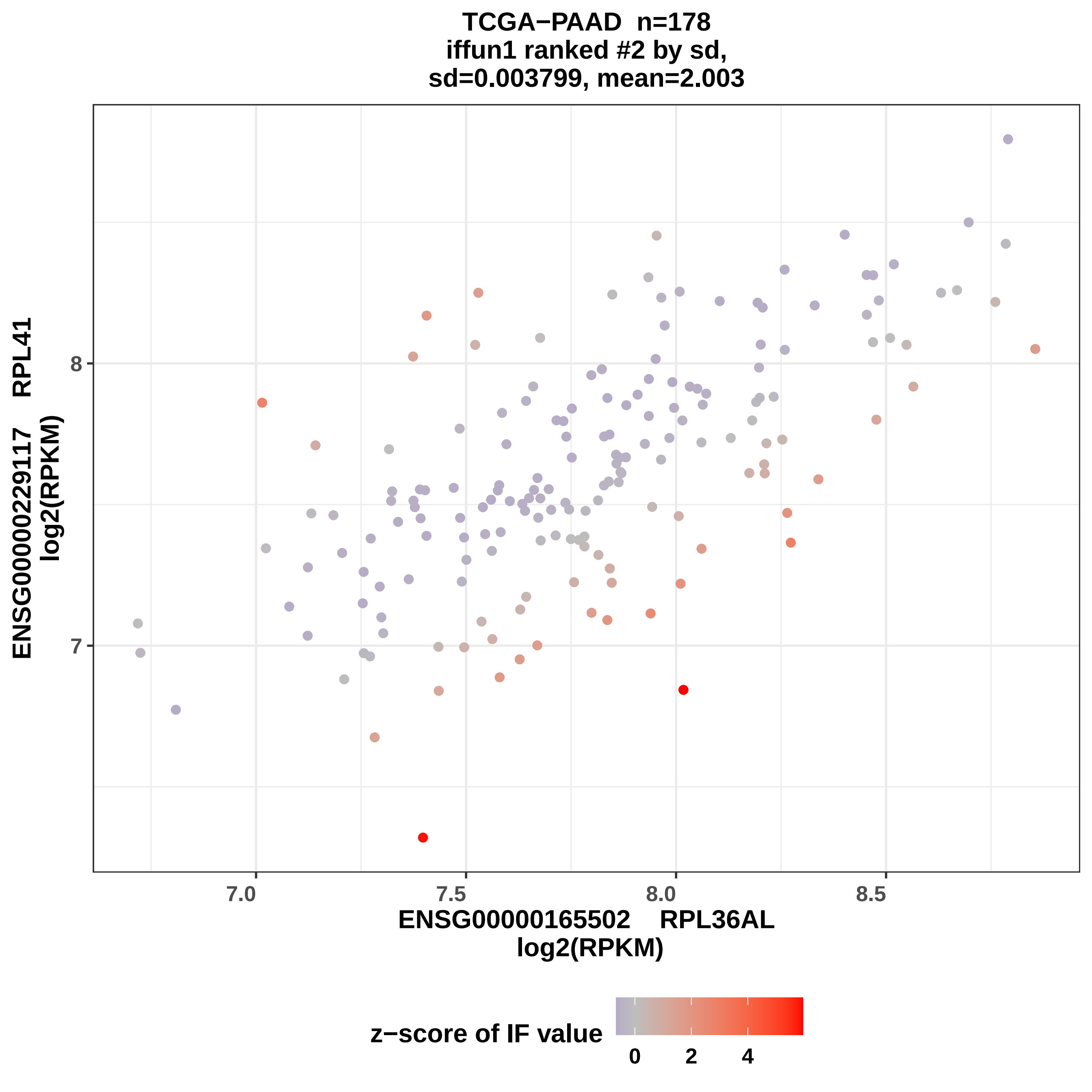

TCGA−PAAD n=178
iffun1 ranked #2 by sd,
sd=0.003799, mean=2.003
ENSG00000229117 RPL41
log2(RPKM)
8
7
7.0
7.5
8.0
8.5
ENSG00000165502 RPL36AL
log2(RPKM)
z−score of IF value
0
2
4

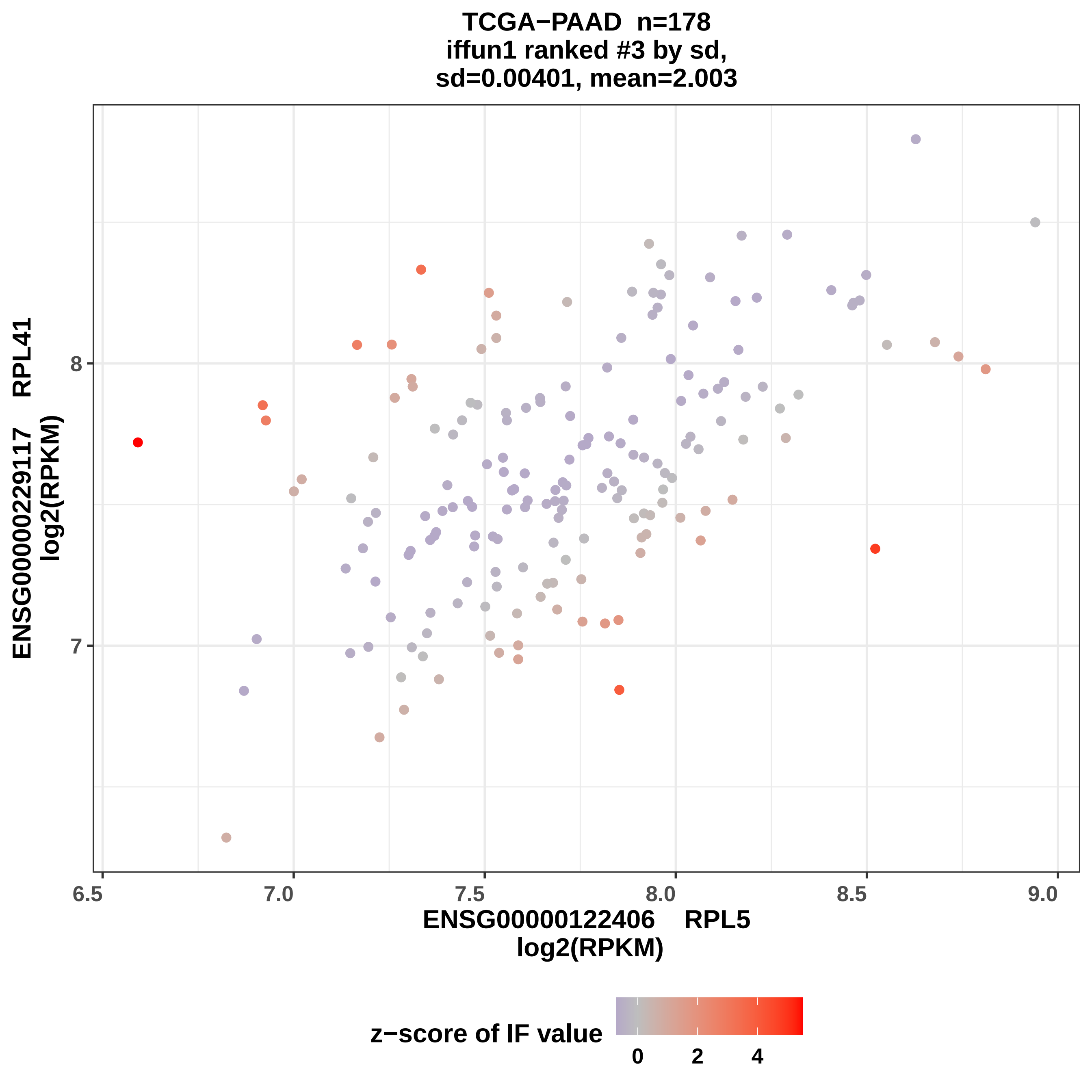

TCGA−PAAD n=178
iffun1 ranked #3 by sd,
sd=0.00401, mean=2.003
ENSG00000229117 RPL41
log2(RPKM)
ENSG00000122406 RPL5
log2(RPKM)
z−score of IF value
0
2
4
6.5
7.0
7.5
8.0
8.5
9.0
7
8

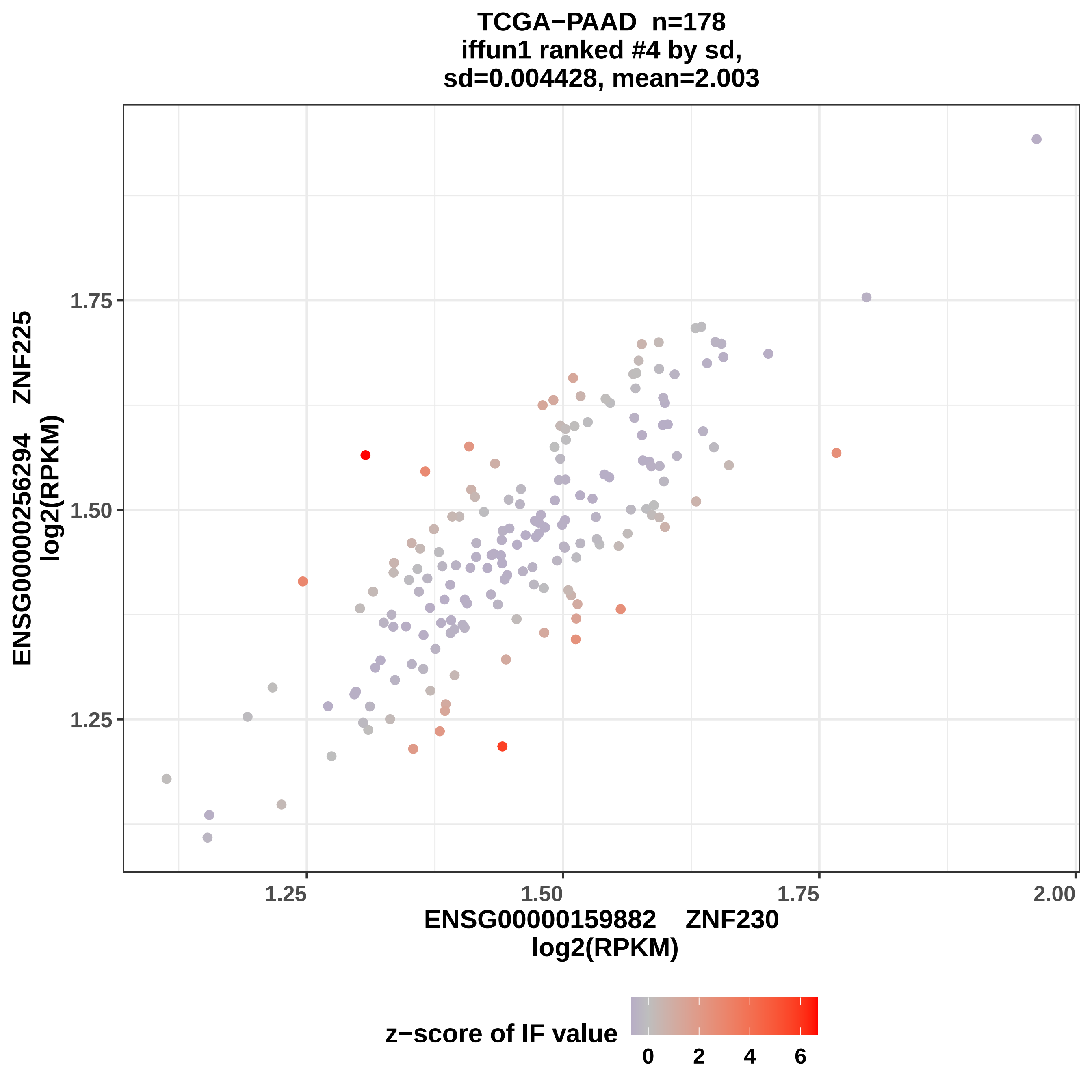

TCGA−PAAD n=178
iffun1 ranked #4 by sd,
sd=0.004428, mean=2.003
ENSG00000256294 ZNF225
log2(RPKM)
1.75
1.50
1.25
1.25
1.50
1.75
2.00
ENSG00000159882 ZNF230
log2(RPKM)
z−score of IF value
0
2
4
6

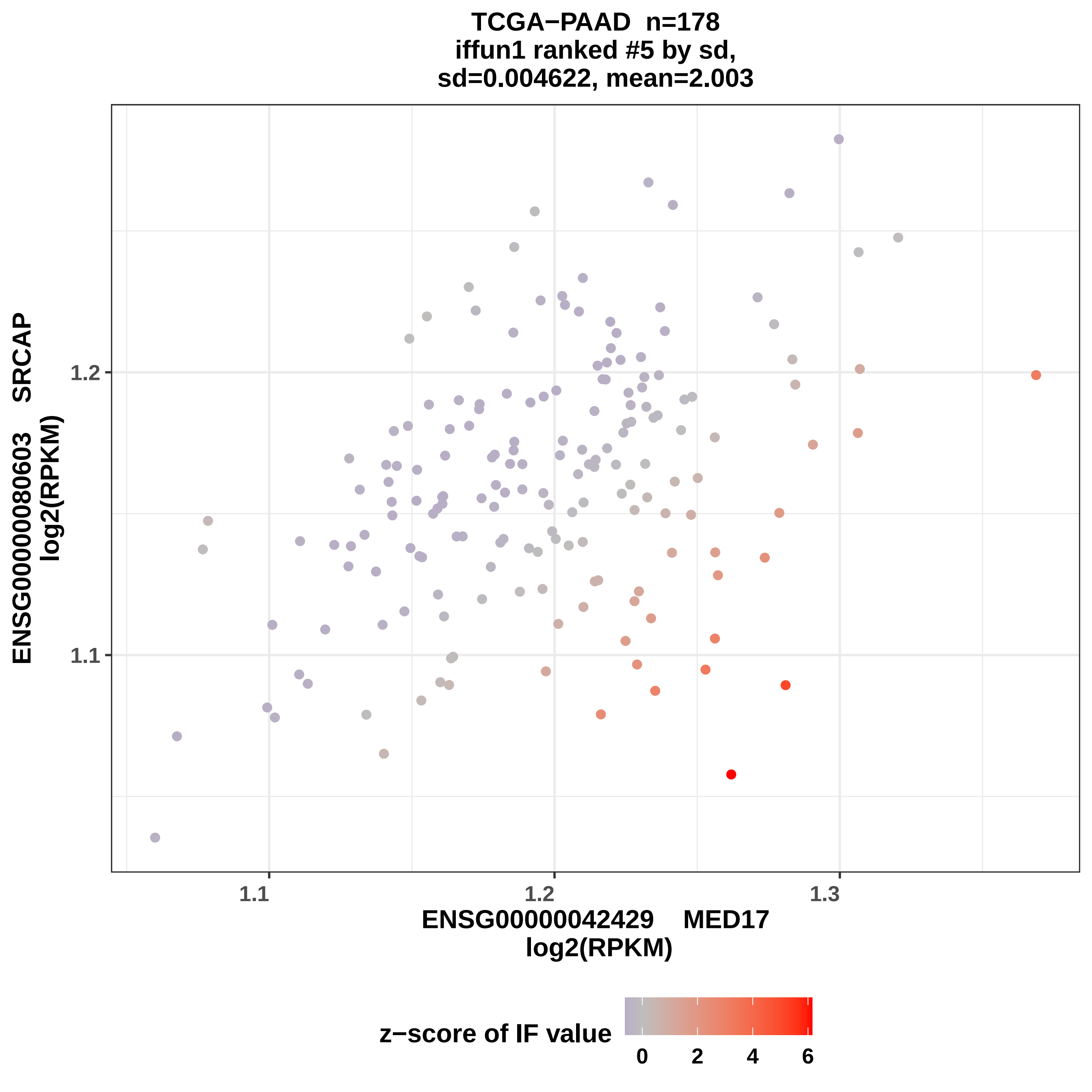

TCGA−PAAD n=178
iffun1 ranked #5 by sd,
sd=0.004622, mean=2.003
ENSG00000080603 SRCAP
log2(RPKM)
1.2
1.1
1.1
1.2
1.3
ENSG00000042429 MED17
log2(RPKM)
z−score of IF value
0
2
4
6

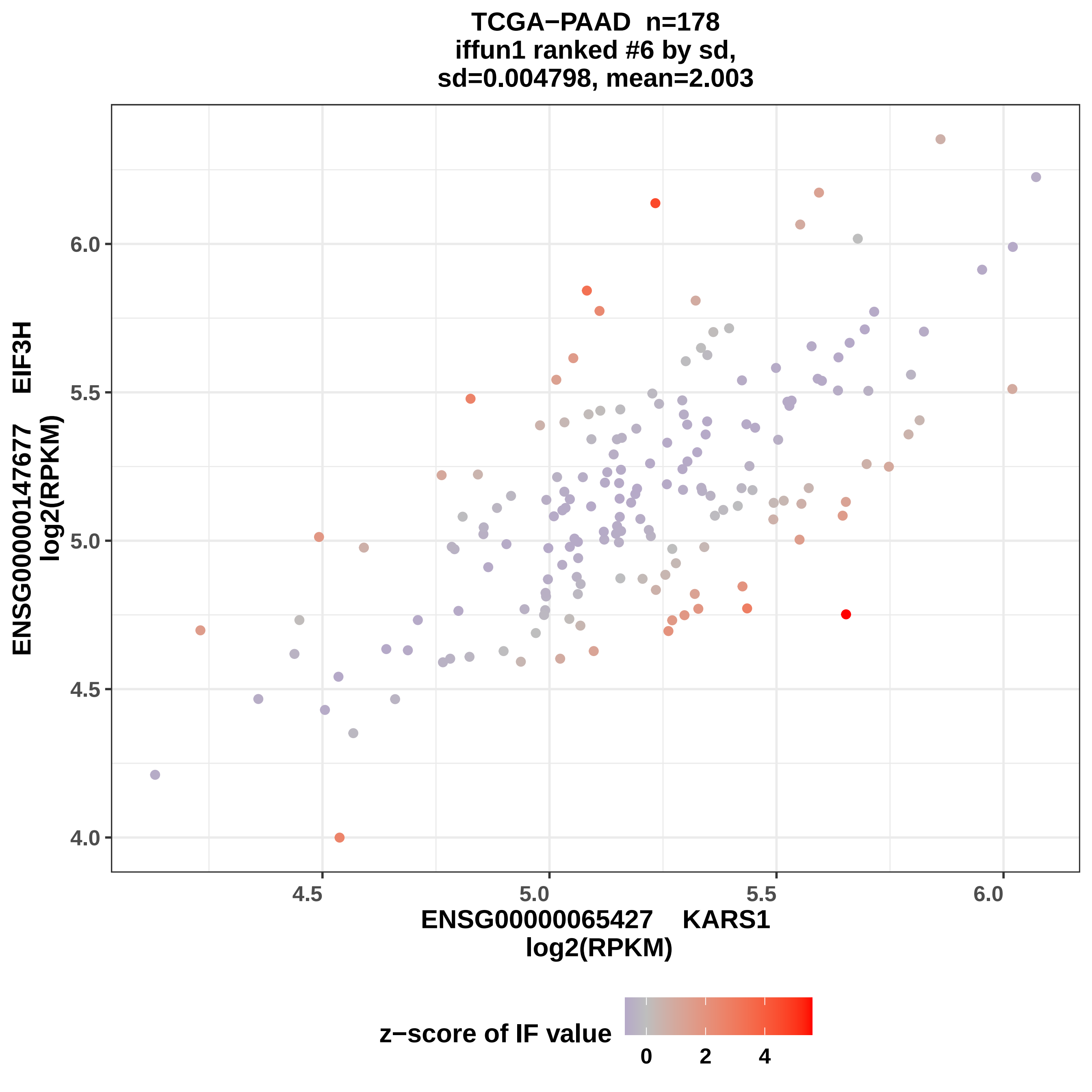
TCGA−PAAD n=178
iffun1 ranked #6 by sd,
sd=0.004798, mean=2.003
ENSG00000147677 EIF3H
log2(RPKM)
6.0
5.5
5.0
4.5
4.0
4.5
5.0
5.5
6.0
ENSG00000065427 KARS1
log2(RPKM)
z−score of IF value
0
2
4

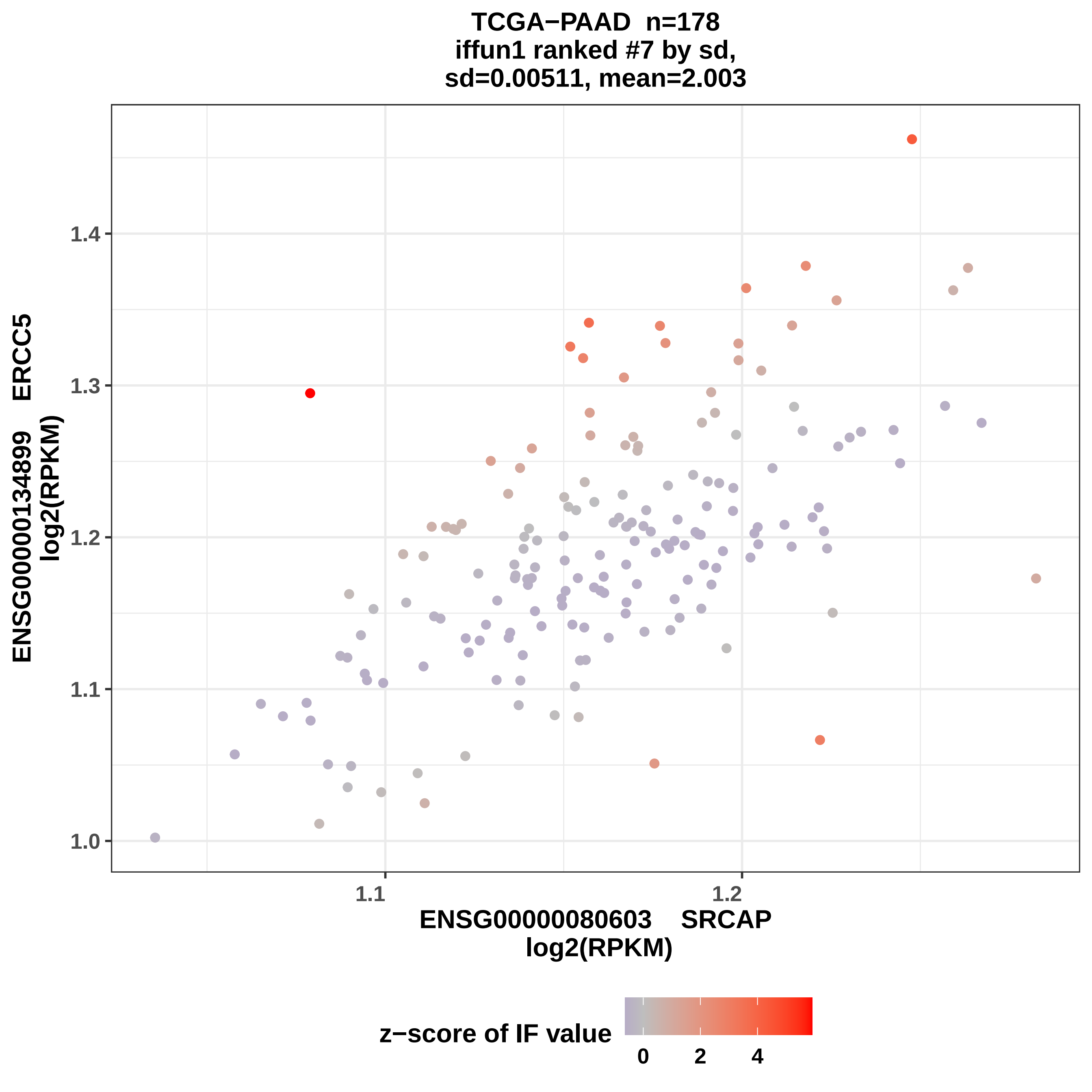

TCGA−PAAD n=178
iffun1 ranked #7 by sd,
sd=0.00511, mean=2.003
ENSG00000134899 ERCC5
log2(RPKM)
1.0
1.1
1.2
1.3
1.4
1.1
1.2
ENSG00000080603 SRCAP
log2(RPKM)
z−score of IF value
0
2
4

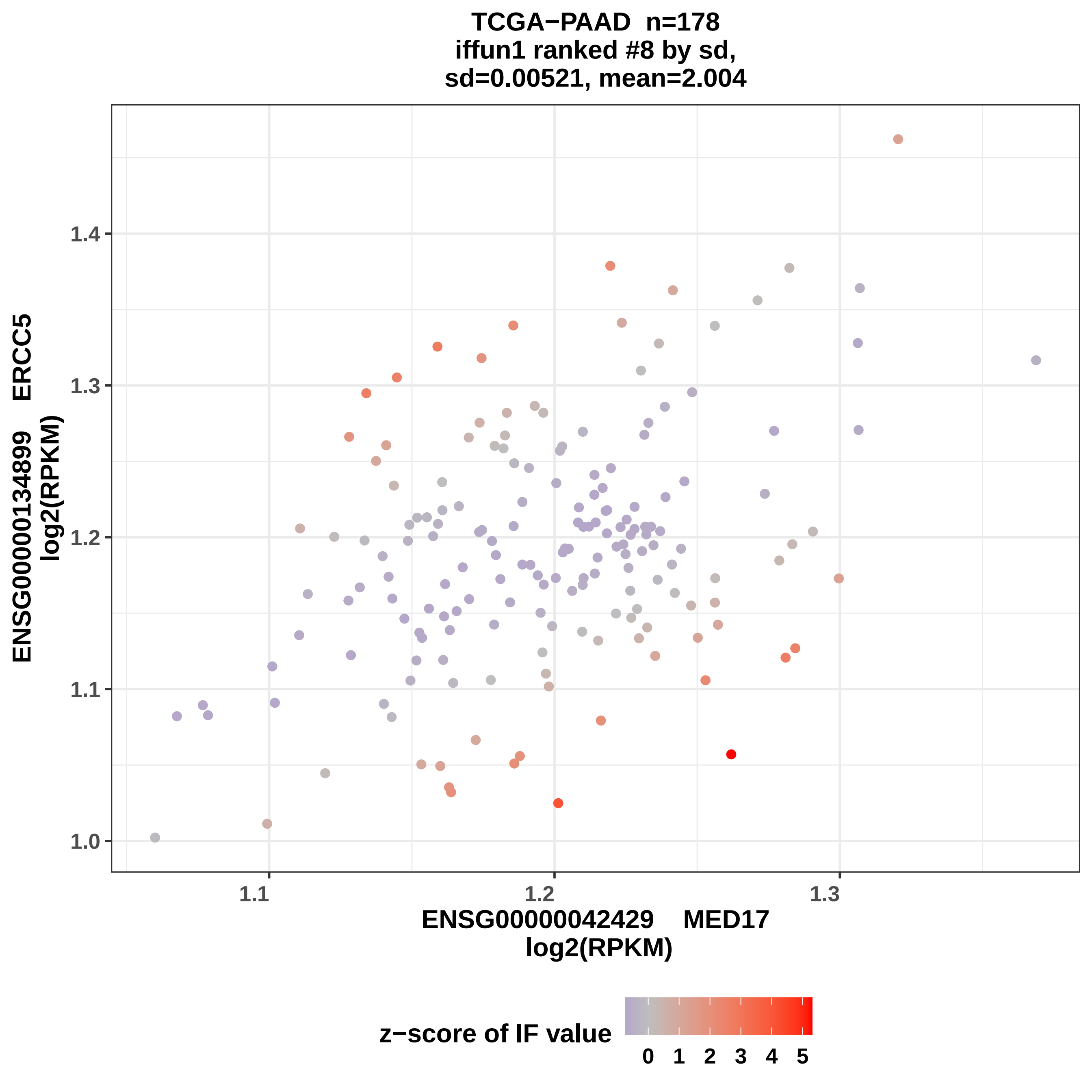
TCGA−PAAD n=178
iffun1 ranked #8 by sd,
sd=0.00521, mean=2.004
ENSG00000134899 ERCC5
log2(RPKM)
1.4
1.3
1.2
1.1
1.0
1.1
1.2
1.3
ENSG00000042429 MED17
log2(RPKM)
z−score of IF value
0 1 2 3 4 5

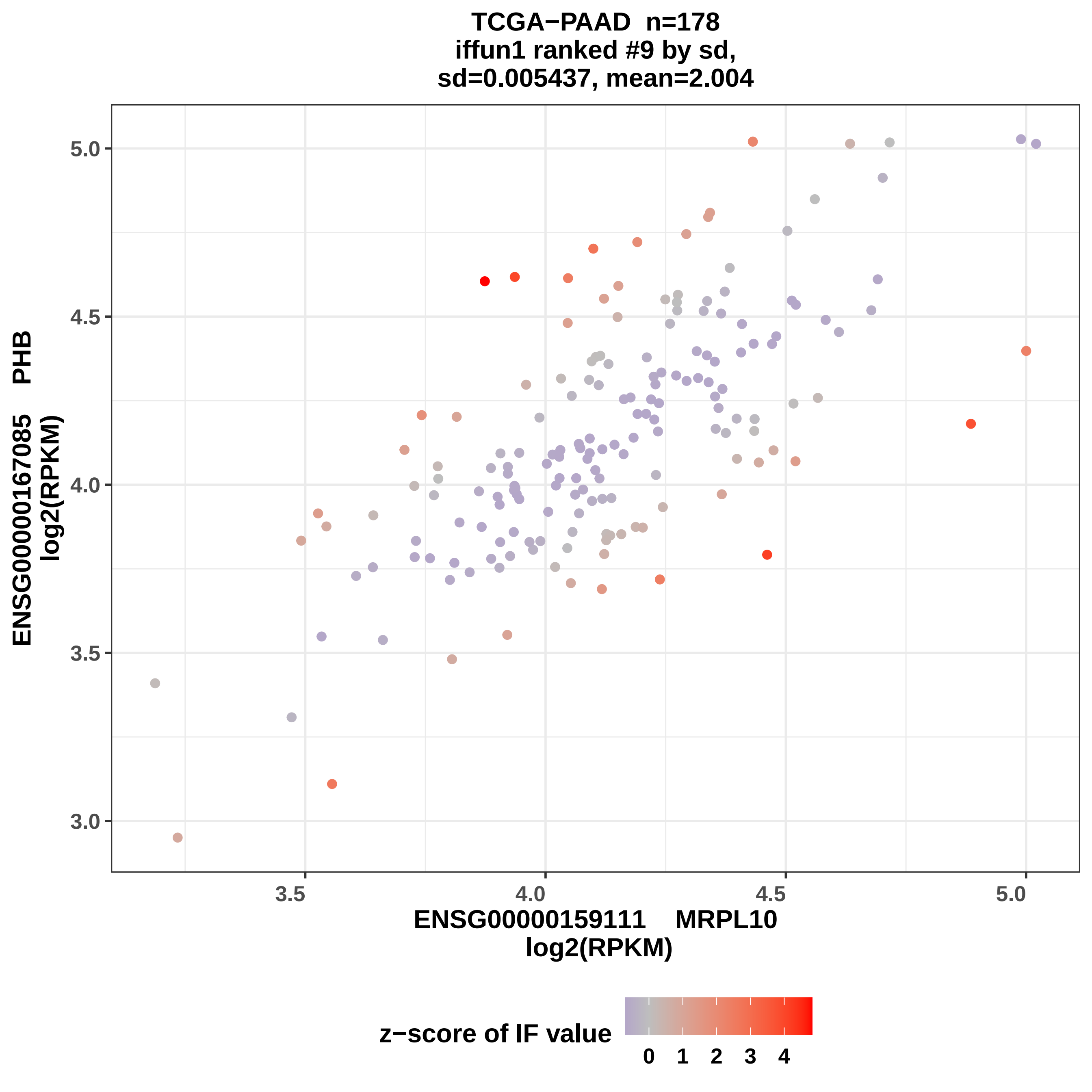
TCGA−PAAD n=178
iffun1 ranked #9 by sd,
sd=0.005437, mean=2.004
ENSG00000167085 PHB
log2(RPKM)
ENSG00000159111 MRPL10
log2(RPKM)
5.0
4.5
4.0
3.5
3.0
3.5
4.0
4.5
5.0
z−score of IF value
0 1 2 3 4

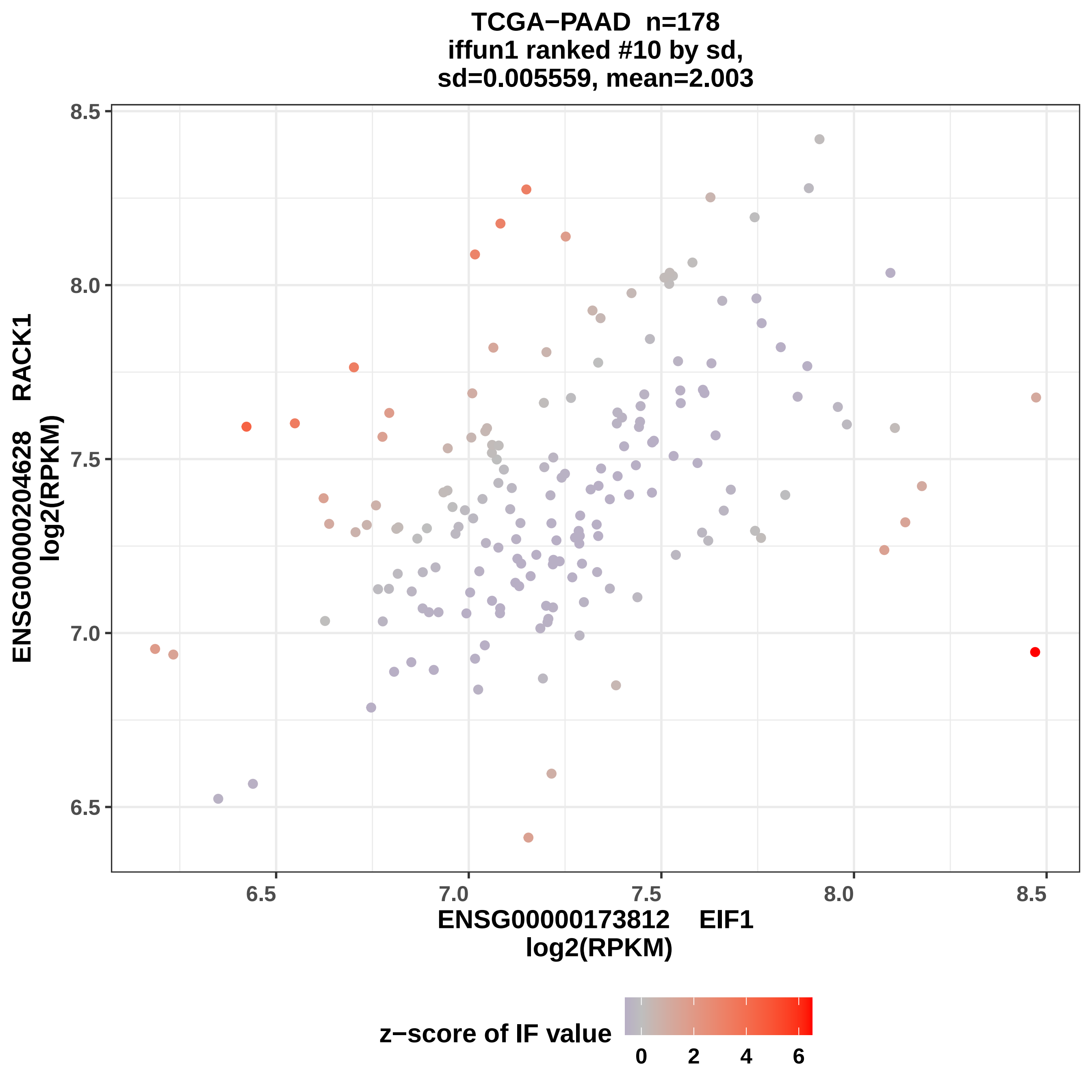

TCGA−PAAD n=178
iffun1 ranked #10 by sd,
sd=0.005559, mean=2.003
ENSG00000204628 RACK1
log2(RPKM)
ENSG00000173812 EIF1
log2(RPKM)
8.5
8.0
7.5
7.0
6.5
6.5
7.0
7.5
8.0
8.5
z−score of IF value
0
2
4
6

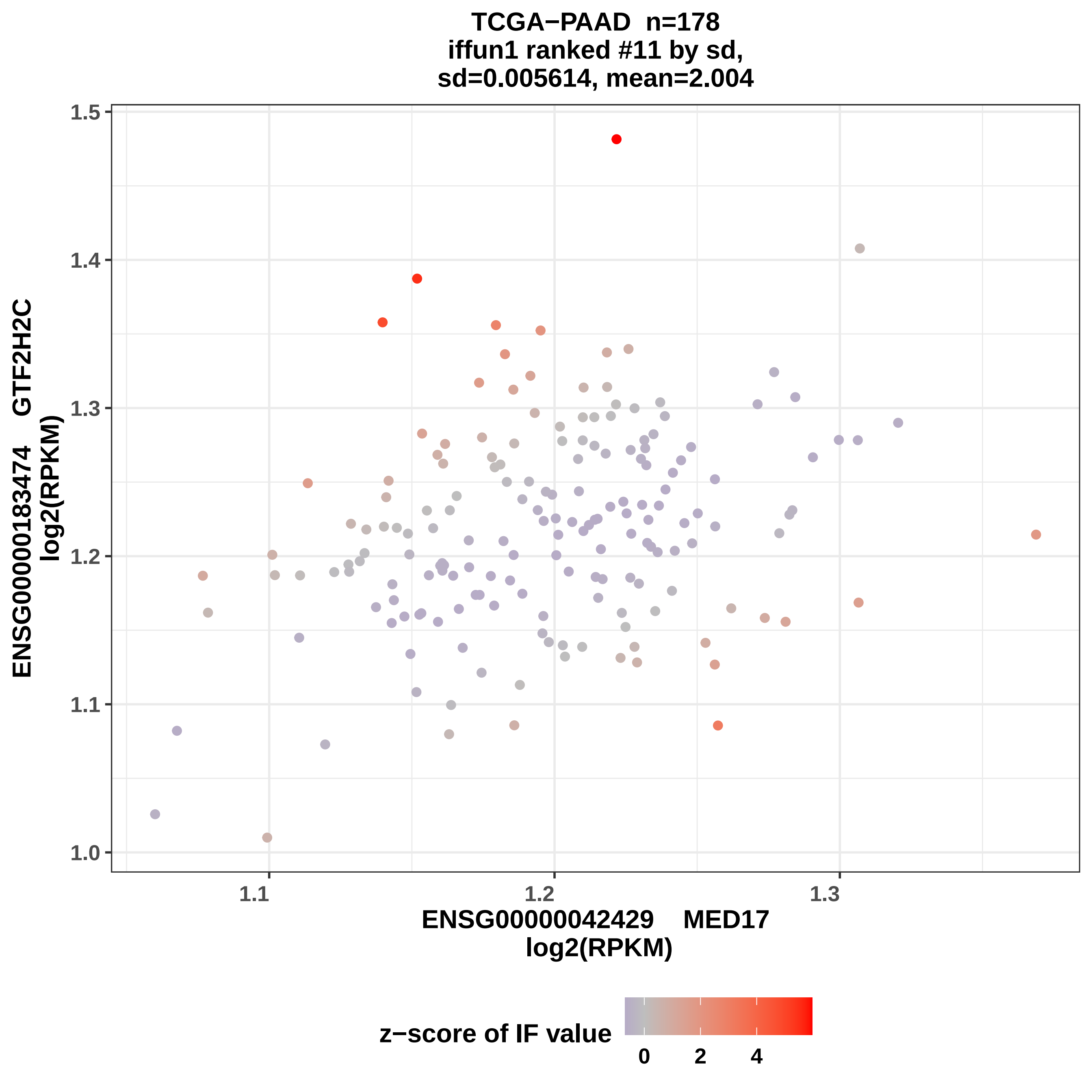
TCGA−PAAD n=178
iffun1 ranked #11 by sd,
sd=0.005614, mean=2.004
ENSG00000183474 GTF2H2C
log2(RPKM)
ENSG00000042429 MED17
log2(RPKM)
z−score of IF value
1.0
1.1
1.2
1.3
1.4
1.5
0
2
4

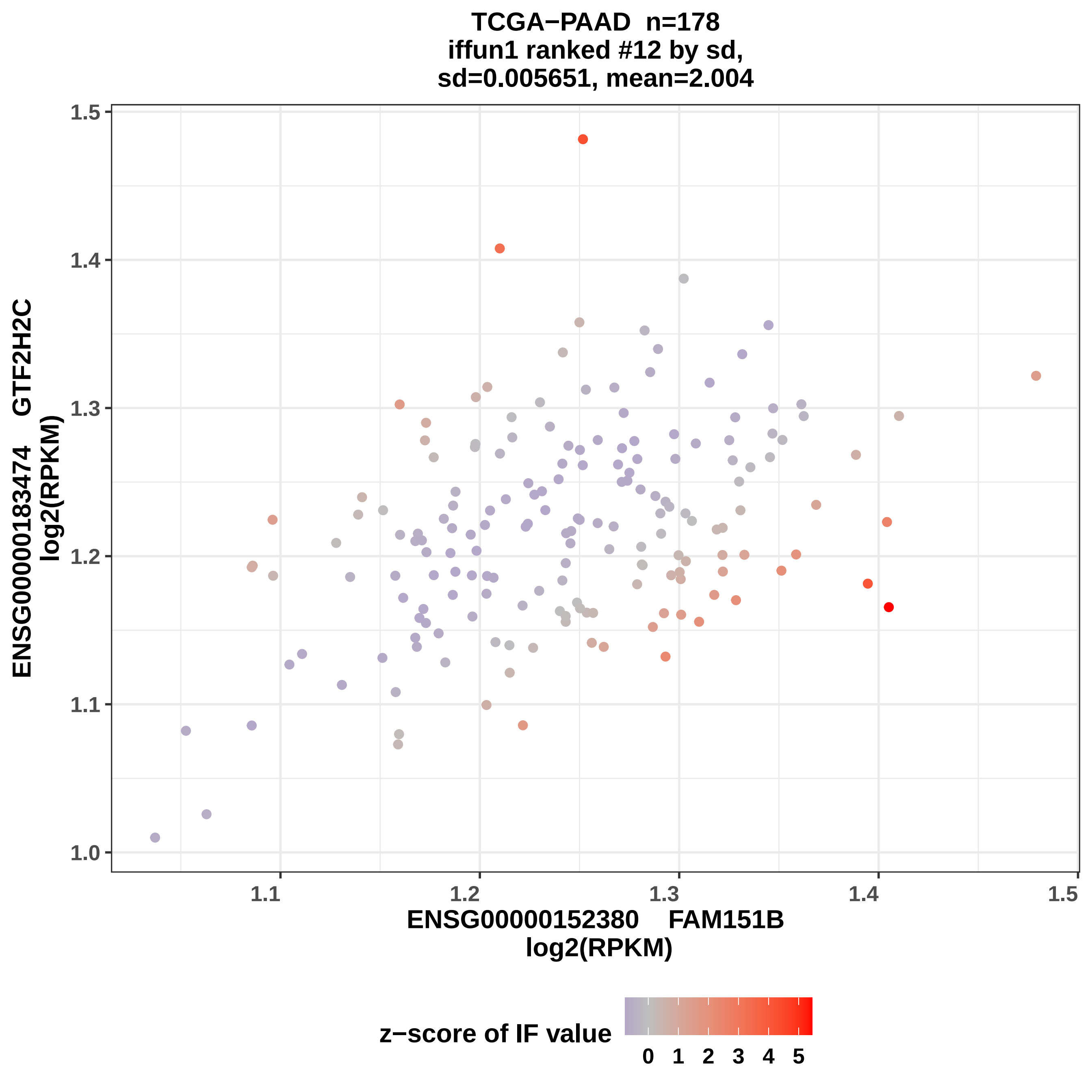

TCGA−PAAD n=178
iffun1 ranked #12 by sd,
sd=0.005651, mean=2.004
ENSG00000183474 GTF2H2C
log2(RPKM)
1.0
1.1
1.2
1.3
1.4
1.5
ENSG00000152380 FAM151B
log2(RPKM)
z−score of IF value
0 1 2 3 4 5

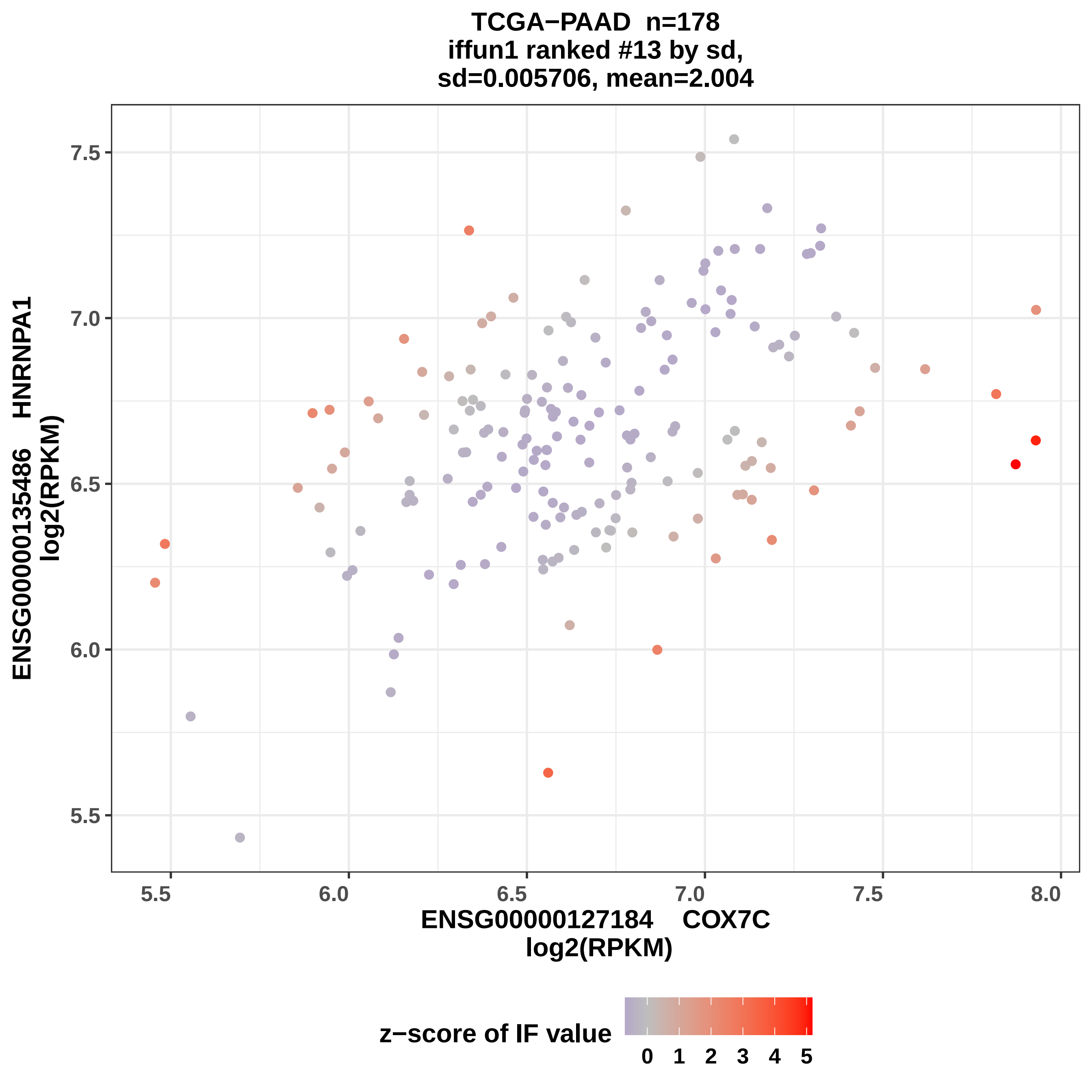
TCGA−PAAD n=178
iffun1 ranked #13 by sd,
sd=0.005706, mean=2.004
ENSG00000135486 HNRNPA1
log2(RPKM)
7.5
7.0
6.5
6.0
5.5
5.5
6.0
6.5
7.0
7.5
8.0
ENSG00000127184 COX7C
log2(RPKM)
z−score of IF value
0
1
2
3
4
5

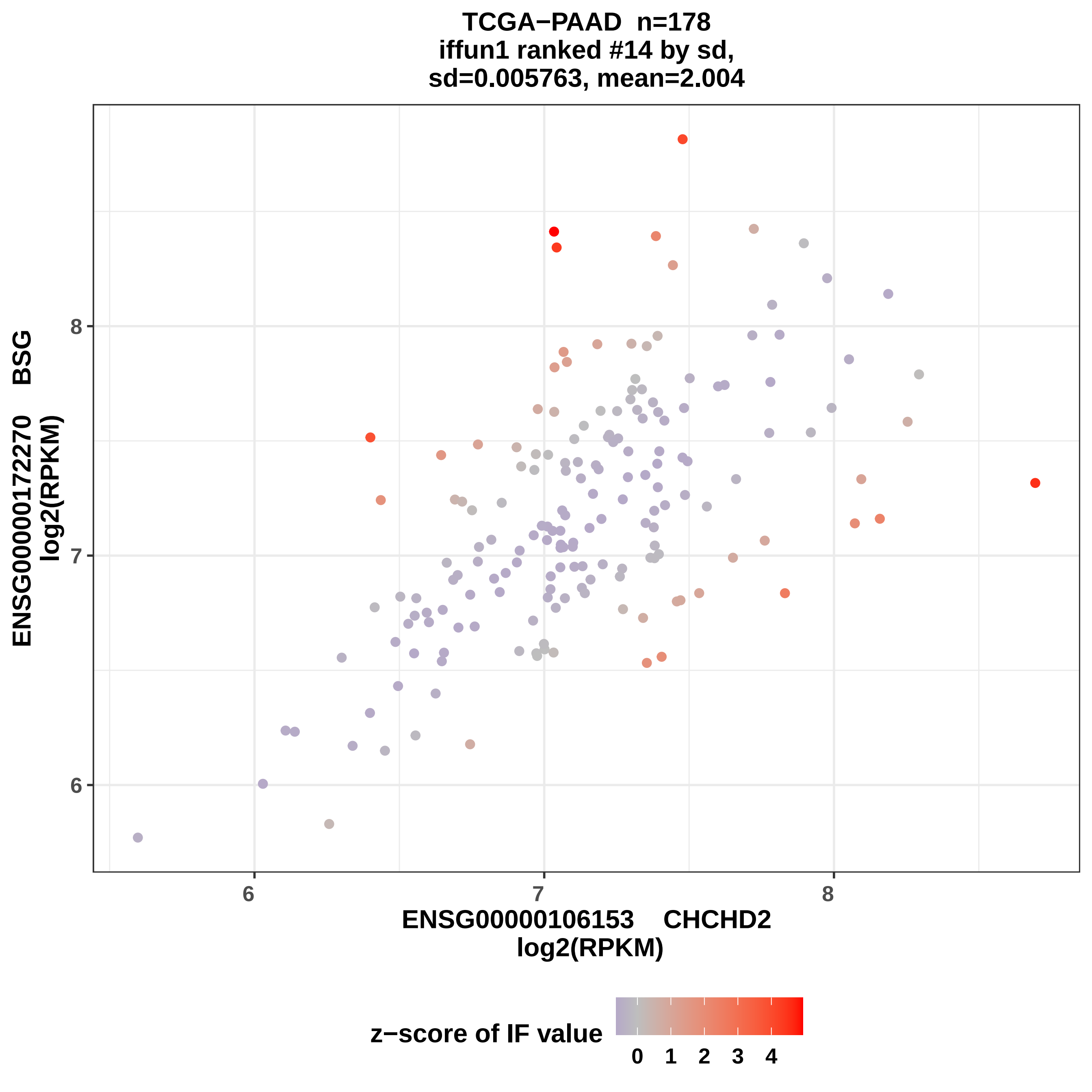
TCGA−PAAD n=178
iffun1 ranked #14 by sd,
sd=0.005763, mean=2.004
ENSG00000172270 BSG
log2(RPKM)
8
7
6
ENSG00000106153 CHCHD2
log2(RPKM)
z−score of IF value
0 1 2 3 4

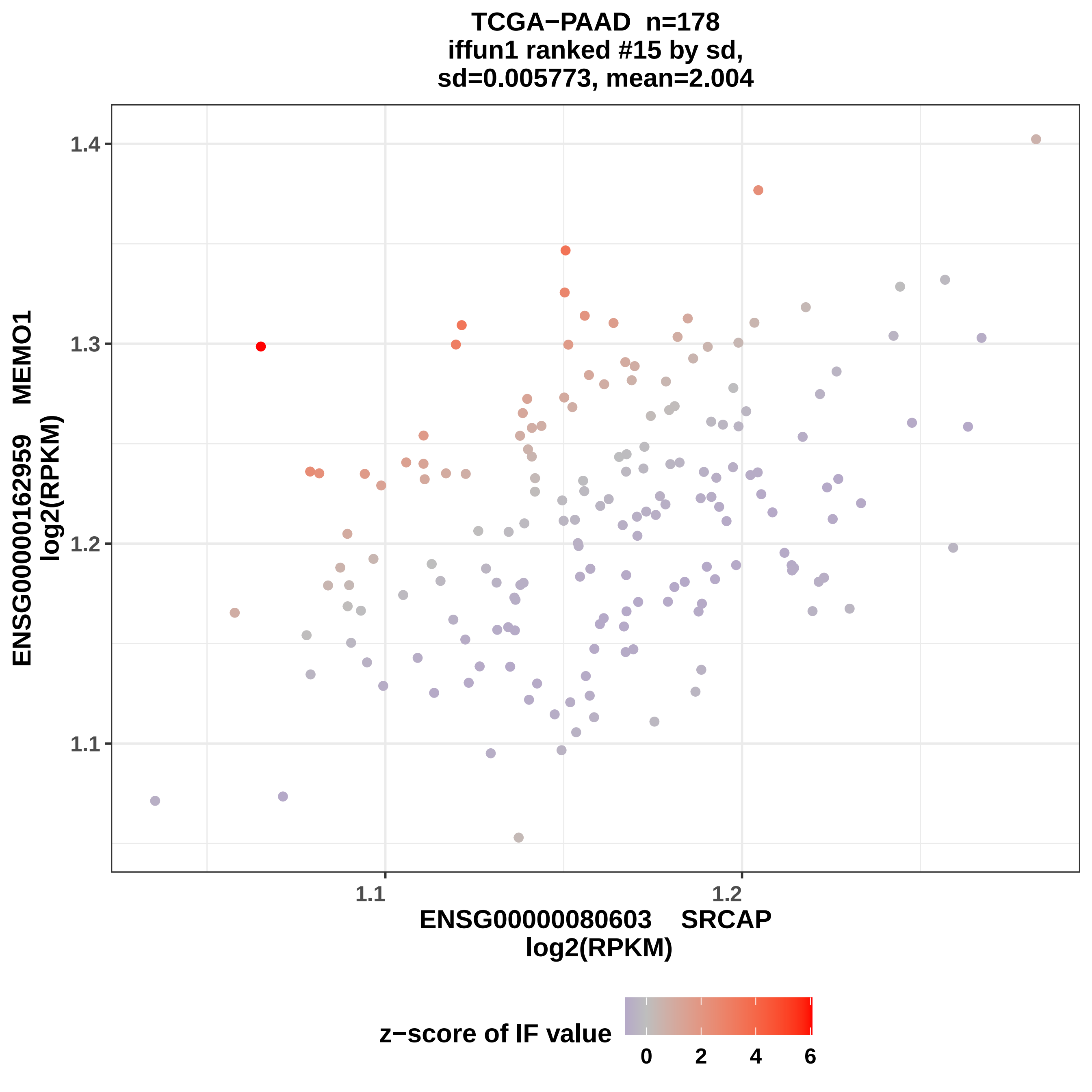

TCGA−PAAD n=178
iffun1 ranked #15 by sd,
sd=0.005773, mean=2.004
ENSG00000162959 MEMO1
log2(RPKM)
1.4
1.3
1.2
1.1
1.1
1.2
ENSG00000080603 SRCAP
log2(RPKM)
z−score of IF value
0
2
4
6

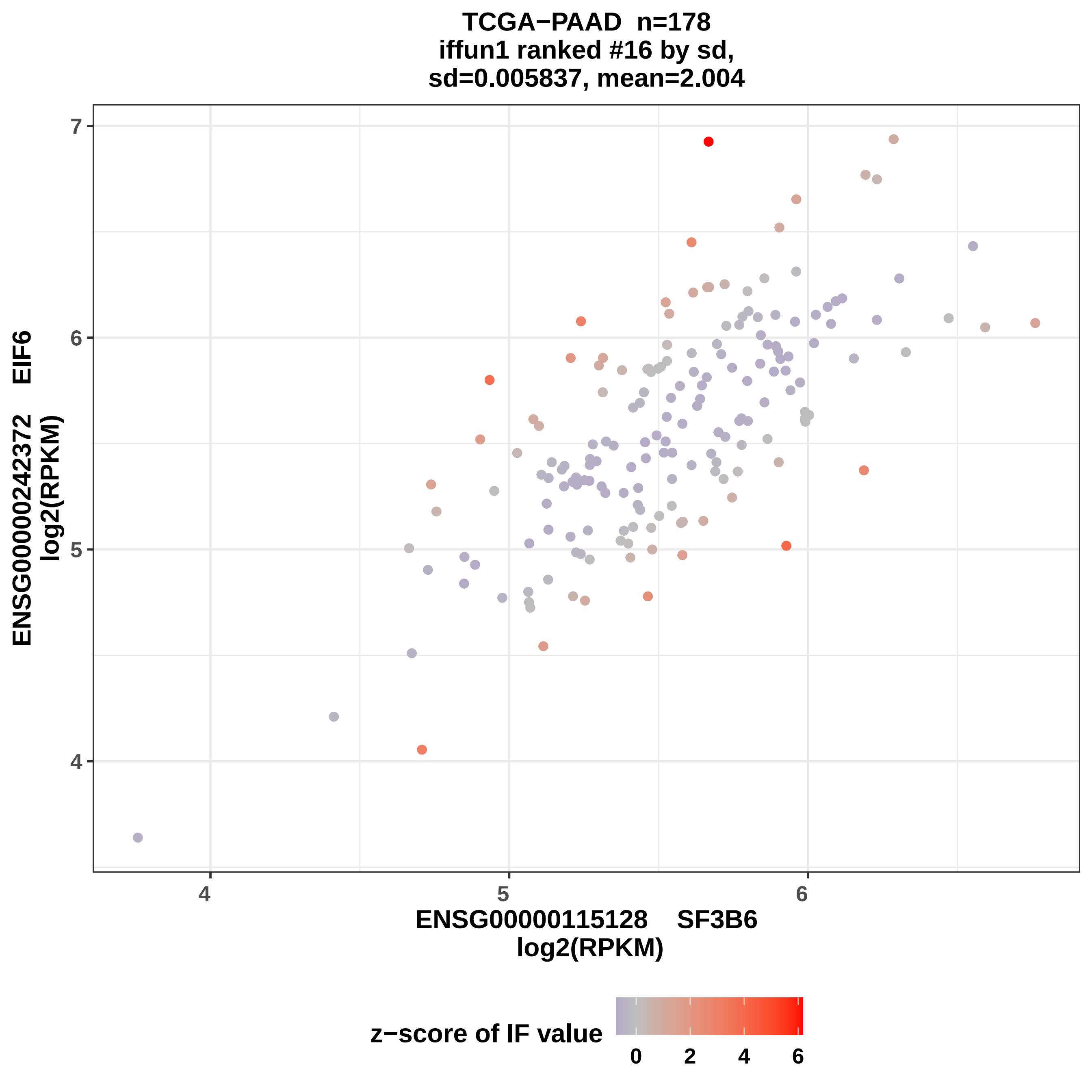

TCGA−PAAD n=178
iffun1 ranked #16 by sd,
sd=0.005837, mean=2.004
ENSG00000242372 EIF6
log2(RPKM)
ENSG00000115128 SF3B6
log2(RPKM)
z−score of IF value
4
5
6
7
0
2
4
6

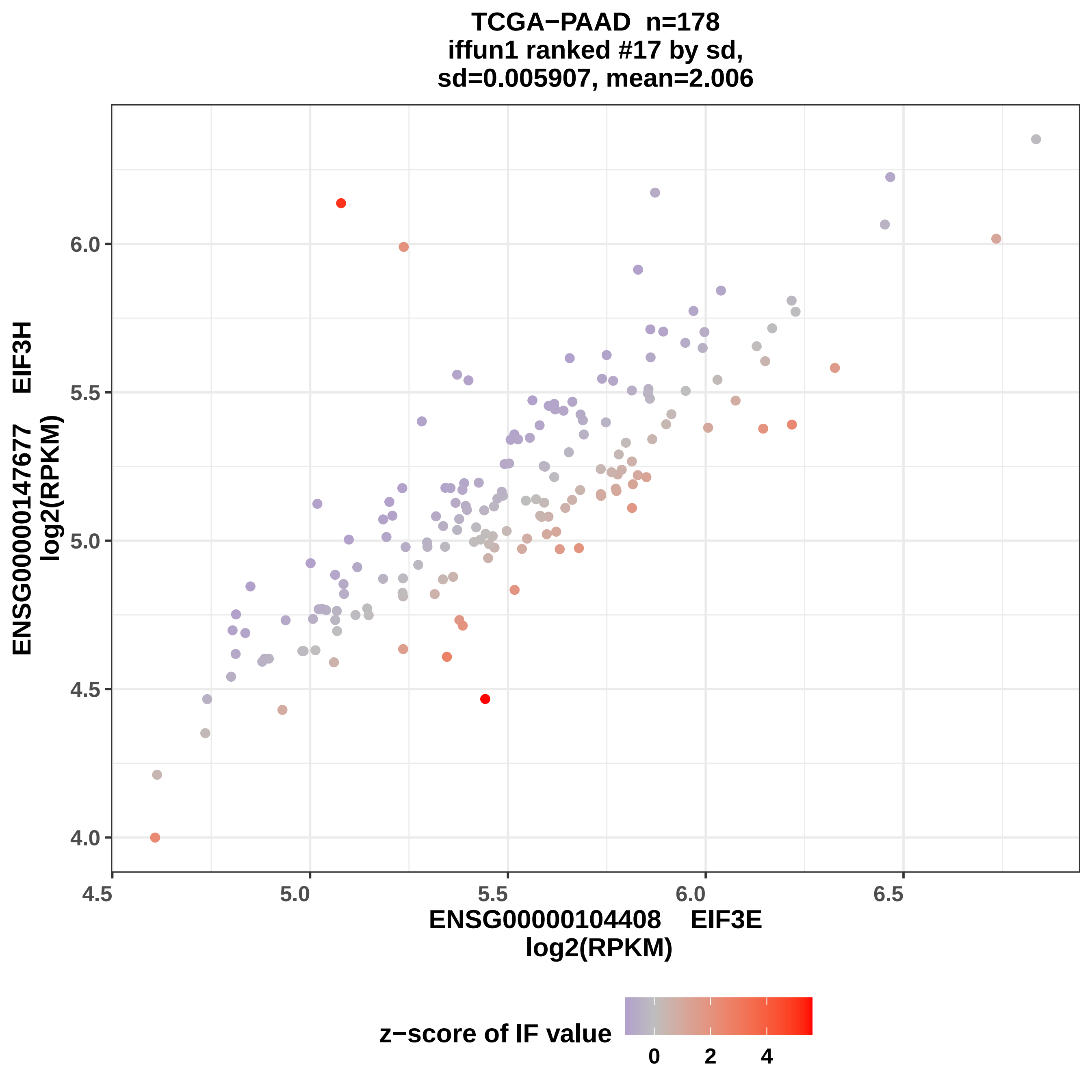

TCGA−PAAD n=178
iffun1 ranked #17 by sd,
sd=0.005907, mean=2.006
ENSG00000147677 EIF3H
log2(RPKM)
ENSG00000104408 EIF3E
log2(RPKM)
4.0
4.5
5.0
5.5
6.0
6.5
z−score of IF value
0
2
4

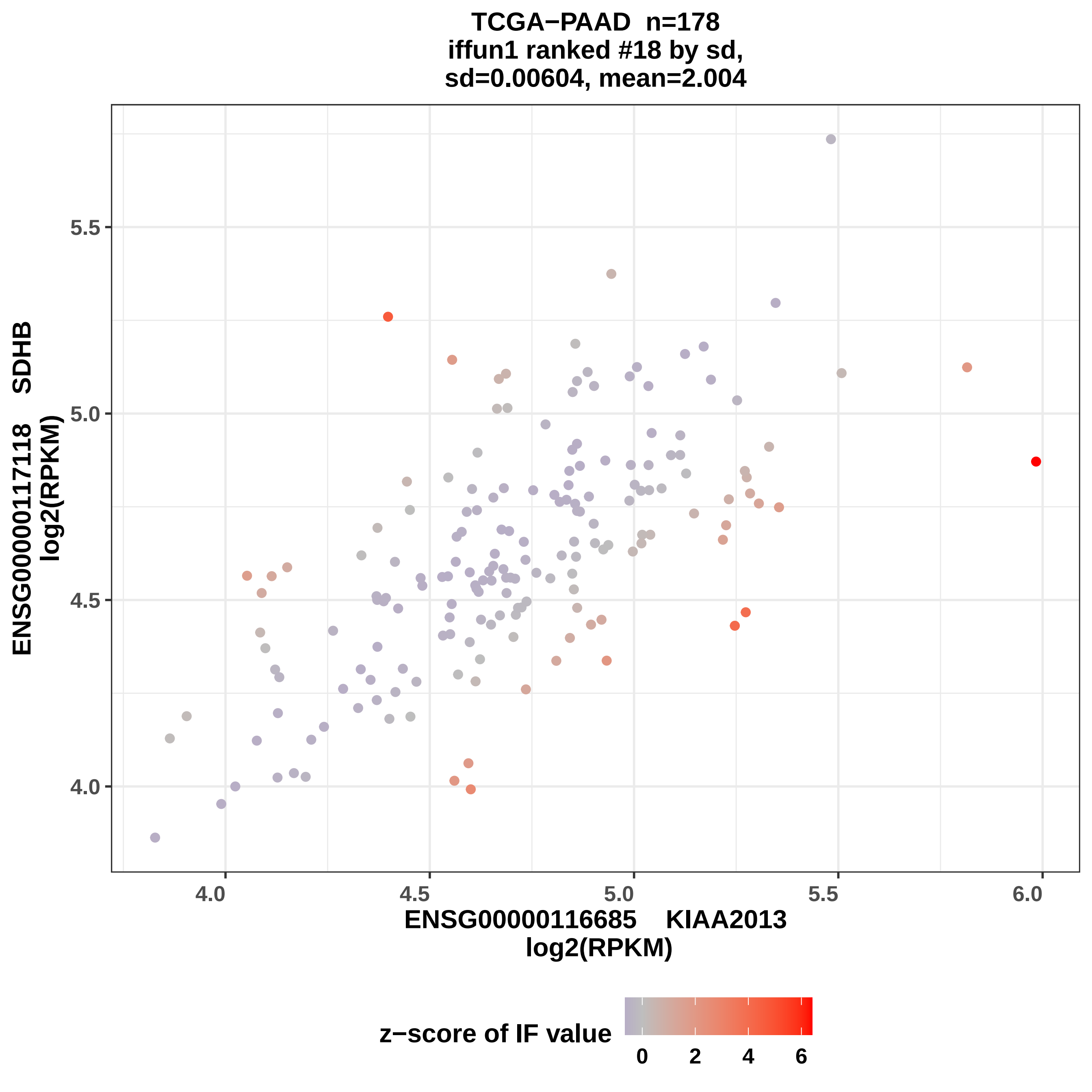

TCGA−PAAD n=178
iffun1 ranked #18 by sd,
sd=0.00604, mean=2.004
ENSG00000117118 SDHB
log2(RPKM)
5.5
5.0
4.5
4.0
4.0
4.5
5.0
5.5
6.0
ENSG00000116685 KIAA2013
log2(RPKM)
z−score of IF value
0
2
4
6

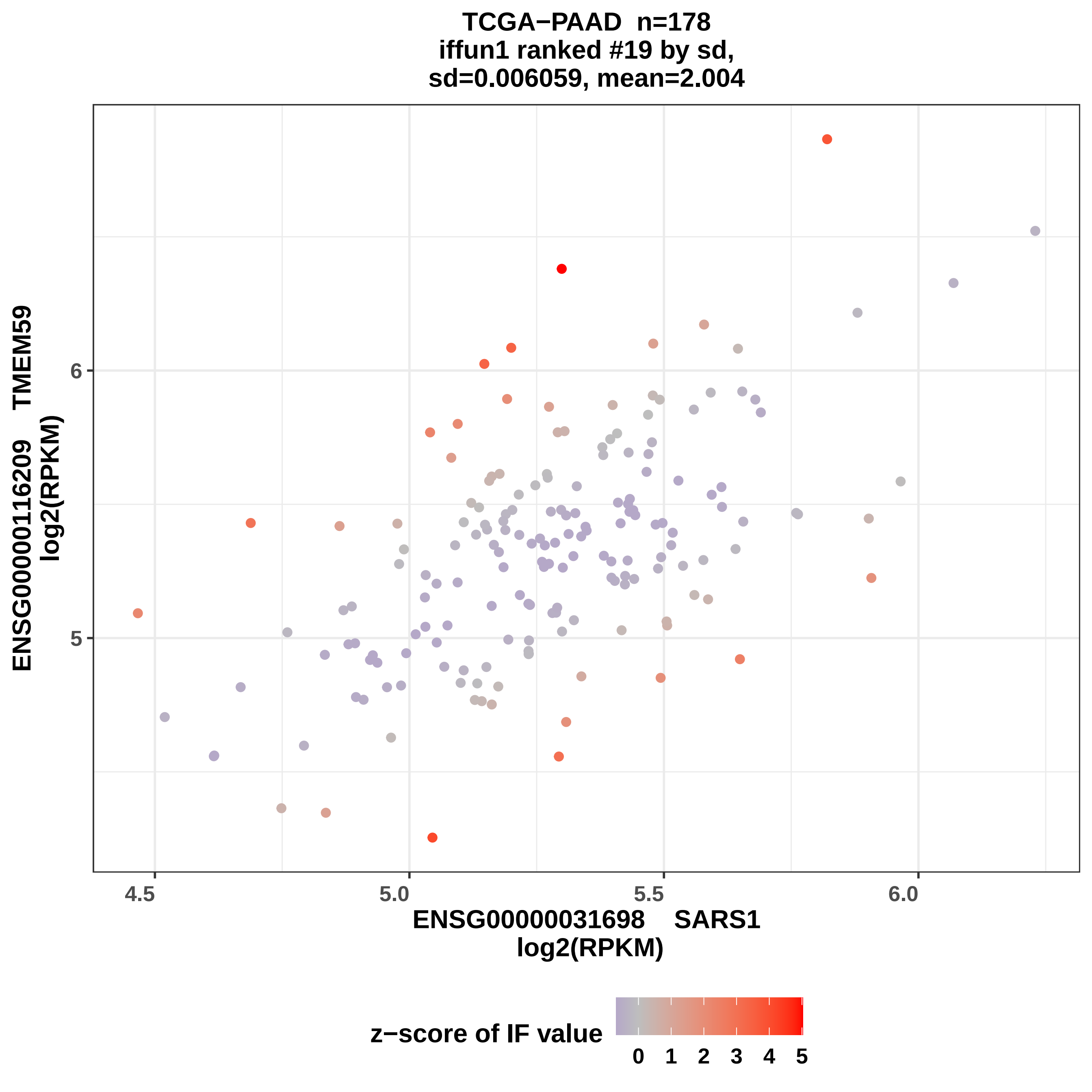
TCGA−PAAD n=178
iffun1 ranked #19 by sd,
sd=0.006059, mean=2.004
ENSG00000116209 TMEM59
log2(RPKM)
6
5
4.5
5.0
5.5
6.0
ENSG00000031698 SARS1
log2(RPKM)
z−score of IF value
0
1
2
3
4
5

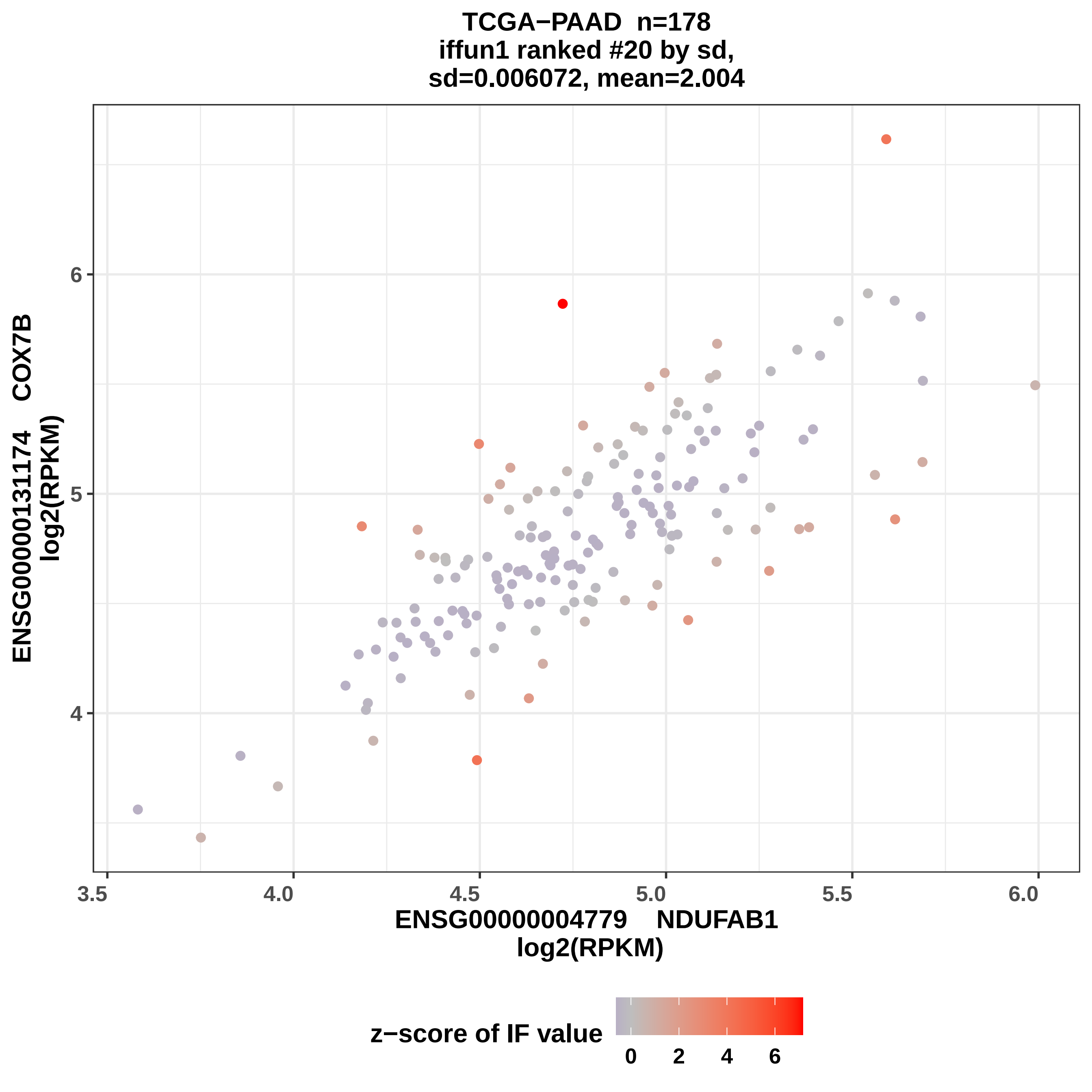

TCGA−PAAD n=178
iffun1 ranked #20 by sd,
sd=0.006072, mean=2.004
ENSG00000131174 COX7B
log2(RPKM)
ENSG00000004779 NDUFAB1
log2(RPKM)
3.5
4.0
4.5
5.0
5.5
6.0
4
5
6
z−score of IF value
0
2
4
6

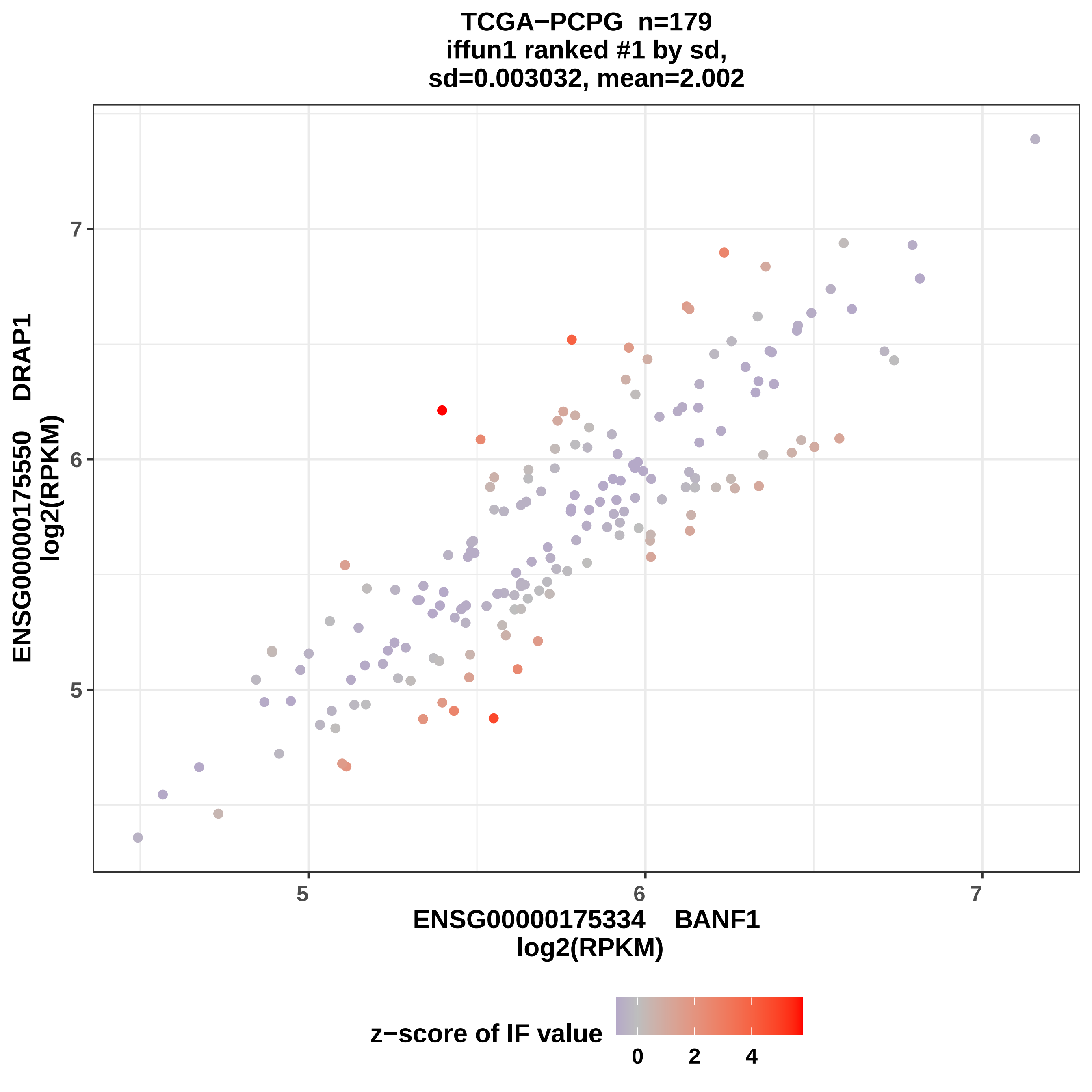
TCGA−PCPG n=179
iffun1 ranked #1 by sd,
sd=0.003032, mean=2.002
ENSG00000175550 DRAP1
log2(RPKM)
7
6
5
5
6
7
ENSG00000175334 BANF1
log2(RPKM)
z−score of IF value
0
2
4

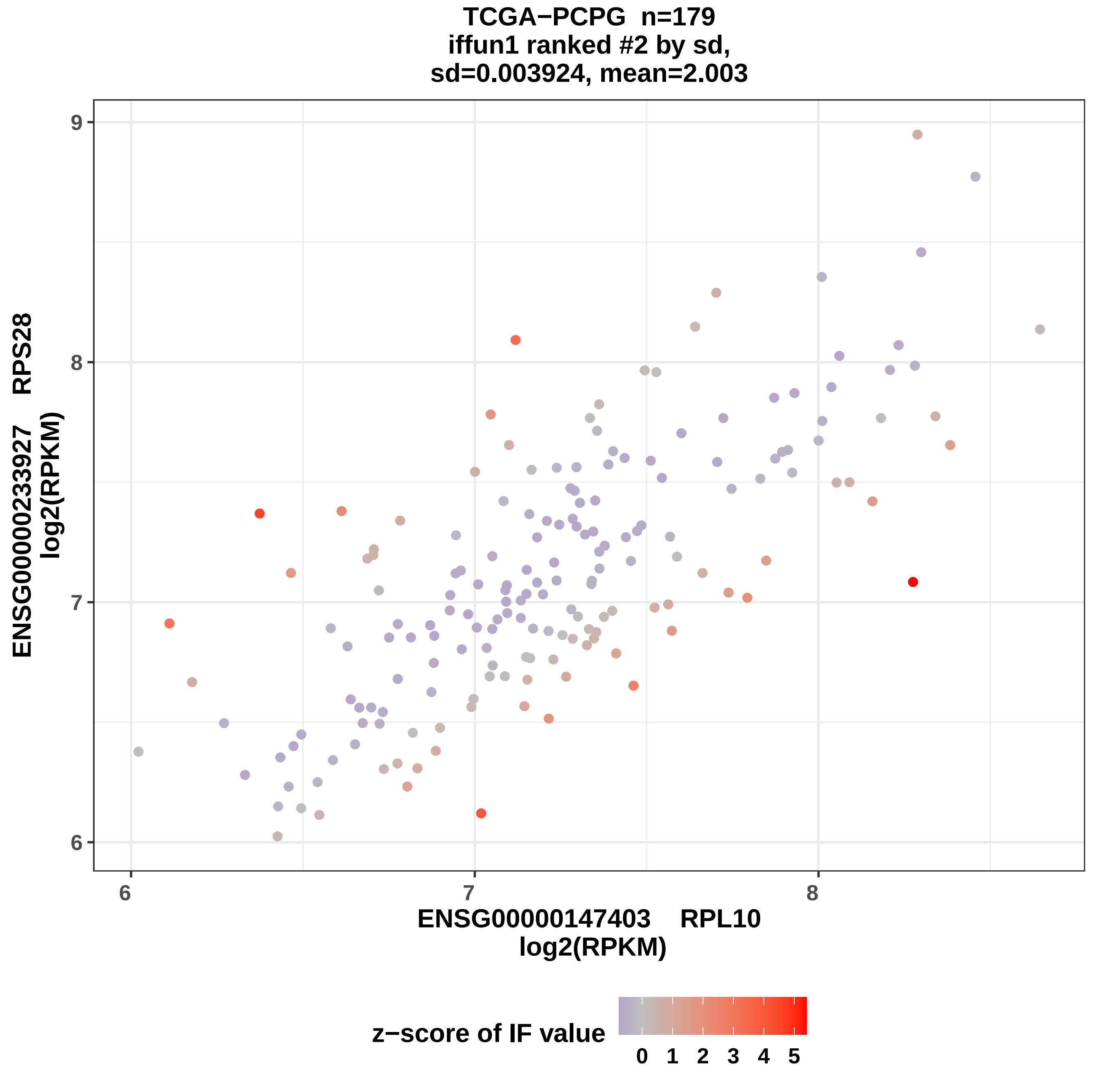
TCGA−PCPG n=179
iffun1 ranked #2 by sd,
sd=0.003924, mean=2.003
ENSG00000233927 RPS28
log2(RPKM)
ENSG00000147403 RPL10
log2(RPKM)
6
7
8
9
z−score of IF value
0 1 2 3 4 5

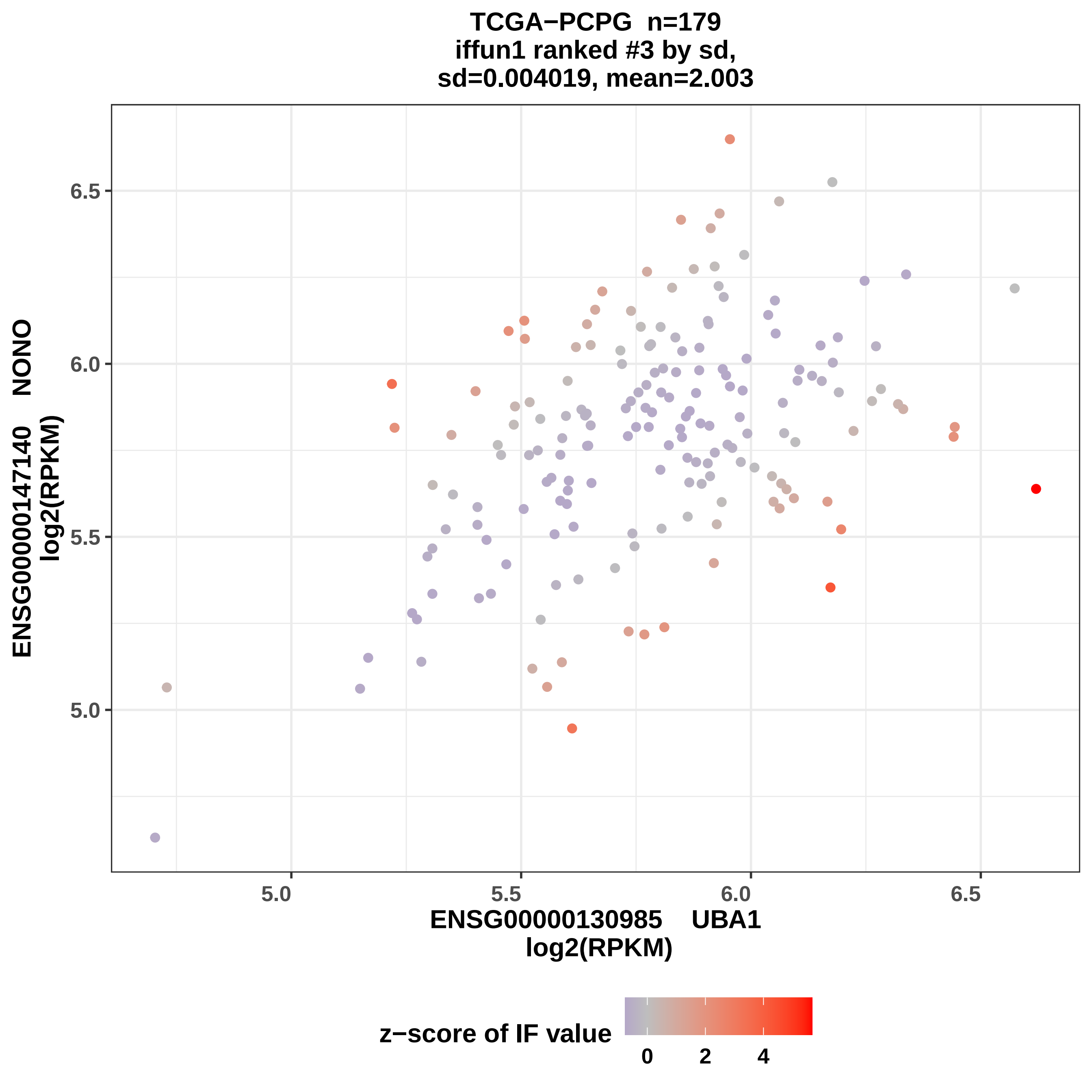
TCGA−PCPG n=179
iffun1 ranked #3 by sd,
sd=0.004019, mean=2.003
ENSG00000147140 NONO
log2(RPKM)
6.5
6.0
5.5
5.0
5.0
5.5
6.0
6.5
ENSG00000130985 UBA1
log2(RPKM)
z−score of IF value
0
2
4

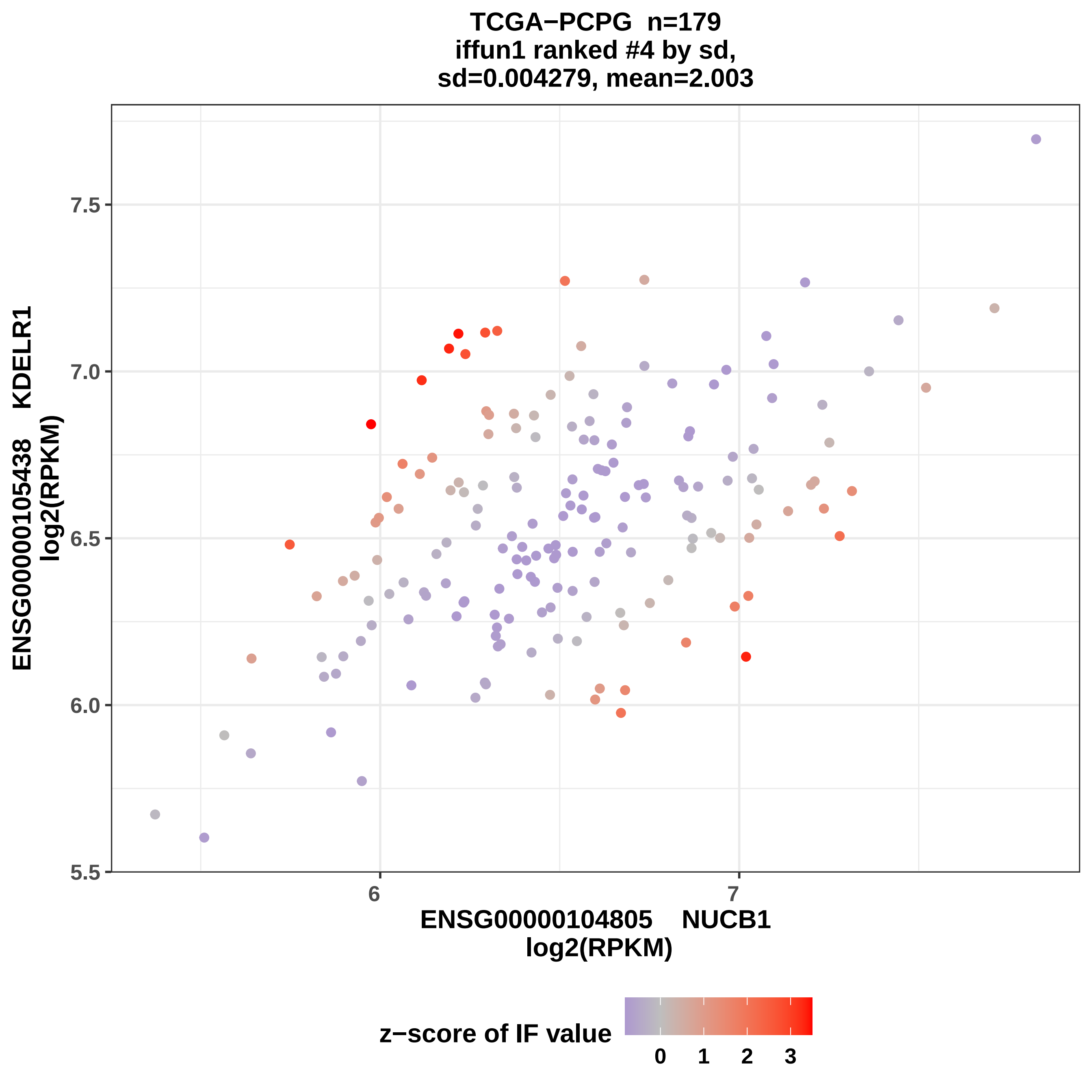

TCGA−PCPG n=179
iffun1 ranked #4 by sd,
sd=0.004279, mean=2.003
ENSG00000105438 KDELR1
log2(RPKM)
7.5
7.0
6.5
6.0
5.5
6
7
ENSG00000104805 NUCB1
log2(RPKM)
z−score of IF value
0
1
2
3

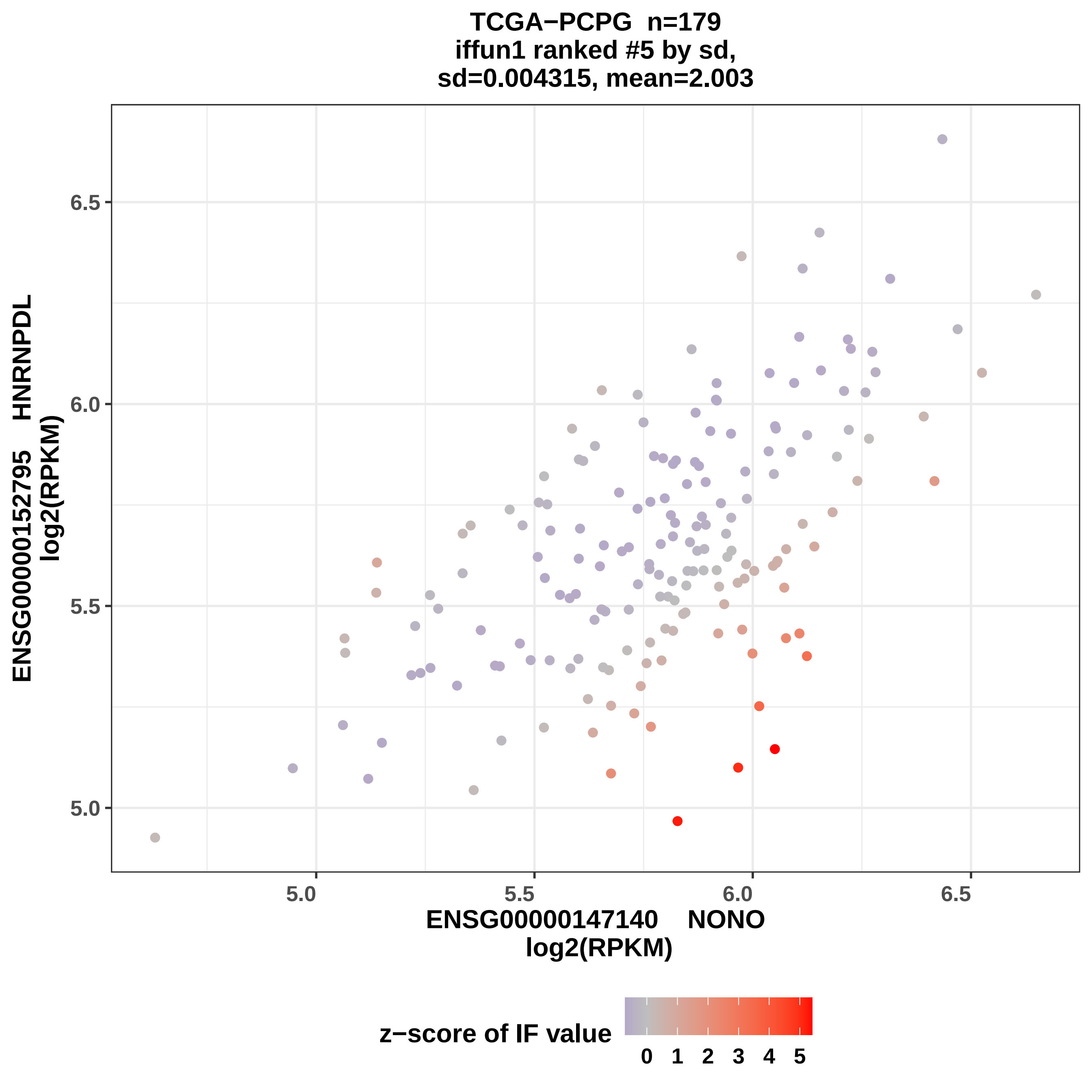

TCGA−PCPG n=179
iffun1 ranked #5 by sd,
sd=0.004315, mean=2.003
ENSG00000152795 HNRNPDL
log2(RPKM)
6.5
6.0
5.5
5.0
5.0
5.5
6.0
6.5
ENSG00000147140 NONO
log2(RPKM)
z−score of IF value
0
1
2
3
4
5

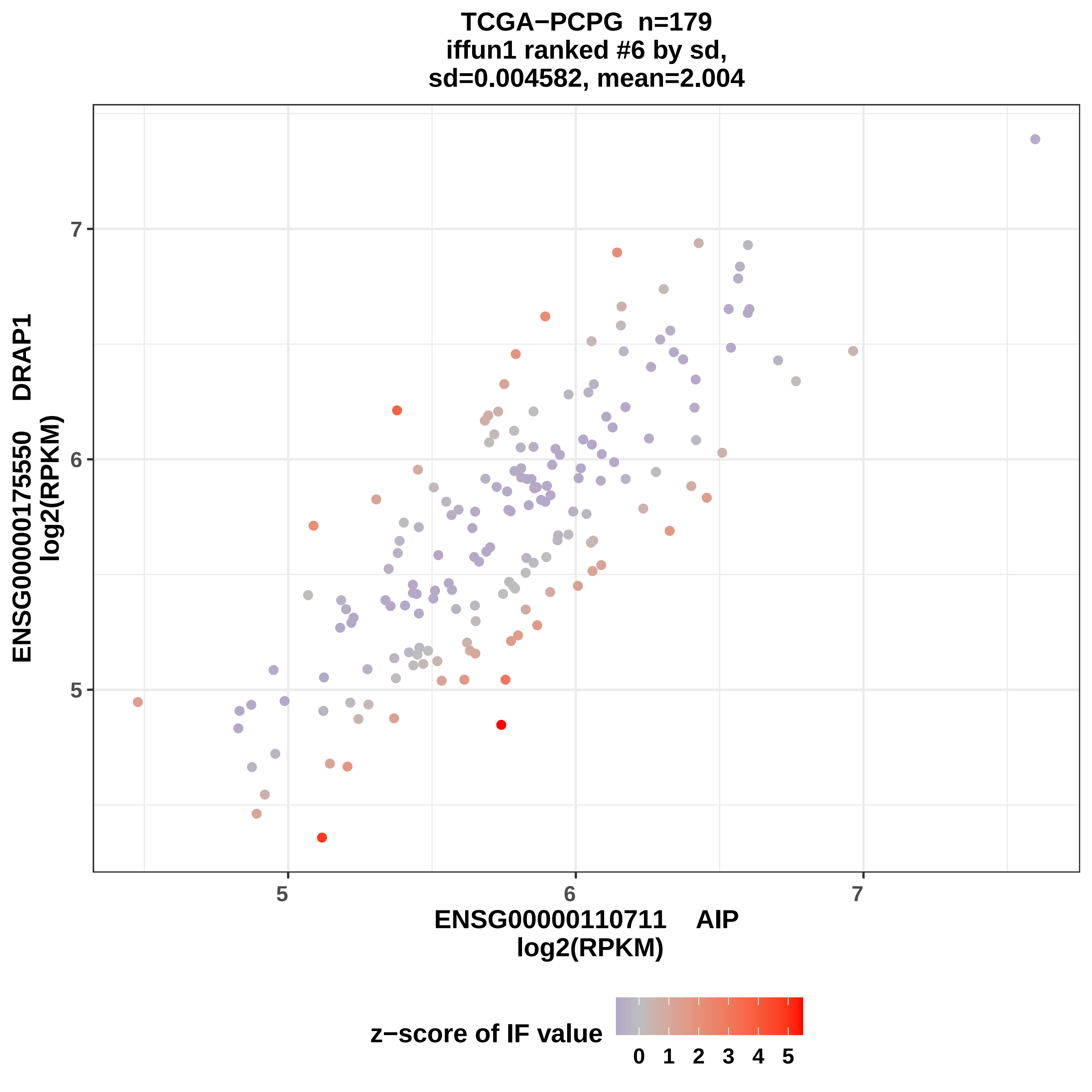

TCGA−PCPG n=179
iffun1 ranked #6 by sd,
sd=0.004582, mean=2.004
ENSG00000175550 DRAP1
log2(RPKM)
7
6
5
5
6
7
ENSG00000110711 AIP
log2(RPKM)
z−score of IF value
0 1 2 3 4 5

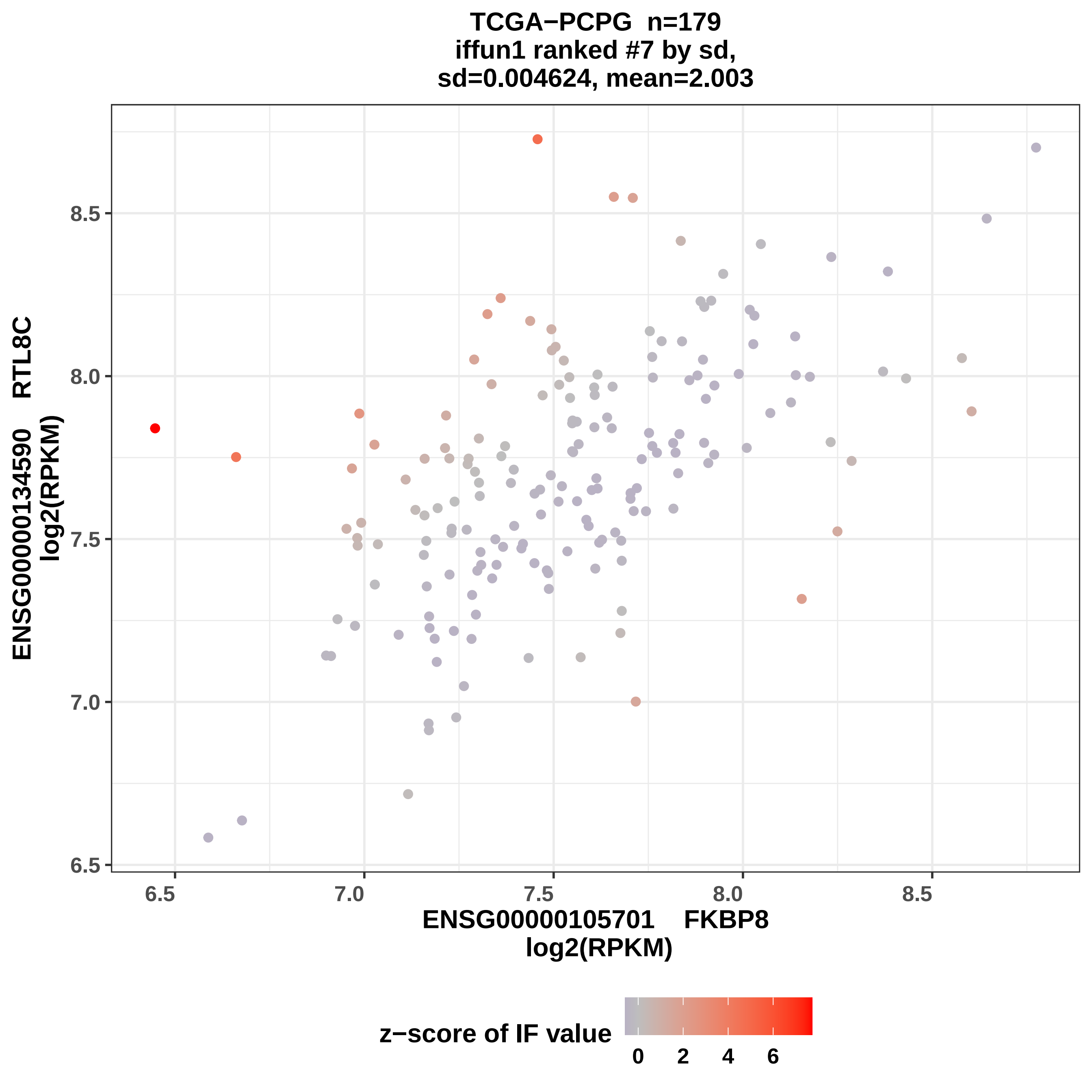

TCGA−PCPG n=179
iffun1 ranked #7 by sd,
sd=0.004624, mean=2.003
ENSG00000134590 RTL8C
log2(RPKM)
8.5
8.0
7.5
7.0
6.5
6.5
7.0
7.5
8.0
8.5
ENSG00000105701 FKBP8
log2(RPKM)
z−score of IF value
0
2
4
6

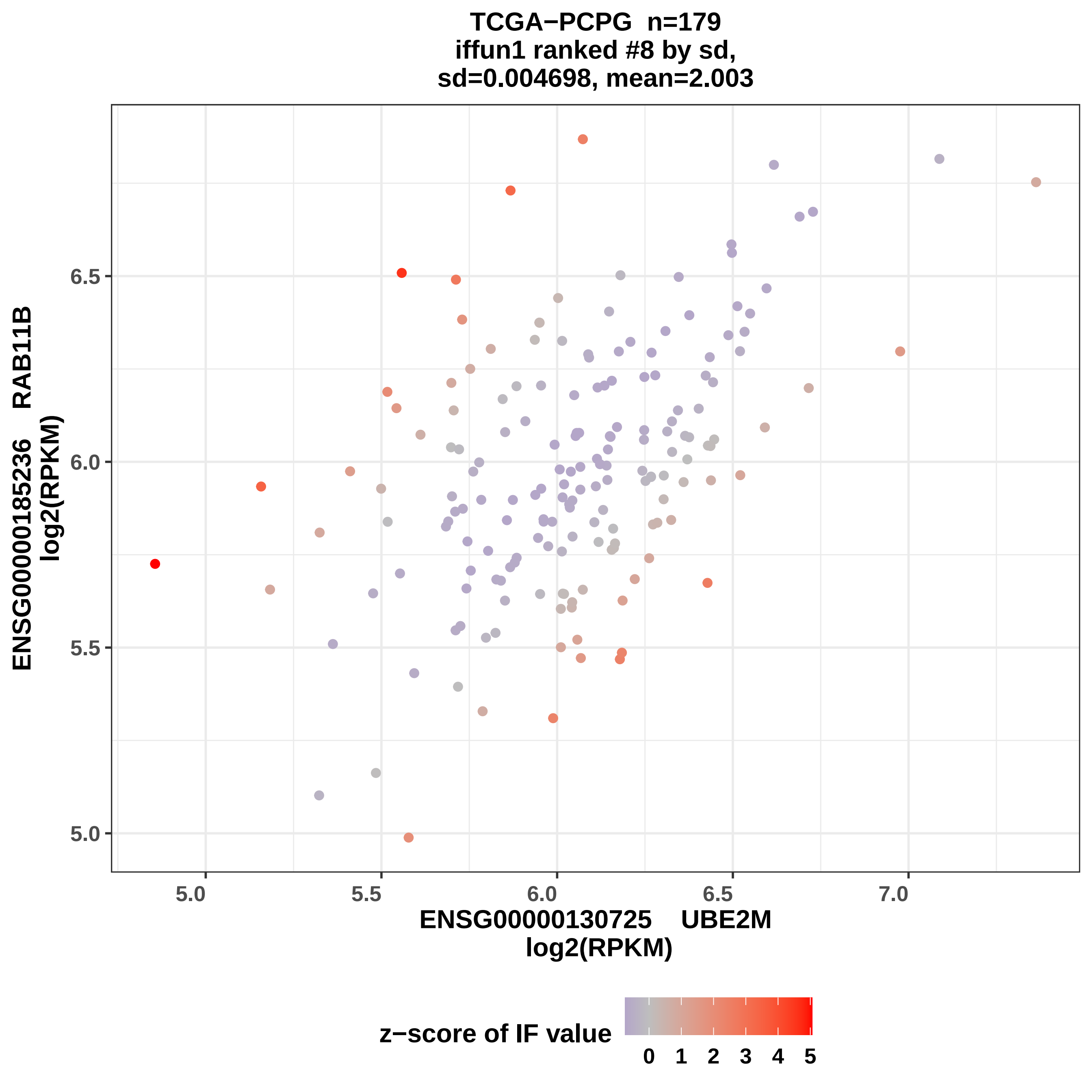

TCGA−PCPG n=179
iffun1 ranked #8 by sd,
sd=0.004698, mean=2.003
ENSG00000185236 RAB11B
log2(RPKM)
6.5
6.0
5.5
5.0
7.0
ENSG00000130725 UBE2M
log2(RPKM)
z−score of IF value
0 1 2 3 4 5

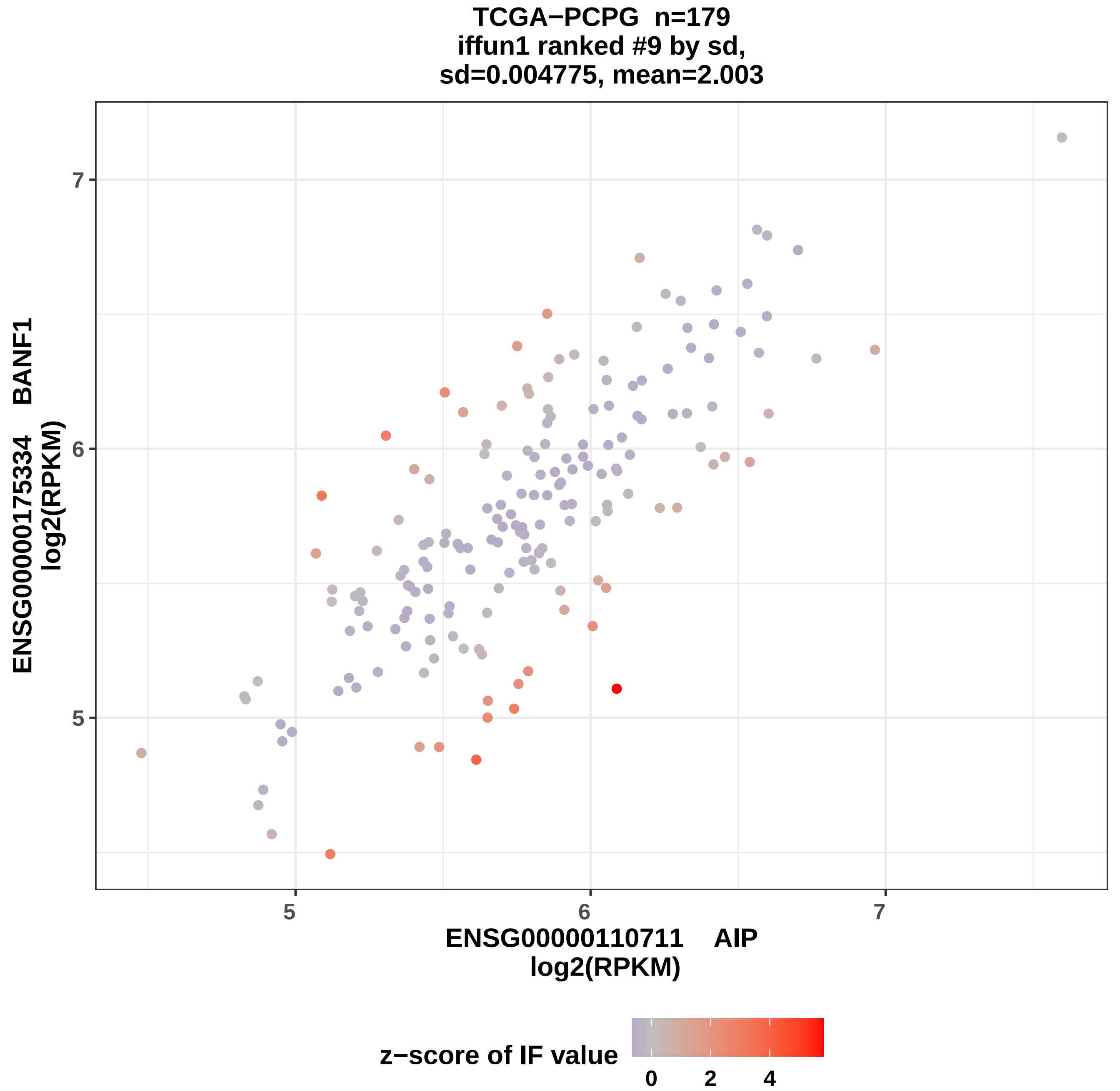

TCGA−PCPG n=179
iffun1 ranked #9 by sd,
sd=0.004775, mean=2.003
ENSG00000175334 BANF1
log2(RPKM)
7
6
5
5
6
7
ENSG00000110711 AIP
log2(RPKM)
z−score of IF value
0
2
4

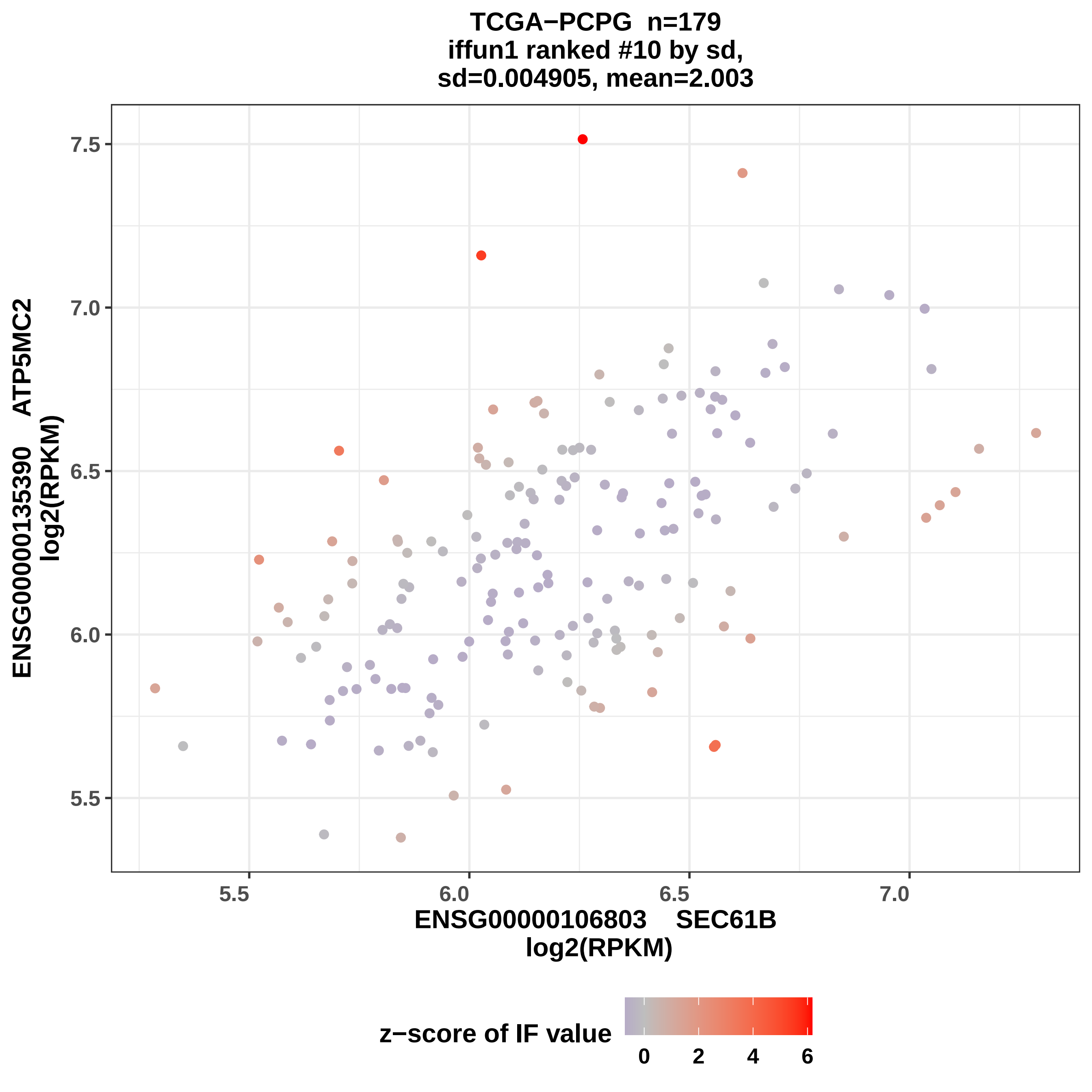

TCGA−PCPG n=179
iffun1 ranked #10 by sd,
sd=0.004905, mean=2.003
ENSG00000135390 ATP5MC2
log2(RPKM)
7.5
7.0
6.5
6.0
5.5
5.5
6.0
6.5
7.0
ENSG00000106803 SEC61B
log2(RPKM)
z−score of IF value
0
2
4
6

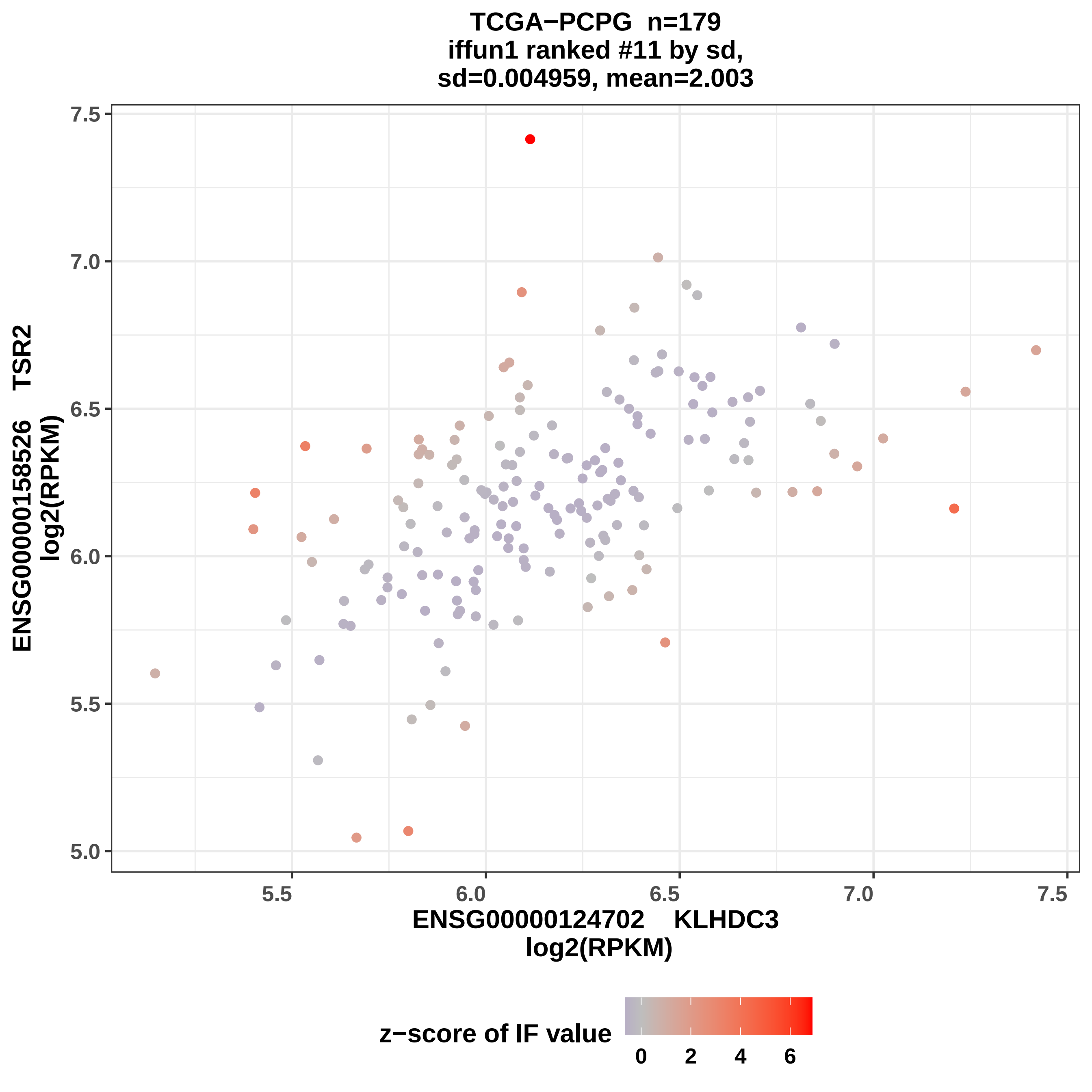

TCGA−PCPG n=179
iffun1 ranked #11 by sd,
sd=0.004959, mean=2.003
ENSG00000158526 TSR2
log2(RPKM)
ENSG00000124702 KLHDC3
log2(RPKM)
7.5
7.0
6.5
6.0
5.5
5.0
5.5
6.0
6.5
7.0
7.5
z−score of IF value
0
2
4
6

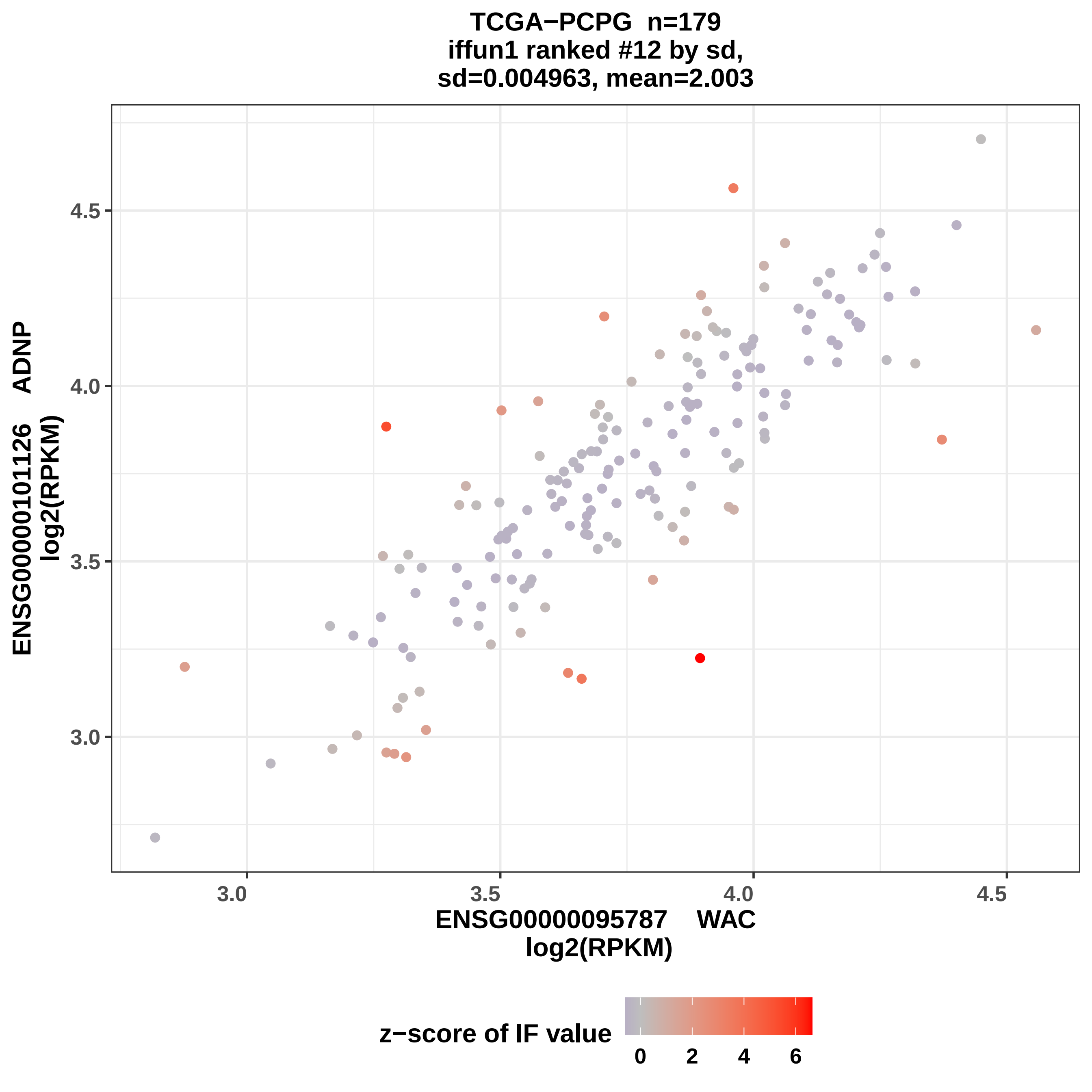
TCGA−PCPG n=179
iffun1 ranked #12 by sd,
sd=0.004963, mean=2.003
ENSG00000101126 ADNP
log2(RPKM)
4.5
4.0
3.5
3.0
3.0
3.5
4.0
4.5
ENSG00000095787 WAC
log2(RPKM)
z−score of IF value
0
2
4
6

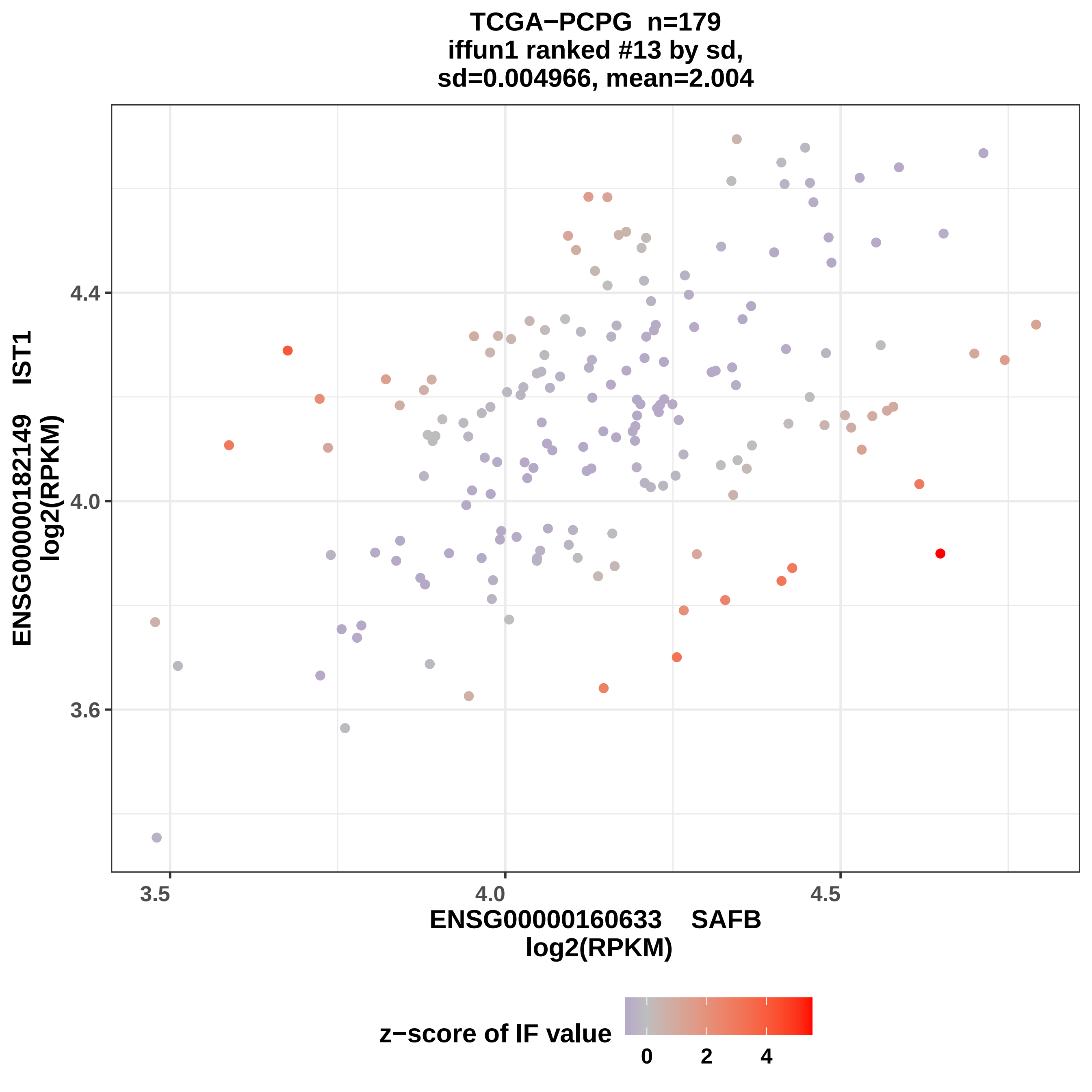
TCGA−PCPG n=179
iffun1 ranked #13 by sd,
sd=0.004966, mean=2.004
ENSG00000182149 IST1
log2(RPKM)
4.4
4.0
3.6
3.5
4.0
4.5
ENSG00000160633 SAFB
log2(RPKM)
z−score of IF value
0
2
4

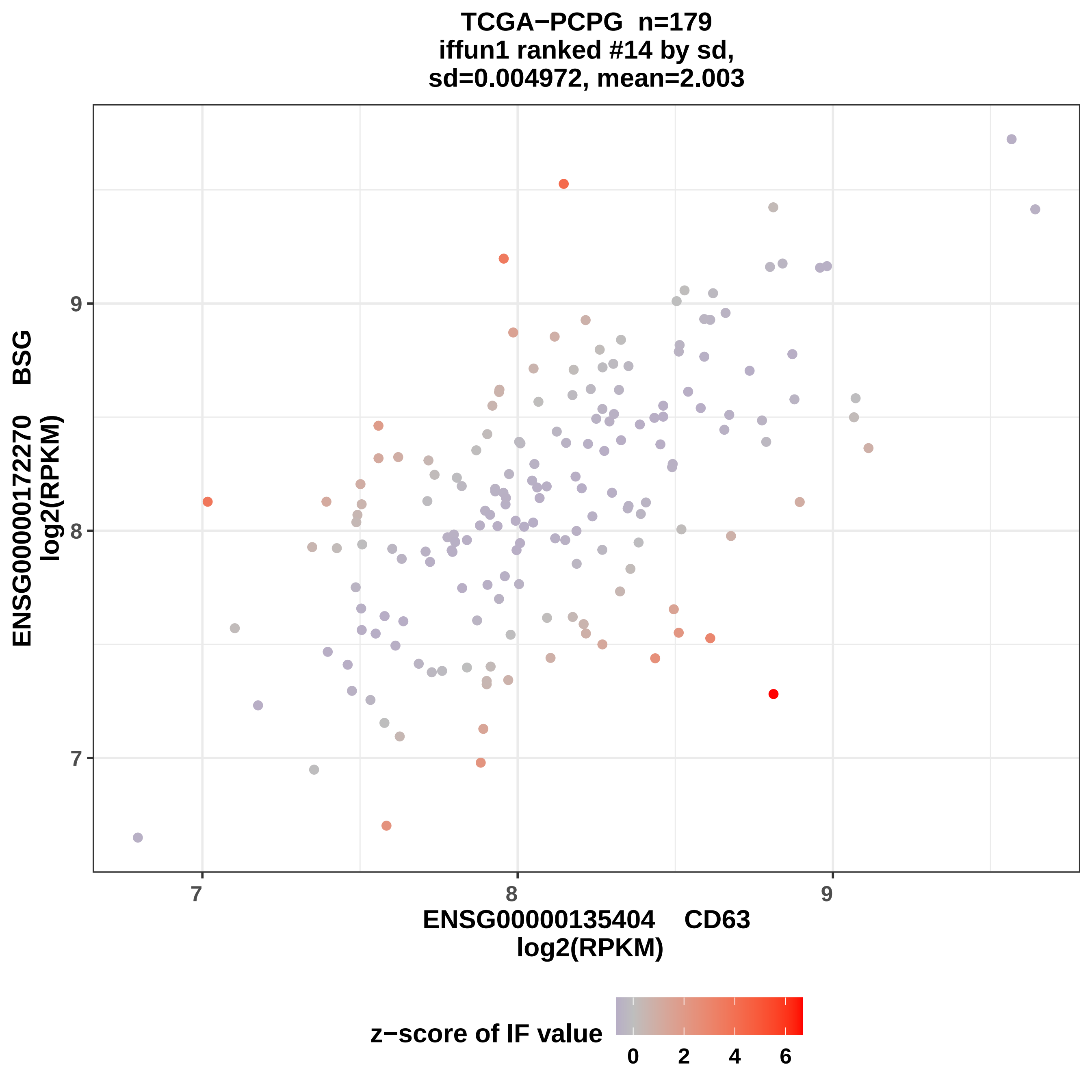
TCGA−PCPG n=179
iffun1 ranked #14 by sd,
sd=0.004972, mean=2.003
ENSG00000172270 BSG
log2(RPKM)
ENSG00000135404 CD63
log2(RPKM)
7
8
9
7
8
9
z−score of IF value
0
2
4
6

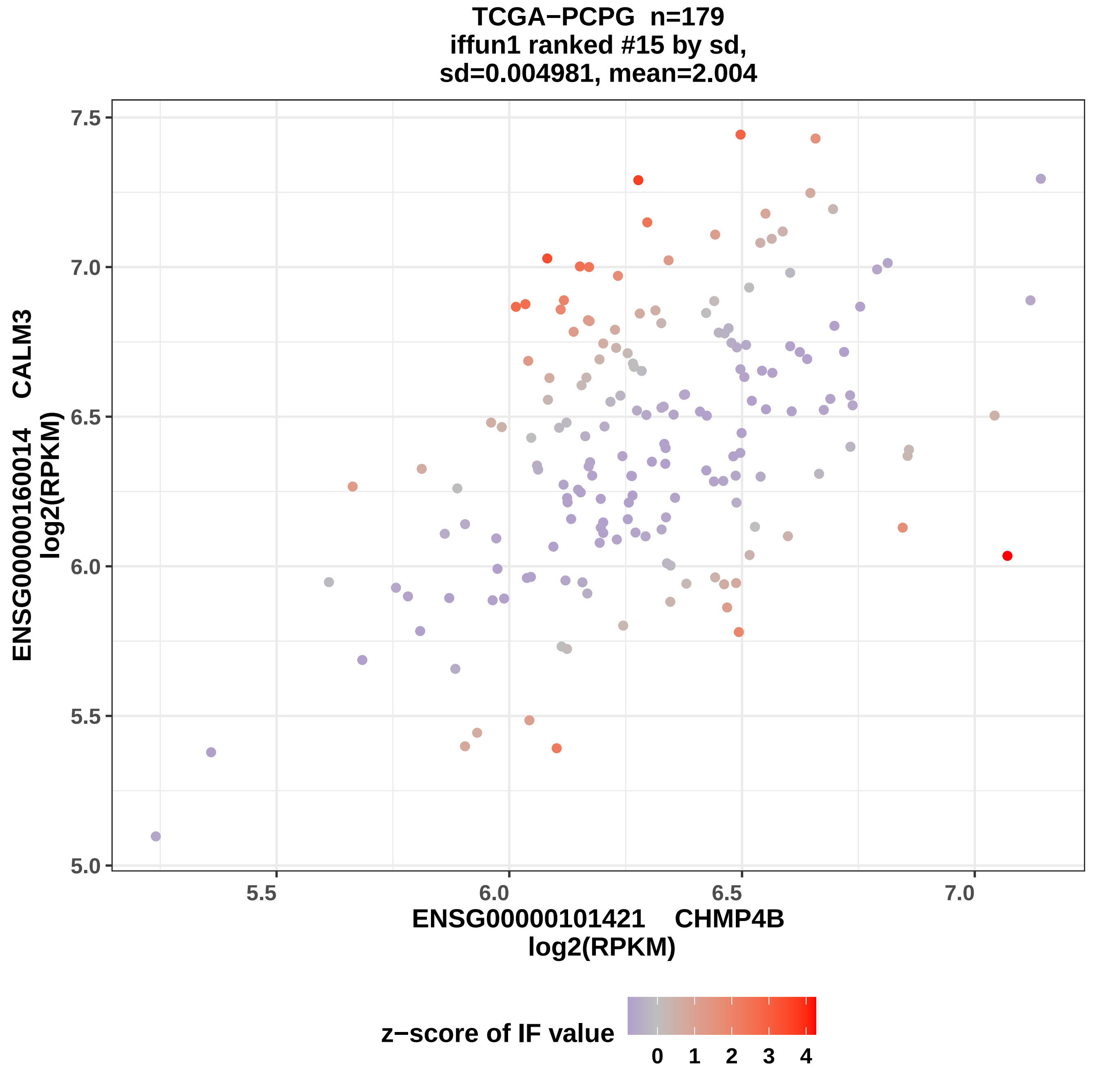

TCGA−PCPG n=179
iffun1 ranked #15 by sd,
sd=0.004981, mean=2.004
ENSG00000160014 CALM3
log2(RPKM)
ENSG00000101421 CHMP4B
log2(RPKM)
7.5
7.0
6.5
6.0
5.5
5.0
5.5
6.0
6.5
7.0
z−score of IF value
0
1
2
3
4

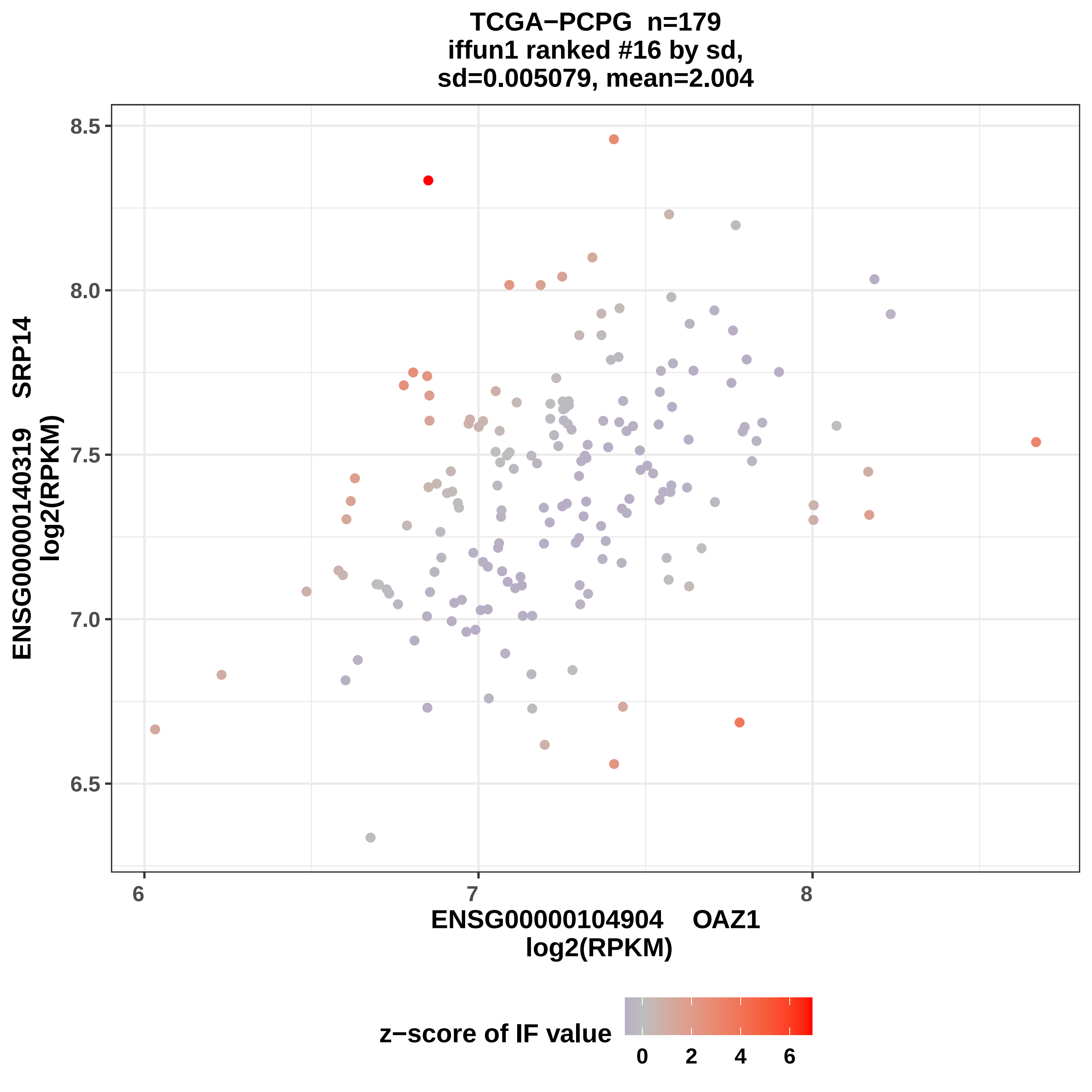
TCGA−PCPG n=179
iffun1 ranked #16 by sd,
sd=0.005079, mean=2.004
ENSG00000140319 SRP14
log2(RPKM)
8.5
8.0
7.5
7.0
6.5
6
7
8
ENSG00000104904 OAZ1
log2(RPKM)
z−score of IF value
0
2
4
6

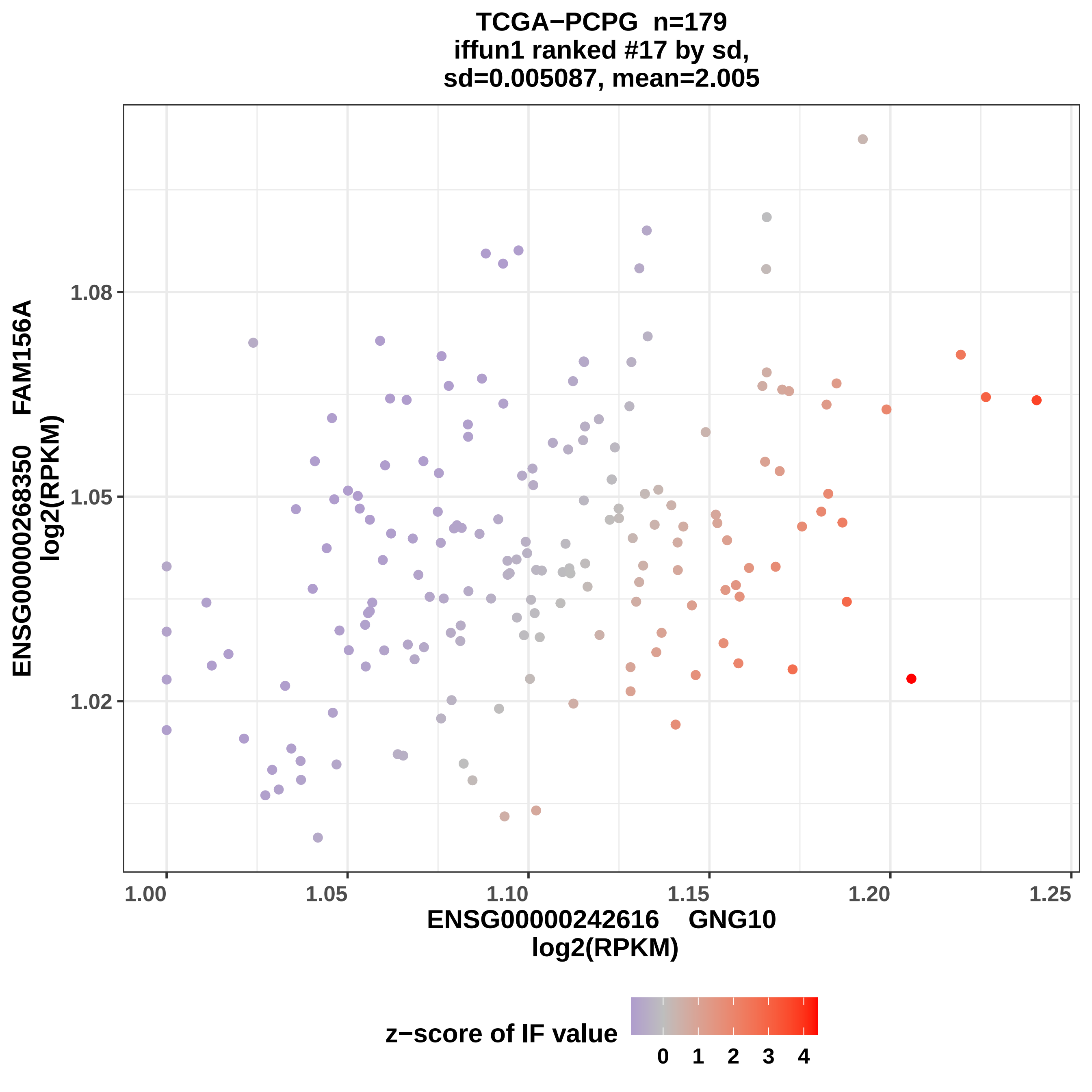
TCGA−PCPG n=179
iffun1 ranked #17 by sd,
sd=0.005087, mean=2.005
ENSG00000268350 FAM156A
log2(RPKM)
1.08
1.05
1.02
1.00
1.05
1.10
1.15
1.20
1.25
ENSG00000242616 GNG10
log2(RPKM)
z−score of IF value
0
1
2
3
4

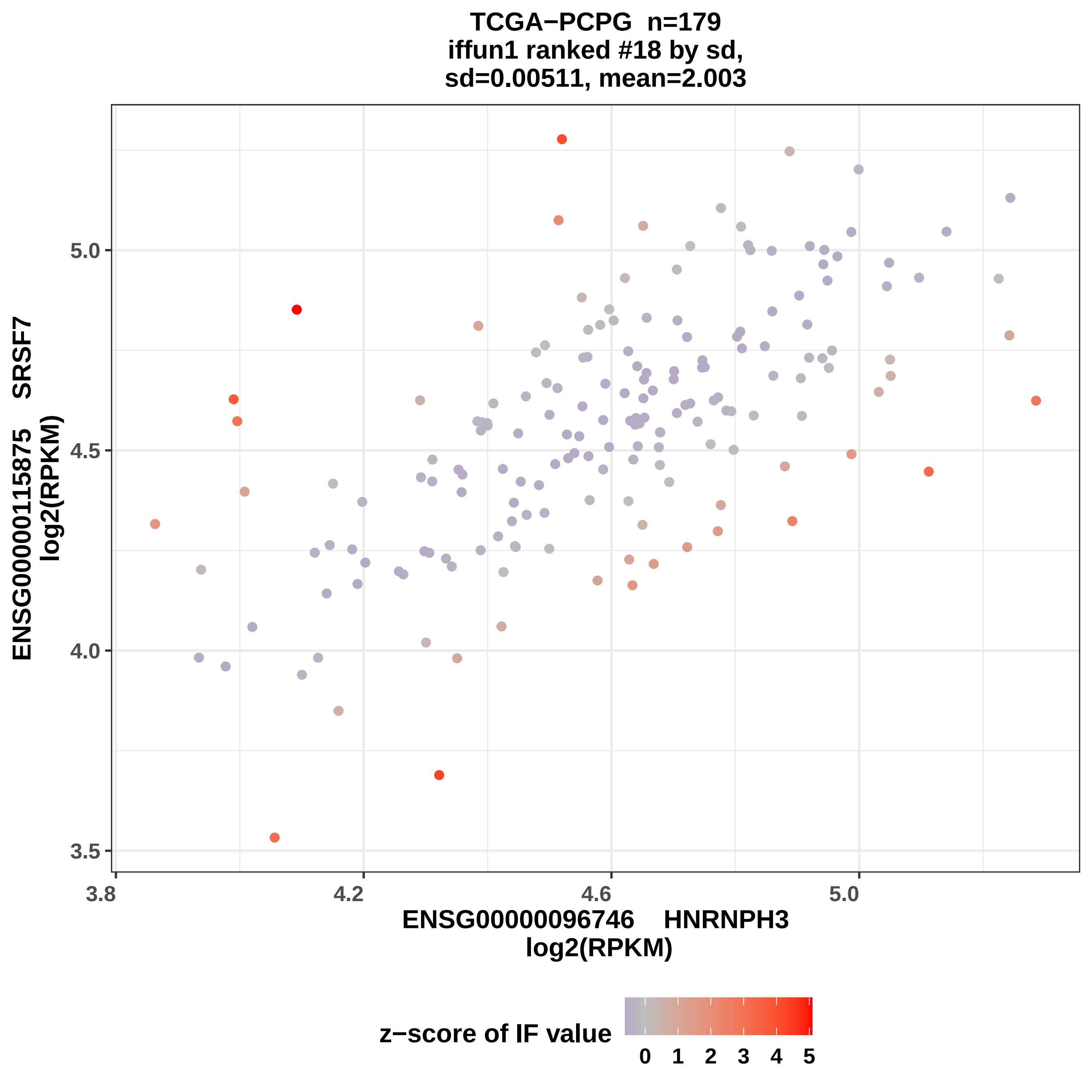
TCGA−PCPG n=179
iffun1 ranked #18 by sd,
sd=0.00511, mean=2.003
ENSG00000115875 SRSF7
log2(RPKM)
5.0
4.5
4.0
3.5
3.8
4.2
4.6
5.0
ENSG00000096746 HNRNPH3
log2(RPKM)
z−score of IF value
0
1
2
3
4
5

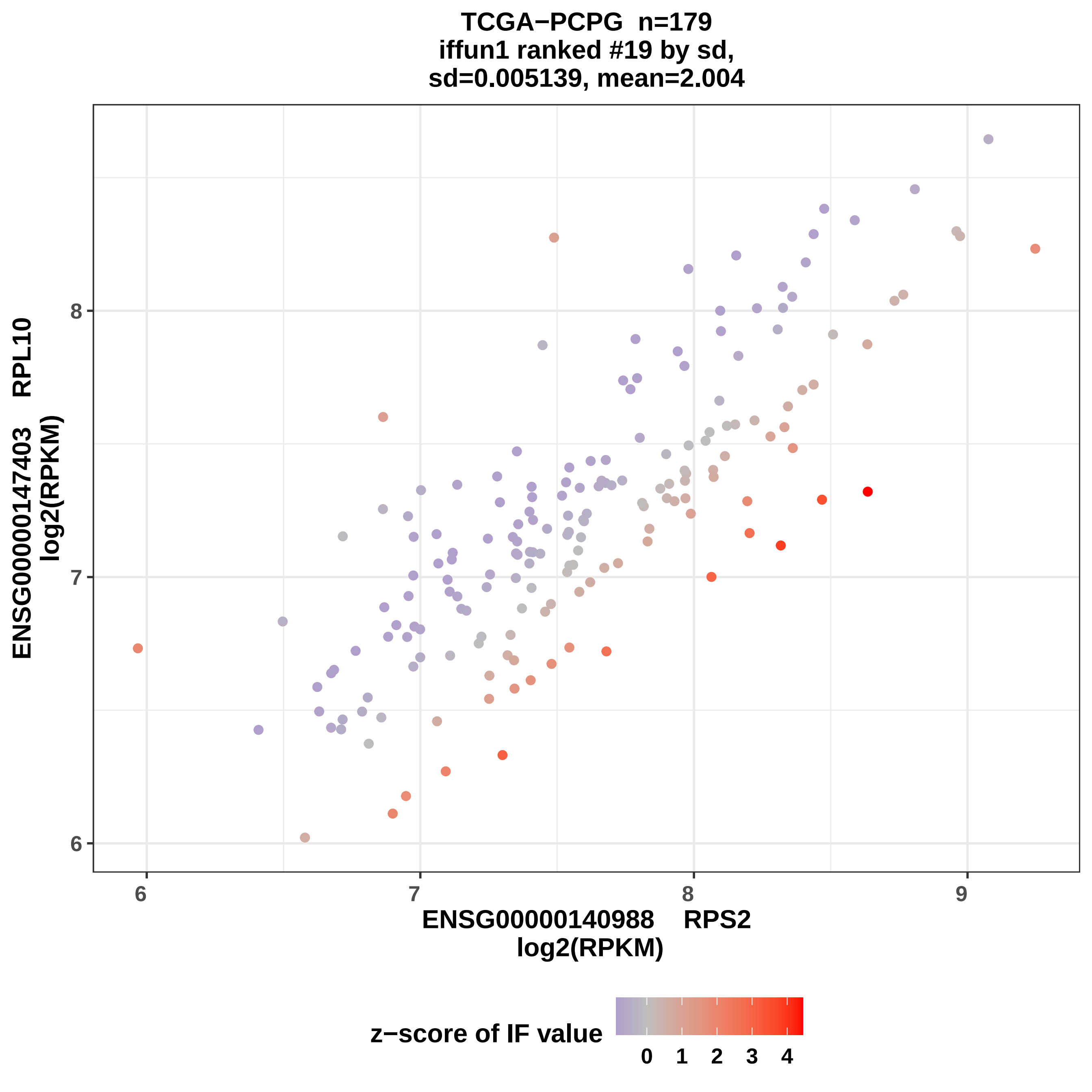

TCGA−PCPG n=179
iffun1 ranked #19 by sd,
sd=0.005139, mean=2.004
ENSG00000147403 RPL10
log2(RPKM)
ENSG00000140988 RPS2
log2(RPKM)
6
7
8
9
z−score of IF value
0
1
2
3
4

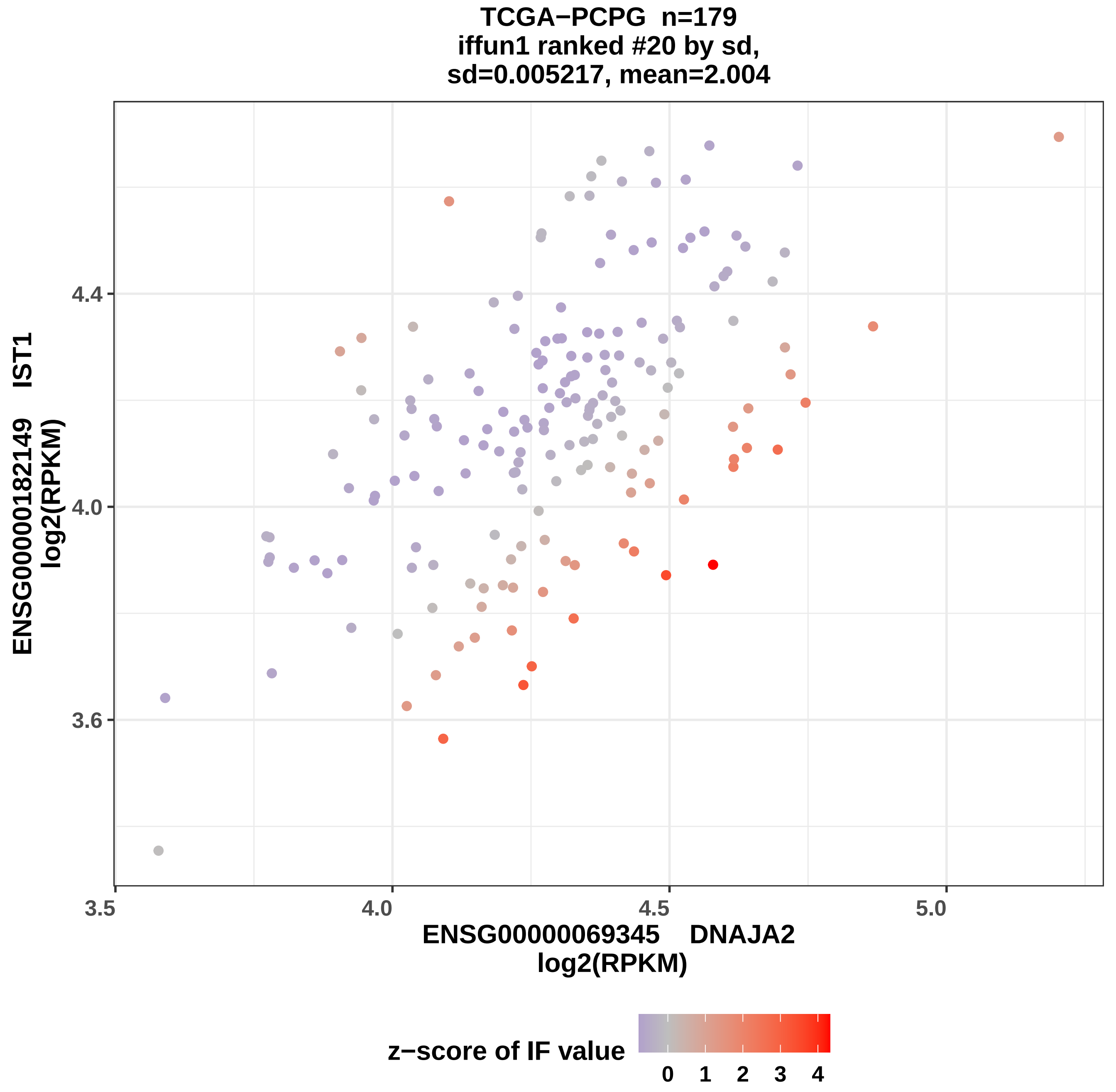
TCGA−PCPG n=179
iffun1 ranked #20 by sd,
sd=0.005217, mean=2.004
ENSG00000182149 IST1
log2(RPKM)
4.4
4.0
3.6
3.5
4.0
4.5
5.0
ENSG00000069345 DNAJA2
log2(RPKM)
z−score of IF value
0
1
2
3
4

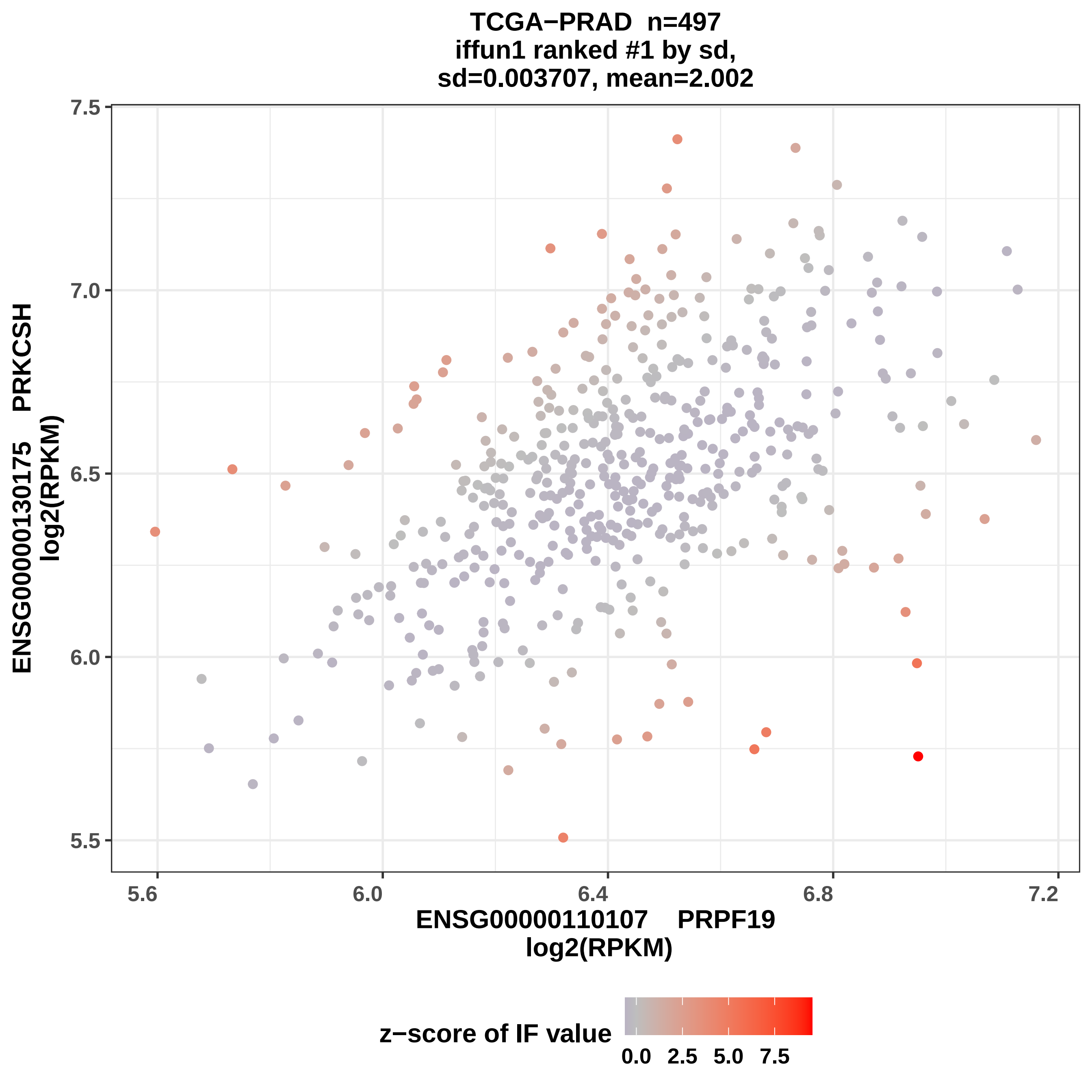

TCGA−PRAD n=497
iffun1 ranked #1 by sd,
sd=0.003707, mean=2.002
ENSG00000130175 PRKCSH
log2(RPKM)
7.5
7.0
6.5
6.0
5.5
5.6
6.0
6.4
6.8
7.2
ENSG00000110107 PRPF19
log2(RPKM)
z−score of IF value
0.0
2.5
5.0
7.5

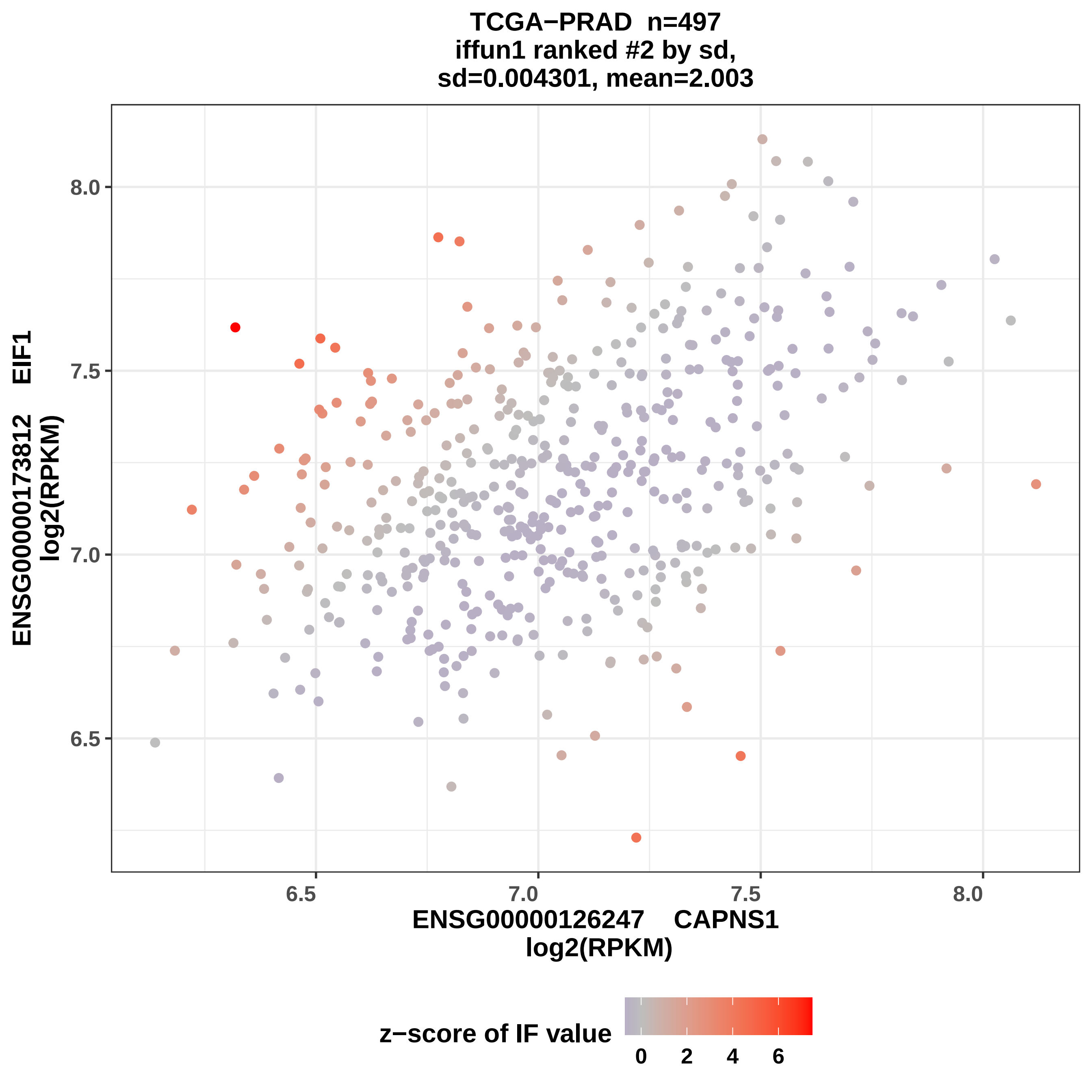

TCGA−PRAD n=497
iffun1 ranked #2 by sd,
sd=0.004301, mean=2.003
ENSG00000173812 EIF1
log2(RPKM)
8.0
7.5
7.0
6.5
6.5
7.0
7.5
8.0
ENSG00000126247 CAPNS1
log2(RPKM)
z−score of IF value
0
2
4
6

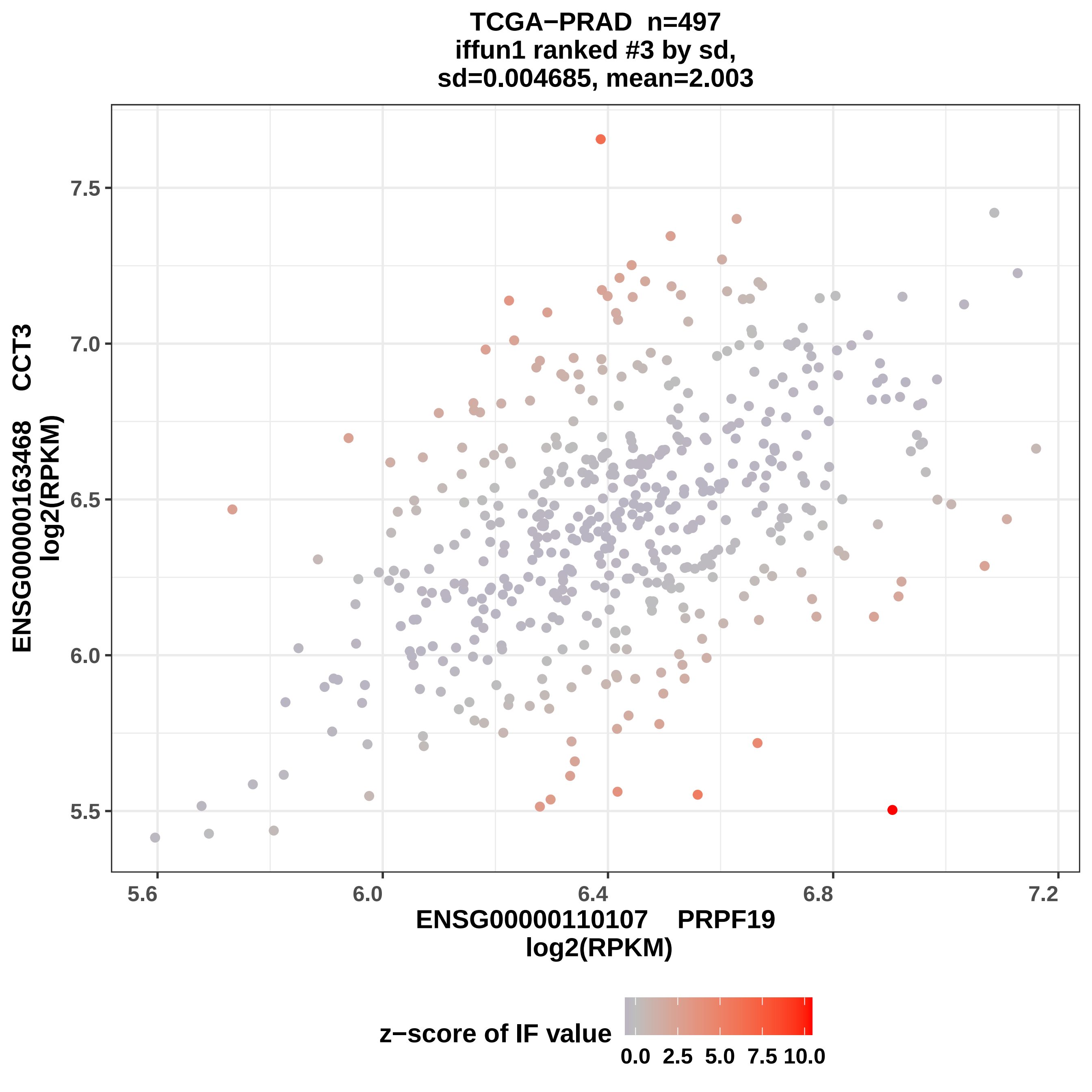
TCGA−PRAD n=497
iffun1 ranked #3 by sd,
sd=0.004685, mean=2.003
ENSG00000163468 CCT3
log2(RPKM)
ENSG00000110107 PRPF19
log2(RPKM)
7.5
7.0
6.5
6.0
5.5
5.6
6.0
6.4
6.8
7.2
z−score of IF value
0.0 2.5 5.0 7.5 10.0

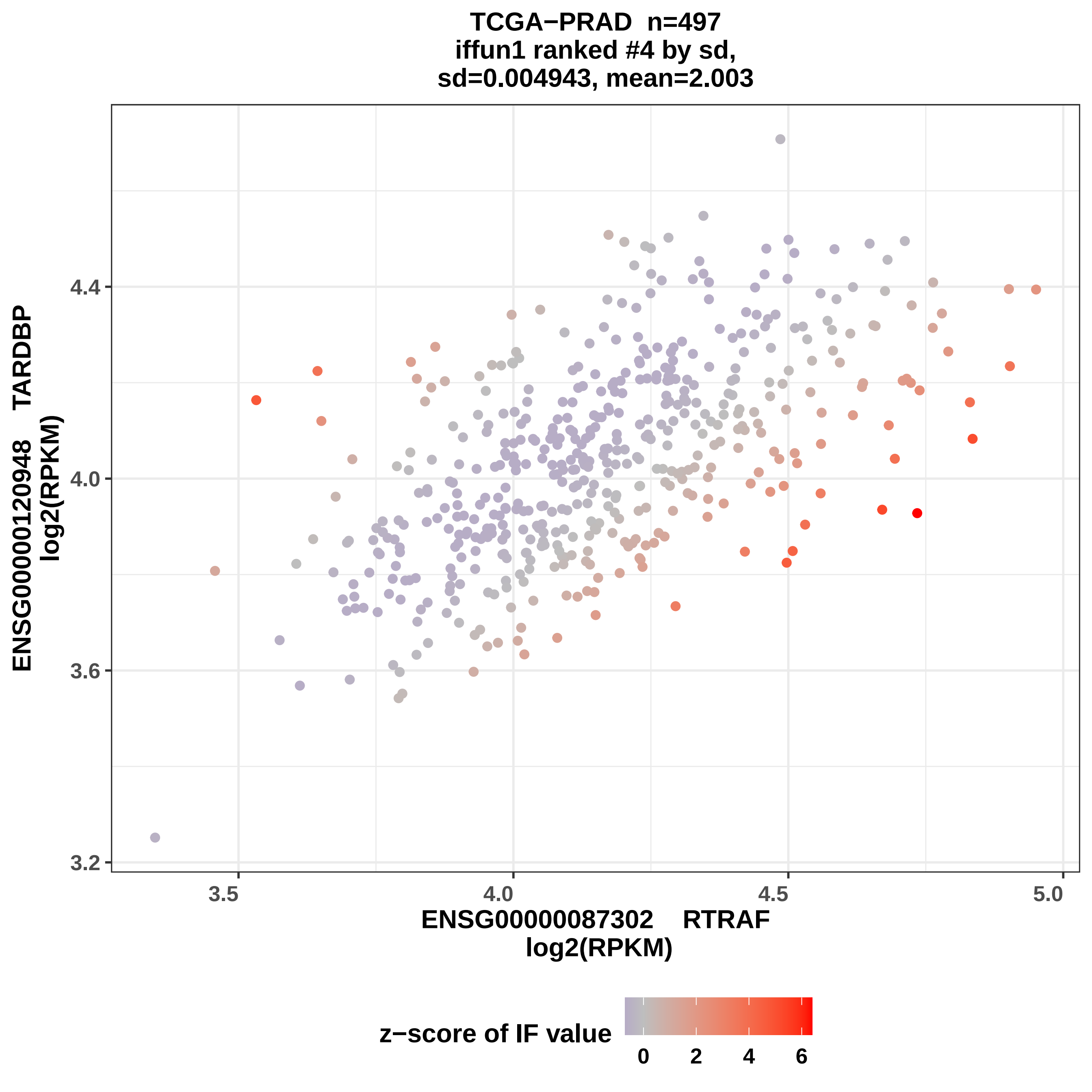
TCGA−PRAD n=497
iffun1 ranked #4 by sd,
sd=0.004943, mean=2.003
ENSG00000120948 TARDBP
log2(RPKM)
4.4
4.0
3.6
3.2
3.5
4.0
4.5
5.0
ENSG00000087302 RTRAF
log2(RPKM)
z−score of IF value
0
2
4
6

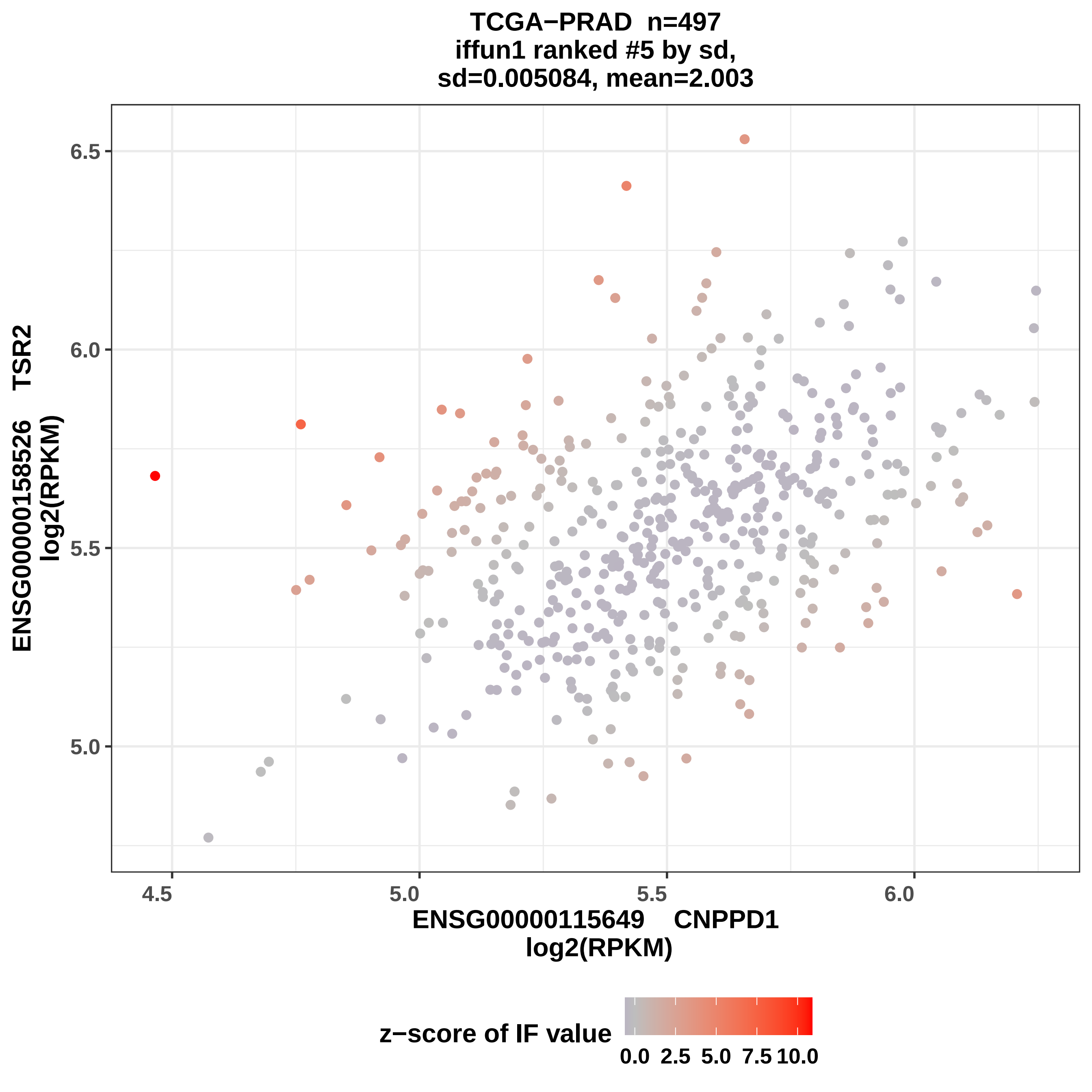
TCGA−PRAD n=497
iffun1 ranked #5 by sd,
sd=0.005084, mean=2.003
ENSG00000158526 TSR2
log2(RPKM)
6.5
6.0
5.5
5.0
4.5
5.0
5.5
6.0
ENSG00000115649 CNPPD1
log2(RPKM)
z−score of IF value
0.0
2.5
5.0
7.5
10.0

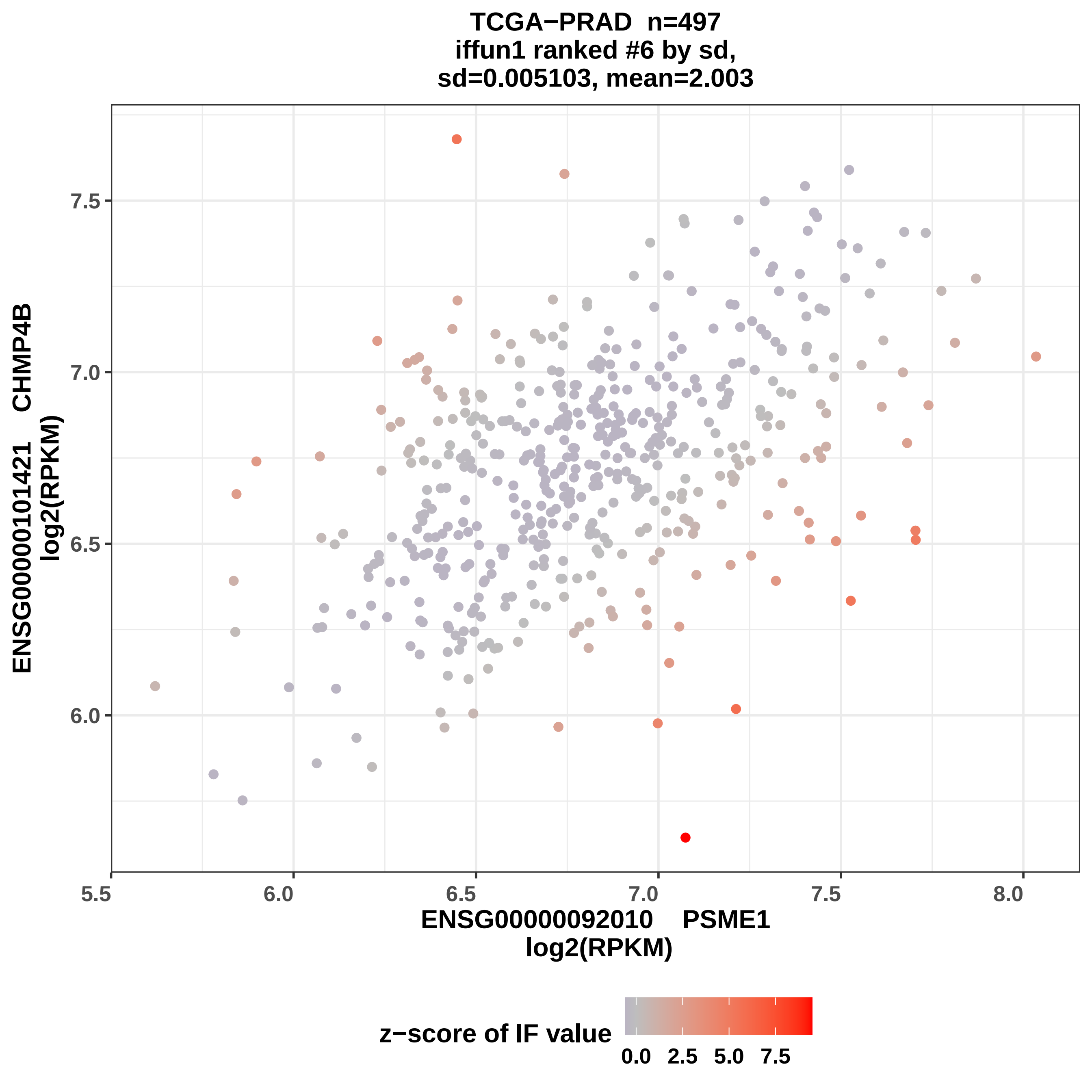

TCGA−PRAD n=497
iffun1 ranked #6 by sd,
sd=0.005103, mean=2.003
ENSG00000101421 CHMP4B
log2(RPKM)
7.5
7.0
6.5
6.0
5.5
6.0
6.5
7.0
7.5
8.0
ENSG00000092010 PSME1
log2(RPKM)
z−score of IF value
0.0
2.5
5.0
7.5

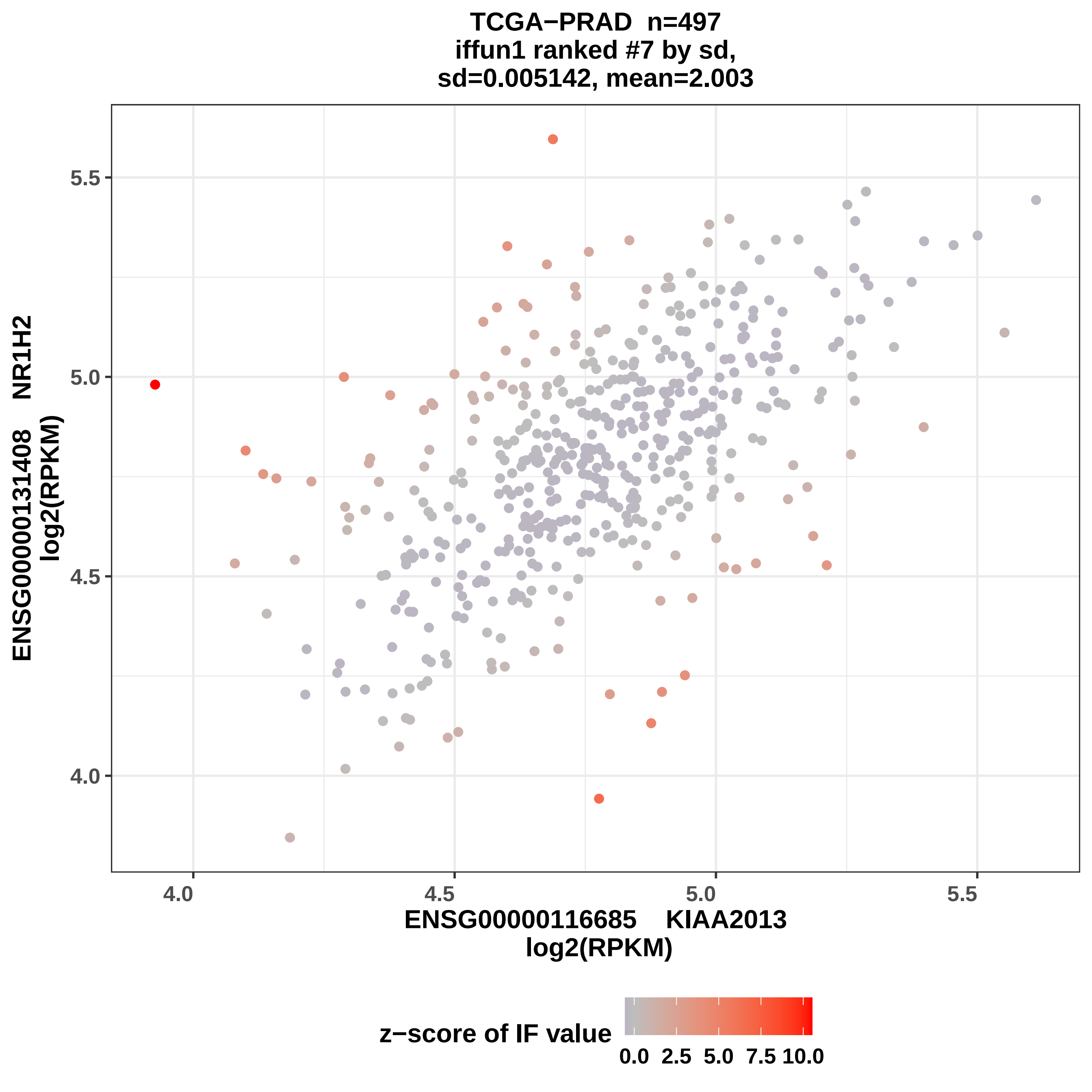
TCGA−PRAD n=497
iffun1 ranked #7 by sd,
sd=0.005142, mean=2.003
ENSG00000131408 NR1H2
log2(RPKM)
ENSG0000116685 KIAA2013
log2(RPKM)
5.5
5.0
4.5
4.0
4.0
4.5
5.0
5.5
z−score of IF value
0.0
2.5
5.0
7.5
10.0

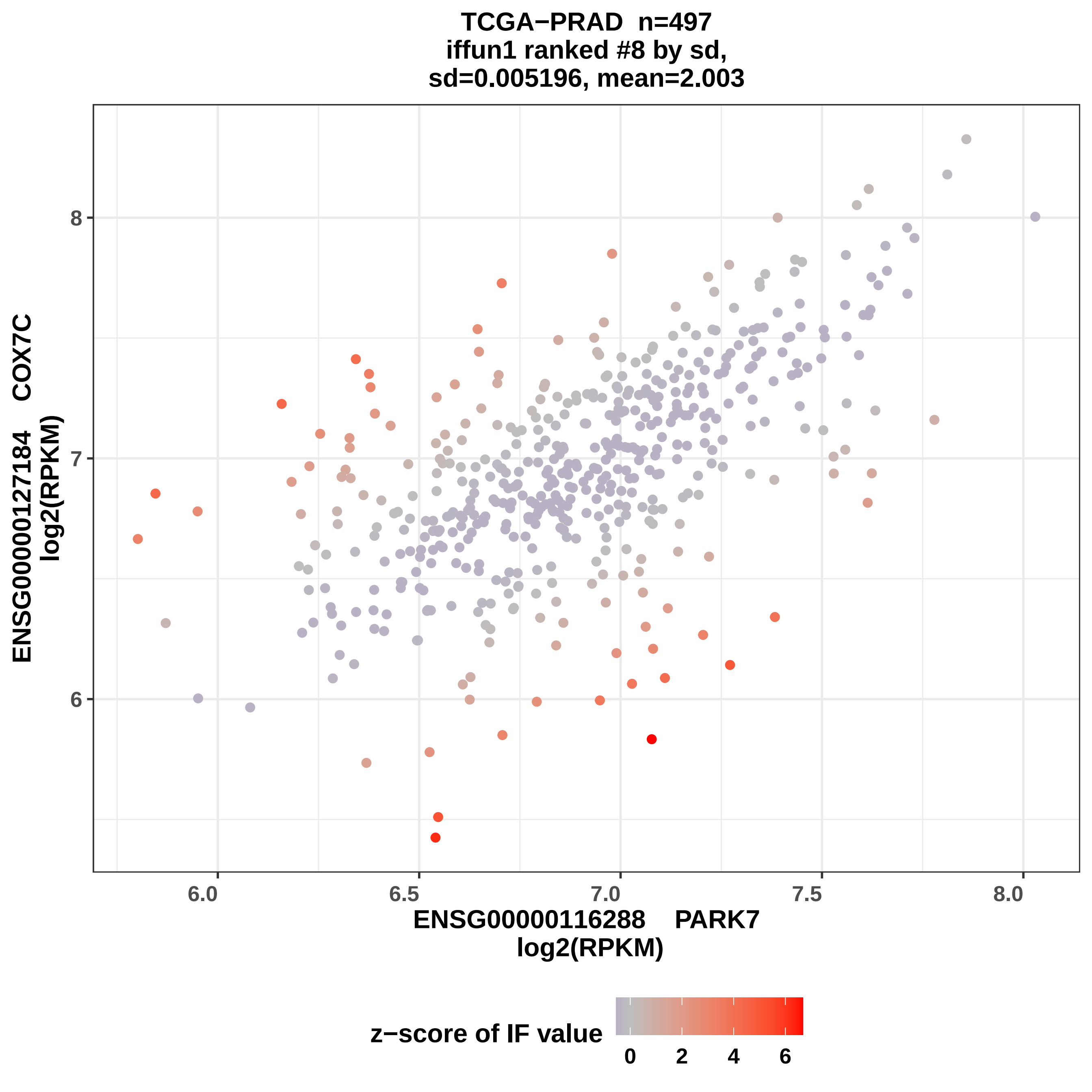
TCGA−PRAD n=497
iffun1 ranked #8 by sd,
sd=0.005196, mean=2.003
ENSG00000127184 COX7C
log2(RPKM)
8
7
6
6.0
6.5
7.0
7.5
8.0
ENSG00000116288 PARK7
log2(RPKM)
z−score of IF value
0
2
4
6

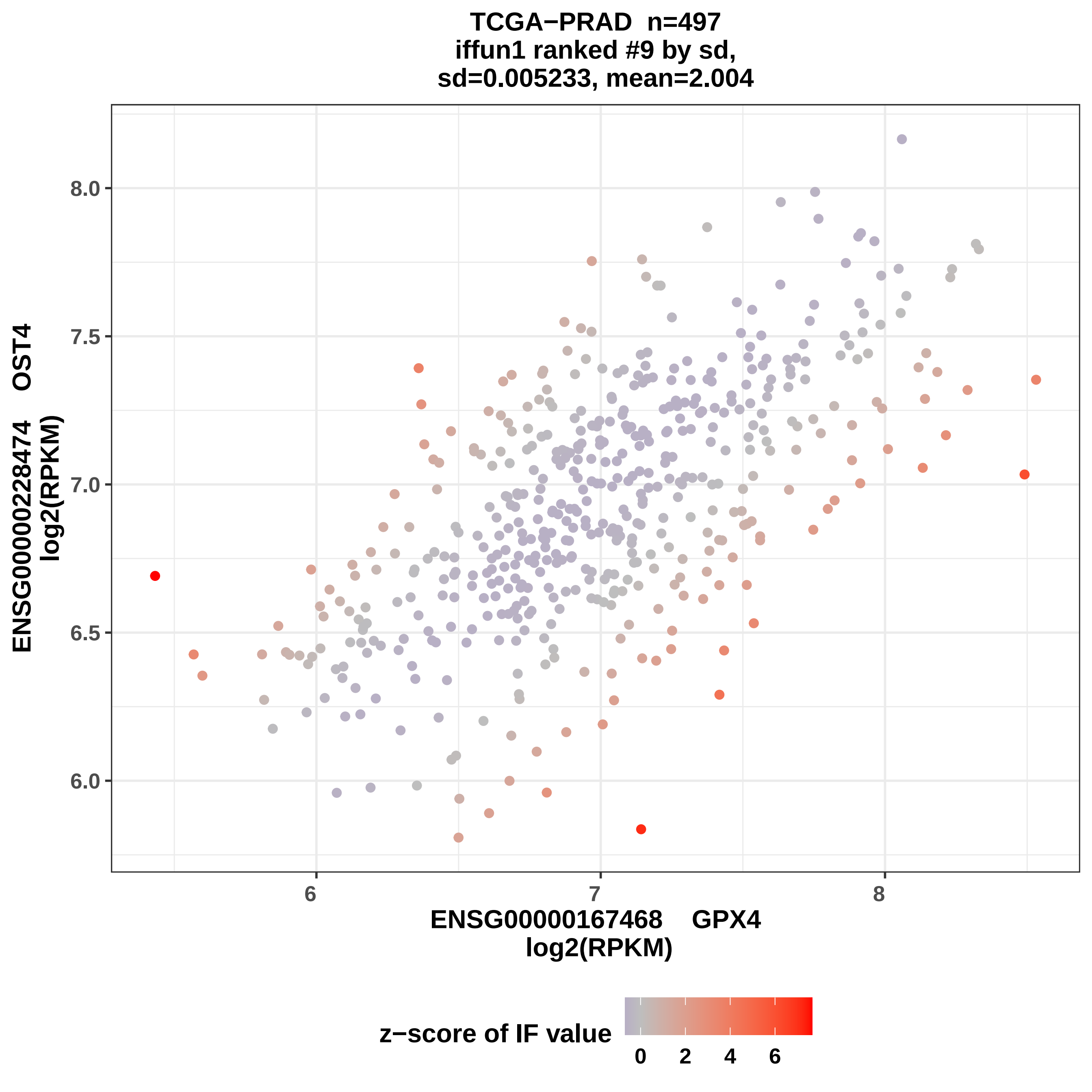

TCGA−PRAD n=497
iffun1 ranked #9 by sd,
sd=0.005233, mean=2.004
ENSG00000228474 OST4
log2(RPKM)
8.0
7.5
7.0
6.5
6.0
6
7
8
ENSG00000167468 GPX4
log2(RPKM)
z−score of IF value
0
2
4
6

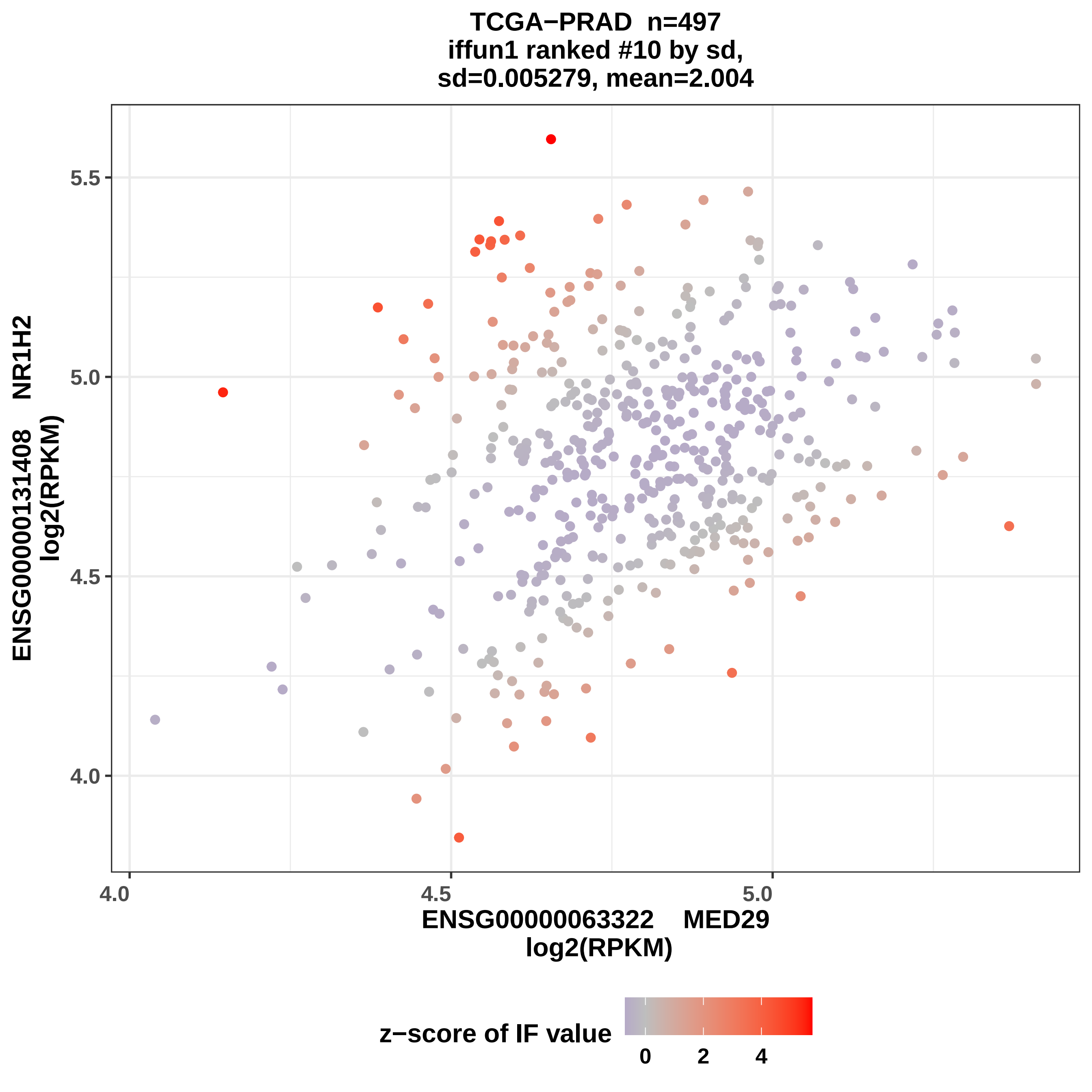

TCGA−PRAD n=497
iffun1 ranked #10 by sd,
sd=0.005279, mean=2.004
ENSG00000131408 NR1H2
log2(RPKM)
5.5
5.0
4.5
4.0
4.0
4.5
5.0
ENSG00000063322 MED29
log2(RPKM)
z−score of IF value
0
2
4

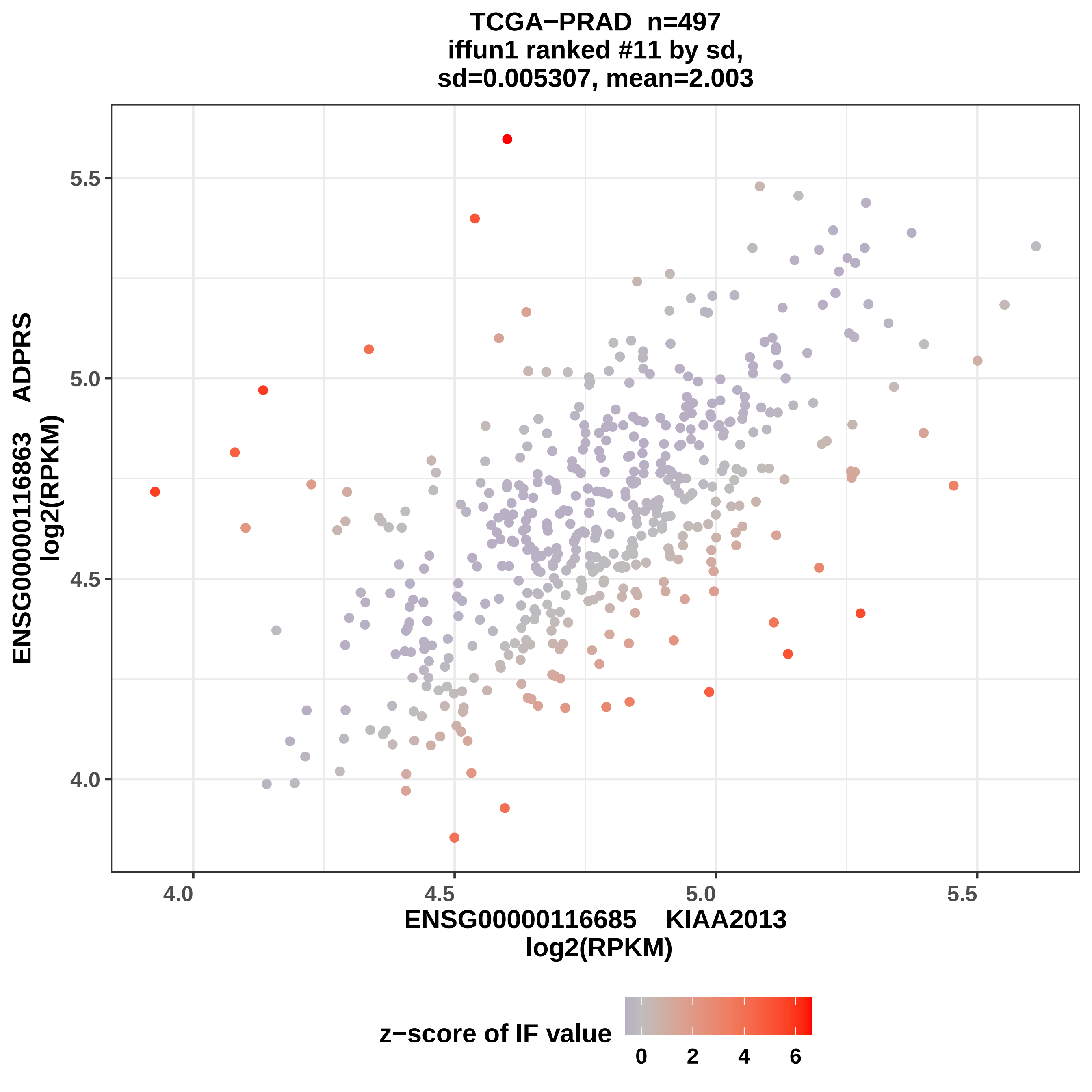

TCGA−PRAD n=497
iffun1 ranked #11 by sd,
sd=0.005307, mean=2.003
ENSG00000116863 ADPRS
log2(RPKM)
5.5
5.0
4.5
4.0
4.0
4.5
5.0
5.5
ENSG0000116685 KIAA2013
log2(RPKM)
z−score of IF value
0
2
4
6

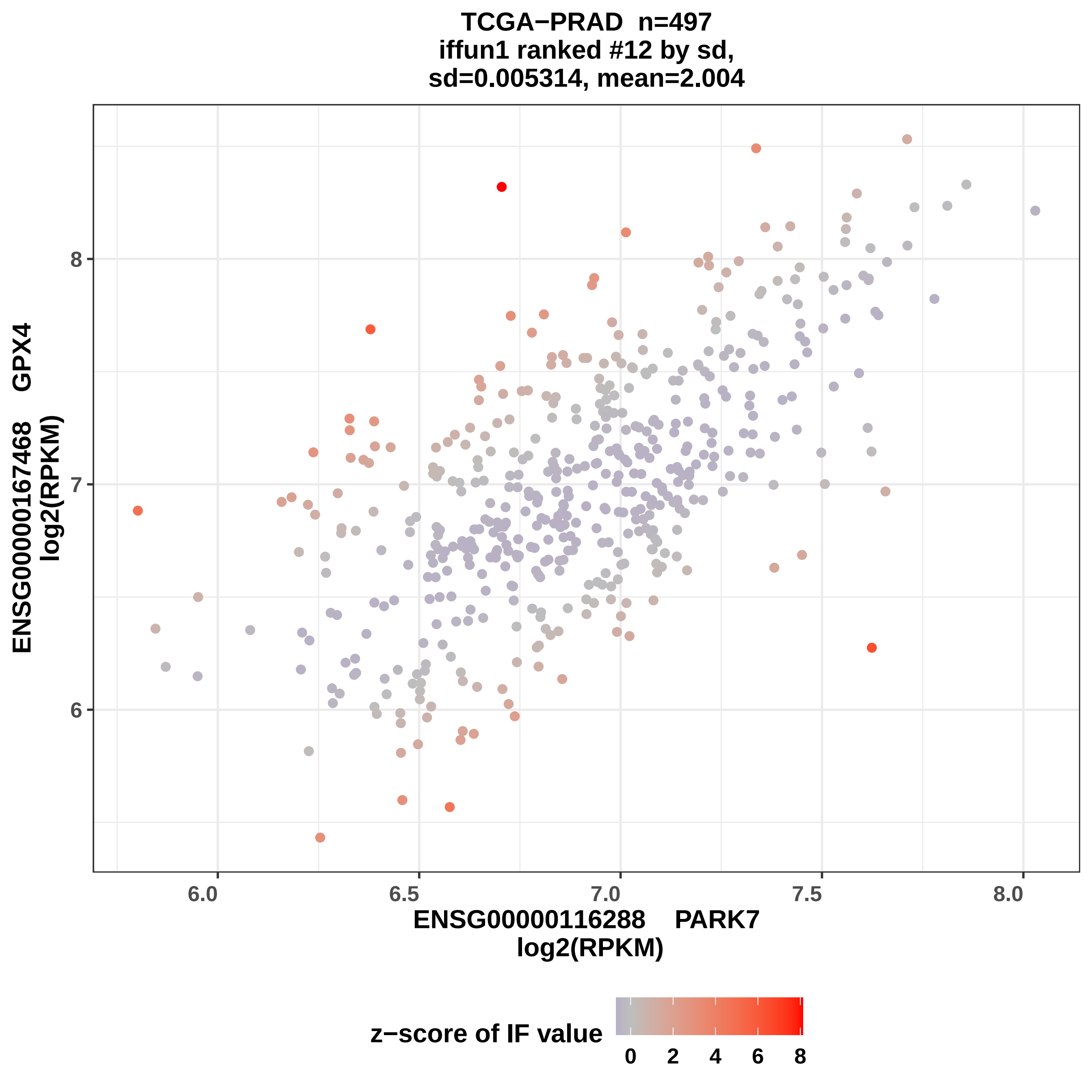
TCGA−PRAD n=497
iffun1 ranked #12 by sd,
sd=0.005314, mean=2.004
ENSG00000167468 GPX4
log2(RPKM)
8
7
6
6.0
6.5
7.0
7.5
8.0
ENSG00000116288 PARK7
log2(RPKM)
z−score of IF value
0
2
4
6
8

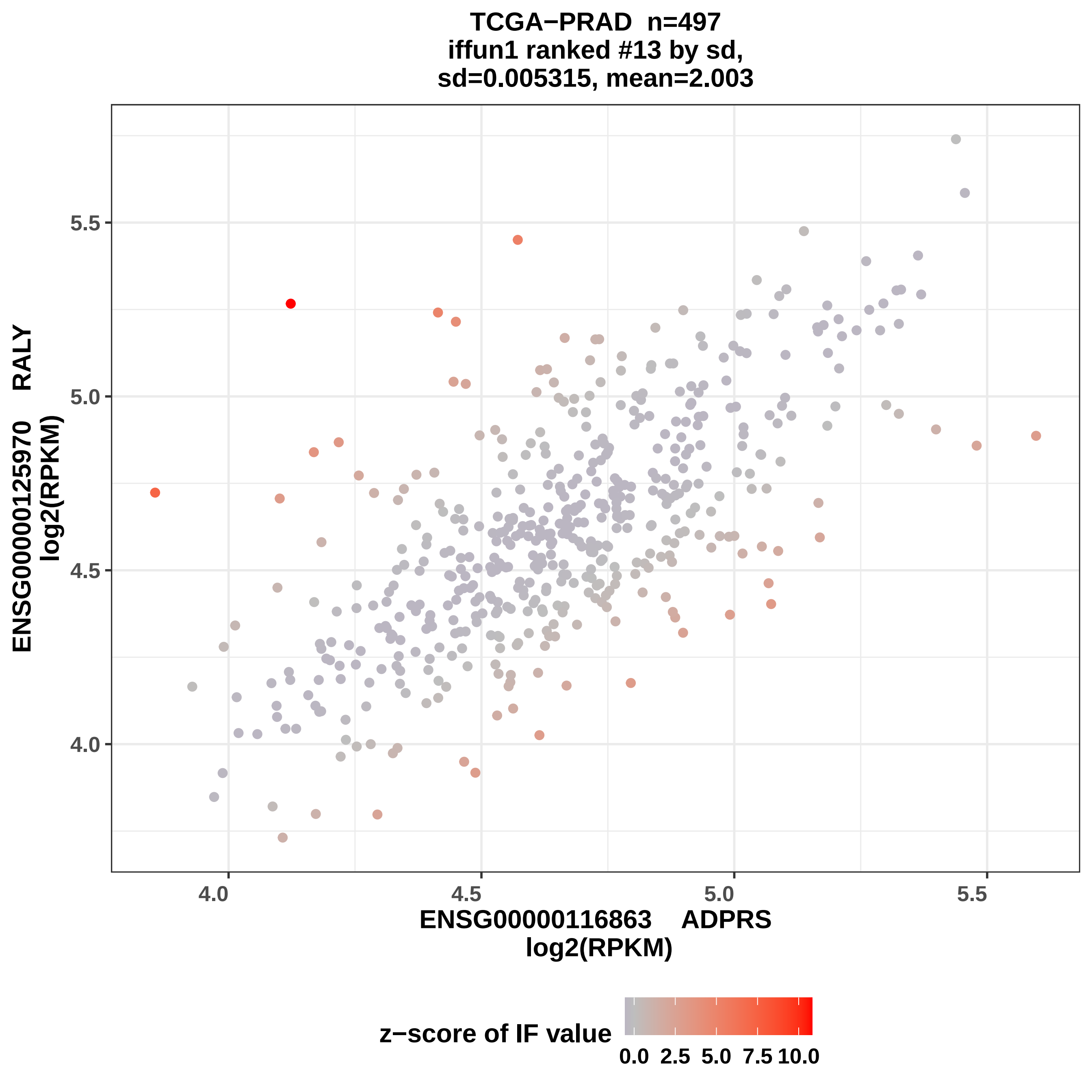
TCGA−PRAD n=497
iffun1 ranked #13 by sd,
sd=0.005315, mean=2.003
ENSG00000125970 RALY
log2(RPKM)
5.5
5.0
4.5
4.0
4.0
4.5
5.0
5.5
ENSG00000116863 ADPRS
log2(RPKM)
z−score of IF value
0.0 2.5 5.0 7.5 10.0

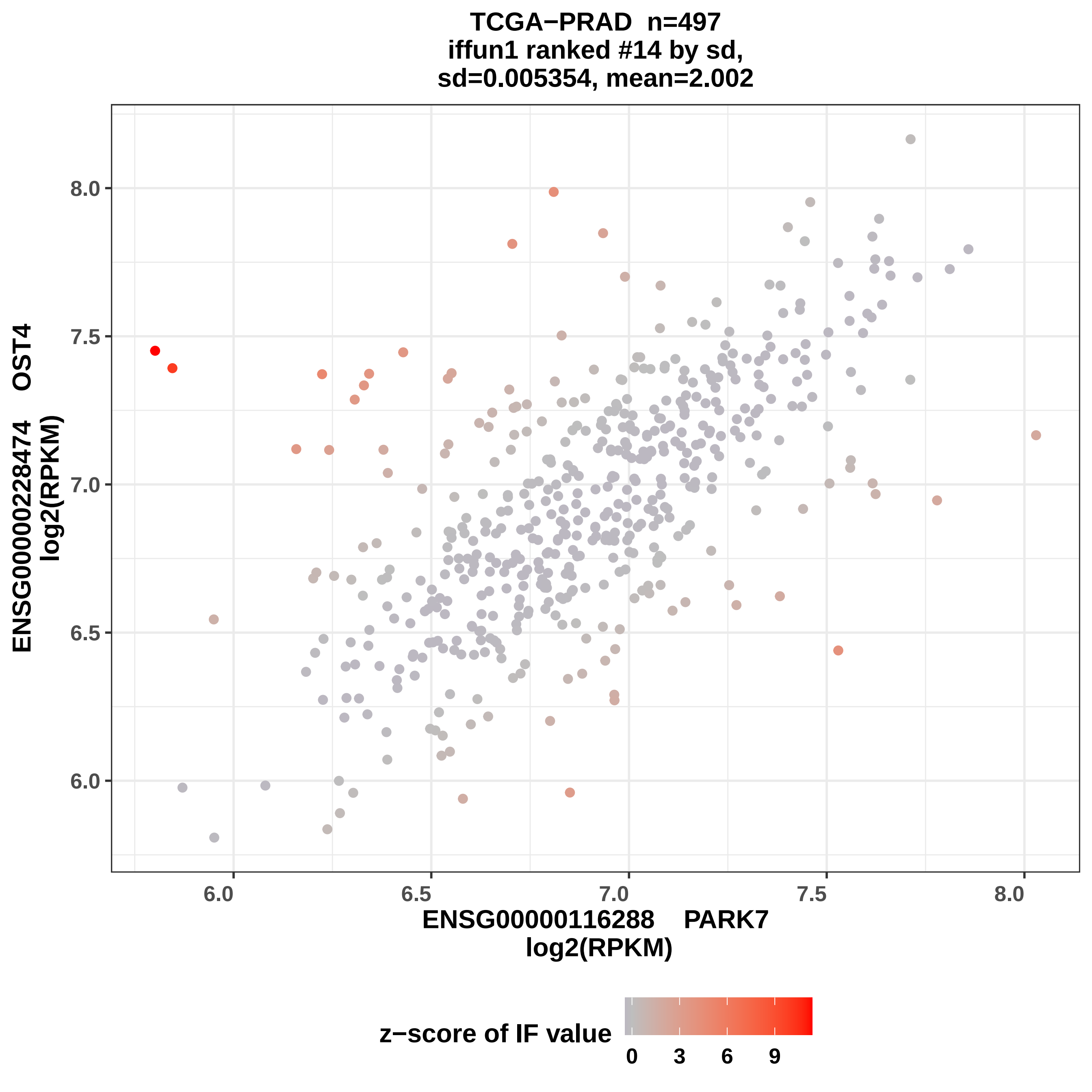
TCGA−PRAD n=497
iffun1 ranked #14 by sd,
sd=0.005354, mean=2.002
ENSG00000228474 OST4
log2(RPKM)
ENSG00000116288 PARK7
log2(RPKM)
8.0
7.5
7.0
6.5
6.0
6.0
6.5
7.0
7.5
8.0
z−score of IF value
0
3
6
9

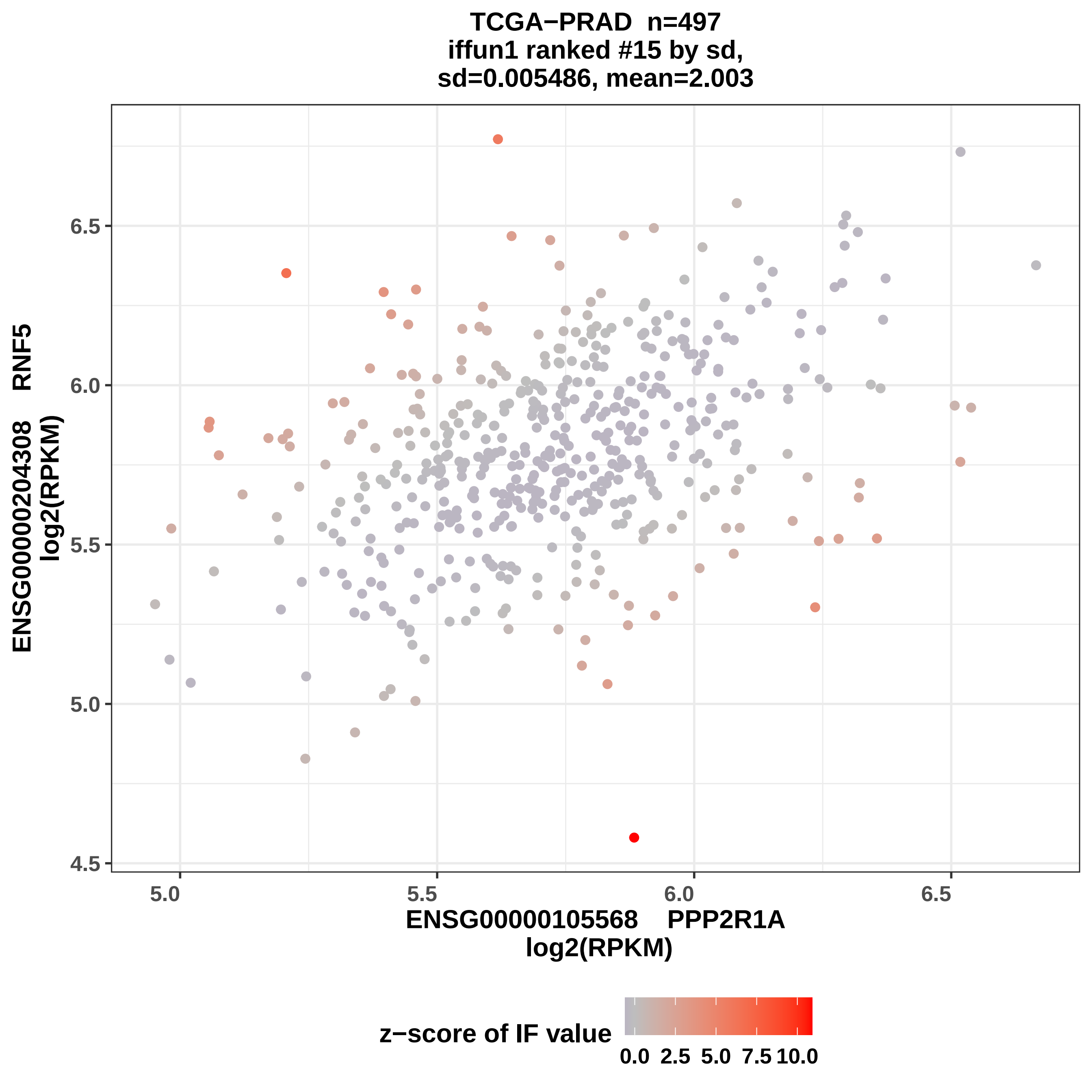

TCGA−PRAD n=497
iffun1 ranked #15 by sd,
sd=0.005486, mean=2.003
ENSG00000204308 RNF5
log2(RPKM)
6.5
6.0
5.5
5.0
4.5
5.0
5.5
6.0
6.5
ENSG00000105568 PPP2R1A
log2(RPKM)
z−score of IF value
0.0 2.5 5.0 7.5 10.0

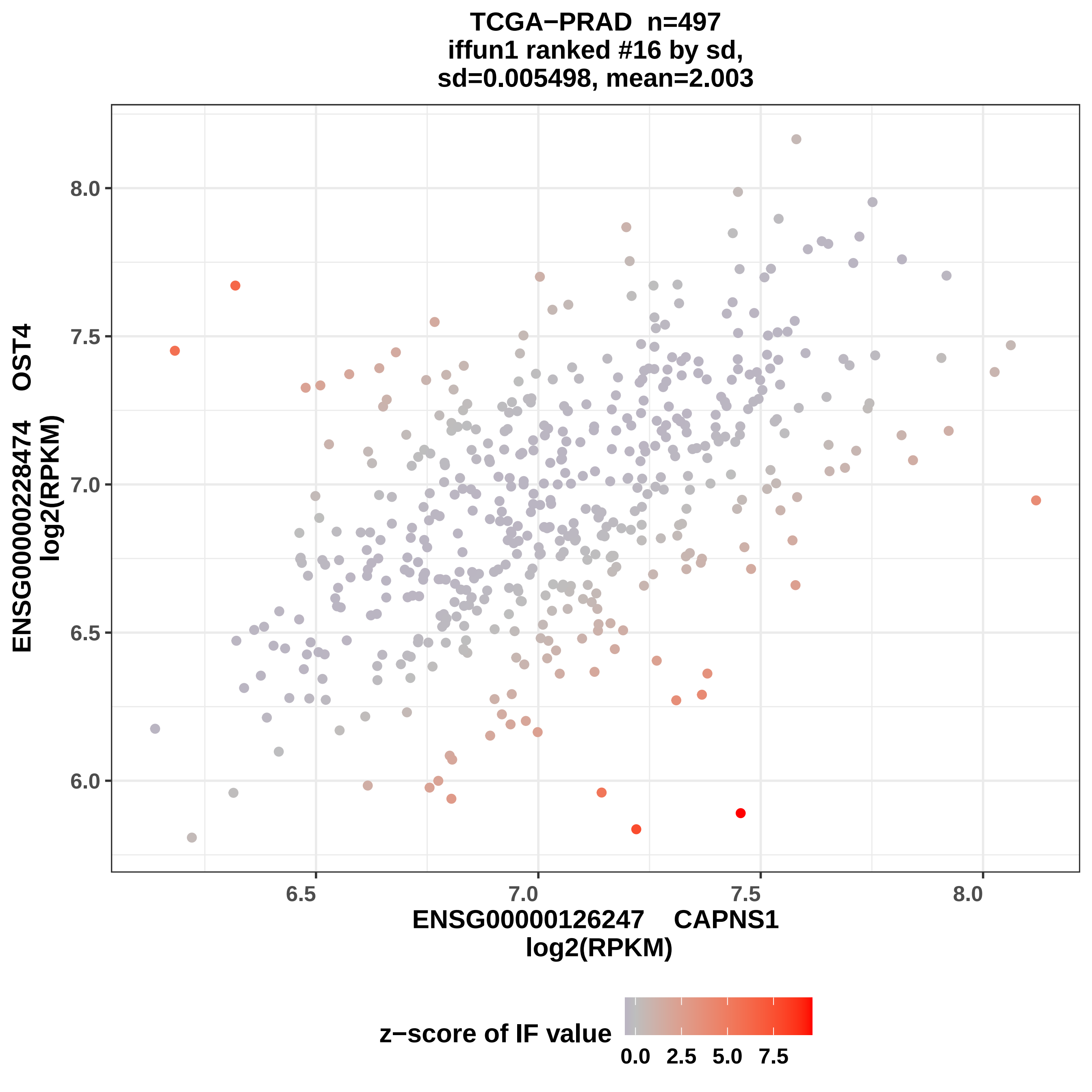

TCGA−PRAD n=497
iffun1 ranked #16 by sd,
sd=0.005498, mean=2.003
ENSG00000228474 OST4
log2(RPKM)
8.0
7.5
7.0
6.5
6.0
6.5
7.0
7.5
8.0
ENSG00000126247 CAPNS1
log2(RPKM)
z−score of IF value
0.0
2.5
5.0
7.5

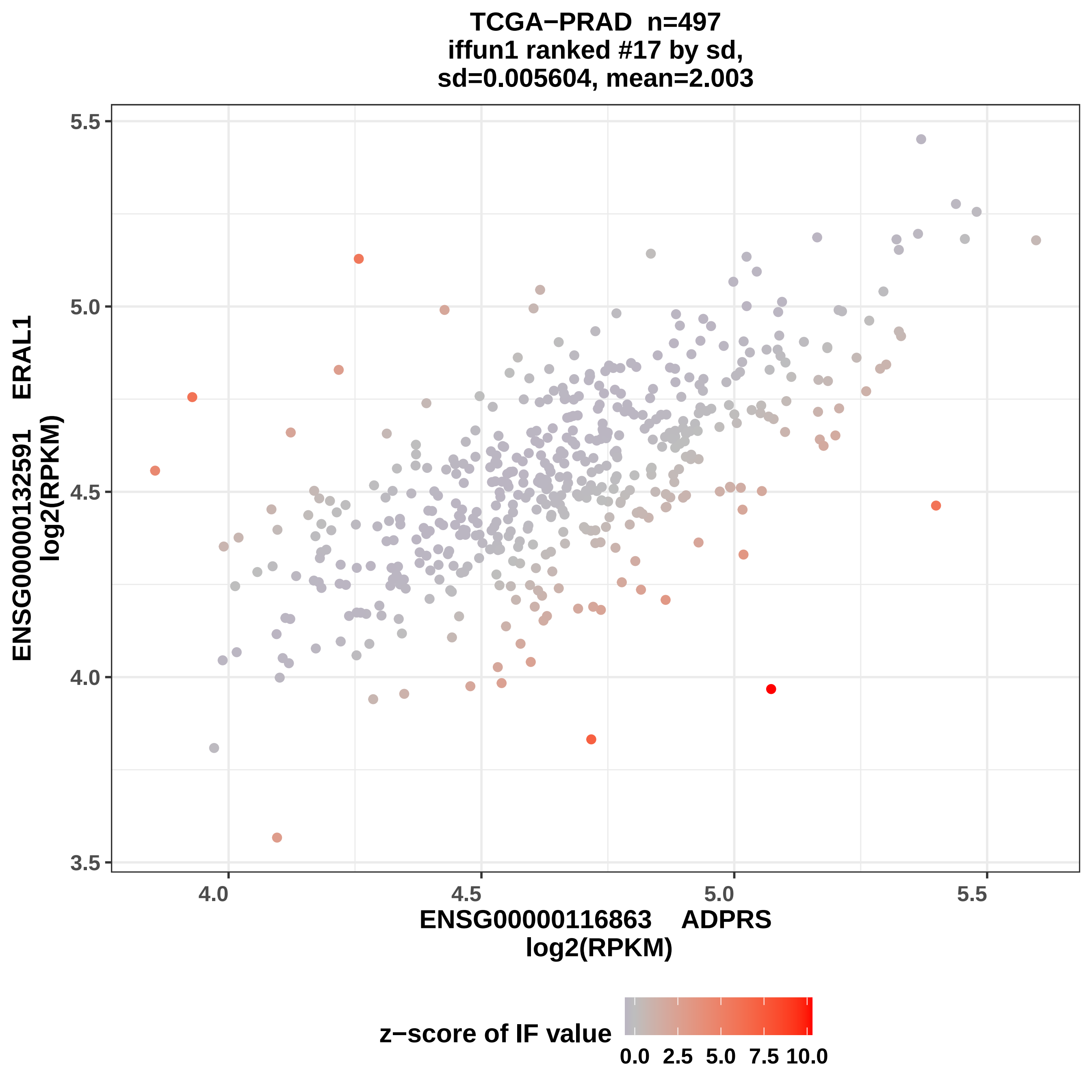

TCGA−PRAD n=497
iffun1 ranked #17 by sd,
sd=0.005604, mean=2.003
ENSG00000132591 ERAL1
log2(RPKM)
ENSG00000116863 ADPRS
log2(RPKM)
3.5
4.0
4.5
5.0
5.5
4.0
4.5
5.0
5.5
z−score of IF value
0.0
2.5
5.0
7.5
10.0

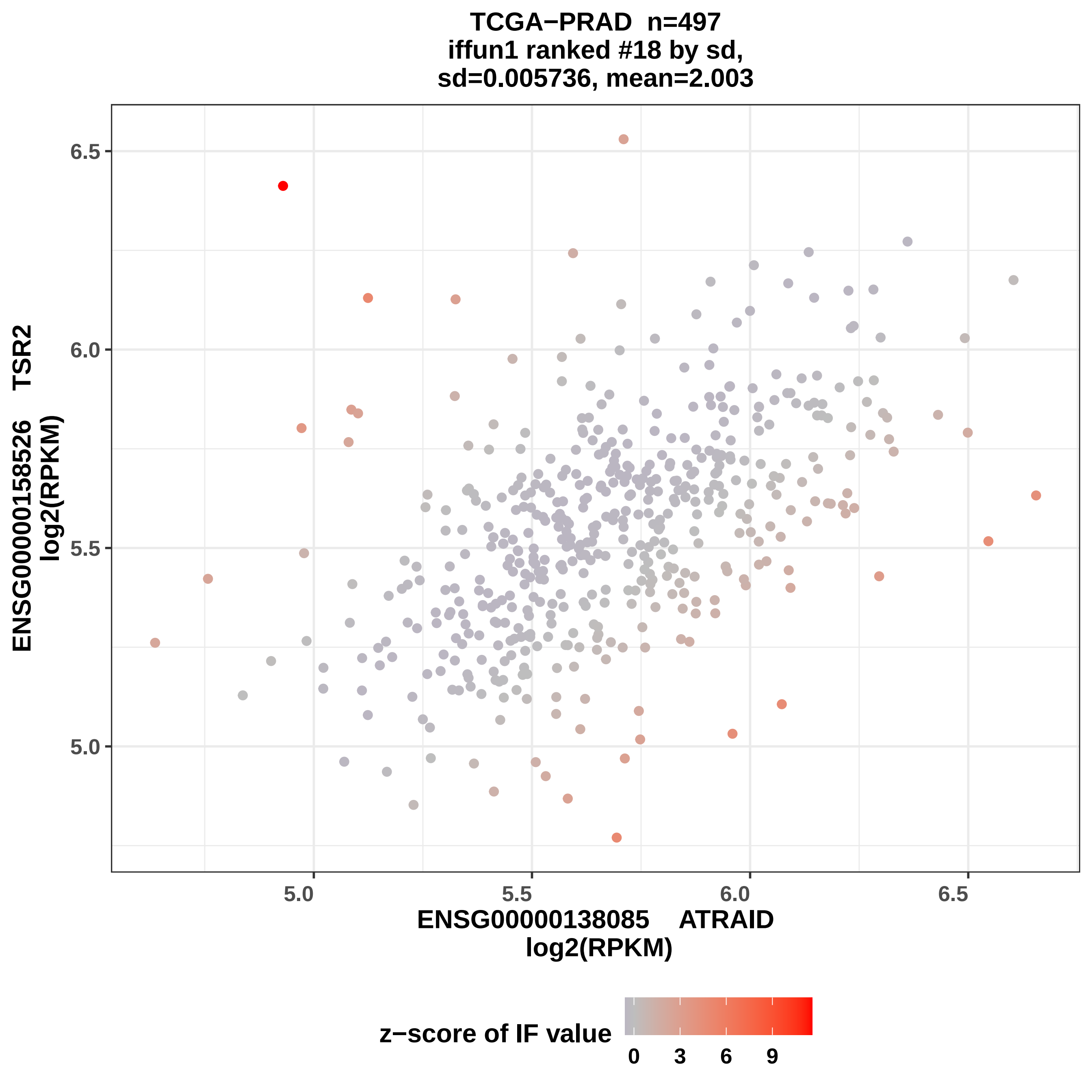
TCGA−PRAD n=497
iffun1 ranked #18 by sd,
sd=0.005736, mean=2.003
ENSG00000158526 TSR2
log2(RPKM)
6.5
6.0
5.5
5.0
ENSG00000138085 ATRAID
log2(RPKM)
z−score of IF value
0
3
6
9

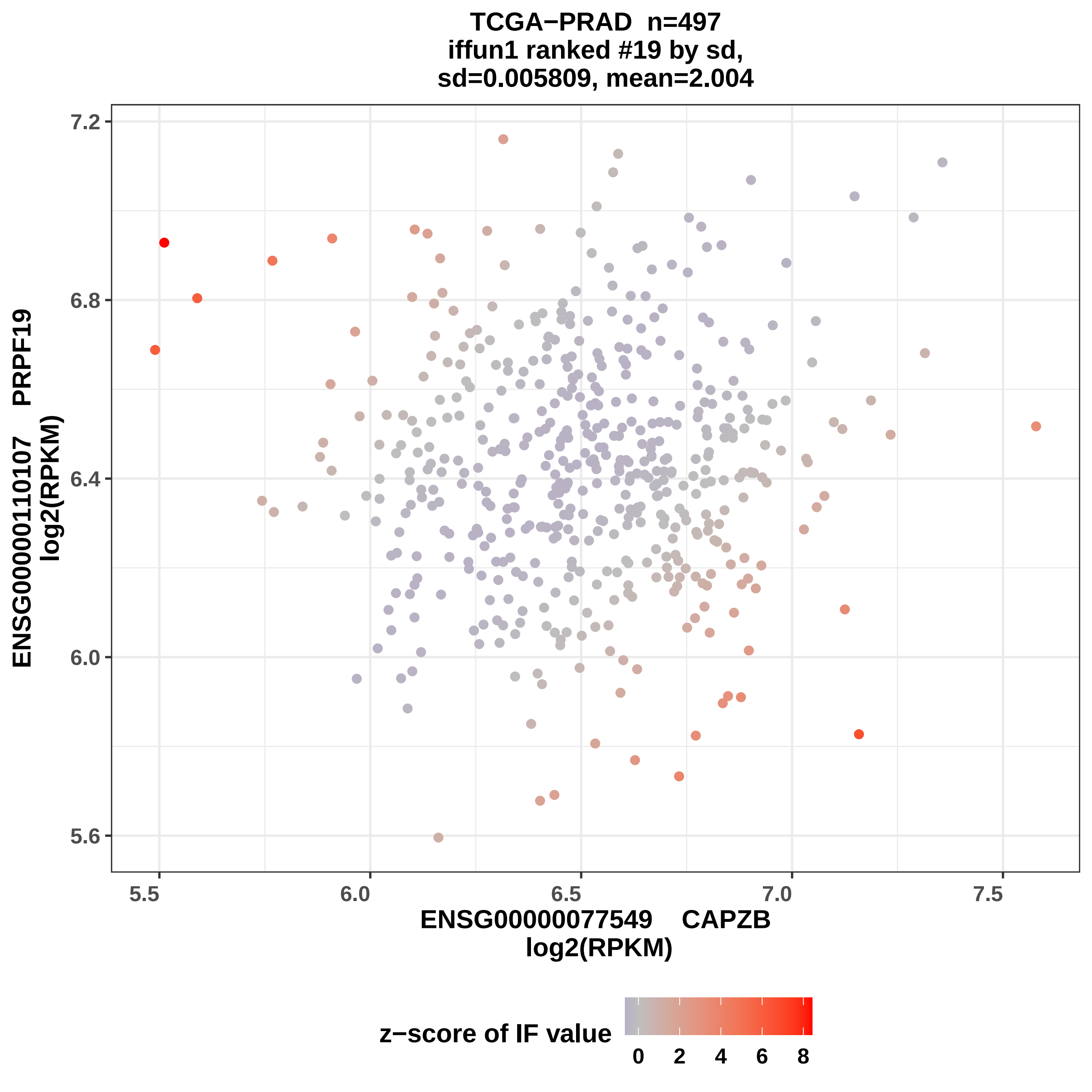

TCGA−PRAD n=497
iffun1 ranked #19 by sd,
sd=0.005809, mean=2.004
ENSG00000110107 PRPF19
log2(RPKM)
7.2
6.8
6.4
6.0
5.6
5.5
6.0
6.5
7.0
7.5
ENSG00000077549 CAPZB
log2(RPKM)
z−score of IF value
0
2
4
6
8

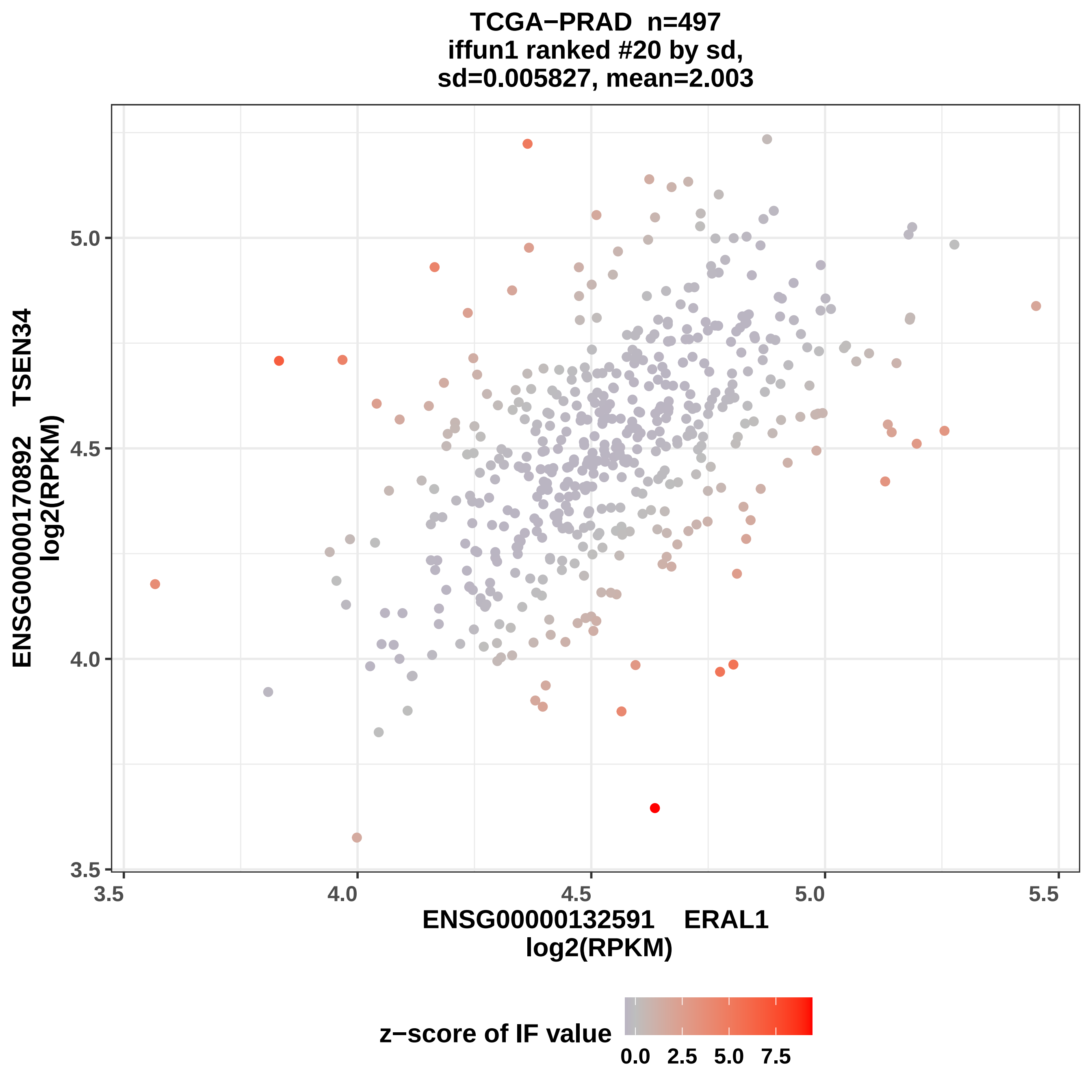
TCGA−PRAD n=497
iffun1 ranked #20 by sd,
sd=0.005827, mean=2.003
ENSG00000170892 TSEN34
log2(RPKM)
ENSG00000132591 ERAL1
log2(RPKM)
3.5
4.0
4.5
5.0
5.5
z−score of IF value
0.0 2.5 5.0 7.5

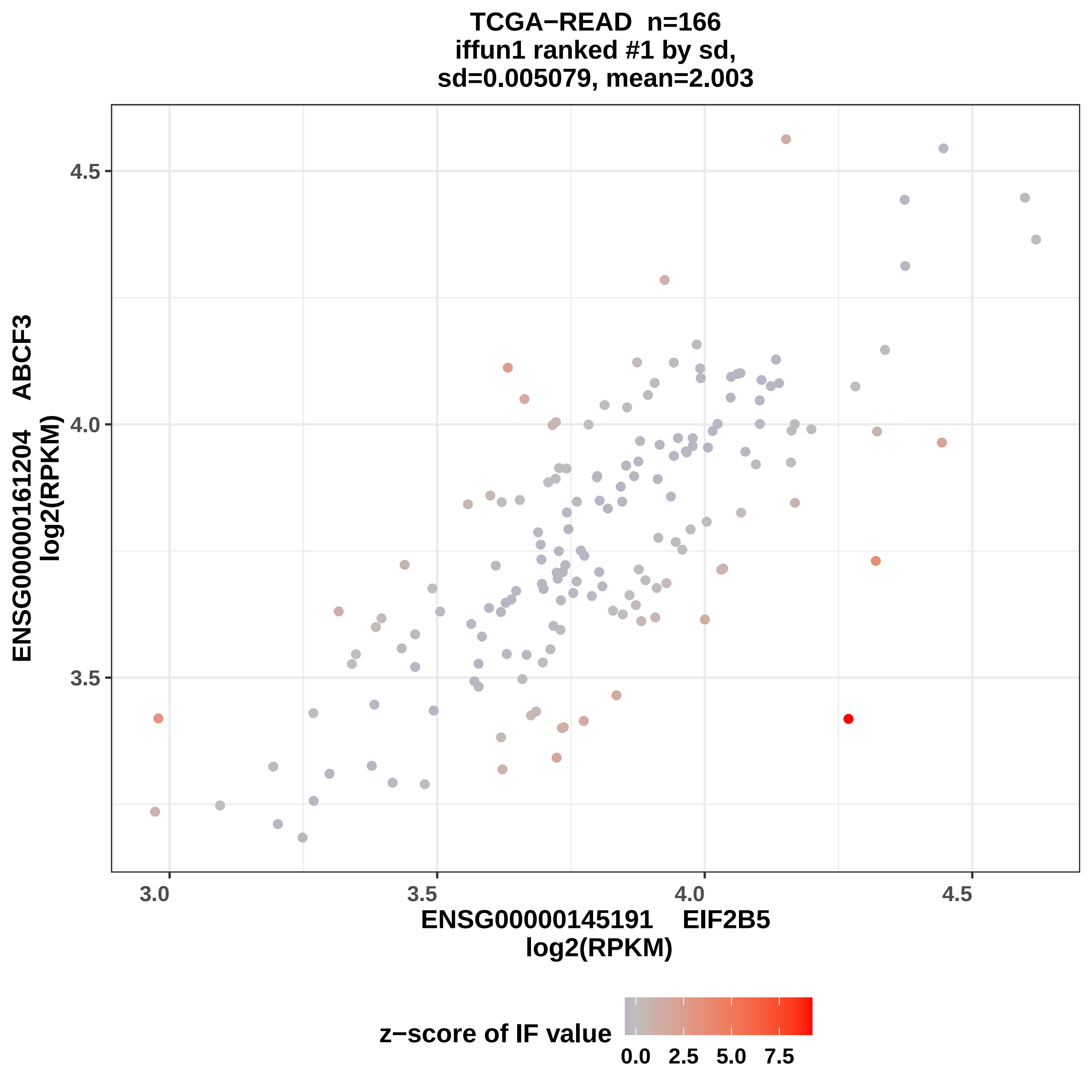

TCGA−READ n=166
iffun1 ranked #1 by sd,
sd=0.005079, mean=2.003
ENSG00000161204 ABCF3
log2(RPKM)
4.5
4.0
3.5
3.0
3.5
4.0
4.5
ENSG00000145191 EIF2B5
log2(RPKM)
z−score of IF value
0.0
2.5
5.0
7.5

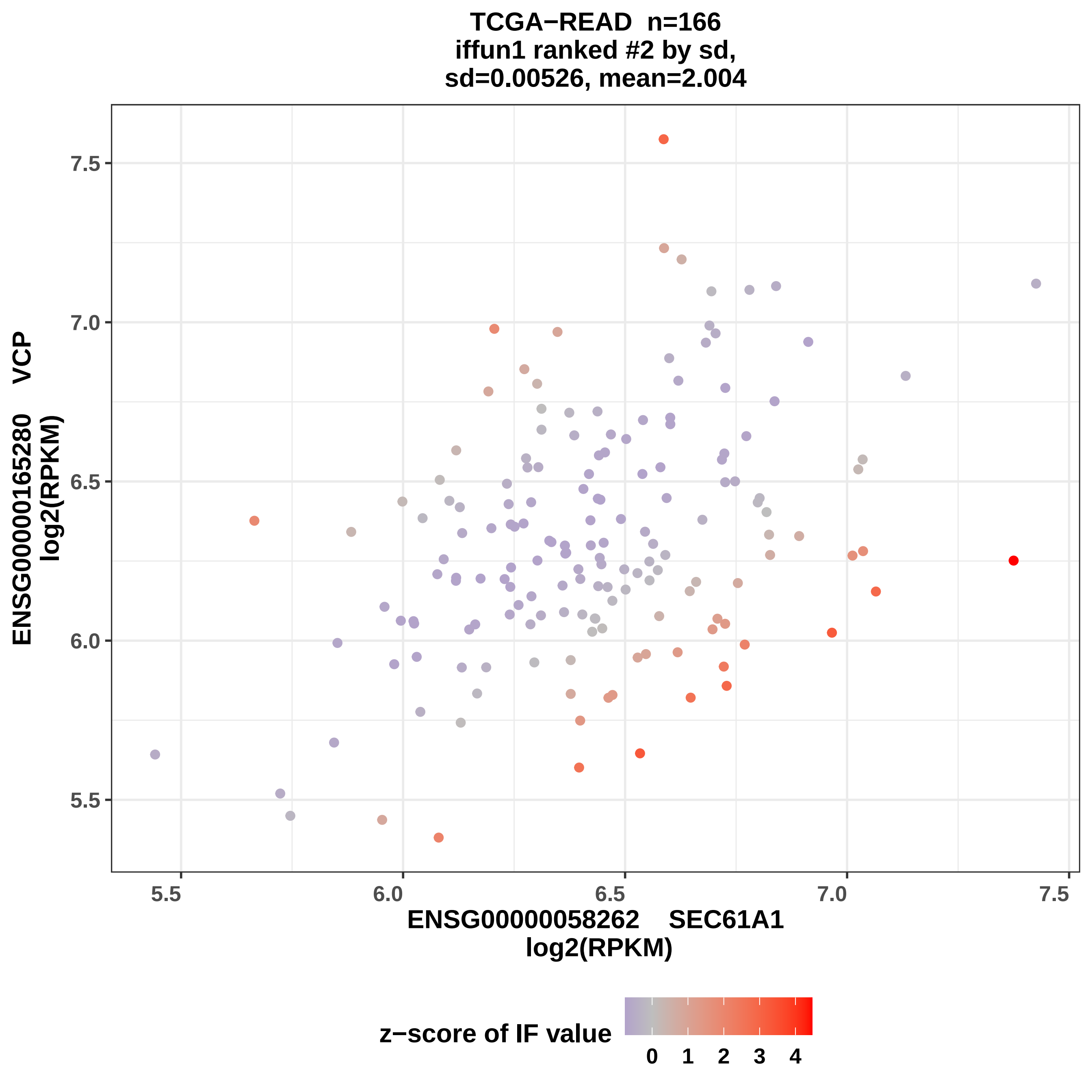

TCGA−READ n=166
iffun1 ranked #2 by sd,
sd=0.00526, mean=2.004
ENSG00000165280 VCP
log2(RPKM)
7.5
7.0
6.5
6.0
5.5
5.5
6.0
6.5
7.0
7.5
ENSG00000058262 SEC61A1
log2(RPKM)
z−score of IF value
0
1
2
3
4

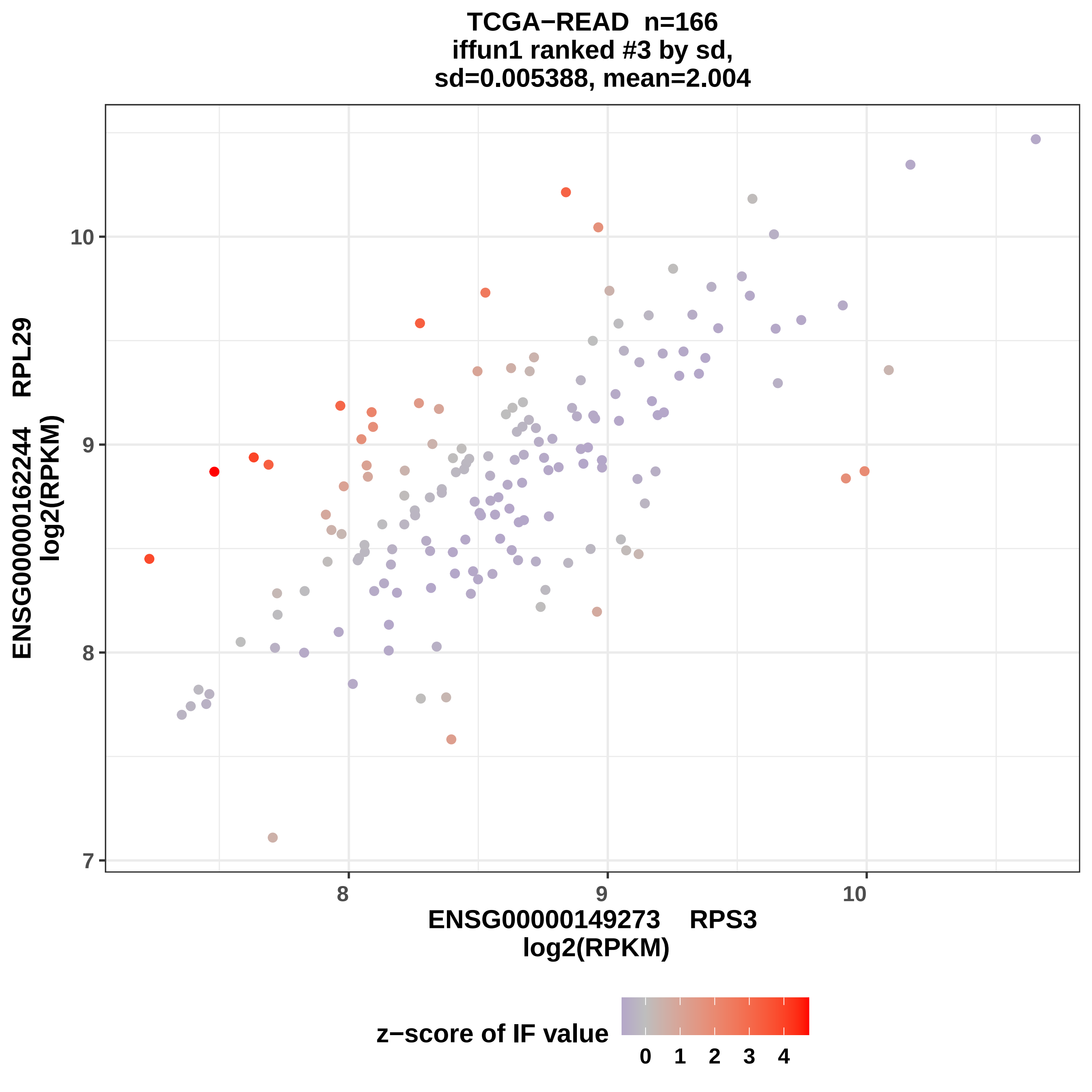

TCGA−READ n=166
iffun1 ranked #3 by sd,
sd=0.005388, mean=2.004
ENSG00000162244 RPL29
log2(RPKM)
ENSG00000149273 RPS3
log2(RPKM)
7
8
9
10
8
9
10
z−score of IF value
0
1
2
3
4

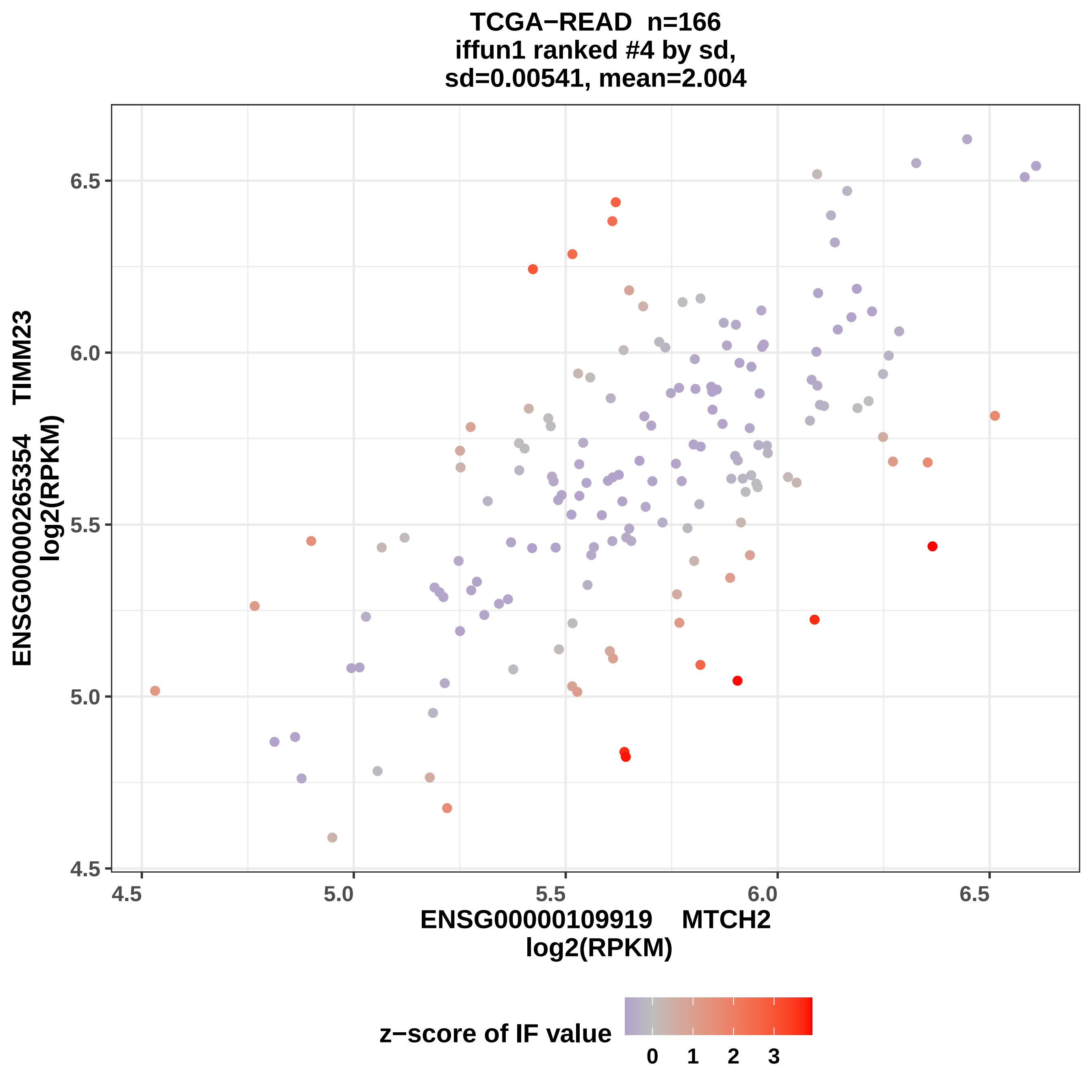
TCGA−READ n=166
iffun1 ranked #4 by sd,
sd=0.00541, mean=2.004
ENSG00000265354 TIMM23
log2(RPKM)
ENSG00000109919 MTCH2
log2(RPKM)
6.5
6.0
5.5
5.0
4.5
4.5
5.0
5.5
6.0
6.5
z−score of IF value
0
1
2
3

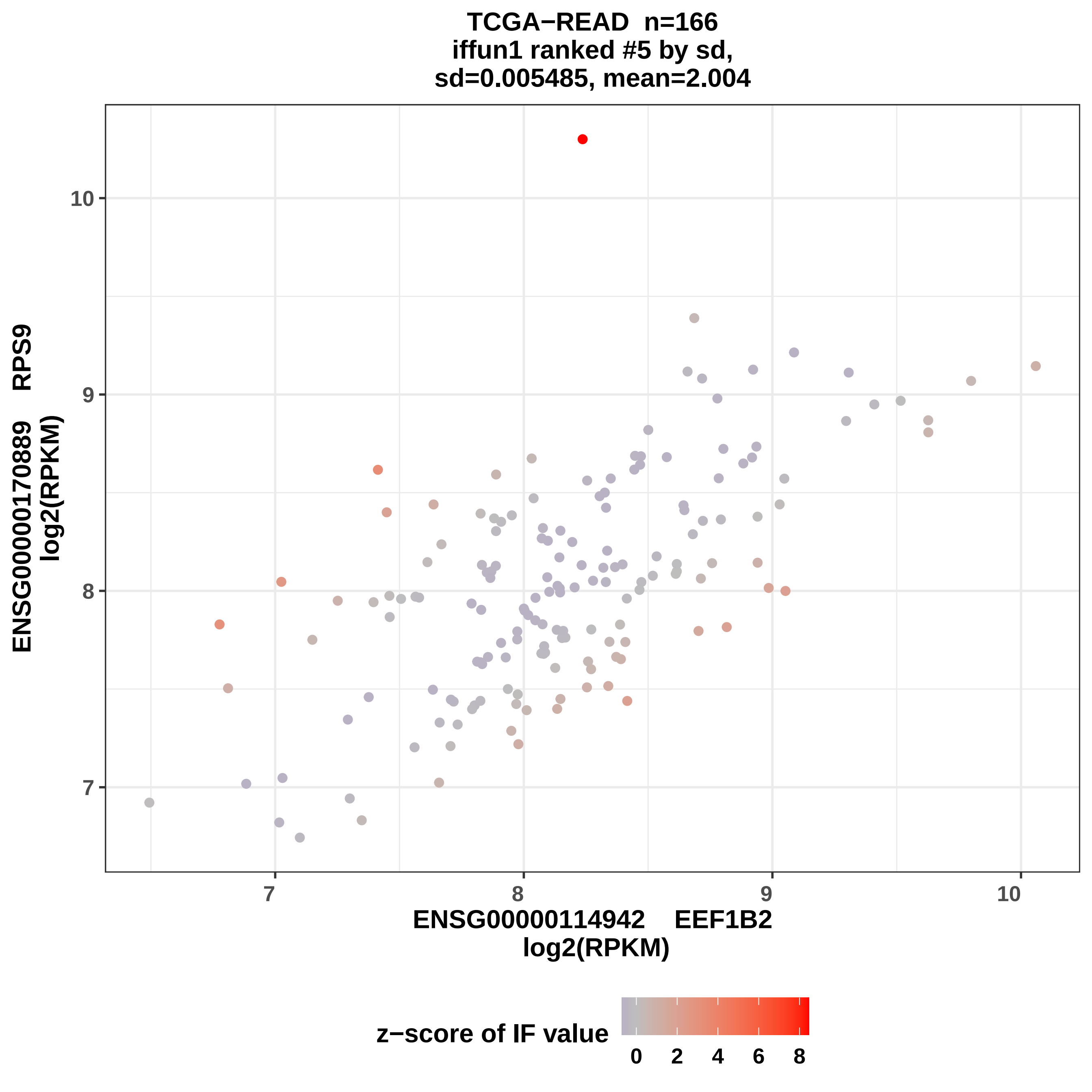

TCGA−READ n=166
iffun1 ranked #5 by sd,
sd=0.005485, mean=2.004
ENSG00000170889 RPS9
log2(RPKM)
10
9
8
7
ENSG00000114942 EEF1B2
log2(RPKM)
z−score of IF value
0
2
4
6
8

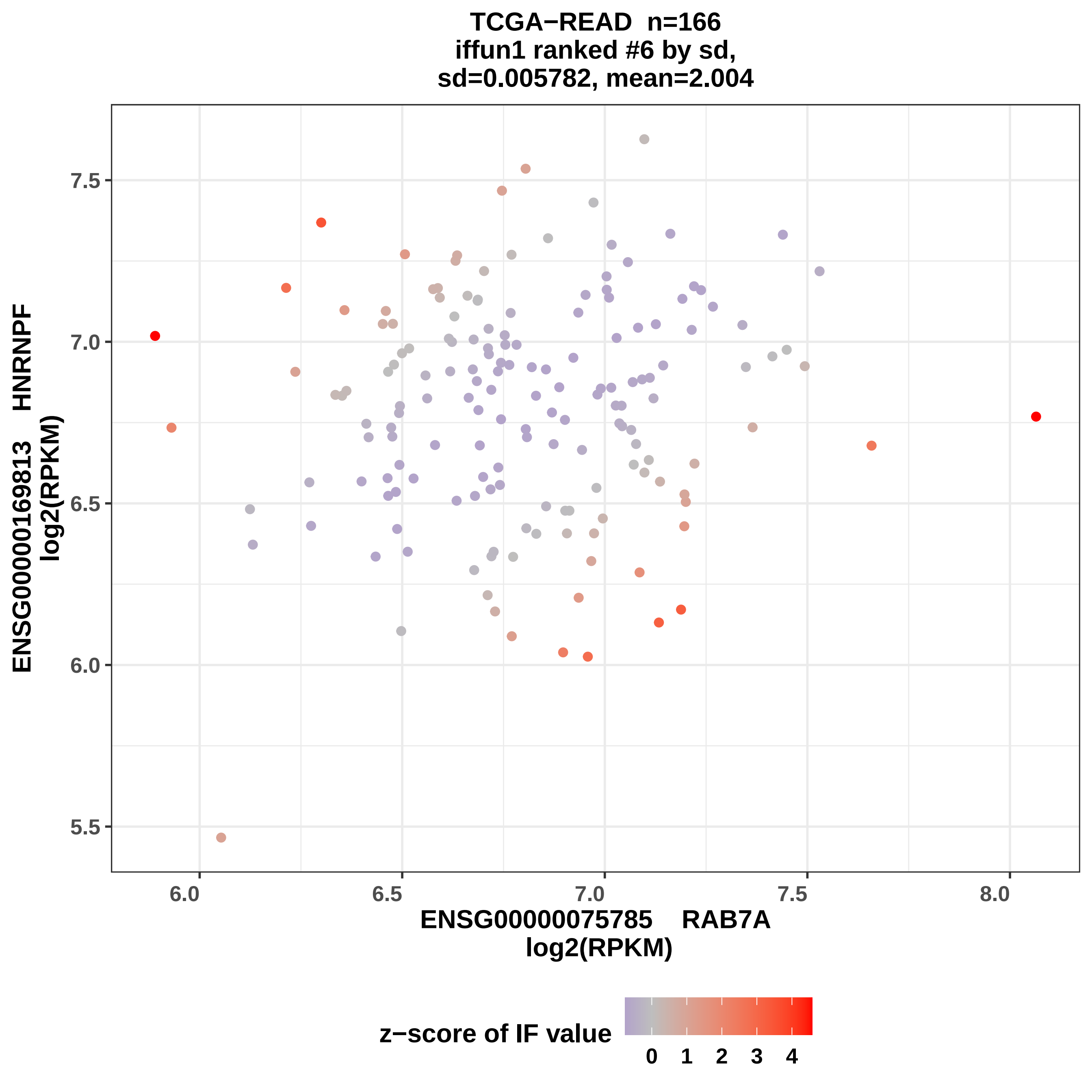

TCGA−READ n=166
iffun1 ranked #6 by sd,
sd=0.005782, mean=2.004
ENSG00000169813 HNRNPF
log2(RPKM)
7.5
7.0
6.5
6.0
5.5
6.0
6.5
7.0
7.5
8.0
ENSG00000075785 RAB7A
log2(RPKM)
z−score of IF value
0 1 2 3 4

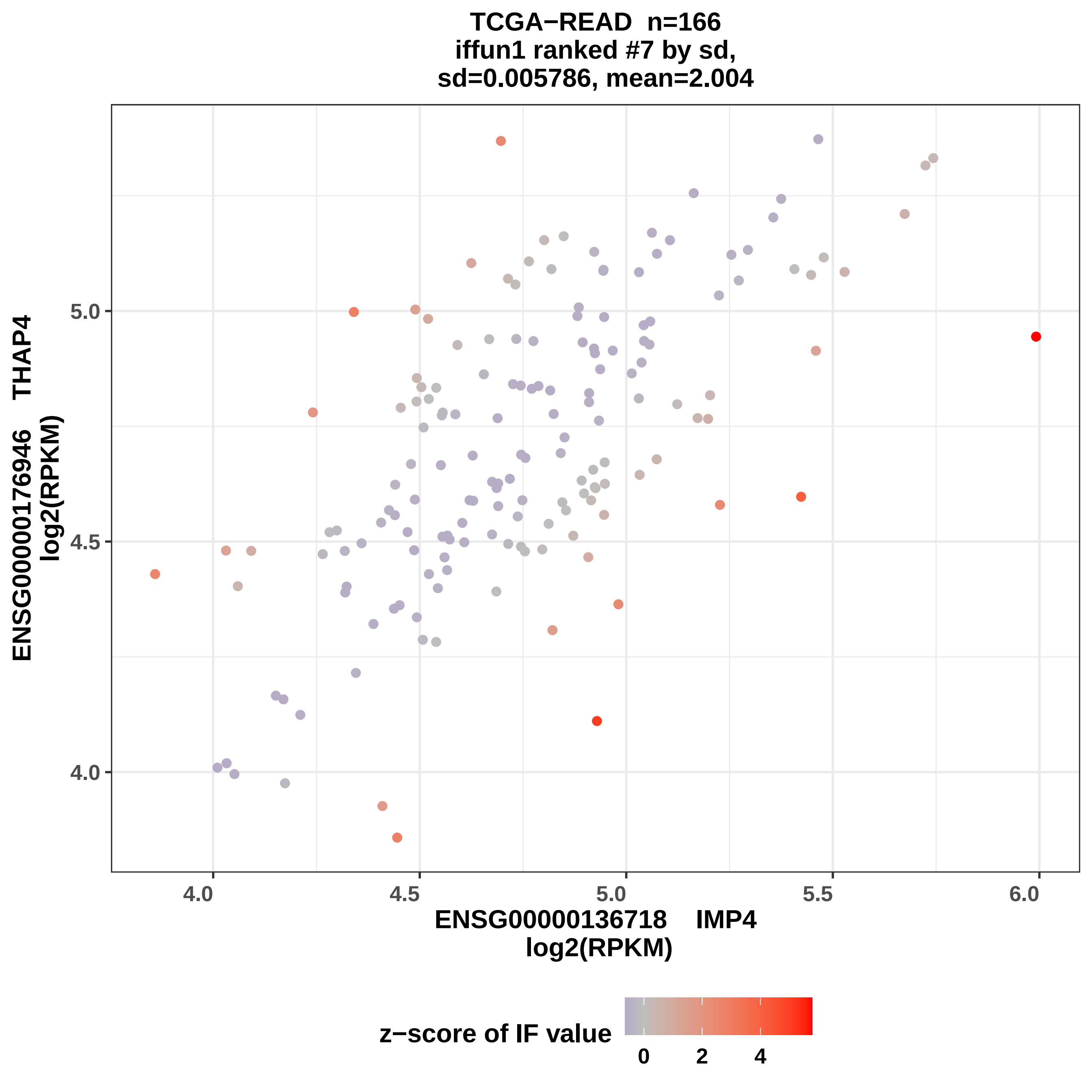
TCGA−READ n=166
iffun1 ranked #7 by sd,
sd=0.005786, mean=2.004
ENSG00000176946 THAP4
log2(RPKM)
ENSG00000136718 IMP4
log2(RPKM)
4.0
4.5
5.0
5.5
6.0
z−score of IF value
0
2
4

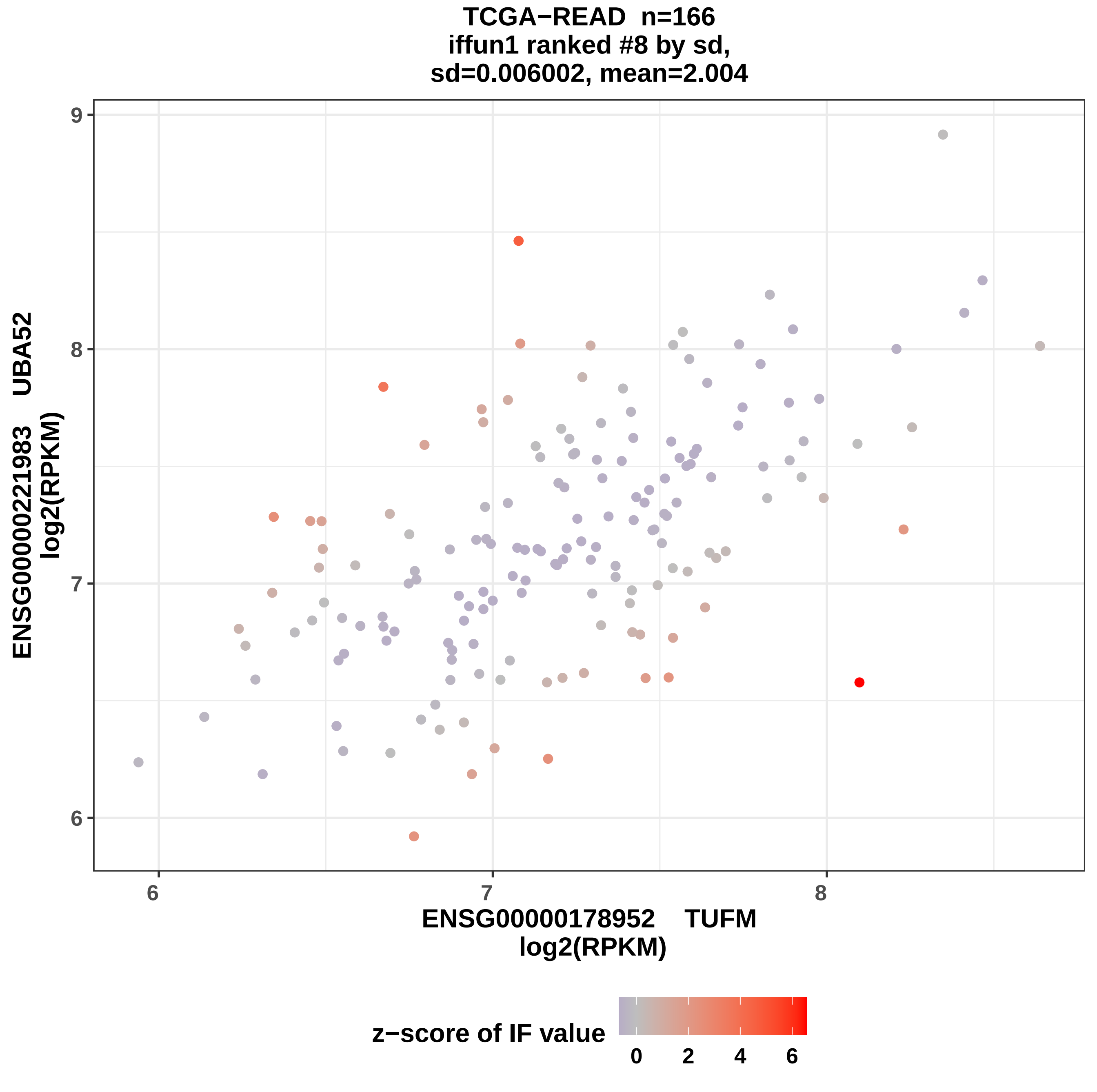
TCGA−READ n=166
iffun1 ranked #8 by sd,
sd=0.006002, mean=2.004
ENSG00000221983 UBA52
log2(RPKM)
6
7
8
9
ENSG00000178952 TUFM
log2(RPKM)
z−score of IF value
0
2
4
6

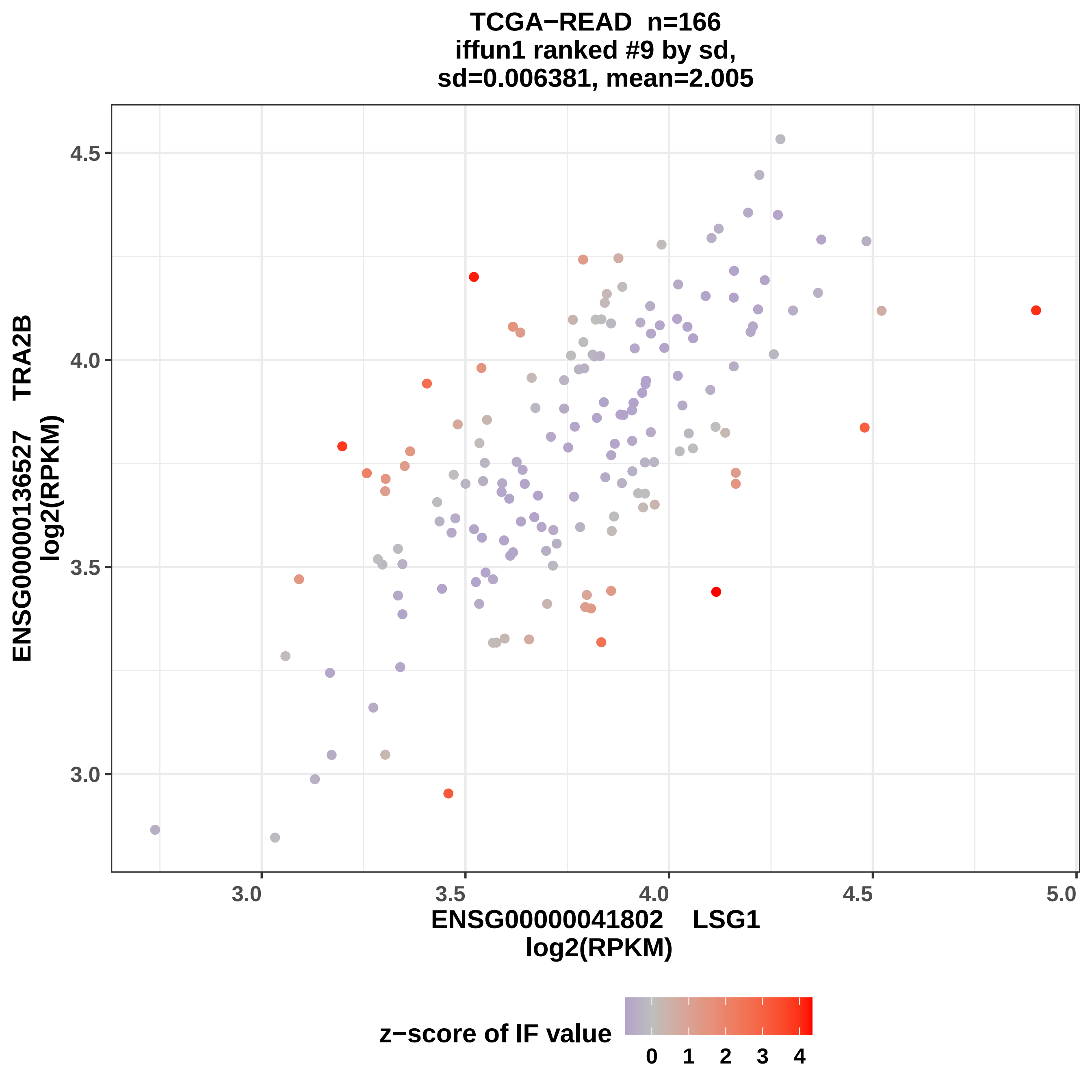

TCGA−READ n=166
iffun1 ranked #9 by sd,
sd=0.006381, mean=2.005
ENSG00000136527 TRA2B
log2(RPKM)
4.5
4.0
3.5
3.0
2.5
3.0
3.5
4.0
4.5
5.0
ENSG00000041802 LSG1
log2(RPKM)
z−score of IF value
0
1
2
3
4

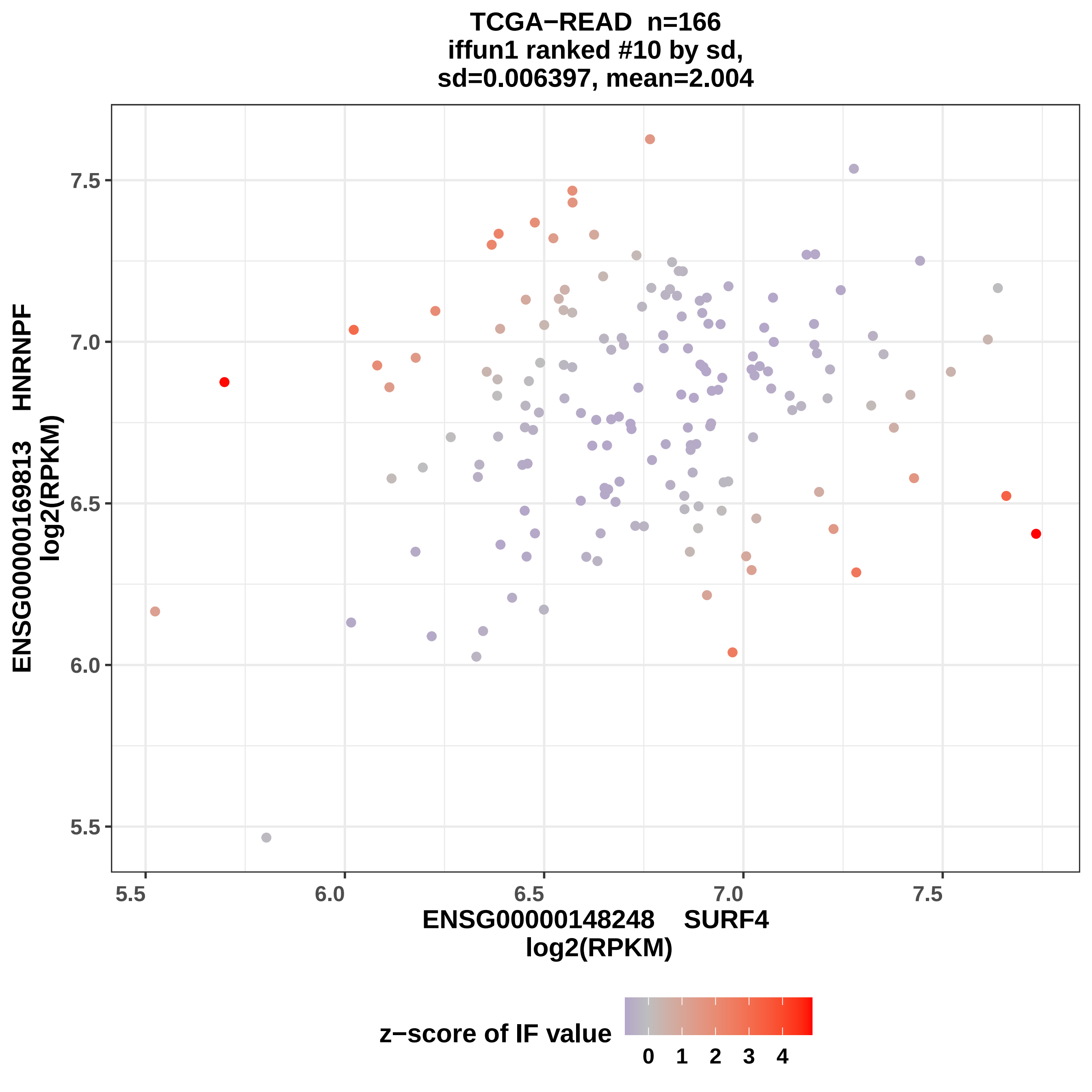

TCGA−READ n=166
iffun1 ranked #10 by sd,
sd=0.006397, mean=2.004
ENSG00000169813 HNRNPF
log2(RPKM)
ENSG00000148248 SURF4
log2(RPKM)
z−score of IF value
0 1 2 3 4
5.5
6.0
6.5
7.0
7.5

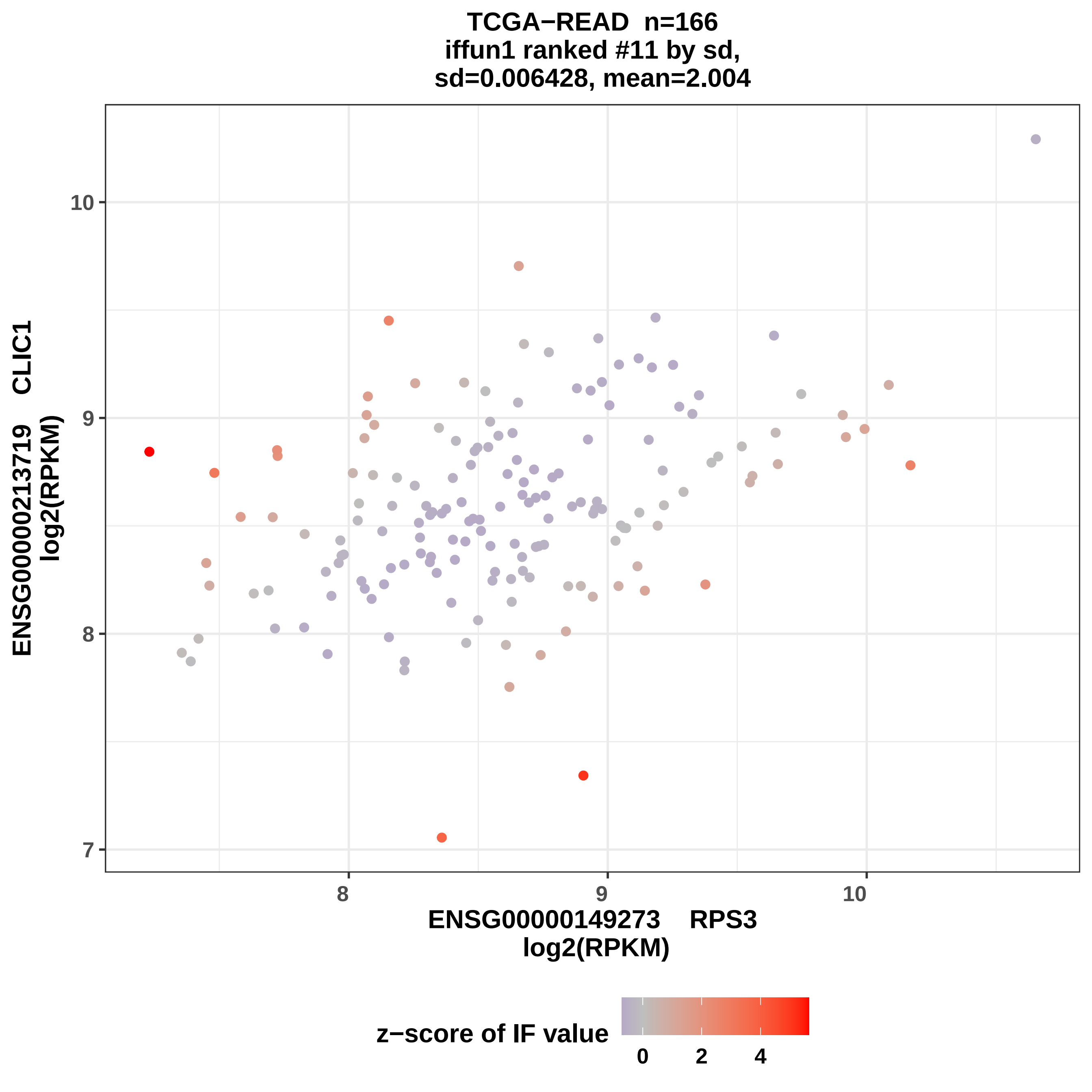

TCGA−READ n=166
iffun1 ranked #11 by sd,
sd=0.006428, mean=2.004
ENSG00000213719 CLIC1
log2(RPKM)
10
9
8
7
8
9
10
ENSG00000149273 RPS3
log2(RPKM)
z−score of IF value
0
2
4

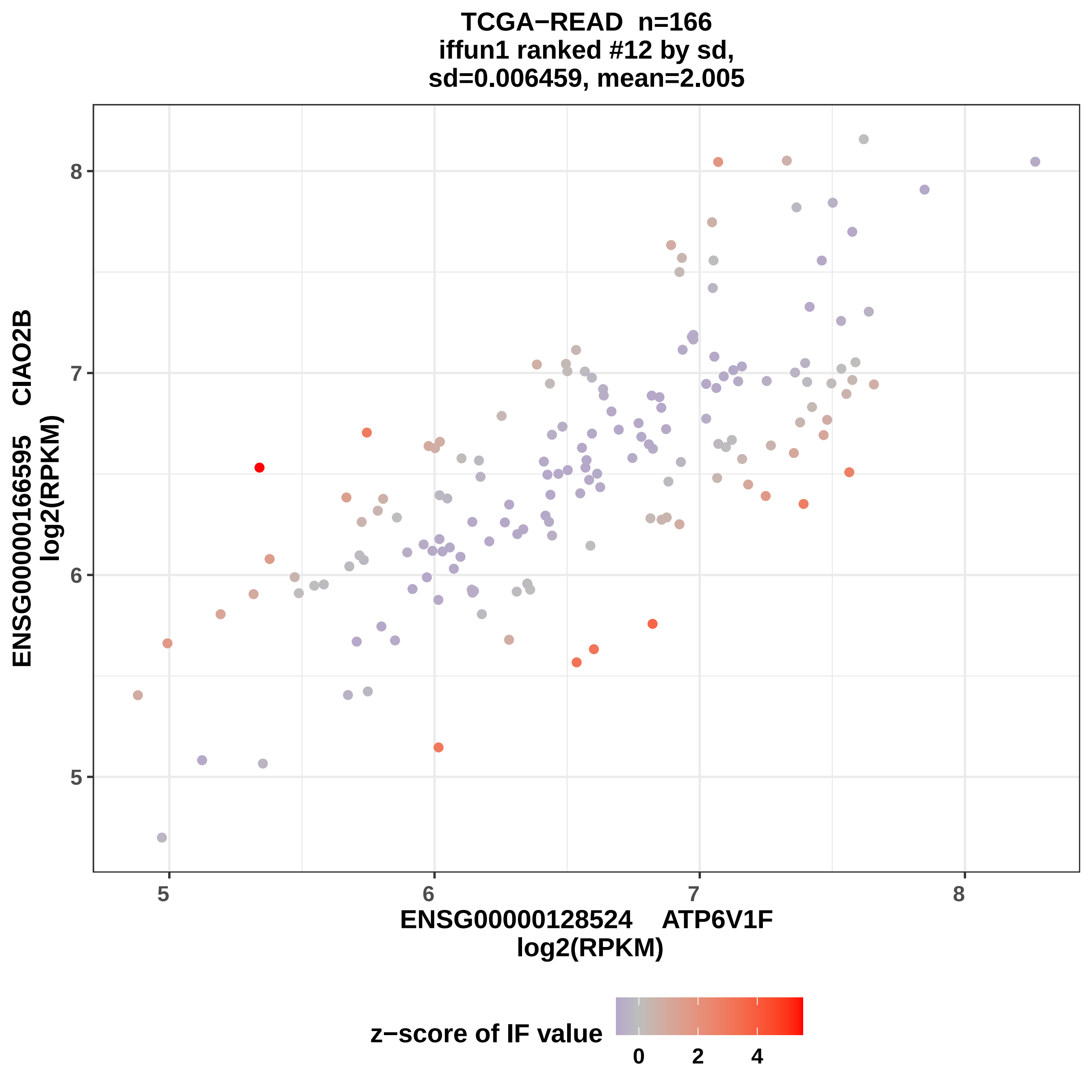
TCGA−READ n=166
iffun1 ranked #12 by sd,
sd=0.006459, mean=2.005
ENSG00000166595 CIAO2B
log2(RPKM)
ENSG00000128524 ATP6V1F
log2(RPKM)
5
6
7
8
z−score of IF value
0
2
4

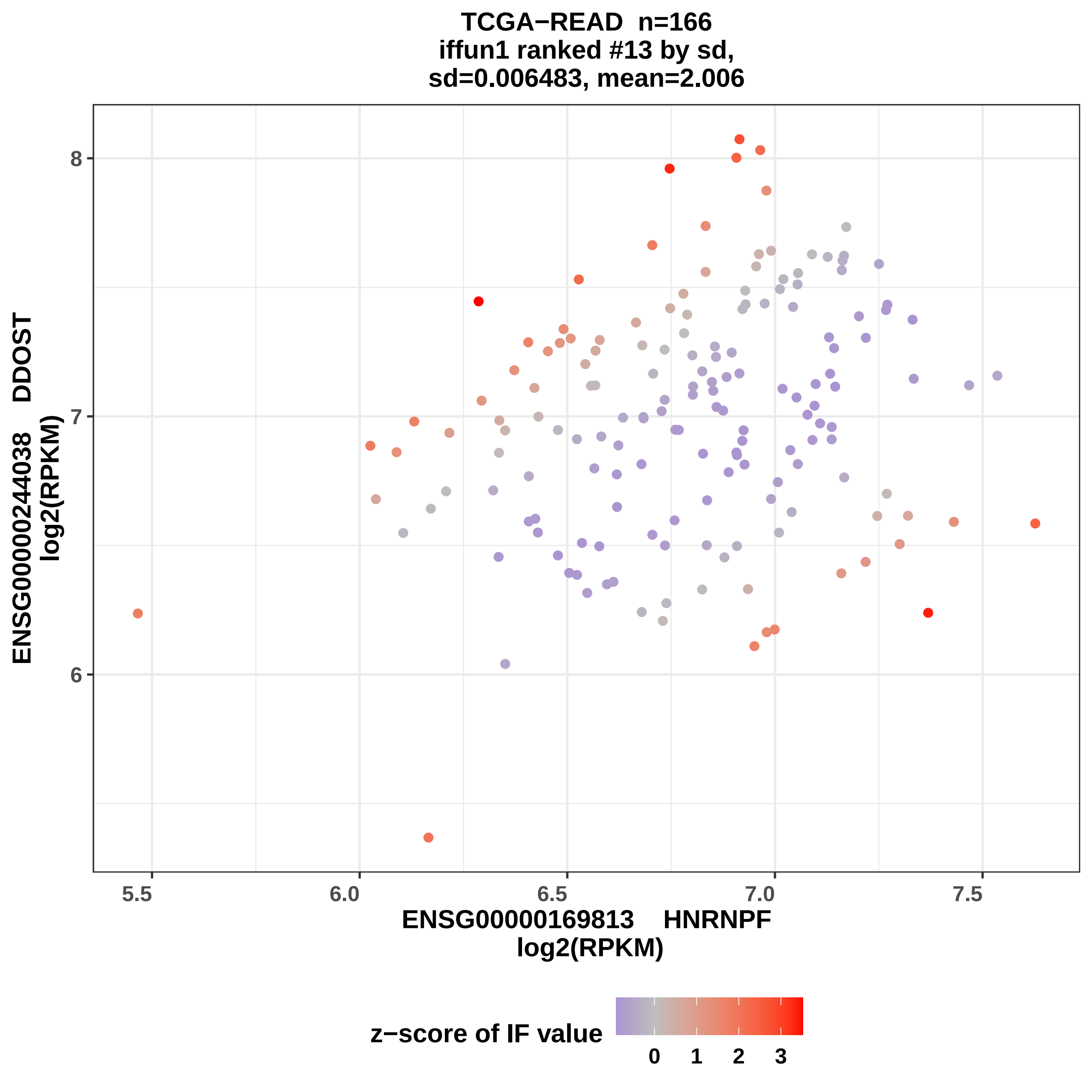
TCGA−READ n=166
iffun1 ranked #13 by sd,
sd=0.006483, mean=2.006
ENSG00000244038 DDOST
log2(RPKM)
8
7
6
5.5
6.0
6.5
7.0
7.5
ENSG00000169813 HNRNPF
log2(RPKM)
z−score of IF value
0
1
2
3

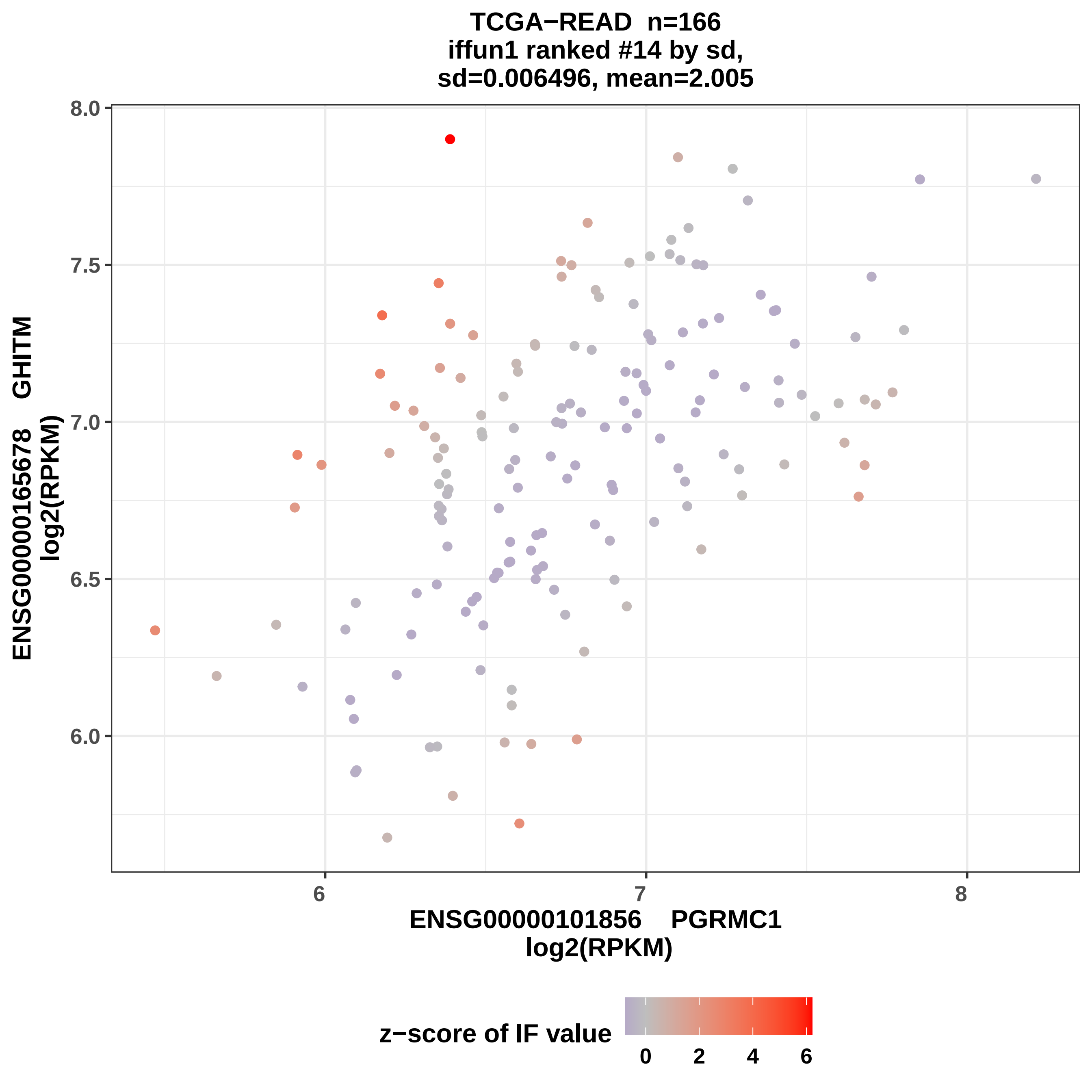

TCGA−READ n=166
iffun1 ranked #14 by sd,
sd=0.006496, mean=2.005
ENSG00000165678 GHITM
log2(RPKM)
ENSG00000101856 PGRMC1
log2(RPKM)
8.0
7.5
7.0
6.5
6.0
6
7
8
z−score of IF value
0
2
4
6

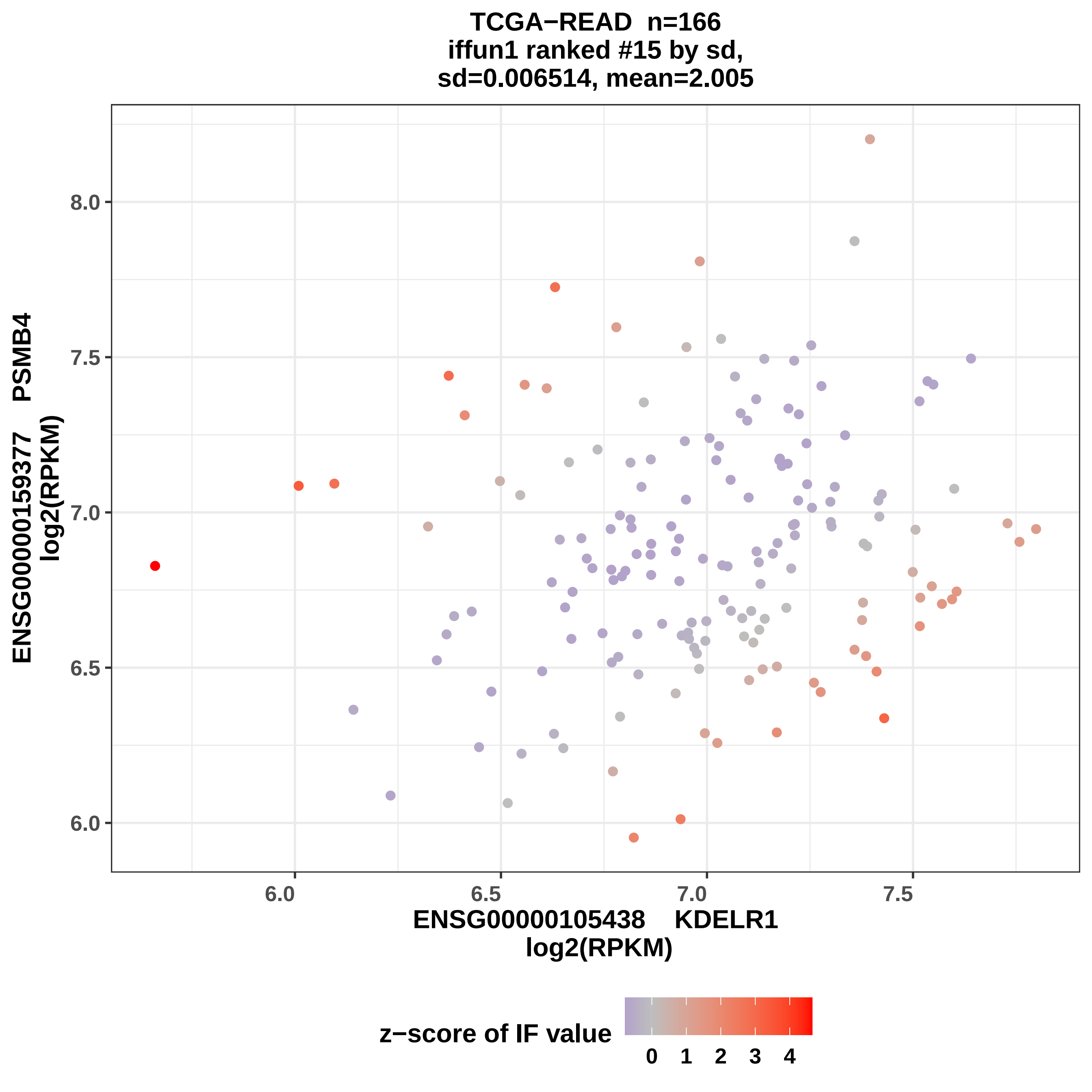

TCGA−READ n=166
iffun1 ranked #15 by sd,
sd=0.006514, mean=2.005
ENSG00000159377 PSMB4
log2(RPKM)
8.0
7.5
7.0
6.5
6.0
6.0
6.5
7.0
7.5
ENSG00000105438 KDELR1
log2(RPKM)
z−score of IF value
0
1
2
3
4

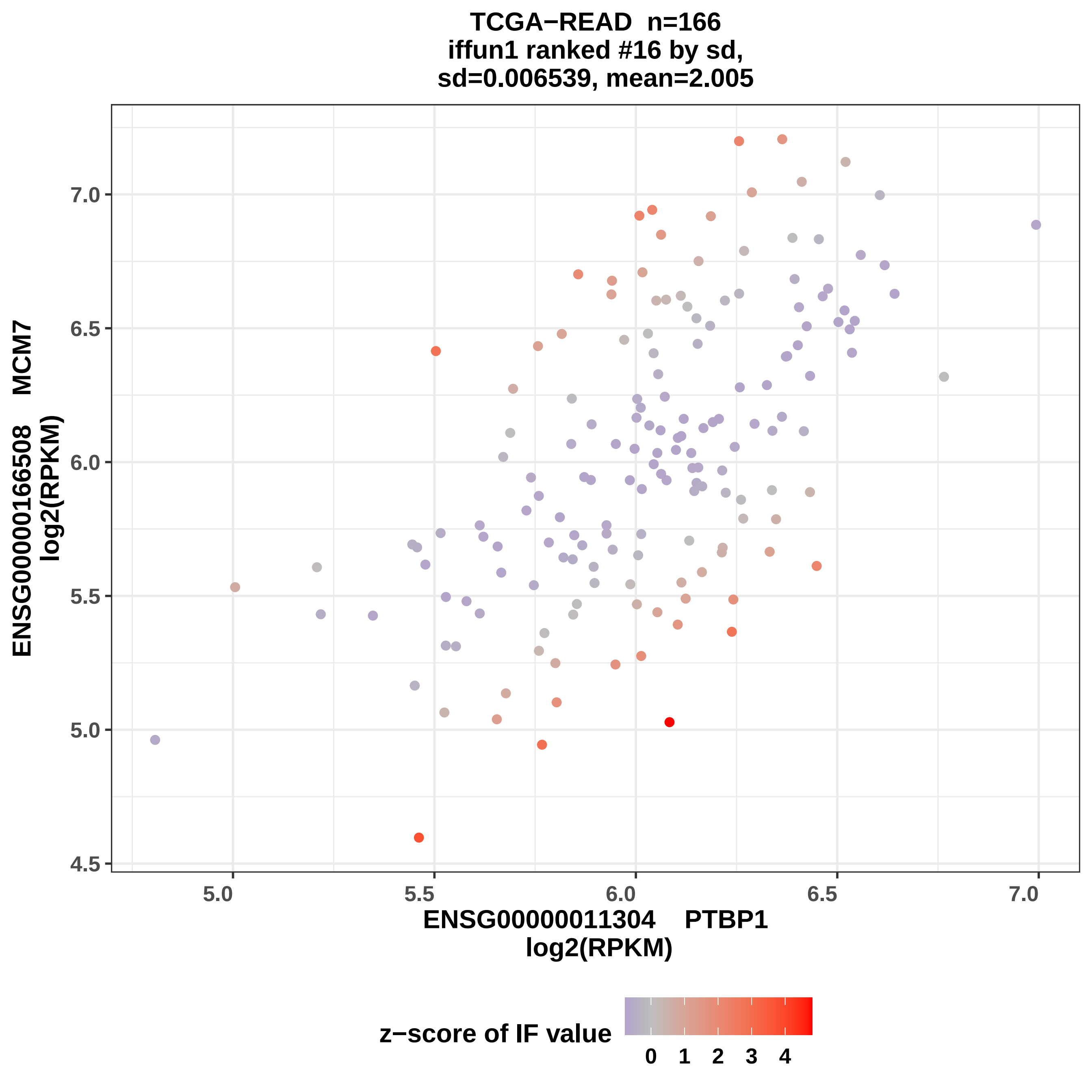
TCGA−READ n=166
iffun1 ranked #16 by sd,
sd=0.006539, mean=2.005
ENSG00000166508 MCM7
log2(RPKM)
7.0
6.5
6.0
5.5
5.0
4.5
5.0
5.5
6.0
6.5
7.0
ENSG00000011304 PTBP1
log2(RPKM)
z−score of IF value
0 1 2 3 4

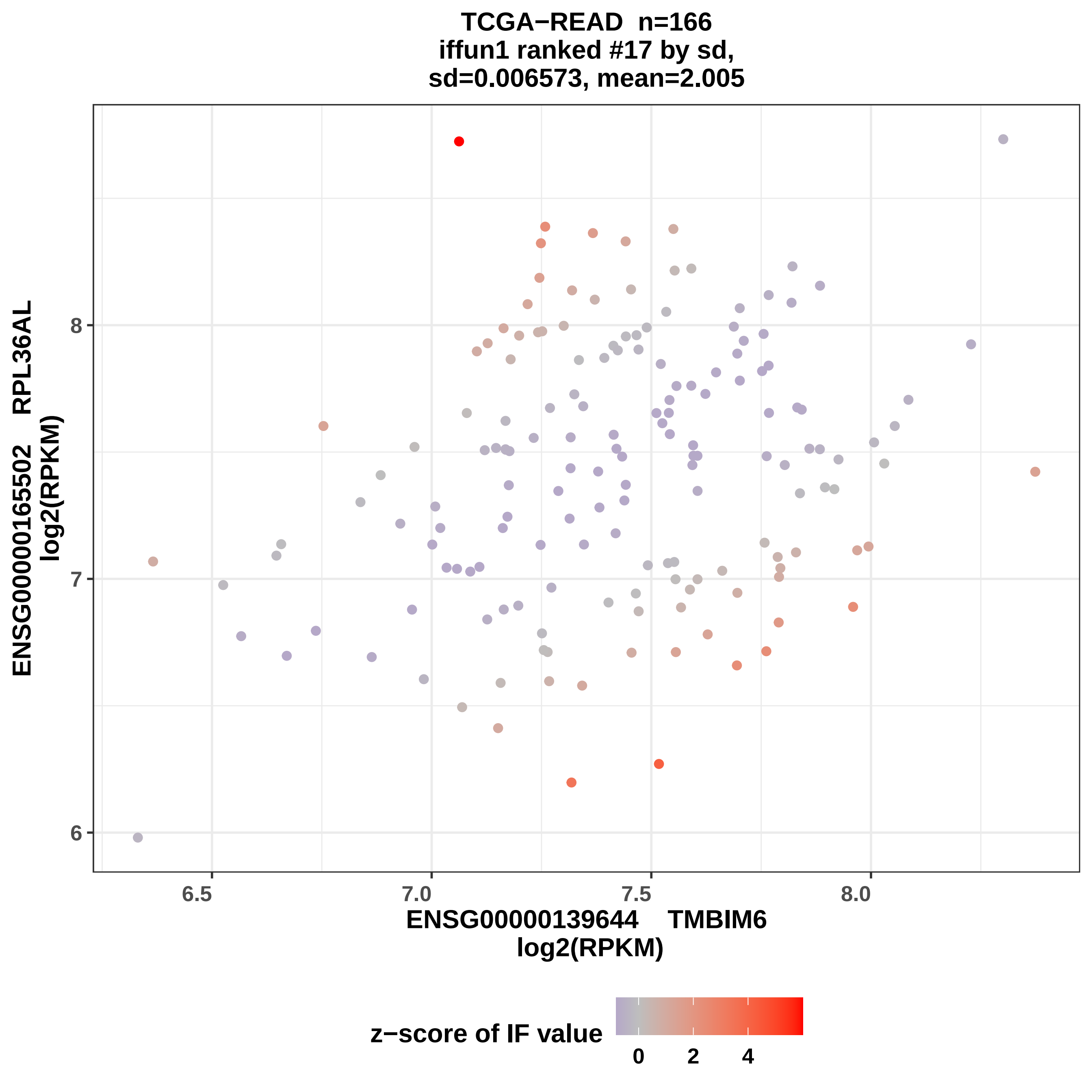

TCGA−READ n=166
iffun1 ranked #17 by sd,
sd=0.006573, mean=2.005
ENSG00000165502 RPL36AL
log2(RPKM)
8
7
6
6.5
7.0
7.5
8.0
ENSG00000139644 TMBIM6
log2(RPKM)
z−score of IF value
0
2
4

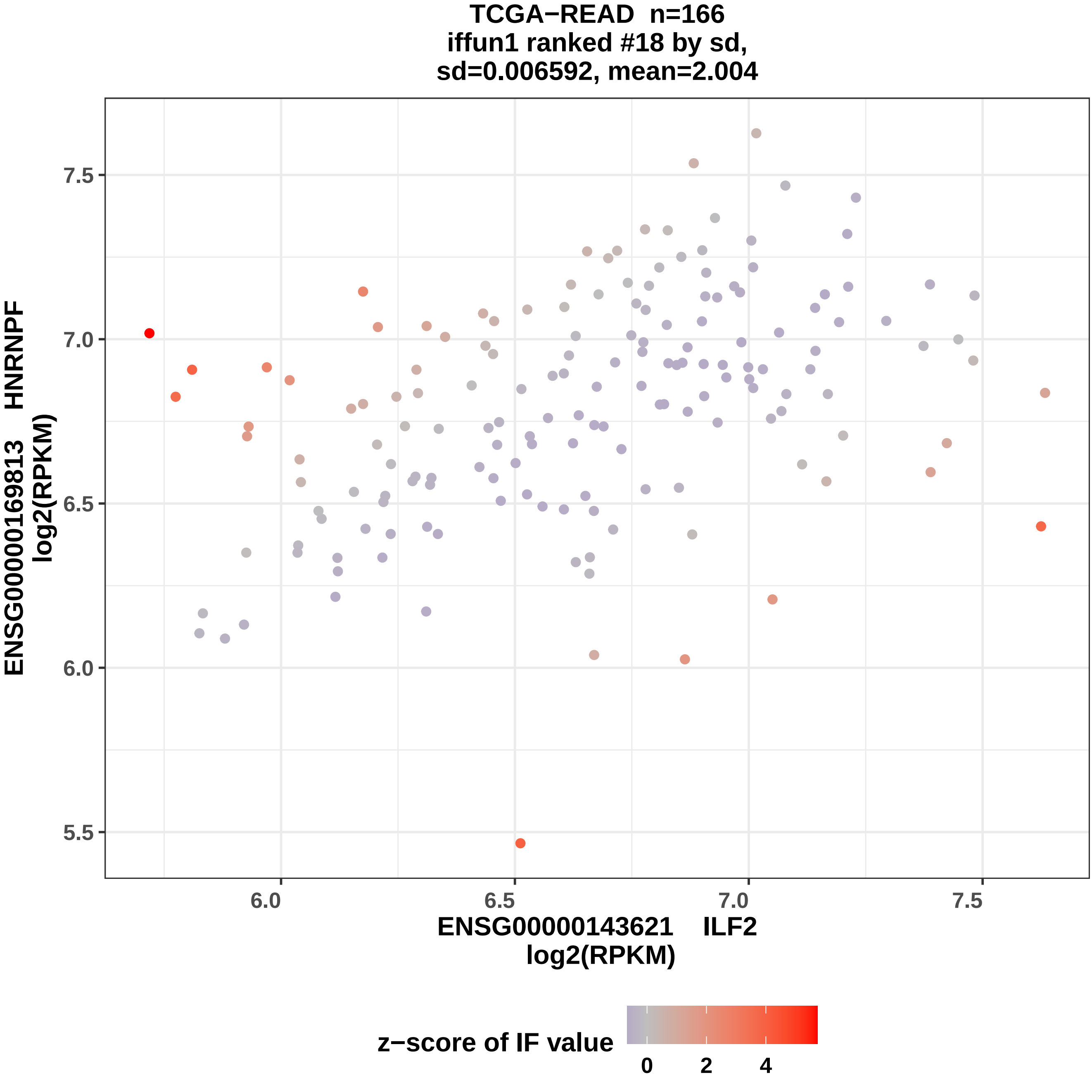

TCGA−READ n=166
iffun1 ranked #18 by sd,
sd=0.006592, mean=2.004
ENSG00000169813 HNRNPF
log2(RPKM)
7.5
7.0
6.5
6.0
5.5
6.0
6.5
7.0
7.5
ENSG00000143621 ILF2
log2(RPKM)
z−score of IF value
0
2
4

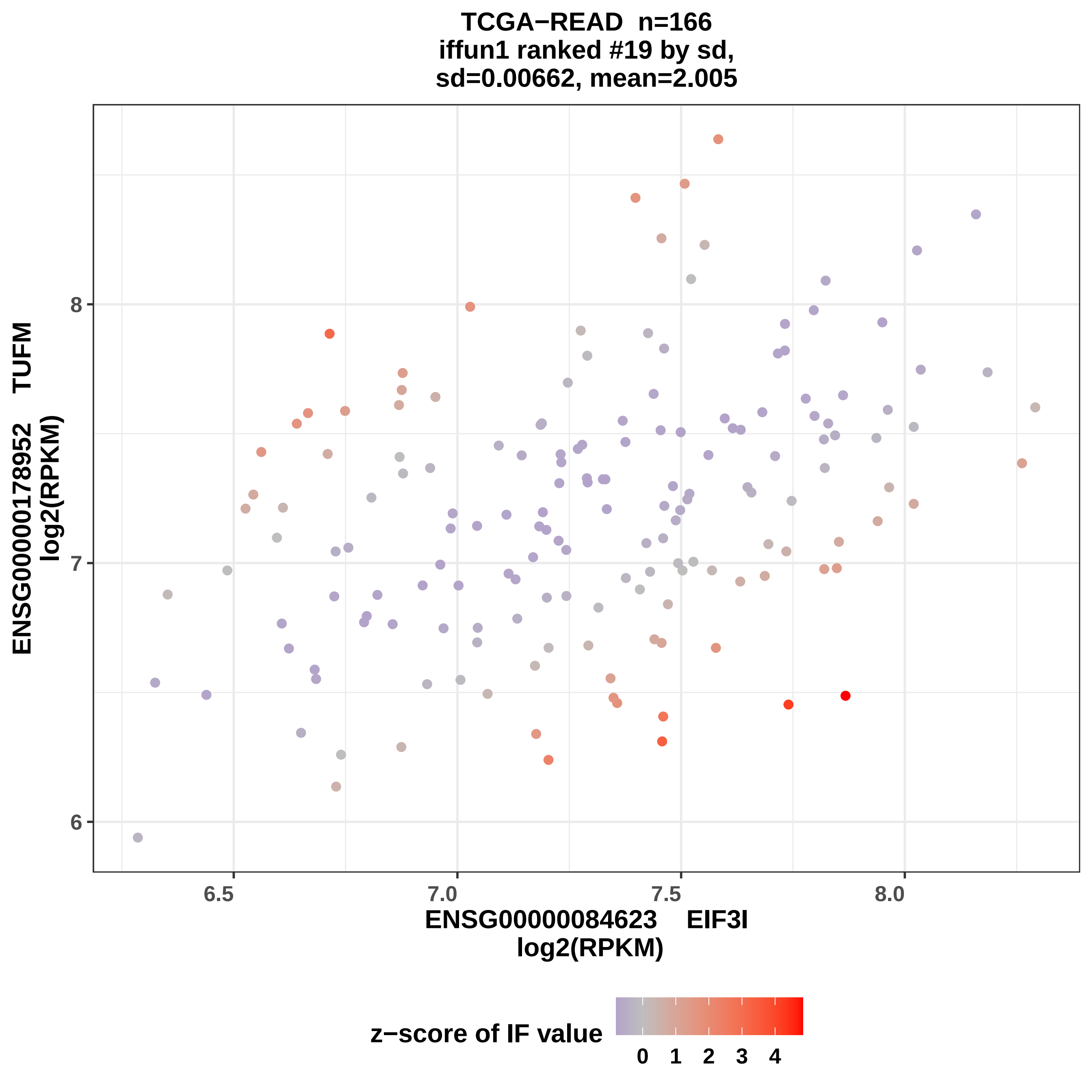
TCGA−READ n=166
iffun1 ranked #19 by sd,
sd=0.00662, mean=2.005
ENSG00000178952 TUFM
log2(RPKM)
8
7
6
6.5
7.0
7.5
8.0
ENSG00000084623 EIF3I
log2(RPKM)
z−score of IF value
0 1 2 3 4

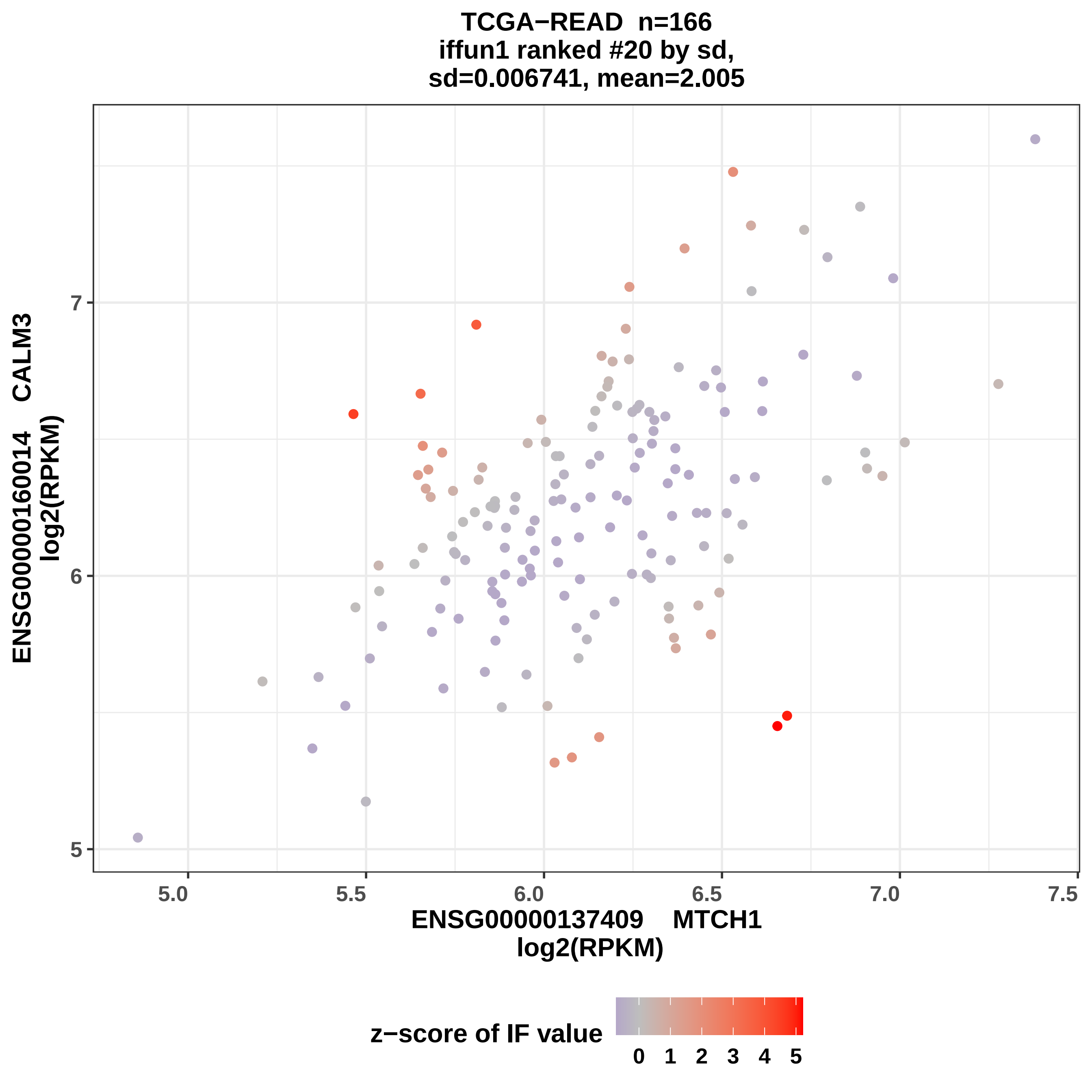
TCGA−READ n=166
iffun1 ranked #20 by sd,
sd=0.006741, mean=2.005
ENSG00000160014 CALM3
log2(RPKM)
ENSG00000137409 MTCH1
log2(RPKM)
5
6
7
5.0
5.5
6.0
6.5
7.0
7.5
z−score of IF value
0
1
2
3
4
5

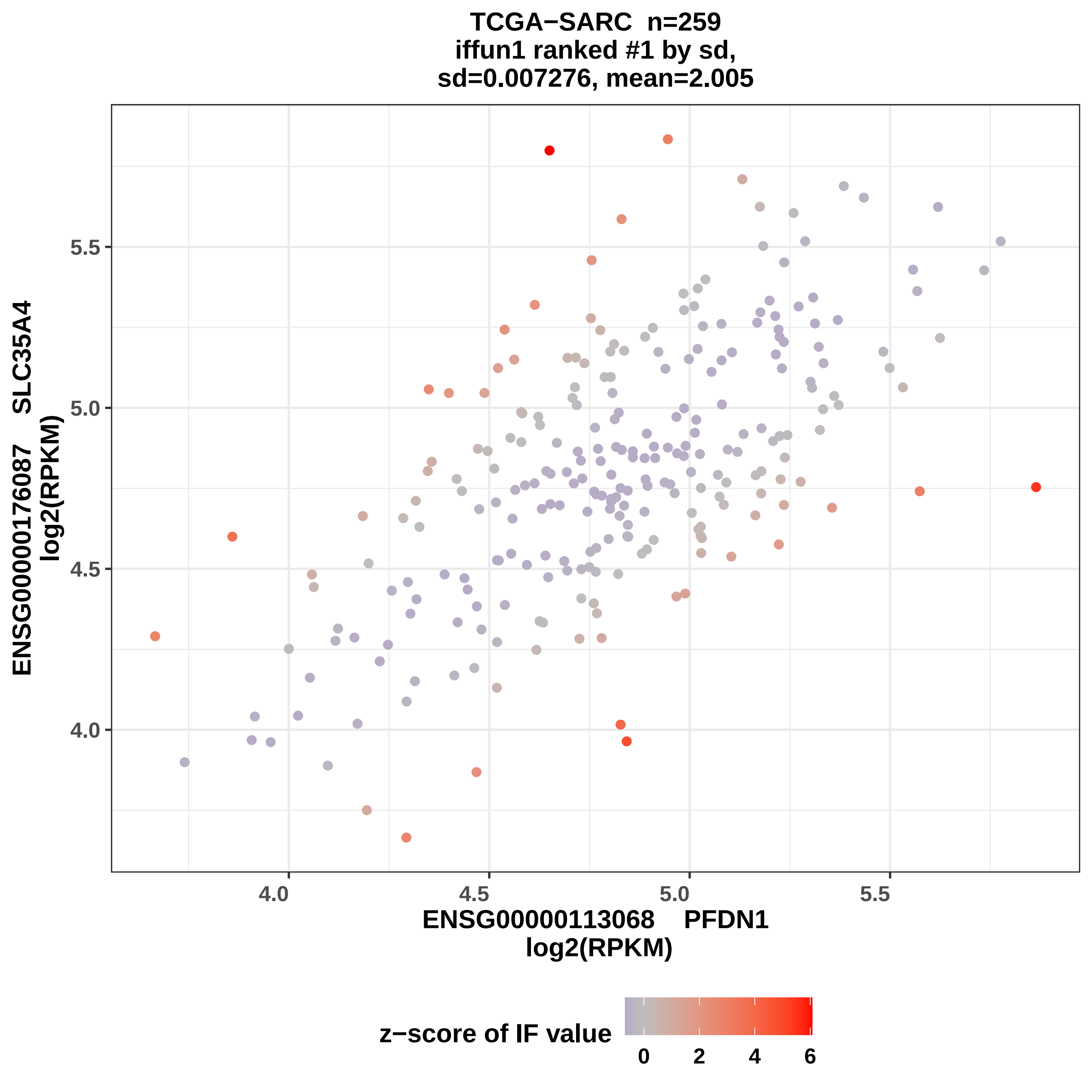
TCGA−SARC n=259
iffun1 ranked #1 by sd,
sd=0.007276, mean=2.005
ENSG00000176087 SLC35A4
log2(RPKM)
5.5
5.0
4.5
4.0
4.0
4.5
5.0
5.5
ENSG00000113068 PFDN1
log2(RPKM)
z−score of IF value
0
2
4
6

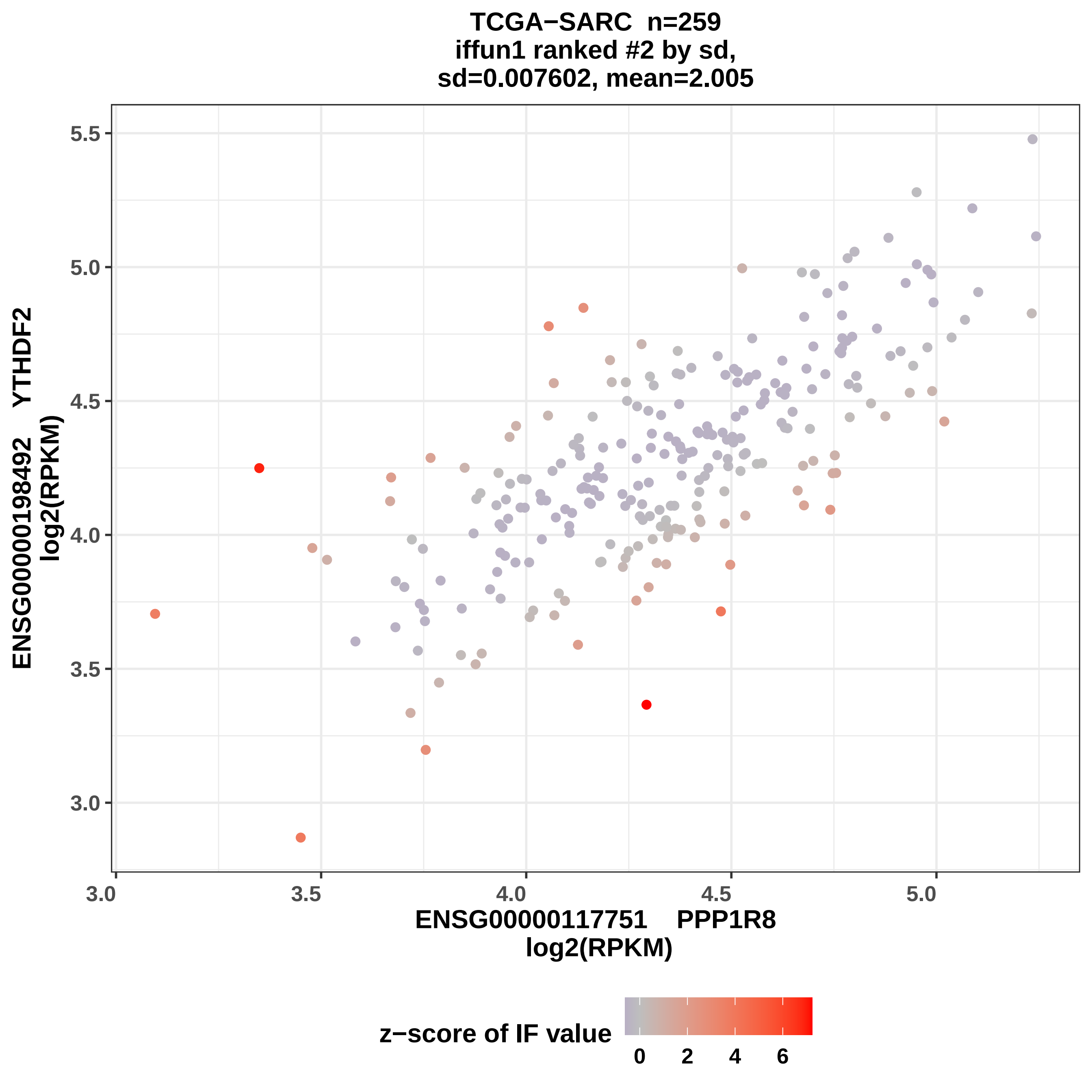

TCGA−SARC n=259
iffun1 ranked #2 by sd,
sd=0.007602, mean=2.005
5.5
5.0
4.5
4.0
3.5
3.0
ENSG00000198492 YTHDF2
log2(RPKM)
3.0
3.5
4.0
4.5
5.0
ENSG00000117751 PPP1R8
log2(RPKM)
z−score of IF value
0
2
4
6

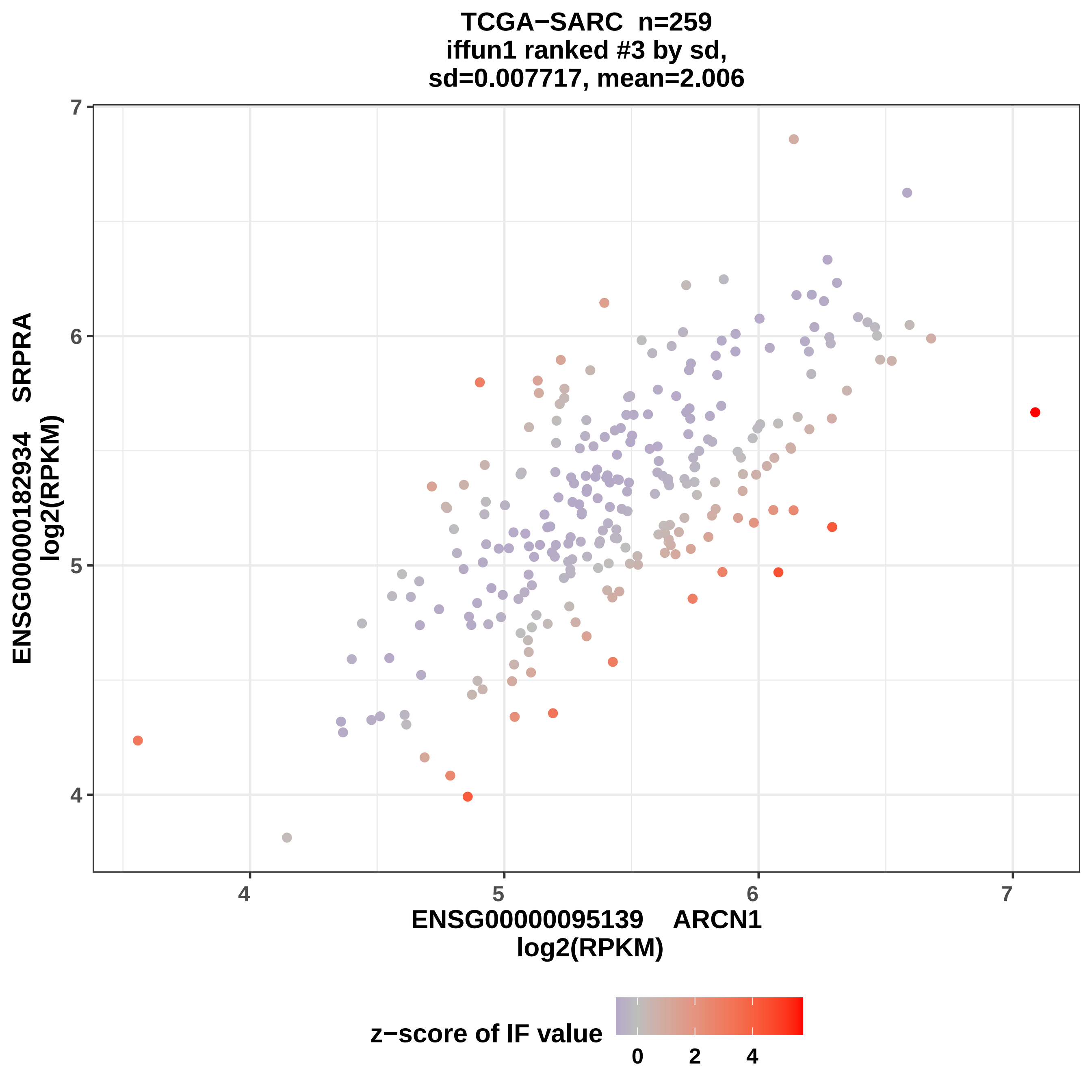

TCGA−SARC n=259
iffun1 ranked #3 by sd,
sd=0.007717, mean=2.006
ENSG00000182934 SRPRA
log2(RPKM)
ENSG00000095139 ARCN1
log2(RPKM)
z−score of IF value
0
2
4

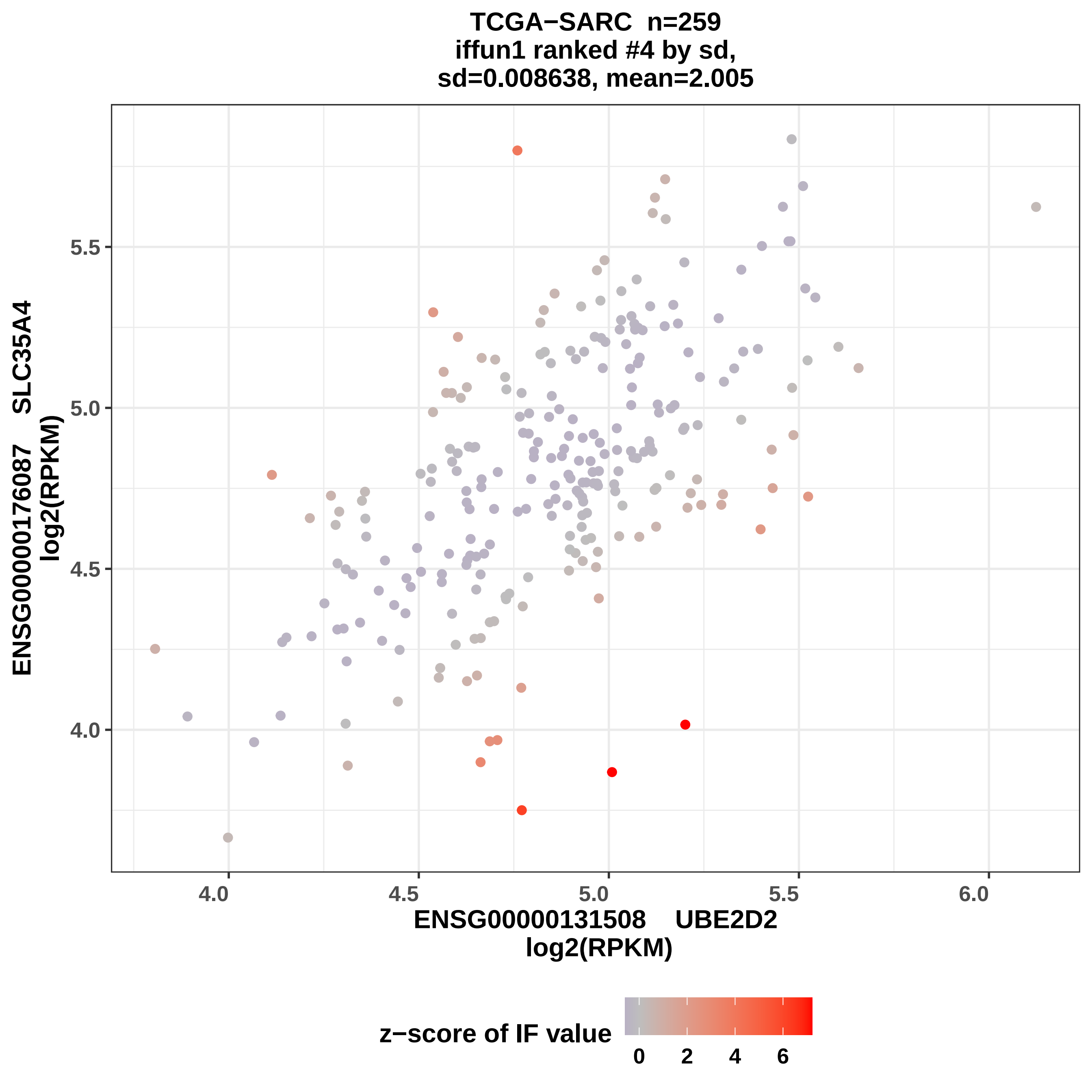
TCGA−SARC n=259
iffun1 ranked #4 by sd,
sd=0.008638, mean=2.005
ENSG00000176087 SLC35A4
log2(RPKM)
5.5
5.0
4.5
4.0
4.0
4.5
5.0
5.5
6.0
ENSG00000131508 UBE2D2
log2(RPKM)
z−score of IF value
0
2
4
6

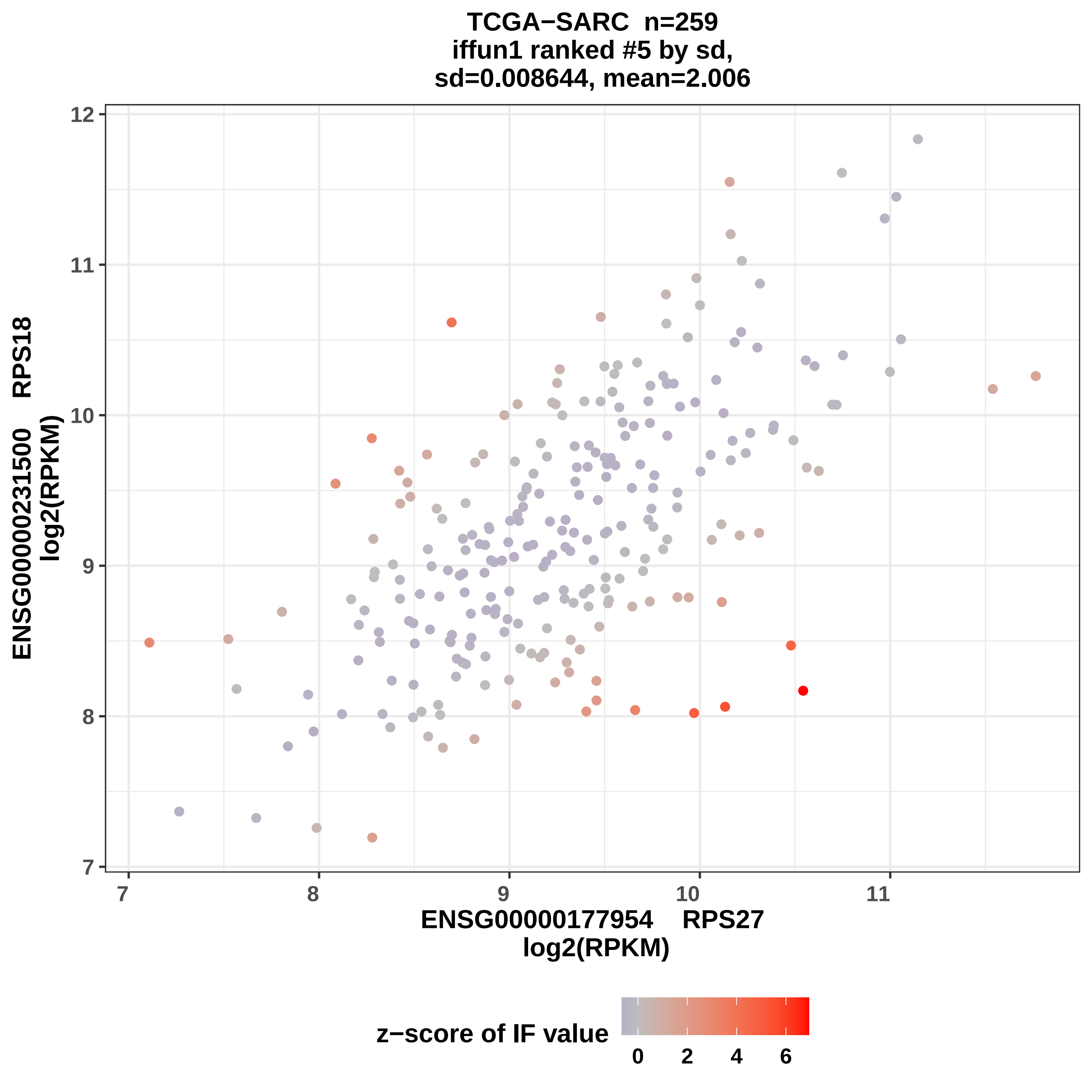
TCGA−SARC n=259
iffun1 ranked #5 by sd,
sd=0.008644, mean=2.006
ENSG00000231500 RPS18
log2(RPKM)
12
11
10
9
8
7
7
8
9
10
11
ENSG00000177954 RPS27
log2(RPKM)
z−score of IF value
0
2
4
6

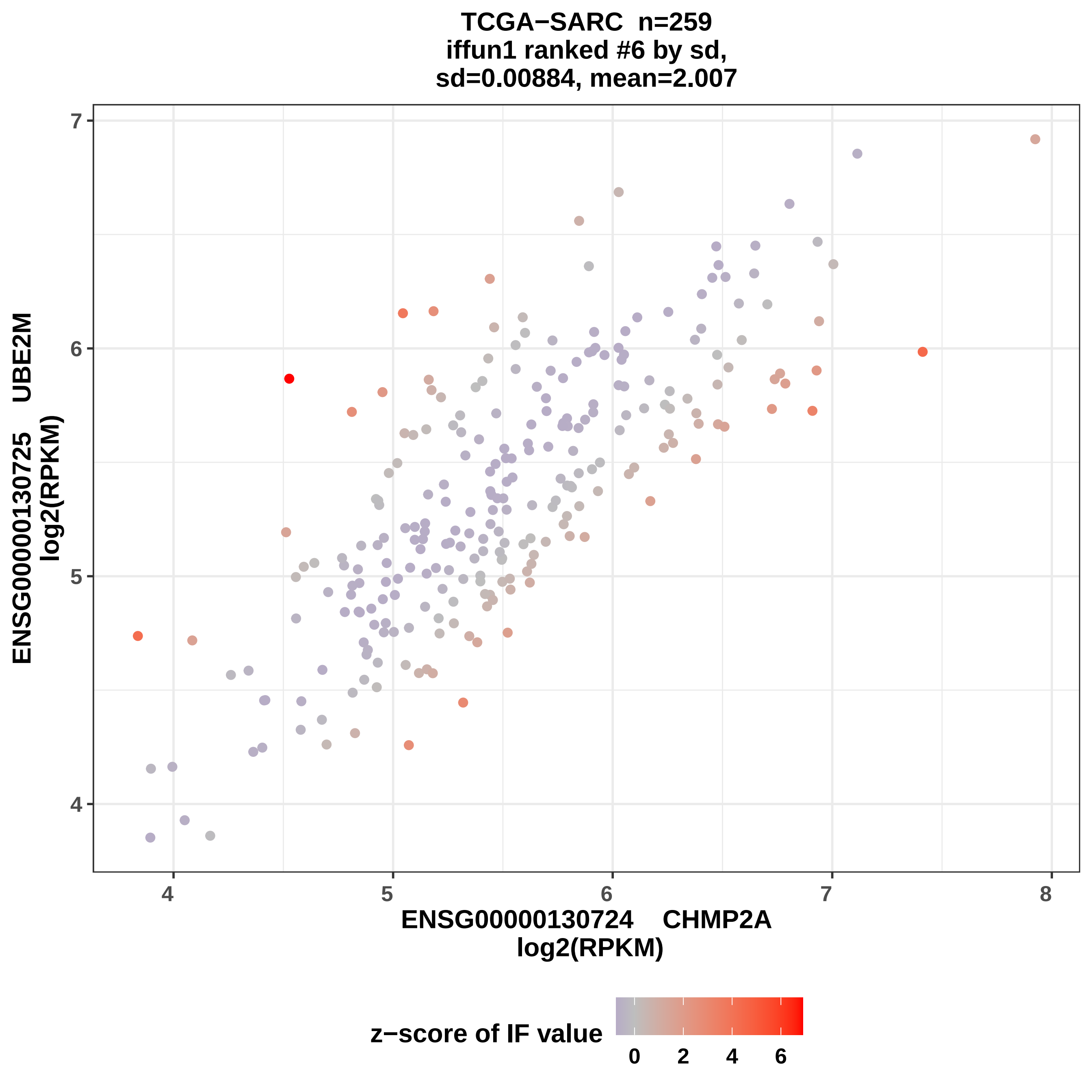
TCGA−SARC n=259
iffun1 ranked #6 by sd,
sd=0.00884, mean=2.007
ENSG00000130725 UBE2M
log2(RPKM)
7
6
5
4
4
5
6
7
8
ENSG00000130724 CHMP2A
log2(RPKM)
z−score of IF value
0
2
4
6

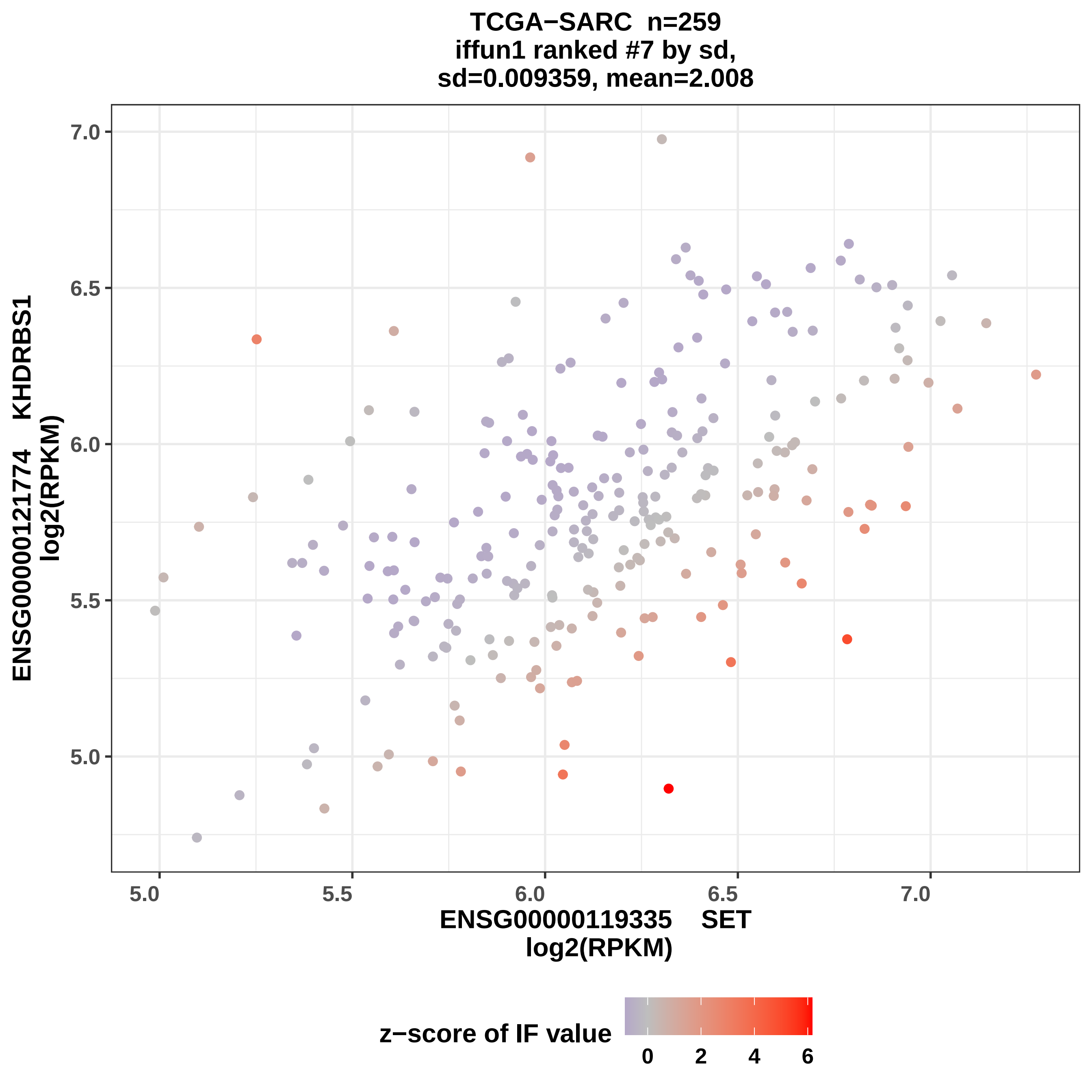
TCGA−SARC n=259
iffun1 ranked #7 by sd,
sd=0.009359, mean=2.008
ENSG00000121774 KHDRBS1
log2(RPKM)
7.0
6.5
6.0
5.5
5.0
5.0
5.5
6.0
6.5
7.0
ENSG00000119335 SET
log2(RPKM)
z−score of IF value
0
2
4
6

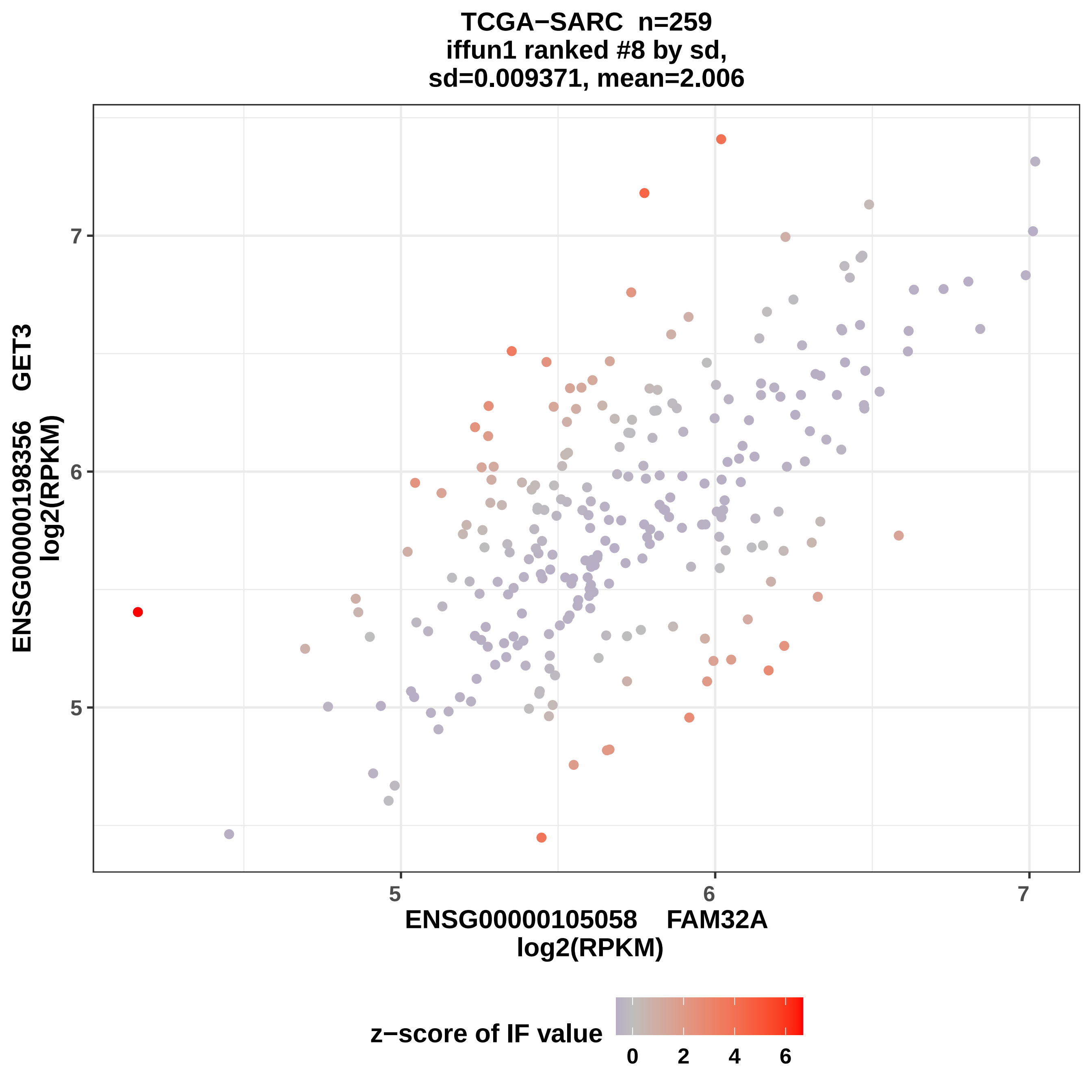

TCGA−SARC n=259
iffun1 ranked #8 by sd,
sd=0.009371, mean=2.006
ENSG00000198356 GET3
log2(RPKM)
ENSG00000105058 FAM32A
log2(RPKM)
5
6
7
5
6
7
z−score of IF value
0
2
4
6

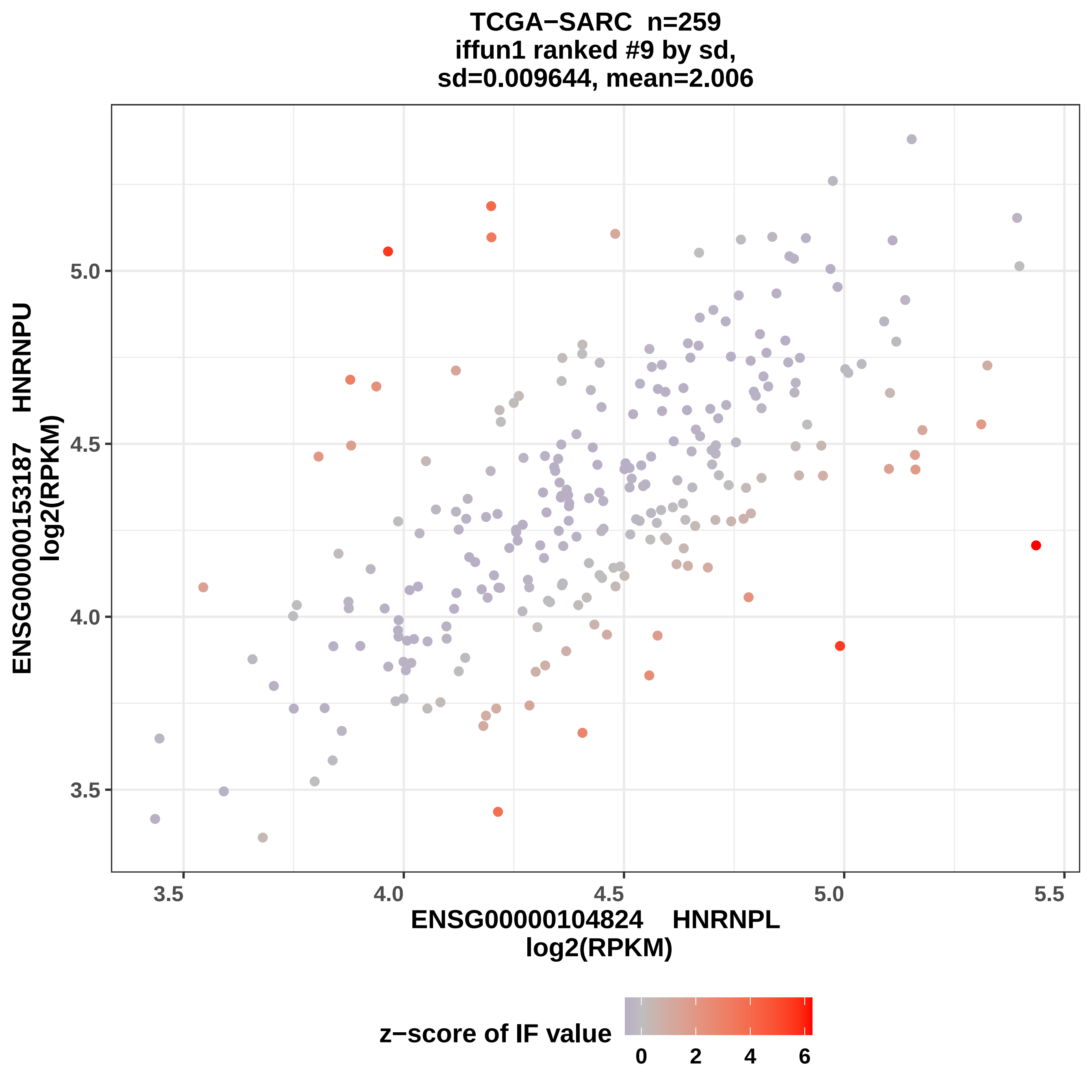
TCGA−SARC n=259
iffun1 ranked #9 by sd,
sd=0.009644, mean=2.006
ENSG00000153187 HNRNPU
log2(RPKM)
5.0
4.5
4.0
3.5
3.5
4.0
4.5
5.0
5.5
ENSG00000104824 HNRNPL
log2(RPKM)
z−score of IF value
0
2
4
6

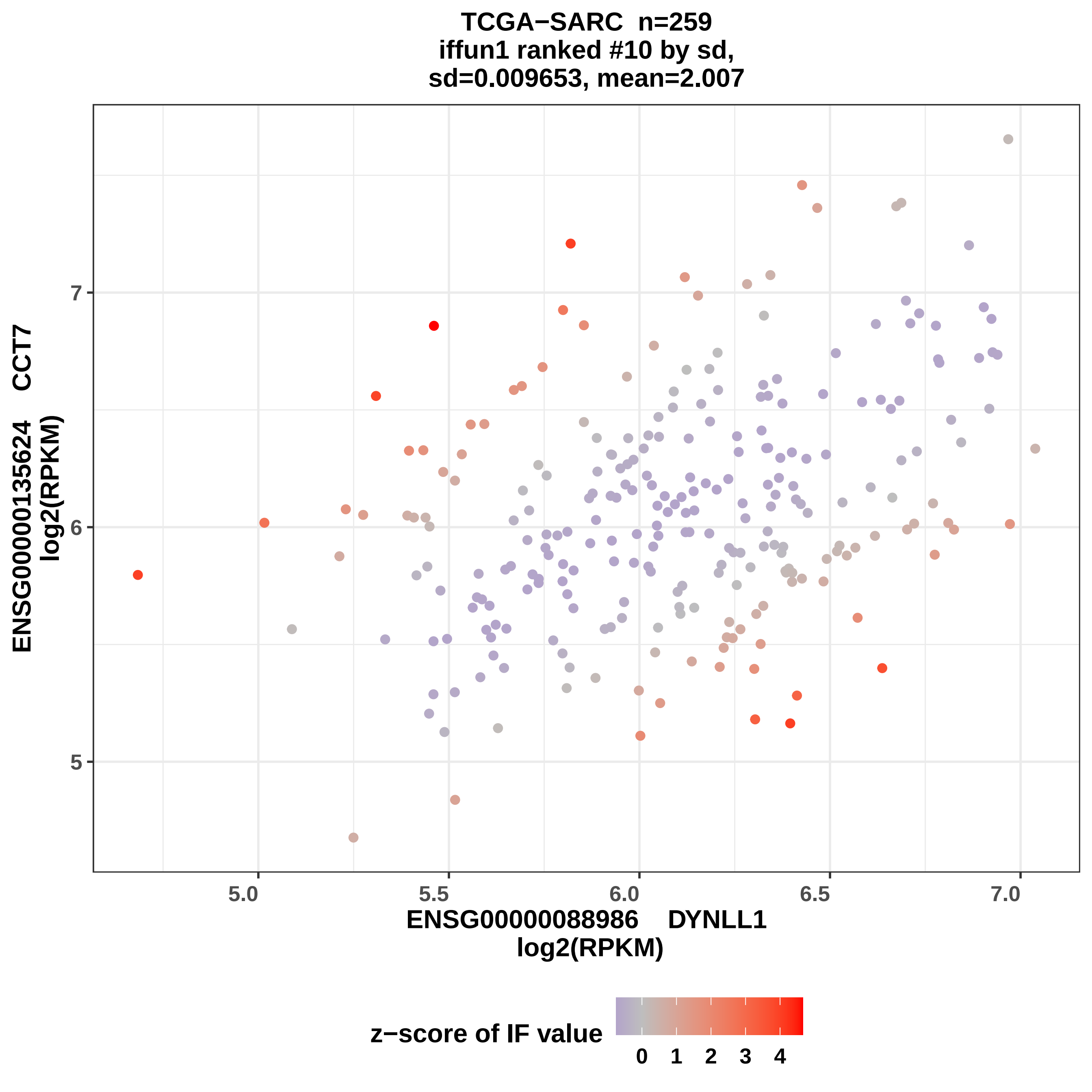
TCGA−SARC n=259
iffun1 ranked #10 by sd,
sd=0.009653, mean=2.007
ENSG00000135624 CCT7
log2(RPKM)
7
6
5
5.0
5.5
6.0
6.5
7.0
ENSG00000088986 DYNLL1
log2(RPKM)
z−score of IF value
0 1 2 3 4

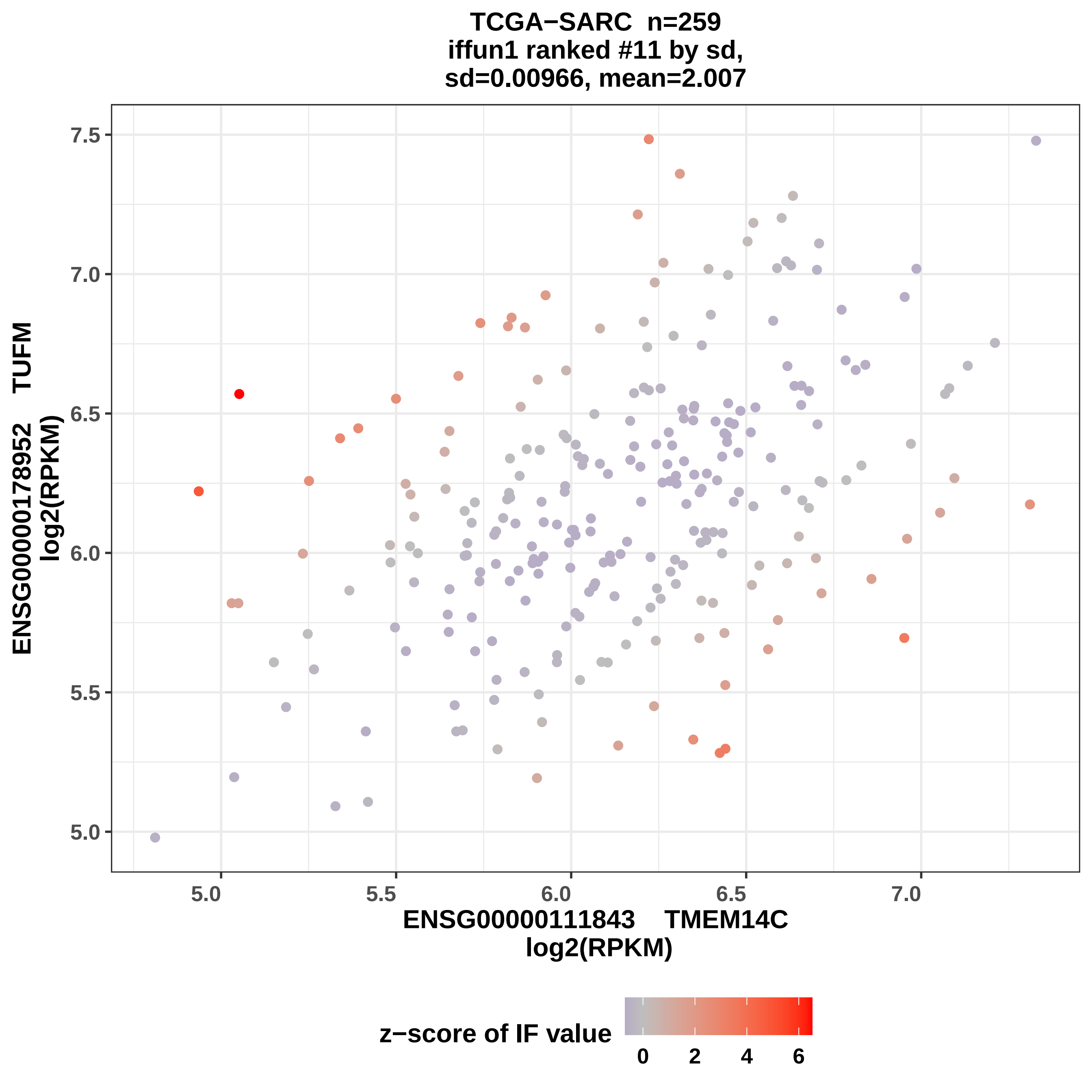

TCGA−SARC n=259
iffun1 ranked #11 by sd,
sd=0.00966, mean=2.007
ENSG00000178952 TUFM
log2(RPKM)
7.5
7.0
6.5
6.0
5.5
5.0
5.0
5.5
6.0
6.5
7.0
ENSG00000111843 TMEM14C
log2(RPKM)
z−score of IF value
0
2
4
6

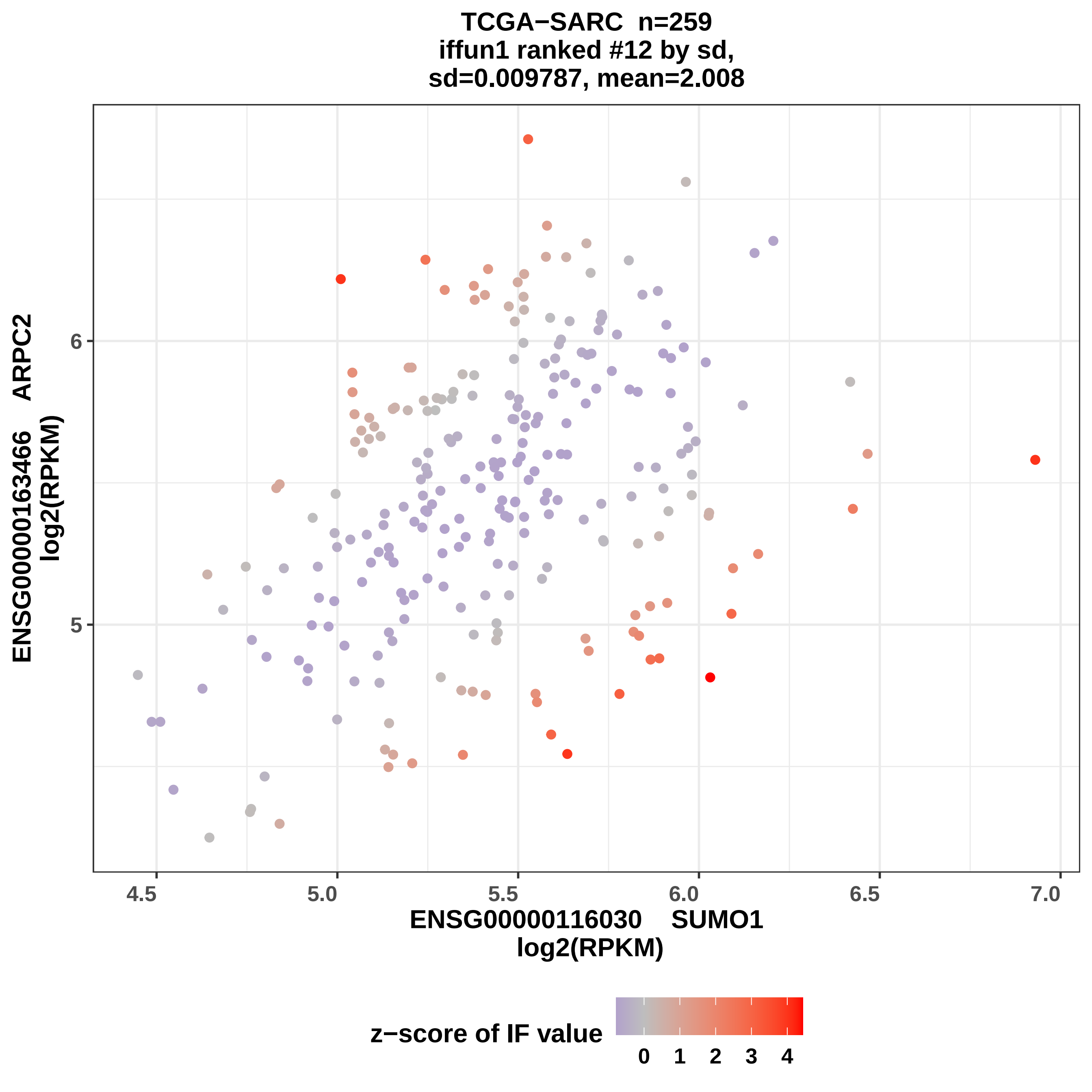

TCGA−SARC n=259
iffun1 ranked #12 by sd,
sd=0.009787, mean=2.008
ENSG00000163466 ARPC2
log2(RPKM)
6
5
4.5
5.0
5.5
6.0
6.5
7.0
ENSG00000116030 SUMO1
log2(RPKM)
z−score of IF value
0
1
2
3
4

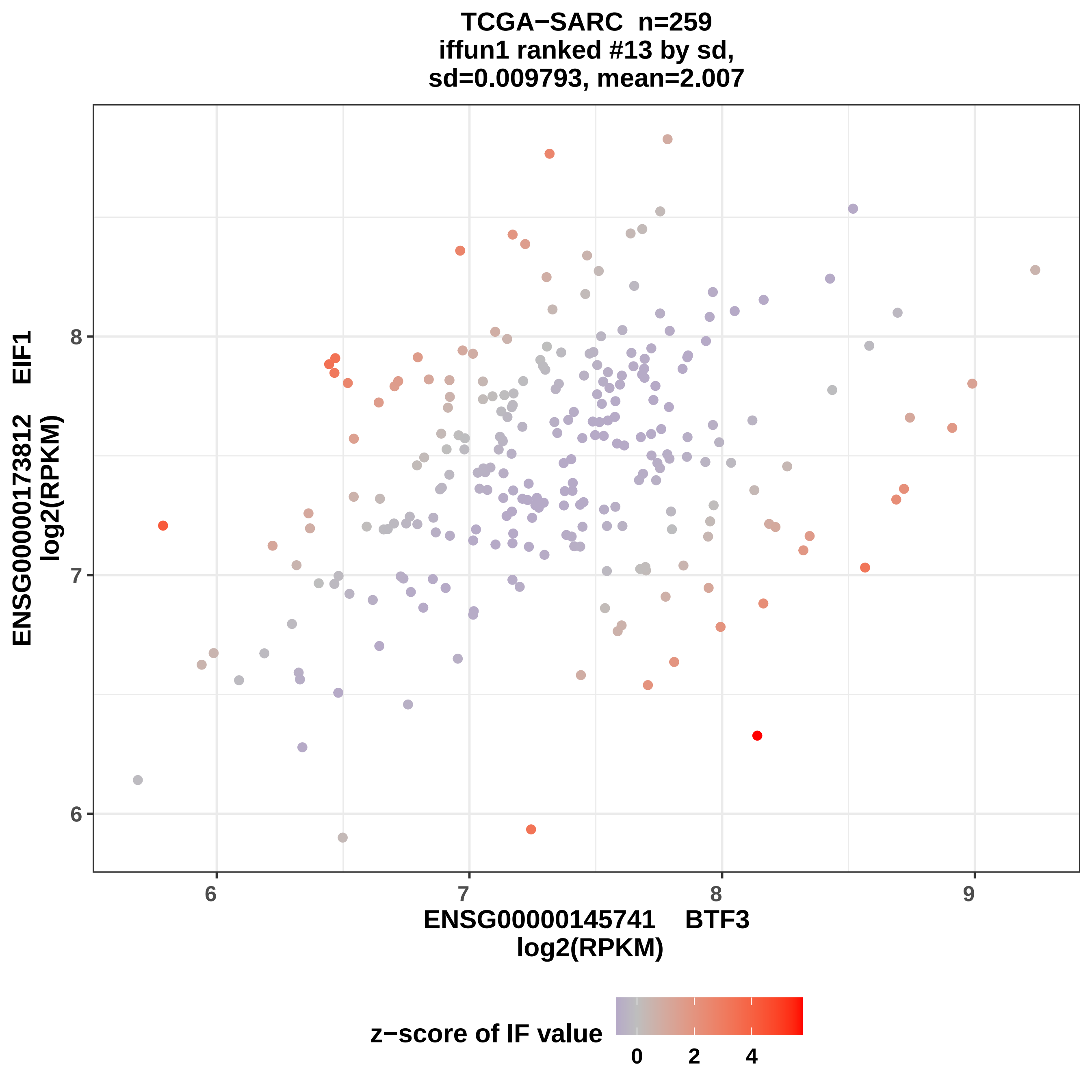
TCGA−SARC n=259
iffun1 ranked #13 by sd,
sd=0.009793, mean=2.007
ENSG00000173812 EIF1
log2(RPKM)
ENSG00000145741 BTF3
log2(RPKM)
6
7
8
9
z−score of IF value
0
2
4

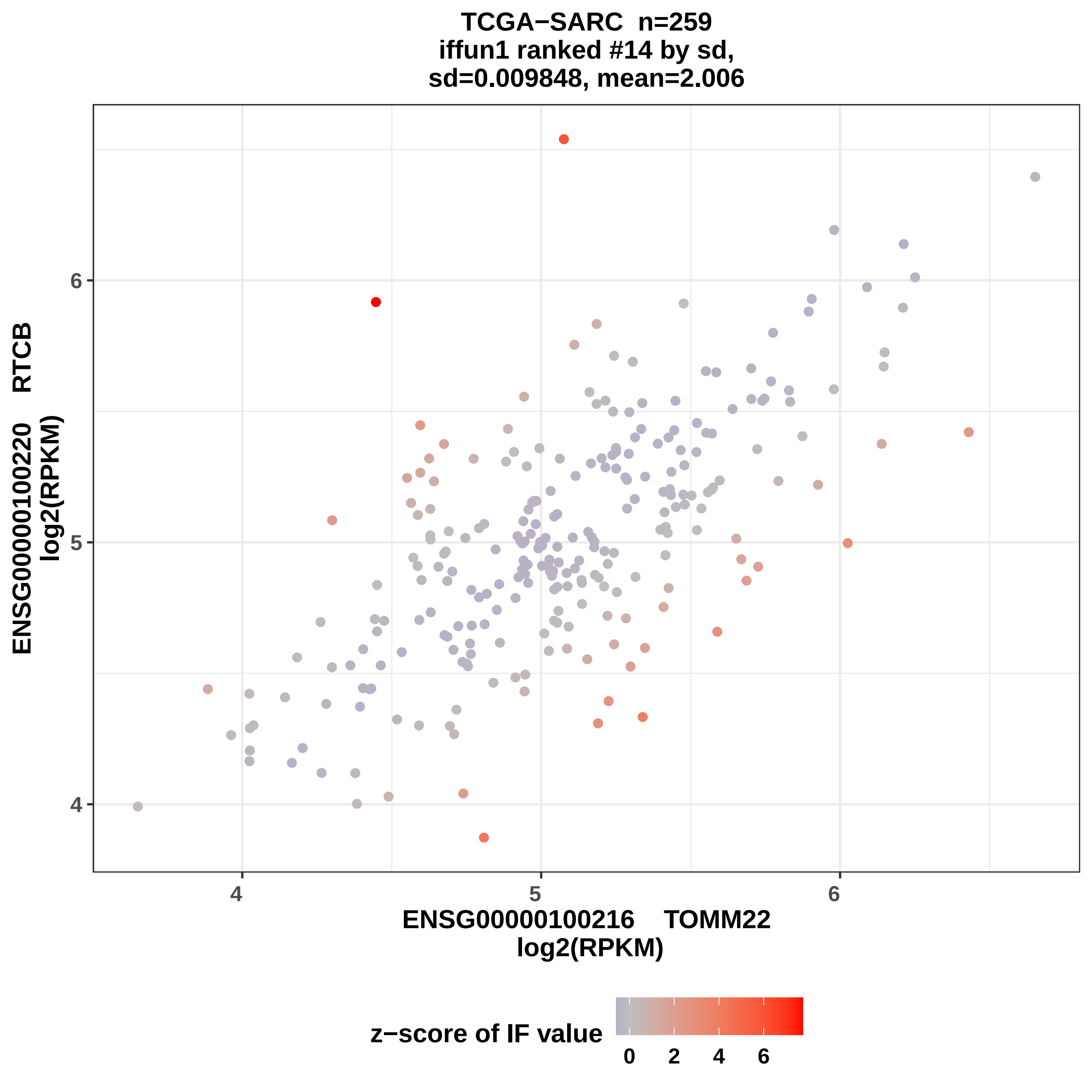

TCGA−SARC n=259
iffun1 ranked #14 by sd,
sd=0.009848, mean=2.006
ENSG00000100220 RTCB
log2(RPKM)
ENSG00000100216 TOMM22
log2(RPKM)
4
5
6
4
5
6
z−score of IF value
0
2
4
6

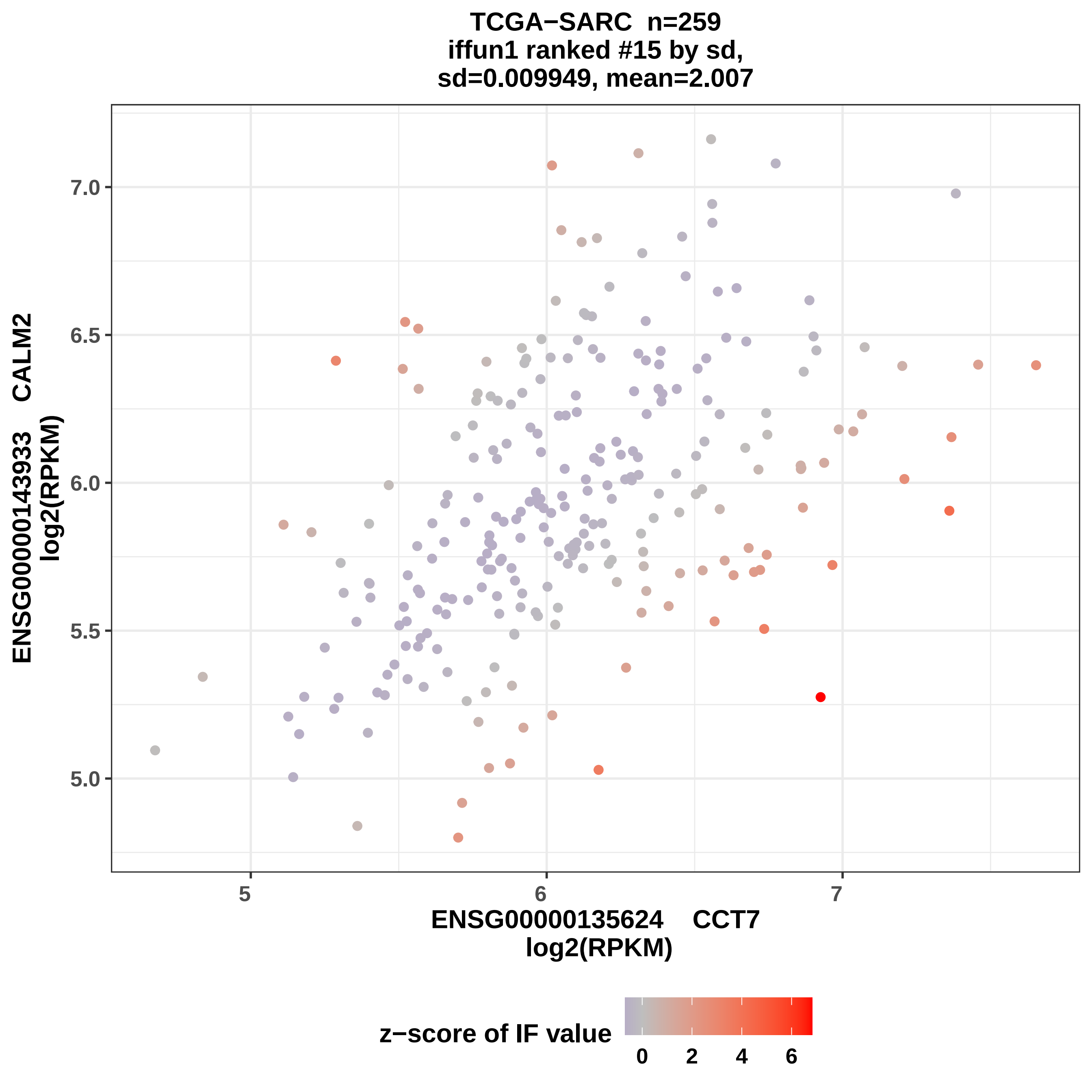

TCGA−SARC n=259
iffun1 ranked #15 by sd,
sd=0.009949, mean=2.007
ENSG00000143933 CALM2
log2(RPKM)
7.0
6.5
6.0
5.5
5.0
5
6
7
ENSG00000135624 CCT7
log2(RPKM)
z−score of IF value
0
2
4
6

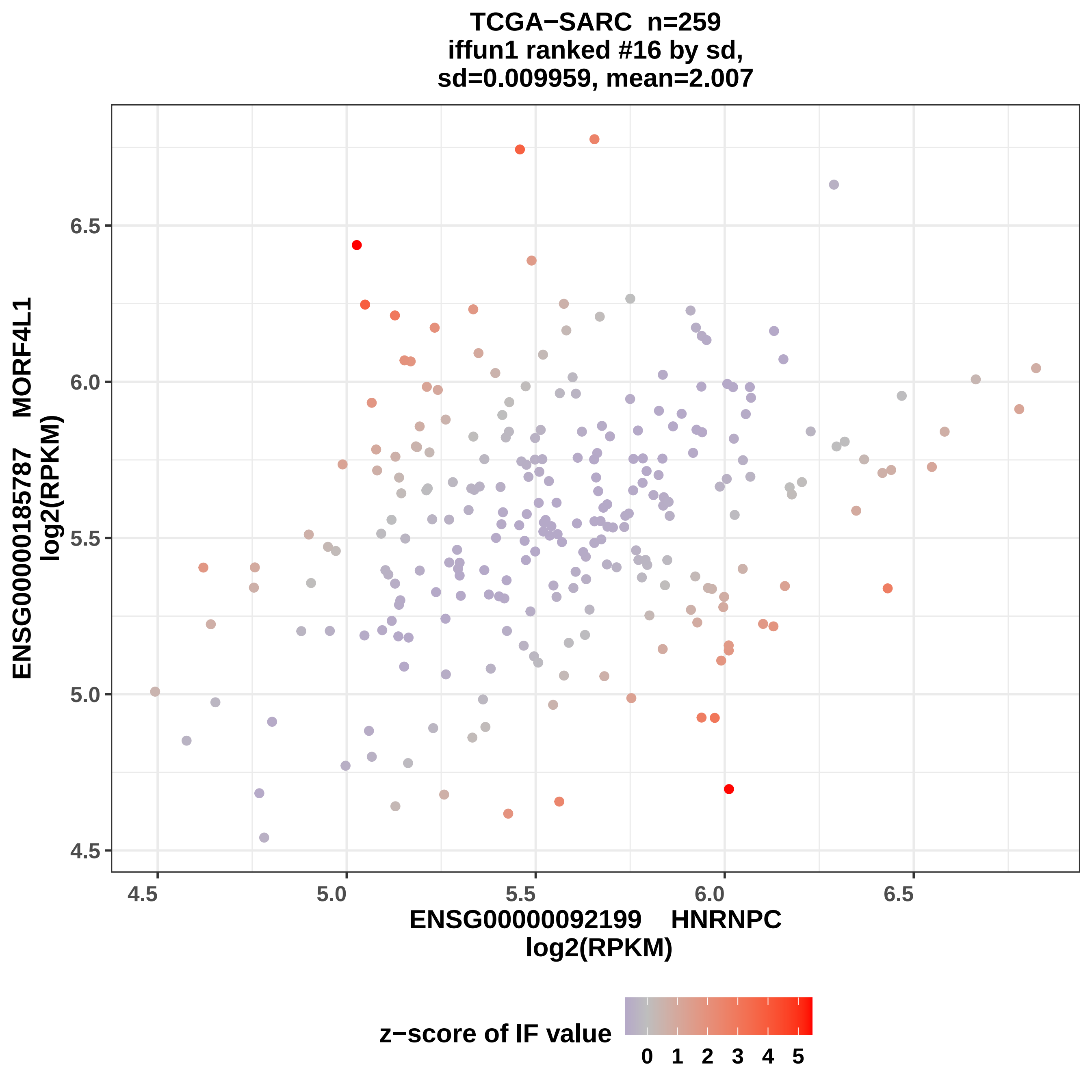
TCGA−SARC n=259
iffun1 ranked #16 by sd,
sd=0.009959, mean=2.007
ENSG00000185787 MORF4L1
log2(RPKM)
6.5
6.0
5.5
5.0
4.5
4.5
5.0
5.5
6.0
6.5
ENSG00000092199 HNRNPC
log2(RPKM)
z−score of IF value
0 1 2 3 4 5

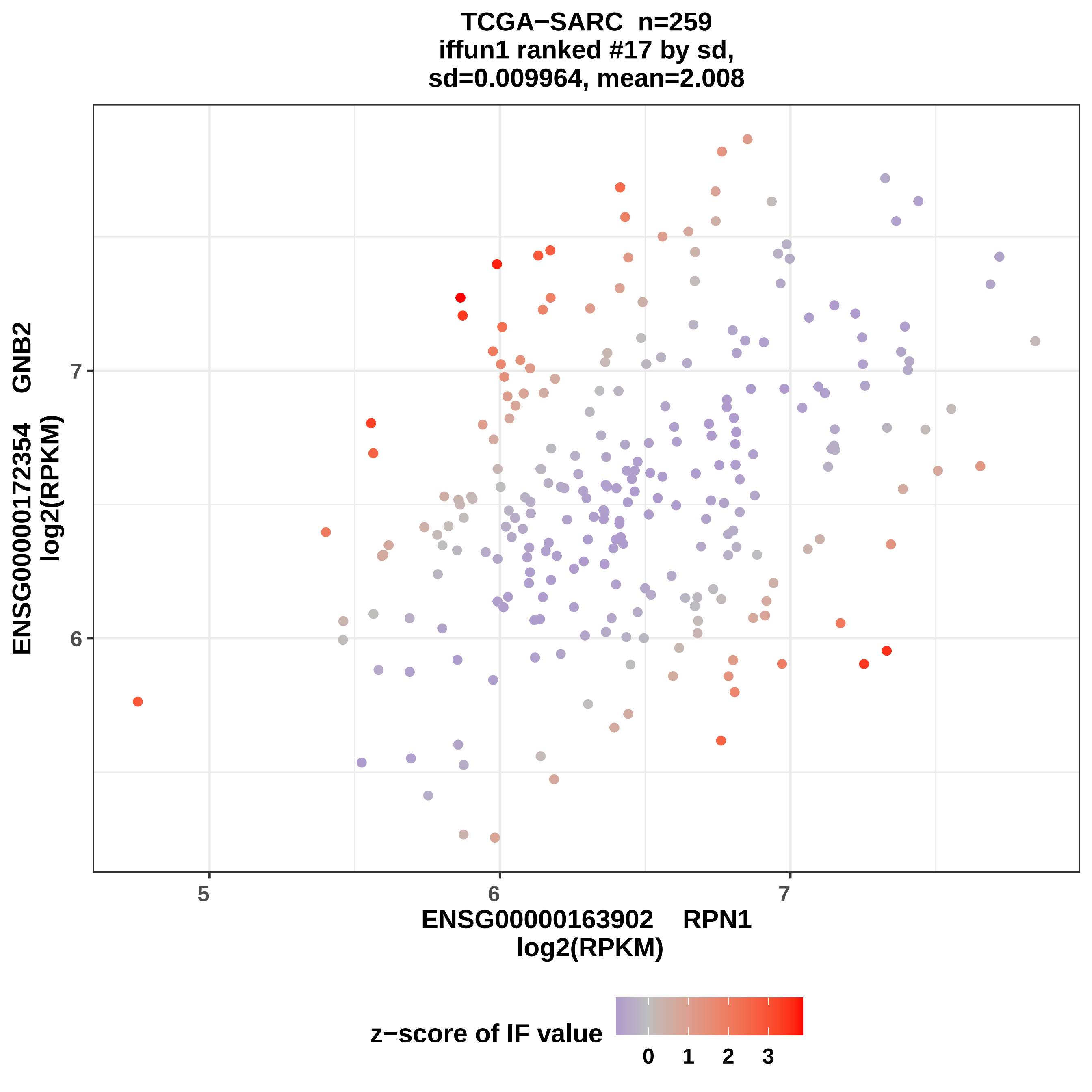
TCGA−SARC n=259
iffun1 ranked #17 by sd,
sd=0.009964, mean=2.008
ENSG00000172354 GNB2
log2(RPKM)
7
6
5
6
7
ENSG00000163902 RPN1
log2(RPKM)
z−score of IF value
0
1
2
3

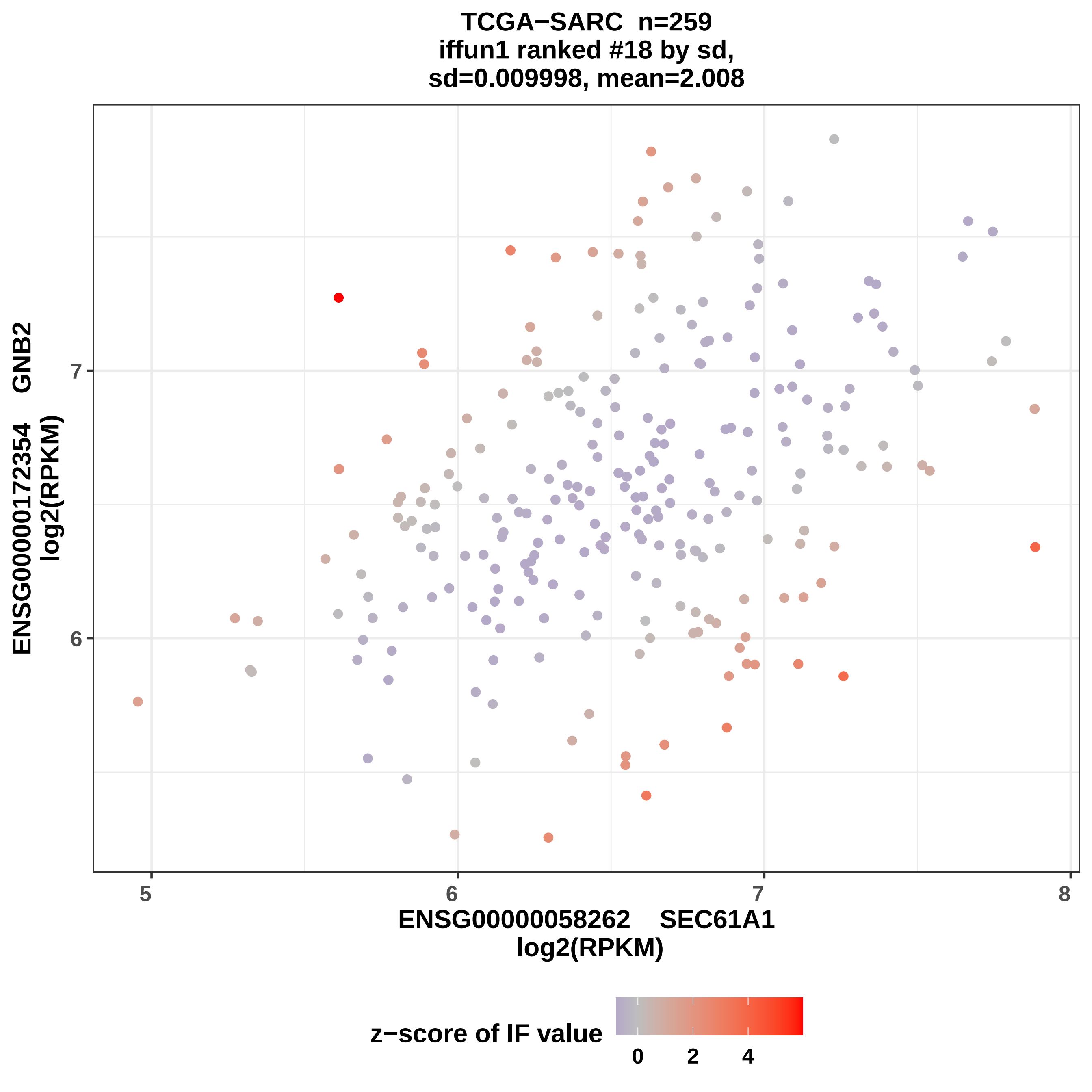

TCGA−SARC n=259
iffun1 ranked #18 by sd,
sd=0.009998, mean=2.008
ENSG00000172354 GNB2
log2(RPKM)
ENSG00000058262 SEC61A1
log2(RPKM)
5
6
7
8
6
7
z−score of IF value
0
2
4

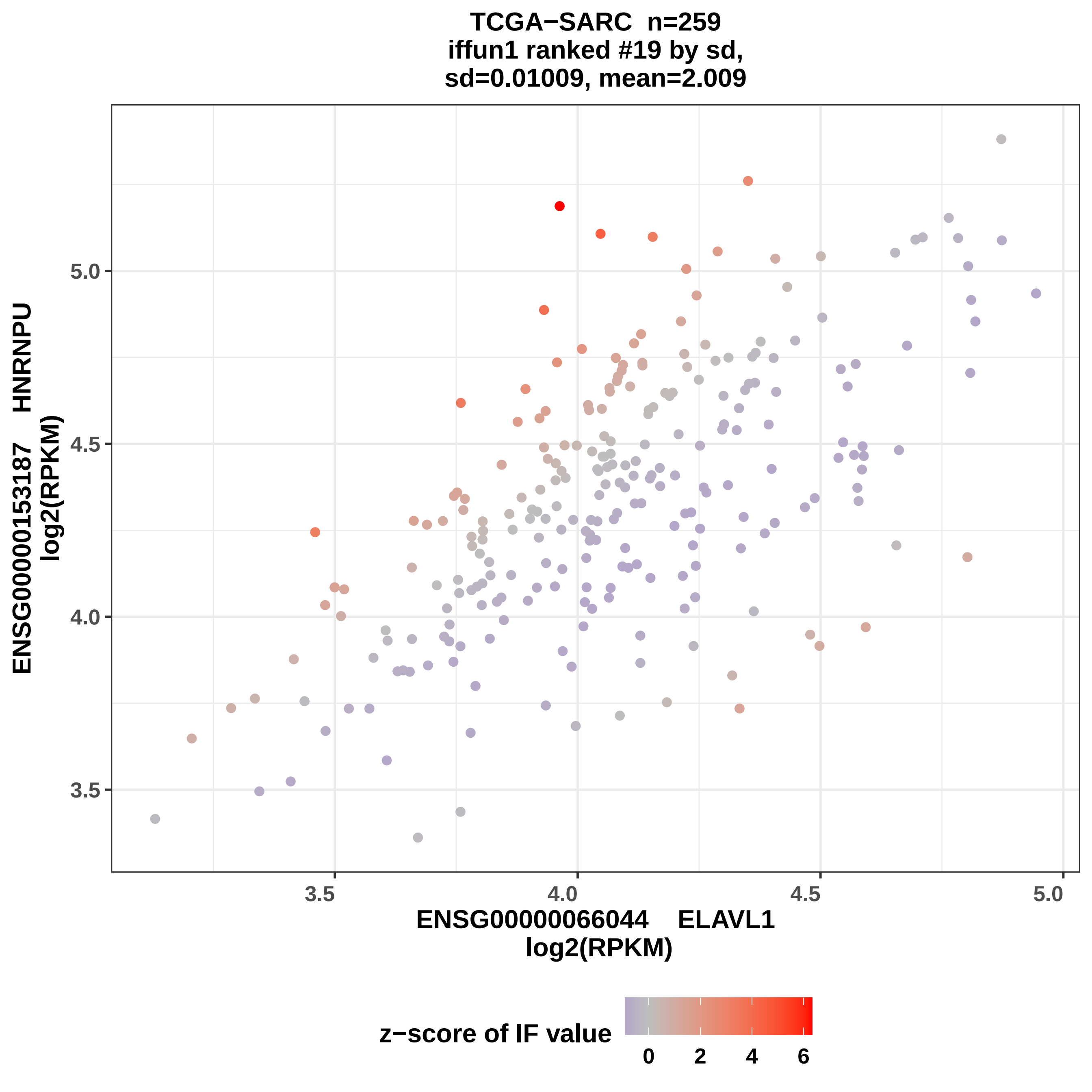

TCGA−SARC n=259
iffun1 ranked #19 by sd,
sd=0.01009, mean=2.009
ENSG00000153187 HNRNPU
log2(RPKM)
5.0
4.5
4.0
3.5
3.5
4.0
4.5
5.0
ENSG00000066044 ELAVL1
log2(RPKM)
z−score of IF value
0
2
4
6

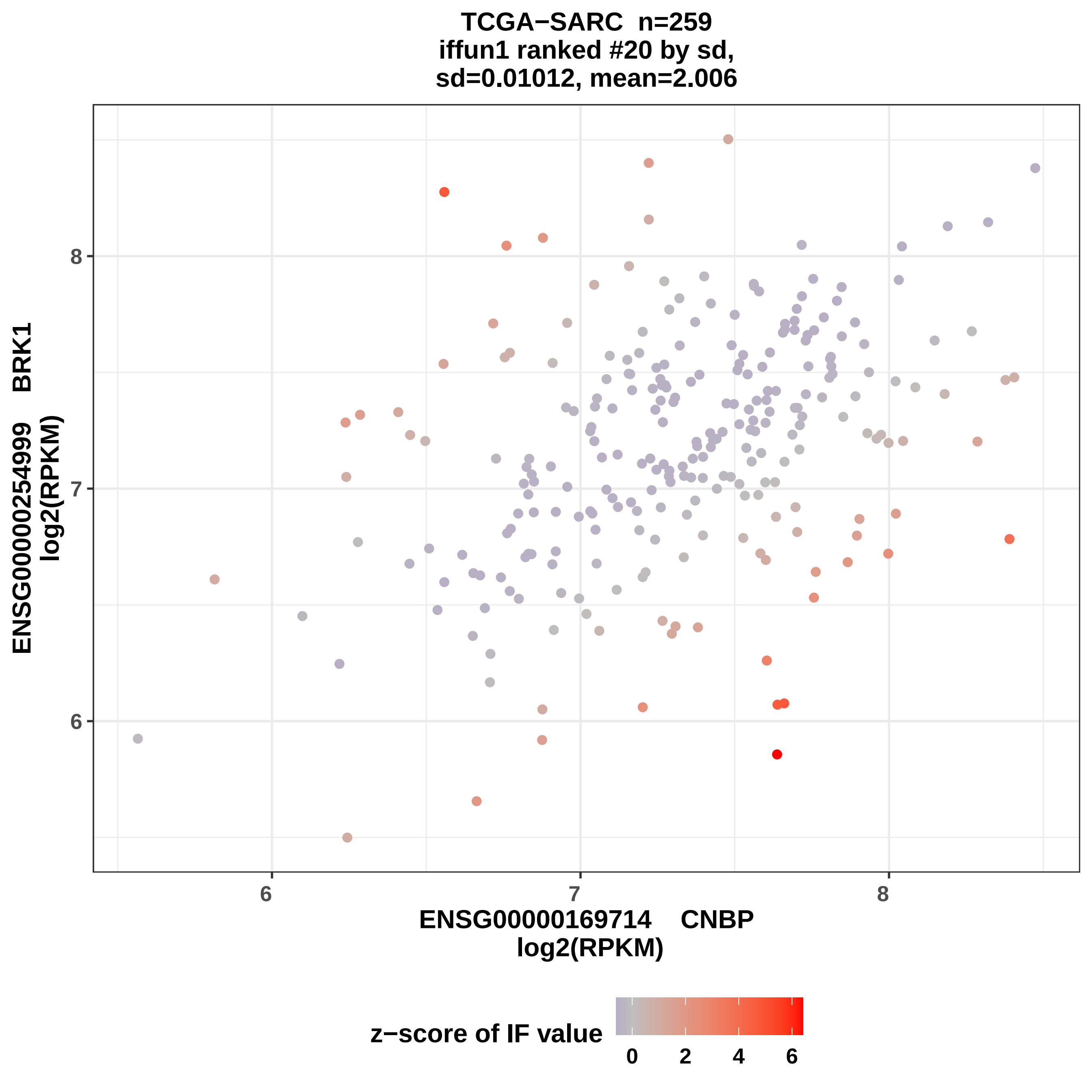
TCGA−SARC n=259
iffun1 ranked #20 by sd,
sd=0.01012, mean=2.006
ENSG00000254999 BRK1
log2(RPKM)
ENSG00000169714 CNBP
log2(RPKM)
6
7
8
6
7
8
z−score of IF value
0
2
4
6

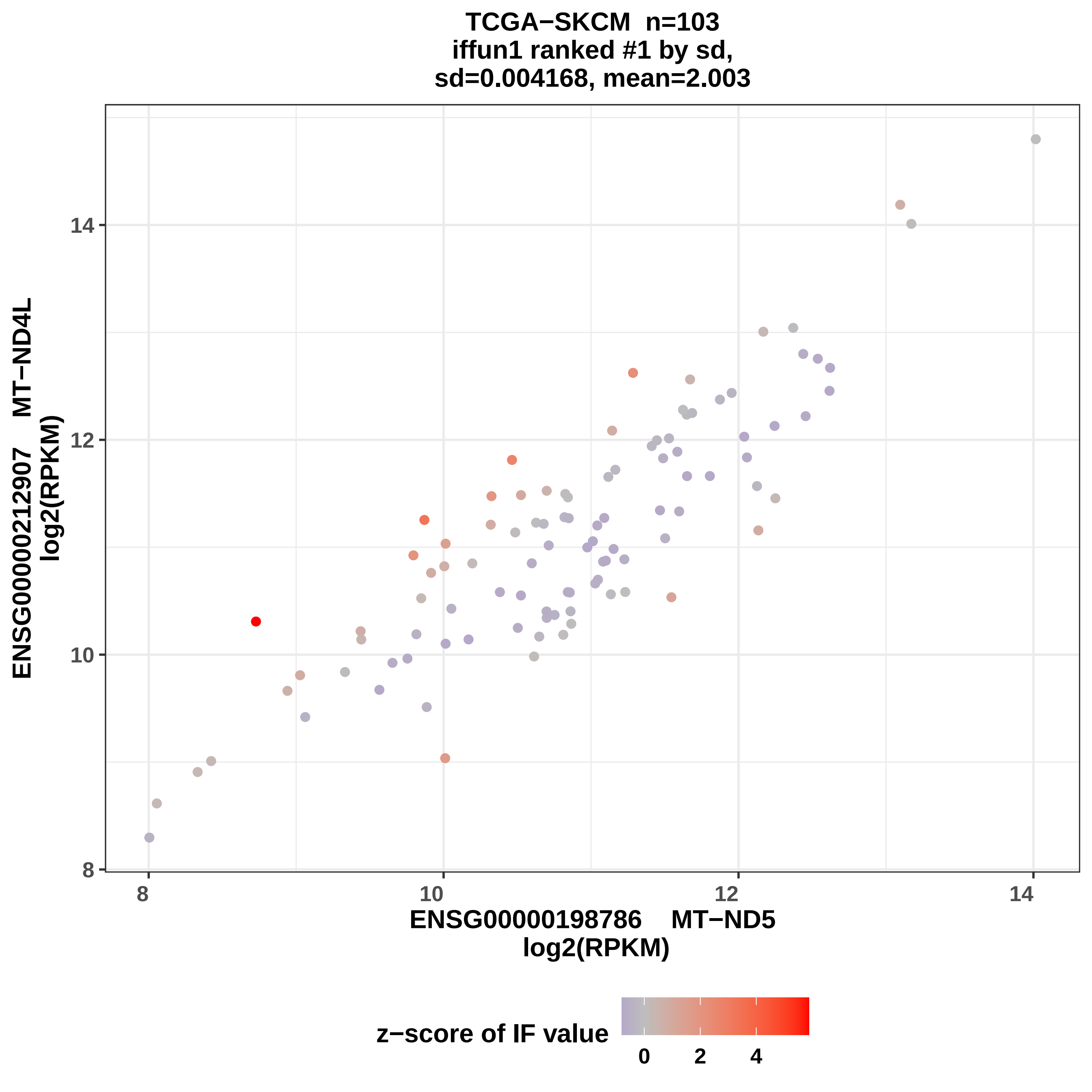

TCGA−SKCM n=103
iffun1 ranked #1 by sd,
sd=0.004168, mean=2.003
ENSG00000212907 MT−ND4L
log2(RPKM)
8
10
12
14
ENSG00000198786 MT−ND5
log2(RPKM)
z−score of IF value
0
2
4

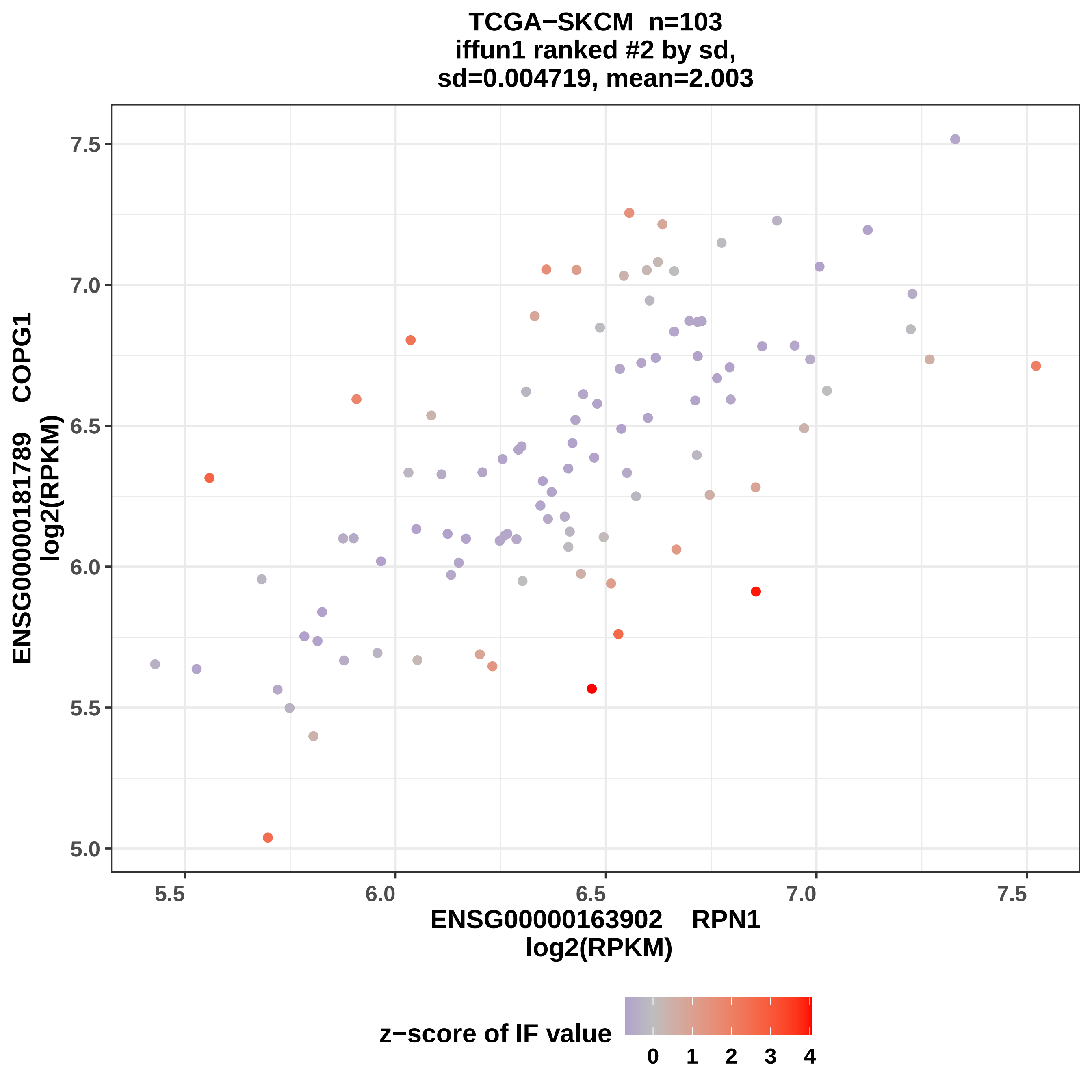

TCGA−SKCM n=103
iffun1 ranked #2 by sd,
sd=0.004719, mean=2.003
ENSG00000181789 COPG1
log2(RPKM)
ENSG00000163902 RPN1
log2(RPKM)
5.0
5.5
6.0
6.5
7.0
7.5
z−score of IF value
0
1
2
3
4

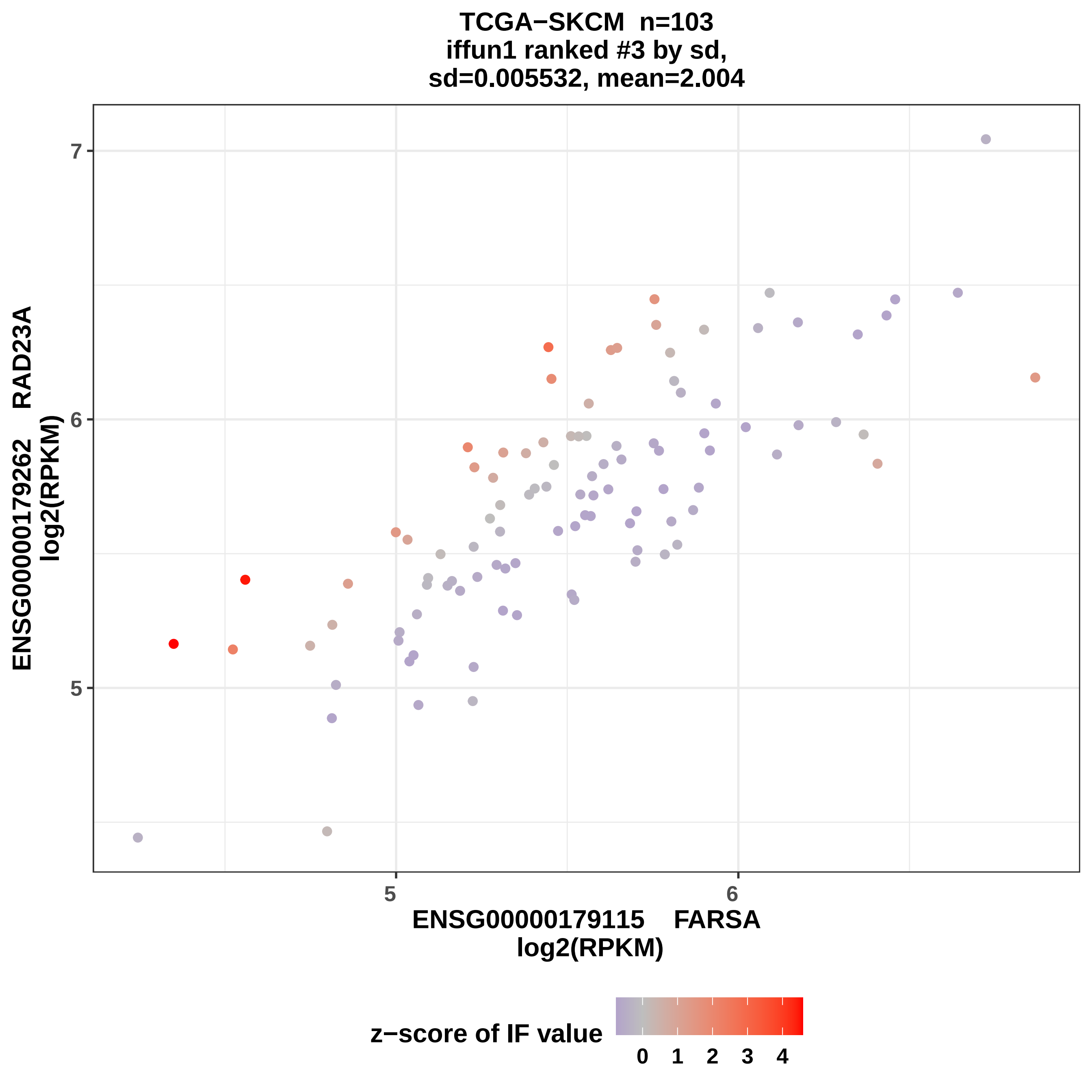
TCGA−SKCM n=103
iffun1 ranked #3 by sd,
sd=0.005532, mean=2.004
7
6
5
5
6
ENSG00000179115 FARSA
log2(RPKM)
ENSG00000179262 RAD23A
log2(RPKM)
z−score of IF value
0
1
2
3
4

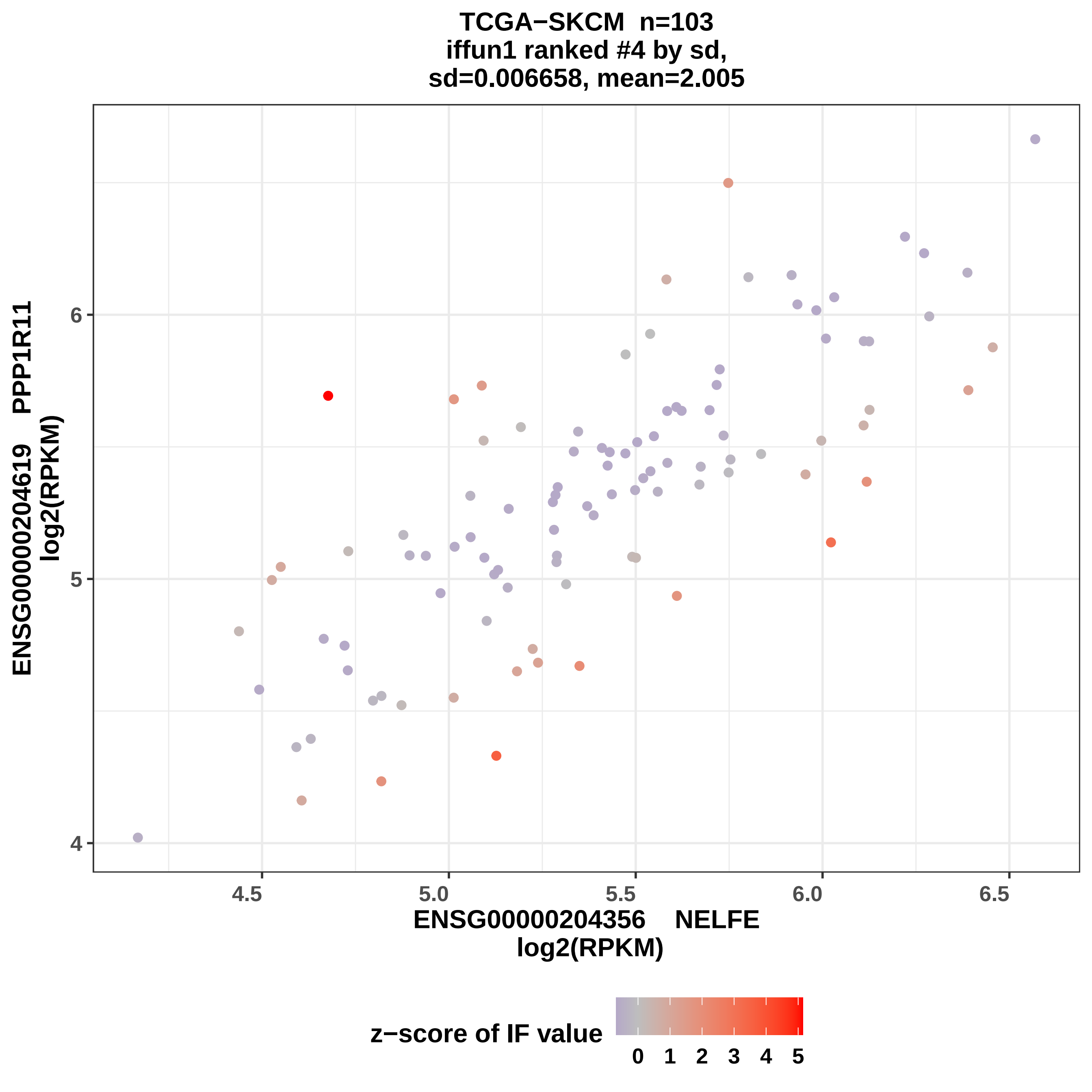
TCGA−SKCM n=103
iffun1 ranked #4 by sd,
sd=0.006658, mean=2.005
ENSG00000204619 PPP1R11
log2(RPKM)
6
5
4
4.5
5.0
5.5
6.0
6.5
ENSG00000204356 NELFE
log2(RPKM)
z−score of IF value
0 1 2 3 4 5

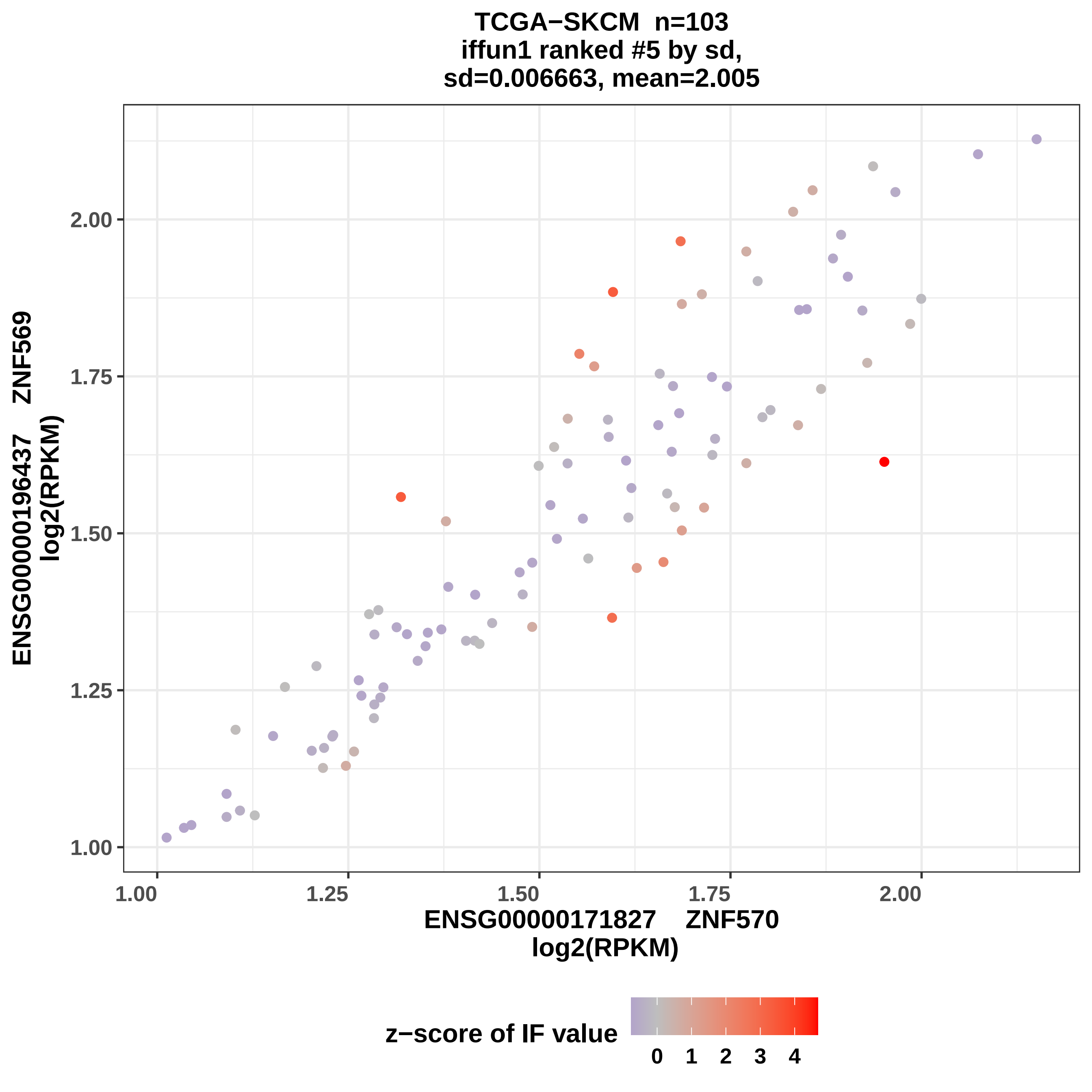

TCGA−SKCM n=103
iffun1 ranked #5 by sd,
sd=0.006663, mean=2.005
ENSG00000196437 ZNF569
log2(RPKM)
1.00
1.25
1.50
1.75
2.00
ENSG00000171827 ZNF570
log2(RPKM)
z−score of IF value
0
1
2
3
4

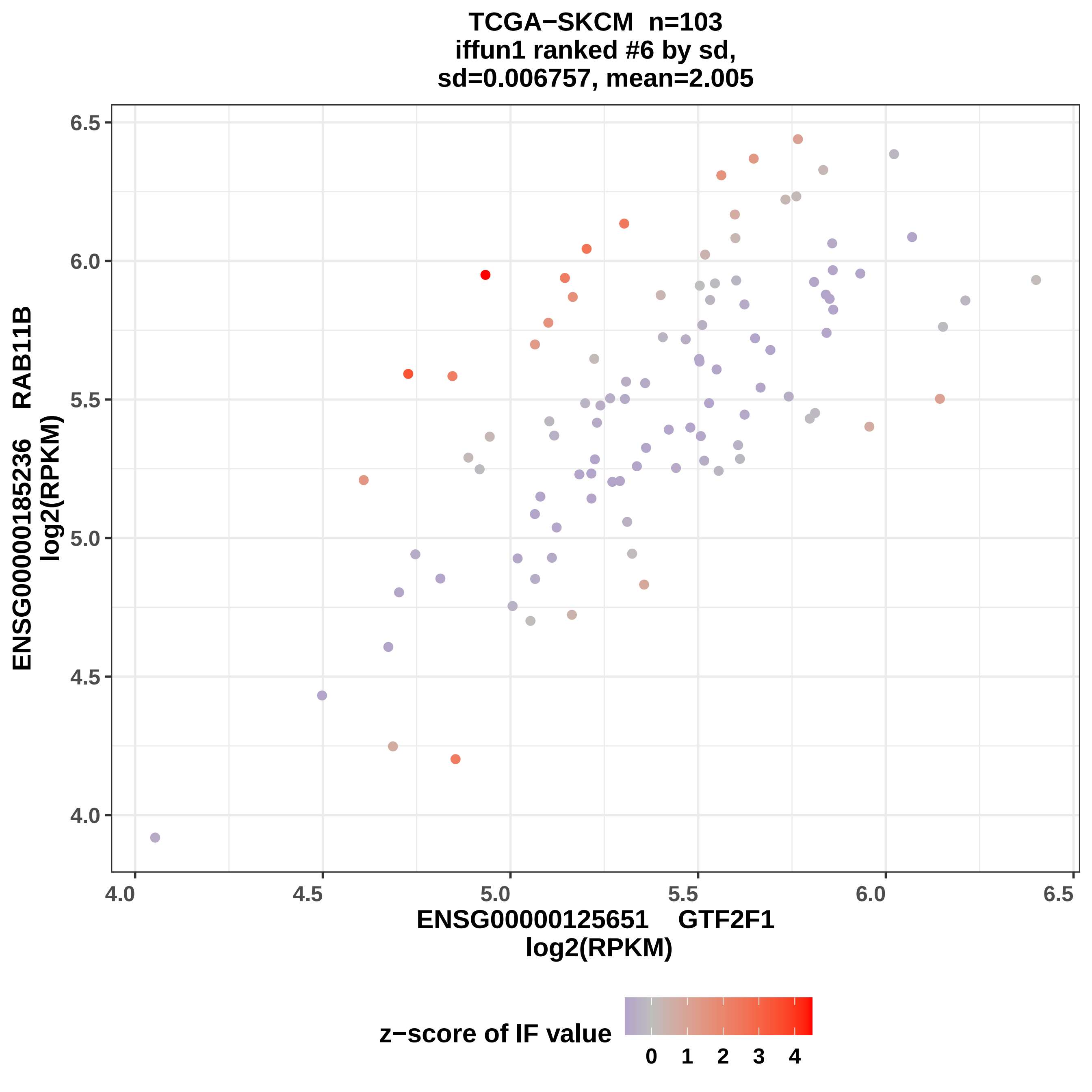
TCGA−SKCM n=103
iffun1 ranked #6 by sd,
sd=0.006757, mean=2.005
ENSG00000185236 RAB11B
log2(RPKM)
6.5
6.0
5.5
5.0
4.5
4.0
4.0
4.5
5.0
5.5
6.0
6.5
ENSG00000125651 GTF2F1
log2(RPKM)
z−score of IF value
0
1
2
3
4

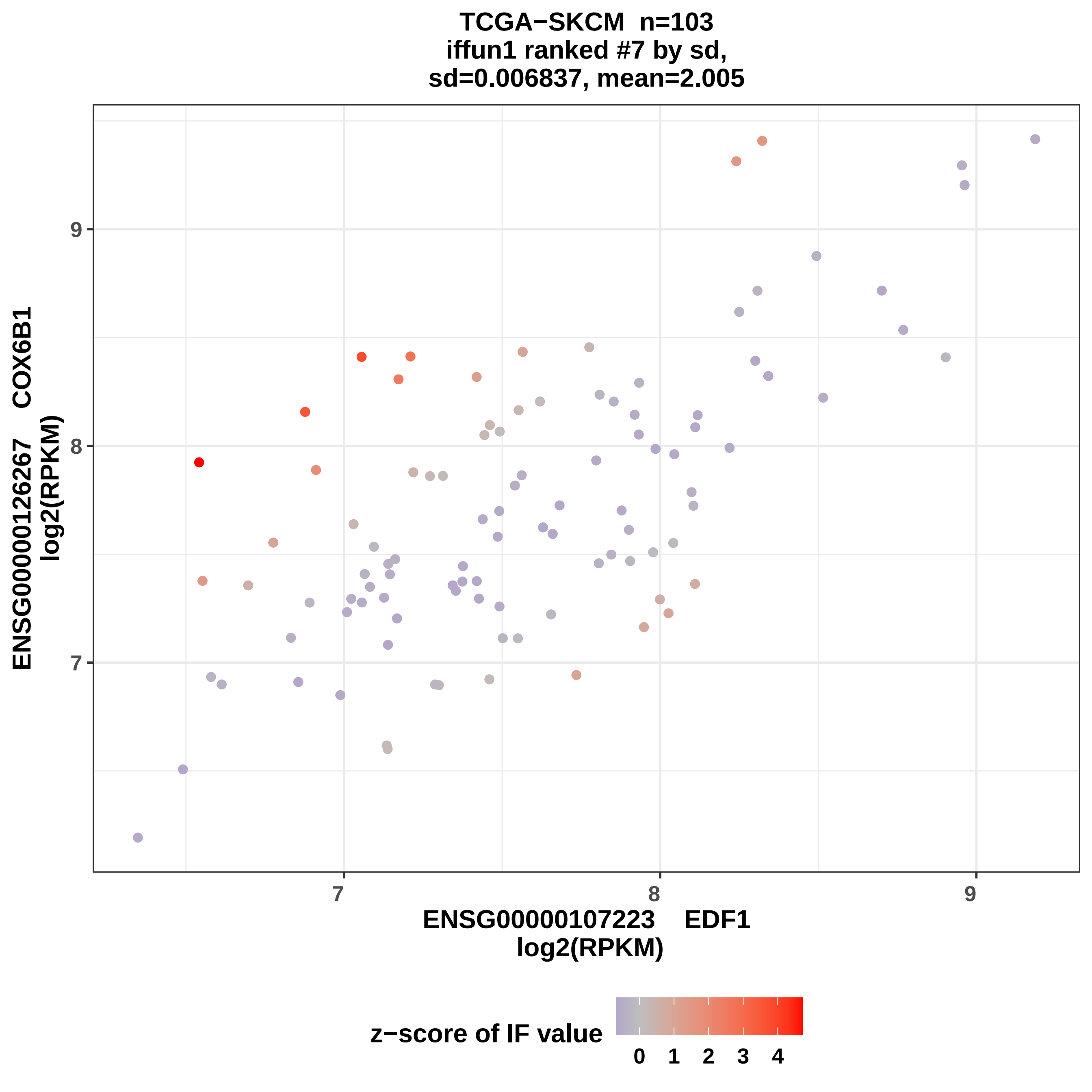
TCGA−SKCM n=103
iffun1 ranked #7 by sd,
sd=0.006837, mean=2.005
ENSG00000126267 COX6B1
log2(RPKM)
9
8
7
7
8
9
ENSG00000107223 EDF1
log2(RPKM)
z−score of IF value
0
1
2
3
4

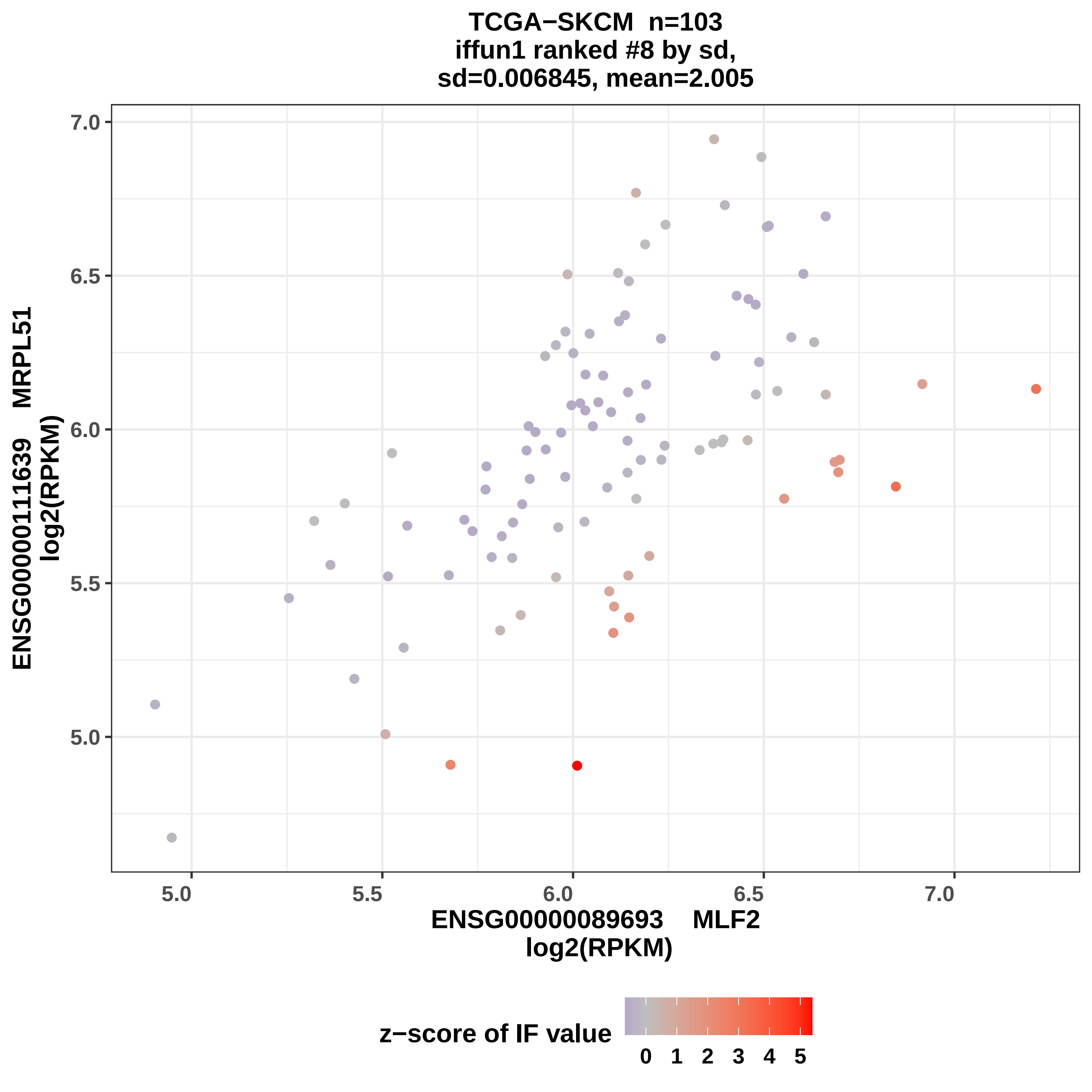
TCGA−SKCM n=103
iffun1 ranked #8 by sd,
sd=0.006845, mean=2.005
ENSG00000111639 MRPL51
log2(RPKM)
7.0
6.5
6.0
5.5
5.0
5.0
5.5
6.0
6.5
7.0
ENSG00000089693 MLF2
log2(RPKM)
z−score of IF value
0 1 2 3 4 5

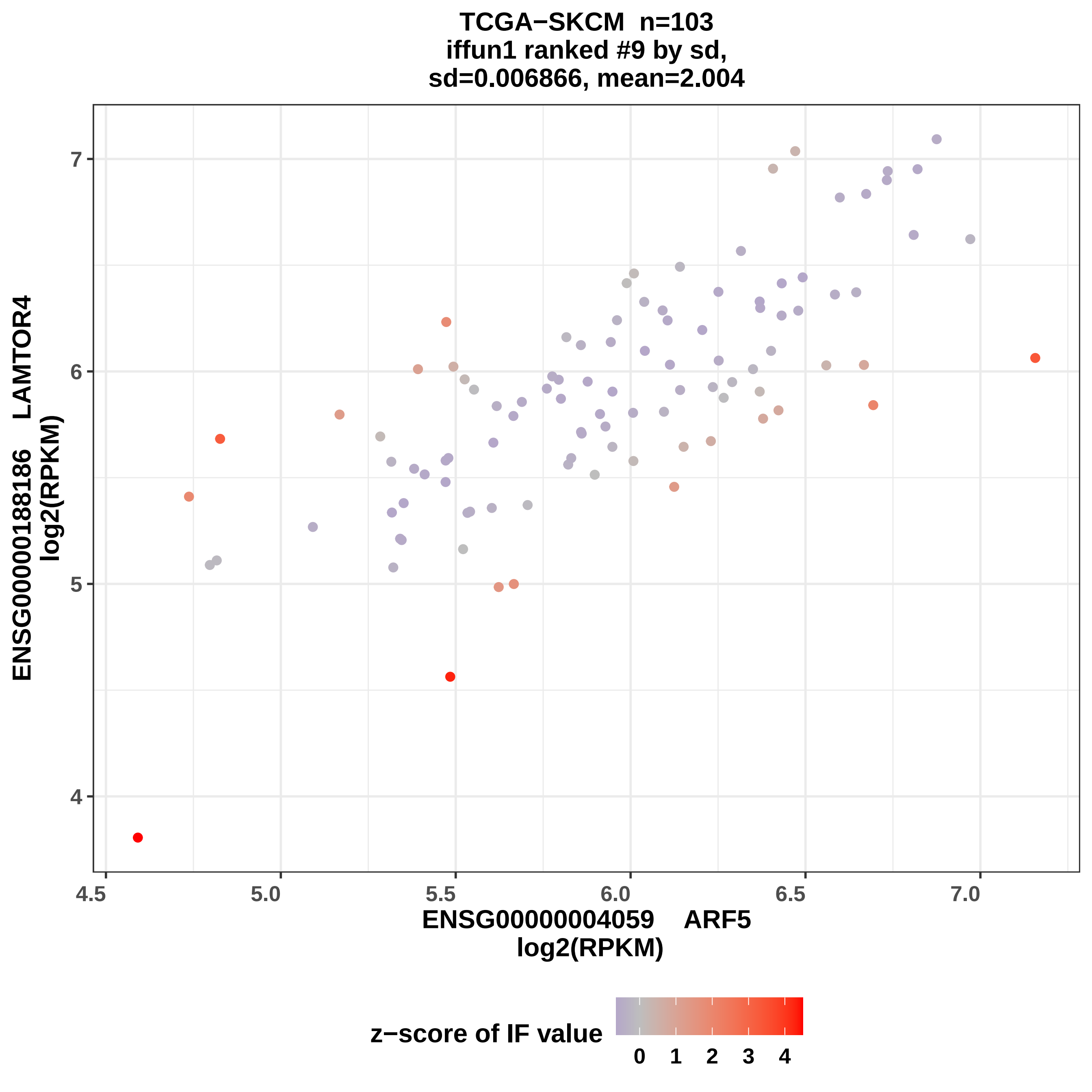
TCGA−SKCM n=103
iffun1 ranked #9 by sd,
sd=0.006866, mean=2.004
ENSG00000188186 LAMTOR4
log2(RPKM)
ENSG00000004059 ARF5
log2(RPKM)
4
5
6
7
4.5
5.0
5.5
6.0
6.5
7.0
z−score of IF value
0
1
2
3
4

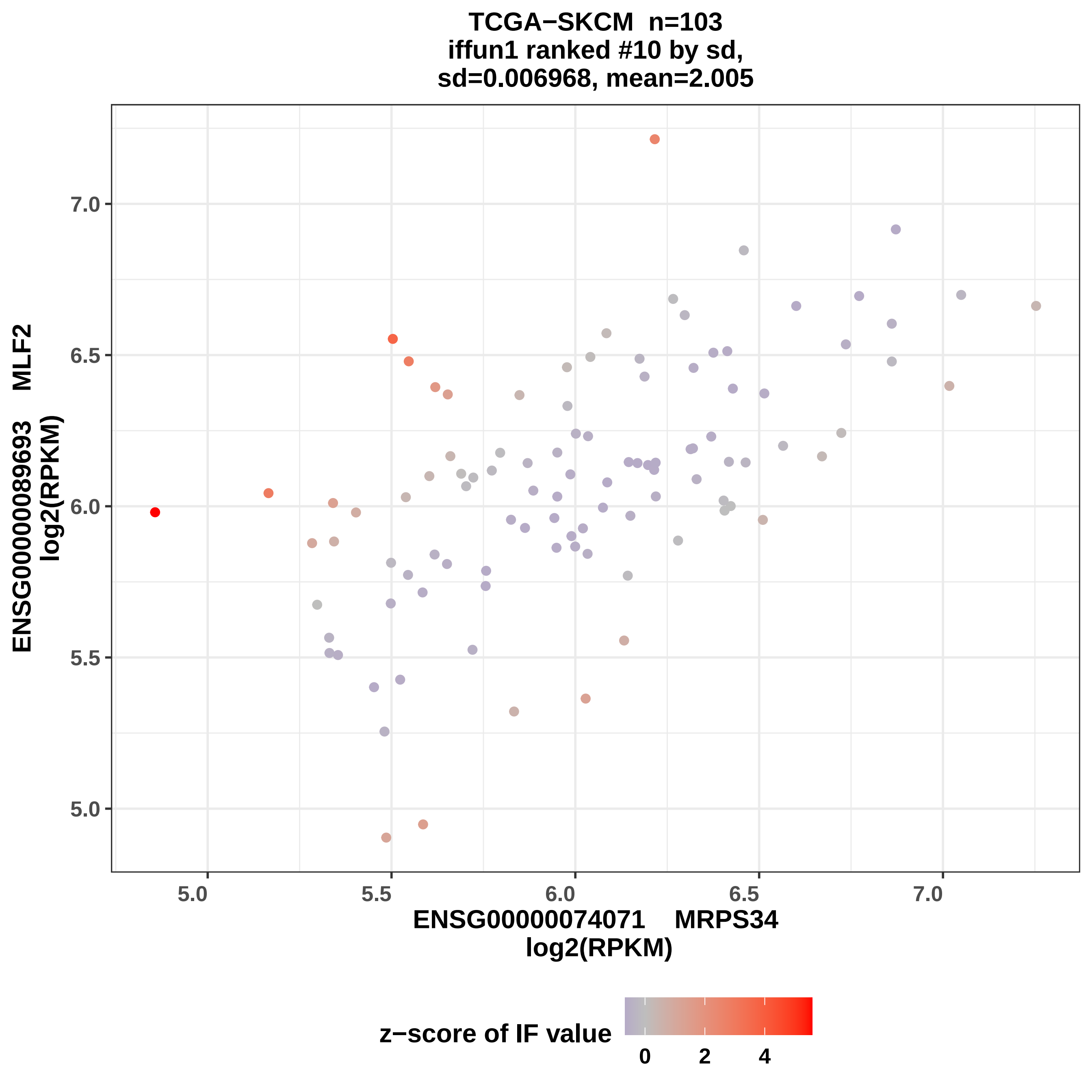

TCGA−SKCM n=103
iffun1 ranked #10 by sd,
sd=0.006968, mean=2.005
ENSG00000089693 MLF2
log2(RPKM)
7.0
6.5
6.0
5.5
5.0
5.0
5.5
6.0
6.5
7.0
ENSG00000074071 MRPS34
log2(RPKM)
z−score of IF value
0
2
4

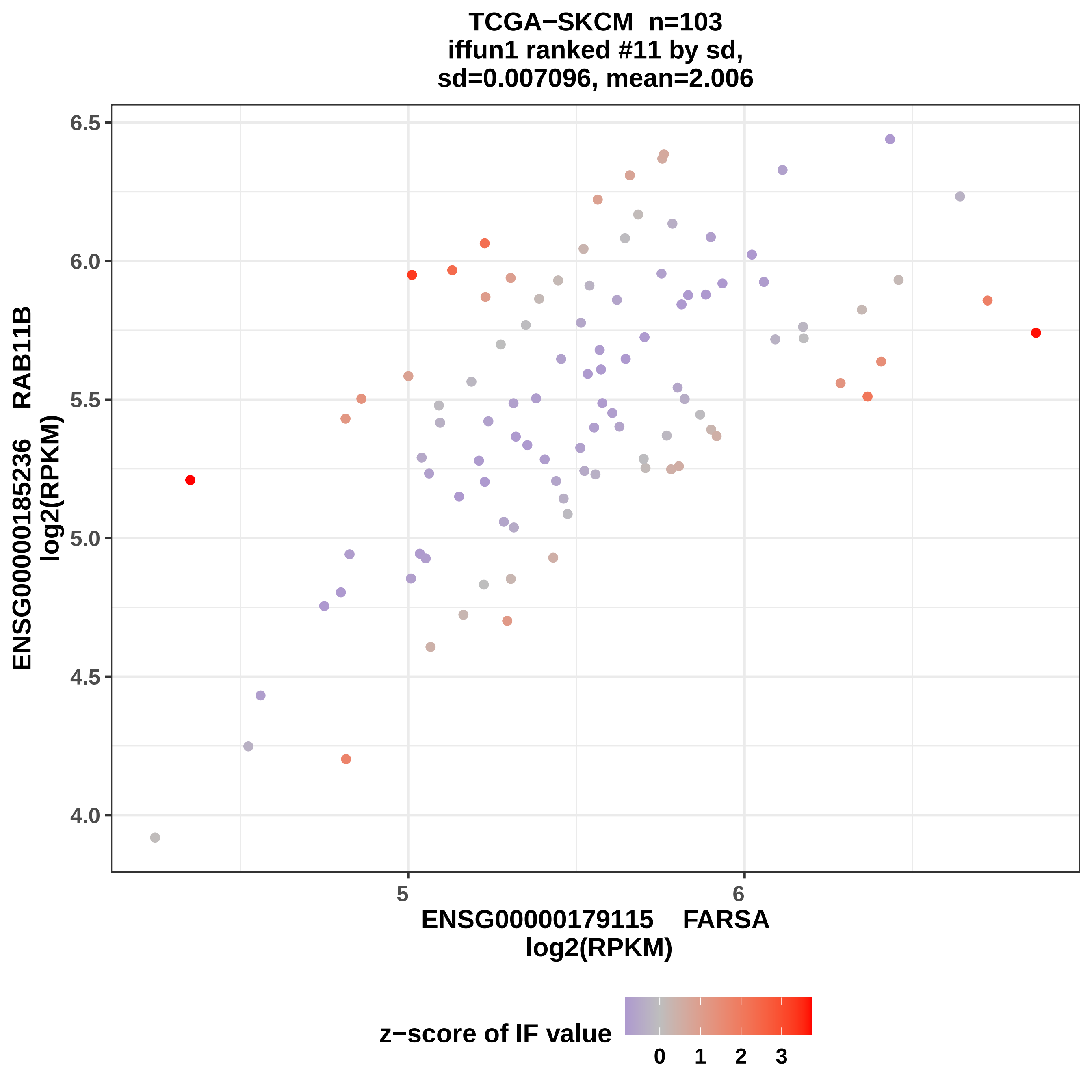
TCGA−SKCM n=103
iffun1 ranked #11 by sd,
sd=0.007096, mean=2.006
ENSG00000185236 RAB11B
log2(RPKM)
6.5
6.0
5.5
5.0
4.5
4.0
5
6
ENSG00000179115 FARSA
log2(RPKM)
z−score of IF value
0
1
2
3

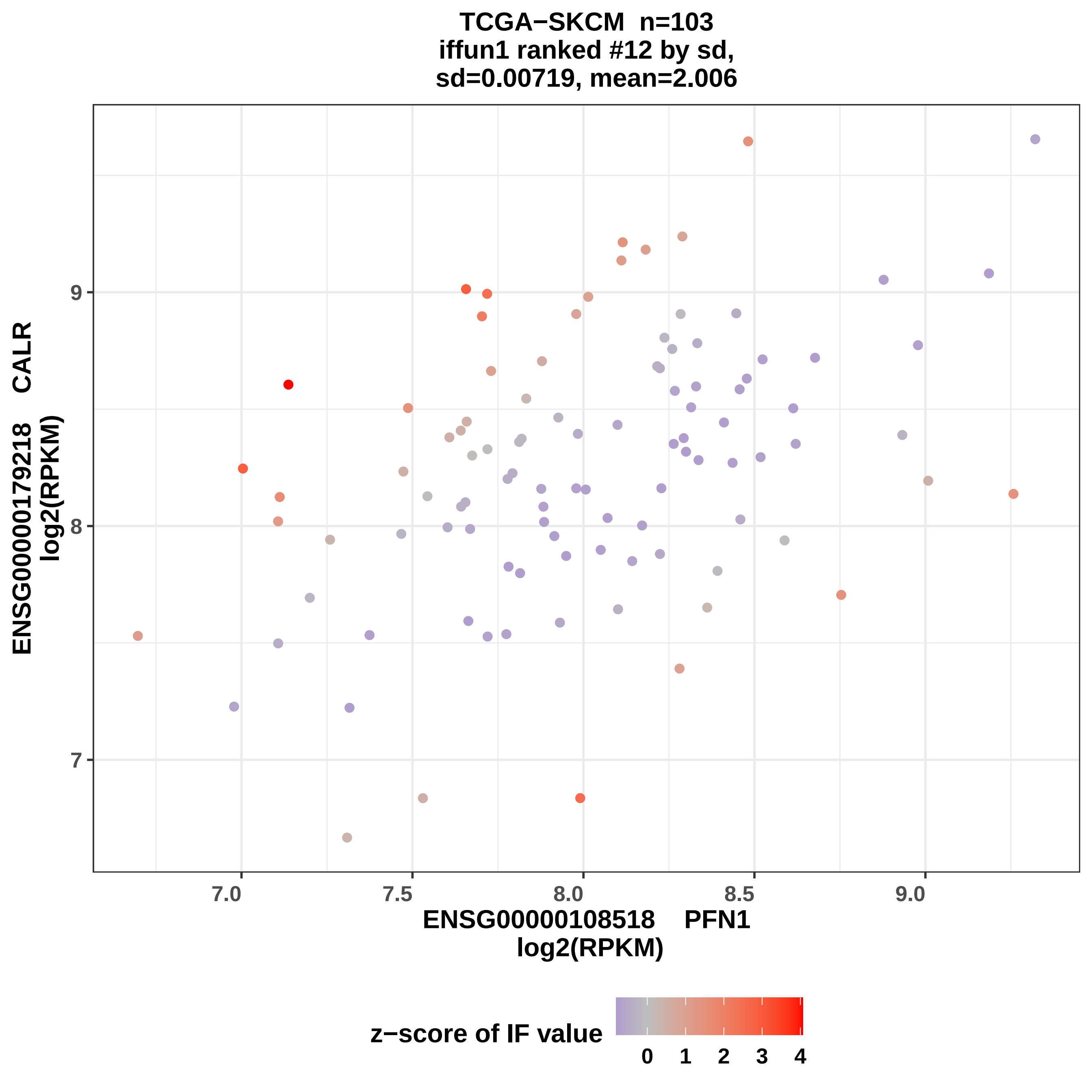

TCGA−SKCM n=103
iffun1 ranked #12 by sd,
sd=0.00719, mean=2.006
ENSG00000179218 CALR
log2(RPKM)
9
8
7
7.0
7.5
8.0
8.5
9.0
ENSG00000108518 PFN1
log2(RPKM)
z−score of IF value
0
1
2
3
4

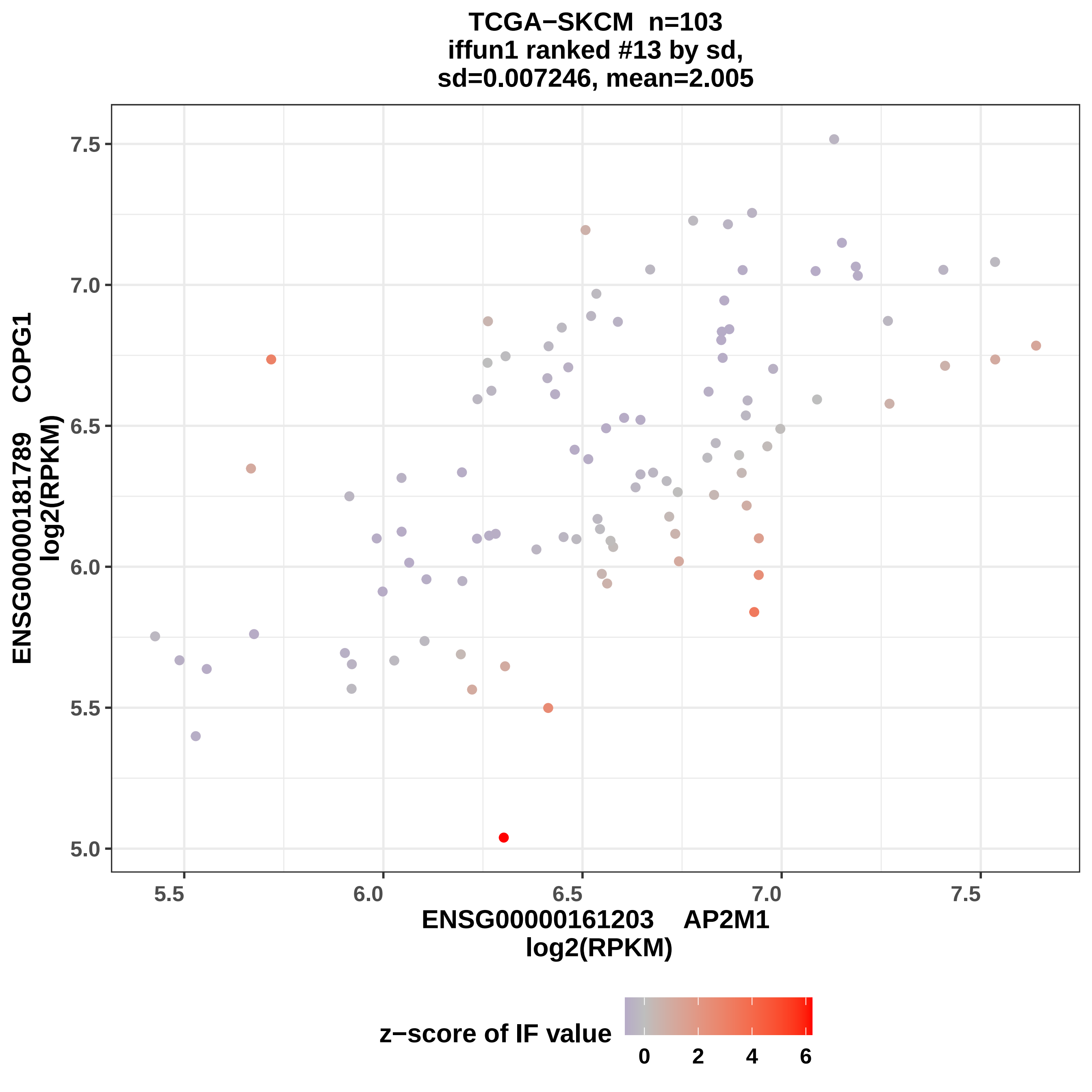

TCGA−SKCM n=103
iffun1 ranked #13 by sd,
sd=0.007246, mean=2.005
ENSG00000181789 COPG1
log2(RPKM)
7.5
7.0
6.5
6.0
5.5
5.0
5.5
6.0
6.5
7.0
7.5
ENSG00000161203 AP2M1
log2(RPKM)
z−score of IF value
0
2
4
6

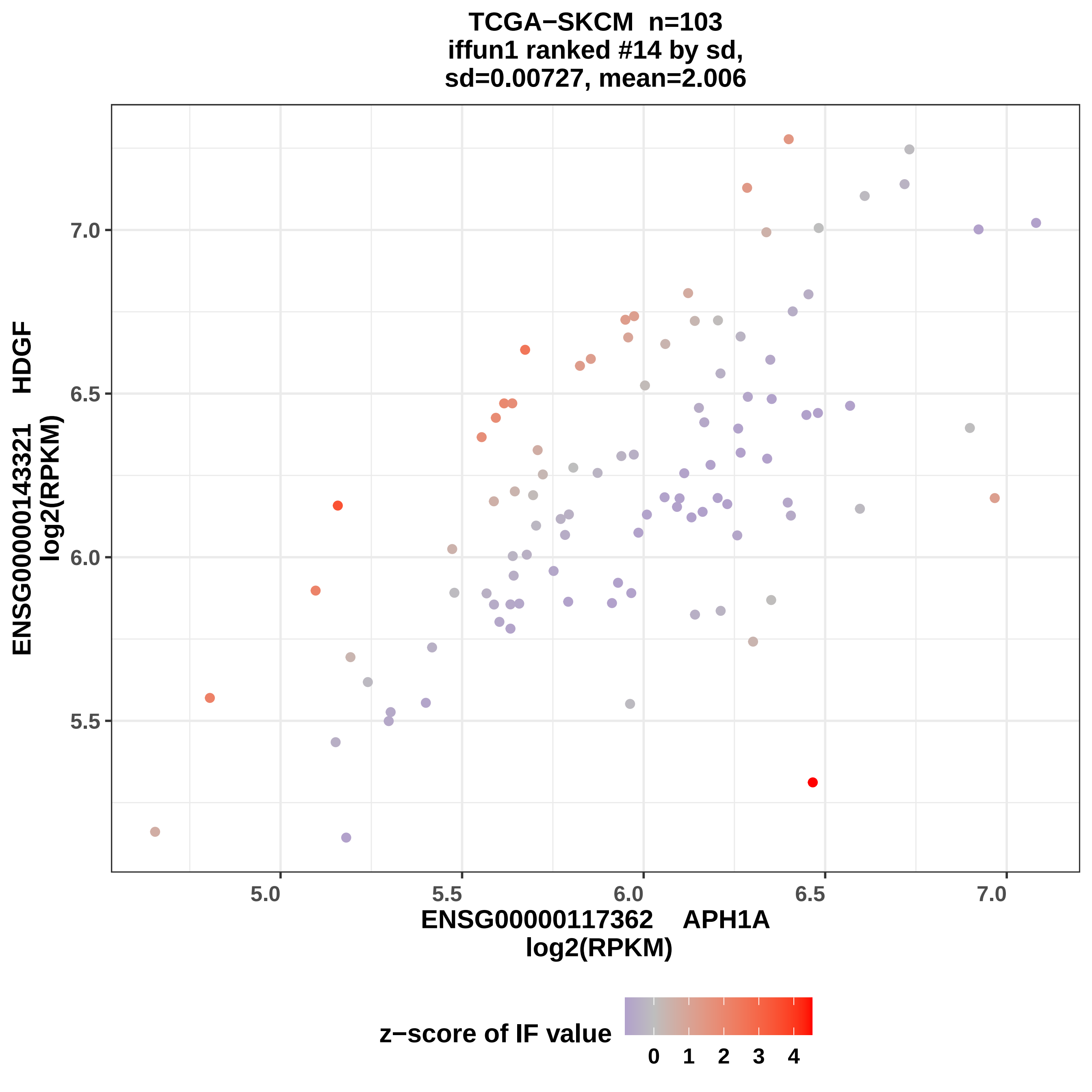

TCGA−SKCM n=103
iffun1 ranked #14 by sd,
sd=0.00727, mean=2.006
7.0
6.5
6.0
5.5
5.0
5.5
6.0
6.5
7.0
ENSG00000143321 HDGF
log2(RPKM)
ENSG00000117362 APH1A
log2(RPKM)
z−score of IF value
0
1
2
3
4

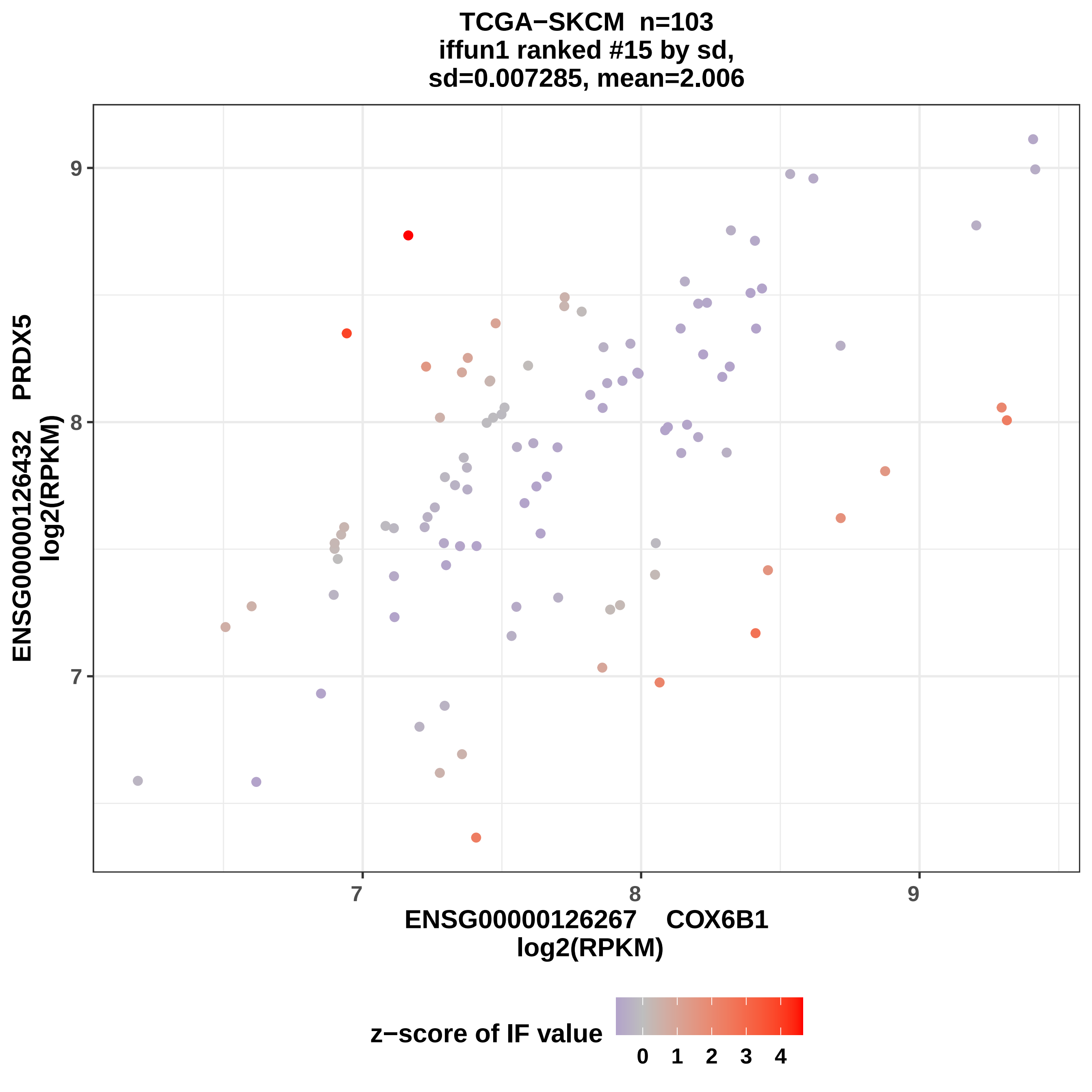
TCGA−SKCM n=103
iffun1 ranked #15 by sd,
sd=0.007285, mean=2.006
ENSG00000126432 PRDX5
log2(RPKM)
ENSG00000126267 COX6B1
log2(RPKM)
9
8
7
7
8
9
z−score of IF value
0
1
2
3
4

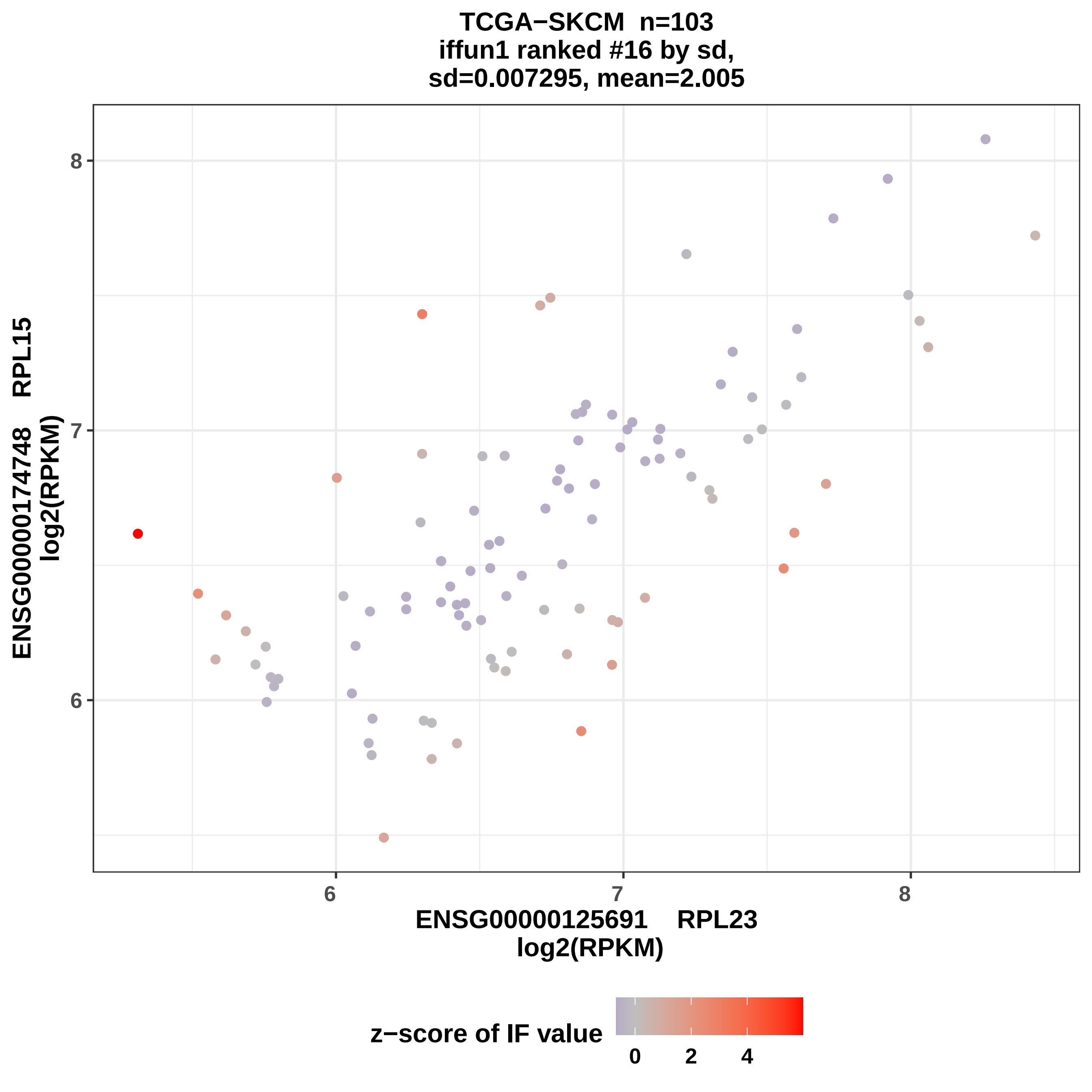

TCGA−SKCM n=103
iffun1 ranked #16 by sd,
sd=0.007295, mean=2.005
ENSG00000174748 RPL15
log2(RPKM)
8
7
6
6
7
8
ENSG00000125691 RPL23
log2(RPKM)
z−score of IF value
0
2
4

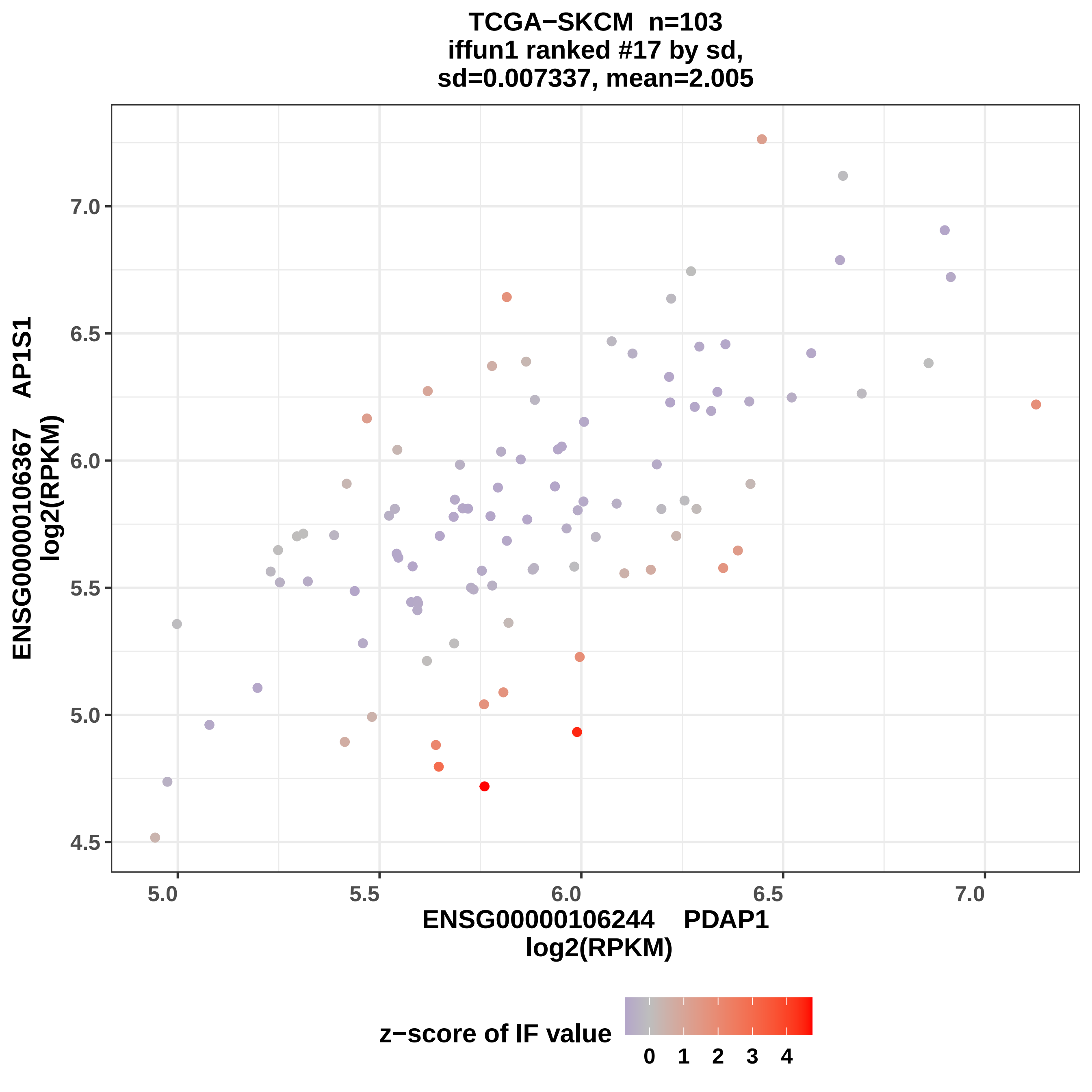

TCGA−SKCM n=103
iffun1 ranked #17 by sd,
sd=0.007337, mean=2.005
ENSG00000106367 AP1S1
log2(RPKM)
7.0
6.5
6.0
5.5
5.0
4.5
5.0
5.5
6.0
6.5
7.0
ENSG00000106244 PDAP1
log2(RPKM)
z−score of IF value
0 1 2 3 4

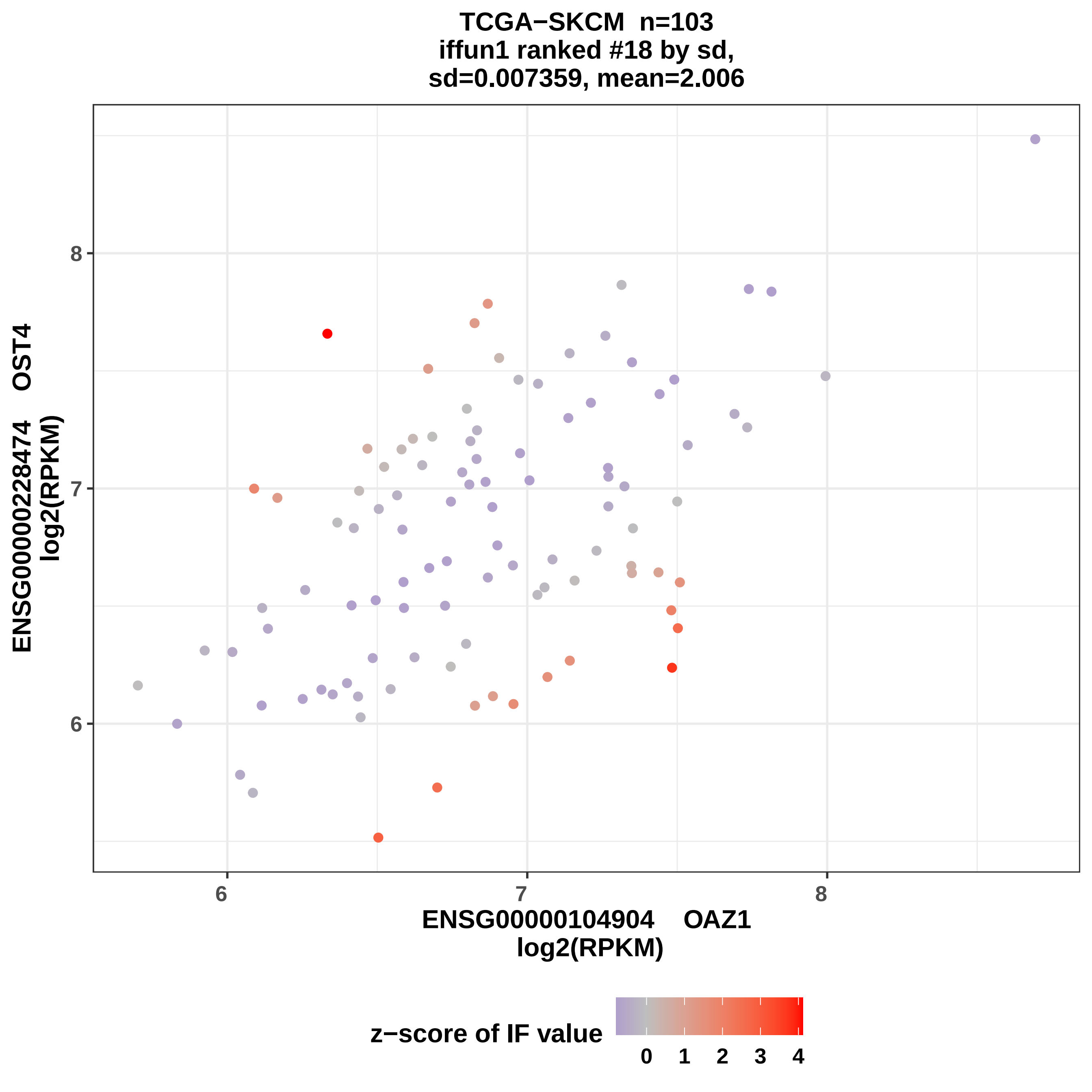
TCGA−SKCM n=103
iffun1 ranked #18 by sd,
sd=0.007359, mean=2.006
ENSG00000228474 OST4
log2(RPKM)
8
7
6
6
7
8
ENSG00000104904 OAZ1
log2(RPKM)
z−score of IF value
0 1 2 3 4

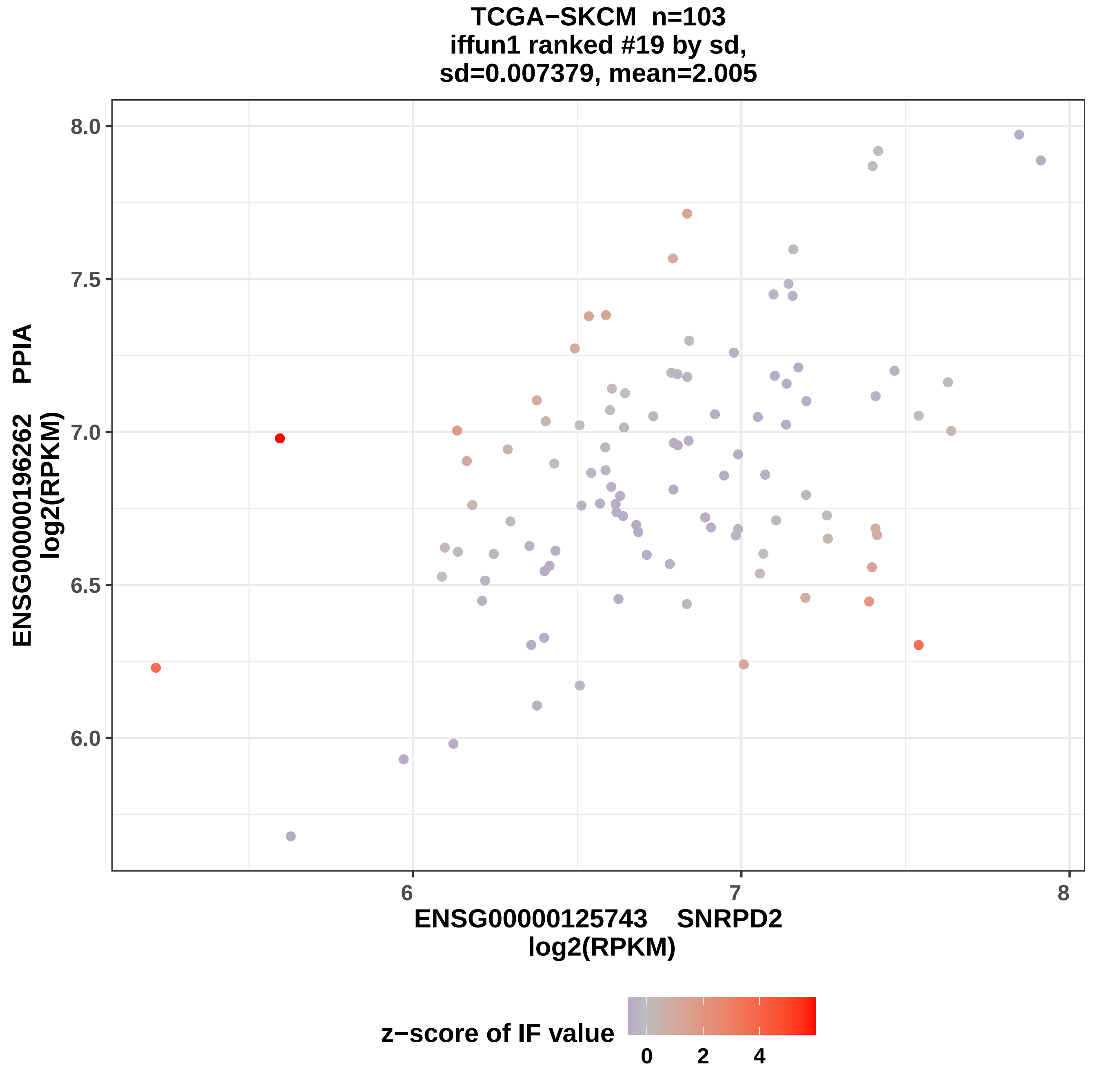
TCGA−SKCM n=103
iffun1 ranked #19 by sd,
sd=0.007379, mean=2.005
ENSG00000196262 PPIA
log2(RPKM)
8.0
7.5
7.0
6.5
6.0
6
7
8
ENSG00000125743 SNRPD2
log2(RPKM)
z−score of IF value
0
2
4

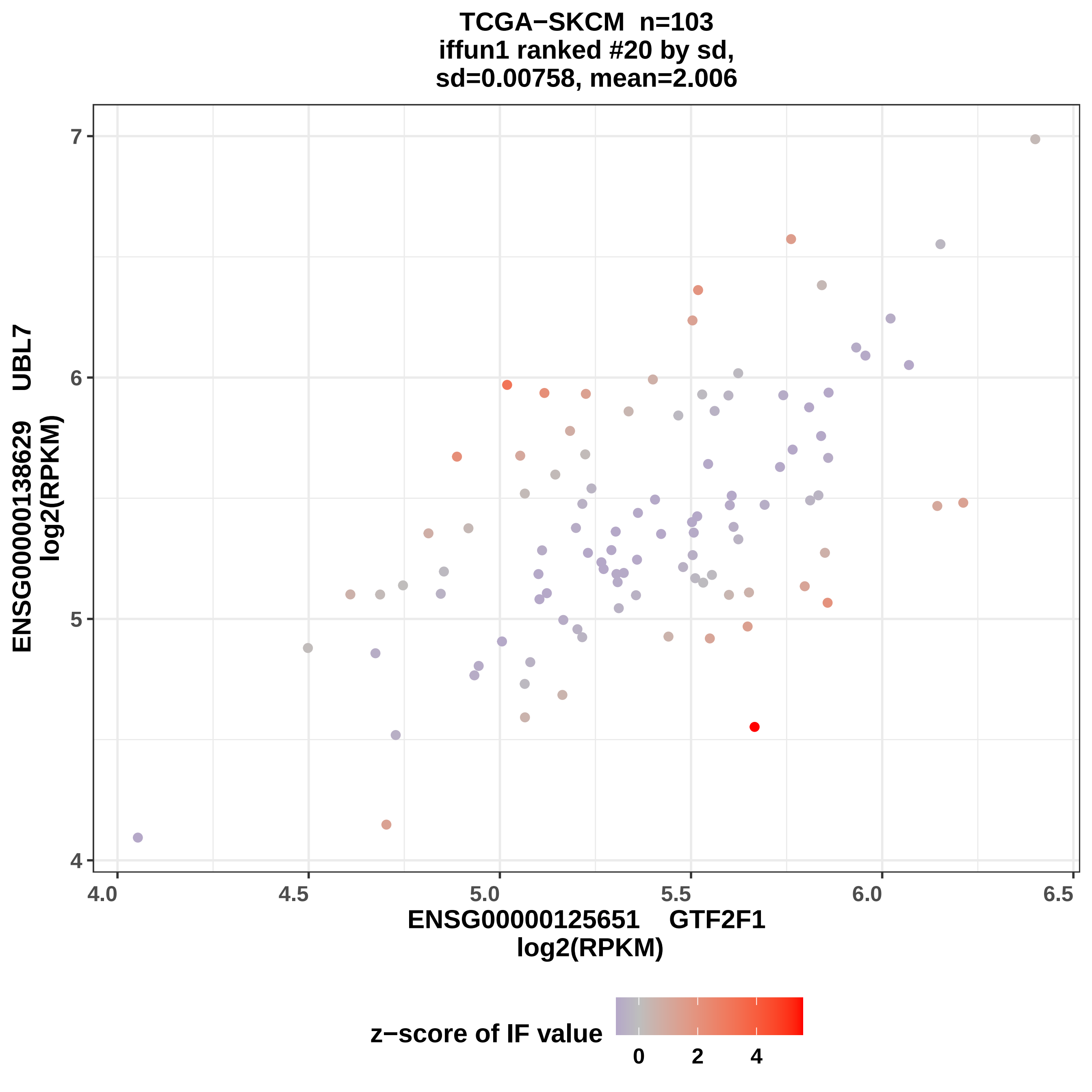
TCGA−SKCM n=103
iffun1 ranked #20 by sd,
sd=0.00758, mean=2.006
ENSG00000138629 UBL7
log2(RPKM)
ENSG00000125651 GTF2F1
log2(RPKM)
z−score of IF value
7
6
5
4
4.0
4.5
5.0
5.5
6.0
6.5
0
2
4

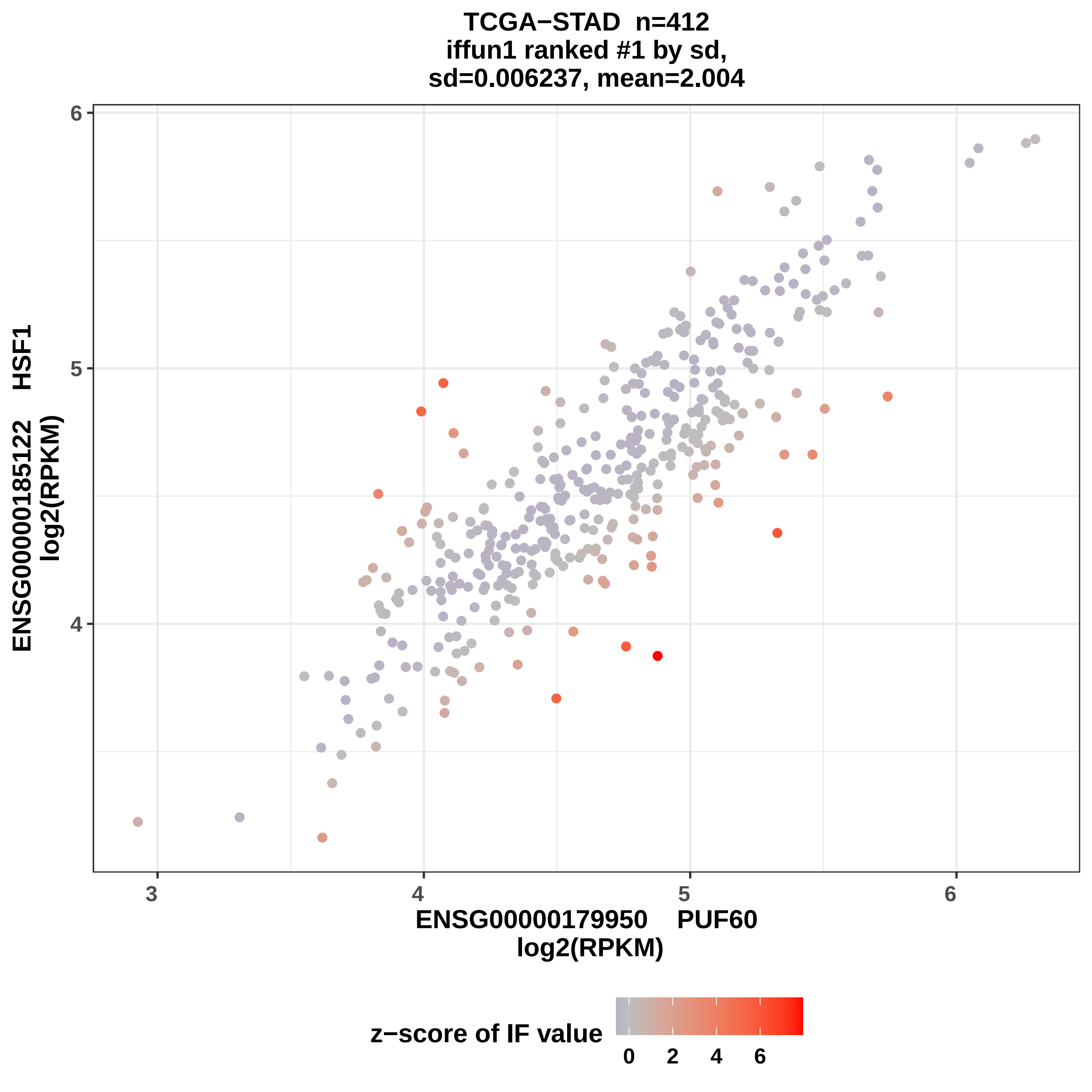

TCGA−STAD n=412
iffun1 ranked #1 by sd,
sd=0.006237, mean=2.004
ENSG00000185122 HSF1
log2(RPKM)
6
5
4
3
4
5
6
ENSG00000179950 PUF60
log2(RPKM)
z−score of IF value
0
2
4
6

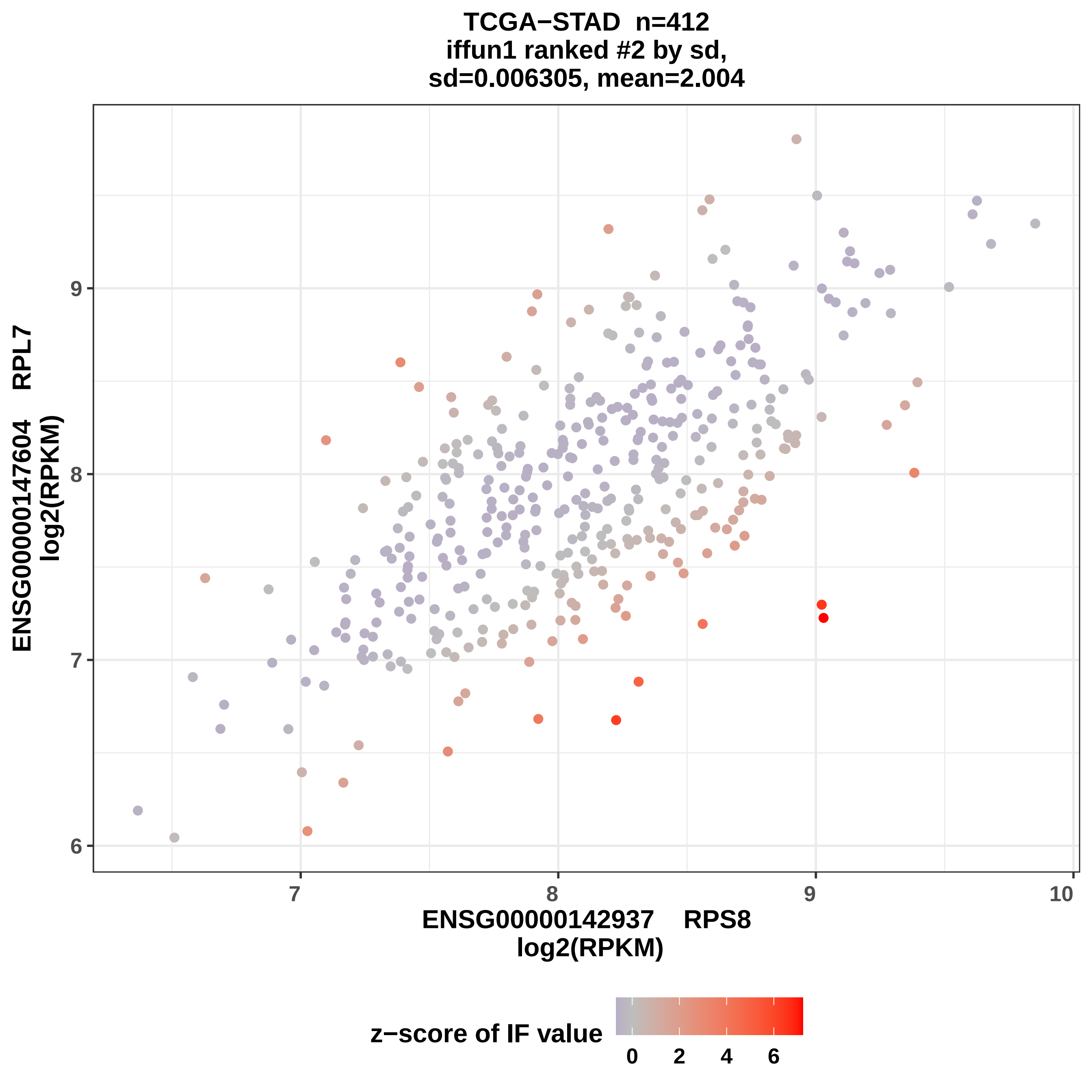
TCGA−STAD n=412
iffun1 ranked #2 by sd,
sd=0.006305, mean=2.004
ENSG00000147604 RPL7
log2(RPKM)
9
8
7
6
7
8
9
10
ENSG00000142937 RPS8
log2(RPKM)
z−score of IF value
0
2
4
6

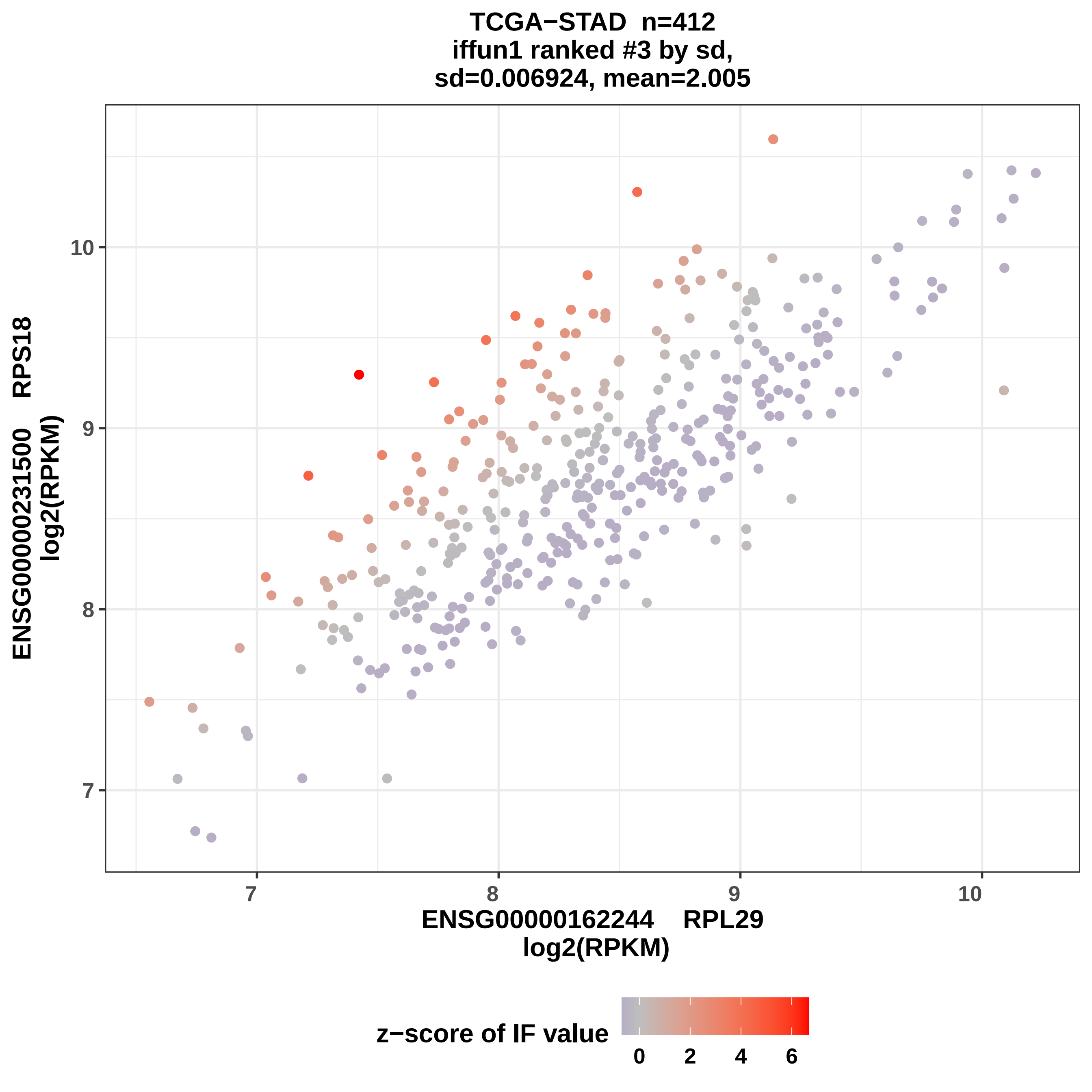
TCGA−STAD n=412
iffun1 ranked #3 by sd,
sd=0.006924, mean=2.005
ENSG00000231500 RPS18
log2(RPKM)
ENSG00000162244 RPL29
log2(RPKM)
7
8
9
10
z−score of IF value
0
2
4
6

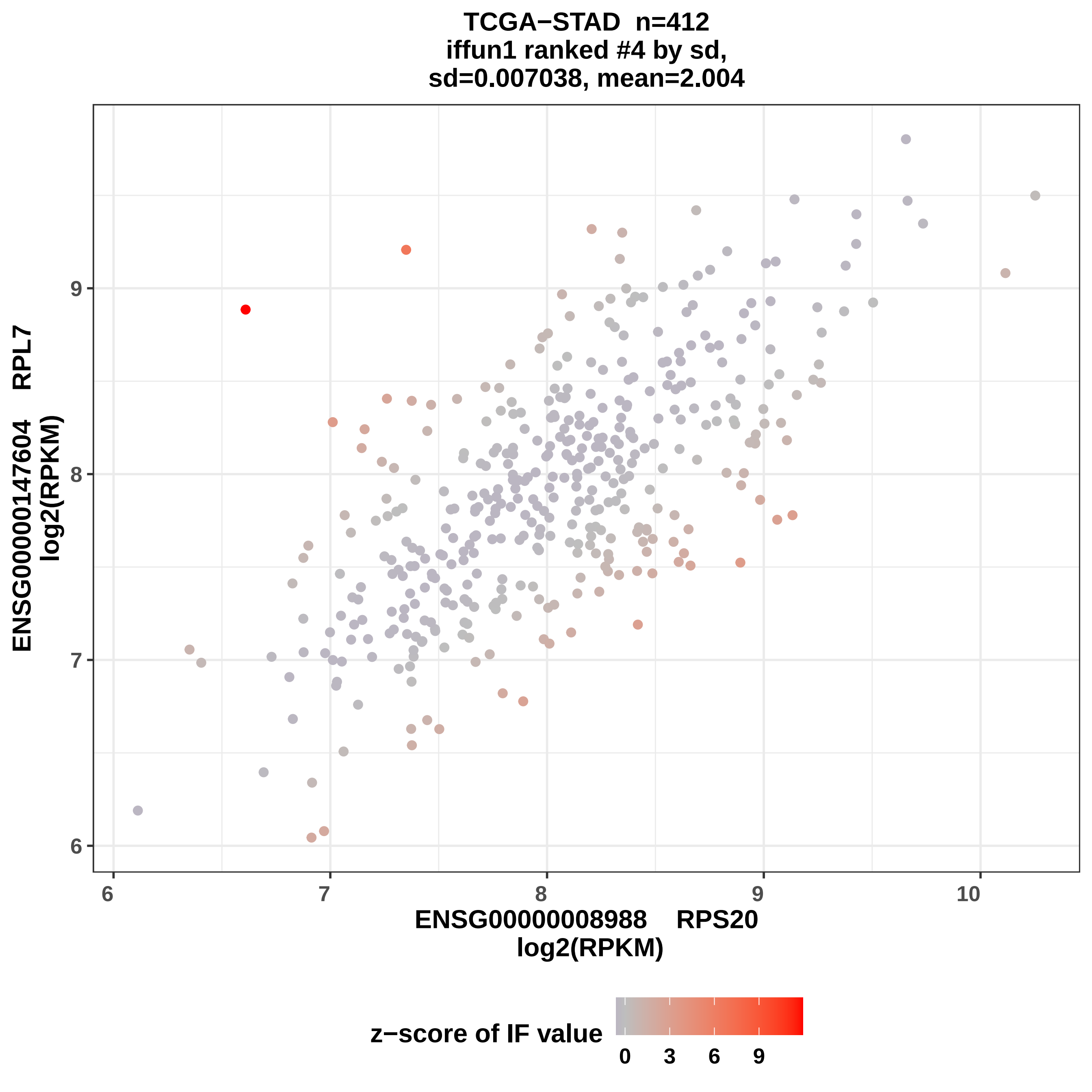

TCGA−STAD n=412
iffun1 ranked #4 by sd,
sd=0.007038, mean=2.004
ENSG00000147604 RPL7
log2(RPKM)
ENSG00000008988 RPS20
log2(RPKM)
6
7
8
9
10
z−score of IF value
0
3
6
9

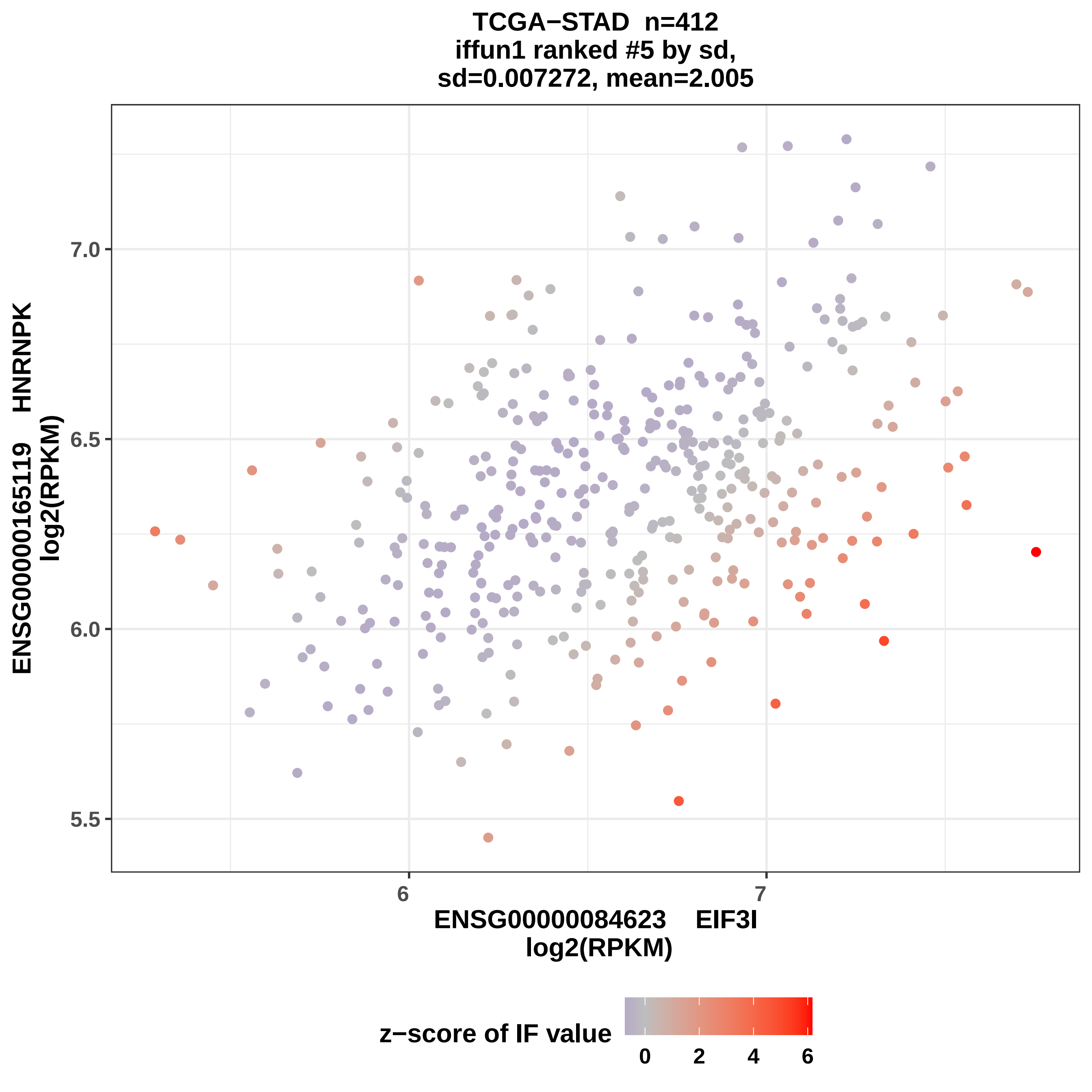
TCGA−STAD n=412
iffun1 ranked #5 by sd,
sd=0.007272, mean=2.005
ENSG00000165119 HNRNPK
log2(RPKM)
ENSG00000084623 EIF3I
log2(RPKM)
7.0
6.5
6.0
5.5
6
7
z−score of IF value
0
2
4
6

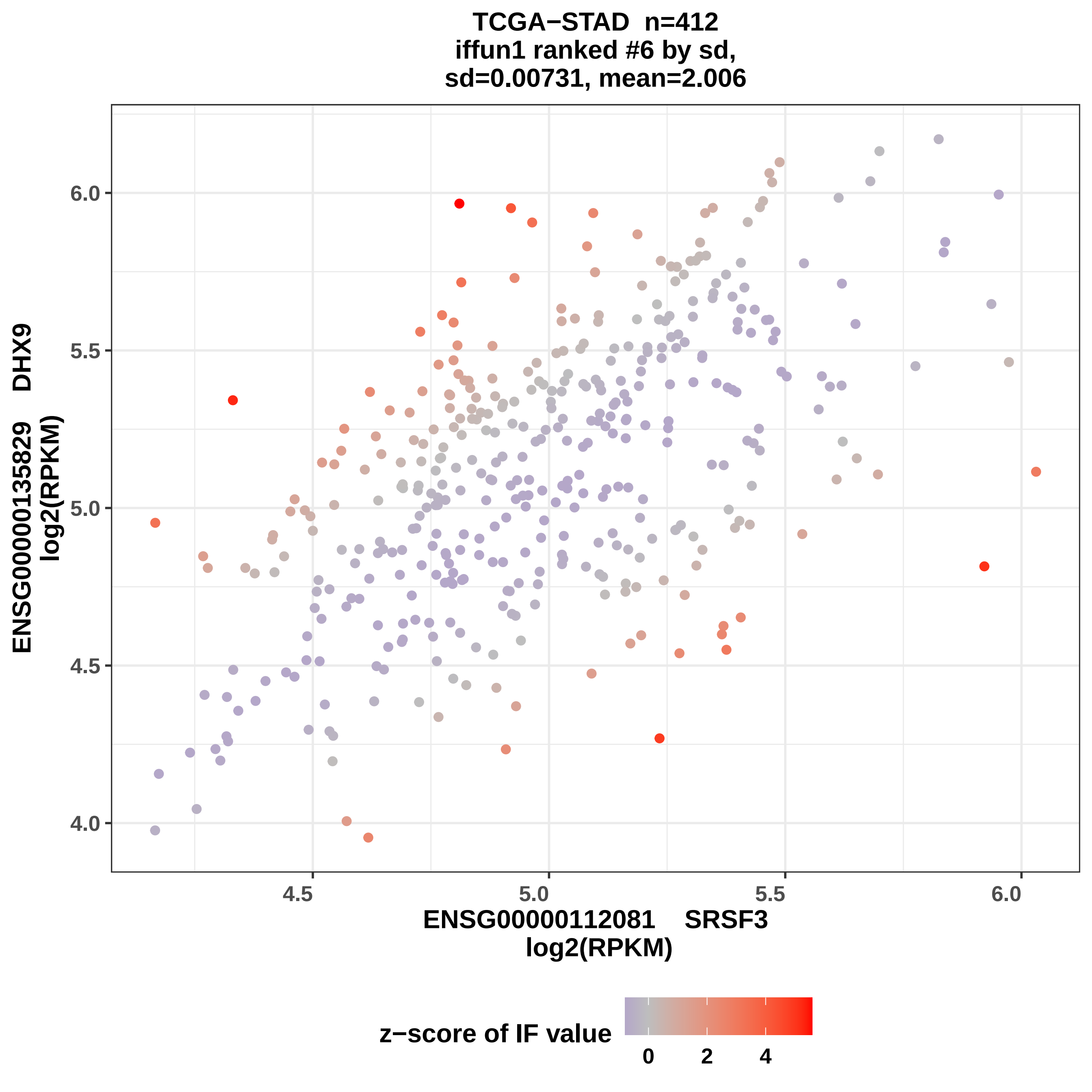
TCGA−STAD n=412
iffun1 ranked #6 by sd,
sd=0.00731, mean=2.006
ENSG00000135829 DHX9
log2(RPKM)
6.0
5.5
5.0
4.5
4.0
4.5
5.0
5.5
6.0
ENSG00000112081 SRSF3
log2(RPKM)
z−score of IF value
0
2
4

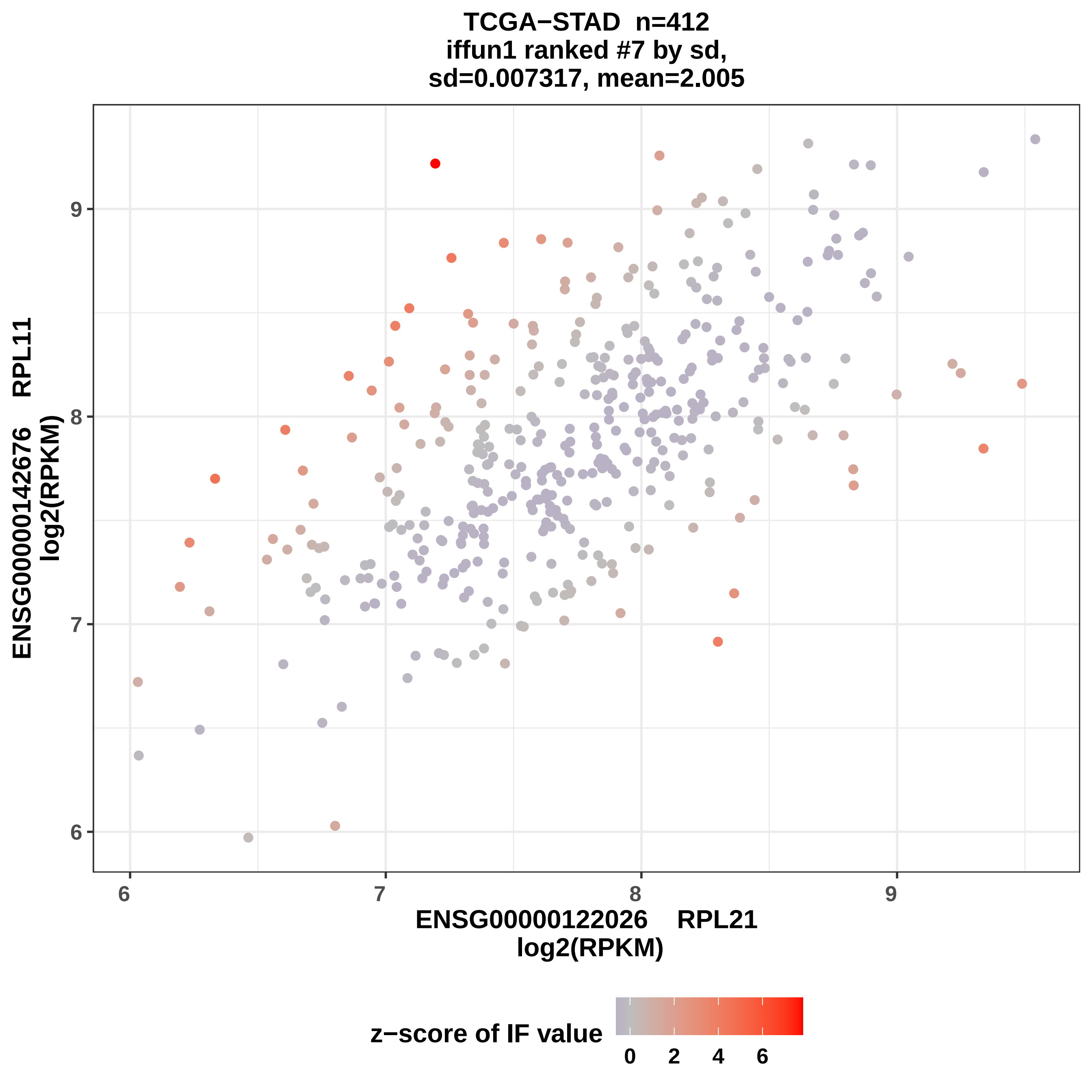
TCGA−STAD n=412
iffun1 ranked #7 by sd,
sd=0.007317, mean=2.005
ENSG00000142676 RPL11
log2(RPKM)
ENSG00000122026 RPL21
log2(RPKM)
6
7
8
9
6
7
8
9
z−score of IF value
0
2
4
6

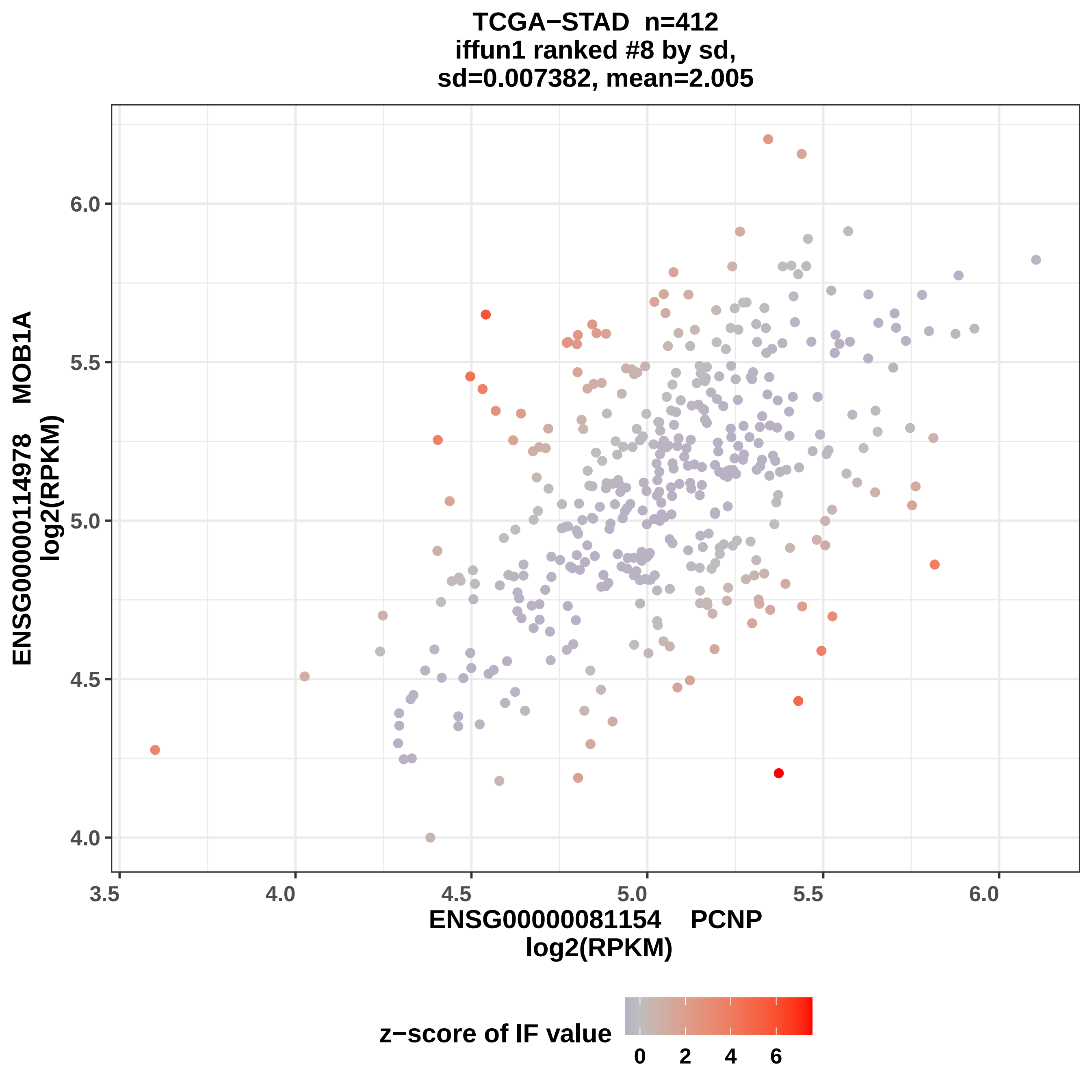
TCGA−STAD n=412
iffun1 ranked #8 by sd,
sd=0.007382, mean=2.005
ENSG00000114978 MOB1A
log2(RPKM)
ENSG00000081154 PCNP
log2(RPKM)
z−score of IF value
0 2 4 6
6.0
5.5
5.0
4.5
4.0
3.5
4.0
4.5
5.0
5.5
6.0

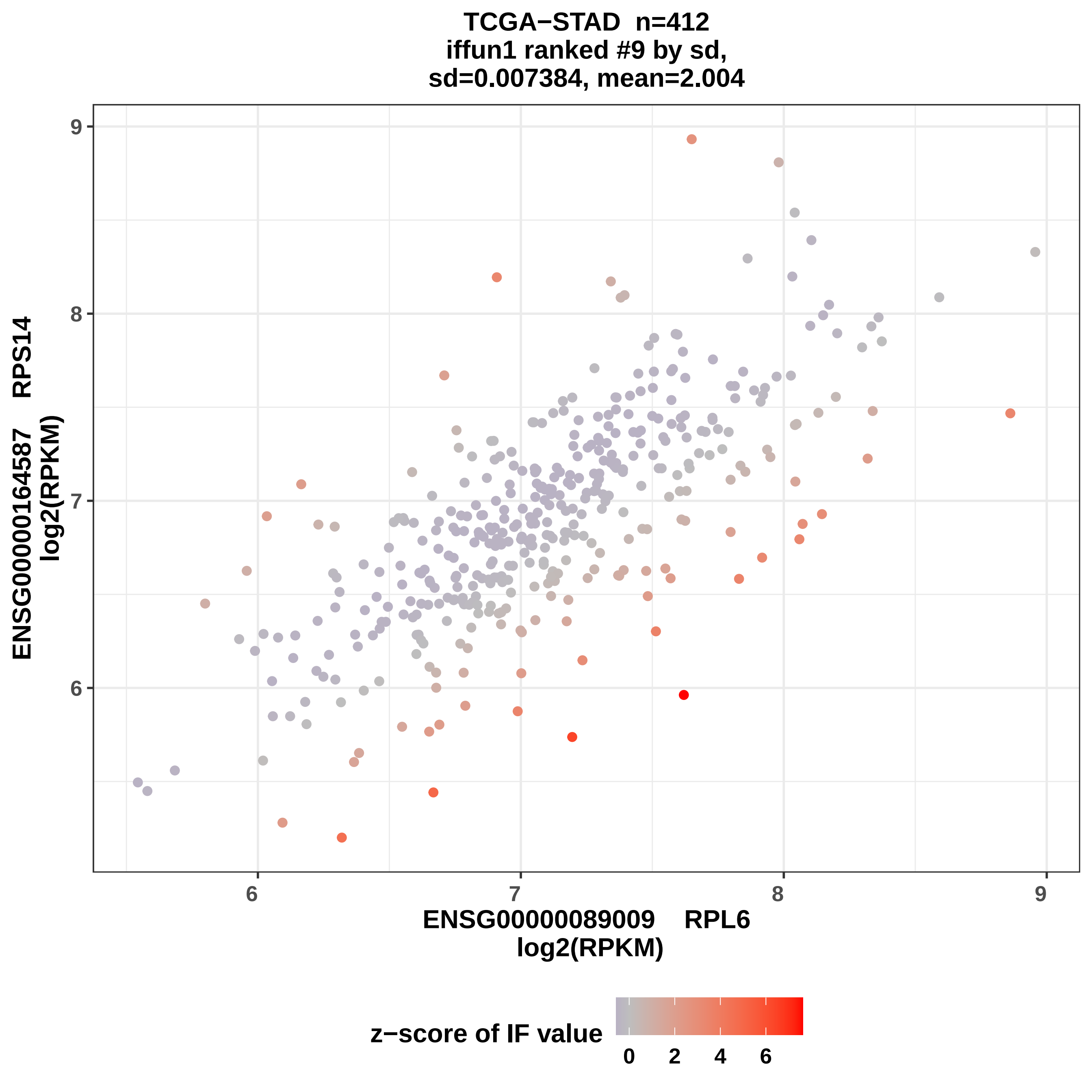

TCGA−STAD n=412
iffun1 ranked #9 by sd,
sd=0.007384, mean=2.004
9
8
7
6
ENSG00000164587 RPS14
log2(RPKM)
6
7
8
9
ENSG00000089009 RPL6
log2(RPKM)
z−score of IF value
0
2
4
6

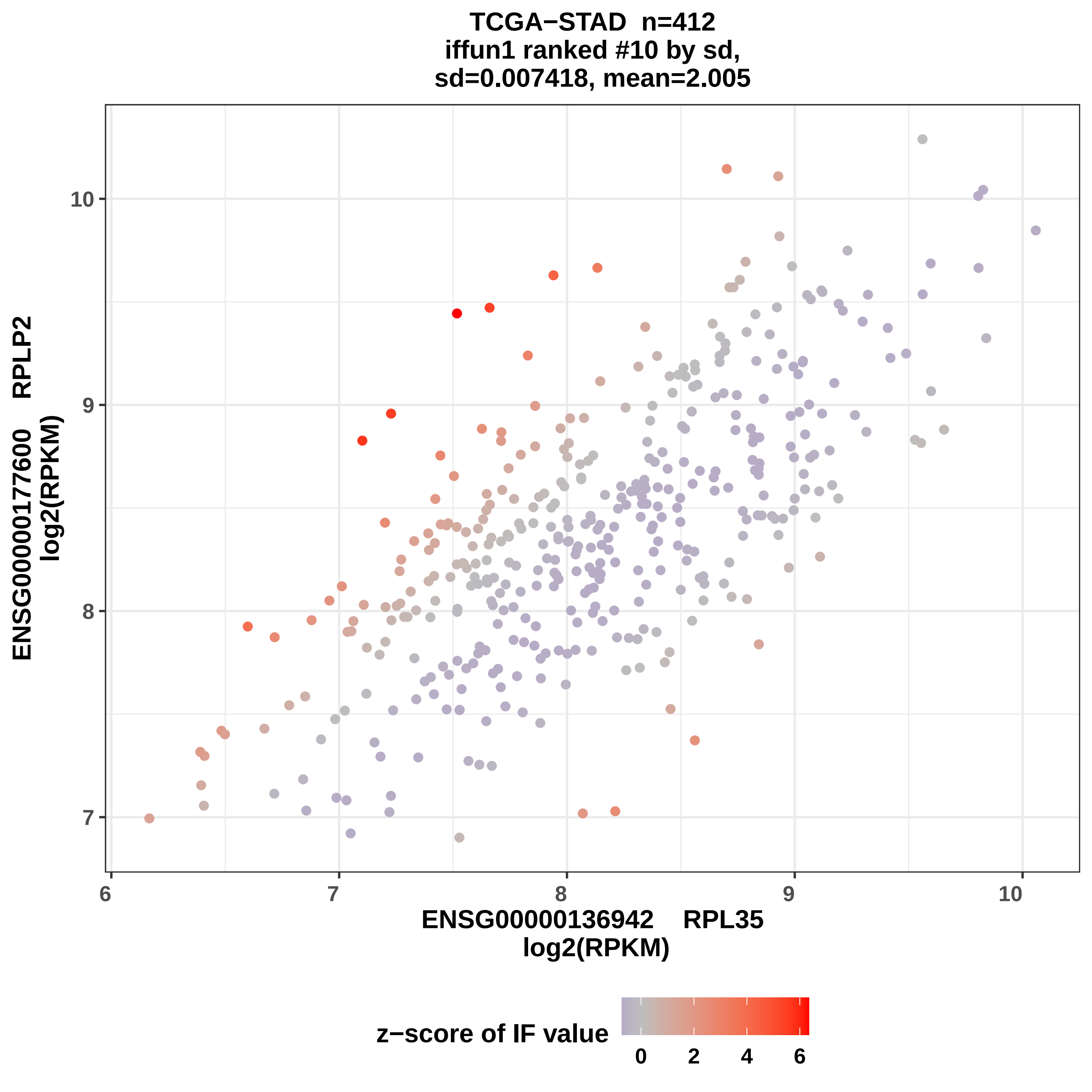

TCGA−STAD n=412
iffun1 ranked #10 by sd,
sd=0.007418, mean=2.005
ENSG00000177600 RPLP2
log2(RPKM)
ENSG00000136942 RPL35
log2(RPKM)
z−score of IF value
10
9
8
7
6
7
8
9
10
0
2
4
6

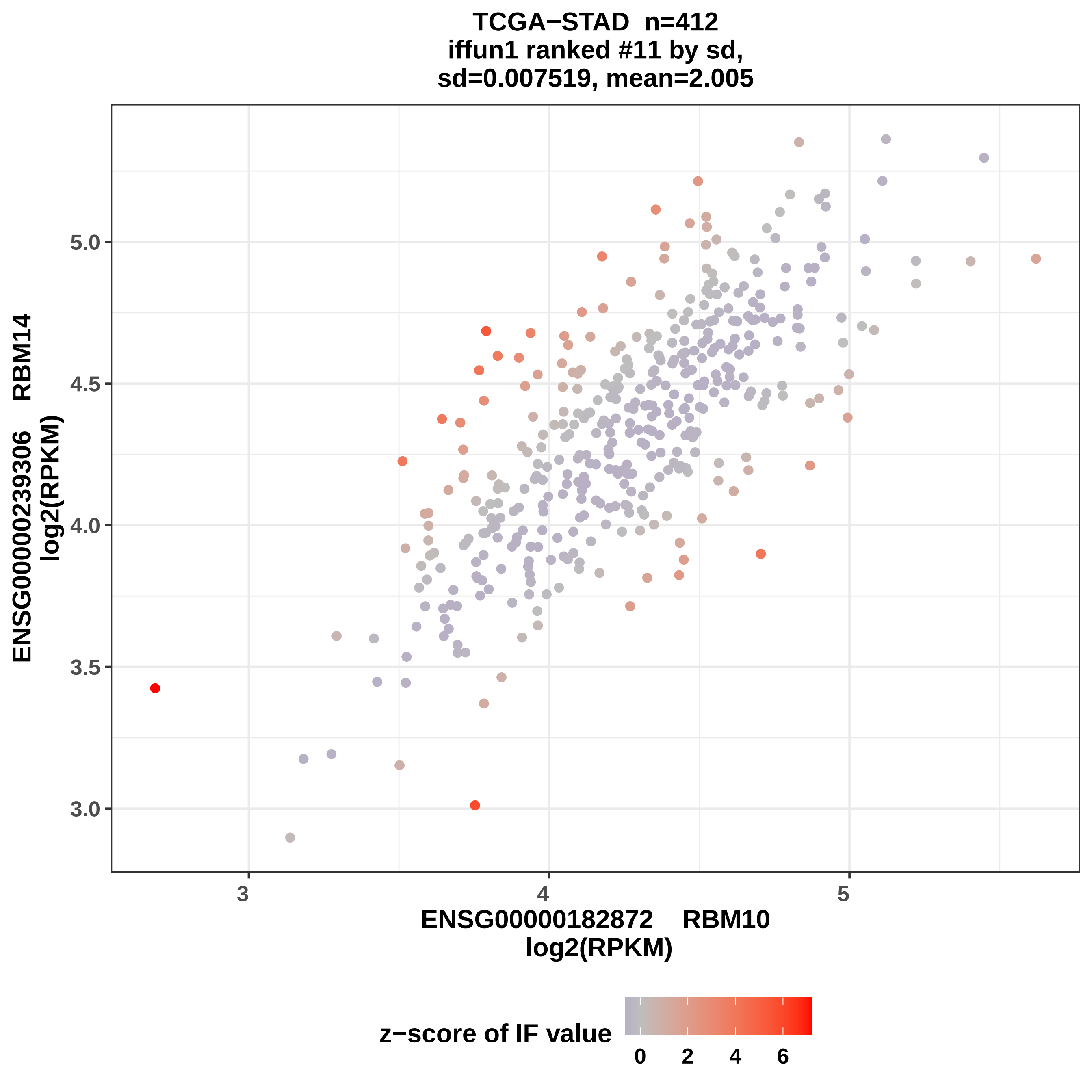
TCGA−STAD n=412
iffun1 ranked #11 by sd,
sd=0.007519, mean=2.005
ENSG00000239306 RBM14
log2(RPKM)
ENSG00000182872 RBM10
log2(RPKM)
z−score of IF value
3.0
3.5
4.0
4.5
5.0
3
4
5
0
2
4
6

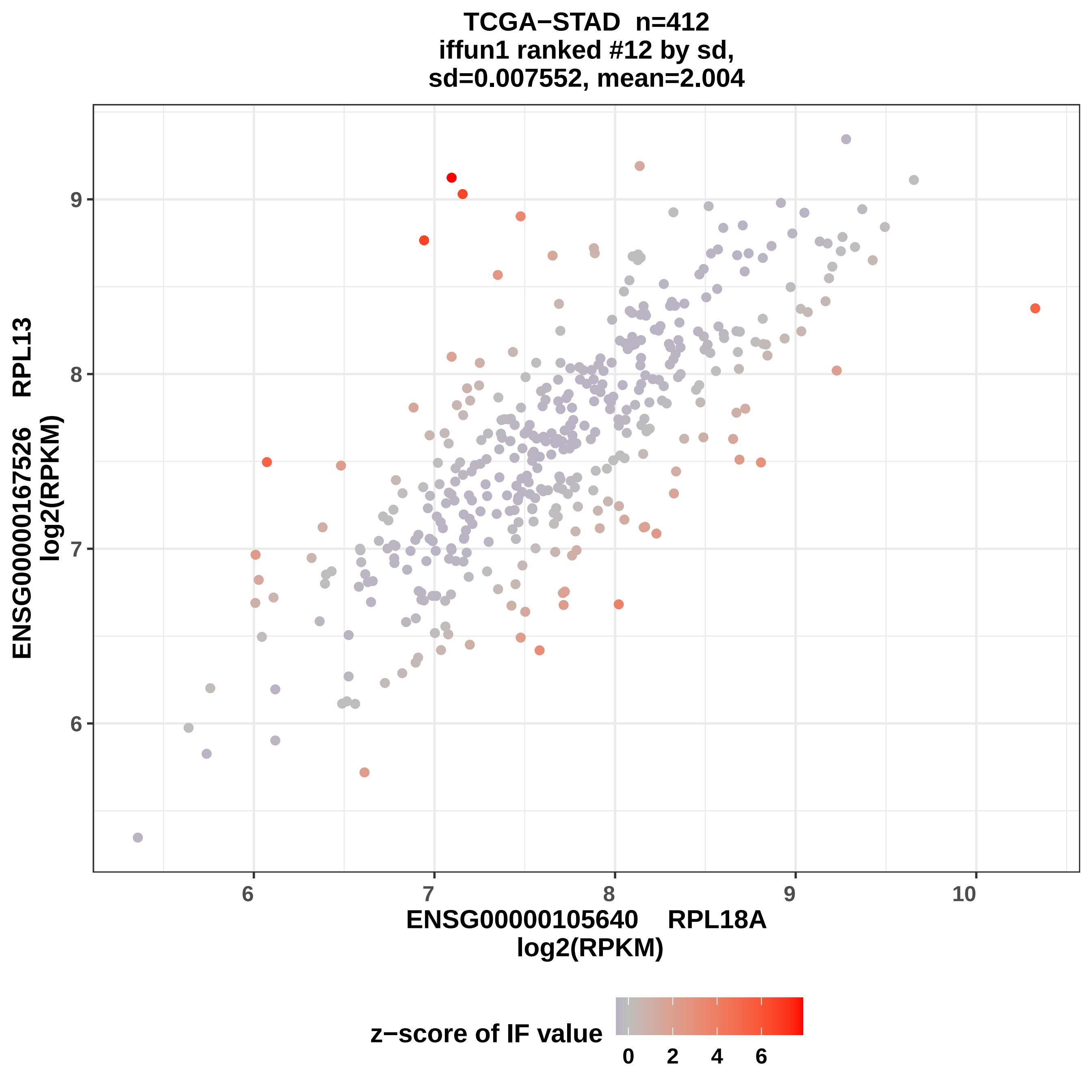
TCGA−STAD n=412
iffun1 ranked #12 by sd,
sd=0.007552, mean=2.004
ENSG00000167526 RPL13
log2(RPKM)
ENSG00000105640 RPL18A
log2(RPKM)
z−score of IF value
0
2
4
6
6
7
8
9
10

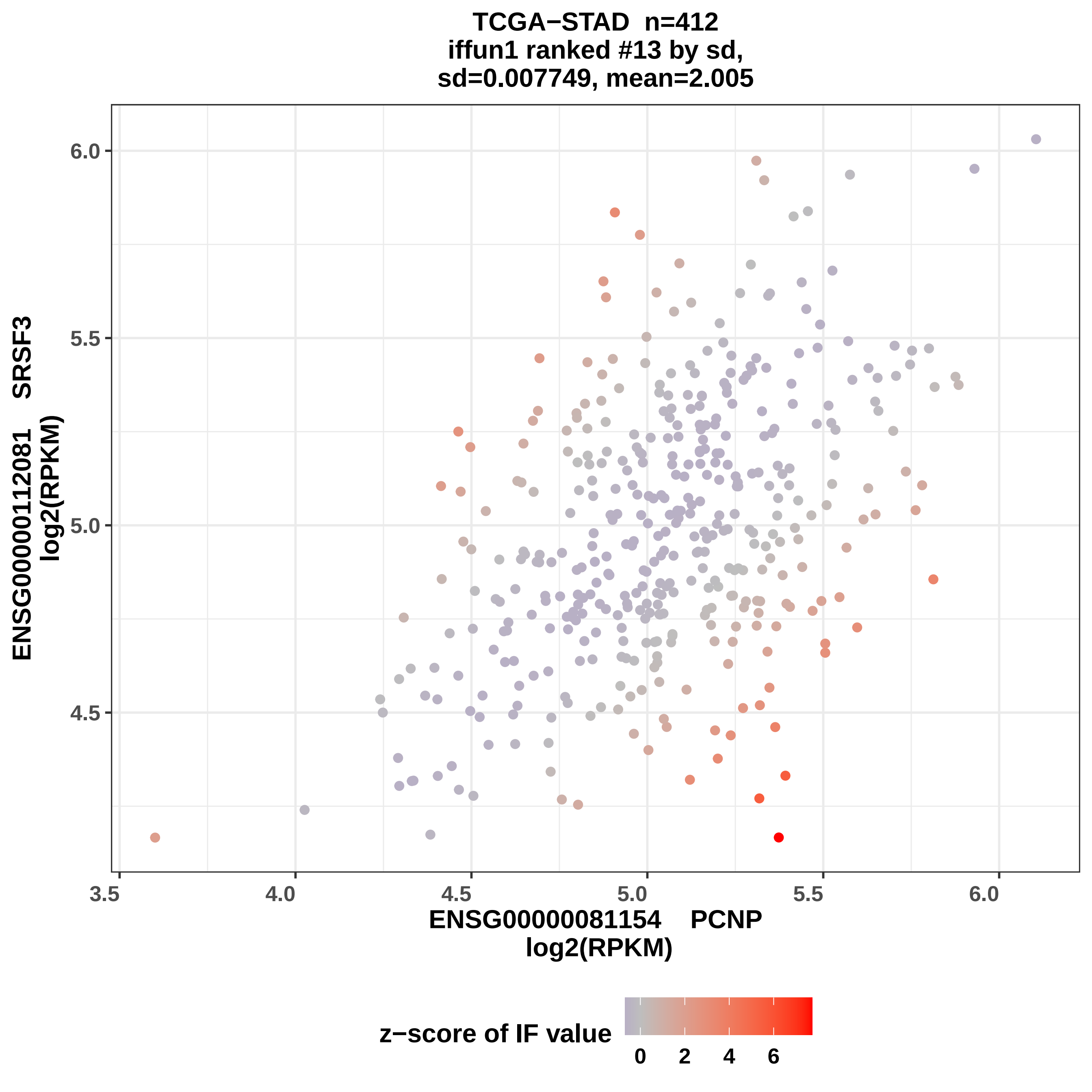
TCGA−STAD n=412
iffun1 ranked #13 by sd,
sd=0.007749, mean=2.005
ENSG00000112081 SRSF3
log2(RPKM)
6.0
5.5
5.0
4.5
3.5
4.0
4.5
5.0
5.5
6.0
ENSG00000081154 PCNP
log2(RPKM)
z−score of IF value
0
2
4
6

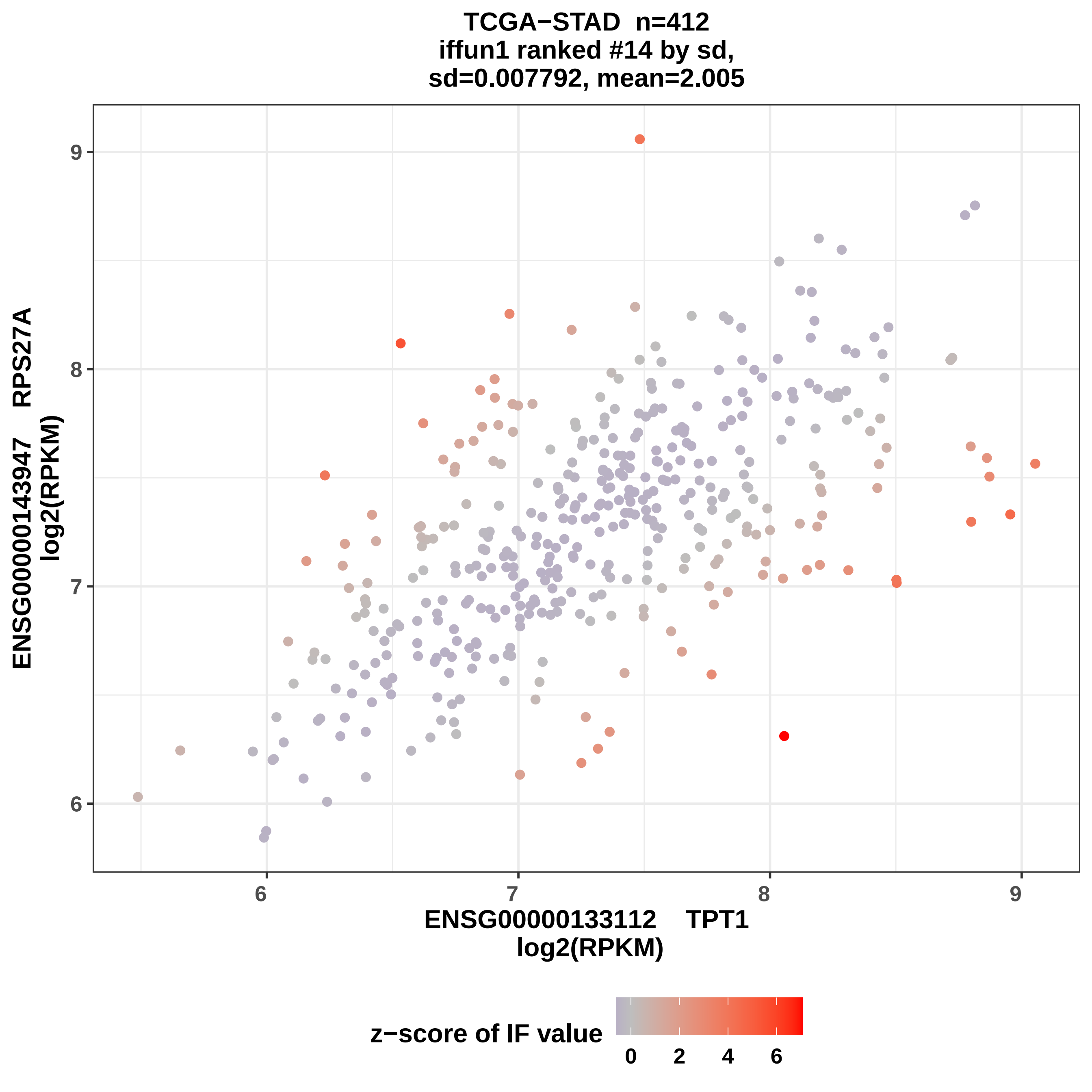
TCGA−STAD n=412
iffun1 ranked #14 by sd,
sd=0.007792, mean=2.005
ENSG00000143947 RPS27A
log2(RPKM)
ENSG00000133112 TPT1
log2(RPKM)
6
7
8
9
z−score of IF value
0
2
4
6

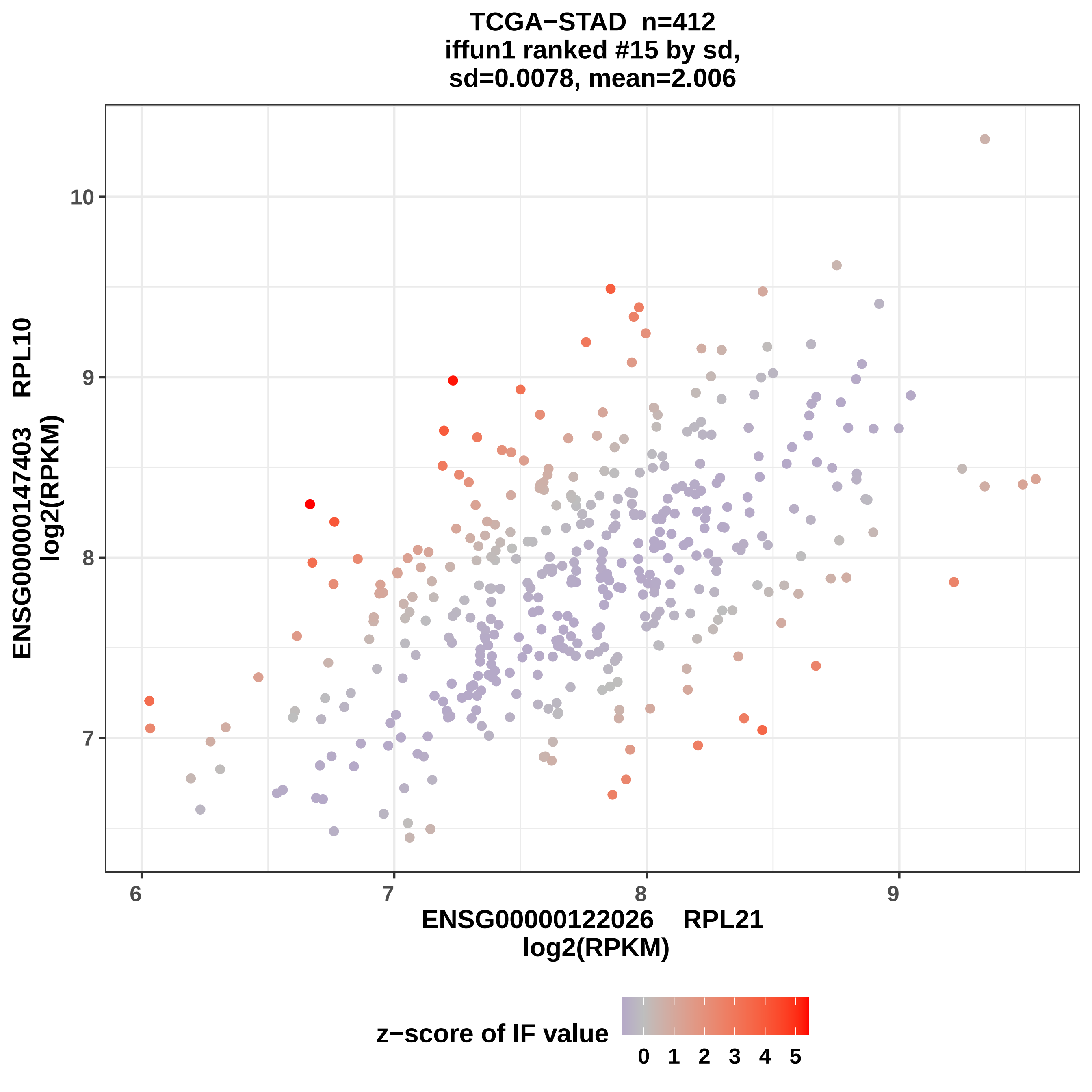
TCGA−STAD n=412
iffun1 ranked #15 by sd,
sd=0.0078, mean=2.006
ENSG00000147403 RPL10
log2(RPKM)
10
9
8
7
6
7
8
9
ENSG00000122026 RPL21
log2(RPKM)
z−score of IF value
0 1 2 3 4 5

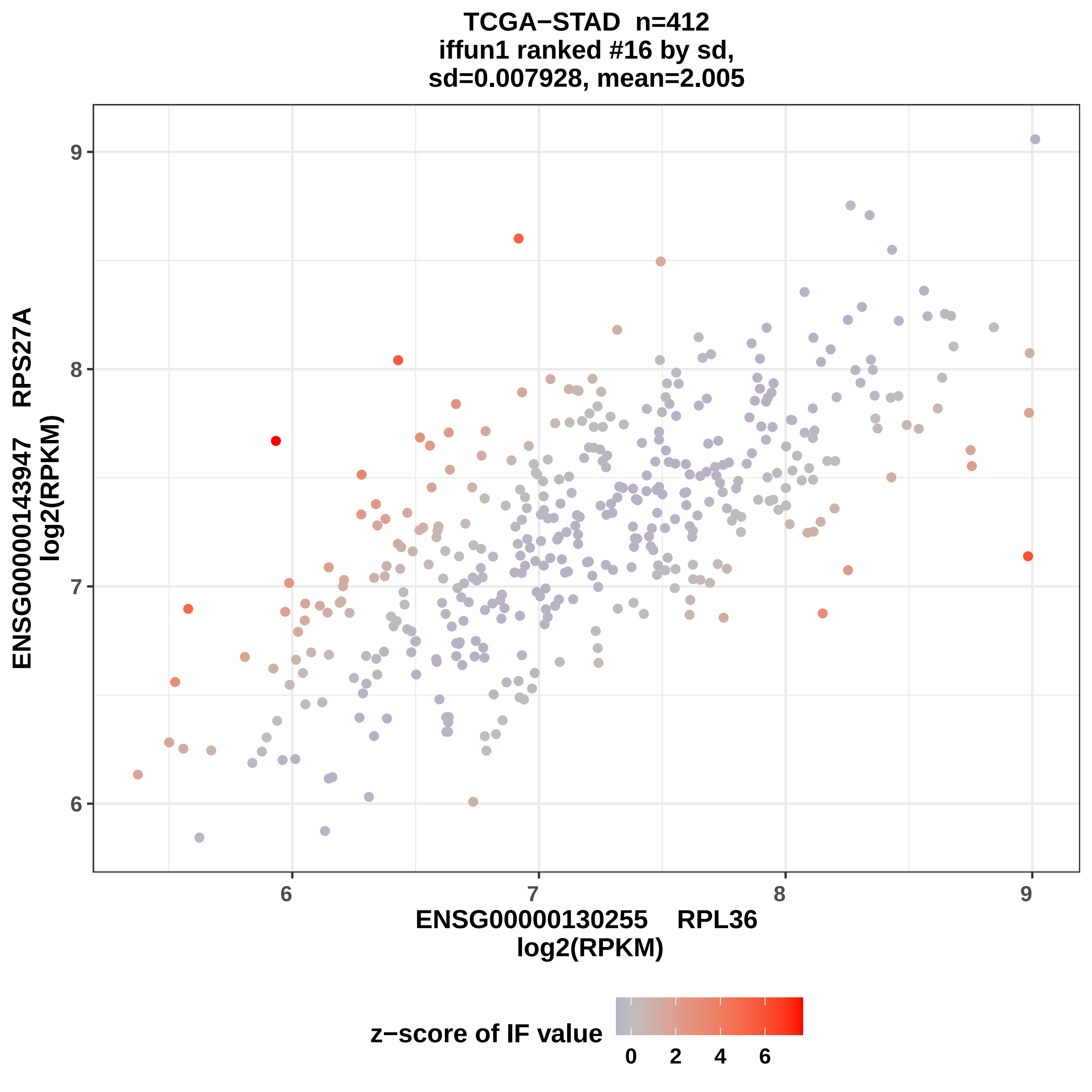

TCGA−STAD n=412
iffun1 ranked #16 by sd,
sd=0.007928, mean=2.005
ENSG00000143947 RPS27A
log2(RPKM)
ENSG00000130255 RPL36
log2(RPKM)
z−score of IF value
0
2
4
6

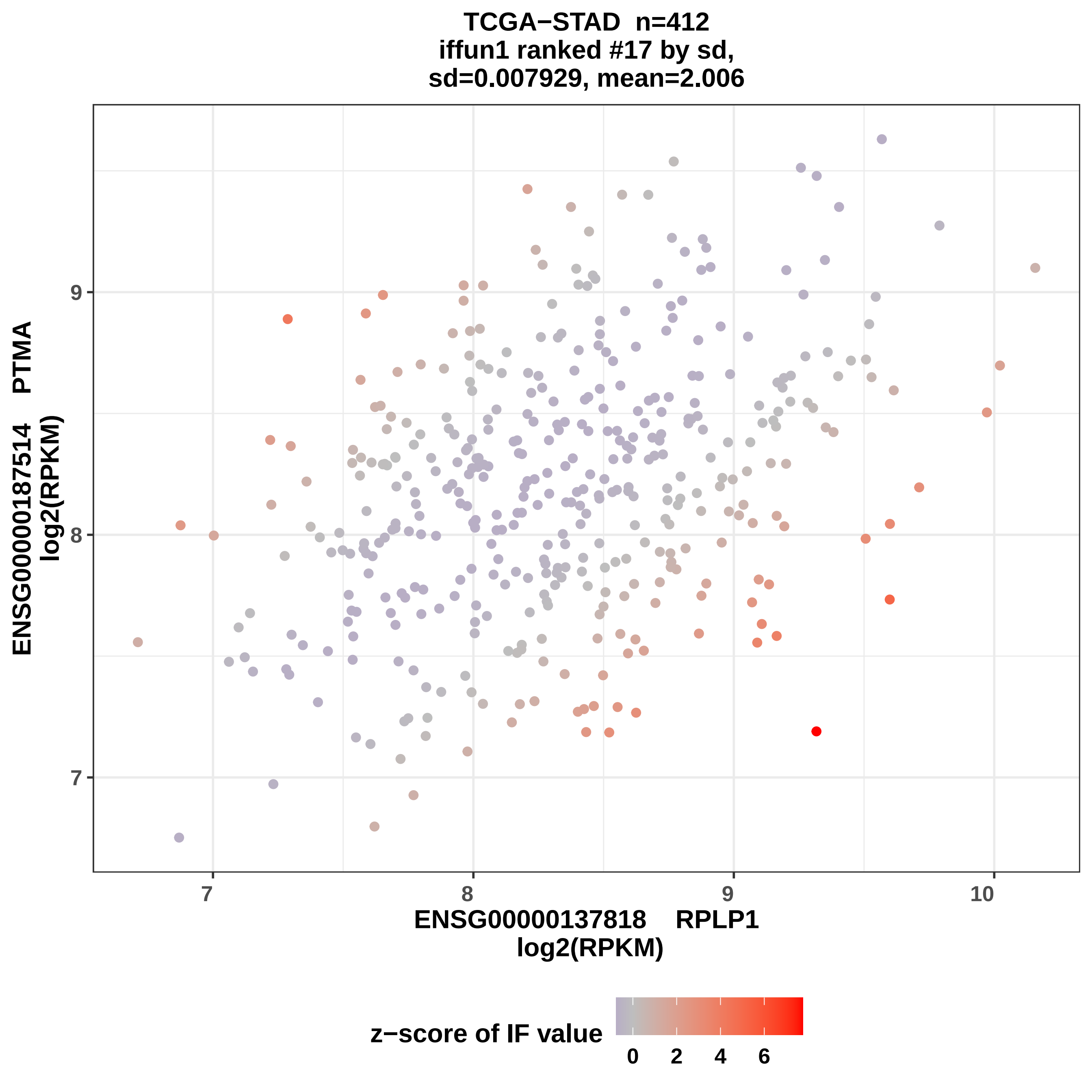
TCGA−STAD n=412
iffun1 ranked #17 by sd,
sd=0.007929, mean=2.006
ENSG00000187514 PTMA
log2(RPKM)
9
8
7
7
8
9
10
ENSG00000137818 RPLP1
log2(RPKM)
z−score of IF value
0
2
4
6

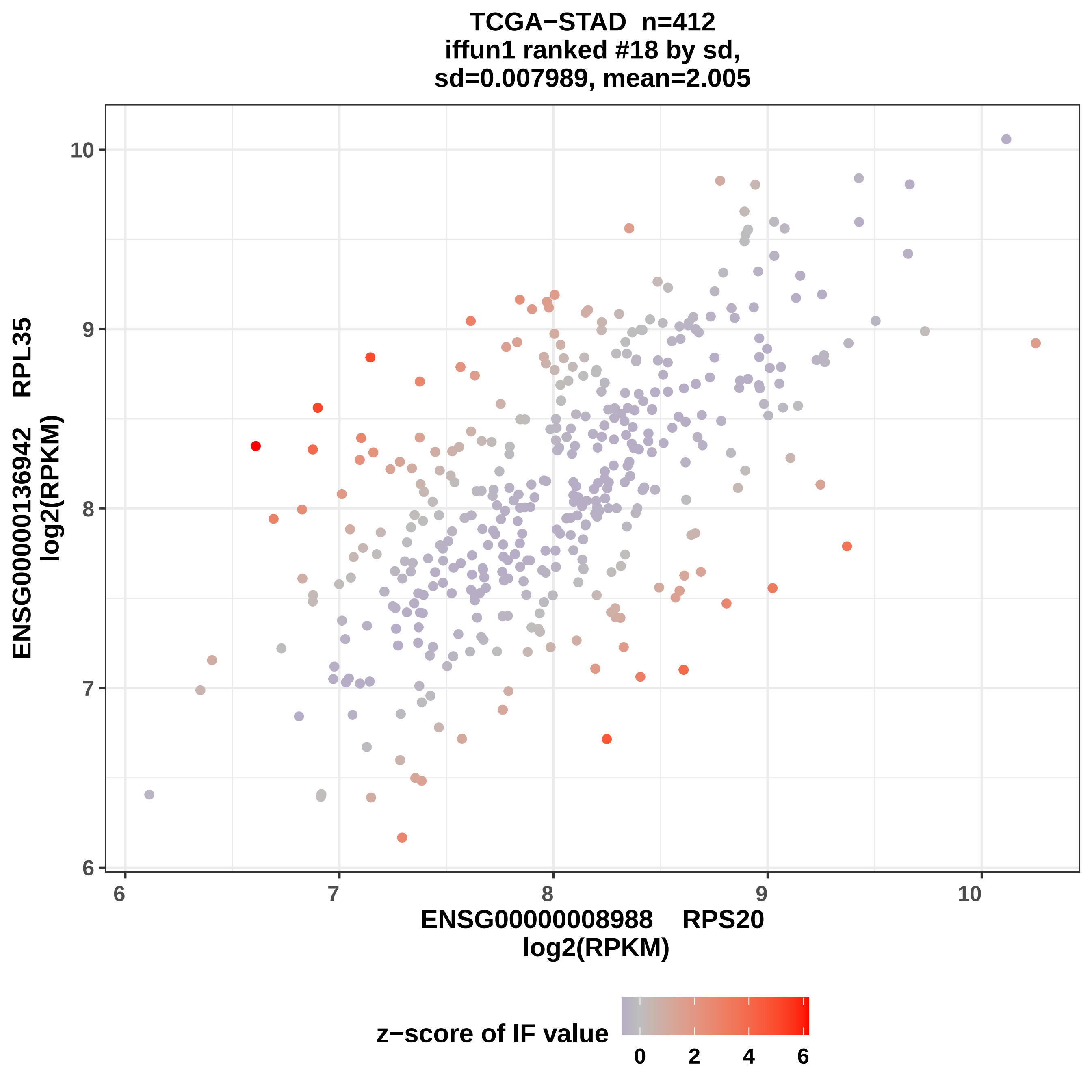

TCGA−STAD n=412
iffun1 ranked #18 by sd,
sd=0.007989, mean=2.005
ENSG00000136942 RPL35
log2(RPKM)
10
9
8
7
6
ENSG00000008988 RPS20
log2(RPKM)
z−score of IF value
0
2
4
6

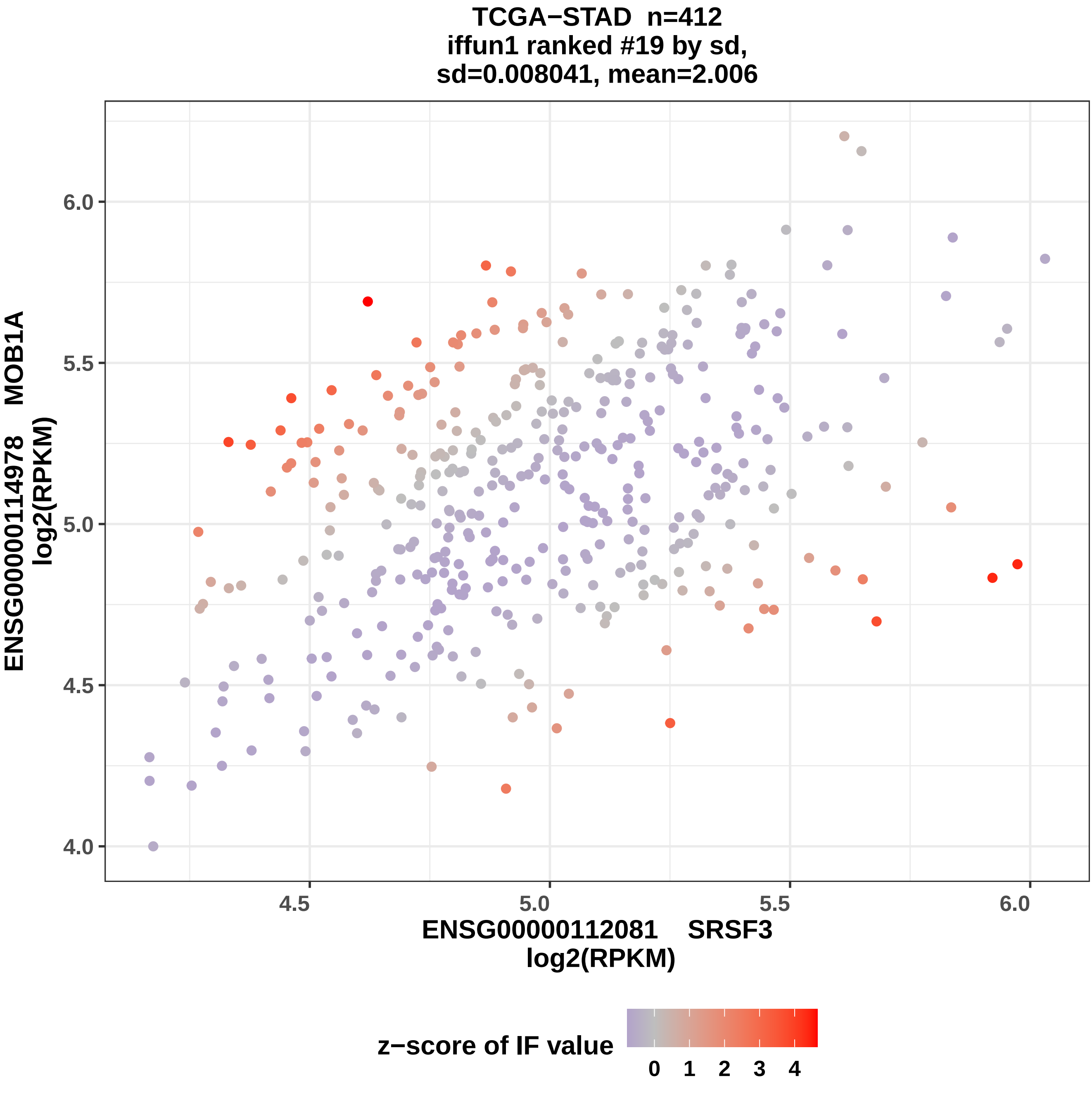

TCGA−STAD n=412
iffun1 ranked #19 by sd,
sd=0.008041, mean=2.006
ENSG00000114978 MOB1A
log2(RPKM)
6.0
5.5
5.0
4.5
4.0
4.5
5.0
5.5
6.0
ENSG00000112081 SRSF3
log2(RPKM)
z−score of IF value
0
1
2
3
4

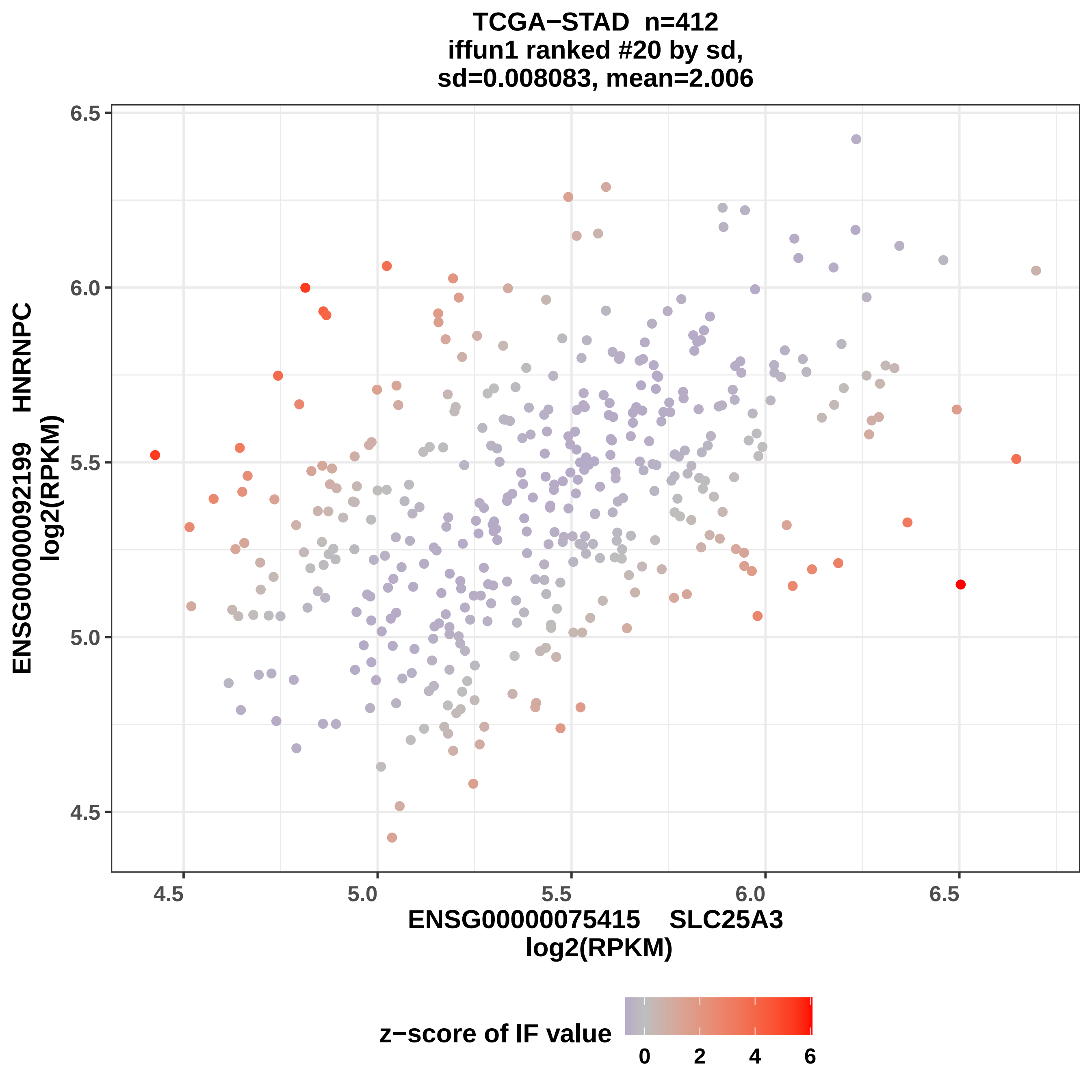
TCGA−STAD n=412
iffun1 ranked #20 by sd,
sd=0.008083, mean=2.006
ENSG00000092199 HNRNPC
log2(RPKM)
6.5
6.0
5.5
5.0
4.5
4.5
5.0
5.5
6.0
6.5
ENSG00000075415 SLC25A3
log2(RPKM)
z−score of IF value
0
2
4
6

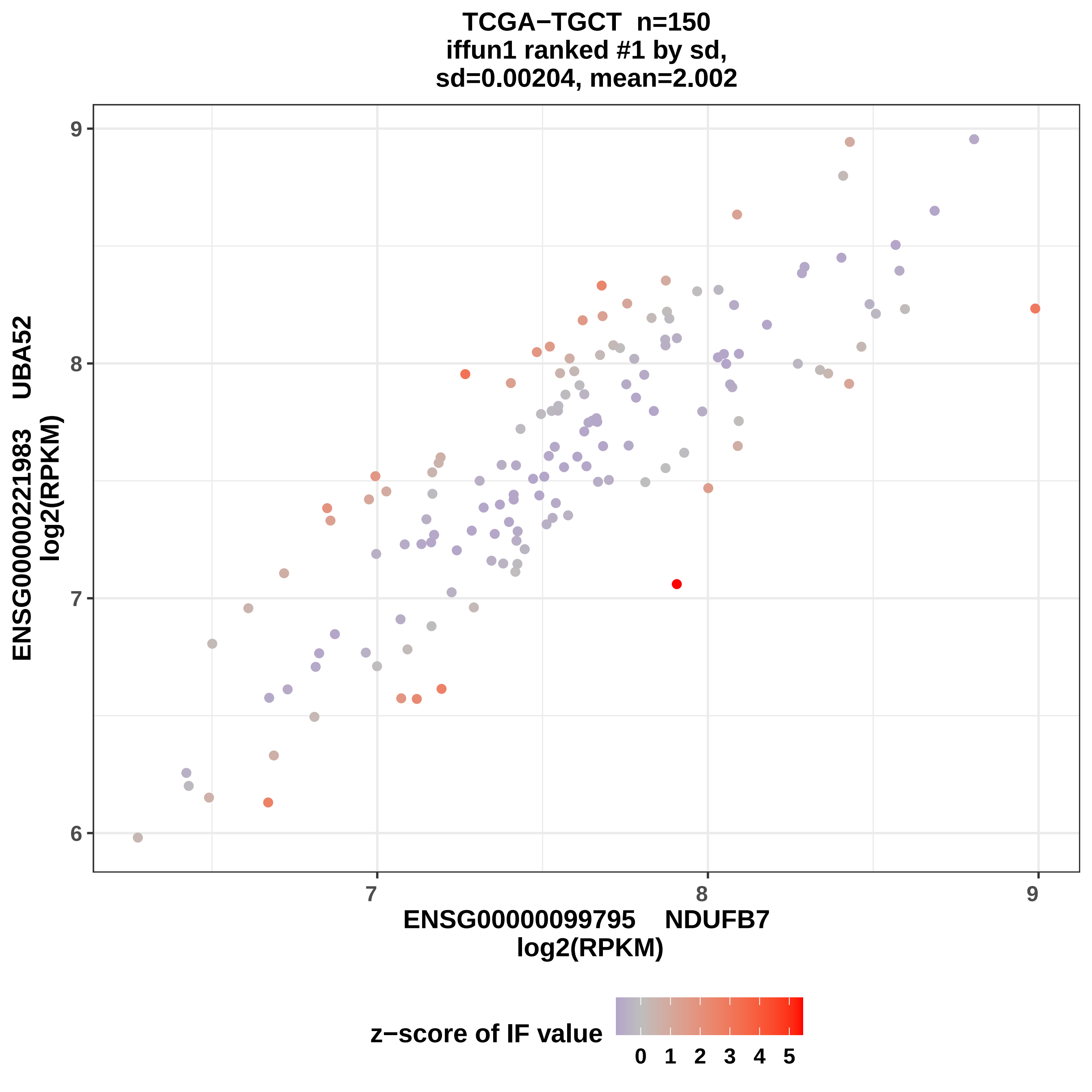
TCGA−TGCT n=150
iffun1 ranked #1 by sd,
sd=0.00204, mean=2.002
ENSG00000221983 UBA52
log2(RPKM)
ENSG00000099795 NDUFB7
log2(RPKM)
z−score of IF value
0 1 2 3 4 5
6
7
8
9
7
8
9

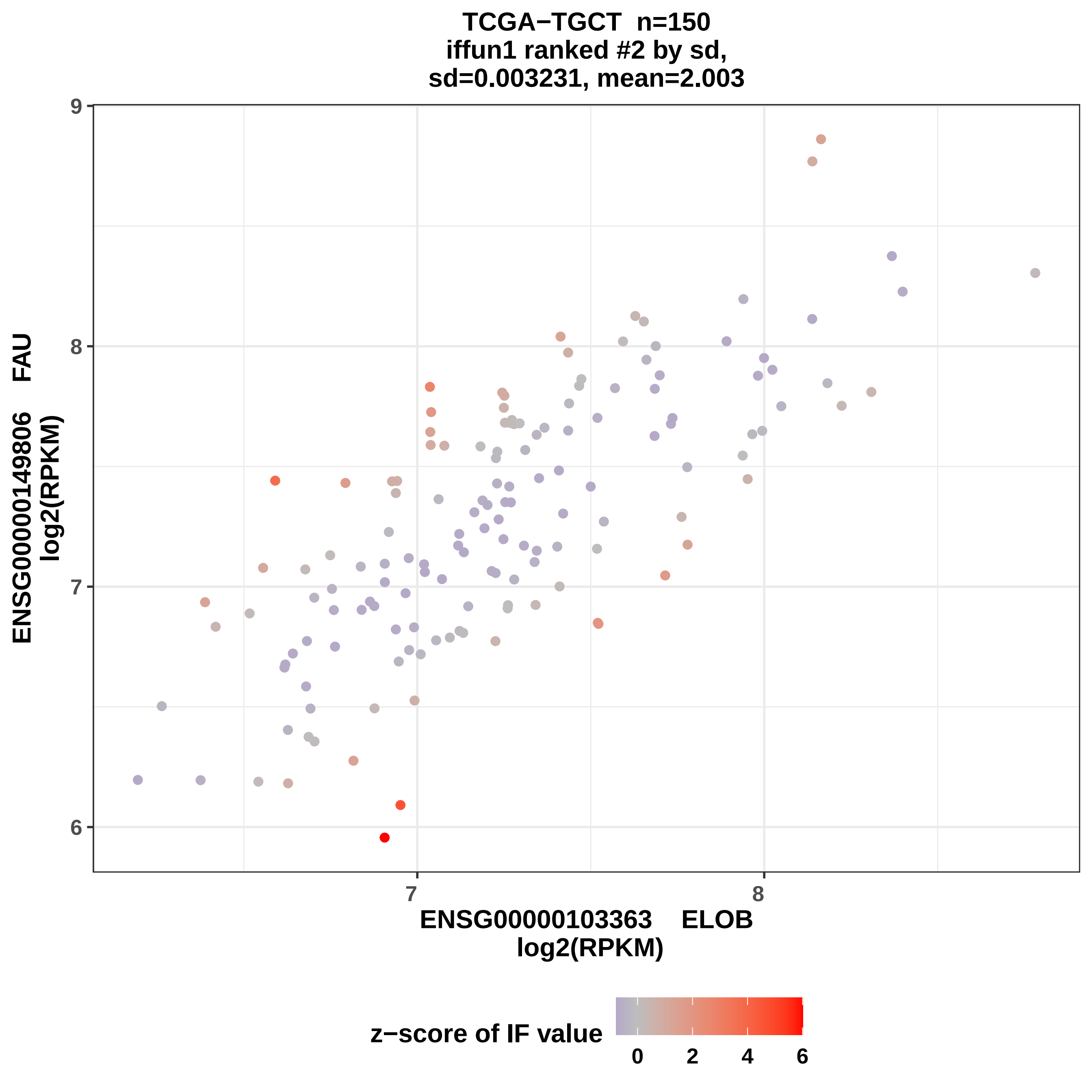
TCGA−TGCT n=150
iffun1 ranked #2 by sd,
sd=0.003231, mean=2.003
ENSG00000149806 FAU
log2(RPKM)
ENSG00000103363 ELOB
log2(RPKM)
z−score of IF value
9
8
7
6
7
8
0
2
4
6

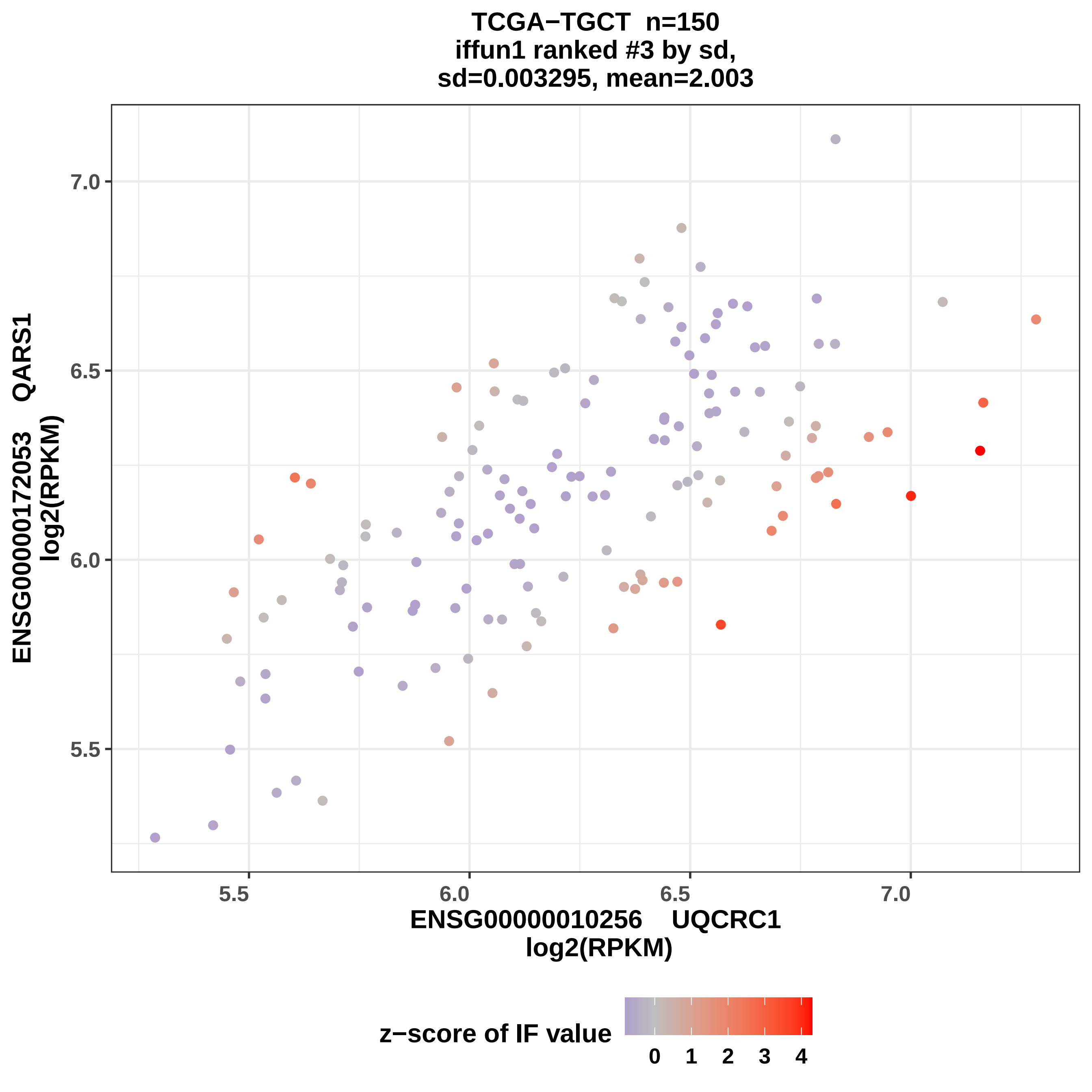
TCGA−TGCT n=150
iffun1 ranked #3 by sd,
sd=0.003295, mean=2.003
ENSG00000172053 QARS1
log2(RPKM)
7.0
6.5
6.0
5.5
5.5
6.0
6.5
7.0
ENSG00000010256 UQCRC1
log2(RPKM)
z−score of IF value
0
1
2
3
4

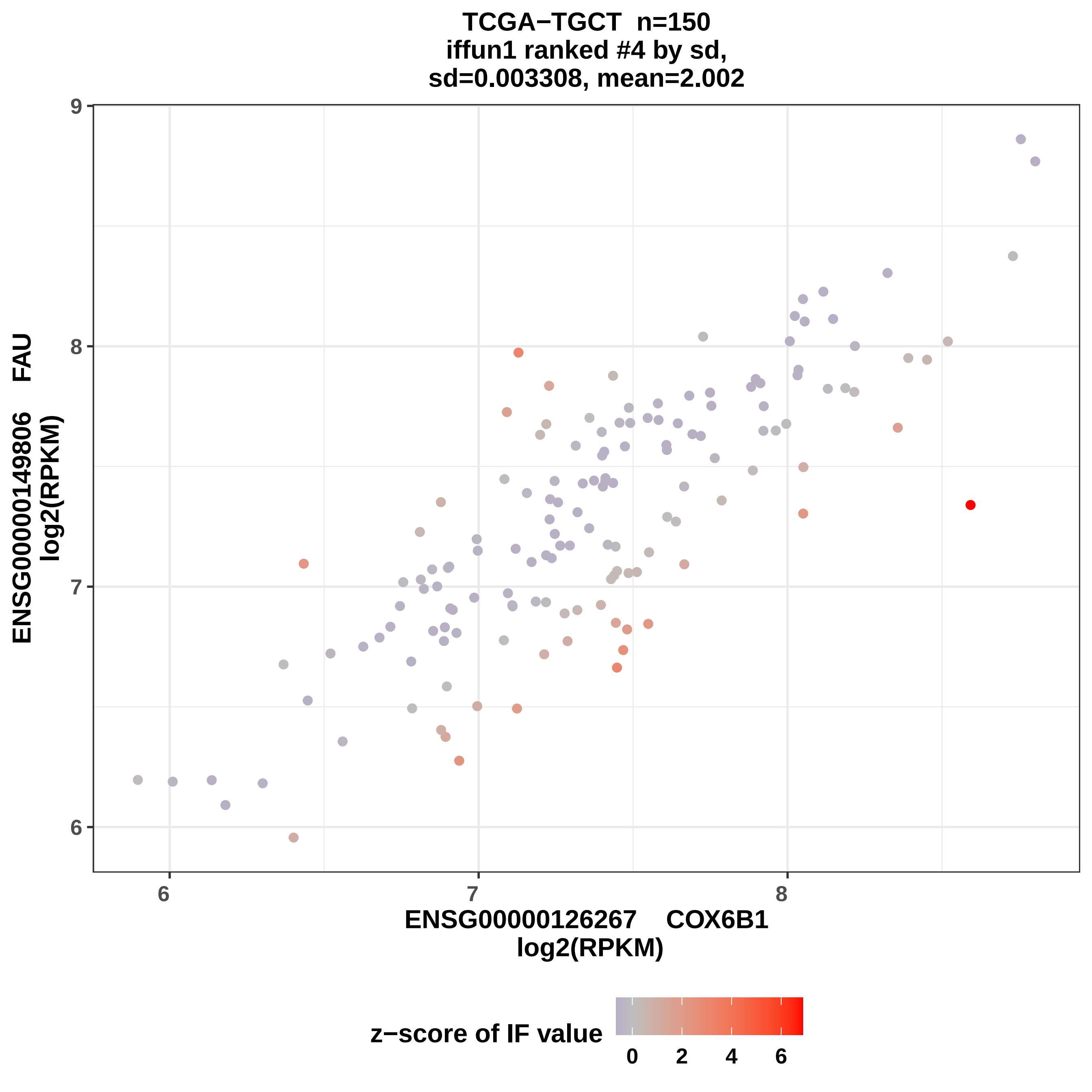
TCGA−TGCT n=150
iffun1 ranked #4 by sd,
sd=0.003308, mean=2.002
ENSG00000149806 FAU
log2(RPKM)
ENSG00000126267 COX6B1
log2(RPKM)
z−score of IF value
6
7
8
9
6
7
8
0
2
4
6

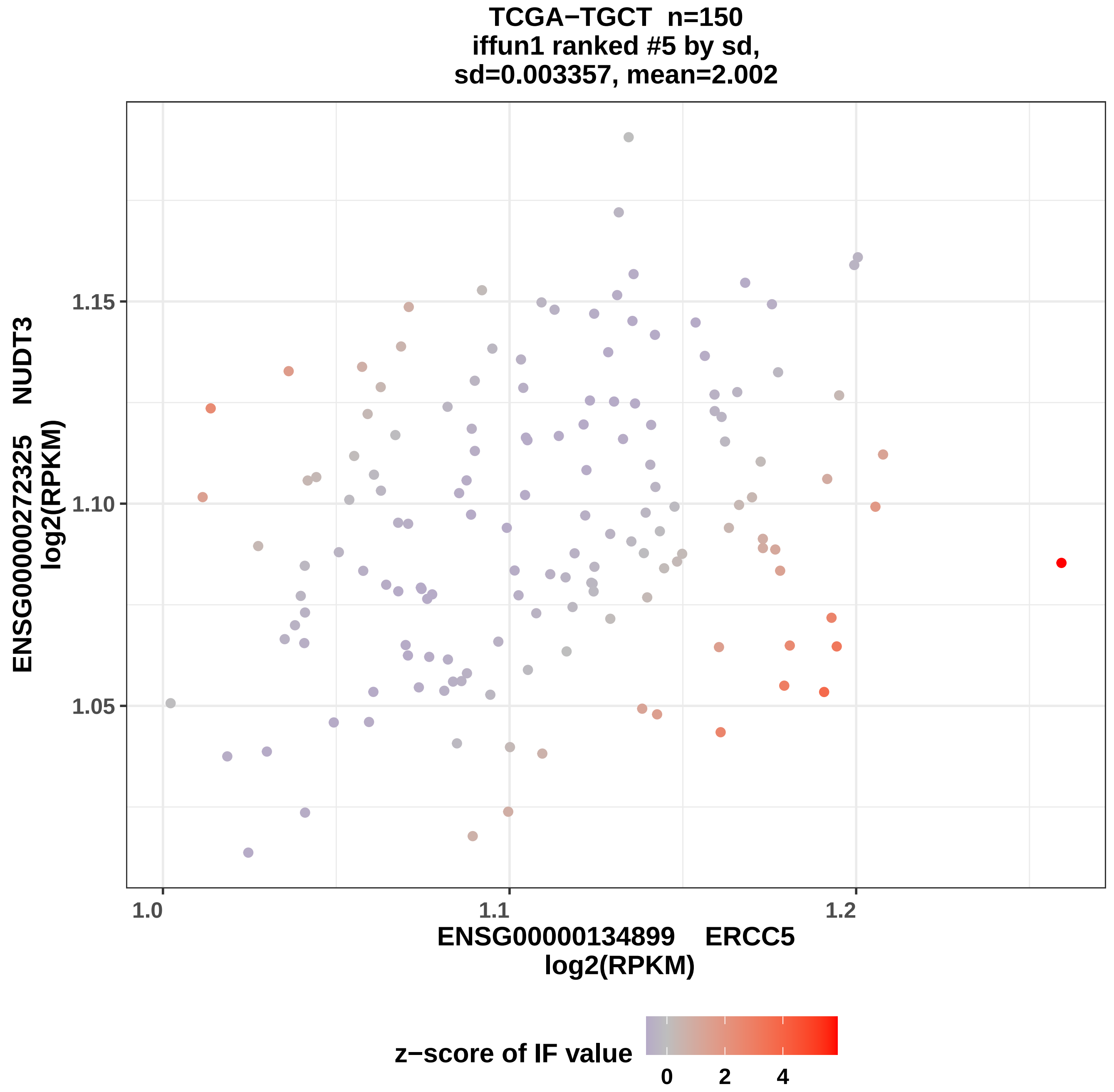

TCGA−TGCT n=150
iffun1 ranked #5 by sd,
sd=0.003357, mean=2.002
ENSG00000272325 NUDT3
log2(RPKM)
1.15
1.10
1.05
1.0
1.1
1.2
ENSG00000134899 ERCC5
log2(RPKM)
z−score of IF value
0
2
4

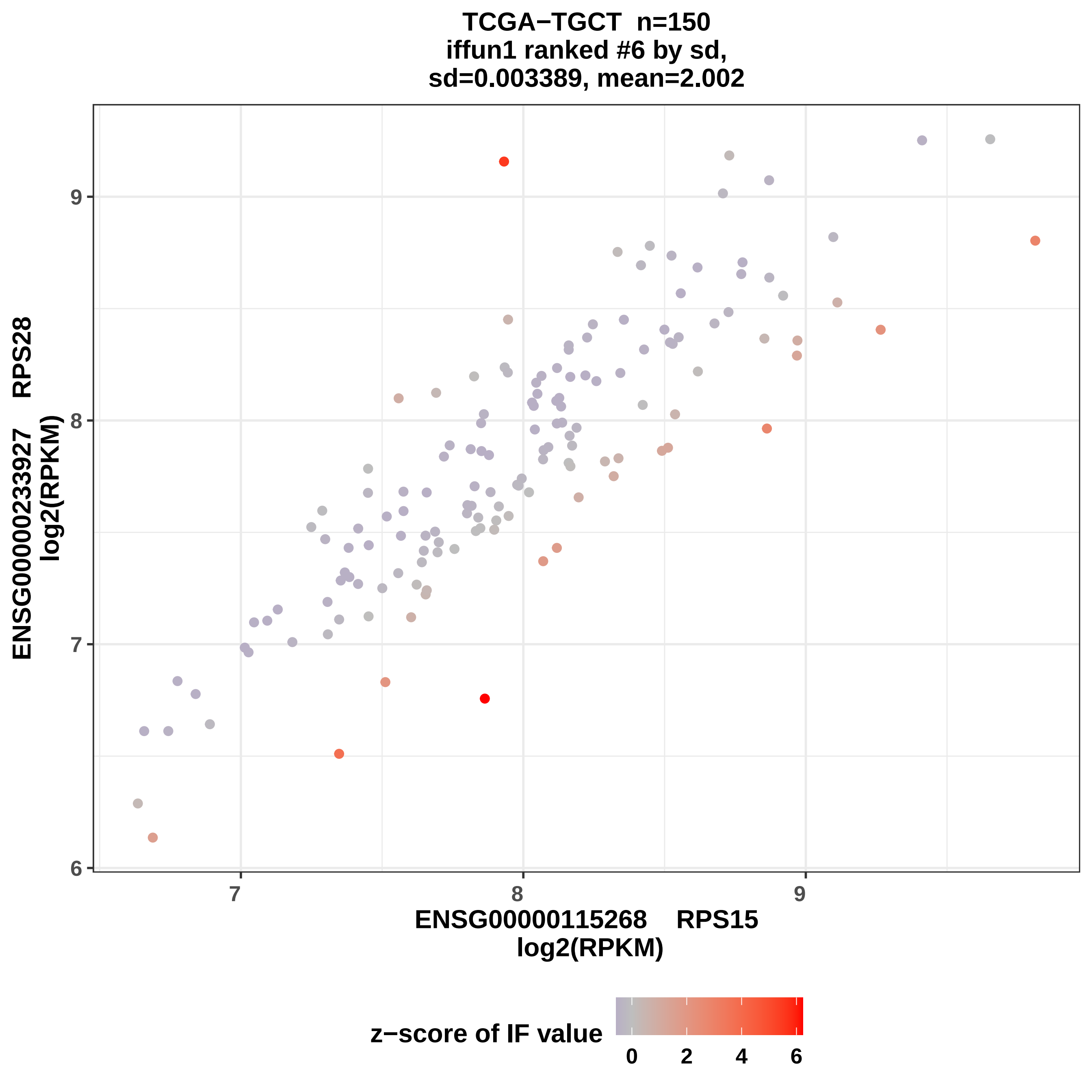
TCGA−TGCT n=150
iffun1 ranked #6 by sd,
sd=0.003389, mean=2.002
ENSG00000233927 RPS28
log2(RPKM)
6
7
8
9
ENSG00000115268 RPS15
log2(RPKM)
7
8
9
z−score of IF value
0
2
4
6

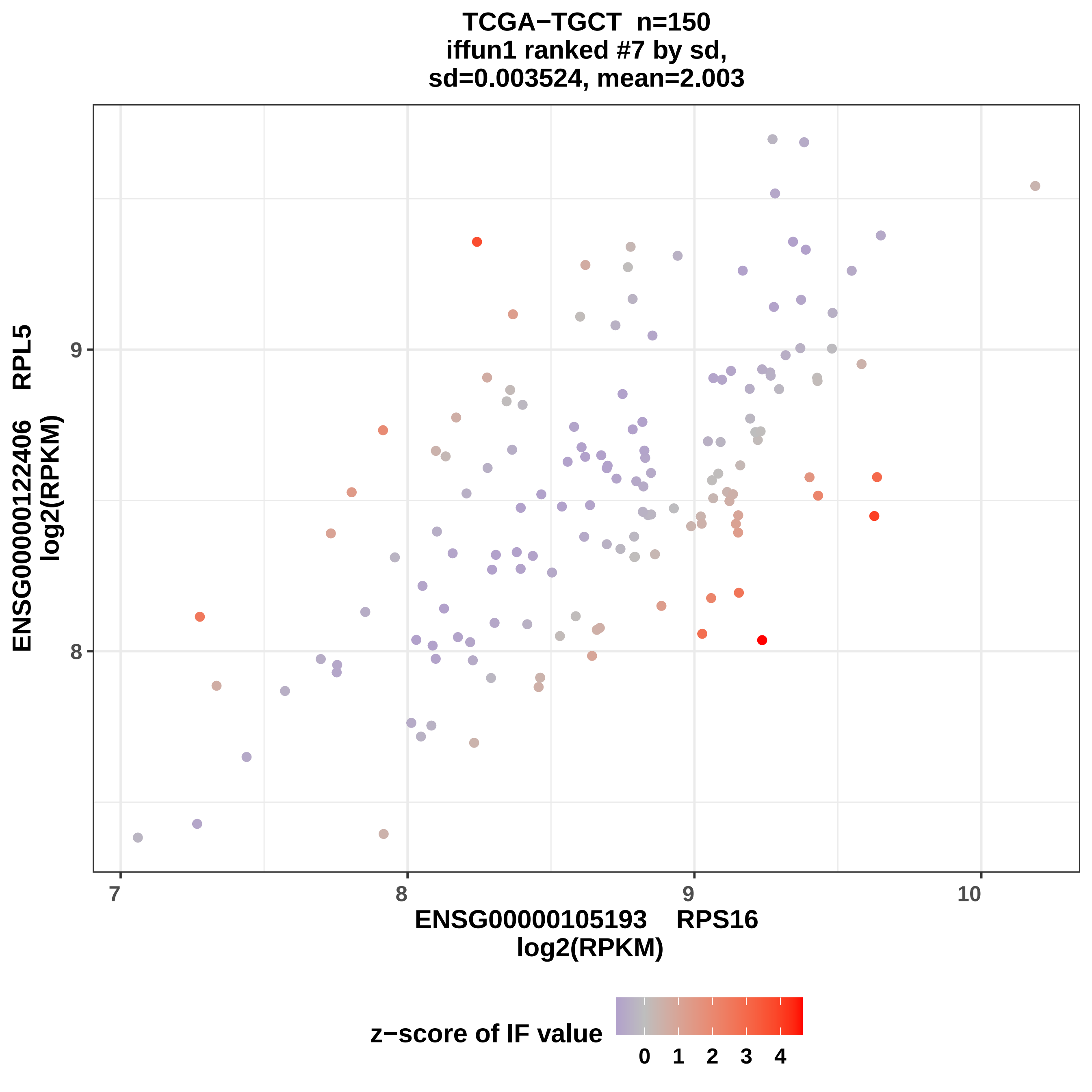
TCGA−TGCT n=150
iffun1 ranked #7 by sd,
sd=0.003524, mean=2.003
ENSG00000122406 RPL5
log2(RPKM)
9
8
7
8
9
10
ENSG00000105193 RPS16
log2(RPKM)
z−score of IF value
0
1
2
3
4

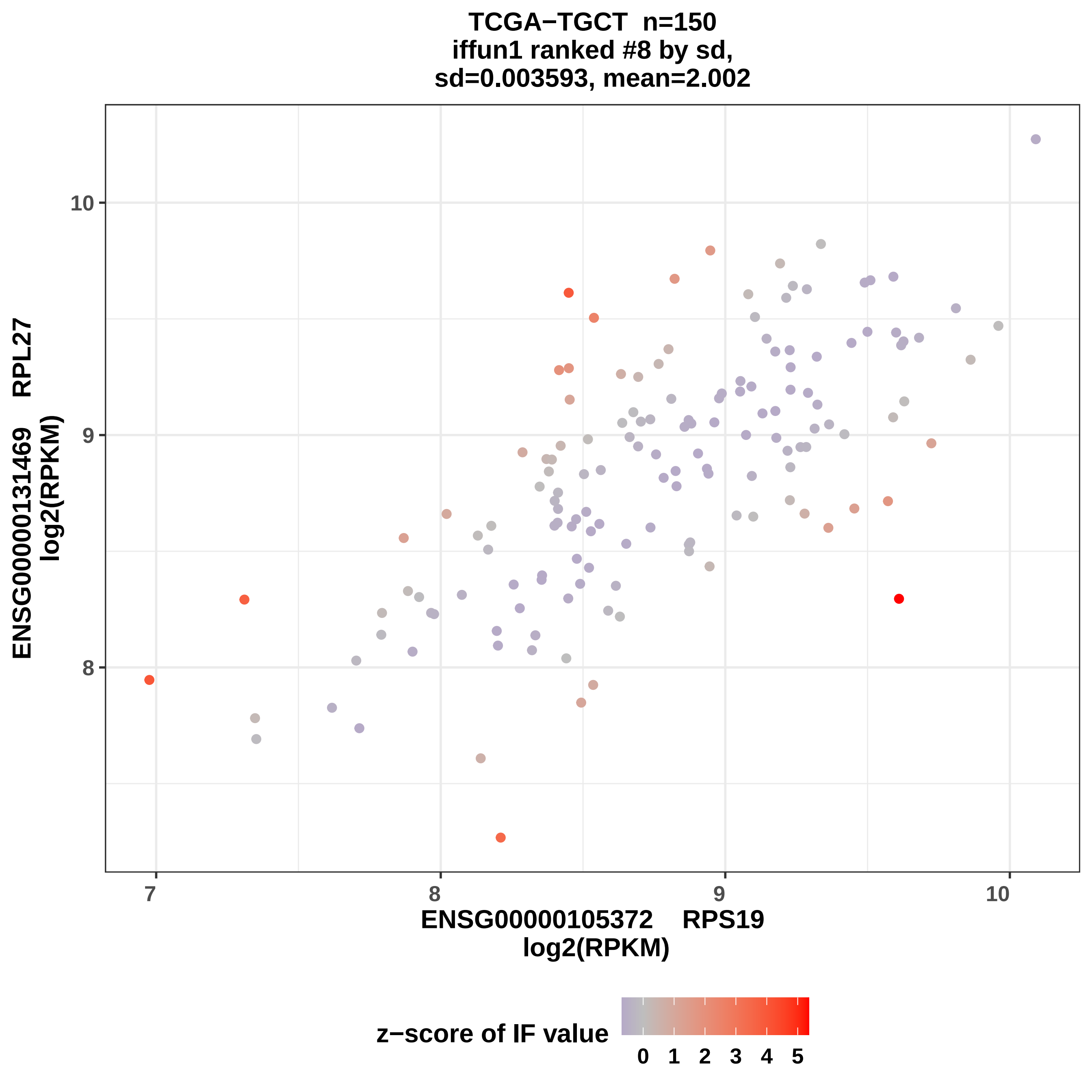
TCGA−TGCT n=150
iffun1 ranked #8 by sd,
sd=0.003593, mean=2.002
ENSG00000131469 RPL27
log2(RPKM)
ENSG00000105372 RPS19
log2(RPKM)
z−score of IF value
0 1 2 3 4 5
7
8
9
10
8
9
10

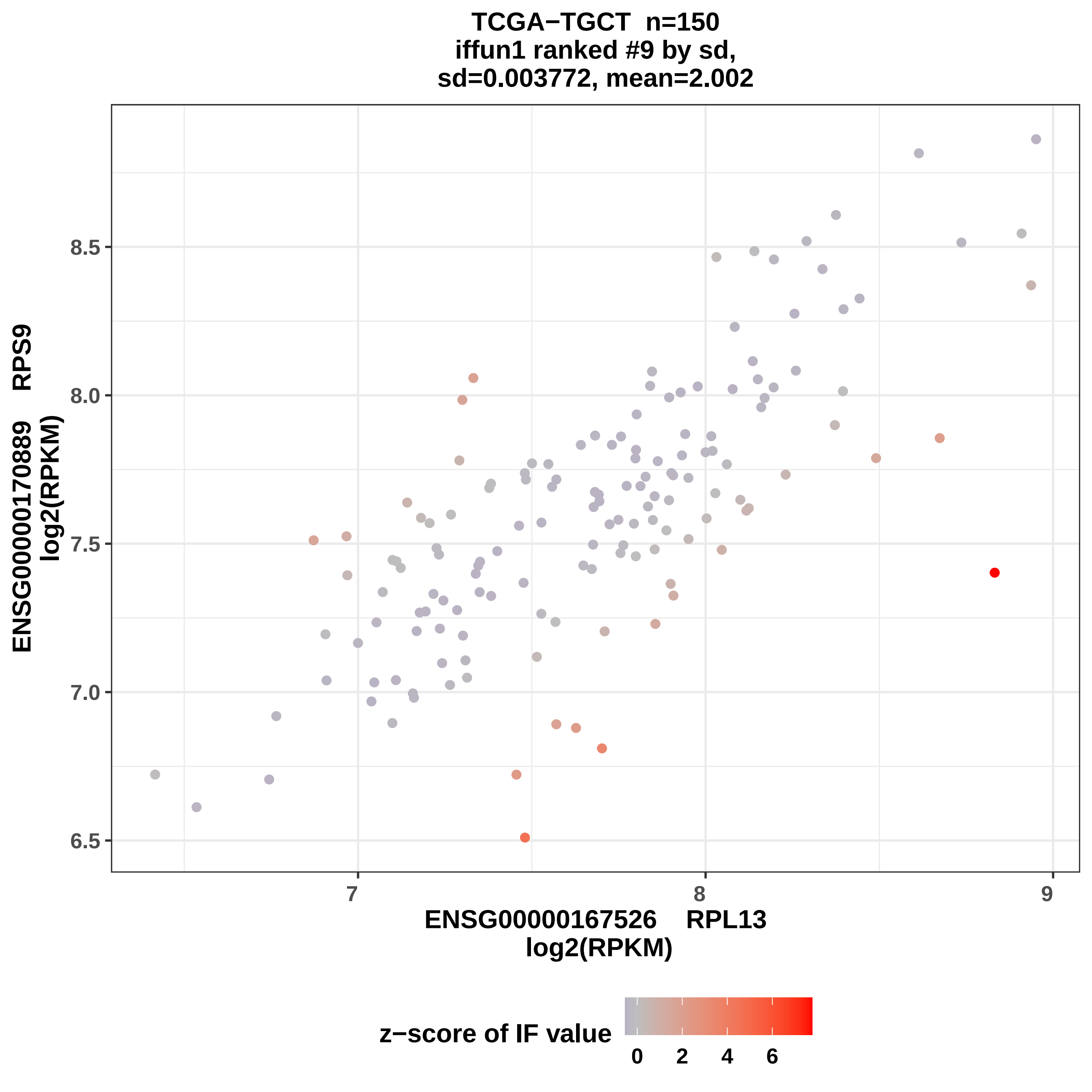
TCGA−TGCT n=150
iffun1 ranked #9 by sd,
sd=0.003772, mean=2.002
ENSG00000170889 RPS9
log2(RPKM)
8.5
8.0
7.5
7.0
6.5
7
8
9
ENSG00000167526 RPL13
log2(RPKM)
z−score of IF value
0
2
4
6

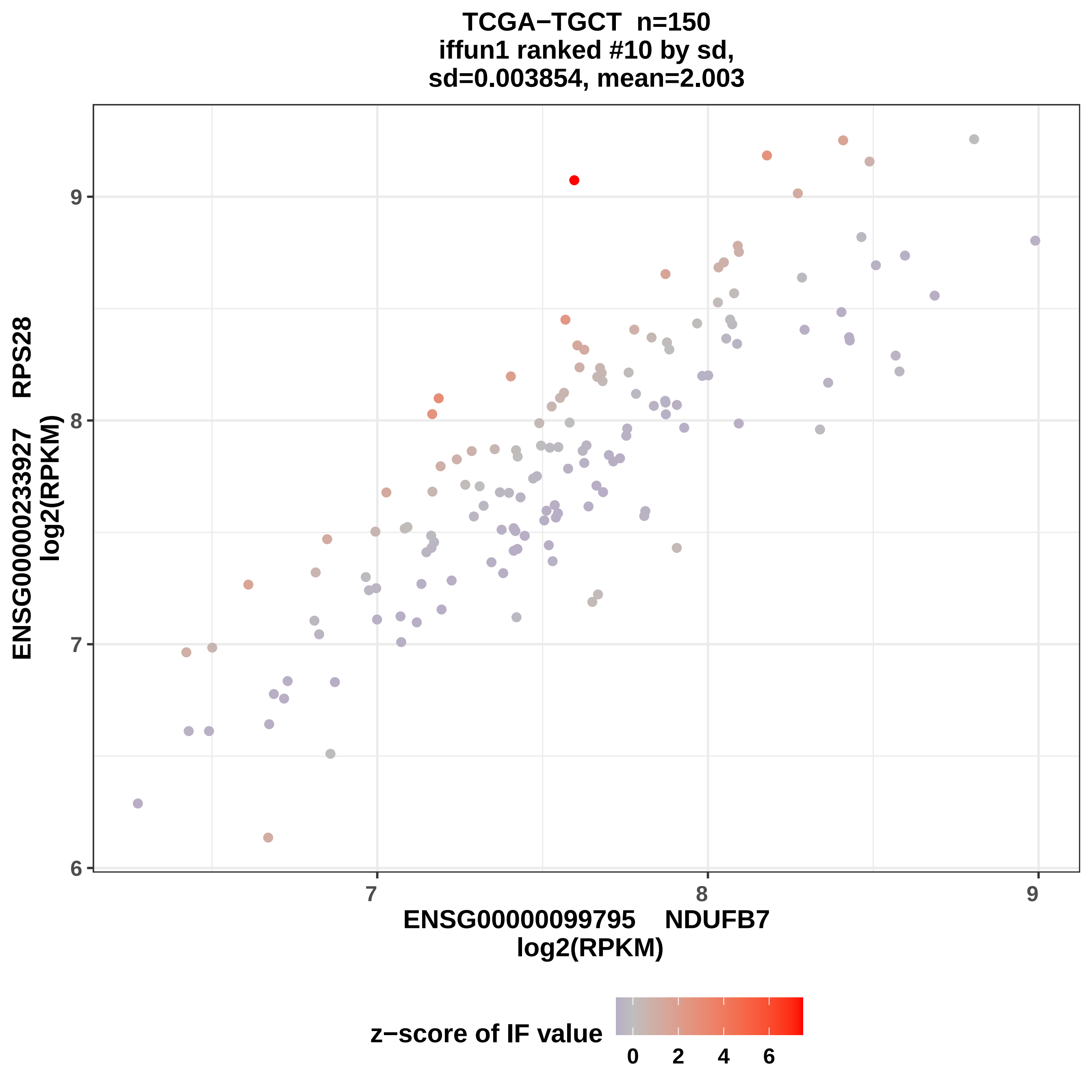
TCGA−TGCT n=150
iffun1 ranked #10 by sd,
sd=0.003854, mean=2.003
ENSG00000233927 RPS28
log2(RPKM)
ENSG00000099795 NDUFB7
log2(RPKM)
6
7
8
9
7
8
9
z−score of IF value
0
2
4
6

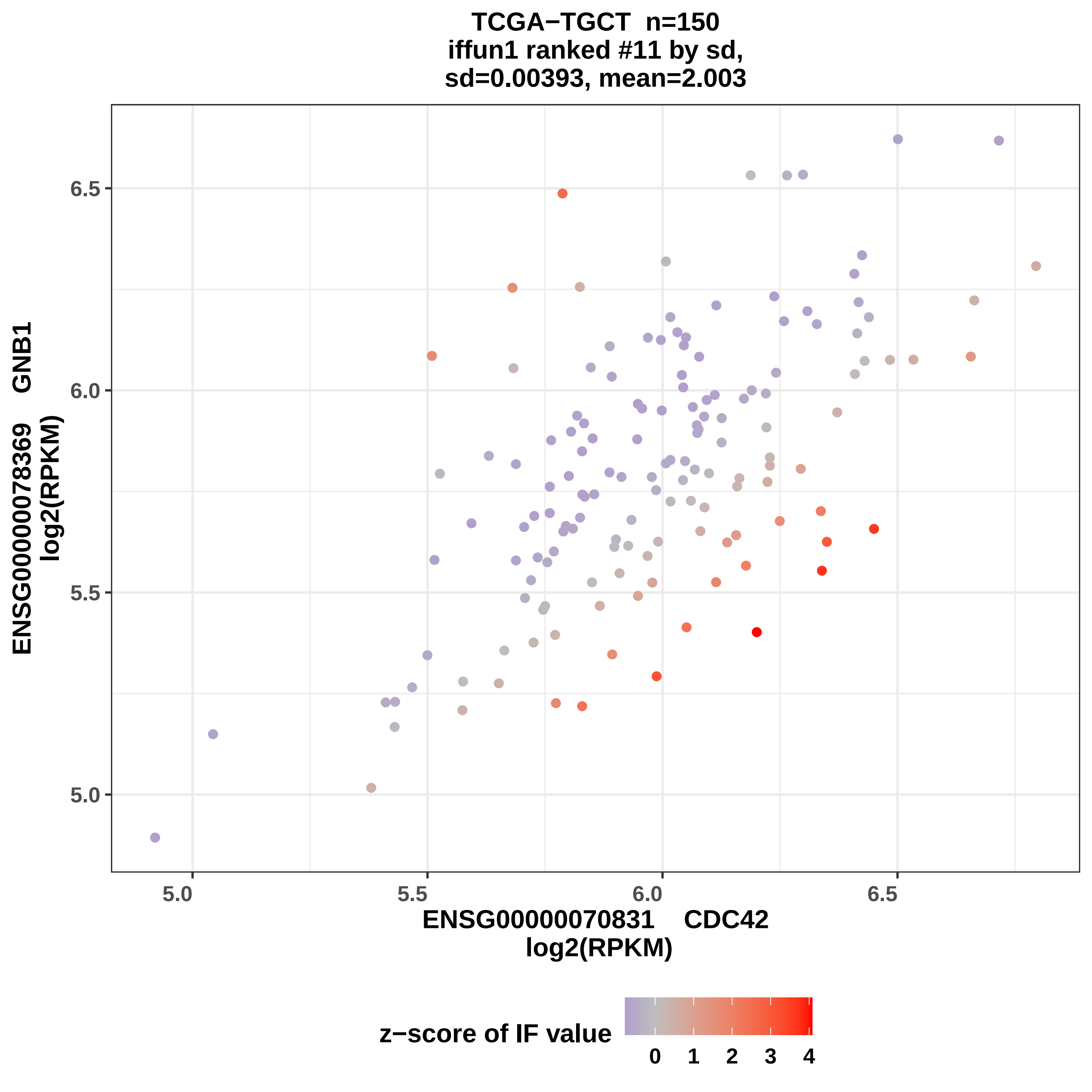
TCGA−TGCT n=150
iffun1 ranked #11 by sd,
sd=0.00393, mean=2.003
ENSG00000078369 GNB1
log2(RPKM)
6.5
6.0
5.5
5.0
5.0
5.5
6.0
6.5
ENSG00000070831 CDC42
log2(RPKM)
z−score of IF value
0
1
2
3
4

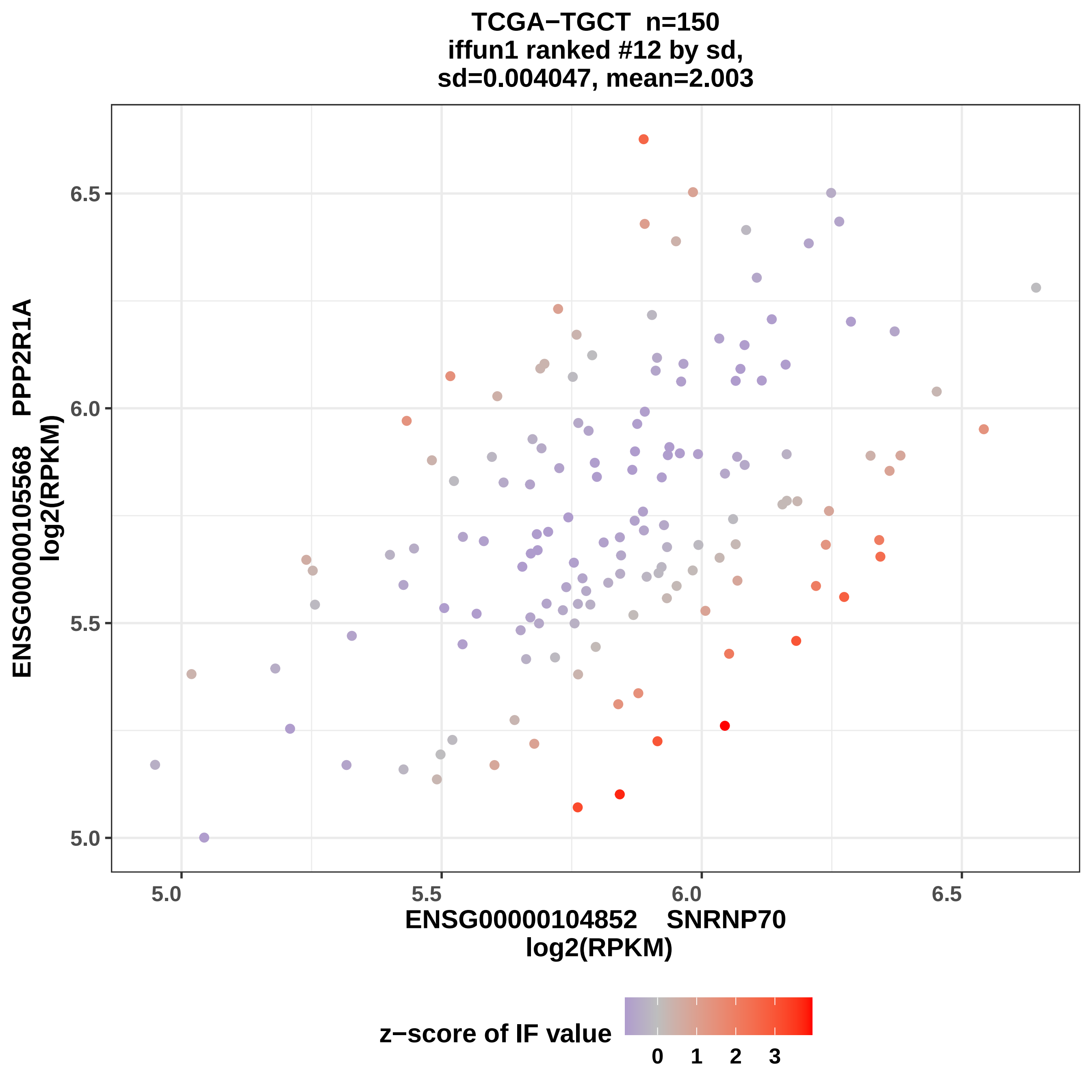

TCGA−TGCT n=150
iffun1 ranked #12 by sd,
sd=0.004047, mean=2.003
ENSG00000105568 PPP2R1A
log2(RPKM)
6.5
6.0
5.5
5.0
5.0
5.5
6.0
6.5
ENSG00000104852 SNRNP70
log2(RPKM)
z−score of IF value
0
1
2
3

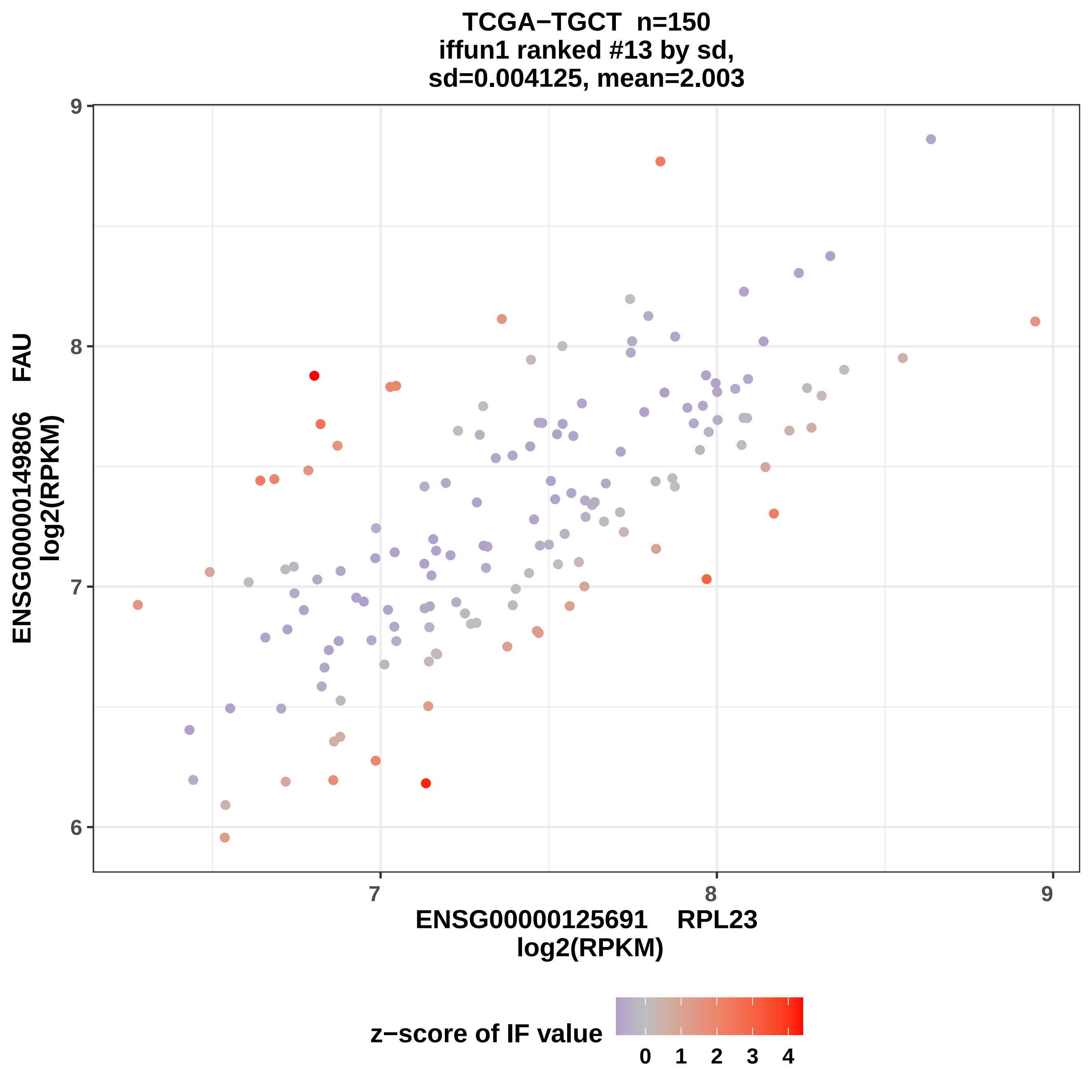

TCGA−TGCT n=150
iffun1 ranked #13 by sd,
sd=0.004125, mean=2.003
ENSG00000149806 FAU
log2(RPKM)
ENSG00000125691 RPL23
log2(RPKM)
z−score of IF value
6
7
8
9
0
1
2
3
4

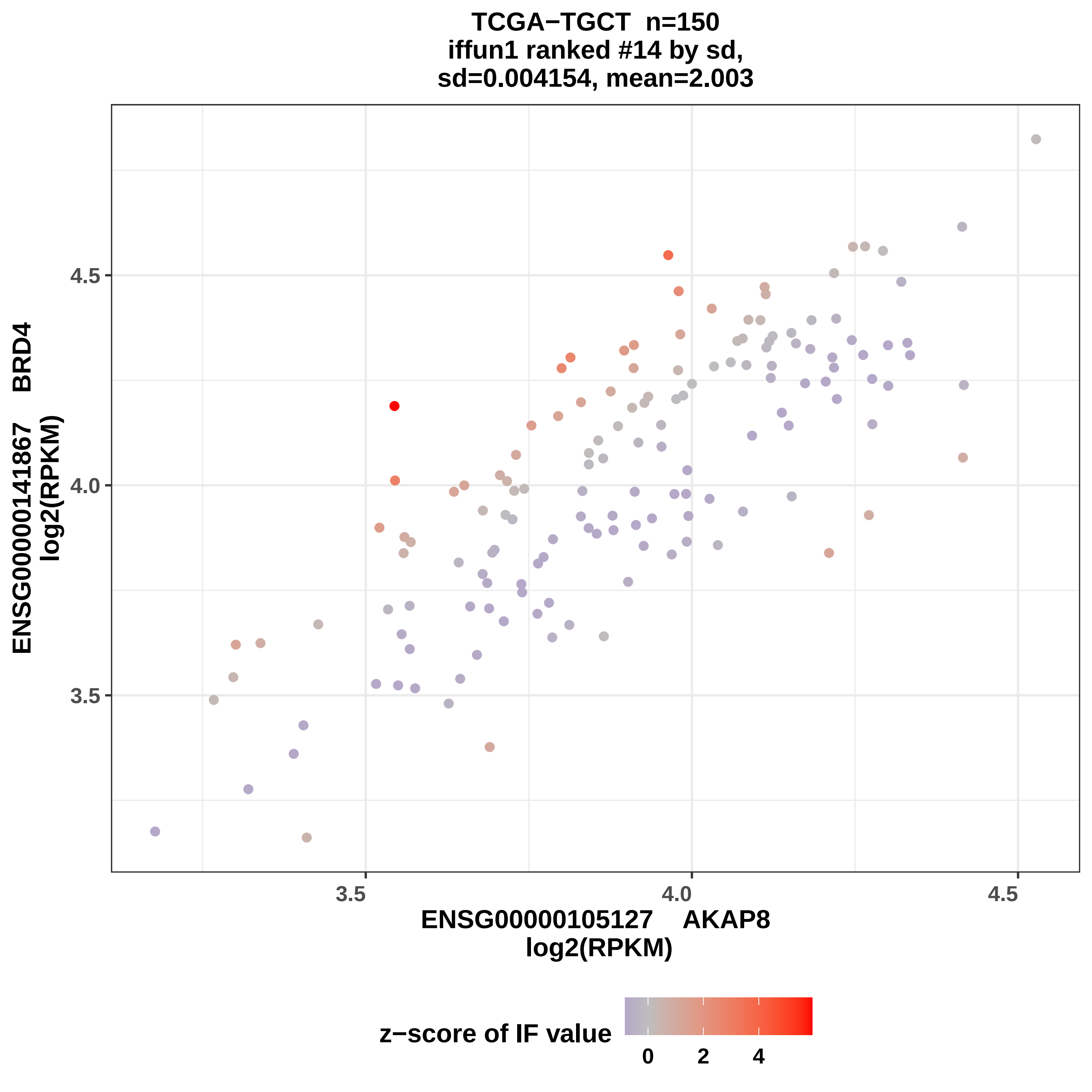
TCGA−TGCT n=150
iffun1 ranked #14 by sd,
sd=0.004154, mean=2.003
ENSG00000141867 BRD4
log2(RPKM)
4.5
4.0
3.5
3.5
4.0
4.5
ENSG00000105127 AKAP8
log2(RPKM)
z−score of IF value
0
2
4

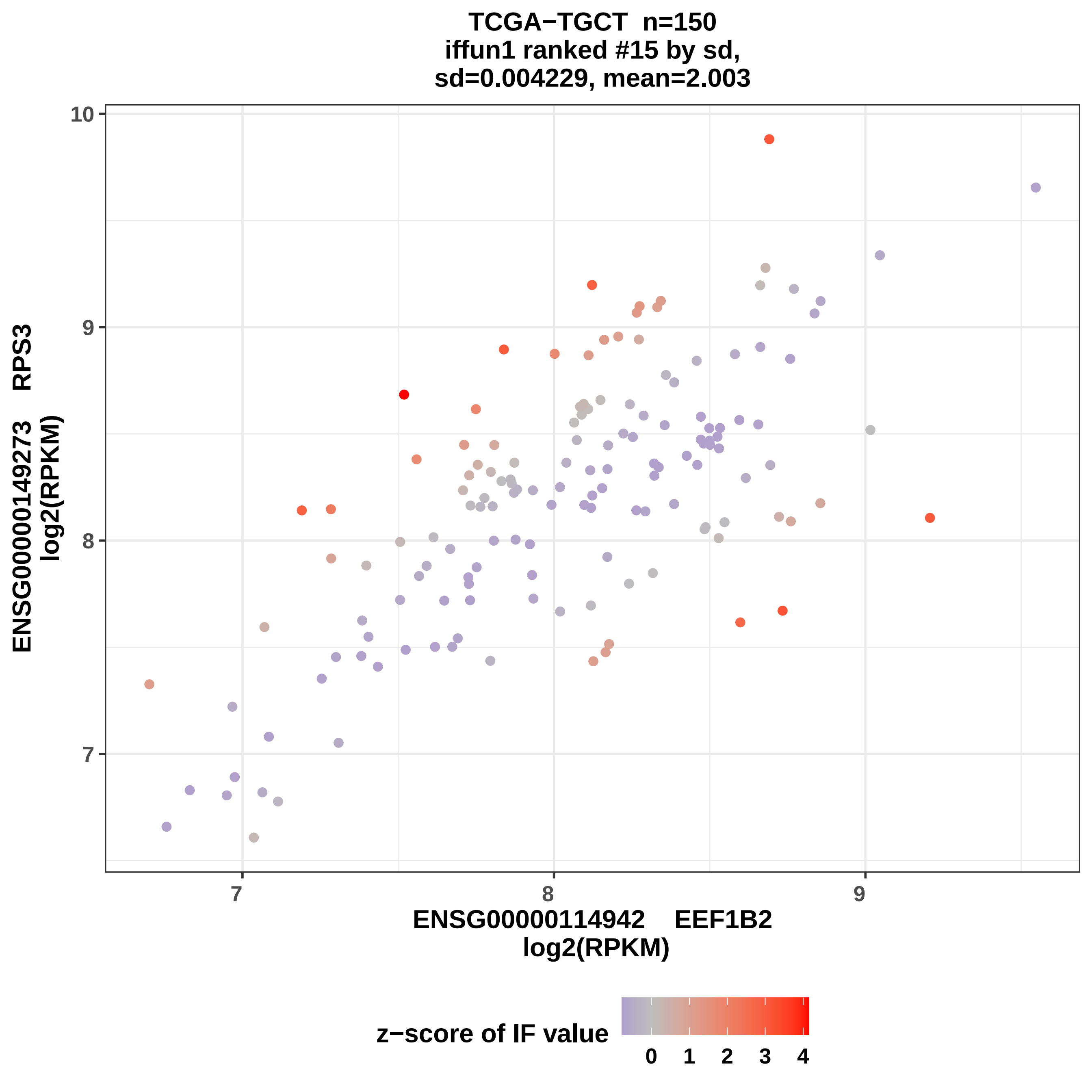

TCGA−TGCT n=150
iffun1 ranked #15 by sd,
sd=0.004229, mean=2.003
ENSG00000149273 RPS3
log2(RPKM)
10
9
8
7
ENSG00000114942 EEF1B2
log2(RPKM)
z−score of IF value
0
1
2
3
4

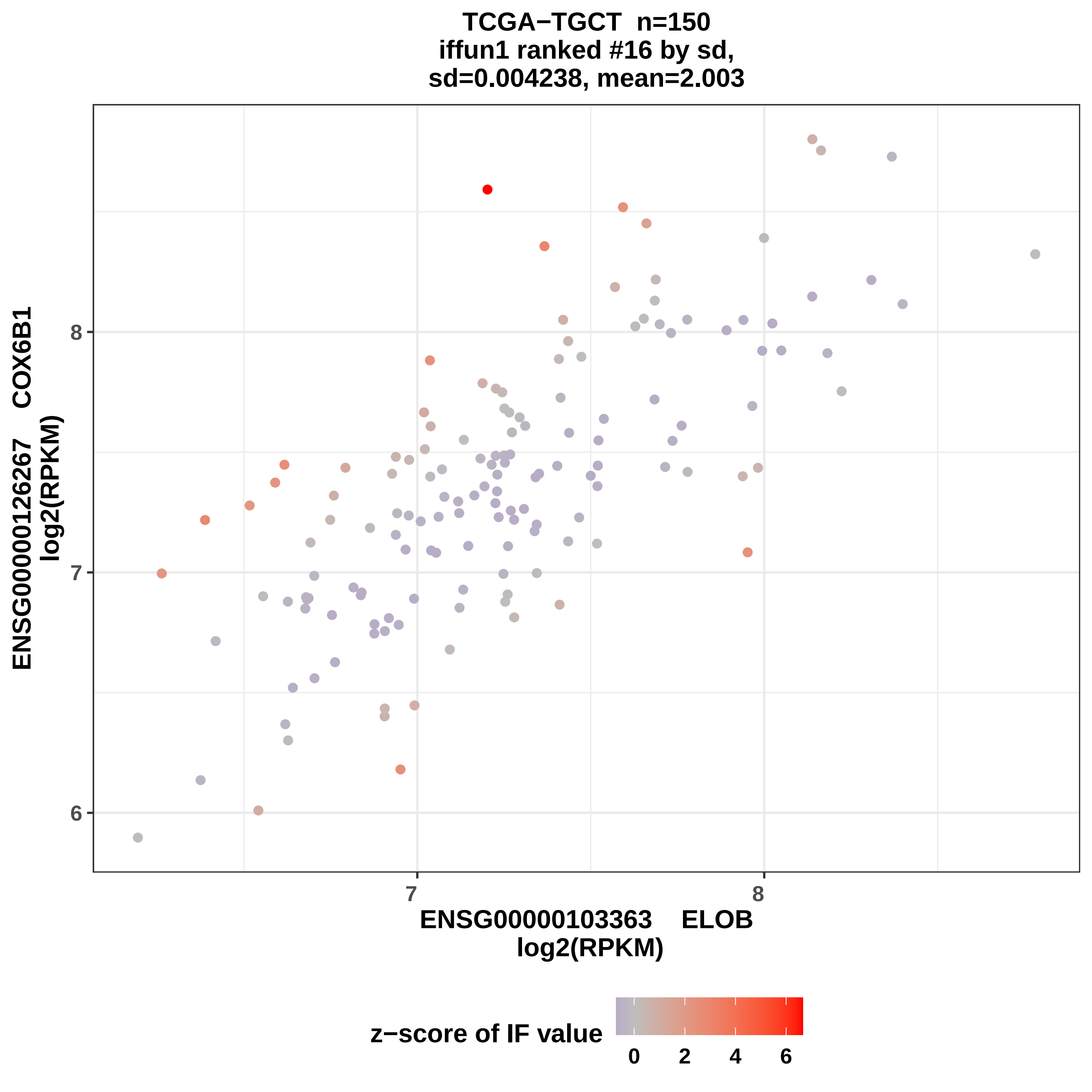

TCGA−TGCT n=150
iffun1 ranked #16 by sd,
sd=0.004238, mean=2.003
ENSG00000126267 COX6B1
log2(RPKM)
8
7
6
7
8
ENSG00000103363 ELOB
log2(RPKM)
z−score of IF value
0
2
4
6

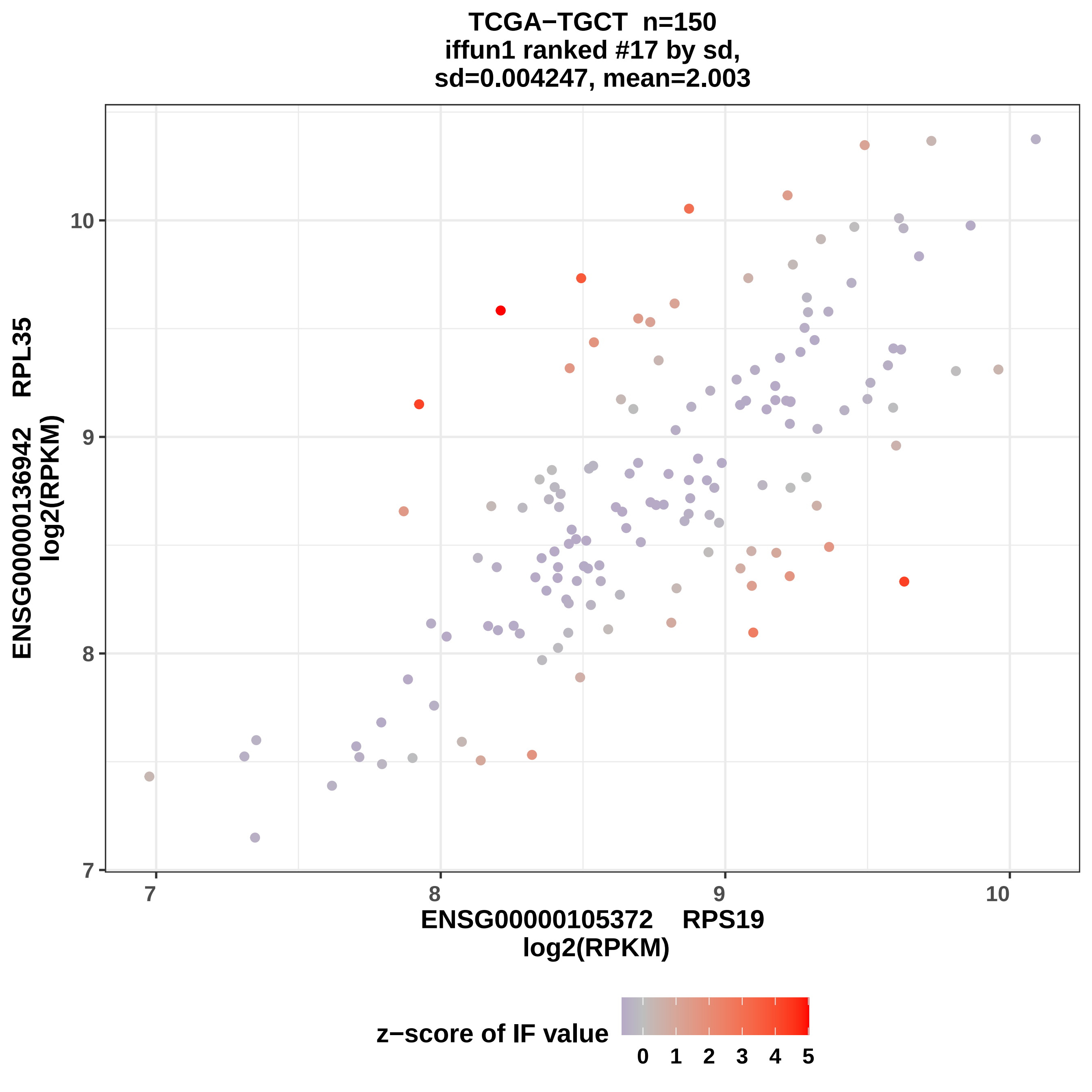

TCGA−TGCT n=150
iffun1 ranked #17 by sd,
sd=0.004247, mean=2.003
ENSG00000136942 RPL35
log2(RPKM)
ENSG00000105372 RPS19
log2(RPKM)
7
8
9
10
z−score of IF value
0 1 2 3 4 5

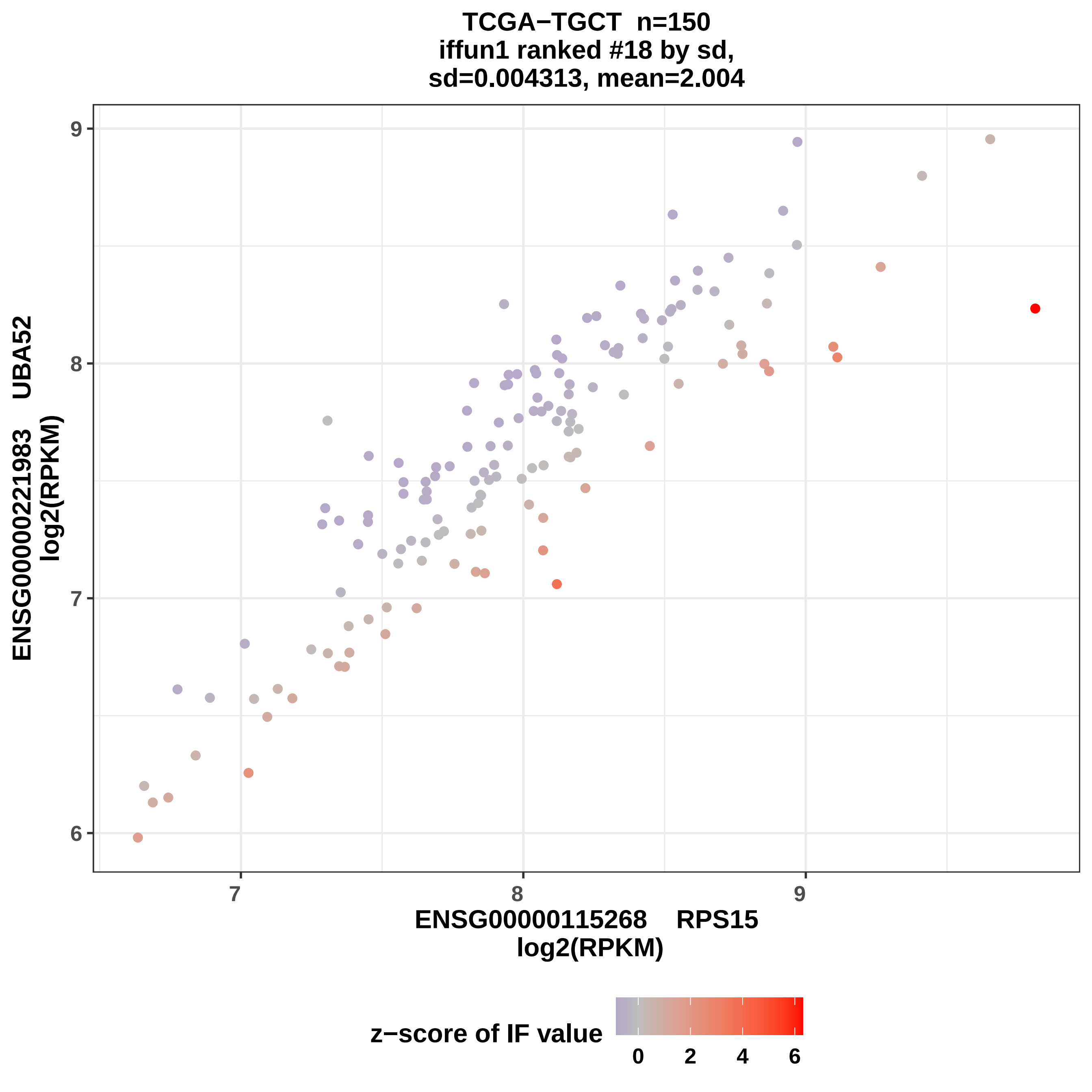

TCGA−TGCT n=150
iffun1 ranked #18 by sd,
sd=0.004313, mean=2.004
ENSG00000221983 UBA52
log2(RPKM)
ENSG00000115268 RPS15
log2(RPKM)
z−score of IF value
9
8
7
6
7
8
9
0
2
4
6

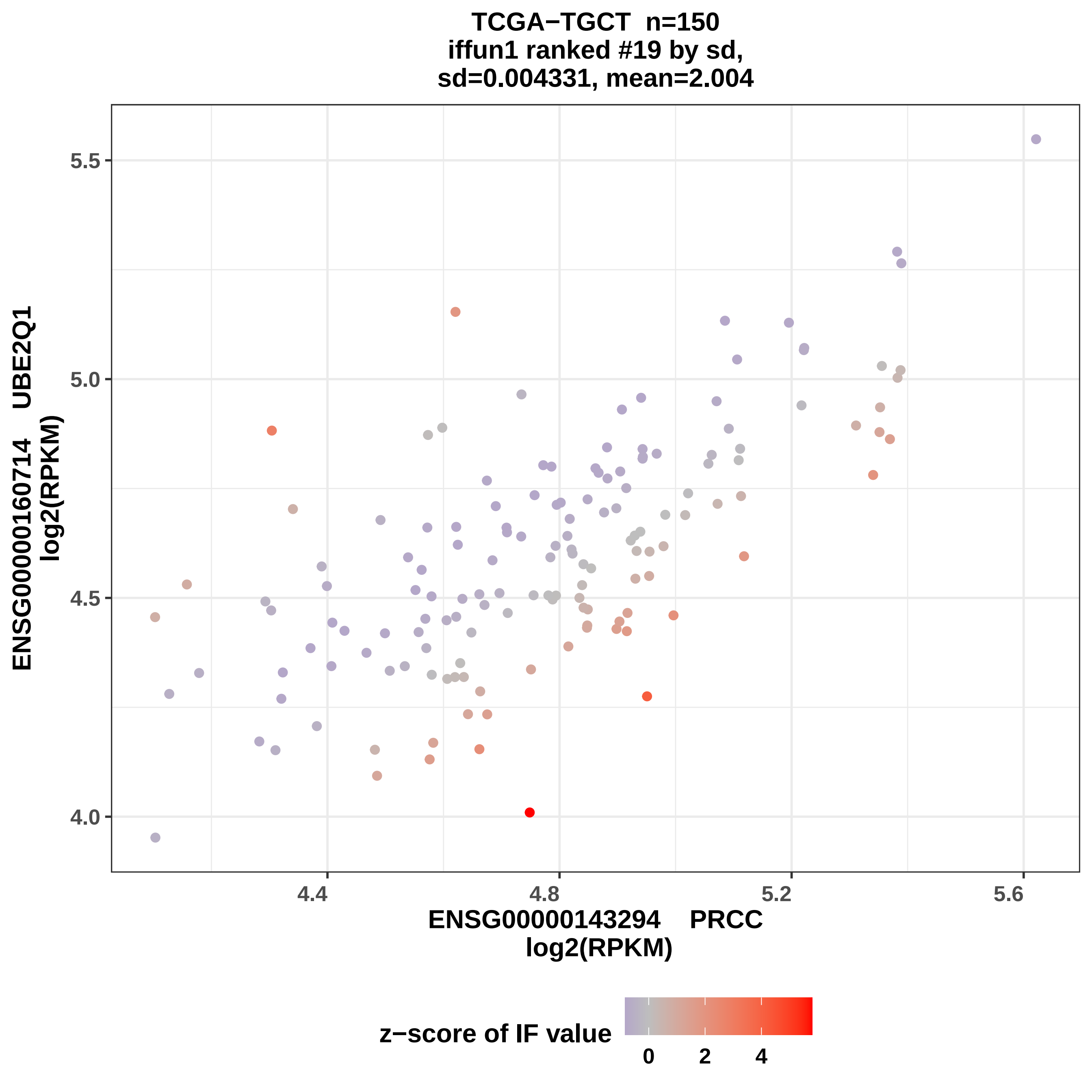

TCGA−TGCT n=150
iffun1 ranked #19 by sd,
sd=0.004331, mean=2.004
ENSG00000160714 UBE2Q1
log2(RPKM)
5.5
5.0
4.5
4.0
4.4
4.8
5.2
5.6
ENSG00000143294 PRCC
log2(RPKM)
z−score of IF value
0
2
4

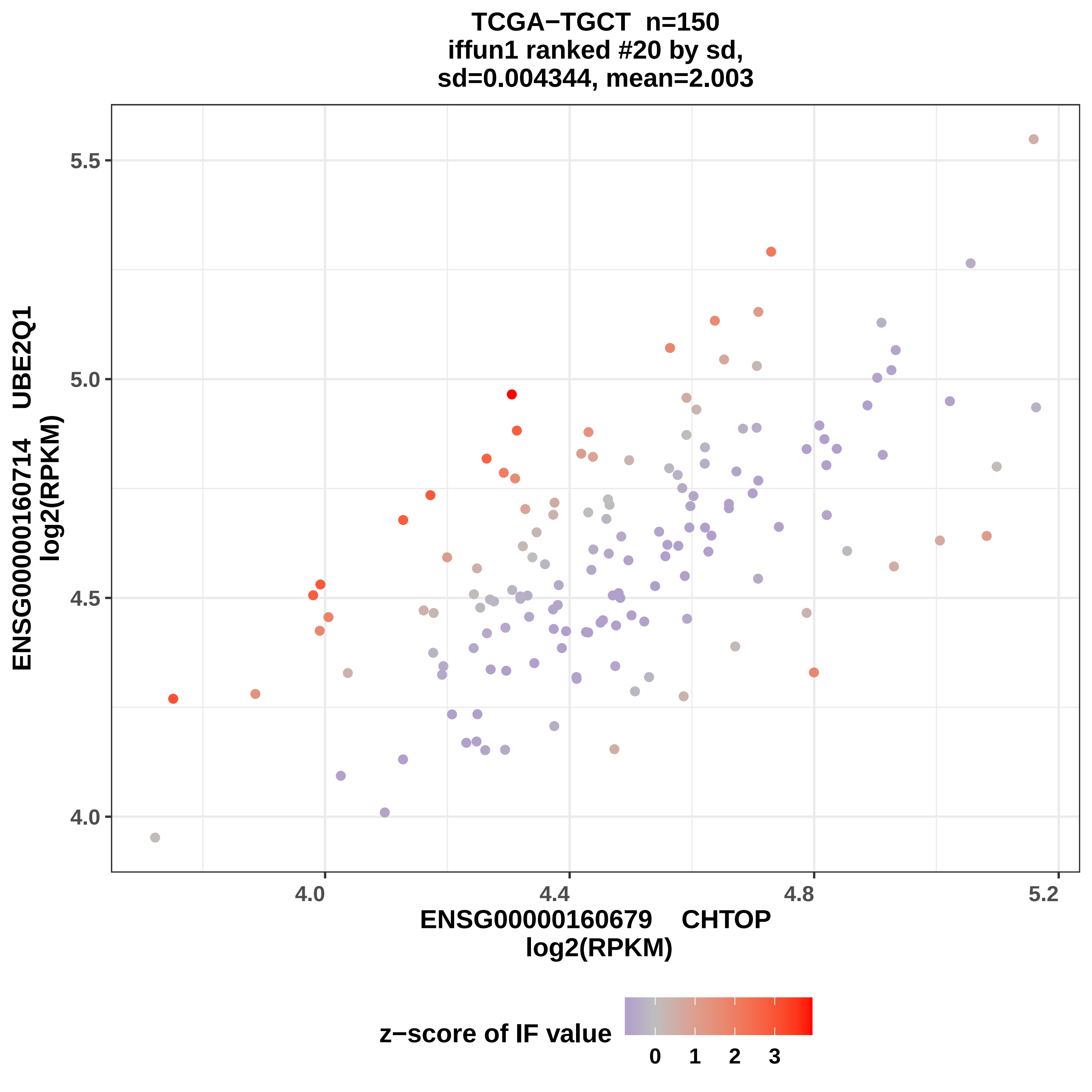
TCGA−TGCT n=150
iffun1 ranked #20 by sd,
sd=0.004344, mean=2.003
ENSG00000160714 UBE2Q1
log2(RPKM)
5.5
5.0
4.5
4.0
4.0
4.4
4.8
5.2
ENSG00000160679 CHTOP
log2(RPKM)
z−score of IF value
0
1
2
3

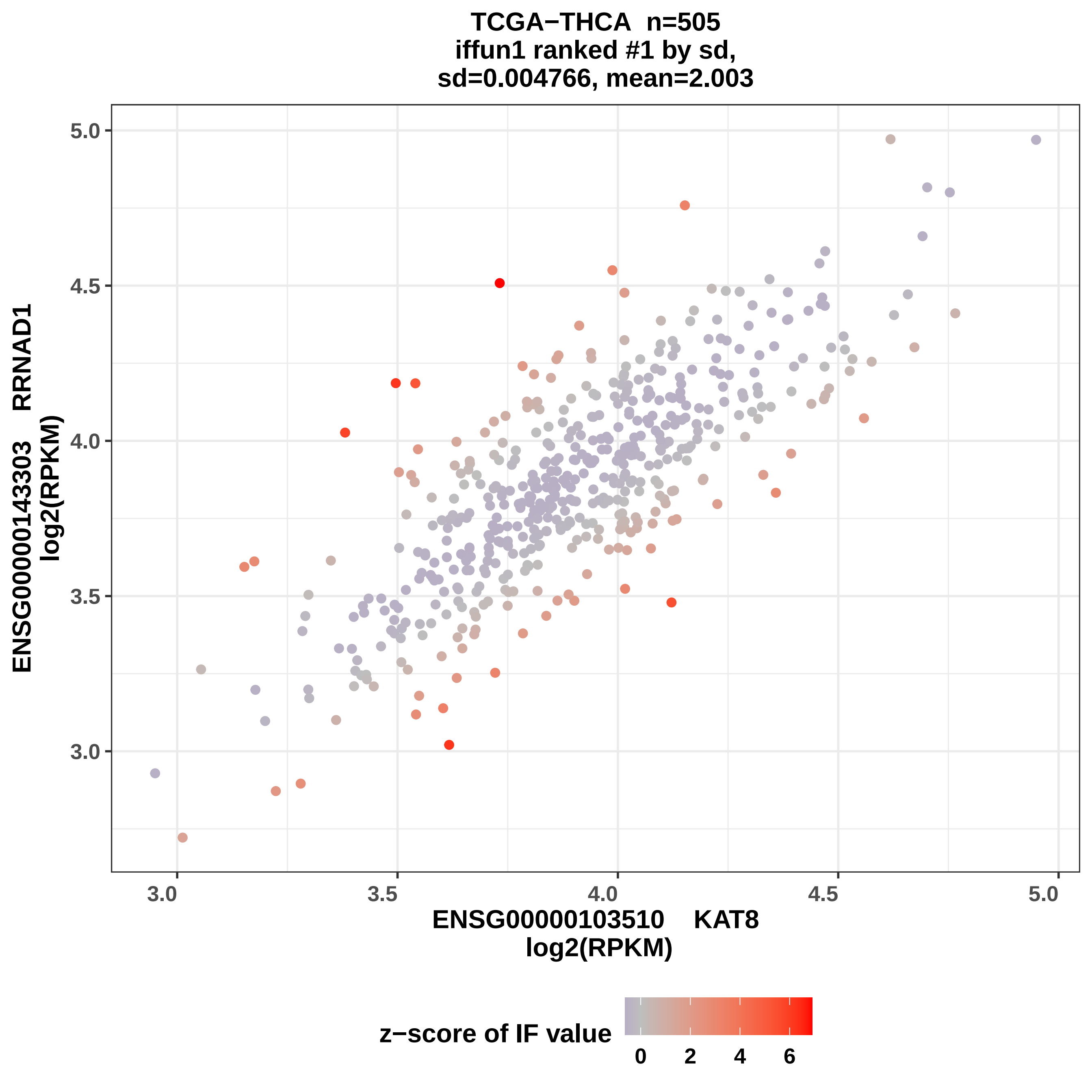
TCGA−THCA n=505
iffun1 ranked #1 by sd,
sd=0.004766, mean=2.003
ENSG00000143303 RRNAD1
log2(RPKM)
5.0
4.5
4.0
3.5
3.0
3.0
3.5
4.0
4.5
5.0
ENSG00000103510 KAT8
log2(RPKM)
z−score of IF value
0
2
4
6

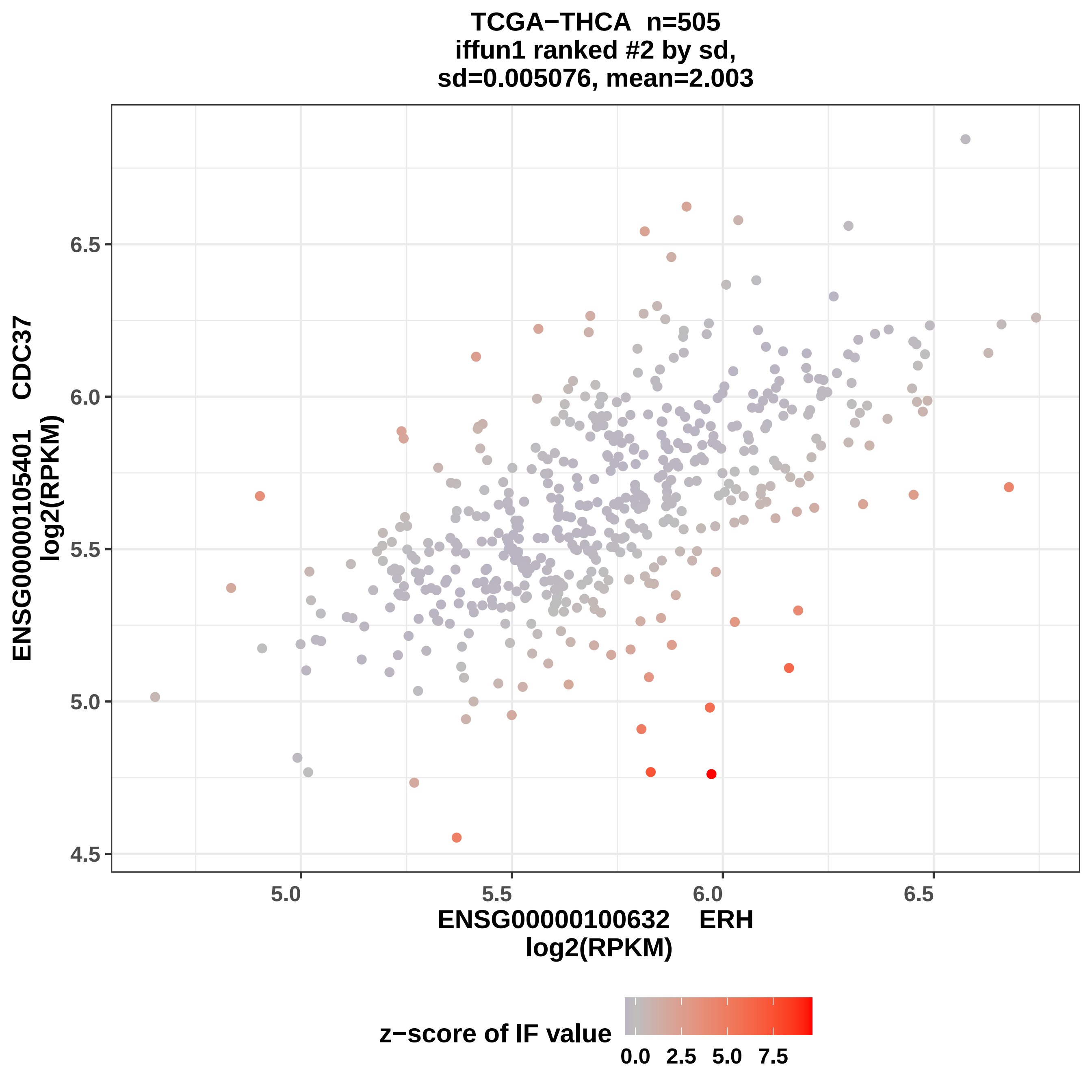

TCGA−THCA n=505
iffun1 ranked #2 by sd,
sd=0.005076, mean=2.003
ENSG00000105401 CDC37
log2(RPKM)
6.5
6.0
5.5
5.0
4.5
5.0
5.5
6.0
6.5
ENSG00000100632 ERH
log2(RPKM)
z−score of IF value
0.0
2.5
5.0
7.5

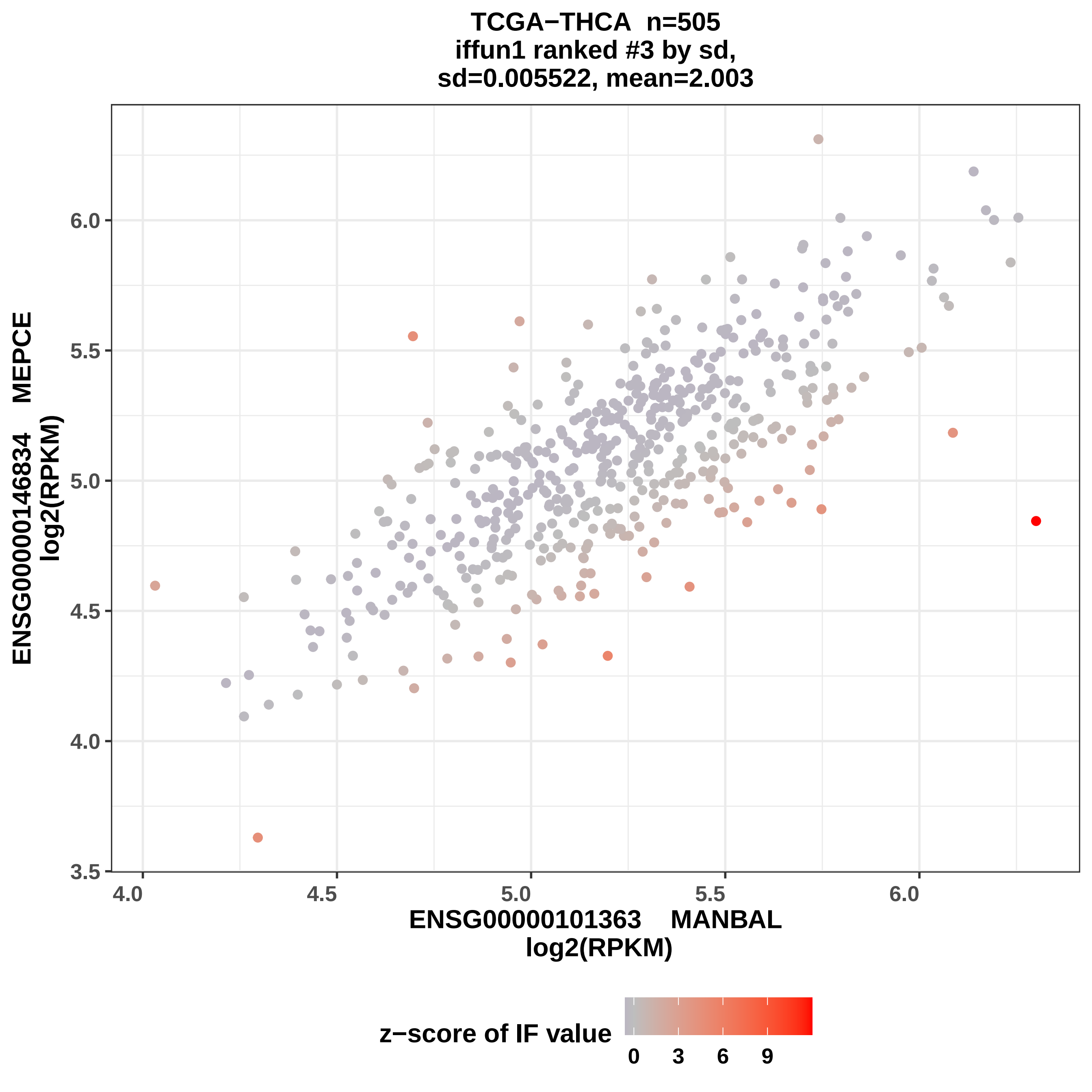
TCGA−THCA n=505
iffun1 ranked #3 by sd,
sd=0.005522, mean=2.003
ENSG00000146834 MEPCE
log2(RPKM)
ENSG00000101363 MANBAL
log2(RPKM)
6.0
5.5
5.0
4.5
4.0
3.5
4.0
4.5
5.0
5.5
6.0
z−score of IF value
0
3
6
9

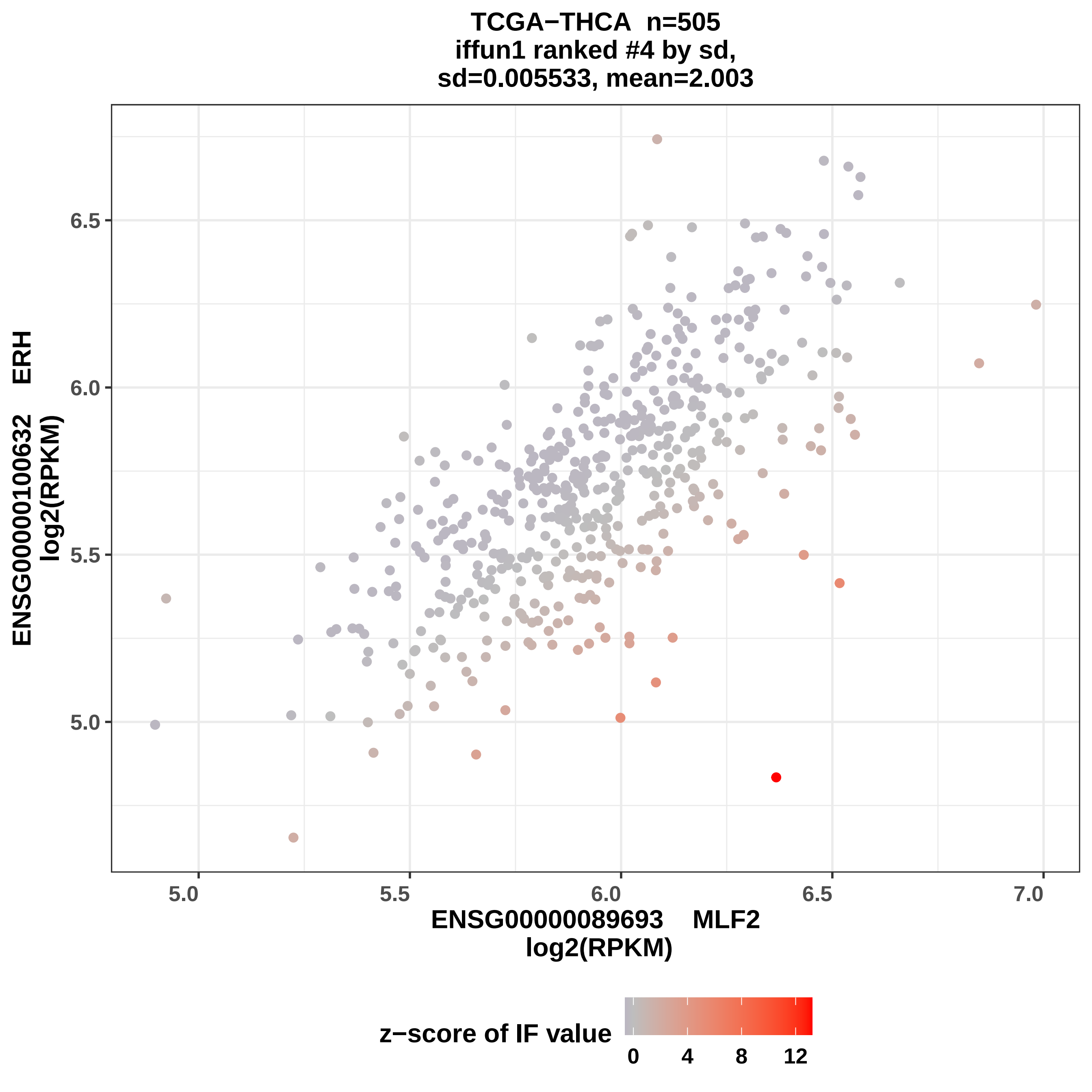
TCGA−THCA n=505
iffun1 ranked #4 by sd,
sd=0.005533, mean=2.003
ENSG00000100632 ERH
log2(RPKM)
6.5
6.0
5.5
5.0
5.0
5.5
6.0
6.5
7.0
ENSG00000089693 MLF2
log2(RPKM)
z−score of IF value
0
4
8
12

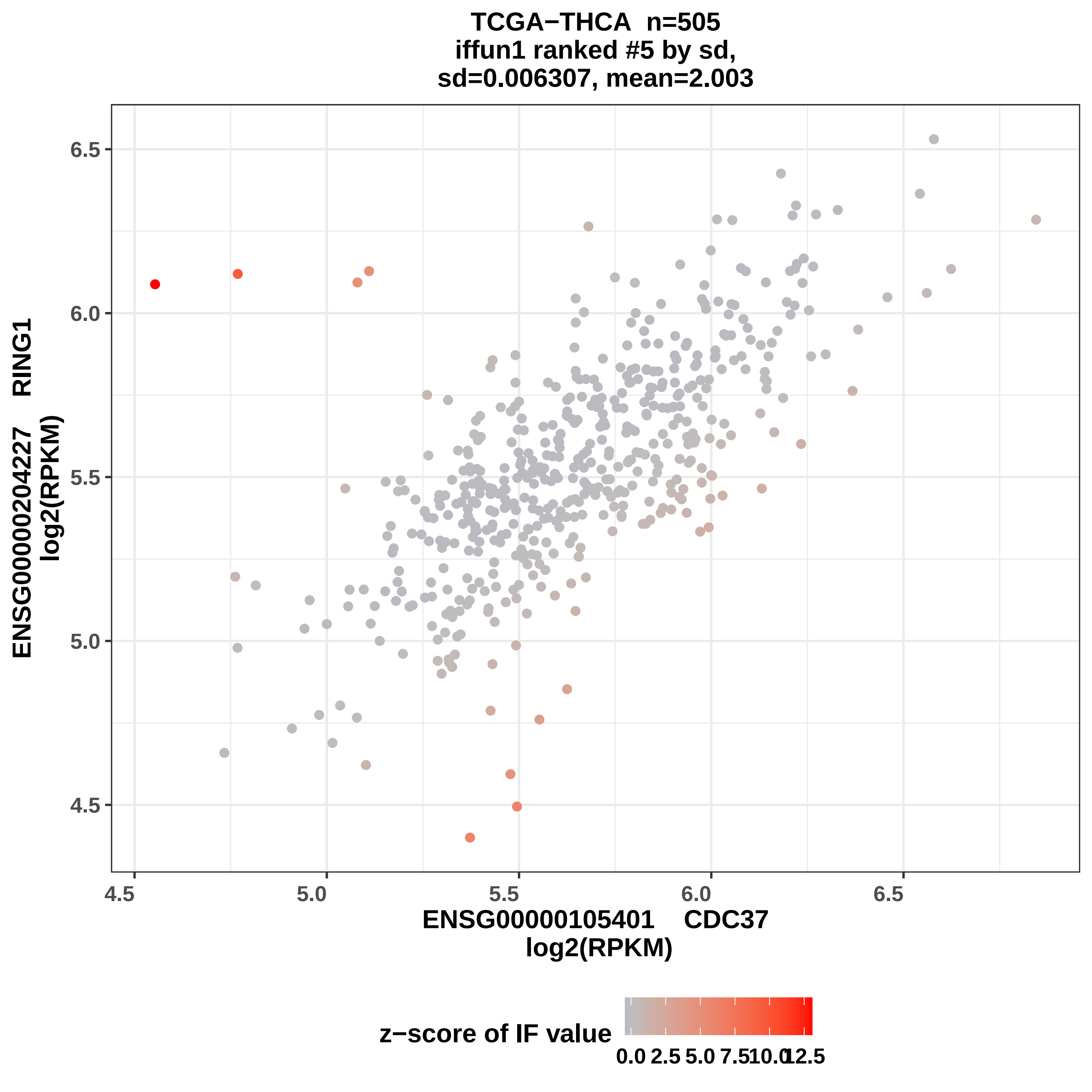

TCGA−THCA n=505
iffun1 ranked #5 by sd,
sd=0.006307, mean=2.003
ENSG00000204227 RING1
log2(RPKM)
6.5
6.0
5.5
5.0
4.5
4.5
5.0
5.5
6.0
6.5
ENSG00000105401 CDC37
log2(RPKM)
z−score of IF value
0.0
2.5
5.0
7.5
10.0
12.5

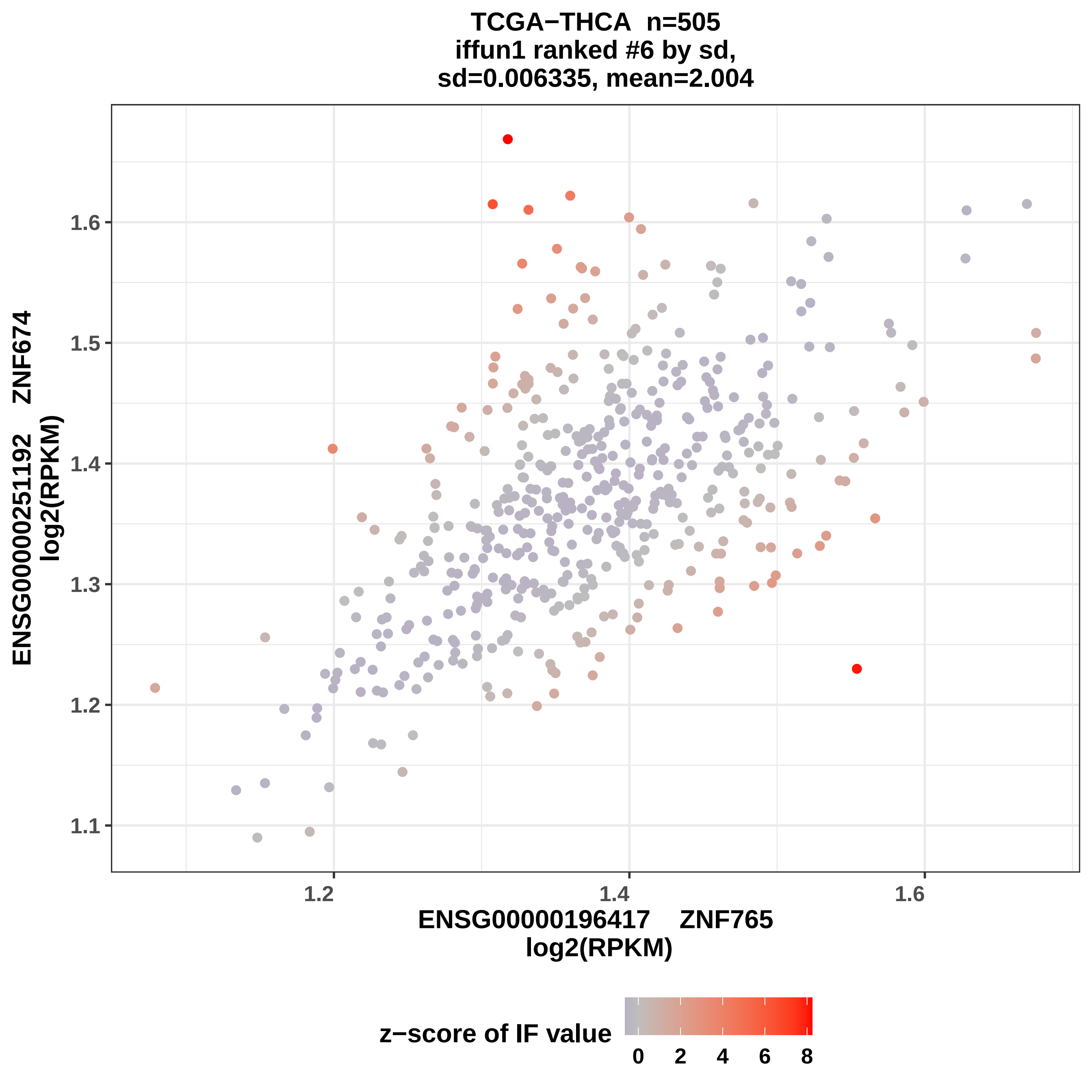
TCGA−THCA n=505
iffun1 ranked #6 by sd,
sd=0.006335, mean=2.004
ENSG00000251192 ZNF674
log2(RPKM)
1.1
1.2
1.3
1.4
1.5
1.6
1.2
1.4
1.6
ENSG00000196417 ZNF765
log2(RPKM)
z−score of IF value
0
2
4
6
8

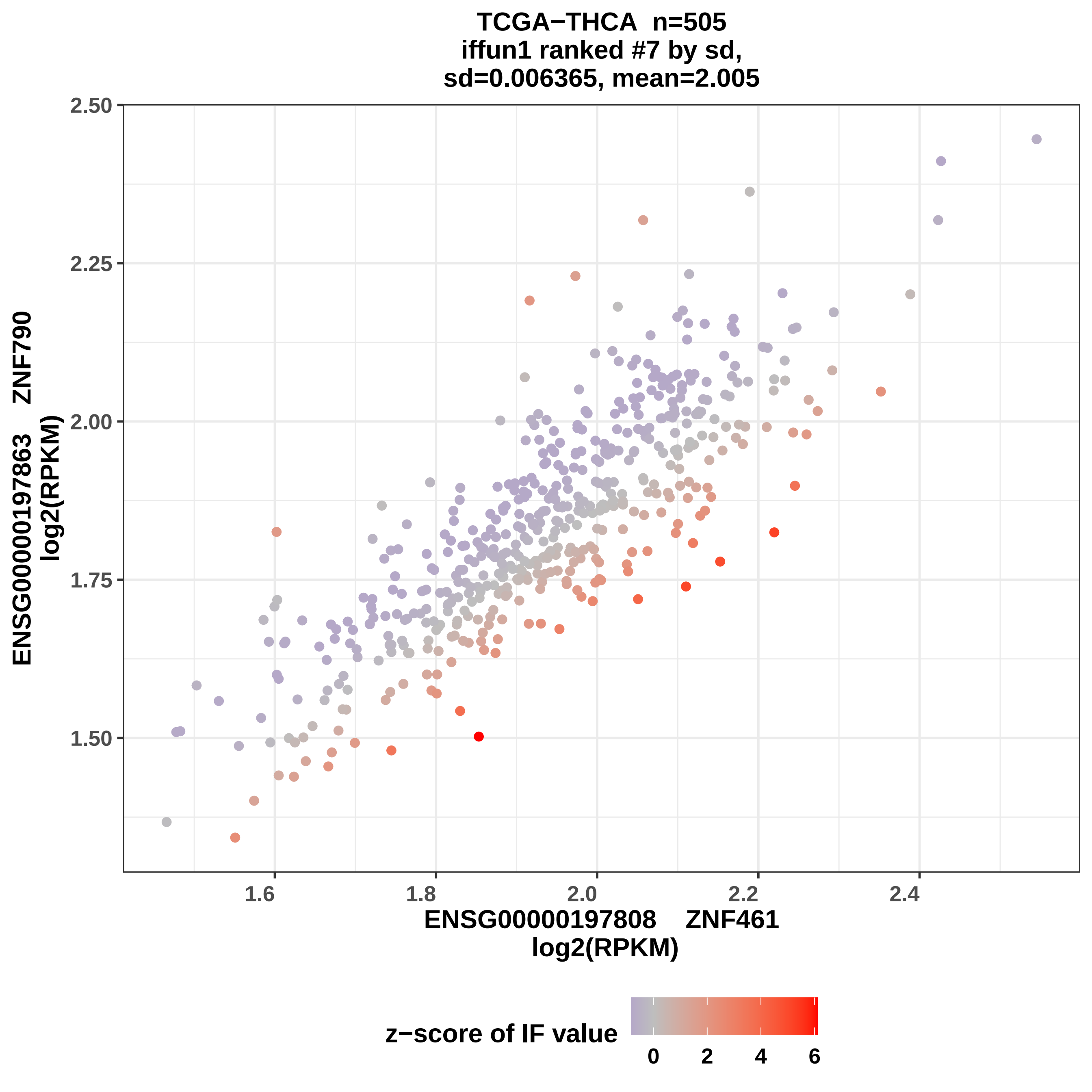

TCGA−THCA n=505
iffun1 ranked #7 by sd,
sd=0.006365, mean=2.005
2.50
2.25
2.00
1.75
1.50
ENSG00000197863 ZNF790
log2(RPKM)
1.6
1.8
2.0
2.2
2.4
ENSG00000197808 ZNF461
log2(RPKM)
z−score of IF value
0
2
4
6

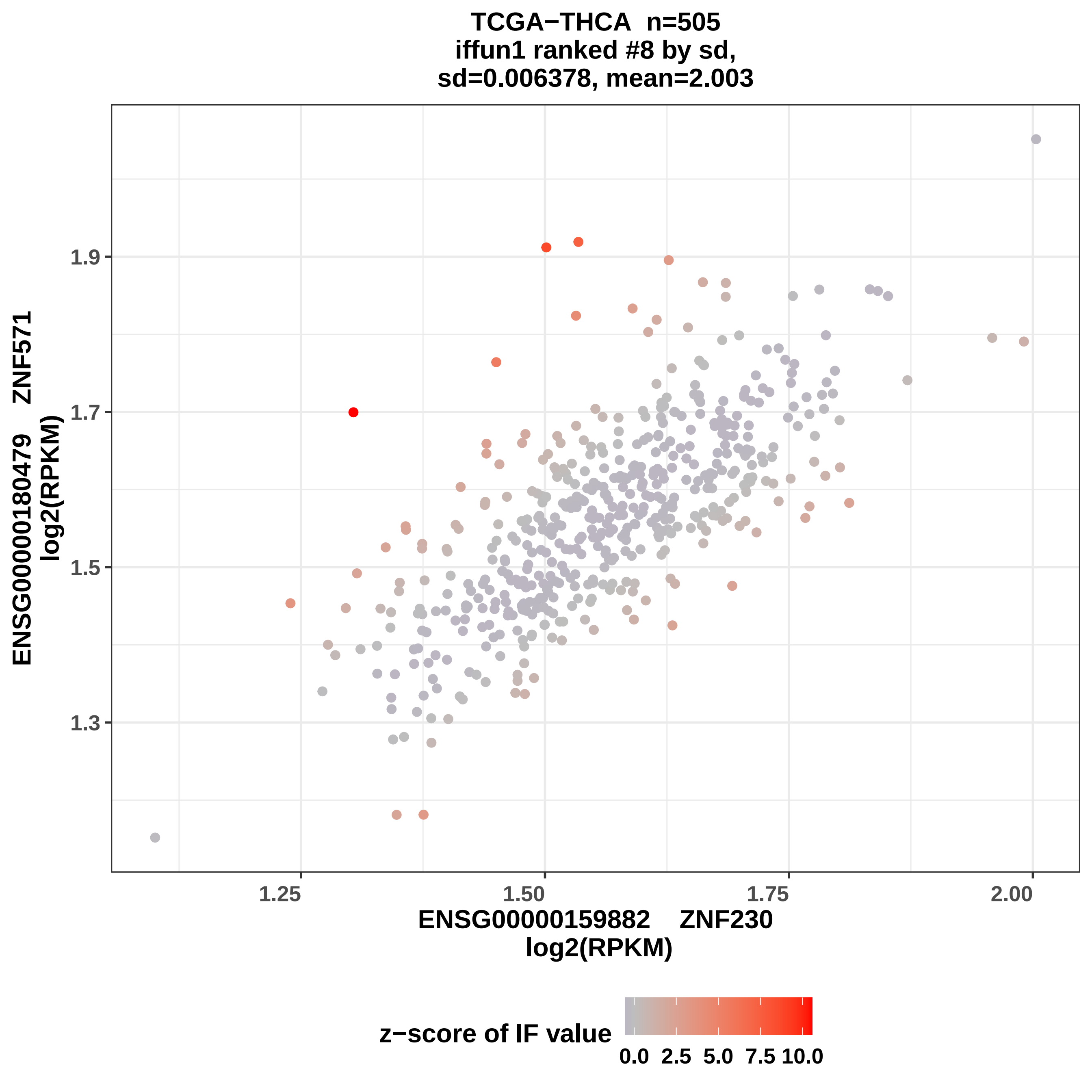
TCGA−THCA n=505
iffun1 ranked #8 by sd,
sd=0.006378, mean=2.003
ENSG00000180479 ZNF571
log2(RPKM)
1.9
1.7
1.5
1.3
1.25
1.50
1.75
2.00
ENSG00000159882 ZNF230
log2(RPKM)
z−score of IF value
0.0
2.5
5.0
7.5
10.0

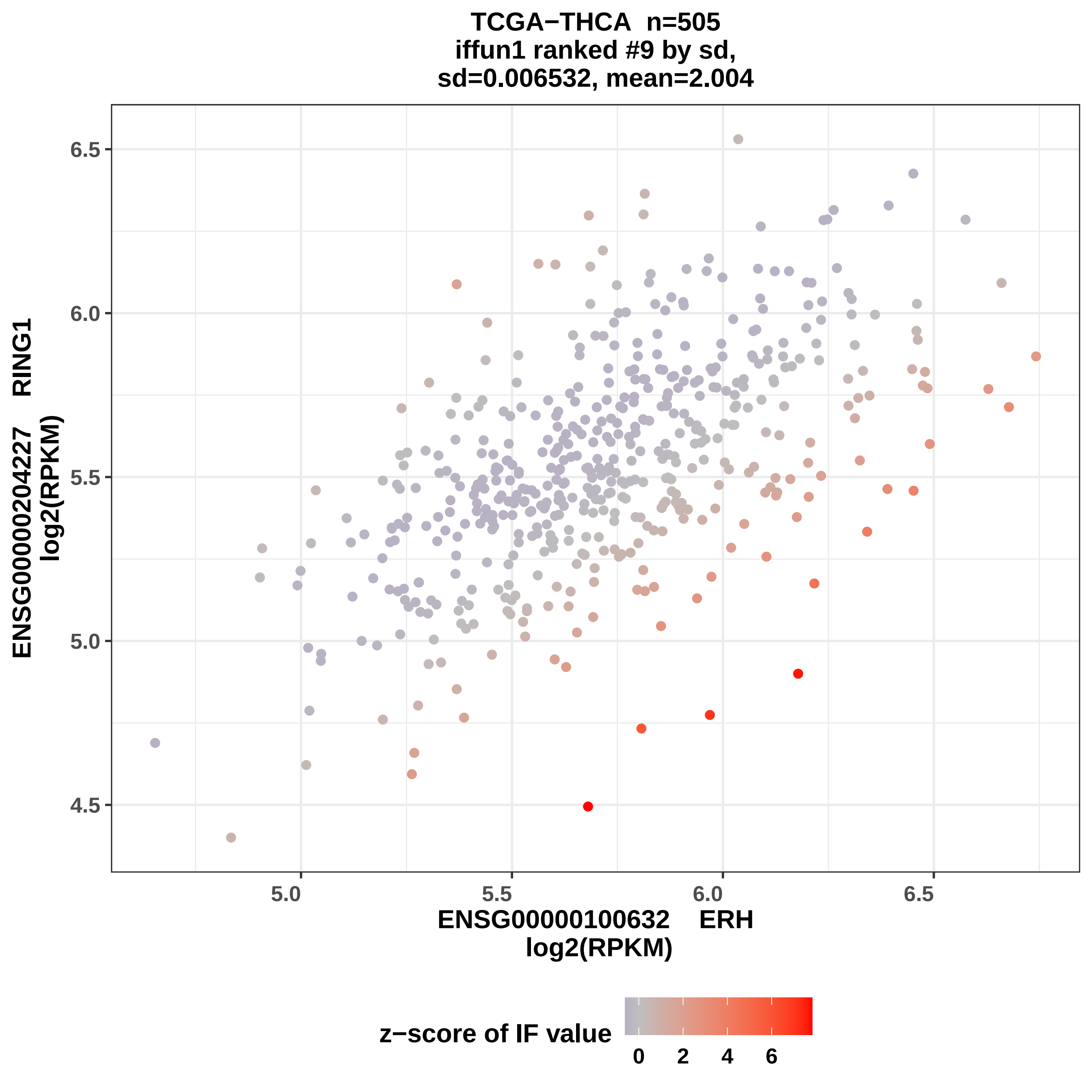
TCGA−THCA n=505
iffun1 ranked #9 by sd,
sd=0.006532, mean=2.004
ENSG00000204227 RING1
log2(RPKM)
6.5
6.0
5.5
5.0
4.5
5.0
5.5
6.0
6.5
ENSG00000100632 ERH
log2(RPKM)
z−score of IF value
0
2
4
6

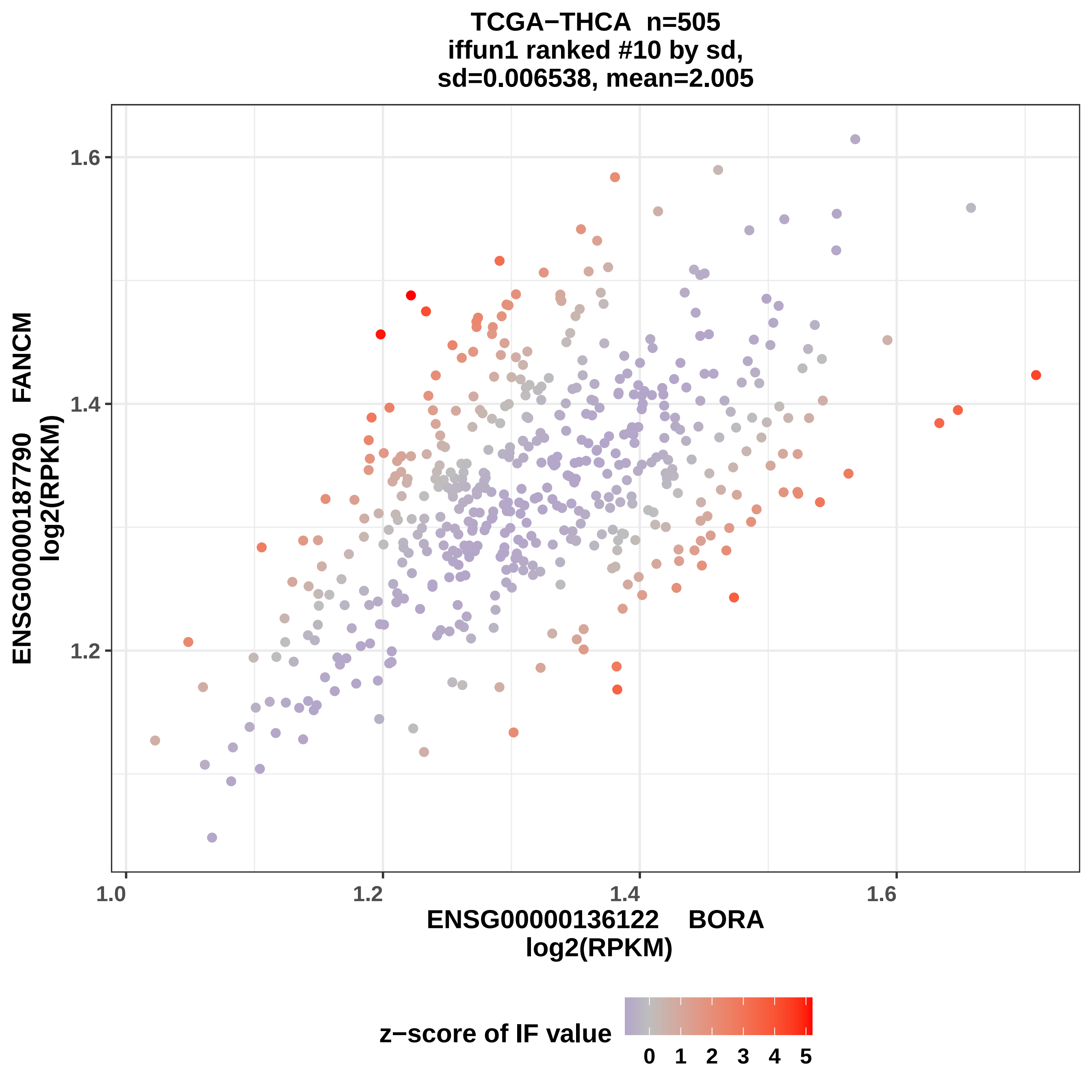

TCGA−THCA n=505
iffun1 ranked #10 by sd,
sd=0.006538, mean=2.005
ENSG00000187790 FANCM
log2(RPKM)
1.6
1.4
1.2
1.0
1.2
1.4
1.6
ENSG00000136122 BORA
log2(RPKM)
z−score of IF value
0 1 2 3 4 5

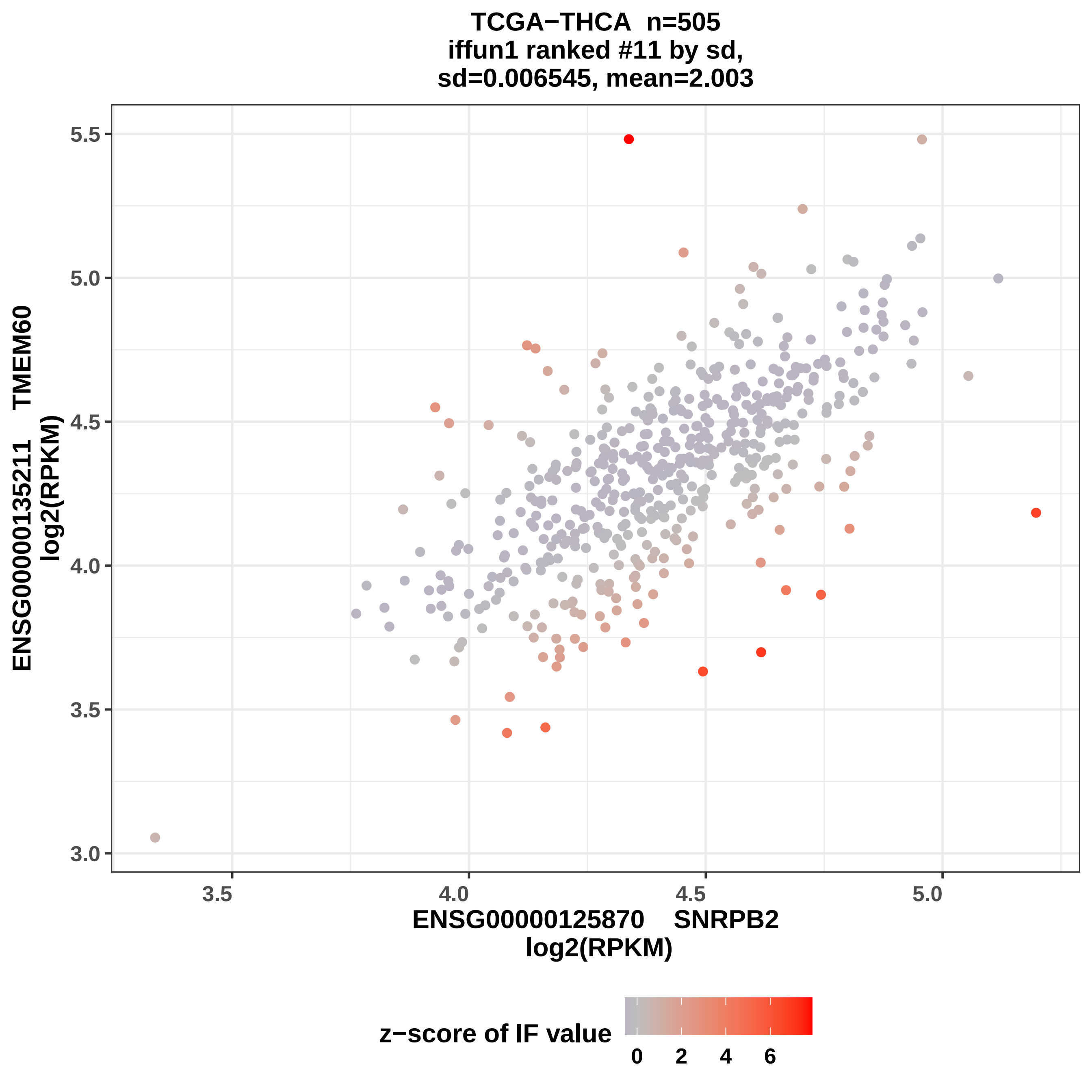

TCGA−THCA n=505
iffun1 ranked #11 by sd,
sd=0.006545, mean=2.003
ENSG00000135211 TMEM60
log2(RPKM)
5.5
5.0
4.5
4.0
3.5
3.0
3.5
4.0
4.5
5.0
ENSG00000125870 SNRPB2
log2(RPKM)
z−score of IF value
0
2
4
6

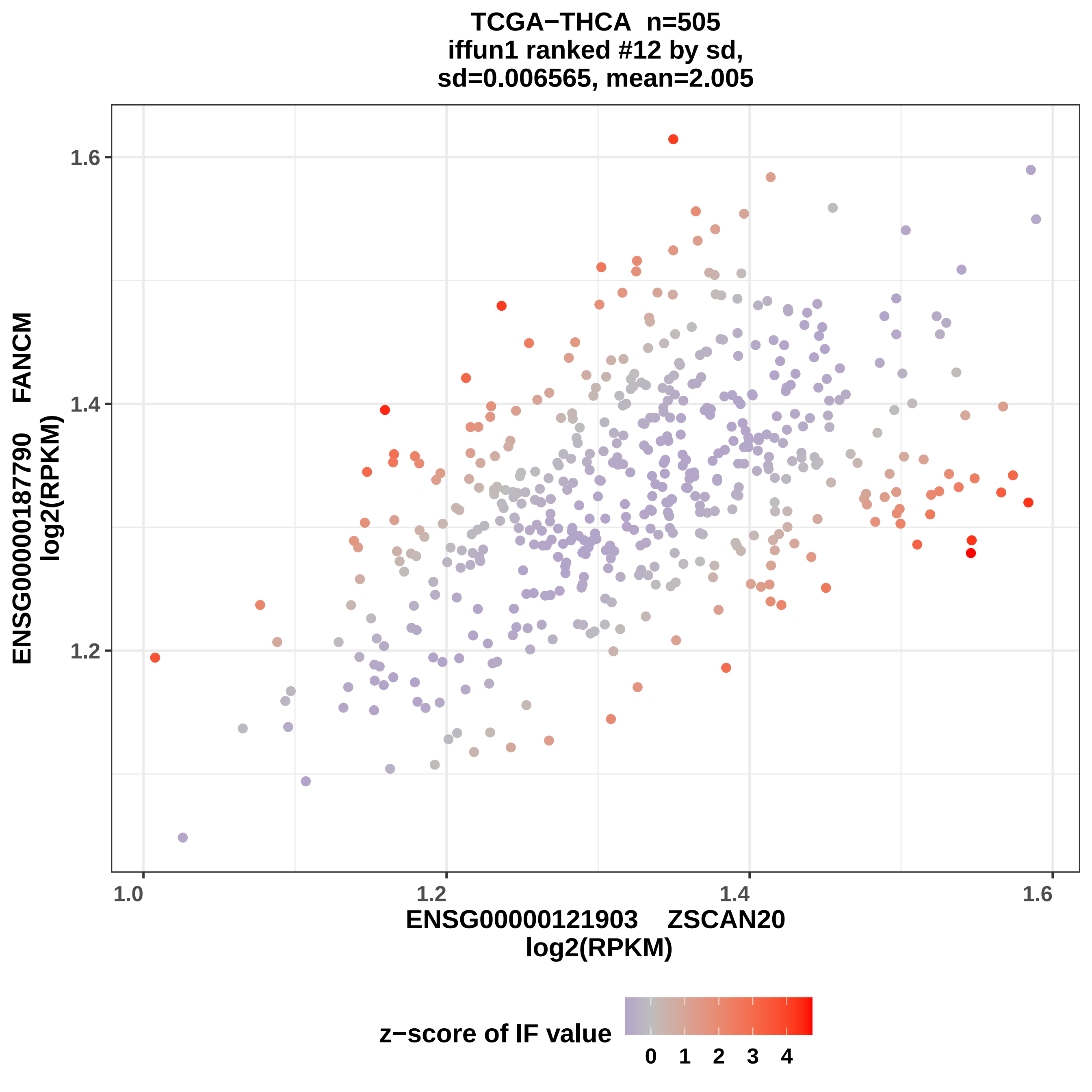

TCGA−THCA n=505
iffun1 ranked #12 by sd,
sd=0.006565, mean=2.005
ENSG00000187790 FANCM
log2(RPKM)
1.6
1.4
1.2
1.0
1.2
1.4
1.6
ENSG00000121903 ZSCAN20
log2(RPKM)
z−score of IF value
0 1 2 3 4

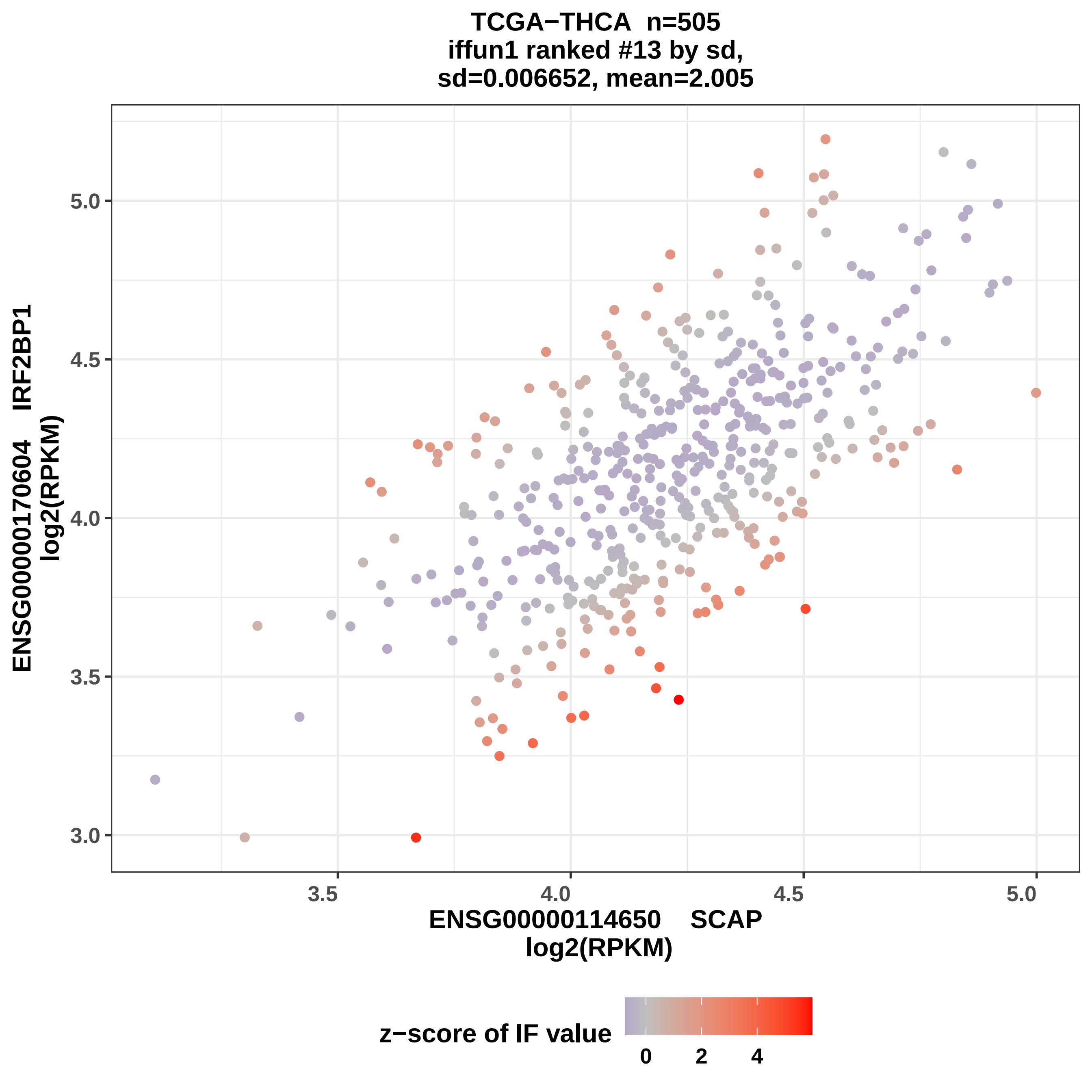

TCGA−THCA n=505
iffun1 ranked #13 by sd,
sd=0.006652, mean=2.005
ENSG00000170604 IRF2BP1
log2(RPKM)
5.0
4.5
4.0
3.5
3.0
3.5
4.0
4.5
5.0
ENSG00000114650 SCAP
log2(RPKM)
z−score of IF value
0
2
4

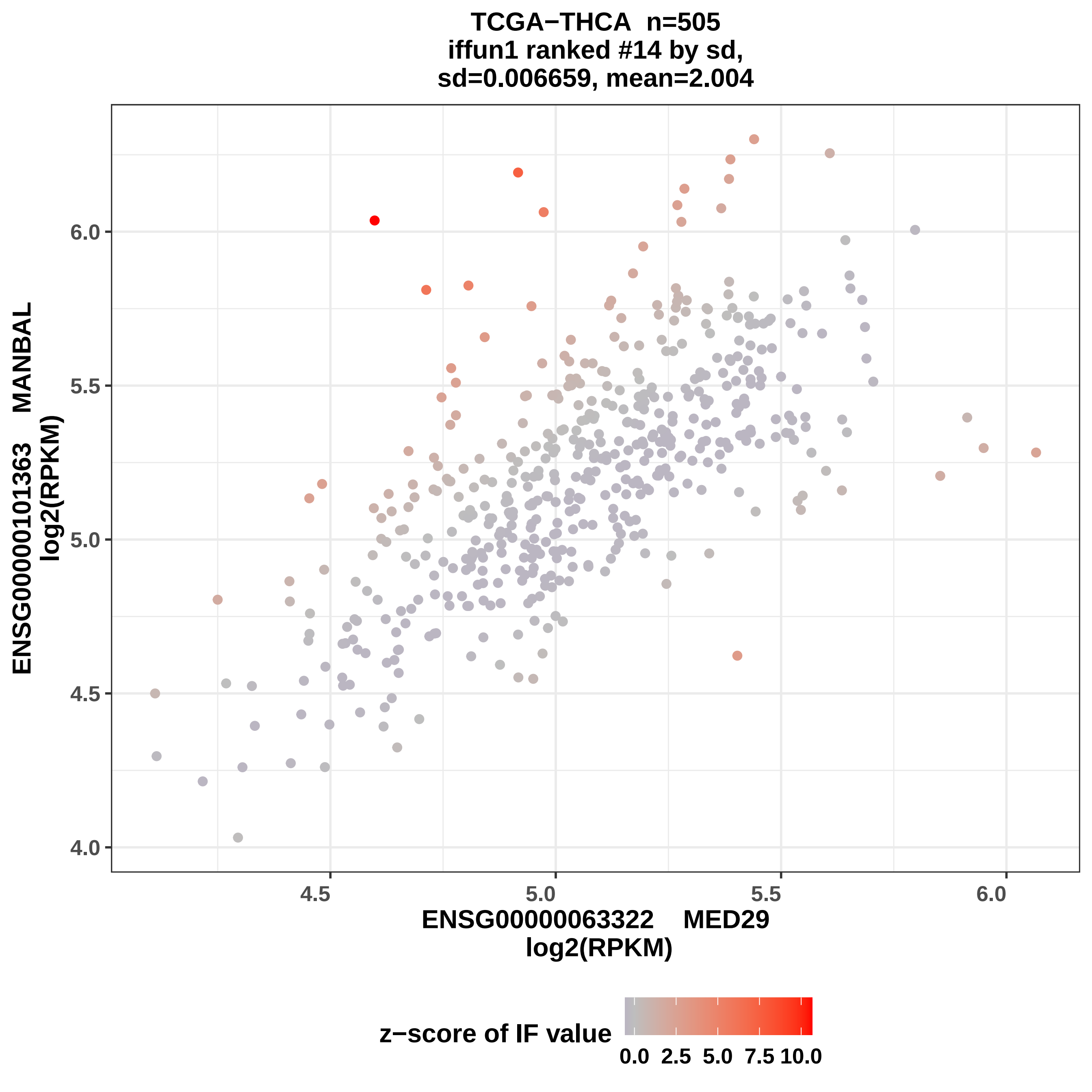
TCGA−THCA n=505
iffun1 ranked #14 by sd,
sd=0.006659, mean=2.004
ENSG00000101363 MANBAL
log2(RPKM)
6.0
5.5
5.0
4.5
4.0
4.5
5.0
5.5
6.0
ENSG00000063322 MED29
log2(RPKM)
z−score of IF value
0.0 2.5 5.0 7.5 10.0

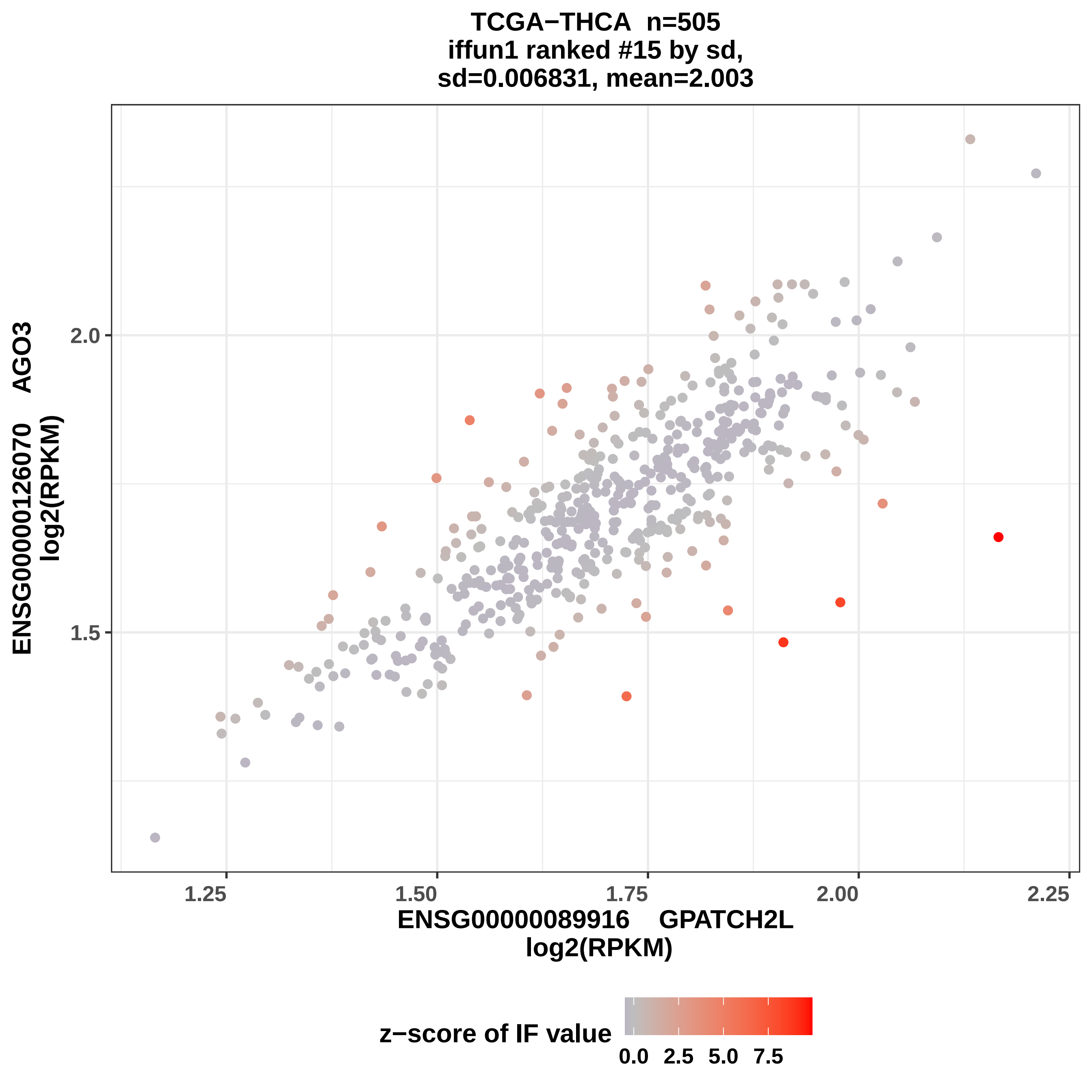

TCGA−THCA n=505
iffun1 ranked #15 by sd,
sd=0.006831, mean=2.003
ENSG00000126070 AGO3
log2(RPKM)
2.0
1.5
1.25
1.50
1.75
2.00
2.25
ENSG00000089916 GPATCH2L
log2(RPKM)
z−score of IF value
0.0
2.5
5.0
7.5

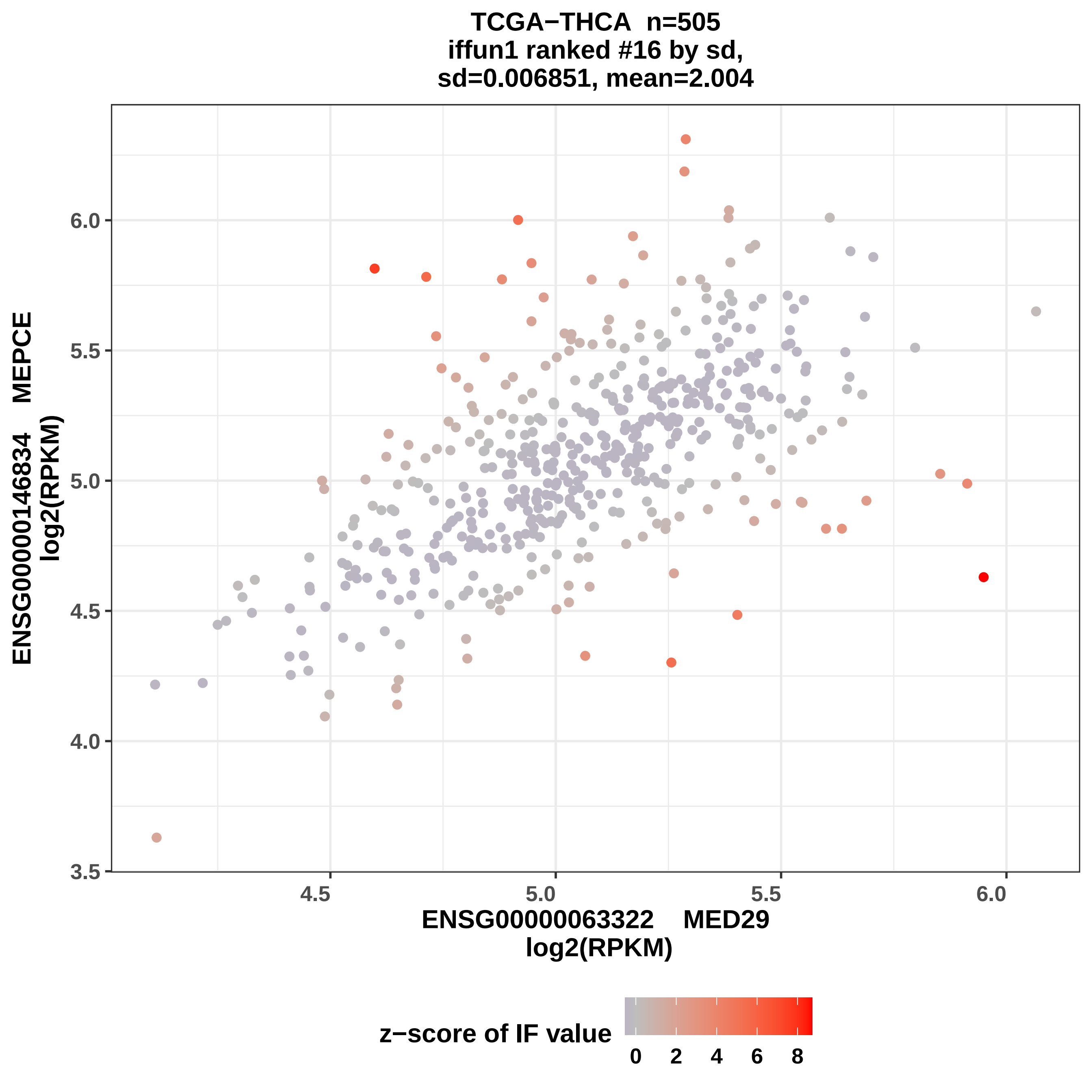
TCGA−THCA n=505
iffun1 ranked #16 by sd,
sd=0.006851, mean=2.004
ENSG00000146834 MEPCE
log2(RPKM)
3.5
4.0
4.5
5.0
5.5
6.0
4.5
5.0
5.5
6.0
ENSG00000063322 MED29
log2(RPKM)
z−score of IF value
0
2
4
6
8

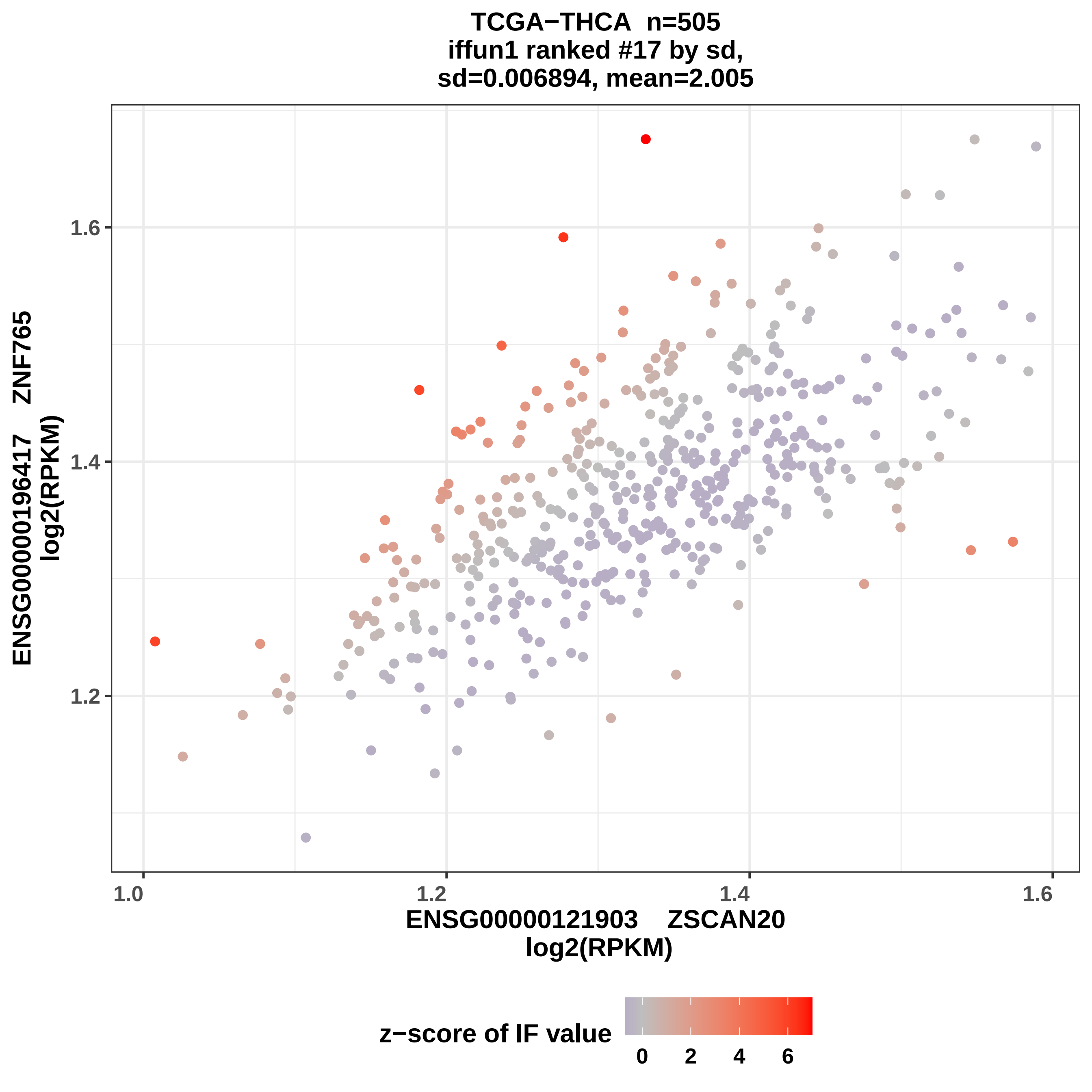

TCGA−THCA n=505
iffun1 ranked #17 by sd,
sd=0.006894, mean=2.005
ENSG00000196417 ZNF765
log2(RPKM)
1.6
1.4
1.2
1.0
1.2
1.4
1.6
ENSG00000121903 ZSCAN20
log2(RPKM)
z−score of IF value
0
2
4
6

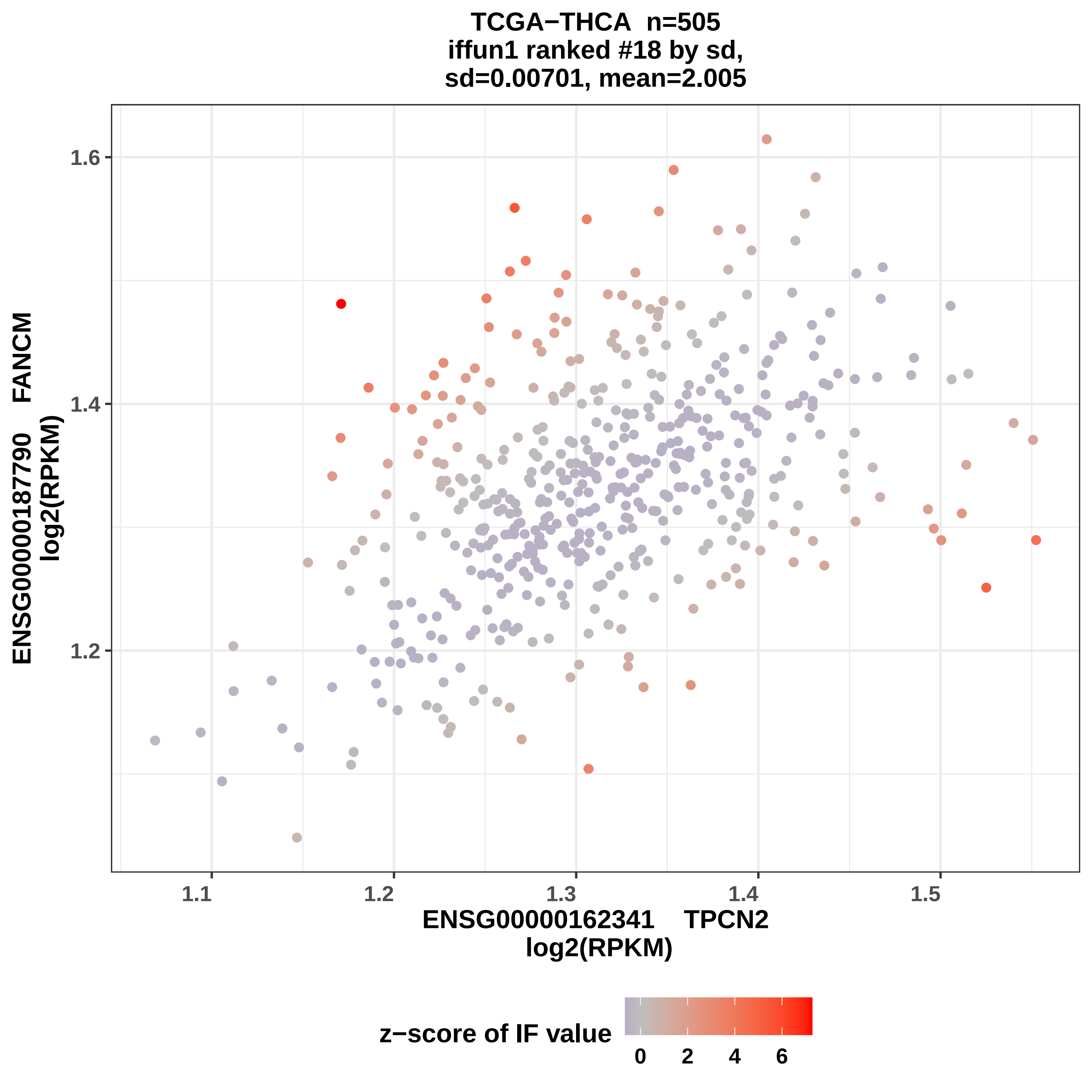
TCGA−THCA n=505
iffun1 ranked #18 by sd,
sd=0.00701, mean=2.005
1.6
1.4
1.2
ENSG00000187790 FANCM
log2(RPKM)
1.1
1.2
1.3
1.4
1.5
ENSG00000162341 TPCN2
log2(RPKM)
z−score of IF value
0
2
4
6

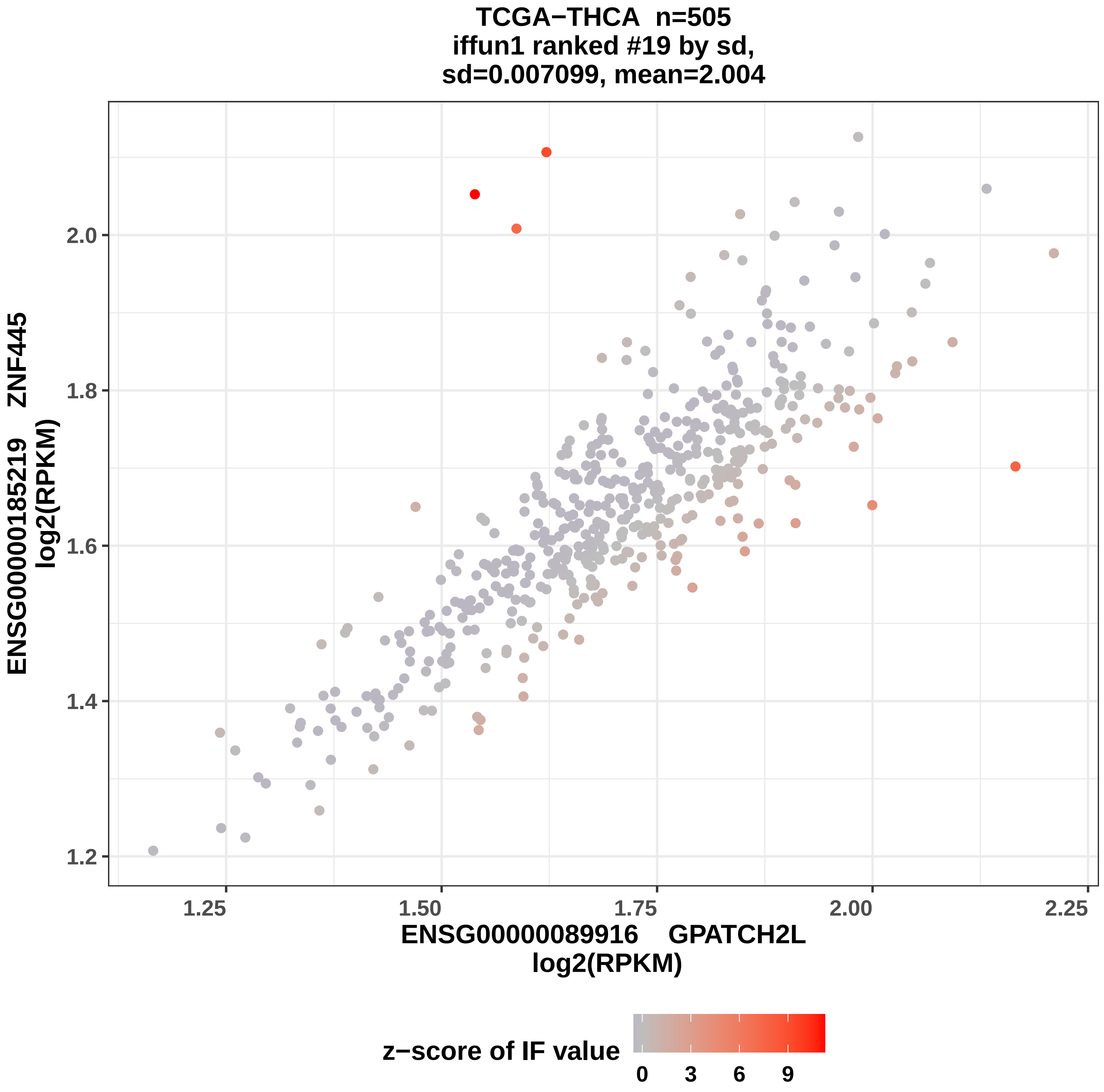
TCGA−THCA n=505
iffun1 ranked #19 by sd,
sd=0.007099, mean=2.004
ENSG00000185219 ZNF445
log2(RPKM)
2.0
1.8
1.6
1.4
1.2
1.25
1.50
1.75
2.00
2.25
ENSG00000089916 GPATCH2L
log2(RPKM)
z−score of IF value
0
3
6
9

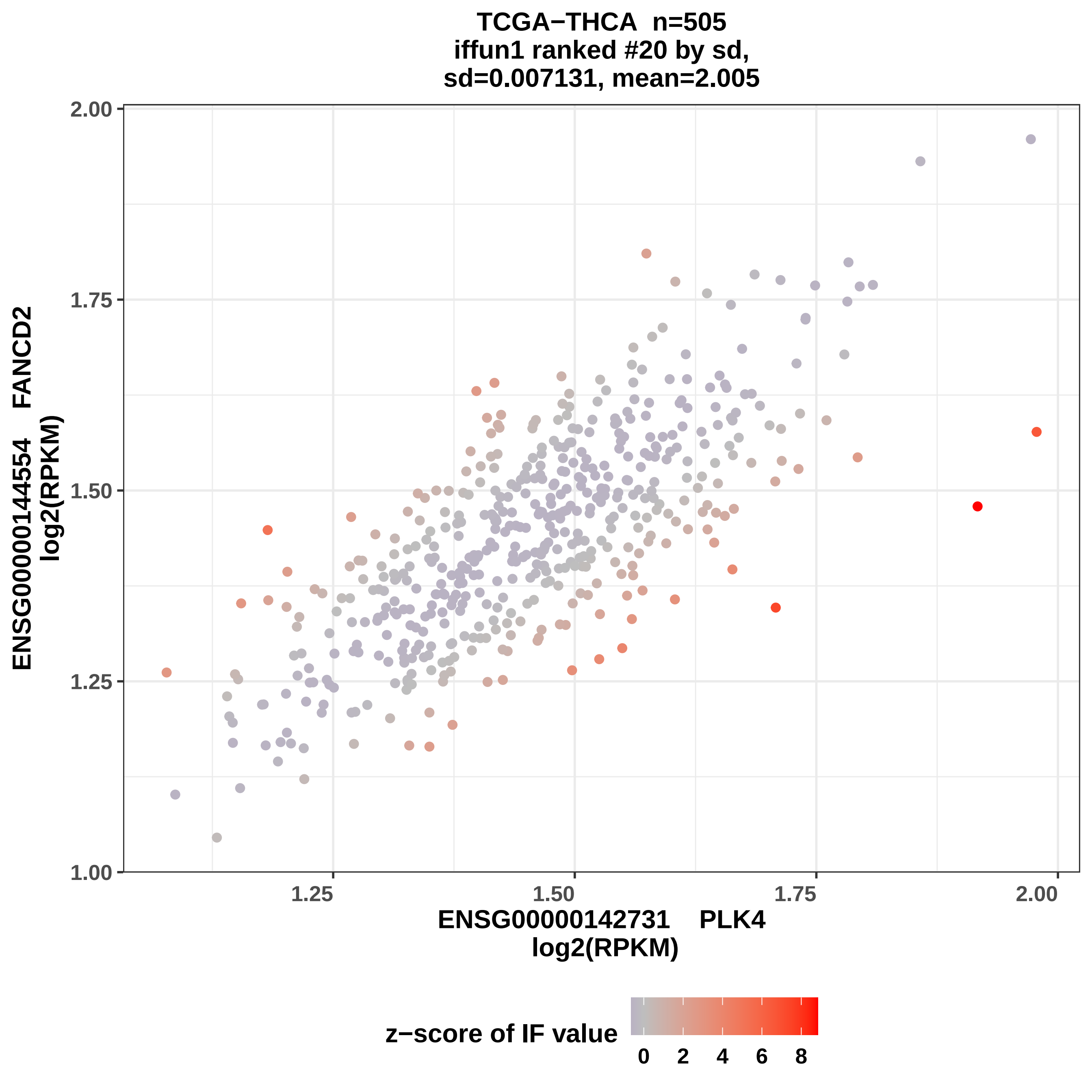
TCGA−THCA n=505
iffun1 ranked #20 by sd,
sd=0.007131, mean=2.005
ENSG00000144554 FANCD2
log2(RPKM)
2.00
1.75
1.50
1.25
1.00
1.25
1.50
1.75
2.00
ENSG00000142731 PLK4
log2(RPKM)
z−score of IF value
0
2
4
6
8

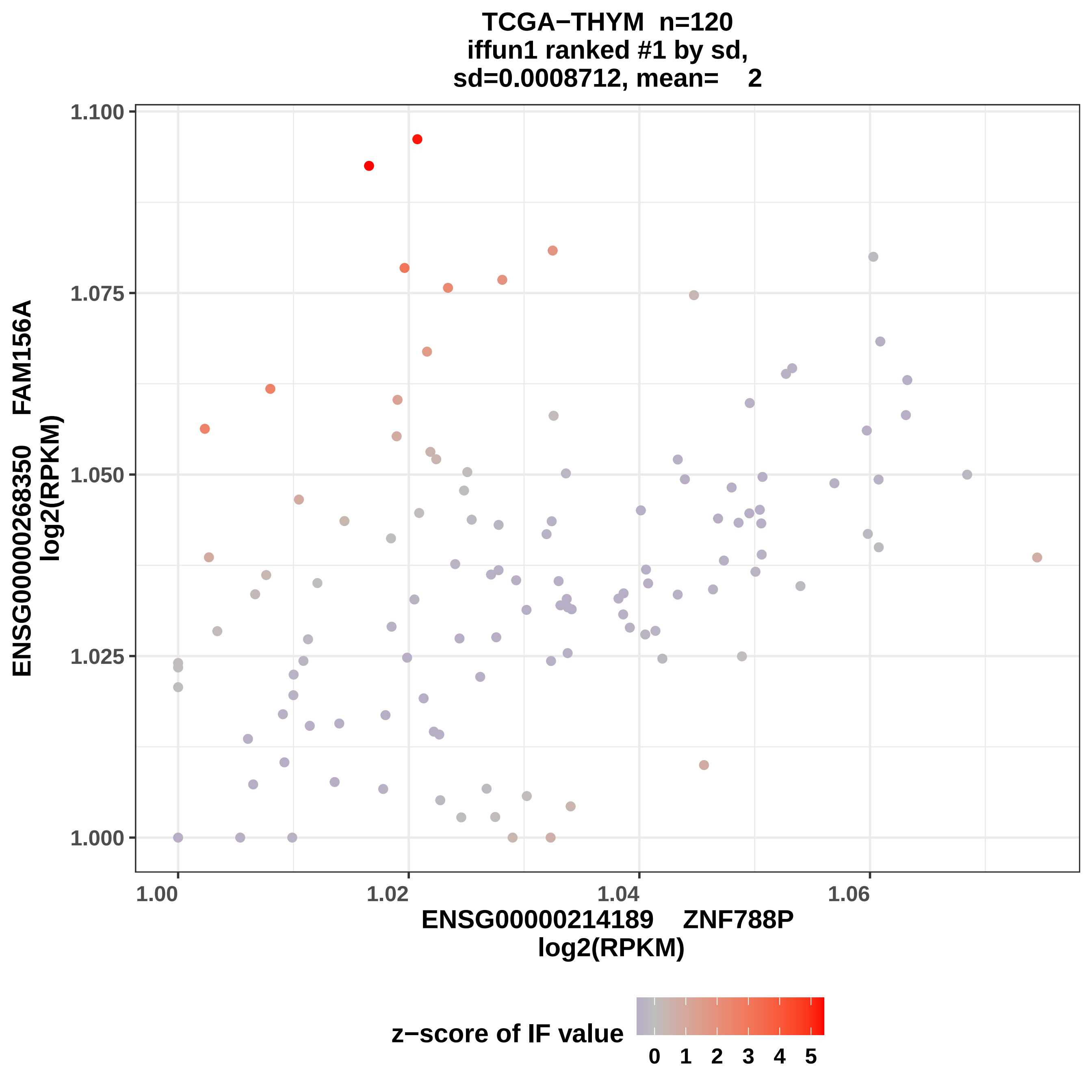
TCGA−THYM n=120
iffun1 ranked #1 by sd,
sd=0.0008712, mean= 2
ENSG00000268350 FAM156A
log2(RPKM)
1.100
1.075
1.050
1.025
1.000
1.00
1.02
1.04
1.06
ENSG00000214189 ZNF788P
log2(RPKM)
z−score of IF value
0 1 2 3 4 5

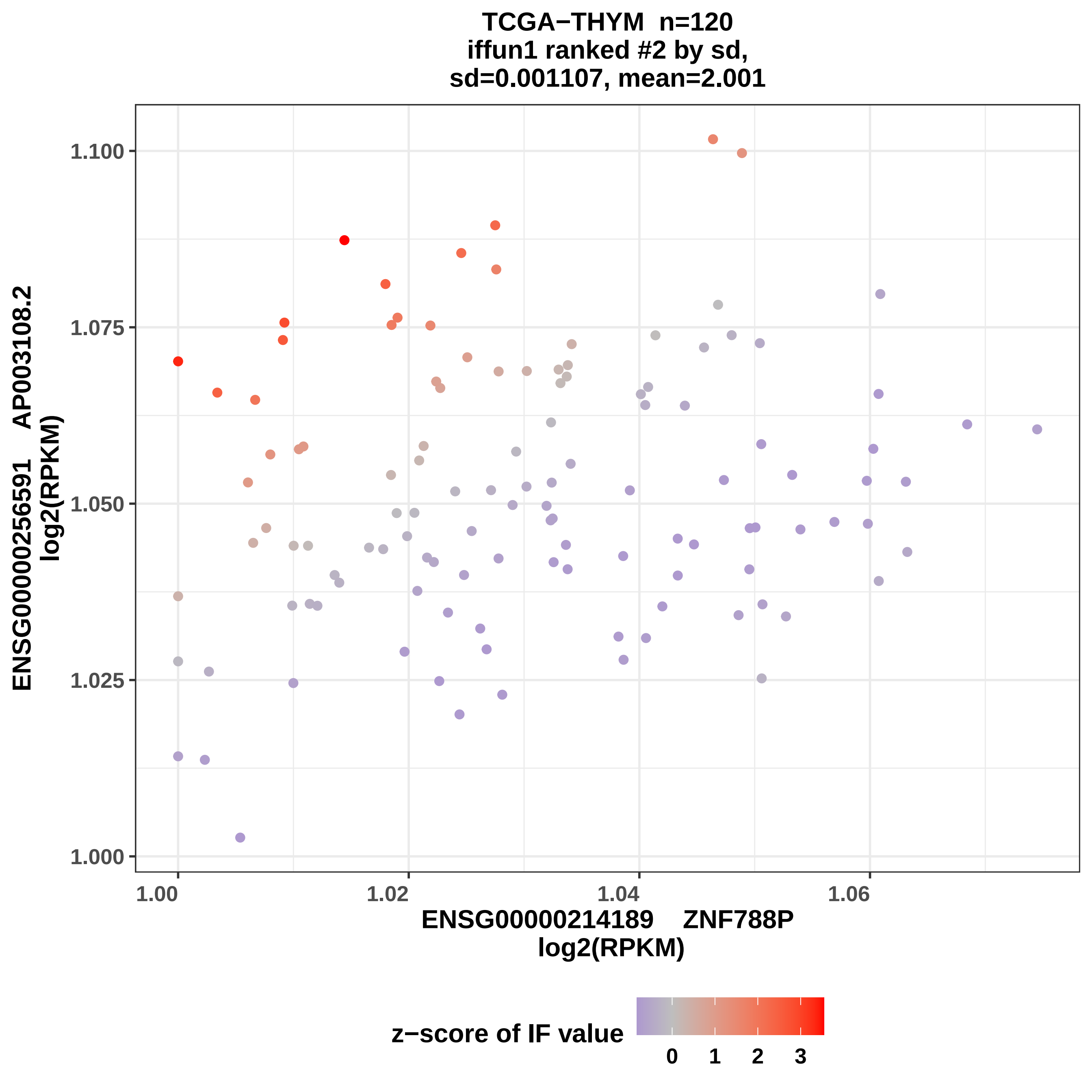

TCGA−THYM n=120
iffun1 ranked #2 by sd,
sd=0.001107, mean=2.001
ENSG00000256591 AP003108.2
log2(RPKM)
1.100
1.075
1.050
1.025
1.000
1.00
1.02
1.04
1.06
ENSG00000214189 ZNF788P
log2(RPKM)
z−score of IF value
0
1
2
3

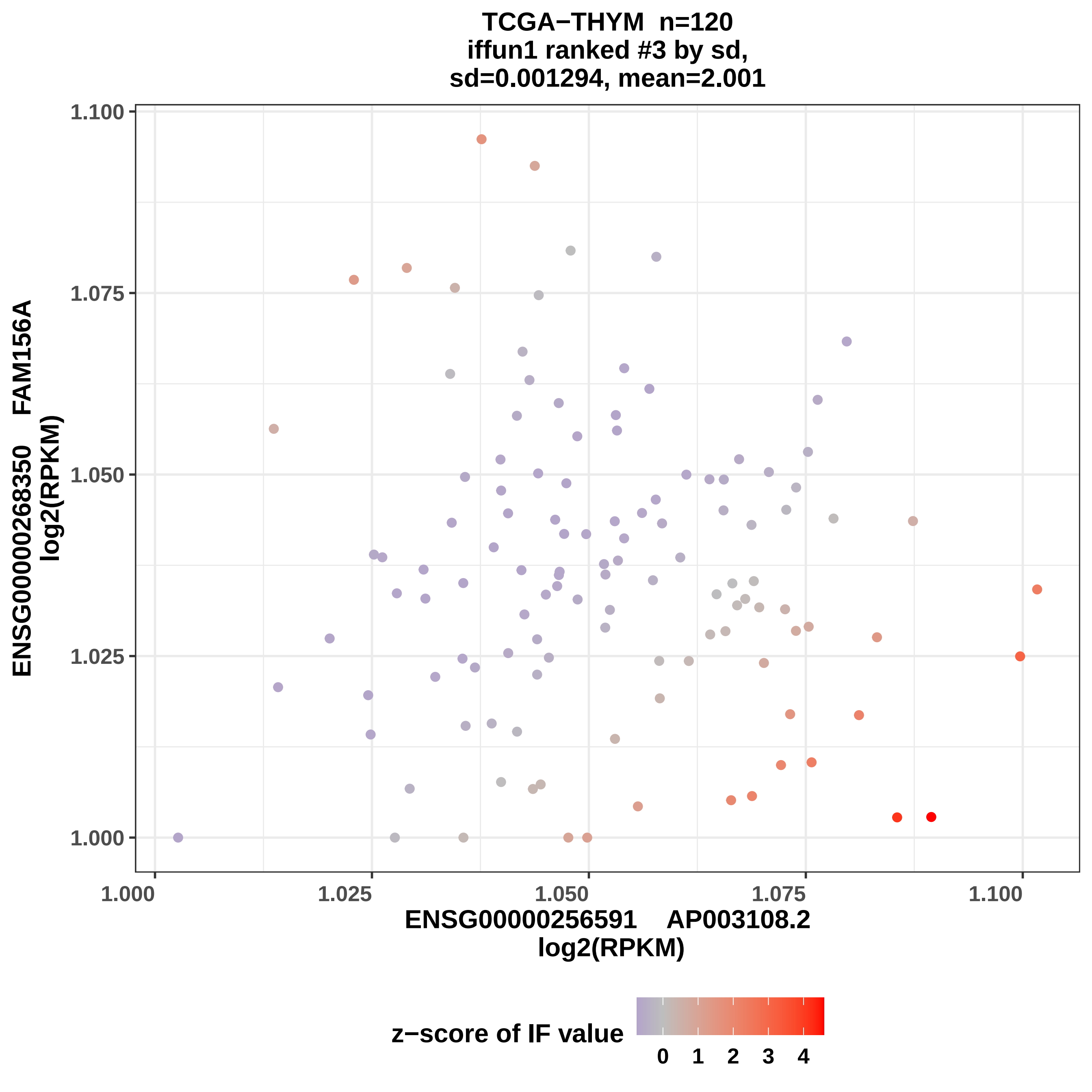

TCGA−THYM n=120
iffun1 ranked #3 by sd,
sd=0.001294, mean=2.001
ENSG00000268350 FAM156A
log2(RPKM)
1.100
1.075
1.050
1.025
1.000
1.000
1.025
1.050
1.075
1.100
ENSG00000256591 AP003108.2
log2(RPKM)
z−score of IF value
0
1
2
3
4

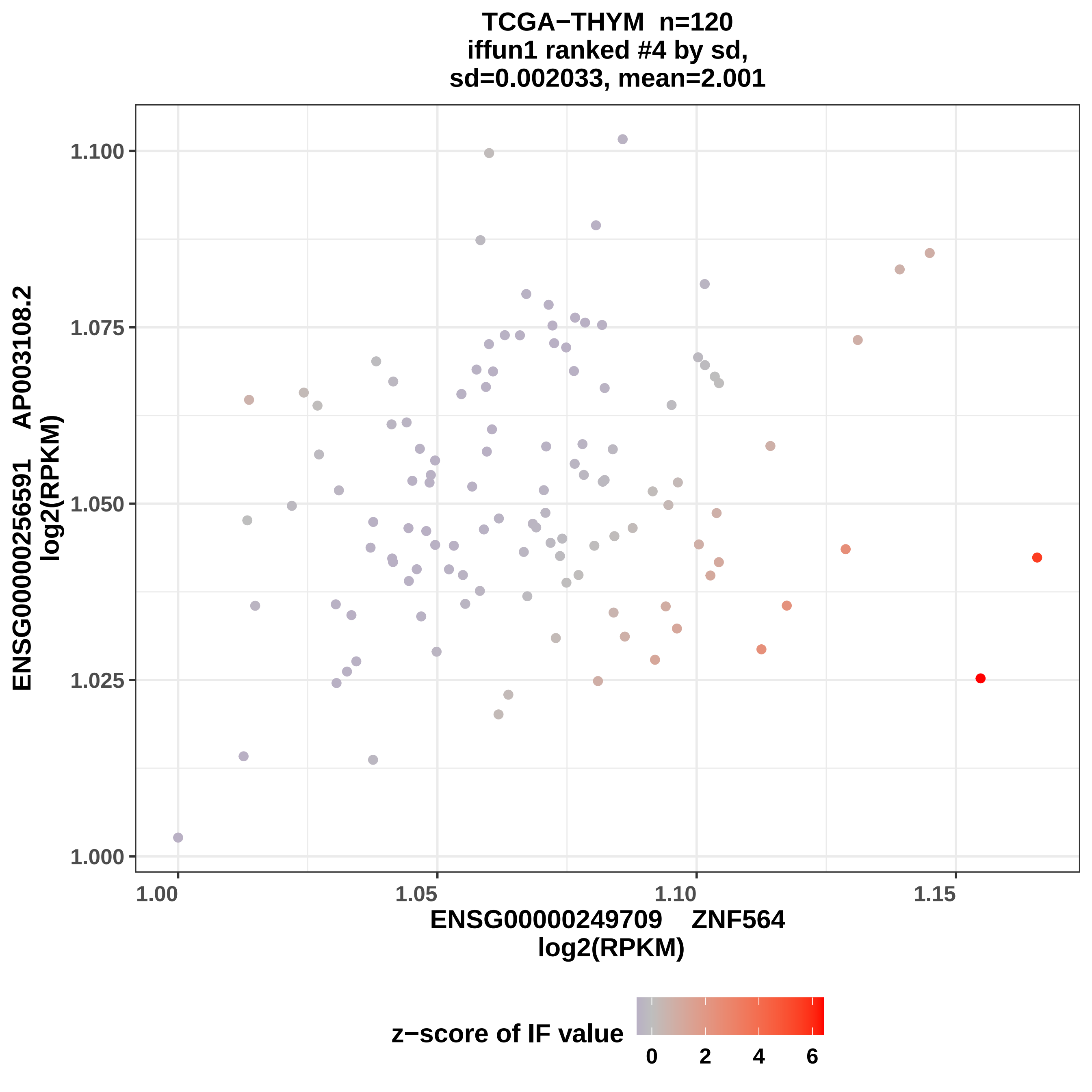
TCGA−THYM n=120
iffun1 ranked #4 by sd,
sd=0.002033, mean=2.001
ENSG00000256591 AP003108.2
log2(RPKM)
1.100
1.075
1.050
1.025
1.000
1.00
1.05
1.10
1.15
ENSG00000249709 ZNF564
log2(RPKM)
z−score of IF value
0
2
4
6

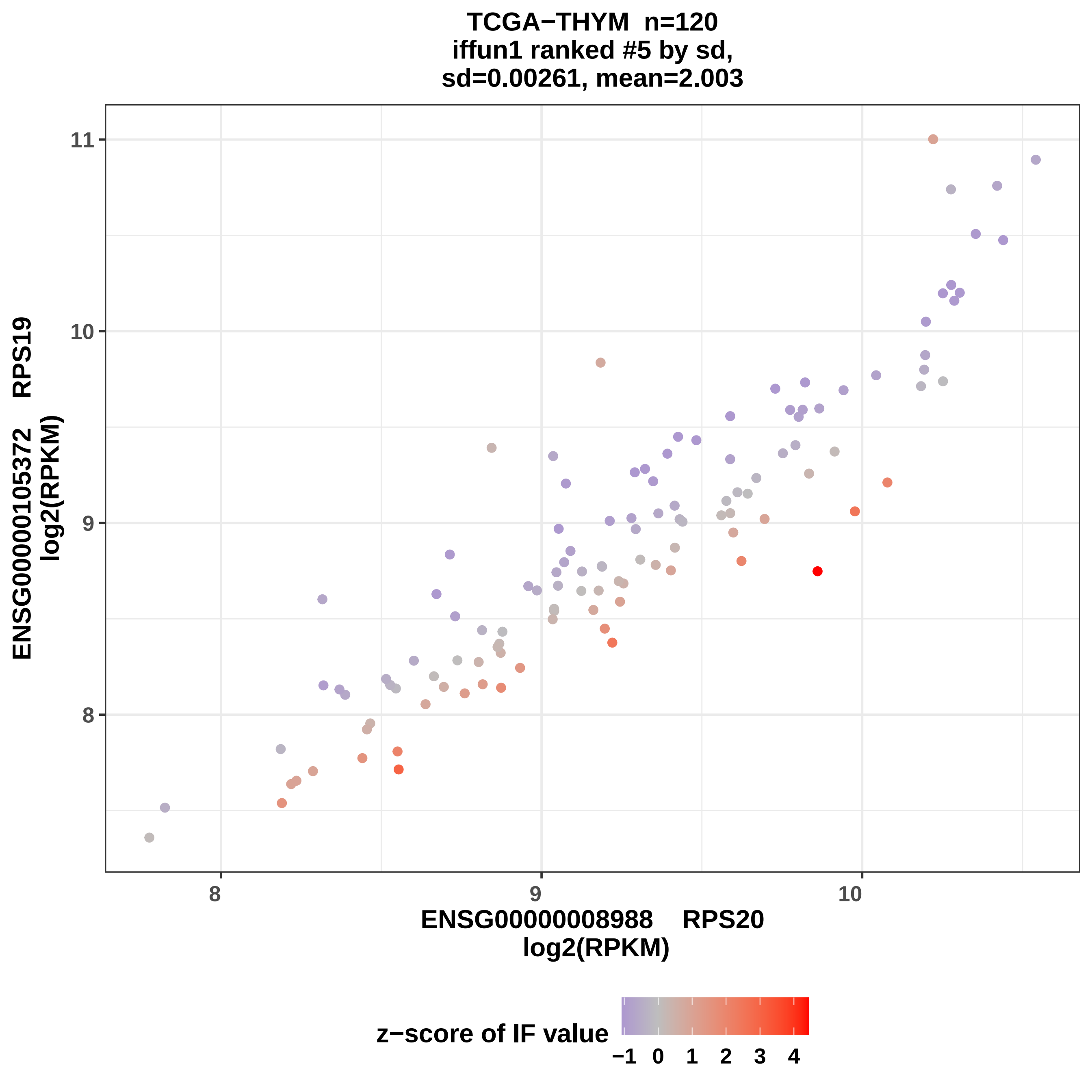

TCGA−THYM n=120
iffun1 ranked #5 by sd,
sd=0.00261, mean=2.003
ENSG00000105372 RPS19
log2(RPKM)
ENSG00000008988 RPS20
log2(RPKM)
11
10
9
8
8
9
10
z−score of IF value
−1
0
1
2
3
4

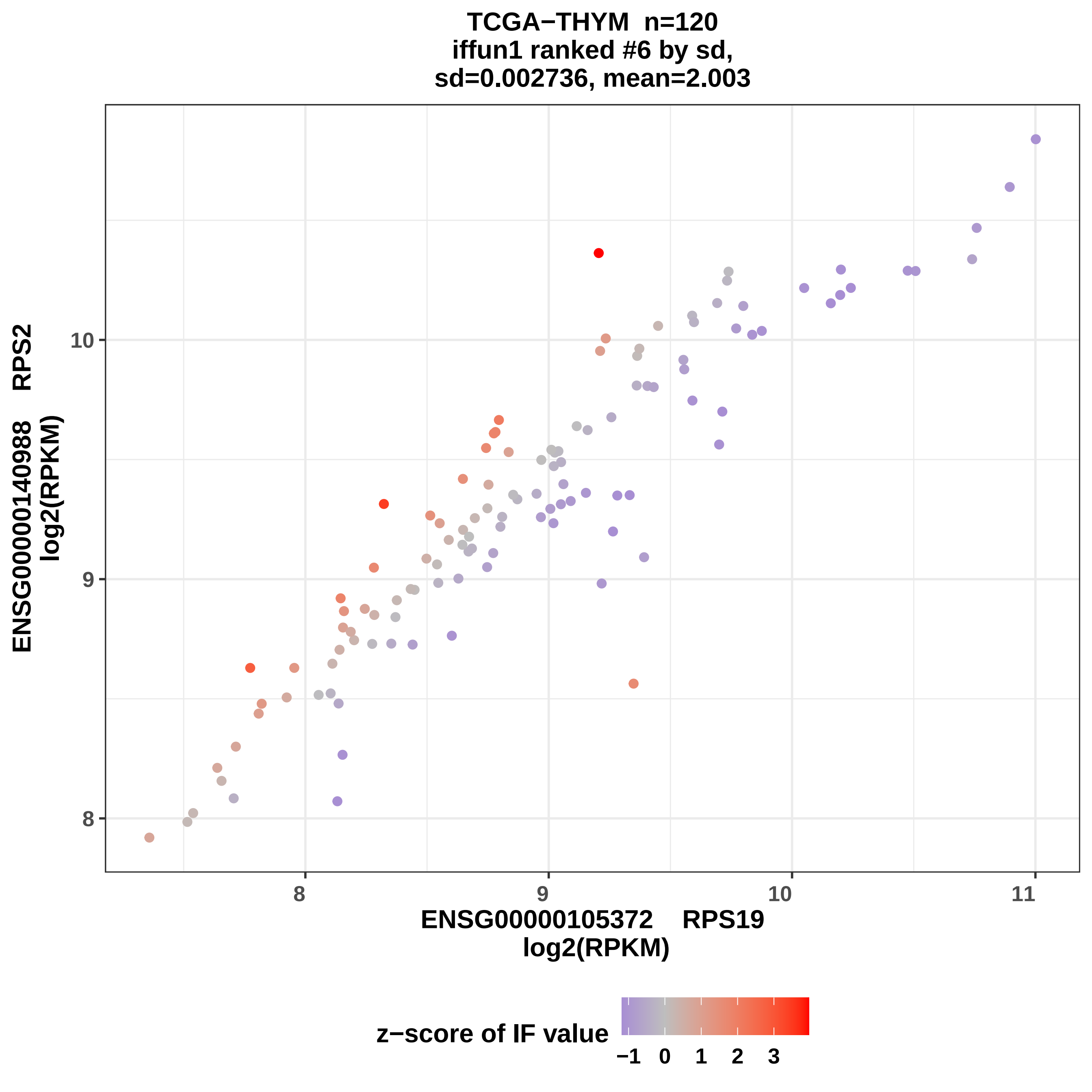

TCGA−THYM n=120
iffun1 ranked #6 by sd,
sd=0.002736, mean=2.003
ENSG00000140988 RPS2
log2(RPKM)
10
9
8
8
9
10
11
ENSG00000105372 RPS19
log2(RPKM)
z−score of IF value
−1 0 1 2 3

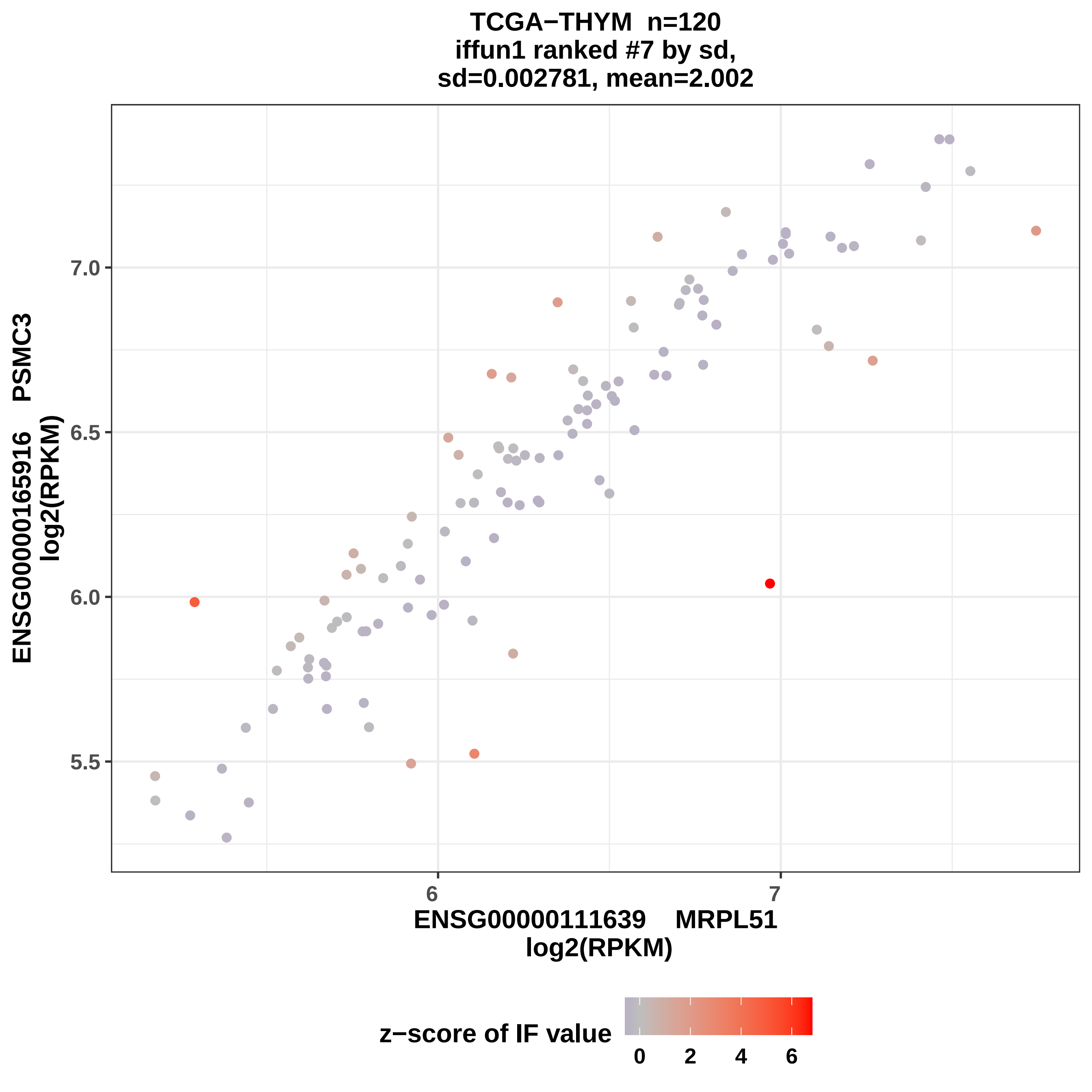
TCGA−THYM n=120
iffun1 ranked #7 by sd,
sd=0.002781, mean=2.002
ENSG00000165916 PSMC3
log2(RPKM)
7.0
6.5
6.0
5.5
6
7
ENSG00000111639 MRPL51
log2(RPKM)
z−score of IF value
0
2
4
6

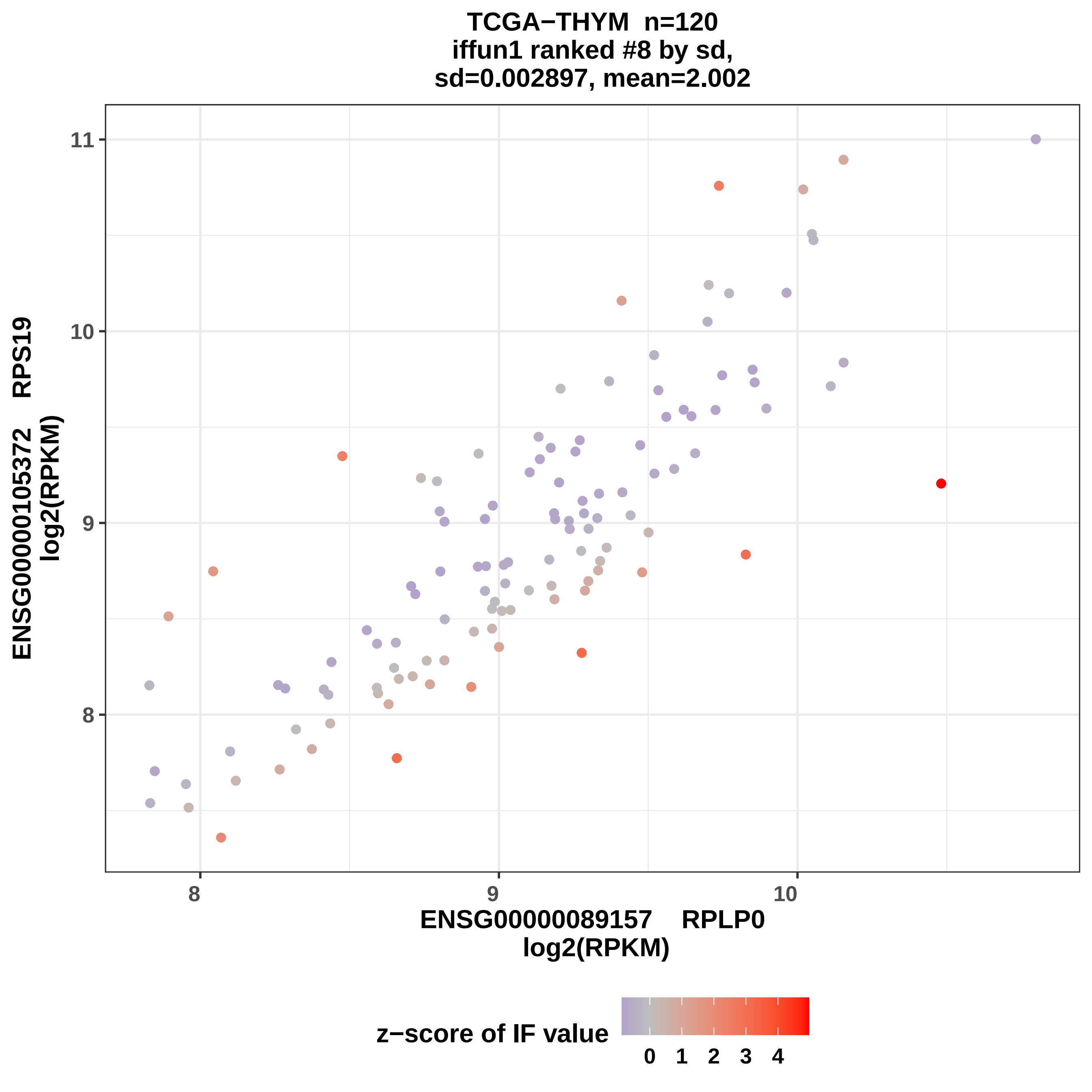

TCGA−THYM n=120
iffun1 ranked #8 by sd,
sd=0.002897, mean=2.002
11
10
9
8
ENSG00000105372 RPS19
log2(RPKM)
8
9
10
ENSG00000089157 RPLP0
log2(RPKM)
z−score of IF value
0
1
2
3
4

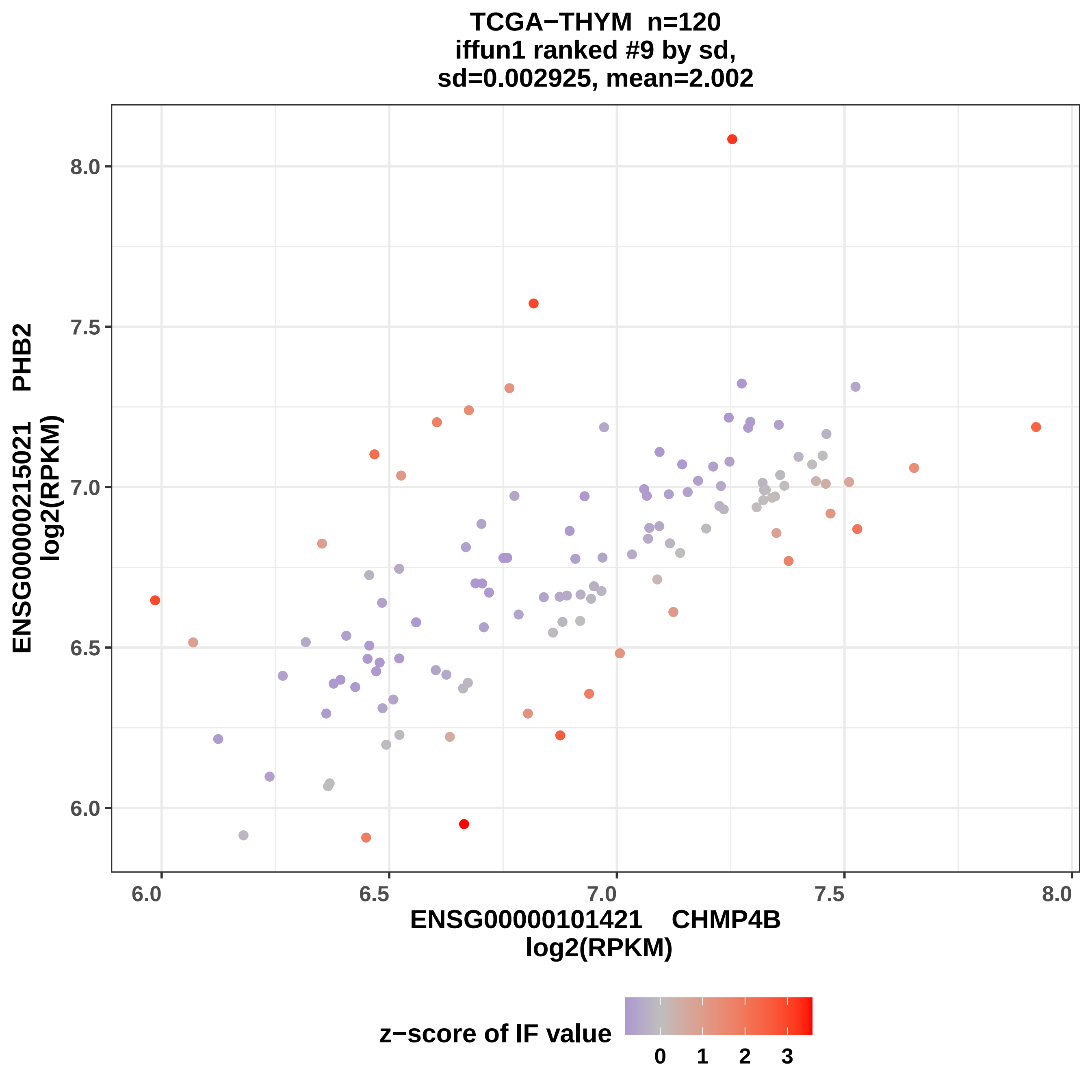
TCGA−THYM n=120
iffun1 ranked #9 by sd,
sd=0.002925, mean=2.002
ENSG00000215021 PHB2
log2(RPKM)
8.0
7.5
7.0
6.5
6.0
6.0
6.5
7.0
7.5
8.0
ENSG00000101421 CHMP4B
log2(RPKM)
z−score of IF value
0
1
2
3

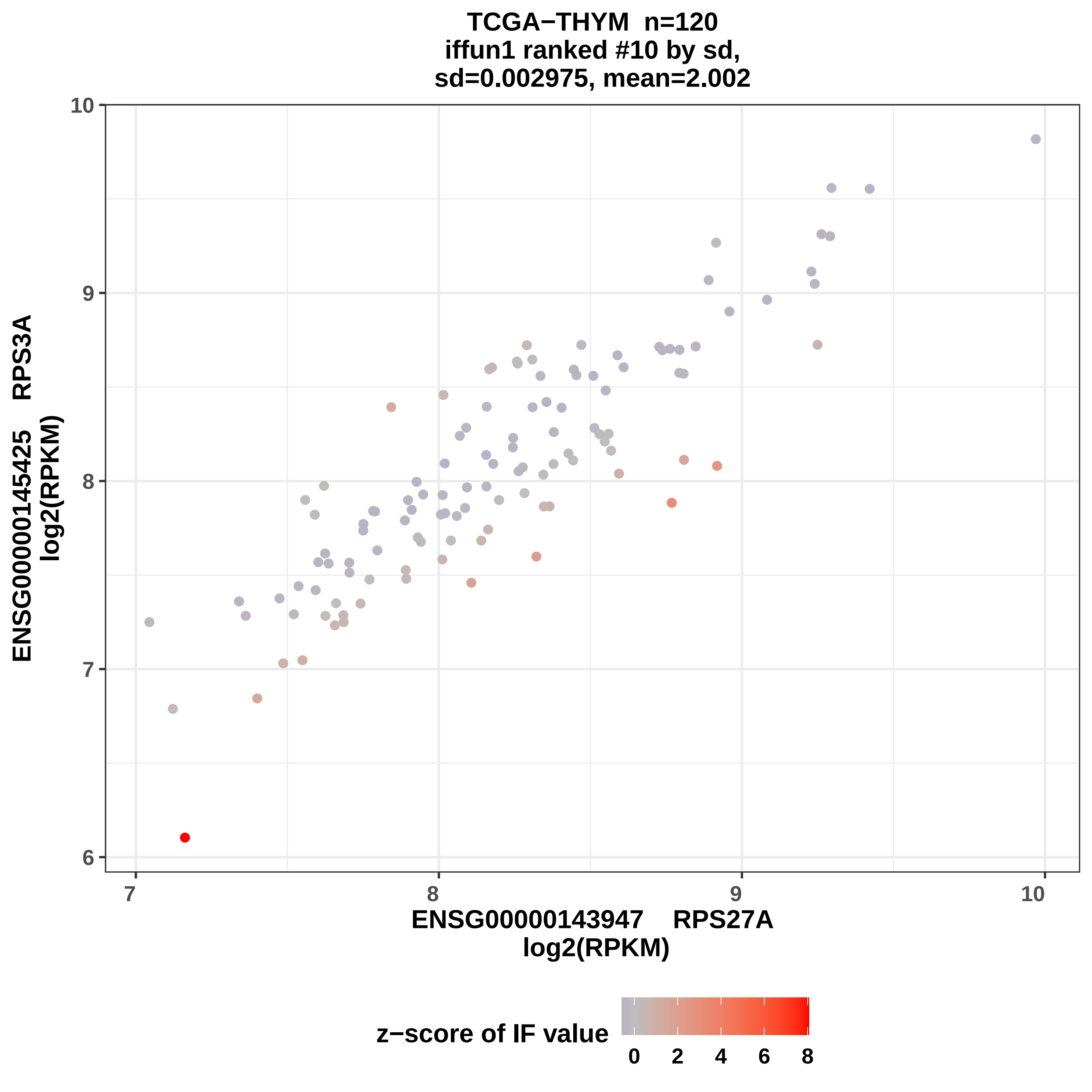

TCGA−THYM n=120
iffun1 ranked #10 by sd,
sd=0.002975, mean=2.002
ENSG00000145425 RPS3A
log2(RPKM)
10
9
8
7
6
7
8
9
10
ENSG00000143947 RPS27A
log2(RPKM)
z−score of IF value
0
2
4
6
8

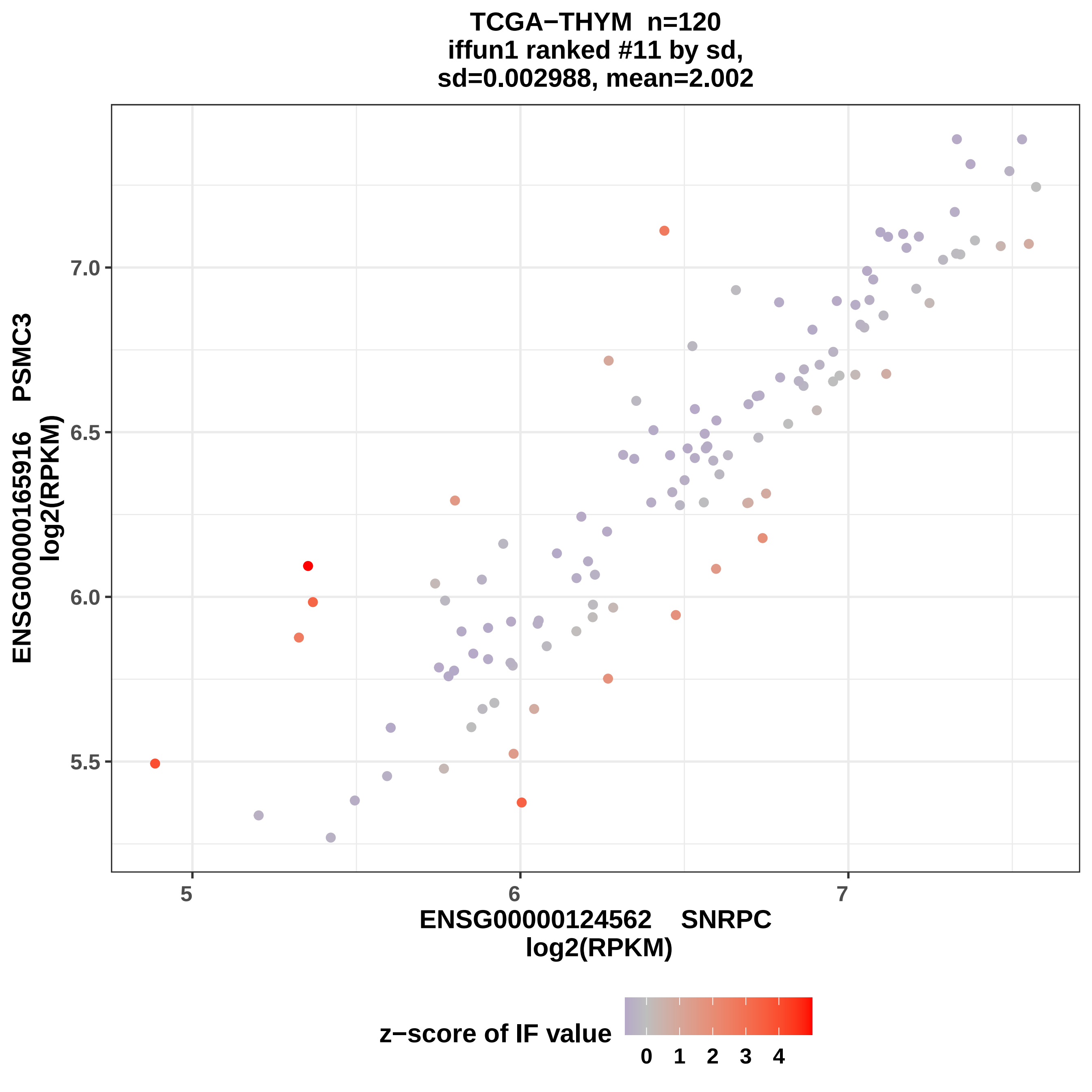
TCGA−THYM n=120
iffun1 ranked #11 by sd,
sd=0.002988, mean=2.002
ENSG00000165916 PSMC3
log2(RPKM)
7.0
6.5
6.0
5.5
5
6
7
ENSG00000124562 SNRPC
log2(RPKM)
z−score of IF value
0 1 2 3 4

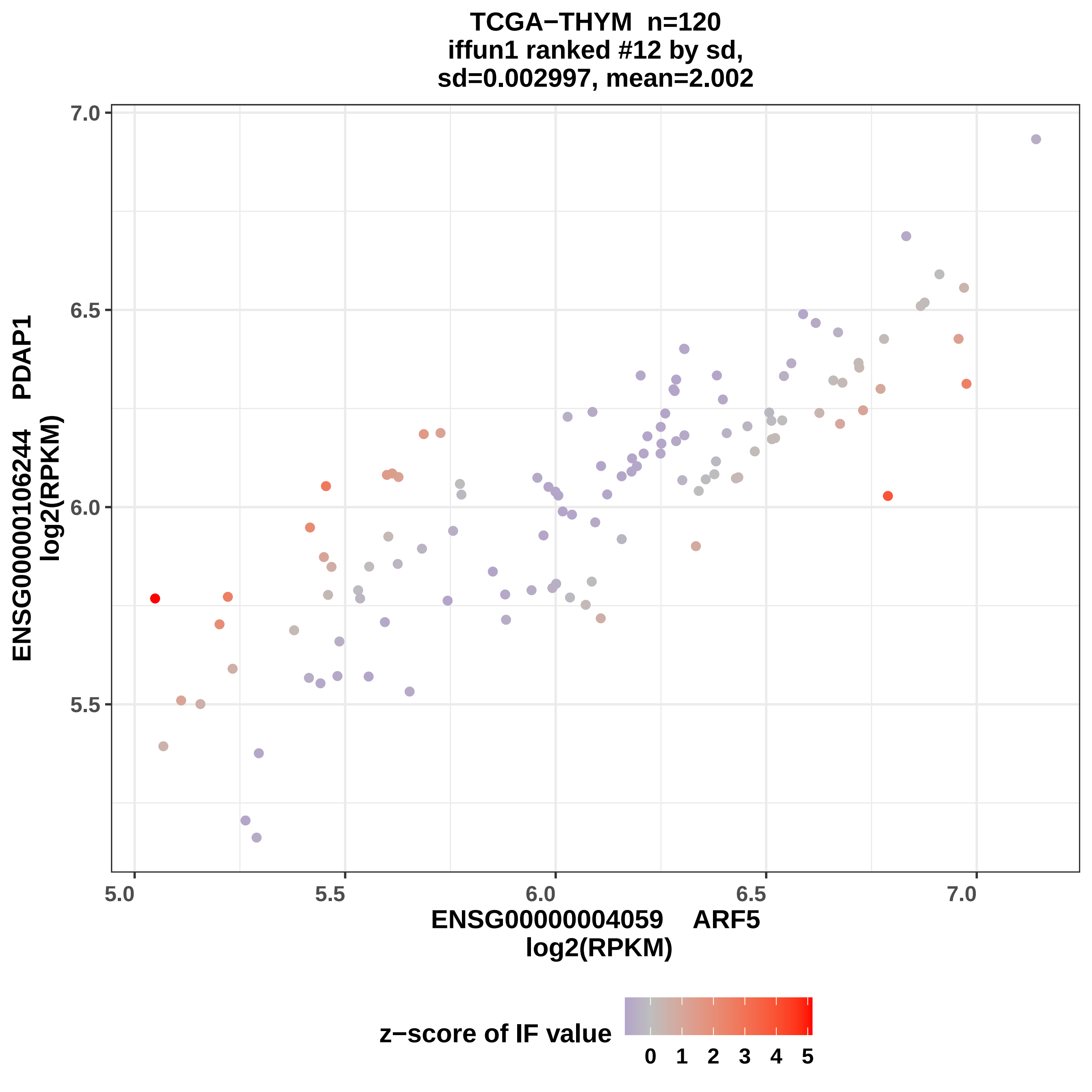
TCGA−THYM n=120
iffun1 ranked #12 by sd,
sd=0.002997, mean=2.002
7.0
6.5
6.0
5.5
ENSG00000106244 PDAP1
log2(RPKM)
5.0
5.5
6.0
6.5
7.0
ENSG00000004059 ARF5
log2(RPKM)
z−score of IF value
0
1
2
3
4
5

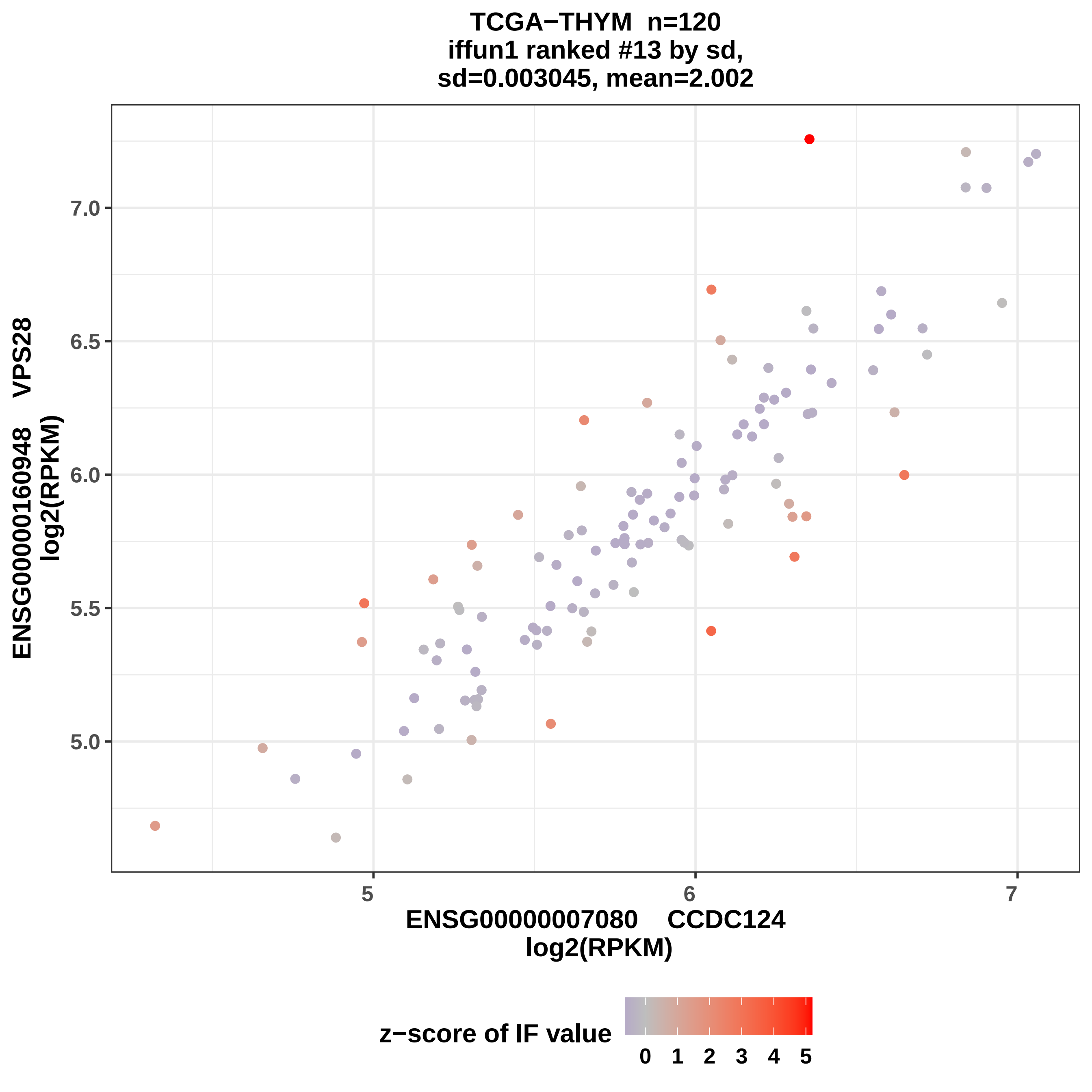
TCGA−THYM n=120
iffun1 ranked #13 by sd,
sd=0.003045, mean=2.002
ENSG00000160948 VPS28
log2(RPKM)
7.0
6.5
6.0
5.5
5.0
5
6
7
ENSG00000007080 CCDC124
log2(RPKM)
z−score of IF value
0
1
2
3
4
5

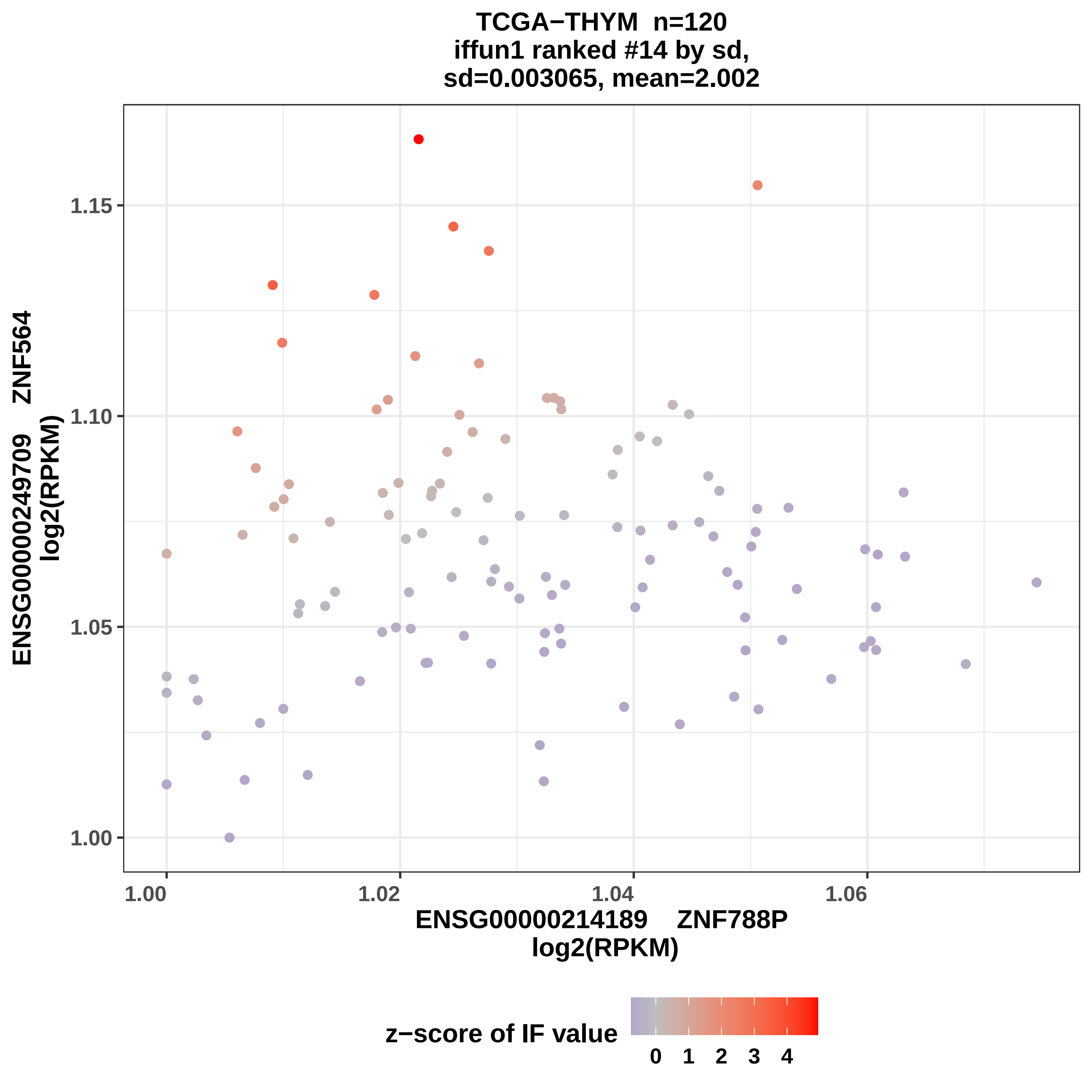

TCGA−THYM n=120
iffun1 ranked #14 by sd,
sd=0.003065, mean=2.002
ENSG00000249709 ZNF564
log2(RPKM)
1.15
1.10
1.05
1.00
1.00
1.02
1.04
1.06
ENSG00000214189 ZNF788P
log2(RPKM)
z−score of IF value
0 1 2 3 4

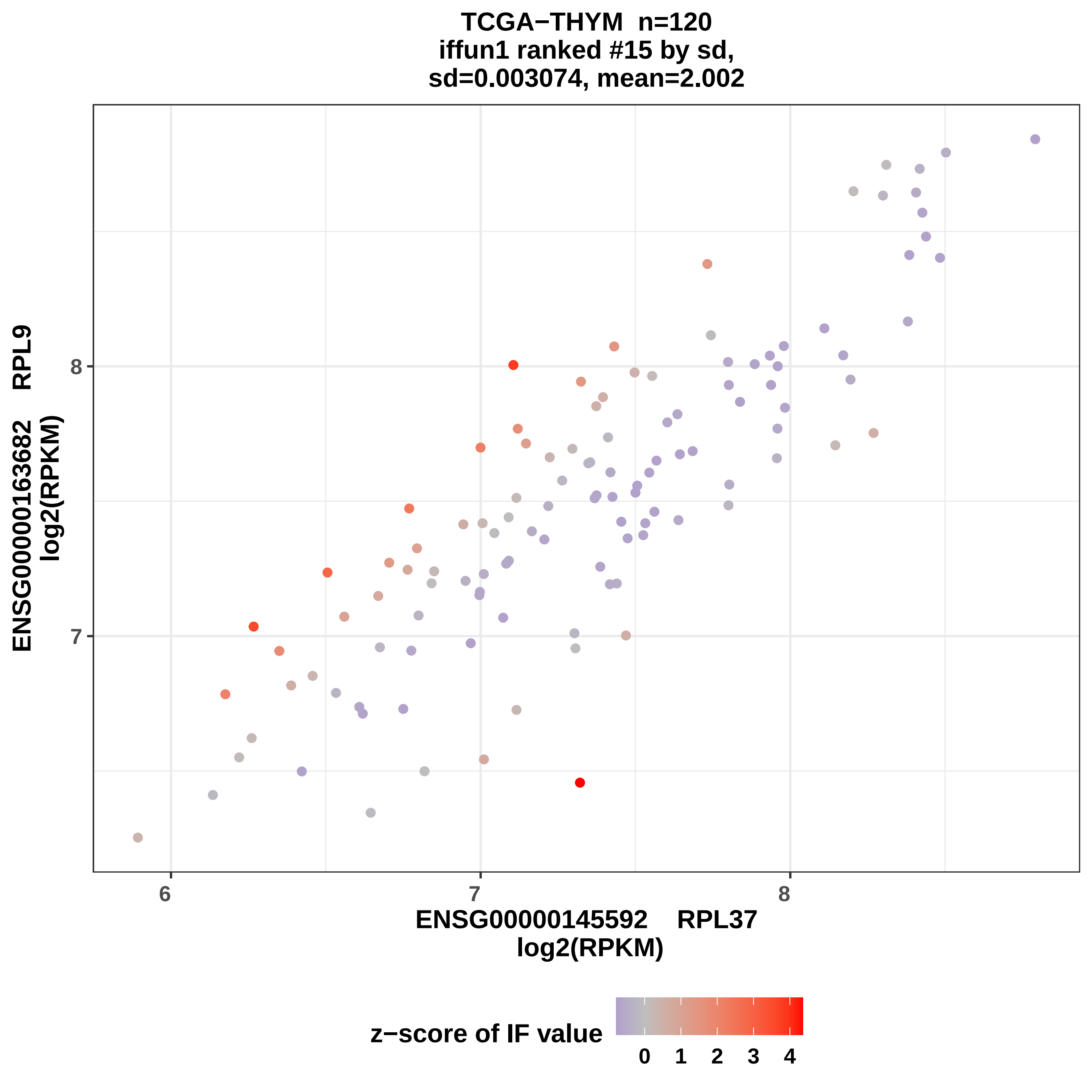

TCGA−THYM n=120
iffun1 ranked #15 by sd,
sd=0.003074, mean=2.002
ENSG00000163682 RPL9
log2(RPKM)
ENSG00000145592 RPL37
log2(RPKM)
z−score of IF value
0
1
2
3
4
6
7
8
7
8

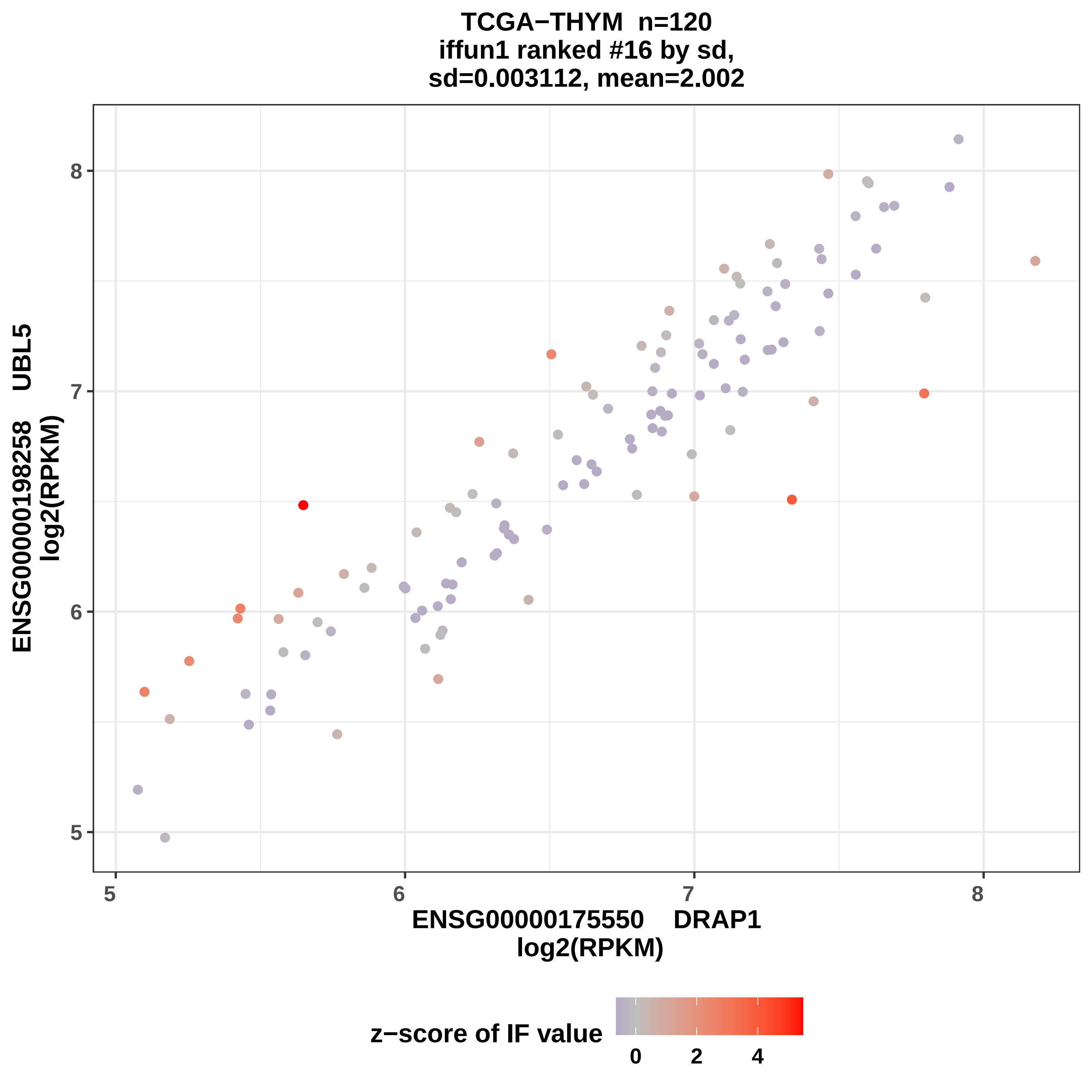
TCGA−THYM n=120
iffun1 ranked #16 by sd,
sd=0.003112, mean=2.002
ENSG00000198258 UBL5
log2(RPKM)
8
7
6
5
5
6
7
8
ENSG0000175550 DRAP1
log2(RPKM)
z−score of IF value
0
2
4

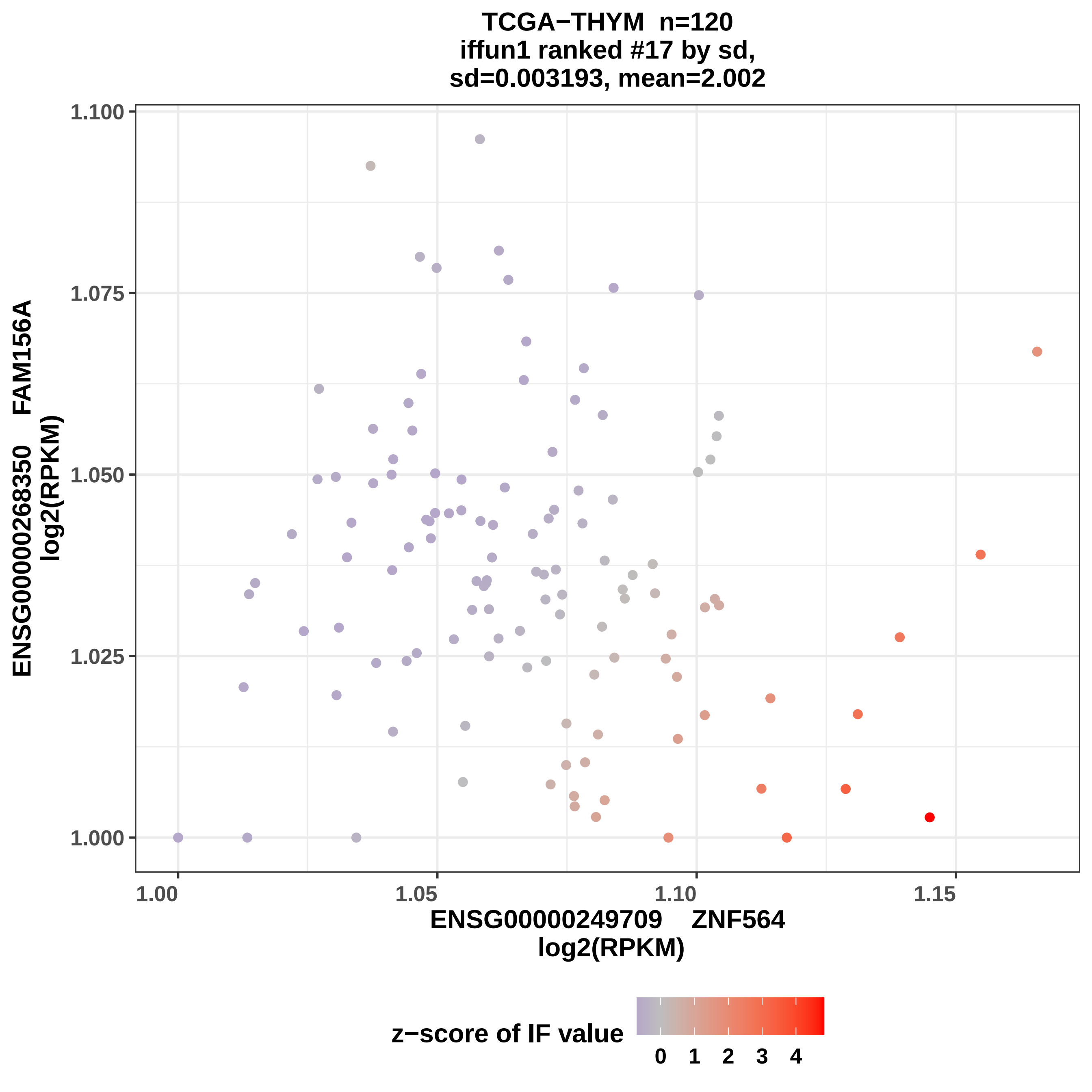

TCGA−THYM n=120
iffun1 ranked #17 by sd,
sd=0.003193, mean=2.002
ENSG00000268350 FAM156A
log2(RPKM)
1.100
1.075
1.050
1.025
1.000
1.00
1.05
1.10
1.15
ENSG00000249709 ZNF564
log2(RPKM)
z−score of IF value
0 1 2 3 4

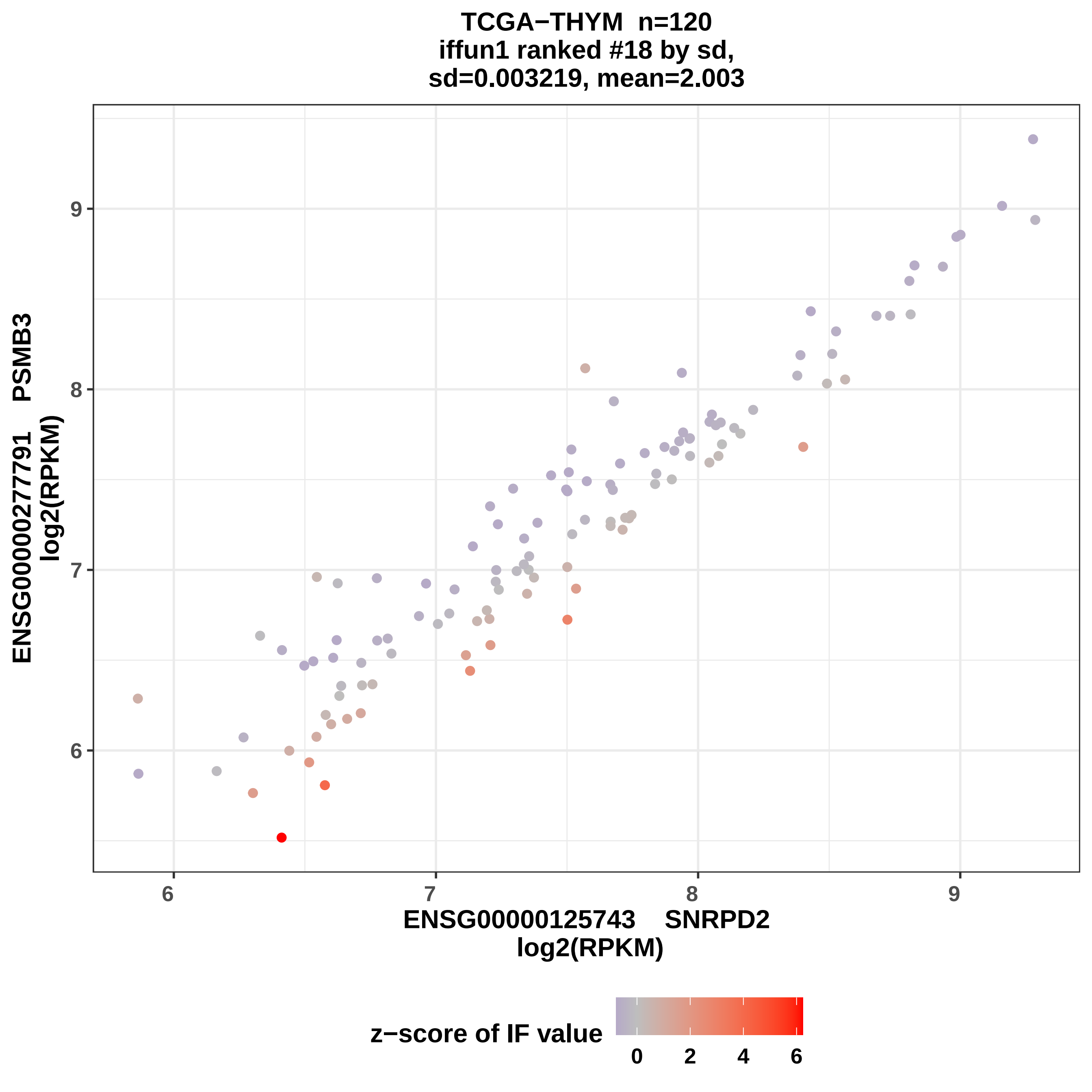

TCGA−THYM n=120
iffun1 ranked #18 by sd,
sd=0.003219, mean=2.003
ENSG00000277791 PSMB3
log2(RPKM)
ENSG00000125743 SNRPD2
log2(RPKM)
6
7
8
9
z−score of IF value
0
2
4
6

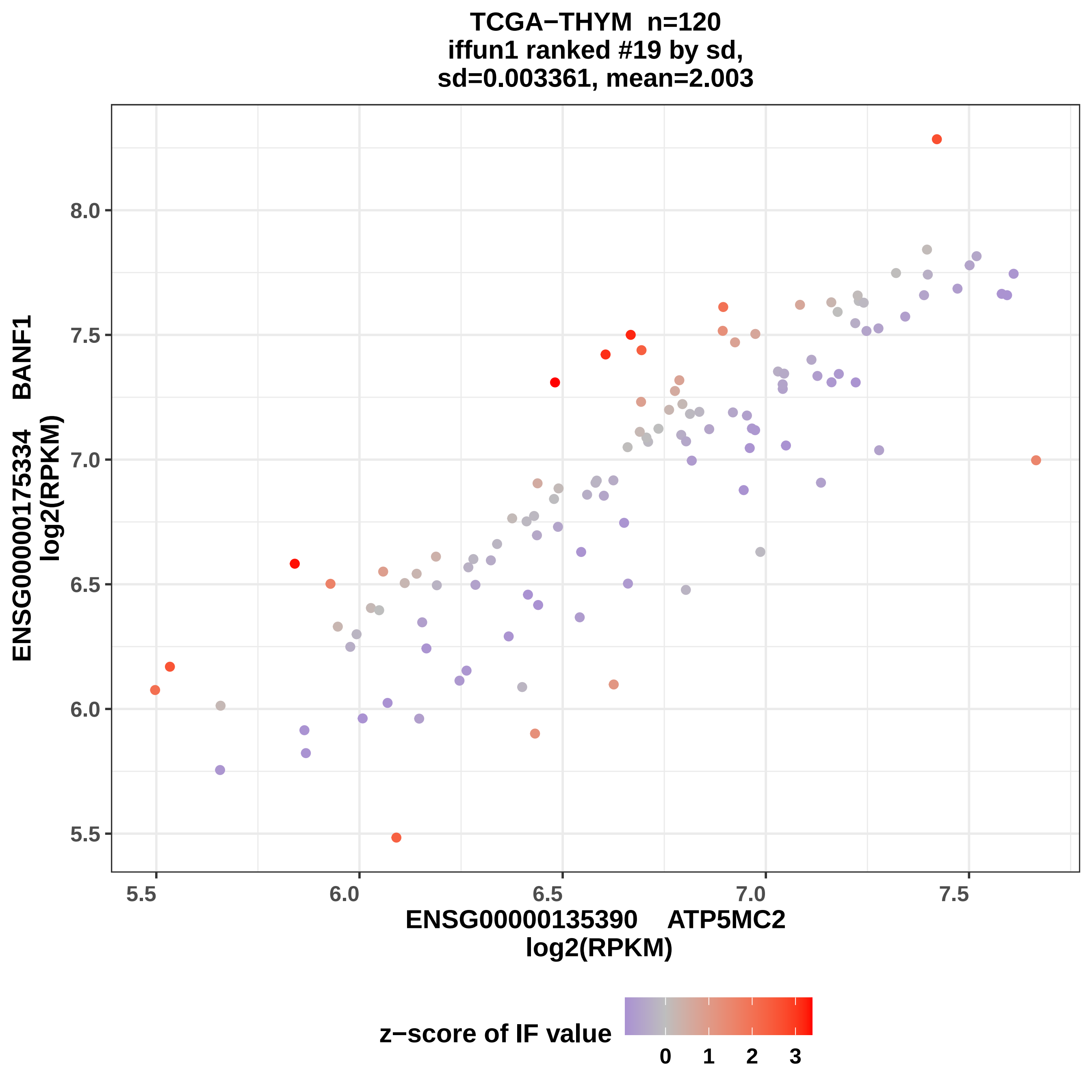
TCGA−THYM n=120
iffun1 ranked #19 by sd,
sd=0.003361, mean=2.003
ENSG00000175334 BANF1
log2(RPKM)
8.0
7.5
7.0
6.5
6.0
5.5
5.5
6.0
6.5
7.0
7.5
ENSG00000135390 ATP5MC2
log2(RPKM)
z−score of IF value
0
1
2
3

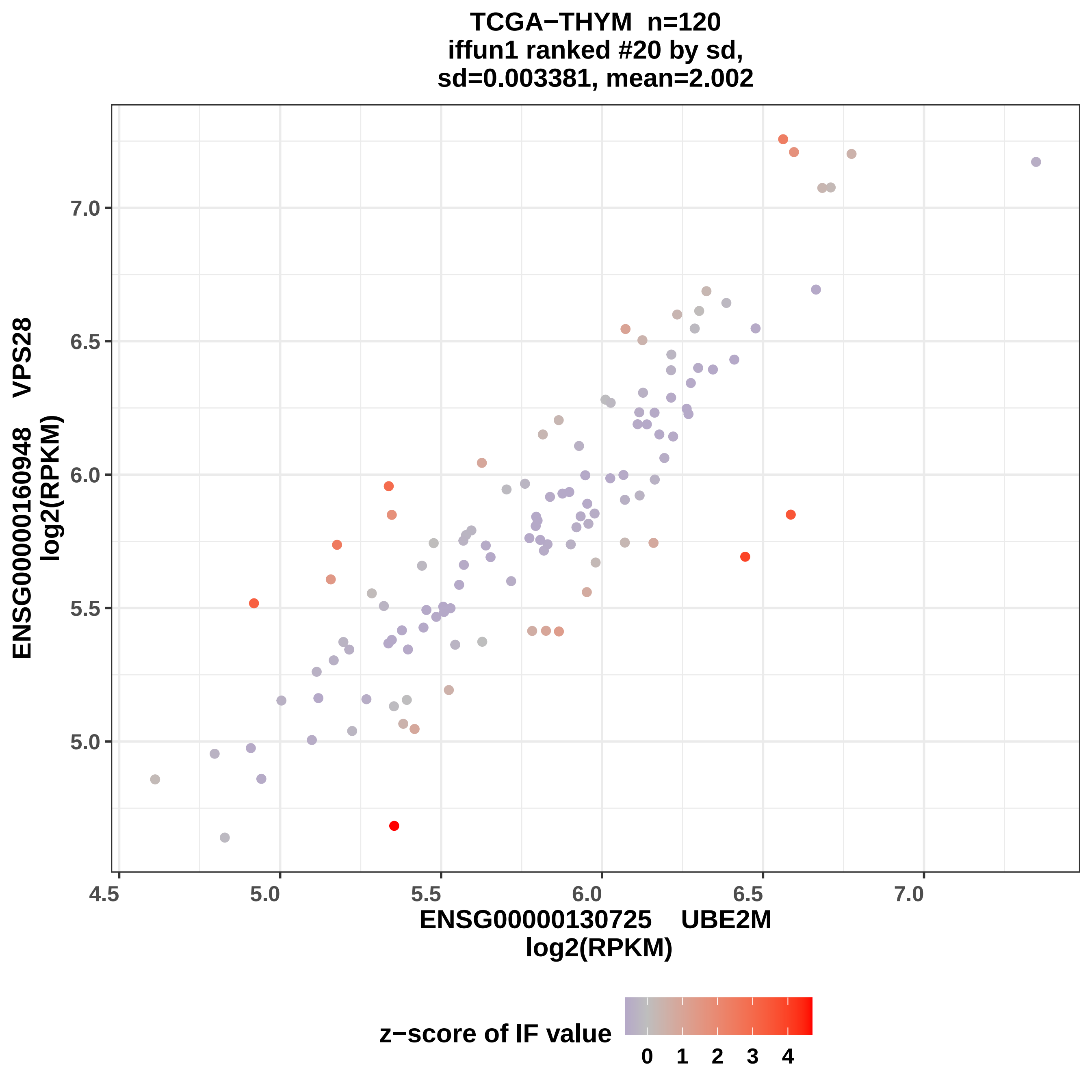

TCGA−THYM n=120
iffun1 ranked #20 by sd,
sd=0.003381, mean=2.002
ENSG00000160948 VPS28
log2(RPKM)
7.0
6.5
6.0
5.5
5.0
4.5
5.0
5.5
6.0
6.5
7.0
ENSG00000130725 UBE2M
log2(RPKM)
z−score of IF value
0
1
2
3
4

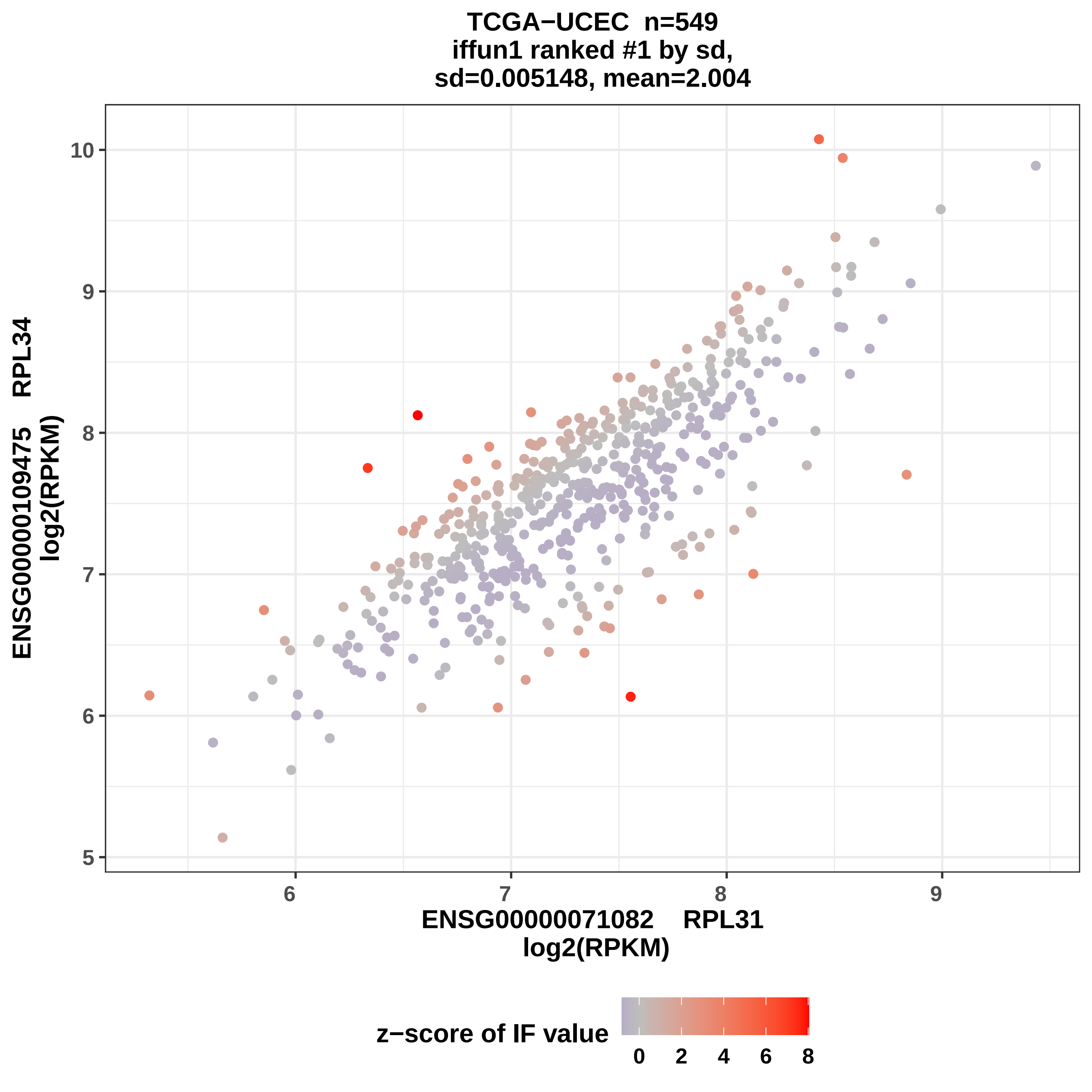

TCGA−UCEC n=549
iffun1 ranked #1 by sd,
sd=0.005148, mean=2.004
ENSG00000109475 RPL34
log2(RPKM)
ENSG00000071082 RPL31
log2(RPKM)
z−score of IF value
10
9
8
7
6
5
6
7
8
9
0
2
4
6
8

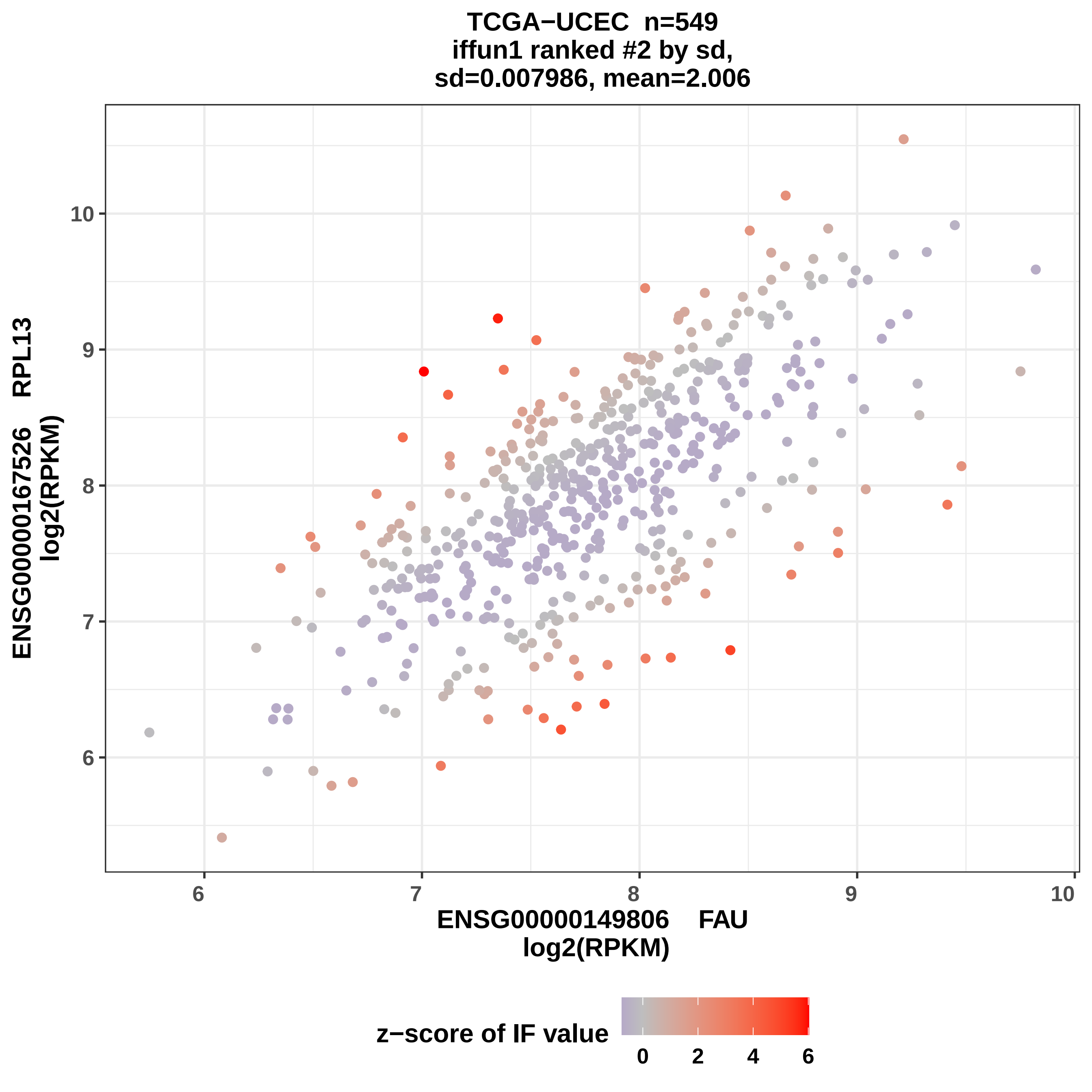
TCGA−UCEC n=549
iffun1 ranked #2 by sd,
sd=0.007986, mean=2.006
ENSG00000167526 RPL13
log2(RPKM)
ENSG00000149806 FAU
log2(RPKM)
z−score of IF value
0
2
4
6

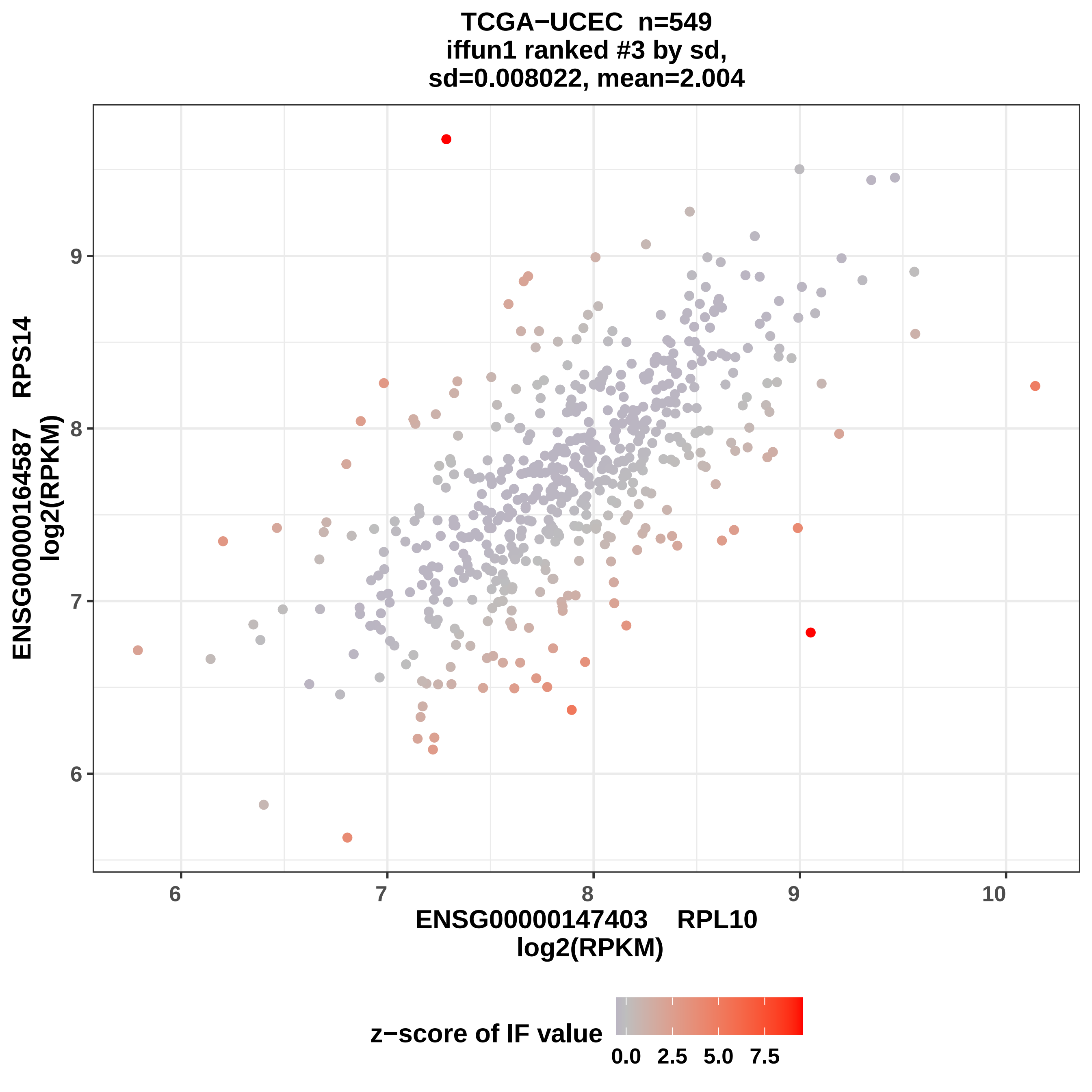
TCGA−UCEC n=549
iffun1 ranked #3 by sd,
sd=0.008022, mean=2.004
ENSG00000164587 RPS14
log2(RPKM)
ENSG00000147403 RPL10
log2(RPKM)
6
7
8
9
10
z−score of IF value
0.0
2.5
5.0
7.5

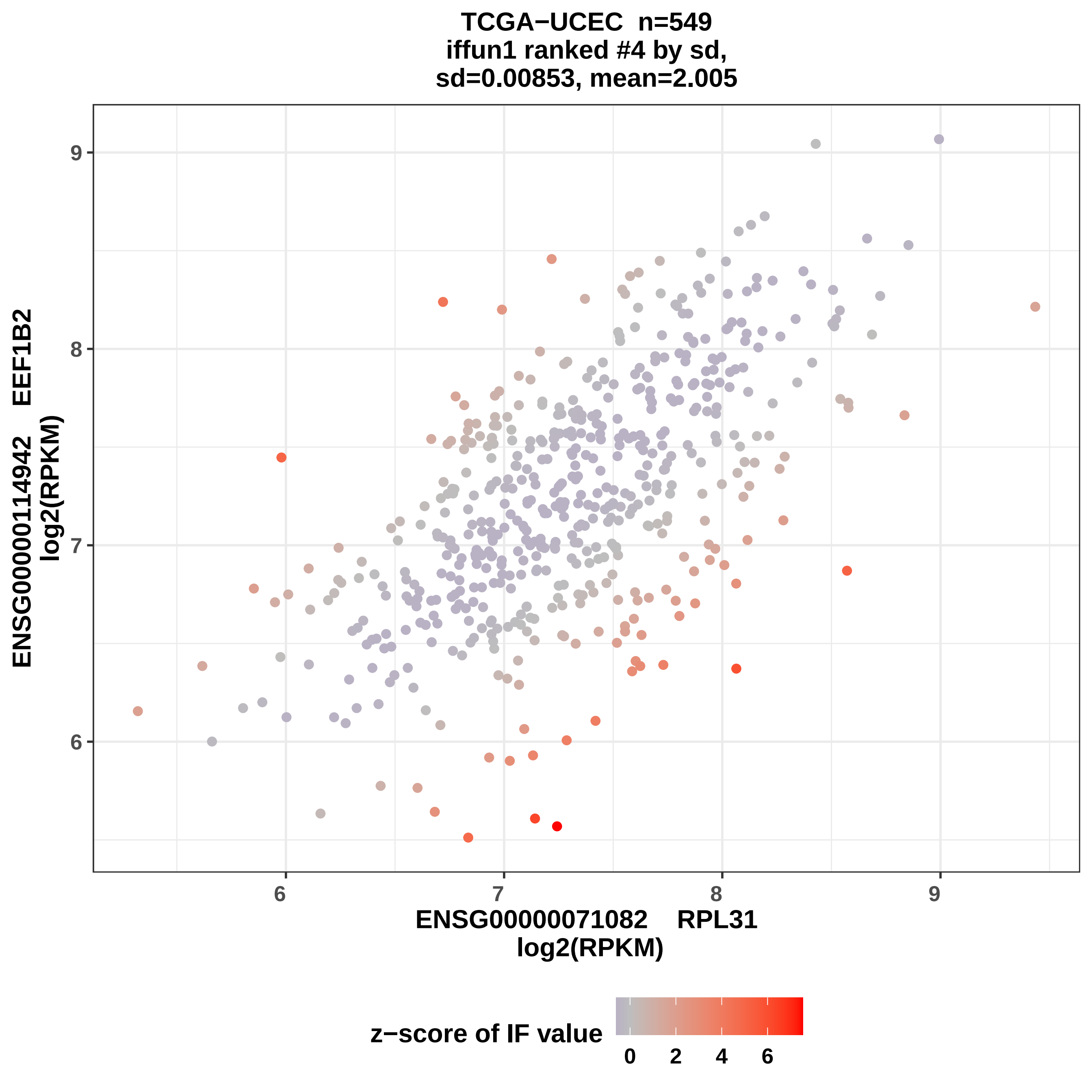
TCGA−UCEC n=549
iffun1 ranked #4 by sd,
sd=0.00853, mean=2.005
ENSG00000114942 EEF1B2
log2(RPKM)
ENSG00000071082 RPL31
log2(RPKM)
6
7
8
9
z−score of IF value
0
2
4
6

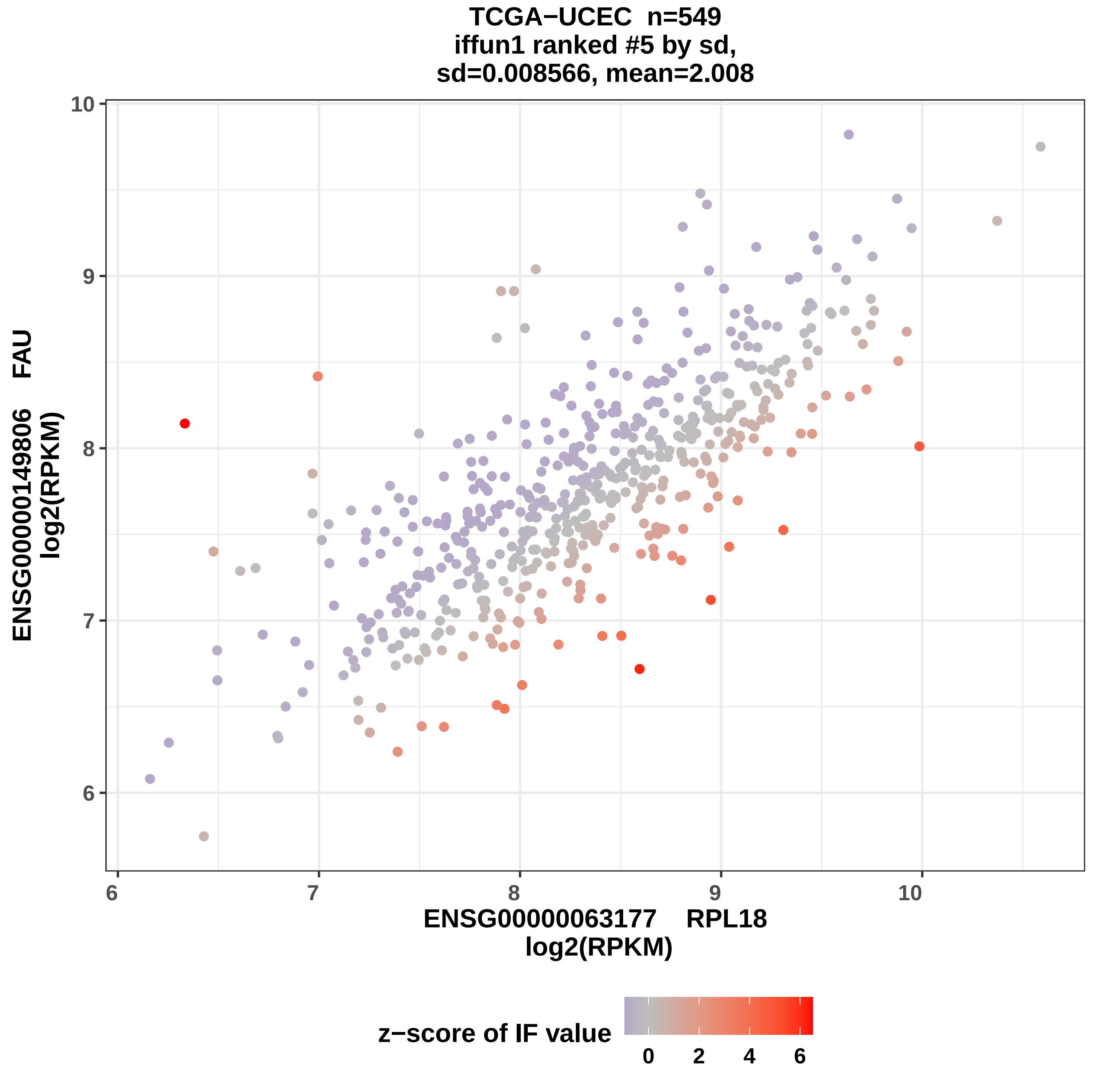
TCGA−UCEC n=549
iffun1 ranked #5 by sd,
sd=0.008566, mean=2.008
ENSG00000149806 FAU
log2(RPKM)
ENSG00000063177 RPL18
log2(RPKM)
6
7
8
9
10
6
7
8
9
10
z−score of IF value
0
2
4
6

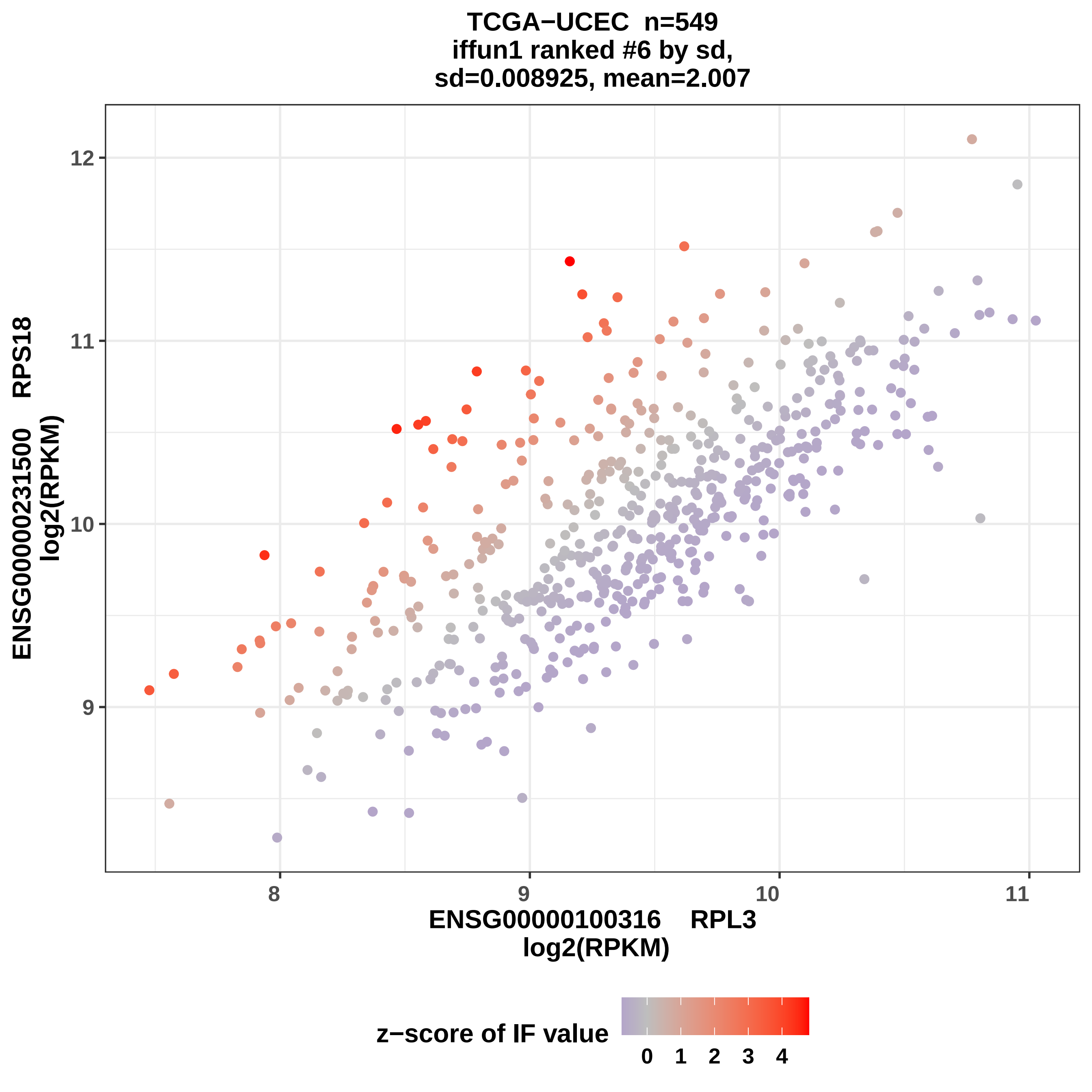
TCGA−UCEC n=549
iffun1 ranked #6 by sd,
sd=0.008925, mean=2.007
ENSG00000231500 RPS18
log2(RPKM)
12
11
10
9
8
9
10
11
ENSG00000100316 RPL3
log2(RPKM)
z−score of IF value
0
1
2
3
4

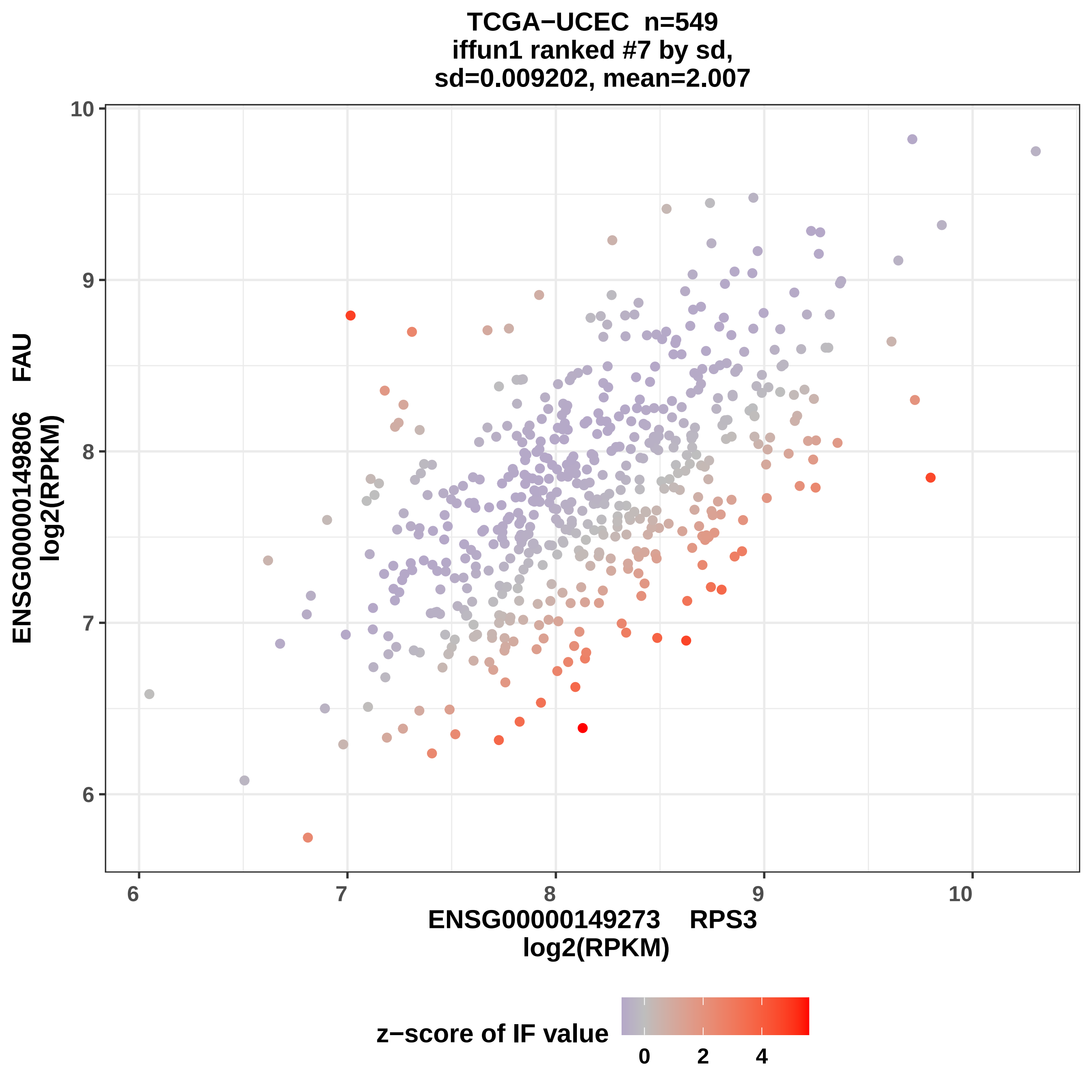
TCGA−UCEC n=549
iffun1 ranked #7 by sd,
sd=0.009202, mean=2.007
ENSG00000149806 FAU
log2(RPKM)
ENSG00000149273 RPS3
log2(RPKM)
6
7
8
9
10
z−score of IF value
0
2
4

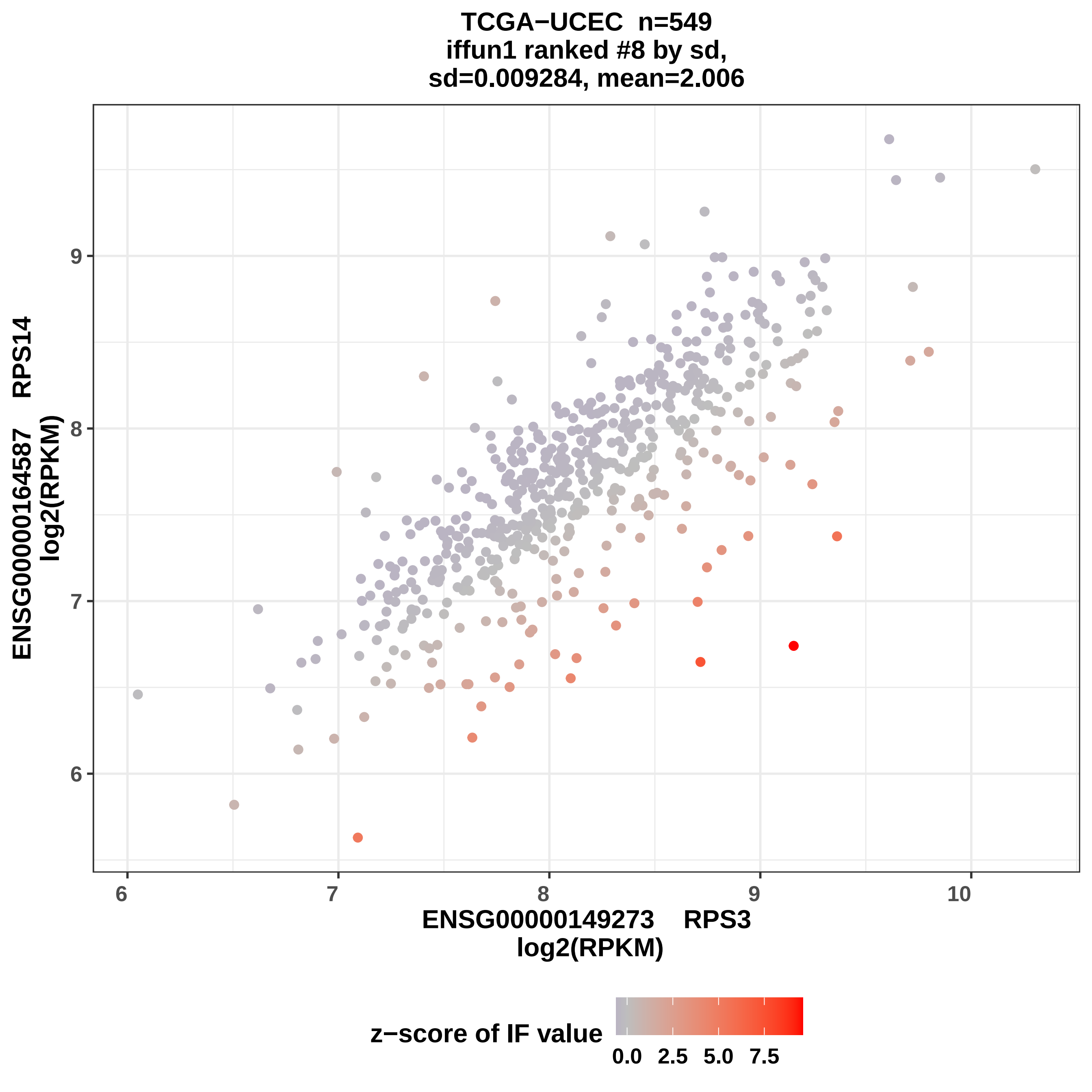
TCGA−UCEC n=549
iffun1 ranked #8 by sd,
sd=0.009284, mean=2.006
ENSG00000164587 RPS14
log2(RPKM)
ENSG00000149273 RPS3
log2(RPKM)
6
7
8
9
10
z−score of IF value
0.0
2.5
5.0
7.5

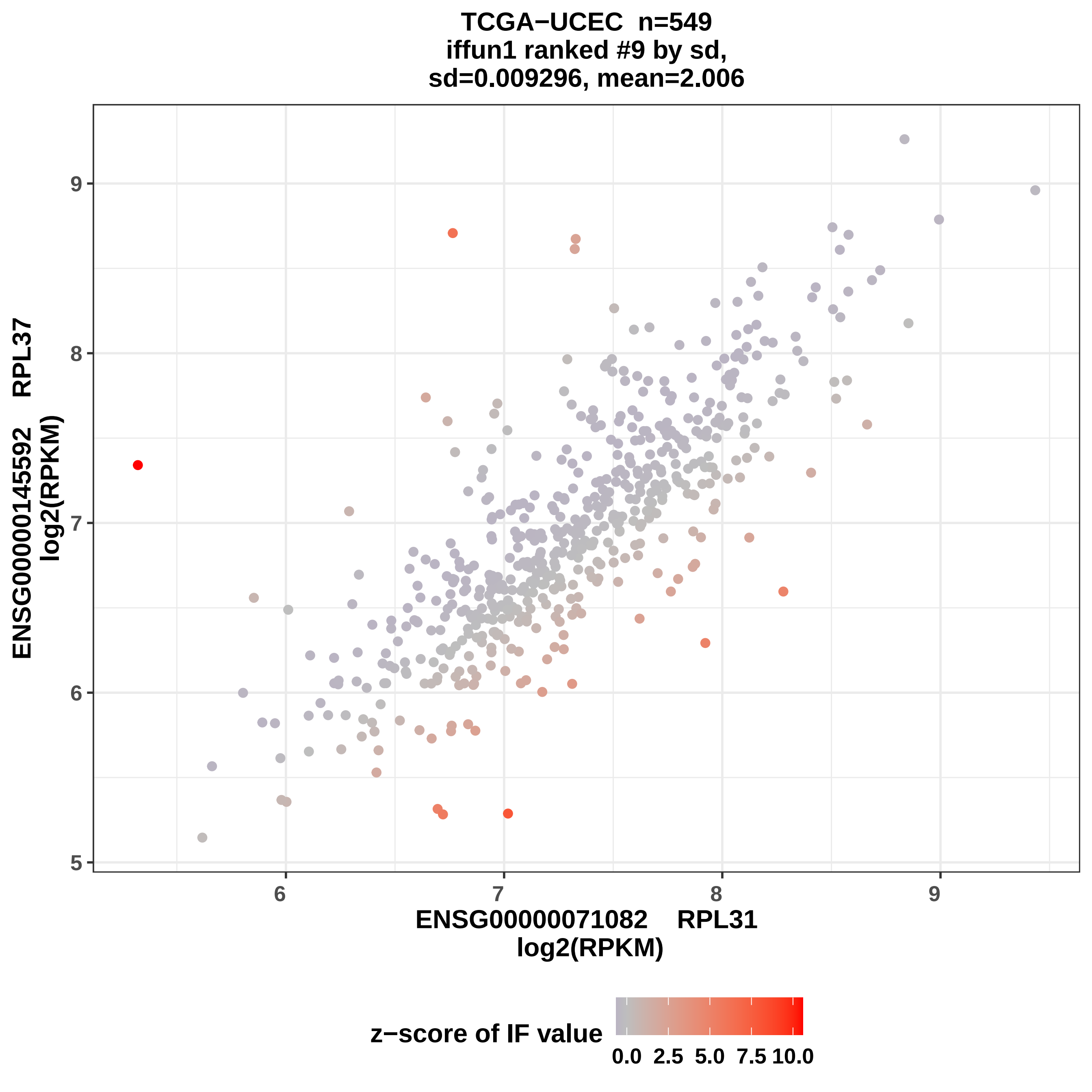
TCGA−UCEC n=549
iffun1 ranked #9 by sd,
sd=0.009296, mean=2.006
ENSG00000145592 RPL37
log2(RPKM)
5
6
7
8
9
6
7
8
9
ENSG00000071082 RPL31
log2(RPKM)
z−score of IF value
0.0 2.5 5.0 7.5 10.0

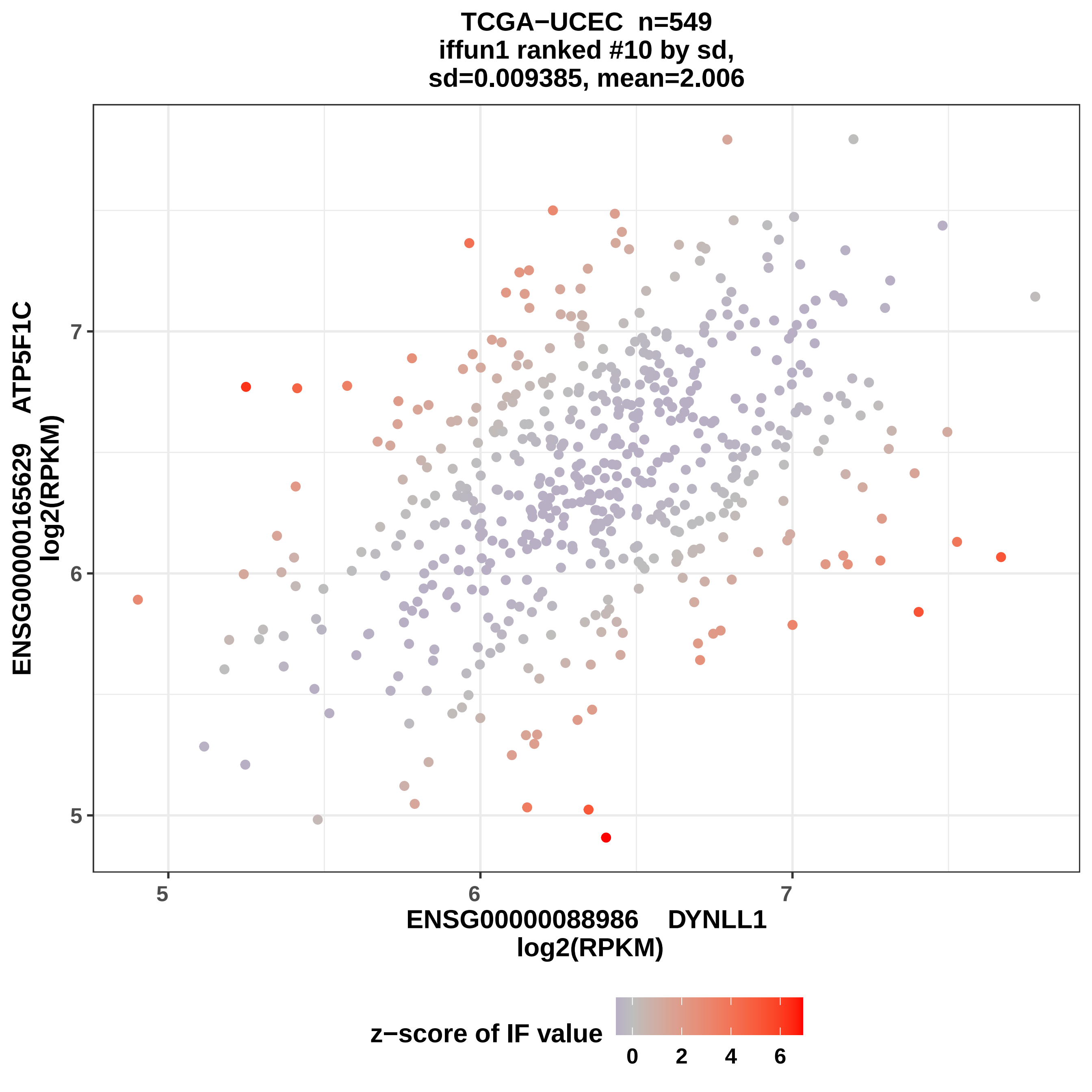

TCGA−UCEC n=549
iffun1 ranked #10 by sd,
sd=0.009385, mean=2.006
ENSG00000165629 ATP5F1C
log2(RPKM)
7
6
5
5
6
7
ENSG00000088986 DYNLL1
log2(RPKM)
z−score of IF value
0
2
4
6

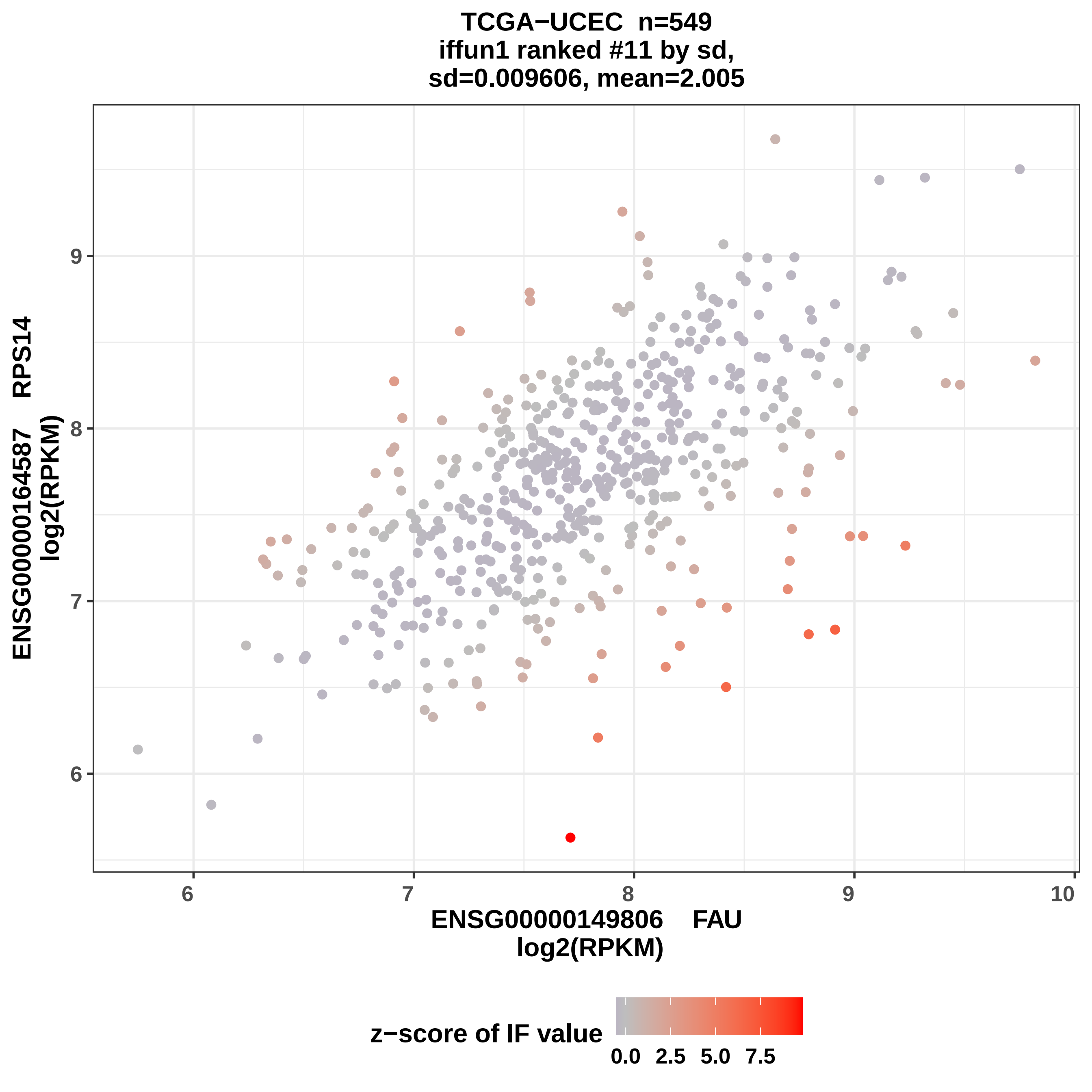
TCGA−UCEC n=549
iffun1 ranked #11 by sd,
sd=0.009606, mean=2.005
ENSG00000164587 RPS14
log2(RPKM)
ENSG00000149806 FAU
log2(RPKM)
z−score of IF value
0.0 2.5 5.0 7.5
6
7
8
9
10

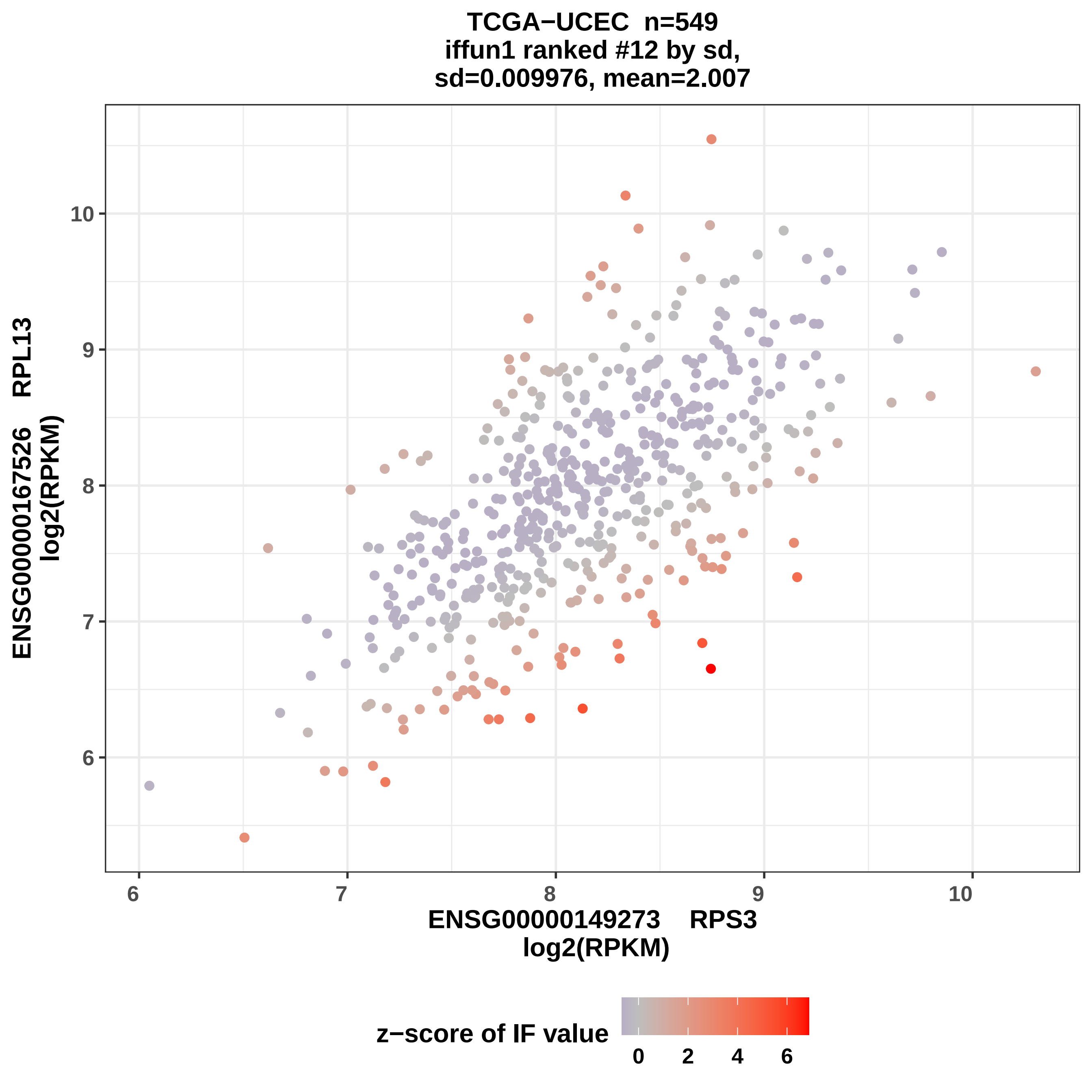

TCGA−UCEC n=549
iffun1 ranked #12 by sd,
sd=0.009976, mean=2.007
ENSG00000167526 RPL13
log2(RPKM)
10
9
8
7
6
6
7
8
9
10
ENSG00000149273 RPS3
log2(RPKM)
z−score of IF value
0
2
4
6

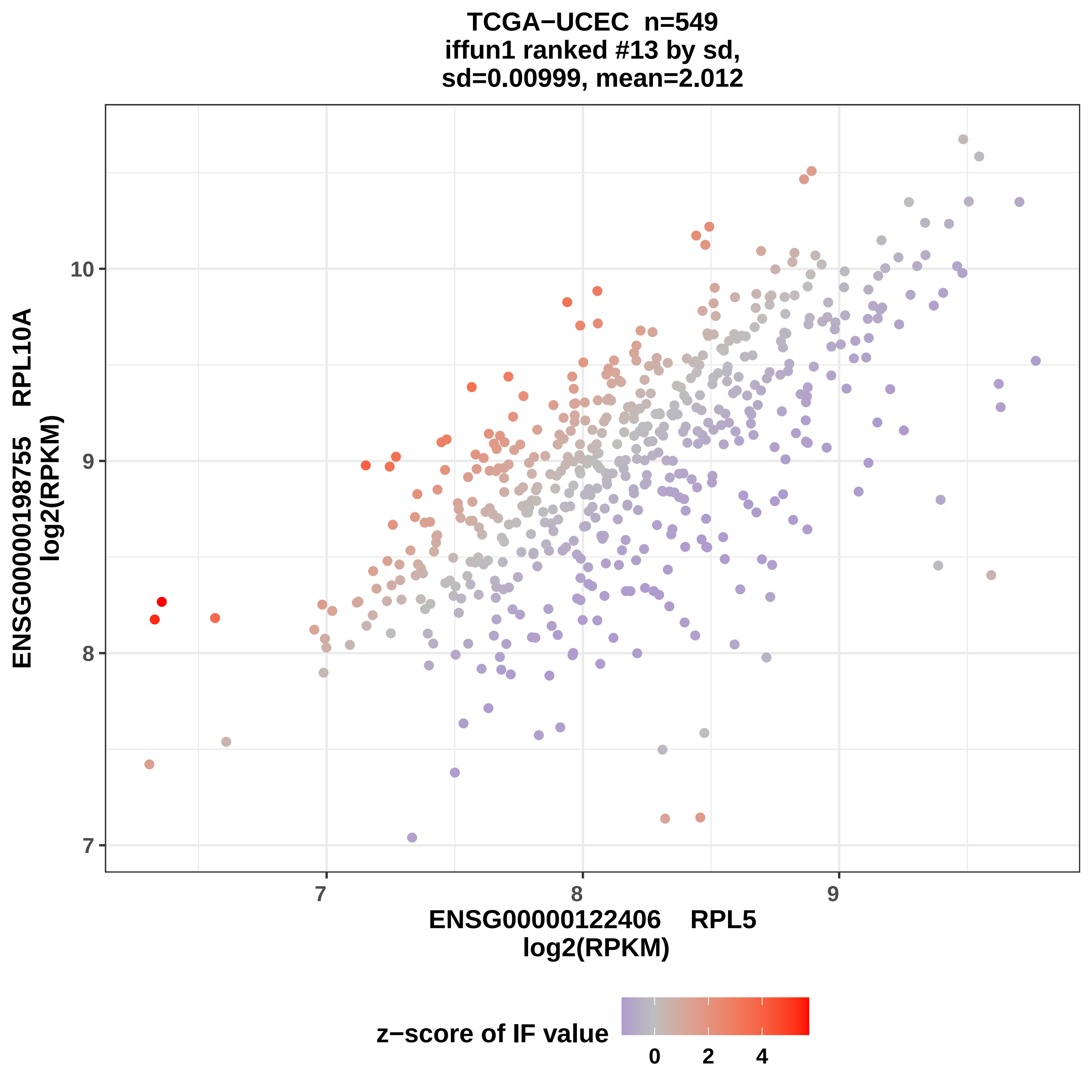

TCGA−UCEC n=549
iffun1 ranked #13 by sd,
sd=0.00999, mean=2.012
ENSG00000198755 RPL10A
log2(RPKM)
10
9
8
7
ENSG00000122406 RPL5
log2(RPKM)
z−score of IF value
0
2
4

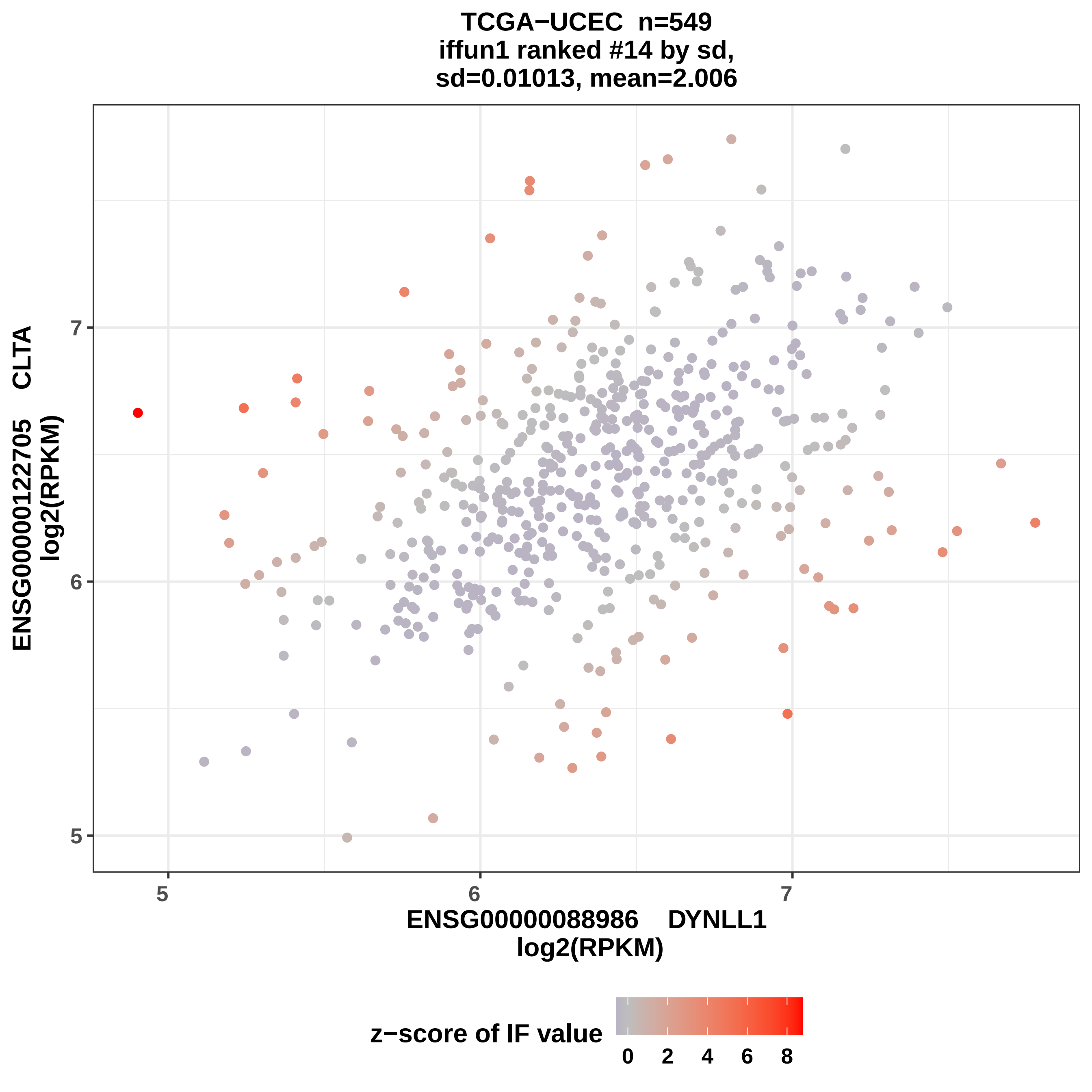

TCGA−UCEC n=549
iffun1 ranked #14 by sd,
sd=0.01013, mean=2.006
ENSG00000122705 CLTA
log2(RPKM)
ENSG00000088986 DYNLL1
log2(RPKM)
5
6
7
z−score of IF value
0
2
4
6
8

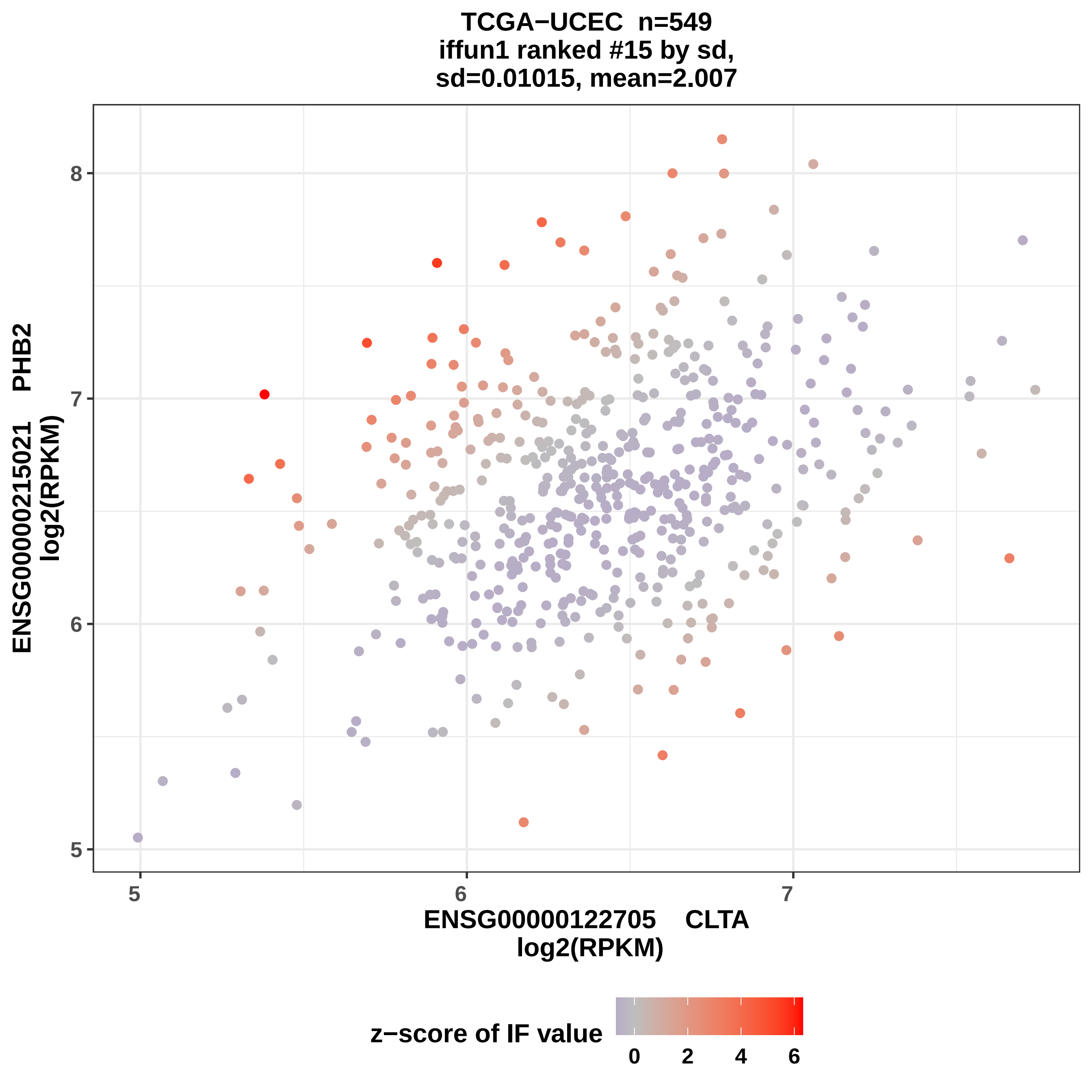

TCGA−UCEC n=549
iffun1 ranked #15 by sd,
sd=0.01015, mean=2.007
ENSG00000215021 PHB2
log2(RPKM)
8
7
6
5
5
6
7
ENSG00000122705 CLTA
log2(RPKM)
z−score of IF value
0
2
4
6

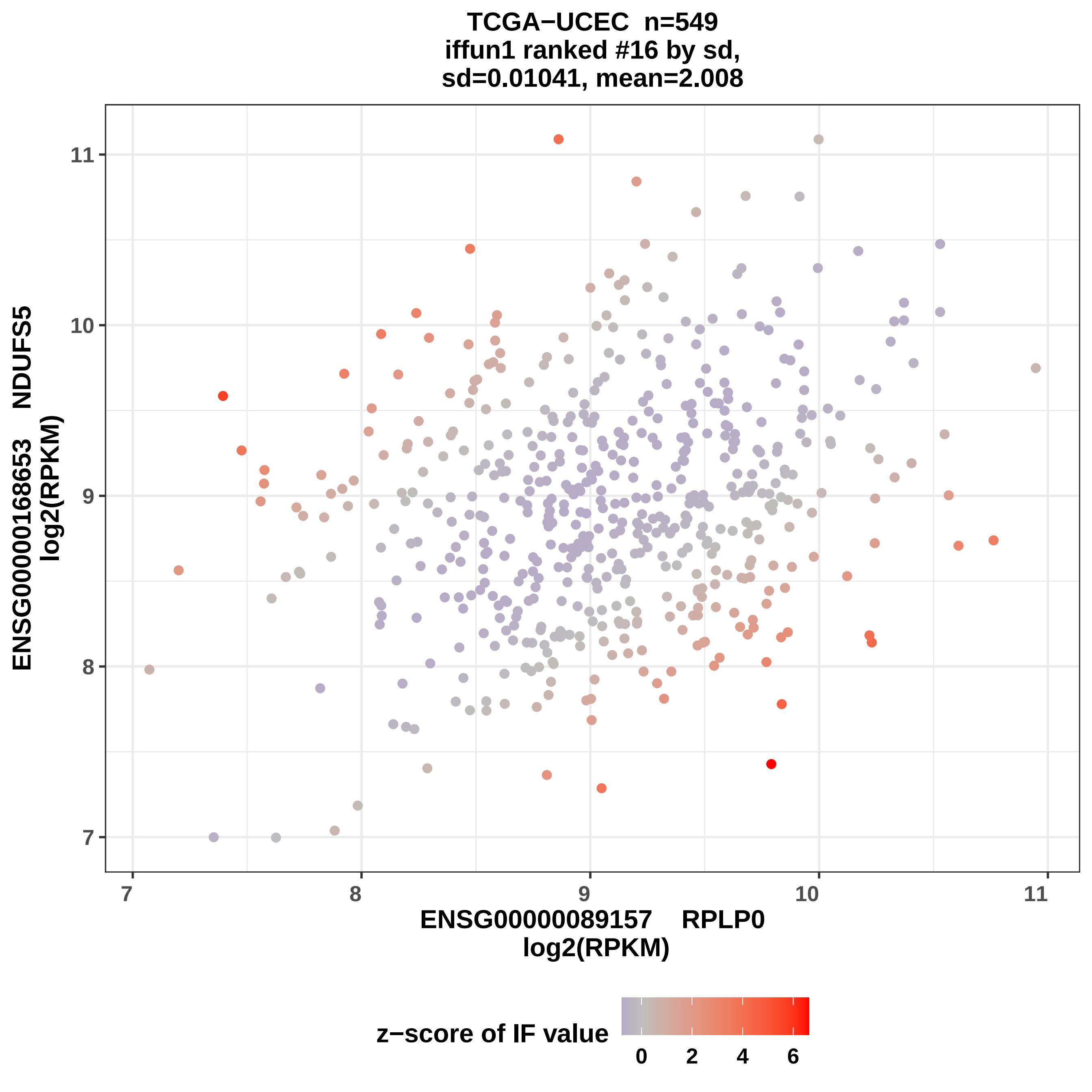
TCGA−UCEC n=549
iffun1 ranked #16 by sd,
sd=0.01041, mean=2.008
ENSG00000168653 NDUFS5
log2(RPKM)
ENSG00000089157 RPLP0
log2(RPKM)
z−score of IF value
7
8
9
10
11
0
2
4
6

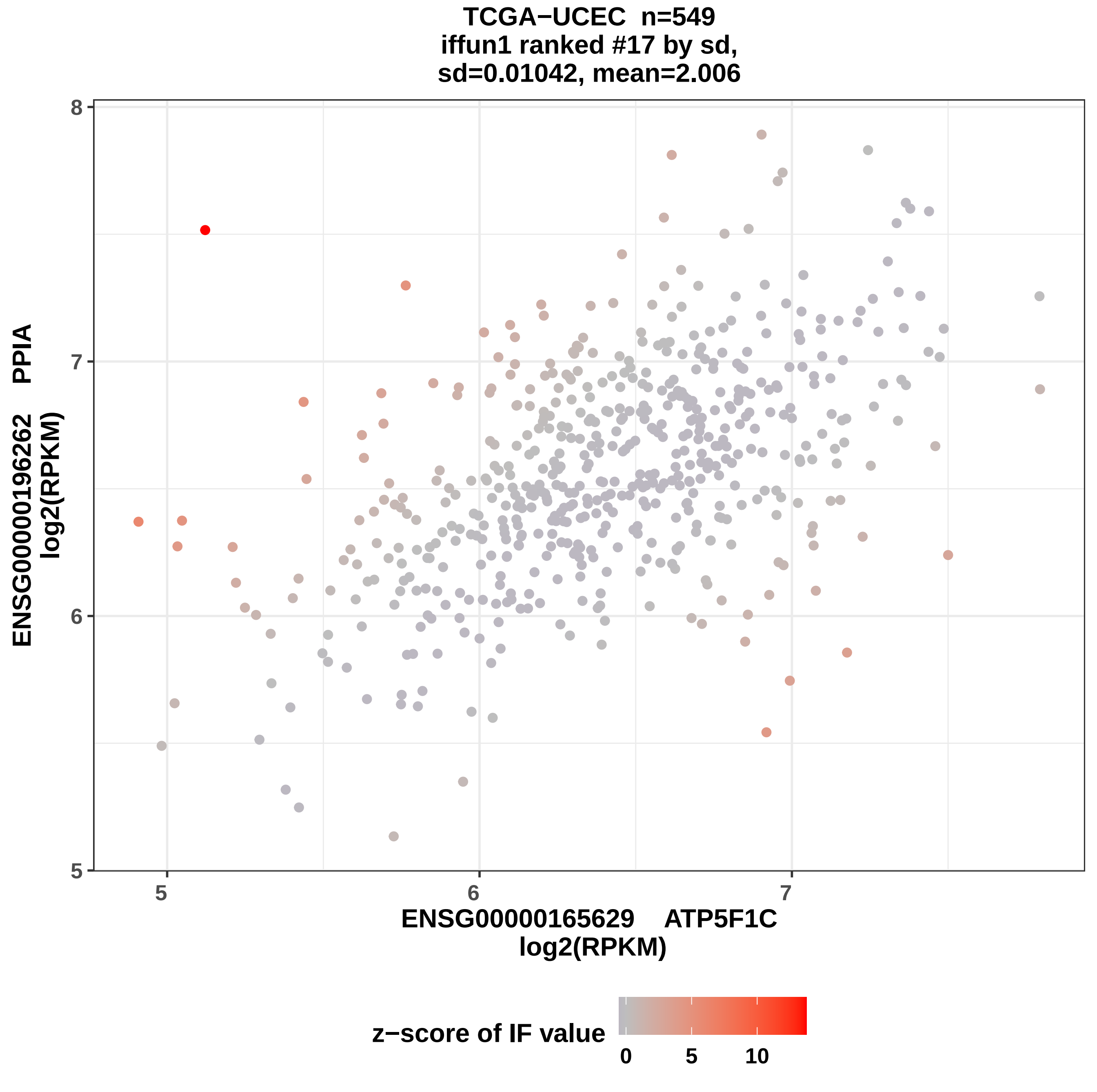
TCGA−UCEC n=549
iffun1 ranked #17 by sd,
sd=0.01042, mean=2.006
ENSG0000196262 PPIA
log2(RPKM)
8
7
6
5
5
6
7
ENSG0000165629 ATP5F1C
log2(RPKM)
z−score of IF value
0
5
10

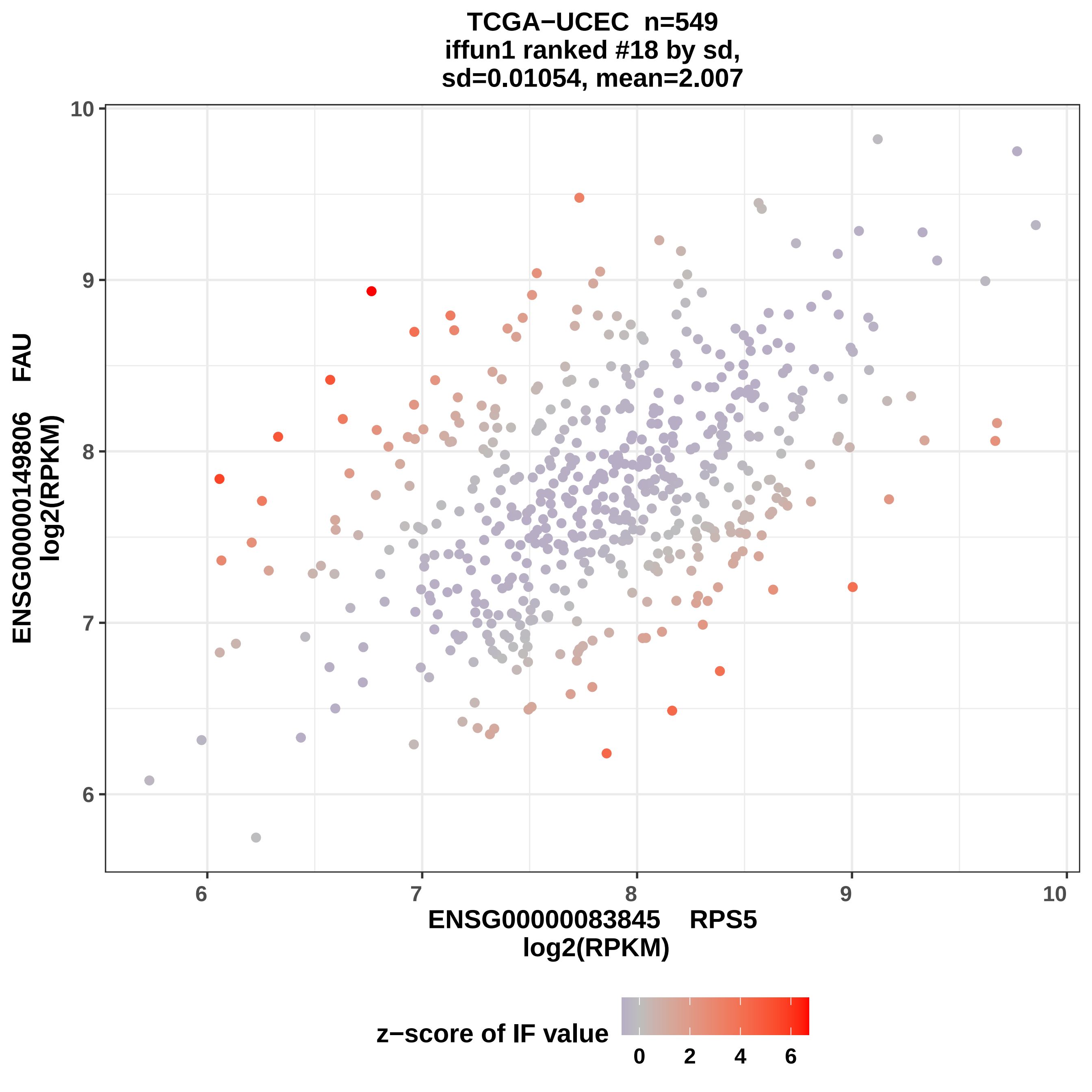
TCGA−UCEC n=549
iffun1 ranked #18 by sd,
sd=0.01054, mean=2.007
ENSG00000149806 FAU
log2(RPKM)
ENSG00000083845 RPS5
log2(RPKM)
z−score of IF value
0 2 4 6
6 7 8 9 10
6 7 8 9 10

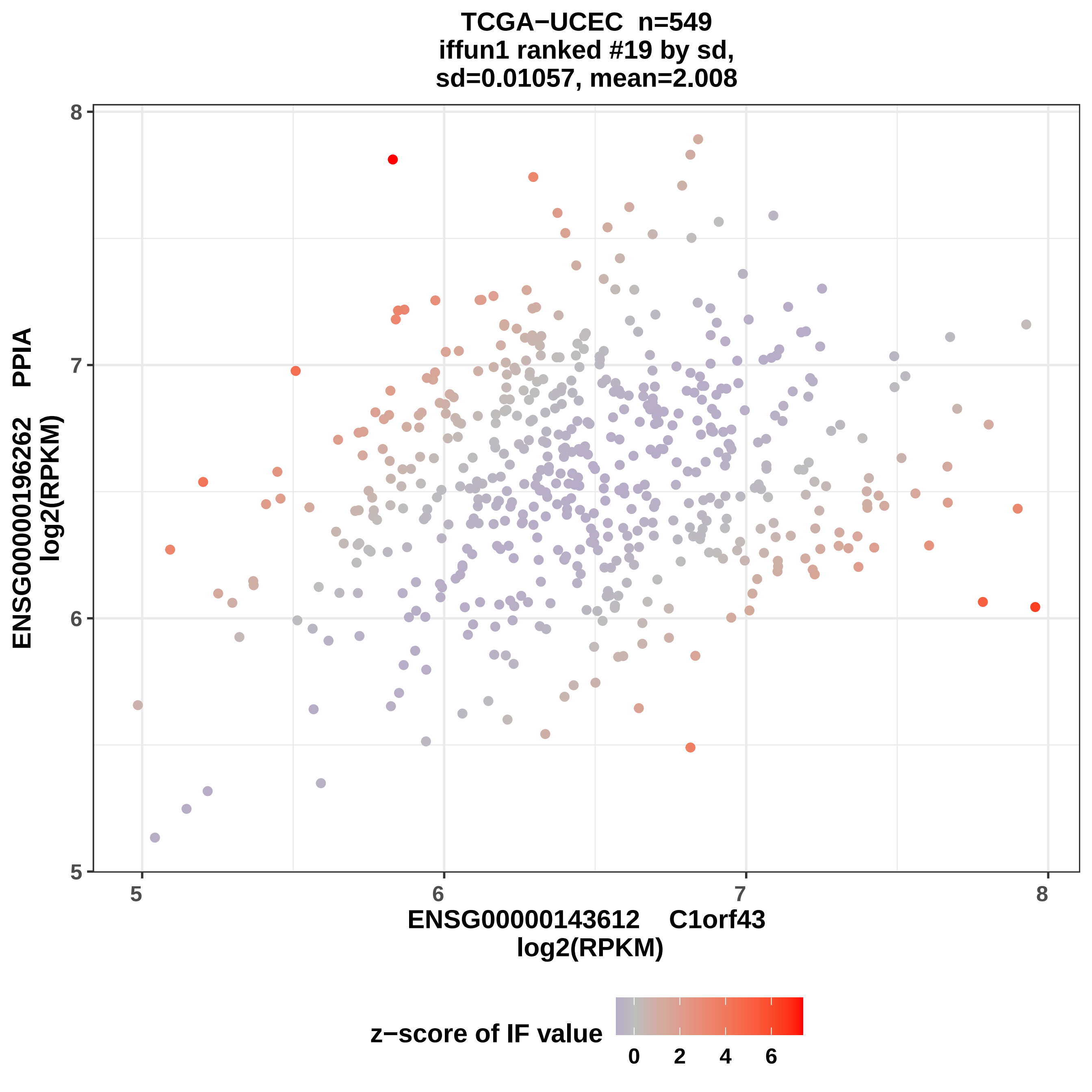
TCGA−UCEC n=549
iffun1 ranked #19 by sd,
sd=0.01057, mean=2.008
ENSG00000196262 PPIA
log2(RPKM)
8
7
6
5
5
6
7
8
ENSG00000143612 C1orf43
log2(RPKM)
z−score of IF value
0
2
4
6

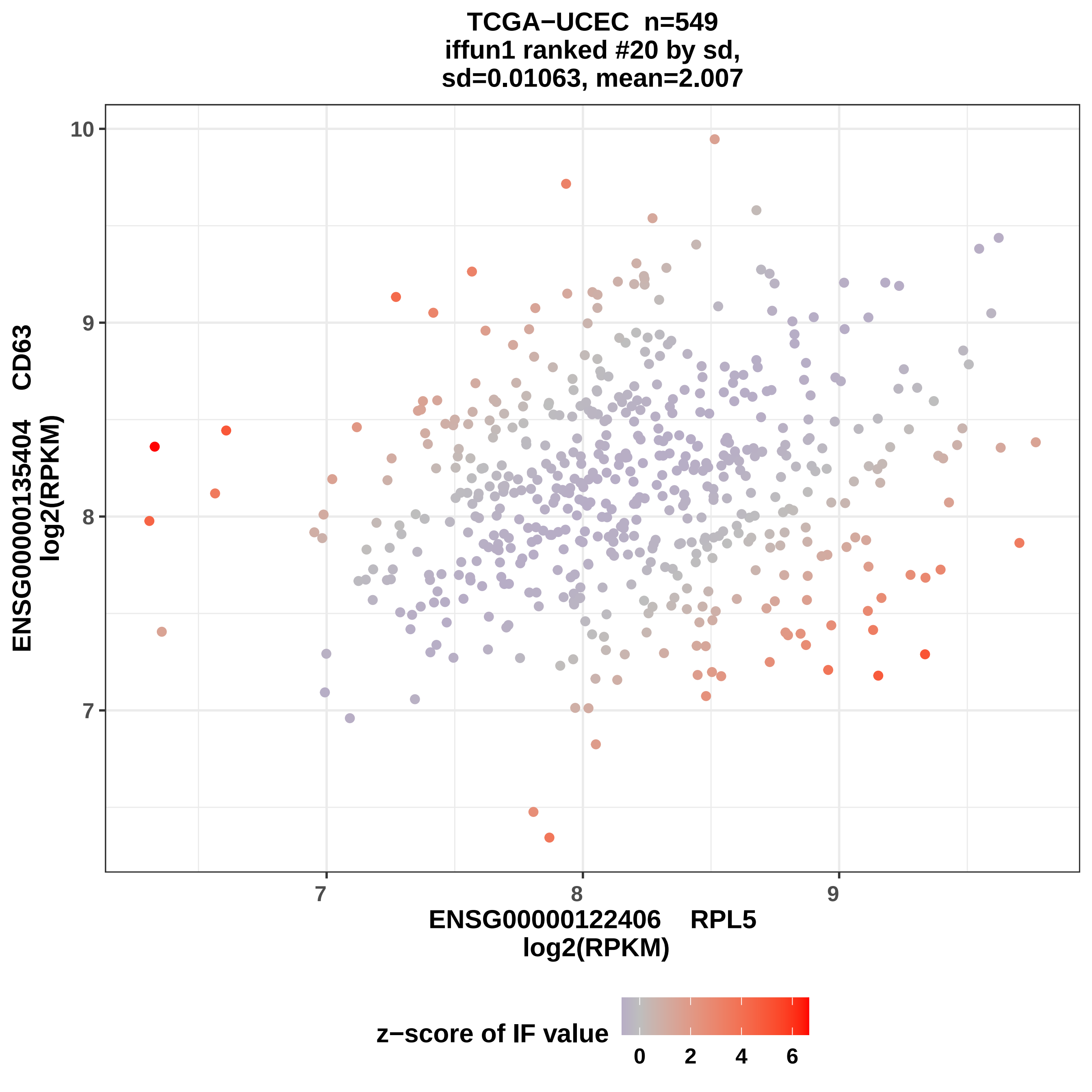
TCGA−UCEC n=549
iffun1 ranked #20 by sd,
sd=0.01063, mean=2.007
10
9
8
7
ENSG00000135404 CD63
log2(RPKM)
7
8
9
ENSG00000122406 RPL5
log2(RPKM)
z−score of IF value
0
2
4
6
